# The *Zymoseptoria tritici* ORFeome: a functional genomics community resource

**DOI:** 10.1101/582205

**Authors:** Yogesh Chaudhari, Timothy C. Cairns, Yaadwinder Sidhu, Victoria Attah, Graham Thomas, Michael Csukai, Nicholas J. Talbot, David J. Studholme, Ken Haynes

## Abstract

Libraries of protein-encoding sequences can be generated by identification of open reading frames (ORFs) from a genome of choice that are then assembled into collections of plasmids termed ORFeome libraries. These represent powerful resources to facilitate functional genomic characterization of genes and their encoded products. Here, we report the generation of an ORFeome for *Zymoseptoria tritici*, which causes the most serious disease of wheat in temperate regions of the world. We screened the genome of strain IP0323 for high confidence gene models, identifying 4075 candidates from 10,933 predicted genes. These were amplified from genomic DNA, cloned into the Gateway^®^ Entry Vector pDONR207, and sequenced, providing a total of 3022 quality-controlled plasmids. The ORFeome includes genes predicted to encode effectors (n = 410) and secondary metabolite biosynthetic proteins (n = 171), in addition to genes residing at dispensable chromosomes (n= 122), or those that are preferentially expressed during plant infection (n = 527). The ORFeome plasmid library is compatible with our previously developed suite of Gateway^®^ Destination vectors, which have various combinations of promoters, selection markers, and epitope tags. The *Z. tritici* ORFeome constitutes a powerful resource for functional genomics, and offers unparalleled opportunities to understand the biology of *Z. tritici*.

## Introduction

Fungal pathogens kill more people per year than malaria, and result in crop destruction or post-harvest spoilage that destroys enough food to feed approximately 10% of the population (1, 2). Technological advances in fungal genomics, transcriptomics, proteomics, metabolomics, bioinformatics, and network analyses, however, now enable pathogenic fungi to be studied as integrated systems, providing unparalleled opportunities to understand their biology (3, 4). Functional genomic approaches, which define the function and interactions of genes and their encoded products at a genome or near-genome level, are increasingly used to dissect host pathogen interactions, virulence factors, drug resistance, and infectious growth during fungal disease (5–9). However, a significant constraint to conducting functional genomic experiments are high reagent and labour costs, due to the necessity to study thousands, or tens of thousands of genes from a given fungal pathogen.

In order to obviate this challenge, community accessible libraries have been developed, which consist of hundreds or thousands of either individual genes, null mutant or over-expression strains, which ultimately enable facile and high throughput experimentation by the end user at a minimal expense (10–17). ORFeomes are collections of open reading frames (ORFs) that are encoded in a library of plasmid vectors. These resources have been generated for several model organisms, including humans, *Escherichia coli, Caenorhabditis elegans*, and *Arabidopsis thaliana* (18–23). ORFeomes have also been developed for the fungal kingdom, including fission yeast (18), budding yeast (24) and, most recently, the human pathogenic yeast *Candida albicans* (8). Usually, ORFeomes are compatible with the Gateway^®^ cloning technology (Invitrogen), which enables rapid and high throughput recombinase-based transfer of an ORF coding sequence to generate expression vectors (25, 26). Community access to hundreds or thousands of such plasmids in a single library enables highly flexible generation of expression vectors for high-throughput functional genomic experiments.

The filamentous ascomycete fungus *Zymoseptoria tritici* (previously *Mycosphaerella graminicola*) (27) causes Septoria tritici blotch, an important foliar disease of wheat (28). *Z. tritici* is a significant threat to international food security, and even with access to some resistant wheat cultivars and frequent fungicide applications, the estimated average yield losses due to this pathogen are still 10% (28). Even more strikingly, approximately 70% of agricultural fungicides in Europe are deployed to control just this single disease (29), which likely drives triazole resistance in fungal pathogens of humans as well as plants (30). These challenges are compounded by very high levels of genome plasticity and gene flow between populations of *Z. tritici* (31, 32). While recent efforts have characterized the underlying cellular biology and infectious growth (33, 34), transcriptionally deployed secondary metabolite loci (35, 36), and components of the secreted effector arsenal [37–40], the vast majority of genes and encoded proteins remain uncharacterized in the laboratory (27).

Recently, there has been a community-wide effort to develop numerous tools, techniques, and resources for *Z. tritici*. This research toolkit includes mutants in the non-homologous end joining pathway for highly efficient gene targeting (41), optimization of conditional expression systems (42), a range of fluorescent translational gene fusion for sub-cellular localization studies (43, 44), optimized virulence assays (45), and a suite of Gateway^®^ Destination vectors (46, 47). These Gateway^®^ destination vectors have been validated using a pilot Gateway^®^ Entry library to generate 32 over-expression mutants, demonstrating the role of a fungal specific transcription factor for *in vitro* hyphal growth (48).

In this study, we report the generation of an improved functional genomics community resource to supplement these tools, by generating a Gateway^®^ compatible *Z. tritici* ORFeome, which to our knowledge is the first such library for a plant infecting fungus. This library is compatible with numerous Gateway^®^ destination vectors that have multiple functionality in *Z. tritici*, including numerous selection markers, epitope tags, and promoters (26, 47). For ORFeome construction, we firstly screened the IP0323 reference genome for high confidence gene models, yielding 4075 candidate ORFs from a possible 10,933 predicted genes. These were PCR amplified from genomic DNA and cloned into the Gateway^®^ Entry vector pDONR207. Quality of ORF sequences was verified by a combination of Sanger and Illumina sequencing, yielding 3022 plasmids that passed quality control checks with 100% sequence verification.

The *Z. tritici* ORFeome described in this study is freely available to the research community. This resource can be rapidly utilized to interrogate the broadest aspects of *Z. tritici* biology, including identification of novel drug targets, mechanisms of drug detoxification and resistance, and pathogen virulence factors, which may ultimately enable development of new disease control strategies.

## 2. Material and Methods

### 2.1 Strains used in this study

*E. coli* One Shot^®^ *ccdB* Survival™ 2 T1R were used for propagation of pDONR207 (Invitrogen, UK) and destination vectors. All Gateway^®^Entry and expression vectors were propagated in *Escherichia coli* DH5α (Invitrogen, UK).

### 2.2 Plasmids used in this study

For generation of Gateway^®^Entry vectors, this study utilized the Gateway^®^Donor vector pDONR207 (Invitrogen, UK) which contains a gentamicin resistance gene for selection in *E. coli*. This plasmid also contains a *ccdB* gene flanked by *attP* sequences for Gateway^®^ mediated recombination using the BP reaction.

### 2.3 Construction of the *Z. tritici* ORFeome

Generation of Gateway^®^ Entry vectors was conducted, as described previously (48). For PCR amplification of each gene of interest, forward primers were designed to include the *attB1* site (ggggacaagtttgtacaaaaaagcaggcttg and the first 20 bp of the gene, and reverse primers to include the *attB2* site (ggggaccactttgtacaagaaagctgggtc) and the last 20 bp of the gene. The stop codon was excluded to enable c-terminal epitope tagging by the end user. Primers were synthesised by Sigma-Aldrich, UK, and are listed in Supplementary File S1. PCRs were conducted using Phusion^®^ High-Fidelity DNA Polymerase (NEB, UK) with a 65 °C primer annealing temperature, an extension of 30 seconds/kb, using *Z. tritici* IP0323 genomic DNA as template. PCR amplicons of predicted sizes were confirmed by gel electrophoresis, PEG purified, and suspended in 10 μl TE buffer (40 mM TRIS base, 20 mM glacial acetic acid, 0.1 mM EDTA, pH8). For construction of Gateway^®^Entry vectors, 150 ng of pDONR207 was mixed with 2.5 μl of purified PCR product, 0.5 μl of Gateway^®^ BP Clonase™ with TE buffer added to a total volume of 10 μl. Reactions were incubated at 25 °C for 12-24h, then treated with Proteinase K (Invitrogen, UK) following the manufacturer’s instructions. *E. coli* strain DH5α was transformed with 5 μl of each reaction mixture. LB supplemented with gentamicin (50 μg/ml) was subsequently used to select transformants, which were grown over-night in LB medium with selection, and plasmids extracted using Plasmid Mini Kit (Qiagen, UK). Plasmids were indexed and stored in 96 well plates and at −20 °C. All Entry vectors are detailed in Supplementary File 1.

### 2.4 Sanger and Illumina sequencing for quality control of Gateway^®^ Entry vectors

In order to confirm replacement of the *ccdB* gene with the ORF encoding sequence, and to confirm high fidelity PCR amplification, a total of 688 Gateway^®^Entry vector were randomly selected and Sanger Sequenced (Eurofins, UK) using primer GOXF (tcgcgttaacgctagcatgga). Quality control reactions are given in Supplementary File S2. A second quality control experiment was conducted using two rounds of Illumina HiSeq 2500 sequencing of 3396 pooled Gateway Entry vectors. Sequencing data are summarized in Supplementary Table S1 and S2. Raw sequencing data are available at the Sequencing Read Archive (49) (accession SRX1267196 and SRX1265386).

## 3. Results

Interrogation of the 10,933 predicted genes in the *Z. tritici* IP0323 reference genome (29) returned 4075 high confidence gene models that had unambiguous start and stop codons (data not shown). This analysis was conducted prior to RNA-seq analysis and comparative genomic analyses (50), which have since improved *Z. tritici* gene models. High-confidence gene models were complemented with a total of 1345 priority ORFs that had been requested during consultation with members of the *Z. tritici* research community. These latter ORFs were included in the ORFeome construction project even if they failed our gene model quality control (primers for all PCR amplifications are included in Supplementary Table S1). An overview of the ORFeome construction project is shown in Figure 1A. PCR amplification utilized genomic DNA as template, which was chosen over cDNA, in order to maintain alternative splice variants during downstream ORFeome expression in *Z. tritici*. Additionally, in order to enable c-terminal epitope-tagging of encoded ORFs using a variety of Destination vectors (26), primers were designed to omit the native stop codon. If expression of the encoded ORFs with a 3’ stop codon is desired, we have developed numerous Destination vectors for this purpose (46).

**Figure 1:**
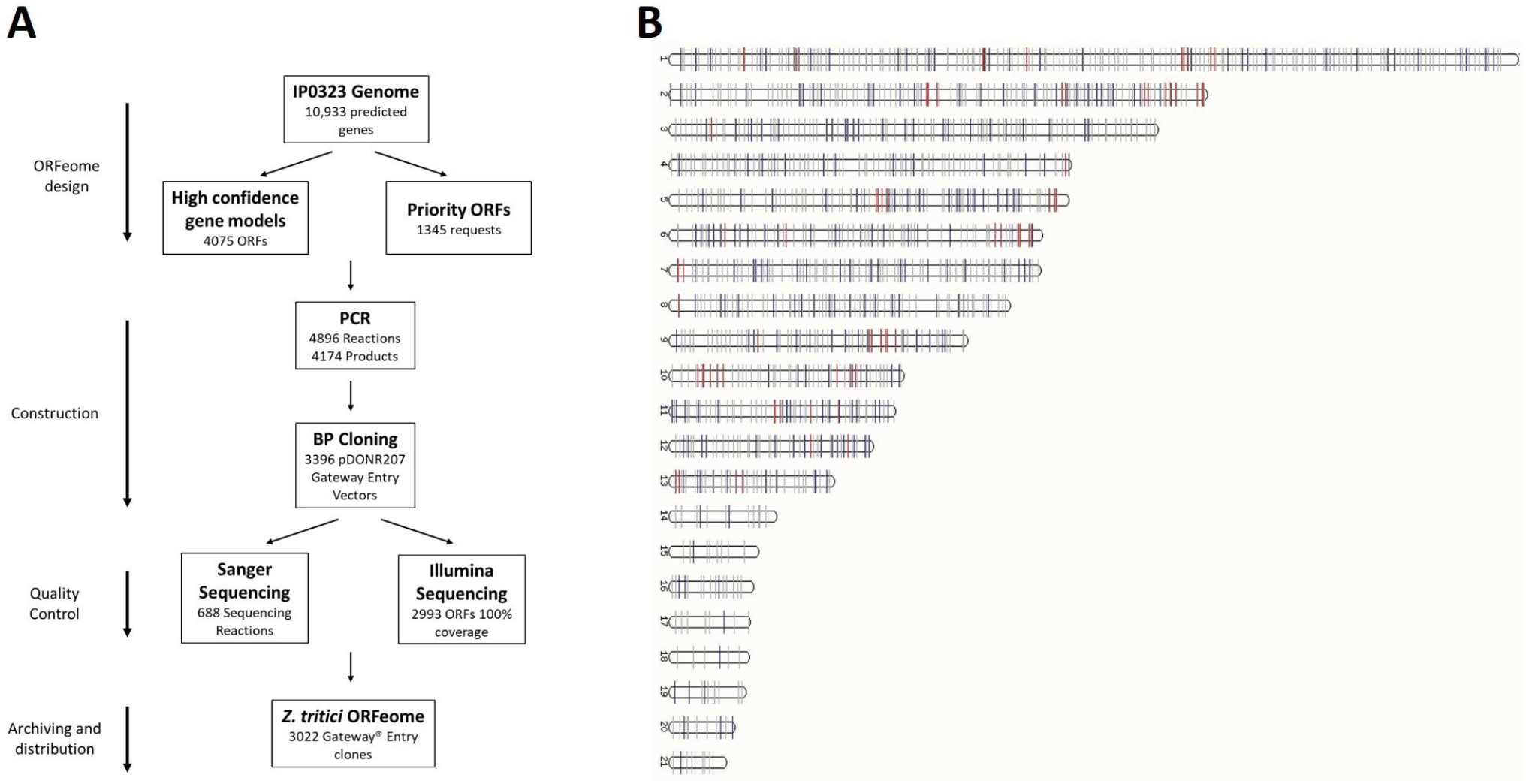
Schematic workflow for ORFeome design, generation, and quality control (**A**). Note that while 374 Entry vectors did not pass quality control, these plasmids are listed in Supplementary Table 2 and can still be distributed. The 3022 ORFs passing quality control were plotted as a function of chromosomal locus (**B**). ORFs are shown as vertical light grey lines, with genes encoding predicted effectors and secondary metabolite biosynthetic genes highlighted in blue and red, respectively. Chromosome numbers are depicted with chromosome 1-13 being core and 14-21 dispensable.

A total of 4896 PCR reactions were conducted in 51 × 96 well plates, which yielded 4174 amplicons of the predicted molecular weight as determined by gel electrophoresis (data not shown). A total of 3396 of these genes were successfully cloned into pDONR207 using the Gateway^®^ BP reaction, as determined by bacterial growth on selection agar, for which plasmids were extracted (Figure 1A and Supplementary File S2). Over 650 Gateway Entry plasmids were randomly selected for sequence verification using Sanger sequencing, with 99.5% passing quality control (Supplementary File S1 and S2). A second quality control experiment was conducted, in which all 3396 ORFs were pooled and sequenced using an Illumina HiSeq 2500 (Supplementary File S1). When combined with Sanger sequencing experiments, a total of 3022 *Z. tritici* ORFs passed quality control checks (Figure 1A). Genes that are represented in the quality controlled *Z. tritici* ORFeome (n = 3022) are plotted as a function of chromosomal locus (Figure 1B), and cover both core and accessory chromosomes (Table 1). All 3396 plasmids are available to end users (Supplementary File S2) with the caveat that plasmids which failed our quality control need be sequence verified by the end user. A summary of ORFeome coverage for 3022 quality controlled ORFs amongst predicted effector encoding genes, secondary metabolite biosynthetic genes, various other functional groups, chromosomal loci, and differentially expressed genes during infection (35) is provided in Table 1. These data indicate that the ORFeome will be applicable for functional genomic experiments to test diverse hypotheses regarding, for example, gene function, expression, or genomic location.

**Table 1.** Summary information for the *Z. tritici* ORFeome.

|  | Predicted category | IP0323 Genome (no. genes) | ORFeome<sup>1</sup> (no. genes) | Coverage in ORFeome (% of genome total) | Reference |
| --- | --- | --- | --- | --- | --- |
| **Total no genes** |  | 10933 | 3022 | 27.6 | (29) |
| **Secreted proteins<sup>2</sup>** | Signal Peptide | 909 | 538 | 59.1 | (53, 54) |
|  | Effector P | 1438 | 410 | 28.5 |  |
| **Chromosomal loci** | Core chromosome | 10278 | 2900 | 28.2 | (29, 36) |
|  | Accessory chromosome | 654 | 122 | 18.6 |  |
|  | Subtelomeric<sup>3</sup> | 2501 | 644 | 25.7 |  |
|  | Secondary metabolite cluster | 682 | 171 | 25.0 |  |
| **Differentially expressed during infection<sup>4</sup>** | 1 DPI | 626 | 194 | 30.9 | (35) |
|  | 4 DPI | 769 | 256 | 33.2 |  |
|  | 9 DPI | 812 | 271 | 33.3 |  |
|  | 14 DPI | 637 | 206 | 32.3 |  |
|  | 21 DPI | 718 | 203 | 28.2 |  |
| **Exemplar GO Terms** | DNA Binding (GO:0003677) | 491 | 76 | 15.4 | (53) |
|  | GTPase Activity (GO:0003924) | 82 | 12 | 14.6 |  |
|  | Protein kinase activity (GO:0004672) | 301 | 53 | 17.6 |  |
|  | Transmembrane transport (GO:0055085) | 950 | 145 | 15.2 |  |

<sup>1</sup> ORFs cloned into pDONR207 that passed quality control (n = 3022, Supplementary File S1 and S2) were assigned various functional categories and are reported both as number of genes, and as a percentage of the predicted total for the IP0323 genome.
<sup>2</sup> Putative effector-encoding genes were predicted from amino acid coding sequences using default parameters in the Effector P prediction algorithm (54). ORFs encoding predicted signal peptides and exemplar GO terms were retrieved using the Ensemble Biomart pipeline (53).
<sup>3</sup> Subtelomeric genes were defined as those residing within 300 kb of the chromosome end. Genes predicted to reside in secondary metabolite biosynthetic gene clusters were identified from AntiSMASH and SMURF predictions (36).
<sup>4</sup> Differentially expressed genes were defined from transcriptional profiling by Rudd and co-workers, with the number of genes significantly upregulated *in planta* at various days post infection (DPI) relative to *in vitro* growth on Czapek-Dox media reported (35). The total predicted number of genes in the IP0323 genome belonging to each functional category is also shown.
ORFeome clones for each functional category are given in Supplemental Dataset S1 and S2.

## 4. Discussion

We have generated an ORFeome library of *Z. tritici* that covers high confidence gene models from the reference genome isolate IP0323 (29) for functional genomic analyses. The ORFeome contains 3022 sequence verified clones. Genes represented in this library are putatively involved in a diverse range of processes, including secreted proteins and putative effectors, biosynthesis of secondary metabolite toxins, drug detoxification, signal sensing and transduction, regulation of gene expression, amongst many others (see Table 1, Supplementary File 1 and 2), and will therefore facilitate functional genomic experiments for diverse aspects of *Z. tritici* biology.

Our strategy prioritised high-confidence gene models for ORFeome construction over a genome-wide cloning approach. While this has resulted in a partial ORFeome for *Z. tritici* IP0323, we believe that a focus on accurate gene models will avoid large-scale future updates of this resource. For example, the first *C. elegans* ORFeome (51) underwent various revisions and additions due to improved gene model predictions (20). More importantly, high confidence gene models will likely limit expression of incorrect ORFs during cost and labour-intensive experiments by end users.

ORFeomes for several model organisms are amplified intron-free sequences from cDNA libraries (20, 21). In contrast, we amplified ORF sequences from genomic DNA in order to maintain the opportunity to generate alternative splice variants (e.g. due to intron skipping) in subsequent *Z. tritici* over-expression or localisation experiments. While the extent of alternative splicing in fungi is not comprehensively determined, an estimated 6.1% of *Z. tritici* genes have splice variants (52). Alternative splicing is thought to predominantly occur for genes required for virulence, multicellularity, and dimorphic switching (52). Our ORFeome will therefore facilitate the study of splice variants that may have critical impacts on infection, or, alternatively, encode promising drug targets in this pathogen.

We have previously demonstrated that the application of a pilot (n = 32) collection of putative DNA-binding protein encoding genes in Entry vectors can enable medium-throughput gene functional analyses in *Z. tritici* (48). The ORFeome generated in this study will drastically increase the throughput of these capabilities for the research community. We predict that functional genomics in *Z. tritici* will enable systems-level understanding of diverse range of processes, including but not limited to growth and development, sensing and signal transduction, virulence, host-pathogen interactions, toxin biosynthesis, drug resistance, and chemical-genetic interactions. Such advances may ultimately lead to novel fungicide development and development of novel resistant wheat cultivars.

## Data Availability

All reagents generated during the current study will be made available. Plasmid requests should be sent to. Raw sequencing data are available at the Sequencing Read Archive (accession SRX1267196 and SRX1265386).

## Supplementary Data

**Supplementary File S1: List of genes included in the *Z. tritici* IPO323 ORFeome project.** Listed for each ORF are: PCR primers used to amplify sequence, KOG/GO terms, and quality control information from Illumia/Sanger sequencing.

**Supplementary File S2: *In silico* characterisation of the ORFeome library.** Listed for each ORFs are: presence at various genomic loci (dispensable chromosome, putative secondary metabolite cluster, subtelomeric locus), transcriptional deployment during infection, and various exemplar GO categories (e.g. DNA binding/GTPase, etc).7

### Funding

This work was funded by a BBSRC BBR grant (<u>BB/I025956/1</u>) to KH and collaborators and a BBSRC CASE studentship (BB/J500793/1), supported by Syngenta UK to YS.

