## Supplementary File S1 for "The *Zymoseptoria tritici* ORFeome: a functional genomics community resource"

### List of genes included in the *Zymoseptoria tritici* IPO323 ORFeome project

Last updated: 25/09/2018

##### Plate #1

| Plate number | PCR primers | Gene model | KOGs | GO terms | QC: coverage by Illumina resequencing | QC: mismatches revealed by Illumina resequencing | QC: alignment against Sanger sequence reads |
| --- | --- | --- | --- | --- | --- | --- | --- |
| 1 | ggggacaagtttgtacaaaaaagcaggcttgatgCCTCACAAGCGAAAAGCGGC, ggggaccactttgtacaagaaagctgggtcACCCTCCGCTCCTCTCCTGG | Mycgr3\|32474  JGI  Ensembl | KOG0133 Deoxyribodipyrimidine photolyase/cryptochrome; | DNA photolyase activity; DNA repair;  deoxyribodipyrimidine photo-lyase activity; | 100 % |  | ``` 1	ATGCCTCACAAGCGAAAAGCGGCCTCGCCACAACAACTTTCGCCTTTCAAGAAACAACATGTGTCTCCGGACAAGTCGAATGGCGCCAGTGCTACCGATCAATATCAAAACAACAGTGGCTTACAGCCCAATCTTCAGCCTCACTTCTACCCACCCGAAATCTCATCCTCGCGATGTGCCCAATATGCCAATGGCGAGCTCTCCAAACCGATCGATGTCCTCGAAACAGCTCTAGCCTCCACCTCGACTCAACGACAGGCCATTGCAGGAGGCCGAGCAGTTCTTCA-TTGGTTCAAGCGAGATCTGC	307 	\|\|\|\|\|\|\|\|\|\|\|\|\|\|\|\|\|\|\|\|\|\|\|\|\|\|\|\|\|\|\|\|\|\|\|\|\|\|\|\|\|\|\|\|\|\|\|\|\|\|\|\|\|\|\|\|\|\|\|\|\|\|\|\|\|\|\|\|\|\|\|\|\|\|\|\|\|\|\|\|\|\|\|\|\|\|\|\|\|\|\|\|\|\|\|\|\|\|\|\|\|\|\|\|\|\|\|\|\|\|\|\|\|\|\|\|\|\|\|\|\|\|\|\|\|\|\|\|\|\|\|\|\|\|\|\|\|\|\|\|\|\|\|\|\|\|\|\|\|\|\|\|\|\|\|\|\|\|\|\|\|\|\|\|\|\|\|\|\|\|\|\|\|\|\|\|\|\|\|\|\|\|\|\|\|\|\|\|\|\|\|\|\|\|\|\|\|\|\|\|\|\|\|\|\|\|\|\|\|\|\|\|\|\|\|\|\|\|\|\|\|\|\|\|\|\|\|\|\|\|\|\|\|\|\|\|\|\|\|\|\|\|\|\|\|\|\|\|\|\|\|\|\|\|\|\|\|\|\|\|\|\|\|\|\|\|\|\|\|\|\|\|\|\|\|\| \|\|\| \|\|\|\|\|\| \|\|\|\|\|\|\|  \|\| \| \|\|\|\|\|\| 	ATGCCTCACAAGCGAAAAGCGGCCTCGCCACAACAACTTTCGCCTTTCAAGAAACAACATGTGTCTCCGGACAAGTCGAATGGCGCCAGTGCTACCGATCAATATCAAAACAACAGTGGCTTACAGCCCAATCTTCAGCCTCACTTCTACCCACCCGAAATCTCATCCTCGCGATGTGCCCAATATGCCAATGGCGAGCTCTCCAAACCGATCGATGTCCTCGAAACAGCTCTAGCCTCCACCTCGACTCAACGACAGGCCATTGCAGGAGGCCGANCAGNTCTTCATTTGGTTCNGGCNANATCTGC ``` |
| 1 | ggggacaagtttgtacaaaaaagcaggcttgatgATCAAGCTGAAGCTCGGCAA, ggggaccactttgtacaagaaagctgggtcACCCTCCTCACTATCCTCCC | Mycgr3\|33410  JGI  Ensembl | KOG4011 Transcription initiation factor TFIID, subunit TAF7; | exo-alpha-sialidase activity; general RNA polymerase II transcription factor activity;  regulation of transcription;  transcription factor TFIID complex; | 100 % | chr\_1:4469573 C->T; |  |
| 1 | ggggacaagtttgtacaaaaaagcaggcttgatgCTCCTCTGTCCGTCTCCCTC, ggggaccactttgtacaagaaagctgggtcATTCTCCCGCCATTCCCGCG | Mycgr3\|33562  JGI  Ensembl | KOG2906 RNA polymerase III subunit C11; | DNA-directed RNA polymerase I activity; DNA-directed RNA polymerase II activity;  DNA-directed RNA polymerase III activity;  DNA-directed RNA polymerase activity;  RNA polymerase activity;  nucleic acid binding;  regulation of transcription;  transcription regulator activity;  zinc ion binding; | 4.71014 % |  | ``` 1	ATGCTCCTCTGTCCGTCTCCCTCTCTCCTTCACCACCAAGACCCTCAAACTAACCTTCCATTCTCCTGCCAAAAAGTCTGCCCGTCCTGCTCCAACTCCCTCATCATCACCACCGTCCCGGCCGCACACTGCACGCCTCTCGACCGCGCCGACGCCGGCAAAAACCGCTTCGAATGCCGCACCTGTCCTTATCAAATGGTCCTCGACCGAAGATACTACGAACGGAAATCCATGACGCTGAAACCCGTCGAGGACATCTTGGGAGGCGCGGAGAGTTGGAAGAATGTGGATAAGACGGAGG	301 	\|\|\|\|\|\|\|\|\|\|\|\|\|\|\|\|\|\|\|\|\|\|\|\|\|\|\|\|\|\|\|\|\|\|\|\| \|\|\|\|\|\|\|\|\|\|\|\|\|\|\|\|\|\|\|\| \|\|\|\|\|\|\|\|\|\|\|\|\|\|\|\|\|\|\|\|\|\|\|\|\|\|\|\|\|\|\|\|\|\|\|\|\|\|\|\|\|\|\|\|\|\|\|\|\|\|\|\|\|\|\|\|\|\|\| \|\|\|\|\|\|\|\|\|\|\|\|\|\|\|\|\|\|\|\|\|\|\|\|\|\|\|\|\|\|\|\|\|\|\|\|\|\|\|\|\|\|\|\|\|\|\|\|\|\|\|\|\|\|\|\|\|\|\|\|\|\|\|\|\|\|\|\|\|\|\|\|\|\|\|\|\|\|\|\|\|\|\|\|\|\|\|\|\|\|\|\|\|\|\|\|\|\|\|\|\|\|\|\|\|\|\|\|\|\|\|\|\|\|\|\|\|\|\|\|\|\|\|\|\|\|\|\|\|\|\|\|\|\|\|\|\|\|\|\|\|\|\|\|\|\|\|\|\|\|\|\|\|\|\|\|\|\|\|\|\|\|\|\|\|\|\|\|\|\|\|\|\|\|\|\|\|\|\|\|\|\|\| 	ATGCTCCTCTGTCCGTCTCCCTCTCTCCTTCACCACNAAGACCCTCAAACTAACCTTNCATTCTCCTGCCAAAAAGTCTGCCCGTCCTGCTCCAACTCCCTCATCATCACCACCGTCGCGGCCGCACACTGCACGCCTCTCGACCGCGCCGACGCCGGCAAAAACCGCTTCGAATGCCGCACCTGTCCTTATCAAATGGTCCTCGACCGAAGATACTACGAACGGAAATCCATGACGCTGAAACCCGTCGAGGACATCTTGGGAGGCGCGGAGAGTTGGAAGAATGTGGATAAGACGGAGG ``` |
| 1 | ggggacaagtttgtacaaaaaagcaggcttgatgCGAGCAGCACGGATCACTCG, ggggaccactttgtacaagaaagctgggtcTGCGCTTCTCCCAATATCTT | Mycgr3\|34626  JGI  Ensembl | KOG1921 Endonuclease III; | DNA binding; DNA repair;  DNA-(apurinic or apyrimidinic site) lyase activity;  base-excision repair;  endonuclease activity;  intracellular; | 100 % |  | ``` 1	ATGCGAGCAGCACGGATCACTCGCGACACAGCTCGAGTAGCAAAGAAACCCTCACCACCACCCAAAACCTCCCGACCTCGACGATCACTACGAGAATTCGCCTACAACGATGATGAAACGAACGAAGACTCCCCTACTCCCTCCGAGGCAGCCAGCGACTTCTCCAGCCCTCCCTCAAGTCCAGACGAAGCATCTCCCGAACCGACATCCACACCCGCCTCAAGCAAGAAACGAAAGCACGAATCGATTCAACCTGCTTCAACCCGCAAGTCACCTCGAAAGGCTCCGAAGAAAGAGACGCCTACCAAAGCTATCAAGCATGAATACGACCCGAATGTCAAGATACATCCTCCACCAGACTGGGAGACCATTTGGTCACACATCGCAACGATGCGTTCCCTAAACCCTGCGCCAGTCGACAGCATGGGCTGCGAGAACCTTGCAGAAGAGACACGCTCCCCACTCGATCGAAGACTACAGACACTGATCGCGTTGATGCTATCGTCTCAGACGAAAGATCCCGTCACGGCGGCTGCAATGAAGAATCTACAAGACAATTTGAAAGGAGGCTTCAACCTCGAAGGACTGCTGAACGTGGAACCGGACGTATTGAACAACCTGATCGCAAAGGTGGGCTTCCACAATAACAAGACGAAGTACATCAAAGCGACGGCGGAGCTGTTGAGAGATAAGTGGAATGGAGACATTCCCGAGACGATCGAGGGATTGGTGTCGCTACCGGGCGTGGGACCGAAGATGGCGTACTTGACCATGTCGTCGGCCTGGGGTAGGGATGAGG	799 	\|\|\|\|\|\|\|\|\|\|\|\|\|\|\|\|\|\|\|\|\|\|\|\|\|\|\|\|\|\|\|\|\|\|\|\|\|\|\|\|\|\|\|\|\|\|\|\|\|\|\|\|\|\|\|\|\|\|\|\|\|\|\|\|\|\|\|\|\|\|\|\|\|\|\|\|\|\|\|\|\|\|\|\|\|\|\|\|\|\|\|\|\|\|\|\|\|\|\|\|\|\|\|\|\|\|\|\|\|\|\|\|\|\|\|\|\|\|\|\|\|\|\|\|\|\|\|\|\|\|\|\|\|\|\|\|\|\|\|\|\|\|\|\|\|\|\|\|\|\|\|\|\|\|\|\|\|\|\|\|\|\|\|\|\|\|\|\|\|\|\|\|\|\|\|\|\|\|\|\|\|\|\|\|\|\|\|\|\|\|\|\|\|\|\|\|\|\|\|\|\|\|\|\|\|\|\|\|\|\|\|\|\|\|\|\|\|\|\|\|\|\|\|\|\|\|\|\|\|\|\|\|\|\|\|\|\|\|\|\|\|\|\|\|\|\|\|\|\|\|\|\|\|\|\|\|\|\|\|\|\|\|\|\|\|\|\|\|\|\|\|\|\|\|\|\|\|\|\|\|\|\|\|\|\|\|\|\|\|\|\|\|\|\|\|\|\|\|\|\|\|\|\|\|\|\|\|\|\|\|\|\|\|\|\|\|\|\|\|\|\|\|\|\|\|\|\|\|\|\|\|\|\|\|\|\|\|\|\|\|\|\|\|\|\|\|\|\|\|\|\|\|\|\|\|\|\|\|\|\|\|\|\|\|\|\|\|\|\|\|\|\|\|\|\|\|\|\|\|\|\|\|\|\|\|\|\|\|\|\|\|\|\|\|\|\|\|\|\|\|\|\|\|\|\|\|\|\|\|\|\|\|\|\|\|\|\|\|\|\|\|\|\|\|\|\|\|\|\|\|\|\|\|\|\|\|\|\|\|\|\|\|\|\|\|\|\|\|\|\|\|\|\|\|\|\|\|\|\|\|\|\|\|\|\|\|\|\|\|\|\|\|\|\|\|\|\|\|\|\|\|\|\|\|\|\|\|\|\|\|\|\|\|\|\|\|\|\|\|\|\|\|\|\|\|\|\|\|\|\|\|\|\|\|\|\|\|\|\|\|\|\|\|\|\|\|\|\|\|\|\|\|\|\|\|\|\|\|\|\|\|\|\|\|\|\|\|\|\|\|\|\|\|\|\|\|\|\|\|\|\|\|\|\|\|\|\|\|\|\|\|\|\|\|\|\|\|\|\|\|\|\|\|\|\|\|\|\|\|\|\|\|\|\|\|\|\|\|\|\|\|\|\|\|\|\|\|\|\|\|\|\|\|\|\|\|  \|\|\|\|\|\|\|\|\|\|\|\|\|\|\|\|\|\|\|\|\|\| \| \|\| \|\| \|\|\|\|\|\|\|\|\|\|\|\|\|\| \|\|\|\|\|\|\|\|\|\|\|\|\| \|\|\|\|\|\|\| \| \|\|\|\|\|\|\|\|\|\|\|\|\|\|\|\|\| \|\| \|\|\|\| \|\|\|\|\| \|\|\|\|\|\|\| \|\|\|\|\|\| \|\|\|\| \|\|\| \|\|\|\| \|\| \|\|\| \|\|\|\| \|\|\| \| \|\|\|  \|\|\|\| \|\|\| \|\|\|\|\|\| \|\|\|\|  \|\|\|\|\| 	ATGCGAGCAGCACGGATCACTCGCGACACAGCTCGAGTAGCAAAGAAACCCTCACCACCACCCAAAACCTCCCGACCTCGACGATCACTACGAGAATTCGCCTACAACGATGATGAAACGAACGAAGACTCCCCTACTCCCTCCGAGGCAGCCAGCGACTTCTCCAGCCCTCCCTCAAGTCCAGACGAAGCATCTCCCGAACCGACATCCACACCCGCCTCAAGCAAGAAACGAAAGCACGAATCGATTCAACCTGCTTCAACCCGCAAGTCACCTCGAAAGGCTCCGAAGAAAGAGACGCCTACCAAAGCTATCAAGCATGAATACGACCCGAATGTCAAGATACATCCTCCACCAGACTGGGAGACCATTTGGTCACACATCGCAACGATGCGTTCCCTAAACCCTGCGCCAGTCGACAGCATGGGCTGCGAGAACCTTGCAGAAGAGACACGCTCCCCACTCGATCGAAGACTACAGACACTGATCGCGTTGATGCTATCGTCTCAGACGAAAGATCCCGTCACGGCGGCTGCAATGAAGAATCTACAAGACAATTTGAAAGGAGGCTTCAACCTCGAAGGACTGCTGAACGTGGAACCGGACGTATTGAACAN-CTGATCGCAAAGGTGGGCTTCC-CNAT-ACNAGACGAAGTACATC-AAGCGACGGCGGANCTGTTGAAA-ATAAGTGGAATGGAGACNTT-CCGANACGATNGAGGGAT-GGTGTC-CTACNGGG-GTGGAAC-GAA-ATGG-GTANT-GACN-TGTC-TCG-CCTGGG-TAGGN-TGAGG ```      ``` 89	TAC-GAGAATTCG-CCTACAACGATGATGAAACGAACGAAGACTCCCCTACTCCCTCCGAGGCAGCCAGCGACTTCTCCAG-CC-CTCCCTCAAGTCCAGACGAAGCATCTCCCGA-ACCGACATCC-ACACCCGCCTCAAGCAAGAAACGAAAGCACGAATCGATTCAACCTGCTTCAACCCGCAAGTCACCTCGAAAGGCTCCGAAGAAAGAGACGCCTACCAAAGCTATCAAGCATGAATACGACCCGAATGTCAAGATACA-TCCTCCACCAGAC-TGGG-AGACCATTTG-GTCACACATCG-CAAC-GATGCGTTCCCTAAACCCTGCGC-CAGTCGACAG-CATGGGCTG-CGAGA-ACCTTGCAGAAGAGACAC-GCTC-CCCACT-CGAT-CGAAGACTA-CAGACACT-GATC-GC-GTTGAT-GCTATCGTCT-CAGACGA-AAGATCCC-GTC-ACGGC-GGCTGCA-ATGAA-GAATC-TACAAGACAA-TTTGAAAGG-AGG-CTT-CAACCTCG-AAGGACTGCTGAACGTGGAACC-GGAC	606 	\|\|\| \|\|\|\|\|\|\|\|\|  \| \| \|\| \|\|\|\|\| \|\| \| \|  \|\|\|\|\|\|\|\|\|\|\|\|\|\|\| \|\|\| \|\| \| \|\| \|\|\|   \|\| \|\|\|\|\|\|\|\| \|\| \|\| \|\|\|\|  \|\|\|\|\|\|\| \|\|   \| \|\| \|\|\|\| \|\|\|\|\|\|\|\|\|\|  \|   \|\| \|\|\|\|\|\| \|\|\|\|\|  \|\|\| \|\| \|\| \| \|\|\|\|\|\|\| \|\|\|\|\|\|\|\| \| \|\|   \|\|\|\|\| \|\|\|\|\|\|\|\|\| \|\|\|\|\|\| \|\|\|\|\|\|\|\|\|\|\|\|\|\|\|\| \|\|\|\|\|\|\|\|\|\|\|\|\|\|\|\|\|\|\|\|\|\|\|\|\|\|\|\|\|\| \| \|\|\| \|\| \|\|\|\|\|\|\|\|\|\| \|  \|\|\|\| \|\|\| \|\| \|\|\| \|\|\|\|\|\|\|\|\| \| \|\| \| \|\|\|\|\|\|\|\|\|\|\|\|\|\|\|\|\|\|\|\|\|\|\| \|\|\|\|\|\|\| \|  \|\|\|\|\|\|\|\|\| \| \|\|\| \|\|\|\|\|\| \| \|\|\|\|\|\|    \|\|\|\| \|\|\|\|\|  \|\|\|\| \|\|\|\|\|\|\|\|\| \|  \|\| \|  \|\|\|\| \|\| \|\|\|\|\|\| \|\|\|   \|\|   \|\|\|\|\|\|\| \|\| \|\| \|  \|\|\| \| \|\|\| \|\|\|\|\|   \|\|\|\|\| \|\|\|\|\| \|  \|  \|\| \| \|\|\|\|\| \|\|\| \| \| \|\|\| \|\| \|\|\|\|  \|\|\|\|\|  \|\|\|\|\|\|\| \|\|\|\|\|\|\| \|\|\|\| 	TACAGAGAATTCGNNCGAAAANGATGANGANAAGNNCGAAGACTCCCCTACACCCNCCNANGCNGCCNNNGANTTCTCCAGNCCGCT-CCTCNNGTCCAGANGANCNANCTNCCGACACCGACATCCNGCGNTCGNCTCAAGNAAGAANNGAANGCGCGNAGCGATTCANCCTGCTTCNANCCNGTAGTCANCTCGAAAGGNTCCGAANAAAGAGACGCCTACCANAGCTATCAAGCATGAATACGACCCGAATGTNANGATNCATTCCTCCACCANAANTGGGTAGANCANTTGNGTCACACATNGNCANCAGATGCGTTCCCTAAACCCTGCGCTCAGTCGANAATCATGGGCTGACTAGANACCTTGNANAAGAGANTNNGCTCTCCCACANCGATNCGAAGACTANCNTACNCCNGATCCGCAGTTGATNGCTNNNGTNNNCAGACGATAANATTCNNGTCNANGGCAGGCTGTNNATGAAAGAATCCTNNATNACNANTTTGANAGGGANGNCTTTCANCCTCNAAAGGAGNGCTGAACNTGGAACCCGGAC ``` |
| 1 | ggggacaagtttgtacaaaaaagcaggcttgatgAGCGACGATGAGCACTCACA, ggggaccactttgtacaagaaagctgggtcGGTCAAGTTCCAAGACAACT | Mycgr3\|34632  JGI  Ensembl |  |  | 100 % |  | ``` 1	ATGAGCGACGATGAGCACTCACATCGAGTGAAGAAGCGGCGCATCTCCCGTGCGTGCGACTACTGCCACAAACGAAGCAAGCGATGCAGAGCCAGTCT-GAACGACTC-CCGCTGCCAAAGCTGCGTTTCTTTCGATCAGC-CATGCACTT-TCAATCGGCCCCAAAAGCGGAGAGGAGCGCC	179 	\|\|\|\| \|\|\|\|\|\|\|\|\| \|\|\|\|\|\|\|\|\|\|\|\|\|\|\|\|\|\|\|\|\|\| \|\|\|\|\|\|\|\|\|\|\|\|\|\|\|\|\|\| \|\| \|\|\| \|\|\| \|\|\|\|\|\|\| \| \|\|\|\|\|\|\|\|\|\|\|\|\|\|\|\|\| \|\| \|\| \|\|\|\|   \|\|\|\|\|\|\|\|\|\|\|\| \|\|\| \|\|\|\| \|\|\|\|\|\|\| \|\| \| \|\|\| \| \| \|\|\|\|\|\|\|\|\|\| \|\| \|\|\|\|\|\|\|\|\|\|\|\| \|\|\|\| 	ATGANCGACGATGANCACTCACATCGAGTGAAGAAGCNGCGCATCTCCCGTGCGTGNGANTACCGCCCCAAACGANGNAAGCGATGCAGAGCCAGNCTTGANCGACANNCCGCTGCCAAAGNTGCNTTTCGTTCGATCNGCACNTGCNCATNTCAATCGGCCNCANAAGCGGAGAGGANCGCC ``` |
| 1 | ggggacaagtttgtacaaaaaagcaggcttgatgCCGTTGATGCGAATCCCCTA, ggggaccactttgtacaagaaagctgggtcTAATGATGCCCACGGCGCTG | Mycgr3\|34913  JGI  Ensembl | KOG2683 Sirtuin 4 and related class II sirtuins (SIR2 family); | (3,5-dichlorophenylurea)acetate amidohydrolase activity; DNA binding;  N-acetylgalactosamine-6-phosphate deacetylase activity;  N2-acetyl-L-lysine deacetylase activity;  NAD binding;  NAD-dependent histone deacetylase activity;  O-succinylbenzoate synthase activity;  UDP-3-O-[3-hydroxymyristoyl] N-acetylglucosamine deacetylase activity;  chromatin silencing;  chromatin silencing complex;  histone deacetylase activity;  indoleacetamide hydrolase activity;  iprodione amidohydrolase activity;  protein N-terminal asparagine amidohydrolase activity;  protein amino acid deacetylation;  regulation of transcription, DNA-dependent;  zinc ion binding; | 0 % |  | ``` 1	ATGCCGTTGATGCGAATCCCCTACACACAACCCCTCCCAAGTCCCACCATCCTCCCCTCCACAGCATCAACCATCCCCGGCGCCGTCTCAGCACTGCGCAACTTCCTCACCCAAAAGTCGACCTCGAACAACAAGACCCTCATCCTCTCCGGCGCCGGCATCTCCGTCGCCTCCGGTCTCGCCGACTACCGCGGCACCAATGGCACATACGTCCTCAACAAGACCTACCGACCCATCTACTTTCACGAATTCAGCGCCAGCCACGAAGCGCGAAAACGATACTGGGCGCGCTCCTTCCTGGGCTGGACGACCCTCCACCGCAGTCGTCCCAACCCCGCTCATCACGCTGTGAAACGACTCGGGGAGTTGGGA	372 	\|\|\|\|\|\|\|\|\|\|\|\|\|\|\|\|\|\|\|\|\|\|\|\|\|\|\|\|\|\|\|\|\|\|\|\|\|\|\|\|\|\|\|\|\|\|\|\|\|\|\|\|\|\|\|\|\|\|\|\|\|\|\|\|\|\|\|\|\|\|\|\|\|\|\|\|\|\|\|\|\|\|\|\|\|\|\|\|\|\|\|\|\|\|\|\|\|\|\|\|\|\|\|\|\|\|\|\|\|\|\|\|\|\|\|\|\|\|\|\|\|\|\|\|\|\|\|\|\|\|\|\|\|\|\|\|\|\|\|\|\|\|\|\|\|\|\|\|\|\|\|\|\|\|\|\|\|\|\|\|\|\|\|\|\|\|\|\|\|\|\|\|\|\|\|\|\|\|\|\|\|\|\|\|\|\|\|\|\|\|\|\|\|\|\|\|\|\|\|\|\|\|\|\|\|\|\|\|\|\|\|\|\|\|\|\|\|\|\|\|\|\|\|\|\|\|\|\|\|\|\|\|\|\|\|\|\|\|\|\|\|\|\|\|\|\|\|\|\|\|\|\|\|\|\|\|\|\|\|\|\|\|\|\|\|\|\|\|\|\|\|\|\|\|\|\|\|\|\|\|\|\|\|\|\|\|\|\|\|\|\|\|\|\|\|\|\|\|\|\|\|\|\|\|\|\|\|\|\|\|\|\|\|\|\|\|\|\|\|\|\|\|\|\|\|\|\|\|\|\|\|\|\|\|\|\|\|\|\|\|\|\|\|\|\|\|\|\|\|\|\|\|\|\| \|\|\|\|\|\|\|\|\|\|\|\|\|\|\|\|\| 	ATGCCGTTGATGCGAATCCCCTACACACAACCCCTCCCAAGTCCCACCATCCTCCCCTCCACAGCATCAACCATCCCCGGCGCCGTCTCAGCACTGCGCAACTTCCTCACCCAAAAGTCGACCTCGAACAACAAGACCCTCATCCTCTCCGGCGCCGGCATCTCCGTCGCCTCCGGTCTCGCCGACTACCGCGGCACCAATGGCACATACGTCCTCAACAAGACCTACCGACCCATCTACTTTCACGAATTCAGCGCCAGCCACGAAGCGCGAAAACGATACTGGGCGCGCTCCTTCCTGGGCTGGACGACCCTCCACCGCAGTCGTCCCAACCCCGCTCATCACGCTGTGAAANGACTCGGGGAGTTGGGA ``` |
| 1 | ggggacaagtttgtacaaaaaagcaggcttgatgAAATTCCGCTCCCTCAAGAG, ggggaccactttgtacaagaaagctgggtcCCGAGCCTCCTCTCTCTTCC | Mycgr3\|37858  JGI  Ensembl | KOG2793 Putative N2,N2-dimethylguanosine tRNA methyltransferase; | DNA binding; | 100 % |  | ``` 1	ATGAAATTCCGCTCCCTCAAGAGTCCCTTCAACATCACCAACGATGCCACAACCAACTCCAAACTCCTCATTCACATCTCCGAAAAGGAAGGCGTGAAAGCCGAAAACCTCTCCCTGGCTACATGGGGTAGTTCCTACGTCTTGGCCAACCACCTCCATCGATGGAAAGATACCCAGCCCATCCACACCGCTCGGGAGAACACATCCAACTCCATCGCTGTACTTGAACTCGGCGCGGGAACAGGACTCGTCGGTCTCTCCGCCGCAGCCATATGGCAACTCCCTGTGATCCTCACAGATCTGGCCCCGATCGTCCCTGGGTTGGCACAGAATATCAACTCGAATCGAGCGTTATTGGCGGAGAAAGGAACTACCGCGGCGTGCGGTTGCTTAGACTGGACGCAGCCGGAGGAACTACTCCTTTCCAATCCTGTTCTTGCTGCATCGGAGCCACAGATCGTGCAATCTTCGCGTAAAGCGCACATTATCCTCGCTGCTGATGTCGTGTACAGCGAGGACCATCCCGAGTTGTTGTTGAAGACAATTCTGGCGTGGTTGGCGCCTGGGCCGGATGCGAGGGTTGTGATTTGCTATCCTGTGAGAATTGCATATATCGATCATATTAGAGACTTATGGGAGCGATTCGAAGCGGCGGGGTTGTTGTGTG	667 	\|\|\|\|\|\|\|\|\|\|\|\|\|\|\|\|\|\|\|\|\|\|\|\|\|\|\|\|\|\|\|\|\|\|\|\|\|\|\|\|\|\|\|\|\|\|\|\|\|\|\|\|\|\|\|\|\|\|\|\|\|\|\|\|\|\|\|\|\|\|\|\|\|\|\|\|\|\|\|\|\|\|\|\|\|\|\|\|\|\|\|\|\|\|\|\|\|\|\|\|\|\|\|\|\|\|\|\|\|\|\|\|\|\|\|\|\|\|\|\|\|\|\|\|\|\|\|\|\|\|\|\|\|\|\|\|\|\|\|\|\|\|\|\|\|\|\|\|\|\|\|\|\|\|\|\|\|\|\|\|\|\|\|\|\|\|\|\|\|\|\|\|\|\|\|\|\|\|\|\|\|\|\|\|\|\|\|\|\|\|\|\|\|\|\|\|\|\|\|\|\|\|\|\|\|\|\|\|\|\|\|\|\|\|\|\|\|\|\|\|\|\|\|\|\|\|\|\|\|\|\|\|\|\|\|\|\|\|\|\|\|\|\|\|\|\|\|\|\|\|\|\|\|\|\|\|\|\|\|\|\|\|\|\|\|\|\|\|\|\|\|\|\|\|\|\|\|\|\|\|\|\|\|\|\|\|\|\|\|\|\|\|\|\|\|\|\|\|\|\|\|\|\|\|\|\|\|\|\|\|\|\|\|\|\|\|\|\|\|\|\|\|\|\|\|\|\|\|\|\|\|\|\|\|\|\|\|\|\|\|\|\|\|\|\|\|\|\|\|\|\|\|\|\|\|\|\|\|\|\|\|\|\|\|\|\|\|\|\|\|\|\|\|\|\|\|\|\|\|\|\|\|\|\|\|\|\|\|\|\|\|\|\|\|\|\|\|\|\|\|\|\|\|\|\|\|\|\|\|\|\|\|\|\|\|\|\|\|\|\|\|\|\|\|\|\|\|\|\|\|\|\|\|\|\|\|\|\|\|\|\|\|\|\|\|\|\|\|\|\|\|\|\|\|\|\|\|\|\|\|\|\|\|\|\|\|\|\|\|\|\|\|\|\|\|\|\|\|\|\|\|\|\|\|\|\|\|\|\|\|\|\|\|\|\|\|\|\|\|\|\|\|\|\|\|\|\|\|\|\|\|\|\|\|\|\|\|\|\|\|\|\|\|\|\|\|\|\|\|\|\|\|\|\|\|\|\|\|\|\|\|\|\|\|\|\|\|\|\|\|\|\|\|\|\|\|\|\|\|\|\|\|\|\|\|\|\|\|\|\|\|\|\|\|\| \|\|\|\|\|\|\|\|\|\|\|\|\|\|\|\|\|\|\|\|\|\| \|\|\|\|\|\|\|\|\|\|\|\|\|\|\|\|\|\|\|\|\|\|\|\|\|\| \|\|\|\|\|\|\|\|\|\|\|\| \|\|\|\|\|\| \|\| \|\|\|\|\|\|\|\|\|\|\|\|\|\|\|\|\|\| 	ATGAAATTCCGCTCCCTCAAGAGTCCCTTCAACATCACCAACGATGCCACAACCAACTCCAAACTCCTCATTCACATCTCCGAAAAGGAAGGCGTGAAAGCCGAAAACCTCTCCCTGGCTACATGGGGTAGTTCCTACGTCTTGGCCAACCACCTCCATCGATGGAAAGATACCCAGCCCATCCACACCGCTCGGGAGAACACATCCAACTCCATCGCTGTACTTGAACTCGGCGCGGGAACAGGACTCGTCGGTCTCTCCGCCGCAGCCATATGGCAACTCCCTGTGATCCTCACAGATCTGGCCCCGATCGTCCCTGGGTTGGCACAGAATATCAACTCGAATCGAGCGTTATTGGCGGAGAAAGGAACTACCGCGGCGTGCGGTTGCTTAGACTGGACGCAGCCGGAGGAACTACTCCTTTCCAATCCTGTTCTTGCTGCATCGGAGCCACAGATCGTGCAATCTTCGCGTAAAGCGCACATTATCCTCGCTGCTGATGTCGTGTACAGCGAGGACCATCCCGAGTTGTTGTTGAAGACAATTCTGGCGTGGTTGGCGCCTGGGCCGGATGCNAGGGTTGTGATTTGCTATCCTGNGAGAATTGCATATATCGATCATATTANAGACTTATGGGANCGATTC-AANCGGCGGGGTTGTTGTGTG ```      ``` 288	GATCCTCACAG-ATCTGGCCCCGATCGTCCCTGGGTTGGCACAGAATATCAACTCGAATCGAG-CGTTATTGGCGGAGAAAGGAACTACCGCGGCGTGCGGTTGCTTAGACTGGACGCAGCCGGAGGAACTACTCCTTTCCAATCCTGTTCTTGCTGCATCGGAGCCACAGATCGTGCAATCTTCGCGTAAAGCGCACATTATCCTCGCTGCTGATGTCGTGTACAGCGAGGACCATCCCGAGTTG-T-TGTTGAAGACAATTCTGGCGTGGTTGG-CGCCTGG-GCCGGATGCGAGGGTTGTGATTTGCTATCCTGT-GAGAATTGCATATATCGATCATATTAGAGACTTATGGGAGCGATTC	645 	\|\|\|\|\|\|\|\| \|\| \|\|\| \|\| \|   \|\|\|\| \|\|\| \|\|\|  \|\|  \|\| \|\|\|\|\|\| \|\|\|\| \|\|\|\| \|\|   \|\|\|\|\|\|\|  \|\| \|\|\|\|\|\|\|\|\|\|\|\|\|  \|\| \|\|\|\|\| \|\|\|\|\|\| \|\|\|\|\|\|  \|\|\|  \|\|\| \|\|\|\|\|\|\|\|\|\|\|\|\|\|\|\|\|\|\|\|\|\|\|\|\|\|\|\|\|\|\|\|\|\|\|\|\|\| \|\|\|\|   \|\|\|\|\|\|\|\|\|\|\|\|   \| \|\|\|     \|\|\| \|\|\|\|\|\|\| \|\|\|\|\|\|\|\|\|\|\|\|\|\|\|\|  \|   \|\|\|\|\|\|\|\|\|\|\|\|\|\|\| \| \| \|\|\|\|   \|\|\|\|\|\|\|\|\|\| \|\| \|\|\|  \|\|\|\|\|\|\| \|\|\| \|\|\|\|\|    \| \|\|\|\|\|\|\|\|\|\|\|\|\|\|\|\|\|\| \|\|\|\|\|\|\|\|\|\|\|\|\|\|\|\|\|\|\|\|\|\|\|\|\|\|\|\|\|\|\|\|\|\|\|\|\|\|\| \| \|\|\|\| 	GATCCTCA-AGTATCAGGACGACATCGGCCCGGGGGAGGTCCATAATATCGACTCTAATC-AGTATTTATTGGACGATAAAGGAACTACCGTCGCTTGCGGCTGCTTATACTGGAATCAGGAGGACGAACTACTCCTTTCCAATCCTGTTCTTGCTGCATCGGACCCACTTTTCGTGCAATCTTTCTGAAAACATGTCATAATCCTCGATGCTGATGTCGTGTACGTCTCCGACCATCCCGAGTTGCTCTTTTGAT--CAATTCTGGCATG-TTGATCGCCTGGAGCCCGATGCCTTTGCTGTGATTTGCTATCCTGTCGAGAATTGCATATATCGATCATATTAGAGACTTATGGGACCCATTC ``` |
| 1 | ggggacaagtttgtacaaaaaagcaggcttgatgACCGTCTACGCTAACGACAC, ggggaccactttgtacaagaaagctgggtcCGTGGTATAAGGATTATACA | Mycgr3\|38718  JGI  Ensembl | KOG2533 Permease of the major facilitator superfamily; | DNA binding; DNA methylation;  N-methyltransferase activity;  integral to membrane;  transport;  transporter activity; | 100 % |  | ``` 1	ATGACCGTCTACG-CTAACGACACTCCCTCTAACAAGTCCAATCCAGAGAAGAACGCCGCCAGCGTGGACCACCTCAATGAGACCAGCGTTCCCTACATCGTTCCAGATAAGCCCATCACACCTGAACTCCTCGCCGCAGTCCCGGGAGGCGTCAAAGCTTCCGAGCTCTTCGAGGATGCCGCTCCCATCCTCGCTCTTAACCTTCCGGATTGGCGTGCCACCGAGAAGAAGCTCG	235 	\|\|\|\|\|\|\|\|\|\|\|\|\| \|\|\|\|\|\|\|\|\|\|\|\|\|\|\|\|\|\|\|\|\|\|\|\|\|\|\|\|\|\|\|\|\|\|\|\|\|\|\|\|\|\|\|\|\|\|\|\|\|\|\|\|\|\|\|\|\|\|\|\|\|\|\|\|\|\|\|\|\|\|\|\|\|\|\|\|\|\|\|\|\|\|\|\|\|\|\|\|\|\|\|\|\|\|\|\|\|\|\|\|\|\|\|\|\|\|\|\|\|\|\|\|\|\|\|\|\|\|\|\|\|\|\|\|\|\|\|\|\|\|\|\|\|\|\|\|\|\|\|\|\|\|\|\|\|\|\|\|\|\|\|\|\|\|\|\|\|\|\|\|\|\|\|\|\|\|\|\|\|\|\|\|\|\|\|\|\|\|\|\|\|\|\|\|\|\|\|\|\|\|\|\|\|\|\|\|\|\|\|\|\|\|\|\|\|\|\|\|\|\|\|\|\|\|\|\|\|\|\|\|\|\| 	ATGACCGTCTACGCCTAACGACACTCCCTCTAACAAGTCCAATCCAGAGAAGAACGCCGCCAGCGTGGACCACCTCAATGAGACCAGCGTTCCCTACATCGTTCCAGATAAGCCCATCACACCTGAACTCCTCGCCGCAGTCCCGGGAGGCGTCAAAGCTTCCGAGCTCTTCGAGGATGCCGCTCCCATCCTCGCTCTTAACCTTCCGGATTGGCGTGCCACCGAGAAGAAGCTCG ```      ``` 235	GCACAAGCTCAACTCGCGGAACCGTCAATGGGAGAGTCTCTGAACTTCGAGTCAGGTGGA	294 	\|\|\|\|\|\|\|\|\|\|\|\|\|\|\|\|\|\|\|\|\|\|\|\|\|\|\|\|\|\|\|\|\|\|\|\|\|\|\|\|\|\|\|\|\|\|\|\| \|\|\|\|  \|\|\|\|\| 	GCACAAGCTCAACTCGCGGAACCGTCAATGGGAGAGTCTCTGAACTTCNAGTCNNGTGGA ``` |
| 1 | ggggacaagtttgtacaaaaaagcaggcttgatgGCATCGGTCGGACCAGCATT, ggggaccactttgtacaagaaagctgggtcGATTACCCTCTTCCTCTTCC | Mycgr3\|39111  JGI  Ensembl |  |  | 100 % | chr\_3:2128176 C->A; | ``` 1	ATGGCATCGGTCGGACCAGCATTGAGGAGAGACCCCTTCGCCAGCGATTCACGAGGCTTCGGCAGTCCGAAGGCGAAGAAGAACATTCCCATCCTCCCGTCGACATTGGTCGGAAGCGGTGGCGATCCGATGGAAGTGAGCCAACCTCAGGTCTCTCCGATGGGTCCGCCCACGAGTAGCCCGACAGCCGGAGAAAGAGCAGGAGAGCAGCAACAGGCACACAGCGAGACGCCGAGCACAAATGGAGGGTCACAACAACCTGTTGGAGCCGCTGCTGCTGCTCAACAGCCCAAGATCGTTCAGACAGCGTTCATACATAAGCTCTATAA	329 	\|\|\|\|\|\|\|\|\|\|\|\|\|\|\|\|\|\|\|\|\|\|\|\|\|\|\|\|\|\|\|\|\|\|\|\|\|\|\|\|\|\|\|\|\|\|\|\|\|\|\|\|\|\|\|\|\|\|\|\|\|\|\|\|\|\|\|\|\|\|\|\|\|\|\|\|\|\|\|\|\|\|\|\|\|\|\|\|\|\|\|\|\|\|\|\|\|\|\|\|\|\|\|\|\|\|\|\|\|\|\|\|\|\|\|\|\|\|\|\|\|\|\|\|\|\|\|\|\|\|\|\|\|\|\|\|\|\|\|\|\|\|\|\|\|\|\|\|\|\|\|\|\|\|\|\|\|\|\|\|\|\|\|\|\|\|\|\|\|\|\|\|\|\|\|\|\|\|\|\|\|\|\|\|\|\|\|\|\|\|\|\|\|\|\|\|\|\|\|\|\|\|\|\|\|\|\|\|\| \|\|\|\|\|\|\|\|\|\|\|\|\|\|\|\|\|\|\|\|\|\|\|\|\|\|\|\|\|\|\|\|\|\|\|\|\|\|\|\|\|\|\|\|\|\| \|\|\|\|\|\|\|\|\|\|\|\|\|\|\|\|\|\|\|\|\|\|\|\|\|\|\|\|\|\|\|\|\|\|\|\|\|\|\|\|\|\|\|\|  \|\|\| \|\|\|\|\|\|\|\|\|\|\|\|  \|\|\|\|\|\|\|\| 	ATGGCATCGGTCGGACCAGCATTGAGGAGAGACCCCTTCGCCAGCGATTCACGAGGCTTCGGCAGTCCGAAGGCGAAGAAGAACATTCCCATCCTCCCGTCGACATTGGTCGGAAGCGGTGGCGATCCGATGGAAGTGAGCCAACCTCAGGTCTCTCCGATGGGTCCGCCCACGAGTAGCCCGACAGCCGGAGAAAGAGCAGGAGAGCANCAACAGGCACACAGCGAGACGCCGAGCACAAATGGAGGGTCACAACNACCTGTTGGAGCCGCTGCTGCTGCTCAACAGCCCAAGATCGTTCNNACANCGTTCATACATANNCTCTATAA ``` |
| 1 | ggggacaagtttgtacaaaaaagcaggcttgatgGCGACAAACGGCCACACCGC, ggggaccactttgtacaagaaagctgggtcGTCCTGGGCCATGGCAGTAT | Mycgr3\|39277  JGI  Ensembl | KOG3334 Transcription initiation factor TFIID, subunit TAF9 (also component of histone acetyltransferase SAGA); | transcription factor TFIID complex; transcription initiation; | 0 % |  |  |
| 1 | ggggacaagtttgtacaaaaaagcaggcttgatgGCGACACCACTGATAGCATG, ggggaccactttgtacaagaaagctgggtcAGCAGGCTCATCACCCGCCG | Mycgr3\|40139  JGI  Ensembl | KOG2845 Activating signal cointegrator 1; | nucleus; regulation of transcription;  transcription regulator activity;  zinc ion binding; | 100 % |  | ``` 1	ATGGCGACACCACTGATAGCATGGGCACTTCCACGCCTTCAGCAAATTCTGCCACTGGACGATGAGTCGCTGAAGCAAGTCATCGCCTACGCCGAAACATTACCCAAAGAGGCTGCGGCAGAACATTTAAAGAACCTCCTCGGCGACGATGCGAAAAGTCTGGAATTCATTTCGAGCTTCAATCTACGGCGAGGAAATGTACCGGCGTCGACGACGAGTCTCGCATCGAATGGTGGGAATGCGGCGGCGTCATCGTCAGAAGCACCAGCACGCAAACCGAGACAGCCGAAGAAGAAACCTAATATTCATGCATTGCCCGCGAGGAGAGTGGAAGGGAGTG	340 	\|\|\|\|\|\|\|\|\|\|\|\|\|\|\|\|\|\|\|\|\|\|\|\|\|\|\|\|\|\|\|\|\|\|\|\|\|\|\|\|\|\|\|\|\|\|\|\|\|\|\|\|\|\|\|\|\|\|\|\|\|\|\|\|\|\|\|\|\|\|\|\|\|\|\|\|\|\|\|\|\|\|\|\|\|\|\|\|\|\|\|\|\|\|\|\|\|\|\|\|\|\|\|\|\|\|\|\|\|\|\|\|\|\|\|\|\|\|\|\|\|\|\|\|\|\|\|\|\|\|\|\|\|\|\|\|\|\|\|\|\|\|\|\|\|\|\|\|\|\|\|\|\|\|\|\|\|\|\|\|\|\|\|\|\|\|\|\|\|\|\|\|\|\|\|\|\|\|\|\|\|\|\|\|\|\|\|\|\|\|\|\|\|\|\|\|\|\|\|\|\|\|\|\|\|\|\|\|\|\|\|\|\|\|\|\|\|\|\|\|\|\|\|\|\|\|\|\|\|\|\|\|\|\|\|\|\|\|\|\|\|\|\|\|\|\|\|\|\|\|\|\|\|\|\|\|\|\|\|\|\|\|\|\|\|\|\|\|\|\|\|\|\|\|\|\|\|\|\|\|\|\|\|\|\|\|\|\|\|\|\|\|\|\|\|\|\|\|\|\|\|\|\|\|\|\|\|\|\|\|\|\|\|\|\|\|\|\|\|\|\|\|\|\|\| \|\|\|\|\|\|\|\|\|\|\|\|\|\| 	ATGGCGACACCACTGATAGCATGGGCACTTCCACGCCTTCAGCAAATTCTGCCACTGGACGATGAGTCGCTGAAGCAAGTCATCGCCTACGCCGAAACATTACCCAAAGAGGCTGCGGCAGAACATTTAAAGAACCTCCTCGGCGACGATGCGAAAAGTCTGGAATTCATTTCGAGCTTCAATCTACGGCGAGGAAATGTACCGGCGTCGACGACGAGTCTCGCATCGAATGGTGGGAATGCGGCGGCGTCATCGTCAGAAGCACCAGCACGCAAACCGAGACAGCCGAAGAAGAAACCTAATATTCATGCATTGCCCGCGAGGANAGTGGAAGGGAGTG ``` |
| 1 | ggggacaagtttgtacaaaaaagcaggcttgatgGCGAGCAGGAGACAACCTCT, ggggaccactttgtacaagaaagctgggtcGCGGATCTTCTTGTGAGGAC | Mycgr3\|41155  JGI  Ensembl | KOG4139 Protein kinase essential for the initiation of DNA replication; | nucleic acid binding; zinc ion binding; | 100 % |  |  |
| 1 | ggggacaagtttgtacaaaaaagcaggcttgatgAGTTCATCTGCCTATCGCGA, ggggaccactttgtacaagaaagctgggtcTGCAAGTGTGCGTGCTTTGC | Mycgr3\|42301  JGI  Ensembl |  |  | 100 % |  | ``` 1	ATGAGTTCATCTGCCTATCGCGAGATAGGCAAAGGACAGTGCGGCAGCGTCTGGGCCGCGGATGTGCCCGGTGAAGCCTGCGCACTGAAACGTCAACATGCCGGAGGTGATGGCGAACACAGCCGATCAGTGATCAAAGACTCCGTAATGCCCCAACGCACTGAGCGTAGTTTCAACGAGTGTCGGACCAGCCTCGAGCTTACGAATCTCAACATTCCGAGATATCATCAGCTTGTGCTCAAGGAAGACGCTCAATGGTGGGCTACTCGCAGAGGCAGATTCCCCAGTGATTTGGAAGCGTGTAATACGCTCATCAGTGAGCGAATACCACCTTTCGAGGGCCCTGTACGGATGAAGATCATCGAGTTATTCTGCCCGGACAAGATCAAGGATGTGGTCATCAACAGCCGCAAAGATGAAGATTGTATGGTGCGGCCTTACCTGGGACGACGAGCTCGCGACTCACGCCCGAGTCCCTTCTTTACGCTCAGGAACAAGGCCCTGACGATTGATCAAATGGAGAATCTCAAGCTCCCAGCTGTCGATTATGCGTGTGCCATGGCGGACGCGCTCGCATTCATGTTCTGGGAGGCC	594 	\|\|\|\|\|\|\|\|\|\|\|\|\|\|\|\|\|\|\|\|\|\|\|\|\|\|\|\|\|\|\|\|\|\|\|\|\|\|\|\|\|\|\|\|\|\|\|\|\|\|\|\|\|\|\|\|\|\|\|\|\|\|\|\|\|\|\|\|\|\|\|\|\|\|\|\|\|\|\|\|\|\|\|\|\|\|\|\|\|\|\|\|\|\|\|\|\|\|\|\|\|\|\|\|\|\|\|\|\|\|\|\|\|\|\|\|\|\|\|\|\|\|\|\|\|\|\|\|\|\|\|\|\|\|\|\|\|\|\|\|\|\|\|\|\|\|\|\|\|\|\|\|\|\|\|\|\|\|\|\|\|\|\|\|\|\|\|\|\|\|\|\|\|\|\|\|\|\|\|\|\|\|\|\|\|\|\|\|\|\|\|\|\|\|\|\|\|\|\|\|\|\|\|\|\|\|\|\|\|\|\|\|\|\|\|\|\|\|\|\|\|\|\|\|\|\|\|\|\|\|\|\|\|\|\|\|\|\|\|\|\|\|\|\|\|\|\|\|\|\|\|\|\|\|\|\|\|\|\|\|\|\|\|\|\|\|\|\|\|\|\|\|\|\|\|\|\|\|\|\|\|\|\|\|\|\|\|\|\|\|\|\|\|\|\|\|\|\|\|\|\|\|\|\|\|\|\|\|\|\|\|\|\|\|\|\|\|\|\|\|\|\|\|\|\|\|\|\|\|\|\|\|\|\|\|\|\|\|\|\|\|\|\|\|\|\|\|\|\|\|\|\|\|\|\|\|\|\|\|\|\|\|\|\|\|\|\|\|\|\|\|\|\|\|\|\|\|\|\|\|\|\|\|\|\|\|\|\|\|\|\|\|\|\|\|\|\|\|\|\|\|\|\|\|\|\|\|\|\|\|\|\|\|\|\|\|\|\|\|\|\|\|\|\|\|\|\|\|\|\|\|\|\|\|\|\|\|\|\|\|\|\|\|\|\|\|\|\|\|\|\| \|\|\|\|\|\|\|\|\|\|\|\|\|\|\|\|\|\|\|\|\|\|\|\|\|\|\|\|\|\|\|\|\| \|\|\|\| \|\|\|\|\|\|\|\|\|\|\|\|\|\|\| \|\|\|\|\|\|\| \|\|\|\|\|\| \|\|\|\|\|\|\|\|\|\|\|\|\|\|  \|\|\|\|\| \|\|\|\|\|\|\| \|\|\|\|\|\| \|\|\|  \|\|  \| \|\|\| \| \|\|\| \|\|\|\|\|\|\|\|\|\| \|\|\| 	ATGAGTTCATCTGCCTATCGCGAGATAGGCAAAGGACAGTGCGGCAGCGTCTGGGCCGCGGATGTGCCCGGTGAAGCCTGCGCACTGAAACGTCAACATGCCGGAGGTGATGGCGAACACAGCCGATCAGTGATCAAAGACTCCGTAATGCCCCAACGCACTGAGCGTAGTTTCAACGAGTGTCGGACCAGCCTCGAGCTTACGAATCTCAACATTCCGAGATATCATCAGCTTGTGCTCAAGGAAGACGCTCAATGGTGGGCTACTCGCAGAGGCAGATTCCCCAGTGATTTGGAAGCGTGTAATACGCTCATCAGTGAGCGAATACCACCTTTCGAGGGCCCTGTACGGATGAAGATCATCGAGTTATTCTGCCCGGACAAGATCAAGGATGTGGTCATCAACAGCCGCAAAGATGAAGATTGTATGGTGCGGCCTTACCTGGGACGACNAGCTCGCGACTCACGCCCGAGTCCCTTCTTTACNCTCANGAACAAGGCCCTGACNATTGATCNAATGGANAATCTCAAGCTCCCNNCTGTCNATTATGCNTGTGCCNTGGNNGANNCNCTCNCNTTCNTGTTCTGGGANGCC ``` |
| 1 | ggggacaagtttgtacaaaaaagcaggcttgatgGCAGACAGCCAAGCCAAACC, ggggaccactttgtacaagaaagctgggtcATGCGAAAAGCGAAAGTCCA | Mycgr3\|42896  JGI  Ensembl | KOG2533 Permease of the major facilitator superfamily; | DNA binding; | 100 % |  | ``` 1	ATGGCAGACAGCCAAGCCAAACCATCCTTCGCAGAGACGATTGCTCCCACTGCAGGGCTGAAAGGGAAAGACTTCGACTCGACTGAGGCTTCGCCCGAGGGCACTCCATCGATACGCTCCGAAGAGCTTGTCGTGGGCGGATATGGCTCGGATCATCATCATGTCTTCGCGAAGCCTGAAGTCGCAGAATACTGGCGTGGAGTATACGAAGCAGCACAGTACGAAGGGCGGCACAGATTCGACCCAGACGTCACATGGTCCGCCGAAGAGGAGAAGAAACTCCTGCGCAAG	291 	\|\|\|\|\|\|\|\|\|\|\|\|\|\|\|\|\|\|\|\|\|\|\|\|\|\|\|\|\|\|\|\|\|\|\|\|\|\|\|\|\|\|\|\|\|\|\|\|\|\|\|\|\|\|\|\|\|\|\|\|\|\|\|\|\|\|\|\|\|\|\|\|\|\|\|\|\|\|\|\|\|\|\|\|\|\|\|\|\|\|\|\|\|\|\|\|\|\|\|\|\|\|\|\|\|\|\|\|\|\|\|\|\|\|\|\|\|\|\|\|\|\|\|\|\|\|\|\|\|\|\|\|\|\|\|\|\|\|\|\|\|\|\|\|\|\|\|\|\|\|\|\|\|\|\|\|\|\|\|\|\|\|\|\|\|\|\|\|\|\|\|\|\|\|\|\|\|\|\|\|\|\|\|\|\|\|\|\|\|\|\|\|\|\|\|\|\|\|\|\|\|\|\|\|\|\|\|\|\|\|\|\|\|\|\|\|\|\|\|\|\|\|\|\|\|\|\|\|\|\|\|\|\|\|\|\|\|\|\|\|\|\|\|\|\|\|\|\|\|\|\|\|\|\|\|\|\|\|\|\|\|\|\|\|\|\|\|\|\|\|\|\|\|\|\|\|\|\|\|\|\|\|\|\|\|\|\|\|\|\|\| 	ATGGCAGACAGCCAAGCCAAACCATCCTTCGCAGAGACGATTGCTCCCACTGCAGGGCTGAAAGGGAAAGACTTCGACTCGACTGAGGCTTCGCCCGAGGGCACTCCATCGATACGCTCCGAAGAGCTTGTCGTGGGCGGATATGGCTCGGATCATCATCATGTCTTCGCGAAGCCTGAAGTCGCAGAATACTGGCGTGGAGTATACGAAGCAGCACAGTACGAAGGGCGGCACAGATTCGACCCAGACGTCACATGGTCCGCCGAAGAGGAGAAGAAACTCCTGCGCAAG ``` |
| 1 | ggggacaagtttgtacaaaaaagcaggcttgatgCCTCACGCACTCTCTCCCGA, ggggaccactttgtacaagaaagctgggtcAAACTCCATCCCATCCTCAT | Mycgr3\|44260  JGI  Ensembl | KOG2851 Eukaryotic-type DNA primase, catalytic (small) subunit; | CTP:2,3-di-O-geranylgeranyl-sn-glycero-1-phosphate cytidyltransferase; DNA primase activity;  DNA replication, synthesis of RNA primer;  DNA-directed DNA polymerase activity;  DNA/RNA helicase activity;  RNA guanylyltransferase activity;  RNA helicase activity;  mannose-phosphate guanylyltransferase activity;  phospholactate guanylyltransferase activity;  sulfate adenylyltransferase activity;  tRNA guanylyltransferase activity;  telomerase activity; | 100 % |  | ``` 1	ATGCCTCACGCACTCTCTCCCGAGCATGCTCCTGAGAGCAGTCCCGTGGACACGATACTCCCGGATGCTTCTGAAAGTCAAGATGTCGACATGTCCGACAGTCCGAACGACACAGCTCGGAGTGCCACGAAAGGACTAGAAGACATGTTTGACGATGAAGACCCTAACGATCCGTATTCTTCAAGCGCAGCGACTCGCGAGTCCAACAGCTCTCAGGGCAAGCCAGATTCCGCACCCAAAGGCAACTTCTCCGAGCCCGAAACACTACGACAATTCTATCAGCGACTGTGTCCTTTTCGACACCTCTTTCAATGGCTCAATCACTCCGCGACACCGCAGCCCGACTTCCAGAACCGCGAATTCTCCTTCTGGCTCCCTGGCAGCACAGTGATTCGATACCTCTCGTTCCCATCTG	415 	\|\|\|\|\|\|\|\|\|\|\|\|\|\|\|\|\|\|\|\|\|\|\|\|\|\|\|\|\|\|\|\|\|\|\|\|\|\|\|\|\|\|\|\|\|\|\|\|\|\|\|\|\|\|\|\|\|\|\|\|\|\|\|\|\|\|\|\|\|\|\|\|\|\|\|\|\|\|\|\|\|\|\|\|\|\|\|\|\|\|\|\|\|\|\|\|\|\|\|\|\|\|\|\|\|\|\|\|\|\|\|\|\|\|\|\|\|\|\|\|\|\|\|\|\|\|\|\|\|\|\|\|\|\|\|\|\|\|\|\|\|\|\|\|\|\|\|\|\|\|\|\|\|\|\|\|\|\|\|\|\|\|\|\|\|\|\|\|\|\|\|\|\|\|\|\|\|\|\|\|\|\|\|\|\|\|\|\|\|\|\|\|\|\|\|\|\|\|\|\|\|\|\|\|\|\|\|\|\|\|\|\|\|\|\|\|\|\|\|\|\|\|\|\|\|\|\|\|\|\|\|\|\|\|\|\|\|\|\|\|\|\|\|\|\|\|\|\|\|\|\|\|\|\|\|\|\|\|\|\|\|\|\|\|\|\|\|\|\|\|\|\|\|\|\|\|\|\|\|\|\|\|\|\|\|\|\|\|\|\|\|\|\|\|\|\|\|\|\|\|\|\|\|\|\|\|\|\|\|\|\|\|\|\|\|\|\|\|\|\|\|\|\|\|\|\|\|\|\|\|\|\|\|\|\|\|\|\| \|\|\|\|\|\|\|\|\|\|\|\|\|\|\|\|\|\|\|\|\|\|\|\|\|\|\|\|\|\|\|\|\|\|\|\|\|\|\|\|\|   \|   \|\|\|\|\|\|\| \| \|\|\|\|\|\|\|\| \|\|\|\|\|\|\|\|\| 	ATGCCTCACGCACTCTCTCCCGAGCATGCTCCTGAGAGCAGTCCCGTGGACACGATACTCCCGGATGCTTCTGAAAGTCAAGATGTCGACATGTCCGACAGTCCGAACGACACAGCTCGGAGTGCCACGAAAGGACTAGAAGACATGTTTGACGATGAAGACCCTAACGATCCGTATTCTTCAAGCGCAGCGACTCGCGAGTCCAACAGCTCTCAGGGCAAGCCAGATTCCGCACCCAAAGGCAACTTCTCCGAGCCCGAAACACTACGACAATTCTATCAGCGACTGTGTCCTTTTCGACACCTCTTTCAATGGCTCAATCACTCCGCGACACCGCANCCCGACTTCCAGAACCGCGAATTCTCCTTCTGGCTCCCTGGNNNCNNNGTGATTCNANACCTCTCGNTCCCATCTG ``` |
| 1 | ggggacaagtttgtacaaaaaagcaggcttgatgTCTCCACCAGCTGCCATGTT, ggggaccactttgtacaagaaagctgggtcAAAAGCATTCTGCTTCTTCC | Mycgr3\|44262  JGI  Ensembl | KOG2960 Protein involved in thiamine biosynthesis and DNA damage tolerance; | thiamin biosynthetic process; | 100 % |  | ``` 1	ATGTCTCCACCAGCTGCCATGTTCCAGGAGCCCGCCTCCATCCCAGTGATGCCCACCAAAGGCACCAAGCCGACAGCCACCATCGAACAGATGGCCAACAACTGGTCCTCGTTCAAGTTCGCTCCCATCCGCGAGTCTCAGGTGTCCCGGGCCATGACTCGCCGCTACTTCAGCGACCTGGACACCTACGCCGAGTCCGACATTGTCATCGTCGGTGCTGGAAGCTGCGGTCTCTCCGCCGCCTACTGTCTCGCCAAAGCCCGACCAGACCTCAAGATCGCCATCATCGAAGCCGGCGTGGCTCCGGGCGGAGGTGCCTGGTTGGGAGGCCAGCTGTTCAGCGCCATGGTCATGAGGAAGCCTGCCGATGCTTTCTTGCGAGAGATCGGTGTACCGTACGAGGACGAAGGTCCCGACTCGAACTTTGTGGTCGTCAAGCACGCTGCTCTGTTCACCAGCACTGTGCTATCCAAAGTGCTGCAATTCGACAACGTCAAGCTCTTCAACGCCACCACGGTCGAGGACCTCATCACCCGTGCCGATGACCAGGGAAAGCTTCGCATCGCTGGCGTGGTCACCAACTGGACTCTCGTCAGCATGCACCA	605 	\|\|\|\|\|\|\|\|\|\|\|\|\|\|\|\|\|\|\|\|\|\|\|\|\|\|\|\|\|\|\|\|\|\|\|\|\|\|\|\|\|\|\|\|\|\|\|\|\|\|\|\|\|\|\|\|\|\|\|\|\|\|\|\|\|\|\|\|\|\|\|\|\|\|\|\|\|\|\|\|\|\|\|\|\|\|\|\|\|\|\|\|\|\|\|\|\|\|\|\|\|\|\|\|\|\|\|\|\|\|\|\|\|\|\|\|\|\|\|\|\|\|\|\|\|\|\|\|\|\|\|\|\|\|\|\|\|\|\|\|\|\|\|\|\|\|\|\|\|\|\|\|\|\|\|\|\|\|\|\|\|\|\|\|\|\|\|\|\|\|\|\|\|\|\|\|\|\|\|\|\|\|\|\|\|\|\|\|\|\|\|\|\|\|\|\|\|\|\|\|\|\|\|\|\|\|\|\|\|\|\|\|\|\|\|\|\|\|\|\|\|\|\|\|\|\|\|\|\|\|\|\|\|\|\|\|\|\|\|\|\|\|\|\|\|\|\|\|\|\|\|\|\|\|\|\|\|\|\|\|\|\|\|\|\|\|\|\|\|\|\|\|\|\|\|\|\|\|\|\|\|\|\|\|\|\|\|\|\|\|\|\|\|\|\|\|\|\|\|\|\|\|\|\|\|\|\|\|\|\|\|\|\|\|\|\|\|\|\|\|\|\|\|\|\|\|\|\|\|\|\|\|\|\|\|\|\|\|\|\|\|\|\|\|\|\|\|\|\|\|\|\|\|\|\|\|\|\|\|\|\|\|\|\|\|\|\|\|\|\|\|\|\|\|\|\|\|\|\|\|\|\|\|\|\|\|\|\|\|\|\|\|\|\|\|\|\|\|\|\|\|\|\|\|\|\|\|\|\|\|\|\|\|\|\|\|\|\|\|\|\|\|\|\|\|\|\|\|\|\|\|\|\|\|\|\|\|\|\|\|\|\|\|\|\|\|\|\|\|\|\|\|\|\|\|\|\|\|\|\|\|\|\|\|\|\|\|\|\|\|\|\|\|\|\|\|\|\|\|\|\|\|\|\|\|\|\|\|\|\|\|\|\|\|\|\|\|\|\|\|\|\|\|\|\|\|\|\|\|\|\|\|\|\|\|\|\|\|\|\|\|\|\|\|\|\|\|\|\|\|\|\|\|\|\|\|\|\|\|\|\|\|\|\|\|\|\|\| \|\|\|\|\|\|\|\|\|\| \|\|\|\|\|\|\|\|\|\|\|\|\|\|\|\|\| \|\|\|\|\|\|\|\|\|\|\|\|\|\|\|\|\| \|\|\|\|\|\|\|\|\| 	ATGTCTCCACCAGCTGCCATGTTCCAGGAGCCCGCCTCCATCCCAGTGATGCCCACCAAAGGCACCAAGCCGACAGCCACCATCGAACAGATGGCCAACAACTGGTCCTCGTTCAAGTTCGCTCCCATCCGCGAGTCTCAGGTGTCCCGGGCCATGACTCGCCGCTACTTCAGCGACCTGGACACCTACGCCGAGTCCGACATTGTCATCGTCGGTGCTGGAAGCTGCGGTCTCTCCGCCGCCTACTGTCTCGCCAAAGCCCGACCAGACCTCAAGATCGCCATCATCGAAGCCGGCGTGGCTCCGGGCGGAGGTGCCTGGTTGGGAGGCCAGCTGTTCAGCGCCATGGTCATGAGGAAGCCTGCCGATGCTTTCTTGCGAGAGATCGGTGTACCGTACGAGGACGAAGGTCCCGACTCGAACTTTGTGGTCGTCAAGCACGCTGCTCTGTTCACCAGCACTGTGCTATCCAAAGTGCTGCAATTCGACAACGTCAAGCTCTTCAACGCCACCACGGTCGAGGACCTCATCACCCGTGCCGATGACCANGGAAAGCTTCNCATCGCTGGCGTGGTCA-CAACTGGACTCTCGTCANCATGCACCA ``` |
| 1 | ggggacaagtttgtacaaaaaagcaggcttgatgTCCGCTGATGAGGTGCAGGT, ggggaccactttgtacaagaaagctgggtcTGCTAGGAACTCACCGTCCA | Mycgr3\|44516  JGI  Ensembl |  |  | 100 % |  | ``` 1	ATGTCCGCTGATGAGGTGCAGGTCAAGTGCGGCGAGGAAAAGCCGGAATGCTTGAACTGTCAGAGAAATGGCAGCTCTGAGGCGTGCGATTACAGCATCAGGTTGAATTGGGGTGGACGGAGCAAGAAAGACAATGATGGAGCACTGGGCGAGGCAGGCGGCTTCACATTTGTCTCTACACCGGGTATCTCGACTGGACACAAAGGCGCCCGGTCTGTCCACGAGCATG	229 	\|\|\|\|\|\|\|\|\|\|\|\|\|\|\|\|\|\|\|\|\|\|\|\|\|\|\|\|\|\|\|\|\|\|\|\|\|\|\|\|\|\|\|\|\|\|\|\|\|\|\|\|\|\|\|\|\|\|\|\|\|\|\|\|\|\|\|\|\|\|\|\|\|\|\|\|\|\|\|\|\|\|\|\|\|\|\|\|\|\|\|\|\|\|\|\|\|\|\|\|\|\|\|\|\|\|\|\|\|\|\|\|\|\|\|\|\|\|\|\|\|\|\|\|\|\|\|\|\|\|\|\|\|\|\|\|\|\|\|\|\|\|\|\|\|\|\|\|\|\|\|\|\|\|\|\|\|\|\|\|\|\|\|\|\|\|\|\|\|\|\|\|\|\|\|\|\|\|\|\|\|\|\|\|\|\|\|\|\|\|\|\|\|\|\|\|\|\|\|\|\|\|\|\|\|\|\|\|\|\|\|\|\|\|\|\|\|\|\|\|\|\|\|\|\|\|\|\|\| 	ATGTCCGCTGATGAGGTGCAGGTCAAGTGCGGCGAGGAAAAGCCGGAATGCTTGAACTGTCAGAGAAATGGCAGCTCTGAGGCGTGCGATTACAGCATCAGGTTGAATTGGGGTGGACGGAGCAAGAAAGACAATGATGGAGCACTGGGCGAGGCAGGCGGCTTCACATTTGTCTCTACACCGGGTATCTCGACTGGACACAAAGGCGCCCGGTCTGTCCACGAGCATG ``` |
| 1 | ggggacaagtttgtacaaaaaagcaggcttgatgGCCATAACGAATGGAGCGAG, ggggaccactttgtacaagaaagctgggtcTCCACTCTCGAAGATCTCGA | Mycgr3\|45479  JGI  Ensembl | KOG2732 DNA polymerase delta, regulatory subunit 55; | DNA polymerase activity; DNA-directed DNA polymerase activity; | 0 % |  |  |
| 1 | ggggacaagtttgtacaaaaaagcaggcttgatgGCAGACCTCAAATCCTTTGA, ggggaccactttgtacaagaaagctgggtcGTCGAAAAGAGCAATGAGCG | Mycgr3\|47390  JGI  Ensembl | KOG1620 Inositol polyphosphate multikinase, component of the ARGR transcription regulatory complex; | inositol polyphosphate multikinase activity; inositol trisphosphate 3-kinase activity; | 100 % |  | ``` 1	ATGGCAGACCTCAAATCCTTTGATCAAACGAAGCTCAAGGCATTCGAGAACGCGGCTGCTGGACACGATGGCGTACTTCTGGACGAATCAGGTGAACTCGTCATCAAACCATGCACAGCGGAAGAAGTCGACTTCTACCAAGAAAGCATCTCACAACACCCCAAATTCTACGAGTTGATGCCAACTTTCATGGGCACCCTCACTCTCGGAGCACCAGCCCAGGTCCTAGACACCACCGCCGCACAGGAGGAACTCACACAAGAGCAAAAAGACCAAACTCTACTCCACGGCAAAGCCTTGACCACCGAGACCGCCATAGTACTGGAGAATCTCGAGCATGGCTTCAAGCAACCCAATGTCCTGGATCTCAAACTCGGCTCCCGTTTGTGGGCAGAAGGAACCAAGCCTGAGAAAGCAGATCGACTCGACAAGGTCTCTGCCGAAACCACAAGTGGCTCTCTCAACTTCCGCATCGCTGGTATGAAAGTCTGGAATGGCCAAGGCTTTGACATCTACGACAAGTTCTATGGACGCAAGTTTAGCAAAGACAATGTCAAGGATGGCTTCGCGGCATTCTTCAGCAGTCTTTCAGCCGGTGTGCCGAAAGATGATGCCCGTTTGCTGCTC	627 	\|\|\|\|\|\|\|\|\|\|\|\|\|\|\|\|\|\|\|\|\|\|\|\|\|\|\|\|\|\|\|\|\|\|\|\|\|\|\|\|\|\|\|\|\|\|\|\|\|\|\|\|\|\|\|\|\|\|\|\|\|\|\|\|\|\|\|\|\|\|\|\|\|\|\|\|\|\|\|\|\|\|\|\|\|\|\|\|\|\|\|\|\|\|\|\|\|\|\|\|\|\|\|\|\|\|\|\|\|\|\|\|\|\|\|\|\|\|\|\|\|\|\|\|\|\|\|\|\|\|\|\|\|\|\|\|\|\|\|\|\|\|\|\|\|\|\|\|\|\|\|\|\|\|\|\|\|\|\|\|\|\|\|\|\|\|\|\|\|\|\|\|\|\|\|\|\|\|\|\|\|\|\|\|\|\|\|\|\|\|\|\|\|\|\|\|\|\|\|\|\|\|\|\|\|\|\|\|\|\|\|\|\|\|\|\|\|\|\|\|\|\|\|\|\|\|\|\|\|\|\|\|\|\|\|\|\|\|\|\|\|\|\|\|\|\|\|\|\|\|\|\|\|\|\|\|\|\|\|\|\|\|\|\|\|\|\|\|\|\|\|\|\|\|\|\|\|\|\|\|\|\|\|\|\|\|\|\|\|\|\|\|\|\|\|\|\|\|\|\|\|\|\|\|\|\|\|\|\|\|\|\|\|\|\|\|\|\|\|\|\|\|\|\|\|\|\|\|\|\|\|\|\|\|\|\|\|\|\|\|\|\|\|\|\|\|\|\|\|\|\|\|\|\|\|\|\|\|\|\|\|\|\|\|\|\|\|\|\|\|\|\|\|\|\|\|\|\|\|\|\|\|\|\|\|\|\|\|\|\|\|\|\|\|\|\|\|\|\|\|\|\|\|\|\|\|\|\|\|\|\|\|\|\|\|\|\|\|\|\|\|\|\|\|\|\|\|\|\|\|\|\|\|\|\|\|\|\|\|\|\|\|\|\|\|\|\|\|\|\|\|\|\|\|\|\|\|\|\|\|\|\|\|\|\|\|\|\|\|\|\|\|\|\|\|\|\|\|\|\|\|\|\|\|\|\|\|\|\|\|\|\|\|\|\|\|\|\|\|\|\|\|\|\|\|\|\|\|\|\|\|\|\|\|\|\|\|\|\|\|\|\|\|\|\|\|\|\|\|\|\|\|\|\|\|\|\|\|\|\|\|\|\|\|\|\|\|\|\|\|\|\|\|\|\|\|\|\|\|\|\|\|\|\|\|\|\| \|\|\|\|\|\|\|\|\|\|\|  \|\| \|\|\|\|\|\|  \|\|\|\| \|\|\|\|\| \|\|\| \|\|\|\|\|\|\|\|\|\|\|\|\|\|\|\|\|\|\|\| 	ATGGCAGACCTCAAATCCTTTGATCAAACGAAGCTCAAGGCATTCGAGAACGCGGCTGCTGGACACGATGGCGTACTTCTGGACGAATCAGGTGAACTCGTCATCAAACCATGCACAGCGGAAGAAGTCGACTTCTACCAAGAAAGCATCTCACAACACCCCAAATTCTACGAGTTGATGCCAACTTTCATGGGCACCCTCACTCTCGGAGCACCAGCCCAGGTCCTAGACACCACCGCCGCACAGGAGGAACTCACACAAGAGCAAAAAGACCAAACTCTACTCCACGGCAAAGCCTTGACCACCGAGACCGCCATAGTACTGGAGAATCTCGAGCATGGCTTCAAGCAACCCAATGTCCTGGATCTCAAACTCGGCTCCCGTTTGTGGGCAGAAGGAACCAAGCCTGAGAAAGCAGATCGACTCGACAAGGTCTCTGCCGAAACCACAAGTGGCTCTCTCAACTTCCGCATCGCTGGTATGAAAGTCTGGAATGGCCAAGGCTTTGACATCTACGACAAGTTCTATGGACGCAAGTTTAGCAAAGACAATGTCAAGGATGGCTTCNCGGCATTCTTCN-CANTCTTTCNNCCGGNGTGCCNAAANATGATGCCCGTTTGCTGCTC ``` |
| 1 | ggggacaagtttgtacaaaaaagcaggcttgatgCCTCCCAAACGCCGCTCCAA, ggggaccactttgtacaagaaagctgggtcTGTTTCGATAAAGCGAGAGC | Mycgr3\|47547  JGI  Ensembl |  |  | 100 % |  | ``` 277	GAGTCTGAAGCGAGGAAGGTTTCTGCGACGCAGATCAACAAATACTGGCGAGCAAAGGAGGCGGAGCGAAAGGCGCCGCGGGTACATCAGCAGGACTTGAGCACGCATGAGAAGATTCTGAGGGAGTGGGATATGAGTGGGAACTATGGG	426 	\|\|\|\|\|\|\|\|\|\|\|\|\|\|\|\|\|\|\|\|\|\|\|\|\|\|\|\|\|\|\|\|\|\|\|\|\|\|\|\|\|\|\|\|\|\|\|\|\|\|\|\|\|\|\|\|\|\|\|\|\|\|\|\|\|\|\|\|\|\|\|\|\|\|\|\|\|\|\|\|\|\|\|\|\|\|\|\|\|\|\|\|\|\|\|\|\|\|\|\|\|\|\|\|\|\|\|\|\|\|\|\|\|\|\|\|\|\|\|\|\|\|\|\|\|\|\|\|\|\|\|\|\|\|\|\|\|\|\|\|\|\|\|\|\|\|\|\|\|\| 	GAGTCTGAAGCGAGGAAGGTTTCTGCGACGCAGATCAACAAATACTGGCGAGCAAAGGAGGCGGAGCGAAAGGCGCCGCGGGTACATCAGCAGGACTTGAGCACGCATGAGAAGATTCTGAGGGAGTGGGATATGAGTGGGAACTATGGG ```      ``` 1	ATGCCTCCCAAACGCCGCTCCAACACACCGCGCACGACACAGCGTCAACAGGGAACACTCTCCTTCCACGGCAAGAGTAAAGTCACAAAACCGACCTCTCCTCGCGGCGCAAAGTCAAAGAAAGATCCCGAGCAGTACGAGCAACTACTGGAGAAAGATATCAAGCCCGAGGTCGAAGAACCCACCACCGCAGAGAAAGCAATCACAGAACAAGTCAAAGCTGAAGTCGCAACACTCGAAGATCCTCTCGATACCACAATCAGCAAGACCGAAGATG	277 	\|\|\|\|\|\|\|\|\|\|\|\|\|\|\|\|\|\|\|\|\|\|\|\|\|\|\|\|\|\|\|\|\|\|\|\|\|\|\|\|\|\|\|\|\|\|\|\|\|\|\|\|\|\|\|\|\|\|\|\|\|\|\|\|\|\|\|\|\|\|\|\|\|\|\|\|\|\|\|\|\|\|\|\|\|\|\|\|\|\|\|\|\|\|\|\|\|\|\|\|\|\|\|\|\|\|\|\|\|\|\|\|\|\|\|\|\|\|\|\|\|\|\|\|\|\|\|\|\|\|\|\|\|\|\|\|\|\|\|\|\|\|\|\|\|\|\|\|\|\|\|\|\|\|\|\|\|\|\|\|\|\|\|\|\|\|\|\|\|\|\|\|\|\|\|\|\|\|\|\|\|\|\|\|\|\|\|\|\|\|\|\|\|\|\|\|\|\|\|\|\|\|\|\|\|\|\|\|\|\|\|\|\|\|\|\|\|\|\|\|\|\|\|\|\|\|\|\|\|\|\|\|\|\|\|\|\|\|\|\|\|\|\|\|\|\|\|\|\|\|\|\|\|\|\|\|\|\|\|\|\|\|\|\|\|\|\|\|\|\|\|\|\|\|\|\|\| 	ATGCCTCCCAAACGCCGCTCCAACACACCGCGCACGACACAGCGTCAACAGGGAACACTCTCCTTCCACGGCAAGAGTAAAGTCACAAAACCGACCTCTCCTCGCGGCGCAAAGTCAAAGAAAGATCCCGAGCAGTACGAGCAACTACTGGAGAAAGATATCAAGCCCGAGGTCGAAGAACCCACCACCGCAGAGAAAGCAATCACAGAACAAGTCAAAGCTGAAGTCGCAACACTCGAAGATCCTCTCGATACCACAATCAGCAAGACCGAAGATG ``` |
| 1 | ggggacaagtttgtacaaaaaagcaggcttgatgTCCGACGCACAGCTCTACGA, ggggaccactttgtacaagaaagctgggtcCTTCTTGAGCAGCATGTAGA | Mycgr3\|47713  JGI  Ensembl | KOG3400 RNA polymerase subunit 8; | DNA-directed RNA polymerase I activity; DNA-directed RNA polymerase II activity;  DNA-directed RNA polymerase III activity;  DNA-directed RNA polymerase activity;  RNA polymerase activity;  transcription; | 0 % |  | ``` 3	GTCCGACGCACAGCTCTACGAAGAGGCCTTCAACGTCAACACCCTCCTCGACCAAGCCTACGACCGCGTCCATCGAGTCATCGGCACAAGCCAGGACTCGTCCACATCCTACACGCTCGACATCAATTGCGAACTCTACCCACTCGTCGCAGGAGAGACATTCTCCATGCTGCTGGCTTCGACGTTGAACTTGGATGGTACGAAAGACGAGGGTCAGGGATGGAGAGGGAAGTCGACGGGCTCGACACTGGCGGATTTGTGGGATTATGTCTGCTATGGGAAAGTGTACAAGGTGGAAGATCCGGAGGATGGGGATAAC	321 	\|\|\|\|\|\|\|\|\|\|\| \| \|\|\|\|\|\|\|\|\|\|\|\|\|\|\|\|\|\|\|\|\|\|\|\|\|\|\|\|\|\|\|\|\|\|\|\|\|\|\|\|\|\|\|\|\|\|\|\|\|\|\|\|\|\|\|\|\|\|\|\|\|\|\|\|\|\|\|\|\|\|\|\|\|\|\|\|\|\|\|\|\|\|\|\|\|\|\|\|\|\|\|\|\|\|\|\|\|\|\|\|\|\|\|\|\|\|\|\|\|\|\|\|\|\|\|\|\|\|\|\|\|\|\|\|\|\|\|\|\|\|\|\|\|\|\|\|\|\|\|\|\|\|\|\|\|\|\|\|\|\|\|\|\|\|\|\|\|\|\|\|\|\|\|\|\|\|\|\|\|\|\|\|\|\|\|\|\|\|\|\|\|\|\|\|\|\|\|\|\|\|\|\|\|\|\|\|\|\|\|\|\|\|\|\|\|\|\|\|\|\|\|\|\|\|\|\|\|\|\|\|\|\|\|\|\|\|\|\|\|\|\|\|\|\|\|\|\|\|\|\|\|\|\|\|\|\|\|\|\|\|\|\|\|\|\|\|\|\|\|\|\|\|\|\|\|\|\|\|\|\|\|\|\|\|\|\|\|\|\|\|\|\|\|\|\|\|\|\|\|\|\|\|\|\|\|\|\|\|\|\|\|\|\|\|\| 	GTCCGACGCACNGNTCTACGAAGAGGCCTTCAACGTCAACACCCTCCTCGACCAAGCCTACGACCGCGTCCATCGAGTCATCGGCACAAGCCAGGACTCGTCCACATCCTACACGCTCGACATCAATTGCGAACTCTACCCACTCGTCGCAGGAGAGACATTCTCCATGCTGCTGGCTTCGACGTTGAACTTGGATGGTACGAAAGACGAGGGTCAGGGATGGAGAGGGAAGTCGACGGGCTCGACACTGGCGGATTTGTGGGATTATGTCTGCTATGGGAAAGTGTACAAGGTGGAAGATCCGGAGGATGGGGATAAC ``` |
| 1 | ggggacaagtttgtacaaaaaagcaggcttgatgCCTCCAAGAGCAGCTGCAAA, ggggaccactttgtacaagaaagctgggtcCATGACGAGCACATTCGCCA | Mycgr3\|51169  JGI  Ensembl |  |  | 0 % |  |  |
| 1 | ggggacaagtttgtacaaaaaagcaggcttgatgGCGGGCAACATCAATCACAC, ggggaccactttgtacaagaaagctgggtcCAACGTATACCGAAACTGCT | Mycgr3\|51346  JGI  Ensembl | KOG2533 Permease of the major facilitator superfamily; | DNA binding; integral to membrane;  transport;  transporter activity; | 0 % |  | ``` 1	ATGGCGGGCAACATCAATCACACAACGACAGGTTCGGAAATTCATCACCATGAGAAAGCCAACCTCCAACAAATCCCGACCAATACGAGCGACGTCAACGACCTCGAGAAAACCGTCCAAGTCGACACGCTACACACCGACGAAGCGATGAAAGTCCTCGCAGTCTACAACGGTCCTCTCGAATGGACACCTCAAGAGGAGAAAAAGCTGCTTCGGAAGGTGGATTTCAAGTTGTTGAGCATCCTTGTCCTGACATATGCATTGCAATATTACGATAAAGCGATGTTATCGCAAGCGG	298 	\|\|\|\|\|\|\|\|\|\|\|\|\|\|\|\|\|\|\|\|\|\|\|\|\|\|\|\|\|\|\|\|\|\|\|\|\|\|\|\|\|\|\|\|\|\|\|\|\|\|\|\|\|\|\|\|\|\|\|\|\|\|\|\|\|\|\|\|\|\|\|\|\|\|\|\|\|\|\|\|\|\|\|\|\|\|\|\|\|\|\|\|\|\|\|\|\|\|\|\|\|\|\|\|\|\|\|\|\|\|\|\|\|\|\|\|\|\|\|\|\|\|\|\|\|\|\|\|\|\|\|\|\|\|\|\|\|\|\|\|\|\|\|\|\|\|\|\|\|\|\|\|\|\|\|\|\|\|\|\|\|\|\|\|\|\|\|\|\|\|\|\|\|\|\|\|\|\|\|\|\|\|\|\|\|\|\|\|\|\|\|\|\|\|\|\|\|\|\|\|\|\|\|\|\|\|\|\|\|\|\|\|\|\|\|\|\|\|\|\|\|\|\|\|\|\|\|\|\|\|\|\|\|\|\|\|\|\|\|\|\|\|\|\|\|\|\|\|\|\|\|\|\|\|\|\|\|\|\|\|\|\|\|\|\|\|\|\|\|\|\|\|\|\|\|\|\|\|\|\|\|\|\|\|\|\|\|\|\|\|\|\|\|\|\|\|\|\| 	ATGGCGGGCAACATCAATCACACAACGACAGGTTCGGAAATTCATCACCATGAGAAAGCCAACCTCCAACAAATCCCGACCAATACGAGCGACGTCAACGACCTCGAGAAAACCGTCCAAGTCGACACGCTACACACCGACGAAGCGATGAAAGTCCTCGCAGTCTACAACGGTCCTCTCGAATGGACACCTCAAGAGGAGAAAAAGCTGCTTCGGAAGGTGGATTTCAAGTTGTTGAGCATCCTTGTCCTGACATATGCATTGCAATATTACGATAAAGCGATGTTATCGCAAGCGG ```      ``` 297	GGCCCTCTTCGGCCTTCGAGACGACCTCAAACTGCGAACGGGAAACCGCTACTCCATGTCCTCCGCCATCTTCTACCTAGGCTTCATCTGCGGCGCCTATCCCGCCATCGTCCTCTCCCAAAAATATCCCATCGAGCGAGTCATGTTCGGCATCGTGATAGCCTGGGGAGCATG	470 	\|\|\|\|\|\|\|\|\|\|\|\|\|\|\|\|\|\|\|\|\|\|\|\|\|\|\|\|\|\|\|\|\|\|\|\|\|\|\|\|\|\|\|\|\|\|\|\|\|\|\|\|\|\|\|\|\|\|\|\|\|\|\|\|\|\|\|\|\|\|\|\|\|\|\|\|\|\|\|\|\|\|\|\|\|\|\|\|\|\|\|\|\|\|\|\|\|\|\|\|\|\|\|\|\|\|\|\|\|\|\|\|\|\|\|\|\|\|\|\|\|\|\|\|\|\|\|\|\|\|\|\|\|\|\|\|\|\|\|\|\|\|\|\|\|\|\|\|\|\|\|\|\|\|\|\|\|\|\|\|\|\|\|\|\|\|\|\|\|\|\|\|\|\| 	GGCCCTCTTCGGCCTTCGAGACGACCTCAAACTGCGAACGGGAAACCGCTACTCCATGTCCTCCGCCATCTTCTACCTAGGCTTCATCTGCGGCGCCTATCCCGCCATCGTCCTCTCCCAAAAATATCCCATCGAGCGAGTCATGTTCGGCATCGTGATAGCCTGGGGAGCATG ``` |
| 1 | ggggacaagtttgtacaaaaaagcaggcttgatgTCTGCCGCCGCGAAAGAGGA, ggggaccactttgtacaagaaagctgggtcAACCTTCAACCACTGCTCAA | Mycgr3\|64857  JGI  Ensembl | KOG3233 RNA polymerase III, subunit C34; | DNA binding; DNA-directed RNA polymerase I activity;  DNA-directed RNA polymerase II activity;  DNA-directed RNA polymerase III activity;  DNA-directed RNA polymerase activity;  RNA polymerase activity;  nucleus;  transcription; | 100 % | chr\_1:123793 T->C; | ``` 147	ATGTCTGCCGCCGCGAAAGAGGAGGACGAGGAAAAAGTGTCCGTCCCCAATGCCAGCGACGCTCTCTATGTCATCATTGCCGCAGCCTCGGTAGACGAAAACGGCGCACGCAGCAAAGTCTTCCAACAAGATGAGCTCCTCCAGCTGGGCAGCATCGCAAACACGCGAGAGCTCATGCCACTGGTACAATTCCTCGCCCAAAAAGGACTCTTCCGCACAGTAAAAGTCGGCGGAAAGCTAGGATGGAGCGTGCGACCACGAGAAGCAGCACGACAAATCGTCAACCTCGACCGAGATGAGAAGACAATCTACGAGATCATCGAAGAAGCACATACCACTGGAATCTGGACACGCGATCTGAAGAAGAAGACGAGCATCGCCCAAAATGTCGTACAGAAGGCGCTGTCGAAGATGGAAAAGGGCAACCTCATCAAATCCGTCAAAAGCGTCAAAGCACCCGCCCAGCGAACATACATGCTCAGCCACCTCTCTCCTAGCGAAGACGTAACAGGAAACAGCTTCTTCGACGCCGGCGATCTCGATGAATCCTTTCGCGACGAACTCCTC	713 	\|\|\|\|\|\|\|\|\|\|\|\|\|\|\|\|\|\|\|\|\|\|\|\|\|\|\|\|\|\|\|\|\|\|\|\|\|\|\|\|\|\|\|\|\|\|\|\|\|\|\|\|\|\|\|\|\|\|\|\|\|\|\|\|\|\|\|\|\|\|\|\|\|\|\|\|\|\|\|\|\|\|\|\|\|\|\|\|\|\|\|\|\|\|\|\|\|\|\|\|\|\|\|\|\|\|\|\|\|\|\|\|\|\|\|\|\|\|\|\|\|\|\|\|\|\|\|\|\|\|\|\|\|\|\|\|\|\|\|\|\|\|\|\|\|\|\|\|\|\|\|\|\|\|\|\|\|\|\|\|\|\|\|\|\|\|\|\|\|\|\|\|\|\|\|\|\|\|\|\|\|\|\|\|\|\|\|\|\|\|\|\|\|\|\|\|\|\|\|\|\|\|\|\|\|\|\|\|\|\|\|\|\|\|\|\|\|\|\|\|\|\|\|\|\|\|\|\|\|\|\|\|\|\|\|\|\|\|\|\|\|\|\|\|\|\|\|\|\|\|\|\|\|\|\|\|\|\|\|\|\|\|\|\|\|\|\|\|\|\|\|\|\|\|\|\|\|\|\|\|\|\|\|\|\|\|\|\|\|\|\|\|\|\|\|\|\|\|\|\|\|\|\|\|\|\|\|\|\|\|\|\|\|\|\|\|\|\|\|\|\|\|\|\|\|\|\|\|\|\|\|\|\|\|\|\|\|\|\|\|\|\|\|\|\|\|\|\|\|\|\|\|\|\|\|\|\|\|\|\|\|\|\|\|\|\|\|\|\|\|\|\|\|\|\|\|\|\|\|\|\|\|\|\|\|\|\|\|\|\|\|\|\|\|\|\|\|\|\|\|\|\|\|\|\|\|\|\|\|\| \|\|\|\|\|\|\|\|\|\|\|\|\|\|\|\|\|\|\|\|\|\|\|\|\|\|\|\|\|\| \|\|\|\|\|\|\|\| \|\|\|\|\|\|\|\|\|\|\|\|\|\|\|\|\|\|\|\|\|\|\| \|\|\|\|\|\| \|\|\|\|\|\|\|\|\|\|\|\|\|\| \|\|\|\| \|\| \|\|\|\|\| \|\|\|\|\|\|\|\|\|\|\|\|\|\| \|\|\|\|\|\|\|\| \|\| \|\| \|\|\|\|\|\|\|\|\|\|\|\| \|\|\|\| \|\| \|\|\|\|\| 	ATGTCTGCCGCCGCGAAAGAGGAGGACGAGGAAAAAGTGTCCGTCCCCAATGCCAGCGACGCTCTCTATGTCATCATTGCCGCAGCCTCGGTAGACGAAAACGGCGCACGCAGCAAAGTCTTCCAACAAGATGAGCTCCTCCAGCTGGGCAGCATCGCAAACACGCGAGAGCTCATGCCACTGGTACAATTCCTCGCCCAAAAAGGACTCTTCCGCACAGTAAAAGTCGGCGGAAAGCTAGGATGGAGCGTGCGACCACGAGAAGCAGCACGACAAATCGTCAACCTCGACCGAGATGAGAAGACAATCTACGAGATCATCGAAGAAGCACATACCACTGGAATCTGGACACGCGATCTGAAGAAGAAGACGAGCATCGCCCAAAATGTCGTACAGAAGGCGCTGTCGAANATGGAAAAGGGCAACCTCATCAAATCCGTCNAAAGCGTCNAAGCACCCGCCCAGCGAACATACNTGCTCANCCACCTCTCTCCTANCGAANACNTAACANGAAACAGCTTCTTCNACGCCGGCNATNTCNATGAATCCTTTCNCGACNAANTCCTC ``` |
| 1 | ggggacaagtttgtacaaaaaagcaggcttgatgAGACTGCTGCTCCTGGACTC, ggggaccactttgtacaagaaagctgggtcCTCGGCCTCCTTCCGCCTAC | Mycgr3\|66397  JGI  Ensembl | KOG2915 tRNA(1-methyladenosine) methyltransferase, subunit GCD14; | DNA-directed DNA polymerase activity; nucleic acid binding;  nucleobase, nucleoside, nucleotide and nucleic acid metabolic process;  nucleotide binding; | 3.70072 % |  |  |
| 1 | ggggacaagtttgtacaaaaaagcaggcttgatgTTCTTCGTCAAGAATATGGA, ggggaccactttgtacaagaaagctgggtcCCCAAGGTAATCCTCGCGAA | Mycgr3\|66842  JGI  Ensembl | KOG3298 DNA-directed RNA polymerase subunit E'; | DNA-directed RNA polymerase I activity; DNA-directed RNA polymerase II activity;  DNA-directed RNA polymerase III activity;  DNA-directed RNA polymerase activity;  RNA binding;  RNA polymerase activity;  transcription; | 100 % |  | ``` 201	ATGTTCTTCGTCAAGAATATGGACCGCGAGGTGACGATCCACCCTTCCTACTTCAACAGCGATGCGGAGAAAATCGTGCTGGACAGACTGCGCGCAGATATCGAAGGGACAATCGAGGGAACACTCATGGTGATTCGGATCGTGGATGTGGAGGATATTAGCGAGCCCAAGCTGCAGCCGGGCACTGGGATGGCGGTCTACCACTTGCACTACCGCGCGATTGTTTGGAGGCCTTTTCGTGGGGAGGTGGTGGATGGATTG	461 	\|\|\|\|\|\|\|\|\|\|\|\|\|\|\|\|\|\|\|\|\|\|\|\|\|\|\|\|\|\|\|\|\|\|\|\|\|\|\|\|\|\|\|\|\|\|\|\|\|\|\|\|\|\|\|\|\|\|\|\|\|\|\|\|\|\|\|\|\|\|\|\|\|\|\|\|\|\|\|\|\|\|\|\|\|\|\|\|\|\|\|\|\|\|\|\|\|\|\|\|\|\|\|\|\|\|\|\|\|\|\|\|\|\|\|\|\|\|\|\|\|\|\|\|\|\|\|\|\|\|\|\|\|\|\|\|\|\|\|\|\|\|\|\|\|\|\|\|\|\|\|\|\|\|\|\|\|\|\|\|\|\|\|\|\|\|\|\|\|\|\|\|\|\|\|\|\|\|\|\|\|\|\|\|\|\|\|\|\|\|\|\|\|\|\|\|\|\|\|\|\|\|\|\|\|\|\|\|\|\|\|\|\|\|\|\|\|\|\|\|\|\|\|\|\|\|\|\|\|\|\|\|\|\|\|\|\|\|\|\|\|\|\|\|\|\|\|\|\|\|\|\|\|\|\|\|\|\|\|\|\| 	ATGTTCTTCGTCAAGAATATGGACCGCGAGGTGACGATCCACCCTTCCTACTTCAACAGCGATGCGGAGAAAATCGTGCTGGACAGACTGCGCGCAGATATCGAAGGGACAATCGAGGGAACACTCATGGTGATTCGGATCGTGGATGTGGAGGATATTAGCGAGCCCAAGCTGCAGCCGGGCACTGGGATGGCGGTCTACCACTTGCACTACCGCGCGATTGTTTGGAGGCCTTTTCGTGGGGAGGTGGTGGATGGATTG ```      ``` 201	ATGTTCTTCGTCAAGAATATGGACCGCGAGGTGACGATCCACCCTTCCTACTTCAACAGCGATGCGGAGAAAATCGTGCTGGACAGACTGCGCGCAGATATCGAAGGGACAATCGAGGGAACACTCATGGTGATTCGGATCGTGGATGTGGAGGATATTAGCGAGCCCAAGCTGCAGCCGGGCACTGGGATGGCGGTCTACCACTTGCACTACCGCGCGATTGTTTGGAGGCCTTTTCGTGGGGAGGTGGTGGATGGATTGGTCAGCAGTGTGGTGAACAATGGCTTCTTCGTGGACGTGGGCGCATTGAGCGTGTTTGTGAGCAAAGCGATGA	534 	\|\|\|\|\|\|\|\|\|\|\|\|\|\|\|\|\|\|\|\|\|\|\|\|\|\|\|\|\|\|\|\|\|\|\|\|\|\|\|\|\|\|\|\|\|\|\|\|\|\|\|\|\|\|\|\|\|\|\|\|\|\|\|\|\|\|\|\|\|\|\|\|\|\|\|\|\|\|\|\|\|\|\|\|\|\|\|\|\|\|\|\|\|\|\|\|\|\|\|\|\|\|\|\|\|\|\|\|\|\|\|\|\|\|\|\|\|\|\|\|\|\|\|\|\|\|\|\|\|\|\|\|\|\|\|\|\|\|\|\|\|\|\|\|\|\|\|\|\|\|\|\|\|\|\|\|\|\|\|\|\|\|\|\|\|\|\|\|\|\|\|\|\|\|\|\|\|\|\|\|\|\|\|\|\|\|\|\|\|\|\|\|\|\|\|\|\|\|\|\|\|\|\|\|\|\|\|\|\|\|\|\|\|\|\|\|\|\|\|\|\|\|\|\|\|\|\|\|\|\|\|\|\|\|\|\|\|\|\|\|\|\|\|\|\|\|\|\|\|\|\|\|\|\|\|\|\|\|\|\|\|\|\|\|\|\|\|\|\|\|\|\|\|\|\|\|\|\|\|\|\|\|\|\|\|\|\|\|\|\|\|\|\|\|\|\|\|\|\|\|\|\|\|\|\|\|\|\|\|\|\|\|\|\|\|\|\|\|\|\|\|\|\|\|\|\|\|\|\|\| \|\|\| 	ATGTTCTTCGTCAAGAATATGGACCGCGAGGTGACGATCCACCCTTCCTACTTCAACAGCGATGCGGAGAAAATCGTGCTGGACAGACTGCGCGCAGATATCGAAGGGACAATCGAGGGAACACTCATGGTGATTCGGATCGTGGATGTGGAGGATATTAGCGAGCCCAAGCTGCAGCCGGGCACTGGGATGGCGGTCTACCACTTGCACTACCGCGCGATTGTTTGGAGGCCTTTTCGTGGGGAGGTGGTGGATGGATTGGTCAGCAGTGTGGTGAACAATGGCTTCTTCGTGGACGTGGGCGCATTGAGCGTGTTTGTGAGCAAAGCGGTGA ``` |
| 1 | ggggacaagtttgtacaaaaaagcaggcttgatgGTTGGACCTTCCGAAAATGC, ggggaccactttgtacaagaaagctgggtcGATCGGCTCAAAGGAACGTG | Mycgr3\|67658  JGI  Ensembl |  |  | 100 % |  | ``` 373	ATGGTTGGACCTTCCGAAAATGCAGGACCTTTGCTCATGGGATTGTCCTGGACAGAAGCCACGATTGCCACGATTGTCGTCGGACTTCGAGCGTACTGCGCATTGAGAGTTGTGCGAAAAGCGGAGTGGGACTTGATATGGGCTCTTCCGGCATGGCTCTGCAATTTCACCTCCCAAACCATCCTCACCATCAGCGTGGCGGCGTACGGCTACGGAAAACATCTAGCGGATATCTCGGA	611 	\|\|\|\|\|\|\|\|\|\|\|\|\|\|\|\|\|\|\|\|\|\|\|\|\|\|\|\|\|\|\|\|\|\|\|\|\|\|\|\|\|\|\|\|\|\|\|\|\|\|\|\|\|\|\|\|\|\|\|\|\|\|\|\|\|\|\|\|\|\|\|\|\|\|\|\|\|\|\|\|\|\|\|\|\|\|\|\|\|\|\|\|\|\|\|\|\|\|\|\|\|\|\|\|\|\|\|\|\|\|\|\|\|\|\|\|\|\|\|\|\|\|\|\|\|\|\|\|\|\|\|\|\|\|\|\|\|\|\|\|\|\|\|\|\|\|\|\|\|\|\|\|\|\|\|\|\|\|\|\|\|\|\|\|\|\|\|\|\|\|\|\|\|\|\|\|  \|\|\|\|\|\|\|\|\|\|\|\|\|\|\| \|\|\|\|\|\|\|\|\|\|\|\|\|\|\|\|\|\|\| \|\|\|\|\|\|\| \|\|\| \|\|\|\|\|\|\|\| \|\|\|\| 	ATGGTTGGACCTTCCGAAAATGCAGGACCTTTGCTCATGGGATTGTCCTGGACAGAAGCCACGATTGCCACGATTGTCGTCGGACTTCGAGCGTACTGCGCATTGAGAGTTGTGCGAAAAGCGGAGTGGGACTTGATATGGGCTCTTCCGGCATGGCTCTGCAATTTCACCTCCCANNCCATCCTCACCATCANCGTGGCGGCGTACGGCTACNGAAAACAACTANCGGATATCNCGGA ``` |
| 1 | ggggacaagtttgtacaaaaaagcaggcttgatgTCGTCTCCGTCCGCCACCAC, ggggaccactttgtacaagaaagctgggtcCACATTCGCACCACCACCAC | Mycgr3\|69317  JGI  Ensembl | KOG3471 RNA polymerase II transcription initiation/nucleotide excision repair factor TFIIH, subunit TFB2; | DNA repair; RNA polymerase II transcription factor activity;  nucleus;  regulation of transcription, DNA-dependent; | 1.5551 % |  |  |
| 1 | ggggacaagtttgtacaaaaaagcaggcttgatgTCACCATCGGGACGAACATC, ggggaccactttgtacaagaaagctgggtcGCCTTCGCCAATCTCGCGCA | Mycgr3\|69693  JGI  Ensembl |  |  | 0 % |  |  |
| 1 | ggggacaagtttgtacaaaaaagcaggcttgatgACCGACCTCGCTCATTCTCT, ggggaccactttgtacaagaaagctgggtcGACATCCTGGAATTCAAGCT | Mycgr3\|69983  JGI  Ensembl | KOG2593 Transcription initiation factor IIE, alpha subunit; | RNA polymerase II transcription factor activity; transcription factor TFIIE complex;  transcription initiation from RNA polymerase II promoter; | 100 % |  | ``` 89	ATGACCGACCTCGCTCATTCTCTGATCCGGACCGTCGTGCGCGCATTCTACCAAACCGAACACATAATCGCCATAGATGCGCTCGTACTCCACTCGACCCTCCAGGATAGCGACCTTGCGCAGCTCCTCGGCGTGCAGACCAAAGGTCTTCACAAGACAATAGGACGTCTTCGTGAAGATGGACTGGTCACAGTACAAAACCGAGCTGAGCGACGCACAGATGGCACGGGAGGATATTATGGAGGTGCGCCGCAGGCCGGCAAAGAACGAGTCACTCATCGACAATGGTACTATCTCAACTTTCATCGCGCCATCGACAGCATCAAGTATCGCATGTATCGGCTGAACAAGCACATTGAAAGTTTGGGAATGCCGACGACGGAGAGGAAGGATCTCTCTTGTCCAAGGTGCAAAAGCCAGTACGCAGAGATTGACGTTATGGACAATATTTCGCCAACTACCGGAGAGTTCCTCTGCCACATGTGCGGGCACGTGCTGGATCCTGTGGAGGAAGATGAGCTTGCGAACGAGAATCAGAGCATGAAACGGCTGAACAATCAGCTGGAGAAATTGCTCAAGCTTATGCAGAAGATCGACGCTACCGATGTTCCCGAGAATGATTTCGAAATGGCTCTGAGCAAGATGAAACCGATTCAGAAATCAGACACGAACCCTGGCTCCCGCACTGAGATCGTGGACCTGCCGAATAGAAATCTGCAGAGCACCAAAGGTCTCGTACTG	829 	\|\|\|\|\|\|\|\|\|\|\|\|\|\|\|\|\|\|\|\|\|\|\|\|\|\|\|\|\|\|\|\|\|\|\|\|\|\|\|\|\|\|\|\|\|\|\|\|\|\|\|\|\|\|\|\|\|\|\|\|\|\|\|\|\|\|\|\|\|\|\|\|\|\|\|\|\|\|\|\|\|\|\|\|\|\|\|\|\|\|\|\|\|\|\|\|\|\|\|\|\|\|\|\|\|\|\|\|\|\|\|\|\|\|\|\|\|\|\|\|\|\|\|\|\|\|\|\|\|\|\|\|\|\|\|\|\|\|\|\|\|\|\|\|\|\|\|\|\|\|\|\|\|\|\|\|\|\|\|\|\|\|\|\|\|\|\|\|\|\|\|\|\|\|\|\|\|\|\|\|\|\|\|\|\|\|\|\|\|\|\|\|\|\|\|\|\|\|\|\|\|\|\|\|\|\|\|\|\|\|\|\|\|\|\|\|\|\|\|\|\|\|\|\|\|\|\|\|\|\|\|\|\|\|\|\|\|\|\|\|\|\|\|\|\|\|\|\|\|\|\|\|\|\|\|\|\|\|\|\|\|\|\|\|\|\|\|\|\|\|\|\|\|\|\|\|\|\|\|\|\|\|\|\|\|\|\|\|\|\|\|\|\|\|\|\|\|\|\|\|\|\|\|\|\|\|\|\|\|\|\|\|\|\|\|\|\|\|\|\|\|\|\|\|\|\|\|\|\|\|\|\|\|\|\|\|\|\|\|\|\|\|\|\|\|\|\|\|\|\|\|\|\|\|\|\|\|\|\|\|\|\|\|\|\|\|\|\|\|\|\|\|\|\|\|\|\|\|\|\|\|\|\|\|\|\|\|\|\|\|\|\|\|\|\|\|\|\|\|\|\|\|\|\|\|\|\|\|\|\|\|\|\|\|\|\|\|\|\|\|\|\|\|\|\|\|\|\|\|\|\|\|\|\|\|\|\|\|\|\|\|\|\|\|\|\|\|\|\|\|\|\|\|\|\|\|\|\|\|\|\|\|\|\|\|\|\|\|\|\|\|\|\|\|\|\|\|\|\|\|\|\|\|\|\|\|\|\|\|\|\|\|\|\|\|\|\|\|\|\|\|\|\|\|\|\|\|\|\|\|\|\|\|\|\|\|\|\|\|\|\|\|\|\|\|\|\|\|\|\| \|\|\|\|\| \|\|\|\|\|\|\|\|\|\|\|\|\|\|\|\|\|\|\|\|\|\|\| \|\|\|\|\| \|\|\|\|\|\|\|\|\| \|\|\|\|\|\|\|\|\|\| \|\| \|\|\|\|\|\|\|\|\|\|\|\|\|\|\|\|\|\|\|\|\|\|\| \|\|\|\|\|\|\|\|\|  \|\|\|\|\|\|\|\|\|\|\| \|\|  \|\|\|\|\|\|\|\|\|\|\|\|\|  \|\|\|\|\|\| \|\| \| \|  \|\|\|\|\|\|\|\|\|\|\|\|\|\|\|\|\| \|\|  \|\|\|\|\|\|\|\|\|\| \|\| \| \|\|\|\| \|\|\|  \| \|\| \|\|\|\|\|\|\|\|\|\| \|\|\|\|\| 	ATGACCGACCTCGCTCATTCTCTGATCCGGACCGTCGTGCGCGCATTCTACCAAACCGAACACATAATCGCCATAGATGCGCTCGTACTCCACTCGACCCTCCAGGATAGCGACCTTGCGCAGCTCCTCGGCGTGCAGACCAAAGGTCTTCACAAGACAATAGGACGTCTTCGTGAAGATGGACTGGTCACAGTACAAAACCGAGCTGAGCGACGCACAGATGGCACGGGAGGATATTATGGAGGTGCGCCGCAGGCCGGCAAAGAACGAGTCACTCATCGACAATGGTACTATCTCAACTTTCATCGCGCCATCGACAGCATCAAGTATCGCATGTATCGGCTGAACAAGCACATTGAAAGTTTGGGAATGCCGACGACGGAGAGGAAGGATCTCTCTTGTCCAAGGTGCAAAAGCCAGTACGCAGAGATTGACGTTATGGACAATATTTCGCCAACTACCGGAGAGTTCCTCTGCCACATGTGCGGGCACGTGCTGGATCCTGTGGAGGAAGATGAGCTTGCGAACGANAATCANAGCATGAAACGGCTGAACAATCANCTGGANAAATTGCTC-AGCTTATGCANAANATCGACGCTACCGATGTTCCCGANAATGATTTCN-AATGGCTCTGANCAN-ATGAAACCGATTCNNAAATCANAC-CNAN-CCTGGCTCCCGCACTGA-ATNNTGGACCTGCCNAANA-AAATNTGCN-ANCA-CAAAGGTCTCNTACTG ``` |
| 1 | ggggacaagtttgtacaaaaaagcaggcttgatgGCAACTTCTGCGCGTCGCAA, ggggaccactttgtacaagaaagctgggtcCGAAGCAGCAGCAGCCGTCT | Mycgr3\|70181  JGI  Ensembl | KOG0519 Sensory transduction histidine kinase; | regulation of transcription, DNA-dependent; two-component response regulator activity;  two-component signal transduction system (phosphorelay); | 2.26372 % |  |  |
| 1 | ggggacaagtttgtacaaaaaagcaggcttgatgGGTAGCCACCGCCATAAGCT, ggggaccactttgtacaagaaagctgggtcATCCTGAAACAGCAATCCCG | Mycgr3\|71099  JGI  Ensembl | KOG0012 DNA damage inducible protein; | carbohydrate metabolic process; hydrolase activity, hydrolyzing O-glycosyl compounds; | 100 % |  | ``` 1	ATGGGTAGCCACCGCCATAAGCTCACCGAGCTCGTTCGCGAGGCAACTTCCTCCGTGATTGCTCACTCAGAACGACTGTCCGGCCACCGCGATAAGAATCGCACGCCAGCTTCTGCGGGAAGAGTAGTGGTCTATCACCAAACACATCATCAACCGAATGATGGTCCGCCAGTGTCCCTGCTGCCTTTGATCAGCAATAACACAGGAGTCACGCATGTGATCATTGCCGCGATTCACCTCAATGACCCAGCCGGTAACATTACTCTCAATGATCATCCGCCAGATCATGAACGCTTCAACACGCTGTGGGGCGAGGTGGCCTGGCTTCAAGCAAGTGGTGTGAAATGTATGGGACTGCTCGGTGGCGCTGCGCAAGGATCTTTCGGCCGATTAGATGGCGAAGATGTTGCGCGCTTTGAATCGTA	425 	\|\|\|\|\|\|\|\|\|\|\|\|\|\|\|\|\|\|\|\|\|\|\|\|\|\|\|\|\|\|\|\|\|\|\|\|\|\|\|\|\|\|\|\|\|\|\|\|\|\|\|\|\|\|\|\|\|\|\|\|\|\|\|\|\|\|\|\|\|\|\|\|\|\|\|\|\|\|\|\|\|\|\|\|\|\|\|\|\|\|\|\|\|\|\|\|\|\|\|\|\|\|\|\|\|\|\|\|\|\|\|\|\|\|\|\|\|\|\|\|\|\|\|\|\|\|\|\|\|\|\|\|\|\|\|\|\|\|\|\|\|\|\|\|\|\|\|\|\|\|\|\|\|\|\|\|\|\|\|\|\|\|\|\|\|\|\|\|\|\|\|\|\|\|\|\|\|\|\|\|\|\|\|\|\|\|\|\|\|\|\|\|\|\|\|\|\|\|\|\|\|\|\|\|\|\|\|\|\|\|\|\|\|\|\|\|\|\|\|\|\|\|\|\|\|\|\|\|\|\|\|\|\|\|\|\|\|\|\|\|\|\|\|\|\|\|\|\|\|\|\|\|\|\|\|\|\|\|\|\|\|\|\|\|\|\|\|\|\|\|\|\|\|\|\|\|\|\|\|\|\|\|\|\|\|\|\|\|\|\|\|\|\|\|\|\|\|\|\|\|\|\|\|\|\|\|\|\|\|\|\|\|\|\|\|\|\|\|\|\|\|\|\|\|\|\|\|\|\|\|\|\|\|\|\|\|\|\|\|\|\|\|\|\|\|\|\|\|\|\|\|\|\|\|\|\|\|\|\|\|\|\|\|\|\|\|\|\|\|\|\|\|\|\|\|\|\|\|\|\|\|\|\|\|\|\|\|\|\|\|\|\|\| \|\|\|\|\|\|\|\|\|\|\|\|\|\|\|\|\|\|\|\|\|\|\|\|\|\|\|\|\|\|\| 	ATGGGTAGCCACCGCCATAAGCTCACCGAGCTCGTTCGCGAGGCAACTTCCTCCGTGATTGCTCACTCAGAACGACTGTCCGGCCACCGCGATAAGAATCGCACGCCAGCTTCTGCGGGAAGAGTAGTGGTCTATCACCAAACACATCATCAACCGAATGATGGTCCGCCAGTGTCCCTGCTGCCTTTGATCAGCAATAACACAGGAGTCACGCATGTGATCATTGCCGCGATTCACCTCAATGACCCAGCCGGTAACATTACTCTCAATGATCATCCGCCAGATCATGAACGCTTCAACACGCTGTGGGGCGAGGTGGCCTGGCTTCAAGCAAGTGGTGTGAAATGTATGGGACTGCTCGGTGGCGCTGCGCAAGGATCTTTCGGCCGATTANATGGCGAAGATGTTGCGCGCTTTGAATCGTA ``` |
| 1 | ggggacaagtttgtacaaaaaagcaggcttgatgTTGGCAATACCATCCGCGTC, ggggaccactttgtacaagaaagctgggtcCTCCACCGAAGACTGAGATG | Mycgr3\|71246  JGI  Ensembl | KOG2267 Eukaryotic-type DNA primase, large subunit; | CTP:2,3-di-O-geranylgeranyl-sn-glycero-1-phosphate cytidyltransferase; DNA primase activity;  DNA replication, synthesis of RNA primer;  DNA-directed DNA polymerase activity;  DNA/RNA helicase activity;  RNA guanylyltransferase activity;  RNA helicase activity;  alpha DNA polymerase:primase complex;  mannose-phosphate guanylyltransferase activity;  phospholactate guanylyltransferase activity;  sulfate adenylyltransferase activity;  tRNA guanylyltransferase activity;  telomerase activity; | 100 % |  | ``` 1	ATGTTGGCAATACCATCCGCGTCGCGCATCGACGCGAAGCGACGGGCGGTGGTCAATCCCAAGCGCCAGCAATTCCAACAGGCCGCATTCGCAGAGCAGGACTATGCACATCGCATGAACTTTTACATCATCCCACCAACAGGCGACATCACACTCGAAGAATTCGAAGAATGGGGCATCGCACGTCTCAAAGTGCTCGCGGAGCTCGAGGCTTGTCAATTCCGGAATAAGACGCCCACCGAGACGGAGGAGTACATGCGACCGATTCTGGCCAAGCACCTGCCGCTCGCGAGTAACAGCTCTCGCAGCAGTGAGGTGAACTTGGAGCGGAAACGAGATCATTACAGTCATTGGACTCTTCGACTAGCATTCTCGAGCACACAGGACCTACGACAGCGATTCTCGCGATTGGAAGCACAGTTGTTCAAGCTGAGGATACAGCAGGATGATCTGCGGGAGAGGAAGTCTTTCATCGAGTCGCTGCCGATGTC	491 	\|\|\|\|\|\|\|\|\|\|\|\|\|\|\|\|\|\|\|\|\|\|\|\|\|\|\|\|\|\|\|\|\|\|\|\|\|\|\|\|\|\|\|\|\|\|\|\|\|\|\|\|\|\|\|\|\|\|\|\|\|\|\|\|\|\|\|\|\|\|\|\|\|\|\|\|\|\|\|\|\|\|\|\|\|\|\|\|\|\|\|\|\|\|\|\|\|\|\|\|\|\|\|\|\|\|\|\|\|\|\|\|\|\|\|\|\|\|\|\|\|\|\|\|\|\|\|\|\|\|\|\|\|\|\|\|\|\|\|\|\|\|\|\|\|\|\|\|\|\|\|\|\|\|\|\|\|\|\|\|\|\|\|\|\|\|\|\|\|\|\|\|\|\|\|\|\|\|\|\|\|\|\|\|\|\|\|\|\|\|\|\|\|\|\|\|\|\|\|\|\|\|\|\|\|\|\|\|\|\|\|\|\|\|\|\|\|\|\|\|\|\|\|\|\|\|\|\|\|\|\|\|\|\|\|\|\|\|\|\|\|\|\|\|\|\|\|\|\|\|\|\|\|\|\|\|\|\|\|\|\|\|\|\|\|\|\|\|\|\|\|\|\|\|\|\|\|\|\|\|\|\|\|\|\|\|\|\|\|\|\|\|\|\|\|\|\|\|\|\|\|\|\|\|\|\|\|\|\|\|\|\|\|\|\|\|\|\|\|\|\|\|\|\|\|\|\|\|\|\|\|\|\|\|\|\|\|\|\|\|\|\|\|\|\|\|\|\|\|\|\|\|\|\|\|\|\|\|\|\|\|\|\|\|\|\|\|\|\|\|\|\|\|\|\|\|\|\|\|\|\|\|\|\|\|\|\|\|\|\|\|\|\|\|\|\|\|\|\|\|\|\|\|\|\|\|\|\|\|\|\|\|\|\|\|\|\|\|\|\|\|\|\|\|\|\|\|\|\|\|\|\|\|\|\|\|\|\|\|\|\|\|\| \|\|\|\|\|\|\|\|\|\|\|\|\|\|\|\|\|\|\|\| \|\|\|\|\|\| \|\| \| \|\| \|\|\|\|\| \|\|\|\|\| 	ATGTTGGCAATACCATCCGCGTCGCGCATCGACGCGAAGCGACGGGCGGTGGTCAATCCCAAGCGCCAGCAATTCCAACAGGCCGCATTCGCAGAGCAGGACTATGCACATCGCATGAACTTTTACATCATCCCACCAACAGGCGACATCACACTCGAAGAATTCGAAGAATGGGGCATCGCACGTCTCAAAGTGCTCGCGGAGCTCGAGGCTTGTCAATTCCGGAATAAGACGCCCACCGAGACGGAGGAGTACATGCGACCGATTCTGGCCAAGCACCTGCCGCTCGCGAGTAACAGCTCTCGCAGCAGTGAGGTGAACTTGGAGCGGAAACGAGATCATTACAGTCATTGGACTCTTCGACTAGCATTCTCGAGCACACAGGACCTACGACAGCGATTCTCGCGATTGGAAGCACAGTTGTTCAAGCTGAGGATACAGCANGATGATCTGCGGGAGAGGAANTCTTTCNTCNANTCNCTGCCNATGTC ``` |
| 1 | ggggacaagtttgtacaaaaaagcaggcttgatgAATCCTTGCGCCAACTGCGC, ggggaccactttgtacaagaaagctgggtcTCGTCTCCTGTCATGATACT | Mycgr3\|72596  JGI  Ensembl |  |  | 100 % |  | ``` 367	GATGAATCCTTGCGCCAACTGCGCTGCATCGAACTCCCAGTGCATCTATGGATCCGAGCCATTGAC	432 	\|\|\|\|\|\|\|\|\|\|\|\|\|\|\|\|\|\|\|\|\|\|\|\|\|\|\|\|\|\|\|\|\|\|\|\|\|\|\|\|\|\|\|\|\|\|\|\|\|\|\|\|\|\|\|\|\|\|\|\|\|\|\|\|\|\| 	GATGAATCCTTGCGCCAACTGCGCTGCATCGAACTCCCAGTGCATCTATGGATCCGAGCCATTGAC ``` |
| 1 | ggggacaagtttgtacaaaaaagcaggcttgatgGTTGCGTGTCCGATCTGCAA, ggggaccactttgtacaagaaagctgggtcATCGCCGTCCGGATTGAGCA | Mycgr3\|74087  JGI  Ensembl | KOG2028 ATPase related to the helicase subunit of the Holliday junction resolvase; | ATP binding; DNA binding;  DNA polymerase activity;  DNA repair;  DNA-directed DNA polymerase activity;  nucleoside-triphosphatase activity;  nucleotide binding; | 100 % |  | ``` 1	ATGGTTGCGTGTCCGATCTGCAACAAGCCCGTCAAGGAAGCCGACATCAACCGCCATATCGACTCGGAATGCAAAAACTTTATCGATGAGCCTTCGCCGCCAGCAACACAGGGCACGCAGTCGCAAGCCAATGGAGCGAAGGCGAAGAGCGCGCTCAGCAAGCCCGCTTCGGGCTTCTTCACGCCGGCGCCGAAGAGAAATGTAAGCTTCAGACTCGACGAGGCTTCGTCGCCTGCACCCGCGTTGCCGAGCACCCAGAAGCCTGC	266 	\|\|\|\|\|\|\|\|\|\|\|\|\|\|\|\|\|\|\|\|\|\|\|\|\|\|\|\|\|\|\|\|\|\|\|\|\|\|\|\|\|\|\|\|\|\|\|\|\|\|\|\|\|\|\|\|\|\|\|\|\|\|\|\|\|\|\|\|\|\|\|\|\|\|\|\|\|\|\|\|\|\|\|\|\|\|\|\|\|\|\|\|\|\|\|\|\|\|\|\|\|\|\|\|\|\|\|\|\|\|\|\|\|\|\|\|\|\|\|\|\|\|\|\|\|\|\|\|\|\|\|\|\|\|\|\|\|\|\|\|\|\|\|\|\|\|\|\|\|\|\|\|\|\|\|\|\|\|\|\|\|\|\|\|\|\|\|\|\|\|\|\|\|\|\|\|\|\|\|\|\|\|\|\|\|\|\|\|\|\|\|\|\|\|\|\|\|\|\|\|\|\|\|\|\|\|\|\|\|\|  \|\|\|\| \|\|\|\|\|\|\|\|\|\| \|\| \|\|\|\|\| \|\|\|\|\|\|\|\|\|\|\| \| \|\|\|\|\|  \|\| \|\|\|\|\| 	ATGGTTGCGTGTCCGATCTGCAACAAGCCCGTCAAGGAAGCCGACATCAACCGCCATATCGACTCGGAATGCAAAAACTTTATCGATGAGCCTTCGCCGCCAGCAACACAGGGCACGCAGTCGCAAGCCAATGGAGCGAAGGCGAAGAGCGCGCTCAGCAAGCCCGCTTCGGGCTTCTTCACGCCGGCGCCGAAGAGAAATGTAAGCTTCNNACTCNACGAGGCTTCNTCNCCTGCNCCCGCGTTGCCNANCACCCNNAANCCTGC ``` |
| 1 | ggggacaagtttgtacaaaaaagcaggcttgatgGAAGATCTTCGGTTCCGACC, ggggaccactttgtacaagaaagctgggtcATTACATGTGGGACAATGCG | Mycgr3\|74232  JGI  Ensembl | KOG1432 Predicted DNA repair exonuclease SIA1; | hydrolase activity; | 0 % |  |  |
| 1 | ggggacaagtttgtacaaaaaagcaggcttgatgGCGCGAACAAAGCAAATATC, ggggaccactttgtacaagaaagctgggtcACCCAAACCACCCCAAGCGC | Mycgr3\|74593  JGI  Ensembl | KOG1745 Histones H3 and H4; | DNA binding; nucleosome; nucleosome assembly;  nucleus; | 100 % |  | ``` 213	ATGGCGCGAACAAAGCAAATATCACGAGGAGGACGGCAGAGCACAGGGGGAAAGTCGGCGGGCAAAACGACACCAGGCAGAGGAGGTGCGAGAGGAGGAGCACGAGGAGGCAAGGCACCGAGATCGAGCACAGGGTCAATGGGAGG	358 	\|\|\|\|\|\|\|\|\|\|\|\|\|\|\|\|\|\|\|\|\|\|\|\|\|\|\|\|\|\|\|\|\|\|\|\|\|\|\|\|\|\|\|\|\|\|\|\|\|\|\|\|\|\|\|\|\|\|\|\|\|\|\|\|\|\|\|\|\|\|\|\|\|\|\|\|\|\|\|\|\|\|\|\|\|\|\|\|\|\|\|\|\|\|\|\|\|\|\|\|\|\|\|\|\|\|\|\|\|\|\|\|\|\|\|\|\|\|\|\|\|\|\|\|\|\|\|\|\|\|\|\|\|\|\|\|\|\|\|\|\|\|\|\|\|\| 	ATGGCGCGAACAAAGCAAATATCACGAGGAGGACGGCAGAGCACAGGGGGAAAGTCGGCGGGCAAAACGACACCAGGCAGAGGAGGTGCGAGAGGAGGAGCACGAGGAGGCAAGGCACCGAGATCGAGCACAGGGTCAATGGGAGG ```      ``` 356	AGGCGTAAAGAAACCGCATCGCTACAAGGCTGGGACCGTTGCTCTTCGCGAGATCCGACGATATCAAAAAAGCACAGAGCTACTGATCGCAAAGCTGCCCTTCTCACGTCTGGTACGAG	474 	\|\|\|\|\|\|\|\|\|\|\|\|\|\|\|\|\|\|\|\|\|\|\|\|\|\|\|\|\|\|\|\|\|\|\|\|\|\|\|\|\|\|\|\|\|\|\|\|\|\|\|\|\|\|\|\|\|\|\|\|\|\|\|\|\|\|\|\|\|\|\|\|\|\|\|\|\|\|\|\|\|\|\|\|\|\|\|\|\|\|\|\|\|\|\|\|\|\|\|\|\|\|\|\|\|\|\|\|\|\|\|\|\|\|\| \|\|\| 	AGGCGTAAAGAAACCGCATCGCTACAAGGCTGGGACCGTTGCTCTTCGCGAGATCCGACGATATCAAAAAAGCACAGAGCTACTGATCGCAAAGCTGCCCTTCTCACGTCTGGTAAGAG ```      ``` 467	GGTACGAGAAGTATGCATGAACGTAGCGCCTGCCTCGTCAGAAGTTTCGCGCTGGCAATCACAAGCCATCCAAGCCCTGCAGGAAGCAGCGGAGGCTTTCCTCGTGCATCTGTTCGAAGACTCTAATCTCTGCGCAATCCATGCGAAACGAGTAACGATCATGCAGAAGGATATTCAACTTGCAAGGCGGATACGCGGCGCTTGGGGTGGTTTGGGT	683 	\|\|\|\|\|\|\|\|\|\|\|\|\|\|\|\|\|\|\|\|\|\|\|\|\|\|\|\|\|\|\|\|\|\|\|\|\|\|\|\|\|\|\|\|\|\|\|\|\|\|\|\|\|\|\|\|\|\|\|\|\|\|\|\|\|\|\|\|\|\|\|\|\|\|\|\|\|\|\|\|\|\|\|\|\|\|\|\|\|\|\|\|\|\|\|\|\|\|\|\|\|\|\|\|\|\|\|\|\|\|\|\|\|\|\|\|\|\|\|\|\|\|\|\|\|\|\|\|\|\|\|\|\|\|\|\|\|\|\|\|\|\|\|\|\|\|\|\|\|\|\|\|\|\|\|\|\|\|\|\|\|\|\|\|\|\|\|\|\|\|\|\|\|\|\|\|\|\|\|\|\|\|\|\|\|\|\|\|\|\|\|\|\|\|\|\|\|\|\|\|\|\|\|\|\|\|\|\|\|\|\|\|\|\|\|\|\| 	GGTACGAGAAGTATGCATGAACGTAGCGCCTGCCTCGTCAGAAGTTTCGCGCTGGCAATCACAAGCCATCCAAGCCCTGCAGGAAGCAGCGGAGGCTTTCCTCGTGCATCTGTTCGAAGACTCTAATCTCTGCGCAATCCATGCGAAACGAGTAACGATCATGCAGAAGGATATTCAACTTGCAAGGCGGATACGCGGCGCTTGGGGTGGTTTGGGT ``` |
| 1 | ggggacaagtttgtacaaaaaagcaggcttgatgCAAAATCCTGTAATGCTACC, ggggaccactttgtacaagaaagctgggtcACAACTCCCCACAAAGAAAT | Mycgr3\|75077  JGI  Ensembl |  |  | 100 % |  | ``` 358	ATGCAAAATCCT-GTAATGCTACCCACCAAGACCATCGCCACCATCAACTCGACCGGTCGGCAGGCCGCATCCTTCATCCGCGCCGCGTCTGCCGTCGGCTGGAACGTCCGCGCCCAAGTCCGCAAAAACGAAGGCCTCGTCGCCGAAGAACTCGGCCTCCTCCCGAACGTCGAGCTCATCGAAGGAGATCTCGTCGGACCGGAACGAAATGCTCTCCTCGCGAAACTCTTCACGGGCGCGAAGATTGCGTTCATCAACACGACGCATTGGGGCGATGAGGTGGCGATAGGGAAGGCCTGTGCGGACGCCGCGAAGAAGGCGGGGATCTCGCATTATGTCTATTCGAGCATGCCGGATCATTCGTCGTTTGGAGAGGATTGGAGGGCGTTG	747 	\|\|\|\|\|\|\|\|\|\|\|\| \|\|\|\|\|\|\|\|\|\|\|\|\|\|\|\|\|\|\|\|\|\|\|\|\|\|\|\|\|\|\|\|\|\|\|\|\|\|\|\|\|\|\|\|\|\|\|\|\|\|\|\|\|\|\|\|\|\|\|\|\|\|\|\|\|\|\|\|\|\|\|\|\|\|\|\|\|\|\|\|\|\|\|\|\|\|\|\|\|\|\|\|\|\|\|\|\|\|\|\|\|\|\|\|\|\|\|\|\|\|\|\|\|\|\|\|\|\|\|\|\|\|\|\|\|\|\|\|\|\|\|\|\|\|\|\|\|\|\|\|\|\|\|\|\|\|\|\|\|\|\|\|\|\|\|\|\|\|\|\|\|\|\|\|\|\|\|\|\|\|\|\|\|\|\|\|\|\|\|\|\|\|\|\|\|\|\|\|\|\|\|\|\|\|\|\|\|\|\|\|\|\|\|\|\|\|\|\|\|\|\|\|\|\|\|\|\|\|\|\|\|\|\|\|\|\|\|\|\|\|\|\|\|\|\|\|\|\|\|\|\|\|\|\|\|\|\|\|\|\|\|\|\|\|\|\|\|\|\|\|\|\|\|\|\|\|\|\|\|\|\|\|\|\|\|\|\|\|\|\|\|\|\|\|\|\|\|\|\|\|\|\|\|\|\|\|\|\|\|\|\|\|\|\|\|\|\|\|\|\|\|\|\|\|\|\|\|\|\|\|\|\|\|\|\|\|\|\|\|\|\|\|\|\|\|\|\|\|\|\|\|\|\|\|\|\|\|\|\|\|\|\|\|\|\|\|\|\|\|\| \|\|\|\|\|\|\|\|\|\|\|\| \|\|\|\| 	ATGCAAAATCCTGGTAATGCTACCCACCAAGACCATCGCCACCATCAACTCGACCGGTCGGCAGGCCGCATCCTTCATCCGCGCCGCGTCTGCCGTCGGCTGGAACGTCCGCGCCCAAGTCCGCAAAAACGAAGGCCTCGTCGCCGAAGAACTCGGCCTCCTCCCGAACGTCGAGCTCATCGAAGGAGATCTCGTCGGACCGGAACGAAATGCTCTCCTCGCGAAACTCTTCACGGGCGCGAAGATTGCGTTCATCAACACGACGCATTGGGGCGATGAGGTGGCGATAGGGAAGGCCTGTGCGGACGCCGCGAAGAAGGCGGGGATCTCGCATTATGTCTATTCGAGCATGCCGGATCATTCGTCGTTTGGANAGGATTGGAGGGNGTTG ``` |
| 1 | ggggacaagtttgtacaaaaaagcaggcttgatgTCAGTCTGGAATCCCGAGAA, ggggaccactttgtacaagaaagctgggtcGATCGTAAGATCCTTCTCCA | Mycgr3\|75105  JGI  Ensembl | KOG2549 Transcription initiation factor TFIID, subunit TAF6 (also component of histone acetyltransferase SAGA); | RNA polymerase II transcription factor activity; nucleus; regulation of transcription factor activity;  transcription initiation;  transcription initiation factor activity; | 100 % |  | ``` 133	ATGTCAGTCTGGAATCCCGAGAATGTCACAGACGTGGCAGAGTCCGTCGGCATCGCTTCACTCAACCGCGATGTGGCAGAGCACCTAGCAAGAGACGTGGAATACCGAATCAGTCAGGTTCTGGAGGAAGCGCTGAAGTTCATGCGCCATGCGAAGCGGACAACACTCACAACACAAGACATTTCACAAGCGCTACGGCTACTGGAGGTAGAGCCATTGTATGGATACGAGTCGACACGGCCGCTGCGATTTGGCGAGGCATCAATAGGTCCTGGCCAGCCGTTGTTCTACGTGGAAGACGAGGAGGTGGATTTCGAGAAGCTCATCAATGCGCCGCTACCGAAGGTTCCACGTGAAATCAGCTTCACTGCGCACTGGCTGGCGGTGGAAGGTGTACAACCCAGCATACCACAAAATCCAACACAAGCAGATCAGAGGAACCAGGAGCTGGCCGCGAAAGGACAAGGCGCAAACACCCTCGCAGCTGTCAGCGGCAACGACAACGTGGCCGTCAAGCCTCTGGTCAAGCACGTCTTGTCGAAAGAACTTCAGCTCTACTTCGAAAGAATATGCTCGGCGATCCTG	717 	\|\|\|\|\|\|\|\|\|\|\|\|\|\|\|\|\|\|\|\|\|\|\|\|\|\|\|\|\|\|\|\|\|\|\|\|\|\|\|\|\|\|\|\|\|\|\|\|\|\|\|\|\|\|\|\|\|\|\|\|\|\|\|\|\|\|\|\|\|\|\|\|\|\|\|\|\|\|\|\|\|\|\|\|\|\|\|\|\|\|\|\|\|\|\|\|\|\|\|\|\|\|\|\|\|\|\|\|\|\|\|\|\|\|\|\|\|\|\|\|\|\|\|\|\|\|\|\|\|\|\|\|\|\|\|\|\|\|\|\|\|\|\|\|\|\|\|\|\|\|\|\|\|\|\|\|\|\|\|\|\|\|\|\|\|\|\|\|\|\|\|\|\|\|\|\|\|\|\|\|\|\|\|\|\|\|\|\|\|\|\|\|\|\|\|\|\|\|\|\|\|\|\|\|\|\|\|\|\|\|\|\|\|\|\|\|\|\|\|\|\|\|\|\|\|\|\|\|\|\|\|\|\|\|\|\|\|\|\|\|\|\|\|\|\|\|\|\|\|\|\|\|\|\|\|\|\|\|\|\|\|\|\|\|\|\|\|\|\|\|\|\|\|\|\|\|\|\|\|\|\|\|\|\|\|\|\|\|\|\|\|\|\|\|\|\|\|\|\|\|\|\|\|\|\|\|\|\|\|\|\|\|\|\|\|\|\|\|\|\|\|\|\|\|\|\|\|\|\|\|\|\|\|\|\|\|\|\|\|\|\|\|\|\|\|\|\|\|\|\|\|\|\|\|\|\|\|\|\|\|\|\|\|\|\|\|\|\|\|\|\|\|\|\|\|\|\|\|\|\|\|\|\|\|\|\|\|\|\|\|\|\|\|\|\|\|\|\|\|\|\|\|\| \|\|\|\|\|\|\|\|\|\|\|\|\|\|\|\|\|\|\|\|\|\|\|\|\|\|\|\|\|\| \|\|\|\|\|\|\|\|\|\|\|\|\|\|\|\|\|\|\|\|\|\|\|\|\|\|\|\|\|\|\|\|\|\|\|\| \|\|\|\|\|\|\|\| \|\| \|\|\|\|\|\| \|\|\|\|\|\|\| \|\| \|\| \|\|\|\|\|\|\|\| \|\|\|\|\|\|\|\|\|\|\|\|  \| \|\|\|\|\|\|\|\|\| \|\|\| \|\|\|\|\|\|  \|\|\|\|\|\|\|\|\|  \|\| \|\|\|\|\|\|\|\|\|\|\|\| \|\| \|\|\| 	ATGTCAGTCTGGAATCCCGAGAATGTCACAGACGTGGCAGAGTCCGTCGGCATCGCTTCACTCAACCGCGATGTGGCAGAGCACCTAGCAAGAGACGTGGAATACCGAATCAGTCAGGTTCTGGAGGAAGCGCTGAAGTTCATGCGCCATGCGAAGCGGACAACACTCACAACACAAGACATTTCACAAGCGCTACGGCTACTGGAGGTAGAGCCATTGTATGGATACGAGTCGACACGGCCGCTGCGATTTGGCGAGGCATCAATAGGTCCTGGCCAGCCGTTGTTCTACGTGGAAGACGAGGAGGTGGATTTCGAGAAGCTCATCAATGCGCCGCTACCGAAGGTTCCACGTGAAATCAGCTTCACTGCGCACTGGCTGGCGGTGGAAGGTGTACAACCCANCATACCACAAAATCCAACACAAGCAGATCANAGGAACCAGGAGCTGGCCGCGAAAGGACAAGGCGCANACACCCTCNCANCTGTCANCGGCAACNACNACNTGGCCGTCNAGCCTCTGGTCAN-CNCGTCTTGTCNAAANAACTTCNNCTCTACTTCNNAANAATATGCTCGGCNATNCTG ``` |
| 1 | ggggacaagtttgtacaaaaaagcaggcttgatgAACGCCGTCGATGCCACCGA, ggggaccactttgtacaagaaagctgggtcCGCAGGAAGAGGAGTACCCG | Mycgr3\|75897  JGI  Ensembl | KOG2487 RNA polymerase II transcription initiation/nucleotide excision repair factor TFIIH, subunit TFB4; | DNA repair; core TFIIH complex;  general RNA polymerase II transcription factor activity;  regulation of transcription, DNA-dependent; | 3.54059 % |  |  |
| 1 | ggggacaagtttgtacaaaaaagcaggcttgatgAGCTCTCCGATCTTGTGGGA, ggggaccactttgtacaagaaagctgggtcAGCTGACTGCGATTCATCCG | Mycgr3\|75907  JGI  Ensembl | KOG2356 Transcriptional activator, adenine-specific DNA methyltransferase; | methylation; methyltransferase activity;  nucleic acid binding;  nucleobase, nucleoside, nucleotide and nucleic acid metabolic process;  nucleus; | 0 % |  |  |
| 1 | ggggacaagtttgtacaaaaaagcaggcttgatgGGCGTCAATGAGAAAAGCCC, ggggaccactttgtacaagaaagctgggtcAGTAATATACCTAAACCCCG | Mycgr3\|76558  JGI  Ensembl | KOG2533 Permease of the major facilitator superfamily; | DNA binding; integral to membrane;  transport;  transporter activity; | 100 % |  | ``` 269	GTGCTTCCTCGACCGCTCCAACATCGGCAATGCCCGAATCCAAGGCATGTCCCGAGACCTCAAACTCGACATCAAGTACCGCTTCAACTGGGCCCTCTCAGTCTTCTACATCGTCTACCTCCTCGTCGAAGTACCCAGCAACATCCTCCTCAAGAAAGTCGGACCACGATTCTACATCCCAGCATTGGTGGTAGGCTTTGGCTTC	473 	\|\|\|\|\|\|\|\|\|\|\|\|\|\|\|\|\|\|\|\|\|\|\|\|\|\|\|\|\|\|\|\|\|\|\|\|\|\|\|\|\|\|\|\|\|\|\|\|\|\|\|\|\|\|\|\|\|\|\|\|\|\|\|\|\|\|\|\|\|\|\|\|\|\|\|\|\|\|\|\|\|\|\|\|\|\|\|\|\|\|\|\|\|\|\|\|\|\|\|\|\|\|\|\|\|\|\|\|\|\|\|\|\|\|\|\|\|\|\|\|\|\|\|\|\|\|\|\|\|\|\|\|\|\|\|\|\|\|\|\|\|\|\|\|\|\|\|\|\|\|\|\|\|\|\|\|\|\|\|\|\|\|\|\|\|\|\|\|\|\|\|\|\|\|\|\|\|\|\|\|\| \|\|\|\|\|\|\|\|\|\|\| \|\|\|\|\|\|\| \|\|\| 	GTGCTTCCTCGACCGCTCCAACATCGGCAATGCCCGAATCCAAGGCATGTCCCGAGACCTCAAACTCGACATCAAGTACCGCTTCAACTGGGCCCTCTCAGTCTTCTACATCGTCTACCTCCTCGTCGAAGTACCCAGCAACATCCTCCTCAAGAAAGTCGGACCACGATTCTACATCCCANCATTGGTGGTANGCTTTGGNTTC ```      ``` 93	ATGGGC-GTCAATGAGAAAAGCCCGTCCATAGGGCAAGAAGGGCGCTCGCCTTCCTCTCACAGCGATTCTCTCCACTCCGATCCTCTGGCCGCAACATGGACCGCCGAAGAAGAAACCCGCGTGCGTCGCAAACTGGACCTTCAAATCGTGCCGATGGTCACAGTGCTCTACTTGATGTG	271 	\|\|\|\|\|\| \|\|\|\|\|\|\|\|\|\|\|\|\|\|\|\|\|\|\|\|\|\|\|\|\|\|\|\|\|\|\|\|\|\|\|\|\|\|\|\|\|\|\|\|\|\|\|\|\|\|\|\|\|\|\|\|\|\|\|\|\|\|\|\|\|\|\|\|\|\|\|\|\|\|\|\|\|\|\|\|\|\|\|\|\|\|\|\|\|\|\|\|\|\|\|\|\|\|\|\|\|\|\|\|\|\|\|\|\|\|\|\|\|\|\|\|\|\|\|\|\|\|\|\|\|\|\|\|\|\|\|\|\|\|\|\|\|\|\|\|\|\|\|\|\|\|\|\|\|\|\|\|\|\|\|\|\|\|\|\|\|\|\|\|\|\|\|\|\|\|\|\|\| 	ATGGGCGGTCAATGAGAAAAGCCCGTCCATAGGGCAAGAAGGGCGCTCGCCTTCCTCTCACAGCGATTCTCTCCACTCCGATCCTCTGGCCGCAACATGGACCGCCGAAGAAGAAACCCGCGTGCGTCGCAAACTGGACCTTCAAATCGTGCCGATGGTCACAGTGCTCTACTTGATGTG ``` |
| 1 | ggggacaagtttgtacaaaaaagcaggcttgatgTCACTCAAGCGCAAAGCAGT, ggggaccactttgtacaagaaagctgggtcCACTCCAGCATCAGCCGGCG | Mycgr3\|76703  JGI  Ensembl | KOG2994 Uracil DNA glycosylase; | DNA N-glycosylase activity; DNA repair;  DNA-3-methyladenine glycosylase III activity;  base-excision repair;  purine-specific mismatch base pair DNA N-glycosylase activity;  single-strand selective uracil DNA N-glycosylase activity;  uracil DNA N-glycosylase activity; | 100 % |  |  |
| 1 | ggggacaagtttgtacaaaaaagcaggcttgatgGCTTCCGCCGACGTGATCAA, ggggaccactttgtacaagaaagctgggtcCTTGTAGAAGTCGAACTTGT | Mycgr3\|76782  JGI  Ensembl | KOG0727 26S proteasome regulatory complex, ATPase RPT3; | ATP binding; ATP-dependent DNA helicase activity;  ATP-dependent RNA helicase activity;  ATP-dependent helicase activity;  ATPase activity;  ATPase activity, coupled;  ATPase activity, coupled to transmembrane movement of ions;  ATPase activity, coupled to transmembrane movement of substances;  ATPase activity, uncoupled;  DNA-dependent ATPase activity;  RNA-dependent ATPase activity;  cytoplasm;  hydrolase activity;  nucleoside-triphosphatase activity;  nucleotide binding;  nucleus;  protein catabolic process;  protein-transmembrane transporting ATPase activity;  single-stranded DNA-dependent ATP-dependent DNA helicase activity; | 100 % |  | ``` 117	GATGGCTTCCGCCGACGTGATCAACCCGGCCAACACACCAGCGCCGCATCTCCAACGACAAATCCCAAACTCCGCCATCACCAACATCGACACCGTTGAAGGTCTTGCAGACTCCGACGACCAATACGCCACATACAAGAAGTTACAGCGACAACTGGAGTACATCAAGCTACAGGAGGAGTACATCAAAGATGAGCAGCGAAGCTTGAAGCGAGAGTTGGTGCGCGCACAGGAGGAGATCAAGCGGATACAAAGTGTGCCGCTGGTCATCGGGCAGTTCATGGAGGCTATCGATCAGAACACTGGGATTGTGCA	431 	\|\|\|\|\|\|\|\|\|\|\|\|\| \|\| \|\|\|\|\|\|\|\|\|\|\|\|\|\|\|\|\|\|\|\|\|\|\|\|\|\|\|\|\|\|\|\|\|\|\|\|\|\|\|\|\|\|\|\|\|\|\|\|\|\|\|\|\|\|\|\|\|\|\|\|\|\|\|\|\|\|\|\|\|\|\|\|\|\|\|\|\|\|\|\|\|\|\|\|\|\|\|\|\|\|\|\|\|\|\|\|\|\|\|\|\|\|\|\|\|\|\|\|\|\|\|\|\|\|\|\|\|\|\|\|\|\|\|\|\|\|\|\|\|\|\|\|\|\|\|\|\|\|\|\|\|\|\|\|\|\|\|\|\|\|\|\|\|\|\|\|\|\|\|\|\|\|\|\|\|\|\|\|\|\|\|\|\|\|\|\|\|\|\|\|\|\|\|\|\|\|\|\|\|\|\|\|\|\|\|\|\|\|\|\|\|\|\|\|\|\|\|\|\|\|\|\|\|\|\|\|\|\|\|\|\|\|\|\|\|\|\|\|\|\|\|\|\|\|\|\|\|\|\|\|\|\|\|\|\|\|\|\|\|\|\|\|\|\|\|\|\|\|\|\|\|\|\|\|\|\|\|\| \|\|\|\|\|\|\|\|\|\|  \|\|  \|\|\|\|\|\| \|\| \|\|\| 	GATGGCTTCCGCCNACNTGATCAACCCGGCCAACACACCAGCGCCGCATCTCCAACGACAAATCCCAAACTCCGCCATCACCAACATCGACACCGTTGAAGGTCTTGCAGACTCCGACGACCAATACGCCACATACAAGAAGTTACAGCGACAACTGGAGTACATCAAGCTACAGGAGGAGTACATCAAAGATGAGCAGCGAAGCTTGAAGCGAGAGTTGGTGCGCGCACAGGAGGAGATCAAGCGGATACAAAGTGTGCCGCTGGTCATCGGGCAGTTCATGGANGCTATCGATCNNAANNCTGGGANTGNGCA ``` |
| 1 | ggggacaagtttgtacaaaaaagcaggcttgatgGCCTCTCCGGTGACAGCAGA, ggggaccactttgtacaagaaagctgggtcCATATTATGCGAATCCCTTC | Mycgr3\|77753  JGI  Ensembl | KOG1216 von Willebrand factor and related coagulation proteins; | nucleus; regulation of transcription, DNA-dependent;  transcription factor activity;  zinc ion binding; | 38.13869 % |  | ``` 1194	GGCCTCTCCTCCGCCGAAGACCGCCTGCACAAACTCCTCCGCGCCAAATACGACGCCGGAATGCTCAAACCCTTCAACTACGTCCGCGGCTACGCTCGCTTGTCAACCTACATGTCCGCGCACCTCACCCGGGACTCCGCGCTGCGCATCCTCCGTCAACTCGAACGGTTTCGTCCGAAATTCCGCGAGGCGATGCAGAGTTTGACTGACATGCAATTGGTTTTGGTGGAGATGTGGTTCGAAAGGAGTTTGATGGACTACGATCGAGTATTTGCGAGCATGGCGATTCCAGCGTGTTGTTGGAGA	1499 	\|\|\|\|\|\|\|\|\|\|\|\|\|\|\|\|\|\|\|\|\|\|\|\|\|\|\|\|\|\|\|\|\|\|\|\|\|\|\|\|\|\|\|\|\|\|\|\|\|\|\|\|\|\|\|\|\|\|\|\|\|\|\|\|\|\|\|\|\|\|\|\|\|\|\|\|\|\|\|\|\|\|\|\|\|\|\|\|\|\|\|\|\|\|\|\|\|\|\|\|\|\|\|\|\|\|\|\|\|\|\|\|\|\|\|\|\|\|\|\|\|\|\|\|\|\|\|\|\|\|\|\|\|\|\|\|\|\|\|\|\|\|\|\|\|\|\|\|\|\|\|\|\|\|\|\|\|\|\|\|\|\|\|\|\|\|\|\|\|\|\|\|\|\|\|\|\|\|\|\|\|\|\|\|\|\|\|\|\|\|\|\|\|\|\|\|\|\|\|\|\|\|\|\|\|\|\|\|\|\|\|\|\|\|\|\|\|\|\|\|\|\|\|\|\|\|\|\|\|\|\|\|\|\|\|\|\|\|\|\|\|\|\|\|\|\|\|\|\|\|\|\|\|\|\|\|\|\|\|\|\|\|\|\|\|\|\|\|\|\|\|\|\|\|\|\|\|\|\|\|\|\|\|\|\|\|\|\|\|\| \|\|\|\|\|\|\|\|\|\|\|\|\|\|\| 	GGCCTCTCCTCCGCCGAAGACCGCCTGCACAAACTCCTCCGCGCCAAATACGACGCCGGAATGCTCAAACCCTTCAACTACGTCCGCGGCTACGCTCGCTTGTCAACCTACATGTCCGCGCACCTCACCCGGGACTCCGCGCTGCGCATCCTCCGTCAACTCGAACGGTTTCGTCCGAAATTCCGCGAGGCGATGCAGAGTTTGACTGACATGCAATTGGTTTTGGTGGAGATGTGGTTCGAAAGGAGTTTGATGGACTACGATCGAGTATTTGCGAGCATGGCGATTCCNGCGTGTTGTTGGAGA ``` |
| 1 | ggggacaagtttgtacaaaaaagcaggcttgatgGACTCCTACGCGCAATACCA, ggggaccactttgtacaagaaagctgggtcACCATCATCATCCTCATCGT | Mycgr3\|83804  JGI  Ensembl | KOG1561 CCAAT-binding factor, subunit B (HAP2); | ATP-dependent DNA helicase activity; ATP-dependent RNA helicase activity;  ATP-dependent helicase activity;  ATPase activity;  ATPase activity, coupled;  ATPase activity, coupled to transmembrane movement of ions;  ATPase activity, coupled to transmembrane movement of substances;  ATPase activity, uncoupled;  DNA-dependent ATPase activity;  RNA-dependent ATPase activity;  nucleus;  protein-transmembrane transporting ATPase activity;  regulation of transcription, DNA-dependent;  single-stranded DNA-dependent ATP-dependent DNA helicase activity;  transcription factor activity; | 100 % |  | ``` 82	GTCCAAC-AG-CCCAA-CTACGCGCA-ACACC-AACAACCGCCGCAACTGGG-ACAGCACAA-CATGATGCCACCGCAGATGGGCTACAATCCTGCCTATGGCATG-CAGCCGCAGTATGGTGTATCG-CCGTCGCAG	210 	\|\|\|\|\|\|\| \|\| \|\|\|\|\|  \|\| \|\|\|\|\|  \| \|\| \|  \|\| \|\|\| \|\|\|\| \|\|\|\| \|\| \|\|\|\|\|\| \|\|\|\|\|\|\|\|\|\|\|\|\|\|\|\|\|\|\|\|\|\|\|\|\|\|\|\| \|\|\|\|\| \|\|\|\|\|\|\|\| \|\|\|\|\|\|\|\|\|\|\|\|\|\|\|\|\|\|\|\|\| \|\|\|\|\|\|\|\|\| 	GTCCAACGAGTCCCAANGTANGCGCANNCNCCNAGTAAGCGCNGCAANTGGGTACNGCACAACCATGATGCCACCGCAGATGGGCTACAATNCTGCCAATGGCATGCCAGCCGCAGTATGGTGTATCGGCCGTCGCAG ```      ``` 367	ATGCCACAGAGAAGAATGTCGCACACGATCAACAGCCCTGCTGTCCCGTCGCAACAACCACCTATGGGCGTACCGCGCCCTCCCACACAGCAGCAGCAAGCCGCCGCACCACCACCACAGGCCGCACAGCAATCGCCGGAAGGACCGCCGGCCGGCGCAAACGAGGAATCCCCCCTCTA	545 	\|\|\|\|\|\|\|\|\|\| \|\| \|\|\|\|\|\|\|\|\|\|\|\|\|\|\|\|\|\|\|\|\|\|\|\|\|\|\|\|\|\|\|\|\|\|\|\| \|\|\|\|\|\|\|\|\|\|\|\|\|\|\|\|\|\|\|\|\|\|\|\|\|\|\|\|\|\|\|\|\|\| \| \|  \| \|\| \| \|\|\|\|\|\|\|\|\|\| \|\| \|\|\|\|\| \|\|\|  \| \| \|  \|\|\|\| \|\|\|\|\|\|\|\|\|\|\| \|\|\|\|\|\|\|\|  \|\|\|\|  \| \|\|\|\|\|\|\|\|\| \|\|\|\| 	ATGCCACAGANAANAATGTCGCACACGATCAACAGCCCTGCTGTCCCGTCNCAACAACCACCTATGGGCGTACCGCGCCCTCCCANANANNANCANCNAGCCGCCGCANCANCACCANAGGNNGNANANNAATCNCCGGAAGGACCNCCGGCCGGNNCAAANNANGAATCCCCC-TCTA ```      ``` 1	ATGGACT-CCTACGCGCAATACCACCCGCAGCAGCAGTCTGCATACCCACATCCTCACAATGGTGCGCCGCAACAAAGTCCCGTCCAACAGCCCAACTACGCGCAACACCAACAACCGCCGCAACTGGGACAGCACAACATGATGCCACCGCAGATGGGCTACAATCCTGCCTATGGCATGCAGCCGCAGTATGGTGTATC-GCCGTCGCAGGCCGCTGCCATGGCCACAGCCGCGGCCTCAGGTTACCCGGCCTACTCGATGCCCGACTCATCAATGCCTGGCTCACTGTCGCAGACTTCGCCGCGCATGAAGACTGATGGATCTTTGCAACGACCTCCGCAGTCACCACGACAACCTCAGATGAAC	366 	\|\|\|\|\|\|\| \|\|\|\|\|\|\|\|\|\|\|\|\|\|\|\|\|\|\|\|\|\|\|\|\|\|\|\|\|\|\|\|\|\|\|\|\|\|\|\|\|\|\|\|\|\|\|\|\|\|\|\|\|\|\|\|\|\|\|\|\|\|\|\|\|\|\|\|\|\|\|\|\|\|\|\|\|\|\|\|\|\|\|\|\|\|\|\|\|\|\|\|\|\|\|\|\|\|\|\|\|\|\|\|\|\|\|\|\|\|\|\|\|\|\|\|\|\|\|\|\|\|\|\|\|\|\|\|\|\|\|\|\|\|\|\|\|\|\|\|\|\|\|\|\|\|\|\|\|\|\|\|\|\|\|\|\|\|\|\|\|\|\|\|\|\|\|\|\|\|\|\|\|\|\|\|\|\|\|\|\|\|\|\|\|\|\|\|\|\|\|\|\| \|\|\|\|\|\|\|\|\|\|\|\|\|\|\|\|\|\|\|\|\|\|\|\|\|\|\|\|\|\|\|\|\|\|\|\|\|\|\|\|\|\|\|\|\|\|\|\|\|\|\|\|\|\|\|\|\|\|\|\|\|\|\|\|\|\|\|\|\|\|\|\|\|\|\|\|\|\|\|\|\|\|\|\|\|\|\|\|\|\|\|\|\|\|\|\|\|\|\|\|\|\|\|\|\|\|\|\|\|\|\|\|\|\|\|\|\|\|\|\|\|\|\|\|\|\|\|\|\|\|\|\|\|\|\|\|\|\|\|\|\|\|\|\|\|\|\|\|\|\|\|\|\|\|\|\|\|\|\|\|\|\|\|\|\|\| 	ATGGACTCCCTACGCGCAATACCACCCGCAGCAGCAGTCTGCATACCCACATCCTCACAATGGTGCGCCGCAACAAAGTCCCGTCCAACAGCCCAACTACGCGCAACACCAACAACCGCCGCAACTGGGACAGCACAACATGATGCCACCGCAGATGGGCTACAATCCTGCCTATGGCATGCAGCCGCAGTATGGTGTATCGGCCGTCGCAGGCCGCTGCCATGGCCACAGCCGCGGCCTCAGGTTACCCGGCCTACTCGATGCCCGACTCATCAATGCCTGGCTCACTGTCGCAGACTTCGCCGCGCATGAAGACTGATGGATCTTTGCAACGACCTCCGCAGTCACCACGACAACCTCAGATGAAC ``` |
| 1 | ggggacaagtttgtacaaaaaagcaggcttgatgGCGCTCCTCGTAGACAAACA, ggggaccactttgtacaagaaagctgggtcATCATCGAAGTCGTCGAGGT | Mycgr3\|85700  JGI  Ensembl | KOG2035 Replication factor C, subunit RFC3; | DNA polymerase activity; DNA-directed DNA polymerase activity;  nucleoside-triphosphatase activity;  nucleotide binding; | 100 % |  | ``` 1	ATGGCGCTCCTCGTAGACAAACACC-GTCCACGGACATTGGACTCCCTCTCGTACCATCCCGAACTCTCAGACCGACTGCGAGCTCTGGCGGCCAGCGGCGACTTTCCCCATCTCCTCATCTATGGCCCTTCCGGAGCAGGCAAGAAGACACGCATTTCGGCCACCCTACGAGCTCTCTACGGTCCAGGAGTCGAGAAAATCAAGATCGACAGTCGCGTGTTCCAGACGACCAGCAATCGCAAGCTCGAATTCAACATCGTCTCATCAAACTACCACCTCGAAATCACTCCCTCAGATGTCGGCAACTACGATCGTGTTGTCATTCAGGATCTGCTCAAAGAGGTGGCACAGACACAGCAGGTGGATGCAGCAGCGAAACAACGCTTCAAGGTCGTCCTCATCAACGAGGCGGACCACTTGACTCGAGATGCTCAGGCGGCGCTGAGGCGAACAATGGAGAAATACAGCCCCAATTTGCGATTGATCCTGGTCGCGAACAGTACAAGTAATATCATTGCTCCTATTCGTTCGCGATGTCTGTTGGTCAGAGTGTCCGCCCCTTCAGAGGAGGAGATTTGCAAAGTGCTGGCAAAGGTGGGCAAGGATGAGCGATACCAGGCATGCGAGACGCTGGAGAAGAGAATCGCCCGGGACAGCAAGCG	662 	\|\|\|\|\|\|\|\|\|\|\|\|\|\|\|\|\|\|\|\|\|\|\|\|\| \|\|\|\|\|\|\|\|\|\|\|\|\|\|\|\|\|\|\|\|\|\|\|\|\|\|\|\|\|\|\|\|\|\|\|\|\|\|\|\|\|\|\|\|\|\|\|\|\|\|\|\|\|\|\|\|\|\|\|\|\|\|\|\|\|\|\|\|\|\|\|\|\|\|\|\|\|\|\|\|\|\|\|\|\|\|\|\|\|\|\|\|\|\|\|\|\|\|\|\|\|\|\|\|\|\|\|\|\|\|\|\|\|\|\|\|\|\|\|\|\|\|\|\|\|\|\|\|\|\|\|\|\|\|\|\|\|\|\|\|\|\|\|\|\|\|\|\|\|\|\|\|\|\|\|\|\|\|\|\|\|\|\|\|\|\|\|\|\|\|\|\|\|\|\|\|\|\|\|\|\|\|\|\|\|\|\|\|\|\|\|\|\|\|\|\|\|\|\|\|\|\|\|\|\|\|\|\|\|\|\|\|\|\|\|\|\|\|\|\|\|\|\|\|\|\|\|\|\|\|\|\|\|\|\|\|\|\|\|\|\|\|\|\|\|\|\|\|\|\|\|\|\|\|\|\|\|\|\|\|\|\|\|\|\|\|\|\|\|\|\|\|\|\|\|\|\|\|\|\|\|\|\|\|\|\|\|\|\|\|\|\|\|\|\|\|\|\|\|\|\|\|\|\|\|\|\|\|\|\|\|\|\|\|\|\|\|\|\|\|\|\|\|\|\|\|\|\|\|\|\|\|\|\|\|\|\|\|\|\|\|\|\|\|\|\|\|\|\|\|\|\|\|\|\|\|\|\|\|\|\|\|\|\|\|\|\|\|\|\|\|\|\|\|\|\|\|\|\|\|\|\|\|\|\|\|\|\|\|\|\|\|\|\|\|\|\|\|\|\|\|\|\|\|\|\|\|\|\|\|\|\|\|\|\|\|\|\|\|\|\|\|\|\|\|\|\|\|\|\|\|\|\|\|\|\|\|\|\|\|\|\|\|\|\|\|\|\|\|\|\|\|\|\|\|\|\|\|\|\|\|\|\|\|\|\|\|\|\|\|\|\|\|\|\|\|\|\|\|\|\|\|\|\|\|\|\|\|\|\|\|\|\|\|\|\|\|\|\|\|\|\|\|\|\|\|\|\|\|\|\|\|\|\|\|\|\|\|\|\|\|\| \|\|\|\|\|\|\|\|\|\|\|\|\|\|\|\| \|\|\|\| \|\| \|\|\|\|\|\|\|\|\|\|\|\|\|\|\|\|\|\|\|  \|\|\|\|\|\|\|\|\|\|\|\|\|\| \|\|\|\|\|\|\|\|  \|\|\|\|\| \| \|  \|\|\|\|\| \|\| \| \|\|\|\| \|\| \|\|\|\| \| \|\|\|\|\|\| 	ATGGCGCTCCTCGTAGACAAACACCGGTCCACGGACATTGGACTCCCTCTCGTACCATCCCGAACTCTCAGACCGACTGCGAGCTCTGGCGGCCAGCGGCGACTTTCCCCATCTCCTCATCTATGGCCCTTCCGGAGCAGGCAAGAAGACACGCATTTCGGCCACCCTACGAGCTCTCTACGGTCCAGGAGTCGAGAAAATCAAGATCGACAGTCGCGTGTTCCAGACGACCAGCAATCGCAAGCTCGAATTCAACATCGTCTCATCAAACTACCACCTCGAAATCACTCCCTCAGATGTCGGCAACTACGATCGTGTTGTCATTCAGGATCTGCTCAAAGAGGTGGCACAGACACAGCAGGTGGATGCAGCAGCGAAACAACGCTTCAAGGTCGTCCTCATCAACGAGGCGGACCACTTGACTCGAGATGCTCAGGCGGCGCTGAGGCGAACAATGGAGAAATACAGCCCCAATTTGCGATTGATCCTGGTCGCGAACAGTACAAGTAATATCATTGCTCCTATTCGTTCGCGATGTCTGTTGGTCANAGTGTCCGCCCCTTCANAGGANGANATTTGCAAAGTGCTGGCAAN-GTGGGCAAGGATGANCGATACCANNCATGCNA-ANNCTGGAAAAAA-AATCNCC-GGGA-ANCAAGCG ``` |
| 1 | ggggacaagtttgtacaaaaaagcaggcttgatgTCGAAAGCAGAGCGAGCCAC, ggggaccactttgtacaagaaagctgggtcAACCTCCAACGCTGCTGTAG | Mycgr3\|85924  JGI  Ensembl | KOG2409 KRR1-interacting protein involved in 40S ribosome biogenesis; | DNA topoisomerase type I activity; | 100 % | chr\_5:98255 G->A; | ``` 1	ATGTCGAAAGCAGAGCGAGCCACAAAAAGGCAGAAGTTCCTCGAAGAGGAGTCCGACGAGAGCGATGCCGGAGTTCCCCTCCTCCAAGTCAACGAGGAGTACGCCCGACGATTCGAGCACAACAAAAAGCGAGAAGAGCTATCGCGGCTGGAAGAAAAGTTCAAGGATAACAAGAGGAAGCGAGATGACGACAGCGAGAATGACTCCTCCTCCGGCTCCGACGAGGACGATGTCGCCGAACTAGCGACCGCCGACCTGGACGAGGAGATCTTT	273 	\|\|\|\|\|\|\|\|\|\|\|\|\|\|\|\|\|\|\|\|\|\|\|\|\|\|\|\|\|\|\|\|\|\|\|\|\|\|\|\|\|\|\|\|\|\|\|\|\|\|\|\|\|\|\|\|\|\|\|\|\|\|\|\|\|\|\|\|\|\|\|\|\|\|\|\|\|\|\|\|\|\|\|\|\|\|\|\|\|\|\|\|\|\|\|\|\|\|\|\|\|\|\|\|\|\|\|\|\|\|\|\|\|\|\|\|\|\|\|\|\|\|\|\|\|\|\|\|\|\|\|\|\|\|\|\|\|\|\|\|\|\|\|\|\|\|\|\|\|\|\|\|\|\|\|\|\|\|\|\|\|\|\|\|\|\|\|\|\|\|\|\|\|\|\|\|\|\|\|\|\|\|\|\|\|\|\|\|\|\|\|\|\|\|\|\|\|\|\|\|\|\|\|\|\|\|\|\|\|\|\|\|\|\|\|\|\|\|\|\|\|\|\|\|\|\|\|\|\|\|\|\|\|\| \|\|\|\|\|\|\|\| \|\|\|\|\|\|\|\|\|\|\|\|\|\|\|\|\| \| \|\| \|\|\|\|\|\| 	ATGTCGAAAGCAGAGCGAGCCACAAAAAGGCAGAAGTTCCTCGAAGAGGAGTCCGACGAGAGCGATGCCGGAGTTCCCCTCCTCCAAGTCAACGAGGAGTACGCCCGACGATTCGAGCACAACAAAAAGCGAGAAGAGCTATCGCGGCTGGAAGAAAAGTTCAAGGATAACAAGAGGAAGCGAGATGACGACAGCGAGAATGACTCCTCCTCCGGCTCCGACGAGGACGATGTCNCCGAACTANCGACCGCCGACCTGGACNANGANATCTTT ``` |
| 1 | ggggacaagtttgtacaaaaaagcaggcttgatgGACGACGGCTACGATGTGCC, ggggaccactttgtacaagaaagctgggtcTGAGCCCACACTGGTGGAGC | Mycgr3\|87533  JGI  Ensembl | KOG0287 Postreplication repair protein RAD18; | DNA repair; damaged DNA binding;  nucleic acid binding;  nucleus;  protein binding;  zinc ion binding; | 0 % |  | ``` 1	ATGGACGACGGCTACGATGTGCCTGACGCCACAGACTGGCTCGAGACGCCTCTCAAGGACTTTGCCAGCCTAGAAACCGCACTTCACTGCCAAATATGCAAGGAGTTCTACGACACACCGATGATCACATCATGCAACCATACTTTCTGTTCGAAGTGCATACGAACCTCACTTTCTGCGGATGGGAAGTGTCCAGCGTGTCGAGGTCCAGATCAAGCAAGCAAGTTGCGCAATAATTGGGCGGTGCAAGAGGTGGTGTCGACATTCATCACAGCAAGGCCGGCAGCTTTACAGGTGGCGAGGAAGGCACTGGAAGAGGAAGGCCAGAAGAAGTCTGGGAAGCGAAAACGACCAGTCGTCCTGGACTCAGAAGATCTCGTCGAGGTAGAGGGCCCGGCTCGAGCAACAAGAAGCAAGAGCAGGCGCATTGCGGCATCACAAGAATCACATACGGAGGCTATCGAGATTGAGGACAGCGAGGATGGGGAAGAGGACTACAGATCCGGCACGCCGGCGGACGATGGCTTAGTCGCCTGCCCGCTCGAATGTGG	551 	\|\|\|\|\|\|\|\|\|\|\|\|\|\|\|\|\|\|\|\|\|\|\|\|\|\|\|\|\|\|\|\|\|\|\|\|\|\|\|\|\|\|\|\|\|\|\|\|\|\|\|\|\|\|\|\|\|\|\|\|\|\|\|\|\|\|\|\|\|\|\|\|\|\|\|\|\|\|\|\|\|\|\|\|\|\|\|\|\|\|\|\|\|\|\|\|\|\|\|\|\|\|\|\|\|\|\|\|\|\|\|\|\|\|\|\|\|\|\|\|\|\|\|\|\|\|\|\|\|\|\|\|\|\|\|\|\|\|\|\|\|\|\|\|\|\|\|\|\|\|\|\|\|\|\|\|\|\|\|\|\|\|\|\|\|\|\|\|\|\|\|\|\|\|\|\|\|\|\|\|\|\|\|\|\|\|\|\|\|\|\|\|\|\|\|\|\|\|\|\|\|\|\|\|\|\|\|\|\|\|\|\|\|\|\|\|\|\|\|\|\|\|\|\|\|\|\|\|\|\|\|\|\|\|\|\|\|\|\|\|\|\|\|\|\|\|\|\|\|\|\|\|\|\|\|\|\|\|\|\|\|\|\|\|\|\|\|\|\|\|\|\|\|\|\|\|\|\|\|\|\|\|\|\|\|\|\|\|\|\|\|\|\|\|\|\|\|\|\|\|\|\|\|\|\|\|\|\|\|\|\|\|\|\|\|\|\|\|\|\|\|\|\|\|\|\|\|\|\|\|\|\|\|\|\|\|\|\|\|\|\|\|\|\|\|\|\|\|\|\|\|\|\|\|\|\|\|\|\|\|\|\|\|\|\|\|\|\|\| \|\|\|\|\|\|\|\|\|\|\|\|\|\|\|\|\| \|\|\|\|\|\|\|\|\|\|\|\|\|\|\|\|\|\|\|\|\|\|\|\|\|\|\|\|\|\|\|\|\|\|\|\|\|\|\|\|\|\|\|\|\|\|\|\|\|\|\|\|\|\|\|\|\|\|\|\|\|\|\|\|\|\|\|\|\|\|\|\|\|\|\|\|\|\|\|\|\|\| \|\|\|\|\|\|\|\|\|\|\|\|\|\|\|\|\|\| \| \|\|\|\|\|\|  \|\|\|\| \|\| \| \|\|\|\|\| \|\|\| \|\|\|\|\|\|\|\| \|\| \|\|\|\|\|\|\| \|\|\| \|\|\|\|\|\|\| 	ATGGACGACGGCTACGATGTGCCTGACGCCACAGACTGGCTCGAGACGCCTCTCAAGGACTTTGCCAGCCTAGAAACCGCACTTCACTGCCAAATATGCAAGGAGTTCTACGACACACCGATGATCACATCATGCAACCATACTTTCTGTTCGAAGTGCATACGAACCTCACTTTCTGCGGATGGGAAGTGTCCAGCGTGTCGAGGTCCAGATCAAGCAAGCAAGTTGCGCAATAATTGGGCGGTGCAAGAGGTGGTGTCGACATTCATCACAGCAAGGCCGGCAGCTTTACAGGTGGCGAGGAAGGCACTGGAAGAGGAAGGCCAGAAGAAGTCTGGGAAGCGAAAACGACCAGTCGTCCTGGACTCANAAGATCTCGTCGAGGTANAGGGCCCGGCTCGAGCAACAAGAAGCAAGAGCAGGCGCATTGCGGCATCACAAGAATCACATACGGAGGCTATCGAGATTGANGACAGCGAGGATGGGGAANANGACTACNNATCCNGCNCNCCGGCNGACNATGGCTTANTCNCCTGCCCNCTCNAATGTGG ``` |
| 1 | ggggacaagtttgtacaaaaaagcaggcttgatgCCGCATTTGCGCATCCCTTT, ggggaccactttgtacaagaaagctgggtcGCGAGCATCTTGCGGCCCAG | Mycgr3\|88443  JGI  Ensembl |  |  | 2.01613 % |  |  |
| 1 | ggggacaagtttgtacaaaaaagcaggcttgatgCTCAAGAACATGAGCGAATT, ggggaccactttgtacaagaaagctgggtcGTCTTTCGCAGCTTCTGTCT | Mycgr3\|89100  JGI  Ensembl | KOG2859 DNA repair protein, member of the recA/RAD51 family; |  | 100 % |  | ``` 1	ATGCTCAAGAACATGAGCGAATTTCGGCAGTCCAATATGCCGCCAGTGTTGGAGCTCATATCACTGGTCTCAGGAGGCGGCAAGACACACCTCCTCTACCATCTCACAGCGATTGCAATCATACCCAAGGCTCGAGGTGGGAGACAGTCCGCCGTGGTGATCATCGACACAGACGGTAGGTTCTCCGTTCCTCGTCTCGCAGAACAAGTCCAACTTCTCCTCGCCCAACACTCTGCCCAAAATCCCGCCAATGCTTCTCCATCACCTCCCTCTGATATGGAAGAGACCATCCACGCCGCGCTGCGACACGTCCACATCTTCCATCCGCATTCCCTTGCCTCCACGACCGCCACGCTCCGTTCTCTCCCAACCTACCTCTTCAACCCTACAGCCCACCACTCCTTCGACCGCGCCGTCGCCTTCATCGCTCTCGACTCCGCCTCAGCATTCTACTGGCAATACCGCTCGGAAGCCGACGACGCCGCACTGGTCCCGCCCTCGGGTATCGCGGGCAGTAACAAATCAGCACGAGCCTACACCGATCTGCTGTCCGCTTTCAGGCACTCCTCCAAAGTCCTGAATGCACCTTTAATCCTCACGAATCGACACCTCGGTGCCCTGAAAGATCCCAACTCCATGGGTCCTTCTTCTCGGTCCTTTCGTCCATCGCTGC-CTCCT	678 	\|\|\|\|\|\|\|\|\|\|\|\|\|\|\|\|\|\|\|\|\|\|\|\|\|\|\|\|\|\|\|\|\|\|\|\|\|\|\|\|\|\|\|\|\|\|\|\|\|\|\|\|\|\|\|\|\|\|\|\|\|\|\|\|\|\|\|\|\|\|\|\|\|\|\|\|\|\|\|\|\|\|\|\|\|\|\|\|\|\|\|\|\|\|\|\|\|\|\|\|\|\|\|\|\|\|\|\|\|\|\|\|\|\|\|\|\|\|\|\|\|\|\|\|\|\|\|\|\|\|\|\|\|\|\|\|\|\|\|\|\|\|\|\|\|\|\|\|\|\|\|\|\|\|\|\|\|\|\|\|\|\|\|\|\|\|\|\|\|\|\|\|\|\|\|\|\|\|\|\|\|\|\|\|\|\|\|\|\|\|\|\|\|\|\|\|\|\|\|\|\|\|\|\|\|\|\|\|\|\|\|\|\|\|\|\|\|\|\|\|\|\|\|\|\|\|\|\|\|\|\|\|\|\|\|\|\|\|\|\|\|\|\|\|\|\|\|\|\|\|\|\|\|\|\|\|\|\|\|\|\|\|\|\|\|\|\|\|\|\|\|\|\|\|\|\|\|\|\|\|\|\|\|\|\|\|\|\|\|\|\|\|\|\|\|\|\|\|\|\|\|\|\|\|\|\|\|\|\|\|\|\|\|\|\|\|\|\|\|\|\|\|\|\|\|\|\|\|\|\|\|\|\|\|\|\|\|\|\|\|\|\|\|\|\|\|\|\|\|\|\|\|\|\|\|\|\|\|\|\|\|\|\|\|\|\|\|\|\|\|\|\|\|\|\|\|\|\|\|\|\|\|\|\|\|\|\|\|\|\|\|\|\|\|\|\|\|\|\|\|\|\|\|\|\|\|\|\|\|\|\|\|\|\|\|\|\|\|\|\|\|\|\|\|\|\|\|\|\|\|\|\|\|\|\|\|\|\|\|\|\|\|\|\|\|\|\|\|\|\|\|\|\|\|\|\|\|\|\|\|\|\|\|\|\|\|\|\|\|\|\|\|\|\|\|\|\|\|\|\|\|\|\|\|\|\|\|\|\|\|\|\|\|\|\|\|\|\|\|\|\|\|\|\|\|\|\|\|\|\|\|\|\|\|\|\|\|\|\|\|\|\|\|\|\| \|\|\|\|\|\|\|\|\|\|\|\|\|\|\|\|\|\|\|\|\|\|\|\|\|\|\|\|\|\|\|\|\|\|\|\|\|\|\|\|\|\|\|\|\|\|\|\|\|\|\|\|\|\|\|\|\|\|\|\|\|\|\|\|\|\|\|\|\|\|\|\|\|\|\|\|\|\|\|\|\|\|\| \|\|\|\|\|\|\|\|\|\|\| \|\|\|\|\| \|\|\|\|\|\|\|\|\|\|\|\|\|\|\|\|\|\|\|\|\|\|\|\|\|\|\|\| \|\| \| \|\|\|\|\|\|\| \|\|\| \|\|\|\|\| 	ATGCTCAAGAACATGAGCGAATTTCGGCAGTCCAATATGCCGCCAGTGTTGGAGCTCATATCACTGGTCTCAGGAGGCGGCAAGACACACCTCCTCTACCATCTCACAGCGATTGCAATCATACCCAAGGCTCGAGGTGGGAGACAGTCCGCCGTGGTGATCATCGACACAGACGGTAGGTTCTCCGTTCCTCGTCTCGCAGAACAAGTCCAACTTCTCCTCGCCCAACACTCTGCCCAAAATCCCGCCAATGCTTCTCCATCACCTCCCTCTGATATGGAAGAGACCATCCACGCCGCGCTGCGACACGTCCACATCTTCCATCCGCATTCCCTTGCCTCCACGACCGCCACGCTCCGTTCTCTCCCAACCTACCTCTTCAACCCTACAGCCCACCACTCCTTCGACCGCGCCGTCGCCTTCATCGCTCTCGACTCCGCCTCAGCATTCTACTGGCAATACCGCTCGGAAGCCGACGACGCCGCACTGGTCCCGCCCTCGGGTATCGCGGGCAGTAACAAATCANCACGAGCCTACACCGATCTGCTGTCCGCTTTCAGGCACTCCTCCAAAGTCCTGAATGCACCTTTAATCCTCACGAATCGACACNTCGGTGCCCTG-AAGATNCCAACTCCATGGGTCCTTCTTCTCGGTCNTT-C-TCCATCGNTGCTCTCCT ``` |
| 1 | ggggacaagtttgtacaaaaaagcaggcttgatgCTTGGACGGATGTTGTTGCG, ggggaccactttgtacaagaaagctgggtcAGCAGCTGAGGCCAGCGGAA | Mycgr3\|89562  JGI  Ensembl | KOG4475 FOG: Immunoglobin and related proteins; | DNA binding; regulation of transcription, DNA-dependent; | 0 % |  |  |
| 1 | ggggacaagtttgtacaaaaaagcaggcttgatgGCGTCTCCGGCTACTGCTGC, ggggaccactttgtacaagaaagctgggtcTCGCCCGGCACTATAGTTGT | Mycgr3\|89727  JGI  Ensembl | KOG2483 Upstream transcription factor 2/L-myc-2 protein; | nucleus; regulation of transcription;  transcription regulator activity; | 100 % |  | ``` 161	ACAACTCGGCGAACAGTGGCCTCCAGGTTCACACGCTCCTCAATCACCCGGACGAGTCACCTTCTCGAAGCTCGATCCCGAACACACCATCATCTGCGAGGATTCCGGCCTC	272 	\|\|\|\|\|\|\|\|\|\|\|\|\|\|\|\|\|\|\|\|\|\|\|\|\| \|\|\|\|\|\|\|\|\|\|\|\|\|\|\|\|\|\|\|\| \|\|\|\|\|\| \| \|\| \|\|\|\|\|\|\| \|\| \|\|\|\|\|\| \|\| \|\| \|   \|\|\|\| \|\|\|\|  \|\|\|\|\|\|\| \|\| \|\|\| 	ACAACTCGGCGAACAGTGGCCTCCANGTTCACACGCTCCTCAATCANCCGGACNANTCNCCTTCTCNAANCTCGATNCCNAANANNNCATCNTCTGNNAGGATTCNGGNCTC ```      ``` 1	ATGGCGTCTCCGGCTACTGCTGCTCAGCAGCACCAAGGACTTCCCTCCATCACGACCCTCACCAATGGCCTGCGACCATCCGCTCCCATCTCGCCCGATCAGCAATCAACGTCCGACACCACCCGTGATTCTGGCACATGGCCGCAACCACAAAATAAGC	160 	\|\|\|\|\|\|\|\|\|\|\|\|\|\|\|\|\|\|\|\|\|\|\|\|\|\|\|\|\|\|\|\|\|\|\|\|\|\|\|\|\|\|\|\|\|\|\|\|\|\|\|\|\|\|\|\|\|\|\|\|\|\|\|\|\|\|\|\|\|\|\|\|\|\|\|\|\|\|\|\|\|\|\|\|\|\|\|\|\|\|\|\|\|\|\|\|\|\|\|\|\|\|\|\|\|\|\|\|\|\|\|\|\|\|\|\|\|\|\|\|\|\|\|\|\|\|\|\|\|\|\|\|\|\|\|\|\|\|\|\|\|\|\|\|\|\|\|\|\|\|\|\|\|\|\|\|\|\|\|\| 	ATGGCGTCTCCGGCTACTGCTGCTCAGCAGCACCAAGGACTTCCCTCCATCACGACCCTCACCAATGGCCTGCGACCATCCGCTCCCATCTCGCCCGATCAGCAATCAACGTCCGACACCACCCGTGATTCTGGCACATGGCCGCAACCACAAAATAAGC ``` |
| 1 | ggggacaagtttgtacaaaaaagcaggcttgatgCTCGCCGGAAGCACGACGGT, ggggaccactttgtacaagaaagctgggtcGAACTGCTGCAATTGCGGTC | Mycgr3\|89934  JGI  Ensembl |  |  | 5.34031 % | chr\_1:4647535 C->A; chr\_1:4647577 A->C; |  |
| 1 | ggggacaagtttgtacaaaaaagcaggcttgatgATCTACCACAACTGCCCTGT, ggggaccactttgtacaagaaagctgggtcCAGACCGCTTGTCCAGAAAA | Mycgr3\|90899  JGI  Ensembl | KOG2992 Nucleolar GTPase/ATPase p130; | DNA binding; cell redox homeostasis;  nucleosome;  nucleosome assembly;  nucleus; | 0 % |  |  |
| 1 | ggggacaagtttgtacaaaaaagcaggcttgatgGCAAACATGAACACCGACGC, ggggaccactttgtacaagaaagctgggtcCAATGCTTCGAGCGACGGCC | Mycgr3\|91241  JGI  Ensembl |  |  | 100 % |  | ``` 1	ATGGCAAACATGAACACCGACGCAGAGGATGAAATCAAACCCCTTCGACAAGCGAATAGAGCTTACCGGCGGCAGCTTGCGGAGACGAACGATGCCCTCAATTACACCAGACGGCAATTGGCATCTGCGCATGGTCTCATCGATCGCCTCAGACACCAAGCCTCAGAGCTCGACACTGTTCCCTCGTCACGAATCGGAGCCGAAGGGAGAATGTTGGCAATAATTGGCCCTAGACTCAAGTTCCTCTCTCCAGACGATGTGGGTTACCCCGACTGCAAGGACGACCCTGTGGACTGGACCGACAACGAGGCATGGCTTCAAGGCTTACGACGACCGTCATCGCCAGACGAAATGGAAGTACTGGACAGAAAGCACACGTATCAAGTTCAGGGAATCAGCGACGTCTGGCCAAACCCGGGCTACCCATGGCTTCGTGGGCAGACACTGCCGCGGCCGCAAACTGAAGCCGACCAGGCTGCTGATGAAGTGAACGAACGCGCAGCAACAGCTGGACAATCAGCTCAGCCGGCAATTTCAACATCTCTCGAACCTCAAAGCAATCATATCAATGAAAGGTCAGCAAAACAAACAGCTTCAGTGGCGGTCGGCAAAGACGGAGAAGCCTAC	627 	\|\|\|\|\|\|\|\|\|\|\|\|\|\|\|\|\|\|\|\|\|\|\|\|\|\|\|\|\|\|\|\|\|\|\|\|\|\|\|\|\|\|\|\|\|\|\|\|\|\|\|\|\|\|\|\|\|\|\|\|\|\|\|\|\|\|\|\|\|\|\|\|\|\|\|\|\|\|\|\|\|\|\|\|\|\|\|\|\|\|\|\|\|\|\|\|\|\|\|\|\|\|\|\|\|\|\|\|\|\|\|\|\|\|\|\|\|\|\|\|\|\|\|\|\|\|\|\|\|\|\|\|\|\|\|\|\|\|\|\|\|\|\|\|\|\|\|\|\|\|\|\|\|\|\|\|\|\|\|\|\|\|\|\|\|\|\|\|\|\|\|\|\|\|\|\|\|\|\|\|\|\|\|\|\|\|\|\|\|\|\|\|\|\|\|\|\|\|\|\|\|\|\|\|\|\|\|\|\|\|\|\|\|\|\|\|\|\|\|\|\|\|\|\|\|\|\|\|\|\|\|\|\|\|\|\|\|\|\|\|\|\|\|\|\|\|\|\|\|\|\|\|\|\|\|\|\|\|\|\|\|\|\|\|\|\|\|\|\| \|\|\|\|\|\|\|\|\|\|\|\|\|\|\|\|\|\|\|\|\|\|\|\|\|\|\|\|\|\|\|\| \|\|\|\|\|\|\|\|\|\|\|\|\|\|\|\|\|\|\|\|\|\|\|\|\|\|\|\|\|\|\|\|\|\|\|\|\|\|\|\|\|\|\|\|\|\|\|\|\|\|\|\|\|\|\|\|\|\|\|\|\|\|\|\|\|\|\|\|\|\|\|\|\|\|\|\|\|\|\|\|\|\|\|\|\|\|\|\|\|\|\|\|\|\|\|\|\|\|\|\|\|\|\|\|\|\|\|\|\|\|\|\|\|\|\|\|\|\|\|\|\|\|\|\|\|\|\|\|\|\|\|\|\|\|\|\|\|\|\|\|\|\|\|\|\|\|\|\|\|\|\|\|\|\|\|\|\|\|\|\|\|\| \|\|\|\|\|\|\| \|\|\|\|\|\|\|\|\|\|\|\| \|\|\|\|\|\|\|\|\|\|\|\|\|\| \|\|\|\|\|\|\|\|\|\|\|\|\|\|\|\|\| \|\|\|\| \|\|\|\|\| \|\|\|\|\| \|\|\|\|\|\|\|\|\|\|\|\|\|\|\|\|\| \| \|\|\|\|\| \|\|\|\|\| \|\|\|\|\|\|\|\|\|  \| \|\|\|\| \|\|\|  \|\|\|\|  \|\|\|\|\|\|\|\| \|\| \|\| \|\|\| \| \|\| \|\|\|\|\| 	ATGGCAAACATGAACACCGACGCAGAGGATGAAATCAAACCCCTTCGACAAGCGAATAGAGCTTACCGGCGGCAGCTTGCGGAGACGAACGATGCCCTCAATTACACCAGACGGCAATTGGCATCTGCGCATGGTCTCATCGATCGCCTCAGACACCAAGCCTCAGAGCTCGACACTGTTCCCTCGTCACGAATCGGAGCCGAAGGGAGAATGTTGGCAATAATTGGCCCTAGACTCAAGTTCCTCTCTCCAGACGATGTGGGTTACCCNGACTGCAAGGACGACCCTGTGGACTGGACCGAGAACGAGGCATGGCTTCAAGGCTTACGACGACCGTCATCGCCAGACGAAATGGAAGTACTGGACAGAAAGCACACGTATCAAGTTCAGGGAATCAGCGACGTCTGGCCAAACCCGGGCTACCCATGGCTTCGTGGGCAGACACTGCCGCGGCCGCAAACTGAANCCGACCANGCTGCTGATGAANTGAACGAACGCGCANCAACAGCTGGACAATCANCTCANCCGGCNATTTCNACATCTCTCGAACCTCANANCAATCNTATCANTGAAAGGTCNNCNAAACNAACN-CTTCNNTGGCGGTCNGCNAANACG-ANAANCCTAC ``` |
| 1 | ggggacaagtttgtacaaaaaagcaggcttgatgACCGCAGTCCGCTCTGCAGG, ggggaccactttgtacaagaaagctgggtcCCTCTGCGCCATCCACCGTT | Mycgr3\|91275  JGI  Ensembl |  |  | 100 % |  | ``` 1	ATGACCGCAGTCCGCTCTGCAGGCGTGCACTACTTTCAGCCGCCGTTTGATCAGATTCAGATGTCCGACCACTACGCCAACGATGCATGGCGCCGCTCATATCCCCAGCAGATGCCGCTGCCATCGAATAACTGGCACGATGGTTCCAACAACATGCCCGTTATGACCCACGGCATGCTCGAGACCGACCTTCGCATGTTTCAACAATCAGAAGCAGGAACCTACAACACGCCCTTCACCAGCGGTCAGCAACGTATCTCGCATGCGCTCGTCTCGCCGCGCCTCGCATTTCCACCTGATGCTGCATGGGCTTCCGACATGGAACTGGGCACGCCATCAACCGGCACATTCCCATCAAACATCGACGAAGGCTACACCATTCGACATACCTCCAACCACCTCACCCTACCATCCACCCACGGACATGACCCCTCCGACACGAACAGCCCTCGCAGCTCCTTCGGCGACCAATCACCCTCATCCCAATATGTCCAACCCATCTTCAAGTCCGCACCAAAACGTCCCGGCATCGCTCGCGCAGTCACAGCACCCGACACAATCCCTCAACGACCACTCCACTCCCCACCAATCAAACGCTCCCCTTCCGACGACGGAGAAGACGACGACTACTTTCCCTCCGCCTCCGACCCAAAAACCTCCTCTCGCAACCGCAAACGTGCACGAATCCCACACACAGCCGTGGAGAGGCGATATAGAGAGAATCTGAACGCACACCTCGATCGTTTGCGGCAGACGGTGCCCGAGCTG	768 	\|\|\|\|\|\|\|\|\|\|\|\|\|\|\|\|\|\|\|\|\|\|\|\|\|\|\|\|\|\|\|\|\|\|\|\|\|\|\|\|\|\|\|\|\|\|\|\|\|\|\|\|\|\|\|\|\|\|\|\|\|\|\|\|\|\|\|\|\|\|\|\|\|\|\|\|\|\|\|\|\|\|\|\|\|\|\|\|\|\|\|\|\|\|\|\|\|\|\|\|\|\|\|\|\|\|\|\|\|\|\|\|\|\|\|\|\|\|\|\|\|\|\|\|\|\|\|\|\|\|\|\|\|\|\|\|\|\|\|\|\|\|\|\|\|\|\|\|\|\|\|\|\|\|\|\|\|\|\|\|\|\|\|\|\|\|\|\|\|\|\|\|\|\|\|\|\|\|\|\|\|\|\|\|\|\|\|\|\|\|\|\|\|\|\|\|\|\|\|\|\|\|\|\|\|\|\|\|\|\|\|\|\|\|\|\|\|\|\|\|\|\|\|\|\|\|\|\|\|\|\|\|\|\|\|\|\|\|\|\|\|\|\|\|\|\|\|\|\|\|\|\|\|\|\|\|\|\|\|\|\|\|\|\|\|\|\|\|\|\|\|\|\|\|\|\|\|\|\|\|\|\|\|\|\|\|\|\|\|\|\|\|\|\|\|\|\|\|\|\|\|\|\|\|\|\|\|\|\|\|\|\|\|\|\|\|\|\|\|\|\|\|\|\|\|\|\|\|\|\|\|\|\|\|\|\|\|\|\|\|\|\|\|\|\|\|\|\|\|\|\|\|\|\|\|\|\|\|\|\|\|\|\|\|\|\|\|\|\|\|\|\|\|\|\|\|\|\|\|\|\|\|\|\|\|\|\|\|\|\|\|\|\|\|\|\|\|\|\|\|\|\|\|\|\|\|\|\|\|\|\|\|\|\|\|\|\|\|\|\|\|\|\|\|\|\|\|\|\|\|\|\|\|\|\|\|\|\|\|\|\|\|\|\|\|\|\|\|\|\|\|\|\|\|\|\|\|\|\|\|\|\|\|\|\|\|\|\|\|\|\|\|\|\|\|\|\|\|\|\|\|\|\|\|\|\|\|\|\|\|\|\|\|\|\|\|\|\|\|\|\|\|\|\|\|\|\|\|\|\|\|\|\|\|\|\|\|\|\|\|\|\|\|\|\|\|\|\|\|\|\|\|\|\|\|\|\|\|\|\|\|\|\|\|\|\|\|\|\|\|\|\|\|\|\|\|\|\|\|\|\|\|\|\|\|\|\|\|\|\|\|\|\|\|\|\|\|\|\|\|\|\|\|\|\|\|\|\|\|\|\|\|\|\|\|\|\|\|\|\|\|\|\|\|\|\|\|\|\|\|\|\|\|\|\|\|\|\|\|\|\|\|\|\|\|\|\|\|\|\|\|\|\|\|\|\|\|\|\|\|\|\|\|\|\|\|\| \|\|\|\|\|\|\|\|\|\|\|\|\|\|\|\|\|\|\|\|\|\|\|\| \|\|\|\|\|\|\|\|\|\|\|\|\|\|\|\|\|\|\|\|\|\|\|\|\|\|\|\|\|\|\|\|\|\|\|\|\|\|\|\|\|\| \|\|\| \|\|\|\|\|\|\|\|\|\|\| \|\|\|\|\|\|\|\|\|\|\| \|\|\|\|\|\|\|\| \| \|\|\|\|\|\| \|\| \|\|\| 	ATGACCGCAGTCCGCTCTGCAGGCGTGCACTACTTTCAGCCGCCGTTTGATCAGATTCAGATGTCCGACCACTACGCCAACGATGCATGGCGCCGCTCATATCCCCAGCAGATGCCGCTGCCATCGAATAACTGGCACGATGGTTCCAACAACATGCCCGTTATGACCCACGGCATGCTCGAGACCGACCTTCGCATGTTTCAACAATCAGAAGCAGGAACCTACAACACGCCCTTCACCAGCGGTCAGCAACGTATCTCGCATGCGCTCGTCTCGCCGCGCCTCGCATTTCCACCTGATGCTGCATGGGCTTCCGACATGGAACTGGGCACGCCATCAACCGGCACATTCCCATCAAACATCGACGAAGGCTACACCATTCGACATACCTCCAACCACCTCACCCTACCATCCACCCACGGACATGACCCCTCCGACACGAACAGCCCTCGCAGCTCCTTCGGCGACCAATCACCCTCATCCCAATATGTCCAACCCATCTTCAAGTCCGCACCAAAACGTCCCGGCATCGCTCGCGCAGTCACAGCACCCGACACAATCCCTCAACGACCACTCCACTCCCCACCAATCAAACGCTCCCCTTCCGACGACGGAGAAGACGACGACTACTTTCCCTCCGCCTCCGA-CCAAAAACCTCCTCTCGCAACCGC-AACGTGCACGAATCCCACACACAGCCGTGGAGAGGCGATATANAGA-AATCTGAACGCNCACCTCGATCG-TTGCGGCAAANGGTGCC-GAACTG ``` |
| 1 | ggggacaagtttgtacaaaaaagcaggcttgatgGAGAGCTTAACGTTGAACGA, ggggaccactttgtacaagaaagctgggtcCATGTCGCAGTCGCGGCTTC | Mycgr3\|91768  JGI  Ensembl | KOG0773 Transcription factor MEIS1 and related HOX domain proteins; | nucleus; regulation of transcription, DNA-dependent;  sequence-specific DNA binding;  transcription factor activity; | 100 % |  | ``` 1	ATGGAGAGCTTAACGTTGAACGAGGGACACTCCTTCCGGCGATACCAGGATCGACCCCAATCGAAGGACCAGCGCCATCGCCCTTCTCGTCTGCCAACTCCCGACACTTCTCGAAGCAGCAACGACAGTCAGAGCCCACCCGAGAGTGGCTCACTTGACTTTCCAACTGTCACGAAGTTTGGACTCGCAGGATTGCCTGACGGCGTTGGTCAACAAAGGCGAGACAGTCGGTATTCTGCCGGCCAATATTCACCACCTTACCGACCAGACGACATGCAAAGACCTTCTTTGCCTCCAATCAAGACGGT	308 	\|\|\|\|\|\|\|\|\|\|\|\|\|\|\|\|\|\|\|\|\|\|\|\|\|\|\|\|\|\|\|\|\|\|\|\|\|\|\|\|\|\|\|\|\|\|\|\|\|\|\|\|\|\|\|\|\|\|\|\|\|\|\|\|\|\|\|\|\|\|\|\|\|\|\|\|\|\|\|\|\|\|\|\|\|\|\|\|\|\|\|\|\|\|\|\|\|\|\|\|\|\|\|\|\|\|\|\|\|\|\|\|\|\|\|\|\|\|\|\|\|\|\|\|\|\|\|\|\|\|\|\|\|\|\|\|\|\|\|\|\|\|\|\|\|\|\|\|\|\|\|\|\|\|\|\|\|\|\|\|\|\|\|\|\|\|\|\|\|\|\|\|\|\|\|\|\|\|\|\|\|\|\|\|\|\|\|\|\|\|\|\|\|\|\|\|\|\|\|\|\|\|\|\|\|\|\|\|\|\|\|\|\|\|\|\|\|\|\|\|\|\|\|\|\|\|\|\|\|\|\|\|\|\|\|\|\|\|\|\|\|\|\|\|\|\|\|\|\|\|\|\|\|\|\|\|\|\|\|\|\|\|\|\|\|\|\|\|\|\|\|\|\|\|\|\|\|\|\|\|\|\|\|\|\|\|\|\|\|\|\|\|\|\|\|\|\|\|\|\|\|\|\|\|\|\|\|\| 	ATGGAGAGCTTAACGTTGAACGAGGGACACTCCTTCCGGCGATACCAGGATCGACCCCAATCGAAGGACCAGCGCCATCGCCCTTCTCGTCTGCCAACTCCCGACACTTCTCGAAGCAGCAACGACAGTCAGAGCCCACCCGAGAGTGGCTCACTTGACTTTCCAACTGTCACGAAGTTTGGACTCGCAGGATTGCCTGACGGCGTTGGTCAACAAAGGCGAGACAGTCGGTATTCTGCCGGCCAATATTCACCACCTTACCGACCAGACGACATGCAAAGACCTTCTTTGCCTCCAATCAAGACGGT ```      ``` 306	GGTCCTCGGCGACAACATAGCAAGTCCACCGACGACGCCTAGCCCACATGGAACGCCCATGCAACCGTCTCCGCATGAGCCATTGTACACCGCACTGGCTTACAAGCCTTCGTCTCTCTACCCGAACAAGAAGCAACGGACAGAGCCCTTTCCAAGTCTGGGACCACAATATGCAACCGCGAACAGGTCCGGCTCT	501 	\|\|\|\|\|\|\|\|\|\|\|\|\|\|\|\|\|\|\| \|\|\|\|\|\|\|\|\|\|\|\|\|\|\|\|\|\|\|\|\|\|\|\|\|\|\|\|\|\|\|\|\|\|\|\|\|\|\|\|\|\|\|\|\|\|\|\|\|\|\|\|\|\|\|\|\|\|\|\|\|\|\|\|\|\|\|\|\|\|\|\|\|\|\|\|\|\|\|\|\|\|\|\|\|\|\|\|\|\|\|\|\|\|\|\|\|\|\|\|\|\|\|\|\|\|\|\|\|\|\|\|\|\|\|\|\|\|\|\|\|\|\|\|\|\|\|\|\|\|\|\|\|\|\|\|\|\|\|\|\|\|\|\|\|\|\|\|\|\|\|\|\|\|\|\|\|\|\|\|\|\|\|\|\|\|\|\|\|\|\|\| \|\|\| 	GGTCCTCGGCGACAACATANCAAGTCCACCGACGACGCCTAGCCCACATGGAACGCCCATGCAACCGTCTCCGCATGAGCCATTGTACACCGCACTGGCTTACAAGCCTTCGTCTCTCTACCCGAACAAGAAGCAACGGACAGAGCCCTTTCCAAGTCTGGGACCACAATATGCAACCGCGAACAGGTCCGGNTCT ``` |
| 1 | ggggacaagtttgtacaaaaaagcaggcttgatgAGCGTCGAGCTGGACCTCGT, ggggaccactttgtacaagaaagctgggtcTTTGCGAGGCCCCTCGATCT | Mycgr3\|92358  JGI  Ensembl | KOG1216 von Willebrand factor and related coagulation proteins; | DNA polymerase activity; DNA-directed DNA polymerase activity; | 18.85099 % |  |  |
| 1 | ggggacaagtttgtacaaaaaagcaggcttgatgGCTCACCCGGCACAGAACCA, ggggaccactttgtacaagaaagctgggtcGTCTCTGCGTTGATACTGCA | Mycgr3\|92880  JGI  Ensembl | KOG4786 Ubinuclein, nuclear protein interacting with cellular and viral transcription factors; |  | 100 % |  | ``` 1	ATGGCTCACCCGGCACAGAACCAGCTGTTCACGCCGCCGGTAGTCTTTGTGGCAGTTCTACCCGCCGGGGCTATCGCCACCGACGCCGTTGGAATGGTCGCCCGTTTCCAATGTATGTCTTCCGAAGAGAGCAGTAAAAGCGTAAAATTCCAGGCGGCAGACGAGAAGATTGTGACGGCAGTTCAGAATGATGGACGATCTTTCATCAGATGGAATAGTCCGAGCAACGGAAACAGACCTGCTATGGTGGAAACCCGCGACGCTGAAATGGAGTCCCTACATGGCCTTGCAAAGCAGAAACTACCCAAGTCGAACCCAAGCCTGGAAGCCGTATTGCCCAAAATGTACGCAAAAGACAGCACGTACGCGGACGATTTAACATGGGTGATCGTGTTCACCCCCATTCAGGCGCGCCGGGCTTTTCCCCACGGCTACACAGACGCTGGCAATCTCACCATCCTCGGGGTTCTTGCCATCTTTCTTCCGTACAAGTGTGGTCGAGACCAGATTTTGAGATCTTTCCACTTCGGATTCTGCGAGGCCGCTAGCGGAATGCTGGAGTCTCTCGCCAACGGGCATGA	581 	\|\|\|\|\|\|\|\|\|\|\|\|\|\|\|\|\|\|\|\|\|\|\|\|\|\|\|\|\|\|\|\|\|\|\|\|\|\|\|\|\|\|\|\|\|\|\|\|\|\|\|\|\|\|\|\|\|\|\|\|\|\|\|\|\|\|\|\|\|\|\|\|\|\|\|\|\|\|\|\|\|\|\|\|\|\|\|\|\|\|\|\|\|\|\|\|\|\|\|\|\|\|\|\|\|\|\|\|\|\|\|\|\|\|\|\|\|\|\|\|\|\|\|\|\|\|\|\|\|\|\|\|\|\|\|\|\|\|\|\|\|\|\|\|\|\|\|\|\|\|\|\|\|\|\|\|\|\|\|\|\|\|\|\|\|\|\|\|\|\|\|\|\|\|\|\|\|\|\|\|\|\|\|\|\|\|\|\|\|\|\|\|\|\|\|\|\|\|\|\|\|\|\|\|\|\|\|\|\|\|\|\|\|\|\|\|\|\|\|\|\|\|\|\|\|\|\|\|\|\|\|\|\|\|\|\|\|\|\|\|\|\|\|\|\|\|\|\|\|\|\|\|\|\|\|\|\|\|\|\|\|\|\|\|\|\|\|\|\|\|\|\|\|\|\|\|\|\|\|\|\|\|\|\|\|\|\|\|\|\|\|\|\|\|\|\|\|\|\|\|\|\|\|\|\|\|\|\|\|\|\|\|\|\|\|\|\|\|\|\|\|\|\|\|\|\|\|\|\|\|\|\|\|\|\|\|\|\|\|\|\|\|\|\|\|\|\|\|\|\|\|\|\|\|\|\|\|\|\|\|\|\|\|\|\|\|\|\|\|\|\|\|\|\|\|\|\|\|\|\|\|\|\|\|\|\|\|\|\|\|\|\|\|\|\|\|\|\|\|\|\|\|\|\|\|\|\|\|\|\|\|\|\|\|\|\|\|\|\|\|\|\|\|\|\|\|\|\|\|\|\|\|\|\|\|\|\|\|\|\|\|\|\|\|\|\|\|\|\|\|\|\|\|\|\|\|\|\|\|\|\|\|\|\|\|\|\|\|\|\|\|\|\|\|\|\|\|\|\|\|\|\|\|\|\|\|\|\|\|\|\| \|\|\|\|\|\|\|\|\|\|\|\|\|\| \|\|\|\|\|\|\| \|\|\|\|\|\|\|\|\|\|\|\|\| \|\|\|\|\|\|\|\| \| \|\|\|\|\|\|\| \| \|\|\|\|\|\|\|\|\|\| \|\|\|\|\|\| \|\|\| \| \|\| \|\|\|\| 	ATGGCTCACCCGGCACAGAACCAGCTGTTCACGCCGCCGGTAGTCTTTGTGGCAGTTCTACCCGCCGGGGCTATCGCCACCGACGCCGTTGGAATGGTCGCCCGTTTCCAATGTATGTCTTCCGAAGAGAGCAGTAAAAGCGTAAAATTCCAGGCGGCAGACGAGAAGATTGTGACGGCAGTTCAGAATGATGGACGATCTTTCATCAGATGGAATAGTCCGAGCAACGGAAACAGACCTGCTATGGTGGAAACCCGCGACGCTGAAATGGAGTCCCTACATGGCCTTGCAAAGCAGAAACTACCCAAGTCGAACCCAAGCCTGGAAGCCGTATTGCCCAAAATGTACGCAAAAGACAGCACGTACGCGGACGATTTAACATGGGTGATCGTGTTCACCCCCATTCAGGCGCGCCGGGCTTTTCCCCACGGCTACACAGACGCTGGCAATCTCACCATCCTCGGGGTTCTTGCCATCTTTCTTCCGTACAANTGTGGTCGAGACCANATTTTGANATCTTTCCACTTCNGATTCTGCNANGCCGCTANCNGAATGCTGGANTCTCTCNCCA-CNGGNATGA ``` |
| 1 | ggggacaagtttgtacaaaaaagcaggcttgatgGAGCCACCAAGCAAAAGACG, ggggaccactttgtacaagaaagctgggtcCCCCTTCCTCTTCTTCCCCA | Mycgr3\|92934  JGI  Ensembl | KOG0340 ATP-dependent RNA helicase; | 8-oxo-7,8-dihydroguanine triphosphatase activity; ATP binding;  ATP-dependent 5'-3' DNA helicase activity;  ATP-dependent helicase activity;  UDP-2,3-diacylglucosamine hydrolase activity;  bis(5'-nucleosyl)-tetraphosphatase activity;  dATP pyrophosphohydrolase activity;  diacylglycerol pyrophosphate phosphatase activity;  dihydroneopterin monophosphate phosphatase activity;  dihydroneopterin triphosphate pyrophosphohydrolase activity;  helicase activity;  nucleic acid binding;  pyrophosphatase activity;  thiamin-pyrophosphatase activity; | 12.98873 % | chr\_5:95699 C->T; |  |
| 1 | ggggacaagtttgtacaaaaaagcaggcttgatgGCTGTTCGAACTTCAGCAGC, ggggaccactttgtacaagaaagctgggtcGTCTTTCGCGGTAATATCCA | Mycgr3\|94275  JGI  Ensembl |  |  | 0 % |  |  |
| 1 | ggggacaagtttgtacaaaaaagcaggcttgatgCCCATCAAATCCACCGGCTG, ggggaccactttgtacaagaaagctgggtcGCCCACAGCTGGAGGCGAAG | Mycgr3\|94536  JGI  Ensembl |  |  | 100 % |  | ``` 1	ATGCCCATCAAATCCACCGGCTGCGCAACCTGTCGCAAACGCAAAATCCGCTGCGACGAGACCCGCCCAAAATGCAACCGCTGCAAAACCCACGGCGTGCCTTGTCCAGGCTATCGTACACCCGCACCCGGCGAGGTCAAGTTCCACGATGAGACGCACGGGATCGTAATGCAATACGGCGGAGGAAGGGGAGCAAGTCAAGGTCGACGACTATCCGCCGACAGCGGGAAGGGTTCGTCGATCATTTCATGGTCTTCCTCTCCGGACGTGCTC	273 	\|\|\|\|\|\|\|\|\|\|\|\|\|\|\|\|\|\|\|\|\|\|\|\|\|\|\|\|\|\|\|\|\|\|\|\|\|\|\|\|\|\|\|\|\|\|\|\|\|\|\|\|\|\|\|\|\|\|\|\|\|\|\|\|\|\|\|\|\|\|\|\|\|\|\|\|\|\|\|\|\|\|\|\|\|\|\|\|\|\|\|\|\|\|\|\|\|\|\|\|\|\|\|\|\|\|\|\|\|\|\|\|\|\|\|\|\|\|\|\|\|\|\|\|\|\|\|\|\|\|\|\|\|\|\|\|\|\|\|\|\|\|\|\|\|\|\|\|\|\|\|\|\|\|\|\|\|\|\|\|\|\|\|\|\|\|\|\|\|\|\|\|\|\|\|\|\|\|\|\|\|\|\|\|\|\|\|\|\|\|\|\|\|\|\|\|\|\|\|\|\|\|\|\|\|\|\|\|\|\|\|\|\|\|\|\|\|\|\|\|\|\|\|\|\|\|\|\|\|\|\|\|\|\|\|\|\|\|\|\|\|\|\|\|\|\|\|\|\|\|\|\|\|\|\|\|\|\|\|\|\|\|\|\|\|\|\|\|\|\|\|\|\| 	ATGCCCATCAAATCCACCGGCTGCGCAACCTGTCGCAAACGCAAAATCCGCTGCGACGAGACCCGCCCAAAATGCAACCGCTGCAAAACCCACGGCGTGCCTTGTCCAGGCTATCGTACACCCGCACCCGGCGAGGTCAAGTTCCACGATGAGACGCACGGGATCGTAATGCAATACGGCGGAGGAAGGGGAGCAAGTCAAGGTCGACGACTATCCGCCGACAGCGGGAAGGGTTCGTCGATCATTTCATGGTCTTCCTCTCCGGACGTGCTC ``` |
| 1 | ggggacaagtttgtacaaaaaagcaggcttgatgGCAGGTCGCGAAAATATGGC, ggggaccactttgtacaagaaagctgggtcCAAGTCTCTCAGATTCTCCA | Mycgr3\|94704  JGI  Ensembl | KOG0260 RNA polymerase II, large subunit; | DNA binding; DNA-directed RNA polymerase II, core complex;  transcription from RNA polymerase II promoter; | 100 % |  | ``` 1	ATGGCAGGTCGCGAAAATATGGCAGCAGGGCAGTACTTGAGCACGCCGGTCCCTACACTTGTCGACGTTCCATCCGTTCTCATCGCCACATTCACGGAATCAAGCCAGCAAGCAACAGCCAGCACATCCACACTACACTCAGAACGAACATACCCCAAAGCTACGACGCTCCCACAATCGGACAACATGCGTTTCAACAACATCAACAACACTGCCAACGACGCCGCAGCCATGCCCACCTCCGCCGCTCCACAGGGCGAGACATTCCGCTTTCCGACACCCTTCCAGATCGACCCCGTCCGCGACAACAACTTTCTCCTGGCCATGAACTGGGCCGATCCAGAGCTCCGTGCCGCTGTGATTCGGTCGTTCGAACGAACTCACCTCGCTCTTGTCAACAACGAACCTCATCCAGGCGGTGAGATCTATGGTCTCAGGCAGGAAATTGAGGACCTGTTCAGCGGCCAGGCGGACAGAGATACCGATAAGGAGATCGCCAAGAGGGTCGAGAGGGTAGAAGCTGAGCGTCTGGAAAGGGCCGCCGAGGAGGGAGCCCGGCGTGCTGAGGCGGAGGAAGCTCAACCTACCGAAGTGGACGAAGAAGCTTTCGCACTGGCGAACCCGACGTCACCAACCTACGCTCCTCGCTCACCTTCTCCGCTGGAGGATCCGTGGACGCCCAGATACGACCCAATTTCACCAGGAACTCCACTTCATCCGTACAATCACCAGCAGTCTC	739 	\|\|\|\|\|\|\|\|\|\|\|\|\|\|\|\|\|\|\|\|\|\|\|\|\|\|\|\|\|\|\|\|\|\|\|\|\|\|\|\|\|\|\|\|\|\|\|\|\|\|\|\|\|\|\|\|\|\|\|\|\|\|\|\|\|\|\|\|\|\|\|\|\|\|\|\|\|\|\|\|\|\|\|\|\|\|\|\|\|\|\|\|\|\|\|\|\|\|\|\|\|\|\|\|\|\|\|\|\|\|\|\|\|\|\|\|\|\|\|\|\|\|\|\|\|\|\|\|\|\|\|\|\|\|\|\|\|\|\|\|\|\|\|\|\|\|\|\|\|\|\|\|\|\|\|\|\|\|\|\|\|\|\|\|\|\|\|\|\|\|\|\|\|\|\|\|\|\|\|\|\|\|\|\|\|\|\|\|\|\|\|\|\|\|\|\|\|\|\|\|\|\|\|\|\|\|\|\|\|\|\|\|\|\|\|\|\|\|\|\|\|\|\|\|\|\|\|\|\|\|\|\|\|\|\|\|\|\|\|\|\|\|\|\|\|\|\|\|\|\|\|\|\|\|\|\|\|\|\|\|\|\|\|\|\|\|\|\|\|\|\|\|\|\|\|\|\|\|\|\|\|\|\|\|\|\|\|\|\|\|\|\|\|\|\|\|\|\|\|\|\|\|\|\|\|\|\|\|\|\|\|\|\|\|\|\|\|\|\|\|\|\|\|\|\|\|\|\|\|\|\|\|\|\|\|\|\|\|\|\|\|\|\|\|\|\|\|\|\|\|\|\|\|\|\|\|\|\|\|\|\|\|\|\|\|\|\|\|\|\|\|\|\|\|\|\|\|\|\|\|\|\|\|\|\|\|\|\|\|\|\|\|\|\|\|\|\|\|\|\|\|\|\|\|\|\|\|\|\|\|\|\|\|\|\|\|\|\|\|\|\|\|\|\|\|\|\|\|\|\|\|\|\|\|\|\|\|\|\|\|\|\|\|\|\|\|\|\|\|\|\|\|\|\|\|\|\|\|\|\|\|\|\|\|\|\|\|\|\|\|\|\|\|\|\|\|\|\|\|\|\|\|\|\|\|\|\|\|\|\|\|\|\|\|\|\|\|\|\|\|\|\|\|\|\|\|\|\|\|\|\|\|\|\|\|\|\|\|\|\|\|\|\|\| \|\|\|\|\|\|\|\|\|\|\|\|\|\|\|\|\|\|\|\|\|\|\|\|\|\|\|\|\|\|\|\|\|\|\|\|\|\|\|\|\| \|\|\|\|\| \|\|\|\|\|\|\|\|\|\|\|\|\|\|\|\|\|\|\|\|\|\|\|\|\|\|\| \|\|\|\|\|\|\|\|\|\|\|\|\|\|\|\|\|\|\| \|\|\|\|\|\|\|\|\|\|\|\|\|\|\|\|\| \|\|\|\| \|\|\| \| \| \|\|\|\|\|\|\|\|\|\|\|\|\|\|\|\|\|\|\|\| \|\|\|\|\| \|\|\| \|\|\|\|\|\| \|\|\| \|\|\|\| \| \|\|\|\|\| \|\| \|\|\|  \|\|\| \|\|\| \| \|\|\|\| \| \|\|\|\|\|\|\| 	ATGGCAGGTCGCGAAAATATGGCAGCAGGGCAGTACTTGAGCACGCCGGTCCCTACACTTGTCGACGTTCCATCCGTTCTCATCGCCACATTCACGGAATCAAGCCAGCAAGCAACAGCCAGCACATCCACACTACACTCAGAACGAACATACCCCAAAGCTACGACGCTCCCACAATCGGACAACATGCGTTTCAACAACATCAACAACACTGCCAACGACGCCGCAGCCATGCCCACCTCCGCCGCTCCACAGGGCGAGACATTCCGCTTTCCGACACCCTTCCAGATCGACCCCGTCCGCGACAACAACTTTCTCCTGGCCATGAACTGGGCCGATCCAGAGCTCCGTGCCGCTGTGATTCGGTCGTTCGAACGAACTCACCTCGCTCTTGTCAACAACGAACCTCATCCAGGCGGTGAGATCTATGGTCTCAGGCAGGAAATTGAGGACCTGTTCAGCGGCCAGGCGGACAGAGATACCGATAAGGAGATCGCCAAGAGGGTCGAGAGGGTAGAAGCTGANCGTCTGGAAAGGGCCGCCGAGGAGGGAGCCCGGCGTGCTGANGCGGANGAAGCTCAACCTACCGAAGTGGACGAANAAGCTTTCGCACTGGCGAA-CCGACGTCACCAACCTANGCTCNTCGNT-ANCTTCTCCGCTGGAGGATCCGNGGACG-CCANATACGANCCA-TTTCNC-AGGAANTC-ACTN-ATC-GTANA-TCAC-ANCAGTCTC ``` |
| 1 | ggggacaagtttgtacaaaaaagcaggcttgatgGGCGACGACGACTACGAGCA, ggggaccactttgtacaagaaagctgggtcCTTCTTCTCTTCTTCCTCGC | Mycgr3\|94893  JGI  Ensembl | KOG3227 Calcium-responsive transcription coactivator; |  | 100 % |  |  |
| 1 | ggggacaagtttgtacaaaaaagcaggcttgatgCCGCCCAAAAAGAAAGCCCG, ggggaccactttgtacaagaaagctgggtcCTTCCCTCTCCCTCTCCCTC | Mycgr3\|96434  JGI  Ensembl | KOG0123 Polyadenylate-binding protein (RRM superfamily); | DNA binding; | 100 % |  | ``` 1	ATGCCGCCCAAAAAGAAAGCCCGTCATTCCCGCGCCGCCACCGCGACAGCCTCACCATCTCCGCCCCCCGCACCTGTCCGCTCGACCTCCCGCTCACGCTCCCCATCACGAACAACCTCACCCGACCCATGGACCGATGCCGAAGAAGCCGCCCTCTTCCGCGCAATACTACGACATAAACCGACAGGCTTTCACAAGCACTTCCGACTTCTTTCACTTCACGCCTGGCTGCTGGAGAACCACCACATCCATCCACACAATGTGCACACGCGACCGCCGGGCATTTGGAAGAAGTTGGACAGTCTGTATGATCTGAAAGCGCTGGATGCGAGGGAGGATGCGAGGCAACTGGATAAGTTGGTGAATCCTTTGCTGGAGGGTGGCGGGAGAGGTG	394 	\|\|\|\|\|\|\|\|\|\|\|\|\|\|\|\|\|\|\|\|\|\|\|\|\|\|\|\|\|\|\|\|\|\|\|\|\|\|\|\|\|\|\|\|\|\|\|\|\|\|\|\|\|\|\|\|\|\|\|\|\|\|\|\|\|\|\|\|\|\|\|\|\|\|\|\|\|\|\|\|\|\|\|\|\|\|\|\|\|\|\|\|\|\|\|\|\|\|\|\|\|\|\|\|\|\|\|\|\|\|\|\|\|\|\|\|\|\|\|\|\|\|\|\|\|\|\|\|\|\|\|\|\|\|\|\|\|\|\|\|\|\|\|\|\|\|\|\|\|\|\|\|\|\|\|\|\|\|\|\|\|\|\|\|\|\|\|\|\|\|\|\|\|\|\|\|\|\|\|\|\|\|\|\|\|\|\|\|\|\|\|\|\|\|\|\|\|\|\|\|\|\|\|\|\|\|\|\|\|\|\|\|\|\|\|\|\|\|\|\|\|\|\|\|\|\|\|\|\|\|\|\|\|\|\|\|\|\|\|\|\|\|\|\|\|\|\|\|\|\|\|\|\|\|\|\|\|\|\|\|\|\|\|\|\|\|\|\|\|\|\|\|\|\|\|\|\|\|\|\|\|\|\|\|\|\|\|\|\|\|\|\|\|\|\|\|\|\|\|\|\|\|\|\|\|\|\|\|\|\|\|\|\|\|\|\|\|\|\|\|\|\|\|\|\|\|\|\|\|\|\|\|\|\|\|\|\|\|\|\|\|\|\|\|\|\|\|\|\|\|\|\|\|\|\|\|\|\|\|\|\|\|\|\|\|\|\|\|\|\|\|\|\|\|\|\|\|\|\|\|\|\|\|\|\|\|\|\| \|\|\|\|\| 	ATGCCGCCCAAAAAGAAAGCCCGTCATTCCCGCGCCGCCACCGCGACAGCCTCACCATCTCCGCCCCCCGCACCTGTCCGCTCGACCTCCCGCTCACGCTCCCCATCACGAACAACCTCACCCGACCCATGGACCGATGCCGAAGAAGCCGCCCTCTTCCGCGCAATACTACGACATAAACCGACAGGCTTTCACAAGCACTTCCGACTTCTTTCACTTCACGCCTGGCTGCTGGAGAACCACCACATCCATCCACACAATGTGCACACGCGACCGCCGGGCATTTGGAAGAAGTTGGACAGTCTGTATGATCTGAAAGCGCTGGATGCGAGGGAGGATGCGAGGCAACTGGATAAGTTGGTGAATCCTTTGCTGGAGGGTGGCGGGANAGGTG ``` |
| 1 | ggggacaagtttgtacaaaaaagcaggcttgatgCCATCGGCGAAACAAACCGC, ggggaccactttgtacaagaaagctgggtcGCGACGATCATCATGGGCCC | Mycgr3\|96672  JGI  Ensembl | KOG1279 Chromatin remodeling factor subunit and related transcription factors; |  | 0 % |  |  |
| 1 | ggggacaagtttgtacaaaaaagcaggcttgatgTCCGACGCCGCCTCGCAGGC, ggggaccactttgtacaagaaagctgggtcCCAGCCATCCTCAGCCTCAG | Mycgr3\|97328  JGI  Ensembl | KOG0335 ATP-dependent RNA helicase; | 8-oxo-7,8-dihydroguanine triphosphatase activity; ATP binding;  ATP-dependent 5'-3' DNA helicase activity;  ATP-dependent helicase activity;  UDP-2,3-diacylglucosamine hydrolase activity;  bis(5'-nucleosyl)-tetraphosphatase activity;  dATP pyrophosphohydrolase activity;  diacylglycerol pyrophosphate phosphatase activity;  dihydroneopterin monophosphate phosphatase activity;  dihydroneopterin triphosphate pyrophosphohydrolase activity;  helicase activity;  nucleic acid binding;  pyrophosphatase activity;  thiamin-pyrophosphatase activity; | 0.99895 % |  |  |
| 1 | ggggacaagtttgtacaaaaaagcaggcttgatgCCATGCAGGATGGCGTCCGG, ggggaccactttgtacaagaaagctgggtcACTCAGCAGCTCTTCCAATG | Mycgr3\|97465  JGI  Ensembl |  |  | 100 % |  |  |
| 1 | ggggacaagtttgtacaaaaaagcaggcttgatgCCGCTATGCCCACGGTTCTA, ggggaccactttgtacaagaaagctgggtcTTGCGGTCGGTTGCGCAGGT | Mycgr3\|97741  JGI  Ensembl | KOG4141 DNA repair and recombination protein RAD52/RAD22; |  | 100 % | chr\_16:324952 G->A; | ``` 1	ATGCCG-CTATGCCCAC-GGTTCTATCGCGAAATGCAAACGTCAAGGCTGCAGGTGGAAGAACAATTCGGCTTCTCTGCCATGCCATTCCTTGGAAGGACCGCGGTCACGATCCAGGGACAGTCGCCTGCAGTCGATGTCGACGACCAGCACAAGCTCATCATGACTCACTGCGCTGCATCACCCACGGCAACAGCGAAGG	199 	\|\|\|\|\|\| \|\|\|\|\|\|\|\|\|\| \|\|\|\|\|\|\|\|\|\|\|\|\|\|\|\|\|\| \|\|\|\|\|\|\|\|\|\|\|\|\|\|\|\|\|\|\|\|\|\|\|\|\|\|\|\| \|\|\|\|\|\|\|\|\|\|\|\|\|\|\|\|\|\|\|\|\|\|\|\|\|\|\|\|\|\|\|\|\|\|\|\|\|\|\|\|\|\|\|\|\|\|\|\|\|\|\|\|\|\|\|\|\|\| \|\|\|\|\|\| \|\| \|\|\|\|\|\|\|  \|\|\|\| \|\|\|\|\|\|\|\|\|\|\|\|\|\|\|\| \|\|\| \|\|\|\|\|\|\|\|\| \|\| \| \| \|\| \| \| \|\|\|\|\|\|\| 	ATGCCGCCTATGCCCACGGGTTCTATCGCGAAATGCNAACGTCAAGGCTGCAGGTGGAAGAACAANTCGGCTTCTCTGCCATGCCATTCCTTGGAAGGACCGCGGTCACGATCCAGGGACAGTCNCCTGCANTCNATGTCGANNACCANCACAAGCTCATCATGAGTCANTGCGCTGCANCANCNANGGNAGCNGCGAAGG ```      ``` 208	TTGGTGCAGCTCTTCTCCTGGGCGGACTTCTTCTCTCACATCG	250 	\|\|\|\|\|\|\|\| \|\|\|\|\|\|\|\|\|\|\|\|\|\|\|\|\|\|\|\|\|\|\|\|\|\|\|\|\|\|\|\|\|\| 	TTGGTGCAACTCTTCTCCTGGGCGGACTTCTTCTCTCACATCG ``` |
| 1 | ggggacaagtttgtacaaaaaagcaggcttgatgAGTCCCAACCGCCGGAACTC, ggggaccactttgtacaagaaagctgggtcGTGAAGGTTCTTCGATATAA | Mycgr3\|97915  JGI  Ensembl |  |  | 100 % |  | ``` 1	ATGAGTCCCAACCGCCGGAACTCGGATCTCAGCCAAGCTTCCTCGACGCCATTAGAAAGGGCGCGTCACGAAGCGGCCAGCAACAACGATTTCGGAACCGGATACGACTCCGACCCCGACGACCTCTTCCTCGTTCACCCAGCGACTGTCTTAATGGTTGACAAGAAGAAGAAGGCCGCTGCGCCACGGAACGTGTTCACAACAGAAGAAGAGGATCAGGTCATACTCGACAACTTCAAACAATACCCTAAGGGGATGAACATTCCATTCACGAATCGATACGAAATGATGAAGAATTCTCTGCTAGGCAGATCAGGAGAAACAACTCGAAGACGATGGCAGCGCCACCTTGACCCTAATCGCAACGGAAAAGAGATTCAGGCAAAAGAAGAACGAGACTCGAAGCGACAGCGATTCCTGCAACACCTTGCTAAGTCTCACCTTAAGGATCCCGACATGGGTCTGGATGATC	472 	\|\|\|\|\|\|\|\|\|\|\|\|\|\|\|\|\|\|\|\|\|\|\|\|\|\|\| \|\|\|\|\|\|\|\|\|\|\|\|\|\|\|\|\|\|\|\|\|\|\|\|\|\| \|\|\|\|\|\|\|\|\|\|\|\| \|\|\|\|\|\|\|\|\|\|\|\|\|\|\|\|\|\|\|\|\|\|\|\|\|\|\|\|\|\|\|\|\|\|\|\|\|\|\|\|\|\|\|\|\|\|\|\|\|\|\|\|\|\|\|\|\|\|\|\|\|\|\|\|\|\|\|\|\|\|\|\|\|\|\|\|\|\|\|\|\|\|\|\|\|\|\|\|\|\|\|\|\|\|\|\|\|\|\|\|\|\|\|\|\|\|\|\|\|\|\|\|\|\|\|\|\|\|\|\|\|\|\|\|\|\|\|\|\|\|\|\|\|\|\|\|\|\|\|\|\|\|\|\|\|\|\|\|\|\|\|\|\|\|\|\|\|\|\|\|\|\|\|\|\|\|\|\|\|\|\|\|\|\|\|\|\|\|\|\|\|\|\|\|\|\|\|\|\|\|\|\|\|\|\|\|\|\|\|\|\|\|\|\|\|\|\|\|\|\|\|\|\|\|\|\|\|\|\|\|\|\|\|\|\|\|\|\|\|\|\|\|\|\|\|\|\|\|\|\|\|\|\|\|\|\|\|\|\|\|\|\|\|\|\|\|\|\|\|\|\|\|\|\|\|\|\|\|\|\|\|\|\|\|\|\|\|\|\|\|\|\|\|\|\|\|\|\|\|\|\|\|\|\|\|\|\|\|\|\|\|\|\|\|\|\|\|\|\|\|\|\|\|\|\| \|\|\|\|\|\|\|\|\|\|\|\|\|\|\|\|\|\|\|\|\|\|\|\|\|\|\|\|\|\|\|\|\|\|\|\|\|\|\|\| \|\|\|\|\|\|\|\|\| \|\|\|\|\|\|\|\|\|\|\|\|\|\|\|\|\|\|\|\|\| \|\|\|\|\|\|\|\|\|\|\|\|\|\|\| 	ATGAGTCCCAACCGCCGGAACTCGGATNTCAGCCAAGCTTCCTCGACGCCATTAAAAAGGGCGCGTCNCGAAGCGGCCAGCAACAACGATTTCGGAACCGGATACGACTCCGACCCCGACGACCTCTTCCTCGTTCACCCAGCGACTGTCTTAATGGTTGACAAGAAGAAGAAGGCCGCTGCGCCACGGAACGTGTTCACAACAGAAGAAGAGGATCAGGTCATACTCGACAACTTCAAACAATACCCTAAGGGGATGAACATTCCATTCACGAATCGATACGAAATGATGAAGAATTCTCTGCTAGGCAGATCAGGAGAAACAACTCGAAGACGATGGCAGCGCCACCTTGACCCTAATCGCAACGGAAAAGAGATTCAGGCNAAAGAAGAACGAGACTCGAAGCGACAGCGATTCCTGCAACNCCTTGCTAANTCTCACCTTAAGGATCCCGACNTGGGTCTGGATGATC ```      ``` 1	ATGAGTCCCAACCGCCGGAACTCGGATCTCAGCCAAGCTTCCTCGACGCCATTAGAAAGGGCGCGTCACGAAGCGGCCAGCAACAACGATTTCGGAACCGGATACGACTCCGACCCCGACGACCTCTTCCTCGTTCACCCAGCGACTGTCTTAATGGTTGACAAGAAGAAGAAGGCCGCTGCGCCACGGAACGTGTTCACAACAGAAGAAGAGGATCAGGTCATACTCGACAACTTCAAACAATACCCTAAGGGGATGAACATTCCATTCACGAATCGATACGAAATGATGAAGAATTCTCTGCTAGGCAGATCAGGAGAAACAACTCGAAGACGATGGCAGCGCCACCTTGACCCTAATCGCAACGGAAAAGAGATTCAGGCAAAAGAAGAACGAGACTCGAAGCGACAGCGATTCCTGCAACACCTTGCTAAGTCTCACCTTAAGGATCCCGACATGGGTCTGGATGATCGTGTTAAGACAGTGGAGGAATCCACCGGCGATGCAACGAATCGAAGCGCCATCGTTGCCGGCATCAAACATCTTCGAGCAGAAGGCATTATATCGAAGAACCTTCAC	579 	\|\|\|\|\|\|\|\|\|\|\|\|\|\|\|\|\|\|\|\|\|\|\|\|\|\|\|\|\|\|\|\|\|\|\|\|\|\|\|\|\|\|\|\|\|\|\|\|\|\|\|\|\|\|\|\|\|\|\|\|\|\|\|\|\|\|\|\|\|\|\|\|\|\|\|\|\|\|\|\|\|\|\|\|\|\|\|\|\|\|\|\|\|\|\|\|\|\|\|\|\|\|\|\|\|\|\|\|\|\|\|\|\|\|\|\|\|\|\|\|\|\|\|\|\|\|\|\|\|\|\|\|\|\|\|\|\|\|\|\|\|\|\|\|\|\|\|\|\|\|\|\|\|\|\|\|\|\|\|\|\|\|\|\|\|\|\|\|\|\|\|\|\|\|\|\|\|\|\|\|\|\|\|\|\|\|\|\|\|\|\|\|\|\|\|\|\|\|\|\|\|\|\|\|\|\|\|\|\|\|\|\|\|\|\|\|\|\|\|\|\|\|\|\|\|\|\|\|\|\|\|\|\|\|\|\|\|\|\|\|\|\|\|\|\|\|\|\|\|\|\|\|\|\|\|\|\|\|\|\|\|\|\|\|\|\|\|\|\|\|\|\|\|\|\|\|\|\|\|\|\|\|\|\|\|\|\|\|\|\|\|\|\|\|\|\|\|\|\|\|\|\|\|\|\|\|\|\|\|\|\|\|\|\|\|\|\|\|\|\|\|\|\|\|\|\|\|\|\|\|\|\|\|\|\|\|\|\|\|\|\|\|\|\|\|\|\|\|\|\|\|\|\|\|\|\|\|\|\|\|\|\|\|\|\|\|\|\|\|\|\|\|\|\|\|\|\|\|\|\|\|\|\|\|\|\|\|\|\|\|\|\|\|\|\|\|\|\|\|\|\|\|\|\|\|\|\|\|\|\|\|\|\|\|\|\|\|\|\|\|\|\|\|\|\|\|\|\|\|\|\|\|\|\|\|\|\|\|\|\|\|\|\|\|\|\|\|\|\|\|\|\|\|\|\|\|\|\|\|\|\|\|\|\|\|\|\|\|\|\|\|\|\|\|\|\|\|\|\|\|\|\|\|\|\|\|\|\|\|\|\|\|\|\|\|\|\|\|\|\|\|\|\|\|\|\|\|\|\|\|\|\|\|\|\|\|\|\|\|\|\|\|\|\|\|\|\|\|\|\|\|\|\|\|\|\|\|\|\|\|\|\|\|\|\|\|\|\|\|\|\|\|\|\|\|\|\|\|\|\|\|\|\|\|\|\|\|\|\|\|\|\|\|\|\|\|\|\|\| 	ATGAGTCCCAACCGCCGGAACTCGGATCTCAGCCAAGCTTCCTCGACGCCATTAGAAAGGGCGCGTCACGAAGCGGCCAGCAACAACGATTTCGGAACCGGATACGACTCCGACCCCGACGACCTCTTCCTCGTTCACCCAGCGACTGTCTTAATGGTTGACAAGAAGAAGAAGGCCGCTGCGCCACGGAACGTGTTCACAACAGAAGAAGAGGATCAGGTCATACTCGACAACTTCAAACAATACCCTAAGGGGATGAACATTCCATTCACGAATCGATACGAAATGATGAAGAATTCTCTGCTAGGCAGATCAGGAGAAACAACTCGAAGACGATGGCAGCGCCACCTTGACCCTAATCGCAACGGAAAAGAGATTCAGGCAAAAGAAGAACGAGACTCGAAGCGACAGCGATTCCTGCAACACCTTGCTAAGTCTCACCTTAAGGATCCCGACATGGGTCTGGATGATCGTGTTAAGACAGTGGAGGAATCCACCGGCGATGCAACGAATCGAAGCGCCATCGTTGCCGGCATCAAACATCTTCGAGCAGAAGGCATTATATCGAAGAACCTTCAC ``` |
| 1 | ggggacaagtttgtacaaaaaagcaggcttgatgGAGGCATGTAGCTGGCCGCC, ggggaccactttgtacaagaaagctgggtcCTTAAACCCCCCACCGCTCT | Mycgr3\|97929  JGI  Ensembl |  |  | 0 % |  |  |
| 1 | ggggacaagtttgtacaaaaaagcaggcttgatgGCGCTATCCACCGGGTGCTC, ggggaccactttgtacaagaaagctgggtcTTGGTGTTGGCGTTCTCCTT | Mycgr3\|98054  JGI  Ensembl | KOG1167 Serine/threonine protein kinase of the CDC7 subfamily involved in DNA synthesis, repair and recombination; |  | 100 % |  | ``` 172	ATTGACGACTCCGTGCTACATGACATGGACCA	203 	\|\|\|\|\|\|\|\|\|\|\|\|\|\|\|\|\|\|\|\|\|\|\|\|\|\|\|\|\|\|\|\| 	ATTGACGACTCCGTGCTACATGACATGGACCA ```      ``` 205	GACGAAGAGGAAGAAGAAGAAATTGACGACTCCGTGCTACATGACATGGACCA	257 	\|\|\|\|\| \|\|\|\|\| \|\|\|\|\|\|\|\| \|\|\|\|\|\|\|\|\|\|\|\|\|\|\|\|\|\|\|\|\|\|\|\|\|\|\|\|\|\|\|\| 	GACGACGAGGAGGAAGAAGACATTGACGACTCCGTGCTACATGACATGGACCA ```      ``` 53	AAGGCATGAGGTACCGCGAGAACAACAACACAG-AGGACGATACCGAAGAAATGGAAGAGAGCCGGGGACCATTGGACGAAAGCGGTGGTACCATGTCTGACGACGAGGAGGAAGAAGACATTGACGACTCCGTGCTACATGACATGGACCACGACGAAGAGGAAGAAGAAGAAATTGACGACTCCGTGCTACATGACATGGACCAATCCGAGCAGACCTTTCAGAACATCAGCCAACGATATCGCCTGGTTCACCGCATTGGCGAAGGCACCTTCTCGACGGTCT	337 	\|\|\|\|\| \|\|\|\|\|  \|\| \|\|\| \| \|\| \|\| \| \|\|\| \|\|\|\|\|\|\|\| \|\|\|\|\|\| \|\|\|\|\|\|\|\|\|\|\|\|\|\|\|\|\|\|\| \|\|\| \|\|\|\|\|\|\|\|\|\|\|\|\|\|\|\|\|\|\|\|\|\|\|\|\|\|\|\|\|\|\|\|\|\|\|\|\|\|\|\|\|\|\|\|\|\|\|\|\|\|\|\|\|\|\|\|\|\|\|\|\|\|\|\|\|\|\|\|\|\|\|\|\|\|\|\|\|\|\|\|\|\|\|\|\| \|\|\|\|\| \|\| \|\| \|\|\|\|\|\|\|\|\|\|\|\|\|\|\|\|\|\|\|\|\|\|\|\|\|\|\|\|\|\|\|\|\|\|\|\|\|\|\|\|\|\|\| \|\|\|\|\|\|\|\| \|\| \|\|\|\| \|\| \|\|\|\|\|\|\|\|\| \|\|\|\|\|\|\| \|\|\|\|\|\|\|\| \|\|\|\| \|\|\|\|\|\|\|\| \|\|\|\| \|\|\|\| 	AAGGCNTGAGGNCCCNCGANACCACCACCCCAGAAGGACGATCCCGAAG-AATGGAAGAGAGCCGGGGAACATNGGACGAAAGCGGTGGTACCATGTCTGACGACGAGGAGGAAGAAGACATTGACGACTCCGTGCTACATGACATGGACCACGACGAANAGGAANAANAANAAATTGACGACTCCGTGCTACATGACATGGACCAATCCGAGCANACCTTTCANAANATCANCCNACGATATCGNCTGGTTCNCCGCATTGCCGAANGCACCTTCNCGACNGTCT ``` |
| 1 | ggggacaagtttgtacaaaaaagcaggcttgatgGCAGAACTCGATGACGAGGA, ggggaccactttgtacaagaaagctgggtcCCATTCTTCCCCAACATCGG | Mycgr3\|98091  JGI  Ensembl | KOG3696 Aspartyl beta-hydroxylase; | ATP-dependent DNA helicase activity; ATP-dependent RNA helicase activity;  ATP-dependent helicase activity;  ATPase activity;  ATPase activity, coupled;  ATPase activity, coupled to transmembrane movement of ions;  ATPase activity, coupled to transmembrane movement of substances;  ATPase activity, uncoupled;  DNA-dependent ATPase activity;  RNA-dependent ATPase activity;  protein-transmembrane transporting ATPase activity;  single-stranded DNA-dependent ATP-dependent DNA helicase activity; | 100 % |  | ``` 312	CTTGGATGATGTCGATGAGGTCGACGGTTTCGAGTATGAGGAGGACGATGGCTTGGATGATGTCGATGAGGTCGATGGTTTCGAGTATGAGGAGGACGATGGCTTCGATGATGA-G--GAGGACGACGGCTTC	441 	\|\|\|\|\|\|\|\|\|\|\|\|\|\|\|\|\|\|\|\|\|\|\|\|\|\|\|\|\|\|\|\|\|\|\|\|\| \|\|\|\|\|\|\|\|\|\|\|\|\|\|\|\|\|\|\|\|\|\|\|\|\|\|\|\|\|\|\|\|\|\|\|\|\| \|\|\|\|\|\|\| \|\|\|\|\|\|\|\|\|\|\|\|\|\|\|\|\|\|\|\|\| \|\|\|\|\|\|\|  \|  \|\|\|\| \|\|\|\|\|\| \|\|\| 	CTTGGATGATGTCGATGAGGTCGACGGTTTCGAGTATAAGGAGGACGATGGCTTGGATGATGTCGATGAGGTCGACGGTTTCGGGTATGAGGAGGACGATGGCTTGGATGATGTCGATGAGGTCGACGGTTTC ```      ``` 57	CTTGGATGATGTCGATGAGGTCGACGGTTTCGAGTATAAGGAGGACGATGGCTTGGATGATGTCGATGAGGTCGACGGTTTCGGGTATGAGGAGGACGATGGCTTGGATGATGTCGATGAGGTCGACGGTTTC	189 	\|\|\|\|\|\|\|\|\|\|\|\|\|\|\|\|\|\|\|\|\|\|\|\|\|\|\|\|\|\|\|\| \|\|\|\| \|\|\|\|\|\|\|\|\|\|\|\|\|\|\|\|\|\|\|\|\|\|\|\|\|\|\|\|\|\|\|\|\|\|\|\|\|\|\|\|\|\|\|\|\|\|\|\|\|\|\|\|\|\|\|\|\|\|\|\|\|\|\|\|\|\|\|\|\|\|\|\|\|\|\|\|\| \|\|\|\| \|\|\| \|\| \|\|\|\|\| 	CTTGGATGATGTCGATGAGGTCGACGGTTTCGGGTATGAGGAGGACGATGGCTTGGATGATGTCGATGAGGTCGACGGTTTCGGGTATGAGGAGGACGATGGCTTGGATGATGTCNATGANGTCNACNGTTTC ```      ``` 210	CTTGGATGATGTCGATGAGGTCGACGGTTTCGAGTATGAGGAGGACGATGGCTTGGATGATGTCGATGAGGTCGACGGTTTCGAGTATGAGGAGGACGATGGCTTGGATGATGTCGATGAGGTCGACGGTTTCGAGTATGAGGAGGACGATGGCTTGGATGATGTCGATGAGGTCGATGGTTTCGAGTATGAGGAGGACGATGGCT	415 	\|\|\|\|\|\|\|\|\|\|\|\|\|\|\|\|\|\|\|\|\|\|\|\|\|\|\|\|\|\|\|\|\|\|\|\|\| \|\|\|\|\|\|\|\|\|\|\|\|\|\|\|\|\|\|\|\|\|\|\|\|\|\|\|\|\|\|\|\|\|\|\|\|\|\|\|\|\|\|\|\|\| \|\|\|\|\|\|\|\|\|\|\|\|\|\|\|\|\|\|\|\|\|\|\|\|\|\|\|\|\|\|\|\|\|\|\|\|\|\|\|\|\|\|\|\|\|\|\|\|\|\| \|\|\|\|\|\|\|\|\|\|\|\|\|\|\|\|\|\|\|\|\|\|\|\|\|\|\|\|\|\|\| \|\|\|\| \|\|\| \|  \|\|\|\|\| \|\| \| \|\|  \|\|\|\|\| \| \|\|\|\| 	CTTGGATGATGTCGATGAGGTCGACGGTTTCGAGTATAAGGAGGACGATGGCTTGGATGATGTCGATGAGGTCGACGGTTTCGGGTATGAGGAGGACGATGGCTTGGATGATGTCGATGAGGTCGACGGTTTCGGGTATGAGGAGGACGATGGCTTGGATGATGTCNATGANGTCNACNGTTTCNAGNANGANNAGGACNANGGCT ```      ``` 108	CTTGGATGATGTCGATGAGGTCGACGGTTTCGGGTATGAGGAGGACGATGGCTTGGATGATGTCGATGAGGTCGACGGTTTCGGGTATGAGGAGGACGATGGCTTGGATGATGTCGATGAGGTCGACGGTTTCGAGTATGAGGAGGACGATGGCTTGGATGATGTCGATGAGGTCGACGGTTTCGAGTATGAGGAGGACGATGGCT	313 	\|\|\|\|\|\|\|\|\|\|\|\|\|\|\|\|\|\|\|\|\|\|\|\|\|\|\|\|\|\|\|\| \|\|\|\| \|\|\|\|\|\|\|\|\|\|\|\|\|\|\|\|\|\|\|\|\|\|\|\|\|\|\|\|\|\|\|\|\|\|\|\|\|\|\|\|\|\|\|\|\|\|\|\|\|\|\|\|\|\|\|\|\|\|\|\|\|\|\|\|\|\|\|\|\|\|\|\|\|\|\|\|\|\|\|\|\|\|\|\|\|\|\|\|\|\|\|\|\|\|\|\| \|\|\|\|\|\|\|\|\|\|\|\|\|\|\|\|\|\|\|\|\|\|\|\|\|\|\|\|\|\|\| \|\|\|\| \|\|\| \|\| \|\|\|\|\| \|\| \| \|\|  \|\|\|\|\| \| \|\|\|\| 	CTTGGATGATGTCGATGAGGTCGACGGTTTCGAGTATAAGGAGGACGATGGCTTGGATGATGTCGATGAGGTCGACGGTTTCGGGTATGAGGAGGACGATGGCTTGGATGATGTCGATGAGGTCGACGGTTTCGGGTATGAGGAGGACGATGGCTTGGATGATGTCNATGANGTCNACNGTTTCNAGNANGANNAGGACNANGGCT ```      ``` 159	CTTGGATGATGTCGATGAGGTCGACGGTTTCGGGTATGAGGAGGACGATGGCTTGGATGATGTCGATGAGGTCGACGGTTTCGAGTATGAGGAGGACGATGGCTTGGATGATGTCGATGAGGTCGACGGTTTCGAGTATGAGGAGGACGATGGCTTGGATGATGTCGATGAGGTCGACGGTTTCGAGTATGAGGAGGACGATGGCT	364 	\|\|\|\|\|\|\|\|\|\|\|\|\|\|\|\|\|\|\|\|\|\|\|\|\|\|\|\|\|\|\|\| \|\|\|\| \|\|\|\|\|\|\|\|\|\|\|\|\|\|\|\|\|\|\|\|\|\|\|\|\|\|\|\|\|\|\|\|\|\|\|\|\|\|\|\|\|\|\|\|\| \|\|\|\|\|\|\|\|\|\|\|\|\|\|\|\|\|\|\|\|\|\|\|\|\|\|\|\|\|\|\|\|\|\|\|\|\|\|\|\|\|\|\|\|\|\|\|\|\|\| \|\|\|\|\|\|\|\|\|\|\|\|\|\|\|\|\|\|\|\|\|\|\|\|\|\|\|\|\|\|\| \|\|\|\| \|\|\| \|\| \|\|\|\|\| \|\| \| \|\|  \|\|\|\|\| \| \|\|\|\| 	CTTGGATGATGTCGATGAGGTCGACGGTTTCGAGTATAAGGAGGACGATGGCTTGGATGATGTCGATGAGGTCGACGGTTTCGGGTATGAGGAGGACGATGGCTTGGATGATGTCGATGAGGTCGACGGTTTCGGGTATGAGGAGGACGATGGCTTGGATGATGTCNATGANGTCNACNGTTTCNAGNANGANNAGGACNANGGCT ```      ``` 261	CTTGGATGATGTCGATGAGGTCGACGGTTTCGAGTATGAGGAGGACGATGGCTTGGATGATGTCGATGAGGTCGACGGTTTCGAGTATGAGGAGGACGATGGCTTGGATGATGTCGATGAGGTCGATGGTTTCGAGTATGAGGAGGACGATGGCTTCGATGATG	424 	\|\|\|\|\|\|\|\|\|\|\|\|\|\|\|\|\|\|\|\|\|\|\|\|\|\|\|\|\|\|\|\|\|\|\|\|\| \|\|\|\|\|\|\|\|\|\|\|\|\|\|\|\|\|\|\|\|\|\|\|\|\|\|\|\|\|\|\|\|\|\|\|\|\|\|\|\|\|\|\|\|\| \|\|\|\|\|\|\|\|\|\|\|\|\|\|\|\|\|\|\|\|\|\|\|\|\|\|\|\|\|\|\|\|\|\|\|\|\|\|\|\|\|\| \|\|\|\|\|\|\| \|\|\|\|\|\|\|\|\|\|\|\|\|\|\|\|\|\|\|\|\| \|\|\|\|\|\|\| 	CTTGGATGATGTCGATGAGGTCGACGGTTTCGAGTATAAGGAGGACGATGGCTTGGATGATGTCGATGAGGTCGACGGTTTCGGGTATGAGGAGGACGATGGCTTGGATGATGTCGATGAGGTCGACGGTTTCGGGTATGAGGAGGACGATGGCTTGGATGATG ```      ``` 57	CTTGGATGATGTCGATGAGGTCGACGGTTTCGAGTATAAGGAGGACGATGGCTTGGATGATGTCGATGAGGTCGACGGTTTC	138 	\|\|\|\|\|\|\|\|\|\|\|\|\|\|\|\|\|\|\|\|\|\|\|\|\|\|\|\|\|\|\|\| \|\|\|\| \|\|\|\|\|\|\|\|\|\|\|\|\|\|\|\|\|\|\|\|\|\|\|\|\|\| \|\|\|\| \|\|\| \|\| \|\|\|\|\| 	CTTGGATGATGTCGATGAGGTCGACGGTTTCGGGTATGAGGAGGACGATGGCTTGGATGATGTCNATGANGTCNACNGTTTC ```      ``` 1	ATGGCAGAACTCGATGACGAGGAGCTGGCTGACATGGGATGCTTTGCCTTGCGTCCCTTGGATGATGTCGATGAGGTCGACGGTTTCGAGTATAAGGAGGACGATGGCTTGGATGATGTCGATGAGGTCGACGGTTTCGGGTATGAGGAGGACGATGGCTTGGATGATGTCGATGAGGTCGACGGTTTCGGGTATGAGGAGGACGATGGCTTGGATGATGTCGATGAGGTCGACGGTTTCGAGTATGAGGAGGACGATGGCT	262 	\|\|\|\|\|\|\|\|\|\|\|\| \|\|\|\|\|\|\|\|\|\|\|\|\|\|\|\|\|\|\|\|\|\|\|\|\|\|\|\|\|\|\|\|\|\|\|\|\|\|\|\|\|\|\|\|\|\|\|\|\|\|\|\|\|\|\|\|\|\|\|\|\|\|\|\|\|\|\|\|\|\|\|\|\|\|\|\|\|\|\|\|\|\|\|\|\|\|\|\|\|\|\|\|\|\|\|\|\|\|\|\|\|\|\|\|\|\|\|\|\|\|\|\|\|\|\|\|\|\|\|\|\|\|\|\|\|\|\|\|\|\|\|\|\|\|\|\|\|\|\|\|\|\|\|\|\|\|\|\|\|\|\|\|\|\|\|\|\|\|\|\|\|\|\|\|\|\|\|\|\|\|\|\|\|\|\|\|\|\|\|\|\|\|\|\|\|\|\|\|\|\|\|\|\|\|\|\|\|\|\|\|\|\|\|\|\|\|\|\|\| \|\|\|\| \|\|\| \|\| \|\|\|\|\| \|\| \| \|\|  \|\|\|\|\| \| \|\|\|\| 	ATGGCAGAACTCNATGACGAGGAGCTGGCTGACATGGGATGCTTTGCCTTGCGTCCCTTGGATGATGTCGATGAGGTCGACGGTTTCGAGTATAAGGAGGACGATGGCTTGGATGATGTCGATGAGGTCGACGGTTTCGGGTATGAGGAGGACGATGGCTTGGATGATGTCGATGAGGTCGACGGTTTCGGGTATGAGGAGGACGATGGCTTGGATGATGTCNATGANGTCNACNGTTTCNAGNANGANNAGGACNANGGCT ``` |
| 1 | ggggacaagtttgtacaaaaaagcaggcttgatgGCCACGAACGGTGCGGCTGC, ggggaccactttgtacaagaaagctgggtcTGAGATGTACTCCGACGCCC | Mycgr3\|98372  JGI  Ensembl | KOG1942 DNA helicase, TBP-interacting protein; | 8-oxo-7,8-dihydroguanine triphosphatase activity; ATP binding;  ATP-dependent 5'-3' DNA helicase activity;  DNA helicase activity;  UDP-2,3-diacylglucosamine hydrolase activity;  bis(5'-nucleosyl)-tetraphosphatase activity;  dATP pyrophosphohydrolase activity;  diacylglycerol pyrophosphate phosphatase activity;  dihydroneopterin monophosphate phosphatase activity;  dihydroneopterin triphosphate pyrophosphohydrolase activity;  nucleoside-triphosphatase activity;  nucleotide binding;  nucleus;  pyrophosphatase activity;  thiamin-pyrophosphatase activity; | 100 % |  | ``` 41	ATGGCCACGAACGGTGCGGCTGCGCCTCCAGCAGCTCCAACGACCGTCAAGATCCAGGAAGTCCAACAGACCAACTCGCGCGAAAACCGAACGGCAGCTCATTCACACATCAAAGGCCTTGGTCTACGCTCAGATGGCTATGCCGATACAAACGCGCAGGGATTCATCGGCCAAACCGCTGCGCGCGAAGCATGCGGCGTAGTTGTGGACTTGATTCGGGCGAAGAAGATGGCTGGAAAAGCTATCCTGCTCGCTGGTGGACCTGGAACTGGAAAGACGGCATTGGCACTTGCGGTTAGTCAGGAGCTTGGAACCAAGGTCCCGTTCTGTCCGATGACCGGCAGTGAAGTGTACAGCGCCGAGGTGAAGAAGACGGAGGCATTGATGGAGAACTTCAGAAGAGCCATCGGTCTGCGGGTGCAGGAGCGGAAAGAGGTGTACGAAGGTGAAGTGGTGGAACTC	502 	\|\|\|\|\|\|\|\|\|\|\|\|\|\|\|\|\|\|\|\|\|\|\|\|\|\|\|\|\|\|\|\|\|\|\|\|\|\|\|\|\|\|\|\|\|\|\|\|\|\|\|\|\|\|\|\|\|\|\|\|\|\|\|\|\|\|\|\|\|\|\|\|\|\|\|\|\|\|\|\|\|\|\|\|\|\|\|\|\|\|\|\|\|\|\|\|\|\|\|\|\|\|\|\|\|\|\|\|\|\|\|\|\|\|\|\|\|\|\|\|\|\|\|\|\|\|\|\|\|\|\|\|\|\|\|\|\|\|\|\|\|\|\|\|\|\|\|\|\|\|\|\|\|\|\|\|\|\|\|\|\|\|\|\|\|\|\|\|\|\|\|\|\|\|\|\|\|\|\|\|\|\|\|\|\|\|\|\|\|\|\|\|\|\|\|\|\|\|\|\|\|\|\|\|\|\|\|\|\|\|\|\|\|\|\|\|\|\|\|\|\|\|\|\|\|\|\|\|\|\|\|\|\|\|\|\|\|\|\|\|\|\|\|\|\|\|\|\|\|\|\|\|\|\|\|\|\|\|\|\|\|\|\|\|\|\|\|\|\|\|\|\|\|\|\|\|\|\|\|\|\|\|\|\|\|\|\|\|\|\|\|\|\|\|\|\|\|\|\|\|\|\|\|\|\|\|\|\|\|\|\|\|\|\|\|\|\|\|\|\|\|\|\|\|\|\|\|\|\|\|\|\|\|\|\|\|\|\|\|\|\|\|\|\|\|\|\|\|\|\|\|\|\|\|\|\|\|\|\|\|\|\|\|\|\|\|\|\| \|\| \|\|\|\|\|\|\|\|\|\|\|\|\|\|\|\|\| \|\|\|\|\|\|\|\|\|\| \|\|\|\|\|\|\|\|\|\|\|\|\|\|\|\|\|\|\|\|\| \|\| \|\|\|\|\|\|\|\|\|\|\|\|\|\|\| \|\| \|\|\|\|\| \|\| \|\| \|\|\|\|\| 	ATGGCCACGAACGGTGCGGCTGCGCCTCCAGCAGCTCCAACGACCGTCAAGATCCAGGAAGTCCAACAGACCAACTCGCGCGAAAACCGAACGGCAGCTCATTCACACATCAAAGGCCTTGGTCTACGCTCAGATGGCTATGCCGATACAAACGCGCAGGGATTCATCGGCCAAACCGCTGCGCGCGAAGCATGCGGCGTAGTTGTGGACTTGATTCGGGCGAAGAAGATGGCTGGAAAAGCTATCCTGCTCGCTGGTGGACCTGGAACTGGAAAGACGGCATTGGCACTTGCGGTTAGTCAGGAGCTTGGAACCAAGGTCCCGTTCTGTCCGATGACCGGCAGTGAAGTGTACAGCGCCGAGGTGAANAANACGGAGGCATTGATGGANAACTTCAGAANAGCCATCGGTCTGCGGGTGCANGANCGGAAAGAGGTGTACNAANGTGAANTGNTGNAACTC ``` |
| 1 | ggggacaagtttgtacaaaaaagcaggcttgatgGCCCTGGAGACAATGCCAGT, ggggaccactttgtacaagaaagctgggtcTCCAAACATCTGCCGAACAT | Mycgr3\|99953  JGI  Ensembl | KOG2731 DNA alkylation damage repair protein; |  | 100 % | chr\_4:1743638 G->T; | ``` 163	AGTCTGAGAAGAAAGGTAGGTCTTTTCATTTATCCCTCTCTGTTCCCGCCACAAATCCAGCGAGAATTGCTCTCGAGACTGCTGCATCGAGACTTGACAAATCCCCAGCACATGACGAATGTCCACACGCATTACAATGTTCAATACCCATCGAAGAGTGAGGACGAGGACGCATCGTTCTTCGACGAGTCTTGCCGCAGTACTGTCTTCGAACCACACGATCCTGCCCTTCACAAGCCCATCTCAACGGAGCAATTCCTGAACAAGAAGCTCCGCTGGACTACACTGGGAG	454 	\|\|\|\|\|\|\|\|\|\|\|\|\|\|\|\|\|\|\|\|\|\|\|\|\|\|\|\|\|\|\|\|\|\|\|\|\|\|\|\|\|\|\|\|\|\|\|\|\|\|\|\|\|\|\|\|\|\|\|\|\|\|\|\|\|\|\|\|\|\|\|\|\|\|\|\|\|\|\|\|\|\|\|\|\|\|\|\|\|\|\|\|\|\|\|\|\|\|\|\|\|\|\|\|\|\|\|\|\|\|\|\|\|\|\|\|\|\|\|\|\|\|\|\|\|\|\|\|\|\|\|\|\|\|\|\|\|\|\|\|\|\|\|\|\|\|\|\|\|\|\|\|\|\|\| \|\|\|\|\|\|\|\|\|\|\| \|\|\|\|\|\|\|\|\|\|\|\|\|\|\| \|\|\|\|\|\|\|\|\|\|\|\|\|\|\| \|\|\|\|\|\|\|\|\|\| \|\|\|\|\|\|\|\|\|\|\|\|\|\|\|\|\|\|\|\|\|\|\|\|\| \|\|\|\|\|\|\|\|\|\|\|\|\|\| \|\|\|\|\|\|\|\|\|\|\|\|\|\| \|\| \|\|\|\|\|\|\|\|\|\|\|\|\|\| \|\|\|\|\|\|\| 	AGTCTGAGAAGAAAGGTAGGTCTTTTCATTTATCCCTCTCTGTTCCCGCCACAAATCCAGCGAGAATTGCTCTCGAGACTGCTGCATCGAGACTTGACAAATCCCCAGCACATGACGAATGTCCACACGCATTACAATGTTCAATACCCATCGAANAGTGAGGACGANGACGCATCGTTCTTCNACGAGTCTTGCCGCANTACTGTCTTCNAACCACACGATCCTGCCCTTCACAANCCCATCTCAACGGANCAATTCCTGAACAANAANCTCCGCTGGACTAC-CTGGGAG ```      ``` 1	ATGGCCCTGGAGACAATGCCAGTGCCGTTGGACGCGCACGCAAAGCCTCCGTCAAGCCTTCGCGATGTCTACAAATATTACCACAAACTCAAGCCGACCGCCCTGGGAAGTCATCCAGGCCTGATCCAATTTGACGGCTGTTCGAGCAGCAGTACCAGCCAC	162 	\|\|\|\|\|\|\|\|\|\|\|\|\|\|\|\|\|\|\|\|\|\|\|\|\|\|\|\|\|\|\|\|\|\|\|\|\|\|\|\|\|\|\|\|\|\|\|\|\|\|\|\|\|\|\|\|\|\|\|\|\|\|\|\|\|\|\|\|\|\|\|\|\|\|\|\|\|\|\|\|\|\|\|\|\|\|\|\|\|\|\|\|\|\|\|\|\|\|\|\|\|\|\|\|\|\|\|\|\|\|\|\|\|\|\|\|\|\|\|\|\|\|\|\|\|\|\|\|\|\|\|\|\|\|\|\|\|\|\|\|\|\|\|\|\|\|\|\|\|\|\|\|\|\|\|\|\|\|\|\|\|\| 	ATGGCCCTGGAGACAATGCCAGTGCCGTTGGACGCGCACGCAAAGCCTCCGTCAAGCCTTCGCGATGTCTACAAATATTACCACAAACTCAAGCCGACCGCCCTGGGAAGTCATCCAGGCCTGATCCAATTTGACGGCTGTTCGAGCAGCAGTACCAGCCAC ``` |
| 1 | ggggacaagtttgtacaaaaaagcaggcttgatgGGCAACGAAGGCAGCTCTCC, ggggaccactttgtacaagaaagctgggtcCCACTGGCCACGCAGATCCC | Mycgr3\|100618  JGI  Ensembl | KOG2682 NAD-dependent histone deacetylases and class I sirtuins (SIR2 family); | (3,5-dichlorophenylurea)acetate amidohydrolase activity; DNA binding;  N-acetylgalactosamine-6-phosphate deacetylase activity;  N2-acetyl-L-lysine deacetylase activity;  NAD binding;  NAD-dependent histone deacetylase activity;  O-succinylbenzoate synthase activity;  UDP-3-O-[3-hydroxymyristoyl] N-acetylglucosamine deacetylase activity;  chromatin silencing;  chromatin silencing complex;  histone deacetylase activity;  indoleacetamide hydrolase activity;  iprodione amidohydrolase activity;  protein N-terminal asparagine amidohydrolase activity;  protein amino acid deacetylation;  regulation of transcription, DNA-dependent;  zinc ion binding; | 100 % |  | ``` 1	ATGGGCAACGAAGGCAGCTCTCCCATCGACGAGTCGATCCCGCCATCCACCCTCTCCGGCCGCACAATCGATGCGCTAGCATCATACATCAAATCCGGCCACTGCAAAAAAGTGGTCGTGATGACCGGCGCCGGAATCTCCACCTCAGCAGGCATTCCCGACTTTCGCTCCCCAGAAACAGGACTCTACGCCAACCTCGCCCGCCTCAATCTCCCCTACGCAGAAGCGGTTTTCGACATCTCCTACTTCCGTGAGAAACCAGAGCCATTCTACACCCTTGCCCACGAACTCTATCCGGGCAAATTCCGCCCGACGATCACACATTCATTCATCAAACTCCTCCATGACAAGGGCCTACTGCTCAAACACTTCACGCAAAACATAGACACTCTGGATCGTGAAGCTGGCGTCCCGGGTGATCTCATCATCGAAGCTCACGGCTCGTTCGCACGGCAGAGCTGCATCGAGTGCAAGGCCTCATATCCGGACGACGACATCCGTGAACACATCAACGCAAAGACTATACCACGGTGCTATGAGTGTAAGGGCCTGGTCAAGCCGGAGATTGTGTTCTTCGGCGAGCAATTACCAGCCGAATTCTTCGCCGCTCGAGGACTGCCTGCAGAAGCGGATTTGTGTATTGTCATGG	649 	\|\|\|\|\|\|\|\|\|\|\|\|\|\|\|\|\|\|\|\|\|\|\|\|\|\|\|\|\|\|\|\|\|\|\|\|\|\|\|\|\|\|\|\|\|\|\|\|\|\|\|\|\|\|\|\|\|\|\|\|\|\|\|\|\|\|\|\|\|\|\|\|\|\|\|\|\|\|\|\|\|\|\|\|\|\|\|\|\|\|\|\|\|\|\|\|\|\|\|\|\|\|\|\|\|\|\|\|\|\|\|\|\|\|\|\|\|\|\|\|\|\|\|\|\|\|\|\|\|\|\|\|\|\|\|\|\|\|\|\|\|\|\|\|\|\|\|\|\|\|\|\|\|\|\|\|\|\|\|\|\|\|\|\|\|\|\|\|\|\|\|\|\|\|\|\|\|\|\|\|\|\|\|\|\|\|\|\|\|\|\|\|\|\|\|\|\|\|\|\|\|\|\|\|\|\|\|\|\|\|\|\|\|\|\|\|\|\|\|\|\|\|\|\|\|\|\|\|\|\|\|\|\|\|\|\|\|\|\|\|\|\|\|\|\|\|\|\|\|\|\|\|\|\|\|\|\|\|\|\|\|\|\|\|\|\|\|\|\|\|\|\|\|\|\|\|\|\|\|\|\|\|\|\|\|\|\|\|\|\|\|\|\|\|\|\|\|\|\|\|\|\|\|\|\|\|\|\|\|\|\|\|\|\|\|\|\|\|\|\|\|\|\|\|\|\|\|\|\|\|\|\|\|\|\|\|\|\|\|\|\|\|\|\|\|\|\|\|\|\|\|\|\|\|\|\|\|\|\|\|\|\|\|\|\|\|\|\|\|\|\|\|\|\|\|\|\|\|\|\|\|\|\|\|\|\|\|\|\|\|\|\|\|\|\|\|\|\|\|\|\|\|\|\|\|\|\|\|\|\|\|\|\|\|\|\|\|\|\|\|\|\|\|\|\|\|\|\|\|\|\|\|\|\|\|\|\|\|\|\|\|\|\|\|\|\|\|\|\|\|\|\|\|\|\|\|\|\|\|\|\|\|\|\|\|\|\|\|\|\|\|\|\|\|\|\|\|\|\|\|\|\|\|\|\|\|\|\|\|\|\|\|\|\|\|\|\|\|\|\|\|\|\|\|\|\|\|\|\|\|\|\|\|\|\|\|\|\|\|\|\|\|\|\|\|\|\|\|\|\|\|\|\|\|\|\|\|\|\|\|\|\|\|\|\|\|\|\|\|\|\|\|\|\|\|\|\|\|\|\|\|\|\| \|\|\|\|\|\|\|\|\|\|\|\|\|\|\|\|\|\|\|\|\|\|\|\|\|\|\| \|\|\|\|\|\|\|\|\|\|\| \|\|\|\|\|\|\|\|\|\|\|\|\|\| \|\|\|\|  \|\| \|\|\|\|\|\|\|\|\|\|\|\|\|\|\|\|\| \|\|\| 	ATGGGCAACGAAGGCAGCTCTCCCATCGACGAGTCGATCCCGCCATCCACCCTCTCCGGCCGCACAATCGATGCGCTAGCATCATACATCAAATCCGGCCACTGCAAAAAAGTGGTCGTGATGACCGGCGCCGGAATCTCCACCTCAGCAGGCATTCCCGACTTTCGCTCCCCAGAAACAGGACTCTACGCCAACCTCGCCCGCCTCAATCTCCCCTACGCAGAAGCGGTTTTCGACATCTCCTACTTCCGTGAGAAACCAGAGCCATTCTACACCCTTGCCCACGAACTCTATCCGGGCAAATTCCGCCCGACGATCACACATTCATTCATCAAACTCCTCCATGACAAGGGCCTACTGCTCAAACACTTCACGCAAAACATAGACACTCTGGATCGTGAAGCTGGCGTCCCGGGTGATCTCATCATCGAAGCTCACGGCTCGTTCGCACGGCAGAGCTGCATCGAGTGCAAGGCCTCATATCCGGACGACGACATCCGTGAACACATCAACGCAAAGACTATACCACGGTGCTATGAGTGTAAGGGCCTGGTCAAGCCGGANATTGTGTTCTTCGGCGAGCAATTACCANCCGAATTCTTCNCCGCTCGAGGACTGNCTGCNNAA-CGGATTTGTGTATTGTCNTGG ``` |
| 1 | ggggacaagtttgtacaaaaaagcaggcttgatgGCTGCCGAGCTCGCGAGTAA, ggggaccactttgtacaagaaagctgggtcTGCCTGAGCCGCAGGCGGAC | Mycgr3\|101609  JGI  Ensembl | KOG0326 ATP-dependent RNA helicase; | 8-oxo-7,8-dihydroguanine triphosphatase activity; ATP binding;  ATP-dependent 5'-3' DNA helicase activity;  ATP-dependent helicase activity;  UDP-2,3-diacylglucosamine hydrolase activity;  bis(5'-nucleosyl)-tetraphosphatase activity;  dATP pyrophosphohydrolase activity;  diacylglycerol pyrophosphate phosphatase activity;  dihydroneopterin monophosphate phosphatase activity;  dihydroneopterin triphosphate pyrophosphohydrolase activity;  helicase activity;  nucleic acid binding;  pyrophosphatase activity;  thiamin-pyrophosphatase activity; | 0 % |  |  |
| 1 | ggggacaagtttgtacaaaaaagcaggcttgatgGCGGAACTCCACGAGGCACT, ggggaccactttgtacaagaaagctgggtcATTGCTCTTCTTCGAATTCG | Mycgr3\|107209  JGI  Ensembl | KOG4786 Ubinuclein, nuclear protein interacting with cellular and viral transcription factors; |  | 100 % |  |  |
| 1 | ggggacaagtttgtacaaaaaagcaggcttgatgCTGATCTCCACGCTGTCCGG, ggggaccactttgtacaagaaagctgggtcTCCAGCCGAATCATACTTCG | Mycgr3\|107320  JGI  Ensembl |  |  | 100 % | chr\_1:3342485 G->T; |  |
| 1 | ggggacaagtttgtacaaaaaagcaggcttgatgCAACGAGGTCGCTTTACCAG, ggggaccactttgtacaagaaagctgggtcCAACTCATCATCCACACTCT | Mycgr3\|107437  JGI  Ensembl | KOG1956 DNA topoisomerase III alpha; | hydrogen ion transmembrane transporter activity; proton-transporting two-sector ATPase complex;  zinc ion binding; | 100 % |  | ``` 1	ATGCAACGAGGTCGCTTTACCAGCAGCAACCGTGGAGGGGAATGGGCAGGACGAGGGAAGTCCCGAGGACGTGGTCGTGGACGAGGAGGATTTGGCAAAAGCACACAGCCAAAAGGCCTCTTCGAGAACGGATCCTGGCTGTGTAATTGTACACCGCGACTACCCGCCGTGCGCTTTCAGGTAAAGAAAGAGGGCGCGAACAAAGGTCGTTTCTTCTACACATGTCAGGAAACGGAGCCGAAACGCTGTGACTTTTTCCTCTGGAACGAAGACGCGCAGCCAAGAGAGGCGGCAGCGGTGTTGA	304 	\|\|\|\|\|\|\|\|\|\|\|\|\|\|\|\|\|\|\|\|\|\|\|\|\|\|\|\|\|\|\|\|\|\|\|\|\|\|\|\|\|\|\|\|\|\|\|\|\|\|\|\|\|\|\|\|\|\|\|\|\|\|\|\|\|\|\|\|\|\|\|\|\|\|\|\|\|\|\|\|\|\|\|\|\|\|\|\|\|\|\|\|\|\|\|\|\|\|\|\|\|\|\|\|\|\|\|\|\|\| \|\|\|\|\|\|\|\|\|\|\|\|\|\|\|\|\|\|\|\|\|\|\|\|\|\|\|\|\|\|\|\|\|\|\|\|\|\|\|\|\|\|\|\|\|\|\|\|\|\|\|\|\|\|\|\|\|\|\|\|\|\|\|\|\|\|\|\|\|\|\|\|\|\|\|\|\|\|\|\|\|\|\|\|\|\|\|\|\|\|\|\|\|\|\|\|\|\|\|\|\|\|\|\|\|\|\|\|\|\|\|\|\|\|\|\| \|\|\|\|\|\|\|\|\|\|\| \|\|\|\|\|\|\|\|\|\|\|\|\|\|\|\|\|\|\|\|\|\|\|\|\|\|\| \|\| \|\| \|\|\|\| \|\| \| \| \| \|\|\| \|  \|\|\|\|\|\|\|\|\| 	ATGCAACGAGGTCGCTTTACCAGCAGCAACCGTGGAGGGGAATGGGCAGGACGAGGGAAGTCCCGAGGACGTGGTCGTGGACGAGGAGGATTTGGCAAAAGCACACAGCCNAAAGGCCTCTTCGAGAACGGATCCTGGCTGTGTAATTGTACACCGCGACTACCCGCCGTGCGCTTTCAGGTAAAGAAAGAGGGCGCGAACAAAGGTCGTTTCTTCTACACATGTCANGAAACGGAGCCNAAACGCTGTGACTTTTTCCTCTGGAACNAANACNCGCANCCNANANANGCGNCNNCGGTGTTGA ``` |
| 1 | ggggacaagtttgtacaaaaaagcaggcttgatgTCTTACTCACACATGGCGCC, ggggaccactttgtacaagaaagctgggtcCGACAACTTGTTCAGGGCCT | Mycgr3\|108026  JGI  Ensembl | KOG3227 Calcium-responsive transcription coactivator; |  | 100 % |  |  |
| 1 | ggggacaagtttgtacaaaaaagcaggcttgatgGCACGCTCTCCCTCCGATGC, ggggaccactttgtacaagaaagctgggtcCCGTCTCCCAAGCCCAAACC | Mycgr3\|108713  JGI  Ensembl |  |  | 0 % |  | ``` 68	ATGGCACGCTCTCCCTCCGATGCCACGCGCTTTACCTCGACAGGACCCTACGTCTCCAATAAACCCGCAGGCACAACATCCTTCACATCCAACCCAACCGCACCATCCCCTTCGCAAGCCACAAACCTCAAATTCGGCCCCTCCGCGCCGGCCAACGAAACGCCTCAACAAAAAATCGCCCGTCTCCGCGCCGCCGCCGCCCTCGCCAAAGCCCCCCAAGAAACATCATGGGACAAGACCGTCCGGCGAGGCCGCATCTGGGCCGACCGCGCCCACCGCTTCACAGCTCTCGGCCTTATCGGCTTCACAGTCATCAGCGCGGCGGTGGCGACGGCAGG-AATCACGGATATGATGATGCACAATCGAA-GACGGAGGAATGAATGGTT	453 	\|\|\|\|\|\|\|\|\|\|\|\|\|\|\|\| \|\|\|\|\|\|\|\|\|\|\|\|\|\|\|\|\|\|\|\|\|\|\|\|\|\|\|\|\|\|\|\|\|\|\|\|\|\|\|\|\|\|\|\|\|\|\|\|\|\|\|\|\|\|\|\|\|\|\|\|\|\|\|\|\|\|\|\|\|\|\|\|\|\|\|\|\|\|\|\|\|\|\|\|\|\|\|\|\|\|\|\|\|\|\|\|\|\|\|\|\|\|\|\|\|\|\|\|\|\|\|\|\|\|\|\|\|\|\|\|\|\|\|\|\|\|\|\|\|\|\|\|\|\|\|\|\|\|\|\|\|\|\|\|\|\|\|\|\|\|\|\|\|\|\|\|\|\|\|\|\|\|\|\|\|\|\|\|\|\|\|\|\|\|\|\|\|\|\|\|\|\|\|\|\|\|\|\|\|\|\|\|\|\|\|\|\|\|\|\|\|\|\|\|\|\|\|\|\|\|\|\|\|\|\|\|\|\|\|\|\|\|\|\|\|\|\|\|\|\|\|\|\|\|\|\|\|\|\|\|\|\|\|\|\|\|\|\|\|\|\|\|\|\|\|\|\|\|\|\|\|\|\|\|\|\|\|\|\|\|\|\|\|\|\|\|\|\|\|\|\|\|\|\|\|\|\|\|\|\|\|\|\|\|\|\|\|\|\|\|\|\|\|\|\|\|\|\|\|\|\|\|\|\|\|\|\|  \|\| \|\|\|\|\|\|\|\|\|\|\|\|\|\|\|\|\|\|\|\|\|\|\|\|\|\|\| \| \|\|\|\|\|\|\|\|\|\|\|\|\|\|\|\|\|\|\| 	ATGGCACGCTCTCCCTNCGATGCCACGCGCTTTACCTCGACAGGACCCTACGTCTCCAATAAACCCGCAGGCACAACATCCTTCACATCCAACCCAACCGCACCATCCCCTTCGCAAGCCACAAACCTCAAATTCGGCCCCTCCGCGCCGGCCAACGAAACGCCTCAACAAAAAATCGCCCGTCTCCGCGCCGCCGCCGCCCTCGCCAAAGCCCCCCAAGAAACATCATGGGACAAGACCGTCCGGCGAGGCCGCATCTGGGCCGACCGCGCCCACCGCTTCACAGCTCTCGGCCTTATCGGCTTCACAGTCATCAGCGCGGCGGTGGCGACGGNNGGTAATCACGGATATGATGATGCACAATCGNAGGACGGAGGAATGAATGGTT ```      ``` 68	ATGGCACGCTCTCCCTCCGATGCCACGCGCTTTACCTCGACAGGACCCTACGTCTCCAATAAACCCGCAGGCACAACATCCTTCACATCCAACCCAACCGCACCATCCCCTTCGCAAGCCACAAACCTCAAATTCGGCCCCTCCGCGCCGGCCAACGAAACGCCTCAACAAAAAATCGCCCGTCTCCGCGCCGCCGCCGCCCTCGCCAAAGCCCCCCAAGAAACATCATGGGACAAGACCGTCCGGCGAGGCCGCATCTGGGCCGACCGCGCCCACCGCTTCACAGCTCTCGGCCTTATCGGCTTCACAGTCATCAGCGCGGCGGTGGCGACGGCAGGAATCACGGATATGATGATGCACAATCGAAGACGGAGGAATGAATGGTTCGCGGAGAAGAGTGCGGAGGCGACGCGGAATATGTTGCTTGCGATTCAAGCGGTAGAGAGGGGCGAGGCGACGGAGGATCAGGTCTTGTTGATTAATCGGGCGAGGGTCGCGGAGCAGGTTGCCGAGGAGAAGAAGAATAAGCCTGGGATTTTTAAGAGGGCTAAAGGATGGTTATATGGGGGTTTGAGTGA-GGAGGAGAAGAGAGGTGGGAGATTAGGGGTTGTTGGTGCTGTGGAAGAGGCGGTAGAGGAGAGGAGGGATGGTGGGACGGGAGGCGTGTTACAGGCGGTCGAGGAGAAGAT-GGAGTCTTCGCGGAGGAGAGAGATGGAAACGTTGGCGGAGAGTATCAATCCGGCGGGAGGATCGTTGGATCAGCGGGCACAGCTTTTGGTCGATTCGGCGTCAAATGTCACGTCAAGTTGGTTTGGGCTTGGGAGACGG	895 	\|\|\|\|\|\|\|\|\|\|\|\|\|\|\|\|\|\|\|\|\|\|\|\|\|\|\|\|\|\|\|\|\|\|\|\|\|\|\|\|\|\|\|\|\|\|\|\|\|\|\|\|\|\|\|\|\|\|\|\|\|\|\|\|\|\|\|\|\|\|\|\|\|\|\|\|\|\|\|\|\|\|\|\|\|\|\|\|\|\|\|\|\|\|\|\|\|\|\|\|\|\|\|\|\|\|\|\|\|\|\|\|\|\|\|\|\|\|\|\|\|\|\|\|\|\|\|\|\|\|\|\|\|\|\|\|\|\|\|\|\|\|\|\|\|\|\|\|\|\|\|\|\|\|\|\|\|\|\|\|\|\|\|\|\|\|\|\|\|\|\|\|\|\|\|\|\|\|\|\|\|\|\|\|\|\|\|\|\|\|\|\|\|\|\|\|\|\|\|\|\|\|\|\|\|\|\|\|\|\|\|\|\|\|\|\|\|\|\|\|\|\|\|\|\|\|\|\|\|\|\|\|\|\|\|\|\|\|\|\|\|\|\|\|\|\|\|\|\|\|\|\|\|\|\|\|\|\|\|\|\|\|\|\|\|\|\|\|\|\|\|\|\|\|\|\|\|\|\|\|\|\|\|\|\|\|\|\|\|\|\|\|\|\|\|\|\|\|\|\|\|\|\|\|\|\|\|\|\|\|\|\|\|\|\|\|\|\|\|\|\|\|\|\|\|\|\|\|\|\|\|\|\|\|\|\|\|\|\|\|\|\|\|\|\|\|\|\|\|\|\|\|\|\|\|\|\|\|\|\|\|\|\|\|\|\|\|\|\|\|\|\|\|\|\|\|\|\|\|\|\|\|\|\|\|\|\|\|\|\|\|\|\|\|\|\|\|\|\|\|\|\|\|\|\|\|\|\|\|\|\|\|\|\|\|\|\|\|\|\|\|\|\|\|\|\|\|\|\|\|\|\|\|\|\|\|\|\|\|\|\|\|\|\|\|\|\|\|\|\|\|\|\|\|\|\|\|\|\|\|\|\|\|\|\|\|\|\|\|\|\|\|\|\|\|\|\|\|\|\|\|\|\|\|\|\|\|\|\|\|\|\|\|\|\|\|\|\|\|\|\|\|\|\|\|\|\|\|\|\|\|\|\|\|\|\|\|\|\|\|\|\|\|\|\|\|\|\|\|\|\|\|\|\|\|\|\|\|\|\|\|\|\|\|\|\|\|\|\|\|\|\|\|\|\|\|\|\|\|\|\|\|\|\|\|\|\|\|\|\|\|\|\|\|\|\|\|\| \|\|\|\|\|\|\|\|\|\|\|\|\|\|\|\|\|\|\|\|\|\|\|\|\|\|\|\|\|\|\|\|\|\|\|\|\|\|\|\|\|\|\|\|\|\|\|\|\|\|\|\|\|\|\|\|\|\|\|\|\|\|\|\|\|\|\|\|\|\|\|\|\|\|\|\|\|\|\|\|\|\|\|\|\|\|\|\|\|\|\|\|\|\|\|\|\|\|\|\|\|\|\|\|\|\|\|\|\|\|\| \|\|\|\|\|\|\|\|\|\|\|\|\|\|\|\|\|\|\|\|\|\|\|\|\|\|\|\|\|\|\|\|\|\|\|\|\|\|\|\|\|\|\|\|\|\|\|\|\|\|\|\|\|\|\|\|\|\|\|\|\|\|\|\|\|\|\|\|\|\|\|\|\|\|\|\|\|\|\|\|\|\|\|\|\|\|\|\|\|\|\|\|\|\|\|\|\|\|\|\|\|\|\|\|\|\|\|\|\|\|\|\|\|\| \|\|\|\|\|\|\|\|\|\|\|\|\|\|\|\|\| \|\|\|\|\|\| 	ATGGCACGCTCTCCCTCCGATGCCACGCGCTTTACCTCGACAGGACCCTACGTCTCCAATAAACCCGCAGGCACAACATCCTTCACATCCAACCCAACCGCACCATCCCCTTCGCAAGCCACAAACCTCAAATTCGGCCCCTCCGCGCCGGCCAACGAAACGCCTCAACAAAAAATCGCCCGTCTCCGCGCCGCCGCCGCCCTCGCCAAAGCCCCCCAAGAAACATCATGGGACAAGACCGTCCGGCGAGGCCGCATCTGGGCCGACCGCGCCCACCGCTTCACAGCTCTCGGCCTTATCGGCTTCACAGTCATCAGCGCGGCGGTGGCGACGGCAGGAATCACGGATATGATGATGCACAATCGAAGACGGAGGAATGAATGGTTCGCGGAGAAGAGTGCGGAGGCGACGCGGAATATGTTGCTTGCGATTCAAGCGGTAGAGAGGGGCGAGGCGACGGAGGATCAGGTCTTGTTGATTAATCGGGCGAGGGTCGCGGAGCAGGTTGCCGAGGAGAAGAAGAATAAGCCTGGGATTTTTAAGAGGGCTAAAGGATGGTTATATGGGGGTTTGAGTGAGGGAGGAGAAGAGAGGTGGGAGATTAGGGGTTGTTGGTGCTGTGGAAGAGGCGGTAGAGGAGAGGAGGGATGGTGGGACGGGAGGCGTGTTACAGGCGGTCGAGGAGAAGATGGGAGTCTTCGCGGAGGAGAGAGATGGAAACGTTGGCGGAGAGTATCAATCCGGCGGGAGGATCGTTGGATCAGCGGGCACAGCTTTTGGTCGATTCGGCGTCAAATGTCACGTC-AGTTGGTTTGGGCTTGG-AGACGG ``` |
| 1 | ggggacaagtttgtacaaaaaagcaggcttgatgGATCCATCCCAGCTGGGCGC, ggggaccactttgtacaagaaagctgggtcTCCATCGTCGTCCCGGACTT | Mycgr3\|109102  JGI  Ensembl | KOG0342 ATP-dependent RNA helicase pitchoune; | ATP binding; ATP-dependent DNA helicase activity;  ATP-dependent RNA helicase activity;  ATP-dependent helicase activity;  ATPase activity;  ATPase activity, coupled;  ATPase activity, coupled to transmembrane movement of ions;  ATPase activity, coupled to transmembrane movement of substances;  ATPase activity, uncoupled;  DNA-dependent ATPase activity;  RNA-dependent ATPase activity;  helicase activity;  nucleic acid binding;  protein-transmembrane transporting ATPase activity;  single-stranded DNA-dependent ATP-dependent DNA helicase activity; | 100 % | chr\_4:1329080 G->A; |  |
| 1 | ggggacaagtttgtacaaaaaagcaggcttgatgACCGCGGAACCGTCCTTCAC, ggggaccactttgtacaagaaagctgggtcTATTTGACCTTTGATGTCGG | Mycgr3\|109254  JGI  Ensembl | KOG1861 Leucine permease transcriptional regulator; |  | 100 % |  | ``` 158	ATGACCGCGGAACCGTCCTTCACGCAAGTTGCGGCGAGGCGAACGCTCGCCAAAGACCCACTGCCGCCGCAGGAAACACCCAAGAAGAAGAAGGAATATCCTCCTTCGCTTCTGGATTATGTGACGCGGGCTTTTTCCGAGTTCAACAAACGCAAAGACGAGCCCGCATTCTCCGGCATTCTCGATGCCGATATTCGCGCGGCACTCTCCAAAGTGATCATGGAAGTGGATGACAGCCCGATCATATATGAGCGTGATTGGAAGGCATTCCCGCTACCTCACCAGATCCTTCTTGAAGCCCGG	460 	\|\|\|\|\|\|\|\|\|\|\|\|\|\|\|\|\|\|\|\|\|\|\|\|\|\|\|\|\|\|\|\|\|\|\|\|\|\|\|\|\|\|\|\|\|\|\|\|\|\|\|\|\|\|\|\|\|\|\|\|\|\|\|\|\|\|\|\|\|\|\|\|\|\|\|\|\|\|\|\|\|\|\|\|\|\|\|\|\|\|\|\|\|\|\|\|\|\|\|\|\|\|\|\|\|\|\|\|\|\|\|\|\|\|\|\|\|\|\|\|\|\|\|\|\|\|\|\|\|\|\|\|\|\|\|\|\|\|\|\|\|\|\|\|\|\|\|\|\|\|\|\|\|\|\|\|\|\|\|\|\|\|\|\|\|\|\|\|\|\|\|\|\|\|\|\|\|\|\|\|\|\|\|\|\|\|\|\|\|\|\|\|\|\|\|\|\|\|\|\|\|\|\|\|\|\|\|\|\|\|\|\|\|\|\|\|\|\|\|\|\|\|\|\|\|\|\|\|\|\|\|\|\|\|\|\|\|\|\|\|\|\|\|\|\|\|\|\|\|\|\| \|\|\|\|\|\|\|\| \|\| \|  \|\|\| \|\|\|\|\|\|\|\|\|  \|\| \|\|\|\|\|\|\| \|\|\|\| \|\|\|\|\| 	ATGACCGCGGAACCGTCCTTCACGCAAGTTGCGGCGAGGCGAACGCTCGCCAAAGACCCACTGCCGCCGCAGGAAACACCCAAGAAGAAGAAGGAATATCCTCCTTCGCTTCTGGATTATGTGACGCGGGCTTTTTCCGAGTTCAACAAACGCAAAGACGAGCCCGCATTCTCCGGCATTCTCGATGCCGATATTCGCGCGGCACTCTCCAAAGTGATCATGGAAGTGGATGACAGCCCGATCATATATGANCGTGATTGNAANGNNTTCNCGCTACCTCNNCANATCCTTCNTGAANCCCGG ``` |
| 1 | ggggacaagtttgtacaaaaaagcaggcttgatgGCGACAGTTTGGACGACGCC, ggggaccactttgtacaagaaagctgggtcCGCCGAACTCACAGCCGATA | Mycgr3\|109438  JGI  Ensembl | KOG2588 Predicted DNA-binding protein; | nucleus; regulation of transcription;  transcription regulator activity; | 100 % |  | ``` 185	CAGCCGATCATTCACCAATATCGCCGTTCAACTCGTTGGAAATTTCCAAGGAAGAGTCCACATCCTCGATCTCTGACTGTGTCGCGCCGTCCACAGCGGAGTACTGGAACAGAGGGACTGGTCTCACCCATCCCCACCACCCCAGCTCGCCGGGACTGAGCCGACCATCAGCAGGGCCCCGACGTTGTAGCGACCCAACCCCTAACATCCTCTCCCCGCCTTCACAGGGTTACTTTCCTCACTTTCCTCACTACCCCAACAACGGCGGCGTTGCGGTCGATCCTCAGACCAATCCATCCGGGCCTCGATCCGCGCGCTCGCTACCGATCAGCTTCCCCGATCACTCCGCACGCACTTCGCCTGGACCTCGCCGCACTTCGGCCCATCACACAGTGCTGA	583 	\|\|\|\|\|\|\|\|\|\|\|\|\|\|\|\|\|\|\|\|\|\|\|\|\|\|\|\|\|\|\|\|\|\|\|\|\|\|\|\|\|\|\|\|\|\|\|\|\|\|\|\|\|\|\|\|\|\|\|\|\|\|\|\|\|\|\|\|\|\|\|\|\|\|\|\|\|\|\|\|\|\|\|\|\|\|\|\|\|\|\|\|\|\|\|\|\|\|\|\|\|\|\|\|\|\|\|\|\|\|\|\|\|\|\|\|\|\|\|\|\|\|\|\|\|\|\|\|\|\|\|\|\|\|\|\|\|\|\|\|\|\|\|\|\|\|\|\|\|\|\|\|\|\|\|\|\|\|\|\|\|\|\|\|\|\|\|\|\|\| \|\|\|\|\|\|\|\|\|\|\|\|\|\|\|\|\|\| \|\|\|\|\|\|\|\|\|\|\|\|\|\|\|\|\|\|\|\|\|\|\|\|\|\|\|\|\|\|\|\|\|\|\|\|\|\|\|\|\|\|\|\|\|\|\|\|\|\|\|\|\|\|\|\|\|\|\|\|\|\|\|\|\|\|\|\|\|\|\|\|\|\|\|\|\|\|\|\|\|\|\|\|\|\|\|\|\|\|\|\|\|\|\|\| \|\|\|\|\|\|\|\|\|\|\|\|\|\|\|\|\|\|\|\|\|\|\|\|\|\|\|\|\|\|\|\|\|\|\|\|\|\|\|\|\| \|\|\|\|\|\|\|\| \|\|\| \| \| \|\|\|\| \|\|\| \|\|\| \| \|\| \|\| \|\|\|\|\|\|\|\| \|\|\|\|\|\|\| \|\|\|  \| \|\|\|\|\|\|\|\| 	CAGCCGATCATTCACCAATATCGCCGTTCAACTCGTTGGAAATTTCCAAGGAAGAGTCCACATCCTCGATCTCTGACTGTGTCGCGCCGTCCACAGCGGAGTACTGGAACAGAGGGACTGGTCTCACCCATCCCCACCACCCCAGCTCGCCGGGACTGAGCCGACCATCANCAGGGCCCCGACGTTGTANCGACCCAACCCCTAACATCCTCTCCCCGCCTTCACAGGGTTACTTTCCTCACTTTCCTCACTACCCCAACAACGGCGGCGTTGCGGTCGATCCTCAAACCAATCCATCCGGGCCTCGATCCGCGCGCTCGCTACCGATNAGCTTCCC-GATNANT-CGCANGCANTTCNC-TGNAC-TCGCCGCA-TTCGGCC-ATCN-A-AGTGCTGA ```      ``` 1	ATGGCGACAGTTTGGACGACGCCGATCTACAATGCGGACCTCATTCTCAACCAGGGCGCCATTCCTTCAGGACCACTGCATATGATCGATTCATCATGGCAACACAACTCCCGGAGGGAAGACATTCAGCACGCTCCGCAAGACGCATACACGCGCACCTCCCACCATTATCACCTCCCATCGCCAG	187 	\|\|\|\|\|\|\|\|\|\|\|\|\|\|\|\|\|\|\|\|\|\|\|\|\|\|\|\|\|\|\|\|\|\|\|\|\|\|\|\|\|\|\|\|\|\|\|\|\|\|\|\|\|\|\|\|\|\|\|\|\|\|\|\|\|\|\|\|\|\|\|\|\|\|\|\|\|\|\|\|\|\|\|\|\|\|\|\|\|\|\|\|\|\|\|\|\|\|\|\|\|\|\|\|\|\|\|\|\|\|\|\|\|\|\|\|\|\|\|\|\|\|\|\|\|\|\|\|\|\|\|\|\|\|\|\|\|\|\|\|\|\|\|\|\|\|\|\|\|\|\|\|\|\|\|\|\|\|\|\|\|\|\|\|\|\|\|\|\|\|\|\|\|\|\|\|\|\|\|\|\|\|\|\|\|\|\| 	ATGGCGACAGTTTGGACGACGCCGATCTACAATGCGGACCTCATTCTCAACCAGGGCGCCATTCCTTCAGGACCACTGCATATGATCGATTCATCATGGCAACACAACTCCCGGAGGGAAGACATTCAGCACGCTCCGCAAGACGCATACACGCGCACCTCCCACCATTATCACCTCCCATCGCCAG ``` |
| 1 | ggggacaagtttgtacaaaaaagcaggcttgatgAATGCAGAAGCTGGACTGGG, ggggaccactttgtacaagaaagctgggtcATCGTCCTCAATCGGCGTGT | Mycgr3\|109489  JGI  Ensembl |  |  | 0.9308 % |  |  |
| 1 | ggggacaagtttgtacaaaaaagcaggcttgatgCCAACTTCGAAACCAGCACC, ggggaccactttgtacaagaaagctgggtcCTTCTTCACTTTCTTCGGCG | Mycgr3\|109968  JGI  Ensembl | KOG2821 RNA polymerase II transcription elongation factor Elongin/SIII, subunit elongin A; | integral to membrane; nucleus; regulation of transcription; | 100 % |  | ``` 218	ATGCCAACTTCGAAACCAGCACCAATTTCACAACCACTACCGACTCCGAATCCAACACCCACATTGTTCGAAATGGCTATCCGAACCCTCCAACGCAACGTCGATGCCATTACAGACATCGCCGACATGCCGTACGAAGTCGCTGAAGGTTTTCTTAAATCTATTCACCAACCGCAACAACTTCGCGCGCTGGAAGTCAAGTGTCCGCAGATTGCCGACGCCGACGCCGATCTCTGGAAGAAGTTCATCGCTCGCGACATCGCCGATTACAAGAAGAAGTGGATTGAGCCGAAGAATCCACGCAGCTGGTGGAAAGTCTACCGCAAGCTCACGAAGGACCAGGAGCTCTTGCAATCCGCACAAGAAGATGCCCTCCGAAAGAGCCTTGTCAGCGCTTCTGTCTCGAAGACTGAACACTCTGCTACCTTTGTCCCATCTGTCGTGCCATTCGCTGGTACGATCACCGACGCTCGTGTCTGCGCTGATGCTCGCGCTCGTGCTCGCACCGCCAACAATGGAGGTCG-CCCATGGCAGGCTTCAGCGAGCAAGAACGGTCTC	775 	\|\|\|\|\|\|\|\|\|\|\|\|\|\|\|\|\|\|\|\|\|\|\|\|\|\|\|\|\|\|\|\|\|\|\|\|\|\|\|\|\|\|\|\|\|\|\|\|\|\|\|\|\|\|\|\|\|\|\|\|\|\|\|\|\|\|\|\|\|\|\|\|\|\|\|\|\|\|\|\|\|\|\|\|\|\|\|\|\|\|\|\|\|\|\|\|\|\|\|\|\|\|\|\|\|\|\|\|\|\|\|\|\|\|\|\|\|\|\|\|\|\|\|\|\|\|\|\|\|\|\|\|\|\|\|\|\|\|\|\|\|\|\|\|\|\|\|\|\|\|\|\|\|\|\|\|\|\|\|\|\|\|\|\|\|\|\|\|\|\|\|\|\|\|\|\|\|\|\|\|\|\|\|\|\|\|\|\|\|\|\|\|\|\|\|\|\|\|\|\|\|\|\|\|\|\|\|\|\|\|\|\|\|\|\|\|\|\|\|\|\|\|\|\|\|\|\|\|\|\|\|\|\|\|\|\|\|\|\|\|\|\|\|\|\|\|\|\|\|\|\|\|\|\|\|\|\|\|\|\|\|\|\|\|\|\|\|\|\|\|\|\|\|\|\|\|\|\|\|\|\|\|\|\|\|\|\|\|\|\|\|\|\|\|\|\|\|\|\|\|\|\|\|\|\|\|\|\|\|\|\|\|\|\|\|\|\|\|\|\|\|\|\|\|\|\|\|\|\|\|\|\|\|\|\|\|\|\|\|\|\|\|\|\|\|\|\|\|\|\|\|\|\|\|\|\|\|\|\|\|\|\|\|\|\|\|\|\|\|\|\|\|\|\|\|\|\|\|\|\|\|\|\|\|\|\|\|\|\|\|\|\|\|\|\|\|\|\|\|\|\|\|\|\|\|\|\|\|\|\|\|\|\|\|\|\|\|\|\|\|\|\|\|\|\|\|\|\|\|\|\|\|\|\|\|\|\|\|\|\|\|\|\|\|\|\|\|\|\|\|\|\|\|\|\|\|\|\|\|\|\|\|\|\|\|\|\|\|\|\|\|\|\|\|\|\|\|\|\|\|\|\|\|\|\|\|\|\|\|\|\|\|\|\|\|\|\|\|\|\|\|\|\|\|\|\|\|\|\|\| \| \|\|\|\|\|\|\|\|\|\|\| \|\|\|\|\|\|\| \|\|\| \|\|\|  \|\|\|\|  \| \|\| \|\|\|\|\|\| 	ATGCCAACTTCGAAACCAGCACCAATTTCACAACCACTACCGACTCCGAATCCAACACCCACATTGTTCGAAATGGCTATCCGAACCCTCCAACGCAACGTCGATGCCATTACAGACATCGCCGACATGCCGTACGAAGTCGCTGAAGGTTTTCTTAAATCTATTCACCAACCGCAACAACTTCGCGCGCTGGAAGTCAAGTGTCCGCAGATTGCCGACGCCGACGCCGATCTCTGGAAGAAGTTCATCGCTCGCGACATCGCCGATTACAAGAAGAAGTGGATTGAGCCGAAGAATCCACGCAGCTGGTGGAAAGTCTACCGCAAGCTCACGAAGGACCAGGAGCTCTTGCAATCCGCACAAGAAGATGCCCTCCGAAAGAGCCTTGTCAGCGCTTCTGTCTCGAAGACTGAACACTCTGCTACCTTTGTCCCATCTGTCGTGCCATTCGCTGGTACGATCACCGACGCTCGTGTCTGCGCTGATGCTCGCGCTCGTGCTCGCACCGCCNANAATGGAGGTCGCCCCATGGNAGGNTTCNNCGAGNNANAANGGTCTC ``` |
| 1 | ggggacaagtttgtacaaaaaagcaggcttgatgAGCGTGCCCAAAATCTCCCT, ggggaccactttgtacaagaaagctgggtcCCTCCGACGATCCCTATCTC | Mycgr3\|110121  JGI  Ensembl | KOG1874 KEKE-like motif-containing transcription regulator (Rlr1)/suppressor of sin4; | intracellular; nucleic acid binding; | 100 % |  | ``` 1	ATGAGCGTGCCCAAAATCTCCCTCTCTCTGGGAGCCGGGCGAAAAGCTTCAGCGCCCACAAATGGCGTCAAGCGCCCGCGCGCAGCACTGCACGAAGACGACGATGATCACGAAGAGCTTGGTCAAACCGAGACGGTGTCGCACTTCGACAAGACTGCAGGCGGTGCAGTCGATGCCAGTCGCAAAAAGGAAGACACGG-GCCCCATGATTATTCCGAAGCAAACGAATAGAGACTGGAAGGAAGCGGCCAATC	253 	\|\|\|\|\|\|\|\|\|\|\|\| \|\|\|\|\|\|\|\|\|\|\|\|\|\|\|\|\|\|\|\|\|\|\|\|\|\|\|\|\|\|\|\| \|\|\|\|  \|\|\|\|\| \| \|\|\|\|\|  \|\|  \|\|\|\| \|\|\|  \|\|     \|\|\| \|\|\|\| \|\| \|\|\|\|\|\|\|\|\|\|\|\|\|\| \| \|\|\|\|  \|\|\|\|\|\|\|\| \| \|\|\|\|\|\| \|\|\|\|\|\|\|\|  \|\|\| \|\|\|\| \|\|\|\| \|\|  \|\| \|\|\|\|  \|\|\|       \|\| \|\|\|\|\|\|\|  \|\| \|\|\|\|\|\|\|\|\|\|\|\|\|\|\|\| \|\| \|\| \|\|\|  \|\|\|\|\|\|\| \| \|\|\|\|\|\|\|\|\|\|\|\|\| 	ATGAGCGTGCCCNAAATCTCCCTCTCTCTGGGAGCCGGGCGAAAANCTTCTTCGCCCNCNAATGGNNTCNNGCGCNCGCNNGCNNNGNTGCNCGAATACNACGATGATCACGAANANCTTGNNCAAACCGANANGGTGTCTCACTTCGANNAGANTGCANGCGGNGCNTTCTATGCNNGTCTNNTTNNGGNAGACACGANGCNCCATGATTATTCCGAAACATACNAATNNAGACTGGNANGAAGCGGCCAATC ```      ``` 1	ATGAGCGTGCCCAAAATCTCCCTCTCTCTGGGAGCCGGGCGAAAAGCTTCAGCGCCCACAAATGGCGTCAAGCGCCCGCGCGCAGCACTGCACGAAGACGACGATGATCACGAAGAGCTTGGTCAAACCGAGACGGTGTCGCACTTCGACAAGACTGCAGGCGGTGCAGTCGATGCCAGTCGCAAAAAGGAAGACACGGGCCCCATGATTATTCCGAAGCAAACGAATAGAGACTGGAAGGAAGCGGCCAATCGGCGCAAACGACAGAAGAGTGGGCTACCTGAAGGCGCGGGTCGTGGACAGGAAGGTCTGGATCAGAGGATGGCAGAAGTCGAGGCGGCGGTGGAGGCTGCGAAGCCGGGCTTTGGACTGAACACATACAACTCGAAGGCTACGGAGACGAATGAAGAGGCCGTGCTTGGGGACGCGGATGCTGCGTCGAAAGCTATGGACGCCGTAGAGAAGTCGCAACAGCTTCCGCCGGAACCTGACGCGGAGGGCGTTTCGCTTC	511 	\|\|\|\|\|\|\|\|\|\|\|\|\|\|\|\|\|\|\|\|\|\|\|\|\|\|\|\|\|\|\|\|\|\|\|\|\|\|\|\|\|\|\|\|\|\|\|\|\|\|\|\|\|\|\|\|\|\|\|\|\|\|\|\|\|\|\|\|\|\|\|\|\|\|\|\|\|\|\|\|\|\|\|\|\|\|\|\|\|\|\|\|\|\|\|\|\|\|\|\|\|\|\|\|\|\|\|\|\|\|\|\|\|\|\|\|\|\|\|\|\|\|\|\|\|\|\|\|\|\|\|\|\|\|\|\|\|\|\|\|\|\|\|\|\|\|\|\|\|\|\|\|\|\|\|\|\|\|\|\|\|\|\|\|\|\|\|\|\|\|\|\|\|\|\|\|\|\|\|\|\|\|\|\|\|\|\|\|\|\|\|\|\|\|\|\|\|\|\|\|\|\|\|\|\|\|\|\|\|\|\|\|\|\|\|\|\|\|\|\|\|\|\|\|\|\|\|\|\|\|\|\|\|\|\|\|\|\|\|\|\|\|\|\|\|\|\|\|\|\|\|\|\|\|\|\|\|\|\|\|\|\|\|\|\|\|\|\|\|\|\|\|\|\|\|\|\|\|\|\|\|\|\|\|\|\|\|\|\|\|\|\|\|\|\|\|\|\|\|\|\|\|\|\|\|\|\|\|\|\|\|\|\|\|\|\|\|\|\|\|\|\|\|\|\|\|\|\|\|\|\|\|\|\|\|\|\|\|\|\|\|\|\|\|\|\|\|\|\|\|\|\|\|\|\|\|\|\|\|\|\|\|\|\|\|\|\|\|\|\|\|\|\|\|\|\|\|\|\|\|\|\|\|\|\|\|\|\|\|\|\|\|\|\|\|\|\|\|\|\|\|\|\|\|\|\|\|\| \|\|\|\|\|\|\|\|\|\|\|\|\|\|\|\|\|\|\|\|\|\|\|\|\|\|\|\|\|\|\| \|\|\|\|\|\|\|\|\|\|\|\|\|\|\|\|\|\| \| \|\| \|\| \|\|\|\|\| \|\|\|\|\|\|\|\|\|\|\|\|\|\|\|\|\|\| \|\|\|\| \|\|\|\|\|\|\|\| \|\|\|\| 	ATGAGCGTGCCCAAAATCTCCCTCTCTCTGGGAGCCGGGCGAAAAGCTTCAGCGCCCACAAATGGCGTCAAGCGCCCGCGCGCAGCACTGCACGAAGACGACGATGATCACGAAGAGCTTGGTCAAACCGAGACGGTGTCGCACTTCGACAAGACTGCAGGCGGTGCAGTCGATGCCAGTCGCAAAAAGGAAGACACGGGCCCCATGATTATTCCGAAGCAAACGAATAGAGACTGGAAGGAAGCGGCCAATCGGCGCAAACGACAGAAGAGTGGGCTACCTGAAGGCGCGGGTCGTGGACAGGAAGGTCTGGATCAGAGGATGGCAGAAGTCGAGGCGGCGGTGGAGGCTGCGAAGCCGGGCTTTGGACTGAACACATACAACTCGAAGGCTACGGAGACGAATGAANAGGCCGTGCTTGGGGACGCGGATGCTGCGTCNAAAGCTATGGACGCCGTANANAANTCNCAACANCTTCCGCCGGAACCTGACNCGGANGGCGTTTCNCTTC ``` |
| 1 | ggggacaagtttgtacaaaaaagcaggcttgatgTCCGACTACGAAGACGACAT, ggggaccactttgtacaagaaagctgggtcTGAGATTGAGCAGAGTGTGT | Mycgr3\|110326  JGI  Ensembl | KOG0990 Replication factor C, subunit RFC5; | ATP binding; DNA polymerase activity;  DNA-directed DNA polymerase activity;  nucleoside-triphosphatase activity;  nucleotide binding; | 100 % | chr\_8:257013 T->\*; | ``` 1	ATGTCCGACTACGAAGACGACATGGACGTCGACATGGAGTCAACCTCCAAACCCTCCGACGACATCACCTTCTCCTCTTCCACCACCGCCAAAGGCAAACGCAGCGCCGCGAACCTCCCCATCGAAGCCGAAGACACCCTCCCGTGGGTCGAGAAGTACCGCCCCAGCTCCCTCGACGACGTCCACGGCCACCAAGACATCCTCGCCACGATCAACAAATTCGTCGACTCCAACCGCCTACCCCATCTCCTCCTCTACGGCCCACCAGGAACAGGAAAGACCTCAACTGTCCTCGCTCTCGCACGACGCATCTACGGCAACAAAAACATGCGACAAATGGTGCTAGAATTGAACGCTTCCGACGACAGAGGTATCGACGTGGTCCGGGAACAAATTAAAACTTTCAGCTCGACGAAACAGATCTTCGCCGGCAGTTTCGACAAGTCAGCCCAGCAAAGCACGATTGCGAATTACAAGCTCATCATCCTCGACGAGGCGGACGCCATGACCAGCACGGCGCAGATGGCGTTGCGGCGGATCATGGAGAAGTACACGGCCAATACGCGGTTCTGCATCATTGCGAATTACACACATAAGCTCAGTCCGGCGTTGTTGTCGCGGTGTACGAGATTTCGATTCTCGCCGTTGAAAGAGGTGGATATCAGACAATTGGTGGACAAGGTCGTGCGGGAGGAGGATGTGAAT	705 	\|\|\|\|\|\|\|\|\| \|\|\|\|\|\|\|\|\|\|\|\|\|\|\|\|\|\|\|\|\|\|\|\|\|\|\|\|\|\|\|\|\|\|\|\|\|\|\|\|\|\|\|\|\|\|\|\|\|\|\|\|\|\|\|\|\|\|\|\|\|\|\|\|\|\|\|\|\|\|\|\|\|\|\|\|\|\|\|\|\|\|\|\|\|\|\|\|\|\|\|\|\|\|\|\|\|\|\|\|\|\|\|\|\|\|\|\|\|\|\|\|\|\|\|\|\|\|\|\|\|\|\|\|\|\|\|\|\|\|\|\|\|\|\|\|\|\|\|\|\|\|\|\|\|\|\|\|\|\|\|\|\|\|\|\|\|\|\|\|\|\|\|\|\|\|\|\|\|\|\|\|\|\|\|\|\|\|\|\|\|\|\|\|\|\|\|\|\|\|\|\|\|\|\|\|\|\|\|\|\|\|\|\|\|\|\|\|\|\|\|\|\|\|\|\|\|\|\|\|\|\|\|\|\|\|\|\|\|\|\|\|\|\|\|\|\|\|\|\|\|\|\|\|\|\|\|\|\|\|\|\|\|\|\|\|\|\|\|\|\|\|\|\|\|\|\|\|\|\|\|\|\|\|\|\|\|\|\|\|\|\|\|\|\|\|\|\|\|\|\|\|\|\|\|\|\|\|\|\|\|\|\|\|\|\|\|\|\|\|\|\|\|\|\|\|\|\|\|\|\|\|\|\|\|\|\|\|\|\|\|\|\|\|\|\|\|\|\|\|\|\|\|\|\|\|\|\|\|\|\|\|\|\|\|\|\|\|\|\|\|\|\|\|\|\|\|\|\|\|\|\|\|\|\|\|\|\|\|\|\|\|\|\|\|\|\|\|\|\|\|\|\|\|\|\|\|\|\|\|\|\|\|\|\|\|\|\|\|\|\|\|\|\|\|\|\|\|\|\|\|\|\|\|\|\|\|\|\|\|\|\|\|\|\|\|\|\|\|\|\|\|\|\|\|\|\|\|\|\|\|\|\|\|\|\|\|\|\|\|\|\|\|\|\|\|\|\|\|\|\|\|\|\|\|\|\|\|\|\|\|\|\|\|\|\|\|\|\|\|\|\|\|\|\|\|\|\|\|\|\|\|\|\|\|\|\|\|\|\|\|\|\|\|\|\|\|\|\|\|\|\|\|\|\|\|\|\|\|\|\|\|\|\|\|\|\|\|\|\|\|\|\|\|\|\|\|\|\|\|\|\|\|\|\|\|\|\|\|\|\|\|\|\|\|\|\|\|\|\|\|\|\|\|\|\|\|\|\|\|\|\|\|\|\|\|\|\|\|\|\|\|\|\|\|\|\|\|\|\|\|\|\|\|\|\|\| \|\|\|\|\|\|\|\|\|\| \|\|\|\|\|\|\|\|\|\|\|\|\|\| \|\|\|\|\|\|\|\|\|\|\|\|\|\|\|\|\|\|\|\| \|\|\|\|\| \|\|\|\|\|\|\|\| \|\|\|\| \|\|\| \|\|\| \|\|\| \|\|\|\|\|\|\|\| 	ATGTCCGAC-ACGAAGACGACATGGACGTCGACATGGAGTCAACCTCCAAACCCTCCGACGACATCACCTTCTCCTCTTCCACCACCGCCAAAGGCAAACGCAGCGCCGCGAACCTCCCCATCGAAGCCGAAGACACCCTCCCGTGGGTCGAGAAGTACCGCCCCAGCTCCCTCGACGACGTCCACGGCCACCAAGACATCCTCGCCACGATCAACAAATTCGTCGACTCCAACCGCCTACCCCATCTCCTCCTCTACGGCCCACCAGGAACAGGAAAGACCTCAACTGTCCTCGCTCTCGCACGACGCATCTACGGCAACAAAAACATGCGACAAATGGTGCTAGAATTGAACGCTTCCGACGACAGAGGTATCGACGTGGTCCGGGAACAAATTAAAACTTTCAGCTCGACGAAACAGATCTTCGCCGGCAGTTTCGACAAGTCAGCCCAGCAAAGCACGATTGCGAATTACAAGCTCATCATCCTCGACGAGGCGGACGCCATGACCAGCACGGCGCAGATGGCGTTGCGGCGGATCATGGAGAAGTACACGGCCAATACGCGGTTCTGCATCATTGCGAATTACACACATAAGCTCAGTCCGGCGTTGTTGTCNCGGTGTACGANATTTCGATTCTCGCNGTTGAAAGAGGTGGATATCANACAATNGGTGGACA-GGTCNTGC-GGANGAGNATGTGAAT ``` |
| 1 | ggggacaagtttgtacaaaaaagcaggcttgatgCCTTCGGCCACGAAGCACAA, ggggaccactttgtacaagaaagctgggtcCCCCTTGCCCAGACTCGGAC | Mycgr3\|110410  JGI  Ensembl | KOG2496 Cdk activating kinase (CAK)/RNA polymerase II transcription initiation/nucleotide excision repair factor TFIIH/TFIIK, cyclin H subunit; |  | 100 % |  | ``` 1	ATGCCTTCGGCCACGAAGCACAACCTTACCGAGGACGACATATATCGCACATCAACGCAGTATCGCTTCTGGTCATTCTCTCCCGAGCAATTGTCCGCACGCCGCAAAGAAACCCACCGACTCGCCCTCCAACGAGCAGCACAATACACTCAAAACGATGCATCCACCGAACATCTCACCCTCGAAGAGGAACTTCGCCTCGTAACCCGATATGCCGAACTCCTCCGCACTACCTCCGACCACCTGAAATGGCCCGCCAACATCAAGTCAACCGCCATCCAATACCTCAGACGATTCTACTTGAGCAATTCGTGCATGACATATCCTCCGAAAGAAATATACAAGACAGTGCTCTTTCTCGCGAGCAAAACTGAGGCCTTCCATATCACACTGAGCAAATTCTGCTCCATGATCAGTGCGGATCCAGAGGCTGTTCTGGCGCCAGAGTACAAAGTGATGCAAGCGCTGCGATTCACACTGGACGTACGGCAGCCATTTCGGGGACTGAAGGGAGTGCTGATGGAGTTGATCAACCTGGTGGATGAGGAAAGA	552 	\|\|\|\|\|\|\|\|\|\|\|\|\|\|\|\|\|\|\|\|\|\|\|\|\|\|\|\|\|\|\|\|\|\|\|\|\|\|\|\|\|\|\|\|\|\|\|\|\|\|\|\|\|\|\|\|\|\|\|\|\|\|\|\|\|\|\|\|\|\|\|\|\|\|\|\|\|\|\|\|\|\|\|\|\|\|\|\|\|\|\|\|\|\|\|\|\|\|\|\|\|\|\|\|\|\|\|\|\|\|\|\|\|\|\|\|\|\|\|\|\|\|\|\|\|\|\|\|\|\|\|\|\|\|\|\|\|\|\|\|\|\|\|\|\|\|\|\|\|\|\|\|\|\|\|\|\|\|\|\|\|\|\|\|\|\|\|\|\|\|\|\|\|\|\|\|\|\|\|\|\|\|\|\|\|\|\|\|\|\|\|\|\|\|\|\|\|\|\|\|\|\|\|\|\|\|\|\|\|\|\|\|\|\|\|\|\|\|\|\|\|\|\|\|\|\|\|\|\|\|\|\|\|\|\|\|\|\|\|\|\|\|\|\|\|\|\|\|\|\|\|\|\|\|\|\|\|\|\|\|\|\|\|\|\|\|\|\|\|\|\|\|\|\|\|\|\|\|\|\|\|\|\|\|\|\|\|\|\|\|\|\|\|\|\|\|\|\|\|\|\|\|\|\|\|\|\|\|\|\|\|\|\|\|\|\|\|\|\|\|\|\|\|\|\|\|\|\|\|\|\|\|\|\|\|\|\|\|\|\|\|\|\|\|\|\|\|\|\|\|\|\|\|\|\|\|\|\|\|\|\|\|\|\|\|\|\|\|\|\|\|\|\|\|\|\|\|\|\|\|\|\|\|\|\|\|\|\|\|\|\|\|\|\|\|\|\|\|\|\|\|\|\|\|\|\|\|\|\|\|\|\|\|\|\|\|\|\|\|\|\|\|\|\|\|\|\|\|\|\|\|\|\|\|\|\|\|\|\|\|\|\|\|\|\|\|\|\|\|\|\|\|\|\|\|\|\|\|\|\|\|\|\|\|\|\|\|\|\|\|\|\|\|\|\|\|\|\|\|\|\|\|\|\|\|\|\|\|\|\|\|\|\|\|\|\|\|\|\|\|\|\|\|\|\|\|\|\|\|\|\|\|\|\|\|\|\|\|\|\|\|\|\|\|\|\|\|\|\|\|\| \|\|\|\|\|\| \|\|\|\|\|\|\| \|\|\|\|\| 	ATGCCTTCGGCCACGAAGCACAACCTTACCGAGGACGACATATATCGCACATCAACGCAGTATCGCTTCTGGTCATTCTCTCCCGAGCAATTGTCCGCACGCCGCAAAGAAACCCACCGACTCGCCCTCCAACGAGCAGCACAATACACTCAAAACGATGCATCCACCGAACATCTCACCCTCGAAGAGGAACTTCGCCTCGTAACCCGATATGCCGAACTCCTCCGCACTACCTCCGACCACCTGAAATGGCCCGCCAACATCAAGTCAACCGCCATCCAATACCTCAGACGATTCTACTTGAGCAATTCGTGCATGACATATCCTCCGAAAGAAATATACAAGACAGTGCTCTTTCTCGCGAGCAAAACTGAGGCCTTCCATATCACACTGAGCAAATTCTGCTCCATGATCAGTGCGGATCCAGAGGCTGTTCTGGCGCCAGAGTACAAAGTGATGCAAGCGCTGCGATTCACACTGGACGTACGGCAGCCATTTCGGGGACTGAAGGGAGTGCTGATGGAGTTGATC-ACCTGGNGGATGAG-AAAGA ``` |
| 1 | ggggacaagtttgtacaaaaaagcaggcttgatgTTCCGAACTCCGAATGCCTC, ggggaccactttgtacaagaaagctgggtcACTCAGCAGCGCGGGCTCGA | Mycgr3\|110483  JGI  Ensembl |  |  | 100 % | chr\_8:1258114 C->T; | ``` 1	ATGTTCCGAACTCCGAATGCCTCTGCATATATCATCAGAGGCACTCTGCCATCGTGGCTCTCTCTCCTCCATCGACTCGTTCCGGGAAACCGCAATGGTCTTGGAATCTACTATACGCACGGTACCCGTCAAACGTCGAAATGGAAAGCTTCAGAGCT	158 	\|\|\|\|\|\|\|\|\|\|\|\|\|\|\|\|\|\|\|\|\|\|\|\|\|\|\|\|\|\|\|\|\|\|\|\|\|\|\|\|\|\|\|\|\|\|\|\|\|\|\|\|\|\|\|\|\|\|\|\|\|\|\|\|\|\|\|\|\|\|\|\|\|\|\|\|\|\|\|\|\|\|\|\|\|\|\|\|\|\|\|\|\|\|\|\|\|\|\|\|\|\|\|\|\|\|\|\|\|\|\|\|\|\|\|\|\|\|\|\|\|\|\|\|\|\|\|\|\|\| \|\|\|\|\|\| \|\| \|\|\|\|\| \|\|\|\|\|\|\|\|\|\|\| 	ATGTTCCGAACTCCGAATGCCTCTGCATATATCATCAGAGGCACTCTGCCATCGTGGCTCTCTCTCCTCCATCGACTCGTTCCGGGAAACCGCAATGGTCTTGGAATCTACTATACGCACGGTACCCGTCNAACGTCNAANTGGAANGCTTCAGAGCT ``` |
| 1 | ggggacaagtttgtacaaaaaagcaggcttgatgGCACATGGCACGGTGGATGC, ggggaccactttgtacaagaaagctgggtcAGAAGTCGGTAAAGTCCACG | Mycgr3\|110870  JGI  Ensembl | KOG1601 GATA-4/5/6 transcription factors; | nucleus; regulation of transcription, DNA-dependent;  sequence-specific DNA binding;  transcription factor activity;  zinc ion binding; | 100 % |  | ``` 350	ATGGCACATGGCACGGTGGATGCCAACCGGATGGGCGGGATGAGACCACTTTCACCGAGAATACCAGCCCTGCTCCAGCGACAGCCATCACAGCAAGAATTGGACGTCGCCCGGCATCTGGTCGAGCACTCTCAGAGCACCCATCATTCAGTATACGACCCGCAATTATCCGTGGAGCCATCACAAAATGCATCACGGATGATGGACCAGAGTTATGCTTCCACAGCTCCCAGCTCGGAATACAACATGAGCGTGACTCCAGCTCAGCAGCCATCTCCGGGTCCTTCAACCTCTGGATCAGTCTCCAGCACGCGACGTTCTCCGACTGCAGGTGCAGCGCCCGCCGGTCAGATGTGCAG	708 	\|\|\|\|\|\|\|\|\|\|\|\|\|\|\|\|\|\|\|\|\|\|\|\|\|\|\|\|\|\|\|\|\|\|\|\|\|\|\|\|\|\|\|\|\|\|\|\|\|\|\|\|\|\|\|\|\|\|\|\|\|\|\|\|\|\|\|\|\|\|\|\|\|\|\|\|\|\|\|\|\|\|\|\|\|\|\|\|\|\|\|\|\|\|\|\|\|\|\|\|\|\|\|\|\|\|\|\|\|\|\|\|\|\|\|\|\|\|\|\|\|\|\|\|\|\|\|\|\|\|\|\|\|\|\|\|\|\|\|\|\|\|\|\|\|\|\|\|\|\|\|\|\|\|\|\|\|\|\|\|\|\|\|\|\|\|\|\|\|\|\|\|\|\|\|\|\|\|\|\|\|\|\|\|\|\|\|\|\|\|\|\|\|\|\|\|\|\|\|\|\|\|\|\|\|\|\|\|\|\|\|\|\|\|\|\|\|\|\|\|\|\|\|\|\|\|\|\|\|\|\|\|\|\|\|\|\|\|\|\|\|\|\|\|\|\|\|\|\|\|\|\|\|\|\|\|\|\|\|\|\|\|\|\|\|\|\|\|\|\|\|\|\|\|\|\|\|\|\|\|\|\|\|\|\|\|\|\|\|\|\|\|\|\|\|\|\|\|\|\|\|\|\|\|\|\|\|\|\|\|\|\|\|\|\|\|\|\|\|\|\|\|\|\|\|\|\|\|\|\|\|\|\|\|\|\|\|\|\|\|\|\|\|\|\|\|\|\|\|\|\|\|\|\|\|\|\|\|\| 	ATGGCACATGGCACGGTGGATGCCAACCGGATGGGCGGGATGAGACCACTTTCACCGAGAATACCAGCCCTGCTCCAGCGACAGCCATCACAGCAAGAATTGGACGTCGCCCGGCATCTGGTCGAGCACTCTCAGAGCACCCATCATTCAGTATACGACCCGCAATTATCCGTGGAGCCATCACAAAATGCATCACGGATGATGGACCAGAGTTATGCTTCCACAGCTCCCAGCTCGGAATACAACATGAGCGTGACTCCAGCTCAGCAGCCATCTCCGGGTCCTTCAACCTCTGGATCAGTCTCCAGCACGCGACGTTCTCCGACTGCAGGTGCAGCGCCCGCCGGTCAGATGTGCAG ```      ``` 706	CAGCAACTGCGGTACAACGAAAACGCCGCTGTGGAGGCGCTCTCCAGCCGGCGCAGTGATCTGCAATGCGTGTGGTCTATACTACAAAGCACGCAACCAGATGCGCCCGGTCGGCATGAAGCGAGGTGCGAGTTCACAGCAAGGCGAAGATGCTCGAGATGATCGACAGCCATCACCTCAACGTGGAGGATCGACGTATGTCACCACACAAAAAGCCATCACTGGAACGTGTCCCGGCGGTGGTCGTTGCAACGGCACTGGTGGTCAGGAAGGTTGTGGAGGTTGTC	992 	\|\|\|\|\|\|\|\|\|\|\|\|\|\|\|\|\|\|\|\|\|\|\|\|\|\|\|\|\|\|\|\|\|\|\|\|\|\|\|\|\|\|\|\|\|\|\|\|\|\|\|\|\|\|\|\|\|\|\|\|\|\|\|\|\|\|\|\|\|\|\|\|\|\|\|\|\|\|\|\|\|\|\|\|\|\|\|\|\|\|\|\|\|\|\|\|\|\|\|\|\|\|\|\|\|\|\|\|\|\|\|\|\|\|\|\|\|\|\|\|\|\|\|\|\|\|\|\|\|\|\|\|\|\|\|\|\|\|\|\|\|\|\|\|\|\|\|\|\|\|\|\|\|\|\|\|\|\|\|\|\|\|\|\|\|\|\|\|\|\|\|\|\|\|\|\|\|\|\|\|\|\|\|\|\|\|\| \|\|\|\|\|\|\|\|\|\|\|\|\|\|\|\|\|\|\|\|\|\|\|\|\|\|\|\|\|\|\|\|\|\|\|\|\|\|\|\|\|\|\|\|\|\|\|\|\|\|\|\|\|\|\|\|\|\|\|\|\|\|\|\|\|\|\|\|\|\|\|\|\|\|\|\|\|\|\|\|\|\|\| \|\|\|\|\|\|\|\|\|\|\|\|\|\|\| 	CAGCAACTGCGGTACAACGAAAACGCCGCTGTGGAGGCGCTCTCCAGCCGGCGCAGTGATCTGCAATGCGTGTGGTCTATACTACAAAGCACGCAACCAGATGCGCCCGGTCGGCATGAAGCGAGGTGCGAGTTCACAGCAAGGCGAAGATGCTCGAGATGATCGACAGCCATCACCTCAACGTGGANGATCGACGTATGTCACCACACAAAAAGCCATCACTGGAACGTGTCCCGGCGGTGGTCGTTGCAACGGCACTGGTGGTCAGGAA-GTTGTGGAGGTTGTC ``` |
| 1 | ggggacaagtttgtacaaaaaagcaggcttgatgCTCCAATACGCCGCTTCCAA, ggggaccactttgtacaagaaagctgggtcCGACGAGCCATTCGTCTCCG | Mycgr3\|110922  JGI  Ensembl | KOG1601 GATA-4/5/6 transcription factors; | nucleus; regulation of transcription, DNA-dependent;  sequence-specific DNA binding;  transcription factor activity;  zinc ion binding; | 100 % |  | ``` 158	CTTCGATGCTCCAATACGCCGCTTCCAACGCGCCTCAGACGTTCTCACGACGATCATCAGTCTTTGCAGACCCGCCCTCAAGATCCACAGACAAGGCATTGTCCAGCTACGATAGCCATACTGGCTACAACATACCGCGACCTCCATCAACGCTGTCGCAAGCCACAACGAGCAGTGAACAACATCGGAGCAAATCGTTCAGTAGTCTGACTGGCTCGCTGGACTCATCGGCAGCGTTGTCACAGTTGGCGATGTTGGCAGCGCAGGCTCCAGCGGCGGAGATGAATACAAACAATGGGAGCAGTGATAG	467 	\|\|\| \|\|\|\|\|\|\|\|\|\|\|\|\|\|\|\|\|\|\|\|\|\|\|\|\|\|\|\|\|\|\|\|\|\|\|\|\|\|\|\|\|\|\|\|\|\|\|\|\|\|\|\|\|\|\|\|\|\|\|\|\|\|\|\|\|\|\|\|\|\|\|\|\|\|\|\|\|\|\|\|\|\|\|\|\|\|\|\|\|\|\|\|\|\|\|\|\|\|\|\|\|\|\|\|\|\|\|\|\|\|\|\|\|\|\|\|\|\|\|\|\|\|\|\|\|\|\|\|\|\|\|\|\|\|\|\|\|\|\|\|\|\|\|\|\|\|\|\|\|\|\|\|\|\|\|\|\|\|\|\|\|\|\|\|\|\|\|\|\|\|\|\|\|\|\|\|\|\|\|\|\|\|\|\|\|\|\|\|\|\|\|\|\|\|\|\|\|\|\|\|\|\|\|\|\|\|\|\|\|\|\|\|\|\|\|\|\|\|\|\|\|\|\|\|\|\|\|\|\|\|\|\|\|\|\|\|\|\|\|\|\|\|\|\|\|\|\|\|\|\|\|\|\|\|\|\|\|\|\|\|\|\|\|\|\|\|\|\|\|\|\|\|\|\|\|\|\|\|\|\|\|\|\|\|\|\|\|\|\|\|\|\|\|\|\|\|\|\|\|\|\|\|\|\|\|\| 	CTT-GATGCTCCAATACGCCGCTTCCAACGCGCCTCAGACGTTCTCACGACGATCATCAGTCTTTGCAGACCCGCCCTCAAGATCCACAGACAAGGCATTGTCCAGCTACGATAGCCATACTGGCTACAACATACCGCGACCTCCATCAACGCTGTCGCAAGCCACAACGAGCAGTGAACAACATCGGAGCAAATCGTTCAGTAGTCTGACTGGCTCGCTGGACTCATCGGCAGCGTTGTCACAGTTGGCGATGTTGGCAGCGCAGGCTCCAGCGGCGGAGATGAATACAAACAATGGGAGCAGTGATAG ``` |
| 1 | ggggacaagtttgtacaaaaaagcaggcttgatgTCACACCGCGGCCAACTCCA, ggggaccactttgtacaagaaagctgggtcTGATGAAACCTGCACGAATT | Mycgr3\|111251  JGI  Ensembl |  |  | 100 % |  | ``` 298	GCGAGCGCAAAAAGTCCAACGACAAGGTCTACCCGAAAGGCTACGTCGAGATGCTTGAGCAGCAGCAGGCTCAGCTGGTTTCTGGTCTCCAAGAAATTTACAGGCGCATGCAAGCCGCCGGAGCCTGGACGGGATCTACGCTCAGCGAGAGCGACAACGGCCAGCCACTTACCCACGACATCCTGAAAGCCCTTGGTCTGCTGGAGTCGAAGAACGACGGATCTGGCGATGTGGAAATGTTCGAAGAGGACCCTTCGCAATTGCAGTCTCGTTTGATCTCTGAAGGCGCCGGCTTTGCTGC	598 	\|\|\|\|\|\|\|\|\|\|\|\|\|\|\|\|\|\|\|\|\|\|\|\|\|\|\|\|\|\|\|\|\|\|\|\|\|\|\|\|\|\|\|\|\|\|\|\|\|\|\|\|\|\|\|\|\|\|\|\|\|\|\|\|\|\|\|\|\|\|\|\|\|\|\|\|\|\|\|\|\|\|\|\|\|\|\|\|\|\|\|\|\|\|\|\|\|\|\|\|\|\|\|\|\|\|\|\|\|\|\|\|\|\|\|\|\|\|\|\|\|\|\|\|\|\|\|\|\|\|\|\|\|\|\|\|\|\|\|\|\|\|\|\| \|\|\|\|\|\|\|\|\|\|\|\|\|\|\|\|\|\|\|\|\|\|\|\|\|\|\|\|\|\|\|\|\|\|\|\|\|\|\|\|\|\|\|\|\|\|\|\|\|\|\|\|\|\|\|\|\|\|\|\|\|\|\| \|\| \|\|\|\|\|\|\|\|\|\|\|\|\|\|\| \|\|\|\|\|\|\|\|\|\|\|\|\|\| \|\| \| \|\|\|\|\|\|\|\| \|\|\|\|\|\|\|\| \|\|\|\| \|\|\|\|\|\|\|\|\|\|\|\|\| \|\| \|\| \|\|\|\|\|\|\|\|\|\| 	GCGAGCGCAAAAAGTCCAACGACAAGGTCTACCCGAAAGGCTACGTCGAGATGCTTGAGCAGCAGCAGGCTCAGCTGGTTTCTGGTCTCCAAGAAATTTACAGGCGCATGCAAGCCGCCGGAGCCTGGACGGGATCTACGCTCANCGAGAGCGACAACGGCCAGCCACTTACCCACGACATCCTGAAAGCCCTTGGTCTGCTGGAGTCNAANAACGACGGATCTGGCNATGTGGAAATGTTCNAANANGACCCTTCNCAATTGCANTCTCNTTTGATCTCTGAANGCNCCNGCTTTGCTGC ```      ``` 48	ATGTCACACCGCGGCCAACTCCATTGCATGAACACGGTCTCGTACCACTCACACACAATGGCACCCAGCGCACATTCACCCGCGACAAAGCTGGGTCACATCTCTCGAACTGCTTCGCAAGATTCAACGACCGATGGCATTCGGAAGCGTGTCTGCAAGGCTTGTGATCGATGCCGTCTCAAGAAGAGCAAGTGCGATGGTTCAAGCCCTTGCTCGAGATGTCGGACCGACAACGCCATCTGTGTTTTTGG	298 	\|\|\|\|\|\|\|\|\|\|\|\|\|\|\|\|\|\|\|\|\|\|\|\|\|\|\|\|\|\|\|\|\|\|\|\|\|\|\|\|\|\|\|\|\|\|\|\|\|\|\|\|\|\|\|\|\|\|\|\|\|\|\|\|\|\|\|\|\|\|\|\|\|\|\|\|\|\|\|\|\|\|\|\|\|\|\|\|\|\|\|\|\|\|\|\|\|\|\|\|\|\|\|\|\|\|\|\|\|\|\|\|\|\|\|\|\|\|\|\|\|\|\|\|\|\|\|\|\|\|\|\|\|\|\|\|\|\|\|\|\|\|\|\|\|\|\|\|\|\|\|\|\|\|\|\|\|\|\|\|\|\|\|\|\|\|\|\|\|\|\|\|\|\|\|\|\|\|\|\|\|\|\|\|\|\|\|\|\|\|\|\|\|\|\|\|\|\|\|\|\|\|\|\|\|\|\|\|\|\|\|\|\|\|\|\|\|\|\|\|\|\|\|\|\|\|\|\|\|\|\|\|\|\|\|\|\|\|\|\|\|\|\|\|\|\|\|\|\|\|\| 	ATGTCACACCGCGGCCAACTCCATTGCATGAACACGGTCTCGTACCACTCACACACAATGGCACCCAGCGCACATTCACCCGCGACAAAGCTGGGTCACATCTCTCGAACTGCTTCGCAAGATTCAACGACCGATGGCATTCGGAAGCGTGTCTGCAAGGCTTGTGATCGATGCCGTCTCAAGAAGAGCAAGTGCGATGGTTCAAGCCCTTGCTCGAGATGTCGGACCGACAACGCCATCTGTGTTTTTGG ``` |
| 2 | ggggacaagtttgtacaaaaaagcaggcttgatgTCTGAGAACAACGGATCCCC, ggggaccactttgtacaagaaagctgggtcGCAGGATCCGCCGATCTGCT | Mycgr3\|31905  JGI  Ensembl | KOG1769 Ubiquitin-like proteins; | protein modification process; | 0 % |  |  |
| 2 | ggggacaagtttgtacaaaaaagcaggcttgatgGCATCTACTGATCCGCAAAA, ggggaccactttgtacaagaaagctgggtcGTCAATCGACATAGCCGCCT | Mycgr3\|33900  JGI  Ensembl |  |  | 100 % |  | ``` 1	ATGGCATCTACTGATCCGCAAAACCGCAAGCCGGAGCCGTCCCGACTTCTCCTCCTCTCCCAGGTTCATCTCCACGCCCCAGGCAGCAAAATTCGCTTCCTAGGCTGCGTGCATTCCTATCGGATCTCGACTGGCACGCTGGTCATTCAAGAGAAGTATGCTGCGGCTCCATCTGCTGTTCCGACCGTACTGGCGAGTATCGACAATGTCCTGGAAGACGTCGATCACGAAATGCTGCAGACAGGAGCATGGGTCAATGTCATAGGTTATATTAGGGCTGGAAAAGGGAAGGAGGATGTCCCGAGAGTGGAAGTTTTGATGATCTGGCCGGCTGGTGCGGTCAAGGTGGATGACTATCACTCTGCGGCGAGGAGCTATCAGGCGGCTATGTCGATTGAC	399 	\|\|\|\|\|\|\|\|\|\|\|\|\|\|\|\|\|\|\|\|\|\|\|\|\|\|\|\|\|\|\|\|\|\|\|\|\|\|\|\|\|\|\|\|\|\|\|\|\|\|\|\|\|\|\|\|\|\|\|\|\|\|\|\|\|\|\|\|\|\|\|\|\|\|\|\|\|\|\|\|\|\|\|\|\|\|\|\|\|\|\|\|\|\|\|\|\|\|\|\|\|\|\|\|\|\|\|\|\|\|\|\|\|\|\|\|\|\|\|\|\|\|\|\|\|\|\|\|\|\|\|\|\|\|\|\|\|\|\|\|\|\|\|\|\|\|\|\|\|\|\|\|\|\|\|\|\|\|\|\|\|\|\|\|\|\|\|\|\|\|\|\|\|\|\|\|\|\|\|\|\|\|\|\|\|\|\|\|\|\|\|\|\|\|\|\|\|\|\|\|\|\|\|\|\|\|\|\|\|\|\|\|\|\|\|\|\|\|\|\|\|\|\|\|\|\|\|\|\|\|\|\|\|\|\|\|\|\|\|\|\|\|\|\|\|\|\|\|\|\|\|\|\|\|\|\|\|\|\|\|\|\|\|\|\|\|\|\|\|\|\|\|\|\|\|\|\|\|\|\|\|\|\|\|\|\|\|\|\|\|\|\|\|\|\|\|\|\|\|\|\|\|\|\|\|\|\|\|\|\|\|\|\|\|\|\|\|\|\|\|\|\|\|\|\|\|\|\|\|\|\|\|\|\|\|\|\|\|\|\|\|\|\|\| \|\|\|\|\|\|\|\|\|\|\|\|\|\|\|\|\|\|\|\|\|\|\|\|\| \|\|\|\|\|\|\|\|\|\|\|\|\|\|\|\|\|\|\|\|\|\|\|\|\|\|\|\| 	ATGGCATCTACTGATCCGCAAAACCGCAAGCCGGAGCCGTCCCGACTTCTCCTCCTCTCCCAGGTTCATCTCCACGCCCCAGGCAGCAAAATTCGCTTCCTAGGCTGCGTGCATTCCTATCGGATCTCGACTGGCACGCTGGTCATTCAAGAGAAGTATGCTGCGGCTCCATCTGCTGTTCCGACCGTACTGGCGAGTATCGACAATGTCCTGGAAGACGTCGATCACGAAATGCTGCAGACAGGAGCATGGGTCAATGTCATAGGTTATATTAGGGCTGGAAAAGGGAAGGAGGATGTCCCGAGAGTGGAAGTTTTGATGATCTGGCCGGCTGGTGCGGTCAANGTGGATGACTATCACTCTGCGGCGANGAGCTATCAGGCGGCTATGTCGATTGAC ``` |
| 2 | ggggacaagtttgtacaaaaaagcaggcttgatgGCCGCATTGGCAAAGCAAAG, ggggaccactttgtacaagaaagctgggtcCAACCCCACTGCCTGTTTGA | Mycgr3\|35055  JGI  Ensembl |  |  | 100 % |  | ``` 203	AGTTCTGCGACACCGCTGACAACGTCTGTCCCGGCGGACAAAAGAGCGGTGCTGGCTGGCATCTGAGCTATGGTAGTGACGTGCAAGCTGCGGCGGCTTTCATCAAACAGGCAGTGGGGTTG	324 	\|\|\|\|\|\|\|\|\|\|\|\|\|\|\|\|\|\|\|\|\|\|\|\|\|\|\|\|\|\|\|\|\|\|\|\|\|\|\|\|\|\|\|\|\|\|\|\|\|\|\|\|\|\|\|\|\|\|\|\|\|\|\|\|\|\|\|\|\|\|\|\|\|\|\|\|\|\|\|\|\|\|\|\|\|\|\|\|\|\|\|\|\|\|\|\|\|\|\|\|\|\|\|\|\|\|\|\|\|\|\|\|\|\|\|\|\|\|\|\|\|\| 	AGTTCTGCGACACCGCTGACAACGTCTGTCCCGGCGGACAAAAGAGCGGTGCTGGCTGGCATCTGAGCTATGGTAGTGACGTGCAAGCTGCGGCGGCTTTCATCAAACAGGCAGTGGGGTTG ```      ``` 1	ATGGCCGCATTGGCAAAGCAAAGGTTGGGCAACTGTCCCGACACCAAGCTCGTCCTCAGCGGCTACTCACAAGGAGGATTCGTCTGTCACAACGCTCTCAGCGCTCAGGGTCTGGACGGTAGCAAGGTCGCTGCGGTTGTCTGCTTCGGCGACCCATTAAACGGTCAACCATTCCAGGGCGTTGACCCGTCCCGAGTCGTAA	202 	\|\|\|\|\|\|\|\|\|\|\|\| \| \|\|\|\|\|\|\|\|\|\|\|\|\|\|\|\|\|\|\|\|\|\|\|\|\|\|\|\|\|\|\|\|\|\|\|\|\|\|\|\|\|\|\|\|\|\|\|\|\|\|\|\|\|\|\|\|\|\|\|\|\|\|\|\|\|\|\|\|\|\|\|\|\|\|\|\|\|\|\|\|\|\|\|\|\|\|\|\|\|\|\|\|\|\|\|\|\|\|\|\|\|\|\|\|\|\|\|\|\|\|\|\|\|\|\|\|\|\|\|\|\|\|\|\|\|\|\|\|\|\|\|\|\|\|\|\|\|\|\|\|\|\|\|\|\|\|\|\|\|\|\|\|\|\|\|\|\|\|\|\|\|\|\|\|\|\|\|\|\|\|\|\|\|\|\|\|\|\|\|\|\|\|\|\|\|\|\| 	ATGGCCGCATTGCCTAAGCAAAGGTTGGGCAACTGTCCCGACACCAAGCTCGTCCTCAGCGGCTACTCACAAGGAGGATTCGTCTGTCACAACGCTCTCAGCGCTCAGGGTCTGGACGGTAGCAAGGTCGCTGCGGTTGTCTGCTTCGGCGACCCATTAAACGGTCAACCATTCCAGGGCGTTGACCCGTCCCGAGTCGTAA ``` |
| 2 | ggggacaagtttgtacaaaaaagcaggcttgatgGCAAAAGGAAGCTGCATGTG, ggggaccactttgtacaagaaagctgggtcACCTACCTCAACCTTGATAT | Mycgr3\|39016  JGI  Ensembl |  |  | 100 % |  | ``` 1	ATGGCAAAAGGAAGCTGCATGTGCGGCCAAATCGGCTACGAATACACCGGCGAACCCGCCACAACAGCGCTGTGCCATTG	80 	\|\|\|\|\|\|\|\|\|\|\|\|\|\|\|\|\|\|\|\|\|\|\|\|\|\|\|\|\|\|\|\|\|\|\|\|\|\|\|\|\|\|\|\|\|\|\|\|\|\|\|\|\|\|\|\|\|\|\|\|\|\|\|\|\|\|\|\|\|\|\|\|\|\|\|\|\|\|\|\| 	ATGGCAAAAGGAAGCTGCATGTGCGGCCAAATCGGCTACGAATACACCGGCGAACCCGCCACAACAGCGCTGTGCCATTG ```      ``` 80	GCACCGACTGTCAAAAGTGGACCGGCGCAGCCTACACTTCCAACGTCGTCGTCCCACGCACTTCCTTCAAAGTCACCAAGGGCAGCCCCAAGGACTATGATGCTGTGGGCGCATCCGGCAAGATCAACAAGCACTGGTTCTGCGGAACTTGTGGCAGTAGCTTGTATACCGAGCTGGAGATCATGCCGGATGTGACGTGCGTCAAGTCGGGTGGTTTGGACGGCGGTGCTGCGAACCATGATATCAAGGTTGAGGTAGGT	339 	\|\|\|\|\|\|\|\|\|\|\|\|\|\|\|\|\|\|\|\|\|\|\|\|\|\|\|\|\|\|\|\|\|\|\|\|\|\|\|\|\|\|\|\|\|\|\|\|\|\|\|\|\|\|\|\|\|\|\|\|\|\|\|\|\|\|\|\|\|\|\|\|\|\|\|\|\|\|\|\|\|\|\|\|\|\|\|\|\|\|\|\|\|\|\|\|\|\|\|\|\|\|\|\|\|\|\|\|\|\|\|\|\|\|\|\|\|\|\|\|\|\|\|\|\|\|\|\|\|\|\|\|\|\|\|\|\|\|\|\|\|\|\|\|\|\|\|\|\|\|\|\|\|\|\|\|\|\|\|\|\|\|\|\|\|\|\|\|\|\|\|\|\|\|\|\|\|\|\|\|\|\|\|\|\|\|\|\|\|\|\|\|\|\|\|\|\|\|\|\|\|\|\|\|\|\|\|\|\|\|\|\|\|\|\|\|\|\|\|\|\|\|\|\|\|\|\|\|\|\|\|\|\|\|\|\|\|\|\|\|\|\|\|\|\|\|\|\|\|\|\|\|\|\|\|\|\|\|\|\| 	GCACCGACTGTCAAAAGTGGACCGGCGCAGCCTACACTTCCAACGTCGTCGTCCCACGCACTTCCTTCAAAGTCACCAAGGGCAGCCCCAAGGACTATGATGCTGTGGGCGCATCCGGCAAGATCAACAAGCACTGGTTCTGCGGAACTTGTGGCAGTAGCTTGTATACCGAGCTGGAGATCATGCCGGATGTGACGTGCGTCAAGTCGGGTGGTTTGGACGGCGGTGCTGCGAACCATGATATCAAGGTTGAGGTAGGT ``` |
| 2 | ggggacaagtttgtacaaaaaagcaggcttgatgTCATACAAGGTCTACCGCGT, ggggaccactttgtacaagaaagctgggtcAGTTGGTGACAATTGGGGAA | Mycgr3\|39787  JGI  Ensembl |  |  | 100 % |  | ``` 1	ATGTCATACAAGGTCTACCGCGTCGCCTATGCGGGTATGCCGCGTGACCACCATGCGATTAACGTGGAGCAGAACGATGATGAGTCCGGCTGGCTCTTCCAGGTCACAGGCGACATCCAGAACGGGATGAGTCACGACGACAAAGCGACAAAGCGGCAAAGAAGCCGGACGAGTCGGCTAGCTACCAGGGCAAAGACTTCCTTGGGAAAGTCACGGTTGCCAACTTTGACCAGATCAAAGGCATATGCTGCTCCATTGCGCCCCCCAAGAAGCAGTTCGACTCGAATCCCATTCTTCCGATCCTGCCTGGCTACCGCAAAGAGGaaaaaaaaaGACACAAAACATACAACACTACGGATTCCCCAATTGTCACCAACT	378 	\|\|\|\|\|\|\|\|\|\|\|\|\|\|\|\|\|\|\|\|\|\|\|\|\|\|\|\|\|\|\|\|\|\|\|\|\|\|\|\|\|\|\|\|\|\|\|\|\|\|\|\|\|\|\|\|\|\|\|\|\|\|\|\|\|\|\|\|\|\|\|\|\|\|\|\|\|\|\|\|\|\|\|\|\|\|\|\|\|\|\|\|\|\|\|\|\|\|\|\|\|\|\|\|\|\|\|\|\|\|\|\|\|\|\|\|\|\|\|\|\|\|\|\|\|\|\|\|\|\|\|\|\|\|\|\|\|\|\|\|\|\|\|\|\|\|\|\|\|\|\|\|\|\|\|\|\|\|\|\|\|\|\|\|\|\|\|\|\|\|\|\|\|\|\|\|\|\|\|\|\|\|\|\|\|\|\|\|\|\|\|\|\|\|\|\|\|\|\|\|\|\|\|\|\|\|\|\|\|\|\|\|\|\|\|\|\|\|\|\|\|\|\|\|\|\|\|\|\|\|\|\|\|\|\|\|\|\|\|\|\|\|\|\|\|\|\|\|\|\|\|\|\|\|\|\|\|\|\|\|\|\|\|\|\|\|\|\|\|\|\|\|\|\|\|\|\|\|\|\|\|\|\|\|\|\|\|\|\|\|\|\|\|\|\|\|\|\|\|\|\|\|\|\|\|\|\|\|\|\|\|\|\|\|\|\|\|\|\|\|\|\|\|\|\|\|\|\|\|\|\|\|\|\|\|\|\|\|\|\|\|\|\|\|\|\|\|\|\|\|\|\|\|\|\|\|\|\|\|\|\|\|\|\|\|\|\|\|\|\|\|\|\|\|\|\|\|\| 	ATGTCATACAAGGTCTACCGCGTCGCCTATGCGGGTATGCCGCGTGACCACCATGCGATTAACGTGGAGCAGAACGATGATGAGTCCGGCTGGCTCTTCCAGGTCACAGGCGACATCCAGAACGGGATGAGTCACGACGACAAAGCGACAAAGCGGCAAAGAAGCCGGACGAGTCGGCTAGCTACCAGGGCAAAGACTTCCTTGGGAAAGTCACGGTTGCCAACTTTGACCAGATCAAAGGCATATGCTGCTCCATTGCGCCCCCCAAGAAGCAGTTCGACTCGAATCCCATTCTTCCGATCCTGCCTGGCTACCGCAAAGAGGAAAAAAAAAGACACAAAACATACAACACTACGGATTCCCCAATTGTCACCAACT ``` |
| 2 | ggggacaagtttgtacaaaaaagcaggcttgatgCTCCACATAGCTGCTCGGCA, ggggaccactttgtacaagaaagctgggtcTTTGCGTCCAGCGCAAAGCG | Mycgr3\|43592  JGI  Ensembl | KOG4177 Ankyrin; |  | 0 % |  |  |
| 2 | ggggacaagtttgtacaaaaaagcaggcttgatgCTTTCAGCCGCGCAAAGGGC, ggggaccactttgtacaagaaagctgggtcAACACGGTCTAGCGTCTCGA | Mycgr3\|43924  JGI  Ensembl |  |  | 100 % | chr\_6:2175375 G->A; chr\_6:2175130 A->G; chr\_6:2175074 T->C; chr\_6:2175116 G->A; chr\_6:2175380 G->A; chr\_6:2175110 G->A; chr\_6:2175084 G->A; chr\_6:2175198 G->A; chr\_6:2175118 G->A; chr\_6:2175143 G->A; chr\_6:2175075 A->G; chr\_6:2175179 G->A; | ``` 1	ATGCTTTCAGCCGCGCAAAGGGCGATTTATAATAAGGTGCTTAAGCACGCCAGGGCGCAAATTGTTACTAGACAAGTTGTCGCGCCCTTGCTGCTTAACATTAATAGCAAAGCTGGTACTAGCAAATCGTTTATTATCGAGGTAATTTCGGCGCATCTCCAGGCTCTTATCTCGAATTACGAGGATGTTATCTTCTGCGCTGCGCCTACCGGCGCCGCCTCGTATAGCATTGCCGGCTTAACGCTACATACCTTACTACGGCTACCTACGAAAGGAGAATTCCACCTACTAGGAGTAAGACAGCTCGCACACCTACAGAGAAAGCTTCGTAGGGTGCTCTACATTATTATTAATAAGAAGTCGATAGTAGGCCTCGAGACGCTAGACCGTGTT	393 	\|\|\|\|\|\|\|\|\|\|\|\| \|\|\|\|\|\|\|\|\|\|\|\|\|\|\|\|\|\|\|\|\|\|\|\|\|\|\|\|\|\|\|\| \|\|\|\| \|\|\|\|\|\|\|\|\|\|\|\|\|\|\|\|\|\|\|\|\|\|\|\|\|\|\|\|\|\|\|\|\|\|\|\|\|\|\|\|\|\|\|\|\|\|\|\|\|\|\|\|\|\|\|\|\|\|\|\|\|\|\|\|\|\|\|\|\|\|\|\|\|\|\|\|\|\|\|\|\|\|\|\|\|\|\|\|\|\|\|\|\|\|\|\|\|\|\|\|\|\|\|\|\|\|\|\|\|\|\|\|\|\|\|\|\|\|\|\|\|\|\|\|\|\|\|\|\|\|\|\|\|\|\|\|\|\|\|\|\|\|\|\|\|\|\|\|\|\|\|\|\|\|\|\|\|\|\|\|\|\|\|\|\|\|\|\|\|\|\|\|\|\|\|\| \|\|\|\|\|\|\|\|\|\|\|\|\|\|\|\|\|\| \|\|\|\|\|\|\|\|\|\|\|\|\|\|\|\|\|\|\|\|\|\|\|\|\|\|\|\|\|\|\|\|\|\|\| \|\|\|\|\|\|\|\|\|\|\|\| \|\|\|\|\|\|\|\|\|\|\| \| \|\|\|\|\| \|\|\|\|\|\|\|\|\|\|\|\|\|\|\|\|\|\|\|\|\|\|\|\|\| \|\|\|\|\|\|\|\|  \|\|\|\|\|\|\|\|\|\|\|\|\|\|\|\|\|\|\|\|\|\|\|\|\|\|\|\|\|\|\|\|\|\|\|\|\|\|\|\|\| 	ATGCTTTCAGCCNCGCAAAGGGCGATTTATAATAAGGTGCTTAAGTACGCTAGGGCGCAAATTGTTACTAGACAAGTTGTCGCGCCCTTGCTGCTTAACATTAATAGCAAAGCTGGTACTAGCAAATCGTTTATTATCGAGGTAATTTCGGCGCATCTCCAGGCTCTTATCTCGAATTACGAGGATGTTATCTTCTGCGCTGCGCCTACCGGCGCCGCCTCGTATAGTATTGCCGGCTTAACGCTATATACCTTACTACGGCTACCTACGAAAGGAGAATTCTACCTACTAGGAGCAAGACAGCTCGTATACCTATAGAGAAAGCTTCGTAGGGTGCTCTATATTATTATCGATAAGAAGTCGATAGTAGGCCTCGAGACGCTAGACCGTGTT ``` |
| 2 | ggggacaagtttgtacaaaaaagcaggcttgatgCCCCCCGGAATACCAGGGGG, ggggaccactttgtacaagaaagctgggtcGAGGAAGTAAAAGTCGTAAC | Mycgr3\|44591  JGI  Ensembl |  |  | 0 % |  | ``` 1	ATGCCCCCCGGAATACCAGGGGGCGCAATGTGCGTTCAAAGATTCGATGATTCACTGAATTCTGCAATTCACATTACTTATCGCATTTCGCTGCGTTCTTCATCGATGCCAGAACCAAGAGATCCGTTGTTGAAAGTTTTGTTTCGATTTACTCGTAAACTCCGACGCAGAGATGCAGTGTTTTTTGGTGGTCCTCCGGGG	201 	\|\|\|\|\|\|\|\|\|\|\|\| \| \|\|\|\|\|\|\|\|\|\|\|\|\|\|\|\|\|\|\|\|\|\|\|\|\|\|\|\|\|\|\|\|\|\|\|\|\|\|\|\|\|\|\|\|\|\|\|\|\|\|\|\|\|\|\|\|\|\|\|\|\|\|\|\|\|\|\|\|\|\|\|\|\|\|\|\|\|\|\|\|\|\|\|\|\|\|\|\|\|\|\|\|\|\|\|\|\|\|\|\|\|\|\|\|\|\|\|\|\|\|\|\|\|\|\|\|\|\|\|\|\|\|\|\|\|\|\|\|\|\|\|\|\|\|\|\|\|\|\|\|\|\|\|\|\|\|\|\|\|\|\|\|\|\|\|\|\|\|\|\|\|\|\|\|\|\|\|\|\|\|\|\|\|\|\|\|\|\|\|\|\|\|\|\|\|\| 	ATGCCCCCCGGACTNCCAGGGGGCGCAATGTGCGTTCAAAGATTCGATGATTCACTGAATTCTGCAATTCACATTACTTATCGCATTTCGCTGCGTTCTTCATCGATGCCAGAACCAAGAGATCCGTTGTTGAAAGTTTTGTTTCGATTTACTCGTAAACTCCGACGCAGAGATGCAGTGTTTTTTGGTGGTCCTCCGGGG ```      ``` 1	ATGCCCCCCGGAATACCAGGGGGCGCAATGTGCGTTCAAAGATTCGATGATTCACTGAATTCTGCAATTCACATTACTTATCGCATTTCGCTGCGTTCTTCATCGATGCCAGAACCAAGAGATCCGTTGTTGAAAGTTTTGTTTCGATTTACTCGTAAACTCCGACGCAGAGATGCAGTGTTTTTTGGTGGTCCTCCGGGGGCGCCCGGCAGGGTCGCCCCCGAAGCAACGGGATGTGTTCACAAAGGGTTGGAGGTCGGACCCGGAGGCCCTCGCTCGGTAATGATCCCTCCGCAGGTTCACCTACGGAGACCTTGTTACGACTTTTACTTCCTC	336 	\|\|\|\|\|\|\|\|\|\|\|\|\|\|\|\|\|\|\|\|\|\|\|\|\|\|\|\|\|\|\|\|\|\|\|\|\|\|\|\|\|\|\|\|\|\|\|\|\|\|\|\|\|\|\|\|\|\|\|\|\|\|\|\|\|\|\|\|\|\|\|\|\|\|\|\|\|\|\|\|\|\|\|\|\|\|\|\|\|\|\|\|\|\|\|\|\|\|\|\|\|\|\|\|\|\|\|\|\|\|\|\|\|\|\|\|\|\|\|\|\|\|\|\|\|\|\|\|\|\|\|\|\|\|\|\|\|\|\|\|\|\|\|\|\|\|\|\|\|\|\|\|\|\|\|\|\|\|\|\|\|\|\|\|\|\|\|\|\|\|\|\|\|\|\|\|\|\|\|\|\|\|\|\|\|\|\|\|\|\|\|\|\|\|\|\|\|\|\|\|\|\|\|\|\|\|\|\|\|\|\|\|\|\|\|\|\|\|\|\|\|\|\|\|\|\|\|\|\|\|\|\|\|\|\|\|\|\|\|\|\|\|\|\|\|\|\|\|\|\|\|\|\|\|\|\|\|\|\|\|\|\|\|\|\|\|\|\|\|\|\|\|\|\|\|\|\|\|\|\|\|\|\|\|\|\|\|\|\|\|\|\|\|\|\|\|\|\|\|\|\|\|\|\|\|\|\|\|\|\|\|\|\|\|\|\|\|\|\|\|\|\|\|\|\|\|\|\|\|\|\|\|\|\|\|\| 	ATGCCCCCCGGAATACCAGGGGGCGCAATGTGCGTTCAAAGATTCGATGATTCACTGAATTCTGCAATTCACATTACTTATCGCATTTCGCTGCGTTCTTCATCGATGCCAGAACCAAGAGATCCGTTGTTGAAAGTTTTGTTTCGATTTACTCGTAAACTCCGACGCAGAGATGCAGTGTTTTTTGGTGGTCCTCCGGGGGCGCCCGGCAGGGTCGCCCCCGAAGCAACGGGATGTGTTCACAAAGGGTTGGAGGTCGGACCCGGAGGCCCTCGCTCGGTAATGATCCCTCCGCAGGTTCACCTACGGAGACCTTGTTACGACTTTTACTTCCTC ``` |
| 2 | ggggacaagtttgtacaaaaaagcaggcttgatgCCCCCCGGAATACCAGGGGG, ggggaccactttgtacaagaaagctgggtcGAGGAAGTAAAAGTCGTAAC | Mycgr3\|45560  JGI  Ensembl |  |  | 0 % |  | ``` 1	ATGCCCCCCGGAATACCAGGGGGCGCAATGTGCGTTCAAAGATTCGATGATTCACTGAATTCTGCAATTCACATTACTTATCGCATTTCGCTGCGTTCTTCATCGATGCCAGAACCAAGAGATCCGTTGTTGAAAGTTTTGTTTCGATTTACTCGTAAACTCCGACGCAGAGATGCAGTGTTTTTTGGTGGTCCTCCGGGG	201 	\|\|\|\|\|\|\|\|\|\|\|\| \| \|\|\|\|\|\|\|\|\|\|\|\|\|\|\|\|\|\|\|\|\|\|\|\|\|\|\|\|\|\|\|\|\|\|\|\|\|\|\|\|\|\|\|\|\|\|\|\|\|\|\|\|\|\|\|\|\|\|\|\|\|\|\|\|\|\|\|\|\|\|\|\|\|\|\|\|\|\|\|\|\|\|\|\|\|\|\|\|\|\|\|\|\|\|\|\|\|\|\|\|\|\|\|\|\|\|\|\|\|\|\|\|\|\|\|\|\|\|\|\|\|\|\|\|\|\|\|\|\|\|\|\|\|\|\|\|\|\|\|\|\|\|\|\|\|\|\|\|\|\|\|\|\|\|\|\|\|\|\|\|\|\|\|\|\|\|\|\|\|\|\|\|\|\|\|\|\|\|\|\|\|\|\|\|\|\| 	ATGCCCCCCGGACTNCCAGGGGGCGCAATGTGCGTTCAAAGATTCGATGATTCACTGAATTCTGCAATTCACATTACTTATCGCATTTCGCTGCGTTCTTCATCGATGCCAGAACCAAGAGATCCGTTGTTGAAAGTTTTGTTTCGATTTACTCGTAAACTCCGACGCAGAGATGCAGTGTTTTTTGGTGGTCCTCCGGGG ```      ``` 1	ATGCCCCCCGGAATACCAGGGGGCGCAATGTGCGTTCAAAGATTCGATGATTCACTGAATTCTGCAATTCACATTACTTATCGCATTTCGCTGCGTTCTTCATCGATGCCAGAACCAAGAGATCCGTTGTTGAAAGTTTTGTTTCGATTTACTCGTAAACTCCGACGCAGAGATGCAGTGTTTTTTGGTGGTCCTCCGGGGGCGCCCGGCAGGGTCGCCCCCGAAGCAACGGGATGTGTTCACAAAGGGTTGGAGGTCGGACCCGGAGGCCCTCGCTCGGTAATGATCCCTCCGCAGGTTCACCTACGGAGACCTTGTTACGACTTTTACTTCCTC	336 	\|\|\|\|\|\|\|\|\|\|\|\|\|\|\|\|\|\|\|\|\|\|\|\|\|\|\|\|\|\|\|\|\|\|\|\|\|\|\|\|\|\|\|\|\|\|\|\|\|\|\|\|\|\|\|\|\|\|\|\|\|\|\|\|\|\|\|\|\|\|\|\|\|\|\|\|\|\|\|\|\|\|\|\|\|\|\|\|\|\|\|\|\|\|\|\|\|\|\|\|\|\|\|\|\|\|\|\|\|\|\|\|\|\|\|\|\|\|\|\|\|\|\|\|\|\|\|\|\|\|\|\|\|\|\|\|\|\|\|\|\|\|\|\|\|\|\|\|\|\|\|\|\|\|\|\|\|\|\|\|\|\|\|\|\|\|\|\|\|\|\|\|\|\|\|\|\|\|\|\|\|\|\|\|\|\|\|\|\|\|\|\|\|\|\|\|\|\|\|\|\|\|\|\|\|\|\|\|\|\|\|\|\|\|\|\|\|\|\|\|\|\|\|\|\|\|\|\|\|\|\|\|\|\|\|\|\|\|\|\|\|\|\|\|\|\|\|\|\|\|\|\|\|\|\|\|\|\|\|\|\|\|\|\|\|\|\|\|\|\|\|\|\|\|\|\|\|\|\|\|\|\|\|\|\|\|\|\|\|\|\|\|\|\|\|\|\|\|\|\|\|\|\|\|\|\|\|\|\|\|\|\|\|\|\|\|\|\|\|\|\|\|\|\|\|\|\|\|\|\|\|\|\|\|\|\| 	ATGCCCCCCGGAATACCAGGGGGCGCAATGTGCGTTCAAAGATTCGATGATTCACTGAATTCTGCAATTCACATTACTTATCGCATTTCGCTGCGTTCTTCATCGATGCCAGAACCAAGAGATCCGTTGTTGAAAGTTTTGTTTCGATTTACTCGTAAACTCCGACGCAGAGATGCAGTGTTTTTTGGTGGTCCTCCGGGGGCGCCCGGCAGGGTCGCCCCCGAAGCAACGGGATGTGTTCACAAAGGGTTGGAGGTCGGACCCGGAGGCCCTCGCTCGGTAATGATCCCTCCGCAGGTTCACCTACGGAGACCTTGTTACGACTTTTACTTCCTC ``` |
| 2 | ggggacaagtttgtacaaaaaagcaggcttgatgTCTTACCATGGAAGCTGCTT, ggggaccactttgtacaagaaagctgggtcCAGCGTTCGCACTGGATTCC | Mycgr3\|46212  JGI  Ensembl |  |  | 0 % |  | ``` 1	ATGTCTTACCATGGAAGCTGCTTCTGTCAAAAAGTCAAAATCGAGTATTGCAGCGAGACCATTACAGCTGTAAGTAATTTAGTGAACTCCGGACGTTCATATGCTAACGTATACAAGGGACTCTGCCACTGCCTCGACTGCCGCAAACTCACCGCAAACCCCTACTCCCTCTGCTTCGTCGCCAGATTCTCCGAGCTGAGAGTCACCGGTGCGGCCAAAGAACTGGTCAAGACGGCGGACAGTGGAAATGTGGTGAAGAATTACTTTTGTGATGATTGTGGTATGGACTGCAGGAATCCAGTGCGAACGCTG	312 	\|\|\|\|\|\|\|\|\|\|\|\|\|\|\|\|\|\|\|\|\|\|\|\|\|\|\|\|\|\|\|\|\|\|\|\|\|\|\|\|\|\|\|\|\|\|\|\|\|\|\|\|\|\|\|\|\|\|\|\|\|\|\|\|\|\|\|\|\|\|\|\|\|\|\|\|\|\|\|\|\|\|\|\|\|\|\|\|\|\|\|\|\|\|\|\|\|\|\|\|\|\|\|\|\|\|\|\|\|\|\|\|\|\|\|\|\|\|\|\|\|\|\|\|\|\|\|\|\|\|\|\|\|\|\|\|\|\|\|\|\|\|\|\|\|\|\|\|\|\|\|\|\|\|\|\|\|\|\|\|\|\|\|\|\|\|\|\|\|\|\|\|\|\|\|\|\|\|\|\|\|\|\|\|\|\|\|\|\|\|\|\|\|\|\|\|\|\|\|\|\|\|\|\|\|\|\|\|\|\|\|\|\|\|\|\|\|\|\|\|\|\|\|\|\|\|\|\|\|\|\|\|\|\|\|\|\|\|\|\|\|\|\|\|\|\|\|\|\|\|\|\|\|\|\|\|\|\|\|\|\|\|\|\|\|\|\|\|\|\|\|\|\|\|\|\|\|\|\|\|\|\|\|\|\|\|\|\|\|\|\|\|\|\|\|\|\|\|\|\|\|\|\|\|\|\|\|\|\|\|\|\| 	ATGTCTTACCATGGAAGCTGCTTCTGTCAAAAAGTCAAAATCGAGTATTGCAGCGAGACCATTACAGCTGTAAGTAATTTAGTGAACTCCGGACGTTCATATGCTAACGTATACAAGGGACTCTGCCACTGCCTCGACTGCCGCAAACTCACCGCAAACCCCTACTCCCTCTGCTTCGTCGCCAGATTCTCCGAGCTGAGAGTCACCGGTGCGGCCAAAGAACTGGTCAAGACGGCGGACAGTGGAAATGTGGTGAAGAATTACTTTTGTGATGATTGTGGTATGGACTGCAGGAATCCAGTGCGAACGCTG ``` |
| 2 | ggggacaagtttgtacaaaaaagcaggcttgatgCAGTTCTCTACCATCACCGC, ggggaccactttgtacaagaaagctgggtcGTTGGTGCCCTCTGGCTCCT | Mycgr3\|48129  JGI  Ensembl |  |  | 0 % |  | ``` 160	GTCGGTGCCCGCATCGACAACGCTCAGCACTTTGTCAACCTGTGTGCCGCCAAGGGCCAGCGAGCTCGCTGCTGCGCCATCCCAGTC	246 	\|\|\|\|\|\|\|\|\|\|\|\|\|\|\|\|\|\|\|\|\|\|\|\|\|\|\|\|\|\|\|\|\|\|\|\|\|\|\|\|\|\|\|\|\|\|\|\|\|\|\|\|\|\|\|\|\|\|\|\|\|\|\|\|\|\|\|\|\|\|\|\|\|\|\|\|\|\|\|\|\|\|\|\|\|\|\| 	GTCGGTGCCCGCATCGACAACGCTCAGCACTTTGTCAACCTGTGTGCCGCCAAGGGCCAGCGAGCTCGCTGCTGCGCCATCCCAGTC ```      ``` 247	CTCGGACAGGCTCTCCTCTGCCAGGAGCCAGAGGGCACCAAC	288 	\|\|\|\|\|\|\|\|\|\|\|\|\|\|\|\|\|\|\|\|\|\|\|\|\|\|\|\|\|\|\|\|\|\|\|\|\|\|\|\|\|\| 	CTCGGACAGGCTCTCCTCTGCCAGGAGCCAGAGGGCACCAAC ```      ``` 1	ATGCAGTTCTCTACCATCACCGCCGTCTTCTTCGCCTCCCTCGCCGTCGCCAGCCCGACTTACGGCGGCAAGTACGAGCCCTGCGGCTCTGCCCTCTACAGCCAGGCTCAGTGCTGTGCTACCGACGTCCTGGAGGTCGCCGGTTTGGACTGCGATGGTG	160 	\|\|\|\|\|\|\|\|\|\|\|\|\|\|\|\|\|\|\|\|\|\|\|\|\|\|\|\|\|\|\|\|\|\|\|\|\|\|\|\|\|\|\|\|\|\|\|\|\|\|\|\|\|\|\|\|\|\|\|\|\|\|\|\|\|\|\|\|\|\|\|\|\|\|\|\|\|\|\|\|\|\|\|\|\|\|\|\|\|\|\|\|\|\|\|\|\|\|\|\|\|\|\|\|\|\|\|\|\|\|\|\|\|\|\|\|\|\|\|\|\|\|\|\|\|\|\|\|\|\|\|\|\|\|\|\|\|\|\|\|\|\|\|\|\|\|\|\|\|\|\|\|\|\|\|\|\|\|\|\| 	ATGCAGTTCTCTACCATCACCGCCGTCTTCTTCGCCTCCCTCGCCGTCGCCAGCCCGACTTACGGCGGCAAGTACGAGCCCTGCGGCTCTGCCCTCTACAGCCAGGCTCAGTGCTGTGCTACCGACGTCCTGGAGGTCGCCGGTTTGGACTGCGATGGTG ``` |
| 2 | ggggacaagtttgtacaaaaaagcaggcttgatgCCGGCCATTCGCGGACCCAA, ggggaccactttgtacaagaaagctgggtcTCGCACGTCTTCATTGTCCG | Mycgr3\|49475  JGI  Ensembl | KOG3408 U1-like Zn-finger-containing protein, probabl erole in RNA processing/splicing; | intracellular; nucleic acid binding;  zinc ion binding; | 100 % |  | ``` 1	ATGCCGGCCATTCGCGGACCCAAGAGCAAGTCGAAGACGAGACGACACACTCGTGACCTCGATCAGGTCCACGCCGACACCCAATCGCAAAAGCACCTCCAGCAGTACAAGGACACAAAAGCTCCGGAGGATCTTCCAGGTTTCGGACAATTCTACTGCACTGAATGCGCAAAATGGTTCGAGTCGGAGCTGAACTACGGCAAGCACATCAAGGGAAAGCCGCACAAGAAGCGTGTGAAGCAGATGAAGGAAGAACCGTACAGTCAGAAGGAGGCTGAAGCTGCGGTGGGCTTGACGACGGACAACGGCAGGAGAGCAGTGGAGACAGACATGATCGATGCGTCGGACAATGAAGACGTGCGA	363 	\|\|\|\|\|\|\|\|\|\|\|\|\|   \|\|\|\|\|\|\|\|\|\|\|\|\|\|\|\|\|\|\|\|\|\|\|\|\|\|\|\|\|\|\|\|\|\|\|\|\|\|\|\|\|\|\|\|\|\|\|\|\|\|\|\|\|\|\|\|\|\|\|\|\|\|\|\|\|\|\|\|\|\|\|\|\|\|\|\|\|\|\|\|\|\|\|\|\|\|\|\|\|\|\|\|\|\|\|\|\|\|\|\|\|\|\|\|\|\|\|\|\|\|\|\|\|\|\|\|\|\|\|\|\|\|\|\|\|\|\|\|\|\|\|\|\|\|\|\|\|\|\|\|\|\|\|\|\|\|\|\|\|\|\|\|\|\|\|\|\|\|\|\|\|\|\|\|\|\|\|\|\|\|\|\|\|\|\|\|\|\|\|\|\|\|\|\|\|\|\|\|\|\|\|\|\|\|\|\|\|\|\|\|\|\|\|\|\|\|\|\|\|\|\|\|\|\|\|\|\|\|\|\|\|\|\|\|\|\|\|\|\|\|\|\|\|\|\|\|\|\|\|\|\|\|\|\|\|\|\|\|\|\|\|\|\|\|\|\|\|\|\|\|\|\|\|\|\|\|\|\|\|\|\|\|\|\|\|\|\|\|\|\|\|\|\|\|\|\|\|\|\|\|\|\|\|\|\|\|\|\|\|\|\|\|\|\|\|\|\|\|\|\|\|\|\|\|\|\|\|\|\|\|\|\|\|\|\|\|\|\|\|\|\|\|\|\|\|\|\|\|\|\|\|\|\|\|\|\|\| 	ATGCCGGCCATTCNNNGACCCAAGAGCAAGTCGAAGACGAGACGACACACTCGTGACCTCGATCAGGTCCACGCCGACACCCAATCGCAAAAGCACCTCCAGCAGTACAAGGACACAAAAGCTCCGGAGGATCTTCCAGGTTTCGGACAATTCTACTGCACTGAATGCGCAAAATGGTTCGAGTCGGAGCTGAACTACGGCAAGCACATCAAGGGAAAGCCGCACAAGAAGCGTGTGAAGCAGATGAAGGAAGAACCGTACAGTCAGAAGGAGGCTGAAGCTGCGGTGGGCTTGACGACGGACAACGGCAGGAGAGCAGTGGAGACAGACATGATCGATGCGTCGGACAATGAAGACGTGCGA ``` |
| 2 | ggggacaagtttgtacaaaaaagcaggcttgatgAGCGCAGAACCGGAGAACGC, ggggaccactttgtacaagaaagctgggtcGCACTGCACTCTCAGTGACC | Mycgr3\|74163  JGI  Ensembl | KOG1783 Small nuclear ribonucleoprotein F; | mRNA metabolic process; ribonucleoprotein complex; | 100 % |  | ``` 1	ATGAGCGCAGAACCGGAGAACGCCCCAAACAGTGGCGAGGAGGCCCGCGATCCAAGTGGCTTCTTGTCGGAAATCATCGGCGCTCCTGTGACCGTTAAGCTAAACTCAGGCGTGGTCTACAAAG	124 	\|\|\|\|\|\|\|\|\|\|\|\|\|\|  \|\|\|\|\|\|\|\|\|\|\|\|\|\|\|\|\|\|\|\|\|\|\|\|\|\|\|\|\|\|\|\|\|\|\|\|\|\|\|\|\|\|\|\|\|\|\|\|\|\|\|\|\|\|\|\|\|\|\|\|\|\|\|\|\|\|\|\|\|\|\|\|\|\|\|\|\|\|\|\|\|\|\|\|\|\|\|\|\|\|\|\|\|\|\|\|\|\|\|\|\|\|\|\|\|\|\|\| 	ATGAGCGCAGAACCNNAGAACGCCCCAAACAGTGGCGAGGAGGCCCGCGATCCAAGTGGCTTCTTGTCGGAAATCATCGGCGCTCCTGTGACCGTTAAGCTAAACTCAGGCGTGGTCTACAAAG ```      ``` 119	ACAA-AGTAGGCGAGCTGCAATCAGTCGATGGATACATGAACATCGCGTTAGAGGACACGAAGGAGATGGTGGATGGCAAGCAAAGACGGAATTACGGCGATGCATTCGTGAGAGGCAACAACGGTAGGTGCAAATTCTTGTGCTGTTGCGATTCGTGGTCACTGAGAGTGCAGTGC	294 	\|\|\|\| \|\|\|\|\|\|\|\|\|\|\|\|\|\|\|\|\|\|\|\|\|\|\|\|\|\|\|\|\|\|\|\|\|\|\|\|\|\|\|\|\|\|\|\|\|\|\|\|\|\|\|\|\|\|\|\|\|\|\|\|\|\|\|\|\|\|\|\|\|\|\|\|\|\|\|\|\|\|\|\|\|\|\|\|\|\|\|\|\|\|\|\|\|\|\|\|\|\|\|\|\|\|\|\|\|\|\|\|\|\|\|\|\|\|\|\|\|\|\|\|\|\|\|\|\|\|\|\|\|\|\|\|\|\|\|\|\|\|\|\|\|\|\|\|\|\|\|\|\|\|\|\|\|\| \|\|\|\|\|\|\|\|\|\|\|\|\|\|\|\|\| 	ACAAGAGTAGGCGAGCTGCAATCAGTCGATGGATACATGAACATCGCGTTAGAGGACACGAAGGAGATGGTGGATGGCAAGCAAAGACGGAATTACGGCGATGCATTCGTGAGAGGCAACAACGGTAGGTGCAAATTCTTGTGCTGTTGCGATTCGTGGNCACTGAGAGTGCAGTGC ``` |
| 2 | ggggacaagtttgtacaaaaaagcaggcttgatgATAGGAAGAGCCGACATCGA, ggggaccactttgtacaagaaagctgggtcCTTAATCAATGAAGCGGAAC | Mycgr3\|74252  JGI  Ensembl |  |  | 0 % |  | ``` 1	ATGATAGGAAGAGCCGACATCGAAGGATCAAAAAGCAACGTCGCTATGAACGCTTGGCTGCCACAAGCCAGTTATCCCTGTGGTAACTTTTCTGGCACCTCTAGCCTCAAATTTCGAGGGACTAAAGGATCGATAGGCCACACTTTCATGGTTTGTATTCACACTGAAAATCAAAATCAAGGGGACTTTTGCCCTTTTGCTCTACTGGAGATTTCTGTTCTCCATGAGTCCCCCTTAGGACACCTGCGTTATGGTTTAACAGATGTGCCGCCCCAGCCAAACTCCCCACCTGACAATGTCATCAACCCGGATCGGCCCGCGAAGGACCTTAACGCCAGAAAATGGTCGGTGAAGACCAGTTCCGCTTCATTGATTAAG	378 	\|\|\|\|\|\|\|\|\|\|\|\|\|\|\|\|\|\|\|\|\|\|\|\|\|\|\|\|\|\|\|\|\|\|\|\|\|\|\|\|\|\|\|\|\|\|\|\|\|\|\|\|\|\|\|\|\|\|\|\|\|\|\|\|\|\|\|\|\|\|\|\|\|\|\|\|\|\|\|\|\|\|\|\|\|\|\|\|\|\|\|\|\|\|\|\|\|\|\|\|\|\|\|\|\|\|\|\|\|\|\|\|\|\|\|\|\|\|\|\|\|\|\|\|\|\|\|\|\|\|\|\|\|\|\|\|\|\|\|\|\|\|\|\|\|\|\|\|\|\|\|\|\|\|\|\|\|\|\|\|\|\|\|\|\|\|\|\|\|\|\|\|\|\|\|\|\|\|\|\|\|\|\|\|\|\|\|\|\|\|\|\|\|\|\|\|\|\|\|\|\|\|\|\|\|\|\|\|\|\|\|\|\|\|\|\|\|\|\|\|\|\|\|\|\|\|\|\|\|\|\|\|\|\|\|\|\|\|\|\|\|\|\|\|\|\|\|\|\|\|\|\|\|\|\|\|\|\|\|\|\|\|\|\|\|\|\|\|\|\|\|\|\|\|\|\|\|\|\|\|\|\|\|\|\|\|\|\|\|\|\|\|\|\|\|\|\|\|\|\|\|\|\|\|\|\|\|\|\|\|\|\|\|\|\|\|\|\|\|\|\|\|\|\|\|\|\|\|\|\|\|\|\|\|\|\|\|\|\|\|\|\|\|\|\|\|\|\|\|\|\|\|\|\|\|\|\|\|\|\|\|\|\|\|\|\|\|\|\|\|\|\|\|\|\|\|\|\| 	ATGATAGGAAGAGCCGACATCGAAGGATCAAAAAGCAACGTCGCTATGAACGCTTGGCTGCCACAAGCCAGTTATCCCTGTGGTAACTTTTCTGGCACCTCTAGCCTCAAATTTCGAGGGACTAAAGGATCGATAGGCCACACTTTCATGGTTTGTATTCACACTGAAAATCAAAATCAAGGGGACTTTTGCCCTTTTGCTCTACTGGAGATTTCTGTTCTCCATGAGTCCCCCTTAGGACACCTGCGTTATGGTTTAACAGATGTGCCGCCCCAGCCAAACTCCCCACCTGACAATGTCATCAACCCGGATCGGCCCGCGAAGGACCTTAACGCCAGAAAATGGTCGGTGAAGACCAGTTCCGCTTCATTGATTAAG ``` |
| 2 | ggggacaagtttgtacaaaaaagcaggcttgatgGCGCATCAAAGGCTGTCTAT, ggggaccactttgtacaagaaagctgggtcCCGGAGTCGATTGTTGGATG | Mycgr3\|83337  JGI  Ensembl |  |  | 0 % |  |  |
| 2 | ggggacaagtttgtacaaaaaagcaggcttgatgTCTGTCCTCGACGCCCTCAA, ggggaccactttgtacaagaaagctgggtcCTGCTCAGTCTCGAAGTAGT | Mycgr3\|83768  JGI  Ensembl | KOG3406 40S ribosomal protein S12; | intracellular; ribosome; structural constituent of ribosome;  translation; | 0 % |  | ``` 1	ATGTCTGTCCTCGACGCCCTCAAGGGAGTCCTCAAGATCTCCTTGATCCACGACGGACTCGCCCGCGGTCTCCGCGAGGCCAGCAAGGCCCTCGATCGTCGCCAGGCTCACATGTGCGTGCTCAACGAGGCGTGCGAGGAGGAGGCATACAAGAAGTTGGTTGTCGCTCTGTGCGGCGAGCACAAGATTCCTCTCATCAAGGTTCCGGATGGCAAGCAGCTCGGCGAGTGGGCGGGTCTC	240 	\|\|\|\|\|\|\|\|\|\|\|\|\|\|\|\|\|\|\|\|\|\|\|\|\|\|\|\|\|\|\|\|\|\|\|\|\|\|\|\|\|\|\|\|\|\|\|\|\|\|\|\|\|\|\|\|\|\|\|\|\|\|\|\|\|\|\|\|\|\|\|\|\|\|\|\|\|\|\|\|\|\|\|\|\|\|\|\|\|\|\|\|\|\|\|\|\|\|\|\|\|\|\|\|\|\|\|\|\|\|\|\|\|\|\|\|\|\|\|\|\|\|\|\|\|\|\|\|\|\|\|\|\|\|\|\|\|\|\|\|\|\|\|\|\|\| \|\|\|\|\|\|\|\|\|\|\|\|\|\|\|\|\|\|\|\|\|\|\|\|\|\|\|\|\|\|\|\|\|\|\|\|\|\|\|\|\|\|\|\|\|\|\|\|\|\|\|\|\|\|\|\|\|\|\|\|\|\|\|\|\|\|\|\|\|\|\|\|\|\|\|\| \| \|  \|\|\|\|\|\|\|\|\|\|\| 	ATGTCTGTCCTCGACGCCCTCAAGGGAGTCCTCAAGATCTCCTTGATCCACGACGGACTCGCCCGCGGTCTCCGCGAGGCCAGCAAGGCCCTCGATCGTCGCCAGGCTCACATGTGCGTGCTCAACGAGGCGTGCGAGGAGGAGGCNTACAAGAAGTTGGTTGTCGCTCTGTGCGGCGAGCACAAGATTCCTCTCATCAAGGTTCCGGATGGCAAGCAGCTCGNCNANNGGGCGGGTCTC ``` |
| 2 | ggggacaagtttgtacaaaaaagcaggcttgatgCCGACCCCCGAATCCGAGTC, ggggaccactttgtacaagaaagctgggtcCTCCGTCTTCGGAACGTAGT | Mycgr3\|84244  JGI  Ensembl | KOG4009 NADH-ubiquinone oxidoreductase, subunit NDUFB10/PDSW; |  | 0 % |  | ``` 1	ATGCCGACCCCCGAATCCGAGTCCTTCAAAGCCTCCAAACCGACCGTCCCGCCGACCTTCGAAGGCGTCGACTACGACGACAACAAAGCCCTCAAAGCCGCGCAGGACTCCATCATCCGCGAGCAATGGGTGCAGAGCATGATGGCGCGGCTGATCCGCGAGGAGATGGGGAAGTGCTATTATCGGGAGGGGGTGAATCATTTGG-AGAAGTGTG	214 	\|\|\|\|\|\|\|\|\|\|\|\|\|\|\|\|\|\|\|\|\|\|\|\|\|\|\|\|\|\|\|\|\|\|\|\|\|\|\|\|\|\|\|\|\|\|\|\|\|\|\|\|\|\|\|\|\|\|\|\|\|\|\|\|\|\|\|\|\|\|\|\|\|\|\|\|\|\|\|\|\|\|\|\|\|\|\|\|\|\|\|\|\|\|\|\|\|\|\|\|\|\|\|\|\|\|\|\|\|\|\|\|\|\|\|\|\|\|\|\|\|\|\|\|\|\|\|\|\|\|\|\|\|\|\|\|\|\|\|\|\|\|\|\|\|\|\|\|\|\|\|\|\|\|\|\|\|\|\|\|\|\|\|\|\|\|\|\|\|\|\|\|\|\|\|\|\|\|\|\|\|\|\|\|\|\|\|\|\|\|\|\|\|\|\|\| \|\| \|\|\|\|  \|\|\|\| \|\|\|\| 	ATGCCGACCCCCGAATCCGAGTCCTTCAAAGCCTCCAAACCGACCGTCCCGCCGACCTTCGAAGGCGTCGACTACGACGACAACAAAGCCCTCAAAGCCGCGCAGGACTCCATCATCCGCGAGCAATGGGTGCAGAGCATGATGGCGCGGCTGATCCGCGAGGAGATGGGGAAGTGCTATTATCGGGAGGGGGTGANTCNTTTGACAGAAATGTG ``` |
| 2 | ggggacaagtttgtacaaaaaagcaggcttgatgCCAGAAACACGCCGCATCGG, ggggaccactttgtacaagaaagctgggtcCTCGCATCTGAAAACCTCCT | Mycgr3\|86169  JGI  Ensembl |  |  | 100 % |  | ``` 1	ATGCCAGAAACACGCCGCATCGGACAGATCATCCGTCTCAAGCCCGAATGTTTGCAAGCCTACAAAGACTGCCATGCTGCGGCTTGGCCAGCTGTCCTCAAGCAAATCAAAGATTGCAATATCTCCGACTACTCAATCTACTTCGATGACCGGAGCTACACGCTATTTGCAACAATGAAATGGACTGGCACCGACTTCGAGGCAGACATGGAGAGCATGAAGGCCAACCCCGAAGTGAGACGTTGGTGGGAAATGACGGACAGTATGCAGGAAACGCTGGTGGACGGCAGTACTGGAAGCACGGATTCGAAGGGCTGGTGGAAAGATCTCGAGGAGGTTTTCAGATGCGAG	351 	\|\|\|\|\|\|\|\|\|\|\|\|\| \|\| \|\|\|\|\|\|\|\|\|\|\|\|\|\|\|\|\|\|\|\|\|\|\|\|\|\|\|\|\|\|\|\|\|\|\|\|\|\|\|\|\|\|\|\|\|\|\|\|\|\|\|\|\|\|\|\|\|\|\|\|\|\|\|\|\|\|\|\|\|\|\|\|\|\|\|\|\|\|\|\|\|\|\|\|\|\|\|\|\|\|\|\|\|\|\|\|\|\|\|\|\|\|\|\|\|\|\|\|\|\|\|\|\|\|\|\|\|\|\|\|\|\|\|\|\|\|\|\|\|\|\|\|\|\|\|\|\|\|\|\|\|\|\|\|\|\|\|\|\|\|\|\|\|\|\|\|\|\|\|\|\|\|\|\|\|\|\|\|\|\|\|\|\|\|\|\|\|\|\|\|\|\|\|\|\|\|\|\|\|\|\|\|\|\|\|\|\|\|\|\|\|\|\|\|\|\|\|\|\|\|\|\|\|\|\|\|\|\|\|\|\|\|\|\|\|\|\|\|\|\|\|\|\|\|\|\|\|\|\|\|\|\|\|\|\|\|\|\|\|\|\|\|\|\|\|\|\|\|\|\|\|\|\|\|\|\|\|\|\|\|\|\|\|\|\|\|\|\|\|\|\|\|\|\|\|\|\|\|\|\|\|\|\|\|\|\|\|\|\|\|\|\|\|\|\|\|\|\|\|\|\|\|\|\|\|\|\|\|\|\|\|\|\|\|\|\|\|\|\|\|\|\|\|\| 	ATGCCAGAAACACNCCNCATCGGACAGATCATCCGTCTCAAGCCCGAATGTTTGCAAGCCTACAAAGACTGCCATGCTGCGGCTTGGCCAGCTGTCCTCAAGCAAATCAAAGATTGCAATATCTCCGACTACTCAATCTACTTCGATGACCGGAGCTACACGCTATTTGCAACAATGAAATGGACTGGCACCGACTTCGAGGCAGACATGGAGAGCATGAAGGCCAACCCCGAAGTGAGACGTTGGTGGGAAATGACGGACAGTATGCAGGAAACGCTGGTGGACGGCAGTACTGGAAGCACGGATTCGAAGGGCTGGTGGAAAGATCTCGAGGAGGTTTTCAGATGCGAG ``` |
| 2 | ggggacaagtttgtacaaaaaagcaggcttgatgACTTCAATCAACGTCTGCAA, ggggaccactttgtacaagaaagctgggtcCTCCCATTTCTTGGTCACGA | Mycgr3\|87428  JGI  Ensembl |  |  | 100 % |  | ``` 1	ATGACTTCAATCAACGTCTGCAAGAACCGAAGCCGCTCAAACTCAACCATGGAAGTGCCAGACGACGAGCGACGCCAATCCACCGACAATCTGCGCCGCTCGACAACCACTGCCAAAACAACCACTGCCAAAACCATCGATCCCGCCGAGAACGCAAAGTACTGCTCCTTCTACAGGCGCCTCAGCCGGGAAGTCAACGAGAAGATTCAAGAAGCCGCGTACAACGACGATGTCACAGTGGAGCAAGTGAAGCAAGAGTCGAAGAGGAAGACCCCTCCGCCGGTCTTGGATGGACCGGAATTGCACGATTTCAGAAAGTTCAGGAAGGAGGACTTTGAGAAGTTCAAGGCTTCGAAGCAGGTGTATAAGGGCGCGGAGGTCGTGACCAAGAAATGGGAG	399 	\|\|\|\|\|\|\|\|\|\|\|\|\|\|\|\|\|\|\|\|\|\|\|\|\|\|\|\|\|\|\|\|\|\|\|\|\|\|\|\|\|\|\|\|\|\|\|\|\|\|\|\|\|\|\|\|\|\|\|\|\|\|\|\|\|\|\|\|\|\|\|\|\|\|\|\|\|\|\|\|\|\|\|\|\|\|\|\|\|\|\|\|\|\|\|\|\|\|\|\|\|\|\|\|\|\|\|\|\|\|\|\|\|\|\|\|\|\|\|\|\|\|\|\|\|\|\|\|\|\|\|\|\|\|\|\|\|\|\|\|\|\|\|\|\|\|\|\|\|\|\|\|\|\|\|\|\|\|\|\|\|\|\|\|\|\|\|\|\|\|\|\|\|\|\|\|\|\|\|\|\|\|\|\|\|\|\|\|\|\|\|\|\|\|\|\|\|\|\|\|\|\|\|\|\|\|\|\|\|\|\|\|\|\|\|\|\|\|\|\|\|\|\|\|\|\|\|\|\|\|\|\|\|\|\|\|\|\|\|\|\|\|\|\|\|\|\|\|\|\|\|\|\|\|\|\|\|\|\|\|\|\|\|\|\|\|\|\|\|\|\|\|\|\|\|\|\|\|\|\|\|\|\|\|\|\|\|\|\|\|\|\|\|\|\|\|\|\|\|\|\|\|\|\|\|\|\|\|\|\|\|\|\|\|\|\|\|\|\|\|\|\|\|\|\|\|\|\|\|\|\|\|\|\|\|\|\|\|\|\|\|\|\|\|\|\|\|\|\|\|\|\|\|\|\|\|\|\|\|\|\|\|\|\|\|\|\|\|\|\|\|\|\|\|\|\|\|\|\|\|\|\|\|\|\|\|\|\|\|\|\|\|\|\|\|\|\|\|\| 	ATGACTTCAATCAACGTCTGCAAGAACCGAAGCCGCTCAAACTCAACCATGGAAGTGCCAGACGACGAGCGACGCCAATCCACCGACAATCTGCGCCGCTCGACAACCACTGCCAAAACAACCACTGCCAAAACCATCGATCCCGCCGAGAACGCAAAGTACTGCTCCTTCTACAGGCGCCTCAGCCGGGAAGTCAACGAGAAGATTCAAGAAGCCGCGTACAACGACGATGTCACAGTGGAGCAAGTGAAGCAAGAGTCGAAGAGGAAGACCCCTCCGCCGGTCTTGGATGGACCGGAATTGCACGATTTCAGAAAGTTCAGGAAGGAGGACTTTGAGAAGTTCAAGGCTTCGAAGCAGGTGTATAAGGGCGCGGAGGTCGTGACCAAGAAATGGGAG ``` |
| 2 | ggggacaagtttgtacaaaaaagcaggcttgatgATTGCATTGAGAGGCGGAGT, ggggaccactttgtacaagaaagctgggtcATGCTTCGGCCTCACATCTG | Mycgr3\|87477  JGI  Ensembl |  |  | 0 % |  |  |
| 2 | ggggacaagtttgtacaaaaaagcaggcttgatgAACACGGACGTTCCACTCAT, ggggaccactttgtacaagaaagctgggtcCGAGCATGATCCCAATCCCA | Mycgr3\|88624  JGI  Ensembl |  |  | 100 % |  | ``` 198	ACCTCCTTCCGCGCGCCTCCCGACCACGGCGTCTTCCGACCTCGTACCCGTCCAGGTCGAGCGCCCGAAACCGAAGCACAATGTGGACGTGGGATTGGGATCATGCTCG	306 	\|\|\|\|\|\|\|\|\|\|\|\|\|\|\|\|\|\|\|\|\|\|\|\|\|\|\|\|\|\|\|\|\|\|\|\|\|\|\|\|\|\|\|\|\|\|\|\|\|\|\|\|\|\|\|\|\|\|\|\|\|\|\|\|\|\|\|\|\|\|\|\|\|\|\|\|\|\|\|\|\|\|\|\|\|\|\|\|\|\|\|\|\|\|\|\|\|\|\|\|\|\|\|\|\|\|\|\|\| 	ACCTCCTTCCGCGCGCCTCCCGACCACGGCGTCTTCCGACCTCGTACCCGTCCAGGTCGAGCGCCCGAAACCGAAGCACAATGTGGACGTGGGATTGGGATCATGCTCG ```      ``` 1	ATGAACACGGACG-TTCCACTCATCTCTCCCGCGCCAATCACCTTCACGCAGACCTCCATCCCTCCCCCTTCGCCCTTCGCCCGAGACTCAGCAAGGACATTCCTCGCGGCGAACATGATTCACTCCATCGATAACAGCAGCGCCATCTCTCCCACCGCCATCTTCGCCATGAGCCACAACACCGAGAAGATCAAGTC	197 	\|\|\|\|\|\|\|\|\|\|\|\|\| \|\|\|\|\|\|\|\|\|\|\|\|\|\|\|\|\|\|\|\|\|\|\|\|\|\|\|\|\|\|\|\|\|\|\|\|\|\|\|\|\|\|\|\|\|\|\|\|\|\|\|\|\|\|\|\|\|\|\|\|\|\|\|\|\|\|\|\|\|\|\|\|\|\|\|\|\|\|\|\|\|\|\|\|\|\|\|\|\|\|\|\|\|\|\|\|\|\|\|\|\|\|\|\|\|\|\|\|\|\|\|\|\|\|\|\|\|\|\|\|\|\|\|\|\|\|\|\|\|\|\|\|\|\|\|\|\|\|\|\|\|\|\|\|\|\|\|\|\|\|\|\|\|\|\|\|\|\|\|\|\|\|\|\|\|\|\|\|\|\|\|\|\|\|\|\|\|\|\|\|\|\|\|\| 	ATGAACACGGACGCTTCCACTCATCTCTCCCGCGCCAATCACCTTCACGCAGACCTCCATCCCTCCCCCTTCGCCCTTCGCCCGAGACTCAGCAAGGACATTCCTCGCGGCGAACATGATTCACTCCATCGATAACAGCAGCGCCATCTCTCCCACCGCCATCTTCGCCATGAGCCACAACACCGAGAAGATCAAGTC ``` |
| 2 | ggggacaagtttgtacaaaaaagcaggcttgatgTCGAACGCAAACCGCCCTCA, ggggaccactttgtacaagaaagctgggtcTTTCCCAACGGTAGGCCCAA | Mycgr3\|88663  JGI  Ensembl |  |  | 0 % |  | ``` 1	ATGTCGAACGCAAACCGCCCTCACATCCTCCCAAAGCTCAACTTCAAGAACTTCAAGAACTTCCTGCTCTCGCAGACTGCTGGCAACAGCTCTGCGCAGGAAGCGAACGACAAGCGAATGGAACAGATGTCTGCGGACGAGAAGAAATCTGGTGGAGACGAGGGAAAGGAGGAGGAATGGACTTTGGTCGACTCGGAGAAAGAGAGGATTGCAAAGGAGTTGAAGGCGGAGGCAGAGGTACGGACGGAAAATGCGGTGTTGGATGAGATCGAGGAGAGCTTTGGGCCTACCGTTGGGAAA	300 	\|\|\|\|\|\|\|\|\|\|\|\|\| \|\|\|\|\|\|\|\|\|\|\|\|\|\|\|\|\|\|\|\|\|\|\|\|\|\|\|\|\|\|\|\|\|\|\|\|\|\|\|\|\|\|\|\|\|\|\|\|\|\|\|\|\|\|\|\|\|\|\|\|\|\|\|\|\|\|\|\|\|\|\|\|\|\|\|\|\|\|\|\|\|\|\|\|\|\|\|\|\|\|\|\|\|\|\|\|\|\|\|\|\|\|\|\|\|\|\|\|\|\|\|\|\|\|\|\|\|\|\|\|\|\|\|\|\|\|\|\|\|\|\|\|\|\|\|\|\|\|\|\|\|\|\|\|\|\|\|\|\|\|\|\|\|\|\|\|\|\|\|\|\|\|\|\|\|\|\|\|\|\|\|\|\|\|\|\|\|\|\|\|\|\|\| \|\|\|\|\|\|\|\|\|\|\|\|\|\|\|\|\|\|\|\|\|\|\|\|\|\|\|\|\|\|\|\|\|\|\|\|\|\|\|\|\|\|\|\|\|\|\|\|\|\|\|\|\|\|\|\|\|\|\|\|\|\|\|\|\|\|\|\|\|\|\|\|\|\|\|\|\|\|\|\|\|\|\|\|\|\|\|\|\|\|\|\|\|\|\|\|\|\|\|\|\|\| 	ATGTCGAACGCAANCCGCCCTCACATCCTCCCAAAGCTCAACTTCAAGAACTTCAAGAACTTCCTGCTCTCGCAGACTGCTGGCAACAGCTCTGCGCAGGAAGCGAACGACAAGCGAATGGAACAGATGTCTGCGGACGAGAAGAAATCTGGTGGAGACGAGGGAAAGGAGGAGGAATGGACTTTGGTCGACTCGGANAAAGAGAGGATTGCAAAGGAGTTGAAGGCGGAGGCAGAGGTACGGACGGAAAATGCGGTGTTGGATGAGATCGAGGAGAGCTTTGGGCCTACCGTTGGGAAA ``` |
| 2 | ggggacaagtttgtacaaaaaagcaggcttgatgGCAGGTCAGGTTGTGAGCAT, ggggaccactttgtacaagaaagctgggtcCAGAGCTAGTTTCGCTACTC | Mycgr3\|88845  JGI  Ensembl |  |  | 100 % |  | ``` 164	CCATCCACAAGACCTTGATGAACAATACGCGCCTGCTCGCATTCGACCTTCGCGCAATCGTTCTTTGCCCTCCGACATGTCCGCCTATGCGCCTCAGATCCTCGGCCGCTCAACACCCTCATCGAGTAGCGAAACTAG-CTCTG	306 	\|\|\|\|\|\|\|\|\|\|\|\|\|\|\|\|\|\|\|\|\|\|\|\|\|\|\|\|\|\|\|\|\|\|\|\|\|\|\|\|\|\|\|\|\|\|\|\|\|\|\|\|\|\|\|\|\|\|\|\|\|\|\|\|\|\|\|\|\|\|\|\|\|\|\|\|\|\|\|\|\|\|\|\|\|\|\|\|\|\|\|\|\|\|\|\|\|\|\|\|\|\|\|\|\|\|\|\|\|\|\|\|\|\|\|\|\|\|\|\|\|\|\|\|\|\|\|\|\|\|\|\|\|\|\|\|\|  \|\|\|\|\| 	CCATCCACAAGACCTTGATGAACAATACGCGCCTGCTCGCATTCGACCTTCGCGCAATCGTTCTTTGCCCTCCGACATGTCCGCCTATGCGCCTCAGATCCTCGGCCGCTCAACACCCTCATCGAGTAGCGAAACTANNCTCTG ```      ``` 1	ATGGCAGGTCAGGTTGTGAGCATCCTGGTCCTTGTTGCTCAACCGCAGCCTACCGTCAGCCGAAATTTGACGCGCTTCCGCGACTTCGTGTTCGATGAGAAGTGGGCTCAGCACGGCTGGTACATGAATGCATGGATGGACGGTCTCATGAAGGCGCTGCTGC	163 	\|\|\|\|\|\|\|\|\|\|\|\|\|\|\|\|\|\|\|\|\|\|\|\|\|\|\|\|\|\|\|\|\|\|\|\|\|\|\|\|\|\|\|\|\|\|\|\|\|\|\|\|\|\|\|\|\|\|\|\|\|\|\|\|\|\|\|\|\|\|\|\|\|\|\|\|\|\|\|\|\|\|\|\|\|\|\|\|\|\|\|\|\|\|\|\|\|\|\|\|\|\|\|\|\|\|\|\|\|\|\|\|\|\|\|\|\|\|\|\|\|\|\|\|\|\|\|\|\|\|\|\|\|\|\|\|\|\|\|\|\|\|\|\|\|\|\|\|\|\|\|\|\|\|\|\|\|\|\|\|\|\|\| 	ATGGCAGGTCAGGTTGTGAGCATCCTGGTCCTTGTTGCTCAACCGCAGCCTACCGTCAGCCGAAATTTGACGCGCTTCCGCGACTTCGTGTTCGATGAGAAGTGGGCTCAGCACGGCTGGTACATGAATGCATGGATGGACGGTCTCATGAAGGCGCTGCTGC ``` |
| 2 | ggggacaagtttgtacaaaaaagcaggcttgatgGAGGACAGAGCCGATGCTGC, ggggaccactttgtacaagaaagctgggtcTGGACAACCGTGAACTGCAG | Mycgr3\|89417  JGI  Ensembl |  |  | 100 % |  | ``` 1	ATGGAGGACAGAGCCGATGCTGCGCTGGCCCGTCACAACGGAGGCATCGCTCCACGCAACGACTCCGAACGCTATCGCGCGGCACAAGCAGCGAGCGTGGTCACTGTGGACTCTGTCCTCATGATCACAGACATGCTGCGGAGGAGAGACGAGCTTGAGGCTACAGCTGACGTGCCATTACTGTGGTCAGAAAGTGCGGATGCGGTGTTGAACGAGTGGAGGAAATTCTTGCACCAACATAAAGGGAAGACCCTTCTTGCGAAAGATCTGCTCGAGGGAATGTATGAGAAGTTTCGGCAGACACTCGTCAACCAGGGTTACTTGTCTCATCATGTGGCTGTTCCAGAGCGATTTGATCGATTCGCCAGGCACATTGCTTTCGCTGCAGTTCACGGTTGTCCA	402 	\|\|\|\|\|\|\|\|\|\|\|\|\|\|\|\|\|\|\|\|\|\|\|\|\|\|\|\|\|\|\|\|\|\|\|\|\|\|\|\|\|\|\|\|\|\|\|\|\|\|\|\|\|\|\|\|\|\|\|\|\|\|\|\|\|\|\|\|\|\|\|\|\|\|\|\|\|\|\|\|\|\|\|\|\|\|\|\|\|\|\|\|\|\|\|\|\|\|\|\|\|\|\|\|\|\|\|\|\|\|\|\|\|\|\|\|\|\|\|\|\|\|\|\|\|\|\|\|\|\|\|\|\|\|\|\|\|\|\|\|\|\|\|\|\|\|\|\|\|\|\|\|\|\|\|\|\|\|\|\|\|\|\|\|\|\|\|\|\|\|\|\|\|\|\|\|\|\|\|\|\|\|\|\|\|\|\|\|\|\|\|\|\|\|\|\|\|\|\|\|\|\|\|\|\|\|\|\|\|\|\|\|\|\|\|\|\|\|\|\|\|\|\|\|\|\|\|\|\|\|\|\|\|\|\|\|\|\|\|\|\|\|\|\|\|\|\|\|\|\|\|\|\|\|\|\|\|\|\|\|\|\|\|\|\|\|\|\|\|\|\|\|\|\|\|\|\|\|\|\|\|\|\|\|\|\|\|\|\|\|\|\|\|\|\|\|\|\|\|\|\|\|\|\|\|\|\|\|\|\|\|\|\|\|\|\|\|\|\|\|\|\|\|\|\|\|\|\|\|\|\|\|\|\|\|\|\|\|\|\|\|\|\|\|\|\|\|\|\|\|\|\|\|\|\|\|\|\|\|\|\|\|\|\|\|\|\|\|\|\|\|\|\|\|\|\|\|\|\|\|\|\|\|\|\|\|\|\|\|\|\|\|\|\|\|\|\|\|\|\|\|\| 	ATGGAGGACAGAGCCGATGCTGCGCTGGCCCGTCACAACGGAGGCATCGCTCCACGCAACGACTCCGAACGCTATCGCGCGGCACAAGCAGCGAGCGTGGTCACTGTGGACTCTGTCCTCATGATCACAGACATGCTGCGGAGGAGAGACGAGCTTGAGGCTACAGCTGACGTGCCATTACTGTGGTCAGAAAGTGCGGATGCGGTGTTGAACGAGTGGAGGAAATTCTTGCACCAACATAAAGGGAAGACCCTTCTTGCGAAAGATCTGCTCGAGGGAATGTATGAGAAGTTTCGGCAGACACTCGTCAACCAGGGTTACTTGTCTCATCATGTGGCTGTTCCAGAGCGATTTGATCGATTCGCCAGGCACATTGCTTTCGCTGCAGTTCACGGTTGTCCA ``` |
| 2 | ggggacaagtttgtacaaaaaagcaggcttgatgCCATTCTCACACTCCCAGAC, ggggaccactttgtacaagaaagctgggtcCCCGCTCCGACCACATACTT | Mycgr3\|90143  JGI  Ensembl |  |  | 100 % |  | ``` 1	ATGCCATTCTCACACTCCCAGACCATGCGCATGATCGCCTACACCTTTGCAGTCGTGCTGACCGGCTTCGGCATCAACGCCATGTCGAACCCTGTCTCTGCACTGTCGTTCTTCGAGCTCGAATACCCGAGTCTTGCTTCGGAGAGAAAGGTCGTTGATGTCCTGCTGTTGGTCTACGGTGCCCGAGACATATTCATGGGCTTCATGCTATTCGTCACGGCGTTCTTCGGCAGCCCCAAGGCAACTGGCTGGACCTTCATCGGGTGTTGGAGCCATGGCAGCTGTGGACGGCGTGGTCTGCAAGTATGTGGTCGGAGCGGG	321 	\|\|\|\|\|\|\|\|\|\|\|\|\|\|\|\|\|\|\|\|\|\|\|\|\|\|\|\|\|\|\|\|\|\|\|\|\|\|\|\|\|\|\|\|\|\|\|\|\|\|\|\|\|\|\|\|\|\|\|\|\|\|\|\|\|\|\|\|\|\|\|\|\|\|\|\|\|\|\|\|\|\|\|\|\|\|\|\|\|\|\|\|\|\|\|\|\|\|\|\|\|\|\|\|\|\|\|\|\|\|\|\|\|\|\|\|\|\|\|\|\|\|\|\|\|\|\|\|\|\|\|\|\|\|\|\|\|\|\|\|\|\|\|\|\|\|\|\|\|\|\|\|\|\|\|\|\|\|\|\|\|\|\|\|\|\|\|\|\|\|\|\|\|\|\|\|\|\|\|\|\|\|\|\|\|\|\|\|\|\|\|\|\|\|\|\|\|\|\|\|\|\|\|\|\|\|\|\|\|\|\|\|\|\|\|\|\|\|\|\|\|\|\|\|\|\|\|\|\|\|\|\|\|\|\|\|\|\|\|\|\|\|\|\|\|\|\|\|\|\|\|\|\|\|\|\|\|\|\|\|\|\|\|\|\|\|\|\|\|\|\|\|\|\|\|\|\|\|\|\|\|\|\|\|\|\|\|\|\|\|\|\|\|\|\|\|\|\|\|\|\|\|\|\|\|\|\|\|\|\|\|\|\|\|\|\|\|\|\|\|\| 	ATGCCATTCTCACACTCCCAGACCATGCGCATGATCGCCTACACCTTTGCAGTCGTGCTGACCGGCTTCGGCATCAACGCCATGTCGAACCCTGTCTCTGCACTGTCGTTCTTCGAGCTCGAATACCCGAGTCTTGCTTCGGAGAGAAAGGTCGTTGATGTCCTGCTGTTGGTCTACGGTGCCCGAGACATATTCATGGGCTTCATGCTATTCGTCACGGCGTTCTTCGGCAGCCCCAAGGCAACTGGCTGGACCTTCATCGGGTGTTGGAGCCATGGCAGCTGTGGACGGCGTGGTCTGCAAGTATGTGGTCGGAGCGGG ``` |
| 2 | ggggacaagtttgtacaaaaaagcaggcttgatgACGGAGCTCGAGAGCCGAAT, ggggaccactttgtacaagaaagctgggtcACCCCGCTCTGTACTCCCTC | Mycgr3\|90311  JGI  Ensembl |  |  | 100 % |  | ``` 1	ATGACGGAGCTCGAGAGCCGAATCTGCCGCGGCGATCCAAAGCAGCAGAGCTCTACGCACACTCCGCCTCGAGCTGCATCCAAGAAGTGCGGCACAAAAGCAGCCTTCCGAACCGGGACAGTCCCGCGATACGGCGTAGAAGAGCTTCCCATAACAGCGACTGATCTGACGAGGCCAGTCGCAATCATGGCATCGAACACGCAGTTCACGGCTTCACTTCCGACTCTGCGCCTACCATTGGACAACAACAGCGACTCGTCTGGACGGACACAGCTATACGATGTACTATCTCAGCCACGAGAGATACTGGATGCAGTCGACCAATGGCTCGAGTTGGACTCGACAGATGGCTTCGCTATGCGAGGGAGTACAGAGCGGGGT	381 	\|\|\|\|\|\|\|\|\|\|\|\|\|\|\|\|\|\|\|\|\|\|\|\|\|\|\|\|\|\|\|\|\|\|\|\|\|\|\|\|\|\|\|\|\|\|\|\|\|\|\|\|\|\|\|\|\|\|\|\|\|\|\|\|\|\|\|\|\|\|\|\|\|\|\|\|\|\|\|\|\|\|\|\|\|\|\|\|\|\|\|\|\|\|\|\|\|\|\|\|\|\|\|\|\|\|\|\|\|\|\|\|\|\|\|\|\|\|\|\|\|\|\|\|\|\|\|\|\|\|\|\|\|\|\|\|\|\|\|\|\|\|\|\|\|\|\|\|\|\|\|\|\|\|\|\|\|\|\|\|\|\|\|\|\|\|\|\|\|\|\|\|\|\|\|\|\|\|\|\|\|\|\|\|\|\|\|\|\|\|\|\|\|\|\|\|\|\|\|\|\|\|\|\|\|\|\|\|\|\|\|\|\|\|\|\|\|\|\|\|\|\|\|\|\|\|\|\|\|\|\|\|\|\|\|\|\|\|\|\|\|\|\|\|\|\|\|\|\|\|\|\|\|\|\|\|\|\|\|\|\|\|\|\|\|\|\|\|\|\|\|\|\|\|\|\|\|\|\|\|\|\|\|\|\|\|\|\|\|\|\|\|\|\|\|\|\|\|\|\|\|\|\|\|\|\|\|\|\|\|\|\|\|\|\|\|\|\|\|\|\|\|\|\|\|\|\|\|\|\|\|\|\|\|\|\|\|\|\|\|\|\|\|\|\|\|\|\|\|\|\|\|\|\|\|\|\|\|\|\|\|\|\|\|\|\|\|\|\|\|\|\|\|\| \|\|\|\|\|\| 	ATGACGGAGCTCGAGAGCCGAATCTGCCGCGGCGATCCAAAGCAGCAGAGCTCTACGCACACTCCGCCTCGAGCTGCATCCAAGAAGTGCGGCACAAAAGCAGCCTTCCGAACCGGGACAGTCCCGCGATACGGCGTAGAAGAGCTTCCCATAACAGCGACTGATCTGACGAGGCCAGTCGCAATCATGGCATCGAACACGCAGTTCACGGCTTCACTTCCGACTCTGCGCCTACCATTGGACAACAACAGCGACTCGTCTGGACGGACACAGCTATACGATGTACTATCTCAGCCACGAGAGATACTGGATGCAGTCGACCAATGGCTCGAGTTGGACTCGACAGATGGCTTCGCTATGCGAGGGAGTACAGANCGGGGT ``` |
| 2 | ggggacaagtttgtacaaaaaagcaggcttgatgATCGGAGACAAGTCGGACCA, ggggaccactttgtacaagaaagctgggtcGCCAGTATCGCCGCGACATT | Mycgr3\|90566  JGI  Ensembl |  |  | 100 % |  | ``` 1	ATGATCGGAGACAAGTCGGACCACTCGGAACGTGAGGAGAATGGCGAGAGCGTCCACGATGGAGACGACAAGAGATCCTTTGATGCTGCTGAAGACGAGCAAAGGCCTCGTACTCCACCGCCGGCCTACCGACATCTAGAGCGCGCGCCATCTTACGCTACGCAAGACACGACGGCAGATCAGACCTTTAAGCGTGAAGAGGTGACCAGCTCGCAATCGACCTCGGCCGGCTTTGCGTCCGCAGCCAGGATTAATGCTGTAATGGGCCCTCCTCCGCCAAAGGAGGAGAAGAAGACCAGCCGATGGAAGCGCTGGCAGCAAGACAGACAGGAAATGCGAGAGCTCGGTGCGCCAAAACACGACAGCGATAAGACCTGGAAATGTCGCGGCGATACTGGC	399 	\|\|\|\|\|\|\|\|\|\|\|\|\|\|\|\|\|\|\|\|\|\|\|\|\|\|\|\|\|\|\|\|\|\|\|\|\|\|\|\|\|\|\|\|\|\|\|\|\|\|\|\|\|\|\|\|\|\|\|\|\|\|\|\|\|\|\|\|\|\|\|\|\|\|\|\|\|\|\|\|\|\|\|\|\|\|\|\|\|\|\|\|\|\|\|\|\|\|\|\|\|\|\|\|\|\|\|\|\|\|\|\|\|\|\|\|\|\|\|\|\|\|\|\|\|\|\|\|\|\|\|\|\|\|\|\|\|\|\|\|\|\|\|\|\|\|\|\|\|\|\|\|\|\|\|\|\|\|\|\|\|\|\|\|\|\|\|\|\|\|\|\|\|\|\|\|\|\|\|\|\|\|\|\|\|\|\|\|\|\|\|\|\|\|\|\|\|\|\|\|\|\|\|\|\|\|\|\|\|\|\|\|\|\|\|\|\|\|\|\|\|\|\|\|\|\|\|\|\|\|\|\|\|\|\|\|\|\|\|\|\|\|\|\|\|\|\|\|\|\|\|\|\|\|\|\|\|\|\|\|\|\|\|\|\|\|\|\|\|\|\|\|\|\|\|\|\|\|\|\|\|\|\|\|\|\|\|\|\|\|\|\|\|\|\|\|\|\|\|\|\|\|\|\|\|\|\|\|\|\|\|\|\|\|\|\|\|\|\|\|\|\|\|\|\|\|\|\|\|\|\|\|\|\|\|\| \|\|\|\|\|\|\|\|\|\|\|\|\|\|\|\| \|\|\|\|\|\|\|\|\|\|\|\|\|\|\|\|\|\|\|\|\|\|\|\|\|\|\|\|\|\|\| \|\|\|\|\|\|\|\|\|\|\|\|\| 	ATGATCGGAGACAAGTCGGACCACTCGGAACGTGAGGAGAATGGCGAGAGCGTCCACGATGGAGACGACAAGAGATCCTTTGATGCTGCTGAAGACGAGCAAAGGCCTCGTACTCCACCGCCGGCCTACCGACATCTAGAGCGCGCGCCATCTTACGCTACGCAAGACACGACGGCAGATCAGACCTTTAAGCGTGAAGAGGTGACCAGCTCGCAATCGACCTCGGCCGGCTTTGCGTCCGCAGCCAGGATTAATGCTGTAATGGGCCCTCCTCCGCCAAAGGAGGAGAAGAAGACCAGCCGATGGAAGCGCTGGCAGCAAGACAGACAGGAAATGNGAGAGCTCGGTGCGCCNAAACACGACAGCGATAAGACCTGGAAATGTCTCGGCGATACTGGC ``` |
| 2 | ggggacaagtttgtacaaaaaagcaggcttgatgCTTCTCCCTCTCTACCTCCT, ggggaccactttgtacaagaaagctgggtcCTCCACTTTCTTCCGAGGTG | Mycgr3\|90809  JGI  Ensembl |  |  | 100 % |  | ``` 1	ATGCTTCTCCCTCTCTACCTCCTCCCTGTCCTCCTCACCCTTACCTCAGCCCGCCAAACCCGCTGCGTCTGCCAAATACCCGAAGGTTCCGACGACTGCAGAGTCGACATCGACACTACGCTGGATGATGTGGATTGCGGCGAGCACTGGGTCAAAAATGCTTGCTCCGGCTCCACTGCTCCGTTCGTGGGCGTCAAGTGTCGCATTTCGAAGGGCTTGCTATCGAGTACTGTGATGAGAATCATCATAGCGATTGTCCAAAGGCAGACGAGGGAGGGGGCTTGTCCACCTCGGAAGAAAGTGGAG	306 	\|\|\|\|\|\|\|\|\|\|\|\|\|\|\|\|\|\|\|\|\|\|\|\|\|\|\|\|\|\|\|\|\|\|\|\|\|\|\|\|\|\|\|\|\|\|\|\|\|\|\|\|\|\|\|\|\|\|\|\|\|\|\|\|\|\|\|\|\|\|\|\|\|\|\|\|\|\|\|\|\|\|\|\|\|\|\|\|\|\|\|\|\|\|\|\|\|\|\|\|\|\|\|\|\|\|\|\|\|\|\|\|\|\|\|\|\|\|\|\|\|\|\|\|\|\|\|\|\|\|\|\|\|\|\|\|\|\|\|\|\|\|\|\|\|\|\|\|\|\|\|\|\|\|\|\|\|\|\|\|\|\|\|\|\|\|\|\|\|\|\|\|\|\|\|\|\|\|\|\|\|\|\|\|\|\|\|\|\|\|\|\|\|\|\|\|\|\|\|\|\|\|\|\|\|\|\|\|\|\|\|\|\|\|\|\|\|\|\|\|\|\|\|\|\|\|\|\|\|\|\|\|\|\|\|\|\|\|\|\|\|\|\|\|\|\|\|\|\|\|\|\|\|\|\|\|\|\|\|\|\|\|\|\|\|\|\|\|\|\|\|\|\|\|\|\|\|\|\|\|\|\|\|\|\|\|\|\|\|\|\|\|\|\|\|\|\|\|\|\|\|\|\|\|\|\| 	ATGCTTCTCCCTCTCTACCTCCTCCCTGTCCTCCTCACCCTTACCTCAGCCCGCCAAACCCGCTGCGTCTGCCAAATACCCGAAGGTTCCGACGACTGCAGAGTCGACATCGACACTACGCTGGATGATGTGGATTGCGGCGAGCACTGGGTCAAAAATGCTTGCTCCGGCTCCACTGCTCCGTTCGTGGGCGTCAAGTGTCGCATTTCGAAGGGCTTGCTATCGAGTACTGTGATGAGAATCATCATAGCGATTGTCCAAAGGCAGACGAGGGAGGGGGCTTGTCCACCTCGGAAGAAAGTGGAG ``` |
| 2 | ggggacaagtttgtacaaaaaagcaggcttgatgGATCAAAGCGCAGGTGTCAC, ggggaccactttgtacaagaaagctgggtcCCTCTCCACCTCGTCCAGCG | Mycgr3\|91216  JGI  Ensembl |  |  | 100 % |  | ``` 1	ATGGATCAAAGCGCAGGTGTCACGACTGTACTCCCTCTCGAGCACCTCGCCCGGAAGGGAGTCCGTACCGCTCGACCGAACCACGTATCTGAATTTGTATACTGCGATACATCAAAGCAGCACACCCTGGCTCTGGACTTCTACCCAGTCGATCGATGGCAAGTCGATCCGGTCTGGTTGCAAGGCGGGCCCAAGAGAGACGAAATTCGCGAAAGCGCACACGGCGTCTTTCGCGAGCATGGATGGCCTAGTCTAGCCGCATACCGGAAGAGCGAATGTCTGCAAGCGCTCAAGAAGGCGCTGGACGAGGTGGAGAGG	318 	\|\|\|\|\|\|\|\|\|\|\|\| \|\| \|\|\|\|\|\|\|\|\|\|\|\|\|\|\|\|\|\|\|\|\|\|\|\|\|\|\|\|\|\|\|\|\|\|\|\|\|\|\|\|\|\|\|\|\|\|\|\|\|\|\|\|\|\|\|\|\|\|\|\|\|\|\|\|\|\|\|\|\|\|\|\|\|\|\|\|\|\|\|\|\|\|\|\|\|\|\|\|\|\|\|\|\|\|\|\|\|\|\|\|\|\|\|\|\|\|\|\|\|\|\|\|\|\|\|\|\|\|\|\|\|\|\|\|\|\|\|\|\|\|\|\|\|\|\|\|\|\|\|\|\|\|\|\|\|\|\|\|\|\|\|\|\|\|\|\|\|\|\|\|\|\|\|\|\|\|\|\|\|\|\|\|\|\|\|\|\|\|\|\|\|\|\|\|\|\|\|\|\|\|\|\|\|\|\|\|\|\|\|\|\|\|\|\|\|\|\|\|\|\|\|\|\|\|\|\|\|\|\|\|\|\|\|\|\|\|\|\|\|\|\|\|\|\|\|\|\|\|\|\|\|\|\|\|\|\|\|\|\|\|\|\|\|\|\|\|\|\|\|\|\|\|\|\|\|\|\|\|\|\|\|\|\|\|\|\|\|\|\|\|\|\|\|\|\|\|\|\|\|\|\|\|\|\|\|\|\|\|\|\|\|\| 	ATGGATCAAAGCNCATGTGTCACGACTGTACTCCCTCTCGAGCACCTCGCCCGGAAGGGAGTCCGTACCGCTCGACCGAACCACGTATCTGAATTTGTATACTGCGATACATCAAAGCAGCACACCCTGGCTCTGGACTTCTACCCAGTCGATCGATGGCAAGTCGATCCGGTCTGGTTGCAAGGCGGGCCCAAGAGAGACGAAATTCGCGAAAGCGCACACGGCGTCTTTCGCGAGCATGGATGGCCTAGTCTAGCCGCATACCGGAAGAGCGAATGTCTGCAAGCGCTCAAGAAGGCGCTGGACGAGGTGGAGAGG ``` |
| 2 | ggggacaagtttgtacaaaaaagcaggcttgatgGTCACTCCCTTCACGACCTT, ggggaccactttgtacaagaaagctgggtcCATCTCCTGCCCGTTGGCGG | Mycgr3\|91290  JGI  Ensembl |  |  | 100 % |  | ``` 1	ATGGTCACTCCCTTCACGACCTTCTCGCCGCGGGAGCACCGGGACACCAGCGACACGGACTACTCCATCCTTGCCAACTTTGGTGCTGGCTGCTGGCTCGTCCTGCCTAAGCTCCAGCTCCGCGTCCACCTCCAGCCCTACGACCTCGTGATCTTCCAGACGAACAGCTTAACCCACGCCACGGCGGCTGTCGACGGCGATTGCGAGGCGGATCAGCGCTGGAGCCTCTCGTACTATCAGCGGAAGGCGGTCCGGAACGACTGCCTTGGCGCCTCGCCGACCTACGCCGCCAACGGGCAGGAGATG	306 	\|\|\|\|\|\|\|\|\|\|\|\|\|\|\|\|\|\|\|\|\|\|\|\|\|\|\|\|\|\|\|\|\|\|\|\|\|\|\|\|\|\|\|\|\|\|\|\|\|\|\|\|\|\|\|\|\|\|\|\|\|\|\|\|\|\|\|\|\|\|\|\|\|\|\|\|\|\|\|\|\|\|\|\|\|\|\|\|\|\|\|\|\|\|\|\|\|\|\|\|\|\|\|\|\|\|\|\|\|\|\|\|\|\|\|\|\|\|\|\|\|\|\|\|\|\|\|\|\|\|\|\|\|\|\|\|\|\|\|\|\|\|\|\|\|\|\|\|\|\|\|\|\|\|\|\|\|\|\|\|\|\|\|\|\|\|\|\|\|\|\|\|\|\|\|\|\|\|\|\|\|\|\|\|\|\|\|\|\|\|\|\|\|\|\|\|\|\|\|\|\|\|\|\|\|\|\|\|\|\|\|\|\|\|\|\|\|\|\|\|\|\|\|\|\|\|\|\|\|\|\|\|\|\|\|\|\|\|\|\|\|\|\|\|\|\|\|\|\|\|\|\|\|\|\|\|\|\|\|\|\|\|\|\|\|\|\|\|\|\|\|\|\|\|\|\|\|\|\|\|\|\|\|\|\|\|\|\|\|\|\|\|\|\|\|\|\|\|\|\|\|\|\|\|\|\| 	ATGGTCACTCCCTTCACGACCTTCTCGCCGCGGGAGCACCGGGACACCAGCGACACGGACTACTCCATCCTTGCCAACTTTGGTGCTGGCTGCTGGCTCGTCCTGCCTAAGCTCCAGCTCCGCGTCCACCTCCAGCCCTACGACCTCGTGATCTTCCAGACGAACAGCTTAACCCACGCCACGGCGGCTGTCGACGGCGATTGCGAGGCGGATCAGCGCTGGAGCCTCTCGTACTATCAGCGGAAGGCGGTCCGGAACGACTGCCTTGGCGCCTCGCCGACCTACGCCGCCAACGGGCAGGAGATG ```      ``` 1	ATGGTCAC-TCCCTTCACGACCTTCTCGCCGCGGGAGCACCGGGACACCAGCGACACGGACTACTCCATCCTTGCCAACTTTGGTGCTGGCTGCTGGCTCGTCCTGCCTAAGCTCCAGCTCCGCGTCCACCTCCAGCCCTACGACCTCGTGATCTTCCAGACGAACAGCTTAACCCACGCCACGGCGGCTGTCGACGGCGATTGCGAGGCGGATCAGCGCTGGAG	224 	\|\|\|\|\|\|\|\| \|\|\|\|\|\|\|\|\|\|\|\|\|\|\|\|\|\|\|\|\|\|\|\|\|\|\|\|\|\|\|\|\|\|\|\|\|\|\|\|\|\|\|\|\|\|\|\|\|\|\|\|\|\|\|\|\|\|\|\|\|\|\|\|\|\|\|\|\|\|\|\|\|\|\|\|\|\|\|\|\|\|\|\|\|\|\|\|\|\|\|\|\|\|\|\|\|\|\|\|\|\|\|\|\|\|\|\|\|\|\|\|\|\|\|\|\|\|\|\|\|\|\|\|\|\|\|\|\|\|\|\|\|\|\|\|\|\|\|\|\|\|\|\|\|\|\|\|\|\|\|\|\|\|\|\|\|\|\|\|\|\|\|\|\|\|\|\|\|\|\|\|\|\|\|\|\|\|\|\|\|\| \|\|\|\|\|\| \|\|\|\|\|  \| \|\|\|\|\|\|\|\|\| \| \|\|\|\|\| 	ATGGTCACATCCCTTCACGACCTTCTCGCCGCGGGAGCACCGGGACACCAGCGACACGGACTACTCCATCCTTGCCAACTTTGGTGCTGGCTGCTGGCTCGTCCTGCCTAAGCTCCAGCTCCGCGTCCACCTCCAGCCCTACGACCTCGTGATCTTCCAGACGAACAGCTTAACCCACGCCACGGCGGCTGNCGACGGAGATTGNNANGCGGATCAGNGATGGAG ```      ``` 1	ATGGTCACTCCCTTCACGACCTTCTCGCCGCGGGAGCACCGGGACACCAGCGACACGGACTACTCCATCCTTGCCAACTTTGGTGCTGGCTGCTGGCTCGTCCTGCCTAAGCTCCAGCTCCGCGTCCACCTCCAGCCCTACGACCTCGTGATCTTCCAGACGAACAGCTTAACCCACGCCACGGCGGCTGTCGACGGCGATTGCGAGGCGGATCAGCGCTGGAGCCTCTCGTACTATCAGCGGAAGGCGGTCCGGAACGACTGCCTTGGCGCCTCGCCGACCTACGCCGCCAACGGGCAGGAGATG	306 	\|\|\|\|\|\|\|\|\|\|\|\|\|\|\|\|\|\|\|\|\|\|\|\|\|\|\|\|\|\|\|\|\|\|\|\|\|\|\|\|\|\|\|\|\|\|\|\|\|\|\|\|\|\|\|\|\|\|\|\|\|\|\|\|\|\|\|\|\|\|\| \|\|\|\|\|\|\|\|\|\|\|\|\|\| \|\|\|\|\|\|\|\|\|\|\|\|\|\|\|\|\|\|\|\|\| \|\|\|\|\|\|\|\|\|\|\|\|\|\|\|\|\|\|\|\|\|\|\|\|\|\|\|\|\|\|\|\|\|\|\|\|\|\|\|\| \|\|\|\|\|\|\|\|\|\|\|\|\|\|\|\|\|\|\|\| \|\|\|\|\|\|\|\|\|\|\| \|\|\|\|\| \|\|\|\|\|\|\|\|\|\|\|\|\|\|\|\|\|\|\|\|\|\|\|\|\|\|\|\|\|\|\|\|\|\|\|\|\|\|\|\|\|\|\|\|\|\|\|\|\|\|\|\|\|\|\|\|\|\|\|\|\|\|\|\|\|\|\|\|\|\|\|\|\|\|\|\|\|\|\|\|\|\|\|\|\|\|\|\|\|\|\|\|\|\|\|\|\|\|\|\|\|\|\|\|\|\|\|\|\|\|\|\|\|\|\|\|\| 	ATGGTCACTCCCTTCACGACCTTCTCGCCGCGGGAGCACCGGGACACCAGCGACACGGACTACTCCATCCTCGCCAACTTTGGTGCCGGCTGCTGGCTCGTCCTGCCTGAGCTCCAGCTCCGCGTCCACCTCCAGCCCTACGACCTCGTAATCTTCCAGACGAACAGCTTGACCCACGCCACAGCGGCCGTCGACGGCGATTGCGAGGCGGATCAGCGCTGGAGCCTCTCGTACTATCAGCGGAAGGCGGTCCGGAACGACTGCCTTGGCGCCTCGCCGACCTACGCCGCCAACGGGCAGGAGATG ``` |
| 2 | ggggacaagtttgtacaaaaaagcaggcttgatgGAGATGTGCTTTCCAGTGGT, ggggaccactttgtacaagaaagctgggtcTGCAGCATGCACGCACGGTT | Mycgr3\|91551  JGI  Ensembl |  |  | 100 % |  | ``` 1	ATGGAGATGTGCTTTCCAGTGGTGTCCACTTTTCCCACCTTCACTCCGAGGCTCGCAGTACTCCACCCTTGTTGGGCAGAACAAGCAGAGTTCAGTCCAATACTCACGAAAGGGTCTGAGAACAACCGCATTCTCTCCGCTTCAACCATTCCATTCCTCAACACCATGCCTCTTCACCAACTCATCTCCTCCGACCAGGTTCCTTCTCTAACCTTCAGGCAAGGGAAGAAGCAGCGGCGATCAACCTCACAGTTGTACAACATCTCCACCATCCGCGCGGGCGATCTCCCACCCTTTCAACCGTGCGTGCATGCTGCA	318 	\|\|\|\|\|\|\|\|\|\|\|\|\|\|\|\|\|\|\|\|\|\|\|\|\|\|\|\|\|\|\|\|\|\|\|\|\|\|\|\|\|\|\|\|\|\|\|\|\|\|\|\|\|\|\|\|\|\|\|\|\|\|\|\|\|\|\|\|\|\|\|\|\|\|\|\|\|\|\|\|\|\|\|\|\|\|\|\|\|\|\|\|\|\|\|\|\|\|\|\|\|\|\|\|\|\|\|\|\|\|\|\|\|\|\|\|\|\|\|\|\|\|\|\|\|\|\|\|\|\|\|\|\|\|\|\|\|\|\|\|\|\|\|\|\|\|\|\|\|\|\|\|\|\|\|\|\|\|\|\|\|\|\|\|\|\|\|\|\|\|\|\|\|\|\|\|\|\|\|\|\|\|\|\|\|\|\|\|\|\|\|\|\|\|\|\|\|\|\|\|\|\|\|\|\|\|\|\|\|\|\|\|\|\|\|\|\|\|\|\|\|\|\|\|\|\|\|\|\|\|\|\|\|\|\|\|\|\|\|\|\|\|\|\|\|\|\|\|\|\|\|\|\|\|\|\|\|\|\|\|\|\|\|\|\|\|\|\|\|\|\|\|\|\|\|\|\|\|\|\|\|\|\|\|\|\|\|\|\|\|\|\|\|\|\|\|\|\|\|\|\|\|\|\|\|\|\|\|\|\|\|\|\|\|\|\|\|\| 	ATGGAGATGTGCTTTCCAGTGGTGTCCACTTTTCCCACCTTCACTCCGAGGCTCGCAGTACTCCACCCTTGTTGGGCAGAACAAGCAGAGTTCAGTCCAATACTCACGAAAGGGTCTGAGAACAACCGCATTCTCTCCGCTTCAACCATTCCATTCCTCAACACCATGCCTCTTCACCAACTCATCTCCTCCGACCAGGTTCCTTCTCTAACCTTCAGGCAAGGGAAGAAGCAGCGGCGATCAACCTCACAGTTGTACAACATCTCCACCATCCGCGCGGGCGATCTCCCACCCTTTCAACCGTGCGTGCATGCTGCA ``` |
| 2 | ggggacaagtttgtacaaaaaagcaggcttgatgTCTCCCCACGCCAGCCAAGC, ggggaccactttgtacaagaaagctgggtcCTTCTTGCCCTTCCCCATCT | Mycgr3\|91610  JGI  Ensembl |  |  | 100 % |  | ``` 1	ATGTCTCCCCACGCCAGCCAAGCCCGCTCCGTCTACCGCCGTCTCCTCCGCGAACTCCCCGCTCGCACACCCTCCCTCCTCGCCAACCCATCGCCAATGCAAAAGCACATCCGCGCCGACTTCTCCGCATCGACAGACTCCGCGTCACTTCAACATCAAGCGACAAAGCCGGTGGAACGACGACTGGAAGAGGCAGAGGAATACGTCAAGTATCTGGCTGGTCAGCGGATGTACACGACTTTGATCGAGAGGTACAATCCGGGCATGAACATGACGGAGGAGGATCGTGTCAGGTTGACGGCGAGGAGGGTGGGAATGGATTTACCGATTGAAGTTCTCAATCAGATGGGGAAGGGCAAGAAG	363 	\|\|\|\|\|\|\|\|\|\|\|\|\|\|\|\|\|\|\|\|\|\|\|\|\|\|\|\|\|\|\|\|\|\|\|\|\|\|\|\|\|\|\|\|\|\|\|\|\|\|\|\|\|\|\|\|\|\|\|\|\|\|\|\|\|\|\|\|\|\|\|\|\|\|\|\|\|\|\|\|\|\|\|\|\|\|\|\|\|\|\|\|\|\|\|\|\|\|\|\|\|\|\|\|\|\|\|\|\|\|\|\|\|\|\|\|\|\|\|\|\|\|\|\|\|\|\|\|\|\|\|\|\|\|\|\|\|\|\|\|\|\|\|\|\|\|\|\|\|\|\|\|\|\|\|\|\|\|\|\|\|\|\|\|\|\|\|\|\|\|\|\|\|\|\|\|\|\|\|\|\|\|\|\|\|\|\|\|\|\|\|\|\|\|\|\|\|\|\|\|\|\|\|\|\|\|\|\|\|\|\|\|\|\|\|\|\|\|\|\|\|\|\|\|\|\|\|\|\|\|\|\|\|\|\|\|\|\|\|\|\|\|\|\|\|\|\|\|\|\|\|\|\|\|\|\|\|\|\|\|\|\|\|\|\|\|\|\|\|\|\|\|\|\|\|\|\|\|\|\|\|\|\|\|\|\|\|\|\|\|\|\|\|\|\|\|\|\|\|\|\|\|\|\|\|\|\|\|\|\|\|\|\|\|\|\|\|\|\|\|\|\|\|\|\|\|\|\|\|\|\|\|\|\|\|\|\|\|\|\|\|\|\|\|\|\|\|\|\|\|\|\|\|\|\|\|\|\|\|\|\|\|\| 	ATGTCTCCCCACGCCAGCCAAGCCCGCTCCGTCTACCGCCGTCTCCTCCGCGAACTCCCCGCTCGCACACCCTCCCTCCTCGCCAACCCATCGCCAATGCAAAAGCACATCCGCGCCGACTTCTCCGCATCGACAGACTCCGCGTCACTTCAACATCAAGCGACAAAGCCGGTGGAACGACGACTGGAAGAGGCAGAGGAATACGTCAAGTATCTGGCTGGTCAGCGGATGTACACGACTTTGATCGAGAGGTACAATCCGGGCATGAACATGACGGAGGAGGATCGTGTCAGGTTGACGGCGAGGAGGGTGGGAATGGATTTACCGATTGAAGTTCTCAATCAGATGGGGAAGGGCAAGAAG ``` |
| 2 | ggggacaagtttgtacaaaaaagcaggcttgatgAGATGCTTCTATGAGCGGCA, ggggaccactttgtacaagaaagctgggtcGTGCAGCAGCGCGAGCATTG | Mycgr3\|91804  JGI  Ensembl |  |  | 0 % |  | ``` 1	ATGAGATGCTTCTATGAGCGGCACTATTCTGACGACGATGAGGACAGTTCATTTGACGCATATGGTCTGCCTCGCGAGGACGGCGAAACGGCCTCCAGCTCGGGAAGCGAGCCCCAGTCTCAACAGCGAGGCAAGATCGAGAAGCTGCCACTGCCAGATGAAATCTGCAATGCAATTGATGGTATCTGCGAGGCCACATGGAGCAAATGGGCGGCTTCAGGACCACTCACCGGACCATTCGCTGATCCTCAGCTGTCAATCGACGACTCTGGACCCATCAGACTTCCGCTGAGCGAGACCGTCCGCACATCGGGTTCGAGGCCTGATTACGGGCCTTCGAAGCAGACCTTCCCTGAAGGCATGCTCTTGATTTCGCAATGCCGCTCACCATACCCGGCAATGCTCGCGCTGCTGCAC	417 	\|\|\|\|\|\|\|\|\|\|\|\|\|\|\|\|\|\|\|\|\|\|\|\|\|\|\|\|\|\|\|\|\|\|\|\|\|\|\|\|\|\|\|\|\|\|\|\|\|\|\|\|\|\|\|\|\|\|\|\|\|\|\|\|\|\|\|\|\|\|\|\|\|\|\|\|\|\|\|\|\|\|\|\|\|\|\|\|\|\|\|\|\|\|\|\|\|\|\|\|\|\|\|\|\|\|\|\|\|\|\|\|\|\|\|\|\|\|\|\|\|\|\|\|\|\|\|\|\|\|\|\|\|\|\|\|\|\|\|\|\|\|\|\|\|\|\|\|\|\|\|\|\|\|\|\|\|\|\|\|\|\|\|\|\|\|\|\|\|\|\|\|\|\|\|\|\|\|\|\|\|\|\|\|\|\|\|\|\|\|\|\|\|\|\|\|\|\|\|\|\|\|\|\|\|\|\|\|\|\|\|\|\|\|\|\|\|\|\|\|\|\|\|\|\|\|\|\|\|\|\|\|\|\|\|\|\|\|\|\|\|\|\|\|\|\|\|\|\|\|\|\|\|\|\|\|\|\|\|\|\|\|\|\|\|\|\|\|\|\|\|\|\|\|\|\|\|\|\|\|\|\|\|\|\|\|\|\|\|\|\|\|\|\|\|\|\|\|\|\|\|\|\|\|\|\|\|\|\|\|\|\|\|\|\|\|\|\|\|\|\|\|\|\|\|\|\|\|\|\|\|\|\|\|\|\|\|\|\|\|\|\|\|\|\|\|\|\|\|\|\|\|\|\|\|\|\|\|\|\|\|\|\|\|\|\|\|\|\|\|\|\|\|\|\|\|\|\|\|\|\|\|\|\|\|\|\|\|\|\|\|\|\|\|\|\|\|\| \|\|\|\|\|\|\|\|\|\|\|\|\|\|\|\|\|\| 	ATGAGATGCTTCTATGAGCGGCACTATTCTGACGACGATGAGGACAGTTCATTTGACGCATATGGTCTGCCTCGCGAGGACGGCGAAACGGCCTCCAGCTCGGGAAGCGAGCCCCAGTCTCAACAGCGAGGCAAGATCGAGAAGCTGCCACTGCCAGATGAAATCTGCAATGCAATTGATGGTATCTGCGAGGCCACATGGAGCAAATGGGCGGCTTCAGGACCACTCACCGGACCATTCGCTGATCCTCAGCTGTCAATCGACGACTCTGGACCCATCAGACTTCCGCTGAGCGAGACCGTCCGCACATCGGGTTCGAGGCCTGATTACGGGCCTTCGAAGCAGACCTTCCCTGAAGGCATGCTCTTGATTTCGCAATGCCGCTCACCATACCCGGCNATGCTCGCGCTGCTGCAC ``` |
| 2 | ggggacaagtttgtacaaaaaagcaggcttgatgGCTTACGAGACCTATCGCTA, ggggaccactttgtacaagaaagctgggtcAATTAGAAATCGTCGAGAAA | Mycgr3\|91834  JGI  Ensembl |  |  | 100 % |  | ``` 1	ATGGCTTACGAGACCTATCGCTACGGCACACCGCGCTTCATGGACGATAGCGTCTATCGCTCCGGGTCCTCCGTATCTTCATCAGGCTCAATCACAAGAGATCCTCGCAACCCACGGACACCCTACATCTCTTCACCGCGACCCGAATACCGCCAAATCGCCGTGGACGAACACGGAATGGCCATCGATCCTCGTCGACAATACGAGGACAGCTTGTATCAACCGCACGGCCAAATGTCTCGATCATATCGCGATGAACAACGACGACGGCAAGCACGCCTAAAACTCCAAGGTTTTTCTCGACGATTTCTAATT	315 	\|\|\|\|\|\|\|\|\|\|\|\|\|\|\|\|\|\|\|\|\|\|\|\|\|\|\|\|\|\|\|\|\|\|\|\|\|\|\|\|\|\|\|\|\|\|\|\|\|\|\|\|\|\|\|\|\|\|\|\|\|\|\|\|\|\|\|\|\|\|\|\|\|\|\|\|\|\|\|\|\|\|\|\|\|\|\|\|\|\|\|\|\|\|\|\|\|\|\|\|\|\|\|\|\|\|\|\|\|\|\|\|\|\|\|\|\|\|\|\|\|\|\|\|\|\|\|\|\|\|\|\|\|\|\|\|\|\|\|\|\|\|\|\|\|\|\|\|\|\|\|\|\|\|\|\|\|\|\|\|\|\|\|\|\|\|\|\|\|\|\|\|\|\|\|\|\|\|\|\|\|\|\|\|\|\|\|\|\|\|\|\|\|\|\|\|\|\|\|\|\|\|\|\|\|\|\|\|\|\|\|\|\|\|\|\|\|\|\|\|\|\|\|\|\|\|\|\|\|\|\|\|\|\|\|\|\|\|\|\|\|\|\|\|\|\|\|\|\|\|\|\|\|\|\|\|\|\|\|\|\|\|\|\|\|\|\|\|\|\|\|\|\|\|\|\|\|\|\|\|\|\|\|\|\|\|\|\|\|\|\|\|\|\|\|\|\|\|\|\|\|\|\|\|\|\|\|\|\|\|\|\|\|\|\| 	ATGGCTTACGAGACCTATCGCTACGGCACACCGCGCTTCATGGACGATAGCGTCTATCGCTCCGGGTCCTCCGTATCTTCATCAGGCTCAATCACAAGAGATCCTCGCAACCCACGGACACCCTACATCTCTTCACCGCGACCCGAATACCGCCAAATCGCCGTGGACGAACACGGAATGGCCATCGATCCTCGTCGACAATACGAGGACAGCTTGTATCAACCGCACGGCCAAATGTCTCGATCATATCGCGATGAACAACGACGACGGCAAGCACGCCTAAAACTCCAAGGTTTTTCTCGACGATTTCTAATT ``` |
| 2 | ggggacaagtttgtacaaaaaagcaggcttgatgTATAATCTCCAAACATGCCG, ggggaccactttgtacaagaaagctgggtcAAACCCGAGGGCGGAGTGGG | Mycgr3\|91932  JGI  Ensembl |  |  | 100 % |  | ``` 1	ATGTATAATCTCCAAACATGCCGACCATCAATCGCAACAATCGACAGAATTGGCGAACGAAACACAACCATCCAAGAATGGCAAGCAAGCGTGGAAGTCGGAATGCAATCCGTTTGCAAAGCGCCATGTCCCACACAATTCATCTCGTATCCAATCCGGCAGTCGGAGAAGCAAGAGGTCCAAAACGACCAATCCATTATCGTACTCATCATCCACAAGTCGATCGAGCCATTGATCGTCTGCCTCTTCCTCCTCAACATTTCCCTGAGCATGTCCTTCCTGGTCCCCATCTTCCAGCCCTTGCACATCCCAACTTCCCACTCCGCCCTCGGGTTT	336 	\|\|\|\|\|\|\|\|\|\|\|\|\|\|\|\|\|\|\|\|\|\|\|\|\|\|\|\|\|\|\|\|\|\|\|\|\|\|\|\|\|\|\|\|\|\|\|\|\|\|\|\|\|\|\|\|\|\|\|\|\|\|\|\|\|\|\|\|\|\|\|\|\|\|\|\|\|\|\|\|\|\|\|\|\|\|\|\|\|\|\|\|\|\|\|\|\|\|\|\|\|\|\|\|\|\|\|\|\|\|\|\|\|\|\|\|\|\|\|\|\|\|\|\|\|\|\|\|\|\|\|\|\|\|\|\|\|\|\|\|\|\|\|\|\|\|\|\|\|\|\|\|\|\|\|\|\|\|\|\|\|\|\|\|\|\|\|\|\|\|\|\|\|\|\|\|\|\|\|\|\|\|\|\|\|\|\|\|\|\|\|\|\|\|\|\|\|\|\|\|\|\|\|\|\|\|\|\|\|\|\|\|\|\|\|\|\|\|\|\|\|\|\|\|\|\|\|\|\|\|\|\|\|\|\|\|\|\|\|\|\|\|\|\|\|\|\|\|\|\|\|\|\|\|\|\|\|\|\|\|\|\|\|\|\|\|\|\|\|\|\|\|\|\|\|\|\|\|\|\|\|\|\|\|\|\|\|\|\|\|\|\|\|\|\|\|\|\|\|\|\|\|\|\|\|\|\|\|\|\|\|\|\|\|\|\|\|\|\|\|\|\|\|\|\|\|\|\|\|\|\|\|\|\|\|\| 	ATGTATAATCTCCAAACATGCCGACCATCAATCGCAACAATCGACAGAATTGGCGAACGAAACACAACCATCCAAGAATGGCAAGCAAGCGTGGAAGTCGGAATGCAATCCGTTTGCAAAGCGCCATGTCCCACACAATTCATCTCGTATCCAATCCGGCAGTCGGAGAAGCAAGAGGTCCAAAACGACCAATCCATTATCGTACTCATCATCCACAAGTCGATCGAGCCATTGATCGTCTGCCTCTTCCTCCTCAACATTTCCCTGAGCATGTCCTTCCTGGTCCCCATCTTCCAGCCCTTGCACATCCCAACTTCCCACTCCGCCCTCGGGTTT ``` |
| 2 | ggggacaagtttgtacaaaaaagcaggcttgatgGCAGCGTTTCCATTCGGCGA, ggggaccactttgtacaagaaagctgggtcCTTCATGTCTCCATCTAAAC | Mycgr3\|92439  JGI  Ensembl | KOG1290 Serine/threonine protein kinase; | ATP binding; protein amino acid phosphorylation;  protein kinase activity; | 0 % |  |  |
| 2 | ggggacaagtttgtacaaaaaagcaggcttgatgCAATCAACTCTCCTAACCAC, ggggaccactttgtacaagaaagctgggtcGGTGAGCATGGCTTCGCGAT | Mycgr3\|92473  JGI  Ensembl |  |  | 100 % |  | ``` 1	ATGCAATCAACTCTCCTAACCACCTGGCCCGCTTACATCCTCTTCCAAGTCTGGAATTGGCAATACATACTCAGCGGCATCCTCTCAATCATCCTCCCAATCATCCTCATTGTCTACAACGATATGAACTCGCGCCTCACACGTATCCTTCTGAGAACTGAGCCAACGATATTCTACATAATCACGTTCTTCGTCAACTTCCTCATGCACTCCGGCCTCCGTGTACCTATTCCCTACACCAACGGAACGGAGCTGACACTATTCCCACAATTCGTGCCAAATACCTGCGCGCAGATGGAAGAGCTTACCGGGAGGCGGGACACGTGGCCATGCCATCGCGAAGCCATGCTCACC	354 	\|\|\|\|\|\|\|\|\|\|\|\|\|\|\|\|\|\|\|\|\|\|\|\|\|\|\|\|\|\|\|\|\|\|\|\|\|\|\|\|\|\|\|\|\|\|\|\|\|\|\|\|\|\|\|\|\|\|\|\|\|\|\|\|\|\|\|\|\|\|\|\|\|\|\|\|\|\|\|\|\|\|\|\|\|\|\|\|\|\|\|\|\|\|\|\|\|\|\|\|\|\|\|\|\|\|\|\|\|\|\|\|\|\|\|\|\|\|\|\|\|\|\|\|\|\|\|\|\|\|\|\|\|\|\|\|\|\|\|\|\|\|\|\|\|\|\|\|\|\|\|\|\|\|\|\|\|\|\|\|\|\|\|\|\|\|\|\|\|\|\|\|\|\|\|\|\|\|\|\|\|\|\|\|\|\|\|\|\|\|\|\|\|\|\|\|\|\|\|\|\|\|\|\|\|\|\|\|\|\|\|\|\|\|\|\|\|\|\|\|\|\|\|\|\|\|\|\|\|\|\|\|\|\|\|\|\|\|\|\|\|\|\|\|\|\|\|\|\|\|\|\|\|\|\|\|\|\|\|\|\|\|\|\|\|\|\|\|\|\|\|\|\|\|\|\|\|\|\|\|\|\|\|\|\|\|\|\|\|\|\|\|\|\|\|\|\|\|\|\|\|\|\|\|\|\|\|\|\|\|\|\|\|\|\|\|\|\|\|\|\|\|\|\|\|\|\|\|\|\|\|\|\|\|\|\|\|\| \|\|\|\|\|\|\|\|\|\|\|\|\|\|\| 	ATGCAATCAACTCTCCTAACCACCTGGCCCGCTTACATCCTCTTCCAAGTCTGGAATTGGCAATACATACTCAGCGGCATCCTCTCAATCATCCTCCCAATCATCCTCATTGTCTACAACGATATGAACTCGCGCCTCACACGTATCCTTCTGAGAACTGAGCCAACGATATTCTACATAATCACGTTCTTCGTCAACTTCCTCATGCACTCCGGCCTCCGTGTACCTATTCCCTACACCAACGGAACGGAGCTGACACTATTCCCACAATTCGTGCCAAATACCTGCGCGCAGATGGAAGAGCTTACCGGGAGGCGGGACACGTGGCCATGCCATCGNGAAGCCATGCTCACC ``` |
| 2 | ggggacaagtttgtacaaaaaagcaggcttgatgTGCACAGACTTATCGTATGT, ggggaccactttgtacaagaaagctgggtcTGGGTCTCCATGATGGGCCG | Mycgr3\|92672  JGI  Ensembl |  |  | 0 % |  |  |
| 2 | ggggacaagtttgtacaaaaaagcaggcttgatgGCGGGTGACGAGAAGACCAA, ggggaccactttgtacaagaaagctgggtcTTTCTCCATCTTAACCACTT | Mycgr3\|92737  JGI  Ensembl |  |  | 0 % |  | ``` 1	ATGGCGGGTGACGAGAAGACCAAGGTCGCCGTGAACGGAGCGACGTCCGGCTACAATCCATCGACACAGGACAATATCACCATGGCTCTCCTCCAGAACGGCGGCATCAAGCGCATACAGAGCACTTTCCAACAACGCCTAGACGAAGCAGGCTGGTCGCAGGCGCTCCGCGACTACGTAGAGAAGCTGTTCCGAAGCGGAGAGGCCCAAACTTACGATGAGGCTCTCAAGATTGTTATGCAGCACATTGCGCTCCAGGGAGACGAGACGAAGTCGAATGGGAACAATGGCTCGTCGGCGCCCGACTTGACTATCCCGAGAGATGCTGCGAAGGATGCTGCGGCGACAGTGAAGAAGGAGTTGGCGCAAGTGGTTAAGATGGAGAAA	387 	\|\|\|\|\|\|\|\|\|\|\|\|\|\|\|\|\|\|\|\|\|\|\|\|\|\|\|\|\|\|\|\|\|\|\|\|\|\|\|\|\|\|\|\|\|\|\|\|\|\|\|\|\|\|\|\|\|\|\|\|\|\|\|\|\|\|\|\|\|\|\|\|\|\|\|\|\|\|\|\|\|\|\|\|\|\|\|\|\|\|\|\|\|\|\|\|\|\|\|\|\|\|\|\|\|\|\|\|\|\|\|\|\|\|\|\|\|\|\|\|\|\|\|\|\|\|\|\|\|\|\|\|\|\|\|\|\|\|\|\|\|\|\|\|\|\|\|\|\|\|\|\|\|\|\|\|\|\|\|\|\|\|\|\|\|\|\|\|\|\|\|\|\|\|\|\|\|\|\|\|\|\|\|\|\|\|\|\|\|\|\|\|\|\|\|\|\|\|\|\|\|\|\|\|\|\|\|\|\|\|\|\|\|\|\|\|\|\|\|\|\|\|\|\|\|\|\|\|\|\|\|\|\|\|\|\|\|\|\|\|\|\|\|\|\|\|\|\|\|\|\|\|\|\|\|\|\|\|\|\|\|\|\|\|\|\|\|\|\|\|\|\|\|\|\|\|\|\|\|\|\|\|\|\|\|\|\|\|\|\|\|\|\|\|\|\|\|\|\|\|\|\|\|\|\|\|\|\|\|\|\|\|\|\|\|\|\|\|\|\|\|\|\|\|\|\|\|\|\|\|\|\|\|\|\|\|\|\|\|\|\|\|\|\|\|\|\|\|\|\|\|\|\|\|\|\|\|\|\|\|\|\|\|\|\|\|\|\|\|\|\|\|\|\|\|\|\|\|\|\|\|\|\|\|\|\|\| 	ATGGCGGGTGACGAGAAGACCAAGGTCGCCGTGAACGGAGCGACGTCCGGCTACAATCCATCGACACAGGACAATATCACCATGGCTCTCCTCCAGAACGGCGGCATCAAGCGCATACAGAGCACTTTCCAACAACGCCTAGACGAAGCAGGCTGGTCGCAGGCGCTCCGCGACTACGTAGAGAAGCTGTTCCGAAGCGGAGAGGCCCAAACTTACGATGAGGCTCTCAAGATTGTTATGCAGCACATTGCGCTCCAGGGAGACGAGACGAAGTCGAATGGGAACAATGGCTCGTCGGCGCCCGACTTGACTATCCCGAGAGATGCTGCGAAGGATGCTGCGGCGACAGTGAAGAAGGAGTTGGCGCAAGTGGTTAAGATGGAGAAA ``` |
| 2 | ggggacaagtttgtacaaaaaagcaggcttgatgATCGGCTTCCCCAGGGTGAA, ggggaccactttgtacaagaaagctgggtcTCCGTCAGATCTGCGATCAG | Mycgr3\|92846  JGI  Ensembl |  |  | 0 % |  |  |
| 2 | ggggacaagtttgtacaaaaaagcaggcttgatgTCTTACGAACAACATGATGC, ggggaccactttgtacaagaaagctgggtcGATGCCAGACGCGCTGTGCA | Mycgr3\|92892  JGI  Ensembl |  |  | 100 % |  | ``` 1	ATGTCTTACGAACAACATGATGCGATGGACAAGACGGGCACGCTCTGGAGTCTGCGCCTCATGGAATCAGAGCTAAAGAAGCGGCTTGTGCAAGCCTTGGCGAGTGAGGGAAACAGCTACGCTGAGAGGACAGAGGCCATCATCGGCGAACACACAGCCGACAGCAACGCGGGCGCAGGTTTGAACGAGCAAGCCACAGCGGAGATCCGTCGGAGAATGGAAGCTCTTTCGATCGATCCTAAAGCCAAATGTCACTTCCGGGGGGCAGTCTTGCTCAAGCTATGCATGATCTCAGCGTGGAAGCTGCAGACACTGAGAGCTAACCATCTGCACAGCGCGTCTGGCATC	348 	\|\|\|\|\|\|\|\|\|\|\|\|\|\|\|\|\|\|\|\|\|\|\|\|\|\|\|\|\|\|\|\|\|\|\|\|\|\|\|\|\|\|\|\|\|\|\|\|\|\|\|\|\|\|\|\|\|\|\|\|\|\|\|\|\|\|\|\|\|\|\|\|\|\|\|\|\|\|\|\|\|\|\|\|\|\|\|\|\|\|\|\|\|\|\|\|\|\|\|\|\|\|\|\|\|\|\|\|\|\|\|\|\|\|\|\|\|\|\|\|\|\|\|\|\|\|\|\|\|\|\|\|\|\|\|\|\|\|\|\|\|\|\|\|\|\|\|\|\|\|\|\|\|\|\|\|\|\|\|\|\|\|\|\|\|\|\|\|\|\|\|\|\|\|\|\|\|\|\|\|\|\|\|\|\|\|\|\|\|\|\|\|\|\|\|\|\|\|\|\|\|\|\|\|\|\|\|\|\|\|\|\|\|\|\|\|\|\|\|\|\|\|\|\|\|\|\|\|\|\|\|\|\|\|\|\|\|\|\|\|\|\|\|\|\|\|\|\|\|\|\|\|\|\|\|\|\|\|\|\|\|\|\|\|\|\|\|\|\|\|\|\|\|\|\|\|\|\|\|\|\|\|\|\|\|\|\|\|\|\|\|\|\|\|\|\|\|\|\|\|\|\|\|\|\|\|\|\|\|\|\|\|\|\|\|\|\|\|\|\|\|\|\|\|\|\|\|\|\|\|\|\|\|\|\|\|\|\|\|\|\|\|\|\|\|\|\|\| 	ATGTCTTACGAACAACATGATGCGATGGACAAGACGGGCACGCTCTGGAGTCTGCGCCTCATGGAATCAGAGCTAAAGAAGCGGCTTGTGCAAGCCTTGGCGAGTGAGGGAAACAGCTACGCTGAGAGGACAGAGGCCATCATCGGCGAACACACAGCCGACAGCAACGCGGGCGCAGGTTTGAACGAGCAAGCCACAGCGGAGATCCGTCGGAGAATGGAAGCTCTTTCGATCGATCCTAAAGCCAAATGTCACTTCCGGGGGGCAGTCTTGCTCAAGCTATGCATGATCTCAGCGTGGAAGCTGCAGACACTGAGAGCTAACCATCTGCACAGCGCGTCTGGCATC ``` |
| 2 | ggggacaagtttgtacaaaaaagcaggcttgatgGATGAAGGGAATACCAATTC, ggggaccactttgtacaagaaagctgggtcTCGCGGACGATGGGTTTCCT | Mycgr3\|93287  JGI  Ensembl |  |  | 0 % |  | ``` 1	ATGGATGAAGGGAATACCAATTCGGCTCACGGCCTTGCTGGAATGCCGAAAGCAAAAGGCCTCGGGCAGAGTAGGAAATGGAAAGCCATGAAGGATCTCAAGAAGACAGACCCAGACCTGGCCCAGCACTTAGCAGAGGACGCTGAAGCCTCCAGCAAAATTTCGGCTTTGCGGGAGCAACAAGCAGAAGGAAAGGACGCAATGCATGCTCATCAACCGGAAGCCAGTCAACCGGTAGTGGAGAAGCCGCGCATCGAGCTTGCGTCGCAGTACGACAGTTCCGCACGTCAGGTCCTGCGCGATGAGTCGCAAGAACCACGCAAGAGTAAGGCAACAGCAGACCGCGCTGATGACGCTGCTAGTCACGACGTAGATCATGCTACGGCCGCTAGCAGCGAGGAAACCCATCGTCCGCGA	417 	\|\|\|\|\|\|\|\|\|\|\|\|\|\|\|\|\|\|\|\|\|\|\|\|\|\|\|\|\|\|\|\|\|\|\|\|\|\|\|\|\|\|\|\|\|\|\|\|\|\|\|\|\|\|\|\|\|\|\|\|\|\|\|\|\|\|\|\|\|\|\|\|\|\|\|\|\|\|\|\|\|\|\|\|\|\|\|\|\|\|\|\|\|\|\|\|\|\|\|\|\|\|\|\|\|\|\|\|\|\|\|\|\|\|\|\|\|\|\|\|\|\|\|\|\|\|\|\|\|\|\|\|\|\|\|\|\|\|\|\|\|\|\|\|\|\|\|\|\|\|\|\|\|\|\|\|\|\|\|\|\|\|\|\|\|\|\|\|\|\|\|\|\|\|\|\|\|\|\|\|\|\|\|\|\|\|\|\|\|\|\|\|\|\|\|\|\|\|\|\|\|\|\|\|\|\|\|\|\|\|\|\|\|\|\|\|\|\|\|\|\|\|\|\|\|\|\|\|\|\|\|\|\|\|\|\|\|\|\|\|\|\|\|\|\|\|\|\|\|\|\|\|\|\|\|\|\|\|\|\|\|\|\|\|\|\|\|\|\|\|\|\|\|\|\|\|\|\|\|\|\|\|\|\|\|\|\|\|\|\|\|\|\|\|\|\|\|\|\|\|\|\|\|\|\|\|\|\|\|\|\|\|\|\|\|\|\|\|\|\|\|\|\|\|\|\|\|\|\|\|\|\|\|\|\|\|\|\|\|\|\|\|\|\|\|\|\|\|\|\|\|\|\|\|\|\|\|\|\|\|\|\|\|\|\|\|\|\|\|\|\|\|\|\|\|\|\|\|\|\|\|\|\| \|\|\|\|\|\|\|\|\|\| \|\|\|\|\|\|\|\|\|\|\|\|\|\|\|\|\|\|\|\|\|\| 	ATGGATGAAGGGAATACCAATTCGGCTCACGGCCTTGCTGGAATGCCGAAAGCAAAAGGCCTCGGGCAGAGTAGGAAATGGAAAGCCATGAAGGATCTCAAGAAGACAGACCCAGACCTGGCCCAGCACTTAGCAGAGGACGCTGAAGCCTCCAGCAAAATTTCGGCTTTGCGGGAGCAACAAGCAGAAGGAAAGGACGCAATGCATGCTCATCAACCGGAAGCCAGTCAACCGGTAGTGGAGAAGCCGCGCATCGAGCTTGCGTCGCAGTACGACAGTTCCGCACGTCAGGTCCTGCGCGATGAGTCGCAAGAACCACGCAAGAGTAAGGCAACAGCAGACCGCGCTGATGACGCTGCTAGTCACGACGTAGATCATGCTACNGCCGCTAGCANCGAGGAAACCCATCGTCCGCGA ``` |
| 2 | ggggacaagtttgtacaaaaaagcaggcttgatgTCGTTGATCGACGACGGCAG, ggggaccactttgtacaagaaagctgggtcACCCTGCTCATCACTGGCAA | Mycgr3\|93312  JGI  Ensembl |  |  | 0 % |  |  |
| 2 | ggggacaagtttgtacaaaaaagcaggcttgatgCCCAACCCCACCCTCCTCAA, ggggaccactttgtacaagaaagctgggtcCTGCCTCCACAAAAACCCCC | Mycgr3\|93790  JGI  Ensembl |  |  | 100 % |  | ``` 1	ATGCCCAACCCCACCCTCCTCAAAACCCACTTCATCATCGACGAAGACAACAGCACCCGCCACACCGAAATCCGCACCGTCCTCCTCAACCGCACCTCCACCCCGCTCTTCTACATCTGCACCCGCGACGGCCGCCACCCCCTCGACCGCAACACCAAAATCCGCACCGCCAGACCCACCAGATCGGGAAGCAAGTGGGGCACTCACGATCCGGGACCTTCGGTGGCGGGCACGCGTCTCTGGTGCGATACGGAGGAGATCCGGGCGATTGCGAAGAGTCATCCGGATGAGAGGCATCGGACGACGTGGACGGATGTTAAAAATGGGAGGGATTGGAGGGTGAAACATCTGGTGGCGGCGAGGGAGTTTAGGTGGGAGGTTgggggga	388 	\|\|\|\|\|\|\|\|\|\|\|\|\|\|\|\|\|\|\|\|\|\|\|\|\|\|\|\|\|\|\|\|\|\|\|\|\|\|\|\|\|\|\|\|\|\|\|\|\|\|\|\|\|\|\|\|\|\|\|\|\|\|\|\|\|\|\|\|\|\|\|\|\|\|\|\|\|\|\|\|\|\|\|\|\|\|\|\|\|\|\|\|\|\|\|\|\|\|\|\|\|\|\|\|\|\|\|\|\|\|\|\|\|\|\|\|\|\|\|\|\|\|\|\|\|\|\|\|\|\|\|\|\|\|\|\|\|\|\|\|\|\|\|\|\|\|\|\|\|\|\|\|\|\|\|\|\|\|\|\|\|\|\|\|\|\|\|\|\|\|\|\|\|\|\|\|\|\|\|\|\|\|\|\|\|\|\|\|\|\|\|\|\|\|\|\|\|\|\|\|\|\|\|\|\|\|\|\|\|\|\|\|\|\|\|\|\|\|\|\|\|\|\|\|\|\|\|\|\|\|\|\|\|\|\|\|\|\|\|\|\|\|\|\|\|\|\|\|\|\|\|\|\|\|\|\|\|\|\|\|\|\|\|\|\|\|\|\|\|\|\|\|\|\|\|\|\|\|\|\|\|\|\|\|\|\|\|\|\|\|\|\|\|\|\|\|\|\|\|\|\|\|\|\|\|\|\|\|\|\|\|\|\|\|\|\|\|\|\|\|\|\|\|\|\|\|\|\|\|\|\|\|\|\|\|\|\|\|\|\|\|\|\|\|\|\|\|\|\|\|\|\|\|\|\|\|\|\|\|\|\|\|\|\|\|\|\|\|\|\|\|\|\|\|\|\|\|\|\|\|\|\|\|\|\|\|\|\| 	ATGCCCAACCCCACCCTCCTCAAAACCCACTTCATCATCGACGAAGACAACAGCACCCGCCACACCGAAATCCGCACCGTCCTCCTCAACCGCACCTCCACCCCGCTCTTCTACATCTGCACCCGCGACGGCCGCCACCCCCTCGACCGCAACACCAAAATCCGCACCGCCAGACCCACCAGATCGGGAAGCAAGTGGGGCACTCACGATCCGGGACCTTCGGTGGCGGGCACGCGTCTCTGGTGCGATACGGAGGAGATCCGGGCGATTGCGAAGAGTCATCCGGATGAGAGGCATCGGACGACGTGGACGGATGTTAAAAATGGGAGGGATTGGAGGGTGAAACATCTGGTGGCGGCGAGGGAGTTTAGGTGGGAGGTTGGGGGGA ``` |
| 2 | ggggacaagtttgtacaaaaaagcaggcttgatgCACTTCCTCCTCCCAACAAC, ggggaccactttgtacaagaaagctgggtcAAATCCTGACGTCGGTGCAT | Mycgr3\|93838  JGI  Ensembl |  |  | 100 % |  | ``` 1	ATGCACTTCCTCCTCCCAACAACACTCCTGCTCACCCTCTCCCTCGCCGCCCCGACCCCCACCGGCTGGATCCCCGGCGCATACTGCAACCTGACCTACTTCGGCAACGGCGGCGTTTGGATCGCACGGATCGACCAGCCATTCAACGACGGAGCCGGCTGCAAAGCCGTCAAGGCAACTCTGAAGAAGTCCATCAAGCTCTCCAACAAGTGGTCCTGCACGGACGATGGCGCTGGCATGACAAGACTCAAGTTCGCCACGAACTTCAAGAATGACAAGGTGAATCACGCGCTGAAGGAGGCGTATCCGATGTATGATTTCGTCGGGGGAAATGTCTGTGCGCCTTATGCACCGACGTCAGGATTT	366 	\|\|\|\|\|\|\|\|\|\|\|\|\|\|\|\|\|\|\|\|\|\|\|\|\|\|\|\|\|\|\|\|\|\|\|\|\|\|\|\|\|\|\|\|\|\|\|\|\|\|\|\|\|\|\|\|\|\|\|\|\|\|\|\|\|\|\|\|\|\|\|\|\|\|\|\|\|\|\|\|\|\|\|\|\|\|\|\|\|\|\|\|\|\|\|\|\|\|\|\|\|\|\|\|\|\|\|\|\|\|\|\|\|\|\|\|\|\|\|\|\|\|\|\|\|\|\|\|\|\|\|\|\|\|\|\|\|\|\|\|\|\|\|\|\|\|\|\|\|\|\|\|\|\|\|\|\|\|\|\|\|\|\|\|\|\|\|\|\|\|\|\|\|\|\|\|\|\|\|\|\|\|\|\|\|\|\|\|\|\|\|\|\|\|\|\|\|\|\|\|\|\|\|\|\|\|\|\|\|\|\|\|\|\|\|\|\|\|\|\|\|\|\|\|\|\|\|\|\|\|\|\|\|\|\|\|\|\|\|\|\|\|\|\|\|\|\|\|\|\|\|\|\|\|\|\|\|\|\|\|\|\|\|\|\|\|\|\|\|\|\|\|\|\|\|\|\|\|\|\|\|\|\|\|\|\|\|\|\|\|\|\|\|\|\|\|\|\|\|\|\|\|\|\|\|\|\|\|\|\|\|\|\|\|\|\|\|\|\|\|\|\|\|\|\|\|\|\|\|\|\|\|\|\|\|\|\|\|\|\|\|\|\|\|\|\|\|\|\|\|\|\|\|\|\|\|\|\|\|\|\|\|\|\|\|\| 	ATGCACTTCCTCCTCCCAACAACACTCCTGCTCACCCTCTCCCTCGCCGCCCCGACCCCCACCGGCTGGATCCCCGGCGCATACTGCAACCTGACCTACTTCGGCAACGGCGGCGTTTGGATCGCACGGATCGACCAGCCATTCAACGACGGAGCCGGCTGCAAAGCCGTCAAGGCAACTCTGAAGAAGTCCATCAAGCTCTCCAACAAGTGGTCCTGCACGGACGATGGCGCTGGCATGACAAGACTCAAGTTCGCCACGAACTTCAAGAATGACAAGGTGAATCACGCGCTGAAGGAGGCGTATCCGATGTATGATTTCGTCGGGGGAAATGTCTGTGCGCCTTATGCACCGACGTCAGGATTT ``` |
| 2 | ggggacaagtttgtacaaaaaagcaggcttgatgCGCGACGAGAAGACCGAAGT, ggggaccactttgtacaagaaagctgggtcATCTCTGAACCTCAGTCCAC | Mycgr3\|94064  JGI  Ensembl |  |  | 100 % |  | ``` 1	ATGCGCGACGAGAAGACCGAAGTCCAGTTCTCGTTACTGTCCGGTGAGGAGGACGAATGCCTTACCGAGAGACCGCACAAGCGTTCTCTAAGATTCTACCTTTGGGGGATCGTGCCTGCTGTGATCATTTACACTCTCGCGGTAGTGGCTATAACTTCCAGCGCTTTCGATGCCAGCTGCGCTGGTGGACCTCGCATTGTGCCCAATATTCTTCAAGACTCCATACGATATCAGCCACAGACCTGGCAGGACCACGACATGATCACTCATCCCTACTTCGGTCACCCGACCGCTGACTTGGACCGTCGATGGAAGGACCTTCTGAAATGTGAGCTGCCGAGCTCGGCCCATTGCGAGCGAGCCCGCGGTGGGGAGGGCTCGAAG-CGAGGTGGACTGAGGTTCAGAGAT	408 	\|\|\|\|\|\|\|\|\|\|\|  \| \|\|\|\|\|\|\|\|\|\|\|\|\|\|\|\|\|\|\|\|\|\|\|\|\|\|\|\|\|\|\|\|\|\|\|\|\|\|\|\|\|\|\|\|\|\|\|\|\|\|\|\|\|\|\|\|\|\|\|\|\|\|\|\|\|\|\|\|\|\|\|\|\|\|\|\|\|\|\|\|\|\|\|\|\|\|\|\|\|\|\|\|\|\|\|\|\|\|\|\|\|\|\|\|\|\|\|\|\|\|\|\|\|\|\|\|\|\|\|\|\|\|\|\|\|\|\|\|\|\|\|\|\|\|\|\|\|\|\|\|\|\|\|\|\|\|\|\|\|\|\|\|\|\|\|\|\|\|\|\|\|\|\|\|\|\|\|\|\|\|\|\|\|\|\|\|\|\|\|\|\|\|\|\|\|\|\|\|\|\|\|\|\|\|\|\|\|\|\|\|\|\|\|\|\|\|\|\|\|\|\|\|\|\|\|\|\|\|\|\|\|\|\|\|\|\|\|\|\|\|\|\|\|\|\|\|\|\|\|\|\|\|\|\|\|\|\|\|\|\|\|\|\|\|\|\|\|\|\|\|\|\|\|\|\|\|\|\|\|\|\|\|\|\|\|\|\|\|\|\|\|\|\|\|\|\|\|\|\|\|\|\|\|\|\|\|\|\|\| \|\|\|\|\|\|\|\|\|\|\|\|\|\|\|\|\|\|\|\|\|\|\|\|\|\|\|\|\|\|\|\|\|\|\|\|\|\|\|\|\|\|\|\|\|\|\|\|\|\|\|\|\|\|\|\|\|\|\|\|\|\|\|\|\| \|\|  \|\|\|\|\|\|\|\|\|\|\|\|\| \|\|\|\|\|\|\|\|\|\| 	ATGCGCGACGANNATACCGAAGTCCAGTTCTCGTTACTGTCCGGTGAGGAGGACGAATGCCTTACCGAGAGACCGCACAAGCGTTCTCTAAGATTCTACCTTTGGGGGATCGTGCCTGCTGTGATCATTTACACTCTCGCGGTAGTGGCTATAACTTCCAGCGCTTTCGATGCCAGCTGCGCTGGTGGACCTCGCATTGTGCCCAATATTCTTCAAGACTCCATACGATATCAGCCACAGACCTGGCAGGACCACGACATGATCACTCATCCCTACTTCGGTCACCCGACCGCTGACTTGGACCGTCGATGGAANGACCTTCTGAAATGTGAGCTGCCGAGCTCGGCCCATTGCGAGCGAGCCCGCGGTGGGGAGGGCTCNAAACCGAGGTGGACTGAAGTTCAGAGAT ``` |
| 2 | ggggacaagtttgtacaaaaaagcaggcttgatgCATGCCTCTTCTACCGCAGT, ggggaccactttgtacaagaaagctgggtcGCGTACCAGGCTAGATGCCC | Mycgr3\|94089  JGI  Ensembl |  |  | 100 % |  | ``` 1	ATGCATGCCTCTTCTACCGCAGTCCTCGCCTTGGGCATGTTCGGTTTCGCCACCGCCGATTTTTACATCTACGCCCAGACGAGCTATGGCCTTCAGGACGGTGGAGGAGCCGCAACTCGTGCCGACGGCTTCAAGTTCTACAACGGTCCTCCGTCCTGTGACGATGTGAATAGCTCCATCTATCAGCACGCCGTCGACGACGCCAGCTACTCCGGCGTACGGTGCAAGGGCTGCGTGAACGCAGGCGGGAGCGCCAACCCCAAGGCCAGCAGGGCGCCAACTCTGCTGGAATGGAACAACGAAATAGGACACTTCAGTGAGTGCAGATCAAAATTTATCTTGCCGTCTCGTGGACTTGTGCTGATCGGATGGGCATCTAGCCTGGTACGC	390 	\|\|\|\|\|\|\|\|\|\|\|\|\|\|\|\|\|\|\|\|\|\|\|\|\|\|\|\|\|\|\|\|\|\|\|\|\|\|\|\|\|\|\|\|\|\|\|\|\|\|\|\|\|\|\|\|\|\|\|\|\|\|\|\|\|\|\|\|\|\|\|\|\|\|\|\|\|\|\|\|\|\|\|\|\|\|\|\|\|\|\|\|\|\|\|\|\|\|\|\|\|\|\|\|\|\|\|\|\|\|\|\|\|\|\|\|\|\|\|\|\|\|\|\|\|\|\|\|\|\|\|\|\|\|\|\|\|\|\|\|\|\|\|\|\|\|\|\|\|\|\|\|\|\|\|\|\|\|\|\|\|\|\|\|\|\|\|\|\|\|\|\|\|\|\|\|\|\|\|\|\|\|\|\|\|\|\|\|\|\|\|\|\|\|\|\|\|\|\|\|\|\|\|\|\|\|\|\|\|\|\|\|\|\|\|\|\|\|\|\|\|\|\|\|\|\|\|\|\|\|\|\|\|\|\|\|\|\|\|\|\|\|\|\|\|\|\|\|\|\|\|\|\|\|\|\|\|\|\|\|\|\|\|\|\|\|\|\|\|\|\|\|\|\|\|\|\|\|\|\|\|\|\|\|\|\|\|\|\|\|\|\|\|\|\|\|\|\|\|\|\|\|\|\|\|\|\|\|\|\|\|\|\|\|\|\|\|\|\|\|\|\|\|\|\|\|\|\|\|\|\|\|\|\|\|\|\|\|\|\|\|\|\|\|\|\|\|\|\|\|\|\|\|\|\|\|\|\|\|\|\|\|\|\|\|\|\|\|\|\|\|\|\|\|\|\|\|\|\|\|\|\|\|\|\|\|\|\|\|\| 	ATGCATGCCTCTTCTACCGCAGTCCTCGCCTTGGGCATGTTCGGTTTCGCCACCGCCGATTTTTACATCTACGCCCAGACGAGCTATGGCCTTCAGGACGGTGGAGGAGCCGCAACTCGTGCCGACGGCTTCAAGTTCTACAACGGTCCTCCGTCCTGTGACGATGTGAATAGCTCCATCTATCAGCACGCCGTCGACGACGCCAGCTACTCCGGCGTACGGTGCAAGGGCTGCGTGAACGCAGGCGGGAGCGCCAACCCCAAGGCCAGCAGGGCGCCAACTCTGCTGGAATGGAACAACGAAATAGGACACTTCAGTGAGTGCAGATCAAAATTTATCTTGCCGTCTCGTGGACTTGTGCTGATCGGATGGGCATCTAGCCTGGTACGC ``` |
| 2 | ggggacaagtttgtacaaaaaagcaggcttgatgACCCACCCATTCATCCTGCT, ggggaccactttgtacaagaaagctgggtcGAGCGTGTCGCCCTCGATGA | Mycgr3\|94091  JGI  Ensembl |  |  | 100 % |  | ``` 1	ATGACCCACCCATTCATCCTGCTTTGCCAGTCCAAACTCTCCCTGATCATGGACATCCCCACAGCCGACGACTTTCTCGCCAGCAAGGACGTGATTAGCGACGATGACCCCGTGAACAACCGATCAAGGACTGCAACGGTCAAGAGGATCGGACCGTACGTTATGAAATACAACGGACGAACGTGTCTCCAGGAAGCCGACAACCTAGCGTACATCGCCGCTCACACCAAAATTCCGGTGCCAAAGTGCTACGCCTCCGGTATCGTGCCTAACAGCGAGTTAAACGGGAAGTATCTTATTATGGACTTCATCGAGGGCGACACGCTC	327 	\|\|\|\|\|\|\|\|\|\|\|\|\|\|\|\|\|\|\|\|\|\|\|\|\|\|\|\|\|\|\|\|\|\|\|\|\|\|\|\|\|\|\|\|\|\|\|\|\|\|\|\|\|\|\|\|\|\|\|\|\|\|\|\|\|\|\|\|\|\|\|\|\|\|\|\|\|\|\|\|\|\|\|\|\|\|\|\|\|\|\|\|\|\|\|\|\|\|\|\|\|\|\|\|\|\|\|\|\|\|\|\|\|\|\|\|\|\|\|\|\|\|\|\|\|\|\|\|\|\|\|\|\|\|\|\|\|\|\|\|\|\|\|\|\|\|\|\|\|\|\|\|\|\|\|\|\|\|\|\|\|\|\|\|\|\|\|\|\|\|\|\|\|\|\|\|\|\|\|\|\|\|\|\|\|\|\|\|\|\|\|\|\|\|\|\|\|\|\|\|\|\|\|\|\|\|\|\|\|\|\|\|\|\|\|\|\|\|\|\|\|\|\|\|\|\|\|\|\|\|\|\|\|\|\|\|\|\|\|\|\|\|\|\|\|\|\|\|\|\|\|\|\|\|\|\|\|\|\|\|\|\|\|\|\|\|\|\|\|\|\|\|\|\|\|\|\|\|\|\|\|\|\|\|\|\|\|\|\|\|\|\|\|\|\|\|\|\|\|\|\|\|\|\|\|\|\|\|\|\|\|\|\|\|\|\|\|\|\|\|\|\|\|\|\|\|\| 	ATGACCCACCCATTCATCCTGCTTTGCCAGTCCAAACTCTCCCTGATCATGGACATCCCCACAGCCGACGACTTTCTCGCCAGCAAGGACGTGATTAGCGACGATGACCCCGTGAACAACCGATCAAGGACTGCAACGGTCAAGAGGATCGGACCGTACGTTATGAAATACAACGGACGAACGTGTCTCCAGGAAGCCGACAACCTAGCGTACATCGCCGCTCACACCAAAATTCCGGTGCCAAAGTGCTACGCCTCCGGTATCGTGCCTAACAGCGAGTTAAACGGGAAGTATCTTATTATGGACTTCATCGAGGGCGACACGCTC ``` |
| 2 | ggggacaagtttgtacaaaaaagcaggcttgatgAAAATCCGACTTCACCTCCT, ggggaccactttgtacaagaaagctgggtcATGCGACGGATGGATGGGAT | Mycgr3\|94131  JGI  Ensembl |  |  | 100 % |  | ``` 1	ATGAAAATCCGACTTCACCTCCTTCCGCCCTTCTCCGCCCAGGAGTTCTCCAAAAGCTCCTATTCACGACCCCTCCACCCCCTCCTTCACCTCACACTCCCCAATAACCCCCTTCCGACCCGCAAGCCGCTCAAACATCCCCAGCGTATGCTCCCAAACCTTCTCCGCCAGTCCCTCCTCCCTCATCGCACGCATGCTCTTCCCACCAGCGACTGTCTCCCCATTCCAATGGACCAGATCCGCTCCGCCGCCGACGAAGCCATTCGAACCAACCAACCGCTACTCTTTGTGGCGCTGCATTTCCATCAACTTTCTCCCTCCCACTTCACACTCGTCGATGGAGGAGCAGTGCATAGCGGGATGGCCGACGCGGCTGGTGGGGAGTTACATCCCATCCATCCGTCGCAT	408 	\|\|\|\|\|\|\|\|\|\|\|\|\|\|\|\|\|\|\|\|\|\|\|\|\|\|\|\|\|\|\|\|\|\|\|\|\|\|\|\|\|\|\|\|\|\|\|\|\|\|\|\|\|\|\|\|\|\|\|\|\|\|\|\|\|\|\|\|\|\|\|\|\|\|\|\|\|\|\|\|\|\|\|\|\|\|\|\|\|\|\|\|\|\|\|\|\|\|\|\|\|\|\|\|\|\|\|\|\|\|\|\|\|\|\|\|\|\|\|\|\|\|\|\|\|\|\|\|\|\|\|\|\|\|\|\|\|\|\|\|\|\|\|\|\|\|\|\|\|\|\|\|\|\|\|\|\|\|\|\|\|\|\|\|\|\|\|\|\|\|\|\|\|\|\|\|\|\|\|\|\|\|\|\|\|\|\|\|\|\|\|\|\|\|\|\|\|\|\|\|\|\|\|\|\|\|\|\|\|\|\|\|\|\|\|\|\|\|\|\|\|\|\|\|\|\|\|\|\|\|\|\|\|\|\|\|\|\|\|\|\|\|\|\|\|\|\|\|\|\|\|\|\|\|\|\|\|\|\|\|\|\|\|\|\|\|\|\|\|\|\|\|\|\|\|\|\|\|\|\|\|\|\|\|\|\|\|\|\|\|\|\|\|\|\|\|\|\|\|\|\|\|\|\|\|\|\|\|\|\|\|\|\|\|\|\|\|\|\|\|\|\|\|\|\|\|\|\|\|\|\|\|\|\|\|\|\|\|\|\|\|\|\|\|\|\|\|\|\|\|\|\|\|\|\|\|\|\|\|\|\|\|\|\|\|\|\|\|\|\|\|\|\|\|\|\|\|\|\|\|\|\|\|\|\|\|\|\|\|\|\|\|\|\|\|\|\|\|\|\|\|\|\|\|\|\|\|\| 	ATGAAAATCCGACTTCACCTCCTTCCGCCCTTCTCCGCCCAGGAGTTCTCCAAAAGCTCCTATTCACGACCCCTCCACCCCCTCCTTCACCTCACACTCCCCAATAACCCCCTTCCGACCCGCAAGCCGCTCAAACATCCCCAGCGTATGCTCCCAAACCTTCTCCGCCAGTCCCTCCTCCCTCATCGCACGCATGCTCTTCCCACCAGCGACTGTCTCCCCATTCCAATGGACCAGATCCGCTCCGCCGCCGACGAAGCCATTCGAACCAACCAACCGCTACTCTTTGTGGCGCTGCATTTCCATCAACTTTCTCCCTCCCACTTCACACTCGTCGATGGAGGAGCAGTGCATAGCGGGATGGCCGACGCGGCTGGTGGGGAGTTACATCCCATCCATCCGTCGCAT ``` |
| 2 | ggggacaagtttgtacaaaaaagcaggcttgatgCCTCCCCGAAACGAAGCAAG, ggggaccactttgtacaagaaagctgggtcTCTACCAGCCGCTCTACGCG | Mycgr3\|94291  JGI  Ensembl |  |  | 0 % |  | ``` 1	ATGCCTCCCCGAAACGAAGCAAGCTACATCCGTACCCGCGCCGAGCTCCAATACCTCATCGACGACCAAGTCAACACAAGCCAACGCCAGCTCGTCCGCCGCATCGACATCGTCCTCGCCAAGCTTCGCGAACCTGGACTCACGAAAGAGTATCGAGCCCTGGGAGCAAGAACGCTCCGCTCCCTGTACGAGGACTTGGAGTATGCGAATGAGAGGATTGTGGCGCTgcgggcggagctggttgagagagagagggctgtggcggagttcgaggagagggagaggcgggagcggcgggatcacgaggagagggtgaggcggcagcgcgtggcggaggagagggaggtggagttgaggagga	361 	\|\|\|\|\|\|\|\|\|\|\|\|\|\|\|\|\|\|\|\|\|\|\|\|\|\|\|\|\|\|\|\|\|\|\|\|\|\|\|\|\|\|\|\|\|\|\|\|\|\|\|\|\|\|\|\|\|\|\|\|\|\|\|\|\|\|\|\|\|\|\|\|\|\|\|\|\|\|\|\|\|\|\|\|\|\|\|\|\|\|\|\|\|\|\|\|\|\|\|\|\|\|\|\|\|\|\|\|\|\|\|\|\|\|\|\|\|\|\|\|\|\|\|\|\|\|\|\|\|\|\|\|\|\|\|\|\|\|\|\|\|\|\|\|\|\|\|\|\|\|\|\|\|\|\|\|\|\|\|\|\|\|\|\|\|\|\|\|\|\|\|\|\|\|\|\|\|\|\|\|\|\|\|\|\|\|\|\|\|\|\|\|\|\|\|\|\|\|\|\|\|\|\|\|\|\|\|\|\|\|\|\|\|\|\|\|\|\|\|\|\|\|\|\|\|\|\|\|\|\|\|\|\|\|\|\|\|\|\|\|\|\|\|\|\|\|\|\|\|\|\|\|\|\|\|\|\|\|\|\|\|\|\|\|\|\|\|\|\|\|\|\|\|\|\|\|\|\|\|\|\|\|\|\|\|\|\|\|\|\|\|\|\|\|\|\|\|\|\|\|\|\|\|\|\|\|\|\|\|\|\|\|\|\|\|\|\|\|\|\|\|\|\|\|\|\|\|\|\|\|\|\|\|\|\|\|\|\|\|\|\|\|\|\|\|\|\|\|\|\|\|\|\|\|\|\|\|\|\|\|\| 	ATGCCTCCCCGAAACGAAGCAAGCTACATCCGTACCCGCGCCGAGCTCCAATACCTCATCGACGACCAAGTCAACACAAGCCAACGCCAGCTCGTCCGCCGCATCGACATCGTCCTCGCCAAGCTTCGCGAACCTGGACTCACGAAAGAGTATCGAGCCCTGGGAGCAAGAACGCTCCGCTCCCTGTACGAGGACTTGGAGTATGCGAATGAGAGGATTGTGGCGCTGCGGGCGGAGCTGGTTGAGAGAGAGAGGGCTGTGGCGGAGTTCGAGGAGAGGGAGAGGCGGGAGCGGCGGGATCACGAGGAGAGGGTGAGGCGGCAGCGCGTGGCGGAGGAGAGGGAGGTGGAGTTGAGGAGGA ``` |
| 2 | ggggacaagtttgtacaaaaaagcaggcttgatgCCACCCAATACCTCGCAACG, ggggaccactttgtacaagaaagctgggtcTCCTCGCTCGAATGGCCTTG | Mycgr3\|94397  JGI  Ensembl |  |  | 100 % |  | ``` 1	ATGCCACCCAATACCTCGCAACGGGATCGTCCTTCAACCTGGCAAGGCGGCCTGATCATTCTGGCATGCGTCTTGCTCACCCAATTCGGCGATCACTTCACCGGCACGCCTCTTCCCCAGCGACTCACAAGCCTATACATGTTCGCCGCCTGCTTCTACTGCATCCGCGCCGGACAGGATCCCGCACTGACGCCTTTGGACCGTTTCGAGGTGTTCGTCATCATTGCCACGGGAATCGCTGCTCAGCTTGTATGCCTGCATACGGGTAGCGTCTGGATGGTACATCTCCACCACTTCGTGGCCACAGTCGTCCTGTACATTGG-CGCAGCA-AGGCCATTCGAGC-GAGGA	348 	\|\|\|\|\|\|\|\|\|\|\|\|\|\|\|\|\|\|\|\|\|\|\|\|\|\|\|\|\|\|\|\|\|\|\|\|\|\|\|\|\|\|\|\|\|\|\|\|\|\|\|\|\|\|\|\|\|\|\|\|\|\|\|\|\|\|\|\|\|\|\|\|\|\|\|\|\|\|\|\|\|\|\|\|\|\|\|\|\|\|\|\|\|\|\|\|\|\|\|\|\|\|\|\|\|\|\|\|\|\|\|\|\|\|\|\|\|\|\|\|\|\|\|\|\|\|\|\|\|\|\|\|\|\|\|\|\|\|\|\|\|\|\|\|\|\|\|\|\|\|\|\|\|\|\|\|\|\|\|\|\|\|\|\|\|\|\|\|\|\|\|\|\|\|\|\|\|\|\|\|\|\|\|\|\|\|\|\|\|\|\|\|\|\|\|\|\|\|\|\|\|\|\|\|\|\|\|\|\|\|\|\|\|\|\|\|\|\|\|\|\|\|\|\|\|\|\|\|\|\|\|\|\|\|\|\|\|\|\|\|\|\|\|\|\| \|\|\|\|\|\|\|\|\|\|\|\|\|\| \|\|\|\|\|\|\|\|\|\|\|\|\|\|\|\|\|\|\| \|\|\|\|\|\|\|\|\|\|\|\|\|\|\|\| \|\|\|\|\|\|\|\| \|\|\|\|\|  \|\|\| \|\|\|\|\| \| \|\| \|  \|\|\| \|\|\|\|\|\|\|\|  \|\|\|\|\| 	ATGCCACCCAATACCTCGCAACGGGATCGTCCTTCAACCTGGCAAGGCGGCCTGATCATTCTGGCATGCGTCTTGCTCACCCAATTCGGCGATCACTTCACCGGCACGCCTCTTCCCCAGCGACTCACAAGCCTATACATGTTCGCCGCCTGCTTCTACTGCATCCGCGCCGGACAGGATCCCGCACTGACGCCTTTGGACCGTTTCGAGGTGTTCGTCATCATTGCCACGGGAATCGCTGCTCANCTTGTATGCCTGCANACGGGTAGCGTCTGGATGGNACATCTCCACCACTTCNTGGCCACANTCGTCNNGTANATTGGNCNCANCTCAGGNCATTCGAGANGAGGA ``` |
| 2 | ggggacaagtttgtacaaaaaagcaggcttgatgTACAGCTCTACAGCCGTCTA, ggggaccactttgtacaagaaagctgggtcCAGACCATTGCAAATCCCAG | Mycgr3\|94408  JGI  Ensembl |  |  | 100 % |  | ``` 1	ATGTACAGCTCTACAGCCGTCTACCTAGCACTAGCAGCATCCACCCTCATCTCAGCATCGGCACAATGGGCATCAACAACCACCTTTACATTCGACGGCCAAGCCCTTCCAACTGGCCTCATCATATCGACAGACACAATCGAAGATCTCGCAGACGCCGACTCTACAGCTAAATTCAACCACCAATTCAACGCTTCCAACGTGCAAATCAGCGACGGCTTTCTACATCTCACCGTGCCAGGCGGTCAGACTGAGAGTCCAATCCAGTGTGGAGAGGTGTCGACGGAGTTTGAGGTCATGTACGCGAGTGTTTGTACGTACGCCATCTTGACCGGTGAAGCTGGGATTTGCAATGGTCTG	360 	\|\|\|\|\|\|\|\|\|\|\|\|\|\|\|\|\|\|\|\|\|\|\|\|\|\|\|\|\|\|\|\|\|\|\|\|\|\|\|\|\|\|\|\|\|\|\|\|\|\|\|\|\|\|\|\|\|\|\|\|\|\|\|\|\|\|\|\|\|\|\|\|\|\|\|\|\|\|\|\|\|\|\|\|\|\|\|\|\|\|\|\|\|\|\|\|\|\|\|\|\|\|\|\|\|\|\|\|\|\|\|\|\|\|\|\|\|\|\|\|\|\|\|\|\|\|\|\|\|\|\|\|\|\|\|\|\|\|\|\|\|\|\|\|\|\|\|\|\|\|\|\|\|\|\|\|\|\|\|\|\|\|\|\|\|\|\|\|\|\|\|\|\|\|\|\|\|\|\|\|\|\|\|\|\|\|\|\|\|\|\|\|\|\|\|\|\|\|\|\|\|\|\|\|\|\|\|\|\|\|\|\|\|\|\|\|\|\|\|\|\|\|\|\|\|\|\|\|\|\|\|\|\|\|\|\|\|\|\|\|\|\|\|\|\|\|\|\|\|\|\|\|\|\|\|\|\|\|\|\|\|\|\|\|\|\|\|\|\|\|\|\|\|\|\|\|\|\|\|\|\|\|\|\|\|\|\|\|\|\|\|\|\|\|\|\|\|\|\|\|\|\|\|\|\|\|\|\|\|\|\|\|\|\|\|\|\|\|\|\|\|\|\|\|\|\|\|\|\|\|\|\|\|\|\|\|\|\|\|\|\|\|\|\|\|\|\|\|\|\|\|\|\|\|\|\|\|\|\|\| 	ATGTACAGCTCTACAGCCGTCTACCTAGCACTAGCAGCATCCACCCTCATCTCAGCATCGGCACAATGGGCATCAACAACCACCTTTACATTCGACGGCCAAGCCCTTCCAACTGGCCTCATCATATCGACAGACACAATCGAAGATCTCGCAGACGCCGACTCTACAGCTAAATTCAACCACCAATTCAACGCTTCCAACGTGCAAATCAGCGACGGCTTTCTACATCTCACCGTGCCAGGCGGTCAGACTGAGAGTCCAATCCAGTGTGGAGAGGTGTCGACGGAGTTTGAGGTCATGTACGCGAGTGTTTGTACGTACGCCATCTTGACCGGTGAAGCTGGGATTTGCAATGGTCTG ``` |
| 2 | ggggacaagtttgtacaaaaaagcaggcttgatgGCGTCCAACAATACATCCGC, ggggaccactttgtacaagaaagctgggtcAAGATGGAGGCAGAGTTGAG | Mycgr3\|94585  JGI  Ensembl |  |  | 0 % |  | ``` 1	ATGGCGTCCAACAATACATCCGCGCAGGTCACAACTCATGATACAG	46 	\|\|\|\|\|\|\|\|\|\|\|\|\|\|\|\|\|\|\|\|\|\|\|\|\|\|\|\|\|\|\|\|\|\|\|\|\|\|\|\|\|\|\|\|\|\| 	ATGGCGTCCAACAATACATCCGCGCAGGTCACAACTCATGATACAG ```      ``` 44	CAGACCACCACGTCCACGTCGCCAAGACTCTCAGTCCTCTATTCGTCGACGATCCACCCGTCAGGTTCATACTCTCCTCACTATCCGAAGAAGCACGTCTCGCCTACATGGACGACTACTTCGGCACTCTGGTCAAAGCCGCAGCTCTCAATTCCGCGGAATTTGTGGAAGCAAATGACTGGGATTGCACCGCCGTCTGGCTACCTCCCGGGAAGAGGATAGATAATTCTTGGACGTGGTGGTCGGCAGGGTTTGTGGGATGTCTGTGGAAGTTGGGA	321 	\|\|\|\|\|\|\|\|\|\|\|\|\|\|\|\|\|\|\|\|\|\|\|\|\|\|\|\|\|\|\|\|\|\|\|\|\|\|\|\|\|\|\|\|\|\|\|\|\|\|\|\|\|\|\|\|\|\|\|\|\|\|\|\|\|\|\|\|\|\|\|\|\|\|\|\|\|\|\|\|\|\|\|\|\|\|\|\|\|\|\|\|\|\|\|\|\|\|\|\|\|\|\|\|\|\|\|\|\|\|\|\|\|\|\|\|\|\|\|\|\|\|\|\|\|\|\|\|\|\|\|\|\|\|\|\|\|\|\|\|\|\|\|\|\|\|\|\|\|\|\|\|\|\|\|\|\|\|\|\|\|\|\|\|\|\|\|\|\|\|\|\|\|\|\|\|\|\|\|\|\|\|\|\|\|\|\|\|\|\|\|\|\|\|\|\|\|\|\|\|\|\|\|\|\|\|\|\|\|\|\|\|\|\|\|\|\|\|\|\|\|\|\|\|\|\|\|\|\|\|\|\|\|\|\|\|\|\|\|\|\|\|\|\|\|\|\|\|\|\|\|\|\|\|\|\|\|\|\|\|\|\|\|\|\|\|\|\|\|\|\| \| \|\|\|\| 	CAGACCACCACGTCCACGTCGCCAAGACTCTCAGTCCTCTATTCGTCGACGATCCACCCGTCAGGTTCATACTCTCCTCACTATCCGAAGAAGCACGTCTCGCCTACATGGACGACTACTTCGGCACTCTGGTCAAAGCCGCAGCTCTCAATTCCGCGGAATTTGTGGAAGCAAATGACTGGGATTGCACCGCCGTCTGGCTACCTCCCGGGAAGAGGATAGATAATTCTTGGACGTGGTGGTCGGCAGGGTTTGTGGGATGTCTGTGGAATTNGGGA ``` |
| 2 | ggggacaagtttgtacaaaaaagcaggcttgatgGGTTCAACGTCCTCACAGCC, ggggaccactttgtacaagaaagctgggtcAAACTGAGTTCCTCCATGCC | Mycgr3\|94738  JGI  Ensembl |  |  | 100 % |  | ``` 1	ATGGGTTCAACGTCCTCACAGCCAGCCCCGAGCCCTCAGGGCAGCCGCAGATATCTGCACGCCCAAGCTGCTAGGTCTATCATCGCAGAGGAAGACGACGCTACCGATGCCGCTGAACGCCTGGCCCGAGAAGCCATCGACTACTTTGGAGAAAGCGAATGGTACACAGCTCGAAGTGAAGTCGACCTTTCCCTTCAGCACTTGCGCGACACACCGGGCACATACGACAATCCCTACGATATCAACGGCCACGTTCACGCCGCTTTGCGAGGTTGTGCCAAGCAAGCCGTTCCTGTTCGGAACACGATCGACTTCCTGACGAGGGATGCTGTGCTGGGACTTTCTAGCCACCTTAGGTACCATGGGCATGGAGGAACTCAGTTTTGA	387 	\|\|\|\|\|\|\|\|\|\|\|\|\|\|\|\|\|\|\|\|\|\|\|\|\|\|\|\|\|\|\|\|\|\|\|\|\|\|\|\|\|\|\|\|\|\|\|\|\|\|\|\|\|\|\|\|\|\|\|\|\|\|\|\|\|\|\|\|\|\|\|\|\|\|\|\|\|\|\|\|\|\|\|\|\|\|\|\|\|\|\|\|\|\|\|\|\|\|\|\|\|\|\|\|\|\|\|\|\|\|\|\|\|\|\|\|\|\|\|\|\|\|\|\|\|\|\|\|\|\|\|\|\|\|\|\|\|\|\|\|\|\|\|\|\|\|\|\|\|\|\|\|\|\|\|\|\|\|\|\|\|\|\|\|\|\|\|\|\|\|\|\|\|\|\|\|\|\|\|\|\|\|\|\|\|\|\|\|\|\|\|\|\|\|\|\|\|\|\|\|\|\|\|\|\|\|\|\|\|\|\|\|\|\|\|\|\|\|\|\|\|\|\|\|\|\|\|\|\|\|\|\|\|\|\|\|\|\|\|\|\|\|\|\|\|\|\|\|\|\|\|\|\|\|\|\|\|\|\|\|\|\|\|\|\|\|\|\|\|\|\|\|\|\|\|\|\|\|\|\|\|\|\|\|\|\|\|\|\|\|\|\|\|\|\|\|\|\|\|\|\|\|\|\|\|\|\|\|\|\|\|\|\|\|\|\|\|\|\|\|\|\|\|\|\|\|\|\|\|\|\|\|\|\|\|\|\|\|\|\|\|\|\|\|\|\|\|\|\|\|\|\|\|\|\|\|\|\|\|\|\|\|\|\|\|\|\| \|\|\|\|\|\|\|\|\|\|\|\| \|\|\|\|\|\| 	ATGGGTTCAACGTCCTCACAGCCAGCCCCGAGCCCTCAGGGCAGCCGCAGATATCTGCACGCCCAAGCTGCTAGGTCTATCATCGCAGAGGAAGACGACGCTACCGATGCCGCTGAACGCCTGGCCCGAGAAGCCATCGACTACTTTGGAGAAAGCGAATGGTACACAGCTCGAAGTGAAGTCGACCTTTCCCTTCAGCACTTGCGCGACACACCGGGCACATACGACAATCCCTACGATATCAACGGCCACGTTCACGCCGCTTTGCGAGGTTGTGCCAAGCAAGCCGTTCCTGTTCGGAACACGATCGACTTCCTGACGAGGGATGCTGTGCTGGGACTTTCTAGCCACCTTAGGTACCATGGGCNTGGAGGAACTCA-TTTTGA ``` |
| 2 | ggggacaagtttgtacaaaaaagcaggcttgatgTCACAAACAAGACTCCGCGA, ggggaccactttgtacaagaaagctgggtcCATCTCGATCTGCGCCAAGA | Mycgr3\|94739  JGI  Ensembl |  |  | 100 % |  | ``` 1	ATGTCACAAACAAGACTCCGCGACCTAGACGACTACTACGTCCGCGTCGGCTATCAGCACCTGGCACTGTCGTTCGAGATCATGCGTCGGCGGTCTCGACTCAATGGCCTTCCGGACGAATCAATGCCCGAAACACTACCCGAAGGCATGCTACAAGCCGAAAGAGAGCCCGAATTTCTCCACGAGCAAGAGCTTGTTCAGGACTATATTACGAATCTCAGAGAAGGGAATGAATCAGCGTCGAGCGGCCTGGAGCCACGCGTCGCCATATTACTCGCGATGTTTGTTTACGAACCCGAGGAGGAGCGCGAACTGTCCGCGGCTTTGCTTCAAGTCGCGGACGACTTCTTGGAGGAGTATCTGCTGATGAGGGAGGAGATCTTGGCGCAGATCGAGATG	399 	\|\|\|\|\|\|\|\|\|\|\|\|\|\|\|\|\|\|\|\|\|\|\|\|\|\|\|\|\|\|\|\|\|\|\|\|\|\|\|\|\|\|\|\|\|\|\|\|\|\|\|\|\|\|\|\|\|\|\|\|\|\|\|\|\|\|\|\|\|\|\|\|\|\|\|\|\|\|\|\|\|\|\|\|\|\|\|\|\|\|\|\|\|\|\|\|\|\|\|\|\|\|\|\|\|\|\|\|\|\|\|\|\|\|\|\|\|\|\|\|\|\|\|\|\|\|\|\|\|\|\|\|\|\|\|\|\|\|\|\|\|\|\|\|\|\|\|\|\|\|\|\|\|\|\|\|\|\|\|\|\|\|\|\|\|\|\|\|\|\|\|\|\|\|\|\|\|\|\|\|\|\|\|\|\|\|\|\|\|\|\|\|\|\|\|\|\|\|\|\|\|\|\|\|\|\|\|\|\|\|\|\|\|\|\|\|\|\|\|\|\|\|\|\|\|\|\|\|\|\|\|\|\|\|\|\|\|\|\|\|\|\|\|\|\|\|\|\|\|\|\|\|\|\|\|\|\|\|\|\|\|\|\|\|\|\|\|\|\|\|\|\|\|\|\|\|\|\|\|\|\|\|\|\|\|\|\|\|\|\|\|\|\|\|\|\|\|\|\|\|\|\|\|\|\|\|\|\|\|\|\|\|\|\|\|\|\|\|\|\|\|\|\|\|\|\|\|\|\|\|\|\|\|\|\|\|\|\|\|\|\|\|\|\|\|\|\|\|\|\|\|\|\|\|\|\|\|\|\|\|\|\|\|\|\|\|\|\|\|\|\|\|\|\|\|\|\|\|\|\|\|\|\|\|\|\|\|\|\| \|\|\|\|\|\|\|\|\| 	ATGTCACAAACAAGACTCCGCGACCTAGACGACTACTACGTCCGCGTCGGCTATCAGCACCTGGCACTGTCGTTCGAGATCATGCGTCGGCGGTCTCGACTCAATGGCCTTCCGGACGAATCAATGCCCGAAACACTACCCGAAGGCATGCTACAAGCCGAAAGAGAGCCCGAATTTCTCCACGAGCAAGAGCTTGTTCAGGACTATATTACGAATCTCAGAGAAGGGAATGAATCAGCGTCGAGCGGCCTGGAGCCACGCGTCGCCATATTACTCGCGATGTTTGTTTACGAACCCGAGGAGGAGCGCGAACTGTCCGCGGCTTTGCTTCAAGTCGCGGACGACTTCTTGGAGGAGTATCTGCTGATGAGGGAGGAGATCTTGGCGCANATCGAGATG ``` |
| 2 | ggggacaagtttgtacaaaaaagcaggcttgatgTTCCTCCCACTCGCAGCCAG, ggggaccactttgtacaagaaagctgggtcGCTTCCGACTCCAAGTCCTT | Mycgr3\|94751  JGI  Ensembl |  |  | 100 % |  | ``` 1	ATGTTCCTCCCACTCGCAGCCAGCGTGGCCTCCGCCTTTACCAAAACAGCGCTACTTATCATCGAAATCATCAGCTTCTCCCTCGCCACAACGAAGCTCCTCTGCATCATAGCCCTTCTTCTCCTTGGACCTTTGATCTATCTCCACCGCCGTAACATTATCAAAGCCACGGAACGCCGCATCGAGAGCACCATCCTCCGTCGCCAACAAGAAGGCTCGAAGCCGCTAGACTTGGAACGCCTGCGCATTCTCATCAGGGACCAGGAGATTCGGGATCGGGATGCGGGCTTGGAGGATAGTGCGAGGATAGACAGGAGAGCTCAAGGACTTGGAGTCGGAAGC	342 	\|\|\|\|\|\|\|\|\|\|\|\|\|\|\|\|\|\|\|\|\|\|\|\|\|\|\|\|\|\|\|\|\|\|\|\|\|\|\|\|\|\|\|\|\|\|\|\|\|\|\|\|\|\|\|\|\|\|\|\|\|\|\|\|\|\|\|\|\|\|\|\|\|\|\|\|\|\|\|\|\|\|\|\|\|\|\|\|\|\|\|\|\|\|\|\|\|\|\|\|\|\|\|\|\|\|\|\|\|\|\|\|\|\|\|\|\|\|\|\|\|\|\|\|\|\|\|\|\|\|\|\|\|\|\|\|\|\|\|\|\|\|\|\|\|\|\|\|\|\|\|\|\|\|\|\|\|\|\|\|\|\|\|\|\|\|\|\|\|\|\|\|\|\|\|\|\|\|\|\|\|\|\|\|\|\|\|\|\|\|\|\|\|\|\|\|\|\|\|\|\|\|\|\|\|\|\|\|\|\|\|\|\|\|\|\|\|\|\|\|\|\|\|\|\|\|\|\|\|\|\|\|\|\|\|\|\|\|\|\|\|\|\|\|\|\|\|\|\|\|\|\|\|\|\|\|\|\|\|\|\|\|\|\|\|\|\|\|\|\|\|\|\|\|\|\|\|\|\|\|\|\|\|\|\|\|\|\|\|\|\|\|\|\|\|\|\|\|\|\|\|\|\|\|\|\|\|\|\|\|\|\|\|\|\|\|\|\|\|\|\|\|\|\|\|\|\|\|\|\|\|\|\|\|\|\|\|\|\|\|\|\| 	ATGTTCCTCCCACTCGCAGCCAGCGTGGCCTCCGCCTTTACCAAAACAGCGCTACTTATCATCGAAATCATCAGCTTCTCCCTCGCCACAACGAAGCTCCTCTGCATCATAGCCCTTCTTCTCCTTGGACCTTTGATCTATCTCCACCGCCGTAACATTATCAAAGCCACGGAACGCCGCATCGAGAGCACCATCCTCCGTCGCCAACAAGAAGGCTCGAAGCCGCTAGACTTGGAACGCCTGCGCATTCTCATCAGGGACCAGGAGATTCGGGATCGGGATGCGGGCTTGGAGGATAGTGCGAGGATAGACAGGAGAGCTCAAGGACTTGGAGTCGGAAGC ``` |
| 2 | ggggacaagtttgtacaaaaaagcaggcttgatgCTCCTGGTCTCGGCCAGAGG, ggggaccactttgtacaagaaagctgggtcCTCGTCCTCGTCCTGCTCGT | Mycgr3\|94812  JGI  Ensembl |  |  | 100 % |  | ``` 1	ATGCTCCTGGTCTC-GGCCAGAGGGACACTATACTTCGCCATCAATCGCCTGGACCTACGCACCTTCCAACCTCCTAACCCGGAGCTCGATTTCGCCTCTCTCGTCGCTCAGCAGGTTGTATGGGCCATTCGTCGTCCGCACATCGTCCAGCCCAACGTCTCCATGACTCCGACTCCCTACTTGACGGTCATCAACGCGAAGGGACTCTACGACTTGGGCGTGTTGTTCGCTGTTCAGGTCGAGAAGGCAACGACGCTGAAGGAATCGTGGCACCACAATCCGCTACGCTTTGGGACGACGAAGCATCAGACTGCGGTGGAGCAGCAGCGCTTTGCTGAGCAGGAGG-ATGATGAC	354 	\|\|\|\|\|\|\|\|\|\|\|\| \|   \|\|\|\|\|\|\|\|\|\|\|\|\|\|\|\|\|\|\|\|\|\|\|\|\|\|\|\|\|\|\|\|\|\|\|\|\|\|\|\|\|\|\|\|\|\|\|\|\|\|\|\|\|\|\|\|\|\|\|\|\|\|\|\|\|\|\|\|\|\|\|\|\|\|\|\|\|\|\|\|\|\|\|\|\|\|\|\|\|\|\|\|\|\|\|\|\|\|\|\|\|\|\|\|\|\|\|\|\|\|\|\|\|\|\|\|\|\|\|\|\|\|\|\|\|\|\|\|\|\|\|\|\|\|\|\|\|\|\|\|\|\|\|\| \|\|\|\|\|\|\|\|\|\|\|\|\|\|\|\|\|\|\|\|\|\|\|\|\|\|\|\|\|\|\|\|\|\|\|\|\|\|\|\|\|\|\|\|\|\|\|\|\|\|\|\|\|\|\|\|\|\|\|\|\|\|\|\|\|\|\|\|\|\|\|\|\|\|\|\|\|\|\|\|\|\|\|\|\|\|\|\|\|\|\|\|\|\|\|\|\|\|\| \|\|\|\|\|\|\|\|\|\|\|\|\|\|\|\|\|\|\|\|\|\|\|\|\|\|\|\|\|\|\|\|\|\|\|\|\|\|\|\|\|\|\|\|\|\|\| \|\|\|\| \|\|\|\|\|\|\|\|\|\| \|\|\|\| \|\|\|\|\|\|\|\|\|\|\| \|\| \| \| \|\|\|\|\|\| 	ATGCTCCTGGTCNCTTNCCAGAGGGACACTATACTTCGCCATCAATCGCCTGGACCTACGCACCTTCCAACCTCCTAACCCGGAGCTCGATTTCGCCTCTCTCGTCGCTCAGCAGGTTGTATGGGCCATTCGTCGTCCGCACATCGTCCAGCCCAACGTCTNCATGACTCCGACTCCCTACTTGACGGTCATCAACGCGAAGGGACTCTACGACTTGGGCGTGTTGTTCGCTGTTCAGGTCGAGAAGGCAACGACGCTGAAAGAATCGTGGCACCACAATCCGCTACGCTTTGGGACGACGAAGCATCANACTGNGGTGGAGCAGNAGCGNTTTGCTGAGCANGANGNANGATGAC ``` |
| 2 | ggggacaagtttgtacaaaaaagcaggcttgatgGTCAAAGTCACTCTTCTCGG, ggggaccactttgtacaagaaagctgggtcGAAGCAGAAATCGTCGTCAC | Mycgr3\|95035  JGI  Ensembl |  |  | 100 % |  | ``` 1	ATGGTCAAAGTCA-CTCTTCTCGGTCTGCTGTTCAGCGCCGTCGCACTCGTGGAAGCAATGCCAAAGCAGAGTCCGCCAGAGCGGAGTCCAAACGAGCAGTCGGTCAAG	108 	\|\|\|\|\|\|\|\|\|\|\|\|\| \|\|\|\|\|\|\|\|\|\|\|\|\|\|\|\|\|\|\|\|\|\|\|\|\|\|\|\|\|\|\|\|\|\|\|\|\|\|\|\|\|\|\|\|\|\|\|\|\|\|\|\|\|\|\|\|\|\|\|\|\|\|\|\|\|\|\|\|\|\|\|\|\|\|\|\|\|\|\|\|\|\|\|\|\|\|\|\|\|\|\| \|\|\| 	ATGGTCAAAGTCACCTCTTCTCGGTCTGCTGTTCAGCGCCGTCGCACTCGTGGAAGCAATGCCAAAGCAGAGTCCGCCAGAGCGGAGTCCAAACGAGCAGTCGGT-AAG ```      ``` 102	GGTCAAGATGTTTAGCAAGCGCATATACCAGAGCGCGTGTCACAAATCGTACAAATGCGACCATCAATCCGACTGCGCTGGACCAGAGAAAATGGACATCGTGCCCGACGGGTGCCCGAAGTGCTATGCAAAATTTCTGTGCGGCGCCGACGGGAAAACCATACCGAATACGCTTCAGGGCTACTGGTGCCTGGGGAGTGACGACGATTTCTGCTTC	318 	\|\|\|\|\|\|\|\|\|\|\|\|\|\|\|\|\|\|\|\|\|\|\|\|\|\|\|\|\|\|\|\|\|\|\|\|\|\|\|\|\|\|\|\|\|\|\|\|\|\|\|\|\|\|\|\|\|\|\|\|\|\|\|\|\|\|\|\|\|\|\|\|\|\|\|\|\|\|\|\|\|\|\|\|\|\|\|\|\|\|\|\|\|\|\|\|\|\|\|\|\|\|\|\|\|\|\|\|\|\|\|\|\|\|\|\|\|\|\|\|\|\|\|\|\|\|\|\|\|\|\|\|\|\|\|\|\|\|\|\|\|\|\|\|\|\|\|\|\|\|\|\|\|\|\|\|\|\|\|\|\|\|\|\|\|\|\|\|\|\|\|\|\|\|\|\|\|\|\|\|\|\|\|\|\|\|\|\|\|\|\|\|\|\|\|\|\|\|\|\|\|\|\|\|\|\|\|\|\|\|\|\|\|\|\|\|\| 	GGTCAAGATGTTTAGCAAGCGCATATACCAGAGCGCGTGTCACAAATCGTACAAATGCGACCATCAATCCGACTGCGCTGGACCAGAGAAAATGGACATCGTGCCCGACGGGTGCCCGAAGTGCTATGCAAAATTTCTGTGCGGCGCCGACGGGAAAACCATACCGAATACGCTTCAGGGCTACTGGTGCCTGGGGAGTGACGACGATTTCTGCTTC ``` |
| 2 | ggggacaagtttgtacaaaaaagcaggcttgatgCTCCGTCAACTCTTCATCAT, ggggaccactttgtacaagaaagctgggtcAGCCGCGGGTCCGGGAAGTG | Mycgr3\|95211  JGI  Ensembl |  |  | 100 % |  | ``` 1	ATGCTCCGTCAACTCTTCATCATCGCATCACTCTGGCTTGGGGCCTACGGCGACAACAACTGTAACTCGGACTGCGTCTGCACTTACTCCAACAACGGCATCCAGTGCATCGCCCAGATCACCGGAATTCACAGGCATGATTGCAAAGGCTATTGCCACGACCGCGCCTGCGGCGGTGTCGACAACAACCGACAATCATGCCGCCTCTGGAACAACGGCTACTGCGACGATAATGGCGGAGCGTACACCAGGGGACGTGGCGGCTACTATGTCGGCTATTGCAACGAGAGAGCTGAGTACAACTCGTGCAATCACGACTGTCCTGGTCATGCGTCACCTGCGGAGCTGTTTGCACTTCCCGGACCCGCGGCT	372 	\|\|\|\|\|\|\|\|\|\|\|\|\|\|\|\|\|\|\|\|\|\|\|\|\|\|\|\|\|\|\|\|\|\|\|\|\|\|\|\|\|\|\|\|\|\|\|\|\|\|\|\|\|\|\|\|\|\|\|\|\|\|\|\|\|\|\|\|\|\|\|\|\|\|\|\|\|\|\|\|\|\|\|\|\|\|\|\|\|\|\|\|\|\|\|\|\|\|\|\|\|\|\|\|\|\|\|\|\|\|\|\|\|\|\|\|\|\|\|\|\|\|\|\|\|\|\|\|\|\|\|\|\|\|\|\|\|\|\|\|\|\|\|\|\|\|\|\|\|\|\|\|\|\|\|\|\|\|\|\|\|\|\|\|\|\|\|\|\|\|\|\|\|\|\|\|\|\|\|\|\|\|\|\|\|\|\|\|\|\|\|\|\|\|\|\|\|\|\|\|\|\|\|\|\|\|\|\|\|\|\|\|\|\|\|\|\|\|\|\|\|\|\|\|\|\|\|\|\|\|\|\|\|\|\|\|\|\|\|\|\|\|\|\|\|\|\|\|\|\|\|\|\|\|\|\|\|\|\|\|\|\|\|\|\|\|\|\|\|\|\|\|\|\|\|\|\|\|\|\|\|\|\|\|\|\|\|\|\|\|\|\|\|\|\|\|\|\|\|\|\|\|\|\|\|\|\|\|\|\|\|\|\|\|\|\|\|\|\|\|\|\|\|\|\|\|\|\|\|\|\|\|\|\|\|\|\|\|\|\|\|\|\|\|\|\|\|\|\|\|\|\|\|\|\|\|\|\|\|\|\|\|\|\|\|\|\|\|\|\|\|\| 	ATGCTCCGTCAACTCTTCATCATCGCATCACTCTGGCTTGGGGCCTACGGCGACAACAACTGTAACTCGGACTGCGTCTGCACTTACTCCAACAACGGCATCCAGTGCATCGCCCAGATCACCGGAATTCACAGGCATGATTGCAAAGGCTATTGCCACGACCGCGCCTGCGGCGGTGTCGACAACAACCGACAATCATGCCGCCTCTGGAACAACGGCTACTGCGACGATAATGGCGGAGCGTACACCAGGGGACGTGGCGGCTACTATGTCGGCTATTGCAACGAGAGAGCTGAGTACAACTCGTGCAATCACGACTGTCCTGGTCATGCGTCACCTGCGGAGCTGTTTGCACTTCCCGGACCCGCGGCT ``` |
| 2 | ggggacaagtttgtacaaaaaagcaggcttgatgTCTTCCACCCCAGCAGCCAA, ggggaccactttgtacaagaaagctgggtcTTCCTCCTTCTCTCCCATCT | Mycgr3\|95225  JGI  Ensembl |  |  | 100 % |  |  |
| 2 | ggggacaagtttgtacaaaaaagcaggcttgatgCCCTCCTCCTCAACCCACAC, ggggaccactttgtacaagaaagctgggtcCCCACCCACCCTCCTCCACC | Mycgr3\|95348  JGI  Ensembl |  |  | 0 % |  |  |
| 2 | ggggacaagtttgtacaaaaaagcaggcttgatgAAGCTTTCAATCACCGGCGC, ggggaccactttgtacaagaaagctgggtcATCATAAACGCAAGCCAATT | Mycgr3\|95697  JGI  Ensembl |  |  | 100 % |  | ``` 1	ATGAAGCTTTCAATCACCGGCGCAATTTTTGGCCTTGGTCTGCTTACTCCCGTCGTCATGGGCTACCTCCATGTTCATTACCATGTCTGCGCTCGGTCAGAGTCCCACCAGTTCTATGACCAGTGTGACTTCAAAAACGCCGGCGCCATCTGCGTACAACTCTCCCACACCGCCTTTCCCAATCTGTATACTAATGCTGGCTCTCAGGCTACCGGTATAACCAGTGTTGACGAGCTACACTGCGACACAAACTATCCCAACTTTCCGGACGAGTGGCACATCGCCAAGTGGACGGACCCAAGGAAAAGCTGTACGGTGAAAGATGTTCATGGCGGGCCGGATGTTGCTGGCCAATTGGCTTGCGTTTATGAT	372 	\|\|\|\|\|\|\|\|\|\|\|\|\|\|\|\|\|\|\|\|\|\|\|\|\|\|\|\|\|\|\|\|\|\|\|\|\|\|\|\|\|\|\|\|\|\|\|\|\|\|\|\|\|\|\|\|\|\|\|\|\|\|\|\|\|\|\|\|\|\|\|\|\|\|\|\|\|\|\|\|\|\|\|\|\|\|\|\|\|\|\|\|\|\|\|\|\|\|\|\|\|\|\|\|\|\|\|\|\|\|\|\|\|\|\|\|\|\|\|\|\|\|\|\|\|\|\|\|\|\|\|\|\|\|\|\|\|\|\|\|\|\|\|\|\|\|\|\|\|\|\|\|\|\|\|\|\|\|\|\|\|\|\|\|\|\|\|\|\|\|\|\|\|\|\|\|\|\|\|\|\|\|\|\|\|\|\|\|\|\|\|\|\|\|\|\|\|\|\|\|\|\|\|\|\|\|\|\|\|\|\|\|\|\|\|\|\|\|\|\|\|\|\|\|\|\|\|\|\|\|\|\|\|\|\|\|\|\|\|\|\|\|\|\|\|\|\|\|\|\|\|\|\|\|\|\|\|\|\|\|\|\|\|\|\|\|\|\|\|\|\|\|\|\|\|\|\|\|\|\|\|\|\|\|\|\|\|\|\|\|\|\|\|\|\|\|\|\|\|\|\|\|\|\|\|\|\|\|\|\|\|\|\|\|\|\|\|\|\|\|\|\|\|\|\|\|\|\|\|\|\|\|\|\|\|\|\|\|\|\|\|\|\|\|\|\|\|\|\|\|\|\|\|\|\|\|\|\|\|\|\|\|\|\|\|\|\|\|\|\|\|\| 	ATGAAGCTTTCAATCACCGGCGCAATTTTTGGCCTTGGTCTGCTTACTCCCGTCGTCATGGGCTACCTCCATGTTCATTACCATGTCTGCGCTCGGTCAGAGTCCCACCAGTTCTATGACCAGTGTGACTTCAAAAACGCCGGCGCCATCTGCGTACAACTCTCCCACACCGCCTTTCCCAATCTGTATACTAATGCTGGCTCTCAGGCTACCGGTATAACCAGTGTTGACGAGCTACACTGCGACACAAACTATCCCAACTTTCCGGACGAGTGGCACATCGCCAAGTGGACGGACCCAAGGAAAAGCTGTACGGTGAAAGATGTTCATGGCGGGCCGGATGTTGCTGGCCAATTGGCTTGCGTTTATGAT ``` |
| 2 | ggggacaagtttgtacaaaaaagcaggcttgatgTCGTTCAATTCTGCAACCAG, ggggaccactttgtacaagaaagctgggtcGGTCTTGCGCTTGCTGGAAG | Mycgr3\|95947  JGI  Ensembl |  |  | 100 % |  | ``` 1	ATGTCGTTCAATTCTGCAACCAGCCGCGACATCACCACGGTCGATCCTCTCTCCACCTCAACATCAAGTCGATCGATCGCTATCGCTATCGCCACGTCTCCCTCCTGCTCCTCATCTCGAAGAGAGAGCTTCAACGCTCCGTCCTCCATCTCAAATCCATACTCATCCTGCGCCTACTCGAGCTGGCAATCTTCATCCACTGGCCACACGAGCGCATATGTCAGCGATGAGGACCTGTGGGGCAACGACGATGAAGCATATCTGAGTGAGCCACCGCCGCCACCGCGCTCGGCTGAAG-TTTGGCTTGCCCGACCTTTACTGCCT	324 	\|\|\|\|\|\|\|\|\|\|\|\|\|\|\|\|\|\|\|\|\|\|\|\|\|\|\|\|\|\|\|\|\|\|\|\|\|\|\|\|\|\|\|\|\|\|\|\|\|\|\|\|\|\|\|\|\|\|\|\|\|\|\|\|\|\|\|\|\|\|\|\|\|\|\|\|\|\|\|\|\|\|\|\|\|\|\|\|\|\|\|\|\|\|\|\|\|\|\|\|\|\|\|\|\|\|\|\|\|\|\|\|\|\|\|\|\|\|\|\|\|\|\|\|\|\|\|\|\|\|\|\|\|\|\|\|\|\|\|\|\|\|\|\|\|\|\|\|\|\|\|\|\|\|\|\|\|\|\|\|\|\|\|\|\|\|\|\|\|\|\|\|\|\|\|\|\|\|\|\|\|\|\|\|\|\|\|\|\|\|\|\|\|\|\|\|\|\|\|\|\|\|\|\|\|\|\|\|\|\|\|\|\|\|\|\|\|\|\|\|\|\|\|\|\|\|\|\|\|\|\|\|\|\|\|\|\|\|\|\|\|\| \|\|\|\|\|\|\|\|\|\|\|\|\|\|\|\|\|\|\|\|\|\|\|\|\|\|\|\|\|\|\|\|\|\| \| \|\|\|\|\|\|\|\|\|\|\|\|\|\|\|\|\|\| \|\|\|\|\|\|\|\|\|\|\|\|\|\|\|\| \|\|\|\|\|\|\|\|\| 	ATGTCGTTCAATTCTGCAACCAGCCGCGACATCACCACGGTCGATCCTCTCTCCACCTCAACATCAAGTCGATCGATCGCTATCGCTATCGCCACGTCTCCCTCCTGCTCCTCATCTCGAAGAGAGAGCTTCAACGCTCCGTCCTCCATCTCAAATCCATACTCATCCTGCGCCTACTCGAGCTGGCAATCTTCATCCACTGGCCACACGAGCGCATATGTCAGCGATGAGGACCTGTGGGGNAACGACGATGAAGCATATCTGAGTGAGCCACCGCTGTCACCGCGCTCGGCTGAAGATTTGGCTTGCCCGACCNTTACTGCCT ``` |
| 2 | ggggacaagtttgtacaaaaaagcaggcttgatgTCTCTCGGTTCAGCCATGTA, ggggaccactttgtacaagaaagctgggtcCTTCGAAGAATACCCCTGCC | Mycgr3\|96157  JGI  Ensembl |  |  | 100 % |  | ``` 49	AAACCCTCCGAGATCCATTCCTCCGCCTCCCACGACCATCATTCTTCGGCCAACGATGCTCAGAAACCACCACTTCAGCGCAACACAACCAACCACTACGAGCAAGCTGTCGCAGCAGCCGGCCAAACTGCCGCCGGAAGCGGAACCGAGGAGAGACCTGCTCTCGACCGACAGCAGAGCTGGAAGATGAGTGACTTGAAGCGCGAGCATGCTGAGAAGATGCTCAGTGTTGAGCCAAAGGGGCAGGGGTATTCTTCGAAG	309 	\|\|\|\|\|\|\|\|\|\|\|\|\|\|\|\|\|\|\|\|\|\|\|\|\|\|\|\|\|\|\|\|\|\|\|\|\|\|\|\|\|\|\|\|\|\|\|\|\|\|\|\|\|\|\|\|\|\|\|\|\|\|\|\|\|\|\|\|\|\|\|\|\|\|\|\|\|\|\|\|\|\|\|\|\|\|\|\|\|\|\|\|\|\|\|\|\|\|\|\|\|\|\|\|\|\|\|\|\|\|\|\|\|\|\|\|\|\|\|\|\|\|\|\|\|\|\|\|\|\|\|\|\|\|\|\|\|\|\|\|\|\|\|\|\|\|\|\|\|\|\|\|\|\|\|\|\|\|\|\|\|\|\|\|\|\|\|\|\|\|\|\|\|\|\|\|\|\|\|\|\|\|\|\|\|\|\|\|\|\|\|\|\|\|\|\|\|\|\|\|\|\|\|\|\|\|\|\|\|\|\|\|\|\|\|\|\|\|\|\|\|\|\|\|\|\|\|\|\|\|\|\|\|\|\|\|\|\|\|\|\|\|\|\|\|\|\|\|\|\|\|\|\|\|\|\|\|\|\|\|\| 	AAACCCTCCGAGATCCATTCCTCCGCCTCCCACGACCATCATTCTTCGGCCAACGATGCTCAGAAACCACCACTTCAGCGCAACACAACCAACCACTACGAGCAAGCTGTCGCAGCAGCCGGCCAAACTGCCGCCGGAAGCGGAACCGAGGAGAGACCTGCTCTCGACCGACAGCAGAGCTGGAAGATGAGTGACTTGAAGCGCGAGCATGCTGAGAAGATGCTCAGTGTTGAGCCAAAGGGGCAGGGGTATTCTTCGAAG ```      ``` 1	ATGTCTCTCGGTTCAGCCATGTACGACTACCAACGAGGCGGCACCACA	48 	\|\|\|\|\|\|\|\|\|\|\|\|\|\|\|\|\|\|\|\|\|\|\|\|\|\|\|\|\|\|\|\|\|\|\|\|\|\|\|\|\|\|\|\|\|\|\|\| 	ATGTCTCTCGGTTCAGCCATGTACGACTACCAACGAGGCGGCACCACA ``` |
| 2 | ggggacaagtttgtacaaaaaagcaggcttgatgGCCGGCTACCATCCCTCCAC, ggggaccactttgtacaagaaagctgggtcCGGCCTCTTCCCCCTGATGG | Mycgr3\|96410  JGI  Ensembl |  |  | 100 % |  | ``` 1	ATGGCCGGCTACCATCCCTCCACCCCCTGGCCCTCAAACCTGGCAGAATTCGACCTCGAAGCCGACATTCTCGTCGAGCAATCCTTCGACTCCCGCACCCCTCAACTGTCTCCAGCTGCTTACGATGACCGAAGCTTCGGTCCGGTGATGGGAACGTTACCGAAAGCCGTTGCTGGGGAGGAGAGTCAGCCGGCGGATGAGGTTGAGAATGCCCGTGATGATCCTTTTGTGACTTTGCCTTTGACGGGGGAGCGGCCGAGTAGAGCATCGGACCCGGTGGAGGATGGGGAGGCGAAGGTGAAGGCTGTGGATGCGGCTCCGATGAGAGCAAAATTGAAGGATGGGAATGAGGGGACTACGATGAGGTTACCCATCAGGGGGAAGAGGCCG	390 	\|\|\|\|\|\|\|\|\|\|\|\|\|\|\|\|\|\|\|\|\|\|\|\|\|\|\|\|\|\|\|\|\|\|\|\|\|\|\|\|\|\|\|\|\|\|\|\|\|\|\|\|\|\|\|\|\|\|\|\|\|\|\|\|\|\|\|\|\|\|\|\|\|\|\|\|\|\|\|\|\|\|\|\|\|\|\|\|\|\|\|\|\|\|\|\|\|\|\|\|\|\|\|\|\|\|\|\|\|\|\|\|\|\|\|\|\|\|\|\|\|\|\|\|\|\|\|\|\|\|\|\|\|\|\|\|\|\|\|\|\|\|\|\|\|\|\|\|\|\|\|\|\|\|\|\|\|\|\|\|\|\|\|\|\|\|\|\|\|\|\|\|\|\|\|\|\|\|\|\|\|\|\|\|\|\|\|\|\|\|\|\|\|\|\|\|\|\|\|\|\|\|\|\|\|\|\|\|\|\|\|\|\|\|\|\|\|\|\|\|\|\|\|\|\|\|\|\|\|\|\|\|\|\|\|\|\|\|\|\|\|\|\|\|\|\|\|\|\|\|\|\|\|\|\|\|\|\|\|\|\|\|\|\|\|\|\|\|\|\|\|\|\|\|\|\|\|\|\|\|\|\|\|\|\|\|\|\|\|\|\|\|\|\|\|\|\|\|\|\|\|\|\|\|\|\|\|\|\|\|\|\|\|\|\|\|\|\|\|\|\|\|\|\|\|\|\|\|\|\|\|\|\|\|\|\|\|\|\|\|\|\|\|\|\|\|\|\|\|\|\|\|\|\|\|\|\|\|\|\|\|\|\|\|\|\|\|\|\|\|\|\|\|\|\|\|\|\|\|\|\|\|\|\|\|\|\|\|\|\| 	ATGGCCGGCTACCATCCCTCCACCCCCTGGCCCTCAAACCTGGCAGAATTCGACCTCGAAGCCGACATTCTCGTCGAGCAATCCTTCGACTCCCGCACCCCTCAACTGTCTCCAGCTGCTTACGATGACCGAAGCTTCGGTCCGGTGATGGGAACGTTACCGAAAGCCGTTGCTGGGGAGGAGAGTCAGCCGGCGGATGAGGTTGAGAATGCCCGTGATGATCCTTTTGTGACTTTGCCTTTGACGGGGGAGCGGCCGAGTAGAGCATCGGACCCGGTGGAGGATGGGGAGGCGAAGGTGAAGGCTGTGGATGCGGCTCCGATGAGAGCAAAATTGAAGGATGGGAATGAGGGGACTACGATGAGGTTACCCATCAGGGGGAAGAGGCCG ``` |
| 2 | ggggacaagtttgtacaaaaaagcaggcttgatgCCGCGAGAGCATTGGGAGAG, ggggaccactttgtacaagaaagctgggtcTGTGCGTCTTCCCGTGAGCC | Mycgr3\|96691  JGI  Ensembl |  |  | 0 % |  | ``` 1	ATGCCGCGAGAGCATTGGGAGAGGAAGGTCTTCCAGGACCACCATGAGACAAGCCGCATCCTTCGCAACACGAAGTCCTGGCTGGTCGATGGAGAGCAAGGCAGAGAGACTGCGCGGATAGGATTGATCAAGAAGCTCTCATCGAAGGCGAAGCCAAACGTGAAGCCTGGAGTCCAGCGGTGGAACAACACATTTGGAAACCGTGTCGGACGAGGAGGAGTGCAACGAACGACAGACGCAGCGCACTGCGAGTCCCTGCGAGGCGCCCCGAGCGGGGTTTTGCCCATCCCGGAGCGTGGCAAGAGAAGGAGAAAGCAACTTTCAGATCATACCAGACGCGGCAAGCGAGGAGCCTTCATGAGTTCGAATCGTCCTGAGAGGAGATATAGAGGGCTCACGGGAAGACGCACA	411 	\|\|\|\|\|\|\|\|\|\|\|\|\| \|\|\|\|\|\|\|\|\|\|\|\|\|\|\|\|\|\|\|\|\|\|\|\|\|\|\|\|\|\|\|\|\|\|\|\|\|\|\|\|\|\|\|\|\|\|\|\|\|\|\|\|\|\|\|\|\|\|\|\|\|\|\|\|\|\|\|\|\|\|\|\|\|\|\|\|\|\|\|\|\|\|\|\|\|\|\|\|\|\|\|\|\|\|\|\|\|\|\|\|\|\|\|\|\|\|\|\|\|\|\|\|\|\|\|\|\|\|\|\|\|\|\|\|\|\|\|\|\|\|\|\|\|\|\|\|\|\|\|\|\|\|\|\|\|\|\|\|\|\|\|\|\|\|\|\|\|\|\|\|\|\|\|\|\|\|\|\|\|\|\|\|\|\|\|\|\|\|\|\|\|\|\|\|\|\|\|\|\|\|\|\|\|\|\|\|\|\|\|\|\|\|\|\|\|\|\|\|\|\|\|\|\|\|\|\|\|\|\|\|\|\|\|\|\|\|\|\|\|\|\|\|\|\|\|\|\|\|\|\|\|\|\|\|\|\|\|\|\|\|\|\|\|\|\|\|\|\|\|\|\|\|\|\|\|\|\|\|\|\|\|\|\|\|\|\|\|\|\|\|\|\|\|\|\|\|\|\|\|\|\|\|\|\|\|\|\|\|\|\|\|\|\|\|\|\|\|\|\|\|\|\|\|\|\|\|\|\|\|\|\|\|\|\|\|\|\|\|\|\|\|\|\|\|\|\|\|\|\|\|\|\|\|\|\|\|\|\|\|\|\|\|\|\|\|\|\|\|\|\|\|\|\|\|\|\|\|\|\|\|\|\|\|\|\|\|\|\|\|\|\|\|\|\|\|\|\|\|\|\|\|\|\|\|\|\|\| 	ATGCCGCGAGAGCNTTGGGAGAGGAAGGTCTTCCAGGACCACCATGAGACAAGCCGCATCCTTCGCAACACGAAGTCCTGGCTGGTCGATGGAGAGCAAGGCAGAGAGACTGCGCGGATAGGATTGATCAAGAAGCTCTCATCGAAGGCGAAGCCAAACGTGAAGCCTGGAGTCCAGCGGTGGAACAACACATTTGGAAACCGTGTCGGACGAGGAGGAGTGCAACGAACGACAGACGCAGCGCACTGCGAGTCCCTGCGAGGCGCCCCGAGCGGGGTTTTGCCCATCCCGGAGCGTGGCAAGAGAAGGAGAAAGCAACTTTCAGATCATACCAGACGCGGCAAGCGAGGAGCCTTCATGAGTTCGAATCGTCCTGAGAGGAGATATAGAGGGCTCACGGGAAGACGCACA ``` |
| 2 | ggggacaagtttgtacaaaaaagcaggcttgatgCCCACGAGTCTGTCTCCTGA, ggggaccactttgtacaagaaagctgggtcCTCATCATCTAAAACCATCC | Mycgr3\|96728  JGI  Ensembl |  |  | 100 % |  | ``` 1	ATGCCCACGAGTCTGTCTCCTGACGTCGAGCAAACACAAGAGCCGTTCGCGGCCACCGAGGAGCTGTACGGGCCACCCACAAAAGAAGATGAGATGGACAAGACCTGGGCGCAGAACGACTACAACACTCGATCTTGGGACATCCTTATGAGCGACATGTACCCCACCATGCGCTCGAACGCCCTGAGATCCGACGCTTATCACAAGAAGTGGGCCCTGATCTTAGAGGAGCTCTCGTCGACCGACACCAAGCCGGAGGTTGAGGCTCGATTGGACGAGATGAGTGGCAGCTTGCACAGGATGGTTTTAGATGATGAG	318 	\|\|\|\|\|\|\|\|\|\|\|\|\|\|\|\|\|\|\|\|\|\|\|\|\|\|\|\|\|\|\|\|\|\|\|\|\|\|\|\|\|\|\|\|\|\|\|\|\|\|\|\|\|\|\|\|\|\|\|\|\|\|\|\|\|\|\|\|\|\|\|\|\|\|\|\|\|\|\|\|\|\|\|\|\|\|\|\|\|\|\|\|\|\|\|\|\|\|\|\|\|\|\|\|\|\|\|\|\|\|\|\|\|\|\|\|\|\|\|\|\|\|\|\|\|\|\|\|\|\|\|\|\|\|\|\|\|\|\|\|\|\|\|\|\|\|\|\|\|\|\|\|\|\|\|\|\|\|\|\|\|\|\|\|\|\|\|\|\|\|\|\|\|\|\|\|\|\|\|\|\|\|\|\|\|\|\|\|\|\|\|\|\|\|\|\|\|\|\|\|\|\|\|\|\|\|\|\|\|\|\|\|\|\|\|\|\|\|\|\|\|\|\|\|\|\|\|\|\|\|\|\|\|\|\|\|\|\|\|\|\|\|\|\|\|\|\|\|\|\|\|\|\|\|\|\|\|\|\|\|\|\|\|\|\|\|\|\|\|\|\|\|\|\|\|\|\|\|\|\|\|\|\|\|\|\|\|\|\|\|\|\|\|\|\|\|\|\|\|\|\|\|\|\|\|\|\|\|\|\|\|\|\|\|\|\|\|\| 	ATGCCCACGAGTCTGTCTCCTGACGTCGAGCAAACACAAGAGCCGTTCGCGGCCACCGAGGAGCTGTACGGGCCACCCACAAAAGAAGATGAGATGGACAAGACCTGGGCGCAGAACGACTACAACACTCGATCTTGGGACATCCTTATGAGCGACATGTACCCCACCATGCGCTCGAACGCCCTGAGATCCGACGCTTATCACAAGAAGTGGGCCCTGATCTTAGAGGAGCTCTCGTCGACCGACACCAAGCCGGAGGTTGAGGCTCGATTGGACGAGATGAGTGGCAGCTTGCACAGGATGGTTTTAGATGATGAG ``` |
| 2 | ggggacaagtttgtacaaaaaagcaggcttgatgCACCTCTCGGCTTTTCTGGC, ggggaccactttgtacaagaaagctgggtcACAAGCATCAGGGTTGGGCA | Mycgr3\|97031  JGI  Ensembl |  |  | 100 % |  | ``` 1	ATGCACCTCTCGGCTTTTCTGGCGACCGTGCTCGGCTTCCAGTTGATTCCCACCATGGTGCTCGCCAAGGACAACCCGCACCAGCATTGCCAATGCATCTATCGCGACACTGGACTCGTGGACGAACGCGCGACAGCACTCGGATGTGACGACCTCGACCACGCAAACATGCACGGCGGTGGAGCCTCGCCGAACTGCGCAGCTGAGGATTTCTGGGGCATCAGGAACGACTGCTGGCAATGCAAGTGCGACCAGTATGTCTCGAACGGGCGTGCATTGGGTTCGAACTGCTGGAACGCTGATGGAGGACGGATCAACATCCACTGCGGCGGTGTGCCC-AACCCTGATGCTTG	353 	\|\|\|\|\|\|\|\|\|\|\|\|\|\|\|\|\|\|\|\|\|\|\|\|\|\|\|\|\|\|\|\|\|\|\|\|\|\|\|\|\|\|\|\|\|\|\|\|\|\|\|\|\|\|\|\|\|\|\|\|\|\|\|\|\|\|\|\|\|\|\|\|\|\|\|\|\|\|\|\|\|\|\|\|\|\|\|\|\|\|\|\|\|\|\|\|\|\|\|\|\|\|\|\|\|\|\|\|\|\|\|\|\|\|\|\|\|\|\|\|\|\|\|\|\|\|\|\|\|\|\|\|\|\|\|\|\|\|\|\|\|\|\|\|\|\|\|\|\|\|\|\|\|\|\|\|\|\|\|\|\|\|\|\|\|\|\|\|\|\|\|\|\|\|\|\|\|\|\|\|\|\|\|\|\|\|\|\|\|\|\|\|\|\|\|\|\|\|\|\|\|\|\|\|\|\|\|\|\|\|\|\|\|\|\|\|\|\|\|\|\|\|\|\|\|\|\|\|\|\|\|\|\|\|\|\|\|\|\|\|\|\|\|\|\|\|\|\|\|\|\|\|\|\|\|\|\|\|\|\|\|\|\|\|\|\|\|\|\|\|\|\|\|\|\|\|\|\|\|\|\|\|\|\|\|\|\|\|\|\|\|\|\|\|\|\|\|\|\|\|\|\|\|\|\|\| \|\|\|\|\|\|\|\|\| \|\|\|\|\|\|\|\|\|\| \|\|\|\|\|\|\|\| \|  \|\|\|\|\|\|\|\|\| \|\|\|\| 	ATGCACCTCTCGGCTTTTCTGGCGACCGTGCTCGGCTTCCAGTTGATTCCCACCATGGTGCTCGCCAAGGACAACCCGCACCAGCATTGCCAATGCATCTATCGCGACACTGGACTCGTGGACGAACGCGCGACAGCACTCGGATGTGACGACCTCGACCACGCAAACATGCACGGCGGTGGAGCCTCGCCGAACTGCGCAGCTGAGGATTTCTGGGGCATCAGGAACGACTGCTGGCAATGCAAGTGCGACCAGTATGTCTCGAACGGGCGTGCATTGGGTTCGAACTGCTGGAACGCTGATGGANGACGGATCANCATCCACTGCNGCGGTGTGNCANAACCCTGATNCTTG ```      ``` 1	ATGCACCTCTCGGCTTTTCTGGCGACCGTGCTCGGCTTCCAGTTGATTCCCACCATGGTGCTCGCCAAGGACAACCCGCACCAGCATTGCCAATGCATCTATCGCGACACTGGACTCGTGGACGAACGCGCGACAGCACTCGGATGTGACGACCTCGACCACGCAAACATGCACGGCGGTGGAGCCTCGCCGAACTGCGCAGCTGAGGATTTCTGGGGCATCAGGAACGACTGCTGGCAATGCAAGTGCGACCAGTATGTCTCGAACGGGCGTGCATTGGGTTCGAACTGCTGGAACGCTGATGGAGGACGGATCAACATCCACTGCGGCGGTGTGCCCAACCCTGATGCTT	352 	\|\|\|\|\|\|\|\|\|\|\|\|\|\|\|\|\|\|\|\|\|\|\|\|\|\|\|\|\|\|\|\|\|\|\|\|\|\|\|\|\|\|\|\|\|\|\|\|\|\|\|\|\|\|\|\|\|\|\|\|\|\|\|\|\|\|\|\|\|\|\|\|\|\|\|\|\|\|\|\|\|\|\|\|\|\|\|\|\|\|\|\|\|\|\|\|\|\|\|\|\|\|\|\|\|\|\|\|\|\|\|\|\|\|\|\|\|\|\|\|\|\|\|\|\|\|\|\|\|\|\|\|\|\|\|\|\|\|\|\|\|\|\|\|\|\|\|\|\|\|\|\|\|\|\|\|\|\|\|\|\|\|\|\|\|\|\|\|\|\|\|\|\|\|\|\|\|\|\|\|\|\|\|\|\|\|\|\|\|\|\|\|\|\|\|\|\|\|\|\|\|\|\|\|\|\|\|\|\|\|\|\|\|\|\|\|\|\|\|\|\|\|\|\|\|\|\|\|\|\|\|\|\|\|\|\|\|\|\|\|\|\|\|\|\|\|\|\|\|\|\|\|\|\|\|\|\|\|\|\|\|\|\|\|\|\|\|\|\|\|\|\|\|\|\|\|\|\|\|\|\|\|\|\|\|\|\|\|\|\|\|\|\|\|\|\|\|\|\|\|\|\|\|\|\|\|\|\|\|\|\|\|\|\|\|\|\|\|\|\|\|\|\|\|\|\|\|\|\|\|\|\|\|\|\|\|\|\|\|\|\| \|\|\|\|\|\|\|\|\|\| 	ATGCACCTCTCGGCTTTTCTGGCGACCGTGCTCGGCTTCCAGTTGATTCCCACCATGGTGCTCGCCAAGGACAACCCGCACCAGCATTGCCAATGCATCTATCGCGACACTGGACTCGTGGACGAACGCGCGACAGCACTCGGATGTGACGACCTCGACCACGCAAACATGCACGGCGGTGGAGCCTCGCCGAACTGCGCAGCTGAGGATTTCTGGGGCATCAGGAACGACTGCTGGCAATGCAAGTGCGACCAGTATGTCTCGAACGGGCGTGCATTGGGTTCGAACTGCTGGAACGCTGATGGAGGACGGATCAACATCCACTGCGGCGGTGTGCCCAANCCTGATGCTT ``` |
| 2 | ggggacaagtttgtacaaaaaagcaggcttgatgCTGCTCGCACTCTTCATCGC, ggggaccactttgtacaagaaagctgggtcGTAACCAAAAACGTAGCCTG | Mycgr3\|97178  JGI  Ensembl |  |  | 0 % |  | ``` 1	ATGCTGCTCGCACTCTTCATCGCCCTGCCATACTTCGGCATCTTCATTACCTACGATCCCTTGGCCGTATTCCACTCGGAACAGCTTCTACAACTCGCTCTCCTCGATATCATCTTCACCCTCAGCATCTGCTTCTTTTACACTCTGAGTTTGCCCACGATCTCCTTTTCCAGTCTCGGTCGAAGTCTCGCCGCCTTCGACTTCTTTCTCGCGGTCTTCATTTGGGCTATGTTAATCAACAGACTTCTCATCCATGAGTTTCATAGCTCCGGCTCGACTGTGATTATCGCGAACATCGCCAGCACCCTCGCTGCTTCCCTTGTTGCCAAGATTGCAGGCTACGTTTTTGGTTAC	354 	\|\|\|\|\|\|\|\|\|\|\|\|\|\|\|\|\|\|\|\|\|\|\|\|\|\|\|\|\|\|\|\|\|\|\|\|\|\|\|\|\|\|\|\|\|\|\|\|\|\|\|\|\|\|\|\|\|\|\|\|\|\|\|\|\|\|\|\|\|\|\|\|\|\|\|\|\|\|\|\|\|\|\|\|\|\|\|\|\|\|\|\|\|\|\|\|\|\|\|\|\|\|\|\|\|\|\|\|\|\|\|\|\|\|\|\|\|\|\|\|\|\|\|\|\|\|\|\|\|\|\|\|\|\|\|\|\|\|\|\|\|\|\|\|\|\|\|\|\|\|\|\|\|\|\|\|\|\|\|\|\|\|\|\|\|\|\|\|\|\|\|\|\|\|\|\|\|\|\|\|\|\|\|\|\|\|\|\|\|\|\|\|\|\|\|\|\|\|\|\|\|\|\|\|\|\|\|\|\|\|\|\|\|\|\|\|\|\|\|\|\|\|\|\|\|\|\|\|\|\|\|\|\|\|\|\|\|\|\|\|\|\|\|\|\|\|\|\|\|\|\|\|\|\|\|\|\|\|\|\|\|\|\|\|\|\|\|\|\|\|\|\|\|\|\|\|\|\|\|\|\|\|\|\|\|\|\|\|\|\|\|\|\|\|\|\|\|\|\|\|\|\|\|\|\|\|\|\|\|\|\|\|\|\|\|\|\|\|\|\|\|\|\|\|\|\|\|\|\|\|\|\|\|\|\|\|\|\|\|\|\|\|\|\|\|\|\|\|\|\|\|\|\|\| 	ATGCTGCTCGCACTCTTCATCGCCCTGCCATACTTCGGCATCTTCATTACCTACGATCCCTTGGCCGTATTCCACTCGGAACAGCTTCTACAACTCGCTCTCCTCGATATCATCTTCACCCTCAGCATCTGCTTCTTTTACACTCTGAGTTTGCCCACGATCTCCTTTTCCAGTCTCGGTCGAAGTCTCGCCGCCTTCGACTTCTTTCTCGCGGTCTTCATTTGGGCTATGTTAATCAACAGACTTCTCATCCATGAGTTTCATAGCTCCGGCTCGACTGTGATTATCGCGAACATCGCCAGCACCCTCGCTGCTTCCCTTGTTGCCAAGATTGCAGGCTACGTTTTTGGTTAC ``` |
| 2 | ggggacaagtttgtacaaaaaagcaggcttgatgGAGCACAATACCTCGAGCAC, ggggaccactttgtacaagaaagctgggtcCTGTCCGTCCTTCGCTGTTG | Mycgr3\|97384  JGI  Ensembl |  |  | 100 % |  |  |
| 2 | ggggacaagtttgtacaaaaaagcaggcttgatgATCTCCTTTGCCGGTCACTT, ggggaccactttgtacaagaaagctgggtcCGTACTGAAATCGAAGTTGT | Mycgr3\|97444  JGI  Ensembl |  |  | 100 % |  | ``` 1	ATGATCTCCTTTGCCGGTCACTTCCTCCTAGAGGTCTGCATGAAGTATCGAGACCAACTCAGTGTCAATGTCGACGAGAATCTCCGCAGAGTGAGCGCTGCCCTGGCGATCTTCGCTCGGCTACCGGTTTTGCCGCAACATTCGGTCTCGCGAGTCACTTCAAGTCTGATGAGGAGGTTGAATGAATGCACGGCAAGTCTGGGCATCGAAGGTCTTCTGACCGAGAGTCCGTTCGCAAACCTGAACGCCTTCACCAGCACAAGTGGCCATGGCAATCTGGAAGGCGGCAACACCACGACCTCCGTGTTTGAATTGGAAATGGATAATGGCTCGTTGCCCAACGATTTTCTACATGCCGACTTCAGCGATATGGCCTTTCCGAATCACAACTTCGATTTCAGTACG	405 	\|\|\|\|\|\|\|\|\|\|\|\|\|\|\|\|\|\|\|\|\|\|\|\|\|\|\|\|\|\|\|\|\|\|\|\|\|\|\|\|\|\|\|\|\|\|\|\|\|\|\|\|\|\|\|\|\|\|\|\|\|\|\|\|\|\|\|\|\|\|\|\|\|\|\|\|\|\|\|\|\|\|\|\|\|\|\|\|\|\|\|\|\|\|\|\|\|\|\|\|\|\|\|\|\|\|\|\|\|\|\|\|\|\|\|\|\|\|\|\|\|\|\|\|\|\|\|\|\|\|\|\|\|\|\|\|\|\|\|\|\|\|\|\|\|\|\|\|\|\|\|\|\|\|\|\|\|\|\|\|\|\|\|\|\|\|\|\|\|\|\|\|\|\|\|\|\|\|\|\|\|\|\|\|\|\|\|\|\|\|\|\|\|\|\|\|\|\|\|\|\|\|\|\|\|\|\|\|\|\|\|\|\|\|\|\|\|\|\|\|\|\|\|\|\|\|\|\|\|\|\|\|\|\|\|\|\|\|\|\|\|\|\|\|\|\|\|\|\|\|\|\|\|\|\|\|\|\|\|\|\|\|\|\|\|\|\|\|\|\|\|\|\|\|\|\|\|\|\|\|\|\|\|\|\|\|\|\|\|\|\|\|\|\|\|\|\|\|\|\|\|\|\|\|\|\|\|\|\|\|\|\|\|\|\|\|\|\|\|\|\|\|\|\|\|\|\|\|\|\|\|\|\|\|\|\|\|\|\|\|\|\|\|\|\|\|\|\|\|\|\|\|\|\|\|\|\|\|\|\|\|\|\|\|\|\|\|\|\|\|\|\|\|\|\|\|\|\|\|\|\|\|\|\|\|\|\|\|\|\|\|\|\|\|\|\|\|\|\|\|\|\|\|\|\| 	ATGATCTCCTTTGCCGGTCACTTCCTCCTAGAGGTCTGCATGAAGTATCGAGACCAACTCAGTGTCAATGTCGACGAGAATCTCCGCAGAGTGAGCGCTGCCCTGGCGATCTTCGCTCGGCTACCGGTTTTGCCGCAACATTCGGTCTCGCGAGTCACTTCAAGTCTGATGAGGAGGTTGAATGAATGCACGGCAAGTCTGGGCATCGAAGGTCTTCTGACCGAGAGTCCGTTCGCAAACCTGAACGCCTTCACCAGCACAAGTGGCCATGGCAATCTGGAAGGCGGCAACACCACGACCTCCGTGTTTGAATTGGAAATGGATAATGGCTCGTTGCCCAACGATTTTCTACATGCCGACTTCAGCGATATGGCCTTTCCGAATCACAACTTCGATTTCAGTACG ``` |
| 2 | ggggacaagtttgtacaaaaaagcaggcttgatgAAGCTCGCCGCCGTCATCTC, ggggaccactttgtacaagaaagctgggtcGCAAGATCCGGGCCCATTGT | Mycgr3\|97449  JGI  Ensembl |  |  | 100 % |  | ``` 1	ATGAAGCTCGCCGCCGTCATCTCTACCGTCCTTCTTGGCGCTGTCCCTGTTCTCGCAGCATACCGTCCAGGCACTGGACCACCAGGCAAATGGTTTCCAGGAGTGAGGGCGTGGTGCGAGGGGTTGCCCGACGGCGGTGGTGATCCTATCGGAGACGCATGCAGGGGAGTGAAAATCGACACGGT	185 	\|\|\|\|\|\|\|\|\|\|\|\|\|\|\|\|\|\|\|\|\|\|\|\|\|\|\|\|\|\|\|\|\|\|\|\|\|\|\|\|\|\|\|\|\|\|\|\|\|\|\|\|\|\|\|\|\|\|\|\|\|\|\|\|\|\|\|\|\|\|\|\|\|\|\|\|\|\|\|\|\|\|\|\|\|\|\|\|\|\|\|\|\|\|\|\|\|\|\|\|\|\|\|\|\|\|\|\|\|\|\|\|\|\|\|\|\|\|\|\|\|\|\|\|\|\|\|\|\|\|\|\|\|\|\|\|\|\|\|\|\|\|\|\|\|\|\|\|\|\|\|\|\|\|\|\|\|\|\|\|\|\|\|\|\|\|\|\|\|\|\|\|\|\|\|\|\|\|\|\|\|\|\|\|\| 	ATGAAGCTCGCCGCCGTCATCTCTACCGTCCTTCTTGGCGCTGTCCCTGTTCTCGCAGCATACCGTCCAGGCACTGGACCACCAGGCAAATGGTTTCCAGGAGTGAGGGCGTGGTGCGAGGGGTTGCCCGACGGCGGTGGTGATCCTATCGGAGACGCATGCAGGGGAGTGAAAATCGACACGGT ```      ``` 183	GGTTCCTGGTGTGAAAGGAACCTGTTGCTTGAGGAACGAGCGGGACTCCAACTTCATGAACTACCTCTGCAATTCATACGACGGAGGCACAAACGGTCACG-TTTACCGCTCCGACAATGGGCCCGGATCTTGC	315 	\|\|\|\|\|\|\|\|\|\|\|\|\|\|\|\|\|\|\|\|\|\|\|\|\|\|\|\|\|\|\|\|\|\|\|\|\|\|\|\|\|\|\|\|\|\|\|\|\|\|\|\|\|\|\|\|\|\|\|\|\|\|\|\|\|\|\|\|\|\|\|\|\|\|\|\|\|\|\|\|\|\|\|\|\|\|\|\|\|\|\|\|\|\|\|\|\|\|\|\|\| \|\|\|\|\|\|\|\|\|\|\|\|\|\|\|\|\|\|\|\|\|\|\|\|\|\|\|\|\|\|\|\| 	GGTTCCTGGTGTGAAAGGAACCTGTTGCTTGAGGAACGAGCGGGACTCCAACTTCATGAACTACCTCTGCAATTCATACGACGGAGGCACAAACGGTCACGTTTTACCGCTCCGACAATGGGCCCGGATCTTGC ```      ``` 183	GGTTCCTGGTGTGAAAGGAACCTGTTGCTTGAGGAACGAGCGGGACTCCAACTTCATGAACTACCTCTGCAATTCATACGACGGAGGCACAAACGGTCACGTTTACCGCTCCGACAATGGGCCCGGATCTT	313 	\|\|\|\|\|\|\|\|\|\|\|\|\|\|\|\|\|\|\|\|\|\|\|\|\|\|\|\|\|\|\|\|\|\|\|\|\|\|\|\|\|\|\|\|\|\|\|\|\|\|\|\|\|\|\|\|\|\|\|\|\|\|\|\|\|\|\|\|\|\|\|\|\|\|\|\|\|\|\|\|\|\|\|\|\|\|\|\|\|\|\|\|\|\|\|\|\|\|\|\|\|\|\|\|\|\|\|\|\|\|\|\|\|\|\|\|\|\|\|\|\|\|\|\|\|\|\|\|\|\|\| 	GGTTCCTGGTGTGAAAGGAACCTGTTGCTTGAGGAACGAGCGGGACTCCAACTTCATGAACTACCTCTGCAATTCATACGACGGAGGCACAAACGGTCACGTTTACCGCTCCGACAATGGGCCCGGATCTT ``` |
| 2 | ggggacaagtttgtacaaaaaagcaggcttgatgGAGCAACCCAAGCCCTCGAC, ggggaccactttgtacaagaaagctgggtcCGACCTTCTCTGCCAGGTGC | Mycgr3\|97543  JGI  Ensembl |  |  | 100 % |  | ``` 1	ATGGAGCAACCCAAGCCCTCGACCGCGCTGTTTGATGTCTTCTTAAGACTCCGCCCTTCCGCTGCCATCGACAAGGAGCGATTTCTGGACGTCGAGCCGGGTGGCCTCTCAGCCCCAACCCACATCACGATCAAGTCTCCGACGGGCGATACAAGAAAGAGAGCAGTTGAGAGATTCGCATTTACGAGAGTGTTCGAGGAGAATGCTGGACAGCGGAGTGTTCGAGGAGAATGCTGGACAGCGCGAACTATTCGAGAACACGGGAGCCGTGGATCTCATTC	281 	\|\|\|\|\|\|\|\|\|\|\|\|\|\|\|\|\|\|\|\|\|\|\|\|\|\|\|\|\|\|\|\|\|\|\|\|\|\|\|\|\|\|\|\|\|\|\|\|\|\|\|\|\|\|\|\|\|\|\|\|\|\|\|\|\|\|\|\|\|\|\|\|\|\|\|\|\|\|\|\|\|\|\|\|\|\|\|\|\|\|\|\|\|\|\|\|\|\|\|\|\|\|\|\|\|\|\|\|\|\|\|\|\|\|\|\|\|\|\|\|\|\|\|\|\|\|\|\|\|\|\|\|\|\|\|\|\|\|\|\|\|\|\|\|\|\|\|\|\|\|\|\|\|\|\|\|\|\|\|\|\|\|\|\|\|\|\|\|\|\|\|\|\|\|\|\|\|\|\|\|\|\|\|\|\|\|\|\|\|\|\|\|\|\|\|\|\|\|\|\|\|\|\|\|\|\|\|\|\|\|\|\|\|\|\|\|\|\|\|\|\|\|\| \|\|\|\|\|\|\|\|\|\|\|\|\|\|\|\|\|\|\|\|\|\|\|\|\|\|\|\|\| \|\|\|\|\|\|\|\|\|\|\|\|\|\|\|\|\|\|\|\|\|\|\|\|\|\|\| 	ATGGAGCAACCCAAGCCCTCGACCGCGCTGTTTGATGTCTTCTTAAGACTCCGCCCTTCCGCTGCCATCGACAAGGAGCGATTTCTGGACGTCGAGCCGGGTGGCCTCTCAGCCCCAACCCACATCACGATCAAGTCTCCGACGGGCGATACAAGAAAGAGAGCAGTTGAGAGATTCGCATTTACGAGAGTGTTCGAGGAGAATGCTGGACAGCGGAGTGTTCNAGGAGAATGCTGGACAGCGCGAACTATTCTAGAACACGGGAGCCGTGGATCTCATTC ``` |
| 2 | ggggacaagtttgtacaaaaaagcaggcttgatgCTCTCTGTCATGCTTCACTC, ggggaccactttgtacaagaaagctgggtcCTTCCCGAGCCACTGTGTAC | Mycgr3\|97579  JGI  Ensembl |  |  | 100 % |  | ``` 1	ATGCTCTCT-GTCATGCTTCACTCCACCAAGAGCCACAACTGCCTGACTCCCGCTGCAGCGACAATGGCAGGTAGCGAGCTTGTGCTTTTCGAACCTAGCGTCTACTGTCTTGCAACTACTCGTAAAGGAGCTCAGTGCAAGAACCTTGTTAAGGAAATAGACCGAATCTCTGCAACGAGACTCATACACAACTCATACGGCGGCTTCGAAAATAATGCTCTTGACGACGACGTGCAACCCGGTATCATCTTCGGTGATGGCTCGAGGAATAGCTGGGAAAATGCGGCGGATTTCCGTGTATATGTCTTTGACGGCAGCGAGGCTCCCTTCTGTCGAGGCATCTTCGAGATTCGTACACAGTGGCTCGGGAAG	372 	\|\|\|\|\|\|\|\|\| \|\|\|\|\|\|\|\|\|\|\|\|\|\|\|\|\|\|\|\|\|\|\|\|\|\|\|\|\|\|\|\|\|\|\|\|\|\|\|\|\|\|\|\|\|\|\|\|\|\|\|\|\|\|\|\|\|\|\|\|\|\|\|\|\|\|\|\|\|\|\|\|\|\|\|\|\|\|\|\|\|\|\|\|\|\|\|\|\|\|\|\|\|\|\|\|\|\|\|\|\|\|\|\|\|\|\|\|\|\|\|\|\|\|\|\|\|\|\|\|\|\|\|\|\|\|\|\|\|\|\|\|\|\|\|\|\|\|\|\|\|\|\|\|\|\|\|\|\|\|\|\|\|\|\|\|\|\|\|\|\|\|\|\|\|\|\|\|\|\|\|\|\|\|\|\|\|\|\|\|\|\|\|\|\|\|\|\|\|\|\|\|\|\|\|\|\|\|\|\|\|\|\|\|\|\|\|\|\|\|\|\|\|\|\|\|\|\|\|\|\|\|\|\|\|\|\|\|\|\|\|\|\|\|\|\|\|\|\|\|\|\|\|\|\|\|\|\|\|\|\|\|\|\|\|\|\|\|\|\|\|\|\|\|\|\|\|\|\|\|\|\|\|\|\|\|\|\|\|\|\|\|\|\|\|\|\|\|\|\|\|\|\|\|\|\|\|\|\|\|\|\|\|\|\|\|\|\|\|\|\|\|\|\|\|\|\|\|\|\|\|\|\|\|\|\|\|\|\|\|\|\|\|\|\|\|\|\|\|\|\|\|\|\|\|\|\|\|\|\|\|\|\|\|\|\|\|\|\|\|\|\|\| 	ATGCTCTCTGGTCATGCTTCACTCCACCAAGAGCCACAACTGCCTGACTCCCGCTGCAGCGACAATGGCAGGTAGCGAGCTTGTGCTTTTCGAACCTAGCGTCTACTGTCTTGCAACTACTCGTAAAGGAGCTCAGTGCAAGAACCTTGTTAAGGAAATAGACCGAATCTCTGCAACGAGACTCATACACAACTCATACGGCGGCTTCGAAAATAATGCTCTTGACGACGACGTGCAACCCGGTATCATCTTCGGTGATGGCTCGAGGAATAGCTGGGAAAATGCGGCGGATTTCCGTGTATATGTCTTTGACGGCAGCGAGGCTCCCTTCTGTCGAGGCATCTTCGAGATTCGTACACAGTGGCTCGGGAAG ``` |
| 2 | ggggacaagtttgtacaaaaaagcaggcttgatgACCGATGTACTCGACTATCT, ggggaccactttgtacaagaaagctgggtcATTGGGGCCAGCTTTTGCAT | Mycgr3\|97628  JGI  Ensembl |  |  | 0 % |  |  |
| 2 | ggggacaagtttgtacaaaaaagcaggcttgatgCCAGCTGGTTCGAACCTCCG, ggggaccactttgtacaagaaagctgggtcCTTCATCACGTAAGTTACGG | Mycgr3\|97740  JGI  Ensembl |  |  | 0 % |  |  |
| 2 | ggggacaagtttgtacaaaaaagcaggcttgatgTTCCACCTGGACGCGCGTAT, ggggaccactttgtacaagaaagctgggtcGACGAGCTCTTGCCCATTCT | Mycgr3\|97770  JGI  Ensembl |  |  | 0 % |  |  |
| 2 | ggggacaagtttgtacaaaaaagcaggcttgatgTCCGACGTACGCGCCGCGCG, ggggaccactttgtacaagaaagctgggtcAAGGAGGGAGGGAGCTGGAC | Mycgr3\|97817  JGI  Ensembl |  |  | 0 % |  |  |
| 2 | ggggacaagtttgtacaaaaaagcaggcttgatgCAGTTGGGCAGTCAGCAAGC, ggggaccactttgtacaagaaagctgggtcAATCAGGTATGAGGTTGCAA | Mycgr3\|97919  JGI  Ensembl |  |  | 100 % |  | ``` 1	ATGCAGTTGGGCAGTCAGCAAGCTCCCCTCAATCTCGATGCGGGCTCGGCGAGTCCAAGTGAGGTCGTGTCTCGCGCTGTCTCCGATGAGGAAGACAGCGAGAGCGATGACGGTGTGATGTCTCCCTCGATCCGGAAAAGTATTGGTGCAGATATGCGAGAGCCTGTCTCCAAGGATGAGGTCCAGCAAGACGAGGATCAGAATGTTGGTAGTGCAAAGATGGAAGAGTCTGTTTTCGAGGTCAAGCTCCAGCAAGATGATCTCGCATCGGATGAAAATTTCGAGAGCAACATTGCAACCTCATACCTGATT	312 	\|\|\|\|\|\|\|\|\|\|\|\|\|\|\|\|\|\|\|\|\|\|\|\|\|\|\|\|\|\|\|\|\|\|\|\|\|\|\|\|\|\|\|\|\|\|\|\|\|\|\|\|\|\|\|\|\|\|\|\|\|\|\|\|\|\|\|\|\|\|\|\|\|\|\|\|\|\|\|\|\|\|\|\|\|\|\|\|\|\|\|\|\|\|\|\|\|\|\|\|\|\|\|\|\|\|\|\|\|\|\|\|\|\|\|\|\|\|\|\|\|\|\|\|\|\|\|\|\|\|\|\|\|\|\|\|\|\|\|\|\|\|\|\|\|\|\|\|\|\|\|\|\|\|\|\|\|\|\|\|\|\|\|\|\|\|\|\|\|\|\|\|\|\|\|\|\|\|\|\|\|\|\|\|\|\|\|\|\|\|\|\|\|\|\|\|\|\|\|\|\|\|\|\|\|\|\|\|\|\|\|\|\|\|\|\|\|\|\|\|\|\|\|\|\|\|\|\|\|\|\|\|\|\|\|\|\|\|\|\|\|\|\|\|\|\|\|\|\|\|\|\|\|\|\|\|\|\|\|\|\|\|\|\|\|\|\|\|\|\|\|\|\|\|\|\|\|\|\|\|\|\|\|\|\|\|\|\|\|\|\|\|\|\|\|\|\|\|\|\|\|\|\|\|\|\|\|\|\|\|\|\| 	ATGCAGTTGGGCAGTCAGCAAGCTCCCCTCAATCTCGATGCGGGCTCGGCGAGTCCAAGTGAGGTCGTGTCTCGCGCTGTCTCCGATGAGGAAGACAGCGAGAGCGATGACGGTGTGATGTCTCCCTCGATCCGGAAAAGTATTGGTGCAGATATGCGAGAGCCTGTCTCCAAGGATGAGGTCCAGCAAGACGAGGATCAGAATGTTGGTAGTGCAAAGATGGAAGAGTCTGTTTTCGAGGTCAAGCTCCAGCAAGATGATCTCGCATCGGATGAAAATTTCGAGAGCAACATTGCAACCTCATACCTGATT ``` |
| 2 | ggggacaagtttgtacaaaaaagcaggcttgatgACCTTCGGGGGCCACTTTCT, ggggaccactttgtacaagaaagctgggtcTCTCTTGCTTCCACGGCGCG | Mycgr3\|97963  JGI  Ensembl |  |  | 100 % |  | ``` 236	AGCGTATCGATCCACGCTTGACCAGAAATTGTGTTCTGAAGCCCTTCTCGCGCCGTGGAAGCAAGAGATA	305 	\|\|\|\|\|\|\|\|\|\|\|\|\|\|\|\|\|\|\|\|\|\|\|\| \|\|\|\|\|\|\|\|\|\|\| \|\|\|\| \|\|\|\|\|\|\|\|\|\|\|\|\|\|\|\|\|\| \|\|\|\|\|\|\|\|\| 	AGCGTATCGATCCACGCTTGACCANAAATTGTGTTCCGAAGNCCTTCTCGCGCCGTGGAANCAAGAGATA ```      ``` 1	ATGACCTTCGGGGGCCACTTTCTTCCTCCCAATCAAATTCACCGCTTCGCTTCTGTGTGTGAGTGGCAGCTCGCCAATCCAGACTCCGCCGCGTTGAACGAAGACCCGATCAAGACCGTGCGGATGTATCTGTTGCTTACCGCAAAGCATCTACGCAATAAGAGAAATAAAAAGGAGGATTGGGCGGAGGAGTTCGGAGGGGAACAAATCGTCCGTCTGTTTGACCAGACGCTCGAGCGTA	241 	\|\|\|\|\|\|\|\|\|\|\|\|\|\|\|\|\|\|\|\|\|\|\|\|\|\|\|\|\|\|\|\|\|\|\|\|\|\|\|\|\|\|\|\|\|\|\|\|\|\|\|\|\|\|\|\|\|\|\|\|\|\|\|\|\|\|\|\|\|\|\|\|\|\|\|\|\|\|\|\|\|\|\|\|\|\|\|\|\|\|\|\|\|\|\|\|\|\|\|\|\|\|\|\|\|\|\|\|\|\|\|\|\|\|\|\|\|\|\|\|\|\|\|\|\|\|\|\|\|\|\|\|\|\|\|\|\|\|\|\|\|\|\|\|\|\|\|\|\|\|\|\|\|\|\|\|\|\|\|\|\|\|\|\|\|\|\|\|\|\|\|\|\|\|\|\|\|\|\|\|\|\|\| \|\|\|\|\|\|\|\|\|\|\|\|\|\|\|\|\|\|\|\|\|\|\|\|\|\|\|\|\|\|\|\|\|\|\|\|\|\|\|\|\|\|\|\|\|\|\|\|\|\|\|\|\| \|\|\| 	ATGACCTTCGGGGGCCACTTTCTTCCTCCCAATCAAATTCACCGCTTCGCTTCTGTGTGTGAGTGGCAGCTCGCCAATCCAGACTCCGCCGCGTTGAACGAAGACCCGATCAAGACCGTGCGGATGTATCTGTTGCTTACCGCAAAGCATCTACGCAATAAGAGAAATAAAAAGGAGGATTGGACGGAGGAGTTCGGAGGGGAACAAATCGTCCGTCTGTTTGACCAGACGCTCGAG-GTA ```      ``` 236	AGCGTATCGATCCACGCTTGACCAGAAATTGTGTTCTGAAGCCCTTCTCGCGCCGTGGAAGCAAGAGA	303 	\|\|\|\|\|\|\|\|\|\|\|\|\|\|\|\|\|\|\|\|\|\|\|\| \|\|\| \|\|\|\|\|\|\|\|\|\|\|\|\|\|\|\|\|\|\|\|\|\|\|\|\|\|\|\|\|\|\|\|\|\|\|\|\|\|\| 	AGCGTATCGATCCACGCTTGACCANAAANTGTGTTCTGAAGCCCTTCTCGCGCCGTGGAAGCAAGAGA ```      ``` 1	ATGACCTTCGGGGGCCACTTTCTTCCTCCCAATCAAATTCACCGCTTCGCTTCTGTGTGTGAGTGGCAGCTCGCCAATCCAGACTCCGCCGCGTTGAACGAAGACCCGATCAAGACCGTGCGGATGTATCTGTTGCTTACCGCAAAGCATCTACGCAATAAGAGAAATAAAAAGGAGGATTGGGCGGAGGAGTTCGGAGGGGAACAAATCGTCCGTCTGTTTGACCAGACGCTCGAGCGTA	241 	\|\|\|\|\|\|\|\|\|\|\|\|\|\|\|\|\|\|\|\|\|\|\|\|\|\|\|\|\|\|\|\|\|\|\|\|\|\|\|\|\|\|\|\|\|\|\|\|\|\|\|\|\|\|\|\|\|\|\|\|\|\|\|\|\|\|\|\|\|\|\|\|\|\|\|\|\|\|\|\|\|\|\|\|\|\|\|\|\|\|\|\|\|\|\|\|\|\|\|\|\|\| \|\|\|\|\|\|\|\|\|\|\|\|\|\|\|\|\|\|\|\|\|\|\|\|\|\|\|\|\|\|\|\|\|\|\|\|\|\|\|\|\|\|\|\|\|\|\|\|\|\|\| \|\|\|\|\|  \|\|\|\|\|\|\|\|\|\|\|\|\|\|\|\|\|\|\|\|\|\|\|\|\|\|\|\|\|\|\|\|\|\|\|\|\|\|\|\|\|\|\|\|\|\|\|\|\|\|\|\|\|\|\|\|\|\|\|\|\|\|\|\|\| \|\|\|\|\|\|\|\|\| \|\|\| 	ATGACCTTCGGGGGCCACTTTCTTCCTCCCAATCAAATTCACCGCTTCGCTTCTGTGTGTGAGTGGCAGCTCGCCAATCCAGACTCCGCCGCGTTGAACGAANACCCGATCAAGACCGTGCGGATGTATCTGTTGCTTACCGCAAAGCATCTACNCAATANCAGAAATAAAAAGGAGGATTGGGCGGAGGAGTTCGGAGGGGAACAAATCGTCCGTCTGTTTGACCAAACGCTCGAG-GTA ```      ``` 1	ATGACCTTCGGGGGCCACTTTCTTCCTCCCAATCAAATTCACCGC-TTCGCTTCTGTGTGTGAGTGGCAGCTCGCCAATCCAGACTCCGCCGCGTTGAACGAAGACCCGATCAAGACCGTGCGGATGTATCTGTTGCTTACCGCAAAGCATCTACGCA-ATAAGAGAAATAAAAAGG-AGGATTGGGCGGAGGAGT-TCGGAGGGGAA-CAAATCGTCCG	215 	\|\|\|\|\|\|\|\|\|\|\|\|\|\|\|\|\|\|\|\|\|\|\|\|\|\|\|\|\|\|\|\|\|\|\|\|\|\|\|\|\|\|\|\|\| \|\|\|\|\|\|\|\|\|\|\|\|\|\| \|\|\|\|\|\|\|\|\|\|\|\|\|\|\|\|\|\|\|\|\|\|\|\|\|\|\|\|\|\|\|\|\|\|\|\|\|\|\|\|\|\|\|\|\|\|\|\|\|\|\| \|\|\|\|\|\|\|\|\|\|\|\|\|\|\|\|\|\| \|\|\|\|\|\|\|\|\|\|\|\|  \|\|\|\|\|\|\|\|\| \|\| \|\| \| \|\|\| \|\|\|\| \|\|\|\| \|\|\|\|\|\|\|\|\|\|\|\|\|  \| \| \| \|\|\|\|\|\| \|\| \| \|\| \|\|\|\|\|\| 	ATGACCTTCGGGGGCCACTTTCTTCCTCCCAATCAAATTCACCGCTTTCGCTTCTGTGTGNGAGTGGCAGCTCGCCAATCCAGACTCCGCCGCGTTGAACGAAGACCCGATCNAGACCGTGCGGATGTATCNGTTGCTTACCGCNNAGCATCTACNCATATTANAGATATAANAAGGTAGGATTGGGCGGANNACTGTNGGAGGGNAAACNAANCGTCCG ``` |
| 2 | ggggacaagtttgtacaaaaaagcaggcttgatgCCGGCCAATCCTCGAGCAGA, ggggaccactttgtacaagaaagctgggtcCCTGGCCATCTTCTTTTCGG | Mycgr3\|97990  JGI  Ensembl |  |  | 100 % |  | ``` 271	CTGTCCAAGATATGTCACATTGCCGCCTCGTTCGCGTCCGAAAAGAAGATGGCCAGG	327 	\|\|\|\|\|\|\|\|\|\|\|\|\|\|\|\|\|\|\|\|\|\|\|\|\|\|\|\|\|\|\|\|\|\|\|\|\|\|\|\|\|\|\|\|\|\|\|\|\|\|\|\|\|\|\|\|\| 	CTGTCCAAGATATGTCACATTGCCGCCTCGTTCGCGTCCGAAAAGAAGATGGCCAGG ```      ``` 1	ATGCCGGCCAATCCTCGAGCAGATGACCATGTCGTTCGCGGTGATCTACTGCAGATTGAGTTTCCGCATAGTCAGGGAGAGTTGCAACTCCGTCAAGCTCCACCGAGCGAGCCGAAAAATGCTACGGGTTTCACAGTCTCCGGAGTCAAAGGCAAAGGATTGCGCTGGGAGATGGAGGATAAAGTCTACTTTACGAATCCTCAACGCCGGGCGGGAATGTACGACGGATACAGGAACATTTCAGCGAGGACGATGTTCGAGCCTTTCACT	270 	\|\|\|\|\|\|\|\|\|\|\|\|\|\|\|\|\|\|\|\|\|\|\|\|\|\|\|\|\|\|\|\|\|\|\|\|\|\|\|\|\|\|\|\|\|\|\|\|\|\|\|\|\|\|\|\|\|\|\|\|\|\|\|\|\|\|\|\|\|\|\|\|\|\|\|\|\|\|\|\|\|\|\|\|\|\|\|\|\|\|\|\|\|\|\|\|\|\|\|\|\|\|\|\|\|\|\|\|\|\|\|\|\|\|\|\|\|\|\|\|\|\|\|\|\|\|\|\|\|\|\|\|\|\|\|\|\|\|\|\|\|\|\|\|\|\|\|\|\|\|\|\|\|\|\|\|\|\|\|\|\|\|\|\|\|\|\|\|\|\|\|\|\|\|\|\|\|\|\|\|\|\|\|\|\|\|\|\|\|\|\|\|\|\|\|\|\|\|\|\|\|\|\|\|\|\|\|\|\|\|\|\|\|\|\|\|\|\|\|\|\|\|\|\|\|\|\|\|\|\|\|\|\|\|\|\|\|\|\|\|\|\|\|\|\|\|\|\|\|\|\|\|\|\|\|\|\|\|\|\|\|\|\|\|\|\|\|\|\|\| 	ATGCCGGCCAATCCTCGAGCAGATGACCATGTCGTTCGCGGTGATCTACTGCAGATTGAGTTTCCGCATAGTCAGGGAGAGTTGCAACTCCGTCAAGCTCCACCGAGCGAGCCGAAAAATGCTACGGGTTTCACAGTCTCCGGAGTCAAAGGCAAAGGATTGCGCTGGGAGATGGAGGATAAAGTCTACTTTACGAATCCTCAACGCCGGGCGGGAATGTACGACGGATACAGGAACATTTCAGCGAGGACGATGTTCGAGCCTTTCACT ``` |
| 2 | ggggacaagtttgtacaaaaaagcaggcttgatgCAAGCCATCTACTCCCAGCT, ggggaccactttgtacaagaaagctgggtcCGTCTTGTCCAACATCAACT | Mycgr3\|98022  JGI  Ensembl |  |  | 100 % |  | ``` 1	ATGCAAGCCATCTACTCCCAGCTTTACGAATACTCTATTCAGGATGCGCACTCGCGACAAAATTATCGTTCCGAGTCGCAATACATTGTCATTCCTTCCATCGGATCCTGCATCTCGACATCCTCGCCGCCGAGGAGGAATCCATCAAGGAGTAAGCCAGATGACACATCCGACCCCATCGTCTCGTCCATGCTCGCCATCGGGCTCATCTCTTCCGTGAGCTTTGCAAAATCGTGGTCGAGCCTCGCAGTCTCCTCAGGCATTGCCGAGGGCGGCAGTGGCAGCCAACTTAGCAGTGGTAGATGGAGAGGCGTCAAAATCTCCCAGCTGTACTCAGCGCCGGGCGGAACGCTCAGGGTCAGCGAGAAGGAGGAAGTACGGCTCGAGGAGTTGATGTTGGACAAGACG	408 	\|\|\|\|\|\|\|\|\|\|\|\|\|\|\|\|\|\|\|\|\|\|\|\|\|\|\|\|\|\|\|\|\|\|\|\|\|\|\|\|\|\|\|\|\|\|\|\|\|\|\|\|\|\|\|\|\|\|\|\|\|\|\|\|\|\|\|\|\|\|\|\|\|\|\|\|\|\|\|\|\|\|\|\|\|\|\|\|\|\|\|\|\|\|\|\|\|\|\|\|\|\|\|\|\|\|\|\|\|\|\|\|\|\|\|\|\|\|\|\|\|\|\|\|\|\|\|\|\|\|\|\|\|\|\|\|\|\|\|\|\|\|\|\|\|\|\|\|\|\|\|\|\|\|\|\|\|\|\|\|\|\|\|\|\|\|\|\|\|\|\|\|\|\|\|\|\|\|\|\|\|\|\|\|\|\|\|\|\|\|\|\|\|\|\|\|\|\|\|\|\|\|\|\|\|\|\|\|\|\|\|\|\|\|\|\|\|\|\|\|\|\|\|\|\|\|\|\|\|\|\|\|\|\|\|\|\|\|\|\|\|\|\|\|\|\|\|\|\|\|\|\|\|\|\|\|\|\|\|\|\|\|\|\|\|\|\|\|\|\|\|\|\|\|\|\|\|\|\|\|\|\|\|\|\|\|\|\|\|\|\|\|\|\|\|\|\|\|\|\|\|\|\|\|\|\|\|\|\|\|\|\|\|\|\|\|\|\|\|\|\|\|\|\|\|\|\|\|\|\|\|\|\|\|\|\|\|\|\|\|\|\|\|\|\|\|\|\|\|\|\|\|\|\|\|\|\|\|\|\|\|\|\|\|\|\|\|\|\|\|\|\|\|\|\|\|\|\|\|\|\|\|\|\|\|\|\|\|\|\|\|\|\|\|\|\|\|\|\|\|\|\|\|\|\|\|\|\| 	ATGCAAGCCATCTACTCCCAGCTTTACGAATACTCTATTCAGGATGCGCACTCGCGACAAAATTATCGTTCCGAGTCGCAATACATTGTCATTCCTTCCATCGGATCCTGCATCTCGACATCCTCGCCGCCGAGGAGGAATCCATCAAGGAGTAAGCCAGATGACACATCCGACCCCATCGTCTCGTCCATGCTCGCCATCGGGCTCATCTCTTCCGTGAGCTTTGCAAAATCGTGGTCGAGCCTCGCAGTCTCCTCAGGCATTGCCGAGGGCGGCAGTGGCAGCCAACTTAGCAGTGGTAGATGGAGAGGCGTCAAAATCTCCCAGCTGTACTCAGCGCCGGGCGGAACGCTCAGGGTCAGCGAGAAGGAGGAAGTACGGCTCGAGGAGTTGATGTTGGACAAGACG ``` |
| 2 | ggggacaagtttgtacaaaaaagcaggcttgatgGGATCAGAGGACACTGGTGC, ggggaccactttgtacaagaaagctgggtcCCCCCCTTGACGACCGACGT | Mycgr3\|98035  JGI  Ensembl |  |  | 100 % |  | ``` 1	ATGGGATCAGAGGAC-ACTGGTGCAGGCGATAAAGAGTTCGACGAGGACATTCCACCTTCGCCAGCGACTATCCGCAGCCAGGTCTCGAACTGCCCTGAAGAGCCCAAGAAGAATGGAGAAGGACGTCTTCAATTGATTCTCATCAGACCAGCTGCCATCATTAACAGTAGCATACTATCCAAGCGCGAGCTCCGTAGCGTCGCCGACCGGGACGATGGCGAACACGAGGATGACAACCTCTCCGCCTTCGGTTCCAGAGCTGAGGGAACGAACGAGGAAGGTCAGGGCGGGAAGAGGGCATACGTCGGTCGTCAAGGGGGG	321 	\|\|\|\|\|\|\|\|\|\|\|\| \|\|  \|\|\|\|\|\|\|\|\|\|\|\|\|\|\|\|\|\|\|\|\|\|\|\|\|\|\|\|\|\|\|\|\|\|\|\|\|\|\|\|\|\|\|\|\|\|\|\|\|\|\|\|\|\|\|\|\|\|\|\|\|\|\|\|\|\|\|\|\|\|\|\|\|\|\|\|\|\|\|\|\|\|\|\|\|\|\|\|\|\|\|\|\|\|\|\|\|\|\|\|\|\|\|\|\|\|\|\|\|\|\|\|\|\|\|\|\|\|\|\|\|\|\|\|\|\|\|\|\|\|\|\|\|\|\|\|\|\|\|\|\|\|\|\|\|\|\|\|\|\|\|\|\|\|\|\|\|\|\|\|\|\|\|\|\|\|\|\|\|\|\|\|\|\|\|\|\|\|\|\|\|\|\|\|\|\|\|\|\|\|\|\|\|\|\|\|\|\|\|\|\|\|\|\|\|\|\|\|\|\|\|\|\|\|\|\|\|\|\|\|\|\|\|\|\|\|\|\|\|\|\|\|\|\|\|\|\|\|\|\|\|\|\|\|\|\|\|\|\|\|\|\|\|\|\|\|\|\|\|\|\|\|\|\|\|\|\|\|\|\|\|\|\|\|\|\|\|\|\|\|\|\|\|\|\|\|\|\|\|\|\|\|\|\|\|\|\|\|\|\|\|\|\|\|\| 	ATGGGATCAGAGNACTTCTGGTGCAGGCGATAAAGAGTTCGACGAGGACATTCCACCTTCGCCAGCGACTATCCGCAGCCAGGTCTCGAACTGCCCTGAAGAGCCCAAGAAGAATGGAGAAGGACGTCTTCAATTGATTCTCATCAGACCAGCTGCCATCATTAACAGTAGCATACTATCCAAGCGCGAGCTCCGTAGCGTCGCCGACCGGGACGATGGCGAACACGAGGATGACAACCTCTCCGCCTTCGGTTCCAGAGCTGAGGGAACGAACGAGGAAGGTCAGGGCGGGAAGAGGGCATACGTCGGTCGTCAAGGGGGG ``` |
| 2 | ggggacaagtttgtacaaaaaagcaggcttgatgCCTCGAGACTGCATCCCGAC, ggggaccactttgtacaagaaagctgggtcCCGAGCGACCCTCTGTTCCT | Mycgr3\|98043  JGI  Ensembl |  |  | 0 % |  |  |
| 2 | ggggacaagtttgtacaaaaaagcaggcttgatgTTGGTGGTCCTGGCGACGGT, ggggaccactttgtacaagaaagctgggtcAGTGCGCTGCACCGGCTTTA | Mycgr3\|98116  JGI  Ensembl |  |  | 0 % |  |  |
| 2 | ggggacaagtttgtacaaaaaagcaggcttgatgAAAAAGGGAGTGGAATTTTG, ggggaccactttgtacaagaaagctgggtcTGGCCTCTCGGCGTTTTCGT | Mycgr3\|102326  JGI  Ensembl |  |  | 0 % |  |  |
| 2 | ggggacaagtttgtacaaaaaagcaggcttgatgCAACAAAGCGAGCGAATACC, ggggaccactttgtacaagaaagctgggtcTAAACCGCGCGCGAACCGCC | Mycgr3\|102794  JGI  Ensembl |  |  | 0 % |  |  |
| 2 | ggggacaagtttgtacaaaaaagcaggcttgatgCTTTCATGCTTTCCGACCAG, ggggaccactttgtacaagaaagctgggtcTGAGGGTGTGGTCGCGGCGG | Mycgr3\|103004  JGI  Ensembl |  |  | 0 % |  |  |
| 2 | ggggacaagtttgtacaaaaaagcaggcttgatgTCTTCCCGTCGAGTCCGGCG, ggggaccactttgtacaagaaagctgggtcGACTTCTCCCCACATCTTCT | Mycgr3\|105691  JGI  Ensembl |  |  | 100 % |  |  |
| 2 | ggggacaagtttgtacaaaaaagcaggcttgatgCTGGAGTACGGCCATTACTA, ggggaccactttgtacaagaaagctgggtcCATCTCCGTCTCGTCTCCAT | Mycgr3\|106705  JGI  Ensembl |  |  | 100 % |  |  |
| 3 | ggggacaagtttgtacaaaaaagcaggcttgatgTTGCCGCACACGAAGAACAC, ggggaccactttgtacaagaaagctgggtcCTGCGACTTCGCAGCCCTCA | Mycgr3\|36271  JGI  Ensembl | KOG0873 C-4 sterol methyl oxidase; | C-4 methylsterol oxidase activity; catalytic activity;  metabolic process; | 100 % |  | ``` 240	AAAAATCCCCTCCGCAAAGGAACAATGGGACTGCGCCATGCTCGTCCTCCTCTCACACTTCACCGTTGAACTCCCCCAAATCTGGCTCTTCCACCCCATGGCTCAATATTTCGGTCTAGCTACATCTGTTCCGTTCCCTCCGTGGTGGAAGATGGCCTACCAAATCGCCATCTTCTTCGTCATGGAAGACACCTGGCACTACTGGATGCACCGCGCCATGCACTGGGGTCCTCTGTACAAGAACATTCACAAGATCCACCACCAATACTCTGCACCTTTCGGACTGGCTGCTGAGTACGCCTCGCCGATCGAAGTCATGGTATTGGGACTCGGAACCGTCGGAAGCCCGATTTTGTGGTGTGCCGTCACCAAGGACTTGCACATCTTGACCATGTACATCTGGATTGTTTGCCGAGTGTTTCAGGCTATCGACGCGCATTCTGGCTACGAGTTTCCGTGGAGTCTCCACCATATTCTGCCTTTCTGGGCCGGTGCGGAGCACCACGACACGCATCACGAGAAGTTCATCGGAAACTACTCGAGCTCGTTCAGGTGGTGGGATTACGTCTTGGACACCGAGAGTGGACCGGAGGCATCGAAGAGGAGGAGAGAGGCCAAGCTGGCGAAGCTGAGGGCTGCGAAGTCGCAG	888 	\|\|\|\|\|\|\|\|\| \|\|  \|\|\|\|\|\|\|\|\|\|\| \|\|\|\| \|   \|\|\|\|  \|  \|\| \|\| \| \|\|\|\|\| \|\|\|\|\|\|\|\|\|\|\|\|\|\|\|\|\|\|\|\|\|\|\|\|\|\|\|\|\|\|\|\|\|\|\|\|\|\|\|\|\|\|\|\|\|\|\|\|\|\|\|\|\|\|\|\|\|\|\|\|\| \|\|\|\|\|\|\|\|\|\|\|\|\|\|\|\|\|\|\|\|\|\|\|\|\|\|\|\|\|\|\|\|\|\|\|\|\|\|\|\|\|\|\|\|\|\|\|\|\|\|\|\|\|\| \|\|\|\|\|\|\|\|\|\|\|\|\|\|\|\|\|\|\|\|\|\|\|\|\|\|\|\|\|\|\|\|\|\|\|\|\|\|\|\|\|\|\|\|\|\|\|\|\|\|\|\|\|\|\|\|\|\|\|\|\|\|\|\|\|\|\|\|\|\|\|\|\|\|\|\|\|\|\|\|\|\|\|\|\|\|\|\|\|\|\|\|\|\|\|\|\|\|\|\|\|\|\|\|\|\|\|\|\|\|\|\|\|\|\|\|\|\|\|\|\|\|\|\|\|\|\|\|\|\|\|\|\|\|\|\|\|\|\|\|\|\|\|\|\|\|\|\|\|\|\|\|\|\|\|\|\|\|\|\|\|\|\|\|\|\|\|\|\|\|\|\|\|\|\|\|\|\|\|\|\|\|\|\|\|\|\|\|\|\|\|\|\|\|\|\|\|\|\|\|\|\|\|\|\|\|\|\|\|\|\|\|\|\|\|\|\|\|\|\|\|\|\|\|\|\|\|\|\|\|\|\|\|\|\|\|\|\|\|\|\|\|\|\|\|\|\|\|\|\|\|\|\|\|\|\|\|\|\|\|\|\|\|\|\|\|\|\|\|\|\|\|\|\|\|\|\|\|\|\|\|\|\|\|\|\|\|\|\|\|\|\|\|\|\|\|\|\|\|\|\|\|\|\|\|\|\|\|\|\|\|\|\|\|\|\|\|\|\|\|\|\|\|\|\|\|\|\|\|\|\|\|\|\|\|\|\|\|\|\|\|\|\|\|\|\|\|\|\|\|\|\|\|\|\|\|\|\|\|\|\|\|\|\|\|\|\|\|\|\|\|\|\|\|\|\|\|\|\|\|\|\|\|\|\|\|\|\|\|\|\|\|\|\|\|\|\|\|\|\|\|\|\|\|\|\|\|\|\|\|\|\|\|\|\|\|\|\|\|\|\|\|\|\|\|\|\|\|\|\|\|\|\|\|\|\|\|\|\|\|\|\|\|\|\|\|\|\|\|\|\|\|\|\|\|\|\|\|\|\|\|\|\|\|\|\|\|\|\|\|\|\|\|\| 	AAAAATCCCNTCNNCAAAGGAACAANGGGANTNNNCCATNNTNNTCNTCNTNTCACANTTCACCGTTGAACTCCCCCAAATCTGGCTCTTCCACCCCATGGCTCAATATTTCGGTCTAGNTACATCTGTTCCGTTCCCTCCGTGGTGGAAGATGGCCTACCAAATCGCCATCTTNTTCGTCATGGAAGACACCTGGCACTACTGGATGCACCGCGCCATGCACTGGGGTCCTCTGTACAAGAACATTCACAAGATCCACCACCAATACTCTGCACCTTTCGGACTGGCTGCTGAGTACGCCTCGCCGATCGAAGTCATGGTATTGGGACTCGGAACCGTCGGAAGCCCGATTTTGTGGTGTGCCGTCACCAAGGACTTGCACATCTTGACCATGTACATCTGGATTGTTTGCCGAGTGTTTCAGGCTATCGACGCGCATTCTGGCTACGAGTTTCCGTGGAGTCTCCACCATATTCTGCCTTTCTGGGCCGGTGCGGAGCACCACGACACGCATCACGAGAAGTTCATCGGAAACTACTCGAGCTCGTTCAGGTGGTGGGATTACGTCTTGGACACCGAGAGTGGACCGGAGGCATCGAAGAGGAGGAGAGAGGCCAAGCTGGCGAAGCTGAGGGCTGCGAAGTCGCAG ``` |
| 3 | ggggacaagtttgtacaaaaaagcaggcttgatgCTAACACGAGCGGTCAACAG, ggggaccactttgtacaagaaagctgggtcGTATCCAACCTTGACCAGGT | Mycgr3\|37911  JGI  Ensembl | KOG1755 Profilin; | actin binding; actin cytoskeleton;  actin cytoskeleton organization and biogenesis;  cytoskeleton organization and biogenesis; | 0 % |  |  |
| 3 | ggggacaagtttgtacaaaaaagcaggcttgatgCGTTTCTCCCTCCTCTCCTC, ggggaccactttgtacaagaaagctgggtcGCTTGTGCCCGAACACCCAA | Mycgr3\|38105  JGI  Ensembl |  |  | 100 % |  | ``` 10	TCCCTCC-TCTCCTCCGCCCTCCTC-TTCGCCACAG-CCGCCT-TCTCCGCCCCCGTCGCTGAGCCTGCAGAGCTCGAGATCCGCCAGCAGGCCACCTACTGTGGCAACCAATACTATTCCGCCTCGCAAGTCTCCGCGGCCGTCAACAAGGGCTACAACTACTACGCCAACGG-CCAACAAGTCG-GGTCGGGCAACTACCCACACCAGT-ACAACAATCG-TGAGGGGTTCAGCTTTGCCGTCAGC-GGACCGTACCAGGAGTTCCCG-ATCTTGGCCAGTGGGTCG-ACGTACTCTGGTGG	302 	\|\|\|\| \|\| \|\|\|\|\|\|\| \|\|\|\|\|\| \|\| \|\|\|\|\|\|\|\|\|\|  \|\| \|\| \|\|\|\| \| \|\|\|\|\|\|\|\|\|\|\|\|\|\|\|\|\| \|\|\|\|\|\|\|\|\|\|\|\|\|\|\|\|\|\|\|\|\|\|\|\|\|\|\|\| \|\|\|\|\|\|\|\|\|\|\|\|\| \|\|\|\|\|\|\|\|\| \|\|\|\|\| \|\|\|\|  \|\|\|\|\|\|\|\|\|\|\|\|\|\| \| \|\|\|\|\|\|\|\|\|\|\|\|\|\|\|\|\|\|\|\|\| \| \|\|\|\|\|\|\|\|\|\|\| \|\|\|\|\|\|\|\|\|\|\|\|\| \|\|\|\|\| \| \|\| \|\|\|\|\|\| \|\|  \|\| \|\|\|\| \|\|\|\|\|\|\|\|\|\|\|  \|    \|\|\|\|\|\|\|\|\|\|\| \|\| \|\|\|\|\|\| \|\|\|\|\|\|\|\|\|\|\|\|\|\|\|\|\|\| \|\|  \|\|\|\|\|\|\|\|\|\| 	TCCCNCCNTCTCCTCTGCCCTCNTCGTTCGCCACAGTNCGNCTNTCTCNGNCCCCGTCGCTGAGCCTGNAGAGCTCGAGATCCGCCAGCAGGCCACCNACTGTGGCAACCANTACTATTCCNCCTCGTAAGTNNCCGCGGCCGTCAACNANGGCTACAACTACTACGCCAACNGACCAACAAGTCGTGGTCGGGCAACTANCCACANC-GTNACAACANTCNATGNGGGGNTCAGCTTTGCCNNCTNTNGGACCGTACCACGANTTCCCGAATCTTGGCCAGTGGGTCGAACNNACTCTGGTGG ```      ``` 1	ATGCGTTTCTCCCTCCTCTCCTCCGCCCTCCTCTTCGCCACAGCCGCCTTCTCCGCCCCCGTCGCTGAGCCTGCAGAGCTCGAGATCCGCCAGCAGGCCACCTACTGTGGCAACCAATACTATTCCGCCTCGCAAGTCTCCGCGGCCGTCAACAAGGGCTACAACTACTACGCCAACGGCCAACAAGTCGGGTCGGGCAACTACCCACACCAGTACAACAATCGTGAGGGGTTCAGCTTTGCCGTCAGCGGACCGTACCAGGAGTTCCCGATCTTGGCCAGTGGGTCGACGTACTCTGGTGGT	303 	\|\|\|\|\|\|\|\|\|\|\|\|\|\|\|\|\| \|\|\|\|\|\|\|\|\|\|\|\|\|\| \|\|\|\|\|\|\|\|\|\|\|\|\|\|\|\|\|\|\|\|\|\|\|\|\|\|\|\|\|\|\|\|\|\|\|\|\|\|\|\|\|\|\|\|\|\|\|\|\|\|\|\|\|\|\|\|\|\|\|\|\|\|\|\|\|\|\|\|\|\|\|\|\|\|\|\|\|\|\|\|\|\|\|\|\|\|\|\|\|\|\|\|\|\|\|\|\|\|\|\|\|\|\|\|\|\|\|\|\|\|\|\|\|\|\|\|\|\|\|\|\|\|\|\|\|\|\|\|\|\|\|\|\|\|\|\|\|\|\|\|\|\|\|\|\|\|\|\|\|\|\|\|\|\|\|\|\|\|\|\|\|\|\|\|\|\|\|\|\|\|\|\|\|\|\|\|\|\|\|\|\|\|\|\|\|\|\|\|\|\|\|\|\|\|\|\|\|\|\|\|\|\|\|\|\|\|\|\|\|\|\|\|\|\|\|\|\|\|\|\|\|\|\|\|\|\|\|\|\|\|\|\|\|\|\|\|\|\|\|\|\|\|\|\|\|\|\|\|\|\|\|\|\|\|\|\|\|\|\|\|\|\|\|\|\|\|\|\|\|\| 	ATGCGTTTCTCCCTCCTNTCCTCCGCCCTCCTNTTCGCCACAGCCGCCTTCTCCGCCCCCGTCGCTGAGCCTGCAGAGCTCGAGATCCGCCAGCAGGCCACCTACTGTGGCAACCAATACTATTCCGCCTCGCAAGTCTCCGCGGCCGTCAACAAGGGCTACAACTACTACGCCAACGGCCAACAAGTCGGGTCGGGCAACTACCCACACCAGTACAACAATCGTGAGGGGTTCAGCTTTGCCGTCAGCGGACCGTACCAGGAGTTCCCGATCTTGGCCAGTGGGTCGACGTACTCTGGTGGT ```      ``` 1	ATGCGTTTCTCCCTCCTCTCCTCCGCCCTCCTCTTCGCCACAGCCGCCTTCTCCGCCCCCGTCGCTGAGCCTGCAGAGCTCGAGATCCGCCAGCAGGCCACCTACTGTGGCAACCAATACTATTCCGCCTCGCAAGTCTCCGCGGCCGTCAACAAGGGCTACAACTACTACGCCAACGGCCAACAAGTCGGGTCGGGCAACTACCCACACCAGTACAACAATCGTGAGGGGTTCAGCTTTGCCGTCAGCGGACCGTACCAGGAGTTCCCGATCTTGGCCAGTGGGTCGACGTACTCTGGTGGT	303 	\|\|\|\|\|\|\|\|\|\|\|\|\|\|\|\|\|\|\|\|\|\|\|\|\|\|\| \|\|\|\|\|\|\|\|\|\|\|\|\|\|\|\|\|\|\|\|\|\|\|\|\|\|\| \|\|\|\|\|\|\|\|\|\|\|\|\|\|\|\|\|\|\|\|\|\|\|\|\|\|\|\|\|\|\|\|\|\|\|\|\|\|\|\|\|\|\|\|\|\|\|\|\|\|\|\|\|\|\|\|\|\|\|\|\|\|\|\|\|\|\|\|\|\|\|\|\|\|\|\|\|\|\|\|\|\|\|\|\|\|\|\|\|\|\|\|\|\|\|\|\|\|\|\|\|\|\|\|\|\|\|\|\|\|\|\|\|\|\|\|\|\|\|\|\|\|\|\|\|\|\|\|\|\|\|\|\|\|\|\|\|\|\|\|\|\|\|\|\|\|\|\|\|\| \|\|\|\|  \|\|\|\|\|\|\|\|\|\|\|\|\|\|\|\|\|\|\|\|\|\|\|\|\|\|\|\|\|\|\|\|\|\|\|\|\|\|\|\|\|\|\|\|\|\|\|\|\|\|\|\|\|\|\|\|\|\|\|\|\|\|\|\|\|\|\|\|\|\|\|\|\|\|\|\|\|\|\|\|\|\|\|\|\|\|\|\|\|\| 	ATGCGTTTCTCCCTCCTCTCCTCCGCCNTCCTCTTCGCCACAGCCGCCTTCTCCGNCCCCGTCGCTGAGCCTGCAGAGCTCGAGATCCGCCAGCAGGCCACCTACTGTGGCAACCAATACTATTCCGCCTCGCAAGTCTCCGCGGCCGTCAACAAGGGCTACAACTACTACGCCAACGGCCAACAAGTCGGGTCGGGCAACTACCCNCACCNTTACAACAATCGTGAGGGGTTCAGCTTTGCCGTCAGCGGACCGTACCAGGAGTTCCCGATCTTGGCCAGTGGGTCGACGTACTCTGGTGGT ```      ``` 1	ATGCGTTTCTCCCTCCTCTCCTCCGCCCTCCTCTTCGCCACAGCCGCCTTCTCCGCCCCCGTCGCTGAGCCTGCAGAGCTCGAGATCCGCCAGCAGGCCACCTACTGTGGCAACCAATACTATTCCGCCTCGCAAGTCTCCGCGGCCGTCAACAAGGGCTACAACTACTACGCCAACGGCCAACAAGTCGGGTCGGGCAACTACCCACACCAGTACAACAATCGTGAGGGGTTCAGCTTTGCCGTCAGCGGACCGTACCAGGAGTTCCCGATCTTGGCCAGTGGGTCGACGTACTCTGGTGGT	303 	\|\|\|\|\|\|\|\|\|\|\|\|\|\|\|\|\|\|\|\|\|\|\|\|\|\|\|\|\|\|\|\|\|\|\|\|\|\|\|\|\|\|\|\|\|\|\|\|\|\|\|\|\|\|\|\|\|\|\|\|\|\|\|\|\|\|\|\|\|\|\|\|\|\|\|\|\|\|\|\|\|\|\|\|\|\|\|\|\|\|\|\|\|\|\|\|\|\|\|\|\|\|\|\|\|\|\|\|\|\|\|\|\|\|\|\|\|\|\|\|\|\|\|\|\|\|\|\|\|\|\|\|\|\|\|\|\|\|\|\|\|\|\|\|\|\|\|\|\|\|\|\|\|\|\|\|\|\|\|\|\|\|\|\|\|\|\|\|\|\|\|\|\|\|\|\|\|\|\|\|\|\|\|\|\|\|\|\|\|\|\|\|\|\|\|\|\|\|\|\|\|\|\|\|\|\|\|\|\|\|\|\|\|\|\|\|\|\|\|\|\|\|\|\|\|\|\|\|\|\|\|\|\|\|\|\|\|\|\|\|\|\|\|\|\|\|\|\|\|\|\|\|\|\|\|\|\|\|\|\|\|\|\|\|\|\|\|\|\|\|\|\|\|\|\|\|\|\|\|\|\|\|\|\|\|\|\|\|\|\|\|\|\|\|\|\|\|\|\|\|\|\|\| 	ATGCGTTTCTCCCTCCTCTCCTCCGCCCTCCTCTTCGCCACAGCCGCCTTCTCCGCCCCCGTCGCTGAGCCTGCAGAGCTCGAGATCCGCCAGCAGGCCACCTACTGTGGCAACCAATACTATTCCGCCTCGCAAGTCTCCGCGGCCGTCAACAAGGGCTACAACTACTACGCCAACGGCCAACAAGTCGGGTCGGGCAACTACCCACACCAGTACAACAATCGTGAGGGGTTCAGCTTTGCCGTCAGCGGACCGTACCAGGAGTTCCCGATCTTGGCCAGTGGGTCGACGTACTCTGGTGGT ```      ``` 301	GGTTCTCCTGGACCGGACCGTGTTGTGTTCAACACGAGGGGACAGTGGGGTGGTACCATCACTCACACTGGCGCGAGCAACAACAACTTTGTTGGGTGTTCGGGCACAAGC	411 	\|\|\|\|\|\|\|\|\|\|\|\|\|\|\|\|\|\|\|\|\|\|\|\|\|\|\|\|\|\|\|\|\|\|\|\|\|\|\|\|\|\|\|\|\|\|\|\|\|\|\|\|\|\|\|\|\|\|\|\|\|\|\|\|\|\|\|\|\|\|\|\|\|\|\|\|\|\|\|\|\|\|\|\|\|\|\|\|\|\|\|\|\|\|\|\|\|\|\|\|\|\|\|\|\|\|\|\|\|\|\| 	GGTTCTCCTGGACCGGACCGTGTTGTGTTCAACACGAGGGGACAGTGGGGTGGTACCATCACTCACACTGGCGCGAGCAACAACAACTTTGTTGGGTGTTCGGGCACAAGC ``` |
| 3 | ggggacaagtttgtacaaaaaagcaggcttgatgCGCTCCTTCACGGCTTTGCT, ggggaccactttgtacaagaaagctgggtcAGCAGGTGCAGAGCAGTTGG | Mycgr3\|41315  JGI  Ensembl |  |  | 100 % |  | ``` 1	ATGCGCTCCTTCACGGCTTTGCTGGCCTCCAGCTCCCTTTGCGCTGTTCTCGCGCTCGCTGCTCCCGAAGG-AAGTGTCCGACGATGGGGCACTGTCAAGCGTTCGAGTGAGAAGGTCGTTCGTAATGATGGTTGCATGACAGAAGAGCAGGCCCAGCAGGTCGCCGACAATTTCAAGGAGCTGATCGCCGCCTACTCTGACGAACTTGCCAACGCGGTCCTCACCGAGGACTTTGTCGACTACTCTGACAGTGTCATCGAGCTCATCAACAACGGATGTCCAAACTCGCCC	291 	\|\|\|\|\| \|\|\|\|\|\|\| \|\|  \|\|\| \|\|\| \|\| \|\|\|\|\| \|\| \|\|\|\|\| \|\|\|\| \|\|\|\|\|\|\|\|\| \|\|\|\|\|\|\|\|\|\|\|\| \|\|\|\|\|\|\|\|\|\|\|\|\|\|\|\|\|\|\|\|\|\|\|\|\|\|\|\|\|\|\|\| \|\|\|\|\|\|\|\|\|\|\|\|\|\|\|\| \|\|\|\|\|\|\|\|\|\|\|\|\|\|\|\|\|\|\|\|\|\|\|\|\|\|\|\|\|\|\|\|\|\|\|\|\|\|\|\|\|\|\|\|\|\|\|\|\|\|\|\|\|\|\|\|\|\|\|\|\|\|\|\|\|\|\|\|\|\| \|\|\|\|\|\|\|\|\|\|\|\|\|\|\|\|\|\|\|\|\|\|\|\|\|\|\|\|\|\|\|\|\|\|\|\|\|\|\|\|\|\|\|\|\|\|\|\|\|\|\|\|\|\|\|\|\|\|\|\|\|\|\|\|\|\|\|\|\|\|\|\|\|\|\|\|\|\|\|\|\|\|\|\|\|\|\|\|\|\|\|\|\|\|\|\|\|\|\| 	ATGCGNTCCTTCANGG-NTTGNTGG-CTNCAGCT-CCNTTGCGNTGTTNTCGCGCTCGNTGCTCCCGAAGGGAAGTGTCCGACGATGGGGCACTGTCAAGCGTTNGAGTGAGAAGGTCGTTNGTAATGATGGTTGCATGACAGAAGAGCAGGCCCAGCAGGTCGCCGACAATTTCAAGGAGCTGATCGCCGCNTACTCTGACGAACTTGCCAACGCGGTCCTCACCGAGGACTTTGTCGACTACTCTGACAGTGTCATCGAGCTCATCAACAACGGATGTCCAAACTCGCCC ```      ``` 292	CTCGGGACTGCCACCTTCTCCAGCCGAGCGGACTTTGAAGCCGGCCAGAGCGCTCAGCCTCCGATCCCGTTCGAGCAGTTGAACATCTGG	381 	\|\|\|\|\|\|\|\|\|\|\|\|\|\|\|\|\|\|\|\|\|\|\|\|\|\|\|\|\|\|\|\|\|\|\|\|\|\|\|\|\|\|\|\|\|\|\|\|\|\|\|\|\|\|\|\|\|\|\|\|\|\|\|\|\|\|\|\|\|\|\|\|\|\|\|\|\|\|\|\|\|\|\|\|\|\|\|\|\|\| 	CTCGGGACTGCCACCTTCTCCAGCCGAGCGGACTTTGAAGCCGGCCAGAGCGCTCAGCCTCCGATCCCGTTCGAGCAGTTGAACATCTGG ```      ``` 380	GGCACACCTGCGACGCCGTCACTCTGCGCTGGAAGTCCGCTGGGCCCGGTCAGGAGCCGCAGCAAGTCACCGGCATCATCGTGATGGAGACTGAGTGTGGCGGTGATGGTACTCGACTCATCAACACCGTCTACTCCGAGTTCAACAGTGGTGCCTGGATGGCCAATCTCGGAATCTTCACGCCTACCAACTGCTCTGCACCTGCT	585 	\|\|\|\|\|\|\|\|\|\|\|\|\|\|\|\|\|\|\|\|\|\|\|\|\|\|\|\|\|\|\|\|\|\|\|\|\|\|\|\|\|\|\|\|\|\|\|\|\|\|\|\|\|\|\|\|\|\|\|\|\|\|\|\|\|\|\|\|\|\|\|\|\|\|\|\|\|\|\|\|\|\|\|\|\|\|\|\|\|\|\|\|\|\|\|\|\|\|\|\|\|\|\|\|\|\|\|\|\|\|\|\|\|\|\|\|\|\|\|\|\|\|\|\|\|\|\|\|\|\|\|\|\|\|\|\|\|\|\|\|\|\|\|\|\|\|\|\|\|\|\|\|\|\|\|\|\|\|\|\|\|\|\|\|\|\|\|\|\|\|\|\|\|\|\|\|\|\|\|\|\|\|\|\|\|\|\|\|\|\|\|\|\|\|\|\|\|\|\|\|\|\|\|\|\|\| 	GGCACACCTGCGACGCCGTCACTCTGCGCTGGAAGTCCGCTGGGCCCGGTCAGGAGCCGCAGCAAGTCACCGGCATCATCGTGATGGAGACTGAGTGTGGCGGTGATGGTACTCGACTCATCAACACCGTCTACTCCGAGTTCAACAGTGGTGCCTGGATGGCCAATCTCGGAATCTTCACGCCTACCAACTGCTCTGCACCTGCT ``` |
| 3 | ggggacaagtttgtacaaaaaagcaggcttgatgTCTTCACCCAGGAACCTCCG, ggggaccactttgtacaagaaagctgggtcTTCATCTTCACCTTTTAACC | Mycgr3\|43835  JGI  Ensembl | KOG3277 Uncharacterized conserved protein; |  | 100 % |  | ``` 252	AACCTCCACCCAACAACAATCCGACGCCGACGCCATCGCGCTCCGCAAAGCCATATCCCCGGCCTACCAACTCTGGTTCACATGCAAAAAGTGCCTCGAACGATCTGGCCACACCATCTCTAAGCAAGCCTACCACTTCGGTACTTGCGTCATCAACTGTCCGAAATGTAAATCCCAGCACCTGATATCGGACAATTTGAAGATTTTCAACGACAAGAGTATGACAATGGAGGATATCGCGAGGGAACATGGACAGCTGTTGAGGAAGGGCAGGTTAGGTGCGGATGGAGATGTGGAGTTTTACGACGACAAGGCGACGGATATTGTGGAGGAGGCGGCGAAGAAGGAACAGGAGAGGTTGGAGAGGTTAAAAGGTGAAGATGAA	636 	\|\|\|\|\|\|\|\|\|\|\|\|\|\|\|\|\|\|\|\|\|\|\|\|\|\|\|\|\|\|\|\|\|\|\|\|\|\|\|\|\|\|\|\|\|\|\|\|\|\|\|\|\|\|\|\|\|\|\|\|\|\|\|\|\|\|\|\|\|\|\|\|\|\|\|\|\|\|\|\|\|\|\|\|\|\|\|\|\|\|\|\|\|\|\|\|\|\|\|\|\|\|\|\|\|\|\|\|\|\|\|\|\|\|\|\|\|\|\|\|\|\|\|\|\|\|\|\|\|\|\|\|\|\|\|\|\|\|\|\|\|\|\|\|\|\|\|\|\|\|\|\|\|\|\|\|\|\|\|\|\|\|\|\|\|\|\|\|\|\|\|\|\|\|\|\|\|\|\|\|\|\|\|\|\|\|\|\|\|\|\|\|\|\|\|\|\|\|\|\|\|\|\|\|\|\|\|\|\|\|\|\|\|\|\|\|\|\|\|\|\|\|\|\|\|\|\|\|\|\|\|\|\|\|\|\|\|\|\|\|\|\|\|\|\|\|\|\|\|\|\|\|\|\|\|\|\|\|\|\|\|\|\|\|\|\|\|\|\|\|\|\|\|\|\|\|\|\|\|\|\|\|\|\|\|\|\|\|\|\|\|\|\|\|\|\|\|\|\|\|\|\|\|\|\|\|\|\|\|\|\|\|\|\|\|\|\|\|\|\|\|\|\|\|\|\|\|\|\|\|\|\|\|\|\|\|\|\|\|\|\|\|\|\|\|\|\|\|\|\|\|\|\|\|\|\|\|\|\|\|\|\|\|\|\|\|\|\|\|\|\|\|\|\|\|\|\|\|\|\|\|\|\|\|\| 	AACCTCCACCCAACAACAATCCGACGCCGACGCCATCGCGCTCCGCAAAGCCATATCCCCGGCCTACCAACTCTGGTTCACATGCAAAAAGTGCCTCGAACGATCTGGCCACACCATCTCTAAGCAAGCCTACCACTTCGGTACTTGCGTCATCAACTGTCCGAAATGTAAATCCCAGCACCTGATATCGGACAATTTGAAGATTTTCAACGACAAGAGTATGACAATGGAGGATATCGCGAGGGAACATGGACAGCTGTTGAGGAAGGGCAGGTTAGGTGCGGATGGAGATGTGGAGTTTTACGACGACAAGGCGACGGATATTGTGGAGGAGGCGGCGAAGAAGGAACAGGAGAGGTTGGAGAGGTTAAAAGGTGAAGATGAA ```      ``` 1	ATGTCTTCACCCAGGAACCTCCGCCGCTTTCAGCGTTCATTTGAGCAAGTCTTCAGGACGTATCGCTCCACTGCAAGGGTACAAAACCAACCTGCCATCCGCGCATTCTCCATCCCTTCCCGCCAACAAACCATCGCATCATCACCTGTGAACAAAACCACCCGCCGCCACAACTCCACCGACTCAGAACCTCAAACCCGGGCTCCACGGCCGCTCACAGACCGCACTCAAGAAGGTCAAACCCCTTCAGAAACCTCCACCCAACAACAATCCGACGCCGACGCCATCGCGCTCCGCAAAGCCATATCCCCGGCCTACCAACTCTGGTTCACATGCAAAAAGTGCCTCGAACGATCTGGCCACACCATCTCTAAGCAAGCCTACCACTTCGGTACTTGCGTCATCAACTGTCCGAAATGTAAATCCCAGCACCTGATATCGGACAATTTGAAGATTTTCAACGACAAGAGTATGACAATGGAGGATATCGCGAGGGAACATGGACAGCTGTTGAGGAAGGGCAGGTTAGGTGCGGATGGAGATGTGGAGTTTTACGACGACAAGGCGACGGATATTGTGGAGGAGGCGGCGAAGAAGGAACAGGAGAGGTTGGAGAGGTTAAAAGGTGAAGATGAA	636 	\|\|\|\|\|\|\|\|\|\|\|\|\|\|\|\|\|\|\|\|\|\|\|\|\|\|\|\|\|\|\|\|\|\|\|\|\|\|\|\|\|\|\|\|\|\|\|\|\|\|\|\|\|\|\|\|\|\|\|\|\|\|\|\|\|\|\|\|\|\|\|\|\|\|\|\|\|\|\|\|\|\|\|\|\|\|\|\|\|\|\|\|\|\|\|\|\|\|\|\|\|\|\|\|\|\|\|\|\|\|\|\|\|\|\|\|\|\|\|\|\|\|\|\|\|\|\|\|\|\|\|\|\|\|\|\|\|\|\|\|\|\|\|\|\|\|\|\|\|\|\|\|\|\|\|\|\|\|\|\|\|\|\|\|\|\|\|\|\|\|\|\|\|\|\|\|\|\|\|\|\|\|\|\|\|\|\|\|\|\|\|\|\|\|\|\|\|\|\|\|\|\|\|\|\|\|\|\|\|\|\|\|\|\|\|\|\|\|\|\|\|\|\|\|\|\|\|\|\|\|\|\|\|\|\|\|\|\|\|\|\|\|\|\|\|\|\|\|\|\|\|\|\|\|\|\|\|\|\|\|\|\|\|\|\|\|\|\|\|\|\|\|\|\|\|\|\|\|\|\|\|\|\|\|\|\|\|\|\|\|\|\|\|\|\|\|\|\|\|\|\|\|\|\|\|\|\|\|\|\|\|\|\|\|\|\|\|\|\|\|\|\|\|\|\|\|\|\|\|\|\|\|\|\|\|\|\|\|\|\|\|\|\|\|\|\|\|\|\|\|\|\|\|\|\|\|\|\|\|\|\|\|\|\|\|\|\|\|\|\|\|\|\|\|\|\|\|\|\|\|\|\|\|\|\|\|\|\|\|\|\|\|\|\|\|\|\|\|\|\|\|\|\|\|\|\|\|\|\|\|\|\|\|\|\|\|\|\|\|\|\|\|\|\|\|\|\|\|\|\|\|\|\|\|\|\|\|\|\|\|\|\|\|\|\|\|\|\|\|\|\|\|\|\|\|\|\|\|\|\|\|\|\|\|\|\|\|\|\|\|\|\|\|\|\|\|\|\|\|\|\|\|\|\|\|\|\|\|\|\|\|\|\|\|\|\|\|\|\|\|\|\|\|\|\|\|\|\|\|\|\|\|\|\|\|\|\|\|\|\|\|\|\|\|\|\|\|\|\|\|\|\|\|\|\|\|\|\|\|\|\|\|\|\|\|\|\|\|\|\|\|\|\|\|\|\|\|\|\|\|\|\|\|\|\|\|\|\|\|\|\|\|\|\|\|\|\|\|\|\|\|\|\|\|\|\|\|\|\|\|\|\|\| \|\|\|\|\|\|\|\|\|\|\| \|\|\|\|\|\|\|\|\|\|\|\|\|  \|\|\|\|\|\|\|\|\| \|\|\|\|\| 	ATGTCTTCACCCAGGAACCTCCGCCGCTTTCAGCGTTCATTTGAGCAAGTCTTCAGGACGTATCGCTCCACTGCAAGGGTACAAAACCAACCTGCCATCCGCGCATTCTCCATCCCTTCCCGCCAACAAACCATCGCATCATCACCTGTGAACAAAACCACCCGCCGCCACAACTCCACCGACTCAGAACCTCAAACCCGGGCTCCACGGCCGCTCACAGACCGCACTCAAGAAGGTCAAACCCCTTCAGAAACCTCCACCCAACAACAATCCGACGCCGACGCCATCGCGCTCCGCAAAGCCATATCCCCGGCCTACCAACTCTGGTTCACATGCAAAAAGTGCCTCGAACGATCTGGCCACACCATCTCTAAGCAAGCCTACCACTTCGGTACTTGCGTCATCAACTGTCCGAAATGTAAATCCCAGCACCTGATATCGGACAATTTGAAGATTTTCAACGACAAGAGTATGACAATGGAGGATATCGCGAGGGAACATGGACAGCTGTTGAGGAAGGGCAGGTTAGGTGCGGATGGAGATGTGGAGTTTTACGACGACAAGGCGACGGATATTGTGGAGGAGGCGGCGAANAAGGAACAGGANAGGTTGGAGAGGTN-AAAGGTGAANATGAA ``` |
| 3 | ggggacaagtttgtacaaaaaagcaggcttgatgAAGCTCCTCACACTCAACTT, ggggaccactttgtacaagaaagctgggtcAACCAGATGGCTCGGCAGGA | Mycgr3\|45617  JGI  Ensembl | KOG1088 Uncharacterized conserved protein; |  | 100 % |  | ``` 1	ATGAAGCTCCTCACACTCAACTTCCTCACATGCGCTCGGAAGTCATGCAAATCCTCCCCCGCCGCGTTTCCTCTACATCCCAAAGAAGCCGAACTCGAACAAGTTGAGCTCGATCTAAATCCGCTATTCCTCACGAACATGCTGCCGAGACTGGACTGGGCAGCGATGAAAACTTTGGCACAAGAA	186 	\|\|\|\|\|\|\|\|\|\|\|\|\|\|\|\|\|\|\|\|\|\|\|\|\|\|\|\|\|\|\|\|\|\|\|\|\|\|\|\|\|\|\|\|\|\|\|\|\|\|\|\|\|\|\|\|\|\|\|\|\|\|\|\|\|\|\|\|\|\|\|\|\|\|\|\|\|\|\|\|\|\|\|\|\|\|\|\|\|\|\|\|\|\|\|\|\|\|\|\|\|\|\|\|\|\|\|\|\|\|\|\|\|\|\|\|\|\|\|\|\|\|\|\|\|\|\|\|\|\|\|\|\|\|\|\|\|\|\|\|\|\|\|\|\|\|\|\|\|\|\|\|\|\|\|\|\|\|\|\|\|\|\|\|\|\|\|\|\|\|\|\|\|\|\|\|\|\|\|\|\|\|\|\|\|\| 	ATGAAGCTCCTCACACTCAACTTCCTCACATGCGCTCGGAAGTCATGCAAATCCTCCCCCGCCGCGTTTCCTCTACATCCCAAAGAAGCCGAACTCGAACAAGTTGAGCTCGATCTAAATCCGCTATTCCTCACGAACATGCTGCCGAGACTGGACTGGGCAGCGATGAAAACTTTGGCACAAGAA ```      ``` 187	CTTGGTCTGCCCAATCTTCCTGCTGAGACTCCCGCCGCCGAGGAGATGGAGTCCGAGGACGGCGAACCTTCGCAAACGCTCAAGGACCTGCATACACTGCTGATGGAGACGAGTATTGCATCGGGAAAGTTGGTTTGCGGAAACTGTGAGCATGAGTATGCTGTGAAGGAGGGCATCGCGAACTTTCTCCTGCCGAGCCATCTGGTT	393 	\|\|\|\|\|\|\|\|\|\|\|\|\|\|\|\|\|\|\|\|\|\|\|\|\|\|\|\|\|\|\|\|\|\|\|\|\|\|\|\|\|\|\|\|\|\|\|\|\|\|\|\|\|\|\|\|\|\|\|\|\|\|\|\|\|\|\|\|\|\|\|\|\|\|\|\|\|\|\|\|\|\|\|\|\|\|\|\|\|\|\|\|\|\|\|\|\|\|\|\|\|\|\|\|\|\|\|\|\|\|\|\|\|\|\|\|\|\|\|\|\|\|\|\|\|\|\|\|\|\|\|\|\|\|\|\|\|\|\|\|\|\|\|\|\|\|\|\|\|\|\|\|\|\|\|\|\|\|\|\|\|\|\|\|\|\|\|\|\|\|\|\|\|\|\|\|\|\|\|\|\|\|\|\|\|\|\|\|\|\|\|\|\|\|\|\|\|\|\|\|\|\|\|\|\|\|\| 	CTTGGTCTGCCCAATCTTCCTGCTGAGACTCCCGCCGCCGAGGAGATGGAGTCCGAGGACGGCGAACCTTCGCAAACGCTCAAGGACCTGCATACACTGCTGATGGAGACGAGTATTGCATCGGGAAAGTTGGTTTGCGGAAACTGTGAGCATGAGTATGCTGTGAAGGAGGGCATCGCGAACTTTCTCCTGCCGAGCCATCTGGTT ```      ``` 3	GAAGCTCCTCACACTCAACTTCCTCACATGCGCTCGGAAGTCATGCAAATCCTCCCCCGCCGCGTTTCCTCTACATCCCAAAGAAGCCGAACTCGAACAAGTTGAGCTCGATCTAAATCCGCTATTCCTCACGAACATGCTGCCGAGACTGGACTGGGCAGCGATGAAAACTTTGGCACAAGAA	186 	\|\|\|\|\|\|\| \|\|\|\|\| \|\|\|\| \|\|\|\|\|\|\|\|\|\|\|\|\|\|\| \|\|\|\|\|\|\|\|\|\|\|\|\|\|\|\|\|\|\|\|\|\|\|\|\|\|\|\|\|\|\|\|\|\|\|\|\|\|\|\|\|\|\|\|\|\|\|\|\|\|\|\|\|\|\|\|\|\|\|\|\|\|\|\|\|\|\|\|\|\|\|\|\|\|\|\|\|\|\|\|\|\|\|\|\|\|\|\|\|\|\|\|\|\|\|\|\|\|\|\|\|\|\|\|\|\|\|\|\|\|\|\|\|\|\|\|\|\|\|\|\|\|\|\|\|\|\|\|\|\|\|\|\|\|\|\|\|\|\|\|\|\|\|\|\|\|\|\|\| 	GAAGCTCNTCACANTCAANTTCCTCACATGCGCTNGGAAGTCATGCAAATCCTCCCCCGCCGCGTTTCCTCTACATCCCAAAGAAGCCGAACTCGAACAAGTTGAGCTCGATCTAAATCCGCTATTCCTCACGAACATGCTGCCGAGACTGGACTGGGCAGCGATGAAAACTTTGGCACAAGAA ``` |
| 3 | ggggacaagtttgtacaaaaaagcaggcttgatgCCAAAACGCTCAGCTCCTCA, ggggaccactttgtacaagaaagctgggtcAGACGGAGGAGACGACTTCT | Mycgr3\|46959  JGI  Ensembl | KOG1817 Ribonuclease; | RNA binding; RNA processing;  ribonuclease III activity; | 0 % |  |  |
| 3 | ggggacaagtttgtacaaaaaagcaggcttgatgTCGCTATACTCCTTCTACAT, ggggaccactttgtacaagaaagctgggtcGCACGACTCCCCAAACCACA | Mycgr3\|47014  JGI  Ensembl | KOG3368 Transport protein particle (TRAPP) complex subunit; | ER to Golgi vesicle-mediated transport; cis-Golgi network; | 100 % |  | ``` 38	CCGAATGCATCTACTCCAAACGCTGGACCTCTccacccaccaccacccccatccccacagaacccaacaaacccagcagccaaggcctcccccccTCCGCCGCAGCAACCGCCAAACTCGCCTCCAAAAACCAAGACGTCCAAAAACTCATCTTCGGCACCATCTTCTCTCTCCGCCGCATGGCCCGCCAACTCGGCGGTCCCTCCGATCAATTCCTCTCCTACCGTACGGGAGAGTACAAACTGCATTACTTCGAGACTCCGACCCAACTCAAACTCGTGCTGCTGACGGATACGCGGGTGGGCAATATGCGCACCGTGTTGCATCAGATTTGGGCCACACTGTATGTCGAGTATGTGGTCAAGTCGCCGCTCGCTCCCGTCGAGCATAAGGGTGGGGTCGGGGTGGGGAATGAATTGTTTGAgggggggTTGGAGAGCTTTATGG	484 	\|\|\|\|\|\|\|\|\|\|\|\|\|\|\|\|\|\|\|\|\|\|\|\|\|\|\|\|\|\|\|\|\|\|\|\|\|\|\|\|\|\|\|\|\|\|\|\|\|\|\|\|\|\|\|\|\|\|\|\|\|\|\|\|\|\|\|\|\|\|\|\|\|\|\|\|\|\|\|\|\|\|\|\|\|\|\|\|\|\|\|\|\|\|\|\|\|\|\|\|\|\|\|\|\|\|\|\|\|\|\|\|\|\|\|\|\|\|\|\|\|\|\|\|\|\|\|\|\|\|\|\|\|\|\|\|\|\|\|\|\|\|\|\|\|\|\|\|\|\|\|\|\|\|\|\|\|\|\|\|\|\|\|\|\|\|\|\|\|\|\|\|\|\|\|\|\|\|\|\|\|\|\|\|\|\|\|\|\|\|\|\|\|\|\|\|\|\|\|\|\|\|\|\|\|\|\|\|\|\|\|\|\|\|\|\|\|\|\|\|\|\|\|\|\|\|\|\|\|\|\|\|\|\|\|\|\|\|\|\|\|\|\|\|\|\|\|\|\|\|\|\|\|\|\|\|\|\|\|\|\|\|\|\|\|\|\|\|\|\|\|\|\|\|\|\|\|\|\|\|\|\|\|\|\|\|\|\|\|\|\|\|\|\|\|\|\|\|\|\|\|\|\|\|\|\|\|\|\|\|\|\|\|\|\|\|\|\|\|\|\|\|\|\|\|\|\|\|\|\|\|\|\|\|\|\|\|\|\|\|\|\|\|\|\|\|\|\|\|\|\|\|\|\|\|\|\|\|\|\|\|\|\|\|\|\|\|\|\|\|\|\|\|\|\|\|\|\|\|\|\|\|\|\|\|\|\|\|\|\|\|\|\|\|\|\|\|\|\|\|\|\|\|\|\|\|\|\|\|\|\|\|\|\|\|\|\|\|\|\|\|\|\|\|\|\|\|\|\|\|\|\|\|\|\|\| \|\| \|\|\| \|\|\| 	CCGAATGCATCTACTCCAAACGCTGGACCTCTCCACCCACCACCACCCCCATCCCCACAGAACCCAACAAACCCAGCAGCCAAGGCCTCCCCCCCTCCGCCGCAGCAACCGCCAAACTCGCCTCCAAAAACCAAGACGTCCAAAAACTCATCTTCGGCACCATCTTCTCTCTCCGCCGCATGGCCCGCCAACTCGGCGGTCCCTCCGATCAATTCCTCTCCTACCGTACGGGAGAGTACAAACTGCATTACTTCGAGACTCCGACCCAACTCAAACTCGTGCTGCTGACGGATACGCGGGTGGGCAATATGCGCACCGTGTTGCATCAGATTTGGGCCACACTGTATGTCGAGTATGTGGTCAAGTCGCCGCTCGCTCCCGTCGAGCATAAGGGTGGGGTCGGGGTGGGGAATGAATTGTTTGAGGGGGGGTTGGANAGNTTTNTGG ```      ``` 1	ATGTCGCTATACTCCTTCTACATCTTCGACCGACACA	37 	\|\|\|\|\|\|\|\|\|\|\|\|\|\|\|\|\|\|\|\|\|\|\|\|\|\|\|\|\|\|\|\|\|\|\|\|\| 	ATGTCGCTATACTCCTTCTACATCTTCGACCGACACA ```      ``` 78	caccacccccatccccacagaacccaacaaacccagcagccaaggcctcccccccTCCGCCGCAGCAACCGCCAAACTCGCCTCCAAAAACCAAGACGTCCAAAAACTCATCTTCGGCACCATCTTCTCTCTCCGCCGCATGGCCCGCCAACTCGGCGGTCCCTCCGATCAATTCCTCTCCTACCGTACGGGAGAGTACAAACTGCATTACTTCGAGACTCCGACCCAACTCAAACTCGTGCTGCTGACGGATACGCGGGTGGGCAATATGCGCACCGTGTTGCATCAGATTTGGGCCACACTGTATGTCGAGTATGTGGTCAAGTCGCCGCTCGCTCCCGTCGAGCATAAGGGTGGGGTCGGGGTGGGGAATGAATTGTTTGAgggggggTTGGAGAGCTTTATGGTGAGTTTGTCTTGTGAACTTCTGTGGTTTGGGGAGTCGTGC	525 	\|\|\|\|\|\|\|\|\|\|\|\|\|\|\|\|\|\|\|\|\|\|\|\|\|\|\|\|\|\|\|\|\|\|\|\|\|\|\|\|\|\|\|\|\|\|\|\|\|\|\|\|\|\|\|\|\|\|\|\|\|\|\|\|\|\|\|\|\|\|\|\|\|\|\|\|\|\|\|\|\|\|\|\|\|\|\|\|\|\|\|\|\|\|\|\|\|\|\|\|\|\|\|\|\|\|\|\|\|\|\|\|\|\|\|\|\|\|\|\|\|\|\|\|\|\|\|\|\|\|\|\|\|\|\|\|\|\|\|\|\|\|\|\|\|\|\|\|\|\|\|\|\|\|\|\|\|\|\|\|\|\|\|\|\|\|\|\|\|\|\|\|\|\|\|\|\|\|\|\|\|\|\|\|\|\|\|\|\|\|\|\|\|\|\|\|\|\|\|\|\|\|\|\|\|\|\|\|\|\|\|\|\|\|\|\|\|\|\|\|\|\|\|\|\|\|\|\|\|\|\|\|\|\|\|\|\|\|\|\|\|\|\|\|\|\|\|\|\|\|\|\|\|\|\|\|\|\|\|\|\|\|\|\|\|\|\|\|\|\|\|\|\|\|\|\|\|\|\|\|\|\|\|\|\|\|\|\|\|\|\|\|\|\|\|\|\|\|\|\|\|\|\|\|\|\|\|\|\|\|\|\|\|\|\|\|\|\|\|\|\|\|\|\|\|\|\|\|\|\|\|\|\|\|\|\|\|\|\|\|\|\|\|\|\|\|\|\|\|\|\|\|\|\|\|\|\|\|\|\|\|\|\|\|\|\|\|\|\|\|\|\|\|\|\|\|\|\|\|\|\|\|\|\|\|\|\|\|\|\|\|\|\|\|\|\|\|\|\|\|\|\|\|\|\|\|\|\|\|\|\|\|\|\|\|\|\|\|\|\|\|\|\|\|\|\|\|\|\|\|\|\|\|\|\|\|\|\|\|\|\|\|\|\|\|\|\|\| 	CACCACCCCCATCCCCACAGAACCCAACAAACCCAGCAGCCAAGGCCTCCCCCCCTCCGCCGCAGCAACCGCCAAACTCGCCTCCAAAAACCAAGACGTCCAAAAACTCATCTTCGGCACCATCTTCTCTCTCCGCCGCATGGCCCGCCAACTCGGCGGTCCCTCCGATCAATTCCTCTCCTACCGTACGGGAGAGTACAAACTGCATTACTTCGAGACTCCGACCCAACTCAAACTCGTGCTGCTGACGGATACGCGGGTGGGCAATATGCGCACCGTGTTGCATCAGATTTGGGCCACACTGTATGTCGAGTATGTGGTCAAGTCGCCGCTCGCTCCCGTCGAGCATAAGGGTGGGGTCGGGGTGGGGAATGAATTGTTTGAGGGGGGGTTGGAGAGCTTTATGGTGAGTTTGTCTTGTGAACTTCTGTGGTTTGGGGAGTCGTGC ``` |
| 3 | ggggacaagtttgtacaaaaaagcaggcttgatgTCTTCCGGCACGTCTCGAGG, ggggaccactttgtacaagaaagctgggtcCCACGGTTGAATGTATCTCA | Mycgr3\|47065  JGI  Ensembl | KOG3327 Thymidylate kinase/adenylate kinase; | ATP binding; dTDP biosynthetic process;  dTTP biosynthetic process;  thymidylate kinase activity; | 100 % |  | ``` 1	ATGTCTTCCGGCACGTCTCGAGGCCATTTGATCGT-CTTCGAAGGTCTCGATCGCTCCGGCAAGAGCACTCAATGCGCACGACTGGTAGATCACCTCCGCTCTCAAGGCAAAGAAGTCTTCCATCAACGCTTTCCGGACCGGACGACTGTGATCGGACAGATGATCAACAGCTATCTCAGTGGGACATCATCGCAGGAGGACCATGCCATTCATCTCTTGTTTTCAGCAAACCGCTGGGAGGCCTCCGAATCGATAAGACAGCACATTGAAAGGGGAACTACTGTGGTCGTTGATCGATATTACTACTCTGGTTGTGTCTACTCTGCAGCCAAGCAGAATCCTGCCATGGACCTTGCGTGGTGTCGGCATCCTGAGATTGGCTTGCCACGTCCCGACTTGTGTCTGTTTCTGGACATCGCTGCTGAGGAGGCTGCTAAACGCGGTGGCTTTGGCACCGAACGATACGAGAAACAGGAACTTCAGGACAGAGTGCGAGACTTGTTTGCCGAGTTGCAAACTCATCCGGACGAGGCTGAAGACATCGTCGTGATTGATGCTGGTCAGAGTGTCGAGACAGTAGAGCAAAGCATTCACGACGCAGTAGGACGCCGGCTCGATGGAGAGCATGCTGTCGGCCCAGTGAGATACATTCAACCGTGG	660 	\|\|\|\|\|\|\|\|\|\|\|\|\|\|\|\| \|\|\|\|\|\|\|\|\|\|\|\|\|\|\|\|\|\|  \|\| \|\|\|\|\|\|\|\|\|\|\|\|\|\| \|\|\|\|\|\|\|\|\|\|\|\|\|\|\|\|\|\|\|\|\|\|\|\|\|\|\|\|\|\|\|\|\|\|\|\|\|\|\|\|\|\|\|\|\|\|\|\|\|\|\|\|\|\|\|\|\|\|\|\|\|\|\|\|\|\|\|\|\|\|\|\|\|\|\|\|\|\|\|\|\|\|\|\|\|\|\|\|\|\|\|\|\|\|\|\|\|\|\|\|\|\|\|\|\|\|\|\|\|\|\|\|\|\|\|\|\|\|\|\|\|\|\|\|\|\|\|\|\|\|\|\|\|\|\|\|\|\|\|\|\|\|\|\|\|\|\|\|\|\|\|\|\|\|\|\|\|\|\|\|\|\|\|\|\|\|\|\|\|\|\|\|\|\|\|\|\|\|\|\|\|\|\|\|\|\|\|\|\|\|\|\|\|\|\|\|\|\|\|\|\|\|\|\|\|\|\|\|\|\|\|\|\|\|\|\|\|\|\|\|\|\|\|\|\|\|\|\|\|\|\|\|\|\|\|\|\|\|\|\|\|\|\|\|\|\|\|\|\|\|\|\|\|\|\|\|\|\|\|\|\|\|\|\|\|\|\|\|\|\|\|\|\|\|\|\|\|\|\|\|\|\|\|\|\|\|\|\|\|\|\|\|\|\|\|\|\|\|\|\|\|\|\|\|\|\|\|\|\|\|\|\|\|\|\|\|\|\|\|\|\|\|\|\|\|\|\|\|\|\|\|\|\|\|\|\|\|\|\|\|\|\|\|\|\|\|\|\|\|\|\|\|\|\|\|\|\|\|\|\|\|\|\|\|\|\|\|\|\|\|\|\|\|\|\|\|\|\|\|\|\|\|\|\|\|\|\|\|\|\|\|\|\|\|\|\|\|\|\|\|\|\|\|\|\|\|\|\|\|\|\|\|\|\|\|\|\|\|\|\|\|\|\|\|\|\|\|\|\|\|\|\|\|\|\|\|\|\|\|\|\|\|\|\|\|\|\|\|\|\|\|\|\|\|\|\|\|\|\|\|\|\|\|\|\|\|\|\|\|\|\|\|\|\|\|\|\|\|\|\|\|\|\|\|\|\|\|\|\|\|\|\|\|\|\|\|\|\|\|\|\|\|\|\|\|\|\|\|\|\|\|\|\|\|\|\|\|\|\|\|\|\|\|\|\|\|\|\|\|\|\|\|\|\|\|\|\|\|\|\|\|\|\|\|\|\|\|\|\|\|\|\|\|\|\|\|\|\|\|\|\|\|\|\|\|\|\|\|\|\|\|\|\|\|\|\|\|\|\|\|\|\|\|\|\|\|\|\|\|\|\|\|\|\|\|\|\|\|\|\|\|\|\|\|\|\| 	ATGTCTTCCGGCACGTNTCGAGGCCATTTGATCGTNTTTNGAAGGTCTCGATCGNTCCGGCAAGAGCACTCAATGCGCACGACTGGTAGATCACCTCCGCTCTCAAGGCAAAGAAGTCTTCCATCAACGCTTTCCGGACCGGACGACTGTGATCGGACAGATGATCAACAGCTATCTCAGTGGGACATCATCGCAGGAGGACCATGCCATTCATCTCTTGTTTTCAGCAAACCGCTGGGAGGCCTCCGAATCGATAAGACAGCACATTGAAAGGGGAACTACTGTGGTCGTTGATCGATATTACTACTCTGGTTGTGTCTACTCTGCAGCCAAGCAGAATCCTGCCATGGACCTTGCGTGGTGTCGGCATCCTGAGATTGGCTTGCCACGTCCCGACTTGTGTCTGTTTCTGGACATCGCTGCTGAGGAGGCTGCTAAACGCGGTGGCTTTGGCACCGAACGATACGAGAAACAGGAACTTCAGGACAGAGTGCGAGACTTGTTTGCCGAGTTGCAAACTCATCCGGACGAGGCTGAAGACATCGTCGTGATTGATGCTGGTCAGAGTGTCGAGACAGTAGAGCAAAGCATTCACGACGCAGTAGGACGCCGGCTCGATGGAGAGCATGCTGTCGGCCCAGTGAGATACATTCAACCGTGG ```      ``` 1	ATGTCTTCCGGCACGTCTCGAGGCCATTTGATCGTCTTCGAAGGTCTCGATCGCTCCGGCAAGAGCACTCAATGCGCACGACTGGTAGATCACCTCCGCTCTCAAGGCAAAGAAGTCTTCCATCAACGCTTTCCGGACCGGACGACTGTGATCGGACAGATGATCAACAGCTATCTCAGTGGGACATCATCGCAGGAGGACCATGCCATTCATCTCTTGTTTTCAGCAAACCGCTGGGAGGCCTCCGAATCGATAAGACAGCACATTGAAAGGGGAACTACTGTGGTCGTTGATCGATATTACTACTCTGGTTGTGTCTACTC	323 	\|\|\|\|\|\|\|\|\|\|\|\|\|\|\|\|\|\|\|\|\|\|\|\|\|\|\|\|\|\|\|\|\|\|\|\|\|\|\|\|\|\|\|\|\|\|\|\|\|\|\|\|\|\|\|\|\|\|\|\|\|\|\|\|\|\|\|\|\|\|\|\|\|\|\|\|\|\|\|\|\|\|\|\|\|\|\|\|\|\|\|\|\|\|\|\|\|\|\|\|\|\|\|\|\|\|\|\|\|\|\|\|\|\|\|\|\|\|\|\|\|\|\|\|\|\|\|\|\|\|\|\|\|\|\|\|\|\|\|\|\|\|\|\|\|\|\|\|\|\|\|\|\|\|\|\|\|\|\|\|\|\|\|\|\|\|\|\|\|\|\|\|\|\|\|\|\|\|\|\|\|\|\|\|\|\|\|\|\|\|\|\|\|\|\|\|\|\|\|\|\|\|\|\|\|\|\|\|\|\|\|\|\|\|\|\|\|\|\|\|\|\|\|\|\|\|\|\|\|\|\|\|\|\|\|\|\|\|\|\|\|\|\|\|\|\|\|\|\|\|\|\|\|\|\|\|\|\|\|\|\|\|\|\|\|\|\|\|\|\|\|\|\|\|\|\|\|\|\|\|\|\|\|\|\|\|\|\|\|\|\|\|\|\|\|\|\|\|\|\|\|\|\|\|\|\|\|\|\|\|\|\|\|\|\|\| \| \|\|\|\| 	ATGTCTTCCGGCACGTCTCGAGGCCATTTGATCGTCTTCGAAGGTCTCGATCGCTCCGGCAAGAGCACTCAATGCGCACGACTGGTAGATCACCTCCGCTCTCAAGGCAAAGAAGTCTTCCATCAACGCTTTCCGGACCGGACGACTGTGATCGGACAGATGATCAACAGCTATCTCAGTGGGACATCATCGCAGGAGGACCATGCCATTCATCTCTTGTTTTCAGCAAACCGCTGGGAGGCCTCCGAATCGATAAGACAGCACATTGAAAGGGGAACTACTGTGGTCGTTGATCGATATTACTACTCTGGTTGTGNCNACTC ``` |
| 3 | ggggacaagtttgtacaaaaaagcaggcttgatgGGCTCCGTCGTCATCCCTCA, ggggaccactttgtacaagaaagctgggtcGTACCGGTACCTCGTCGAGT | Mycgr3\|47771  JGI  Ensembl | KOG3414 Component of the U4/U6.U5 snRNP/mitosis protein DIM1; | mitosis; spliceosome; | 0 % |  |  |
| 3 | ggggacaagtttgtacaaaaaagcaggcttgatgGTGACCAGCACTCAACTTGG, ggggaccactttgtacaagaaagctgggtcAACCAGTACACCAATGTACT | Mycgr3\|50695  JGI  Ensembl | KOG3339 Predicted glycosyltransferase; | N-acetylglucosaminyldiphosphodolichol N-acetylglucosaminyltransferase activity; | 100 % |  | ``` 34	TTCCTCTTCCTCCTCCTCATTGCGGCCAGTTCCACGATCCATCGACTGCGAACAGTCTTGAGCCCGAATCGCTTCAGAGCCGTCCGCCGCCGACATGATACCTCAGCGCCGACACACTTGATGATAGTGCTCGGGAGTGGAGGGCACACTGCTGAAATGATCGCCATGCTCAGCCGAGCTCTGCACGAGAAGGAGTCGGCAATACGGATGGACTGGGAGGACTTTTCCCATCGTACATGGGTGGTGAGCTCAGGAGATGGGATATCAGCAGATCGAGCGCGGCAGTTTGAGGACATGACTGCCAAGGCAGAGACTGCTTCTGCCGGCACATATGCCATATCCACGGTTCCGCGCGCGCGAGAGATCCATCAGAGCGCTATGACTGCGCCTATCTCCTGTCTACGCTGTCTTGTCGCATGTCTGACTGCTCTCATGCTATCTTCCGACGCCTCAAGAGACTTCCCGGATGTCATCCTTTGCAATGGTCCTGCTACAGCAACAATCCTAATCTTCACCTCGATCCTGCTCCGGTTCTTCGACATCAAAGGCTGCAATTCACGCGGAAAGATGCGCACGGTTTATATTGAATCATGGGCTAGGGTGAAGAAGTTGAGTCTGTCCGGCAAACTGCTGTCCAGGATCGTTGATCGCTTTCTGGTTCAATGGCCTCAACTGGAAGCGGCGACTGGTGGGAGAGGACAGTACATTGGTGTACTGGTT	753 	\|\|\|\|\| \|\|\|\|\|\| \|\|\|\|\|\|\| \| \|\|\|\|\|\|\|\|\|\|\|\|\|\|\|\|\|\|\|\|\|\|\|\|\|\|\|\|\|\|\|\|\| \|\|\|\|\|\|\|\|\|\|\|\|\|\|\|\|\|\|\|\|\|\|\|\|\|\|\|\|\|\|\|\|\|\|\|\|\|\|\|\|\|\|\|\| \|\|\|\|\|\|\|\|\|\|\|\|\|\|\|\|\|\|\|\|\|\|\|\|\|\|\|\|\|\|\|\|\|\|\|\|\|\|\|\|\|\|\|\|\|\|\|\|\|\|\|\|\|\|\|\|\|\|\|\|\|\|\|\|\|\|\|\|\|\|\|\|\|\|\|\|\|\|\|\|\|\|\|\|\|\|\|\|\|\|\|\|\|\|\|\|\|\|\|\|\|\|\|\|\|\|\|\|\|\|\|\|\|\|\|\|\|\|\|\|\|\|\|\|\|\|\|\|\|\|\|\|\|\|\|\|\|\|\|\|\|\|\|\|\|\|\|\|\|\|\|\|\|\|\|\|\|\|\|\|\|\|\|\|\|\|\|\|\|\|\|\|\|\|\|\|\|\|\|\|\|\|\|\|\|\|\|\|\|\|\|\|\|\|\|\|\|\|\|\|\|\|\|\|\|\|\|\|\|\|\|\|\|\|\|\|\|\|\|\|\|\|\|\|\|\|\|\|\|\|\|\|\|\|\|\|\|\|\|\|\|\|\|\|\|\|\|\|\|\|\|\|\|\|\|\|\|\|\|\|\|\|\|\|\|\|\|\|\|\|\|\|\|\|\|\|\|\|\|\|\|\|\|\|\|\|\|\|\|\|\|\|\|\|\|\|\|\|\|\|\|\|\|\|\|\|\|\|\|\|\|\|\|\|\|\|\|\|\|\|\|\|\|\|\|\|\|\|\|\|\|\|\|\|\|\|\|\|\|\|\|\|\|\|\|\|\|\|\|\|\|\|\|\|\|\|\|\|\|\|\|\|\|\|\|\|\|\|\|\|\|\|\|\|\|\|\|\|\|\|\|\|\|\|\|\|\|\|\|\|\|\|\|\|\|\|\|\|\|\|\|\|\|\|\|\|\|\|\|\|\|\|\|\|\|\|\|\|\|\|\|\|\|\|\|\|\|\|\|\|\|\|\|\|\|\|\|\|\|\|\|\|\|\|\|\|\|\|\|\|\|\|\|\|\|\|\|\|\|\|\|\|\|\|\|\|\|\|\|\|\|\|\|\|\|\|\|\|\|\|\|\|\|\|\|\|\|\|\|\|\|\|\|\|\|\|\|\|\|\|\|\|\|\|\|\|\|\|\|\|\|\|\|\|\|\|\|\|\|\|\|\|\|\|\|\|\|\|\|\|\|\|\|\|\|\|\|\|\|\|\|\|\|\|\|\|\|\|\|\|\|\|\|\|\|\|\|\|\|\|\|\|\|\|\|\|\|\|\|\|\|\|\|\|\|\|\|\|\|\|\|\|\|\|\|\|\|\|\|\|\|\|\|\|\|\|\|\|\|\|\|\|\|\|\|\|\|\|\|\|\|\|\|\|\|\|\|\| 	TTCCTTTTCCTCNTCCTCATNGGGGCCAGTTCCACGATCCATCGACTGCGAACAGTTTTGAGCCCGAATCGCTTCAGAGCCGTCCGCCGCCGACATGATACNTCAGCGCCGACACACTTGATGATAGTGCTCGGGAGTGGAGGGCACACTGCTGAAATGATCGCCATGCTCAGCCGAGCTCTGCACGAGAAGGAGTCGGCAATACGGATGGACTGGGAGGACTTTTCCCATCGTACATGGGTGGTGAGCTCAGGAGATGGGATATCAGCAGATCGAGCGCGGCAGTTTGAGGACATGACTGCCAAGGCAGAGACTGCTTCTGCCGGCACATATGCCATATCCACGGTTCCGCGCGCGCGAGAGATCCATCAGAGCGCTATGACTGCGCCTATCTCCTGTCTACGCTGTCTTGTCGCATGTCTGACTGCTCTCATGCTATCTTCCGACGCCTCAAGAGACTTCCCGGATGTCATCCTTTGCAATGGTCCTGCTACAGCAACAATCCTAATCTTCACCTCGATCCTGCTCCGGTTCTTCGACATCAAAGGCTGCAATTCACGCGGAAAGATGCGCACGGTTTATATTGAATCATGGGCTAGGGTGAAGAAGTTGAGTCTGTCCGGCAAACTGCTGTCCAGGATCGTTGATCGCTTTCTGGTTCAATGGCCTCAACTGGAAGCGGCGACTGGTGGGAGAGGACAGTACATTGGTGTACTGGTT ```      ``` 1	ATGGTGACCAGCACTCAACTTGGACTCCTCTCTTTCCTCTTCCTCCTCCTCATTGCGGCCAGTTCCACGATCCATCGACTGCGAACAGTCTTGAGCCCGAATCGCTTCAGAGCCGTCCGCCGCCGACATGATACCTCAGCGCCGACACACTTGATGATAGTGCTCGGGAGTGGAGGGCACACTGCTGAAATGATCGCCATGCTCAGCCGAGCTCTGCACGAGAAGGAGTCGGCAATACGGATGGACTGGGAGGACTTTTCCCATCGTACATGGGTGGTGAGCTCAGGAGATGGGATATCAGCAGATCGAGCGCGGCAGTTTGAGGACATGACTGCCAAGGCAGAGACTGCTTCTGCCGGCACATATGCCATATCCACGGTTCCGCGCGCGCGAGAGATCCATCAGAGCGCTATGACTGCGCCTATCTCCTGTCTACGCTGTCTTGTCGCATGTCTGACTGCTCTCATGCTATCTTCCGACGCCTCAAGAGACTTCCCGGATGTCATCCTTTGCAATGGTCCTGCTACAGCAACAATCCTAATCTTCACCTCGATCCTGCTCCGGTTCTTCGACATCAAAGGCTGCAATTCACGCGGAAAGATGCGCACGGTTTATATTGAATCATGGGCTAGGGTGAAGAAGTTGAGTCTGTCCGGCAAACTGCTGTCCAGGATCGTTGATCGCTTTCTGGTTCAATGGCCTCAACTGGAAGCGGCGACTGGTGGGAGAGGACAGTACATTGGTGTACTGGTT	753 	\|\|\|\|\|\|\|\|\|\|\|\|\|\|\|\|\|\|\|\|\|\|\|\|\|\|\|\|\|\|\|\|\|\|\|\|\|\|\|\|\|\|\|\|\|\|\|\|\|\|\|\|\|\|\|\|\|\|\|\|\|\|\|\|\|\|\|\|\|\|\|\|\|\|\|\|\|\|\|\|\|\|\|\|\|\|\|\|\|\|\|\|\|\|\|\|\|\|\|\|\|\|\|\|\|\|\|\|\|\|\|\|\|\|\|\|\|\|\|\|\|\|\|\|\|\|\|\|\|\|\|\|\|\|\|\|\|\|\|\|\|\|\|\|\|\|\|\|\|\|\|\|\|\|\|\|\|\|\|\|\|\|\|\|\|\|\|\|\|\|\|\|\|\|\|\|\|\|\|\|\|\|\|\|\|\|\|\|\|\|\|\|\|\|\|\|\|\|\|\|\|\|\|\|\|\|\|\|\|\|\|\|\|\|\|\|\|\|\|\|\|\|\|\|\|\|\|\|\|\|\|\|\|\|\|\|\|\|\|\|\|\|\|\|\|\|\|\|\|\|\|\|\|\|\|\|\|\|\|\|\|\|\|\|\|\|\|\|\|\|\|\|\|\|\|\|\|\|\|\|\|\|\|\|\|\|\|\|\|\|\|\|\|\|\|\|\|\|\|\|\|\|\|\|\|\|\|\|\|\|\|\|\|\|\|\|\|\|\|\|\|\|\|\|\|\|\|\|\|\|\|\|\|\|\|\|\|\|\|\|\|\|\|\|\|\|\|\|\|\|\|\|\|\|\|\|\|\|\|\|\|\|\|\|\|\|\|\|\|\|\|\|\|\|\|\|\|\|\|\|\|\|\|\|\|\|\|\|\|\|\|\|\|\|\|\|\|\|\|\|\|\|\|\|\|\|\|\|\|\|\|\|\|\|\|\|\|\|\|\|\|\|\|\|\|\|\|\|\|\|\|\|\|\|\|\|\|\|\|\|\|\|\|\|\|\|\|\|\|\|\|\|\|\|\|\|\|\|\|\|\|\|\|\|\|\|\|\|\|\|\|\|\|\|\|\|\|\|\|\|\|\|\|\|\|\|\|\|\|\|\|\|\|\|\|\|\|\|\|\|\|\|\|\|\|\|\|\|\|\|\|\|\|\|\|\|\|\|\|\|\|\|\|\|\|\|\|\|\|\|\|\|\|\|\|\|\|\|\|\|\|\|\|\|\|\|\|\|\|\|\|\|\|\|\|\|\|\|\|\|\|\|\|\|\|\|\|\|\|\|\|\|\|\|\|\|\|\|\|\|\|\|\|\|\|\|\|\|\|\|\|\|\|\|\|\|\|\|\|\|\|\|\|\|\|\|\|\|\|\|\|\|\|\|\|\|\|\|\|\|\|\|\|\|\|\|\|\|\|\|\|\|\|\|\|\|\|\|\|\|\|\|\|\|\|\|\|\|\|\|\|\|\|\|\|\|\|\|\|\|\|\|\|\|\|\|\|\|\|\|\|\|\|\|\|\|\|\|\|\|\|\|\|\|\|\|\|\|\|\|\|\|\|\|\|\|\|\|\|\|\|\|\|\|\|\|\|\|\|\|\|\|\|\|\|\|\|\|\|\|\|\|\|\|\|\|\|\|\|\|\|\|\|\|\|\|\|\|\| \|\|\|\|\|\|\|\|\|\|\|\|\| 	ATGGTGACCAGCACTCAACTTGGACTCCTCTCTTTCCTCTTCCTCCTCCTCATTGCGGCCAGTTCCACGATCCATCGACTGCGAACAGTCTTGAGCCCGAATCGCTTCAGAGCCGTCCGCCGCCGACATGATACCTCAGCGCCGACACACTTGATGATAGTGCTCGGGAGTGGAGGGCACACTGCTGAAATGATCGCCATGCTCAGCCGAGCTCTGCACGAGAAGGAGTCGGCAATACGGATGGACTGGGAGGACTTTTCCCATCGTACATGGGTGGTGAGCTCAGGAGATGGGATATCAGCAGATCGAGCGCGGCAGTTTGAGGACATGACTGCCAAGGCAGAGACTGCTTCTGCCGGCACATATGCCATATCCACGGTTCCGCGCGCGCGAGAGATCCATCAGAGCGCTATGACTGCGCCTATCTCCTGTCTACGCTGTCTTGTCGCATGTCTGACTGCTCTCATGCTATCTTCCGACGCCTCAAGAGACTTCCCGGATGTCATCCTTTGCAATGGTCCTGCTACAGCAACAATCCTAATCTTCACCTCGATCCTGCTCCGGTTCTTCGACATCAAAGGCTGCAATTCACGCGGAAAGATGCGCACGGTTTATATTGAATCATGGGCTAGGGTGAAGAAGTTGAGTCTGTCCGGCAAACTGCTGTCCAGGATCGTTGATCGCTTTCTGGTTCAATGGCCTCAACTGGAAGCGGCGACTGGTGGGAGAGGACAGTACA-TGGTGTACTGGTT ``` |
| 3 | ggggacaagtttgtacaaaaaagcaggcttgatgAAGCTTTCACTTCTTGCTGC, ggggaccactttgtacaagaaagctgggtcGAGGCCACTAAACACCGACA | Mycgr3\|51381  JGI  Ensembl |  |  | 100 % |  | ``` 382	GACACCATGAACAGCTTCGCAGCTGCTGGCATCTCCATCTCCCTCGTGTCGATTGGCAACGAAATCAGCAACGGCTTCTTATTTCCCCTCGGGAAACTGGGAACGACGACTGGAAACTACAACTGCGCTCGACTTCTCCATTCCGCTTCTTCTGGCATCAGAGATTCCAAAATCTCAGGAGCCAAGATCCTCATCCACGTTGCCAACGGATGGAACTTCGCCACGCAGAAGTACTTCTACGACACTGTCCTTGGTGCAGGACCTTTGACTTCTGCCGACTACGACGTTCAGGCTGTCTCCTACTACCCGATGTATGGCAGTGGTGCCACACTCGCCAGTTTGAAGTCCTCCTTGAGCCAAATGTCTTCCAAATATGGCAAAAGTGTGATGGTTGTTGAGACTAACTGGCCCGTCAGCTGTCCAAATCCAAGTTCGGCTTTCCCCTCGGACCTTTCAAGCATTCCCTTCTCAGCTGCAGGACAAGCAACCTTTTTGAAGGATGTGGCGGGCGTGGCGGGCTCGGGTCTGATGTATTGGGAGCCGGCATGGCTGAAGAATGCAGGACTTGGGACCAGTTGTGATGACAATCTGATGTTCGATCAGACGGGAACGGCGCGGGGCTCTTTGTCGGTGTTTAGTGGCCTC	1026 	\|\|\| \|\|\| \|\|\|\|\|\| \|\| \|\|\|\|  \|  \|\|\|\|\| \|\|\|\|\| \|\|\|\|\| \| \|\| \|\|\| \|\|\|\|\| \|\|\|\|\|\|\|\|\|\|\|\|\|\|\|\|\| \|\|\|\|\|\|\|\|\|\|\|\|\|\|\|\|\|\|\|\|\|\|\|\|\|\|\|\|\|\|\|\|\|\|\|\|\|\| \|\|\|\|\| \|\|\|\|\|\| \|\|\|\|\|\|\|\|\|\|\|\|\|\|\|\|\|\|\|\|\|\|\|\|\|\|\|\|\|\|\|\|\|\|\|\|\|\|\|\|\|\|\|\|\|\|\|\|\|\|\|\|\|\|\|\|\|\|\| \|\|\|\|\|\|\|\|\|\|\|\|\|\|\|\|\|\|\|\|\|\|\|\|\|\|\|\|\|\|\|\|\|\|\|\|\|\|\|\|\|\|\|\|\|\|\|\|\|\|\|\|\|\|\|\|\|\|\|\|\|\|\|\|\|\|\|\|\|\|\|\|\|\|\|\|\|\|\|\|\|\|\|\|\|\|\|\|\|\|\|\|\|\|\|\|\|\|\|\|\|\|\|\|\|\|\|\|\|\|\|\|\|\|\|\|\|\|\|\|\|\|\|\|\|\|\|\|\|\|\|\|\|\|\|\|\|\|\|\|\|\|\|\|\|\|\|\|\|\|\|\|\|\|\|\|\|\|\|\|\|\|\|\|\|\|\|\|\|\|\|\|\|\|\|\|\|\|\|\|\|\|\|\|\|\|\|\|\|\|\|\|\|\|\|\|\|\|\|\|\|\|\|\|\|\|\|\|\|\|\|\|\|\|\|\|\|\|\|\|\|\|\|\|\|\|\|\|\|\|\|\|\|\|\|\|\|\|\|\|\|\|\|\|\|\|\|\|\|\|\|\|\|\|\|\|\|\|\|\|\|\|\|\|\|\|\|\|\|\|\|\|\|\|\|\|\|\|\|\|\|\|\|\|\|\|\|\|\|\|\|\|\|\|\|\|\|\|\|\|\|\|\|\|\|\|\|\|\|\|\|\|\|\|\|\|\|\|\|\|\|\|\|\|\|\|\|\|\|\|\|\|\|\|\|\|\|\|\|\|\|\|\|\|\|\|\|\|\|\|\|\|\|\|\|\|\|\|\|\|\|\|\|\|\|\|\|\|\|\|\|\|\|\|\|\|\|\|\|\|\|\|\|\|\|\|\|\|\|\|\|\|\|\|\|\|\|\|\|\|\|\|\|\|\|\|\|\|\|\|\|\|\|\|\|\|\|\|\|\|\|\|\|\|\|\|\|\|\|\|\|\|\|\|\|\|\|\|\|\|\|\|\|\|\|\|\|\|\|\|\|\|\|\|\| 	GACNCCA-GAACAGNTTNGCAGNNGNNGGCATNTCCATNTCCCTNGNGTNGATNGGCAANGAAATCAGCAACGGCTTNTTATTTCCCCTCGGGAAACTGGGAACGACGACTGGAAANTACAANTGCGCTNGACTTCTCCATTCCGCTTCTTCTGGCATCAGAGATTCCAAAATCTCAGGAGCCAAGATCNTCATCCACGTTGCCAACGGATGGAACTTCGCCACGCAGAAGTACTTCTACGACACTGTCCTTGGTGCAGGACCTTTGACTTCTGCCGACTACGACGTTCAGGCTGTCTCCTACTACCCGATGTATGGCAGTGGTGCCACACTCGCCAGTTTGAAGTCCTCCTTGAGCCAAATGTCTTCCAAATATGGCAAAAGTGTGATGGTTGTTGAGACTAACTGGCCCGTCAGCTGTCCAAATCCAAGTTCGGCTTTCCCCTCGGACCTTTCAAGCATTCCCTTCTCAGCTGCAGGACAAGCAACCTTTTTGAAGGATGTGGCGGGCGTGGCGGGCTCGGGTCTGATGTATTGGGAGCCGGCATGGCTGAAGAATGCAGGACTTGGGACCAGTTGTGATGACAATCTGATGTTCGATCAGACGGGAACGGCGCGGGGCTCTTTGTCGGTGTTTAGTGGCCTC ``` |
| 3 | ggggacaagtttgtacaaaaaagcaggcttgatgTCTTTCTCTCTCTTCGGTGG, ggggaccactttgtacaagaaagctgggtcATTGAAGATTCCCGAACCTC | Mycgr3\|52569  JGI  Ensembl | KOG0082 G-protein alpha subunit (small G protein superfamily); | mitochondrial intermembrane space protein transporter complex; protein import into mitochondrial inner membrane;  protein targeting to mitochondrion; | 100 % |  | ``` 192	ATTAGTCGACACATGCACAAAGAAGTGCATTCCCAACCAGTACCGCGAAGGCGAACTCAACAAGGGCGAATCCGTCTGCCTCGACCGCTGCGTATCAAAGTTCTTCGAGGTCAACATCAAAGTGTCGGAGAAGATGCAAGGAGAGGCGGCACAAAGGCAGGGAGGTTCGGGAATCTTCAAT	372 	\|\|\|\|\|\|\|\|\|\|\|\|\|\|\|\|\|\|\|\|\|\|\|\|\|\|\|\|\|\|\|\|\|\|\|\|\|\|\|\|\|\|\|\|\|\|\|\|\|\|\|\|\|\|\|\|\|\|\|\|\|\|\|\|\|\|\|\|\|\|\|\|\|\|\|\|\|\|\|\|\|\|\|\|\|\|\|\|\|\|\|\|\|\|\|\|\|\|\|\|\|\|\|\|\|\|\|\|\|\|\|\|\|\|\|\|\|\|\|\|\|\|\|\|\|\|\|\|\|\|\|\|\|\|\|\|\|\|\|\|\|\|\|\|\|\|\|\|\|\|\|\|\|\|\|\|\|\|\|\|\|\|\|\|\|\|\|\|\|\|\|\|\|\|\|\|\|\|\|\|\| 	ATTAGTCGACACATGCACAAAGAAGTGCATTCCCAACCAGTACCGCGAAGGCGAACTCAACAAGGGCGAATCCGTCTGCCTCGACCGCTGCGTATCAAAGTTCTTCGAGGTCAACATCAAAGTGTCGGAGAAGATGCAAGGAGAGGCGGCACAAAGGCAGGGAGGTTCGGGAATCTTCAAT ```      ``` 88	ATGTCTTTCTCTCTCTTCGGTGGCGGACGCCCGCAGCCCTCATCCGCGGAAAAGATCGCAGCAGCAGAAGCCGAGATTGAGATGGTATCCAACATGTTCAACCA	191 	\|\|\|\|\|\|\|\|\|\|\|\|\|\|\|\|\|\|\|\|\|\|\|\|\|\|\|\|\|\|\|\|\|\|\|\|\|\|\|\|\|\|\|\|\|\|\|\|\|\|\|\|\|\|\|\|\|\|\|\|\|\|\|\|\|\|\|\|\|\|\|\|\|\|\|\|\|\|\|\|\|\|\|\|\|\|\|\|\|\|\|\|\|\|\|\|\|\|\|\|\|\|\|\| 	ATGTCTTTCTCTCTCTTCGGTGGCGGACGCCCGCAGCCCTCATCCGCGGAAAAGATCGCAGCAGCAGAAGCCGAGATTGAGATGGTATCCAACATGTTCAACCA ``` |
| 3 | ggggacaagtttgtacaaaaaagcaggcttgatgCGTCCTCTAACCGACACCGA, ggggaccactttgtacaagaaagctgggtcACCAGCAAACAGTGTATCTT | Mycgr3\|56835  JGI  Ensembl | KOG3492 Ribosome biogenesis protein NIP7; | RNA binding; | 100 % |  | ``` 100	CAACCTCCTCACCGACACCACAACCGACCCAAACTCCAAAACTCCCGACCGCTACGTCTTCCGCGTACAAAAAGATCGCGTCTACTACATCCGCGAATCGCTCGCCAACCTCGCCACCTCCGTCGCACGGGACTCTCTCCTTTCCCTCGGAACATGTCTAGGAAAATTCACCAAGACGGGCAAGTTCCGCTTGCACATCACGGCGTTGGATGTCATTGCGCCGCATGCGAGGTATAAGGTGTGGGTGAAGGCGAATGGAGAGATGCCGTTTTTGTATGGTGGACATGTGGTCAAGGCGCATGTGGGACGGTGGAGTGAGGATGCACCGGAGCATGCGGGAGTGATTGTTATGAGTATGAATGATACGCCTTTGG	473 	\|\|\|\| \|\| \|\|\|\| \|\|\|\|\|\|\|\|\|\|\|\|\|\|\|\|\|\|\|\|\|\|\|\|\|\|\|\|\|\|\|\|\|\|\|\|\|\|\|\|\|\|\|\|\|\|\|\|\|\|\|\|\|\|\|\|\|\|\|\|\|\|\|\|\|\|\|\|\|\|\|\|\|\|\|\|\|\|\|\|\|\|\|\|\|\|\|\|\|\|\|\|\|\|\|\|\|\|\|\|\|\|\|\|\|\|\|\|\|\|\|\|\|\|\|\|\|\|\|\|\|\|\|\|\|\|\|\|\|\|\|\|\|\|\|\|\|\|\|\|\|\|\|\|\|\|\|\|\|\|\|\|\|\|\|\|\|\|\|\|\|\|\|\|\|\|\|\|\|\|\|\|\|\|\|\|\|\|\|\|\|\|\|\|\|\|\|\|\|\|\|\|\|\|\|\|\|\|\|\|\|\|\|\|\|\|\|\|\|\|\|\|\|\|\|\|\|\|\|\|\|\|\|\|\|\|\|\|\|\|\|\|\|\|\|\|\|\|\|\|\|\|\|\|\|\|\|\|\|\|\|\|\|\|\|\|\|\|\|\|\|\|\|\|\|\|\|\|\|\|\|\|\|\|\|\|\|\|\|\|\|\|\|\|\|\|\|\|\|\|\|\|\|\|\|\|\|\|\|\|\|\|\|\|\|\|\|\|\|\|\|\|\|\|\|\|\|\|\|\|\|\|\|\|\|\|\|\|\|\|\|\|\|\|\|\|\|\|\|\|\|\|\|\|\|\|\|\|\|\|\|\|\|\|\|\|\|\|\|\|\| 	CAACNTCNTCACNGACACCACAACCGACCCAAACTCCAAAACTCCCGACCGCTACGTCTTCCGCGTACAAAAAGATCGCGTCTACTACATCCGCGAATCGCTCGCCAACCTCGCCACCTCCGTCGCACGGGACTCTCTCCTTTCCCTCGGAACATGTCTAGGAAAATTCACCAAGACGGGCAAGTTCCGCTTGCACATCACGGCGTTGGATGTCATTGCGCCGCATGCGAGGTATAAGGTGTGGGTGAAGGCGAATGGAGAGATGCCGTTTTTGTATGGTGGACATGTGGTCAAGGCGCATGTGGGACGGTGGAGTGAGGATGCACCGGAGCATGCGGGAGTGATTGTTATGAGTATGAATGATACGCCTTTGG ```      ``` 472	GGGATTCGGCGTCACAGCGAGATCAACAGCAGAGGCACGAAAACTGGATCCGACTGGAATCGTCACCTTTCGACAAGCAGATGTTGGCGAGTACCTGCGTGACGAAGATACACTGTTTGCTGGT	595 	\|\|\|\|\|\|\|\|\|\|\|\|\|\|\|\|\|\|\|\|\|\|\|\|\|\|\|\|\|\|\|\|\|\|\|\|\|\|\|\|\|\|\|\|\|\|\|\|\|\|\|\|\|\|\|\|\|\|\|\|\|\|\|\|\|\|\|\|\|\|\|\|\|\|\|\|\|\|\|\|\|\|\|\|\|\|\|\|\|\|\|\|\|\|\|\|\|\|\|\|\|\|\|\|\|\|\|\|\|\|\|\|\|\|\|\|\|\|\|\|\|\|\|\| 	GGGATTCGGCGTCACAGCGAGATCAACAGCAGAGGCACGAAAACTGGATCCGACTGGAATCGTCACCTTTCGACAAGCAGATGTTGGCGAGTACCTGCGTGACGAAGATACACTGTTTGCTGGT ``` |
| 3 | ggggacaagtttgtacaaaaaagcaggcttgatgAGCTCCACCATGACTGTGAC, ggggaccactttgtacaagaaagctgggtcCATGCGACGGATCTCAGTCT | Mycgr3\|62428  JGI  Ensembl | KOG0142 Isopentenyl pyrophosphate:dimethylallyl pyrophosphate isomerase; | hydrolase activity; isopentenyl-diphosphate delta-isomerase activity;  isoprenoid biosynthetic process; | 100 % |  | ``` 216	GGATAAGGACGACGCACCAATTGGCAGTGCGAGCAAGAAAGTATGCCATCTCATGGAGAACATCAACAAGGGCTTGCTCCACCGCGCATTCTCCGTCTTCCTCTTCGACTCGCAAGATCGCCTCCTCCTGCAACAACGCGCTTCAGAAAAGATCACATTTCCCGACCTGTGGACAAACACATGCTGCTCTCACCCGCTAGGCATTCCCGGCGAGACTGGTGCGACCCTCGAGGCTGCTGTGGCTGGTGTCAAGCGAGCAGCACAACGGAAGCTTGATCAAGAGCTTGGGATCAAGAAGGAGCAGGTGCCGATCGAAAAGTTCGACTTCCTCACACGGATACACTACCTCGCACCCAGCGATGGAAAGTGGGGCGAGCACGAGATTGATTACATTCTGTTTATCAAGGCGGATGTGGACTTGAACATCAATCCGAACGAGGTGCAGGCGACCAAGTACGTGTCGGCAGATGAGCTGAAGGATATGTTCAAGGATGATTCGTTGAAGTTCACTCCGTGGTTCAAGCTCATTTGCCAGAGCATGCTATTCGAGTGGTGGCAGAATCTGAATGGAGGACTGGACAAATACAAGGGCGAGACTGAGATCCGTCGCATG	828 	\|\|\|\|\|\|\|\|  \| \|\|\| \|\|\|\|\|\|\| \|\|\|\|\|\|\|\|\|\|\|\|\|\|\|\|\|\|\|\| \|\|\|\| \|\|\|\|\|\|\|\|\|\|\|\|\|\|\| \|\|\|\|\|\|\|\|\|\| \|\|\| \|\|\| \|\|\|\|\| \|\|\| \| \|\|\| \| \|\| \|\|\|\| \|\|\|\|\|\|\|\|\|\| \|\|\|\|\| \|\|\|\|\|\|\|\|\|\|\|\|\|\|\|\|\|\|\|\|\|\|\|\|\|\|\|\|\|\|\|\|\|\|\|\|\|\|\|\|\|\|\|\|\|\|\|\|\|\|\|\|\|\|\|\|\|\|\|\|\|\|\|\|\|\|\|\|\|\|\|\|\|\|\|\|\|\|\|\|\|\|\|\|\|\|\|\|\|\|\|\|\|\|\|\|\|\|\|\|\|\|\|\|\|\|\|\|\|\|\|\|\|\|\|\|\|\|\|\|\|\|\|\|\|\|\|\|\|\|\|\|\|\|\|\|\|\|\|\|\|\|\|\|\|\|\|\|\|\|\|\|\|\|\|\|\|\|\|\|\|\|\|\|\|\|\|\|\|\|\|\|\|\|\|\|\|\|\|\|\|\|\|\|\|\|\|\|\|\|\|\|\|\|\|\|\|\|\|\|\|\|\|\|\|\|\|\|\|\|\|\|\|\|\|\|\|\|\|\|\|\|\|\|\|\|\|\|\|\|\|\|\|\|\|\|\|\|\|\|\|\|\|\|\|\|\|\|\|\|\|\|\|\|\|\|\|\|\|\|\|\|\|\|\|\|\|\|\|\|\|\|\|\|\|\|\|\|\|\|\|\|\|\|\|\|\|\|\|\|\|\|\|\|\|\|\|\|\|\|\|\|\|\|\|\|\|\|\|\|\|\|\|\|\|\|\|\|\|\|\|\|\|\|\|\|\|\|\|\|\|\|\|\|\|\|\|\|\|\|\|\|\|\|\|\|\|\|\|\|\|\|\|\|\|\|\|\|\|\|\|\|\|\|\|\|\|\|\|\|\|\|\|\|\|\|\|\|\|\|\|\|\|\|\|\|\|\|\|\|\|\|\|\|\|\|\|\|\|\|\|\|\|\|\|\|\|\|\|\|\|\|\|\|\|\|\|\|\|\|\|\|\|\|\|\|\|\|\|\|\|\|\|\|\|\|\|\|\|\|\|\|\|\|\|\|\|\|\|\|\|\|\|\|\|\|\|\|\|\|\|\|\|\|\|\|\|\|\|\|\|\|\|\|\|\|\|\|\|\|\|\|\|\|\| 	GGATAAGGNNGNCGCNCCAATTGNCAGTGCGAGCAAGAAAGTATNCCATNTCATGGAGAACATCANCAAGGGCTTGNTCCNCCGNGCATTNTCCNTNTTCNTNTTNGACTNGCAAGATCGCNTCCTCNTGCAACAACGCGCTTCAGAAAAGATCACATTTCCCGACCTGTGGACAAACACATGCTGCTCTCACCCGCTAGGCATTCCCGGCGAGACTGGTGCGACCCTCGAGGCTGCTGTGGCTGGTGTCAAGCGAGCAGCACAACGGAAGCTTGATCAAGAGCTTGGGATCAAGAAGGAGCAGGTGCCGATCGAAAAGTTCGACTTCCTCACACGGATACACTACCTCGCACCCAGCGATGGAAAGTGGGGCGAGCACGAGATTGATTACATTCTGTTTATCAAGGCGGATGTGGACTTGAACATCAATCCGAACGAGGTGCAGGCGACCAAGTACGTGTCGGCAGATGAGCTGAAGGATATGTTCAAGGATGATTCGTTGAAGTTCACTCCGTGGTTCAAGCTCATTTGCCAGAGCATGCTATTCGAGTGGTGGCAGAATCTGAATGGAGGACTGGACAAATACAAGGGCGAGACTGAGATCCGTCGCATG ```      ``` 37	ATGAGCTCCACCATGACTGTGACAGAGACCGAACCGACATACACCGAAAGAATCCTCCAGCTATTTCCCGAGATCAACACGAGCCTCGCGACCCAGAACCGGTCAGCAACTCAAGACCAAGATCTCGCTGGCTACGATGAAGAGCAGATTCGATTAATGGACGAGGTGTGCATTGTGCTGGATAAGGACGACGCACCAATTGGCAGTGCGAGCAAGAAAGTATGCCATCTCATGGAGAACATCAACAAGGGCTTGCTCCACCGCGCATTCTCCGTCTTCCTCTTCGACTCGCAAGATCGCCTCCTCCTGCAACAACGCGCTTCAGAAAAGATCACATTTCCCGACCTGTGGACAAACACATGCTGCTCTCACCCGCTAGGCATTCCCGGCGAGACTGGTGCGACCCTCGAGGCTGCTGTGGCTGGTGTCAAGCGAGCAGCACAACGGAAGCTTGATCAAGAGCTTGGGATCAAGAAGGAGCAGGTGCCGATCGAAAAGTTCGACTTCCTCACACGGATACACTACCTCGCACCCAGCGATGGAAAGTGGGGCGAGCACGAGATTGATTACATTCTGTTTATCAAGGCGGATGTGGACTTGAACATCAATCCGAACGAGGTGCAGGCGACCAAGTACGTGTCGGCAGATGAGCTGAAGGATATGTTCAAGGATGATTCGTTGAAGTTCACTCCGTGGTTCAAGCTCATTTGCCAGAGCATGCTATTCGAGTGGTGGCAGAATCTGAATGGAGGACTGGACAAATAC-AAGGGCGAGACTGAGATCCGTCGCATG	828 	\|\|\|\|\|\|\|\|\|\|\|\|\|\|\|\|\|\|\|\|\|\|\|\|\|\|\|\|\|\|\|\|\|\|\|\|\|\|\|\|\|\|\|\|\|\|\|\|\|\|\|\|\|\|\|\|\|\|\|\|\|\|\|\|\|\|\|\|\|\|\|\|\|\|\|\|\|\|\|\|\|\|\|\|\|\|\|\|\|\|\|\|\|\|\|\|\|\|\|\|\|\|\|\|\|\|\|\|\|\|\|\|\|\|\|\|\|\|\|\|\|\|\|\|\|\|\|\|\|\|\|\|\|\|\|\|\|\|\|\|\|\|\|\|\|\|\|\|\|\|\|\|\|\|\|\|\|\|\|\|\|\|\|\|\|\|\|\|\|\|\|\|\|\|\|\|\|\|\|\|\|\|\|\|\|\|\|\|\|\|\|\|\|\|\|\|\|\|\|\|\|\|\|\|\|\|\|\|\|\|\|\|\|\|\|\|\|\|\|\|\|\|\|\|\|\|\|\|\|\|\|\|\|\|\|\|\|\|\|\|\|\|\|\|\|\|\|\|\|\|\|\|\|\|\|\|\|\|\|\|\|\|\|\|\|\|\|\|\|\|\|\|\|\|\|\|\|\|\|\|\|\|\|\|\|\|\|\|\|\|\|\|\|\|\|\|\|\|\|\|\|\|\|\|\|\|\|\|\|\|\|\|\|\|\|\|\|\|\|\|\|\|\|\|\|\|\|\|\|\|\|\|\|\|\|\|\|\|\|\|\|\|\|\|\|\|\|\|\|\|\|\|\|\|\|\|\|\|\|\|\|\|\|\|\|\|\|\|\|\|\|\|\|\|\|\|\|\|\|\|\|\|\|\|\|\|\|\|\|\|\|\|\|\|\|\|\|\|\|\|\|\|\|\|\|\|\|\|\|\|\|\|\|\|\|\|\|\|\|\|\|\|\|\|\|\|\|\|\|\|\|\|\|\|\|\|\|\|\|\|\|\|\|\|\|\|\|\|\|\|\|\|\|\|\|\|\|\|\|\|\|\|\|\|\|\|\|\|\|\|\|\|\|\|\|\|\|\|\|\|\|\|\|\|\|\|\|\|\|\|\|\|\|\|\|\|\|\|\|\|\|\|\|\|\|\|\|\|\|\|\|\|\|\|\|\|\|\|\|\|\|\|\|\|\|\|\|\|\|\|\|\|\|\|\|\|\|\|\|\|\|\|\|\|\|\|\|\|\|\|\|\|\|\|\|\|\|\|\|\|\|\|\|\|\|\|\|\|\|\|\|\|\|\|\|\|\|\|\|\|\|\|\|\|\|\|\|\|\|\|\|\|\|\|\|\|\|\|\|\|\|\|\|\|\|\|\|\|\|\|\|\|\|\|\|\|\|\|\|\|\|\|\|\|\|\|\|\|\|\|\|\|\|\|\|\|\|\|\|\|\|\|\|\|\|\|\|\|\|\|\|\|\|\|\|\|\|\|\|\|\|\|\|\|\|\|\|\|\|\|\|\|\|\|\|\|\|\|\|\|\|\|\|\|\|\|\|\|\|\|\|\|\|\|\|\|\|\|\|\|\|\|\|\|\|\|\|\|\|\|\|\|\|\|\|\|\|\|\|\|\|\|\|\|\|\|\|\|\|\|\|\|\|\|\|\|\|\|\|\|\|\|\|\|\|\|\|\|\|\|\|\|\|\|\|\|\|\|\|\|\|\|\|\|\| \|\|\|\|\|\|\|\|\|\|\|\|\|\|\|\|\|\|\|\|\|\|\|\|\|\|\| 	ATGAGCTCCACCATGACTGTGACAGAGACCGAACCGACATACACCGAAAGAATCCTCCAGCTATTTCCCGAGATCAACACGAGCCTCGCGACCCAGAACCGGTCAGCAACTCAAGACCAAGATCTCGCTGGCTACGATGAAGAGCAGATTCGATTAATGGACGAGGTGTGCATTGTGCTGGATAAGGACGACGCACCAATTGGCAGTGCGAGCAAGAAAGTATGCCATCTCATGGAGAACATCAACAAGGGCTTGCTCCACCGCGCATTCTCCGTCTTCCTCTTCGACTCGCAAGATCGCCTCCTCCTGCAACAACGCGCTTCAGAAAAGATCACATTTCCCGACCTGTGGACAAACACATGCTGCTCTCACCCGCTAGGCATTCCCGGCGAGACTGGTGCGACCCTCGAGGCTGCTGTGGCTGGTGTCAAGCGAGCAGCACAACGGAAGCTTGATCAAGAGCTTGGGATCAAGAAGGAGCAGGTGCCGATCGAAAAGTTCGACTTCCTCACACGGATACACTACCTCGCACCCAGCGATGGAAAGTGGGGCGAGCACGAGATTGATTACATTCTGTTTATCAAGGCGGATGTGGACTTGAACATCAATCCGAACGAGGTGCAGGCGACCAAGTACGTGTCGGCAGATGAGCTGAAGGATATGTTCAAGGATGATTCGTTGAAGTTCACTCCGTGGTTCAAGCTCATTTGCCAGAGCATGCTATTCGAGTGGTGGCAGAATCTGAATGGAGGACTGGACAAATACAAAGGGCGAGACTGAGATCCGTCGCATG ``` |
| 3 | ggggacaagtttgtacaaaaaagcaggcttgatgGCTCGCTCACCGCCCACACG, ggggaccactttgtacaagaaagctgggtcCTCCGGCCTGAACTTCCCTC | Mycgr3\|65192  JGI  Ensembl | KOG4008 rRNA processing protein RRP7; |  | 100 % |  | ``` 250	GGCGGCGCGCGCGTGGAAAGTGTAGCATTTGAAGGTGCCAGAGTCGGAAAAGCGATCAATGCGCCGGTAGCGCCAGTACAGCAAGGCAAGAAGCGGAAACGTGGAGCAGATGGAGAGGACGGTGCACAACTACCAGAAGAAGTCGGAGCGTTGCCGGAGGTCTGGGATCGAGAATTGCACCGCAGTGGAGGAACGGCCATTGTGACGTTCGTGGACAGAGTCAGCGCGGAAATGGCTTTGAAGGAGGTCAAGAGAGCTATCAAGAACAAGAGCGTTATCGTATGGACAAGTGAGAAAGGTGTTGCGCCATCTCTTGGCTCAGCAAGATACCGCACACATCATACTCTGCGATACCCGGATCATGCGGTTTTGCAGGCATCAGTGGACAACTTCATGGCAGCGTTCACAGCGCAGGAGGAGGCGAAAGCGAAGGCACTGGCGAAATTACGAGCAGAGCCGGACGAAGATGGGTTCATCACTGTTACCCGCGGAGGACGGGTCGGAGCAGGCAGAGAAGACGAGGCGAAAGCGAAAGAGGCAGAGTTGGCGAAGAGGGACAAGAATCGAATCAAGGAGGACTTCTACCGATTCCAGGTGCGAGAGAAGAAGAAAGAGCAGGCGCAGGATCTGGTCAAGGGCTTCCAGGAGGATCAAAGGAGGGTAGAGGAGATGAAGCGGAGAAGAGGGAAGTTCAGGCCGGAG	951 	\|\|\|\|\|\|\|\|\|\|\|\|   \|\|\|\|\|\|\|\|\| \|  \|\|\|\|\| \|\| \|\|\|\|\|\|\|\|\|\|\|\|\|\|\|\|\|\|\|\|\|\|\|\|\|\|\|\|\|\|\|\|\|\|\|\|\|\|\|\|\|\|\|\|\|\|\|\| \|\|\|\|\|\|\|\|\|\|\|\|\|\|\|\|\|\|\|\|\|\|\|\|\|\|\|\|\|\|\|\|\|\|\|\|\|\|\|\|\|\|\|\|\|\|\|\|\|\|\|\|\|\|\|\|\|\|\|\|\|\|\|\|\|\|\|\|\|\|\|\|\|\|\|\|\|\|\|\|\|\|\|\|\|\|\|\|\|\|\|\|\|\|\|\|\|\|\|\|\|\|\|\|\|\|\|\|\|\|\|\|\|\|\|\|\|\|\|\|\|\|\|\|\|\|\|\|\|\|\|\|\|\|\|\|\|\|\|\|\|\|\|\|\|\|\|\|\|\|\|\|\|\|\|\|\|\|\|\|\|\|\|\|\|\|\|\|\|\|\|\|\|\|\|\|\|\|\|\|\|\|\|\|\|\|\|\|\|\|\|\|\|\|\|\|\|\|\|\|\|\|\|\|\|\|\|\|\|\|\|\|\|\|\|\|\|\|\|\|\|\|\|\|\|\|\|\|\|\|\|\|\|\|\|\|\|\|\|\|\|\|\|\|\|\|\|\|\|\|\|\|\|\|\|\|\|\|\|\|\|\|\|\|\|\|\|\|\|\|\|\|\|\|\|\|\|\|\|\|\|\|\|\|\|\|\|\|\|\|\|\|\|\|\|\|\|\|\|\|\|\|\|\|\|\|\|\|\|\|\|\|\|\|\|\|\|\|\|\|\|\|\|\|\|\|\|\|\|\|\|\|\|\|\|\|\|\|\|\|\|\|\|\|\|\|\|\|\|\|\|\|\|\|\|\|\|\|\|\|\|\|\|\|\|\|\|\|\|\|\|\|\|\|\|\|\|\|\|\|\|\|\|\|\|\|\|\|\|\|\|\|\|\|\|\|\|\|\|\|\|\|\|\|\|\|\|\|\|\|\|\|\|\|\|\|\|\|\|\|\|\|\|\|\|\|\|\|\|\|\|\|\|\|\|\|\|\|\|\|\|\|\|\|\|\|\|\|\|\|\|\|\|\|\|\|\|\|\|\|\|\|\|\|\|\|\|\|\|\|\|\|\|\|\|\|\|\|\|\|\|\|\|\|\|\|\|\|\|\|\|\|\|\|\|\|\|\|\|\|\|\|\|\|\|\|\|\|\|\|\|\|\|\|\|\|\|\|\|\|\|\|\|\|\|\|\|\|\|\|\|\|\|\|\|\|\|\|\|\|\|\|\|\|\|\|\|\|\|\|\|\|\|\|\|\|\|\|\|\|\|\|\|\|\|\|\|\|\|\|\|\|\|\|\|\|\|\|\|\|\|\|\|\|\|\|\|\|\|\|\|\|\|\|\|\|\|\|\|\|\|\|\|\|\|\|\|\|\|\|\|\|\|\|\|\| 	GGCGGCGCGCGC-NNGAAAGTGTANC-NTTGAA-GTNCCAGAGTCGGAAAAGCGATCAATGCGCCGGTAGCGCCAGTACAGCAAGNCAAGAAGCGGAAACGTGGAGCAGATGGAGAGGACGGTGCACAACTACCAGAAGAAGTCGGAGCGTTGCCGGAGGTCTGGGATCGAGAATTGCACCGCAGTGGAGGAACGGCCATTGTGACGTTCGTGGACAGAGTCAGCGCGGAAATGGCTTTGAAGGAGGTCAAGAGAGCTATCAAGAACAAGAGCGTTATCGTATGGACAAGTGAGAAAGGTGTTGCGCCATCTCTTGGCTCAGCAAGATACCGCACACATCATACTCTGCGATACCCGGATCATGCGGTTTTGCAGGCATCAGTGGACAACTTCATGGCAGCGTTCACAGCGCAGGAGGAGGCGAAAGCGAAGGCACTGGCGAAATTACGAGCAGAGCCGGACGAAGATGGGTTCATCACTGTTACCCGCGGAGGACGGGTCGGAGCAGGCAGAGAAGACGAGGCGAAAGCGAAAGAGGCAGAGTTGGCGAAGAGGGACAAGAATCGAATCAAGGAGGACTTCTACCGATTCCAGGTGCGAGAGAAGAAGAAAGAGCAGGCGCAGGATCTGGTCAAGGGCTTCCAGGAGGATCAAAGGAGGGTAGAGGAGATGAAGCGGAGAAGAGGGAAGTTCAGGCCGGAG ``` |
| 3 | ggggacaagtttgtacaaaaaagcaggcttgatgACTACAACATCAGGCAAGAC, ggggaccactttgtacaagaaagctgggtcATCCAGAAAGCCCATTTCAT | Mycgr3\|65316  JGI  Ensembl | KOG3349 Predicted glycosyltransferase; | N-acetylglucosaminyldiphosphodolichol N-acetylglucosaminyltransferase activity; carbohydrate binding;  carbohydrate metabolic process;  lipid glycosylation;  transferase activity, transferring hexosyl groups; | 100 % |  | ``` 1	ATGACTACAACATCAGGCAAGACTTGCTTTGTCACAATCGGCGCGACAGCTTCCTTTGTCGGTCTCATCAAAGCCGCACTCGCACCGGATTTCTTGGCTGCGTTGAAGTCGCATGGCTACACACATTTACTTGTCCAGTACGGTCAAGATGGCAAAGCGCTCTTCGACTCCTGCATTGGATCGCTAGATCAAATCGACATTCACATCAGCGGCTTTGACTTGGACAAGGCTGGACTTGC	239 	\|\|\|\|\|\|\|\|\|\|\|\|\|\|\|\|\|\|\|\|\|\|\|\|\|\|\|\|\|\|\|\|\|\|\|\|\|\|\|\|\|\|\|\|\|\|\|\|\|\|\|\|\|\|\|\|\|\|\|\|\|\|\|\|\|\|\|\|\|\|\|\|\|\|\|\|\|\|\|\|\|\|\|\|\|\|\|\|\|\|\|\|\|\|\|\|\|\|\|\|\|\|\|\|\|\|\|\|\|\|\|\|\|\|\|\|\|\|\|\|\|\|\|\|\|\|\|\|\|\|\|\|\|\|\|\|\|\|\|\|\|\|\|\|\|\|\|\|\|\|\|\|\|\|\|\|\|\|\|\|\|\|\|\|\|\|\|\|\|\|\|\|\|\|\|\|\|\|\|\|\|\|\|\|\|\|\|\|\|\|\|\|\|\|\|\|\|\|\|\|\|\|\|\|\|\|\|\| \|\|\|\|\|\|\|\|\| \|\|\|\|\|\| \| \|\|\|\| \| \|\|\|\| 	ATGACTACAACATCAGGCAAGACTTGCTTTGTCACAATCGGCGCGACAGCTTCCTTTGTCGGTCTCATCAAAGCCGCACTCGCACCGGATTTCTTGGCTGCGTTGAAGTCGCATGGCTACACACATTTACTTGTCCAGTACGGTCAAGATGGCAAAGCGCTCTTCGACTCCTGCATTGGATCGCTAGATCAAATCGACATTCACATCANCGGCTTTGAATTGGACCANGCTGNANTTGC ```      ``` 1	ATGACTACAACATCAGGCAAGACTTG-CTTTGTCACAATCGGCGCGACAGCTTCCTTTGTCGGTCTCATCAAAGCCGCACTCGCACCGGATTTCTTGGCTGCGTTGAAGTCGCATGGCTACACACATTTACTTGTCCAGTACGGTCAAGATGGCAAAGCGCTCTTCGACTCCTGCATTGGATCGCTAGATCAAATCGACATTCACATCAGCGGCTTTGACTTGGACAAGGCTGGACTTGCGGAGTACATGCTTCAAGCCAAGGGCGAAAGGAATACTGGCCGTCAAGAAGGAGTGGTAGTCAGTCACGCTGG	311 	\|\|\|\| \| \|\|\|\|\|\|\|\|\|\|\|\|\|\|\|\|\|\|\|  \|\|\|\|\|\|\|\|\|\|\| \|\|\|\| \|\|\|\|\|\|\|\|\| \|\|\|\|\|\|\|\|\| \|\|\|\|\|\|\|\|\|\|\|\|\|\|\|\|\|\|\|\|\|\|\|\|\|\|\|\|\|\|\|\|\| \|\|\|\|\|\|\|\|\|\|\|\|\|\|\|\|\|\|\|\|\|\|\|\|\|\|\|\|\|\|\|\|\|\|\|\|\|\|\|\|\|\|\|\|\|\|\|\|\|\|\|\|\|\|\|\|\|\|\|\|\|\|\|\|\|\|\|\|\|\|\|\|\|\|\|\|\|\|\|\|\|\|\|\|\|\|\|\|\|\|\|\|\|\|\|\|\|\|\|\|\|\|\|\|\|\|\|\|\|\|\|\|\|\|\|\|\|\|\|\|\|\|\|\|\|\|\|\|\|\|\|\|\|\|\|\|\|\|\|\|\|\|\|\|\|\|\|\|\|\|\|\|\|\|\|\|\|\|\|\|\|\|\|\|\|\|\|\|\|\|\|\|\|\|\|\|\|\|\|\|\|\|\|\|\|\|\|\|\|\|\|\|\|\|\|\|\|\|\|\|\|\|\|\|\|\|\|\|\|\|\|\|\| 	ATGANTNCAACATCAGGCAAGACTTGNTTTTGTCACAATNGGCGNGACAGCTTCNTTTGTCGGTNTCATCAAAGCCGCACTCGCACCGGATTTCTTGGNTGCGTTGAAGTCGCATGGCTACACACATTTACTTGTCCAGTACGGTCAAGATGGCAAAGCGCTCTTCGACTCCTGCATTGGATCGCTAGATCAAATCGACATTCACATCAGCGGCTTTGACTTGGACAAGGCTGGACTTGCGGAGTACATGCTTCAAGCCAAGGGCGAAAGGAATACTGGCCGTCAAGAAGGAGTGGTAGTCAGTCACGCTGG ```      ``` 310	GGCTCCGGCACGATTTTGGATGCTCTTCGCATCTCTGTGCCGATCATCGTAGTTCCAAATTCCGAGCTGCTGGACAATCATCAAGTCGAGCTCGCTGAAGCGCTTGCCGAGCAGGAATATGTCGTGCATGGTAGCCTCCATGCGCTCGGCCAAGCCTTAAACGAGGCAGCACAGTTGGGCAAGAAGAGCAAATCCTGGCCACCACCTAACAGCGGGTTTCATAGAGAAGCTCGAGGCCTGAAAGGCGTGCTAGACGATGAAATGGGCTTTCTGGAT	585 	\|\|\|\|\|\|\|\|\|\|\|\|\|\|\|\|\|\|\|\|\|\|\|\|\|\|\|\|\|\|\|\|\|\|\|\|\|\|\|\|\|\|\|\|\|\|\|\|\|\|\|\|\|\|\|\|\|\|\|\|\|\|\|\|\|\|\|\|\|\|\|\|\|\|\|\|\|\|\|\|\|\|\|\|\|\|\|\|\|\|\|\|\|\|\|\|\|\|\|\|\|\|\|\|\|\|\|\|\|\|\|\|\|\|\|\|\|\|\|\|\|\|\|\|\|\|\|\|\|\|\|\|\|\|\|\|\|\|\|\|\|\|\|\|\|\|\|\|\|\|\|\|\|\|\|\|\|\|\|\|\|\|\|\|\|\|\|\|\|\|\|\|\|\|\|\|\|\|\|\|\|\|\|\|\|\|\|\|\|\|\|\|\|\|\|\|\|\|\|\|\|\|\|\|\|\|\|\|\|\|\|\|\|\|\|\|\|\|\|\|\|\|\|\|\|\|\|\|\|\|\|\|\|\|\|\|\|\|\|\|\|\|\|\|\|\|\|\|\|\|\|\|\|\|\|\|\|\|\|\|\|\|\|\|\|\|\|\|\|\|\|\|\|\|\|\| 	GGCTCCGGCACGATTTTGGATGCTCTTCGCATCTCTGTGCCGATCATCGTAGTTCCAAATTCCGAGCTGCTGGACAATCATCAAGTCGAGCTCGCTGAAGCGCTTGCCGAGCAGGAATATGTCGTGCATGGTAGCCTCCATGCGCTCGGCCAAGCCTTAAACGAGGCAGCACAGTTGGGCAAGAAGAGCAAATCCTGGCCACCACCTAACAGCGGGTTTCATAGAGAAGCTCGAGGCCTGAAAGGCGTGCTAGACGATGAAATGGGCTTTCTGGAT ``` |
| 3 | ggggacaagtttgtacaaaaaagcaggcttgatgCTGTCAAGAACACTACGCAG, ggggaccactttgtacaagaaagctgggtcATCCACTTCGCCACTTTCAA | Mycgr3\|66192  JGI  Ensembl | KOG0453 Aconitase/homoaconitase (aconitase superfamily); |  | 100 % |  | ``` 530	TTCTCATATGTCGACGATCGGATGTTTGTGTGCAGGGCAGTGGTGACTGGTGTCGAAAGCGAGCCAATGCGAATCAAGGAGAAGACCAAGCGGAGACAACGGCACGTCAAGAAGGTCAAGAGCAAGCATCGGTACACGATTATCAAGATCAAGGAACTGCGTGTGAAGAGTGTGGAAGAGATTGAAAGTGGCGAAGTGGAT	730 	\|\|\|\|\|\|\|\|\|\|\|\|\|\|\|\|\|\|\|\|\|\|\|\|\|\|\|\|\|\|\|\|\|\|\|\|\|\|\|\|\|\|\|\|\|\|\|\|\|\|\|\|\|\|\|\|\|\|\|\|\|\|\|\|\|\|\|\|\|\|\|\|\|\|\|\|\|\|\|\|\|\|\|\|\|\|\|\|\|\|\|\|\|\|\|\|\|\|\|\|\|\|\|\|\|\|\|\|\|\|\|\|\|\|\|\|\|\|\|\|\|\|\|\|\|\|\|\|\|\|\|\|\|\|\|\|\|\|\|\|\|\|\|\|\|\|\|\|\|\|\|\|\|\|\|\|\|\|\|\|\|\|\|\|\|\|\|\|\|\|\|\|\|\|\|\|\|\|\|\|\|\|\|\|\|\|\|\|\|\|\|\|\|\|\|\|\|\|\|\|\| 	TTCTCATATGTCGACGATCGGATGTTTGTGTGCAGGGCAGTGGTGACTGGTGTCGAAAGCGAGCCAATGCGAATCAAGGAGAAGACCAAGCGGAGACAACGGCACGTCAAGAAGGTCAAGAGCAAGCATCGGTACACGATTATCAAGATCAAGGAACTGCGTGTGAAGAGTGTGGAAGAGATTGAAAGTGGCGAAGTGGAT ```      ``` 209	CAAAATGCCGAACACCTCGAGTTACCTGCTCCCCACTCCCAGTCTTCGGGCCAAACTTCTTCACCTCCACAACTCTCCGAGTCCGTTCGCGAACTCCTACCACTTCTGCGGTCACAAGGCCCTCACTACATGACCGTTCACATCCATGGTAACCCATACCTCATCACAGAAGGAGATACTGTACGGCTGCCTTTCCTGATGCAAGGCGTCGAACCGGGAGATGTCATACGCCTCAACCGGGCGATCAACTTGGGAAGCAGAGACTATGCGTTGAAAGCACCAGCGGCGTCACCGAAGATGAAGAGTCCGACAACAGCCACC	529 	\|\|\|\|\|\| \| \|\| \| \|\|\| \|\| \|\| \|\|\|\| \|\|\|\|\|\|\|\|\|\|\|\|\|\| \|\| \|\|\|\|\|\|\|\| \|\| \|\|\|\|\|\|\|\|\|\|\|\|\|\|\|\|\|\|\|\|\|\|\|\|\|\|\|\|\|\|\|\|\|\|\|\|\|\|\|\|\|\|\|\|\|\| \|\|\|\|\|\|\|\|\|\|\|\|\|\|\|\|\|\|\|\|\|\|\|\|\|\|\|\|\|\|\|\|\|\|\|\|\|\|\|\|\|\|\|\|\|\|\|\|\|\|\|\|\|\|\|\|\|\|\|\|\|\|\|\|\|\|\|\|\|\|\|\|\|\|\|\|\|\|\|\|\|\|\|\|\|\|\|\|\|\|\|\|\|\|\|\|\|\|\|\|\|\|\|\|\|\|\|\|\|\|\|\|\|\|\|\|\|\|\|\|\|\|\|\|\|\|\|\|\|\|\|\|\|\|\|\|\|\|\|\|\|\|\|\|\|\|\|\|\|\|\|\|\|\|\|\|\|\|\|\|\|\|\|\|\|\|\|\|\|\|\|\|\|\|\|\|\|\|\|\|\|\|\|\|\|\|\|\|\|\|\|\|\|\|\|\|\|\|\|\|\|\|\|\|\|\|\|\|\|\|\|\|\|\|\| 	CAAAATNCNGANCNCCTNGANTTNCCTGNTCCCCACTCCCAGTNTTNGGGCCAAANTTNTTCACCTCCACAACTCTCCGAGTCCGTTCGCGAACTCCTACCACTTNTGCGGTCACAAGGCCCTCACTACATGACCGTTCACATCCATGGTAACCCATACCTCATCACAGAAGGAGATACTGTACGGCTGCCTTTCCTGATGCAAGGCGTCGAACCGGGAGATGTCATACGCCTCAACCGGGCGATCAACTTGGGAAGCAGAGACTATGCGTTGAAAGCACCAGCGGCGTCACCGAAGATGAAGAGTCCGACAACAGCCACC ``` |
| 3 | ggggacaagtttgtacaaaaaagcaggcttgatgATCCGACGACAGGCGCGACA, ggggaccactttgtacaagaaagctgggtcCAGATAGTCCTTCTTCTTGC | Mycgr3\|66663  JGI  Ensembl | KOG2781 U3 small nucleolar ribonucleoprotein (snoRNP) component; |  | 100 % |  | ``` 227	GAATACAGTCATCTCTCCGGAGTCGTTGATCCGCGCGTTTTGGTCACCACCTCGCGGGACCCCAGCAGTCGGCTGGCCTCATTCGCCAAAGAAATTCGTCTGCTCCTCCCCACCAGCATTCGTCTGAACCGCGGAAATCTCGTACTCCCCAACCTTGTTGGCAGCGCAAAGTCCAGCGGCCTGTCCGATCTCATACTCCTCCACGAACATCGAGGAACACCTACCGCGATGACCGTGTCACACTTTCCTCATGGACCGACCGCATCCTTCAGCTTGCACAACGTTGTCCTGCGTCACGACATCCCCAATGCTTCCCGAGGCACCGTCAGCGAAAGCTACCCGCATCTGATATTTGAAGGCTTCACGACGCCGTTGGGACGGCGTGTGGTGAAGATTTTGCAACACCTGTTCCCACCACGAGAGGGTGGAACAAAGCTGGGTTCAAGAGTCGTGACATTCAAGAATATTGAAGACAGTATTGAAGTGAGGCACCACGTTTTTGTCAAGACGGCGTACCAGAGCGTGGAGCTGGCCGAAGTTGGACCACGAATGACGATGCGTTTGTTTGAGATTCGACAAGGCACGGCAGAGAATAAGGATGGTGACGTGGAGTGGCATATGAATCAGTATACTCGGACCAGCAAGAAGAAGGACTATCTG	886 	\|\|\|\| \|\|\|\|\|\|\| \| \|\|\|\|\|\|\|\| \|\|\|\|\|\|\|\|\|\|\| \|\|\|\| \|\|\|\|\|\|\|\|\|\|\| \|\|\|\|\|\|\|\|\|\|\|\|\|\|\|\|\|\|\| \|\|\|\| \|\|\|\|\|\|\|\|\|\|\|\|\|\|\|\|\|\| \|\| \|\|\|\|\|\|\|\|\|\|\|\|\|\|\|\|\|\|\|\|\|\|\| \|\|\|\|\|\|\|\|\|\|\|\|\|\|\|\|\|\|\|\|\|\|\|\|\|\|\|\|\|\|\|\|\|\|\|\|\|\|\|\|\|\|\|\|\|\|\|\|\|\|\|\|\|\|\|\|\|\|\|\|\|\|\|\|\|\|\|\|\|\|\|\|\|\|\|\|\|\|\|\|\|\|\|\|\|\|\|\|\|\|\|\|\|\|\|\|\|\|\|\|\|\|\|\|\|\|\|\|\|\|\|\|\|\|\|\|\|\|\|\|\|\|\|\|\|\|\|\|\|\|\|\|\|\|\|\|\|\|\|\|\|\|\|\|\|\|\|\|\|\|\|\|\|\|\|\|\|\|\|\|\|\|\|\|\|\|\|\|\|\|\|\|\|\|\|\|\|\|\|\|\|\|\|\|\|\|\|\|\|\|\|\|\|\|\|\|\|\|\|\|\|\|\|\|\|\|\|\|\|\|\|\|\|\|\|\|\|\|\|\|\|\|\|\|\|\|\|\|\|\|\|\|\|\|\|\|\|\|\|\|\|\|\|\|\|\|\|\|\|\|\|\|\|\|\|\|\|\|\|\|\|\|\|\|\|\|\|\|\|\|\|\|\|\|\|\|\|\|\|\|\|\|\|\|\|\|\|\|\|\|\|\|\|\|\|\|\|\|\|\|\|\|\|\|\|\|\|\|\|\|\|\|\|\|\|\|\|\|\|\|\|\|\|\|\|\|\|\|\|\|\|\|\|\|\|\|\|\|\|\|\|\|\|\|\|\|\|\|\|\|\|\|\|\|\|\|\|\|\|\|\|\|\|\|\|\|\|\|\|\|\|\|\|\|\|\|\|\|\|\|\|\|\|\|\|\|\|\|\|\|\|\|\|\|\|\|\|\|\|\|\|\|\|\|\|\|\|\|\|\|\|\|\|\|\|\|\|\|\|\|\|\|\|\|\|\|\|\|\|\|\|\|\|\|\|\|\|\|\|\|\|\|\|\|\|\|\|\|\|\|\|\|\|\|\|\|\|\|\|\|\|\|\|\|\|\|\|\|\|\|\|\|\|\|\|\|\|\|\|\|\|\|\|\|\|\|\|\|\|\|\|\|\|\|\|\|\|\|\|\|\|\|\|\|\|\|\|\|\|\|\|\|\|\|\|\|\|\|\|\|\|\|\|\|\|\|\|\|\|\|\|\|\|\|\|\| 	GAATNCAGTCATNTNTCCGGAGTNGTTGATCCGCGNGTTT-GGTCACCACCTNGCGGGACCCCAGCAGTCGGNTGGCNTCATTCGCCAAAGAAATTNGTNTGCTCCTCCCCACCAGCATTCGTNTGAACCGCGGAAATCTCGTACTCCCCAACCTTGTTGGCAGCGCAAAGTCCAGCGGCCTGTCCGATCTCATACTCCTCCACGAACATCGAGGAACACCTACCGCGATGACCGTGTCACACTTTCCTCATGGACCGACCGCATCCTTCAGCTTGCACAACGTTGTCCTGCGTCACGACATCCCCAATGCTTCCCGAGGCACCGTCAGCGAAAGCTACCCGCATCTGATATTTGAAGGCTTCACGACGCCGTTGGGACGGCGTGTGGTGAAGATTTTGCAACACCTGTTCCCACCACGAGAGGGTGGAACAAAGCTGGGTTCAAGAGTCGTGACATTCAAGAATATTGAAGACAGTATTGAAGTGAGGCACCACGTTTTTGTCAAGACGGCGTACCAGAGCGTGGAGCTGGCCGAAGTTGGACCACGAATGACGATGCGTTTGTTTGAGATTCGACAAGGCACGGCAGAGAATAAGGATGGTGACGTGGAGTGGCATATGAATCAGTATACTCGGACCAGCAAGAAGAAGGACTATCTG ``` |
| 3 | ggggacaagtttgtacaaaaaagcaggcttgatgGTCGTCTTCGCCCTTTTGAT, ggggaccactttgtacaagaaagctgggtcCGACGGTCTGACCAGATACG | Mycgr3\|66828  JGI  Ensembl | KOG3369 Transport protein particle (TRAPP) complex subunit; | ER to Golgi vesicle-mediated transport; cis-Golgi network; | 100 % |  | ``` 37	GGCGGCCTAGTCTACCACCGCACCTTCGCCCCAGGTCTCCAGAAGATCGACTCAAACGACTACTTGATCCTAGCCGGCACATTTCACGGCGTGCACGCCATCAGTCGAAGCATCAACCCGGTACCTCCTCTCCCTCAACCGCCGGGCAACCGCAAACCTCAGACCACGGGCATCGAAAGTCTGGAGAGCAGCCACTTCCGATTGACGTGTTTCCAAACGCCGACGGGCGTGAAGTTCCTACTGATCACAAGTCCGGAACAGCCAAATACGGAATTGGTCGTGCGGAGGTGCTATGAGATTTATGGCGACTTCGTGATGAAGAATCCGTTCTATAATCTGGAGATGCCAATTCGGGTGGAGAAGTTTGATCGTGCGCTGGGGTCGTATCTGGTCAGACC-GTCG	438 	\|\|\|\|\|\|\|\|\|\|\|\|\|\|\|\|\|\|\|\|\|\|\|\|\|\|\|\|\|\|\|\|\|\|\|\|\|\|\|\|\|\|\|\|\|\|\|\|\|\|\|\|\|\|\|\|\|\|\|\|\|\|\|\|\|\|\|\|\|\|\|\|\|\|\|\|\|\|\|\|\|\|\|\|\|\|\|\|\|\|\|\|\|\|\|\|\|\|\|\|\|\|\|\|\|\|\|\|\|\|\|\|\|\|\|\|\|\|\|\|\|\|\|\|\|\|\|\|\|\|\|\|\|\|\|\|\|\|\|\|\|\|\|\|\|\|\|\|\|\|\|\|\|\|\|\|\|\|\|\|\|\|\|\|\|\|\|\|\|\|\|\|\|\|\|\|\|\|\|\|\|\|\|\|\|\|\|\|\|\|\|\|\|\|\|\|\|\|\|\|\|\|\|\|\|\|\|\|\|\|\|\|\|\|\|\|\|\|\|\|\|\|\|\|\|\|\|\|\|\|\|\|\|\|\|\|\|\|\|\|\|\|\|\|\|\|\|\|\|\|\|\|\|\|\|\|\|\|\|\|\|\|\|\|\|\|\|\|\|\|\|\|\|\|\|\|\|\|\|\|\|\|\|\|\|\|\|\|\|\|\|\|\|\|\|\|\|\|\|\|\|\|\|\|\|\|\|\|\|\|\|\|\|\|\|\|\|\|\|\|\|\|\|\|\|\|\|\|\|\|\|\|\|\|\|\|\|\|\|\|\|\|\|\|\|\|\|\|\|\|\|\|\|\|\|\|\|\|\|\|\|\|\|\|\|\|\|\|\|\|\|\|\|\|\|\|\|\|\|\|\|\|\|\|\|\|\|\|\|\|\|\|\|\|\|\|\|\| \|\|\|\| 	GGCGGCCTAGTCTACCACCGCACCTTCGCCCCAGGTCTCCAGAAGATCGACTCAAACGACTACTTGATCCTAGCCGGCACATTTCACGGCGTGCACGCCATCAGTCGAAGCATCAACCCGGTACCTCCTCTCCCTCAACCGCCGGGCAACCGCAAACCTCAGACCACGGGCATCGAAAGTCTGGAGAGCAGCCACTTCCGATTGACGTGTTTCCAAACGCCGACGGGCGTGAAGTTCCTACTGATCACAAGTCCGGAACAGCCAAATACGGAATTGGTCGTGCGGAGGTGCTATGAGATTTATGGCGACTTCGTGATGAAGAATCCGTTCTATAATCTGGAGATGCCAATTCGGGTGGAGAAGTTTGATCGTGCGCTGGGGTCGTATCTGGTCAGACCCGTCG ``` |
| 3 | ggggacaagtttgtacaaaaaagcaggcttgatgGCACCGCCAGCGATTATCGC, ggggaccactttgtacaagaaagctgggtcCCGCCGCTTTTCAACAGCTT | Mycgr3\|66977  JGI  Ensembl | KOG3111 D-ribulose-5-phosphate 3-epimerase; | carbohydrate metabolic process; ribulose-phosphate 3-epimerase activity; | 0 % |  |  |
| 3 | ggggacaagtttgtacaaaaaagcaggcttgatgTCGCAAGCTCACGACTTGCA, ggggaccactttgtacaagaaagctgggtcGCTTGCAGCCTCGACTTCTT | Mycgr3\|67149  JGI  Ensembl |  |  | 100 % |  | ``` 510	GAGATGACTTCTTCACGGCGTTGAAGAAGAAGCTGAGCAAGGCGGCGGATGACACGAGGCTACAGGGTACGGCGCTGGGAAGGCCAAATGAGTTCAAGGAGGTGGTGGTTGAAGGTGGTCCCGTGCAGCACGTGAGAGATTGGAAGCCGAAGTTGGAGGAGAGATCTGTGAATGTTCAGGCACAGAACACGGCGGAGAGATCAAAAGAACCACAGGACGGACTGGATACAGTGGACGATATCGCAAAGGACTCCGCTATGGAAGAAGTCGAGGCTGCAAGC	790 	\|\|\|\|\|\|\|\|\|\|\|\|\|\|\|\|\|\|\|\|\|\|\|\|\|\|\|\|\|\|\|\|\|\|\|\|\|\|\|\|\|\|\|\|\|\|\|\|\|\|\|\|\|\|\|\|\|\|\|\|\|\|\|\|\|\|\|\|\|\|\|\|\|\|\|\|\|\|\|\|\|\|\|\|\|\|\|\|\|\|\|\|\|\|\|\|\|\|\|\|\|\|\|\|\|\|\|\|\|\|\|\|\|\|\|\|\|\|\|\|\|\|\|\|\|\|\|\|\|\|\|\|\|\|\|\|\|\|\|\|\|\|\|\|\|\|\|\|\|\|\|\|\|\|\|\|\|\|\|\|\|\|\|\|\|\|\|\|\|\|\|\|\|\|\|\|\|\|\|\|\|\|\|\|\|\|\|\|\|\|\|\|\|\|\|\|\|\|\|\|\|\|\|\|\|\|\|\|\|\|\|\|\|\|\|\|\|\|\|\|\|\|\|\|\|\|\|\|\|\|\|\|\|\|\|\|\|\|\|\|\|\|\|\|\|\|\|\|\|\|\|\|\|\|\|\|\|\|\|\|\|\|\|\|\|\|\|\|\|\|\|\|\|\|\|\|\|\|\|\|\| 	GAGATGACTTCTTCACGGCGTTGAAGAAGAAGCTGAGCAAGGCGGCGGATGACACGAGGCTACAGGGTACGGCGCTGGGAAGGCCAAATGAGTTCAAGGAGGTGGTGGTTGAAGGTGGTCCCGTGCAGCACGTGAGAGATTGGAAGCCGAAGTTGGAGGAGAGATCTGTGAATGTTCAGGCACAGAACACGGCGGAGAGATCAAAAGAACCACAGGACGGACTGGATACAGTGGACGATATCGCAAAGGACTCCGCTATGGAAGAAGTCGAGGCTGCAAGC ```      ``` 419	GACACGAAGGACGTGCGAGAGTTTACGAGATGGTACGATTCTCCTCAGTACCGCGAGATCAAGCGTGTGGCGGGCATCATACCGGACGATAG	510 	\|\|\|\|\|\|\|\|\|\|\|\|\|\|\|\|\|\|\|\|\|\|\|\|\|\|\|\|\|\|\|\|\|\|\|\|\|\|\|\|\|\|\|\|\|\|\|\|\|\|\|\|\|\|\|\|\|\|\|\|\|\|\|\|\|\|\|\|\|\|\|\|\|\|\|\|\|\|\|\|\|\|\|\|\|\|\|\|\|\|\|\| 	GACACGAAGGACGTGCGAGAGTTTACGAGATGGTACGATTCTCCTCAGTACCGCGAGATCAAGCGTGTGGCGGGCATCATACCGGACGATAG ```      ``` 238	CGAGCGTCACCTCCTCCTTCTCGCCTCCACCACAGCACAACATGCCTGGTCCAGCCATAATGACCCGGTCCCGACCATCACCGACCTGCGCATGGCGCTGTCCGACTGTGGTGTCCTAATACCCTACTCCGACCCCTCAGAAGAAGCCTGGAAAGAGCGCATGCGCAAGCCCTTGTCAGAAG	419 	\|\|\|\| \|\|\|\|\|\|\|\| \| \|\|\| \| \|\| \|\|\|\|\|\|\|\|\|\|\|\|\|\|\|\|\|\|\|\| \|\|\|\|\|\|\|\|\|\|\|\|\|\|\|\|\|\|\|\|\| \|\|\|\|\|\|\|\|\|\|\|\|\|\|\|\|\|\|\|\|\|\|\|\|\|\|\|\|\|\|\|\|\|\|\|\|\|\|\|\|\|\|\|\|\|\|\|\|\|\|\|\|\|\|\|\|\|\|\|\|\|\|\|\|\|\|\|\|\|\|\|\|\|\|\|\|\|\|\|\|\|\|\|\|\|\|\|\|\|\|\|\|\|\|\|\|\|\|\|\|\|\|\|\|\|\|\|\|\|\|\|\|\|\| 	CGAGNGTCACCTCNTNCTTNTNGCNTCCACCACAGCACAACATGCNTGGTCCAGCCATAATGACCCGNTCCCGACCATCACCGACCTGCGCATGGCGCTGTCCGACTGTGGTGTCCTAATACCCTACTCCGACCCCTCAGAAGAAGCCTGGAAAGAGCGCATGCGCAAGCCCTTGTCAGAAG ``` |
| 3 | ggggacaagtttgtacaaaaaagcaggcttgatgCTCTCCGCTCTGGCCTCCCT, ggggaccactttgtacaagaaagctgggtcTTCTCGTTGAAGCAACACCC | Mycgr3\|67720  JGI  Ensembl | KOG3342 Signal peptidase I; | membrane; peptidase activity;  proteolysis;  signal peptidase I activity;  signal peptide processing; | 100 % |  | ``` 387	GGAGCGAACCTCTCCAACTCCTCACGAAAGGAGACAACAACGCTGCAGACGACACGCAGCTATACGCCCGCGGCCAAAGCTACCTGAATCGCAAGGACGACGTCGTCGGGAGTGTGTTTGGGTTCATACCTTTTGTTGGCTACGTTACAATCCTGCTCAGCGAGTACCCATGGCTGAAAACAGCAATGTTGGTATTCATGGGCTTAGGGGTGTTGCTTCAACGAGAA	613 	\|\|\|\|\|\|\|\|\|\|\|\|\|\|\|\|\|\|\|\|\|\|\|\|\|\|\|\|\|\|\|\|\|\|\|\|\|\|\|\|\|\|\|\|\|\|\|\|\|\|\|\|\|\|\|\|\|\|\|\|\|\|\|\|\|\|\|\|\|\|\|\|\|\|\|\|\|\|\|\|\|\|\|\|\|\|\|\|\|\|\|\|\|\|\|\|\|\|\|\|\|\|\|\|\|\|\|\|\|\|\|\|\|\|\|\|\|\|\|\|\|\|\|\|\|\|\|\|\|\|\|\|\|\|\|\|\|\|\|\|\|\|\|\|\|\|\|\|\|\|\|\|\|\|\|\|\|\|\|\|\|\|\|\|\|\|\|\|\|\|\|\|\|\|\|\|\|\|\|\|\|\|\|\|\|\|\|\|\|\|\|\|\|\|\|\|\|\|\|\|\|\|\|\|\|\|\|\|\|\|\|\|\|\|\|\|\|\|\|\|\|\|\|\|\|\|\| 	GGAGCGAACCTCTCCAACTCCTCACGAAAGGAGACAACAACGCTGCAGACGACACGCAGCTATACGCCCGCGGCCAAAGCTACCTGAATCGCAAGGACGACGTCGTCGGGAGTGTGTTTGGGTTCATACCTTTTGTTGGCTACGTTACAATCCTGCTCAGCGAGTACCCATGGCTGAAAACAGCAATGTTGGTATTCATGGGCTTAGGGGTGTTGCTTCAACGAGAA ```      ``` 86	ATGCTCTCCGCTCTGGCCTCCCTGGGCAGCCAGCCACGCCAATTGGCCGCATCGGTCCTCAACTTCGCACTGGTGCTGTCAACTGCTTTCATGCTCTGGAAAGGACTCTCAGTCATAACAGACTCTTCATCGCCCATCGTTGTGGTACTTTCAGGCAGTATGGAGCCGGCGTTTCAAAGAGGAGACTTGCTATTCCTCTGGAATCGCGGCATGGATACACAAGTGGGCGAGATTGTGGTCTACAACGTTCGCGGCAAGGACATTCCGATTGTGCACAGAGTCGTGAGGAGATTCGGCGGAGGG	388 	\|\|\|\|\|\|\|\|\|\|\|\|\|\|\|\|\|\|\|\|\|\|\|\|\|\|\|\|\|\|\|\|\|\|\|\|\|\|\|\|\|\|\|\|\|\|\|\|\|\|\|\|\|\|\|\|\|\|\|\|\|\|\|\|\|\|\|\|\|\|\|\|\|\|\|\|\|\|\|\|\|\|\|\|\|\|\|\|\|\|\|\|\|\|\|\|\|\|\|\|\|\|\|\|\|\|\|\|\|\|\|\|\|\|\|\|\|\|\|\|\|\|\|\|\|\|\|\|\|\|\|\|\|\|\|\|\|\|\|\|\|\|\|\|\|\|\|\|\|\|\|\|\|\|\|\|\|\|\|\|\|\|\|\|\|\|\|\|\|\|\|\|\|\|\|\|\|\|\|\|\|\|\|\|\|\|\|\|\|\|\|\|\|\|\|\|\|\|\|\|\|\|\|\|\|\|\|\|\|\|\|\|\|\|\|\|\|\|\|\|\|\|\|\|\|\|\|\|\|\|\|\|\|\|\|\|\|\|\|\|\|\|\|\|\|\|\|\|\|\|\|\|\|\|\|\|\|\|\|\|\|\|\|\|\|\|\|\|\|\|\|\|\|\|\|\|\|\|\|\|\|\|\|\|\|\|\|\|\|\|\|\|\|\|\|\|\|\|\|\|\|\|\| 	ATGCTCTCCGCTCTGGCCTCCCTGGGCAGCCAGCCACGCCAATTGGCCGCATCGGTCCTCAACTTCGCACTGGTGCTGTCAACTGCTTTCATGCTCTGGAAAGGACTCTCAGTCATAACAGACTCTTCATCGCCCATCGTTGTGGTACTTTCAGGCAGTATGGAGCCGGCGTTTCAAAGAGGAGACTTGCTATTCCTCTGGAATCGCGGCATGGATACACAAGTGGGCGAGATTGTGGTCTACAACGTTCGCGGCAAGGACATTCCGATTGTGCACAGAGTCGTGAGGAGATTCGGCGGAGGG ``` |
| 3 | ggggacaagtttgtacaaaaaagcaggcttgatgAAGACCTCCATTTTGACCCT, ggggaccactttgtacaagaaagctgggtcGTGACCACCCTTCCCACCGG | Mycgr3\|67799  JGI  Ensembl |  |  | 60.1626 % |  |  |
| 3 | ggggacaagtttgtacaaaaaagcaggcttgatgGCCGACGCCGAGCAGAAGGT, ggggaccactttgtacaagaaagctgggtcCTCCGTCTTGCTGAAGAGGA | Mycgr3\|68026  JGI  Ensembl | KOG0864 Ran-binding protein RANBP1 and related RanBD domain proteins; | intracellular transport; | 0 % |  | ``` 602	GTGGTTCCGGACATGAAGCTTTCACCAAACGTGGGATCCGACCGCAGCTGGGTGTGGAACGCTGCCGCAGATGTTTCCGAGGGCGAGCCAGAGGCTTCCACTCTTGCTATTCGCTTCGCCAACGCTGAGA	731 	\|\|\|\|\|\|\|\|\|\|\|\|\|\|\|\|\|\|\|\|\|\|\|\|\|\|\|\|\|\|\|\|\|\|\|\|\|\|\|\|\|\|\|\|\|\|\|\|\|\|\|\|\|\|\|\|\|\|\|\|\|\|\|\|\|\|\|\|\|\|\|\|\|\|\|\|\|\|\|\|\|\|\|\|\|\|\|\|\|\|\|\|\|\|\|\|\|\|\|\|\|\|\|\|\|\|\|\|\|\|\|\|\|\|\|\|\|\|\|\|\|\|\|\|\|\|\|\|\|\| 	GTGGTTCCGGACATGAAGCTTTCACCAAACGTGGGATCCGACCGCAGCTGGGTGTGGAACGCTGCCGCAGATGTTTCCGAGGGCGAGCCAGAGGCTTCCACTCTTGCTATTCGCTTCGCCAACGCTGAGA ```      ``` 146	ACCAAGCCGGAAGAGAGCAAGACCGTCACGGAAAAGGCTTCCGAGACGGCTT-CCGCAGTAGCTGGCAACGTGTTTGGCATGTTCGGAGGTGGCCCTAAGAAA-GAGGCGAAGAAGGACGATGACGATGCGGAGAATGACCGCTCCGG-AAGTGCCAAGGCGCAGAAGGACAAGGAGAACGCGGACAAGGATGACGATGAGAAGGCCGATGAAGAGGAGGCAGATGTGCACTTCGAGCCGGTCGTGCATCTCACCGAGAAGGTTGATACGAAGACGAACGAGGAGGCCGAAGAGCAGACTTTCAAGATGCGGGCGAAGCTCTTCAAGTTCGACCGTGAGAGCAGGGAGTGGAAGGAGCGCGGCACTGGTGATGTGAGACTTCTCAAGCACAAGGAGAACAGCAAGACGAGATTGGTGATGCGACGAGACAAGACTTTGAAGGTCTGCGCCAACCACTACG	602 	\|\|\| \|\|\| \|\| \|\| \|\| \|\|\|\|  \|\|\|\| \|\|\| \|\|\| \|\|\|\|\|\|\| \|\|\|\|\|\|\|\| \|  \|\|\|\|\|\|\|\|\|\|\|\|\|\|\|\|\|\|\|\|\|\|\|\|\|\|\|\|\|\|\|\|\|\|\|\|\|\|\| \|\|\|\|\|\|\| \|\|\|\|\|\|\|\|\|\|\|\|\|\|\|\|\|\|\|\|\|\|\|\|\|\|\|\|\|\|\|\|\|\|\|\|\|\|\|\|\|\|\|\| \|\|\|\|\|\|\|\|\|\|\|\|\|\|\|\|\|\|\|\|\|\|\|\|\|\|\|\|\|\|\|\|\|\|\|\|\|\|\|\|\|\|\|\|\|\|\|\|\|\|\|\|\|\|\|\|\|\|\|\|\|\|\|\|\|\|\|\|\|\|\|\|\|\|\|\|\|\|\|\|\|\|\|\|\|\|\|\|\|\|\|\|\|\|\|\|\|\|\|\|\|\|\|\|\|\|\|\|\|\|\|\|\|\|\|\|\|\|\|\|\|\|\|\|\|\|\|\|\|\|\|\|\|\|\|\|\|\|\|\|\|\|\|\|\|\|\|\|\|\|\|\|\|\|\|\|\|\|\|\|\|\|\|\|\|\|\|\|\|\|\|\|\|\|\|\|\|\|\|\|\|\|\|\|\|\|\|\|\|\|\|\|\|\|\|\|\|\|\|\|\|\|\|\|\|\|\|\|\|\|\|\|\|\|\|\|\|\|\|\|\|\|\|\|\|\|\|\|\|\|\|\|\|\|\|\|\|\|\|\|\|\|\|\|\|\|\|\|\|\|\|\|\|\|\|\|\|\|\|\|\|\|\|\|\|\|\|\|\|\|\|\|\|\|\|\|\|\|\|\|\|\|\|\|\|\|\|\|\|\|\|\|\|\|\|\|\|\|\|\|\|\|\|\|\|\|\|\|\|\|\| 	ACC-AGCNGG-AGNGANCAAG-NCGTCNCGG-AAANGCTTCCGNGACGGCTTNCNNCAGTAGCTGGCAACGTGTTTGGCATGTTCGGAGGTGGCCNTAAGAAAAGAGGCGAAGAAGGACGATGACGATGCGGAGAATGACCGCTCCGGGAAGTGCCAAGGCGCAGAAGGACAAGGAGAACGCGGACAAGGATGACGATGAGAAGGCCGATGAAGAGGAGGCAGATGTGCACTTCGAGCCGGTCGTGCATCTCACCGAGAAGGTTGATACGAAGACGAACGAGGAGGCCGAAGAGCAGACTTTCAAGATGCGGGCGAAGCTCTTCAAGTTCGACCGTGAGAGCAGGGAGTGGAAGGAGCGCGGCACTGGTGATGTGAGACTTCTCAAGCACAAGGAGAACAGCAAGACGAGATTGGTGATGCGACGAGACAAGACTTTGAAGGTCTGCGCCAACCACTACG ```      ``` 732	ACGCCAACCTCTTCAAGGAAGCCTTCATCAAGGCGCAGCAGGAGAATGAGGTCCTCTTCAGCAAGACGGAG	802 	\|\|\|\|\|\|\|\|\|\|\|\|\|\|\|\|\|\|\|\|\|\|\|\|\|\|\|\|\|\|\|\|\|\|\|\|\|\|\|\|\|\|\|\|\|\|\|\|\|\|\|\|\|\|\|\|\|\|\|\|\|\|\|\|\|\|\|\|\|\|\| 	ACGCCAACCTCTTCAAGGAAGCCTTCATCAAGGCGCAGCAGGAGAATGAGGTCCTCTTCAGCAAGACGGAG ``` |
| 3 | ggggacaagtttgtacaaaaaagcaggcttgatgGCAGCGAGCACTCAGACGAC, ggggaccactttgtacaagaaagctgggtcCGCACCACCAGACCTTCCAT | Mycgr3\|68633  JGI  Ensembl |  |  | 100 % |  | ``` 277	AGCAAAATGAGATTGACAATGGCATCGTGAAGCTGGAGAGTTTGCTGAACGCGACGGTGGACAAGGACTTTGACAAATTCGAGATCTACACGCTGCGAAACATTCTCGCTGTCGGACACGATGAGGAGGATTTGGCGAACTGGGTACGACTGGACCACTATCAAGGCGTAGACATCGCCGGCGCGGAGAACGTCCCGTCGCCTGAGGAAGTCCAGCTTCAGCGGCGCAAGCTCAACGAGACCGCGAAACTGAACACGATGCTCAAGGCAGAGGAAGCGAAGAATGCCGCGATTCTGTCACAACTGACAACGCTGATCGGAGCCGAAGCGAAGTCGCAAGAAGGCGACGAGAGCGAATCACCCTTCGCATTCCTCAACTCTTCCGCCCAAACTGCCAACGCACAGCAACTCACCCAGGATACGCAATACGCTGTGAACCAGATTCCAGCATTACGCCGACTTTTGGACCAATTGAAAGATGCGATGCGGACGATACCTAATACACGGCAAATGCGACTTGACGATGAAGACTCCATTGAGGGCAGGAGAAGGAGATATCTACACAATCAAAGCAGGAGGGCTCTGGAGAGGAAGGGCATCGATGTGGAGGCTTCTGCTGGTGCCTCGATCGCTACTGGGAGGAAAGTGGGCCGCGATGAGCTGGAGGGTATCGAGGCTGTTGCGCAGGCGCTTGCAGGGGCTGAGAATGGAAGGTCTGGTGGTGCG	1001 	\|\|\|\|\|\|\|\|\|\|\|\|\|\|\| \| \|\|\|\|\| \|\|\|\|\|\|\|\| \|\|\|\|\|\|  \|\|  \|\|\|\|\|\| \| \|\|\|\|\|\|\|\|\|\|\|\|\|\|\|\|\|\|\|\|\|\|\|\|\|\|\|\|\|\|\| \|\|\|\|\|\| \|\|\|\|\|\|\|\|\|\|\|\|\|\|\|\|\|\|\|\|\|\|\|\|\|\|\|\|\|\|\|\|\|\|\|\|\|\|\|\|\|\|\|\|\|\|\|\|\|\|\|\|\|\|\|\|\|\|\|\|\|\|\|\|\|\|\|\|\|\|\|\|\|\|\|\|\|\|\|\|\|\|\|\|\|\|\|\|\|\|\|\|\|\|\|\|\|\|\|\|\|\|\|\|\|\|\|\|\|\|\|\|\|\|\|\|\|\|\|\|\|\|\|\|\|\|\|\|\|\|\|\|\|\|\|\|\|\|\|\|\|\|\|\|\|\|\|\|\|\|\|\|\|\|\|\|\|\|\|\|\|\|\|\|\|\|\|\|\|\|\|\|\|\|\|\|\|\|\|\|\|\|\|\|\|\|\|\|\|\|\|\|\|\|\|\|\|\|\|\|\|\|\|\|\|\|\|\|\|\|\|\|\|\|\|\|\|\|\|\|\|\|\|\|\|\|\|\|\|\|\|\|\|\|\|\|\|\|\|\|\|\|\|\|\|\|\|\|\|\|\|\|\|\|\|\|\|\|\|\|\|\|\|\|\|\|\|\|\|\|\|\|\|\|\|\|\|\|\|\|\|\|\|\|\|\|\|\|\|\|\|\|\|\|\|\|\|\|\|\|\|\|\|\|\|\|\|\|\|\|\|\|\|\|\|\|\|\|\|\|\|\|\|\|\|\|\|\|\|\|\|\|\|\|\|\|\|\|\|\|\|\|\|\|\|\|\|\|\|\|\|\|\|\|\|\|\|\|\|\|\|\|\|\|\|\|\|\|\|\|\|\|\|\|\|\|\|\|\|\|\|\|\|\|\|\|\|\|\|\|\|\|\|\|\|\|\|\|\|\|\|\|\|\|\|\|\|\|\|\|\|\|\|\|\|\|\|\|\|\|\|\|\|\|\|\|\|\|\|\|\|\|\|\|\|\|\|\|\|\|\|\|\|\|\|\|\|\|\|\|\|\|\|\|\|\|\|\|\|\|\|\|\|\|\|\|\|\|\|\|\|\|\|\|\|\|\|\|\|\|\|\|\|\|\|\|\|\|\|\|\|\|\|\|\|\|\|\|\|\|\|\|\|\|\|\|\|\|\|\|\|\|\|\|\|\|\|\|\|\|\|\|\|\|\|\|\|\|\|\|\|\|\|\|\|\|\|\|\|\|\|\|\|\|\|\|\|\|\|\|\|\|\|\|\|\|\|\|\|\|\|\|\|\|\|\|\|\|\|\|\|\|\|\|\|\|\|\|\|\|\|\|\|\|\|\|\|\|\|\|\|\|\|\|\|\|\|\|\|\|\|\|\|\|\|\|\|\|\|\|\|\|\|\|\|\|\|\|\|\|\|\|\|\|\|\|\|\|\|\|\|\| 	AGCAAAATGAGATTGNCCATGGCNTCGTGAAGNTGGAGA-NTTNNTGAACGNGNCGGTGGACAAGGACTTTGACAAATTCGAGATNTACACGNTGCGAAACATTCTCGCTGTCGGACACGATGAGGAGGATTTGGCGAACTGGGTACGACTGGACCACTATCAAGGCGTAGACATCGCCGGCGCGGAGAACGTCCCGTCGCCTGAGGAAGTCCAGCTTCAGCGGCGCAAGCTCAACGAGACCGCGAAACTGAACACGATGCTCAAGGCAGAGGAAGCGAAGAATGCCGCGATTCTGTCACAACTGACAACGCTGATCGGAGCCGAAGCGAAGTCGCAAGAAGGCGACGAGAGCGAATCACCCTTCGCATTCCTCAACTCTTCCGCCCAAACTGCCAACGCACAGCAACTCACCCAGGATACGCAATACGCTGTGAACCAGATTCCAGCATTACGCCGACTTTTGGACCAATTGAAAGATGCGATGCGGACGATACCTAATACACGGCAAATGCGACTTGACGATGAAGACTCCATTGAGGGCAGGAGAAGGAGATATCTACACAATCAAAGCAGGAGGGCTCTGGAGAGGAAGGGCATCGATGTGGAGGCTTCTGCTGGTGCCTCGATCGCTACTGGGAGGAAAGTGGGCCGCGATGAGCTGGAGGGTATCGAGGCTGTTGCGCAGGCGCTTGCAGGGGCTGAGAATGGAAGGTCTGGTGGTGCG ```      ``` 39	ATGGCAGCGAGCACTCAGACGACGACTGCGCTGCTGACAGAGCACTTCCGCTACACTCCACTGACCCTTCTGGACGATATCATCAATACCGTGAACGAACTCGTCTTCCGCGCGGTGAATGCGATCGAAGAAGGATTTGGAGGGACAACAGCAGAACAGCTCGGCTTCACATTGGACGATAGTGTCGCCAAAGCTTTGCCCAGTGAAGCTGCTAGACGAGATGCGCTTGCAGAGTTGAAGCAAAATGAGATTGACAATGGCATCGTGAAGCTGGAGAGTTTGCTGAACGCGACGGTGGACAAGGACTTTGACAAATTCGAGATCTACACGCTGCGAAACATTCTCGCTGTCGGACACGATGAGGAGGATTTGGCGAACTGGGTACGACTGGACCACTATCAAGGCGTAGACATCGCCGGCGCGGAGAACGTCCCGTCGCCTGAGGAAGTCCAGCTTCAGCGGCGCAAGCTCAACGAGACCGCGAAACTGAACACGATGCTCAAGGCAGAGGAAGCGAAGAATGCCGCGATTCTGTCACAACTGACAACGCTGATCGGAGCCGAAGCGAAGTCGCAAGAAGGCGACGAGAGCGAATCACCCTTCGCATTCCTCAACTCTTCCGCCCAAACTGCCAACGCACAGCAACTCACCCAGGATACGCAATACGCTGTGAACCAGATTCCAGCATTACGCCGACTTTTGGACCAATTGAAAGATGCGATGCGGACGATACCTAATACACGGCAAATGCGACTTGACGATGAAGACTCCATTGAGGGCAGGAGAAGGAGATATCTACACAATCAAAGCAGGAGGGCTCTGGAGAGGAAGGGCATCGATGTGGAGGCTTCTGCTGGTGCCTCGATCGCTACTGGGAGGAAAGTGGGCCGCGATGAGCTGGAGGGTATCGAGGCTGTTGCGCAGGCGCTTGCAGGGGCTGAGAATGGAAGGTCTGGTGGTGCG	1001 	\|\|\|\|\|\|\|\|\|\|\|\|\|\|\|\|\|\|\|\|\|\|\|\|\|\|\|\|\|\|\|\|\|\|\|\|\|\|\|\|\|\|\|\|\|\|\|\|\|\|\|\|\|\|\|\|\|\|\|\|\|\|\|\|\|\|\|\|\|\|\|\|\|\|\|\|\|\|\|\|\|\|\|\|\|\|\|\|\|\|\|\|\|\|\|\|\|\|\|\|\|\|\|\|\|\|\|\|\|\|\|\|\|\|\|\|\|\|\|\|\|\|\|\|\|\|\|\|\|\|\|\|\|\|\|\|\|\|\|\|\|\|\|\|\|\|\|\|\|\|\|\|\|\|\|\|\|\|\|\|\|\|\|\|\|\|\|\|\|\|\|\|\|\|\|\|\|\|\|\|\|\|\|\|\|\|\|\|\|\|\|\|\|\|\|\|\|\|\|\|\|\|\|\|\|\|\|\|\|\|\|\|\|\|\|\|\|\|\|\|\|\|\|\|\|\|\|\|\|\|\|\|\|\|\|\|\|\|\|\|\|\|\|\|\|\|\|\|\|\|\|\|\|\|\|\|\|\|\|\|\|\|\|\|\|\|\|\|\|\|\|\|\|\|\|\|\|\|\|\|\|\|\|\|\|\|\|\|\|\|\|\|\|\|\|\|\|\|\|\|\|\|\|\|\|\|\|\|\|\|\|\|\|\|\|\|\|\|\|\|\|\|\|\|\|\|\|\|\|\|\|\|\|\|\|\|\|\|\|\|\|\|\|\|\|\|\|\|\|\|\|\|\|\|\|\|\|\|\|\|\|\|\|\|\|\|\|\|\|\|\|\|\|\|\|\|\|\|\|\|\|\|\|\|\|\|\|\|\|\|\|\|\|\|\|\|\|\|\|\|\|\|\|\|\|\|\|\|\|\|\|\|\|\|\|\|\|\|\|\|\|\|\|\|\|\|\|\|\|\|\|\|\|\|\|\|\|\|\|\|\|\|\|\|\|\|\|\|\|\|\|\|\|\|\|\|\|\|\|\|\|\|\|\|\|\|\|\|\|\|\|\|\|\|\|\|\|\|\|\|\|\|\|\|\|\|\|\|\|\|\|\|\|\|\|\|\|\|\|\|\|\|\|\|\|\|\|\|\|\|\|\|\|\|\|\|\|\|\|\|\|\|\|\|\|\|\|\|\|\|\|\|\|\|\|\|\|\|\|\|\|\|\|\|\|\|\|\|\|\|\|\|\|\|\|\|\|\|\|\|\|\|\|\|\|\|\|\|\|\|\|\|\|\|\|\|\|\|\|\|\|\|\|\|\|\|\|\|\|\|\|\|\|\|\|\|\|\|\|\|\|\|\|\|\|\|\|\|\|\|\|\|\|\|\|\|\|\|\|\|\|\|\|\|\|\|\|\|\|\|\|\|\|\|\|\|\|\|\|\|\|\|\|\|\|\|\|\|\|\|\|\|\|\|\|\|\|\|\|\|\|\|\|\|\|\|\|\|\|\|\|\|\|\|\|\|\|\|\|\|\|\|\|\|\|\|\|\|\|\|\|\|\|\|\|\|\|\|\|\|\|\|\|\|\|\|\|\|\|\|\|\|\|\|\|\|\|\|\|\|\|\|\|\|\|\|\|\|\|\|\|\|\|\|\|\|\|\|\|\|\|\|\|\|\|\|\|\|\|\|\|\|\|\|\|\|\|\|\|\|\|\|\|\|\|\|\|\|\|\|\|\|\|\|\|\|\|\|\|\|\|\|\|\|\|\|\|\|\|\|\|\|\|\|\|\|\|\|\|\|\|\|\|\|\|\|\|\|\|\|\|\|\|\|\|\|\|\|\|\|\|\|\|\|\|\|\|\|\|\|\|\|\|\|\|\|\|\|\|\|\|\|\|\|\|\|\|\|\|\|\|\|\|\|\|\|\|\|\|\|\|\|\|\|\|\|\|\|\|\|\|\|\|\|\|\|\|\|\|\|\|\|\|\|\|\|\|\|\|\|\|\|\|\|\|\|\|\|\|\|\|\|\|\|\|\|\|\|\|\|\|\|\|\|\|\|\|\|\|\|\|\|\|\| \|\|\|\|\|\|\|\|\|\|\| \|\|\|\|\|\| \|\| \| \| \|\|\|\|\|\|\| \|\|\|\| 	ATGGCAGCGAGCACTCAGACGACGACTGCGCTGCTGACAGAGCACTTCCGCTACACTCCACTGACCCTTCTGGACGATATCATCAATACCGTGAACGAACTCGTCTTCCGCGCGGTGAATGCGATCGAAGAAGGATTTGGAGGGACAACAGCAGAACAGCTCGGCTTCACATTGGACGATAGTGTCGCCAAAGCTTTGCCCAGTGAAGCTGCTAGACGAGATGCGCTTGCAGAGTTGAAGCAAAATGAGATTGACAATGGCATCGTGAAGCTGGAGAGTTTGCTGAACGCGACGGTGGACAAGGACTTTGACAAATTCGAGATCTACACGCTGCGAAACATTCTCGCTGTCGGACACGATGAGGAGGATTTGGCGAACTGGGTACGACTGGACCACTATCAAGGCGTAGACATCGCCGGCGCGGAGAACGTCCCGTCGCCTGAGGAAGTCCAGCTTCAGCGGCGCAAGCTCAACGAGACCGCGAAACTGAACACGATGCTCAAGGCAGAGGAAGCGAAGAATGCCGCGATTCTGTCACAACTGACAACGCTGATCGGAGCCGAAGCGAAGTCGCAAGAAGGCGACGAGAGCGAATCACCCTTCGCATTCCTCAACTCTTCCGCCCAAACTGCCAACGCACAGCAACTCACCCAGGATACGCAATACGCTGTGAACCAGATTCCAGCATTACGCCGACTTTTGGACCAATTGAAAGATGCGATGCGGACGATACCTAATACACGGCAAATGCGACTTGACGATGAAGACTCCATTGAGGGCAGGAGAAGGAGATATCTACACAATCAAAGCAGGAGGGCTCTGGAGAGGAAGGGCATCGATGTGGAGGCTTCTGCTGGTGCCTCGATCGCTACTGGGAGGAAAGTGGGCCGCGATGAGCTGGAGGGTATCGAGGCTGTTGCGCAG-CGCTTGCAGGG-CTGAGA-TG-A-G-TCTGGTG-TGCG ``` |
| 3 | ggggacaagtttgtacaaaaaagcaggcttgatgGCGCTCCCTGGACAGATTCT, ggggaccactttgtacaagaaagctgggtcAACCGGCTTCGCCACCTTCC | Mycgr3\|69638  JGI  Ensembl | KOG3409 Exosomal 3'-5' exoribonuclease complex, subunit ski4 (Csl4); | 3'-5'-exoribonuclease activity; 5'-3' exoribonuclease activity;  RNA binding; | 100 % |  | ``` 15	ATGGCGCTCCCTGGACAGATTCTCGGACCAACAAGTGCCAACAGTCTCGGCACAGGAACTCACATCCAAGGCACCGATCTCTGCGCAAGCATTTCCGGTCCCATCCTCTCCCAACCTTCAACAAGCAAAACCTCAAAACTTCCCACCGTTTCCATCGCCCGCACGACAGGTACTCTCCTTCCCGAAGTTGGAACCGTCATCCTAGGCAAAATCACACGCACCGGCCCACGACAAGCCAACGTCACGATCCTGGCCCTCGGTTCCGCAGGCGAACACGTCTGTGGCGATGCCTTTCCCGCTCTCATCCGACAGCAAGACATTCGCGCAACGGAGATCGACAAGGTGAAAGTCGCGGAGAGTTTTCGGGTTGGGGATGTGGTGCGTGCGACGGTGATTAGTCTAGGAGATGAGAGGAGCTATTACTTGACGACCGCGAGGAATGAGCTGGGCGTTGTGATGGCGCAGAGTGAGTGGGGAAATCAGATGTATCCCATCTCGTGGAAGGAGTTTCAGGATCCGGAGACGGGATTGAGGGAGCTGAGGAA	559 	\|\|\|\|\|\|\|\|\|\|\|\|\|\|\|\|\|\|\|\|\|\|\|\|\|\|\|\|\|\|\|\|\|\|\|\|\|\|\|\|\|\|\|\|\|\|\|\|\|\|\|\|\|\|\|\|\|\|\|\|\|\|\|\|\|\|\|\|\|\|\|\|\|\|\|\|\|\|\|\|\|\|\|\|\|\|\|\|\|\|\|\|\|\|\|\|\|\|\|\|\|\|\|\|\|\|\|\|\|\|\|\|\|\|\|\|\|\|\|\|\|\|\|\|\|\|\|\|\|\|\|\|\|\|\|\|\|\|\|\|\|\|\|\|\|\|\|\|\|\|\|\|\|\|\|\|\|\|\|\|\|\|\|\|\|\|\|\|\|\|\|\|\|\|\|\|\|\|\|\|\|\|\|\|\|\|\|\|\|\|\|\|\|\|\|\|\|\|\|\|\|\|\|\|\|\|\|\|\|\|\|\|\|\|\|\|\|\|\|\|\|\|\|\|\|\|\|\|\|\|\|\|\|\|\|\|\|\|\|\|\|\|\|\|\|\|\|\|\|\|\|\|\|\|\|\|\|\|\|\|\|\|\|\|\|\|\|\|\|\|\|\|\|\|\|\|\|\|\|\|\|\|\|\|\|\|\|\|\|\|\|\|\|\|\|\|\|\|\|\|\|\|\|\|\|\|\|\|\|\|\|\|\|\|\|\|\|\|\|\|\|\|\|\|\|\|\|\|\|\|\|\|\|\|\|\|\|\|\|\|\|\|\|\|\|\|\|\|\|\|\|\|\|\|\|\|\|\|\|\|\|\|\|\|\|\|\|\|\|\|\|\|\|\|\|\|\|\|\|\|\|\|\|\|\|\|\|\|\|\|\|\|\|\|\|\|\|\|\|\|\|\|\|\|\|\|\|\|\|\|\|\|\|\|\|\|\|\|\|\|\|\|\|\|\|\|\|\|\|\|\|\|\|\|\|\|\|\|\|\|\|\|\|\|\|\|\|\|\|\|\|\|\|\|\|\|\|\|\|\|\|\|\|\| \|\|\|\|\|\|\|\|\|\|\|\|\|\|\|\|\| \|\|\|\|\|\|\|\|\|\|\|\|\|\|\|\|\|\|\|\|\|\|\|\|\|\|\|\|\|\|\|\|\|\|\|\|\|\| \|\|\|\|\|\|\|\|\|\| \|\|\| \|\|\|\| \|\|\| 	ATGGCGCTCCCTGGACAGATTCTCGGACCAACAAGTGCCAACAGTCTCGGCACAGGAACTCACATCCAAGGCACCGATCTCTGCGCAAGCATTTCCGGTCCCATCCTCTCCCAACCTTCAACAAGCAAAACCTCAAAACTTCCCACCGTTTCCATCGCCCGCACGACAGGTACTCTCCTTCCCGAAGTTGGAACCGTCATCCTAGGCAAAATCACACGCACCGGCCCACGACAAGCCAACGTCACGATCCTGGCCCTCGGTTCCGCAGGCGAACACGTCTGTGGCGATGCCTTTCCCGCTCTCATCCGACAGCAAGACATTCGCGCAACGGAGATCGACAAGGTGAAAGTCGCGGAGAGTTTTCGGGTTGGGGATGTGGTGCGTGCGACGGTGATTAGTCTAGGAGATGAGAGGAGCTATTACTTGACGACCGCGAGGAATGAGCTGGGCGTTGTGATGGCGCANAGTGAGTGGGGAAATCANATGTATCCCATCTCGTGGAAGGAGTTTCAGGATCCGGANACGGGATTGANGGANCTGANGAA ```      ``` 114	CCCATCCTCTCCCAACCTTCAACAAGCAAAACCTCAAAACTTCCCACCGTTTCCATCGCCCGCACGACAGGTACTCTCCTTCCCGAAGTTGGAACCGTCATCCTAGGCAAAATCACACGCACCGGCCCACGACAAGCCAACGTCACGATCCTGGCCCTCGGTTCCGCAGGCGAACACGTCTGTGGCGATGCCTTTCCCGCTCTCATCCGACAGCAAGACATTCGCGCAACGGAGATCGACAAGGTGAAAGTCGCGGAGAGTTTTCGGGTTGGGGATGTGGTGCGTGCGACGGTGATTAGTCTAGGAGATGAGAGGAGCTATTACTTGACGACCGCGAGGAATGAGCTGGGCGTTGTGATGGCGCAGAGTGAGTGGGGAAATCAGATGTATCCCATCTCGTGGAAGGAGTTTCAGGATCCGGAGACGGGATTGAGGGAGCTGAGGAAGGTGGCGAAGCCGGTT	575 	\|\|\|\|\|\| \|\|\|\|\|\|\|\|\| \|\|\|\|\|\|\|\|\|\|\|\|\|\|\|\|\|\|\|\|\|\|\|\|\|\|\|\|\|\|\|\|\|\|\|\|\|\|\|\|\|\|\|\|\|\|\|\|\|\|\|\|\|\|\|\|\|\|\|\|\|\|\|\|\|\|\|\|\|\|\|\|\|\|\|\|\|\|\|\|\|\|\|\|\|\|\|\|\|\|\|\|\|\|\|\|\|\|\|\|\|\|\|\|\|\|\|\|\|\|\|\|\|\|\|\|\|\|\|\|\|\|\|\|\|\|\|\|\|\|\|\|\|\|\|\|\|\|\|\|\|\|\|\|\|\|\|\|\|\|\|\|\|\|\|\|\|\|\|\|\|\|\|\|\|\|\|\|\|\|\|\|\|\|\|\|\|\|\|\|\|\|\|\|\|\|\|\|\|\|\|\|\|\|\|\|\|\|\|\|\|\|\|\|\|\|\|\|\|\|\|\|\|\|\|\|\|\|\|\|\|\|\|\|\|\|\|\|\|\|\|\|\|\|\|\|\|\|\|\|\|\|\|\|\|\|\|\|\|\|\|\|\|\|\|\|\|\|\|\|\|\|\|\|\|\|\|\|\|\|\|\|\|\|\|\|\|\|\|\|\|\|\|\|\|\|\|\|\|\|\|\|\|\|\|\|\|\|\|\|\|\|\|\|\|\|\|\|\|\|\|\|\|\|\|\|\|\|\|\|\|\|\|\|\|\|\|\|\|\|\|\|\|\|\|\|\|\|\|\|\|\|\|\|\|\|\|\|\|\|\|\|\|\|\|\|\|\|\|\|\|\|\|\|\|\|\|\|\|\|\|\|\|\|\|\|\|\|\|\|\|\|\|\|\|\|\|\|\|\|\|\|\|\|\|\|\|\|\|\|\|\|\|\|\|\|\|\|\|\|\|\|\|\|\|\|\|\|\|\|\|\|\|\|\|\|\|\|\|\|\|\|\|\|\|\|\|\|\|\|\|\|\|\|\| 	CCCATCNTCTCCCAACNTTCAACAAGCAAAACCTCAAAACTTCCCACCGTTTCCATCGCCCGCACGACAGGTACTCTCCTTCCCGAAGTTGGAACCGTCATCCTAGGCAAAATCACACGCACCGGCCCACGACAAGCCAACGTCACGATCCTGGCCCTCGGTTCCGCAGGCGAACACGTCTGTGGCGATGCCTTTCCCGCTCTCATCCGACAGCAAGACATTCGCGCAACGGAGATCGACAAGGTGAAAGTCGCGGAGAGTTTTCGGGTTGGGGATGTGGTGCGTGCGACGGTGATTAGTCTAGGAGATGAGAGGAGCTATTACTTGACGACCGCGAGGAATGAGCTGGGCGTTGTGATGGCGCAGAGTGAGTGGGGAAATCAGATGTATCCCATCTCGTGGAAGGAGTTTCAGGATCCGGAGACGGGATTGAGGGAGCTGAGGAAGGTGGCGAAGCCGGTT ``` |
| 3 | ggggacaagtttgtacaaaaaagcaggcttgatgTCCTACTTCTCCCCTGATGC, ggggaccactttgtacaagaaagctgggtcATCGCGACGGCTGTTGACAT | Mycgr3\|69673  JGI  Ensembl | KOG1106 Uncharacterized conserved protein; |  | 100 % |  | ``` 333	TCCCCTGATGCCATCCTGACGGATGCGCAGAAGGCGCCCTGCGAATTCGGTCTCGCCGTTCCGCAGCTCACACCTCTCAACAATGGATCTGCAATCGACAAGGGCACCAAATTGGAACTGCCTTTGTGGATAGCAGAAATGCTGGCTGTGTCGAAACCTGCGGACACGGCTCTGGCGACGCTGGATATGCCCGCACCGCTCGGCCATCGTGTACTGAATGCATTACGAGCGGATCCGAAGTCTGTTGATGTCAGAGCGCAGGCGCAGTGGTTTTATGGGTTGGGCGAGAGGATGCTCGAATTGTTCGAGGATGAGGAGGTTGCGGAGGTGTTGAACGAT	671 	\|\|\|\|\|\|\|\|\|\|\|\|\|\|\|\|\|\|\|\|\|\|\|\|\|\|\|\|\|\|\|\|\|\|\|\|\|\|\|\|\| \|\|\|\|\|\|\|\|\|\|\|\|\|\|\|\|\|\|\|\|\|\|\|\|\|\|\|\|\|\|\|\|\|\|\|\|\|\|\|\|\|\|\|\|\|\|\|\|\|\|\|\|\|\|\|\|\|\|\|\|\|\|\|\|\|\|\|\|\|\|\|\|\|\|\|\|\|\|\|\|\|\|\|\|\|\|\|\|\|\|\|\|\|\|\|\|\|\|\|\|\|\|\|\|\|\|\|\|\|\|\|\|\|\|\|\|\|\|\|\|\|\|\|\|\|\|\|\|\|\|\|\|\|\|\|\|\|\|\|\|\|\|\|\|\|\|\|\|\|\|\|\|\|\|\|\|\|\|\|\|\|\|\|\|\|\|\|\|\|\|\|\|\|\|\|\|\|\|\|\|\|\|\|\|\|\|\|\|\|\|\|\|\|\|\|\|\|\|\|\|\|\|\|\|\|\|\|\|\|\|\|\|\|\|\|\|\|\|\|\|\|\|\|\|\|\|\|\|\|\|\|\|\|\|\|\|\|\|\|\|\|\|\|\|\|\|\|\|\|\|\|\|\|\|\|\|\|\|\|\|\|\|\|\|\|\|\|\|\|\|\|\|\|\|\|\|\|\|\|\|\|\|\|\|\|\|\|\|\|\|\|\|\|\|\|\|\| 	TCCCCTGATGCCATCCTGACGGATGCGCAGAAGGCGCCCTGNGAATTCGGTCTCGCCGTTCCGCAGCTCACACCTCTCAACAATGGATCTGCAATCGACAAGGGCACCAAATTGGAACTGCCTTTGTGGATAGCAGAAATGCTGGCTGTGTCGAAACCTGCGGACACGGCTCTGGCGACGCTGGATATGCCCGCACCGCTCGGCCATCGTGTACTGAATGCATTACGAGCGGATCCGAAGTCTGTTGATGTCAGAGCGCAGGCGCAGTGGTTTTATGGGTTGGGCGAGAGGATGCTCGAATTGTTCGAGGATGAGGAGGTTGCGGAGGTGTTGAACGAT ```      ``` 672	ACATTCAAGCAAAGATCACTGGAGATTGCAGACAAGGCGCAGAATACCCGTAATGTACAGCAAGGCAGTGATAACGAGTTCATGAGTGGACTGGATGAGACGGAAAGGCAAT	783 	\|\|\|\|\|\|\|\|\|\|\|\|\|\|\|\|\|\|\|\|\|\|\|\|\|\|\|\|\|\|\|\|\|\|\|\|\|\|\|\|\|\|\|\|\|\|\|\|\|\|\|\|\|\|\|\|\|\|\|\|\|\|\|\|\|\|\|\|\|\|\|\|\|\|\|\|\|\|\|\|\|\|\|\|\|\|\|\|\|\|\|\|\|\|\|\|\|\|\|\|\|\|\|\|\|\|\|\|\|\|\|\| 	ACATTCAAGCAAAGATCACTGGAGATTGCAGACAAGGCGCAGAATACCCGTAATGTACAGCAAGGCAGTGATAACGAGTTCATGAGTGGACTGGATGAGACGGAAAGGCAAT ```      ``` 784	TATTCCGTGCAGCACACGATGGCAGTCATGCCGTGAGGAAGTGGCTAGATGTCAACAGCCGTCGCGAT	851 	\|\|\|\|\|\|\|\|\|\|\|\|\|\|\|\|\|\|\|\|\|\|\|\|\|\|\|\|\|\|\|\|\|\|\|\|\|\|\|\|\|\|\|\|\|\|\|\|\|\|\|\|\|\|\|\|\|\|\|\|\|\|\|\|\|\|\|\| 	TATTCCGTGCAGCACACGATGGCAGTCATGCCGTGAGGAAGTGGCTAGATGTCAACAGCCGTCGCGAT ``` |
| 3 | ggggacaagtttgtacaaaaaagcaggcttgatgGCAACACATCAACCCTTTCC, ggggaccactttgtacaagaaagctgggtcCCCTCCCATACAGATCCCGA | Mycgr3\|70288  JGI  Ensembl | KOG0028 Ca2+-binding protein (centrin/caltractin), EF-Hand superfamily protein; | calcium ion binding; | 65.61403 % |  | ``` 264	CCGCAGCAATCGAACCCTGCGGGACTCGGAGCAACACAACAACAACGCGAAGCGCAGCGCTTAGAACGAGAAAGACATGAGCGTGCGGAGCGGGAACGGCGAGAAGCTGAAGAGAGGGAAACGCTGGAGCAGTTGTCGGAGGAGCAAAGGGAGGAGATAAATGAAGCT	431 	\|\|\|\|\| \| \|\| \|\|\|\|\|\|\|\|\|\|\|\|\| \| \|\|\|\|\|\|\|\|\|\|\|\|\|\|\|\| \|\|\| \|\|\|\|\|\|\|\|\|\|\| \|\|\|\|\|\|\|\|\|\|\|\|\|\|\|\|\|\|\|\|\|\| \|\|\|\|\|\|\|\|\|\|\|\|\|\|\|\|\|\|\|\|\|\|\|\|\|\|\|\|\|\|\|\|\|\|\|\|\|\|\|\|\|\|\|\|\|\|\|\|\|\|\|\|\|\|\|\|\|\|\|\|\|\|\|\|\|\|\|\|\|\|\|\|\|\|\|\|\|\|\|\|\|\|\|\|\| 	CCGCANCCATNGAACCCTGCGGGANTNGGAGCAACACAACAACNACGNGAAGCGCAGCGNTTAGAACGAGAAAGACATGAGCNTGCGGAGCGGGAACGGCGAGAAGCTGAAGAGAGGGAAACGCTGGAGCAGTTGTCGGAGGAGCAAAGGGAGGAGATAAATGAAGCT ```      ``` 432	TTCTCGCTCTTCGACCTCGACAAAGACTCCCACATCGACTACCACGAACTCAAAGTCGCGCTGAAAGCCCTCGGCTTCGAAGTCCCCAAATCCGAACTCCTGCAAATCCTCCAATCCCACGGCGTGCCCGCTTCCGCACTGCAGACCTCAGCAAGTACGTCTAAAGCGCCCGTACCACAACAACCGACATTCTCAGGCCCTTCGCGACTACTGCTCAGTCATGCGGCGTTCACCTCTGTGGCAGCACAGCGAATATTGGCGAGAGACCCGAGGGAGGAGATCTTGAGGGCTTTTGAGCTATTCGATGCAGATGGCAAGGGCAGAATCGAGTTGCAGGACTTGAGGAGGGTCGCGAGGGAGTTGGGCGAGGGACTGCAGGAGGAGGAACTGGCGGCGATGATTGAGGAGTTTGACATTAGAGGTGAAGGAGGGATTGGTAGAGATGAGTTTGTCGGGATCTGTATGGGAGGG	902 	\|\|\|\|\|\|\|\|\|\|\|\|\|\|\|\|\|\|\|\|\|\|\|\|\|\|\|\|\|\|\|\|\|\|\|\|\|\|\|\|\|\|\|\|\|\|\|\|\|\|\|\|\|\|\|\|\|\|\|\|\|\|\|\|\|\|\|\|\|\|\|\|\|\|\|\|\|\|\|\|\|\|\|\|\|\|\|\|\|\|\|\|\|\|\|\|\|\|\|\|\|\|\|\|\|\|\|\|\|\|\|\|\|\|\|\|\|\|\|\|\|\|\|\|\|\|\|\|\|\|\|\|\|\|\|\|\|\|\|\|\|\|\|\|\|\|\|\|\|\|\|\|\|\|\|\|\|\|\|\|\|\|\|\|\|\|\|\|\|\|\|\|\|\|\|\|\|\|\|\|\|\|\|\|\|\|\|\|\|\|\|\|\|\|\|\|\|\|\|\|\|\|\|\|\|\|\|\|\|\|\|\|\|\|\|\|\|\|\|\|\|\|\|\|\|\|\|\|\|\|\|\|\|\|\|\|\|\|\|\|\|\|\|\|\|\|\|\|\|\|\|\|\|\|\|\|\|\|\|\|\|\|\|\|\|\|\|\|\|\|\|\|\|\|\|\|\|\|\|\|\|\|\|\|\|\|\|\|\|\|\|\|\|\|\|\|\|\|\|\|\|\|\|\|\|\|\|\|\|\|\|\|\|\|\|\|\|\|\|\|\|\|\|\|\|\|\|\|\|\|\|\|\|\|\|\|\|\|\|\|\|\|\|\|\|\|\|\|\|\|\|\|\|\|\|\|\|\|\|\|\|\|\|\|\|\|\|\|\|\|\|\|\|\|\|\|\|\|\|\|\|\|\|\|\|\|\|\|\|\|\|\|\|\|\|\|\|\|\|\|\|\|\|\|\|\|\|\|\|\|\|\|\|\|\|\|\|\|\|\|\|\|\|\|\|\|\|\|\|\|\|\|\|\|\|\|\|\|\|\|\|\|\|\|\|\|\|\|\|\|\|\|\|\|\|\|\|\|\|\|\|\|\|\|\|\|\|\|\|\|\| 	TTCTCGCTCTTCGACCTCGACAAAGACTCCCACATCGACTACCACGAACTCAAAGTCGCGCTGAAAGCCCTCGGCTTCGAAGTCCCCAAATCCGAACTCCTGCAAATCCTCCAATCCCACGGCGTGCCCGCTTCCGCACTGCAGACCTCAGCAAGTACGTCTAAAGCGCCCGTACCACAACAACCGACATTCTCAGGCCCTTCGCGACTACTGCTCAGTCATGCGGCGTTCACCTCTGTGGCAGCACAGCGAATATTGGCGAGAGACCCGAGGGAGGAGATCTTGAGGGCTTTTGAGCTATTCGATGCAGATGGCAAGGGCAGAATCGAGTTGCAGGACTTGAGGAGGGTCGCGAGGGAGTTGGGCGAGGGACTGCAGGAGGAGGAACTGGCGGCGATGATTGAGGAGTTTGACATTAGAGGTGAAGGAGGGATTGGTAGAGATGAGTTTGTCGGGATCTGTATGGGAGGG ``` |
| 3 | ggggacaagtttgtacaaaaaagcaggcttgatgGCGCATCAGGCGGACGCAAA, ggggaccactttgtacaagaaagctgggtcGTCTGCGTCGCTCATCTCAC | Mycgr3\|70421  JGI  Ensembl | KOG2889 Predicted PRP38-like splicing factor; |  | 0 % |  |  |
| 3 | ggggacaagtttgtacaaaaaagcaggcttgatgGCCAACACCGAGTTCGGCTA, ggggaccactttgtacaagaaagctgggtcCTCATCATAGTCCTCCCCTC | Mycgr3\|70961  JGI  Ensembl | KOG2721 Uncharacterized conserved protein; |  | 0 % |  |  |
| 3 | ggggacaagtttgtacaaaaaagcaggcttgatgGACGCCGCAAAGGACAACGC, ggggaccactttgtacaagaaagctgggtcCTGCTCGCACGAAGACAGCA | Mycgr3\|72125  JGI  Ensembl | KOG3381 Uncharacterized conserved protein; |  | 100 % |  | ``` 125	CACAGGGCATCTTCGATGCTCTAGTACCCGCATATCCCAGTCTGGCAGATCCTGCCTCGCCACCAGAAGTGCTCTCCGACCTGTCCGACTCTGCCTCCGAAGACGAACGAGAGGAAATCGACGAGCAGGAAATCTACGATCTCATCTCCACCATCACCGATCCCGAGCATCCGCTCACCCTCGGCTCGCTCGGCGTCGTGAACCTGGACGACATCAAGATCTTCCCACCTTCCTCGCCGCGGTCGCGCATCAGCTCTGTGCGGGTGTTGATCACTCCCACGACCACTGCGTGCTCTTTGACGACGGTCATCGGATTGGGCGTCAAGGTCCGCCTGGTCAATGCCCTGCCACCGCGCTTTCGAATCGATGTACGCATCAAGGAAGGCACTTCTTCATCTGCCGACGAGGCCAACAAGCAGCTGGGCGACAAGGAGCGTGTCGCTGCCGCCATGGAGAACCGCAATCTGATCAACATGGTGAATCACATGCTGTCTTCGTGCGAGCAG	630 	\|\|\|\|\|\|\|\|\|\| \|\| \|\|\|\| \| \|\|\|\|\|\|\| \|\|\|\|\|\| \|\|\|\|\| \|\|\|\|\|\|\|\|\| \|\|\|\|\| \|\|\|\|\|\|\|\|\|\|\|\|\| \| \|\| \|\|\| \|\|\|\| \| \|\| \|\|\|\|\|\| \|\|\|\|\|\|\|\| \|\|\|\|\|\|\|\|\|\|\| \|\|\|\|\|\|\|\|\|\|\|\|\|\| \|\| \|\|\|\|\|\|\|\|\|\|\|\|\|\|\|\|\|\|\|\|\|\|\|\|\|\|\|\|\|\|\|\|\|\|\|\|\|\|\|\|\|\|\|\|\|\|\|\|\|\|\|\|\|\|\|\|\|\|\|\|\|\|\|\|\|\|\|\|\|\|\|\|\|\|\|\|\|\|\|\|\|\|\|\|\|\|\|\|\|\|\|\|\|\|\|\|\|\|\|\|\|\|\|\|\|\|\|\|\|\|\|\|\|\|\|\|\|\|\|\|\|\|\|\|\|\|\|\|\|\|\|\|\|\|\|\|\|\|\|\|\|\|\|\|\|\|\|\|\|\|\|\|\|\|\|\|\|\|\|\|\|\|\|\|\|\|\|\|\|\|\|\|\|\|\|\|\|\|\|\|\|\|\|\|\|\|\|\|\|\|\|\|\|\|\|\|\|\|\|\|\|\|\|\|\|\|\|\|\|\|\|\|\|\|\|\|\|\|\|\|\|\|\|\|\|\|\|\|\|\|\|\|\|\|\|\|\|\|\|\|\|\|\|\|\|\|\|\|\|\|\|\|\|\|\|\|\|\|\|\|\|\|\|\|\|\|\|\|\|\|\|\|\|\|\|\|\|\|\|\|\|\|\|\|\|\|\|\|\|\|\|\|\|\|\|\|\|\|\|\|\|\|\|\|\|\|\|\|\|\|\|\|\|\|\|\|\|\|\|\|\|\|\|\|\|\|\|\|\|\|\|\|\|\|\|\|\|\|\|\|\|\|\|\|\|\|\|\|\|\|\|\|\|\|\|\|\|\|\|\|\|\|\|\|\|\|\|\|\| 	CACAGGGCATNTTNGATGNTNTAGTACCNGCATATNCCAGTNTGGCAGATCNTGCCTNGCCACCAGAAGTGNTNTCNGACNTGTCNGNCTNTGCCTCNGAAGACGANCGAGAGGAAATNGACGAGCAGGAAATNTANGATCTCATCTCCACCATCACCGATCCCGAGCATCCGCTCACCCTCGGCTCGCTCGGCGTCGTGAACCTGGACGACATCAAGATCTTCCCACCTTCCTCGCCGCGGTCGCGCATCAGCTCTGTGCGGGTGTTGATCACTCCCACGACCACTGCGTGCTCTTTGACGACGGTCATCGGATTGGGCGTCAAGGTCCGCCTGGTCAATGCCCTGCCACCGCGCTTTCGAATCGATGTACGCATCAAGGAAGGCACTTCTTCATCTGCCGACGAGGCCAACAAGCAGCTGGGCGACAAGGAGCGTGTCGCTGCCGCCATGGAGAACCGCAATCTGATCAACATGGTGAATCACATGCTGTCTTCGTGCGAGCAG ``` |
| 3 | ggggacaagtttgtacaaaaaagcaggcttgatgCTGCTTTACTACTCGCTCGC, ggggaccactttgtacaagaaagctgggtcGAGTCGGATCCCGAGCCGTC | Mycgr3\|72169  JGI  Ensembl |  |  | 0 % |  |  |
| 3 | ggggacaagtttgtacaaaaaagcaggcttgatgCATTTCTCCCACATCTTGCC, ggggaccactttgtacaagaaagctgggtcCTGCGCAATCTTGTAATCGT | Mycgr3\|73448  JGI  Ensembl |  |  | 100 % |  | ``` 217	AGCATCGAGGTCCGCGCCATTCCGACCAGTTGGCGCATTACCGGCCAGGCCTTCAATG	274 	\|\|\|\|\|\|\|\|\|\|\|\|\|\|\|\|\|\|\|\|\|\|\|\|\|\|\|\|\|\|\|\|\|\|\|\|\|\|\|\|\|\|\|\|\|\|\|\|\|\|\|\|\|\|\|\|\|\| 	AGCATCGAGGTCCGCGCCATTCCGACCAGTTGGCGCATTACCGGCCAGGCCTTCAATG ```      ``` 274	GGAGGTGATTGCAGAAACGTCCTCTACGTAGTTCAGTCCAACGGCCGGGAGAACATCTGTCTCGGCAACAGTGACTACTCCGGCGGCTCCTATATCTTCCAAAACTTGAGGCGCTCCATCGACGCCGCTCCTCAGGAAGCCTGTCCTGCATCGGGCTGCAACGTCCGTAGGGGCAACGAGATGGTCTTCGAGGGTGGCCCGAGTTACAACATGTCTGCGCTTGATGAGTCGGACTATGATGAATTG	519 	\|\|\|\|\|\|\|\|\|\|\|\|\|\|\|\|\|\|\|\|\|\|\| \|\|\|\|\|\|\|\|\|\|\|\|\|\|\|\|\|\|\|\|\|\|\|\|\|\|\|\|\|\|\|\|\|\|\|\|\|\|\|\|\|\|\|\|\|\|\|\|\|\|\|\|\|\|\|\|\|\|\|\|\|\|\|\|\|\|\|\|\|\|\|\|\|\|\|\|\|\|\|\|\|\|\|\|\|\|\|\|\|\|\|\|\|\|\|\|\|\|\|\|\|\|\|\|\|\|\|\|\|\|\|\|\|\|\|\|\|\|\|\|\|\|\|\|\|\|\|\|\|\|\|\|\|\|\|\|\|\|\|\|\|\|\|\|\|\|\|\|\|\|\|\|\|\|\|\|\|\|\|\|\|\|\|\|\|\|\|\|\|\|\|\|\|\|\|\|\|\|\|\|\|\|\|\|\|\|\|\|\|\|\|\|\|\|\|\|\|\|\|\|\|\|\|\|\|\|\|\|\|\|\|\|\|\|\|\|\|\|\|\|\|\| 	GGAGGTGATTGCAGAAACGTCCTNTACGTAGTTCAGTCCAACGGCCGGGAGAACATCTGTCTCGGCAACAGTGACTACTCCGGCGGCTCCTATATCTTCCAAAACTTGAGGCGCTCCATCGACGCCGCTCCTCAGGAAGCCTGTCCTGCATCGGGCTGCAACGTCCGTAGGGGCAACGAGATGGTCTTCGAGGGTGGCCCGAGTTACAACATGTCTGCGCTTGATGAGTCGGACTATGATGAATTG ```      ``` 274	GGAGGTGATTGCAGAAACGTCCTCTACGTAGTTCAGTCCAACGGCCGGGAGAACATCTGTCTCGGCAACAGTGACTACTCCGGCGGCTCCTATATCTTCCAAAACTTGAGGCGCTCCATCGACGCCGCTCCTCAGGAAGCCTGTCCTGCATCGGGCTGCAACGTCCGTAGGGGCAACGAGATGGTCTTCGAGGGTGGCCCGAGTTACAACATGTCTGCGCTTGATGAGTCGGACTATGATGAATTG	519 	\|\|\|\|\|\|\|\|\|\|\|\|\|\|\|\|\|\|\|\|\|\|\|\|\|\|\|\|\|\|\|\|\|\|\|\|\|\|\|\|\|\|\|\|\|\|\|\|\|\|\|\|\|\|\|\|\|\|\|\|\|\|\|\|\|\|\|\|\|\|\|\|\|\|\|\|\|\|\|\|\|\|\|\|\|\|\|\|\|\|\|\|\|\|\|\|\|\|\|\|\|\|\|\|\|\|\|\|\|\|\|\|\|\|\|\|\|\|\|\|\|\|\|\|\|\|\|\|\|\|\|\|\|\|\|\|\|\|\|\|\|\|\|\|\|\|\|\|\|\|\|\|\|\|\|\|\|\|\|\|\|\|\|\|\|\|\|\|\|\|\|\|\|\|\|\|\|\|\|\|\|\|\|\|\|\|\|\|\|\|\|\|\|\|\|\|\|\|\|\|\|\|\|\|\|\|\|\|\|\|\|\|\|\|\|\|\|\|\|\|\|\|\|\|\|\|\|\|\|\|\|\|\|\|\|\|\|\|\|\|\|\|\|\|\|\| 	GGAGGTGATTGCAGAAACGTCCTCTACGTAGTTCAGTCCAACGGCCGGGAGAACATCTGTCTCGGCAACAGTGACTACTCCGGCGGCTCCTATATCTTCCAAAACTTGAGGCGCTCCATCGACGCCGCTCCTCAGGAAGCCTGTCCTGCATCGGGCTGCAACGTCCGTAGGGGCAACGAGATGGTCTTCGAGGGTGGCCCGAGTTACAACATGTCTGCGCTTGATGAGTCGGACTATGATGAATTG ```      ``` 519	GCTCAGCATCTTCGAGACTGGAGCTCATTGGACCGAGTTTCCGGCCAAATTCGACGATTACAAGATTGCGCAG	591 	\|\|\|\|\|\|\|\|\|\|\|\|\|\|\|\|\|\|\|\|\|\|\|\|\|\|\|\|\|\|\|\|\|\|\|\|\|\|\|\|\|\|\|\|\|\|\|\|\|\|\|\|\|\|\|\|\|\|\|\|\|\|\|\|\|\|\|\|\|\|\|\|\| 	GCTCAGCATCTTCGAGACTGGAGCTCATTGGACCGAGTTTCCGGCCAAATTCGACGATTACAAGATTGCGCAG ``` |
| 3 | ggggacaagtttgtacaaaaaagcaggcttgatgGTGCGCAAATTGAAGCATCA, ggggaccactttgtacaagaaagctgggtcTAGCAGGTCAAAGTCGTCCA | Mycgr3\|73498  JGI  Ensembl | KOG4655 U3 small nucleolar ribonucleoprotein (snoRNP) component; | RNA binding; | 0 % |  |  |
| 3 | ggggacaagtttgtacaaaaaagcaggcttgatgGCCCCTCCAACCAGGATCAC, ggggaccactttgtacaagaaagctgggtcGTCCCTCGTCGTCCTGATCT | Mycgr3\|73624  JGI  Ensembl | KOG3076 5'-phosphoribosylglycinamide formyltransferase; | 'de novo' IMP biosynthetic process; biosynthetic process;  hydroxymethyl-, formyl- and related transferase activity;  phosphoribosylglycinamide formyltransferase activity; | 100 % | chr\_7:80985 G->A; | ``` 83	GAATCTCCAAGCTCTCATCGACGCCTCGAAAACATCCTCGCTGTCGAATACCACCTTTGTTCGTGTCATATCCGACCGCAAATCTGCCTACGGCCTGATTCGTGCCTCCAATGCCAACATCCCAACAAGCTACCACTCAATCCTGCCATACAAGAAGTCGCATCCCGATACCTCTTCCAACCCGACGTTCAATGCAGCTCGCGAAGCATACGATGCAGATCTCGCAAAGCTTGTGCTCGCCGATGAGCCACAATTAGTCGTTTGCGCTGGCTTCATGCGGATCCTAACTCCCTCTTTCCTTAACCCACTCGCAATGGCAAATGTGCCGATCATCAATCTTCACCCATCGCTGCACGGCGACTTGATCGGAGCGGGATGCATTGAACGAGCATGGGAGGAGTTCCAACAAGGCAGGCGAAAGAAGACGGGAATCATGATACACTATGTCATCGCGGAAGTCGATATGGGAGAACCTATCCTGCAGAAAGAGGTCGACATCGAGGGCTGTACTTCGCTCGTAGACCTGCAGGCGAGGATTCATGAGGCGGAGCATGTGCTCATCGTTGATGGCGCAAGCACTGTTGTCGAGAAGATCAGGACGACGAGGGAC	692 	\|\|\|\| \| \|\|\|\| \| \|\|\|\| \|\|  \|\|\| \|\|\|\|\|\|\|\|\|\|\| \|\|\|\|\| \|\|\|\|\|\|\|\|\|\|\|\|\|\|\|\| \|\| \|\|\|\|\|\|\|\|\|\|\|\|\|\|\|\|\|\| \|\|\| \|\|\|\|\|\| \|\|\|\|\| \|\|\|\|\|\|\|\|\|\|\|\|\|\|\|\|\|\|\|\|\|\|\|\|\|\|\|\|\|\|\|\|\|\|\|\|\|\|\|\|\|\|\|\|\|\|\|\|\|\|\|\|\|\|\|\|\|\|\|\|\|\|\|\|\|\|\|\|\|\|\|\|\|\|\|\|\|\|\|\|\|\|\|\|\|\|\|\|\|\|\|\|\|\|\|\|\|\|\|\|\|\|\|\|\|\|\|\|\|\|\|\|\|\|\|\|\|\|\|\|\|\|\|\|\|\|\|\|\|\|\|\|\|\|\|\|\|\|\|\|\|\|\|\|\|\|\|\|\|\|\|\|\|\|\|\|\|\|\|\|\|\|\|\|\|\|\|\|\|\|\|\|\|\|\|\|\|\|\|\|\|\|\|\|\|\|\|\|\|\|\|\|\|\|\|\|\|\|\|\|\|\|\|\|\|\|\|\|\|\|\|\|\|\|\|\|\|\|\|\|\|\|\|\|\|\|\|\|\|\|\|\|\|\|\|\|\|\|\|\|\|\|\|\|\|\|\|\|\|\|\|\|\|\|\|\|\|\|\|\|\|\|\|\|\|\|\|\|\|\|\|\|\|\|\|\|\|\|\|\|\|\|\|\|\|\|\|\|\|\|\|\|\|\|\|\|\|\|\|\|\|\|\|\|\|\|\|\|\|\|\|\|\|\|\|\|\|\|\|\|\|\|\|\|\|\|\|\|\|\|\|\|\|\|\|\|\|\|\|\|\|\|\|\|\|\|\|\|\|\|\|\|\|\|\|\|\|\|\|\|\|\|\|\|\|\|\|\|\|\|\|\|\|\|\|\|\|\|\|\|\|\|\|\|\|\|\|\|\|\|\|\|\|\|\|\|\|\|\|\|\|\|\|\|\|\|\|\|\|\|\|\|\|\|\|\|\|\|\|\|\|\|\|\|\|\|\|\|\|\|\|\|\|\|\|\|\|\|\|\|\|\|\|\|\|\|\|\|\|\|\|\|\|\|\|\|\|\|\|\|\|\|\|\|\|\|\|\|\|\|\|\|\|\|\|\|\|\|\|\|\|\|\|\|\|\|\|\|\|\|\|\|\|\|\|\|\|\|\|\|\|\|\|\|\|\|\|\|\| 	GAATNTNCAAGNTNTCATNGANNCCTNGAAAACATCCTNGCTGTNGAATACCACCTTTGTTNGTATCATATCCGACCGCAAATNTGCNTACGGCNTGATTNGTGCCTCCAATGCCAACATCCCAACAAGCTACCACTCAATCCTGCCATACAAGAAGTCGCATCCCGATACCTCTTCCAACCCGACGTTCAATGCAGCTCGCGAAGCATACGATGCAGATCTCGCAAAGCTTGTGCTCGCCGATGAGCCACAATTAGTCGTTTGCGCTGGCTTCATGCGGATCCTAACTCCCTCTTTCCTTAACCCACTCGCAATGGCAAATGTGCCGATCATCAATCTTCACCCATCGCTGCACGGCGACTTGATCGGAGCGGGATGCATTGAACGAGCATGGGAGGAGTTCCAACAAGGCAGGCGAAAGAAGACGGGAATCATGATACACTATGTCATCGCGGAAGTCGATATGGGAGAACCTATCCTGCAGAAAGAGGTCGACATCGAGGGCTGTACTTCGCTCGTAGACCTGCAGGCGAGGATTCATGAGGCGGAGCATGTGCTCATCGTTGATGGCGCAAGCACTGTTGTCGAGAAGATCAGGACGACGAGGGAC ``` |
| 3 | ggggacaagtttgtacaaaaaagcaggcttgatgTCCGACGACGATGCATTCCC, ggggaccactttgtacaagaaagctgggtcCGCAGCAGCACCCTCCACTC | Mycgr3\|73812  JGI  Ensembl | KOG0530 Protein farnesyltransferase, alpha subunit/protein geranylgeranyltransferase type I, alpha subunit; | protein amino acid prenylation; protein prenyltransferase activity; | 100 % |  | ``` 252	TGGTTGTACCGTGCGTCTATACTCTTCGCGCTAGGGTCGGATCTGAGAGAGGAATTGGAGTGGTTGAGTGACATCGCGTTGTCGCATCAGAAGAACTACC-AGATTTGGCATCATCGAAATCTGATCGTTGATAAACTGGGGAGTGCGGATGGGGAAGGGGAGTTTGTGGAGAGGATGTTTGAGTTGGACGGGAAGAATTATCATGTTTGGAGTTATCGGCAGTGGCTTGTCAAGAGGTTTGGATTGTGGGAGGGACATGGGGAGATGGAGTTTGTGGAGAGGATGTTGAGTAAAGATGGGAGGAATAATTCGGCGTGGAATCATCGGTGGTTTGTGGTCAATGGGAGGGAAGACGAGGGAGTGGCTGGCGTCAAAGGGGACGAGGAGGTGAGGAGGACGGAGCTGAGGTTCGCGATGGACAAGATAAGGGAGATACCCGGGAATCAGAGTGCGTGGAGCTATTTGAGGGGAGTGGTGAGGGTTGGAGGGATGAGGTTGAAGGAACTGAAGGGGTTTGTGGAGGAGTTTGCGAGTCTGGGAGAAGAGGATGGCGTGAGGAGTAGTCATGCGTTGGATATGCTGGCGGATGTGTTGGCGGAGGAGGATGGAGAGGGGAAGAAGAAGGCTGCGGAGGCGTTGGATCTGTTGGCGAAGAAGTATGATCCAATCAGGAAGAACTATTGGGAGTATCGGAAGGGGTTGTTGGGAGTGGAGGGTGCTGCTGCG	977 	\|\|\|\|\| \| \|\|\|\|\|\|\|\|\|\| \|\|\|\| \|\|\| \| \|\|\|\|\| \|\|\|\|\|\| \|\|\|\|\|\|\|\|\|\|\|\|\|\|\|\|\|\|\| \|\| \|\|\|\|\|\|\|\|\| \|\|\|\|\|\|\| \| \|\|\|\|\|\|\|\|\|\|\|\|\|\| \|\|\| \|\| \|\|\|\|\|\|\|\|\|\|\|\|\|\|\|\|\|\|\|\|\|\|\|\|\|\|\| \|\|\|\|\| \| \|\|\|\|\|\|\|\|\|\|\|\|\|\|\|\|\|\|\|\|\|\|\|\|\|\|\|\|\|\|\|\|\|\|\|\|\|\|\|\|\|\|\|\|\|\| \|\|\|\|\|\|\|\|\|\|\|\|\|\|\|\|\|\|\|\|\|\|\|\|\|\|\|\| \|\|\|\|\|\|\|\|\|\|\|\|\|\|\|\|\|\|\|\|\|\|\|\|\|\|\|\|\|\|\|\|\|\|\|\|\|\|\|\|\|\|\|\|\|\|\|\|\|\|\|\|\|\|\|\|\|\|\|\|\|\|\|\|\|\|\|\|\|\|\|\|\|\|\|\|\|\|\|\|\|\|\|\|\|\|\|\|\|\|\|\|\|\|\|\|\|\|\|\|\|\|\|\|\|\|\|\|\|\|\|\|\|\|\|\|\|\|\|\|\|\|\|\|\|\|\|\|\|\|\|\|\|\|\|\|\|\|\|\|\|\|\|\|\|\|\|\|\|\|\|\|\|\|\|\|\|\|\|\|\|\|\|\|\|\|\|\|\|\|\|\|\|\|\|\|\|\|\|\|\|\|\|\|\|\|\|\|\|\|\|\|\|\|\|\|\|\|\|\|\|\|\|\|\|\|\|\|\|\|\|\|\|\|\|\|\|\|\|\|\|\|\|\|\|\|\|\|\|\|\|\|\|\|\|\|\|\|\|\|\|\|\|\|\|\|\|\|\|\|\|\|\|\|\|\|\|\|\|\|\|\|\|\|\|\|\|\|\|\|\|\|\|\|\|\|\|\|\|\|\|\|\|\|\|\|\|\|\|\|\|\|\|\|\|\|\|\|\|\|\|\|\|\|\|\|\|\|\|\|\|\|\|\|\|\|\|\|\|\|\|\|\|\|\|\|\|\|\|\|\|\|\|\|\|\|\|\|\|\|\|\|\|\|\|\|\|\|\|\|\|\|\|\|\|\|\|\|\|\|\|\|\|\|\|\|\|\|\|\|\|\|\|\|\|\|\|\|\|\|\|\|\|\|\|\|\|\|\|\|\|\|\|\|\|\|\|\|\|\|\|\|\|\|\|\|\|\|\|\|\|\|\|\|\|\|\|\|\|\|\|\|\|\|\|\|\|\|\|\|\|\|\|\|\|\|\|\|\|\|\|\|\|\|\|\|\|\|\|\|\|\|\|\|\|\|\|\|\|\|\|\|\|\|\|\|\|\|\|\|\|\|\|\|\|\|\|\|\|\|\|\|\|\|\|\|\|\|\|\|\|\|\|\|\|\|\|\|\|\|\|\|\|\|\|\|\| 	TGGTTTTNCCGTGCGTCTTTACTNTTCNCNCTAGGNTCGGATNTGAGAGAGGAATTGGAGTGNTTNAGTGACATCCCGTTGTCNCNTCAGAAGAACTACCCAGA-TTNGCATCATCGAAATCTGATCGTTGATAANCTGGG-ANTGCGGATGGGGAAGGGGAGTTTGTGGAGAGGATGTTTGAGTTGGACNGGAAGAATTATCATGTTTGGAGTTATCGCCAGTGGCTTGTCAAGAGGTTTGGATTGTGGGAGGGACATGGGGAGATGGAGTTTGTGGAGAGGATGTTGAGTAAAGATGGGAGGAATAATTCGGCGTGGAATCATCGGTGGTTTGTGGTCAATGGGAGGGAAGACGAGGGAGTGGCTGGCGTCAAAGGGGACGAGGAGGTGAGGAGGACGGAGCTGAGGTTCGCGATGGACAAGATAAGGGAGATACCCGGGAATCAGAGTGCGTGGAGCTATTTGAGGGGAGTGGTGAGGGTTGGAGGGATGAGGTTGAAGGAACTGAAGGGGTTTGTGGAGGAGTTTGCGAGTCTGGGAGAAGAGGATGGCGTGAGGAGTAGTCATGCGTTGGATATGCTGGCGGATGTGTTGGCGGAGGAGGATGGAGAGGGGAAGAAGAAGGCTGCGGAGGCGTTGGATCTGTTGGCGAAGAAGTATGATCCAATCAGGAAGAACTATTGGGAGTATCGGAAGGGGTTGTTGGGAGTGGAGGGTGCTGCTGCG ``` |
| 3 | ggggacaagtttgtacaaaaaagcaggcttgatgGCCAGCGGCTACGACCGCGC, ggggaccactttgtacaagaaagctgggtcCTCCGCGCCGCCAGCAGGTC | Mycgr3\|74348  JGI  Ensembl | KOG0183 20S proteasome, regulatory subunit alpha type PSMA7/PRE6; | endopeptidase activity; proteasome core complex;  proteasome endopeptidase activity;  threonine endopeptidase activity;  ubiquitin-dependent protein catabolic process; | 100 % |  | ``` 371	GAAGCGCAATCACATCGTCTCACTGTGGAGGATCCCGTCAGCATTGAGTACATCACCAAATATGTGGCCGGCGTGCAACAGCGATACACCCAGAGTGGAGGAGTGCGGCCGTTTGGCATCTCCACGCTCATCGTGGGTTTCGATCCTGGCAGCAAGGAGGCCAGACTATACCAGACCGAGCCATCAGGCATCTACTCCGCATGG	574 	\|\|\|\| \|\|\|\|\|\|\|\|\|\| \|\|\|\|\|\|\|\| \|\|\|\|\|\|\|\|\|\|\|\|\|\|\|\|\|\|\|\|\|\|\|\|\|\|\|\|\|\|\|\|\|\|\|\|\|\|\|\|\|\|\|\|\|\|\|\|\|\|\|\|\|\|\|\|\|\|\|\|\|\|\|\|\|\|\|\|\|\|\|\|\|\|\|\|\|\|\|\|\|\| \|\|\|\|\|\|\|\|\|\|\|\|\|\|\|\|\|\|\|\|\|\|\|\|\|\|\|\|\|\|\|\|\|\|\|\|\|\|\|\|\|\|\|\|\|\|\|\|\|\|\|\|\|\|\|\|\|\|\|\|\|\|\|\|\|\|\|\|\|\|\|\|\|\|\|\|\|\|\|\|\|\|\|\|\|\|\|\|\|\|\|\|\|\|\|\| 	GAAGNGCAATCACATNGTCTCACT-TGGAGGATCCCGTCAGCATTGAGTACATCACCAAATATGTGGCCGGCGTGCAACAGCGATACACCCAGAGTGGAGGAGTGCGNCCGTTTGGCATCTCCACGCTCATCGTGGGTTTCGATCCTGGCAGCAAGGAGGCCAGACTATACCAGACCGAGCCATCAGGCATCTACTCCGCATGG ```      ``` 573	GGAAAGCCAACGCCATCGGCCGCTCCTCCAAAACCGTCCGCGAATTCCTCGAACGCAATCACAAAGACGCCATGCCTCGCGACGACACAATCGAGCTGACCATCAAGTCCCTCCTTGAAGTCGTGCAGACCGGAGCCAAGAACATCGAGATCGCCATCATGGCACCAGGCAAGGGCGTGGAGATGCTGGCGGGTGAGGAGATTGAGAAGATTGTGGAGAAGATCAACACGGAGAAGGATGCTGCTGCGGAGGCGAACGCGAGCCGGAGAGGAGGTGCTAGAGGTGCTGCGGCGGCGGGTGATGCGAGCAGTGCGGATGGTGCGGCGCAAGTGCTGGCAAACAGACCTGCTGGCGGCGCGGAG	934 	\|\|\|\|\|\|\|\|\|\|\|\|\|\|\|\|\|\|\|\|\|\|\|\|\|\|\|\|\|\|\|\|\|\|\|\|\|\|\|\|\|\|\|\|\|\|\|\|\|\|\|\|\|\|\|\|\|\|\|\|\|\|\|\|\|\|\|\|\|\|\|\|\|\|\|\|\|\|\|\|\|\|\|\|\|\|\|\|\|\|\|\|\|\|\|\|\|\|\|\|\|\|\|\|\|\|\|\|\|\|\|\|\|\|\|\|\|\|\|\|\|\|\|\|\|\|\|\|\|\|\|\|\|\|\|\|\|\|\|\|\|\|\|\|\|\|\|\|\|\|\|\|\|\|\|\|\|\|\|\|\|\|\|\|\|\|\|\|\|\|\|\|\|\|\|\|\|\|\|\|\|\|\|\|\|\|\|\|\|\|\|\|\|\|\|\|\|\|\|\|\|\|\|\|\|\|\|\|\|\|\|\|\|\|\|\|\|\|\|\|\|\|\|\|\|\|\|\|\|\|\|\|\|\|\| \|\|\|\|\|\|\|\|\|\|\|\|\|\|\|\|\|\|\|\|\|\|\|\|\|\|\|\|\|\|\|\|\|\|\|\|\|\|\|\|\|\|\|\|\|\|\|\|\|\|\|\|\|\|\|\|\|\|\|\|\|\|\|\|\|\|\|\|\|\|\|\|\|\|\|\|\|\|\|\|\|\|\|\|\|\|\|\|\|\|\|\|\|\|\|\|\|\|\|\|\|\|\|\|\|\|\|\|\|\|\|\|\|\|\|\|\|\|\|\|\|\|\|\|\|\| 	GGAAAGCCAACGCCATCGGCCGCTCCTCCAAAACCGTCCGCGAATTCCTCGAACGCAATCACAAAGACGCCATGCCTCGCGACGACACAATCGAGCTGACCATCAAGTCCCTCCTTGAAGTCGTGCAGACCGGAGCCAAGAACATCGAGATCGCCATCATGGCACCAGGCAAGGGCGTGGAGATGCTGGCGGGTGAGGAGATTGAGAAGATTGTGGAGAAGATCAACACGGAGAAAGATGCTGCTGCGGAGGCGAACGCGAGCCGGAGAGGAGGTGCTAGAGGTGCTGCGGCGGCGGGTGATGCGAGCAGTGCGGATGGTGCGGCGCAAGTGCTGGCAAACAGACCTGCTGGCGGCGCGGAG ```      ``` 573	GGAAAGCCAACGCCATCGGCCGCTCCTCCAAAACCGTCCGCGAATTCCTCGAACGCAATCACAAAGACGCCATGCCTCGCGACGACACAATCGAGCTGACCATCAAGTCCCTCCTTGAAGTCGTGCAGACCGGAGCCAAGAACATCGAGATCGCCATCATGGCACCAGGCAAGGGCGTGGAGATGCTGGCGGGTGAGGAGATTGAGAAGATTGTGGAGAAGATCAACACGGAGAAGGATGCTGCTGCGGAGGCGAACGCGAGCCGGAGAGGAGGTGCTAGAGGTGCTGCGGCGGCGGGTGATGCGAGCAGTGCGGATGGTGCGGCGCAAGTGCTGGCAAACAGACCTGCTGGCGGCGCGGAG	934 	\|\|\|\|\|\|\|\|\|\|\|\|\|\|\|\|\|\|\|\|\|\|\|\|\|\|\|\|\|\|\|\|\|\|\|\|\|\|\|\|\|\|\|\|\|\|\|\|\|\|\|\|\|\|\|\|\|\|\|\|\|\|\|\|\|\|\|\|\|\|\|\|\|\|\|\|\|\|\|\|\|\|\|\|\|\|\|\|\|\|\|\|\|\|\|\|\|\|\|\|\|\|\|\|\|\|\|\|\|\|\|\|\|\|\|\|\|\|\|\|\|\|\|\|\|\|\|\|\|\|\|\|\|\|\|\|\|\|\|\|\|\|\|\|\|\|\|\|\|\|\|\|\|\|\|\|\|\|\|\|\|\|\|\|\|\|\|\|\|\|\|\|\|\|\|\|\|\|\|\|\|\|\|\|\|\|\|\|\|\|\|\|\|\|\|\|\|\|\|\|\|\|\|\|\|\|\|\|\|\|\|\|\|\|\|\|\|\|\|\|\|\|\|\|\|\|\|\|\|\|\|\|\|\|\|\|\|\|\|\|\|\|\|\|\|\|\|\|\|\|\|\|\|\|\|\|\|\|\|\|\|\|\|\|\|\|\|\|\|\|\|\|\|\|\|\|\|\|\|\|\|\|\|\|\|\|\|\|\|\|\|\|\|\|\|\|\|\|\|\|\|\|\|\|\|\|\|\|\|\|\|\|\|\|\|\|\|\|\|\|\|\|\|\|\|\|\|\|\|\|\|\|\|\|\|\|\|\|\|\|\|\|\|\|\|\|\|\|\|\|\|\|\|\|\|\|\|\|\|\|\|\| 	GGAAAGCCAACGCCATCGGCCGCTCCTCCAAAACCGTCCGCGAATTCCTCGAACGCAATCACAAAGACGCCATGCCTCGCGACGACACAATCGAGCTGACCATCAAGTCCCTCCTTGAAGTCGTGCAGACCGGAGCCAAGAACATCGAGATCGCCATCATGGCACCAGGCAAGGGCGTGGAGATGCTGGCGGGTGAGGAGATTGAGAAGATTGTGGAGAAGATCAACACGGAGAAGGATGCTGCTGCGGAGGCGAACGCGAGCCGGAGAGGAGGTGCTAGAGGTGCTGCGGCGGCGGGTGATGCGAGCAGTGCGGATGGTGCGGCGCAAGTGCTGGCAAACAGACCTGCTGGCGGCGCGGAG ```      ``` 95	ATGGCCAGCGGCTACGACCGCGCCCTCTCCGTCTTCTCCCCCGATGGCCACGTCTTTCAGGTCGAATATGCTCTCGAAGCCGTCAAGCGCGGCACCTGCGCCGTAGCAGTCAAGGGCGCCGACGTCGTCGTTCTAGGCTGCGAGAAGCGATCCGCCCTCAAACTCCAAGACACCCGCATCACTCCTTCCAAGATCAGCATGGTCGACACCCACGTCTGCCTCGCCTTCGCCGGTCTCAACGCCGATGCACGCATTCTCGTCGACAAAGCTCGTCTCGAAGCGCAATCACATCGTCTCACTGTGGAGGATCCCGTCAGCATTGAGTACATCACCAAATATGTGGCCGGCGTGCAACAGCGATACACCCAGAGTGGAGGAGTGCGGCCGTTTGGCATCTCCACGCTCATCGTGGGTTTCGATCCTGGCAGCAAGGAGGCCAGACTATACCAGACCGAGCCATCAGGCATCTACTCCGCATGG	574 	\|\|\|\|\|\|\|\|\|\|\|\|\|\|\|\|\|\|\|\|\|\|\|\|\|\|\|\|\|\|\|\|\|\|\|\|\|\|\|\|\|\|\|\|\|\|\|\|\|\|\|\|\|\|\|\|\|\|\|\|\|\|\|\|\|\|\|\|\|\|\|\|\|\|\|\|\|\|\|\|\|\|\|\|\|\|\|\|\|\|\|\|\|\|\|\|\|\|\|\|\|\|\|\|\|\|\|\|\|\|\|\|\|\|\|\|\|\|\|\|\|\|\|\|\|\|\|\|\|\|\|\|\|\|\|\|\|\|\|\|\|\|\|\|\|\|\|\|\|\|\|\|\|\|\|\|\|\|\|\|\|\|\|\|\|\|\|\|\|\|\|\|\|\|\|\|\|\|\|\|\|\|\|\|\|\|\|\|\|\|\|\|\|\|\|\|\|\|\|\|\|\|\|\|\|\|\|\|\|\|\|\|\|\|\|\|\|\|\|\|\|\|\|\|\|\|\|\|\|\|\|\|\|\|\|\|\|\|\|\|\|\|\|\|\|\|\|\|\|\|\|\|\|\|\|\|\|\|\|\|\|\|\|\|\|\|\|\|\|\|\|\|\|\|\|\|\|\|\|\|\|\|\|\|\|\|\|\|\|\|\|\|\|\|\|\|\|\|\|\|\|\|\|\|\|\|\|\|\|\|\|\|\|\|\|\|\|\|\|\|\|\|\|\|\|\|\|\|\|\|\|\|\|\|\|\|\|\|\|\|\|\|\|\|\|\|\|\|\|\|\|\|\|\|\|\|\|\|\|\|\|\|\|\|\|\|\|\|\|\|\|\|\|\|\|\|\|\|\|\|\|\|\|\|\|\|\|\|\|\|\|\|\|\|\|\|\|\|\|\|\|\|\|\|\|\|\|\|\|\|\|\|\|\|\|\|\|\|\|\|\|\|\|\|\|\|\|\|\|\|\|\|\|\|\|\|\|\|\|\|\|\|\|\|\|\|\|\|\|\|\|\|\|\|\|\|\|\|\|\|\|\|\|\|\|\|\|\|\|\|\|\|\|\|\|\|\|\|\|\| 	ATGGCCAGCGGCTACGACCGCGCCCTCTCCGTCTTCTCCCCCGATGGCCACGTCTTTCAGGTCGAATATGCTCTCGAAGCCGTCAAGCGCGGCACCTGCGCCGTAGCAGTCAAGGGCGCCGACGTCGTCGTTCTAGGCTGCGAGAAGCGATCCGCCCTCAAACTCCAAGACACCCGCATCACTCCTTCCAAGATCAGCATGGTCGACACCCACGTCTGCCTCGCCTTCGCCGGTCTCAACGCCGATGCACGCATTCTCGTCGACAAAGCTCGTCTCGAAGCGCAATCACATCGTCTCACTGTGGAGGATCCCGTCAGCATTGAGTACATCACCAAATATGTGGCCGGCGTGCAACAGCGATACACCCAGAGTGGAGGAGTGCGGCCGTTTGGCATCTCCACGCTCATCGTGGGTTTCGATCCTGGCAGCAAGGAGGCCAGACTATACCAGACCGAGCCATCAGGCATCTACTCCGCATGG ``` |
| 3 | ggggacaagtttgtacaaaaaagcaggcttgatgCAGTCCAGGACATTATTGCG, ggggaccactttgtacaagaaagctgggtcACCCTTCTTCTCTTTCTCTG | Mycgr3\|74842  JGI  Ensembl | KOG3309 Ferredoxin; | 2 iron, 2 sulfur cluster binding; electron carrier activity;  electron transport;  iron-sulfur cluster binding; | 100 % |  | ``` 504	GGAGGAACCATCAGACGATGAGAACGACATGCTTGACCTTGCTTTCGGACTTACAGAGACGAGTCGATTGGGATGCCAGGTCGTGATGAGTAAAGAAATCGATGGACTTGTTGTAAAGCTGCCTAGCATGACGAGGAATTTGCAGGCCTCGGACTTTGCAGAGAAAGAGAAGAAGGGT	681 	\|\|\|\|\|\|\|\|\|\|\|\|\|\|\|\|\|\|\|\|\|\|\|\|\|\|\|\|\|\|\|\|\|\|\|\|\|\|\|\|\|\|\|\|\|\|\|\|\|\|\|\|\|\|\|\|\|\|\|\|\|\|\|\|\|\|\|\|\|\|\|\|\|\|\|\|\|\|\|\|\|\|\|\|\|\|\|\|\|\|\|\|\|\|\|\|\|\|\|\|\|\|\|\|\|\|\|\|\|\|\|\|\|\|\|\|\|\|\|\|\|\|\|\|\|\|\|\|\|\|\|\|\|\|\|\|\|\|\|\|\|\|\|\|\|\|\|\|\|\|\|\|\|\|\|\|\|\|\|\|\|\|\|\|\|\|\|\|\|\|\|\|\|\|\|\|\|\| 	GGAGGAACCATCAGACGATGAGAACGACATGCTTGACCTTGCTTTCGGACTTACAGAGACGAGTCGATTGGGATGCCAGGTCGTGATGAGTAAAGAAATCGATGGACTTGTTGTAAAGCTGCCTAGCATGACGAGGAATTTGCAGGCCTCGGACTTTGCAGAGAAAGAGAAGAAGGGT ```      ``` 34	ATGCAGTCCAGGACATTATTGCGATCGCAATGCCTGGCACTCTCACGGACGGCGACCACGAGTCTGCGAACGGCTTCACGAGACCATGTGACCAATGCTAGTCGGCAGCTGAGGTCCAGTCCATCCACATTTCCATCGCAAAGAGCATACGCTTCGAGTACGCGGAGTGGATTCCCTCTTATACCGGCCCGATGCTCGCATCGTTCATACACCTCCTCGCCTCATCGGACATTCCACACGACCTCTGTGCGACAGCACGGCGACCTCACGCCACCTAAGCCGGGCGAGGAAAGAAAAGTGACATTTATCGACAAGGATGGGCAAGAATCGACATTCGAGGTCGCCGATGGCGACAACTTGTTGGACATCGCACAAGCAAATGATTTGGAAATGGAGGGTGCATGCGGCGGGAGCTGTGCGTGCAGTACGTGTCATGTCATTGTGAAGGACGAGGACATATATGACAAGATGG	505 	\|\|\|\|\|\|\|\|\|\| \| \|\|\|\|  \|\|\|\|\|\|\|\|\|\|\|\|\|\|\|\|\|\|\|\|\|\| \|\|\|\|\|\|\|\|\|\|\|\|\|\|\|\|\|\|\|\|\| \|\|\|\|\|\|\|\|\|\|\|\|\|\|\|\|\|\|\|\|\|\|\|\|\|\|\|\|\|\|\|\|\|\|\|\|\|\|\|\|\|\|\|\|\|\|\|\|\|\|\|\|\|\|\|\|\|\|\|\|\|\|\|\|\|\|\|\|\|\|\|\|\|\|\|\|\|\|\|\|\|\|\|\|\|\|\|\|\|\|\|\|\|\|\|\|\|\|\|\|\|\|\|\|\|\|\|\|\|\|\|\|\|\|\|\|\|\|\|\|\|\|\|\|\|\|\|\|\|\|\|\|\|\|\|\|\|\|\|\|\|\|\|\|\|\|\|\|\|\|\|\|\|\|\|\|\|\|\|\|\|\|\|\|\|\|\|\|\|\|\|\|\|\|\|\|\|\|\|\|\|\|\|\|\|\|\|\|\|\|\|\|\|\|\|\|\|\|\|\|\|\|\|\|\|\|\|\|\|\|\|\|\|\|\|\|\|\|\|\|\|\|\|\|\|\|\|\|\|\|\|\|\|\|\|\|\|\|\|\|\|\|\|\|\|\|\|\|\|\|\|\|\|\|\|\|\|\|\|\|\|\|\|\|\|\|\|\|\|\|\|\|\|\|\|\|\|\|\|\|\|\|\|\|\|\|\|\|\|\|\|\|\|\|\|\|\|\|\|\|\|\|\|\|\|\|\|\|\|\|\|\|\|\|\|\|\|\|\|\|\|\|\|\|\|\|\|\|\|\|\|\|\|\|\|\|\|\|\|\|\|\|\|\|\|\|\|\|\|\|\|\|\|\|\|\|\|\|\|\|\|\|\|\|\|\|\|\|\|\|\|\|\|\|\|\|\|\|\|\|\|\|\|\|\|\|\|\|\|\|\|\|\|\|\|\|\|\|\|\|\|\|\|\|\|\|\|\| 	ATGCAGTCCA-GNCATT-NTGCGATCGCAATGCCTGGCACTTTCACGGACGGCGACCACGAGTNTGCGAACGGCTTCACGAGACCATGTGACCAATGCTAGTCGGCAGCTGAGGTCCAGTCCATCCACATTTCCATCGCAAAGAGCATACGCTTCGAGTACGCGGAGTGGATTCCCTCTTATACCGGCCCGATGCTCGCATCGTTCATACACCTCCTCGCCTCATCGGACATTCCACACGACCTCTGTGCGACAGCACGGCGACCTCACGCCACCTAAGCCGGGCGAGGAAAGAAAAGTGACATTTATCGACAAGGATGGGCAAGAATCGACATTCGAGGTCGCCGATGGCGACAACTTGTTGGACATCGCACAAGCAAATGATTTGGAAATGGAGGGTGCATGCGGCGGGAGCTGTGCGTGCAGTACGTGTCATGTCATTGTGAAGGACGAGGACATATATGACAAGATGG ``` |
| 3 | ggggacaagtttgtacaaaaaagcaggcttgatgTCTGAAAGAAAGGTCCTCTC, ggggaccactttgtacaagaaagctgggtcCACTAGGGACGGCTTCTTCT | Mycgr3\|74966  JGI  Ensembl | KOG2989 Uncharacterized conserved protein; |  | 100 % |  | ``` 738	CTTTCAAGCGCCAGCCCAAGAAGAAGAAGGATTTCACCTCCGCGCTCGGAATCAAGAAGAAGCCGTCCCTAGTG	811 	\|\|\|\|\|\|\|\|\|\|\|\|\|\|\|\|\|\|\|\|\|\|\|\|\|\|\|\|\|\|\|\|\|\|\|\|\|\|\|\|\|\|\|\|\|\|\|\|\|\|\|\|\|\|\|\|\|\|\|\|\|\|\|\|\|\|\|\|\|\|\|\|\|\| 	CTTTCAAGCGCCAGCCCAAGAAGAAGAAGGATTTCACCTCCGCGCTCGGAATCAAGAAGAAGCCGTCCCTAGTG ```      ``` 97	GAAGCGAGGCCCTAAGC-AGGTCGGACCAAAAGTGCAGACAGTCCGCCTGATGGCACCTTTCAGCATGCGATGCACATCCTGCGGCGAATATGTATACAAAGGACGCAAGTTCAATGCGCGCAAGGAGACGACCGACGAGAAATACTACGCCATCACGATTTTCCGCTTCTACATTCGGTGTACGAGATGTTCGGCCGAGATCACCTTTAAGACGGACCCGAAGAACATGGACTATACGTGTGAGAAAGGTGCGAAGCGAAATTTTGAGCCATGGCGAGAAGCGAAATTGGCggaggagacggaggaggagcgactggacaggatcgcaaaggaggaagaggagcgagATGCGATGAAGGATCTGGAAAAGAAAACGCTGGATGCGAAGACGGAGATGGCCATTGCGGATGCGCTGGACGATGTGAGACACAGAAACGCCGCCCGAGAAAGGATAGGCAAGGATGGTGAAATCAGCACAGTACCCGTGGCGGACAAGCGGGATGAGGAAGCTGAACAGATCGAGAGGGAGATTGAGGAACAGGCTCGACTGGCATTTCAAAGTGCTACTGGAGACAAGGTGCGAAGACTGGGAGAACAATTAGGAGACGATTTGCTGCTGGACGAGCCAATCATCGAGACAA	737 	\|\|\|\|\|\|\|\|\|\|\| \|\|\|\|\| \|\| \| \|\|\|\|\|\|\|\|\|\|\|\|\|\|\|\|\|\| \|\|\| \| \|\|\|\| \|\| \|\|\|\|\|\|\|\|\|\|\|\| \|\|\|\|\|\|\|\|\|\|\|\|\|\|\|\| \|\|\|\|\|\|\|\|\|\|\|\|\|\|\|\|\|\|\|\|\|\|\|\|\|\|\|\|\|\|\|\|\|\|\|\|\|\|\|\|\|\|\|\|\|\|\|\|\|\|\|\|\|\|\|\|\|\|\| \|\|\|\|\|\|\|\|\|\|\|\|\|\|\|\|\|\|\|\|\|\|\| \|\|\|\|\|\|\|\|\|\|\|\|\|\|\|\|\|\|\|\|\|\|\|\|\|\|\|\|\|\|\|\|\|\|\|\|\|\|\|\|\|\|\|\|\|\|\|\|\|\|\|\|\|\|\|\|\|\|\|\|\|\|\|\|\|\|\|\|\|\|\|\|\|\|\|\|\|\|\|\|\|\|\|\|\|\|\|\|\|\|\|\|\|\|\|\|\|\|\|\|\|\|\|\|\|\|\|\|\|\|\|\|\|\|\|\|\|\|\|\|\|\|\|\|\|\|\|\|\|\|\|\|\|\|\|\|\|\|\|\|\|\|\|\|\|\|\|\|\|\|\|\|\|\|\|\|\|\|\|\|\|\|\|\|\|\|\|\|\|\|\|\|\|\|\|\|\|\|\|\|\|\|\|\|\|\|\|\|\|\|\|\|\|\|\|\|\|\|\|\|\|\|\|\|\|\|\|\|\|\|\|\|\|\|\|\|\|\|\|\|\|\|\|\|\|\|\|\|\|\|\|\|\|\|\|\|\|\|\|\|\|\|\|\|\|\|\|\|\|\|\|\|\|\|\|\|\|\|\|\|\|\|\|\|\|\|\|\|\|\|\|\|\|\|\|\|\|\|\|\|\|\|\|\|\|\|\|\|\|\|\|\|\|\|\|\|\|\|\|\|\|\|\|\|\|\|\|\|\|\|\|\|\|\|\|\|\|\|\|\|\|\|\|\|\|\|\|\|\|\|\|\|\|\|\|\|\|\|\|\|\|\|\|\|\|\|\|\|\|\|\|\|\|\|\|\|\|\|\|\|\|\|\|\|\|\|\|\|\|\|\|\|\|\|\|\|\|\|\|\|\|\|\|\|\|\|\|\|\|\|\|\|\|\|\|\|\|\|\|\|\|\|\|\|\|\|\|\|\|\|\|\|\|\|\|\|\|\|\|\|\|\|\|\|\|\|\|\|\|\|\|\|\|\|\|\|\|\|\|\|\|\|\|\|\|\|\|\|\|\|\|\|\|\|\|\|\|\|\|\|\|\|\|\|\|\|\|\|\|\|\|\| 	GAAGCGAGGCCNTAAGCCAGNT-GGACCAAAAGTGCAGACANTCCNCNTGATNGCNCCTTTCAGCATGNGATGCACATCCTGCGGNGAATATGTATACAAAGGACGCAAGTTCAATGCGCGCAAGGAGACGACCGACGAGAAATANTACGCCATCACGATTTTCCGCTTNTACATTCGGTGTACGAGATGTTCGGCCGAGATCACCTTTAAGACGGACCCGAAGAACATGGACTATACGTGTGAGAAAGGTGCGAAGCGAAATTTTGAGCCATGGCGAGAAGCGAAATTGGCGGAGGAGACGGAGGAGGAGCGACTGGACAGGATCGCAAAGGAGGAAGAGGAGCGAGATGCGATGAAGGATCTGGAAAAGAAAACGCTGGATGCGAAGACGGAGATGGCCATTGCGGATGCGCTGGACGATGTGAGACACAGAAACGCCGCCCGAGAAAGGATAGGCAAGGATGGTGAAATCAGCACAGTACCCGTGGCGGACAAGCGGGATGAGGAAGCTGAACAGATCGAGAGGGAGATTGAGGAACAGGCTCGACTGGCATTTCAAAGTGCTACTGGAGACAAGGTGCGAAGACTGGGAGAACAATTAGGAGACGATTTGCTGCTGGACGAGCCAATCATCGAGACAA ``` |
| 3 | ggggacaagtttgtacaaaaaagcaggcttgatgTCGCTCCAGGATATCAAGCA, ggggaccactttgtacaagaaagctgggtcTCGTTCGATAACAGTCGTGA | Mycgr3\|75204  JGI  Ensembl | KOG3022 Predicted ATPase, nucleotide-binding; | ATP binding; | 0 % |  |  |
| 3 | ggggacaagtttgtacaaaaaagcaggcttgatgTCCGCTCCCGACCTCACAGA, ggggaccactttgtacaagaaagctgggtcCTCAAGAGCAGTGTACATGT | Mycgr3\|75239  JGI  Ensembl |  |  | 100 % |  | ``` 110	ATGTCCGCTCCCGACCTCACAGAATCACAAACCACATTTCCCTCCGACAGCGAGATCTACGATGCCAACCACTCAAATTCCAACGACTCGGATGAACCGGCGTCCTCAAGCTCAAGTCCGATGATCCTCTATTCACCGCCGACGTTCTGGAGTCTGATGAGGGGTGCCGCCATCAATCTTCTACTACCATTTGTCAACGGATTGATGCTGGGATTTGGGGAGTTAATTGCACATGAGGCGGCGTACAGGTTAGGTTGGGGCAAGACGAAGGT	381 	\|\|\|\|\|\|\|\|\|\|\|\|\|\|\|\|\|\|\|\|\|\|\|\|\|\|\|\|\|\|\|\|\|\|\|\|\|\|\|\|\|\|\|\|\|\|\|\|\|\|\|\|\|\|\|\|\|\|\|\|\|\|\|\|\|\|\|\|\|\|\|\|\|\|\|\|\|\|\|\|\|\|\|\|\|\|\|\|\|\|\|\|\|\|\|\|\|\|\|\|\|\|\|\|\|\|\|\|\|\|\|\|\|\|\|\|\|\|\|\|\|\|\|\|\|\|\|\|\|\|\|\|\|\|\|\|\|\|\|\|\|\|\|\|\|\|\|\|\|\|\|\|\|\|\|\|\|\|\|\|\|\|\|\|\|\|\|\|\|\|\|\|\|\|\|\|\|\|\|\|\|\|\|\|\|\|\|\|\|\|\|\|\|\|\|\|\|\|\|\|\|\|\|\|\|\|\|\|\|\|\|\|\|\|\|\|\|\|\|\|\|\|\|\|\|\|\|\|\|\|\|\|\|\|\|\|\|\|\|\|\|\|\|\|\|\|\|\|\|\|\|\|\|\|\|\|\|\|\|\|\|\|\|\|\|\|\|\|\|\|\|\| 	ATGTCCGCTCCCGACCTCACAGAATCACAAACCACATTTCCCTCCGACAGCGAGATCTACGATGCCAACCACTCAAATTCCAACGACTCGGATGAACCGGCGTCCTCAAGCTCAAGTCCGATGATCCTCTATTCACCGCCGACGTTCTGGAGTCTGATGAGGGGTGCCGCCATCAATCTTCTACTACCATTTGTCAACGGATTGATGCTGGGATTTGGGGAGTTAATTGCACATGAGGCGGCGTACAGGTTAGGTTGGGGCAAGACGAAGGT ```      ``` 378	AGGTCTTCCCCTCGCACCGAACGCACTCATTAGGCCCCGGGATCGAAGTGAAGGATGATCCAGTCGAGCGAACACGACGAGACGGACGGGAGTTGGACATGTACACTGCTCTTGAG	493 	\|\|\|\|\|\|\|\|\|\|\|\|\|\|\|\|\|\|\|\|\|\|\|\|\|\|\|\|\|\|\|\|\|\|\|\|\|\|\|\|\|\|\|\|\|\|\|\|\|\|\|\|\|\|\|\|\|\|\|\|\|\|\|\|\|\|\|\|\|\|\|\|\|\|\|\|\|\|\|\|\|\|\|\|\|\|\|\|\|\|\|\|\|\|\|\|\|\|\|\|\|\|\|\|\|\|\|\|\|\|\|\|\|\|\|\| 	AGGTCTTCCCCTCGCACCGAACGCACTCATTAGGCCCCGGGATCGAAGTGAAGGATGATCCAGTCGAGCGAACACGACGAGACGGACGGGAGTTGGACATGTACACTGCTCTTGAG ``` |
| 3 | ggggacaagtttgtacaaaaaagcaggcttgatgCACTTCGCCTCTCTCTTCCT, ggggaccactttgtacaagaaagctgggtcACCCAACAATCCTCCTGACG | Mycgr3\|77055  JGI  Ensembl |  |  | 3.79965 % |  |  |
| 3 | ggggacaagtttgtacaaaaaagcaggcttgatgGGTCGTCTTCACTCCAACGG, ggggaccactttgtacaagaaagctgggtcTGCGACCATGGTGCTGGCGG | Mycgr3\|80995  JGI  Ensembl | KOG0400 40S ribosomal protein S13; | intracellular; ribosome; structural constituent of ribosome;  translation; | 100 % |  | ``` 339	AGGCCGTCGCCGTCCGCAAGCATCTCGAGCGCAACCGCAAGGACAAGGACAGCAAGTTCCGCCTCATTCTCATCGAGTCCCGCATCCACCGCCTCTCCCGCTACTACAAGACCGTCGGTGTTCTCCCACCCACCTGGAGATACGAGTCTGCCACCGCCAGCACC-ATGGTCGCA	511 	\|\|\|\|\|\|\|\|\|\|\|\|\|\|\|\|\|\|\|\|\|\|\|\|\|\|\|\|\|\|\|\|\|\|\|\|\|\|\|\|\|\|\|\|\|\|\|\|\|\|\|\|\|\|\|\|\|\|\|\|\|\|\|\|\|\|\|\|\|\|\|\|\|\|\|\|\|\|\|\|\|\|\|\|\|\|\|\|\|\|\|\|\|\|\|\|\|\|\|\|\|\|\|\|\|\|\|\|\|\|\|\|\|\|\|\|\|\|\|\|\|\|\|\|\|\|\|\|\|\|\|\|\|\|\|\|\|\|\|\|\|\|\|\|\|\|\|\|\|\|\|\|\|\|\|\|\|\|\|\|\|\|\|\| \|\|\|\|\|\|\|\|\| 	AGGCCGTCGCCGTCCGCAAGCATCTCGAGCGCAACCGCAAGGACAAGGACAGCAAGTTCCGCCTCATTCTCATCGAGTCCCGCATCCACCGCCTCTCCCGCTACTACAAGACCGTCGGTGTTCTCCCACCCACCTGGAGATACGAGTCTGCCACCGCCAGCACCCATGGTCGCA ```      ``` 339	AGGCCGTCGCCGTCCGCAAGCATCTCGAGCGCAACCGCAAGGACAAGGACAGCAAGTTCCGCCTCATTCTCATCGAGTCCCGCATCCACCGCCTCTCCCGCTACTACAAGACCGTCGGTGTTCTCCCACCCACCTGGAGATACGAGTCTGCCACCGCCAGCACCATGGTCGCA	511 	\|\|\|\|\|\|\|\|\|\|\|\|\|\|\|\|\|\|\|\|\|\|\|\|\|\|\|\|\|\|\|\|\|\|\|\|\|\|\|\|\|\|\|\|\|\|\|\|\|\|\|\|\|\|\|\|\|\|\|\|\|\|\|\|\|\|\|\|\|\|\|\|\|\|\|\|\|\|\|\|\|\|\|\|\|\|\|\|\|\|\|\|\|\|\|\|\|\|\|\|\|\|\|\|\|\|\|\|\|\|\|\|\|\|\|\|\|\|\|\|\|\|\|\|\|\|\|\|\|\|\|\|\|\|\|\|\|\|\|\|\|\|\|\|\|\|\|\|\|\|\|\|\|\|\|\|\|\|\|\|\|\|\|\|\|\|\|\|\|\|\|\|\| 	AGGCCGTCGCCGTCCGCAAGCATCTCGAGCGCAACCGCAAGGACAAGGACAGCAAGTTCCGCCTCATTCTCATCGAGTCCCGCATCCACCGCCTCTCCCGCTACTACAAGACCGTCGGTGTTCTCCCACCCACCTGGAGATACGAGTCTGCCACCGCCAGCACCATGGTCGCA ```      ``` 260	GGTAACAAGATTCTCCGCATCCTCAAGGGCAACGG	294 	\|\|\|\|\|\|\|\|\|\|\|\|\|\|\|\|\|\|\|\|\|\|\|\|\|\|\|\|\|\|\|\|\|\|\| 	GGTAACAAGATTCTCCGCATCCTCAAGGGCAACGG ```      ``` 59	ATGGGTCGTCTTCACTCCAACGGAAAGGGAATCTCTTCCTCCGCCATCCCATACTCGCGCACCGCCCCGGCATGGCTCAAGACCACCCCGGAGCAGGTCGTTGATCAGATTTGCAAGCTCGCAAAGAAGGGTGCCACTCCTTCGCAGATTGGTGTCGTCCTCCGTGACAGCCACGGTGTTGCCCAGGTCAAGATTGTGACTGGT	262 	\|\|\|\|\|\|\|\|\|\|\|\|\|\|\|\|\|\|\|\|\|\|\|\|\|\|\|\|\|\|\|\|\|\|\|\|\|\|\|\|\|\|\|\|\|\|\|\|\|\|\|\|\|\|\|\|\|\|\|\|\|\|\|\|\|\|\|\|\|\|\|\|\|\|\|\|\|\|\|\|\|\|\|\|\|\|\|\|\|\|\|\|\|\|\|\|\|\|\|\|\|\|\|\|\|\|\|\|\|\|\|\|\|\|\|\|\|\|\|\|\|\|\|\|\|\|\|\|\|\|\|\|\|\|\|\|\|\|\|\|\|\|\|\|\|\|\|\|\|\|\|\|\|\|\|\|\|\|\|\|\|\|\|\|\|\|\|\|\|\|\|\|\|\|\|\|\|\|\|\|\|\|\|\|\|\|\|\|\|\|\|\|\|\|\|\|\|\|\|\|\|\|\|\| 	ATGGGTCGTCTTCACTCCAACGGAAAGGGAATCTCTTCCTCCGCCATCCCATACTCGCGCACCGCCCCGGCATGGCTCAAGACCACCCCGGAGCAGGTCGTTGATCAGATTTGCAAGCTCGCAAAGAAGGGTGCCACTCCTTCGCAGATTGGTGTCGTCCTCCGTGACAGCCACGGTGTTGCCCAGGTCAAGATTGTGACTGGT ```      ``` 290	AAC-GGCCTCGCACCAGAGATTCCAGAGGACCTCTACATGCTCATCAAGAAGG	341 	\|\|\| \|\|\|\|\|\|\|\|\|\|\|\|\|\|\|\|\|\|\|\|\|\|\|\|\|\|\|\|\|\|\|\|\|\|\|\|\|\|\|\|\|\|\|\|\|\|\|\|\| 	AACAGGCCTCGCACCAGAGATTCCAGAGGACCTCTACATGCTCATCAAGAAGG ```      ``` 339	AGGCCGTCGCCGTCCGCAAGCATCTCGAGCGCAACCGCAAGGAC-AAGGACAGCAAGTTCCGCCTCATTCTCATCGAGTCCCGCATCCACCGCCTCTCCCGCTACTACAAGACCGTCGGTGTTCTCCCACCCACCTGGAGATACGAGTCTGCCACCGCCAGCACCATGGTCGCA	511 	\|\|\|\|\|\|\|\|\|\|\|\|\|\|\|\|\|\|\|\|\|\|\|\|\|\|\|\|\|\|\|\|\|\|\|\|\|\|\|\|\|\|\|\| \|\|\|\|\|\|\|\|\|\|\|\|\|\|\|\|\|\|\|\|\|\|\|\|\|\|\|\|\|\|\|\|\|\|\|\|\|\|\|\|\|\|\|\|\|\|\|\|\|\|\|\|\|\|\|\|\|\|\|\|\|\|\|\|\|\|\|\|\|\|\|\|\|\|\|\|\|\|\|\|\|\|\|\|\|\|\|\|\|\|\|\|\|\|\|\|\|\|\|\|\|\|\|\|\|\|\|\|\|\|\|\|\|\|\|\|\|\|\|\|\|\|\|\|\|\|\|\|\| 	AGGCCGTCGCCGTCCGCAAGCATCTCGAGCGCAACCGCAAGGACAAAGGACAGCAAGTTCCGCCTCATTCTCATCGAGTCCCGCATCCACCGCCTCTCCCGCTACTACAAGACCGTCGGTGTTCTCCCACCCACCTGGAGATACGAGTCTGCCACCGCCAGCACCATGGTCGCA ``` |
| 3 | ggggacaagtttgtacaaaaaagcaggcttgatgAGTGCCCTCTACAATTCCGC, ggggaccactttgtacaagaaagctgggtcTCGTAACCACTTCAACACAA | Mycgr3\|82382  JGI  Ensembl | KOG3251 Golgi SNAP receptor complex member; | intracellular protein transport; membrane; | 100 % |  | ``` 79	CCTGCGGCAATCAAAATCCATCCGTGCCGACCTCGACATCCTCAGTTCCCAACCCTCCTCCTCCGCCGCTCTCCAAGGCCAACTCACCGCAGCCATAACCTCCTTCTCCCGCACTCTAGACGACTACCAACGCAGCATAAACGACGAACTCGTCCCTGACAAAGCCACCAAAGGCCAAGAACGACTGACGAATTTCCGCTCCGAACTCCTCGACTTTCGCTCTCGATTCGCAGAACTCAAACACACCCAGCAGGAACAAGCATATGCGCAGAATCGATCGGAACTACTCGGGCGGAGACCGCACGCTGCGAATACACCGGAAAACCCATATGCTGATGCTGCGATTGCGAGTGGCGCGGGACACGGCGAAAATGTGAATCCGCTGTTTCGCACGAGTAATCCGAATTTCGTGCCGGGACAGCAGGGGAATAGCTACTCGGCATACTCGTCGCCGTATGGACTGGGAGCGGATGAAGCGAGGGAGAGCCATGCGTTGAGAGAGAACTCGTTCTTTGGAAGCGCAAATCAGACGCTGGATGAATACTTGGAGCGAGGGAGGGCGGTGTTGGGAGACTTGGGAGATCAGAGGGAGATGTTGAAGGGAACACAGAAGAAGTTGTATTCCATTGGGAGCACGCTGGGAATTTCGGGAGACACGATAAGGATGGTGGAGAGGAGAGCGAAGCAGGACAAGTCGATATTCTGGGGTGGTGTGGTGGTCTTCATCATTTTCGTGTACTTTGTGTTGAAGTGGTTACGA	838 	\|\|\|\|\|\|\|  \|\|\| \|\|\|\|\|\|\|\|\|\|\|\|\|\|\|\|\|\|\|\| \|\|\|\|\| \|\|\|\|\|\|\| \|\|\|\|\|\|\|\| \|\| \|\| \|\|\|\|\| \| \|\|\|\|\|\|\|\|\|\|\|\|\|\|\|\|\|\|\|\|\|\|\|\|\|\|\|\|\|\|\|\|\|\|\|\|\|\|\|\|\|\|\|\|\|\|\|\|\|\|\|\|\|\|\|\|\|\|\|\|\|\|\|\|\|\|\|\|\|\|\|\|\|\|\|\|\|\|\|\|\|\|\|\|\|\|\|\|\|\|\|\|\|\|\|\|\|\|\|\|\|\|\|\|\|\|\|\|\|\|\|\|\|\|\|\|\|\|\|\|\|\|\|\|\|\|\|\|\|\|\|\|\|\|\|\|\|\|\|\|\|\|\|\|\|\|\|\|\|\|\|\|\|\|\|\|\|\|\|\|\|\|\|\|\|\|\|\|\|\|\|\|\|\|\|\|\|\|\|\|\|\|\|\|\|\|\|\|\|\|\|\|\|\|\|\|\|\|\|\|\|\|\|\|\|\|\|\|\|\|\|\|\|\|\|\|\|\|\|\|\|\|\|\|\|\|\|\|\|\|\|\|\|\|\|\|\|\|\|\|\|\|\|\|\|\|\|\|\|\|\|\|\|\|\|\|\|\|\|\|\|\|\|\|\|\|\|\|\|\|\|\|\|\|\|\|\|\|\|\|\|\|\|\|\|\|\|\|\|\|\|\|\|\|\|\|\|\|\|\|\|\|\|\|\|\|\|\|\|\|\|\|\|\|\|\|\|\|\|\|\|\|\|\|\|\|\|\|\|\|\|\|\|\|\|\|\|\|\|\|\|\|\|\|\|\|\|\|\|\|\|\|\|\|\|\|\|\|\|\|\|\|\|\|\|\|\|\|\|\|\|\|\|\|\|\|\|\|\|\|\|\|\|\|\|\|\|\|\|\|\|\|\|\|\|\|\|\|\|\|\|\|\|\|\|\|\|\|\|\|\|\|\|\|\|\|\|\|\|\|\|\|\|\|\|\|\|\|\|\|\|\|\|\|\|\|\|\|\|\|\|\|\|\|\|\|\|\|\|\|\|\|\|\|\|\|\|\|\|\|\|\|\|\|\|\|\|\|\|\|\|\|\|\|\|\|\|\|\|\|\|\|\|\|\|\|\|\|\|\|\|\|\|\|\|\|\|\|\|\|\|\|\|\|\|\|\|\|\|\|\|\|\|\|\|\|\|\|\|\|\|\|\|\|\|\|\|\|\|\|\|\|\|\|\|\|\|\|\|\|\|\|\|\|\|\|\|\|\|\|\|\|\|\|\|\|\|\|\|\|\|\|\|\|\|\|\|\|\|\|\|\|\|\|\|\|\|\|\|\|\|\|\|\|\|\|\|\|\|\|\|\|\|\|\|\|\|\|\|\|\|\|\|\|\|\|\|\|\|\|\|\|\|\|\|\|\|\|\|\|\|\|\|\|\|\|\|\|\|\|\|\|\|\|\|\|\|\|\|\|\|\|\|\|\|\|\|\|\|\|\|\|\|\|\|\|\|\|\|\|\|\|\|\|\|\|\|\|\|\|\|\|\|\|\|\|\|\|\|\|\|\|\|\|\|\|\|\|\| 	CCTGCGGNCATC-AAATCCATCCGTGCCGACCTNGACATNCTCAGTTNCCAACCCTTCTTCTNCGCCGNTTTCCAAGGCCAACTCACCGCAGCCATAACCTCCTTCTCCCGCACTCTAGACGACTACCAACGCAGCATAAACGACGAACTCGTCCCTGACAAAGCCACCAAAGGCCAAGAACGACTGACGAATTTCCGCTCCGAACTCCTCGACTTTCGCTCTCGATTCGCAGAACTCAAACACACCCAGCAGGAACAAGCATATGCGCAGAATCGATCGGAACTACTCGGGCGGAGACCGCACGCTGCGAATACACCGGAAAACCCATATGCTGATGCTGCGATTGCGAGTGGCGCGGGACACGGCGAAAATGTGAATCCGCTGTTTCGCACGAGTAATCCGAATTTCGTGCCGGGACAGCAGGGGAATAGCTACTCGGCATACTCGTCGCCGTATGGACTGGGAGCGGATGAAGCGAGGGAGAGCCATGCGTTGAGAGAGAACTCGTTCTTTGGAAGCGCAAATCAGACGCTGGATGAATACTTGGAGCGAGGGAGGGCGGTGTTGGGAGACTTGGGAGATCAGAGGGAGATGTTGAAGGGAACACAGAAGAAGTTGTATTCCATTGGGAGCACGCTGGGAATTTCGGGAGACACGATAAGGATGGTGGAGAGGAGAGCGAAGCAGGACAAGTCGATATTCTGGGGTGGTGTGGTGGTCTTCATCATTTTCGTGTACTTTGTGTTGAAGTGGTTACGA ``` |
| 3 | ggggacaagtttgtacaaaaaagcaggcttgatgGCGGATCGCAAAGGCGCATC, ggggaccactttgtacaagaaagctgggtcACAGATGACTCCATCGTTCT | Mycgr3\|83502  JGI  Ensembl | KOG4727 U1-like Zn-finger protein; | intracellular; nucleic acid binding;  zinc ion binding; | 100 % |  | ``` 1	ATGGCGGATCGCAAAGGCGCATCGTCGAGTAATGACACATCTTTCCGCAAGACTTGGAATCGCGATGAATACGCCGCAAAAGCAAGCGAACGCGATCAAAAGATCAAGGAAGAGGGCAAAGCTCGATACGAAGCTGCGCTTGAAGGGAAGAAATACCATCGCCGAGCCTCGACACCACCAGACACGCGCGATACAGAAGCACGCAAGGCAAGGTTGAATGTCGCAGATCAGGTGGGGAAGACACAACTTGTAGTAGCGGGCGGACAAGGAAAGCGAGGCAAGAGTGCCGGATTCTACTGCGATGCTTGCGACCTCACGTTCAAGGACAACCTGCAATTTGTGGAGCATCTGAACAGCAAGCAACATTTGATCGCTACGGGAGAGACTGGCGAGGTGCGCAGAGCCACACTAGAGGATGTCAAGGAGCGATTCGAGTATCTCAAGCGGAAGCGAGACGAGGAGAAAGCTAGGGACGCAGTTGATCTGACCACAAGAATTGAAGTCGCGAAAGAGGAAATGGACAAGGAGCGTGAAGAGAAGAGGCGGAAGCGCAATGAGAAACGCAGGAAGACTAAGGATGGGATGGGCGTGCCGGAGGTCAAGCTCGAGAACGATGGAGTCATCTGT	627 	\|\|\|\|\|\|\|\|\|\|\|\|\|\|\|\|\|\|\|\|\|\|\|\|\|\|\|\|\|\|\|\|\|\|\|\|\|\|\|\|\|\|\|\|\|\|\|\|\|\|\|\|\|\|\|\|\|\|\|\|\|\|\|\|\|\|\|\|\|\|\|\|\|\|\|\|\|\|\|\|\|\|\|\|\|\|\|\|\|\|\|\|\|\|\|\|\|\|\|\|\|\|\|\|\|\|\|\|\|\|\|\|\|\|\|\|\|\|\|\|\|\|\|\|\|\|\|\|\|\|\|\|\|\|\|\|\|\|\|\|\|\|\|\|\|\|\|\|\|\|\|\|\|\|\|\|\|\|\|\|\|\|\|\|\|\|\|\|\|\|\|\|\|\|\|\|\|\|\|\|\|\|\|\|\|\|\|\|\|\|\|\|\|\|\|\|\|\|\|\|\|\|\|\|\|\|\|\|\|\|\|\|\|\|\|\|\|\|\|\|\|\|\|\|\|\|\|\|\|\|\|\|\|\|\|\|\|\|\|\|\|\|\|\|\|\|\|\|\|\|\|\|\|\|\|\|\|\|\|\|\|\|\|\|\|\|\|\|\|\|\|\|\|\|\|\|\|\|\|\|\|\|\|\|\|\|\|\|\|\|\|\|\|\|\|\|\|\|\|\|\|\|\|\|\|\|\|\|\|\|\|\|\|\|\|\|\|\|\|\|\|\|\|\|\|\|\|\|\|\|\|\|\|\|\|\|\|\|\|\|\|\|\|\|\|\|\|\|\|\|\|\|\|\|\|\|\|\|\|\|\|\|\|\|\|\|\|\|\|\|\|\|\|\|\|\|\|\|\|\|\|\|\|\|\|\|\|\|\|\|\|\|\|\|\|\|\|\|\|\|\|\|\|\|\|\|\|\|\|\|\|\|\|\|\|\|\|\|\|\|\|\|\|\|\|\|\|\|\|\|\|\|\|\|\|\|\|\|\|\|\|\|\|\|\|\|\|\|\|\|\|\|\|\|\|\|\|\|\|\|\|\|\|\|\|\|\|\|\|\|\|\|\|\|\|\|\|\|\|\|\|\|\|\|\|\|\|\|\|\|\|\|\|\|\|\|\|\|\|\|\|\|\|\|\|\|\|\|\|\|\|\|\|\|\|\|\|\|\|\|\|\|\|\|\|\|\|\|\|\|\|\|\|\|\|\|\|\|\|\|\|\|\|\|\|\|\|\|\|\|\|\|\|\|\|\|\|\|\|\|\|\|\|\|\|\|\|\|\|\|\|\|\|\|\|\|\|\|\|\|\|\|\|\|\|\|\|\|\|\|\|\|\|\|\|\| \|\|\|\|\|\|\|\|\|\|\|\|\|\|\|\|\|\|\|\|\|\|\|\|\|\|\|\|\|\| 	ATGGCGGATCGCAAAGGCGCATCGTCGAGTAATGACACATCTTTCCGCAAGACTTGGAATCGCGATGAATACGCCGCAAAAGCAAGCGAACGCGATCAAAAGATCAAGGAAGAGGGCAAAGCTCGATACGAAGCTGCGCTTGAAGGGAAGAAATACCATCGCCGAGCCTCGACACCACCAGACACGCGCGATACAGAAGCACGCAAGGCAAGGTTGAATGTCGCAGATCAGGTGGGGAAGACACAACTTGTAGTAGCGGGCGGACAAGGAAAGCGAGGCAAGAGTGCCGGATTCTACTGCGATGCTTGCGACCTCACGTTCAAGGACAACCTGCAATTTGTGGAGCATCTGAACAGCAAGCAACATTTGATCGCTACGGGAGAGACTGGCGAGGTGCGCAGAGCCACACTAGAGGATGTCAAGGAGCGATTCGAGTATCTCAAGCGGAAGCGAGACGAGGAGAAAGCTAGGGACGCAGTTGATCTGACCACAAGAATTGAAGTCGCGAAAGAGGAAATGGACAAGGAGCGTGAAGAGAAGAGGCGGAAGCGCAATGAGAAACGCAGGAAGACTAAGGATGGGATGGGCGTGCCGGANGTCAAGCTCGAGAACGATGGAGTCATCTGT ```      ``` 1	ATGGCGGATCGCAAAGGCGCATCGTCGAGTAATGACACATCTTTCCGCAAGACTTGGAATCGCGATGAATACGCCGCAAAAGCAAGCGAACGCGATCAAAAGATCAAGGAAGAGGGCAAAGCTCGATACGAAGCTGCGCTTGAAGGGAAGAAATACCATCGCCGAGCCTCGACACCACCAGACACGCGCGATACAGAAGCACGCAAGGCAAGGTTGAATGTCGCAGATCAGGTGGGGAAGACACAACTTGTAGTAGCGGGCGGACAAGGAAAGCGAGGCAAGAGTGCCGGATTCTACTGCGATGCTTGCGACCTCACGTTCAAGGACAACCTGCAATTTGTGGAGCATCTGAACAGCAAGCAACATTTGATCGCTACGGGAGAGACTGGCGAGGTGCGCAGAGCCACACTAGAGGATGTCAAGGAGCGATTCGAGTATCTCAAGCGGAAGCGAGACGAGGAGAAAGCTAGGGACGCAGTTGATCTGACCACAAGAATTGAAGTCGCGAAAGAGGAAATGGACAAGGAGCGTGAAGAGAAGAGGCGGAAGCGCAATGAGAAACGCAGGAAGACTAAGGATGGGATGGGCGTGCCGGAGGTCAAGCTCGAGAACGATGGAGTCATCTGT	627 	\|\|\| \|\|\|\|\|\|\|\|\|\|\|\|\|\|\|\| \|\|\|\|\|\|\|\|\|\|\|\|\|\|\|\|\|\|\|\|\|\|\|\|\|\|\|\|\|\|\|\|\|\|\|\|\|\|\|\|\|\|\|\|\|\|\|\|\|\|\|\|\|\|\|\|\|\|\|\|\|\|\|\|\|\|\|\|\|\|\|\|\|\|\|\|\|\|\|\|\|\|\|\|\|\|\|\|\|\|\|\|\|\|\|\|\|\|\|\|\|\|\|\|\|\|\|\|\|\|\|\|\|\|\|\|\|\|\|\|\|\|\|\|\|\|\|\|\|\|\|\|\|\|\|\|\|\|\|\|\|\|\|\|\|\|\|\|\|\|\|\|\|\|\|\|\|\|\|\|\|\|\|\|\|\|\|\|\|\|\|\|\|\|\|\|\|\|\|\|\|\|\|\|\|\|\|\|\|\|\|\|\|\|\|\|\|\|\|\|\|\|\|\|\|\|\|\|\|\|\|\|\|\|\|\|\|\|\|\|\|\|\|\|\|\|\|\|\|\|\|\|\|\|\|\|\|\|\|\|\|\|\|\|\|\|\|\|\|\|\|\|\|\|\|\|\|\|\|\|\|\|\|\|\|\|\|\|\|\|\|\|\|\|\|\|\|\|\|\|\|\|\|\|\|\|\|\|\|\|\|\|\|\|\|\|\|\|\|\|\|\|\|\|\|\|\|\|\|\|\|\|\|\|\|\|\|\|\|\|\|\|\|\|\|\|\|\|\|\|\|\|\|\|\|\|\|\|\|\|\|\|\|\|\|\|\|\|\|\|\|\|\|\|\|\|\|\|\|\|\|\|\|\|\|\|\|\|\|\|\|\|\|\|\|\|\|\|\|\|\|\|\|\|\|\|\|\|\|\|\|\|\|\|\|\|\|\|\|\|\|\|\|\|\|\|\|\|\|\|\|\|\|\|\|\|\|\|\|\|\|\|\|\|\|\|\|\|\|\|\|\|\|\|\|\|\|\|\|\|\|\|\|\|\|\|\|\|\|\|\|\|\|\|\|\|\|\|\|\|\|\|\|\|\|\|\|\|\|\|\|\|\|\|\|\|\|\|\|\|\|\|\|\|\|\|\|\|\|\|\|\|\|\|\|\|\|\|\|\|\|\|\|\|\|\|\|\|\|\|\|\|\|\|\|\|\|\|\|\|\|\|\|\|\|\|\|\|\|\|\|\|\|\|\|\|\|\|\|\|\|\|\|\|\|\|\|\|\|\|\|\|\|\|\|\|\|\|\|\|\|\|\|\|\|\|\|\|\|\|\|\|\|\|\|\|\|\|\|\|\|\|\|\|\|\|\|\|\|\|\|\|\|\|\|\|\|\|\|\|\|\|\|\|\|\| 	ATGNCGGATCGCAAAGGCGCNTCGTCGAGTAATGACACATCTTTCCGCAAGACTTGGAATCGCGATGAATACGCCGCAAAAGCAAGCGAACGCGATCAAAAGATCAAGGAAGAGGGCAAAGCTCGATACGAAGCTGCGCTTGAAGGGAAGAAATACCATCGCCGAGCCTCGACACCACCAGACACGCGCGATACAGAAGCACGCAAGGCAAGGTTGAATGTCGCAGATCAGGTGGGGAAGACACAACTTGTAGTAGCGGGCGGACAAGGAAAGCGAGGCAAGAGTGCCGGATTCTACTGCGATGCTTGCGACCTCACGTTCAAGGACAACCTGCAATTTGTGGAGCATCTGAACAGCAAGCAACATTTGATCGCTACGGGAGAGACTGGCGAGGTGCGCAGAGCCACACTAGAGGATGTCAAGGAGCGATTCGAGTATCTCAAGCGGAAGCGAGACGAGGAGAAAGCTAGGGACGCAGTTGATCTGACCACAAGAATTGAAGTCGCGAAAGAGGAAATGGACAAGGAGCGTGAAGAGAAGAGGCGGAAGCGCAATGAGAAACGCAGGAAGACTAAGGATGGGATGGGCGTGCCGGAGGTCAAGCTCGAGAACGATGGAGTCATCTGT ``` |
| 3 | ggggacaagtttgtacaaaaaagcaggcttgatgGCCTCCGTCTACAAATCCCT, ggggaccactttgtacaagaaagctgggtcAGCAAAAAGCACCCTCTCAT | Mycgr3\|87034  JGI  Ensembl | KOG2971 RNA-binding protein required for biogenesis of the ribosomal 60S subunit; |  | 0 % |  |  |
| 3 | ggggacaagtttgtacaaaaaagcaggcttgatgCCCGCCCCTTACAAACTCAT, ggggaccactttgtacaagaaagctgggtcCTCCAACTGCCTCCCTTTCC | Mycgr3\|88000  JGI  Ensembl | KOG0235 Phosphoglycerate mutase; | 2,3-bisphosphoglycerate-dependent phosphoglycerate mutase activity; 2,3-bisphosphoglycerate-independent phosphoglycerate mutase activity;  catalytic activity;  glycolysis;  intramolecular transferase activity, phosphotransferases;  metabolic process;  phosphoglycerate mutase activity; | 100 % |  | ``` 86	TCCCACTCAACGAACAAGGAATCAAAGAAGCCACTCAAGCCGGGAAACTGCTCAAAAAACACAACTACCTCCCCGAAATCCTCTTCACCTCCCTCCTCCGCCGCGCCATCGTCACCGCTGACCTCGCTCTCGACGAAGCTGACAGACTCTGGATCCCCGTCAAGCGATCCTGGAAACTCAACGAACGCCACTACGGCGGTCTGCAGGGTCTCGACAAAGCTGAAGCCGCTGAACGAGAAGGTGCAGAACAAGTCCATATCTGGCGGAGGTCGTTTGATGTTCGGCCTCCTGCTACGAGTGAGGAGTATCGGAAGTCAGAGGATGAAGGTGGGAGGTATGCTTCGCAAGGGATTGAAGTGCCGGCTACGGAGTGCTTGAAAGATGTGCTTGTGAGGGTTCTGCCGTATTGGACGGAGGATATTGTGGGTGAGTTGAAGAAGGGGAGGACGGTGTTGATCGCGGCGCATGGGAATAGTTTGAGGGCGTTGATGAAGGAGCTGGAGGGGATTAGTGATGAGGATATTCCGGGCGTGGAGATTCCGACGGGGGTGCCGATTGTTTATGAGTTGGATGAGGAGTTCAAGGGGAAAGGGAGGCAGTTGGAG	690 	\|\|\|\| \|\|\|\| \| \| \|\|\| \|\| \|\|\|\|  \|\| \|\|\|\|\|\|\|\| \|\|\|\|\|\|\|\|\|\| \|\|\|\|\|\|\|\|\|\|\|\|\|\|\|\|\|\|\|\|\|\|\|\|\|\|\|\|\|\|\|\|\|\|\|\|\|\|\|\|\|\|\|\|\| \|\|\|\|\|\|\|\|\|\|\|\|\|\|\|\|\|\|\|\|\|\|\|\|\|\|\|\|\|\|\|\|\|\|\|\|\|\|\|\|\|\|\|\|\|\|\|\|\|\|\|\|\|\|\|\|\|\|\|\|\|\|\|\|\|\|\|\|\|\|\|\|\|\|\|\|\|\|\|\|\|\|\|\|\|\|\|\|\|\|\|\|\|\|\|\|\|\|\|\|\|\|\|\|\|\|\|\|\|\|\|\|\|\|\|\|\|\|\|\|\|\|\|\|\|\|\|\|\|\|\|\|\|\|\|\|\|\|\|\|\|\|\|\|\|\|\|\|\|\|\|\|\|\|\|\|\|\|\|\|\|\|\|\|\|\|\|\|\|\|\|\|\|\|\|\|\|\|\|\|\|\|\|\|\|\|\|\|\|\|\|\|\|\|\|\|\|\|\|\|\|\|\|\|\|\|\|\|\|\|\|\|\|\|\|\|\|\|\|\|\|\|\|\|\|\|\|\|\|\|\|\|\|\|\|\|\|\|\|\|\|\|\|\|\|\|\|\|\|\|\|\|\|\|\|\|\|\|\|\|\|\|\|\|\|\|\|\|\|\|\|\|\|\|\|\|\|\|\|\|\|\|\|\|\|\|\|\|\|\|\|\|\|\|\|\|\|\|\|\|\|\|\|\|\|\|\|\|\|\|\|\|\|\|\|\|\|\|\|\|\|\|\|\|\|\|\|\|\|\|\|\|\|\|\|\|\|\|\|\|\|\|\|\|\|\|\|\|\|\|\|\|\|\|\|\|\|\|\|\|\|\|\|\|\|\|\|\|\|\|\|\|\|\|\|\|\|\|\|\|\|\|\|\|\|\|\|\|\|\|\|\|\|\|\|\|\|\|\|\|\|\|\|\|\|\|\|\|\|\|\|\|\|\|\|\|\|\|\|\|\|\|\|\|\|\|\|\|\|\|\|\|\|\|\|\|\|\|\|\|\|\|\|\|\|\|\|\|\|\|\|\|\|\|\|\|\|\|\|\|\|\|\|\|\|\|\|\|\|\|\|\|\|\|\|\|\|\|\|\|\|\|\|\|\|\|\|\|\|\|\|\|\|\|\|\|\|\|\|\|\|\|\|\|\|\|\|\|\| 	TCCCNCTCANCNANCAANGA-TCAANNAANCCACTCAANCCGGGAAACTNCTCAAAAAACACAACTACCTCCCCGAAATCCTCTTCACCTCCCTCNTCCGCCGCGCCATCGTCACCGCTGACCTCGCTCTCGACGAAGCTGACAGACTCTGGATCCCCGTCAAGCGATCCTGGAAACTCAACGAACGCCACTACGGCGGTCTGCAGGGTCTCGACAAAGCTGAAGCCGCTGAACGAGAAGGTGCAGAACAAGTCCATATCTGGCGGAGGTCGTTTGATGTTCGGCCTCCTGCTACGAGTGAGGAGTATCGGAAGTCAGAGGATGAAGGTGGGAGGTATGCTTCGCAAGGGATTGAAGTGCCGGCTACGGAGTGCTTGAAAGATGTGCTTGTGAGGGTTCTGCCGTATTGGACGGAGGATATTGTGGGTGAGTTGAAGAAGGGGAGGACGGTGTTGATCGCGGCGCATGGGAATAGTTTGAGGGCGTTGATGAAGGAGCTGGAGGGGATTAGTGATGAGGATATTCCGGGCGTGGAGATTCCGACGGGGGTGCCGATTGTTTATGAGTTGGATGAGGAGTTCAAGGGGAAAGGGAGGCAGTTGGAG ``` |
| 3 | ggggacaagtttgtacaaaaaagcaggcttgatgAGGCTCGCGCTATTGCTGTT, ggggaccactttgtacaagaaagctgggtcGTCAAGCTGAGCCTCGTCCA | Mycgr3\|88451  JGI  Ensembl |  |  | 0 % |  |  |
| 3 | ggggacaagtttgtacaaaaaagcaggcttgatgGCGGCAGACGAGACAGAGAT, ggggaccactttgtacaagaaagctgggtcACTGATCTTCATCTTCCTCC | Mycgr3\|89342  JGI  Ensembl | KOG3144 Ethanolamine-P-transferase GPI11/PIG-F, involved in glycosylphosphatidylinositol anchor biosynthesis; | GPI anchor biosynthetic process; endoplasmic reticulum membrane;  integral to membrane; | 100 % |  | ``` 223	CCACCACCTCCCAAATCCAAACCAGGCCAAAAGAAACTGCCCAAACCAGTGCCAGATCTCGGGTCAAAGCTAGTG	297 	\|\|\|\|\|\|\| \|\|\|\|\|\|\|\|\|\|\|\|\|\|\|\|\|\|\|\|\|\|\|\|\|\|\|\|\|\|\|\|\|\|\|\|\|\|\|\|\|\|\|\|\|\|\|\|\|\|\|\|\|\|\|\|\|\|\|\|\|\|\|\|\|\|\| 	CCACCACNTCCCAAATCCAAACCAGGCCAAAAGAAACTGCCCAAACCAGTGCCAGATCTCGGGTCAAAGCTAGTG ```      ``` 297	GCCCGCGTTCCTCTCTTTCGTGCTGACGCTCACACTATCCGCTCCCGTCCTCTACATAGCCGCCATCCTGTTCGGCGCGCCGCTCATCTCGCATCACCTCCACACGCTTCTCCTAGCCGTCCACCTCGCCCTCCTCACGACTCCACCACTCTTCTACGTCCACGGTCTTCACGCGCCCACATGGCTAGCGCTCGCGAGTTTGCAACAGCCGGTCGACGAGACATATGGAATGGCGCTGGGCGCGGTCGTAGGAGCTTGGGTGGGAGCTGTGCCAATTCCGCTGGACTGGGATCGCGAGTGGCAGAAGTGGCCGGTTACGATTATTTGTGGGATGTATGGAGGTGCTGTGCTGGGCAAGGTGGCGGGTGGCTGGCTCTTAAGAGGGAGGAAGATGAAGATCAGT	699 	\|\|\|\|\|\|\|\|\|\|\|\|\|\|\|\|\|\|\|\|\|\|\|\|\|\|\|\|\|\|\|\|\|\|\|\|\|\|\|\|\|\|\|\|\|\|\|\|\|\|\|\|\|\|\|\|\|\|\|\|\|\|\|\|\|\|\|\|\|\|\|\|\|\|\|\|\|\|\|\|\|\|\|\|\|\|\|\|\|\|\|\|\|\|\|\|\|\|\|\|\|\|\|\|\|\|\|\|\|\|\|\|\|\|\|\|\|\|\|\|\|\|\|\|\|\|\|\|\|\|\|\|\|\|\|\|\|\|\|\|\|\|\|\|\|\|\|\|\|\|\|\|\|\|\|\|\|\|\|\|\|\|\|\|\|\|\|\|\|\|\|\|\|\|\|\|\|\|\|\|\|\|\|\|\|\|\|\|\|\|\|\|\|\|\|\|\|\|\|\|\|\|\|\|\|\|\|\|\|\|\|\|\|\|\|\|\|\|\|\|\|\|\|\|\|\|\|\|\|\|\|\|\|\|\|\|\|\|\|\|\|\|\|\|\|\|\|\|\|\|\|\|\|\|\|\|\|\|\|\|\|\|\|\|\|\|\|\|\|\|\|\|\|\|\|\|\|\|\|\|\|\|\|\|\|\|\|\|\|\|\|\|\|\|\|\|\|\|\|\|\|\|\|\|\|\|\|\|\|\|\|\|\|\|\|\|\|\|\|\|\|\|\|\|\|\|\|\|\|\|\|\|\|\|\|\|\|\|\|\|\|\|\|\|\|\|\|\|\|\|\|\|\|\|\|\|\|\|\|\|\|\|\|\|\|\|\|\|\|\|\|\|\|\|\|\|\|\|\|\|\|\|\|\|\|\|\|\|\|\|\|\|\|\|\|\|\|\|\|\|\|\|\| 	GCCCGCGTTCCTCTCTTTCGTGCTGACGCTCACACTATCCGCTCCCGTCCTCTACATAGCCGCCATCCTGTTCGGCGCGCCGCTCATCTCGCATCACCTCCACACGCTTCTCCTAGCCGTCCACCTCGCCCTCCTCACGACTCCACCACTCTTCTACGTCCACGGTCTTCACGCGCCCACATGGCTAGCGCTCGCGAGTTTGCAACAGCCGGTCGACGAGACATATGGAATGGCGCTGGGCGCGGTCGTAGGAGCTTGGGTGGGAGCTGTGCCAATTCCGCTGGACTGGGATCGCGAGTGGCAGAAGTGGCCGGTTACGATTATTTGTGGGATGTATGGAGGTGCTGTGCTGGGCAAGGTGGCGGGTGGCTGGCTCTTAAGAGGGAGGAAGATGAAGATCAGT ``` |
| 3 | ggggacaagtttgtacaaaaaagcaggcttgatgCGTCCACGAACTCTCCGAGC, ggggaccactttgtacaagaaagctgggtcATGCTCATTGTTTCCACCGC | Mycgr3\|89454  JGI  Ensembl | KOG3192 Mitochondrial J-type chaperone; | chaperone binding; heat shock protein binding;  protein folding; | 100 % |  | ``` 158	CATTCAACATTACACCTTCCCTCCGACAACAACAGTCGCGCGCGCAGCAGCACGAAGCCGCCTCCGAAACGTCCGCCAGACCCACGCAGACCCATTACTCCCTCTTCCCCAGCACCTTTCCCGCTGGGCCACCACCAGCTTCACCCTTCACGCCTTCCCTTCCAGCCCTCCGGCGTGAATTTCTCACTCTCCAACAAAAATCTCACCCGGACCTCTTCCCTCCCGCAGAGAAACCCCGCGCCGAAGCCTTGTCCATGCTCATCAACGAGGCGTATCGCACATTGCAAGATCCCTTGCGACGGGCGGAGTACCTCTTATCGCTGCAAGGCGTAGACCTCGACGATGAGTCCGCAAAGCTGGGCGACCAAGCGGCTGGTGAGGGAGGGGAGCTTTTGATGGAGGTCATGGAAGCGAGAGAGGCGGTCGACGAGGTTGAGGACGAGGCTGGGTTGGAGGCGTTAAAGAGCGACAATGAAGAGAGGATGAAGGGGAGTGTGGCGGGGTTGGAAGCCGCTTTTGCGGAAGGTAATTTGGAGGCGGCGAGGAGGGAGGCCGTCAGGCTGCGCTATTGGAAGAACATCGAGGAGAGTATACGCGGCTGGGAGAAGGGTCACGGCGGTGGAAACAATGAGCAT	792 	\|\|\|\|\|\|\|\|\|\|\| \|\|\| \|\|\|\| \|\| \|\|\|\|\|\|\|\|\|\| \|\|\|\|\|\|\|\|\|\|\|\|\|\|\|\|\| \|\|\|\|\|\|\|\|\|\|\|\|\|\|\|\|\| \|\|\|\|\|\|\|\|\|\|\|\|\|\|\|\|\|\|\|\|\|\|\|\|\|\|\|\|\|\|\|\|\|\|\|\|\|\|\|\|\|\|\|\|\|\|\|\|\|\|\|\| \|\|\|\|\|\|\|\|\|\|\|\|\|\|\|\|\|\|\|\|\|\|\|\|\|\|\|\|\|\|\|\|\|\|\|\|\|\|\|\|\|\|\|\|\|\|\|\|\|\|\|\|\|\|\|\|\|\|\|\|\|\|\|\|\|\|\|\|\|\|\|\|\|\|\|\|\|\|\|\|\|\|\|\|\|\|\|\|\|\|\|\|\|\|\|\|\|\|\|\|\|\|\|\|\|\|\|\|\|\|\|\|\|\|\|\|\|\|\|\|\|\|\|\|\|\|\|\|\|\|\|\|\|\|\|\|\|\|\|\|\|\|\|\|\|\|\|\|\|\|\|\|\|\|\|\|\|\|\|\|\|\|\|\|\|\|\|\|\|\|\|\|\|\|\|\|\|\|\|\|\|\|\|\|\|\|\|\|\|\|\|\|\|\|\|\|\|\|\|\|\|\|\|\|\|\|\|\|\|\|\|\|\|\|\|\|\|\|\|\|\|\|\|\|\|\|\|\|\|\|\|\|\|\|\|\|\|\|\|\|\|\|\|\|\|\|\|\|\|\|\|\|\|\|\|\|\|\|\|\|\|\|\|\|\|\|\|\|\|\|\|\|\|\|\|\|\|\|\|\|\|\|\|\|\|\|\|\|\|\|\|\|\|\|\|\|\|\|\|\|\|\|\|\|\|\|\|\|\|\|\|\|\|\|\|\|\|\|\|\|\|\|\|\|\|\|\|\|\|\|\|\|\|\|\|\|\|\|\|\|\|\|\|\|\|\|\|\|\|\|\|\|\|\|\|\|\|\|\|\|\|\|\|\|\|\|\|\|\|\|\|\|\|\|\|\|\|\|\|\|\|\|\|\|\|\|\|\|\|\|\|\|\|\|\|\|\|\|\|\|\|\|\|\|\|\|\|\|\|\|\|\|\|\|\|\|\|\|\|\|\|\|\|\|\|\|\|\|\|\|\|\|\|\|\|\|\|\|\|\|\|\|\|\|\|\|\|\|\|\|\|\|\|\|\|\|\|\|\|\|\|\|\|\|\|\|\|\|\|\|\|\|\|\|\|\|\|\|\|\|\|\|\|\|\|\|\|\|\|\|\|\|\|\|\|\|\|\|\|\|\|\|\|\|\|\|\|\|\|\|\| 	CATTCAACATTNCACNTTCCTTCNGACAACAACANTCGCGCGCGCAGCAGCA-GAAGCCGCCTCCGAAACNTCCGCCAGACCCACGCAGACCCATTACTCCCTCTTCCCCAGCACCTTTCCCGNTGGGCCACCACCAGCTTCACCCTTCACGCCTTCCCTTCCAGCCCTCCGGCGTGAATTTCTCACTCTCCAACAAAAATCTCACCCGGACCTCTTCCCTCCCGCAGAGAAACCCCGCGCCGAAGCCTTGTCCATGCTCATCAACGAGGCGTATCGCACATTGCAAGATCCCTTGCGACGGGCGGAGTACCTCTTATCGCTGCAAGGCGTAGACCTCGACGATGAGTCCGCAAAGCTGGGCGACCAAGCGGCTGGTGAGGGAGGGGAGCTTTTGATGGAGGTCATGGAAGCGAGAGAGGCGGTCGACGAGGTTGAGGACGAGGCTGGGTTGGAGGCGTTAAAGAGCGACAATGAAGAGAGGATGAAGGGGAGTGTGGCGGGGTTGGAAGCCGCTTTTGCGGAAGGTAATTTGGAGGCGGCGAGGAGGGAGGCCGTCAGGCTGCGCTATTGGAAGAACATCGAGGAGAGTATACGCGGCTGGGAGAAGGGTCACGGCGGTGGAAACAATGAGCAT ``` |
| 3 | ggggacaagtttgtacaaaaaagcaggcttgatgCACTTCAATCCTCTCCTCGC, ggggaccactttgtacaagaaagctgggtcGGCAGACGACGCCGAGCCGT | Mycgr3\|89705  JGI  Ensembl |  |  | 100 % |  | ``` 133	TACACGTTCAGCATCGCAGGCCCGAAGTCTGCCGGAAAGAATGTACCTAGCTTCAATGCCAAGTGCGCGGCCCAAATCCCAGGATTGGACTCTACATACCGCCCTTGCACGCTGGGTCCGAATTCCGACGGAAGTCCACGTCCTTCAGCCTTTGATGCATACTTCTTCCTCGACAAGAGCGACGCCCGCTACAATTCGCTGTCAGTCCG	341 	\|\|\|\| \|\|\| \|\|\|\|\| \|\|\|\|\|\|\| \|\|\|\|\|\|\|\|\|\|\|\|\|\| \|\|\|\|\|\|\|\| \| \|\|\|\|\|\|\|\| \|\|\|\|\|\|\|\|\|\|\|\|\|\|\|\|\|\|\|\|\|\|\|\|\|\|\|\|\|\|\|\|\| \|\|\|\|\|\|\|\|\|\|\|\|\|\|\|\|\|\|\| \|\|\|\|\|\|\|\|\|\|\|\|\|\|\|\|\|\|\|\|\|\|\|\|\|\|\|\|\|\| \|\|\|\|\|\|\|\|\|\|\|\|\|\|\|\|\|\|\|\|\|\|\|\|\|\|\|\|\|\|\|\|\|\|\|\|\|\|\|\|\|\|\|\|\|\|\|\|\|\|\|\|\|\|\|\|\|\|\|\|\|\|\|\|\|\| 	TACAAGTTNAGCATNGCAGGCCNGAAGTCTGCCGGAANGAATGTACNTNGCTTCAATNCCAAGTGCGCGGCCCAAATCCCAGGATTGGACTTTACATACCGCCCTTGCACGNTGGGTCCGAATTCCGACGGAAGTCCACGTCNTTCAGCCTTTGATGCATACTTCTTCCTCGACAAGAGCGACGCCCGCTACAATTCGCTGTCAGTCCG ```      ``` 1	ATGCACTTCAATCCTCTCCTCGCCCTCGCATTCGCATCCTTCTCCTCGGCGGCCGCCGTAGCGAGAGACGACAAGCTCGTCTGGACCGTCAGCGGTCTCTCCGCGACATGCAGCAGCGACACGAGCTGCTCCTACACGTTCAGCATCGCAGGCCCGAAGTCTGCCGGAAAGAATGTACCTAGCTTCAATGCCAAGTGCGCGGCCCAAATCCCAGGATTGGACTCTACATACCGCCCTTGCACGCTGGGTCCGAATTCCGACGGAAGTCCACGTCCTTCAGCCTTTGATGCATACTTCTTCCTCGACAAGAGCGACGCCCGCTACAATTCGCTGTCAGTCCG	341 	\|\|\|\|\| \|\|\|\| \| \|\|\|\|\|\|\|\|\|\|\|\|\|\|\|\|\|\|\|\|\|\|\|\|\|\|\|\|\|\|\|\|\|\|\|\|\|\|\|\|\|\|\|\|\|\|\|\|\|\|\|\|\|\|\|\|\|\|\|\|\|\|\|\|\|\|\|\|\|\|\|\|\|\|\|\|\|\|\|\|\|\|\|\|\|\|\|\|\|\|\|\|\|\|\|\|\|\|\|\|\|\|\|\|\|\|\|\|\|\|\|\|\|\|\|\|\|\|\|\|\|\|\|\|\|\|\|\|\|\|\|\|\|\|\|\|\|\|\|\|\|\|\|\|\|\|\|\|\|\|\|\|\|\|\|\|\|\|\|\|\|\|\|\|\|\|\|\|\|\|\|\|\|\|\|\|\|\|\|\|\|\|\|\|\|\|\|\|\|\|\|\|\|\|\|\|\|\|\|\|\|\|\|\|\|\|\|\|\|\|\|\|\|\|\|\|\|\|\|\|\|\|\|\|\|\|\|\|\|\|\|\|\|\|\|\|\|\|\|\|\|\|\|\|\|\|\|\|\|\|\|\|\|\|\|\|\|\|\|\|\|\|\|\|\|\|\|\|\|\|\|\|\|\|\|\|\|\|\|\|\|\|\|\|\|\|\|\|\|\|\|\|\|\|\|\|\|\|\|\|\|\|\|\|\|\|\|\|\|\|\|\|\|\|\|\|\|\|\|\|\|\|\|\|\|\|\|\| 	ATGCAGTTCACT-CTCTCCTCGCCCTCGCATTCGCATCCTTCTCCTCGGCGGCCGCCGTAGCGAGAGACGACAAGCTCGTCTGGACCGTCAGCGGTCTCTCCGCGACATGCAGCAGCGACACGAGCTGCTCCTACACGTTCAGCATCGCAGGCCCGAAGTCTGCCGGAAAGAATGTACCTAGCTTCAATGCCAAGTGCGCGGCCCAAATCCCAGGATTGGACTCTACATACCGCCCTTGCACGCTGGGTCCGAATTCCGACGGAAGTCCACGTCCTTCAGCCTTTGATGCATACTTCTTCCTCGACAAGAGCGACGCCCGCTACAATTCGCTGTCAGTCCG ```      ``` 1	ATGCACTTCAATCCTCTCCTCGCCCTCGCATTCGCATCCTTCTCCTCGGCGGCCGCCGTAGCGAGAGACGACAAGCTCGTCTGGACCGTCAGCGGTCTCTCCGCGACATGCAGCAGCGACACGAGCTGCTCCTACACGTTCAGCATCGCAGGCCCGAAGTCTGCCGGAAAGAATGTACCTAGCTTCAATGCCAAGTGCGCGGCCCAAATCCCAGGATTGGACTCTACATACCGCCCTTGCACGCTGGGTCCGAATTCCGACGGAAGTCCACGTCCTTCAGCCTTTGATGCATACTTCTTCCTCGACAAGAGCGACGCCCGCTACAATTCGCTGTCAGTCCG	341 	\|\|\|\|\|\|\|\|\|\|\|\|\|\|\|\|\|\|\|\|\|\|\|\|\|\|\|\|\|\|\|\|\|\|\|\|\|\|\|\|\|\|\|\|\|\|\|\|\|\|\|\|\|\|\|\|\|\|\|\|\|\|\|\|\|\|\|\|\|\|\|\|\|\|\|\|\|\|\|\|\|\|\|\|\|\|\|\|\|\|\|\|\|\|\|\|\|\|\|\|\|\|\|\|\|\|\|\|\|\|\|\|\|\|\|\|\|\|\|\|\|\|\|\|\|\|\|\|\|\|\|\|\|\|\|\|\|\|\|\|\|\|\|\|\|\|\|\|\|\|\|\|\|\|\|\|\|\|\|\|\|\|\|\|\|\|\|\|\|\|\|\|\|\|\|\|\|\|\|\|\|\|\|\|\|\|\|\|\|\|\|\|\|\|\|\|\|\|\|\|\|\|\|\|\|\|\|\|\|\|\|\|\|\|\|\|\|\|\|\|\|\|\|\|\|\|\|\|\|\|\|\|\|\|\|\|\|\|\|\|\|\|\|\|\|\|\|\|\|\|\|\|\|\|\|\|\|\|\|\|\|\|\|\|\|\|\|\|\|\|\|\|\|\|\|\|\|\|\|\|\|\|\|\|\|\|\|\|\|\|\|\|\|\|\|\|\|\|\|\|\|\|\|\|\|\|\|\|\|\|\|\|\|\|\|\|\|\|\|\|\|\|\|\|\|\|\|\|\|\|\|\|\|\|\|\|\|\|\|\|\| 	ATGCACTTCAATCCTCTCCTCGCCCTCGCATTCGCATCCTTCTCCTCGGCGGCCGCCGTAGCGAGAGACGACAAGCTCGTCTGGACCGTCAGCGGTCTCTCCGCGACATGCAGCAGCGACACGAGCTGCTCCTACACGTTCAGCATCGCAGGCCCGAAGTCTGCCGGAAAGAATGTACCTAGCTTCAATGCCAAGTGCGCGGCCCAAATCCCAGGATTGGACTCTACATACCGCCCTTGCACGCTGGGTCCGAATTCCGACGGAAGTCCACGTCCTTCAGCCTTTGATGCATACTTCTTCCTCGACAAGAGCGACGCCCGCTACAATTCGCTGTCAGTCCG ```      ``` 24	CCTCG-CATTCGCATCC-TTCTCC-TCGGCGGCC-GCCGTAGCGAGAGACGACAAGCTCGTCTGGACCG-TCAGCGGTCTCTCCG-CGACATGCAGCAGCGACACGAGCTGCTCCTACACGTTCAGCATCGCAGGCCCGAAGTCTGCCGGAAAGAATGTACCTAGCTTCAATGCCAAGTGCGCGGCCCAAATCCCAGGATTGGACTCTACATACCGCCCTTGCACGCTGGGTCCGAATTCCGACGGAAGTCCACGTCCTTCA-GCCTTTGATGCATACTTCTTCCTCGACAAGAGCGACGCCCGCTACAATTCGCTG-TCAGTCCGC	342 	\|\|\|\|\|  \|\|\|\|\|\|\|\|\|\| \|\|\|\|\|\| \|\|\|\|  \|\|\| \| \|\| \|\|\| \|\|\|\|\| \|\|\|\|\|\| \|\|\|\| \|\|\|\|\| \| \|\|\|\|\|\|\| \|\|\|\|\|\|\|  \|\|\|\|\|\| \|\|  \|\| \|\|\|  \|\|\|\|\|\|\| \|\|\|\|\|\|\|\| \|\|\|  \|\|\| \| \|\|\|\| \|\|\|\|\|\|\| \|\|\|\| \| \|\|\|\|\|\|\|\| \| \|\|\|\|\|\|\|\|\|\|\|\|\|\|\| \| \| \| \|\|\|\|\|\|\| \|\|\|\|\|\|\|\|\|\|\|\|\|\|\|\|\|\|\|\|\| \|\|\|\|\|\| \|\|\|\|\|\| \|\| \|\|\|\|\|\|\| \| \|\|\|\|  \| \| \|\|\|\|\|\|\|\| \| \|\|\|\|\|\|\|\|\|\|\|\|\|\|\|\|\|\|\|\|\|\| \|\|\|\|\|\|\|\|\|\|  \|\|\| \|\| \|\|\|\|\|\|\|\|\|\| \|\|\|\| \|\|\|\| 	CCTCGAGATTCGCATCCTTTCTCCATCGGNNGCCGGNCGAAGCNAGAGANGACAAGNTCGTNTGGACNGTTCAGCGGNCTCTCCGTNGACATGAAGNTGCNACANAAGCTGCTNCTACACGTNCAGNNTCGNAAGCCCTAAGTCTGNCGGAGANAATGTACCNANCTTCAATGCCAAGTGNGNGNCTCAAATCCNAGGATTGGACTCTACATACCGNCCTTGCNCGCTGGNTCNGAATTCCTAAGGAANNCNANGTCCTTCAGGTCTTTGATGCATACTTCTTCCTCNACAAGAGCGANNCCCNCTNCAATTCGCTGNTCAGNCCGC ```      ``` 341	GCCTAAACAAGAAGACCTTTGAGATGTTCCCATTTGTGGACGGCTCGGCGTCGTCTGCC	399 	\|\|\|\|\|\|\|\|\|\|\|\|\|\|\|\|\|\|\|\|\|\|\|\|\|\|\|\|\|\|\|\|\|\|\|\|\|\|\|\|\|\|\|\|\|\|\|\|\|\|\|\|\|\|\|\|\|\|\| 	GCCTAAACAAGAAGACCTTTGAGATGTTCCCATTTGTGGACGGCTCGGCGTCGTCTGCC ```      ``` 1	ATGCACTTCAATCCTCTCCTCGCCCTCGCATTCGCATCCTTCTCCTCGGCGGCCGCCGTAGCGAGAGACGACAAGCTCGTCTGGACCGTCAGCGGTCTCTCCGCGACATGCAGCAGCGACACGAGCTGCTCCTACACGTTCAGCATCGCAGGCCCGAAGTCTGCCGGAAAGAATGTACCTAGCTTCAATGCCAAGTGCGCGGCCCAAATCCCAGGATTGGACTCTACATACCGCCCTTGCACGCTGGGTCCGAATTCCGACGGAAGTCCACGTCCTTCAGCCTTTGATGCATACTTCTTCCTCGACAAGAGCGACGCCCGCTACAATTCGCTGTCAGTCCG	341 	\|\|\|\|\| \|\|\|\| \|\| \| \|\|\|\|\|\|\|\|\|\| \|\|\|\|\| \|\|\|\|\|\|\|\| \|\|\|\|\|\|\|\|\|\|\|\|\|\|\|\|\|\|\|\|\|\|\|\|\|\|\|\|\|\|\|\|\|\|\|\|\|\|\|\|\|\|\|\|\|\|\|\|\|\|\|\|\|\|\|\|\|\|\|\|\|\|\|\|\|\|\|\|\|\|\|\|\|\|\|\|\|\|\|\|\|\|\|\|\|\|\|\|\|\|\|\|\|\|\|\|\|\|\|\|\|\|\|\|\|\|\|\|\|\|\|\|\|\|\|\|\|\|\|\|\|\|\|\|\|\|\|\|\|\|\|\|\|\|\|\|\|\|\|\|\|\|\|\|\|\|\|\|\|\|\|\|\|\|\|\|\|\|\|\|\|\|\|\|\|\|\|\|\|\|\|\|\|\|\|\|\|\|\|\|\|\|\|\|\|\|\|\|\|\|\|\|\|\|\|\|\|\|\|\|\|\|\|\|\|\|\|\|\|\|\|\|\|\|\|\|\|\|\|\|\|\|\|\|\|\|\|\|\|\|\|\|\|\|\|\|\|\|\|\|\|\|\|\|\|\|\|\|\|\|\|\|\|\|\|\|\|\|\|\|\|\|\|\|\|\|\|\|\|\|\|\|\|\|\|\|\|\|\|\|\|\|\|\|\|\|\|\|\|\|\|\|\|\|\|\|\|\|\| 	ATGCAGTTCACTC-TNTCCTCGCCCTNGCATTNGCATCCTTNTCCTCGGCGGCCGCCGTAGCGAGAGACGACAAGCTCGTCTGGACCGTCAGCGGTCTCTCCGCGACATGCAGCAGCGACACGAGCTGCTCCTACACGTTCAGCATCGCAGGCCCGAAGTCTGCCGGAAAGAATGTACCTAGCTTCAATGCCAAGTGCGCGGCCCAAATCCCAGGATTGGACTCTACATACCGCCCTTGCACGCTGGGTCCGAATTCCGACGGAAGTCCACGTCCTTCAGCCTTTGATGCATACTTCTTCCTCGACAAGAGCGACGCCCGCTACAATTCGCTGTCAGTCCG ```      ``` 1	ATGCACTTCAATCCTCTCCTCGCCCTCGCATTCGCATCCTTCTCCTCGGCGGCCGCCGTAGCGAGAGACGACAAGCTCGTCTGGACCGTCAGCGGTCTCTCCGCGACATGCAGCAGCGACACGAGCTGCTCCTACACGTTCAGCATCGCAGGCCCGAAGTCTGCCGGAAAGAATGTACCTAGCTTCAATGCCAAGTGCGCGGCCCAAATCCCAGGATTGGACTCTACATACCGCCCTTGCACGCTGGG-TCCGAATTCCGACGGAAGTCCACGTCCTTCAGCCTTTGATGCATACTTCTTCCTCGACAAGAGCGACGCCCGCTACAAT-TCGCTGTC	335 	\|\|\|\|\|\|\|\|\|\|\|\|\|\|\|\|\|\|\|\|\|\|\|\|\|\|\|\|\|\|\|\|\|\|\|\|\|\|\|\|\|\|\|\|\|\|\|\|\|\|\|\|\|\|\|\|\|\|\|\|\|\|\|\|\|\|\|\|\|\|\|\|\|\| \|\|\|\|\|\|\|\|\|\|\|\|\| \|\|\|\|\|\|\|\|\|\|\|\|\|\|\|\|\|\|\|\|\|\|\|\|\|\|\|\|\|\|\|\|\|\|\|\|\|\|\|\|\|\|\|\|\|\|\|\|\|\|\|\|\|\|\|\|\|\|\|\|\|\|\|\|\|\|\|\|\|\|\|\|\|\|\|\|\|\|\|\|\|\|\|\|\|\|\|\|\|\|\|\|\| \|\|\|\|\|\|\|\|\|\|\|\|\|\|\|\|\|\|\|\|\|\|\|\|\|\|\|\|\|\|\|\|\|\|\|\|\|\|\|\|\|\|\|\|\|\|\|\|\|\|\|\|\|\|\|\|\|\|\|\|\|\|\|\|\| \|\|\|\|\|\|\|\|\|\|\|\|\|\|\|\|\|\|\|\|\|\|\|\|\|\|\|\|\|\|\| \|\|\|\|\|\|\|\|\|\|\|\|\|\|\|\|\|\|\|\|\|\|\|\|\|\| \| \| \|\|\|\| \|\|\|\|\|\|\|\|\|\|\| \|\| \|\|\|\|\| 	ATGCACTTCAATCCTCTCCTCGCCCTCGCATTCGCATCCTTCTCCTCGGCGGCCGCCGTAGCGAGAGACGACAANCTCGTCTGGACCGNCAGCGGTCTCTCCGCGACATGCAGCAGCGACACGAGCTGCTCCTACACGTTCAGCATCGCAGGCCCGAAGTCTGCCGGAAAGAATGTACCTAGNTTCAATGCCAAGTGCGCGGCCCAAATCCCAGGATTGGACTCTACATACCGCCCTTGCACGCTGGGTTCCGAATTCCGACGGAAGTCCACGTCCTTCAACCTTTGATGCATACTTCTTCCTCGACNANANCGACNCCCGCTACAATNTCNCTGTC ``` |
| 3 | ggggacaagtttgtacaaaaaagcaggcttgatgCAGATCAAGATGCAGCTCCT, ggggaccactttgtacaagaaagctgggtcCTGCTGGGCTTCCTCTTCCG | Mycgr3\|90776  JGI  Ensembl | KOG1808 AAA ATPase containing von Willebrand factor type A (vWA) domain; |  | 81.20205 % | chr\_2:1922858 G->T; chr\_2:1922819 A->G; chr\_2:1922828 A->C; chr\_2:1922840 T->C; chr\_2:1922885 C->T; chr\_2:1922888 G->A; chr\_2:1923107 C->T; chr\_2:1922806 T->C; chr\_2:1923110 T->C; chr\_2:1922912 T->C; | ``` 1	ATGCAGATCAAGATGCAGCTCCTCTCTCTGGTACTGATCCCGCTCATCGTTGATGCAGCTGCCATTC	67 	\|\|\|\|\|\|\|\|\|\|\|\|\|\|\|\|\|\|\|\|\|\|\|\|\|\|\|\|\|\|\|\|\|\|\|\|\|\|\|\|\|\|\|\|\|\|\|\|\|\|\|\|\|\|\|\|\|\|\|\|\|\|\|\|\|\|\| 	ATGCAGATCAAGATGCAGCTCCTCTCTCTGGTACTGATCCCGCTCATCGTTGATGCAGCTGCCATTC ```      ``` 200	AGGCCAAGCGAGATGAGGCATACGACTACCCTCGAGACAGCGGCTACTCGTTCATCACCAAGCCGGAAGAGGAGACCGAGCAGAAGGTCAAGGCCAAGCGAGATGAGGCATACGACTACCCTCGAGACAGCGGCTACTCGTTCATCACCAAGCCGGAAGAGGAGACTAAGGACAAGGCCAAGGCCAAGCGAGATGAGGCATACGACTACCCTCGAGACAGCGGATACTCGTTCATCACCAAGCCGGAAGAGGAGACCGAGCAG	462 	\|\|\|\|\|\|\|\|\|\|\|\|\|\|\|\|\|\|\|\|\|\|\|\|\|\|\|\| \|\| \|\|\|\|\|\|\|\|\|\|\|\|\|\|\|\|\|\|\|\|\|\|\|\|\|\|\|\|\|\|\|\|\|\|\|\|\|\|\|\|\|\|\|\|\|\|\|\|\|\|\|\|\|\|\| \|\|\|\|\|\|\|\|\|\|\|\| \|\|\|\|\|\|\|\| \|\|\|\|\|\|\|\|\|\|\| \|\|\|\|\|\|\|\|\|\|\|\|\|\|\|\|\| \|\|\|\|\|\|\|\|\|\|\|\|\|\|\|\|\|\|\|\|\|\|\|\|\|\|\| \|  \|\|\| \|   \|\| \|\|\|\|\|\|\|\|\|\|\| \|\|\|\|\|\|\|\|\|\|\|\|\|\|\|\|\|\|\|\| \|\|\|\|\|\|\|\| \|\|\|\|\| \|\|\|\|\|\|\|\|\|\|\|\|\|\|\|\|\|\|\|\|\|\|\|  \|\| \|\|\|\|\| 	AGGCCAAGCGAGATGAGGCATACGACTATCCCCGAGACAGCGGCTACTCGTTCATCACCAAGCCGGAAGAGGAGACCGAGCAGAAGGCCAAGGCCAAGCGGGATGAGGCCTACGACTACCCCCGAGACAGCGGCTACTCTTTCATCACCAAGCCGGAAGAGGAGACTGA--ACA-GA--AA-GCCAAGCGAGACGAGGCATACGACTACCCTCGCGACAGCGGCTACTCCTTCATCACCAAGCCGGAAGAGGAAGCCCAGCAG ```      ``` 372	CAAGGCCAAGGCCAAGCGAGATGAGGCATACGACTACCCTCGAGACAGCGGATACTCGTTCATCACCAAGCCGGAAGAGGAGACCGAGCAGAAGGTCAAGGCCAAGCGAGATGAGGCATACGACTACCCTCGAGACAGCGGCTACTCGTTCATCACCAAGCCGGAAGAGGAGACCGAGCAGAAGGCCAAGGCCAAGCGGGATGAGGCCTACGACTACCCCCGAGACAGCGGCTACTCTTTCATCACCAAGCCGGAAGAGGA	632 	\|\|\| \|\|\| \|\|\|\|\|\|\|\|\|\|\|\|\|\|\|\|\|\|\|\|\|\|\|\|\|\|\|\| \|\| \|\|\|\|\|\|\|\|\|\|\| \|\|\|\|\|\|\|\|\|\|\|\|\|\|\|\|\|\|\|\|\|\|\|\|\|\|\|\|\|\|\|\|\|\|\|\|\|\|\|\|\|\|\| \|\|\|\|\|\|\|\|\|\|\|\| \|\|\|\|\|\|\|\| \|\|\|\|\|\|\|\|\|\|\| \|\|\|\|\|\|\|\|\|\|\|\|\|\|\|\|\| \|\|\|\|\|\|\|\|\|\|\|\|\|\|\|\|\|\|\|\|\|\|\|\|\|\| \|\| \|\|\|\|\|    \|  \|\|\|\|\|\|\|\| \|\| \|\|\|\|\| \|\|\|\|\|\|\|\|\|\|\| \|\| \|\|\|\|\|\|\|\|\|\|\|\|\|\| \|\|\|\|\|\|\|\|\|\|\|\|\|\|\|\|\|\|\|\|\|\|\| 	CAATGCCGAGGCCAAGCGAGATGAGGCATACGACTATCCCCGAGACAGCGGCTACTCGTTCATCACCAAGCCGGAAGAGGAGACCGAGCAGAAGGCCAAGGCCAAGCGGGATGAGGCCTACGACTACCCCCGAGACAGCGGCTACTCTTTCATCACCAAGCCGGAAGAGGAGACTGAACAGAA----A--GCCAAGCGAGACGAGGCATACGACTACCCTCGCGACAGCGGCTACTCCTTCATCACCAAGCCGGAAGAGGA ```      ``` 470	AGGCCAAGCGAGATGAGGCATACGACTACCCTCGAGACAGCGGCTACTCGTTCATCACCAAGCCGGAAGAGGAGACCGAGCAGAAGGCCAAGGCCAAGCGGGATGAGGCCTACGACTACCCCCGAGACAGCGGCTACTCTTTCATCACCAAGCCGGAAGAGGAGACTGAACAGAAAGCCAAGCGAGACGAGGCATACGACTACCCTCGCGACAGCGGCTACTCCTTCATCACCAAGCCGGAAGAGGAAGCCCAGCAG	726 	\|\|\|\|\|\|\|\|\|\|\|\|\|\|\|\|\|\|\|\|\|\|\|\|\|\|\|\| \|\| \|\|\|\|\|\|\|\|\|\|\|\|\|\|\|\|\|\|\|\|\|\|\|\|\|\|\|\|\|\|\|\|\|\|\|\|\|\|\|\|\|\|\|\|\|\|\|\|\|\|\|\|\|\|\|\|\|\|\|\|\|\|\|\|\|\|\|\|\|\|\|\|\|\|\|\|\|\|\|\|\|\|\|\|\|\|\|\|\|\|\|\|\|\|\|\|\|\|\|\|\|\|\|\|\|\|\|\|\|\|\|\|\|\|\|\|\|\|\|\|\|\|\|\|\|\|\|\|\|\|\|\|\|\|\|\|\|\|\|\|\|\|\|\|\|\|\|\|\|\|\|\|\|\|\|\|\|\|\|\|\|\|\|\|\|\|\|\|\|\|\|\|\|\|\|\|\|\|\|\|\|\|\|\|\|\|\|\|\|\|\|\|\|\|\|\|\|\|\|\|\|\|\|\|\|\|\|\|\|\|\|\|\|\|\|\|\|\|\|\|\|\|\|\|\| 	AGGCCAAGCGAGATGAGGCATACGACTATCCCCGAGACAGCGGCTACTCGTTCATCACCAAGCCGGAAGAGGAGACCGAGCAGAAGGCCAAGGCCAAGCGGGATGAGGCCTACGACTACCCCCGAGACAGCGGCTACTCTTTCATCACCAAGCCGGAAGAGGAGACTGAACAGAAAGCCAAGCGAGACGAGGCATACGACTACCCTCGCGACAGCGGCTACTCCTTCATCACCAAGCCGGAAGAGGAAGCCCAGCAG ```      ``` 68	CAAAGCCTGTGGAAAGTCGCGCTCTCCAAGGAAGCAATGCCGAGGCCAAGCGAGATGAGGCATACGACTATCCCCGAGACAGCGGCTACTCGTTCATCACCAAGCCGGAAGAGGAGACCGAGCAGAAGGTCAAGGCCAAGCGAGATGAGGCATACGACTACCCTCGAGACAGCGGCTACTCGTTCATCACCAAGCCGGAAGAGGAGACCGAGCAGAAGGTCAAGGCCAAGCGAGATGAGGCATACGACTACCCTCGAGACAGCGGCTACTCGTTCATCACCAAGCCGGAAGAGGA	362 	\|\|\|\|\|\|\|\|\|\|\|\|\|\|\|\|\|\|\|\|\|\|\|\|\|\|\|\|\|\|\|\|\|\|\|\|\|\|\|\|\|\|\|\|\|\|\|\|\|\|\|\|\|\|\|\|\|\|\|\|\|\|\|\|\|\|\|\|\|\|\|\|\|\|\|\|\|\|\|\|\|\|\|\|\|\|\|\|\|\|\|\|\|\|\|\|\|\|\|\|\|\|\|\|\|\|\|\|\|\|\|\|\|\|\|\|\|\|\|\|\|\|\|\|\|\|\|\|\| \|\|\|\|\|\|\|\|\|\|\|\| \|\|\|\|\|\|\|\| \|\|\|\|\|\|\|\|\|\|\| \|\|\|\|\|\|\|\|\|\|\|\|\|\|\|\|\| \|\|\|\|\|\|\|\|\|\|\|\|\|\|\|\|\|\|\|\|\|\|\|\|\|\| \|\| \|\|\|\|\|    \|  \|\|\|\|\|\|\|\|\|\|\| \|\|\|\|\|\|\|\|\|\|\|\|\|\|\|\|\|\|\|\| \|\|\|\|\|\|\|\|\|\|\|\|\|\| \|\|\|\|\|\|\|\|\|\|\|\|\|\|\|\|\|\|\|\|\|\|\| 	CAAAGCCTGTGGAAAGTCGCGCTCTCCAAGGAAGCAATGCCGAGGCCAAGCGAGATGAGGCATACGACTATCCCCGAGACAGCGGCTACTCGTTCATCACCAAGCCGGAAGAGGAGACCGAGCAGAAGGCCAAGGCCAAGCGGGATGAGGCCTACGACTACCCCCGAGACAGCGGCTACTCTTTCATCACCAAGCCGGAAGAGGAGACTGAACAGAA----A--GCCAAGCGAGACGAGGCATACGACTACCCTCGCGACAGCGGCTACTCCTTCATCACCAAGCCGGAAGAGGA ```      ``` 290	AGGCCAAGCGAGATGAGGCATACGACTACCCTCGAGACAGCGGCTACTCGTTCATCACCAAGCCGGAAGAGGAGACTAAGGACAAGGCCAAGGCCAAGCGAGATGAGGCATACGACTACCCTCGAGACAGCGGATACTCGTTCATCACCAAGCCGGAAGAGGAGACCGAGCAGAAGGTCAAGGCCAAGCGAGATGAGGCATACGACTACCCTCGAGACAGCGGCTACTCGTTCATCACCAAGCCGGAAGAGGAGACCGAGCAG	552 	\|\|\|\|\|\|\|\|\|\|\|\|\|\|\|\|\|\|\|\|\|\|\|\|\|\|\|\| \|\| \|\|\|\|\|\|\|\|\|\|\|\|\|\|\|\|\|\|\|\|\|\|\|\|\|\|\|\|\|\|\|\|\|\|\|\|\|\|\|\|\|\|\|\|  \|\| \| \|\|\|\|\|\|\|\|\|\|\|\|\|\|\|\|\| \|\|\|\|\|\|\|\| \|\|\|\|\|\|\|\|\|\|\| \|\|\|\|\|\|\|\|\|\|\| \|\|\|\|\| \|\|\|\|\|\|\|\|\|\|\|\|\|\|\|\|\|\|\|\|\|\|\|\|\|\| \|\| \|\|\|\|\|    \|  \|\|\|\|\|\|\|\|\|\|\| \|\|\|\|\|\|\|\|\|\|\|\|\|\|\|\|\|\|\|\| \|\|\|\|\|\|\|\|\|\|\|\|\|\| \|\|\|\|\|\|\|\|\|\|\|\|\|\|\|\|\|\|\|\|\|\|\|  \|\| \|\|\|\|\| 	AGGCCAAGCGAGATGAGGCATACGACTATCCCCGAGACAGCGGCTACTCGTTCATCACCAAGCCGGAAGAGGAGACCGAGCAGAAGGCCAAGGCCAAGCGGGATGAGGCCTACGACTACCCCCGAGACAGCGGCTACTCTTTCATCACCAAGCCGGAAGAGGAGACTGAACAGAA----A--GCCAAGCGAGACGAGGCATACGACTACCCTCGCGACAGCGGCTACTCCTTCATCACCAAGCCGGAAGAGGAAGCCCAGCAG ``` |
| 3 | ggggacaagtttgtacaaaaaagcaggcttgatgCCGTCGATCAAGACTCTAGC, ggggaccactttgtacaagaaagctgggtcCCTCGACACCAAGTCCGTGG | Mycgr3\|91702  JGI  Ensembl |  |  | 0 % |  |  |
| 3 | ggggacaagtttgtacaaaaaagcaggcttgatgCTCACTCTCACCCTACTTGC, ggggaccactttgtacaagaaagctgggtcCCACCCTATCATGGCGCATA | Mycgr3\|91885  JGI  Ensembl |  |  | 100 % |  | ``` 1	ATGCTCACTCTCACCCTACTTGCGCTTCTCCTCCCGACAACCCTTTGCCAGACCTTCTCACTCGATATGGCCTCCGATCCCACGCGCCAATCTCTCTCCGCTTCCCTCATGAACATAACCTTCTCCAACCCCTCCGACGTCTGCAGCAGTCCCAACACTACCATCGATAGCTCCAGCCTTTCCATTGCCATCCGGACACTGCCGCAGTCTCGGACGTGCTTCGACTTTGCGAAAATTTTCCGCAATCCGGAGAAGACGTATACTTCCAAAGGTGCAGATTGCAAGGGTGACGGATCCTGCGGGGTCAATTATACCGTGGTCTCACCTTCCGACTCGTCTGTATACAGCTCGACAGCGAATTACAGTCAGGTATTCTATACGCAAGTTCAGAACCCTCTCGCTGGAATTGACTCGAACAGAGCCAACTCGGGAGAAGCTGGACGGCTCGAGATGGTCGCATATCGTGCGCAGGGCTGCAGAGGTCGGAAGGCTAGCTACGAATGGAACTGTCTTGAGCAGGACGGAAGGTGTGATTCGATGCCATGGCCGGTGAGAAGTTTCAGCGTGGAGATGACGAAGGCGGGAGATGTGGGAGTGGGGAGTTGTCGAGCTGCAAAGTTGAACGGAGTGGATAGGAAGAGGCCGGGAGTGGGTTTCGCTGTGGTGATCTTTGCAACGATATGCGCCATGATAGGGTGG	699 	\|\|\|\|\|\|\| \| \|\|\|\|\|\|\|\|\| \|\|\|\| \|\| \|\|\|\|\|\|\|\|\|\| \|\|\|\|\|\|\|\|\|\|\|\|\|\|\|\|\| \|\|\|\|\|\|\|\|\|\|\|\|\|\|\|\|\|\|\|\|\|\|\|\|\|\|\|\|\|\|\|\|\|\|\|\|\|\| \|\|\|\|\|\|\|\|\| \|\|\|\|\|\|\|\|\|\|\|\|\|\|\|\|\|\|\|\|\|\|\|\|\| \|\|\|\|\|\|\|\|\|\|\|\|\|\|\|\|\|\|\|\|\|\|\|\|\|\|\|\|\|\|\|\|\|\|\|\|\|\|\|\|\|\|\|\|\|\|\|\|\|\|\|\|\|\|\|\|\|\|\|\|\|\|\|\|\|\|\|\|\|\|\|\|\|\|\|\|\|\|\|\|\|\|\|\|\|\|\|\|\|\|\|\|\|\|\|\|\|\|\|\|\|\|\|\|\|\|\|\|\|\|\|\|\|\|\|\|\|\|\|\|\|\|\|\|\|\|\|\|\|\|\|\|\|\|\|\|\|\|\|\|\|\|\|\|\|\|\|\|\|\|\|\|\|\|\|\|\|\|\|\|\|\|\|\|\|\|\|\|\|\|\|\|\|\|\|\|\|\|\|\|\|\|\|\|\|\|\|\|\|\|\|\|\|\|\|\|\|\|\|\|\|\|\|\|\|\|\|\|\|\|\|\|\|\|\|\|\|\|\|\|\|\|\|\|\|\|\|\|\|\|\|\|\|\|\|\|\|\|\|\|\|\|\|\|\|\|\|\|\|\|\|\|\|\|\|\|\|\|\|\|\|\|\|\|\|\|\|\|\|\|\|\|\|\|\|\|\|\|\|\|\|\|\|\|\|\|\|\|\|\|\|\|\|\|\|\|\|\|\|\|\|\|\|\|\|\|\|\|\|\|\|\|\|\|\|\|\|\|\|\|\|\|\|\|\|\|\|\|\|\|\|\|\|\|\|\|\|\|\|\|\|\|\|\|\|\|\|\|\|\|\|\|\|\|\|\|\|\|\|\|\|\|\|\|\|\|\|\|\|\|\|\|\|\|\|\|\|\|\|\|\|\|\|\|\|\|\|\|\|\|\|\|\|\|\|\|\|\|\|\|\|\|\|\|\|\|\|\|\|\|\|\|\|\|\|\|\|\|\|\|\|\|\|\|\|\|\|\|\|\|\|\|\|\|\|\|\|\|\|\|\|\|\|\|\|\|\|\|\|\|\|\|\|\|\|\|\|\|\|\|\|\|\|\|\|\|\|\|\|\|\|\|\|\|\|\|\|\|\|\|\|\|\|\|\|\|\|\|\|\|\|\|\|\|\|\|\|\|\|\|\|\|\|\|\|\|\|\|\|\|\|\|\|\|\|\|\|\|\|\|\|\|\|\|\|\|\|\|\|\|\|\|\|\|\|\|\|\|\|\|\|\|\|\|\|\|\|\|\|\|\|\|\|\|\|\|\|\|\|\|\|\|\|\|\|\|\| 	ATGCTCA-TNTCACCCTAC-TGCGNTTNTCCTCCCGACNACCCTTTGCCAGACCTTNTCACTCGATATGGCCTCCGATCCCACGCGCCAATCTCTTTCCGCTTCCNTCATGAACATAACCTTCTCCAACCCNTCCGACGTCTGCAGCAGTCCCAACACTACCATCGATAGCTCCAGCCTTTCCATTGCCATCCGGACACTGCCGCAGTCTCGGACGTGCTTCGACTTTGCGAAAATTTTCCGCAATCCGGAGAAGACGTATACTTCCAAAGGTGCAGATTGCAAGGGTGACGGATCCTGCGGGGTCAATTATACCGTGGTCTCACCTTCCGACTCGTCTGTATACAGCTCGACAGCGAATTACAGTCAGGTATTCTATACGCAAGTTCAGAACCCTCTCGCTGGAATTGACTCGAACAGAGCCAACTCGGGAGAAGCTGGACGGCTCGAGATGGTCGCATATCGTGCGCAGGGCTGCAGAGGTCGGAAGGCTAGCTACGAATGGAACTGTCTTGAGCAGGACGGAAGGTGTGATTCGATGCCATGGCCGGTGAGAAGTTTCAGCGTGGAGATGACGAAGGCGGGAGATGTGGGAGTGGGGAGTTGTCGAGCTGCAAAGTTGAACGGAGTGGATAGGAAGAGGCCGGGAGTGGGTTTCGCTGTGGTGATCTTTGCAACGATATGCGCCATGATAGGGTGG ``` |
| 3 | ggggacaagtttgtacaaaaaagcaggcttgatgCATTTCTCACACGCCATTCT, ggggaccactttgtacaagaaagctgggtcGGAAGGAGGCCCAGAAGGCG | Mycgr3\|93075  JGI  Ensembl |  |  | 100 % |  | ``` 1	ATGCATTTCTCACACGCCATTCTTTTTCTGGCTTCTGCCCTTCCAGCTGTGGTGCTCGCCGACACCACAGGAAGTCGAATCGCGGTCCCAATCGCAGCCGGTCCGGGAAACACCAAGCGCAGCTTCTCGCCATTGCTCCGCATGCTTCGCAGAGACGACCCTCAACCTGCGTCTTGCACAGCTCCTGACGATACTCCATGCCTTCCAAAGGGTGTTGCACCTCACACGCCCCACACCTGCAACATTCCGCTCACGAACTCTCCCGCTTACATCGGACTTGACGCGGTCCGCCGTAACGGGTATCCTTCTTGCAGCTCCTCAAATACCTGCTCATGCTCTG	340 	\|\|\|\|\|\|\|\|\|\|\|\|\|\|\|\|\|\|\|\|\|\|\|\|\|\|\|\|\|\|\|\|\|\|\|\|\|\|\|\|\|\|\|\|\|\|\|\|\|\|\|\|\|\|\|\|\|\|\|\|\|\|\|\|\|\|\|\|\|\|\|\|\|\|\|\|\|\|\|\|\|\|\|\|\|\|\|\|\|\|\|\|\|\|\|\|\|\|\|\|\|\|\|\|\|\|\|\|\|\|\|\|\|\|\|\|\|\|\|\|\|\|\|\|\|\|\|\|\|\|\|\|\|\|\|\|\|\|\|\|\|\|\|\|\|\|\|\|\|\|\|\|\|\|\|\|\|\|\|\|\|\|\|\|\|\|\|\|\|\|\|\|\|\|\|\|\|\|\|\|\|\|\|\|\|\|\|\|\|\|\|\|\|\|\|\|\|\|\|\|\|\|\|\|\|\|\|\|\|\|\|\|\|\|\|\|\|\|\|\|\|\|\|\|\|\|\|\|\|\|\|\|\|\|\|\|\|\|\|\|\|\|\|\|\|\|\|\|\|\|\|\|\|\|\|\|\|\|\|\|\|\|\|\|\|\|\|\|\|\|\|\|\|\|\|\|\|\|\|\|\|\|\|\|\|\|\|\|\|\|\|\|\|\|\|\|\|\|\|\|\|\|\|\|\|\|\|\|\|\|\|\|\|\|\|\|\|\|\|\|\|\|\|\|\|\|\|\|\|\|\|\|\|\|\|\|\|\|\|\| 	ATGCATTTCTCACACGCCATTCTTTTTCTGGCTTCTGCCCTTCCAGCTGTGGTGCTCGCCGACACCACAGGAAGTCGAATCGCGGTCCCAATCGCAGCCGGTCCGGGAAACACCAAGCGCAGCTTCTCGCCATTGCTCCGCATGCTTCGCAGAGACGACCCTCAACCTGCGTCTTGCACAGCTCCTGACGATACTCCATGCCTTCCAAAGGGTGTTGCACCTCACACGCCCCACACCTGCAACATTCCGCTCACGAACTCTCCCGCTTACATCGGACTTGACGCGGTCCGCCGTAACGGGTATCCTTCTTGCAGCTCCTCAAATACCTGCTCATGCTCTG ```      ``` 89	CAATCGCAGCCGGTCCGGGAAACACCAAGCGCAGCTTCTCGCCATTGCTCCGCATGCTTCGCAGAGACGACCCTCAACCTGCGTCTTGCACAGCTCCTGACGATACTCCATGCCTTCCAAAGGGTGTTGCACCTCACACGCCCCACACCTGCAACATTCCGCTCACGAACTCTCCCGCTTACATCGGACTTGACGCGGTCCGCCGTAACGGGTATCCTTCTTGCAGCTCCTCAAATACCTGCTCATGCTCTG	340 	\|\|\|\| \|\|\| \| \|\|\|\| \|\|\|\|\|\|\|\|\|\|\|\|\| \|\|\|\| \|\| \| \|\|\|\|\| \| \|  \|\|\|\|\| \|\| \|\|\|\|\|\|\| \|\|\|\|\|\|\|\|\|\| \|\| \|\| \|\|\|\|\|\|\|\| \|\|\|\|\|\|\|\|\|\|\| \|\|\|\|\|\|\|\|\|\|\|\|\|\|\|\|\|\|\|\|\|\|\|\|\|\| \|\|\|\|\|\|\|\|\|\|\|\|\|\|\| \|\|\|\|\|\|\|\|\|\|\|\|\|\|\|\|\|\|\|\|\|\|\|\|\|\|\|\|\|\|\|\|\|\|\|\|\|\|\|\|\|\|\|\|\|\|\|\|\|\|\|\|\|\|\|\|\|\|\|\|\|\|\|\|\|\|\|\|\|\|\|\|\|\|\|\|\|\|\|\|\|\|\|\|\|\|\|\|\|\|\|\|\|\|\|\|\|\|\|\|\|\|\| 	CAATNGCANCNGGTCNGGGAAACACCAAGNGCAGNTTNTNGCCAT-GNTNNGCATGNTTNGCAGAGANGACCCTCAACNTGNGTNTTGCACAGNTCCTGACGATANTCCATGCCTTCCAAAGGGTGTTGCACNTCACACGCCCCACACNTGCAACATTCCGCTCACGAACTCTCCCGCTTACATCGGACTTGACGCGGTCCGCCGTAACGGGTATCCTTCTTGCAGCTCCTCAAATACCTGCTCATGCTCTG ```      ``` 340	GCCATCGGTGTGCCGCATGATGTTCTCATCGAGATCTGCAGAGAAATCATCTCACGCCAGGATCCCTCGCTCCACGTCGTCGACGCTTGCGCAAGCGGTCCCACTGGTGAGGACGGTCACTGCGACCTGGCAGGCCCAACTTACCCGGACTTCTCGGGCACCGACGTCACGGTGTACAACGGCTGGGTGTGCAAGAACGGAAAATATGTTGGCGAGCACACGCCTTCTGGGCCTCCTTCC	579 	\|\|\|\|\|\|\|\|\|\|\|\|\|\|\|\|\|\|\|\|\|\|\|\|\|\|\|\|\|\|\|\|\|\|\|\|\|\|\|\|\|\|\|\|\|\|\|\|\|\|\|\|\|\|\|\|\|\|\|\|\|\|\|\|\|\|\|\|\|\|\|\|\|\|\|\|\|\|\|\|\|\|\|\|\|\|\|\|\|\|\|\|\|\|\|\|\|\|\|\|\|\|\|\|\|\|\|\|\|\|\|\|\|\|\|\|\|\|\|\|\|\|\|\|\|\|\|\|\|\|\|\|\|\|\|\|\|\|\|\|\|\|\|\|\|\|\|\|\|\|\|\|\|\|\|\|\|\|\|\|\|\|\|\|\|\|\|\|\|\|\|\|\|\|\|\|\|\|\|\|\|\|\|\|\|\|\|\|\|\|\|\|\|\|\|\|\|\|\|\|\|\|\|\|\|\|\|\|\|\|\|\|\|\|\|\|\|\|\|\|\|\|\|\|\|\|\|\|\|\|\|\|\|\|\|\|\|\|\|\| 	GCCATCGGTGTGCCGCATGATGTTCTCATCGAGATCTGCAGAGAAATCATCTCACGCCAGGATCCCTCGCTCCACGTCGTCGACGCTTGCGCAAGCGGTCCCACTGGTGAGGACGGTCACTGCGACCTGGCAGGCCCAACTTACCCGGACTTCTCGGGCACCGACGTCACGGTGTACAACGGCTGGGTGTGCAAGAACGGAAAATATGTTGGCGAGCACACGCCTTCTGGGCCTCCTTCC ```      ``` 340	GCCATCGGTGTGCCGCATGATGTTCTCATCGAGATCTGCAGAGAAATCATCTCACGCCAGGATCCCTCGCTCCACGTCGTCGACGCTTGCGCAAGCGGTCCCACTGGTGAGGACGGTCACTGCGACCTGGCAGGCCCAACTTACCCGGACTTCTCGGGCACCGACGTCACGGTGTACAACGGCTGGGTGTGCAAGAACGGAAAATATGTTGGCGAGCACACGCCTTCTGGGCCTCCTTCC	579 	\|\|\|\|\|\|\|\|\|\|\|\|\|\|\|\|\|\|\|\|\|\|\|\|\|\|\|\|\|\|\|\|\|\|\|\|\|\|\|\|\|\|\|\|\|\|\|\|\|\|\|\|\|\|\|\|\|\|\|\|\|\|\|\|\|\|\|\|\|\|\|\|\|\|\|\|\|\|\|\|\|\|\|\|\|\|\|\|\|\|\|\|\|\|\|\|\|\|\|\|\|\|\|\|\|\|\|\|\|\|\|\|\|\|\|\|\|\|\|\|\|\|\|\|\|\|\|\|\|\|\|\|\|\|\|\|\|\|\|\|\|\|\|\|\|\|\|\|\|\|\|\|\|\|\|\|\|\|\|\|\|\|\|\|\|\|\|\|\|\|\|\|\|\|\|\|\|\|\|\|\|\|\|\|\|\|\|\|\|\|\|\|\|\| \|\|\|\|\|\|\|\|\|\|\|\|\|\|\|\|\|\| \|  \| \|  \|\|\|\|\|\|\|\|\| \|\|\|\|\|\|\|\| 	GCCATCGGTGTGCCGCATGATGTTCTCATCGAGATCTGCAGAGAAATCATCTCACGCCAGGATCCCTCGCTCCACGTCGTCGACGCTTGCGCAAGCGGTCCCACTGGTGAGGACGGTCACTGCGACCTGGCAGGCCCAACTTACCCGGACTTCTCGGGCACCGACGTCACGGTGTACAACGGCTGGGTGTGCAANAACGGAAAATATGTTGGCNANNA-ANNCCTTCTGGGNCTCCTTCC ``` |
| 3 | ggggacaagtttgtacaaaaaagcaggcttgatgTCGCAGTTGCCGCTTCTCGC, ggggaccactttgtacaagaaagctgggtcCGATAAGTCCTCGCCATATC | Mycgr3\|93371  JGI  Ensembl |  |  | 100 % |  | ``` 237	CGGTCGACTCTACACCATTGCGTCTTCGAAAGCCTGGGATTTCCTTGGGCCCCCATCATCCACCACCGATCAATACTGGGACGATTTATCTGCAGAAGGCCACGAGATCATAATGGTTCGAAGCCAGGAGCTTGAAGCAGCTGGATACAACGCCTCCAAGTACTTTCAAGGTCCAACATCTTGGGGTTTCGGTAAAGACATGTATCCAGTCCAAGTGGACGTCTTCCACCAGATAC	472 	\|\|\|\| \|\| \| \|\|\| \|\|\|\|\|\| \|\| \|\| \|\|\|\|\|\| \|\|\|\|\|\|\|\|\|\|\|\|\|\|\|\|\|\|\|\|\|\|\|\|\|\|\|\|\|\|\|\|\|\|\|\|\|\|\|\|\|\|\|\|\|\|\|\|\|\|\|\|\|\|\|\|\|\|\|\|\|\|\|\|\|\|\|\|\|\|\|\|\|\|\|\|\|\|\|\|\|\|\|\|\|\|\|\|\|\|\|\|\|\|\|\|\|\|\|\|\|\|\|\|\|\|\|\|\|\|\|\|\|\|\|\|\|\|\|\|\|\|\|\|\|\|\|\|\|\|\|\|\|\|\|\|\|\|\|\|\|\|\|\|\|\|\|\|\|\|\|\|\|\|\|\|\|\|\|\|\|\|\|\|\|\|\|\|\|\|\|\|\|\|\|\|\|\|\|\|\|\|\|\|\|\|\|\|\|\|\|\|\|\|\|\|\|\|\|\|\|\| 	CGGTNGANTNTACNCCATTGNGTNTTNGAAAGCNTGGGATTTCCTTGGGCCCCCATCATCCACCACCGATCAATACTGGGACGATTTATCTGCAGAAGGCCACGAGATCATAATGGTTCGAAGCCAGGAGCTTGAAGCAGCTGGATACAACGCCTCCAAGTACTTTCAAGGTCCAACATCTTGGGGTTTCGGTAAAGACATGTATCCAGTCCAAGTGGACGTCTTCCACCAGATAC ```      ``` 473	ACTGTCTCGATCAGCTTCGCAAACAGATGTATGCAGATCACTACTACCCTCGAGATCAAGAGGCAGACTGGCAACACCTGGGTCATTGCTTGCACATCCTGCTTCAGAACATACAGTGCCACGCAGATACCAATCTGATCCCTCATCGCTGGGTTGAGAACAGCTCAAGGCCATATGCGCAGTTCAGCATTGACAAGCAGTGCCGCAACTTCCATGCGTTAAAGAGATGGAACAAGAAGCACAGTGTACTAGTTTCCAACGATATGTGGGCCTCTGCGAAGCCACACGCAGATTCTTACGTGTGGGAAGGATATGGCGAGGACTTATCG	801 	\|\|\|\|\|\|\|\|\|\|\|\|\|\|\|\|\|\|\|\|\|\|\|\|\|\|\|\|\|\|\|\|\|\|\|\|\|\|\|\|\|\|\|\|\|\|\|\|\|\|\|\|\|\|\|\|\|\|\|\|\|\|\|\|\|\|\|\|\|\|\|\|\|\|\|\|\|\|\|\|\|\|\|\|\|\|\|\|\|\|\|\|\|\|\|\|\|\|\|\|\|\|\|\|\|\|\|\|\|\|\|\|\|\|\|\|\|\|\|\|\|\|\|\|\|\|\|\|\|\|\|\|\|\|\|\|\|\|\|\|\|\|\|\|\|\|\|\|\|\|\|\|\|\|\|\|\|\|\|\|\|\|\|\|\|\|\|\|\|\|\|\|\|\|\|\|\|\|\|\|\|\|\|\|\|\|\|\|\|\|\|\|\|\|\|\|\|\|\|\|\|\|\|\|\|\|\|\|\|\|\|\|\|\|\|\|\|\|\|\|\|\|\|\|\|\|\|\|\|\|\|\|\|\|\|\|\|\|\|\|\|\|\|\|\|\|\|\|\|\|\|\|\|\|\|\|\|\|\|\|\|\|\|\|\|\|\|\|\|\|\|\|\|\|\|\|\|\|\|\|\|\|\|\|\|\|\|\|\|\|\|\|\|\|\|\|\|\|\|\|\|\|\|\|\|\|\|\|\|\|\|\|\|\|\|\|\|\|\|\|\|\|\|\|\|\|\|\|\| 	ACTGTCTCGATCAGCTTCGCAAACAGATGTATGCAGATCACTACTACCCTCGAGATCAAGAGGCAGACTGGCAACACCTGGGTCATTGCTTGCACATCCTGCTTCAGAACATACAGTGCCACGCAGATACCAATCTGATCCCTCATCGCTGGGTTGAGAACAGCTCAAGGCCATATGCGCAGTTCAGCATTGACAAGCAGTGCCGCAACTTCCATGCGTTAAAGAGATGGAACAAGAAGCACAGTGTACTAGTTTCCAACGATATGTGGGCCTCTGCGAAGCCACACGCAGATTCTTACGTGTGGGAAGGATATGGCGAGGACTTATCG ``` |
| 3 | ggggacaagtttgtacaaaaaagcaggcttgatgAGATTCATCGCTGCCACTAC, ggggaccactttgtacaagaaagctgggtcGGCATCCTTTGCAACTACGC | Mycgr3\|95478  JGI  Ensembl |  |  | 100 % |  | ``` 289	CGGTCGAAACTTCAGGAGCGGAAGAAGCGCGGCGAAATCGCGAACAATCCCAATCCAGTCTACACGGTACCTGAAGTCGAACGGTCGAAACTTCAGGAGCGGAAGAAGCGCGGCGAAATCGCGAACAATCCCAATCCAGTCTACACGGTACCTGAAGTCGAACGGTCGAAACTTCAGGAGCGGAAGAAGCGCGGCGAAATCGCGAACAATCCCAATCCAGTCTACACGGTACCTGAAGTCGAACGGTCGAGACTTCAGGAGCGGAAGAAGCGCGGCGAAATCGCGAACAATCCTAATCCAGTCTACCCTGGACCTG	604 	\|\|\|\| \| \|\|\|\|\| \|\|\|\|\| \|\|\|\|\|\|\|\|\| \|\|\| \|\|\|\|\| \| \|\|\|\|\|\|\|\|\|\|\|\|\|\|\|\|\|\| \|\|\|\|\|\|\|\|\|\| \|\|\|\|\|\|\|\|\| \|\|\|\| \|\|\|\| \|\|\|\|\|\|\|\|\|\|\|\|\|\|\|\|\|\|\|\|\|\| \|\|\|\|\|\|\|\|\|\|\|\|\|\|\|\|\|\|\| \|\|\|\|\|\| \|\|\|\|\|\|\|\|\|\|\|\|\|\|\|\|\|\|\|\|\|\|\|\|\|\|\|\|\|\|\|\|\|\|\|\|\|\|\|\|\|\|\|\|\|\|\|\|\|\|\|\|\|\|\|\|\|\|\|\|\|\|\|\|\|\|\|\|\|\|\|\|\|\|\|\|\|\|\|\|\|\|\|\|\|\|\|\|\|\|\|\|\|\|\|\|\|\|\|\|\|\|\|\|\|\|\|\|\| \|\|\|\|\|\|\|\|\|\|\|\|\|\|\|\|\|\|\|\|\|\|\|\|\|\|\|\|\|\|\|\|\|\|\|\|\|\|\|\|\|\| \|\|\|\|\|\|\|\|\|\|\|\| \| \| \|\|\|\|\| 	CGGTTGGAACTTGAGGAGNGGAAGAAGCNCGGNGAAATNGNGAACAATCCCAATCCAGTNTACACGGTACNTGAAGTCGANCGGTNGAAANTTCAGGAGCGGAAGAAGCGCGGNGAAATCGCGAACAATCCCA-TCCAGTNTACACGGTACCTGAAGTCGAACGGTCGAAACTTCAGGAGCGGAAGAAGCGCGGCGAAATCGCGAACAATCCCAATCCAGTCTACACGGTACCTGAAGTCGAACGGTCGAAACTTCAGGAGCGGAAGAAGCGCGGCGAAATCGCGAACAATCCCAATCCAGTCTACACGGTACCTG ```      ``` 98	ATCCAGTCTACCTAGGAGCTAGCATTAAGCGGTCGGAACTTGAGGAGCGGAAGAAGCGCGGCGAAATCGCGAACAATCCCAATCCAGTCTACACGGTACCTGAAGTCGAACGGTCGAAACTTCAGGAGCGGAAGAAGCGCGGCGAAATCGCGAACAATCCCAATCCAGTCTACACGGTACCTGAAGTCGAACGGTCGAAACTTCAGGAGCGGAAGAAGCGCGGCGAAATCGCGAACAATCCCAATCCAGTCTACACGGTACCTGAAGTCGAACGGTCGAAACTTCAGGAGCGGAAGAAGCGCGGCGAAATCGCGAACAATCCCAATCCAGTCTACACGGTACCTGAAGTCGAACGGTCGAAACTTCAGGAGCGGAAGAAGCGCGGCGAAATCGCGAACAATCCCAATCCAGTCTACACGGTACCTGAAGTCGAACGGTCGAGACTTCAGGAGCGGAAGAAGCGCGGCGAAATCGCGAACAATCCTAATCCAGTCTACCCTGGACCTGGCGTGGAAAGAGCAAAGCTCGAGGAGCATGGCAAAAGCCCAACCGCCGACGAAACGATCGCGAAGCGCCAAACTCCAAAGTCGAGCGTAGTTGCAAAGGATGCC	708 	\|\|\|\|\|\|\| \|  \|\| \|\|\|\| \|\|\|\|\|\|\|\|\|\|\|\|\|\| \|\|\|\|\|\|\|\|\|\|\|\|\| \|\|\|\|\|\|\|\|\| \|\|\| \|\|\|\|\| \| \|\|\|\|\|\|\|\|\|\|\|\|\|\|\|\|\|\| \|\|\|\|\|\|\|\|\|\| \|\|\|\|\|\|\|\|\| \|\|\|\| \|\|\|\| \|\|\|\|\|\|\|\|\|\|\|\|\|\|\|\|\|\|\|\|\|\| \|\|\|\|\|\|\|\|\|\|\|\|\|\|\|\|\|\|\| \|\|\|\|\|\| \|\|\|\|\|\|\|\|\|\|\|\|\|\|\|\|\|\|\|\|\|\|\|\|\|\|\|\|\|\|\|\|\|\|\|\|\|\|\|\|\|\|\|\|\|\|\|\|\|\|\|\|\|\|\|\|\|\|\|\|\|\|\|\|\|\|\|\|\|\|\|\|\|\|\|\|\|\|\|\|\|\|\|\|\|\|\|\|\|\|\|\|\|\|\|\|\|\|\|\|\|\|\|\|\|\|\|\|\|\|\|\|\|\|\|\|\|\|\|\|\|\|\|\|\|\|\|\|\|\|\|\|\|\|\|\|\|\|\|\|\|\|\|\|\|\|\|\|\|\|\|\|\|\|\|\|\|\|\|\|\|\|\|\|\|\|\|\|\|\|\|\|\|\|\|\|\|\|\|\|\|\|\|\|\|\|\|\|\|\|\|\|\|\|\|\|\|\|\|\|\|\|\|\|\|\|\|\|\|\|\|\|\|\|\|\|\|\|\|\|\|\|\|\|\|\|\|\|\|\|\|\|\|\|\|\|\|\|\|\|\|\|\|\|\|\|\|\|\|\|\|\|\|\|\|\|\|\|\|\|\|\|\|\|\|\|\|\|\|\|\|\|\|\|\|\|\|\|\|\|\|\|\|\|\|\|\|\|\|\|\|\|\|\|\|\|\|\|\|\|\|\|\|\|\|\|\|\|\|\|\|\|\|\|\|\|\|\|\|\|\|\|\|\|\|\|\|\|\|\|\|\|\|\|\|\|\|\|\|\|\|\|\|\|\|\|\|\|\|\|\|\|\|\|\|\|\|\|\|\|\|\|\|\|\|\|\|\|\|\|\|\|\|\|\|\|\|\|\|\|\|\|\|\|\|\|\|\|\|\|\|\|\|\|\|\|\|\|\|\|\|\|\|\|\|\|\|\|\|\|\|\|\|\|\|\|\|\|\|\|\|\|\|\|\|\|\|\|\|\|\|\|\|\|\|\|\|\|\|\|\| 	ATCCAGTNT-NCTNGGAG-TAGCATTAAGCGGTTGGAACTTGAGGAGNGGAAGAAGCNCGGNGAAATNGNGAACAATCCCAATCCAGTNTACACGGTACNTGAAGTCGANCGGTNGAAANTTCAGGAGCGGAAGAAGCGCGGNGAAATCGCGAACAATCCCA-TCCAGTNTACACGGTACCTGAAGTCGAACGGTCGAAACTTCAGGAGCGGAAGAAGCGCGGCGAAATCGCGAACAATCCCAATCCAGTCTACACGGTACCTGAAGTCGAACGGTCGAAACTTCAGGAGCGGAAGAAGCGCGGCGAAATCGCGAACAATCCCAATCCAGTCTACACGGTACCTGAAGTCGAACGGTCGAAACTTCAGGAGCGGAAGAAGCGCGGCGAAATCGCGAACAATCCCAATCCAGTCTACACGGTACCTGAAGTCGAACGGTCGAGACTTCAGGAGCGGAAGAAGCGCGGCGAAATCGCGAACAATCCTAATCCAGTCTACCCTGGACCTGGCGTGGAAAGAGCAAAGCTCGAGGAGCATGGCAAAAGCCCAACCGCCGACGAAACGATCGCGAAGCGCCAAACTCCAAAGTCGAGCGTAGTTGCAAAGGATGCC ```      ``` 87	GAACAATCCCGATCCAGTCTACCTAGG-AGCTAGCA-TTAAGCGGTCGGAACTTGAGGAGCGGAAGAAGCGCGGCGAAATCGCGAACAATCCCAATCCAGTCTACACGGTACCTGAAGTCGAACGGTCGAAACTTCAGGAGCGGAAGAAGCGCGGCGAAATCGCGAACAATCCCAATCCAGTCTACACGGTACCTG	280 	\|\|\|\|\|\|\|\|\|\| \|\|\|\|\|\|\|\|\|\|\|   \|\| \| \|\| \| \| \|  \| \|\|\|\|\|\| \|\|\|\|\| \|\|\|\|\|\|\|\|\|\|\|\|\|\|\|\|\|\|\|\|\|\|\|\|\|\|\|\|\|\|\|\|\|\|\|\|\|\|\|\|\|\|\|\|\|\|\|\|\|\|\|\|\|\|\|\|\|\|\|\|\|\|\|\|\|\|\|\|\|\|\|\|\|\|\| \|\|\|\|\|\|\|\|\|\|\|\|\|\|\|\|\|\|\|\|\|\|\|\|\|\|\|\|\|\|\|\|\|\|\|\|\|\|\|\|\|\| \|\|\|\|\|\|\|\|\|\|\|\| \| \| \|\|\|\|\| 	GAACAATCCCAATCCAGTCTAC-ACGGTACCT-GAAGTCGAACGGTCGAAACTTCAGGAGCGGAAGAAGCGCGGCGAAATCGCGAACAATCCCAATCCAGTCTACACGGTACCTGAAGTCGAACGGTCGAGACTTCAGGAGCGGAAGAAGCGCGGCGAAATCGCGAACAATCCTAATCCAGTCTACCCTGGACCTG ```      ``` 87	GAACAATCCCGATCCAGTCTACCTAGG-AGCTAGCA-TTAAGCGGTCGGAACTTGAGGAGCGGAAGAAGCGCGGCGAAATCGCGAACAATCCCAATCCAGTCTACACGGTACCTGAAGTCGAACGGTCGAAACTTCAGGAGCGGAAGAAGCGCGGCGAAATCGCGAACAATCCCAATCCAGTCTACACGGTACCTGAAGTCGAACGGTCGAAACTTCAGGAGCGGAAGAAGCGCGGCGAAATCGCGAACAATCCCAATCCAGTCTACACGGTACCTG	361 	\|\|\|\|\|\|\|\|\|\| \|\|\|\|\|\|\|\|\|\|\|   \|\| \| \|\| \| \| \|  \| \|\|\|\|\|\| \|\|\|\|\| \|\|\|\|\|\|\|\|\|\|\|\|\|\|\|\|\|\|\|\|\|\|\|\|\|\|\|\|\|\|\|\|\|\|\|\|\|\|\|\|\|\|\|\|\|\|\|\|\|\|\|\|\|\|\|\|\|\|\|\|\|\|\|\|\|\|\|\|\|\|\|\|\|\|\|\|\|\|\|\|\|\|\|\|\|\|\|\|\|\|\|\|\|\|\|\|\|\|\|\|\|\|\|\|\|\|\|\|\|\|\|\|\|\|\|\|\|\|\|\|\|\|\|\|\|\|\|\|\|\|\|\|\|\|\|\|\|\|\|\|\|\|\|\|\|\|\|\|\|\|\|\|\|\|\|\| \|\|\|\|\|\|\|\|\|\|\|\|\|\|\|\|\|\|\|\|\|\|\|\|\|\|\|\|\|\|\|\|\|\|\|\|\|\|\|\|\|\| \|\|\|\|\|\|\|\|\|\|\|\| \| \| \|\|\|\|\| 	GAACAATCCCAATCCAGTCTAC-ACGGTACCT-GAAGTCGAACGGTCGAAACTTCAGGAGCGGAAGAAGCGCGGCGAAATCGCGAACAATCCCAATCCAGTCTACACGGTACCTGAAGTCGAACGGTCGAAACTTCAGGAGCGGAAGAAGCGCGGCGAAATCGCGAACAATCCCAATCCAGTCTACACGGTACCTGAAGTCGAACGGTCGAGACTTCAGGAGCGGAAGAAGCGCGGCGAAATCGCGAACAATCCTAATCCAGTCTACCCTGGACCTG ```      ``` 208	CGGTCGAAACTTCAGGAGCGGAAGAAGCGCGGCGAAATCGCGAACAATCCCAATCCAGTCTACACGGTACCTGAAGTCGAACGGTCGAAACTTCAGGAGCGGAAGAAGCGCGGCGAAATCGCGAACAATCCCAATCCAGTCTACACGGTACCTGAAGTCGAACGGTCGAAACTTCAGGAGCGGAAGAAGCGCGGCGAAATCGCGAACAATCCCAATCCAGTCTACACGGTACCTGAAGTCGAACGGTCGAAACTTCAGGAGCGGAAGAAGCGCGGCGAAATCGCGAACAATCCCAATCCAGTCTACACGGTACCTGAAGTCGAACGGTCGAGACTTCAGGAGCGGAAGAAGCGCGGCGAAATCGCGAACAATCCTAATCCAGTCTACCCTGGACCTG	604 	\|\|\|\| \| \|\|\|\|\| \|\|\|\|\| \|\|\|\|\|\|\|\|\| \|\|\| \|\|\|\|\| \| \|\|\|\|\|\|\|\|\|\|\|\|\|\|\|\|\|\| \|\|\|\|\|\|\|\|\|\| \|\|\|\|\|\|\|\|\| \|\|\|\| \|\|\|\| \|\|\|\|\|\|\|\|\|\|\|\|\|\|\|\|\|\|\|\|\|\| \|\|\|\|\|\|\|\|\|\|\|\|\|\|\|\|\|\|\| \|\|\|\|\|\| \|\|\|\|\|\|\|\|\|\|\|\|\|\|\|\|\|\|\|\|\|\|\|\|\|\|\|\|\|\|\|\|\|\|\|\|\|\|\|\|\|\|\|\|\|\|\|\|\|\|\|\|\|\|\|\|\|\|\|\|\|\|\|\|\|\|\|\|\|\|\|\|\|\|\|\|\|\|\|\|\|\|\|\|\|\|\|\|\|\|\|\|\|\|\|\|\|\|\|\|\|\|\|\|\|\|\|\|\|\|\|\|\|\|\|\|\|\|\|\|\|\|\|\|\|\|\|\|\|\|\|\|\|\|\|\|\|\|\|\|\|\|\|\|\|\|\|\|\|\|\|\|\|\|\|\|\|\|\|\|\|\|\|\|\|\|\|\|\|\|\|\|\|\|\|\|\|\|\|\|\|\|\|\|\|\|\|\|\|\| \|\|\|\|\|\|\|\|\|\|\|\|\|\|\|\|\|\|\|\|\|\|\|\|\|\|\|\|\|\|\|\|\|\|\|\|\|\|\|\|\|\| \|\|\|\|\|\|\|\|\|\|\|\| \| \| \|\|\|\|\| 	CGGTTGGAACTTGAGGAGNGGAAGAAGCNCGGNGAAATNGNGAACAATCCCAATCCAGTNTACACGGTACNTGAAGTCGANCGGTNGAAANTTCAGGAGCGGAAGAAGCGCGGNGAAATCGCGAACAATCCCA-TCCAGTNTACACGGTACCTGAAGTCGAACGGTCGAAACTTCAGGAGCGGAAGAAGCGCGGCGAAATCGCGAACAATCCCAATCCAGTCTACACGGTACCTGAAGTCGAACGGTCGAAACTTCAGGAGCGGAAGAAGCGCGGCGAAATCGCGAACAATCCCAATCCAGTCTACACGGTACCTGAAGTCGAACGGTCGAAACTTCAGGAGCGGAAGAAGCGCGGCGAAATCGCGAACAATCCCAATCCAGTCTACACGGTACCTG ```      ``` 127	CGGTCGGAACTTGAGGAGCGGAAGAAGCGCGGCGAAATCGCGAACAATCCCAATCCAGTCTACACGGTACCTGAAGTCGAACGGTCGAAACTTCAGGAGCGGAAGAAGCGCGGCGAAATCGCGAACAATCCCAATCCAGTCTACACGGTACCTGAAGTCGAACGGTCGAAACTTCAGGAGCGGAAGAAGCGCGGCGAAATCGCGAACAATCCCAATCCAGTCTACACGGTACCTGAAGTCGAACGGTCGAAACTTCAGGAGCGGAAGAAGCGCGGCGAAATCGCGAACAATCCCAATCCAGTCTACACGGTACCTGAAGTCGAACGGTCGAAACTTCAGGAGCGGAAGAAGCGCGGCGAAATCGCGAACAATCCCAATCCAGTCTACACGGTACCTG	523 	\|\|\|\| \| \|\| \|\| \|\|\|\|\|\|\|\|\|\|\|\|\|\|\|\|\|\|\| \|\|\|\|\|\|\|\|\|\|\|\|\|\|\|\|\|\|\| \|\|\|\|\|\| \|\|\|\|\|\|\|\|\|\|\|\|\|\|\|\|\|\|\|\|\|\|\|\|\|\|\|\|\|\|\|\|\|\|\|\|\|\|\|\|\|\|\|\|\|\|\|\|\|\|\|\|\|\|\|\|\|\|\|\|\|\|\|\|\|\|\|\|\|\|\|\|\|\|\|\|\|\|\|\|\|\|\|\|\|\|\|\|\|\|\|\|\|\|\|\|\|\|\|\|\|\|\|\|\|\|\|\|\|\|\|\|\|\|\|\|\|\|\|\|\|\|\|\|\|\|\|\|\|\|\|\|\|\|\|\|\|\|\|\|\|\|\|\|\|\|\|\|\|\|\|\|\|\|\|\|\|\|\|\|\|\|\|\|\|\|\|\|\|\|\|\|\|\|\|\|\|\|\|\|\|\|\|\|\|\|\|\|\|\|\|\|\|\|\|\|\|\|\|\|\|\|\|\|\|\|\|\|\|\|\|\|\|\|\|\|\|\|\|\|\|\|\|\|\|\|\|\|\|\|\|\|\|\|\|\|\|\|\|\|\|\|\|\|\|\|\|\|\|\|\|\|\|\|\|\|\|\|\|\|\|\|\|\|\|\|\|\|\|\|\| \|\|\|\|\|\|\|\|\|\|\|\|\|\|\|\|\|\|\|\|\|\|\|\|\|\|\|\|\|\|\|\|\|\|\|\|\|\|\|\|\|\| \|\|\|\|\|\|\|\|\|\|\|\| \| \| \|\|\|\|\| 	CGGTNGAAANTTCAGGAGCGGAAGAAGCGCGGNGAAATCGCGAACAATCCCA-TCCAGTNTACACGGTACCTGAAGTCGAACGGTCGAAACTTCAGGAGCGGAAGAAGCGCGGCGAAATCGCGAACAATCCCAATCCAGTCTACACGGTACCTGAAGTCGAACGGTCGAAACTTCAGGAGCGGAAGAAGCGCGGCGAAATCGCGAACAATCCCAATCCAGTCTACACGGTACCTGAAGTCGAACGGTCGAAACTTCAGGAGCGGAAGAAGCGCGGCGAAATCGCGAACAATCCCAATCCAGTCTACACGGTACCTGAAGTCGAACGGTCGAGACTTCAGGAGCGGAAGAAGCGCGGCGAAATCGCGAACAATCCTAATCCAGTCTACCCTGGACCTG ```      ``` 87	GAACAATCCCGATCCAGTCTACCTAGG-AGCTAGCA-TTAAGCGGTCGGAACTTGAGGAGCGGAAGAAGCGCGGCGAAATCGCGAACAATCCCAATCCAGTCTACACGGTACCTGAAGTCGAACGGTCGAAACTTCAGGAGCGGAAGAAGCGCGGCGAAATCGCGAACAATCCCAATCCAGTCTACACGGTACCTGAAGTCGAACGGTCGAAACTTCAGGAGCGGAAGAAGCGCGGCGAAATCGCGAACAATCCCAATCCAGTCTACACGGTACCTGAAGTCGAACGGTCGAAACTTCAGGAGCGGAAGAAGCGCGGCGAAATCGCGAACAATCCCAATCCAGTCTACACGGTACCTG	442 	\|\|\|\|\|\|\|\|\|\| \|\|\|\|\|\|\| \|\| \|  \|\| \| \|\| \| \| \|  \| \|\|\|\|\|\| \|\|\|\|\| \|\|\|\|\|\|\|\|\|\|\|\|\|\|\|\|\|\|\|\|\|\|\|\|\|\|\|\|\|\|\|\|\|\|\|\|\|\|\|\|\|\|\|\|\|\|\|\|\|\|\|\|\|\|\|\|\|\|\|\|\|\|\|\|\|\|\|\|\|\|\|\|\|\|\|\|\|\|\|\|\|\|\|\|\|\|\|\|\|\|\|\|\|\|\|\|\|\|\|\|\|\|\|\|\|\|\|\|\|\|\|\|\|\|\|\|\|\|\|\|\|\|\|\|\|\|\|\|\|\|\|\|\|\|\|\|\|\|\|\|\|\|\|\|\|\|\|\|\|\|\|\|\|\|\|\|\|\|\|\|\|\|\|\|\|\|\|\|\|\|\|\|\|\|\|\|\|\|\|\|\|\|\|\|\|\|\|\|\|\|\|\|\|\|\|\|\|\|\|\|\|\|\|\|\|\|\|\|\|\|\|\|\|\|\|\|\|\|\|\|\|\|\|\|\|\|\|\|\|\|\|\|\|\|\|\|\| \|\|\|\|\|\|\|\|\|\|\|\|\|\|\|\|\|\|\|\|\|\|\|\|\|\|\|\|\|\|\|\|\|\|\|\|\|\|\|\|\|\| \|\|\|\|\|\|\|\|\|\|\|\| \| \| \|\|\|\|\| 	GAACAATCCC-ATCCAGTNTA-CACGGTACCT-GAAGTCGAACGGTCGAAACTTCAGGAGCGGAAGAAGCGCGGCGAAATCGCGAACAATCCCAATCCAGTCTACACGGTACCTGAAGTCGAACGGTCGAAACTTCAGGAGCGGAAGAAGCGCGGCGAAATCGCGAACAATCCCAATCCAGTCTACACGGTACCTGAAGTCGAACGGTCGAAACTTCAGGAGCGGAAGAAGCGCGGCGAAATCGCGAACAATCCCAATCCAGTCTACACGGTACCTGAAGTCGAACGGTCGAGACTTCAGGAGCGGAAGAAGCGCGGCGAAATCGCGAACAATCCTAATCCAGTCTACCCTGGACCTG ```      ``` 370	CGGTCGAAACTTCAGGAGCGGAAGAAGCGCGGCGAAATCGCGAACAATCCCAATCCAGTCTACACGGTACCTGAAGTCGAACGGTCGAAACTTCAGGAGCGGAAGAAGCGCGGCGAAATCGCGAACAATCCCAATCCAGTCTACACGGTACCTGAAGTCGAACGGTCGAGACTTCAGGAGCGGAAGAAGCGCGGCGAAATCGCGAACAATCCTAATCCAGTCTACCCTGGACCTG	604 	\|\|\|\| \| \|\|\|\|\| \|\|\|\|\| \|\|\|\|\|\|\|\|\| \|\|\| \|\|\|\|\| \| \|\|\|\|\|\|\|\|\|\|\|\|\|\|\|\|\|\| \|\|\|\|\|\|\|\|\|\| \|\|\|\|\|\|\|\|\| \|\|\|\| \|\|\|\| \|\|\|\|\|\|\|\|\|\|\|\|\|\|\|\|\|\|\|\|\|\| \|\|\|\|\|\|\|\|\|\|\|\|\|\|\|\|\|\|\| \|\|\|\|\|\| \|\|\|\|\|\|\|\|\|\|\|\|\|\|\|\|\|\|\|\|\|\|\|\|\|\|\|\| \|\|\|\|\|\|\|\|\|\|\|\|\|\|\|\|\|\|\|\|\|\|\|\|\|\|\|\|\|\|\|\|\|\|\|\|\|\|\|\|\|\| \|\|\|\|\|\|\|\|\|\|\|\| \| \| \|\|\|\|\| 	CGGTTGGAACTTGAGGAGNGGAAGAAGCNCGGNGAAATNGNGAACAATCCCAATCCAGTNTACACGGTACNTGAAGTCGANCGGTNGAAANTTCAGGAGCGGAAGAAGCGCGGNGAAATCGCGAACAATCCCA-TCCAGTNTACACGGTACCTGAAGTCGAACGGTCGAAACTTCAGGAGCGGAAGAAGCGCGGCGAAATCGCGAACAATCCCAATCCAGTCTACACGGTACCTG ```      ``` 451	CGGTCGAAACTTCAGGAGCGGAAGAAGCGCGGCGAAATCGCGAACAATCCCAATCCAGTCTACACGGTACCTGAAGTCGAACGGTCGAGACTTCAGGAGCGGAAGAAGCGCGGCGAAATCGCGAACAATCCTAATCCAGTCTACCCTGGACCTG	604 	\|\|\|\| \| \|\|\|\|\| \|\|\|\|\| \|\|\|\|\|\|\|\|\| \|\|\| \|\|\|\|\| \| \|\|\|\|\|\|\|\|\|\|\|\|\|\|\|\|\|\| \|\|\|\|\|\|\|\|\|\| \|\|\|\|\|\|\|\|\| \|\|\|\| \|\| \| \|\|\|\|\|\|\|\|\|\|\|\|\|\|\|\|\|\|\|\|\|\| \|\|\|\|\|\|\|\|\|\|\|\|\|\|\|\|\| \| \|\|\|\|\|\| \|\|\| \| \| \|\|\|\|\| 	CGGTTGGAACTTGAGGAGNGGAAGAAGCNCGGNGAAATNGNGAACAATCCCAATCCAGTNTACACGGTACNTGAAGTCGANCGGTNGAAANTTCAGGAGCGGAAGAAGCGCGGNGAAATCGCGAACAATCCCA-TCCAGTNTACACGGTACCTG ```      ``` 87	GAACAATCCCGATCCAGTCTACCTAGG-AGCTAGCA-TTAAGCGGTCGGAACTTGAGGAGCGGAAGAAGCGCGGCGAAATCGCGAACAATCCCAATCCAGTCTACACGGTACCTG	199 	\|\|\|\|\|\|\|\|\|\| \|\|\|\|\|\|\|\|\|\|\|   \|\| \| \|\| \| \| \|  \| \|\|\|\|\|\|  \|\|\|\| \|\|\|\|\|\|\|\|\|\|\|\|\|\|\|\|\|\|\|\|\|\|\|\|\|\|\|\|\|\|\|\|\|\|\|\|\| \|\|\|\|\|\|\|\|\|\|\|\| \| \| \|\|\|\|\| 	GAACAATCCCAATCCAGTCTAC-ACGGTACCT-GAAGTCGAACGGTCGAGACTTCAGGAGCGGAAGAAGCGCGGCGAAATCGCGAACAATCCTAATCCAGTCTACCCTGGACCTG ``` |
| 3 | ggggacaagtttgtacaaaaaagcaggcttgatgCAGTTCACTCTCTCCCTCCT, ggggaccactttgtacaagaaagctgggtcAGCGCCAGTGTTCGGACCCG | Mycgr3\|95537  JGI  Ensembl |  |  | 3.51852 % |  |  |
| 3 | ggggacaagtttgtacaaaaaagcaggcttgatgCAGATCACGAACACCTTCCT, ggggaccactttgtacaagaaagctgggtcGCACGTGATCGATATCGAAG | Mycgr3\|95672  JGI  Ensembl |  |  | 0 % |  |  |
| 3 | ggggacaagtttgtacaaaaaagcaggcttgatgCAATATTCCGCCATCACCCT, ggggaccactttgtacaagaaagctgggtcGACATAGAGGCAGCTGATAC | Mycgr3\|95831  JGI  Ensembl |  |  | 100 % |  | ``` 155	GCATCTTCATCGGTGGTCCCCAATTCGCTCAATGCGTGAACAACAAGCGTCCAAACTCCGCCACCTTCGCCAGATACTCCGACGACACCATCTCACTATGGCACACCGAGAATCCAACTCAACAACTCTGGGTGAGCCCCTCTTCCTCAGGCTCAGGTGAAGGAGTCGTTGGATACACCACTGGAAATGGTGCTACACCTCATCACGACTTGCGAAGCCCCTTCAAGATCGACAAGAATGGACGCCTGACATTCAACGGTGTTGGAGCCAAAGCTTGCCCGGGTACGGGCAACAGATACCTTCTCTATTTCACCAACCGCCAATTGGAGAAGGGATGCGTCGACGTCACGCTCCAAACTTTCAAGACTCCTGCTAGTATCAGCTGCCTCTATGTC	549 	\|\|\|\| \|\|\|\|\| \|\| \|\|\|\|\| \|\| \|\| \| \|\|\|\|\|\| \|\|\|\| \|\|\|\|\|\|\|\|\|\|\|\|\|\|  \|\| \|\|\|\|\| \|\|\|\|\|\|\|\|\|\|\| \|\|\|\|\| \|\|\|\|\|\|\|\|\|\|\|\|\|\|\|\|\|\|\|\|\|\|\|\|\|\|\|\|\|\|\|\|\|\|\|\|\|\|\|\|\|\|\|\|\|\|\|\|\|\|\|\|\|\|\|\|\|\|\|\|\|\|\|\|\|\|\|\|\|\|\|\|\|\|\|\|\|\|\|\|\|\|\|\|\|\|\|\|\|\|\|\|\|\|\|\|\|\|\|\|\|\|\|\|\|\|\|\|\|\|\|\|\|\|\|\|\|\|\|\|\|\|\|\|\|\|\|\|\|\|\|\|\|\|\|\|\|\|\|\|\|\|\|\|\|\|\|\|\|\|\|\|\|\|\|\|\|\|\|\|\|\|\|\|\|\|\|\|\|\|\|\|\|\|\|\|\|\|\|\|\|\|\|\|\|\|\|\|\|\|\|\|\|\|\|\|\|\|\|\|\|\|\|\|\|\|\|\|\|\|\|\|\|\|\|\|\|\|\|\|\|\|\|\|\|\|\|\|\|\|\|\|\|\|\|\|\|\|\|\|\|\|\|\|\|\|\|\|\|\|\|\|\|\|\|\|\|\|\|\|\|\|\|\|\|\|\|\|\|\|\|\|\|\|\|\|\|\|\|\|\|\|\|\|\|\|\|\|\|\|\|\|\|\|\|\|\|\|\|\|\|\|\|\|\|\|\|\|\|\|\|\| 	GCATNTTCATNGGNGGTCCNCANTTNGNTCAATGNGTGANCAACAAGCGTCCAANNTCNGCCACNTTCGCCAGATANTCCGANGACACCATCTCACTATGGCACACCGAGAATCCAACTCAACAACTCTGGGTGAGCCCCTCTTCCTCAGGCTCAGGTGAAGGAGTCGTTGGATACACCACTGGAAATGGTGCTACACCTCATCACGACTTGCGAAGCCCCTTCAAGATCGACAAGAATGGACGCCTGACATTCAACGGTGTTGGAGCCAAAGCTTGCCCGGGTACGGGCAACAGATACCTTCTCTATTTCACCAACCGCCAATTGGAGAAGGGATGCGTCGACGTCACGCTCCAAACTTTCAAGACTCCTGCTAGTATCAGCTGCCTCTATGTC ```      ``` 1	ATGCAATATTCCGCCATCACCCTCATGACTTTGGTCTCCTCTGTCCTGGCCGTTCCAACTACACAGTCCAATGTGCCCGGCTTCAACGAGAAATTTGGAATCAAGGCCCACGGCAAAGGCATCAACGGCATCAACGTTGTAGCCACTGGCGACCGCATCTTCATCGGTGGTCCCCAATTCGCTCAATGCGTGAACAACAAGCGTCCAAACTCCGCCACCTTCGCCAGATACTCCGACGACACCATCTCACTATGGCACACCGAGAATCCAACTCAACAACTCTGGGTGAGCCCCTCTTCCTCAGGCTCAGGTGAAGGAGTCGTTGGATACACCACTGGAAATGGTGCTACACCTCATCACGACTTGCGAAGCCCCTTCAAGATCGACAAGAATGGACGCCTGACATTCAACGGTGTTGGAGCCAAAGCTTGCCCGGGTACGGGCAACAGATACCTTCTCTATTTCACCAACCGCCAATTGGAGAAGGGATGCGTCGACGTCACGCTCCAAACTTTCAAGACTCCTGCTAGTATCAGCTGCCTCTATGTC	549 	\|\|\|\|\|\|\|\|\|\|\|\|\|\|\|\|\|\|\|\|\|\|\|\|\|\|\|\|\|\|\|\|\|\|\|\|\|\|\|\|\|\|\|\|\|\|\|\|\|\|\|\|\|\|\|\|\|\|\|\|\|\|\|\|\|\|\|\|\|\|\|\|\|\|\|\|\|\|\|\|\|\|\|\|\|\|\|\|\|\|\|\|\|\|\|\|\|\|\|\|\|\|\|\|\|\|\|\|\|\|\|\|\|\|\|\|\|\|\|\|\|\|\|\|\|\|\|\|\|\|\|\|\|\|\|\|\|\|\|\|\|\|\|\|\|\|\|\|\|\|\|\|\|\|\|\|\|\|\|\|\|\|\|\|\|\|\|\|\|\|\|\|\|\|\|\|\|\|\|\|\|\|\|\|\|\|\|\|\|\|\|\|\|\|\|\|\|\|\|\|\|\|\|\|\|\|\|\|\|\|\|\|\|\|\|\|\|\|\|\|\|\|\|\|\|\|\|\|\|\|\|\|\|\|\|\|\|\|\|\|\|\|\|\|\|\|\|\|\|\|\|\|\|\|\|\|\|\|\|\|\|\|\|\|\|\|\|\|\|\|\|\|\|\|\|\|\|\|\|\|\|\|\|\|\|\|\|\|\|\|\|\|\|\|\|\|\|\|\|\|\|\|\|\|\|\|\|\|\|\|\|\|\|\|\|\|\|\|\|\|\|\|\|\|\|\|\|\|\|\|\|\|\|\|\|\|\|\|\|\|\|\|\|\|\|\|\|\|\|\|\|\|\|\|\|\|\|\|\|\|\|\|\|\|\|\|\|\|\|\|\|\|\|\|\|\|\|\|\|\|\|\|\|\|\|\|\|\|\|\|\|\|\|\|\|\|\|\|\|\|\|\|\|\|\|\|\|\|\|\|\|\|\|\|\|\|\|\|\|\|\|\|\|\|\|\|\|\|\|\|\|\|\|\|\|\|\|\|\|\|\|\|\|\|\|\|\|\|\|\|\|\|\|\|\|\|\|\|\|\|\|\|\|\|\|\|\|\|\|\|\|\|\|\|\|\|\|\|\|\|\|\|\|\|\|\|\|\|\|\|\|\|\|\|\|\|\|\|\|\|\|\|\|\|\|\|\|\|\|\|\|\|\|\|\|\|\|\|\|\|\|\|\|\|\|\|\|\|\|\|\|\|\|\|\|\|\|\|\|\|\|\|\|\|\|\|\|\|\| 	ATGCAATATTCCGCCATCACCCTCATGACTTTGGTCTCCTCTGTCCTGGCCGTTCCAACTACACAGTCCAATGTGCCCGGCTTCAACGAGAAATTTGGAATCAAGGCCCACGGCAAAGGCATCAACGGCATCAACGTTGTAGCCACTGGCGACCGCATCTTCATCGGTGGTCCCCAATTCGCTCAATGCGTGAACAACAAGCGTCCAAACTCCGCCACCTTCGCCAGATACTCCGACGACACCATCTCACTATGGCACACCGAGAATCCAACTCAACAACTCTGGGTGAGCCCCTCTTCCTCAGGCTCAGGTGAAGGAGTCGTTGGATACACCACTGGAAATGGTGCTACACCTCATCACGACTTGCGAAGCCCCTTCAAGATCGACAAGAATGGACGCCTGACATTCAACGGTGTTGGAGCCAAAGCTTGCCCGGGTACGGGCAACAGATACCTTCTCTATTTCACCAACCGCCAATTGGAGAAGGGATGCGTCGACGTCACGCTCCAAACTTTCAAGACTCCTGCTAGTATCAGCTGCCTCTATGTC ``` |
| 3 | ggggacaagtttgtacaaaaaagcaggcttgatgCAGTTCATCATCCTCGCCCT, ggggaccactttgtacaagaaagctgggtcGTTGGACTTGACCTCCTCGC | Mycgr3\|96543  JGI  Ensembl | KOG3070 Predicted RNA-binding protein containing PIN domain and invovled in translation or RNA processing; | cell communication; cell wall; extracellular region; | 100 % |  | ``` 1	ATGCAGTTCATCATCCTCGCCCTTGCTGCCCTCGCGGCTGCCGCTCCTGGCGGCGCTCCCAGCTACGGTGGCAGTGGCAGTGGCATGGCCGGTGGCATGGCCGGTGGCCACGGAGGTAGCGGCGGCTACTCTGGCGGCTCCAACAACGGTGGCTCCAACGGTGGTTCCAACGGCGGCTCCTCCAACGGTGGTTCCAACGGTGGTTCCAACGGAGGCAAATTCCAATGCTCTGCGCCCCTCCAGAGCAGCCCACAGTGCTGCGCTGTCAACGCGCTCGGGGTTGCTTCCCTTCCCTGCAATGCCC	304 	\|\|\|\|\|\|\|\|\|\|\|\|\|\|\|\|\|\|\|\|\|\|\|\|\|\|\|\|\|\|\|\|\|\|\|\|\|\|\|\|\|\|\|\|\|\|\|\|\|\|\|\|\|\|\|\|\|\|\|\|\|\|\|\|\|\|\|\|\|\|\|\|\|\|\|\|\|\|\|\|\|\|\|\|\|\|\|\|\|\|\|\|\|\|\|\|\|\|\|\|\|\|\|\|\|\|\|\|\|\|\|\|\|\|\|\|\|\|\|\|\|\|\|\|\|\|\|\|\|\|\|\|\|\|\|\|\|\|\|\|\|\|\|\|\|\|\|\|\|\|\|\|\|\|\|\|\|\|\|\|\|\|\|\|\|\|\|\|\|\|\|\|\|\|\|\|\|\|\|\|\|\|\|\|\|\|\|\|\|\|\|\|\|\|\|\|\|\|\|\|\|\|\|\|\|\|\|\|\|\|\|\|\|\|\|\|\|\|\|\|\|\|\|\|\|\|\|\|\|\|\|\|\|\|\|\|\|\|\|\|\|\|\|\|\|\|\|\|\|\|\|\|\|\|\|\|\|\|\|\|\|\|\|\|\|\|\|\|\|\|\|\|\|\|\|\|\|\|\|\|\|\|\|\|\|\|\|\|\|\|\|\|\|\|\|\|\|\|\|\|\|\|\|\| 	ATGCAGTTCATCATCCTCGCCCTTGCTGCCCTCGCGGCTGCCGCTCCTGGCGGCGCTCCCAGCTACGGTGGCAGTGGCAGTGGCATGGCCGGTGGCATGGCCGGTGGCCACGGAGGTAGCGGCGGCTACTCTGGCGGCTCCAACAACGGTGGCTCCAACGGTGGTTCCAACGGCGGCTCCTCCAACGGTGGTTCCAACGGTGGTTCCAACGGAGGCAAATTCCAATGCTCTGCGCCCCTCCAGAGCAGCCCACAGTGCTGCGCTGTCAACGCGCTCGGGGTTGCTTCCCTTCCCTGCAATGCCC ```      ``` 391	CTCGGTGCCGGCGTCGCCTGCGAGGAGGTCAAGTCCAAC	429 	\|\|\|\|\|\|\|\|\|\|\|\|\|\|\|\|\|\|\|\|\|\|\|\|\|\|\|\|\|\|\|\|\|\|\|\|\|\|\| 	CTCGGTGCCGGCGTCGCCTGCGAGGAGGTCAAGTCCAAC ```      ``` 1	ATGCAGTTCATCATCCTCGCCCTTGCTGCCCTCGCGGCTGCCGCTCCTGGCGGCGCTCCCAGCTACGGTGGCAGTGGCAGTGGCATGGCCGGTGGCATGGCCGGTGGCCACGGAGGTAGCGGCGGCTACTCTGGCGGCTCCAACAACGGTGGCTCCAACGGTGGTTCCAACGGCGGCTCCTCCAACGGTGGTTCCAACGGTGGTTCCAACGGAGGCAAATTCCAATGCTCTGCGCCCCTCCAGAGCAGCCCACAGTGCTGCGCTGTCAACGCGCTCGGGGTTGCTTCCCTTCCCTGCAATGCCC	304 	\|\|\|\|\|\|\|\|\|\|\|\|\|\|\|\|\|\|\|\|\|\|\|\|\|\|\|\|\|\|\|\|\|\|\|\|\|\|\|\|\|\|\|\|\|\|\|\|\|\|\|\|\|\|\|\|\|\|\|\|\|\| \|\|\|\|\|\|\|\|\|\|\|\|\|\|\|\|\|\|\|\|\|\|\|\|\|\|\|\|\|\|\|\|\|\|\|\|\|\|\|\|\|\|\|\|\|\|\|\|\|\|\|\|\|\|\|\|\|\|\|\|\|\|\|\|\|\|\|\|\|\|\|\|\|\|\|\|\|\|\|\|\|\|\|\|\|\|\|\|\|\|\|\|\|\|\|\|\|\|\|\|\|\|\|\|\|\|\|\|\|\|\|\|\|\|\|\|\|\|\|\|\|\|\|\|\|\|\|\|\|\|\|\|\|\|\|\|\|\|\|\|\|\|\|\|\|\|\|\|\|\|\|\|\|\|\|\|\|\|\|\|\|\|\|\|\|\|\|\|\|\|\|\|\|\|\|\|\|\|\|\|\|\|\|\|\|\|\|\|\|\|\|\|\|\|\|\|\|\|\|\|\|\|\|\|\|\|\|\|\|\|\|\|\|\|\|\|\|\|\|\|\|\|\|\|\|\|\|\|\|\|\|\|\|\|\|\|\|\|\|\|\| 	ATGCAGTTCATCATCCTCGCCCTTGCTGCCCTCGCGGCTGCCGCTCCTGGCGGCGCTCCCAGNTACGGTGGCAGTGGCAGTGGCATGGCCGGTGGCATGGCCGGTGGCCACGGAGGTAGCGGCGGCTACTCTGGCGGCTCCAACAACGGTGGCTCCAACGGTGGTTCCAACGGCGGCTCCTCCAACGGTGGTTCCAACGGTGGTTCCAACGGAGGCAAATTCCAATGCTCTGCGCCCCTCCAGAGCAGCCCACAGTGCTGCGCTGTCAACGCGCTCGGGGTTGCTTCCCTTCCCTGCAATGCCC ```      ``` 305	CCACTCGCGACATCGAATCGAGGGAGGACTTTGTCAAGTACTGCGGCCAGTCTGGCGCCACCGCTCAGTGCTGCGTCTTGCCTGCT	390 	\|\|\|\|\|\|\|\|\|\|\|\|\|\|\|\|\|\|\|\|\|\|\|\|\|\|\|\|\|\|\|\|\|\|\|\|\|\|\|\|\|\|\|\|\|\|\|\|\|\|\|\|\|\|\|\|\|\|\|\|\|\|\|\|\|\|\|\|\|\|\|\|\|\|\|\|\|\|\|\|\|\|\|\|\|\| 	CCACTCGCGACATCGAATCGAGGGAGGACTTTGTCAAGTACTGCGGCCAGTCTGGCGCCACCGCTCAGTGCTGCGTCTTGCCTGCT ``` |
| 3 | ggggacaagtttgtacaaaaaagcaggcttgatgCAGTCCATCATCGCCACTTC, ggggaccactttgtacaagaaagctgggtcAGTGCCATCGTTGGTCAGGG | Mycgr3\|96868  JGI  Ensembl |  |  | 100 % |  | ``` 128	CCTCGG-TGTC-TTTCGACATCACTCCGTACGATCCTGCTACCACTTCTGGGTCCGG-CGGTGTCTACCACTGTACGCAATACGACGTGGTCACCGGAGAGCTGGTTGCCAATTTTGAGGAGATGACGACGTACAGTTGCGGCCCGGGGTCTCCGTTCAGCTTCTCGTACTCCGAAAAGGATGGACCCTTCAAGTACAGGCTGTACATGTGGCAAGAAGCGACTCCGCAGACTCCCACGCGTTGGGCTTTTACACCGATCCGGGCCCCCGCATGCAACACCGCCGATGGGCATCAGATCTGCTACGCTCCGGACAGCTTGATCACCCTGACCAACGATGGCACT	468 	\|\|\|\|\|\| \|\|\|\| \|\|  \|\|\|\|\|\| \|\|\| \|\|\|\|\|\|\|\|\|\|\| \|\|\|\|\|\|\|\| \|\|\|\|\|\|\|\|\| \|\|\|\|\|\| \| \|\|\|\|\|\|\|\|\|\|\|\|\|\|\|\| \|\|\|\|\|\|\|\| \|\|\| \|\|\|\|\|\|\|\|\|\|\|\|\|\|\|\|\|\|\|\|\|\|\|\|\|\|\|\|\| \|\|\|\|\|\|\|\|\|\|\|\|\|\|\|\|\|\|\|\|\|\|\|\|\|\|\|\|\|\|\|\|\|\|\|\|\|\|\|\|\|\|\|\|\|\|\|\|\|\|\|\|\|\|\|\|\|\|\|\|\|\|\|\|\|\|\|\|\|\|\|\|\|\|\|\|\|\|\|\|\|\|\|\|\|\|\|\|\|\|\|\|\|\|\|\|\|\|\|\|\|\|\|\|\|\|\|\|\|\|\|\|\|\|\|\|\|\|\|\|\|\|\|\|\|\|\|\|\|\|\|\|\|\|\|\|\|\|\|\|\|\|\|\|\|\|\|\|\|\|\|\|\|\|\|\|\|\|\|\|\|\|\|\|\|\|\|\|\|\|\|\|\|\| \|\|\|\|\|\|\|\|\|\|\|\|\|\|\|\|\|\|\|\|\|\|\|\|\|\|\|\|\|\|\|\|\|\|\|\|\|\|\|\|\|\| 	CCTCGGTTGTCNTTAGGACATCNCTCNGTACGATCCTGATACCACTTNTGGGTCCGGNCGGTGTATNCCACTGTACGCAATACTACGTGGTCNCCGCAGAGCTGGTTGCCAATTTTGAGGAGATGATGACGTACAGTTGCGGCCCGGGGTCTCCGTTCAGCTTCTCGTACTCCGAAAAGGATGGACCCTTCAAGTACAGGCTGTACATGTGGCAAGAAGCGACTCCGCAGACTCCCACGCGTTGGGCTTTTACACCGATCCGGGCCCCCGCATGCAACACCGCCGATGGGCATCAGATCTGNTACGCTCCGGACAGCTTGATCACCCTGACCAACGATGGCACT ```      ``` 1	ATGCAGTCCATCATCGCCACTTCCATTCTGGCATTCTTCGCCAGCTCAGCCATCGCCGCTCTTCCGGACAGAAGCTCCTACAACATCGAGAACTTCTCCGAGACCAAGTCGCTCGACAACGACATTACCTCGGTGTCTTTCGACATCACTCCGTACGATCCTGCTACCACTTCTGGGTCCGGCGGTGTCTACCACTGTACGCAATACGACGTGGTCACCGGAGAGCTGGTTGCCAATTTTGAGGA	245 	\|\|\|\|\|\|\|\|\|\|\|\|\|\|\|\|\|\|\|\|\|\|\|\|\|\|\|\|\|\|\|\|\|\|\|\|\|\|\|\|\|\|\|\|\|\|\|\|\|\|\|\|\|\|\|\|\|\|\|\|\|\|\|\|\|\|\|\|\|\|\|\|\|\|\|\|\|\|\|\|\|\|\|\|\|\|\|\|\|\|\|\|\|\|\|\|\|\|\|\|\|\|\|\|\|\|\|\|\|\|\|\|\|\|\|\|\|\|\|\|\|\|\|\|\|\|\|\|\|\|\|\|\|\|\|\|\|\|\|\|\|\|\|\|\|\|\|\|\|\|\|\|\|\|\|\|\|\|\|\|\|\|\|\|\|\|\|\|\|\|\|\|\|\|\|\|\|\|\|\|\|\|\|\|\|\|\|\|\| \|\|\|\|\|\|\| \|\|\|\| \|\|\|\|\|\|\|\|\|\|\|\|\|\|\|\|\|\|\| \|\|\|\|\|\|\| \|\|\|  \|\|\|\|\|\|\|\|\| 	ATGCAGTCCATCATCGCCACTTCCATTCTGGCATTCTTCGCCAGCTCAGCCATCGCCGCTCTTCCGGACAGAAGCTCCTACAACATCGAGAACTTCTCCGAGACCAAGTCGCTCGACAACGACATTACCTCGGTGTCTTTCGACATCACTCCGTACGATCCTGCTACCACTTCTGGGTCCGGCGGTGTCNACCACTGNACGCTATACGACGTGGTCACCGGANAGCTGGTAGCCGNTTTTGAGGA ``` |
| 3 | ggggacaagtttgtacaaaaaagcaggcttgatgAAGTTCTTCACGATCCTCTC, ggggaccactttgtacaagaaagctgggtcTGCCTTATAAGCAACAAGGC | Mycgr3\|97262  JGI  Ensembl |  |  | 0 % |  | ``` 1	ATGAAGTTCTTCACGATCCTCTCGTTCCTTCTTCCTCTCTGCCTGGCCCTGGTCCTTCCCAGAAGTATCACCGATGGATCCGATCAGGCCGTTGCAGCCATTGAAGCGACCTTCGAATTCGTCTTCGAGAATGGAGGTGCGGGGGCCACGGAGGAGGCTGAGGTTCTTGTTGAACCTCCTCCCGGCGTAGGTCGGGCTGGCGCAGCAGGCTACGCACCATCAGGCACATCAATTACGGGCCACAAACCTATGGACGGGGCCACCAAGTTCTTTTTCAGGAAATACGACGAATTTACTGGCTTTATAGGAGAGGGAGAGTTCATTTACAGCGAAGCTGAACACCGAATCACGGAGTTTCACGCGAATAGCGGCAGAATTGAACGAGCTGGAATTGTGATCAAACAACTCTATCCCAAGCTGCGCCTTGTTGCTTATAAGGCA	441 	\|\|\|\|\|\|\|\|\|\|\|\|\|\|\|\|\|\|\|\|\|\|\|\|\|\|\|\|\|\|\|\|\|\|\|\|\|\|\|\|\|\|\|\|\|\|\|\|\|\|\|\|\|\|\|\|\|\|\|\|\|\|\|\|\|\|\|\|\|\|\|\|\|\|\|\|\|\|\|\|\|\|\|\|\|\|\|\|\|\|\|\|\|\|\|\|\|\|\|\|\|\|\|\|\|\|\|\|\|\|\|\|\|\|\|\|\|\|\|\|\|\|\|\|\|\|\|\|\|\|\|\|\|\|\|\|\|\|\|\|\|\|\|\|\|\|\|\|\|\|\|\|\|\|\|\|\|\|\|\|\|\|\|\|\|\|\|\|\|\|\|\|\|\|\|\|\|\|\|\|\|\|\|\|\|\|\|\|\|\|\|\|\|\|\|\|\|\|\|\|\|\|\|\|\|\|\|\|\|\|\|\|\|\|\|\|\|\|\|\|\|\|\|\|\|\|\|\|\|\|\|\|\|\|\|\|\|\|\|\|\|\|\|\|\|\|\|\|\|\|\|\|\|\|\|\|\|\|\|\|\|\|\|\|\|\|\|\|\|\|\|\|\|\|\|\|\|\|\|\|\|\|\|\|\|\|\|\|\|\|\|\|\|\|\|\|\|\|\|\|\|\|\|\|\|\|\|\|\|\|\|\|\|\|\|\|\|\|\|\|\|\|\|\|\|\|\|\|\|\|\|\|\|\|\|\|\|\|\|\|\|\|\|\|\|\|\|\|\|\|\|\|\|\|\|\|\|\|\|\|\|\|\|\|\|\|\|\|\|\|\|\|\|\|\|\|\|\|\|\|\|\|\|\|\|\|\|\|\|\|\|\|\|\|\|\|\|\|\|\|\|\|\|\|\|\|\|\|\|\|\|\|\|\|\|\|\|\|\|\|\|\|\|\|\|\|\|\|\|\|\|\|\|\|\|\|\|\|\|\|\| 	ATGAAGTTCTTCACGATCCTCTCGTTCCTTCTTCCTCTCTGCCTGGCCCTGGTCCTTCCCAGAAGTATCACCGATGGATCCGATCAGGCCGTTGCAGCCATTGAAGCGACCTTCGAATTCGTCTTCGAGAATGGAGGTGCGGGGGCCACGGAGGAGGCTGAGGTTCTTGTTGAACCTCCTCCCGGCGTAGGTCGGGCTGGCGCAGCAGGCTACGCACCATCAGGCACATCAATTACGGGCCACAAACCTATGGACGGGGCCACCAAGTTCTTTTTCAGGAAATACGACGAATTTACTGGCTTTATAGGAGAGGGAGAGTTCATTTACAGCGAAGCTGAACACCGAATCACGGAGTTTCACGCGAATAGCGGCAGAATTGAACGAGCTGGAATTGTGATCAAACAACTCTATCCCAAGCTGCGCCTTGTTGCTTATAAGGCA ```      ``` 3	GAAGTTCTTCACGATCCTCTCGTTCCTTCTTCCT-CTCTGCCTGGCCCTGGTCCTTCCCAGAAGTATCACCGATGGATCCGATCAGGCCGTTGCAGCCATTGAAGCGACCTTCGAATTCGTCTTCGAGAATGGAGGTGCGGGGGCCACGGAGGAGGCTGAGGTTCTTGTTGAACCTCCTCCCGGCGTAGGTCGGGCTGGCGCAGCAGGCTACGCACCATCAGGCACATCAATTACGGGCCACAAACCTATGGACGGGGCCACCAAGTTCTTTTTCAGGAAATACGACGAATTTACTGGCTTTATAGGAGAGGGAGAGTTCATTTACAGCGAAGCTGAACACCGAATCACGGAGTTTCACGCGAATAGCGGCAGAATTGAACGAGCTGGAATTGTGATCAAACAACTCTATCCCAAGCTGCGCCTTGTTGCTTATAAGGCA	441 	\|\|\|\|\|\| \|\|\|\| \|\|\|\| \| \|\|\|\|\|\| \|\|\|\|\|\|\|\|  \| \|\|\|\|\|\|\|\|\| \|\|\|\|\|\|\|\|\|\|\|\|\|\|\|\|\|\|\|\| \|\|\|\|\|\|\|\|\|\|\|\|\|\|\|\|\| \|\|\|\|\|\|\|\|\|\|\|\|\|\|\|\|\|\|\|\|\|\|\|\|\|\|\|\|\|\|\|\|\|\|\|\|\|\|\|\|\|\|\|\|\|\|\|\|\|\|\|\|\|\|\|\|\|\|\|\|\|\|\|\|\|\|\|\|\|\|\|\|\|\|\|\|\|\|\|\|\|\|\|\|\|\|\|\|\|\|\|\|\|\|\|\|\|\|\|\|\|\|\|\|\|\|\|\|\|\|\|\|\|\|\|\|\|\|\|\|\|\|\|\|\|\|\|\|\|\|\|\|\|\|\|\|\|\|\|\|\|\|\|\|\|\|\|\|\|\|\|\|\|\|\|\|\|\|\|\|\|\|\|\|\|\|\|\|\|\|\|\|\|\|\|\|\|\|\|\|\|\|\|\|\|\|\|\|\|\|\|\|\|\|\|\|\|\|\|\|\|\|\|\|\|\|\|\|\|\|\|\|\|\|\|\|\|\|\|\|\|\|\|\|\|\|\|\|\|\|\|\|\|\|\|\|\|\|\|\|\|\|\|\|\|\|\|\|\|\|\|\|\|\|\|\|\|\|\|\|\|\|\|\|\|\|\|\|\|\|\|\|\|\|\|\|\|\|\|\|\|\|\|\|\|\|\|\|\|\|\|\|\|\|\|\|\|\|\|\|\|\|\|\|\|\|\|\|\|\|\|\|\|\|\|\|\|\|\|\|\|\|\|\|\|\|\|\|\|\|\|\|\|\|\|\|\|\|\|\|\|\|\|\|\|\|\|\|\|\|\|\|\| 	GAAGTTNTTCANGATCNTNTCGTTCNTTCTTCCTNTTTTGCCTGGCCNTGGTCCTTCCCAGAAGTATCNCCGATGGATCCGATCAGNCCGTTGCAGCCATTGAAGCGACCTTCGAATTCGTCTTCGAGAATGGAGGTGCGGGGGCCACGGAGGAGGCTGAGGTTCTTGTTGAACCTCCTCCCGGCGTAGGTCGGGCTGGCGCAGCAGGCTACGCACCATCAGGCACATCAATTACGGGCCACAAACCTATGGACGGGGCCACCAAGTTCTTTTTCAGGAAATACGACGAATTTACTGGCTTTATAGGAGAGGGAGAGTTCATTTACAGCGAAGCTGAACACCGAATCACGGAGTTTCACGCGAATAGCGGCAGAATTGAACGAGCTGGAATTGTGATCAAACAACTCTATCCCAAGCTGCGCCTTGTTGCTTATAAGGCA ``` |
| 3 | ggggacaagtttgtacaaaaaagcaggcttgatgCAGCCCTCGACGTTCTTCCA, ggggaccactttgtacaagaaagctgggtcGAGTCCCAACCCAACTTGTC | Mycgr3\|97283  JGI  Ensembl |  |  | 100 % |  | ``` 84	CAACGTCTACAAGGAAGAACACGTCTACGACATCCTCCAGCCCGGTCAATGGTCCTCGCCCTTCGAAGTCACCTACAACACAGGCTGGCTTACCACAGGCCGAGACTGGTGGCTCGTGCTTGGGCACAACGACAACGACCCGATCGGCAAGAACTCCATGTCTCAATTCTACACCGACCCGGAGAACTTCCAAGACATCATCAACTGGCTCGAGAGTGCGGCGCCGACTTTGATCGGCGCTGCGCTGACAGCGGCTGAGATCCTTCAACCTGAGCTCGCGGCCGTTACAGCGGTGGCTGGAGTTGCGGCGGAAGCTCTGTGTACCGCGATGCTCAACGATGCCGCCACGGCCGGATACAAGCAGCACATCTTGACGGACGAAGATAAGGGCAAAGTCTTGCAGATCTGGATCAATAACGACGGCACCGTCAACTTCATCTCTCCATCGGGACGCTCGGACACGGAGTATGAGACTAGACAAGTTGGGTTGGGACTC	579 	\|\|\|\|\|\| \|\|\|\|\|\|\|\|\|\|\|\|\|\| \|\| \|\| \|\|\|\|\|\|\|\|\|\|\|\|\|\|\|\|\|\|\|\|\|\|\|\|\|\|\|\|\|\|\|\|\|\|\| \|\|\|\|\|\|\|\|\|\|\|\|\|\|\|\|\|\|\|\|\|\|\|\|\|\|\|\|\|\|\|\|\|\|\|\|\|\|\|\|\|\|\|\|\|\|\|\|\|\|\|\|\|\|\|\|\|\|\|\|\|\|\|\|\|\|\|\|\|\|\|\|\|\|\|\|\|\|\|\|\|\|\|\|\|\|\|\|\|\|\|\|\|\|\|\|\|\|\|\|\|\|\|\|\|\|\|\|\|\|\|\|\|\|\|\|\|\|\|\|\|\|\|\|\|\|\|\|\|\|\|\|\|\|\|\|\|\|\|\|\|\|\|\|\|\|\|\|\|\|\|\|\|\|\|\|\|\|\|\|\|\|\|\|\|\|\|\|\|\|\|\|\|\|\|\|\|\|\|\|\|\|\|\|\|\|\|\|\|\|\|\|\|\|\|\|\|\|\|\|\|\|\|\|\|\|\|\|\|\|\|\|\|\|\|\|\|\|\|\|\|\|\|\|\|\|\|\|\|\|\|\|\|\|\|\|\|\|\|\|\|\|\|\|\|\|\|\|\|\|\|\|\|\|\|\|\|\|\|\|\|\|\|\|\|\|\|\|\|\|\|\|\|\|\|\|\|\|\|\|\|\|\|\|\|\|\|\|\|\|\|\|\|\|\|\|\|\|\|\|\|\|\|\|\|\|\|\|\|\|\|\|\|\|\|\|\|\|\|\|\|\|\|\|\|\|\|\|\|\|\|\|\|\|\|\|\|\|\|\|\|\|\|\|\|\|\|\|\|\|\|\|\|\|\|\|\|\|\|\|\|\|\|\|\|\|\|\|\|\|\|\|\|\|\|\|\|\|\|\|\|\|\|\|\|\|\|\|\|\|\|\|\|\|\|\|\|\|\|\|\|\|\|\|\|\|\|\|\|\|\|\|\|\|\|\|\|\|\|\|\|\|\|\| \|\|\|\|\|\|\| 	CAACGTNTACAAGGAAGAACANGTNTANGACATCCTCCAGCCCGGTCAATGGTCCTCGCCCTTNGAAGTCACCTACAACACAGGCTGGCTTACCACAGGCCGAGACTGGTGGCTCGTGCTTGGGCACAACGACAACGACCCGATCGGCAAGAACTCCATGTCTCAATTCTACACCGACCCGGAGAACTTCCAAGACATCATCAACTGGCTCGAGAGTGCGGCGCCGACTTTGATCGGCGCTGCGCTGACAGCGGCTGAGATCCTTCAACCTGAGCTCGCGGCCGTTACAGCGGTGGCTGGAGTTGCGGCGGAAGCTCTGTGTACCGCGATGCTCAACGATGCCGCCACGGCCGGATACAAGCAGCACATCTTGACGGACGAAGATAAGGGCAAAGTCTTGCAGATCTGGATCAATAACGACGGCACCGTCAACTTCATCTCTCCATCGGGACGCTCGGACACGGAGTATGAGACTAGACAAGTTGGGT-GGGACTC ```      ``` 1	ATGCAGCCCTCGACGTTCTTCCAGTCGCTCTTTGCCTTCACTCTCCTCTTCCCGCTCGC	59 	\|\|\|\|\|\|\|\|\|\|\|\|\|\|\|\|\|\|\|\|\|\|\|\|\|\|\|\|\|\|\|\|\|\|\|\|\|\|\|\|\|\|\|\|\|\|\|\|\|\|\|\| \|\|\|\|\|\| 	ATGCAGCCCTCGACGTTCTTCCAGTCGCTCTTTGCCTTCACTCTCCTCTTCCNGCTCGC ``` |
| 3 | ggggacaagtttgtacaaaaaagcaggcttgatgTCTTTCACCGTCAATACCTT, ggggaccactttgtacaagaaagctgggtcCAACTTCAACACACCCGCCT | Mycgr3\|97371  JGI  Ensembl | KOG0619 FOG: Leucine rich repeat; | protein binding; | 100 % |  | ``` 242	TTTCCCACCTCCACCTCTCCGGACCATCCAGCCACCCTCTGGGAATTAACTTCCTGCCCTACCTTGAAGACTTGACCTCCTTCACCGCCAGCGAGAACACCTTCACCGGAGAGCTGGCGGACATCCCTCTCCCGTCGAAATTGTCTACCATCGACATCTCCTTCAACCAACTCACCGGCTCCATCCCGGATCCAGCAGTGTGGGAACTCGACTCCCTTCAAAGTCTCGACTTGGGGTATAATCGCCTTTCCGGCAATCTGCCCAGCACCATCTTTGCGCAGAAGGATCTGCACCTTCTGTTCGTGCCGAATAACAAGCTCGGCGGCGCGATTCCAGAGTTTGCGCATACGGCGCTGAGGGTTGTCTCGCTGGCATCGTGTTCGTTCACGGGTGCAATCCCGAAGAGTATTGGCAAGGCGAAGAACTTGGAGGAGGTGTACTTTGATGGCAATGCTGGGCTGAACGGAAAGGTCCCGGAGGAGCTGTGCAGCTTGCCGAAGTTGAGGTCCGCGGATCTGACAAAGACGGGTGTGGATAAGAGTAGATTGGGACCGGCTTGTCAGAAGGCGCAGAAGGCGGGTGTGTTGAAGTTG	834 	\|\|\| \|\|\|\| \|\|\|\|\| \| \|\| \|\|\|\|\|\|\|\|\|\|\|\|\|\|\|\|\|\| \|\|\|\|\|\|\|\|\|\|\|\|\|\|\|\|\|\|\|\|\|\|\|\|\|\|\|\|\|\|\|\|\|\|\|\|\|\|\|\|\|\|\|\|\|\|\|\|\|\|\|\|\|\|\|\|\|\|\|\|\|\|\|\|\|\|\|\|\|\|\|\|\|\|\|\|\|\|\|\|\|\|\|\|\|\|\|\|\|\|\|\|\|\|\|\|\|\|\|\|\|\|\|\|\|\|\|\|\|\|\|\|\|\|\|\|\|\|\|\|\|\|\|\|\|\|\|\|\|\|\|\|\|\|\|\|\|\|\|\|\|\|\|\|\|\|\|\|\|\|\|\|\|\|\|\|\|\|\|\|\|\|\|\|\|\|\|\|\|\|\|\|\|\|\|\|\|\|\|\|\|\|\|\|\|\|\|\|\|\|\|\|\|\|\|\|\|\|\|\|\|\|\|\|\|\|\|\|\|\|\|\|\|\|\|\|\|\|\|\|\|\|\|\|\|\|\|\|\|\|\|\|\|\|\|\|\|\|\|\|\|\|\|\|\|\|\|\|\|\|\|\|\|\|\|\|\|\|\|\|\|\|\|\|\|\|\|\|\|\|\|\|\|\|\|\|\|\|\|\|\|\|\|\|\|\|\|\|\|\|\|\|\|\|\|\|\|\|\|\|\|\|\|\|\|\|\|\|\|\|\|\|\|\|\|\|\|\|\|\|\|\|\|\|\|\|\|\|\|\|\|\|\|\|\|\|\|\|\|\|\|\|\|\|\|\|\|\|\|\|\|\|\|\|\|\|\|\|\|\|\|\|\|\|\|\|\|\|\|\|\|\|\|\|\|\|\|\|\|\|\|\|\|\|\|\|\|\|\|\|\|\|\|\|\|\|\|\|\|\|\|\|\|\|\|\|\|\|\|\|\|\|\|\|\|\|\|\|\|\|\|\|\|\|\|\|\|\|\|\|\|\|\|\|\|\|\|\|\|\|\|\|\|\|\|\|\|\|\|\|\|\|\|\|\|\|\|\|\|\|\|\|\|\|\|\|\|\|\|\|\|\|\|\|\|\|\|\|\|\|\|\|\|\|\|\|\|\|\|\|\|\|\|\|\|\|\|\|\|\|\|\|\|\|\|\|\|\|\|\|\|\|\|\|\|\|\|\|\|\|\|\|\|\|\|\|\|\|\|\|\|\|\|\|\|\|\|\|\|\|\|\|\|\|\|\|\|\|\|\|\|\|\|\| 	TTTNCCACNTCCACNTNTCNGGACCATCCAGCCACCCTNTGGGAATTAACTTCCTGCCCTACCTTGAAGACTTGACCTCCTTCACCGCCAGCGAGAACACCTTCACCGGAGAGCTGGCGGACATCCCTCTCCCGTCGAAATTGTCTACCATCGACATCTCCTTCAACCAACTCACCGGCTCCATCCCGGATCCAGCAGTGTGGGAACTCGACTCCCTTCAAAGTCTCGACTTGGGGTATAATCGCCTTTCCGGCAATCTGCCCAGCACCATCTTTGCGCAGAAGGATCTGCACCTTCTGTTCGTGCCGAATAACAAGCTCGGCGGCGCGATTCCAGAGTTTGCGCATACGGCGCTGAGGGTTGTCTCGCTGGCATCGTGTTCGTTCACGGGTGCAATCCCGAAGAGTATTGGCAAGGCGAAGAACTTGGAGGAGGTGTACTTTGATGGCAATGCTGGGCTGAACGGAAAGGTCCCGGAGGAGCTGTGCAGCTTGCCGAAGTTGAGGTCCGCGGATCTGACAAAGACGGGTGTGGATAAGAGTAGATTGGGACCGGCTTGTCAGAAGGCGCAGAAGGCGGGTGTGTTGAAGTTG ``` |
| 3 | ggggacaagtttgtacaaaaaagcaggcttgatgCACCTGTCCCTTCACGCCAC, ggggaccactttgtacaagaaagctgggtcGGGACAAGGAGCGCCAGTGT | Mycgr3\|97500  JGI  Ensembl |  |  | 100 % |  | ``` 19	GCCAC-TATGAAGACCTCT-GC-TCTCATTCTCGCCATCACCGC-GCTTTATAGCTCC-GGC-GTGCTGGCAGACC-ACCAGG-AACCCCTCGATGCTA-GTGCCTACGTCTGCATCCGCAATAATAAGG-TTATGGGC-GAGG-ACG-GCTGCTACAAGAGAAACGG-CGTGGACCGTATCACTCACTGCCACAATGCCG	205 	\|\|\|\|\| \|\| \|\|\|\|\|\|\|\|\|\| \|\| \|\|\|\|\|\|\|   \|\|\|\| \|\|  \|\| \|\| \|\|\|\|\|\| \|\|\|  \|\| \|\|\|  \|\| \| \|\|\| \| \|\|\|\| \|\|\| \|\|\|\| \| \|\|\|\| \|\|\|\|\|\|\|\|\|\|\|\|\|\|\|\|\|\|\|\| \|\|\|\|\|\|\|\|  \|\|\|\|\|\|\|\| \|\|\|  \|\|\| \|\| \|\|\|\| \|\|\|\| \|\| \|\|\| \|\|\|\|   \|\| \|\|\|\| \| \| \|\| \|\|\|\|\| \|\|\|\| 	GCCACGTANGAAGACCTCTTGCATCTCATTGGGGCCANCANNGCAGCNTTATAGNTCCNCGCNGTGGNGGGANACCTANCAGGCAACNCCTCNAGGCTAGGTGCCTACGTCTGCATCCGCNATAATAAGCNTTATGGGCAGAGNCACGAGCGGCTANAAGAAAANCGGTCGTGNTTCGGATCANTGAGTGNCACAANGCCG ```      ``` 1	ATGCACCTGTCCCTTCACGCCACTATGAAGACCTCTGCTCTCATTCTCGCCATCACCGCGCTTTATAGCTCCGGCGTGCTGGCAGACCACCAGGAACCCCTCGATGCTAGTGCCTACGTCTGCATCCGCAATAATAAGGTTATGGGCGAGGACGGCTGCTACAAGAGAAACGGCGTGGACCGTATCACTCACTGCCACAATGCCGGCAACGGTCACTATCACTACTCAGAGTGCCGC	237 	\|\|\|\|\|\|\|\|\|\|\|\|\|\|\|\|\|\|\|\|\|\|\|\|\|\|\|\|\|\|\|\|\|\|\|\|\|\|\|\|\|\|\|\|\|\|\|\|\|\|\|\|\|\|\|\|\|\|\|\|\|\|\|\|\|\|\|\|\|\|\|\|\|\|\|\|\|\|\|\|\|\|\|\|\|\|\|\|\|\|\|\|\|\|\|\|\|\|\|\|\|\|\|\|\|\|\|\|\|\|\|\|\|\|\|\|\|\|\|\|\|\|\|\|\|\|\|\|\|\|\|\|\|\|\|\|\|\|\|\|\|\|\|\|\|\|\|\|\|\|\|\|\|\|\|\|\|\|\|\|\|\|\|\|\|\|\|\|\|\|\|\|\|\|\|\|\|\|\|\|\|\|\|\|\|\|\|\|\|\|\|\|\|\|\|\|\|\|\|\|\|\|\|\|\|\|\|\|\|\|\|\|\|\|\|\|\|\|\|\|\|\|\|\|\|\|\|\|\|\|\|\|\|\|\|\|\| 	ATGCACCTGTCCCTTCACGCCACTATGAAGACCTCTGCTCTCATTCTCGCCATCACCGCGCTTTATAGCTCCGGCGTGCTGGCAGACCACCAGGAACCCCTCGATGCTAGTGCCTACGTCTGCATCCGCAATAATAAGGTTATGGGCGAGGACGGCTGCTACAAGAGAAACGGCGTGGACCGTATCACTCACTGCCACAATGCCGGCAACGGTCACTATCACTACTCAGAGTGCCGC ```      ``` 238	ACTATCTGTTGCGTCGTCGCTAGTCGCAGCGGTGAAGAGTTCATAGACGTAACGCAACACTGCACCGGAGTGGGTGGAGTGCTTACGTCCTACAACGACCTGCCCGGAAACAAGAAATACCACTGTCCACTGGTCAACACCTGTGTCCCGGATAACACTGGCGCTCCTTGTCCC	411 	\|\|\|\|\|\|\|\|\|\|\|\|\|\|\|\|\|\|\|\|\|\|\|\|\|\|\|\|\|\|\|\|\|\|\|\|\|\|\|\|\|\|\|\|\|\|\|\|\|\|\|\|\|\|\|\|\|\|\|\|\|\|\|\|\|\|\|\|\|\|\|\|\|\|\|\|\|\|\|\|\|\|\|\|\|\|\|\|\|\|\|\|\|\|\|\|\|\|\|\|\|\|\|\|\|\|\|\|\|\|\|\|\|\|\|\|\|\|\|\|\|\|\|\|\|\|\|\|\|\|\|\|\|\|\|\|\|\|\|\|\|\|\|\|\|\|\|\|\|\|\|\|\|\|\|\|\|\|\|\|\|\|\|\|\|\|\|\|\|\|\|\|\|\| 	ACTATCTGTTGCGTCGTCGCTAGTCGCAGCGGTGAAGAGTTCATAGACGTAACGCAACACTGCACCGGAGTGGGTGGAGTGCTTACGTCCTACAACGACCTGCCCGGAAACAAGAAATACCACTGTCCACTGGTCAACACCTGTGTCCCGGATAACACTGGCGCTCCTTGTCCC ```      ``` 1	ATGCACCTGTCCCTTCACGCCACTATGAAGACCTCTGCTCTCATTCTCGCCATCACCGCGCTTTATAGCTCCGGCGTGCTGGCAGACCACCAGGAACCCCTCGATGCTAGTGCCTACGTCTGCATCCGCAATAATAAGGTTATGGGCGAGGACGGCTGCTACAAGAGAAACGGCGTGGACCGTATCACTCACTGCCACAATGCCGGCAACGGTCACTATCACTACTCAGAGTGCCGC	237 	\|\|\|\|\|\|\|\|\|\|\|\|\|\|\|\|\|\|\|\|\|\|\|\|\|\|\|\|\|\|\|\|\|\| \|\|\|\| \|\|\|\|\| \| \|\|\|\|\|\|\|\|\|\|\|\|\|\|\|\|\|\|\|\|\|\|\|\|\|\|\|\|\|\|\|\|\|\|\|\|\|\|\|\|\|\|\|\|\|\|\|\|\|\|\|\|\|\|\|\|\|\|\|\|\|\|\|\|\|\|\|\|\|\|\|\|\|\|\|\|\|\|\|\|\|\|\|\|\|\|\|\|\|\|\|\|\|\|\|\|\|\|\|\|\|\|\|\|\|\|\|\|\|\|\|\|\|\|\|\|\|\|\|\|\|\|\|\|\|\|\|\|\|\|\|\|\|\|\|\|\|\|\|\|\|\|\|\|\|\|\|\|\|\|\|\|\|\|\|\|\|\|\|\|\|\|\|\|\|\|\|\|\|\|\|\|\|\|\|\|\|\|\|\|\|\|\|\|\|\|\|\|\| 	ATGCACCTGTCCCTTCACGCCACTATGAAGACCTNTGCTNTCATTNTNGCCATCACCGCGCTTTATAGCTCCGGCGTGCTGGCAGACCACCAGGAACCCCTCGATGCTAGTGCCTACGTCTGCATCCGCAATAATAAGGTTATGGGCGAGGACGGCTGCTACAAGAGAAACGGCGTGGACCGTATCACTCACTGCCACAATGCCGGCAACGGTCACTATCACTACTCAGAGTGCCGC ``` |
| 3 | ggggacaagtttgtacaaaaaagcaggcttgatgTTGAACCACCATCGCTTCCA, ggggaccactttgtacaagaaagctgggtcACATGCAGGGAATTGGTCGA | Mycgr3\|97526  JGI  Ensembl |  |  | 100 % |  | ``` 360	GATCCCACCGATCGATGCTCACCCGGGTGGGCCTGCTCAGCCCGCTGTTGCCCCCGGGTCAGATCCCAAGTCGACGTGTCGCGAGATCCTTTCCAAGGGAGGCGACACCGGTATCGGCATCGCGAATGGATGCTGGATCCCCTACGAGGGAACCCGAGTTCTGCAATGCATGCTCACGTGCAACAACTCCCCCTCCTGCAGCCCGGGCGACGACCCGAACGAGGTTCACAAATACCTCGACCAATTCCCTGCATGT	615 	\|\|\|\|\|\|\|\|\|\|\|\|\|\|\|\|\|\|\|\|\|\|\|\|\|\|\|\|\|\|\|\|\|\|\|\|\|\|\|\|\|\|\|\|\|\|\|\|\|\|\|\|\|\|\|\|\|\|\|\|\|\|\|\|\|\|\|\|\|\|\|\|\|\|\|\|\|\|\|\|\|\|\|\|\|\|\|\|\|\|\|\|\|\|\|\|\|\|\|\|\|\|\|\|\|\|\|\|\|\|\|\|\|\|\|\|\|\|\|\|\|\|\|\|\|\|\|\|\|\|\|\|\|\|\|\|\|\|\|\|\|\|\|\|\|\|\|\|\|\|\|\|\|\|\|\|\|\|\|\|\|\|\|\|\|\|\|\|\|\|\|\|\|\|\|\|\|\|\|\|\|\|\|\|\|\|\|\|\|\|\|\|\|\|\|\|\|\|\|\|\|\|\|\|\|\|\|\|\|\|\|\|\|\|\|\|\|\|\|\|\|\|\|\|\|\|\|\|\|\|\|\|\|\|\|\|\|\|\|\|\|\|\|\|\|\|\|\|\|\|\|\|\|\|\|\| 	GATCCCACCGATCGATGCTCACCCGGGTGGGCCTGCTCAGCCCGCTGTTGCCCCCGGGTCAGATCCCAAGTCGACGTGTCGCGAGATCCTTTCCAAGGGAGGCGACACCGGTATCGGCATCGCGAATGGATGCTGGATCCCCTACGAGGGAACCCGAGTTCTGCAATGCATGCTCACGTGCAACAACTCCCCCTCCTGCAGCCCGGGCGACGACCCGAACGAGGTTCACAAATACCTCGACCAATTCCCTGCATGT ```      ``` 175	TTCCAGAAACCTCTGGGTGTCGGATGCCAACGCGACTCGGAATGCCAGAGAGGCCCAAACCACAGGTCCATCTACCCATACACCAATGGCAAGGACTGGATGTCTATCAACTACGACGTCATCCTGACCCAAGGCACCGGAATCTGCGACACCGATCACCAGGGAGGCACCGTGTGCAACGCTGTG	360 	\|\|\|\|\|\|\|\|\|\| \| \|\|\|\|\|\|\| \|\|\|\|\|\|\|\|\|\|\| \|\|\|\| \|\|\|\|\| \|\|\|\|\|\|\|\|\|\|\|\|\|\|\|\|\|\|\|\|\|\|\|\|\|\|\|\|\|\|\|\|\|\|\|\|\|\|\|\|\|\|\|\|\|\|\|\|\|\|\|\|\|\|\|\|\|\|\| \|\|\|\|\|\| \|\|\|\|\|\|\|\|\|\|\|\|\|\|\|\|\|\|\|\|\|\|\|\|\|\|\|\|\|\|\|\|\|\|\|\|\|\|\|\|\|\|\|\|\|\|\|\|\|\|\|\|\|\|\|\|\|\|\|\|\|\|\|\|\|\|\|\|\|\|\|\|\|\|\| 	TTCCAGAAACNTNTGGGTGTNGGATGCCAACGNGACTNGGAATNCCAGAGAGGCCCAAACCACAGGTCCATCTACCCATACACCAATGGCAAGGACTGGATGTNTATCAANTACGACGTCATCCTGACCCAAGGCACCGGAATCTGCGACACCGATCACCAGGGAGGCACCGTGTGCAACGCTGTG ``` |
| 3 | ggggacaagtttgtacaaaaaagcaggcttgatgGCGAACCCGCTCGACACCGA, ggggaccactttgtacaagaaagctgggtcTCTAAACTTGCTGATAATGA | Mycgr3\|98375  JGI  Ensembl | KOG1666 V-SNARE; | intracellular protein transport; membrane; | 100 % |  | ``` 115	GCCGGCTCTGAGCTCTTCAGCAACTACGAAGCGGAACTCAAGCTAGTACAAGCAGACCTATCACAGAAACTCGATCAGATTCCTGAGCTGTCCGGCGAACAGCGAAAGGCAGCTGTTTCGCAGGCAGATCGCGCGCTTGAGGAGGCCAAGGAGCTAATAGAGTCCATGCGTTTGGAGAAGCAGAACATTCCACAAGCACTCAAGATCAAGGTCAACCAACGCTTCCGCAATTACCAGACCGACGTGGATGCTGCAGGACGGAAACTCAAAGGCATGCAGGACGATAGATCAGCGCTCTTCGGCAAGCGATACACCGACAATCCACAGGATGAGCAGCTCGAGCAGAGGCAGCAGCTTCTGGGAGGCACGGAGAGGTTAGAGCGGTCGAGCGGCAGGCTGCGGGAGAGTCAGCGCATCGCGAACGAGACGGAGGATATCGGTCGGAACACATTGGGAGATCTCGCGAGGCAGAGGGAGACGATTGAGCACACCAGGACTACGTTGCTGCAGAGCGAGGGTTACACGGACAGGAGCAATAAGACCCTCAAGGGCATGGCACGAAG	677 	\|\|\|\|\| \| \|\|\|\| \| \|\|\|\|\|\|\|\| \|  \|\|\|\| \|\|\|\| \|\|\|\|\| \|\|\|\| \|\|\|\|\|\|\|\|\| \|\|\|\|\|\|\|\|\|\|\|\|\| \|\|\|\|\|\|\|\|\|\| \|\|\|\|\|\| \|\|\|\|\|\|\|\|\|\|\|\|\|\|\|\|\|\|\|\|\|\|\|\|\|\|\|\|\|\|\|\|\|\|\|\|\|\|\|\|\|\|\|\|\|\|\|\|\|\|\|\|\|\|\|\|\|\|\|\|\|\|\| \|\|\|\|\|\|\|\|\|\|\|\|\|\|\|\|\|\|\|\|\|\|\|\|\|\|\|\|\|\|\|\|\|\|\|\|\|\|\|\|\|\|\|\|\|\|\|\|\|\|\|\|\|\|\|\|\|\|\|\|\|\|\|\|\|\|\|\|\|\|\|\|\|\|\|\|\|\|\|\|\|\|\|\|\|\|\|\|\|\|\|\|\|\|\|\|\|\|\|\|\|\|\|\|\|\|\|\|\|\|\|\|\|\|\|\|\|\|\|\|\|\|\|\|\|\|\|\|\|\|\|\|\|\|\|\|\|\|\|\|\|\|\|\|\|\|\|\|\|\|\|\|\|\|\|\|\|\|\|\|\|\|\|\|\|\|\|\|\|\|\|\|\|\|\|\|\|\|\|\|\|\|\|\|\|\|\|\|\|\|\|\|\|\|\|\|\|\|\|\|\|\|\|\|\|\|\|\|\|\|\|\|\|\|\|\|\|\|\|\|\|\|\|\|\|\|\|\|\|\|\|\|\|\|\|\|\|\|\|\|\|\|\|\|\|\|\|\|\|\|\|\|\|\|\|\|\|\|\|\|\|\|\|\|\|\|\|\|\|\|\|\|\|\|\|\|\|\|\|\|\|\|\|\|\|\|\|\|\|\|\|\|\|\|\|\|\|\|\|\|\|\|\|\|\|\|\|\|\|\|\|\|\|\|\|\|\|\|\|\|\|\|\|\|\|\|\|\|\|\|\|\|\|\|\|\|\|\|\|\|\|\|\|\|\|\|\|\|\|\|\|\|\|\|\|\|\|\|\|\|\|\|\|\|\|\|\|\|\|\|\|\|\|\|\|\|\|\|\|\|\|\|\|\|\|\|\|\|\|\|\|\|\|\|\|\|\|\|\|\|\|\|\|\|\|\|\|\|\| 	GCCGGNTNTGAGTTNTTCAGCAANTNNGAAGNGGAANTCAAGNTAGTNCAAGCAGACNTATCACAGAAACTNGATCAGATTCNTGAGCTNTCCGGCGAACAGCGAAAGGCAGCTGTTTCGCAGGCAGATCGCGCGCTTGAGGAGGCCAAGGAGNTAATAGAGTCCATGCGTTTGGAGAAGCAGAACATTCCACAAGCACTCAAGATCAAGGTCAACCAACGCTTCCGCAATTACCAGACCGACGTGGATGCTGCAGGACGGAAACTCAAAGGCATGCAGGACGATAGATCAGCGCTCTTCGGCAAGCGATACACCGACAATCCACAGGATGAGCAGCTCGAGCAGAGGCAGCAGCTTCTGGGAGGCACGGAGAGGTTAGAGCGGTCGAGCGGCAGGCTGCGGGAGAGTCAGCGCATCGCGAACGAGACGGAGGATATCGGTCGGAACACATTGGGAGATCTCGCGAGGCAGAGGGAGACGATTGAGCACACCAGGACTACGTTGCTGCAGAGCGAGGGTTACACGGACAGGAGCAATAAGACCCTCAAGGGCATGGCACGAAG ```      ``` 673	CGA-AGAATGGCAACAAACAAGATCATCACAGTGGCTATCATTGCGGTTTTGGTGATCTTGATCCTCGCTGTCATTATCAGCAAGTTTAGA	762 	\|\|\| \|\|\|\|\|\|\|\|\|\|\|\|\|\|\|\|\|\|\|\|\|\|\|\|\|\|\|\|\|\|\|\|\|\|\|\|\|\|\|\|\|\|\|\|\|\|\|\|\|\|\|\|\|\|\|\|\|\|\|\|\|\|\|\|\|\|\|\|\|\|\|\|\|\|\|\|\|\|\|\|\|\|\|\|\|\|\| 	CGACAGAATGGCAACAAACAAGATCATCACAGTGGCTATCATTGCGGTTTTGGTGATCTTGATCCTCGCTGTCATTATCAGCAAGTTTAGA ``` |
| 3 | ggggacaagtttgtacaaaaaagcaggcttgatgGCCGCCGAACAGCGCAAGCT, ggggaccactttgtacaagaaagctgggtcCCATCGGGGACGACCTCTTC | Mycgr3\|98404  JGI  Ensembl | KOG0796 Spliceosome subunit; |  | 100 % |  | ``` 154	AACACAAAGCAAGATCTCGGCTCTTGCGACCGCATGCACCTACCCAACCTGCGCGAGGAATATCTGGCCGCCACGGACGATCAGCGACGAGATTGGGGATTCGAATTCGACTACCAACGAGATATCGGAAAATATGTCAGCGAGTGCGACAGGCGCATAGAAGGTGCTCAGCGCAGACTCGAGAAGACACCCGAGGAGATACGACAGACGAATGCGCTGCTCAAGCAGATCAGTGACCTGGGCCGAACGATCGAGGCAGGCCTGATTGAGGTGGAGATCATGTGCGAGCTCGGAGCGGTCAATCTTGGTGTCCAGGAATTTCACAATTTGAAACTGAAAAAGGTTCAGAAGGATGAGCGCGAGCGCGAGCTGAAGGCGCTATCAGACACGTCTGGGCCATCAGGTCATCAAAAGCTACAAGTCTGCGACGTGTGCGGCGCATACCTGTCGCGACTGGACAACGATCGACGTTTGGCGGATCACTTCTTCGGCAAGATGCACCTGGGATACGCGCAAATGCGAAAGGAGTATGACAGGCTGGCGAAAGAGCTCAAGGGAAGGCCGCCGCCACGACGAGACGAGCCTGTGGAAGAGAGAGGCGGATATGATGATGGCGGATAtggaggaggatacggaggaagaaatgggggaggctatggaggtggtcgaggagggggtggaggtggagggtatgggcgtagaggtggagatgggggaggcttcggtggaggCGGAGGAAGAGGTCGTCCCCGATGG	909 	\|\|\| \|\|\|\| \|\|\| \|\| \| \|\| \| \|\|\| \|\|\|\| \|\|\|\|\|\|\|\|\|\|\|\|\|\|\|\|\|\| \|\|\|\|\|\|\|\|\|\| \|\|\|\|\|\|\| \|\|\|\|\|\|\|\|\|\|\|\|\|\|\|\|\|\|\|\|\|\|\|\|\|\|\|\|\|\|\|\|\|\|\|\|\|\|\|\|\|\|\|\|\|\|\|\|\|\|\|\|\|\|\|\|\|\|\|\|\|\|\|\|\| \|\|\|\|\|\|\|\|\|\|\|\|\|\|\|\|\|\|\|\|\|\|\|\|\|\|\|\|\|\|\|\|\|\|\|\|\|\|\|\|\|\|\|\|\|\|\|\|\|\|\|\|\|\|\|\|\|\|\|\|\|\|\|\|\|\|\|\|\|\|\|\|\|\|\|\|\|\|\|\|\|\|\|\|\|\|\|\|\|\|\|\|\|\|\|\|\|\|\|\|\|\|\|\|\|\|\|\|\|\|\|\|\|\|\|\|\|\|\|\|\|\|\|\|\|\|\|\|\|\|\|\|\|\|\|\|\|\|\|\|\|\|\|\|\|\|\|\|\|\|\|\|\|\|\|\|\|\|\|\|\|\|\|\|\|\|\|\|\|\|\|\|\|\|\|\|\|\|\|\|\|\|\|\|\|\|\|\|\|\|\|\|\|\|\|\|\|\|\|\|\|\|\|\|\|\|\|\|\|\|\|\|\|\|\|\|\|\|\|\|\|\|\|\|\|\|\|\|\|\|\|\|\|\|\|\|\|\|\|\|\|\|\|\|\|\|\|\|\|\|\|\|\|\|\|\|\|\|\|\|\|\|\|\|\|\|\|\|\|\|\|\|\|\|\|\|\|\|\|\|\|\|\|\|\|\|\|\|\|\|\|\|\|\|\|\|\|\|\|\|\|\|\|\|\|\|\|\|\|\|\|\|\|\|\|\|\|\|\|\|\|\|\|\|\|\|\|\|\|\|\|\|\|\|\|\|\|\|\|\|\|\|\|\|\|\|\|\|\|\|\|\|\|\|\|\|\|\|\|\|\|\|\|\|\|\|\|\|\|\|\|\|\|\|\|\|\|\|\|\|\|\|\|\|\|\|\|\|\|\|\|\|\|\|\|\|\|\|\|\|\|\|\|\|\|\|\|\|\|\|\|\|\|\|\|\|\|\|\|\|\|\|\|\|\|\|\|\|\|\|\|\|\|\|\|\|\|\|\|\|\|\|\|\|\|\|\|\|\|\|\|\|\|\|\|\|\|\|\|\|\|\|\|\|\|\|\|\|\|\|\|\|\|\|\|\|\|\|\|\|\|\|\|\|\|\|\|\|\|\|\|\|\|\|\|\|\|\|\|\|\|\|\|\|\|\|\|\|\|\|\|\|\|\|\|\|\|\|\|\|\|\|\|\|\|\|\|\|\|\|\|\|\|\|\|\|\|\|\|\|\|\|\|\|\|\|\|\|\|\|\|\|\|\|\|\|\|\|\|\|\|\|\|\|\|\|\|\|\|\|\|\|\|\|\|\|\|\|\|\|\|\|\|\|\|\|\|\|\|\|\|\|\|\|\|\|\|\|\|\|\|\|\|\|\|\|\|\|\|\|\|\|\|\|\|\|\|\|\|\| 	AACNCAAANCAANATTTNGGTTTTTGNGACCNCATGCACCTACCCAACCTNCGCGAGGAATNTCTGGCCNCCACGGACGATCAGCGACGAGATTGGGGATTCGAATTCGACTACCAACGAGATATCGGAAAATATNTCAGCGAGTGCGACAGGCGCATAGAAGGTGCTCAGCGCAGACTCGAGAAGACACCCGAGGAGATACGACAGACGAATGCGCTGCTCAAGCAGATCAGTGACCTGGGCCGAACGATCGAGGCAGGCCTGATTGAGGTGGAGATCATGTGCGAGCTCGGAGCGGTCAATCTTGGTGTCCAGGAATTTCACAATTTGAAACTGAAAAAGGTTCAGAAGGATGAGCGCGAGCGCGAGCTGAAGGCGCTATCAGACACGTCTGGGCCATCAGGTCATCAAAAGCTACAAGTCTGCGACGTGTGCGGCGCATACCTGTCGCGACTGGACAACGATCGACGTTTGGCGGATCACTTCTTCGGCAAGATGCACCTGGGATACGCGCAAATGCGAAAGGAGTATGACAGGCTGGCGAAAGAGCTCAAGGGAAGGCCGCCGCCACGACGAGACGAGCCTGTGGAAGAGAGAGGCGGATATGATGATGGCGGATATGGAGGAGGATACGGAGGAAGAAATGGGGGAGGCTATGGAGGTGGTCGAGGAGGGGGTGGAGGTGGAGGGTATGGGCGTAGAGGTGGAGATGGGGGAGGCTTCGGTGGAGGCGGAGGAAGAGGTCGTCCCCGATGG ``` |
| 3 | ggggacaagtttgtacaaaaaagcaggcttgatgTCGAGTAAACGAGGCGGTCG, ggggaccactttgtacaagaaagctgggtcTGAGTTGTCGTTGCGTTTCG | Mycgr3\|98638  JGI  Ensembl | KOG2278 RNA:NAD 2'-phosphotransferase TPT1; | tRNA splicing; transferase activity, transferring phosphorus-containing groups; | 100 % |  | ``` 363	CCCCACAGATATTCCTACACAAACGCCGTTACCCTCCGAAGATCCGAAAGACTACCTCATTCGTGCAAATCAAGGCCACAGTCTGAAGGTGGAATCAGAAGGGCTCCTAAAACCCATCACGCCGGAGACCGCACCGAAAAACGTCGTCCACGGAACCACGCACTCAGCCTGGCCGCTCATTGTCGCTTCTGGAGGTCTCAAACCGATGGGTCGAAACCACGCACACTTTGCTTCTGGACTCCCGGCAGGCTTCAAGTCGCTGGTAGAAGAAGACAGTGCAGCGACGGATGCGGCGCCGGTGATCTCGGGCATGCGCAAGTCTTCTACTGTATTGATGTTTCTGGACATCGCCAAGGCGATGGAGGCGGGCGTGAACTTTTGGCTGAGCGACAACGGTGTCATTCTGAGTGAGGGGAATGAGAACGGGGTCATTCCTCTGGAAGTCTTCCGGCGTGTGGAGGATCGGACGGGGGAAGGAGTCTTGTTGGAGGATGGCATGATTGTCAAGGAAGCGCCTGAGAAGTGGAGTGCGAAACGCAACGACAACTCA	912 	\|\|\|\| \|\|\|\|\|\|\|\|\| \| \| \|\|\| \| \|\|\|\|\| \|\|\|\|\| \|\|\|\|\|\|\| \|\|\|\|\|\| \|\|\| \|\|\|\|\|\|\|\|\|\|\|\|\|\|\|\|\|\|\|\|\|\|\|\|\|\| \|\|\|\|\|\|\|\|\|\|\|\|\|\|\|\|\|\|\|\|\|\|\|\|\|\|\|\|\|\|\|\|\|\|\|\|\|\|\|\|\|\|\|\|\|\|\|\|\|\|\|\|\|\|\|\|\|\|\|\|\|\|\|\|\|\|\|\|\|\|\|\|\|\|\|\|\|\|\|\|\|\|\|\|\|\|\|\|\|\|\|\|\|\|\|\|\|\|\|\|\|\|\|\|\|\|\|\|\|\|\|\|\|\|\|\|\|\|\|\|\|\|\|\|\|\|\|\|\|\|\|\|\|\|\|\|\|\|\|\|\|\|\|\|\|\|\|\|\|\|\|\|\|\|\|\|\|\|\|\|\|\|\|\|\|\|\|\|\|\|\|\|\|\|\|\|\|\|\|\|\|\|\|\|\|\|\|\|\|\|\|\|\|\|\|\|\|\|\|\|\|\|\|\|\|\|\|\|\|\|\|\|\|\|\|\|\|\|\|\|\|\|\|\|\|\|\|\|\|\|\|\|\|\|\|\|\|\|\|\|\|\|\|\|\|\|\|\|\|\|\|\|\|\|\|\|\|\|\|\|\|\|\|\|\|\|\|\|\|\|\|\|\|\|\|\|\|\|\|\|\|\|\|\|\|\|\|\|\|\|\|\|\|\|\|\|\|\|\|\|\|\|\|\|\|\|\|\|\|\|\|\|\|\|\|\|\|\|\|\|\|\|\|\|\|\|\|\|\|\|\|\|\|\|\|\|\|\|\|\|\|\|\|\|\|\|\|\|\|\|\|\|\|\|\|\|\|\|\|\|\|\|\|\|\|\|\|\|\|\|\|\|\|\|\|\|\|\|\|\|\|\|\|\|\|\|\|\|\|\|\|\|\|\|\|\|\|\|\|\|\|\|\|\|\|\|\|\|\|\|\|\|\|\|\|\|\|\|\|\|\|\|\|\|\|\|\|\|\|\|\|\|\|\|\|\|\|\|\|\|\|\|\|\|\|\|\|\|\|\|\|\|\|\|\|\|\|\|\|\|\|\|\|\|\|\|\| 	CCCCNCAGATATTCNTNCNCAANCNCCGTTNCCCTCNGAAGATCNGAAAGANTACNTCATTCGTGCAAATCAAGGCCACAGTNTGAAGGTGGAATCAGAAGGGCTCCTAAAACCCATCACGCCGGAGACCGCACCGAAAAACGTCGTCCACGGAACCACGCACTCAGCCTGGCCGCTCATTGTCGCTTCTGGAGGTCTCAAACCGATGGGTCGAAACCACGCACACTTTGCTTCTGGACTCCCGGCAGGCTTCAAGTCGCTGGTAGAAGAAGACAGTGCAGCGACGGATGCGGCGCCGGTGATCTCGGGCATGCGCAAGTCTTCTACTGTATTGATGTTTCTGGACATCGCCAAGGCGATGGAGGCGGGCGTGAACTTTTGGCTGAGCGACAACGGTGTCATTCTGAGTGAGGGGAATGAGAACGGGGTCATTCCTCTGGAAGTCTTCCGGCGTGTGGAGGATCGGACGGGGGAAGGAGTCTTGTTGGAGGATGGCATGATTGTCAAGGAAGCGCCTGAGAAGTGGAGTGCGAAACGCAACGACAACTCA ``` |
| 3 | ggggacaagtttgtacaaaaaagcaggcttgatgTCCGACGATCCTTTCCTAGC, ggggaccactttgtacaagaaagctgggtcAAGGAATAGTACGAACATTA | Mycgr3\|100899  JGI  Ensembl | KOG3202 SNARE protein TLG1/Syntaxin 6; |  | 100 % |  | ``` 761	GACAAATACTCCAGCTGCTGCATCTCGATGCTGATCATTGTCCTCATCATCCTGCTAATGTTCGTACTATTCCTT	835 	\|\|\|\|\|\|\|\|\|\|\|\|\|\|\|\|\|\|\|\|\|\|\|\|\|\|\|\|\|\|\|\|\|\|\|\|\|\|\|\|\|\|\|\|\|\|\|\|\|\|\|\|\|\|\|\|\|\|\|\|\|\|\|\|\|\|\|\|\|\|\|\|\|\|\| 	GACAAATACTCCAGCTGCTGCATCTCGATGCTGATCATTGTCCTCATCATCCTGCTAATGTTCGTACTATTCCTT ```      ``` 137	GATCCTTTCCTAGCGGCTCAGAGCGATGTTCTGTCCCTCCTCCAACAATCTCGACCACTCCTCTCTTCCTACCTCCGAATCCGATCGTCAGCGTCGAGCTCTAATTCTCCAGAACTCGCCGAAGCTCGATCCGAACTCGAAAGCACCCTCACGGATATAAGCACTGATCTCGAAGATCTCGTCGAGAGTGTCCGCGCTGTTGAAGGCGATCCCTATAAATATGGACTTGATGTCGCAGAAGTTGGGCGGAGGCGGAAGCTTGTGGCCGACGTTGGTCGAGAAGTGGAGGAGATGCGGCAGAAGCTGAACGAGACTGTGATCGCGGCAGACCGTAAAGCTGCGCTGGCACATCCAGATGCCTTTGCTACAGACCCTGGCGAGGATGAGGACGACTACGGCGCGTGGGAAGAGCAGCGACAGATGGAGATGATGCATGAGCAAGACGAGGCTCTCGATGGCGTCTTCCAGACAGTGGGCAATCTGCGTGCACAGGCAGATACGATGGGACGAGAACTGGAAGAGCAGGCAGAAATGCTCGAGGATACGGAGAACATCACAGATCGAGTGGGATCCAAGCTTGGAGTTGGCATGAAGAAGATCAGATATGTCATTGAGAAGAATGAGG	761 	\|\|\|\|\| \|\|\|\| \|  \|\| \|\|\| \|\| \|\|\| \|\| \| \|\|\| \|\| \|\|\|\|\|\| \|\| \| \|\|\|\|\| \| \|\| \| \|\|\|\|\|\|\| \|\|\|\|\|\|\|\|\|\|\|\|\|\|\|\|\|\|\|\|\|\|\|\|\|\|\| \|\|\|\|\| \|\|\|\|\|\|\|\|\|\|\|\|\|\|\|\|\|\|\|\|\|\|\|\|\|\|\|\|\|\|\|\|\|\|\|\|\|\|\|\|\|\|\|\|\|\|\|\|\|\|\|\|\|\|\|\|\|\|\|\|\|\|\|\|\|\|\|\|\|\|\|\|\|\|\|\|\|\|\|\|\|\|\|\|\|\|\|\|\|\|\|\|\|\|\|\|\|\|\|\|\|\|\|\|\|\|\|\|\|\|\|\|\|\|\|\|\|\|\|\|\|\|\|\|\|\|\|\|\|\|\|\|\|\|\|\|\|\|\|\|\|\|\|\|\|\|\|\|\|\|\|\|\|\|\|\|\|\|\|\|\|\|\|\|\|\|\|\|\|\|\|\|\|\|\|\|\|\|\|\|\|\|\|\|\|\|\|\|\|\|\|\|\|\|\|\|\|\|\|\|\|\|\|\|\|\|\|\|\|\|\|\|\|\|\|\|\|\|\|\|\|\|\|\|\|\|\|\|\|\|\|\|\|\|\|\|\|\|\|\|\|\|\|\|\|\|\|\|\|\|\|\|\|\|\|\|\|\|\|\|\|\|\|\|\|\|\|\|\|\|\|\|\|\|\|\|\|\|\|\|\|\|\|\|\|\|\|\|\|\|\|\|\|\|\|\|\|\|\|\|\|\|\|\|\|\|\|\|\|\|\|\|\|\|\|\|\|\|\|\|\|\|\|\|\|\|\|\|\|\|\|\|\|\|\|\|\|\|\|\|\|\|\|\|\|\|\|\|\|\|\|\|\|\|\|\|\|\|\|\|\|\|\|\|\|\|\|\|\|\|\|\|\|\|\|\|\|\|\|\|\|\|\|\|\|\|\|\|\|\|\|\|\|\|\|\|\|\|\|\|\|\|\|\|\|\|\|\|\|\|\|\|\|\|\|\|\|\|\|\|\|\|\|\|\|\|\|\|\|\|\|\|\|\|\|\|\|\|\|\|\|\|\|\|\|\|\|\|\|\|\|\|\|\|\|\|\|\|\|\|\|\|\|\|\|\|\|\|\|\|\|\|\|\|\|\|\|\|\|\|\|\|\|\|\|\|\|\|\|\|\|\|\|\|\|\|\|\|\|\|\|\|\|\|\|\|\|\|\|\|\|\|\|\|\|\|\|\| 	GATCC-TTCCNA-NGGNTCANAGNGATNTTNTNTCC-TCNTCCAACNATTTNGACCANTTCTTTTTTCCTACNTCCGAATCCGATCGTCAGCGTCGAGCTNTAATTNTCCAGAACTCGCCGAAGCTCGATCCGAACTCGAAAGCACCCTCACGGATATAAGCACTGATCTCGAAGATCTCGTCGAGAGTGTCCGCGCTGTTGAAGGCGATCCCTATAAATATGGACTTGATGTCGCAGAAGTTGGGCGGAGGCGGAAGCTTGTGGCCGACGTTGGTCGAGAAGTGGAGGAGATGCGGCAGAAGCTGAACGAGACTGTGATCGCGGCAGACCGTAAAGCTGCGCTGGCACATCCAGATGCCTTTGCTACAGACCCTGGCGAGGATGAGGACGACTACGGCGCGTGGGAAGAGCAGCGACAGATGGAGATGATGCATGAGCAAGACGAGGCTCTCGATGGCGTCTTCCAGACAGTGGGCAATCTGCGTGCACAGGCAGATACGATGGGACGAGAACTGGAAGAGCAGGCAGAAATGCTCGAGGATACGGAGAACATCACAGATCGAGTGGGATCCAAGCTTGGAGTTGGCATGAAGAAGATCAGATATGTCATTGAGAAGAATGAGG ``` |
| 3 | ggggacaagtttgtacaaaaaagcaggcttgatgCTGCAACAGAGATTGTTGCG, ggggaccactttgtacaagaaagctgggtcTGAGTTCTTCACCACGCCAA | Mycgr3\|102569  JGI  Ensembl | KOG3003 Molecular chaperone of the GrpE family; | adenyl-nucleotide exchange factor activity; chaperone binding;  protein folding;  protein homodimerization activity; | 0 % |  |  |
| 3 | ggggacaagtttgtacaaaaaagcaggcttgatgGACATTAATGCTATCATTAC, ggggaccactttgtacaagaaagctgggtcATCCGCCTCAACATCCTCCC | Mycgr3\|103274  JGI  Ensembl |  |  | 0 % |  | ``` 49	ATGGACATTAATGCTATCATTACATGTGGTCTCTTCTCTTTGACCTGGGCTCATGAGCTTGCAGCTGTTGATGCGTCG-TTCCGTGTACTGTTCAATATGCCACGCA	154 	\|\|\|\|\|\|\|\|\|\|\|\|\|\|\|\|\|\|\|\|\|\|\|\|\|\|\|\|\|\|\|\|\|\|\|\|\|\|\|\|\|\|\|\|\|\|\|\|\|\|\|\|\|\|\|\|\|\|\|\|\|\|\|\|\|\|\|\|\|\|\|\|\|\|\|\|\|\| \|\|\|\| \|\| \|  \|\|\|\|\|  \|\|\|\|\|\|\|\|\|\| 	ATGGACATTAATGCTATCATTACATGTGGTCTCTTCTCTTTGACCTGGGCTCATGAGCTTGCAGCTGTTGATGCGTCGNTTCCNTGNANNGTTCANNATGCCACGCA ```      ``` 49	ATGGACATTAATGCTATCATTACATGTGGTCTCTTCTCTTTGACCTGGGCTCATGAGCTTGCAGCTGTTGATGCGTCGTTCCGTGTACTGTTCAATATGCCACGCAGAGCGGAGAGCACAAAAATCCATCACCAGCGCAGTATTTCGTTGCAGTCGAGCGGCATCGACACGTCAACGCTCTGGTTCATTCACTTCCTTCACGACTTTCCAGCCGCCTTGCTTGACCCTCGCGCCGCTGCACCCAATTCGCCATCTCACGACAGTTCGCTAAGCACCACTATGCCTCAATCATCTCCCAATACCGCCTCCGCCGCGCCCATAGCTGTACTCTCCGCCTGCTCCTCGGCCTTCTTACGGCGATACTCATCCAGATACGCCTGCGCGGCATTGCCTTCGATGGCGGCAAAGTTACCTGTGGCGCTCTTCTTTCTTCTACCTGCGCTGTTCGCAGAAGCGGAGCCGAGGGTGGGGAAGGACGAGGATGAGGAAATGTCGATGGATGATTGGCGGGAGGATGTTGAGGCGGAT	576 	\|\|\|\|\|\|\|\|\|\|\|\|\|\|\|\|\|\|\|\|\|\|\|\|\|\|\|\|\|\|\|\| \|\|\|\|\|\|\|\|\|\|\|\|\|\|\|\| \|\|\|\|\|\|\|\|\|\|\|\|\|\|\|\|\|\|\|\|\|\|\|\|\|\|\|\|\|\|\|\|\|\|\|\|\|\|\|\|\|\|\|\|\|\|\|\|\|\|\|\|\|\|\|\|\|\|\|\|\|\|\|\|\|\|\|\|\|\|\|\|\|\|\|\|\|\|\|\|\|\|\|\|\|\|\|\|\|\|\|\|\|\|\|\|\|\|\|\|\|\|\|\|\|\|\|\|\|\|\|\|\|\| \|\|\|\|\|\|\|\|\|\|\|\|\|\| \|\|\|\|\|\|\|\|\|\|\|\|\|\|\|\|\|\|\|\|\|\|\|\|\|\|\|\|\|\|\|\|\|\|\|\|\|\|\|\|\|\|\|\|\|\|\|\|\|\|\|\|\|\|\|\|\|\|\|\|\|\|\|\|\|\|\|\|\|\|\|\|\|\|\|\|\|\|\|\|\|\|\|\|\|\|\|\|\|\|\|\|\|\|\|\|\|\|\|\|\|\|\|\|\|\|\|\|\|\|\|\|\|\|\|\|\|\|\|\|\|\|\|\|\|\|\|\|\|\|\|\|\|\|\|\|\|\|\|\|\|\|\|\|\|\|\|\|\|\|\|\|\|\|\|\|\|\|\|\|\|\|\|\|\|\|\|\|\|\|\|\|\|\|\|\|\|\|\|\|\|\|\|\|\|\|\|\|\|\|\|\|\|\|\|\|\|\|\|\|\|\|\|\|\|\|\|\|\|\|\|\|\|\|\|\|\|\|\|\|\|\|\|\|\|\|\|\|\|\|\|\|\|\|\|\|\|\|\|\|\|\|\|\|\|\|\|\|\|\|\|\|\|\|\|\|\|\|\|\|\|\|\|\|\|\|\|\|\|\|\|\|\|\|\|\|\|\|\|\|\|\|\|\|\|\|\|\|\|\|\|\|\|\|\|\|\|\|\|\|\|\|\|\|\|\|\|\|\|\|\|\|\|\|\|\|\|\|\|\|\|\|\|\|\|\|\|\|\|\|\|\|\|\|\|\|\|\|\|\|\|\|\|\|\|\|\|\| 	ATGGACATTAATGCTATCATTACATGTGGTCTNTTCTCTTTGACCTGGGNTCATGAGCTTGCAGCTGTTGATGCGTCGTTCCGTGTACTGTTCAATATGCCACGCAGAGCGGAGAGCACAAAAATCCATCACCAGCGCAGTATTTCGTTGCAGTCGAGCGGCATNGACACGTCAACGCTNTGGTTCATTCACTTCCTTCACGACTTTCCAGCCGCCTTGCTTGACCCTCGCGCCGCTGCACCCAATTCGCCATCTCACGACAGTTCGCTAAGCACCACTATGCCTCAATCATCTCCCAATACCGCCTCCGCCGCGCCCATAGCTGTACTCTCCGCCTGCTCCTCGGCCTTCTTACGGCGATACTCATCCAGATACGCCTGCGCGGCATTGCCTTCGATGGCGGCAAAGTTACCTGTGGCGCTCTTCTTTCTTCTACCTGCGCTGTTCGCAGAAGCGGAGCCGAGGGTGGGGAAGGACGAGGATGAGGAAATGTCGATGGATGATTGGCGGGAGGATGTTGAGGCGGAT ``` |
| 3 | ggggacaagtttgtacaaaaaagcaggcttgatgGTCCAACTCGAAGAAGTGCC, ggggaccactttgtacaagaaagctgggtcCAACGCAGGCTTTGCGCCGG | Mycgr3\|103356  JGI  Ensembl | KOG4111 Translocase of outer mitochondrial membrane complex, subunit TOM22; | P-P-bond-hydrolysis-driven protein transmembrane transporter activity; intracellular protein transport;  mitochondrial outer membrane; | 100 % |  | ``` 486	AGCTTCTTACTGCTGGCGCGCCAACCGGCGCAAAGCCTGCGTTG	529 	\|\|\|\|\|\|\|\|\|\|\|\|\|\|\|\|\|\|\|\|\|\|\|\|\|\|\|\|\|\|\|\|\|\|\|\|\|\|\|\|\|\|\|\| 	AGCTTCTTACTGCTGGCGCGCCAACCGGCGCAAAGCCTGCGTTG ```      ``` 197	GAATCAGAGATATCAGACGTGGACGAAGACGACGCTTTCGACGAGACCATCCTCGACCGAGTTTACGCCCTCAAAGACATCGTCCCGCCCACCTACCGCAAGCGCATCGCCAGCGCCGCCGACACCGGCTACGCATGGATATCATCGGGTCTGAGCTTCAGCGGAAAGACGTTGTGGGTTGTGAGCACGAGCGCGCTGCTTTTGGGTGTGCCGTGGGCGTTGGCGTTCAGCGAGGAGCAACAGTACCAGGAGATGGAGAGGGAGATGCGCATGCAGCAGAGTGCGAATGAG	487 	\|\|\|\|\|\|\|\|\|\|\|\|\|\|\|\|\|\|\|\|\|\|\|\|\|\|\|\|\|\|\|\|\|\|\|\|\|\|\|\|\|\|\|\|\|\|\|\|\|\|\|\|\|\|\|\|\|\|\|\|\|\|\|\|\|\|\|\|\|\|\|\|\|\|\|\|\|\|\|\|\|\|\|\|\|\|\|\|\|\|\|\|\|\|\|\|\|\|\|\|\|\|\|\|\|\|\|\|\|\|\|\|\|\|\|\|\|\|\|\|\|\|\|\|\|\|\|\|\|\|\|\|\|\|\|\|\|\|\|\|\|\|\|\|\|\|\|\|\|\|\|\|\|\|\|\|\|\|\|\|\|\|\|\|\|\|\|\|\|\|\|\|\|\|\|\|\|\|\|\|\|\|\|\|\|\|\|\|\|\|\|\|\|\|\|\|\|\|\|\|\|\|\|\|\|\|\|\|\|\|\|\|\|\|\|\|\|\|\|\|\|\|\|\|\|\|\|\|\|\|\|\|\|\|\|\|\|\|\|\|\|\|\|\|\|\|\|\|\|\|\|\|\|\|\|\|\|\|\|\|\|\|\|\|\|\|\|\|\|\|\|\|\|\|\|\|\|\|\|\|\|\|\|\|\|\|\|\|\|\|\| 	GAATCAGAGATATCAGACGTGGACGAAGACGACGCTTTCGACGAGACCATCCTCGACCGAGTTTACGCCCTCAAAGACATCGTCCCGCCCACCTACCGCAAGCGCATCGCCAGCGCCGCCGACACCGGCTACGCATGGATATCATCGGGTCTGAGCTTCAGCGGAAAGACGTTGTGGGTTGTGAGCACGAGCGCGCTGCTTTTGGGTGTGCCGTGGGCGTTGGCGTTCAGCGAGGAGCAACAGTACCAGGAGATGGAGAGGGAGATGCGCATGCAGCAGAGTGCGAATGAG ```      ``` 104	ATGGTCCAACTCGAAGAAGTGCCCGATGAGGAGCTCTTCGCGCAACAGTCAGGACCAAAGGCAGAGGATGAGGAAGAATGGGACACAAGTGACG	197 	\|\|\|\|\|\|\|\|\|\|\|\|\|\|\|\|\|\|\|\|\|\|\|\|\|\|\|\|\|\|\|\|\|\|\|\|\|\|\|\|\|\|\|\|\|\|\|\|\|\|\|\|\|\|\|\|\|\|\|\|\|\|\|\|\|\|\|\|\|\|\|\|\|\|\|\|\|\|\|\|\|\|\|\|\|\|\|\|\|\|\|\|\|\| 	ATGGTCCAACTCGAAGAAGTGCCCGATGAGGAGCTCTTCGCGCAACAGTCAGGACCAAAGGCAGAGGATGAGGAAGAATGGGACACAAGTGACG ``` |
| 3 | ggggacaagtttgtacaaaaaagcaggcttgatgCACTTCCAAACCATCTTCGC, ggggaccactttgtacaagaaagctgggtcGTTGTGAGGAGGAAGCGTCA | Mycgr3\|104404  JGI  Ensembl |  |  | 100 % |  | ``` 317	GTGCCTTCAGCATCGCTGTCATTCCAGGTCAGCTAAATGGCAACGGCGTCCTGGGAGGTTTCGCCGACTTCGCGGATCTGTCGCAAAATGCCTTGTGGGCTGCGCAAAACGCTGGCGCGGCCACGGGTGGTTGGCTGCACCCAAGAGTTGACTCGTCTGGTACCCTGCCATGCGACCAGCCACCAGCAAACGGCGGCCACGACAACCAGGTTTACGTCCAGTTCATCGTTGCCAGCTCGGAGTCCAACCCGCCTCAGCTGACGCTTCCTCCTCACAAC	594 	\|\|\|\|\|\|\|\|\|\|\|\|\|\|\|\|\|\|\|\|\|\|\|\|\|\|\|\|\|\|\|\|\|\|\|\|\|\|\|\|\|\|\|\|\|\|\|\|\|\|\|\|\|\|\|\|\|\|\|\|\|\|\|\|\|\|\|\|\|\|\|\|\|\|\|\|\|\|\|\|\|\|\|\|\|\|\|\|\|\|\|\|\|\|\|\|\|\|\|\|\|\|\|\|\|\|\|\|\|\|\|\|\|\|\|\|\|\|\|\|\|\|\|\|\|\|\|\|\|\|\|\|\|\|\|\|\|\|\|\|\|\|\|\|\|\|\|\|\|\|\|\|\|\|\|\|\|\|\|\|\|\|\|\|\|\|\|\|\|\|\|\|\|\|\|\|\|\|\|\|\|\|\|\|\|\|\|\|\|\|\|\|\|\|\|\|\|\|\|\|\|\|\|\|\|\|\|\|\|\|\|\|\|\|\|\|\|\|\|\|\|\|\|\|\|\|\|\|\|\|\|\|\|\|\|\|\|\|\|\|\|\|\|\|\|\|\|\|\|\|\|\|\|\|\|\|\|\|\|\|\|\|\|\|\|\|\|\|\|\|\|\|\|\|\|\|\|\| 	GTGCCTTCAGCATCGCTGTCATTCCAGGTCAGCTAAATGGCAACGGCGTCCTGGGAGGTTTCGCCGACTTCGCGGATCTGTCGCAAAATGCCTTGTGGGCTGCGCAAAACGCTGGCGCGGCCACGGGTGGTTGGCTGCACCCAAGAGTTGACTCGTCTGGTACCCTGCCATGCGACCAGCCACCAGCAAACGGCGGCCACGACAACCAGGTTTACGTCCAGTTCATCGTTGCCAGCTCGGAGTCCAACCCGCCTCAGCTGACGCTTCCTCCTCACAAC ```      ``` 57	GATG-CACTTCCAAACCATCTTCGCCGCCGGCCTCCTGCAAGCCGCGGCTGTGTCGGCCGTCCACTACCTCACTCCTGACAACGCTGGCCGCAACCAGGGAGCCAACTATTGTGGCCAATACGCACCCATGGACGACCAATTTGGCCTCAAGTCATGCTCCCAGGAAGACGTCCAGCTTCTCATCAACTCTTTCTTGGCAACTCCAGACGCCTACATCTCCATCGGTACCAGCTGGGCGCGCGCCGGCACTGCGGGTGCTTGT	318 	\|\|\|\| \|\| \|\|\|\|\|\|\|\|\|\|\| \|\| \|\|\|\|\|\|\|\|\| \|\| \|\|\|\|\|\|\|\|\|\|\|\| \|\|\|\|\|\|\|\|\|\|\|\|\|\|\| \|\|\|\|\|\|\|\|\|\|\|\|\|\|\|\|\|\|\|\|\|\|\|\|\|\|\|\|\|\|\|\|\|\|\|\|\|\|\|\|\|\|\|\|\|\|\|\|\|\|\|\|\|\|\|\|\|\|\|\|\|\|\|\|\|\|\|\|\|\|\|\|\|\|\|\|\|\|\|\|\|\|\|\|\|\|\|\|\|\|\|\|\|\|\|\|\|\|\|\|\|\|\|\|\|\|\|\|\|\|\|\|\|\|\|\|\|\|\|\|\|\|\|\|\|\|\|\|\|\|\|\|\|\|\|\|\|\|\|\|\|\|\|\|\|\|\|\|\|\|\|\|\|\|\|\|\|\|\|\|\|\|\|\|\|\|\|\|\|\|\|\|\|\|\|\|\|\|\|\|\|\|\|\|\|\|\|\|\|\|\|\|\|\|\|\|\|\| 	GATGNCANTTCCAAACCATNTTNGCCGCCGGCNTCNTGCAAGCCGCGGNTGTGTCGGCCGTCCANTACCTCACTCCTGACAACGCTGGCCGCAACCAGGGAGCCAACTATTGTGGCCAATACGCACCCATGGACGACCAATTTGGCCTCAAGTCATGCTCCCAGGAAGACGTCCAGCTTCTCATCAACTCTTTCTTGGCAACTCCAGACGCCTACATCTCCATCGGTACCAGCTGGGCGCGCGCCGGCACTGCGGGTGCTTGT ``` |
| 3 | ggggacaagtttgtacaaaaaagcaggcttgatgGGCGATCGACGCGGCGGCGG, ggggaccactttgtacaagaaagctgggtcTGTCAAACTTTGAATCAGAC | Mycgr3\|104520  JGI  Ensembl | KOG1774 Small nuclear ribonucleoprotein E; | mRNA metabolic process; ribonucleoprotein complex; | 100 % |  | ``` 379	GGCCAGATTCTGCTCAAAGGAGACAATGTCTGTCTGATTCAAAGTTTGACA	429 	\|\|\|\|\|\|\|\|\|\|\|\|\|\|\|\|\|\|\|\|\|\|\|\|\|\|\|\|\|\|\|\|\|\|\|\|\|\|\|\|\|\|\|\|\|\|\|\|\|\|\| 	GGCCAGATTCTGCTCAAAGGAGACAATGTCTGTCTGATTCAAAGTTTGACA ```      ``` 133	ATGGGCGATCGACGCGGCGGCGGCCAGCGCAAGGTCCTCCTCCCTCCAATCAACTTCATCTTCAAGTTGCTGCAACAACACAGCACGGTGCAGATCTGGTTATACGAGCAGCTTGGAATCCGTATCGAGGGCAAGATTCGCGGATTTGACGAATTCATGAATTTGGTGATCGACGAGGCGGTGGAGGTAAAGCAGCCAGGCAAAGGACAAACCGAGGCAGATGTGAAGGATCAGAGGCGGGAGTTGGG	380 	\|\|\|\|\|\|\|\|\|\|\|\|\|\|\|\|\|\|\|\|\|\|\|\|\|\|\|\|\|\|\|\|\|\|\|\|\|\|\|\|\|\|\|\|\|\|\|\|\|\|\|\|\|\|\|\|\|\|\|\|\|\|\|\|\|\|\|\|\|\|\|\|\|\|\|\|\|\|\|\|\|\|\|\|\|\|\|\|\|\|\|\|\|\|\|\|\|\|\|\|\|\|\|\|\|\|\|\|\|\|\|\|\|\|\|\|\|\|\|\|\|\|\|\|\|\|\|\|\|\|\|\|\|\|\|\|\|\|\|\|\|\|\|\|\|\|\|\|\|\|\|\|\|\|\|\|\|\|\|\|\|\|\|\|\|\|\|\|\|\|\|\|\|\|\|\|\|\|\|\|\|\|\|\|\|\|\|\|\|\|\|\|\|\|\|\|\|\|\|\|\|\|\|\|\|\|\|\|\|\|\|\|\|\|\|\|\|\|\|\|\|\|\|\|\|\|\|\|\|\|\|\|\|\|\|\|\|\|\|\|\|\|\|\|\|\|\|\| 	ATGGGCGATCGACGCGGCGGCGGCCAGCGCAAGGTCCTCCTCCCTCCAATCAACTTCATCTTCAAGTTGCTGCAACAACACAGCACGGTGCAGATCTGGTTATACGAGCAGCTTGGAATCCGTATCGAGGGCAAGATTCGCGGATTTGACGAATTCATGAATTTGGTGATCGACGAGGCGGTGGAGGTAAAGCAGCCAGGCAAAGGACAAACCGAGGCAGATGTGAAGGATCAGAGGCGGGAGTTGGG ``` |
| 3 | ggggacaagtttgtacaaaaaagcaggcttgatgCAGTTCTCCACCACTGTCCT, ggggaccactttgtacaagaaagctgggtcAAGCTGGGCAGGAGTGTAGC | Mycgr3\|104697  JGI  Ensembl |  |  | 100 % |  | ``` 100	ATGCAGTTCTCCACCACTGTCCTCGCCATCTTGGCCACCGCCGGCACCTCCTTCGCCGCCCCGGCTCCTGTCCCGGATGTCGCAGTCTCCAAGATGGTCTCAAACGCCCAATGGACGATCCGGGACTTCAAGCGTCAATGCGCCAACGGCTCTTGCAGCTTCACTTACGCGATCGACAACGGCGCCACCGTCACCGGCTGCGCCTACTCCGTCGCGGCAAACGGCGGCAAGTCGGCCGACCAGACCGACTACAGCAACATCAAGTGCGGCGACTACACCATCGGCAGCGGCTGGAGCGGCCAGTTCGGCCCCGGCAACGGTTTCACGACCCTGTCTGTTGTCGACAGCTCTCGCAAGATCATCTACCCGTCTTACAGCGACGCGACCATCGCCAACGCTGGCAATGGCCCTGTCAAGCCCGACCAGAGCTACACTCCTGCCCAGCTT	546 	\|\|\|\|\|\|\|\| \|\|\|\|\|\|\|\|\|\|\|\|\|\| \|\|\|\|\| \|\|\|\|\|\|\|\|\|\|\|\|\|\|\|\|\|\|\|\| \|\| \|\|\|\|\|\|\|\|\|\|\|\|\|\|\|\|\|\|\|\|\|\|\|\|\|\|\|\|\|\|\|\|\|\|\|\|\|\|\|\|\|\|\|\|\|\|\|\|\|\|\|\|\|\|\|\|\|\|\|\|\|\|\|\|\|\|\|\|\|\|\|\|\|\|\|\|\|\|\|\|\|\|\|\|\|\|\|\|\|\|\|\|\|\|\|\|\|\|\|\|\|\|\|\|\|\|\|\|\|\|\|\|\|\|\|\|\|\|\|\|\|\|\|\|\|\|\|\|\|\|\|\|\|\|\|\|\|\|\|\|\|\|\|\|\|\|\|\|\|\|\|\|\|\|\|\|\|\|\|\|\|\|\|\|\|\|\|\|\|\|\|\|\|\|\|\|\|\|\|\|\|\|\|\|\|\|\|\|\|\|\|\|\|\|\|\|\|\|\|\|\|\|\|\|\|\|\|\|\|\|\|\|\|\|\|\|\|\|\|\|\|\|\|\|\|\|\|\|\|\|\|\|\|\|\|\|\|\|\|\|\|\|\|\|\|\|\|\|\|\|\|\|\|\|\|\|\|\|\|\|\|\|\|\|\|\|\|\|\|\|\|\|\|\|\|\|\|\|\|\|\|\|\|\|\|\|\|\|\|\|\|\|\|\|\|\|\|\|\|\|\|\|\|\|\|\|\|\|\|\|\|\|\|\|\|\|\|\|\|\|\|\|\|\|\|\|\|\|\|\|\|\|\|\|\|\|\|\|\|\|\|\|\|\|\|\|\|\|\|\|\|\|\|\|\|\|\|\|\|\|\|\|\|\|\|\|\|\|\|\|\|\|\|\|\|\|\|\|\|\|\|\|\|\|\|\|\|\|\|\|\|\|\| 	ATGCAGTTNTCCACCACTGTCCTNGCCATNTTGGCCACCGCCGGCACCTCNTTNGCCGCCCCGGCTCCTGTCCCGGATGTCGCAGTCTCCAAGATGGTCTCAAACGCCCAATGGACGATCCGGGACTTCAAGCGTCAATGCGCCAACGGCTCTTGCAGCTTCACTTACGCGATCGACAACGGCGCCACCGTCACCGGCTGCGCCTACTCCGTCGCGGCAAACGGCGGCAAGTCGGCCGACCAGACCGACTACAGCAACATCAAGTGCGGCGACTACACCATCGGCAGCGGCTGGAGCGGCCAGTTCGGCCCCGGCAACGGTTTCACGACCCTGTCTGTTGTCGACAGCTCTCGCAAGATCATCTACCCGTCTTACAGCGACGCGACCATCGCCAACGCTGGCAATGGCCCTGTCAAGCCCGACCAGAGCTACACTCCTGCCCAGCTT ``` |
| 3 | ggggacaagtttgtacaaaaaagcaggcttgatgAAGATCTTCACTCTCCTCTC, ggggaccactttgtacaagaaagctgggtcTGAAGTCTTGACCTTGGTAA | Mycgr3\|104754  JGI  Ensembl |  |  | 0 % |  | ``` 204	ATTGAAGCGACCTTCGAATTCGTCTTCGAAGAGGCAGGTGGGCAGGGGGCTACGGAGGAGGCTGAGGTTATTATC-A-CGAC-TGAAGGCAATACCGGTCGAAACGCCGCCGCAGCAGGCTACGCACCATCAGGCACATCAATCACAGGATCCAAAT-T-TGAGAAATTTGGCACGCGA-TTCTTCTTCAGAAAATTCGACGAATTCACTGGCTTT	413 	\|\|\|\|\|\|\|\|\|\|\|\|\|\|\|\|\|\|\|\|\|\|\|\|\|\|\|\|\|  \| \| \|\|\|\|  \|\| \|\|\|\|\|\| \|\|\|\|\|\|\|\|\|\|\|\|\|\|\|\|\|\| \|\| \|  \| \|  \| \|   \|\|\|  \|\|  \|\|\|\|\|    \|\| \| \|\|\|\|\|\|\|\|\|\|\|\|\|\|\|\|\|\|\|\|\|\|\|\|\|\|\|\|\|\|\|\|\| \|\| \|\|   \|\|\|\|  \| \|\| \|\|     \| \|\|\| \| \| \|\|\|\|\| \|\|\|\|\| \|\|\|\| \|\|\|\|\|\|\|\|\| \|\|\|\|\|\|\|\|\| 	ATTGAAGCGACCTTCGAATTCGTCTTCGAGAATGGAGGT--GC-GGGGGCCACGGAGGAGGCTGAGGTTCTTGTTGAACCTCCTCCCGGCG-TA--GGTCGG---GCTGGCGCAGCAGGCTACGCACCATCAGGCACATCAATTACGGGCCACAAACCTATG-GACGG-GGCCAC-CAAGTTCTTTTTCAGGAAATACGACGAATTTACTGGCTTT ``` |
| 3 | ggggacaagtttgtacaaaaaagcaggcttgatgGCACCCAATATGAGTCTCTT, ggggaccactttgtacaagaaagctgggtcAAGTCCTTGCCGTAATCTCT | Mycgr3\|105015  JGI  Ensembl | KOG3343 Vesicle coat complex COPI, zeta subunit; |  | 100 % |  | ``` 98	TCCTCCTCTCCACCGACGACAACTCGCGCATCCTCGCGAAATACTACAGTCCACCACACATCCCGCAAGGCGCTCAGGGCAACAACTATCCCGGCGCGAATCCATACCCGACCGTGAAAGAGCAGAAAGCATTCGAGAAGGGATTGTTGGAGAAGACGGCCAAGCAGACGAGCGATGTCATCCTCTACGACAACCGTGTGGTGGTGTTCAAGATGGAGCAGGATGTCATGATGTATGTGGTTGGAGGCGCGGAGGAGAATGAGATTATGCTGTACAATGTGGTTTTGGCGTTGAGGGATAGTTTGACGATCTTGCTGAA	416 	\|\|\|\|\| \| \|\|\|\|\| \|\| \|\|\|\|\|\|\|\|\| \|\|\|\|\|\|\|\|\|\|\|\|\|\|\|\| \|\|\|\|\|\|\|\|\|\|\|\|\|\|\|\|\|\|\|\|\|\|\|\|\|\|\|\|\|\|\|\|\|\|\|\|\|\|\|\|\|\|\|\|\|\|\|\|\|\|\|\|\|\|\|\|\|\|\|\|\|\|\|\|\|\|\|\|\|\|\|\|\|\|\|\|\|\|\|\|\|\|\|\|\|\|\|\|\|\|\|\|\|\|\|\|\|\|\|\|\|\|\|\|\|\|\|\|\|\|\|\|\|\|\|\|\|\|\|\|\|\|\|\|\|\|\|\|\|\|\|\|\|\|\|\|\|\|\|\|\|\|\|\|\|\|\|\|\|\|\|\|\|\|\|\|\|\|\|\|\|\|\|\|\|\|\|\|\|\|\|\|\|\|\|\|\|\|\|\|\|\|\|\|\|\|\|\|\|\|\|\|\|\|\|\|\|\|\|\|\|\|\|\|\|\|\|\|\|\|\|\|\|\|\|\|\|\|\|\|\|\|\|\|\|\|\|\|\|\|\|\|\|\|\|\|\|\|\|\|\|\|\|\|\|\|\|\|\|\|\|\|\|\|\|\|\|\|\|\|\|\|\|\|\|\|\|\|\|\|\|\|\|\|\| 	TCCTCNTNTCCACNGANGACAACTCGNGCATCCTCGCGAAATANTACAGTCCACCACACATCCCGCAAGGCGCTCAGGGCAACAACTATCCCGGCGCGAATCCATACCCGACCGTGAAAGAGCAGAAAGCATTCGAGAAGGGATTGTTGGAGAAGACGGCCAAGCAGACGAGCGATGTCATCCTCTACGACAACCGTGTGGTGGTGTTCAAGATGGAGCAGGATGTCATGATGTATGTGGTTGGAGGCGCGGAGGAGAATGAGATTATGCTGTACAATGTGGTTTTGGCGTTGAGGGATAGTTTGACGATCTTGCTGAA ```      ``` 417	AAACTCCGTGGACAAGCGCACCGTGATTGAAAACTACGACCTGGCCTCCCTCTGCATCGACGAGATTGTCGACGACGGCATCATCCTCGAGACCGACCCAGTCGTCATCGCAAGTCGTGTCAGCCGTCCTCCACAACAGGATATGCCCAACATGCAAGGTATTGATCTCAGCGAGGAGGGATTGTTGAAAATCTACCAATTCGGAAAGCAAAAGTTGGGCGAGAGATTACGGCAAGGACTT	657 	\|\|\|\|\|\|\|\|\|\|\|\|\|\|\|\|\|\|\|\|\|\|\|\|\|\|\|\|\|\|\|\|\|\|\|\|\|\|\|\|\|\|\|\|\|\|\|\|\|\|\|\|\|\|\|\|\|\|\|\|\|\|\|\|\|\|\|\|\|\|\|\|\|\|\|\|\|\|\|\|\|\|\|\|\|\|\|\|\|\|\|\|\|\|\|\|\|\|\|\|\|\|\|\|\|\|\|\|\|\|\|\|\|\|\|\|\|\|\|\|\|\|\|\|\|\|\|\|\|\|\|\|\|\|\|\|\|\|\|\|\|\|\|\|\|\|\|\|\|\|\|\|\|\|\|\|\|\|\|\|\|\|\|\|\|\|\|\|\|\|\|\|\|\|\|\|\|\|\|\|\|\|\|\|\|\|\|\|\|\|\|\|\|\|\|\|\|\|\|\|\|\|\|\|\|\|\|\|\|\|\|\|\|\|\|\|\|\|\|\|\|\|\|\|\|\|\|\|\|\|\|\|\|\|\|\|\|\|\|\|\| 	AAACTCCGTGGACAAGCGCACCGTGATTGAAAACTACGACCTGGCCTCCCTCTGCATCGACGAGATTGTCGACGACGGCATCATCCTCGAGACCGACCCAGTCGTCATCGCAAGTCGTGTCAGCCGTCCTCCACAACAGGATATGCCCAACATGCAAGGTATTGATCTCAGCGAGGAGGGATTGTTGAAAATCTACCAATTCGGAAAGCAAAAGTTGGGCGAGAGATTACGGCAAGGACTT ``` |
| 3 | ggggacaagtttgtacaaaaaagcaggcttgatgAAGTCCGCTTGCGCCATCCT, ggggaccactttgtacaagaaagctgggtcCTCGTACTGCCAAGCAGCGG | Mycgr3\|105677  JGI  Ensembl |  |  | 0 % |  |  |
| 3 | ggggacaagtttgtacaaaaaagcaggcttgatgCTGAAGACATCACGCATCTT, ggggaccactttgtacaagaaagctgggtcTTTCGGCTTCGGCTTTCCAA | Mycgr3\|106251  JGI  Ensembl | KOG4526 Predicted membrane protein; |  | 100 % |  | ``` 151	GAAGACATCACGCATCTTCCAGCTCTTTCGCCAAGTGCCGACCACCCAAGAAGCTCTGCTCAGAGCCTCACGACGGACAGAACTCACAGGCCTGCGACTACCGTCGAAAACGATACCGCAACGGCGAGGTCTCACCACAACCACCATCCGATCCGCAAGACCAGCTCTTTCATCAACATGGCGAGCTTCACTATTCCGCCGCGCGCGACAGACACGATACAACAGCACGAATCCCACCGGTCCGAAGATCGATGAGACGAAATTGTCCATGTCGGAACGCATGAGGGCGATGTCGAAGAAATATGGCTGGACGGTGGTGGGGATATACCTGGGACTGTCGGCGATTGATTTTCCATTCTGTTACTTGGCTGTACGATGGTTTGGCACTGAGCGGATAGCAAAGGTCGAACACGCGATCATGGATTACTTCTGGACGACACTGGAAAAGGTTGCGCCGGGATTGATGGCGCGGAGGGAGACCAAAGAGGCGCTGGATGCTGCTGAGGAGGCTACAAGGGATGCGGATGCTGCTGTTGCGGAGAAGGCGAAGCATGAGAATGCTAGTA	716 	\|\|\|\|\|\|\|\|\|\| \|\| \| \|\|\|\|\|\| \| \|\|\|  \|\| \|\|\|\|\|\|\|\|\|\|\|\|\|\|\|\|\|\|\|\| \| \|\| \|\|\|\|\|  \|\|\|\|\|\|\|\| \|\|\|\|\|\|\|\|\|\| \|\|\|\|\|\|\|\| \|\| \|\|\|\|\|\|\|\|\|\|\|\|\|\|\|\|\|\|\|\|\|\|\|\|\|\|\|\|\|\|\|\|\|\|\|\|\|\|\|\|\|\|\|\|\|\|\|\|\|\|\|\|\|\|\|\|\|\|\|\|\|\|\|\|\|\|\|\|\|\|\|\|\|\|\|\|\|\|\|\|\|\|\|\|\|\|\|\|\|\|\|\|\|\|\|\|\|\|\|\|\|\|\|\|\|\|\|\|\|\|\|\|\|\|\|\|\|\|\|\|\|\|\|\|\|\|\|\|\|\|\|\|\|\|\|\|\|\|\|\|\|\|\|\|\|\|\|\|\|\|\|\|\|\|\|\|\|\|\|\|\|\|\|\|\|\|\|\|\|\|\|\|\|\|\|\|\|\|\|\|\|\|\|\|\|\|\|\|\|\|\|\|\|\|\|\|\|\|\|\|\|\|\|\|\|\|\|\|\|\|\|\|\|\|\|\|\|\|\|\|\|\|\|\|\|\|\|\|\|\|\|\|\|\|\|\|\|\|\|\|\|\|\|\|\|\|\|\|\|\|\|\|\|\|\|\|\|\|\|\|\|\|\|\|\|\|\|\|\|\|\|\|\|\|\|\|\|\|\|\|\|\|\|\|\|\|\|\|\|\|\|\|\|\|\|\|\|\|\|\|\|\|\|\|\|\|\|\|\|\|\|\|\|\|\|\|\|\|\|\|\|\|\|\|\|\|\|\|\|\|\|\|\|\|\|\|\|\|\|\|\|\|\|\|\|\|\|\|\|\|\|\|\|\|\|\|\|\|\|\|\|\|\|\|\|\|\|\|\|\|\|\|\|\|\|\|\|\|\|\|\|\|\|\|\|\|\|\|\|\|\|\|\|\|\|\|\|\|\|\|\|\|\|\|\|\|\|\|\|\|\|\|\|\|\|\|\|\|\|\|\|\|\|\|\|\|\|\|\|\|\|\|\|\|\|\|\|\|\|\|\|\|\|\|\|\|\|\|\|\|\|\|\|\|\|\|\|\|\|\|\|\|\|\|\|\|\|\| 	GAAGACATCA-GCNT-TTCCAG-TTTTTNNCC-AGTGCCGACCACCCAAGAAGNT-TGNTCAGA-NCTCACGAC-GACAGAACTCNCAGGCCTGNGANTACCGTCGAAAACGATACCGCAACGGCGAGGTCTCACCACAACCACCATCCGATCCGCAAGACCAGCTCTTTCATCAACATGGCGAGCTTCACTATTCCGCCGCGCGCGACAGACACGATACAACAGCACGAATCCCACCGGTCCGAAGATCGATGAGACGAAATTGTCCATGTCGGAACGCATGAGGGCGATGTCGAAGAAATATGGCTGGACGGTGGTGGGGATATACCTGGGACTGTCGGCGATTGATTTTCCATTCTGTTACTTGGCTGTACGATGGTTTGGCACTGAGCGGATAGCAAAGGTCGAACACGCGATCATGGATTACTTCTGGACGACACTGGAAAAGGTTGCGCCGGGATTGATGGCGCGGAGGGAGACCAAAGAGGCGCTGGATGCTGCTGAGGAGGCTACAAGGGATGCGGATGCTGCTGTTGCGGAGAAGGCGAAGCATGAGAATGCTAGTA ```      ``` 146	ATGCTGAAGACATCACGCATCTTCCAGCTCTTTCGCCAAGTGCCGACCACCCAAGAAGCTCTGCTCAGAGCCTCACGACGGACAGAACTCACAGGCCTGCGACTACCGTCGAAAACGATACCGCAACGGCGAGGTCTCACCACAACCACCATCCGATCCGCAAGACCAGCTCTTTCATCAACATGGCGAGCTTCACTATTCCGCCGCGCGCGACAGACACGATACAACAGCACGAATCCCACCGGTCCGAAGATCGATGAGACGAAATTGTCCATGTCGGAACGCATGAGGGCGATGTCGAAGAAATATGGCTGGACGGTGGTGGGGATATACCTGGGACTGTCGGCGATTGATTTTCCATTCTGTTACTTGGCTGTACGATGGTTTGGCACTGAGCGGATAGCAAAGGTCGAACACGCGATCATGGATTACTTCTGGACGACACTGGAAAAGGTTGCGCCGGGATTGATGGCGCGGAGGGAGACCAAAGAGGCGCTGGATGCTGCTGAGGAGGCTACAAGGGATGCGGATGCTGCTGTTGCGGAGAAGGCGAAGCATGAGAATGCTAGTA	716 	\|\|\|\|\|\|\|\|\|\|\|\|\|\|\|\|\|\|\|\|\|\|\|\|\|\|\|\|\|\|\|\|\|\|\|\|\|\|\|\|\|\|\|\|\|\|\|\|\|\|\|\|\|\|\|\|\|\|\|\|\|\|\|\|\|\|\|\|\|\|\|\|\|\|\|\|\|\|\|\|\|\|\|\|\|\|\|\|\|\|\|\|\|\|\|\|\|\|\|\|\|\|\|\|\|\|\|\|\|\|\|\|\|\|\|\|\|\|\|\|\|\|\|\|\|\|\|\|\|\|\|\|\|\|\|\|\|\|\|\|\|\|\|\|\|\|\|\|\|\|\|\|\|\|\|\|\|\|\|\|\|\|\|\|\|\|\|\|\|\|\|\|\|\|\|\|\|\|\|\|\|\|\|\|\|\|\|\|\|\|\|\|\|\|\|\|\|\|\|\|\|\|\|\|\|\|\|\|\|\|\|\|\|\|\|\|\|\|\|\|\|\|\|\|\|\|\|\|\|\|\|\|\|\|\|\|\|\|\|\|\|\|\|\|\|\|\|\|\|\|\|\|\|\|\|\|\|\|\|\|\|\|\|\|\|\|\|\|\|\|\|\|\|\|\|\|\|\|\|\|\|\|\|\|\|\|\|\|\|\|\|\|\|\|\|\|\|\|\|\|\|\|\|\|\|\|\|\|\|\|\|\|\|\|\|\|\|\|\|\|\|\|\|\|\|\|\|\|\|\|\|\|\|\|\|\|\|\|\|\|\|\|\|\|\|\|\|\|\|\|\|\|\|\|\|\|\|\|\|\|\|\|\|\|\|\|\|\|\|\|\|\|\|\|\|\|\|\|\|\|\|\|\|\|\|\|\|\|\|\|\|\|\|\|\|\|\|\|\|\|\|\|\|\|\|\|\|\|\|\|\|\|\|\|\|\|\|\|\|\|\|\|\|\|\|\|\|\|\|\|\|\|\|\|\|\|\|\|\|\|\|\|\|\|\|\|\|\|\|\|\|\|\|\|\|\|\|\|\|\|\|\|\|\|\|\|\|\|\|\|\|\|\|\|\|\|\|\|\|\|\|\|\|\|\|\|\|\|\|\|\|\|\|\|\|\|\|\|\|\|\|\|\|\|\|\|\|\|\|\|\|\|\|\|\|\|\|\|\|\|\|\|\|\|\|\|\|\|\|\|\|\|\|\|\|\|\|\|\|\|\|\|\|\|\|\|\|\|\|\|\|\|\|\|\|\|\|\|\|\|\|\|\|\|\|\|\|\|\|\|\| 	ATGCTGAAGACATCACGCATCTTCCAGCTCTTTCGCCAAGTGCCGACCACCCAAGAAGCTCTGCTCAGAGCCTCACGACGGACAGAACTCACAGGCCTGCGACTACCGTCGAAAACGATACCGCAACGGCGAGGTCTCACCACAACCACCATCCGATCCGCAAGACCAGCTCTTTCATCAACATGGCGAGCTTCACTATTCCGCCGCGCGCGACAGACACGATACAACAGCACGAATCCCACCGGTCCGAAGATCGATGAGACGAAATTGTCCATGTCGGAACGCATGAGGGCGATGTCGAAGAAATATGGCTGGACGGTGGTGGGGATATACCTGGGACTGTCGGCGATTGATTTTCCATTCTGTTACTTGGCTGTACGATGGTTTGGCACTGAGCGGATAGCAAAGGTCGAACACGCGATCATGGATTACTTCTGGACGACACTGGAAAAGGTTGCGCCGGGATTGATGGCGCGGAGGGAGACCAAAGAGGCGCTGGATGCTGCTGAGGAGGCTACAAGGGATGCGGATGCTGCTGTTGCGGAGAAGGCGAAGCATGAGAATGCTAGTA ```      ``` 714	GTATCTGGACCCAATTGCTCTTCGCCTACGGAGTTCACAAGTCCCTCATCTTCATCCGAGTGCCGCTCACGGTTGCCATCACGCCGAAAGTCGTCAAGACATTGCGGAAATGGGGTTATCAGATTGGAAAGCCGAAGCCGAAA	856 	\|\|\|\|\|\|\|\|\|\|\|\|\|\|\|\|\|\|\|\|\|\|\|\|\|\|\|\|\|\|\|\|\|\|\|\|\|\|\|\|\|\|\|\|\|\|\|\|\|\|\|\|\|\|\|\|\|\|\|\|\|\|\|\|\|\|\|\|\|\|\|\|\|\|\|\|\|\|\|\|\|\|\|\|\|\|\|\|\|\|\|\|\|\|\|\|\|\|\|\|\|\|\|\|\|\|\|\|\|\|\|\|\|\|\|\|\|\|\|\|\|\|\|\|\|\|\|\|\|\|\|\|\|\|\|\|\|\|\|\|\|\|\| 	GTATCTGGACCCAATTGCTCTTCGCCTACGGAGTTCACAAGTCCCTCATCTTCATCCGAGTGCCGCTCACGGTTGCCATCACGCCGAAAGTCGTCAAGACATTGCGGAAATGGGGTTATCAGATTGGAAAGCCGAAGCCGAAA ``` |
| 3 | ggggacaagtttgtacaaaaaagcaggcttgatgCTCAGAGACCTGCCACCCTT, ggggaccactttgtacaagaaagctgggtcGCCTTCCGGGAGCAGGTCTT | Mycgr3\|106497  JGI  Ensembl | KOG4774 Uncharacterized conserved protein; |  | 0 % |  |  |
| 3 | ggggacaagtttgtacaaaaaagcaggcttgatgTATGGCGAGCAAGGCAACAA, ggggaccactttgtacaagaaagctgggtcGCTCAGTCGTTGCAAATAGC | Mycgr3\|107715  JGI  Ensembl | KOG3303 Predicted alpha-helical protein, potentially involved in replication/repair; |  | 100 % |  | ``` 45	CATCCAAAACCTGCCACACCTACCACCATACGCAACGACGACAATCCGTGCGGTGACAAAAGAAGTCCGCGAGCTGGACCGCGACGCGACCGCGCTTCTGGCACCATACAGACTCGGAAGCACTCAAGGAAGCAATGGATCTCAGCCTCAATCATCGTTCAACCCGGCAGACGACCCGGCTTTGGCCTGCACACTCCTGGTCAACCACCTCTCCACCCGCAGGAACAAGCGATGCCTAATGGCATACCACAAAACCCGCACGGACAAGCTGGAAGCCATGTGCTGGGAGGGTGTCGATGTCTCTGATTTGCAGCAAGCTTCCGCCTCGACTACCGCTGGTGGACAGAGTGGACCGACGAGCAGTCTGAGCGCAGAGGAAGAACAGTACGTTCATCAGTACAGCGACCTTCTCGCTGCGTACAAGGGTCAGTGGACCGACATCGACTTGACTGGAAGTCTAGAGCCTCCGAAAGATATCTTCATCGATGTGCGGGTGCTGAAGGATGCGGGTGAAATTCAGACGGAGTACGGGTCGATCACTTTGACGAAGAACAGCCAGTTCTACGTTCGTCAAGCTGATGTCGAGCGGCTCATTCATCAAGGCTATTTGCAACGACTGAGC	666 	\|\|\|\|\|\|\|\|\|\|\|\|\|\|\|\|\|\|\|\|\|\|\|\|\|\|\|\|\|\|\|\|\|\|\| \|\| \|\|\|\|\|\|\|\|\|\|\|\|\|\|\|\|\|\|\|\|\|\|\|\|\|\|\|\|\|\|\|\|\|\|\|\|\|\|\|\|\|\|\|\|\|\|\|\|\|\|\|\|\|\|\|\|\|\|\|\|\|\|\|\|\|\|\|\|\|\|\|\|\|\|\|\|\|\|\|\|\|\|\|\|\|\|\|\|\|\|\|\|\|\|\|\|\|\|\|\|\|\|\|\|\|\|\|\|\|\|\|\|\|\|\|\|\|\|\|\|\|\|\|\|\|\|\|\|\|\|\|\|\|\|\|\|\|\|\|\|\|\|\|\|\|\|\|\|\|\|\|\|\|\|\|\|\|\|\|\|\|\|\|\|\|\|\|\|\|\|\|\|\|\|\|\|\|\|\|\|\|\|\|\|\|\|\|\|\|\|\|\|\|\|\|\|\|\|\|\|\|\|\|\|\|\|\|\|\|\|\|\|\|\|\|\|\|\|\|\|\|\|\|\|\|\|\|\|\|\|\|\|\|\|\|\|\|\|\|\|\|\|\|\|\|\|\|\|\|\|\|\|\|\|\|\|\|\|\|\|\|\|\|\|\|\|\|\|\|\|\|\|\|\|\|\|\|\|\|\|\|\|\|\|\|\|\|\|\|\|\|\|\|\|\|\|\|\|\|\|\|\|\|\|\|\|\|\|\|\|\|\|\|\|\|\|\|\|\|\|\|\|\|\|\|\|\|\|\|\|\|\|\|\|\|\|\|\|\|\|\|\|\|\|\|\|\|\|\|\|\|\|\|\|\|\|\|\|\|\|\|\|\|\|\|\|\|\|\|\|\|\|\|\|\|\|\|\|\|\|\|\|\|\|\|\|\|\|\|\|\|\|\|\|\|\|\|\|\|\|\|\|\|\|\|\|\|\|\|\|\|\|\|\|\|\|\|\|\|\|\|\|\|\|\|\|\|\|\|\|\|\|\|\|\|\|\|\|\|\|\|\|\|\|\|\|\|\|\|\|\|\|\|\|\|\|\|\|\|\|\|\|\|\|\|\|\|\|\|\|\|\|\|\|\|\|\|\|\|\|\|\|\|\|\|\|\|\|\|\|\|\|\|\|\|\|\|\|\|\|\|\|\|\|\|\|\|\|\|\|\|\|\|\|\|\|\|\|\|\|\|\|\|\|\|\|\|\|\|\|\|\|\|\|\|\|\|\|\|\|\|\|\|\|\|\|\|\|\|\|\|\|\|\|\|\|\|\|\|\|\|\|\|\|\|\|\|\|\|\|\|\|\|\|\|\|\|\|\|\|\|\|\| 	CATCCAAAACCTGCCACACCTACCACCATACGCAANGANGACAATCCGTGCGGTGACAAAAGAAGTCCGCGAGCTGGACCGCGACGCGACCGCGCTTCTGGCACCATACAGACTCGGAAGCACTCAAGGAAGCAATGGATCTCAGCCTCAATCATCGTTCAACCCGGCAGACGACCCGGCTTTGGCCTGCACACTCCTGGTCAACCACCTCTCCACCCGCAGGAACAAGCGATGCCTAATGGCATACCACAAAACCCGCACGGACAAGCTGGAAGCCATGTGCTGGGAGGGTGTCGATGTCTCTGATTTGCAGCAAGCTTCCGCCTCGACTACCGCTGGTGGACAGAGTGGACCGACGAGCAGTCTGAGCGCAGAGGAAGAACAGTACGTTCATCAGTACAGCGACCTTCTCGCTGCGTACAAGGGTCAGTGGACCGACATCGACTTGACTGGAAGTCTAGAGCCTCCGAAAGATATCTTCATCGATGTGCGGGTGCTGAAGGATGCGGGTGAAATTCAGACGGAGTACGGGTCGATCACTTTGACGAAGAACAGCCAGTTCTACGTTCGTCAAGCTGATGTCGAGCGGCTCATTCATCAAGGCTATTTGCAACGACTGAGC ```      ``` 1	ATGTATGGCGAGCAAGGCAACAAGCTCGTGGTCGATGCGAAGCGCATCCAAAACCTGCCACACCTACCACCATACGCAACGACGACAATCCGTGCGGTGACAAAAGAAGTCCGCGAGCTGGACCGCGACGCGACCGCGCTTCTGGCACCATACAGACTC-GGAAGCACTCAAGGAAGCAATGGATCTCAGCCTCAATCATC	200 	\|\|\|\|\|\|\|\|\|\|\|\|\|\|\|\|\|\|\|\|\|\|\|\|\|\|\|\|\|\|\|\|\|\|\|\|\|\|\|\|\|\|\|\|\|\|\|\|\|\|\|\|\|\|\|\|\|\|\|\|\|\|\|\|\|\|\|\|\|\|\|\|\|\|\|\|\|\|\|\|\|\|\|\|\|\|\|\|\|\|\|\|\|\|\|\|\|\|\|\|\|\|\|\|\|\|\|\|\|\|\|\|\|\|\|\|\|\|\|\|\|\|\|\|\|\|\|\|\| \|\|\|\|\|\|\|\|\|\|\|\|\|\|\|\|\|\|\|\|\|\|\|\|\|\|\|\|\| \|\|\|\|\|\|\|\|\|\|\|\|\|\|\|\| \|\|\|\|\|\|\|\|\|\|\|  \|\|\|\| \| \|\|\|\| 	ATGTATGGCGAGCAAGGCAACAAGCTCGTGGTCGATGCGAAGCGCATCCAAAACCTGCCACACCTACCACCATACGCAACGACGACAATCCGTGCGGTGACAAAAGAAGTCCGCGAGCTGGACCGCGACNCGACCGCGCTTCTGGCACCATACAGACTCAGGAAGCACTCAAGGAAACAATGGATCTCNACCTCNANCATC ``` |
| 3 | ggggacaagtttgtacaaaaaagcaggcttgatgCCTGAATCGAAGTCGAGATG, ggggaccactttgtacaagaaagctgggtcCGGCCACTTTGGAAGTTTCG | Mycgr3\|108384  JGI  Ensembl | KOG3242 Oligoribonuclease (3'->5' exoribonuclease); | T/G mismatch-specific endonuclease activity; exonuclease activity;  intracellular;  nuclease activity;  phosphoric ester hydrolase activity; | 100 % |  | ``` 26	ATGCCTGAATCGAAGTCGAGATGGCCGCTGGTGTGGATCGACTGCGAGATGACTGGCCTCAACGACACGAAGGACAGCATCATGTCCCTAGCATGCTTCATCACGGACTACGATCTCAACGTGCTTGACGAGACTGGCTACGAAGCCGTCATTCACCACTCACAGGAACAGCTGGATGCCATGGGAGAGTGGTGCACGAGCCACCATGGCTCTTCGGGATTGACAAAGGCGTGTCTGGACTCGTCAACAACAGCAGAGCAAGCTGCCGAAGAACTCCTTCAGTACATTCAGAAGTACGCTCCAGAGAGGAAGAAAGCGCTTCTGGCTGGCAACACTGTACATGCCGATAAGATGTTCCTCGCTAAGGAGCCCTGGAGGAAAGTTATTTCGCACCTGCATCACAGGATTCTGGATGTATCGGCGATCAAAGAAGCAGCAAAGCGATGGGCACCCGAGGAATTGATCAAGCAAAGTCCGTCGAAACAAGGACTACACGAGGCGAAAGCCGATATTCTGGAGAGTATTGAGGAGGCACGATTCTA	567 	\|\|\|\|\|\|\|\|\|\|\|\|\|\|\|\|\|\|\|\|\|\|\|\|\|\|\|\|\|\|\|\|\|\|\|\|\|\|\|\|\|\|\|\|\|\|\|\|\|\|\|\|\|\|\|\|\|\|\|\|\|\|\|\|\|\|\|\|\|\|\|\|\|\|\|\|\|\|\|\|\|\|\|\|\|\|\|\|\|\|\|\|\|\|\|\|\|\|\|\|\|\|\|\|\|\|\|\|\|\|\|\|\|\|\|\|\|\|\|\|\|\|\|\|\|\|\|\|\|\|\|\|\|\|\|\|\|\|\|\|\|\|\|\|\|\|\|\|\|\|\|\|\|\|\|\|\|\|\|\|\|\|\|\|\|\|\|\|\|\|\|\|\|\|\|\|\|\|\|\|\|\|\|\|\|\|\|\|\|\|\|\|\|\|\|\|\|\|\|\|\|\|\|\|\|\|\|\|\|\|\|\|\|\|\|\|\|\|\|\|\|\|\|\|\|\|\|\|\|\|\|\|\|\|\|\|\|\|\|\|\|\|\|\|\|\|\|\|\|\|\|\|\|\|\|\|\|\|\|\|\|\|\|\|\|\|\|\|\|\|\|\|\|\|\|\|\|\|\|\|\|\|\|\|\|\|\|\|\|\|\|\|\|\|\|\|\|\|\|\|\|\|\|\|\|\|\|\|\|\|\|\|\|\|\|\|\|\|\|\|\|\|\|\|\|\|\|\|\|\|\|\|\|\|\|\|\|\|\|\|\|\|\|\|\|\|\|\|\|\|\|\|\|\|\|\|\|\|\|\|\|\|\|\|\|\|\|\|\|\|\|\|\|\|\|\|\|\|\|\|\|\|\|\|\|\|\|\|\|\|\|\|\|\|\|\|\|\|\|\|\|\|\|\|\|\|\|\|\|\|\|\|\|\|\|\|\|\|\|\|\|\|\|\|\|\|\|\|\|\|\|\|\|\|\| \|\| \|\|\|\|\|\|\|\|\|\|\|\|\|\|\|\| \|\|\|\|\|\|\|\|\|\|\| \|\|\|\| \|\|\|\|\|\| \|\|\|\| \|\|\|\|\|\|\|\|\|\| \| \|\| \|\|\| \|\|\|\|\|\|\|\|\|\|\|\|\| \|\|\| \|\|\|  \|\| \|\| \| \|\|\|\|\|\| 	ATGCCTGAATCGAAGTCGAGATGGCCGCTGGTGTGGATCGACTGCGAGATGACTGGCCTCAACGACACGAAGGACAGCATCATGTCCCTAGCATGCTTCATCACGGACTACGATCTCAACGTGCTTGACGAGACTGGCTACGAAGCCGTCATTCACCACTCACAGGAACAGCTGGATGCCATGGGAGAGTGGTGCACGAGCCACCATGGCTCTTCGGGATTGACAAAGGCGTGTCTGGACTCGTCAACAACAGCAGAGCAAGCTGCCGAAGAACTCCTTCAGTACATTCAGAAGTACGCTCCAGAGAGGAAGAAAGCGCTTCTGGCTGGCAACACTGTACATGCCGATAAGATGTTCCTCGCTAAGGAGCCCTGGAGGAAAGTTATTTCGCACCTGCATCACAGGATTCTGGATGTATCGGCGATCAAAGAAGCANCANAGCGATGGGCACCCGANGAATTGATCAANCAAANTCCGTCNAAACNAGGACTACACNANGCNAAANCCGATATTCTGGANAGTTTTGNNGANGCNC-ATTCTA ```      ``` 26	ATGCCTGAATCGAAGTCGAGATGGCCGCTGGTGTGGATCGACTGCGAGATGACTGGCCTCAACGACACGAAGGACAGCATCATGTCCCTAGCATGCTTCATCACGGACTACGATCTCAACGTGCTTGACGAGACTGGCTACGAAGCCGTCATTCACCACTCACAGGAACAGCTGGATGCCATGGGAGAGTGGTGCACGAGCCACCATGGCTCTTCGGGATTGACAAAGGCGTGTCTGGACTCGTCAACAACAGCAGAGCAAGCTGCCGAAGAACTCCTTCAGTACATTCAGAAGTACGCTCCAGAGAGGAAGAAAGCGCTTCTGGCTGGCAACACTGTACATGCCGATAAGATGTTCCTCGCTAAGGAGCCCTGGAGGAAAGTTATTTCGCACCTGCATCACAGGATTCTGGATGTATCGGCGATCAAAGAAGCAGCAAAGCGATGGGCACCCGAGGAATTGATCAAGCAAAGTCCGTCGAAACAAGGACTACACGAGGCGAAAGCCGATATTCTGGAGAGTATTGAGGAGGCACGATTCTACAAGCGAGTGTTCTTTCAAAACCCTGCCGAGGTTGAGAGTGAGCAGTCGAAACTTCCAAAGTGGCCG	634 	\|\|\|\|\|\|\|\|\|\| \|\|\|\|\|\|\|\|\|\|\|\|\|\|\|\|\|\|\|\|\|\|\|\|\|\|\|\|\|\| \|\|\|\|\|\|\|\|\|\|\|\|\|\|\|\|\|\|\|\|\|\|\|\|\|\|\|\|\|\|\|\|\|\|\|\|\|\|\|\|\|\|\|\|\|\|\|\|\|\|\|\|\|\|\|\|\|\|\|\|\|\|\|\|\| \|\|\|\|\|\|\|\|\|\|\|\|\|\|\|\|\|\|\|\|\|\|\|\|\|\|\|\|\|\|\|\|\|\|\|\|\|\|\|\|\|\|\|\|\|\|\|\|\|\|\|\|\|\|\|\|\|\|\|\|\|\|\|\|\|\|\|\|\|\|\|\|\|\|\|\|\|\|\|\|\|\|\|\|\|\|\|\|\|\|\|\|\|\|\|\|\|\|\|\|\|\|\|\|\|\|\|\|\|\|\|\|\|\|\|\|\|\|\|\|\|\|\|\|\|\|\|\|\|\|\|\|\|\|\|\|\|\|\|\|\|\|\|\|\|\|\|\|\|\|\|\|\|\|\|\|\|\|\|\|\|\|\|\|\|\|\|\|\|\|\|\|\|\|\|\|\|\|\|\|\|\|\|\|\|\|\|\|\|\|\|\|\|\|\|\|\|\|\|\|\|\|\|\|\|\|\|\|\|\|\|\|\|\|\|\|\|\|\|\|\|\|\|\|\|\|\|\|\|\|\|\|\|\|\|\|\|\|\|\|\|\|\|\|\|\|\|\|\|\|\|\|\|\|\|\|\|\|\|\|\|\|\|\|\|\|\|\|\|\|\|\|\|\|\|\|\|\|\|\|\|\|\|\|\|\|\|\|\|\|\|\|\|\|\|\|\|\|\|\|\|\|\|\|\|\|\|\|\|\|\|\|\|\|\|\|\|\|\|\|\|\|\|\|\|\|\|\|\|\|\|\|\|\|\|\|\|\|\|\|\|\|\|\|\|\|\|\|\|\|\|\|\|\|\|\|\|\|\|\|\|\|\|\|\|\|\|\|\|\|\|\|\|\|\|\|\|\|\|\|\|\|\|\|\|\|\|\|\|\|\|\|\|\|\|\|\|\|\|\|\|\|\|\|\|\|\|\|\|\|\|\|\|\|\|\|\|\|\|\|\|\|\|\|\|\|\|\|\|\|\|\|\|\|\|\|\|\|\|\|\|\|\|\|\|\|\|\|\|\|\|\|\|\|\|\|\|\|\|\|\|\|\|\|\|\|\|\|\|\|\|\|\|\|\|\|\|\|\|\|\|\|\|\|\|\|\|\|\|\|\|\|\|\|\|\|\|\|\|\|\| 	ATGCCTGAATNGAAGTCGAGATGGCCGCTGGTGTGGATCGANTGCGAGATGACTGGCCTCAACGACACGAAGGACAGCATCATGTCCCTAGCATGCTTCATCACGGANTACGATCTCAACGTGCTTGACGAGACTGGCTACGAAGCCGTCATTCACCACTCACAGGAACAGCTGGATGCCATGGGAGAGTGGTGCACGAGCCACCATGGCTCTTCGGGATTGACAAAGGCGTGTCTGGACTCGTCAACAACAGCAGAGCAAGCTGCCGAAGAACTCCTTCAGTACATTCAGAAGTACGCTCCAGAGAGGAAGAAAGCGCTTCTGGCTGGCAACACTGTACATGCCGATAAGATGTTCCTCGCTAAGGAGCCCTGGAGGAAAGTTATTTCGCACCTGCATCACAGGATTCTGGATGTATCGGCGATCAAAGAAGCAGCAAAGCGATGGGCACCCGAGGAATTGATCAAGCAAAGTCCGTCGAAACAAGGACTACACGAGGCGAAAGCCGATATTCTGGAGAGTATTGAGGAGGCACGATTCTACAAGCGAGTGTTCTTTCAAAACCCTGCCGAGGTTGAGAGTGAGCAGTCGAAACTTCCAAAGTGGCCG ``` |
| 3 | ggggacaagtttgtacaaaaaagcaggcttgatgCCCAGCAAAATCGTTCTAGC, ggggaccactttgtacaagaaagctgggtcACAGTTGTGTCCACAATTCG | Mycgr3\|108482  JGI  Ensembl |  |  | 100 % |  | ``` 110	ATGCCCAGCAAAATCGTTCTAGCCCATCTGTTCGGGGCGGTTTCCGTCGTCCTCGCAAGAACGTCGTTTCACAAAGAGCGGTGCTGCCTCTGGAGCGATACACTGAATAGATGGTATGGATTGCCGTCGGCCACCCAACAGGCGTGCAACTTCATCGACCATTCCGACTACGAGTCCGACGATGAGGGTTGGTGCCACGCTAAGGGCGGCTACGGCATCGTAGACGAATGCTTCAGCTGCCTGTGCTGGAACGGTGCGAATGCTCACTCCTGTGGCGGTGGGGGAAAGTGGGGCGGCGCAACGACGAATTGTGGACACAACTGT	433 	\|\|\|\|\|\|\|\|\|\|\|\|\|\|\|\|\|\|\|\|\|\|\|\|\|\|\|\|\|\|\|\|\|\|\|\|\|\|\|\|\|\|\|\|\|\|\|\|\|\|\|\|\|\|\|\|\|\|\|\|\|\|\|\|\|\|\|\|\|\|\|\|\|\|\|\|\|\|\|\|\|\|\|\|\|\|\|\|\|\|\|\|\|\|\|\|\|\|\|\|\|\|\|\|\|\|\|\|\|\|\|\|\|\|\|\|\|\|\|\|\|\|\|\|\|\|\|\|\|\|\|\|\|\|\|\|\|\|\|\|\|\|\|\|\|\|\|\|\|\|\|\|\|\|\|\|\|\|\|\|\|\|\|\|\|\|\|\|\|\|\|\|\|\|\|\|\|\|\|\|\|\|\|\|\|\|\|\|\|\|\|\|\|\|\|\|\|\|\|\|\|\|\|\|\|\|\|\|\|\|\|\|\|\|\|\|\|\|\|\|\|\|\|\|\|\|\|\|\|\|\|\|\|\|\|\|\|\|\|\|\|\|\|\|\|\|\|\|\|\|\|\|\|\|\|\|\|\|\|\|\|\|\|\|\|\|\|\|\|\|\|\|\|\|\|\|\|\|\|\|\|\|\|\|\|\|\|\|\|\|\|\|\|\|\|\|\|\|\|\|\|\|\|\|\|\|\|\|\|\|\|\|\|\|\|\|\|\|\|\|\|\|\|\| 	ATGCCCAGCAAAATCGTTCTAGCCCATCTGTTCGGGGCGGTTTCCGTCGTCCTCGCAAGAACGTCGTTTCACAAAGAGCGGTGCTGCCTCTGGAGCGATACACTGAATAGATGGTATGGATTGCCGTCGGCCACCCAACAGGCGTGCAACTTCATCGACCATTCCGACTACGAGTCCGACGATGAGGGTTGGTGCCACGCTAAGGGCGGCTACGGCATCGTAGACGAATGCTTCAGCTGCCTGTGCTGGAACGGTGCGAATGCTCACTCCTGTGGCGGTGGGGGAAAGTGGGGCGGCGCAACGACGAATTGTGGACACAACTGT ``` |
| 3 | ggggacaagtttgtacaaaaaagcaggcttgatgGCCCTCCAAGGATCGCTCGC, ggggaccactttgtacaagaaagctgggtcTCTCTTGTTGTAGTCCTTGA | Mycgr3\|108767  JGI  Ensembl | KOG3095 Transcription initiation factor IIE, beta subunit; |  | 100 % |  | ``` 423	GCCCAGAAGAACATCCCTTTCATCCGAACAGCCCTCGCTGGTAATGAGCGCGTGCAGTTCGTCTCAAAGAGTGAAAGCGGCAACGGCCATGATAGCTTCCGCTACCGTCCACAGCACCCTGTCACCAACGCGGAGGAACTCAAGAACTACCTCGCACGCCAAATTACAGCCAGAGGAATTCCCGTGAAGGAACTGAAAGACGGCTGGCCAGGCTGCATTCCCGCCATCGACGCGCTTGAAAAGGAAGGTGCAATTCTGGTAACACGATTGACCAAGGACAACACCCCGAAGATGGTCTGGCCCGACTCCCCGTCCTACCACGTGCACATCGACGACGACTTTCGCGAGTTCTGGAACAAGACGAAGTTGCCTCCGACGGAGACGGAGATCAGGACTGAGCTGGAGAAAGCTGGGCTCACGCCTACGAGTCAGGTGAAGGAGACGAACAAGACGGATAACAAGAAGAAGGATCGCAAGCGGCCGAACCGCAGGGGAGGCATGAAGACGACCAATTCGCACATGACGGGCATTCTCAAGGACTACAACAAGAGA	974 	\|\|\|\| \|\|\|\|\|\|\|\|\|\|\|\|\|\|\|\|\|\|\|\|\|\| \|\|\|\|\|\|\|\|\| \|\|\|\|\|\|\|\|\|\|\|\|\|\|\|\|\|\|\|\|\| \|\| \|\|\|\|\|\|\|\|\|\|\|\|\|\|\|\|\|\|\|\|\|\|\|\|\|\|\|\|\|\|\|\| \|\|\|\|\| \| \|\|\|\|\|\|\|\|\|\|\|\|\|\|\|\|\|\|\|\|\|\|\|\|\|\|\|\|\|\|\|\|\|\|\|\|\|\|\|\|\|\|\|\|\|\|\|\|\|\|\|\|\|\|\|\|\|\|\|\|\|\|\|\|\|\|\|\|\|\|\|\|\|\|\|\|\|\|\|\|\|\|\|\|\|\|\|\|\|\|\|\|\|\|\|\|\|\|\|\|\|\|\|\|\|\|\|\|\|\|\|\|\|\|\|\|\|\|\|\|\|\|\|\|\|\|\|\|\|\|\|\|\|\|\|\|\|\|\|\|\|\|\|\|\|\|\|\|\|\|\|\|\|\|\|\|\|\|\|\|\|\|\|\|\|\|\|\|\|\|\|\|\|\|\|\|\|\|\|\|\|\|\|\|\|\|\|\|\|\|\|\|\|\|\|\|\|\|\|\|\|\|\|\|\|\|\|\|\|\|\|\|\|\|\|\|\|\|\|\|\|\|\|\|\|\|\|\|\|\|\|\|\|\|\|\|\|\|\|\|\|\|\|\|\|\|\|\|\|\|\|\|\|\|\|\|\|\|\|\|\|\|\|\|\|\|\|\|\|\|\|\|\|\|\|\|\|\|\|\|\|\|\|\|\|\|\|\|\|\|\|\|\|\|\|\|\|\|\|\|\|\|\|\|\|\|\|\|\|\|\|\|\|\|\|\|\|\|\|\|\|\|\|\|\|\|\|\|\|\|\|\|\|\|\|\|\|\|\|\|\|\|\|\|\|\|\|\|\|\|\|\|\|\|\|\|\|\|\|\|\|\|\|\|\|\|\|\|\|\|\|\|\|\|\|\|\|\|\|\|\|\|\|\|\|\|\|\|\|\|\|\|\|\|\|\|\|\|\|\|\|\|\|\|\|\|\|\|\|\|\|\|\|\|\|\|\|\|\|\|\|\|\|\|\|\|\|\|\|\|\|\|\|\|\|\|\|\|\|\|\|\|\|\|\|\|\|\| 	GCCCNGAAGAACATCCCTTTCATCCGANCAGCCCTCGNTGGTAATGAGCGCGTGCAGTTNGTNTCAAAGAGTGAAAGCGGCAACGGCCATGATAGNTTCCGNTNCCGTCCACAGCACCCTGTCACCAACGCGGAGGAACTCAAGAACTACCTCGCACGCCAAATTACAGCCAGAGGAATTCCCGTGAAGGAACTGAAAGACGGCTGGCCAGGCTGCATTCCCGCCATCGACGCGCTTGAAAAGGAAGGTGCAATTCTGGTAACACGATTGACCAAGGACAACACCCCGAAGATGGTCTGGCCCGACTCCCCGTCCTACCACGTGCACATCGACGACGACTTTCGCGAGTTCTGGAACAAGACGAAGTTGCCTCCGACGGAGACGGAGATCAGGACTGAGCTGGAGAAAGCTGGGCTCACGCCTACGAGTCAGGTGAAGGAGACGAACAAGACGGATAACAAGAAGAAGGATCGCAAGCGGCCGAACCGCAGGGGAGGCATGAAGACGACCAATTCGCACATGACGGGCATTCTCAAGGACTACAACAAGAGA ``` |
| 3 | ggggacaagtttgtacaaaaaagcaggcttgatgCGCTCAATAGCGCAGACAGC, ggggaccactttgtacaagaaagctgggtcTTGTGTTTCCGCTCCCGCCG | Mycgr3\|108774  JGI  Ensembl |  |  | 22.86282 % |  | ``` 307	TTCAAAATTCCCGACTTCCAGCGCATGGTGTTGAAGTTTCCTTTTGTCGATCGTCTCAACGACGGCTACTCTTGCTTCCAGTACTCCGCGCCACTGATCCTCAGTTTCGACAACCTGATCGCTCTCGCGGGATCGAAAGCTTACGGTCCAACCTTTGCTGGCTCCTTCGATCCGCCGTGCGATGGATACGCCGACGATGGCCAGGGCGCTACAGTTCTGAGCGCATATGTCGTACCTCCGGGTCAGCATCTTGGCTCTAAGCCGGCTTTCGAAGGCAGATTCAAAGCTGCCTGCAGTAGTGTCCCATACGACCAGAAAGT	626 	\|\|\| \|\|\| \| \|\|\|\| \|\|\|\|\|\| \|\|\|\|\|\|\|\|\| \|\|\|\| \|\|\|\| \|\|\|\|\| \|\|\| \|\| \|\|\|\| \|\|\|\|\| \|\| \| \|\|\| \|\|\|\|\|\|\|\| \|\|\|\|\|\|\|\|\| \|\|\|\|\|\|\|\|\|\|\|\|\| \|\|\|\|\|\|\|\|\|\|\|\|\|\|\|\|\|\|\|\|\|\|\|\|\|\|\|\|\|\|\|\|\|\|\|\|\|\|\|\|\|\|\|\|\|\|\|\|\|\|\|\|\|\|\|\|\|\|\|\|\|\|\|\|\|\|\|\|\|\|\|\|\|\|\|\|\|\|\|\|\|\|\|\|\|\|\|\|\|\|\|\|\|\|\|\|\|\|\|\|\|\|\|\|\|\|\|\|\|\|\|\|\|\|\|\|\|\|\|\|\|\|\|\|\|\|\|\|\|\|\|\|\|\|\|\|\|\|\|\|\|\|\|\|\|\|\|\| \|\|\|\|\|\|\|\|\|\|\|\|\|\|\|\|\|\|\|\|\|\|\|\|\|\|\|\|\|\|\|\|\|\|\|\|\|\|\|\|\|\|\|\|\|\|\|\|\|\|\|\|\|\|\|\|\|\|\|\|\|\|\| 	TTC-AAANTNCCGA-TTCCAGNGCATGGTGT-GAAG-TTCC-TTTGTNGATNGTNTCAANGACGGNTANTNTTGNTTCCAGTANTCCGCGCCANTGATCCTCAGTTTNGACAACCTGATCGCTCTCGCGGGATCGAAAGCTTACGGTCCAACCTTTGCTGGCTCCTTCGATCCGCCGTGCGATGGATACGCCGACGATGGCCAGGGCGCTACAGTTCTGAGCGCATATGTCGTACCTCCGGGTCAGCATCTTGGCTNTAAGCCGGCTTTCGAAGGCAGATTCAAAGCTGCCTGCAGTAGTGTCCCATACGACCAGAAAGT ```      ``` 1	ATGCGCTCAATAGCGCAGACAGCAGCGCTGGCAGCAGCCTTGTTCCACCTCACCGCCGCAGCGCTCACCTGCAAGACCCATAGCCAGCCGAACCCAATCGCAAAGCAGTATCCAAAAGATGTCACCGGCACCATCAACGGCACCACGGCCATCGTGCCCATTCCATACGACGTAGCCCGCAGCATCATTCCCTCGGAGTACGGCATCCTCCGTGACGCCTACGAAAAGCTCATGCCAGATCTTCCCAAAGACATGTACCCGGTCCACTACTCCGGACTGCTCGACCACGGTCTCGGGCTGGGCGACTTCAAAATTCCCGACTTCCAGCGCATGGTGTTGAAGTTTCCTTTTGTCGATCGTCTCAACGACGGCTACTCTTGCTTCCAGTACTCCGCGCCACTGATCCTCAGTTTCGACAACCTGATCGCTCTCGCGGGATCGAAAGCTTACGGTCCAACCTTTGCTGGCTCCTTCGATCCGCCGTGCGATGGATACGCCGACGATGGCCAGGGCGCTACAGTTCTGAGCGCATATGTCGTACCTCCGGGTCAGCATCTTGGCTCTAAGCCGGCTTTCGAAGGCAGATTCAAAGCTGCCTGCAGTAGTGTCCCATACGACCAGAAAGT	626 	\|\|\|\|\|\|\|\|\|\|\|\|\|\|\|\|\|\|\|\|\|\|\|\|\|\|\|\|\|\|\|\|\|\|\|\|\|\|\|\|\|\|\|\|\|\|\|\|\|\|\|\|\|\|\|\|\|\|\|\|\|\|\|\|\|\|\|\|\|\|\|\|\|\|\|\|\|\|\|\|\|\|\|\|\|\|\|\|\|\|\|\|\|\|\|\|\|\|\|\|\|\|\|\|\|\|\|\|\|\|\|\|\|\|\|\|\|\|\|\|\|\|\|\|\|\|\|\|\|\|\|\|\|\|\|\|\|\|\|\|\|\|\|\|\|\|\|\|\|\|\|\|\|\|\|\|\|\|\|\|\|\|\|\|\|\|\|\|\|\|\|\|\|\|\|\|\|\|\|\|\|\|\|\|\|\|\|\|\|\|\|\|\|\|\|\|\|\|\|\|\|\|\|\|\|\|\|\|\|\|\|\|\|\|\|\|\|\|\|\|\|\|\|\|\|\|\|\|\|\|\|\|\|\|\|\|\|\|\|\|\|\|\|\|\|\|\|\|\|\|\|\|\|\|\|\|\|\|\|\|\|\|\|\|\|\|\|\|\|\|\|\|\|\|\|\|\|\|\|\|\|\|\|\|\|\|\|\|\|\|\|\|\|\|\|\|\|\|\|\|\|\|\|\|\|\|\|\|\|\|\|\|\|\|\|\|\|\|\|\|\|\|\|\|\|\|\|\|\|\|\|\|\|\|\|\|\|\|\|\|\|\|\|\|\|\|\|\|\|\|\|\|\|\|\|\|\|\|\|\|\|\|\|\|\|\|\|\|\|\|\|\|\|\|\|\|\|\|\|\|\|\|\|\|\|\|\|\|\|\|\|\|\|\|\|\|\|\|\|\|\|\|\|\|\|\|\|\|\|\|\|\|\|\|\|\|\|\|\|\|\|\|\|\|\|\|\|\|\|\|\|\|\|\|\|\|\|\|\|\|\|\|\|\|\|\|\|\|\|\|\|\|\|\|\|\|\|\|\|\|\|\|\|\|\|\|\|\|\|\|\|\|\|\|\|\|\|\|\|\|\|\|\|\|\|\|\|\|\|\|\|\|\|\|\|\|\|\|\|\|\|\|\|\|\|\|\|\|\|\|\|\|\|\|\|\|\|\|\|\|\|\|\|\|\|\|\|\|\|\|\|\|\|\|\|\|\|\|\|\|\|\|\|\|\|\|\|\|\|\|\|\|\|\|\|\|\|\|\|\|\|\|\|\|\|\|\|\|\|\|\|\|\|\|\|\|\|\|\|\|\|\|\|\|\|\|\|\|\|\|\|\|\|\|\|\|\|\|\|\|\|\|\|\|\|\|\|\|\|\|\|\|\|\|\|\|\|\|\|\|\|\|\|\|\|\| 	ATGCGCTCAATAGCGCAGACAGCAGCGCTGGCAGCAGCCTTGTTCCACCTCACCGCCGCAGCGCTCACCTGCAAGACCCATAGCCAGCCGAACCCAATCGCAAAGCAGTATCCAAAAGATGTCACCGGCACCATCAACGGCACCACGGCCATCGTGCCCATTCCATACGACGTAGCCCGCAGCATCATTCCCTCGGAGTACGGCATCCTCCGTGACGCCTACGAAAAGCTCATGCCAGATCTTCCCAAAGACATGTACCCGGTCCACTACTCCGGACTGCTCGACCACGGTCTCGGGCTGGGCGACTTCAAAATTCCCGACTTCCAGCGCATGGTGTTGAAGTTTCCTTTTGTCGATCGTCTCAACGACGGCTACTCTTGCTTCCAGTACTCCGCGCCACTGATCCTCAGTTTCGACAACCTGATCGCTCTCGCGGGATCGAAAGCTTACGGTCCAACCTTTGCTGGCTCCTTCGATCCGCCGTGCGATGGATACGCCGACGATGGCCAGGGCGCTACAGTTCTGAGCGCATATGTCGTACCTCCGGGTCAGCATCTTGGCTCTAAGCCGGCTTTCGAAGGCAGATTCAAAGCTGCCTGCAGTAGTGTCCCATACGACCAGAAAGT ```      ``` 627	CTTCGTGAACATCACGAACCAGCCCATCTTCGGCAGCGGCAAGCCTGTGTGCGACAACTACATCACACTGTACAACACCAGCGTGACGCAA-GGCGAGTTCGCTCCGGTGGCCGTAGAAGGAACGGTCAAAGTCGGTCCGCCATTCTATCCCCAGGACGCAACACTTGAGGCATTTGGCTACCGGATGGACAACGCATTCGTGGAGAGGAATGATGTTCCTTGCGAAAAGCTCAAAGGCTACGCGGGTACTGGTCCAGGCGACTCGGCGGGAGCGGAAAC-ACAA	909 	\|\|\|\|\|\|\|\|\|\|\|\|\|\|\|\|\|\|\|\|\|\|\|\|\|\|\|\|\|\|\|\|\|\|\|\|\|\|\|\|\|\|\|\|\|\|\|\|\|\|\|\|\|\|\|\|\|\|\|\|\|\|\|\|\|\|\|\|\|\|\|\|\|\|\|\|\|\|\|\|\|\|\|\|\|\|\|\|\|\|\| \|\|\|\|\|\|\|\|\|\|\|\|\|\|\|\|\|\|\|\|\|\|\|\|\|\|\| \|\|\|\|\|\|\|\|\|\|\|\|\|\|\|\|\|\|\|\|\|\|\|\|\|\|\|\|\|\|\|\|\|\|\|\|\|\|\|\|\|\|\|\|\|\|\|\|\| \|\|\|\|\|\|\|\|\|\|\|\|\|\|\|\|\|\|\|\|\| \|\|\|\|\|\|\|\|\|\|\|\|\|\|\|\|\|\| \|\|\|\|\|\|\|\| \|\|\|\|\|\|\|\| \|\|\|\|\|\|\| \|\|\|\|\|\|\|\|\|\|\|\|\|\|\| \| \|\|\|\|\|\|\|\|\|\|\|\|\|\|\|\|\|\|\|\|\|\|\|\|\| \|\|\|\| 	CTTCGTGAACATCACGAACCAGCCCATCTTCGGCAGCGGCAAGCCTGTGTGCGACAACTACATCACACTGTACAACACCAGCGTGACGCAAGGGCGAGTTCGCTCCGGTGGCCGTAGAAAGAACGGTCAAAGTCGGTCCGCCATTCTATCCCCAGGACGCAACACTTGA-GCATTTGGCTACCGGATGGAC-ACGCATTCGTGGAGAGGA-TGATGTTC-TTGCGAAA-GCTCAAA-GCTACGCGGGTACTG-T-CAGGCGACTCGGCGGGAGCGGAAACCACAA ```      ``` 627	CTTCGTGAACATCACGAACCAGCCCATCTTCGGCAGCGGCAAGCCTGTGTGCGACAACTACATCACACTGTACAACACCAGCGTGACGCAAGGCGAGTTCGCTCCGGTGGCCGTAGAAGGAACGGTCAAAGTCGGTCCGCCATTCTATCCCCAGGACGCAACACTTGAGGCATTTGGCTACCGGATGGACAACGCATTCGTGGAGAGGAATGATGTTCCTTGCGAAAAGCTCAAAGGCTACGCGGGTACTGGTCCAGGCGACTCGGCGGGAGCGGAAACACAA	909 	\|\|\|\|\|\|\|\|\|\|\|\|\|\|\|\|\|\|\|\|\|\|\|\|\|\|\|\|\|\|\|\|\|\|\|\|\|\|\|\|\|\|\|\|\|\|\|\|\|\|\|\|\|\|\|\|\|\|\|\|\|\|\|\|\|\|\|\|\|\|\|\|\|\|\|\|\|\|\|\|\|\|\|\|\|\|\|\|\|\|\|\|\|\|\|\|\|\|\|\|\|\|\|\|\|\|\|\|\|\|\|\|\|\|\|\|\|\|\|\|\|\|\|\|\|\|\|\|\|\|\|\|\|\|\|\|\|\|\|\|\|\|\|\|\|\|\|\|\|\|\|\|\|\|\|\|\|\|\|\|\|\|\|\|\|\|\|\|\|\|\|\|\|\|\|\|\|\|\|\|\|\|\|\|\|\|\|\|\|\|\|\|\|\|\|\|\|\|\|\|\|\|\|\|\|\|\|\|\|\|\|\|\|\|\|\|\|\|\|\|\|\|\|\|\|\|\|\|\|\|\|\|\|\|\|\|\|\|\|\|\|\|\|\|\|\|\|\|\|\|\|\|\|\|\|\|\|\|\|\|\|\|\|\|\|\|\|\|\|\|\|\|\|\|\|\|\|\|\|\|\|\|\| 	CTTCGTGAACATCACGAACCAGCCCATCTTCGGCAGCGGCAAGCCTGTGTGCGACAACTACATCACACTGTACAACACCAGCGTGACGCAAGGCGAGTTCGCTCCGGTGGCCGTAGAAGGAACGGTCAAAGTCGGTCCGCCATTCTATCCCCAGGACGCAACACTTGAGGCATTTGGCTACCGGATGGACAACGCATTCGTGGAGAGGAATGATGTTCCTTGCGAAAAGCTCAAAGGCTACGCGGGTACTGGTCCAGGCGACTCGGCGGGAGCGGAAACACAA ``` |
| 3 | ggggacaagtttgtacaaaaaagcaggcttgatgGGCGTGTGGGACATTTTCAG, ggggaccactttgtacaagaaagctgggtcACCCTCGATCAAGACTTTGC | Mycgr3\|108883  JGI  Ensembl | KOG3324 Mitochondrial import inner membrane translocase, subunit TIM23; | mitochondrial inner membrane; protein transport;  protein transporter activity; | 100 % |  | ``` 52	ATGGGCGTGTGGGACATTTTCAGCGGCAAATCCACCCAGACCGCAACCGCATCCGACTCCTCCTTCGACGCCTCCGCCGCGGCGGCCAGCAGCAGTCCTCTCCCTCCGACTTTCGTTACACCCACCGCCGACGATGTCGGGTCATTCATAAGCACACCCGGCGCATTCGATCCATCAGCACTTCACCCGCTCGCGGGCTTAAATCAGGACACCCTCGACTACCTGACCCTCGACGACGATGTGCTCACAGATCTACCCGGCGGAAAGTCATTATTACCATCACGAGGATGGTCAGACGATCTTTGTTATGGAACAGGAGTGACATATCTCGCGGCGCTGTCAGTGGGAGGAGCATGGGGTTTGACAGAAGGATTGAGCCGGTTGCCGGCCACAGCACCACCGAAGTTGCGCCTCAACTCGGCGCTGAACGCCATCACACGGAGAGGACCATTCTTGGGCAACTCGGCGGGCGTGGTCGCGATGATGTACAATGGGATAAACAGCACGTTGGGTTACTACCGAGGGAAGCATGATTCGTTCAACAGTGTGGCAGCGGGTGGCATCAGTGGAATGATCTTCAGGAGCACGCACGGTGTGAGGCCGATGATGATCTCTGGAGGTATTGTGGCATCCGTTGCAGGCGCATGGGCTGT	704 	\|\|\|\|\|\| \| \| \|\|\|\|\| \|\|\|\|\|\|\|\|\|\| \|\|\|\|\|\|\|\|\|\|\|\|\|\|\|\|\|\|\|\|\|\|\|\|\|\|\|\|\|\| \|\|\|\|\|\| \|\|\|\|\| \|\|\|\|\|\|\|\|\|\|\|\|\|\|\|\|\|\|\|\|\|\|\|\|\|\|\|\|\|\|\| \|\|\|\|\|\|\|\|\|\|\|\|\|\|\|\|\|\|\|\|\|\|\|\|\|\|\|\|\|\|\|\|\|\|\|\|\|\|\|\|\|\|\|\|\|\|\|\|\|\|\|\|\|\|\|\|\|\|\|\|\|\|\|\|\|\|\|\|\|\|\|\|\|\|\|\|\|\|\|\|\|\|\|\|\|\|\|\|\|\|\|\|\|\|\|\|\|\|\|\|\|\|\|\|\|\|\|\|\|\|\|\|\|\|\|\|\|\|\|\|\|\|\|\|\|\|\|\|\|\|\|\|\|\|\|\|\|\|\|\|\|\|\|\|\|\|\|\|\|\|\|\|\|\|\|\|\|\|\|\|\|\|\|\|\|\|\|\|\|\|\|\|\|\|\|\|\|\|\|\|\|\|\|\|\|\|\|\|\|\|\|\|\|\|\|\|\|\|\|\|\|\|\|\|\|\|\|\|\|\|\|\|\|\|\|\|\|\|\|\|\|\|\|\|\|\|\|\|\|\|\|\|\|\|\|\|\|\|\|\|\|\|\|\|\|\|\|\|\|\|\|\|\|\|\|\|\|\|\|\|\|\|\|\|\|\|\|\|\|\|\|\|\|\|\|\|\|\|\|\|\|\|\|\|\|\|\|\|\|\|\|\|\|\|\|\|\|\|\|\|\|\|\|\|\|\|\|\|\|\|\|\|\|\|\|\|\|\|\|\|\|\|\|\|\|\|\|\|\|\|\|\|\|\|\|\|\|\|\|\|\|\|\|\|\|\|\|\|\|\|\|\|\|\|\|\|\|\|\|\|\|\|\|\|\|\|\|\|\|\|\|\|\|\|\|\|\|\|\|\|\|\|\|\|\|\|\|\|\|\|\|\|\|\|\|\|\|\|\|\|\|\|\|\|\|\|\|\|\|\|\|\|\|\|\|\|\|\|\|\|\|\|\|\|\|\|\|\|\|\|\|\|\|\|\|\|\|\|\|\|\|\|\|\|\|\|\|\|\|\|\|\|\|\|\|\|\|\|\|\|\|\|\|\|\|\|\|\|\|\|\|\|\|\|\|\|\|\|\|\|\|\|\|\|\|\|\|\|\|\|\|\|\|\|\|\|\|\|\|\|\|\|\|\|\|\|\|\|\|\|\|\|\|\|\|\|\|\|\|\|\|\|\|\|\|\|\|\|\|\|\|\|\|\|\|\|\|\|\|\|\|\|\|\|\|\|\|\|\| 	ATGGGCNT-T-GGACA-TTTCAGCGGC-AATCCACCCAGACCGCAACCGCATCCGACTNCTCCTTNGACGCNTCCGCCGCGGCGGCCAGCAGCAGTCCTCTCCNTCCGACTTTCGTTACACCCACCGCCGACGATGTCGGGTCATTCATAAGCACACCCGGCGCATTCGATCCATCAGCACTTCACCCGCTCGCGGGCTTAAATCAGGACACCCTCGACTACCTGACCCTCGACGACGATGTGCTCACAGATCTACCCGGCGGAAAGTCATTATTACCATCACGAGGATGGTCAGACGATCTTTGTTATGGAACAGGAGTGACATATCTCGCGGCGCTGTCAGTGGGAGGAGCATGGGGTTTGACAGAAGGATTGAGCCGGTTGCCGGCCACAGCACCACCGAAGTTGCGCCTCAACTCGGCGCTGAACGCCATCACACGGAGAGGACCATTCTTGGGCAACTCGGCGGGCGTGGTCGCGATGATGTACAATGGGATAAACAGCACGTTGGGTTACTACCGAGGGAAGCATGATTCGTTCAACAGTGTGGCAGCGGGTGGCATCAGTGGAATGATCTTCAGGAGCACGCACGGTGTGAGGCCGATGATGATCTCTGGAGGTATTGTGGCATCCGTTGCAGGCGCATGGGCTGT ``` |
| 3 | ggggacaagtttgtacaaaaaagcaggcttgatgCAGTTCACAACTCTGTCCAT, ggggaccactttgtacaagaaagctgggtcCTTCTTGGTACCGTAGTGGT | Mycgr3\|109137  JGI  Ensembl |  |  | 100 % |  | ``` 121	CCGTGACTCCTTCTGGCACGTCACCGACATGCAGGCCAACTGCCGCGACGAGGCCACCTCCCAGGAGTGCTTTTACCTCTTCAACATCCACGGCGCCAAGGACGGCAAACTCCCCGGCTTCGAAGCTCAATGCCGGACGCAGTCCCTCCCGGGCGACAGGAACAAGCGCACGTGCTACATCCAGCCCGGGCACGCCGACCAGGTCGTGCAGGAGGTGTCTGCTTACCTCAAGTTCGACCCGCGCCACGACAACACGGAGGACAATGAGTTGTATGTGGAGCTGAAGTTCACGTCTGGG	418 	\|\|\|\|\|\| \| \|\|\| \|\|\|\|\| \|\|\|\|\| \|\|\|\|\|\|\|\|\|\|\|\|\|\| \|\|\|\|\| \|\|\|\|\|\|\|\|\|\|\|\|\|\|\|\|\|\|\|\|\|\|\|\|\|\|\|\|\|\|\|\| \|\|\|\|\|\|\|\|\|\|\|\|\|\|\|\|\|\|\|\|\|\|\|\|\|\|\|\|\|\|\|\|\|\|\|\|\|\|\|\|\|\|\|\|\|\|\|\|\|\|\|\|\|\|\|\|\|\|\|\|\|\|\|\|\|\|\|\|\|\|\|\|\|\|\|\|\|\|\|\|\|\|\|\|\|\|\|\|\|\|\|\|\|\|\|\|\|\|\|\|\|\|\|\|\|\|\|\|\|\|\|\|\|\|\|\|\|\|\|\|\|\|\|\|\|\|\|\|\|\|\|\|\|\|\|\|\|\|\|\|\|\|\|\|\|\|\|\|\|\|\|\|\|\|\|\|\|\|\|\|\|\|\|\|\|\|\|\|\|\|\|\|\|\|\|\|\|\|\|\|\|\|\|\|\|\|\|\|\|\|\|\|\|\|\|\|\|\|\|\|\|\|\|\|\|\|\|\|\|\|\|\|\|\|\|\|\|\|\| 	CCGTGANT-CTTNTGGCANGTCACNGACATGCAGGCCAANTGCCGNGACGAGGCCACCTCCCAGGAGTGCTTTTACCTNTTCAACATCCACGGCGCCAAGGACGGCAAACTCCCCGGCTTCGAAGCTCAATGCCGGACGCAGTCCCTCCCGGGCGACAGGAACAAGCGCACGTGCTACATCCAGCCCGGGCACGCCGACCAGGTCGTGCAGGAGGTGTCTGCTTACCTCAAGTTCGACCCGCGCCACGACAACACGGAGGACAATGAGTTGTATGTGGAGCTGAAGTTCACGTCTGGG ```      ``` 417	GGGACAGCAACGAGGTTCAGACTTGGCAGGCCAACAGGATCACGACCTACGACAACTTCGTCACCACGAAGGACTTCGATCTCATCCCCACGTACCACTACGGTACCAAGAAG	529 	\|\|\|\|\|\|\|\|\|\|\|\|\|\|\|\|\|\|\|\|\|\|\|\|\|\|\|\|\|\|\|\|\|\|\|\|\|\|\|\|\|\|\|\|\|\|\|\|\|\|\|\|\|\|\|\|\|\|\|\|\|\|\|\|\|\|\|\|\|\|\|\|\|\|\|\|\|\|\|\|\|\|\|\|\|\|\|\|\|\|\|\|\|\|\|\|\|\|\|\|\|\|\|\|\|\|\|\|\|\|\|\|\| 	GGGACAGCAACGAGGTTCAGACTTGGCAGGCCAACAGGATCACGACCTACGACAACTTCGTCACCACGAAGGACTTCGATCTCATCCCCACGTACCACTACGGTACCAAGAAG ```      ``` 68	ATGCAGTTCACAACTCTG-TCCATCTTCACGACCC-TCACGGC-CCTCGCCGCCGCCCGTGACTCCTTCTGG	136 	\|\|\|\|\|\|\|\|\|\|\|\|\|\|\|\|\|\| \|\|\|\|\|\|\|\|\|\|\|\|\|\|\|\| \|\|\|\|\|\|\| \|\| \|\|\|\|\|\|\|\|\|\|\|\|\|\|\|\|\|\|\|\|\|\|\|\|\| 	ATGCAGTTCACAACTCTGNTCCATCTTCACGACCCNTCACGGCGCCNCGCCGCCGCCCGTGACTCCTTCTGG ``` |
| 3 | ggggacaagtttgtacaaaaaagcaggcttgatgTGGTCTCTACCCAAGATTGG, ggggaccactttgtacaagaaagctgggtcTCGCTGCTCTTCGATACTGT | Mycgr3\|110052  JGI  Ensembl |  |  | 100 % |  | ``` 436	ATGTGGTCTCTACCCAAGATTGGACTGTTGGCTCCGGCCTACGTGCTGGTTTTCCCCGCCTACACAGCAGGCGCATGCCTCGACCATTGGCATGGAACCGACCACCCTGCCAAGCCTTTCGGCCACGCCGCAGTCAAGCGCCACGAGGCCGCGATGATCCAAAAGCGCGACATCATTAGTTACATTCGACTCGCGCTTACCCTCCGATTGATCGCAGAAGGATTCCAGGGTGCTTGTGACAACCATTTCGGACTCCTTGCGCGAGGAGATGGCTCGGGTGGTCTGGATCTGACAGTCGGACCCACGGGCAATGCCGGAGCCAACTGGTTGGACACGACGATTGTCATCGATCCGGACTCCGCAAGTGATGATGCACCCAGGAGAATCGGGTATTTAACTCATGCTCAGTACAACGACGCGGATTCTGTCACTGCCCACTACAGCGCGGACGAAACTCAACACTATATGGAGGCGAATGGTTACCCCCAGAATCATGAGTTTTTCATCGAGATCCAATACAGTATCGAAGAGCAGCGA	972 	\|\|\|\|\|\|\|\|\| \|\|\|\|\|\|\|\|\|\|\|\|\|\|\|\|\|\|\|\|\|\|\|\|\|\|\|\| \|\|\|\|\|\| \|\|\|\|\|\|\|\|\|\|\|\|\| \|\|\|\|\|\|\|\|\|\|\|\|\|\|\|\|\|\|\|\|\|\|\|\|\|\|\|\|\|\|\|\|\|\|\|\|\|\|\|\|\|\|\|\|\|\|\|\|\|\|\|\|\|\|\|\|\|\|\|\|\|\|\|\|\|\|\|\|\|\|\|\|\|\|\|\|\|\|\|\|\|\|\|\|\|\|\|\|\|\|\|\|\|\|\|\|\|\|\|\|\|\|\|\|\|\|\|\|\|\|\|\|\|\|\|\|\|\|\|\|\|\|\|\|\|\|\|\|\|\|\|\|\|\|\|\|\|\|\|\|\|\|\|\|\|\|\|\|\|\|\|\|\|\|\|\|\|\|\|\|\|\|\|\|\|\|\|\|\|\|\|\|\|\|\|\|\|\|\|\|\|\|\|\|\|\|\|\|\|\|\|\|\|\|\|\|\|\|\|\|\|\|\|\|\|\|\|\|\|\|\|\|\|\|\|\|\|\|\|\|\|\|\|\|\|\|\|\|\|\|\|\|\|\|\|\|\|\|\|\|\|\|\|\|\|\|\|\|\|\|\|\|\|\|\|\|\|\|\|\|\|\|\|\|\|\|\|\|\|\|\|\|\|\|\|\|\|\|\|\|\|\|\|\|\|\|\|\|\|\|\|\|\|\|\|\|\|\|\|\|\|\|\|\|\|\|\|\|\|\|\|\|\|\|\|\|\|\|\|\|\|\|\|\|\|\|\|\|\|\|\|\|\|\|\|\|\|\|\|\|\|\|\|\|\|\|\|\|\|\|\|\|\|\|\|\|\|\|\|\|\|\|\|\|\|\|\|\|\|\|\|\|\|\|\|\|\|\|\|\|\|\|\|\|\|\|\|\|\|\|\|\|\|\|\|\|\|\|\|\|\|\|\|\|\|\|\|\|\|\|\|\|\|\|\|\|\|\|\|\|\|\|\|\|\|\|\|\|\|\|\|\|\|\|\|\|\|\|\|\|\|\|\|\|\|\|\|\|\|\|\|\|\|\|\|\|\|\|\|\|\|\|\|\|\|\|\|\|\|\|\|\|\|\|\|\|\| 	ATGTGGTCTNTACCCAAGATTGGACTGTTGGCTCCGGCNTACGTGNTGGTTTTCCCCGCNTACACAGCAGGCGCATGCCTCGACCATTGGCATGGAACCGACCACCCTGCCAAGCCTTTCGGCCACGCCGCAGTCAAGCGCCACGAGGCCGCGATGATCCAAAAGCGCGACATCATTAGTTACATTCGACTCGCGCTTACCCTCCGATTGATCGCAGAAGGATTCCAGGGTGCTTGTGACAACCATTTCGGACTCCTTGCGCGAGGAGATGGCTCGGGTGGTCTGGATCTGACAGTCGGACCCACGGGCAATGCCGGAGCCAACTGGTTGGACACGACGATTGTCATCGATCCGGACTCCGCAAGTGATGATGCACCCAGGAGAATCGGGTATTTAACTCATGCTCAGTACAACGACGCGGATTCTGTCACTGCCCACTACAGCGCGGACGAAACTCAACACTATATGGAGGCGAATGGTTACCCCCAGAATCATGAGTTTTTCATCGAGATCCAATACAGTATCGAAGAGCAGCGA ``` |
| 3 | ggggacaagtttgtacaaaaaagcaggcttgatgCAGTTCTCCATCTTCTCCCT, ggggaccactttgtacaagaaagctgggtcAGCCTTCACACAGGCCCAGT | Mycgr3\|110386  JGI  Ensembl |  |  | 100 % |  | ``` 43	ATGCAGTTCTCCATCTTCTCCCTTGCTACTCTTGTAGCCATAGCATCCGCTACTCCCTTTGCCATGCCCGAAGCCTACGCCGGCACTCTTGACACTACTTCTTCCTCTCTCTCGGCCCGTCAGTCCAACTGCACTGAATACTGCTCCAACAGCGCCGGCTGCGTCTGT	210 	\|\|\|\|\|\|\|\|\|\|\|\|\|\|\|\|\|\|\|\|\|\|\|\|\|\|\|\|\|\|\|\|\|\|\|\|\|\|\|\|\|\|\|\|\|  \|\|\|\|\|\|\|\|\|\|\|\|\|\|\|\|\|\|\|\|\|\|\|\|\|\|\|\|\|\|\|\|\|\|\|\|\|\|\|\|\|\|\|\|\|\|\|\|\|\|\|\|\|\|\|\|\|\|\|\|\|\|\|\|\|\|\|\|\|\|\|\|\|\|\|\|\|\|\|\|\|\|\|\|\|\|\|\|\|\|\|\|\|\|\|\|\|\|\|\|\|\|\|\|\|\|\|\|\|\|\|\|\|\|\|\|\| \|\|\| 	ATGCAGTTCTCCATCTTCTCCCTTGCTACTCTTGTAGCCATAGCANNCGCTACTCCCTTTGCCATGCCCGAAGCCTACGCCGGCACTCTTGACACTACTTCTTCCTCTCTCTCGGCCCGTCAGTCCAACTGCACTGAATACTGCTCCAACAGCGCCGGCTGCGTATGT ```      ``` 205	GTCTGTACTGTCCGTCCCTCCGACTGCACCGCCTTCTACACCGTCCAGTCAGGCGACACCTGCCTCTCCATCGCCAAAGTCTTCGGCAACTTCACCGTCACCCAATTCTATCGCTGGAACCCTTCCGTCGGACAAACCTGCTTTGGACTGCAAGCCTACGTGCCTGTCTGCATCGACACTCCGTGGTATGAGTTTACTCCACCCGTTCAGCCAGAGTTCGGCACGAAGTACACGCCTGAGCAGACGCCTGTGCCGGTCATGCCGGGGATTGTGGATGGGTGTGAGCTGTATGAGTTGGTTGAGCCCGGGAAGAGAGTTGAGGAGTTGGCGGCGGAGAATGGATTCGCGGTGGAGAAGTTCGCTGAGTGGAATGGGAATAGTACGACTGCTTGGGCGACTTACTGGGCCTGTGTGAAGGCT	624 	\|\|\|\|\|\|\|\|\|\|\|\|\|\|\|\|\|\|\|\|\|\|\|\|\|\|\|\|\|\|\|\|\|\|\|\|\|\|\|\|\|\|\|\|\|\|\|\|\|\|\|\|\|\|\|\|\|\|\|\|\|\|\|\|\|\|\|\|\|\|\|\|\|\|\|\|\|\|\|\|\|\|\|\|\|\|\|\|\|\|\|\|\|\|\|\|\|\|\|\|\|\|\|\|\|\|\|\|\|\|\|\|\|\|\|\|\|\|\|\|\|\|\|\|\|\|\|\|\|\|\|\|\|\|\|\|\|\|\|\|\|\|\|\|\|\|\|\|\|\|\|\|\|\|\|\|\|\|\|\|\|\|\|\|\|\|\|\|\|\|\|\|\|\|\|\|\|\|\|\|\|\|\|\|\|\|\|\|\|\|\|\|\|\|\|\|\|\|\|\|\|\|\|\|\|\|\|\|\|\|\|\|\|\|\|\|\|\|\|\|\|\|\|\|\|\|\|\|\|\|\|\|\|\|\|\|\|\|\|\|\|\|\|\|\|\|\|\|\|\|\|\|\|\|\|\|\|\|\|\|\|\|\|\|\|\|\|\|\|\|\|\|\|\|\|\|\|\|\|\|\|\|\|\|\|\|\|\|\|\|\|\|\|\|\|\|\|\|\|\|\|\|\|\|\|\|\|\|\|\|\|\|\|\|\|\|\|\|\|\|\|\|\|\|\|\|\|\|\|\|\|\|\|\|\|\|\|\|\|\|\|\|\|\|\|\|\|\|\|\|\|\|\|\|\|\|\|\|\|\|\|\|\|\|\|\|\|\|\|\|\|\|\|\|\|\|\|\|\|\|\|\|\|\|\|\|\|\|\|\|\|\|\|\|\|\|\|\|\|\|\|\|\|\|\|\|\|\|\|\|\|\|\|\|\|\|\|\|\|\| 	GTCTGTACTGTCCGTCCCTCCGACTGCACCGCCTTCTACACCGTCCAGTCAGGCGACACCTGCCTCTCCATCGCCAAAGTCTTCGGCAACTTCACCGTCACCCAATTCTATCGCTGGAACCCTTCCGTCGGACAAACCTGCTTTGGACTGCAAGCCTACGTGCCTGTCTGCATCGACACTCCGTGGTATGAGTTTACTCCACCCGTTCAGCCAGAGTTCGGCACGAAGTACACGCCTGAGCAGACGCCTGTGCCGGTCATGCCGGGGATTGTGGATGGGTGTGAGCTGTATGAGTTGGTTGAGCCCGGGAAGAGAGTTGAGGAGTTGGCGGCGGAGAATGGATTCGCGGTGGAGAAGTTCGCTGAGTGGAATGGGAATAGTACGACTGCTTGGGCGACTTACTGGGCCTGTGTGAAGGCT ```      ``` 43	ATGCAGTTCTCCATCTTCTCCCTTGCTACTCTTGTAGCCATAGCATCCGCTACTCCCTTTGCCATGCCCGAAGCCTACGCCGGCACTCTTGACACTACTTCTTCCTCTCTCTCGGCCCGTCAGTCCAACTGCACTGAATACTGCTCCAACAGCGCCGGCTGCGTCTGT	210 	\|\|\|\|\|\|\|\| \|\|\|\|\| \|\| \|\|\|\|\|\|\| \|\| \| \| \|\|\|\|\|\|\|\|\|\|\|\|\|\|\|\| \|\|\|\|\|\|\|\|\|\|\|\|\|\|\|\|\|\|\|\|\|\|\|\| \|\|\|\|\|\|\|\|\|\|\|\|\|\|\|\|\|\|\| \|\|\|\|\|\|\|\|\|\|\|\|\|\|\|\|\|\|\|\|\|\|\|\|\|\|\|\|\|\|\|\|\|\|\|\|\|\|\|\|\|\|\|\|\|\|\|\|\|\|\|\|\|\|\|\|\|\|\|\|\|\|\|\|\|\|\|\|\| \|\|\| 	ATGCAGTTNTCCATNTTNTCCCTTGNTANTNTNGTAGCCATAGCATCCGNTACTCCCTTTGCCATGCCCGAAGCNTACGCCGGCACTCTTGACANTACTTCTTCCTCTCTCTCGGCCCGTCAGTCCAACTGCACTGAATACTGCTCCAACAGCGCCGGCTGCGTATGT ``` |
| 3 | ggggacaagtttgtacaaaaaagcaggcttgatgCAGCGCATTCTCAGCCTCGC, ggggaccactttgtacaagaaagctgggtcACCTCCTCCTCCTTCTCCCC | Mycgr3\|110887  JGI  Ensembl |  |  | 0 % |  |  |
| 3 | ggggacaagtttgtacaaaaaagcaggcttgatgCAGTTCACTCAGGCTTCCGC, ggggaccactttgtacaagaaagctgggtcAGAGCTCACTTGCCAGGTGA | Mycgr3\|111636  JGI  Ensembl |  |  | 31.86583 % |  | ``` 76	CCAGCCA-ACGGCAACCATGCAGGCACCGACAACGGCAACAACCACTGCGGAGATACAACCATACCTCAGCCGATCATCGGCCAAGGCCCGCCTGTCGACAGGAACGACTGCCAAGAACTCAAGAACGCGTTCGCCAATGCTCACACCGACTACTACGTCACTGGACCCGGCCAGTTGATCCTTGCTCACGGGACCTGCAAATTTTATGCCACGGTCGATAGGGGTCAAACGGCCATCCTTGGAGGTGACGATGCCAGTGATCTCATTGGGACCGCCTTCAGTTACTGGGTCGATGGTCAATCCGGCTCTAAGGGAACCTATATCACCGGCCAGAACGCTGAAGCTCCTTGCGACAACGCCAACGGCCAGACCCAGGTCACCGTCACCTGGCAAGTGAGCTCT	477 	\|\|\|\|\|\|\|   \|\|\|\| \|\|\| \|\|\|\|\|\| \| \| \|\|   \|\|\|\|\|\|\|\|\|\| \|\|\|\|\|\|\|\|\|\|\|\|\|\|\|\|\|\|\|\|\|\|\|\|\|\|\|\|\|\|\|\|\|\|\|\|\|\|\|\|\|\|\|\|\|\|\|\|\|\|\|\|\|\|\|\|\|\|\|\|\|\|\|\|\|\|\|\|\|\|\|\|\|\|\|\|\|\|\|\|\|\|\|\|\|\|\|\|\|\|\|\|\|\|\|\|\|\|\|\|\|\|\|\|\|\|\| \|\|\|\|\|\|\|\|\|\|\|\|\|\|\|\|\|\|\|\|\|\|\|\|\|\|\|\|\|\|\|\|\|\|\|\|\|\|\|\|\|\|\|\|\|\|\|\|\|\|\|\|\|\|\|\|\|\|\|\|\|\|\|\|\|\|\|\|\|\|\|\|\|\|\|\|\|\|\|\|\|\|\|\|\|\|\|\|\|\|\|\|\|\|\|\|\|\|\|\|\|\|\|\|\|\|\|\|\|\|\|\|\|\|\|\|\|\|\|\|\|\|\|\|\|\|\|\|\|\|\|\|\|\|\|\|\|\|\|\|\|\|\|\|\|\|\|\|\|\|\|\|\|\| \|\|\|\|\|\|\|\|\|\|\|\|\|\|\|\|\|\|\|\|\|\|\|\|\|\|\|\|\|\|\|\|\|\|\|\|\|\|\|\|\|\|\|\|\|\|\|\|\|\|\|\|\|\|\|\|\|\|\|\|\|\|\|\|\|\|\|\|\|\|\|\|\|\|\|\|\|\|\|\|\|\|\|\|\|\|\|\|\|\|\|\|\|\| 	CCAGCCANGGGGCANCCANGCAGGCNCNGNCANNNGCAACAACCANTGCGGAGATACAACCATACCTCAGCCGATCATCGGCCAAGGCCCGCCTGTCGACAGGAACGACTGCCAAGAACTCAAGAACGCGTTCGCCAATGCTCACACCGACTANTACGTCACTGGACCCGGCCAGTTGATCCTTGCTCACGGGACCTGCAAATTTTATGCCACGGTCGATAGGGGTCAAACGGCCATCCTTGGAGGTGACGATGCCAGTGATCTCATTGGGACCGCCTTCAGTTACTGGGTCGATGGTCAATCCGGCTNTAAGGGAACCTATATCACCGGCCAGAACGCTGAAGCTCCTTGCGACAACGCCAACGGCCAGACCCAGGTCACCGTCACCTGGCAAGTGAGCTCT ``` |
| 4 | ggggacaagtttgtacaaaaaagcaggcttgatgGTGTCCGTCCACGAGGAGGA, ggggaccactttgtacaagaaagctgggtcTCGCCCGAATCTAAACTCCG | Mycgr3\|33116  JGI  Ensembl | KOG2818 Predicted undecaprenyl diphosphate synthase; | metabolic process; transferase activity; | 100 % | chr\_1:3046759 C->T; chr\_1:3046151 G->A; chr\_1:3046081 C->T; |  |
| 4 | ggggacaagtttgtacaaaaaagcaggcttgatgGCAGATGAAGACATGTGGCG, ggggaccactttgtacaagaaagctgggtcCTCTGACAAAGTCCGATAGG | Mycgr3\|35337  JGI  Ensembl | KOG2078 tRNA modification enzyme; |  | 100 % | chr\_1:3947665 T->C; chr\_1:3947997 A->G; chr\_1:3947393 T->C; |  |
| 4 | ggggacaagtttgtacaaaaaagcaggcttgatgCCGAAGGCCACGAGATCCAA, ggggaccactttgtacaagaaagctgggtcAACATCCATCATAGTATCAT | Mycgr3\|37123  JGI  Ensembl | KOG3871 Cell adhesion complex protein bystin; |  | 0 % |  |  |
| 4 | ggggacaagtttgtacaaaaaagcaggcttgatgGCACAGTCTGCCGAGCAGTC, ggggaccactttgtacaagaaagctgggtcAGCAGGCGGCATGAAACTCG | Mycgr3\|39191  JGI  Ensembl | KOG2647 Predicted Dolichyl-phosphate-mannose-protein mannosyltransferase; |  | 100 % | chr\_3:1139089 T->C; chr\_3:1139628 G->A; chr\_3:1139494 C->T; chr\_3:1139606 A->G; |  |
| 4 | ggggacaagtttgtacaaaaaagcaggcttgatgCGACCAGACGGACCTCGCGA, ggggaccactttgtacaagaaagctgggtcGCTAGCAATCTCATCTTTGT | Mycgr3\|40195  JGI  Ensembl | KOG3110 Riboflavin kinase; | FMN adenylyltransferase activity; riboflavin biosynthetic process;  riboflavin kinase activity; | 4.15861 % |  |  |
| 4 | ggggacaagtttgtacaaaaaagcaggcttgatgTCCTCCACCAAGGAAGACAA, ggggaccactttgtacaagaaagctgggtcCTCAGGCTGCCGATTTGTCG | Mycgr3\|40511  JGI  Ensembl | KOG1472 Histone acetyltransferase SAGA/ADA, catalytic subunit PCAF/GCN5 and related proteins; | N-acetyltransferase activity; histone acetyltransferase activity;  metabolic process; | 100 % | chr\_4:573645 A->G; chr\_4:574103 T->C; chr\_4:574236 A->G; chr\_4:574016 T->A; chr\_4:573422 A->G; chr\_4:574310 A->G; chr\_4:574289 T->C; | ``` 1	ATGTCCTCCACCAAGGAAGACAAGCGCAAAGCCGAAGACGAAATCGAGCACGATCCTAAGCGACTCAAGCCCTCAGTATCGCCTCCACCAAGCGACGACACACCCACCATCCGACACGAGCTCTTCCCCGAGAAGCCCGCCGTGACAGAAGAGCGCAACGGCGACATCTCATTCCGCGTGGTCAACAATGACGGCCAGAACTCCTCCTTCATCATCCTCACAGGCCTGAAGTGCATCTTCCAGAAGCAATTACCCAAGATGCCAAAGGACTACATCGCGCGATTGGTATACGATCGCACGCACTTGAGCATCGCAATCGTGAAGAAGCCTCCAGCAGGAAGCTTTGCCGAAAGCTCTGGTCTTCCCGGAGAAGTCGTTGGAGGCATTACATATCGACCATTCAAGGGTCGCCAGTTCGCGGAGATTGTCTTCTGTGCTATCAGCAGCGACCAGCAAGTCAAAGGATACGGCGCACATCTCATGAGCCACCTCAAAGACTACGTCAAGGCCACGAGCGATGTCATGTACTTCCTCACCTACGCCGACAACTACGCTATCGGGTACTTTAAGAAGCAAGGATTCACAAAGGAGATCACACTGGACAAACCGAAATGGATGGGCTACATCAAGGATTACGAGGGAGGAACAATCATGCAGTGTTCCATGCTGCCGAAGATTAGGTATCTCGAATCAGGTCGCATGCTGCTGAAGCAAAAGGCCGCAGTTCACGCCAAAATTCGTGCTGTCAGCAAAAGTTACGAGATACACCAGCCGCCGTCAGAATGGCGCAAAGTGAAACCTGGACAACCGCTGCCGGAGATCGATCCTCTGACCATCGATGCGATCAAGGCTACAGGGTGGTCGCCAGACATGGACGTTCTCGCACGGCAGCCTCGTCGTAATCCTTCACACAGCTTACTCCTCGGTTTGCTGTCCGCACTCCAGACAAGTTCTTCTGCGTGGCCTTTCCTGCAGCCTGTTAACGGTGAAGAAGTGCACGACTATTACGACGTGATCAAAGAGCCCATGGATCTGAGTACGATGGAGAGTAAGCTTGAGAAGGATCAGTACG-AGAATGTGGAGGACTTT-GTCA	1093 	\|\|\|\|\|\|\|\|\|\|\|\|\|\|\|\|\|\|\|\|\|\|\|\|\|\|\|\|\|\|\|\|\|\|\|\|\|\|\|\|\|\|\|\|\|\|\|\|\|\|\|\|\|\|\|\|\|\|\|\|\|\|\|\|\|\|\|\|\|\|\|\|\|\|\|\|\|\|\|\|\|\|\|\|\|\|\|\|\|\|\|\|\|\|\|\|\|\|\|\|\|\|\|\|\|\|\|\|\|\|\|\|\|\|\|\|\|\|\|\|\|\|\|\|\|\|\|\|\|\|\|\|\|\|\|\|\|\|\|\|\|\|\|\|\|\|\|\|\|\|\|\|\|\|\|\|\|\|\|\|\|\|\|\|\|\|\|\|\|\|\|\|\|\|\|\|\|\|\|\|\|\|\|\|\|\|\|\|\|\|\|\|\|\|\|\|\|\|\|\|\|\|\|\|\|\|\|\|\|\|\|\|\|\|\|\|\|\|\|\|\|\|\|\|\|\|\|\|\|\|\|\|\|\|\|\|\|\|\|\|\|\|\|\|\|\|\|\|\|\|\|\|\|\|\|\|\|\|\|\|\|\|\|\|\|\|\|\|\|\|\|\|\|\|\|\|\|\|\|\|\|\|\|\|\|\|\|\|\|\|\|\|\|\|\|\|\|\|\|\|\|\|\|\|\|\|\|\|\|\|\|\|\|\|\|\|\|\|\|\|\| \|\|\|\|\|\|\|\|\|\|\|\|\|\|\|\|\|\|\|\|\|\|\|\|\|\|\|\|\|\|\|\|\|\|\|\|\|\|\|\|\|\|\|\|\|\|\|\|\|\|\|\|\|\|\|\|\|\|\|\|\|\|\|\|\|\|\|\|\|\|\|\|\|\|\|\|\|\|\|\|\|\|\|\|\|\|\|\|\|\|\|\|\|\|\|\|\|\|\|\|\|\|\|\|\|\|\|\|\|\|\|\|\|\|\|\|\|\|\|\|\|\|\|\|\|\|\|\|\|\|\|\|\|\|\|\|\|\|\|\|\|\|\|\|\|\|\|\|\|\|\|\|\|\|\|\|\|\|\|\|\|\|\|\|\|\|\|\|\|\|\|\|\|\|\|\|\|\|\|\|\|\|\|\|\|\|\|\|\|\|\|\|\|\|\|\|\|\|\|\|\|\|\|\|\|\|\|\|\|\|\|\|\|\|\|\|\|\|\|\|\|\| \|\|\|\|\|\|\|\|\|\|\|\|\|\|\|\|\|\|\|\|\|\|\|\|\|\|\|\|\|\|\|\|\|\|\|\|\|\|\|\|\|\|\|\|\|\|\|\|\|\|\|\|\|\|\|\|\|\|\|\|\|\|\|\|\|\|\|\|\|\|\|\|\|\|\|\|\|\|\|\|\|\|\|\|\|\|\|\|\|\|\|\|\|\|\|\|\|\|\|\|\|\|\|\|\|\|\|\|\|\|\|\|\|\|\|\|\|\|\|\|\|\|\|\|\|\|\|\|\|\|\|\|\|\|\|\|\|\|\|\|\|\|\|\|\|\|\|\|\|\|\|\|\|\|\|\|\|\|\|\|\|\|\|\|\|\|\|\|\|\|\|\|\|\|\|\|\|\|\|\|\|\|\|\|\|\|\|\|\|\|\|\|\|\|\|\|\|\|\|\|\|\|\|\|\|\|\|\|\|\|\|\|\|\|\|\|\|\|\|\|\|\|\|\|\|\|\|\|\|\|\|\|\|\|\|\|\|\|\|\|\|\|\|\|\|\|\|\|\|\|\|\|\|\|\|\|\| \|\|\|\|\|\|\|\|\|\|\|\|\|\|\|\|\|\|\|\|\|\|\|\|\|\|\|\|\|\|\|\|\|\|\|\|\|\|\|\|\|\|\|\|\|\|\|\|\|\|\|\|\|\|\|\|\|\|\|\|\|\|\|\|\|\|\|\|\|\|\|\|\|\|\|\|\|\|\|\|\|\|\|\|\|\|\|\|\|\|\|\|\|\|\|\|\|\|\|\|\|\|\|\|\|\|\|\|\|\|\|\| \|\|\|\| \|\|\|\|\|\|\|\|\|\|\|\|\|\|\|\|\|\|\|\|\| \|\|\|\|\| \|\| \| \|\|\|\|\|\|\|\|\|\|\|\|\|\|\| \|\|\|\|\|\| \|\|\| \| \|\|\|\|\|\| \|\| \|\|\|\|\|\|\|\|\| \|\|\|\|\|\|\|\|\|\|\|\|\|\|\|  \|\|\|\|\| \|\|\| \|\|\|\|\|\| \|\|\|\|\|\|\| \| \|\|\|\| \|\|\| \|\| \| \| \|\|\|\|\|\|\|\|\| \|\|\|  \|\|\|\|\|\|\|\|\|\|\|\| \|\|\|\| 	ATGTCCTCCACCAAGGAAGACAAGCGCAAAGCCGAAGACGAAATCGAGCACGATCCTAAGCGACTCAAGCCCTCAGTATCGCCTCCACCAAGCGACGACACACCCACCATCCGACACGAGCTCTTCCCCGAGAAGCCCGCCGTGACAGAAGAGCGCAACGGCGACATCTCATTCCGCGTGGTCAACAATGACGGCCAGAACTCCTCCTTCATCATCCTCACAGGCCTGAAGTGCATCTTCCAGAAGCAATTACCCAAGATGCCAAAGGACTACATCGCGCGATTGGTATACGATCGCACGCACTTGAGCATCGCAATCGTGGAGAAGCCTCCAGCAGGAAGCTTTGCCGAAAGCTCTGGTCTTCCCGGAGAAGTCGTTGGAGGCATTACATATCGACCATTCAAGGGTCGCCAGTTCGCGGAGATTGTCTTCTGTGCTATCAGCAGCGACCAGCAAGTCAAAGGATACGGCGCACATCTCATGAGCCACCTCAAAGACTACGTCAAGGCCACGAGCGATGTCATGTACTTCCTCACCTACGCCGGCAACTACGCTATCGGGTACTTTAAGAAGCAAGGATTCACAAAGGAGATCACACTGGACAAACCGAAATGGATGGGCTACATCAAGGATTACGAGGGAGGAACAATCATGCAGTGTTCCATGCTGCCGAAGATTAGGTATCTCGAATCAGGTCGCATGCTGCTGAAGCAAAAGGCCGCAGTTCACGCCAAAATTCGTGCTGTCAGCAAAAGTTACGAGATACACCAGCCGCCGTCAGAATGGCGCAAAGTGAAACCTGTACAACCGCTGCCGGAGATCGATCCTCTGACCATCGATGCGATCAAGGCTACAGGGTGGTCGCCAGACATGGACGTTCTCGCACGGCAGCCTCGTCGTAATCCTTCACACAGCATACT-CTCGGTTTGCTGTCCGCACTC-AGACA-GT-C-TCTGCGTGGCCTTTC-TGCAGC-TGT-A-CGGTGA-GA-GTGCACGAC-ATTACGACGTGATCA--GAGCC-ATG-ATCTGA-TACGATG-A-AGTA-GCT-GA-A-G-ATCAGTACGGAGATGGTGGAGGACTTTTGTCA ``` |
| 4 | ggggacaagtttgtacaaaaaagcaggcttgatgGTCAAGGAAACCAAGTTCTA, ggggaccactttgtacaagaaagctgggtcCTGGCTAGCACACTGCACCC | Mycgr3\|42499  JGI  Ensembl | KOG0712 Molecular chaperone (DnaJ superfamily); | heat shock protein binding; protein folding;  unfolded protein binding; | 100 % | chr\_5:2553744 C->T; |  |
| 4 | ggggacaagtttgtacaaaaaagcaggcttgatgGAGACCAAGATACTGGCCGT, ggggaccactttgtacaagaaagctgggtcACTCTCGACACGAATTTCAG | Mycgr3\|43124  JGI  Ensembl | KOG3051 RNA binding/translational regulation protein of the SUA5 family; |  | 100 % |  |  |
| 4 | ggggacaagtttgtacaaaaaagcaggcttgatgGCCTCACATACCACTGAAAC, ggggaccactttgtacaagaaagctgggtcACTCGTCTTACCCACAGTCT | Mycgr3\|45907  JGI  Ensembl | KOG2924 Deoxyhypusine synthase; | deoxyhypusine synthase activity; peptidyl-lysine modification to hypusine;  spermidine catabolic process to deoxyhypusine, using deoxyhypusine synthase; | 100 % | chr\_8:1665135 A->G; |  |
| 4 | ggggacaagtttgtacaaaaaagcaggcttgatgAGCCTGCCAAATGGCGACAT, ggggaccactttgtacaagaaagctgggtcATCTCGACTCCTATTCGCCA | Mycgr3\|46254  JGI  Ensembl | KOG4685 tRNA splicing endonuclease SEN2; | tRNA splicing; tRNA-intron endonuclease activity;  tRNA-intron endonuclease complex; | 100 % | chr\_8:1418933 T->C; chr\_8:1418940 C->T; chr\_8:1419462 T->C; chr\_8:1418328 C->T; chr\_8:1418293 G->A; chr\_8:1418612 A->G; chr\_8:1419632 C->A; |  |
| 4 | ggggacaagtttgtacaaaaaagcaggcttgatgGCGACAACAAAACCCATACA, ggggaccactttgtacaagaaagctgggtcTCTCTTCTTCGAGTTCACAC | Mycgr3\|46438  JGI  Ensembl | KOG2463 Predicted RNA-binding protein Nob1p involved in 26S proteasome assembly; |  | 0 % |  |  |
| 4 | ggggacaagtttgtacaaaaaagcaggcttgatgTCCGGCGCCCGCCACTGGGA, ggggaccactttgtacaagaaagctgggtcCCTCTGACCACCAGGCGGGA | Mycgr3\|46623  JGI  Ensembl | KOG0131 Splicing factor 3b, subunit 4; | nucleic acid binding; | 3.80952 % |  |  |
| 4 | ggggacaagtttgtacaaaaaagcaggcttgatgGCCTCCCAATTTGCCCTCCG, ggggaccactttgtacaagaaagctgggtcCTTCGTCCACTCTTTCGCCT | Mycgr3\|46814  JGI  Ensembl | KOG1309 Suppressor of G2 allele of skp1; |  | 100 % |  |  |
| 4 | ggggacaagtttgtacaaaaaagcaggcttgatgCAGAAACTCATCGCCCGCAC, ggggaccactttgtacaagaaagctgggtcTGCTTTCTCTTGCCCAACCG | Mycgr3\|50825  JGI  Ensembl | KOG1708 Mitochondrial/chloroplast ribosomal protein L24; | intracellular; ribosome; structural constituent of ribosome;  translation; | 100 % | chr\_12:817440 T->C; chr\_12:817141 C->T; |  |
| 4 | ggggacaagtttgtacaaaaaagcaggcttgatgTCAAACCCACGCATAGAAGA, ggggaccactttgtacaagaaagctgggtcCCGTTTCTTCGGAGCCTTGA | Mycgr3\|55177  JGI  Ensembl | KOG3198 Signal recognition particle, subunit Srp19; | 7S RNA binding; SRP-dependent cotranslational protein targeting to membrane;  signal recognition particle; | 100 % |  |  |
| 4 | ggggacaagtttgtacaaaaaagcaggcttgatgTCTGAGCTGGTGGACACGCT, ggggaccactttgtacaagaaagctgggtcGTTCTTCAGGCCAATCTTGC | Mycgr3\|55692  JGI  Ensembl | KOG3078 Adenylate kinase; | ATP binding; adenylate kinase activity;  nucleobase, nucleoside, nucleotide and nucleic acid metabolic process;  nucleobase, nucleoside, nucleotide kinase activity;  nucleotide kinase activity;  phosphotransferase activity, phosphate group as acceptor; | 100 % |  |  |
| 4 | ggggacaagtttgtacaaaaaagcaggcttgatgGCTCGCAACGCAGAGAAAGC, ggggaccactttgtacaagaaagctgggtcAAGCTTGTCCAACAACTTGC | Mycgr3\|57452  JGI  Ensembl | KOG3068 mRNA splicing factor; |  | 100 % | chr\_3:3380539 T->C; chr\_3:3380941 T->C; chr\_3:3380703 C->A; |  |
| 4 | ggggacaagtttgtacaaaaaagcaggcttgatgGCAGAAAAGATCCTCCAGAA, ggggaccactttgtacaagaaagctgggtcCTCATCGTCCTCCTCCAGAT | Mycgr3\|61494  JGI  Ensembl | KOG0425 Ubiquitin-protein ligase; | negative regulation of ubiquitin-protein ligase activity; positive regulation of ubiquitin-protein ligase activity;  post-translational protein modification;  regulation of protein metabolic process;  regulation of ubiquitin-protein ligase activity;  regulation of ubiquitin-protein ligase activity during meiotic cell cycle;  small conjugating protein ligase activity;  ubiquitin-protein ligase activity; | 100 % | chr\_8:990614 C->T; |  |
| 4 | ggggacaagtttgtacaaaaaagcaggcttgatgGCCACCGCAACGGCGCTCCC, ggggaccactttgtacaagaaagctgggtcCAATTGACCCTTCTTCTGCA | Mycgr3\|65258  JGI  Ensembl | KOG3117 Protein involved in rRNA processing; |  | 5.78898 % |  |  |
| 4 | ggggacaagtttgtacaaaaaagcaggcttgatgGCCTCCACAGAAGCTGTCGG, ggggaccactttgtacaagaaagctgggtcTGCCTCCTTTGTCCGATCAG | Mycgr3\|65482  JGI  Ensembl | KOG2145 Cytoplasmic tryptophanyl-tRNA synthetase; | ATP binding; aminoacyl-tRNA ligase activity;  cytoplasm;  nucleotide binding;  tRNA aminoacylation for protein translation;  translation;  tryptophan-tRNA ligase activity;  tryptophanyl-tRNA aminoacylation; | 0 % |  |  |
| 4 | ggggacaagtttgtacaaaaaagcaggcttgatgGCGCCCGCTCTGCAAATCTC, ggggaccactttgtacaagaaagctgggtcGTGTATCTTCTTGATACGAT | Mycgr3\|65974  JGI  Ensembl | KOG3202 SNARE protein TLG1/Syntaxin 6; | cell communication; phosphoinositide binding;  protein binding; | 1.97664 % |  |  |
| 4 | ggggacaagtttgtacaaaaaagcaggcttgatgGCAAATGTGTTCAGCCTCAA, ggggaccactttgtacaagaaagctgggtcTTTGATCTCCGTCTTGCCCA | Mycgr3\|66253  JGI  Ensembl | KOG1909 Ran GTPase-activating protein; | protein binding; | 100 % | chr\_1:3048478 T->C; chr\_1:3048292 T->C; chr\_1:3048583 C->T; chr\_1:3048444 A->G; chr\_1:3048460 T->C; |  |
| 4 | ggggacaagtttgtacaaaaaagcaggcttgatgGCTTCAGACGAGCCGCAGAT, ggggaccactttgtacaagaaagctgggtcCTGTATCTTTTTGCCCTTCA | Mycgr3\|67122  JGI  Ensembl | KOG1430 C-3 sterol dehydrogenase/3-beta-hydroxysteroid dehydrogenase and related dehydrogenases; | 3-beta-hydroxy-delta5-steroid dehydrogenase activity; C-3 sterol dehydrogenase (C-4 sterol decarboxylase) activity;  steroid biosynthetic process;  sterol-4-alpha-carboxylate 3-dehydrogenase (decarboxylating) activity; | 100 % | chr\_1:4738052 T->C; chr\_1:4737246 C->T; chr\_1:4737521 T->C; chr\_1:4737269 T->C; chr\_1:4738142 A->G; chr\_1:4737230 C->A; chr\_1:4737293 A->G; |  |
| 4 | ggggacaagtttgtacaaaaaagcaggcttgatgCTTCCCGTCCGCACGATCAA, ggggaccactttgtacaagaaagctgggtcAGCCTCGATCTTTGCCTTCT | Mycgr3\|67666  JGI  Ensembl | KOG3031 Protein required for biogenesis of the ribosomal 60S subunit; |  | 100 % | chr\_1:5992542 A->T; chr\_1:5993401 G->A; |  |
| 4 | ggggacaagtttgtacaaaaaagcaggcttgatgTCGAACTACGGCTACCAATC, ggggaccactttgtacaagaaagctgggtcCGCACCGGTAGCCGCTTTCA | Mycgr3\|67821  JGI  Ensembl | KOG3103 Rab GTPase interacting factor, Golgi membrane protein; |  | 100 % | chr\_2:567078 C->T; chr\_2:567375 T->A; chr\_2:566740 A->G; |  |
| 4 | ggggacaagtttgtacaaaaaagcaggcttgatgGCTCTGGAACGAGGTGCTCG, ggggaccactttgtacaagaaagctgggtcTCTGTACCTCTCCTCCTTCA | Mycgr3\|68166  JGI  Ensembl | KOG3346 Phosphatidylethanolamine binding protein; |  | 100 % | chr\_2:1371528 T->C; chr\_2:1371867 G->A; chr\_2:1372582 A->G; chr\_2:1371974 A->G; chr\_2:1371452 G->A; chr\_2:1372314 C->T; chr\_2:1371787 A->G; chr\_2:1371475 A->G; chr\_2:1372174 A->G; |  |
| 4 | ggggacaagtttgtacaaaaaagcaggcttgatgGCCCCAGGAGACCTTCAACC, ggggaccactttgtacaagaaagctgggtcCGCAGCGTCCGGATCTGGTA | Mycgr3\|68937  JGI  Ensembl | KOG1446 Histone H3 (Lys4) methyltransferase complex and RNA cleavage factor II complex, subunit SWD2; |  | 100 % | chr\_2:2931759 C->T; chr\_2:2931651 A->G; |  |
| 4 | ggggacaagtttgtacaaaaaagcaggcttgatgGCCTCCAAATCAATATGTAC, ggggaccactttgtacaagaaagctgggtcCTCCGTCGTCTTTCCGGTAT | Mycgr3\|69123  JGI  Ensembl |  |  | 100 % | chr\_2:3535973 A->G; chr\_2:3537221 G->A; chr\_2:3536866 T->C; |  |
| 4 | ggggacaagtttgtacaaaaaagcaggcttgatgCCGGTCGACTACAGCAAATG, ggggaccactttgtacaagaaagctgggtcGTCCATTCCCGGGTCGTCAA | Mycgr3\|69545  JGI  Ensembl | KOG2260 Cell division cycle 37 protein, CDC37; |  | 100 % | chr\_3:1094458 G->A; chr\_3:1094350 A->G; chr\_3:1094220 C->T; chr\_3:1094907 T->C; chr\_3:1094689 G->A; |  |
| 4 | ggggacaagtttgtacaaaaaagcaggcttgatgGACTCCTCCGATGAAGACTC, ggggaccactttgtacaagaaagctgggtcCCCATCCATCAACCTCGTCC | Mycgr3\|70424  JGI  Ensembl |  |  | 100 % | chr\_3:3123054 A->G; chr\_3:3123817 T->C; chr\_3:3124107 T->A; chr\_3:3124101 T->C; |  |
| 4 | ggggacaagtttgtacaaaaaagcaggcttgatgTTCTACGACCTTAACATCCC, ggggaccactttgtacaagaaagctgggtcTGTTTGGTGTTCCGTAGCTC | Mycgr3\|72059  JGI  Ensembl | KOG2363 Protein subunit of nuclear ribonuclease P (RNase P); | ribonuclease P activity; ribonuclease activity;  tRNA processing; | 100 % | chr\_5:1184356 T->G; |  |
| 4 | ggggacaagtttgtacaaaaaagcaggcttgatgCAGCCCCTGAGCGGCACAGA, ggggaccactttgtacaagaaagctgggtcTCTATTGTCCGCAGTAAACA | Mycgr3\|72420  JGI  Ensembl | KOG2788 Glycosyltransferase; | UDP-N-acetylglucosamine-dolichyl-phosphate N-acetylglucosaminephosphotransferase activity; integral to membrane;  phospho-N-acetylmuramoyl-pentapeptide-transferase activity; | 100 % |  |  |
| 4 | ggggacaagtttgtacaaaaaagcaggcttgatgCCAGCGCCGATCTTGAACTG, ggggaccactttgtacaagaaagctgggtcTCGAGCATTCTTTGGATCAG | Mycgr3\|72630  JGI  Ensembl | KOG2751 Beclin-like protein; | autophagy; | 3.01783 % |  |  |
| 4 | ggggacaagtttgtacaaaaaagcaggcttgatgTCGCAACCCAGCACAGGCGA, ggggaccactttgtacaagaaagctgggtcATTCTTGCCTCTCCTCTTTT | Mycgr3\|72790  JGI  Ensembl | KOG0551 Hsp90 co-chaperone CNS1 (contains TPR repeats); | CTD phosphatase activity; calcium-dependent protein serine/threonine phosphatase activity;  calcium-dependent protein serine/threonine phosphatase regulator activity;  magnesium-dependent protein serine/threonine phosphatase activity;  magnesium-dependent protein serine/threonine phosphatase complex;  myosin phosphatase activity;  myosin phosphatase complex;  myosin phosphatase regulator activity;  phosphoprotein phosphatase activity; | 100 % | chr\_6:387027 T->C; chr\_6:387394 G->A; chr\_6:386788 A->T; chr\_6:386514 G->A; |  |
| 4 | ggggacaagtttgtacaaaaaagcaggcttgatgCCGAAATCCGCTCGAAAGCC, ggggaccactttgtacaagaaagctgggtcAGCAAAATGTATGCCCTCTT | Mycgr3\|73499  JGI  Ensembl | KOG0820 Ribosomal RNA adenine dimethylase; | 1-phenanthrol methyltransferase activity; 2-polyprenyl-6-methoxy-1,4-benzoquinone methyltransferase activity;  C-methyltransferase activity;  C-terminal protein carboxyl methyltransferase activity;  DNA-methyltransferase activity;  N-methyltransferase activity;  O-methyltransferase activity;  RNA methyltransferase activity;  S-methyltransferase activity;  arginine N-methyltransferase activity;  cobalt-precorrin-3 C17-methyltransferase activity;  cobalt-precorrin-5B C1-methyltransferase activity;  cobalt-precorrin-6B C5-methyltransferase activity;  cobalt-precorrin-7 C15-methyltransferase activity;  demethylmenaquinone methyltransferase activity;  dimethyladenosine transferase activity;  dimethylamine methyltransferase activity;  hydroxyneurosporene-O-methyltransferase activity;  lysine N-methyltransferase activity;  mRNA methyltransferase activity;  methanol-specific methylcobalamin:coenzyme M methyltransferase activity;  methylamine-specific methylcobalamin:coenzyme M methyltransferase activity;  methylarsonite methyltransferase activity;  monomethylamine methyltransferase activity;  protein methyltransferase activity;  protein-arginine N-methyltransferase activity;  protein-arginine N5-methyltransferase activity;  protein-leucine O-methyltransferase activity;  protein-lysine N-methyltransferase activity;  rRNA (adenine) methyltransferase activity;  rRNA (adenine-N6,N6-)-dimethyltransferase activity;  rRNA (cytosine) methyltransferase activity;  rRNA (cytosine-C5-967)-methyltransferase activity;  rRNA (guanine) methyltransferase activity;  rRNA (uridine) methyltransferase activity;  rRNA (uridine-2'-O-)-methyltransferase activity;  rRNA methyltransferase activity;  rRNA modification;  rRNA processing;  selenocysteine methyltransferase activity;  site-specific DNA-methyltransferase (cytosine-specific) activity;  tRNA (adenine)-methyltransferase activity;  tRNA (adenine-57, 58-N(1)-) methyltransferase activity;  tRNA (cytosine)-methyltransferase activity;  tRNA (guanine) methyltransferase activity;  tRNA (guanosine) methyltransferase activity;  tRNA (uracil) methyltransferase activity;  tRNA (uridine) methyltransferase activity;  tRNA methyltransferase activity;  trimethylamine methyltransferase activity; | 100 % | chr\_6:2357263 A->G; chr\_6:2356445 A->G; |  |
| 4 | ggggacaagtttgtacaaaaaagcaggcttgatgAAGATCAAAGCGCTCACACG, ggggaccactttgtacaagaaagctgggtcCTGCTCGGTCGCTAAAATCA | Mycgr3\|73576  JGI  Ensembl | KOG0268 Sof1-like rRNA processing protein (contains WD40 repeats); |  | 100 % | chr\_6:2598789 C->A; chr\_6:2597833 T->C; |  |
| 4 | ggggacaagtttgtacaaaaaagcaggcttgatgGCCGCAGAGGACAACGCAAA, ggggaccactttgtacaagaaagctgggtcCTCCAGCATGCTCGCCTGGT | Mycgr3\|73579  JGI  Ensembl | KOG1463 26S proteasome regulatory complex, subunit RPN6/PSMD11; | proteasome endopeptidase activity; threonine endopeptidase activity; | 100 % | chr\_6:2600526 A->G; chr\_6:2600596 A->G; chr\_6:2600550 G->A; chr\_6:2600251 T->C; chr\_6:2600637 G->A; |  |
| 4 | ggggacaagtttgtacaaaaaagcaggcttgatgGCGACTTTAGACTCCGCCAA, ggggaccactttgtacaagaaagctgggtcCGGTCTCCAATACCTCCATC | Mycgr3\|73816  JGI  Ensembl | KOG1511 Mevalonate kinase MVK/ERG12; | ATP binding; cytoplasm; isoprenoid biosynthetic process;  kinase activity;  metabolic process;  mevalonate kinase activity;  phosphorylation;  phosphotransferase activity, alcohol group as acceptor; | 100 % | chr\_7:630848 G->T; chr\_7:630101 A->G; chr\_7:630629 G->A; chr\_7:629969 C->T; chr\_7:629840 T->A; |  |
| 4 | ggggacaagtttgtacaaaaaagcaggcttgatgGCGGCACAGACTGGAGCAAG, ggggaccactttgtacaagaaagctgggtcAGCCGGCGACTGATTAATCT | Mycgr3\|73948  JGI  Ensembl | KOG0313 Microtubule binding protein YTM1 (contains WD40 repeats); | 1-alkyl-2-acetylglycerophosphocholine esterase activity; | 100 % |  |  |
| 4 | ggggacaagtttgtacaaaaaagcaggcttgatgTCGCTCGCTGCAGGGCCTGG, ggggaccactttgtacaagaaagctgggtcCCTGCCAAGCACTTTGTCGG | Mycgr3\|74097  JGI  Ensembl | KOG0366 Protein geranylgeranyltransferase type II, beta subunit; | Rab-protein geranylgeranyltransferase activity; catalytic activity; | 100 % | chr\_7:1367757 T->C; chr\_7:1368267 G->A; chr\_7:1367588 A->T; |  |
| 4 | ggggacaagtttgtacaaaaaagcaggcttgatgGCTCTTCGACTCGCGACGAG, ggggaccactttgtacaagaaagctgggtcTCCCGTGAAAGCCATGCTCT | Mycgr3\|74146  JGI  Ensembl | KOG3049 Succinate dehydrogenase, Fe-S protein subunit; | electron carrier activity; electron transport;  iron-sulfur cluster binding;  oxidoreductase activity;  succinate dehydrogenase (ubiquinone) activity;  tricarboxylic acid cycle; | 100 % | chr\_7:1491909 C->T; chr\_7:1492286 G->A; chr\_7:1491689 A->G; chr\_7:1491534 G->\*; |  |
| 4 | ggggacaagtttgtacaaaaaagcaggcttgatgTCGGCACCCACAGAGCACGT, ggggaccactttgtacaagaaagctgggtcTTCGAGAGGCGCTGCTGCGG | Mycgr3\|74650  JGI  Ensembl | KOG2014 SMT3/SUMO-activating complex, AOS1/RAD31 component; | catalytic activity; negative regulation of ubiquitin-protein ligase activity;  positive regulation of ubiquitin-protein ligase activity;  protein modification process;  regulation of ubiquitin-protein ligase activity;  regulation of ubiquitin-protein ligase activity during meiotic cell cycle;  small protein activating enzyme activity;  ubiquitin cycle;  ubiquitin-protein ligase activity; | 100 % | chr\_8:977188 A->G; chr\_8:976974 T->C; chr\_8:977290 T->C; chr\_8:976731 T->G; chr\_8:977566 T->C; chr\_8:977868 G->C; |  |
| 4 | ggggacaagtttgtacaaaaaagcaggcttgatgGCCGAAGAACAACGCAGTCT, ggggaccactttgtacaagaaagctgggtcTCCCCGATTGATGGTGTTCC | Mycgr3\|75819  JGI  Ensembl | KOG2830 Protein phosphatase 2A-associated protein; | regulation of signal transduction; response to biotic stimulus; | 100 % | chr\_9:1665934 T->C; chr\_9:1665789 A->T; | ``` 1	ATGGCCGAAGAACAACGCAGTCTGCGAGCAGTATACGAAGATGCCGAATCAAAACGAAACACCATCGAGAGCAGCTTCGACAGCAACTCCTCCGACTTTCAGGACAATCTTCTCGCCGCCATTAAGCTCTACGAACAGTGTCTGCAAATAGCCGACCAGATCTCTCTCTTCAGTCCCAATGAGTCACTCGAAGACATCAGCTCCGTCGATCTGCAATACCTTCTCCTCAACTACCGCATCGCCGAGCTCATCCTGCGAATAAACTCGCGAGAGCAACGAAAGGCCAACATCGAACGAGGACAGCGGAGCTATGAGAAGTTTCTCAAGCAGCTGGACAACTACGATATACTCACAAAAGATGATGCAAGGTTACTGGAACAATACCTCGACTCGCCGACCACATTCTCGGTCGCCTCAACCGCAGATGCTGCAGCAAGACGAGACACAAAGATCAAGCGTTTCAAGGAGGAGAAAGCACTAAGGCAAAAGTTGGAATATCTTCGCAGGAATCCGCTGGCTTTACAAAACGACGATACCGCGTATCGGGACCTGCAACTCACGAATGTGGCATTTGCAACGCACCAGTCCTTCGCCGCGTTGGAGTCGATCGCACAGGAGCTGCATATCCTGTCCCTCGCTCCTCCACCTCAAGCTCCGTCATCACAACCTCAAGTCCTAGATGCTCGTGAACGGAATGGCCGTGGGGATGGCTACAGCGAACGTCTCGATGCACCGATGTCCCACCTTTCCGCCGGCATGAAGGGTCCTCTGCTTGACAAAAGTGGCAAGCCACTGAGGCCGTTCACGTTGACAGGGAAGAGAACAGAGTTTCGAGAAGGTGTCTTCCGGCCGGACCA-TAGTTTGCCCACGATGAGCATAGACGAATACCTGGAGGAGGAGAGGAA-GCGTGGAGGCAT-G-ATAGA-TGGTG-G-AGGACCGCAAAGCCAGTTGACACCGC-AGGT-CGACGAGGATG-ATTATGATTTGGCGGACAAGGCGACAATGAAGGCAAGAGATTGG	1014 	\|\|\|\|\|\|\|\|\|\|\|\|\|\|\|\|\|\|\|\|\|\|\|\|\|\|\|\|\|\|\|\|\|\| \|\|\|\|\|\|\|\|\|\|\|\|\|\|\|\|\|\|\|\|\|\|\|\|\|\|\|\|\|\|\|\|\|\|\|\|\|\|\|\|\|\|\|\|\|\|\|\|\|\|\|\|\|\|\|\|\|\|\|\|\|\|\|\|\|\|\|\|\|\|\|\|\|\|\|\|\|\|\|\|\|\|\|\|\|\|\|\|\|\|\|\|\|\|\|\|\|\|\|\|\|\|\|\|\|\|\|\|\|\|\|\|\|\|\|\|\|\|\|\|\|\|\|\|\|\|\|\|\|\|\|\|\|\|\|\|\|\|\|\|\|\|\|\| \|\|\|\|\|\|\|\|\|\|\|\|\|\|\|\|\|\|\|\|\|\|\|\|\|\|\|\|\|\|\|\|\|\|\|\|\|\|\|\|\|\|\|\|\|\|\|\|\|\|\|\|\|\|\|\|\|\|\|\|\|\|\|\|\|\|\|\|\|\|\|\|\|\|\|\|\|\|\|\|\|\|\|\|\|\|\|\|\|\|\|\|\|\|\|\|\|\|\|\|\|\|\|\|\|\|\|\|\|\|\|\|\|\|\|\|\|\|\|\|\|\|\|\|\|\|\|\|\|\|\|\|\|\|\|\|\|\|\|\|\|\|\|\|\|\|\|\|\|\|\|\|\|\|\|\|\|\|\|\|\|\|\|\|\|\|\|\|\|\|\|\|\|\|\|\|\|\|\|\|\|\|\|\|\|\|\|\|\|\|\|\|\|\|\|\|\|\|\|\|\|\|\|\|\|\|\|\|\|\|\|\|\|\|\|\|\|\|\|\|\|\|\|\|\|\|\|\|\|\|\|\|\|\|\|\|\|\|\|\|\|\|\|\|\|\|\|\|\|\|\|\|\|\|\|\|\|\|\|\|\|\|\|\|\|\|\|\|\|\|\|\|\|\|\|\|\|\|\|\|\|\|\|\|\|\|\|\|\|\|\|\|\|\|\|\|\|\|\|\|\|\|\|\|\|\|\|\|\|\|\|\|\|\|\|\|\|\|\|\|\|\|\|\|\|\|\|\|\|\|\|\|\|\|\|\|\|\|\|\|\|\|\|\|\|\|\|\|\|\|\|\|\|\|\|\|\|\|\|\|\|\|\|\|\|\|\|\|\|\|\|\|\|\|\|\|\|\|\|\|\|\|\|\|\|\|\|\|\|\|\|\|\|\|\|\|\|\|\|\|\|\|\|\|\|\|\|\|\|\|\|\|\|\|\|\|\|\|\|\|\|\|\|\|\|\|\|\|\|\|\|\|\|\|\|\|\|\|\|\|\|\|\|\|\|\|\|\|\|\|\|\|\|\|\|\|\|\|\|\|\|\|\|\|\|\|\|\|\|\|\|\|\|\|\|\|\|\|\|\|\|\|\|\|\|\|\|\|\|\|\|\|\|\|\|\|\|\|\|\|\|\|\|\|\|\|\|\|\|\|\|\|\|\|\|\|\|\|\|\|\|\|\|\|\|\|\|\|\|\|\|\|\|\|\|\|\|\|\|\|\|\|\|\|\|\|\|\|\|\|\|\|\|\|\|\|\|\|\|\|\|\|\|\|\|\|\|\|\|\|\|\|\|\|\|\|\|\|\|\|\|\|\|\|\|\|\|\|\|\|\|\|\|\|\|\|\|\|\|\|\|\|\|\|\|\|\|\|\|\|\|\|\|\|\|\|\|\|\|\|\|\|\|\|\|\|\|\|\|\|\|\|\|\|\|\|\|\|\|\|\| \|\|\|\|\|\|\|\|\|\|\|\|\|\|\| \|\|\|\|\|\|\|\|\|\|\|\|\|\|\|\|\|\|\| \|\|\|\|\|\|\|\|\|\|\|\|\|\|\|\|\|\|\| \|\|\|\|\|\|\|\|\|\|\|\|\| \|\|\|\|\| \|\| \| \|\|\| \| \|\|\|\|\| \|\| \| \| \|\|\| \| \|\|\|\|\| \| \|\| \|\|\|  \|\|\|\|\|  \|\|  \|\| \|\|\|\| \|\| \| \|\|\|\|\|\|\|\|\|\|\| \|\|\|\| \|\|\|\|\|\| \|\| \|\| \|\|\|\| \|\|\|\|\|\|\|\|\|\|\|\|\|\|\| \|\|\|\|\| 	ATGGCCGAAGAACAACGCAGTCTGCGAGCAGTATTCGAAGATGCCGAATCAAAACGAAACACCATCGAGAGCAGCTTCGACAGCAACTCCTCCGACTTTCAGGACAATCTTCTCGCCGCCATTAAGCTCTACGAACAGTGTCTGCAAATAGCCGACCAGATCTCTCTCTTCAGTCCCAACGAGTCACTCGAAGACATCAGCTCCGTCGATCTGCAATACCTTCTCCTCAACTACCGCATCGCCGAGCTCATCCTGCGAATAAACTCGCGAGAGCAACGAAAGGCCAACATCGAACGAGGACAGCGGAGCTATGAGAAGTTTCTCAAGCAGCTGGACAACTACGATATACTCACAAAAGATGATGCAAGGTTACTGGAACAATACCTCGACTCGCCGACCACATTCTCGGTCGCCTCAACCGCAGATGCTGCAGCAAGACGAGACACAAAGATCAAGCGTTTCAAGGAGGAGAAAGCACTAAGGCAAAAGTTGGAATATCTTCGCAGGAATCCGCTGGCTTTACAAAACGACGATACCGCGTATCGGGACCTGCAACTCACGAATGTGGCATTTGCAACGCACCAGTCCTTCGCCGCGTTGGAGTCGATCGCACAGGAGCTGCATATCCTGTCCCTCGCTCCTCCACCTCAAGCTCCGTCATCACAACCTCAAGTCCTAGATGCTCGTGAACGGAATGGCCGTGGGGATGGCTACAGCGAACGTCTCGATGCACCGATGTCCCACCTTTCCGCCGGCATGAAGGGTCCTCTGCTTGACAAAAGTGGCAAGCCACTGAGGCCGTTCACGTTGACAGGGAAGAG-ACAGAGTTTCGAGAA-GTGTCTTCCGGCCGGACCATTAGTTTGCCCACGATGAGC-TAGACGAATACCTTGAGGAAGAAAAGAACGTGTGGAAGCCTTGTATAAAGTGGTGAGGAGCACCCAAAAGCATGTA-AC-CCGCGAGCTACGACGAGGATGTATTA-GATTTGCCG-AC-AGGCTACAATGAAGGCAAGA-ATTGG ``` |
| 4 | ggggacaagtttgtacaaaaaagcaggcttgatgGAGAACGAGAAGATCTCGGA, ggggaccactttgtacaagaaagctgggtcAGAAGCCCAAACGTCCTGTC | Mycgr3\|76790  JGI  Ensembl | KOG2908 26S proteasome regulatory complex, subunit RPN9/PSMD13; | proteasome endopeptidase activity; threonine endopeptidase activity; | 100 % | chr\_11:565438 T->C; chr\_11:565834 T->C; chr\_11:566296 G->A; chr\_11:565572 T->C; chr\_11:565196 A->G; chr\_11:566250 A->G; chr\_11:565175 A->G; |  |
| 4 | ggggacaagtttgtacaaaaaagcaggcttgatgGCTTCCTCCTCCTCCGCCGA, ggggaccactttgtacaagaaagctgggtcACAAAGATGCTCCTCCGTCT | Mycgr3\|77157  JGI  Ensembl | KOG2833 Mevalonate pyrophosphate decarboxylase; | ATP binding; diphosphomevalonate decarboxylase activity;  isoprenoid biosynthetic process;  kinase activity;  phosphorylation; | 100 % | chr\_12:55311 T->C; chr\_12:54904 T->C; |  |
| 4 | ggggacaagtttgtacaaaaaagcaggcttgatgAACAGTGTTCTGCCCGTCGA, ggggaccactttgtacaagaaagctgggtcCCACCCGCTACTAGGTCCGA | Mycgr3\|77431  JGI  Ensembl |  |  | 100 % | chr\_12:773321 G->A; chr\_12:772713 G->T; chr\_12:773184 T->C; chr\_12:772603 A->G; chr\_12:773291 A->G; |  |
| 4 | ggggacaagtttgtacaaaaaagcaggcttgatgTCCTTCTCCACCCTCGTCAA, ggggaccactttgtacaagaaagctgggtcCGTATCCAACCACTCCAAAA | Mycgr3\|77532  JGI  Ensembl | KOG2794 Delta-aminolevulinic acid dehydratase; | metal ion binding; porphobilinogen synthase activity;  tetrapyrrole biosynthetic process; | 0 % |  |  |
| 4 | ggggacaagtttgtacaaaaaagcaggcttgatgGTGTCTGCGCTGCCCAGAAC, ggggaccactttgtacaagaaagctgggtcTACGCCTGCACTTTCACCAA | Mycgr3\|77701  JGI  Ensembl | KOG0757 Mitochondrial carrier protein - Rim2p/Mrs12p; | binding; membrane; mitochondrial inner membrane;  transport;  transporter activity; | 100 % |  |  |
| 4 | ggggacaagtttgtacaaaaaagcaggcttgatgCCGCGATTTCTCACTCCGGC, ggggaccactttgtacaagaaagctgggtcAACCCTGCGCCGTGGAAAAG | Mycgr3\|77839  JGI  Ensembl | KOG4322 Anaphase-promoting complex (APC), subunit 5; |  | 100 % | chr\_13:615596 C->T; chr\_13:615869 A->T; chr\_13:616293 A->T; chr\_13:615281 T->C; |  |
| 4 | ggggacaagtttgtacaaaaaagcaggcttgatgCCGACCACCACCAACAACAT, ggggaccactttgtacaagaaagctgggtcCCCATGCATCACCACTCCCA | Mycgr3\|86699  JGI  Ensembl | KOG2976 Protein involved in autophagy and nutrient starvation; | autophagy; cytoplasm; | 100 % | chr\_7:68434 G->T; chr\_7:68239 A->G; |  |
| 4 | ggggacaagtttgtacaaaaaagcaggcttgatgACCGACTCGTCACGACCCTC, ggggaccactttgtacaagaaagctgggtcCTTCTTCACCGTGTCCTCCA | Mycgr3\|86952  JGI  Ensembl | KOG2915 tRNA(1-methyladenosine) methyltransferase, subunit GCD14; |  | 100 % | chr\_7:1637125 G->A; chr\_7:1636911 A->T; chr\_7:1637307 A->G; chr\_7:1636272 A->G; chr\_7:1636723 A->G; chr\_7:1636373 A->G; |  |
| 4 | ggggacaagtttgtacaaaaaagcaggcttgatgACAAACCAACTCCGCATCCT, ggggaccactttgtacaagaaagctgggtcTCGCAGCGCTCGGTAGGCGA | Mycgr3\|87000  JGI  Ensembl | KOG0237 Glycinamide ribonucleotide synthetase (GARS)/Aminoimidazole ribonucleotide synthetase (AIRS); | phosphoribosylamine-glycine ligase activity; purine base biosynthetic process; | 100 % | chr\_8:85386 T->C; chr\_8:85636 C->T; |  |
| 4 | ggggacaagtttgtacaaaaaagcaggcttgatgCCTCCTGCACCGAAACGAAG, ggggaccactttgtacaagaaagctgggtcCGCCTCCACCTTTCCCCCAC | Mycgr3\|87313  JGI  Ensembl | KOG2974 Uncharacterized conserved protein; |  | 100 % | chr\_8:1956645 T->C; chr\_8:1956824 A->G; chr\_8:1956509 A->T; chr\_8:1956869 T->A; chr\_8:1955942 T->C; |  |
| 4 | ggggacaagtttgtacaaaaaagcaggcttgatgGCAGGAAAACCTGTGACCGC, ggggaccactttgtacaagaaagctgggtcCTTCAACTCCCAGTCCTTGT | Mycgr3\|87527  JGI  Ensembl | KOG2823 Cellular protein (glioma tumor suppressor candidate region gene 2); |  | 100 % | chr\_9:1222031 T->C; chr\_9:1222851 G->A; chr\_9:1222677 A->G; |  |
| 4 | ggggacaagtttgtacaaaaaagcaggcttgatgGCCGCTGATGGGGACCTCAA, ggggaccactttgtacaagaaagctgggtcCGCGGTCTTCTCTTCTGCTT | Mycgr3\|87823  JGI  Ensembl | KOG0691 Molecular chaperone (DnaJ superfamily); | heat shock protein binding; | 100 % | chr\_10:961025 C->T; chr\_10:961163 T->C; chr\_10:960601 T->C; |  |
| 4 | ggggacaagtttgtacaaaaaagcaggcttgatgGAGTACTTTGCGAACAATCC, ggggaccactttgtacaagaaagctgggtcAGAACGAGTCTTTCCAGCTT | Mycgr3\|88465  JGI  Ensembl | KOG2728 Uncharacterized conserved protein with similarity to phosphopantothenoylcysteine synthetase/decarboxylase; | phosphopantothenate--cysteine ligase activity; | 0 % |  |  |
| 4 | ggggacaagtttgtacaaaaaagcaggcttgatgGCCATCTCACGCAAACGAAA, ggggaccactttgtacaagaaagctgggtcAGTCCAGACATCTCTCCACC | Mycgr3\|89109  JGI  Ensembl |  |  | 100 % | chr\_1:1926652 A->G; chr\_1:1926079 T->C; chr\_1:1926799 T->C; chr\_1:1926506 T->C; chr\_1:1926455 A->G; |  |
| 4 | ggggacaagtttgtacaaaaaagcaggcttgatgTCCTCGATAGGGAGTGGAAG, ggggaccactttgtacaagaaagctgggtcCTCGTCGTCTGCTTCCCCTT | Mycgr3\|89327  JGI  Ensembl |  |  | 2.82051 % |  |  |
| 4 | ggggacaagtttgtacaaaaaagcaggcttgatgAAGATCACCGCTCTACTTTT, ggggaccactttgtacaagaaagctgggtcCCTCTCCACCATAAAGATCT | Mycgr3\|90089  JGI  Ensembl |  |  | 100 % | chr\_1:5229424 A->G; chr\_1:5229354 C->T; chr\_1:5229342 T->A; |  |
| 4 | ggggacaagtttgtacaaaaaagcaggcttgatgGCCCAACAACTGCAGTCGCT, ggggaccactttgtacaagaaagctgggtcAGTGTGAGTTGTTGGAGCAA | Mycgr3\|91615  JGI  Ensembl | KOG0945 Alpha-aminoadipic semialdehyde dehydrogenase-phosphopantetheinyl transferase; |  | 100 % | chr\_3:1390316 G->A; chr\_3:1390849 G->T; chr\_3:1390817 A->G; chr\_3:1390111 T->C; chr\_3:1390492 T->C; chr\_3:1391029 T->C; chr\_3:1391017 A->T; | ``` 1	ATGGCCCAACAACTGCAGTCGCTGTCAGACAGCCTGCCGAGTGTCTCGACTCTGGTCAATGGAATAACCAACGCCTTCCACAATGCCACATCTCTGGTCAGTGGCGCTCCGCACGCTGGCGAAATTACCTGCTGGCTGATCGATACTCGAAAGCTGTGGCCTGGAACCAAAATCTGGGATGCTCCTGGCGCAGCAGAAGCCATGGGTTTGATTTCTGTGGCCGAGCAGAATGCCATCTCGACGAAGATGTTCATTCAAGACGCACGCATGAGCCTTGCAAGTGCACTGATCAAGCGACTCTATATCTCCAGAACCCTGGACATCCCCTGGAAGGACGTGATTCTTGCTAGGAAAGGTGATGCACAGCATGGAAAGCCATGCGCGGTAGATGTGGCCGGCAAGCCCATCGAGGGCATCGACTTCAACGTGTCACACCAGGCAGGACTGGTGACCTTGATCGGATGGAACGGCAAGCTGCGACATTCGTACAGCTCTGAAGGTATCATCGAGGGATCTGTCAATACTCCAGCCGCCGAAGACTTTGACACGAATGTCATGGTGGGCACTGACATCGTTTGCGTACACGAACGAAACGATGACCGCACCATAGAAGAGGAAGGTCTGGAGAGCTGGGTCGACACCTTCGACTTTGTCTTCAGCGATGAAGAGCGATGGAGCATGAAGTTTGACGTC	693 	\|\|\|\|\|\|\|\|\|\|\|\|\|\|\|\|\|\|\|\|\|\|\|\|\|\|\|\|\|\|\|\|\|\|\|\|\|\|\|\|\|\|\|\|\|\|\|\|\|\|\|\|\|\|\|\|\|\|\|\|\|\|\|\|\|\|\|\|\|\|\|\|\|\|\|\|\|\|\|\|\|\|\|\|\|\|\|\|\|\|\|\|\|\|\|\|\|\|\|\|\|\|\|\|\|\|\|\|\|\|\|\|\|\|\|\|\|\|\|\|\|\|\|\|\|\|\|\|\|\|\|\|\|\|\|\|\|\|\|\|\|\|\|\| \|\|\|\|\|\|\|\|\|\|\| \|\|\|\|\|\|\|\|\|\|\|\|\|\|\|\|\|\|\|\|\|\|\|\|\|\|\|\|\|\|\|\|\|\|\|\|\|\|\|\|\|\|\|\|\|\|\|\|\|\|\|\|\|\|\|\|\|\|\|\|\|\|\|\|\|\|\|\|\|\|\|\|\|\|\|\|\|\|\|\|\|\|\|\|\|\|\|\|\|\|\|\|\|\|\|\|\|\|\|\|\|\|\|\|\|\|\|\|\|\|\|\|\|\|\|\|\|\|\|\|\|\|\|\|\|\|\|\|\|\|\|\|\|\|\|\|\|\|\|\|\|\|\|\|\|\|\|\|\|\|\|\|\|\|\|\|\|\|\|\|\|\|\|\|\|\|\| \|\|\|\|\|\|\|\|\|\|\|\|\|\|\|\|\|\|\|\|\|\|\|\|\|\|\|\|\|\|\| \|\|\|\|\|\|\|\|\|\|\|\|\|\|\|\|\|\|\|\|\|\|\|\| \|\|\|\|\| \|\|\|\|\|\|\|\|\|\|\|\|\|\|\|\|\|\|\|\|\|\|\|\|\|\|\|\|\|\|\|\|\|\|\|\|\|\|\|\|\|\|\|\|\|\|\|\|\|\|\|\|\|\|\|\|\|\|\|\|\|\|\|\|\|\|\|\|\|\|\|\|\|\|\|\|\|\|\|\|\|\|\|\|\|\|\|\|\|\| \|\|\|\|\|\|\|\|\|\|\|\|\|\|\|\|\|\|\|\|\|\|\|\|\|\|\|\|\|\|\|\|\|\|\|\|\|\|\|\|\|\|\|\|\|\|\|\|\|\|\|\|\|\|\| \|\|\|\|\|\|\|\|\|\|\|\|\|\|\|\|\|\|\|\|\|\|\|\|\|\|\|\|\|\|\|\|\|\|\|\|\|\|\|\|\|\|\|\|\|\|\|\|\|\| \|\|\|\|\|\|\|\|\|\|\|\|\|\|\|\|\| \|\|\|\|\|\|\|\|\|\|\|\|\|\|\|\|\|\|\|\|\|\|\|\|\|\|\|\|\|\|\|\|\|\|\|\|\|\|\|\|\|\|\|\|\|\|\|\|\|\|\|\|\|\|\|\|\|\|\|\|\|\| \| \|\|\|\|\|\|\|\|\|\|\|\| \|\|\|\|\|\|\|\|\|\|\| 	ATGGCCCAACAACTGCAGTCGCTGTCAGACAGCCTGCCGAGTGTCTCGACTCTGGTCAATGGAATAACCAACGCCTTCCACAATGCCACATCTCTGGTCAGTGGCGCTCCGCACGCTGGCGAAATTACCTGCTGGCTGATCGATGCTCGAAAGCTGAGGCCTGGAACCAAAATCTGGGATGCTCCTGGCGCAGCAGAAGCCATGGGTTTGATTTCTGTGGCCGAGCAGAATGCCATCTCGACGAAGATGTTCATTCAAGACGCACGCATGAGCCTTGCAAGTGCACTGATCAAGCGACTCTATATCTCCAGAACCCTGGACATCACCTGGAAGGACGTGATTCTTGCTAGGAAAGGCGATGCACAGCATGGAAAGCCATGCTCGGTAAATGTGGCCGGCAAGCCCATCGAGGGCATCGACTTCAACGTGTCACACCAGGCAGGACTGGTGACCTTGATCGGATGGAACGGCAAGCTGCCACATTCGTACAGCTCTGAAGGTATCATCGAGGGATCTGTCAATACTCCAGCCGCCTAAGACTTTGACACGAATGTCATGGTGGGCACTGACATCGTTTGCGTACACTAACGAAACGATGACCGCCCCATAGAAGAGGAAGGTCTGGAGAGCTGGGTCGACACCTTCGACTTTGTCTTCAGCGATGAAAA-CGATGGAGCATGGAGTTTGACGTC ``` |
| 4 | ggggacaagtttgtacaaaaaagcaggcttgatgAAAGGCCTTTTCACGTCCGC, ggggaccactttgtacaagaaagctgggtcATAAACACTCCCCGTCCAAT | Mycgr3\|92097  JGI  Ensembl |  |  | 0 % |  |  |
| 4 | ggggacaagtttgtacaaaaaagcaggcttgatgAAGCTCAATATCGGGAGCAA, ggggaccactttgtacaagaaagctgggtcTGCGGACGCTGCTTCTTTGA | Mycgr3\|92792  JGI  Ensembl |  |  | 100 % | chr\_4:2292594 G->A; chr\_4:2292791 A->G; chr\_4:2292890 T->C; |  |
| 4 | ggggacaagtttgtacaaaaaagcaggcttgatgATGTCCCTCAAAGCATTGGC, ggggaccactttgtacaagaaagctgggtcCCGAATCTTGTTCCCTCCAT | Mycgr3\|93433  JGI  Ensembl |  |  | 100 % | chr\_5:2094408 G->A; chr\_5:2095218 T->C; chr\_5:2095041 C->G; chr\_5:2094281 A->G; chr\_5:2094855 A->T; chr\_5:2094755 G->A; chr\_5:2094476 A->G; chr\_5:2094911 T->C; |  |
| 4 | ggggacaagtttgtacaaaaaagcaggcttgatgGCATCGACCACAGGAAAGCG, ggggaccactttgtacaagaaagctgggtcCTCGGCATCATCCTCAGAGC | Mycgr3\|94337  JGI  Ensembl | KOG0294 WD40 repeat-containing protein; |  | 100 % | chr\_7:553780 A->G; |  |
| 4 | ggggacaagtttgtacaaaaaagcaggcttgatgGACAAGGACCGATTCCCCAT, ggggaccactttgtacaagaaagctgggtcAAACCCGCCGCCCTTAGCAT | Mycgr3\|95216  JGI  Ensembl | KOG1869 Splicing coactivator SRm160/300, subunit SRm300; |  | 100 % | chr\_8:1302505 T->C; chr\_8:1302117 T->C; chr\_8:1301322 C->G; |  |
| 4 | ggggacaagtttgtacaaaaaagcaggcttgatgGTCAAGCGTCGCCGCGAACG, ggggaccactttgtacaagaaagctgggtcCTCGATCGCAGTGGCAGCAG | Mycgr3\|97302  JGI  Ensembl | KOG1181 FOG: Low-complexity; |  | 100 % | chr\_13:281710 A->T; chr\_13:282048 G->A; chr\_13:282063 G->A; chr\_13:281880 T->G; chr\_13:282103 T->C; chr\_13:281773 T->C; |  |
| 4 | ggggacaagtttgtacaaaaaagcaggcttgatgTCGCTGCCGAGCGACTACAC, ggggaccactttgtacaagaaagctgggtcCCTGCCCGCGGCGTTTCGCG | Mycgr3\|98311  JGI  Ensembl | KOG1113 cAMP-dependent protein kinase types I and II, regulatory subunit; | (S)-lactate 2-kinase activity; ATP-dependent polynucleotide kinase activity;  GTP-dependent polynucleotide kinase activity;  LPPG:FO 2-phospho-L-lactate transferase activity;  cAMP-dependent protein kinase complex;  cAMP-dependent protein kinase regulator activity;  cobinamide kinase activity;  cyclin-dependent protein kinase activating kinase regulator activity;  cyclin-dependent protein kinase regulator activity;  cytidine kinase activity;  eukaryotic elongation factor-2 kinase activator activity;  eukaryotic elongation factor-2 kinase regulator activity;  glycerate 2-kinase activity;  inositol tetrakisphosphate 2-kinase activity;  phenol kinase activity;  phosphatidylinositol phosphate kinase activity;  phosphofructokinase activity;  phosphoinositide 3-kinase activity;  phosphorylase kinase regulator activity;  polynucleotide kinase activity;  protein amino acid phosphorylation;  signal transduction; | 100 % | chr\_1:1707248 T->C; chr\_1:1707669 T->C; chr\_1:1707860 T->C; |  |
| 4 | ggggacaagtttgtacaaaaaagcaggcttgatgGCCGACGTAGCAGCGCCCGT, ggggaccactttgtacaagaaagctgggtcATCCTCATCCTCACTGACCG | Mycgr3\|99535  JGI  Ensembl | KOG1813 Predicted E3 ubiquitin ligase; | nucleic acid binding; protein binding;  zinc ion binding; | 100 % | chr\_3:1626944 A->G; chr\_3:1626926 T->C; chr\_3:1626612 C->T; |  |
| 4 | ggggacaagtttgtacaaaaaagcaggcttgatgGCGTTGGACACCTTCTTCAA, ggggaccactttgtacaagaaagctgggtcCTTCCACAACTTACTCAAAC | Mycgr3\|99610  JGI  Ensembl | KOG0728 26S proteasome regulatory complex, ATPase RPT6; | ATP binding; cytoplasm; hydrolase activity;  nucleoside-triphosphatase activity;  nucleotide binding;  nucleus;  protein catabolic process; | 100 % | chr\_3:2206740 G->\*; chr\_3:2206244 T->C; chr\_3:2206591 A->G; chr\_3:2205589 C->A; |  |
| 4 | ggggacaagtttgtacaaaaaagcaggcttgatgCAGAACCTCATCCACACCTT, ggggaccactttgtacaagaaagctgggtcGACACTCATGGTGAAGTCGT | Mycgr3\|99699  JGI  Ensembl | KOG2981 Protein involved in autophagocytosis during starvation; |  | 100 % | chr\_3:2858281 T->C; chr\_3:2857778 A->G; chr\_3:2858662 A->G; chr\_3:2857882 T->C; |  |
| 4 | ggggacaagtttgtacaaaaaagcaggcttgatgTCTACCTCCACATCCGCCAT, ggggaccactttgtacaagaaagctgggtcGTGCTGCGCTCCTCCAACAC | Mycgr3\|99758  JGI  Ensembl | KOG2655 Septin family protein (P-loop GTPase); | GTP binding; cell cycle; | 100 % | chr\_3:3370644 T->A; chr\_3:3369869 G->A; |  |
| 4 | ggggacaagtttgtacaaaaaagcaggcttgatgACCCAATTCGCACCCCTCAA, ggggaccactttgtacaagaaagctgggtcGCTACCTGATATCCGGTGGA | Mycgr3\|100065  JGI  Ensembl | KOG2872 Uroporphyrinogen decarboxylase; | porphyrin biosynthetic process; uroporphyrinogen decarboxylase activity; | 100 % |  |  |
| 4 | ggggacaagtttgtacaaaaaagcaggcttgatgGCGACGAACCGTAATCGACG, ggggaccactttgtacaagaaagctgggtcCATCTCTCCGAACAGCCTCG | Mycgr3\|100874  JGI  Ensembl | KOG0418 Ubiquitin-protein ligase; | negative regulation of ubiquitin-protein ligase activity; positive regulation of ubiquitin-protein ligase activity;  post-translational protein modification;  regulation of protein metabolic process;  regulation of ubiquitin-protein ligase activity;  regulation of ubiquitin-protein ligase activity during meiotic cell cycle;  small conjugating protein ligase activity;  ubiquitin-protein ligase activity; | 5.62092 % |  |  |
| 4 | ggggacaagtttgtacaaaaaagcaggcttgatgGCCTCCGAAACCAACGGCAC, ggggaccactttgtacaagaaagctgggtcCGATGACGTACCCTCTCCTT | Mycgr3\|101162  JGI  Ensembl | KOG2703 C4-type Zn-finger protein; | protein binding; zinc ion binding; | 100 % | chr\_9:580148 T->A; chr\_9:579859 A->G; chr\_9:580027 T->C; chr\_9:580077 G->A; chr\_9:580338 G->A; |  |
| 4 | ggggacaagtttgtacaaaaaagcaggcttgatgTCGCTCACCAACTGTCGCTT, ggggaccactttgtacaagaaagctgggtcTGCAACCACGTTATCTTCGC | Mycgr3\|101353  JGI  Ensembl | KOG2916 Translation initiation factor 2, alpha subunit (eIF-2alpha); | RNA binding; eukaryotic translation initiation factor 2 complex;  translation;  translation initiation factor activity; | 100 % | chr\_10:242383 G->A; chr\_10:242648 A->G; chr\_10:241790 A->G; chr\_10:241781 A->G; |  |
| 4 | ggggacaagtttgtacaaaaaagcaggcttgatgCTTCGATTCACGCAACTTTT, ggggaccactttgtacaagaaagctgggtcAAGCTCGCTCTTCGCATCGT | Mycgr3\|101365  JGI  Ensembl | KOG0191 Thioredoxin/protein disulfide isomerase; | cell redox homeostasis; isomerase activity;  protein disulfide isomerase activity; | 100 % | chr\_10:347459 C->T; | ``` 534	ACTGCAAGAGCCTCGCCCCAATCTGGGAGACGGTCGCCGCCGACTTCGCCTCCGAGCCCTCCGTTCTTATCGCCAAGGTCGACGCCGATGCACCCAATGGCAAGAAAACCGCTGAGCGCTACGAGGTCAGGTCCTACCCAACAATTCTGTACTTCCCCAAGGGTTCGAAAGACGCAGTACCATACACGGGCGGACGTACCGAGGCCGATTTGGTGACCTTCATGAACGAAAAGGCTGGAACTTTCCGCTCTCCAGGCGGTGGTCTCAACGCTCTGGCCGGTGTCATCCCATCGCTCGACGAGGCCGTTGCGGCCTTGAAGACCGGCGGCGAGAAGGCGTACAAGGAGCTGGCCAAGCAAG-CCGGCGCGCTGCAGGACAAGACCGCTGAGTACTACGCCAAGGTTGCGAGCAAGGCTGAAAACAACGCGGGA-TACCTCGAGAAGGAATACACAAGGTTGCAGAACCTGATCGGCAAGGGCAACCTTGCGCCGGAGAAGCTGGACGATCTGA-TCAGCCG	1050 	\|\|\|\|\|\|\|\|\|\|\|\|\|\|\|\|\|\|\|\|\|\|\|\|\|\|\|\|\|\|\|\|\|\|\|\|\|\|\|\|\|\|\|\|\|\|\|\|\|\|\|\|\|\|\|\|\|\|\|\|\|\|\|\|\|\|\|\|\|\|\|\|\|\|\|\|\|\|\|\|\|\|\|\|\|\|\|\|\|\|\|\|\|\|\|\|\|\|\|\|\|\|\|\|\|\|\|\|\|\|\|\|\|\|\|\|\|\|\|\|\|\|\|\|\|\|\|\|\|\|\|\|\|\|\|\|\|\|\|\|\|\|\|\|\|\|\|\|\|\|\|\|\|\|\|\|\|\|\|\|\|\|\|\|\|\|\|\|\|\|\|\|\|\|\|\|\|\|\|\|\|\|\|\|\|\|\|\|\|\|\|\|\|\|\|\|\|\|\|\|\|\|\|\|\|\|\|\|\|\|\|\|\|\|\|\|\|\|\|\|\|\|\|\|\|\|\|\|\|\|\|\|\|\|\|\|\|\|\|\|\|\|\|\|\|\|\|\|\|\|\|\|\|\|\|\|\|\|\|\|\|\|\|\|\|\|\|\|\|\|\|\|\|\|\|\|\|\|\|\|\|\|\|\|\|\|\|\|\|\|\|\|\|\|\|\|\|\|\|\|\|\|\|\|\|\|\|\|\|\|\|\|\|\|\|\|\|\|\|\|\|\|\|\|\|\|\|\|\|\|\|\|\|\|\|\|\|\|\|\|\|\|\|\|\|\|\|\|\|\|\|\|\|\|\|\|\|\|\|\| \|\|\|\|\|\|\|\|\|\|\|\|\|\| \|\| \|\|\|\|\|\|\|\|\|\|\|\|\|\|\|\|\|\|\|\|\| \| \| \|\|\|\|\|\|\| \| \|\|\|\|\|  \|\| \|\|\|\|\|\|\| \|\|\|\|\|\|\|\|\|\| \| \| \|\|\| \|\| \| \| \|\|\|\|\| \| \|\|\|\|\|\| \|\| \|\|\|\|\| \| \| \|\|\|\|\|\|\|\| \|\|\|\|\|\|\|\|\|\|\|\|\|\|\|\| \|\|\|\|\|\|\| 	ACTGCAAGAGCCTCGCCCCAATCTGGGAGACGGTCGCCGCCGACTTCGCCTCCGAGCCCTCCGTTCTTATCGCCAAGGTCGACGCCGATGCACCCAATGGCAAGAAAACCGCTGAGCGCTACGAGGTCAGGTCCTACCCAACAATTCTGTACTTCCCCAAGGGTTCGAAAGACGCAGTACCATACACGGGCGGACGTACCGAGGCCGATTTGGTGACCTTCATGAACGAAAAGGCTGGAACTTTCCGCTCTCCAGGCGGTGGTCTCAACGCTCTGGCCGGTGTCATCCCATCGCTCGACGAGGCCGTTGCGGCCTTGAAGACCGGCGGCGAGAAGGCGTACAAGGAGCTGGCCAAGCAAGCCCGGCGCGCTGCAG-AC-AGACCGCTGAGTACTACGCCA-G-T-GCGAGCA-G-CTGAA--CA-CGCGGGAATACCTCGAGA-G-A-TAC-CA-G-T-GCAGA-C-TGATCG-CA-GGGCA-C-TGGCGCCGGA-AAGCTGGACGATCTGAATCAGCCG ```      ``` 17	ATGCTTCGATTCACGCAACTTTTCACTGCGGCGCTCGTCGTAGTCAGCACCTCTGCTGCCGCTGTCGTCGACCTCACGCCCTCAAACTTCGACTCCATCGTGCTCAAGTCCGGCAAGCCAGCGCTTGTCGAGTTCTTTGCACCATGGTGCGGCCACTGCAAGACTCTCGCTCCCATCTACGAAGAGCTCGCGGGTTCCTTCGCGTCTTCCACAGACAAAATCACCATCGCCAAAGTCGACGCCGACGAGCACAAGTCGCTGGGAACGAAATACGGAATCAAGGGATTTCCCACGATCAAGTACTTTGATGGCAGCGGGAAGTCGGAGCCAGAGGATTACAAGAAGGGCCGGGATATCGACAGCTTGACTGAGTTCATCACCGAGAAGATCGGCGTCAAGCCAAAGGGTGCGAAGAAGGTTGCATCGTCAGTCGAGATGTTGAACGACAGCACTTTTGACAAGCAGATTGGCGGTGACATGGATGCCATTGTCGCTTTCACAGCGCCGTGGTGTGGAC	533 	\|\|\|\|\|\|\|\|\|\|\|\|\|\|\|\|\|\|\|\|\|\|\|\|\|\|\|\|\|\|\|\|\|\|\|\|\|\|\|\|\|\|\|\|\|\|\|\|\|\|\|\|\|\|\|\|\|\|\|\|\|\|\|\|\|\|\|\|\|\|\|\|\|\|\|\|\|\|\|\|\|\|\|\|\|\|\|\|\|\|\|\|\|\|\|\|\|\|\|\|\|\|\|\|\|\|\|\|\|\|\|\|\|\|\|\|\|\|\|\|\|\|\|\|\|\|\|\|\|\|\|\|\|\|\|\|\|\|\|\|\|\|\|\|\|\|\|\|\|\|\|\|\|\|\|\|\|\|\|\|\|\|\|\|\|\|\|\|\|\|\|\|\|\|\|\|\|\|\|\|\|\|\|\|\|\|\|\|\|\|\|\|\|\|\|\|\|\|\|\|\|\|\|\|\|\|\|\|\|\|\|\|\|\|\|\|\|\|\|\|\|\|\|\|\|\|\|\|\|\|\|\|\|\|\|\|\|\|\|\|\|\|\|\|\|\|\|\|\|\|\|\|\|\| \|\|\|\|\|\|\|\|\|\|\|\|\|\|\|\|\|\|\|\|\|\|\|\|\|\|\|\|\|\|\|\|\|\|\|\|\|\|\|\|\|\|\|\|\|\|\|\|\|\|\|\|\|\|\|\|\|\|\|\|\|\|\|\|\|\|\|\|\|\|\|\|\|\|\|\|\|\|\|\|\|\|\|\|\|\|\|\|\|\|\|\|\|\|\|\|\|\|\|\|\|\|\|\|\|\|\|\|\|\|\|\|\|\|\|\|\|\|\|\|\|\|\|\|\|\|\|\|\|\|\|\|\|\|\|\|\|\|\|\|\|\|\|\|\|\|\|\|\|\|\|\|\|\|\|\|\|\|\|\|\|\|\|\|\|\|\|\|\|\|\|\|\|\|\|\|\|\|\|\|\|\|\|\|\|\|\|\|\|\|\|\|\|\|\|\|\|\|\|\|\|\|\|\|\|\|\|\|\|\|\|\|\|\|\|\|\|\|\|\|\|\|\|\|\|\|\|\|\|\|\|\|\|\|\|\|\|\|\|\|\|\|\|\|\|\|\|\|\|\|\|\|\|\|\|\|\|\|\|\|\|\| 	ATGCTTCGATTCACGCAACTTTTCACTGCGGCGCTCGTCGTAGTCAGCACCTCTGCTGCCGCTGTCGTCGACCTCACGCCCTCAAACTTCGACTCCATCGTGCTCAAGTCCGGCAAGCCAGCGCTTGTCGAGTTCTTTGCACCATGGTGCGGCCACTGCAAGACTCTCGCTCCCATCTACGAAGAGCTCGCGGGTTCCTTCGCGTCTTCCACAGACAAAATCACCATCGCCAAAGTCGACGCCGACGAGCACAAATCGCTGGGAACGAAATACGGAATCAAGGGATTTCCCACGATCAAGTACTTTGATGGCAGCGGGAAGTCGGAGCCAGAGGATTACAAGAAGGGCCGGGATATCGACAGCTTGACTGAGTTCATCACCGAGAAGATCGGCGTCAAGCCAAAGGGTGCGAAGAAGGTTGCATCGTCAGTCGAGATGTTGAACGACAGCACTTTTGACAAGCAGATTGGCGGTGACATGGATGCCATTGTCGCTTTCACAGCGCCGTGGTGTGGAC ``` |
| 4 | ggggacaagtttgtacaaaaaagcaggcttgatgGCCTCCGACGAGATCGTCTG, ggggaccactttgtacaagaaagctgggtcTTTCGCAAGCAACATCTCCC | Mycgr3\|101478  JGI  Ensembl | KOG3064 RNA-binding nuclear protein (MAK16) containing a distinct C4 Zn-finger; |  | 100 % |  |  |
| 4 | ggggacaagtttgtacaaaaaagcaggcttgatgCTCGCGGAGCACTTCATCGC, ggggaccactttgtacaagaaagctgggtcGTCCGAATCGTCCTCGCTTG | Mycgr3\|101861  JGI  Ensembl | KOG0646 WD40 repeat protein; |  | 100 % | chr\_13:424636 C->T; chr\_13:423860 T->C; chr\_13:423812 G->A; |  |
| 4 | ggggacaagtttgtacaaaaaagcaggcttgatgACACCGGCACGCTTCCTCCT, ggggaccactttgtacaagaaagctgggtcCCTGTCGAAGTTCTTCAAAT | Mycgr3\|102131  JGI  Ensembl | KOG3326 Uncharacterized conserved protein; |  | 100 % | chr\_1:1353450 C->T; chr\_1:1354240 T->C; chr\_1:1354360 A->G; chr\_1:1353487 A->T; | ``` 458	ATCAAAGCCGGAAAAGAGGCACACTAGAGTCCGATTTGCTCCTATCCACATTCGCCGACGAGAACCTTGGTGCCATGACAAAGGCGCAGCTGGAGCAATACGACTCTTTCCTTGACGAGAATGACTGGGACATCTACTATTGGGTTACACAGGAGCCAGTTCCTACTTCGATGGCATACGCTGAGGGATCGGGGCCGGATATGGCATCGCCGGAAGCGCAGGGCAAGGATGTTGCGGAGCAGAAGCCGGACACACAAGGCAAGAGAGTGCGCGCGCCTGGAACCGGTGAATGGGCTCAGACTGTTGGTACTTTCAAGCCTGCATACAGACCCGTCCCTCAGCGATGGAAGAACAGCGAGATTCTGTCGCTGTTGAGGAAGCATGTGATTGACCGCAGCGCCGGCAGTGTGCACATCGCTGGAAAGGACACATCGAATATGGATGGCGAGAGGAAGGGCGGCGGACTTGGCTTTATGCCTGATTTGAAGAACTTCGACAGG	957 	\|\|\|\|\|\|\|\|\|\|\|\|\|\|\|\|\|\|\|\|\|\|\|\|\|\|\|\|\|\|\|\|\|\|\|\|\|\|\|\|\|\|\|\|\|\|\|\|\|\|\|\|\|\|\|\|\|\|\|\|\|\|\|\|\|\|\|\|\|\|\|\|\|\|\|\|\|\|\|\|\|\|\|\|\|\|\|\|\|\|\|\|\|\|\|\|\|\|\|\|\|\|\|\|\|\|\|\|\|\|\|\|\|\|\|\|\|\|\|\|\|\|\|\|\|\|\|\|\|\|\|\|\|\|\|\|\|\|\|\|\|\|\|\|\|\|\|\|\|\|\|\|\|\|\|\|\|\|\|\|\|\|\|\|\|\|\|\|\|\|\|\|\|\|\|\|\|\|\|\|\|\|\|\|\|\|\|\|\|\|\|\|\|\|\|\|\|\|\|\|\|\|\|\|\|\|\|\|\|\|\|\|\|\|\|\|\|\|\|\|\|\|\|\|\|\|\|\|\|\|\|\|\|\|\|\|\|\|\|\|\|\|\|\|\|\|\|\|\|\|\|\|\|\|\|\|\|\|\|\|\|\|\|\|\|\|\|\|\|\|\|\|\|\|\|\|\|\|\|\|\|\|\|\|\|\|\|\|\|\|\|\|\|\|\|\|\|\|\|\|\|\|\|\|\|\|\|\|\|\|\|\|\|\|\|\|\|\|\|\|\|\|\|\|\|\|\|\|\|\|\|\|\|\|\|\|\|\|\|\|\|\|\|\|\|\|\|\|\|\| \|\|\|\|\|\|\|\|\|\|\|\|\|\|\|\|\|\|\|\|\|\|\|\|\|\|\|\|\|\|\|\|\|\|\|\|\|\|\|\|\|\|\|\|\|\|\|\|\|\|\|\|\|\|\|\|\|\|\|\|\|\|\| \|\|\|\|\|\|\|\|\|\|\|\|\|\|\|\|\|\|\| \|\|\|\|\|\|\|\|\|\|\|\|\|\|\|\| \|\|\|\| \| \|\|\|\|\|\|\|\|\|\|\|\|\|\|\|\|\|\| \|\|\|\|\|\|\|\| \|\| \|\| \|\|\|\|\|\|\| 	ATCAAAGCCGGAAAAGAGGCACACTAGAGTCCGATTTGCTCCTATCCACATTCGCCGACGAGAACCTTGGTGCCATGACAAAGGCGCAGCTGGAGCAATACGACTCTTTCCTTGACGAGAATGACTGGGACATCTACTATTGGGTTACACAGGAGCCAGTTCCTACTTCGATGGCATACGCTGAGGGATCGGGGCCGGATATGGCATCGCCGGAAGCGCAGGGCAAGGATGTTGCGGAGCAGAAGCCGGACACACAAGGCAAGAGAGTGCGCGCGCCTGGAACCGGTGAATGGGCTCAGACTGTTGGTACTTTCAAGCCTGCATACAGACCCGTCCCTCAGCGATGGAAG-ACAGCGAGATTCTGTCGCTGTTGAGGAAGCATGTGATTGACCGCAGCGCCGGCAGTGTGCACA-CGCTGGAAAGGACACATCG-ATATGGATGGCGAGAGAAAGG-C-GCGGACTTGGCTTTATGC-TGATTTGA-GA-CT-CGACAGG ```      ``` 16	ATGACACCGGCACGCTTCCTCCTCCGATCCAGCCAGTTCGCCGTGCGCAGCGTGCGACCCTTCGGCACCACATCTCGCTTGGCCGCAGTGCCAACCAACGACAGGCACAATGACACTGCAGAGACATACCGCAAGTTCATGAAGGAGAAGCCATTGAACCCGCATTTGACCAACTCCACATCGACAATCGCCAACGAAATGCCCAACATCGGCGAGGATGCGGCTCCGCCAGAGCTCATCACAAGCGTGGACTCCAAGTTCGTGCCCAAGGACTCCAAGCCTGAAAACACCGAGAGGATGACCGGCGGTACACAATCTGGATCGCCAAAGTCTGGACCCAACGCAGATCTCGATGTAGGTGAAATCGAGGGCGGCACGTTCAAGGTTGAGCCTATTCGTCGAACTGGCGAAGACGTCAACACTACTCGTGCACGTCTACTGT	457 	\|\|\|\|\|\|\|\|\|\|\|\|\|\|\|\|\|\|\|\|\|\|\|\|\|\|\|\|\|\|\|\|\|\|\|\|\| \|\|\|\|\|\|\|\|\|\|\|\|\|\|\|\|\|\|\|\|\|\|\|\|\|\|\|\|\|\|\|\|\|\|\|\|\|\|\|\|\|\|\|\|\|\|\|\|\|\|\|\|\|\|\|\|\|\|\|\|\|\|\|\|\|\|\|\|\|\|\|\|\|\|\|\|\|\|\|\|\|\|\|\|\|\|\|\|\|\|\|\|\|\|\|\|\|\|\|\|\|\|\|\|\|\|\|\|\|\|\|\|\|\|\|\|\|\|\| \|\|\|\|\|\|\|\|\|\|\|\|\|\|\|\|\|\|\|\|\|\|\|\|\|\|\|\|\|\|\|\|\|\|\|\|\|\|\|\|\|\|\|\|\|\|\|\|\|\|\|\|\|\|\|\|\|\|\|\|\|\|\|\|\|\|\|\|\|\|\|\|\|\|\|\|\|\|\|\|\|\|\|\|\|\|\|\|\|\|\|\|\|\|\|\|\|\|\|\|\|\|\|\|\|\|\|\|\|\|\|\|\|\|\|\|\|\|\|\|\|\|\|\|\|\|\|\|\|\|\|\|\|\|\|\|\|\|\|\|\|\|\|\|\|\|\|\|\|\|\|\|\|\|\|\|\|\|\|\|\|\|\|\|\|\|\|\|\|\|\|\|\|\|\|\|\|\|\|\|\|\|\|\|\|\|\|\|\|\|\|\|\|\|\|\|\|\|\|\|\|\|\|\|\|\|\|\|\|\|\|\|\|\|\|\|\|\|\|\|\|\|\|\|\|\|\|\|\|\|\|\|\|\|\|\|\|\|\|\|\|\|\|\|\|\|\|\|\|\|\|\|\|\|\|\|\|\|\|\|\|\|\|\|\|\|\|\|\|\|\|\|\|\|\|\|\|\|\|\|\|\|\|\| 	ATGACACCGGCACGCTTCCTCCTCCGATCCAGCCAGTCCGCCGTGCGCAGCGTGCGACCCTTCGGCACCACATCTCGCTTGGCCGCAGTGCCAACCAACGACAGGCACAATGACACTGCAGAGACATACCGCAAGTTCATGAAGGAGAAGCCATTGAGCCCGCATTTGACCAACTCCACATCGACAATCGCCAACGAAATGCCCAACATCGGCGAGGATGCGGCTCCGCCAGAGCTCATCACAAGCGTGGACTCCAAGTTCGTGCCCAAGGACTCCAAGCCTGAAAACACCGAGAGGATGACCGGCGGTACACAATCTGGATCGCCAAAGTCTGGACCCAACGCAGATCTCGATGTAGGTGAAATCGAGGGCGGCACGTTCAAGGTTGAGCCTATTCGTCGAACTGGCGAAGACGTCAACACTACTCGTGCACGTCTACTGT ``` |
| 4 | ggggacaagtttgtacaaaaaagcaggcttgatgGCTGAAGTCGCAAAGAACCC, ggggaccactttgtacaagaaagctgggtcGACGGTGAACTTGGCCTTCG | Mycgr3\|102384  JGI  Ensembl | KOG1670 Translation initiation factor 4F, cap-binding subunit (eIF-4E) and related cap-binding proteins; | RNA binding; cytoplasm; translation initiation factor activity;  translational initiation; | 100 % | chr\_1:2618738 A->G; chr\_1:2619092 T->C; |  |
| 4 | ggggacaagtttgtacaaaaaagcaggcttgatgAGATACTCCTTCGTCTCCGC, ggggaccactttgtacaagaaagctgggtcCAGTCCCCTGCCGGAAGGCA | Mycgr3\|102996  JGI  Ensembl |  |  | 100 % | chr\_1:5898389 C->G; |  |
| 4 | ggggacaagtttgtacaaaaaagcaggcttgatgAAGTTCTCTCAGCTCAGCAC, ggggaccactttgtacaagaaagctgggtcGCATCTCCACGAGGAGATGT | Mycgr3\|104000  JGI  Ensembl |  |  | 100 % |  |  |
| 4 | ggggacaagtttgtacaaaaaagcaggcttgatgCCGAAATTCTTCTGCGACTA, ggggaccactttgtacaagaaagctgggtcCTGCCCAGGCGGTCCACCGT | Mycgr3\|104068  JGI  Ensembl | KOG3454 U1 snRNP-specific protein C; | nucleic acid binding; nucleus; zinc ion binding; | 100 % |  |  |
| 4 | ggggacaagtttgtacaaaaaagcaggcttgatgCACTCCAAAGTCGTGATCAT, ggggaccactttgtacaagaaagctgggtcCAACAATGGGTTTGACCTGT | Mycgr3\|104491  JGI  Ensembl | KOG0404 Thioredoxin reductase; | FAD binding; cytoplasm; electron transport;  oxidoreductase activity;  removal of superoxide radicals;  thioredoxin-disulfide reductase activity; | 100 % | chr\_5:1209836 A->T; |  |
| 4 | ggggacaagtttgtacaaaaaagcaggcttgatgGCTCCCCAACCCGTATACAT, ggggaccactttgtacaagaaagctgggtcATCAAGACGCTCGATCACAA | Mycgr3\|104498  JGI  Ensembl | KOG1390 Acetyl-CoA acetyltransferase; |  | 100 % | chr\_5:1261135 C->T; chr\_5:1261324 T->C; chr\_5:1261229 G->A; |  |
| 4 | ggggacaagtttgtacaaaaaagcaggcttgatgGCGGCCATGTTCAACTCGGC, ggggaccactttgtacaagaaagctgggtcATCCTTCTTCAACTCCGAAT | Mycgr3\|105119  JGI  Ensembl | KOG0179 20S proteasome, regulatory subunit beta type PSMB1/PRE7; | proteasome core complex; proteasome endopeptidase activity;  threonine endopeptidase activity;  ubiquitin-dependent protein catabolic process; | 100 % |  |  |
| 4 | ggggacaagtttgtacaaaaaagcaggcttgatgGGCGACAACTTTCTCCACCT, ggggaccactttgtacaagaaagctgggtcCTGGTCGCCGCCGCGCCGAC | Mycgr3\|106350  JGI  Ensembl | KOG2975 Translation initiation factor 3, subunit f (eIF-3f); | proteasome endopeptidase activity; threonine endopeptidase activity; | 100 % | chr\_12:735892 A->G; |  |
| 4 | ggggacaagtttgtacaaaaaagcaggcttgatgTCTGTCCAACGCCCTTCGGC, ggggaccactttgtacaagaaagctgggtcCGTCCTCAACTCCGCAAAGA | Mycgr3\|108597  JGI  Ensembl | KOG1298 Squalene monooxygenase; | metabolic process; oxidoreductase activity;  squalene monooxygenase activity; | 100 % | chr\_3:1625420 A->G; chr\_3:1625514 T->A; |  |
| 4 | ggggacaagtttgtacaaaaaagcaggcttgatgAATGCAAACGAGCTGTCAAA, ggggaccactttgtacaagaaagctgggtcCAATTCATCTCTCTTGACTT | Mycgr3\|109553  JGI  Ensembl | KOG0161 Myosin class II heavy chain; | endoplasmic reticulum; | 3.49702 % |  |  |
| 4 | ggggacaagtttgtacaaaaaagcaggcttgatgGCGGAAAACGAAGTCAAACA, ggggaccactttgtacaagaaagctgggtcATCGAACTGCTGCTCCGGGC | Mycgr3\|109914  JGI  Ensembl | KOG3151 26S proteasome regulatory complex, subunit RPN12/PSMD8; | proteasome regulatory particle; proteolysis; | 100 % | chr\_6:1680741 C->T; chr\_6:1680842 C->T; |  |
| 4 | ggggacaagtttgtacaaaaaagcaggcttgatgGACACCCTCGTGGCTCAATA, ggggaccactttgtacaagaaagctgggtcATCCAAGTTCGCCGTGACGT | Mycgr3\|109970  JGI  Ensembl | KOG0175 20S proteasome, regulatory subunit beta type PSMB5/PSMB8/PRE2; | endopeptidase activity; proteasome core complex;  proteasome endopeptidase activity;  threonine endopeptidase activity;  ubiquitin-dependent protein catabolic process; | 100 % | chr\_6:1978072 C->T; chr\_6:1977892 T->A; chr\_6:1977751 T->C; chr\_6:1977682 A->G; chr\_6:1978392 T->A; chr\_6:1978186 C->T; |  |
| 4 | ggggacaagtttgtacaaaaaagcaggcttgatgCAGTACCTCACCAACACCCT, ggggaccactttgtacaagaaagctgggtcAGCAGAGCCAACGTCCATCA | Mycgr3\|109991  JGI  Ensembl |  |  | 100 % | chr\_6:2301482 A->G; chr\_6:2301638 A->G; chr\_6:2301522 A->T; chr\_6:2301563 A->G; |  |
| 4 | ggggacaagtttgtacaaaaaagcaggcttgatgGCGCACCGGAATCCCCTCAA, ggggaccactttgtacaagaaagctgggtcCAAGCTGAATCTGGTCCGGA | Mycgr3\|110502  JGI  Ensembl | KOG2780 Ribosome biogenesis protein RPF1, contains IMP4 domain; |  | 100 % | chr\_8:1347893 C->T; chr\_8:1347801 G->A; chr\_8:1348625 G->C; chr\_8:1348064 T->C; chr\_8:1347878 T->C; |  |
| 4 | ggggacaagtttgtacaaaaaagcaggcttgatgTCCACACCAATCGGCACCCC, ggggaccactttgtacaagaaagctgggtcAGCCTCCTTCTCCCTCGCCT | Mycgr3\|110683  JGI  Ensembl | KOG3075 Ribose 5-phosphate isomerase; | pentose-phosphate shunt, non-oxidative branch; ribose-5-phosphate isomerase activity; | 100 % | chr\_9:482364 T->C; chr\_9:482696 G->A; chr\_9:482335 T->C; chr\_9:482742 A->G; |  |
| 4 | ggggacaagtttgtacaaaaaagcaggcttgatgTCGACTATAAAACCCCTCGC, ggggaccactttgtacaagaaagctgggtcAGACGGTCTCATTGCTGCAC | Mycgr3\|111432  JGI  Ensembl | KOG0645 WD40 repeat protein; | lanosterol synthase activity; | 100 % | chr\_12:404978 A->G; chr\_12:405477 C->T; chr\_12:405634 A->T; chr\_12:405264 A->G; chr\_12:404738 T->A; chr\_12:405104 T->C; chr\_12:404775 T->C; chr\_12:405271 T->C; chr\_12:404398 C->T; |  |
| 5 | ggggacaagtttgtacaaaaaagcaggcttgatgCCACTCAGAAAGAAGAGGAC, ggggaccactttgtacaagaaagctgggtcGCCTCCGAATCCAAGCCCTC | Mycgr3\|31277  JGI  Ensembl | KOG1514 Origin recognition complex, subunit 1, and related proteins; | ATP binding; DNA binding;  nucleoside-triphosphatase activity;  nucleotide binding; | 0 % |  |  |
| 5 | ggggacaagtttgtacaaaaaagcaggcttgatgTCTATACATCCGTCGCGGCA, ggggaccactttgtacaagaaagctgggtcGAGTCCTTCCATGTCGTCTC | Mycgr3\|31393  JGI  Ensembl | KOG0272 U4/U6 small nuclear ribonucleoprotein Prp4 (contains WD40 repeats); | RNA splicing; | 100 % |  | ``` 61	AACGAGTCTGCAGCCCTCGACACCAACTACCAATTACCTTCCTCTGGCGGCGGACCTTCTCAAAATGCAGCGATTGTATTCGACGAGTTTGCGCGAAAGCGAAGAGCTGCGGCACTGGCTGTACCGACGGACGATCTACGAGTAAGGACGGAGCTGAGAGCACGCGGCGAACCCATCACATTATTCGGCGA	251 	\|\|\| \|\|\|\|\|\|\|\|\|\|\|\|\|\|\|\|\|\|\|\|\|\|\|\|\|\|\|\|\|\|\|\|\|\|\|\|\|\|\|\|\|\|\|\|\|\|\|\|\|\|\|\|\|\|\|\|\|\|\|\|\|\|\|\|\|\|\|\|\|\|\|\|\|\|\|\|\|\|\|\|\|\|\|\|\|\|\|\|\|\|\|\|\|\|\|\|\|\|\| \|\|\|\|\|\|\|\|\|\|\|\|\|\|\|\|\|\|\|\|\|\|\|\|\|\|\|\|\|\|\|\|\|\|\| \|\|\|\|\| \|\|\|\|\|\| \|\|\|\| \| \| \| \| \|  \|\|\|\|\|\|\|\|\|\| \|\|\| \|\| \|\|\|\| 	AAC-AGTCTGCAGCCCTCGACACCAACTACCAATTACCTTCCTCTGGCGGCGGACCTTCTCAAAATGCAGCGATTGTATTCGACGAGTTTGCGCGAAAGCGAANAGCTGCGGCACTGGCTGTACCGACGGACGATCTACNAGTAANGACGGANCTGANANCNCNCNGNNAACCCATCAC-TTANTCNGCGA ```      ``` 1	ATGTCTATACATCCGTCGCGGCAGGCGTACGTGGAGGAAGAGTCACAGCAGCAACAGTCCAACGAG	66 	\|\|\|\|\|\|\|\|\|\|\|\|\|\|\|\|\|\|\|\|\|\|\|\|\|\|\|\|\|\|\|\|\|\|\|\|\|\|\|\|\|\|\|\|\|\|\|\|\|\|\|\|\|\|\|\|\|\|\|\|\|\|\|\|\|\| 	ATGTCTATACATCCGTCGCGGCAGGCGTACGTGGAGGAAGAGTCACAGCAGCAACAGTCCAACGAG ``` |
| 5 | ggggacaagtttgtacaaaaaagcaggcttgatgGGACCCGCAAAGAAGAAGCA, ggggaccactttgtacaagaaagctgggtcCTTTCCACCCTTGCGATTCT | Mycgr3\|31414  JGI  Ensembl | KOG0127 Nucleolar protein fibrillarin NOP77 (RRM superfamily); | nucleic acid binding; | 0.8726 % |  |  |
| 5 | ggggacaagtttgtacaaaaaagcaggcttgatgTGGTGCACAAAATTATGGCT, ggggaccactttgtacaagaaagctgggtcTGGAAGGCTGAGGAAAAATT | Mycgr3\|38910  JGI  Ensembl |  |  | 100 % |  |  |
| 5 | ggggacaagtttgtacaaaaaagcaggcttgatgCCTGGCGCTACCCGTCGCGG, ggggaccactttgtacaagaaagctgggtcTGCGACAGGCACGGCATTGA | Mycgr3\|42271  JGI  Ensembl | KOG3763 mRNA export factor TAP/MEX67; | DNA binding; intracellular;  nucleosome;  nucleosome assembly;  nucleus;  protein binding;  transport; | 1.03842 % |  |  |
| 5 | ggggacaagtttgtacaaaaaagcaggcttgatgATCACAAGAAGGGCGTCTTC, ggggaccactttgtacaagaaagctgggtcCAACGCTCCTCCCACTAACG | Mycgr3\|45693  JGI  Ensembl | KOG3964 Phosphatidylglycerolphosphate synthase; | CDP-diacylglycerol-serine O-phosphatidyltransferase activity; catalytic activity;  metabolic process; | 100 % |  | ``` 1	ATGATCACAAGAAGGGCGTCTTCATGCCTGCGGCTGGTCAAGCAAAGCACATGCAAGCGGTCCTCTAATAGCCATGGCTTCAGCCAGAAACGATTCGCCAGCAGTGCATCTACTCCTGCAACAGCTTCGAGCACAACACCATCTGCAGCATCTCAAATCGGCCTCATAACGGGAGAGCTTGATAAGCTTTCGCCTCGCTTCGATGTTTCCGCCGACTCGATCGAGATCTTGAAGTCTCCTGCAGAGTTCTATCAAACGCTCAAG	264 	\|\|\|\|\|\|\|\|\|\|\|\|\|\|\|\|\|\|\|\|\|\|\|\|\|\|\|\|\|\|\|\|\|\|\|\|\|\|\|\|\|\|\|\|\|\|\|\|\|\|\|\|\|\|\|\|\|\|\|\|\|\|\|\|\|\|\|\|\|\|\|\|\|\|\|\|\|\|\|\|\|\|\|\|\|\|\|\|\|\|\|\|\|\|\|\|\|\|\|\|\|\|\|\|\|\|\|\|\|\|\|\|\|\|\|\|\|\|\|\|\|\|\|\|\|\|\|\|\|\|\|\|\|\|\|\|\|\|\|\|\|\|\|\|\|\|\|\|\|\|\|\|\|\|\|\|\|\|\|\|\|\|\|\|\|\|\|\|\|\|\|\|\|\|\|\|\|\|\|\|\|\|\|\|\|\|\|\|\|\|\|\|\|\|\|\|\|\|\|\|\|\|\|\|\|\|\|\|\|\|\|\|\|\|\|\|\|\|\|\|\|\|\|\|\|\|\|\|\|\|\|\|\|\|\|\|\|\|\|\|\|\|\|\|\|\|\|\|\|\|\|\|\|\|\|\|\|\|\|\|\|\|\|\| 	ATGATCACAAGAAGGGCGTCTTCATGCCTGCGGCTGGTCAAGCAAAGCACATGCAAGCGGTCCTCTAATAGCCATGGCTTCAGCCAGAAACGATTCGCCAGCAGTGCATCTACTCCTGCAACAGCTTCGAGCACAACACCATCTGCAGCATCTCAAATCGGCCTCATAACGGGAGAGCTTGATAAGCTTTCGCCTCGCTTCGATGTTTCCGCCGACTCGATCGAGATCTTGAAGTCTCCTGCAGAGTTCTATCAAACGCTCAAG ```      ``` 263	AGACGAAGATCAGCAAAGCACAGAAGAGGATCTACCTTTCAACGCTGTACATTGGGAAGAGTGAACATGAGCTG	336 	\|\|\|\|\|\|\|\|\|\|\|\|\|\|\|\|\|\|\|\|\|\|\|\|\|\|\|\|\|\|\|\|\|\|\|\|\|\|\|\|\|\|\|\|\|\|\|\|\|\|\|\|\|\|\|\|\|\|\|\|\|\|\|\|\|\|\|\|\|\|\|\|\|\| 	AGACGAAGATCAGCAAAGCACAGAAGAGGATCTACCTTTCAACGCTGTACATTGGGAAGAGTGAACATGAGCTG ```      ``` 336	GATCGACACCATTCGCGCTGCACTCAAAGCGAATCCGAACCTGAAAGTCTCGCTTCTAACAGACTATCTGAGAGGAACCCGCGAGGCTCCAAATCCGTCATGTGCAAGTTTGCTTGCTTCCCTGGTGTCCGACTTCGGACCGGATCGAGTTGATGTCCGGATGTACCACACCCCGAACCTGACCGGCACTCGCAAGGCGATTCTTCCGAAGAGAATCAACGAAGGCTGGGGTTTACAACACATGAAGCTCTATGGTATCGATGATGAGATCATCATGTCTGGGGCAAATCTGTCGGATGACTACTTCACGAACCGTCAAGATCGATACCACGTGTTCAAGTCCGCGGAGATCACAGAGTATTTTTCGAAACTGTA-TCGTACGGTGGCCAGCCTAAGCTATCGGGTGAG	743 	\|\|\|\|\|\|\|\|\|\|\|\|\|\|\|\|\|\|\|\|\|\|\|\|\|\|\|\|\|\|\|\|\|\|\|\|\|\|\|\|\|\|\|\|\|\|\|\|\|\|\|\|\|\|\|\|\|\|\|\|\|\|\|\|\|\|\|\|\|\|\|\|\|\|\|\|\|\|\|\|\|\|\|\|\|\|\|\|\|\|\|\|\|\|\|\|\|\|\|\|\|\|\|\|\|\|\|\|\|\|\|\|\|\|\|\|\|\|\|\|\|\|\|\|\|\|\|\|\|\|\|\|\|\|\|\|\|\|\|\|\|\|\|\|\|\|\|\|\|\|\|\|\|\|\|\|\|\|\|\|\|\|\|\|\|\|\|\|\|\|\|\|\|\|\|\|\|\|\|\|\|\|\|\|\|\|\|\|\|\|\|\|\|\|\|\|\|\|\|\|\|\|\|\|\|\|\|\|\|\| \|\|\|\|\|\|\|\|\|\|\|\|\|\|\|\|\|\|\|\|\|\|\|\|\|\|\|\|\|\|\|\|\|\|\|\|\|\|\|\|\|\|\|\|\|\|\|\|\|\|\|\|\|\|\|\|\|\|\|\|\|\|\|\|\|\|\|\|\|\|\|\|\|\| \|\|\|\|\|\|\|\|\|\|\|\|\|\|\|\|\|\|\| \|\|\|\|\|\|\|\|\|\|\|\|\|\|\|\|\|\|\|\|\|\|\|\|\|\|\|\|\|\|\|\|\|\|\|\|\| \|\|\|\| \|\|\|\|\|\| \|\|\|\|\|\|\|\| \| \|\| \|\|\|\|  \|\|\|\|\|\|\|\| \|\| \| \|\| \|\|  \|\|\|\|\|\|\|\| \|\|\| 	GATCGACACCATTCGCGCTGCACTCAAAGCGAATCCGAACCTGAAAGTCTCGCTTCTAACAGACTATCTGAGAGGAACCCGCGAGGCTCCAAATCCGTCATGTGCAAGTTTGCTTGCTTCCCTGGTGTCCGACTTCGGACCGGATCGAGTTGATGTCCGGATGTACCACACCCCGAACCTGACCGGCACTCGCAAGGCGATTCTTCCGAAAAGAATCAACGAAGGCTGGGGTTTACAACACATGAAGCTCTATGGTATCGATGATGAGATCATCATGTCTGGGGC-AATCTGTCGGATGACTACTNCACGAACCGTCAAGATCGATACCACGTGTTCAAGTCCCCGGANATCACAAAGTATTTTNCNAA-CTGTTNTCGTACGG-GGNCNGCNTAN-CTATCGGGNGAG ``` |
| 5 | ggggacaagtttgtacaaaaaagcaggcttgatgTCTTGGGACGAGGAGAAACA, ggggaccactttgtacaagaaagctgggtcTCGTCCGAAAAATTTATACT | Mycgr3\|50627  JGI  Ensembl | KOG2466 Uridine permease/thiamine transporter/allantoin transport; | membrane; nucleobase transmembrane transporter activity;  nucleobase, nucleoside, nucleotide and nucleic acid transport; | 100 % | chr\_12:1036375 C->T; | ``` 1	ATGTCTTGGGACGAGGAGAAACAACATGGCCTCGCTGAAGAGATCATTACCGAGGCACCTTCACAATTGCCACAACGACCTCTAACAACACAAGAGAAACTCGAAGCCACCCGATTCGACTCGAGGACAATCCACTCCCGAGACGCGTCTTCAAGCAACAATGGCTTGGAGTCTCACATTCAACCACATACCAGCATCCTCTCCCGTCTACGCAACATCGAAGCAACGCTCGATCGAAGATTAGGCGTCGAATCCGAAGCCATCGACCGCAAGTTGCCCGAAGACCGCAGACCCCAACCATGGCACGCCCAACTGAACATGGCCTTGATCTGGACCTCCGGAACCATGAACATCAGCTGTTTCGCGACGGGATTCATCGGCTGGCAATTCGGTCTCAGTCTCAGGCAGTCGATCTTGATCACCGTCTTCGGCTCGTTGCTCGGTGGCGCCGTGTCTGGCTTCTGTGCGACGATGGGTGCGCCGACCGGTTTGCGACAGATTTCGATATCGCGGTATAGCTTTGGGTGGTACCCGAATAAGATCGTCGCGGCGTTGAATACGATTCAGCAGTTGGGTTGGAGTGCGGCGGGGTGTATCACCGGTGGACTTGCGTTGACGGCAGTCTCGAACGGGAGGGTCTCGGTTGTGGTGGGGATCATCATCATCGCTACTTGCAGTCTGGTGATTAGCTTCATCGGACTGCGAGCGATTCTGGTGTATGA	722 	\|\|\|\|\|\|\|\|\|\|\|\|\|\|\|\|\|\|\|\|\|\|\|\|\|\|\|\|\|\|\|\|\|\|\|\|\|\|\|\|\|\|\|\|\|\|\|\|\|\|\|\|\|\|\|\|\|\|\|\|\|\|\|\|\|\|\|\|\|\|\|\|\|\|\|\|\|\|\|\|\|\|\|\|\|\|\|\|\|\|\|\|\|\|\|\|\|\|\|\|\|\|\|\|\|\|\|\|\|\|\|\|\|\|\|\|\|\|\|\|\|\|\|\|\|\|\|\|\|\|\|\|\|\|\|\|\|\|\|\|\|\|\|\|\|\|\|\|\|\|\|\|\|\|\|\|\|\|\|\|\|\|\|\|\|\|\|\|\|\|\|\|\|\|\|\|\|\|\|\|\|\|\|\|\|\|\|\|\|\|\|\|\|\|\|\|\|\|\|\|\|\|\|\|\|\|\|\|\|\|\|\|\|\|\|\|\|\|\|\|\|\|\|\|\|\|\|\|\|\|\|\|\|\|\|\|\|\|\|\|\|\|\|\|\|\|\|\|\|\|\|\|\|\|\|\|\|\|\|\|\|\|\|\|\|\|\|\|\|\|\|\|\|\|\|\|\|\|\|\|\|\|\|\|\|\|\|\|\|\|\|\|\|\|\|\|\|\|\|\|\|\|\|\|\|\|\|\|\|\|\|\|\|\|\|\|\|\|\|\|\|\|\|\|\|\|\|\|\|\|\|\|\|\|\|\|\|\|\|\|\|\|\|\|\|\|\|\|\|\|\|\|\|\|\|\|\|\|\|\|\|\|\|\|\|\|\|\|\|\|\|\|\|\|\|\|\|\|\|\|\|\|\|\|\|\|\|\|\|\|\|\|\|\|\|\|\|\|\|\|\|\|\|\|\|\|\|\|\|\|\|\|\|\|\|\|\|\|\|\|\|\|\|\|\|\|\|\|\|\|\|\|\|\|\|\|\|\|\|\|\|\|\|\|\|\|\|\|\|\|\|\|\|\|\|\|\|\|\|\|\|\|\|\|\|\|\|\|\|\|\|\|\|\|\|\|\|\|\|\|\|\|\|\|\|\|\|\|\|\|\|\|\|\|\|\|\|\|\|\|\|\|\|\|\|\|\|\|\|\|\|\|\|\|\|\|\|\|\|\|\|\|\|\|\|\|\|\|\|\|\|\|\|\|\|\|\|\|\|\|\|\|\|\|\|\|\|\|\|\|\|\|\|\|\|\|\|\|\|\|\|\|\|\|\|\|\|\|\|\|\|\|\|\|\|\|\|\|\|\|\|\|\|\|\|\|\|\|\|\|\|\| \|\|\|\|\|\|\|\|\|\|\|\|\|\|\|\|\|\|\|\|\|\|\|\|\|\|\|\|\|\|\|\|\|\|\|\|\|\|\|\|\|\|\|\|\|\|\|\|\|\|\|\|\|\|\|\|\|\|\|\|\|\|\|\|\|\|\|\|\|\|\|\|\|\|\|\|\|\|\|\|\|\|\|\|\|\|\|\|\|\|\|\|\|\|\| \|\|\|\|\|\|\|\|\|\| \|\|\| \|\|\|\|\|\|\|\|\|\|\|\|\|\|\|\|\|\| 	ATGTCTTGGGACGAGGAGAAACAACATGGCCTCGCTGAAGAGATCATTACCGAGGCACCTTCACAATTGCCACAACGACCTCTAACAACACAAGAGAAACTCGAAGCCACCCGATTCGACTCGAGGACAATCCACTCCCGAGACGCGTCTTCAAGCAACAATGGCTTGGAGTCTCACATTCAACCACATACCAGCATCCTCTCCCGTCTACGCAACATCGAAGCAACGCTCGATCGAAGATTAGGCGTCGAATCCGAAGCCATCGACCGCAAGTTGCCCGAAGACCGCAGACCCCAACCATGGCACGCCCAACTGAACATGGCCTTGATCTGGACCTCCGGAACCATGAACATCAGCTGTTTCGCGACGGGATTCATCGGCTGGCAATTCGGTCTCAGTCTCAGGCAGTCGATCTTGATCACCGTCTTCGGCTCGTTGCTCGGTGGCGCCGTGTCTGGCTTCTGTGCGACGATGGGTGCGCCGACCGGTTTGCGACAGATTTCGATATCGCGGTATAGCTTTGGGTGGTACCCGAATAAGATCGTCGCGGCGTTGAATACGATTCAGCAGTTGGGTTGGAGTGCGGCGGGGTATATCACCGGTGGACTTGCGTTGACGGCAGTCTCGAACGGGAGGGTCTCGGTTGTGGTGGGGATCATCATCATCGCTACTTGCAGTCTGGTGATTANCTTCATCGGA-TGCNAGCGATTCTGGTGTATGA ``` |
| 5 | ggggacaagtttgtacaaaaaagcaggcttgatgTCGGAGCAGCAAAAGCCAGA, ggggaccactttgtacaagaaagctgggtcCATGCCCTCATTTGCCAAGC | Mycgr3\|55269  JGI  Ensembl | KOG1467 Translation initiation factor 2B, delta subunit (eIF-2Bdelta/GCD2); | cellular metabolic process; | 100 % |  | ``` 380	ATGTCGGAGCAGCAAAAGCCAGAGCAGAAGCCCGCCGCGGCAGCCACACCCGCAGCTGCAGCTGCG-TCTACGTCTA-CAGAGGAAAAG-ATTTCTCC-TGCGGAAGCAAA-GAAGA	491 	\|\|\|\|\|\|\|\|\|\|\|\|\|\|\|\|\|\|\|\|\|\|\|\|\|\|\|\|\|\|\|\|\|\|\|\|\|\|\|\|\|\|\|\|\|\|\|\|\|\|\|\|\|\|\|\|\|\|\|\|\|\|\|\|\|\| \|\|\|\|\|\|\|\|\|\| \|\|\|\| \|\|\|\|\|  \|\|\|\|\|\| \| \|\|\|\|\|\|\|\| \|\|  \|\|\|\|\| 	ATGTCGGAGCAGCAAAAGCCAGAGCAGAAGCCCGCCGCGGCAGCCACACCCGCAGCTGCAGCTGCGCTCTACGTCTANCAGACGAAAAACATTTCTTCNTGCGGAAGNAANNGAAGA ``` |
| 5 | ggggacaagtttgtacaaaaaagcaggcttgatgCCGTCCCGCGAAGATGTCTA, ggggaccactttgtacaagaaagctgggtcAGCCTTCGCGAAATCTTGCA | Mycgr3\|55454  JGI  Ensembl | KOG2790 Phosphoserine aminotransferase; | L-serine biosynthetic process; metabolic process;  phosphoserine transaminase activity; | 0 % |  |  |
| 5 | ggggacaagtttgtacaaaaaagcaggcttgatgAGCAGCTGCAGAGCGCTCTC, ggggaccactttgtacaagaaagctgggtcCCCCAGAAATCTCCTGATTT | Mycgr3\|57150  JGI  Ensembl |  |  | 0 % |  |  |
| 5 | ggggacaagtttgtacaaaaaagcaggcttgatgGGTGACTCTCTCGTCCACCA, ggggaccactttgtacaagaaagctgggtcAGCATTCTGATCCAACCCCA | Mycgr3\|57630  JGI  Ensembl | KOG2323 Pyruvate kinase; | glycolysis; magnesium ion binding;  potassium ion binding;  pyruvate kinase activity; | 100 % |  | ``` 168	GGCCCCAAGACCAACTCCGCCGAAAAGATCAACATGCTTCGCACCGCCGGCCTCAACGTCGTCCGCATGAACTTCTCCCACGGCTCCTACGAATACCACCAGTCCGTCATCGACAATGCCCGCCAAGCAGAGAAAGAGCAGGAAGGTCGTTCCGTCGCCATCGCCCTCGACACCAAGGGTCCCGAGATCCGCACGGGCAACACCCCCGGAGACGAGGATATTCCCATCTCTGCGGGCACCGAGATCAACATCACCACCGACGACAAATATGCCACCGCCAGTGACGCTCAGAACATGTATGTCGATTACAAGAACATTACAAAAGTCATCGAGGCCGGCCGCACAATCTTTGTCGACGATGGAGTCCTCGCATTCGAGGTGCTCGAGGTGGTCGATGATAAGACTCTCCG	577 	\|\|\|\|\|\|\|\|\|\|\|\|\|\|\|\|\|\|\|\|\|\|\|\|\|\|\|\|\|\|\|\|\|\|\|\|\|\|\|\|\|\|\|\|\|\|\|\|\|\|\|\|\|\|\|\|\|\|\|\|\|\|\|\|\|\|\|\|\|\|\|\|\|\|\|\|\|\|\|\|\|\|\|\|\|\|\|\|\|\|\|\|\|\|\|\|\|\|\|\|\|\|\|\|\|\|\|\|\|\|\|\|\|\|\|\|\|\|\|\|\|\|\|\|\|\|\|\|\|\|\|\|\|\|\|\|\|\|\|\|\|\|\|\|\|\|\|\|\|\|\|\|\|\|\|\|\|\|\|\|\|\|\|\|\|\|\|\|\|\|\|\|\|\|\|\|\|\|\|\|\|\|\|\|\|\|\|\|\|\|\|\|\|\|\|\|\|\|\|\|\|\|\|\|\|\|\|\|\|\|\|\|\|\|\|\|\|\|\|\|\|\|\|\|\|\|\|\|\|\|\|\|\|\|\|\|\|\|\|\|\|\|\|\|\|\|\|\|\|\|\|\|\|\|\|\|\|\|\|\|\|\|\|\|\|\|\|\|\|\|\|\|\|\|\|\|\|\|\|\|\|\|\|\|\|\|\|\|\|\| \|\|\|\|\|\|\|\|\|\|\|\|\|\|\|\|\|\|\|\|\|\|\|\|\|\|\|\|\|\|\|\|\|\|\|\|\|\|\|\|\|\|\|\|\|\|\|\|\|\|\|\|\|\|\|\|\|\|\|\|\|\|\|\|\|\|\|\|\|\|\|\|\|\|\|\|\|\|\|\|\|\|\|\| \|\|\|\|\|\|\|\|\|\|\|\|\|\|\|\|\| \|\|\|\|\|\|\| \| \|\|\|\|\|\| 	GGCCCCAAGACCAACTCCGCCGAAAAGATCAACATGCTTCGCACCGCCGGCCTCAACGTCGTCCGCATGAACTTCTCCCACGGCTCCTACGAATACCACCAGTCCGTCATCGACAATGCCCGCCAAGCAGAGAAAGAGCAGGAAGGTCGTTCCGTCGCCATCGCCCTCGACACCAAGGGTCCCGAGATCCGCACGGGCAACACCCCCGGAGACGAGGATATTCCCATCTCTGCGGGCACCGAGATCAACATCACCACCGACGACAAATATGCCACCGCCAGTGACGCTCANAACATGTATGTCGATTACAAGAACATTACAAAAGTCATCGAGGCCGGCCGCACAATCTTTGTCGACGATGGAGTCCTCGCATTCNAGGTGCTCGAGGTGGTCNATGATAANANTCTCCG ```      ``` 48	ATGGGTGACTCTCTCGTCCACCAGAGCTATGGCCGCACCCGTGTGCAATGGATCGCCAGCCTCAACACCGAGTACCGCCCTCCGACCCAGTTCCGCCGCACCAGCATCATCTGCACCATCGG	169 	\|\|\|\|\|\|\|\|\|\|\|\|\|\|\|\|\|\|\|\|\|\|\|\|\|\|\|\|\|\|\|\|\|\|\|\|\|\|\|\|\|\|\|\|\|\|\|\|\|\|\|\|\|\|\|\|\|\|\|\|\|\|\|\|\|\|\|\|\|\|\|\|\|\|\|\|\|\|\|\|\|\|\|\|\|\|\|\|\|\|\|\|\|\|\|\|\|\|\|\|\|\|\|\|\|\|\|\|\|\|\|\|\|\|\|\|\|\|\|\|\|\| 	ATGGGTGACTCTCTCGTCCACCAGAGCTATGGCCGCACCCGTGTGCAATGGATCGCCAGCCTCAACACCGAGTACCGCCCTCCGACCCAGTTCCGCCGCACCAGCATCATCTGCACCATCGG ``` |
| 5 | ggggacaagtttgtacaaaaaagcaggcttgatgCTCGACATCAACGACTTCAT, ggggaccactttgtacaagaaagctgggtcTGACTTGACCTCCTCCGCCT | Mycgr3\|61156  JGI  Ensembl | KOG2509 Seryl-tRNA synthetase; | ATP binding; aminoacyl-tRNA ligase activity;  cytoplasm;  nucleotide binding;  serine-tRNA ligase activity;  seryl-tRNA aminoacylation;  tRNA aminoacylation for protein translation;  translation; | 100 % |  | ``` 64	ATGCTCGACATCAACGACTTCATCAAGGAACGCGGCGGTGACCCGGCCAAAATCAAGGAATCGCAGCGTCGCCGCCATGCACCGGAAGAGGTCGTAGACGAGGTCATCGCATTATTCGAGGACCACCGCA-AAACTCACTATGCCGCCGCAACGGAGATGGGAGCCA-AGCTCAACAAGGTGCAAAAGGACATGGGCCTGAACAAGA	268 	\|\|\|\|\|\|\|\|\|\|\|\|\|\|\|\|\|\|\|\|\|\|\|\|\|\|\|\|\|\|\|\|\|\|\|\|\|\|\|\|\|\|\|\|\|\|\|\|\|\|\|\|\|\|\|\|\|\|\|\|\|\|\|\|\|\|\|\|\|\|\|\|\|\|\|\|\|\|\|\|\|\|\|\|\|\|\|\|\|\|\|\|\|\|\|\| \|\|\|\| \|\|\|\|\|\|\|\|\|\|\|\|\|\|\|\|\|\|\|\|\|\| \|\|\|\|  \|\|\| \|\|\|\|\|\|\|\|\|\|\|\|\| \| \|\|\|\|\|\|\|\|\|\|\| \|\|\|  \| \|\|\| \|\|  \|\|\|\|\|\| \|\|  \| \|\|\| \|\|\| \| \|\|\|\|\|\| 	ATGCTCGACATCAACGACTTCATCAAGGAACGCGGCGGTGACCCGGCCAAAATCAAGGAATCGCAGCGTCGCCGCCATGCACCGGAAGAGGTCGTANACGANGTCATCGCATTATTCGAGGACCTCCGCCTAAANTCACTATGCCGCCNCNACGGAGATGGGNGCCCCANCTCNACGCGGTGCATAANNATATGNGCCNGNACAAGA ``` |
| 5 | ggggacaagtttgtacaaaaaagcaggcttgatgGCGCCGTCTTTGGAGGAACC, ggggaccactttgtacaagaaagctgggtcATCCTCTGCCAGTATTTGAT | Mycgr3\|62484  JGI  Ensembl | KOG3022 Predicted ATPase, nucleotide-binding; |  | 0 % |  |  |
| 5 | ggggacaagtttgtacaaaaaagcaggcttgatgACGGAACTCACGATCCCGCT, ggggaccactttgtacaagaaagctgggtcCTCAGCACCCTCCTCTTCCC | Mycgr3\|65107  JGI  Ensembl | KOG4533 Uncharacterized conserved protein; |  | 100 % |  | ``` 1	ATGACGGAACTCACGATCCCGCTCCCCTCCTCCGCCTCTCCCACCTTCACCCATCTCATCCTAGTATGCGGCCACGCGACCTACATTGGTGCCACAACTCCCTCCTCCGCGTCCCTCGATGAACAGAACTGGTCTCTTCAATCCTTCCAGCGCTCCAATCCTACAACTCACAAACCCGGCGAGCACCATACATTCCTATCTCATATCCACTCCGCCGCCTCTCTCGCTGCTTCAACGCCGTCCTCCCTGCTCATCTTCTCCGGTGGGCAGACCACGCCCACCGAACCATCTAACCCTGATGAATTGACCGAAGCTCAGTCACACGCAAGACTACTTCAAATGCACATCTCCACTCTCCCTGCTTCATCTCAGGAAAACCTCACCTACGCCTCTGAGAACTCGGCGACGGATTCGTACCAAAATCTCCTCTTCTCTATTCTTCTCTTTCATCACCTGACACACGCATACCCTGCGCACGTAACGGTGATCACCCACGCTTTCAAAACTCCCCGGTTCCTACATCTCCACGGCCCAGCGATCAAATTTCCCGCTCATCGCCTGCGAGTGCTGGGTATCAATCCACCTTTCACTGCACAAGAAGCAG-AGGAGACGGAGATCGGGGAAAGGGAGAGAGGCTATGCGCCTTGGAAGGAGGATTTGTATGGTGTTGGAGAGGTGCTGGGCCGGAAGAGGAGGGAGAGAGGGTGGGACGATGAGAGGGTCAAAGATGGATGGATGGATGAGTTGTCGGACGAAGTGAGGGAGCTGCTGGAGTGGAAAGGTGGGAGCACGGGGAGAGAGATCTTTCCTA	811 	\|\|\|\|\|\|\|\|\|\|\|\|\|\|\|\|\|\|\|\|\|\|\|\|\|\|\|\|\|\|\|\|\|\|\|\|\|\|\|\|\|\|\|\|\|\|\|\|\|\|\|\|\|\|\|\|\|\|\|\|\|\|\|\|\|\|\|\|\|\|\|\|\|\|\|\|\|\|\|\|\|\|\|\|\|\|\|\|\|\|\|\|\|\|\|\|\|\|\|\|\|\|\|\|\|\|\|\|\|\|\|\|\|\|\|\|\|\|\|\|\|\|\|\|\|\|\|\|\|\|\|\|\|\|\|\|\|\|\|\|\|\|\|\|\|\|\|\|\|\|\|\|\|\|\|\|\|\|\|\|\|\|\|\|\|\|\|\|\|\|\|\|\|\|\|\|\|\|\|\|\|\|\|\|\|\|\|\|\|\|\|\|\|\|\|\|\|\|\|\|\|\|\|\|\|\|\|\|\|\|\|\|\|\|\|\|\|\|\|\|\|\|\|\|\|\|\|\|\|\|\|\|\|\|\|\|\|\|\|\|\|\|\|\|\|\|\|\|\|\|\|\|\|\|\|\|\|\|\|\|\|\|\|\|\|\|\|\|\|\|\|\|\|\|\|\|\|\|\|\|\|\|\|\|\|\|\|\|\|\|\|\|\|\|\|\|\|\|\|\|\|\|\|\|\|\|\|\|\|\|\|\|\|\|\|\|\|\|\|\|\|\|\|\|\|\|\|\|\|\|\|\|\|\|\|\|\|\|\|\|\|\|\|\|\|\|\|\|\|\|\|\|\|\|\|\|\|\|\|\|\|\|\|\|\|\|\|\|\|\|\|\|\|\|\|\|\|\|\|\|\|\|\|\|\|\|\|\|\|\|\|\|\|\|\|\|\|\|\|\|\|\|\|\|\|\|\|\|\|\|\|\|\|\|\|\|\|\|\|\|\|\|\|\|\|\|\|\|\|\|\|\|\|\|\|\|\|\|\|\|\|\|\|\|\|\|\|\|\|\|\|\|\|\|\|\|\|\|\|\|\|\|\|\|\|\|\|\|\|\|\|\|\|\|\|\|\|\|\|\|\|\|\|\|\|\|\|\|\|\|\|\|\|\|\|\|\|\|\|\|\|\|\|\|\|\|\|\|\|\|\|\|\|\|\|\|\|\|\|\|\|\|\|\|\|\|\|\|\|\|\|\|\|\|\|\|\|\|\|\|\|\|\|\|\|\|\|\|\|\|\|\|\|\|\|\|\|\|\|\|\|\|\|\|\|\|\|\|\|\|\|\|\|\|\|\|\|\|\|\|\|\|\|\|\|\|\|\|\|\|\|\|\|\|\|\|\|\|\|\|\|\|\|\| \|\|\|\|\|\|\|\|\|\|\|\|\|\|\|\|\|\|\|\|\|\|\|\|\|\|\|\|\|\|\|\|\|\|\|\|\|\|\|\|\|\|\|\|\|\|\|\|\|\|\|\|\|\|\|\|\|\|\|\|\|\|\|\|\|\|\|\|\|\|\|\|\|\|\|\|\|\|\|\|\|\|\|\|\| \| \|\|\|\|\|\|\|\|\|\|\|\|\|\|\|\|\|\|\|\|\|\|\|\| \| \|\|\|\| \|\|\|\|\|\|\|\|\|\|\|\|\|\|\|\|\|\|\|\|\|\|\|\|\|\|\|\|\|\|\|\| \|\|\|\|\|\|\|\|\|\|\|\|\|\|\|\|\|\|\|\|\|\|\|\|\|\|\|\|\|\|\|\|  \|\|\|\|\|\|\|\|\|\|\|  \|\|\|\|\|\|\| 	ATGACGGAACTCACGATCCCGCTCCCCTCCTCCGCCTCTCCCACCTTCACCCATCTCATCCTAGTATGCGGCCACGCGACCTACATTGGTGCCACAACTCCCTCCTCCGCGTCCCTCGATGAACAGAACTGGTCTCTTCAATCCTTCCAGCGCTCCAATCCTACAACTCACAAACCCGGCGAGCACCATACATTCCTATCTCATATCCACTCCGCCGCCTCTCTCGCTGCTTCAACGCCGTCCTCCCTGCTCATCTTCTCCGGTGGGCAGACCACGCCCACCGAACCATCTAACCCTGATGAATTGACCGAAGCTCAGTCACACGCAAGACTACTTCAAATGCACATCTCCACTCTCCCTGCTTCATCTCAGGAAAACCTCACCTACGCCTCTGAGAACTCGGCGACGGATTCGTACCAAAATCTCCTCTTCTCTATTCTTCTCTTTCATCACCTGACACACGCATACCCTGCGCACGTAACGGTGATCACCCACGCTTTCAAAACTCCCCGGTTCCTACATCTCCACGGCCCAGCGATCAAATTTCCCGCTCATCGCCTGCGAGTGCTGGGTATCAATCCACCTTTCACTGCACAAGAAGCAGAAGGAGACGGAGATCGGGGAAAGGGAGAGAGGCTATGCGCCTTGGAAGGAGGATTTGTATGGTGTTGGAGAGGTGCTGGGCCGGAAAAAGAGGGAGAGAGGGTGGGACGATGANANGGTC-AAGATGGATGGATGGATGAGTTGTCGGACGAATTGAGGGAGCTGCTGGAGTGGAAAGGTGGGAGCNGGGGGAGAGAGAN-TTTCCTA ``` |
| 5 | ggggacaagtttgtacaaaaaagcaggcttgatgGCCATCTCAACGCGACTGGC, ggggaccactttgtacaagaaagctgggtcCCATTCCGCAACCTCACCCG | Mycgr3\|65235  JGI  Ensembl | KOG1231 Proteins containing the FAD binding domain; | D-lactate dehydrogenase (cytochrome) activity; FAD binding;  catalytic activity;  oxidoreductase activity; | 0 % |  |  |
| 5 | ggggacaagtttgtacaaaaaagcaggcttgatgTCGTACGATTACATTGTCAA, ggggaccactttgtacaagaaagctgggtcTCTCCTCCTCTGGCCACCAC | Mycgr3\|65342  JGI  Ensembl | KOG0342 ATP-dependent RNA helicase pitchoune; | 8-oxo-7,8-dihydroguanine triphosphatase activity; ATP binding;  ATP-dependent 5'-3' DNA helicase activity;  ATP-dependent helicase activity;  UDP-2,3-diacylglucosamine hydrolase activity;  bis(5'-nucleosyl)-tetraphosphatase activity;  dATP pyrophosphohydrolase activity;  diacylglycerol pyrophosphate phosphatase activity;  dihydroneopterin monophosphate phosphatase activity;  dihydroneopterin triphosphate pyrophosphohydrolase activity;  helicase activity;  nucleic acid binding;  pyrophosphatase activity;  thiamin-pyrophosphatase activity; | 0 % |  |  |
| 5 | ggggacaagtttgtacaaaaaagcaggcttgatgGATCCGTACAAGCTCTTGTC, ggggaccactttgtacaagaaagctgggtcATCGAAGCCACCAAAGTGTG | Mycgr3\|65543  JGI  Ensembl | KOG0344 ATP-dependent RNA helicase; | 8-oxo-7,8-dihydroguanine triphosphatase activity; ATP binding;  ATP-dependent 5'-3' DNA helicase activity;  ATP-dependent helicase activity;  UDP-2,3-diacylglucosamine hydrolase activity;  bis(5'-nucleosyl)-tetraphosphatase activity;  dATP pyrophosphohydrolase activity;  diacylglycerol pyrophosphate phosphatase activity;  dihydroneopterin monophosphate phosphatase activity;  dihydroneopterin triphosphate pyrophosphohydrolase activity;  helicase activity;  nucleic acid binding;  pyrophosphatase activity;  thiamin-pyrophosphatase activity; | 0 % |  |  |
| 5 | ggggacaagtttgtacaaaaaagcaggcttgatgCCTTCAGCCTCGAGCATCGC, ggggaccactttgtacaagaaagctgggtcCAATTCGCCACGACTGGCAT | Mycgr3\|65850  JGI  Ensembl | KOG1459 Squalene synthetase; | biosynthetic process; farnesyl-diphosphate farnesyltransferase activity;  integral to membrane;  lipid biosynthetic process;  transferase activity; | 100 % |  | ``` 36	ATGCCTTCAGCCTCGAGCATCGCCTACTACCTTGTCCACGTTTCAGAACTTCGCGCCATCATCCAATGGAAAGTATGGCATAACCCCGTACACGAGCGTGACGAGAGCAAAGAGTCGCACGACCTCAAAGAATGCTTCCGATTCCTGCAACTCACAAGCCGATCCTTTGCCGCCGTCATCCAAGAGCTTCATCCCGAGCTCCTCGTACCCGTATGCCTTTTCTACCTCATCCTTCGCGGACTGGACACGGTCGAGGATGACATGACCATCAGCATCGCGGCGAAAGAGCCATTGCTACGCGACTTTCACAACCACATTGAAGATGAGGCATGGAATTTTGACGGCAACGGCCCGAACGAAAAGGATCGCGAGCTGCTGGTCAAGTTTGACTGTGTGACGAGGGAATTCAATAAGTGCAAAGATGCGTATCGGGCCATCATTAAGGATATCACAAAGCGCATGGGCAATGGCATGGCTGATTACGCCAAAAACGCGGATCACCTGGCCAACGGTGTGCAGACGATCAAGGACTACGAGCTGTACTGCCACTATGTGGCTGGCCTTGTCGGAGAAGGGCTCACGCGGTTGTTCGTCGAGGCGAAGCTTGCCAATCCTGCGCTTCTGCAAAGGCCAGAGTTGATGGAGTCCATGGGTCAATTACTGCAACAGACCAACATCATTCGCGACATCAGGGAGGACCACGACGACAAACGGTATTTCTGGCCCAAGGAGATCTGGTCAAAGCACGTCGAGAAGTTCAACGATCTCTTCGAGCCTCAAAATCGAGAAAAGGCATTGCAATGC	839 	\|\|\|\|\|\|\|\|\|\|\|\|\|\|\|\|\|\|\|\|\|\|\|\|\|\|\|\|\|\|\|\|\|\|\|\|\|\|\|\|\|\|\|\|\|\|\|\|\|\|\|\|\|\|\|\|\|\|\|\|\|\|\|\|\|\|\|\|\|\|\|\|\|\|\|\|\|\|\|\|\|\|\|\|\|\|\|\|\|\|\|\|\|\|\|\|\|\|\|\|\|\|\|\|\|\|\|\|\|\|\|\|\|\|\|\|\|\|\|\|\|\|\|\|\|\|\|\|\|\|\|\|\|\|\|\|\|\|\|\|\|\|\|\|\|\|\|\|\|\|\|\|\|\|\|\|\|\|\|\|\|\|\|\|\|\|\|\|\|\|\|\|\|\|\|\|\|\|\|\|\|\|\|\|\|\|\|\|\|\|\|\|\|\|\|\|\|\|\|\|\|\|\|\|\|\|\|\|\|\|\|\|\|\|\|\|\|\|\|\|\|\|\|\|\|\|\|\|\|\|\|\|\|\|\|\|\|\|\|\|\|\|\|\|\|\|\|\|\|\|\|\|\|\|\|\|\|\|\|\|\|\|\|\|\|\|\|\|\|\|\|\|\|\|\|\|\|\|\|\|\|\|\|\|\|\|\|\|\|\|\|\|\|\|\|\|\|\|\|\|\|\|\|\|\|\|\|\|\|\|\|\|\|\|\|\|\|\|\|\|\|\|\|\|\|\|\|\|\|\|\|\|\|\|\|\|\|\|\|\|\|\|\|\|\|\|\|\|\|\|\|\|\|\|\|\|\|\|\|\|\|\|\|\|\|\|\|\|\|\|\|\|\|\|\|\|\|\|\|\|\|\|\|\|\|\|\|\|\|\|\|\|\|\|\|\|\|\|\|\|\|\|\|\|\|\|\|\|\|\|\|\|\|\|\|\|\|\|\|\|\|\|\|\|\|\|\|\|\|\|\|\|\|\|\|\|\|\|\|\|\|\|\|\|\|\|\|\|\|\|\|\|\|\|\|\|\|\|\|\|\|\|\|\|\|\|\|\|\|\|\|\|\|\|\|\|\|\|\|\|\|\|\|\|\|\|\|\|\|\|\|\|\|\|\|\|\|\|\|\|\|\|\|\|\|\|\|\|\|\|\|\|\|\|\|\|\|\|\|\|\|\|\|\|\|\|\|\|\|\|\|\|\|\|\|\|\|\|\|\|\|\|\|\|\|\|\|\|\|\|\|\|\|\|\|\|\|\|\|\|\|\|\|\|\|\|\|\|\|\|\|\|\|\|\|\|\|\|\|\|\|\|\|\|\|\|\|\|\|\|\|\|\|\|\|\|\|\|\|\|\|\|\|\|\|\|\|\|\|\|\|\|\|\|\|\|\|\|\|\|\|\|\|\|\|\|\|\|\|\|\|\|\|\|\|\|\|\|\|\|\|\|\|\|\|\|\|\|\|\|\|\|\|\|\|\|\|\|\|\|\|\|\|\|\|\|\|\|\|\|\|\|\|\|\|\|\|\|\|\|\|\|\|\|\|\|\|\|\|\|\|\|\|\|\|\|\|\|\|\|\|\|\|\|\|\|\|\| \|\|\|\|\|\|\|\|\|\|\|\|\|\|\|\|\|\| \|\|\| \|\|\|\|\|\|\|\|\|\|  \| \|\|\|\| \|\|\|\| \|\|\| \|\|\|\|\|\|\|\|\|\|\| \| \| \|\|\|\|\| \|\| \| \|\|\| \|  \|\|\|\|\|\| \|\|\| 	ATGCCTTCAGCCTCGAGCATCGCCTACTACCTTGTCCACGTTTCAGAACTTCGCGCCATCATCCAATGGAAAGTATGGCATAACCCCGTACACGAGCGTGACGAGAGCAAAGAGTCGCACGACCTCAAAGAATGCTTCCGATTCCTGCAACTCACAAGCCGATCCTTTGCCGCCGTCATCCAAGAGCTTCATCCCGAGCTCCTCGTACCCGTATGCCTTTTCTACCTCATCCTTCGCGGACTGGACACGGTCGAGGATGACATGACCATCAGCATCGCGGCGAAAGAGCCATTGCTACGCGACTTTCACAACCACATTGAAGATGAGGCATGGAATTTTGACGGCAACGGCCCGAACGAAAAGGATCGCGAGCTGCTGGTCAAGTTTGACTGTGTGACGAGGGAATTCAATAAGTGCAAAGATGCGTATCGGGCCATCATTAAGGATATCACAAAGCGCATGGGCAATGGCATGGCTGATTACGCCAAAAACGCGGATCACCTGGCCAACGGTGTGCAGACGATCAAGGACTACGAGCTGTACTGCCACTATGTGGCTGGCCTTGTCGGAGAAGGGCTCACGCGGTTGTTCGTCGAGGCGAAGCTTGCCAATCCTGCGCTTCTGCAAAGGCCAGAGTTGATGGAGTCCATGGGTCAATTACTGCAACAGACCAACATCATTCGCGACATCAGGGAGGACCACGACGACNAACGGTATTTCTGGCCCANGGANATCTGGTCAAN-CNCGTCNAGAANTTC-ACGATCTCTTCNANC-TCAAA-TCNANAAANGN-ATTGCA-TGC ``` |
| 5 | ggggacaagtttgtacaaaaaagcaggcttgatgGCAGACTTCGACTCATTCAT, ggggaccactttgtacaagaaagctgggtcCGACTTCACCCTGTAGAACT | Mycgr3\|66686  JGI  Ensembl | KOG0922 DEAH-box RNA helicase; | ATP binding; ATP-dependent DNA helicase activity;  ATP-dependent RNA helicase activity;  ATP-dependent helicase activity;  ATPase activity;  ATPase activity, coupled;  ATPase activity, coupled to transmembrane movement of ions;  ATPase activity, coupled to transmembrane movement of substances;  ATPase activity, uncoupled;  DNA-dependent ATPase activity;  RNA-dependent ATPase activity;  helicase activity;  nucleic acid binding;  protein-transmembrane transporting ATPase activity;  single-stranded DNA-dependent ATP-dependent DNA helicase activity; | 100 % |  | ``` 72	ATGGCAGACTTCGACTCATTCATCCCGGCATTGCACAAGCCCTCGTCGTTGCTTCCCATAGCACGGCATCGGGACGCATTACTCTATACCATCGAGAAGCTTCCGGTCACAGTCATCATCGGTCAGACTGGCAGCGGAAAGACCACTCAACTTCCTCAATATATGGA	238 	\|\|\|\|\|\|\|\|\|\|\|\|\|\|\|\|\|\|\|\|\|\|\|\|\|\|\|\|\|\|\|\|\|\|\|\|\|\|\|\|\|\|\|\|\|\|\|\|\|\|\|\|\|\|\|\|\|\|\|\|\|\|\|\|\|\|\|\|\|\|\|\|\|\|\|\|\|\|\|\|\|\|\|\|\|\|\|\|\|\|\|\|\|\|\|\|\|\|\|\|\|\|\|\|\|\|\|\|\|\|\|\|\|\|\|\|\|\|\|\|\|\| \|\|\|\|\|\|\|\|\|\| \|\|\|\|\|\| \| \|\|\| \|\|\|\|\|\|\|\| \|\|\|\|\|\|\| \|\|\| 	ATGGCAGACTTCGACTCATTCATCCCGGCATTGCACAAGCCCTCGTCGTTGCTTCCCATAGCACGGCATCGGGACGCATTACTCTATACCATCGAGAAGCTTCCGGTCACAGTCATCATCGGNCAGACTGGCANCGGAAANA-CACCCAACTTCCACAATATANGGA ``` |
| 5 | ggggacaagtttgtacaaaaaagcaggcttgatgGCGGAAGTTCTCCAGGCGTC, ggggaccactttgtacaagaaagctgggtcAAACTTCTGCCTGCCGAGAG | Mycgr3\|66791  JGI  Ensembl | KOG0285 Pleiotropic regulator 1; | 1-alkyl-2-acetylglycerophosphocholine esterase activity; | 0 % |  |  |
| 5 | ggggacaagtttgtacaaaaaagcaggcttgatgTCCGCCATCCTCTCCGCCGA, ggggaccactttgtacaagaaagctgggtcCCATCCACCACCCGCAGCAT | Mycgr3\|66934  JGI  Ensembl | KOG2439 Nuclear architecture related protein; | ferredoxin hydrogenase activity; ferredoxin hydrogenase complex; | 100 % |  | ``` 63	ATGTCCGCCATCCTCTCCGCCGACGATCTGAATGACTTCATCAATCCCGGTGTCGCGTGCATTAAGCCGGTCGAGACTCTACCCAAGCAGCAACCTCCGGAGGACTCGGGT-GCATACGAGGTAACTACGG-AGGA	196 	\|\|\|\|\|\|\|\|\|\|\|\|\|\|\|\|\|\|\|\|\|\|\|\|\|\|\|\|\|\|\|\|\|\|\|\|\|\|\|\|\|\|\|\|\|\|\|\|\|\|\|\|\|\|\|\|\|\|\|\|\|\|\|\|\|\|\|\|\|\|\|\|\|\|\|\|\|\|\|\|\|\|\|\|\|\|\|\|\|\|\|\|\|\|\|\|\|\|\|\|\|\|\|\| \|\|\|\|\|\| \|\|\|\|\|\|\|\|\|\| \|\|\|\|\|\|\|\| \|\|\|\| 	ATGTCCGCCATCCTCTCCGCCGACGATCTGAATGACTTCATCAATCCCGGTGTCGCGTGCATTAAGCCGGTCGAGACTCTACCCAAGCAGCAACCTCCGGAGGANTCGGGTGGCATACGAGGGAACTACGGNAGGA ``` |
| 5 | ggggacaagtttgtacaaaaaagcaggcttgatgGCGACCATCGCGGATGAGCT, ggggaccactttgtacaagaaagctgggtcTCCATCCTGCTTCTGCTTCT | Mycgr3\|67218  JGI  Ensembl | KOG2574 mRNA splicing factor PRP31; |  | 100 % |  |  |
| 5 | ggggacaagtttgtacaaaaaagcaggcttgatgTCACATCTCGCACCAAACGC, ggggaccactttgtacaagaaagctgggtcGTGCTGCGACCACTCAATAG | Mycgr3\|67552  JGI  Ensembl | KOG1549 Cysteine desulfurase NFS1; | cysteine desulfurase activity; metabolic process; | 0 % |  |  |
| 5 | ggggacaagtttgtacaaaaaagcaggcttgatgGCTCCCACAAAACGTCTCAA, ggggaccactttgtacaagaaagctgggtcGACCTTCCGCTTTACCAAAC | Mycgr3\|67807  JGI  Ensembl | KOG1536 Biotin holocarboxylase synthetase/biotin-protein ligase; | biotin-[acetyl-CoA-carboxylase] ligase activity; catalytic activity;  protein modification process; | 0 % |  |  |
| 5 | ggggacaagtttgtacaaaaaagcaggcttgatgGCGGCCTCAATAGAACGCCT, ggggaccactttgtacaagaaagctgggtcTGATCCCTCTCCGGGCTCCA | Mycgr3\|69324  JGI  Ensembl | KOG2201 Pantothenate kinase PanK and related proteins; | ATP binding; coenzyme A biosynthetic process;  pantothenate kinase activity; | 0 % |  |  |
| 5 | ggggacaagtttgtacaaaaaagcaggcttgatgAAGTCGTTTCTCGGCTTGAC, ggggaccactttgtacaagaaagctgggtcCTCGGACTTGAGCTGACCAA | Mycgr3\|70312  JGI  Ensembl | KOG1153 Subtilisin-related protease/Vacuolar protease B; | cerevisin activity; identical protein binding;  negative regulation of catalytic activity;  proteolysis;  subtilase activity; | 0 % |  |  |
| 5 | ggggacaagtttgtacaaaaaagcaggcttgatgTCACATAACACAGATCCGAG, ggggaccactttgtacaagaaagctgggtcCTTCTTGCCGCGACTACTTG | Mycgr3\|70324  JGI  Ensembl | KOG2270 Serine/threonine protein kinase involved in cell cycle control; | 3-phosphoinositide-dependent protein kinase activity; AMP-activated protein kinase activity;  ATP binding;  DNA-dependent protein kinase activity;  GTP-dependent protein kinase activity;  JUN kinase activity;  JUN kinase kinase activity;  JUN kinase kinase kinase activity;  Janus kinase activity;  MAP kinase 1 activity;  MAP kinase 2 activity;  MAP kinase kinase activity;  MAP kinase kinase kinase kinase activity;  MAP/ERK kinase kinase activity;  MP kinase activity;  NF-kappaB-inducing kinase activity;  SAP kinase activity;  atypical protein kinase C activity;  calcium-dependent protein kinase C activity;  casein kinase I activity;  casein kinase activity;  cyclic nucleotide-dependent protein kinase activity;  cyclin-dependent protein kinase activating kinase activity;  eukaryotic translation initiation factor 2alpha kinase activity;  galactosyltransferase-associated kinase activity;  glycogen synthase kinase 3 activity;  protein amino acid phosphorylation;  protein serine/threonine kinase activity;  protein-tyrosine kinase activity;  receptor signaling protein serine/threonine kinase activity;  ribosomal protein S6 kinase activity;  transmembrane receptor protein kinase activity;  transmembrane receptor protein serine/threonine kinase activity; | 0 % |  |  |
| 5 | ggggacaagtttgtacaaaaaagcaggcttgatgCCTCCTACACGAGCGAGCGC, ggggaccactttgtacaagaaagctgggtcTGCACCCGCACTCCCCGCCT | Mycgr3\|70456  JGI  Ensembl | KOG2074 RNA polymerase II transcription initiation/nucleotide excision repair factor TFIIH, subunit TFB1; |  | 0 % |  |  |
| 5 | ggggacaagtttgtacaaaaaagcaggcttgatgGCCGACCAAGAAATGCAAAA, ggggaccactttgtacaagaaagctgggtcCGACCTCTCCATATCAAACA | Mycgr3\|70475  JGI  Ensembl | KOG0296 Angio-associated migratory cell protein (contains WD40 repeats); | calcium ion binding; | 0 % |  |  |
| 5 | ggggacaagtttgtacaaaaaagcaggcttgatgCAACGCCCGGCCTACGCAGT, ggggaccactttgtacaagaaagctgggtcCTTCTTCTTGCTGGCCTTTC | Mycgr3\|71377  JGI  Ensembl | KOG3094 Predicted membrane protein; |  | 0 % |  |  |
| 5 | ggggacaagtttgtacaaaaaagcaggcttgatgGACGAAATCGCACCGGAGTA, ggggaccactttgtacaagaaagctgggtcCGCCTCGTACTCCAACCCTT | Mycgr3\|71522  JGI  Ensembl | KOG1439 RAB proteins geranylgeranyltransferase component A (RAB escort protein); | Rab GDP-dissociation inhibitor activity; protein transport;  regulation of GTPase activity; | 0 % |  |  |
| 5 | ggggacaagtttgtacaaaaaagcaggcttgatgGGTTTCGACTCTACCCAGAC, ggggaccactttgtacaagaaagctgggtcCCACACCCATCCGAGATACT | Mycgr3\|71649  JGI  Ensembl | KOG1286 Amino acid transporters; | amino acid transmembrane transporter activity; amino acid transport;  integral to membrane;  membrane;  transport; | 100 % |  | ``` 361	GTTCCCCACCGCCGGCTCCTTCCCCCACTGGGCCACCCGCTTCATCGACCCCTCCGTTGGCTTCTCCCTCGCCATCTCCTACGGCTACTGCTACACGATCGCCATCGCCTCCGAAGCCTCCGCCTCCGCCATTATCGTTTCCTACTGGACTTCCCTTACCCCAGCCGTCGTCATCACCGTGTCCCTCGCTCTCATCCTCTTCATCAACCTCATGGACGTCCGCTGGTTCGGTGAATCCGAAGTCATTGCGGGATCGATCAAAGTCCTCTGCTTCCTTGG	639 	\|\|\|\|\|\|\|\|\|\|\|\|\|\|\|\|\|\|\|\|\|\|\|\|\|\|\|\|\|\|\|\|\|\|\|\|\|\|\|\|\|\|\|\|\|\|\|\|\|\|\|\|\|\|\|\|\|\|\|\|\|\|\|\|\|\|\|\|\|\|\|\|\|\|\|\|\|\|\|\|\|\|\|\|\|\|\|\|\|\|\|\|\|\|\|\|\|\|\|\|\|\|\|\|\|\|\|\|\|\|\|\|\|\|\|\|\|\|\|\|\|\|\|\|\|\|\|\|\|\|\|\|\|\|\|\|\|\|\|\|\|\|\|\|\|\|\|\|\|\|\|\|\|\|\|\|\|\|\|\|\|\|\| \|\|\|\|\|\|\|\|\|\|\|\|\|\|\|\|\|\|\|\|\|\|\|\|\|\|\|\|\|\|\|\|\|\|\|\|\|\|\|\|\|\|\|\|\|\|\|\|\|\|\|\|\|\|\|\|\|\|\|\|\|\|\|\|\|\|\|\|\|\|\|\|\|\| \|\| \|\| \|\|\|\|\|\|\|\|\|\| \|\| \|\|\| \|\|\|\|\|\|\|\|\|\|\|\|\|\|\|\| 	GTTCCCCACCGCCGGCTCCTTCCCCCACTGGGCCACCCGCTTCATCGACCCCTCCGTTGGCTTCTCCCTCGCCATCTCCTACGGCTACTGCTACACGATCGCCATCGCCTCCGAAGCCTCCGCCTCCGCCATTATCGTTTCCTACTGGACTTCCCTTACCCCANCCGTCGTCATCACCGTGTCCCTCGCTCTCATCCTCTTCATCAACCTCATGGACGTCCGCTGGTTCGGTGAATCCNAANTCNTTGCGGGATCNATNAAANTCCTCTGCTTCCTTGG ```      ``` 40	GATGGGTTTCGACTCTACCCAGACCAAGTCGCACGCTGGCGGACTGCATCCTGGCAGCGACTCGGACGCTCAGTCTTTGCATCATGATGGGACTATTGAAGAACCTCAGATCAACGTCGGCGGACATGGAGGGACGCAACGGAGACTGCGTGACTACCAGGTTACCATGATTGGGTTTTCGTCGGGGATTGGAACCG	236 	\|\|\|\|\|\|\|\|\|\|\|\|\|\|\|\|\|\|\|\|\|\|\|\|\|\|\|\|\|\|\|\|\|\|\|\|\|\|\|\|\|\|\|\|\|\|\|\|\|\|\|\|\|\|\|\|\|\|\|\|\|\|\|\|\|\|\|\|\|\|\|\|\|\|\|\|\|\|\|\|\|\|\|\|\|\|\|\|\|\|\|\|\|\|\|\|\|\|\|\|\|\|\|\|\|\|\|\|\|\|\|\|\|\|\|\|\|\|\|\|\|\|\|\|\|\|\|\|\|\|\|\|\|\|\|\|\|\|\|\|\|\|\|\|\|\|\|\|\|\|\|\|\|\|\|\|\|\|\|\|\|\|\|\|\|\|\|\|\|\|\|\|\|\|\|\|\|\|\|\|\|\|\|\|\|\|\|\|\|\|\|\|\|\|\|\|\| 	GATGGGTTTCGACTCTACCCAGACCAAGTCGCACGCTGGCGGACTGCATCCTGGCAGCGACTCGGACGCTCAGTCTTTGCATCATGATGGGACTATTGAAGAACCTCAGATCAACGTCGGCGGACATGGAGGGACGCAACGGAGACTGCGTGACTACCAGGTTACCATGATTGGGTTTTCGTCGGGGATTGGAACCG ```      ``` 236	GGTCTCTTCATCGGCACAGGCTCAGCATACGCCCAAGCCGGTCCAGCAGGTCTCCTCCTCGCGTACCTCATCGTCGGAGCAATCCTCTGGTGTGTGATGCAAAGCATCGCCGAGTTGGCCACATTG	361 	\|\|\|\|\|\|\|\|\|\|\|\|\|\|\|\|\|\|\|\|\|\|\|\|\|\|\|\|\|\|\|\|\|\|\|\|\|\|\|\|\|\|\|\|\|\|\|\|\|\|\|\|\|\|\|\|\|\|\|\|\|\|\|\|\|\|\|\|\|\|\|\|\|\|\|\|\|\|\|\|\|\|\|\|\|\|\|\|\|\|\|\|\|\|\|\|\|\|\|\|\|\|\|\|\|\|\|\|\|\|\|\|\|\|\|\|\|\|\|\|\|\|\|\|\|\| 	GGTCTCTTCATCGGCACAGGCTCAGCATACGCCCAAGCCGGTCCAGCAGGTCTCCTCCTCGCGTACCTCATCGTCGGAGCAATCCTCTGGTGTGTGATGCAAAGCATCGCCGAGTTGGCCACATTG ``` |
| 5 | ggggacaagtttgtacaaaaaagcaggcttgatgGGATCTGGCAAGAAAGAAGG, ggggaccactttgtacaagaaagctgggtcCGAAGGAGCCTCCTCATCGC | Mycgr3\|71923  JGI  Ensembl | KOG2423 Nucleolar GTPase; | GTP binding; intracellular; | 100 % |  | ``` 46	ATGGGATCTGGCAAGAAAGAAGGCGCCCGCCGGGAACGG-CAGGGCAAGACTGGAGATGGCATG	108 	\|\|\|\|\|\|\|\|\|\|\|\|\|\|\|\|\|\|\|\|\|\|\|\|\|\|\|\|\|\|\|\|\|\|\|\|\|\|\| \|\| \|\|\|\|\|\|\|\|\|\|\|\| \|\|\|\|\|\|\|\| 	ATGGGATCTGGCAAGAAAGAAGGCGCCCGCCGGGAACGGCCANGGCAAGACTGGANATGGCATG ``` |
| 5 | ggggacaagtttgtacaaaaaagcaggcttgatgAGGTTATTACAATGGGCCCT, ggggaccactttgtacaagaaagctgggtcCGCTCGCTTCTGCTTCCCAA | Mycgr3\|73380  JGI  Ensembl | KOG2291 Oligosaccharyltransferase, alpha subunit (ribophorin I); | dolichyl-diphosphooligosaccharide-protein glycotransferase activity; endoplasmic reticulum;  integral to membrane;  protein amino acid glycosylation; | 100 % |  |  |
| 5 | ggggacaagtttgtacaaaaaagcaggcttgatgAACGGCAAACGACCGCACGA, ggggaccactttgtacaagaaagctgggtcCTTGCTCCGCGCCAGCGTCC | Mycgr3\|73598  JGI  Ensembl | KOG2769 Putative u4/u6 small nuclear ribonucleoprotein; |  | 100 % |  | ``` 5	GATGAACGGCAAACGACCGCACGAGACGAATGGCGATGATGGCGATGCCAAGCGGCAGCGCGCTCTGGATGGCTCACCTTTGCCGGTCAAATCAGCGGATCATGAAATGTCAGAGCTTGACAGGAAGAAGGCGGCGACCCAGGCGAAGATTGCTGCCATGAAGCAAGCGTTAGCAGCGAAAAAGGCCGGTGCCGATGCTGCAGCGGCTCCTCCACCGCCGCCTCCACCCGCCAACGATGTCGATGCCCAGAAAGCTCGAGCCGCGCAGAAGATCCAGGAGCTCAAGGCGAAGCTTGCGGCGAAGAAGCTGGAGAATGGTGCTTCTCCTGTCAACGGATCATTTGCAAGCCCGCCACCTCCTCCTCAACAGTCCGAGCAGGCACTCCGAATCGCAGAAGCTCGTGCAAAGGCTGCAGAACTGGCAGGGAAGCGCGGTCCCAGCGCAGCGCCCACACCTCCTCC	466 	\|\|\|\|\|\|\|\|\|\|\|\|\|\|\|\|\|\|\|\|\|\|\|\|\|\|\|\|\|\|\|\|\|\|\|\|\|\|\|\|\|\|\|\|\|\|\|\|\|\|\|\|\|\|\|\|\|\|\|\|\|\|\|\|\|\|\|\|\|\|\|\|\|\|\|\|\|\|\|\|\|\|\|\|\|\|\|\|\|\|\|\|\|\|\|\|\|\|\|\|\|\|\|\|\|\|\|\|\|\|\|\|\|\|\|\|\|\|\|\|\|\|\|\|\|\|\|\|\|\|\|\|\|\|\|\|\|\|\|\|\|\|\|\|\|\|\|\|\|\|\|\|\|\|\|\|\|\|\|\|\|\|\|\|\|\|\|\|\|\|\|\|\|\|\|\|\|\|\|\|\|\|\|\|\|\|\|\|\|\|\|\|\|\|\|\|\|\|\|\|\|\|\|\|\|\|\|\|\|\|\|\|\|\|\|\|\|\|\|\|\|\|\|\|\|\|\|\|\|\|\|\|\|\|\|\|\|\|\|\|\|\|\|\|\|\|\|\|\|\|\|\|\|\|\|\|\|\|\|\|\|\|\|\|\|\|\|\|\|\|\|\|\|\|\|\|\|\|\|\|\|\|\|\|\|\|\|\|\|\|\|\|\|\|\|\|\|\|\|\|\|\|\|\|\|\|\|\|\|\|\|\|\|\|\|\|\|\|\|\|\|\|\|\|\|\|\|\|\|\|\|\|\|\|\|\|\|\|\|\|\|\|\|\|\|\|\|\|\|\|\|\|\|\|\|\|\|\|\|\|\|\|\|\|\|\|\|\|\|\|\|\|\|\|\|\|\|\|\|\|\|\|\|\|\|\|\|\|\|\|\|\|\|\| \|\| \|\|\|\|\|\|\|\|\|\|\|\|\|\|\|\|\| \|\|\|\|\|\|\|  \|\|\|\| \| \|\|\|\|\| \|\| \| \|\| \|\| \|\|\|\| \|\|\|\|\|\|\|\| 	GATGAACGGCAAACGACCGCACGAGACGAATGGCGATGATGGCGATGCCAAGCGGCAGCGCGCTCTGGATGGCTCACCTTTGCCGGTCAAATCAGCGGATCATGAAATGTCAGAGCTTGACAGGAAGAAGGCGGCGACCCAGGCGAAGATTGCTGCCATGAAGCAAGCGTTAGCAGCGAAAAAGGCCGGTGCCGATGCTGCAGCGGCTCCTCCACCGCCGCCTCCACCCGCCAACGATGTCGATGCCCAGAAAGCTCGAGCCGCGCAGAAGATCCAGGAGCTCAAGGCGAAGCTTGCGGCGAAGAAGCTGGAGAATGGTGCTTCTCCTGTCAACGGATCATTTGCAAGCCCGCCACCTCCTCCTCAACAGTCCGAGCAGGCACTCCGAATCGCANAANCTCGTGCAAAGGCTGCANAACTGGCNNGGAANCNCGGTCNCANCNCANCGNCCACNCCTCCTCC ``` |
| 5 | ggggacaagtttgtacaaaaaagcaggcttgatgGGAGTCGGACGTCGCATGGG, ggggaccactttgtacaagaaagctgggtcTCTCGTCAGACTCTGTACAG | Mycgr3\|74360  JGI  Ensembl | KOG1122 tRNA and rRNA cytosine-C5-methylase (nucleolar protein NOL1/NOP2); |  | 0 % |  |  |
| 5 | ggggacaagtttgtacaaaaaagcaggcttgatgGTACTCGCAAAGAGCAAGAA, ggggaccactttgtacaagaaagctgggtcATCGTCGTAATGCGCCAAAC | Mycgr3\|74700  JGI  Ensembl | KOG1424 Predicted GTP-binding protein MMR1; | GTP binding; intracellular; | 100 % |  | ``` 5	ATGGTACTCGCAAAGAGCAAGAACGCCGTCGGGCTGGGAAACAGCCTGATGAACGACCGCTTCGGTCGCGGCAAGAACAGCAACATGCATCGTGCAAACCAAGCCGGCGCAATCAAGCGCAAGGATCCTCTCGGGAACGAATATGTCACGAACGCAGCGCAGGAGGCGGCTTGGGTGAAGATGCG-CAGTGTGACGGAGCAGGGCGCGCTGGATGAA	220 	\|\|\|\|\|\|\|\|\|\|\|\|\|\|\|\|\|\|\|\|\|\|\|\|\|\|\|\|\|\|\|\|\|\|\|\|\|\|\|\|\|\|\|\|\|\|\|\|\|\|\|\|\|\|\|\|\|\|\|\|\|\|\|\|\|\|\|\|\|\|\|\|\|\|\|\|\|\|\|\|\|\|\|\|\|\|\|\|\|\|\|\|\|\|\|\|\|\|\|\|\|\|\|\|\|\|\|\|\|\|\|\|\|\|\|\|\|\|\|\|\|\|\|\|\|\|\|\|\|\|\|\|\|\|\|\|\|\|\|\|\|\|\|\|\|\|\|\|\|\|\|\|\|\|\|\|\|\|\|\|\|\|\|\|\|\|\|\|\|\|\|\|\| \|\|\|\|\|\|\|\|\|\|  \|\| \|\|\| \|\|\|\|\| \|\| \| \| \|\| \|\| \|\|\|\|\| 	ATGGTACTCGCAAAGAGCAAGAACGCCGTCGGGCTGGGAAACAGCCTGATGAACGACCGCTTCGGTCGCGGCAAGAACAGCAACATGCATCGTGCAAACCAAGCCGGCGCAATCAAGCGCAAGGATCCTCTCGGGAACGAATATGTCACGAACGCAGCGCAGGAGGCGGCTTGNGTGAAGATGCACCATTGTNACGGANCANGACTCGNTGAATGAA ```      ``` 5	ATGGTACTCGCAAAGAGCAAGAACGCCGTCGGGCTGGGAAACAGCCTGATGAACGACCGCTTCGGTCGCGGCAAGAACAGCAACATGCATCGTGCAAACCAAGCCGGCGCAATCAAGCGCAAGGATCCTCTCGGGAACGAATATGTCACGAACGCAGCGCAGGAGGCGGCTTGGGTGAAGATGC	188 	\|\|\|\|\|\|\|\|\|\|\|\|\|\|\|\|\|\|\|\|\|\|\|\|\|\|\|\|\|\|\|\|\|\|\|\|\|\|\|\|\|\|\|\|\|\|\|\|\|\|\|\|\|\|\|\|\|\|\|\|\|\|\|\|\|\|\|\|\|\|\|\|\|\|\|\|\|\|\|\|\|\|\|\|\|\|\|\|\|\|\|\|\|\|\|\|\|\|\|\|\|\|\|\|\|\|\|\|\|\|\|\|\|\|\|\|\|\|\|\|\|\|\|\|\|\|\|\|\|\|\|\|\|\|\|\|\|\|\|\|\|\|\|\|\|\|\|\|\|\|\|\|\|\|\|\| \|\|\|\| \|\|\|\|\|\|\|\|\|\|\|\|\|\|\|\|\| \|\|\|\| 	ATGGTACTCGCAAAGAGCAAGAACGCCGTCGGGCTGGGAAACAGCCTGATGAACGACCGCTTCGGTCGCGGCAAGAACAGCAACATGCATCGTGCAAACCAAGCCGGCGCAATCAAGCGCAAGGATCCTCTCGGGAACGAATATGTCACGAACGCANCGCATGAGGCGGCTTGGGTGAAAATGC ``` |
| 5 | ggggacaagtttgtacaaaaaagcaggcttgatgGCGCTCGTCAAGAAGCAGGC, ggggaccactttgtacaagaaagctgggtcATCACTGTCGTCGCTATCAG | Mycgr3\|74715  JGI  Ensembl | KOG2529 Pseudouridine synthase; | RNA binding; pseudouridylate synthase activity;  tRNA pseudouridine synthesis;  tRNA-pseudouridine synthase activity; | 0 % |  |  |
| 5 | ggggacaagtttgtacaaaaaagcaggcttgatgGTTCTCCAAGATCTCGGGCG, ggggaccactttgtacaagaaagctgggtcCCTCTTCCCACCTCCACCCA | Mycgr3\|75124  JGI  Ensembl | KOG0780 Signal recognition particle, subunit Srp54; | 7S RNA binding; GTP binding;  SRP-dependent cotranslational protein targeting to membrane;  membrane;  nucleoside-triphosphatase activity;  nucleotide binding;  signal recognition particle; | 100 % |  | ``` 1	ATGGTTCTCCAAGATCTCGGGCGACGTATCAACGCCGCCGTCAACGACCTCACACGATCTCAGACCCTCGACGAGAAGGCCTTCGATGGCATGATCAAGGAGATCTCCAACGCCCTGGTAGAAGCCGACGTCTCAATAAAGCTCGTCTCGACCCTACGCAAGACCATCAAAACCTCCGTCAAGTTCTCAGAACTCCCACCCTCCGCCAACAAGAGACGCATAATACAAAAAGCCGTGTTCGACGCTCTCGTCGACATAGTAGATCCACATCAAAAACCATACGAACCTCGAAAAGGCAAATCAAATGTCATCATGTTCGTCGGATTACAAGGTGCCGGCAAGACCACAACCTGCACAAAACTCGCCCGATGGTATCAAGCACGAGGCTTCAAAACCTGCCTCGTCTGCGCCGACACATTCCGAGCAGGAGCCTTCGATCAACTCAAACAAAACGCCACAAAAGCCAAAATTCCCTACTACGGCTCCCTCACACAAACAGACCCCGTAGTCGTCGCCCGCGATGGAGTCAACCAATTCAAGAAGGAACGCTTCGAGATCATCATCGTTGACACCTCCGGTCGCCACCACCAAGAAGACGCTCTCTTCGCCGAAATGACCGAGATCCAATCCGCCGTGCAACCCCA	644 	\|\|\|\|\|\|\|\|\|\|\|\|\|\|\|\|\|\|\|\|\|\|\|\|\|\|\|\|\|\|\|\|\|\|\|\|\|\|\|\|\|\|\|\|\|\|\|\|\|\|\|\|\|\|\|\|\|\|\|\|\|\|\|\|\|\|\|\|\|\|\|\|\|\|\|\|\|\|\|\|\|\|\|\|\|\|\|\|\|\|\|\|\|\|\|\|\|\|\|\|\|\|\|\|\|\|\|\|\|\|\|\|\|\|\|\|\|\|\|\|\|\|\|\|\|\|\|\|\|\|\|\|\|\|\|\|\|\|\|\|\|\|\|\|\|\|\|\|\|\|\|\|\|\|\|\|\|\|\|\|\|\|\|\|\|\|\|\|\|\|\|\|\|\|\|\|\|\|\|\|\|\|\|\|\|\|\|\|\|\|\|\|\|\|\|\|\|\|\|\|\|\|\|\|\|\|\|\|\|\|\|\|\|\|\|\|\|\|\|\|\|\|\|\|\|\|\|\|\|\|\|\|\|\|\|\|\|\|\|\|\|\|\|\|\|\|\|\|\|\|\|\|\|\|\|\|\|\|\|\|\|\|\|\|\|\|\|\|\|\|\|\|\|\|\|\|\|\|\|\|\|\|\|\|\|\|\|\|\|\|\|\|\|\|\|\|\|\|\|\|\|\|\|\|\|\|\|\|\|\|\|\|\|\|\|\|\|\|\|\|\|\|\|\|\|\|\|\|\|\|\|\|\|\|\|\|\|\|\|\|\|\|\|\|\|\|\|\|\|\|\|\|\|\|\|\|\|\|\|\|\|\|\|\|\|\|\|\|\|\|\|\|\|\|\|\|\|\|\|\|\|\|\|\|\|\|\|\|\|\|\|\|\|\|\|\|\|\|\|\|\|\|\|\|\|\|\|\|\|\|\|\|\|\|\|\|\|\|\|\|\|\|\|\|\|\|\|\|\|\|\|\|\|\|\|\|\|\|\|\|\|\|\|\|\|\|\|\|\|\|\|\|\|\|\|\|\|\|\|\|\|\|\|\|\|\|\|\|\|\|\|\|\|\|\|\|\|\|\|\|\|\|\|\|\|\|\|\|\|\|\|\|\|\|\|\|\|\|\|\|\|\|\|\|\|\|\|\|\|\|\|\|\|\|\|\|\|\|\|\|\|\|\|\|\|\|\|\| \|\|\| \|\|\|\| \|\|\|\|\|\|\|\|\|\|\|\|\|\|\|\|\|\|\|\|\|\|\|\|\|\|\|\|\|\|\|\|\|\|\|\|\|\|\|\|\|\|\|\|\| \|\|\|\|\|   \|\| \|\|\|\|\|\|\|\|\|\|\|  \| \|\|\|\|\|\|\|\|\|\| \|\|\|\|  \|\|\|\|\|\|\|\|\|\|  \|\|\|\|\| 	ATGGTTCTCCAAGATCTCGGGCGACGTATCAACGCCGCCGTCAACGACCTCACACGATCTCAGACCCTCGACGAGAAGGCCTTCGATGGCATGATCAAGGAGATCTCCAACGCCCTGGTAGAAGCCGACGTCTCAATAAAGCTCGTCTCGACCCTACGCAAGACCATCAAAACCTCCGTCAAGTTCTCAGAACTCCCACCCTCCGCCAACAAGAGACGCATAATACAAAAAGCCGTGTTCGACGCTCTCGTCGACATAGTAGATCCACATCAAAAACCATACGAACCTCGAAAAGGCAAATCAAATGTCATCATGTTCGTCGGATTACAAGGTGCCGGCAAGACCACAACCTGCACAAAACTCGCCCGATGGTATCAAGCACGAGGCTTCAAAACCTGCCTCGTCTGCGCCGACACATTCCGAGCAGGAGCCTTCGATCAACTCAAACAAAACGCCACAAAAGCCAAAATTCCCTACTACGGCTCCCTCACACAAACAGACCCCGTAGTCGTCGCCCGCGATGGAGTCNACC-ATTCNAGAAGGAACGCTTCGAGATCATCATCGTTGACACCTCCGGTCGCCNCCACCNNNAANACGCTCTCTTCNNCNAAATGACCGANATCCNNTCCGCCGTGCN-CCCCA ``` |
| 5 | ggggacaagtttgtacaaaaaagcaggcttgatgTCTCAGTCGACCGCATCCTC, ggggaccactttgtacaagaaagctgggtcCTCCTCTTCATCGCTACTCA | Mycgr3\|75156  JGI  Ensembl | KOG2030 Predicted RNA-binding protein; |  | 0.93583 % |  |  |
| 5 | ggggacaagtttgtacaaaaaagcaggcttgatgCAGCGCGCTTTCACATCCGC, ggggaccactttgtacaagaaagctgggtcGAAGCCCATGCCGCCCATGC | Mycgr3\|75170  JGI  Ensembl | KOG0356 Mitochondrial chaperonin, Cpn60/Hsp60p; | ATP binding; cellular protein metabolic process;  chaperonin ATPase activity;  chaperonin ATPase complex;  protein binding; | 0 % |  |  |
| 5 | ggggacaagtttgtacaaaaaagcaggcttgatgCAGCGTCGCGAGCGAGGAGT, ggggaccactttgtacaagaaagctgggtcGTCTAGCAAAATTGCACCGT | Mycgr3\|75543  JGI  Ensembl | KOG2013 SMT3/SUMO-activating complex, catalytic component UBA2; | catalytic activity; negative regulation of ubiquitin-protein ligase activity;  positive regulation of ubiquitin-protein ligase activity;  protein modification process;  regulation of ubiquitin-protein ligase activity;  regulation of ubiquitin-protein ligase activity during meiotic cell cycle;  small protein activating enzyme activity;  ubiquitin cycle;  ubiquitin-protein ligase activity; | 0 % |  |  |
| 5 | ggggacaagtttgtacaaaaaagcaggcttgatgGCCTCCCTCGACACGCTGGA, ggggaccactttgtacaagaaagctgggtcCGACCAGACATCCTCCTGGT | Mycgr3\|75805  JGI  Ensembl | KOG1158 NADP/FAD dependent oxidoreductase; | FMN binding; NADPH-hemoprotein reductase activity;  electron carrier activity;  electron transport;  iron ion binding;  oxidoreductase activity; | 100 % |  | ``` 47	ATGGCCTCCCTCGACACGCTGGACATCGTCGTCCTGGCCATCCTCCTCCTCGGAACCATCGCATACTTCACCAAAGGCACATACTGGGCAGTCCCACCGAAGGACCCGTACGCGTCTCTCACAAATGGATCCGCCAAGGCAGGCAAGAGCAGGGACATTCTTGAAAAGCTGGAAGAGACGGGCAAGAACTGTGTCATCTTCTACGGCTCTCAGACAGGAACGGCCGAAGATTATGCTTCTCGCCTCGCCAAGGAGGGTTCATCGCGTTTTGGATTGAAGACCATGA-CCGCCG-ATTTGGAAGAGTACGATTA	357 	\|\|\|\|\|\|\|\|\|\|\|\|\|\|\|\|\|\|\|\|\|\|\|\|\|\|\|\|\|\|\|\|\|\|\|\|\|\|\|\|\|\|\|\|\|\|\|\|\|\|\|\|\|\|\|\|\|\|\|\|\|\|\|\|\|\|\|\|\|\|\|\|\|\|\|\|\|\|\|\|\|\|\|\|\|\|\|\|\|\|\|\|\|\|\|\|\|\|\|\|\|\|\|\|\|\|\|\|\|\|\|\|\|\|\|\|\|\|\|\|\|\|\|\|\|\|\|\|\|\|\|\|\|\|\|\|\|\|\|\|\|\|\|\|\|\|\|\|\|\|\|\|\|\|\|\|\|\|\|\|\|\|\|\|\|\|\|\|\|\|\|\|\|\|\|\|\|\|\|\|\|\|\|\|\|\|\|\|\|\|\|\|\|\|\|\|\|\|\|\|\|\|\|\|\|\|\|\|\|\|\|\|\|\|\|\|\|\|\|\|\|\|\|\|\|\|\|\|\|\|\|\|\|\|\|\|\|\|\|\|\|\|\|\|\|\|\|\|\|\|\|\|\|\|\|\|\|\|\|\|\|\|\| \|\|\|\|\|\|\|\|\|\|\|\|\|\| \|\| \|\|\|\| \|\|\|\|\|  \|\|\|\|\| \|\| \| \|\|\| \|\|\|\| 	ATGGCCTCCCTCGACACGCTGGACATCGTCGTCCTGGCCATCCTCCTCCTCGGAACCATCGCATACTTCACCAAAGGCACATACTGGGCAGTCCCACCGAAGGACCCGTACGCGTCTCTCACAAATGGATCCGCCAAGGCAGGCAAGAGCAGGGACATTCTTGAAAAGCTGGAAGAGACGGGCAAGAACTGTGTCATCTTCTACGGCTCTCAGACAGGAACGGCCGAAGATTATGCTTCTCGCCTCGCCAAGGAGGGTTCATCNCGTTTTGGATTGAANACNATGAACCGCCNAATTTGAAATANTACNATTA ``` |
| 5 | ggggacaagtttgtacaaaaaagcaggcttgatgGCCCAAACGGACCAAAAGAA, ggggaccactttgtacaagaaagctgggtcGGCCTGACCAAGCTTCTTCC | Mycgr3\|76149  JGI  Ensembl | KOG2533 Permease of the major facilitator superfamily; |  | 0 % |  |  |
| 5 | ggggacaagtttgtacaaaaaagcaggcttgatgGCGGACGCTCCAGAGGAGCC, ggggaccactttgtacaagaaagctgggtcCGCCTTCTTGCCCGCATCCT | Mycgr3\|76205  JGI  Ensembl | KOG1148 Glutaminyl-tRNA synthetase; | ATP binding; aminoacyl-tRNA ligase activity;  cytoplasm;  glutamine-tRNA ligase activity;  glutaminyl-tRNA aminoacylation;  glutamyl-tRNA aminoacylation;  nucleotide binding;  tRNA aminoacylation for protein translation;  translation; | 0 % |  |  |
| 5 | ggggacaagtttgtacaaaaaagcaggcttgatgTTATCAAGGACGGGCTACAT, ggggaccactttgtacaagaaagctgggtcTGAGTCCAAGATCTTCAGCA | Mycgr3\|76370  JGI  Ensembl | KOG1800 Ferredoxin/adrenodoxin reductase; | NADPH-adrenodoxin reductase activity; electron transport;  ferredoxin-NADP+ reductase activity; | 100 % |  | ``` 1	ATGTTATCAAGGACGGGCTACATCTGTCGGAGCTGCCAACGCTCACAGAAACATGTCACCGTCACGGGGACTCGACTGCTAGGGGACAATGTCCGCTACTACAGCACACGACCACGGCCGCTGCATCTTGCCATCATCGGCTCTGGTCCGGCTGGCTTCTACTCCGCATACAGACTATTGAAAAGTCTACCAGATGCGCGCGTCGACATGTACGAGGGACTGCCCTCGCCATATGGACTTGTACGATTTGGTATCGCTCCAGATCACCCAGAAGCaaaa	279 	\|\|\|\|\|\|\|\|\|\|\|\|\|\|\|\|\|\|\|\|\|\|\|\|\|\|\|\|\|\|\|\|\|\|\|\|\|\|\|\|\|\|\|\|\|\|\|\|\|\|\|\|\|\|\|\|\|\|\|\|\|\|\|\|\|\|\|\|\|\|\|\|\|\|\|\|\|\|\|\|\|\|\|\|\|\|\|\|\|\|\|\|\|\|\|\|\|\|\|\|\|\|\|\|\|\|\|\|\|\|\|\|\|\|\|\|\|\|\|\|\|\|\|\|\|\|\|\|\|\|\|\|\|\|\|\|\|\|\|\|\|\|\|\|\|\|\|\|\|\|\|\|\|\|\|\|\|\|\|\|\|\|\|\|\|\|\|\|\|\|\|\|\|\|\|\|\|\|\|\|\|\|\|\|\|\|\|\|\|\|\|\|\|\|\|\|\|\|\|\|\|\|\|\|\|\|\|\|\|\|\|\|\|\|\|\|\|\|\|\|\|\|\|\|\|\|\|\|\|\|\|\|\|\|\|\|\|\|\|\|\|\|\|\|\|\|\|\|\|\|\|\|\|\|\|\|\|\|\|\|\|\|\|\|\|\|\|\|\|\|\|\|\|\|\|\|\|\|\| 	ATGTTATCAAGGACGGGCTACATCTGTCGGAGCTGCCAACGCTCACAGAAACATGTCACCGTCACGGGGACTCGACTGCTAGGGGACAATGTCCGCTACTACAGCACACGACCACGGCCGCTGCATCTTGCCATCATCGGCTCTGGTCCGGCTGGCTTCTACTCCGCATACAGACTATTGAAAAGTCTACCAGATGCGCGCGTCGACATGTACGAGGGACTGCCCTCGCCATATGGACTTGTACGATTTGGTATCGCTCCAGATCACCCAGAAGCAAAA ```      ``` 1	ATGTTATCAAGGACGGGCTACAT-CTGTCGGAGCTGCCAACG-CTCACAGAAACATGTCACCGTCACGGGGACTCGACTGCTA-GGGGACAATGTCCGCTACTACAGCACACGACCACGGCCGCTGCATCTTGCCATCATCGGCTCTGGTCCGGCTGGCTTCTACTCCGCATACAGACTATTGAAAAGTCTACCA-GATGCGCGCGTCGACATGTACGAGGGACTGCCCTCGCCATATGGACTTGTACGATTT-GGTATCG-CTCCAGATCACCCAGAAGCa	276 	\|\|\|\|\|\|\|\|\|\|\|\|\|\| \| \|\|\|\|\|\| \|\|\| \|\|\|\|\|\|\|\|\|\|\|\|\|\|  \|\|\|\|\|\|\|\|\|\|\|\|\|\|\|\|\|\|\| \|\|\|\|\|\|\|\|\|\|\|\|\|\|\|\|\|\|\|  \|\|\|\|\|\|\|\|\|\|\| \| \|\| \|\|\|\|\| \|\|\|\|\|\|\|\|\|\|\|\|\|\|\|\|\|\|\|\|\|\|\|\|\|\|\|\|\|\|\|\|\|\|\|\|\|\|\|\|\| \|\|\|\|\|\|\|\| \|\|\|\|\|\|\|\|\| \|\|\|\|\|\|\|\|\|\|\|\|\|\|\|\|\|\|\|\|\|\|\|\|\|\| \|\|\|\| \|\|\|\|\|\|\|\|\|\| \|\|\|\|\|\|\|\|\|\|\|\|\|\|\|\|\|\|\|\|\|\|\|\|\|\|\|\|\|\|\|\|\|\|\|\|\|\|\|\|  \|\|\|\|\|\|\| \|\|\|\|\|\|\|\|\|\|\|\|\|\|\|\|\|\|\|\| 	ATGTTATCAAGGACNGNCTACATCCTGGCGGAGCTGCCAACGNGTCACAGAAACATGTCACCGNCACGGGGACTCGACTGCTANAGGGACAATGTCNGNTANTACAGNACACGACCACGGCCGCTGCATCTTGCCATCATCGGCTCTGGNCCGGCTGGNTTCTACTCCNCATACAGACTATTGAAAAGTCTACCAGGATGGGCGCGTCGACNTGTACGAGGGACTGCCCTCGCCATATGGACTTGTACGATTNCGGTATCGACTCCAGATCACCCAGAAGCA ```      ``` 280	aaaTGCTCCGAGACTCTAGTTGAAGTAGCGAAATATCCCGGCTTCAACTACATCGGCAACACGCCCATAGGCGATGGACCGTCCCAACTACCTCTCTCTTTGATAGCTCCCCACTACGATGCAATGCTCTTCGCGTATGGCGCCTCGAAGGACAAGAGATTGGGTTTGCCTGGCGAAGATTTCAAAGGCGTACACTCGGCAAGAGCATTTGTAGGATGGTACAATGGGTTGCCGGAATATGCAGATCTGCAACCAGACCTCGAATCAGGAGATACTGCCGTGGTCATTGGACAAGGAAATGTAGCATTGGATGTGACGCGCATTCTCCTCGCGCCACTGGATGAGCTGAGAAAGACGGATATTTCAGAGCATGCTGTGGAAGCTCTCAGCAAAAGCAGGATTCGCGACCTGAAAATTCTCGGACGTAGAGGACCTCTGCAAGCTCCATATACGATCAAA	738 	\|\|\|\|\|\|\|\|\|\|\|\|\|\|\|\|\|\|\|\|\|\|\|\|\|\|\|\|\|\|\|\|\|\|\|\|\|\|\|\|\|\|\|\|\|\|\|\|\|\|\|\|\|\|\|\|\|\|\|\|\|\|\|\|\|\|\|\|\|\|\|\|\|\|\|\|\|\|\|\|\|\|\|\|\|\|\|\|\|\|\|\|\|\|\|\|\|\|\|\|\|\|\|\|\|\|\|\|\|\|\|\|\|\|\|\|\|\|\|\|\|\|\|\|\|\|\|\|\|\|\|\|\|\|\|\|\|\|\|\|\|\|\|\|\|\|\|\|\|\|\|\|\|\|\|\|\|\|\|\|\|\|\|\|\|\|\|\|\|\|\|\|\|\|\|\|\|\|\|\|\|\|\|\|\|\|\|\|\|\|\|\|\|\|\|\|\|\|\|\|\|\|\|\|\|\|\|\|\|\|\|\|\|\|\|\|\|\|\|\|\|\|\|\|\|\|\|\|\|\|\|\|\|\|\|\|\|\|\|\|\|\|\|\|\|\|\|\|\|\|\|\|\|\|\|\|\|\|\|\|\|\|\|\|\|\|\|\|\|\|\|\|\|\|\|\|\|\|\|\|\|\|\|\|\|\|\|\|\|\|\|\|\|\|\|\|\|\|\|\|\|\|\|\|\|\|\|\|\|\|\|\|\|\|\|\|\|\|\|\|\|\|\|\|\|\|\|\|\|\|\|\|\|\|\|\|\|\|\|\|\|\|\|\|\|\|\|\|\|\|\|\|\|\|\|\|\|\|\|\|\|\|\|\|\|\|\|\|\|\|\|\|\|\|\|\|\|\|\|\|\|\|\|\|\|\|\|\|\|\|\|\|\|\|\|\|\|\|\|\|\| \|\|\|\|\|\|\|\|\| \|\|\|\|\|\|\|\|\|\|\|\|\|\|\|\|\|\|\|\|\| \|\|\|\|\|\|\|\|\|\|\|\|\|\|\|\|\|\|\|\|\|\|\|\|\| 	AAATGCTCCGAGACTCTAGTTGAAGTAGCGAAATATCCCGGCTTCAACTACATCGGCAACACGCCCATAGGCGATGGACCGTCCCAACTACCTCTCTCTTTGATAGCTCCCCACTACGATGCAATGCTCTTCGCGTATGGCGCCTCGAAGGACAAGAGATTGGGTTTGCCTGGCGAAGATTTCAAAGGCGTACACTCGGCAAGAGCATTTGTAGGATGGTACAATGGGTTGCCGGAATATGCAGATCTGCAACCAGACCTCGAATCAGGAGATACTGCCGTGGTCATTGGACAAGGAAATGTAGCATTGGATGTGACGCGCATTCTCCTCGCGCCACTGGATGAGCTGAGAAAGACGGATATTTCAGAGCATGCTGTGGAAGCTCTCAGCAAAAGCAGGATNCGCGACCTGNAAATTCTCGGACGTAGAGGACNTCTGCAAGCTCCATATACGATCAAA ``` |
| 5 | ggggacaagtttgtacaaaaaagcaggcttgatgTCGGTCGACTACCTCCTCCA, ggggaccactttgtacaagaaagctgggtcTTCCGCCTTGCTCTTCTTCT | Mycgr3\|77007  JGI  Ensembl | KOG2573 Ribosome biogenesis protein - Nop56p/Sik1p; |  | 100 % |  | ``` 97	ATGTCGGTCGACTACCTCCTCCACGAGTCCTCTGTGGGCTACGCCATTTTCTCCGTCAAGCTCCCATCTGATACCGTTGGTGCTCGCCTGAAGGAAGTACAAGATGCGCAGACCGATCTCGCTCGATTCGGAAAGATGGTGCAGCTGGTCTCCTTTGCACCTTTCCAGGGCGCCGGACAAGCATTGGAGAACGCAAACGACGTCTCTGAGGGCATCATGAGCGACTACTTGCGCACTGTGCTGGAGGCCAACCTTCCCAAAGCGAGCAAGAAGAACAAGATCACTATTGGTGTCGGCGACAAGGGTCTTGCAGGAAGCGTCAAGGCTGGATTCACTGGCATTGAGTGCGAGACTCCGGAAACCAGCGAGCTGGCCGCCGACCTTCTCAGAGGACTGCGTCTGCACGCCGAGAAGCTGCTCAAGAACTTGCAGACGGGCGATATCTCTCGCGCTCAGCTTGGTCTTGGTCACGCCTACTCCCGCGCTAAGGTCAAGTTCAGCGTGCAGAAGAACGACAACCACATCATCCAGGCTATCGCAACGATTGATCACCTCGACAAGGCTGTCAACACTTTCTCCATGCGTGTTCGCGAATGGTACGGCTGGCATTTCCCGGAGCTCATCAGGATTGTGAGCGAGAACCAGAAATACGCACAATGCGCACTTTACATCGGTGATAAAAAGACTTTGACGGAGGACAGCCTACACGATTTGGCCAAAATCGTGGATGA	827 	\|\|\|\|\|\|\|\|\|\|\|\|\|\|\|\|\|\|\|\|\|\|\|\|\|\|\|\|\|\|\|\|\|\|\|\|\|\|\|\|\|\|\|\|\|\|\|\|\|\|\|\|\|\|\|\|\|\|\|\|\|\|\|\|\|\|\|\|\|\|\|\|\|\|\|\|\|\|\|\|\|\|\|\|\|\|\|\|\|\|\|\|\|\|\|\|\|\|\|\|\|\|\|\|\|\|\|\|\|\|\|\|\|\|\|\|\|\|\|\|\|\|\|\|\|\|\|\|\|\|\|\|\|\|\|\|\|\|\|\|\|\|\|\|\|\|\|\|\|\|\|\|\|\|\|\|\|\|\|\|\|\|\|\|\|\|\|\|\|\|\|\|\|\|\|\|\|\|\|\|\|\|\|\|\|\|\|\|\|\|\|\|\|\|\|\|\|\|\|\|\|\|\|\|\|\|\|\|\|\|\|\|\|\|\|\|\|\|\|\|\|\|\|\|\|\|\|\|\|\|\|\|\|\|\|\|\|\|\|\|\|\|\|\|\|\|\|\|\|\|\|\|\|\|\|\|\|\|\|\|\|\|\|\|\|\|\|\|\|\|\|\|\|\|\|\|\|\|\|\|\|\|\|\|\|\|\|\|\|\|\|\|\|\|\|\|\|\|\|\|\|\|\|\|\|\|\|\|\|\|\|\|\|\|\|\|\|\|\|\|\|\|\|\|\|\|\|\|\|\|\|\|\|\|\|\|\|\|\|\|\|\|\|\|\|\|\|\|\|\|\|\|\|\|\|\|\|\|\|\|\|\|\|\|\|\|\|\|\|\|\|\|\|\|\|\|\|\|\|\|\|\|\|\|\|\|\|\|\|\|\|\|\|\|\|\|\|\|\|\|\|\|\|\|\|\|\|\|\|\|\|\|\|\|\|\|\|\|\|\|\|\|\|\|\|\|\|\|\|\|\|\|\|\|\|\|\|\|\|\|\|\|\|\|\|\|\|\|\|\|\|\|\|\|\|\|\|\|\|\|\|\|\|\|\|\|\|\|\|\|\|\|\|\|\|\|\|\|\|\|\|\|\|\|\|\|\|\|\|\|\|\|\|\|\|\|\|\|\|\|\|\|\|\|\|\| \|\|\|\|\|\|\|\|\|\|\|\|\|\|\|\|\|\|\|\|\|\|\|\|\|\|\|\|\|\|\|\|\|\|\|\|\|\|\|\|\|\|\|\|\|\|\|\|\|\|\|\|\|\|\|\|\|\|\|\|\|\|\|\|\|\|\|\|\|\|\|\|\|\|\|\|\|\|\|\|\|\|\|\|\|\|\|\|\|\|\|\|\|\|\|\|\|\|\|\|\|\|\|\|\|\|\|\|\|\|\|\|\|\|\|\|\|\|\|\|\|\|\|\|\|\|\|\|\|\|\| \|\|\|\|\| \|\|\|\|\|\| \|\|\|\|\|\|\|\|\|\|\|\|\|\|\|\|\|\|\|\|\|\|\|\|\|\|\|\|\|\|  \|\|\|\|\|\|\|\|\|\|\| \|\| \| \|\|\|\|\| \| \|\|\|\|\|\|\|\|\|\|\| \|\| \|\| \|\|\|\| 	ATGTCGGTCGACTACCTCCTCCACGAGTCCTCTGTGGGCTACGCCATTTTCTCCGTCAAGCTCCCATCTGATACCGTTGGTGCTCGCCTGAAGGAAGTACAAGATGCGCAGACCGATCTCGCTCGATTCGGAAAGATGGTGCAGCTGGTCTCCTTTGCACCTTTCCAGGGCGCCGGACAAGCATTGGAGAACGCAAACGACGTCTCTGAGGGCATCATGAGCGACTACTTGCGCACTGTGCTGGAGGCCAACCTTCCCAAAGCGAGCAAGAAGAACAAGATCACTATTGGTGTCGGCGACAAGGGTCTTGCAGGAAGCGTCAAGGCTGGATTCACTGGCATTGAGTGCGAGACTCCGGAAACCAGCGAGCTGGCCGCCGACCTTCTCAGAGGACTGCGTCTGCACGCCGAGAAGCTGCTCAAGAACTTGCAGACGGGCGATATCTCTCGCGCTCAGCTTGGTCTTGGTCACGCCTACTCCCGCGCTAAGGTCAAGTTCAGCGTGCANAAGAACGACAACCACATCATCCAGGCTATCGCAACGATTGATCACCTCGACAAGGCTGTCAACACTTTCTCCATGCGTGTTCGCGAATGGTACGGCTGGCATTTCCCGGAGCTCATCAGGATTGTGAGCGANAACCANAAATACNCACAATGCGCACTTTACATCGGTGATAAAAN-ACTTTGACGGANGANA-CCTACNCNATTTGGCCAAA-TCNTGNATGA ``` |
| 5 | ggggacaagtttgtacaaaaaagcaggcttgatgTCAGCTGCACAACTCCTCAA, ggggaccactttgtacaagaaagctgggtcTTCCCCATCTTCCAACCCAG | Mycgr3\|77111  JGI  Ensembl | KOG0359 Chaperonin complex component, TCP-1 zeta subunit (CCT6); | ATP binding; cellular protein metabolic process;  protein binding; | 100 % |  | ``` 11	ATGTCAGCTGCACAACTCCTCAACCCCAAGGCGGAGTCGCGACGCCGCGGTGAGGCTCTCAGAGTCAACATTAATGCTGGAGAAGGCCTGAAAGATGTTCTCTCCTCCAACTTGGGTCCAACGGGCACCCTCAAAATGCTGGTCGATGGCGCAGGAGGCATCAAGCTCACAAAGGATGGCAGCGTGCTGTTGAAAGAGATGCAAATTCAGAACCCCACAGCTGTCATGATTGCCCGCGCAGCGACTGCACAAGACGAGATTTGCGGAGATGGCACAACGAGTGTGGTTATGCTGGTGGGAGAACTGCTGAAGCAAGCAGATCGCTACATCAGTGAGGGACTACACCCAAGAGTCATCACAGATGGTTACGAGGTGGCCAAGAACGAGACACTACGATTCCTGGACTCCTTCAAACTCCCGAAAGAAGTCGACCGCGAACTCCTCCTCTCCGTCGCCCGCACATCGCTCTCCACCAAGATCAACTCCACCCTAGCCGAGCAACTCACACCCGATATCGTCGATGCCGTCCTCGCCATCTATGAAGCTCCCGCAAAGCCCGACCTGCACATGATCGAGATCATGACCATGCAGCACCGCACAGCAGCAGACACCCAGCTCATCCGTGGTCTCGCATTAGACCACGGCGCTCGCCATCCCGATATGCCCAAAGACGTCAAGAATGCCTTCGTCCTCACCCTCAACGTCAGCTTGGAATACGAAAAGAGCGAGATCAACAGTGGATTCTACTACTCCAGCGCAGAACAGCGGGACAAACTGGTGGAGAGTGAGCGTCGATTCGTGGACGACAAGCTGCGCAAGATTGTC	835 	\|\|\|\|\|\|\|\|\|\|\|\|\|\|\|\|\|\|\|\|\|\|\|\|\|\|\|\|\|\|\|\|\|\|\|\|\|\|\|\|\|\|\|\|\|\|\|\|\|\|\|\|\|\|\|\|\|\|\|\|\|\|\|\|\|\|\|\|\|\|\|\|\|\|\|\|\|\|\|\|\|\|\|\|\|\|\|\|\|\|\|\|\|\|\|\|\|\|\|\|\|\|\|\|\|\|\|\|\|\|\|\|\|\|\|\|\|\|\|\|\|\|\|\|\|\|\|\|\|\|\|\|\|\|\|\|\|\|\|\|\|\|\|\|\|\|\|\|\|\|\|\|\|\|\|\|\|\|\|\|\|\|\|\|\|\|\|\|\|\|\|\|\|\|\|\|\|\|\|\|\|\|\|\|\|\|\|\|\|\|\|\|\|\|\|\|\|\|\|\|\|\|\|\|\|\|\|\|\|\|\|\|\|\|\|\|\|\|\|\|\|\|\|\|\|\|\|\|\|\|\|\|\|\|\|\|\|\|\|\|\|\|\|\|\|\|\|\|\|\|\|\|\|\|\|\|\|\|\|\|\|\|\|\|\|\|\|\|\|\|\|\|\|\|\|\|\|\|\|\|\|\|\|\|\|\|\|\|\|\|\|\|\|\|\|\|\|\|\|\|\|\|\|\|\|\|\|\|\|\|\|\|\|\|\|\|\|\|\|\|\|\|\|\|\|\|\|\|\|\|\|\|\|\|\|\|\|\|\|\|\|\|\|\|\|\|\|\|\|\|\|\|\|\|\|\|\|\|\|\|\|\|\|\|\|\|\|\|\|\|\|\|\|\|\|\|\|\|\|\|\|\|\|\|\|\|\|\|\|\|\|\|\|\|\|\|\|\|\|\|\|\|\|\|\|\|\|\|\|\|\|\|\|\|\|\|\|\|\|\|\|\|\|\|\|\|\|\|\|\|\|\|\|\|\|\|\|\|\|\|\|\|\|\|\|\|\|\|\|\|\|\|\|\|\|\|\|\|\|\|\|\|\|\|\|\|\|\|\|\|\|\|\|\|\|\|\|\|\|\|\|\|\|\|\|\|\|\|\|\|\|\|\|\|\|\|\|\|\|\|\|\|\|\|\|\|\|\|\|\|\|\|\|\|\|\|\|\|\|\|\|\|\|\|\|\|\|\|\|\|\|\|\|\|\|\|\|\|\|\|\|\|\|\|\|\|\|\|\|\|\|\|\|\|\|\|\|\|\|\|\|\|\|\|\|\|\|\|\|\|\|\|\|\|\|\|\|\|\|\|\|\|\|\|\|\|\|\|\|\|\|\|\|\|\|\|\|\|\|\|\|\|\|\|\|\|\|\|\|\|\|\|\|\|\|\|\|\|\|\|\|\|\|\|\|\|\|\|\|\|\|\|\|\|\|\|\|\|\|\|\|\|\|\|\|\|\|\|\|\|\|\|\|\|\|\|\|\|\|\|\|\|\|\|\|\|\|\|\|\|\|\|\|\|\|\|\|\|\|\|\|\|\|\|\|\|\|\|\|\|\|\|\|\|\|\|\|\|\|\|\|\|\|\|\|\|\|\|\|\|\|\|\|\|\|\|\|\|\|\|\| \|\|\|\|\|\|\|\|\|\|\| \|\|\|\|\|\|\| \|\|\|\|\|\|\|\|\|\|\|\|\|\|\|\|\| \|\|\|\|\|\|\|\| \|\|\|\|  \|\|\|\|\|\|\| \| \|\|\|\|\|\|\|\|\|\|\| \| \|\|\|\|\| \| \|\| \|\|\|\| \|\|\| \|\|\|\|\|\| 	ATGTCAGCTGCACAACTCCTCAACCCCAAGGCGGAGTCGCGACGCCGCGGTGAGGCTCTCAGAGTCAACATTAATGCTGGAGAAGGCCTGAAAGATGTTCTCTCCTCCAACTTGGGTCCAACGGGCACCCTCAAAATGCTGGTCGATGGCGCAGGAGGCATCAAGCTCACAAAGGATGGCAGCGTGCTGTTGAAAGAGATGCAAATTCAGAACCCCACAGCTGTCATGATTGCCCGCGCAGCGACTGCACAAGACGAGATTTGCGGAGATGGCACAACGAGTGTGGTTATGCTGGTGGGAGAACTGCTGAAGCAAGCAGATCGCTACATCAGTGAGGGACTACACCCAAGAGTCATCACAGATGGTTACGAGGTGGCCAAGAACGAGACACTACGATTCCTGGACTCCTTCAAACTCCCGAAAGAAGTCGACCGCGAACTCCTCCTCTCCGTCGCCCGCACATCGCTCTCCACCAAGATCAACTCCACCCTAGCCGAGCAACTCACACCCGATATCGTCGATGCCGTCCTCGCCATCTATGAAGCTCCCGCAAAGCCCGACCTGCACATGATCGAGATCATGACCATGCAGCACCGCACAGCAGCAGACACCCAGCTCATCCGTGGTCTCGCATTAGACCACGGCGCTCGCCATCCCGATATGCCCAAAGACGTCAAGAATGCCTTCGTCCTCACCCTCAACGTCAGCTTGGAATACGAAA-GAGCGAGATCA-CAGTGGA-TCTACTACTCCAGCGCANAACAGCGGNACAAN-TGGTGGA-ANTGAGCGTCGAT-CNTGGACAA-AANCTGCCCAA-ATTGTC ``` |
| 5 | ggggacaagtttgtacaaaaaagcaggcttgatgGAGAGCCGGGAAGAAGAGCG, ggggaccactttgtacaagaaagctgggtcCTCCCATCCCAGCCCATAAC | Mycgr3\|77140  JGI  Ensembl | KOG2997 F-box protein FBX9; | phosphopantetheine binding; | 0 % |  |  |
| 5 | ggggacaagtttgtacaaaaaagcaggcttgatgAACGGTGCAAAGAAGGAAGT, ggggaccactttgtacaagaaagctgggtcCGAGGTCTGCAACGCAGCCT | Mycgr3\|77226  JGI  Ensembl | KOG1247 Methionyl-tRNA synthetase; | ATP binding; aminoacyl-tRNA ligase activity;  cytoplasm;  methionine-tRNA ligase activity;  methionyl-tRNA aminoacylation;  nucleotide binding;  tRNA aminoacylation for protein translation;  translation; | 100 % |  | ``` 117	ATGAACGGTGCAAAGAAGGAAGTCATCCTGCCGGTGGAGGGCAAGAAGAACGTGTTGATCACTTCCGCATTGCCATACGTCAACAATGTACCTCATTTGGGAAACGTAATTGGCAGCGTGCTCTCGGCAGACGTCTTCTCCCGCTACAGCAAAGCACGAGGTCTTCCCACATTGTTCATATGCGGCACAGACGAATATGGCACAGCGACTGAGACGAAAGCGATCGAAGAGGGCATCACACCGCAAGAGCTCTGCGACAAATACAACAAGCTCCACAAGGAAGTCTACGACTGGTTCGAGATTGACTTTGACCACTTCGGGCGCACGCCGACCCAGCAACAGACGGATATTGCGCAGGATATCTTTCTGAAGCTATACAAGAACGGATACCTCGAGGA	514 	\|\|\|\|\|\|\|\|\|\|\|\|\|\|\|\|\|\|\|\|\|\|\|\|\|\|\|\|\|\|\|\|\|\|\|\|\|\|\|\|\|\|\|\|\|\|\|\|\|\|\|\|\|\|\|\|\|\|\|\|\|\|\|\|\|\|\|\|\|\|\|\|\|\|\|\|\|\|\|\|\|\|\|\|\|\|\|\|\|\|\|\|\|\|\|\|\|\|\|\|\|\|\|\|\|\|\|\|\|\|\|\|\|\|\|\|\|\|\|\|\|\|\|\|\|\|\|\|\|\|\|\|\|\|\|\|\|\|\|\|\|\|\|\|\|\|\|\|\|\|\|\|\|\|\|\|\|\|\|\|\|\|\|\|\|\|\|\|\|\|\|\|\|\|\|\|\|\|\|\|\|\|\|\|\|\|\|\|\|\|\|\|\|\|\|\|\|\|\|\|\|\|\|\|\|\|\|\|\|\|\|\|\|\|\|\|\|\|\|\|\|\|\|\|\|\|\|\|\|\|\|\|\|\|\|\|\|\|\|\|\|\|\|\|\|\|\|\|\|\|\|\|\|\|\|\|\|\|\|\|\|\|\|\|\|\|\|\|\|\|\|\|\|\|\|\|\|\|\|\|\|\|\|\|\|\|\|\|\|\|\|\|\|\|\|\|\|\|\|\|\|\|\|\|\|\|\|\|\|\|\|\|\|\|\|\|\|\|\|\|\|\|\|\|\|\|\|\|\|\|\|\|\|\|\| \|\|\|\|\| \|\|\|\|\|\|\|\|\|\|\| \|\| \|\|\|\|\|\|\| \|\|\|\|\|  \|\|\|\|\|\| \| \|\|\| \|\| \|\|\|\|\| \|\|\|\| 	ATGAACGGTGCAAAGAAGGAAGTCATCCTGCCGGTGGAGGGCAAGAAGAACGTGTTGATCACTTCCGCATTGCCATACGTCAACAATGTACCTCATTTGGGAAACGTAATTGGCAGCGTGCTCTCGGCAGACGTCTTCTCCCGCTACAGCAAAGCACGAGGTCTTCCCACATTGTTCATATGCGGCACAGACGAATATGGCACAGCGACTGAGACGAAAGCGATCGAAGAGGGCATCACACCGCAAGAGCTCTGCGACAAATACAACAAGCTCCACAAGGAAGTCTACGACTGGTTCGAGATTGACTTTGACCACTTCGGGCGCACGCCGACCCANCAACANACGGATATTGCNCANGATATCTGTCTGANNCTATACNANAACNGANACCTCNAGGA ``` |
| 5 | ggggacaagtttgtacaaaaaagcaggcttgatgCCTTCGTTGGATCCTTCTTT, ggggaccactttgtacaagaaagctgggtcATCCCAATGTCTCCTGATGA | Mycgr3\|77760  JGI  Ensembl | KOG3059 N-acetylglucosaminyltransferase complex, subunit PIG-C/GPI2, required for phosphatidylinositol biosynthesis; | GPI anchor biosynthetic process; integral to membrane;  phosphatidylinositol N-acetylglucosaminyltransferase activity; | 100 % |  |  |
| 5 | ggggacaagtttgtacaaaaaagcaggcttgatgGCGAAGAAACGCAAGGCTCC, ggggaccactttgtacaagaaagctgggtcCAGTCTCGTGCTCTTCACCA | Mycgr3\|83972  JGI  Ensembl | KOG3118 Disrupter of silencing SAS10; | hydrogen ion transmembrane transporter activity; proton-transporting two-sector ATPase complex;  rRNA processing;  small subunit processome; | 0 % |  |  |
| 5 | ggggacaagtttgtacaaaaaagcaggcttgatgTCTTCCGTCAAGAGACGCAA, ggggaccactttgtacaagaaagctgggtcCGCCGTCGACTTTGAGGAGG | Mycgr3\|84402  JGI  Ensembl | KOG0330 ATP-dependent RNA helicase; | ATP binding; ATP-dependent helicase activity;  helicase activity;  nucleic acid binding; | 9.22935 % |  |  |
| 5 | ggggacaagtttgtacaaaaaagcaggcttgatgCTCTGGGTGCCATTTTGGGC, ggggaccactttgtacaagaaagctgggtcCGGAAGGCTCAGATACAACT | Mycgr3\|84465  JGI  Ensembl |  |  | 100 % |  |  |
| 5 | ggggacaagtttgtacaaaaaagcaggcttgatgCGAATATCGGGCCTGCACGC, ggggaccactttgtacaagaaagctgggtcAAGCTCTTCTCGCGTGGGAA | Mycgr3\|86118  JGI  Ensembl | KOG2608 Endoplasmic reticulum membrane-associated oxidoreductin involved in disulfide bond formation; | FAD binding; electron carrier activity;  endoplasmic reticulum;  endoplasmic reticulum membrane;  oxidoreductase activity, acting on sulfur group of donors, disulfide as acceptor; protein binding;  protein thiol-disulfide exchange;  protein-methionine-R-oxide reductase activity; | 0 % |  |  |
| 5 | ggggacaagtttgtacaaaaaagcaggcttgatgTTGGTCCATTCGCTCCTCAT, ggggaccactttgtacaagaaagctgggtcCTTGTTGTAGAAAGCAGCGT | Mycgr3\|89162  JGI  Ensembl |  |  | 0 % |  |  |
| 5 | ggggacaagtttgtacaaaaaagcaggcttgatgGAGGCCGCCGCAACGAAGAG, ggggaccactttgtacaagaaagctgggtcGACCGCAACTCCAGCACCCT | Mycgr3\|89374  JGI  Ensembl | KOG1721 FOG: Zn-finger; | cell redox homeostasis; intracellular;  nucleic acid binding;  zinc ion binding; | 0 % |  |  |
| 5 | ggggacaagtttgtacaaaaaagcaggcttgatgAAGCGAAAATTGAACGAGCG, ggggaccactttgtacaagaaagctgggtcCTTCTTTCCCCTCCCCTTGG | Mycgr3\|92212  JGI  Ensembl | KOG0346 RNA helicase; | 8-oxo-7,8-dihydroguanine triphosphatase activity; ATP binding;  ATP-dependent 5'-3' DNA helicase activity;  ATP-dependent helicase activity;  UDP-2,3-diacylglucosamine hydrolase activity;  bis(5'-nucleosyl)-tetraphosphatase activity;  dATP pyrophosphohydrolase activity;  diacylglycerol pyrophosphate phosphatase activity;  dihydroneopterin monophosphate phosphatase activity;  dihydroneopterin triphosphate pyrophosphohydrolase activity;  helicase activity;  nucleic acid binding;  pyrophosphatase activity;  thiamin-pyrophosphatase activity; | 24.16413 % |  |  |
| 5 | ggggacaagtttgtacaaaaaagcaggcttgatgAAGCTCTACATCCACGCCAT, ggggaccactttgtacaagaaagctgggtcTCCGGAGATTTTGTAAGAAG | Mycgr3\|93000  JGI  Ensembl |  |  | 0 % |  |  |
| 5 | ggggacaagtttgtacaaaaaagcaggcttgatgGTCTTCCACGACGCCGAGGA, ggggaccactttgtacaagaaagctgggtcTTGTGCTGAGTCGACCAAGG | Mycgr3\|93675  JGI  Ensembl | KOG4132 Uroporphyrinogen III synthase UROS/HEM4; | tetrapyrrole biosynthetic process; uroporphyrinogen-III synthase activity; | 0 % |  |  |
| 5 | ggggacaagtttgtacaaaaaagcaggcttgatgCCGCCTGGACTGATCAGTTC, ggggaccactttgtacaagaaagctgggtcATCCGCCTCCACGGCAACAT | Mycgr3\|95409  JGI  Ensembl | KOG2009 Transcription initiation factor TFIIIB, Bdp1 subunit; | DNA binding; | 0 % |  |  |
| 5 | ggggacaagtttgtacaaaaaagcaggcttgatgGAGTCCTCACGAGGGCCTCC, ggggaccactttgtacaagaaagctgggtcTGCGTCCCCATTCCCAGCAG | Mycgr3\|98811  JGI  Ensembl | KOG1915 Cell cycle control protein (crooked neck); | RNA processing; intracellular; | 100 % |  | ``` 1	ATGGAGTCCTCACGAGGGCCTCCACGGG-TCAAGAACAAGGCCGCCGCGCCTCAGCAGATCTCCGCCGAGCAGCTTCTCCGAGAAGCCGTCGACCGACAGGAGCCCGGTCTCGCCGCGCCTACACAGCGATTCGCGGATCTCGAGGAACTTCATGAGTACCAGGGCCGGAAACG	173 	\|\|\|\|\|\|\|\|\|\|\|\|\|\|\|\|\|\|\|\|\|\|\|\|\|\|\|\| \|\|\|\|\|\|\|\|\|\|\|\|\|\|\|\|\|\|\|\|\|\|\|\|\|\|\|\|\|\|\|\|\|\|\|\|\|\|\|\|\|\|\|\|\|\|\|\|\|\|\|\|\|\|\|\|\|\|\|\|\|\|\|\|\|\|\|\|\|\|\|\|\|\|\|\|\|\|\|\|\|\|\|\|\|\| \|\|\|\|\|\|\|\|\|\|\|\|\|\|\|\|\| \|\|\|\|\|\|\|\|\|\|\|\|\|\|\|\|\|\| \|\|\|   \|\|\| \|\|\|\|\|\| \|\|\|\| 	ATGGAGTCCTCACGAGGGCCTCCACGGGATCAAGAACAAGGCCGCCGCGCCTCAGCAGATCTCCGCCGAGCAGCTTCTCCGAGAAGCCGTCGACCGACAGGAGCCCGGTCTCGCCNCGCCTACACAGCGATTCNCGGATCTCGAGGAACTTCNTGANNNCCANGGCCGGNAACG ``` |
| 5 | ggggacaagtttgtacaaaaaagcaggcttgatgTCGAACGAATACAACTACGA, ggggaccactttgtacaagaaagctgggtcCGAGAACCACCCCTTCTTCT | Mycgr3\|99310  JGI  Ensembl | KOG0721 Molecular chaperone (DnaJ superfamily); | heat shock protein binding; | 0 % |  |  |
| 5 | ggggacaagtttgtacaaaaaagcaggcttgatgCCGGGCGTGATCGCCGAACC, ggggaccactttgtacaagaaagctgggtcTCTCCGTCCACCTCCCCTAC | Mycgr3\|99349  JGI  Ensembl | KOG2330 Splicing factor 3b, subunit 2; | mRNA processing; nucleus; | 100 % |  | ``` 98	AGGTCTCCGCAGTACCACCGCAGCCAGATGTCGCCGAGAAGCCGGAAGAGCCTGCACAAGCAGCAGCGCCCACGACCGATGCGGCAGACGATGAAACCACGATAATCGAGATCGACGACGACCTGCCCAAAGATGATGCGCTTTTCTCACAGTACGCCGGTGTGCTAGCAAGGTTCCAGAAGACCGACGAGGACGCCAAAGAGAACGAGCCAGAAGCGAAGCCGGAGGTATACTATGATGATGACGACAATGTTCAGTCAGAGGACGAGGAGGAGGCTAC	377 	\|\|\|\|\|\|\|\|\|\|\|\|\|\|\|\|\|\|\|\|\|\|\|\|\|\|\|\|\|\|\|\|\|\|\|\|\|\|\|\|\|\|\|\|\|\|\|\|\|\|\|\|\|\|\|\|\|\|\|\|\|\|\|\|\|\|\|\|\|\|\|\|\|\|\|\|\|\|\|\|\|\|\|\|\|\|\|\|\|\|\|\|\|\|\|\|\|\|\|\|\|\|\|\|\|\|\|\|\|\|\|\|\|\|\|\|\|\|\|\|\|\|\|\|\|\|\|\|\|\|\|\|\|\|\|\|\|\|\|\|\|\|\|\|\|\|\|\|\|\|\|\|\|\|\|\|\|\|\|\|\|\|\|\|\|\|\|\|\|\|\|\|\|\|\|\|\|\|\|\|\|\|\|\|\|\|\|\|\|\|\|\|\|\|\|\|\|\|\|\|\|\|\|\|\|\|\|\|\|\|\|\| \|\|\|\|\|\|\|\|\|\|\|\|\|\|\|\|\|\|\|\|\|\|\|\|\|\|\|\|\|\|\|\|\|\|\| \|\|\|\|\|\|\|\|\|\|  \|\|\|\|\| \|\|\|\| \|\| \|\|\|\|\| 	AGGTCTCCGCAGTACCACCGCAGCCAGATGTCGCCGAGAAGCCGGAAGAGCCTGCACAAGCAGCAGCGCCCACGACCGATGCGGCAGACGATGAAACCACGATAATCGAGATCGACGACGACCTGCCCAAAGATGATGCGCTTTTCTCACAGTACGCCGGTGTGCTAGCAAGGTTCCAGAAGACCGACGAGGACGCCAAAGAGAACGAGCCANAAGCGAAGCCGGAGGTATACTATGATGATGACGACNATGTTCAGTCNNAGGACNAGGANGANGCTAC ```      ``` 1	ATGCCGGGCGTGATCGCCGAACCTCAATCTGGTGCGAAGGCTCGCAAGCCAACGAAGAATGAGCAGCGTCGCGCAAAGAAGAAGCAACAGAAGCAAGAGGT	101 	\|\|\|\|\|\|\|\|\|\|\|\|\|\|\|\|\|\|\|\|\|\|\|\|\|\|\|\|\|\|\|\|\|\|\|\|\|\|\|\|\|\|\|\|\|\|\|\|\|\|\|\|\|\|\|\|\|\|\|\|\|\|\|\|\|\|\|\|\|\|\|\|\|\|\|\|\|\|\|\|\|\|\|\|\|\|\|\|\|\|\|\|\|\|\|\|\|\|\|\|\| 	ATGCCGGGCGTGATCGCCGAACCTCAATCTGGTGCGAAGGCTCGCAAGCCAACGAAGAATGAGCAGCGTCGCGCAAAGAAGAAGCAACAGAAGCAAGAGGT ``` |
| 5 | ggggacaagtttgtacaaaaaagcaggcttgatgCCGAAGAATCTGAAGCTCAA, ggggaccactttgtacaagaaagctgggtcAGCCCGCGTGCGCTTACTCT | Mycgr3\|99446  JGI  Ensembl | KOG3080 Nucleolar protein-like/EBNA1-binding protein; |  | 100 % |  | ``` 15	ATGCCGAAGAATCTGAAGCTCAAGGAGGCTCTCAATCGGCACAAGGGCGTCGACCACAAGCTAGAGCGCCAAAAGAAATTGCAGAAAGAGGCGGAGAAGAAGAAGCGTTCGAAGAAGGGCAGCAGC	140 	\|\|\|\|\|\|\|\|\|\|\|\|\|\|\|\|\|\|\|\|\|\|\|\|\|\|\|\|\|\|\|\|\|\|\|\|\|\|\|\|\|\|\|\|\|\|\|\|\|\|\| \|\|\|\|\|\|\|\|\|\|\|\|\|\|\|\|\|\| \|\|\|\|\|\|\|\|\|\|\|\|\|\|\|\|\|\|\|\|\|\|\|\| \|\| \|\|\|\|\| \|\|\|\|\| \|\|\|\|\| \|\|\|\| \|\|\|\| 	ATGCCGAAGAATCTGAAGCTCAAGGAGGCTCTCAATCGGCACAAGGGCGTCNACCACAAGCTAGAGCGCCNAAAGAAATTGCAGAAAGAGGCGGANAANAAGAACCGTTCCAAGAANGGCACCAGC ``` |
| 5 | ggggacaagtttgtacaaaaaagcaggcttgatgTCAAGCCTCGCCGCGCTCCT, ggggaccactttgtacaagaaagctgggtcCCACTTTCCACCCTTCCCCT | Mycgr3\|99562  JGI  Ensembl | KOG2376 Signal recognition particle, subunit Srp72; |  | 0 % |  |  |
| 5 | ggggacaagtttgtacaaaaaagcaggcttgatgCCGACCATTGCGGTGGATAA, ggggaccactttgtacaagaaagctgggtcCAGGAACACTTCGAGGTTGA | Mycgr3\|99865  JGI  Ensembl | KOG2472 Phenylalanyl-tRNA synthetase beta subunit; | ATP binding; RNA binding;  cytoplasm;  nucleotide binding;  phenylalanine-tRNA ligase activity;  phenylalanine-tRNA ligase complex;  phenylalanyl-tRNA aminoacylation;  translation; | 100 % |  | ``` 204	ATGCCGACCATTGCGGTGGATAAGGAGGACCTATATAAGGCATTGGGCAAGGAATACACCAAGGATGAGTTCGATGAGCTGTGCTTCGAGTTCGGCATTGAGCTTGACGAGGACACCTCCGAGCACAAGCGACCGATCGTAGATGGCGTGGA	355 	\|\|\|\|\|\|\|\|\|\|\|\|\|\|\|\|\|\|\|\|\|\|\|\|\|\|\|\|\|\|\|\|\|\|\|\|\|\|\|\|\|\|\|\|\|\|\|\|\|\|\|\|\|\|\|\|\|\|\|\|\|\|\|\|\|\|\|\|\|\|\|\|\|\|\|\|\|\|\|\|\|\|\|\|\|\|\|\|\|\|\|\|\|\|\|\|\|\|\|\|\|\|\|\|\|\|\|\|\|\|\|\|\|\|\|\|\|\|\|\|\|\|\|\|\|\|\|\|\|\|\|\|\|\|\|\|\|\|\|\|\| \|\|\| \|\|\|\|\|\| 	ATGCCGACCATTGCGGTGGATAAGGAGGACCTATATAAGGCATTGGGCAAGGAATACACCAAGGATGAGTTCGATGAGCTGTGCTTCGAGTTCGGCATTGAGCTTGACGAGGACACCTCCGAGCACAAGCGACCGATCGTAAATGNCGTGGA ``` |
| 5 | ggggacaagtttgtacaaaaaagcaggcttgatgCCCTCCATCCTCACAGAAGA, ggggaccactttgtacaagaaagctgggtcCCACTCATCATCACTCTTCT | Mycgr3\|100083  JGI  Ensembl | KOG3671 Actin regulatory protein (Wiskott-Aldrich syndrome protein); |  | 0 % |  |  |
| 5 | ggggacaagtttgtacaaaaaagcaggcttgatgGCGGGCACATCTGTCGAGGC, ggggaccactttgtacaagaaagctgggtcAGTGACGATTCCCTCACTTT | Mycgr3\|100443  JGI  Ensembl | KOG3702 Nuclear polyadenylated RNA binding protein; |  | 0 % |  |  |
| 5 | ggggacaagtttgtacaaaaaagcaggcttgatgGCCGCGCCCAACGGCGACTC, ggggaccactttgtacaagaaagctgggtcCTTCTTATACTTTTCGTGCA | Mycgr3\|101867  JGI  Ensembl | KOG0007 Splicing factor 3a, subunit 1; | RNA binding; RNA processing; | 100 % |  | ``` 132	GCGCCCGCCGACGACGACCTCGCCCACAAAGTCCCAGCCGGCACCGTCCTTCCCCCCAAAGTTGTGCGCAGCAAGATCGAAAAAGTCGCTTCATACGTCGCCCGCGCCGGCGAGAAATTCGAAGACTCGGTCCGACAAAAGAATGCCGGCACAAACCAGACCACTTTCCTCGAACCGGACGATCCATACTTCCCTTACTACAAATGGCGCGTGGGCGAGAT-AAAAGCTGGTCGAGGTCACGATCCGGATAATAATGCTGCGGCGAAGACGGAGGTCACATTCC-AGGGACGAGA-GTCGAGGGCGC-CGAAGCCGCCAGCGGAG	452 	\|\|\|\|\|\|\|\|\|\|\| \|\|\|\|\|\|\|\|\|\|\|\|\|\|\|\|\|\|\| \|\|\|\|\| \|\|\| \|\|\|\| \|\|\| \|\|\| \|\|\|\|\|\| \|\| \|\|\|\|\|\|\|\|\|\|\|\| \|\|\|\|\|\|\|\|\|\|\| \|\|\| \|\|  \| \|\| \|\|\|\|\|\|\|\|\|\|\|\|\|\|  \|\|\|\|\|\|\| \|\|\| \|\|\|\|\|\|\|\|\|\|\|\|\|\|\|\|\|\|\|\|\|\|\|\|\|\|\|\|\|\|\|\|\|\|  \|\| \|\|\|\|\|\|\|\|\|\|\|\|\|\|\|\| \|\| \|\| \|\| \|\| \|\|\|\|\| \|\|\|\|\|\| \|\|\|\|\|\|\|\|  \|\|\|\|\|\| \|\| \|\|\|\|\|\|\|\|\|\|\| \|\|\|\|\| \|\| \|\| \|  \| \|\|\|\|\|\|\|\|\|\|\|\|\|\| \|\|\|\|\|\| \| \|\|\|\|\| \|\|\|\| \| \|\|\|\|\|\|\|\|  \|\|\|\| \| \| \|\|\| \|\|\|\| 	GCGCCCGCCGANGACGACCTCGCCCACAAAGACCCAGNCGGNACCGNCCTACCCNCCAAAGATGNGCGCAGCAAGATNGAAAAAGTCGCNTCANACNNCNCCNGCGCCGGCGAGAAAGGCGAAGACNCGGNCCGACAAAAGAATGCCGGCACAAACCAGACCACTNNCCACGAACCGGACGATCCAAACATCNCTAACNACAAANGGCGCGNGGGCGAGAANAAAAGCNGGACGAGGTCACGANCCGGACAAAAAGGAAGNGGCGAAGACGGAGGNCACATTACNAGGGANGAGAAGACGAGGGCGNACGAANCNGACAGAGGAG ```      ``` 105	ATGGCCGCGCCCAACGGCG-ACTCCGCCGCG-CCCGCCGACGACGACCTCGCCCACAAAGTCCCAGCCGGCACCG-TCCTTCCCCCCAAAG-TTGTGCGCAGCAAG-ATCGAAAAAGTCGC-TTCATACGTCGCCCGCGCCGGCGAGAAATTCGAAGACTCGGTCCGACAAAAGAATGCCGGCACAAACCA-GACCACTTTCCTCGAACCGGACGATCCATACTTCCCTTACTACAAATGGCGCGTGGGCGAGATAAAAGCTGGTCGAGGTCACGAT-CCGGATAATAATGCTGCGGCGAAGACGGAGGTCACATTCCAGGG-ACGAGAG-TCGAGGGCGCCGAAGC-CGCC-AGCGG-AGT-TCAAGT-TCAGCGCGCGC-ATGCC-GAATATT-TCGGCGCAG-GAT-TTGG-AGATTGTCAAGGT-GACGGCGC-T-GTATGCGGC-GAAGAACGGGAGG-AGTTGGGTCACGG-CG-TTGAGTCAACGGG-AAGGAGGCA-ATTAT	584 	\|\|\|\|\|\|\|\|\| \|\|\| \| \| \| \|\|\|\|\|\|\|\|\|\|\| \|\|\|\|\|\|\|\|\|\|\|\|\|\|\|\|\|\|\|\|\|\|\| \|\|\|\|\|\|\|\|\|\|\|\|\|\|\| \|\|\| \|\|\|\|\|\|\|\|\|\|\|\|\|\|\| \|\|\|\|\|\|\|\|\| \|\|\|\| \|\|\|\|\|\|\|\|\|\|\|\|\|\| \|\|\| \|\| \|\|\|\|\|\|\|\|\|\|\|\|\|\|\|\|\|\|\|\|\| \|\|\|\|\|\|\|\|\|\|\|\|\|\|\|\|\|\| \|\|\|\|\|\|\|\|\|\|\|\|\|\|\|\|\|\|\|\|\|  \|\|\|\|\|\|\|\|\|\|\|\|\|\|\|\|\|\|\|\|\|\|\|\|\|\|\|\|\|\|\|\|\|\| \|\|\|\|\|\|\|\|\|\|\|\|\|\|\|\|\|\|\|\|\|\|\|\|\|\|\|\|\|\|\|\|\|\|\|\| \|\|\|\|\|\|\|\|\|\|\|\| \|\|\|\|\|\|\|\|\|\|\|\|\| \|\| \|\| \|\|\|\|\| \|\|\|\|\|\|\|\|\|\|\|\|\| \|\|\|\| \|\|\|\|\|\|\| \|\|\|\|\|\|\| \|\|\|  \|   \|\|   \|\|\|\|  \|\|\|    \|\|\| \|\|\|\| \|\|\| \|  \|\|\|\|  \|\|\|\|\|\|\| \|\|\|\|\|\|\|   \|\|\| \|\|\|\| \|\|\|\|\|\|\|\|\| \|\|  \|\|\| \|  \| \| \|\|\|\| \| \|\| \|\|\| \|\|\|\|\|\|\|\|\| \|\| \|\|\|\|\|\|\|  \| \|\| \|\| \|\|\|\|\|\|\|\|\|\| \|\|\|\| \|\|\|  \|\|\|\|\| 	ATGGCCGCGNCCACCNGTGTACTCCGCCGCGTCCCGCCGACGACGACCTCGCCCANAAAGTCCCAGCCGGCNCCGTTCCTTCCCCCCAAAGCTTGTGCGCANCAAGCATCGAAAAAGTCGCNTTCNTANGTCGCCCGCGCCGGCGAGAAANTCGAAGACTCGGTCCGACNAAAGAATGCCGGCACAAACCANAACCACTTTCCTCGAACCGGACGATCCATACTTCCNTTACTACAAATGGCGCGTGGGCGAGATAAAAGCTGGNCGAGGTCACGATCCCGGATAATAATGNTGNGGNGAAGANGGAGGTCACATTCNAGGGNACGAGAGTTCGAGGGTGCCTNATTNCGNTNAGCGNNAGTNNTNAGTNTCAGAGCGNGTNATGCTNGAATATTNTCGGCGCCNTGATNTTGGCAGATTGTCANGGNNGACNGNCCNTNGTATNCNGCNGAAAAACGGGAGGAAGNTGGGTCATNGNCGNTTNAGTCAACGGGNAAGGNGGCCNATTAT ```      ``` 105	ATGGCCGCGCCCAACGGCGACTCCGCCGCGCCCGCCGACGACGACCTCGCCCACAAAGTCCCAGCCGGCACCGTCCTTCCCCCCAAAGTTGTGCGCAGCAAGATCGAAAAAGTCGCTTCATACGTCGCCCGCGCCGGCGAGAAATTCGAAGACTCGGTCCGACAAAAGAATGCCGGCACAAACCAGACCACTTTCCTCGAACCGGACGATCCATACTTCCCTTACTACAAATGGCGCGTGGGCGAGATAAAAGCTGGTCGAGGTCACGATCCGGATAATAATGCTGCGGCGAAGACGGAGGTCACATTCCAGGGACGAGAGTCGAGGGCGCCGAAGCCGCCAGCGGAGTTCAAGTTCAGCGCGCGCATGCCGAATATTTCGGCGCAGGATTTGGAGATTGTCAAGGTGACGGCGCTGTATGCGGCGAAGAACGGGAGGAGTTGGGTCACGGCGTTGAGTCAACGGGAAGGAGGCAATTATCAATTTGATTTCTTGAGGCCACAGCATTCGTTGAATATGTTCTATACGAAGCTGGTGGATCAGTACAAAGACTTGTTGGAAGGGGAGACGGCGGATGGTGGGAAGCCGCAGAGAGAGAGGATTGCGGATCTGGAGGCGAATGTCGGCGACCGATTTCGCATGCTGGAGAGGGCGAAGAAGAGGGCCGAGTGGTTGAAGTGGCAGGAGAAGCAGAAAGTCGAGACGGAGCAGAAGGAGGAGAAGGACCGCATTGCGTTTGCGGAGATTGATTGGCATGACTTCACGGTCGTGGCTACGGTTGTGTTTGAC	893 	\|\|\|\|\|\|\|\|\|\|\|\|\|\|\|\|\|\|\|\|\|\|\|\|\|\|\|\|\|\|\|\|\|\|\|\|\|\|\|\|\|\|\|\|\|\|\|\|\|\|\|\|\|\|\|\|\|\|\|\|\|\|\|\|\|\|\|\|\|\|\|\|\|\|\|\|\|\|\|\|\|\|\|\|\|\|\|\|\|\|\|\|\|\|\|\|\|\|\|\|\|\|\|\|\|\|\|\|\|\|\|\|\|\|\|\|\|\|\|\|\|\|\|\|\|\|\|\|\|\|\|\|\|\|\|\|\|\|\|\|\|\|\|\|\|\|\|\|\|\|\|\|\|\|\|\|\|\|\|\|\|\|\|\|\|\|\|\|\|\|\|\|\|\|\|\|\|\|\|\|\|\|\|\|\|\|\|\|\|\|\|\|\|\|\|\|\|\|\|\|\|\|\|\|\|\|\|\|\|\|\|\|\|\|\|\|\|\|\|\|\|\|\|\|\|\|\|\|\|\|\|\|\|\|\|\|\|\|\|\|\|\|\|\|\|\|\|\|\|\|\|\|\|\|\|\|\|\|\|\|\|\|\|\|\|\|\|\|\|\|\|\|\|\|\|\|\|\|\|\|\|\|\|\|\|\|\|\|\|\|\|\|\|\|\|\|\|\|\|\|\|\|\|\|\|\|\|\|\|\|\|\|\|\|\|\|\|\|\|\|\|\|\|\|\|\|\|\|\|\|\|\|\|\|\|\|\|\|\|\|\|\|\|\|\|\|\|\|\|\|\|\|\|\|\|\|\|\|\|\|\|\|\|\|\|\|\|\|\|\|\|\|\|\|\|\|\|\|\|\|\|\|\|\|\|\|\|\|\|\|\|\|\|\|\|\|\|\|\|\|\|\|\|\|\|\|\|\|\|\|\|\|\|\|\|\|\|\|\|\|\|\|\|\|\|\|\|\|\|\|\|\|\|\|\|\|\|\|\|\|\|\|\|\|\|\|\|\|\|\|\|\|\|\|\|\|\|\|\|\|\|\|\|\|\|\|\|\|\|\|\|\|\|\|\|\|\|\|\|\|\|\|\|\|\|\|\|\|\|\|\|\|\|\|\|\|\|\|\|\|\|\|\|\|\|\|\|\|\|\|\|\|\|\|\|\|\|\|\|\|\|\|\|\|\|\|\|\|\|\|\|\|\|\|\|\|\|\|\|\|\|\|\|\|\|\|\|\|\|\|\|\|\|\|\|\|\|\|\|\|\|\|\|\|\|\|\|\|\|\|\|\|\|\|\|\|\|\|\|\|\|\|\|\|\|\|\|\|\|\|\|\|\|\|\|\|\|\|\|\|\|\|\|\|\|\|\|\|\|\|\|\|\|\|\|\|\|\|\|\|\|\|\|\|\|\|\|\|\|\|\|\|\|\|\|\|\|\|\|\|\|\|\|\|\|\|\|\|\|\|\|\|\|\|\|\| \|\| \| \|\|\|\|\|\|\|\|\|\|\|\|\|\|\|\|\|\|\|\|\|\|\|\|\|\|\|\|\|\|\|\|\|\|\|\|\|\|\|\|\|\|\|\|\|\|\|\|\|\|\|\|\|\|\|\|\|\|\|\|\|\|\|\|\|\|\|\|\|\|\|\|\|\|\|\|\|\|\|\|\|\|\| \|\|\|\|\|\|\|\|\|\|\|\|\|\| \| \|\|\|\|\|\|\|\|\|\|\|\|\|\| \| \|\|\|\|\|\|\|\|\| 	ATGGCCGCGCCCAACGGCGACTCCGCCGCGCCCGCCGACGACGACCTCGCCCACAAAGTCCCAGCCGGCACCGTCCTTCCCCCCAAAGTTGTGCGCAGCAAGATCGAAAAAGTCGCTTCATACGTCGCCCGCGCCGGCGAGAAATTCGAAGACTCGGTCCGACAAAAGAATGCCGGCACAAACCAGACCACTTTCCTCGAACCGGACGATCCATACTTCCCTTACTACAAATGGCGCGTGGGCGAGATAAAAGCTGGTCGAGGTCACGATCCGGATAATAATGCTGCGGCGAAGACGGAGGTCACATTCCAGGGACGAGAGTCGAGGGCGCCGAAGCCGCCAGCGGAGTTCAAGTTCAGCGCGCGCATGCCGAATATTTCGGCGCAGGATTTGGAGATTGTCAAGGTGACGGCGCTGTATGCGGCGAAGAACGGGAGGAGTTGGGTCACGGCGTTGAGTCAACGGGAAGGAGGCAATTATCAATTTGATTTCTTGAGGCCACAGCATTCGTTGAATATGTTCTATACGAAGCTGGTGGATCAGTACAAAGACTTGTTGGAAGGGGAGACGGCGGATGGTGGGAAGCCGCAGAGAGAGAGGATTGCGGATCTGGAGGCGAATGTCGGCGACCGATTTCGCATGCTGGAGAGGGCGAAAAAAA-GGCCGAGTGGTTGAAGTGGCAGGAGAAGCAGAAAGTCGAGACGGAGCAGAAGGAGGAGAAGGACCGCATTGCGTTTGCGGAGA-TGATTGGCATGACTNCNCGGTCGTGGCTACGNT-GTGTTTGAC ``` |
| 5 | ggggacaagtttgtacaaaaaagcaggcttgatgGCCGACAACAATCTCAACCG, ggggaccactttgtacaagaaagctgggtcATGCTGATGCTGATCCCGTG | Mycgr3\|102416  JGI  Ensembl | KOG0909 Peptide:N-glycanase; |  | 13.6059 % | chr\_1:2742717 C->G; |  |
| 5 | ggggacaagtttgtacaaaaaagcaggcttgatgGCACCCAAAGACTTCGCGGC, ggggaccactttgtacaagaaagctgggtcAGCACCTTGACCACTTCGAG | Mycgr3\|103034  JGI  Ensembl | KOG2537 Phosphoglucomutase/phosphomannomutase; | carbohydrate metabolic process; intramolecular transferase activity, phosphotransferases; | 100 % |  |  |
| 5 | ggggacaagtttgtacaaaaaagcaggcttgatgGCGCAAGGCCTGTCTCCGGG, ggggaccactttgtacaagaaagctgggtcAGAAGGCTTGGGCAACTTCC | Mycgr3\|103549  JGI  Ensembl | KOG1597 Transcription initiation factor TFIIB; | regulation of transcription, DNA-dependent; transcription factor complex;  transcription initiation;  transcription regulator activity;  zinc ion binding; | 0 % |  |  |
| 5 | ggggacaagtttgtacaaaaaagcaggcttgatgGCAACTGCATCAGTATTCAC, ggggaccactttgtacaagaaagctgggtcGCGAATGGTAGTAGCCTTGG | Mycgr3\|104090  JGI  Ensembl | KOG0305 Anaphase promoting complex, Cdc20, Cdh1, and Ama1 subunits; |  | 0 % |  |  |
| 5 | ggggacaagtttgtacaaaaaagcaggcttgatgGAAGTCCTCCTCCGCCAGTC, ggggaccactttgtacaagaaagctgggtcAGCCGCCGCAGCAACAGCAA | Mycgr3\|105601  JGI  Ensembl | KOG1360 5-aminolevulinate synthase; | 5-aminolevulinate synthase activity; biosynthetic process;  metabolic process;  pyridoxal phosphate binding;  tetrapyrrole biosynthetic process;  transferase activity;  transferase activity, transferring nitrogenous groups; | 2.19017 % |  |  |
| 5 | ggggacaagtttgtacaaaaaagcaggcttgatgCCGACCGCCGCACGAGACAT, ggggaccactttgtacaagaaagctgggtcGTACAGCACCCTCGTCCGGA | Mycgr3\|105609  JGI  Ensembl | KOG0068 D-3-phosphoglycerate dehydrogenase, D-isomer-specific 2-hydroxy acid dehydrogenase superfamily; | NAD binding; cofactor binding;  metabolic process;  oxidoreductase activity, acting on the CH-OH group of donors, NAD or NADP as acceptor; phosphoglycerate dehydrogenase activity; | 0 % |  |  |
| 5 | ggggacaagtttgtacaaaaaagcaggcttgatgGACGCAGTCAAGGGCACCAT, ggggaccactttgtacaagaaagctgggtcGTCCTCCTTCTCAATCACAG | Mycgr3\|106864  JGI  Ensembl | KOG2388 UDP-N-acetylglucosamine pyrophosphorylase; | UDP-N-acetylglucosamine diphosphorylase activity; metabolic process;  nucleotidyltransferase activity; | 0 % |  |  |
| 5 | ggggacaagtttgtacaaaaaagcaggcttgatgGCCAACTCCACAAGCTTACA, ggggaccactttgtacaagaaagctgggtcCTCCTTCTTTTCACCTGTCA | Mycgr3\|106974  JGI  Ensembl | KOG1216 von Willebrand factor and related coagulation proteins; |  | 27.45744 % |  | ``` 1	ATGGCCAACTCCACAAGCTTACACCAACCCCTTCCCTCCTACCTACCATCCACCCTTCCTCATCAGCAGCCTCGACTCGTCTACTTCATCACTCGTCCGGGCAATGTTCCTGTGGCACTCATTCCTGTTGACGAACTGCCACTCTCCGTCAAGCTCGTCCACGCATCCAGAGTCTTGACTCCGGATCAGACGTATGGCATGCACTATGTTGGAGCGGCACCAGTCAG-CGGTGGTC	235 	\|\|\|\|\|\|\|\|\|\|\|\|\|\|\|\|\|\|\|\|\|\|\|\|\|\|\|\|\|\|\|\|\|\|\|\|\|\|\|\|\|\|\|\|\|\|\|\|\|\|\|\|\|\|\|\|\|\|\|\|\|\|\|\|\|\|\|\|\|\|\|\|\|\|\|\|\|\|\|\|\|\|\|\|\|\|\|\|\|\|\|\|\|\|\|\|\|\|\|\|\|\|\|\|\|\|\|\|\|\|\|\|\|\|\|\|\|\|\|\|\|\|\|\|\|\|\|\|\|\|\|\|\|\|\|\|\|\|\|\|\|\|\|\|\|\|\|\|\|\|\|\|\|\|\|\|\|\|\|\|\|\|\|\|\|\|\|\|\|\|\|\|\|\|\|\|\|\|\|\|\|\|\|\|\|\|\|\|\|\| \|\|\|\|\|\|\|\|\|\|\|\|\|\|\|\|\|\|\|\|\|\|\|\|\| \|\|\| \| \|\| \| \| \|\|\|\|\|\| 	ATGGCCAACTCCACAAGCTTACACCAACCCCTTCCCTCCTACCTACCATCCACCCTTCCTCATCAGCAGCCTCGACTCGTCTACTTCATCACTCGTCCGGGCAATGTTCCTGTGGCACTCATTCCTGTTGACGAACTGCCACTCTCCGTCAAGCTCGTCCACGCATCCAGAGTCTTGACTCCGGATCAGANGTATGGCATGCACTATGTTGGAGCGNCACNANTCTGACNGTGGTC ``` |
| 5 | ggggacaagtttgtacaaaaaagcaggcttgatgGGCGACCCGCAATACCTCAG, ggggaccactttgtacaagaaagctgggtcACTCCGTTCGGAACCGCGCA | Mycgr3\|107306  JGI  Ensembl | KOG0687 26S proteasome regulatory complex, subunit RPN7/PSMD6; | proteasome endopeptidase activity; threonine endopeptidase activity; | 100 % |  | ``` 92	ATGGG-CGACCCG-CAATAC-CTCAGC-TTCCCAA-CG-CTGG-CA-CTCG-CGCA-GAATGT-C-TTCACCG-TCGCCTC-TCCATCA-ACATCA-TC-CT-CCACGAGGCAGTCCGCACTCACTGCCTTGCAAGATGACATC-CGCGAGCACAAGATGGCGCCTCTCTACGCATATCTGGCTCATCCCCAAACGGGC-AAGCTGAATTCTCCAGGTGAAGGCGGCGCCTCTCTTTCACCCACAAACACCCGTGCTTCGCCTCCATCTCCTGGCCATCTCCTTCGAAGGACG-AGCAGCATAAACGCTC-CATCAATCG-TCGGCGTACTGGGTGGTAAGACAGA-CACCACTGTGACA-CTACCATGGGATGAAGCACTCT	449 	\|\|\|\|\|  \|\|\|\|\|\| \|\|\|\|\|\|  \|\|\|\|\| \|\|\| \|\|\| \|\| \|\|\|\|  \| \|\|\|\|   \|\|  \|\|\|\|\| \| \|\|\|\| \|\| \|\|\|\|\|\|\| \|\|\|\|\|\|\| \|  \|\|\| \|\| \|\|     \|\|\|\| \|\| \|\|\|\|\|\|\|\|\|\| \| \|\|\|\|\|\|\|\|\|\|\|\|\|\|\|\| \|\|\|\|\|\|\|\|\|\| \|\|\|\|\|\|\|\|\|\|\|\|\|\|\|\|\|\|\|\|\|\|\|\|\|\|\|\|\|\|\|\|\|\|\|\|\|\| \|\|\|  \|\|\|\|\|\|\|\|\|\|\|\| \|\|\|\|\|\|\|\|\|\|\|\|\|\|\|\|\|\|\|\|\|\|\|\|\|\|\|\| \|\|\|\|\|\|\|\|\|\|\|\|\|\|\|\|\|\|\|\|\|\|\|\|\|\|\|\|\|\|\|\|\| \|\|\|\|\|\|\|\|\|\|\|\|\|\|\|\|  \|\|\|\|\| \|\|\|\|\|\|\|\|\|  \|\|\|\|\| \|   \|\| \|\|\|\| \|\|\|\|\| \|\|\|\|\|\|\|\|\| \| \| \|\|\| \|\|\|\|    \|\|\| \|\|\|\|\|\|\| \|\| \|\|\|\|\|\|\| 	ATGGGTGGACCCGNCAATACNGTCAGCTTTCNCAANCGTCTGGANACCTCGNNACANTAATGTACTTTCANCGNTCGCCTCTTCCATCAGANNTCATTCNCTNGNGGGAGGNAGACCGCACTCACAGNCTTGCAAGATGACATCNCGCGAGCACAGGATGGCGCCTCTCTACGCATATCTGGCTCATCCCCAAANGGGNNAAGCTGAATTCTTCAGGTGAAGGCGGCGCCTCTCTTTCACCNACAAACACCCGTGCTTCGCCTCCATCTCCTGGCAATCTCCTTCGAAGGACNTAGCAGTATAAACGCTATCATCANTNTNTCNGCGTNCTGGGNGGTAAGACANANCNCCANTGTGGNNNCTANCATGGGANGANGCACTCT ```      ``` 119	CCAACGCTGGCACTCGCG-CAGAATGTCTTCAC-CGTCG-CCTCTCCA-TCAACATCATCCTCCACGAGGCAGTCCGCACTCACTGCCTTGCAAGATGACATCCGCGAGCACAAGATGGCGCCTCTCTACGCATATCTGGCTCATCCCCAAACGGGCAAGCTGAATTCTCCAGGTGAAGGCGGCGCCTCTCTTTCACCCACAAACACCCGTGCTTCGCCT-CC-ATCTCCTGGCCATCTCCTTCGAAGGACGAGCAGCATAAACGCTCCATCAATCGTCGGCG-TACTGGGTGGTAAGACAGACACCA-CTGT	423 	\|\|\|\|\|\|\|\|\|\| \|\|\| \|\|\| \|\|\|\|\|\|\|\|\| \|\| \|  \|\|\|\|  \|\|\|\|\|\|\| \| \|\| \|\|\|\|\|\|\|\|\|\|\|\|\|\|\|\| \|\| \| \|\|\|\|\|\|\| \|\|\|\|\|\|\| \|\|\|\| \|\| \|\|\|\|\|\|\|\|\|\|\|\|\|\|\|\|\|\|\| \|\|\|\|\|\|\|\|\|\|\|\|\|\|\|\|\| \|\|\|\| \|\|\| \|\|\|\|\|\|\|\|\|\|\|\|\|\|\|\|\|\|\|\|\|\| \|\|\|\| \|\| \|\|\|\|\|\|\| \| \|\|\|\|\|\|\|\|\|  \| \|\|\|\|\|\|\| \|\|\|\|\|\|\|\|    \|\|   \|\|\|\|\|\|\|  \|\|\| \|\|\|   \|\|\|\|\| \|\|\|\| \| \|\|\|\|\|\|\|\|\|\|\| \|\| \|\|\| \|\|\|\|\|\| \|  \|\|\|\|\|\|\|\|\|\|\|\| \|\|\|\|\|\|\|\| \|\|\|\| 	CCAACGCTGGNACTNGCGACAGAATGTCNTCNCTNGTCGNNCTCTCCANTNAANATCATCCTCCACGAGGNAGACNGCACTCANTGCCTTGNAAGANGATATCCGCGAGCACAAGATGGNGCCTCTCTACGCATATCNGGCTNATCNCCAAACGGGCAAGCTGAATTCTNCAGGNGANGGCGGCGNCNCTCTTTCACANANAAACACCNGTGCTTCGNNNACCNGACTCCTGGNNATCNCCTNNNAAGGANGAGCNGTATAAACGCTCCNTCNATCNTCGGCGNTNNTGGGTGGTAAGANAGACACCAACTGT ```      ``` 92	ATGGGCGACCCGCAATACCTCAGCTTCCCAACGCTGGCACTCGCGCAGAATGTCTTCACCGTCGCCTCTCCATCAACATCATCCTCCACGAGGCAGTCCGCACTCACTGCCTTGCAAGATGACATCCGCGAGCACAAGATGGCGCCTCTCTACGCATATCTGGCTCATCCCCAAACGGGCAAGCTGAATTCTCCAGGTGAAGGCGGCGCCTCTCTTTCACCCACAAACACCCGTGCTTCGCCTCCATCTCCTGGCCATCTCCTTCGAAGGACGAGCAGCATAAACGCTCCATCAATCGTCGGCGTACTGGGTGGTAAGACAGACACCACTGTGACACTACCATGGGATGAAGCACTCTACGAGCAATTACGGGCGGACAACGAGAAAGAGCTCGCGGCGATACAAAAGGAGGAAGACGACGCGGTGGAGCAGGCAGGTGAGACGGAGATTCAGACAGCGAAGGGAAAGCGCGCAGAGTTCTACGCGCGGGTCGGTGACAAGGATAAAGCATTGGAGGAGTTTGAGAAGCTACTGGAGAAGAGCGCCATCCTCGGGACAAAGATCGACATCGTGCTTGCTATTATCCGCATCGGACTCTTCTTCGACGACAAGCTGCTGGTGAAGAAGAACGTCGACCGCGCTCAGCAGCTGGTCGAGAGTGGCGGAGACTGGGACCGCCGCAATCGCCTCAAGGCCTACAATGGTCTGCATCTCATCACCATCCGTGCCTACAACCTCGCAGCACCGCTTCTTCTCGACTCGCTCTCCACCTTCACATCCACTGAACTGTGCTCATACAGCAACCTTGTCGTCTATGCCACCCTCGCTGGATCCGTCAGTCTAC	935 	\|\|\|\|\|\|\|\|\|\|\|\|\|\|\|\|\|\|\|\|\|\|\|\|\|\|\|\|\|\|\|\|\|\|\|\|\|\|\|\|\|\|\|\|\|\|\|\|\|\|\|\|\|\|\|\|\|\|\|\|\|\|\|\|\|\|\|\|\|\|\|\|\|\|\|\|\|\|\|\|\|\|\|\|\|\|\|\|\|\|\|\|\|\|\|\|\|\|\|\|\|\|\|\|\|\|\|\|\|\|\|\|\|\|\|\|\|\|\|\|\|\|\|\|\|\|\|\|\|\|\|\|\|\|\|\|\|\|\|\|\|\|\|\|\|\|\|\|\|\|\|\|\|\|\|\|\|\|\|\|\|\|\|\|\|\|\|\|\|\|\|\|\|\|\|\|\|\|\|\|\|\|\|\|\|\|\|\|\|\|\|\|\|\|\|\|\|\|\|\|\|\|\|\|\|\|\|\|\|\|\|\|\|\|\|\|\|\|\|\|\|\|\|\|\|\|\|\|\|\|\|\|\|\|\|\|\|\|\|\|\|\|\|\|\|\|\|\|\|\|\|\|\|\|\|\|\|\|\|\|\|\|\|\|\|\|\|\|\|\|\|\|\|\|\|\|\|\|\|\|\|\|\|\|\|\|\|\|\|\|\|\|\|\|\|\|\|\|\|\|\|\|\|\|\|\|\|\|\|\|\|\|\|\|\|\|\|\|\|\|\|\|\|\|\|\|\|\|\|\|\|\|\|\|\|\|\|\|\|\|\|\|\|\|\|\|\|\|\|\|\|\|\|\|\|\|\|\|\|\|\|\|\|\|\|\|\|\|\|\|\|\|\|\|\|\|\|\|\|\|\|\|\|\|\|\|\|\|\|\|\|\|\|\|\|\|\|\|\|\|\|\|\|\|\|\|\|\|\|\|\|\|\|\|\|\|\|\|\|\|\|\|\|\|\|\|\|\|\|\|\|\|\|\|\|\|\|\|\|\|\|\|\|\|\|\|\|\|\|\|\|\|\|\|\|\|\|\|\|\|\|\|\|\|\|\|\|\|\|\|\|\|\|\|\|\|\|\|\|\|\|\|\|\|\|\|\|\|\|\|\|\|\|\|\|\|\|\|\|\|\|\|\|\|\|\|\|\|\|\|\|\|\|\|\|\|\|\|\|\|\|\|\|\|\|\|\|\|\|\|\|\|\|\|\|\|\|\|\|\|\|\|\|\|\|\|\|\|\|\|\|\|\|\|\|\|\|\|\|\|\|\|\|\|\|\|\|\|\|\|\|\|\|\|\|\|\|\|\|\|\|\|\|\|\|\|\|\|\|\|\|\|\|\|\|\|\|\|\|\|\|\|\|\|\|\|\|\|\|\|\|\|\|\|\|\|\|\|\|\|\|\|\|\|\|\|\|\|\|\|\|\|\|\|\|\|\|\|\|\|\|\|\|\|\|\|\|\|\|\|\|\|\|\|\|\|\|\|\|\|\|\|\|\|\|\|\|\|\|\|\|\|\|\|\|\|\|\|\|\|\|\| \|\|\|\| \|\|\|  \|\|\|\|\|\|\|\|\|\|\|\|\|\|\|\|\|\|\|\|\|\|\|\|\|\| \|\|\|\|\|\|\|\| \|\|\| \|\|\|\|\|\|\|\|\|\|\|\|\|\|\|\|\|\|\|\|\|\|\|\|\|\|\|\|\| \| \|\| \|\| \|\|\|\| \|\|\| \|\|\|\|\|  \|\| \|  \|\|\|\|\|\|\|\|\|\|\|  \|\|\|\|\|\|\|\|\|\|\|\|\|\|\|\| \|\| \|\|\| \|\|\| \|\| \|\|\| \|\|\| 	ATGGGCGACCCGCAATACCTCAGCTTCCCAACGCTGGCACTCGCGCAGAATGTCTTCACCGTCGCCTCTCCATCAACATCATCCTCCACGAGGCAGTCCGCACTCACTGCCTTGCAAGATGACATCCGCGAGCACAAGATGGCGCCTCTCTACGCATATCTGGCTCATCCCCAAACGGGCAAGCTGAATTCTCCAGGTGAAGGCGGCGCCTCTCTTTCACCCACAAACACCCGTGCTTCGCCTCCATCTCCTGGCCATCTCCTTCGAAGGACGAGCAGCATAAACGCTCCATCAATCGTCGGCGTACTGGGTGGTAAGACAGACACCACTGTGACACTACCATGGGATGAAGCACTCTACGAGCAATTACGGGCGGACAACGAGAAAGAGCTCGCGGCGATACAAAAGGAGGAAGACGACGCGGTGGAGCAGGCAGGTGAGACGGAGATTCAGACAGCGAAGGGAAAGCGCGCAGAGTTCTACGCGCGGGTCGGTGACAAGGATAAAGCATTGGAGGAGTTTGAGAAGCTACTGGAGAAGAGCGCCATCCTCGGGACAAAGATCGACATCGTGCTTGCTATTATCCGCATCGGACTCTTCTTCGACGACAAGCTGCTGGTGAAGAAGAACGTCGACCGCGCTCAGCAGCTGGTCGAGAGTGGCGGAGACTGGGACCGCCGCANTCGCNTCAN-GCCTACAATGGTCTGCATCTCATCACNATCCGTGCNTAC-ACCTCGCAGCACCGCTTCTTCTCGACTCGNTNTC-AC-TTCA-ATC-ACTGAN-TGGGN-CATACAGCAACN-TGTCGTCTATGCCACC-TCNCTGNATC-GT-AGTNTAC ``` |
| 5 | ggggacaagtttgtacaaaaaagcaggcttgatgAGAGCGTCTCCAAGCCGATT, ggggaccactttgtacaagaaagctgggtcTTCCTTCAACGTCAACTTCT | Mycgr3\|107981  JGI  Ensembl | KOG1919 RNA pseudouridylate synthases; | RNA binding; pseudouridine synthase activity;  pseudouridine synthesis;  pseudouridylate synthase activity; | 0 % |  |  |
| 5 | ggggacaagtttgtacaaaaaagcaggcttgatgACGTTCCGCGGGAACGAGTC, ggggaccactttgtacaagaaagctgggtcCTTCCACCCCACCTTCTTCG | Mycgr3\|108402  JGI  Ensembl | KOG4503 Uncharacterized conserved membrane protein; |  | 0 % |  |  |
| 5 | ggggacaagtttgtacaaaaaagcaggcttgatgGCCGACCAGAACAACCCACC, ggggaccactttgtacaagaaagctgggtcTCTCCTGCCCAGATAACTCC | Mycgr3\|108919  JGI  Ensembl | KOG2196 Nuclear porin; | ATP binding; aminoacyl-tRNA ligase activity;  cytoplasm;  nuclear pore;  nucleotide binding;  structural constituent of nuclear pore;  tRNA aminoacylation for protein translation;  translation; | 100 % |  |  |
| 5 | ggggacaagtttgtacaaaaaagcaggcttgatgACTTCTATGATCACAGCGAC, ggggaccactttgtacaagaaagctgggtcGTTCTCGTCCATCGACTCCC | Mycgr3\|109282  JGI  Ensembl | KOG0270 WD40 repeat-containing protein; |  | 100 % |  | ``` 13	TATG-ATCACAGC-GACCGC-ATGGGTCCCTCGAGGCCATGCGGCGGCCTTTCCCACGAAG-TACCAAATGGACGACGACGAATACGCTCGCATTGCGAAACTGGCGAAGCTCGAGCTCGACGATGCAAAGGTAGATTTGGAAAAGGCGAGGGATGCGTTGAACAAAGTGAACATGGGAGATACTTCAGAAGGAGCTGAAGACGATGACAAGGACGAGGAGGACTCGGACAACGACAGCGACGACGAGGCCGCAAAGGACGACGATCTTGCGGAATACGACATGGAGCACTACGACGACGAACCGAACGTT-CCCGGCGAGGAGGATGACGAACCTGCGGGCGGTGTTTT-CAGCAACATCAAA-GACCTT-GCATATCACACAGACAACAAC-GACGA-TCCATACCTAA-CGATCCCTGAAG-GAGAGC-TATCAG-ACGACG-AAGAGCGAGAAGAGCTGCAAATCC-TGC-CCACC-GA-CAGTG-TTCTGCTATCGGCAAG-AATAGAAG-ACGAGGT-CG-CACA-TCTGGA-AGT-ATACGTCTAC-GAAG-ATG-GTGCGGACAACTT-GTACGTTCACC-ACG-ATGTCATGCTG-CCCGCC-GTCCC-ACTTTGCGTAG-AATGGGTCGGAA-CAAAA-GT-CGGACAAAACGCCGCCTCGAGCGATGGAGGGAACTTCGCCGCCGT-GGGGACAATGGATCCGGACATTGAGCTGTGGGATCTGGATATCGTGGACTGTATGTACCCAAACGCAGTCCTC	742 	\|\|\|\| \|\|\|  \|\|\| \|\|\|\|\|\|  \|\|\|\|\|\|\|\|\|\|\|\|\|\| \|\|\|\|\|\|\|\|\|\|\|\|\|\|\|\|\|\|\|\| \|\|\| \|\|\|\|\|\|\|\|\|\|\|\| \|\|\|\|\|\|\|\|\|\|\|\|\|\|\|\|\|\|\|\|\|\|\|\|\|\|\|\|\|\|\|\|\|\|\|\|\|\|\|\|\|\|\|\|\|\|\|\|\|\|\|\|\|\|\|\|\|\|\|\|\|\|\|\|\|\|\|\|\|\|\|\|\|\|\|\|\|\|\|\|\|\|\|\|\| \|\|\|\|\|\|\|\|\|\|\|\|\|\|\|\|\|\|\|\|\|\|\|\|\|\|\|\|\|\|\|\|\|\|\|\|\|\|\|\|\|\|\|\|\|\|\|\|\|\|\|\|\|\|\|\|\|\|\|\|\|\|\|\|\|\|\|\|\|\|\|\|\|\|\| \|\|\|\|\|\|\|\|\|\|\|\|\|\|\|\|\|  \|\|\|\|\|\|\|\|\|\|\|\|\|\|\|\|\|\|\|\|\|\|\|\|\|\|\|\|\|\|\|\|\|\|\|\|\|\|\|\|\|\|\|\|\|\|\|\|\|\|\|\|\|\|\| \|\|\|\|\|\|\|\|\|\|\| \|\|\|\|\|\|\|\|\|\|\|\|\|  \|\|\|\|\|\|\|\|\| \| \|\|\|\|\|\|\|\|\|\|\|\|\| \|\|\|\|\|\| \|\| \|\|\|\|\|\|\|\|\| \|\| \|\|  \| \|\|\|\|  \|\|\|\|\|\|\|\|\|   \|\|\|\|\|\|\|\|\|\|\|  \|\|\|\|\|  \|\|\|\|\|  \|\|\|\|\|  \|\|\|\|\|\|\|\|\|\|\|\|\|\|\|\|  \| \|\|\|\| \|\|\| \|\|\|\|\| \|\|   \|\|\| \|\|\|\|\|\|\|\|\| \|\|\| \|\| \|\|\|\| \|\|\| \|\|\|\| \|\| \|  \|\|\|\| \|\|\|\|\|  \| \| \|\| \| \|\|\| \| \|\|   \|\|\| \|\|\|\|\|\|\|\|\|\|\|\|\| \|\|\| \|\|\| \|\|\| \|\|\| \|\|\|\|\|  \|\|\|\| \|\|\|\|\|\| \| \|\|  \|\|\|\|\|\|\| \|\|\| \|\|\|\|\|\|\|\|\|\| \| \|\| \|  \|\| \|\|\|\|\|\|\|  \|\|\|\| \|\|\|\|    \| \|\|\|\|\|\|\|\|   \|\| \|\|\| \|    \|\|\| \|\|  \|\|\|\|\|\|\|\|\|\| \|\|\|\|\|\| \|\|\|\|\|\| \| \|\|\|   \|  \| \|\|\|\| \|\|\|\| \|\|\|\|\| \|\|\|\|\|\|\|\|\|\| 	TATGCATCNNAGCGGACCGCNCTGGGTCCCTCGAGGNCATGCGGCGGCCTTTCCCACTAAGATACCAAATGGACNACGACGAATACGCTCGCATTGCGAAACTGGCGAAGCTCGAGCTCGACGATGCAAAGGTAGATTTGGAAAAGGCGAGGGATGCGTTNAACAAAGTGAACATGGGAGATACTTCAGAAGGAGCTGAAGACGATGACAAGGACGAGGAGGACTCGGACAACGACNGCGACGACGAGGCCGCANNGGACGACGATCTTGCGGAATACGACATGGAGCACTACGACGACGAACCGAACGTTNCCCGGCGAGGANGATGACGAACCTGNNGGCGGTGTTCTNCAGCAACATCAAAAGACCTTGGCNTATCACACANACNACNNCCGACGTNTCCATACCTTNNCGATCCCTGAANNGAGAGGNTATCANCACGACNAAAGAGCGAGAAGAGCTNNANATCCCTGCTCCACCNGAANCGTGTTTCTGCTATNGGCNAGGAATATAAGGACGAAGTTCNTCACANTCTGGTNANTNATTCNTCTCCCGANNNATGCGTGCGGACAACTTTGTAAGTTNACCCACGGATGTCNAGCTGNCCCGCCCGNCCNNACTTTGCNTAGNAATGGGTCGGGAGCAGANNGTCCGGACAAGNCGCCNCCTCTGNNGNTGGAGGGANTATC-CCGNCNNNNGGGGCATGGGATCCGGACCTTGAGCNGTGGGANCNGGANNNCTNGTACTGGATGT-CCCAA-CGCAGTCCTC ```      ``` 5	ATGACTTCTATGATCACAGCGACCGCATGGGTCCCTCGAGGCCATGCGGCGGCCTTTCCCACGAAGTACCAAATGGACGACGACGAATACGCTCGCATTGCGAAACTGGCGAAGCTCGAGCTCGACGATGCAAAGGTAGATTTGGAAAAGGCGAGGGATGCGTTGAACAAAGTGAACATGGGAGATACTTCAGAAGGAGCTGAAGACGATGACAAGGACGAGGAGGACTCGGACAACGACAGCGACGACGAGGCCGCAAAGGACGACGATCTTGCGGAATACGACATGGAGCACTACGACGACGAACCGAACGTTCCCGGCGAGGAGGATGACGAACCTGCGGGCGGTGTTTTCAGCAACATCAAAGACCTTGCATATCACACAGACAACAACGACGATCCATACCTAACGATCCCTGAAGGAGAGCTATCAGACGACGAAGAGCGAGAAGAGCTGCAAATCCTGCCCACCGACAGTGTTCTGCTATCGGCAAGAATAGAAGACGAGGTCGCACATCTGGAAGTATACGTCTACGAAGATGGTGCGGACAACTTGTACGTTCACCACGATGTCATGCTGCCCGCCGTCCCACTTTGCGTAGAATGGGTCGGAACAAAAGTCGGACAAAACGCCGCCTCGAGCGATGGAGGGAACTTCGCCGCCGTGGGGACAATGGATCCGGACATTGAGCTGTGGGATCTGGATATCGTGGACTGTATGTACCCAAACGCAGTCCTCGGACAAAACTCGCAAGGCGGAGGTATGGACATGCCCGAACCAGTGCTCAAGAAAAAGAAGAAAAAGACCAAGAAGTCAAACGACGCCT	830 	\|\|\|\|\|\|\|\|\|\|\|\|\|\|\|\|\| \|\|\|\|\|\|\|\|\|\|\|\|\|\|\|\|\|\|\|\|\|\|\|\|\|\|\|\|\|\|\|\|\|\|\|\|\|\|\|\|\|\|\|\|\|\|\|\|\|\|\|\|\|\|\|\|\|\|\|\|\|\|\|\|\|\|\|\|\|\|\|\|\|\|\|\|\|\|\|\|\|\|\|\|\|\|\|\|\|\|\|\|\|\|\|\|\|\|\|\|\|\|\|\|\|\|\|\|\|\|\|\|\|\|\|\|\|\|\|\|\|\|\|\|\|\|\|\|\|\|\|\|\|\|\|\|\|\|\|\|\|\|\|\|\|\|\|\|\|\|\|\|\|\|\|\|\|\|\|\|\|\|\|\|\|\|\|\|\|\|\|\|\|\|\|\|\|\|\|\|\|\|\|\|\|\|\|\|\|\|\|\|\|\|\|\|\|\|\|\|\|\|\|\|\|\|\|\|\|\|\|\|\|\|\|\|\|\|\|\|\|\|\|\|\|\|\|\|\|\|\|\|\|\|\|\|\|\|\|\|\|\|\|\|\|\|\|\|\|\|\|\|\|\|\|\|\|\|\|\|\|\|\|\|\|\|\|\|\|\|\|\|\|\|\|\|\|\|\|\|\|\|\|\|\|\|\|\|\|\|\|\|\|\|\|\|\|\|\|\|\|\|\|\|\|\|\|\|\|\|\|\|\|\|\|\|\|\|\|\|\|\|\|\|\|\|\|\|\|\|\|\|\|\|\|\|\|\|\|\|\|\|\|\|\|\|\|\|\|\|\|\|\|\|\|\|\|\|\|\|\|\|\|\|\|\|\|\|\|\|\|\|\|\|\|\|\|\|\|\|\|\|\|\|\|\|\|\|\|\|\|\|\|\|\|\|\|\|\|\|\|\|\|\|\|\|\|\|\|\|\|\|\|\|\|\|\|\|\|\|\|\|\|\|\|\|\|\|\|\|\|\|\|\|\|\|\|\|\|\|\|\|\|\|\|\|\|\|\|\|\|\|\|\|\|\|\|\|\|\|\|\|\|\|\|\|\|\|\|\|\|\|\|\|\|\|\|\|\|\|\|\|\|\|\|\|\|\|\|\|\|\|\|\|\|\|\|\|\|\|\|\|\|\|\|\|\|\|\|\|\|\|\|\|\|\|\|\|\|\|\|\|\|\|\|\|\|\|\|\|\|\|\|\|\|\|\|\|\|\|\|\|\|\|\|\|\|\|\|\|\|\|\|\|\|\|\|\|\|\|\|\|\|\|\|\|\|\|\|\|\|\|\|\|\|\|\|\|\|\|\|\|\|\|\|\|\|\|\|\|\|\|\|\|\|\|\|\|\|\|\|\|\|\|\|\|\|\|\|\|\|\|\|\|\|\|\|\|\|\|\|\|\|\|\|\|\|\|\|\|\|\|\|\|\|\|\|\|\|\|\|\|\|\|\|\|\|\|\|\|\|\|\|\|\|\|\|\|\|\|\|\|\|\|\|\|\|\|\|\|\|\|\|\|\|\|\|\|\|\|\|\|\|\|\|\|\|\|\|\|\|\|\|\|\|\|\|\|\|\|\|\|\|\|\|\|\|\|\|\|\|\|\|\|\|\|\|\|\|\|\|\|\|\| \|\|\|\|\|\|\|\|\| \|\|\|\|\|\|\|\|\|\|\|\|\|\|\|\|\|\|\|\|  \|\|\|\|\|\|\|\|\|\|\|\| \|\|\|\|\| \|\| \|\|\|\|\| \|\|\| \| \|\| \|\|\| \| \|\|\|\| \|\|\| 	ATGACTTCTATGATCACNGCGACCGCATGGGTCCCTCGAGGCCATGCGGCGGCCTTTCCCACGAAGTACCAAATGGACGACGACGAATACGCTCGCATTGCGAAACTGGCGAAGCTCGAGCTCGACGATGCAAAGGTAGATTTGGAAAAGGCGAGGGATGCGTTGAACAAAGTGAACATGGGAGATACTTCAGAAGGAGCTGAAGACGATGACAAGGACGAGGAGGACTCGGACAACGACAGCGACGACGAGGCCGCAAAGGACGACGATCTTGCGGAATACGACATGGAGCACTACGACGACGAACCGAACGTTCCCGGCGAGGAGGATGACGAACCTGCGGGCGGTGTTTTCAGCAACATCAAAGACCTTGCATATCACACAGACAACAACGACGATCCATACCTAACGATCCCTGAAGGAGAGCTATCAGACGACGAAGAGCGAGAAGAGCTGCAAATCCTGCCCACCGACAGTGTTCTGCTATCGGCAAGAATAGAAGACGAGGTCGCACATCTGGAAGTATACGTCTACGAAGATGGTGCGGACAACTTGTACGTTCACCACGATGTCATGCTGCCCGCCGTCCCACTTTGCGTAGAATGGGTCGGAACAAAAGTCGGACAAAACGCCGCCTCGAGCGATGGAGGGAACTTCGCCGCCGTGGGGACAATGGATCCGGACATTGAGCTGTGGGATCTGGATATCGTGGACTGTATGTACCCAAACGCAGTCCTCGGAC-AAACTCGCANGGCGGAGGTATGGACATGCCN-AACCAGTGCTCA-GAAAA-GA-GAAAA-GACNA-GA-GTCNA-CGACNCCT ```      ``` 120	TCGAGCTCGACGATGCAAAGGTAGATTTGGAAAAGGCGAGGGATGCGTTGAACAAAGTGAACATGGGAGATACTTCAGAAGGAGCTGAAGACGATGACAAGGACGAGGAGGACTCGGACAACGACAGCGACGACGAGGCCGCAAAGGACG-ACGATCTTGCGGAATACGACATGGAGCACTACGACGACGAACCGAACGTTCCCGGCGAGGAGGATGACGAACCTGCGGGCGGTGTTTTCAGCAAC-ATCAAAGACCTTGCATATCACACAGACAACA-ACGACGATCC-ATACCTA-ACGAT-CCCTG-AAGG-AGAGC-TATCAGACGACGAAGAGCGAGAA-GAGCT-GCAAAT-CCTGCCCACC-GACAGTGTT-CTGCTATCGGCAAG-A-ATAG-AAG-ACGAGG-T-CGCACA-TCTGGAAGTATACGTCTACG-AAGATGGTGCG-GACAACTT-GTA-CGTTCACCACGATGT-CATGCTGCCCG-CCGTCCCACTTTGCGTAGAA-TGGGTCGGAACAAAAG	623 	\|\|\| \|\| \|\|\|\|\|\|\|\|\|\|\|\|\|\|\|\|\|\|\|\|\|\| \|\|\|\|\|\|\|\|\|\|\|\|\|\|\|\|\|\|\|\|\|\|\|\| \|\|\|\| \|\|\|\|\|\|\|\|\|\|\|\|\|\|\|\|\|\|\|\|\|\|\|\|\|\|\|\|\| \|\|\|\|\|\|\|\|  \|\|\|\| \|\|\|\|\|\|\|\|\|\| \|\|\| \| \|\|\| \|\|\|\|\|\|\|\| \|\|\|\|\|\|\|\|\|\|\|\|\|\|\| \|\|\|\|\|\|\|\|\|\|\|\|\|\|\|\|\|\|\|\|\|\|\|\|\| \|\|\|\|\|\|\|\|\| \|\|\|\|\|\|\|\|\|\|\|\|\|\|\|\|\|\|\|\|\|\|\|\|\|\|\|\|\|\|\| \|\|\| \|\|\|\|\|\| \|\|\|\|\|\|\|\|\| \|\|\|\|\|\| \|\|\|\|\|\|\|\|\|\|\|\|\| \|\|\|\|\|\|\|\|\|\| \|\| \|\|  \|\|\|\|\|\|\|\|\|  \|\|\|\|\|\|  \|\| \|\| \|\|\|\|\| \|\|\|\| \|\|\|\|  \| \|\| \|\| \|\|\|\|\|\| \| \| \|\|\|\| \|\|\|\|\| \|\|\| \|\| \|\|\|\|\|\| \|\|\| \|\|\|\|\|\|\|\|\| \|\|\|\|\| \|\| \|\|\| \| \| \|\|\|  \|\|  \|\|\|\|\|\| \| \| \| \|\| \|\| \|\|\|\|\|\|\|\|\|\|\|\|\|\|\|\|  \|\| \|\|\|\|\|\|\|  \|\| \|\|\|\|\| \| \| \| \| \|\|\| \|  \| \|\| \|\|\|\|  \|\|\|\|  \|\|\|\| \| \|\|\|\|\|\|\| \|  \|\| \|\|\|\|  \|\|\|\|\|\|\|\|\|\| 	TCGNGCGCGACGATGCAAAGGTAGATTTGNAAAAGGCGAGGGATGCGTTGAACANAGTGNACATGGGAGATACTTCAGAAGGAGCTGAANACGATGACN-GGACNAGGAGGACTCNGACTANGACTGCGACGACTAGGCCGCAAAGGACGTACGATCTTGCGGAATACGACATGGANCACTACGACNACGAACCGAACGTTCCCGGCGAGGAGGATGANGAANCTGCGGNCGGTGTTTTNAGCAACNATCAAAGACCTTGTATATCACACATACCACNNACGACGATCNNATACCTTNACNATTCCCTGCAAGGCAGAGTNTNTCNGATGACGAANANCNAGAAAGAGCTTGCATATNCCTGCCNACCCGACAGTGTTNCTGCTNTCNGCANGTANATANCAATCACGAGGNTNCTC-CATTCNGGAAGTATACGTCTACNTAANATGGTGCCNGANAACTTNGNANCNTNCACNANNANGTACATGNNGCCCCCCCGTNCTACTTTGCCTTNAACTGGGCNGGAACAAAAG ```      ``` 10	TTCTATGA-TCACAGCGACCGCATGGG-TCCCTCGAGGCCA-TGCGGCGGCCTTTCCCACGAAGTACCAAATGGACGACGACGAATACGCTCGCATTGCGAAACTGGCGAAGCTCGAGCTCGACGATGCAAAGGTAGATTTGGAAAAGGCGAGGGATGCGTTGAACAAAGTGAACATGGGAGATACTTCAGAAGGAGCTGAAGACGATGACAAGGACGAGGAGGACTCGGACAACGACAGCGACGACGAGGCCGCAAAGGACGACG-ATCTTGCGGAATACGACATGGAGCA-CTACGACGACGAACCGAACGTTCC-CGGCGAGGAGGATGACG-AACCTGCGGGCGGTGTTTTC-AGCAACATCAAAGACCTTGCATATCACA-CAG-ACAACAACGACGATCCATACCTAACGATCCCTGAA-GG-A-GAG-CTATCAGACGACGAAGAGC-GAGAAGAGCTGCAAAT-CCTG-CCCACCGACAGTGTT-CTGCTATCG-GCAAGAATAGAAGACGAGGTCG-CACATC-TGGAAGTATACGTCTACGAAGATGGTGC-GGACAACTTGTACGTTCACCACGATGTCATGCTGCCC-GCCGTC	592 	\|\|\|\| \|\|\| \|\|  \|\|\|\|\|\|\|\|\|\|\|\|\|\| \|\|\|\| \|\|\|\|\|\|\|\| \|\|\|\|\|\|\|\| \|\|\|\|\|\|\|\|\|\|\|\|\| \|\|\| \|\|\|\| \| \|\|\|\|\|\|\|\|\| \|\|\|\| \|\| \|\|\|\|\|\|\|\|\|\|\|\|\|\|  \|\|\|\|\|\|\|\|\|\|\|\|\|\|\|\|\|\|\|\| \|\|\|\|\| \|\|\|\|\|\|\|\|\|\|\|\|\|\|\|\|\|\|\|\|\|\|\|\|\| \|\|\|\| \|\|\|\| \|\| \|\|\| \|\|\|\|\|\|\|\|\|\|\|\|\|\|\|\|\|\|\|\|\|  \|\|\|\|\|\|\|\|\|\|\|\|\|\|\|\|\|\|\|\|\|\|\|\|\|\|\|\|\| \|\|\|\|\| \|\|\|\|\|\|\|\|\|\|\|\| \|\| \|\|\|\|\|\|\|\|\|\|\| \|\|\|\|\|\|\|\|\|\| \|\|\|\|\|  \|\|\|\|\|\|\| \|\| \|\|\|\| \|\|\|\|\| \|\|\|\|\|\|\|\|\|\| \|\|\|\|\|\|\|\|\| \|\| \|\|\|\| \|\|\|\|\|\|\|\|\|\| \|\|\|\|\|\|\|\|  \|\| \|\|\|\|\|\|\| \|\|\|\|\|\|\|\| \|\|\|\|\|\|\|\| \|\|\| \|\| \|\|\|\|\|\|\|\|\|\|\|\|\| \|\|\|\|\|\|\|\|\|\|\| \|  \|   \|\| \| \|\|\| \|\|\|\|\|\| \|\|  \| \|\| \|\|  \|\|\|\|\|\|\|\|\|\|\| \|\|   \|\|\|\| \|\|\|\| \|\|\|\| \| \|\|\| \| \|\|   \|\| \|\|\|\|\|\|\|\| \|\|\|\|\|\|\|\| \|\|   \|\|\|\|\|  \|\|\| \| \|\|\|\|\|\|\|\|\| \| \|\| \|\|\|\|\|\|\| \|\| \|\|\|\|\| \|\|\|\| \|\|\|  \| \|\|\|\|\| \|\| \|\| \|\|\|\| \| \|\|\|\| 	TTCTNTGACTCTNAGCGACCGCATGGGCTCCCNCGAGGCCANTGCGGCGGNCTTTCCCACGAAGAACCNAATGNANGACGACGAANACGCNCGNATTGCGAAACTGGCNNAGCTCGAGCTCGACGATGCANAGGTACATTTGGAAAAGGCGAGGGATGCGTTNAACANAGTGNACNTGGNAGATACTTCAGAAGGAGCTGANNACGATGACAAGGACGAGGAGGACTCGGACNACGACNGCGACGACGAGGNCGNAAAGGACGACGTATCTTGCGGANTACGANTTGGAGCACCTNCGACCACGAATCGAACGTTCCNCGGCGAGGANGANGACGAAACCTGCGGGNGGTGTTTTTNAGNAACATCANAGACCTTGTATATCACANCAGCACTACAACGACGATCCTTACCTAACGATTCNNGGNNGGGAAGAGCCTATCATACTNCNAANAGTNGAGAAGAGCTGNAACGNCCTGCCCCANCGACNGNGTTTCNGCNNNCGNGCAAGAATNGAAGACGANGTTNTCACATNNTGGNANTATACGTCTNCNAANATGGTGCNGGTCAACTNGTACNTTCNNCNCGATGNCAAGCNGCCCCGTCGTC ```      ``` 25	GACCGCATGGGTCCCTCGAGGCCATGCGGCGGCCTTTCCCACGAAGTACCAAATGGACGACGACGAATACGCTCGCATTGCGAAACTGGCGAAGCTCGAGCTCGACGATGCAAAGGTAGATTTGGAAAAGGCGAGGGATGCGTTGAACAAAGTGAACATGGGAGATACTTCAGAAGGAGCTGAAGACGATGACAAGGACGAGGAGGACTCGGACAACGACAGCGACGACGAGGCCGCAAAGGACGACGATCTTGCGGAATACGACATGGAGCACTACGACGACGAAC-CGAACGTTCCCGGCGAGG-AGGATGACGAA-CCTGCGGGCGGTGTTTTCAGCAACATCAAAGACCTTG-CATATCACA-CAG-ACAACAACGACGATCCATACC-TAAC-GATCCCTGAAGG-AGAGCTATC-AGACGACGAAGAGCGAG-AAGAG	457 	\|\|\| \|\|\|\|\|\|\|\|\|\|\|\|\|\|\|\|\|\|\|\|\|\|\|\|\|\|\|\| \|\|\|\|\|\|\|\|\| \| \| \|\|\|\|\|\|\|\|\|\|\|\|\|\|\|\|\|\|\|\|\|\|\| \|\|\|\|\|\|\|\|\|\|  \|\|\|\|\|\|\|\| \|\|\|\|\|\|\|\|\|\|\|\|\|\| \|\|\|\|\|\|\|\|\|\|\|\|\|\|\|\|\|\|\|\|\| \|\|\|\|\|\|\|\|\| \|\|\|\|\|\|\|\|\|\|\|\|\|\|\| \|\|\|\|\|\|\|\|\|\|\|\|\| \|\|\|\|\|\|\|\|\|\|\|\|\|\|\|\|\|\| \|\|\|\|\|\|\|\|\|\|\|\|\|\|\|\|\|\|\|\|\|\|\|\|\|\|\|\|\|\|\|\|\|\|\|\|\|\|\|\|\|\|\| \|\|\| \|\|\|\|\|\|\|\|\|\|\|\|\|\|\|\|\|\|\|\|\|\|\|\|\|\|\|\|\|\|\|\|\|\|\|\|\|\|\|\|\|\|\|\|\|\|\|\|\|\|  \|\|\|\|\|\|\|\|\|\|\|\|\|\|\|\|\|\| \|\|\|\|\|\|\|\|\|\|\| \|\|\| \| \|\|\|\|\|\|\| \|\|\|\|\|\| \|\|\|\|\|\|\|\|\|\|\| \| \|\| \|\|\|\|\|\|\|\|  \| \| \|\|\|\|  \|\|\|\|\|\|\|\|\|\|\|\|\|\|\| \|\|\|  \|\|\|\|\|\|\|\|\|\|\|\| \| \|\| \| \|\| \|\|\|\|\|\|\| \|\| \|\|\|\| \| \|\|\|\|\| 	GACAGCATGGGTCCCTCGAGGCCATGCGGCGGGCTTTCCCACNANGNACCAAATGGACGACGACGAATACNCTCGCATTGCNNAACTGGCGNAGCTCGAGCTCGACNATGCAAAGGTAGATTTGGAAANGGCGAGGGANGCGTTGAACAAAGTGNACATGGGAGATACNTCAGAAGGAGCTGAAGACNATGACAAGGACGAGGAGGACTCGGACAACGACAGCGACGACGACGCCNCAAAGGACGACGATCTTGCGGAATACGACATGGAGCACTACGACGACGAAANCGAACGTTCCCGGCGAGGAAGGATGACGAACCCTNCNGGCGGTGATTTCAGNAACATCAAAGANCNTGGCATATCACNNCNGTACAANGACGACGATCCATACCATAANNGATCCCTGAAGGTANAGNTNTCTAGACGACCAANAGCGNGGAAGAG ```      ``` 140	GTAGATTTGGAAAAGGCGAGGGATGCGTTGAACAAAGTGAACATGGGAGATACTTCAGAAGGAGCTGAAGACGATGACAAGGACGAGGAGGACTCGGACAACG-ACAGCGACGACGAGGCCGCAAAGGACGACGATCTTGC-GGAATACGACATGGAGCACTACGACGACGAACCGAACGTTCCCGGCGAGGAGGATGACGAACCTGCG-GGCGGTGTTTTCAGCAACATCAAAGACCTTGCATATCACACAGACAACAA-CGACGATCCATACC-TAACGATCCCTGAAGGAGAGCTATCAG-ACGACG-AAGAGCGAGAA-GAGCTGCAAATCCTGCCC-ACCGACAGTGTTCTGCTAT-C-GGCAAGAATAGAAGACGAGG-TCGCACATCTGGAAGTATACGTCTA-CGAA-GATGGTGCGG-ACAACTTG-TACG-TTCACC-ACGATGTC-ATGCTGCC-CGCCGTCCCAC-TTTGCGTAGAATGGGTCGGAACAAAAGTCGGACAAAACGCCGCCTCGAGCGATGGAGGGAACTTCGCCGCCGTGGGGACAATGGATCCGGACATTGAGCTGTGGGATCTGGATATCGTGGACTGTATGTACCCAAACGCAGTCCTCGGACAAA	749 	\|\|\|\|\|  \|\|\|\| \|\|\|\|\|\|\|\|\|\|\|\|\|\|\|\|\|\|\|\|\|\|\|\|\|\|\|\|\|\|\|\|\|\|\|\|\|\|\|\|\|\|\|\|\|\|\|\|\|\|\|\|\|\|\|\|\|\|\|\|\|\|\|\|\|\|\|\|\|\|\|\|\|\|\|\|\|\|\|\|\|\|\|\|\|\| \|\|\|\| \|\|\|\|\|\|\|\|\|\|\|\|\|\|\|\|\|\|\|\|\|\|\|\|\|\|\|\|\|\|\|\|\|\|\|\|\| \|\|\|\|\|\|\|\|\|\|\|\|\|\|\|\|\|\|\|\|\|\|\|\|\| \|\|\|\|\|\|\|\|\|\|\|\|\| \|\|\|\|\|\|\|\|\|\|\|\|\|\| \|\|\|\|\|\|\|\|\|\|   \|\|\|\|\| \|\|\|\| \|\|\| \|\|\|\|\|\| \|\|\|\|\|\|\|\|\| \|\|\|\|\|\|\|\|\|\| \|\|\|\|\|\|\| \|\|\|\|\|\|\|\|\|\|\|\|\|\| \|\|\| \|\|\|\| \|\|\|\|\|\|\|\|\|\| \|\|\|\|\|\|\| \|\|\|\|\|\| \|\| \|\|\|\|\|\|\|\| \|\|\|\|\|\|\| \|\|  \|\|\|\|\|\| \|\|\|\|\| \|\|\|\| \|\|\|\|\|\|\|  \| \|\|\| \|\|\|\| \|\|\|\|\|\|\|\|\|\|\| \|\|\|\|\|\|\|\|\|\|\|\|\|\|\|\|\|\|\|\|\|\|\|\|\| \| \|\| \|\| \|\| \|\|\|\| \|\|\|\| \|\|\| \|\|\|  \|\|\|\| \| \|\|\|\|\|\|\|\| \|\|\| \|\|\|\| \|\|\|\|\| \|\|\|\|  \|\|\|\|\|\|\|\| \|\| \|\|\|\|\|\|\|\|\| \|\|\|\|\|\|\| \|\| \| \|  \|\|\| \|\|\|\| \| \|\|\| \|\|\| \|\|\|\|\| \|\| \|\|\|\|\|\|\|\| \|\|\| \|\|\|\|\|\|\|\|\|\|\|\| \|\|\|\|\|\|\|\|\|\|\|\|\|\| \|\|\|\|\|\|  \|\|\|\| \| \|\| \|  \| \| \| \|\|\| \|\|\|\|\|\|\|\|\|\|\|\|\|\|\| 	GTAGAGCTGGAGAAGGCGAGGGATGCGTTGAACAAAGTGAACATGGGAGATACTTCAGAAGGAGCTGAAGACGATGACAAGGACGAGGAGGACTCGGANAACGNACAGCGACGACGAGGCCGCAAAGGACGACGATCTTGCGGGAATACGACATGGAGCACTACGACNACGAACCGAACGTNCCCGGCGAGGAGGANGACGAACCTGNNNGGCGGNGTTTNCAGNAACATCNAAGACCTTGNATATCACACANACAACAACCGACGATCCATACCTTAANGATCNCTGAAGGAGANCTATCAGTACGACGNAANAGCGAGAAAGAGCTGCNAANTCTGCCCTACCGANAGTGATCTGCTAANCGGGCNAGAACAGAAGACGAGGTTCGCACATCTGGAAGTATACGTCTAACNAANGAGGGNGCGGGACAANTTGNTACNNTTCANCNACGATGTCTATGNTGCCNCGCCGCCCCANNTTTGCGTANAANGGGTCGGAANAAAAGTCNGAAANANNGCCNCCTCNANCGANGGANGGAACNTCACCGCCGTGNGGAAAATGGATCCGGANATTGAGCTGTGGGA-CTGGATNCCGTGNANTGAANNTCCNC-AACNCAGTCCTCGGACAAA ``` |
| 5 | ggggacaagtttgtacaaaaaagcaggcttgatgGCGGACGCAGACAAGAAGAC, ggggaccactttgtacaagaaagctgggtcAGGCTTACACCTGCCCATAT | Mycgr3\|109954  JGI  Ensembl | KOG0555 Asparaginyl-tRNA synthetase; | ATP binding; aminoacyl-tRNA ligase activity;  asparagine-tRNA ligase activity;  asparaginyl-tRNA aminoacylation;  aspartate-tRNA ligase activity;  aspartyl-tRNA aminoacylation;  cytoplasm;  nucleic acid binding;  nucleotide binding;  tRNA aminoacylation for protein translation;  translation; | 100 % |  |  |
| 5 | ggggacaagtttgtacaaaaaagcaggcttgatgTCGTCCTTCTTCACTCTTCC, ggggaccactttgtacaagaaagctgggtcATCGAAACCTTCGAAGCCGT | Mycgr3\|110058  JGI  Ensembl | KOG0299 U3 snoRNP-associated protein (contains WD40 repeats); |  | 100 % |  | ``` 34	ATGTCGTCCTTCTTCACTCTTCCCGCCTCACAGCGGAAGAGGAAGCGACCCGAAAATAACTCCACCGCATCACGATCAAAATCACAGAAAGAAGGCAAATCTCGCCCGTCACGAGACGAATCCATTTCCGGCAGCGATGTGTCCGACGATGATGTTCCCCGCGACGATGGACCAGCGCAATCCTCCTCTGACGAAGACGAGGAGTTTTTCGACGAAGATCCAGCCGCCAAACGAATTCGCCTAGCCGAGCAATATCTCGCCAATACACGAAAGGAAGTTGAGGAAGAAGTCGGATTCGATGCGAAAGATGTGGATCAGGAGAATTTGCGACGCCGCATGGGTGACAGATTGAAGGAGGATACTGCCGAGACGAAGGGAAAGCTGTACCGCTGGATTGCTTCGGACTTCGACTGGAGTAAAGCAACGAGATCACAATTTCGACACCACCAAAAGCCACTGACTGGCGTCGCTGTTGCCGCTGGCTTCGCATACACAGTCGGCAAGGATTCAATCCTCGTCAAGTGGAGATTGGCTCCCCCTCCCGCCGCCGACGACCCTCCGAACCGATCTCGAAAGCCCACGAAACTCCTCGTCTCCCAACCTTTCGGCAAACCGACCGCCAATGCAAATGCAATCCGCCACACTTCCGCAGTCCGATGTATAGCTGCTTCACAAGACGGCAAATTCGTTGCCACCGGCGCGACAGATTCATCCCTGATCGTATGGGAAGCTCAAACACTCAAGCCGCTCAAACTCTTCCCTCACCACCGCGATGCCGTCCTCTCTCTCTCCTTTCGTCGCGGCACAAATCAGCTCTTCTCCGCCTCTCGCGATCGTA	871 	\|\|\|\|\|\|\|\|\|\|\|\|\|\|\|\|\|\|\|\|\|\|\|\|\|\|\|\|\|\|\|\|\|\|\|\|\|\|\|\|\|\|\|\|\|\|\|\|\|\|\|\|\|\|\|\|\|\|\|\|\|\|\|\|\|\|\|\|\|\|\|\|\|\|\|\|\|\|\|\|\|\|\|\|\|\|\|\|\|\|\|\|\|\|\|\|\|\|\|\|\|\|\|\|\|\|\|\|\|\|\|\|\|\|\|\|\|\|\|\|\|\|\|\|\|\|\|\|\|\|\|\|\|\|\|\|\|\|\|\|\|\|\|\|\|\|\|\|\|\|\|\|\|\|\|\|\|\|\|\|\|\|\|\|\|\|\|\|\|\|\|\|\|\|\|\|\|\|\|\|\|\|\|\|\|\|\|\|\|\|\|\|\|\|\|\|\|\|\|\|\|\|\|\|\|\|\|\|\|\|\|\|\|\|\|\|\|\|\|\|\|\|\|\|\|\|\|\|\|\|\|\|\|\|\|\|\|\|\|\|\|\|\|\|\|\|\|\|\|\|\|\|\|\|\|\|\|\|\|\|\|\|\|\|\|\|\|\|\|\|\|\|\|\|\|\|\|\|\|\|\|\|\|\|\|\|\|\|\|\|\|\|\|\|\|\|\|\|\|\|\|\|\|\|\|\|\|\|\|\|\|\|\|\|\|\|\|\|\|\|\|\|\|\|\|\|\|\|\|\|\|\|\|\|\|\|\|\|\|\|\|\|\|\|\|\|\|\|\|\|\|\|\|\|\|\|\|\|\|\|\|\|\|\|\|\|\|\|\|\|\|\|\|\|\|\|\|\|\|\|\|\|\|\|\|\|\|\|\|\|\|\|\|\|\|\|\|\|\|\|\|\|\|\|\|\|\|\|\|\|\|\|\|\|\|\|\|\|\|\|\|\|\|\|\|\|\|\|\|\|\|\|\|\|\|\|\|\|\|\|\|\|\|\|\|\|\|\|\|\|\|\|\|\|\|\|\|\|\|\|\|\|\|\|\|\|\|\|\|\|\|\|\|\|\|\|\|\|\|\|\|\|\|\|\|\|\|\|\|\|\|\|\|\|\|\|\|\|\|\|\|\|\|\|\|\|\|\|\|\|\|\|\|\|\|\|\|\|\|\|\|\|\|\|\|\|\|\|\|\|\|\|\|\|\|\|\|\|\|\|\|\|\|\|\|\|\|\|\|\|\|\|\|\|\|\|\|\|\|\|\|\|\|\|\|\|\|\|\|\|\|\|\|\|\|\|\|\|\|\|\|\|\|\|\|\|\|\|\|\|\|\|\|\|\|\|\|\|\|\|\|\|\|\|\|\|\|\|\|\|\|\|\|\|\|\|\|\|\|\|\|\|\|\|\|\|\|\|\|\|\|\|\|\|\|\|\|\|\|\|\|\|\|\|\|\|\|\|\|\|\|\|\|\|\|\|\|\|\|\|\|\|\|\|\|\|\|\|\|\|\|\|\|\|\|\|\|\|\|\|\| \|\|\|\|\|\|\|\|\|\|\|\|\| \|\|\|\|\|\|\|\|\|\|\|\|\|\|\|\| \|\|\|\|\|\|\|\|\|\|\|\|\|\|\|\|\|\|\|\|\|\|\|\|\|\|\|\|\|\|\|\| \|\|\|\|\|\| \|\|\|\|\|\|\| \|\|\| \|\|\|\|\| \|\|\|\|\|\|\|\|\|\|\|\|\|\|\|\|\|\|\|\| \|\| \|\|\|\|\|\|\|\|\| \|\|\| \|\|\|\|\|\|\|\|\| \| \|\|\|\|\|\|\|\|\| \|\|\| \|\|\| 	ATGTCGTCCTTCTTCACTCTTCCCGCCTCACAGCGGAAGAGGAAGCGACCCGAAAATAACTCCACCGCATCACGATCAAAATCACAGAAAGAAGGCAAATCTCGCCCGTCACGAGACGAATCCATTTCCGGCAGCGATGTGTCCGACGATGATGTTCCCCGCGACGATGGACCAGCGCAATCCTCCTCTGACGAAGACGAGGAGTTTTTCGACGAAGATCCAGCCGCCAAACGAATTCGCCTAGCCGAGCAATATCTCGCCAATACACGAAAGGAAGTTGAGGAAGAAGTCGGATTCGATGCGAAAGATGTGGATCAGGAGAATTTGCGACGCCGCATGGGTGACAGATTGAAGGAGGATACTGCCGAGACGAAGGGAAAGCTGTACCGCTGGATTGCTTCGGACTTCGACTGGAGTAAAGCAACGAGATCACAATTTCGACACCACCAAAAGCCACTGACTGGCGTCGCTGTTGCCGCTGGCTTCGCATACACAGTCGGCAAGGATTCAATCCTCGTCAAGTGGAGATTGGCTCCCCCTCCCGCCGCCGACGACCCTCCGAACCGATCTCGAAAGCCCACGAAACTCCTCGTCTCCCAACCTTTCGGCAAACCGACCGCCAATGCAAATGCAATCCGCCACACTTCCGCAGTCCGATGTATAGCTGCTTCACAAGACGGC-AATTCGTTGCCACNGGCGCGACAGATTCAT-CCTGATCGTATGGGAAGCTCAAACACTCAAGCNGCTCAA-CTCTTCC-TCA-CACCGNGATGCCGTCCTCTCTCTCTCNTT-CGTCGCGGC-CAA-TCAGCTCTT-T-CGCCTCTCGNGAT-GTA ``` |
| 5 | ggggacaagtttgtacaaaaaagcaggcttgatgGCCACGAACGGCGACTACCC, ggggaccactttgtacaagaaagctgggtcGCTCGACTGTGGGTCGAGCG | Mycgr3\|110080  JGI  Ensembl | KOG0466 Translation initiation factor 2, gamma subunit (eIF-2gamma; | GTP binding; GTPase activity;  protein-synthesizing GTPase activity;  translation; | 0 % |  |  |
| 5 | ggggacaagtttgtacaaaaaagcaggcttgatgGGTCTCCTCCAGGAAGTCCT, ggggaccactttgtacaagaaagctgggtcGTTCTTCTCCTCCTTCTCCT | Mycgr3\|110231  JGI  Ensembl | KOG0684 Cytochrome P450; | electron transport; heme binding;  iron ion binding;  monooxygenase activity;  sterol 14-demethylase activity; | 100 % |  | ``` 1	ATGGGTCTCCTCCAGGAAGTCCTCGCGCAGTTCGACGCGCAATTCGGCCAGACCAGCCTCTGGAAACTTGTCGGACTTGGATTCCTCGCCTTCAGCACGCTCGCCATCCTCCTCAATGTCCTCTCACAACTTCTCTTCCGTGGCAAGTTGTCCGATCCGCCACTCGTATTCCACTGGGTGCCCTTCATCGGAAGCACCATCACCTACGGCATCGACCCATACAAGTTCTTCTTCTCCTGTCGGGAAAAGTATGGAGATGTCTTTACATTCATCCTGCTGGGAAAGAAGACGACGGTGTGCTTGGGCACCAAGGGCAATGATTTTATTTTGAATGGAAAACTGAAGGACGTCAACGCGGAGGAGATATACAGCCCGCTGACCACTCCTGTCTTTGGCAAGGATG	403 	\|\|\|\|\|\|\|\|\|\|\|\|\|\|\|\|\|\|\|\|\|\|\|\|\|\|\|\|\|\|\|\|\|\|\|\|\|\|\|\|\|\|\|\|\|\|\|\|\|\|\|\|\|\|\|\|\|\|\|\|\|\|\|\|\|\|\|\|\|\|\|\|\|\|\|\|\|\|\|\|\|\|\|\|\|\|\|\|\|\|\|\|\|\|\|\|\|\|\|\|\|\|\|\|\|\|\|\|\|\|\|\|\|\|\|\|\|\|\|\|\|\|\|\|\|\|\|\|\|\|\|\|\|\|\|\|\|\|\|\|\|\|\|\|\|\|\|\|\|\|\|\|\|\|\|\|\|\|\|\|\|\|\|\|\|\|\|\|\|\|\|\|\|\|\|\|\|\|\|\|\|\|\|\|\|\|\|\|\|\|\|\|\|\|\|\|\|\|\|\|\|\|\|\|\|\|\|\|\|\|\|\|\|\|\|\|\|\|\|\|\|\|\|\|\|\|\|\|\|\|\|\|\|\|\|\|\|\|\|\|\|\|\|\|\|\|\|\|\|\|\|\|\|\|\|\|\|\|\|\|\|\|\|\|\|\|\|\|\|\|\|\|\|\|\|\|\|\|\|\|\|\|\|\|\|\|\|\|\|\|\|\|\|\|\|\|\|\|\|\|\|\|\|\|\|\|\|\|\|\|\|\|\|\|\|\|\|\|\|\|\|\|\|\|\|\|\|\|\|\|\|\|\|\|\|\|\|\|\|\|\|\|\|\|\|\|\|\|\|\|\|\|\|\|\|\|\|\|\| \|\| \|\|\|\|\|\|\|\|\|\|\|\|\|\|\|\|\|\|\|\|\|\|\|\|\|\|\|\|\|\|\|\| \|\| \|\|\|\| 	ATGGGTCTCCTCCAGGAAGTCCTCGCGCAGTTCGACGCGCAATTCGGCCAGACCAGCCTCTGGAAACTTGTCGGACTTGGATTCCTCGCCTTCAGCACGCTCGCCATCCTCCTCAATGTCCTCTCACAACTTCTCTTCCGTGGCAAGTTGTCCGATCCGCCACTCGTATTCCACTGGGTGCCCTTCATCGGAAGCACCATCACCTACGGCATCGACCCATACAAGTTCTTCTTCTCCTGTCGGGAAAAGTATGGAGATGTCTTTACATTCATCCTGCTGGGAAAGAAGACGACGGTGTGCTTGGGCACCAAGGGCAATGATTTTATTTTGAATGGAAAACTGAAGGACGTCAACGCGGANGANATATACAGCCCGCTGACCACTCCTGTCTTTGGNAANGATG ``` |
| 5 | ggggacaagtttgtacaaaaaagcaggcttgatgGCTCTCCCAGGACTGTCTCT, ggggaccactttgtacaagaaagctgggtcAGCCACCAAATCCGCCACAG | Mycgr3\|110776  JGI  Ensembl | KOG2749 mRNA cleavage and polyadenylation factor IA/II complex, subunit CLP1; |  | 100 % |  | ``` 195	CCCAGG-CTTGAATCTCCCACAGACATCCTCCACCCCCTTGGCCGATTCAAAAAACACCATACAAGCATCAAGATTAGAGGACCTACCA-TCGCGGACAGAATGGCGATTCGAGGTAGCGTTCTCTCAACACTACAG-CATCAAGCTGGAGTCCGGCCATGCGGAAATGTTTGGTGTTGAGCTCGCCCTCAAACAAGTCTACACGTTCTCCGGATTCAAGGGCGCAATCTTCACATGG-CAAGGCTGTCAGCTCGAGGTGACGGGAAATGCAGA-GAGTGAATACATTGGACAAGAAACCGACTACGCAGTGGAGTGGCTCAATGTGCACGGCATGCTTGATACGCTTCGATCACAGAACCCAACAGATGGCC-CAAGAGTGCTTGTGGTCGGCCCTGACTTTGTTG-GCAAGACCAGTCTGGTGCG-AACTCTTGCAGCGTGGGAGGCACGCAGC-GGATACACGCCCACAGTGCTGAATCTTGATCCTCGAGAAGGCATTCTGGCACCTCCGAGTAGTCTGACAGCGG-TCACTGTT-GATTCGCAGT-TGG-AAGTAGATAGTGGATATGGCA-TTCCG-TCTTCGTCGGGCCCC-ACACTTTCACCTGTGA-GAACGCCGCTGA	805 	\|\|\|\|\|\| \|\|\|\|\|\| \|\|\|\|\|\|\|\| \|\|\|\|\|\|\|\| \|\|\|\|\|\|\|\|\|\| \|\|\|\|\|\|\|\|\|\|\|\| \|\|\|\|\|\|\|\|\|\|\|\|\|\|\|\|\|\|\|\|\|\|\|\|\|\|\|\|\|\|\|\|\| \|\|\|\|\|\|\|\|\|\|\|\|\|\|\|\|\|\|\|\|\|\|\|\|\|\|\|\| \|\|\|\|\|\|\|\|\|\|\|\|\|\|\|\|\|\|  \|\|\|\|\|\|\|\|\|\|\|\|\|\|\|\|\|\|\|\|\|\|\|\|\|\|\|\|\|\|\|\|\|\|\|\|\| \|\|\|\|\|\|\|\|\|\|\|\|\|\|\|\|\|\|\|\|\|\|\|\|\|\|\|\|\|\|\|\|\|\|\|\|\| \|\|\|\|\|\|\| \|\|\|\|\|\|\|\|\|\|\|\|\|\|\| \|\|\|\|\|\|\|\| \|\|\|\|\|\|\|\|\|\|\|\|\|  \|\|\|\|\|\|\|\|\| \| \|\|\|\|\|\|\|\|\| \|\|\|\| \|\|\|\|\|\|\|\|\|\|\|\|\|\|\|\|\|\|\|\| \|\|\|\|\|\|\|\|\| \|\| \|\|\|\|\|\|\|\|\|\|\|\|\|\|\|\|\|\|\| \|\|\|\| \|\|\|\|\|\|\|\|\|\|\| \| \|\|\|\|\|\|\|\| \| \| \|\|\| \|\|\| \|\|\|\|\|\|\|\|\|  \| \|\|\|\|\|\|\| \|\| \|\|\|\|\|\| \|\|\| \|\|\|\| \| \| \|\|\|\|\| \|\|\|\|\|\|\| \|\|\|\| \|\|\| \|\|    \|\|  \| \|\| \|\|\|\|\|\|\|\|\| \| \|\| \| \|\|\| \|\|\|  \|\|\|\|\|\| \| \| \| \| \| \| \|\| \| \|\|\|\|\|\|\|\| \|\|\|\|\| \| \|\|\|\| \| \|\|\|\|\|\|\|\|   \|\|\| \|\|\|\|\|  \|\|\|\|\|\|\| \| \|\| \| \|\|\|\|\| \|\|\| \|\| \|\|\|\|  \|\| \| \|\|\|\|\| \|\|\|\|\| \|\| \|\|\|\|\|\|\| \|\|\|\| 	CCCAGGNCTTGAANCTCCCACANACATCCTCNACCCCCTTGGTCGATTCAAAAAATACCATACAAGCATCAAGATTAGAGGACCTACCANTCGCGGACAGAATGGCGATTCGAGGTAGNGTTCTCTCAACACTACAGNNATCAAGCTGGAGTCCGGCCATGCGGAAATGTTTGGTGNTGAGCTCGCCCTCAAACAAGTCTACACGTTCTCCGGANTCAAGGGNGCAATCTTCACATGGTCAAGGCTGNCAGCTCGAGGTGANNGGAAATGCANAGGAGTGAATANATTGNACAAGAAACCGACTACGCAGCGGAGTGGCTNAAAGTGCACGGCATGCTTGATANGCTTNGATCACAGAACNCNACAGATGGNCTCNAGANTGCNTGTGGTCGGANCNGACTTTGATGNGCAAGANCAGNCTGGNGNGAAACTCNTGCAGCGNGGGANGCANGCNTTNGGCAAAACNCCCACAGTGNTNAAACNTGANCCTNCAGAAGGNANTNTNGNANCNCCNATTAGTCTGANAGCGGGTAACTGGTNGATTCGCANACTGGCAAGTAANTAGTGGAAAGGGNANTTCCGNTCTNCGACGGGGTCCNANACTTTAACCTGGGANGAACGCCNCTGA ```      ``` 172	ATGGCTCTCCCAGGACTGTCTCTCCCAGGCTTGAATCTCCCACAGACATCCTCCACCCCCTTGGCCGATTCAAAAAACACCATACAAGCATCAAGATTAGAGGACCTACCATCGCGGACAGAATGGCGATTCGAGGTAGCGTTCTCTCAACACTACAGCATCAAGCTGGAGTCCGGCCATGCGGAAATGTTTGGTGTTGAGCTCGCCCTCAAACAAGTCTACACGTTCTCCGGATTCAAGGGCGCAATCTTCACATGGCAAGGCTGTCAGCTCGAGGTGACGGGAAATGCAGAGAGTGAATACATTGGACAAGAAACCGACTACGCAGTGGAGTGGCTCAATGTGCACGGCATGCTTGATACGCTTCGATCACAGAACCCAACAGATGGCCCAAGAGTGCTTGTGGTCGGCCCTGACTTTGTTGGCAAGACCAGTCTGGTGCGAACTCTTGCAGCGTGGGAGGCACGCAGCGGATACACGCCCACAGTGCTGAATCTTGATCCTCGAGAAGGCATTCTGGCACCTCCGAGTAGTCTGACAGCGGTCACTGTTGATTCGCAGTTGGAAGTAGATAGTGGATATGGCATTCCGTCTTCGTCGGGCCCCACACTTTCACCTGTGAGAACGCCGCTGATCTATCACTTTCCCTACGCTTCTCCACTCGAAAAGCCTGAAGTCTACAAAGCAGTAGTCACTCGGATGGCGCTCAGTGTGATGAACAAGTTGGAGGAAAATCAGGCGACAAAACGGGGTGGCATCATTGTCGACACACCGGGCACACTGAATGATCCACGGAGCAACTAC	975 	\|\|\|\|\|\|\|\|\|\|\|\|\|\|\|\|\|\|\|\|\|\|\|\|\|\|\|\|\|\|\|\|\|\|\|\|\|\|\|\|\|\|\|\|\|\|\|\|\|\|\|\|\|\|\|\|\|\|\|\|\|\|\|\|\|\|\|\|\|\|\|\|\|\|\|\|\|\|\|\|\|\|\|\|\|\|\|\|\|\|\|\|\|\|\|\|\|\|\|\|\|\|\|\|\|\|\|\|\|\|\|\|\|\|\|\|\|\|\|\|\|\|\|\|\|\|\|\|\|\|\|\|\|\|\|\|\|\|\|\|\|\|\|\|\|\|\|\|\|\|\|\|\|\|\|\|\|\|\|\|\|\|\|\|\|\|\|\|\|\|\|\|\|\|\|\|\|\|\|\|\|\|\|\|\|\|\|\|\|\|\|\|\|\|\|\|\|\|\|\|\|\|\|\|\|\|\|\|\|\|\|\|\|\|\|\|\|\|\|\|\|\|\|\|\|\|\|\|\|\|\|\|\|\|\|\|\|\|\|\|\|\|\|\|\|\|\|\|\|\|\|\|\|\|\|\|\|\|\|\|\|\|\|\|\|\|\|\|\|\|\|\|\|\|\|\|\|\|\|\|\|\|\|\|\|\|\|\|\|\|\|\|\|\|\|\|\|\|\|\|\|\|\|\|\|\|\|\|\|\|\|\|\|\|\|\|\|\|\|\|\|\|\|\|\|\|\|\|\|\|\|\|\|\|\|\|\|\|\|\|\|\|\|\|\|\|\|\|\|\|\|\|\|\|\|\|\|\|\|\|\|\|\|\|\|\|\|\|\|\|\|\|\|\|\|\|\|\|\|\|\|\|\|\|\|\|\|\|\|\|\|\|\|\|\|\|\|\|\|\|\|\|\|\|\|\|\|\|\|\|\|\|\|\|\|\|\|\|\|\|\|\|\|\|\|\|\|\|\|\|\|\|\|\|\|\|\|\|\|\|\|\|\|\|\|\|\|\|\|\|\|\|\|\|\|\|\|\|\|\|\|\|\|\|\|\|\|\|\|\|\|\|\|\|\|\|\|\|\|\|\|\|\|\|\|\|\|\|\|\|\|\|\|\|\|\|\|\|\|\|\|\|\|\|\|\|\|\|\|\|\|\|\|\|\|\|\|\|\|\|\|\|\|\|\|\|\|\|\|\|\|\|\|\|\|\|\|\|\|\|\|\|\|\|\|\|\|\|\|\|\|\|\|\|\|\|\|\|\|\|\|\|\|\|\|\|\|\|\|\|\|\|\|\|\|\|\|\|\|\|\|\|\|\|\|\|\|\|\|\|\|\|\|\|\|\|\|\|\|\|\|\|\|\|\|\|\|\|\|\|\|\|\|\|\|\|\|\|\|\|\|\|\|\|\|\|\|\|\|\|\|\|\|\|\|\|\|\|\|\|\|\|\|\|\|\|\|\|\|\|\|\|\|\|\|\|\|\|\|\|\|\|\|\|\|\|\|\|\|\|\|\|\|\|\|\|\|\|\|\|\|\|\|\|\|\|\|\|\|\|\|\|\|\|\|\|\|\|\|\|\|\|\|\|\|\|\|\|\|\|\|\|\|\|\|\|\|\|\|\|\|\|\|\| \|\|\|\|\| \|\|\|\|\|\|\|\|\|\|\|\|\|\|\| \|\|\|\|\| \|\|\|\|\|\|\|\|\|\|\|\|\|\|\| \|\|\|\|\|\|  \|\| \|\|\|\|\|\|\|\|\|\|\| \|\|\|\|\| \| \|\|\|\| 	ATGGCTCTCCCAGGACTGTCTCTCCCAGGCTTGAATCTCCCACAGACATCCTCCACCCCCTTGGCCGATTCAAAAAACACCATACAAGCATCAAGATTAGAGGACCTACCATCGCGGACAGAATGGCGATTCGAGGTAGCGTTCTCTCAACACTACAGCATCAAGCTGGAGTCCGGCCATGCGGAAATGTTTGGTGTTGAGCTCGCCCTCAAACAAGTCTACACGTTCTCCGGATTCAAGGGCGCAATCTTCACATGGCAAGGCTGTCAGCTCGAGGTGACGGGAAATGCAGAGAGTGAATACATTGGACAAGAAACCGACTACGCAGTGGAGTGGCTCAATGTGCACGGCATGCTTGATACGCTTCGATCACAGAACCCAACAGATGGCCCAAGAGTGCTTGTGGTCGGCCCTGACTTTGTTGGCAAGACCAGTCTGGTGCGAACTCTTGCAGCGTGGGAGGCACGCAGCGGATACACGCCCACAGTGCTGAATCTTGATCCTCGAGAAGGCATTCTGGCACCTCCGAGTAGTCTGACAGCGGTCACTGTTGATTCGCAGTTGGAAGTAGATAGTGGATATGGCATTCCGTCTTCGTCGGGCCCCACACTTTCACCTGTGAGAACGCCGCTGATCTATCACTTTCCCTACGCTTCTCCACTCGAAAAGCCTGAAGTCTACAAAGCAGTAGTCACTCGGATGGCGCTCAGTGTGATGAACAAGTNGGAGG-AAATCAGGCGACAAA-CGGGG-GGCATCATTGTCGAC-CACCGGN-AC-CTGAATGATCC-CGGAGNA-CTAC ```      ``` 172	ATGGCTCTCCCAGGACTGTCTCTCCCAGGCTTGAATCTCCCACAGACATCCTCCACCCCCTTGGCCGATTCAAAAAACACCATACAAGCATCAAGATTAGAGGACCTACCATCGCGGACAGAATGGCGATTCGAGGTAGCGTTCTCTCAACACTACAGCATCAAGCTGGAGTCCGGCCATGCGGAAATGTTTGGTGTTGAGCTCGCCCTCAAACAAGTCT-ACACGTTCTCC-GGATTCAAGGGCGCAATCTTCACATGG-CAAGGCTGTCAGCTCG-AGGTGACGGGAAATGCAGA-GAGTGAATACATTGG-ACAAG-AAACCGACTACGCAGTGGAGTGGCTC-AATGTGC-ACGGCA-TGCTTGATACGCTTCGATCACAG-AACCCAACAGATGGCCCAA-GAGTGCTTG-TGGTCGGCCCTGACTTT-GTTG-GCAAGACCA-GTCTGGTGCGAACTCTTGCAG-CGT-GGGAGGCACGC	639 	\|\|\|\|\|\|\|\|\|\|\|\| \|\| \|\|\| \|\| \|\|\|  \|\|\|\|\|\|\|  \| \| \|\|\|\|\|\|\|\|\|\|\|\| \|\|\| \|\| \|\|\|\|\|\|\|\|\|\|\|\|\|\| \|\|\|\|\|\|\|\|\|\|\|\|\| \| \|\|\|\|\|\|\|\|\|\|\|\|\|\|\|\|\| \|\| \|\|\|\|\|\|\|\|\|\|\|\|\|\|\|\|\|\|\| \|\|\|\|\|\|\|\|\|\|\|\|\|\|\|\|\|\| \|\|\|\|\|\|\|\|\|\| \|\|\|\|\|\|\|\|\|\|\|\|\|\|\|\|\|\|\|\|\|\|\|\|\|\|\|\|\|\|\| \| \|\|\|\|\|\|\|\|\| \|\|\|\|\|\|\| \|\|\|\|\| \|\|\|\|\|\|\|\|\|\|\| \|\|\| \|\|\|\|\|\|\|\|\|\|\|\|\|\|\|\|\|\|\|\|\|\|  \|\|\|\|\| \|\|\|\|\|\|\| \|\| \|\| \|\|\|\| \| \|\|\|\|\|\|\| \| \|\|\|\|\|\|\|\|\|  \|\|\|  \|\|\|\|  \|\|\|\|\|\|\|\|\|\|\|\|\|\| \|\|\|\|\|\|\|\|\|\|\| \|\| \|\|\|\| \|  \|\|  \|\|\|\|\|\|\|\|\|\|   \|\|  \| \|    \|\|\|\| \|\|\|\|\|\|\| \| \|\|\|\| \|\| \|\|\|\| \| \|\|  \|\|\|\|\|\|\|\|\|\|\|\|\| \| \|\| \|\|\|\| \| \|\| \| \|\|\|\|\|\|\|\|\|\|\|\|\|\| \|\|\|  \|\|  \|\|\|\|\|\| \|\|\|\| 	ATGGCTCTCCCACGANTGTNTCNCCCCNGCTTGAAGNTNCNACAGACATCCTCNACCNCCNTGGCCGATTCAAAANACACCATACAAGCNTNAAGATTAGAGGACCTACNATNGCGGACAGAATGGCGATTCNAGGTAGCGTTCTCTCAACNCTACAGCATCNAGCTGGAGTCCGGCCATGCGGAAATGTTTGGNGNTGAGCTCGCNCTCAAACNAGTCTNACACGTTCTCCGGGANTCAAGGGCGCAATCTTCACATGNACAAGGNTGTCAGCNCGCAGNTGACCGNAAATGCACAGGAGTGAATANNTTGTNACAANNAAACCGACTACGCANTGGAGTGGCTCNAAAGTGCCANNGCNNTGCTTGATACNNGTCNGTGANNTNAACCNAACAGATNGNCCAAAGANTGCTNGNTGNCCGGCCCTGACTTTNGNTGNGCAANANCANGGCTGGTGCGAACTCTNGCAATCGNCGGGAGGNACGC ``` |
| 5 | ggggacaagtttgtacaaaaaagcaggcttgatgGCGGGAATTTTCGAGCAGTC, ggggaccactttgtacaagaaagctgggtcGTGCCCATGTCCATCGTCTC | Mycgr3\|110785  JGI  Ensembl | KOG0360 Chaperonin complex component, TCP-1 alpha subunit (CCT1); | ATP binding; cellular protein metabolic process;  protein binding;  protein folding;  unfolded protein binding; | 100 % |  | ``` 114	ATGGCGGGAATTTTCGAGCAGTCGCGCAATCAAGGCACCTTGTTTCTGGGAGGTCAGAAGATTAGCGGCGCAGATATTCGCGACCAGAATGTACTCGCTACACAAGCCATTGCCAACATCGTGAAGTCATCATTCGGGCCTAGCGGTCTGGATAAGATGATGGTGGATGACATCGGCGAGGTGACAGTCACAAACGATGGAGCCACGATCCTCAGTCTGCTCAACGTCGAGCATCCAGCTGGAAAGATTCTTGTGGACCTCGCTCAGCAACAAGACAAGGAAGTCGGAGATGGAACAACCAGTGTGGTGCTCATCGCTGCGGAATTACTACGAAGAGCAAACGAGCTGGTCAAGATGCGCATACACCCCACCACAATCATCACAGGCTACAGACTGGCGCTGCGCGAGGCTGTCCGATATATGAACGAGAATGTGAGCACAAAGGTGGAGACTCTGGGCCGGGAAAGCTTGGTCAACATTGCAAAGACGAGCATGTCTAGCAAAATCATCGGTGGTGACTCCGAGTTCTTCTCCA	648 	\|\|\|\|\|\|\|\|\|\|\|\|\|\|\|\|\|\|\|\|\|\|\|\|\|\|\|\|\|\|\|\|\|\|\|\|\|\|\|\|\|\|\|\|\|\|\|\|\|\|\|\|\|\|\|\|\|\|\|\|\|\|\|\|\|\|\|\|\|\|\|\|\|\|\|\|\|\|\|\|\|\|\|\|\|\|\|\|\|\|\|\|\|\|\|\|\|\|\|\|\|\|\|\|\|\|\|\|\|\|\|\|\|\|\|\|\|\|\|\|\|\|\|\|\|\|\|\|\|\|\|\|\|\|\|\|\|\|\|\|\|\|\|\|\|\|\|\|\|\|\|\|\|\|\|\|\|\|\|\|\|\|\|\|\|\|\|\|\|\|\|\|\|\|\|\|\|\|\|\|\|\|\|\|\|\|\|\|\|\|\|\|\|\|\|\|\|\|\|\|\|\|\|\|\|\|\|\|\|\|\|\|\|\|\|\|\|\|\|\|\|\|\|\|\|\|\|\|\|\|\|\|\|\|\|\|\|\|\|\|\|\|\|\|\|\|\|\|\|\|\|\|\|\|\|\|\|\|\|\|\|\|\|\|\|\|\|\|\|\|\|\|\|\|\|\|\|\|\|\|\|\|\|\|\|\|\|\|\|\|\|\|\|\|\|\|\|\|\|\|\|\|\|\|\|\|\|\|\|\|\|\|\|\|\|\|\|\|\|\|\|\|\|\|\|\|\|\|\|\|\|\|\|\|\|\|\|\|\|\|\|\|\|\|\|\|\|\|\|\|\|\|\|\|\|\|\|\|\|\|\|\|\|\|\|\|\|\|\|\|\|\|\|\|\|\|\|\|\|\|\|\|\|\|\|\|\|\|\|\|\|\|\|\|\|\|\|\|\|\|\|\|\|\|\|\|\|\|\|\|\|\|\|\|\|\|\|\|\|\|\|\|\|\|\|\|\|\| \|\|\|\|\|\|\| \|\|\| \| \|\|\|\|\|\|\|\|\|\|\|\|\|\|\|\|\|\|\|\|\|\|\| \|\|\|\|\|\|\|\|\|\| \|\|\|\|\|\|\| \|\| \| \|\|\|\|\|\|\|\| \| \|\|\|\|\| \|\|   \|\| \|\|\|\|\|\| \| \|\|\| \|\| \|\|\| 	ATGGCGGGAATTTTCGAGCAGTCGCGCAATCAAGGCACCTTGTTTCTGGGAGGTCAGAAGATTAGCGGCGCAGATATTCGCGACCAGAATGTACTCGCTACACAAGCCATTGCCAACATCGTGAAGTCATCATTCGGGCCTAGCGGTCTGGATAAGATGATGGTGGATGACATCGGCGAGGTGACAGTCACAAACGATGGAGCCACGATCCTCAGTCTGCTCAACGTCGAGCATCCAGCTGGAAAGATTCTTGTGGACCTCGCTCAGCAACAAGACAAGGAAGTCGGAGATGGAACAACCAGTGTGGTGCTCATCGCTGCGGAATTACTACGAAGAGCAAACGAGCTGGTCAAGATGCGCATACACCCCACCACAATCATCACAGGCTACAGACTGGCGCTGCGCGAGGCTGTCCGATATATGAACGANAATGTGANCACNANGGTGGAGACTCTGGGCCGGGAAANCTTGGTCAACNTTGCAAANACNANCATGTCTANC-AAATCNTCNNNGGNGACTCCNANTTCNTCNCCA ``` |
| 5 | ggggacaagtttgtacaaaaaagcaggcttgatgCCCGCAATCATGGAAGAGGA, ggggaccactttgtacaagaaagctgggtcCCGCGAACTGCTGAACATGA | Mycgr3\|110944  JGI  Ensembl | KOG2983 Uncharacterized conserved protein; |  | 1.03653 % |  |  |
| 5 | ggggacaagtttgtacaaaaaagcaggcttgatgACGAGACCTAAGAAGGACGT, ggggaccactttgtacaagaaagctgggtcTAGCGTTTTCCGCGCTACGT | Mycgr3\|110994  JGI  Ensembl | KOG1440 CDP-diacylglycerol synthase; | membrane; phosphatidate cytidylyltransferase activity;  phospholipid biosynthetic process; | 100 % |  | ``` 210	ATGACGAGACCTAAGAAGGACGTGAAGTTTCAGCACCGGGAGCAAAGGCGCATGAGCAAGAGCAGCAGCATGTCAGACTTCAACAACGACTCGCCAAATGAGGAGCGAAGCTCGTCCAGAAGCAATGGAGCTAAGAGTCCTACGCGTAACGGGTTCGTCGGAAAGGATGAGAGGCCCAAGGCAGAGAGGACCGATACCACAGCGAGTGGATGGGCTACGGAAAATGAGGACGAGAAG	446 	\|\|\|\|\|\|\|\|\|\|\|\|\|\|\|\|\|\|\|\|\|\|\|\|\|\|\|\|\|\|\|\|\|\|\|\|\|\|\|\|\|\|\|\|\|\|\|\|\|\|\|\|\|\|\|\|\|\|\|\|\|\|\|\|\|\|\|\|\|\|\|\|\|\|\|\|\|\|\|\|\|\|\|\|\|\|\|\|\|\|\|\|\|\|\|\|\|\|\|\|\|\|\|\|\|\|\|\|\|\|\|\|\|\|\|\|\|\|\|\|\|\|\|\|\|\|\|\|\|\|\|\|\|\|\|\|\|\|\|\|\|\|\|\|\|\|\|\|\|\|\|\|\|\|\|\|\|\|\|\|\|\|\|\|\|\|\|\|\|\|\|\|\|\|\|\|\|\|\|\|\|\|\|\|\|\|\|\|\|\|\|\|\|\|\|\|\|\|\|\|\|\|\|\|\|\|\|\|\|\|\|\|\|\|\|\|\|\|\|\|\|\|\|\|\|\|\|\|\|\|\|\|\|\|\|\|\| 	ATGACGAGACCTAAGAAGGACGTGAAGTTTCAGCACCGGGAGCAAAGGCGCATGAGCAAGAGCAGCAGCATGTCAGACTTCAACAACGACTCGCCAAATGAGGAGCGAAGCTCGTCCAGAAGCAATGGAGCTAAGAGTCCTACGCGTAACGGGTTCGTCGGAAAGGATGAGAGGCCCAAGGCAGAGAGGACCGATACCACAGCGAGTGGATGGGCTACGGAAAATGAGGACGAGAAG ```      ``` 445	AGTCGTTCATCACCCGCGCCATCTGGACGTTCGCCATGATCGGCATCTTCTTCGGATCCATGTTCGCAGGCCATATCTACGTGATAATCATCGTGACGGCTGTGCAGATCATCTCGTTCAAGGAGGTCATTGCCATTTCCAACGTGCCCAGCAGAGCCAGATCGCTTCGTTTCACGAAAACGCTAAACTGGTACTTTCTCGGAACGACAATGTACTTTCTCTATGGCGAGAGTGTCATCTACTACTTCAAGCACATTGTGCTTATCGACCGCGTGCTGCTACCCTTTGCGACCCACCACAGGTTCATCAGCTTTATGCTCTACGTGATGGG	775 	\|\|\|\|\|\|\|\|\|\|\|\|\|\|\|\|\|\|\|\|\|\|\|\|\|\|\|\|\|\|\|\|\|\|\|\|\|\|\|\|\|\|\|\|\|\|\|\|\|\|\|\|\|\|\|\|\|\|\|\|\|\|\|\|\|\|\|\|\|\|\|\|\|\|\|\|\|\|\|\|\|\|\|\|\|\|\|\|\|\|\|\|\|\|\|\|\|\|\|\|\|\|\|\|\|\|\|\|\|\|\|\|\|\|\|\|\|\|\|\|\|\|\|\|\|\|\|\|\|\|\|\|\|\|\|\|\|\|\|\|\|\|\|\|\|\|\|\|\|\|\|\|\|\|\|\|\|\|\|\|\|\|\|\|\|\|\|\|\|\|\|\|\|\|\|\|\|\|\|\|\|\|\|\|\|\|\|\|\|\|\|\|\|\|\|\|\|\|\|\|\|\|\|\|\|\|\|\|\|\|\|\|\|\|\|\|\|\|\|\|\|\|\|\|\|\|\|\|\|\|\|\|\|\|\|\|\|\|\|\|\|\|\|\|\|\|\|\| \|\|\|\|\|\|\|\|\|\|\|\|\|\|\|\|\|\|\|\|\|\|\|\|\|\|\|\|\|\|\|\|\|\|\|\|\|\|\|\|\|\|\|\|\|\|\|\|\|\|\|\|\|\|\|\|\|\|\|\| \|\|\|\|\|\|\|\|\|\|\|\|\|\|\|\|\|\|\|\|\| 	AGTCGTTCATCACCCGCGCCATCTGGACGTTCGCCATGATCGGCATCTTCTTCGGATCCATGTTCGCAGGCCATATCTACGTGATAATCATCGTGACGGCTGTGCAGATCATCTCGTTCAAGGAGGTCATTGCCATTTCCAACGTGCCCAGCAGAGCCAGATCGCTTCGTTTCACGAAAACGCTAAACTGGTACTTTCTCGGAACGACAATGTACTTTCTCTATGGCGAGAGTGTCATCTACTACTTC-AGCACATTGTGCTTATCGACCGCGTGCTGCTACCCTTTGCGACCCACCACAGGTTCATCANCTTTATGCTCTACGTGATGGG ``` |
| 5 | ggggacaagtttgtacaaaaaagcaggcttgatgGCGACAACGGTACCTCTCGA, ggggaccactttgtacaagaaagctgggtcAAACGGAGGGATCGCGCCGC | Mycgr3\|111073  JGI  Ensembl | KOG3296 Translocase of outer mitochondrial membrane complex, subunit TOM40; | anion transport; mitochondrial outer membrane;  voltage-gated anion channel activity; | 100 % |  | ``` 127	ATGGCGACAACGGTACCTCTCGAAAAAACTGCGGTGCCAGTCCCTGTCGAAGTCGAGTTCCCTCCGCAAGATGTAACACTGCCGTCAGTGCGAACAGGGGGATTCCTCGACACCGTCCTCGACACATATCAGAGCTTCCAGGCACGGAGAGATGCGCTCGGTCTCAGCAACCCAGGCACAGTCGATCAAATAGCAAAGGAGGTCCAGCGCGATGTCTTCCTCACCAACCAGATATTCTCTGGTTTGCGCGCGGAGCTCAACAAGACCTTCTCCATCCTGCCACTCTTCCAGATCAGCCATGCCTTCTCGAGCGGATCCCAGATGCTCTCGCCTTACACATTCCTCGCCTTGTATGGCACCAACAAT	492 	\|\|\|\|\|\|\|\|\|\|\|\|\|\|\|\|\|\|\|\|\|\|\|\|\|\|\|\|\|\|\|\|\|\|\|\|\|\|\|\|\|\|\|\|\|\|\|\|\|\|\|\|\|\|\|\|\|\|\|\|\|\|\|\|\|\|\|\|\|\|\|\|\|\|\|\|\|\|\|\|\|\|\|\|\|\|\|\|\|\|\|\|\|\|\|\|\|\|\|\|\|\|\|\|\|\|\|\|\|\|\|\|\|\|\|\|\|\|\|\|\|\|\|\|\|\|\|\|\|\|\|\|\|\|\|\|\|\|\|\|\|\|\|\|\|\|\|\|\|\|\|\|\|\|\|\|\|\|\|\|\|\|\|\|\|\|\|\|\|\|\|\|\|\|\|\|\|\|\|\|\|\|\|\|\|\|\|\|\|\|\|\|\|\|\|\|\|\|\|\|\|\|\|\|\|\|\|\|\|\|\|\|\|\|\|\|\|\|\|\|\|\|\|\|\|\|\|\|\|\|\|\|\|\|\|\|\|\|\|\|\|\|\|\|\|\|\|\|\|\|\|\|\|\|\|\|\|\|\|\|\|\|\|\|\|\|\|\|\|\|\|\|\|\|\|\|\|\|\|\|\|\|\|\|\|\|\|\|\|\|\|\|\|\|\|\|\|\|\|\|\|\|\|\|\|\|\|\|\|\|\|\|\|\|\|\|\|\|\|\|\|\|\|\|\|\|\|\|\|\|\|\|\|\|\|\|\|\|\|\|\|\|\|\|\|\|\|\|\|\|\|\|\|\|\|\|\|\|\|\|\|\|\|\|\|\| 	ATGGCGACAACGGTACCTCTCGAAAAAACTGCGGTGCCAGTCCCTGTCGAAGTCGAGTTCCCTCCGCAAGATGTAACACTGCCGTCAGTGCGAACAGGGGGATTCCTCGACACCGTCCTCGACACATATCAGAGCTTCCAGGCACGGAGAGATGCGCTCGGTCTCAGCAACCCAGGCACAGTCGATCAAATAGCAAAGGAGGTCCAGCGCGATGTCTTCCTCACCAACCAGATATTCTCTGGTTTGCGCGCGGAGCTCAACAAGACCTTCTCCATCCTGCCACTCTTCCAGATCAGCCATGCCTTCTCGAGCGGATCCCAGATGCTCTCGCCTTACACATTCCTCGCCTTGTATGGCACCAACAAT ```      ``` 493	ATGCTCTGCCAAGCTCAGCTCGACTCCGACGGTTCCCTCAACGCTCGATTTAATGGCCGCATCACAAGCCGTCTCATCAACAAACTCGCGGTGCAGATTCAGCCTGGCTCGATGGCTGGACCAGGTGGCGCGCAAGTATCTCTCGAACAGGACTACACTGGCGACGACTTCACCGCATCTCTCAAGAGCATCAATCCGAGTGTGTTGGAAGGTGGATTGACGGG-AATGTTCATCGGATCATACCTCCAGAGCGTCACTCCCAGGTTATCACTCGGAATCGAGGGTGTATTGCAAAAGCCAGGTGGTGGAATGGGCCCGGAGGTGGCCATGAGCTACGCAGGACGATACAAGGGCGATGACTGGATCGCCAGCGCGCAACTTCTGACTCAGGGTGGCATGCAAGTGA	898 	\|\|\|\|\|\|\|\|\|\|\|\|\|\|\|\|\|\|\|\|\|\|\|\|\|\|\|\|\|\|\|\|\|\|\|\|\|\|\|\|\|\|\|\|\|\|\|\|\|\|\|\|\|\|\|\|\|\|\|\|\|\|\|\|\|\|\|\|\|\|\|\|\|\|\|\|\|\|\|\|\|\|\|\|\|\|\|\|\|\|\|\|\|\|\|\|\|\|\|\|\|\|\|\|\|\|\|\|\|\|\|\|\|\|\|\|\|\|\|\|\|\|\|\|\|\|\|\|\|\|\|\|\|\|\|\|\|\|\|\|\|\|\|\|\|\|\|\|\|\|\|\|\|\|\|\|\|\|\|\|\|\|\|\|\|\|\|\|\|\|\|\|\|\|\|\|\|\|\|\|\|\|\|\|\|\|\|\|\|\|\|\|\|\|\|\|\|\|\|\|\|\|\|\|\|\|\|\|\|\|\|\|\|\|\|\|\|\|\|\|\|\|\|\| \|\|\|\|\|\|\|\|\|\|\|\|\|\|\|\|\|\|\|\|\|\|\|\|\|\|\|\|\|\|\|\|\|\|\|\|\|\|\|\|\|\|\|\|\|\|\|\|\|\|\|\|\|\|\|\|\|\|\|\|\|\|\|\|\| \|\| \|\|\|\|\|\|\|\|\|\|\|\|\|\|\|\|\|\|\|\|\|\|\|\|\|\|\|\|\|\|\|\|\|\|\|\|\|\|\|\|\|\|\|\|\|\|\|\|\|\|\|\|\|\|\|\|\|\|\|\|\|\|\|\|\|\|\|\|\|\|\|\|\|\|\| \|\|\|\|\|\|\|\|  \|\|\|\|\|\|\|\|\|\|\|\|\| \|\| \|\|\|\|\|\|\|\|\|\| 	ATGCTCTGCCAAGCTCAGCTCGACTCCGACGGTTCCCTCAACGCTCGATTTAATGGCCGCATCACAAGCCGTCTCATCAACAAACTCGCGGTGCAGATTCAGCCTGGCTCGATGGCTGGACCAGGTGGCGCGCAAGTATCTCTCGAACAGGACTACACTGGCGACGACTTCACCGCATCTCTCAAGAGCATCAATCCGAGTGTGTTGGAAGGTGGATTGACGGGNAATGTTCATCGGATCATACCTCCAGAGCGTCACTCCCAGGTTATCACTCGGAATCGAGGGTGTATNGC-AAAGCCAGGTGGTGGAATGGGCCCGGAGGTGGCCATGAGCTACGCAGGACGATACAAGGGCGATGACTGGATCGCNAGCGCGCAN-TTCTGACTCAGGG-GGNATGCAAGTGA ``` |
| 5 | ggggacaagtttgtacaaaaaagcaggcttgatgTCCCCATCATCATCCCAACT, ggggaccactttgtacaagaaagctgggtcCCGCCTTTCTCCGCCGCCTC | Mycgr3\|111700  JGI  Ensembl | KOG1187 Serine/threonine protein kinase; |  | 1.36452 % |  |  |
| 5 | ggggacaagtttgtacaaaaaagcaggcttgatgGCGGCCCTCCGACTACCCTC, ggggaccactttgtacaagaaagctgggtcTCGGTCGGATCGGGAAGCGG | Mycgr3\|111717  JGI  Ensembl | KOG2672 Lipoate synthase; | catalytic activity; iron-sulfur cluster binding;  lipoate biosynthetic process;  lipoate synthase activity; | 1.04396 % |  |  |
| 6 | ggggacaagtttgtacaaaaaagcaggcttgatgCATTGTCCTAATACGAGTAT, ggggaccactttgtacaagaaagctgggtcGTACTCGAAATCATCGTCCA | Mycgr3\|31749  JGI  Ensembl | KOG0967 ATP-dependent DNA ligase I; | ATP binding; DNA ligase (ATP) activity;  DNA recombination;  DNA repair;  DNA replication; | 0 % |  |  |
| 6 | ggggacaagtttgtacaaaaaagcaggcttgatgGTCCGCGACGGCGACACGGC, ggggaccactttgtacaagaaagctgggtcGCTTTTCCGTCCCAAAACTT | Mycgr3\|32716  JGI  Ensembl | KOG2676 Uncharacterized conserved protein; |  | 0.58923 % |  |  |
| 6 | ggggacaagtttgtacaaaaaagcaggcttgatgGCTCCAGTCCCGCACTCCCA, ggggaccactttgtacaagaaagctgggtcCTCCACCTCCATCAACCTCG | Mycgr3\|32817  JGI  Ensembl | KOG1980 Uncharacterized conserved protein; |  | 0 % |  |  |
| 6 | ggggacaagtttgtacaaaaaagcaggcttgatgCCCGCAGTCGTAGCACCACC, ggggaccactttgtacaagaaagctgggtcCCACAACGCACTGTCCTTAT | Mycgr3\|35747  JGI  Ensembl | KOG0969 DNA polymerase delta, catalytic subunit; | DNA binding; DNA polymerase activity;  DNA replication;  DNA-directed DNA polymerase activity;  membrane;  nucleic acid binding;  nucleobase, nucleoside, nucleotide and nucleic acid metabolic process;  nucleotide binding;  nucleus;  transport;  transporter activity; | 0 % |  |  |
| 6 | ggggacaagtttgtacaaaaaagcaggcttgatgCCTCCTCCACGTTCACGTCA, ggggaccactttgtacaagaaagctgggtcTTCATACTCCTCACCCTCAT | Mycgr3\|37171  JGI  Ensembl | KOG1598 Transcription initiation factor TFIIIB, Brf1 subunit; | regulation of transcription, DNA-dependent; transcription factor complex;  transcription initiation;  transcription regulator activity;  zinc ion binding; | 0 % |  |  |
| 6 | ggggacaagtttgtacaaaaaagcaggcttgatgGCCGACCGATATCGAAACAC, ggggaccactttgtacaagaaagctgggtcGCCTCCAACAATCTCGCACA | Mycgr3\|40742  JGI  Ensembl | KOG0588 Serine/threonine protein kinase; | 3-phosphoinositide-dependent protein kinase activity; AMP-activated protein kinase activity;  ATP binding;  DNA-dependent protein kinase activity;  GTP-dependent protein kinase activity;  JUN kinase activity;  JUN kinase kinase activity;  JUN kinase kinase kinase activity;  Janus kinase activity;  MAP kinase 1 activity;  MAP kinase 2 activity;  MAP kinase kinase activity;  MAP kinase kinase kinase kinase activity;  MAP/ERK kinase kinase activity;  MP kinase activity;  NF-kappaB-inducing kinase activity;  SAP kinase activity;  atypical protein kinase C activity;  calcium-dependent protein kinase C activity;  casein kinase I activity;  casein kinase activity;  cyclic nucleotide-dependent protein kinase activity;  cyclin-dependent protein kinase activating kinase activity;  eukaryotic translation initiation factor 2alpha kinase activity;  galactosyltransferase-associated kinase activity;  glycogen synthase kinase 3 activity;  protein amino acid phosphorylation;  protein kinase activity;  protein serine/threonine kinase activity;  protein-tyrosine kinase activity;  receptor signaling protein serine/threonine kinase activity;  ribosomal protein S6 kinase activity;  transmembrane receptor protein kinase activity;  transmembrane receptor protein serine/threonine kinase activity; | 0 % |  |  |
| 6 | ggggacaagtttgtacaaaaaagcaggcttgatgTCGTCGCCAACGAGTCGACG, ggggaccactttgtacaagaaagctgggtcGGCAACGCCCGTCAACCTTC | Mycgr3\|43403  JGI  Ensembl | KOG0478 DNA replication licensing factor, MCM4 component; | ATP binding; ATP-dependent DNA helicase activity;  ATP-dependent RNA helicase activity;  ATP-dependent helicase activity;  ATPase activity;  ATPase activity, coupled;  ATPase activity, coupled to transmembrane movement of ions;  ATPase activity, coupled to transmembrane movement of substances;  ATPase activity, uncoupled;  DNA binding;  DNA replication initiation;  DNA-dependent ATPase activity;  RNA-dependent ATPase activity;  nucleus;  protein-transmembrane transporting ATPase activity;  single-stranded DNA-dependent ATP-dependent DNA helicase activity; | 0.66667 % |  |  |
| 6 | ggggacaagtttgtacaaaaaagcaggcttgatgAAGTTCTTCATCGAGGACCT, ggggaccactttgtacaagaaagctgggtcAGCATCCAACTCCATCATCG | Mycgr3\|43883  JGI  Ensembl | KOG1131 RNA polymerase II transcription initiation/nucleotide excision repair factor TFIIH, 5'-3' helicase subunit RAD3; | 8-oxo-7,8-dihydroguanine triphosphatase activity; ATP binding;  ATP-dependent 5'-3' DNA helicase activity;  ATP-dependent DNA helicase activity;  ATP-dependent helicase activity;  DNA binding;  UDP-2,3-diacylglucosamine hydrolase activity;  bis(5'-nucleosyl)-tetraphosphatase activity;  cytoplasm;  dATP pyrophosphohydrolase activity;  diacylglycerol pyrophosphate phosphatase activity;  dihydroneopterin monophosphate phosphatase activity;  dihydroneopterin triphosphate pyrophosphohydrolase activity;  hydrolase activity, acting on acid anhydrides, in phosphorus-containing anhydrides; inorganic diphosphatase activity;  magnesium ion binding;  nucleic acid binding;  nucleobase, nucleoside, nucleotide and nucleic acid metabolic process;  nucleotide-excision repair;  nucleus;  phosphate metabolic process;  pyrophosphatase activity;  thiamin-pyrophosphatase activity; | 0 % |  |  |
| 6 | ggggacaagtttgtacaaaaaagcaggcttgatgCGCCTGTGGACCCCCGAAAG, ggggaccactttgtacaagaaagctgggtcGTTCCCCTTCTTCTCAGCCG | Mycgr3\|44229  JGI  Ensembl | KOG0323 TFIIF-interacting CTD phosphatases, including NLI-interacting factor; | CTD phosphatase activity; calcium-dependent protein serine/threonine phosphatase activity;  calcium-dependent protein serine/threonine phosphatase regulator activity;  intracellular;  magnesium-dependent protein serine/threonine phosphatase activity;  magnesium-dependent protein serine/threonine phosphatase complex;  myosin phosphatase activity;  myosin phosphatase complex;  myosin phosphatase regulator activity;  phosphoprotein phosphatase activity; | 0 % |  |  |
| 6 | ggggacaagtttgtacaaaaaagcaggcttgatgAGCGAGTTCAACATCAGTGA, ggggaccactttgtacaagaaagctgggtcAGCCCCGGTCTCCACGACCT | Mycgr3\|44387  JGI  Ensembl | KOG0414 Chromosome condensation complex Condensin, subunit D2; | chromosome condensation; mitosis; nucleus; | 0 % |  |  |
| 6 | ggggacaagtttgtacaaaaaagcaggcttgatgTCCGCCGACGTGATAACGCC, ggggaccactttgtacaagaaagctgggtcTCTCCTGCCTCCTCCCCTAA | Mycgr3\|44713  JGI  Ensembl | KOG2163 Centromere/kinetochore protein zw10 involved in mitotic chromosome segregation; |  | 0 % |  |  |
| 6 | ggggacaagtttgtacaaaaaagcaggcttgatgGCTTCCTCGTCGTTCAGAGA, ggggaccactttgtacaagaaagctgggtcATCGCCCATCCACCCAGTCA | Mycgr3\|54282  JGI  Ensembl | KOG2887 Membrane protein involved in ER to Golgi transport; |  | 0 % |  |  |
| 6 | ggggacaagtttgtacaaaaaagcaggcttgatgTCGATATCAGTATACATTGG, ggggaccactttgtacaagaaagctgggtcTGGGCCCCAGAGGAGGTCGT | Mycgr3\|54658  JGI  Ensembl | KOG4674 Uncharacterized conserved coiled-coil protein; |  | 0 % |  |  |
| 6 | ggggacaagtttgtacaaaaaagcaggcttgatgTCTGGAGAGCCAGACCACGC, ggggaccactttgtacaagaaagctgggtcTTCGTAAAGATCCTCCTCGG | Mycgr3\|55128  JGI  Ensembl | KOG0730 AAA+-type ATPase; | ATP binding; ATP-dependent DNA helicase activity;  ATP-dependent RNA helicase activity;  ATP-dependent helicase activity;  ATPase activity;  ATPase activity, coupled;  ATPase activity, coupled to transmembrane movement of ions;  ATPase activity, coupled to transmembrane movement of substances;  ATPase activity, uncoupled;  DNA-dependent ATPase activity;  RNA-dependent ATPase activity;  hydrolase activity;  nucleoside-triphosphatase activity;  nucleotide binding;  protein-transmembrane transporting ATPase activity;  single-stranded DNA-dependent ATP-dependent DNA helicase activity; | 0 % |  |  |
| 6 | ggggacaagtttgtacaaaaaagcaggcttgatgGCGTCGGCCTCACCCTCATC, ggggaccactttgtacaagaaagctgggtcGTCGCCGAGCACACTTTCCT | Mycgr3\|55591  JGI  Ensembl | KOG2011 Sister chromatid cohesion complex Cohesin, subunit STAG/IRR1/SCC3; | DNA ligase (ATP) activity; | 0 % |  |  |
| 6 | ggggacaagtttgtacaaaaaagcaggcttgatgGCCCAATACCAGCCTCAGCA, ggggaccactttgtacaagaaagctgggtcCTGCTTGGGATGCTGCGGAG | Mycgr3\|57099  JGI  Ensembl | KOG4305 RhoGEF GTPase; | Rho guanyl-nucleotide exchange factor activity; intracellular;  intracellular signaling cascade;  regulation of Rho protein signal transduction;  small GTPase regulator activity; | 0 % |  |  |
| 6 | ggggacaagtttgtacaaaaaagcaggcttgatgGTTTCCGCTTCCAAGGCCGC, ggggaccactttgtacaagaaagctgggtcGACACCTCCAGCTGCCAACA | Mycgr3\|61825  JGI  Ensembl | KOG0062 ATPase component of ABC transporters with duplicated ATPase domains/Translation elongation factor EF-3b; | ATP binding; ATPase activity;  nucleoside-triphosphatase activity;  nucleotide binding; | 0 % |  |  |
| 6 | ggggacaagtttgtacaaaaaagcaggcttgatgGCCGACAATCGCTTCAGTGC, ggggaccactttgtacaagaaagctgggtcCCTCTTCTTCTTCTGACTCT | Mycgr3\|62083  JGI  Ensembl | KOG2318 Uncharacterized conserved protein; |  | 0 % |  |  |
| 6 | ggggacaagtttgtacaaaaaagcaggcttgatgGACTACGAGGCCCTCAAGGA, ggggaccactttgtacaagaaagctgggtcGCCAGTACCGCTGACAGCCA | Mycgr3\|64280  JGI  Ensembl | KOG1986 Vesicle coat complex COPII, subunit SEC23; | COPII vesicle coat; ER to Golgi vesicle-mediated transport;  intracellular protein transport;  protein binding;  zinc ion binding; | 0 % |  |  |
| 6 | ggggacaagtttgtacaaaaaagcaggcttgatgGCGGACGAAGTGAGCATCGA, ggggaccactttgtacaagaaagctgggtcACGCTTTTTGCCCTTCGACT | Mycgr3\|64524  JGI  Ensembl | KOG1189 Global transcriptional regulator, cell division control protein; | DNA ligase (ATP) activity; | 0 % |  |  |
| 6 | ggggacaagtttgtacaaaaaagcaggcttgatgCTCGGCTCCTCTCTCATACC, ggggaccactttgtacaagaaagctgggtcCCGCACACTAGTCATCCCCA | Mycgr3\|64888  JGI  Ensembl | KOG2480 3-hydroxy-3-methylglutaryl-CoA (HMG-CoA) reductase; | NADP binding; coenzyme A metabolic process;  coenzyme binding;  hydroxymethylglutaryl-CoA reductase (NADPH) activity;  integral to membrane;  isoprenoid biosynthetic process; | 0.7743 % |  |  |
| 6 | ggggacaagtttgtacaaaaaagcaggcttgatgGCCTCGACACCTCACCACCC, ggggaccactttgtacaagaaagctgggtcCCTCCTCACGGAACTCGCCC | Mycgr3\|65473  JGI  Ensembl | KOG2000 Gamma-tubulin complex, DGRIP91/SPC98 component; | collagenase activity; matrilysin-2 activity;  microtubule cytoskeleton organization and biogenesis;  microtubule organizing center;  prenyl protein specific endopeptidase activity;  spindle pole;  stromelysin 3 activity; | 0 % |  |  |
| 6 | ggggacaagtttgtacaaaaaagcaggcttgatgTCCACCCACCGCTCACAGGC, ggggaccactttgtacaagaaagctgggtcTAGTGCATTCAATACAGCCT | Mycgr3\|66246  JGI  Ensembl | KOG3745 Exocyst subunit - Sec10p; | cytoplasm; exocytosis; vesicle docking; | 0 % |  |  |
| 6 | ggggacaagtttgtacaaaaaagcaggcttgatgGTCCGCCTCCGCGAGATCCA, ggggaccactttgtacaagaaagctgggtcTGTCTTCGTGGCGCGCCCAA | Mycgr3\|66801  JGI  Ensembl | KOG0307 Vesicle coat complex COPII, subunit SEC31; | triacylglycerol lipase activity; | 0 % |  |  |
| 6 | ggggacaagtttgtacaaaaaagcaggcttgatgGCGACTTTCCTCGAGACTTC, ggggaccactttgtacaagaaagctgggtcAACCAATTCGCCAATCTTGC | Mycgr3\|67108  JGI  Ensembl | KOG1058 Vesicle coat complex COPI, beta subunit; | intracellular protein transport; membrane coat;  protein binding;  protein complex assembly;  vesicle-mediated transport; | 0 % |  |  |
| 6 | ggggacaagtttgtacaaaaaagcaggcttgatgACCTCTCTCAAGCGGTCGCA, ggggaccactttgtacaagaaagctgggtcAGCCAACTGCAGCTTAATCT | Mycgr3\|67423  JGI  Ensembl | KOG2102 Exosomal 3'-5' exoribonuclease complex, subunit Rrp44/Dis3; | 3'-5'-exoribonuclease activity; 5'-3' exoribonuclease activity;  RNA binding;  ribonuclease activity; | 0 % |  |  |
| 6 | ggggacaagtttgtacaaaaaagcaggcttgatgGCGTATCAGTCAGAAGCACC, ggggaccactttgtacaagaaagctgggtcCCTGGAAAGCACCCCCTTCA | Mycgr3\|67486  JGI  Ensembl |  |  | 28.17073 % |  |  |
| 6 | ggggacaagtttgtacaaaaaagcaggcttgatgCGATCACGATGTCGACACCA, ggggaccactttgtacaagaaagctgggtcCGCTCTCCCTACACCCGCAG | Mycgr3\|68179  JGI  Ensembl | KOG3657 Mitochondrial DNA polymerase gamma, catalytic subunit; | DNA binding; DNA polymerase activity;  DNA replication;  DNA-directed DNA polymerase activity;  gamma DNA polymerase complex; | 0.54457 % |  |  |
| 6 | ggggacaagtttgtacaaaaaagcaggcttgatgGCTATCGACACCGCCCTACG, ggggaccactttgtacaagaaagctgggtcGCATCGCACGTCATAGACAC | Mycgr3\|68349  JGI  Ensembl | KOG1835 Uncharacterized conserved protein; | cysteine-type endopeptidase activity; proteolysis; | 0 % |  |  |
| 6 | ggggacaagtttgtacaaaaaagcaggcttgatgGATGTCAACGCCGTCCTCAC, ggggaccactttgtacaagaaagctgggtcCCACCGACGGCGTTGATTTT | Mycgr3\|69066  JGI  Ensembl | KOG1241 Karyopherin (importin) beta 1; | 1-phosphatidylinositol-3-kinase activity; cytoplasm; intracellular protein transport;  nuclear pore;  nucleus;  phosphatidylinositol 3-kinase, class I, catalyst activity;  phosphatidylinositol 3-kinase, class I, regulator activity;  protein import into nucleus, docking;  protein transporter activity; | 0 % |  |  |
| 6 | ggggacaagtttgtacaaaaaagcaggcttgatgGAGACCAAAGAGGGGACCAC, ggggaccactttgtacaagaaagctgggtcCCCCTCCGCCCACCCAAAAA | Mycgr3\|69132  JGI  Ensembl | KOG2126 Glycosylphosphatidylinositol anchor synthesis protein; | catalytic activity; | 10.32953 % |  |  |
| 6 | ggggacaagtttgtacaaaaaagcaggcttgatgTCTCTCCTTATGAATGAGGA, ggggaccactttgtacaagaaagctgggtcCTCTCCTATTCCGAGTCCAA | Mycgr3\|69542  JGI  Ensembl | KOG3739 Stress-activated MAP kinase-interacting protein, Sin1p; |  | 0 % |  |  |
| 6 | ggggacaagtttgtacaaaaaagcaggcttgatgTACATCAAACAGATCATCAT, ggggaccactttgtacaagaaagctgggtcCTTTCCACTGATCTGACCCT | Mycgr3\|69561  JGI  Ensembl | KOG0964 Structural maintenance of chromosome protein 3 (sister chromatid cohesion complex Cohesin, subunit SMC3); | ATP binding; ATPase activity;  DNA metabolic process;  chromosome;  chromosome organization and biogenesis;  protein binding; | 0 % |  |  |
| 6 | ggggacaagtttgtacaaaaaagcaggcttgatgTCGCAGCTTGGACAGTCCGC, ggggaccactttgtacaagaaagctgggtcATACGCTCCCAAGTCCTGCC | Mycgr3\|70159  JGI  Ensembl | KOG2168 Cullins; | nuclear pore; transport; | 0 % |  |  |
| 6 | ggggacaagtttgtacaaaaaagcaggcttgatgGCATCAGAATCACGCCATGT, ggggaccactttgtacaagaaagctgggtcAGCCGCATCTCCAATATCTC | Mycgr3\|70427  JGI  Ensembl | KOG0274 Cdc4 and related F-box and WD-40 proteins; |  | 0 % |  |  |
| 6 | ggggacaagtttgtacaaaaaagcaggcttgatgGCGCAAAAGAAGTCCACGAA, ggggaccactttgtacaagaaagctgggtcCTCGCTCTCATCTTCCTCGT | Mycgr3\|70984  JGI  Ensembl | KOG2256 Predicted protein involved in nuclear export of pre-ribosomes; | DNA ligase (ATP) activity; | 0 % |  |  |
| 6 | ggggacaagtttgtacaaaaaagcaggcttgatgAGCAACATCCTCGATCAAAA, ggggaccactttgtacaagaaagctgggtcATCATACTGTGGCCCTTGAT | Mycgr3\|71868  JGI  Ensembl | KOG1999 RNA polymerase II transcription elongation factor DSIF/SUPT5H/SPT5; |  | 0.65833 % |  |  |
| 6 | ggggacaagtttgtacaaaaaagcaggcttgatgGACGACCTTTACGACGAGTT, ggggaccactttgtacaagaaagctgggtcAGCCCCATCCAACATGCCCG | Mycgr3\|72117  JGI  Ensembl | KOG0468 U5 snRNP-specific protein; | GTP binding; GTPase activity;  protein-synthesizing GTPase activity;  translation; | 0 % |  |  |
| 6 | ggggacaagtttgtacaaaaaagcaggcttgatgCCAAAGAAAGCCATCGATGC, ggggaccactttgtacaagaaagctgggtcTCTACCACTCTTCTTCCCTC | Mycgr3\|72396  JGI  Ensembl | KOG2036 Predicted P-loop ATPase fused to an acetyltransferase; |  | 0 % |  |  |
| 6 | ggggacaagtttgtacaaaaaagcaggcttgatgTCCGGTCGCAAGGACTTTCT, ggggaccactttgtacaagaaagctgggtcCTCCAAGCCTCTAGCCACAA | Mycgr3\|72776  JGI  Ensembl | KOG0495 HAT repeat protein; | RNA processing; intracellular;  nuclear mRNA splicing, via spliceosome;  nucleus; | 0.67664 % |  |  |
| 6 | ggggacaagtttgtacaaaaaagcaggcttgatgGCGCCCACATCGAAAGCCAC, ggggaccactttgtacaagaaagctgggtcAGGCCCGACCTGGAACTTCA | Mycgr3\|73125  JGI  Ensembl | KOG0216 RNA polymerase I, second largest subunit; | DNA binding; DNA-directed RNA polymerase I activity;  DNA-directed RNA polymerase II activity;  DNA-directed RNA polymerase III activity;  DNA-directed RNA polymerase activity;  RNA polymerase activity;  nucleus;  transcription; | 0 % |  |  |
| 6 | ggggacaagtttgtacaaaaaagcaggcttgatgTCTACAAAGCCATTGAAGGA, ggggaccactttgtacaagaaagctgggtcTAACCTTACGCCCTTGACCA | Mycgr3\|73738  JGI  Ensembl | KOG2468 Dolichol kinase; | membrane; phosphatidate cytidylyltransferase activity;  phospholipid biosynthetic process; | 0 % |  |  |
| 6 | ggggacaagtttgtacaaaaaagcaggcttgatgGACGATGCGGAATACGGCAA, ggggaccactttgtacaagaaagctgggtcTGAGTACTCATCCCCCATAT | Mycgr3\|74015  JGI  Ensembl | KOG0482 DNA replication licensing factor, MCM7 component; | ATP binding; ATP-dependent DNA helicase activity;  ATP-dependent RNA helicase activity;  ATP-dependent helicase activity;  ATPase activity;  ATPase activity, coupled;  ATPase activity, coupled to transmembrane movement of ions;  ATPase activity, coupled to transmembrane movement of substances;  ATPase activity, uncoupled;  DNA binding;  DNA replication initiation;  DNA-dependent ATPase activity;  RNA-dependent ATPase activity;  nucleoside-triphosphatase activity;  nucleotide binding;  nucleus;  protein-transmembrane transporting ATPase activity;  single-stranded DNA-dependent ATP-dependent DNA helicase activity; | 0 % |  |  |
| 6 | ggggacaagtttgtacaaaaaagcaggcttgatgCCTCGTGTACGCGCGCCGAC, ggggaccactttgtacaagaaagctgggtcATCCCCACCCCCACCCTCTG | Mycgr3\|74061  JGI  Ensembl | KOG2328 Chromosome condensation complex Condensin, subunit H; | mitosis; mitotic cell cycle;  nucleus; | 0.79848 % |  |  |
| 6 | ggggacaagtttgtacaaaaaagcaggcttgatgTCACAAGCAGGCGCCAACTC, ggggaccactttgtacaagaaagctgggtcAGCAAGGTAGCCGAGATCGC | Mycgr3\|76524  JGI  Ensembl | KOG2166 Cullins; | cullin-RING ubiquitin ligase complex; ubiquitin protein ligase binding;  ubiquitin-dependent protein catabolic process; | 3.47863 % |  |  |
| 6 | ggggacaagtttgtacaaaaaagcaggcttgatgCATCCGGTCTCGATCTTGAC, ggggaccactttgtacaagaaagctgggtcCCTCCGCTCACTCTTCCTCC | Mycgr3\|76905  JGI  Ensembl | KOG1183 N-acetylglucosaminyltransferase complex, subunit PIG-Q/GPI1, required for phosphatidylinositol biosynthesis; | GPI anchor biosynthetic process; integral to membrane;  phosphatidylinositol N-acetylglucosaminyltransferase activity; | 0 % |  |  |
| 6 | ggggacaagtttgtacaaaaaagcaggcttgatgGAATCCTCACGCATATTTGT, ggggaccactttgtacaagaaagctgggtcAGCATCGTCGTCCTCGGCCT | Mycgr3\|85600  JGI  Ensembl | KOG0110 RNA-binding protein (RRM superfamily); | nucleic acid binding; | 0 % |  |  |
| 6 | ggggacaagtttgtacaaaaaagcaggcttgatgGCCACCGATACGACCGCATA, ggggaccactttgtacaagaaagctgggtcTCCAATCCAAGCCTCCTTTA | Mycgr3\|86683  JGI  Ensembl | KOG3818 DNA polymerase epsilon, subunit B; | DNA binding; DNA polymerase activity;  DNA replication;  DNA-directed DNA polymerase activity;  nucleus; | 4.85933 % |  |  |
| 6 | ggggacaagtttgtacaaaaaagcaggcttgatgAACGAAGATGCGGCCATTCA, ggggaccactttgtacaagaaagctgggtcAACGAAGTCGGCAGAGTACG | Mycgr3\|87605  JGI  Ensembl | KOG0694 Serine/threonine protein kinase; | ATP binding; calcium-dependent protein kinase C activity;  calcium-independent protein kinase C activity;  intracellular;  intracellular signaling cascade;  protein amino acid phosphorylation;  protein kinase C activity;  protein kinase activity;  protein serine/threonine kinase activity;  protein-tyrosine kinase activity;  signal transduction; | 0 % |  |  |
| 6 | ggggacaagtttgtacaaaaaagcaggcttgatgTTGTTATGCCCATTATGGCA, ggggaccactttgtacaagaaagctgggtcGCGCCTCCCGTTTGTCGCCG | Mycgr3\|89011  JGI  Ensembl | KOG1856 Transcription elongation factor SPT6; | protein binding; | 0.4537 % |  |  |
| 6 | ggggacaagtttgtacaaaaaagcaggcttgatgAAGCGAAAGATCGGAGCGTT, ggggaccactttgtacaagaaagctgggtcATTACCCGCGTTACCTCTCT | Mycgr3\|90616  JGI  Ensembl | KOG2229 Protein required for actin cytoskeleton organization and cell cycle progression; | FK506-sensitive peptidyl-prolyl cis-trans isomerase; cyclophilin; cyclophilin-type peptidyl-prolyl cis-trans isomerase activity;  peptidyl-prolyl cis-trans isomerase activity; | 0 % |  |  |
| 6 | ggggacaagtttgtacaaaaaagcaggcttgatgCCTCCCACGCGAGCCTCCAA, ggggaccactttgtacaagaaagctgggtcGATCTCCCTACACCACCGTC | Mycgr3\|93387  JGI  Ensembl | KOG2228 Origin recognition complex, subunit 4; |  | 0 % |  |  |
| 6 | ggggacaagtttgtacaaaaaagcaggcttgatgGTACGCTCATACCGGAAGTA, ggggaccactttgtacaagaaagctgggtcCGCCATACTAGCGAAAGCTC | Mycgr3\|93412  JGI  Ensembl | KOG0306 WD40-repeat-containing subunit of the 18S rRNA processing complex; | rRNA processing; small subunit processome; | 0 % |  |  |
| 6 | ggggacaagtttgtacaaaaaagcaggcttgatgGCGGTCGATGCACCCGTATC, ggggaccactttgtacaagaaagctgggtcGCCCTCCCTGCCCCCCACAA | Mycgr3\|93631  JGI  Ensembl | KOG0412 Golgi transport complex COD1 protein; |  | 0 % |  |  |
| 6 | ggggacaagtttgtacaaaaaagcaggcttgatgAATTCTTCACAAGCTGTGGC, ggggaccactttgtacaagaaagctgggtcGTCCTCCATCTCAACATCAC | Mycgr3\|94196  JGI  Ensembl | KOG0262 RNA polymerase I, large subunit; | DNA binding; DNA-directed RNA polymerase I activity;  DNA-directed RNA polymerase II activity;  DNA-directed RNA polymerase III activity;  DNA-directed RNA polymerase activity;  RNA polymerase activity;  transcription; | 0 % |  |  |
| 6 | ggggacaagtttgtacaaaaaagcaggcttgatgGCCAACTCGAACGTGGGACA, ggggaccactttgtacaagaaagctgggtcAATCACCAACCACTCCTCCC | Mycgr3\|96617  JGI  Ensembl | KOG2076 RNA polymerase III transcription factor TFIIIC; |  | 0 % |  |  |
| 6 | ggggacaagtttgtacaaaaaagcaggcttgatgCTACAGAAGGCCAAAGCACC, ggggaccactttgtacaagaaagctgggtcGTCGCTCTCATCGTCGTCGA | Mycgr3\|98287  JGI  Ensembl | KOG0355 DNA topoisomerase type II; | ATP binding; DNA binding;  DNA metabolic process;  DNA topoisomerase (ATP-hydrolyzing) activity;  DNA topological change;  chromosome;  nucleus; | 0 % |  |  |
| 6 | ggggacaagtttgtacaaaaaagcaggcttgatgCCGGGCGCGGCGTCTCCCAT, ggggaccactttgtacaagaaagctgggtcATATCCCATGCCATCATTCA | Mycgr3\|98356  JGI  Ensembl | KOG4124 Putative transcriptional repressor regulating G2/M transition; | intracellular; nucleic acid binding;  zinc ion binding; | 0 % |  |  |
| 6 | ggggacaagtttgtacaaaaaagcaggcttgatgGATATCCACCGATCACGTTT, ggggaccactttgtacaagaaagctgggtcCTTCGTCTCTTCGGCCTCCC | Mycgr3\|98483  JGI  Ensembl | KOG2048 WD40 repeat protein; |  | 0 % |  |  |
| 6 | ggggacaagtttgtacaaaaaagcaggcttgatgGCAGAAGGACGCATCGCCAA, ggggaccactttgtacaagaaagctgggtcAACCCTAGCCGCCCTCAAAT | Mycgr3\|99387  JGI  Ensembl | KOG1539 WD repeat protein; | rRNA processing; small subunit processome; | 0 % |  |  |
| 6 | ggggacaagtttgtacaaaaaagcaggcttgatgCCGAAAATCCTCTCATGTCA, ggggaccactttgtacaagaaagctgggtcACGAATAGCAGCCTGAAGCT | Mycgr3\|99512  JGI  Ensembl |  |  | 0 % |  |  |
| 6 | ggggacaagtttgtacaaaaaagcaggcttgatgGCGTCCAAGGCTGACGCTCT, ggggaccactttgtacaagaaagctgggtcCCCCAACCTCTTGAGAGCAA | Mycgr3\|99627  JGI  Ensembl | KOG0970 DNA polymerase alpha, catalytic subunit; | DNA binding; DNA polymerase activity;  DNA replication;  DNA-directed DNA polymerase activity;  nucleic acid binding;  nucleobase, nucleoside, nucleotide and nucleic acid metabolic process;  nucleotide binding;  nucleus; | 0 % |  |  |
| 6 | ggggacaagtttgtacaaaaaagcaggcttgatgCCGCCCTCCCAACTCAAGCG, ggggaccactttgtacaagaaagctgggtcCCGCCCCGCCTTCTTCTTCT | Mycgr3\|100069  JGI  Ensembl | KOG2147 Nucleolar protein involved in 40S ribosome biogenesis; |  | 0 % |  |  |
| 6 | ggggacaagtttgtacaaaaaagcaggcttgatgGCACAGCAGAATACGCCACG, ggggaccactttgtacaagaaagctgggtcGACAAGATCCCATTTCGCCA | Mycgr3\|100162  JGI  Ensembl | KOG3322 Ribonucleases P/MRP protein subunit; | ribonuclease P activity; tRNA 5'-leader removal; | 0.78563 % |  |  |
| 6 | ggggacaagtttgtacaaaaaagcaggcttgatgGCGTTGAACCACGCTTCCCA, ggggaccactttgtacaagaaagctgggtcCAACTTCATCTCCTCGAACC | Mycgr3\|100193  JGI  Ensembl | KOG0432 Valyl-tRNA synthetase; | ATP binding; aminoacyl-tRNA ligase activity;  cytoplasm;  nucleotide binding;  tRNA aminoacylation for protein translation;  translation;  valine-tRNA ligase activity;  valyl-tRNA aminoacylation; | 0 % |  |  |
| 6 | ggggacaagtttgtacaaaaaagcaggcttgatgGCGCGACTCGGGCACAGGGG, ggggaccactttgtacaagaaagctgggtcCCTTCGCTTTTTTCCACCTT | Mycgr3\|100344  JGI  Ensembl | KOG2124 Glycosylphosphatidylinositol anchor synthesis protein; |  | 0 % |  |  |
| 6 | ggggacaagtttgtacaaaaaagcaggcttgatgGAGATGATGTCCTCACGGGT, ggggaccactttgtacaagaaagctgggtcAATACCAACTCCTGGCGGCC | Mycgr3\|100550  JGI  Ensembl | KOG2173 Integral membrane protein; |  | 0.93423 % |  |  |
| 6 | ggggacaagtttgtacaaaaaagcaggcttgatgGCGCCGCCCACAAATGCCCG, ggggaccactttgtacaagaaagctgggtcCCCAAGCAGCCTTGCAGCCG | Mycgr3\|101180  JGI  Ensembl | KOG0343 RNA Helicase; | 8-oxo-7,8-dihydroguanine triphosphatase activity; ATP binding;  ATP-dependent 5'-3' DNA helicase activity;  ATP-dependent helicase activity;  UDP-2,3-diacylglucosamine hydrolase activity;  bis(5'-nucleosyl)-tetraphosphatase activity;  dATP pyrophosphohydrolase activity;  diacylglycerol pyrophosphate phosphatase activity;  dihydroneopterin monophosphate phosphatase activity;  dihydroneopterin triphosphate pyrophosphohydrolase activity;  helicase activity;  nucleic acid binding;  pyrophosphatase activity;  thiamin-pyrophosphatase activity; | 0.78093 % |  |  |
| 6 | ggggacaagtttgtacaaaaaagcaggcttgatgGGCAAAAAGGACCACCAGTA, ggggaccactttgtacaagaaagctgggtcTTCCAAATTCTCCTTAAACG | Mycgr3\|101192  JGI  Ensembl | KOG1386 Nucleoside phosphatase; | adenosine-diphosphatase activity; apyrase activity;  hydrolase activity; | 0 % |  |  |
| 6 | ggggacaagtttgtacaaaaaagcaggcttgatgAATGGACTCGCTGGACAACC, ggggaccactttgtacaagaaagctgggtcCAAGAAGCCGGACTCTGACA | Mycgr3\|101362  JGI  Ensembl | KOG3417 Ras1 guanine nucleotide exchange factor; | guanyl-nucleotide exchange factor activity; intracellular;  regulation of small GTPase mediated signal transduction;  small GTPase mediated signal transduction; | 0 % |  |  |
| 6 | ggggacaagtttgtacaaaaaagcaggcttgatgGCCGGCGAAGATCAAGCGCA, ggggaccactttgtacaagaaagctgggtcAGCCCTCTTTCTCCTCTTCC | Mycgr3\|101428  JGI  Ensembl | KOG1951 GTP-binding protein AARP2 involved in 40S ribosome biogenesis; |  | 0 % |  |  |
| 6 | ggggacaagtttgtacaaaaaagcaggcttgatgTCCAAAATCGAGAAGCTCTC, ggggaccactttgtacaagaaagctgggtcCAACACCTCCGAAATGCTCT | Mycgr3\|106885  JGI  Ensembl | KOG0962 DNA repair protein RAD50, ABC-type ATPase/SMC superfamily; | ATP binding; ATPase activity;  DNA repair;  Mre11 complex; | 0 % |  |  |
| 6 | ggggacaagtttgtacaaaaaagcaggcttgatgGCTTCCGGGCCTTCAACAGC, ggggaccactttgtacaagaaagctgggtcCACCGAATCTGTCACCTTGG | Mycgr3\|107174  JGI  Ensembl | KOG2710 Rho GTPase-activating protein; | intracellular; signal transduction; | 0 % |  |  |
| 6 | ggggacaagtttgtacaaaaaagcaggcttgatgCTCGGGTTCTCACCCAGCCT, ggggaccactttgtacaagaaagctgggtcGAGCTGCTCGGCAATCGCCA | Mycgr3\|107212  JGI  Ensembl | KOG0433 Isoleucyl-tRNA synthetase; | ATP binding; aminoacyl-tRNA ligase activity;  catalytic activity;  cytoplasm;  isoleucine-tRNA ligase activity;  isoleucyl-tRNA aminoacylation;  metabolic process;  nucleotide binding;  oxidoreductase activity;  tRNA aminoacylation for protein translation;  translation; | 0 % |  |  |
| 6 | ggggacaagtttgtacaaaaaagcaggcttgatgCCTCCACCACCGCATACGCG, ggggaccactttgtacaagaaagctgggtcTCTCCGCCCTGGTGGACGGT | Mycgr3\|107402  JGI  Ensembl | KOG2072 Translation initiation factor 3, subunit a (eIF-3a); |  | 0 % |  |  |
| 6 | ggggacaagtttgtacaaaaaagcaggcttgatgGCTGAGCGGTTCAGAAGCGA, ggggaccactttgtacaagaaagctgggtcGTTACAATGTACGACAATCG | Mycgr3\|107460  JGI  Ensembl |  |  | 0 % |  |  |
| 6 | ggggacaagtttgtacaaaaaagcaggcttgatgTCTGAAGGCGACGTGCAGCC, ggggaccactttgtacaagaaagctgggtcCGCGAAGATGTCAGAGATCA | Mycgr3\|107618  JGI  Ensembl | KOG1175 Acyl-CoA synthetase; | acetate-CoA ligase activity; catalytic activity;  metabolic process; | 0 % |  |  |
| 6 | ggggacaagtttgtacaaaaaagcaggcttgatgGCCCCGTCCGCAACAATGGA, ggggaccactttgtacaagaaagctgggtcATCATCCTCATCAGAGAACA | Mycgr3\|108146  JGI  Ensembl | KOG0062 ATPase component of ABC transporters with duplicated ATPase domains/Translation elongation factor EF-3b; | ATP binding; ATPase activity;  nucleoside-triphosphatase activity;  nucleotide binding; | 4.85589 % |  |  |
| 6 | ggggacaagtttgtacaaaaaagcaggcttgatgCCGTCCATATCAACCCTGCT, ggggaccactttgtacaagaaagctgggtcGCTGGCCACCAGCGTATCGA | Mycgr3\|108192  JGI  Ensembl |  |  | 0 % |  |  |
| 6 | ggggacaagtttgtacaaaaaagcaggcttgatgGCGGATGAGGCGGAGGAGTT, ggggaccactttgtacaagaaagctgggtcCTGTCCTCTTCCAATATTCT | Mycgr3\|108665  JGI  Ensembl | KOG1820 Microtubule-associated protein; |  | 0 % |  |  |
| 6 | ggggacaagtttgtacaaaaaagcaggcttgatgCAATTATGCGTGACTTCTCC, ggggaccactttgtacaagaaagctgggtcGTCCTGCTCACATGGTGGTT | Mycgr3\|108706  JGI  Ensembl | KOG4540 Putative lipase essential for disintegration of autophagic bodies inside the vacuole; | catalytic activity; triacylglycerol lipase activity; | 0.80611 % |  |  |
| 6 | ggggacaagtttgtacaaaaaagcaggcttgatgTCTCGATTCTTTCACGGCAG, ggggaccactttgtacaagaaagctgggtcCTGCGCTCGGATGGCATTGC | Mycgr3\|108717  JGI  Ensembl | KOG1076 Translation initiation factor 3, subunit c (eIF-3c); | eukaryotic translation initiation factor 3 complex; translation initiation factor activity;  translational initiation; | 0 % |  |  |
| 6 | ggggacaagtttgtacaaaaaagcaggcttgatgGCAAGCAGAGCCTCTCTCGA, ggggaccactttgtacaagaaagctgggtcTTCCGGCGACCTGCCTAAGG | Mycgr3\|109027  JGI  Ensembl |  |  | 0 % |  |  |
| 6 | ggggacaagtttgtacaaaaaagcaggcttgatgTATCTCCCACGCCAGCAGAT, ggggaccactttgtacaagaaagctgggtcCTTCGAGGCGAGATTGATGA | Mycgr3\|109152  JGI  Ensembl | KOG2475 CDC45 (cell division cycle 45)-like protein; | DNA replication initiation; | 0 % |  |  |
| 6 | ggggacaagtttgtacaaaaaagcaggcttgatgGCGGCCTACCCGTCCTCCAC, ggggaccactttgtacaagaaagctgggtcAGGCACAACCAAAACCCCAC | Mycgr3\|109911  JGI  Ensembl | KOG3257 Mitochondrial/chloroplast ribosomal protein L11; | Kdo transferase activity; catalytic activity;  intracellular;  membrane;  regulation of telomerase activity;  regulation of transferase activity;  ribosome;  structural constituent of ribosome;  translation; | 14.12145 % |  |  |
| 6 | ggggacaagtttgtacaaaaaagcaggcttgatgTTCAACTTCACGGCGTTGCT, ggggaccactttgtacaagaaagctgggtcAGGCTTGTGCACCACCAACC | Mycgr3\|109995  JGI  Ensembl | KOG1135 mRNA cleavage and polyadenylation factor II complex, subunit CFT2 (CPSF subunit); |  | 0 % |  |  |
| 6 | ggggacaagtttgtacaaaaaagcaggcttgatgGCGACCGAAAAAGCCCAGCC, ggggaccactttgtacaagaaagctgggtcCTCCACCTCGACAGCCTGCG | Mycgr3\|110152  JGI  Ensembl | KOG2753 Uncharacterized conserved protein, contains PCI domain; | nucleus; ribosome biogenesis and assembly; | 0 % |  |  |
| 6 | ggggacaagtttgtacaaaaaagcaggcttgatgCCCATCGCTTTGGGCCTCGA, ggggaccactttgtacaagaaagctgggtcCTCGTCCTCCATCTTGACGT | Mycgr3\|110412  JGI  Ensembl | KOG2708 Predicted metalloprotease with chaperone activity (RNAse H/HSP70 fold); | O-sialoglycoprotein endopeptidase activity; collagenase activity;  endopeptidase activity;  matrilysin-2 activity;  prenyl protein specific endopeptidase activity;  proteolysis;  stromelysin 3 activity;  zinc ion binding; | 0 % |  |  |
| 6 | ggggacaagtttgtacaaaaaagcaggcttgatgTGGCGCAGGATCTATCTCGC, ggggaccactttgtacaagaaagctgggtcCATTTTCTTGCCCGTCGGAA | Mycgr3\|110450  JGI  Ensembl | KOG1147 Glutamyl-tRNA synthetase; | ATP binding; aminoacyl-tRNA ligase activity;  cytoplasm;  glutamate-tRNA ligase activity;  glutamate-tRNA ligase complex;  glutamyl-tRNA aminoacylation;  nucleotide binding;  tRNA aminoacylation for protein translation;  translation; | 0 % |  |  |
| 6 | ggggacaagtttgtacaaaaaagcaggcttgatgGCGCCCAAACGGAAAGCGGC, ggggaccactttgtacaagaaagctgggtcCTCCAGCAAACCCTGCTGCT | Mycgr3\|110466  JGI  Ensembl | KOG1123 RNA polymerase II transcription initiation/nucleotide excision repair factor TFIIH, 3'-5' helicase subunit SSL2; | 8-oxo-7,8-dihydroguanine triphosphatase activity; ATP binding;  ATP-dependent 5'-3' DNA helicase activity;  ATP-dependent DNA helicase activity;  ATP-dependent helicase activity;  DNA binding;  UDP-2,3-diacylglucosamine hydrolase activity;  bis(5'-nucleosyl)-tetraphosphatase activity;  dATP pyrophosphohydrolase activity;  diacylglycerol pyrophosphate phosphatase activity;  dihydroneopterin monophosphate phosphatase activity;  dihydroneopterin triphosphate pyrophosphohydrolase activity;  helicase activity;  hydrolase activity;  nucleic acid binding;  nucleotide-excision repair;  nucleus;  pyrophosphatase activity;  thiamin-pyrophosphatase activity; | 0 % |  |  |
| 6 | ggggacaagtttgtacaaaaaagcaggcttgatgGCAGACTCTCTCGTGACATA, ggggaccactttgtacaagaaagctgggtcCCCCTTCAGATCTTCCGTGA | Mycgr3\|111035  JGI  Ensembl | KOG1128 Uncharacterized conserved protein, contains TPR repeats; |  | 0 % |  |  |
| 6 | ggggacaagtttgtacaaaaaagcaggcttgatgAACGACCACGACGACACCCC, ggggaccactttgtacaagaaagctgggtcCCGCCCCCTATCCCCAAAAT | Mycgr3\|111161  JGI  Ensembl | KOG0118 FOG: RRM domain; | nucleic acid binding; | 0.88823 % |  |  |
| 6 | ggggacaagtttgtacaaaaaagcaggcttgatgCCCAAAGAGGGCCTCAAGCG, ggggaccactttgtacaagaaagctgggtcCAACCCCATCCTCTCCCCCG | Mycgr3\|111327  JGI  Ensembl | KOG0629 Glutamate decarboxylase and related proteins; | RNA binding; carboxy-lyase activity;  carboxylic acid metabolic process;  pyridoxal phosphate binding; | 0 % |  |  |
| 6 | ggggacaagtttgtacaaaaaagcaggcttgatgGCGCTCGTCCAGTCCACCAA, ggggaccactttgtacaagaaagctgggtcTCTTCTTGAATATCCGCCGG | Mycgr3\|111506  JGI  Ensembl | KOG0619 FOG: Leucine rich repeat; | protein binding; | 0.6563 % |  |  |
| 6 | ggggacaagtttgtacaaaaaagcaggcttgatgGCGATGAGTATCGAGGAGCT, ggggaccactttgtacaagaaagctgggtcCTCGATTTCATGGTAGGAAT | Mycgr3\|111551  JGI  Ensembl | KOG2020 Nuclear transport receptor CRM1/MSN5 (importin beta superfamily); | cytoplasm; intracellular protein transport;  nuclear pore;  nucleus;  protein import into nucleus, docking;  protein transporter activity; | 0 % |  |  |
| 6 | ggggacaagtttgtacaaaaaagcaggcttgatgGACGAGCTATTCGATGTCTT, ggggaccactttgtacaagaaagctgggtcGCGAACATCGAGGGCATGAG | Mycgr3\|111559  JGI  Ensembl | KOG0948 Nuclear exosomal RNA helicase MTR4, DEAD-box superfamily; | 8-oxo-7,8-dihydroguanine triphosphatase activity; ATP binding;  ATP-dependent 5'-3' DNA helicase activity;  ATP-dependent helicase activity;  UDP-2,3-diacylglucosamine hydrolase activity;  bis(5'-nucleosyl)-tetraphosphatase activity;  dATP pyrophosphohydrolase activity;  diacylglycerol pyrophosphate phosphatase activity;  dihydroneopterin monophosphate phosphatase activity;  dihydroneopterin triphosphate pyrophosphohydrolase activity;  helicase activity;  nucleic acid binding;  pyrophosphatase activity;  thiamin-pyrophosphatase activity; | 0.61118 % |  |  |
| 7 | ggggacaagtttgtacaaaaaagcaggcttgatgGCGCCCAAAGGCAAAGGCGG, ggggaccactttgtacaagaaagctgggtcCTTTCTCCCCTCGACCATAA | Mycgr3\|39964  JGI  Ensembl | KOG3258 Parvulin-like peptidyl-prolyl cis-trans isomerase; | FK506-sensitive peptidyl-prolyl cis-trans isomerase; cyclophilin; cyclophilin-type peptidyl-prolyl cis-trans isomerase activity;  isomerase activity;  peptidyl-prolyl cis-trans isomerase activity; | 4.71311 % |  |  |
| 7 | ggggacaagtttgtacaaaaaagcaggcttgatgTCTCCACGAAAGATCCACCT, ggggaccactttgtacaagaaagctgggtcCGCACCCATACTTCCCGTCT | Mycgr3\|40001  JGI  Ensembl | KOG4754 Predicted phosphoglycerate mutase; |  | 100 % |  | ``` 1	ATGTCTCCACGAAAGATCCACCTCCTCCGCCACGCTCAAGCATACCACAACGTCCCACCGTGCAACTACAACATTCCGGATCCCACCTTGACTGAATTGGGACTCGAACAGACGTCGACTTTCGCCCGCGAGTTCCCACATCACGGCAATGTGTCGGTCATCCTCTCCTCGCCCATGACGAGAACTCTCGAGACGGCGCTTCTATGTTTTCCTCAGGAA	219 	\|\|\|\|\|\|\|\|\|\|\|\|\|\|\|\|\|\|\|\|\|\|\|\|\|\|\|\|\|\|\|\|\|\|\|\|\|\|\|\|\|\|\|\|\|\|\|\|\|\|\|\|\|\|\|\|\|\|\|\|\|\|\|\|\|\|\|\|\|\|\|\|\|\|\|\|\|\|\|\|\|\|\|\|\|\|\|\|\|\|\|\|\|\|\|\|\|\|\|\|\|\|\|\|\|\|\|\|\|\|\|\|\|\|\|\|\|\|\|\|\|\|\|\|\|\|\|\|\|\|\|\|\|\|\|\|\|\|\|\|\|\|\|\|\|\|\|\|\|\|\|\|\|\|\|\|\|\|\|\|\|\|\|\|\|\|\|\|\|\|\|\|\|\|\|\|\|\|\|\|\|\|\|\|\|\|\|\|\|\|\|\|\|\|\|\|\|\|\|\|\|\|\|\|\|\|\|\|\|\|\|\|\|\|\|\|\|\|\| 	ATGTCTCCACGAAAGATCCACCTCCTCCGCCACGCTCAAGCATACCACAACGTCCCACCGTGCAACTACAACATTCCGGATCCCACCTTGACTGAATTGGGACTCGAACAGACGTCGACTTTCGCCCGCGAGTTCCCACATCACGGCAATGTGTCGGTCATCCTCTCCTCGCCCATGACGAGAACTCTCGAGACGGCGCTTCTATGTTTTCCTCAGGAA ```      ``` 220	CACGAACATCGACACCATCACAAAGGACTGAAGATATACGCCGTTCCTGAGCTTCAAGAGACGTCCGATCACCCCTGCGACACCGGCTCGAGTCGAGCAGACTTGGAGAAGGTGTTTGATGGAAAGAACATCGATTTCAGCGCGTTGAAAGAAGGCTGGCAGCTGAAGACGGGAAGTATGGGTGCG	405 	\|\|\|\|\|\|\|\|\|\|\|\|\|\|\|\|\|\|\|\|\|\|\|\|\|\|\|\|\|\|\|\|\|\|\|\|\|\|\|\|\|\|\|\|\|\|\|\|\|\|\|\|\|\|\|\|\|\|\|\|\|\|\|\|\|\|\|\|\|\|\|\|\|\|\|\|\|\|\|\|\|\|\|\|\|\|\|\|\|\|\|\|\|\|\|\|\|\|\|\|\|\|\|\|\|\|\|\|\|\|\|\|\|\|\|\|\|\|\|\|\|\|\|\|\|\|\|\|\|\|\|\|\|\|\|\|\|\|\|\|\|\|\|\|\|\|\|\|\|\|\|\|\|\|\|\|\|\|\|\|\|\|\|\|\|\|\|\|\|\|\|\|\|\|\|\|\|\|\|\|\|\|\|\|\|\| 	CACGAACATCGACACCATCACAAAGGACTGAAGATATACGCCGTTCCTGAGCTTCAAGAGACGTCCGATCACCCCTGCGACACCGGCTCGAGTCGAGCAGACTTGGAGAAGGTGTTTGATGGAAAGAACATCGATTTCAGCGCGTTGAAAGAAGGCTGGCAGCTGAAGACGGGAAGTATGGGTGCG ``` |
| 7 | ggggacaagtttgtacaaaaaagcaggcttgatgCAGCAACCAACACCACCAGA, ggggaccactttgtacaagaaagctgggtcGGCAGGAGGGGTATCATGAA | Mycgr3\|42373  JGI  Ensembl |  |  | 100 % |  | ``` 358	GATGCGAAGTTCCTTCTAGTTGTGATTCATGATACCCCTCCTGCC	402 	\|\|\|\|\|\|\|\|\|\|\|\|\|\|\|\|\|\|\|\|\|\|\|\|\|\|\|\|\|\|\|\|\|\|\|\|\|\|\|\|\|\|\|\|\| 	GATGCGAAGTTCCTTCTAGTTGTGATTCATGATACCCCTCCTGCC ```      ``` 1	ATGCAGCAACCAACACCACCAGAAGAGACTGCAAATCTCCTTGCGCGTTTGTCACAACGACCAGGAGTTCAAGGCACCCTCATCCTCTCCCGCGACACAGGAGCCATTGTGCGATCATCTGGGCTCATCACAAACGAAGACGGCGAGACTGAGGCCGCGCGACCTACAAGCAGCGATCAAGTAAACGGAGACGGCGAAGTGAAGAAGAAGGGAACAAGAAACGCAGAGGATGTGGCACGGCTGGTCTGGAGATTCGTGCAGAGTGCAGGGACAATGATTGAGGAGCTGAATGGAGATTCGGACGAGGCGAAGCTGCTGAGGATACGGACGAAGAAGAATGAGTTAGTGATTGTGCCGG	358 	\|\|\|\|\|\|\|\|\|\|\|\|\|\|\|\|\|\|\|\|\|\|\|\|\|\|\|\|\|\|\|\|\|\|\|\|\|\|\|\|\|\|\|\|\|\|\|\|\|\|\|\|\|\|\|\|\|\|\|\|\|\|\|\|\|\|\|\|\|\|\|\|\|\|\|\|\|\|\|\|\|\|\|\|\|\|\|\|\|\|\|\|\|\|\|\|\|\|\|\|\|\|\|\|\|\|\|\|\|\|\|\|\|\|\|\|\|\|\|\|\|\|\|\|\|\|\|\|\|\|\|\|\|\|\|\|\|\|\|\|\|\|\|\|\|\|\|\|\|\|\|\|\|\|\|\|\|\|\|\|\|\|\|\|\|\|\|\|\|\|\|\|\|\|\|\|\|\|\|\|\|\|\|\|\|\|\|\|\|\|\|\|\|\|\|\|\|\|\|\|\|\|\|\|\|\|\|\|\|\|\|\|\|\|\|\|\|\|\|\|\|\|\|\|\|\|\|\|\|\|\|\|\|\|\|\|\|\|\|\|\|\|\|\|\|\|\|\|\|\|\|\|\|\|\|\|\|\|\|\|\|\|\|\|\|\|\|\|\|\|\|\|\|\|\|\|\|\|\|\|\|\|\|\|\|\|\|\|\|\|\|\|\|\|\|\|\|\|\|\|\|\|\|\|\|\|\|\|\|\|\|\|\|\|\|\|\|\|\|\|\|\|\|\|\|\|\|\|\|\|\|\|\|\|\|\|\|\|\|\|\|\|\|\|\|\|\|\|\|\|\|\|\|\|\|\|\|\| 	ATGCAGCAACCAACACCACCAGAAGAGACTGCAAATCTCCTTGCGCGTTTGTCACAACGACCAGGAGTTCAAGGCACCCTCATCCTCTCCCGCGACACAGGAGCCATTGTGCGATCATCTGGGCTCATCACAAACGAAGACGGCGAGACTGAGGCCGCGCGACCTACAAGCAGCGATCAAGTAAACGGAGACGGCGAAGTGAAGAAGAAGGGAACAAGAAACGCAGAGGATGTGGCACGGCTGGTCTGGAGATTCGTGCAGAGTGCAGGGACAATGATTGAGGAGCTGAATGGAGATTCGGACGAGGCGAAGCTGCTGAGGATACGGACGAAGAAGAATGAGTTAGTGATTGTGCCGG ``` |
| 7 | ggggacaagtttgtacaaaaaagcaggcttgatgGCCGACGACGCCGTCAAGTC, ggggaccactttgtacaagaaagctgggtcTTCCTTCTTCGGCTCGGAGC | Mycgr3\|43793  JGI  Ensembl | KOG3328 HGG motif-containing thioesterase; |  | 0 % |  |  |
| 7 | ggggacaagtttgtacaaaaaagcaggcttgatgCCAGTTGCTTCCCGACTACT, ggggaccactttgtacaagaaagctgggtcCATGCCTTTTCTAATAGTGC | Mycgr3\|43977  JGI  Ensembl |  |  | 100 % |  | ``` 168	TACACATCTCGCAGAGCTGTCCAAATATCCCTCGGACTTCATACTTGTCGGAGGCCCTTACCTAGGCGGGCCTGTACAAGACGACGCGCCAATGGACATGCTCGGTTCCGTGATACTAGTCCGGGCATCGACCCGGGAAGAGGCTCTCGGAGAGCTACGCAAGGACATATTCGCAAGGAAAGCGGTCTGGGATTTCGATCAAGCCCTCGTCTTGCCTTTTAAGAGCACTATTAGAAAAGGCATG	411 	\|\|\|\|\|\|\|\|\|\|\|\|\|\|\|\|\|\|\|\|\|\|\|\|\|\|\|\|\|\|\|\|\|\|\|\|\|\|\|\|\|\|\|\|\|\|\|\|\|\|\|\|\|\|\|\|\|\|\|\|\|\|\|\|\|\|\|\|\|\|\|\|\|\|\|\|\|\|\|\|\|\|\|\|\|\|\|\|\|\|\|\|\|\|\|\|\|\|\|\|\|\|\|\|\|\|\|\|\|\|\|\|\|\|\|\|\|\|\|\|\|\|\|\|\|\|\|\|\|\|\|\|\|\|\|\|\|\|\|\|\|\|\|\|\|\|\|\|\|\|\|\|\|\|\|\|\|\|\|\|\|\|\|\|\|\|\|\|\|\|\|\|\|\|\|\|\|\|\|\|\|\|\|\|\|\|\|\|\|\|\|\|\|\|\|\|\|\|\|\|\|\|\|\|\|\|\|\|\|\|\|\|\|\|\|\|\|\|\|\|\|\|\|\|\|\|\|\|\|\|\|\|\| \|\|\|\|\|\|\|\|\|\| 	TACACATCTCGCAGAGCTGTCCAAATATCCCTCGGACTTCATACTTGTCGGAGGCCCTTACCTAGGCGGGCCTGTACAAGACGACGCGCCAATGGACATGCTCGGTTCCGTGATACTAGTCCGGGCATCGACCCGGGAAGAGGCTCTCGGAGAGCTACGCAAGGACATATTCGCAAGGAAAGCGGTCTGGGATTTCGATCAAGCCCTCGTCTTGCCTTTTAAGAGCACTATTANAAAAGGCATG ```      ``` 1	ATGCCAGTTGCTTCCCGACTACTTTGGACCCTACCGGCCACAAGGCCAAGGCGTCTGCCCGCTACTGCATCAAGATATGCTGAGGCACTGCGACGGCACAATCACGAGTGGTTGGTCATTCTACCTGATCAGGAAGGAGCTTTACAGCGTCGGGTTCAGCATCGATC	167 	\|\|\|\|\|\|\|\|\|\|\|\|\|\|\|\|\|\|\|\|\|\|\|\|\|\|\|\|\|\|\|\|\|\|\|\|\|\|\|\|\|\|\|\|\|\|\|\|\|\|\|\|\|\|\|\|\|\|\|\|\|\|\|\|\|\|\|\|\|\|\|\|\|\|\|\|\|\|\|\|\|\|\|\|\|\|\|\|\|\|\|\|\|\|\|\|\|\|\|\|\|\|\|\|\|\|\|\|\|\|\|\|\|\|\|\|\|\|\|\|\|\|\|\|\|\|\|\|\|\|\|\|\|\|\|\|\|\|\|\|\|\|\|\|\|\|\|\|\|\|\|\|\|\|\|\|\|\|\|\|\|\|\|\|\|\|\| 	ATGCCAGTTGCTTCCCGACTACTTTGGACCCTACCGGCCACAAGGCCAAGGCGTCTGCCCGCTACTGCATCAAGATATGCTGAGGCACTGCGACGGCACAATCACGAGTGGTTGGTCATTCTACCTGATCAGGAAGGAGCTTTACAGCGTCGGGTTCAGCATCGATC ``` |
| 7 | ggggacaagtttgtacaaaaaagcaggcttgatgGATATCAACGTATCCAGGGG, ggggaccactttgtacaagaaagctgggtcTCTTAACCTAAAAATGCTTA | Mycgr3\|44834  JGI  Ensembl | KOG0271 Notchless-like WD40 repeat-containing protein; |  | 0 % |  |  |
| 7 | ggggacaagtttgtacaaaaaagcaggcttgatgTCCACTCAACAACAAGTCCG, ggggaccactttgtacaagaaagctgggtcCCGCGGACTTGGTATCACGC | Mycgr3\|48799  JGI  Ensembl |  |  | 100 % |  | ``` 1	ATGTCCACTCAACAACAAGTCCGCG-TCGGCATAGCCCTCTTCATCCTCCACCCCTCCTCCACCCCCTCAAACCCAACCTTCCTCCTCGGCCGCCGCAAAGGCTCCCACGGAGCCGGAACCTACGCCCTTCCAGGCGGGCACCTCGAGTTCGGCGAAGATCCCGCTACCTGCGCTGGGCGAGAAGCGATGGAAGAGACAGGACTGGAAGTCGTGGATATCGAGTTCTTTACTGCGACGAACGATTATATGCCTGACGAGGGGAAGCATTATATCACGCTCTGGATGGTTGGACGACGGAAAGATGAGAGTGTAGAGGCGGAGGTGATGGAGAAGGATAAGTGTGAGGGGTGGGAGTGGTGTGC-TTGGGAG-AAGTTTG	376 	\|\|\|\|\|\|\|\|\|\|\|\|\|\|\|\|\|\|\|\| \|\|\|\| \|\|\|\|\|\|\|\|\|\|\|\|\|\|\|\|\|\|\|\|\|\|\|\|\|\|\|\|\|\|\|\|\|\|\|\|\|\|\|\|\|\|\|\|\|\|\|\|\|\|\|\|\|\|\|\| \|\|\|\|\|\|\|\|\|\|\|\|\|\|\| \|\|\|\|\|\|\|\|\|\|\|\|\|\|\|\|\|\|\|\|\|\|\|\|\|\|\|\|\|\|\|\|\|\|\|\|\|\|\|\|\|\|\|\|\|\|\|\|\|\|\|\|\|\|\|\|\|\|\|\|\|\|\|\|\|\|\|\|\|\|\|\|\|\|\|\|\|\|\|\|\|\|\|\|\|\|\|\|\|\|\|\|\|\|\|\|\|\|\|\|\|\|\|\|\|\|\|\|\|\|\|\|\|\|\|\|\|\|\|\|\|\|\|\|\|\|\|\|\|\|\|\|\|\|\|\|\|\|\|\|\|\|\|\|\|\|\|\|\|\|\|\|\|\|\|\|\|\|\|\|\|\|\|\|\|\|\|\|\|\|\|\|\|\|\|\|\|\|\|\|\|\|\|\|\|\|\|\|\|\|\|\|\|\|\|\|\|\|\|\|\|\|\|\|\|\|\|\|\|\|\|\|\|\|\|\|\|\|\|\|\|\|\|\|\|\|\|\|\|\|\|\|\|\|\|\|\|\|\|\|\|\|\|\|\|\|\|\|\|\|\|\|\| \|\|\|\|\|\|\|\|\|\| \|\|\| \|\|  \|\|\|\|\|\|\| 	ATGTCCACTCAACAACAAGTGCGCGTTCGGCATAGCCCTCTTCATCCTCCACCCCTCCTCCACCCCCTCAAACCCAACCTTCGTCCTCGGCCGCCGCAGAGGCTCCCACGGAGCCGGAACCTACGCCCTTCCAGGCGGGCACCTCGAGTTCGGCGAAGATCCCGCTACCTGCGCTGGGCGAGAAGCGATGGAAGAGACAGGACTGGAAGTCGTGGATATCGAGTTCTTTACTGCGACGAACGATTATATGCCTGACGAGGGGAAGCATTATATCACGCTCTGGATGGTTGGACGACGGAAAGATGAGAGTGTAGAGGCGGAGGTGATGGAGAAGGATAAGTGTGAGGGGTGGNAGTGGTGTGCNTTGNGANNAAGTTTG ``` |
| 7 | ggggacaagtttgtacaaaaaagcaggcttgatgGCCGACAAATCTCAGAATCC, ggggaccactttgtacaagaaagctgggtcCAAAGCCCTGGGATCATCCC | Mycgr3\|49175  JGI  Ensembl | KOG0940 Ubiquitin protein ligase RSP5/NEDD4; | protein binding; | 0 % |  |  |
| 7 | ggggacaagtttgtacaaaaaagcaggcttgatgCCGACCTCGGACGAATACGT, ggggaccactttgtacaagaaagctgggtcGCTGGAGTCGATGATCTCTT | Mycgr3\|49268  JGI  Ensembl |  |  | 0 % |  |  |
| 7 | ggggacaagtttgtacaaaaaagcaggcttgatgGCCTACCTTCTCCCAAGCTT, ggggaccactttgtacaagaaagctgggtcCGCGTGATGGTACAGAACAG | Mycgr3\|51082  JGI  Ensembl |  |  | 0 % |  |  |
| 7 | ggggacaagtttgtacaaaaaagcaggcttgatgGCGGAAGTACACACTCAAAT, ggggaccactttgtacaagaaagctgggtcGAAAACTTCTGTGCATTGGA | Mycgr3\|51649  JGI  Ensembl |  |  | 2.0202 % |  | ``` 54	CCGACT-AGACCACATTACATTCGACTCT-GGCCGAGAACTAGACGACCGAAATGTGGAGAGACTATTAGGTATTTTCAAGCTGCAAGGCTGTCAACGTGAGGAACCTGGTCACTCCATTCCTGTCTTAGTCGACAAAGACACTCTTGCCGCCGCGCTCGCGCACGAAGGCTCCCCTCCCCGGATCCTTCACCAAACTTCTGTGGAAGCCGAGCCGTTGCTCTTCCCGAGTGCCATCACATTAACCTCACTCCACGGCAAGCACCGTGTTCATGCAGCACGGAAGTTCCTTCTTCCCGTCGATCAGTGGTGGACGGTGAAGGTCTTCGATACTGGTCAGCAAACATTCCCTCCCGTCAGGCGTTTTCTTACCGAGGTCTCTAGGCCTTCCTCTTTGCGC-ACG-AGAACACCTTCGATACGAG-TACTCCAA-TGCACAGAAGTTTTC	495 	\|\|\|\|\|\| \|\|\|\|\|\|\|\|\|\|\|\|\|\|\|\|\|\|\|\|\|\| \|\|\|\|\|\|\|\|\|\|\|\|\|\|\|\|\|\|\|\|\|\|\|\|\|\|\|\|\|\|\|\|\|\|\|\|\|\|\|\|\|\|\|\|\|\|\|\|\|\|\|\|\|\|\|\|\|\|\|\|\|\|\|\|\|\|\|\|\|\|\|\|\|\|\|\|\|\|\|\|\|\|\|\|\|\|\|\|\|\|\|\|\|\|\|\|\|\|\|\|\|\|\|\|\|\|\|\|\|\|\|\|\|\|\|\|\|\|\|\|\|\|\|\|\|\|\|\|\|\|\|\|\|\|\|\|\|\|\|\|\|\|\|\|\|\|\|\|\|\|\|\|\|\|\|\|\|\|\|\|\|\|\|\|\|\|\|\|\|\|\|\|\|\|\|\|\|\|\|\|\|\|\|\|\|\|\|\|\|\|\|\|\|\|\|\|\|\|\|\|\|\|\|\|\|\|\|\|\|\|\|\|\|\|\|\|\|\|\|\|\|\|\|\|\|\|\|\|\|\|\|\|\|\|\|\|\|\|\|\|\|\|\|\|\|\|\|\|\|\|\|\|\|\|\|\|\|\|\|\|\|\|\|\|\|\|\|\|\|\|\|\|\|\|\|\|\|\|\|\|\|\|\|\|\|\|\|\|\|\|\|\|\|\|\|\|\|\|\|\|\|\|\|\|\|\|\|\|\|\|\|\|\|\|\|\|\|\|\|\|\|\|\|\|\|\|\|\|\|\|\|\|\|\|\|\|\|\|\|\|\|\|\|\|\|\|\|\|\|\|\|\|\|\|\|\|\|\|\|\|\|\| \|\|\|\|\|\| \|\|\| \|\|\|\|\|\|\|\|\|\|\|\|\|  \|\|\|\| \|\|\|\|\|\|\|\| \|\| \|\|\| \|\|\|\|\|\|\|\| 	CCGACTCAGACCACATTACATTCGACTCTAGGCCGAGAACTAGACGACCGAAATGTGGAGAGACTATTAGGTATTTTCAAGCTGCAAGGCTGTCAACGTGAGGAACCTGGTCACTCCATTCCTGTCTTAGTCGACAAAGACACTCTTGCCGCCGCGCTCGCGCACGAAGGCTCCCCTCCCCGGATCCTTCACCAAACTTCTGTGGAAGCCGAGCCGTTGCTCTTCCCGAGTGCCATCACATTAACCTCACTCCACGGCAAGCACCGTGTTCATGCAGCACGGAAGTTCCTTCTTCCCGTCGATCAGTGGTGGACGGTGAAGGTCTTCGATACTGGTCAGCAAACATTCCCTCCCGTCAGGCGTTTTCTTACCGAGGTCTCTAGGCCTTCCTCNTTGCGCCACGAAGAACACCTTCGACNCGAGGTACTCCAANTGNACANAAGTTTTC ``` |
| 7 | ggggacaagtttgtacaaaaaagcaggcttgatgACAGCCCGCCAGCGACGACC, ggggaccactttgtacaagaaagctgggtcCCTGCATTGGATCAAGGGCC | Mycgr3\|51711  JGI  Ensembl | KOG1869 Splicing coactivator SRm160/300, subunit SRm300; |  | 60.04141 % |  | ``` 226	GACAGCCCGCCAGCGACGACCCCCCAGCAAGAACACTCCAGCGATAGCCATACGTCGACAGCCATACGTCGACAGCCTCCAGCGACAACCTCCAACCGGCAACTCTCCAATCGACAACACTCCAGCGACAGCCATACGTCAACAGCCTCCATCGAACTCAGGTCCACTCATGATCTTGTCGTCTGGCTTCCTGTCTCTCAGTTCACCGCCACGCACAGCTCGACGGGAGCCAGCACTTGGCCCTTGATCCAATGCAGG	483 	\|\|\|\|\|\|\|\|\|\|\|\|\|  \|\|\|\|\|\|\|\|\|\|\|\|\|\|\|\|\|\|\|\|\|\|\|\|\|\|\|\|\|\|\|\|\|\|\|\|\|\|\|\|\|\|\|\|\|\|\|\|\|\|\|\|\|\|\|\|\|\|\|\|\|\|\|\|\|\|\|\|\|\|\|\|\|\|\|\|\|\|\|\|\|\|\|\|\|\|\|\|\|\|\|\|\|\|\|\|\|\|\|\|\|\|\|\|\|\|\|\|\|\|\|\|\|\|\|\|\|\|\|\|\|\|\|\|\|\|\|\|\|\|\|\|\|\|\|\|\|\|\|\|\|\|\|\|\|\|\|\|\|\|\|\|\|\|\|\|\|\|\|\|\|\|\|\|\|\|\|\|\|\|\|\|\|\|\|\|\|\|\|\|\|\|\|\|\|\|\|\|\|\|\|\|\|\|\|\|\|\|\|\|\|\|\|\|\|\|\|\|\|\|\|\|\|\|\|\|\|\|\|\|\|\|\|\|\|\|\|\|\|\|\|\|\|\|\|\|\|\|\|\|\|\|\| 	GACAGCCCGCCAGNNACGACCCCCCAGCAAGAACACTCCAGCGATAGCCATACGTCGACAGCCATACGTCGACAGCCTCCAGCGACAACCTCCAACCGGCAACTCTCCAATCGACAACACTCCAGCGACAGCCATACGTCAACAGCCTCCATCGAACTCAGGTCCACTCATGATCTTGTCGTCTGGCTTCCTGTCTCTCAGTTCACCGCCACGCACAGCTCGACGGGAGCCAGCACTTGGCCCTTGATCCAATGCAGG ``` |
| 7 | ggggacaagtttgtacaaaaaagcaggcttgatgCTCTTCACACGCGCTCTGCT, ggggaccactttgtacaagaaagctgggtcCGAAACGGTGATGCTCGCAG | Mycgr3\|60105  JGI  Ensembl |  |  | 100 % |  | ``` 36	ATGCTCTTCACACGCGCTCTGCTCGGCCTCGCCGCCGCTACGTACTCTCTGGCCGCTCCCATCACCGAGGCCTCTCCCAACCCGGACTCGGCCACCCACCTCTTCGCTCGCGAGAGCCCGGGCACTGGCACGAACAATGGCTACTACTACTCCTTCTGGACCGACGGTGGAGGCCAAGTGAACTACAACAATGGCCCGGGCGGCTCGTACACGGTCTCGTGGAGCAACGTTGGGAACTTCGTTGCGGGCAAGGGTTGGGCTACAGGGTCTGCTAG	310 	\|\|\|\|\|\|\|\|\|\|\|\|\|\|\|\|\|\|\|\|\|\|\|\|\|\|\|\|\|\|\|\|\|\|\|\|\|\|\|\|\|\|\|\|\|\|\|\|\|\|\|\|\|\|\|\|\|\|\|\|\|\|\|\|\|\|\|\|\|\|\|\|\|\|\|\|\|\|\|\|\|\|\|\|\|\|\|\|\|\|\|\|\|\|\|\|\|\|\|\|\|\|\|\|\|\|\|\|\|\|\|\|\|\|\|\|\|\|\|\|\|\|\|\|\|\|\|\|\|\|\|\|\|\|\|\|\|\|\|\|\|\|\|\|\|\|\|\|\|\|\|\|\|\|\|\|\|\|\|\|\|\|\|\|\|\|\|\|\|\|\|\|\|\|\|\|\|\|\|\|\|\|\|\|\|\|\|\|\|\|\|\|\|\|\|\|\|\|\|\|\|\|\|\|\|\|\|\|\|\|\|\|\|\|\|\|\|\|\|\|\|\|\|\|\|\|\|\|\|\|\|\|\|\|\|\|\|\|\|\|\|\|\|\|\|\|\|\|\|\|\|\|\|\|\|\|\|\|\|\|\|\|\|\|\|\|\|\|\|\|\|\|\|\|\| 	ATGCTCTTCACACGCGCTCTGCTCGGCCTCGCCGCCGCTACGTACTCTCTGGCCGCTCCCATCACCGAGGCCTCTCCCAACCCGGACTCGGCCACCCACCTCTTCGCTCGCGAGAGCCCGGGCACTGGCACGAACAATGGCTACTACTACTCCTTCTGGACCGACGGTGGAGGCCAAGTGAACTACAACAATGGCCCGGGCGGCTCGTACACGGTCTCGTGGAGCAACGTTGGGAACTTCGTTGCGGGCAAGGGTTGGGCTACAGGGTCTGCTAG ```      ``` 485	TATCCTCCAAACCACCCGCGTCAACCAGCCCTCCATTGACGGCACGCAGACCTTTCAGCAATTCTGGAGCGTGCGCCAGCAGAAGCGCGTCGGCGGCACGGTGACCATGAAGAACCACTTCGACGCTTGGTCGAGGCTGGGCATGAAGCTGGGTTCGTCGCAGAACTACCAGATTTTGGCGACCGAGGGATACCAGAGCAGCGGGTCTGCGAGCATCACCGTT	707 	\|\|\|\|\|\|\|\|\|\|\|\|\|\|\|\|\|\|\|\|\|\|\|\|\|\|\|\|\|\|\|\|\|\|\|\|\|\|\|\|\|\|\|\|\|\|\|\|\|\|\|\|\|\|\|\|\|\|\|\|\|\|\|\|\|\|\|\|\|\|\|\|\|\|\|\|\|\|\|\|\|\|\|\|\|\|\|\|\|\|\|\|\|\|\|\|\|\|\|\|\|\|\|\|\| \|\|\|\|\|\|\|\|\|\|\|\|\|\|\|\|\|\|\|\|\| \|\|\|\|\|\| \|\|\|\|\|\|\|\|\|\|\|\| \|\|\|\|\|\| \|\|\|\| \|\| \|\|\|\|\|\| \| \|\|\|\| \|\|\|\|\|\| \|\|\|\| \|\|\|\| \| \| \|\|\|\|\|\| \|\|\|\|\| \| \|\|\|\|\|\| \|\|\| 	TATCCTCCAAACCACCCGCGTCAACCAGCCCTCCATTGACGGCACGCAGACCTTTCAGCAATTCTGGAGCGTGCGCCAGCAGAAGCGCGTCGGCGGCACGGTGACNATGAAGAACCACTTCGACGCTGGGTCGAAGCTGGGCATGAANCTGGGTNCGTCNCA-AACTAC-ANATTT-GGCGAC-GAGGAATACAA-A-CAGCGG-TCTGCNA-CATCAC-GTT ```      ``` 309	AGAACCATCAACTACTCCGGCTCCTTCAACCCCTCCGGCAACGGCTACATCGCCGCCTACGGCTGGACCCGCAACCCTCTGATCGAATACTACGTCGTCGAATCCTACGGCACCTACAACCCCAGCTCTGCGGCTCAGAAGAAGGGAACCGTCACCACCGACGGCGGCACCTACGA	484 	\|\|\|\|\|\|\|\|\|\|\|\|\|\|\|\|\|\|\|\|\|\|\|\|\|\|\|\|\|\|\|\|\|\|\|\|\|\|\|\|\|\|\|\|\|\|\|\|\|\|\|\|\|\|\|\|\|\|\|\|\|\|\|\|\|\|\|\|\|\|\|\|\|\|\|\|\|\|\|\|\|\|\|\|\|\|\|\|\|\|\|\|\|\|\|\|\|\|\|\|\|\|\|\|\|\|\|\|\|\|\|\|\|\|\|\|\|\|\|\|\|\|\|\|\|\|\|\|\|\|\|\|\|\|\|\|\|\|\|\|\|\|\|\|\|\|\|\|\|\|\|\|\|\|\|\|\|\|\|\|\|\|\|\|\|\|\|\|\|\|\|\|\|\|\|\| 	AGAACCATCAACTACTCCGGCTCCTTCAACCCCTCCGGCAACGGCTACATCGCCGCCTACGGCTGGACCCGCAACCCTCTGATCGAATACTACGTCGTCGAATCCTACGGCACCTACAACCCCAGCTCTGCGGCTCAGAAGAAGGGAACCGTCACCACCGACGGCGGCACCTACGA ``` |
| 7 | ggggacaagtttgtacaaaaaagcaggcttgatgGCATCCAGAACCCTCTTCCT, ggggaccactttgtacaagaaagctgggtcTGGCGCCTTGGTCTTCAACT | Mycgr3\|69516  JGI  Ensembl | KOG3348 BolA (bacterial stress-induced morphogen)-related protein; |  | 100 % |  | ``` 62	ATGGCATCCAGAACCCTCTTCCTGCGGCTTACGCGCGCACCGACAAAACTCTCCAGTCCATCAACACTCCTCCCTCGAATCGCACAACCATCACGAAGACCTCTCCAACGCCTCTCAGTCCGCACTTACTCCGCCTCGCCCTCCACCGAAGTCCAAGCGCCCGACTACCTCAACGAGAGCGAGCGCCAAATTTTCGACAAGCTCAAAACCTCCCTCCAACCCAGCAAGCTCGAAGTTCAAGACATCTCCGGAGGGTGCGGCTCCATGTACGGCTTGGACATTGTGTCGGAGCAGTTTAAGGGGCTGTCGGTCATCAAACAGCATCGGATGGTTAATCAGATTCTGGGAGAGGAGATCAAGAAGTGGCATGGAGTGCAGTTGAAGACCAAGGCGCCA	457 	\|\|\|\|\|\|\|\|\|\|\|\| \|\|\|\|\|\|\|\|\|\|\|\|\|\|\|\|\|\|\|\|\|\|\|\|\|\|\|\|\|\|\|\|\|\|\|\|\|\|\|\|\|\|\|\|\|\|\|\|\|\|\|\|\|\|\|\|\|\|\|\|\|\|\|\|\|\|\|\|\|\|\|\|\|\|\|\|\|\|\|\|\|\|\|\|\|\|\|\|\|\|\|\|\|\|\|\|\|\|\|\|\|\|\|\|\|\|\|\|\|\|\|\|\|\|\|\|\|\|\|\|\|\|\|\|\|\|\|\|\|\|\|\|\|\|\|\|\|\|\|\|\|\|\|\|\|\|\|\|\|\|\|\|\|\|\|\|\|\|\|\|\|\|\|\|\|\|\|\|\|\|\|\|\|\|\|\|\|\|\|\|\|\|\|\|\|\|\|\|\|\|\|\|\|\|\|\|\|\|\|\|\|\|\|\|\|\|\|\|\|\|\|\|\|\|\|\|\|\|\|\|\|\|\|\|\|\|\|\|\|\|\|\|\|\|\|\|\|\|\|\|\|\|\|\|\|\|\|\|\|\|\|\|\|\|\|\|\|\|\|\|\|\|\|\|\|\|\|\|\|\|\|\|\|\|\|\|\|\|\|\|\|\|\|\|\|\|\|\|\|\|\|\|\|\|\|\|\|\|\|\|\|\|\|\|\|\|\|\|\|\|\|\|\|\|\|\|\|\|\|\|\|\|\|\|\|\|\|\|\|\|\|\|\|\|\|\|\|\|\|\|\|\|\|\|\|\|\|\|\|\|\|\|\|\|\|\|\|\|\|\|\|\|\|\|\|\|\|\|\|\|\|\|\|\|\|\|\|\|\|\|\|\|\| 	ATGGCATCCAGANCCCTCTTCCTGCGGCTTACGCGCGCACCGACAAAACTCTCCAGTCCATCAACACTCCTCCCTCGAATCGCACAACCATCACGAAGACCTCTCCAACGCCTCTCAGTCCGCACTTACTCCGCCTCGCCCTCCACCGAAGTCCAAGCGCCCGACTACCTCAACGAGAGCGAGCGCCAAATTTTCGACAAGCTCAAAACCTCCCTCCAACCCAGCAAGCTCGAAGTTCAAGACATCTCCGGAGGGTGCGGCTCCATGTACGGCTTGGACATTGTGTCGGAGCAGTTTAAGGGGCTGTCGGTCATCAAACAGCATCGGATGGTTAATCAGATTCTGGGAGAGGAGATCAAGAAGTGGCATGGAGTGCAGTTGAAGACCAAGGCGCCA ``` |
| 7 | ggggacaagtttgtacaaaaaagcaggcttgatgCAGGGCGAAGTCGCTCCTCG, ggggaccactttgtacaagaaagctgggtcGTCGTCCTTCATTTTCACCG | Mycgr3\|81367  JGI  Ensembl |  |  | 100 % |  | ``` 21	ATGCAGGGCGAAGTCGCTCCTCGAAACGCCCGGAAGCCTGGCGCCCAAGTTGACGATGCGGTCCTCGCGGACGCTACGATCAAATGCACCATGGAAGGCAACTTTGCGAATGGCGAAGTCTCTTCGGAGACATTTACTGTTTCTCTCCGTTCAATTCATGGCGCTGGAATGCATCTTATCTTTCGCGATTGGGCCGACGGCGGCGTCCAGGATCAGCTTCTCGTCGGGAATAGACAAGACATCGTTGCAGACGCGATGTGCACCTTTGGCGAAACCATGCGCAGCGGAATGTACACTTGCAAGTTCGACGCGAGGCTACCGGACGGCGAATGTCTGTTCGCACTGGAAGTGTCGCAGTGGTTACATGGACATGGCGATGGACCGGCGGCGGTGAAAATGAAGGACGAC	428 	\|\|\|\|\|\|\|\|\|\|\|\|\|\|\| \|\|\|\|\|\|\|\|\|\|\|\|\|\|\|\|\|\|\|\|\|\|\|\|\|\|\|\|\|\|\|\|\|\|\|\|\|\|\|\|\|\|\|\|\|\|\|\|\|\|\|\|\|\|\|\|\|\|\|\|\|\|\|\|\|\|\|\|\|\|\|\|\|\|\|\|\|\|\|\|\|\|\|\|\|\|\|\|\|\|\|\|\|\|\|\|\|\|\|\|\|\|\|\|\|\|\|\|\|\|\|\|\|\|\|\|\|\|\|\|\|\|\|\|\|\|\|\|\|\|\|\|\|\|\|\|\|\|\|\|\|\|\|\|\|\|\|\|\|\|\|\|\|\|\|\|\|\|\|\|\|\|\|\|\|\|\|\|\|\|\|\|\|\|\|\|\|\|\|\|\|\|\|\|\|\|\|\|\|\|\|\|\|\|\|\|\|\|\|\|\|\|\|\|\|\|\|\|\|\|\|\|\|\|\|\|\|\|\|\|\|\|\|\|\|\|\|\|\|\|\|\|\|\|\|\|\|\|\|\|\|\|\|\|\|\|\|\|\|\|\|\|\|\|\|\|\|\|\|\|\|\|\|\|\|\|\|\|\|\|\|\|\|\|\|\|\|\|\|\|\|\|\|\|\|\|\|\|\|\|\|\|\|\|\|\|\|\|\|\|\|\|\|\|\|\|\|\|\|\|\|\|\|\|\|\|\|\|\|\|\|\|\|\|\|\|\|\|\|\|\|\|\|\|\|\|\|\|\|\|\|\|\|\|\|\|\|\|\|\|\|\|\|\|\|\|\|\|\|\|\|\|\|\|\|\|\|\|\|\|\|\|\|\|\|\|\|\|\|\|\|\|\|\|\|\|\|\|\|\|\|\| 	ATGCAGGGCGAAGTCNCTCCTCGAAACGCCCGGAAGCCTGGCGCCCAAGTTGACGATGCGGTCCTCGCGGACGCTACGATCAAATGCACCATGGAAGGCAACTTTGCGAATGGCGAAGTCTCTTCGGAGACATTTACTGTTTCTCTCCGTTCAATTCATGGCGCTGGAATGCATCTTATCTTTCGCGATTGGGCCGACGGCGGCGTCCAGGATCAGCTTCTCGTCGGGAATAGACAAGACATCGTTGCAGACGCGATGTGCACCTTTGGCGAAACCATGCGCAGCGGAATGTACACTTGCAAGTTCGACGCGAGGCTACCGGACGGCGAATGTCTGTTCGCACTGGAAGTGTCGCAGTGGTTACATGGACATGGCGATGGACCGGCGGCGGTGAAAATGAAGGACGAC ``` |
| 7 | ggggacaagtttgtacaaaaaagcaggcttgatgTACAGATCCGAGTCGAGATT, ggggaccactttgtacaagaaagctgggtcTTGCGGCCGAGCACTCGTAC | Mycgr3\|85479  JGI  Ensembl | KOG0454 3-isopropylmalate dehydratase (aconitase superfamily); | metabolic process; | 0 % |  |  |
| 7 | ggggacaagtttgtacaaaaaagcaggcttgatgTCACAAGTATCATCCTACCT, ggggaccactttgtacaagaaagctgggtcCCTGACATACCATCCCCATT | Mycgr3\|86065  JGI  Ensembl | KOG3455 Predicted membrane protein; | integral to membrane; | 100 % |  | ``` 1	ATGTCACAAGTATCATCCTACCTCCCCCCGGGAGAAGGCTTCCTACCAAAGTGGCTCCTCTTC	63 	\|\|\|\|\|\|\|\|\|\|\|\|\|\| \|\|\| \| \|\|\|\|\|\|\| \|\|\|\|\|\|\|\|\|\|\|\|\|\|\|\|\|\|\|\|\|\|\|\|\|\|\|\|\|\|\|\|\|\| 	ATGTCACAAGTATCNTCCGAGCTCCCCCNGGGAGAAGGCTTCCTACCAAAGTGGCTCCTCTTC ```      ``` 64	ATCGCAGTAGTCGCCCTCGGCAACAGCATCCAAAGCTATACTTCCCTCCGCGGCACCCGCGAAGTCTACGCCGGCAACAACCCCAGCGCCGTCACTCCTCTATCAGCCCGCACATTCGGAACCTGGACAGGCCTCAGCAGCATTATCCGCCTCTACGCCGCGTACAACATTTCTAACCCGGTGGTCTACCAACTCGCGCTCTGGACATACGGGATCGCATTTGCGCATTTCTTCAGCGAGTGGTTGATCTTCGGTACGGCGAAGTGGGGACGTGGATTGGGAGGGCCGGTGTTTGTGAGTACGGTTACTACGGTGTGGATGGTGGCGCAATGGGGATGGTATGTCAGG	411 	\|\|\|\|\|\|\|\|\|\|\|\|\|\|\|\|\|\|\|\|\|\|\|\|\|\|\|\|\|\|\|\|\|\|\|\|\|\|\|\|\|\|\|\|\|\|\|\|\|\|\|\|\|\|\|\|\|\|\|\|\|\|\|\|\|\|\|\|\|\|\|\|\|\|\|\|\|\|\|\|\|\|\|\|\|\|\|\|\|\|\|\|\|\|\|\|\|\|\|\|\|\|\|\|\|\|\|\|\|\|\|\|\|\|\|\|\|\|\|\|\|\|\|\|\|\|\|\|\|\|\|\|\|\|\|\|\|\|\|\|\|\|\|\|\|\|\|\|\|\|\|\|\|\|\|\|\|\|\|\|\|\|\|\|\|\|\|\|\|\|\|\|\|\|\|\|\|\|\|\|\|\|\|\|\|\|\|\|\|\|\|\|\|\|\|\|\|\|\|\|\|\|\|\|\|\|\|\|\|\|\|\|\|\|\|\|\|\|\|\|\|\|\|\|\|\|\|\|\|\|\|\|\|\|\|\|\|\|\|\|\|\|\|\|\|\|\|\|\|\|\|\|\|\|\|\|\|\|\|\|\|\|\|\|\|\|\|\|\|\|\|\|\|\|\|\|\|\|\|\|\|\|\|\|\|\|\|\|\|\|\|\|\|\|\|\|\|\|\|\|\|\|\|\|\|\|\|\|\|\|\|\|\|\|\|\|\|\|\|\|\|\|\|\|\|\|\|\|\|\|\|\|\|\|\|\|\|\|\|\|\|\|\|\|\|\|\|\| 	ATCGCAGTAGTCGCCCTCGGCAACAGCATCCAAAGCTATACTTCCCTCCGCGGCACCCGCGAAGTCTACGCCGGCAACAACCCCAGCGCCGTCACTCCTCTATCAGCCCGCACATTCGGAACCTGGACAGGCCTCAGCAGCATTATCCGCCTCTACGCCGCGTACAACATTTCTAACCCGGTGGTCTACCAACTCGCGCTCTGGACATACGGGATCGCATTTGCGCATTTCTTCAGCGAGTGGTTGATCTTCGGTACGGCGAAGTGGGGACGTGGATTGGGAGGGCCGGTGTTTGTGAGTACGGTTACTACGGTGTGGATGGTGGCGCAATGGGGATGGTATGTCAGG ``` |
| 7 | ggggacaagtttgtacaaaaaagcaggcttgatgCTCGCCTCAACCATTTTCAC, ggggaccactttgtacaagaaagctgggtcTAGCACACCAAATGCGATCG | Mycgr3\|88743  JGI  Ensembl |  |  | 100 % |  | ``` 1	ATGCTCGCCTCAACCATTTTCACCGCCTGGCTCCTCGCTGCACCAATTGTCAGCGGCTTCACCTTCACAGCACCACCCAAAGGCACCAAGCTGGACCTCTCCCAAAACATCACAATTTCCTGGACCGGCGAAAACGAACAATTCGAAGAACTCGATCTCACGTTCAATGCAGTCGGGTCGAGCAGATTCGGGTATGGCATAGCCTATAACCTCTCTTCCTCCACCGGCTCTTTGATCTGGGATCCAAGGAATGTGACAGAGGCACTGCAGAGCACCAACATATCATTGAGCAAGGGCGAGGATCACTACTTCGAGGCGAAGCTTCACCCGAACGGTACAGATGGGACGCGGAGTGCTGGCGCGGGACTGGAGAGCGAGAAGTACAAGATTACGGGATATCCTTTGATCGGTGCGGCGGCGGCGCTGCAGCCTCAGGTTGGAGCAGCGGTTTTGACGGTGGTGTTCATCTCGATCGCATTTGGTGTGCTA	489 	\|\|\|\|\|\|\|\|\|\|\|\| \|\|\|\|\|\|\|\|\|\|\|\|\|\|\|\|\|\|\|\|\|\|\|\|\|\|\|\|\|\|\|\|\|\|\|\|\|\|\|\|\|\|\|\|\|\|\|\|\|\|\|\|\|\|\|\|\|\|\|\|\|\|\|\|\|\|\|\|\|\|\|\|\|\|\|\|\|\|\|\|\|\|\|\|\|\|\|\|\|\|\|\|\|\|\|\|\|\|\|\|\|\|\|\|\|\|\|\|\|\|\|\|\|\|\|\|\|\|\|\|\|\|\|\|\|\|\|\|\|\|\|\|\|\|\|\|\|\|\|\|\|\|\|\|\|\|\|\|\|\|\|\|\|\|\|\|\|\|\|\|\|\|\|\|\|\|\|\|\|\|\|\|\|\|\|\|\|\|\|\|\|\|\|\|\|\|\|\|\|\|\|\|\|\|\|\|\|\|\|\|\|\|\|\|\|\|\|\|\|\|\|\|\|\|\|\|\|\|\|\|\|\|\|\|\|\|\|\|\|\|\|\|\|\|\|\|\|\|\|\|\|\|\|\|\|\|\|\|\|\|\|\|\|\|\|\|\|\|\|\|\|\|\|\|\|\|\|\|\|\|\|\|\|\|\|\|\|\|\|\|\|\|\|\|\|\|\|\|\|\|\|\|\|\|\|\|\|\|\|\|\|\|\|\|\|\|\|\|\|\|\|\|\|\|\|\|\|\|\|\|\|\|\|\|\|\|\|\|\|\|\|\|\|\|\|\|\|\|\|\|\|\|\|\|\|\|\|\|\|\|\|\|\|\|\|\|\|\|\|\|\|\|\|\|\|\|\|\|\|\|\|\|\|\|\|\|\|\|\|\|\|\|\|\|\|\|\|\|\|\|\|\|\|\|\|\|\|\|\|\|\|\|\|\|\|\|\|\|\|\|\|\|\|\|\|\|\|\|\|\|\|\|\|\|\|\|\|\|\|\|\|\|\|\|\|\|\|\|\|\|\|\|\|\|\|\|\|\|\|\|\|\|\|\|\|\|\|\|\|\|\|\|\|\|\|\|\|\|\|\|\|\|\|\|\|\| 	ATGCTCGCCTCANCCATTTTCACCGCCTGGCTCCTCGCTGCACCAATTGTCAGCGGCTTCACCTTCACAGCACCACCCAAAGGCACCAAGCTGGACCTCTCCCAAAACATCACAATTTCCTGGACCGGCGAAAACGAACAATTCGAAGAACTCGATCTCACGTTCAATGCAGTCGGGTCGAGCAGATTCGGGTATGGCATAGCCTATAACCTCTCTTCCTCCACCGGCTCTTTGATCTGGGATCCAAGGAATGTGACAGAGGCACTGCAGAGCACCAACATATCATTGAGCAAGGGCGAGGATCACTACTTCGAGGCGAAGCTTCACCCGAACGGTACAGATGGGACGCGGAGTGCTGGCGCGGGACTGGAGAGCGAGAAGTACAAGATTACGGGATATCCTTTGATCGGTGCGGCGGCGGCGCTGCAGCCTCAGGTTGGAGCAGCGGTTTTGACGGTGGTGTTCATCTCGATCGCATTTGGTGTGCTA ```      ``` 1	ATGCTCGCCTCAACCATTTTCACCGCCTGGCTCCTCGCTGCACCAATTGTCAGCGGCTTCACCTTCACAGCACCACCCAAAGGCACCAAGCTGGACCTCTCCCAAAACATCACAATTTCCTGGACCGGCGAAAACGAACAATTCGAAGAACTCGATCTCACGTTCAATGCAGTCGGGTCGAGCAGATTCGGGTATGGCATAGCCTATAACCTCTCTTCCTCCACCGGCTCTTTGATCTGGGATCCAAGGAATGTGACAGAGGCACTGCAGAGCACCAACATATCATTGAGCAAGGGCGAGGATCACTACTTCGAGGCGAAGCTTCACCCGAACGGTACAGATGGGACGCGGAGTGCTGGCGCGGGACTGGAGAGCGAGAAGTACAAGATTACGGGATATCCTTTGATCGGTGCGGCGGCGGCGCTGCAGCCTCAGGTTGGAGCAGCGGTTTTGACGGTGGTGTTCATCTCGATCGCATTTGGTGTGCTA	489 	\|\|\|\|\|\|\|\|\|\|\|\|\|\|\|\|\|\|\|\|\|\|\|\|\|\|\|\|\|\|\|\|\|\|\|\|\|\|\|\|\|\|\|\|\|\|\|\|\|\|\|\|\|\|\|\|\|\|\|\|\|\|\|\|\|\|\|\|\|\|\|\|\|\|\|\|\|\|\|\|\|\|\|\|\|\|\|\|\|\|\|\|\|\|\|\|\|\|\|\|\|\|\|\|\|\|\|\|\|\|\|\|\|\|\|\|\|\|\|\|\|\|\|\|\|\|\|\|\|\|\|\|\|\|\|\|\|\|\|\|\|\|\|\|\|\|\|\|\|\|\|\|\|\|\|\|\|\|\|\|\|\|\|\|\|\|\|\|\|\|\|\|\|\|\|\|\|\|\|\|\|\|\|\|\|\|\|\|\|\|\|\|\|\|\|\|\|\|\|\|\|\|\|\|\|\|\|\|\|\|\|\|\|\|\|\|\|\|\|\|\|\|\|\|\|\|\|\|\|\|\|\|\|\|\|\|\|\|\|\|\|\|\|\|\|\|\|\|\|\|\|\|\|\|\|\|\|\|\|\|\|\|\|\|\|\|\|\|\|\|\|\|\|\|\|\|\|\|\|\|\|\|\|\|\|\|\|\|\|\|\|\|\|\|\|\|\|\|\|\|\|\|\|\|\|\|\|\|\|\|\|\|\|\|\|\|\|\|\|\|\|\|\|\|\|\|\|\|\|\|\|\|\|\|\|\|\|\|\|\|\|\|\|\|\|\|\|\|\|\|\|\|\|\|\|\|\|\|\|\|\|\|\|\|\|\|\|\|\|\|\|\|\|\|\|\|\|\|\|\|\|\|\|\|\|\|\|\|\|\|\|\|\|\|\|\|\|\|\|\|\|\|\|\|\|\|\|\|\|\|\|\|\|\|\|\|\|\|\|\|\|\|\|\|\|\|\|\|\|\|\|\|\|\|\|\|\|\|\|\|\|\|\|\|\|\|\|\|\|\|\|\|\|\|\|\|\|\|\|\|\|\|\|\|\|\|\|\|\|\|\|\|\|\|\|\|\|\|\|\|\|\|\|\|\|\|\|\|\| 	ATGCTCGCCTCAACCATTTTCACCGCCTGGCTCCTCGCTGCACCAATTGTCAGCGGCTTCACCTTCACAGCACCACCCAAAGGCACCAAGCTGGACCTCTCCCAAAACATCACAATTTCCTGGACCGGCGAAAACGAACAATTCGAAGAACTCGATCTCACGTTCAATGCAGTCGGGTCGAGCAGATTCGGGTATGGCATAGCCTATAACCTCTCTTCCTCCACCGGCTCTTTGATCTGGGATCCAAGGAATGTGACAGAGGCACTGCAGAGCACCAACATATCATTGAGCAAGGGCGAGGATCACTACTTCGAGGCGAAGCTTCACCCGAACGGTACAGATGGGACGCGGAGTGCTGGCGCGGGACTGGAGAGCGAGAAGTACAAGATTACGGGATATCCTTTGATCGGTGCGGCGGCGGCGCTGCAGCCTCAGGTTGGAGCAGCGGTTTTGACGGTGGTGTTCATCTCGATCGCATTTGGTGTGCTA ``` |
| 7 | ggggacaagtttgtacaaaaaagcaggcttgatgGCCATCTTCAAGAAAAAGAA, ggggaccactttgtacaagaaagctgggtcGGCCGCCGTCTCCGCCGCCT | Mycgr3\|88817  JGI  Ensembl | KOG2812 Uncharacterized conserved protein; |  | 0 % |  |  |
| 7 | ggggacaagtttgtacaaaaaagcaggcttgatgTGCCAGTACTACCACTGGAG, ggggaccactttgtacaagaaagctgggtcGCCGATTTGCAAACGCTCCT | Mycgr3\|89085  JGI  Ensembl |  |  | 0 % |  |  |
| 7 | ggggacaagtttgtacaaaaaagcaggcttgatgTCGGCCACCATCATACTTTG, ggggaccactttgtacaagaaagctgggtcGCTCCAAAACTCGACCTGCT | Mycgr3\|89185  JGI  Ensembl |  |  | 100 % |  | ``` 1	ATGTCGGCCACCATCATACTTTGCAATACGCCAATCTACATCATGACCACAGGCAGCCGTCCTCCAAGCGACCCCATTCCGGCTCGTCGATTCACTACGTCTGCCGCTCTTCACGCCCACCTTCGCTCTCACAAGCGCTTCGTGCTTTCCTGGACCGTCTTTCGACAACACCTGACGGACCACTGTGCTTTTGACTCAGCTACGGATGCTGTCGTCTACGCGAAAGAGCCCGCGTTCGAGGCCAAAATGGTACGTAACGAGGCAGAGCTGAACGAAGCGTTTCGCGAAATGAGAAACGGAAGGGGTGGAGTTGGATTCTGGGTCGTGAGAGGAGATGTGTTCG	343 	\|\|\|\|\|\|\|\|\|\|\|\|\|\|\|\|\|\|\|\|\|\|\|\|\|\|\|\|\|\|\|\|\|\|\|\|\|\|\|\|\|\|\|\|\|\|\|\|\|\|\|\|\|\|\|\|\|\|\|\|\|\|\|\|\|\|\|\|\|\|\|\|\|\|\|\|\|\|\|\|\|\|\|\|\|\|\|\|\|\|\|\|\|\|\|\|\|\|\|\|\|\|\|\|\|\|\|\|\|\|\|\|\|\|\|\|\|\|\|\|\|\|\|\|\|\|\|\|\|\|\|\|\|\|\|\|\|\|\|\|\|\|\|\|\|\|\|\|\|\|\|\|\|\|\|\|\|\|\|\|\|\|\|\|\|\|\|\|\|\|\|\|\|\|\|\|\|\|\|\|\|\|\|\|\|\|\|\|\|\|\|\|\|\|\|\|\|\|\|\|\|\|\|\|\|\|\|\|\|\|\|\|\|\|\|\|\|\|\|\|\|\|\|\|\|\|\|\|\|\|\|\|\|\|\|\|\|\|\|\|\|\|\|\|\|\|\|\|\|\|\|\|\|\|\|\|\|\|\|\|\|\|\|\|\|\|\|\|\|\|\|\|\|\|\|\|\|\|\|\|\|\|\|\|\|\|\|\|\|\|\|\|\|\|\|\|\|\|\|\|\|\|\|\|\|\|\|\|\|\|\|\|\|\|\|\|\|\|\|\|\|\|\|\|\|\|\|\|\|\|\|\|\|\|\|\|\|\|\|\|\|\|\| 	ATGTCGGCCACCATCATACTTTGCAATACGCCAATCTACATCATGACCACAGGCAGCCGTCCTCCAAGCGACCCCATTCCGGCTCGTCGATTCACTACGTCTGCCGCTCTTCACGCCCACCTTCGCTCTCACAAGCGCTTCGTGCTTTCCTGGACCGTCTTTCGACAACACCTGACGGACCACTGTGCTTTTGACTCAGCTACGGATGCTGTCGTCTACGCGAAAGAGCCCGCGTTCGAGGCCAAAATGGTACGTAACGAGGCAGAGCTGAACGAAGCGTTTCGCGAAATGAGAAACGGAAGGGGTGGAGTTGGATTCTGGGTCGTGAGAGGAGATGTGTTCG ```      ``` 343	GCGAATGGATTCTTGGCCTCTGCGAAGCAGGTCGAGTTTTGGAGC	387 	\|\|\|\|\|\|\|\|\|\|\|\|\|\|\|\|\|\|\|\|\|\|\|\|\|\|\|\|\|\|\|\|\|\|\|\|\|\|\|\|\|\|\|\|\| 	GCGAATGGATTCTTGGCCTCTGCGAAGCAGGTCGAGTTTTGGAGC ``` |
| 7 | ggggacaagtttgtacaaaaaagcaggcttgatgCAGTTCATCAGATCCGCTCA, ggggaccactttgtacaagaaagctgggtcCCCTGAAGAGTTGATTTTCT | Mycgr3\|89270  JGI  Ensembl |  |  | 0 % |  |  |
| 7 | ggggacaagtttgtacaaaaaagcaggcttgatgCGACACTTCGAGCCCAAGGA, ggggaccactttgtacaagaaagctgggtcTAACCATGCTCTGCCGTGTC | Mycgr3\|90306  JGI  Ensembl |  |  | 0 % |  |  |
| 7 | ggggacaagtttgtacaaaaaagcaggcttgatgTCTTCCAACTTCACTCCCCA, ggggaccactttgtacaagaaagctgggtcCTCCTCCACCCGACAATACA | Mycgr3\|91033  JGI  Ensembl |  |  | 100 % |  | ``` 1	ATGTCTTCCAACTTCACTCCCCAAAAAGCCTGCTCCGACCGTCCCAACAGCAGCACTACCAGCAGCATGACTGCCGACACCAGAAGAGACGCATATTTCGCCAATCTCTACGTGCTCCCAAGCAGCGACGGCACCAGCTATGCCAACGCCATCGCCACCAGCAGCACCACCTCCTCCACCAGCCAAGAGAGCTTAATGGTGAGATTGGCCTCGCCGGCCCTGTTGAATCACCAAGTGAGGCTATGCGACCTCGGATGTAAATTTTGCGCAGAGGATGAGAAGGAGAGGAGGAAAGCTCGAGGAGTGAGAGGGACGCTGAGGCGGTGGTGGGGAGGTTGGAAGGGTGAGAAGCCAGCATATTGCGGAAAGGAGATTTGGGAGAGGCACACGAAGGAGCAGTGCCGGGTGTATTGTCGGGTGGAGGAG	426 	\|\|\|\|\|\|\|\|\|\|\|\|\|\|\|\|\|\|\|\|\|\|\|\|\|\|\|\|\|\|\|\|\|\|\|\|\|\|\|\|\|\|\|\|\|\|\|\|\|\|\|\|\|\|\|\|\|\|\|\|\|\|\|\|\|\|\|\|\|\|\|\|\|\|\|\|\|\|\|\|\|\|\|\|\|\|\|\|\|\|\|\|\|\|\|\|\|\|\|\|\|\|\|\|\|\|\|\|\|\|\|\|\|\|\|\|\|\|\|\|\|\|\|\|\|\|\|\|\|\|\|\|\|\|\|\|\|\|\|\|\|\|\|\|\|\|\|\|\|\|\|\|\|\|\|\|\|\|\|\|\|\|\|\|\|\|\|\|\|\|\|\|\|\|\|\|\|\|\|\|\|\|\|\|\|\|\|\|\|\|\|\|\|\|\|\|\|\|\|\|\|\|\|\|\|\|\|\|\|\|\|\|\|\|\|\|\|\|\|\|\|\|\|\|\|\|\|\|\|\|\|\|\|\|\|\|\|\|\|\|\|\|\|\|\|\|\|\|\|\|\|\|\|\|\|\|\|\|\|\|\|\|\|\|\|\|\|\|\|\|\|\|\|\|\|\|\|\|\|\|\|\|\|\|\|\|\|\|\|\|\|\|\|\|\|\|\|\|\|\|\|\|\|\|\|\|\|\|\|\|\|\|\|\|\|\|\|\|\|\|\|\|\|\|\|\|\|\|\|\|\|\|\|\|\|\|\|\|\|\|\|\|\|\|\|\|\|\|\|\|\|\|\|\|\|\|\|\|\|\|\|\|\|\|\|\|\|\|\|\|\|\|\|\|\|\|\|\|\|\|\|\|\|\|\|\|\|\|\|\|\|\|\|\|\|\|\|\|\|\|\|\|\|\|\|\|\|\|\|\|\|\|\|\|\|\|\|\|\|\|\|\|\|\|\|\| 	ATGTCTTCCAACTTCACTCCCCAAAAAGCCTGCTCCGACCGTCCCAACAGCAGCACTACCAGCAGCATGACTGCCGACACCAGAAGAGACGCATATTTCGCCAATCTCTACGTGCTCCCAAGCAGCGACGGCACCAGCTATGCCAACGCCATCGCCACCAGCAGCACCACCTCCTCCACCAGCCAAGAGAGCTTAATGGTGAGATTGGCCTCGCCGGCCCTGTTGAATCACCAAGTGAGGCTATGCGACCTCGGATGTAAATTTTGCGCAGAGGATGAGAAGGAGAGGAGGAAAGCTCGAGGAGTGAGAGGGACGCTGAGGCGGTGGTGGGGAGGTTGGAAGGGTGAGAAGCCAGCATATTGCGGAAAGGAGATTTGGGAGAGGCACACGAAGGAGCAGTGCCGGGTGTATTGTCGGGTGGAGGAG ``` |
| 7 | ggggacaagtttgtacaaaaaagcaggcttgatgTTGACACCTGGGTCCTTTTG, ggggaccactttgtacaagaaagctgggtcCTGCGCCTCGTCCCTGATCT | Mycgr3\|91122  JGI  Ensembl |  |  | 100 % |  | ``` 147	AGAGTGGGGGCTGTACTTCGCATGCGCCGCCCGCAAAGGGAAGCTTGGCTTGGACGAGCAAGCTCCACTTCGCGGCTATGGTGGAGCCTTCTTCAACCACGCTGGGTCCATGGCCTTGCTCGTCGGTCCCGTGTATGCGCCTGATGAAACGG-AAGCCAAGCATGTGATGGGATGGA	322 	\|\|\|\|\|\|\|\|\|\|\|\|\|\|\|\|\|\|\|\|\|\|\|\|\|\|\|\|\|\|\|\|\|\|\|\|\|\|\|\|\|\|\|\|\|\|\|\|\|\|\|\|\|\|\|\|\|\|\|\|\|\|\|\|\|\|\|\|\|\|\| \|\|\|\|\|\|\|\|\|\|\|\|\|\|\|\|\|\|\|\|\|\|\|\|\|\|\|\|\|\|\|\|\|\|\|\|\|\|\|\|\|\|\|\|\|\|\|\|\|\|\|\|\|\|\|\|\|\|\|\|\|\|\|\|\|\|\|\| \|\|\|\|\|\|\|\|  \| \|\|\|\|\|\|\|  \|\|\|\|\|\|\|\|\|\|\| \|\|\| 	AGAGTGGGGGCTGTACTTCGCATGCGCCGCCCGCAAAGGGAAGCTTGGCTTGGACGAGCAAGCTCCACTTCNCGGCTATGGTGGAGCCTTCTTCAACCACGCTGGGTCCATGGCCTTGCTCGTCGGTCCCGTGTATGCGCNTGATGAAAANGNAAGCCAANNATGTGATGGGANGGA ```      ``` 1	ATGTTGACACCTGGGTCCTTTTGGGAGGACAAAGGCCACTGCATACTTCTCCGCGATGATCAGACTTTGTGCAACGACATCTTACTACTACTGCAGACTGCTGCAATTCCAGACGCACATCTGGACCAATACCTCACGCAACGAGAAGAGTGGGGGC	157 	\|\|\|\|\|\|\|\|\|\|\|\|\| \|\|\|\|\|\|\|\|\|\|\|\|\|\|\|\|\|\|\|\|\|\|\|\|\|\|\|\|\|\|\|\|\|\|\|\|\|\|\|\|\|\|\|\|\|\|\|\|\|\|\|\|\|\|\|\|\|\|\|\|\|\|\|\|\|\|\|\|\|\|\|\|\|\|\|\|\|\|\|\|\|\|\|\|\|\|\|\|\|\|\|\|\|\|\|\|\|\|\|\|\|\|\|\|\|\|\|\|\|\|\|\|\|\|\|\|\|\|\|\|\|\|\|\|\|\|\|\|\|\|\|\|\|\| \|\|\| \|\|\|\| 	ATGTTGACACCTGCGTCCTTTTGGGAGGACAAAGGCCACTGCATACTTCTCCGCGATGATCAGACTTTGTGCAACGACATCTTACTACTACTGCAGACTGCTGCAATTCCAGACGCACATCTGGACCAATACCTCACGCAACGAGAAG-GTGTGGGC ``` |
| 7 | ggggacaagtttgtacaaaaaagcaggcttgatgTCGGTGGATACTGAAGGTTT, ggggaccactttgtacaagaaagctgggtcTCGCCGACATAACCACTCTA | Mycgr3\|91293  JGI  Ensembl |  |  | 3.0303 % |  |  |
| 7 | ggggacaagtttgtacaaaaaagcaggcttgatgTCAGGCTACGGCTACCCTTC, ggggaccactttgtacaagaaagctgggtcAGCCGGATCCGCTGTTCCGC | Mycgr3\|91694  JGI  Ensembl |  |  | 0 % |  |  |
| 7 | ggggacaagtttgtacaaaaaagcaggcttgatgACACCGAGCATGCAGCAATT, ggggaccactttgtacaagaaagctgggtcGGTTGCGGCATTGGCATGAC | Mycgr3\|91742  JGI  Ensembl |  |  | 0 % |  |  |
| 7 | ggggacaagtttgtacaaaaaagcaggcttgatgTCCGCCCACGGTCGTGCCAT, ggggaccactttgtacaagaaagctgggtcGGGCAGCACTTCCTTCACGC | Mycgr3\|92917  JGI  Ensembl |  |  | 100 % |  | ``` 1	ATGTCCGCCCACGG-TCGTGCCATCATGCAGAACGTCCCGCAGAACAATCCGACTTTCACCAACACGGACGGCCGCACATCACCAGACCTTGCACCTCGCACCGGCCCCGACGCCATCAGCCCCCAGGAAGCCGCCTTCGACGACGCCTTCCGCGCAGTGGTCGACATCGCGCTTGCGAAAGGCTTGCCCAACGAGCTCGCTCTGTGCGAGACTGCCGCCAAGATCAAACAAAACCTGACCACAGCCCTGGCCCAGGTTGACCCGGACCAGGCCAACCGGGCCGAGACTGCCATTCGGGCCATCGTCACGTCCTGGGTGGCCTTGGACAATCCGCCCTCGAATGGCAAGTGGAAGTTCAGGATGACGAAGGAGAAACAGAGTCGCCAGAAGCAGAGTCGCCAGAAATGGGCGGCGGACGCGATTTTCGAGTGCAGTTTCATGAAGAGTTGCGTGAAGGAAGTGCTGCCC	468 	\|\|\|\|\|\|\|\|\|\|\|\|\|\| \|\|\|\|\|\|\|\|\|\|\|\|\|\|\|\|\|\|\|\|\|\|\|\|\|\|\|\|\|\|\|\|\|\|\|\|\|\|\|\|\|\|\|\|\|\|\|\|\|\|\|\|\|\|\|\|\|\|\|\|\|\|\|\|\|\|\|\|\|\|\|\|\|\|\|\|\|\|\|\|\|\|\|\|\|\|\|\|\|\|\|\|\|\|\|\|\|\|\|\|\|\|\|\|\|\|\|\|\|\|\|\|\|\|\|\|\|\|\|\|\|\|\|\|\|\|\|\|\|\|\|\|\|\|\|\|\|\|\|\|\|\|\|\|\|\|\|\|\|\|\|\|\|\|\|\|\|\|\|\|\|\|\|\|\|\|\|\|\|\|\|\|\|\|\|\|\|\|\|\|\|\|\|\|\|\|\|\|\|\|\|\|\|\|\|\|\|\|\|\|\|\|\|\|\|\|\|\|\|\|\|\|\|\|\|\|\|\|\|\|\|\|\|\|\|\|\|\|\|\|\|\|\|\|\|\|\|\|\|\|\|\|\|\|\|\|\|\|\|\|\|\|\|\|\|\|\|\|\|\|\|\|\|\|\|\|\|\|\|\|\|\|\|\|\|\|\|\|\|\|\|\|\|\|\|\|\|\|\|\|\|\|\|\|\|\|\|\|\|\|\|\|\|\|\|\|\|\|\|\|\|\|\|\|\|\|\|\|\|\|\|\|\|\|\|\|\|\|\|\|\|\|\|\|\|\|\|\|\|\|\|\|\|\|\|\|\|\|\|\|\|\|\|\|\|\|\|\|\|\|\|\|\|\|\|\|\|\|\|\|\|\|\|\|\|\|\|\|\|\|\|\|\|\|\|\|\|\|\|\|\|\|\|\|\|\|\|\|\|\|\|\|\|\|\|\|\|\|\|\|\|\|\|\|\|\|\|\|\|\|\|\|\|\|\|\|\|\|\|\|\|\|\|\|\|\|\|\|\|\|\|\|\|\|\|\|\|\|\|\|\|\|\|\| 	ATGTCCGCCCACGGCTCGTGCCATCATGCAGAACGTCCCGCAGAACAATCCGACTTTCACCAACACGGACGGCCGCACATCACCAGACCTTGCACCTCGCACCGGCCCCGACGCCATCAGCCCCCAGGAAGCCGCCTTCGACGACGCCTTCCGCGCAGTGGTCGACATCGCGCTTGCGAAAGGCTTGCCCAACGAGCTCGCTCTGTGCGAGACTGCCGCCAAGATCAAACAAAACCTGACCACAGCCCTGGCCCAGGTTGACCCGGACCAGGCCAACCGGGCCGAGACTGCCATTCGGGCCATCGTCACGTCCTGGGTGGCCTTGGACAATCCGCCCTCGAATGGCAAGTGGAAGTTCAGGATGACGAAGGAGAAACAGAGTCGCCAGAAGCAGAGTCGCCAGAAATGGGCGGCGGACGCGATTTTCGAGTGCAGTTTCATGAAGAGTTGCGTGAAGGAAGTGCTGCCC ``` |
| 7 | ggggacaagtttgtacaaaaaagcaggcttgatgCTTCACTCAAACACGACGAG, ggggaccactttgtacaagaaagctgggtcTTCTGGTCCTGCACAACCGA | Mycgr3\|93029  JGI  Ensembl |  |  | 100 % |  | ``` 1	ATGCTTCACTCAAACACGACGAGGAAACGCGCCCTGAACAAACAAGAAACAGAGCGAGCACTCTACCTCAGCCTCTACGAGCTCATGAATCGCGCCTCCACAGTCGGAGATGAGGAGACGGCTTTCGCGCTGGCGAGCATATTGGCGCAGG	151 	\|\|\|\|\|\|\|\|\|\|\|\|\|\|\|\|\|\|\|\|\|\|\|\|\|\|\|\|\|\|\|\|\|\|\|\|\|\|\|\|\|\|\|\|\|\|\|\|\|\|\|\|\|\|\|\|\|\|\|\|\|\|\|\|\|\|\|\|\|\|\|\|\|\|\|\|\|\|\|\|\|\|\|\|\|\|\|\|\|\|\|\|\|\|\|\|\|\|\|\|\|\|\|\|\|\|\|\|\|\|\|\|\|\|\|\|\|\|\|\|\|\|\|\|\|\|\|\|\|\|\|\|\|\|\|\|\|\|\|\|\|\|\|\|\|\|\|\|\|\|\| 	ATGCTTCACTCAAACACGACGAGGAAACGCGCCCTGAACAAACAAGAAACAGAGCGAGCACTCTACCTCAGCCTCTACGAGCTCATGAATCGCGCCTCCACAGTCGGAGATGAGGAGACGGCTTTCGCGCTGGCGAGCATATTGGCGCAGG ```      ``` 148	CAGGACGAACACTGCCCGATCCTCTACCGCGCCCACGCCCACATGCTCCTCGGACTTTGGCTCGACGCACCGGAAAGCATCGAGCATGCCAAAGAAGCCATCACCATGCTGGAGGAAGACTACTATGATGAGCTCGAAGCTGACGGACGCTTGCAGGACTTCGATCGAGAGCAAGTTGCGCTGGCGAAGAGGATTTTGCGCATGGCGGAGGAGAAGGCGGCGGCTGTTGAGAAGAACGTGGAGGCAGACATCGGTTGTGCAGGACCAGAA	417 	\|\|\|\|\|\|\|\|\|\|\|\|\|\|\|\|\|\|\|\|\|\|\|\|\|\|\|\|\|\|\|\|\|\|\|\|\|\|\|\|\|\|\|\|\|\|\|\|\|\|\|\|\|\|\|\|\|\|\|\|\|\|\|\|\|\|\|\|\|\|\|\|\|\|\|\|\|\|\|\|\|\|\|\|\|\|\|\|\|\|\|\|\|\|\|\|\|\|\|\|\|\|\|\|\|\|\|\|\|\|\|\|\|\|\|\|\|\|\|\|\|\|\|\|\|\|\|\|\|\|\|\|\|\|\|\|\|\|\|\|\|\|\|\|\|\|\|\|\|\|\|\|\|\|\|\|\|\|\|\|\|\|\|\|\|\|\|\|\|\|\|\|\|\|\|\|\|\|\|\|\|\|\|\|\|\|\|\|\|\|\|\|\|\|\|\|\|\|\|\|\|\|\|\|\|\|\|\|\|\|\|\|\|\|\|\|\|\|\|\|\|\|\|\|\|\|\|\|\|\|\|\|\|\|\|\|\|\|\|\|\|\|\|\|\|\|\|\|\|\|\|\|\|\|\|\|\|\|\|\|\|\|\|\|\|\|\|\|\|\| 	CAGGACGAACACTGCCCGATCCTCTACCGCGCCCACGCCCACATGCTCCTCGGACTTTGGCTCGACGCACCGGAAAGCATCGAGCATGCCAAAGAAGCCATCACCATGCTGGAGGAAGACTACTATGATGAGCTCGAAGCTGACGGACGCTTGCAGGACTTCGATCGAGAGCAAGTTGCGCTGGCGAAGAGGATTTTGCGCATGGCGGAGGAGAAGGCGGCGGCTGTTGAGAAGAACGTGGAGGCAGACATCGGTTGTGCAGGACCAGAA ``` |
| 7 | ggggacaagtttgtacaaaaaagcaggcttgatgTTCAACGACTACTGCACCGC, ggggaccactttgtacaagaaagctgggtcACCCAGATTCGAGTTCGGAC | Mycgr3\|93035  JGI  Ensembl |  |  | 100 % |  | ``` 194	GCTACAAAGCAACCATCAGCCAGGTCACGGGTCCCAAGACAGCGGTCACGGAGGATTGTCCCGAGGACATGCAAGCTGGTTGCTGCGATCCGGTATGTGGCCCTGCGATTCCGTTTGAGGTCGTTTGTGGGGCTGAATCGTGGGATGCACAGACTCTTCTTGAACTGACCGGATCGGCACCGTGTACGGGTCCGAACTCGAATCTGGGT	402 	\|\|\|\|\|\|\|\|\|\|\|\|\|\|\|\|\|\|\|\|\|\|\|\|\|\|\|\|\|\|\|\|\|\|\|\|\|\|\|\|\|\|\|\|\|\|\|\|\|\|\|\|\|\|\|\|\|\|\|\|\|\|\|\|\|\|\|\|\|\|\|     \|\|\|\|\|\|\|\|\|\|\|\|\|\|\|\|\|\|\|\|\|\|\|\|\|\|\|\|\|\|\|\|\|\|\|\|\|\|\|\|\|\|\|\|\|\|\|\|\|\|\|\|\|\|\|\|\|\|\|\|\|\|\|\|\|\|\|\|\|\|\|\|\|\|\|\|\|\|\|\|\|\|\|\|\|\|\|\|\|\|\|\|\|\|\|\|\|\|\|\|\|\|\|\|\|\|\|\|\|\|\|\|\|\|\|\|\|\|\|\|\|\|\|\|\|\|\|\|\|\|\|\|\| 	GCTACAAAGCAACCATCAGCCAGGTCACGGGTCCCAAGACAGCGGTCACGGAGGATTGTCCCGAGGACATGNNNNNTGGTTGCTGCGATCCGGTATGTGGCCCTGCGATTCCGTTTGAGGTCGTTTGTGGGGCTGAATCGTGGGATGCACAGACTCTTCTTGAACTGACCGGATCGGCACCGTGTACGGGTCCGAACTCGAATCTGGGT ```      ``` 24	GACAATCAGCCTGCTCCTGGCCCTCCTCGCCGCAACAGGGACTCTGGCAACATTCACCTGTCCCAAACCCGTCAACCCCATTGCCAATAGTCCTTCGAACGCCTGCTGCAAGGAGCTGAGCGATCTACGTGTCAAATTAGGCGGATTCTCACTCTACTATGGCAAGAGCTG	194 	\|\|\|\|\|\|\|\|\|\|\|\|\|\|\|\|\|\|\|\|\|\|\|\|\|\|\|\|\|\|\|\|\|\|\|\|\|\|\|\|\|\|\|\|\|\|\|\|\|\|\|\|\|\|\|\|\|\|\|\|\|\|\|\|\|\|\|\|\|\|\|\|\|\|\|\|\|\|\|\|\|\|\|\|\|\|\|\|\|\|\|\|\|\|\|\|\|\|\|\|\|\|\|\|\|\|\|\|\|\|\|\|\|\|\|\|\|\|\|\|\|\|\|\|\|\|\|\|\|\|\|\|\|\|\|\|\|\|\|\|\|\|\|\|\|\|\|\|\|\|\|\|\|\|\|\|\|\|\|\|\|\|\|\|\|\|\|\|\|\|\| 	GACAATCAGCCTGCTCCTGGCCCTCCTCGCCGCAACAGGGACTCTGGCAACATTCACCTGTCCCAAACCCGTCAACCCCATTGCCAATAGTCCTTCGAACGCCTGCTGCAAGGAGCTGAGCGATCTACGTGTCAAATTAGGCGGATTCTCACTCTACTATGGCAAGAGCTG ``` |
| 7 | ggggacaagtttgtacaaaaaagcaggcttgatgGCCGTCAAAGGTAGCGTGAT, ggggaccactttgtacaagaaagctgggtcCGTCCTGTCTGACCGGAATC | Mycgr3\|93409  JGI  Ensembl |  |  | 0 % |  |  |
| 7 | ggggacaagtttgtacaaaaaagcaggcttgatgGACGTGCCTACTGTCGCCGA, ggggaccactttgtacaagaaagctgggtcAATTCCCAAGTCGAATTGCC | Mycgr3\|93428  JGI  Ensembl |  |  | 0 % |  |  |
| 7 | ggggacaagtttgtacaaaaaagcaggcttgatgCACTTCTTCGCAGCATGTAC, ggggaccactttgtacaagaaagctgggtcACACTGGCAGTACGCCTTTG | Mycgr3\|93442  JGI  Ensembl |  |  | 100 % |  | ``` 1	ATGCACTTCTTCGCAGCATGTACTTGCTTATGGCTGGTGGCCCCGATGGTTTTCGCCGACTTTGCTCTCGACTCTCCAGCTCTCAATGCTCAACAATTCCCGATTGGACTGCCGCAAGACGGCCCTCTCACTCTCACGCGAACGACGACAAGGACGAGATATGACTTCGAGTACCACACCCTCACCATGACAGAAGAAACTTCGATCCCTGCGCAGACGACAGCCGAGCATGCCAATGAAGCGACAGACCCGGGCACGGCCTCGTGCGGCGAAACGGCGTGTGCAGTGTGCAGGTGGACATCCAGCGAGTGCGAATATGGAACAACAGAATG	332 	\|\|\|\|\|\|\|\|\|\|\|\|\|\|\|\|\|\|\|\|\|\|\|\|\|\|\|\|\|\|\|\|\|\|\|\|\|\|\|\|\|\|\|\|\|\|\|\|\|\|\|\|\|\|\|\|\|\|\|\|\|\|\|\|\|\|\|\|\|\|\|\|\|\|\|\|\|\|\|\|\|\|\|\|\|\|\|\|\|\|\|\|\|\|\|\|\|\|\|\|\|\|\|\|\|\|\|\|\|\|\|\|\|\|\|\|\|\|\|\|\|\|\|\|\|\|\|\|\|\|\|\|\|\|\|\|\|\|\|\|\|\|\|\|\|\|\|\|\|\|\|\|\|\|\|\|\|\|\|\|\|\|\|\|\|\|\|\|\|\|\|\|\|\|\|\|\|\|\|\|\|\|\|\|\|\|\|\|\|\|\|\|\|\|\|\|\|\|\|\|\|\|\|\|\|\|\|\|\|\|\|\|\|\|\|\|\|\|\|\|\|\|\|\|\|\|\|\|\|\|\|\|\|\|\|\|\|\|\|\|\|\|\|\|\|\|\|\|\|\|\|\|\|\|\|\|\|\|\|\|\|\|\|\|\|\|\|\|\|\|\|\|\|\|\|\|\|\|\|\|\|\|\|\|\|\|\|\|\|\|\|\|\|\|\|\|\|\|\|\|\|\|\|\|\|\|\|\|\|\|\|\|\|\|\|\|\|\|\|\|\|\|\|\|\|\|\|\|\|\|\|\| 	ATGCACTTCTTCGCAGCATGTACTTGCTTATGGCTGGTGGCCCCGATGGTTTTCGCCGACTTTGCTCTCGACTCTCCAGCTCTCAATGCTCAACAATTCCCGATTGGACTGCCGCAAGACGGCCCTCTCACTCTCACGCGAACGACGACAAGGACGAGATATGACTTCGAGTACCACACCCTCACCATGACAGAAGAAACTTCGATCCCTGCGCAGACGACAGCCGAGCATGCCAATGAAGCGACAGACCCGGGCACGGCCTCGTGCGGCGAAACGGCGTGTGCAGTGTGCAGGTGGACATCCAGCGAGTGCGAATATGGAACAACAGAATG ```      ``` 332	GCCATGAATGCGACGCAAAGGCGTACTGCCAGTGT	366 	\|\|\|\|\|\|\|\|\|\|\|\|\|\|\|\|\|\|\|\|\|\|\|\|\|\|\|\|\|\|\|\|\|\|\| 	GCCATGAATGCGACGCAAAGGCGTACTGCCAGTGT ``` |
| 7 | ggggacaagtttgtacaaaaaagcaggcttgatgCCTACCGCCATGGGTCCCTG, ggggaccactttgtacaagaaagctgggtcCCCAAAGATGACATTCCCGA | Mycgr3\|93639  JGI  Ensembl |  |  | 100 % |  | ``` 1	ATGCCTACCGCCATGGGTCCCTGCGTCAACTGCGCCAAAGCCGCCCGCATCAAGTGTGCCCACTGCGACGATGGCGATTCCTTCACCGCCTACTGCGGCACCAAGTGTCTGAAGAAACATCAAGCCAAGCACCAAGAAGTCTGCAAGGAGAAGAAGAAAGTCCAGAGCGCAAAGGCACTCGTTCCTTCCGAGCACGCAACAAAGGCGACCGTCCCGTCCGAGGCGCACGCAAACGACGATGACGGCGGCTGCGCAGCTTGTGGGAAGGCTCAGACTACTCTCACATGCCTGCGCTGCACTGACGGCATCGTCCACTACTGCAACGAGAACTGCCACGACGCCTACATCAAGAAGCACAAGTTGGTGTGCGTCGCTCAGACGTTGCGCAAGCAGGCGGTAGGGGAGGGAGGGCATGCAATTCTCGACAATCCAGCATGGGATGGAGTCTTCGGGAATGTCATCTTTGGG	468 	\|\|\|\|\|\|\|\|\|\|\|\|\|\|\|\|\|\|\|\|\|\|\|\|\|\|\|\|\|\|\|\|\|\|\|\|\|\|\|\|\|\|\|\|\|\|\|\|\|\|\|\|\|\|\|\|\|\|\|\|\|\|\|\|\|\|\|\|\|\|\|\|\|\|\|\|\|\|\|\|\|\|\|\|\|\|\|\|\|\|\|\|\|\|\|\|\|\|\|\|\|\|\|\|\|\|\|\|\|\|\|\|\|\|\|\|\|\|\|\|\|\|\|\|\|\|\|\|\|\|\|\|\|\|\|\|\|\|\|\|\|\|\|\|\|\|\|\|\|\|\|\|\|\|\|\|\|\|\|\|\|\|\|\|\|\|\|\|\|\|\|\|\|\|\|\|\|\|\|\|\|\|\|\|\|\|\|\|\|\|\|\|\|\|\|\|\|\|\|\|\|\|\|\|\|\|\|\|\|\|\|\|\|\|\|\|\|\|\|\|\|\|\|\|\|\|\|\|\|\|\|\|\|\|\|\|\|\|\|\|\|\|\|\|\|\|\|\|\|\|\|\|\|\|\|\|\|\|\|\|\|\|\|\|\|\|\|\|\|\|\|\|\|\|\|\|\|\|\|\|\|\|\|\|\|\|\|\|\|\|\|\|\|\|\|\|\|\|\|\|\|\|\|\|\|\|\|\|\|\|\|\|\|\|\|\|\|\|\|\|\|\|\|\|\|\|\|\|\|\|\|\|\|\|\|\|\|\|\|\|\|\|\|\|\|\|\|\|\|\|\|\|\|\|\|\|\|\|\|\|\|\|\|\|\|\|\|\|\|\|\|\|\|\|\|\|\|\|\|\|\|\|\|\|\|\|\|\|\|\|\|\|\|\|\|\|\|\|\|\|\|\|\|\|\|\|\|\|\|\|\|\|\|\|\|\|\|\|\|\|\|\|\|\|\|\|\|\|\|\|\|\|\|\|\|\|\|\|\|\|\|\|\|\|\|\|\|\|\|\|\|\|\|\|\|\|\|\|\|\|\|\|\|\|\|\|\|\| 	ATGCCTACCGCCATGGGTCCCTGCGTCAACTGCGCCAAAGCCGCCCGCATCAAGTGTGCCCACTGCGACGATGGCGATTCCTTCACCGCCTACTGCGGCACCAAGTGTCTGAAGAAACATCAAGCCAAGCACCAAGAAGTCTGCAAGGAGAAGAAGAAAGTCCAGAGCGCAAAGGCACTCGTTCCTTCCGAGCACGCAACAAAGGCGACCGTCCCGTCCGAGGCGCACGCAAACGACGATGACGGCGGCTGCGCAGCTTGTGGGAAGGCTCAGACTACTCTCACATGCCTGCGCTGCACTGACGGCATCGTCCACTACTGCAACGAGAACTGCCACGACGCCTACATCAAGAAGCACAAGTTGGTGTGCGTCGCTCAGACGTTGCGCAAGCAGGCGGTAGGGGAGGGAGGGCATGCAATTCTCGACAATCCAGCATGGGATGGAGTCTTCGGGAATGTCATCTTTGGG ``` |
| 7 | ggggacaagtttgtacaaaaaagcaggcttgatgAATTCCTCCAACTGGCTCTA, ggggaccactttgtacaagaaagctgggtcTCCAATCCTCTTCTTTTTCG | Mycgr3\|93679  JGI  Ensembl |  |  | 100 % |  | ``` 1	ATGAATTCCTCCAACTGGCTCTACGCCGGCCCCTCCTCCTCAATCCCCGACCTCTCCACCTCCAACACCCCCACCTCCGACACCAAATTCACCTCCGCCGACTCCTCCAGCCCACCACCCTGCCTCATCCTCTCTCTCGCAGCCTCTCCACCCGCTGCCCTGACACCCTCAGCCGCCTCCCTCACAATCGCCCCCGAACCACAACTCCTCATCTTCCGGTACCGCTCCAAGATCCACGCGATCGACCATTCCTGTCCTCATCAGTCGTACCCTCTGTCCGATGGCAGTATCTATGACATTGAAGACTTTGGAATCGTGCTGTCGGCGGGAATCGCGTGTCCGAAGCATGGGTGGACGTTTGATCTGTTCACGGGGGAGAGTGATCGGGGCGGGTATAAATTGGGAGTGTGGGATGTTGAGGTGAGAGGGGAGGAAGTCTGGGTTAGGAGGAGGCCGAAAAAGAAGAGGATTGG	473 	\|\|\|\|\|\|\|\|\|\|\|\|\|\|\|\|\|\|\|\|\|\|\|\|\|\|\|\|\|\|\|\|\|\|\|\|\|\|\|\|\|\|\|\|\|\|\|\|\|\|\|\|\|\|\|\|\|\|\|\|\|\|\|\|\|\|\|\|\|\|\|\|\|\|\|\|\|\|\|\|\|\|\|\|\|\|\|\|\|\|\|\|\|\|\|\|\|\|\|\|\|\|\|\|\|\|\|\|\|\|\|\|\|\|\|\|\|\|\|\|\|\|\|\|\|\|\|\|\|\|\|\|\|\|\|\|\|\|\|\|\|\|\|\|\|\|\|\|\|\|\|\|\|\|\|\|\|\|\|\|\|\|\|\|\|\|\|\|\|\|\|\|\|\|\|\|\|\|\|\|\|\|\|\|\|\|\|\|\|\|\|\|\|\|\|\|\|\|\|\|\|\|\|\|\|\|\|\|\|\|\|\|\|\|\|\|\|\|\|\|\|\|\|\|\|\|\|\|\|\|\|\|\|\|\|\|\|\|\|\|\|\|\|\|\|\|\|\|\|\|\|\|\|\|\|\|\|\|\|\|\|\|\|\|\|\|\|\|\|\|\|\|\|\|\|\|\|\|\|\|\|\|\|\|\|\|\|\|\|\|\|\|\|\|\|\|\|\|\|\|\|\|\|\|\|\|\|\|\|\|\|\|\|\|\|\|\|\|\|\|\|\|\|\|\|\|\|\|\|\|\|\|\|\|\|\|\|\|\|\|\|\|\|\|\|\|\|\|\|\|\|\|\|\|\|\|\|\|\|\|\|\|\|\|\|\|\|\|\|\|\|\|\|\|\|\|\|\|\|\|\|\|\|\|\|\|\|\|\|\|\|\|\|\|\|\|\|\|\|\|\|\|\|\|\|\|\|\|\|\|\|\|\|\|\|\|\|\|\|\|\|\|\|\| \|\|\|\|\|\| \|\|\| \|\|\|\|\|\|  \| \|\|\|\|\|\|\|\|\|\|\|\| \| \|\|\|\|\|\|\|\| \|\|\| 	ATGAATTCCTCCAACTGGCTCTACGCCGGCCCCTCCTCCTCAATCCCCGACCTCTCCACCTCCAACACCCCCACCTCCGACACCAAATTCACCTCCGCCGACTCCTCCAGCCCACCACCCTGCCTCATCCTCTCTCTCGCAGCCTCTCCACCCGCTGCCCTGACACCCTCAGCCGCCTCCCTCACAATCGCCCCCGAACCACAACTCCTCATCTTCCGGTACCGCTCCAAGATCCACGCGATCGACCATTCCTGTCCTCATCAGTCGTACCCTCTGTCCGATGGCAGTATCTATGACATTGAAGACTTTGGAATCGTGCTGTCGGCGGGAATCGCGTGTCCGAAGCATGGGTGGACGTTTGATCTGTTCACGGGGGAGAGTGATCGGGGCGGGTATAAATTGGGAGTGTGGGATGTTGAGGTGANAGGGGANGAACTCTGGGNNATGAGGAGGCCGAANANGAAGAGGANTGG ``` |
| 7 | ggggacaagtttgtacaaaaaagcaggcttgatgAGCGACTTCCAAGAACTAAG, ggggaccactttgtacaagaaagctgggtcAGTAGAAGATGGGTTGCGGA | Mycgr3\|93730  JGI  Ensembl |  |  | 0 % |  |  |
| 7 | ggggacaagtttgtacaaaaaagcaggcttgatgTCGGACTATGAAGAAGAAGA, ggggaccactttgtacaagaaagctgggtcAGGAGCCCGCAGCCTAGCCC | Mycgr3\|94075  JGI  Ensembl |  |  | 100 % |  | ``` 1	ATGTCGGACTATGAAGAAGAAGATGAGCTCGGCATCGATCCCGCGATCGCGGCAGCGATGGGCTTCACAGGTTTCGGCGCCAAACCCAGCAAGAAGCGAAAATTCGACCGCACCGACGGCTTCGTCGACCCTTCCATCTCGGCCACATCATCGTTACCTCAAGGCAAAGGAGCGAATAACACACCGATAGGCGAGCGGAAGTCTGGCAAATCAAAGACGCCAGCACCTGGTTCAGACGTGGGTATGAACGCAGTGGAGAATGGAGGCGCGCAGGTGCCGATCCGATCGCAGCCGCAGAGCGATGGAAACGCGGAGTCAAATGCGACGAGTGTGACAGCTGCCAGTACTGGTCCGCCGAGTCTGGAAGCTTTGCGATGGGGTGTGAAGAACGACAGAGGGGATCTGGTTGTGTTCATGCCGAGTTTCTTGGAGGATCCTTGGGCTAGGCTGCGGGCTCCT	459 	\|\|\|\|\|\|\|\|\|\|\|\|\|\|\|\|\|\|\|\|\|\|\|\|\|\|\|\|\|\|\|\|\|\|\|\|\|\|\|\|\|\|\|\|\|\|\|\|\|\|\|\|\|\|\|\|\|\|\|\|\|\|\|\|\|\|\|\|\|\|\|\|\|\|\|\|\|\|\|\|\|\|\|\|\|\|\|\|\|\|\|\|\|\|\|\|\|\|\|\|\|\|\|\|\|\|\|\|\|\|\|\|\|\|\|\|\|\|\|\|\|\|\|\|\|\|\|\|\|\|\|\|\|\|\|\|\|\|\|\|\|\|\|\|\|\|\|\|\|\|\|\|\|\|\|\|\|\|\|\|\|\|\|\|\|\|\|\|\|\|\|\|\|\|\|\|\|\|\|\|\|\|\|\|\|\|\|\|\|\|\|\|\|\|\|\|\|\|\|\|\|\|\|\|\|\|\|\|\|\|\|\|\|\|\|\|\|\|\|\|\|\|\|\|\|\|\|\|\|\|\|\|\|\|\|\|\|\|\|\|\|\|\|\|\|\|\|\|\|\|\|\|\|\|\|\|\|\|\|\|\|\|\|\|\|\|\|\|\|\|\|\|\|\|\|\|\|\|\|\|\|\|\|\|\|\|\|\|\|\|\|\|\|\|\|\|\|\|\|\|\|\|\|\|\|\|\|\|\|\|\|\|\|\|\|\|\|\|\|\|\|\|\|\|\|\|\|\|\|\|\|\|\|\|\|\|\|\|\|\|\|\|\|\|\|\|\|\|\|\|\|\|\|\|\|\|\|\|\|\|\|\|\|\|\|\|\|\|\|\|\|\|\|\|\|\|\|\|\|\|\|\|\|\|\|\|\|\|\|\|\|\|\|\|\|\|\|\|\|\|\|\|\|\|\|\|\|\|\|\|\|\|\|\|\|\|\|\|\|\|\|\|\|\|\|\|\|\|\|\|\|\|\|\|\|\|\|\|\|\|\|\|\|\|\|\|\|\|\|\|\|\|\|\|\|\|\|\|\| 	ATGTCGGACTATGAAGAAGAAGATGAGCTCGGCATCGATCCCGCGATCGCGGCAGCGATGGGCTTCACAGGTTTCGGCGCCAAACCCAGCAAGAAGCGAAAATTCGACCGCACCGACGGCTTCGTCGACCCTTCCATCTCGGCCACATCATCGTTACCTCAAGGCAAAGGAGCGAATAACACACCGATAGGCGAGCGGAAGTCTGGCAAATCAAAGACGCCAGCACCTGGTTCAGACGTGGGTATGAACGCAGTGGAGAATGGAGGCGCGCAGGTGCCGATCCGATCGCAGCCGCAGAGCGATGGAAACGCGGAGTCAAATGCGACGAGTGTGACAGCTGCCAGTACTGGTCCGCCGAGTCTGGAAGCTTTGCGATGGGGTGTGAAGAACGACAGAGGGGATCTGGTTGTGTTCATGCCGAGTTTCTTGGAGGATCCTTGGGCTAGGCTGCGGGCTCCT ``` |
| 7 | ggggacaagtttgtacaaaaaagcaggcttgatgGCCTCCAAACTCTGGCAACC, ggggaccactttgtacaagaaagctgggtcGCCCTCCACAGCCTTCCTCC | Mycgr3\|94220  JGI  Ensembl |  |  | 100 % |  | ``` 1	ATGGCCTCCAAACTCTGGCAACCCATCGACCCAACCAAAGCCGCACCCGTCCCAATCCGCCGCAAGAATACCATCCCAGGCACATGGCTCACCCTCACACCCTCGACCACCTCTTCCACCAACATCACCAACGCCGAAGCCGTGTCGCCCACCCTCGACCCCCAATCAGCTTCCTCATCGTTCCTCACCAACGACAACTCCACCGCCAGCGACGGCTCCTCGCGACACTCTTCTTTCACCTCAGCCGGGAACGAGCTGACAAGTCCGTTGAAGAGTGATGTCGAGGTCTCTCCGCGGCACTCTTCGTTTGGCTCGACGGGGAATGAGATGCGGCTTCCGATGAAGAGTAATGTCCAGGCTGGAGTGCATGGAGATGCGGCGGAGAGGAGGGAGGCGAACGTTGCGCTGGGGAGGAAGGCTGTGGAGGGC	429 	\|\|\|\|\|\|\|\|\|\|\|\|\|\|\|\|\|\|\|\|\|\|\|\|\|\|\|\|\|\|\|\|\|\|\|\|\|\|\|\|\|\|\|\|\|\|\|\|\|\|\|\|\|\|\|\|\|\|\|\|\|\|\|\|\|\|\|\|\|\|\|\|\|\|\|\|\|\|\|\|\|\|\|\|\|\|\|\|\|\|\|\|\|\|\|\|\|\|\|\|\|\|\|\|\|\|\|\|\|\|\|\|\|\|\|\|\|\|\|\|\|\|\|\|\|\|\|\|\|\|\|\|\|\|\|\|\|\|\|\|\|\|\|\|\|\|\|\|\|\|\|\|\|\|\|\|\|\|\|\|\|\|\|\|\|\|\|\|\|\|\|\|\|\|\|\|\|\|\|\|\|\|\|\|\|\|\|\|\|\|\|\|\|\|\|\|\|\|\|\|\|\|\|\|\|\|\|\|\|\|\|\|\|\|\|\|\|\|\|\|\|\|\|\|\|\|\|\|\|\|\|\|\|\|\|\|\|\|\|\|\|\|\|\|\|\|\|\|\|\|\|\|\|\|\|\|\|\|\|\|\|\|\|\|\|\|\|\|\|\|\|\|\|\|\|\|\|\|\|\|\|\|\|\|\|\|\|\|\|\|\|\|\|\|\|\|\|\|\|\|\|\|\|\|\|\|\|\|\|\|\|\|\|\|\|\|\|\|\|\|\|\|\|\|\|\|\|\|\|\|\|\|\|\|\|\|\|\|\|\|\|\|\|\|\|\|\|\|\|\|\|\|\|\|\|\|\|\|\|\|\|\|\|\|\|\|\|\|\|\|\|\|\|\|\|\|\|\|\|\|\|\|\|\|\|\|\|\|\|\|\|\|\|\|\|\|\|\|\|\|\|\|\|\|\|\|\|\|\|\|\|\|\|\|\|\|\|\|\|\|\|\|\|\|\|\|\|\|\| 	ATGGCCTCCAAACTCTGGCAACCCATCGACCCAACCAAAGCCGCACCCGTCCCAATCCGCCGCAAGAATACCATCCCAGGCACATGGCTCACCCTCACACCCTCGACCACCTCTTCCACCAACATCACCAACGCCGAAGCCGTGTCGCCCACCCTCGACCCCCAATCAGCTTCCTCATCGTTCCTCACCAACGACAACTCCACCGCCAGCGACGGCTCCTCGCGACACTCTTCTTTCACCTCAGCCGGGAACGAGCTGACAAGTCCGTTGAAGAGTGATGTCGAGGTCTCTCCGCGGCACTCTTCGTTTGGCTCGACGGGGAATGAGATGCGGCTTCCGATGAAGAGTAATGTCCAGGCTGGAGTGCATGGAGATGCGGCGGAGAGGAGGGAGGCGAACGTTGCGCTGGGGAGGAAGGCTGTGGAGGGC ``` |
| 7 | ggggacaagtttgtacaaaaaagcaggcttgatgCCTACTATGCCCAATTTCGA, ggggaccactttgtacaagaaagctgggtcTTTGTCAAATCTACTCTTTT | Mycgr3\|94584  JGI  Ensembl | KOG2959 Transcriptional regulator; |  | 100 % |  | ``` 1	ATGCCTACTATGCCCAATTTCGACATACCTGATTCACCACCTCCACCACCAGTCAACAGCGAAGAGGCCGCAGTACTCGCTGCCACAACGAAGAAGATTGAGCGCTTCCTTGAGTTGAAGAAGAAGGGTGTACACTTCAATGAGCGTCTCCAGAATTCCACCAGCCTACGGAATCCCAGCTTGTTGCCGAAGTTGATGCAGTTTGCGGATATCACAAGGGAGGACAGCTACAGGAGTAGTCTACCGGAAGGACTGGGCATATCGGTCACATGGCCGGAAGAATGTTACGTTGAAGGTCTGCTGCGACAGAACGAGCGGAAACAGAAGAAGAAGCTAGCAGAGAGGGACAAAGTCGACTTTGTGCCGGCTGTCGCGAAAGACGAGGCCAAGGCGCAGGTGCAGAGCGCGGCCAGTGAAGTAGGCAGTGGGAAAAAGAGTAGATTTGACAAA	450 	\|\|\|\|\|\|\|\|\|\|\|\|\|\|\|\|\|\|\|\|\|\|\|\|\|\|\|\|\|\|\|\|\|\|\|\|\|\|\|\|\|\|\|\|\|\|\|\|\|\|\|\|\|\|\|\|\|\|\|\|\|\|\|\|\|\|\|\|\|\|\|\|\|\|\|\|\|\|\|\|\|\|\|\|\|\|\|\|\|\|\|\|\|\|\|\|\|\|\|\|\|\|\|\|\|\|\|\|\|\|\|\|\|\|\|\|\|\|\|\|\|\|\|\|\|\|\|\|\|\|\|\|\|\|\|\|\|\|\|\|\|\|\|\|\|\|\|\|\|\|\|\|\|\|\|\|\|\|\|\|\|\|\|\|\|\|\|\|\|\|\|\|\|\|\|\|\|\|\|\|\|\|\|\|\|\|\|\|\|\|\|\|\|\|\|\|\|\|\|\|\|\|\|\|\|\|\|\|\|\|\|\|\|\|\|\|\|\|\|\|\|\|\|\|\|\|\|\|\|\|\|\|\|\|\|\|\|\|\|\|\|\|\|\|\|\|\|\|\|\|\|\|\|\|\|\|\|\|\|\|\|\|\|\|\|\|\|\|\|\|\|\|\|\|\|\|\|\|\|\|\|\|\|\|\|\|\|\|\|\|\|\|\|\|\|\|\|\|\|\|\|\|\|\|\|\|\|\|\|\|\|\|\|\|\|\|\|\|\|\|\|\|\|\|\|\|\|\|\|\|\|\|\|\|\|\|\|\|\|\|\|\|\|\|\|\|\|\|\|\|\|\|\|\|\|\|\|\|\|\|\|\|\|\|\|\|\|\|\|\|\|\|\|\|\|\|\|\|\|\|\|\|\|\|\|\|\|\|\|\|\|\|\|\|\|\|\|\|\|\|\|\|\|\|\|\|\|\|\|\|\|\|\|\|\|\|\|\|\|\|\|\|\|\|\|\|\|\|\|\|\|\|\|\|\|\|\|\|\|\|\|\|\|\|\|\|\|\|\|\| 	ATGCCTACTATGCCCAATTTCGACATACCTGATTCACCACCTCCACCACCAGTCAACAGCGAAGAGGCCGCAGTACTCGCTGCCACAACGAAGAAGATTGAGCGCTTCCTTGAGTTGAAGAAGAAGGGTGTACACTTCAATGAGCGTCTCCAGAATTCCACCAGCCTACGGAATCCCAGCTTGTTGCCGAAGTTGATGCAGTTTGCGGATATCACAAGGGAGGACAGCTACAGGAGTAGTCTACCGGAAGGACTGGGCATATCGGTCACATGGCCGGAAGAATGTTACGTTGAAGGTCTGCTGCGACAGAACGAGCGGAAACAGAAGAAGAAGCTAGCAGAGAGGGACAAAGTCGACTTTGTGCCGGCTGTCGCGAAAGACGAGGCCAAGGCGCAGGTGCAGAGCGCGGCCAGTGAAGTAGGCAGTGGGAAAAAGAGTAGATTTGACAAA ``` |
| 7 | ggggacaagtttgtacaaaaaagcaggcttgatgTCCGGTCTCAACAAATCCGC, ggggaccactttgtacaagaaagctgggtcGGTCACACGGAGCATCATTC | Mycgr3\|94679  JGI  Ensembl |  |  | 0 % |  |  |
| 7 | ggggacaagtttgtacaaaaaagcaggcttgatgTGGACCAACGCCCTCGGAGG, ggggaccactttgtacaagaaagctgggtcGACCTTCAATTCGTCCCTCA | Mycgr3\|94702  JGI  Ensembl |  |  | 0 % |  |  |
| 7 | ggggacaagtttgtacaaaaaagcaggcttgatgGTCACTTCCCGTTGGGTAGT, ggggaccactttgtacaagaaagctgggtcTGATACAAAACCAGCCCCAG | Mycgr3\|94772  JGI  Ensembl |  |  | 0 % |  |  |
| 7 | ggggacaagtttgtacaaaaaagcaggcttgatgGCTTTCCCCAGCCAGCTCGT, ggggaccactttgtacaagaaagctgggtcTGCCTTCGGCCTCCAGACCT | Mycgr3\|94779  JGI  Ensembl |  |  | 100 % |  | ``` 1	ATGGCTTTCCCCAGCCAGCTCGTCGGCTCAAAACCACAACATCCACTGTCCAGGCCAACCTTCCCCGACGCCGACTCACCACCAGCTGTTCCTGGCAAGGACACCTACCGAACGAAAGCTCTGCCGTCTCCGCCGCCTCGGACACCGACTATGGCATACTTGCCTGTCCCAGCGAGGACTGCTGAAATTCCGTCGAATCCTAGTCCAGAGCGAATACAGGAGCAGGCAAGATCAGCCCAAAACGAGCCGTTTCCTGCCTTTGATTTTTGCTCGGACGGAGGAGCGAGCAGAGAGGATCAGATCGAGCGGATTGATGAGGAAGTGCGGAAGAGCAGCTGGGTGTTGAATGGAGACTTGAATCGAGCGGATCGATGTACCGACTCGCCGTTGGAGGAGGAGGCGAAAGAAGGCGAGGCTGCTGAGAATTCTACGGCGAAGGAGAAGAAGACGAGGGAAAAGTGGTTCCAGGTCTGGAGGCCGAAGGCA	486 	\|\|\|\|\|\|\|\|\|\|\|\|\|\|\|\|\|\|\|\|\|\|\|\|\|\|\|\|\|\|\|\|\|\|\|\|\|\|\|\|\|\|\|\|\|\|\|\|\|\|\|\|\|\|\|\|\|\|\|\|\|\|\|\|\|\|\|\|\|\|\|\|\|\|\|\|\|\|\|\|\|\|\|\|\|\|\|\|\|\|\|\|\|\|\|\|\|\|\|\|\|\|\|\|\|\|\|\|\|\|\|\|\|\|\|\|\|\|\|\|\|\|\|\|\|\|\|\|\|\|\|\|\|\|\|\|\|\|\|\|\|\|\|\|\|\|\|\|\|\|\|\|\|\|\|\|\|\|\|\|\|\|\|\|\|\|\|\|\|\|\|\|\|\|\|\|\|\|\|\|\|\|\|\|\|\|\|\|\|\|\|\|\|\|\|\|\|\|\|\|\|\|\|\|\|\|\|\|\|\|\|\|\|\|\|\|\|\|\|\|\|\|\|\|\|\|\|\|\|\|\|\|\|\|\|\|\|\|\|\|\|\|\|\|\|\|\|\|\|\|\|\|\|\|\|\|\|\|\|\|\|\|\|\|\|\|\|\|\|\|\|\|\|\|\|\|\|\|\|\|\|\|\|\|\|\|\|\|\|\|\|\|\|\|\|\|\|\|\|\|\|\|\|\|\|\|\|\|\|\|\|\|\|\|\|\|\|\|\|\|\|\|\|\|\|\|\|\|\|\|\|\|\|\|\|\|\|\|\|\|\|\|\|\|\|\|\|\|\|\|\|\|\|\|\|\|\|\|\|\|\|\|\|\|\|\|\|\|\|\|\|\|\|\|\|\|\|\|\|\|\|\|\|\|\|\|\|\|\|\|\|\|\|\|\|\|\|\|\|\|\|\|\|\|\|\|\|\|\|\|\|\|\|\|\|\|\|\|\|\|\|\|\|\|\|\|\|\|\|\|\|\|\|\|\|\|\|\|\|\| \|\|\|\|\|\|\|\|\|\|\|\|\|\|\|\|\|\|\|\|\|\|\|\|\|\|\|\|\|\|\|\|\|\|\|\|\|\|\|\|\|\|\|\|\| 	ATGGCTTTCCCCAGCCAGCTCGTCGGCTCAAAACCACAACATCCACTGTCCAGGCCAACCTTCCCCGACGCCGACTCACCACCAGCTGTTCCTGGCAAGGACACCTACCGAACGAAAGCTCTGCCGTCTCCGCCGCCTCGGACACCGACTATGGCATACTTGCCTGTCCCAGCGAGGACTGCTGAAATTCCGTCGAATCCTAGTCCAGAGCGAATACAGGAGCAGGCAAGATCAGCCCAAAACGAGCCGTTTCCTGCCTTTGATTTTTGCTCGGACGGAGGAGCGAGCAGAGAGGATCAGATCGAGCGGATTGATGAGGAAGTGCGGAAGAGCAGCTGGGTGTTGAATGGAGACTTGAATCGAGCGGATCGATGTACCGACTCGCCGTTGGAGGAGGAGGCGAAAGAAGGCGAGGCTGCTGAGAATTCTACGGCGAAGGANAAGAAGACGAGGGAAAAGTGGTTCCAGGTCTGGAGGCCGAAGGCA ``` |
| 7 | ggggacaagtttgtacaaaaaagcaggcttgatgGCAAGAAGTGCGCGATCCTC, ggggaccactttgtacaagaaagctgggtcCTTCTTTCCCGCCTTCTTCT | Mycgr3\|95012  JGI  Ensembl |  |  | 0 % |  |  |
| 7 | ggggacaagtttgtacaaaaaagcaggcttgatgGATCCCAACTCGCCAGACTA, ggggaccactttgtacaagaaagctgggtcACGCTCTCTAATCGGCAGAT | Mycgr3\|95075  JGI  Ensembl |  |  | 0 % |  |  |
| 7 | ggggacaagtttgtacaaaaaagcaggcttgatgAACACCATCATCGACAACCA, ggggaccactttgtacaagaaagctgggtcGTTTCTGTCTGTTTCGCAAC | Mycgr3\|95076  JGI  Ensembl |  |  | 0 % |  |  |
| 7 | ggggacaagtttgtacaaaaaagcaggcttgatgTCTCGCACCAGACCGATCTC, ggggaccactttgtacaagaaagctgggtcCTGCATAGCACAGCAGCCGT | Mycgr3\|95203  JGI  Ensembl |  |  | 100 % |  | ``` 1	ATGTCTCGCACCAGACCGATCTCCTTCGCAACCGGCGGTGTCCACGCCACCGACGCGCCCGTCACCCTCTTCTACCGCGAGCCCGATGACCGAAACTACCACGCCTGCGGCCTATACCACACCCCAGGCCTCCACGCCGTCATCACCGTGGCCCTCGACGCCATGTGCGACACTCCCGGCCGCACCATCTTGCGCACCGACGACGGCTCTGGCGCTGCAGTCTCGGTCGAGGGGCTAGTTCATATCCTCGCCACGGAGACAAAGCTCGGCGGAGGACATCTGGACAGCGTCTACGTGACGGTCTACTCCTCGGATCCGTTTGCTGTTGCCGGTCCGGCGGCGGCCGCGTTGGCTTCGCCGGCTGGCCAGGCGCAGAGAGTGGCCGCGGGAGAGCGGATTAGCTTCAATGAGAAGCACGGCTGCTGTGCTATGCAG	435 	\|\|\|\|\|\|\|\|\|\|\|\|\|\|\|\|\|\|\|\|\|\|\|\|\|\|\|\|\|\|\|\|\|\|\|\|\|\|\|\|\|\|\|\|\|\|\|\|\|\|\|\|\|\|\|\|\|\|\|\|\|\|\|\|\|\|\|\|\|\|\|\|\|\|\|\|\|\|\|\|\|\|\|\|\|\|\|\|\|\|\|\|\|\|\|\|\|\|\|\|\|\|\|\|\|\|\|\|\|\|\|\|\|\|\|\|\|\|\|\|\|\|\|\|\|\|\|\|\|\|\|\|\|\|\|\|\|\|\|\|\|\|\|\|\|\|\|\|\|\|\|\|\|\|\|\|\|\|\|\|\|\|\|\|\|\|\|\|\|\|\|\|\|\|\|\|\|\|\|\|\|\|\|\|\|\|\|\|\|\|\|\|\|\|\|\|\|\|\|\|\|\|\|\|\|\|\|\|\|\|\|\|\|\|\|\|\|\|\|\|\|\|\|\|\|\|\|\|\|\|\|\|\|\|\|\|\|\|\|\|\|\|\|\|\|\|\|\|\|\|\|\|\|\|\|\|\|\|\|\|\|\|\|\|\|\|\|\|\|\|\|\|\|\|\|\|\|\|\|\|\|\|\|\|\|\|\|\|\|\|\|\|\|\|\|\|\|\|\|\|\|\|\|\|\|\|\|\|\|\|\|\|\|\|\|\|\|\|\|\|\|\|\|\|\|\|\|\|\|\|\|\|\|\|\|\|\|\|\|\|\|\|\|\|\|\|\|\|\|\|\|\|\|\|\|\|\|\|\|\|\|\|\|\|\|\|\|\|\|\|\|\|\|\|\|\|\|\|\|\|\|\|\|\|\|\|\|\|\|\|\|\|\|\|\|\|\|\|\|\|\|\|\|\|\|\|\|\|\|\|\|\|\|\|\|\|\|\|\|\|\|\|\|\|\|\|\|\|\|\|\|\|\|\|\| 	ATGTCTCGCACCAGACCGATCTCCTTCGCAACCGGCGGTGTCCACGCCACCGACGCGCCCGTCACCCTCTTCTACCGCGAGCCCGATGACCGAAACTACCACGCCTGCGGCCTATACCACACCCCAGGCCTCCACGCCGTCATCACCGTGGCCCTCGACGCCATGTGCGACACTCCCGGCCGCACCATCTTGCGCACCGACGACGGCTCTGGCGCTGCAGTCTCGGTCGAGGGGCTAGTTCATATCCTCGCCACGGAGACAAAGCTCGGCGGAGGACATCTGGACAGCGTCTACGTGACGGTCTACTCCTCGGATCCGTTTGCTGTTGCCGGTCCGGCGGCGGCCGCGTTGGCTTCGCCGGCTGGCCAGGCGCAGAGAGTGGCCGCGGGAGAGCGGATTAGCTTCAATGAGAAGCACGGCTGCTGTGCTATGCAG ``` |
| 7 | ggggacaagtttgtacaaaaaagcaggcttgatgTCCAGCTCCGACGAAGAAGA, ggggaccactttgtacaagaaagctgggtcCCGCGACCGTCCAGACTCGC | Mycgr3\|95218  JGI  Ensembl |  |  | 100 % |  | ``` 1	ATGTCCAGCTCCGACGAAGAAGACAACAAGTCATGGAATCCTGCCACCGAAACGCATCCCAGCATCGACCTCCCCCTAGCCTTCAAAATCTACTGTGCCAAATCCCGCCTCACCCACTCGGAGTTCTACATCGCACACGAGAAGAACAAAAGACTACACGCCATCTCCGGCGGCTACAAAGGCTGGCTCAAATCCACCGTCGGACTCCACCAAGGTCCCAGCATCGACACCCCAACTCTAGCCAACGCCATCATCAAGCACAAAACAATCGAAGTCACCCTCCCGACTGGCACCACG	297 	\|\|\|\|\|\|\|\|\|\|\|\|\|\|\|\|\|\|\|\|\|\|\|\|\|\|\|\|\|\|\|\|\|\|\|\|\|\|\|\|\|\|\|\|\|\|\|\|\|\|\|\|\|\|\|\|\|\|\|\|\|\|\|\|\|\|\|\|\|\|\|\|\|\|\|\|\|\|\|\|\|\|\|\|\|\|\|\|\|\|\|\|\|\|\|\|\|\|\|\|\|\|\|\|\|\|\|\|\|\|\|\|\|\|\|\|\|\|\|\|\|\|\|\|\|\|\|\|\|\|\|\|\|\|\|\|\|\|\|\|\|\|\|\|\|\|\|\|\|\|\|\|\|\|\|\|\|\|\|\|\|\|\|\|\|\|\|\|\|\|\|\|\|\|\|\|\|\|\|\|\|\|\|\|\|\|\|\|\|\|\|\|\|\|\|\|\|\|\|\|\|\|\|\|\|\|\|\|\|\|\|\|\|\|\|\|\|\|\|\|\|\|\|\|\|\|\|\|\|\|\|\|\|\|\|\|\|\|\|\|\|\|\|\|\|\|\|\|\|\|\|\|\|\|\|\|\|\|\|\|\|\|\|\|\|\|\|\|\|\|\|\|\|\|\|\|\|\|\|\|\|\|\|\|\|\|\|\|\|\|\|\|\|\|\|\|\| 	ATGTCCAGCTCCGACGAAGAAGACAACAAGTCATGGAATCCTGCCACCGAAACGCATCCCAGCATCGACCTCCCCCTAGCCTTCAAAATCTACTGTGCCAAATCCCGCCTCACCCACTCGGAGTTCTACATCGCACACGAGAAGAACAAAAGACTACACGCCATCTCCGGCGGCTACAAAGGCTGGCTCAAATCCACCGTCGGACTCCACCAAGGTCCCAGCATCGACACCCCAACTCTAGCCAACGCCATCATCAAGCACAAAACAATCGAAGTCACCCTCCCGACTGGCACCACG ```      ``` 297	GCCCCGAGGTCGCGAGTCTGGACGGTCGCGG	327 	\|\|\|\|\|\|\|\|\|\|\|\|\|\|\|\|\|\|\|\|\|\|\|\|\|\|\|\|\|\|\| 	GCCCCGAGGTCGCGAGTCTGGACGGTCGCGG ``` |
| 7 | ggggacaagtttgtacaaaaaagcaggcttgatgTCCCGCGCCGCCGAAGGGGC, ggggaccactttgtacaagaaagctgggtcTAAAGATACCAGCCATCCGG | Mycgr3\|96177  JGI  Ensembl |  |  | 83.43949 % |  | ``` 1	ATGTCCCGCGCCGCCGAAGGGGCGGGGAGGGCAAGTCGTGGCGTGAGGCAGGATGAGGTAGAAGTGGAGTTGGAAGGCCAGCGCTGTACTGCCGAGGTGGATGTAGGGCACGTTCTCCGGTCGAGCGAGGGGTACAGGGGAGATGTATTTCCGGAAGAGATGTTCTTCGATGTGGGCGGGGAGAGGAGGGCCGAGTTGCGGGGTGGTGTTGACGGTGAAGCGGCAGATCCAGGATTTTTTGAGGACGGCGAGGGATTTGGGGAAGAGGAGGATTTCTTTGAC	282 	\|\|\|\|\|\|\|\|\|\|\|\| \|\|\|\|\|\|\|\|\|\|\|\|\|\|\|\|\|\|\|\|\|\|\|\|\|\|\|\|\|\|\|\|\|\|\|\|\|\|\|\|\|\|\|\|\|\|\|\|\|\|\|\|\|\|\|\|\|\|\|\|\|\|\|\|\|\|\|\|\|\|\|\|\|\|\|\|\|\|\|\|\|\|\|\|\|\|\|\|\|\|\|\|\|\|\|\|\|\|\|\|\|\|\|\|\|\|\|\|\|\|\|\|\|\|\|\|\|\|\|\|\|\|\|\|\|\|\|\|\|\|\|\|\|\|\|\|\|\|\|\|\|\|\|\|\|\|\|\|\|\|\|\|\|\|\|\|\|\|\|\|\|\|\|\|\|\|\|\|\|\|\|\|\|\|\|\|\|\|\|\|\|\|\|\|\|\|\|\|\|\|\|\|\|\|\|\|\|\|\|\|\|\|\|\|\|\|\|\|\|\|\|\|\|\|\|\|\|\|\|\|\|\|\|\|\|\|\|\|\|\|\|\|\|\|\|\|\|\|\|\|\|\|\|\|\|\|\|\|\|\|\|\|\|\|\|\|\|\|\|\|\|\|\|\|\|\|\|\|\| 	ATGTCCCGCGCCNCCGAAGGGGCGGGGAGGGCAAGTCGTGGCGTGAGGCAGGATGAGGTAGAAGTGGAGTTGGAAGGCCAGCGCTGTACTGCCGAGGTGGATGTAGGGCACGTTCTCCGGTCGAGCGAGGGGTACAGGGGAGATGTATTTCCGGAAGAGATGTTCTTCGATGTGGGCGGGGAGAGGAGGGCCGAGTTGCGGGGTGGTGTTGACGGTGAAGCGGCAGATCCAGGATTTTTTGAGGACGGCGAGGGATTTGGGGAAGAGGAGGATTTCTTTGAC ``` |
| 7 | ggggacaagtttgtacaaaaaagcaggcttgatgGCGTCCACCTCCCACCCAAC, ggggaccactttgtacaagaaagctgggtcATCCTGCGGAATAGCACTAA | Mycgr3\|96240  JGI  Ensembl |  |  | 100 % |  | ``` 1	ATGGCGTCCACCTCCCACCCAACGCAAACCCACCCCTCCGCCCCTCCCGCCGCCCTCTCAAAAGAAGACTCCACCGCCACCACCACCTCCTCCACCTCCACCGACTTCGAAGTCGCGTCGCCGATGAAATTCACCCACACCGGTGACCCGTTGGCCGATCGACCGAGCGTGCAGGATGCGGAACCCAAGCTGTCACGACACATCAGAGCGATTCCCGATGATGGCAACGAGGAGGATGGGAAAGAGAAGGGATTGATGGCGAAGTTGAAGGCGAAACTCCTAGGTTCAAAGGGGAAGGAGCACGCGCATCACGAAGCTCCGACGAGGACGTCGGAAGATGTGGAGGTTGTGGGTACGATGAGTGGGAGTCCGGTTAGGGATGGAGGAGGGTTGGAGAGGCATATTAGTGCTATTCCGCAGGAT	423 	\|\|\|\|\|\|\|\|\|\|\|\|\|\|\|\|\|\|\|\|\|\|\|\|\|\|\|\|\|\|\|\|\|\|\|\|\|\|\|\|\|\|\|\|\|\|\|\|\|\|\|\|\|\|\|\|\|\|\|\|\|\|\|\|\|\|\|\|\|\|\|\|\|\|\|\|\|\|\|\|\|\|\|\|\|\|\|\|\|\|\|\|\|\|\|\|\|\|\|\|\|\|\|\|\|\|\|\|\|\|\|\|\|\|\|\|\|\|\|\|\|\|\|\|\|\|\|\|\|\|\|\|\|\|\|\|\|\|\|\|\|\|\|\|\|\|\|\|\|\|\|\|\|\|\|\|\|\|\|\|\|\|\|\|\|\|\|\|\|\|\|\|\|\|\|\|\|\|\|\|\|\|\|\|\|\|\|\|\|\|\|\|\|\|\|\|\|\|\|\|\|\|\|\|\|\|\|\|\|\|\|\|\|\|\|\|\|\|\|\|\|\|\|\|\|\|\|\|\|\|\|\|\|\|\|\|\|\|\|\|\|\|\|\|\|\|\|\|\|\|\|\|\|\|\|\|\|\|\|\|\|\|\|\|\|\|\|\|\|\|\|\|\|\|\|\|\|\|\|\|\|\|\|\|\|\|\|\|\|\|\|\|\|\|\|\|\|\|\|\|\|\|\|\|\|\|\|\|\|\|\|\|\|\|\|\|\|\|\|\|\|\|\|\|\|\|\|\|\|\|\|\|\|\|\|\|\|\|\|\|\|\|\|\|\|\|\|\|\|\|\|\|\|\|\|\|\|\|\|\|\|\|\|\|\|\|\|\|\|\|\|\|\|\|\|\|\|\|\|\|\|\|\|\|\|\|\|\|\|\|\|\|\|\|\|\|\|\|\|\|\|\|\|\|\|\|\|\|\|\|\|\|\|\|\|\|\|\|\|\|\|\|\| 	ATGGCGTCCACCTCCCACCCAACGCAAACCCACCCCTCCGCCCCTCCCGCCGCCCTCTCAAAAGAAGACTCCACCGCCACCACCACCTCCTCCACCTCCACCGACTTCGAAGTCGCGTCGCCGATGAAATTCACCCACACCGGTGACCCGTTGGCCGATCGACCGAGCGTGCAGGATGCGGAACCCAAGCTGTCACGACACATCAGAGCGATTCCCGATGATGGCAACGAGGAGGATGGGAAAGAGAAGGGATTGATGGCGAAGTTGAAGGCGAAACTCCTAGGTTCAAAGGGGAAGGAGCACGCGCATCACGAAGCTCCGACGAGGACGTCGGAAGATGTGGAGGTTGTGGGTACGATGAGTGGGAGTCCGGTTAGGGATGGAGGAGGGTTGGAGAGGCATATTAGTGCTATTCCGCAGGAT ``` |
| 7 | ggggacaagtttgtacaaaaaagcaggcttgatgGCCACTCTCCGCAACCTCTC, ggggaccactttgtacaagaaagctgggtcCTTCTTCAAGCCAAAGATTT | Mycgr3\|96315  JGI  Ensembl |  |  | 0 % |  |  |
| 7 | ggggacaagtttgtacaaaaaagcaggcttgatgACCTACAAGTACAAACATCT, ggggaccactttgtacaagaaagctgggtcTGGGCCTTCCTTAATACGGT | Mycgr3\|96646  JGI  Ensembl |  |  | 100 % |  | ``` 382	AGTCCAAGCGACCGTATTAAGGAAGGCC	409 	\|\|\|\|\|\|\|\|\|\|\|\|\|\|\|\|\|\|\|\|\|\|\|\|\|\|\|\| 	AGTCCAAGCGACCGTATTAAGGAAGGCC ```      ``` 1	ATGACCTACAAGTACAAACATCTCTTTCTGGAGGCGCAAGACGAGGAGGAGAGCGACGCTACCGAGATGGACAACCCCTACGCCGGTGATCCGCTCATAGCGTGTATTGGATTCCTAGAGCCAGCCCGCGTCGAGTGCCTAACGCATGTCCAAAGCAAGAAGCTGTTGCGCTTGGTAATCGAAGGAGAGTTCCGGAAGATGAAGAAGCTGCCTAAGCTGCACGTCCGCGGCGAGCCAAAGTTCTACGAGAGGATTAGACAGATCGCACCGAGAGTTATAGAATGGGAGGAGAGAGTAGTAAAAGCGGAGAAGGCAGCGGATAAGAAGGGGATTAAGAGGAACAGGAAAGAACACCTCTTACTATGGAACTGTAGGGAGTTAAG	383 	\|\|\|\|\|\|\|\|\|\|\|\|\|\|\|\|\|\|\|\|\|\|\|\|\|\|\|\|\|\|\|\|\|\|\|\|\|\|\|\|\|\|\|\|\|\|\|\|\|\|\|\|\|\|\|\|\|\|\|\|\|\|\|\|\|\|\|\|\|\|\|\|\|\|\|\|\|\|\|\|\|\|\|\|\|\|\|\|\|\|\|\|\|\|\|\|\|\|\|\|\|\|\|\|\|\|\|\|\|\|\|\|\|\|\|\|\|\|\|\|\|\|\|\|\|\|\|\|\|\|\|\|\|\|\|\|\|\|\|\|\|\|\|\|\|\|\|\|\|\|\|\|\|\|\|\|\|\|\|\|\|\|\|\|\|\|\|\|\|\|\|\|\|\|\|\|\|\|\|\|\|\|\|\|\|\|\|\|\|\|\|\|\|\|\|\|\|\|\|\|\|\|\|\|\|\|\|\|\|\|\|\|\|\|\|\|\|\|\|\|\|\|\|\|\|\|\|\|\|\|\|\|\|\|\|\|\|\|\|\|\|\|\|\|\|\|\|\|\|\|\|\|\|\|\|\|\|\|\|\|\|\|\|\|\|\|\|\|\|\|\|\|\|\|\|\|\|\|\|\|\|\|\|\|\|\|\|\|\|\|\|\|\|\|\|\|\|\|\|\|\|\|\|\|\|\|\|\|\|\|\|\|\|\|\|\|\|\|\|\|\|\|\|\|\|\|\|\|\|\|\|\|\|\|\|\|\|\|\|\|\|\|\|\|\|\|\|\|\|\|\|\|\|\|\|\|\|\|\|\|\|\|\|\|\|\|\|\|\|\|\|\|\|\|\|\|\|\|\|\|\|\|\| 	ATGACCTACAAGTACAAACATCTCTTTCTGGAGGCGCAAGACGAGGAGGAGAGCGACGCTACCGAGATGGACAACCCCTACGCCGGTGATCCGCTCATAGCGTGTATTGGATTCCTAGAGCCAGCCCGCGTCGAGTGCCTAACGCATGTCCAAAGCAAGAAGCTGTTGCGCTTGGTAATCGAAGGAGAGTTCCGGAAGATGAAGAAGCTGCCTAAGCTGCACGTCCGCGGCGAGCCAAAGTTCTACGAGAGGATTAGACAGATCGCACCGAGAGTTATAGAATGGGAGGAGAGAGTAGTAAAAGCGGAGAAGGCAGCGGATAAGAAGGGGATTAAGAGGAACAGGAAAGAACACCTCTTACTATGGAACTGTAGGGAGTTAAG ``` |
| 7 | ggggacaagtttgtacaaaaaagcaggcttgatgGCCTTCGGCGAAGATCAAGA, ggggaccactttgtacaagaaagctgggtcAGACAAGTCCATCGCATCCG | Mycgr3\|96754  JGI  Ensembl |  |  | 0 % |  |  |
| 7 | ggggacaagtttgtacaaaaaagcaggcttgatgTCAGCTGGAGATAATCCGTC, ggggaccactttgtacaagaaagctgggtcAATGCACCAGTTATGCGTAT | Mycgr3\|96784  JGI  Ensembl |  |  | 100 % |  | ``` 1	ATGTCAGCTGGAGATAATCCGTCCGCTGTCATGTGCAACTACCGCATGCGCATAGCATGGCTTGGCTCACTTCCTGATGATCAGGTCGTTGCGCCCGGACCGGATGTGACGAGTGGCCGCTCGCCAGTCCACTTGCTTACCATTCAAACCACCCAGCTTCTCGTCGATTATCTGAAATTCTGGCGCATCTACCTCGTTATCGCGTCCAAGATCCCTGACAGGCCTCGCCGGCGTTATACGGACTCGAACAGTGACCAAGCAAAACAGATTGGATACTGGTGGGATTGTGAACGCCCAGACCACGATGTTAATAAAGCAAGGAGCTGGCGTGACAGCTTGGAGAGCTTACCCTCCGAGCTGCGCAAGAAGTCGACCGACTCATGGCGGCCCGCAGAGCTAATACAACGACTTTACGACTTCACGTTCGATACGCATAACTGGTGCATT	447 	\|\|\|\|\|\|\|\|\|\|\|\|\|\|\|\|\|\|\|\|\|\|\|\|\|\|\|\|\|\|\|\|\|\|\|\|\|\|\|\|\|\|\|\|\|\|\|\|\|\|\|\|\|\|\|\|\|\|\|\|\|\|\|\|\|\|\|\|\|\|\|\|\|\|\|\|\|\|\|\|\|\|\|\|\|\|\|\|\|\|\|\|\|\|\|\|\|\|\|\|\|\|\|\|\|\|\|\|\|\|\|\|\|\|\|\|\|\|\|\|\|\|\|\|\|\|\|\|\|\|\|\|\|\|\|\|\|\|\|\|\|\|\|\|\|\|\|\|\|\|\|\|\|\|\|\|\|\|\|\|\|\|\|\|\|\|\|\|\|\|\|\|\|\|\|\|\|\|\|\|\|\|\|\|\|\|\|\|\|\|\|\|\|\|\|\|\|\|\|\|\|\|\|\|\|\|\|\|\|\|\|\|\|\|\|\|\|\|\|\|\|\|\|\|\|\|\|\|\|\|\|\|\|\|\|\|\|\|\|\|\|\|\|\|\|\|\|\|\|\|\|\|\|\|\|\|\|\|\|\|\|\|\|\|\|\|\|\|\|\|\|\|\|\|\|\|\|\|\|\|\|\|\|\|\|\|\|\|\|\|\|\|\|\|\|\|\|\|\|\|\|\|\|\|\|\|\|\|\|\|\|\|\|\|\|\|\|\|\|\|\|\|\|\|\|\|\|\|\|\|\|\|\|\|\|\|\|\|\|\|\|\|\|\|\|\|\|\|\|\|\|\|\|\|\|\|\|\|\|\|\|\|\|\|\|\|\|\|\|\|\|\|\|\|\|\|\|\|\|\|\|\|\|\|\|\|\|\|\|\|\|\|\|\|\|\|\|\|\|\|\|\|\|\|\|\|\|\|\|\|\|\|\|\|\|\|\|\|\|\|\|\|\|\|\|\|\|\|\|\|\|\|\|\|\|\|\|\|\|\|\|\|\|\|\|\|\| 	ATGTCAGCTGGAGATAATCCGTCCGCTGTCATGTGCAACTACCGCATGCGCATAGCATGGCTTGGCTCACTTCCTGATGATCAGGTCGTTGCGCCCGGACCGGATGTGACGAGTGGCCGCTCGCCAGTCCACTTGCTTACCATTCAAACCACCCAGCTTCTCGTCGATTATCTGAAATTCTGGCGCATCTACCTCGTTATCGCGTCCAAGATCCCTGACAGGCCTCGCCGGCGTTATACGGACTCGAACAGTGACCAAGCAAAACAGATTGGATACTGGTGGGATTGTGAACGCCCAGACCACGATGTTAATAAAGCAAGGAGCTGGCGTGACAGCTTGGAGAGCTTACCCTCCGAGCTGCGCAAGAAGTCGACCGACTCATGGCGGCCCGCAGAGCTAATACAACGACTTTACGACTTCACGTTCGATACGCATAACTGGTGCATT ``` |
| 7 | ggggacaagtttgtacaaaaaagcaggcttgatgGGCTGGGTAAAGTTCGCCCT, ggggaccactttgtacaagaaagctgggtcACCCAGACCAGACGAGCACG | Mycgr3\|96813  JGI  Ensembl |  |  | 0 % |  |  |
| 7 | ggggacaagtttgtacaaaaaagcaggcttgatgAACTCCACACACGACTCAAC, ggggaccactttgtacaagaaagctgggtcGCTCTTCAACAGACGCACGA | Mycgr3\|96870  JGI  Ensembl |  |  | 100 % |  | ``` 1	ATGAACTCCACACACGACTCAACCGCCGGCGGTGTCGGACGAGTAGGACACGAACGTATCGGGGAAGAATACCTCACTCGGCTGGGCTACTCGAAGAAAGTCGGGTTTCTGGTAGGATCACATGCTGCAGCAAAGAGATTTCTCTGCGGGACCGATCCTGCCTATCACGATACGCTTTCGGGAGCGAGCAAGAAGTCGCTGGTCTTTCAAGGAGAGCCCATGAGAGGCGACGAGCTTAATGAGTGGGCTGCGAATCCTTGGTGCGACGAGATGTGCCA	278 	\|\|\|\|\|\|\|\|\|\|\|\|\|\|\|\|\|\|\|\|\|\|\|\|\|\|\|\|\|\|\|\|\|\|\|\|\|\|\|\|\|\|\|\|\|\|\|\|\|\|\|\|\|\|\|\|\|\|\|\|\|\|\|\|\|\|\|\|\|\|\|\|\|\|\|\|\|\|\|\|\|\|\|\|\|\|\|\|\|\|\|\|\|\|\|\|\|\|\|\|\|\|\|\|\|\|\|\|\|\|\|\|\|\|\|\|\|\|\|\|\|\|\|\|\|\|\|\|\|\|\|\|\|\|\|\|\|\|\|\|\|\|\|\|\|\|\|\|\|\|\|\|\|\|\|\|\|\|\|\|\|\|\|\|\|\|\|\|\|\|\|\|\|\|\|\|\|\|\|\|\|\|\|\|\|\|\|\|\|\|\|\|\|\|\|\|\|\|\|\|\|\|\|\|\|\|\|\|\|\|\|\|\|\|\|\|\|\|\|\|\|\|\|\|\|\|\|\|\|\|\|\|\|\|\|\|\|\|\|\|\|\|\|\|\|\|\|\|\|\|\|\|\|\|\|\|\|\|\|\|\|\|\|\|\|\|\|\|\|\|\|\|\|\|\|\|\|\| 	ATGAACTCCACACACGACTCAACCGCCGGCGGTGTCGGACGAGTAGGACACGAACGTATCGGGGAAGAATACCTCACTCGGCTGGGCTACTCGAAGAAAGTCGGGTTTCTGGTAGGATCACATGCTGCAGCAAAGAGATTTCTCTGCGGGACCGATCCTGCCTATCACGATACGCTTTCGGGAGCGAGCAAGAAGTCGCTGGTCTTTCAAGGAGAGCCCATGAGAGGCGACGAGCTTAATGAGTGGGCTGCGAATCCTTGGTGCGACGAGATGTGCCA ```      ``` 265	GACGAGATGTGCCA-ACTCCGAAAATGGGACGACGCAGCGAAGGACGTAGGCCTCGAGACCGATCCCGCAAACGCGTACGAGGCGATGATCGTGCGTCTGTTGAAGAGC	372 	\|\|\| \|\|\|\| \|\| \|\| \|\|\|\|\|\|\|\|\|\|\|\|\|\|\|\|\|\|\|\|\|\|\|\|\|\|\|\|\|\|\|\|\|\|\|\|\|\|\|\|\|\|\|\|\|\|\|\|\|\|\|\|\|\|\|\|\|\|\|\|\|\|\|\|\|\|\|\|\|\|\|\|\|\|\|\|\|\|\|\|\|\|\|\|\|\|\|\|\|\|\|\|\|\| 	GAC-AGAT-TGGCAGACTCCGAAAATGGGACGACGCAGCGAAGGACGTAGGCCTCGAGACCGATCCCGCAAACGCGTACGAGGCGATGATCGTGCGTCTGTTGAAGAGC ``` |
| 7 | ggggacaagtttgtacaaaaaagcaggcttgatgCCGGGACCTTCGCCGAACGA, ggggaccactttgtacaagaaagctgggtcTGTGGCTTTCCGCAAAGAGG | Mycgr3\|96984  JGI  Ensembl |  |  | 100 % |  | ``` 1	ATGCCGGGACCTTCGCCGAACGACGCTATGCAAGCATTGGGCCTGCTGTTCGTGCATGCTGAAAACAACGATGAGTTGCACAGCATAGCAAGTGCCTGGCCGAGTCCGCAACCGCTTACCCAACAGATGTTCCTGGTGAATCACGGGAATCTGCAGACGTCTTTCGGCCGAACCGTCGTCCTCGTTACGGTGTGCCTAATTTTCACGCCCGACCATTGCCGCCATGCAGGTCTCGCACTATTGACGCAACACGACATCTTGCTCGCTGTCAATCTTGTGCGCACACGATTGTCGCACGGTCGCACACAGTCTCCATATTCCGCTCTGCCGCCTGCCCTTGGCGCATATTATTGCAAGCACGACCAATTCTCAGCGTCAGGCGTCGGCAAGTCGCTTGGAGCTCAAGGGCCGGCGGATGATCTCATGCTTCGCGCCTCTTTGCGGAAAGCCACA	453 	\|\|\|\|\|\|\|\|\|\|\|\|\|\|\|\|\|\|\|\|\|\|\|\|\|\|\|\|\|\|\|\|\|\|\|\|\|\|\|\|\|\|\|\|\|\|\|\|\|\|\|\|\|\|\|\|\|\|\|\|\|\|\|\|\|\|\|\|\|\|\|\|\|\|\|\|\|\|\|\|\|\|\|\|\|\|\|\|\|\|\|\|\|\|\|\|\|\|\|\|\|\|\|\|\|\|\|\|\|\|\|\|\|\|\|\|\|\|\|\|\|\|\|\|\|\|\|\|\|\|\|\|\|\|\|\|\|\|\|\|\|\|\|\|\|\|\|\|\|\|\|\|\|\|\|\|\|\|\|\|\|\|\|\|\|\|\|\|\|\|\|\|\|\|\|\|\|\|\|\|\|\|\|\|\|\|\|\|\|\|\|\|\|\|\|\|\|\|\|\|\|\|\|\|\|\|\|\|\|\|\|\|\|\|\|\|\|\|\|\|\|\|\|\|\|\|\|\|\|\|\|\|\|\|\|\|\|\|\|\|\|\|\|\|\|\|\|\|\|\|\|\|\|\|\|\|\|\|\|\|\|\|\|\|\|\|\|\|\|\|\|\|\|\|\|\|\|\|\|\|\|\|\|\|\|\|\|\|\|\|\|\|\|\|\|\|\|\|\|\|\|\|\|\|\|\|\|\|\|\|\|\|\|\|\|\|\|\|\|\|\|\|\|\|\|\|\|\|\|\|\|\|\|\|\|\|\|\|\|\|\|\|\|\|\|\|\|\|\|\|\|\|\|\|\|\|\|\|\|\|\|\|\|\|\|\|\|\|\|\|\|\|\|\|\|\|\|\|\|\|\|\|\|\|\|\|\|\|\|\|\|\|\|\|\|\|\|\|\|\|\|\|\|\|\|\|\|\|\|\|\|\|\|\|\|\|\|\|\|\|\|\|\|\|\|\|\|\|\|\|\|\|\|\|\|\|\|\|\|\|\|\|\|\|\|\|\|\|\|\|\|\|\| 	ATGCCGGGACCTTCGCCGAACGACGCTATGCAAGCATTGGGCCTGCTGTTCGTGCATGCTGAAAACAACGATGAGTTGCACAGCATAGCAAGTGCCTGGCCGAGTCCGCAACCGCTTACCCAACAGATGTTCCTGGTGAATCACGGGAATCTGCAGACGTCTTTCGGCCGAACCGTCGTCCTCGTTACGGTGTGCCTAATTTTCACGCCCGACCATTGCCGCCATGCAGGTCTCGCACTATTGACGCAACACGACATCTTGCTCGCTGTCAATCTTGTGCGCACACGATTGTCGCACGGTCGCACACAGTCTCCATATTCCGCTCTGCCGCCTGCCCTTGGCGCATATTATTGCAAGCACGACCAATTCTCAGCGTCAGGCGTCGGCAAGTCGCTTGGAGCTCAAGGGCCGGCGGATGATCTCATGCTTCGCGCCTCTTTGCGGAAAGCCACA ``` |
| 7 | ggggacaagtttgtacaaaaaagcaggcttgatgCATTGCAAACTCCCATCCTT, ggggaccactttgtacaagaaagctgggtcAGCATCGGGACAGTTGAATG | Mycgr3\|97498  JGI  Ensembl |  |  | 0 % |  |  |
| 7 | ggggacaagtttgtacaaaaaagcaggcttgatgGCGAAAGCCATTTCGAAGGT, ggggaccactttgtacaagaaagctgggtcGTCATCCATAATCGAACGGA | Mycgr3\|97541  JGI  Ensembl |  |  | 0 % |  |  |
| 7 | ggggacaagtttgtacaaaaaagcaggcttgatgGCCACCACCAGATTCATCAG, ggggaccactttgtacaagaaagctgggtcCAGCCCTAGTAACGACTCGA | Mycgr3\|97556  JGI  Ensembl |  |  | 0 % |  |  |
| 7 | ggggacaagtttgtacaaaaaagcaggcttgatgCTGCCGGCGGAAGAGCTGCT, ggggaccactttgtacaagaaagctgggtcGCCGCCTCCGAATGGCTTCC | Mycgr3\|97625  JGI  Ensembl |  |  | 0 % |  |  |
| 7 | ggggacaagtttgtacaaaaaagcaggcttgatgTCGATACACGATGTCTCCCA, ggggaccactttgtacaagaaagctgggtcGCGGATAACCGTGCTCGTAG | Mycgr3\|97637  JGI  Ensembl |  |  | 0 % |  | ``` 1	ATGTCGATACACGATGTCTCCCACGAATCGGCCTGCATATTCATTGCACGAAGTCAGTATCCGACTATGGACAACGACTGGATGACGACGGATGACGATGGTGTGCCGACAAAATCGGCGGCACGTCTTGCTTCAAGGCAGGCTGGTCATGCCAGCACAGGCGGATCCTTCGAAACGCTCGGCTCCATTTCCGTCATCCAGGAAGCCAACAACGTCGCCCAAGACACAACAACCTCACAGTCTGATCTCGTCAGGTGGAGACGCACGCATAAGTACGAGCACAATCAGAAACGAAACCGCACCGCCGAAACCGCCGCCAGTAGCCTCCTCACGAGCACCACGCCCAGCCATGACGAAAAGCAAAATCATGAGCGAGCACTGCAGAAAGAGACTCGATCTCGTTGCGTAGTCCAACAAAGCGCACCGACCGCCGACGCGCAGCATGCTACAACTACTGTCATTAAGCCTTCTACGAGCACGGTTATCCGC	489 	\|\|\|\|\|\|\|\|\|\|\|\|\|\|\|\|\|\|\|\|\|\|\|\|\|\|\|\|\|\|\|\|\|\|\|\|\|\|\|\|\|\|\|\|\|\|\|\|\|\|\|\|\|\|\|\|\|\|\|\|\|\|\|\|\|\|\|\|\|\|\|\|\|\|\|\|\|\|\|\|\|\|\|\|\|\|\|\|\|\|\|\|\|\|\|\|\|\|\|\|\|\|\|\|\|\|\|\|\|\|\|\|\|\|\|\|\|\|\|\|\|\|\|\|\|\|\|\|\|\|\|\|\|\|\|\|\|\|\|\|\|\|\|\|\|\|\|\|\|\|\|\|\|\|\|\|\|\|\|\|\|\|\|\|\|\|\|\|\|\|\|\|\|\|\|\|\|\|\|\|\|\|\|\|\|\|\|\|\|\|\|\|\|\|\|\|\|\|\|\|\|\|\|\|\|\|\|\|\|\|\|\|\|\|\|\|\|\|\|\|\|\|\|\|\|\|\|\|\|\|\|\|\|\|\|\|\|\|\|\|\|\|\|\|\|\|\|\|\|\|\|\|\|\|\|\|\|\|\|\|\|\|\|\|\|\|\|\|\|\|\|\|\|\|\|\|\|\|\|\|\|\|\|\|\|\|\|\|\|\|\|\|\|\|\|\|\|\|\|\|\|\|\|\|\|\|\|\|\|\|\|\|\|\|\|\|\|\|\|\|\|\|\|\|\|\|\|\|\|\|\|\|\|\|\|\|\|\|\|\|\|\|\|\|\|\|\|\|\|\|\|\|\|\|\|\|\|\|\|\|\|\|\|\|\|\|\|\|\|\|\|\|\|\|\|\|\|\|\|\|\|\|\|\|\|\|\|\|\|\|\|\|\|\|\|\|\|\|\|\|\|\|\|\|\|\|\|\|\|\|\|\|\|\|\|\|\|\|\|\|\|\|\|\|\|\|\|\|\|\|\|\|\|\|\|\|\|\|\|\|\|\|\|\|\|\|\|\|\|\|\|\|\|\|\|\|\|\|\|\|\|\|\|\|\|\|\|\|\|\|\|\|\|\|\|\|\|\|\|\|\|\|\|\|\|\|\|\|\| 	ATGTCGATACACGATGTCTCCCACGAATCGGCCTGCATATTCATTGCACGAAGTCAGTATCCGACTATGGACAACGACTGGATGACGACGGATGACGATGGTGTGCCGACAAAATCGGCGGCACGTCTTGCTTCAAGGCAGGCTGGTCATGCCAGCACAGGCGGATCCTTCGAAACGCTCGGCTCCATTTCCGTCATCCAGGAAGCCAACAACGTCGCCCAAGACACAACAACCTCACAGTCTGATCTCGTCAGGTGGAGACGCACGCATAAGTACGAGCACAATCAGAAACGAAACCGCACCGCCGAAACCGCCGCCAGTAGCCTCCTCACGAGCACCACGCCCAGCCATGACGAAAAGCAAAATCATGAGCGAGCACTGCAGAAAGAGACTCGATCTCGTTGCGTAGTCCAACAAAGCGCACCGACCGCCGACGCGCAGCATGCTACAACTACTGTCATTAAGCCTTCTACGAGCACGGTTATCCGC ``` |
| 7 | ggggacaagtttgtacaaaaaagcaggcttgatgTTTCGATTCAGTGCGCAGGC, ggggaccactttgtacaagaaagctgggtcGAATCGTGGAGGTGTCAGAA | Mycgr3\|97659  JGI  Ensembl |  |  | 100 % |  | ``` 391	ACGATCCTCACACCTCTACGATCCTCACACCTCCCCGATTCTGACACCTCCCCGATTCTGACACCTCCACGAT	463 	\|\|\|\| \|\|\|\|\|\|\|\|\|\|\|\|\|\|\|\|\|\|\|\|\|\|\|\|\|\|\|\|\| \|\|\|\| \|\| \|\|\|\|\|\|\|\| \|\|\|\|\|\|  \|\|\| \|\|\| \|\|\|\|\| 	ACGA-CCTCACACCTCTACGATCCTCACACCTCCACGATCCTCACACCTCCACGATTCGCACATCTCGACGAT ```      ``` 1	ATGTTTCGATTCAGTGCGCAGGCGCGCGGAGGGGAGGAGAAGATCGTTCTCGTCGAGGCCGTGTCGATGCAGGGAAATTTCGTCAGATTGATGTGTTCGCGTATGTCGCAGGAAGAGGTCCGTGTTGCTGGCGTCGTGGATGCGTATGAGATGGCCGTGGTCGTAGTTCTACGTCAGTCACTTCGTCTTGAAATTCCCGAGCTCCGTCCGGTCTGCAGAGGAGATGTCCCGAGCCGTCTTCTCCAACGATTCGAGAAGCCGAACTTTACCCGGCCCCGCACGCCTTTCGATctcacccttcacgacctcacaccttcacgatcacgacctcacacctctacgatcctcacacctccacgatcctcacacctccacgatTCGCACATCTCGACGATCCTCACACCTCTACGATCCTCACACCTCCCCGATTCTGACACCTCCCCGATTCTGACACCTCCACGATTC	465 	\|\|\|\|\|\|\|\|\|\|\|\|\| \|\|\|\|\|\|\|\|\|\|\|\|\|\|\|\|\|\|\|\|\|\|\|\|\|\|\|\|\|\|\|\|\|\|\|\|\|\|\|\|\|\|\|\|\|\|\|\|\|\|\|\|\|\|\|\|\|\|\|\|\|\|\|\|\|\|\|\|\|\|\|\|\|\|\|\|\|\|\|\|\|\|\|\|\|\|\|\|\|\|\|\|\|\|\|\|\|\|\|\|\|\|\|\|\|\|\|\|\|\|\|\|\|\|\|\|\|\|\|\|\|\|\|\|\|\|\|\|\|\|\|\|\|\|\|\|\|\|\|\|\|\|\|\|\|\|\|\|\|\|\|\|\|\|\|\|\|\|\|\|\|\|\|\|\|\|\|\|\|\|\|\|\|\|\|\|\|\|\|\|\|\|\|\|\|\|\|\|\|\|\|\|\|\|\|\|\|\|\|\|\|\|\|\|\|\|\|\|\|\|\|\|\|\|\|\|\|\|\|\|\|\|\|\|\|\|\|\|\|\|\|\|\|\|\|\|\|\|\|\|\|\|\|\|\|\|\|\|\|\|\|\|\|\|\|\|\|\|\|\|\|\|\|\|\|\|\|\|\|\|\|\|\|\|\|\|\|\|\|\|\|\|\|\|\|\|\|\|\|\|\|\|\|\|\|\|\|\|\|\|\|\|\|\|\|\|\|\|\|\|\|\|\|\|\|\|\|\|\|\|\|\|\|\|\|\|\|\|\|\|\|\|\|\|\|\|\|\|\|\|\|\|\|\|\|\|\|\|\|\|\|\|\|\|\|\|\|\|\|\|\|\|\|\|\|\|\|\|\|\|\|\|\|\|\|\|\|\|\|\|\|\|\|\|\|\|\|\|\|\|\|\|\|\|\|\|\|\|\|\|\|\|\|\|\|\|\|\|\|\|\|\|\|\|\|\|\|\|\|\|\|\|\|\|\|\|\|\|\|\|\|\|\|\|\|\|\|\|\|\|\|\|\|\|\|\|\|\|\|\|\| 	ATGTTTCGATTCANTGCGCAGGCGCGCGGAGGGGAGGAGAAGATCGTTCTCGTCGAGGCCGTGTCGATGCAGGGAAATTTCGTCAGATTGATGTGTTCGCGTATGTCGCAGGAAGAGGTCCGTGTTGCTGGCGTCGTGGATGCGTATGAGATGGCCGTGGTCGTAGTTCTACGTCAGTCACTTCGTCTTGAAATTCCCGAGCTCCGTCCGGTCTGCAGAGGAGATGTCCCGAGCCGTCTTCTCCAACGATTCGAGAAGCCGAACTTTACCCGGCCCCGCACGCCTTTCGATCTCACCCTTCACGACCTCACACCTTCACGATCACGACCTCACACCTCTACGATCCTCACACCTCCACGATCCTCACACCTCCACGATTCGCACATCTCGACGATCCTCACACCTCTACGATCCTCACACCTCCCCGATTCTGACACCTCCCCGATTCTGACACCTCCACGATTC ``` |
| 7 | ggggacaagtttgtacaaaaaagcaggcttgatgGCTAACATCGCTGTGAAGCG, ggggaccactttgtacaagaaagctgggtcATCTTGCGTGACTGGATTCA | Mycgr3\|97774  JGI  Ensembl |  |  | 100 % |  | ``` 1	ATGGCTAACATCGCTGTGAAGCGAGGGCATCTTCCCCTTCTCAACCGCATACAGCAGCTTAAGGCGAGTCCTTTACCTTTCCCTCCGCTACCGCCGGGAGGCACGAACCAACAACAGCTCGAATCTAGCAGGCCGAACCAGGGACAGGTTGTCCTCGAATACGACGACTCACCAAGCCGGACGTGGAGAAAACAATCACCCTGCCTTAGTCGAATAAGTCCGGTCTCTCTCTCTAGCGAATCGAGCTCCTCTACCTCCGACCTTCACACCTACCATTCCACCGCAAGCTCCCCGAGTCTTCGCAGTCGAGACGCAGTTGACATGGAACGACCGCTCACACCAATCGACAGCGACGGCGACAACGGGAAAGTCGTGATCCGAGTGCGACGCTTCATCAAGCGAGACTTTGAGAAACAATCACTATGCATCATTCGCATGAATCCAGTCACGCAAGAT	456 	\|\|\|\|\|\|\|\|\|\|\|\|\|\|\|\|\|\|\|\|\|\|\|\|\|\|\|\|\|\|\|\|\|\|\|\|\|\|\|\|\|\|\|\|\|\|\|\|\|\|\|\|\|\|\|\|\|\|\|\|\|\|\|\|\|\|\|\|\|\|\|\|\|\|\|\|\|\|\|\|\|\|\|\|\|\|\|\|\|\|\|\|\|\|\|\|\|\|\|\|\|\|\|\|\|\|\|\|\|\|\|\|\|\|\|\|\|\|\|\|\|\|\|\|\|\|\|\|\|\|\|\|\|\|\|\|\|\|\|\|\|\|\|\|\|\|\|\|\|\|\|\|\|\|\|\|\|\|\|\|\|\|\|\|\|\|\|\|\|\|\|\|\|\|\|\|\|\|\|\|\|\|\|\|\|\|\|\|\|\|\|\|\|\|\|\|\|\|\|\|\|\|\|\|\|\|\|\|\|\|\|\|\|\|\|\|\|\|\|\|\|\|\|\|\|\|\|\|\|\|\|\|\|\|\|\|\|\|\|\|\|\|\|\|\|\|\|\|\|\|\|\|\|\|\|\|\|\|\|\|\|\|\|\|\|\|\|\|\|\|\|\|\|\|\|\|\|\|\|\|\|\|\|\|\|\|\|\|\|\|\|\|\|\|\|\|\|\|\|\|\|\|\|\|\|\|\|\|\|\|\|\|\|\|\|\|\|\|\|\|\|\|\|\|\|\|\|\|\|\|\|\|\|\|\|\|\|\|\|\|\|\|\|\|\|\|\|\|\|\|\|\|\|\|\|\|\|\|\|\|\|\|\|\|\|\|\|\|\|\|\|\|\|\|\|\|\|\|\|\|\|\|\|\|\|\|\|\|\|\|\|\|\|\|\|\|\|\|\|\|\|\|\|\|\|\|\|\|\|\|\|\|\|\|\|\|\|\|\|\|\|\|\|\|\|\|\|\|\|\|\|\|\|\|\|\|\|\|\|\|\|\|\|\|\|\|\|\|\|\|\|\|\|\|\|\| 	ATGGCTAACATCGCTGTGAAGCGAGGGCATCTTCCCCTTCTCAACCGCATACAGCAGCTTAAGGCGAGTCCTTTACCTTTCCCTCCGCTACCGCCGGGAGGCACGAACCAACAACAGCTCGAATCTAGCAGGCCGAACCAGGGACAGGTTGTCCTCGAATACGACGACTCACCAAGCCGGACGTGGAGAAAACAATCACCCTGCCTTAGTCGAATAAGTCCGGTCTCTCTCTCTAGCGAATCGAGCTCCTCTACCTCCGACCTTCACACCTACCATTCCACCGCAAGCTCCCCGAGTCTTCGCAGTCGAGACGCAGTTGACATGGAACGACCGCTCACACCAATCGACAGCGACGGCGACAACGGGAAAGTCGTGATCCGAGTGCGACGCTTCATCAAGCGAGACTTTGAGAAACAATCACTATGCATCATTCGCATGAATCCAGTCACGCAAGAT ``` |
| 7 | ggggacaagtttgtacaaaaaagcaggcttgatgCAACTGTCGATTGGACTGGA, ggggaccactttgtacaagaaagctgggtcGTAGCGGGCGTGCGGGCCTC | Mycgr3\|97802  JGI  Ensembl |  |  | 0 % |  |  |
| 7 | ggggacaagtttgtacaaaaaagcaggcttgatgGCCGAACCTCCATCGCCGAC, ggggaccactttgtacaagaaagctgggtcGGCGCCGTCCTTCCTTCGAA | Mycgr3\|97807  JGI  Ensembl |  |  | 0 % |  |  |
| 7 | ggggacaagtttgtacaaaaaagcaggcttgatgTTGGAGCAATTGGACCTGGA, ggggaccactttgtacaagaaagctgggtcATTCGATTTCCGAACTTCCG | Mycgr3\|97959  JGI  Ensembl |  |  | 100 % |  | ``` 1	ATGTTGGAGCAATTGGACCTGGATACACCTGCCACCCCGTTACACTCGGAAGCTATTGTCAGCCGAAAGCGCAAACGACAGGAACAGGATCAAGCCCAGGACGTGGTTGCTGAAGGCGATTTCGACAGTGATTCTGGCAGTGCAATGGACGCTGAGATTGGTCGACGTGGTCGAAGCCCTCTCCTGGATCGATCCTCGACGCTATGGGAACTTGTGGGATCGCATCACGATGGTATTGAATCTGACAGGCCTAATCCTCCTTCCGACTCTGAAAGAGCAGACCAATACTCACCTATCCAGATCGACGTATTTGACGATAATCCGCTGGACGGCGGATCTGAACATGCGAGCAACAACACCTCATACGACACGCACCGCCTTCCGGAACGCAAAGCCAAAGACCCACCGGCACTCGGCACACGAGCCCTAAAGCGGAGACGCGACGAAGCGCCAAAGCGGCCAAGTCCGGAAGTTCGGAAATCGAAT	486 	\|\|\|\|\|\|\|\|\|\|\|\|\|\|\|\|\|\|\|\|\|\|\|\|\|\|\|\|\|\|\|\|\|\|\|\|\|\|\|\|\|\|\|\|\|\|\|\|\|\|\|\|\|\|\|\|\|\|\|\|\|\|\|\|\|\|\|\|\|\|\|\|\|\|\|\|\|\|\|\|\|\|\|\|\|\|\|\|\|\|\|\|\|\|\|\|\|\|\|\|\|\|\|\|\|\|\|\|\|\|\|\|\|\|\|\|\|\|\|\|\|\|\|\|\|\|\|\|\|\|\|\|\|\|\|\|\|\|\|\|\|\|\|\|\|\|\|\|\|\|\|\|\|\|\|\|\|\|\|\|\|\|\|\|\|\|\|\|\|\|\|\|\|\|\|\|\|\|\|\|\|\|\|\|\|\|\|\|\|\|\|\|\|\|\|\|\|\|\|\|\|\|\|\|\|\|\|\|\|\|\|\|\|\|\|\|\|\|\|\|\|\|\|\|\|\|\|\|\|\|\|\|\|\|\|\|\|\|\|\|\|\|\|\|\|\|\|\|\|\|\|\|\|\|\|\|\|\|\|\|\|\|\|\|\|\|\|\|\|\|\|\|\|\|\|\|\|\|\|\|\|\|\|\|\|\|\|\|\|\|\|\|\|\|\|\|\|\|\|\|\|\|\|\|\|\|\|\|\|\|\|\|\|\|\|\|\|\|\|\|\|\|\|\|\|\|\|\|\|\|\|\|\|\|\|\|\|\|\|\|\|\|\|\|\|\|\|\|\|\|\|\|\|\|\|\|\|\|\|\|\|\|\|\|\|\|\|\|\|\|\|\|\|\|\|\|\|\|\|\|\|\|\|\|\|\|\|\|\|\|\|\|\|\|\|\|\|\|\|\|\|\|\|\|\|\|\|\|\|\|\|\|\|\|\|\|\|\|\|\|\|\|\|\|\|\|\|\|\|\|\|\|\|\|\|\|\|\|\|\|\|\|\|\|\|\|\|\|\|\|\|\|\|\|\|\|\|\|\|\|\|\|\|\|\|\|\|\|\|\|\|\|\|\|\|\|\|\|\|\|\|\|\|\|\|\| 	ATGTTGGAGCAATTGGACCTGGATACACCTGCCACCCCGTTACACTCGGAAGCTATTGTCAGCCGAAAGCGCAAACGACAGGAACAGGATCAAGCCCAGGACGTGGTTGCTGAAGGCGATTTCGACAGTGATTCTGGCAGTGCAATGGACGCTGAGATTGGTCGACGTGGTCGAAGCCCTCTCCTGGATCGATCCTCGACGCTATGGGAACTTGTGGGATCGCATCACGATGGTATTGAATCTGACAGGCCTAATCCTCCTTCCGACTCTGAAAGAGCAGACCAATACTCACCTATCCAGATCGACGTATTTGACGATAATCCGCTGGACGGCGGATCTGAACATGCGAGCAACAACACCTCATACGACACGCACCGCCTTCCGGAACGCAAAGCCAAAGACCCACCGGCACTCGGCACACGAGCCCTAAAGCGGAGACGCGACGAAGCGCCAAAGCGGCCAAGTCCGGAAGTTCGGAAATCGAAT ``` |
| 7 | ggggacaagtttgtacaaaaaagcaggcttgatgGCCGAGAAACGGGCTGAAGC, ggggaccactttgtacaagaaagctgggtcTTCGGGTGAGCGAGGGTTGG | Mycgr3\|97994  JGI  Ensembl |  |  | 100 % |  | ``` 1	ATGGCCGAGAAACGGGCTGAAGCCGCCGCTATGATACAGAGGGCACATAACAAGGCTGCTGCTGAGGTCACAGGCCAGAAGCTGGGTGACAGCCAAAACAGCGATCATCGAGGTTATATCGGCGGTTCGATCTCCCTATCCGACATCATCCACTTCCGATGCCGCGACTACCACTGCAAGTGCCGTACCGGTGTAATGCATCTCGGTCGATTCTTGGCTCCTGTACACGATCAGCGTACGGTTCGTAAGCACGCCGGCGAGCGTTGTCTTGTGTTTGTTAAGATTCAGCGACTGGTCGATAAAGCGGATGTCCCAGGAGGTCTGCAAGCCGGGAACGAAGTCCTTGTCCGCGTGGAGGATGAAAATGTCGGACCCAGTCGATATATACTCGCCTATGGCAGTGCGAATGGTCCCAACCCTCGCTCACCCGAA	432 	\|\|\|\|\|\|\|\|\|\|\|\|\|\|\|\|\|\|\|\|\|\|\|\|\|\|\|\|\|\|\|\|\|\|\|\|\|\|\|\|\|\|\|\|\|\|\|\|\|\|\|\|\|\|\|\|\|\|\|\|\|\|\|\|\|\|\|\|\|\|\|\|\|\|\|\|\|\|\|\|\|\|\|\|\|\|\|\|\|\|\|\|\|\|\|\|\|\|\|\|\|\|\|\|\|\|\|\|\|\|\|\|\|\|\|\|\|\|\|\|\|\|\|\|\|\|\|\|\|\|\|\|\|\|\|\|\|\|\|\|\|\|\|\|\|\|\|\|\|\|\|\|\|\|\|\|\|\|\|\|\|\|\|\|\|\|\|\|\|\|\|\|\|\|\|\|\|\|\|\|\|\|\|\|\|\|\|\|\|\|\|\|\|\|\|\|\|\|\|\|\|\|\|\|\|\|\|\|\|\|\|\|\|\|\|\|\|\|\|\|\|\|\|\|\|\|\|\|\|\|\|\|\|\|\|\|\|\|\|\|\|\|\|\|\|\|\|\|\|\|\|\|\|\|\|\|\|\|\|\|\|\|\|\|\|\|\|\|\|\|\|\|\|\|\|\|\|\|\|\|\|\|\|\|\|\|\|\|\|\|\|\|\|\|\|\|\|\|\|\|\|\|\|\|\|\|\|\|\|\|\|\|\|\|\|\|\|\|\|\|\|\|\|\|\|\|\|\|\|\|\|\|\|\|\|\|\|\|\|\|\|\|\|\|\|\|\|\|\|\|\|\|\|\|\|\|\|\|\|\|\|\|\|\|\|\|\|\|\|\|\|\|\|\|\|\|\|\|\|\|\|\|\|\|\|\|\|\|\|\|\|\|\|\|\|\|\|\|\|\|\|\|\|\|\|\|\|\|\|\|\|\|\|\|\|\|\|\|\|\|\|\|\|\|\|\|\|\|\|\|\|\| 	ATGGCCGAGAAACGGGCTGAAGCCGCCGCTATGATACAGAGGGCACATAACAAGGCTGCTGCTGAGGTCACAGGCCAGAAGCTGGGTGACAGCCAAAACAGCGATCATCGAGGTTATATCGGCGGTTCGATCTCCCTATCCGACATCATCCACTTCCGATGCCGCGACTACCACTGCAAGTGCCGTACCGGTGTAATGCATCTCGGTCGATTCTTGGCTCCTGTACACGATCAGCGTACGGTTCGTAAGCACGCCGGCGAGCGTTGTCTTGTGTTTGTTAAGATTCAGCGACTGGTCGATAAAGCGGATGTCCCAGGAGGTCTGCAAGCCGGGAACGAAGTCCTTGTCCGCGTGGAGGATGAAAATGTCGGACCCAGTCGATATATACTCGCCTATGGCAGTGCGAATGGTCCCAACCCTCGCTCACCCGAA ``` |
| 7 | ggggacaagtttgtacaaaaaagcaggcttgatgAACGGCTACACGACCGGACA, ggggaccactttgtacaagaaagctgggtcGACCTTCCGGGGCAATGTTC | Mycgr3\|98020  JGI  Ensembl |  |  | 0 % |  |  |
| 7 | ggggacaagtttgtacaaaaaagcaggcttgatgTATCTTCGCATACGGACAGA, ggggaccactttgtacaagaaagctgggtcGGGTGAGGGTGAAAACGGCT | Mycgr3\|98110  JGI  Ensembl | KOG0245 Kinesin-like protein; | ATP binding; microtubule associated complex;  microtubule motor activity;  microtubule-based movement; | 100 % |  | ``` 1	ATGTATCTTCGCATACGGACAGACAGGAAGTGGAAAGTCGTACATTATTGTAGGTACACCGGACGAGCCCGGCCTCATTCCACGCACAGTCAAGGCCTCTTCCAACATGTCAAGGCCTCTTCCAACGAGTCGAATCGGATAACTTACAGCATTTACGTTCCCTACATCGAGAACTACAACGAGCGCGACAAAGACCTGCTCACCTATCGCACCAATCCATTGACCTACCTCAACATCCGCAAATCCGAATTTGACGGCGTCTATGTACAGAACCCCACCGATGAGCCCGTCAAGTCTTACGACGACATTCTCCACCTTGAGAAGATGGGCGACATCAACCGCACAACCGCTTCCACCAAAATGAACGACATCTCTTCCCGCTCCCACGCCCCCTCCGCTCCTCCCGCTCCCAAGCCGTTTTCACCCTCACCC	432 	\|\|\|\|\|\|\|\|\|\|\|\|\|\|\|\|\|\|\|\|\|\|\|\|\|\|\|\|\|\|\|\|\|\|\|\|\|\|\|\|\|\|\|\|\|\|\|\|\|\|\|\|\|\|\|\|\|\|\|\|\|\|\|\|\|\|\|\|\|\|\|\|\|\|\|\|\|\|\|\|\|\|\|\|\|\|\|\|\|\|\|\|\|\|\|\|\|\|\|\|\|\|\|\|\|\|\|\|\|\|\|\|\|\|\|\|\|\|\|\|\|\|\|\|\|\|\|\|\|\|\|\|\|\|\|\|\|\|\|\|\|\|\|\|\|\|\|\|\|\|\|\|\|\|\|\|\|\|\|\|\|\|\|\|\|\|\|\|\|\|\|\|\|\|\|\|\|\|\|\|\|\|\|\|\|\|\|\|\|\|\|\|\|\|\|\|\|\|\|\|\|\|\|\|\|\|\|\|\|\|\|\|\|\|\|\|\|\|\|\|\|\|\|\|\|\|\|\|\|\|\|\|\|\|\|\|\|\|\|\|\|\|\|\|\|\|\|\|\|\|\|\|\|\|\|\|\|\|\|\|\|\|\|\|\|\|\|\|\|\|\|\|\|\|\|\|\|\|\|\|\|\|\|\|\|\|\|\|\|\|\|\|\|\|\|\|\|\|\|\|\|\|\|\|\|\|\|\|\|\|\|\|\|\|\|\|\|\|\|\|\|\|\|\|\|\|\|\|\|\|\|\|\|\|\|\|\|\|\|\|\|\|\|\|\|\|\|\|\|\|\|\|\|\|\|\|\|\|\|\|\|\|\|\|\|\|\|\|\|\|\|\|\|\|\|\|\|\|\|\|\|\|\|\|\|\|\|\|\|\|\|\|\|\|\|\|\|\|\|\|\|\|\|\|\|\|\|\|\|\|\|\|\|\|\|\|\|\|\|\|\|\|\|\|\|\|\|\|\|\|\|\| 	ATGTATCTTCGCATACGGACAGACAGGAAGTGGAAAGTCGTACATTATTGTAGGTACACCGGACGAGCCCGGCCTCATTCCACGCACAGTCAAGGCCTCTTCCAACATGTCAAGGCCTCTTCCAACGAGTCGAATCGGATAACTTACAGCATTTACGTTCCCTACATCGAGAACTACAACGAGCGCGACAAAGACCTGCTCACCTATCGCACCAATCCATTGACCTACCTCAACATCCGCAAATCCGAATTTGACGGCGTCTATGTACAGAACCCCACCGATGAGCCCGTCAAGTCTTACGACGACATTCTCCACCTTGAGAAGATGGGCGACATCAACCGCACAACCGCTTCCACCAAAATGAACGACATCTCTTCCCGCTCCCACGCCCCCTCCGCTCCTCCCGCTCCCAAGCCGTTTTCACCCTCACCC ``` |
| 7 | ggggacaagtttgtacaaaaaagcaggcttgatgTCAATTGAAATGATCTTTGC, ggggaccactttgtacaagaaagctgggtcGGGCATCTGTATGATAGACT | Mycgr3\|101080  JGI  Ensembl |  |  | 100 % |  | ``` 12	GATGTCAATTGAAATGATCTTTGCTTCCGCCGTTCGTCTGCCTCATATCCCGCACCCGCCTCGGCCTCTACCAATCGATTGCGGCACAGAAGGCACAATCACCAGACTCAAAATGACACTCACCCCACTCCGTATGAGCTTCGCGCATCCGTCGAAGCTCAGCACCAGGCGCATCCATATGCTCAATGTCTTCAAGTGCTGTAGAAAACGCATCCTGCGCCTCGAAGTGGTCAAGCTGAGCCCTTTCGTAATCTTCCAAATCCCGCAGGAGCACAGACTCAAGTCTATCATACAGATGCCC	312 	\|\|\|\|\|\|\|\|\|\|\|\|\|\|\|\|\|\|\|\|\|\|\|\|\|\|\|\|\|\|\|\|\|\|\|\|\|\|\|\|\|\|\|\|\|\|\|\|\|\|\|\|\|\|\|\|\|\|\|\|\|\|\|\|\|\|\|\|\|\|\|\|\|\|\|\|\|\|\|\|\|\|\|\|\|\|\|\|\|\|\|\|\|\|\|\|\|\|\|\|\|\|\|\|\|\|\|\|\|\|\|\|\|\|\|\|\|\|\|\|\|\|\|\|\|\|\|\|\|\|\|\|\|\|\|\|\|\|\|\|\|\|\|\|\|\|\|\|\|\|\|\|\|\|\|\|\|\|\|\|\|\|\|\|\|\|\|\|\|\|\|\|\|\|\|\|\|\|\|\|\|\|\|\|\|\|\|\|\| \| \|\|\|\|\|\|\|\|\|\|\|\|\|\|\|\|\|\|\|\|\|\|\|\|\|\|\|\|\|\|\|\|\|\|\|\|\|\|\|\|\|\|\|\|\|\|\|\|\|\|\|\|\|\|\|\|\|\|\|\|\|\|\|\|\|\|\|\|\|\|\|\|\|\|\|\|\|\|\|\|\|\|\|\|\|\|\|\|\|\|\|\|\|\|\|\|\|\|\|\|\|\|\|\|\|\|\|\|\| 	GATGTCAATTGAAATGATCTTTGCTTCCGCCGTTCGTCTGCCTCATATCCCGCACCCGCCTCGGCCTCTACCAATCGATTGCGGCACAGAAGGCACAATCACCAGACTCAAAATGACACTCACCCCACTCCGTATGAGCTTCGCGCATCCGTCGAAGCTCAGCACCAGGCGCATCCATATGCTCAATGTNTGCAAGTGCTGTAGAAAACGCATCCTGCGCCTCGAAGTGGTCAAGCTGAGCCCTTTCGTAATCTTCCAAATCCCGCAGGAGCACAGACTCAAGTCTATCATACAGATGCCC ``` |
| 7 | ggggacaagtttgtacaaaaaagcaggcttgatgGCATTGAAGTCCAACCTTTT, ggggaccactttgtacaagaaagctgggtcGAGGACGAGGATACCACCCA | Mycgr3\|102204  JGI  Ensembl |  |  | 100 % |  | ``` 49	TTTGGGATGTTTCGAGTCTTCCAGATCCGATCACTGTCATATTGGAAAGATTGTACATGGTGCTTCAGCAAGAGATAGTCGTCGTGGGCAGCAGCTCAGCCCGATTAGGCTCAATGCGGCGGATGACTTGCTGCTCGCAGGGCTCGCTCCATTGGATTCCCTGGCTCAACGCCTAGGAGCAACTCTCCACGAAGATGTTCCAGTACCTCTGGCGGCTGCAATGGTTCTCCTCCACCAATGTCCGGAGGAGCAATTTTTAGCTCACTTCCGGATGAGCCAAGGTTCGATGCGGTATTACTGCAGGCGCCTTTCGACAGGTCCCTTTTTGGGTGGTATCCTCGTCCTC	394 	\|\|\|\|\|\|\|\|\|\|\|\|\|\|\|\|\|\|\|\|\|\|\|\|\|\|\|\|\|\|\|\|\|\|\|\|\|\|\|\|\|\|\|\|\|\|\|\|\|\|\|\|\|\|\|\|\|\|\|\|\|\|\|\|\|\|\|\|\|\|\|\|\|\|\|\|\|\|\|\|\|\|\|\|\|\|\|\|\|\|\|\|\|\|\|\|\|\|\|\|\|\|\|\|\|\|\|\|\|\|\|\|\|\|\|\|\|\|\|\|\|\|\|\|\|\|\|\|\|\|\|\|\|\|\|\|\|\|\|\|\|\|\|\|\|\|\|\|\|\|\|\|\|\|\|\|\|\|\|\|\|\|\|\|\|\|\|\|\|\|\|\|\|\|\|\|\|\|\|\|\|\|\|\|\|\|\|\|\|\|\|\|\|\|\|\|\|\|\|\|\|\|\|\|\|\|\|\|\|\|\|\|\|\|\|\|\|\|\|\|\|\|\|\|\|\|\|\|\|\|\|\|\|\|\|\|\|\|\|\|\|\|\|\|\|\|\|\|\|\|\|\|\|\|\|\|\|\|\|\|\|\|\|\|\|\|\|\|\|\|\|\|\|\|\|\|\|\|\|\|\|\|\|\|\|\|\|\|\|\|\|\|\|\|\|\|\|\|\|\|\|\|\|\|\|\|\| \|\|\|\|\|\|\|\|\|\|\|\|\|\|\|\|\|\|\|\|\|\|\|\|\|\|\|\|\|\|\|\|\|\|\|\|\|\| 	TTTGGGATGTTTCGAGTCTTCCAGATCCGATCACTGTCATATTGGAAAGATTGTACATGGTGCTTCAGCAAGAGATAGTCGTCGTGGGCAGCAGCTCAGCCCGATTAGGCTCAATGCGGCGGATGACTTGCTGCTCGCAGGGCTCGCTCCATTGGATTCCCTGGCTCAACGCCTAGGAGCAACTCTCCACGAAGATGTTCCAGTACCTCTGGCGGCTGCAATGGTTCTCCTCCACCAATGTCCGGAGGAGCAATTTTTAGCTCACTTCCGGATGAGCCAAGGTTCGATGCGGTATTACTGCAGGCGCNTTTCGACAGGTCCCTTTTTGGGTGGTATCCTCGTCCTC ``` |
| 7 | ggggacaagtttgtacaaaaaagcaggcttgatgCCAGCTCATCAGCTGGACAA, ggggaccactttgtacaagaaagctgggtcAACACGGCTTGGGGATTGCG | Mycgr3\|102437  JGI  Ensembl |  |  | 0 % |  |  |
| 7 | ggggacaagtttgtacaaaaaagcaggcttgatgTCGAAGACCTGCTCGTGCTG, ggggaccactttgtacaagaaagctgggtcCTCGAGACACACCGACCAGT | Mycgr3\|102916  JGI  Ensembl |  |  | 0 % |  |  |
| 7 | ggggacaagtttgtacaaaaaagcaggcttgatgGCCAACCCTGTTGATGATCG, ggggaccactttgtacaagaaagctgggtcACGGACAGGAGTCTGCGGCA | Mycgr3\|103089  JGI  Ensembl |  |  | 100 % |  | ``` 267	ATGGCCAACCCTGTTGATGATCGGGAGCGAGAACCCGGCACAGCCCACGCTGCTCGACTCCAGGAGCTTGGACATGTTCCGAACAGCTCATCAGATGGTTTCCTCGAAACCATTGCCAGCGTGAGCATGTTCATACAGGACACCGCTGTCGAATGTCTGGTCCTGTGTGATTTCTGGTTGCC	448 	\|\|\|\|\|\|\|\|\|\|\| \|\|\|\|\|\|\|\|\|\|\|\|\|\|\|\|\|\|\|\|\|\|\|\|\|\|\|\|\|\|\|\|\|\|\|\|\|\|\|\|\|\|\|\|\|\|\|\|\|\|\|\|\|\|\|\|\|\|\|\|\|\|\|\|\|\|\|\|\|\|\|\|\|\|\|\|\|\|\|\|\| \|\|\|\|\|\|\|\|\|\|\|\|\|\|\|\|\|\|\|\|\| \|\|\|\|\| \|\|\|\| \|\|\|\|\|\|\|\|\|\| \|\|\|\|\|\| \|\|\|\|\|\|\|\| \|\|\|\|\|\|\|\|\|\|\|\|\| \|\|\|\|\|\| \| \|\|\|\|\| 	ATGGCCAACCCNGTTGATGATCGGGAGCGAGAACCCGGCACAGCCCACGCTGCTCGACTCCAGGAGCTTGGACATGTTCCGAACAGCTCATCANATGGTTTCCTCGAAACCATTGNCAGCGNGAGCNTGTTCATACANGACACCNCTGTCGAANGTCTGGTCCTGTGNGATTTCCGNTTGCC ``` |
| 7 | ggggacaagtttgtacaaaaaagcaggcttgatgATCCTCGCTTTGGCTTCGTC, ggggaccactttgtacaagaaagctgggtcTAAGTCTTGTCGTAGGTGCA | Mycgr3\|103254  JGI  Ensembl |  |  | 0 % |  |  |
| 7 | ggggacaagtttgtacaaaaaagcaggcttgatgGACCTGTCTCTTGACACGGA, ggggaccactttgtacaagaaagctgggtcCCAACGATGAGAGCCAATGC | Mycgr3\|103434  JGI  Ensembl |  |  | 100 % |  |  |
| 7 | ggggacaagtttgtacaaaaaagcaggcttgatgCTCTCAGCGATGACAGCGCT, ggggaccactttgtacaagaaagctgggtcAATGGGCCCTTTAGGCAAGG | Mycgr3\|103713  JGI  Ensembl |  |  | 0 % |  | ``` 57	GATGCTCTCAGCGATGACAGCGCTCTCATGGCTGCTGCTGACGAATATTCTGGCCACCCCTGCAGCAGCGACTTATCAGTCACCAGGATGTTGCTCCAAACTGCCCTACGTTGCCCTCAAGCCACTCTTCAAAGTCCCCGGCGTTCAGTCCTACTGCTCTTCAAAATACCCGCTCGCTGTCGTCACCAGCACCATCACCAACCCAGCCGCTACCAGAACAACTATCGTTCAAACCGTCACCGCAACGACAACAGTCGACACTTCGGTTCAGCAAGGCTCCTGTACAATTTCCTTGCCTAAAGGGCCCATTTGAC	370 	\|\|\|\|\|\|\|\|\|\|\|\|\|\|\|\|\|\|\|\|\|\|\|\|\|\|\|\|\|\|\|\|\|\|\|\|\|\|\|\|\|\|\|\|\|\|\|\|\|\|\|\|\|\|\|\|\|\|\|\|\|\|\|\|\|\|\|\|\|\|\|\|\|\|\|\|\|\|\|\|\|\|\|\|\|\|\|\|\|\|\|\|\|\|\|\|\|\|\|\|\|\|\|\|\|\|\|\|\|\|\|\|\|\|\|\|\|\|\|\|\|\|\|\|\|\|\|\|\|\|\|\|\|\|\|\|\|\|\|\|\|\|\|\|\|\|\|\|\|\|\|\|\|\|\|\|\|\|\|\|\|\|\|\|\|\|\|\|\|\|\|\|\|\|\|\|\|\|\|\|\|\|\|\|\|\|\|\|\|\|\|\|\|\|\|\|\|\|\|\|\|\|\|\|\|\|\|\|\|\|\|\|\|\|\|\|\|\|\|\|\|\|\|\|\|\|\|\|\|\|\|\|\|\|\|\|\|\|\|\|\|\|\|\|\|\|\|\|\|\|\|\|\|\|\|\|\|\|\|\|\|\|\|\|\|\|\|\|\|\|\|\|\|\|\|\|\|\|\|\|\|\|\|\|\|\|\|\|\|\|\|\|\|\|\|\|\|\|\|\|\|\|\|\|\|\|\|\|\|\| \|\|\| 	GATGCTCTCAGCGATGACAGCGCTCTCATGGCTGCTGCTGACGAATATTCTGGCCACCCCTGCAGCAGCGACTTATCAGTCACCAGGATGTTGCTCCAAACTGCCCTACGTTGCCCTCAAGCCACTCTTCAAAGTCCCCGGCGTTCAGTCCTACTGCTCTTCAAAATACCCGCTCGCTGTCGTCACCAGCACCATCACCAACCCAGCCGCTACCAGAACAACTATCGTTCAAACCGTCACCGCAACGACAACAGTCGACACTTCGGTTCAGCAAGGCTCCTGTACAATTTCCTTGCCTAAAGGGCCCATT-GAC ``` |
| 7 | ggggacaagtttgtacaaaaaagcaggcttgatgAAACATTGCAGACAAAACGA, ggggaccactttgtacaagaaagctgggtcTGCCATGATGCGAATCATGC | Mycgr3\|104086  JGI  Ensembl |  |  | 0 % |  |  |
| 7 | ggggacaagtttgtacaaaaaagcaggcttgatgCTCTCCGTCGCCGTCGTGCA, ggggaccactttgtacaagaaagctgggtcGGAGAACGTGATGGACAAAA | Mycgr3\|104118  JGI  Ensembl |  |  | 100 % |  | ``` 129	GATGCTCTCCGTCGCCGTCGTGCACCACGTCCTCACTGTCCTCATCATCATCCTCATACCCACCCTGGTACTCAAGCACGTCAGCCAAGGTAGGCTTGAACATGCCATCAATTTCATCCATCAGTGCGTTCGCTCTCAGCTCGTCCGCTTTTTGGACGACTTCGTCCGCTTGCTTCGCCTTTGCTTCACTTTCAAGTATATCCTCTTCGTCACTCCAGGTCTAACTGCCAGTGCGCGTGCGGGCAATGTCACGAGCGTCTTGATATTCCTCATCTGGATTTTCCTCTTCTGCCTCTTCATCTGCTCGATTTTGTCCATCACGTTCTCCTGAC	460 	\|\|\|\|\|\|\|\|\|\|\|\|\|\|\|\|\|\|\|\|\|\|\|\|\|\|\|\|\|\|\|\|\|\|\|\|\|\|\|\|\|\|\|\|\|\|\|\|\|\|\|\|\|\|\|\|\|\|\|\|\|\|\|\|\|\|\|\|\|\|\|\|\|\|\|\|\|\|\|\|\|\|\|\|\|\|\|\|\|\|\|\|\|\|\|\|\|\|\|\|\|\|\|\|\|\|\|\|\|\|\|\|\|\|\|\|\|\|\|\|\|\|\|\|\|\|\|\|\|\|\|\|\|\|\|\|\|\|\|\|\|\|\|\|\|\|\|\|\|\|\|\|\|\|\|\|\|\|\|\|\|\|\|\|\|\|\|\|\|\|\|\|\|\|\|\|\|\|\|\|\|\|\|\|\|\|\|\|\|\|\|\|\|\|\|\|\|\|\|\|\|\|\|\|\|\|\|\|\|\|\|\|\|\|\|\|\|\|\|\|\|\|\|\|\|\|\|\|\|\|\|\|\|\|\|\|\|\|\|\|\|\|\|\|\|\|\|\|\|\|\|\|\|\|\|\|\|\|\|\|\|\|\|\|\|\|\|\|\|\|\|\|\|\|\|\|\|\|\|\|\|\|\|\|\|\|\|\|\|\|\|\|\|\|\|\|\|\|\|\|\|\|\|\|\|\|\|\|\|\|\|\|\|\|\|\|\|\|\|\|\|\|\|\|\|\|\|\| \|\|\| 	GATGCTCTCCGTCGCCGTCGTGCACCACGTCCTCACTGTCCTCATCATCATCCTCATACCCACCCTGGTACTCAAGCACGTCAGCCAAGGTAGGCTTGAACATGCCATCAATTTCATCCATCAGTGCGTTCGCTCTCAGCTCGTCCGCTTTTTGGACGACTTCGTCCGCTTGCTTCGCCTTTGCTTCACTTTCAAGTATATCCTCTTCGTCACTCCAGGTCTAACTGCCAGTGCGCGTGCGGGCAATGTCACGAGCGTCTTGATATTCCTCATCTGGATTTTCCTCTTCTGCCTCTTCATCTGCTCGATTTTGTCCATCACGTTCTCC-GAC ``` |
| 7 | ggggacaagtttgtacaaaaaagcaggcttgatgGTGTTCCATGCGATCAATCA, ggggaccactttgtacaagaaagctgggtcCCGAAGCATTGCATTATCAG | Mycgr3\|104278  JGI  Ensembl |  |  | 0 % |  |  |
| 7 | ggggacaagtttgtacaaaaaagcaggcttgatgCCGTCCGAAGAGGAAAAGCT, ggggaccactttgtacaagaaagctgggtcGCGCTTGGGGAAGTCTTCTT | Mycgr3\|105104  JGI  Ensembl |  |  | 0 % |  |  |
| 7 | ggggacaagtttgtacaaaaaagcaggcttgatgCACTCTCCAATGCAATTCGC, ggggaccactttgtacaagaaagctgggtcGTATCCGTAGCCTGGTGGTC | Mycgr3\|105419  JGI  Ensembl | KOG1216 von Willebrand factor and related coagulation proteins; |  | 100 % |  | ``` 403	CCACCGTCAAGTCCACCACGGCCTACGTCCCCAAGACCATCACGTCCATCGTCACCAAGACCACGACCAAAGATGTCCCCTTCACCACGGTCATTACTGGCGATAAGCCCAAGACCATCATCGTCACCAAGACCATCCCGCAAACCACTCAGACCGTCAGCAAATCGACCACCAAGATCCCCAAGCCCACCACTGAATACAAGACGACGGTCATCACCACCGTCGTTCCCGTCGTCAAAACCGGCTACTCCGAAAGCGTAGTCACCGTCACCAAGCCGGTGACCAAGACCATTACGACTACCGAGACCTCGATCTCGACCGGCAAGGGCGAGACCACCTCGACTGGTACCAAGAAGACGACCGCCACGGAGACCAAGGCGGAGACCTACACCACTTCCACCAAGAGCGTCGGCAGCACGACTGTGGTTACCAAGAAGCCATTCACCACCACCAAGGT	859 	\|\|\|\|\|\|\|\|\|\|\|\|\|\|\|\|\|\|\|\|\|\|\|\|\|\|\|\|\|\|\|\|\|\|\|\|\|\|\|\|\|\|\|\|\|\|\|\|\|\|\|\|\|\|\|\|\|\|\|\|\|\|\|\|\|\|\|\|\|\|\|\|\|\|\|\|\|\|\|\|\|\|\|\|\|\|\|\|\|\|\|\|\|\|\|\|\|\|\|\|\|\|\|\|\|\|\|\|\|\|\|\|\|\|\|\|\|\|\|\|\|\|\|\|\|\|\|\|\|\|\|\|\|\|\|\|\|\|\|\|\|\|\|\|\|\|\|\|\|\|\|\|\|\|\|\|\|\|\|\|\|\|\|\|\|\|\|\|\|\|\|\|\|\|\|\|\|\|\|\|\|\|\|\|\|\|\|\|\|\|\|\|\|\|\|\|\|\|\|\|\|\|\|\|\|\|\|\|\|\|\|\|\|\|\|\|\|\|\|\|\|\|\|\|\|\|\|\|\|\|\|\|\|\|\|\|\|\|\|\|\|\|\|\|\|\|\|\|\|\|\|\|\|\|\|\|\|\|\|\|\|\|\|\|\|\|\|\|\|\|\|\|\|\|\|\|\|\|\|\|\|\|\|\|\|\|\|\|\|\|\|\|\|\|\|\|\|\|\|\|\|\|\|\|\|\|\|\|\|\|\|\|\|\|\|\|\|\|\|\|\|\|\|\|\|\|\|\|\|\|\|\|\| \|\|\|\|\|\|\|\|\|\|\|\|\|\|\| \|\|\|\|\|\|\|\|\|\|\| \|\|\|\|\|\|\|\|\|\|\| \|\|\|\|\|\|\|\|\|\|\| \|\|\|\|\| \|\|\| \| \|\| \| \|\|\|\|\|\|\|\|  \|\|\|  \|\| \|\|\|\|\|\| \|\| \|\|\|\| \|\| \|\|\| \|\| \|\| \|\| \|\|\|\|\| 	CCACCGTCAAGTCCACCACGGCCTACGTCCCCAAGACCATCACGTCCATCGTCACCAAGACCACGACCAAAGATGTCCCCTTCACCACGGTCATTACTGGCGATAAGCCCAAGACCATCATCGTCACCAAGACCATCCCGCAAACCACTCAGACCGTCAGCAAATCGACCACCAAGATCCCCAAGCCCACCACTGAATACAAGACGACGGTCATCACCACCGTCGTTCCCGTCGTCAAAACCGGCTACTCCGAAAGCGTAGTCACCGTCACCAAGCCGGTGACCAAGACCATTACGACTACCGAGACCTCGATCTCGACCGGCAAGGGCGAGA-CACCTCGACTGGTAC-AAGAAGACGAC-GCCACGGAGAC-AAGGCGGAGAC-TACAC-ACTNC-ACNA-GAGCGTCGN-AGCN-GA-TGTGGT-AC-AAGA-GC-ATT-AC-AC-AC-AAGGT ```      ``` 95	GATGCACTCTCCAATGCAATTCGCGGCTCTGGCCATGTTCGCCCTCAGTGGCATGGTCTCTGCTGTGCCTGGCAAGTCCAGTACTACCGTCACGCCCCCATCTTATGAGACTACCACTCCTCCTCCACCTCCTCCACCATATGGCGGTAGCACTACTACCTCGTCTACGAAGAAGAccactaccacgtccaagtccagctccacgaccaagtccacctccacgaccaagtccagctccacgaccaagtccacctccaccTGCGCCGTGAGCACCAGCATTTCCACTTCCGTTGGAGAGTCTCCCTACA	402 	\|\|\|\|\|\|\|\|\|\|\|\|\|\|\|\|\|\|\|\|\|\|\|\|\|\|\|\|\|\|\|\|\|\|\|\|\|\|\|\|\|\|\|\|\|\|\|\|\|\|\|\|\|\|\|\|\|\|\|\|\|\|\|\|\|\|\|\|\|\|\|\|\|\|\|\|\|\|\|\|\|\|\|\|\|\|\|\|\|\|\|\|\|\|\|\|\|\|\|\|\|\|\|\|\|\|\|\|\|\|\|\|\|\|\|\|\|\|\|\|\|\|\|\|\|\|\|\|\|\|\|\|\|\|\|\|\|\|\|\|\|\|\|\|\|\|\|\|\|\|\|\|\|\|\|\|\|\|\|\|\|\|\|\|\|\|\|\|\|\|\|\|\|\|\|\|\|\|\|\|\|\|\|\|\|\|\|\|\|\|\|\|\|\|\|\|\|\|\|\|\|\|\|\|\|\|\|\|\|\|\|\|\|\|\|\|\|\|\|\|\|\|\|\|\|\|\|\|\|\|\|\|\|\|\|\|\|\|\|\|\|\|\|\|\|\|\|\|\|\|\|\|\|\|\|\|\|\|\|\|\|\|\|\|\|\|\|\|\|\|\|\|\|\|\|\|\|\|\|\|\|\|\|\|\|\|\|\|\|\|\|\|\|\|\|\|\|\|\|\|\|\|\|\|\|\|\|\| 	GATGCACTCTCCAATGCAATTCGCGGCTCTGGCCATGTTCGCCCTCAGTGGCATGGTCTCTGCTGTGCCTGGCAAGTCCAGTACTACCGTCACGCCCCCATCTTATGAGACTACCACTCCTCCTCCACCTCCTCCACCATATGGCGGTAGCACTACTACCTCGTCTACGAAGAAGACCACTACCACGTCCAAGTCCAGCTCCACGACCAAGTCCACCTCCACGACCAAGTCCAGCTCCACGACCAAGTCCACCTCCACCTGCGCCGTGAGCACCAGCATTTCCACTTCCGTTGGAGAGTCTCCCTACA ``` |
| 7 | ggggacaagtttgtacaaaaaagcaggcttgatgCTTGCCGACGCGGACGCGAT, ggggaccactttgtacaagaaagctgggtcTCTGACAGCACTTCTCGCCC | Mycgr3\|105632  JGI  Ensembl |  |  | 100 % |  | ``` 179	ATGCTTGCCGACGCGGACGCGATGGGCTGTCCGAGACGGTTAGCTGAGGCGGAGGTAAGTGAGCGCAGCATGGCCGCAAGGGTGGATACGCAAGATGCGCAGTGCGGGAAGTCGGGCTCGGTTGAGACTTTGATCATGAGAGCTCGCCAGGTCTGGTCGCCTTCGATGCGTGCAAGCGGGATGTTCGAGCCGTTCGTGAAGGACGCGGCAGCGACACTGAGGGCGAGAAGTGCTGTC	415 	\|\|\|\|\|\|\|\|\|\|\|\| \| \|\|\|\|\|\|\|\|\|\|\|\|\|\|\|\|\|\|\|\|\|\|\|\|\|\|\|\|\|\|\|\|\|\|\|\|\|\|\|\|\|\|\|\|\|\|\|\|\|\|\|\|\|\|\|\|\|\|\|\|\|\|\|\|\|\|\|\|\|\|\|\|\|\|\|\|\|\|\|\|\|\|\|\|\|\|\|\|\|\|\|\|\|\|\|\|\|\|\|\|\|\|\|\|\|\|\|\|\|\|\|\|\|\|\|\|\|\|\|\|\|\|\|\|\|\|\|\|\|\|\|\|\|\|\|\|\|\|\|\|\|\|\|\|\|\|\|\|\|\|\|\|\|\|\|\|\|\|\|\|\|\|\|\|\|\|\|\|\|\|\| \|\|\|\|\|\|\|\|\|\|\|\|\|\|\|\|\|\|\|\|\|\|\|\|\|\|\|\|\|\|\|\|\| \|\|\|\|\| \|\| \|\|\| \|\|\| 	ATGCTTGCCGACNCNGACGCGATGGGCTGTCCGAGACGGTTAGCTGAGGCGGAGGTAAGTGAGCGCAGCATGGCCGCAAGGGTGGATACGCAAGATGCGCAGTGCGGGAAGTCGGGCTCGGTTGAGACTTTGATCATGAGAGCTCGCCAGGTCTGGTCGCCTTCGATGCGTGCAAGCGGGATGTTCNAGCCGTTCGTGAAGGACGCGGCAGCGACACTGANGGCGANAANTGCNGTC ``` |
| 7 | ggggacaagtttgtacaaaaaagcaggcttgatgATCCCATGCGAAAAGCATGA, ggggaccactttgtacaagaaagctgggtcCGGATCCTGCTCCCCGCCGA | Mycgr3\|105802  JGI  Ensembl |  |  | 0 % |  |  |
| 7 | ggggacaagtttgtacaaaaaagcaggcttgatgGATACTATGCAGGGCCAACA, ggggaccactttgtacaagaaagctgggtcCCTGTGATTGTCATGTCCCA | Mycgr3\|106585  JGI  Ensembl |  |  | 100 % |  | ``` 84	GATGGATACTATGCAGGGCCAACAGCGGCGCAGTCTGCGTGCGTGCGGTGGCAAGGACAACACAACACAACACGGCACGAGTCCGCAAGGACTTCTCCGGCACAAACGAGGGAGGGAGTCTCCACCGCGGCAGACTCGGCGTGCTATGGAGGGAGGACGAAGCGGCAAAAGTCGGCGGCCCAGTGTGAGGGCAGGCAAGGCAAGGTGGGATTTGGGACATGACAATCACAGG	315 	\|\|\|\|\|\|\|\|\|\|\|\|\|\| \|\|\|\|\|\|\|\|\|\|\|\|\|\|\|\|\|\|\|\|\|\|\|\|\|\|\|\|\|\|\|\|\|\|\|\|\|\|\|\|\|\|\|\|\|\|\|\|\|\|\|\|\|\|\|\|\|\|\|\|\|\|\|\|\|\|\|\|\|\|\|\|\|\|\|\|\|\|\|\|\|\|\|\|\|\|\|\|\|\|\|\|\|\|\|\|\|\|\|\|\|\|\|\|\|\|\|\|\|\|\|\|\|\|\|\|\|\|\|\|\|\|\|\|\|\|\|\|\|\|\|\|\|\|\|\|\|\|\|\|\|\|\|\|\|\|\|\|\|\|\|\|\|\|\|\|\|\|\|\|\|\|\|\|\|\|\|\|\|\|\|\|\|\|\|\|\|\|\|\|\|\|\|\|\|\|\|\|\|\|\|\|\|\|\|\|\|\|\|\|\|\|\|\|\|\|\|\|\|\|\|\|\|\|\|\|\| 	GATGGATACTATGCNGGGCCAACAGCGGCGCAGTCTGCGTGCGTGCGGTGGCAAGGACAACACAACACAACACGGCACGAGTCCGCAAGGACTTCTCCGGCACAAACGAGGGAGGGAGTCTCCACCGCGGCAGACTCGGCGTGCTATGGAGGGAGGACGAAGCGGCAAAAGTCGGCGGCCCAGTGTGAGGGCAGGCAAGGCAAGGTGGGATTTGGGACATGACAATCACAGG ``` |
| 7 | ggggacaagtttgtacaaaaaagcaggcttgatgACTCTGATTGTGAAAGCAGG, ggggaccactttgtacaagaaagctgggtcTCTTACTGCTCCTGTCAACA | Mycgr3\|106588  JGI  Ensembl |  |  | 0 % |  |  |
| 7 | ggggacaagtttgtacaaaaaagcaggcttgatgTACGAGCAACTGGCAAGGCC, ggggaccactttgtacaagaaagctgggtcGTCGAAACGCAGCTGTAGTC | Mycgr3\|106690  JGI  Ensembl |  |  | 100 % |  | ``` 25	GATGTACGAGCAACTGGCAAGGCCCGGCTATACACCCACCGCGTCTTCGACCACTCGATTTACAGAGCTTGCGGCCCATTACGAACCCGCTTTGTCGATAATTCACCCTCACGGAGCACGCACGTTGATCCTCAACGGCGAAGACGTACTGGCCGCGTTCTGCAAGCAGACGGCTTCACGGCGCCCTTCTAAGAGATGGGAAAGGATGTGTCCAAGACAGGATACAGAGGCTGAGTGTGATGTCACAAACCACCCTCAAACCTCTTTGCTGCGGGCCATGGAAAGCGGCAGAGCTGTCAATGGGGCTGCGGGTGGACTACAGCTGCGTTTCGAC	358 	\|\|\|\|\|\|\|\|\|\|\|\|\|\|\|\|\|\|\|\|\|\|\|\|\|\|\|\|\|\|\|\|\|\|\|\|\|\|\|\|\|\|\|\|\|\|\|\|\|\|\|\|\|\|\|\|\|\|\|\|\|\|\|\|\|\|\|\|\|\|\|\|\|\|\|\|\|\|\|\|\|\|\|\|\|\|\|\|\|\|\|\|\|\|\|\|\|\|\|\|\|\|\|\|\|\|\|\|\|\|\|\|\|\|\|\|\|\|\|\|\|\|\|\|\|\|\|\|\|\|\|\|\|\|\|\|\|\|\|\|\|\|\|\|\|\|\|\|\|\|\|\|\|\|\|\|\|\|\|\|\|\|\|\|\|\|\|\|\|\|\|\|\|\|\|\|\|\|\|\|\|\|\|\|\|\|\|\|\|\|\|\|\|\|\|\|\|\|\|\|\|\|\|\|\|\|\|\|\|\|\|\|\|\|\|\|\|\|\|\|\|\|\|\|\|\|\|\|\|\|\|\|\|\|\|\|\|\|\|\|\|\|\|\|\|\|\|\|\|\|\|\|\|\|\|\|\|\|\|\|\|\|\|\|\|\|\|\|\|\|\|\|\|\|\|\|\|\|\|\|\|\|\|\|\|\|\|\|\|\|\|\|\|\|\|\|\|\|\|\|\|\|\|\|\|\|\|\|\|\|\|\|\|\|\|\|\|\|\|\|\|\|\|\|\|\|\|\|\|\|\|\|\|\| 	GATGTACGAGCAACTGGCAAGGCCCGGCTATACACCCACCGCGTCTTCGACCACTCGATTTACAGAGCTTGCGGCCCATTACGAACCCGCTTTGTCGATAATTCACCCTCACGGAGCACGCACGTTGATCCTCAACGGCGAAGACGTACTGGCCGCGTTCTGCAAGCAGACGGCTTCACGGCGCCCTTCTAAGAGATGGGAAAGGATGTGTCCAAGACAGGATACAGAGGCTGAGTGTGATGTCACAAACCACCCTCAAACCTCTTTGCTGCGGGCCATGGAAAGCGGCAGAGCTGTCAATGGGGCTGCGGGTGGACTACAGCTGCGTTTCGAC ``` |
| 7 | ggggacaagtttgtacaaaaaagcaggcttgatgCAAACCACTTGGAGGTCAGT, ggggaccactttgtacaagaaagctgggtcCACAACTCCATACACAACCA | Mycgr3\|106743  JGI  Ensembl |  |  | 0 % |  |  |
| 7 | ggggacaagtttgtacaaaaaagcaggcttgatgGAATTCATCGTCACTACCAA, ggggaccactttgtacaagaaagctgggtcGGCGGGACGACGATTGGCGG | Mycgr3\|106800  JGI  Ensembl |  |  | 0 % |  |  |
| 7 | ggggacaagtttgtacaaaaaagcaggcttgatgATCCGTTTTTGGACCCTCTT, ggggaccactttgtacaagaaagctgggtcAGCGATAGAGAGCTCTTCCG | Mycgr3\|111789  JGI  Ensembl |  |  | 0 % |  | ``` 1	ATGATCCGTTTTTGGACCCTCTTACAAACCCGCCTTCGCCTTCGGATCCGCTTGCATATCCTACGTCGAACCCCACCTCTATCGCATCCACCACCACCACAATCGCCGCCACCGCCAAATTCGGTGGATCGAGCACGCACCGAACTCACAGTGCATCTCGACGATAAGCACGCTCTCGACTGCTACGATGTGCTATTGCGCGCATATCGCGCCTCGCCGCTATATGAGCCGGACAGTAAGGAGGATCCTTACGCTAGGATCGAGATTCTCGAAGCAACACTCGCACATTTGGACACCGCAGTCTGCGGGCGAGCGGAAGAGCTCTCTATCGCT	333 	\|\|\|\|\|\|\|\|\|\|\|\|\|\|\|\|\|\|\|\|\|\|\|\|\|\|\|\|\|\|\|\|\|\|\|\|\|\|\|\|\|\|\|\|\|\|\|\|\|\|\|\|\|\|\|\|\|\|\|\|\|\|\|\|\|\|\|\|\|\|\|\|\|\|\|\|\|\|\|\|\|\|\|\|\|\|\|\|\|\|\|\|\|\|\|\|\|\|\|\|\|\|\|\|\|\|\|\|\|\|\|\|\|\|\|\|\|\|\|\|\|\|\|\|\|\|\|\|\|\|\|\|\|\|\|\|\|\|\|\|\|\|\|\|\|\|\|\|\|\|\|\|\|\|\|\|\|\|\|\|\|\|\|\|\|\|\|\|\|\|\|\|\|\|\|\|\|\|\|\|\|\|\|\|\|\|\|\|\|\|\|\|\|\|\|\|\|\|\|\|\|\|\|\|\|\|\|\|\|\|\|\|\|\|\|\|\|\|\|\|\|\|\|\|\|\|\|\|\|\|\|\|\|\|\|\|\|\|\|\|\|\|\|\|\|\|\|\|\|\|\|\|\|\|\|\|\|\|\|\|\|\|\|\|\|\|\|\|\|\|\|\|\|\|\|\|\|\|\|\|\|\|\|\|\|\|\|\|\|\|\|\|\|\|\|\|\|\|\|\|\|\|\|\|\|\|\|\|\|\|\|\|\|\|\|\|\|\|\|\|\|\|\|\|\|\|\|\|\|\|\|\|\| 	ATGATCCGTTTTTGGACCCTCTTACAAACCCGCCTTCGCCTTCGGATCCGCTTGCATATCCTACGTCGAACCCCACCTCTATCGCATCCACCACCACCACAATCGCCGCCACCGCCAAATTCGGTGGATCGAGCACGCACCGAACTCACAGTGCATCTCGACGATAAGCACGCTCTCGACTGCTACGATGTGCTATTGCGCGCATATCGCGCCTCGCCGCTATATGAGCCGGACAGTAAGGAGGATCCTTACGCTAGGATCGAGATTCTCGAAGCAACACTCGCACATTTGGACACCGCAGTCTGCGGGCGAGCGGAAGAGCTCTCTATCGCT ``` |
| 8 | ggggacaagtttgtacaaaaaagcaggcttgatgGGCAAACTGATCAAGAATCA, ggggaccactttgtacaagaaagctgggtcAGCCTTCTCAGCTCGTCTTC | Mycgr3\|33479  JGI  Ensembl |  |  | 100 % |  |  |
| 8 | ggggacaagtttgtacaaaaaagcaggcttgatgTATGCTGGAGAATCGAGGCC, ggggaccactttgtacaagaaagctgggtcGCCTTCTGGCATCTCGAATC | Mycgr3\|35535  JGI  Ensembl |  |  | 0 % |  |  |
| 8 | ggggacaagtttgtacaaaaaagcaggcttgatgTCCGTCCTCGCCCCCAATGG, ggggaccactttgtacaagaaagctgggtcAAGATGTCGATGGCTTGACG | Mycgr3\|35688  JGI  Ensembl |  |  | 100 % |  |  |
| 8 | ggggacaagtttgtacaaaaaagcaggcttgatgTTCGAAACAACCTTCACCCA, ggggaccactttgtacaagaaagctgggtcAGCCCTCGCCGGACCCTCCT | Mycgr3\|36641  JGI  Ensembl |  |  | 100 % |  |  |
| 8 | ggggacaagtttgtacaaaaaagcaggcttgatgGCTCCCATCACCACCGCCTC, ggggaccactttgtacaagaaagctgggtcCTTCAACACCGTGTTGGCAC | Mycgr3\|37176  JGI  Ensembl | KOG4137 Uncharacterized conserved protein; |  | 100 % |  |  |
| 8 | ggggacaagtttgtacaaaaaagcaggcttgatgTTTGTTCGGAGACTGTACAG, ggggaccactttgtacaagaaagctgggtcGTTCACGCCGGTAGCTTTAG | Mycgr3\|37496  JGI  Ensembl | KOG2726 Mitochondrial polypeptide chain release factor; | translation release factor activity; translational termination; | 100 % |  |  |
| 8 | ggggacaagtttgtacaaaaaagcaggcttgatgGCAGACCTTCAAGACACTGC, ggggaccactttgtacaagaaagctgggtcCCACCATCGCCGTATCCTGG | Mycgr3\|37650  JGI  Ensembl |  |  | 100 % |  |  |
| 8 | ggggacaagtttgtacaaaaaagcaggcttgatgCCTGCCAAACAAGTCGTCCT, ggggaccactttgtacaagaaagctgggtcCGGCATGACCGCGATACACT | Mycgr3\|38865  JGI  Ensembl | KOG2317 Putative translation initiation inhibitor UK114/IBM1; |  | 100 % |  |  |
| 8 | ggggacaagtttgtacaaaaaagcaggcttgatgAAGTTCACCACGACCTTCAC, ggggaccactttgtacaagaaagctgggtcAGCAAGGAACTCCACAGCCC | Mycgr3\|39947  JGI  Ensembl |  |  | 100 % |  |  |
| 8 | ggggacaagtttgtacaaaaaagcaggcttgatgGTTTACACCATGGTCTGGTA, ggggaccactttgtacaagaaagctgggtcACAAGAATAGCCGTCGTCTT | Mycgr3\|41575  JGI  Ensembl | KOG3752 Ribonuclease H; | nucleic acid binding; ribonuclease H activity; | 100 % |  |  |
| 8 | ggggacaagtttgtacaaaaaagcaggcttgatgGGAGAAAATGTACCACCCAA, ggggaccactttgtacaagaaagctgggtcCTCCCTTGAGTACGAAATCT | Mycgr3\|43038  JGI  Ensembl | KOG3856 Uncharacterized conserved protein; |  | 100 % |  |  |
| 8 | ggggacaagtttgtacaaaaaagcaggcttgatgATTCTCAGCGGCCTCGAAAT, ggggaccactttgtacaagaaagctgggtcTAACCCCACTCCTCCTTGCC | Mycgr3\|45051  JGI  Ensembl |  |  | 100 % |  |  |
| 8 | ggggacaagtttgtacaaaaaagcaggcttgatgTCAACCCCAGGAGAAACCTC, ggggaccactttgtacaagaaagctgggtcACCAGCATACTCCCCTTCGC | Mycgr3\|45705  JGI  Ensembl |  |  | 100 % |  |  |
| 8 | ggggacaagtttgtacaaaaaagcaggcttgatgAGTTCCGCCATTGATCGAGC, ggggaccactttgtacaagaaagctgggtcGGGCACCACTCCCAGTGGCT | Mycgr3\|50993  JGI  Ensembl |  |  | 100 % | chr\_13:156960 C->A; |  |
| 8 | ggggacaagtttgtacaaaaaagcaggcttgatgGATCCATCCACAACAACCGC, ggggaccactttgtacaagaaagctgggtcATCCTCCAACGACTGCACAG | Mycgr3\|51197  JGI  Ensembl |  |  | 100 % |  |  |
| 8 | ggggacaagtttgtacaaaaaagcaggcttgatgGGCGTGTTTCAAACGAAGAA, ggggaccactttgtacaagaaagctgggtcCATCACTCCGTCGTGTTCGC | Mycgr3\|56211  JGI  Ensembl |  |  | 100 % |  |  |
| 8 | ggggacaagtttgtacaaaaaagcaggcttgatgGCCGTCCCACGAAGTCGAGT, ggggaccactttgtacaagaaagctgggtcCTTCTTCTTTCCCTTTTGAT | Mycgr3\|61154  JGI  Ensembl | KOG4844 Mitochondrial ribosomal protein S27; |  | 100 % |  |  |
| 8 | ggggacaagtttgtacaaaaaagcaggcttgatgCCCAGCTTTCATTTCCACCT, ggggaccactttgtacaagaaagctgggtcCATCAATCCAAAATGGGCGG | Mycgr3\|79746  JGI  Ensembl |  |  | 100 % |  |  |
| 8 | ggggacaagtttgtacaaaaaagcaggcttgatgGACGCTCGCCCCGCCCTCAT, ggggaccactttgtacaagaaagctgggtcCCGAAACCACCAATTGACCG | Mycgr3\|83976  JGI  Ensembl | KOG2132 Uncharacterized conserved protein, contains JmjC domain; |  | 100 % |  |  |
| 8 | ggggacaagtttgtacaaaaaagcaggcttgatgCGCATTCCGTACGCCCCAAC, ggggaccactttgtacaagaaagctgggtcCTTCTCCTCCGGCATCTTCA | Mycgr3\|86913  JGI  Ensembl |  |  | 100 % |  |  |
| 8 | ggggacaagtttgtacaaaaaagcaggcttgatgCCAGAATACGACGTCCGTAC, ggggaccactttgtacaagaaagctgggtcAGCTTTGCCGTTCGCATCCT | Mycgr3\|87613  JGI  Ensembl |  |  | 0 % |  |  |
| 8 | ggggacaagtttgtacaaaaaagcaggcttgatgGCCGTCGCCGTGGTAGACTG, ggggaccactttgtacaagaaagctgggtcACCAGCGATGCTAATCTTTG | Mycgr3\|88614  JGI  Ensembl |  |  | 100 % |  |  |
| 8 | ggggacaagtttgtacaaaaaagcaggcttgatgTCGTACTACGACCCACCTTC, ggggaccactttgtacaagaaagctgggtcTCGCCGCCCACTGCTCCTCC | Mycgr3\|89015  JGI  Ensembl |  |  | 100 % |  |  |
| 8 | ggggacaagtttgtacaaaaaagcaggcttgatgTCGGGCCAAGGGGGTTACTA, ggggaccactttgtacaagaaagctgggtcTGAAGCAGATAACTCCTGCA | Mycgr3\|89276  JGI  Ensembl | KOG1703 Adaptor protein Enigma and related PDZ-LIM proteins; |  | 100 % |  |  |
| 8 | ggggacaagtttgtacaaaaaagcaggcttgatgAAGTTTCAAGCCAGAACGGA, ggggaccactttgtacaagaaagctgggtcAGGGCCGCGCAGCCTCATGA | Mycgr3\|89618  JGI  Ensembl |  |  | 0 % |  |  |
| 8 | ggggacaagtttgtacaaaaaagcaggcttgatgCGCACCACCTTCGCCACCTT, ggggaccactttgtacaagaaagctgggtcAGCGCAGAACATAGCAGCGC | Mycgr3\|90056  JGI  Ensembl |  |  | 100 % |  |  |
| 8 | ggggacaagtttgtacaaaaaagcaggcttgatgGCATCCCAGCCTCCGAATGA, ggggaccactttgtacaagaaagctgggtcAACCACCAACTTCATCAAAA | Mycgr3\|90119  JGI  Ensembl |  |  | 0 % |  |  |
| 8 | ggggacaagtttgtacaaaaaagcaggcttgatgGCTAAGAAGGTCGACCCTCA, ggggaccactttgtacaagaaagctgggtcTGGTGCGAAGACATCGTTGT | Mycgr3\|90387  JGI  Ensembl | KOG1685 Uncharacterized conserved protein; |  | 0 % |  |  |
| 8 | ggggacaagtttgtacaaaaaagcaggcttgatgAACGCAGTCGAAATGTCTGC, ggggaccactttgtacaagaaagctgggtcAGCACCCGCGAGGAGATGCA | Mycgr3\|91254  JGI  Ensembl |  |  | 100 % |  |  |
| 8 | ggggacaagtttgtacaaaaaagcaggcttgatgCCAACCACGGAAAAGCTCCC, ggggaccactttgtacaagaaagctgggtcACACTCATCCTCATCCACCT | Mycgr3\|91423  JGI  Ensembl |  |  | 100 % |  |  |
| 8 | ggggacaagtttgtacaaaaaagcaggcttgatgTCCTTCCTCCGTCGACTATC, ggggaccactttgtacaagaaagctgggtcGGACGTGGCTTCTGCATAAC | Mycgr3\|91515  JGI  Ensembl |  |  | 100 % |  |  |
| 8 | ggggacaagtttgtacaaaaaagcaggcttgatgGCGTCTCCCTGGCCTGTCCC, ggggaccactttgtacaagaaagctgggtcACAAATCTCTCCAAGCTCCT | Mycgr3\|91593  JGI  Ensembl |  |  | 0 % |  |  |
| 8 | ggggacaagtttgtacaaaaaagcaggcttgatgTCCGGCAACGACGGCAAAAC, ggggaccactttgtacaagaaagctgggtcGGTTCGCGAGCACGCGGCCA | Mycgr3\|91797  JGI  Ensembl |  |  | 100 % |  |  |
| 8 | ggggacaagtttgtacaaaaaagcaggcttgatgTCAGACCCAAGCACCCACCC, ggggaccactttgtacaagaaagctgggtcCGGCCGGAAATTGTGACACA | Mycgr3\|92762  JGI  Ensembl |  |  | 100 % |  |  |
| 8 | ggggacaagtttgtacaaaaaagcaggcttgatgTCAGACATACCGAACAGCAG, ggggaccactttgtacaagaaagctgggtcAAATTCTTCCTCCGCTATGA | Mycgr3\|92832  JGI  Ensembl |  |  | 3.72439 % |  |  |
| 8 | ggggacaagtttgtacaaaaaagcaggcttgatgTCTCTCAACGCAACACGGTC, ggggaccactttgtacaagaaagctgggtcTGACTCTCCATTGATCACCT | Mycgr3\|93469  JGI  Ensembl |  |  | 0 % |  |  |
| 8 | ggggacaagtttgtacaaaaaagcaggcttgatgACGGACGAGCACGATAGATT, ggggaccactttgtacaagaaagctgggtcTCCCAGGTCTGCATATCGTG | Mycgr3\|93755  JGI  Ensembl |  |  | 7.6779 % |  |  |
| 8 | ggggacaagtttgtacaaaaaagcaggcttgatgCTTGAAGAGCAAGACCTCCT, ggggaccactttgtacaagaaagctgggtcCTCCTCTGCCGATCGCGACA | Mycgr3\|94556  JGI  Ensembl |  |  | 0 % |  |  |
| 8 | ggggacaagtttgtacaaaaaagcaggcttgatgGCCATCCAATGGACCAACGA, ggggaccactttgtacaagaaagctgggtcTCGCGGGCGCAAGTTTCTCG | Mycgr3\|94710  JGI  Ensembl |  |  | 100 % |  |  |
| 8 | ggggacaagtttgtacaaaaaagcaggcttgatgACAAAGGAAGAGGAGCAAGA, ggggaccactttgtacaagaaagctgggtcAGTCATACGGTCCAAGTACG | Mycgr3\|94816  JGI  Ensembl |  |  | 100 % |  |  |
| 8 | ggggacaagtttgtacaaaaaagcaggcttgatgACCACCCTCACACCCCTCCA, ggggaccactttgtacaagaaagctgggtcTTCTTCTTCGATGGGCGTGG | Mycgr3\|94925  JGI  Ensembl |  |  | 5.62852 % |  |  |
| 8 | ggggacaagtttgtacaaaaaagcaggcttgatgAAATCCGTCGCCGAGAGCTT, ggggaccactttgtacaagaaagctgggtcCCACCTCCATCCCTTCTCCC | Mycgr3\|95596  JGI  Ensembl |  |  | 100 % |  |  |
| 8 | ggggacaagtttgtacaaaaaagcaggcttgatgGCAACTGACAACGATATTCA, ggggaccactttgtacaagaaagctgggtcCTCATCCATCTCCACCCATC | Mycgr3\|95998  JGI  Ensembl |  |  | 100 % |  |  |
| 8 | ggggacaagtttgtacaaaaaagcaggcttgatgTGCGACACCATCAGCATCCC, ggggaccactttgtacaagaaagctgggtcTTCAATCGTCATACCGCCGA | Mycgr3\|96356  JGI  Ensembl |  |  | 100 % |  |  |
| 8 | ggggacaagtttgtacaaaaaagcaggcttgatgTTGCTCACCTCCATCCTCTC, ggggaccactttgtacaagaaagctgggtcATTACGAAACTCGCAGTCGA | Mycgr3\|96820  JGI  Ensembl |  |  | 100 % |  |  |
| 8 | ggggacaagtttgtacaaaaaagcaggcttgatgTGGACCACCATCACCCGGAC, ggggaccactttgtacaagaaagctgggtcATTGGGATCGCTCACGACCA | Mycgr3\|96844  JGI  Ensembl |  |  | 100 % |  |  |
| 8 | ggggacaagtttgtacaaaaaagcaggcttgatgCGCTTCTTTTCCGTCCTTAC, ggggaccactttgtacaagaaagctgggtcTTCAGTCGCCCTGTAGTAGT | Mycgr3\|96876  JGI  Ensembl |  |  | 100 % |  |  |
| 8 | ggggacaagtttgtacaaaaaagcaggcttgatgAAGACATTCTTTGCCCTCGC, ggggaccactttgtacaagaaagctgggtcTTTGCTACCTCCTCCGAGTA | Mycgr3\|96910  JGI  Ensembl |  |  | 100 % |  |  |
| 8 | ggggacaagtttgtacaaaaaagcaggcttgatgCTGTTCATCCTCTCCGCCAT, ggggaccactttgtacaagaaagctgggtcATTCGTCGGCCAGACTTCCG | Mycgr3\|97163  JGI  Ensembl |  |  | 100 % |  |  |
| 8 | ggggacaagtttgtacaaaaaagcaggcttgatgCTGCTGCTTCATCTAGGGAC, ggggaccactttgtacaagaaagctgggtcGACAGCGAAGTGGTCTTTGT | Mycgr3\|97214  JGI  Ensembl |  |  | 100 % |  |  |
| 8 | ggggacaagtttgtacaaaaaagcaggcttgatgCTCTCCCGTCTCAAAACAAT, ggggaccactttgtacaagaaagctgggtcGCCAAAGGTACTAAGAAACC | Mycgr3\|97477  JGI  Ensembl |  |  | 100 % |  |  |
| 8 | ggggacaagtttgtacaaaaaagcaggcttgatgCCCGACGCAGGACTGAATAA, ggggaccactttgtacaagaaagctgggtcTTGGCGTGAGTGGTCGCGGA | Mycgr3\|97734  JGI  Ensembl |  |  | 0 % |  |  |
| 8 | ggggacaagtttgtacaaaaaagcaggcttgatgATCTTGTACTCATTTCTGGG, ggggaccactttgtacaagaaagctgggtcCCCTTTGCGCTCTCCACCAG | Mycgr3\|97755  JGI  Ensembl |  |  | 0 % |  |  |
| 8 | ggggacaagtttgtacaaaaaagcaggcttgatgCTGTGCATCAGCCCTCGCAC, ggggaccactttgtacaagaaagctgggtcAGTCCTTCTCGGGAGTGGAG | Mycgr3\|97820  JGI  Ensembl | KOG4157 beta-1,6-N-acetylglucosaminyltransferase, contains WSC domain; |  | 100 % |  |  |
| 8 | ggggacaagtttgtacaaaaaagcaggcttgatgAGTCCCTCCAACCGAGAAGT, ggggaccactttgtacaagaaagctgggtcCTGCTTCTTCTCCTCCTTCA | Mycgr3\|97843  JGI  Ensembl |  |  | 100 % |  |  |
| 8 | ggggacaagtttgtacaaaaaagcaggcttgatgCAGCCAGCCAGCGAGCACGA, ggggaccactttgtacaagaaagctgggtcTCCGTCCATTTCCTCCTTAG | Mycgr3\|97845  JGI  Ensembl |  |  | 100 % |  |  |
| 8 | ggggacaagtttgtacaaaaaagcaggcttgatgGAGCCAGATGCATATGCATT, ggggaccactttgtacaagaaagctgggtcCACGCTCAGGTTCCTGCTCC | Mycgr3\|97879  JGI  Ensembl |  |  | 100 % |  |  |
| 8 | ggggacaagtttgtacaaaaaagcaggcttgatgCCGCCGCTTCCAACAACACG, ggggaccactttgtacaagaaagctgggtcCTCGCCAACGAAAACCGCGT | Mycgr3\|97965  JGI  Ensembl |  |  | 100 % |  |  |
| 8 | ggggacaagtttgtacaaaaaagcaggcttgatgGACTTTCTTGATCAGAGTGG, ggggaccactttgtacaagaaagctgggtcTTCTCCTTCTTCATCAAACA | Mycgr3\|97976  JGI  Ensembl |  |  | 0 % |  |  |
| 8 | ggggacaagtttgtacaaaaaagcaggcttgatgACCTCTTCCCCTCGTACTAC, ggggaccactttgtacaagaaagctgggtcGCGGAGGCGGGAAAGACGAG | Mycgr3\|98029  JGI  Ensembl |  |  | 0 % |  |  |
| 8 | ggggacaagtttgtacaaaaaagcaggcttgatgCGAGACCATCGTGGTCAGCA, ggggaccactttgtacaagaaagctgggtcAATCGTATCAAACCAACCAA | Mycgr3\|102033  JGI  Ensembl |  |  | 100 % |  |  |
| 8 | ggggacaagtttgtacaaaaaagcaggcttgatgCAAACCACAGTGCGTTCTCT, ggggaccactttgtacaagaaagctgggtcGAAGTCCGTAACCCTTCCAT | Mycgr3\|102272  JGI  Ensembl |  |  | 100 % |  |  |
| 8 | ggggacaagtttgtacaaaaaagcaggcttgatgAACCGGCCTCGCAATCACCA, ggggaccactttgtacaagaaagctgggtcCGTGAAGTGGTCTTCTACGT | Mycgr3\|102412  JGI  Ensembl |  |  | 100 % |  |  |
| 8 | ggggacaagtttgtacaaaaaagcaggcttgatgCATCAGGCCCTTGCGGCAAA, ggggaccactttgtacaagaaagctgggtcTCGGTCATGTACGTTCGAGC | Mycgr3\|102869  JGI  Ensembl |  |  | 100 % |  |  |
| 8 | ggggacaagtttgtacaaaaaagcaggcttgatgCCTCGTGACGGATCAGGACA, ggggaccactttgtacaagaaagctgggtcCTTCGAGGTGGTCTCGCCGC | Mycgr3\|103030  JGI  Ensembl |  |  | 0 % |  |  |
| 8 | ggggacaagtttgtacaaaaaagcaggcttgatgTCAGATTCAGAAGCTCCCCG, ggggaccactttgtacaagaaagctgggtcATCCTCATCTTCCTCTTCCT | Mycgr3\|103199  JGI  Ensembl |  |  | 100 % |  |  |
| 8 | ggggacaagtttgtacaaaaaagcaggcttgatgGGATCGATCATCGCCGACGG, ggggaccactttgtacaagaaagctgggtcTATATCGTTTGCTTGGAGGA | Mycgr3\|103270  JGI  Ensembl |  |  | 100 % |  |  |
| 8 | ggggacaagtttgtacaaaaaagcaggcttgatgGATATTCCAGACAGCGAGCC, ggggaccactttgtacaagaaagctgggtcCCAACTCTCATCAAGGCTGC | Mycgr3\|103297  JGI  Ensembl |  |  | 10.94891 % |  |  |
| 8 | ggggacaagtttgtacaaaaaagcaggcttgatgTTCCGCAGCTCGCTCGCGCG, ggggaccactttgtacaagaaagctgggtcCGCGACCGTGCTAGGAACAC | Mycgr3\|103360  JGI  Ensembl |  |  | 9.72222 % |  |  |
| 8 | ggggacaagtttgtacaaaaaagcaggcttgatgCCGCTCCTTCCCGCCGACAT, ggggaccactttgtacaagaaagctgggtcCATCGTGCAGCTCGGATCAC | Mycgr3\|103509  JGI  Ensembl |  |  | 100 % |  |  |
| 8 | ggggacaagtttgtacaaaaaagcaggcttgatgAGGCGAACAACGTGGATCAA, ggggaccactttgtacaagaaagctgggtcGCTCGAGCCACACCCAAACG | Mycgr3\|103655  JGI  Ensembl |  |  | 0 % |  |  |
| 8 | ggggacaagtttgtacaaaaaagcaggcttgatgGTCGATCAGGCTACCGCGCC, ggggaccactttgtacaagaaagctgggtcAGACCAGAGCTCTGGGCGGA | Mycgr3\|103951  JGI  Ensembl |  |  | 100 % |  |  |
| 8 | ggggacaagtttgtacaaaaaagcaggcttgatgAGAGATTCTTGTCCATCCGA, ggggaccactttgtacaagaaagctgggtcCCTTCGCTGCCCGCCTTGTC | Mycgr3\|103979  JGI  Ensembl |  |  | 0 % |  |  |
| 8 | ggggacaagtttgtacaaaaaagcaggcttgatgCGGCGCGCATTGTCACATGC, ggggaccactttgtacaagaaagctgggtcGAACAGGTGCAAGAGGCTGT | Mycgr3\|104253  JGI  Ensembl |  |  | 100 % |  |  |
| 8 | ggggacaagtttgtacaaaaaagcaggcttgatgGCAGTGCCAGACGCGTGTCC, ggggaccactttgtacaagaaagctgggtcCTGCTTTGTCTGCGGCTTGT | Mycgr3\|104380  JGI  Ensembl |  |  | 100 % |  |  |
| 8 | ggggacaagtttgtacaaaaaagcaggcttgatgACAACTCAAGCTCAAGCTCC, ggggaccactttgtacaagaaagctgggtcGCGGGCATAGCTATTGACGC | Mycgr3\|104559  JGI  Ensembl |  |  | 100 % |  |  |
| 8 | ggggacaagtttgtacaaaaaagcaggcttgatgGCGCTGCAGCATCGGTCTCT, ggggaccactttgtacaagaaagctgggtcTTTCATCATTTCGTTGCGAC | Mycgr3\|104645  JGI  Ensembl |  |  | 0 % |  |  |
| 8 | ggggacaagtttgtacaaaaaagcaggcttgatgCAAAACTCAACGTCATGCGC, ggggaccactttgtacaagaaagctgggtcGTCCATCGTCCGGGGTAGCC | Mycgr3\|104669  JGI  Ensembl |  |  | 100 % |  |  |
| 8 | ggggacaagtttgtacaaaaaagcaggcttgatgGCCGCGGCCGTCGCAAACCC, ggggaccactttgtacaagaaagctgggtcCTTTGGCTCATCATAAGGAC | Mycgr3\|104840  JGI  Ensembl |  |  | 0 % |  |  |
| 8 | ggggacaagtttgtacaaaaaagcaggcttgatgATTACGCAGTTCTGCAAATA, ggggaccactttgtacaagaaagctgggtcCTCTTCGGCCGCGGGATTTC | Mycgr3\|104874  JGI  Ensembl |  |  | 100 % |  |  |
| 8 | ggggacaagtttgtacaaaaaagcaggcttgatgCGCTTTTCGCAACAACGCTC, ggggaccactttgtacaagaaagctgggtcCCGGCAGATTGGGAAGGATG | Mycgr3\|105578  JGI  Ensembl |  |  | 100 % |  |  |
| 8 | ggggacaagtttgtacaaaaaagcaggcttgatgTCCGTCACACTCAATACCAC, ggggaccactttgtacaagaaagctgggtcTCTCTCATCCGCCAACGGGT | Mycgr3\|105653  JGI  Ensembl | KOG0884 Similar to cyclophilin-type peptidyl-prolyl cis-trans isomerase; | FK506-sensitive peptidyl-prolyl cis-trans isomerase; cyclophilin; cyclophilin-type peptidyl-prolyl cis-trans isomerase activity;  peptidyl-prolyl cis-trans isomerase activity;  protein folding; | 100 % |  |  |
| 8 | ggggacaagtttgtacaaaaaagcaggcttgatgGAAGTGAGGGGGCTTGTCGT, ggggaccactttgtacaagaaagctgggtcTTGCGTCAGCATCAGCACGA | Mycgr3\|105739  JGI  Ensembl |  |  | 100 % |  |  |
| 8 | ggggacaagtttgtacaaaaaagcaggcttgatgCGGACAGCGCACAGTAGTTG, ggggaccactttgtacaagaaagctgggtcAAGCAGCAGCCCAGTCATGG | Mycgr3\|106004  JGI  Ensembl |  |  | 100 % |  |  |
| 8 | ggggacaagtttgtacaaaaaagcaggcttgatgGACCACAGAACAGACTCAAA, ggggaccactttgtacaagaaagctgggtcTAAAGCACAAACAGCCAGAT | Mycgr3\|106205  JGI  Ensembl |  |  | 100 % |  |  |
| 8 | ggggacaagtttgtacaaaaaagcaggcttgatgCCGCTCTTCGAGATTCAACA, ggggaccactttgtacaagaaagctgggtcAAAGATTGTTGTGTTCTTCA | Mycgr3\|106348  JGI  Ensembl |  |  | 100 % |  |  |
| 8 | ggggacaagtttgtacaaaaaagcaggcttgatgCCCTTTGCACTTCAACATCC, ggggaccactttgtacaagaaagctgggtcAGCGCTGGGGAACTCCAGCG | Mycgr3\|106556  JGI  Ensembl |  |  | 100 % |  |  |
| 8 | ggggacaagtttgtacaaaaaagcaggcttgatgCGATTGCAAGATGGTGGAAA, ggggaccactttgtacaagaaagctgggtcCAGCACTCCATCAACATGGA | Mycgr3\|106603  JGI  Ensembl |  |  | 100 % |  |  |
| 8 | ggggacaagtttgtacaaaaaagcaggcttgatgCCGAAAACATGCCAGGAGCT, ggggaccactttgtacaagaaagctgggtcCGTGGGTAGTTGAGGAGGGT | Mycgr3\|106719  JGI  Ensembl |  |  | 100 % |  |  |
| 8 | ggggacaagtttgtacaaaaaagcaggcttgatgGTCTCCACAAAGCCAACTCC, ggggaccactttgtacaagaaagctgggtcACCTTGCTCCAAGCTCACGC | Mycgr3\|106817  JGI  Ensembl |  |  | 100 % |  |  |
| 8 | ggggacaagtttgtacaaaaaagcaggcttgatgCTGCGCTACTTTGGCGAAGT, ggggaccactttgtacaagaaagctgggtcGCGAAATCCACAAGTTTCAT | Mycgr3\|107968  JGI  Ensembl |  |  | 100 % |  |  |
| 9 | ggggacaagtttgtacaaaaaagcaggcttgatgGATCCTGACAGTCCCGCATC, ggggaccactttgtacaagaaagctgggtcTCTGGTCTTCGAATGTACAC | Mycgr3\|31411  JGI  Ensembl |  |  | 100 % |  |  |
| 9 | ggggacaagtttgtacaaaaaagcaggcttgatgTTCGCGTTCAAGACTCTAAC, ggggaccactttgtacaagaaagctgggtcCCACTGCTTCAAAGCCATGA | Mycgr3\|34196  JGI  Ensembl |  |  | 100 % |  |  |
| 9 | ggggacaagtttgtacaaaaaagcaggcttgatgGCTGTCACTCAAGACACCGC, ggggaccactttgtacaagaaagctgggtcTACCATCTCTTCCTCTCTAA | Mycgr3\|38493  JGI  Ensembl |  |  | 100 % |  |  |
| 9 | ggggacaagtttgtacaaaaaagcaggcttgatgCCACCGCCGAGAAACATTTA, ggggaccactttgtacaagaaagctgggtcGATCGGAACGTGATGTCTCA | Mycgr3\|40192  JGI  Ensembl |  |  | 100 % |  |  |
| 9 | ggggacaagtttgtacaaaaaagcaggcttgatgCCTTCGAAAGACATCGAATT, ggggaccactttgtacaagaaagctgggtcTTCAGAGCGACTACAGCTCG | Mycgr3\|41428  JGI  Ensembl |  |  | 100 % |  |  |
| 9 | ggggacaagtttgtacaaaaaagcaggcttgatgATGCAAGCCAGCAAGAAGAT, ggggaccactttgtacaagaaagctgggtcCTCCTCCTTAGCACCACCAC | Mycgr3\|46537  JGI  Ensembl | KOG4554 Protein involved in inorganic phosphate transport; |  | 100 % |  |  |
| 9 | ggggacaagtttgtacaaaaaagcaggcttgatgGTCTGCCTTACCCTGTCACC, ggggaccactttgtacaagaaagctgggtcCTCCTCCCCCTGCAACCTCT | Mycgr3\|47196  JGI  Ensembl | KOG1736 Glia maturation factor beta; | actin binding; intracellular; | 100 % |  |  |
| 9 | ggggacaagtttgtacaaaaaagcaggcttgatgTTCGGACTCATCATCTCCGG, ggggaccactttgtacaagaaagctgggtcGTCTGCTCGGTCCAGGAATG | Mycgr3\|47656  JGI  Ensembl | KOG4067 Uncharacterized conserved protein; |  | 100 % |  |  |
| 9 | ggggacaagtttgtacaaaaaagcaggcttgatgTTTGCTATCAGATCTTCTCG, ggggaccactttgtacaagaaagctgggtcGGATCCATTCTGCCAATCAC | Mycgr3\|49178  JGI  Ensembl |  |  | 100 % |  |  |
| 9 | ggggacaagtttgtacaaaaaagcaggcttgatgGACGGCATGGGCCATATGAG, ggggaccactttgtacaagaaagctgggtcGTGATGCTCGACGTCAATCA | Mycgr3\|70912  JGI  Ensembl | KOG3386 Copper transporter; | copper ion transmembrane transporter activity; copper ion transport;  integral to membrane; | 100 % |  |  |
| 9 | ggggacaagtttgtacaaaaaagcaggcttgatgGCCTCCGTTCCCGCCGAAAT, ggggaccactttgtacaagaaagctgggtcCGAAGGAGGAGCCCAGTCGT | Mycgr3\|71496  JGI  Ensembl |  |  | 100 % |  |  |
| 9 | ggggacaagtttgtacaaaaaagcaggcttgatgTCGGTCACAACAACTGCCAA, ggggaccactttgtacaagaaagctgggtcCCACTCTCCTCCGACCTTGT | Mycgr3\|72695  JGI  Ensembl |  |  | 100 % |  |  |
| 9 | ggggacaagtttgtacaaaaaagcaggcttgatgCCAACGCCACTCGACAATGC, ggggaccactttgtacaagaaagctgggtcGGCATTGGACTTTGGCGGAT | Mycgr3\|73735  JGI  Ensembl |  |  | 100 % |  |  |
| 9 | ggggacaagtttgtacaaaaaagcaggcttgatgTCCGCCCTTCCCCGCTCCCT, ggggaccactttgtacaagaaagctgggtcCTGCTCATCGGTGCTCTGGC | Mycgr3\|76263  JGI  Ensembl | KOG3490 Transcription elongation factor SPT4; | nucleus; positive regulation of transcription;  positive transcription elongation factor activity;  zinc ion binding; | 0 % |  |  |
| 9 | ggggacaagtttgtacaaaaaagcaggcttgatgGCCACCAAACGCATCATAAA, ggggaccactttgtacaagaaagctgggtcTCTTCTCCCCACAGCAAACC | Mycgr3\|76430  JGI  Ensembl | KOG0417 Ubiquitin-protein ligase; | negative regulation of ubiquitin-protein ligase activity; positive regulation of ubiquitin-protein ligase activity;  post-translational protein modification;  regulation of protein metabolic process;  regulation of ubiquitin-protein ligase activity;  regulation of ubiquitin-protein ligase activity during meiotic cell cycle;  small conjugating protein ligase activity;  ubiquitin-protein ligase activity; | 100 % |  |  |
| 9 | ggggacaagtttgtacaaaaaagcaggcttgatgCCAATGCCGTCGAAAGCTCC, ggggaccactttgtacaagaaagctgggtcCACCTCCGGCACATCCGCCT | Mycgr3\|77548  JGI  Ensembl |  |  | 100 % |  |  |
| 9 | ggggacaagtttgtacaaaaaagcaggcttgatgAATACGATTTCATCTGGATT, ggggaccactttgtacaagaaagctgggtcTGCCGCAGCTTCCAGACCTC | Mycgr3\|80635  JGI  Ensembl |  |  | 100 % |  |  |
| 9 | ggggacaagtttgtacaaaaaagcaggcttgatgAATTCGAGCATCGCAAGCTC, ggggaccactttgtacaagaaagctgggtcATTCTCACGAACTGGACTGC | Mycgr3\|83189  JGI  Ensembl |  |  | 100 % |  |  |
| 9 | ggggacaagtttgtacaaaaaagcaggcttgatgTCGGAGAACATCCCAGAGTC, ggggaccactttgtacaagaaagctgggtcGTCATCGTCGTGTATCGTAA | Mycgr3\|83558  JGI  Ensembl | KOG4055 Uncharacterized conserved protein; |  | 100 % |  |  |
| 9 | ggggacaagtttgtacaaaaaagcaggcttgatgGCTCCGATAACTCTGCGCCC, ggggaccactttgtacaagaaagctgggtcATCTTTGCGACTCTCGTCGC | Mycgr3\|85086  JGI  Ensembl |  |  | 100 % |  |  |
| 9 | ggggacaagtttgtacaaaaaagcaggcttgatgACCTCCCAGAACACGTCACG, ggggaccactttgtacaagaaagctgggtcGATCCGGAGCCCGAGATATG | Mycgr3\|88601  JGI  Ensembl |  |  | 100 % |  |  |
| 9 | ggggacaagtttgtacaaaaaagcaggcttgatgCTACCGCCCAAGGTTGCACT, ggggaccactttgtacaagaaagctgggtcTCCTAGCATGACCGTTCCCC | Mycgr3\|88794  JGI  Ensembl |  |  | 100 % |  |  |
| 9 | ggggacaagtttgtacaaaaaagcaggcttgatgTGCTACAAGGAGACGCCCTA, ggggaccactttgtacaagaaagctgggtcTTCGGAGGACGATCTCAGCA | Mycgr3\|88936  JGI  Ensembl | KOG4157 beta-1,6-N-acetylglucosaminyltransferase, contains WSC domain; |  | 100 % |  |  |
| 9 | ggggacaagtttgtacaaaaaagcaggcttgatgAGCCAAAGGCATCTTGAGGT, ggggaccactttgtacaagaaagctgggtcAAACAGTTCCTGCATCCATG | Mycgr3\|89574  JGI  Ensembl |  |  | 100 % |  |  |
| 9 | ggggacaagtttgtacaaaaaagcaggcttgatgACCACCTCGCTGCAGTCTCT, ggggaccactttgtacaagaaagctgggtcCTTCTTCGCCTTTGCGGCAG | Mycgr3\|89752  JGI  Ensembl | KOG3797 Peripheral-type benzodiazepine receptor and related proteins; | integral to membrane; | 100 % |  |  |
| 9 | ggggacaagtttgtacaaaaaagcaggcttgatgGGCGAAGAACACTCAAAACT, ggggaccactttgtacaagaaagctgggtcGACTTCCTCGAACACCCACA | Mycgr3\|90085  JGI  Ensembl |  |  | 100 % |  |  |
| 9 | ggggacaagtttgtacaaaaaagcaggcttgatgGCCGACAGCACCACTGCTCG, ggggaccactttgtacaagaaagctgggtcGTCCATCTTGTTGGAACTTC | Mycgr3\|90830  JGI  Ensembl |  |  | 100 % |  |  |
| 9 | ggggacaagtttgtacaaaaaagcaggcttgatgAACGAGACGGCCCGCATCGA, ggggaccactttgtacaagaaagctgggtcCGCCTTCCCTATGGACTGCT | Mycgr3\|90841  JGI  Ensembl | KOG4293 Predicted membrane protein, contains DoH and Cytochrome b-561/ferric reductase transmembrane domains; |  | 100 % |  |  |
| 9 | ggggacaagtttgtacaaaaaagcaggcttgatgTCGTCCACCAACCACCCCAC, ggggaccactttgtacaagaaagctgggtcCGACTTCTCCTCCTCCTCCT | Mycgr3\|90849  JGI  Ensembl |  |  | 7.26073 % |  |  |
| 9 | ggggacaagtttgtacaaaaaagcaggcttgatgTCAAAGAAGACAGAGGATAG, ggggaccactttgtacaagaaagctgggtcCTTCGCCGTCCCATCGGGCA | Mycgr3\|91009  JGI  Ensembl |  |  | 3.08642 % |  |  |
| 9 | ggggacaagtttgtacaaaaaagcaggcttgatgTCATCCTTCGACCTCTTCGA, ggggaccactttgtacaagaaagctgggtcGCAGGGGGAGGGGAGGTGGG | Mycgr3\|91131  JGI  Ensembl |  |  | 3.25203 % |  |  |
| 9 | ggggacaagtttgtacaaaaaagcaggcttgatgTCCCAGAACACTATGCAGGG, ggggaccactttgtacaagaaagctgggtcTGTGGCAATGTCCCTCTGAA | Mycgr3\|91670  JGI  Ensembl |  |  | 100 % |  |  |
| 9 | ggggacaagtttgtacaaaaaagcaggcttgatgGCCTCAACGGACGCACTGCA, ggggaccactttgtacaagaaagctgggtcGAGCTCGTCCGCATAATATT | Mycgr3\|91684  JGI  Ensembl |  |  | 100 % |  |  |
| 9 | ggggacaagtttgtacaaaaaagcaggcttgatgGAGAATATGTGCGATCCTGT, ggggaccactttgtacaagaaagctgggtcAGTCTCCTTGGAATAGTATC | Mycgr3\|91787  JGI  Ensembl |  |  | 100 % |  |  |
| 9 | ggggacaagtttgtacaaaaaagcaggcttgatgACCCCCAAACCCAAACCCCT, ggggaccactttgtacaagaaagctgggtcGTTGCAAATCTCCCTCTCAC | Mycgr3\|92025  JGI  Ensembl |  |  | 100 % |  |  |
| 9 | ggggacaagtttgtacaaaaaagcaggcttgatgGGTCTTCTACACTCACTCGC, ggggaccactttgtacaagaaagctgggtcAGCAAGCACATTATATCCCA | Mycgr3\|92094  JGI  Ensembl |  |  | 100 % |  |  |
| 9 | ggggacaagtttgtacaaaaaagcaggcttgatgGGCGCCGAACTATCGAGGAC, ggggaccactttgtacaagaaagctgggtcTTGGTGTGGCTTCCTCGCGT | Mycgr3\|92117  JGI  Ensembl |  |  | 100 % |  |  |
| 9 | ggggacaagtttgtacaaaaaagcaggcttgatgCTGAACGCCTGTAGATGGGC, ggggaccactttgtacaagaaagctgggtcGCCTGAGAGCTTGGAAAGCA | Mycgr3\|92766  JGI  Ensembl | KOG0541 Alkyl hydroperoxide reductase/peroxiredoxin; | antioxidant activity; oxidoreductase activity;  peroxiredoxin activity; | 100 % |  |  |
| 9 | ggggacaagtttgtacaaaaaagcaggcttgatgGCCAGTCCAGCCTTTGCCGT, ggggaccactttgtacaagaaagctgggtcGAACGTCACCGGCGGAGAGG | Mycgr3\|92841  JGI  Ensembl |  |  | 100 % |  |  |
| 9 | ggggacaagtttgtacaaaaaagcaggcttgatgGCCAACGTCACCACCAACAT, ggggaccactttgtacaagaaagctgggtcAACCAGCCCAGCCCCTCCCC | Mycgr3\|92889  JGI  Ensembl |  |  | 100 % |  |  |
| 9 | ggggacaagtttgtacaaaaaagcaggcttgatgTCCGATTCCGAGCAGACCAC, ggggaccactttgtacaagaaagctgggtcCTTCCCCTGAAGATCTTCCG | Mycgr3\|93041  JGI  Ensembl |  |  | 100 % |  |  |
| 9 | ggggacaagtttgtacaaaaaagcaggcttgatgTACACCTCCATCACCTTCGT, ggggaccactttgtacaagaaagctgggtcCAACGCCATCATAAGAGCTC | Mycgr3\|93231  JGI  Ensembl | KOG4157 beta-1,6-N-acetylglucosaminyltransferase, contains WSC domain; |  | 100 % |  |  |
| 9 | ggggacaagtttgtacaaaaaagcaggcttgatgAACGCCATCTCAATACCGTG, ggggaccactttgtacaagaaagctgggtcAAAAATGACATCTACCAGGC | Mycgr3\|93245  JGI  Ensembl | KOG1869 Splicing coactivator SRm160/300, subunit SRm300; | intracellular; nucleic acid binding;  zinc ion binding; | 100 % |  |  |
| 9 | ggggacaagtttgtacaaaaaagcaggcttgatgGCCTCGCAGAACTCTCCCCC, ggggaccactttgtacaagaaagctgggtcTGAAGGCACAAAGTAGGCTC | Mycgr3\|93353  JGI  Ensembl | KOG3318 Predicted membrane protein; |  | 100 % |  |  |
| 9 | ggggacaagtttgtacaaaaaagcaggcttgatgTACCCTTTCCTGCTCTCTGC, ggggaccactttgtacaagaaagctgggtcCGCGGACATCTCCCTCATAA | Mycgr3\|93408  JGI  Ensembl | KOG1238 Glucose dehydrogenase/choline dehydrogenase/mandelonitrile lyase (GMC oxidoreductase family); |  | 0 % |  |  |
| 9 | ggggacaagtttgtacaaaaaagcaggcttgatgCAGTTCTCGGAAGCCCTTCT, ggggaccactttgtacaagaaagctgggtcAGAAGGAGGCGAGCCAGGCG | Mycgr3\|93609  JGI  Ensembl |  |  | 100 % |  |  |
| 9 | ggggacaagtttgtacaaaaaagcaggcttgatgTCGAAGAAGGAAATCGATGT, ggggaccactttgtacaagaaagctgggtcGCCCGTGTAGTAGCATTCCA | Mycgr3\|93990  JGI  Ensembl |  |  | 100 % |  |  |
| 9 | ggggacaagtttgtacaaaaaagcaggcttgatgCCTTCAATCTCGACAGTCCT, ggggaccactttgtacaagaaagctgggtcCTCGGCGTAGTTGAGGTAGT | Mycgr3\|94065  JGI  Ensembl | KOG4744 Uncharacterized conserved protein; |  | 44.91393 % |  |  |
| 9 | ggggacaagtttgtacaaaaaagcaggcttgatgGGTCTCTTCGAATCCGACGA, ggggaccactttgtacaagaaagctgggtcCATAAACATCTTCGCAACTT | Mycgr3\|94438  JGI  Ensembl |  |  | 100 % |  |  |
| 9 | ggggacaagtttgtacaaaaaagcaggcttgatgACCAACTCAAACGGCATCCT, ggggaccactttgtacaagaaagctgggtcCGGCTGGCGCAACGCAGCCT | Mycgr3\|94666  JGI  Ensembl | KOG0613 Projectin/twitchin and related proteins; |  | 100 % |  |  |
| 9 | ggggacaagtttgtacaaaaaagcaggcttgatgGATCCGATCGAGATGCTGCA, ggggaccactttgtacaagaaagctgggtcATCCTCCTTCGTCTCCTCTC | Mycgr3\|94680  JGI  Ensembl | KOG4362 Transcriptional regulator BRCA1; | protein binding; zinc ion binding; | 100 % |  |  |
| 9 | ggggacaagtttgtacaaaaaagcaggcttgatgGCCTCCAACGCCGAAGACAT, ggggaccactttgtacaagaaagctgggtcCCATACTGTCTTCTGCCCCA | Mycgr3\|94757  JGI  Ensembl |  |  | 100 % |  |  |
| 9 | ggggacaagtttgtacaaaaaagcaggcttgatgGAGGAGAACACGACAATGAA, ggggaccactttgtacaagaaagctgggtcGAGCTTCTCCCACACCACCA | Mycgr3\|94760  JGI  Ensembl |  |  | 3.1746 % |  |  |
| 9 | ggggacaagtttgtacaaaaaagcaggcttgatgTCTTCCTTCACCAACTCCAA, ggggaccactttgtacaagaaagctgggtcCTTTTTCTCATTATTCCCAT | Mycgr3\|94780  JGI  Ensembl |  |  | 100 % |  |  |
| 9 | ggggacaagtttgtacaaaaaagcaggcttgatgGACTTGGAGAAAGCCAACAC, ggggaccactttgtacaagaaagctgggtcCGGTCTACCTTGCTGTACGC | Mycgr3\|94930  JGI  Ensembl |  |  | 100 % |  |  |
| 9 | ggggacaagtttgtacaaaaaagcaggcttgatgGCCAAGGGTACCGTAGACTG, ggggaccactttgtacaagaaagctgggtcAAGCAAGTCGTCGAAGCCGT | Mycgr3\|95133  JGI  Ensembl |  |  | 100 % |  |  |
| 9 | ggggacaagtttgtacaaaaaagcaggcttgatgTCTGCTCCCTCCGCCTTCGA, ggggaccactttgtacaagaaagctgggtcAACCATGATCTCACCAGCCC | Mycgr3\|95395  JGI  Ensembl | KOG0334 RNA helicase; |  | 100 % |  |  |
| 9 | ggggacaagtttgtacaaaaaagcaggcttgatgCACACCTGGCTCCTTTCCAA, ggggaccactttgtacaagaaagctgggtcCTTCTTGCTCTTTCGTGTTG | Mycgr3\|95526  JGI  Ensembl |  |  | 100 % |  |  |
| 9 | ggggacaagtttgtacaaaaaagcaggcttgatgGCTACCCCCGCCGCGCATCC, ggggaccactttgtacaagaaagctgggtcAGATGCATTATTTGCTGCGA | Mycgr3\|95759  JGI  Ensembl |  |  | 100 % |  |  |
| 9 | ggggacaagtttgtacaaaaaagcaggcttgatgGGCATGAACCCCTTCTTCGG, ggggaccactttgtacaagaaagctgggtcAGTGTTCCTCGTCAACCGCA | Mycgr3\|96134  JGI  Ensembl | KOG1596 Fibrillarin and related nucleolar RNA-binding proteins; |  | 100 % |  |  |
| 9 | ggggacaagtttgtacaaaaaagcaggcttgatgGCGCCAACAAAATCTTACGT, ggggaccactttgtacaagaaagctgggtcGCGAAAGGAGCACGTCAAAC | Mycgr3\|96290  JGI  Ensembl |  |  | 3.5473 % |  |  |
| 9 | ggggacaagtttgtacaaaaaagcaggcttgatgTCCAGAGCGGCGATGCAATC, ggggaccactttgtacaagaaagctgggtcATTGACCTGCTTCTCGCCCA | Mycgr3\|96555  JGI  Ensembl |  |  | 100 % |  |  |
| 9 | ggggacaagtttgtacaaaaaagcaggcttgatgAAACTATCCCTCATCACGGT, ggggaccactttgtacaagaaagctgggtcTTGCGCATTCGTTGCTATTG | Mycgr3\|96838  JGI  Ensembl |  |  | 100 % |  |  |
| 9 | ggggacaagtttgtacaaaaaagcaggcttgatgACCTCCTCGGACGACAAAAC, ggggaccactttgtacaagaaagctgggtcCTCCCTCGGATCCATCCTCA | Mycgr3\|97451  JGI  Ensembl |  |  | 100 % |  |  |
| 9 | ggggacaagtttgtacaaaaaagcaggcttgatgGTTCCACCGCCCGTCGAGCT, ggggaccactttgtacaagaaagctgggtcGAACTGTTTCGCCTCTTCTT | Mycgr3\|97615  JGI  Ensembl |  |  | 100 % |  |  |
| 9 | ggggacaagtttgtacaaaaaagcaggcttgatgGATTTGCAGGCCATGAGCTC, ggggaccactttgtacaagaaagctgggtcCAGGTCTTTATCCTGCAGTG | Mycgr3\|97634  JGI  Ensembl |  |  | 0 % |  |  |
| 9 | ggggacaagtttgtacaaaaaagcaggcttgatgGCTACCACCAACAACGCAAT, ggggaccactttgtacaagaaagctgggtcCAAATGTTCCCAAGTATCAG | Mycgr3\|97724  JGI  Ensembl |  |  | 100 % |  |  |
| 9 | ggggacaagtttgtacaaaaaagcaggcttgatgGTGAGAAGCTGTGGTATGCT, ggggaccactttgtacaagaaagctgggtcCAGCTCGGCGAAGTAGTCCT | Mycgr3\|97752  JGI  Ensembl |  |  | 100 % |  |  |
| 9 | ggggacaagtttgtacaaaaaagcaggcttgatgTCCTCCAAAGATGTCTCCCT, ggggaccactttgtacaagaaagctgggtcTATGGCGATGTGTTGGATAT | Mycgr3\|97903  JGI  Ensembl |  |  | 0 % |  |  |
| 9 | ggggacaagtttgtacaaaaaagcaggcttgatgCCCCGCCTCGTCCCACCCGC, ggggaccactttgtacaagaaagctgggtcCTGCTTCAGTCTCTGCAACC | Mycgr3\|97945  JGI  Ensembl |  |  | 100 % |  |  |
| 9 | ggggacaagtttgtacaaaaaagcaggcttgatgCCTGTCACTGTTCCACCCGC, ggggaccactttgtacaagaaagctgggtcCTCATTCAAAGCCCGCAACA | Mycgr3\|98045  JGI  Ensembl | KOG4293 Predicted membrane protein, contains DoH and Cytochrome b-561/ferric reductase transmembrane domains; |  | 100 % |  |  |
| 9 | ggggacaagtttgtacaaaaaagcaggcttgatgGCAACTCTCAAGGCAACCTC, ggggaccactttgtacaagaaagctgggtcCCTCTTCGCCTTCTTAGCCA | Mycgr3\|99182  JGI  Ensembl |  |  | 100 % |  |  |
| 9 | ggggacaagtttgtacaaaaaagcaggcttgatgGTAGCCATCAAGAATGCCCT, ggggaccactttgtacaagaaagctgggtcTTCCGGCACAGGAAGCACCA | Mycgr3\|101672  JGI  Ensembl |  |  | 0 % |  |  |
| 9 | ggggacaagtttgtacaaaaaagcaggcttgatgCCGCGACAGCAACACATGAA, ggggaccactttgtacaagaaagctgggtcTGCAGGCTTCAGAAGCCGGA | Mycgr3\|102473  JGI  Ensembl |  |  | 100 % |  |  |
| 9 | ggggacaagtttgtacaaaaaagcaggcttgatgGCAGCTTCGCACTCTCCAAA, ggggaccactttgtacaagaaagctgggtcGTGCAGCTGTGGCAGCATCA | Mycgr3\|103029  JGI  Ensembl |  |  | 100 % |  |  |
| 9 | ggggacaagtttgtacaaaaaagcaggcttgatgAAAGGAAACCAATGGCGGAC, ggggaccactttgtacaagaaagctgggtcCGCTGACATCTTAAAGCGAC | Mycgr3\|103481  JGI  Ensembl |  |  | 100 % |  |  |
| 9 | ggggacaagtttgtacaaaaaagcaggcttgatgTCCCGGTTTGATCGAGCGTC, ggggaccactttgtacaagaaagctgggtcGGGGCATGTCGCATCGCAGT | Mycgr3\|104462  JGI  Ensembl |  |  | 100 % |  |  |
| 9 | ggggacaagtttgtacaaaaaagcaggcttgatgATCATTTGGGTATACCTGCT, ggggaccactttgtacaagaaagctgggtcTGTTTCAGACGATAAACTGA | Mycgr3\|105227  JGI  Ensembl |  |  | 100 % |  |  |
| 9 | ggggacaagtttgtacaaaaaagcaggcttgatgAACAAGCGGCATCCTTTGAG, ggggaccactttgtacaagaaagctgggtcGTTGATGATGTCGCTCAAAG | Mycgr3\|105304  JGI  Ensembl |  |  | 21.03175 % |  |  |
| 9 | ggggacaagtttgtacaaaaaagcaggcttgatgTCACACGAGAGGATGGCGAG, ggggaccactttgtacaagaaagctgggtcGTTACGGTAAACAACATCGC | Mycgr3\|106049  JGI  Ensembl |  |  | 100 % |  |  |
| 9 | ggggacaagtttgtacaaaaaagcaggcttgatgCCATCCAAACTCAGCAAGGT, ggggaccactttgtacaagaaagctgggtcGACCGATCCTCTCGGCGGGA | Mycgr3\|106133  JGI  Ensembl |  |  | 100 % |  |  |
| 9 | ggggacaagtttgtacaaaaaagcaggcttgatgTTCCGCCAAACTCTCCTCCG, ggggaccactttgtacaagaaagctgggtcCTTCCTAGCCTCCTTCGCAG | Mycgr3\|106214  JGI  Ensembl |  |  | 0 % |  |  |
| 9 | ggggacaagtttgtacaaaaaagcaggcttgatgGCGCTGTCAAAGAACACGCC, ggggaccactttgtacaagaaagctgggtcCGTTTCCTGTCGAGAGCTTT | Mycgr3\|106242  JGI  Ensembl |  |  | 100 % |  |  |
| 9 | ggggacaagtttgtacaaaaaagcaggcttgatgGCACTCTGGCCGCTCAACTC, ggggaccactttgtacaagaaagctgggtcGACCGGTCCTTCGACCAATT | Mycgr3\|106390  JGI  Ensembl |  |  | 100 % |  |  |
| 9 | ggggacaagtttgtacaaaaaagcaggcttgatgCGAGGACTATCGGACCACTC, ggggaccactttgtacaagaaagctgggtcACACCAGGAACCGTTTTGCC | Mycgr3\|106525  JGI  Ensembl |  |  | 100 % |  |  |
| 9 | ggggacaagtttgtacaaaaaagcaggcttgatgTCACCTAGAAAGGCTAAGTC, ggggaccactttgtacaagaaagctgggtcCAAGGCTGGGCTCGGCGTAG | Mycgr3\|106583  JGI  Ensembl |  |  | 100 % |  |  |
| 9 | ggggacaagtttgtacaaaaaagcaggcttgatgTTTCTGCTTTCAAGATGCAT, ggggaccactttgtacaagaaagctgggtcTGCCCCGTCAACCGCATCTT | Mycgr3\|106657  JGI  Ensembl |  |  | 100 % |  |  |
| 9 | ggggacaagtttgtacaaaaaagcaggcttgatgAAACGGTACCGCTGTAGGTA, ggggaccactttgtacaagaaagctgggtcAACATCTCTTCGACGTGTGG | Mycgr3\|106700  JGI  Ensembl |  |  | 100 % |  |  |
| 9 | ggggacaagtttgtacaaaaaagcaggcttgatgCCAAAAGAAAAGGTTTTCCT, ggggaccactttgtacaagaaagctgggtcAACCGCTGGATGGCCTCGCA | Mycgr3\|109261  JGI  Ensembl |  |  | 100 % |  |  |
| 9 | ggggacaagtttgtacaaaaaagcaggcttgatgAGTTGGGGGTCGAGGAGGGC, ggggaccactttgtacaagaaagctgggtcCGCCGCATTCACACATTGAA | Mycgr3\|109706  JGI  Ensembl |  |  | 100 % |  |  |
| 9 | ggggacaagtttgtacaaaaaagcaggcttgatgGCAAATGCCTCCGCCTTCAA, ggggaccactttgtacaagaaagctgggtcTCGCTTTTCTTTGGGTCTGT | Mycgr3\|110283  JGI  Ensembl |  |  | 100 % |  |  |
| 9 | ggggacaagtttgtacaaaaaagcaggcttgatgGCTTGCAATGCGCCCAAGAC, ggggaccactttgtacaagaaagctgggtcGACCTCCACGGCGGTCGAGA | Mycgr3\|110715  JGI  Ensembl |  |  | 100 % |  |  |
| 10 | ggggacaagtttgtacaaaaaagcaggcttgatgGCGACCTCACTCCTTTCGCA, ggggaccactttgtacaagaaagctgggtcAGCATCTTGTCCCCAAAGCT | Mycgr3\|32848  JGI  Ensembl |  |  | 100 % |  |  |
| 10 | ggggacaagtttgtacaaaaaagcaggcttgatgCCCCGCCGCATCATCTACGG, ggggaccactttgtacaagaaagctgggtcCTGCGGGTCCGGAATCGGTT | Mycgr3\|33778  JGI  Ensembl |  |  | 100 % |  |  |
| 10 | ggggacaagtttgtacaaaaaagcaggcttgatgTCTGAGGAGACGCCGAAAAA, ggggaccactttgtacaagaaagctgggtcGAAGAACGAGTAGTCTCCCC | Mycgr3\|36836  JGI  Ensembl |  |  | 100 % |  |  |
| 10 | ggggacaagtttgtacaaaaaagcaggcttgatgCCCATCGAAGTCAGCTCCCT, ggggaccactttgtacaagaaagctgggtcGGATCTGCCCGCGGATCCCT | Mycgr3\|36885  JGI  Ensembl |  |  | 100 % |  |  |
| 10 | ggggacaagtttgtacaaaaaagcaggcttgatgTACCATCTCGCGCAAGCTCT, ggggaccactttgtacaagaaagctgggtcCCTCATGACAGGTGGTCTGC | Mycgr3\|37446  JGI  Ensembl | KOG0076 GTP-binding ADP-ribosylation factor-like protein yARL3; | GTP binding; intracellular;  intracellular protein transport;  protein transport;  small GTPase mediated signal transduction; | 100 % |  |  |
| 10 | ggggacaagtttgtacaaaaaagcaggcttgatgTCTCACAACTTCGCACCATA, ggggaccactttgtacaagaaagctgggtcCTCCTCATCAACAAACGAGC | Mycgr3\|37539  JGI  Ensembl |  |  | 100 % |  |  |
| 10 | ggggacaagtttgtacaaaaaagcaggcttgatgACCTTTCAAAGGCTGTGGCA, ggggaccactttgtacaagaaagctgggtcGGTGAATGAAACATGAGGCA | Mycgr3\|38519  JGI  Ensembl |  |  | 100 % |  |  |
| 10 | ggggacaagtttgtacaaaaaagcaggcttgatgCAACCTTCTCCGTATCGAGG, ggggaccactttgtacaagaaagctgggtcAATGTGAACTAGTTCAACCA | Mycgr3\|40021  JGI  Ensembl |  |  | 100 % |  |  |
| 10 | ggggacaagtttgtacaaaaaagcaggcttgatgACGTGCAAGATCCATGTTGC, ggggaccactttgtacaagaaagctgggtcCAACCCCACCTGCCCCGACA | Mycgr3\|42015  JGI  Ensembl |  |  | 100 % |  |  |
| 10 | ggggacaagtttgtacaaaaaagcaggcttgatgGTCGAAGTAAGAAGTTACAG, ggggaccactttgtacaagaaagctgggtcAAGCCTTCCCTTGAACACCG | Mycgr3\|44946  JGI  Ensembl |  |  | 100 % |  |  |
| 10 | ggggacaagtttgtacaaaaaagcaggcttgatgTCGAACAAACTCGAGACCCA, ggggaccactttgtacaagaaagctgggtcAATCTTCGCCGCCCTTGGCC | Mycgr3\|46779  JGI  Ensembl |  |  | 100 % |  |  |
| 10 | ggggacaagtttgtacaaaaaagcaggcttgatgTCTTCCCAACCTCGAGTCCG, ggggaccactttgtacaagaaagctgggtcGAACAAGACTCCTTTCGTAC | Mycgr3\|46922  JGI  Ensembl | KOG3143 Imidazoleglycerol-phosphate dehydratase; | histidine biosynthetic process; imidazoleglycerol-phosphate dehydratase activity; | 100 % |  |  |
| 10 | ggggacaagtttgtacaaaaaagcaggcttgatgCCTGTATACCTGGCAGTGCG, ggggaccactttgtacaagaaagctgggtcCTTCGCCTCGTCCGGCTCGG | Mycgr3\|54855  JGI  Ensembl | KOG1110 Putative steroid membrane receptor Hpr6.6/25-Dx; | heme binding; transition metal ion binding; | 100 % |  |  |
| 10 | ggggacaagtttgtacaaaaaagcaggcttgatgGAACAACTCCTCCTCCCTCA, ggggaccactttgtacaagaaagctgggtcAGATCCCTTCAACACCATGC | Mycgr3\|64933  JGI  Ensembl | KOG4066 Cell growth regulatory protein CGR11; |  | 100 % |  |  |
| 10 | ggggacaagtttgtacaaaaaagcaggcttgatgCTGAAGATATGGAGCATGAA, ggggaccactttgtacaagaaagctgggtcCTTCATGACCCGGTCAAAGC | Mycgr3\|65387  JGI  Ensembl | KOG0420 Ubiquitin-protein ligase; | negative regulation of ubiquitin-protein ligase activity; positive regulation of ubiquitin-protein ligase activity;  post-translational protein modification;  regulation of protein metabolic process;  regulation of ubiquitin-protein ligase activity;  regulation of ubiquitin-protein ligase activity during meiotic cell cycle;  small conjugating protein ligase activity;  ubiquitin-protein ligase activity; | 3.45964 % |  |  |
| 10 | ggggacaagtttgtacaaaaaagcaggcttgatgGCCACCATGACGGATGACTA, ggggaccactttgtacaagaaagctgggtcCAACAGCGAGAGTGTGAGAT | Mycgr3\|66540  JGI  Ensembl |  |  | 100 % |  |  |
| 10 | ggggacaagtttgtacaaaaaagcaggcttgatgAGCAAGAACCGCCCTTCCCA, ggggaccactttgtacaagaaagctgggtcCATCTTCGCCCTCTCCTTCG | Mycgr3\|69151  JGI  Ensembl |  |  | 100 % |  |  |
| 10 | ggggacaagtttgtacaaaaaagcaggcttgatgTCCTCCCCTTCCTCCGAAGA, ggggaccactttgtacaagaaagctgggtcCCCCTCCAAAGCCTTCCCAA | Mycgr3\|69157  JGI  Ensembl |  |  | 100 % |  |  |
| 10 | ggggacaagtttgtacaaaaaagcaggcttgatgGGCCGCAACCGCCCCTCCAC, ggggaccactttgtacaagaaagctgggtcTTCACCGTCTTTCGCCAAAC | Mycgr3\|69686  JGI  Ensembl | KOG4549 Magnesium-dependent phosphatase; |  | 100 % |  |  |
| 10 | ggggacaagtttgtacaaaaaagcaggcttgatgGCGCACGATGGCCAGACATC, ggggaccactttgtacaagaaagctgggtcTGCCTCCTTCTCTTGCATCT | Mycgr3\|72236  JGI  Ensembl |  |  | 100 % |  |  |
| 10 | ggggacaagtttgtacaaaaaagcaggcttgatgGACGAACTCAATCTATCAGC, ggggaccactttgtacaagaaagctgggtcTTGCCTCATCTGCTCCAAGT | Mycgr3\|74555  JGI  Ensembl | KOG3489 Mitochondrial import inner membrane translocase, subunit TIM8; | mitochondrial intermembrane space protein transporter complex; protein import into mitochondrial inner membrane;  protein targeting to mitochondrion; | 100 % |  |  |
| 10 | ggggacaagtttgtacaaaaaagcaggcttgatgGTCCACTCCAGCAGTATGAA, ggggaccactttgtacaagaaagctgggtcATGACCGTCGCAGTGATCGC | Mycgr3\|77248  JGI  Ensembl | KOG3386 Copper transporter; | copper ion transmembrane transporter activity; copper ion transport;  integral to membrane; | 100 % |  |  |
| 10 | ggggacaagtttgtacaaaaaagcaggcttgatgTTTCTCTTACGTACCCCTCG, ggggaccactttgtacaagaaagctgggtcATGGCCCTCCTTACTCGGCT | Mycgr3\|79756  JGI  Ensembl | KOG4210 Nuclear localization sequence binding protein; |  | 100 % | chr\_3:200344 G->A; |  |
| 10 | ggggacaagtttgtacaaaaaagcaggcttgatgGTCAAAATCACCGGTTATGA, ggggaccactttgtacaagaaagctgggtcGGAACCATGCTCGGCGCTCA | Mycgr3\|82703  JGI  Ensembl |  |  | 100 % |  |  |
| 10 | ggggacaagtttgtacaaaaaagcaggcttgatgGCACAAGCACCTTCGATTGA, ggggaccactttgtacaagaaagctgggtcATCCAGTCTCCGGCGAGATG | Mycgr3\|84104  JGI  Ensembl | KOG0504 FOG: Ankyrin repeat; |  | 100 % |  |  |
| 10 | ggggacaagtttgtacaaaaaagcaggcttgatgAAGCTCACAATCGCTTCCCT, ggggaccactttgtacaagaaagctgggtcATTGTCCAGGCCCTGGCATT | Mycgr3\|86867  JGI  Ensembl |  |  | 100 % |  |  |
| 10 | ggggacaagtttgtacaaaaaagcaggcttgatgCGCACAGAATGTGCTTCCCT, ggggaccactttgtacaagaaagctgggtcCCGCAATAGGTTTCGAATCG | Mycgr3\|87062  JGI  Ensembl |  |  | 100 % |  |  |
| 10 | ggggacaagtttgtacaaaaaagcaggcttgatgCCCAATATCCACCCAATCCT, ggggaccactttgtacaagaaagctgggtcCTTGGACGACTTCTTCGCCG | Mycgr3\|87443  JGI  Ensembl |  |  | 100 % |  |  |
| 10 | ggggacaagtttgtacaaaaaagcaggcttgatgCGTTCTACTCTGTTTATCTG, ggggaccactttgtacaagaaagctgggtcGTGGTGGTGGTGGAGGAGAA | Mycgr3\|88665  JGI  Ensembl |  |  | 0 % |  |  |
| 10 | ggggacaagtttgtacaaaaaagcaggcttgatgAAGTCTAAAGCATCGATCGA, ggggaccactttgtacaagaaagctgggtcTGACTTGTGACCAGCATGGG | Mycgr3\|90803  JGI  Ensembl |  |  | 100 % |  |  |
| 10 | ggggacaagtttgtacaaaaaagcaggcttgatgGAGGCCTCTCTCCTTGGACT, ggggaccactttgtacaagaaagctgggtcGATTGTTTCTGGCATGCCCA | Mycgr3\|90973  JGI  Ensembl |  |  | 0 % |  |  |
| 10 | ggggacaagtttgtacaaaaaagcaggcttgatgTCTTACCTCGAGCCCACTAC, ggggaccactttgtacaagaaagctgggtcTCCCACCTTCTCCTTCTGGT | Mycgr3\|91263  JGI  Ensembl |  |  | 100 % |  |  |
| 10 | ggggacaagtttgtacaaaaaagcaggcttgatgGCTATCAGCGCTGCAAACAC, ggggaccactttgtacaagaaagctgggtcGTGCTGCCGCTCTCGTTCTC | Mycgr3\|91395  JGI  Ensembl |  |  | 100 % |  |  |
| 10 | ggggacaagtttgtacaaaaaagcaggcttgatgGCGGAAGCAGCAATGGCAGA, ggggaccactttgtacaagaaagctgggtcATCTCCATGCTCACTGCCAT | Mycgr3\|91792  JGI  Ensembl |  |  | 6.07407 % |  |  |
| 10 | ggggacaagtttgtacaaaaaagcaggcttgatgACAAAATCCAACCTCTACGT, ggggaccactttgtacaagaaagctgggtcCACCTCGTTCCTCGGCGCCG | Mycgr3\|92253  JGI  Ensembl |  |  | 0 % |  |  |
| 10 | ggggacaagtttgtacaaaaaagcaggcttgatgCTTGTCCGCACGTCAATGCC, ggggaccactttgtacaagaaagctgggtcCATCACGGCCAATACACCCG | Mycgr3\|92901  JGI  Ensembl |  |  | 100 % |  |  |
| 10 | ggggacaagtttgtacaaaaaagcaggcttgatgGTCGCCTTCAAACTCCTCAC, ggggaccactttgtacaagaaagctgggtcTGCTCGATAACCCCGTGGGC | Mycgr3\|92990  JGI  Ensembl |  |  | 100 % |  |  |
| 10 | ggggacaagtttgtacaaaaaagcaggcttgatgTCTACAGCAACACCTTACTA, ggggaccactttgtacaagaaagctgggtcAAAGTCCACCGGTGCTGGGT | Mycgr3\|93936  JGI  Ensembl |  |  | 100 % |  |  |
| 10 | ggggacaagtttgtacaaaaaagcaggcttgatgGCGCCCAAAATCCACCAAAA, ggggaccactttgtacaagaaagctgggtcAAACCATTTCGACCACATCT | Mycgr3\|94054  JGI  Ensembl | KOG0721 Molecular chaperone (DnaJ superfamily); | heat shock protein binding; | 100 % |  |  |
| 10 | ggggacaagtttgtacaaaaaagcaggcttgatgTCTTCTCGATCCACCCGCCT, ggggaccactttgtacaagaaagctgggtcTTGCCCACCAGCGTTCTCCG | Mycgr3\|94429  JGI  Ensembl |  |  | 3.39506 % |  |  |
| 10 | ggggacaagtttgtacaaaaaagcaggcttgatgGCCACTCCACGCCGCGAGTT, ggggaccactttgtacaagaaagctgggtcGCACAGCAGCTCTTCAAACG | Mycgr3\|94681  JGI  Ensembl |  |  | 100 % |  |  |
| 10 | ggggacaagtttgtacaaaaaagcaggcttgatgAGCCTCACGCGATCCGCCAG, ggggaccactttgtacaagaaagctgggtcGAAAGCAGGCATCTCACCAG | Mycgr3\|94696  JGI  Ensembl |  |  | 100 % |  |  |
| 10 | ggggacaagtttgtacaaaaaagcaggcttgatgTCAAGTCCAGCAGACCCTTC, ggggaccactttgtacaagaaagctgggtcGCGTTGACTGAATTTGCTGA | Mycgr3\|94736  JGI  Ensembl |  |  | 0 % |  |  |
| 10 | ggggacaagtttgtacaaaaaagcaggcttgatgCCTCTCGACCGAGCTCGCTT, ggggaccactttgtacaagaaagctgggtcAACCCACGTAATCAACTTCT | Mycgr3\|94829  JGI  Ensembl |  |  | 100 % |  |  |
| 10 | ggggacaagtttgtacaaaaaagcaggcttgatgACCGGTCTTTCATCCAAAGT, ggggaccactttgtacaagaaagctgggtcATGCTGCTGGTGCGATAGTA | Mycgr3\|95546  JGI  Ensembl |  |  | 0 % |  |  |
| 10 | ggggacaagtttgtacaaaaaagcaggcttgatgCAGCTCCTCCATCTATACGC, ggggaccactttgtacaagaaagctgggtcACACTTCCATTCTCCGAAAC | Mycgr3\|95579  JGI  Ensembl |  |  | 100 % |  |  |
| 10 | ggggacaagtttgtacaaaaaagcaggcttgatgGCCGAGTACGAAGACAAACT, ggggaccactttgtacaagaaagctgggtcCGAGCGCATGTGCTCCGTTG | Mycgr3\|95609  JGI  Ensembl |  |  | 2.77778 % |  |  |
| 10 | ggggacaagtttgtacaaaaaagcaggcttgatgTTCTCCTCCCTCGTAAACTC, ggggaccactttgtacaagaaagctgggtcCCCAAACATCTTCAACATCT | Mycgr3\|95738  JGI  Ensembl |  |  | 49.12281 % |  |  |
| 10 | ggggacaagtttgtacaaaaaagcaggcttgatgGACAAAAACATCTGGACCGA, ggggaccactttgtacaagaaagctgggtcTGCACTTGGGCGCGGGAAGT | Mycgr3\|95852  JGI  Ensembl |  |  | 3.08642 % |  |  |
| 10 | ggggacaagtttgtacaaaaaagcaggcttgatgCACAGCATTATGGCTTCACC, ggggaccactttgtacaagaaagctgggtcGACGTGGAGGATGTCAAGGT | Mycgr3\|95997  JGI  Ensembl |  |  | 100 % |  |  |
| 10 | ggggacaagtttgtacaaaaaagcaggcttgatgGGAAGCCGACGATTAGCCGA, ggggaccactttgtacaagaaagctgggtcGCCTATATATTTGTACGCCC | Mycgr3\|96127  JGI  Ensembl |  |  | 2.93686 % |  |  |
| 10 | ggggacaagtttgtacaaaaaagcaggcttgatgGACCTCCTCCAATCCGTTCG, ggggaccactttgtacaagaaagctgggtcTCGCCGTGCCTGGACCTGCT | Mycgr3\|96292  JGI  Ensembl | KOG4520 Predicted coiled-coil protein; |  | 100 % |  |  |
| 10 | ggggacaagtttgtacaaaaaagcaggcttgatgGATAACACTGCCGACATCAG, ggggaccactttgtacaagaaagctgggtcCTTCTCAAGCTCGTGCGAGT | Mycgr3\|96367  JGI  Ensembl |  |  | 100 % |  |  |
| 10 | ggggacaagtttgtacaaaaaagcaggcttgatgAGCTCCACCAAAGACTACAC, ggggaccactttgtacaagaaagctgggtcCAAAAGCCAATTACTGCGCT | Mycgr3\|96371  JGI  Ensembl |  |  | 100 % |  |  |
| 10 | ggggacaagtttgtacaaaaaagcaggcttgatgGCCACCAACGCCGTCATTGC, ggggaccactttgtacaagaaagctgggtcATGGAGCCAGTTGCTGCCCT | Mycgr3\|96372  JGI  Ensembl |  |  | 100 % |  |  |
| 10 | ggggacaagtttgtacaaaaaagcaggcttgatgCACTTCAAATACCTCGCCGC, ggggaccactttgtacaagaaagctgggtcCTCCTTCTTCCATCCACCAT | Mycgr3\|96384  JGI  Ensembl | KOG1995 Conserved Zn-finger protein; |  | 5.64399 % |  |  |
| 10 | ggggacaagtttgtacaaaaaagcaggcttgatgCTACGTAACAACAACGACAC, ggggaccactttgtacaagaaagctgggtcCGGCGGCGGTGAGTGGGCGG | Mycgr3\|96643  JGI  Ensembl |  |  | 0 % |  |  |
| 10 | ggggacaagtttgtacaaaaaagcaggcttgatgGCTTCGGAACCAAGCGAGTC, ggggaccactttgtacaagaaagctgggtcGGCCTCGAGTGCCTTGCCGG | Mycgr3\|96736  JGI  Ensembl |  |  | 100 % |  |  |
| 10 | ggggacaagtttgtacaaaaaagcaggcttgatgTCGCACATTTCCAGCTTGCT, ggggaccactttgtacaagaaagctgggtcGTCCTGGCGAAATTGATTCC | Mycgr3\|96751  JGI  Ensembl |  |  | 100 % |  |  |
| 10 | ggggacaagtttgtacaaaaaagcaggcttgatgCTTTCCGCCACCAGCACATC, ggggaccactttgtacaagaaagctgggtcCGGCATCGAAAGCGTCACCA | Mycgr3\|96943  JGI  Ensembl |  |  | 2.84858 % |  |  |
| 10 | ggggacaagtttgtacaaaaaagcaggcttgatgGACTCCGTCAATTTTATTAT, ggggaccactttgtacaagaaagctgggtcTACCGTGTCTATATCGTAAT | Mycgr3\|97594  JGI  Ensembl |  |  | 100 % |  |  |
| 10 | ggggacaagtttgtacaaaaaagcaggcttgatgCAACTCCCGCATGCCGACAG, ggggaccactttgtacaagaaagctgggtcGGGAGCGAGCAGCAAGTGGA | Mycgr3\|97653  JGI  Ensembl |  |  | 100 % |  |  |
| 10 | ggggacaagtttgtacaaaaaagcaggcttgatgTCGCGCAAGTCCTCCATTCC, ggggaccactttgtacaagaaagctgggtcACCATCCAGCGGCGACTTGT | Mycgr3\|97663  JGI  Ensembl |  |  | 100 % |  |  |
| 10 | ggggacaagtttgtacaaaaaagcaggcttgatgCAGGGCGAAAGCGGCATTTT, ggggaccactttgtacaagaaagctgggtcGGCTGTGAGCCTCTGGAGGA | Mycgr3\|97716  JGI  Ensembl |  |  | 100 % |  |  |
| 10 | ggggacaagtttgtacaaaaaagcaggcttgatgGACGTGGGCGGCGGAGAGGT, ggggaccactttgtacaagaaagctgggtcGGAGAGGTGACATTGTTCAC | Mycgr3\|97772  JGI  Ensembl |  |  | 100 % |  |  |
| 10 | ggggacaagtttgtacaaaaaagcaggcttgatgGTTGGATTCCAGTGTTCGAA, ggggaccactttgtacaagaaagctgggtcTCGGGAGTTGCATCTTCTGA | Mycgr3\|97867  JGI  Ensembl |  |  | 100 % |  |  |
| 10 | ggggacaagtttgtacaaaaaagcaggcttgatgTCGGCGCCACTTCTGATGGT, ggggaccactttgtacaagaaagctgggtcTTGTGCTTCGTATCCATGCC | Mycgr3\|97970  JGI  Ensembl |  |  | 100 % |  |  |
| 10 | ggggacaagtttgtacaaaaaagcaggcttgatgCAGCAGGCGTTCAAAGGACT, ggggaccactttgtacaagaaagctgggtcAGCCGGCTGGCAGTTCAGCA | Mycgr3\|97995  JGI  Ensembl |  |  | 100 % |  |  |
| 10 | ggggacaagtttgtacaaaaaagcaggcttgatgATCCAGGAGCGGCCCGTCAC, ggggaccactttgtacaagaaagctgggtcGCTCTTTCTGCCATACTTCC | Mycgr3\|98025  JGI  Ensembl |  |  | 2.89017 % |  |  |
| 10 | ggggacaagtttgtacaaaaaagcaggcttgatgCCAGGAGTCACCGAGCCTCT, ggggaccactttgtacaagaaagctgggtcAGCAGTAAAGTCTCGCAGCT | Mycgr3\|98518  JGI  Ensembl | KOG2351 RNA polymerase II, fourth largest subunit; | DNA-directed RNA polymerase I activity; DNA-directed RNA polymerase II activity;  DNA-directed RNA polymerase III activity;  DNA-directed RNA polymerase activity;  RNA polymerase activity;  transcription; | 100 % |  |  |
| 10 | ggggacaagtttgtacaaaaaagcaggcttgatgCCCAAGCCCAAACGAAACAT, ggggaccactttgtacaagaaagctgggtcATCCTCATCCTCGGAGTCGG | Mycgr3\|101301  JGI  Ensembl | KOG2925 Predicted translation initiation factor related to eIF-1A; | RNA binding; translation initiation factor activity;  translational initiation; | 100 % |  |  |
| 10 | ggggacaagtttgtacaaaaaagcaggcttgatgTTCGCCCGCACCCTCCGCCA, ggggaccactttgtacaagaaagctgggtcTGAGTTTCCACCGCTGCCAC | Mycgr3\|102517  JGI  Ensembl | KOG2992 Nucleolar GTPase/ATPase p130; |  | 100 % |  |  |
| 10 | ggggacaagtttgtacaaaaaagcaggcttgatgGCGGAAGCACCGCTCATCAC, ggggaccactttgtacaagaaagctgggtcCGCCCTGCCACCATCCTCTC | Mycgr3\|102741  JGI  Ensembl | KOG3037 Cell membrane glycoprotein; |  | 100 % |  |  |
| 10 | ggggacaagtttgtacaaaaaagcaggcttgatgCTGCCCGGAGGAGGTCTGGG, ggggaccactttgtacaagaaagctgggtcCCACGCACGTAGGTCTAACG | Mycgr3\|103042  JGI  Ensembl |  |  | 100 % |  |  |
| 10 | ggggacaagtttgtacaaaaaagcaggcttgatgGGACAGAACAGGAAGCGAAC, ggggaccactttgtacaagaaagctgggtcGCACATCGAATCGTCATAAT | Mycgr3\|103157  JGI  Ensembl |  |  | 100 % |  |  |
| 10 | ggggacaagtttgtacaaaaaagcaggcttgatgTGGTCTGAGCATGATTACAA, ggggaccactttgtacaagaaagctgggtcGCCAGTGACCCTATGCGGGA | Mycgr3\|103805  JGI  Ensembl |  |  | 100 % |  |  |
| 10 | ggggacaagtttgtacaaaaaagcaggcttgatgCAAGCGAGTAGTTGCTCTGC, ggggaccactttgtacaagaaagctgggtcATTCATGCCTCCCCAAAGTG | Mycgr3\|103942  JGI  Ensembl |  |  | 100 % |  |  |
| 10 | ggggacaagtttgtacaaaaaagcaggcttgatgGGTTCCTCGGCATCCAAAGC, ggggaccactttgtacaagaaagctgggtcGACCATGTTGATCTCACGCT | Mycgr3\|104020  JGI  Ensembl | KOG3751 Growth factor receptor-bound proteins (GRB7, GRB10, GRB14); |  | 100 % |  |  |
| 10 | ggggacaagtttgtacaaaaaagcaggcttgatgGCGCAGAACATCTTCTACTG, ggggaccactttgtacaagaaagctgggtcAAAGCTGTGCGCTCTCAAGT | Mycgr3\|104256  JGI  Ensembl |  |  | 7.03704 % |  |  |
| 10 | ggggacaagtttgtacaaaaaagcaggcttgatgTCCGCTCGCGGCGGCGCCGG, ggggaccactttgtacaagaaagctgggtcCTCAGCGCTGTTGACAAGAT | Mycgr3\|104440  JGI  Ensembl |  |  | 100 % |  |  |
| 10 | ggggacaagtttgtacaaaaaagcaggcttgatgTTTCTCGCGCTGATCCTCCG, ggggaccactttgtacaagaaagctgggtcCGTCTGCAAAGTGAACCCCA | Mycgr3\|104769  JGI  Ensembl |  |  | 100 % |  |  |
| 10 | ggggacaagtttgtacaaaaaagcaggcttgatgGAGCGACATATACAGCTGGC, ggggaccactttgtacaagaaagctgggtcTTTCGGCCACTTCTTCCATC | Mycgr3\|105063  JGI  Ensembl |  |  | 100 % |  |  |
| 10 | ggggacaagtttgtacaaaaaagcaggcttgatgTGCTGCTCCCAACGTCGCCG, ggggaccactttgtacaagaaagctgggtcACGACATTCCCCGCCACTCT | Mycgr3\|105731  JGI  Ensembl |  |  | 100 % |  |  |
| 10 | ggggacaagtttgtacaaaaaagcaggcttgatgAATCTCTTCCCCATGGCGCA, ggggaccactttgtacaagaaagctgggtcAGACTTACCTCCGAACCCCC | Mycgr3\|105914  JGI  Ensembl | KOG1283 Serine carboxypeptidases; |  | 100 % |  |  |
| 10 | ggggacaagtttgtacaaaaaagcaggcttgatgGGCCTTTTCTCCAGCTCTTC, ggggaccactttgtacaagaaagctgggtcATTGATCGGCTTCGCGTTCT | Mycgr3\|106110  JGI  Ensembl |  |  | 100 % |  |  |
| 10 | ggggacaagtttgtacaaaaaagcaggcttgatgGCTCAACCACGCACAACGCT, ggggaccactttgtacaagaaagctgggtcACCCTCCTTCTTCGCCTTTG | Mycgr3\|106408  JGI  Ensembl | KOG4102 Uncharacterized conserved protein; |  | 100 % |  |  |
| 10 | ggggacaagtttgtacaaaaaagcaggcttgatgCGTCTCCCATACATCCCAGA, ggggaccactttgtacaagaaagctgggtcGCTCATGTCCACGCCATACT | Mycgr3\|107354  JGI  Ensembl |  |  | 100 % |  |  |
| 10 | ggggacaagtttgtacaaaaaagcaggcttgatgTCCACAATCACAGCTCCCAC, ggggaccactttgtacaagaaagctgggtcTCGTCCACCTCTTTTCTTAA | Mycgr3\|107883  JGI  Ensembl | KOG4032 Uncharacterized conserved protein; |  | 0 % |  |  |
| 10 | ggggacaagtttgtacaaaaaagcaggcttgatgAGCGTTGTCGCCGGCGGACC, ggggaccactttgtacaagaaagctgggtcACCACCTTCCCTCTCCATTC | Mycgr3\|109211  JGI  Ensembl |  |  | 0 % |  |  |
| 10 | ggggacaagtttgtacaaaaaagcaggcttgatgTCTTCCCAAACCACCTTTGG, ggggaccactttgtacaagaaagctgggtcATTACCAACAGCCTTCGAGC | Mycgr3\|110160  JGI  Ensembl |  |  | 100 % |  |  |
| 10 | ggggacaagtttgtacaaaaaagcaggcttgatgGCGGCAGGCATGAAAGGCTT, ggggaccactttgtacaagaaagctgggtcAGAGCCATGATCCTCCGCCT | Mycgr3\|110333  JGI  Ensembl |  |  | 5.50964 % |  |  |
| 10 | ggggacaagtttgtacaaaaaagcaggcttgatgACAGTATCATCCTATGACCT, ggggaccactttgtacaagaaagctgggtcGTACATCGGGAATCGGGAAC | Mycgr3\|110841  JGI  Ensembl |  |  | 100 % |  |  |
| 10 | ggggacaagtttgtacaaaaaagcaggcttgatgGGACCCAATGGCTTACCTCC, ggggaccactttgtacaagaaagctgggtcATTCGAGTGCCAGCGATTGC | Mycgr3\|111734  JGI  Ensembl | KOG1015 Transcription regulator XNP/ATRX, DEAD-box superfamily; |  | 3.01659 % |  |  |
| 11 | ggggacaagtttgtacaaaaaagcaggcttgatgACAGAGAACAAAGCCATAAT, ggggaccactttgtacaagaaagctgggtcATCCTTCGGCATAGTGAGTT | Mycgr3\|32378  JGI  Ensembl | KOG3308 Uncharacterized protein of the uridine kinase family; | ATP binding; phosphotransferase activity, alcohol group as acceptor;  uridine kinase activity; | 0 % |  |  |
| 11 | ggggacaagtttgtacaaaaaagcaggcttgatgCATAACTCCAACTCTGACAA, ggggaccactttgtacaagaaagctgggtcGGCATCTTCCACCATAACCT | Mycgr3\|33813  JGI  Ensembl | KOG3354 Gluconate kinase; | carbohydrate metabolic process; gluconokinase activity;  kinase activity; | 2.63158 % |  |  |
| 11 | ggggacaagtttgtacaaaaaagcaggcttgatgGCCACCTCCCTCTCCCTCAC, ggggaccactttgtacaagaaagctgggtcCCTAACCCAATTCCTCACCC | Mycgr3\|36961  JGI  Ensembl |  |  | 0 % |  |  |
| 11 | ggggacaagtttgtacaaaaaagcaggcttgatgCCCAAATCCGCCTCCTTCGA, ggggaccactttgtacaagaaagctgggtcGACCTTCTTCATCTTCTCCA | Mycgr3\|39493  JGI  Ensembl |  |  | 0 % |  |  |
| 11 | ggggacaagtttgtacaaaaaagcaggcttgatgGCCCCCCTCCGTACCCTCAG, ggggaccactttgtacaagaaagctgggtcCAACTTCTCACCACCACCCT | Mycgr3\|40375  JGI  Ensembl | KOG2585 Uncharacterized conserved protein; |  | 0 % |  |  |
| 11 | ggggacaagtttgtacaaaaaagcaggcttgatgCCTCTACAACGAACGCAACC, ggggaccactttgtacaagaaagctgggtcCTCCCAATATCCAAGATTTG | Mycgr3\|42623  JGI  Ensembl |  |  | 0 % |  |  |
| 11 | ggggacaagtttgtacaaaaaagcaggcttgatgGCTCTCTCCTTTCTCCGAGC, ggggaccactttgtacaagaaagctgggtcTTTGCTCTTCTTTCCATCAT | Mycgr3\|42649  JGI  Ensembl |  |  | 0 % |  |  |
| 11 | ggggacaagtttgtacaaaaaagcaggcttgatgGCTCATCAAGGAAAGACAGT, ggggaccactttgtacaagaaagctgggtcGTGCCTCATCAACTTACAGG | Mycgr3\|42990  JGI  Ensembl | KOG0725 Reductases with broad range of substrate specificities; | metabolic process; oxidoreductase activity; | 0 % |  |  |
| 11 | ggggacaagtttgtacaaaaaagcaggcttgatgGCCTCATTACGAACCATACT, ggggaccactttgtacaagaaagctgggtcTATCCGCATGCCTTTGCTCC | Mycgr3\|43109  JGI  Ensembl | KOG2969 Uncharacterized conserved protein; |  | 0 % |  |  |
| 11 | ggggacaagtttgtacaaaaaagcaggcttgatgGCAAAACACGTCGTCTTCGA, ggggaccactttgtacaagaaagctgggtcCTCCGCGTGCTTCACCAGCT | Mycgr3\|44448  JGI  Ensembl |  |  | 0 % |  |  |
| 11 | ggggacaagtttgtacaaaaaagcaggcttgatgGAGTCCACGCACGCTCTTGA, ggggaccactttgtacaagaaagctgggtcACCACTGATGCGATACTGAA | Mycgr3\|44786  JGI  Ensembl |  |  | 0 % |  |  |
| 11 | ggggacaagtttgtacaaaaaagcaggcttgatgGCGTCCAAACGTCCGATCTC, ggggaccactttgtacaagaaagctgggtcCCTCGCCTTCCTCTCTGCCG | Mycgr3\|45248  JGI  Ensembl |  |  | 0 % |  |  |
| 11 | ggggacaagtttgtacaaaaaagcaggcttgatgCCCATCCGCCCAGCAACCTA, ggggaccactttgtacaagaaagctgggtcCTTCTCCACATACTCCTCAA | Mycgr3\|49004  JGI  Ensembl |  |  | 0 % |  |  |
| 11 | ggggacaagtttgtacaaaaaagcaggcttgatgAGTTCGTGTCGGATCGAAAG, ggggaccactttgtacaagaaagctgggtcCAGTGTATTCCTCATCTGCC | Mycgr3\|51492  JGI  Ensembl |  |  | 0 % |  |  |
| 11 | ggggacaagtttgtacaaaaaagcaggcttgatgCCAAAGACCATCTTGATATC, ggggaccactttgtacaagaaagctgggtcCCACTCAATGGGCTCGCCGA | Mycgr3\|66113  JGI  Ensembl | KOG1611 Predicted short chain-type dehydrogenase; | F420H2 dehydrogenase activity; malolactic enzyme activity;  metabolic process;  oxidoreductase activity;  regulation of oxidoreductase activity;  sulfur oxygenase reductase activity; | 0 % |  |  |
| 11 | ggggacaagtttgtacaaaaaagcaggcttgatgATCTGCCAAAGATGCCTGCA, ggggaccactttgtacaagaaagctgggtcCCGCATCTCCTTCAAGAAAT | Mycgr3\|66222  JGI  Ensembl | KOG3435 Mitochondrial/chloroplast ribosomal protein L54/L37; |  | 0 % |  |  |
| 11 | ggggacaagtttgtacaaaaaagcaggcttgatgGGGAAGGAAGAGCAGCAGGA, ggggaccactttgtacaagaaagctgggtcTCCTTCCACCGAGAGTTGTC | Mycgr3\|67443  JGI  Ensembl | KOG4018 Uncharacterized conserved protein, contains RWD domain; |  | 0 % |  |  |
| 11 | ggggacaagtttgtacaaaaaagcaggcttgatgGCGAGCGCAGTATCGAACCT, ggggaccactttgtacaagaaagctgggtcCTTCTTTCCGTAGCACACTA | Mycgr3\|68480  JGI  Ensembl | KOG1270 Methyltransferases; |  | 0 % |  |  |
| 11 | ggggacaagtttgtacaaaaaagcaggcttgatgTCCATCTTGGCGCCCCTACG, ggggaccactttgtacaagaaagctgggtcTCGCCCCTCCACCTTCTTCC | Mycgr3\|69353  JGI  Ensembl | KOG4073 Pterin carbinolamine dehydratase PCBD/dimerization cofactor of HNF1; |  | 0 % |  |  |
| 11 | ggggacaagtttgtacaaaaaagcaggcttgatgGCCAATCAGAATGCCACAAC, ggggaccactttgtacaagaaagctgggtcCTTGTAGATCTCATCAAAAG | Mycgr3\|72054  JGI  Ensembl | KOG3314 Ku70-binding protein; | extracellular region; hormone activity;  metallopeptidase activity;  proteolysis;  zinc ion binding; | 0 % |  |  |
| 11 | ggggacaagtttgtacaaaaaagcaggcttgatgGCCGACCTCGACGCAACCCC, ggggaccactttgtacaagaaagctgggtcCGATCGGTTCCACATTCTTG | Mycgr3\|72400  JGI  Ensembl | KOG3195 Uncharacterized membrane protein NPD008/CGI-148; | integral to membrane; | 0 % |  |  |
| 11 | ggggacaagtttgtacaaaaaagcaggcttgatgGCCTCCCTCCTAACCTCCCA, ggggaccactttgtacaagaaagctgggtcCCTCATCAACCCACCCAAGT | Mycgr3\|73067  JGI  Ensembl |  |  | 0 % |  |  |
| 11 | ggggacaagtttgtacaaaaaagcaggcttgatgATATTGAGCTTGCATCCGCA, ggggaccactttgtacaagaaagctgggtcACTACCTAGCTTTGCCTCGA | Mycgr3\|74230  JGI  Ensembl |  |  | 0 % |  |  |
| 11 | ggggacaagtttgtacaaaaaagcaggcttgatgCGCAAAATCCCCACCCACCC, ggggaccactttgtacaagaaagctgggtcCTTCTTCTTCTTATTGACCT | Mycgr3\|74659  JGI  Ensembl | KOG1698 Mitochondrial/chloroplast ribosomal protein L19; | intracellular; ribosome; structural constituent of ribosome;  translation; | 0 % |  |  |
| 11 | ggggacaagtttgtacaaaaaagcaggcttgatgGGCGAAACGGAGAAACCCGC, ggggaccactttgtacaagaaagctgggtcCCCCAAAGTCTTCATCTTCT | Mycgr3\|75437  JGI  Ensembl | KOG1150 Predicted molecular chaperone (DnaJ superfamily); | heat shock protein binding; protein folding;  unfolded protein binding; | 0 % |  |  |
| 11 | ggggacaagtttgtacaaaaaagcaggcttgatgTCATCTCCTCCGCTCAACGT, ggggaccactttgtacaagaaagctgggtcTCTTCCACCAGCAACATCAA | Mycgr3\|75800  JGI  Ensembl | KOG0725 Reductases with broad range of substrate specificities; | 3-oxoacyl-[acyl-carrier-protein] reductase activity; metabolic process;  oxidoreductase activity; | 0 % |  |  |
| 11 | ggggacaagtttgtacaaaaaagcaggcttgatgGCCCAAGCCTACAAACCCGG, ggggaccactttgtacaagaaagctgggtcCCACTTCGCCTCCGGACTCG | Mycgr3\|76134  JGI  Ensembl | KOG3221 Glycolipid transfer protein; |  | 0 % |  |  |
| 11 | ggggacaagtttgtacaaaaaagcaggcttgatgCGCCTCTCAACCCTCGACAT, ggggaccactttgtacaagaaagctgggtcAAGTGGCAGAAATTCCAAGA | Mycgr3\|78893  JGI  Ensembl |  |  | 0 % |  |  |
| 11 | ggggacaagtttgtacaaaaaagcaggcttgatgCAGTTCTCGAGCGGTGTAAA, ggggaccactttgtacaagaaagctgggtcTTTGCCACTTCCGACAACGA | Mycgr3\|80260  JGI  Ensembl |  |  | 0 % |  |  |
| 11 | ggggacaagtttgtacaaaaaagcaggcttgatgCCAATGCTCGACCCCGCAAC, ggggaccactttgtacaagaaagctgggtcTTGAAACTCTCCAAATTCTC | Mycgr3\|87405  JGI  Ensembl | KOG2500 Uncharacterized conserved protein; |  | 0 % |  |  |
| 11 | ggggacaagtttgtacaaaaaagcaggcttgatgCTCCCCACCCCAACAACCTC, ggggaccactttgtacaagaaagctgggtcCTTGTCCCTCCAAATCCTCA | Mycgr3\|87952  JGI  Ensembl | KOG3191 Predicted N6-DNA-methyltransferase; | methylation; methyltransferase activity;  nucleic acid binding;  site-specific DNA-methyltransferase (adenine-specific) activity; | 0 % |  |  |
| 11 | ggggacaagtttgtacaaaaaagcaggcttgatgGTGACGACCGCCTCCAGCGA, ggggaccactttgtacaagaaagctgggtcAACCAGCTTCAGCACTGATG | Mycgr3\|89515  JGI  Ensembl |  |  | 0 % |  |  |
| 11 | ggggacaagtttgtacaaaaaagcaggcttgatgGCTCCCACCGACAAAGTCAC, ggggaccactttgtacaagaaagctgggtcGATCCAAGTCTGCGCAGTCG | Mycgr3\|90590  JGI  Ensembl |  |  | 0 % |  |  |
| 11 | ggggacaagtttgtacaaaaaagcaggcttgatgGTTGACTCATCCCTACCATA, ggggaccactttgtacaagaaagctgggtcGGCCAACATCTTTGTGATCC | Mycgr3\|90696  JGI  Ensembl |  |  | 0 % |  |  |
| 11 | ggggacaagtttgtacaaaaaagcaggcttgatgGCCAGCACTCCGAATCAGCA, ggggaccactttgtacaagaaagctgggtcATTCCCACGGACAAACTCTG | Mycgr3\|91211  JGI  Ensembl |  |  | 0 % |  |  |
| 11 | ggggacaagtttgtacaaaaaagcaggcttgatgGACCAGTCGCCCGCCGCTCC, ggggaccactttgtacaagaaagctgggtcTCCTCGACGTCCTGCTCGAT | Mycgr3\|91789  JGI  Ensembl |  |  | 0 % |  |  |
| 11 | ggggacaagtttgtacaaaaaagcaggcttgatgGCAGACTCAATCGATCCATT, ggggaccactttgtacaagaaagctgggtcATCTGCGCCCTTGTCCCCGT | Mycgr3\|91862  JGI  Ensembl |  |  | 0 % |  |  |
| 11 | ggggacaagtttgtacaaaaaagcaggcttgatgACCATCCCACCCTTCGAAGC, ggggaccactttgtacaagaaagctgggtcCAACATCCCCTTCCCGACCA | Mycgr3\|91870  JGI  Ensembl | KOG4656 Copper chaperone for superoxide dismutase; | carbon-oxygen lyase activity; carbon-oxygen lyase activity, acting on phosphates;  carbon-oxygen lyase activity, acting on polysaccharides;  metal ion binding;  metal ion transport;  superoxide metabolic process; | 25.35948 % |  |  |
| 11 | ggggacaagtttgtacaaaaaagcaggcttgatgTCCGACTCCCACTCTCACGA, ggggaccactttgtacaagaaagctgggtcTGTAGTATGCGTCGGATCAA | Mycgr3\|92057  JGI  Ensembl |  |  | 0 % |  |  |
| 11 | ggggacaagtttgtacaaaaaagcaggcttgatgGCCGCAGCTCTCGTCTTTGG, ggggaccactttgtacaagaaagctgggtcCGCCTTCCACTCATCCCTTC | Mycgr3\|92115  JGI  Ensembl | KOG4039 Serine/threonine kinase TIP30/CC3; |  | 0 % |  |  |
| 11 | ggggacaagtttgtacaaaaaagcaggcttgatgTCGGATTCTCCTCTCAAATC, ggggaccactttgtacaagaaagctgggtcCATGCGCTGCTGTCTCCTCT | Mycgr3\|92174  JGI  Ensembl |  |  | 0 % |  |  |
| 11 | ggggacaagtttgtacaaaaaagcaggcttgatgGCCTCCATAACGCGCCCCAT, ggggaccactttgtacaagaaagctgggtcGCCCCTCCCTTCCTCCCAAA | Mycgr3\|92547  JGI  Ensembl | KOG1187 Serine/threonine protein kinase; |  | 0 % |  |  |
| 11 | ggggacaagtttgtacaaaaaagcaggcttgatgCACTCCGACGAAGAGAGCAG, ggggaccactttgtacaagaaagctgggtcCAGGAAAGCCTTGATGTGAT | Mycgr3\|92813  JGI  Ensembl |  |  | 0 % |  |  |
| 11 | ggggacaagtttgtacaaaaaagcaggcttgatgACAAGCACCGACAGGGCTGC, ggggaccactttgtacaagaaagctgggtcGGTCCGAATATTCTTCCAGC | Mycgr3\|92924  JGI  Ensembl |  |  | 0 % |  |  |
| 11 | ggggacaagtttgtacaaaaaagcaggcttgatgTCACTTCAGACCGACTGCGA, ggggaccactttgtacaagaaagctgggtcACTCGTCTCCGGTTCGCCGA | Mycgr3\|93449  JGI  Ensembl |  |  | 0 % |  |  |
| 11 | ggggacaagtttgtacaaaaaagcaggcttgatgGCCGGCCAGACTTCAATTCC, ggggaccactttgtacaagaaagctgggtcCATCTTCGAAATCTTCGCCA | Mycgr3\|93775  JGI  Ensembl |  |  | 0 % |  |  |
| 11 | ggggacaagtttgtacaaaaaagcaggcttgatgGACGAACTTCTCGAAACCCA, ggggaccactttgtacaagaaagctgggtcTATCGTAAGCCTCGTCGCTG | Mycgr3\|93820  JGI  Ensembl | KOG0663 Protein kinase PITSLRE and related kinases; |  | 0 % |  |  |
| 11 | ggggacaagtttgtacaaaaaagcaggcttgatgTCACAAAAGCGCAAGCGCGA, ggggaccactttgtacaagaaagctgggtcCTTTTTCCCAAAATTCTTCA | Mycgr3\|93943  JGI  Ensembl | KOG3095 Transcription initiation factor IIE, beta subunit; |  | 0 % |  |  |
| 11 | ggggacaagtttgtacaaaaaagcaggcttgatgGATTCCCACCAACAAGACGA, ggggaccactttgtacaagaaagctgggtcCATCACCATCTCGTCATCCC | Mycgr3\|94401  JGI  Ensembl |  |  | 0 % |  |  |
| 11 | ggggacaagtttgtacaaaaaagcaggcttgatgTGTGACTTGAGCGACACCGC, ggggaccactttgtacaagaaagctgggtcCTTCGCCGAGCGCTGCGAGA | Mycgr3\|94628  JGI  Ensembl |  |  | 2.92072 % |  |  |
| 11 | ggggacaagtttgtacaaaaaagcaggcttgatgGCGTCCTACAACCCACGCCC, ggggaccactttgtacaagaaagctgggtcCCAGTGCCGCACCACAATCG | Mycgr3\|94714  JGI  Ensembl |  |  | 0 % |  |  |
| 11 | ggggacaagtttgtacaaaaaagcaggcttgatgGACACCACAAAATCAAGCGC, ggggaccactttgtacaagaaagctgggtcCTTCTCCACCCAGCGAAGCA | Mycgr3\|94767  JGI  Ensembl |  |  | 0 % |  |  |
| 11 | ggggacaagtttgtacaaaaaagcaggcttgatgGCCGCCATAATCGTCAATCT, ggggaccactttgtacaagaaagctgggtcCCCAAAACTCATCATGATGC | Mycgr3\|94796  JGI  Ensembl |  |  | 0 % |  |  |
| 11 | ggggacaagtttgtacaaaaaagcaggcttgatgACAGCTCTAGGAACACTCGA, ggggaccactttgtacaagaaagctgggtcAGGAGTATCCATGATCTCCG | Mycgr3\|94832  JGI  Ensembl |  |  | 0 % |  |  |
| 11 | ggggacaagtttgtacaaaaaagcaggcttgatgTCCACCTCCGAGCACAACAA, ggggaccactttgtacaagaaagctgggtcCCACTCCCTCTGCCCATCCC | Mycgr3\|94910  JGI  Ensembl |  |  | 0 % |  |  |
| 11 | ggggacaagtttgtacaaaaaagcaggcttgatgGCCTTCACTCGCTCCACACT, ggggaccactttgtacaagaaagctgggtcGTGGAGAAGCTCGTACTGCA | Mycgr3\|96080  JGI  Ensembl | KOG2177 Predicted E3 ubiquitin ligase; | protein binding; zinc ion binding; | 0 % |  |  |
| 11 | ggggacaagtttgtacaaaaaagcaggcttgatgACAGCAAACACGGCCAACAT, ggggaccactttgtacaagaaagctgggtcATCGAATCCAGGAACGTCGG | Mycgr3\|96366  JGI  Ensembl |  |  | 100 % |  |  |
| 11 | ggggacaagtttgtacaaaaaagcaggcttgatgGCGGAAGTGCAGCTTGCGAT, ggggaccactttgtacaagaaagctgggtcGTAATGATCGTGGAAGCCGG | Mycgr3\|96527  JGI  Ensembl | KOG0516 Dystonin, GAS (Growth-arrest-specific protein), and related proteins; |  | 0 % |  |  |
| 11 | ggggacaagtttgtacaaaaaagcaggcttgatgCCAGGCAAAACCCTCCTCGT, ggggaccactttgtacaagaaagctgggtcCTCATGAACCTCCTCCTCCC | Mycgr3\|97265  JGI  Ensembl | KOG1201 Hydroxysteroid 17-beta dehydrogenase 11; | 3-oxoacyl-[acyl-carrier-protein] reductase activity; | 0 % |  |  |
| 11 | ggggacaagtttgtacaaaaaagcaggcttgatgGCACCCACATCCATCGAAAA, ggggaccactttgtacaagaaagctgggtcCGATTGTCCTCCTACCTGCT | Mycgr3\|97372  JGI  Ensembl |  |  | 0 % |  |  |
| 11 | ggggacaagtttgtacaaaaaagcaggcttgatgGAACACTCTTCATTCGACCT, ggggaccactttgtacaagaaagctgggtcTGATAGCTGCTCGACCACAG | Mycgr3\|97525  JGI  Ensembl |  |  | 0 % |  |  |
| 11 | ggggacaagtttgtacaaaaaagcaggcttgatgCCCCGCCTCCATCTCATCTT, ggggaccactttgtacaagaaagctgggtcACGGAGCATCGCTCGGAACT | Mycgr3\|97649  JGI  Ensembl |  |  | 0 % |  |  |
| 11 | ggggacaagtttgtacaaaaaagcaggcttgatgCAAGAGCAGAAGGAGGACAT, ggggaccactttgtacaagaaagctgggtcACAGAGTCGCAGATCCGCCC | Mycgr3\|97902  JGI  Ensembl | KOG1474 Transcription initiation factor TFIID, subunit BDF1 and related bromodomain proteins; |  | 0 % |  |  |
| 11 | ggggacaagtttgtacaaaaaagcaggcttgatgGTCGATGCGAATCCCGGCCA, ggggaccactttgtacaagaaagctgggtcGCTTTTCTCGTCCATCTCAT | Mycgr3\|97968  JGI  Ensembl |  |  | 0 % |  |  |
| 11 | ggggacaagtttgtacaaaaaagcaggcttgatgTCGGGAATGTTGGCGTCGAT, ggggaccactttgtacaagaaagctgggtcGTCCAGAGTTCGGCGTCCTT | Mycgr3\|98073  JGI  Ensembl |  |  | 0 % |  |  |
| 11 | ggggacaagtttgtacaaaaaagcaggcttgatgCCCAAATCGAAGCGCTCCCG, ggggaccactttgtacaagaaagctgggtcACCCTCCTCCATAGCGCCGG | Mycgr3\|99074  JGI  Ensembl | KOG0816 Protein involved in mRNA turnover; | intracellular; ribosome biogenesis and assembly; | 0 % |  |  |
| 11 | ggggacaagtttgtacaaaaaagcaggcttgatgATGAGTACGGCGGCGGCGAT, ggggaccactttgtacaagaaagctgggtcCGCCGCCTCCGTTCTCCTCA | Mycgr3\|101983  JGI  Ensembl |  |  | 0 % |  |  |
| 11 | ggggacaagtttgtacaaaaaagcaggcttgatgCCTGGAGAATCCATGGATCG, ggggaccactttgtacaagaaagctgggtcTCTCTGGGACAGGGGCGAAT | Mycgr3\|102155  JGI  Ensembl |  |  | 0 % |  |  |
| 11 | ggggacaagtttgtacaaaaaagcaggcttgatgCCTCCCAAGCCGAAGTCAGG, ggggaccactttgtacaagaaagctgggtcTCCGAAGCTCACTCGTTTTC | Mycgr3\|103315  JGI  Ensembl | KOG4475 FOG: Immunoglobin and related proteins; |  | 3.58974 % |  |  |
| 11 | ggggacaagtttgtacaaaaaagcaggcttgatgTCCACCGCCACCCCTCGAAT, ggggaccactttgtacaagaaagctgggtcAGCAGACAACTTCGCTGCCC | Mycgr3\|103418  JGI  Ensembl |  |  | 0 % |  |  |
| 11 | ggggacaagtttgtacaaaaaagcaggcttgatgTCCGCCAACCGCAACCCAGA, ggggaccactttgtacaagaaagctgggtcATTCCGCTCCGAAGCAACAC | Mycgr3\|103423  JGI  Ensembl |  |  | 0 % |  |  |
| 11 | ggggacaagtttgtacaaaaaagcaggcttgatgGCCTCCCAATACCAAGGTCC, ggggaccactttgtacaagaaagctgggtcCCATATCTTATCATCTCTTT | Mycgr3\|103676  JGI  Ensembl |  |  | 0 % |  |  |
| 11 | ggggacaagtttgtacaaaaaagcaggcttgatgCAGTCCTTCATCCTCTTGGC, ggggaccactttgtacaagaaagctgggtcCTTCGAGCAGACAACGCCGT | Mycgr3\|103792  JGI  Ensembl |  |  | 3.37423 % |  |  |
| 11 | ggggacaagtttgtacaaaaaagcaggcttgatgCACATCGATGACACCCGCGA, ggggaccactttgtacaagaaagctgggtcATCGACCTCCATCGCATCGG | Mycgr3\|104494  JGI  Ensembl |  |  | 0 % |  |  |
| 11 | ggggacaagtttgtacaaaaaagcaggcttgatgCATCATTCCCTCAACTCTAC, ggggaccactttgtacaagaaagctgggtcCAAGCAAACTCTTGTCCGTC | Mycgr3\|105090  JGI  Ensembl |  |  | 0 % |  |  |
| 11 | ggggacaagtttgtacaaaaaagcaggcttgatgTTGTCCTCCCTTCTCAGCCT, ggggaccactttgtacaagaaagctgggtcCGAGTTGCCTCTCAATCCCT | Mycgr3\|105659  JGI  Ensembl |  |  | 0 % |  |  |
| 11 | ggggacaagtttgtacaaaaaagcaggcttgatgTCACGACAGAAGACCGTGGC, ggggaccactttgtacaagaaagctgggtcTTTCCATCGAATCATGTTCT | Mycgr3\|106138  JGI  Ensembl |  |  | 0 % |  |  |
| 11 | ggggacaagtttgtacaaaaaagcaggcttgatgCGTCTCAACTGCTTTCGTGC, ggggaccactttgtacaagaaagctgggtcGCCGATCATGTCAATCTCGT | Mycgr3\|106356  JGI  Ensembl |  |  | 0 % |  |  |
| 11 | ggggacaagtttgtacaaaaaagcaggcttgatgCGGCCACTCGCATTGCATTC, ggggaccactttgtacaagaaagctgggtcACACCAACATCTCCAGAAGA | Mycgr3\|106721  JGI  Ensembl |  |  | 0 % |  |  |
| 11 | ggggacaagtttgtacaaaaaagcaggcttgatgCCTCCCGTGCCCGTGCACAT, ggggaccactttgtacaagaaagctgggtcTGATTCGCGATTGATCCTCT | Mycgr3\|107168  JGI  Ensembl | KOG1187 Serine/threonine protein kinase; |  | 0 % |  |  |
| 11 | ggggacaagtttgtacaaaaaagcaggcttgatgCGCTCCTCAATCTATCTCCT, ggggaccactttgtacaagaaagctgggtcAATGTAAATCGTCCCAGCAA | Mycgr3\|107289  JGI  Ensembl |  |  | 0 % |  |  |
| 11 | ggggacaagtttgtacaaaaaagcaggcttgatgTCAGTCCTCCGATCACGACT, ggggaccactttgtacaagaaagctgggtcTCTCTTCTTATCGGGATCCT | Mycgr3\|107387  JGI  Ensembl |  |  | 0 % |  |  |
| 11 | ggggacaagtttgtacaaaaaagcaggcttgatgGACCACGGCGGGATGGACAT, ggggaccactttgtacaagaaagctgggtcATTACAGCAAAAGGTCAGAT | Mycgr3\|107812  JGI  Ensembl | KOG3386 Copper transporter; | copper ion transmembrane transporter activity; copper ion transport;  integral to membrane; | 0 % |  |  |
| 11 | ggggacaagtttgtacaaaaaagcaggcttgatgTCATCAACCCTGACCCAAAC, ggggaccactttgtacaagaaagctgggtcACCCGAACCACCTCCTCCTC | Mycgr3\|107837  JGI  Ensembl | KOG1283 Serine carboxypeptidases; |  | 0 % |  |  |
| 11 | ggggacaagtttgtacaaaaaagcaggcttgatgGCGAAAACACCTAAGGTCAA, ggggaccactttgtacaagaaagctgggtcCGTAACCTCGTCGACATCCT | Mycgr3\|108001  JGI  Ensembl |  |  | 0 % |  |  |
| 11 | ggggacaagtttgtacaaaaaagcaggcttgatgGCCACTGCAACCGCTCCCAA, ggggaccactttgtacaagaaagctgggtcCATCTTCACTCCCCTGATCT | Mycgr3\|108479  JGI  Ensembl | KOG0406 Glutathione S-transferase; | glutathione transferase activity; | 0 % |  |  |
| 11 | ggggacaagtttgtacaaaaaagcaggcttgatgGCACCCATCCGCCCCGCCAA, ggggaccactttgtacaagaaagctgggtcATCCTTCGCAAAAGCCTTGT | Mycgr3\|110296  JGI  Ensembl |  |  | 0 % |  |  |
| 11 | ggggacaagtttgtacaaaaaagcaggcttgatgCCTTTCATCCACGGAATCGA, ggggaccactttgtacaagaaagctgggtcTGGGTGCTTTGGTGAAGTCG | Mycgr3\|110884  JGI  Ensembl |  |  | 0 % |  |  |
| 11 | ggggacaagtttgtacaaaaaagcaggcttgatgGCCTCCAATGGCGCTACGAA, ggggaccactttgtacaagaaagctgggtcTGTTGGTTTGAAGCATTCCG | Mycgr3\|111064  JGI  Ensembl | KOG3109 Haloacid dehalogenase-like hydrolase; | catalytic activity; metabolic process;  phosphoglycolate phosphatase activity; | 0 % |  |  |
| 11 | ggggacaagtttgtacaaaaaagcaggcttgatgCCCACCACGGAAATCTTCAT, ggggaccactttgtacaagaaagctgggtcCTGCTCCCTCACAAACTCCA | Mycgr3\|111173  JGI  Ensembl |  |  | 0 % |  |  |
| 12 | ggggacaagtttgtacaaaaaagcaggcttgatgTCCAAACCACACATCATTCT, ggggaccactttgtacaagaaagctgggtcCATTCTCTCGCCAGCAAAGC | Mycgr3\|32167  JGI  Ensembl |  |  | 100 % |  |  |
| 12 | ggggacaagtttgtacaaaaaagcaggcttgatgGCCTCTGTGTGCCTGGTCGG, ggggaccactttgtacaagaaagctgggtcAAGATTCCACTCCGTCCTGC | Mycgr3\|33027  JGI  Ensembl | KOG4039 Serine/threonine kinase TIP30/CC3; |  | 100 % |  |  |
| 12 | ggggacaagtttgtacaaaaaagcaggcttgatgGCCACCTCAATCCCCTGGAC, ggggaccactttgtacaagaaagctgggtcCGTAGCCTGACGTACTACGA | Mycgr3\|34317  JGI  Ensembl |  |  | 100 % |  |  |
| 12 | ggggacaagtttgtacaaaaaagcaggcttgatgATCTATATTGATGAAGGTGA, ggggaccactttgtacaagaaagctgggtcAAGCGAGCCTTCGGGCCACT | Mycgr3\|35346  JGI  Ensembl |  |  | 100 % |  |  |
| 12 | ggggacaagtttgtacaaaaaagcaggcttgatgGCGCCAAACATTTCCAAACG, ggggaccactttgtacaagaaagctgggtcAGCCTCCGTCAATTCCACGA | Mycgr3\|35421  JGI  Ensembl |  |  | 100 % |  |  |
| 12 | ggggacaagtttgtacaaaaaagcaggcttgatgACAACCGCCCACAGACCGAC, ggggaccactttgtacaagaaagctgggtcTCGCACATATTTGTCCATGA | Mycgr3\|39745  JGI  Ensembl | KOG3228 Uncharacterized conserved protein; | nuclear mRNA splicing, via spliceosome; spliceosome; | 100 % |  |  |
| 12 | ggggacaagtttgtacaaaaaagcaggcttgatgGCTGACTCGGGTACTACTTC, ggggaccactttgtacaagaaagctgggtcACCTCTGGTCTTGCCAAACA | Mycgr3\|40933  JGI  Ensembl | KOG3200 Uncharacterized conserved protein; |  | 100 % |  |  |
| 12 | ggggacaagtttgtacaaaaaagcaggcttgatgCCTCCGCTTCCTGCAGCACG, ggggaccactttgtacaagaaagctgggtcTGGCTCAGCCAGCTCCTCAA | Mycgr3\|41530  JGI  Ensembl | KOG1643 Triosephosphate isomerase; | metabolic process; triose-phosphate isomerase activity; | 100 % |  |  |
| 12 | ggggacaagtttgtacaaaaaagcaggcttgatgACTTCCGAAAATCCCATCGT, ggggaccactttgtacaagaaagctgggtcCTTGATCTCCGTCCACGGGC | Mycgr3\|42293  JGI  Ensembl |  |  | 100 % |  |  |
| 12 | ggggacaagtttgtacaaaaaagcaggcttgatgAATGCAGATGAGGCGGCGTT, ggggaccactttgtacaagaaagctgggtcAAGGGTCCTTGACATGTACT | Mycgr3\|42625  JGI  Ensembl |  |  | 0 % |  |  |
| 12 | ggggacaagtttgtacaaaaaagcaggcttgatgCATGCTGCATCGCATGTTCT, ggggaccactttgtacaagaaagctgggtcTTCTTCGGTCTTGCAATCCC | Mycgr3\|44154  JGI  Ensembl | KOG1204 Predicted dehydrogenase; | F420H2 dehydrogenase activity; malolactic enzyme activity;  metabolic process;  oxidoreductase activity;  regulation of oxidoreductase activity;  sulfur oxygenase reductase activity; | 100 % |  |  |
| 12 | ggggacaagtttgtacaaaaaagcaggcttgatgGCGCGTGCGGTGCCACCCGC, ggggaccactttgtacaagaaagctgggtcACACTCTTGAGAAAGCGTCA | Mycgr3\|44586  JGI  Ensembl |  |  | 100 % |  |  |
| 12 | ggggacaagtttgtacaaaaaagcaggcttgatgTCGAGCAACAAGGAAGTGGT, ggggaccactttgtacaagaaagctgggtcCCAAGGCTGAATCTCATCCG | Mycgr3\|47551  JGI  Ensembl | KOG1208 Dehydrogenases with different specificities (related to short-chain alcohol dehydrogenases); | metabolic process; oxidoreductase activity; | 100 % |  |  |
| 12 | ggggacaagtttgtacaaaaaagcaggcttgatgTCGACCTCAACTTCCACCCG, ggggaccactttgtacaagaaagctgggtcCTTCAATACGACAATCACCA | Mycgr3\|47666  JGI  Ensembl | KOG3202 SNARE protein TLG1/Syntaxin 6; |  | 0 % |  |  |
| 12 | ggggacaagtttgtacaaaaaagcaggcttgatgGCTCCCAACTCCCGCATCAT, ggggaccactttgtacaagaaagctgggtcCAACTTCCCAACCCCCTCAT | Mycgr3\|49444  JGI  Ensembl | KOG4754 Predicted phosphoglycerate mutase; | catalytic activity; metabolic process; | 100 % |  |  |
| 12 | ggggacaagtttgtacaaaaaagcaggcttgatgCACTACACCGCCGCTCTGCG, ggggaccactttgtacaagaaagctgggtcTATCGAAGCCGACTTACTGT | Mycgr3\|49574  JGI  Ensembl |  |  | 100 % |  |  |
| 12 | ggggacaagtttgtacaaaaaagcaggcttgatgTCGTATTCTCGACCACAATC, ggggaccactttgtacaagaaagctgggtcAAGCAAGACGTTCTCCTGTA | Mycgr3\|50020  JGI  Ensembl |  |  | 100 % |  |  |
| 12 | ggggacaagtttgtacaaaaaagcaggcttgatgGCCGAGCCGCAACCATCGAA, ggggaccactttgtacaagaaagctgggtcGACAGCACTCGAAACCCACG | Mycgr3\|53017  JGI  Ensembl |  |  | 100 % |  |  |
| 12 | ggggacaagtttgtacaaaaaagcaggcttgatgGCGACCATGTCAATCACCAG, ggggaccactttgtacaagaaagctgggtcAGGCGTCGTGGATGGCATTC | Mycgr3\|66529  JGI  Ensembl | KOG3422 Mitochondrial ribosomal protein L16; | intracellular; ribosome; structural constituent of ribosome;  translation; | 0 % |  |  |
| 12 | ggggacaagtttgtacaaaaaagcaggcttgatgGCCACCGAACCGATCATTCC, ggggaccactttgtacaagaaagctgggtcGTCTTCTTTCTTCTTGTCCG | Mycgr3\|68720  JGI  Ensembl |  |  | 100 % |  |  |
| 12 | ggggacaagtttgtacaaaaaagcaggcttgatgTCTCAAGTAAAGCCCTTCGC, ggggaccactttgtacaagaaagctgggtcATGTCCACTCGCCAACATCC | Mycgr3\|71140  JGI  Ensembl | KOG1200 Mitochondrial/plastidial beta-ketoacyl-ACP reductase; | 3-oxoacyl-[acyl-carrier-protein] reductase activity; metabolic process;  oxidoreductase activity; | 100 % |  |  |
| 12 | ggggacaagtttgtacaaaaaagcaggcttgatgGCCGACCACCCACCAACCAC, ggggaccactttgtacaagaaagctgggtcCGCTGCCGGCTTCCTCACGA | Mycgr3\|72253  JGI  Ensembl |  |  | 100 % |  |  |
| 12 | ggggacaagtttgtacaaaaaagcaggcttgatgGAGAAACCACCTGAATACGC, ggggaccactttgtacaagaaagctgggtcCATGACAGCAAAAGTCGCCC | Mycgr3\|74323  JGI  Ensembl |  |  | 100 % |  |  |
| 12 | ggggacaagtttgtacaaaaaagcaggcttgatgGAAGCCCGCCGTCCTCCAGA, ggggaccactttgtacaagaaagctgggtcAAAAATCCCCATCCGTCCAC | Mycgr3\|74422  JGI  Ensembl | KOG4493 Uncharacterized conserved protein; |  | 100 % |  |  |
| 12 | ggggacaagtttgtacaaaaaagcaggcttgatgGACTCCGCCATCGACCTCTC, ggggaccactttgtacaagaaagctgggtcTTCCCACTCCGTCCCCTTGA | Mycgr3\|75755  JGI  Ensembl | KOG0795 Chorismate mutase; | aromatic amino acid family biosynthetic process; chorismate mutase activity; | 100 % |  |  |
| 12 | ggggacaagtttgtacaaaaaagcaggcttgatgGACAACCCCATCCACGAAAT, ggggaccactttgtacaagaaagctgggtcATCGACCTTCTTCACCCCCT | Mycgr3\|76127  JGI  Ensembl |  |  | 76.99724 % |  |  |
| 12 | ggggacaagtttgtacaaaaaagcaggcttgatgTCCTCCTTCACCTCCAGCGG, ggggaccactttgtacaagaaagctgggtcAGAATGTCCAGTCGTCGTCC | Mycgr3\|76146  JGI  Ensembl |  |  | 100 % |  |  |
| 12 | ggggacaagtttgtacaaaaaagcaggcttgatgGCATCTCGCAATGCGCAAGT, ggggaccactttgtacaagaaagctgggtcCACCTCTCTCAGATTCATCG | Mycgr3\|83119  JGI  Ensembl |  |  | 100 % |  |  |
| 12 | ggggacaagtttgtacaaaaaagcaggcttgatgTCGCTTAACACAGCAAAGAA, ggggaccactttgtacaagaaagctgggtcCAATCTACTGTACATGCCCT | Mycgr3\|84211  JGI  Ensembl | KOG1014 17 beta-hydroxysteroid dehydrogenase type 3, HSD17B3; | 2-deoxy-D-gluconate 3-dehydrogenase activity; metabolic process;  oxidoreductase activity; | 100 % |  |  |
| 12 | ggggacaagtttgtacaaaaaagcaggcttgatgTCTCAACCAAAATACGAACT, ggggaccactttgtacaagaaagctgggtcCTCCTGCCTATCCAACTCCG | Mycgr3\|86002  JGI  Ensembl |  |  | 100 % |  |  |
| 12 | ggggacaagtttgtacaaaaaagcaggcttgatgGCACATATCATAATCACAGG, ggggaccactttgtacaagaaagctgggtcCATGCCAGCAAGTCTTCTGA | Mycgr3\|88767  JGI  Ensembl |  |  | 5.00642 % |  |  |
| 12 | ggggacaagtttgtacaaaaaagcaggcttgatgGCAGATGTTGCGAACCGCGC, ggggaccactttgtacaagaaagctgggtcCTGCTCCTCAATGCGCGAAG | Mycgr3\|88990  JGI  Ensembl |  |  | 100 % |  |  |
| 12 | ggggacaagtttgtacaaaaaagcaggcttgatgTCGTTGTTACGATCATCTCG, ggggaccactttgtacaagaaagctgggtcACACCGCCCTTCGAAGAATC | Mycgr3\|89490  JGI  Ensembl |  |  | 100 % |  |  |
| 12 | ggggacaagtttgtacaaaaaagcaggcttgatgGCGACCAACGAGAACAACGA, ggggaccactttgtacaagaaagctgggtcAGTCGTGATCCTCACTCGCT | Mycgr3\|89506  JGI  Ensembl |  |  | 100 % |  |  |
| 12 | ggggacaagtttgtacaaaaaagcaggcttgatgGCCACACGGGAATGGCACAT, ggggaccactttgtacaagaaagctgggtcAGCATCGGCTTGTCGACAGA | Mycgr3\|90565  JGI  Ensembl |  |  | 100 % |  |  |
| 12 | ggggacaagtttgtacaaaaaagcaggcttgatgTCTGCCATCAACAAGGAAAA, ggggaccactttgtacaagaaagctgggtcAGCGCGCAACGCGGGCTCTC | Mycgr3\|90749  JGI  Ensembl |  |  | 100 % |  |  |
| 12 | ggggacaagtttgtacaaaaaagcaggcttgatgAAGCTCCTCTGTCTCCACGG, ggggaccactttgtacaagaaagctgggtcCAGCATCCCCATCATGACCA | Mycgr3\|91126  JGI  Ensembl | KOG2551 Phospholipase/carboxyhydrolase; |  | 100 % |  |  |
| 12 | ggggacaagtttgtacaaaaaagcaggcttgatgTCAAGCAAATACAGCGAGAT, ggggaccactttgtacaagaaagctgggtcCCAAGCCGACGCACCGGTAA | Mycgr3\|91176  JGI  Ensembl |  |  | 0 % |  |  |
| 12 | ggggacaagtttgtacaaaaaagcaggcttgatgGCGCCGCAACCCACCACACC, ggggaccactttgtacaagaaagctgggtcCTCCTCCGTCACGACCGCCA | Mycgr3\|91335  JGI  Ensembl | KOG1415 Ubiquitin C-terminal hydrolase UCHL1; | intracellular; ubiquitin thiolesterase activity;  ubiquitin-dependent protein catabolic process; | 0 % |  |  |
| 12 | ggggacaagtttgtacaaaaaagcaggcttgatgCAATTCTTCAACCCATTCAC, ggggaccactttgtacaagaaagctgggtcCAACTCCGCAAACACCCAGC | Mycgr3\|91346  JGI  Ensembl | KOG1216 von Willebrand factor and related coagulation proteins; |  | 100 % |  |  |
| 12 | ggggacaagtttgtacaaaaaagcaggcttgatgGGGCCCAAAAGAGCGACAGC, ggggaccactttgtacaagaaagctgggtcAACAAAGCCGTACTCCTCGT | Mycgr3\|91380  JGI  Ensembl |  |  | 0 % |  |  |
| 12 | ggggacaagtttgtacaaaaaagcaggcttgatgGACAACACGACAAATCCTCA, ggggaccactttgtacaagaaagctgggtcGCCGGCGGTCTCCAGAATCT | Mycgr3\|91810  JGI  Ensembl |  |  | 100 % |  |  |
| 12 | ggggacaagtttgtacaaaaaagcaggcttgatgTCGGGCTCCTCTGGCAAATC, ggggaccactttgtacaagaaagctgggtcCGATTCGTCCGTCCAATGCT | Mycgr3\|92058  JGI  Ensembl |  |  | 100 % |  |  |
| 12 | ggggacaagtttgtacaaaaaagcaggcttgatgTCCGCTGTCACCGCTGACTT, ggggaccactttgtacaagaaagctgggtcATCGTCAGAGTCTTCTTCGT | Mycgr3\|92845  JGI  Ensembl |  |  | 100 % |  |  |
| 12 | ggggacaagtttgtacaaaaaagcaggcttgatgTCCAAGCCACGAATCGTCAA, ggggaccactttgtacaagaaagctgggtcATGAATCGGCATGAACGGAA | Mycgr3\|93791  JGI  Ensembl |  |  | 2.56082 % |  |  |
| 12 | ggggacaagtttgtacaaaaaagcaggcttgatgTTCCTCAGCATGATCAACCT, ggggaccactttgtacaagaaagctgggtcCTGTATCGGACCGACTGTGT | Mycgr3\|94441  JGI  Ensembl |  |  | 100 % |  |  |
| 12 | ggggacaagtttgtacaaaaaagcaggcttgatgTCGGCACCGAAGCTGTTCCT, ggggaccactttgtacaagaaagctgggtcCTTTGGCTGAGGCCACGAAA | Mycgr3\|94490  JGI  Ensembl | KOG0868 Glutathione S-transferase; | glutathione transferase activity; | 100 % |  |  |
| 12 | ggggacaagtttgtacaaaaaagcaggcttgatgGTCAGTCTCCAGCCGCCACA, ggggaccactttgtacaagaaagctgggtcATCATTCTTGCCGAGAATTT | Mycgr3\|94706  JGI  Ensembl |  |  | 100 % |  |  |
| 12 | ggggacaagtttgtacaaaaaagcaggcttgatgGCAGCATCAACGCAGAACGA, ggggaccactttgtacaagaaagctgggtcGAGCTCTTTCCATTTATCCG | Mycgr3\|94788  JGI  Ensembl |  |  | 100 % |  |  |
| 12 | ggggacaagtttgtacaaaaaagcaggcttgatgTCCGTCAGTCCAGAACATGC, ggggaccactttgtacaagaaagctgggtcATTGTCGTCGTGGGACGAGT | Mycgr3\|94795  JGI  Ensembl |  |  | 0 % |  |  |
| 12 | ggggacaagtttgtacaaaaaagcaggcttgatgGCGAACATCCTCGACGTCCC, ggggaccactttgtacaagaaagctgggtcGGGCATGCCAGCCTCCACCC | Mycgr3\|94811  JGI  Ensembl |  |  | 0 % |  |  |
| 12 | ggggacaagtttgtacaaaaaagcaggcttgatgCCTCATCCAACCTTGCTGAC, ggggaccactttgtacaagaaagctgggtcCGCCGGCCTCTTCCCTCTCC | Mycgr3\|94819  JGI  Ensembl |  |  | 100 % |  |  |
| 12 | ggggacaagtttgtacaaaaaagcaggcttgatgGCAAAGTTCACCATCACCTC, ggggaccactttgtacaagaaagctgggtcTGGCGCCTTGCCCACCCAGC | Mycgr3\|94820  JGI  Ensembl |  |  | 0 % |  |  |
| 12 | ggggacaagtttgtacaaaaaagcaggcttgatgTCTGCCGCCCATCGCATTGT, ggggaccactttgtacaagaaagctgggtcTCGCTTCTTCTCCGCCAGCC | Mycgr3\|94942  JGI  Ensembl |  |  | 100 % |  |  |
| 12 | ggggacaagtttgtacaaaaaagcaggcttgatgGCAGGCAATGGGGGCGACAG, ggggaccactttgtacaagaaagctgggtcGTTGTTTCTCAAGAAAGCGA | Mycgr3\|95512  JGI  Ensembl |  |  | 100 % |  |  |
| 12 | ggggacaagtttgtacaaaaaagcaggcttgatgACATCGACATCACCACAACA, ggggaccactttgtacaagaaagctgggtcATACTCTCCCGGCCGTGCGG | Mycgr3\|95520  JGI  Ensembl |  |  | 100 % |  |  |
| 12 | ggggacaagtttgtacaaaaaagcaggcttgatgGCGACCAAGTTCACACTCTT, ggggaccactttgtacaagaaagctgggtcAGACTTCCCAGCCTCGATCA | Mycgr3\|96055  JGI  Ensembl |  |  | 100 % |  |  |
| 12 | ggggacaagtttgtacaaaaaagcaggcttgatgTCAGATGTCAGATGCAAGGC, ggggaccactttgtacaagaaagctgggtcAGAGAGCAACAACTTCTTCA | Mycgr3\|96644  JGI  Ensembl |  |  | 100 % |  |  |
| 12 | ggggacaagtttgtacaaaaaagcaggcttgatgCAAGACCTGCAACCGGGTAT, ggggaccactttgtacaagaaagctgggtcGTCCAGGTCCATGCGGTCCT | Mycgr3\|96755  JGI  Ensembl |  |  | 100 % |  |  |
| 12 | ggggacaagtttgtacaaaaaagcaggcttgatgTTCGAAGCGGCGGTCGCGGG, ggggaccactttgtacaagaaagctgggtcAGGAGGATGATTGCCATTGA | Mycgr3\|97605  JGI  Ensembl |  |  | 5.01882 % |  |  |
| 12 | ggggacaagtttgtacaaaaaagcaggcttgatgCCGCCTCAAGATCAGTCCAC, ggggaccactttgtacaagaaagctgggtcGAAACTGCCCGTGTCGCGGC | Mycgr3\|97687  JGI  Ensembl |  |  | 100 % |  |  |
| 12 | ggggacaagtttgtacaaaaaagcaggcttgatgCCACCCGACCTCGACTTCCC, ggggaccactttgtacaagaaagctgggtcAGCACCGATATTGCTCGCCT | Mycgr3\|97733  JGI  Ensembl | KOG1245 Chromatin remodeling complex WSTF-ISWI, large subunit (contains heterochromatin localization, PHD and BROMO domains); |  | 100 % |  |  |
| 12 | ggggacaagtttgtacaaaaaagcaggcttgatgAATTGCGGTGTGTACATTAC, ggggaccactttgtacaagaaagctgggtcTGAAGGGAATTGATGTATCC | Mycgr3\|97803  JGI  Ensembl |  |  | 100 % |  |  |
| 12 | ggggacaagtttgtacaaaaaagcaggcttgatgTCTATCAAGCATGCGACCAA, ggggaccactttgtacaagaaagctgggtcCTTCAACCGCTCCATCAGCT | Mycgr3\|97851  JGI  Ensembl | KOG0504 FOG: Ankyrin repeat; |  | 0 % |  |  |
| 12 | ggggacaagtttgtacaaaaaagcaggcttgatgCATCAAAAGTCTGCTCCCGT, ggggaccactttgtacaagaaagctgggtcATGCACAAGCACACGTCGGG | Mycgr3\|98016  JGI  Ensembl |  |  | 100 % |  |  |
| 12 | ggggacaagtttgtacaaaaaagcaggcttgatgCAGAACTTTAGTATGACCGG, ggggaccactttgtacaagaaagctgggtcGCGCCACCACTTGCAGTCTC | Mycgr3\|98076  JGI  Ensembl |  |  | 0 % |  |  |
| 12 | ggggacaagtttgtacaaaaaagcaggcttgatgGTCACATACACCGTCAAGCG, ggggaccactttgtacaagaaagctgggtcAGTCAGACCCGGCAAAGAGG | Mycgr3\|98106  JGI  Ensembl |  |  | 100 % |  |  |
| 12 | ggggacaagtttgtacaaaaaagcaggcttgatgGACATGGCAGACGCGGCCGA, ggggaccactttgtacaagaaagctgggtcTTCCGGGCTCATAACCAGTT | Mycgr3\|98117  JGI  Ensembl |  |  | 100 % |  |  |
| 12 | ggggacaagtttgtacaaaaaagcaggcttgatgGCTGTGACAACAGCACCTTC, ggggaccactttgtacaagaaagctgggtcAGAATTGAAAATCCTCGAGT | Mycgr3\|100955  JGI  Ensembl |  |  | 100 % |  |  |
| 12 | ggggacaagtttgtacaaaaaagcaggcttgatgCCTGAATACGCGACCTACCC, ggggaccactttgtacaagaaagctgggtcTGGATCTCCAACCCAACCTC | Mycgr3\|101874  JGI  Ensembl | KOG0725 Reductases with broad range of substrate specificities; | carbonyl reductase (NADPH) activity; metabolic process;  oxidoreductase activity; | 100 % |  |  |
| 12 | ggggacaagtttgtacaaaaaagcaggcttgatgGAGGTCACCCCATCAACAAG, ggggaccactttgtacaagaaagctgggtcCTGCGAGAGGTCACGCATAT | Mycgr3\|102377  JGI  Ensembl |  |  | 0 % |  |  |
| 12 | ggggacaagtttgtacaaaaaagcaggcttgatgCGCATCAATAGCAACGACTT, ggggaccactttgtacaagaaagctgggtcTGATGGATAGCATTTGTGTT | Mycgr3\|103159  JGI  Ensembl |  |  | 0 % |  |  |
| 12 | ggggacaagtttgtacaaaaaagcaggcttgatgTCAGGACTCAAGGACAAGAT, ggggaccactttgtacaagaaagctgggtcCAGATGTCCCAAAACCGCAT | Mycgr3\|103459  JGI  Ensembl | KOG0541 Alkyl hydroperoxide reductase/peroxiredoxin; | antioxidant activity; oxidoreductase activity;  peroxiredoxin activity; | 100 % |  |  |
| 12 | ggggacaagtttgtacaaaaaagcaggcttgatgTCGCAATACATAGGCAAAAC, ggggaccactttgtacaagaaagctgggtcTTGCGCTGGCGGTGAACTTG | Mycgr3\|103797  JGI  Ensembl | KOG3203 Mitochondrial/chloroplast ribosomal protein L13; | intracellular; ribosome; structural constituent of ribosome;  translation; | 100 % |  |  |
| 12 | ggggacaagtttgtacaaaaaagcaggcttgatgCACACGCCAACACTGCGGTA, ggggaccactttgtacaagaaagctgggtcAGTACAAAGAAACTCTTTCG | Mycgr3\|104117  JGI  Ensembl |  |  | 100 % |  |  |
| 12 | ggggacaagtttgtacaaaaaagcaggcttgatgGATTACATCCTCCACGATCA, ggggaccactttgtacaagaaagctgggtcATAGACAATGTTTCGCCATT | Mycgr3\|104414  JGI  Ensembl |  |  | 100 % |  |  |
| 12 | ggggacaagtttgtacaaaaaagcaggcttgatgTCGAACCCTCGAGCCAAGCC, ggggaccactttgtacaagaaagctgggtcGATGCCCAACGGATTGACAG | Mycgr3\|104515  JGI  Ensembl | KOG1922 Rho GTPase effector BNI1 and related formins; |  | 100 % |  |  |
| 12 | ggggacaagtttgtacaaaaaagcaggcttgatgGCATCGTTCACGCTCGCCTA, ggggaccactttgtacaagaaagctgggtcTTTCACGCTCCGCCTGATCA | Mycgr3\|104703  JGI  Ensembl |  |  | 100 % |  |  |
| 12 | ggggacaagtttgtacaaaaaagcaggcttgatgACCGAGACCGGCCCACCACC, ggggaccactttgtacaagaaagctgggtcGTCCTCCATCGCGTCCTCGT | Mycgr3\|105272  JGI  Ensembl |  |  | 64.17624 % |  |  |
| 12 | ggggacaagtttgtacaaaaaagcaggcttgatgAACCGATCATTACGCGCAGC, ggggaccactttgtacaagaaagctgggtcCTTCACATCCTTGGGCAAGA | Mycgr3\|105511  JGI  Ensembl |  |  | 100 % |  |  |
| 12 | ggggacaagtttgtacaaaaaagcaggcttgatgACTGTCATCAACAAGAACTT, ggggaccactttgtacaagaaagctgggtcAGCAGACTTGCGCTTCTTCT | Mycgr3\|105646  JGI  Ensembl | KOG3401 60S ribosomal protein L26; | intracellular; large ribosomal subunit;  ribosome;  structural constituent of ribosome;  translation; | 0 % |  |  |
| 12 | ggggacaagtttgtacaaaaaagcaggcttgatgCGCAGGACTTCTTGCTGGCG, ggggaccactttgtacaagaaagctgggtcCGTCGAGGGGTTCTGATTTC | Mycgr3\|105805  JGI  Ensembl |  |  | 100 % |  |  |
| 12 | ggggacaagtttgtacaaaaaagcaggcttgatgAACAAGGAGTCCGATGCTGC, ggggaccactttgtacaagaaagctgggtcTTCCCAGCTCTGTACCCGAG | Mycgr3\|106008  JGI  Ensembl |  |  | 100 % |  |  |
| 12 | ggggacaagtttgtacaaaaaagcaggcttgatgGGCGACTTCCTAGCAAACCT, ggggaccactttgtacaagaaagctgggtcTTGCCTGTTGTCCACCTTCT | Mycgr3\|106144  JGI  Ensembl |  |  | 100 % |  |  |
| 12 | ggggacaagtttgtacaaaaaagcaggcttgatgGCTCGAAGGCGACGACCTCC, ggggaccactttgtacaagaaagctgggtcAGCAGGCTCCTTTGGCGCCA | Mycgr3\|106186  JGI  Ensembl | KOG4697 Integral membrane protein involved in transport between the late Golgi and endosome; |  | 2.75762 % |  |  |
| 12 | ggggacaagtttgtacaaaaaagcaggcttgatgCCCATGGCCTACAACAGTCC, ggggaccactttgtacaagaaagctgggtcGTGCCCTCTCCCAGGCGCGA | Mycgr3\|106353  JGI  Ensembl |  |  | 100 % |  |  |
| 12 | ggggacaagtttgtacaaaaaagcaggcttgatgGCTCCCCGCGCCCGCCGCCC, ggggaccactttgtacaagaaagctgggtcAGGCCTGCTCAAATGCGAGT | Mycgr3\|106733  JGI  Ensembl |  |  | 100 % |  |  |
| 12 | ggggacaagtttgtacaaaaaagcaggcttgatgCCCCTCTCCCTCTGGTTCAT, ggggaccactttgtacaagaaagctgggtcAACCAACGCCTTCGTAACCC | Mycgr3\|107673  JGI  Ensembl |  |  | 100 % |  |  |
| 12 | ggggacaagtttgtacaaaaaagcaggcttgatgACTTCACAGCGACCATTCTT, ggggaccactttgtacaagaaagctgggtcTAAGCTCATTTGATCCGTCG | Mycgr3\|107929  JGI  Ensembl |  |  | 100 % |  |  |
| 12 | ggggacaagtttgtacaaaaaagcaggcttgatgTTGCAACGAAGTGCTCGCAG, ggggaccactttgtacaagaaagctgggtcTCTCTGCTCGTAAGGTGCGC | Mycgr3\|110313  JGI  Ensembl | KOG3456 NADH:ubiquinone oxidoreductase, NDUFS6/13 kDa subunit; |  | 100 % |  |  |
| 12 | ggggacaagtttgtacaaaaaagcaggcttgatgGCAAGGCTACGGAGAGGTTC, ggggaccactttgtacaagaaagctgggtcGGCGGTGGAGTCGTCGATCG | Mycgr3\|111231  JGI  Ensembl |  |  | 100 % |  |  |
| 13 | ggggacaagtttgtacaaaaaagcaggcttgatgGGACTGATCGTCCACCACCT, ggggaccactttgtacaagaaagctgggtcCTTATGGCCGGCAAAGCCCT | Mycgr3\|31926  JGI  Ensembl | KOG0867 Glutathione S-transferase; | glutathione transferase activity; | 100 % |  |  |
| 13 | ggggacaagtttgtacaaaaaagcaggcttgatgGGCATCGTACGCACCGCACT, ggggaccactttgtacaagaaagctgggtcTCGGGTGCAGTTGGAGATGG | Mycgr3\|32504  JGI  Ensembl |  |  | 100 % |  |  |
| 13 | ggggacaagtttgtacaaaaaagcaggcttgatgGCGACCTTGATCACGGACAC, ggggaccactttgtacaagaaagctgggtcGTATTCGCAAACTGCTGGAC | Mycgr3\|35104  JGI  Ensembl |  |  | 100 % |  |  |
| 13 | ggggacaagtttgtacaaaaaagcaggcttgatgTCCTCATCCCACGGCGCACT, ggggaccactttgtacaagaaagctgggtcCCCCTGCGTCTTCTTGATAA | Mycgr3\|36827  JGI  Ensembl | KOG1014 17 beta-hydroxysteroid dehydrogenase type 3, HSD17B3; |  | 100 % |  |  |
| 13 | ggggacaagtttgtacaaaaaagcaggcttgatgCGTTTCTCCGCTGTCATCAC, ggggaccactttgtacaagaaagctgggtcGGTCAGTCTGAAGACAATGT | Mycgr3\|38087  JGI  Ensembl |  |  | 100 % |  |  |
| 13 | ggggacaagtttgtacaaaaaagcaggcttgatgAGCTCATCCGAAGAAACTCC, ggggaccactttgtacaagaaagctgggtcTGATATAGTATAAGAGACTT | Mycgr3\|38362  JGI  Ensembl | KOG2105 Predicted metal-dependent hydrolase, contains AlaS domain; | alanine-tRNA ligase activity; | 0 % |  |  |
| 13 | ggggacaagtttgtacaaaaaagcaggcttgatgACTTCCCAGTTTCCTGTCTA, ggggaccactttgtacaagaaagctgggtcCATGAAACCGGTTCCTGTCG | Mycgr3\|39030  JGI  Ensembl | KOG0725 Reductases with broad range of substrate specificities; | 3-oxoacyl-[acyl-carrier-protein] reductase activity; metabolic process;  oxidoreductase activity; | 100 % |  |  |
| 13 | ggggacaagtttgtacaaaaaagcaggcttgatgTATGCCCACCAACCGAGCGG, ggggaccactttgtacaagaaagctgggtcACTCCCCGACCCACCACCCC | Mycgr3\|41022  JGI  Ensembl |  |  | 100 % |  |  |
| 13 | ggggacaagtttgtacaaaaaagcaggcttgatgGTACTGCTCACGACCACGGC, ggggaccactttgtacaagaaagctgggtcCCTGTGTTTTCCCTTCCTGC | Mycgr3\|43297  JGI  Ensembl |  |  | 100 % |  |  |
| 13 | ggggacaagtttgtacaaaaaagcaggcttgatgTCTCTGGGCGACAGCGAAGC, ggggaccactttgtacaagaaagctgggtcGTCAAAGGCGCTGGCGGGGA | Mycgr3\|43304  JGI  Ensembl |  |  | 4.82353 % |  |  |
| 13 | ggggacaagtttgtacaaaaaagcaggcttgatgACAACGGCCTTCCGAGACCC, ggggaccactttgtacaagaaagctgggtcCTTCTCAATCCCCAACTTCT | Mycgr3\|43351  JGI  Ensembl |  |  | 100 % |  |  |
| 13 | ggggacaagtttgtacaaaaaagcaggcttgatgGCACGAGACCTTGCGACACG, ggggaccactttgtacaagaaagctgggtcTGGCTCGTCTGCATACGCTG | Mycgr3\|44375  JGI  Ensembl |  |  | 100 % |  |  |
| 13 | ggggacaagtttgtacaaaaaagcaggcttgatgTACGATGCCGAAAAGGCCAT, ggggaccactttgtacaagaaagctgggtcCTGACCCTCCTTCTTACTCT | Mycgr3\|48694  JGI  Ensembl |  |  | 100 % |  |  |
| 13 | ggggacaagtttgtacaaaaaagcaggcttgatgCCGTCCAATGTCCTCATCGT, ggggaccactttgtacaagaaagctgggtcCCACGGAAGCTGCAGAGGTG | Mycgr3\|49284  JGI  Ensembl | KOG1611 Predicted short chain-type dehydrogenase; | acylglycerone-phosphate reductase activity; metabolic process;  oxidoreductase activity; | 100 % |  |  |
| 13 | ggggacaagtttgtacaaaaaagcaggcttgatgGACGACTGGACAGCGACAGC, ggggaccactttgtacaagaaagctgggtcCCGGTTGTACGGATCGACAA | Mycgr3\|50056  JGI  Ensembl |  |  | 100 % |  |  |
| 13 | ggggacaagtttgtacaaaaaagcaggcttgatgAACGCTGCTACTGACGACGG, ggggaccactttgtacaagaaagctgggtcCAAGTAGCCGGGATGTAACG | Mycgr3\|50563  JGI  Ensembl |  |  | 100 % |  |  |
| 13 | ggggacaagtttgtacaaaaaagcaggcttgatgCTCGCCTCGGGATTCACGAA, ggggaccactttgtacaagaaagctgggtcTGACGCCCCTCTGAACTGAT | Mycgr3\|69677  JGI  Ensembl | KOG4463 Uncharacterized conserved protein; |  | 100 % |  |  |
| 13 | ggggacaagtttgtacaaaaaagcaggcttgatgCCTGGCAACAAGAACGTCGT, ggggaccactttgtacaagaaagctgggtcCGACAGAGCTATGATCCGAT | Mycgr3\|69689  JGI  Ensembl |  |  | 0 % |  |  |
| 13 | ggggacaagtttgtacaaaaaagcaggcttgatgCCATTTCCCTCACTCCCCTC, ggggaccactttgtacaagaaagctgggtcCTCCCGAAATCCTCTCTCCC | Mycgr3\|73372  JGI  Ensembl | KOG3399 Predicted Yippee-type zinc-binding protein; |  | 100 % |  |  |
| 13 | ggggacaagtttgtacaaaaaagcaggcttgatgCCTGCCCCCACCGCTCTCCA, ggggaccactttgtacaagaaagctgggtcGAACCTCTCCTTCATCCTGC | Mycgr3\|73841  JGI  Ensembl | KOG3273 Predicted RNA-binding protein Pno1p interacting with Nob1p and involved in 26S proteasome assembly; | RNA binding; | 100 % |  |  |
| 13 | ggggacaagtttgtacaaaaaagcaggcttgatgGCGGACCCGAATCAGGCGGG, ggggaccactttgtacaagaaagctgggtcAATGGCGTCGGTCGAGCTGA | Mycgr3\|75178  JGI  Ensembl |  |  | 100 % |  |  |
| 13 | ggggacaagtttgtacaaaaaagcaggcttgatgAAGACCCCCTCACCCTCCTT, ggggaccactttgtacaagaaagctgggtcCTTCAACCCCAACGGCTCCA | Mycgr3\|75187  JGI  Ensembl |  |  | 100 % |  |  |
| 13 | ggggacaagtttgtacaaaaaagcaggcttgatgTCTCTCTCCACCAAGGCCGC, ggggaccactttgtacaagaaagctgggtcGCCAATAACCAGAGCACCAT | Mycgr3\|75376  JGI  Ensembl | KOG1671 Ubiquinol cytochrome c reductase, subunit RIP1; | electron transport; membrane; oxidoreductase activity;  ubiquinol-cytochrome-c reductase activity; | 0 % |  |  |
| 13 | ggggacaagtttgtacaaaaaagcaggcttgatgCAACCATCCCTCCGAATCGC, ggggaccactttgtacaagaaagctgggtcGTCCTCCAACAGAAACCTCA | Mycgr3\|75463  JGI  Ensembl | KOG3179 Predicted glutamine synthetase; | catalytic activity; | 100 % |  |  |
| 13 | ggggacaagtttgtacaaaaaagcaggcttgatgGCGTCGACAGCGCAGCCACC, ggggaccactttgtacaagaaagctgggtcCTCCTCGTCCAACTGGAATG | Mycgr3\|76609  JGI  Ensembl | KOG2730 Methylase; |  | 100 % |  |  |
| 13 | ggggacaagtttgtacaaaaaagcaggcttgatgGATCCGAACGACGCCCTCGC, ggggaccactttgtacaagaaagctgggtcAACCTCAAGCATTCCCTTCA | Mycgr3\|77840  JGI  Ensembl | KOG0161 Myosin class II heavy chain; |  | 100 % |  |  |
| 13 | ggggacaagtttgtacaaaaaagcaggcttgatgCAGTTCTCACGCTCTTTCCT, ggggaccactttgtacaagaaagctgggtcGAGTGCCATGGCGGCAAGAG | Mycgr3\|78594  JGI  Ensembl | KOG4157 beta-1,6-N-acetylglucosaminyltransferase, contains WSC domain; |  | 0 % |  |  |
| 13 | ggggacaagtttgtacaaaaaagcaggcttgatgCTCCTCACCCTCGCCCTCTC, ggggaccactttgtacaagaaagctgggtcCATCAGAAACCCAGCCACCA | Mycgr3\|79783  JGI  Ensembl | KOG4157 beta-1,6-N-acetylglucosaminyltransferase, contains WSC domain; |  | 0 % |  |  |
| 13 | ggggacaagtttgtacaaaaaagcaggcttgatgGCTCCCAAAGTCAAGACGAG, ggggaccactttgtacaagaaagctgggtcCCGCTTCACGACCAGATCGC | Mycgr3\|83529  JGI  Ensembl |  |  | 100 % |  |  |
| 13 | ggggacaagtttgtacaaaaaagcaggcttgatgGCTCTGCTGCAAGGCAAAGT, ggggaccactttgtacaagaaagctgggtcCTGCAGATTTACGAACAATC | Mycgr3\|83548  JGI  Ensembl | KOG1200 Mitochondrial/plastidial beta-ketoacyl-ACP reductase; | 3-oxoacyl-[acyl-carrier-protein] reductase activity; metabolic process;  oxidoreductase activity; | 100 % |  |  |
| 13 | ggggacaagtttgtacaaaaaagcaggcttgatgGCTACCACCACGAACACCAG, ggggaccactttgtacaagaaagctgggtcGCCGAAGACCAGAGCTCGCA | Mycgr3\|83566  JGI  Ensembl |  |  | 100 % |  |  |
| 13 | ggggacaagtttgtacaaaaaagcaggcttgatgGCCGACAAAACCCTCGTCCT, ggggaccactttgtacaagaaagctgggtcCCACGGCTCCGCTCCCTCGT | Mycgr3\|85689  JGI  Ensembl |  |  | 100 % |  |  |
| 13 | ggggacaagtttgtacaaaaaagcaggcttgatgTGGAAGAAAGTCCGAATCAC, ggggaccactttgtacaagaaagctgggtcAGCCGCCTTCCTGAGTGTCT | Mycgr3\|88619  JGI  Ensembl |  |  | 100 % |  |  |
| 13 | ggggacaagtttgtacaaaaaagcaggcttgatgTCGAATCAATCCACCGCCAA, ggggaccactttgtacaagaaagctgggtcAGCTGCTAGTGGTACAGCTG | Mycgr3\|88630  JGI  Ensembl |  |  | 100 % |  |  |
| 13 | ggggacaagtttgtacaaaaaagcaggcttgatgGACGAGCTGAAGAAGTCACC, ggggaccactttgtacaagaaagctgggtcGGTGTATTGGGAAGGAAGAA | Mycgr3\|89155  JGI  Ensembl | KOG2533 Permease of the major facilitator superfamily; | integral to membrane; transport; transporter activity; | 0 % |  |  |
| 13 | ggggacaagtttgtacaaaaaagcaggcttgatgGCTGACTCAGACTACTCGCC, ggggaccactttgtacaagaaagctgggtcCTGTGGATCAGGCAGCTGAT | Mycgr3\|89996  JGI  Ensembl |  |  | 100 % |  |  |
| 13 | ggggacaagtttgtacaaaaaagcaggcttgatgGCAGATGTCGATACGAATGA, ggggaccactttgtacaagaaagctgggtcCAACTCTTTCTTTCCAGCAT | Mycgr3\|90609  JGI  Ensembl | KOG1110 Putative steroid membrane receptor Hpr6.6/25-Dx; | heme binding; transition metal ion binding; | 100 % |  |  |
| 13 | ggggacaagtttgtacaaaaaagcaggcttgatgTCTACTTCCATGGCGGCCAC, ggggaccactttgtacaagaaagctgggtcCACTCCTGGCGAAACGTCTC | Mycgr3\|90801  JGI  Ensembl |  |  | 100 % |  |  |
| 13 | ggggacaagtttgtacaaaaaagcaggcttgatgGCGGCCATACAAACGAGAAT, ggggaccactttgtacaagaaagctgggtcGAAGAGCGCGAGTCTCACTG | Mycgr3\|90839  JGI  Ensembl | KOG4297 C-type lectin; |  | 0 % |  |  |
| 13 | ggggacaagtttgtacaaaaaagcaggcttgatgACCACATCCACGACTCAACC, ggggaccactttgtacaagaaagctgggtcCTGATGAGCTTCTTGCCACC | Mycgr3\|91624  JGI  Ensembl |  |  | 100 % |  |  |
| 13 | ggggacaagtttgtacaaaaaagcaggcttgatgTGCAAGTCGTGGACCCTCCG, ggggaccactttgtacaagaaagctgggtcATTGCTGATCGCAAATCCCA | Mycgr3\|91755  JGI  Ensembl |  |  | 100 % |  |  |
| 13 | ggggacaagtttgtacaaaaaagcaggcttgatgTCCGGCTCATCACCATGGAA, ggggaccactttgtacaagaaagctgggtcGCGGTGGATGTTTTGCGTGT | Mycgr3\|92103  JGI  Ensembl |  |  | 100 % |  |  |
| 13 | ggggacaagtttgtacaaaaaagcaggcttgatgAGCAAATCCAAGCGAAGCGG, ggggaccactttgtacaagaaagctgggtcGAACTCCTTGCAGAGGTTCT | Mycgr3\|92520  JGI  Ensembl |  |  | 100 % |  |  |
| 13 | ggggacaagtttgtacaaaaaagcaggcttgatgGAAGCCAGCACGCCGCCTTC, ggggaccactttgtacaagaaagctgggtcCATGCCCGCGAATTCGGCAA | Mycgr3\|92804  JGI  Ensembl |  |  | 0 % |  |  |
| 13 | ggggacaagtttgtacaaaaaagcaggcttgatgTCGGTATCCTCGCCATCCCG, ggggaccactttgtacaagaaagctgggtcCGGTATTCTGGCGCCGCTCA | Mycgr3\|92919  JGI  Ensembl |  |  | 100 % |  |  |
| 13 | ggggacaagtttgtacaaaaaagcaggcttgatgAAGCTGCGTAAGGCGCAGTA, ggggaccactttgtacaagaaagctgggtcCAAGCGTATGAAGTACGCCG | Mycgr3\|92980  JGI  Ensembl |  |  | 100 % |  |  |
| 13 | ggggacaagtttgtacaaaaaagcaggcttgatgTCTCCTCTCATCATCGACGA, ggggaccactttgtacaagaaagctgggtcAGAAAGCATCGCTTGTACTG | Mycgr3\|93128  JGI  Ensembl |  |  | 100 % |  |  |
| 13 | ggggacaagtttgtacaaaaaagcaggcttgatgTCCAGCCAAAAGACATACTT, ggggaccactttgtacaagaaagctgggtcAGCAGGCACGCAGTAAACAC | Mycgr3\|93377  JGI  Ensembl |  |  | 100 % |  |  |
| 13 | ggggacaagtttgtacaaaaaagcaggcttgatgGCCTCCGATTCTGCACCACC, ggggaccactttgtacaagaaagctgggtcGTACGTCGTCCACATCTCGG | Mycgr3\|93594  JGI  Ensembl |  |  | 100 % |  |  |
| 13 | ggggacaagtttgtacaaaaaagcaggcttgatgACGCTCCCCAAGTACGCTCT, ggggaccactttgtacaagaaagctgggtcTAACCTCAGCCCCTCCATCT | Mycgr3\|93695  JGI  Ensembl |  |  | 100 % |  |  |
| 13 | ggggacaagtttgtacaaaaaagcaggcttgatgAAAGTCAACACAAGCACTCT, ggggaccactttgtacaagaaagctgggtcCTCGCCTCTGTCACCATCGC | Mycgr3\|94108  JGI  Ensembl |  |  | 0 % |  |  |
| 13 | ggggacaagtttgtacaaaaaagcaggcttgatgCCGAAGCGCAAGAGAGCCGC, ggggaccactttgtacaagaaagctgggtcTTTCTTCCTCCTCTTTCGCT | Mycgr3\|94286  JGI  Ensembl | KOG3253 Predicted alpha/beta hydrolase; |  | 2.44755 % |  |  |
| 13 | ggggacaagtttgtacaaaaaagcaggcttgatgGTCTGCTCTGGCCTCATTGT, ggggaccactttgtacaagaaagctgggtcGTATCCTGGATCACCAAGAG | Mycgr3\|94396  JGI  Ensembl |  |  | 0 % |  |  |
| 13 | ggggacaagtttgtacaaaaaagcaggcttgatgGCTGAGAACACGGAGGCACA, ggggaccactttgtacaagaaagctgggtcATGTCCCAACAACGAATACC | Mycgr3\|94694  JGI  Ensembl |  |  | 100 % |  |  |
| 13 | ggggacaagtttgtacaaaaaagcaggcttgatgTTGGCCTCGACTGCGGAGTT, ggggaccactttgtacaagaaagctgggtcCTCCTTCAACTCCGGCTTCG | Mycgr3\|94759  JGI  Ensembl |  |  | 100 % |  |  |
| 13 | ggggacaagtttgtacaaaaaagcaggcttgatgCCGGGTGTATGGGAGCCGAC, ggggaccactttgtacaagaaagctgggtcCGTAGTGACCTCCCACGAAG | Mycgr3\|94773  JGI  Ensembl |  |  | 100 % |  |  |
| 13 | ggggacaagtttgtacaaaaaagcaggcttgatgTCGTCTCGCGCCCTGGAAGA, ggggaccactttgtacaagaaagctgggtcCGAGAGCTGCAGCGAGCCGA | Mycgr3\|94781  JGI  Ensembl |  |  | 100 % |  |  |
| 13 | ggggacaagtttgtacaaaaaagcaggcttgatgGACGAGCAGACGTTCCGCCT, ggggaccactttgtacaagaaagctgggtcACGCCTCCCGCGAATGCCTG | Mycgr3\|94793  JGI  Ensembl |  |  | 100 % |  |  |
| 13 | ggggacaagtttgtacaaaaaagcaggcttgatgACGTCGACGGTCAACCCAAT, ggggaccactttgtacaagaaagctgggtcAGTCGGTCGCCCGAAGACGC | Mycgr3\|94797  JGI  Ensembl |  |  | 0 % |  |  |
| 13 | ggggacaagtttgtacaaaaaagcaggcttgatgGGCATCCTCGACATTCCCGC, ggggaccactttgtacaagaaagctgggtcGGGCCCGATCCTATCTCCCG | Mycgr3\|94800  JGI  Ensembl |  |  | 0 % |  |  |
| 13 | ggggacaagtttgtacaaaaaagcaggcttgatgTCCATCTTCCTTCTTCTCGT, ggggaccactttgtacaagaaagctgggtcTCGGCGTGAACTGCGATCGG | Mycgr3\|94850  JGI  Ensembl |  |  | 100 % |  |  |
| 13 | ggggacaagtttgtacaaaaaagcaggcttgatgTCAGCGCCCCCGCCCCCGCC, ggggaccactttgtacaagaaagctgggtcGGTGGCGGCGGAATGCCAGG | Mycgr3\|95024  JGI  Ensembl | KOG1812 Predicted E3 ubiquitin ligase; | zinc ion binding; | 2.42424 % |  |  |
| 13 | ggggacaagtttgtacaaaaaagcaggcttgatgCCGATCCCGCCTGGCCAGAA, ggggaccactttgtacaagaaagctgggtcCATAATCACACAACAAGAGC | Mycgr3\|95055  JGI  Ensembl | KOG1187 Serine/threonine protein kinase; |  | 100 % |  |  |
| 13 | ggggacaagtttgtacaaaaaagcaggcttgatgGCCGGCGCCAAAGTCTTCGC, ggggaccactttgtacaagaaagctgggtcTTTCGATTGCTCTTCAGCCG | Mycgr3\|95325  JGI  Ensembl |  |  | 100 % |  |  |
| 13 | ggggacaagtttgtacaaaaaagcaggcttgatgGACTCACACAAGCGCAAGCA, ggggaccactttgtacaagaaagctgggtcCAGCGTATCAGCATGCGCAA | Mycgr3\|95536  JGI  Ensembl |  |  | 100 % |  |  |
| 13 | ggggacaagtttgtacaaaaaagcaggcttgatgCCGTCCTCCACAGCAGAAAC, ggggaccactttgtacaagaaagctgggtcAGGCCTCCGGTATGCCGGCA | Mycgr3\|95728  JGI  Ensembl |  |  | 100 % |  |  |
| 13 | ggggacaagtttgtacaaaaaagcaggcttgatgGCCAAAACATACCCGTTCAG, ggggaccactttgtacaagaaagctgggtcAAGATCTTCCTCGCCGTCAT | Mycgr3\|95879  JGI  Ensembl | KOG1081 Transcription factor NSD1 and related SET domain proteins; | nucleus; | 100 % |  |  |
| 13 | ggggacaagtttgtacaaaaaagcaggcttgatgACCACCACAGAAAGCCGCCT, ggggaccactttgtacaagaaagctgggtcCGGTAGACCACGCTCGCCAC | Mycgr3\|96284  JGI  Ensembl |  |  | 100 % |  |  |
| 13 | ggggacaagtttgtacaaaaaagcaggcttgatgGTCGCCTCTCAGAGAGCGCC, ggggaccactttgtacaagaaagctgggtcACCAGACTCGTCCGCATACA | Mycgr3\|97003  JGI  Ensembl |  |  | 100 % |  |  |
| 13 | ggggacaagtttgtacaaaaaagcaggcttgatgTCCTCCAAATGGTTGAACCA, ggggaccactttgtacaagaaagctgggtcTGTCTGTTCGTCCTGTAACT | Mycgr3\|97041  JGI  Ensembl |  |  | 100 % |  |  |
| 13 | ggggacaagtttgtacaaaaaagcaggcttgatgAGCCAGTACCCATGGCAGAA, ggggaccactttgtacaagaaagctgggtcACCACTGAACATGCGATCGA | Mycgr3\|97176  JGI  Ensembl |  |  | 100 % |  |  |
| 13 | ggggacaagtttgtacaaaaaagcaggcttgatgTCCTCCTTCTCCCTCCCCCA, ggggaccactttgtacaagaaagctgggtcCATCAACGCCCTCCCCCCAT | Mycgr3\|97474  JGI  Ensembl | KOG0725 Reductases with broad range of substrate specificities; | 3-oxoacyl-[acyl-carrier-protein] reductase activity; metabolic process;  oxidoreductase activity; | 100 % |  |  |
| 13 | ggggacaagtttgtacaaaaaagcaggcttgatgGCGCGCACCAAGACACGATC, ggggaccactttgtacaagaaagctgggtcACGTGGCTCGTTTCCGACTG | Mycgr3\|97693  JGI  Ensembl |  |  | 0 % |  |  |
| 13 | ggggacaagtttgtacaaaaaagcaggcttgatgGCGGCTGGGCCATGGCTGCA, ggggaccactttgtacaagaaagctgggtcGCGCCGCAGCTTCAGTGCAT | Mycgr3\|97881  JGI  Ensembl |  |  | 0 % |  |  |
| 13 | ggggacaagtttgtacaaaaaagcaggcttgatgAACGTTCAGACATTTGCAAA, ggggaccactttgtacaagaaagctgggtcGTTCCCAGAGCCAGGCACAC | Mycgr3\|97921  JGI  Ensembl |  |  | 0 % |  |  |
| 13 | ggggacaagtttgtacaaaaaagcaggcttgatgGACGAGACTCACAGGAACGG, ggggaccactttgtacaagaaagctgggtcAAAGAGCGCATTTGTGTTCG | Mycgr3\|97961  JGI  Ensembl |  |  | 100 % |  |  |
| 13 | ggggacaagtttgtacaaaaaagcaggcttgatgGCAACCTGCTCTCGAGCGCG, ggggaccactttgtacaagaaagctgggtcCCAAGCCCTCGCCGTGCCTC | Mycgr3\|97975  JGI  Ensembl | KOG1181 FOG: Low-complexity; |  | 100 % |  |  |
| 13 | ggggacaagtttgtacaaaaaagcaggcttgatgTCCGCCTCGGACGCCTCGAC, ggggaccactttgtacaagaaagctgggtcTTCATCTTCAAAGACCTCAT | Mycgr3\|98061  JGI  Ensembl |  |  | 2.33918 % |  |  |
| 13 | ggggacaagtttgtacaaaaaagcaggcttgatgCTCATTGATTCGGGAAGAGG, ggggaccactttgtacaagaaagctgggtcCGCCAGGACTTCGTTCCCGG | Mycgr3\|98717  JGI  Ensembl |  |  | 100 % |  |  |
| 13 | ggggacaagtttgtacaaaaaagcaggcttgatgTCTTCCAACCTCAAGCCTCT, ggggaccactttgtacaagaaagctgggtcCATCTCCTTGCCCTTCGGCA | Mycgr3\|100643  JGI  Ensembl | KOG0867 Glutathione S-transferase; | glutathione transferase activity; | 0 % |  |  |
| 13 | ggggacaagtttgtacaaaaaagcaggcttgatgGCCTCCGGCTCCGCCCCTGC, ggggaccactttgtacaagaaagctgggtcTCCTTTGTGAATGCACCGGC | Mycgr3\|101300  JGI  Ensembl | KOG0859 Synaptobrevin/VAMP-like protein; | integral to membrane; vesicle-mediated transport; | 100 % |  |  |
| 13 | ggggacaagtttgtacaaaaaagcaggcttgatgCTGTCCACGTTCCCACGCAA, ggggaccactttgtacaagaaagctgggtcCGCGCTCTCACCGTCAGCTG | Mycgr3\|102552  JGI  Ensembl |  |  | 100 % |  |  |
| 13 | ggggacaagtttgtacaaaaaagcaggcttgatgCACGTCAAAGACCGCACCTT, ggggaccactttgtacaagaaagctgggtcCATCTTTGCCGGCATGCGGA | Mycgr3\|103663  JGI  Ensembl | KOG1199 Short-chain alcohol dehydrogenase/3-hydroxyacyl-CoA dehydrogenase; | 3-hydroxyacyl-CoA dehydrogenase activity; metabolic process;  oxidoreductase activity; | 100 % |  |  |
| 13 | ggggacaagtttgtacaaaaaagcaggcttgatgCGCACATGGGCACAATTGGT, ggggaccactttgtacaagaaagctgggtcCCTGTCTAGCAATGTCAGCG | Mycgr3\|103811  JGI  Ensembl |  |  | 100 % |  | ``` 1	ACCCGACACCGCATCACGCGCTCGGGTGCTCAAGCCATTCAT	42 	\|\|\|\|\|\|\|\|\|\|\|\|\|\|\|\|\|\|\|\|\|\|\|\|\|\|\|\|\|\|\|\|\|\|\|\|\|\|\|\|\|\| 	ACCCGACACCGCATCACGCGCTCGGGTGCTCAAGCCATTCAT ``` |
| 13 | ggggacaagtttgtacaaaaaagcaggcttgatgGCTCAATCATTCCGCCAGCT, ggggaccactttgtacaagaaagctgggtcCTTGATATCCTTGCTCTGCA | Mycgr3\|103912  JGI  Ensembl |  |  | 100 % |  |  |
| 13 | ggggacaagtttgtacaaaaaagcaggcttgatgAGCGCCTACAGCAAGCAGAA, ggggaccactttgtacaagaaagctgggtcGGTCGCTGCGGAAAACTTGG | Mycgr3\|104022  JGI  Ensembl | KOG4259 Putative nucleic acid-binding protein Hcc-1/proliferation associated cytokine-inducible protein, contains SAP domain; | nucleic acid binding; | 100 % |  |  |
| 13 | ggggacaagtttgtacaaaaaagcaggcttgatgAGGGCCACTCTACGCCTCTT, ggggaccactttgtacaagaaagctgggtcCGGCGCCTGAGTCTTCCCAA | Mycgr3\|104644  JGI  Ensembl | KOG3365 NADH:ubiquinone oxidoreductase, NDUFA5/B13 subunit; | electron transport; mitochondrial inner membrane;  oxidoreductase activity, acting on NADH or NADPH; | 100 % |  |  |
| 13 | ggggacaagtttgtacaaaaaagcaggcttgatgCAAAGAATCACATCCACAAC, ggggaccactttgtacaagaaagctgggtcCTCGATGGTGATGCGCTTCG | Mycgr3\|104881  JGI  Ensembl | KOG0710 Molecular chaperone (small heat-shock protein Hsp26/Hsp42); |  | 100 % |  |  |
| 13 | ggggacaagtttgtacaaaaaagcaggcttgatgCCTGAACGATCCGGCTTCCA, ggggaccactttgtacaagaaagctgggtcACGTGGATACGGCGCACGGA | Mycgr3\|105469  JGI  Ensembl |  |  | 100 % |  |  |
| 13 | ggggacaagtttgtacaaaaaagcaggcttgatgGTTTTTAGCTGCTTTGGAAG, ggggaccactttgtacaagaaagctgggtcCAGCCTTGCGACCAGGCCGA | Mycgr3\|105798  JGI  Ensembl |  |  | 0 % |  |  |
| 13 | ggggacaagtttgtacaaaaaagcaggcttgatgAAGTACGGTGTTTGGCTTCT, ggggaccactttgtacaagaaagctgggtcCGAACCTTGCCCGACTCCGA | Mycgr3\|106672  JGI  Ensembl |  |  | 100 % |  |  |
| 13 | ggggacaagtttgtacaaaaaagcaggcttgatgCATCCGAGCCAATGGATGCG, ggggaccactttgtacaagaaagctgggtcCCCTAGATACCTCACAAGTG | Mycgr3\|106928  JGI  Ensembl |  |  | 2.2864 % |  |  |
| 13 | ggggacaagtttgtacaaaaaagcaggcttgatgGACAGAGATCTCGTGCTCCC, ggggaccactttgtacaagaaagctgggtcCACTTCCTGCTTCTCGCCCC | Mycgr3\|107417  JGI  Ensembl |  |  | 0 % |  |  |
| 13 | ggggacaagtttgtacaaaaaagcaggcttgatgTCTCTCAAGCGGAAGCAGGA, ggggaccactttgtacaagaaagctgggtcTGGCGGCTTCTCGCCATCTT | Mycgr3\|107461  JGI  Ensembl | KOG3299 Uncharacterized conserved protein; |  | 100 % |  |  |
| 13 | ggggacaagtttgtacaaaaaagcaggcttgatgTCTGGACCATACGACCAAGG, ggggaccactttgtacaagaaagctgggtcCCAGCGCTGATTGCTGTGCT | Mycgr3\|107982  JGI  Ensembl | KOG1676 K-homology type RNA binding proteins; |  | 2.95858 % |  |  |
| 14 | ggggacaagtttgtacaaaaaagcaggcttgatgTCTCCTCACTCCACGGTGGT, ggggaccactttgtacaagaaagctgggtcTCCTTTCCTCAACATCCAGC | Mycgr3\|34180  JGI  Ensembl |  |  | 100 % |  |  |
| 14 | ggggacaagtttgtacaaaaaagcaggcttgatgCCCTCCTCCCGCCAATACAA, ggggaccactttgtacaagaaagctgggtcTGCCTTGAATTCCCTCTTGA | Mycgr3\|35060  JGI  Ensembl | KOG1752 Glutaredoxin and related proteins; | cell redox homeostasis; electron carrier activity;  protein disulfide oxidoreductase activity; | 100 % |  |  |
| 14 | ggggacaagtttgtacaaaaaagcaggcttgatgGCACTATCAGAAAAGATCTG, ggggaccactttgtacaagaaagctgggtcCCTCCACAGCACACTCTGCT | Mycgr3\|35783  JGI  Ensembl |  |  | 100 % |  |  |
| 14 | ggggacaagtttgtacaaaaaagcaggcttgatgGTCGCCTTTTCCTCGACATC, ggggaccactttgtacaagaaagctgggtcGACTTGCTTCTCTCCAAGCG | Mycgr3\|36430  JGI  Ensembl |  |  | 100 % |  |  |
| 14 | ggggacaagtttgtacaaaaaagcaggcttgatgTCCTCCCCTCCCGCAGAAAC, ggggaccactttgtacaagaaagctgggtcCAACTTCGAAAAGTCCCCAC | Mycgr3\|37766  JGI  Ensembl | KOG1680 Enoyl-CoA hydratase; | catalytic activity; enoyl-CoA hydratase activity;  metabolic process; | 100 % |  |  |
| 14 | ggggacaagtttgtacaaaaaagcaggcttgatgACGAGCTCAAACGACATCTC, ggggaccactttgtacaagaaagctgggtcCTTCAGACCGCCAGCTATGA | Mycgr3\|38460  JGI  Ensembl | KOG0725 Reductases with broad range of substrate specificities; | carbonyl reductase (NADPH) activity; metabolic process;  oxidoreductase activity; | 100 % |  |  |
| 14 | ggggacaagtttgtacaaaaaagcaggcttgatgTCCGTCGACTCAAGTCGTGT, ggggaccactttgtacaagaaagctgggtcAGCAGGATGGACTCCCCCAT | Mycgr3\|39543  JGI  Ensembl | KOG0725 Reductases with broad range of substrate specificities; | metabolic process; oxidoreductase activity; | 100 % |  |  |
| 14 | ggggacaagtttgtacaaaaaagcaggcttgatgATTTTCTCCCGAACCTCAAA, ggggaccactttgtacaagaaagctgggtcAGCATTAATCTGATGCTGCC | Mycgr3\|40317  JGI  Ensembl | KOG0382 Carbonic anhydrase; | carbonate dehydratase activity; one-carbon compound metabolic process;  zinc ion binding; | 0 % |  |  |
| 14 | ggggacaagtttgtacaaaaaagcaggcttgatgTCGGGAATAGGCGCAGAGCT, ggggaccactttgtacaagaaagctgggtcCAGGTACGATCCCCCTGAAA | Mycgr3\|41539  JGI  Ensembl | KOG4188 Uncharacterized conserved protein; |  | 100 % |  |  |
| 14 | ggggacaagtttgtacaaaaaagcaggcttgatgTCTCCATCAGTAACAGTCCC, ggggaccactttgtacaagaaagctgggtcCGCCGCCGCACAATGCCCAC | Mycgr3\|41667  JGI  Ensembl | KOG1200 Mitochondrial/plastidial beta-ketoacyl-ACP reductase; | D-arabinitol 2-dehydrogenase activity; metabolic process;  oxidoreductase activity; | 100 % |  |  |
| 14 | ggggacaagtttgtacaaaaaagcaggcttgatgGCTTCCGGCGCAAAGAAGCT, ggggaccactttgtacaagaaagctgggtcCTTCGCCGTCTCCTGCTGCT | Mycgr3\|41701  JGI  Ensembl |  |  | 2.46361 % |  |  |
| 14 | ggggacaagtttgtacaaaaaagcaggcttgatgCCTCCAACAACAGCACTCTC, ggggaccactttgtacaagaaagctgggtcATCCCTGGCCTCCTTGTCCA | Mycgr3\|42788  JGI  Ensembl |  |  | 100 % |  |  |
| 14 | ggggacaagtttgtacaaaaaagcaggcttgatgCTAACCTTAAACCCAGCCCT, ggggaccactttgtacaagaaagctgggtcAACTACAATCTCCTTCCCAC | Mycgr3\|43486  JGI  Ensembl |  |  | 100 % |  |  |
| 14 | ggggacaagtttgtacaaaaaagcaggcttgatgCAGCAGGCCACGGACGAACT, ggggaccactttgtacaagaaagctgggtcAACTGTTGGATCACCTTTGT | Mycgr3\|44136  JGI  Ensembl |  |  | 100 % |  |  |
| 14 | ggggacaagtttgtacaaaaaagcaggcttgatgGCACCACCAAGCAAGCTCCG, ggggaccactttgtacaagaaagctgggtcATCTTTCTTGACCAGAATCA | Mycgr3\|46885  JGI  Ensembl |  |  | 0 % |  |  |
| 14 | ggggacaagtttgtacaaaaaagcaggcttgatgGCCGATTCCGACTCCGACTC, ggggaccactttgtacaagaaagctgggtcTGTCTTCTTCCCAGCACCCT | Mycgr3\|47396  JGI  Ensembl | KOG4234 TPR repeat-containing protein; |  | 100 % |  |  |
| 14 | ggggacaagtttgtacaaaaaagcaggcttgatgACATTCCCTCTTCCTCCTGC, ggggaccactttgtacaagaaagctgggtcCAGCACTCCAGGAAACTGCT | Mycgr3\|49006  JGI  Ensembl |  |  | 0 % |  |  |
| 14 | ggggacaagtttgtacaaaaaagcaggcttgatgCATCTCCTATCCATCCTTCT, ggggaccactttgtacaagaaagctgggtcCAACTCATCCCTTCCCTGCG | Mycgr3\|49854  JGI  Ensembl |  |  | 0 % |  |  |
| 14 | ggggacaagtttgtacaaaaaagcaggcttgatgGGCCGCTACGACTTTGCCGC, ggggaccactttgtacaagaaagctgggtcAGGATGAAGTGCCACACCAC | Mycgr3\|50818  JGI  Ensembl |  |  | 100 % |  |  |
| 14 | ggggacaagtttgtacaaaaaagcaggcttgatgAATGTCTCCGCAACGGTCAA, ggggaccactttgtacaagaaagctgggtcCTCGAAGTCACTGCGCGGTT | Mycgr3\|55823  JGI  Ensembl | KOG2961 Predicted hydrolase (HAD superfamily); |  | 100 % |  |  |
| 14 | ggggacaagtttgtacaaaaaagcaggcttgatgGGCCTCGCCTACAACATCTA, ggggaccactttgtacaagaaagctgggtcATTCACCTGACACAACAACT | Mycgr3\|58850  JGI  Ensembl | KOG3399 Predicted Yippee-type zinc-binding protein; | FK506-sensitive peptidyl-prolyl cis-trans isomerase; cyclophilin; cyclophilin-type peptidyl-prolyl cis-trans isomerase activity;  peptidyl-prolyl cis-trans isomerase activity; | 100 % |  |  |
| 14 | ggggacaagtttgtacaaaaaagcaggcttgatgCACTCAGAACACTCTCAGAA, ggggaccactttgtacaagaaagctgggtcCTTGCCCGCCGCCTCATCCT | Mycgr3\|64883  JGI  Ensembl |  |  | 100 % |  |  |
| 14 | ggggacaagtttgtacaaaaaagcaggcttgatgTCGACCGCGGCGCGCCGACG, ggggaccactttgtacaagaaagctgggtcATCATCCCAACTCTTCTCCA | Mycgr3\|65747  JGI  Ensembl | KOG0419 Ubiquitin-protein ligase; | negative regulation of ubiquitin-protein ligase activity; positive regulation of ubiquitin-protein ligase activity;  post-translational protein modification;  regulation of protein metabolic process;  regulation of ubiquitin-protein ligase activity;  regulation of ubiquitin-protein ligase activity during meiotic cell cycle;  small conjugating protein ligase activity;  ubiquitin-protein ligase activity; | 0 % |  |  |
| 14 | ggggacaagtttgtacaaaaaagcaggcttgatgTCGGAATACTGGAAGTCGAC, ggggaccactttgtacaagaaagctgggtcCTTCTTGCCTTTCCGTTTCT | Mycgr3\|67069  JGI  Ensembl | KOG0150 Spliceosomal protein FBP21; | nucleic acid binding; nucleus; zinc ion binding; | 0 % |  |  |
| 14 | ggggacaagtttgtacaaaaaagcaggcttgatgTGGGAACCAGTCAGCCTTCC, ggggaccactttgtacaagaaagctgggtcAATACGCATGTCTTTGTCAA | Mycgr3\|70331  JGI  Ensembl |  |  | 100 % |  |  |
| 14 | ggggacaagtttgtacaaaaaagcaggcttgatgGCAACATCTACCTCCGGAAC, ggggaccactttgtacaagaaagctgggtcTAGCTTCGCCTTCCCACTCT | Mycgr3\|70943  JGI  Ensembl | KOG1455 Lysophospholipase; | acylglycerol lipase activity; catalytic activity; | 100 % |  |  |
| 14 | ggggacaagtttgtacaaaaaagcaggcttgatgGCAGCACCTCGGGCGCTTCC, ggggaccactttgtacaagaaagctgggtcTGGTCCACTGCTTCCTCTCT | Mycgr3\|73826  JGI  Ensembl |  |  | 100 % |  |  |
| 14 | ggggacaagtttgtacaaaaaagcaggcttgatgCTCAAGGCGTGGCTGCGCGA, ggggaccactttgtacaagaaagctgggtcTCCTCGCCCATAAAGCCCAG | Mycgr3\|74100  JGI  Ensembl | KOG4269 Rac GTPase-activating protein BCR/ABR; | intracellular; signal transduction; | 62.41611 % |  |  |
| 14 | ggggacaagtttgtacaaaaaagcaggcttgatgTCCTACACCACCACATCTCG, ggggaccactttgtacaagaaagctgggtcGTAAAGGAAGGGCACCACCT | Mycgr3\|74856  JGI  Ensembl | KOG4650 Predicted steroid reductase; | cytoplasm; integral to membrane;  lipid metabolic process;  oxidoreductase activity, acting on the CH-CH group of donors; | 2.23793 % |  |  |
| 14 | ggggacaagtttgtacaaaaaagcaggcttgatgGAGCACCTCCCCGAATCCAA, ggggaccactttgtacaagaaagctgggtcCTCCTTGCTCTTACTCTTCC | Mycgr3\|76119  JGI  Ensembl |  |  | 100 % |  |  |
| 14 | ggggacaagtttgtacaaaaaagcaggcttgatgAAATTCTTCTCCTCCTCCCA, ggggaccactttgtacaagaaagctgggtcCGCCCTCTGCATCTCCCTCT | Mycgr3\|77228  JGI  Ensembl | KOG3336 Predicted member of the intramitochondrial sorting protein family; |  | 100 % |  |  |
| 14 | ggggacaagtttgtacaaaaaagcaggcttgatgCGTTTCACCACCGCCGTCCT, ggggaccactttgtacaagaaagctgggtcAAGCATGCCGATGTGGTCGA | Mycgr3\|81079  JGI  Ensembl | KOG0260 RNA polymerase II, large subunit; |  | 100 % |  |  |
| 14 | ggggacaagtttgtacaaaaaagcaggcttgatgACATCCAAGAAAGTCCTCGT, ggggaccactttgtacaagaaagctgggtcAAACTCCGCTCCGACAAACT | Mycgr3\|84210  JGI  Ensembl |  |  | 100 % |  |  |
| 14 | ggggacaagtttgtacaaaaaagcaggcttgatgGATCTTCGAGGACTCACACC, ggggaccactttgtacaagaaagctgggtcGTTCAATCCACCCGCGCCAA | Mycgr3\|84757  JGI  Ensembl |  |  | 100 % |  |  |
| 14 | ggggacaagtttgtacaaaaaagcaggcttgatgTCTATGTCCGTCGAAGACGC, ggggaccactttgtacaagaaagctgggtcCGGAAGTACATACCCTCCAT | Mycgr3\|84831  JGI  Ensembl | KOG0725 Reductases with broad range of substrate specificities; | carbonyl reductase (NADPH) activity; metabolic process;  oxidoreductase activity; | 100 % |  |  |
| 14 | ggggacaagtttgtacaaaaaagcaggcttgatgACCTCCGCCGCCGCCGACGA, ggggaccactttgtacaagaaagctgggtcACCATGCTCGTCTTCACTCA | Mycgr3\|85860  JGI  Ensembl | KOG1994 Predicted RNA binding protein, contains G-patch and Zn-finger domains; | intracellular; nucleic acid binding; | 2.32558 % |  |  |
| 14 | ggggacaagtttgtacaaaaaagcaggcttgatgTCAACCGAGGAAGCGATCTC, ggggaccactttgtacaagaaagctgggtcAAAGTCCACCCCTCTCCGTA | Mycgr3\|88390  JGI  Ensembl | KOG0118 FOG: RRM domain; | nucleic acid binding; | 0 % |  |  |
| 14 | ggggacaagtttgtacaaaaaagcaggcttgatgGACCGGAACCCGGGCCAAGC, ggggaccactttgtacaagaaagctgggtcGGCCCACTGGACGGCGACAT | Mycgr3\|88732  JGI  Ensembl |  |  | 0 % |  |  |
| 14 | ggggacaagtttgtacaaaaaagcaggcttgatgATCCCCGGCAGAGCCAAAAA, ggggaccactttgtacaagaaagctgggtcAAACCCTTCTTCGCATCGCA | Mycgr3\|89072  JGI  Ensembl |  |  | 100 % |  |  |
| 14 | ggggacaagtttgtacaaaaaagcaggcttgatgACGACCACCGATCTCGAGAA, ggggaccactttgtacaagaaagctgggtcAGCGCAAGCCTCTCCCGGCA | Mycgr3\|89604  JGI  Ensembl | KOG1216 von Willebrand factor and related coagulation proteins; |  | 100 % |  |  |
| 14 | ggggacaagtttgtacaaaaaagcaggcttgatgTCGACAACTCGCTTCGCGAA, ggggaccactttgtacaagaaagctgggtcTCTTCCACCAGTGATATCGA | Mycgr3\|89635  JGI  Ensembl | KOG1200 Mitochondrial/plastidial beta-ketoacyl-ACP reductase; | estradiol 17-beta-dehydrogenase activity; metabolic process;  oxidoreductase activity; | 0 % |  |  |
| 14 | ggggacaagtttgtacaaaaaagcaggcttgatgGCGCGACTCTGGACAAGCTT, ggggaccactttgtacaagaaagctgggtcGCTCGCTGTCGTGTCCAGGG | Mycgr3\|89637  JGI  Ensembl | KOG0161 Myosin class II heavy chain; | 3-phosphoinositide-dependent protein kinase activity; AMP-activated protein kinase activity;  DNA-dependent protein kinase activity;  GTP-dependent protein kinase activity;  JUN kinase activity;  JUN kinase kinase activity;  JUN kinase kinase kinase activity;  Janus kinase activity;  MAP kinase 1 activity;  MAP kinase 2 activity;  MAP kinase kinase activity;  MAP kinase kinase kinase kinase activity;  MAP/ERK kinase kinase activity;  MP kinase activity;  NF-kappaB-inducing kinase activity;  SAP kinase activity;  atypical protein kinase C activity;  calcium-dependent protein kinase C activity;  casein kinase I activity;  casein kinase activity;  cyclic nucleotide-dependent protein kinase activity;  cyclin-dependent protein kinase activating kinase activity;  eukaryotic translation initiation factor 2alpha kinase activity;  galactosyltransferase-associated kinase activity;  glycogen synthase kinase 3 activity;  receptor signaling protein serine/threonine kinase activity;  ribosomal protein S6 kinase activity;  transmembrane receptor protein kinase activity;  transmembrane receptor protein serine/threonine kinase activity; | 0 % |  |  |
| 14 | ggggacaagtttgtacaaaaaagcaggcttgatgGTCACTCCGAAAATGTCTTG, ggggaccactttgtacaagaaagctgggtcTGGCGTTGCATTCGCAGACA | Mycgr3\|89984  JGI  Ensembl | KOG4157 beta-1,6-N-acetylglucosaminyltransferase, contains WSC domain; |  | 100 % |  |  |
| 14 | ggggacaagtttgtacaaaaaagcaggcttgatgGCGAAGAATGGGAAGAGCTT, ggggaccactttgtacaagaaagctgggtcGTCACCCTCCCTCTTCACTC | Mycgr3\|90122  JGI  Ensembl |  |  | 2.2779 % |  |  |
| 14 | ggggacaagtttgtacaaaaaagcaggcttgatgACGTCCGAACTTCACCTCGC, ggggaccactttgtacaagaaagctgggtcACGAGGCATTGATAATGCGC | Mycgr3\|90408  JGI  Ensembl |  |  | 100 % |  |  |
| 14 | ggggacaagtttgtacaaaaaagcaggcttgatgCAGTCCGCCAGCATGGCTTC, ggggaccactttgtacaagaaagctgggtcCCTCGAGCAAACACGACAAT | Mycgr3\|91382  JGI  Ensembl |  |  | 100 % |  |  |
| 14 | ggggacaagtttgtacaaaaaagcaggcttgatgCGCACCACCATAGCGGCTTC, ggggaccactttgtacaagaaagctgggtcACAAACCCCCACCACCGCGA | Mycgr3\|92048  JGI  Ensembl |  |  | 2.14447 % |  |  |
| 14 | ggggacaagtttgtacaaaaaagcaggcttgatgGCGAATCCCACAGCTCAACA, ggggaccactttgtacaagaaagctgggtcTTCCGATGCCATGCCCTCCT | Mycgr3\|92216  JGI  Ensembl |  |  | 100 % |  |  |
| 14 | ggggacaagtttgtacaaaaaagcaggcttgatgGCGTCTCCTTACCAGCAGAC, ggggaccactttgtacaagaaagctgggtcTGCGCCGGTACTGAAAGCGA | Mycgr3\|92308  JGI  Ensembl |  |  | 100 % |  |  |
| 14 | ggggacaagtttgtacaaaaaagcaggcttgatgGCCGACTCAAACGAGGCCAG, ggggaccactttgtacaagaaagctgggtcAGCACCACCTCCCTCGTTTC | Mycgr3\|92485  JGI  Ensembl |  |  | 0 % |  |  |
| 14 | ggggacaagtttgtacaaaaaagcaggcttgatgGCCAACCTGTTCAACCCCGA, ggggaccactttgtacaagaaagctgggtcAGAAGCTTTCATATCAGGCT | Mycgr3\|93087  JGI  Ensembl |  |  | 0 % |  |  |
| 14 | ggggacaagtttgtacaaaaaagcaggcttgatgAAGCACATCCCAGAGCGCCG, ggggaccactttgtacaagaaagctgggtcTCGTGAGCTTAAGCCCTCGT | Mycgr3\|93582  JGI  Ensembl |  |  | 100 % |  |  |
| 14 | ggggacaagtttgtacaaaaaagcaggcttgatgACCGACCTATCAAGCGCAGC, ggggaccactttgtacaagaaagctgggtcAGCGAAAGGCAAGCCGCTCG | Mycgr3\|93586  JGI  Ensembl |  |  | 0 % |  |  |
| 14 | ggggacaagtttgtacaaaaaagcaggcttgatgGTCACACCTCATGGTCAGCG, ggggaccactttgtacaagaaagctgggtcGAGGCAAGAGTGATTGGCCC | Mycgr3\|93882  JGI  Ensembl | KOG2533 Permease of the major facilitator superfamily; |  | 100 % |  |  |
| 14 | ggggacaagtttgtacaaaaaagcaggcttgatgTTGTACATCATCGGCCTGGG, ggggaccactttgtacaagaaagctgggtcTTTTTGGATACCATAGTCTC | Mycgr3\|93945  JGI  Ensembl | KOG3123 Diphthine synthase; | diphthine synthase activity; metabolic process;  methyltransferase activity;  peptidyl-diphthamide biosynthetic process from peptidyl-histidine; | 100 % |  |  |
| 14 | ggggacaagtttgtacaaaaaagcaggcttgatgTCAACTGGCACCATCGGCGT, ggggaccactttgtacaagaaagctgggtcGAGTCCAGCGTCGGTCAGCT | Mycgr3\|94431  JGI  Ensembl |  |  | 100 % |  |  |
| 14 | ggggacaagtttgtacaaaaaagcaggcttgatgACTCCACCCTCTCCGCACGC, ggggaccactttgtacaagaaagctgggtcGCTGCCCGTTTTGTCTTTCT | Mycgr3\|94674  JGI  Ensembl |  |  | 100 % |  |  |
| 14 | ggggacaagtttgtacaaaaaagcaggcttgatgGATGGAATGCGTGACTTGAC, ggggaccactttgtacaagaaagctgggtcCTCCTCGTCGTCGGTATATT | Mycgr3\|94804  JGI  Ensembl |  |  | 100 % |  |  |
| 14 | ggggacaagtttgtacaaaaaagcaggcttgatgTTCCGGATCTGGGATCTCGC, ggggaccactttgtacaagaaagctgggtcCTCGGACTCATCCATGCCCT | Mycgr3\|94805  JGI  Ensembl |  |  | 100 % |  |  |
| 14 | ggggacaagtttgtacaaaaaagcaggcttgatgTCTTCATCAACGGCATACGG, ggggaccactttgtacaagaaagctgggtcATCACGTCTCAGACCTTCGT | Mycgr3\|95079  JGI  Ensembl |  |  | 100 % |  |  |
| 14 | ggggacaagtttgtacaaaaaagcaggcttgatgTCTTCCACCACGCACAACTG, ggggaccactttgtacaagaaagctgggtcGTTCTGCGCTGGCGTGTTGG | Mycgr3\|95080  JGI  Ensembl |  |  | 0 % |  |  |
| 14 | ggggacaagtttgtacaaaaaagcaggcttgatgCCAGTACACAACAATGACCC, ggggaccactttgtacaagaaagctgggtcAACCCTGTCGAGTTCCTCAA | Mycgr3\|95327  JGI  Ensembl | KOG0939 E3 ubiquitin-protein ligase/Putative upstream regulatory element binding protein; | intracellular; protein modification process;  ubiquitin cycle;  ubiquitin-protein ligase activity; | 100 % |  |  |
| 14 | ggggacaagtttgtacaaaaaagcaggcttgatgTCCACCAACACCCAATCCTC, ggggaccactttgtacaagaaagctgggtcTCGAGGCAGATCCTGAAACT | Mycgr3\|95438  JGI  Ensembl | KOG2249 3'-5' exonuclease; | exonuclease activity; intracellular; | 100 % |  |  |
| 14 | ggggacaagtttgtacaaaaaagcaggcttgatgAACGACGACCGGTATTCCCG, ggggaccactttgtacaagaaagctgggtcATATCTCGGACGCCGGAACG | Mycgr3\|96164  JGI  Ensembl | KOG4676 Splicing factor, arginine/serine-rich; |  | 100 % |  |  |
| 14 | ggggacaagtttgtacaaaaaagcaggcttgatgCCACCCAAAGCCTCCCAAAC, ggggaccactttgtacaagaaagctgggtcACTTCCGTTGTTGTTCCGGG | Mycgr3\|96401  JGI  Ensembl |  |  | 100 % |  |  |
| 14 | ggggacaagtttgtacaaaaaagcaggcttgatgGACAGTCTCATGCACACCAG, ggggaccactttgtacaagaaagctgggtcTCTGTAGCGCAGTACATAAT | Mycgr3\|96812  JGI  Ensembl |  |  | 100 % |  |  |
| 14 | ggggacaagtttgtacaaaaaagcaggcttgatgGGTATTATCAAGGCATTGTC, ggggaccactttgtacaagaaagctgggtcTTGTAGTCGTTCCTGCAGTG | Mycgr3\|97057  JGI  Ensembl | KOG2806 Chitinase; | negative regulation of ubiquitin-protein ligase activity; positive regulation of ubiquitin-protein ligase activity;  regulation of ubiquitin-protein ligase activity;  regulation of ubiquitin-protein ligase activity during meiotic cell cycle;  ubiquitin-protein ligase activity; | 100 % |  |  |
| 14 | ggggacaagtttgtacaaaaaagcaggcttgatgGCCGCCCTTCGCAAAACTTT, ggggaccactttgtacaagaaagctgggtcCAGCCACGATTTCGTACGTA | Mycgr3\|97075  JGI  Ensembl |  |  | 100 % |  |  |
| 14 | ggggacaagtttgtacaaaaaagcaggcttgatgGTCAACAAAACAGTGAAGAG, ggggaccactttgtacaagaaagctgggtcCCTCATCACAACCACCCGAT | Mycgr3\|97141  JGI  Ensembl | KOG1216 von Willebrand factor and related coagulation proteins; |  | 2.17155 % |  |  |
| 14 | ggggacaagtttgtacaaaaaagcaggcttgatgGCGCCGCTCTTCCCACGAGT, ggggaccactttgtacaagaaagctgggtcGTCCATGTCGGCGATGTGCT | Mycgr3\|97404  JGI  Ensembl |  |  | 100 % |  |  |
| 14 | ggggacaagtttgtacaaaaaagcaggcttgatgACGGGAAAGCTGAAAGCGAT, ggggaccactttgtacaagaaagctgggtcTGCGTGGCGCGCCGAATCCA | Mycgr3\|97557  JGI  Ensembl |  |  | 0 % |  |  |
| 14 | ggggacaagtttgtacaaaaaagcaggcttgatgGACGTCGAAGACCAAGACTC, ggggaccactttgtacaagaaagctgggtcCTCCTGATCCCACCACCCAC | Mycgr3\|97711  JGI  Ensembl |  |  | 100 % |  |  |
| 14 | ggggacaagtttgtacaaaaaagcaggcttgatgGCGAAGATCGAGAGTGGACC, ggggaccactttgtacaagaaagctgggtcGATCTCTCTTTTCCCCAGGC | Mycgr3\|97765  JGI  Ensembl | KOG2593 Transcription initiation factor IIE, alpha subunit; |  | 100 % |  |  |
| 14 | ggggacaagtttgtacaaaaaagcaggcttgatgGACCTTACGAAGTGGACTAC, ggggaccactttgtacaagaaagctgggtcGCGCTGATCCGCCTCGCAAT | Mycgr3\|97779  JGI  Ensembl |  |  | 100 % |  | ``` 694	GTCACTCCCTTCACAACCTTCTCGCCGCGGGAGCACCGGGACACCAGCGACACGGACTACTCCATCCTCGCCAACTTTGGTGCCGGCTGCTGGCTCGTCCTGCCTGAGCTCCAGCTCCGCGTCCACCTCCAGCCCTACGACCTCGTAATCTTCCAGACGAACAGCTTGACCCACGCCACAGCGGCCGTCGACGGCGATTGCGAGGCGGATCAGCGCT	910 	\|\|\|\|\|\|\|\|\|\|\|\|\|\| \|\|\|\|\|\|\|\|\|\|\|\|\|\|\|\|\|\|\|\|\|\|\|\|\|\|\|\|\|\|\|\|\|\|\|\|\|\|\|\|\|\|\|\|\|\|\|\|\|\|\|\|\| \|\|\|\|\|\|\|\|\|\|\|\|\|\| \|\|\|\|\|\|\|\|\|\|\|\|\|\|\|\|\|\|\|\|\| \|\|\|\|\|\|\|\|\|\|\|\|\|\|\|\|\|\|\|\|\|\|\|\|\|\|\|\|\|\|\|\|\|\|\|\|\|\|\|\| \|\|\|\|\|\|\|\|\|\|\|\|\|\|\|\|\|\|\|\| \|\|\|\|\|\|\|\|\|\|\| \|\|\|\|\| \|\|\|\|\|\|\|\|\|\|\|\|\|\|\|\|\|\|\|\|\|\|\|\|\|\|\|\|\|\|\| 	GTCACTCCCTTCACGACCTTCTCGCCGCGGGAGCACCGGGACACCAGCGACACGGACTACTCCATCCTTGCCAACTTTGGTGCTGGCTGCTGGCTCGTCCTGCCTAAGCTCCAGCTCCGCGTCCACCTCCAGCCCTACGACCTCGTGATCTTCCAGACGAACAGCTTAACCCACGCCACGGCGGCTGTCGACGGCGATTGCGAGGCGGATCAGCGCT ```      ``` 694	GTCAC-TCCCTTCACAACCTTCTCGCCGCGGGAGCACCGGGACACCAGCGACACGGACTACTCCATCCTCGCCAACTTTGGTGCCGGCTGCTGGCTCGTCCTGCCTGAGCTCCAGCTCCGCGTCCACCTCCAGCCCTACGACCTCGTAATCTTCCAGACGAACAGCTTGACCCACGCCACAGCGGCCGTCGACGGCGATTGCGAGGCGGATCAG	906 	\|\|\|\|\| \|\|\|\|\|\|\|\|\| \|\|\|\|\|\|\|\|\|\|\|\|\|\|\|\|\|\|\|\|\|\|\|\|\|\|\|\|\|\|\|\|\|\|\|\|\|\|\|\|\|\|\|\|\|\|\|\|\|\|\|\|\| \|\|\|\|\|\|\|\|\|\|\|\|\|\| \|\|\|\|\|\|\|\|\|\|\|\|\|\|\|\|\|\|\|\|\| \|\|\|\|\|\|\|\|\|\|\|\|\|\|\|\|\|\|\|\|\|\|\|\|\|\|\|\|\|\|\|\|\|\|\|\|\|\|\|\| \|\|\|\|\|\|\|\|\|\|\|\|\|\|\|\|\|\|\|\| \|\|\|\|\|\|\|\|\|\|\| \|\|\|\|\| \| \|\|\|\|\|\| \|\|\|\|\|  \| \|\|\|\|\|\|\|\|\| 	GTCACATCCCTTCACGACCTTCTCGCCGCGGGAGCACCGGGACACCAGCGACACGGACTACTCCATCCTTGCCAACTTTGGTGCTGGCTGCTGGCTCGTCCTGCCTAAGCTCCAGCTCCGCGTCCACCTCCAGCCCTACGACCTCGTGATCTTCCAGACGAACAGCTTAACCCACGCCACGGCGGCTGNCGACGGAGATTGNNANGCGGATCAG ```      ``` 694	GTCACTCCCTTCACAACCTTCTCGCCGCGGGAGCACCGGGACACCAGCGACACGGACTACTCCATCCTCGCCAACTTTGGTGCCGGCTGCTGGCTCGTCCTGCCTGAGCTCCAGCTCCGCGTCCACCTCCAGCCCTACGACCTCGTAATCTTCCAGACGAACAGCTTGACCCACGCCACAGCGGCCGTCGACGGCGATTGCGAGGCGGATCAGCGCT	910 	\|\|\|\|\|\|\|\|\|\|\|\|\|\| \|\|\|\|\|\|\|\|\|\|\|\|\|\|\|\|\|\|\|\|\|\|\|\|\|\|\|\|\|\|\|\|\|\|\|\|\|\|\|\|\|\|\|\|\|\|\|\|\|\|\|\|\|\|\|\|\|\|\|\|\|\|\|\|\|\|\|\|\|\|\|\|\|\|\|\|\|\|\|\|\|\|\|\|\|\|\|\|\|\|\|\|\|\|\|\|\|\|\|\|\|\|\|\|\|\|\|\|\|\|\|\|\|\|\|\|\|\|\|\|\|\|\|\|\|\|\|\|\|\|\|\|\|\|\|\|\|\|\|\|\|\|\|\|\|\|\|\|\|\|\|\|\|\|\|\|\|\|\|\|\|\|\|\|\|\|\|\|\|\|\|\|\|\|\|\|\|\|\|\|\|\|\|\|\|\|\|\|\|\|\|\|\|\|\|\|\|\|\|\|\|\| 	GTCACTCCCTTCACGACCTTCTCGCCGCGGGAGCACCGGGACACCAGCGACACGGACTACTCCATCCTCGCCAACTTTGGTGCCGGCTGCTGGCTCGTCCTGCCTGAGCTCCAGCTCCGCGTCCACCTCCAGCCCTACGACCTCGTAATCTTCCAGACGAACAGCTTGACCCACGCCACAGCGGCCGTCGACGGCGATTGCGAGGCGGATCAGCGCT ``` |
| 14 | ggggacaagtttgtacaaaaaagcaggcttgatgGCGCGCCTTGGTGCACGAGA, ggggaccactttgtacaagaaagctgggtcCCATAGCCGACAAGCGGTGG | Mycgr3\|97957  JGI  Ensembl |  |  | 0 % |  |  |
| 14 | ggggacaagtttgtacaaaaaagcaggcttgatgAACAACCACCGAACTCCCGC, ggggaccactttgtacaagaaagctgggtcTGCTATCAAGTCCTCGACCG | Mycgr3\|98895  JGI  Ensembl |  |  | 100 % |  |  |
| 14 | ggggacaagtttgtacaaaaaagcaggcttgatgCTCTCGTCCTCGATAACTTC, ggggaccactttgtacaagaaagctgggtcGCAGCTGGGAGTTACAATGA | Mycgr3\|102970  JGI  Ensembl |  |  | 100 % |  |  |
| 14 | ggggacaagtttgtacaaaaaagcaggcttgatgTCGTCCAACCGACGCGGCGT, ggggaccactttgtacaagaaagctgggtcTCCTCCATGCATTAGGCTCG | Mycgr3\|103841  JGI  Ensembl | KOG3341 RNA polymerase II transcription factor complex subunit; |  | 100 % |  | ``` 1	ATGTCGTCCAACCGACGCGGCGTAGGCCTCAGTGCCTTCTCCAACGCTGCAATCTCTTCAGATAAATATGCTGCCCACGGGGCCGCACTCCGGACAAGCCATGCCGATGCACTCGCGAACCAACTCTCGGTCTTCCAGGCCGCCCTGCACAACTTCTCCTTGACCCACGCGAAAGACATTCGCAGCAATCCGACCTTTCGAGCCGAATTCGCCCGTATGTGCCATGCTATTGGGGTCGATCCACTTGCGGGAAGCAACGCCAAGGGCAATGCTGCAGCAGATGGCAAGGGTGGTAGTGTTTGGGCAAAGATGCTGGGAACAAGTTTAAACGACTTCTACTTTGAGTTGGGTGTCAGGCTGGTGGAGGTCTGTCGAGAGAGCAGAGCGGAGAATGGAGGTATGATAGCACTTGCGGAAGCTCGGAAGCGAATCGCGAAAGGGCGTGGCCTGGTAGGAGGCGGCATCGAGGTGACGGACGATGATGTCGAGCGAGCATTGGAGAGCCTCGAGCCTCTCGGGGGCATGTTCAAGATCACGAAGCTCGGCAGCACAAAGTTCATCAGATCTGTGCCGAAGGAGCTGAACCAGGATCAATCGAGAGTTCTCGAAGTCATCGACATCATGGGCTTTGTGACTGTTTCAATGCTGCGCGCGAACCTGCAATGGGAGAAGGCTAGAGCGATCGCCGTCATTGACGACATGGTAATGGACGGTCTGTTGTGGGTTGACGAGCAAGGAGAAGAAGTCGAATACTGGAGCCCGAGCCTAATGCATGGAGGA	780 	\|\|\|\|\|\|\|\|\|\|\|\|\|\|\|\|\|\|\|\|\|\|\|\|\|\|\|\|\|\|\|\|\|\|\|\|\|\|\|\|\|\|\|\|\|\|\|\|\|\|\|\|\|\|\|\|\|\|\|\|\|\|\|\|\|\|\|\|\|\|\|\|\|\|\|\|\|\|\|\|\|\|\|\|\|\|\|\|\|\|\|\|\|\|\|\|\|\|\|\|\|\|\|\|\|\|\|\|\|\|\|\|\|\|\|\|\|\|\|\|\|\|\|\|\|\|\|\|\|\|\|\|\|\|\|\|\|\|\|\|\|\|\|\|\|\|\|\|\|\|\|\|\|\|\|\|\|\|\|\|\|\|\|\|\|\|\|\|\|\|\|\|\|\|\|\|\|\|\|\|\|\|\|\|\|\|\|\|\|\|\|\|\|\|\|\|\|\|\|\|\|\|\|\|\|\|\|\|\|\|\|\|\|\|\|\|\|\|\|\|\|\|\|\|\|\|\|\|\|\|\|\|\|\|\|\|\|\|\|\|\|\|\|\|\|\|\|\|\|\|\|\|\|\|\|\|\|\|\|\|\|\|\|\|\|\|\|\|\|\|\|\|\|\|\|\|\|\|\|\|\|\|\|\|\|\|\|\|\|\|\|\|\|\|\|\|\|\|\|\|\|\|\|\|\|\|\|\|\|\|\|\|\|\|\|\|\|\|\|\|\|\|\|\|\|\|\|\|\|\|\|\|\|\|\|\|\|\|\|\|\|\|\|\|\|\|\|\|\|\|\|\|\|\|\|\|\|\|\|\|\|\|\|\|\|\|\|\|\|\|\|\|\|\|\|\|\|\|\|\|\|\|\|\|\|\|\|\|\|\|\|\|\|\|\|\|\|\|\|\|\|\|\|\|\|\|\|\|\|\|\|\|\|\|\|\|\|\|\|\|\|\|\|\|\|\|\|\|\|\|\|\|\|\|\|\|\|\|\|\|\|\|\|\|\|\|\|\|\|\|\|\|\|\|\|\|\|\|\|\|\|\|\|\|\|\|\|\|\|\|\|\|\|\|\|\|\|\|\|\|\|\|\|\|\|\|\|\|\|\|\|\|\|\|\|\|\|\|\|\|\|\|\|\|\|\|\|\|\|\|\|\|\|\|\|\|\|\|\|\|\|\|\|\|\|\|\|\|\|\|\|\|\|\|\|\|\|\|\|\|\|\|\|\|\|\|\|\|\|\|\|\|\|\|\|\|\|\|\|\|\|\|\|\|\|\|\|\|\|\|\|\|\|\|\|\|\|\|\|\|\|\|\|\|\|\|\|\|\|\|\|\|\|\|\|\|\|\|\|\|\|\|\|\|\|\|\|\|\|\|\|\|\|\|\|\|\|\|\|\|\|\|\|\|\|\|\|\|\|\|\|\|\|\|\|\|\|\|\|\|\|\|\|\|\|\|\|\|\|\|\|\|\|\|\|\|\|\|\|\|\|\|\|\|\|\|\|\|\|\|\|\|\|\|\|\|\|\|\|\|\|\|\|\|\|\|\|\|\|\|\|\|\|\|\|\|\|\|\|\|\|\|\|\|\|\|\|\|\|\|\|\|\|\|\|\|\|\|\|\|\| \|\|\|\|\|\|\|\|\|\|\|\|\|\|\|\|\|\|\|\|\|\|\|\|\|\|\|\|\|\|\|\|\|\|\|\|\|\|\|\|\|\|\|\|\|\|\|\|\|\|\|\|\|\|\|\|\|\| 	ATGTCGTCCAACCGACGCGGCGTAGGCCTCAGTGCCTTCTCCAACGCTGCAATCTCTTCAGATAAATATGCTGCCCACGGGGCCGCACTCCGGACAAGCCATGCCGATGCACTCGCGAACCAACTCTCGGTCTTCCAGGCCGCCCTGCACAACTTCTCCTTGACCCACGCGAAAGACATTCGCAGCAATCCGACCTTTCGAGCCGAATTCGCCCGTATGTGCCATGCTATTGGGGTCGATCCACTTGCGGGAAGCAACGCCAAGGGCAATGCTGCAGCAGATGGCAAGGGTGGTAGTGTTTGGGCAAAGATGCTGGGAACAAGTTTAAACGACTTCTACTTTGAGTTGGGTGTCAGGCTGGTGGAGGTCTGTCGAGAGAGCAGAGCGGAGAATGGAGGTATGATAGCACTTGCGGAAGCTCGGAAGCGAATCGCGAAAGGGCGTGGCCTGGTAGGAGGCGGCATCGAGGTGACGGACGATGATGTCGAGCGAGCATTGGAGAGCCTCGAGCCTCTCGGGGGCATGTTCAAGATCACGAAGCTCGGCAGCACAAAGTTCATCAGATCTGTGCCGAAGGAGCTGAACCAGGATCAATCGAGAGTTCTCGAAGTCATCGACATCATGGGCTTTGTGACTGTTTCAATGCTGCGCGCGAACCTGCAATGGGAGAAGGCTAGAGCGATCGCCGTCATTGACGACATGGTAATGGACGGTCTGTTGT-GGTTGACGAGCAAGGAGAAGAAGTCGAATACTGGAGCCCGAGCCTAATGCATGGAGGA ``` |
| 14 | ggggacaagtttgtacaaaaaagcaggcttgatgGCCATACTCGACAAAGCTAT, ggggaccactttgtacaagaaagctgggtcGGCTTTGGCGCGACCAGTGT | Mycgr3\|108540  JGI  Ensembl |  |  | 100 % |  |  |
| 14 | ggggacaagtttgtacaaaaaagcaggcttgatgGATCCTGTCTCTTCCGCCGC, ggggaccactttgtacaagaaagctgggtcCTCCTCAAGTGCACCACACT | Mycgr3\|108619  JGI  Ensembl |  |  | 100 % |  |  |
| 14 | ggggacaagtttgtacaaaaaagcaggcttgatgAGGTCGAGCGGGAGTACCAA, ggggaccactttgtacaagaaagctgggtcATCAGAATCATCCGCCCAAT | Mycgr3\|108829  JGI  Ensembl |  |  | 100 % |  |  |
| 14 | ggggacaagtttgtacaaaaaagcaggcttgatgCGGTCCACCCAGATCTTCAA, ggggaccactttgtacaagaaagctgggtcGAACTTCCCAGCACCCGCAG | Mycgr3\|109000  JGI  Ensembl |  |  | 100 % |  |  |
| 14 | ggggacaagtttgtacaaaaaagcaggcttgatgGCCAAACTCCTCAAGAAGCC, ggggaccactttgtacaagaaagctgggtcCTCCTCATACCGCACGTTAC | Mycgr3\|109139  JGI  Ensembl | KOG0806 Carbon-nitrogen hydrolase; | hydrolase activity, acting on carbon-nitrogen (but not peptide) bonds; nitrogen compound metabolic process; | 100 % |  |  |
| 14 | ggggacaagtttgtacaaaaaagcaggcttgatgCAATCTCAAAGGGTCATGAT, ggggaccactttgtacaagaaagctgggtcGCTTTTCGCCTTGAACGTAT | Mycgr3\|109163  JGI  Ensembl | KOG1579 Homocysteine S-methyltransferase; | homocysteine S-methyltransferase activity; | 100 % |  |  |
| 14 | ggggacaagtttgtacaaaaaagcaggcttgatgGCCTCAACAACACCCCTCAA, ggggaccactttgtacaagaaagctgggtcCTCATCGTAATCATCTAGCT | Mycgr3\|109300  JGI  Ensembl | KOG4780 Uncharacterized conserved protein; |  | 100 % |  |  |
| 14 | ggggacaagtttgtacaaaaaagcaggcttgatgGCTCGACAAATGCACTATGG, ggggaccactttgtacaagaaagctgggtcTGCCTTGACCTTCCCGGCAG | Mycgr3\|109442  JGI  Ensembl |  |  | 100 % |  |  |
| 14 | ggggacaagtttgtacaaaaaagcaggcttgatgGCGTCCCAAGCTCTCCGGGA, ggggaccactttgtacaagaaagctgggtcTCCCTTGTGGCCCCAAGCCG | Mycgr3\|109530  JGI  Ensembl |  |  | 100 % |  |  |
| 14 | ggggacaagtttgtacaaaaaagcaggcttgatgCGATCCAAAAAAGACAACGT, ggggaccactttgtacaagaaagctgggtcACGCCGCGGCGCCATAGATC | Mycgr3\|110440  JGI  Ensembl |  |  | 0 % |  |  |
| 14 | ggggacaagtttgtacaaaaaagcaggcttgatgTCTTTCAGCACATCCAAGCC, ggggaccactttgtacaagaaagctgggtcCCTCCAGGATCCAGCCATCA | Mycgr3\|111522  JGI  Ensembl |  |  | 0 % |  |  |
| 14 | ggggacaagtttgtacaaaaaagcaggcttgatgGTGCTCCCCGACAAAGCCAA, ggggaccactttgtacaagaaagctgggtcATCAACAATCTCACATCCCT | Mycgr3\|111548  JGI  Ensembl |  |  | 2.26537 % |  |  |
| 14 | ggggacaagtttgtacaaaaaagcaggcttgatgCGCAACGATCTCCTTGAAGC, ggggaccactttgtacaagaaagctgggtcAGCCGTCACAGGTGTGTCGT | Mycgr3\|111747  JGI  Ensembl |  |  | 100 % |  |  |
| 15 | ggggacaagtttgtacaaaaaagcaggcttgatgTTGAGCAAACAGGCCATGTG, ggggaccactttgtacaagaaagctgggtcAAATGATGACTGCTCAGGTC | Mycgr3\|33604  JGI  Ensembl | KOG0223 Aquaporin (major intrinsic protein family); | membrane; transport; transporter activity; | 100 % |  |  |
| 15 | ggggacaagtttgtacaaaaaagcaggcttgatgACCCGCGACAACGACCTCCA, ggggaccactttgtacaagaaagctgggtcCACCCACCCGGCATCGACGG | Mycgr3\|35031  JGI  Ensembl | KOG1200 Mitochondrial/plastidial beta-ketoacyl-ACP reductase; | 3-oxoacyl-[acyl-carrier-protein] reductase activity; metabolic process;  oxidoreductase activity; | 100 % |  |  |
| 15 | ggggacaagtttgtacaaaaaagcaggcttgatgGCCGACTCCGACCCGTCGCT, ggggaccactttgtacaagaaagctgggtcCTTCCTCGTCGCCAAAACCA | Mycgr3\|36084  JGI  Ensembl | KOG3010 Methyltransferase; |  | 2.1097 % |  |  |
| 15 | ggggacaagtttgtacaaaaaagcaggcttgatgCAGATCAACTCGTTCCTTGC, ggggaccactttgtacaagaaagctgggtcCACTGCAAAAGAATCAATTA | Mycgr3\|36978  JGI  Ensembl | KOG1437 Fasciclin and related adhesion glycoproteins; |  | 0 % |  |  |
| 15 | ggggacaagtttgtacaaaaaagcaggcttgatgTCCGTCAAATCACGAGTAGC, ggggaccactttgtacaagaaagctgggtcCCACTCGCTGACCTCTTCAT | Mycgr3\|36990  JGI  Ensembl | KOG1208 Dehydrogenases with different specificities (related to short-chain alcohol dehydrogenases); | metabolic process; oxidoreductase activity; | 100 % |  |  |
| 15 | ggggacaagtttgtacaaaaaagcaggcttgatgGGTCTTTTCGACACGCTCCC, ggggaccactttgtacaagaaagctgggtcCATTAGTAGGCCCAGTTTCA | Mycgr3\|37599  JGI  Ensembl | KOG0725 Reductases with broad range of substrate specificities; | 2,3-dihydro-2,3-dihydroxybenzoate dehydrogenase activity; metabolic process;  oxidoreductase activity; | 100 % |  |  |
| 15 | ggggacaagtttgtacaaaaaagcaggcttgatgGCTGCCTCAACAAACCCCTC, ggggaccactttgtacaagaaagctgggtcCTCCTCACCGGCTTTCCTCC | Mycgr3\|37896  JGI  Ensembl |  |  | 0 % |  |  |
| 15 | ggggacaagtttgtacaaaaaagcaggcttgatgGCTCTCACGCTTCAGGACAG, ggggaccactttgtacaagaaagctgggtcCTTCCCGAAGTTCTTCTCCA | Mycgr3\|38394  JGI  Ensembl | KOG4169 15-hydroxyprostaglandin dehydrogenase and related dehydrogenases; | 15-hydroxyprostaglandin dehydrogenase (NAD+) activity; metabolic process;  oxidoreductase activity; | 100 % |  |  |
| 15 | ggggacaagtttgtacaaaaaagcaggcttgatgTCATTGACGGACCTAATCTC, ggggaccactttgtacaagaaagctgggtcGCTTGATCGCCTCTTCAACA | Mycgr3\|39862  JGI  Ensembl | KOG2920 Predicted methyltransferase; |  | 100 % |  |  |
| 15 | ggggacaagtttgtacaaaaaagcaggcttgatgCCTCCAATCTCCTTCAAACC, ggggaccactttgtacaagaaagctgggtcCAACTTCGAAGCCACCCACT | Mycgr3\|42801  JGI  Ensembl | KOG1680 Enoyl-CoA hydratase; | catalytic activity; enoyl-CoA hydratase activity;  metabolic process; | 100 % |  |  |
| 15 | ggggacaagtttgtacaaaaaagcaggcttgatgCCGACCAAAGCGCCCGTAGC, ggggaccactttgtacaagaaagctgggtcACTCTTCTTCCCGAACAACG | Mycgr3\|43828  JGI  Ensembl |  |  | 100 % |  |  |
| 15 | ggggacaagtttgtacaaaaaagcaggcttgatgCCCTCCTCCAACCAATCCTT, ggggaccactttgtacaagaaagctgggtcCTCCCTCGCTCCCCAAACCA | Mycgr3\|47036  JGI  Ensembl | KOG0316 Conserved WD40 repeat-containing protein; |  | 100 % |  |  |
| 15 | ggggacaagtttgtacaaaaaagcaggcttgatgGCTCCCAGCAGAGGATACTC, ggggaccactttgtacaagaaagctgggtcGGCTATCCGACTCTTTATGT | Mycgr3\|48269  JGI  Ensembl | KOG2972 Uncharacterized conserved protein; | glucose-1-phosphate adenylyltransferase activity; | 100 % |  |  |
| 15 | ggggacaagtttgtacaaaaaagcaggcttgatgGGCGCCTGGGCACTTACGCT, ggggaccactttgtacaagaaagctgggtcATTCCCGCAATACCTTCGAT | Mycgr3\|51460  JGI  Ensembl |  |  | 100 % |  |  |
| 15 | ggggacaagtttgtacaaaaaagcaggcttgatgGCCGCAGCCATCATCTCCGG, ggggaccactttgtacaagaaagctgggtcCGCCTTCCCCAACTCCTCAC | Mycgr3\|52044  JGI  Ensembl | KOG3124 Pyrroline-5-carboxylate reductase; | proline biosynthetic process; pyrroline-5-carboxylate reductase activity; | 0 % |  |  |
| 15 | ggggacaagtttgtacaaaaaagcaggcttgatgTCCGCACGATTCGGCCTGCG, ggggaccactttgtacaagaaagctgggtcCTTGGAAACCTTGGCCTCCA | Mycgr3\|52717  JGI  Ensembl | KOG1589 Uncharacterized conserved protein; |  | 100 % |  |  |
| 15 | ggggacaagtttgtacaaaaaagcaggcttgatgTCTCAAATACACGCCATGTC, ggggaccactttgtacaagaaagctgggtcATCGCGGAATTTCCTGTTCT | Mycgr3\|58858  JGI  Ensembl | KOG1664 Vacuolar H+-ATPase V1 sector, subunit E; | hydrogen ion transporting ATP synthase activity, rotational mechanism; hydrogen ion transporting ATPase activity, rotational mechanism;  hydrogen-transporting two-sector ATPase activity;  proton-transporting ATP synthase complex;  proton-transporting two-sector ATPase complex; | 100 % |  |  |
| 15 | ggggacaagtttgtacaaaaaagcaggcttgatgGAAGAGCAAGTAGAACGATT, ggggaccactttgtacaagaaagctgggtcACCAGTCTCCTCCGCAACCA | Mycgr3\|64085  JGI  Ensembl | KOG2702 Predicted panthothenate kinase/uridine kinase-related protein; | pantothenate kinase activity; | 100 % |  |  |
| 15 | ggggacaagtttgtacaaaaaagcaggcttgatgGGTCTCAGAATGGACGACAT, ggggaccactttgtacaagaaagctgggtcCAACGGTAGTAAGTGACAGC | Mycgr3\|65563  JGI  Ensembl | KOG3129 26S proteasome regulatory complex, subunit PSMD9; | protein binding; | 100 % |  |  |
| 15 | ggggacaagtttgtacaaaaaagcaggcttgatgGCACAGGCACTCAACAGCAT, ggggaccactttgtacaagaaagctgggtcCGACCTCAATCCCAGCAAGA | Mycgr3\|66025  JGI  Ensembl | KOG3083 Prohibitin; | membrane; | 100 % |  |  |
| 15 | ggggacaagtttgtacaaaaaagcaggcttgatgGCCGAAGACCGTCCCGCCTC, ggggaccactttgtacaagaaagctgggtcGCTCATCTTCGCGTTCGAAA | Mycgr3\|66328  JGI  Ensembl | KOG2265 Nuclear distribution protein NUDC; |  | 100 % |  |  |
| 15 | ggggacaagtttgtacaaaaaagcaggcttgatgCGCGTCTCAACCTTCGTCCT, ggggaccactttgtacaagaaagctgggtcCATGAAGGTCCTGCTGATGA | Mycgr3\|66795  JGI  Ensembl |  |  | 100 % |  |  |
| 15 | ggggacaagtttgtacaaaaaagcaggcttgatgTCTCTCTTCTCGTTCTCGAC, ggggaccactttgtacaagaaagctgggtcACTGGCCTGGATCTCCTTTG | Mycgr3\|67314  JGI  Ensembl | KOG3063 Membrane coat complex Retromer, subunit VPS26; | retromer complex; vacuolar transport; | 100 % |  |  |
| 15 | ggggacaagtttgtacaaaaaagcaggcttgatgGCGTCCCTTCGTCTCGCAGT, ggggaccactttgtacaagaaagctgggtcCCATATTCCAAACCACTTCT | Mycgr3\|67504  JGI  Ensembl |  |  | 100 % |  |  |
| 15 | ggggacaagtttgtacaaaaaagcaggcttgatgTCCAACGCCTGCGCGACGTG, ggggaccactttgtacaagaaagctgggtcCCTCATGCTGCCACTCCCAC | Mycgr3\|68371  JGI  Ensembl |  |  | 0 % |  |  |
| 15 | ggggacaagtttgtacaaaaaagcaggcttgatgTCTCCCACGACATCCAACCT, ggggaccactttgtacaagaaagctgggtcCTCATCATGAGCAAACCCGA | Mycgr3\|68418  JGI  Ensembl |  |  | 100 % |  |  |
| 15 | ggggacaagtttgtacaaaaaagcaggcttgatgGCCACCCGCGCCGCCCACAA, ggggaccactttgtacaagaaagctgggtcCTGCTGCTGATCCTTCACCA | Mycgr3\|68497  JGI  Ensembl | KOG0894 Ubiquitin-protein ligase; | post-translational protein modification; regulation of protein metabolic process;  small conjugating protein ligase activity; | 100 % |  |  |
| 15 | ggggacaagtttgtacaaaaaagcaggcttgatgAAGCTCCTCTACCCAACTTC, ggggaccactttgtacaagaaagctgggtcAATCATATTCCTCAACTCCT | Mycgr3\|69051  JGI  Ensembl | KOG0069 Glyoxylate/hydroxypyruvate reductase (D-isomer-specific 2-hydroxy acid dehydrogenase superfamily); | cofactor binding; gluconate 2-dehydrogenase activity;  gluconate dehydrogenase activity;  oxidoreductase activity, acting on the CH-OH group of donors, NAD or NADP as acceptor; | 2.02343 % |  |  |
| 15 | ggggacaagtttgtacaaaaaagcaggcttgatgACAGACCCCAGTCGAGAGCG, ggggaccactttgtacaagaaagctgggtcCGTCCCGCTCCCTCTTCCCG | Mycgr3\|69761  JGI  Ensembl |  |  | 100 % |  |  |
| 15 | ggggacaagtttgtacaaaaaagcaggcttgatgTCTCTGCAAAAGCTCTCCGA, ggggaccactttgtacaagaaagctgggtcCGCTTTCCCTGTGCTCCCGA | Mycgr3\|70268  JGI  Ensembl | KOG4153 Fructose 1,6-bisphosphate aldolase; | 2,4-dihydroxyhept-2-ene-1,7-dioate aldolase activity; 4-hydroxy-2-ketopimelate aldolase activity;  D-threonine aldolase activity;  L-allo-threonine aldolase activity;  fructose-bisphosphate aldolase activity;  glycolysis;  hexulose-6-phosphate synthase activity;  zinc ion binding; | 100 % |  |  |
| 15 | ggggacaagtttgtacaaaaaagcaggcttgatgTCGGACTCCCAGGTGACGCT, ggggaccactttgtacaagaaagctgggtcTTTCTTCTTGTCCTTGCTGG | Mycgr3\|70471  JGI  Ensembl | KOG3424 40S ribosomal protein S24; | intracellular; ribosome; structural constituent of ribosome;  translation; | 2.93255 % |  |  |
| 15 | ggggacaagtttgtacaaaaaagcaggcttgatgCCAGACTCTGTCCTTGTGCT, ggggaccactttgtacaagaaagctgggtcGCGATATCTCGAGTGACAGC | Mycgr3\|71002  JGI  Ensembl |  |  | 4.29043 % |  |  |
| 15 | ggggacaagtttgtacaaaaaagcaggcttgatgTCCGCCTCCACGCCTTTCGC, ggggaccactttgtacaagaaagctgggtcCACCCATTGCAATCCTAACC | Mycgr3\|71621  JGI  Ensembl | KOG2631 Class II aldolase/adducin N-terminal domain protein; | metal ion binding; | 100 % |  |  |
| 15 | ggggacaagtttgtacaaaaaagcaggcttgatgACCTCCAAGAAAATCCGCAT, ggggaccactttgtacaagaaagctgggtcATCTCCTGAAGCAGCAACAC | Mycgr3\|71881  JGI  Ensembl | KOG3947 Phosphoesterases; | hydrolase activity; | 100 % |  |  |
| 15 | ggggacaagtttgtacaaaaaagcaggcttgatgCCAGAATCACCACGCCTCGC, ggggaccactttgtacaagaaagctgggtcCTTGTTCTCGAATTTCAATC | Mycgr3\|72036  JGI  Ensembl | KOG0409 Predicted dehydrogenase; | (R)-2-hydroxyglutarate dehydrogenase activity; (R)-2-hydroxyisocaproate dehydrogenase activity;  2-hydroxytetrahydrofuran dehydrogenase activity;  3-keto sterol reductase activity;  3-ketoglucose-reductase activity;  5-exo-hydroxycamphor dehydrogenase activity;  C-3 sterol dehydrogenase (C-4 sterol decarboxylase) activity;  aldo-keto reductase activity;  epoxide dehydrogenase activity;  gluconate dehydrogenase activity;  isocitrate dehydrogenase activity;  mevaldate reductase activity;  pentose-phosphate shunt;  phenylcoumaran benzylic ether reductase activity;  phosphogluconate dehydrogenase (decarboxylating) activity;  steroid dehydrogenase activity;  steroid dehydrogenase activity, acting on the CH-CH group of donors;  steroid dehydrogenase activity, acting on the CH-OH group of donors, NAD or NADP as acceptor; | 100 % |  |  |
| 15 | ggggacaagtttgtacaaaaaagcaggcttgatgAAGGTCGCGGTGCTTCAGTT, ggggaccactttgtacaagaaagctgggtcTCCAGCTTTCTGCTGTAAGA | Mycgr3\|74759  JGI  Ensembl | KOG0806 Carbon-nitrogen hydrolase; | (3,5-dichlorophenylurea)acetate amidohydrolase activity; N-acetylgalactosamine-6-phosphate deacetylase activity;  N2-acetyl-L-lysine deacetylase activity;  O-succinylbenzoate synthase activity;  UDP-3-O-[3-hydroxymyristoyl] N-acetylglucosamine deacetylase activity;  histone deacetylase activity;  hydrolase activity, acting on carbon-nitrogen (but not peptide) bonds;  indoleacetamide hydrolase activity;  iprodione amidohydrolase activity;  nitrogen compound metabolic process;  protein N-terminal asparagine amidohydrolase activity; | 0 % |  |  |
| 15 | ggggacaagtttgtacaaaaaagcaggcttgatgGCTCCAACAATCCTCGTCGT, ggggaccactttgtacaagaaagctgggtcCTGCAGCATCTGCGCCAGAC | Mycgr3\|77151  JGI  Ensembl |  |  | 100 % |  |  |
| 15 | ggggacaagtttgtacaaaaaagcaggcttgatgTCGTCCTCCAACCGCAAGAA, ggggaccactttgtacaagaaagctgggtcTTTGGCCTTCTTAGCTTGCT | Mycgr3\|77714  JGI  Ensembl |  |  | 100 % |  |  |
| 15 | ggggacaagtttgtacaaaaaagcaggcttgatgTCACCCAGCTTCCGCATCCA, ggggaccactttgtacaagaaagctgggtcAGTGGCGCGTTCCCACAGCC | Mycgr3\|85907  JGI  Ensembl | KOG2084 Predicted histone tail methylase containing SET domain; | nucleus; | 100 % |  |  |
| 15 | ggggacaagtttgtacaaaaaagcaggcttgatgCCAAAGCGCTCGCGATCACC, ggggaccactttgtacaagaaagctgggtcCTTCCTCGCCCGCTTGTGCT | Mycgr3\|87660  JGI  Ensembl | KOG0719 Molecular chaperone (DnaJ superfamily); | heat shock protein binding; | 100 % |  |  |
| 15 | ggggacaagtttgtacaaaaaagcaggcttgatgATCCCTCCCATCATCGCTCT, ggggaccactttgtacaagaaagctgggtcCATCGGTGCCTTGATTCGCA | Mycgr3\|89119  JGI  Ensembl | KOG4245 Predicted metal-dependent hydrolase of the TIM-barrel fold; | catalytic activity; metabolic process; | 100 % |  |  |
| 15 | ggggacaagtttgtacaaaaaagcaggcttgatgGCTGAGAACAACGGAGAGCA, ggggaccactttgtacaagaaagctgggtcGGCGGCGCCACTGTCCTCAA | Mycgr3\|89718  JGI  Ensembl | KOG0802 E3 ubiquitin ligase; | protein binding; zinc ion binding; | 0 % |  |  |
| 15 | ggggacaagtttgtacaaaaaagcaggcttgatgGAATATATCAAGGAAGAGCC, ggggaccactttgtacaagaaagctgggtcGAGGAGATCCCATATAGGTT | Mycgr3\|90578  JGI  Ensembl |  |  | 0 % |  |  |
| 15 | ggggacaagtttgtacaaaaaagcaggcttgatgGACACCATGGAAGCCGCATT, ggggaccactttgtacaagaaagctgggtcTCTCGCCTTCTTCTTCTCCT | Mycgr3\|90840  JGI  Ensembl | KOG4297 C-type lectin; |  | 100 % |  |  |
| 15 | ggggacaagtttgtacaaaaaagcaggcttgatgCGTTCCACACTCCTCACCCT, ggggaccactttgtacaagaaagctgggtcAACAACCCATGCCAAAATCC | Mycgr3\|91361  JGI  Ensembl |  |  | 100 % |  |  |
| 15 | ggggacaagtttgtacaaaaaagcaggcttgatgACGAGCATCATCTCATCCAG, ggggaccactttgtacaagaaagctgggtcCTCCCACCCACTATCACCCA | Mycgr3\|92888  JGI  Ensembl |  |  | 100 % |  |  |
| 15 | ggggacaagtttgtacaaaaaagcaggcttgatgTGGGATTGGGTTTACAAAAC, ggggaccactttgtacaagaaagctgggtcCTCATTAACATCCTCATCAA | Mycgr3\|93553  JGI  Ensembl | KOG0672 Halotolerance protein HAL3 (contains flavoprotein domain); |  | 2.16718 % |  |  |
| 15 | ggggacaagtttgtacaaaaaagcaggcttgatgTCTTCTCCTCCTCAACCCAC, ggggaccactttgtacaagaaagctgggtcCCTCACAGCATCCAAAACCC | Mycgr3\|93831  JGI  Ensembl | KOG2305 3-hydroxyacyl-CoA dehydrogenase; | 3-hydroxyacyl-CoA dehydrogenase activity; fatty acid metabolic process;  oxidoreductase activity; | 100 % |  |  |
| 15 | ggggacaagtttgtacaaaaaagcaggcttgatgCAGAAGCTCGACAATGGTAC, ggggaccactttgtacaagaaagctgggtcTCCCCCGCCGGAGGACGCAA | Mycgr3\|94102  JGI  Ensembl |  |  | 100 % |  |  |
| 15 | ggggacaagtttgtacaaaaaagcaggcttgatgGACACCTGCCGCGAACAAAT, ggggaccactttgtacaagaaagctgggtcAAGATCCTTCATGATCCCAG | Mycgr3\|94146  JGI  Ensembl |  |  | 100 % |  |  |
| 15 | ggggacaagtttgtacaaaaaagcaggcttgatgTCTTCCATGAAGACGTTCCT, ggggaccactttgtacaagaaagctgggtcGACTTCTTCCTCCAGCAAGT | Mycgr3\|94173  JGI  Ensembl | KOG2924 Deoxyhypusine synthase; | deoxyhypusine synthase activity; spermidine catabolic process to deoxyhypusine, using deoxyhypusine synthase; | 100 % |  |  |
| 15 | ggggacaagtttgtacaaaaaagcaggcttgatgAGTCAATTTGCCCTCATCAC, ggggaccactttgtacaagaaagctgggtcTCGCCGTGCTCGAAGAAGGT | Mycgr3\|94547  JGI  Ensembl | KOG1209 1-Acyl dihydroxyacetone phosphate reductase and related dehydrogenases; | metabolic process; oxidoreductase activity;  ribitol 2-dehydrogenase activity; | 100 % |  |  |
| 15 | ggggacaagtttgtacaaaaaagcaggcttgatgAAATACCGATCTGGAGCTGT, ggggaccactttgtacaagaaagctgggtcTAGCAAGCAGGCCAAAGCTC | Mycgr3\|94571  JGI  Ensembl | KOG1216 von Willebrand factor and related coagulation proteins; | exo-alpha-sialidase activity; | 4.50161 % |  |  |
| 15 | ggggacaagtttgtacaaaaaagcaggcttgatgCGCAAGTCAGCCCGTGTCCA, ggggaccactttgtacaagaaagctgggtcATCGTCCAGAACGATGACCA | Mycgr3\|94688  JGI  Ensembl |  |  | 2.02343 % |  |  |
| 15 | ggggacaagtttgtacaaaaaagcaggcttgatgTCGGCACCGAAATACTTCAT, ggggaccactttgtacaagaaagctgggtcTTTCTGCTCCGGCACTTCCT | Mycgr3\|94691  JGI  Ensembl |  |  | 100 % |  |  |
| 15 | ggggacaagtttgtacaaaaaagcaggcttgatgGTCCATTCGATCGTCCGCCG, ggggaccactttgtacaagaaagctgggtcACTGGACTGCGGATCCGGTC | Mycgr3\|94708  JGI  Ensembl |  |  | 0 % |  |  |
| 15 | ggggacaagtttgtacaaaaaagcaggcttgatgTCGCAACAGACCGCAGCCGA, ggggaccactttgtacaagaaagctgggtcCATCCAGGCATAACTCATCT | Mycgr3\|94743  JGI  Ensembl |  |  | 100 % |  |  |
| 15 | ggggacaagtttgtacaaaaaagcaggcttgatgGAAGATCTCACCACCACCAC, ggggaccactttgtacaagaaagctgggtcGATATCGCGGTACTCGTTCC | Mycgr3\|94825  JGI  Ensembl |  |  | 100 % |  |  |
| 15 | ggggacaagtttgtacaaaaaagcaggcttgatgGCAGATGATCACAAGAGCAC, ggggaccactttgtacaagaaagctgggtcTGGACCGGCATACTTGTGAC | Mycgr3\|94988  JGI  Ensembl | KOG0143 Iron/ascorbate family oxidoreductases; |  | 0 % |  |  |
| 15 | ggggacaagtttgtacaaaaaagcaggcttgatgAGTGGTCTTACTGCATTGAG, ggggaccactttgtacaagaaagctgggtcTCCTTCTGCTGGTGGAATTG | Mycgr3\|95227  JGI  Ensembl |  |  | 100 % |  |  |
| 15 | ggggacaagtttgtacaaaaaagcaggcttgatgGCGTCCGACCTCTCCGCCCA, ggggaccactttgtacaagaaagctgggtcATCTCTCGGCGTGAACGACG | Mycgr3\|95761  JGI  Ensembl | KOG1307 K+-dependent Ca2+/Na+ exchanger NCKX1 and related proteins; |  | 100 % |  |  |
| 15 | ggggacaagtttgtacaaaaaagcaggcttgatgGTGTTCAACACGCGGGACCG, ggggaccactttgtacaagaaagctgggtcACGTCGCTCCGCATCACCCG | Mycgr3\|96096  JGI  Ensembl |  |  | 4.07524 % |  |  |
| 15 | ggggacaagtttgtacaaaaaagcaggcttgatgTCTGATCCACCAACAACGAT, ggggaccactttgtacaagaaagctgggtcGTCGTAGTCTCCCGACGACG | Mycgr3\|96231  JGI  Ensembl |  |  | 0 % |  |  |
| 15 | ggggacaagtttgtacaaaaaagcaggcttgatgGGATTCCAGCAGCTCTGGGC, ggggaccactttgtacaagaaagctgggtcCTTACAAAAGTAAGGTTTGC | Mycgr3\|96437  JGI  Ensembl |  |  | 0 % |  |  |
| 15 | ggggacaagtttgtacaaaaaagcaggcttgatgTGGCCAGCTCGTTTGACGGA, ggggaccactttgtacaagaaagctgggtcGGCCATGACTCCCGGGTTGC | Mycgr3\|96753  JGI  Ensembl |  |  | 100 % |  |  |
| 15 | ggggacaagtttgtacaaaaaagcaggcttgatgGAAGCAGAGCTCGCGCGCCG, ggggaccactttgtacaagaaagctgggtcTCCTTTGATACGTTTCGTCG | Mycgr3\|97006  JGI  Ensembl |  |  | 100 % |  |  |
| 15 | ggggacaagtttgtacaaaaaagcaggcttgatgAAAGCAAAAGAAGAGCCACG, ggggaccactttgtacaagaaagctgggtcATTACCGAGCAAACTGTCGG | Mycgr3\|97078  JGI  Ensembl |  |  | 100 % | chr\_12:841603 C->A; |  |
| 15 | ggggacaagtttgtacaaaaaagcaggcttgatgCCACGCCCTCACCGCAACTA, ggggaccactttgtacaagaaagctgggtcCACTCCCGTCTTGAGCTGCT | Mycgr3\|97552  JGI  Ensembl | KOG4207 Predicted splicing factor, SR protein superfamily; |  | 0 % |  |  |
| 15 | ggggacaagtttgtacaaaaaagcaggcttgatgATGTGGGAGGGAGCAGGAGG, ggggaccactttgtacaagaaagctgggtcGGGCAAAATCCCCAGTCCAC | Mycgr3\|97651  JGI  Ensembl |  |  | 100 % |  |  |
| 15 | ggggacaagtttgtacaaaaaagcaggcttgatgATCCTCAGTCTTTCATCAAT, ggggaccactttgtacaagaaagctgggtcATTGGACAGCTTTTCCGTGC | Mycgr3\|97660  JGI  Ensembl |  |  | 100 % |  |  |
| 15 | ggggacaagtttgtacaaaaaagcaggcttgatgCCGTCACCAACAGGTCGTGG, ggggaccactttgtacaagaaagctgggtcGACCTCTCCAATGACCTCAG | Mycgr3\|97678  JGI  Ensembl |  |  | 0 % |  |  |
| 15 | ggggacaagtttgtacaaaaaagcaggcttgatgGACACTACTTTCTCCGGTTC, ggggaccactttgtacaagaaagctgggtcCTCAACACCGTCAGTGTCAG | Mycgr3\|97685  JGI  Ensembl |  |  | 100 % |  |  |
| 15 | ggggacaagtttgtacaaaaaagcaggcttgatgCATGGGTACATCGCACAACG, ggggaccactttgtacaagaaagctgggtcACCTGAACCTCGCGGAGGAG | Mycgr3\|97721  JGI  Ensembl |  |  | 100 % |  |  |
| 15 | ggggacaagtttgtacaaaaaagcaggcttgatgGCCACCGTCTACCATAATTT, ggggaccactttgtacaagaaagctgggtcATGATGGCATGAAATCTTGC | Mycgr3\|98027  JGI  Ensembl | KOG0161 Myosin class II heavy chain; |  | 100 % |  |  |
| 15 | ggggacaagtttgtacaaaaaagcaggcttgatgGCCGCCAGCAACCCCTCCAA, ggggaccactttgtacaagaaagctgggtcTGATACGGGAACACTCTTCG | Mycgr3\|98928  JGI  Ensembl | KOG0911 Glutaredoxin-related protein; | cell redox homeostasis; | 100 % |  |  |
| 15 | ggggacaagtttgtacaaaaaagcaggcttgatgCGGCTCTTCCTCCTGCCCGT, ggggaccactttgtacaagaaagctgggtcCTTCGTCTCCTCTTTCGTCT | Mycgr3\|99187  JGI  Ensembl | KOG4539 Uncharacterized conserved protein; |  | 100 % |  |  |
| 15 | ggggacaagtttgtacaaaaaagcaggcttgatgGAAGCCCTCGAACAGACCCT, ggggaccactttgtacaagaaagctgggtcTCCACGACGTCTGGTCGAGG | Mycgr3\|100168  JGI  Ensembl | KOG3211 Predicted endoplasmic reticulum membrane protein Lec35/MPDU1 involved in monosaccharide-P-dolichol utilization; |  | 2.38353 % |  |  |
| 15 | ggggacaagtttgtacaaaaaagcaggcttgatgGCGTTGAAGAGAATAAACAA, ggggaccactttgtacaagaaagctgggtcGATGGCGTACTTTCGAGTCC | Mycgr3\|100608  JGI  Ensembl | KOG0417 Ubiquitin-protein ligase; | negative regulation of ubiquitin-protein ligase activity; positive regulation of ubiquitin-protein ligase activity;  post-translational protein modification;  regulation of protein metabolic process;  regulation of ubiquitin-protein ligase activity;  regulation of ubiquitin-protein ligase activity during meiotic cell cycle;  small conjugating protein ligase activity;  ubiquitin-protein ligase activity; | 100 % |  |  |
| 15 | ggggacaagtttgtacaaaaaagcaggcttgatgTCCGCACCCATCGCCATCAT, ggggaccactttgtacaagaaagctgggtcCCACTTCTCCAACATAGGCC | Mycgr3\|101134  JGI  Ensembl | KOG1014 17 beta-hydroxysteroid dehydrogenase type 3, HSD17B3; | carbonyl reductase (NADPH) activity; metabolic process;  oxidoreductase activity; | 100 % |  |  |
| 15 | ggggacaagtttgtacaaaaaagcaggcttgatgGCAGCTTCAATCGTAGACAT, ggggaccactttgtacaagaaagctgggtcGAACCCCATGAATCCACCAT | Mycgr3\|101325  JGI  Ensembl | KOG1200 Mitochondrial/plastidial beta-ketoacyl-ACP reductase; | 2-deoxy-D-gluconate 3-dehydrogenase activity; metabolic process;  oxidoreductase activity; | 100 % |  |  |
| 15 | ggggacaagtttgtacaaaaaagcaggcttgatgCCAATCACAGAAGGCGCACC, ggggaccactttgtacaagaaagctgggtcGACCTCCGCCGACTCTCCAT | Mycgr3\|101664  JGI  Ensembl | KOG3008 Quinolinate phosphoribosyl transferase; | NAD biosynthetic process; nicotinate-nucleotide diphosphorylase (carboxylating) activity;  pyridine nucleotide biosynthetic process; | 2.20126 % |  |  |
| 15 | ggggacaagtttgtacaaaaaagcaggcttgatgGCACGCAACCGCTCTCCGTC, ggggaccactttgtacaagaaagctgggtcCGAGGACGTCTCAATCTGCA | Mycgr3\|101851  JGI  Ensembl | KOG0189 Phosphoadenosine phosphosulfate reductase; | metabolic process; phosphoadenylyl-sulfate reductase (thioredoxin) activity;  sulfate assimilation, phosphoadenylyl sulfate reduction by phosphoadenylyl-sulfate reductase (thioredoxin); transferase activity; | 100 % |  |  |
| 15 | ggggacaagtttgtacaaaaaagcaggcttgatgGCCTCGATCTTTTCTTCGCT, ggggaccactttgtacaagaaagctgggtcGGATGGCTGCTTCAGTGCAA | Mycgr3\|102570  JGI  Ensembl |  |  | 100 % |  |  |
| 15 | ggggacaagtttgtacaaaaaagcaggcttgatgTCGGGCGTCAAGAAATTCCA, ggggaccactttgtacaagaaagctgggtcGAACACCATCTCGTGCGGCC | Mycgr3\|102740  JGI  Ensembl | KOG1694 60s ribosomal protein L6; | intracellular; ribosome; structural constituent of ribosome;  translation; | 0 % |  |  |
| 15 | ggggacaagtttgtacaaaaaagcaggcttgatgGATTCGTACCAGCAGAATTT, ggggaccactttgtacaagaaagctgggtcCTTGCGTCTTGCAAGCCGTG | Mycgr3\|104366  JGI  Ensembl |  |  | 100 % |  | ``` 408	GCTTGCGAGGAGTGTCGAAAGCGCAAGCAGCGCTGTGACGGCAACGAACCGTGTGCCGCGTGCGAGGAGCAAGGCACCATGTGTCGCTACCGAGACGTGCCACCCGCGAAG	518 	\|\|\|\|\|\|\|\|\|\|\|\|\|\|\|\|\|\|\|\|\|\|\|\|\|\|\|\|\|\|\|\|\|\|\|\|\|\|\|\|\|\|\|\|\|\|\|\|\|\|\|\|\|\|\|\|\|\|\|\|\|\|\|\|\|\|\|\|\|\|\|\|\|\|\|\|\|\|\|\|\|\|\|\|\|\|\|\|\|\|\|\|\|\|\|\|\|\|\|\|\|\|\|\|\|\|\|\|\|\|\| 	GCTTGCGAGGAGTGTCGAAAGCGCAAGCAGCGCTGTGACGGCAACGAACCGTGTGCCGCGTGCGAGGAGCAAGGCACCATGTGTCGCTACCGAGACGTGCCACCCGCGAAG ```      ``` 518	GAAAGATCCTACCATCGACAAGATCCTGGAGATGTCCAAGGAGCACAACAAGAGCCTCGAATCGTTGACCCAAGGTATTAATAGTCTGCAAACGAGCTTGAGAGGCATCGAAGCACGGCTTGCAAGACGCAAG	650 	\|\|\|\|\|\|\|\|\|\|\|\|\|\|\|\|\|\|\|\|\|\|\|\|\|\|\|\|\|\|\|\|\|\|\|\|\|\|\|\|\|\|\|\|\|\|\|\|\|\|\|\|\|\|\|\|\|\|\|\|\|\|\|\|\|\|\|\|\|\|\|\|\|\|\|\|\|\|\|\|\|\|\|\|\|\|\|\|\|\|\|\|\|\|\|\|\|\|\|\|\|\|\|\|\|\|\|\|\|\|\|\|\|\|\|\|\|\|\|\|\|\|\|\|\|\|\|\|\|\|\|\|\| 	GAAAGATCCTACCATCGACAAGATCCTGGAGATGTCCAAGGAGCACAACAAGAGCCTCGAATCGTTGACCCAAGGTATTAATAGTCTGCAAACGAGCTTGAGAGGCATCGAAGCACGGCTTGCAAGACGCAAG ```      ``` 72	ATGGATTCGTACCAGCAGAATTTTGCGCAGCAAGCGGCCTGGTACTACCCTGGAATGCAAGAACCTTTTGTTCCTCCACAGCAATTCTTTTCACATCCCGACCAACCATGGAGCCCAACAGCTCCGACGATGCGCCCTCATCCTGTTCTTGACGTTACAGCAGCGCCTCAAAATATCCCACACGCCACCGAAAGTTTCGGATACTCCGAAAGATCAAGCAGACCCGAAAGTACGCAGCTACCGGGCTCTCTCTCCTCACCCAGCACGCCCGGGCATTGGAGTTCACATGGACATGAGTTGGAGCCGAGAAGGATTCGGATCAAGAGAATCAGAACAG	408 	\|\|\|\|\|\|\|\|\|\|\|\|\|\|\|\|\|\|\|\|\|\|\|\|\|\|\|\|\|\|\|\|\|\|\|\|\|\|\|\|\|\|\|\|\|\|\|\|\|\|\|\|\|\|\|\|\|\|\|\|\|\|\|\|\|\|\|\|\|\|\|\|\|\|\|\|\|\|\|\|\|\|\|\|\|\|\|\|\|\|\|\|\|\|\|\|\|\|\|\|\|\|\|\|\|\|\|\|\|\|\|\|\|\|\|\|\|\|\|\|\|\|\|\|\|\|\|\|\|\|\|\|\|\|\|\|\|\|\|\|\|\|\|\|\|\|\|\|\|\|\|\|\|\|\|\|\|\|\|\|\|\|\|\|\|\|\|\|\|\|\|\|\|\|\|\|\|\|\|\|\|\|\|\|\|\|\|\|\|\|\|\|\|\|\|\|\|\|\|\|\|\|\|\|\|\|\|\|\|\|\|\|\|\|\|\|\|\|\|\|\|\|\|\|\|\|\|\|\|\|\|\|\|\|\|\|\|\|\|\|\|\|\|\|\|\|\|\|\|\|\|\|\|\|\|\|\|\|\|\|\|\|\|\|\|\|\|\|\|\|\|\|\|\|\|\|\|\|\|\|\|\|\|\|\|\|\|\|\|\|\|\|\|\|\|\|\|\|\|\|\|\|\|\|\|\|\|\|\|\|\|\|\|\|\|\|\|\|\|\|\|\|\|\|\|\|\|\|\|\|\|\|\|\|\|\|\| 	ATGGATTCGTACCAGCAGAATTTTGCGCAGCAAGCGGCCTGGTACTACCCTGGAATGCAAGAACCTTTTGTTCCTCCACAGCAATTCTTTTCACATCCCGACCAACCATGGAGCCCAACAGCTCCGACGATGCGCCCTCATCCTGTTCTTGACGTTACAGCAGCGCCTCAAAATATCCCACACGCCACCGAAAGTTTCGGATACTCCGAAAGATCAAGCAGACCCGAAAGTACGCAGCTACCGGGCTCTCTCTCCTCACCCAGCACGCCCGGGCATTGGAGTTCACATGGACATGAGTTGGAGCCGAGAAGGATTCGGATCAAGAGAATCAGAACAG ``` |
| 15 | ggggacaagtttgtacaaaaaagcaggcttgatgCGCGTTTCGACACCTCCGCC, ggggaccactttgtacaagaaagctgggtcACCATTGGTACTCTCCGTCT | Mycgr3\|105616  JGI  Ensembl | KOG0504 FOG: Ankyrin repeat; |  | 100 % |  |  |
| 15 | ggggacaagtttgtacaaaaaagcaggcttgatgGACCATCTCTGCCCAAGACA, ggggaccactttgtacaagaaagctgggtcGTCGGAAGCAAGAGGCGTCG | Mycgr3\|106276  JGI  Ensembl |  |  | 97.90026 % | chr\_12:259650 C->T; |  |
| 15 | ggggacaagtttgtacaaaaaagcaggcttgatgGCTCACAACGGATATCCTCC, ggggaccactttgtacaagaaagctgggtcATTGGGGCATTGGAGACCCG | Mycgr3\|106406  JGI  Ensembl |  |  | 100 % |  | ``` 465	GCTTGCGAAGAGTGTCGGAAAAAGAAGCAGAAGTGTGATGGTGAACAGCCTTGTCAAGCTTGCAAGGAGCAAAACATCGACTGCCAGTATCGAGAGGTCCCACCAACAAA	574 	\|\|\|\|\|\|\|\|\|\|\|\|\|\|\|\|\|\|\|\|\|\|\|\|\|\|\|\|\|\|\|\|\|\|\|\|\|\|\|\|\|\|\|\|\|\|\|\|\|\|\|\|\|\|\|\|\|\|\|\|\|\|\|\|\|\|\|\|\|\|\|\|\|\|\|\|\|\|\|\|\|\|\|\|\|\|\|\|\|\|\|\|\|\|\|\|\|\|\|\|\|\|\|\|\|\|\|\|\|\| 	GCTTGCGAAGAGTGTCGGAAAAAGAAGCAGAAGTGTGATGGTGAACAGCCTTGTCAAGCTTGCAAGGAGCAAAACATCGACTGCCAGTATCGAGAGGTCCCACCAACAAA ```      ``` 575	AAAGGACCACAGTCTAGAGAAGCTTGCGGAATTGATGGAGAACTTCAGCTATCAGCTCGTCTCGCTGAGCCGCCGGATCGACACCATGGGCATGCAACTCAAGCAGAACGAGGACCGAATGGCGATGTTGCCCACTTCCTGTTGGCATTGCGGCTCGGGTCTCCAATGCCCCAAT	749 	\|\|\|\|\|\|\|\|\|\|\|\|\|\|\|\|\|\|\|\|\|\|\|\|\|\|\|\|\|\|\|\|\|\|\|\|\|\|\|\|\|\|\|\|\|\|\|\|\|\|\|\|\|\|\|\|\|\|\|\|\|\|\|\|\|\|\|\|\|\|\|\|\|\|\|\|\|\|\|\|\|\|\|\|\|\|\|\|\|\|\|\|\|\|\|\|\|\|\|\|\|\|\|\|\|\|\|\|\|\|\|\|\|\|\|\|\|\|\|\|\|\|\|\|\|\|\|\|\|\|\|\|\|\|\|\|\|\|\|\|\|\|\|\|\|\|\|\|\|\|\|\|\|\|\|\|\|\|\|\|\|\|\|\|\|\|\|\|\|\|\|\|\|\|\| 	AAAGGACCACAGTCTAGAGAAGCTTGCGGAATTGATGGAGAACTTCAGCTATCAGCTCGTCTCGCTGAGCCGCCGGATCGACACCATGGGCATGCAACTCAAGCAGAACGAGGACCGAATGGCGATGTTGCCCACTTCCTGTTGGCATTGCGGCTCGGGTCTCCAATGCCCCAAT ```      ``` 102	ATGGCTCACAACGGATATCCTCCTCAAGGCCTACCCATGAGTCACCGAGGCACCTACTATCCCACCACTCCTCCTCGACACCCGATGTACACATACAACCACTTTGCCGGCCAAGACCTCACGCCATCATATGAGTATGAGGCGATCACTTTCGGACAACCTGCCGTGCAGTTGTCGACTCGAGCTCTTTCAGAGTCATTTGTCTTTCAAGCAAATGGTTCACCGCAAAGTGCATCGTTTGAGGATCCACCGCCTCGACTGGAGACTCCACGGTCTCAGCATGCGGGCTCGCACAAGACTCCTGCTGTCCCGATCAGTCCAAGTCTTGCCACGCCCCGCGTAAAAGCAAAGCGCACCCCAACTG	465 	\|\|\|\|\|\|\|\|\|\|\|\|\|\|\|\|\|\|\|\|\|\|\|\|\|\|\|\|\|\|\|\|\|\|\|\|\|\|\|\|\|\|\|\|\|\|\|\|\|\|\|\|\|\|\|\|\|\|\|\|\|\|\|\|\|\|\|\|\|\|\|\|\|\|\|\|\|\|\|\|\|\|\|\|\|\|\|\|\|\|\|\|\|\|\|\|\|\|\|\|\|\|\|\|\|\|\|\|\|\|\|\|\|\|\|\|\|\|\|\|\|\|\|\|\|\|\|\|\|\|\|\|\|\|\|\|\|\|\|\|\|\|\|\|\|\|\|\|\|\|\|\|\|\|\|\|\|\|\|\|\|\|\|\|\|\|\|\|\|\|\|\|\|\|\|\|\|\|\|\|\|\|\|\|\|\|\|\|\|\|\|\|\|\|\|\|\|\|\|\|\|\|\|\|\|\|\|\|\|\|\|\|\|\|\|\|\|\|\|\|\|\|\|\|\|\|\|\|\|\|\|\|\|\|\|\|\|\|\|\|\|\|\|\|\|\|\|\|\|\|\|\|\|\|\|\|\|\|\|\|\|\|\|\|\|\|\|\|\|\|\|\|\|\|\|\|\|\|\|\|\|\|\|\|\|\|\|\|\|\|\|\|\|\|\|\|\|\|\|\|\|\|\|\|\|\|\|\|\|\|\|\|\|\|\|\|\|\|\|\|\|\|\|\|\|\|\|\|\|\|\|\|\|\|\|\|\|\|\|\|\|\|\|\|\|\|\|\|\|\|\|\|\|\|\|\|\|\|\|\|\|\|\|\| 	ATGGCTCACAACGGATATCCTCCTCAAGGCCTACCCATGAGTCACCGAGGCACCTACTATCCCACCACTCCTCCTCGACACCCGATGTACACATACAACCACTTTGCCGGCCAAGACCTCACGCCATCATATGAGTATGAGGCGATCACTTTCGGACAACCTGCCGTGCAGTTGTCGACTCGAGCTCTTTCAGAGTCATTTGTCTTTCAAGCAAATGGTTCACCGCAAAGTGCATCGTTTGAGGATCCACCGCCTCGACTGGAGACTCCACGGTCTCAGCATGCGGGCTCGCACAAGACTCCTGCTGTCCCGATCAGTCCAAGTCTTGCCACGCCCCGCGTAAAAGCAAAGCGCACCCCAACTG ``` |
| 15 | ggggacaagtttgtacaaaaaagcaggcttgatgCGACTCCCATGTGCACTCGA, ggggaccactttgtacaagaaagctgggtcAGACTCAGCTCCGGTTGGAA | Mycgr3\|106709  JGI  Ensembl |  |  | 100 % |  |  |
| 15 | ggggacaagtttgtacaaaaaagcaggcttgatgTCGGCCTCCTCCTTGAAGTA, ggggaccactttgtacaagaaagctgggtcTAGCTTTGCGAATTTCGGCG | Mycgr3\|106870  JGI  Ensembl | KOG1681 Enoyl-CoA isomerase; | 3-carboxymuconate cycloisomerase type II activity; C-8 sterol isomerase activity;  catalytic activity;  delta5-delta2,4-dienoyl-CoA isomerase activity;  metabolic process;  trans-2, cis-3 decenoyl-ACP isomerase activity; | 100 % |  |  |
| 15 | ggggacaagtttgtacaaaaaagcaggcttgatgGCCAACGTTCAAGACCGCGG, ggggaccactttgtacaagaaagctgggtcAAGCAACTTCAGATGTCCGG | Mycgr3\|107627  JGI  Ensembl | KOG3282 Uncharacterized conserved protein; | aminoacyl-tRNA hydrolase activity; | 100 % |  |  |
| 15 | ggggacaagtttgtacaaaaaagcaggcttgatgTCGAGCCTTTCTCCGCAAAA, ggggaccactttgtacaagaaagctgggtcCGGTCCTTGCATCACAGCGT | Mycgr3\|108058  JGI  Ensembl | KOG1669 Predicted mRNA cap-binding protein related to eIF-4E; | RNA binding; cytoplasm; translation initiation factor activity;  translational initiation; | 100 % |  |  |
| 15 | ggggacaagtttgtacaaaaaagcaggcttgatgGAGACAGAAAGCTTCACACC, ggggaccactttgtacaagaaagctgggtcGCGAACCGAAACTCTCGGAC | Mycgr3\|108910  JGI  Ensembl | KOG1003 Actin filament-coating protein tropomyosin; |  | 100 % |  |  |
| 15 | ggggacaagtttgtacaaaaaagcaggcttgatgAGCGACGATACACCAGCTCC, ggggaccactttgtacaagaaagctgggtcCTCCACCGAGACCTGCGCCG | Mycgr3\|109402  JGI  Ensembl |  |  | 100 % |  |  |
| 15 | ggggacaagtttgtacaaaaaagcaggcttgatgACCAGCACAAAGACGTGTAT, ggggaccactttgtacaagaaagctgggtcATAGGTCACCGTCCGAGTCG | Mycgr3\|110482  JGI  Ensembl |  |  | 100 % |  |  |
| 16 | ggggacaagtttgtacaaaaaagcaggcttgatgACCATTGCGCCACCACTTCC, ggggaccactttgtacaagaaagctgggtcCTCCTTCCATTCCATCGAGG | Mycgr3\|31787  JGI  Ensembl | KOG1205 Predicted dehydrogenase; | F420H2 dehydrogenase activity; malolactic enzyme activity;  metabolic process;  oxidoreductase activity;  regulation of oxidoreductase activity;  sulfur oxygenase reductase activity; | 0 % |  |  |
| 16 | ggggacaagtttgtacaaaaaagcaggcttgatgCATCCCACCCACACCCTCTT, ggggaccactttgtacaagaaagctgggtcCATGGCCGACCGACCTTTGA | Mycgr3\|33254  JGI  Ensembl |  |  | 0 % |  |  |
| 16 | ggggacaagtttgtacaaaaaagcaggcttgatgGCGACTAGACGAACGACCAA, ggggaccactttgtacaagaaagctgggtcGACGCCAGGCATCCCAATAA | Mycgr3\|33473  JGI  Ensembl | KOG3750 Inositol phospholipid synthesis protein, Scs3p; |  | 0 % |  |  |
| 16 | ggggacaagtttgtacaaaaaagcaggcttgatgCGAGACTTCCTCGAGCAAAG, ggggaccactttgtacaagaaagctgggtcCTTCTTCTTCTGCTGCAGCA | Mycgr3\|33888  JGI  Ensembl | KOG1515 Arylacetamide deacetylase; | 1-oxa-2-oxocycloheptane lactonase activity; 3,4-dihydrocoumarin hydrolase activity;  butyrolactone hydrolase activity;  lipase activity;  phospholipase activity;  sulfolactone hydrolase activity; | 0 % |  |  |
| 16 | ggggacaagtttgtacaaaaaagcaggcttgatgGGGGCTGAACTGAGTGACGA, ggggaccactttgtacaagaaagctgggtcGAATGTATACCTGAAGCCCA | Mycgr3\|35107  JGI  Ensembl | KOG2533 Permease of the major facilitator superfamily; |  | 0 % |  |  |
| 16 | ggggacaagtttgtacaaaaaagcaggcttgatgGGCTGGTTCTGGGCCGATTC, ggggaccactttgtacaagaaagctgggtcCTTCCTCCCCTGCCTCGCCA | Mycgr3\|35508  JGI  Ensembl | KOG3996 Holocytochrome c synthase/heme-lyase; | holocytochrome-c synthase activity; mitochondrion; | 0 % |  |  |
| 16 | ggggacaagtttgtacaaaaaagcaggcttgatgTCCGCGAGCCCTTTCCTTAA, ggggaccactttgtacaagaaagctgggtcTCGGAGACAGAGTAGCGCTC | Mycgr3\|35722  JGI  Ensembl |  |  | 0 % |  |  |
| 16 | ggggacaagtttgtacaaaaaagcaggcttgatgTCCGAAGAGCCCGACGATCG, ggggaccactttgtacaagaaagctgggtcATCGGCACCACCACTGGCAT | Mycgr3\|35819  JGI  Ensembl | KOG4718 Non-SMC (structural maintenance of chromosomes) element 1 protein (NSE1); | protein binding; zinc ion binding; | 0 % |  |  |
| 16 | ggggacaagtttgtacaaaaaagcaggcttgatgATGTCTGAAACCAAGAATAT, ggggaccactttgtacaagaaagctgggtcCAGCTTCGCTTGAACACTTA | Mycgr3\|36735  JGI  Ensembl |  |  | 0 % |  |  |
| 16 | ggggacaagtttgtacaaaaaagcaggcttgatgGGCGCCTGGGCACTCACGCT, ggggaccactttgtacaagaaagctgggtcGGCCTTCGACACTCGGCACT | Mycgr3\|37720  JGI  Ensembl | KOG1710 MYND Zn-finger and ankyrin repeat protein; | zinc ion binding; | 0 % |  |  |
| 16 | ggggacaagtttgtacaaaaaagcaggcttgatgCTCGACTATCCAGCGCCTTT, ggggaccactttgtacaagaaagctgggtcCTTGATCCCTGTGTACGCTT | Mycgr3\|38707  JGI  Ensembl |  |  | 0 % |  |  |
| 16 | ggggacaagtttgtacaaaaaagcaggcttgatgTCGTCTCCTAAGGTTCCCGA, ggggaccactttgtacaagaaagctgggtcTGTGGATTCCAGCAGAGACA | Mycgr3\|39133  JGI  Ensembl |  |  | 2.0979 % |  |  |
| 16 | ggggacaagtttgtacaaaaaagcaggcttgatgACAGCCAGAACACGATACAC, ggggaccactttgtacaagaaagctgggtcATCCCGCTCCGGCAACGCCC | Mycgr3\|39307  JGI  Ensembl |  |  | 0 % |  |  |
| 16 | ggggacaagtttgtacaaaaaagcaggcttgatgTCCTCCCAACCGTCACCCGT, ggggaccactttgtacaagaaagctgggtcAGCCGCACCACCCCTCTGCT | Mycgr3\|42346  JGI  Ensembl | KOG0747 Putative NAD+-dependent epimerases; |  | 0 % |  |  |
| 16 | ggggacaagtttgtacaaaaaagcaggcttgatgGGCAAACTCATGATCTACGG, ggggaccactttgtacaagaaagctgggtcTCGCACCAACCGCGATGCTC | Mycgr3\|43638  JGI  Ensembl | KOG2733 Uncharacterized membrane protein; |  | 0 % |  |  |
| 16 | ggggacaagtttgtacaaaaaagcaggcttgatgGCCAATTACTTTGTCAACCT, ggggaccactttgtacaagaaagctgggtcGCTCTCGTGCGATGGCACTC | Mycgr3\|44018  JGI  Ensembl |  |  | 0 % |  |  |
| 16 | ggggacaagtttgtacaaaaaagcaggcttgatgTCTCAGAAGCTCATTGTCAT, ggggaccactttgtacaagaaagctgggtcTCCGCGTGTCCTTCGGGCCA | Mycgr3\|44529  JGI  Ensembl |  |  | 0 % |  |  |
| 16 | ggggacaagtttgtacaaaaaagcaggcttgatgGCAATCCTCCTAGGAAAACG, ggggaccactttgtacaagaaagctgggtcCCGCCCTCTCCCTCTTCCTC | Mycgr3\|46371  JGI  Ensembl | KOG4210 Nuclear localization sequence binding protein; |  | 0 % |  |  |
| 16 | ggggacaagtttgtacaaaaaagcaggcttgatgGCCACCAACGGAGCACCAAA, ggggaccactttgtacaagaaagctgggtcAGTGCTGTGCTCCACAGGCT | Mycgr3\|46601  JGI  Ensembl |  |  | 0 % |  |  |
| 16 | ggggacaagtttgtacaaaaaagcaggcttgatgGCGGCCAACTACTGGGACTC, ggggaccactttgtacaagaaagctgggtcCTTATTCAAGAGCCCGCCAC | Mycgr3\|47503  JGI  Ensembl | KOG0794 CDK8 kinase-activating protein cyclin C; |  | 2.12766 % |  |  |
| 16 | ggggacaagtttgtacaaaaaagcaggcttgatgTCAAGCCAACACACCAAGGG, ggggaccactttgtacaagaaagctgggtcGCTTTCACGTGGTATGGCAT | Mycgr3\|50779  JGI  Ensembl |  |  | 0 % |  |  |
| 16 | ggggacaagtttgtacaaaaaagcaggcttgatgGACCCCACATTCGCAGCAGC, ggggaccactttgtacaagaaagctgggtcGGTTTTCAACAGCATCGCAG | Mycgr3\|51070  JGI  Ensembl |  |  | 0 % |  |  |
| 16 | ggggacaagtttgtacaaaaaagcaggcttgatgGCCTCAAAAGTCCCACAATA, ggggaccactttgtacaagaaagctgggtcGAACTTGTGGAGCTCCTTTC | Mycgr3\|51378  JGI  Ensembl | KOG0534 NADH-cytochrome b-5 reductase; | cytochrome-b5 reductase activity; electron transport;  oxidoreductase activity; | 0 % |  |  |
| 16 | ggggacaagtttgtacaaaaaagcaggcttgatgAATAGCCTCGACACGCTCCC, ggggaccactttgtacaagaaagctgggtcCAGCTGCAGACCCAGTTGGA | Mycgr3\|59280  JGI  Ensembl | KOG1205 Predicted dehydrogenase; | F420H2 dehydrogenase activity; malolactic enzyme activity;  metabolic process;  oxidoreductase activity;  regulation of oxidoreductase activity;  sulfur oxygenase reductase activity; | 0 % |  |  |
| 16 | ggggacaagtttgtacaaaaaagcaggcttgatgACGACAGCCAATACAACTAA, ggggaccactttgtacaagaaagctgggtcCTCGACGGTAATGCTGAATG | Mycgr3\|61690  JGI  Ensembl |  |  | 0 % |  |  |
| 16 | ggggacaagtttgtacaaaaaagcaggcttgatgGCCGACGCCTCAGCTCCAAA, ggggaccactttgtacaagaaagctgggtcACAAGCACATCCCTCCTTCG | Mycgr3\|66333  JGI  Ensembl | KOG0092 GTPase Rab5/YPT51 and related small G protein superfamily GTPases; | GTP binding; GTPase activity;  intracellular;  intracellular protein transport;  nucleocytoplasmic transport;  protein binding;  protein transport;  signal transduction;  small GTPase mediated signal transduction;  small monomeric GTPase activity; | 0 % |  |  |
| 16 | ggggacaagtttgtacaaaaaagcaggcttgatgCCGTACGAAAGCACCATCAA, ggggaccactttgtacaagaaagctgggtcCGAATTCTCCTTAACGACCT | Mycgr3\|67546  JGI  Ensembl |  |  | 3.09735 % |  |  |
| 16 | ggggacaagtttgtacaaaaaagcaggcttgatgTCTTCCACTCGGCGCGTAGG, ggggaccactttgtacaagaaagctgggtcAACTTTCCTCGGCACTGCCC | Mycgr3\|67812  JGI  Ensembl |  |  | 0 % |  |  |
| 16 | ggggacaagtttgtacaaaaaagcaggcttgatgGATTCCAACGCTATAGACAG, ggggaccactttgtacaagaaagctgggtcCCCAAACCTCTCCCTCGCCA | Mycgr3\|67943  JGI  Ensembl | KOG1973 Chromatin remodeling protein, contains PHD Zn-finger; |  | 0 % |  |  |
| 16 | ggggacaagtttgtacaaaaaagcaggcttgatgCCGCTCGCCGACTACCTCGC, ggggaccactttgtacaagaaagctgggtcCTCATCCATCTGCCAAGCAT | Mycgr3\|67969  JGI  Ensembl | KOG2654 Uncharacterized conserved protein; |  | 2.29541 % |  |  |
| 16 | ggggacaagtttgtacaaaaaagcaggcttgatgGCCGGTCACGAAGCAGCACC, ggggaccactttgtacaagaaagctgggtcCATATCCTGGAAAGGCGCAA | Mycgr3\|67984  JGI  Ensembl |  |  | 0 % |  |  |
| 16 | ggggacaagtttgtacaaaaaagcaggcttgatgACGACCCTCCAATTCCAACA, ggggaccactttgtacaagaaagctgggtcAATGTAGCGCGTCAGCAGAG | Mycgr3\|68377  JGI  Ensembl |  |  | 0 % |  |  |
| 16 | ggggacaagtttgtacaaaaaagcaggcttgatgAAGTTTGCCGAGACCATCAT, ggggaccactttgtacaagaaagctgggtcCTGCTGCAGAGTAAGAAGCG | Mycgr3\|68922  JGI  Ensembl |  |  | 0 % |  |  |
| 16 | ggggacaagtttgtacaaaaaagcaggcttgatgCCGGTCCCACGCGAAGAATA, ggggaccactttgtacaagaaagctgggtcCAGCTTCGCCTTCCTCCCAC | Mycgr3\|69507  JGI  Ensembl | KOG0725 Reductases with broad range of substrate specificities; | 2,4-dienoyl-CoA reductase (NADPH) activity; metabolic process;  oxidoreductase activity; | 0 % |  |  |
| 16 | ggggacaagtttgtacaaaaaagcaggcttgatgCCACGTCCGACAGTCGTCGA, ggggaccactttgtacaagaaagctgggtcTCGAATGAAAGGGATCATGG | Mycgr3\|69522  JGI  Ensembl | KOG2628 Farnesyl cysteine-carboxyl methyltransferase; | C-terminal protein amino acid methylation; integral to membrane;  protein-S-isoprenylcysteine O-methyltransferase activity; | 0 % |  |  |
| 16 | ggggacaagtttgtacaaaaaagcaggcttgatgTCAGTTCAAGTCCCTCGAGC, ggggaccactttgtacaagaaagctgggtcCAACGGCTTGATCCCCTTGT | Mycgr3\|71344  JGI  Ensembl | KOG1540 Ubiquinone biosynthesis methyltransferase COQ5; | 1-phenanthrol methyltransferase activity; 2-polyprenyl-6-methoxy-1,4-benzoquinone methyltransferase activity;  C-methyltransferase activity;  C-terminal protein carboxyl methyltransferase activity;  DNA-methyltransferase activity;  N-methyltransferase activity;  O-methyltransferase activity;  RNA methyltransferase activity;  S-methyltransferase activity;  arginine N-methyltransferase activity;  cobalt-precorrin-3 C17-methyltransferase activity;  cobalt-precorrin-5B C1-methyltransferase activity;  cobalt-precorrin-6B C5-methyltransferase activity;  cobalt-precorrin-7 C15-methyltransferase activity;  demethylmenaquinone methyltransferase activity;  dimethyladenosine transferase activity;  dimethylamine methyltransferase activity;  hydroxyneurosporene-O-methyltransferase activity;  lysine N-methyltransferase activity;  mRNA methyltransferase activity;  methanol-specific methylcobalamin:coenzyme M methyltransferase activity;  methylamine-specific methylcobalamin:coenzyme M methyltransferase activity;  methylarsonite methyltransferase activity;  methyltransferase activity;  monomethylamine methyltransferase activity;  protein methyltransferase activity;  protein-arginine N-methyltransferase activity;  protein-arginine N5-methyltransferase activity;  protein-leucine O-methyltransferase activity;  protein-lysine N-methyltransferase activity;  rRNA (adenine) methyltransferase activity;  rRNA (adenine-N6,N6-)-dimethyltransferase activity;  rRNA (cytosine) methyltransferase activity;  rRNA (cytosine-C5-967)-methyltransferase activity;  rRNA (guanine) methyltransferase activity;  rRNA (uridine) methyltransferase activity;  rRNA (uridine-2'-O-)-methyltransferase activity;  rRNA methyltransferase activity;  selenocysteine methyltransferase activity;  site-specific DNA-methyltransferase (cytosine-specific) activity;  tRNA (adenine)-methyltransferase activity;  tRNA (adenine-57, 58-N(1)-) methyltransferase activity;  tRNA (cytosine)-methyltransferase activity;  tRNA (guanine) methyltransferase activity;  tRNA (guanosine) methyltransferase activity;  tRNA (uracil) methyltransferase activity;  tRNA (uridine) methyltransferase activity;  tRNA methyltransferase activity;  trimethylamine methyltransferase activity; | 0 % |  |  |
| 16 | ggggacaagtttgtacaaaaaagcaggcttgatgAAGTTCACCACCCTCGTCCT, ggggaccactttgtacaagaaagctgggtcACACCTCCTCGTCCAACCAT | Mycgr3\|71724  JGI  Ensembl |  |  | 0 % |  |  |
| 16 | ggggacaagtttgtacaaaaaagcaggcttgatgGATGCTACGGCTGAGCAGAC, ggggaccactttgtacaagaaagctgggtcCAGAGCCGCCAATGAAATAA | Mycgr3\|71790  JGI  Ensembl |  |  | 0 % |  |  |
| 16 | ggggacaagtttgtacaaaaaagcaggcttgatgATCGCACGGTTGTCCCTTGC, ggggaccactttgtacaagaaagctgggtcGTCCCTCTTCCCAATGCCAT | Mycgr3\|72545  JGI  Ensembl |  |  | 0 % |  |  |
| 16 | ggggacaagtttgtacaaaaaagcaggcttgatgCCTTTCTCCCACCACTCCCA, ggggaccactttgtacaagaaagctgggtcAGGCTTGAAGAATGCATGTG | Mycgr3\|74092  JGI  Ensembl |  |  | 0 % |  |  |
| 16 | ggggacaagtttgtacaaaaaagcaggcttgatgGGCATTATCGGTAACGTGGG, ggggaccactttgtacaagaaagctgggtcGTTCTCTTGCTTCGCATATG | Mycgr3\|74217  JGI  Ensembl |  |  | 0 % |  |  |
| 16 | ggggacaagtttgtacaaaaaagcaggcttgatgGCCACTCCAAGCATGCCCAT, ggggaccactttgtacaagaaagctgggtcAGGCTTTGGAGGAATGGGAG | Mycgr3\|76101  JGI  Ensembl | KOG0725 Reductases with broad range of substrate specificities; | metabolic process; oxidoreductase activity; | 0 % |  |  |
| 16 | ggggacaagtttgtacaaaaaagcaggcttgatgACCCGCTCCGCTCTCATCGC, ggggaccactttgtacaagaaagctgggtcAAGCAAAGCGGCGAGTCCGA | Mycgr3\|76591  JGI  Ensembl |  |  | 100 % |  | ``` 34	ATGACCCGCTCCGCTCTCATCGCCGCCGCCATTGCCGGCGCGTGGACCATCACACCGACGCTCGCCGTCCCGCCCGAGGACTTTGGCTTCCCTTCCGCACCGAACGACACGATCCTCGACGTCTCCTTCCGCCAAACACTCAGCGGATACGAAGACGTCGTGCCAGGAAGCCGCTACGGTATCGCCATCGCCGAGAATGAGCCCAATCTCAGTCTCAACACCACCGCCTACCCTCGACTGGACTCGTACCAAGGCGAATACATCGTCATCATGGTCGATCCGGATGCCTCATACCCGGAGAATCCCTCCAACCGCTTCATTCTGCACTGGCTGCGACAAAATCTCAGCTCAGCAGAGAACGCCAATCCTCCACCAGCTGGTCTGCAAATTGTCACCGACACGGCGAAGTTACTAAACAGCTTCACACCCGCAATAGTCCAATATCGCCGTCCTTCGCCACCGACCAACTCCTCCGCTCATCGCTACATCTTGTACGCATTCGAGCAGCCACAGCACTTCCGCCTGCCAGAGCAGTGGTCCGGCTTGAGCGGCAGCAACCGCACGGGATTCAACCTGACGTCGTTCATCAGCGACACAAACCTGGGAATGCCATGCGCGGCAAACTACTTCTACACTTCCAACCAGACTGTTGTACCTGATGACTTTGTCTCTACCAATGGCTCTTCCTACCCAGGTGGTGATGGCACGGCGATCACTTCCGGCGATCCACATGGTCCTGGCTCCTCTGCTCCAAACAACACCCACAGCGCCAACTCGACCATGCCGACTCGCATGCCGAATGCGACAACAAGTGGTGGTGCTGGACCATCGCCGACCTCATCTCCGACGACATCTGCGGAACCTTCCTCGACCTCGGCCTCCA-CTGGTGGAACAGCCATG-TTGACCGGCGC-CGGA----A-GTCTTGTCGCCGC-AGGCCTCG-TCGGACTCGCCGCTTTG	987 	\|\|\|\|\|\|\|\|\|\|\|\|\|\|\|\|\|\|\|\|\|\|\|\|\|\|\|\|\|\|\|\|\|\|\|\|\|\|\|\|\|\|\|\|\|\|\|\|\|\|\|\|\|\|\|\|\|\|\|\|\|\|\|\|\|\|\|\|\|\|\|\|\|\|\|\|\|\|\|\|\|\|\|\|\|\|\|\|\|\|\|\|\|\|\|\|\|\|\|\|\|\|\|\|\|\|\|\|\|\|\|\|\|\|\|\|\|\|\|\|\|\|\|\|\|\|\|\|\|\|\|\|\|\|\|\|\|\|\|\|\|\|\|\|\|\|\|\|\|\|\|\|\|\|\|\|\|\|\|\|\|\|\|\|\|\|\|\|\|\|\|\|\|\|\|\|\|\|\|\|\|\|\|\|\|\|\|\|\|\|\|\|\|\|\|\|\|\|\|\|\|\|\|\|\|\|\|\|\|\|\|\|\|\|\|\|\|\|\|\|\|\|\|\|\|\|\|\|\|\|\|\|\|\|\|\|\|\|\|\|\|\|\|\|\|\|\|\|\|\|\|\|\|\|\|\|\|\|\|\|\|\|\|\|\|\|\|\|\|\|\|\|\|\|\|\|\|\|\|\|\|\|\|\|\|\|\|\|\|\|\|\|\|\|\|\|\|\|\|\|\|\|\|\|\|\|\|\|\|\|\|\|\|\|\|\|\|\|\|\|\|\|\|\|\|\|\|\|\|\|\|\|\|\|\|\|\|\|\|\|\|\|\|\|\|\|\|\|\|\|\|\|\|\|\|\|\|\|\|\|\|\|\|\|\|\|\|\|\|\|\|\|\|\|\|\|\|\|\|\|\|\|\|\|\|\|\|\|\|\|\|\|\|\|\|\|\|\|\|\|\|\|\|\|\|\|\|\|\|\|\|\|\|\|\|\|\|\|\|\|\|\|\|\|\|\|\|\|\|\|\|\|\|\|\|\|\|\|\|\|\|\|\|\|\|\|\|\|\|\|\|\|\|\|\|\|\|\|\|\|\|\|\|\|\|\|\|\|\|\|\|\|\|\|\|\|\|\|\|\|\|\|\|\|\|\|\|\|\|\|\|\|\|\|\|\|\|\|\|\|\|\|\|\|\|\|\|\|\|\|\|\|\|\|\|\|\|\|\|\|\|\|\|\|\|\|\|\|\|\|\|\|\|\|\|\|\|\|\|\|\|\|\|\|\|\|\|\|\|\|\|\|\|\|\|\|\|\|\|\|\|\|\|\|\|\|\|\|\|\|\|\|\|\|\|\|\|\|\|\|\|\|\|\|\|\|\|\|\|\|\|\|\|\|\|\|\|\|\|\|\|\|\|\|\|\|\|\|\|\|\|\|\|\|\|\|\|\|\|\|\|\|\|\|\|\|\|\|\|\|\|\|\|\|\|\|\|\|\|\|\|\|\|\|\|\|\|\|\|\|\|\|\|\|\|\|\|\|\|\|\|\|\|\|\|\|\|\|\|\|\|\|\|\|\|\|\|\|\|\|\|\|\|\|\|\|\|\|\|\|\|\|\|\|\|\|\|\|\|\|\|\|\|\|\|\|\|\|\|\|\|\|\|\|\|\|\|\|\|\|\|\|\|\|\|\|\|\|\|\|\|\|\|\|\|\|\|\|\|\|\|\|\|\|\|\|\|\|\|\|\|\|\|\|\|\|\|\|\|\|\|\|\|\|\|\|\|\|\|\|\|\|\|\|\|\|\|\|\|\|\|\|\|\|\|\|\|\|\|\|\|\|\|\|\|\|\|\|\|\|\|\|\|\|\|\|\|\|\|\|\|\|\|\|\|\|\|\|\|\|\|\|\|\|\|\|\|\|\|\|\|\|\|\|\|\|\|\|\|\|\|\|\|\|\|\|\|\|\|\|\|\|\|\|\|\|\|\|\|\|\|\|\|\|\|\|\|\|\|\|\|\|\|\|\|\|\|\|\|\|\|\|\| \|\|\|\|\|\|\|\|\|\|\| \|\|\|\|\| \|\|\|\|\|\|\| \|\|\| \|\|\|\|    \| \|\|\|\|\| \|\|  \|\|\| \| \|\|\|\|\|\| \|\| \|\|\|\|\| \| \|\|\|\|\|\| 	ATGACCCGCTCCGCTCTCATCGCCGCCGCCATTGCCGGCGCGTGGACCATCACACCGACGCTCGCCGTCCCGCCCGAGGACTTTGGCTTCCCTTCCGCACCGAACGACACGATCCTCGACGTCTCCTTCCGCCAAACACTCAGCGGATACGAAGACGTCGTGCCAGGAAGCCGCTACGGTATCGCCATCGCCGAGAATGAGCCCAATCTCAGTCTCAACACCACCGCCTACCCTCGACTGGACTCGTACCAAGGCGAATACATCGTCATCATGGTCGATCCGGATGCCTCATACCCGGAGAATCCCTCCAACCGCTTCATTCTGCACTGGCTGCGACAAAATCTCAGCTCAGCAGAGAACGCCAATCCTCCACCAGCTGGTCTGCAAATTGTCACCGACACGGCGAAGTTACTAAACAGCTTCACACCCGCAATAGTCCAATATCGCCGTCCTTCGCCACCGACCAACTCCTCCGCTCATCGCTACATCTTGTACGCATTCGAGCAGCCACAGCACTTCCGCCTGCCAGAGCAGTGGTCCGGCTTGAGCGGCAGCAACCGCACGGGATTCAACCTGACGTCGTTCATCAGCGACACAAACCTGGGAATGCCATGCGCGGCAAACTACTTCTACACTTCCAACCAGACTGTTGTACCTGATGACTTTGTCTCTACCAATGGCTCTTCCTACCCAGGTGGTGATGGCACGGCGATCACTTCCGGCGATCCACATGGTCCTGGCTCCTCTGCTCCAAACAACACCCACAGCGCCAACTCGACCATGCCGACTCGCATGCCGAATGCGACAACAAGTGGTGGTGCTGGACCATCGCCGACCTCATCTCCGACGACATCTGCGGAACCTTCCTCGACCTCGGCCTCCACCTGGTGGAACAACCATGCTTGACCGACGCGCGGACGGCATGTCTTCTCCACGCCACGCCTCGATCCGACTCCCTGCTTTG ``` |
| 16 | ggggacaagtttgtacaaaaaagcaggcttgatgCCACGACCGACATATACCAT, ggggaccactttgtacaagaaagctgggtcTGGTTCTCCCAGCATGCTTC | Mycgr3\|85442  JGI  Ensembl |  |  | 0 % |  |  |
| 16 | ggggacaagtttgtacaaaaaagcaggcttgatgCTCTCCCAGAACATCGACAG, ggggaccactttgtacaagaaagctgggtcCTGAACTCCAGGTCTGAGCA | Mycgr3\|86044  JGI  Ensembl | KOG2892 Porphobilinogen deaminase; | hydroxymethylbilane synthase activity; tetrapyrrole biosynthetic process; | 0 % |  |  |
| 16 | ggggacaagtttgtacaaaaaagcaggcttgatgCCGCGCAGAGTAACCAACCC, ggggaccactttgtacaagaaagctgggtcGCTAGGCGGCAAAGACGCAG | Mycgr3\|88242  JGI  Ensembl | KOG2663 Acetolactate synthase, small subunit; | acetolactate synthase activity; amino acid binding;  branched chain family amino acid biosynthetic process;  metabolic process; | 0 % |  |  |
| 16 | ggggacaagtttgtacaaaaaagcaggcttgatgGCGACCCTGGATCCCGCTCT, ggggaccactttgtacaagaaagctgggtcTGATTTTTCATCGCTCCCGT | Mycgr3\|88477  JGI  Ensembl | KOG3021 Predicted kinase; | (S)-lactate 2-kinase activity; ATP-dependent polynucleotide kinase activity;  GTP-dependent polynucleotide kinase activity;  LPPG:FO 2-phospho-L-lactate transferase activity;  cobinamide kinase activity;  cyclin-dependent protein kinase activating kinase regulator activity;  cyclin-dependent protein kinase regulator activity;  cytidine kinase activity;  eukaryotic elongation factor-2 kinase activator activity;  eukaryotic elongation factor-2 kinase regulator activity;  glycerate 2-kinase activity;  inositol tetrakisphosphate 2-kinase activity;  kinase activity;  phenol kinase activity;  phosphatidylinositol phosphate kinase activity;  phosphofructokinase activity;  phosphoinositide 3-kinase activity;  phosphorylase kinase regulator activity;  polynucleotide kinase activity; | 1.98413 % |  |  |
| 16 | ggggacaagtttgtacaaaaaagcaggcttgatgTCGAAACAGGCCAAGAAGGA, ggggaccactttgtacaagaaagctgggtcCTTGTCGCCTTTAACGACCT | Mycgr3\|88629  JGI  Ensembl |  |  | 0 % |  |  |
| 16 | ggggacaagtttgtacaaaaaagcaggcttgatgGCCAACCTCCAATTCCCCTA, ggggaccactttgtacaagaaagctgggtcACTCCTCCCCTCCGCATCCC | Mycgr3\|89653  JGI  Ensembl | KOG4748 Subunit of Golgi mannosyltransferase complex; | 1,2-dihydroxy-phenanthrene glycosyltransferase activity; 1-phenanthrol glycosyltransferase activity;  9-phenanthrol UDP-glucuronosyltransferase activity;  9-phenanthrol glycosyltransferase activity;  O antigen polymerase activity;  UDP-N-acetylglucosamine-peptide N-acetylglucosaminyltransferase activity;  UDP-glucose:glycoprotein glucosyltransferase activity;  acetylgalactosaminyltransferase activity;  acetylglucosaminyltransferase activity;  alpha(1,3)-fucosyltransferase activity;  alpha(1,6)-fucosyltransferase activity;  alpha-1,2-galactosyltransferase activity;  alpha-1,2-mannosyltransferase activity;  alpha-1,3-galactosyltransferase activity;  alpha-1,3-mannosyltransferase activity;  alpha-1,6-mannosyltransferase activity;  beta-1,4-mannosyltransferase activity;  cellulose synthase activity;  dolichyl pyrophosphate Glc1Man9GlcNAc2 alpha-1,3-glucosyltransferase activity; dolichyl pyrophosphate Man7GlcNAc2 alpha-1,3-glucosyltransferase activity;  dolichyl pyrophosphate Man9GlcNAc2 alpha-1,3-glucosyltransferase activity;  dolichyl-phosphate-glucose-glycolipid alpha-glucosyltransferase activity;  fucosyltransferase activity;  galactosyltransferase activity;  glucosyltransferase activity;  glycolipid mannosyltransferase activity;  inositol phosphoceramide synthase activity;  integral to membrane;  lipopolysaccharide-1,6-galactosyltransferase activity;  mannosyltransferase activity;  oligosaccharyl transferase activity;  phenanthrol glycosyltransferase activity;  transferase activity, transferring hexosyl groups; | 0 % |  |  |
| 16 | ggggacaagtttgtacaaaaaagcaggcttgatgTCGCTCGACCTACAAAGCCA, ggggaccactttgtacaagaaagctgggtcAGCCTCTTCTGCACTCTTCC | Mycgr3\|89942  JGI  Ensembl |  |  | 0 % |  |  |
| 16 | ggggacaagtttgtacaaaaaagcaggcttgatgGATGACCCATTCGTATACCC, ggggaccactttgtacaagaaagctgggtcAAGACGAACGACAACGGCAT | Mycgr3\|90579  JGI  Ensembl |  |  | 0 % |  |  |
| 16 | ggggacaagtttgtacaaaaaagcaggcttgatgCCTGCCAAAACGCGGGCGAA, ggggaccactttgtacaagaaagctgggtcCGCGCAGGCGCAGGAGGGAG | Mycgr3\|90773  JGI  Ensembl |  |  | 2.2335 % |  |  |
| 16 | ggggacaagtttgtacaaaaaagcaggcttgatgCGACGCATCACACCGCGGAC, ggggaccactttgtacaagaaagctgggtcCCTGTACCATCCGTTGCCCA | Mycgr3\|90780  JGI  Ensembl | KOG0039 Ferric reductase, NADH/NADPH oxidase and related proteins; | electron transport; membrane; oxidoreductase activity; | 0 % |  |  |
| 16 | ggggacaagtttgtacaaaaaagcaggcttgatgCCGGTGCTTGTCAAACTGAC, ggggaccactttgtacaagaaagctgggtcCTCGGCGAATCTCATGTTCT | Mycgr3\|91042  JGI  Ensembl |  |  | 0 % |  |  |
| 16 | ggggacaagtttgtacaaaaaagcaggcttgatgGCTCGCAGCCGTCCAGCAAT, ggggaccactttgtacaagaaagctgggtcGTGCGGATTAGGCCTCCGAC | Mycgr3\|91727  JGI  Ensembl | KOG1815 Predicted E3 ubiquitin ligase; | zinc ion binding; | 0 % |  |  |
| 16 | ggggacaagtttgtacaaaaaagcaggcttgatgATCGCCATCCGACCCCCACC, ggggaccactttgtacaagaaagctgggtcAAAGATCCTCTTCATCCCCC | Mycgr3\|92053  JGI  Ensembl |  |  | 0 % |  |  |
| 16 | ggggacaagtttgtacaaaaaagcaggcttgatgGAGCGTCCCACCAAGCCAGT, ggggaccactttgtacaagaaagctgggtcCGTGTGAACTGGAGTAGACT | Mycgr3\|93696  JGI  Ensembl |  |  | 0 % |  |  |
| 16 | ggggacaagtttgtacaaaaaagcaggcttgatgGAACGAATCAAGTCAATACT, ggggaccactttgtacaagaaagctgggtcCACACCTTTCTTCAACTCGT | Mycgr3\|93756  JGI  Ensembl |  |  | 0 % |  |  |
| 16 | ggggacaagtttgtacaaaaaagcaggcttgatgCCATCTCTCAACACCCTTAT, ggggaccactttgtacaagaaagctgggtcCACAACAAGCAGGAAAGCAA | Mycgr3\|94299  JGI  Ensembl |  |  | 0 % |  |  |
| 16 | ggggacaagtttgtacaaaaaagcaggcttgatgGCCAGAGTCACACGCAGCAA, ggggaccactttgtacaagaaagctgggtcGCTGCGTCCCATGATCGTGA | Mycgr3\|94341  JGI  Ensembl |  |  | 0 % |  |  |
| 16 | ggggacaagtttgtacaaaaaagcaggcttgatgGCCAACATTGCCAAAGACAC, ggggaccactttgtacaagaaagctgggtcCTCCAACGGACCAAAATGAA | Mycgr3\|94450  JGI  Ensembl | KOG1282 Serine carboxypeptidases (lysosomal cathepsin A); |  | 0 % |  |  |
| 16 | ggggacaagtttgtacaaaaaagcaggcttgatgTCCGCCACAGACCTCGCCGA, ggggaccactttgtacaagaaagctgggtcCTCCCCCAAGAAGAACTCTT | Mycgr3\|94695  JGI  Ensembl | KOG4178 Soluble epoxide hydrolase; | catalytic activity; cis-stilbene-oxide hydrolase activity; | 0 % |  |  |
| 16 | ggggacaagtttgtacaaaaaagcaggcttgatgTCCCAGTCCACGCGTACGGT, ggggaccactttgtacaagaaagctgggtcCGGGATAACAACCCACCACT | Mycgr3\|94792  JGI  Ensembl |  |  | 0 % |  |  |
| 16 | ggggacaagtttgtacaaaaaagcaggcttgatgGTTCCCGTCACTGTCATCAT, ggggaccactttgtacaagaaagctgggtcCTCCTCTTCTCCCTCGGAGT | Mycgr3\|94802  JGI  Ensembl |  |  | 0 % |  |  |
| 16 | ggggacaagtttgtacaaaaaagcaggcttgatgCCGCTAAACTACCTCGCCAT, ggggaccactttgtacaagaaagctgggtcGCCTGTCTTGTCGTCGAGAT | Mycgr3\|94821  JGI  Ensembl |  |  | 0 % |  |  |
| 16 | ggggacaagtttgtacaaaaaagcaggcttgatgCCCAACACCACCCTAATAGT, ggggaccactttgtacaagaaagctgggtcTTCACTCCCCCGACTCTCAC | Mycgr3\|94962  JGI  Ensembl |  |  | 2.15686 % |  |  |
| 16 | ggggacaagtttgtacaaaaaagcaggcttgatgCAGCTCATCAACCTTTTCGC, ggggaccactttgtacaagaaagctgggtcGTTCTTCCTCGCAGAGTCGA | Mycgr3\|95063  JGI  Ensembl |  |  | 32.21545 % |  |  |
| 16 | ggggacaagtttgtacaaaaaagcaggcttgatgGCCTCGTTTCCCAACTTCAG, ggggaccactttgtacaagaaagctgggtcCTTACCAGCAGCACTCGACT | Mycgr3\|95860  JGI  Ensembl |  |  | 2.10843 % |  |  |
| 16 | ggggacaagtttgtacaaaaaagcaggcttgatgGCCAACAACATCGAAGCTGC, ggggaccactttgtacaagaaagctgggtcAGCAGAAGTGGCCGAGCCAG | Mycgr3\|96244  JGI  Ensembl |  |  | 0 % |  |  |
| 16 | ggggacaagtttgtacaaaaaagcaggcttgatgGCTGCCGCTTCTACCGATCA, ggggaccactttgtacaagaaagctgggtcACGACAGAGCGATGCGAAGC | Mycgr3\|96552  JGI  Ensembl |  |  | 0 % |  |  |
| 16 | ggggacaagtttgtacaaaaaagcaggcttgatgGAGAGTCACAATTCGGCCGC, ggggaccactttgtacaagaaagctgggtcGCTGCGATCCTGTGGGTATT | Mycgr3\|96847  JGI  Ensembl |  |  | 0 % |  |  |
| 16 | ggggacaagtttgtacaaaaaagcaggcttgatgACCCAATTCATCATCCAAGG, ggggaccactttgtacaagaaagctgggtcAATGAGCTCGAAGAATCCTC | Mycgr3\|97320  JGI  Ensembl |  |  | 0 % |  |  |
| 16 | ggggacaagtttgtacaaaaaagcaggcttgatgTCACGCTCTTATTGCAACAC, ggggaccactttgtacaagaaagctgggtcGCAGCGCGAACCGCGCCGCG | Mycgr3\|97616  JGI  Ensembl |  |  | 0 % |  |  |
| 16 | ggggacaagtttgtacaaaaaagcaggcttgatgCCGCCTTTCTTCATGGCACC, ggggaccactttgtacaagaaagctgggtcTGATTCGAAAAGTTCGTTGA | Mycgr3\|97674  JGI  Ensembl |  |  | 0 % |  |  |
| 16 | ggggacaagtttgtacaaaaaagcaggcttgatgCCGGATCACGGCCATGTCTA, ggggaccactttgtacaagaaagctgggtcAACACGGAAGAACGCCTCTT | Mycgr3\|97809  JGI  Ensembl | KOG0382 Carbonic anhydrase; |  | 2.18254 % |  |  |
| 16 | ggggacaagtttgtacaaaaaagcaggcttgatgTCCGCCGGAGTGAGGGAAAG, ggggaccactttgtacaagaaagctgggtcGCCTTTATGTCGGTCGGCTG | Mycgr3\|97933  JGI  Ensembl |  |  | 0 % |  |  |
| 16 | ggggacaagtttgtacaaaaaagcaggcttgatgTCGAAGCGACCGCGGACACT, ggggaccactttgtacaagaaagctgggtcTTCGGATTTCCACGCGACCA | Mycgr3\|98514  JGI  Ensembl |  |  | 1.94872 % |  |  |
| 16 | ggggacaagtttgtacaaaaaagcaggcttgatgCCTCCCCGATTACAGCTCCT, ggggaccactttgtacaagaaagctgggtcCCATAATCTGTTCTCGGCCC | Mycgr3\|99059  JGI  Ensembl | KOG0846 Mitochondrial/chloroplast ribosomal protein L15/L10; | intracellular; large ribosomal subunit;  ribosome;  structural constituent of ribosome;  translation; | 0 % |  |  |
| 16 | ggggacaagtttgtacaaaaaagcaggcttgatgGCACCAAAGAAGAAGGTCGA, ggggaccactttgtacaagaaagctgggtcCAGACGACGACCACGGCGAC | Mycgr3\|99159  JGI  Ensembl | KOG0407 40S ribosomal protein S14; | intracellular; ribosome; structural constituent of ribosome;  translation; | 0 % |  |  |
| 16 | ggggacaagtttgtacaaaaaagcaggcttgatgTCATCCAGAAAGAAGATCCT, ggggaccactttgtacaagaaagctgggtcGCACGAGCAGCCTTGCTCAT | Mycgr3\|99493  JGI  Ensembl | KOG0394 Ras-related GTPase; | ATP binding; GTP binding;  GTPase activity;  intracellular;  intracellular protein transport;  nucleocytoplasmic transport;  protein binding;  protein transport;  regulation of transcription, DNA-dependent;  signal transduction;  small GTPase mediated signal transduction;  small monomeric GTPase activity;  transcription factor binding; | 0 % |  |  |
| 16 | ggggacaagtttgtacaaaaaagcaggcttgatgTCCACTAAACCCCTCCCCAA, ggggaccactttgtacaagaaagctgggtcTGATTTGGAGCTCTTGGCCT | Mycgr3\|101009  JGI  Ensembl | KOG3316 Transport protein particle (TRAPP) complex subunit; |  | 0 % |  |  |
| 16 | ggggacaagtttgtacaaaaaagcaggcttgatgGATATCGATGAGCCACATGG, ggggaccactttgtacaagaaagctgggtcGTGTTTCATAGCGGTAGCAT | Mycgr3\|101681  JGI  Ensembl | KOG0223 Aquaporin (major intrinsic protein family); | membrane; transport; transporter activity; | 2.28401 % |  |  |
| 16 | ggggacaagtttgtacaaaaaagcaggcttgatgGTAAAATGCCATCCGCAAAG, ggggaccactttgtacaagaaagctgggtcCATCAACACCTTGGCGAATG | Mycgr3\|103488  JGI  Ensembl |  |  | 0 % |  |  |
| 16 | ggggacaagtttgtacaaaaaagcaggcttgatgTCGTCCAAAATCATTCTCTA, ggggaccactttgtacaagaaagctgggtcCTGATCATTCTCCAACCAAG | Mycgr3\|103524  JGI  Ensembl |  |  | 0 % |  |  |
| 16 | ggggacaagtttgtacaaaaaagcaggcttgatgGTCTCCTCCACACAACAGCA, ggggaccactttgtacaagaaagctgggtcCCCACTCTTCCCAACCATCC | Mycgr3\|104372  JGI  Ensembl |  |  | 0 % |  |  |
| 16 | ggggacaagtttgtacaaaaaagcaggcttgatgCCTCCGAAGCGAAAGACCGC, ggggaccactttgtacaagaaagctgggtcTCCTCTCTTTGCGTTCTTGA | Mycgr3\|105431  JGI  Ensembl | KOG2868 Decapping enzyme complex component DCP1; |  | 0 % |  |  |
| 16 | ggggacaagtttgtacaaaaaagcaggcttgatgAAGTACTCATACGCCGCCCT, ggggaccactttgtacaagaaagctgggtcGTAAGAGAGAGGAGCACCGA | Mycgr3\|106335  JGI  Ensembl |  |  | 0 % |  |  |
| 16 | ggggacaagtttgtacaaaaaagcaggcttgatgGTGGAACTCAGGAAAAGGCC, ggggaccactttgtacaagaaagctgggtcAGCAGCATCGACCTTCTCCG | Mycgr3\|106420  JGI  Ensembl | KOG0855 Alkyl hydroperoxide reductase, thiol specific antioxidant and related enzymes; | antioxidant activity; oxidoreductase activity;  peroxiredoxin activity; | 0 % |  |  |
| 16 | ggggacaagtttgtacaaaaaagcaggcttgatgACTTCCCTCCTCCGCTTCCA, ggggaccactttgtacaagaaagctgggtcGCAAGCCGCCGCGATCATCG | Mycgr3\|107889  JGI  Ensembl | KOG0118 FOG: RRM domain; |  | 2.4173 % |  |  |
| 16 | ggggacaagtttgtacaaaaaagcaggcttgatgGCGTCTTTAATGCGCATCGT, ggggaccactttgtacaagaaagctgggtcAACTGCTTGCTCTCTCAAAA | Mycgr3\|107972  JGI  Ensembl | KOG0401 Translation initiation factor 4F, ribosome/mRNA-bridging subunit (eIF-4G); |  | 2.06972 % |  |  |
| 16 | ggggacaagtttgtacaaaaaagcaggcttgatgGCGACCACCACAATCGCTGC, ggggaccactttgtacaagaaagctgggtcGAACGTGCTCCAGACCATTC | Mycgr3\|108290  JGI  Ensembl | KOG1729 FYVE finger containing protein; |  | 0 % |  |  |
| 16 | ggggacaagtttgtacaaaaaagcaggcttgatgGAGCTTCTTCGCCTTCCAGC, ggggaccactttgtacaagaaagctgggtcCAAAGGACTCTCCAGCCCAC | Mycgr3\|108770  JGI  Ensembl | KOG2486 Predicted GTPase; | GTP binding; | 0 % |  |  |
| 16 | ggggacaagtttgtacaaaaaagcaggcttgatgTCAAAAGCAGGAGAACCCCT, ggggaccactttgtacaagaaagctgggtcAACATACTCCACCTCCCCAT | Mycgr3\|110754  JGI  Ensembl |  |  | 0 % |  |  |
| 16 | ggggacaagtttgtacaaaaaagcaggcttgatgCACGCCCAACGCTCTTTCAC, ggggaccactttgtacaagaaagctgggtcAGCATAGTTTTCAGACGCCA | Mycgr3\|110756  JGI  Ensembl |  |  | 0 % |  |  |
| 17 | ggggacaagtttgtacaaaaaagcaggcttgatgGACCACCACCGTCCGCACCC, ggggaccactttgtacaagaaagctgggtcACGCTCCCGAACCTCACTCC | Mycgr3\|32807  JGI  Ensembl |  |  | 100 % |  |  |
| 17 | ggggacaagtttgtacaaaaaagcaggcttgatgGCACCGACTCGGATCATAGA, ggggaccactttgtacaagaaagctgggtcATTTTGTGCGAGTCGATAAG | Mycgr3\|33800  JGI  Ensembl |  |  | 0 % |  |  |
| 17 | ggggacaagtttgtacaaaaaagcaggcttgatgCTGTTTCAAAGAGCACTACT, ggggaccactttgtacaagaaagctgggtcGACAATGACACCAGGCATCT | Mycgr3\|34033  JGI  Ensembl |  |  | 100 % |  |  |
| 17 | ggggacaagtttgtacaaaaaagcaggcttgatgTGTATCTCTGTCTTCTCAAC, ggggaccactttgtacaagaaagctgggtcCGAACTAGCTGCCCATCCTT | Mycgr3\|36216  JGI  Ensembl | KOG2342 Uncharacterized conserved protein; |  | 100 % |  |  |
| 17 | ggggacaagtttgtacaaaaaagcaggcttgatgTCCAACGGCCAACCCATCCC, ggggaccactttgtacaagaaagctgggtcCAACTTCGGCTTCGCATTGG | Mycgr3\|41694  JGI  Ensembl | KOG1680 Enoyl-CoA hydratase; | catalytic activity; enoyl-CoA hydratase activity;  metabolic process; | 1.86393 % |  |  |
| 17 | ggggacaagtttgtacaaaaaagcaggcttgatgAAAGCCATCAACATCCCCGC, ggggaccactttgtacaagaaagctgggtcTAATCGCCCTGACTCTGGGA | Mycgr3\|41893  JGI  Ensembl | KOG1198 Zinc-binding oxidoreductase; | NADPH:quinone reductase activity; oxidoreductase activity;  zinc ion binding; | 0 % |  |  |
| 17 | ggggacaagtttgtacaaaaaagcaggcttgatgGGCTCAGCAGCACCTGCTCT, ggggaccactttgtacaagaaagctgggtcCTCCGCACCCAACTGCCTCG | Mycgr3\|41898  JGI  Ensembl |  |  | 100 % |  |  |
| 17 | ggggacaagtttgtacaaaaaagcaggcttgatgGTGCTGCATCTGCTGGGCAA, ggggaccactttgtacaagaaagctgggtcCCGCAGCTTAACATTGCTAT | Mycgr3\|44079  JGI  Ensembl | KOG4297 C-type lectin; |  | 100 % |  |  |
| 17 | ggggacaagtttgtacaaaaaagcaggcttgatgGCCGTCTTGTTGGCGCACGG, ggggaccactttgtacaagaaagctgggtcCCAATCGAGTTTCTTGGCTT | Mycgr3\|44611  JGI  Ensembl | KOG2876 Molybdenum cofactor biosynthesis pathway protein; | catalytic activity; iron-sulfur cluster binding; | 100 % |  |  |
| 17 | ggggacaagtttgtacaaaaaagcaggcttgatgGCCGACACCAACACATCCTT, ggggaccactttgtacaagaaagctgggtcTTTTAACTCATTTGCAAGTT | Mycgr3\|44943  JGI  Ensembl | KOG4169 15-hydroxyprostaglandin dehydrogenase and related dehydrogenases; | 15-hydroxyprostaglandin dehydrogenase (NAD+) activity; metabolic process;  oxidoreductase activity; | 100 % |  |  |
| 17 | ggggacaagtttgtacaaaaaagcaggcttgatgGTCGAACCAAACCCTCTCCC, ggggaccactttgtacaagaaagctgggtcCTCCACCGCCTCGATCTTGA | Mycgr3\|48752  JGI  Ensembl |  |  | 100 % |  |  |
| 17 | ggggacaagtttgtacaaaaaagcaggcttgatgCTCGAATACCTGCGATTGAA, ggggaccactttgtacaagaaagctgggtcTCGCCGAGATCCTTGCCATG | Mycgr3\|50884  JGI  Ensembl | KOG1515 Arylacetamide deacetylase; | hormone-sensitive lipase activity; | 0 % |  |  |
| 17 | ggggacaagtttgtacaaaaaagcaggcttgatgTGGCGCATCCCGAAAGACAC, ggggaccactttgtacaagaaagctgggtcATGCCCGTTGTCCCTCATAT | Mycgr3\|63387  JGI  Ensembl |  |  | 100 % |  |  |
| 17 | ggggacaagtttgtacaaaaaagcaggcttgatgGCCGCAAATTGGGGAACACA, ggggaccactttgtacaagaaagctgggtcCAACGAAAGCTTCATCTTGA | Mycgr3\|65361  JGI  Ensembl | KOG1208 Dehydrogenases with different specificities (related to short-chain alcohol dehydrogenases); | 1-phenylethanol dehydrogenase activity; chloral hydrate dehydrogenase activity;  glucose dehydrogenase activity;  hydroxymethylmethylsilanediol oxidase activity;  ketoreductase activity;  metabolic process;  myrtenol dehydrogenase activity;  oxidoreductase activity;  pinocarveol dehydrogenase activity;  versicolorin reductase activity; | 100 % |  |  |
| 17 | ggggacaagtttgtacaaaaaagcaggcttgatgGCTCCCTCACGCACCTCACC, ggggaccactttgtacaagaaagctgggtcCGCCTTCACTGCAGAGTTCA | Mycgr3\|65666  JGI  Ensembl | KOG3290 Peroxisomal phytanoyl-CoA hydroxylase; |  | 100 % |  |  |
| 17 | ggggacaagtttgtacaaaaaagcaggcttgatgTCAGCCAAAGTTCCTCCTTC, ggggaccactttgtacaagaaagctgggtcCAGCGACTCCTCAATGCCAA | Mycgr3\|66153  JGI  Ensembl |  |  | 100 % |  |  |
| 17 | ggggacaagtttgtacaaaaaagcaggcttgatgCCTCCACGAATAAGACTGTC, ggggaccactttgtacaagaaagctgggtcAGCCTCGCTCTTGGACCCCG | Mycgr3\|66422  JGI  Ensembl |  |  | 100 % |  |  |
| 17 | ggggacaagtttgtacaaaaaagcaggcttgatgGACGCCTACTCGACCGGCGC, ggggaccactttgtacaagaaagctgggtcTTCGTCTTCATCGTCATCCT | Mycgr3\|66569  JGI  Ensembl | KOG1609 Protein involved in mRNA turnover and stability; | protein binding; zinc ion binding; | 100 % |  |  |
| 17 | ggggacaagtttgtacaaaaaagcaggcttgatgGAAGCCGAAAATAGGGTCAA, ggggaccactttgtacaagaaagctgggtcTACATCAATATCGACAGGAC | Mycgr3\|67844  JGI  Ensembl |  |  | 0 % |  |  |
| 17 | ggggacaagtttgtacaaaaaagcaggcttgatgGCCCACCTCAACTACGCTCG, ggggaccactttgtacaagaaagctgggtcCAACTTCGAAGCTACCTTCC | Mycgr3\|70045  JGI  Ensembl | KOG1599 Uricase (urate oxidase); | purine base metabolic process; urate oxidase activity; | 100 % |  |  |
| 17 | ggggacaagtttgtacaaaaaagcaggcttgatgAAGATACTCATCACTGGAGC, ggggaccactttgtacaagaaagctgggtcCTTCTTCAAGCTCGCTGCAA | Mycgr3\|70266  JGI  Ensembl | KOG2774 NAD dependent epimerase; | L-threonine 3-dehydrogenase activity; | 2.1164 % |  |  |
| 17 | ggggacaagtttgtacaaaaaagcaggcttgatgGCTATCGACCGACGACCGCA, ggggaccactttgtacaagaaagctgggtcTAACAGCGGCATGTTGAATG | Mycgr3\|70541  JGI  Ensembl |  |  | 100 % |  |  |
| 17 | ggggacaagtttgtacaaaaaagcaggcttgatgGGTGCACTGAGCATCCTCAT, ggggaccactttgtacaagaaagctgggtcATCCATATCCTCCAGACTCC | Mycgr3\|71290  JGI  Ensembl |  |  | 100 % |  |  |
| 17 | ggggacaagtttgtacaaaaaagcaggcttgatgGGCAATCTCGCTATCCTCGG, ggggaccactttgtacaagaaagctgggtcTGCCAACATCTCCCTGATCG | Mycgr3\|71493  JGI  Ensembl |  |  | 100 % |  |  |
| 17 | ggggacaagtttgtacaaaaaagcaggcttgatgCCTCGGGACCTCTCCGCACT, ggggaccactttgtacaagaaagctgggtcATGATTGTCGTACTCGAACT | Mycgr3\|72359  JGI  Ensembl | KOG4144 Arylalkylamine N-acetyltransferase; | N-acetyltransferase activity; aralkylamine N-acetyltransferase activity;  metabolic process; | 100 % |  |  |
| 17 | ggggacaagtttgtacaaaaaagcaggcttgatgGCACCATCTGCGATCACCAC, ggggaccactttgtacaagaaagctgggtcAAGCTTGTGCGCTCCACCCG | Mycgr3\|72416  JGI  Ensembl | KOG3699 Cytoskeletal protein Adducin; | L-fuculose-phosphate aldolase activity; metal ion binding; | 100 % |  |  |
| 17 | ggggacaagtttgtacaaaaaagcaggcttgatgGAAGCTCCCCCGTTCTACTC, ggggaccactttgtacaagaaagctgggtcTATCGAACAATGTGATCCAC | Mycgr3\|72554  JGI  Ensembl |  |  | 100 % |  |  |
| 17 | ggggacaagtttgtacaaaaaagcaggcttgatgACCATCTCCCTCCCTTTCAC, ggggaccactttgtacaagaaagctgggtcCACCTCAACAGTACCCTCCA | Mycgr3\|73076  JGI  Ensembl | KOG3033 Predicted PhzC/PhzF-type epimerase; | biosynthetic process; catalytic activity; | 100 % |  |  |
| 17 | ggggacaagtttgtacaaaaaagcaggcttgatgCTCGCCCGGAGAGCACTGCC, ggggaccactttgtacaagaaagctgggtcACCACAGCTACCAGCACTAT | Mycgr3\|75026  JGI  Ensembl | KOG2382 Predicted alpha/beta hydrolase; | T/G mismatch-specific endonuclease activity; nuclease activity;  phosphoric ester hydrolase activity; | 100 % |  |  |
| 17 | ggggacaagtttgtacaaaaaagcaggcttgatgGCCTCCTCCCCAACCGTCCT, ggggaccactttgtacaagaaagctgggtcAGATGTATATACTCCGGTCG | Mycgr3\|75982  JGI  Ensembl | KOG1371 UDP-glucose 4-epimerase/UDP-sulfoquinovose synthase; |  | 0 % |  |  |
| 17 | ggggacaagtttgtacaaaaaagcaggcttgatgCCCGTCCTCCCTCAACTCAT, ggggaccactttgtacaagaaagctgggtcCTTCTGGTCTCGATTAGAAG | Mycgr3\|76388  JGI  Ensembl |  |  | 100 % |  |  |
| 17 | ggggacaagtttgtacaaaaaagcaggcttgatgTCCTCCTACCTCTCCAACCT, ggggaccactttgtacaagaaagctgggtcTCGTCCATACCTCCTCCCAC | Mycgr3\|76532  JGI  Ensembl |  |  | 100 % | chr\_10:1211349 C->T; |  |
| 17 | ggggacaagtttgtacaaaaaagcaggcttgatgGCTCTACCCCAACGAGCTCG, ggggaccactttgtacaagaaagctgggtcTCTCGGCCTGCTAGACAACG | Mycgr3\|77482  JGI  Ensembl | KOG2625 Uncharacterized conserved protein; |  | 100 % |  |  |
| 17 | ggggacaagtttgtacaaaaaagcaggcttgatgGCGAACGACCTCGACAACCT, ggggaccactttgtacaagaaagctgggtcCTTACAGTTCAACACCGCAC | Mycgr3\|77625  JGI  Ensembl | KOG2689 Predicted ubiquitin regulatory protein; | intracellular; nucleic acid binding;  zinc ion binding; | 100 % |  |  |
| 17 | ggggacaagtttgtacaaaaaagcaggcttgatgCCGAGCACACACGAAGCAGC, ggggaccactttgtacaagaaagctgggtcGCATGATACAACAACCTTGA | Mycgr3\|84764  JGI  Ensembl | KOG1198 Zinc-binding oxidoreductase; | NADPH:quinone reductase activity; oxidoreductase activity;  zinc ion binding; | 100 % |  |  |
| 17 | ggggacaagtttgtacaaaaaagcaggcttgatgCTCACGCTCAGAGCCTATCT, ggggaccactttgtacaagaaagctgggtcGTATCGATAGCAGAGTATTC | Mycgr3\|86183  JGI  Ensembl |  |  | 100 % |  |  |
| 17 | ggggacaagtttgtacaaaaaagcaggcttgatgCCGTCTCAAACCGTCCCCGA, ggggaccactttgtacaagaaagctgggtcCGAAGACTTCAATACCGCAA | Mycgr3\|88783  JGI  Ensembl |  |  | 100 % |  |  |
| 17 | ggggacaagtttgtacaaaaaagcaggcttgatgTCTACCAACGAAGCGGTTCG, ggggaccactttgtacaagaaagctgggtcGTTCAGCTTATGAAGCACCT | Mycgr3\|88803  JGI  Ensembl | KOG0068 D-3-phosphoglycerate dehydrogenase, D-isomer-specific 2-hydroxy acid dehydrogenase superfamily; | cofactor binding; oxidoreductase activity, acting on the CH-OH group of donors, NAD or NADP as acceptor; phosphoglycerate dehydrogenase activity; | 0 % |  |  |
| 17 | ggggacaagtttgtacaaaaaagcaggcttgatgGCTTCAACGCAGCGCTCAGA, ggggaccactttgtacaagaaagctgggtcAACCTCGTGCTGCATGTTTG | Mycgr3\|88858  JGI  Ensembl |  |  | 0 % |  |  |
| 17 | ggggacaagtttgtacaaaaaagcaggcttgatgGCACAAAGCGAATACGACAA, ggggaccactttgtacaagaaagctgggtcGGCGAGGACCTTGGTTGGTC | Mycgr3\|89227  JGI  Ensembl |  |  | 0 % |  |  |
| 17 | ggggacaagtttgtacaaaaaagcaggcttgatgACTTACAACATTCCCGATGA, ggggaccactttgtacaagaaagctgggtcCTCCTCCTCAACAAACTCCC | Mycgr3\|91389  JGI  Ensembl | KOG4409 Predicted hydrolase/acyltransferase (alpha/beta hydrolase superfamily); | aminopeptidase activity; cytoplasm; prolyl aminopeptidase activity;  proteolysis; | 100 % |  |  |
| 17 | ggggacaagtttgtacaaaaaagcaggcttgatgGCCAACTCATCAGAACAAAC, ggggaccactttgtacaagaaagctgggtcTCCTCCCGGTCCACTCCGCC | Mycgr3\|91538  JGI  Ensembl |  |  | 100 % |  |  |
| 17 | ggggacaagtttgtacaaaaaagcaggcttgatgGCCAAACCCATCCCCGAAAC, ggggaccactttgtacaagaaagctgggtcTCGTCCTCGGATGCCGGCAG | Mycgr3\|91646  JGI  Ensembl |  |  | 100 % |  |  |
| 17 | ggggacaagtttgtacaaaaaagcaggcttgatgGAAGATGTGCCGTCGTACCG, ggggaccactttgtacaagaaagctgggtcACCGGCATGCATCCGGCTCC | Mycgr3\|92385  JGI  Ensembl |  |  | 0 % |  |  |
| 17 | ggggacaagtttgtacaaaaaagcaggcttgatgAATCTAGAGCCTGGTGAAGC, ggggaccactttgtacaagaaagctgggtcGTGGTGTCGAATCAGATGTG | Mycgr3\|93271  JGI  Ensembl | KOG2084 Predicted histone tail methylase containing SET domain; | nucleus; | 0 % |  |  |
| 17 | ggggacaagtttgtacaaaaaagcaggcttgatgGTGGACGAGAAAGTTATTGT, ggggaccactttgtacaagaaagctgggtcCTTCCTCGCTGAACGCTGCC | Mycgr3\|93484  JGI  Ensembl |  |  | 100 % |  |  |
| 17 | ggggacaagtttgtacaaaaaagcaggcttgatgGCACCCAAGCCATCGAAACG, ggggaccactttgtacaagaaagctgggtcGTCCTGATCATTCTCTCCGA | Mycgr3\|94703  JGI  Ensembl | KOG2146 Splicing coactivator SRm160/300, subunit SRm160 (contains PWI domain); |  | 0 % |  |  |
| 17 | ggggacaagtttgtacaaaaaagcaggcttgatgGCGAAGCTCAAGCGTATCTC, ggggaccactttgtacaagaaagctgggtcTAGCTCGGACATCTTGGACA | Mycgr3\|94720  JGI  Ensembl |  |  | 100 % |  |  |
| 17 | ggggacaagtttgtacaaaaaagcaggcttgatgTCCCGCAACACCCGCGCTTA, ggggaccactttgtacaagaaagctgggtcCTCCCTCTCGTCCTCCGCCT | Mycgr3\|94744  JGI  Ensembl |  |  | 100 % |  |  |
| 17 | ggggacaagtttgtacaaaaaagcaggcttgatgTCAAACCTCCGCCTCCTAAT, ggggaccactttgtacaagaaagctgggtcTCCCAACACCATCGCCTCCA | Mycgr3\|94830  JGI  Ensembl |  |  | 100 % |  |  |
| 17 | ggggacaagtttgtacaaaaaagcaggcttgatgCCGCGTCCCAGCAATGACAC, ggggaccactttgtacaagaaagctgggtcCTTCGAAGAAGGCCGCTTTC | Mycgr3\|94843  JGI  Ensembl |  |  | 100 % |  |  |
| 17 | ggggacaagtttgtacaaaaaagcaggcttgatgTCCCTCTACCCAACAACCCT, ggggaccactttgtacaagaaagctgggtcCTTTCGTCCTTCGCGCTCGA | Mycgr3\|95221  JGI  Ensembl |  |  | 100 % |  |  |
| 17 | ggggacaagtttgtacaaaaaagcaggcttgatgGCTTCAGCGGCCATGGAATC, ggggaccactttgtacaagaaagctgggtcGTTCGGAGGGTCTGGGAGGG | Mycgr3\|95369  JGI  Ensembl |  |  | 100 % |  |  |
| 17 | ggggacaagtttgtacaaaaaagcaggcttgatgCTTCACCTATCCTGCGCAAG, ggggaccactttgtacaagaaagctgggtcGAGATAGGCGACTTGCGCGG | Mycgr3\|95513  JGI  Ensembl |  |  | 100 % |  |  |
| 17 | ggggacaagtttgtacaaaaaagcaggcttgatgCCAGGCATCTGGGACGCCAC, ggggaccactttgtacaagaaagctgggtcTTGCCCGGTCGCCTGTAGTA | Mycgr3\|95528  JGI  Ensembl |  |  | 0 % |  |  |
| 17 | ggggacaagtttgtacaaaaaagcaggcttgatgTCGTATGTTGCCCAAAATGA, ggggaccactttgtacaagaaagctgggtcGACATCGATAAGTCTGCCTC | Mycgr3\|95610  JGI  Ensembl |  |  | 100 % |  |  |
| 17 | ggggacaagtttgtacaaaaaagcaggcttgatgCCGTCGTACCAATTGCGAAA, ggggaccactttgtacaagaaagctgggtcCTTTCCACCTTCGAGAGCCT | Mycgr3\|95824  JGI  Ensembl | KOG1216 von Willebrand factor and related coagulation proteins; |  | 100 % |  |  |
| 17 | ggggacaagtttgtacaaaaaagcaggcttgatgGCGCTTCACTTCATTATCAT, ggggaccactttgtacaagaaagctgggtcGTGTGGGCGCATCAATGCCT | Mycgr3\|96081  JGI  Ensembl | KOG0297 TNF receptor-associated factor; | protein binding; zinc ion binding; | 0 % |  |  |
| 17 | ggggacaagtttgtacaaaaaagcaggcttgatgGCTATGGACGACTTGCCAAA, ggggaccactttgtacaagaaagctgggtcGAAGAAGAACAAATCTTCCA | Mycgr3\|96279  JGI  Ensembl |  |  | 0 % |  |  |
| 17 | ggggacaagtttgtacaaaaaagcaggcttgatgCAGTTCACCAACCTCATCAC, ggggaccactttgtacaagaaagctgggtcGACGAGGAGGACTGCGGAAC | Mycgr3\|96704  JGI  Ensembl | KOG3599 Ca2+-modulated nonselective cation channel polycystin; |  | 0 % |  |  |
| 17 | ggggacaagtttgtacaaaaaagcaggcttgatgGCGATCCTCGACAGCTTTCC, ggggaccactttgtacaagaaagctgggtcGTCCTCAGTCAAATCGACGA | Mycgr3\|96992  JGI  Ensembl |  |  | 100 % |  |  |
| 17 | ggggacaagtttgtacaaaaaagcaggcttgatgACGGCAAAAGCAACAACAGC, ggggaccactttgtacaagaaagctgggtcCCCTCCTTTCTTAGTATTCC | Mycgr3\|97083  JGI  Ensembl |  |  | 100 % |  |  |
| 17 | ggggacaagtttgtacaaaaaagcaggcttgatgACTCGTGCTCACTTTGATAA, ggggaccactttgtacaagaaagctgggtcGTCGTAGGCCTTGGTCATGC | Mycgr3\|97243  JGI  Ensembl |  |  | 100 % |  |  |
| 17 | ggggacaagtttgtacaaaaaagcaggcttgatgGAGCCGACATATGGCACTAC, ggggaccactttgtacaagaaagctgggtcCAACAGCATATCGACTTCCT | Mycgr3\|97683  JGI  Ensembl | KOG0401 Translation initiation factor 4F, ribosome/mRNA-bridging subunit (eIF-4G); |  | 100 % |  |  |
| 17 | ggggacaagtttgtacaaaaaagcaggcttgatgCCGCCCGCCGCCGATATCAC, ggggaccactttgtacaagaaagctgggtcCGAAGCCCACAACAGGTCCG | Mycgr3\|97696  JGI  Ensembl |  |  | 100 % |  |  |
| 17 | ggggacaagtttgtacaaaaaagcaggcttgatgGGAGGCGCCCTCTCCATGTT, ggggaccactttgtacaagaaagctgggtcAACATGCCGGCGGGTCTTTC | Mycgr3\|98945  JGI  Ensembl | KOG0072 GTP-binding ADP-ribosylation factor-like protein ARL1; | GTP binding; intracellular;  intracellular protein transport;  protein transport;  small GTPase mediated signal transduction; | 0 % |  |  |
| 17 | ggggacaagtttgtacaaaaaagcaggcttgatgGCGCCGAAACCAGCAGAGGT, ggggaccactttgtacaagaaagctgggtcTCCAACGGCTTCCTCCCACT | Mycgr3\|99376  JGI  Ensembl |  |  | 0 % |  |  |
| 17 | ggggacaagtttgtacaaaaaagcaggcttgatgCCAGTCACGACCAACACCGG, ggggaccactttgtacaagaaagctgggtcCTTCTTCCGCAAGACGAGAA | Mycgr3\|100372  JGI  Ensembl | KOG1541 Predicted protein carboxyl methylase; |  | 100 % |  |  |
| 17 | ggggacaagtttgtacaaaaaagcaggcttgatgGCATCAATACCAAATCTCAA, ggggaccactttgtacaagaaagctgggtcCACTCCATCGTCTTTGTTCT | Mycgr3\|100580  JGI  Ensembl | KOG2918 Carboxymethyl transferase; | 1-phenanthrol methyltransferase activity; 2-polyprenyl-6-methoxy-1,4-benzoquinone methyltransferase activity;  C-methyltransferase activity;  C-terminal protein carboxyl methyltransferase activity;  DNA-methyltransferase activity;  N-methyltransferase activity;  O-methyltransferase activity;  RNA methyltransferase activity;  S-methyltransferase activity;  arginine N-methyltransferase activity;  cobalt-precorrin-3 C17-methyltransferase activity;  cobalt-precorrin-5B C1-methyltransferase activity;  cobalt-precorrin-6B C5-methyltransferase activity;  cobalt-precorrin-7 C15-methyltransferase activity;  demethylmenaquinone methyltransferase activity;  dimethyladenosine transferase activity;  dimethylamine methyltransferase activity;  hydroxyneurosporene-O-methyltransferase activity;  lysine N-methyltransferase activity;  mRNA methyltransferase activity;  methanol-specific methylcobalamin:coenzyme M methyltransferase activity;  methylamine-specific methylcobalamin:coenzyme M methyltransferase activity;  methylarsonite methyltransferase activity;  monomethylamine methyltransferase activity;  protein methyltransferase activity;  protein-arginine N-methyltransferase activity;  protein-arginine N5-methyltransferase activity;  protein-leucine O-methyltransferase activity;  protein-lysine N-methyltransferase activity;  rRNA (adenine) methyltransferase activity;  rRNA (adenine-N6,N6-)-dimethyltransferase activity;  rRNA (cytosine) methyltransferase activity;  rRNA (cytosine-C5-967)-methyltransferase activity;  rRNA (guanine) methyltransferase activity;  rRNA (uridine) methyltransferase activity;  rRNA (uridine-2'-O-)-methyltransferase activity;  rRNA methyltransferase activity;  selenocysteine methyltransferase activity;  site-specific DNA-methyltransferase (cytosine-specific) activity;  tRNA (adenine)-methyltransferase activity;  tRNA (adenine-57, 58-N(1)-) methyltransferase activity;  tRNA (cytosine)-methyltransferase activity;  tRNA (guanine) methyltransferase activity;  tRNA (guanosine) methyltransferase activity;  tRNA (uracil) methyltransferase activity;  tRNA (uridine) methyltransferase activity;  tRNA methyltransferase activity;  trimethylamine methyltransferase activity; | 0 % |  |  |
| 17 | ggggacaagtttgtacaaaaaagcaggcttgatgCATGCGATCGCACTCACCTG, ggggaccactttgtacaagaaagctgggtcAGCCACAGCACTGGTATCTT | Mycgr3\|100840  JGI  Ensembl | KOG2913 Predicted membrane protein; |  | 2.03095 % |  |  |
| 17 | ggggacaagtttgtacaaaaaagcaggcttgatgCCTCCCAAGAAAGGCGCCAC, ggggaccactttgtacaagaaagctgggtcTGACCCTCCACCGCCCAATG | Mycgr3\|101221  JGI  Ensembl | KOG1763 Uncharacterized conserved protein, contains CCCH-type Zn-finger; | nucleic acid binding; zinc ion binding; | 0 % |  |  |
| 17 | ggggacaagtttgtacaaaaaagcaggcttgatgCCGTCACTCTTCTCGGTCAT, ggggaccactttgtacaagaaagctgggtcTGATGAGCGACGGCTGAAGA | Mycgr3\|102108  JGI  Ensembl |  |  | 0 % |  |  |
| 17 | ggggacaagtttgtacaaaaaagcaggcttgatgCCTTCGCCACGCTCATCGCC, ggggaccactttgtacaagaaagctgggtcCCGACTGCTGGATCGAGGTG | Mycgr3\|102360  JGI  Ensembl |  |  | 100 % |  |  |
| 17 | ggggacaagtttgtacaaaaaagcaggcttgatgGCTCTGCTCCTTTCACGCCG, ggggaccactttgtacaagaaagctgggtcAGGAGCTCCATACAACCGTA | Mycgr3\|102895  JGI  Ensembl | KOG4781 Uncharacterized conserved protein; |  | 0 % |  |  |
| 17 | ggggacaagtttgtacaaaaaagcaggcttgatgTCTTCCTCAAGTGCGGCCGA, ggggaccactttgtacaagaaagctgggtcCGCCCGCTGCGAACCCTTCC | Mycgr3\|102907  JGI  Ensembl |  |  | 100 % |  |  |
| 17 | ggggacaagtttgtacaaaaaagcaggcttgatgGAAAGTCAGCAGCTCCAAAC, ggggaccactttgtacaagaaagctgggtcTTCAGCCTCCCCGCTTCCGC | Mycgr3\|103603  JGI  Ensembl |  |  | 100 % |  |  |
| 17 | ggggacaagtttgtacaaaaaagcaggcttgatgTTTGCACGACGCGTTGTCGA, ggggaccactttgtacaagaaagctgggtcGAGCCGGCCGTGAGAGATGT | Mycgr3\|103617  JGI  Ensembl |  |  | 100 % |  |  |
| 17 | ggggacaagtttgtacaaaaaagcaggcttgatgCAATTCCTCTCCCTCTTCGC, ggggaccactttgtacaagaaagctgggtcCTGGCCGACACCACCAAGAA | Mycgr3\|104571  JGI  Ensembl | KOG3346 Phosphatidylethanolamine binding protein; |  | 100 % |  |  |
| 17 | ggggacaagtttgtacaaaaaagcaggcttgatgTTCGCCACAAAACTCAACGT, ggggaccactttgtacaagaaagctgggtcGTTGACCTTTGCGGTCCATT | Mycgr3\|105871  JGI  Ensembl |  |  | 100 % |  |  |
| 17 | ggggacaagtttgtacaaaaaagcaggcttgatgACCGACCACACCCCACAACG, ggggaccactttgtacaagaaagctgggtcCGCCGCCTGAGGCTCAACAT | Mycgr3\|106430  JGI  Ensembl |  |  | 100 % |  |  |
| 17 | ggggacaagtttgtacaaaaaagcaggcttgatgCAGTACGGTCCAGTTCCAGA, ggggaccactttgtacaagaaagctgggtcTGCCCACTTTTTGCCCCTCT | Mycgr3\|107028  JGI  Ensembl | KOG0725 Reductases with broad range of substrate specificities; | 15-hydroxyprostaglandin dehydrogenase (NAD+) activity; metabolic process;  oxidoreductase activity; | 100 % |  |  |
| 17 | ggggacaagtttgtacaaaaaagcaggcttgatgGCTATCTCGAATCCCGTCGA, ggggaccactttgtacaagaaagctgggtcGAGCCTCAACTTCAATCGGC | Mycgr3\|107713  JGI  Ensembl |  |  | 100 % |  |  |
| 17 | ggggacaagtttgtacaaaaaagcaggcttgatgACCGACGAAGAGAGCTGGCA, ggggaccactttgtacaagaaagctgggtcCACCACAATATCCTTCTCCA | Mycgr3\|108008  JGI  Ensembl | KOG2375 Protein interacting with poly(A)-binding protein; |  | 100 % |  |  |
| 17 | ggggacaagtttgtacaaaaaagcaggcttgatgCACCGCTACCAGTCGAACAG, ggggaccactttgtacaagaaagctgggtcAAGGCCCTCCATCGCCTTCA | Mycgr3\|108336  JGI  Ensembl |  |  | 100 % |  |  |
| 17 | ggggacaagtttgtacaaaaaagcaggcttgatgTTCACCACCTCCTCGGCGGG, ggggaccactttgtacaagaaagctgggtcCTCCGCTTGGACTGGTGGTG | Mycgr3\|108406  JGI  Ensembl |  |  | 0 % |  |  |
| 17 | ggggacaagtttgtacaaaaaagcaggcttgatgCCCGACCAACCCTCCACCAA, ggggaccactttgtacaagaaagctgggtcATTCCCTCCTCCTTCTTGCT | Mycgr3\|110329  JGI  Ensembl | KOG1887 Ubiquitin carboxyl-terminal hydrolase; |  | 100 % |  |  |
| 17 | ggggacaagtttgtacaaaaaagcaggcttgatgGCCGTCGAACCGCTTGATAC, ggggaccactttgtacaagaaagctgggtcGTGGCCGTAGAAAGCTACGG | Mycgr3\|110395  JGI  Ensembl |  |  | 100 % |  |  |
| 17 | ggggacaagtttgtacaaaaaagcaggcttgatgCCCGCCACCTCATCGACCTA, ggggaccactttgtacaagaaagctgggtcTAGCCACGGCGCAGGCAGAA | Mycgr3\|110914  JGI  Ensembl |  |  | 100 % |  |  |
| 17 | ggggacaagtttgtacaaaaaagcaggcttgatgGTGGCTCCTCTCCGCACAAG, ggggaccactttgtacaagaaagctgggtcCGCCTTGCCCAACTTCTCCG | Mycgr3\|111108  JGI  Ensembl | KOG3954 Electron transfer flavoprotein, alpha subunit; | FAD binding; electron carrier activity;  electron transport; | 0 % |  |  |
| 17 | ggggacaagtttgtacaaaaaagcaggcttgatgGGCACTCGTCATCTCATCCT, ggggaccactttgtacaagaaagctgggtcCGCATACAGAACTTCCGCGT | Mycgr3\|111193  JGI  Ensembl |  |  | 100 % |  |  |
| 17 | ggggacaagtttgtacaaaaaagcaggcttgatgCTCTCCAAGATCACCACCAC, ggggaccactttgtacaagaaagctgggtcAGAAGTGGCCTCCTTGGCCT | Mycgr3\|111676  JGI  Ensembl |  |  | 100 % |  |  |
| 18 | ggggacaagtttgtacaaaaaagcaggcttgatgTCATTCGCACCTCAGACCCC, ggggaccactttgtacaagaaagctgggtcGTCTTCCGCCGGCAGACCGA | Mycgr3\|33694  JGI  Ensembl | KOG1208 Dehydrogenases with different specificities (related to short-chain alcohol dehydrogenases); | (R)-2-hydroxyglutarate dehydrogenase activity; (R)-2-hydroxyisocaproate dehydrogenase activity;  2-hydroxytetrahydrofuran dehydrogenase activity;  3-keto sterol reductase activity;  3-ketoglucose-reductase activity;  5-exo-hydroxycamphor dehydrogenase activity;  C-3 sterol dehydrogenase (C-4 sterol decarboxylase) activity;  aldo-keto reductase activity;  epoxide dehydrogenase activity;  gluconate dehydrogenase activity;  isocitrate dehydrogenase activity;  metabolic process;  mevaldate reductase activity;  oxidoreductase activity;  phenylcoumaran benzylic ether reductase activity;  steroid dehydrogenase activity;  steroid dehydrogenase activity, acting on the CH-CH group of donors;  steroid dehydrogenase activity, acting on the CH-OH group of donors, NAD or NADP as acceptor; | 100 % |  |  |
| 18 | ggggacaagtttgtacaaaaaagcaggcttgatgGCGGGCGGCAGCACGTTCAC, ggggaccactttgtacaagaaagctgggtcCAGTCTACTGGATCGTACCG | Mycgr3\|34023  JGI  Ensembl |  |  | 100 % |  |  |
| 18 | ggggacaagtttgtacaaaaaagcaggcttgatgGGTGCGGCCATACTGAATCA, ggggaccactttgtacaagaaagctgggtcCACTTCCTGGATCTCGTTGA | Mycgr3\|35631  JGI  Ensembl |  |  | 100 % |  |  |
| 18 | ggggacaagtttgtacaaaaaagcaggcttgatgGCAGAAGAAACCAGGACCGT, ggggaccactttgtacaagaaagctgggtcCTTCCTGTCCTGCTGCTTCT | Mycgr3\|37838  JGI  Ensembl |  |  | 3.64444 % |  |  |
| 18 | ggggacaagtttgtacaaaaaagcaggcttgatgGGCGGCGGCGAGCTGAAAGA, ggggaccactttgtacaagaaagctgggtcATACCCCCTCCCCCTCTCCA | Mycgr3\|41081  JGI  Ensembl | KOG0069 Glyoxylate/hydroxypyruvate reductase (D-isomer-specific 2-hydroxy acid dehydrogenase superfamily); | cofactor binding; glyoxylate reductase (NADP) activity;  oxidoreductase activity, acting on the CH-OH group of donors, NAD or NADP as acceptor; | 100 % |  |  |
| 18 | ggggacaagtttgtacaaaaaagcaggcttgatgCCCGAAGCAAGAATATTACC, ggggaccactttgtacaagaaagctgggtcTCGTCGCGACGCTTCTCGCA | Mycgr3\|41107  JGI  Ensembl | KOG0123 Polyadenylate-binding protein (RRM superfamily); | zinc ion binding; | 3.84615 % |  |  |
| 18 | ggggacaagtttgtacaaaaaagcaggcttgatgCACCTCTTCTCGCTGTTCAT, ggggaccactttgtacaagaaagctgggtcATTGGCATTCATGCCATATT | Mycgr3\|43397  JGI  Ensembl |  |  | 0 % |  |  |
| 18 | ggggacaagtttgtacaaaaaagcaggcttgatgAAGGTCAGCAAGATCATCGT, ggggaccactttgtacaagaaagctgggtcTTGGCACACAGTCAGCGATT | Mycgr3\|43539  JGI  Ensembl | KOG2362 Uncharacterized Fe-S protein; | catalytic activity; molybdenum ion binding;  pyridoxal phosphate binding; | 100 % |  |  |
| 18 | ggggacaagtttgtacaaaaaagcaggcttgatgGCTCCAAGCCTGCCCAGAAC, ggggaccactttgtacaagaaagctgggtcGTTGCGCTCGATGTTGTTCT | Mycgr3\|44286  JGI  Ensembl | KOG1198 Zinc-binding oxidoreductase; | NADPH:quinone reductase activity; oxidoreductase activity;  zinc ion binding; | 100 % |  |  |
| 18 | ggggacaagtttgtacaaaaaagcaggcttgatgACAGGAGTCCCGGACAAACC, ggggaccactttgtacaagaaagctgggtcCAACACCTTCAGTCTCTCAG | Mycgr3\|45968  JGI  Ensembl | KOG0535 Sulfite oxidase, molybdopterin-binding component; | electron transport; molybdenum ion binding;  oxidoreductase activity;  sulfite oxidase activity; | 100 % |  |  |
| 18 | ggggacaagtttgtacaaaaaagcaggcttgatgGCCACCACCTCCGATCCCCG, ggggaccactttgtacaagaaagctgggtcCAATTCCACCTCTTTCAGAT | Mycgr3\|46160  JGI  Ensembl | KOG2855 Ribokinase; | D-ribose metabolic process; ribokinase activity; | 0 % |  |  |
| 18 | ggggacaagtttgtacaaaaaagcaggcttgatgGTGTCCGAAACTTGGGATCC, ggggaccactttgtacaagaaagctgggtcAAACACTACCGTACCGTTCA | Mycgr3\|47647  JGI  Ensembl |  |  | 100 % |  |  |
| 18 | ggggacaagtttgtacaaaaaagcaggcttgatgCCTACTCCCGGTCGCGGCGG, ggggaccactttgtacaagaaagctgggtcCAGCTTCTCCCTCCGAACAG | Mycgr3\|48130  JGI  Ensembl | KOG3016 Acyl-CoA thioesterase; | palmitoyl-CoA hydrolase activity; | 100 % |  |  |
| 18 | ggggacaagtttgtacaaaaaagcaggcttgatgTCGATTCAACAGGTCACTCT, ggggaccactttgtacaagaaagctgggtcGCCTTCCAACCTCTTCACAG | Mycgr3\|56299  JGI  Ensembl |  |  | 100 % | chr\_3:672125 A->G; |  |
| 18 | ggggacaagtttgtacaaaaaagcaggcttgatgACGACCTCCACGTTCAAACA, ggggaccactttgtacaagaaagctgggtcATCATAAAACACCAAACACC | Mycgr3\|63510  JGI  Ensembl |  |  | 100 % | chr\_11:334743 G->A; chr\_11:335025 G->A; chr\_11:334969 A->G; |  |
| 18 | ggggacaagtttgtacaaaaaagcaggcttgatgGCGAGCTCAACTCAACTGGG, ggggaccactttgtacaagaaagctgggtcCACCGCCACCTCGTCTCCTT | Mycgr3\|65284  JGI  Ensembl |  |  | 100 % |  |  |
| 18 | ggggacaagtttgtacaaaaaagcaggcttgatgCCTTCAACACCACCACCAGT, ggggaccactttgtacaagaaagctgggtcCTTGCTGTCCTCAGCCTCGA | Mycgr3\|65551  JGI  Ensembl |  |  | 100 % |  |  |
| 18 | ggggacaagtttgtacaaaaaagcaggcttgatgTCGAACTACAACCAGTACTC, ggggaccactttgtacaagaaagctgggtcATTCTTACTCGGCGGGCCCA | Mycgr3\|66031  JGI  Ensembl | KOG0810 SNARE protein Syntaxin 1 and related proteins; | membrane; | 100 % |  |  |
| 18 | ggggacaagtttgtacaaaaaagcaggcttgatgGCTGCGCCTCCGCCATCGTC, ggggaccactttgtacaagaaagctgggtcCAGTAACCCGTTCGCTTGTG | Mycgr3\|66908  JGI  Ensembl | KOG0089 Methylenetetrahydrofolate dehydrogenase/methylenetetrahydrofolate cyclohydrolase; | catalytic activity; folic acid and derivative biosynthetic process;  methylenetetrahydrofolate dehydrogenase (NAD+) activity;  methylenetetrahydrofolate dehydrogenase activity; | 100 % |  |  |
| 18 | ggggacaagtttgtacaaaaaagcaggcttgatgTCCTACCCTCCCCGACTCAA, ggggaccactttgtacaagaaagctgggtcAATCGCGGTATACCCTCCAT | Mycgr3\|66956  JGI  Ensembl | KOG0725 Reductases with broad range of substrate specificities; | 3-oxoacyl-[acyl-carrier-protein] reductase activity; metabolic process;  oxidoreductase activity; | 100 % |  |  |
| 18 | ggggacaagtttgtacaaaaaagcaggcttgatgGTCAAGACACGCCAATGGCA, ggggaccactttgtacaagaaagctgggtcATCGATATGTGTGACCAACT | Mycgr3\|67099  JGI  Ensembl | KOG1196 Predicted NAD-dependent oxidoreductase; | NADPH:quinone reductase activity; oxidoreductase activity;  zinc ion binding; | 3.69702 % |  |  |
| 18 | ggggacaagtttgtacaaaaaagcaggcttgatgTCTTCAAACCTGAGTAATGG, ggggaccactttgtacaagaaagctgggtcCCAAGCGCAATGCCCAGCAT | Mycgr3\|67679  JGI  Ensembl | KOG0725 Reductases with broad range of substrate specificities; | D-arabinitol 2-dehydrogenase activity; metabolic process;  oxidoreductase activity; | 100 % |  |  |
| 18 | ggggacaagtttgtacaaaaaagcaggcttgatgTCAGACGTGCCTCTGCTCGT, ggggaccactttgtacaagaaagctgggtcGAATTCGTCGTCGTCGTCGG | Mycgr3\|68104  JGI  Ensembl | KOG3206 Alpha-tubulin folding cofactor B; |  | 0 % |  |  |
| 18 | ggggacaagtttgtacaaaaaagcaggcttgatgGCTTTCACCCAATTGAAGAC, ggggaccactttgtacaagaaagctgggtcAACCTGCACCGGGAATTGGT | Mycgr3\|68477  JGI  Ensembl |  |  | 0 % |  |  |
| 18 | ggggacaagtttgtacaaaaaagcaggcttgatgTCTTCGCACGCAGGCTCACA, ggggaccactttgtacaagaaagctgggtcCTTCGCCTCTTTGGCCTGCT | Mycgr3\|68555  JGI  Ensembl | KOG4842 Protein involved in sister chromatid separation and/or segregation; | metallopeptidase activity; proteolysis; zinc ion binding; | 3.56506 % |  |  |
| 18 | ggggacaagtttgtacaaaaaagcaggcttgatgGCGACCAAACCCAACAAAAC, ggggaccactttgtacaagaaagctgggtcCAACTTCGCCCTCGCCCTCT | Mycgr3\|69661  JGI  Ensembl | KOG1683 Hydroxyacyl-CoA dehydrogenase/enoyl-CoA hydratase; | catalytic activity; dodecenoyl-CoA delta-isomerase activity;  metabolic process; | 100 % |  |  |
| 18 | ggggacaagtttgtacaaaaaagcaggcttgatgGCGACCAAGCTGTTCGTCAC, ggggaccactttgtacaagaaagctgggtcAGGTCGAGCCTCGCCCGCGG | Mycgr3\|69901  JGI  Ensembl |  |  | 100 % |  |  |
| 18 | ggggacaagtttgtacaaaaaagcaggcttgatgTCCGCCCCACGAATCCTCTC, ggggaccactttgtacaagaaagctgggtcTGACCACCTCGAAAAATCCG | Mycgr3\|69961  JGI  Ensembl | KOG2661 Peptidase family M48; | membrane; metalloendopeptidase activity;  metallopeptidase activity;  proteolysis;  zinc ion binding; | 100 % |  |  |
| 18 | ggggacaagtttgtacaaaaaagcaggcttgatgCCAGTCCGCGTCCTCATCGT, ggggaccactttgtacaagaaagctgggtcAAGCTTACTCTCCTTCGGCT | Mycgr3\|70413  JGI  Ensembl |  |  | 100 % |  |  |
| 18 | ggggacaagtttgtacaaaaaagcaggcttgatgGCATCCAACGAATCCTTCGA, ggggaccactttgtacaagaaagctgggtcGTTCTCCAAAACATACCGAT | Mycgr3\|70497  JGI  Ensembl | KOG0023 Alcohol dehydrogenase, class V; | (R)-2-hydroxyglutarate dehydrogenase activity; (R)-2-hydroxyisocaproate dehydrogenase activity;  2-hydroxytetrahydrofuran dehydrogenase activity;  3-keto sterol reductase activity;  3-ketoglucose-reductase activity;  5-exo-hydroxycamphor dehydrogenase activity;  C-3 sterol dehydrogenase (C-4 sterol decarboxylase) activity;  aldo-keto reductase activity;  epoxide dehydrogenase activity;  gluconate dehydrogenase activity;  isocitrate dehydrogenase activity;  mevaldate reductase activity;  oxidoreductase activity;  phenylcoumaran benzylic ether reductase activity;  steroid dehydrogenase activity;  steroid dehydrogenase activity, acting on the CH-CH group of donors;  steroid dehydrogenase activity, acting on the CH-OH group of donors, NAD or NADP as acceptor; zinc ion binding; | 100 % |  |  |
| 18 | ggggacaagtttgtacaaaaaagcaggcttgatgCGTGCCATCCAAGTCAAAGA, ggggaccactttgtacaagaaagctgggtcCAACTTACTCTCCTTCACCT | Mycgr3\|71113  JGI  Ensembl | KOG1198 Zinc-binding oxidoreductase; | NADPH:quinone reductase activity; oxidoreductase activity;  zinc ion binding; | 0 % |  |  |
| 18 | ggggacaagtttgtacaaaaaagcaggcttgatgAAGACCGAAGTCGGAGTCTC, ggggaccactttgtacaagaaagctgggtcAGTGCCATCCTTTCGCTTCA | Mycgr3\|71170  JGI  Ensembl |  |  | 3.75587 % |  |  |
| 18 | ggggacaagtttgtacaaaaaagcaggcttgatgGCCGACGACGAGATGGATCC, ggggaccactttgtacaagaaagctgggtcCTCCGCCGGGACGAAGGGCT | Mycgr3\|71192  JGI  Ensembl | KOG2825 Putative arsenite-translocating ATPase; | ATP binding; anion transport;  antimonite-transporting ATPase activity;  arsenite transmembrane-transporting ATPase activity;  electron transport;  membrane;  oxidoreductase activity; | 100 % |  |  |
| 18 | ggggacaagtttgtacaaaaaagcaggcttgatgTCTGAAGAGATCACGCCTTT, ggggaccactttgtacaagaaagctgggtcTTTCGTTCTGGTCGAGTCAG | Mycgr3\|72213  JGI  Ensembl | KOG2565 Predicted hydrolases or acyltransferases (alpha/beta hydrolase superfamily); | catalytic activity; cis-stilbene-oxide hydrolase activity;  epoxide hydrolase activity;  membrane;  microsome;  response to toxin; | 100 % | chr\_5:1451825 G->A; chr\_5:1452208 A->G; chr\_5:1452118 A->G; |  |
| 18 | ggggacaagtttgtacaaaaaagcaggcttgatgCATCTCCAGTCTAGCATCCT, ggggaccactttgtacaagaaagctgggtcATTCGCCCTCCCACCAATCT | Mycgr3\|72662  JGI  Ensembl |  |  | 100 % |  |  |
| 18 | ggggacaagtttgtacaaaaaagcaggcttgatgCTGGAGTTCAGGACCGCGGG, ggggaccactttgtacaagaaagctgggtcTGGCCTCATCTGTTCTCGCA | Mycgr3\|73544  JGI  Ensembl | KOG0277 Peroxisomal targeting signal type 2 receptor; |  | 100 % |  |  |
| 18 | ggggacaagtttgtacaaaaaagcaggcttgatgAGCATCGCGAGCGACAAAAG, ggggaccactttgtacaagaaagctgggtcTGATGGGTGATATGCCTTGT | Mycgr3\|73568  JGI  Ensembl |  |  | 100 % |  |  |
| 18 | ggggacaagtttgtacaaaaaagcaggcttgatgGCGGGCCGCTTCTTTTTATC, ggggaccactttgtacaagaaagctgggtcCATAAGTCCTCTCAAGAACA | Mycgr3\|73621  JGI  Ensembl | KOG0503 Asparaginase; | amino acid metabolic process; asparaginase activity;  asparagine metabolic process; | 100 % | chr\_7:74720 A->G; chr\_7:74452 T->G; chr\_7:74441 T->C; |  |
| 18 | ggggacaagtttgtacaaaaaagcaggcttgatgCCCTCCTTCGACTCCGCCAA, ggggaccactttgtacaagaaagctgggtcAGGCCTCAATGAATCTGAGA | Mycgr3\|73993  JGI  Ensembl | KOG1663 O-methyltransferase; | catechol O-methyltransferase activity; | 100 % |  |  |
| 18 | ggggacaagtttgtacaaaaaagcaggcttgatgGGGAGAGCGTGTGCGAGATC, ggggaccactttgtacaagaaagctgggtcCGCTCCGCTCTGCTCAGTAC | Mycgr3\|74145  JGI  Ensembl |  |  | 100 % | chr\_7:1490865 C->T; chr\_7:1490378 A->G; |  |
| 18 | ggggacaagtttgtacaaaaaagcaggcttgatgCTGCCCGTGGTTCGATCCAT, ggggaccactttgtacaagaaagctgggtcTAGCAGACCGACAGCTCTCT | Mycgr3\|74955  JGI  Ensembl | KOG2713 Mitochondrial tryptophanyl-tRNA synthetase; | ATP binding; aminoacyl-tRNA ligase activity;  cytoplasm;  nucleotide binding;  tRNA aminoacylation for protein translation;  translation;  tryptophan-tRNA ligase activity;  tryptophanyl-tRNA aminoacylation; | 100 % |  |  |
| 18 | ggggacaagtttgtacaaaaaagcaggcttgatgGCTTCTCCTACGTTCACAAC, ggggaccactttgtacaagaaagctgggtcCATCTTGCTCGGCACCCACT | Mycgr3\|75576  JGI  Ensembl | KOG1680 Enoyl-CoA hydratase; | catalytic activity; enoyl-CoA hydratase activity;  metabolic process; | 100 % |  |  |
| 18 | ggggacaagtttgtacaaaaaagcaggcttgatgGCAGAAGTACCCCAGGAGGC, ggggaccactttgtacaagaaagctgggtcGCCACTGCCAGTTGGCTCAG | Mycgr3\|75774  JGI  Ensembl | KOG3969 Uncharacterized conserved protein; | GPI-anchor transamidase activity; cyanophycinase activity;  deadenylation-dependent decapping;  hydrolase activity;  protein binding;  regulation of hydrolase activity; | 100 % |  |  |
| 18 | ggggacaagtttgtacaaaaaagcaggcttgatgGCCGACCGAATACCACCCGA, ggggaccactttgtacaagaaagctgggtcGATGAAGTCTCCACCACCGA | Mycgr3\|75839  JGI  Ensembl | KOG4761 Proteasome formation inhibitor PI31; |  | 100 % |  |  |
| 18 | ggggacaagtttgtacaaaaaagcaggcttgatgGCCTCGCAGCCAGAGGCGGA, ggggaccactttgtacaagaaagctgggtcTTGAACAACACAACGGCACC | Mycgr3\|75944  JGI  Ensembl |  |  | 100 % |  |  |
| 18 | ggggacaagtttgtacaaaaaagcaggcttgatgCCTCCGCGACCATCCACCCG, ggggaccactttgtacaagaaagctgggtcAGCCTCAGCAACCTTCTCCC | Mycgr3\|77496  JGI  Ensembl | KOG3141 Mitochondrial/chloroplast ribosomal protein L3; | intracellular; ribosome; structural constituent of ribosome;  translation; | 100 % |  |  |
| 18 | ggggacaagtttgtacaaaaaagcaggcttgatgGCTTTCACTCCGGGACAAGA, ggggaccactttgtacaagaaagctgggtcGCCATATAGTTGAGCAGAGA | Mycgr3\|84377  JGI  Ensembl | KOG0399 Glutamate synthase; | FAD binding; electron transport;  oxidoreductase activity; | 100 % |  |  |
| 18 | ggggacaagtttgtacaaaaaagcaggcttgatgCCACCACCAGCCTCCCTCTC, ggggaccactttgtacaagaaagctgggtcCTCCATCGTCTCCGCCAACA | Mycgr3\|87050  JGI  Ensembl |  |  | 100 % |  |  |
| 18 | ggggacaagtttgtacaaaaaagcaggcttgatgCTTCCCCCCTCCTCATCCTC, ggggaccactttgtacaagaaagctgggtcCCGACTCTTCCACCTCACCC | Mycgr3\|88121  JGI  Ensembl |  |  | 22.81335 % | chr\_11:1335862 C->A; |  |
| 18 | ggggacaagtttgtacaaaaaagcaggcttgatgGATCGCTTCCGCTGGCAGAT, ggggaccactttgtacaagaaagctgggtcAGATGCAGCAGCACTTGGAG | Mycgr3\|89163  JGI  Ensembl |  |  | 100 % |  |  |
| 18 | ggggacaagtttgtacaaaaaagcaggcttgatgAGCACCGCCTGTTCGTTCTT, ggggaccactttgtacaagaaagctgggtcGTGAGTGATTGAAGAGACAG | Mycgr3\|89169  JGI  Ensembl |  |  | 100 % |  |  |
| 18 | ggggacaagtttgtacaaaaaagcaggcttgatgTACTCGTCATATGCTGCTGA, ggggaccactttgtacaagaaagctgggtcCTTGCCGTGTCTCTTGCTCA | Mycgr3\|89241  JGI  Ensembl |  |  | 100 % |  |  |
| 18 | ggggacaagtttgtacaaaaaagcaggcttgatgCGCCGATTCACGAACAAGTT, ggggaccactttgtacaagaaagctgggtcATCCCCTCCCTCCCCAAAAA | Mycgr3\|90294  JGI  Ensembl |  |  | 100 % |  |  |
| 18 | ggggacaagtttgtacaaaaaagcaggcttgatgCCGATATTACGCGCGAGCTC, ggggaccactttgtacaagaaagctgggtcTTTCACGGGCGCGTTGGGAT | Mycgr3\|90968  JGI  Ensembl | KOG2992 Nucleolar GTPase/ATPase p130; |  | 100 % | chr\_2:2564244 A->G; chr\_2:2564461 T->C; |  |
| 18 | ggggacaagtttgtacaaaaaagcaggcttgatgGCGCTCGTCGAATACTCCGA, ggggaccactttgtacaagaaagctgggtcACCCCACCCATATTTACTTG | Mycgr3\|91519  JGI  Ensembl | KOG2992 Nucleolar GTPase/ATPase p130; |  | 100 % |  |  |
| 18 | ggggacaagtttgtacaaaaaagcaggcttgatgTCGCGCACCAGCACAGCGAG, ggggaccactttgtacaagaaagctgggtcTTTGCTTTCCCTTTTCGCCT | Mycgr3\|92432  JGI  Ensembl |  |  | 100 % | chr\_4:922251 T->C; |  |
| 18 | ggggacaagtttgtacaaaaaagcaggcttgatgGCGTACCTGTTCCGAGTCTA, ggggaccactttgtacaagaaagctgggtcGTATCTCCGCTCCCTCCTCT | Mycgr3\|92512  JGI  Ensembl |  |  | 81.19429 % |  |  |
| 18 | ggggacaagtttgtacaaaaaagcaggcttgatgGACGTTGCAATTGCTGTGAA, ggggaccactttgtacaagaaagctgggtcGGAGCCATCTTCCACCTCTG | Mycgr3\|92769  JGI  Ensembl |  |  | 100 % |  |  |
| 18 | ggggacaagtttgtacaaaaaagcaggcttgatgCCGATATGTATTGAATGTCG, ggggaccactttgtacaagaaagctgggtcGCCCAAGCCTAGAGTCTCGA | Mycgr3\|92812  JGI  Ensembl | KOG3134 Predicted membrane protein; |  | 100 % |  |  |
| 18 | ggggacaagtttgtacaaaaaagcaggcttgatgGCGGATGCACGAGAACGACT, ggggaccactttgtacaagaaagctgggtcCTTTTCCCAGAACAACAAGC | Mycgr3\|94552  JGI  Ensembl | KOG3599 Ca2+-modulated nonselective cation channel polycystin; |  | 100 % |  |  |
| 18 | ggggacaagtttgtacaaaaaagcaggcttgatgTCAGACAACCGCCCCAAACC, ggggaccactttgtacaagaaagctgggtcCGCACCACCGGCAGCAACTG | Mycgr3\|94595  JGI  Ensembl | KOG1187 Serine/threonine protein kinase; |  | 100 % |  |  |
| 18 | ggggacaagtttgtacaaaaaagcaggcttgatgGGTTCCCAACAGTCTCATCC, ggggaccactttgtacaagaaagctgggtcCTCTCCACTCTCGCTCGCTG | Mycgr3\|94669  JGI  Ensembl | KOG0800 FOG: Predicted E3 ubiquitin ligase; | protein binding; zinc ion binding; | 100 % |  |  |
| 18 | ggggacaagtttgtacaaaaaagcaggcttgatgTCCACCCACGCCAACATCGT, ggggaccactttgtacaagaaagctgggtcAGTCCCAAGCTTCGGAGGCT | Mycgr3\|94912  JGI  Ensembl | KOG0591 NIMA (never in mitosis)-related G2-specific serine/threonine protein kinase; | 3-phosphoinositide-dependent protein kinase activity; AMP-activated protein kinase activity;  ATP binding;  DNA-dependent protein kinase activity;  GTP-dependent protein kinase activity;  JUN kinase activity;  JUN kinase kinase activity;  JUN kinase kinase kinase activity;  Janus kinase activity;  MAP kinase 1 activity;  MAP kinase 2 activity;  MAP kinase kinase activity;  MAP kinase kinase kinase kinase activity;  MAP/ERK kinase kinase activity;  MP kinase activity;  NF-kappaB-inducing kinase activity;  SAP kinase activity;  atypical protein kinase C activity;  calcium-dependent protein kinase C activity;  casein kinase I activity;  casein kinase activity;  cyclic nucleotide-dependent protein kinase activity;  cyclin-dependent protein kinase activating kinase activity;  eukaryotic translation initiation factor 2alpha kinase activity;  galactosyltransferase-associated kinase activity;  glycogen synthase kinase 3 activity;  protein amino acid phosphorylation;  protein kinase activity;  protein serine/threonine kinase activity;  protein-tyrosine kinase activity;  receptor signaling protein serine/threonine kinase activity;  ribosomal protein S6 kinase activity;  transmembrane receptor protein kinase activity;  transmembrane receptor protein serine/threonine kinase activity; | 100 % |  |  |
| 18 | ggggacaagtttgtacaaaaaagcaggcttgatgTTTCAACGCTTCGACGTGAA, ggggaccactttgtacaagaaagctgggtcGTACTTAAAGACCCGACCCA | Mycgr3\|95098  JGI  Ensembl | KOG1208 Dehydrogenases with different specificities (related to short-chain alcohol dehydrogenases); | (R)-2-hydroxyglutarate dehydrogenase activity; (R)-2-hydroxyisocaproate dehydrogenase activity;  2-hydroxytetrahydrofuran dehydrogenase activity;  3-keto sterol reductase activity;  3-ketoglucose-reductase activity;  5-exo-hydroxycamphor dehydrogenase activity;  C-3 sterol dehydrogenase (C-4 sterol decarboxylase) activity;  aldo-keto reductase activity;  epoxide dehydrogenase activity;  gluconate dehydrogenase activity;  isocitrate dehydrogenase activity;  mevaldate reductase activity;  phenylcoumaran benzylic ether reductase activity;  steroid dehydrogenase activity;  steroid dehydrogenase activity, acting on the CH-CH group of donors;  steroid dehydrogenase activity, acting on the CH-OH group of donors, NAD or NADP as acceptor; | 0 % |  |  |
| 18 | ggggacaagtttgtacaaaaaagcaggcttgatgCACAAAGACAAGTTCCTCAG, ggggaccactttgtacaagaaagctgggtcATTTCGCCACGGCCAAGAGA | Mycgr3\|96726  JGI  Ensembl | KOG1208 Dehydrogenases with different specificities (related to short-chain alcohol dehydrogenases); | (R)-2-hydroxyglutarate dehydrogenase activity; (R)-2-hydroxyisocaproate dehydrogenase activity;  2-hydroxytetrahydrofuran dehydrogenase activity;  3-keto sterol reductase activity;  3-ketoglucose-reductase activity;  5-exo-hydroxycamphor dehydrogenase activity;  C-3 sterol dehydrogenase (C-4 sterol decarboxylase) activity;  aldo-keto reductase activity;  epoxide dehydrogenase activity;  gluconate dehydrogenase activity;  isocitrate dehydrogenase activity;  metabolic process;  mevaldate reductase activity;  oxidoreductase activity;  phenylcoumaran benzylic ether reductase activity;  steroid dehydrogenase activity;  steroid dehydrogenase activity, acting on the CH-CH group of donors;  steroid dehydrogenase activity, acting on the CH-OH group of donors, NAD or NADP as acceptor; | 100 % |  |  |
| 18 | ggggacaagtttgtacaaaaaagcaggcttgatgCAACAGGTCCGGTACAATCC, ggggaccactttgtacaagaaagctgggtcTAGCTCTTCCGACCATGTCG | Mycgr3\|96752  JGI  Ensembl |  |  | 100 % |  |  |
| 18 | ggggacaagtttgtacaaaaaagcaggcttgatgCTCACCCTCGCATTCAGAGC, ggggaccactttgtacaagaaagctgggtcAGGCGACTGAACCGGAGGCC | Mycgr3\|97922  JGI  Ensembl |  |  | 88.15427 % | chr\_19:50356 C->G; chr\_19:50416 C->T; chr\_19:50355 C->T; chr\_19:50346 C->A; chr\_19:50415 A->G; chr\_19:50347 C->T; chr\_19:50360 G->A; chr\_19:50388 T->G; chr\_19:50401 G->A; chr\_19:50362 G->A; chr\_19:50413 T->C; chr\_19:50386 A->G; chr\_19:50389 C->G; chr\_19:50406 C->A; |  |
| 18 | ggggacaagtttgtacaaaaaagcaggcttgatgTCGAGATCAATGGCGCTAGG, ggggaccactttgtacaagaaagctgggtcGTAGGCGATACGTCGAGTGC | Mycgr3\|98034  JGI  Ensembl |  |  | 100 % |  |  |
| 18 | ggggacaagtttgtacaaaaaagcaggcttgatgGCCAGCTACAACGCAATGTC, ggggaccactttgtacaagaaagctgggtcTCTTGATATCATGCCGGAGT | Mycgr3\|98050  JGI  Ensembl |  |  | 100 % |  |  |
| 18 | ggggacaagtttgtacaaaaaagcaggcttgatgGCTCCTTCCGCCCGCAGCCG, ggggaccactttgtacaagaaagctgggtcGGCAAACGCCTTGGAGTACA | Mycgr3\|98275  JGI  Ensembl | KOG0546 HSP90 co-chaperone CPR7/Cyclophilin; | FK506-sensitive peptidyl-prolyl cis-trans isomerase; cyclophilin; cyclophilin-type peptidyl-prolyl cis-trans isomerase activity;  peptidyl-prolyl cis-trans isomerase activity;  protein folding; | 100 % |  |  |
| 18 | ggggacaagtttgtacaaaaaagcaggcttgatgGCAAGCCTTATCGAGCTGGA, ggggaccactttgtacaagaaagctgggtcTATCGTGTATTTCGAGCCAA | Mycgr3\|98581  JGI  Ensembl |  |  | 100 % |  |  |
| 18 | ggggacaagtttgtacaaaaaagcaggcttgatgATCGAAGCGACCATCTCACG, ggggaccactttgtacaagaaagctgggtcGCGCTTGACCCATGCTGGCA | Mycgr3\|99301  JGI  Ensembl | KOG1697 Mitochondrial/chloroplast ribosomal protein S9; | intracellular; ribosome; structural constituent of ribosome;  translation; | 100 % |  |  |
| 18 | ggggacaagtttgtacaaaaaagcaggcttgatgGCCTCCTCCCTTTCGGTCCA, ggggaccactttgtacaagaaagctgggtcTGAAAACGTCCTCTTCTCCC | Mycgr3\|101592  JGI  Ensembl |  |  | 100 % |  |  |
| 18 | ggggacaagtttgtacaaaaaagcaggcttgatgCCGCCCGCCATGTCAAAATT, ggggaccactttgtacaagaaagctgggtcCGGTCGAGGAAGATTCTGCG | Mycgr3\|103615  JGI  Ensembl | KOG3721 Mitochondrial endonuclease; | endonuclease activity; nucleic acid binding; | 100 % |  |  |
| 18 | ggggacaagtttgtacaaaaaagcaggcttgatgACTTCGGTCCTGGTCACCGG, ggggaccactttgtacaagaaagctgggtcCCACGGCGCATCTTTGCCGT | Mycgr3\|104300  JGI  Ensembl | KOG1611 Predicted short chain-type dehydrogenase; | acylglycerone-phosphate reductase activity; metabolic process;  oxidoreductase activity; | 100 % |  |  |
| 18 | ggggacaagtttgtacaaaaaagcaggcttgatgCTGCTCATGTCGTTCGTGTT, ggggaccactttgtacaagaaagctgggtcTTGGCTTGCGGCGGCATGAT | Mycgr3\|105478  JGI  Ensembl |  |  | 0 % |  |  |
| 18 | ggggacaagtttgtacaaaaaagcaggcttgatgTGGGATGGGAAATTGGGTCT, ggggaccactttgtacaagaaagctgggtcTCGGCCGCCTGCCTGTTCCA | Mycgr3\|105598  JGI  Ensembl |  |  | 100 % |  |  |
| 18 | ggggacaagtttgtacaaaaaagcaggcttgatgGCCGACCAACGAGCACACAT, ggggaccactttgtacaagaaagctgggtcTGGCTGTGCTGGCTGCATTT | Mycgr3\|106061  JGI  Ensembl | KOG3478 Prefoldin subunit 6, KE2 family; | prefoldin complex; protein folding;  unfolded protein binding; | 6.82196 % |  |  |
| 18 | ggggacaagtttgtacaaaaaagcaggcttgatgCACTTCTCTCAGCTCCTCGC, ggggaccactttgtacaagaaagctgggtcCGCAATCCCAAACCACTTCA | Mycgr3\|106075  JGI  Ensembl |  |  | 100 % |  |  |
| 18 | ggggacaagtttgtacaaaaaagcaggcttgatgGCCACCAAGTTAGCATTGAC, ggggaccactttgtacaagaaagctgggtcCTTGTTCTTGACGGTAAACT | Mycgr3\|106960  JGI  Ensembl | KOG1201 Hydroxysteroid 17-beta dehydrogenase 11; | acylglycerone-phosphate reductase activity; metabolic process;  oxidoreductase activity; | 100 % |  |  |
| 18 | ggggacaagtttgtacaaaaaagcaggcttgatgTCCTTGGTCAACCTCGCCAA, ggggaccactttgtacaagaaagctgggtcCCACACTCGGCACAACGGCA | Mycgr3\|107068  JGI  Ensembl |  |  | 100 % |  |  |
| 18 | ggggacaagtttgtacaaaaaagcaggcttgatgGCTCGAGACAAGAGGTTGAT, ggggaccactttgtacaagaaagctgggtcTCCATAATTGTCCAGCATTC | Mycgr3\|107566  JGI  Ensembl | KOG1022 Acetylglucosaminyltransferase EXT2/exostosin 2; | glucuronyl-galactosyl-proteoglycan 4-alpha-N-acetylglucosaminyltransferase activity; | 6.54297 % |  |  |
| 18 | ggggacaagtttgtacaaaaaagcaggcttgatgAGTTCCGGAAGCAGTGCAAA, ggggaccactttgtacaagaaagctgggtcAGTTGGTAGCTCAACATGAG | Mycgr3\|107596  JGI  Ensembl | KOG2814 Transcription coactivator complex, P50 component (LigT RNA ligase/phosphodiesterase family); |  | 100 % |  |  |
| 18 | ggggacaagtttgtacaaaaaagcaggcttgatgCGCGCTGCACTCGCGTTGGC, ggggaccactttgtacaagaaagctgggtcAAGGCTGCTCCCACACTTCA | Mycgr3\|107711  JGI  Ensembl |  |  | 100 % |  |  |
| 18 | ggggacaagtttgtacaaaaaagcaggcttgatgAAGCACGAGACCCTCACATC, ggggaccactttgtacaagaaagctgggtcCATCGTACCGCTCTCCGTTG | Mycgr3\|108330  JGI  Ensembl |  |  | 100 % |  |  |
| 18 | ggggacaagtttgtacaaaaaagcaggcttgatgCCATCTCGATACGCCCTCCC, ggggaccactttgtacaagaaagctgggtcGCGAAGCGATCCGTAGCGCC | Mycgr3\|108353  JGI  Ensembl |  |  | 100 % |  |  |
| 18 | ggggacaagtttgtacaaaaaagcaggcttgatgGCGTCCCTCCTCCGCCGCGA, ggggaccactttgtacaagaaagctgggtcCTCCCTCACCGGCGCATTCC | Mycgr3\|108362  JGI  Ensembl |  |  | 3.26797 % |  |  |
| 18 | ggggacaagtttgtacaaaaaagcaggcttgatgTCGTCGCCCGCTTTGCGTCC, ggggaccactttgtacaagaaagctgggtcCCATCTCCTTCCGCGACCTT | Mycgr3\|108530  JGI  Ensembl |  |  | 100 % |  |  |
| 18 | ggggacaagtttgtacaaaaaagcaggcttgatgAAGCTCGCGGTTTCCACCGT, ggggaccactttgtacaagaaagctgggtcAAGCGCCTTGATCGCAGCGT | Mycgr3\|110066  JGI  Ensembl |  |  | 100 % | chr\_7:273698 A->G; chr\_7:273575 T->C; chr\_7:273487 T->C; |  |
| 18 | ggggacaagtttgtacaaaaaagcaggcttgatgGCCAACAATGACGACATGGA, ggggaccactttgtacaagaaagctgggtcACCAGCACGGGCCATGCTTG | Mycgr3\|110625  JGI  Ensembl |  |  | 100 % |  |  |
| 18 | ggggacaagtttgtacaaaaaagcaggcttgatgTCTCTTCTCACGCCACTCCT, ggggaccactttgtacaagaaagctgggtcCATAAATCCTTTAACGAGGT | Mycgr3\|111117  JGI  Ensembl |  |  | 3.75361 % |  |  |
| 18 | ggggacaagtttgtacaaaaaagcaggcttgatgCTCTCTTTCTTCTTGGGTTT, ggggaccactttgtacaagaaagctgggtcAAAGTTACTCGCCGTATTCA | Mycgr3\|111482  JGI  Ensembl | KOG4157 beta-1,6-N-acetylglucosaminyltransferase, contains WSC domain; |  | 100 % |  |  |
| 18 | ggggacaagtttgtacaaaaaagcaggcttgatgGTCCGATCCCTCAGCTCACG, ggggaccactttgtacaagaaagctgggtcCACCATCCCAGCCAAGACGG | Mycgr3\|111647  JGI  Ensembl | KOG2504 Monocarboxylate transporter; | integral to membrane; transport; transporter activity; | 100 % | chr\_13:577331 G->A; chr\_13:577822 C->T; chr\_13:577973 C->T; chr\_13:577467 T->C; |  |
| 19 | ggggacaagtttgtacaaaaaagcaggcttgatgCCTTCATTGCCAGACACACA, ggggaccactttgtacaagaaagctgggtcAACCCGCACCACCAATTTCT | Mycgr3\|32934  JGI  Ensembl | KOG1198 Zinc-binding oxidoreductase; | NADPH:quinone reductase activity; oxidoreductase activity;  zinc ion binding; | 100 % |  |  |
| 19 | ggggacaagtttgtacaaaaaagcaggcttgatgAATGGGTGCCAGGAAATCCT, ggggaccactttgtacaagaaagctgggtcAGAAGACGAAGTGCGGAGCT | Mycgr3\|35311  JGI  Ensembl |  |  | 100 % |  |  |
| 19 | ggggacaagtttgtacaaaaaagcaggcttgatgCGGATACCCTTTGCTCGCCG, ggggaccactttgtacaagaaagctgggtcCGCTGTATAGCTGGCGCCTC | Mycgr3\|36249  JGI  Ensembl | KOG0223 Aquaporin (major intrinsic protein family); | membrane; transport; transporter activity; | 100 % |  |  |
| 19 | ggggacaagtttgtacaaaaaagcaggcttgatgCTCTTGACCCAGTCTCAAGT, ggggaccactttgtacaagaaagctgggtcAGGGCGCCAACTCTCGAGGT | Mycgr3\|37988  JGI  Ensembl |  |  | 1.7452 % |  |  |
| 19 | ggggacaagtttgtacaaaaaagcaggcttgatgTCCGAATTCACAATCCCCTC, ggggaccactttgtacaagaaagctgggtcCCACACCAAAACGCTCTTCA | Mycgr3\|39151  JGI  Ensembl | KOG0022 Alcohol dehydrogenase, class III; | cofactor binding; oxidoreductase activity;  oxidoreductase activity, acting on the CH-OH group of donors, NAD or NADP as acceptor; zinc ion binding; | 0 % |  |  |
| 19 | ggggacaagtttgtacaaaaaagcaggcttgatgTCCGGCGAAAAGCGAGCGGC, ggggaccactttgtacaagaaagctgggtcCCGCGGCAGCTCTCCCAGCA | Mycgr3\|41112  JGI  Ensembl | KOG0265 U5 snRNP-specific protein-like factor and related proteins; |  | 17.83877 % |  |  |
| 19 | ggggacaagtttgtacaaaaaagcaggcttgatgCTTCCTCCAACGTCCACCCC, ggggaccactttgtacaagaaagctgggtcTCTCTCCACGTGCTCGACTC | Mycgr3\|42049  JGI  Ensembl | KOG1228 Uncharacterized conserved protein; |  | 0 % |  |  |
| 19 | ggggacaagtttgtacaaaaaagcaggcttgatgAAGACTTCCCTCCTCGTCGA, ggggaccactttgtacaagaaagctgggtcACAAACGCGGACCTTACCAA | Mycgr3\|42327  JGI  Ensembl |  |  | 100 % |  |  |
| 19 | ggggacaagtttgtacaaaaaagcaggcttgatgGCAGTGCCCTCCTCCTCCGC, ggggaccactttgtacaagaaagctgggtcCGACCTGTCTACCTCTAACC | Mycgr3\|43901  JGI  Ensembl |  |  | 3.43612 % |  |  |
| 19 | ggggacaagtttgtacaaaaaagcaggcttgatgTCGACCATCGATCAAGAAAC, ggggaccactttgtacaagaaagctgggtcTCTCTCCTTCTTGCCGCTTC | Mycgr3\|45223  JGI  Ensembl | KOG1430 C-3 sterol dehydrogenase/3-beta-hydroxysteroid dehydrogenase and related dehydrogenases; | 3-beta-hydroxy-delta5-steroid dehydrogenase activity; C-3 sterol dehydrogenase (C-4 sterol decarboxylase) activity;  steroid biosynthetic process;  sterol-4-alpha-carboxylate 3-dehydrogenase (decarboxylating) activity; | 100 % |  |  |
| 19 | ggggacaagtttgtacaaaaaagcaggcttgatgCAAAACTTGGTATGGGTAGA, ggggaccactttgtacaagaaagctgggtcCGCCGCCAAAATCTCCTTAA | Mycgr3\|45401  JGI  Ensembl |  |  | 100 % |  |  |
| 19 | ggggacaagtttgtacaaaaaagcaggcttgatgATCGTTGTACGTTCAAGTCA, ggggaccactttgtacaagaaagctgggtcTTGCACCATCCGATCTTCAC | Mycgr3\|45762  JGI  Ensembl |  |  | 100 % |  |  |
| 19 | ggggacaagtttgtacaaaaaagcaggcttgatgCGCTCGTTCTTCTGTTTTGC, ggggaccactttgtacaagaaagctgggtcTGCTTTACGCGAACAGACCT | Mycgr3\|47097  JGI  Ensembl | KOG2938 Predicted inosine-uridine preferring nucleoside hydrolase; | purine nucleosidase activity; | 100 % |  |  |
| 19 | ggggacaagtttgtacaaaaaagcaggcttgatgGATCATCAGCTTCTTCTCTA, ggggaccactttgtacaagaaagctgggtcACAAACGTCCGCTCCTCCCT | Mycgr3\|47934  JGI  Ensembl | KOG1515 Arylacetamide deacetylase; | 1-oxa-2-oxocycloheptane lactonase activity; 3,4-dihydrocoumarin hydrolase activity;  butyrolactone hydrolase activity;  lipase activity;  phospholipase activity;  phosphopantetheine binding;  sulfolactone hydrolase activity; | 100 % |  |  |
| 19 | ggggacaagtttgtacaaaaaagcaggcttgatgTCTTCCGATCACAAGGTCCC, ggggaccactttgtacaagaaagctgggtcGAACACAGCCATTGAAGAAG | Mycgr3\|48514  JGI  Ensembl | KOG1205 Predicted dehydrogenase; | cysteine-type endopeptidase activity; estradiol 17-beta-dehydrogenase activity;  metabolic process;  oxidoreductase activity;  proteolysis; | 100 % |  |  |
| 19 | ggggacaagtttgtacaaaaaagcaggcttgatgCCCAAGATCAACGTTGCTCG, ggggaccactttgtacaagaaagctgggtcACCATCAAGTTTTGACGGCT | Mycgr3\|48517  JGI  Ensembl | KOG0024 Sorbitol dehydrogenase; | L-iditol 2-dehydrogenase activity; oxidoreductase activity;  zinc ion binding; | 100 % |  |  |
| 19 | ggggacaagtttgtacaaaaaagcaggcttgatgCCTGATCTTCCTGGAACTTC, ggggaccactttgtacaagaaagctgggtcAGCCTCGAAACCATGCGTAC | Mycgr3\|49627  JGI  Ensembl | KOG0759 Mitochondrial oxoglutarate/malate carrier proteins; | binding; membrane; transport; | 100 % |  |  |
| 19 | ggggacaagtttgtacaaaaaagcaggcttgatgGTTCATTCAATCGTTTTCGC, ggggaccactttgtacaagaaagctgggtcCTTTGAGCATATGTCCACCT | Mycgr3\|50311  JGI  Ensembl |  |  | 100 % | chr\_12:161984 A->T; chr\_12:162699 G->A; |  |
| 19 | ggggacaagtttgtacaaaaaagcaggcttgatgTCACAAATTCTAAGCTTCCT, ggggaccactttgtacaagaaagctgggtcCAGCTTCGCACTGACGCCCC | Mycgr3\|66157  JGI  Ensembl | KOG1208 Dehydrogenases with different specificities (related to short-chain alcohol dehydrogenases); | (R)-2-hydroxyglutarate dehydrogenase activity; (R)-2-hydroxyisocaproate dehydrogenase activity;  2-hydroxytetrahydrofuran dehydrogenase activity;  3-keto sterol reductase activity;  3-ketoglucose-reductase activity;  5-exo-hydroxycamphor dehydrogenase activity;  C-3 sterol dehydrogenase (C-4 sterol decarboxylase) activity;  aldo-keto reductase activity;  epoxide dehydrogenase activity;  gluconate dehydrogenase activity;  isocitrate dehydrogenase activity;  metabolic process;  mevaldate reductase activity;  oxidoreductase activity;  phenylcoumaran benzylic ether reductase activity;  steroid dehydrogenase activity;  steroid dehydrogenase activity, acting on the CH-CH group of donors;  steroid dehydrogenase activity, acting on the CH-OH group of donors, NAD or NADP as acceptor; | 0 % |  |  |
| 19 | ggggacaagtttgtacaaaaaagcaggcttgatgGCCGAGATCAACGGCGAAAT, ggggaccactttgtacaagaaagctgggtcTGAGTTCCCTGACACACCCG | Mycgr3\|66280  JGI  Ensembl | KOG3199 Nicotinamide mononucleotide adenylyl transferase; | NAD biosynthetic process; biosynthetic process;  nicotinamide-nucleotide adenylyltransferase activity;  nucleotidyltransferase activity; | 0 % |  |  |
| 19 | ggggacaagtttgtacaaaaaagcaggcttgatgTCGCCCTCGTATGGCTCTTT, ggggaccactttgtacaagaaagctgggtcCAGCTGATCTTTCGGAAACA | Mycgr3\|68213  JGI  Ensembl |  |  | 100 % |  |  |
| 19 | ggggacaagtttgtacaaaaaagcaggcttgatgACGTCCCTCCTCAAAGGCCT, ggggaccactttgtacaagaaagctgggtcCAACTTCGGGATGAACATGG | Mycgr3\|69411  JGI  Ensembl | KOG4650 Predicted steroid reductase; | cytoplasm; integral to membrane;  lipid metabolic process;  oxidoreductase activity, acting on the CH-CH group of donors; | 100 % |  |  |
| 19 | ggggacaagtttgtacaaaaaagcaggcttgatgGCCGCTCTCACCAGGCTATA, ggggaccactttgtacaagaaagctgggtcCAGTCTTGACGACAATGCCG | Mycgr3\|70263  JGI  Ensembl | KOG0538 Glycolate oxidase; | electron transport; oxidoreductase activity; | 100 % |  |  |
| 19 | ggggacaagtttgtacaaaaaagcaggcttgatgCGCTCGTCTATTACACGACT, ggggaccactttgtacaagaaagctgggtcCTGGCCGCCGCCACCACCAA | Mycgr3\|70703  JGI  Ensembl | KOG2940 Predicted methyltransferase; |  | 100 % |  |  |
| 19 | ggggacaagtttgtacaaaaaagcaggcttgatgTCGTCGAAACCAGGCGTTGT, ggggaccactttgtacaagaaagctgggtcAACTTTGCTGGCCTTCTCCA | Mycgr3\|70724  JGI  Ensembl |  |  | 100 % |  |  |
| 19 | ggggacaagtttgtacaaaaaagcaggcttgatgGCCGAAGAATCTGCATCCAG, ggggaccactttgtacaagaaagctgggtcGGGATTTTCCTGCATGATCC | Mycgr3\|71420  JGI  Ensembl |  |  | 100 % |  |  |
| 19 | ggggacaagtttgtacaaaaaagcaggcttgatgACGAGCCGAACCAGAATTCA, ggggaccactttgtacaagaaagctgggtcCTCATCCGCCACCCAGGCAA | Mycgr3\|72012  JGI  Ensembl |  |  | 100 % |  |  |
| 19 | ggggacaagtttgtacaaaaaagcaggcttgatgCCCGTCTATCTTCTACACGG, ggggaccactttgtacaagaaagctgggtcAGGCTGTACTCTTGACGGGA | Mycgr3\|73025  JGI  Ensembl |  |  | 100 % |  |  |
| 19 | ggggacaagtttgtacaaaaaagcaggcttgatgAATCCTTCGAAGACTTATCA, ggggaccactttgtacaagaaagctgggtcTTTTACAAGACCGCCATAGT | Mycgr3\|75416  JGI  Ensembl | KOG2646 Ribosomal protein S5; | intracellular; ribosome; structural constituent of ribosome;  translation; | 0 % |  |  |
| 19 | ggggacaagtttgtacaaaaaagcaggcttgatgTCTCTCACAGGCGCCATCAA, ggggaccactttgtacaagaaagctgggtcCGGCATTTCATTAACCCTTT | Mycgr3\|76259  JGI  Ensembl |  |  | 100 % |  |  |
| 19 | ggggacaagtttgtacaaaaaagcaggcttgatgCTCCGCACCGCGCCGCTGCT, ggggaccactttgtacaagaaagctgggtcCTCCTTCTTTGGCGGGCTCT | Mycgr3\|76293  JGI  Ensembl | KOG4075 Cytochrome c oxidase, subunit IV/COX5b; | aa3-type cytochrome c oxidase; ba3-type cytochrome c oxidase;  caa3-type cytochrome c oxidase;  cbb3-type cytochrome c oxidase;  cytochrome-c oxidase activity;  electron transport; | 0 % |  |  |
| 19 | ggggacaagtttgtacaaaaaagcaggcttgatgGCATCCCGAACCCTCCGCCC, ggggaccactttgtacaagaaagctgggtcCGCAATCTTCCCCTTATTCA | Mycgr3\|77366  JGI  Ensembl | KOG1504 Ornithine carbamoyltransferase OTC/ARG3; | 'de novo' pyrimidine base biosynthetic process; amino acid binding;  amino acid metabolic process;  aspartate carbamoyltransferase activity;  carboxyl- or carbamoyltransferase activity;  ornithine carbamoyltransferase activity;  ornithine carbamoyltransferase complex; | 100 % |  |  |
| 19 | ggggacaagtttgtacaaaaaagcaggcttgatgCGGTTCCAATTCATTCTCAT, ggggaccactttgtacaagaaagctgggtcCTTGACCCAGCGCTCCATCA | Mycgr3\|77793  JGI  Ensembl |  |  | 0 % |  |  |
| 19 | ggggacaagtttgtacaaaaaagcaggcttgatgGATAAATCCCGCATCCTCTG, ggggaccactttgtacaagaaagctgggtcCCACTCCCGATTCTGCCTCA | Mycgr3\|77886  JGI  Ensembl | KOG1563 Mitochondrial protein Surfeit 1/SURF1/SHY1, required for expression of cytochrome oxidase; |  | 100 % |  |  |
| 19 | ggggacaagtttgtacaaaaaagcaggcttgatgCCTGGCCGAGCGTGCGGTTC, ggggaccactttgtacaagaaagctgggtcAGCACTTGGCCAGGCGAGCA | Mycgr3\|78344  JGI  Ensembl |  |  | 0 % |  |  |
| 19 | ggggacaagtttgtacaaaaaagcaggcttgatgGCCGCCCCGCCTCTACCGCC, ggggaccactttgtacaagaaagctgggtcGTACTGCCCTTGCTGTTGCC | Mycgr3\|81065  JGI  Ensembl |  |  | 100 % |  |  |
| 19 | ggggacaagtttgtacaaaaaagcaggcttgatgACTTCGCCCACACCCATACC, ggggaccactttgtacaagaaagctgggtcTCGACTCTGGGCCTCCGCAC | Mycgr3\|85716  JGI  Ensembl | KOG2938 Predicted inosine-uridine preferring nucleoside hydrolase; | purine nucleosidase activity; | 100 % |  |  |
| 19 | ggggacaagtttgtacaaaaaagcaggcttgatgTCCACACAACCAATCCGGAT, ggggaccactttgtacaagaaagctgggtcCAAGTGCGCGTATCTCGCCG | Mycgr3\|86450  JGI  Ensembl | KOG2741 Dimeric dihydrodiol dehydrogenase; | oxidoreductase activity; | 100 % |  | ``` 1	ATGTCCACACAACCAATCCGGATCGGCTTCATCGGACTCAACAAGACCGACTCCTGGGCCGTTAATGCGCATCTACCATATCTGAAAGACTCCGCCAAGTACAAAATCGTAGCCCTACTGAACAGCTCCAAGGATTCTGCCCAAGCCGCCATCGAATCCCACGGCCTGTCATCCGACACCAAGGCCCATGGCTCAACAGAGGCTTTCGCAGCCGATGACAACATTGATCTCATAGTTTGCAGTGTGAGGGTGGACAGACACTACCCCATACTCAAACCAATCCTCGAAGCCACGAAAGCGAAAGCAGTTCATTGCGAATGGCCGCTCGGGAAGAATCTGGCCGAAGCCGAGGAACTCACTGCTTTGGCGAAGAGCAAAGGTCTCAAGACGATCATCGGTCTGCAAGGTCGTCAATCGCCAACCGGACATGTCGTGAAAAGATTGCTCGACGAAGGAAGAATTGGGAAGCCCTTGAGTGTTACAATCACATTCGCAGGCTACAATTTCGGCGAAGCCGATCTCGAACCTCTAGCATATCTGTCCGACATCGAAATCGGCGGAAACATGGTGACGATCCATTTCTCCCACACCCTGGACGTCATCACCAACGCCGTGGGCCAGCTCGCATCCTACTCCGTAGTCCTCGACACCAAGCGCGAGAAGACTCTCCTTCGGAACAAACCTCACACCTACAAGCCGCAAAGCGAGACCGACGAGCCCGTCCAAATCGTCGGAGAAGTCAAGCGGACTTCGCACGATCAGATCGCCATCCAAGGTCATCTCACGAACGGCGCGGTGTTCTCGTTCCACATGCGAGGCGGCCATCCCTTTCCAGATACACCCGGTCTTGAGTGGCGAGTGTACGGTAAAGCTGGTGAGATCAGGATTACGACGCCTTCTGCGAATTTACACTTCGGCGGGTCGGGGCATAAGATCCAGTTGCACGATCACAAGTCGGGAAGCGTGAAGAA-CATTGAAGTGCCGAAGGATGAGTGGGACGAGAAGGAGCTGCCGATCACCGCGCGTGCTCCTGCGAGGATAT-ATGAGGCGTTTGCTGATGGGAAGGAGGATGGGTACGGGACTTGGGA-GGA-TG-CGGTGGTCAGG	1104 	\|\|\|\|\|\|\|\|\|\|\|\|\|\|\|\|\|\|\|\|\|\|\|\|\|\|\|\|\|\|\|\|\|\|\|\|\|\|\|\|\|\|\|\|\|\|\|\|\|\|\|\|\|\|\|\|\|\|\|\|\|\|\|\|\|\|\|\|\|\|\|\|\|\|\|\|\|\|\|\|\|\|\|\|\|\|\|\|\|\|\|\|\|\|\|\|\|\|\|\|\|\|\|\|\|\|\|\|\|\|\|\|\|\|\|\|\|\|\|\|\|\|\|\|\|\|\|\|\|\|\|\|\|\|\|\|\|\|\|\|\|\|\|\|\|\|\|\|\|\|\|\|\|\|\|\|\|\|\|\|\|\|\|\|\|\|\|\|\|\|\|\|\|\|\|\|\|\|\|\|\|\|\|\|\|\|\|\|\|\|\|\|\|\|\|\|\|\|\|\|\|\|\|\|\|\|\|\|\|\|\|\|\|\|\|\|\|\|\|\|\|\|\|\|\|\|\|\|\|\|\|\|\|\|\|\|\|\|\|\|\|\|\|\|\|\|\|\|\|\|\|\|\|\|\|\|\|\|\|\|\|\|\|\|\|\|\|\|\|\|\|\|\|\|\|\|\|\|\|\|\|\|\|\|\|\|\|\|\|\|\|\|\|\|\|\|\|\|\|\|\|\|\|\|\|\|\|\|\|\|\|\|\|\|\|\|\|\|\|\|\|\|\|\|\|\|\|\|\|\|\|\|\|\|\|\|\|\|\|\|\|\|\|\|\|\|\|\|\|\|\|\|\|\|\|\|\|\|\|\|\|\|\|\|\|\|\|\|\|\|\|\|\|\|\|\|\|\|\|\|\|\|\|\|\|\|\|\|\|\|\|\|\|\|\|\|\|\|\|\|\|\|\|\|\|\|\|\|\|\|\|\|\|\|\|\|\|\|\|\|\|\|\|\|\|\|\|\|\|\|\|\|\|\|\|\|\|\|\|\|\|\|\|\|\|\|\|\|\|\|\|\|\|\|\|\|\|\|\|\|\|\|\|\|\|\|\|\|\|\|\|\|\|\|\|\|\|\|\|\|\|\|\|\|\|\|\|\|\|\|\|\|\|\|\|\|\|\|\|\|\|\|\|\|\|\|\|\|\|\|\|\|\|\|\|\|\|\|\|\|\|\|\|\|\|\|\|\|\|\|\|\|\|\|\|\|\|\|\|\|\|\|\|\|\|\|\|\|\|\|\|\|\|\|\|\|\|\|\|\|\|\|\|\|\|\|\|\|\|\|\|\|\|\|\|\|\|\|\|\|\|\|\|\|\|\|\|\|\|\|\|\|\|\|\|\|\|\|\|\|\|\|\|\|\|\|\|\|\|\|\|\|\|\|\|\|\|\|\|\|\|\|\|\|\|\|\|\|\|\|\|\|\|\|\|\|\|\|\|\|\|\|\|\|\|\|\|\|\|\|\|\|\|\|\|\|\|\|\|\|\|\|\|\|\|\|\|\|\|\|\|\|\|\|\|\|\|\|\|\|\|\|\|\|\|\|\|\|\|\|\|\|\|\|\|\|\|\|\|\|\|\|\|\|\|\|\|\|\|\|\|\|\|\|\|\|\|\|\|\|\|\|\|\|\|\|\|\|\|\|\|\|\|\|\|\|\|\|\|\|\|\|\|\|\|\|\|\|\|\|\|\|\|\|\|\|\|\|\|\|\|\|\|\|\|\|\|\|\|\|\|\|\|\|\|\|\|\|\|\|\|\|\|\|\|\|\|\|\|\|\|\|\|\|\|\|\|\|\|\|\|\|\|\|\|\|\|\|\|\|\|\|\|\|\|\|\|\|\|\|\|\|\|\|\|\|\|\|\|\|\|\|\|\|\|\|\|\|\|\|\|\|\|\|\|\|\|\|\|\|\|\|\|\|\|\|\|\|\|\|\|\|\|\|\|\|\|\|\|\|\|\|\|\|\|\|\|\|\|\|\|\|\|\|\|\|\|\|\|\|\|\|\|\|\|\|\|\|\|\|\|\|\|\|\|\|\|\|\|\|\|\|\|\|\|\|\|\|\|\|\|\|\|\|\|\|\|\|\|\|\|\|\|\|\|\|\|\|\|\|\|\|\|\|\|\|\|\|\|\|\|\| \|\|\|\|\|\|\| \|\|\|\|\|\|\|\|\|\| \|\|\|\|\|\| \|\|\|\|\|\|\| \| \|\|\|\| \|\|\|\|\|\|\|\|\|\|  \|\|\|\|\|\|\|\|\|\|\|\|\|\|  \|\|\|\|\|\|\|\|\| \|\|\|\|\|\|\|\|\|\|\| \|\|\|\|\| \|\|\|\| \|\|\|\|\|\|\|\|\|\| \|\|\|\| \|\| \|\|\|\|\|\|\|\|\|\| \|\|\|\|\| \|\|\| \|\| \|\|\|\|\| \|\|\|\|\| 	ATGTCCACACAACCAATCCGGATCGGCTTCATCGGACTCAACAAGACCGACTCCTGGGCCGTTAATGCGCATCTACCATATCTGAAAGACTCCGCCAAGTACAAAATCGTAGCCCTACTGAACAGCTCCAAGGATTCTGCCCAAGCCGCCATCGAATCCCACGGCCTGTCATCCGACACCAAGGCCCATGGCTCAACAGAGGCTTTCGCAGCCGATGACAACATTGATCTCATAGTTTGCAGTGTGAGGGTGGACAGACACTACCCCATACTCAAACCAATCCTCGAAGCCACGAAAGCGAAAGCAGTTCATTGCGAATGGCCGCTCGGGAAGAATCTGGCCGAAGCCGAGGAACTCACTGCTTTGGCGAAGAGCAAAGGTCTCAAGACGATCATCGGTCTGCAAGGTCGTCAATCGCCAACCGGACATGTCGTGAAAAGATTGCTCGACGAAGGAAGAATTGGGAAGCCCTTGAGTGTTACAATCACATTCGCAGGCTACAATTTCGGCGAAGCCGATCTCGAACCTCTAGCATATCTGTCCGACATCGAAATCGGCGGAAACATGGTGACGATCCATTTCTCCCACACCCTGGACGTCATCACCAACGCCGTGGGCCAGCTCGCATCCTACTCCGTAGTCCTCGACACCAAGCGCGAGAAGACTCTCCTTCGGAACAAACCTCACACCTACAAGCCGCAAAGCGAGACCGACGAGCCCGTCCAAATCGTCGGAGAAGTCAAGCGGACTTCGCACGATCAGATCGCCATCCAAGGTCATCTCACGAACGGCGCGGTGTTCTCGTTCCACATGCGAGGCGGCCATCCCTTTCCAGATACACCCGGTCTTGAGTGGCGAGTGTACGGTAAAGCTGGTGAGATCAGGATTACGACGCCTTCTGCGAATTTACACTTCGGCGGGTCGGGGCATAAGATCCAGTTGCACGATCACA-GTCGGGA-GCGTGAAGAAACATTGA-GTGCCGATG-ATGA-TGGGACGAGA--GAGCTGCCGATCACGCCGCGTGCTC-TGCGAGGATATTATGAG-CGTT-GCTGATGGGA-GGAG-AT-GGTACGGGACCTGGGAAGGAATGGCGGTG-TCAGG ``` |
| 19 | ggggacaagtttgtacaaaaaagcaggcttgatgCTCATAACATCAATCGAATA, ggggaccactttgtacaagaaagctgggtcCCACTCTCGGATCTCGCCTC | Mycgr3\|87695  JGI  Ensembl |  |  | 100 % |  |  |
| 19 | ggggacaagtttgtacaaaaaagcaggcttgatgACGACCTACAACATCGCCTC, ggggaccactttgtacaagaaagctgggtcCTTCTTCCCGAAAACGACCT | Mycgr3\|87717  JGI  Ensembl | KOG0785 Isocitrate dehydrogenase, alpha subunit; | metabolic process; oxidoreductase activity, acting on the CH-OH group of donors, NAD or NADP as acceptor; tartrate dehydrogenase activity; | 0 % |  |  |
| 19 | ggggacaagtttgtacaaaaaagcaggcttgatgCAGAAGCCTACCACAAACTA, ggggaccactttgtacaagaaagctgggtcCGGCTTCGCCTGCTTATTGT | Mycgr3\|88946  JGI  Ensembl | KOG1441 Glucose-6-phosphate/phosphate and phosphoenolpyruvate/phosphate antiporter; |  | 100 % |  |  |
| 19 | ggggacaagtttgtacaaaaaagcaggcttgatgGCAGCGGGCCGGCGACTGCA, ggggaccactttgtacaagaaagctgggtcCGCCCATTCCTTGTCCAGCT | Mycgr3\|89020  JGI  Ensembl |  |  | 100 % |  |  |
| 19 | ggggacaagtttgtacaaaaaagcaggcttgatgAAGGTGACCACTACGAACAT, ggggaccactttgtacaagaaagctgggtcGCGTGAGAGAGCAAAACTTA | Mycgr3\|89811  JGI  Ensembl | KOG3753 Circadian clock protein period; |  | 100 % |  |  |
| 19 | ggggacaagtttgtacaaaaaagcaggcttgatgAGAATTAAGACCACCACCCC, ggggaccactttgtacaagaaagctgggtcATTTGGACGAACATTCAAAT | Mycgr3\|90067  JGI  Ensembl |  |  | 0 % |  |  |
| 19 | ggggacaagtttgtacaaaaaagcaggcttgatgAACACGCACTCCGACCGAGT, ggggaccactttgtacaagaaagctgggtcTGGGTTGTTCGCAATCCAGG | Mycgr3\|90144  JGI  Ensembl |  |  | 100 % |  |  |
| 19 | ggggacaagtttgtacaaaaaagcaggcttgatgGCGCGCAATTTCCAGCTCTT, ggggaccactttgtacaagaaagctgggtcGAACCTGCTGCCGTCGAAAC | Mycgr3\|91170  JGI  Ensembl |  |  | 100 % |  |  |
| 19 | ggggacaagtttgtacaaaaaagcaggcttgatgCTGCTACCCAACAGTTTCTA, ggggaccactttgtacaagaaagctgggtcCCCAATGACCTTGCTCGCAT | Mycgr3\|92039  JGI  Ensembl | KOG3599 Ca2+-modulated nonselective cation channel polycystin; |  | 100 % |  |  |
| 19 | ggggacaagtttgtacaaaaaagcaggcttgatgGCGGAACACATCGAGAAAGC, ggggaccactttgtacaagaaagctgggtcTCCTCGGCAGCGTCGGTCAT | Mycgr3\|92364  JGI  Ensembl |  |  | 100 % |  |  |
| 19 | ggggacaagtttgtacaaaaaagcaggcttgatgGCCCAACTCTCCTCGATCGC, ggggaccactttgtacaagaaagctgggtcCATGCTTCCCAAGCTGAGAA | Mycgr3\|92583  JGI  Ensembl |  |  | 100 % |  | ``` 1	ATGGCCCAACTCTCCTCGATCGCACAGGCGCTTGCCGTGCAGACCTCCTCCGTCGCCATTCCGAATGCACGCAGCACCGCCGCCGACTACTCCGGCACTGCCAACTTCGAAAAGGCCGCCAACGCCTTGCTCACTCCGCCGAGCTCCATCTCTCCTCAGCTTCAAGCCCAGCGCACCCGTGCTGCGGCGTTCTCGCCACCGCCGAACAACATGAGTCAAGATACGGAGATGGAGGATTCTGCTGGCGAATACTCGCCGGATTTGGACAACATCCAGCCGTCGTCGACCGGCATGCCTCTCAGCCTCGGTGCCTTGTCTGATCTGGACAACGCGGTGACCATCACCCCGAGCATGCTGGCTAAGCAGCACCTGCCGGGCATCATGCTCGGTAACGGGCCACGACCCATCCGCTATGTGATGGGCGAGCTATGCCAAACAGTACCTGGCTTCTCGCGCATCGCACCGGCCAAGGCTCGCAGGATTGTGGTCGCGGCGCTCGGCAACCCTGCTGGAGGTGGACCCAACGGAGATGTCGTTTTCTCCAAGACTGGCTGGGGACGATGGGATGCGCACATCAAGGGGTCTTCTCGTGATTCCGCCATCGGCTCCTTCAACGAGGGCAACCTGTCGCCTCCACGCAGTGAGCAGGGCAGCTACGCCATGTCGCACAGTGATTCAGCAATCCACATGCACGCTCAGCATATGAATCAGAAGTACCGAGAAGGCTATTCTGGTGGTAGTTGGACCGCATCTAGCATCCGCGAGGAGGACGAACTCGACATGGATATGGATCCGGTGGAAGACGCCGCTGATAACATGTCTCTGGACGGCGATTCCTCTGACTCCTCGTCAGAAGAGTTGAACGAGGATACCGACGACGAAGATTGGGCTGCCGTTGGTATCGAAGCGTTGCGGAAGGCTTCAATGTCCACTTCCAACGTGACTCCTCGGCAGAACATCCGTGTCGGCAGTATCCCGTACAACGGACCCTGGGCATCGAAGAGCTGGCAGCGACCAACTA-TATCACAAAGTCGATCGTTCCAGTCAGCCTCGGTTCCGAACGGCGCCAACTTCCGAAA	1079 	\|\|\|\|\|\|\|\|\|\|\|\|\|\|\|\|\|\|\|\|\|\|\|\|\|\|\|\|\|\|\|\|\|\|\|\|\|\|\|\|\|\|\|\|\|\|\|\|\|\|\|\|\|\|\|\|\|\|\|\|\|\|\|\|\|\|\|\|\|\|\|\|\|\|\|\|\|\|\|\|\|\|\|\|\|\|\|\|\|\|\|\|\|\|\|\|\|\|\|\|\|\|\|\|\|\|\|\|\|\|\|\|\|\|\|\|\|\|\|\|\|\|\|\|\|\|\|\|\|\|\|\|\|\|\|\|\|\|\|\|\|\|\|\|\|\|\|\|\|\|\|\|\|\|\|\|\|\|\|\|\|\|\|\|\|\|\|\|\|\|\|\|\|\|\|\|\|\|\|\|\|\|\|\|\|\|\|\|\|\|\|\|\|\|\|\|\|\|\|\|\|\|\|\|\|\|\|\|\|\|\|\|\|\|\|\|\|\|\|\|\|\|\|\|\|\|\|\|\|\|\|\|\|\|\|\|\|\|\|\|\|\|\|\|\|\|\|\|\|\|\|\|\|\|\|\|\|\|\|\|\|\|\|\|\|\|\|\|\|\|\|\|\|\|\|\|\|\|\|\|\|\|\|\|\|\|\|\|\|\|\|\|\|\|\|\|\|\|\|\|\|\|\|\|\|\|\|\|\|\|\|\|\|\|\|\|\|\|\|\|\|\|\|\|\|\|\|\|\|\|\|\|\|\|\|\|\|\|\|\|\|\|\|\|\|\|\|\|\|\|\|\|\|\|\|\|\|\|\|\|\|\|\|\|\|\|\|\|\|\|\|\|\|\|\|\|\|\|\|\|\|\|\|\|\|\|\|\|\|\|\|\|\|\|\|\|\|\|\|\|\|\|\|\|\|\|\|\|\|\|\|\|\|\|\|\|\|\|\|\|\|\|\|\|\|\|\|\|\|\|\|\|\|\|\|\|\|\|\|\|\|\|\|\|\|\|\|\|\|\|\|\|\|\|\|\|\|\|\|\|\|\|\|\|\|\|\|\|\|\|\|\|\|\|\|\|\|\|\|\|\|\|\|\|\|\|\|\|\|\|\|\|\|\|\|\|\|\|\|\|\|\|\|\|\|\|\|\|\|\|\|\|\|\|\|\|\|\|\|\|\|\|\|\|\|\|\|\|\|\|\|\|\|\|\|\|\|\|\|\|\|\|\|\|\|\|\|\|\|\|\|\|\|\|\|\|\|\|\|\|\|\|\|\|\|\|\|\|\|\|\|\|\|\|\|\|\|\|\|\|\|\|\|\|\|\|\|\|\|\|\|\|\|\|\|\|\|\|\|\|\|\|\|\|\|\|\|\|\|\|\|\|\|\|\|\|\|\|\|\|\|\|\|\|\|\|\|\|\|\|\|\|\|\|\|\|\|\|\|\|\|\|\|\|\|\|\|\|\|\|\|\|\|\|\|\|\|\|\|\|\|\|\|\|\|\|\|\|\|\|\|\|\|\|\|\|\|\|\|\|\|\|\|\|\|\|\|\|\|\|\|\|\|\|\|\|\|\|\|\|\|\|\|\|\|\|\|\|\|\|\|\|\|\|\|\|\|\|\|\|\|\|\|\|\|\|\|\|\|\|\|\|\|\|\|\|\|\|\|\|\|\|\|\|\|\|\|\|\|\|\|\|\|\|\|\|\|\|\|\|\|\|\|\|\|\|\|\|\|\|\|\|\|\|\|\|\|\|\|\|\|\|\|\|\|\|\|\|\|\|\|\|\|\|\|\|\|\|\|\|\|\|\|\|\|\|\|\|\|\|\|\|\|\|\|\|\|\|\|\|\|\|\|\|\|\|\|\|\|\|\|\|\|\|\|\|\|\|\|\|\|\|\|\|\|\|\|\|\|\|\|\|\|\|\|\|\|\|\|\|\|\|\|\|\|\|\|\|\|\|\|\|\|\|\|\|\|\|\|\|\|\|\|\|\|\| \|\|\|\|\|\|\|\|\|\|\|\|\|\|\|\|\|\|\|\|\|\|\|\|\|\|\|\|\|\|\|\|\|\|\|\| \|\|\|\|\|\|\|\|\|\|\|\|\|\|\|\|\|\|\|\|\|\| \|\|\|\|\|\|\| \|\|\|\|\|\|\|\|\|\|\|\|\|\|\|\|\|\|\|\|\|\|\|\|\|\| \|\|\|\|\|\|\|\|\|\|\|\|\|\|\|\|\|\| \|\|\|\|\|\| \|\|\|\|\|\|\| \| \|\|\| \|\|\|\|\|\|\|\|\|\|\|\|\|\|\|\|\| \|\|\|\|\|\|\|\|\|\|\|\|\|\|\|\|\|\| \|\|\|\|\|\|\|\|\|\|\|\|\|\|\|\|\|\|\|\|\| 	ATGGCCCAACTCTCCTCGATCGCACAGGCGCTTGCCGTGCAGACCTCCTCCGTCGCCATTCCGAATGCACGCAGCACCGCCGCCGACTACTCCGGCACTGCCAACTTCGAAAAGGCCGCCAACGCCTTGCTCACTCCGCCGAGCTCCATCTCTCCTCAGCTTCAAGCCCAGCGCACCCGTGCTGCGGCGTTCTCGCCACCGCCGAACAACATGAGTCAAGATACGGAGATGGAGGATTCTGCTGGCGAATACTCGCCGGATTTGGACAACATCCAGCCGTCGTCGACCGGCATGCCTCTCAGCCTCGGTGCCTTGTCTGATCTGGACAACGCGGTGACCATCACCCCGAGCATGCTGGCTAAGCAGCACCTGCCGGGCATCATGCTCGGTAACGGGCCACGACCCATCCGCTATGTGATGGGCGAGCTATGCCAAACAGTACCTGGCTTCTCGCGCATCGCACCGGCCAAGGCTCGCAGGATTGTGGTCGCGGCGCTCGGCAACCCTGCTGGAGGTGGACCCAACGGAGATGTCGTTTTCTCCAAGACTGGCTGGGGACGATGGGATGCGCACATCAAGGGGTCTTCTCGTGATTCCGCCATCGGCTCCTTCAACGAGGGCAACCTGTCGCCTCCACGCAGTGAGCAGGGCAGCTACGCCATGTCGCACAGTGATTCAGCAATCCACATGCACGCTCAGCATATGAATCAGAAGTACCGAGAAGGCTATTCTGGTGGTAGTTGGACCGCATCTAGCATCCGCGAGGAGGACGAACTCGACATGGATATGGATCCGGTGGAAGACGCCGCTGATAACATGTCTCTGGACGGCGATTCCTCTGACTCCTCGTCAGAAGAGTTGAACGAGGATACCGACGACGAAGATT-GGCTGCCGTTGGTATCGAAGCGTTGCGGAAGGCTTC-ATGTCCACTTCCAACGTGACTC-TCGGCAG-ACATCCGTGTCGGCAGTATCCCGTAC-ACGGACCCTGGGCATCGA-GAGCTG-CAGCGAC-A-CTATTATCACAAAGTCGATCG-TCCAGTCAGCCTCGGTTCGGAACGGCGCCAACTTCCGAAA ``` |
| 19 | ggggacaagtttgtacaaaaaagcaggcttgatgGCTCAGGCTCTTGGTTTCAT, ggggaccactttgtacaagaaagctgggtcGTTCTTCCAATAGGCCTGCG | Mycgr3\|92790  JGI  Ensembl |  |  | 0 % |  |  |
| 19 | ggggacaagtttgtacaaaaaagcaggcttgatgGCTGCAGGCAAGAAGGCGAA, ggggaccactttgtacaagaaagctgggtcGAAGTCGCGACCACGGACAC | Mycgr3\|93337  JGI  Ensembl | KOG2084 Predicted histone tail methylase containing SET domain; | nucleus; | 100 % |  |  |
| 19 | ggggacaagtttgtacaaaaaagcaggcttgatgGCTGAAGCAAAGTTCCGCCT, ggggaccactttgtacaagaaagctgggtcTCGTTTGCGCGGGACTCCCT | Mycgr3\|93617  JGI  Ensembl |  |  | 100 % |  |  |
| 19 | ggggacaagtttgtacaaaaaagcaggcttgatgAAGCGCACCCAGGAAGAAGC, ggggaccactttgtacaagaaagctgggtcTGCACTGCTTGCGATAGCGA | Mycgr3\|94659  JGI  Ensembl |  |  | 0 % |  |  |
| 19 | ggggacaagtttgtacaaaaaagcaggcttgatgTCCAACAATACCATCAACAA, ggggaccactttgtacaagaaagctgggtcTGATCGAAATAATGGGTTCG | Mycgr3\|94877  JGI  Ensembl |  |  | 0 % |  |  |
| 19 | ggggacaagtttgtacaaaaaagcaggcttgatgTCTTCCAATATCAAAGTAAC, ggggaccactttgtacaagaaagctgggtcAATTGAGACAGTCTCGCCGA | Mycgr3\|95030  JGI  Ensembl | KOG3798 Predicted Zn-dependent hydrolase (beta-lactamase superfamily); |  | 100 % | chr\_8:787477 T->C; chr\_8:787867 T->C; chr\_8:787653 C->T; chr\_8:787797 A->G; |  |
| 19 | ggggacaagtttgtacaaaaaagcaggcttgatgGCTCCCGCAATGAGACGTAT, ggggaccactttgtacaagaaagctgggtcGCACCTCTTCTTCGCATTCC | Mycgr3\|95115  JGI  Ensembl |  |  | 100 % |  |  |
| 19 | ggggacaagtttgtacaaaaaagcaggcttgatgGCTTATACCGACATCAAAAA, ggggaccactttgtacaagaaagctgggtcGTACCCATCGTTGATTCGCC | Mycgr3\|95183  JGI  Ensembl |  |  | 0 % |  |  |
| 19 | ggggacaagtttgtacaaaaaagcaggcttgatgTTCAAAGCTATCCTGCTCAG, ggggaccactttgtacaagaaagctgggtcCTTGTTCGACGGTGCCGTAG | Mycgr3\|95636  JGI  Ensembl |  |  | 100 % |  |  |
| 19 | ggggacaagtttgtacaaaaaagcaggcttgatgGCCACGCTCCGCATTTTCTT, ggggaccactttgtacaagaaagctgggtcGGCAATGGCAGCAAAACCGT | Mycgr3\|96011  JGI  Ensembl |  |  | 100 % |  |  |
| 19 | ggggacaagtttgtacaaaaaagcaggcttgatgGCGAGATTGAGATTGAGATT, ggggaccactttgtacaagaaagctgggtcACCGAGCTGGGTTTTTGGGA | Mycgr3\|96142  JGI  Ensembl |  |  | 100 % |  |  |
| 19 | ggggacaagtttgtacaaaaaagcaggcttgatgACTGGCAGAGAAGACGAGGC, ggggaccactttgtacaagaaagctgggtcAAAGTCGACCATTCTTCGTC | Mycgr3\|96168  JGI  Ensembl |  |  | 100 % |  |  |
| 19 | ggggacaagtttgtacaaaaaagcaggcttgatgGCAATCACTGGAGCCGCTCC, ggggaccactttgtacaagaaagctgggtcGGTCAAGTCAATCACTTCCG | Mycgr3\|96993  JGI  Ensembl |  |  | 100 % |  |  |
| 19 | ggggacaagtttgtacaaaaaagcaggcttgatgACGACCCCGTTTCTCCTTCA, ggggaccactttgtacaagaaagctgggtcACACACCGTCAATGAAATGC | Mycgr3\|97333  JGI  Ensembl |  |  | 100 % |  |  |
| 19 | ggggacaagtttgtacaaaaaagcaggcttgatgGAGCACCGCGCCGTGAGCAA, ggggaccactttgtacaagaaagctgggtcCTTGGACGGCTTCTCGTTCT | Mycgr3\|97382  JGI  Ensembl |  |  | 100 % |  |  |
| 19 | ggggacaagtttgtacaaaaaagcaggcttgatgTTCGCCAATCCCCTCGACTC, ggggaccactttgtacaagaaagctgggtcCCTCCTGAGCACCGGCCGCA | Mycgr3\|97679  JGI  Ensembl |  |  | 100 % |  |  |
| 19 | ggggacaagtttgtacaaaaaagcaggcttgatgGCGGCAATGTCAGCTCTGCC, ggggaccactttgtacaagaaagctgggtcGTAGCTCTCTTGGCTTGGCA | Mycgr3\|97729  JGI  Ensembl |  |  | 100 % |  |  |
| 19 | ggggacaagtttgtacaaaaaagcaggcttgatgGCGGAAAACAACGAAGCCGA, ggggaccactttgtacaagaaagctgggtcAATCCATCCGCATCGATCTT | Mycgr3\|97896  JGI  Ensembl |  |  | 100 % |  |  |
| 19 | ggggacaagtttgtacaaaaaagcaggcttgatgACCGGCGAGTACTCTGACGA, ggggaccactttgtacaagaaagctgggtcCGATGGGATGCGAACCGGAG | Mycgr3\|98072  JGI  Ensembl |  |  | 100 % |  |  |
| 19 | ggggacaagtttgtacaaaaaagcaggcttgatgTCCGTCGACATCAACTGGGA, ggggaccactttgtacaagaaagctgggtcAACGAGGAATGTCCAGAAGC | Mycgr3\|98339  JGI  Ensembl |  |  | 0 % |  |  |
| 19 | ggggacaagtttgtacaaaaaagcaggcttgatgGATCCCTCAGAAGTGAACGT, ggggaccactttgtacaagaaagctgggtcATCGACATCCGGCACCGGCA | Mycgr3\|98975  JGI  Ensembl |  |  | 100 % |  |  |
| 19 | ggggacaagtttgtacaaaaaagcaggcttgatgTCAGGCATCAAAGTGGTCTT, ggggaccactttgtacaagaaagctgggtcCTTGTGGTAATTATCGAGCG | Mycgr3\|102471  JGI  Ensembl | KOG1575 Voltage-gated shaker-like K+ channel, subunit beta/KCNAB; | F420H2 dehydrogenase activity; malolactic enzyme activity;  oxidoreductase activity;  regulation of oxidoreductase activity;  sulfur oxygenase reductase activity; | 100 % |  |  |
| 19 | ggggacaagtttgtacaaaaaagcaggcttgatgGCCGACACCGACCTCGAGCT, ggggaccactttgtacaagaaagctgggtcTGCCACTCCTTTCTCCTTGC | Mycgr3\|103058  JGI  Ensembl | KOG0401 Translation initiation factor 4F, ribosome/mRNA-bridging subunit (eIF-4G); |  | 100 % |  |  |
| 19 | ggggacaagtttgtacaaaaaagcaggcttgatgTGTTCCCGCGAGAGTCACGA, ggggaccactttgtacaagaaagctgggtcACGGATGACTGGTTGAGGGG | Mycgr3\|103909  JGI  Ensembl |  |  | 0 % |  |  |
| 19 | ggggacaagtttgtacaaaaaagcaggcttgatgAAGCTCAACATCTCCTTCCC, ggggaccactttgtacaagaaagctgggtcCTTGCGCATGCTGGACGCAC | Mycgr3\|104155  JGI  Ensembl | KOG1646 40S ribosomal protein S6; | intracellular; ribosome; structural constituent of ribosome;  translation; | 100 % |  |  |
| 19 | ggggacaagtttgtacaaaaaagcaggcttgatgGGAGCTTTCTACGAAACGAT, ggggaccactttgtacaagaaagctgggtcAACGCCTGCCATCATCTTTA | Mycgr3\|105954  JGI  Ensembl |  |  | 100 % |  |  |
| 19 | ggggacaagtttgtacaaaaaagcaggcttgatgGCGACTGGAAACAATGACAT, ggggaccactttgtacaagaaagctgggtcATGCGCATGATACTCCTCCA | Mycgr3\|106959  JGI  Ensembl | KOG4157 beta-1,6-N-acetylglucosaminyltransferase, contains WSC domain; |  | 0 % |  |  |
| 19 | ggggacaagtttgtacaaaaaagcaggcttgatgGACTCTCCCGGCTCGCCTCC, ggggaccactttgtacaagaaagctgggtcGCCCTCCACAACCGCCATCC | Mycgr3\|107313  JGI  Ensembl | KOG2992 Nucleolar GTPase/ATPase p130; |  | 100 % |  |  |
| 19 | ggggacaagtttgtacaaaaaagcaggcttgatgGATGCAAAGGCAGCCTCGCC, ggggaccactttgtacaagaaagctgggtcTCCAAACATCGTCCTCAAAC | Mycgr3\|107492  JGI  Ensembl | KOG0758 Mitochondrial carnitine-acylcarnitine carrier protein; | binding; membrane; transport; | 100 % |  |  |
| 19 | ggggacaagtttgtacaaaaaagcaggcttgatgGCGCCCGCTCCTCGCCCTCG, ggggaccactttgtacaagaaagctgggtcATCCTTGTACGCGCCAACCA | Mycgr3\|107688  JGI  Ensembl | KOG1407 WD40 repeat protein; | 1-alkyl-2-acetylglycerophosphocholine esterase activity; heme binding;  transition metal ion binding; | 100 % |  |  |
| 19 | ggggacaagtttgtacaaaaaagcaggcttgatgGCACTCCCGATCCGCACGCG, ggggaccactttgtacaagaaagctgggtcAGCGAGCTGACAAAGATGAA | Mycgr3\|107808  JGI  Ensembl |  |  | 100 % | chr\_2:77514 C->T; chr\_2:77548 A->G; chr\_2:77875 A->G; |  |
| 19 | ggggacaagtttgtacaaaaaagcaggcttgatgATATTCCATACACCTCCGAC, ggggaccactttgtacaagaaagctgggtcCAGACCGACAAATCTCCACA | Mycgr3\|108172  JGI  Ensembl |  |  | 100 % |  |  |
| 19 | ggggacaagtttgtacaaaaaagcaggcttgatgGCTGCCACGAAACAGACTAC, ggggaccactttgtacaagaaagctgggtcAGCAGCCACAATCGGCGCCG | Mycgr3\|109022  JGI  Ensembl |  |  | 3.81062 % |  |  |
| 19 | ggggacaagtttgtacaaaaaagcaggcttgatgCAGTTGAAAGACGAAGTCCA, ggggaccactttgtacaagaaagctgggtcAGTCACCGTTTCATCCCGAC | Mycgr3\|109375  JGI  Ensembl | KOG2084 Predicted histone tail methylase containing SET domain; | nucleus; | 100 % |  |  |
| 19 | ggggacaagtttgtacaaaaaagcaggcttgatgTCCACCGTCCACCACCTCTT, ggggaccactttgtacaagaaagctgggtcCTTCTCCTCTACCACTCCGC | Mycgr3\|109649  JGI  Ensembl |  |  | 100 % |  |  |
| 19 | ggggacaagtttgtacaaaaaagcaggcttgatgCCAACTCAACAAAGGACCCT, ggggaccactttgtacaagaaagctgggtcCCGAGCATGATTCTGGTGCC | Mycgr3\|110455  JGI  Ensembl |  |  | 100 % |  |  |
| 19 | ggggacaagtttgtacaaaaaagcaggcttgatgAGCAACGCAACACTGGCTGC, ggggaccactttgtacaagaaagctgggtcGTACGCATCCTTTCCTCCAA | Mycgr3\|110793  JGI  Ensembl | KOG0409 Predicted dehydrogenase; | 3-hydroxyisobutyrate dehydrogenase activity; valine metabolic process; | 100 % |  |  |
| 19 | ggggacaagtttgtacaaaaaagcaggcttgatgGCCGACCAACAATCCCTCAA, ggggaccactttgtacaagaaagctgggtcTGCCCTCGCAGCACTTCTCT | Mycgr3\|111048  JGI  Ensembl | KOG2160 Armadillo/beta-catenin-like repeat-containing protein; |  | 100 % |  |  |
| 19 | ggggacaagtttgtacaaaaaagcaggcttgatgAAATTCTTCACGACCACGGC, ggggaccactttgtacaagaaagctgggtcGAATTGGCCGGATGGATCAT | Mycgr3\|111203  JGI  Ensembl |  |  | 0 % |  |  |
| 19 | ggggacaagtttgtacaaaaaagcaggcttgatgGCCACCGAGGCGCACTGTGC, ggggaccactttgtacaagaaagctgggtcCGCCGAACCTGTAGCCTCAA | Mycgr3\|111249  JGI  Ensembl | KOG3274 Uncharacterized conserved protein, AMMECR1; |  | 0 % |  |  |
| 19 | ggggacaagtttgtacaaaaaagcaggcttgatgTCTCTCCACACCCCAGCACC, ggggaccactttgtacaagaaagctgggtcAGCATGGACCTCAGTATTCG | Mycgr3\|111569  JGI  Ensembl |  |  | 100 % |  | ``` 474	GGCTTGTGAAGAGTGCCGCAAGAGGAAGCAAAAGTGTGATGGCGAGCACGTATGCCAATCATGCAAGGAGCAAAAGTTGGAGTGCAAATACCGTGAGGTCGTCCCCACAAA	584 	\|\|\|\|\|\|\|\|\|\|\|\|\|\|\|\|\|\|\|\|\|\|\|\|\|\|\|\|\|\|\|\|\|\|\|\|\|\|\|\|\|\|\|\|\|\|\|\|\|\|\|\|\|\|\|\|\|\|\|\|\|\|\|\|\|\|\|\|\|\|\|\|\|\|\|\|\|\|\|\|\|\|\|\|\|\|\|\|\|\|\|\|\|\|\|\|\|\|\|\|\|\|\|\|\|\|\|\|\|\|\| 	GGCTTGTGAAGAGTGCCGCAAGAGGAAGCAAAAGTGTGATGGCGAGCACGTATGCCAATCATGCAAGGAGCAAAAGTTGGAGTGCAAATACCGTGAGGTCGTCCCCACAAA ```      ``` 1	ATGTCTCTCCACACCCCAGCACCAAAGAGCCTCAAGCGCAACTTTCAGCGTATGATGATGGACCACCACTCTCACGGCATGATCCCGATGGAACAACCTCACTACAACGAGTTCCCACCTGGCGTGTTCTACACTCCTCCGCCACCACCGCCTCAACACGTCCTCTTCAGCAACCCATTCCCAGTATCGCCATATGAATACTCCCCAGATGACTACTTCCCATCGGAGTCGACTAGCTACTACAACGCCGACCGCTCTTGCTCAGATCCTTTCATTCAAGAGGTCGATTCGTCCTTCAGCGGCATGGAGAGTGGCAGCTCTTCCTGCAATGTGTCCGTGGGACCAGCAACTCCTCCATCAGCAACATCTCCGCTACACCACCACTACACAGAAGGTCGGCCAACAAAGTTCCCTGCTGTTCCAATCACGCCTAGCCCATCAACTCCTCGTCAAAAAGCCAAGCGCACCGCAACGG	475 	\|\|\|\|\|\|\|\|\|\|\|\|\|\|\|\|\|\|\|\|\|\|\|\|\|\|\|\|\|\|\|\|\|\|\|\|\|\|\|\|\|\|\|\|\|\|\|\|\|\|\|\|\|\|\|\|\|\|\|\|\|\|\|\|\|\|\|\|\|\|\|\|\|\|\|\|\|\|\|\|\|\|\|\|\|\|\|\|\|\|\|\|\|\|\|\|\|\|\|\|\|\|\|\|\|\|\|\|\|\|\|\|\|\|\|\|\|\|\|\|\|\|\|\|\|\|\|\|\|\|\|\|\|\|\|\|\|\|\|\|\|\|\|\|\|\|\|\|\|\|\|\|\|\|\|\|\|\|\|\|\|\|\|\|\|\|\|\|\|\|\|\|\|\|\|\|\|\|\|\|\|\|\|\|\|\|\|\|\|\|\|\|\|\|\|\|\|\|\|\|\|\|\|\|\|\|\|\|\|\|\|\|\|\|\|\|\|\|\|\|\|\|\|\|\|\|\|\|\|\|\|\|\|\|\|\|\|\|\|\|\|\|\|\|\|\|\|\|\|\|\|\|\|\|\|\|\|\|\|\|\|\|\|\|\|\|\|\|\|\|\|\|\|\|\|\|\|\|\|\|\|\|\|\|\|\|\|\|\|\|\|\|\|\|\|\|\|\|\|\|\|\|\|\|\|\|\|\|\|\|\|\|\|\|\|\|\|\|\|\|\|\|\|\|\|\|\|\|\|\|\|\|\|\|\|\|\|\|\|\|\|\|\|\|\|\|\|\|\|\|\|\|\|\|\|\|\|\|\|\|\|\|\|\|\|\|\|\|\|\|\|\|\|\|\|\|\|\|\|\|\|\|\|\|\|\|\|\|\|\|\|\|\|\|\|\|\|\|\|\|\|\|\|\|\|\|\|\|\|\|\|\|\|\|\|\|\|\|\|\|\|\|\|\|\|\|\|\|\|\|\|\|\|\|\|\|\|\|\|\|\|\|\|\|\|\|\|\|\|\|\|\|\|\|\|\|\|\|\|\|\|\|\|\|\|\|\|\|\|\|\|\|\|\|\| 	ATGTCTCTCCACACCCCAGCACCAAAGAGCCTCAAGCGCAACTTTCAGCGTATGATGATGGACCACCACTCTCACGGCATGATCCCGATGGAACAACCTCACTACAACGAGTTCCCACCTGGCGTGTTCTACACTCCTCCGCCACCACCGCCTCAACACGTCCTCTTCAGCAACCCATTCCCAGTATCGCCATATGAATACTCCCCAGATGACTACTTCCCATCGGAGTCGACTAGCTACTACAACGCCGACCGCTCTTGCTCAGATCCTTTCATTCAAGAGGTCGATTCGTCCTTCAGCGGCATGGAGAGTGGCAGCTCTTCCTGCAATGTGTCCGTGGGACCAGCAACTCCTCCATCAGCAACATCTCCGCTACACCACCACTACACAGAAGGTCGGCCAACAAAGTTCCCTGCTGTTCCAATCACGCCTAGCCCATCAACTCCTCGTCAAAAAGCCAAGCGCACCGCAACGG ```      ``` 585	AAAGGACAACTCATTGGAGAAGCTGATGAGCCTCATCGACAGCTTCGGCAAGTCAATGGAGGCTCTCAACCAACGCCTCGATACCATGAGCGCGAGCGTTGTCTCCATCCAGTCTCGGCTCGACTCACAATGTGCATCGACGAATACTGAGGTCCATGCT	744 	\|\|\|\|\|\|\|\|\|\|\|\|\|\|\|\|\|\|\|\|\|\|\|\|\|\|\|\|\|\|\|\|\|\|\|\|\|\|\|\|\|\|\|\|\|\|\|\|\|\|\|\|\|\|\|\|\|\|\|\|\|\|\|\|\|\|\|\|\|\|\|\|\|\|\|\|\|\|\|\|\|\|\|\|\|\|\|\|\|\|\|\|\|\|\|\|\|\|\|\|\|\|\|\|\|\|\|\|\|\|\|\|\|\|\|\|\|\|\|\|\|\|\|\|\|\|\|\|\|\|\|\|\|\|\|\|\|\|\|\|\|\|\|\|\|\|\|\|\|\|\|\|\|\|\|\|\|\|\|\| 	AAAGGACAACTCATTGGAGAAGCTGATGAGCCTCATCGACAGCTTCGGCAAGTCAATGGAGGCTCTCAACCAACGCCTCGATACCATGAGCGCGAGCGTTGTCTCCATCCAGTCTCGGCTCGACTCACAATGTGCATCGACGAATACTGAGGTCCATGCT ``` |
| 19 | ggggacaagtttgtacaaaaaagcaggcttgatgGCCAACGCAATGCAGCCGGC, ggggaccactttgtacaagaaagctgggtcGATCTCCCCATCGACAGAGT | Mycgr3\|111774  JGI  Ensembl |  |  | 100 % |  |  |
| 20 | ggggacaagtttgtacaaaaaagcaggcttgatgCCTGAATTTCCCGCCCACGG, ggggaccactttgtacaagaaagctgggtcTGAACTCGAGAACATATGCT | Mycgr3\|31520  JGI  Ensembl | KOG1284 Bifunctional GTP cyclohydrolase II/3,4-dihydroxy-2butanone-4-phosphate synthase; | GTP cyclohydrolase II activity; riboflavin biosynthetic process; | 100 % |  |  |
| 20 | ggggacaagtttgtacaaaaaagcaggcttgatgGCGAAGTCTACGGAACCTTG, ggggaccactttgtacaagaaagctgggtcTCCACCGCGTCCAACACAAT | Mycgr3\|32274  JGI  Ensembl | KOG1430 C-3 sterol dehydrogenase/3-beta-hydroxysteroid dehydrogenase and related dehydrogenases; | 3-beta-hydroxy-delta5-steroid dehydrogenase activity; steroid biosynthetic process; | 100 % |  |  |
| 20 | ggggacaagtttgtacaaaaaagcaggcttgatgAAGTCCTTCGCCGCCGCAGC, ggggaccactttgtacaagaaagctgggtcAGCACGACGAGTCCTCATCG | Mycgr3\|32500  JGI  Ensembl |  |  | 100 % |  |  |
| 20 | ggggacaagtttgtacaaaaaagcaggcttgatgGCATTAAGACTACTGACCAG, ggggaccactttgtacaagaaagctgggtcCAACTCATCCCTCCCCTTCC | Mycgr3\|33113  JGI  Ensembl | KOG2964 Arginase family protein; | agmatinase activity; arginase activity;  arginine catabolic process;  endoplasmic reticulum;  hydrolase activity, acting on carbon-nitrogen (but not peptide) bonds, in linear amidines; metal ion binding; | 100 % | chr\_1:5930860 T->C; chr\_1:5931350 G->A; chr\_1:5930697 A->G; |  |
| 20 | ggggacaagtttgtacaaaaaagcaggcttgatgGCGCTCGATGAGCCGTCTGC, ggggaccactttgtacaagaaagctgggtcCAGCAAGACATGACTCTCCC | Mycgr3\|33464  JGI  Ensembl |  |  | 100 % |  |  |
| 20 | ggggacaagtttgtacaaaaaagcaggcttgatgCCGCCTCCCCATCGAATGGA, ggggaccactttgtacaagaaagctgggtcAGAATCACCCCACACCTCCC | Mycgr3\|36261  JGI  Ensembl |  |  | 100 % |  |  |
| 20 | ggggacaagtttgtacaaaaaagcaggcttgatgTCATCAGCCGCATCACCATT, ggggaccactttgtacaagaaagctgggtcCGGCAAATCAAACCAATTGC | Mycgr3\|37854  JGI  Ensembl | KOG0198 MEKK and related serine/threonine protein kinases; | 3-phosphoinositide-dependent protein kinase activity; AMP-activated protein kinase activity;  ATP binding;  DNA-dependent protein kinase activity;  GTP-dependent protein kinase activity;  JUN kinase activity;  JUN kinase kinase activity;  JUN kinase kinase kinase activity;  Janus kinase activity;  MAP kinase 1 activity;  MAP kinase 2 activity;  MAP kinase kinase activity;  MAP kinase kinase kinase kinase activity;  MAP/ERK kinase kinase activity;  MP kinase activity;  NF-kappaB-inducing kinase activity;  SAP kinase activity;  atypical protein kinase C activity;  calcium-dependent protein kinase C activity;  casein kinase I activity;  casein kinase activity;  cyclic nucleotide-dependent protein kinase activity;  cyclin-dependent protein kinase activating kinase activity;  eukaryotic translation initiation factor 2alpha kinase activity;  galactosyltransferase-associated kinase activity;  glycogen synthase kinase 3 activity;  protein amino acid phosphorylation;  protein kinase activity;  protein serine/threonine kinase activity;  receptor signaling protein serine/threonine kinase activity;  ribosomal protein S6 kinase activity;  transmembrane receptor protein kinase activity;  transmembrane receptor protein serine/threonine kinase activity; | 100 % |  |  |
| 20 | ggggacaagtttgtacaaaaaagcaggcttgatgGATCCTCAATTGATACTCTT, ggggaccactttgtacaagaaagctgggtcAGGCCGCCTGATCAAGAGAA | Mycgr3\|38483  JGI  Ensembl | KOG2497 Predicted methyltransferase; |  | 0 % |  |  |
| 20 | ggggacaagtttgtacaaaaaagcaggcttgatgGGCTCGATCGAACAAGGAGA, ggggaccactttgtacaagaaagctgggtcTGCCTTCACCCTCATCCCAC | Mycgr3\|38733  JGI  Ensembl | KOG0143 Iron/ascorbate family oxidoreductases; | gibberellin 2-beta-dioxygenase activity; | 100 % |  |  |
| 20 | ggggacaagtttgtacaaaaaagcaggcttgatgGCAGGCGAGGTCCGTCAACC, ggggaccactttgtacaagaaagctgggtcCAACTTCACCCCGGCCCGGT | Mycgr3\|38862  JGI  Ensembl |  |  | 100 % |  |  |
| 20 | ggggacaagtttgtacaaaaaagcaggcttgatgGCCTCGGAAACCCAGACCAG, ggggaccactttgtacaagaaagctgggtcTTGCCTCCACGCTACAACAT | Mycgr3\|41900  JGI  Ensembl | KOG2426 Dihydroxyacetone kinase/glycerone kinase; | glycerol metabolic process; glycerone kinase activity; | 100 % |  |  |
| 20 | ggggacaagtttgtacaaaaaagcaggcttgatgCATGTCACGACACTCCTTCA, ggggaccactttgtacaagaaagctgggtcAAGCTTTAGGCCACCACGCC | Mycgr3\|42064  JGI  Ensembl |  |  | 100 % |  |  |
| 20 | ggggacaagtttgtacaaaaaagcaggcttgatgCTGACCTCCCTCCTCCTAAC, ggggaccactttgtacaagaaagctgggtcAGTGCAGTGCAGCTCGCCGT | Mycgr3\|42164  JGI  Ensembl |  |  | 100 % |  |  |
| 20 | ggggacaagtttgtacaaaaaagcaggcttgatgGCAACATCCTCTTTTCCCGC, ggggaccactttgtacaagaaagctgggtcTGTTTTGGTCTGGTAGAAGA | Mycgr3\|43194  JGI  Ensembl |  |  | 100 % | chr\_6:2479121 T->C; chr\_6:2480131 T->C; |  |
| 20 | ggggacaagtttgtacaaaaaagcaggcttgatgTCATCCTACACCGTCCAGCA, ggggaccactttgtacaagaaagctgggtcCTCCGGCACCACATACTCCC | Mycgr3\|43667  JGI  Ensembl | KOG1250 Threonine/serine dehydratases; | catalytic activity; metabolic process;  pyridoxal phosphate binding; | 100 % |  |  |
| 20 | ggggacaagtttgtacaaaaaagcaggcttgatgTCGGATCACATCTCAGACCG, ggggaccactttgtacaagaaagctgggtcCCCTCTCAAAGGCAAAACGT | Mycgr3\|45644  JGI  Ensembl | KOG0317 Predicted E3 ubiquitin ligase, integral peroxisomal membrane protein; | peroxisomal membrane; peroxisome organization and biogenesis;  protein binding;  zinc ion binding; | 100 % |  |  |
| 20 | ggggacaagtttgtacaaaaaagcaggcttgatgTTCCCACCATCCGAAGGCTA, ggggaccactttgtacaagaaagctgggtcACGCTCGCCCGCCTCCCGAC | Mycgr3\|46378  JGI  Ensembl | KOG2913 Predicted membrane protein; |  | 0 % |  |  |
| 20 | ggggacaagtttgtacaaaaaagcaggcttgatgTCCCAACCCAAAATCATCAT, ggggaccactttgtacaagaaagctgggtcCAGGCTGGACTTTACAGTTC | Mycgr3\|49121  JGI  Ensembl |  |  | 0 % |  |  |
| 20 | ggggacaagtttgtacaaaaaagcaggcttgatgCCAACAGAGTTCATCAGTCT, ggggaccactttgtacaagaaagctgggtcAGTCCCATCCTTACCAACGG | Mycgr3\|49158  JGI  Ensembl |  |  | 100 % |  |  |
| 20 | ggggacaagtttgtacaaaaaagcaggcttgatgCCACAGCAGCCGATCCGGAT, ggggaccactttgtacaagaaagctgggtcGTGGAACTTCCTCATCACCT | Mycgr3\|50077  JGI  Ensembl |  |  | 100 % |  |  |
| 20 | ggggacaagtttgtacaaaaaagcaggcttgatgGGGCTCGAGGCGACGATGAT, ggggaccactttgtacaagaaagctgggtcTGCAGTATCCATGCGATCGT | Mycgr3\|53825  JGI  Ensembl | KOG2884 26S proteasome regulatory complex, subunit RPN10/PSMD4; |  | 100 % |  | ``` 377	GCTCGCACTCAAACATCGCCAGAATAAATCCCAACGTCAACGCATCATCGTGTTCATTTGCTCGCCCATCGCCGACTCTTCAAGCACCCTCACCAAACTTGCCAAGCGGATGAAGAAGAACAACACCAGCATCGACATCATCGCCTTCGGTGATCTTTCGGACGAGAACCTCGACAAGCTGCGCGCGTTCAACGAGGCTGTCAAATCCAACGACGGCTCACATCTTGAAATCATCCCACCGGGCCCGAACCTTCTATCCGACACGATCGTGGCTTCCCCGATATTGGCAGGCGAAGGAGGTGCCGCAACAGCAAATGGTGGTGGCGCCGGC-GGAGATGGCGGCGCAGGCGG-AGACTT-CG-AATTTGGCGTGGATCCTAACATGGACCCTGAGTTGGCACTC-GTACTGCGGATGAGTATGGAAGAGGAACGCGAGCGACAG-GAGCGGGAGAAGAAGGCAAGAGACGAGGCAGAGGGAAAGACAAATCTCGAGAGCGTGCCAGAAGCT-GCCGAGGGAG-AATCAAGCGAGAA-GCAGCCT-CTACTGGATAAGA-GTGGCGAT	932 	\|\|\|\|\|\|\|\|\|\|\|\|\|\|\|\|\|\|\|\|\|\|\|\|\| \|\|\|\|\|\|\|\|\|\|\|\|\|\|\|\|\|\|\|\|\|\|\|\|\|\|\|\|\|\|\|\|\|\|\|\|\|\|\|\|\|\|\|\|\|\|\|\|\|\|\|\|\|\|\|\|\|\|\|\|\|\|\|\|\|\|\|\|\|\|\|\|\|\|\|\|\|\|\|\|\|\|\|\|\|\|\|\|\|\|\|\|\|\|\|\|\|\|\|\|\|\|\|\|\|\|\|\|\|\|\|\|\|\|\|\|\|\|\|\|\|\|\|\|\|\|\|\|\|\|\|\|\|\|\|\|\|\|\|\|\|\|\|\|\|\|\|\|\|\|\|\|\|\|\|\|\|\|\|\|\|\|\|\|\|\|\|\|\|\|\|\|\|\|\|\|\|\|\|\|\|\|\|\|\|\|\|\|\|\|\|\|\|\|\|\|\|\|\|\|\|\|\|\|\|\|\|\|\|\|\|\|\|\|\|\|\|\|\|\|\|\|\|\|\|\|\|\|\|\|\|\|\|\|\|\|\|\|\|\|\|\|\|\|\|\|\|\|\|\|\|\|\|\|\|\|\|\|\|\|\|\|\|\|\|\|\|\|\|\|\|\|\|\|\|\|\|\|\|\|\|\|\|\|\|\|\|\|\|\|\|\|\|\|\|\|\|\|\|\|\|\|\|\|\| \|\|\|\|\|\|\|\|\|\|\|\|\|\|\|\|\|\|\|\| \|\|\|\|\|\| \|\| \|\|\|\|\|\|\|\|\|\|\|\|\|\|\|\|\|\|\|\|\|\|\|\|\|\|\|\|\|\|\|\|\|\|\|\|\|\|\|\|\| \|\|\|\|\|\|\|\|\|\|\|\|\| \|\|\|\|\|\|\|\|\|\|\|\|\|\|\|\| \|\|\|\|\|\|\|\| \|\|\|\|\|\|\|\|\|\|\| \|\|\|\|\|\|\| \| \|\|\|\| \|\|\| \|\|\|\|\|\|\|\|\|\|\|\|  \|\|\|\|\|\|\|\|\|\|\|\|\|\|\| \|\|\|\|\| \|\|\| \|\|\| \|\| \|\|\|\|\| \|\| \|\|\|\| \|\| \|\|\|\| \| \|\|\|\| \| \|\|\|\| \|\|\|\|\|\|\|\| 	GCTCGCACTCAAACATCGCCAGAATGAATCCCAACGTCAACGCATCATCGTGTTCATTTGCTCGCCCATCGCCGACTCTTCAAGCACCCTCACCAAACTTGCCAAGCGGATGAAGAAGAACAACACCAGCATCGACATCATCGCCTTCGGTGATCTTTCGGACGAGAACCTCGACAAGCTGCGCGCGTTCAACGAGGCTGTCAAATCCAACGACGGCTCACATCTTGAAATCATCCCACCGGGCCCGAACCTTCTATCCGACACGATCGTGGCTTCCCCGATATTGGCAGGCGAAGGAGGTGCCGCAACAGCAAATGGTGGTGGCGCCGGCGGGAGATGGCGGCGCAGGCGGGAGACTTTCGAAATTTGGCGTGGATCCTAACATGGACCCTGAGTTGGCACTCGGTACTGCGGATGAATATGGAAGAGGAACGCCAGCGACAGTGAGCGGGAGAATAAGGCAAAACACGACGCATAGGGAAAGACAATCCTCGAGAGCGTGCCAAAAGCTTGCCAAGG-AGCAATCATGC-AGAAAGC-GCCTTCGACTG-ACAAGAAGTGGCGAT ```      ``` 119	TGTCGACAACAGCGAGAGCAGTCGGAACGGAGACTACGTACCATCTCGATGGGAAGCACAAGTAGATGCAGCCAACCTCATCTTCCACTCGAAGACACAAGCAAACCCAGAATCATCAGTCGGTCTCATGAGCATGGGCGGCAGCGGTCCTCAGGTGCTCACAACACTCACTACAAACCCTGGCCAAGTACTGGATGGACTTCATCGAACAAAGGTCAAGGGCGAGTCACATCTTTACACGGGAATCATGATCGCCTCG	377 	\|\|\|\|\|\|\|\|\|\|\|\|\|\|\|\|\|\|\|\|\|\|\|\|\|\|\|\|\|\|\|\|\|\|\|\|\|\|\|\|\|\|\|\|\|\|\|\|\|\|\|\|\|\|\|\|\|\|\|\|\|\|\|\|\|\|\|\|\|\|\|\|\|\|\|\|\|\|\|\|\|\|\| \|\|\|\|\|\|\|\|\|\|\|\|\|\|\|\|\|\|\|\|\|\|\|\|\|\|\|\|\|\|\|\|\|\|\|\|\|\|\|\|\|\|\|\|\|\|\|\|\|\|\|\|\|\|\|\|\|\|\|\|\|\|\|\|\|\|\|\|\|\|\|\|\|\|\|\|\|\|\|\|\|\|\|\|\|\|\|\|\|\|\|\|\|\|\|\|\|\|\|\|\|\|\|\|\|\|\|\|\|\|\|\|\|\|\|\|\|\|\|\|\|\|\|\|\|\|\|\|\|\|\|\|\|\|\|\|\|\|\|\|\|\|\|\|\|\|\|\|\|\|\|\|\|\|\|\|\|\|\|\|\|\|\|\|\|\|\|\|\|\|\|\|\|\|\| 	TGTCGACAACAGCGAGAGCAGTCGGAACGGAGACTACGTACCATCTCGATGGGAAGCACAAGTAGATGCAGCCAACCTCATCTCCCACTCGAAGACACAAGCAAACCCAGAATCATCAGTCGGTCTCATGAGCATGGGCGGCAGCGGTCCTCAGGTGCTCACAACACTCACTACAAACCCTGGCCAAGTACTGGATGGACTTCATCGAACAAAGGTCAAGGGCGAGTCACATCTTTACACGGGAATCATGATCGCCTCG ``` |
| 20 | ggggacaagtttgtacaaaaaagcaggcttgatgCCACTTCCTCGGATAAGTCC, ggggaccactttgtacaagaaagctgggtcTGTCGAAGCCTGCAATATTT | Mycgr3\|53893  JGI  Ensembl | KOG1436 Dihydroorotate dehydrogenase; | 'de novo' pyrimidine base biosynthetic process; dihydroorotate dehydrogenase activity;  dihydroorotate oxidase activity; | 100 % |  |  |
| 20 | ggggacaagtttgtacaaaaaagcaggcttgatgTCGGCTGCCAAGGAGACCGC, ggggaccactttgtacaagaaagctgggtcGTGCTTCGCCGTCACCGACT | Mycgr3\|54914  JGI  Ensembl | KOG0756 Mitochondrial tricarboxylate/dicarboxylate carrier proteins; | binding; membrane; transport; | 100 % |  |  |
| 20 | ggggacaagtttgtacaaaaaagcaggcttgatgGCTTCACAACGACCATCAAA, ggggaccactttgtacaagaaagctgggtcATCCTCGCTCTCGGCAGAAA | Mycgr3\|58228  JGI  Ensembl |  |  | 100 % |  |  |
| 20 | ggggacaagtttgtacaaaaaagcaggcttgatgCCGGATGCGCGACGCATACG, ggggaccactttgtacaagaaagctgggtcCGGCCCCAACCTTTGCGTGC | Mycgr3\|63728  JGI  Ensembl | KOG2632 Rhomboid family proteins; | integral to membrane; | 100 % | chr\_11:947583 A->G; |  |
| 20 | ggggacaagtttgtacaaaaaagcaggcttgatgTCCTCCGCCGAACGACCAAC, ggggaccactttgtacaagaaagctgggtcAGAGCCGAGAGCAGCGTCCT | Mycgr3\|64873  JGI  Ensembl | KOG1712 Adenine phosphoribosyl transferases; | adenine phosphoribosyltransferase activity; adenine salvage;  nucleoside metabolic process;  transferase activity; | 100 % |  |  |
| 20 | ggggacaagtttgtacaaaaaagcaggcttgatgTCGTCAACAGTCAAGCCGCC, ggggaccactttgtacaagaaagctgggtcCCTCCCACTCAAGCGTTCCG | Mycgr3\|66140  JGI  Ensembl | KOG4178 Soluble epoxide hydrolase; | hydrolase activity; | 100 % |  |  |
| 20 | ggggacaagtttgtacaaaaaagcaggcttgatgGGCTGGTTTTGGGATAGCAA, ggggaccactttgtacaagaaagctgggtcCTTGTCCCAGCCCCATAGCC | Mycgr3\|66163  JGI  Ensembl |  |  | 100 % | chr\_1:2899073 A->G; chr\_1:2898467 A->\*; chr\_1:2898686 T->C; |  |
| 20 | ggggacaagtttgtacaaaaaagcaggcttgatgGCTGATGTCGCGGCTGCTGC, ggggaccactttgtacaagaaagctgggtcTCCATCCTTGGCAGTTGCCC | Mycgr3\|66216  JGI  Ensembl | KOG0143 Iron/ascorbate family oxidoreductases; | iron ion binding; oxidoreductase activity; | 1.61702 % |  |  |
| 20 | ggggacaagtttgtacaaaaaagcaggcttgatgAATCTCCTCTACTACTCACT, ggggaccactttgtacaagaaagctgggtcATCCTCCCACTCCTCATTCA | Mycgr3\|66596  JGI  Ensembl | KOG4504 Cation-independent mannose-6-phosphate receptor CI-MPR; |  | 100 % | chr\_1:3724210 A->G; chr\_1:3724871 T->A; |  |
| 20 | ggggacaagtttgtacaaaaaagcaggcttgatgCGTTACATCAACATCCTCGC, ggggaccactttgtacaagaaagctgggtcCGACAAGTATGCCTTGTCAA | Mycgr3\|66866  JGI  Ensembl |  |  | 0 % |  |  |
| 20 | ggggacaagtttgtacaaaaaagcaggcttgatgGCCAACCAACACCCCTCCGA, ggggaccactttgtacaagaaagctgggtcCCCCATCAATTCCTCCACAT | Mycgr3\|67094  JGI  Ensembl | KOG0768 Mitochondrial carrier protein PET8; | binding; intracellular;  membrane;  ribosome;  structural constituent of ribosome;  translation;  transport; | 100 % |  |  |
| 20 | ggggacaagtttgtacaaaaaagcaggcttgatgTCGACCAAATCCTCCAAAGT, ggggaccactttgtacaagaaagctgggtcACTCCAAGGCAAAAGACCCT | Mycgr3\|67921  JGI  Ensembl | KOG0748 Predicted membrane proteins, contain hemolysin III domain; | integral to membrane; | 100 % |  |  |
| 20 | ggggacaagtttgtacaaaaaagcaggcttgatgAGCCGACCGACTTTGATCAA, ggggaccactttgtacaagaaagctgggtcAAGCTTCGAGTCCTGCTTCA | Mycgr3\|68247  JGI  Ensembl | KOG3016 Acyl-CoA thioesterase; | acyl-CoA metabolic process; acyl-CoA thioesterase activity;  palmitoyl-CoA hydrolase activity; | 100 % |  |  |
| 20 | ggggacaagtttgtacaaaaaagcaggcttgatgGTCTCCTTCTGGCCCTTCGG, ggggaccactttgtacaagaaagctgggtcAGCCTTCTGCCTCTGTCGGG | Mycgr3\|69219  JGI  Ensembl | KOG2846 Predicted membrane protein; |  | 100 % |  |  |
| 20 | ggggacaagtttgtacaaaaaagcaggcttgatgCTAGCGAGAGTAGGCCTCGG, ggggaccactttgtacaagaaagctgggtcACCCTCTCCAAGCACGTCCT | Mycgr3\|73148  JGI  Ensembl | KOG4589 Cell division protein FtsJ; | 1-phenanthrol methyltransferase activity; 2-polyprenyl-6-methoxy-1,4-benzoquinone methyltransferase activity;  C-methyltransferase activity;  C-terminal protein carboxyl methyltransferase activity;  DNA-methyltransferase activity;  N-methyltransferase activity;  O-methyltransferase activity;  RNA methyltransferase activity;  S-methyltransferase activity;  arginine N-methyltransferase activity;  cobalt-precorrin-3 C17-methyltransferase activity;  cobalt-precorrin-5B C1-methyltransferase activity;  cobalt-precorrin-6B C5-methyltransferase activity;  cobalt-precorrin-7 C15-methyltransferase activity;  demethylmenaquinone methyltransferase activity;  dimethyladenosine transferase activity;  dimethylamine methyltransferase activity;  hydroxyneurosporene-O-methyltransferase activity;  lysine N-methyltransferase activity;  mRNA methyltransferase activity;  methanol-specific methylcobalamin:coenzyme M methyltransferase activity;  methylamine-specific methylcobalamin:coenzyme M methyltransferase activity;  methylarsonite methyltransferase activity;  monomethylamine methyltransferase activity;  protein methyltransferase activity;  protein-arginine N-methyltransferase activity;  protein-arginine N5-methyltransferase activity;  protein-leucine O-methyltransferase activity;  protein-lysine N-methyltransferase activity;  rRNA (adenine) methyltransferase activity;  rRNA (adenine-N6,N6-)-dimethyltransferase activity;  rRNA (cytosine) methyltransferase activity;  rRNA (cytosine-C5-967)-methyltransferase activity;  rRNA (guanine) methyltransferase activity;  rRNA (uridine) methyltransferase activity;  rRNA (uridine-2'-O-)-methyltransferase activity;  rRNA methyltransferase activity;  selenocysteine methyltransferase activity;  site-specific DNA-methyltransferase (cytosine-specific) activity;  tRNA (adenine)-methyltransferase activity;  tRNA (adenine-57, 58-N(1)-) methyltransferase activity;  tRNA (cytosine)-methyltransferase activity;  tRNA (guanine) methyltransferase activity;  tRNA (guanosine) methyltransferase activity;  tRNA (uracil) methyltransferase activity;  tRNA (uridine) methyltransferase activity;  tRNA methyltransferase activity;  trimethylamine methyltransferase activity; | 100 % |  |  |
| 20 | ggggacaagtttgtacaaaaaagcaggcttgatgCGGTTATTAGCAATAGCAGA, ggggaccactttgtacaagaaagctgggtcCCTCTGCCCAAACCTCCTAA | Mycgr3\|73308  JGI  Ensembl |  |  | 100 % |  |  |
| 20 | ggggacaagtttgtacaaaaaagcaggcttgatgCGTCTCCTCTCCCTCCTGAC, ggggaccactttgtacaagaaagctgggtcAAACGGAGGCAGCCAGAATG | Mycgr3\|73506  JGI  Ensembl | KOG2603 Oligosaccharyltransferase, gamma subunit; | dolichyl-diphosphooligosaccharide-protein glycotransferase activity; | 100 % | chr\_6:2379578 T->C; chr\_6:2378960 A->G; chr\_6:2378950 A->T; chr\_6:2378614 A->G; chr\_6:2379128 T->C; |  |
| 20 | ggggacaagtttgtacaaaaaagcaggcttgatgTTCACATCAAGTGATGCCCG, ggggaccactttgtacaagaaagctgggtcCAGAAGCAACCTTGAATTCG | Mycgr3\|74052  JGI  Ensembl | KOG1097 Adenine deaminase/adenosine deaminase; | adenosine deaminase activity; deaminase activity;  deoxyadenosine deaminase activity;  purine ribonucleoside monophosphate biosynthetic process; | 100 % | chr\_7:1215685 A->G; chr\_7:1216421 C->T; chr\_7:1216361 T->C; chr\_7:1215767 A->G; chr\_7:1216110 T->C; |  |
| 20 | ggggacaagtttgtacaaaaaagcaggcttgatgCCTCGCCCATATCTCAAGAC, ggggaccactttgtacaagaaagctgggtcAAGCTTGCTGCTGACCGACT | Mycgr3\|74181  JGI  Ensembl |  |  | 100 % |  |  |
| 20 | ggggacaagtttgtacaaaaaagcaggcttgatgTTCTTCGATACCCTACCCAA, ggggaccactttgtacaagaaagctgggtcGTAGTCCTCAACAAGAGCCA | Mycgr3\|74379  JGI  Ensembl |  |  | 100 % |  |  |
| 20 | ggggacaagtttgtacaaaaaagcaggcttgatgCGCCTCCTCACTTCTCTCCT, ggggaccactttgtacaagaaagctgggtcTCGCATACCCCACGCATTCG | Mycgr3\|75286  JGI  Ensembl | KOG0538 Glycolate oxidase; | L-lactate dehydrogenase (cytochrome) activity; electron transport;  oxidoreductase activity; | 100 % |  |  |
| 20 | ggggacaagtttgtacaaaaaagcaggcttgatgGCGGGCCCCGATCTCAGCCA, ggggaccactttgtacaagaaagctgggtcCTTCCTCTTCGTCCTCTTCT | Mycgr3\|75621  JGI  Ensembl | KOG2249 3'-5' exonuclease; | exonuclease activity; intracellular;  poly(A)-specific ribonuclease activity; | 3.4687 % |  |  |
| 20 | ggggacaagtttgtacaaaaaagcaggcttgatgCCAAAGATGCTCAGGATAGA, ggggaccactttgtacaagaaagctgggtcGAATGACAAGATGATCTTCT | Mycgr3\|77239  JGI  Ensembl |  |  | 0 % |  |  |
| 20 | ggggacaagtttgtacaaaaaagcaggcttgatgTCCGAAGTGCAGACCAAGAA, ggggaccactttgtacaagaaagctgggtcAGACCACTTGATCACCGGCT | Mycgr3\|77716  JGI  Ensembl | KOG0022 Alcohol dehydrogenase, class III; | alcohol dehydrogenase activity; alcohol dehydrogenase activity, iron-dependent;  alcohol dehydrogenase activity, metal ion-independent;  alcohol dehydrogenase activity, zinc-dependent;  oxidoreductase activity;  zinc ion binding; | 100 % |  |  |
| 20 | ggggacaagtttgtacaaaaaagcaggcttgatgCACCTCCCTCCACCACCGAG, ggggaccactttgtacaagaaagctgggtcCTCGCCGTCCTCCATGAATC | Mycgr3\|84034  JGI  Ensembl |  |  | 100 % |  |  |
| 20 | ggggacaagtttgtacaaaaaagcaggcttgatgCAAGCCGCCCGCTTCCACGC, ggggaccactttgtacaagaaagctgggtcAGTCTGTGCTGCTAAGAAAT | Mycgr3\|86872  JGI  Ensembl | KOG0024 Sorbitol dehydrogenase; | (R,R)-butanediol dehydrogenase activity; NAD binding;  oxidoreductase activity;  oxidoreductase activity, acting on the CH-OH group of donors, NAD or NADP as acceptor; zinc ion binding; | 100 % |  |  |
| 20 | ggggacaagtttgtacaaaaaagcaggcttgatgTCCGCAGTCACTGATGTTCT, ggggaccactttgtacaagaaagctgggtcGAAGTTTTGTGCAATGCCCG | Mycgr3\|88276  JGI  Ensembl | KOG4157 beta-1,6-N-acetylglucosaminyltransferase, contains WSC domain; | exo-alpha-sialidase activity; | 100 % | chr\_12:808900 T->C; chr\_12:809735 G->A; chr\_12:809240 C->T; chr\_12:809308 T->C; chr\_12:809812 A->G; |  |
| 20 | ggggacaagtttgtacaaaaaagcaggcttgatgTCCACCCCCCGCCCAACCTG, ggggaccactttgtacaagaaagctgggtcCGCCCAAGCCCCAATCAAAC | Mycgr3\|88667  JGI  Ensembl | KOG1558 Fe2+/Zn2+ regulated transporter; | acid phosphatase activity; membrane; metal ion transmembrane transporter activity;  metal ion transport; | 100 % |  |  |
| 20 | ggggacaagtttgtacaaaaaagcaggcttgatgCCGTCCGCCAGAGAGCGGTC, ggggaccactttgtacaagaaagctgggtcATCCTCCTCCTCCTCCCCCT | Mycgr3\|88827  JGI  Ensembl |  |  | 0 % |  |  |
| 20 | ggggacaagtttgtacaaaaaagcaggcttgatgGCCCCTCTACTTCGATCTAT, ggggaccactttgtacaagaaagctgggtcGTAGCAATGCACCAGTTGCT | Mycgr3\|88916  JGI  Ensembl |  |  | 0 % |  |  |
| 20 | ggggacaagtttgtacaaaaaagcaggcttgatgTCTTCCCTCTGGTCTTGCGC, ggggaccactttgtacaagaaagctgggtcAAGCTTCTGTGCCTTGCCCT | Mycgr3\|88932  JGI  Ensembl | KOG1721 FOG: Zn-finger; |  | 100 % |  |  |
| 20 | ggggacaagtttgtacaaaaaagcaggcttgatgGCAACGCCTCAGAACGAGGC, ggggaccactttgtacaagaaagctgggtcAGCCGAGCCAAACCCCCTAC | Mycgr3\|89152  JGI  Ensembl | KOG4475 FOG: Immunoglobin and related proteins; |  | 100 % |  |  |
| 20 | ggggacaagtttgtacaaaaaagcaggcttgatgGCGAACGCCGTGGACATGGA, ggggaccactttgtacaagaaagctgggtcGCCAAAGCTTGTGACCGTCC | Mycgr3\|89497  JGI  Ensembl | KOG4307 RNA binding protein RBM12/SWAN; |  | 100 % |  |  |
| 20 | ggggacaagtttgtacaaaaaagcaggcttgatgCTGAGTGTCTTGGAAACCGT, ggggaccactttgtacaagaaagctgggtcGATGCGGTAGCTATGGTGGA | Mycgr3\|90682  JGI  Ensembl | KOG0017 FOG: Transposon-encoded proteins with TYA, reverse transcriptase, integrase domains in various combinations; |  | 100 % |  |  |
| 20 | ggggacaagtttgtacaaaaaagcaggcttgatgCTCTCCCTCTTCCGCACCCT, ggggaccactttgtacaagaaagctgggtcCCGAGAGACAATGCCGGTAG | Mycgr3\|90788  JGI  Ensembl |  |  | 100 % | chr\_2:1974087 C->T; chr\_2:1974160 A->G; chr\_2:1974710 C->T; |  |
| 20 | ggggacaagtttgtacaaaaaagcaggcttgatgAACAATCAACAGTACGCACA, ggggaccactttgtacaagaaagctgggtcTTCCGCTGCAGGAGAATTCG | Mycgr3\|90984  JGI  Ensembl | KOG1721 FOG: Zn-finger; | intracellular; nucleic acid binding;  zinc ion binding; | 100 % |  |  |
| 20 | ggggacaagtttgtacaaaaaagcaggcttgatgAAGGCCAACTTTTTGAAAGG, ggggaccactttgtacaagaaagctgggtcGTCCTCCTCCGTCAGGTCGA | Mycgr3\|91546  JGI  Ensembl |  |  | 100 % | chr\_3:1162194 G->A; chr\_3:1161443 A->G; |  |
| 20 | ggggacaagtttgtacaaaaaagcaggcttgatgGCGAATGGTGCAAGCTGTGT, ggggaccactttgtacaagaaagctgggtcGACGTTCTGCGCCACAGGAG | Mycgr3\|91738  JGI  Ensembl | KOG1815 Predicted E3 ubiquitin ligase; | protein binding; zinc ion binding; | 0 % |  |  |
| 20 | ggggacaagtttgtacaaaaaagcaggcttgatgTATCTCTCCACCCTTGATCT, ggggaccactttgtacaagaaagctgggtcGCTCCGCCGGGAGAATATCG | Mycgr3\|92063  JGI  Ensembl |  |  | 0 % |  |  |
| 20 | ggggacaagtttgtacaaaaaagcaggcttgatgGACTACTCGCAGCATTATCC, ggggaccactttgtacaagaaagctgggtcGCCATGCAGACTGGTGATTG | Mycgr3\|92874  JGI  Ensembl |  |  | 100 % |  |  |
| 20 | ggggacaagtttgtacaaaaaagcaggcttgatgTCTGCAGACACACCGACTGC, ggggaccactttgtacaagaaagctgggtcGAACAGGGGAGCGATGACAC | Mycgr3\|93019  JGI  Ensembl |  |  | 100 % | chr\_5:465638 T->C; chr\_5:465350 T->C; |  |
| 20 | ggggacaagtttgtacaaaaaagcaggcttgatgACACTCGCTGCCATCGACGT, ggggaccactttgtacaagaaagctgggtcGTCGCCATCAGAGCTGTCCT | Mycgr3\|94264  JGI  Ensembl | KOG0670 U4/U6-associated splicing factor PRP4; |  | 100 % |  |  |
| 20 | ggggacaagtttgtacaaaaaagcaggcttgatgCATTTCATCACCAAGCTGCT, ggggaccactttgtacaagaaagctgggtcGAGTCGAGTGCGCAAGAGGA | Mycgr3\|94555  JGI  Ensembl |  |  | 1.67785 % |  |  |
| 20 | ggggacaagtttgtacaaaaaagcaggcttgatgGCGCCCAGAATCCAGCTTCC, ggggaccactttgtacaagaaagctgggtcTGCGGCGGTTTCGTCAAAAG | Mycgr3\|94682  JGI  Ensembl | KOG2177 Predicted E3 ubiquitin ligase; | protein binding; zinc ion binding; | 1.72555 % |  |  |
| 20 | ggggacaagtttgtacaaaaaagcaggcttgatgTCCCACAAGCGCGCGAGCCT, ggggaccactttgtacaagaaagctgggtcGTCCATCTCATTTGACATGT | Mycgr3\|94892  JGI  Ensembl |  |  | 100 % |  |  |
| 20 | ggggacaagtttgtacaaaaaagcaggcttgatgTTCCTCACCACCATTACCCT, ggggaccactttgtacaagaaagctgggtcTCTGTCGCTTTTCGGTCCTT | Mycgr3\|95108  JGI  Ensembl |  |  | 100 % |  |  |
| 20 | ggggacaagtttgtacaaaaaagcaggcttgatgCTTATCACTATCATCAGTAT, ggggaccactttgtacaagaaagctgggtcCTCCTGTTCAACATCCACCA | Mycgr3\|96505  JGI  Ensembl |  |  | 0 % |  |  |
| 20 | ggggacaagtttgtacaaaaaagcaggcttgatgCCTACCAAACGCTACAAGTC, ggggaccactttgtacaagaaagctgggtcGAGCTCTCGCAAGGCCCGCG | Mycgr3\|97455  JGI  Ensembl |  |  | 100 % |  |  |
| 20 | ggggacaagtttgtacaaaaaagcaggcttgatgTCGGGGGAAGACGATACCTT, ggggaccactttgtacaagaaagctgggtcCTCAGGCTTCATTCGCTTCG | Mycgr3\|97502  JGI  Ensembl | KOG0118 FOG: RRM domain; |  | 100 % | chr\_13:968124 T->C; chr\_13:967553 T->C; chr\_13:967796 T->C; chr\_13:968190 A->T; |  |
| 20 | ggggacaagtttgtacaaaaaagcaggcttgatgATGCCAGCCATGGACGAGCG, ggggaccactttgtacaagaaagctgggtcTCGCCCTCCGCATCCGCACA | Mycgr3\|97532  JGI  Ensembl |  |  | 100 % |  |  |
| 20 | ggggacaagtttgtacaaaaaagcaggcttgatgCCGCGTTTCGCGGCCGAAGG, ggggaccactttgtacaagaaagctgggtcCCAGCCCTTCTGAGTAGGCA | Mycgr3\|97669  JGI  Ensembl |  |  | 100 % | chr\_15:394668 T->C; chr\_15:394180 A->G; chr\_15:394226 A->G; |  |
| 20 | ggggacaagtttgtacaaaaaagcaggcttgatgTCGGGACCGTATCTGTCGAG, ggggaccactttgtacaagaaagctgggtcTGCATCTCCAATGATGTTCA | Mycgr3\|97731  JGI  Ensembl |  |  | 100 % |  |  |
| 20 | ggggacaagtttgtacaaaaaagcaggcttgatgCCTCTCGCTCCTTTTGCCGT, ggggaccactttgtacaagaaagctgggtcCTCGCAGAGCGGCACATCAT | Mycgr3\|98417  JGI  Ensembl | KOG0257 Kynurenine aminotransferase, glutamine transaminase K; | 1-aminocyclopropane-1-carboxylate synthase activity; 6-aminohexanoate transaminase activity;  N2-acetyl-L-lysine aminotransferase activity;  acetyldiaminopimelate aminotransferase activity;  aspartate-2-keto-4-methylthiobutyrate transaminase activity;  biosynthetic process;  branched-chain amino acid:2-keto-4-methylthiobutyrate aminotransferase activity; pyridoxal phosphate binding;  transferase activity, transferring nitrogenous groups; | 100 % |  |  |
| 20 | ggggacaagtttgtacaaaaaagcaggcttgatgGCCTCCAACAAACCCTTCAA, ggggaccactttgtacaagaaagctgggtcTACTTCGATGGTCCGGCCTT | Mycgr3\|99108  JGI  Ensembl | KOG2742 Predicted oxidoreductase; | electron transport; metabolic process;  oxidoreductase activity; | 100 % |  |  |
| 20 | ggggacaagtttgtacaaaaaagcaggcttgatgTCTGCTCAAGAGAACGCCGA, ggggaccactttgtacaagaaagctgggtcCTGCACTCCATTTGTCTTCC | Mycgr3\|99130  JGI  Ensembl |  |  | 100 % |  |  |
| 20 | ggggacaagtttgtacaaaaaagcaggcttgatgCCTCAGATACCACGGTCGGC, ggggaccactttgtacaagaaagctgggtcTTGCTTCTTCTCGAAGATGA | Mycgr3\|99255  JGI  Ensembl |  |  | 100 % |  |  |
| 20 | ggggacaagtttgtacaaaaaagcaggcttgatgACCACCACCGCCCACCCATC, ggggaccactttgtacaagaaagctgggtcACCCTTCAAATACTTCTCCG | Mycgr3\|100653  JGI  Ensembl |  |  | 100 % |  |  |
| 20 | ggggacaagtttgtacaaaaaagcaggcttgatgGTCGCAGGATCTTGCCTCAT, ggggaccactttgtacaagaaagctgggtcCTTCCTCTTCAGCGCCTCCA | Mycgr3\|101275  JGI  Ensembl | KOG1371 UDP-glucose 4-epimerase/UDP-sulfoquinovose synthase; | UDP-glucose 4-epimerase activity; catalytic activity;  cellular metabolic process;  coenzyme binding;  galactose metabolic process; | 100 % |  |  |
| 20 | ggggacaagtttgtacaaaaaagcaggcttgatgTCTGCTGATCCGGACTCTGC, ggggaccactttgtacaagaaagctgggtcGCGGTCTTGGCAGTGGAAGC | Mycgr3\|104082  JGI  Ensembl |  |  | 100 % |  |  |
| 20 | ggggacaagtttgtacaaaaaagcaggcttgatgTCGACCATTCCACAGCAATA, ggggaccactttgtacaagaaagctgggtcAACATTCTGCTTCTCACGCT | Mycgr3\|104351  JGI  Ensembl |  |  | 100 % |  |  |
| 20 | ggggacaagtttgtacaaaaaagcaggcttgatgTCATACAACGACGACGAGCG, ggggaccactttgtacaagaaagctgggtcCTTCATGAACTTGCTGGCCA | Mycgr3\|104653  JGI  Ensembl | KOG0118 FOG: RRM domain; |  | 100 % |  |  |
| 20 | ggggacaagtttgtacaaaaaagcaggcttgatgTCCACCGCCAAGTCCCCCCT, ggggaccactttgtacaagaaagctgggtcCTTCAACGTGGTCTTCGCCT | Mycgr3\|105021  JGI  Ensembl | KOG1575 Voltage-gated shaker-like K+ channel, subunit beta/KCNAB; | (R)-2-hydroxyglutarate dehydrogenase activity; (R)-2-hydroxyisocaproate dehydrogenase activity;  2-hydroxytetrahydrofuran dehydrogenase activity;  3-keto sterol reductase activity;  3-ketoglucose-reductase activity;  5-exo-hydroxycamphor dehydrogenase activity;  C-3 sterol dehydrogenase (C-4 sterol decarboxylase) activity;  aldo-keto reductase activity;  epoxide dehydrogenase activity;  gluconate dehydrogenase activity;  isocitrate dehydrogenase activity;  mevaldate reductase activity;  oxidoreductase activity;  phenylcoumaran benzylic ether reductase activity;  steroid dehydrogenase activity;  steroid dehydrogenase activity, acting on the CH-CH group of donors;  steroid dehydrogenase activity, acting on the CH-OH group of donors, NAD or NADP as acceptor; | 100 % |  |  |
| 20 | ggggacaagtttgtacaaaaaagcaggcttgatgTCGAAAACAGTCTCCATCAG, ggggaccactttgtacaagaaagctgggtcTTGCTCATCACCGACCTTCT | Mycgr3\|105124  JGI  Ensembl | KOG0908 Thioredoxin-like protein; | cell redox homeostasis; electron carrier activity;  glycerol ether metabolic process;  protein disulfide oxidoreductase activity;  thioredoxin-disulfide reductase activity; | 100 % |  |  |
| 20 | ggggacaagtttgtacaaaaaagcaggcttgatgGCTCCTGCTATGTACGAGAT, ggggaccactttgtacaagaaagctgggtcTACTTGTGCAGTACGTCTAC | Mycgr3\|107335  JGI  Ensembl | KOG1573 Aldehyde reductase; | carbohydrate metabolic process; hydrolase activity, hydrolyzing O-glycosyl compounds;  inositol oxygenase activity; | 100 % |  |  |
| 20 | ggggacaagtttgtacaaaaaagcaggcttgatgGGTCTCTTGTCGAGCCTTGT, ggggaccactttgtacaagaaagctgggtcCCGCGGCGAGCCAAATTCCA | Mycgr3\|107494  JGI  Ensembl |  |  | 100 % |  |  |
| 20 | ggggacaagtttgtacaaaaaagcaggcttgatgCCAAAGAAGCGCTCCTTCAA, ggggaccactttgtacaagaaagctgggtcGGTCTTGGTCAAGGCCAGAA | Mycgr3\|108514  JGI  Ensembl |  |  | 100 % |  |  |
| 20 | ggggacaagtttgtacaaaaaagcaggcttgatgAAGCTCTCCGCCTCTACGAT, ggggaccactttgtacaagaaagctgggtcCTTCCTCGCCGTCGGCCCAC | Mycgr3\|109278  JGI  Ensembl |  |  | 100 % |  |  |
| 20 | ggggacaagtttgtacaaaaaagcaggcttgatgGCGGCATCGCTCAGTGCGAA, ggggaccactttgtacaagaaagctgggtcGACATCAGTTTGCTTCGTGT | Mycgr3\|109332  JGI  Ensembl |  |  | 100 % | chr\_5:167561 C->T; chr\_5:167590 A->T; chr\_5:168021 A->G; chr\_5:167778 G->A; |  |
| 20 | ggggacaagtttgtacaaaaaagcaggcttgatgATGGCATCCCCTTCAATCTG, ggggaccactttgtacaagaaagctgggtcATCCCGAATACATGACGTCC | Mycgr3\|110421  JGI  Ensembl |  |  | 100 % |  |  |
| 20 | ggggacaagtttgtacaaaaaagcaggcttgatgGCGGACCAGCTCAAGAATGC, ggggaccactttgtacaagaaagctgggtcTCCCAACTTCAACCCACTCA | Mycgr3\|110438  JGI  Ensembl |  |  | 100 % |  |  |
| 20 | ggggacaagtttgtacaaaaaagcaggcttgatgTCCGCTGTACGCCCTTTCGC, ggggaccactttgtacaagaaagctgggtcCTGTCGTTTCTTCTCGACAA | Mycgr3\|111265  JGI  Ensembl |  |  | 3.30722 % |  |  |
| 20 | ggggacaagtttgtacaaaaaagcaggcttgatgGGCAACGCCTCCTCCAAACA, ggggaccactttgtacaagaaagctgggtcCACCCTCCCCGCCGGCGTCC | Mycgr3\|111415  JGI  Ensembl | KOG0161 Myosin class II heavy chain; |  | 100 % | chr\_12:309567 T->C; chr\_12:309112 T->C; chr\_12:308626 T->C; chr\_12:309425 T->C; chr\_12:308640 T->C; |  |
| 20 | ggggacaagtttgtacaaaaaagcaggcttgatgGACTACGACGATACCTCAGG, ggggaccactttgtacaagaaagctgggtcGTCGCTGGGCGTGGAAGATC | Mycgr3\|111645  JGI  Ensembl |  |  | 100 % |  |  |
| 21 | ggggacaagtttgtacaaaaaagcaggcttgatgGCTCAATCTCGTACCAAGAC, ggggaccactttgtacaagaaagctgggtcGATGACTACCACGCAGTGAA | Mycgr3\|32734  JGI  Ensembl | KOG1209 1-Acyl dihydroxyacetone phosphate reductase and related dehydrogenases; | acylglycerone-phosphate reductase activity; metabolic process;  oxidoreductase activity; | 100 % |  |  |
| 21 | ggggacaagtttgtacaaaaaagcaggcttgatgTCGCAACAACAACAAATCCC, ggggaccactttgtacaagaaagctgggtcAGTCCAAGGCGCAGTCCCGA | Mycgr3\|34966  JGI  Ensembl | KOG4453 Predicted ER membrane protein; | membrane; phosphatidate cytidylyltransferase activity;  phospholipid biosynthetic process; | 1.69082 % |  |  |
| 21 | ggggacaagtttgtacaaaaaagcaggcttgatgCCCGCCAAACTCCCCTACCA, ggggaccactttgtacaagaaagctgggtcTGCTCTCCCTGGCCACAACT | Mycgr3\|36801  JGI  Ensembl |  |  | 100 % |  |  |
| 21 | ggggacaagtttgtacaaaaaagcaggcttgatgGAATACAATGATTCGTCTCG, ggggaccactttgtacaagaaagctgggtcCCGAGCTCTCGATGTTGGAG | Mycgr3\|37647  JGI  Ensembl | KOG2510 SWI-SNF chromatin-remodeling complex protein; |  | 100 % |  |  |
| 21 | ggggacaagtttgtacaaaaaagcaggcttgatgGCGAACGAGGAGCGGCCGTA, ggggaccactttgtacaagaaagctgggtcAGTATCTTTGATGCTGCTTC | Mycgr3\|38006  JGI  Ensembl | KOG2847 Phosphate acyltransferase; | acyltransferase activity; metabolic process; | 100 % |  |  |
| 21 | ggggacaagtttgtacaaaaaagcaggcttgatgTCTCAAGATCAACGCCAATA, ggggaccactttgtacaagaaagctgggtcGGGCATGGTCTTGACTTCAG | Mycgr3\|39200  JGI  Ensembl | KOG2733 Uncharacterized membrane protein; |  | 100 % | chr\_3:761443 G->\*; |  |
| 21 | ggggacaagtttgtacaaaaaagcaggcttgatgTCAAAAATCATCCTAGCCAT, ggggaccactttgtacaagaaagctgggtcTTTGCCCCTCAAGCTATCCG | Mycgr3\|39578  JGI  Ensembl |  |  | 3.25397 % |  |  |
| 21 | ggggacaagtttgtacaaaaaagcaggcttgatgACAACAGGCGAGCCAACGGT, ggggaccactttgtacaagaaagctgggtcGCTGGATGTTGCATCTGTGG | Mycgr3\|40719  JGI  Ensembl |  |  | 0 % |  |  |
| 21 | ggggacaagtttgtacaaaaaagcaggcttgatgGGCGCCCTTCGCAACATCGT, ggggaccactttgtacaagaaagctgggtcGTTGCCTCTGAACACCTCTA | Mycgr3\|41304  JGI  Ensembl | KOG1312 DHHC-type Zn-finger proteins; | zinc ion binding; | 100 % |  |  |
| 21 | ggggacaagtttgtacaaaaaagcaggcttgatgACACCATATACTGAGACCAA, ggggaccactttgtacaagaaagctgggtcAACCTCAATCCCCAACCCCC | Mycgr3\|43806  JGI  Ensembl |  |  | 100 % |  |  |
| 21 | ggggacaagtttgtacaaaaaagcaggcttgatgAGGTTACTGCTTGGAAGATG, ggggaccactttgtacaagaaagctgggtcCTTCTTCTCCTCGCCCTTGC | Mycgr3\|45102  JGI  Ensembl | KOG2275 Aminoacylase ACY1 and related metalloexopeptidases; | acetylornithine deacetylase activity; metallopeptidase activity;  proteolysis; | 1.80251 % |  |  |
| 21 | ggggacaagtttgtacaaaaaagcaggcttgatgTCCACCGCAACCAACCCTCC, ggggaccactttgtacaagaaagctgggtcAAGCTTCCTCATCCCCATCT | Mycgr3\|48057  JGI  Ensembl | KOG2614 Kynurenine 3-monooxygenase and related flavoprotein monooxygenases; | metabolic process; oxidoreductase activity; | 100 % |  |  |
| 21 | ggggacaagtttgtacaaaaaagcaggcttgatgCCTCCCATCAGAGTCGGCGT, ggggaccactttgtacaagaaagctgggtcGCTCCACGCCACAGTCAGCC | Mycgr3\|48601  JGI  Ensembl | KOG2742 Predicted oxidoreductase; | electron transport; metabolic process;  oxidoreductase activity; | 100 % |  |  |
| 21 | ggggacaagtttgtacaaaaaagcaggcttgatgATTTTCTTCCACTCCCTCTT, ggggaccactttgtacaagaaagctgggtcCTTTTGAACCTTCGGCAGCA | Mycgr3\|50623  JGI  Ensembl | KOG3159 Lipoate-protein ligase A; | UDP-N-acetylmuramoylalanyl-D-glutamyl-2,6-diaminopimelate-D-alanyl-D-alanine ligase activity; catalytic activity;  coenzyme F420-0 gamma-glutamyl ligase activity;  coenzyme F420-2 alpha-glutamyl ligase activity;  protein modification process;  ribosomal S6-glutamic acid ligase activity; | 100 % |  |  |
| 21 | ggggacaagtttgtacaaaaaagcaggcttgatgGCCTTCCAACCGTACACTCC, ggggaccactttgtacaagaaagctgggtcCGCACGCCTACCAAAGAACC | Mycgr3\|52686  JGI  Ensembl |  |  | 100 % |  |  |
| 21 | ggggacaagtttgtacaaaaaagcaggcttgatgCCTCAGCGAATCCCCGAAGC, ggggaccactttgtacaagaaagctgggtcCCTGCAGCAGTAGCCACCGT | Mycgr3\|64923  JGI  Ensembl | KOG0725 Reductases with broad range of substrate specificities; | carbonyl reductase (NADPH) activity; metabolic process;  oxidoreductase activity; | 100 % |  |  |
| 21 | ggggacaagtttgtacaaaaaagcaggcttgatgGCAGAACCGATGTCCATCTT, ggggaccactttgtacaagaaagctgggtcCTTGGACCCTCCGTCTTGCC | Mycgr3\|64973  JGI  Ensembl | KOG3106 ER lumen protein retaining receptor; | ER retention sequence binding; integral to membrane;  protein retention in ER; | 100 % |  |  |
| 21 | ggggacaagtttgtacaaaaaagcaggcttgatgACTGCGATATTCTCTCAACC, ggggaccactttgtacaagaaagctgggtcCAGATTGATATACAGCGGCA | Mycgr3\|65233  JGI  Ensembl |  |  | 100 % |  |  |
| 21 | ggggacaagtttgtacaaaaaagcaggcttgatgGACTCCCGGGCCGACGTTCA, ggggaccactttgtacaagaaagctgggtcATGCTTTTCACCATCGAAGA | Mycgr3\|65279  JGI  Ensembl |  |  | 100 % |  |  |
| 21 | ggggacaagtttgtacaaaaaagcaggcttgatgGACAGCCTCCAAGAGAAATA, ggggaccactttgtacaagaaagctgggtcCTCCCCATCAGCAAATCCCC | Mycgr3\|67922  JGI  Ensembl | KOG4178 Soluble epoxide hydrolase; | catalytic activity; epoxide hydrolase activity; | 100 % |  |  |
| 21 | ggggacaagtttgtacaaaaaagcaggcttgatgTCTCCTCGGGAAGATGACTG, ggggaccactttgtacaagaaagctgggtcTTCTGCAAGAGGAGTCCCAT | Mycgr3\|68707  JGI  Ensembl | KOG0134 NADH:flavin oxidoreductase/12-oxophytodienoate reductase; | FMN binding; NADPH dehydrogenase activity;  oxidoreductase activity; | 100 % |  |  |
| 21 | ggggacaagtttgtacaaaaaagcaggcttgatgACAGATCTCACAGTGGGTGT, ggggaccactttgtacaagaaagctgggtcGATCGCTTCAACAAGACACC | Mycgr3\|69702  JGI  Ensembl | KOG3210 Imidazoleglycerol-phosphate synthase subunit H-like; |  | 0 % |  |  |
| 21 | ggggacaagtttgtacaaaaaagcaggcttgatgGCGAGACCAACAACCTCAAA, ggggaccactttgtacaagaaagctgggtcCTCAACCTCCAACTTCTCCA | Mycgr3\|69926  JGI  Ensembl | KOG3980 RNA 3'-terminal phosphate cyclase; | RNA-3'-phosphate cyclase activity; | 100 % |  |  |
| 21 | ggggacaagtttgtacaaaaaagcaggcttgatgCCGCCTCAGATCAAGCAAGA, ggggaccactttgtacaagaaagctgggtcCTCCCCAGCCATAGCAGCAT | Mycgr3\|70110  JGI  Ensembl | KOG0153 Predicted RNA-binding protein (RRM superfamily); | nucleic acid binding; | 100 % |  |  |
| 21 | ggggacaagtttgtacaaaaaagcaggcttgatgATCTCCAAGCCATCTCGACG, ggggaccactttgtacaagaaagctgggtcGGTGGAGTACCTCGCAGCGA | Mycgr3\|70396  JGI  Ensembl |  |  | 100 % |  |  |
| 21 | ggggacaagtttgtacaaaaaagcaggcttgatgGTCGCGACACGAAATCACCC, ggggaccactttgtacaagaaagctgggtcCCTGACTTTCTTGCTTTCTC | Mycgr3\|70745  JGI  Ensembl |  |  | 100 % |  |  |
| 21 | ggggacaagtttgtacaaaaaagcaggcttgatgGAGCCCTTCGAATACACTCC, ggggaccactttgtacaagaaagctgggtcCAGATCTGATCTAGCCTCCT | Mycgr3\|71772  JGI  Ensembl | KOG3857 Alcohol dehydrogenase, class IV; | alcohol dehydrogenase activity; alcohol dehydrogenase activity, iron-dependent;  alcohol dehydrogenase activity, metal ion-independent;  alcohol dehydrogenase activity, zinc-dependent;  metabolic process;  metal ion binding;  oxidoreductase activity; | 100 % |  |  |
| 21 | ggggacaagtttgtacaaaaaagcaggcttgatgGTTCTCGAAGCAATCCGCTA, ggggaccactttgtacaagaaagctgggtcCAGCCCGCCAACAGTGCTTG | Mycgr3\|73955  JGI  Ensembl | KOG1468 Predicted translation initiation factor related to eIF-2B alpha/beta/delta subunits (CIG2/IDI2); | cellular biosynthetic process; cellular metabolic process; | 0 % |  |  |
| 21 | ggggacaagtttgtacaaaaaagcaggcttgatgATGCAAGCGGAAACAATATC, ggggaccactttgtacaagaaagctgggtcCTGCTTCGTCGGTCTCCCTG | Mycgr3\|74109  JGI  Ensembl | KOG1198 Zinc-binding oxidoreductase; | NADPH:quinone reductase activity; oxidoreductase activity;  zinc ion binding; | 100 % |  |  |
| 21 | ggggacaagtttgtacaaaaaagcaggcttgatgGTTGCAGACCCGGCAGAAGG, ggggaccactttgtacaagaaagctgggtcAAGATCTACCATAGAAGCAG | Mycgr3\|74152  JGI  Ensembl | KOG1162 Predicted small molecule transporter; | integral to membrane; | 100 % |  |  |
| 21 | ggggacaagtttgtacaaaaaagcaggcttgatgTCCACTCCACGAATTCACGA, ggggaccactttgtacaagaaagctgggtcTTCTTTGGAATGCTCAACAC | Mycgr3\|74162  JGI  Ensembl | KOG3010 Methyltransferase; |  | 100 % |  |  |
| 21 | ggggacaagtttgtacaaaaaagcaggcttgatgCCGAAATCTGTCAATCCTCT, ggggaccactttgtacaagaaagctgggtcCGTTCCACGAACGGCCACAC | Mycgr3\|74207  JGI  Ensembl |  |  | 100 % |  |  |
| 21 | ggggacaagtttgtacaaaaaagcaggcttgatgGCTCAAACTGCTCCCGCTCA, ggggaccactttgtacaagaaagctgggtcCTTCTCCTCTTCATCGTCAA | Mycgr3\|74353  JGI  Ensembl | KOG4041 Protein phosphatase 1, regulatory (inhibitor) subunit PPP1R2; |  | 100 % |  |  |
| 21 | ggggacaagtttgtacaaaaaagcaggcttgatgCGACGAGGTGTAGCCATATT, ggggaccactttgtacaagaaagctgggtcAGGCTCATGGCGATTTATCA | Mycgr3\|74362  JGI  Ensembl |  |  | 100 % |  |  |
| 21 | ggggacaagtttgtacaaaaaagcaggcttgatgCAATTCCACCTCATCACCCT, ggggaccactttgtacaagaaagctgggtcCTCCCCACTCCGATACTGCT | Mycgr3\|76021  JGI  Ensembl |  |  | 100 % |  |  |
| 21 | ggggacaagtttgtacaaaaaagcaggcttgatgTCCGAAGAAGAACCCTTCCC, ggggaccactttgtacaagaaagctgggtcAATCACGCAACTCCACCCAA | Mycgr3\|76564  JGI  Ensembl | KOG1527 Uroporphyrin III methyltransferase; | porphyrin biosynthetic process; | 100 % |  |  |
| 21 | ggggacaagtttgtacaaaaaagcaggcttgatgTCGCACTCATGGCTCATGAA, ggggaccactttgtacaagaaagctgggtcCTTCCGCAGCACAGCCTCGT | Mycgr3\|76594  JGI  Ensembl |  |  | 100 % |  |  |
| 21 | ggggacaagtttgtacaaaaaagcaggcttgatgGACTTCAAGAACCCCAACAA, ggggaccactttgtacaagaaagctgggtcCAGCCTGCTCTTCTCCGGTA | Mycgr3\|76652  JGI  Ensembl | KOG0538 Glycolate oxidase; | L-lactate dehydrogenase (cytochrome) activity; electron transport;  oxidoreductase activity; | 100 % |  |  |
| 21 | ggggacaagtttgtacaaaaaagcaggcttgatgCGTACCACCATCGCAGCCAG, ggggaccactttgtacaagaaagctgggtcAGCTGTCATCACGTCAGTCT | Mycgr3\|76704  JGI  Ensembl |  |  | 100 % |  |  |
| 21 | ggggacaagtttgtacaaaaaagcaggcttgatgCCTGGCCAATACGCCCCCTC, ggggaccactttgtacaagaaagctgggtcCATCTGCCCCTTCAACCACT | Mycgr3\|77854  JGI  Ensembl | KOG1515 Arylacetamide deacetylase; | 1-oxa-2-oxocycloheptane lactonase activity; 3,4-dihydrocoumarin hydrolase activity;  butyrolactone hydrolase activity;  lipase activity;  phospholipase activity;  sulfolactone hydrolase activity; | 100 % |  |  |
| 21 | ggggacaagtttgtacaaaaaagcaggcttgatgGTCGCGACCTATGTCCGACC, ggggaccactttgtacaagaaagctgggtcTCTCATCTCTTGCTGGCCAT | Mycgr3\|84117  JGI  Ensembl | KOG1401 Acetylornithine aminotransferase; | acetylornithine transaminase activity; arginine metabolic process;  pyridoxal phosphate binding;  transaminase activity; | 100 % |  |  |
| 21 | ggggacaagtttgtacaaaaaagcaggcttgatgGAGACGTCTGGTTTCAGCGA, ggggaccactttgtacaagaaagctgggtcGTAACTCCTCGGCAAGCCCA | Mycgr3\|85738  JGI  Ensembl | KOG0139 Short-chain acyl-CoA dehydrogenase; | acyl-CoA dehydrogenase activity; acyl-CoA reductase activity;  electron transport;  oxidoreductase activity, acting on the CH-CH group of donors;  propionyl-CoA dehydrogenase activity;  short-branched-chain-acyl-CoA dehydrogenase activity;  thiol-driven fumarate reductase activity;  very-long-chain-acyl-CoA dehydrogenase activity; | 100 % |  |  |
| 21 | ggggacaagtttgtacaaaaaagcaggcttgatgGCTGCGCCGCAAGCTCAGGT, ggggaccactttgtacaagaaagctgggtcAGCCGGCTGCCCTTTAACTT | Mycgr3\|88104  JGI  Ensembl | KOG0032 Ca2+/calmodulin-dependent protein kinase, EF-Hand protein superfamily; | ATP binding; calmodulin-dependent protein kinase activity;  protein amino acid phosphorylation;  protein kinase activity;  protein serine/threonine kinase activity;  protein-tyrosine kinase activity; | 100 % |  |  |
| 21 | ggggacaagtttgtacaaaaaagcaggcttgatgGTCAGACCAAGCACTTCGCA, ggggaccactttgtacaagaaagctgggtcATACCTCAGCATGCTCTTCC | Mycgr3\|89741  JGI  Ensembl |  |  | 100 % |  |  |
| 21 | ggggacaagtttgtacaaaaaagcaggcttgatgGCGCGGACCAAGAAGATAAA, ggggaccactttgtacaagaaagctgggtcACGCTGACTCTGTTGCTTTG | Mycgr3\|90065  JGI  Ensembl |  |  | 100 % |  |  |
| 21 | ggggacaagtttgtacaaaaaagcaggcttgatgTCGGACGACGACCAGCCCTT, ggggaccactttgtacaagaaagctgggtcATTGCCATTTGCGGGTCGAT | Mycgr3\|90453  JGI  Ensembl |  |  | 0 % |  |  |
| 21 | ggggacaagtttgtacaaaaaagcaggcttgatgGTGCTCAAGTCTCTACTCAT, ggggaccactttgtacaagaaagctgggtcAACCGGCACCACTGTATTCC | Mycgr3\|90532  JGI  Ensembl |  |  | 100 % |  |  |
| 21 | ggggacaagtttgtacaaaaaagcaggcttgatgAAGCTCTCCCTCCTGGGCTC, ggggaccactttgtacaagaaagctgggtcGTAATCCGTGGGCTCTTCGC | Mycgr3\|91242  JGI  Ensembl | KOG4754 Predicted phosphoglycerate mutase; | catalytic activity; metabolic process; | 100 % |  |  |
| 21 | ggggacaagtttgtacaaaaaagcaggcttgatgTCAGCGACTACAGAACGAAT, ggggaccactttgtacaagaaagctgggtcTACAAAGGTGCCGATCAGCC | Mycgr3\|91286  JGI  Ensembl |  |  | 0 % |  |  |
| 21 | ggggacaagtttgtacaaaaaagcaggcttgatgTCGAGCACTCTGGCCGCTGC, ggggaccactttgtacaagaaagctgggtcGCCCAACTTCAAAGCCGTTG | Mycgr3\|91436  JGI  Ensembl |  |  | 100 % |  |  |
| 21 | ggggacaagtttgtacaaaaaagcaggcttgatgTCACCCGAAATGGCAGCCTA, ggggaccactttgtacaagaaagctgggtcTGTGTCAAACGCAATAGCAG | Mycgr3\|91860  JGI  Ensembl |  |  | 100 % |  |  |
| 21 | ggggacaagtttgtacaaaaaagcaggcttgatgCGCTTCATCCAGAAGACAGT, ggggaccactttgtacaagaaagctgggtcGACAGGATGCTCGATGTCGA | Mycgr3\|92715  JGI  Ensembl |  |  | 100 % |  |  |
| 21 | ggggacaagtttgtacaaaaaagcaggcttgatgGACGACCGTAGCCTTGGGCC, ggggaccactttgtacaagaaagctgggtcGTCGTGTCTGCGATCCAAGT | Mycgr3\|93483  JGI  Ensembl | KOG0663 Protein kinase PITSLRE and related kinases; | protein binding; zinc ion binding; | 3.29412 % |  |  |
| 21 | ggggacaagtttgtacaaaaaagcaggcttgatgGCGGATCAGAAGCCCGTGAT, ggggaccactttgtacaagaaagctgggtcCCTGCCGCCTCTCCACGGCC | Mycgr3\|93485  JGI  Ensembl |  |  | 100 % |  |  |
| 21 | ggggacaagtttgtacaaaaaagcaggcttgatgCCGCCATCCAAGAAAGACAA, ggggaccactttgtacaagaaagctgggtcCTTCGTTGCAAACTTCTTCC | Mycgr3\|93921  JGI  Ensembl |  |  | 0 % |  |  |
| 21 | ggggacaagtttgtacaaaaaagcaggcttgatgTCTTCAAAGATCCCATCCGC, ggggaccactttgtacaagaaagctgggtcCTTTTTTGCTTTCTTTGGAC | Mycgr3\|94684  JGI  Ensembl |  |  | 0 % |  |  |
| 21 | ggggacaagtttgtacaaaaaagcaggcttgatgATGGCGCATGCAAGCTTTCC, ggggaccactttgtacaagaaagctgggtcCGCTCCAAATGTAAATCCTC | Mycgr3\|94838  JGI  Ensembl | KOG4456 Inner centromere protein (INCENP), C-terminal domain; |  | 100 % |  |  |
| 21 | ggggacaagtttgtacaaaaaagcaggcttgatgCTCCGAGTTCTGGCATACAT, ggggaccactttgtacaagaaagctgggtcTATGCGGTTGCACGCCGGAA | Mycgr3\|95102  JGI  Ensembl |  |  | 100 % |  |  |
| 21 | ggggacaagtttgtacaaaaaagcaggcttgatgTCCGACATCTCCAATGAACC, ggggaccactttgtacaagaaagctgggtcCAAGCAGCCTCCCTCTCCGA | Mycgr3\|95365  JGI  Ensembl |  |  | 100 % |  |  |
| 21 | ggggacaagtttgtacaaaaaagcaggcttgatgGCTTTCCGGAAACGCAATGT, ggggaccactttgtacaagaaagctgggtcAAATTCTACATCCTTACTTG | Mycgr3\|95776  JGI  Ensembl | KOG3949 RNA polymerase II elongator complex, subunit ELP4; |  | 100 % |  | ``` 1	ATGGCTTTCCGGAAACGCAATGTCGTCGTTGGTAAGCCACCAGTCGCCTCGCCGTCTAGTGAGAGTCCAGCGCCTCTAGGCGTTCGACCTTCGCCATTGACTTCTCACTCCACCACATCAACCGGCACTCCCTCCTTCGATAGCCTGCTCGGCGGGCATGGAGGTCTCGCACTCGGTTACAGTCTGCTCGTATCAGAGAGTGGCACGACAGACTTCTCGGGCACTTTGCTTCGATACTTCGCTGCGGAAGGCATCTGTCAAGAGCACCACGTACATATTGTGGGGGCCGGCGAAGGATGGGTGAAAGAACTTCCAGGCGTGGCAGAGGAGAGAAGTCGAGACAAAGCTGCGGGAAAGAGTGCCTCCGATGAGGAGAAGATGAAGATTGCGTGGAGGTACGAGAGGTTGGGACAAGCGGGAGAAAGAG	427 	\|\|\|\|\|\|\|\|\|\|\|\|\|\|\|\|\|\|\|\|\|\|\|\|\|\|\|\|\|\|\|\|\|\|\|\|\|\|\|\|\|\|\|\|\|\|\|\|\|\|\|\|\|\|\|\|\|\|\|\|\|\|\|\|\|\|\|\|\|\|\|\|\|\|\|\|\|\|\|\|\|\|\|\|\|\|\|\|\|\|\|\|\|\|\|\|\|\|\|\|\|\|\|\|\|\|\|\|\|\|\|\|\|\|\|\|\|\|\|\|\|\|\|\|\|\|\|\|\|\|\|\|\|\|\|\|\|\|\|\|\|\|\|\|\|\|\|\|\|\|\|\|\|\|\|\|\|\|\|\|\|\|\|\|\|\|\|\|\|\|\|\|\|\|\|\|\|\|\|\|\|\|\|\|\|\|\|\|\|\|\|\|\|\|\|\|\|\|\|\|\|\|\|\|\|\|\|\|\|\|\|\|\|\|\|\|\|\|\|\|\|\|\|\|\|\|\|\|\|\|\|\|\|\|\|\|\|\|\|\|\|\|\|\|\|\|\|\|\|\|\|\|\|\|\|\|\|\|\|\|\|\|\|\|\|\|\|\|\|\|\|\|\|\|\|\|\|\|\|\|\|\|\|\|\|\|\|\|\|\|\|\|\|\|\|\|\|\|\|\|\|\|\|\|\|\|\|\|\|\|\|\|\|\|\|\|\|\|\|\|\|\|\|\|\|\|\|\|\|\|\|\|\|\|\|\|\|\|\|\|\|\|\|\|\|\|\|\|\|\|\|\|\|\|\|\|\|\|\|\|\|\|\|\|\|\|\|\|\|\|\|\|\|\|\|\|\|\|\|\|\|\|\|\|\|\|\|\|\|\|\|\|\|\|\|\|\|\|\|\|\|\|\|\|\|\|\|\|\|\|\|\|\|\|\|\|\|\|\|\|\|\|\|\|\|\|\| 	ATGGCTTTCCGGAAACGCAATGTCGTCGTTGGTAAGCCACCAGTCGCCTCGCCGTCTAGTGAGAGTCCAGCGCCTCTAGGCGTTCGACCTTCGCCATTGACTTCTCACTCCACCACATCAACCGGCACTCCCTCCTTCGATAGCCTGCTCGGCGGGCATGGAGGTCTCGCACTCGGTTACAGTCTGCTCGTATCAGAGAGTGGCACGACAGACTTCTCGGGCACTTTGCTTCGATACTTCGCTGCGGAAGGCATCTGTCAAGAGCACCACGTACATATTGTGGGGGCCGGCGAAGGATGGGTGAAAGAACTTCCAGGCGTGGCAGAGGAGAGAAGTCGAGACAAAGCTGCGGGAAAGAGTGCCTCCGATGAGGAGAAGATGAAGATTGCGTGGAGGTACGAGAGGTTGGGACAAGCGGGAGAAAGAG ```      ``` 426	AGCTCTTCCTGATCGAATCGCAGCACCAGCGGCAGACGGTGTCCCAGTACCTTCGCCCTTTTGCCACACGTTTGACCTCACCAAGCGACTGGCAGTACCTACCGGCGCCAAAGTCAATCACATCCCGATATCCTTGACATCAACATTCGAGAACATCCTCCAATCTCTCGCGAAAAGCCTTGCTGCCACTCCTCCAACTAGCATTCATCGACTAGTCGTTCCTTCTCTCCTCAGTCCAGCACTCTACCCTCCAAACGCCAGCCAACCCGACCAAGTGCTAGGCTTCTTTCGCTCGCTACGAGCGCTGCTACGACAGCACCACAATCGCCTAGTCGCCATGATAACCCTACCACTGGAGCTCTACCCTCGCCACTCCGGCCTGGTTCGATGGATGGAGATCCTCAGCGATGGTGTCATCGAGCTCACGCCGTTCCCGCATCTCATGGATTCATCCAGTGCTCTCGCGGAAAGTGGTGGTGCACGTGGTAACGATGAGCAACCGCAAGGCATGGTCAAGG-TGCACAAGCTG	954 	\|\|\|\|\|\|\|\|\|\|\|\|\|\|\|\|\|\|\|\|\|\|\|\|\|\|\|\|\|\|\|\|\|\|\|\|\|\|\|\|\|\|\|\|\|\|\|\|\|\|\|\|\|\|\|\|\|\|\|\|\|\|\|\|\|\|\|\|\|\|\|\|\|\|\|\|\|\|\|\|\|\|\|\|\|\|\|\|\|\|\|\|\|\|\|\|\|\|\|\|\|\|\|\|\|\|\|\|\|\|\|\|\|\|\|\|\|\|\|\|\|\|\|\|\|\|\|\|\|\|\|\|\|\|\|\|\|\|\|\|\|\|\|\|\|\|\|\|\|\|\|\|\|\|\|\|\|\|\|\|\|\|\|\|\|\|\|\|\|\|\|\|\|\|\|\|\|\|\|\|\|\|\|\|\|\|\|\|\|\|\|\|\|\|\|\|\|\|\|\|\|\|\|\|\|\|\|\|\|\|\|\|\|\|\|\|\|\|\|\|\|\|\|\|\|\|\|\|\|\|\|\|\|\|\|\|\|\|\|\|\|\|\|\|\|\|\|\|\|\|\|\|\|\|\|\|\|\|\|\|\|\|\|\|\|\|\|\|\|\|\|\|\|\|\|\|\|\|\|\|\|\|\|\|\|\|\|\|\|\|\|\|\|\|\|\|\|\|\|\|\|\|\|\|\|\|\|\|\|\|\|\|\|\|\|\|\|\|\|\|\|\|\|\|\|\|\|\|\|\|\|\|\|\|\|\|\|\|\|\|\|\|\|\|\|\|\|\|\|\|\|\|\|\|\|\|\|\|\|\|\|\|\|\|\|\|\|\|\|\|\|\|\|\|\|\|\|\|\|\|\|\| \|\|\|\|\|\|\|\|\|\|\|\|\|\|\|\|\|\|\|\|\|\|\|\|\|\|\|\|\|\|\|\|\|\|\|\|\|\|\|\|\|\|\|\|\|\|\|\|\|\|\|\|\|\|\|\|\|\|\|\|\|\| \|\| \|\|\| \|\|\|\|\|\|\|\|\|\|\|\|\| \|\|\|\|\|\| \|\| \|\|\|\|\|\|\|\| \|\| \|\|\|\|\|\|\|\|\| \|\|\|\| \|\|\|\|\|\|\|\|\|\|\|\|\|\| \|\|\|\|\|\| \|\|\|\| 	AGCTCTTCCTGATCGAATCGCAGCACCAGCGGCAGACGGTGTCCCAGTACCTTCGCCCTTTTGCCACACGTTTGACCTCACCAAGCGACTGGCAGTACCTACCGGCGCCAAAGTCAATCACATCCCGATATCCTTGACATCAACATTCGAGAACATCCTCCAATCTCTCGCGAAAAGCCTTGCTGCCACTCCTCCAACTAGCATTCATCGACTAGTCGTTCCTTCTCTCCTCAGTCCAGCACTCTACCCTCCAAACGCCAGCCAACCCGACCAAGTGCTAGGCTTCTTTCGCTCGCTACGAGCGCTGCTACGACAGCACCACAATCGCCTAGTCGCCATGATAACCCTACCACTGGAGCTCTACCCTCGCCACTCCGGCCTGCTTCGATGGATGGAGATCCTCAGCGATGGTGTCATCGAGCTCACGCCGTTCCCGCATCTCATG-AT-CAT-CAGTGCTCTCGCG-AAAGTG-TG-TGCACGTG-TA-CGATGAGCA-CCGC-AGGCATGGTCAAGGGTGCACA-GCTG ``` |
| 21 | ggggacaagtttgtacaaaaaagcaggcttgatgACAGATCAAGATGCAGACTA, ggggaccactttgtacaagaaagctgggtcTGCTCCCGTCCTCATCGCCT | Mycgr3\|95952  JGI  Ensembl |  |  | 1.61812 % |  |  |
| 21 | ggggacaagtttgtacaaaaaagcaggcttgatgGAAAACCAGATGGATCCGCC, ggggaccactttgtacaagaaagctgggtcCTCGGCAGTGAGTCTGAAGA | Mycgr3\|96914  JGI  Ensembl |  |  | 100 % |  |  |
| 21 | ggggacaagtttgtacaaaaaagcaggcttgatgGAGCATGGATTCGTGAGCGA, ggggaccactttgtacaagaaagctgggtcAGCAGACTTATTCTTCGCAG | Mycgr3\|97383  JGI  Ensembl |  |  | 100 % |  |  |
| 21 | ggggacaagtttgtacaaaaaagcaggcttgatgCCCAATAAGGTGGTCGGTGA, ggggaccactttgtacaagaaagctgggtcGAATCGTCTTCCCAACCAAA | Mycgr3\|97840  JGI  Ensembl |  |  | 100 % | chr\_17:495319 C->T; |  |
| 21 | ggggacaagtttgtacaaaaaagcaggcttgatgGAGAAGTCGATAGGGATGGG, ggggaccactttgtacaagaaagctgggtcGTGGTCGCTTCCGGTCGCTC | Mycgr3\|98002  JGI  Ensembl |  |  | 3.04731 % |  |  |
| 21 | ggggacaagtttgtacaaaaaagcaggcttgatgGATCAATCCCGCGCCATCAA, ggggaccactttgtacaagaaagctgggtcTGCCATGCCATGATGCCTGA | Mycgr3\|98284  JGI  Ensembl | KOG3250 COP9 signalosome, subunit CSN7; |  | 100 % |  |  |
| 21 | ggggacaagtttgtacaaaaaagcaggcttgatgGAGTTCATAGTCGGTACCTT, ggggaccactttgtacaagaaagctgggtcCAACCACACTGGAGTCGCAG | Mycgr3\|99485  JGI  Ensembl |  |  | 100 % |  |  |
| 21 | ggggacaagtttgtacaaaaaagcaggcttgatgGTACTAGGCGTTGGTCCACA, ggggaccactttgtacaagaaagctgggtcCATAGTCCCTCTGCGACATG | Mycgr3\|99976  JGI  Ensembl | KOG4157 beta-1,6-N-acetylglucosaminyltransferase, contains WSC domain; |  | 100 % |  |  |
| 21 | ggggacaagtttgtacaaaaaagcaggcttgatgGCCTTCATGCTCCGGCGACC, ggggaccactttgtacaagaaagctgggtcCTTCCTTCGTCCCTGTTGCA | Mycgr3\|102019  JGI  Ensembl | KOG1630 Growth hormone-induced protein and related proteins; |  | 100 % |  |  |
| 21 | ggggacaagtttgtacaaaaaagcaggcttgatgGTCCAAAACAAAGCTCTCAT, ggggaccactttgtacaagaaagctgggtcGTGCTTGCTAACCTCCAAGA | Mycgr3\|102173  JGI  Ensembl | KOG1196 Predicted NAD-dependent oxidoreductase; | 2-alkenal reductase activity; oxidoreductase activity;  zinc ion binding; | 100 % |  |  |
| 21 | ggggacaagtttgtacaaaaaagcaggcttgatgCAGTACGCTATCGTCGCCGC, ggggaccactttgtacaagaaagctgggtcAAGGGCGGCAACGAGGCCGA | Mycgr3\|104730  JGI  Ensembl | KOG1216 von Willebrand factor and related coagulation proteins; |  | 2.9661 % |  |  |
| 21 | ggggacaagtttgtacaaaaaagcaggcttgatgTCAGTCGAACCTGGCAGCGA, ggggaccactttgtacaagaaagctgggtcATGATCAAACAACTCCTCCC | Mycgr3\|105629  JGI  Ensembl |  |  | 100 % |  |  |
| 21 | ggggacaagtttgtacaaaaaagcaggcttgatgGGCAGCATGGGCCTTCCATG, ggggaccactttgtacaagaaagctgggtcGCACCACCCATACTTCTCGT | Mycgr3\|105712  JGI  Ensembl |  |  | 100 % |  |  |
| 21 | ggggacaagtttgtacaaaaaagcaggcttgatgGCTGCTGTTCCATTGCTGAA, ggggaccactttgtacaagaaagctgggtcTGGTACATGATCATAGATAT | Mycgr3\|106332  JGI  Ensembl |  |  | 100 % |  |  |
| 21 | ggggacaagtttgtacaaaaaagcaggcttgatgCCAAAGGCACGCAAATCAGC, ggggaccactttgtacaagaaagctgggtcGGCATGCACATGGACTAGCC | Mycgr3\|106814  JGI  Ensembl |  |  | 100 % |  |  |
| 21 | ggggacaagtttgtacaaaaaagcaggcttgatgAGCAGACCGACCTTGGCGTC, ggggaccactttgtacaagaaagctgggtcGTCCTCCCGAATACACCTCG | Mycgr3\|107064  JGI  Ensembl |  |  | 2.1575 % |  |  |
| 21 | ggggacaagtttgtacaaaaaagcaggcttgatgAGCGCCAGTACGCCTATGTC, ggggaccactttgtacaagaaagctgggtcCTCCTCTTCATCAAGCTTTG | Mycgr3\|108052  JGI  Ensembl |  |  | 1.92503 % |  |  |
| 21 | ggggacaagtttgtacaaaaaagcaggcttgatgGCGCGCGACAAGAAGCGTCA, ggggaccactttgtacaagaaagctgggtcACCCTTGTTCTTCTTGCGCT | Mycgr3\|108195  JGI  Ensembl | KOG4415 Uncharacterized conserved protein; |  | 1.50435 % |  |  |
| 21 | ggggacaagtttgtacaaaaaagcaggcttgatgTCGACTCAGAACCAACCCAT, ggggaccactttgtacaagaaagctgggtcAATACTCATGCCTCTCGCGG | Mycgr3\|108594  JGI  Ensembl |  |  | 100 % |  |  |
| 21 | ggggacaagtttgtacaaaaaagcaggcttgatgTACCTATGTCGAAGCGTCTC, ggggaccactttgtacaagaaagctgggtcCGAGCCTGTGTTCGACACCC | Mycgr3\|108925  JGI  Ensembl | KOG4391 Predicted alpha/beta hydrolase BEM46; |  | 0 % |  |  |
| 21 | ggggacaagtttgtacaaaaaagcaggcttgatgTCGAAACGAAAACGGGACGA, ggggaccactttgtacaagaaagctgggtcGTCGAAGCTAATCTTCTTAC | Mycgr3\|109076  JGI  Ensembl | KOG2992 Nucleolar GTPase/ATPase p130; | phosphoenolpyruvate-dependent sugar phosphotransferase system; sugar:hydrogen symporter activity; | 3.05882 % |  |  |
| 21 | ggggacaagtttgtacaaaaaagcaggcttgatgGCCAAGGTCTCCTTCAATGA, ggggaccactttgtacaagaaagctgggtcATCTTCTCCATCCTCCAACA | Mycgr3\|109240  JGI  Ensembl | KOG2612 Predicted integral membrane protein; |  | 100 % |  |  |
| 21 | ggggacaagtttgtacaaaaaagcaggcttgatgCCGGCCCACGACGATGACCT, ggggaccactttgtacaagaaagctgggtcAGACGCAGCAACCCTCTTCT | Mycgr3\|109310  JGI  Ensembl | KOG1818 Membrane trafficking and cell signaling protein HRS, contains VHS and FYVE domains; |  | 100 % |  |  |
| 21 | ggggacaagtttgtacaaaaaagcaggcttgatgGTCGCAACAATGGACACGAC, ggggaccactttgtacaagaaagctgggtcCCTCAACGGCTCCTCACTCT | Mycgr3\|109668  JGI  Ensembl | KOG4650 Predicted steroid reductase; | cytoplasm; integral to membrane;  lipid metabolic process;  oxidoreductase activity, acting on the CH-CH group of donors; | 100 % |  |  |
| 21 | ggggacaagtttgtacaaaaaagcaggcttgatgACAGAGACACTTCATCCAAC, ggggaccactttgtacaagaaagctgggtcTTCATCCACCGCTACCTTGG | Mycgr3\|109818  JGI  Ensembl |  |  | 100 % |  |  |
| 21 | ggggacaagtttgtacaaaaaagcaggcttgatgAGCAACTATGGCTTCTTCGG, ggggaccactttgtacaagaaagctgggtcTGAGTACTGCCTTCCCTTCA | Mycgr3\|109847  JGI  Ensembl |  |  | 100 % |  |  |
| 21 | ggggacaagtttgtacaaaaaagcaggcttgatgGCGACAACAGCAGCCACTGG, ggggaccactttgtacaagaaagctgggtcGACTGCTGCAGACCTTGCTC | Mycgr3\|109866  JGI  Ensembl | KOG1520 Predicted alkaloid synthase/Surface mucin Hemomucin; |  | 100 % |  |  |
| 21 | ggggacaagtttgtacaaaaaagcaggcttgatgCTCTCTTTCTCACTATCACG, ggggaccactttgtacaagaaagctgggtcCGGCTTCGTCAACCCCAAGA | Mycgr3\|109874  JGI  Ensembl |  |  | 100 % |  |  |
| 21 | ggggacaagtttgtacaaaaaagcaggcttgatgCCCACCCCAATCGAGCTCGC, ggggaccactttgtacaagaaagctgggtcGGCCCTCGCATCCACCACCG | Mycgr3\|110463  JGI  Ensembl |  |  | 100 % |  |  |
| 21 | ggggacaagtttgtacaaaaaagcaggcttgatgTTCGTTGCGCCAACGCTTGC, ggggaccactttgtacaagaaagctgggtcTCCGGCGGCTTCCTTCTCCC | Mycgr3\|110595  JGI  Ensembl |  |  | 100 % |  |  |
| 21 | ggggacaagtttgtacaaaaaagcaggcttgatgTCCACCTTCGAACAAACCAT, ggggaccactttgtacaagaaagctgggtcATCCTCGAAGACTCTCCACT | Mycgr3\|110635  JGI  Ensembl |  |  | 100 % |  |  |
| 21 | ggggacaagtttgtacaaaaaagcaggcttgatgGACAATGTCGTGAAGCCGAA, ggggaccactttgtacaagaaagctgggtcCTCCACAAACACCCCACTAT | Mycgr3\|110851  JGI  Ensembl |  |  | 100 % |  |  |
| 21 | ggggacaagtttgtacaaaaaagcaggcttgatgTCTGTTCAACCAAAGGACAC, ggggaccactttgtacaagaaagctgggtcCTCGACCTCTTCGATCTTCT | Mycgr3\|110875  JGI  Ensembl | KOG3158 HSP90 co-chaperone p23; |  | 100 % |  |  |
| 21 | ggggacaagtttgtacaaaaaagcaggcttgatgTCTGCCTTCCACGGCATTCC, ggggaccactttgtacaagaaagctgggtcTTTCGGGCACGCAGGCGCTT | Mycgr3\|111159  JGI  Ensembl |  |  | 100 % |  |  |
| 21 | ggggacaagtttgtacaaaaaagcaggcttgatgTCTCCTATTAGACGCTCGCA, ggggaccactttgtacaagaaagctgggtcAGACGGCGTGATCTTGCCCT | Mycgr3\|111644  JGI  Ensembl |  |  | 100 % |  |  |
| 22 | ggggacaagtttgtacaaaaaagcaggcttgatgCCGCCAAGTCAGACTGTCAA, ggggaccactttgtacaagaaagctgggtcTTGCGCGGGCATGGAATTCG | Mycgr3\|31477  JGI  Ensembl | KOG1438 Anthranilate phosphoribosyltransferase; | anthranilate phosphoribosyltransferase activity; metabolic process;  transferase activity, transferring glycosyl groups;  tryptophan biosynthetic process; | 100 % |  |  |
| 22 | ggggacaagtttgtacaaaaaagcaggcttgatgGCAGGATCACAGGAGAAAGT, ggggaccactttgtacaagaaagctgggtcCGCCTTTGCGTCCTCCTCCT | Mycgr3\|32540  JGI  Ensembl |  |  | 1.65913 % |  |  |
| 22 | ggggacaagtttgtacaaaaaagcaggcttgatgTCAGCTCCACATCCCACGAC, ggggaccactttgtacaagaaagctgggtcCTGCTCTGCGTCCGGCTGCC | Mycgr3\|32957  JGI  Ensembl | KOG3178 Hydroxyindole-O-methyltransferase and related SAM-dependent methyltransferases; | O-methyltransferase activity; | 100 % |  |  |
| 22 | ggggacaagtttgtacaaaaaagcaggcttgatgTCCGTTTCAACACATCCTCC, ggggaccactttgtacaagaaagctgggtcTTTGGGTCGGTGTGGTGGTT | Mycgr3\|33249  JGI  Ensembl |  |  | 0 % |  |  |
| 22 | ggggacaagtttgtacaaaaaagcaggcttgatgGCCGACCTCTCGACCTGTCC, ggggaccactttgtacaagaaagctgggtcGCCTGCGACATTCTCCGCTG | Mycgr3\|33257  JGI  Ensembl | KOG1251 Serine racemase; | catalytic activity; metabolic process;  pyridoxal phosphate binding; | 100 % |  |  |
| 22 | ggggacaagtttgtacaaaaaagcaggcttgatgGGCGATTCTGGCGTAGCCAT, ggggaccactttgtacaagaaagctgggtcCCTCGCCACAGCTCCGTCGT | Mycgr3\|33363  JGI  Ensembl | KOG1269 SAM-dependent methyltransferases; | sterol 24-C-methyltransferase activity; | 100 % |  |  |
| 22 | ggggacaagtttgtacaaaaaagcaggcttgatgAAATCATTCATTACGTTCGC, ggggaccactttgtacaagaaagctgggtcTCTCCTCATTCCAAACCGCC | Mycgr3\|34374  JGI  Ensembl |  |  | 100 % |  | ``` 1	ATGAAATCATTCATTACGTTCGCCACGCTCTGGGCAGGCTTCACCTCCGCCGCTCCAGTATCAACCCACGTCTCTGGTGGTCAGGAGCACATTGTCATCATCGACAAATCCCACCCAACACCACCAGGAGTCGCCGAAGTCCTCAGGAGACTCGAACTCAATCCACAACACTCCGATGTCCGCCACATCTTCAACAACAGCGCCTTTCAAGGCTTCTCCGCGTCCATGAAATCCCACTGTCTCGACCTCCTCGCCAACATGACCGATGTCTCCATGGTCGAGGCCGCGATCTCCGTCACGCGTGGCGCCTCAGCTCCCTACGATACCCGACCCGATGCACCGTGGGGTCTACAAGGCATCTCGACATCCGCCTCTGTCAGCGGTAGTGCTTCCAGGTTGGACCACACCTATGCATACTCCGACTCCTCGCTCGGCACCGGCGCAGACATCTACATCGTCGACACGGGTGTCTACGCCGAACACAATATCTTCAACGGCCGCGCGAAGATGGTCTGGTCCTACAACCAAGACATGTCGGATAAAGACGGCCACGGCACCCACGTTGCCGGCACAGCAGCAGGCGACATTCTTGGCGTGGCATCCAAGGCTCGCGTATTCGGTGTCAAAGCCCTCGATGCCGATGGTGGTGGGTGGTCGTCCAATGTCGTGGCGGGCATCGACTATGTCATCAAAGCCCACGACGCACGCAAGGCCGCCTCCGACCCTTCATTCGCCGGCTCCGTAATTTCCATGTCGCTAGCTTCCACTGGTCGAGTGGAAGCCATCAGCCAAGCAGTCAATGCCGCCATCAGCGGAGGGATTCACGTCTGTGTCGCAGCAGGCAATAATGGCGAAGACGCCTGCAATTCGTCTCCCGCTTCCTCC-GGAGGCTCCCACGGCGGAGCTATCACAGTCGGCGCAGTGGACATTCGCGGTAAGAAGGCGAAGTTCTCGAACGACGGCGACTGCGTCGATG-TCTATGCTCCGGGTGTGGACATCGTGAGCTCCTGGATCGGAGAGAGGAACATGATCAATTCGCTATCCGGGACATCCATGGCGGCGCC-ACATG-TG-ACTGGGATCATCGCTTATGCCATGGCGAACCGT-ACATTGGCTGAGAACC	1120 	\|\|\|\|\|\|\|\|\|\|\|\|\|\|\|\|\|\|\|\|\|\|\|\|\|\|\|\|\|\|\|\|\|\|\|\|\|\|\|\|\|\|\|\|\|\|\|\|\|\|\|\|\|\|\|\|\|\|\|\|\|\|\|\|\|\|\|\|\|\|\|\|\|\|\|\|\|\|\|\|\|\|\|\|\|\|\|\|\|\|\|\|\|\|\|\|\|\|\|\|\|\|\|\|\|\|\|\|\|\|\|\|\|\|\|\|\|\|\|\|\|\|\|\|\|\|\|\|\|\|\|\|\|\|\|\|\|\|\|\|\|\|\|\|\|\|\|\|\|\|\|\|\|\|\|\|\|\|\|\|\|\|\|\|\|\|\|\|\|\|\|\|\|\|\|\|\|\|\|\|\|\|\|\|\|\|\|\|\|\|\|\|\|\|\|\|\|\|\|\|\|\|\|\|\|\|\|\|\|\|\|\|\|\|\|\|\|\|\|\|\|\|\|\|\|\|\|\|\|\|\|\|\|\|\|\|\|\|\|\|\|\|\|\|\|\|\|\|\|\|\|\|\|\|\|\|\|\|\|\|\|\|\|\|\|\|\|\|\|\|\|\|\|\|\|\|\|\|\|\|\|\|\|\|\|\|\|\|\|\|\|\|\|\|\|\|\|\|\|\|\|\|\|\|\|\|\|\|\|\|\|\|\|\|\|\|\|\|\|\|\|\|\|\|\|\|\|\|\|\|\|\|\|\|\|\|\|\|\|\|\|\|\|\|\|\|\|\|\|\|\|\|\|\|\|\|\|\|\|\|\|\|\|\|\|\|\|\|\|\|\|\|\|\|\|\|\|\|\|\|\|\|\|\|\|\|\|\|\|\|\|\|\|\|\|\|\|\|\|\|\|\|\|\|\|\|\|\|\|\|\|\|\|\|\|\|\|\|\|\|\|\|\|\|\|\|\|\|\|\|\|\|\|\|\|\|\|\|\|\|\|\|\|\|\|\|\|\|\|\|\|\|\|\|\|\|\|\|\|\|\|\|\|\|\|\|\|\|\|\|\|\|\|\|\|\|\|\|\|\|\|\|\|\|\|\|\|\|\|\|\|\|\|\|\|\|\|\|\|\|\|\|\|\|\|\|\|\|\|\|\|\|\|\|\|\|\|\|\|\|\|\|\|\|\|\|\|\|\|\|\|\|\|\|\|\|\|\|\|\|\|\|\|\|\|\|\|\|\|\|\|\|\|\|\|\|\|\|\|\|\|\|\|\|\|\|\|\|\|\|\|\|\|\|\|\|\|\|\|\|\|\|\|\|\|\|\|\|\|\|\|\|\|\|\|\|\|\|\|\|\|\|\|\|\|\|\|\|\|\|\|\|\|\|\|\|\|\|\|\|\|\|\|\|\|\|\|\|\|\|\|\|\|\|\|\|\|\|\|\|\|\|\|\|\|\|\|\|\|\|\|\|\|\|\|\|\|\|\|\|\|\|\|\|\|\|\|\|\|\|\|\|\|\|\|\|\|\|\|\|\|\|\|\|\|\|\|\|\|\|\|\|\|\|\|\|\|\|\|\|\|\|\|\|\|\|\|\|\|\|\|\|\|\|\|\|\|\|\|\|\|\|\|\|\|\|\|\|\|\|\|\|\|\|\|\|\|\|\|\|\|\|\|\|\|\|\|\|\|\|\|\|\|\|\|\|\|\|\|\|\|\|\|\|\|\|\|\|\|\|\|\|\|\|\|\|\|\|\|\|\|\|\|\|\|\|\|\|\|\|\|\|\|\|\|\|\|\|\|\|\|\|\|\|\|\|\|\|\|\|\|\|\|\|\|\|\|\|\|\|\|\|\|\|\|\|\|\|\|\|\|\|\|\|\|\|\|\|\|\|\|\|\|\|\|\|\|\|\|\|\|\|\|\|\|\|\|\|\|\|\|\|\|\|\|\|\|\|\|\|\|\|\|\|\|\|\|\|\|\|\|\|\|\|\| \|\|\|\|\|\|\|\|\|\|\|\|\|\|\|\|\|\|\|\|\|\|\|\|\|\|\|\|\|\|\|\|\|\|\|\|\|\| \|\|\|\|\|\|\|\|\|\|\|\|\|\|\|\|\|\|\|\|\|\|\|\|\|\|\|\|\|\| \|\|\|\|\|\|\|\|\|\|\|\|\|\|\|\|\|\|\|\|\| \|\|\|\|\|\|\|\|\|\|\|\|\|\|\|\|\|\|\|\|\|\|\|\|\|\|\|\|\|\|\| \| \|\|\|\|\|\|\|\|\|\|\|\|\|\| \|\|\|\|\|\|\|\| \| \|\|\|\|\|\|\| \|\|\|\|\|\|\|\| \|\|\|\|\|\|\|\|\|\|\| \|\|\|\|\| \|\| \|\|\|\|\|\|\|\|\|\|\|\|\|\|\| \|\|\|\| \|\|\| \|\|\|   \|\| \|\|\|\|\|\|\|\|\|\|\| \|\|\|\| 	ATGAAATCATTCATTACGTTCGCCACGCTCTGGGCAGGCTTCACCTCCGCCGCTCCAGTATCAACCCACGTCTCTGGTGGTCAGGAGCACATTGTCATCATCGACAAATCCCACCCAACACCACCAGGAGTCGCCGAAGTCCTCAGGAGACTCGAACTCAATCCACAACACTCCGATGTCCGCCACATCTTCAACAACAGCGCCTTTCAAGGCTTCTCCGCGTCCATGAAATCCCACTGTCTCGACCTCCTCGCCAACATGACCGATGTCTCCATGGTCGAGGCCGCGATCTCCGTCACGCGTGGCGCCTCAGCTCCCTACGATACCCGACCCGATGCACCGTGGGGTCTACAAGGCATCTCGACATCCGCCTCTGTCAGCGGTAGTGCTTCCAGGTTGGACCACACCTATGCATACTCCGACTCCTCGCTCGGCACCGGCGCAGACATCTACATCGTCGACACGGGTGTCTACGCCGAACACAATATCTTCAACGGCCGCGCGAAGATGGTCTGGTCCTACAACCAAGACATGTCGGATAAAGACGGCCACGGCACCCACGTTGCCGGCACAGCAGCAGGCGACATTCTTGGCGTGGCATCCAAGGCTCGCGTATTCGGTGTCAAAGCCCTCGATGCCGATGGTGGTGGGTGGTCGTCCAATGTCGTGGCGGGCATCGACTATGTCATCAAAGCCCACGACGCACGCAAGGCCGCCTCCGACCCTTCATTCGCCGGCTCCGTAATTTCCATGTCGCTAGCTTCCACTGGTCGAGTGGAAGCCATCAGCCAAGCAGTCAATGCCGCCATCAGCGGAGGGATTCACGTCTGTGTCGCAGCAGGCAATAATGGCGAAGACGCCTGCAATTCGTCTCCCGCTTCCTCCGGGAGGCTCCCACGGCGGAGCTATCACAGTCGGCGCAGT-GACATTCGCGGTAAGAAGGCGAAGTTCTCG-ACGACGGCGACTGCGTCGATGTTCTATGCTCCGGGTGTGGACATCGTGAGCTC-T-GATCGGAGAGAGGA-CATGATCA-T-CGCTATC-GGGACATC-ATGGCGGCGCCCACATGGTGAACTGGGATCATCGCT-ATGC-ATG-CGAT--GTAACATTGGCTGAAAACC ``` |
| 22 | ggggacaagtttgtacaaaaaagcaggcttgatgGCCAAACGACGGGTCAAGAA, ggggaccactttgtacaagaaagctgggtcCTCTCCATCATCATTATCGA | Mycgr3\|35447  JGI  Ensembl | KOG2963 RNA-binding protein required for 60S ribosomal subunit biogenesis; |  | 100 % |  | ``` 1	ATGGCCAAACGACGGGTCAAGAAGCGAACACATGTTGGCGCACAGATTGCTACTGGCGTCAAGACCGGACATGCGGCCTCGAAACCCGGCACACGCGCACCCAAGAGCATGGTCATTCGGGTTGGAGCCAGCGAGGTCGGTCCCAGCGTCACGCAATTGGTGAAGGATGTCCGTGGTATGATGGAGCCAGACACGGCATCTCGATTGAAAGAGCGACGGAGCAACAAGTTGCGCGACTACACCACCATGGCCGGTCCGCTCGGCGTTACTCATCTCATGCTGTTCAGTCGCAGCAGCACCGGCAACACAAATTTGCGGTTGGCATTGACACCCCGTGGTCCGACATTACATTTCCGCGTGGAGAAGTACAGCTTGTGCAAGGACATCTTCCAAAGCATGAAAAGGCCAAAGAGCGGCGGCACGGAGTACTTGACGGCGCCTTTACTGGTTATGAACAACTTCAACTCCAAGGACGACGACGATGCCACGAACAACACCAAGATACCGAAACATCTCGAATCGCTGGCGACGAGCGTCTTCCAATCCCTTTTTCCACCCATCAATCCGGCCGCAACTCCTTTGAACAGCATCAAGCGTGTATTGCTGCTCGATCGCGAGCCACAGGACCCCAATTCCGACTCTGCAACATACATCTTGAACCTACGCCACTACGCAATCGAAACTCGTACGGCCAAGTCTGTACCTAAACCACTCCGCAGATTGGACGCAGCCGAGAAAAGTGCACATGCTGGGCAAAAACGCAAAAGAGGAGCGCTTCCCAATCTTGGAAAGCTCAATGATGTCTCCGACTATCTCCTCGATCCTCACGCCGCGGACGGGTTCACATCCGGGTCCGAATCAGAGCCAGACACCGACGCTGAAATCGAGATCGAGATGACCAACGCGCGAAAGGTGCAGAGAAACAGATCACAAAAGGAAGGCTCAGATCGG-AAAGAGC-GAGTAAATGTTCAAAAGAAGGCCGTAAAGATGCATGAGCTTGGTCCGAGGATGAGGTTGAGATTGACAAAGGTCGAGGAAGGCGTGTGCAATGGGAAGATCATG-TGGCACGAGTACATACACAAGACGAAGGAGGAGATTAGCAAGATG	1107 	\|\|\|\|\|\|\|\|\|\|\|\|\|\|\|\|\|\|\|\|\|\|\|\|\|\|\|\|\|\|\|\|\|\|\|\|\|\|\|\|\|\|\|\|\|\|\|\|\|\|\|\|\|\|\|\|\|\|\|\|\|\|\|\|\|\|\|\|\|\|\|\|\|\|\|\|\|\|\|\|\|\|\|\|\|\|\|\|\|\|\|\|\|\|\|\|\|\|\|\|\|\|\|\|\|\|\|\|\|\|\|\|\|\|\|\|\|\|\|\|\|\|\|\|\|\|\|\|\|\|\|\|\|\|\|\|\|\|\|\|\|\|\|\|\|\|\|\|\|\|\|\|\|\|\|\|\|\|\|\|\|\|\|\|\|\|\|\|\|\|\|\|\|\|\|\|\|\|\|\|\|\|\|\|\|\|\|\|\|\|\|\|\|\|\|\|\|\|\|\|\|\|\|\|\|\|\|\|\|\|\|\|\|\|\|\|\|\|\|\|\|\|\|\|\|\|\|\|\|\|\|\|\|\|\|\|\|\|\|\|\|\|\|\|\|\|\|\|\|\|\|\|\|\|\|\|\|\|\|\|\|\|\|\|\|\|\|\|\|\|\|\|\|\|\|\|\|\|\|\|\|\|\|\|\|\|\|\|\|\|\|\|\|\|\|\|\|\|\|\|\|\|\|\|\|\|\|\|\|\|\|\|\|\|\|\|\|\|\|\|\|\|\|\|\|\|\|\|\|\|\|\|\|\|\|\|\|\|\|\|\|\|\|\|\|\|\|\|\|\|\|\|\|\|\|\|\|\|\|\|\|\|\|\|\|\|\|\|\|\|\|\|\|\|\|\|\|\|\|\|\|\|\|\|\|\|\|\|\|\|\|\|\|\|\|\|\|\|\|\|\|\|\|\|\|\|\|\|\|\|\|\|\|\|\|\|\|\|\|\|\|\|\|\|\|\|\|\|\|\|\|\|\|\|\|\|\|\|\|\|\|\|\|\|\|\|\|\|\|\|\|\|\|\|\|\|\|\|\|\|\|\|\|\|\|\|\|\|\|\|\|\|\|\|\|\|\|\|\|\|\|\|\|\|\|\|\|\|\|\|\|\|\|\|\|\|\|\|\|\|\|\|\|\|\|\|\|\|\|\|\|\|\|\|\|\|\|\|\|\|\|\|\|\|\|\|\|\|\|\|\|\|\|\|\|\|\|\|\|\|\|\|\|\|\|\|\|\|\|\|\|\|\|\|\|\|\|\|\|\|\|\|\|\|\|\|\|\|\|\|\|\|\|\|\|\|\|\|\|\|\|\|\|\|\|\|\|\|\|\|\|\|\|\|\|\|\|\|\|\|\|\|\|\|\|\|\|\|\|\|\|\|\|\|\|\|\|\|\|\|\|\|\|\|\|\|\|\|\|\|\|\|\|\|\|\|\|\|\|\|\|\|\|\|\|\|\|\|\|\|\|\|\|\|\|\|\|\|\|\|\|\|\|\|\|\|\|\|\|\|\|\|\|\|\|\|\|\|\|\|\|\|\|\|\|\|\|\|\|\|\|\|\|\|\|\|\|\|\|\|\|\|\|\|\|\|\|\|\|\|\|\|\|\|\|\|\|\|\|\|\|\|\|\|\|\|\|\|\|\|\|\|\|\|\|\|\|\|\|\|\|\|\|\|\|\|\|\|\|\|\|\|\|\|\|\|\|\|\|\|\|\|\|\|\|\|\|\|\|\|\|\|\|\|\|\|\|\|\|\|\|\|\|\|\|\|\|\|\|\|\|\|\|\|\|\|\|\|\|\|\|\|\|\|\|\|\|\|\|\|\|\|\|\|\|\|\|\|\|\|\|\|\|\|\|\|\|\|\|\|\|\|\|\|\|\|\|\|\|\|\|\|\|\|\|\|\|\|\|\|\|\|\|\|\|\|\|\|\|\|\|\|\|\|\|\|\|\|\|\|\|\|\|\|\|\|\|\|\|\|\|\|\|\|\|\|\|\|\|\|\|\|\|\|\|\|\|\|\|\|\|\|\|\|\|\|\|\|\|\|\|\|\|\|\|\|\|\|\|\|\|\|\|\|\|\|\|\|\|\|\|\|\|\| \|\| \|\|\|\|\|\|\|\|\|\|\|\|\| \|\|\|\|\|\|\| \|\|\|\|\|\|\|\|\|\|\|\| \|\|\|\|\|\|\|\|\|\|\|\|\|\|\|\|\|\|\|\|\|\|\|\|\|\|\|\|\|\|\|\|\|\|\|\|\|\|\|\|\|\| \| \|\|\|\| \|\|\|\|\|\|\|\|\|\|\|\|\|\|\|\| \|\|\| \|\|\|\|\|\| \|\|\| \|\|\|\|\|\|\|\|\| \|\|\|\| \|\|\|\|\|\|\| \|\|\| \| \|\|\|\|\|\|\|\|\|\|\|\|\|\|\|\|\|\|\|\| \|\|\|\|\| 	ATGGCCAAACGACGGGTCAAGAAGCGAACACATGTTGGCGCACAGATTGCTACTGGCGTCAAGACCGGACATGCGGCCTCGAAACCCGGCACACGCGCACCCAAGAGCATGGTCATTCGGGTTGGAGCCAGCGAGGTCGGTCCCAGCGTCACGCAATTGGTGAAGGATGTCCGTGGTATGATGGAGCCAGACACGGCATCTCGATTGAAAGAGCGACGGAGCAACAAGTTGCGCGACTACACCACCATGGCCGGTCCGCTCGGCGTTACTCATCTCATGCTGTTCAGTCGCAGCAGCACCGGCAACACAAATTTGCGGTTGGCATTGACACCCCGTGGTCCGACATTACATTTCCGCGTGGAGAAGTACAGCTTGTGCAAGGACATCTTCCAAAGCATGAAAAGGCCAAAGAGCGGCGGCACGGAGTACTTGACGGCGCCTTTACTGGTTATGAACAACTTCAACTCCAAGGACGACGACGATGCCACGAACAACACCAAGATACCGAAACATCTCGAATCGCTGGCGACGAGCGTCTTCCAATCCCTTTTTCCACCCATCAATCCGGCCGCAACTCCTTTGAACAGCATCAAGCGTGTATTGCTGCTCGATCGCGAGCCACAGGACCCCAATTCCGACTCTGCAACATACATCTTGAACCTACGCCACTACGCAATCGAAACTCGTACGGCCAAGTCTGTACCTAAACCACTCCGCAGATTGGACGCAGCCGAGAAAAGTGCACATGCTGGGCAAAAACGCAAAAGAGGAGCGCTTCCCAATCTTGGAAAGCTCAATGATGTCTCCGACTATCTCCTCGATCCTCACGCCGCGGACGGGTTCACATCCGGGTCCGAATCAGAGCCAGACACCGACGCTGAAATCGAGATCGAGATGACCAACGCGCGAAAGGTGCAGAGAAACAGATCACAAA-GG-AGGCTCAGATCGGGAAAGAGCGGAGTAAATGTTC-AAAGAAGGCCGTAAAGATGCATGAGCTTGGTCCGAGGATGAG-T-GAGA-TGACAAAGGTCGAGGA-GGC-TGTGCA-TGG-AAGATCATGGTGGC-CGAGTAC-TACCC-AGACGAAGGAGGAGATTAGCCAGATG ``` |
| 22 | ggggacaagtttgtacaaaaaagcaggcttgatgCTGCGAAAGGCGGCTCTCTT, ggggaccactttgtacaagaaagctgggtcACTTCGAGGCATCATCAGTC | Mycgr3\|35494  JGI  Ensembl |  |  | 0 % |  |  |
| 22 | ggggacaagtttgtacaaaaaagcaggcttgatgGCGTCTGCCATTAAAGCCAA, ggggaccactttgtacaagaaagctgggtcGTCAATCTCATCCAGGAAAA | Mycgr3\|39456  JGI  Ensembl |  |  | 100 % |  | ``` 372	AGACAAGATTCCACCTGGTTTCAATACCGCCTTTAAAATTGTCAAGGAGTTCATCGATCCAGGCCTCAGTTGCGACGCCTATGCTGATCAGCCTTGGCTCTATGGGCCTAGTCTCAGCTGTTGGTTTGCTCTCCGCATCGGAGAAAAGGTCAACGAAGGCGAAGACTTTCCCGCACCCGGCGTGATGACTGAGGGAGCAGATGGAACCGGTAAAGATCTCAGAGAGCGACTAGGCCTCCCGGAAAACAACGAAAAGCGACGAAAGTACTTCCTGGACGCAAAGCATCGCCAACCtttttttttCGAAAAAGGTCGATGCTATCAAGGAGACTTCTACAACCCTTACTTGGACTTTGGTAATTTTAGTTTGAAACTGCCAGGTTTTACTTTGAAAGTGATCAAATACGTCGGAGACAAGACGCATTGTCTTCGATACGTGTTCAAGAACCGAGAGACCGGCGACATGTACTTCAACGTCAATTTCAAATTGCTCTGGGGTGAAGAGC-TGCAAAAGGCCCTGACGGCAGACCAGAAAGAGCAGGCAAGCTCTGAAAGAGACTCTGGAATTGGCTTTCAAGATCAAGCCTCAGATGGGGCTACCGACGGCGCATCGAATGGTGTTGCAGC	998 	\|\|\|\|\|\|\|\|\|\|\|\|\|\|\|\|\|\|\|\|\|\|\|\|\|\|\|\|\|\|\|\|\|\|\|\|\|\|\|\|\|\|\|\|\|\|\|\|\|\|\|\|\|\|\|\|\|\|\|\|\|\|\|\|\|\|\|\|\|\|\|\|\|\|\|\|\|\|\|\|\|\|\|\|\|\|\|\|\|\|\|\|\|\|\|\|\|\|\|\|\|\|\|\|\|\|\|\|\|\|\|\|\|\|\|\|\|\|\|\|\|\|\|\|\|\|\|\|\|\|\|\|\|\|\|\|\|\|\|\|\|\|\|\|\|\|\|\|\|\|\|\|\|\|\|\|\|\|\|\|\|\|\|\|\|\|\|\|\|\|\|\|\|\|\|\|\|\|\|\|\|\|\|\|\|\|\|\|\|\|\|\|\|\|\|\|\|\|\|\|\|\|\|\|\|\|\|\|\|\|\|\|\|\|\|\|\|\|\|\|\|\|\|\|\|\|\|\|\|\|\|\|\|\|\|\|\|\|\|\|\|\|\|\|\|\|\|\|\|\|\|\|\|\|\|\|\|\|\|\|\|\|\|\|\|\|\|\|\|\|\|\|\|\|\|\|\|\|\|\|\|\|\|\|\|\|\|\|\|\|\|\|\|\|\|\|\|\|\|\|\|\|\|\|\|\|\|\|\|\| \|\|\|\|\|\|\|\|\|\|\|\|\|\|\|\|\|\|\|\|\|\|\|\|\|\|\|\|\|\|\|\|\|\|\|\|\|\|\|\|\|\|\|\|\|\|\|\|\|\|\|\|\|\|\|\|\|\|\|\|\|\|\|\|\|\|\|\|\|\|\|\|\|\|\|\|\|\|\|\|\|\|\|\|\|\|\|\|\|\|\|\|\|\|\|\|\|\|\|\|\|\|\|\|\|\|\|\|\|\|\|\|\|\|\|\|\|\|\|\|\|\|\|\|\|\|\|\|\|\|\|\|\|\|\|\|\|\|\|\|\|\|\|\|\|\|\|\|\|\|\|\|\|\|\|\|\|\|\|\|\|\|\|\|\|\|\|\|\|\|\|\|\| \|\|\|\|\|\|\|\|\|\|\|\|\|\|\|\|\|\|\|\|\| \|\|\| \|\|\| \|\|\| \|\|\|\|\|\|\|\|\|\|\|\|\|\|\|\|\|\|\|\|\|\|\| \| \|\|\|\|\|\|\|\| \|\|\|\|\|\|\|\|\| \|\|\| \|\|\|\| \|\|\|\|\|\|\|\|\|\| \|\| \|\|\|\|\|\|\|\|  \|\|\|\|  \|\|\|\|\|  \|\|\| \|  \|\|\|\|\| \|\|\|\|\| 	AGACAAGATTCCACCTGGTTTCAATACCGCCTTTAAAATTGTCAAGGAGTTCATCGATCCAGGCCTCAGTTGCGACGCCTATGCTGATCAGCCTTGGCTCTATGGGCCTAGTCTCAGCTGTTGGTTTGCTCTCCGCATCGGAGAAAAGGTCAACGAAGGCGAAGACTTTCCCGCACCCGGCGTGATGACTGAGGGAGCAGATGGAACCGGTAAAGATCTCAGAGAGCGACTAGGCCTCCCGGAAAACAACGAAAAGCGACGAAAGTACTTCCTGGACGCAAAGCATCGCCAACCTTTTTTTTTCGAAAAA-GTCGATGCTATCAAGGAGACTTCTACAACCCTTACTTGGACTTTGGTAATTTTAGTTTGAAACTGCCAGGTTTTACTTTGAAAGTGATCAAATACGTCGGAGACAAGACGCATTGTCTTCGATACGTGTTCAAGAACCGAGAGACCGGCGACATGTACTTCAACGTCAATTTC-AATTGCTCTGGGGTGAAGAGCTTGCCAAACGCCTTGACGGCAGACCAGAAAGAGCAG-C-AGCTCTGACAGAGACTCT-GAACTGGCCTTCAAGATCA-GC-TCAGATGGC-CTACGTACGGCCTATCCATGGGTGTGGCAGC ```      ``` 1	ATGGCGTCTGCCATTAAAGCCAAATTGACGGGAGGATCAGCTGCCAACGACGCTGACCTGCAGGAAGAACGAGACAAGTACAGACTTCTAGTCACAGCCGGTCCGTCCTATGATGAGAAGACGCACAAGACCGTCAAGGTCAATTCCGAGATTCCAACGACAATCGAGAACGAGTTCTTGACAGCGAAGATCAATGTCCGCGTGAAAGGCTTCCGTGGTTTGCCAAACACAAGCCCATCAACATCGAAGGACTACTTCGACGACCCTATGCATGCAAAAGACCAATACTCGCTTGGTTTCAGCTTCGTTCCGAAGCAGGATATTCCCAGTGTAGATACTGTCTGGGGCAACGACTTCGACCATAGCATTAA	371 	\|\|\|\|\|\|\|\|\|\|\|\|\|\|\|\|\|\|\|\|\|\|\|\|\|\|\|\|\|\|\|\|\|\|\|\|\|\|\|\|\|\|\|\|\|\|\|\|\|\|\|\|\|\|\|\|\|\|\|\|\|\|\|\|\|\|\|\|\|\|\|\|\|\|\|\|\|\|\|\|\|\|\|\|\|\|\|\|\|\|\|\|\|\|\|\|\|\|\|\|\|\|\|\|\|\|\|\|\|\|\|\|\|\|\|\|\|\|\|\|\|\|\|\|\|\|\|\|\|\|\|\|\|\|\|\|\|\|\|\|\|\|\|\|\|\|\|\|\|\|\|\|\|\|\|\|\|\|\|\|\|\|\|\|\|\|\|\|\|\|\|\|\|\|\|\|\|\|\|\|\|\|\|\|\|\|\|\|\|\|\|\|\|\|\|\|\|\|\|\|\|\|\|\|\|\|\|\|\|\|\|\|\|\|\|\|\|\|\|\|\|\|\|\|\|\|\|\|\|\|\|\|\|\|\|\|\|\|\|\|\|\|\|\|\|\|\|\|\|\|\|\|\|\|\|\|\|\|\|\|\|\|\|\|\|\|\|\|\|\|\|\|\|\|\|\|\|\|\|\|\|\|\|\|\|\|\|\|\|\|\|\|\|\|\|\|\|\|\|\|\|\|\|\|\|\|\|\|\|\|\|\|\|\|\|\|\|\|\|\|\|\|\|\|\|\|\|\|\|\|\|\|\|\|\|\|\|\|\|\|\|\|\|\|\|\|\|\|\|\|\|\|\|\|\|\|\|\|\|\|\|\|\|\|\|\|\|\|\|\|\| 	ATGGCGTCTGCCATTAAAGCCAAATTGACGGGAGGATCAGCTGCCAACGACGCTGACCTGCAGGAAGAACGAGACAAGTACAGACTTCTAGTCACAGCCGGTCCGTCCTATGATGAGAAGACGCACAAGACCGTCAAGGTCAATTCCGAGATTCCAACGACAATCGAGAACGAGTTCTTGACAGCGAAGATCAATGTCCGCGTGAAAGGCTTCCGTGGTTTGCCAAACACAAGCCCATCAACATCGAAGGACTACTTCGACGACCCTATGCATGCAAAAGACCAATACTCGCTTGGTTTCAGCTTCGTTCCGAAGCAGGATATTCCCAGTGTAGATACTGTCTGGGGCAACGACTTCGACCATAGCATTAA ``` |
| 22 | ggggacaagtttgtacaaaaaagcaggcttgatgGCTTCCAACGACACTCCAAA, ggggaccactttgtacaagaaagctgggtcAACCCCAAGCAACCTACTCA | Mycgr3\|40731  JGI  Ensembl | KOG0786 3-isopropylmalate dehydrogenase; | 3-isopropylmalate dehydrogenase activity; cytoplasm; leucine biosynthetic process;  metabolic process;  oxidoreductase activity, acting on the CH-OH group of donors, NAD or NADP as acceptor; | 0 % |  |  |
| 22 | ggggacaagtttgtacaaaaaagcaggcttgatgGCTCCCATTAACGTCCTCAT, ggggaccactttgtacaagaaagctgggtcCTTCAAAGTCCACACTCCAT | Mycgr3\|41072  JGI  Ensembl |  |  | 100 % |  |  |
| 22 | ggggacaagtttgtacaaaaaagcaggcttgatgGCCAGATTGTATTATCCCCT, ggggaccactttgtacaagaaagctgggtcTGCGAACCACGGATGCTCAA | Mycgr3\|42881  JGI  Ensembl | KOG1290 Serine/threonine protein kinase; | ATP binding; protein amino acid phosphorylation;  protein kinase activity;  protein serine/threonine kinase activity; | 100 % |  |  |
| 22 | ggggacaagtttgtacaaaaaagcaggcttgatgAAGTTCTCTTCCACTGCCGC, ggggaccactttgtacaagaaagctgggtcCGATTTCTGGAAACCAGGGA | Mycgr3\|43487  JGI  Ensembl |  |  | 100 % |  |  |
| 22 | ggggacaagtttgtacaaaaaagcaggcttgatgTCAGCCTCCGGGCCTTTCGT, ggggaccactttgtacaagaaagctgggtcATCCTCAGCATCCCAACCAA | Mycgr3\|46265  JGI  Ensembl | KOG1268 Glucosamine 6-phosphate synthetases, contain amidotransferase and phosphosugar isomerase domains; |  | 100 % | chr\_8:1009977 G->A; chr\_8:1010912 C->T; |  |
| 22 | ggggacaagtttgtacaaaaaagcaggcttgatgGACAGCACGACGAATTCAGC, ggggaccactttgtacaagaaagctgggtcTAATACAGAATCCCTCTTGG | Mycgr3\|48833  JGI  Ensembl |  |  | 100 % |  |  |
| 22 | ggggacaagtttgtacaaaaaagcaggcttgatgCCCGCCGCCCTCACCCCCGA, ggggaccactttgtacaagaaagctgggtcCTTCACCAAAACCCCCCACT | Mycgr3\|50597  JGI  Ensembl | KOG0975 Branched chain aminotransferase BCAT1, pyridoxal phosphate enzymes type IV superfamily; | branched-chain-amino-acid transaminase activity; catalytic activity;  metabolic process; | 100 % |  |  |
| 22 | ggggacaagtttgtacaaaaaagcaggcttgatgCGCCTACGCTCACATGGTAG, ggggaccactttgtacaagaaagctgggtcGAAGCTGGTGTGCAAGCCGT | Mycgr3\|51173  JGI  Ensembl |  |  | 100 % |  |  |
| 22 | ggggacaagtttgtacaaaaaagcaggcttgatgGGTTGCGGTCAGAGTTCAGA, ggggaccactttgtacaagaaagctgggtcTATCAGTCCGCACATGCGCA | Mycgr3\|52873  JGI  Ensembl | KOG0082 G-protein alpha subunit (small G protein superfamily); | G-protein coupled receptor protein signaling pathway; GTP binding;  guanyl nucleotide binding;  signal transducer activity; | 0 % |  |  |
| 22 | ggggacaagtttgtacaaaaaagcaggcttgatgCTCCAACCTCGGATACGGCC, ggggaccactttgtacaagaaagctgggtcGTTCTTCGCTCTCCGCTTGC | Mycgr3\|53366  JGI  Ensembl | KOG0438 Mitochondrial/chloroplast ribosomal protein L2; | RNA binding; intracellular;  large ribosomal subunit;  ribosome;  structural constituent of ribosome;  transferase activity;  translation; | 100 % |  | ``` 79	ATGCTCCAACCTCGGATACGGCCGTCACAATGCCGCTCATTCATCACCTCCTTGAGACGAACATATGCAACACCGGTCCCAAAGACGCCCGTACAGGCAGCCACCACAACTCAGGACACATCCGCCTTCACACCTTCGCGAGCGCCGACGAGAAAGCAGACCGGCTTCCAATTGAAGACTTACACCCCACGGACTCCCGGTACCCGACATCTCAGACGACCCATCAATGACCATCTCTGGAAAGGCGGGCCGTACAAGAAGTTGACATTCCCGAAAATCGGTCATGGAAAGGGTGGACGGAACAACTCAGGCCGTGTGACGATGAGACATCGTGGAGGTGGACACCGACGAAGAATAAGGACGGTGGATTATGTTCGATCAGAGCCTGGAAAGCATTTGGTGGAGCGCATCGAGTACGACCCTAATCGATCTGCTCATCTGGCGCTGGTTGAGAACATGGAGACCAAAAAGAAGAGCTACATCATTGCTGCAGAGGGAATGCGTGCAGGAGATGTGGTCGAGTCATTTCGCTCAGGAATCCCAAAAGAGCTATTGGAGAGCATGGGTGGTGTGGTAGATCCGGGTATTCTTGCCGCCAAGACTGCGTTCAGAGGGAACTGCTTGCCGATGCAGATGGTGCCGCTAGGCACTCAAGTGTACTGTGTTGGATCGGCGGCGGTTCGAGGAGCCGTGTTCTGTCGCAGTGCAGGTACACACGCCACTGTCATTGCAAAGGAAGAG-AAGGGAAAGAGCGTGGCTGATGTCACTGTCAGACTCCAGAGTGGGGAGGTGCGGAAAGTGCACAGAGACGCATGTGCTACTATTGGCGTTGCCAGCAACCCTCATTGGCACTTCAGACAATTGGGCAAGGCTGGTCGAAGCAGGTGGCTGAATATCCGACCTACAGTCCGTGGTGTGGCCATGAACGCGGTCGATCACCCTCACGGTGGTGGTCGTGGTAAGAGTAAGGGTAACGTGCACCCCGTGTCGATCTGGGGCACTC	1081 	\|\|\|\|\|\|\|\|\|\|\|\|\|\|\|\|\|\|\|\|\|\|\|\|\|\|\|\|\|\|\|\|\|\|\|\|\|\|\|\|\|\|\|\|\|\|\|\|\|\|\|\|\|\|\|\|\|\|\|\|\|\|\|\|\|\|\|\|\|\|\|\|\|\|\|\|\|\|\|\|\|\|\|\|\|\|\|\|\|\|\|\|\|\|\|\|\|\|\|\|\|\|\|\|\|\|\|\|\|\|\|\|\|\|\|\|\|\|\|\|\|\|\|\|\|\|\|\|\|\|\|\|\|\|\|\|\|\|\|\|\|\|\|\|\|\|\|\|\|\|\|\|\|\|\|\|\|\|\|\|\|\|\|\|\|\|\|\|\|\|\|\|\|\|\|\|\|\|\|\|\|\|\|\|\|\|\|\|\|\|\|\|\|\|\|\|\|\|\|\|\|\|\|\|\|\|\|\|\|\|\|\|\|\|\|\|\|\|\|\|\|\|\|\|\|\|\|\|\|\|\|\|\|\|\|\|\|\|\|\|\|\|\|\|\|\|\|\|\|\|\|\|\|\|\|\|\|\|\|\|\|\|\|\|\|\|\|\|\|\|\|\|\|\|\|\|\|\|\|\|\|\|\|\|\|\|\|\|\|\|\|\|\|\|\|\|\|\|\|\|\|\|\|\|\|\|\|\|\|\|\|\|\|\|\|\|\|\|\|\|\|\|\|\|\|\|\|\|\|\|\|\|\|\|\|\|\|\|\|\|\|\|\|\|\|\|\|\|\|\|\|\|\|\|\|\|\|\|\|\|\|\|\|\|\|\|\|\|\|\|\|\|\|\|\|\|\|\|\|\|\|\|\|\|\|\|\|\|\|\|\|\|\|\|\|\|\|\|\|\|\|\|\|\|\|\|\|\|\|\|\|\|\|\|\|\|\|\|\|\|\|\|\|\|\|\|\|\|\|\|\|\|\|\|\|\|\|\|\|\|\|\|\|\|\|\|\|\|\|\|\|\|\|\|\|\|\|\|\|\|\|\|\|\|\|\|\|\|\|\|\|\|\|\|\|\|\|\|\|\|\|\|\|\|\|\|\|\|\|\|\|\|\|\|\|\|\|\|\|\|\|\|\|\|\|\|\|\|\|\|\|\|\|\|\|\|\|\|\|\|\|\|\|\|\|\|\|\|\|\|\|\|\|\|\|\|\|\|\|\|\|\|\|\|\|\|\|\|\|\|\|\|\|\|\|\|\|\|\|\|\|\|\|\|\|\|\|\|\|\|\|\|\|\|\|\|\|\|\|\|\|\|\|\|\|\|\|\|\|\|\|\|\|\|\|\|\|\|\|\|\|\|\|\|\|\|\|\|\|\|\|\|\|\|\|\|\|\|\|\|\|\|\|\|\|\|\|\|\|\|\|\|\|\|\|\|\|\|\|\|\|\|\|\|\|\|\|\|\|\|\|\|\|\|\|\|\|\|\|\|\|\|\|\|\|\|\|\|\|\|\|\|\|\|\|\|\|\|\|\|\|\|\|\|\|\|\|\|\|\|\|\|\|\|\|\|\|\|\|\|\|\|\|\|\|\|\|\|\|\|\|\|\|\|\|\|\|\|\|\|\|\|\|\|\|\|\|\|\|\|\|\|\|\|\|\|\|\|\|\|\| \|\|\|\|\|\|\|\|\|\|\|\|\|\|\|\|\|\|\|\|\|\|\|\|\|\|\|\|\|\|\|\|\|\|\|\|\|\|\|\|\|\|\|\|\|\|\|\|\|\|\|\|\|\|\|\|\|\|\|\|\|\|\|\|\|\|\|\|\|\|\|\|\|\|\|\|\|\|\|\|\|\|\|\|\|\|\|\|\|\|\|\|\|\|\|\|\|\|\|\|\|\|\|\|\|\|\|\|\|\|\|\|\|\|\|\|\|\|\|\|\|\|\|\|\|\|\|\|\|\|\|\|\|\|\|\|\|\|\|\|\|\|\|\|\|\|\|\|\|\| \|\|\|\|\|\|\|\|\|\|\|\|\|\|\|\|\|\|\|\|\| \|\|\| \|\|\|\|\|\|\|\|\|\|\|\|\|\|\|\|\|\|\|\|\|\|\|\|\|\|\|\|\|\|\|\|\|\|\|\|\|\|\|\| \|\| \|\|\|\|\| \|  \| \|\|\|\|\|\|\|\|\|\|\|\|\|\|\|\|\|\|\|\|\|\|\|\|\|\|\|\|\|\| 	ATGCTCCAACCTCGGATACGGCCGTCACAATGCCGCTCATTCATCACCTCCTTGAGACGAACATATGCAACACCGGTCCCAAAGACGCCCGTACAGGCAGCCACCACAACTCAGGACACATCCGCCTTCACACCTTCGCGAGCGCCGACGAGAAAGCAGACCGGCTTCCAATTGAAGACTTACACCCCACGGACTCCCGGTACCCGACATCTCAGACGACCCATCAATGACCATCTCTGGAAAGGCGGGCCGTACAAGAAGTTGACATTCCCGAAAATCGGTCATGGAAAGGGTGGACGGAACAACTCAGGCCGTGTGACGATGAGACATCGTGGAGGTGGACACCGACGAAGAATAAGGACGGTGGATTATGTTCGATCAGAGCCTGGAAAGCATTTGGTGGAGCGCATCGAGTACGACCCTAATCGATCTGCTCATCTGGCGCTGGTTGAGAACATGGAGACCAAAAAGAAGAGCTACATCATTGCTGCAGAGGGAATGCGTGCAGGAGATGTGGTCGAGTCATTTCGCTCAGGAATCCCAAAAGAGCTATTGGAGAGCATGGGTGGTGTGGTAGATCCGGGTATTCTTGCCGCCAAGACTGCGTTCAGAGGGAACTGCTTGCCGATGCAGATGGTGCCGCTAGGCACTCAAGTGTACTGTGTTGGATCGGCGGCGGTTCGAGGAGCCGTGTTCTGTCGCAGTGCAGGTACACACGCCACTGTCATTGCAAAGGAAGAGAAAGGGAAAGAGCGTGGCTGATGTCACTGTCAGACTCCAGAGTGGGGAGGTGCGGAAAGTGCACAGAGACGCATGTGCTACTATTGGCGTTGCCAGCAACCCTCATTGGCACTTCAGACAATTGGGCAAGGCTGGTCGAAGCAGGTGGCTG-ATATCCGACCTACAGTCCGTG-TGT-GCCATGAACGCGGTCGATCACCCTCACGGTGGTGGTCGTG-TA-GAGTATG--T-ACGTGCACCCCGTGTCGATCTGGGGCACTC ``` |
| 22 | ggggacaagtttgtacaaaaaagcaggcttgatgCGTGTACGGAATGCCTCTCG, ggggaccactttgtacaagaaagctgggtcCTCAGTTTTCCATTGCTCAA | Mycgr3\|55061  JGI  Ensembl | KOG0562 Predicted hydrolase (HIT family); |  | 100 % |  |  |
| 22 | ggggacaagtttgtacaaaaaagcaggcttgatgTCGCCCGCCGCTACACTCCC, ggggaccactttgtacaagaaagctgggtcCGCCGCCTGAGCGATTGGAA | Mycgr3\|62164  JGI  Ensembl | KOG0567 HEAT repeat-containing protein; |  | 100 % |  |  |
| 22 | ggggacaagtttgtacaaaaaagcaggcttgatgCTCTGTACATACCGGATCGT, ggggaccactttgtacaagaaagctgggtcAAGGCCTTCTTTGGCAAGCA | Mycgr3\|65536  JGI  Ensembl |  |  | 100 % |  | ``` 33	GATGCTCTGTACATACCGGATCGTATCGCGGAGGGTGCTCTGTACATCCAGCGCCGGCAGAAGGCAAACTATAACGCGTCGTTGTCTGCACGCGACCACTTTCCAACGCAGCAAAGAACCGTCAGCCCAGGACCCGCGATTACCGAACGATGTGGACGATCTCATCGAAGACAAATTTGCGGTACTGAAGTCGACGTACGACGTTCCGAAACATCCGATCATCTTGGCCCATGGTCTTCTAGGATTCGATGAACTGCATCTGGCCGGAAATCTCTTGCCGGGCGTTGAATATTGGTATGG	332 	\|\|\|\|\|\|\|\|\|\|\|\|\|\|\|\|\|\|\|\|\|\|\|\|\|\|\|\|\|\|\|\|\|\|\|\|\|\|\|\|\|\|\|\|\|\|\|\|\|\|\|\|\|\|\|\|\|\|\|\|\|\|\|\|\|\|\|\|\|\|\|\|\|\|\|\|\|\|\|\|\|\|\|\|\|\|\|\|\|\|\|\|\|\|\|\|\|\|\|\|\|\|\|\|\|\|\|\|\|\|\|\|\|\|\|\|\|\|\|\|\|\|\|\|\|\|\|\|\|\|\|\|\|\|\|\|\|\|\|\|\|\|\|\|\|\|\|\|\|\|\|\|\|\|\|\|\|\|\|\|\|\|\|\|\|\|\|\|\|\|\|\|\|\|\|\|\|\|\|\|\|\|\|\|\|\|\|\|\|\|\|\|\|\|\|\|\|\|\|\|\|\|\|\|\|\|\|\|\|\|\|\|\|\|\|\|\|\|\|\|\|\|\|\|\|\|\|\|\|\|\|\|\|\|\|\|\|\|\|\|\|\|\|\|\|\|\|\|\|\|\|\|\|\|\|\|\|\|\|\|\|\|\|\|\|\|\|\|\|\|\|\|\|\|\|\|\|\|\|\|\|\|\|\|\|\|\|\|\|\|\|\|\|\|\|\|\|\|\|\| 	GATGCTCTGTACATACCGGATCGTATCGCGGAGGGTGCTCTGTACATCCAGCGCCGGCAGAAGGCAAACTATAACGCGTCGTTGTCTGCACGCGACCACTTTCCAACGCAGCAAAGAACCGTCAGCCCAGGACCCGCGATTACCGAACGATGTGGACGATCTCATCGAAGACAAATTTGCGGTACTGAAGTCGACGTACGACGTTCCGAAACATCCGATCATCTTGGCCCATGGTCTTCTAGGATTCGATGAACTGCATCTGGCCGGAAATCTCTTGCCGGGCGTTGAATATTGGTATGG ```      ``` 474	AGGTCTTGACAGCAGATACATGATATCCCGTCTGCAGCCTCCTAACGTGAAGGTGCTATCACTAACCACCGTCGCCACTCCACATCGCGGATCCTCCTTTGCGGACTTTGTCTTCAAAGGCATCGGCGTCACGAGACTTCCCAAGCTCTACAAAGCCCTCGAATTTGTCGGCGTGGAAACTGGCGCTTTCAAACAATTGACCTGCGAGTACATGGCGGAGTCGTTCAATCCACGCACGCCCGACGACGAAAGCGTGCAGTACTACAGCTACGGAGCCGACTTCCAGCCGCACCTGATGTCCGTGTTTCGCAAAAGTCACAATGT-GATTCAGGATATCGAGGGACCAAACGATGGACTTGTTAGCGTTCAGAGCAGTCGATGGGGTACCTACAAAGGGACACT-GAATCATGTCAACCATCT-GGATTTGATCAACTGGACGAA	914 	\|\|\|\|\|\|\|\|\|\|\|\|\|\|\|\|\|\|\|\|\|\|\|\|\|\|\|\|\|\|\|\|\|\|\|\|\|\|\|\|\|\|\|\|\|\|\|\|\|\|\|\|\|\|\|\|\|\|\|\|\|\|\|\|\|\|\|\|\|\|\|\|\|\|\|\|\|\|\|\|\|\|\|\|\|\|\|\|\|\|\|\|\|\|\|\|\|\|\|\|\|\|\|\|\|\|\|\|\|\|\|\|\|\|\|\|\|\|\|\|\|\|\|\|\|\|\|\|\|\|\|\|\|\|\|\|\|\|\|\|\|\|\|\|\|\|\|\|\|\|\|\|\|\|\|\|\|\|\|\|\|\|\|\|\|\|\|\|\|\|\|\|\|\|\|\|\|\|\|\|\|\|\|\|\|\|\|\|\|\|\|\|\|\|\|\|\|\|\|\|\|\|\|\|\|\|\|\|\|\|\|\|\|\|\|\|\|\|\|\|\|\|\|\|\|\|\|\|\|\|\|\|\|\|\|\|\|\|\|\|\|\|\|\|\|\|\|\|\|\|\|\|\|\|\|\|\|\|\|\|\|\|\|\|\|\|\|\|\|\|\|\|\|\|\|\| \|\|\|\|\| \|\|\|\|\|\|\|\|\|\|\|\|\|\|\|\| \|\|\|\|\|\|\|\|\|\|\|\|\|\|\|\|\|\|\|\| \|\|\| \|\| \|\|\| \|\|\|\|\|\|\|\|\|\|\|\| \|\|\|\|\|\|\|\|\|\|\|\| \|\|\| \|\|\|\|\| \|\| \|\|\|\|\|\|\|\|\|\|\|\|\| \|\|\| \|\|\| \| \|\| \|\|\|\|\| \|\|\|\|\|\|\|\|\|\|\| \| \|\|\|\| \|\|\|\|\|\|\|\|\|\| \|\|\|\|\|\|\|\|\|\| 	AGGTCTTGACAGCAGATACATGATATCCCGTCTGCAGCCTCCTAACGTGAAGGTGCTATCACTAACCACCGTCGCCACTCCACATCGCGGATCCTCCTTTGCGGACTTTGTCTTCAAAGGCATCGGCGTCACGAGACTTCCCAAGCTCTACAAAGCCCTCGAATTTGTCGGCGTGGAAACTGGCGCTTTCAAACAATTGACCTGCGAGTACATGGCGGAGTCGTTCAATCCACGCACGCCCGACGACGAAAGCGTGCAGTACTACAGCTACGGAGC-GACTT-CAGCCGCACCTGATGT-CGTGTTTCGCAAAAGTCACA-TGTTGA-TCAAGATATCGAGGGAACAAACGATGGACCTGT-AGCGTCCA-AGCAGTCGATGGGTTACTTACCACGG-ACACTTGAATCATGTCA-C-ATCTGGGATTTGATCCACTGGACGAA ```      ``` 326	GGTATGGGATCACTCAAGCTCTAGCAGCGAAGAATGTCGAGGTCATCACAGCTTCCGTTCCACCATCTGGGAGCATCGAAGTGCGAGCGGCAAGACTCGCTGAAGTCATCGAGAGGAAGGCGAACGGAAAGGCCGTCAACATCATAGC	473 	\|\|\|\|\|\|\|\|\|\|\|\|\|\|\|\|\|\|\|\|\|\|\|\|\|\|\|\|\|\|\|\|\|\|\|\|\|\|\|\|\|\|\|\|\|\|\|\|\|\|\|\|\|\|\|\|\|\|\|\|\|\|\|\|\|\|\|\|\|\|\|\|\|\|\|\|\|\|\|\|\|\|\|\|\|\|\|\|\|\|\|\|\|\|\|\|\|\|\|\|\|\|\|\|\|\|\|\|\|\|\|\|\|\|\|\|\|\|\|\|\|\|\|\|\|\|\|\|\|\|\|\|\|\|\|\|\|\|\|\|\|\|\|\|\|\|\|\| 	GGTATGGGATCACTCAAGCTCTAGCAGCGAAGAATGTCGAGGTCATCACAGCTTCCGTTCCACCATCTGGGAGCATCGAAGTGCGAGCGGCAAGACTCGCTGAAGTCATCGAGAGGAAGGCGAACGGAAAGGCCGTCAACATCATAGC ``` |
| 22 | ggggacaagtttgtacaaaaaagcaggcttgatgGCGAAACAGATCAAGCAATG, ggggaccactttgtacaagaaagctgggtcCGAAACCTTGACAACGACCT | Mycgr3\|65978  JGI  Ensembl | KOG1198 Zinc-binding oxidoreductase; | NADPH:quinone reductase activity; oxidoreductase activity;  zinc ion binding; | 100 % |  |  |
| 22 | ggggacaagtttgtacaaaaaagcaggcttgatgGTGGTCACCGCTCCGCCCTC, ggggaccactttgtacaagaaagctgggtcATCAGACCTAACGCTGTACG | Mycgr3\|66270  JGI  Ensembl |  |  | 100 % |  |  |
| 22 | ggggacaagtttgtacaaaaaagcaggcttgatgTTCCAATTCCGACTTCCTCA, ggggaccactttgtacaagaaagctgggtcGGCACCGTGAGTCAATCTCT | Mycgr3\|67102  JGI  Ensembl |  |  | 100 % |  |  |
| 22 | ggggacaagtttgtacaaaaaagcaggcttgatgACGGAAACACGCAAGATTGT, ggggaccactttgtacaagaaagctgggtcAGCCTTGACAAACTTCTTTC | Mycgr3\|67645  JGI  Ensembl | KOG2495 NADH-dehydrogenase (ubiquinone); | FAD binding; NADH dehydrogenase activity;  electron transport;  oxidoreductase activity; | 0 % |  |  |
| 22 | ggggacaagtttgtacaaaaaagcaggcttgatgCCTTGCAAACCAGCAGTATG, ggggaccactttgtacaagaaagctgggtcTAACTGCGCCCTTGGTGCCG | Mycgr3\|67847  JGI  Ensembl |  |  | 100 % | chr\_2:597482 G->A; |  |
| 22 | ggggacaagtttgtacaaaaaagcaggcttgatgAGTGAAGCAATCATCCGTCC, ggggaccactttgtacaagaaagctgggtcCGTCGACTTCGGGTGATGAT | Mycgr3\|68530  JGI  Ensembl | KOG0725 Reductases with broad range of substrate specificities; | 2-deoxy-D-gluconate 3-dehydrogenase activity; metabolic process;  oxidoreductase activity; | 100 % |  | ``` 315	GGGTATTGGAGGTGCATCCGGGACGGCAGTCGACGTGGATATGGTCGAATGGGCGAAGAGCATGGAGGTCAACGTAGCATCAATGGTCATGATGTCAAAGTACGCAATTCCTGTAATGAAGAAGAACGATGGGCAATGGCGAGGAAGTATTGTGAATATGGCCTCTGTTGCTGGCATCCGAGGTGGCAATCCTCATCTGCTCTATCCTACGAGTAAAGGCGCTATTGTT	543 	\|\|\|\|\|\|\|\|\|\|\|\|\|\|\|\|\|\|\|\|\|\|\|\|\|\|\|\|\|\|\|\|\|\|\|\|\|\|\|\|\|\|\|\|\|\|\|\|\|\|\|\|\|\|\|\|\|\|\|\|\|\|\|\|\|\|\|\|\|\|\|\|\|\|\|\|\|\|\|\|\|\|\|\|\|\|\|\|\|\|\|\|\|\|\|\|\|\|\|\|\|\|\|\|\|\|\|\|\|\|\|\|\|\|\|\|\|\|\|\|\|\|\|\|\|\|\|\|\|\|\|\|\|\|\|\|\|\|\|\|\|\|\|\|\|\|\|\|\|\|\|\|\|\|\|\|\|\|\|\|\|\|\|\|\|\|\|\|\|\|\|\|\|\|\|\|\|\|\|\|\|\|\|\|\|\|\|\|\|\|\|\|\|\|\|\|\|\|\|\|\|\|\|\|\|\|\|\|\|\|\|\|\|\|\|\|\|\|\|\|\|\|\|\|\|\|\|\|\| 	GGGTATTGGAGGTGCATCCGGGACGGCAGTCGACGTGGATATGGTCGAATGGGCGAAGAGCATGGAGGTCAACGTAGCATCAATGGTCATGATGTCAAAGTACGCAATTCCTGTAATGAAGAAGAACGATGGGCAATGGCGAGGAAGTATTGTGAATATGGCCTCTGTTGCTGGCATCCGAGGTGGCAATCCTCATCTGCTCTATCCTACGAGTAAAGGCGCTATTGTT ```      ``` 544	AACATGACGCGGGCAATGGCCTCGCATCACGCTCCTGATGGGATTCGAGTGAACTGTGTTTGTCCTGGCATGGTCTACACTCCACACATGGGCGGATATGATGGTGGTATGAGCGATGCGGCTCGAGCATCGCGGAAGAACAGGAGTCTTCTGAAGACCGAGGGCAATGGGTGGGATGTCGGCGCTGCGATCAGGTTCTTGGCTGGAGATGAAGCTCGATGG	765 	\|\|\|\|\|\|\|\|\|\|\|\|\|\|\|\|\|\|\|\|\|\|\|\|\|\|\|\|\|\|\|\|\|\|\|\|\|\|\|\|\|\|\|\|\|\|\|\|\|\|\|\|\|\|\|\|\|\|\|\|\|\|\|\|\|\| \|\|\|\|\|\|\|\|\|\|\|\|\|\|\|\|\|\|\|\|\|\|\|\|\|\|\|\|\|\|\|\|\|\|\|\|\|\|\|\|\|\|\|\|\|\|\|\|\|\|\|\|\|\|\|\|\|\|\|\|\|\|\|\|\|\|\|\|\|\|\|\|\|\|\|\|\|\|\|\|\|\|\|\|\|\|\|\|\|\|\|\|\|\|\|\|\|\|\|\|\|\|\|\|\|\|\|\|\|\|\|\|\|\|\|\|\|\|\|\|\|\|\|\|\|\|\|\|\|\|\|\|\|\|\|\|\|\|\|\|\|\|\|\|\|\|\|\|\|\|\|\|\|\|\| 	AACATGACGCGGGCAATGGCCTCGCATCACGCTCCTGATGGGATTCGAGTGAACTGTGTTTGTCCT-GCATGGTCTACACTCCACACATGGGCGGATATGATGGTGGTATGAGCGATGCGGCTCGAGCATCGCGGAAGAACAGGAGTCTTCTGAAGACCGAGGGCAATGGGTGGGATGTCGGCGCTGCGATCAGGTTCTTGGCTGGAGATGAAGCTCGATGG ```      ``` 1	ATGAGTGAAGCAATCATCCGTCCAGTCCGGCACTTTCACAACCGTGTCGCCATTGTGACCGGTGCCGGAGCTCACGGCGATGGCATCGGGAATGGACGTGCTGCGGCCATCTTGCTCGCAGAGGAGGGATGTTCAGTCGTCTGTGTGGACTTGAAAATCGAGTTAGCAGAGAAGACGGTTGAGATGATCAAGGAGGACGGTAAAGGCAAGGCCATGGCCATCAAGGCGAATGTGGCGGACGAAGAAGATTGTAAGAGGGTGGTGGATGAAGCTGTAAAGCAGTGGGGAAGGTTGGATATCCTCGTCAACACCGTGGGTAT	320 	\|\|\|\|\|\|\|\|\|\|\|\|\|\|\|\|\|\|\|\|\|\|\|\|\|\|\|\|\|\|\|\|\|\|\|\|\|\|\|\|\|\|\|\|\|\|\|\|\|\|\|\|\|\|\|\|\|\|\|\|\|\|\|\|\|\|\|\|\|\|\|\|\|\|\|\|\|\|\|\|\|\|\|\|\|\|\|\|\|\|\|\|\|\|\|\|\|\|\|\|\|\|\|\|\|\|\|\|\|\|\|\|\|\|\|\|\|\|\|\|\|\|\|\|\|\|\|\|\|\|\|\|\|\|\|\|\|\|\|\|\|\|\|\|\|\|\|\|\|\|\|\|\|\|\|\|\|\|\|\|\|\|\|\|\|\|\|\|\|\|\|\|\|\|\|\|\|\|\|\|\|\|\|\|\|\|\|\|\|\|\|\|\|\|\|\|\|\|\|\|\|\|\|\|\|\|\|\|\|\|\|\|\|\|\|\|\|\|\|\|\|\|\|\|\|\|\|\|\|\|\|\|\|\|\|\|\|\|\|\|\|\|\|\|\|\|\|\|\|\|\|\|\|\|\|\|\|\|\|\|\|\|\|\|\|\|\|\|\|\|\|\|\|\|\|\|\|\|\|\|\|\|\|\|\|\|\|\|\|\|\|\|\|\|\|\|\|\|\|\|\|\|\|\|\|\|\|\|\|\|\|\|\|\|\|\| \|\|\| 	ATGAGTGAAGCAATCATCCGTCCAGTCCGGCACTTTCACAACCGTGTCGCCATTGTGACCGGTGCCGGAGCTCACGGCGATGGCATCGGGAATGGACGTGCTGCGGCCATCTTGCTCGCAGAGGAGGGATGTTCAGTCGTCTGTGTGGACTTGAAAATCGAGTTAGCAGAGAAGACGGTTGAGATGATCAAGGAGGACGGTAAAGGCAAGGCCATGGCCATCAAGGCGAATGTGGCGGACGAAGAAGATTGTAAGAGGGTGGTGGATGAAGCTGTAAAGCAGTGGGGAAGGTTGGATATCCTCGTCAACACCGTGG-TAT ``` |
| 22 | ggggacaagtttgtacaaaaaagcaggcttgatgGCTACCACCGGAACAGTCAC, ggggaccactttgtacaagaaagctgggtcGATGTACGATACTTTCACGC | Mycgr3\|69603  JGI  Ensembl | KOG3527 Erythrocyte membrane protein 4.1 and related proteins of the ERM family; |  | 100 % |  |  |
| 22 | ggggacaagtttgtacaaaaaagcaggcttgatgATCCTTGGCCAGATAGCATG, ggggaccactttgtacaagaaagctgggtcATCCGCCTTACGCCTGTTCA | Mycgr3\|70297  JGI  Ensembl | KOG1515 Arylacetamide deacetylase; | extracellular region; hormone activity; | 100 % |  |  |
| 22 | ggggacaagtttgtacaaaaaagcaggcttgatgTTCCCTCGTCGCCTTCTCCC, ggggaccactttgtacaagaaagctgggtcCCCCATAGCCTTTCCCGTCC | Mycgr3\|70658  JGI  Ensembl |  |  | 100 % |  |  |
| 22 | ggggacaagtttgtacaaaaaagcaggcttgatgGCACCCGGAGCACTCCTCGT, ggggaccactttgtacaagaaagctgggtcGAGATTGCCACGCAAGTCCT | Mycgr3\|72272  JGI  Ensembl | KOG0638 4-hydroxyphenylpyruvate dioxygenase; | 4-hydroxyphenylpyruvate dioxygenase activity; aromatic amino acid family metabolic process; | 100 % |  |  |
| 22 | ggggacaagtttgtacaaaaaagcaggcttgatgACCTCCAACAATGGCTTCCA, ggggaccactttgtacaagaaagctgggtcCGCCCCCTTGAAAACATCAA | Mycgr3\|72424  JGI  Ensembl | KOG2445 Nuclear pore complex component (sc Seh1); |  | 100 % |  |  |
| 22 | ggggacaagtttgtacaaaaaagcaggcttgatgAAGTCCCTCACCCTCCTCTC, ggggaccactttgtacaagaaagctgggtcCTTCGTCACCGGCCTAGCAC | Mycgr3\|74298  JGI  Ensembl |  |  | 100 % |  | ``` 317	GCGGCCCCTGCCCCGGCCTCAACGTCGCCGCAAACCACGGCTTCCTCGCCCGCGACGGCATAACCACCTTCGCCGAGCTCGTCAACGCCCAACAAAACATCTACAACGTCGGCTACGACCTCTCCCTCCTCCTCGCCTTCCTCGGCCTGCAAGCCGACGGAGACCTCGTAACAACCAAACTCTCCATCGGCTGCGACGCCACAACCCGCACCTCCATCGCGCCCTTACTCACCGGCTCACAACCCGGTCTTGCCGGCCACAACAAATTCGAAGGCGACACCTCCCTCACCCGCAACGACTTCTTCACCGGCGGCGGCGACAACTTCTCTTTCAACCGCACCCTGTTCGACATGATGTCGTCCACCACAGGCGGGAATTTCAACCGCGAAAACCTCGCGGTCTACCGCAAACAGCGCTACGACCAGTCCGTCGCGGAAAACGCCAACTTCTTCTTCGGACCCCTCGCGCTCCTCCTCTTCGGCGCCGCCTCGTTCCTCTACGAACTCATGCCATCGGGATCGAGGAACTACGCTCCCGACCAGGAAACCATGAATTCCTTCTTCATCGACGAACGCTTGCCGTCCAACTGGATCAACCGCGTCGAACCCTACTCCAACGGCGACGTCGTCCGGGAGATTCTGGCCATGTACCTCGAAGCGCCCGTTCTCTTCGGTGGCAACACCGCTCAAGGGAAGTTTT	1015 	\|\|\|\|\|\|\|\|\|\|\|\|\|\|\|\|\|\|\|\|\|\|\|\|\|\|\|\|\|\|\|\|\|\|\|\|\|\|\|\|\|\|\|\|\|\|\|\|\|\|\|\|\|\|\|\|\|\|\|\|\|\|\|\|\|\|\|\|\|\|\|\|\|\|\|\|\|\|\|\|\|\|\|\|\|\|\|\|\|\|\|\|\|\|\|\|\|\|\|\|\|\|\|\|\|\|\|\|\|\|\|\|\|\|\|\|\|\|\|\|\|\|\|\|\|\|\|\|\|\|\|\|\|\|\|\|\|\|\|\|\|\|\|\|\|\|\|\|\|\|\|\|\|\|\|\|\|\|\|\|\|\|\|\|\|\|\|\|\|\|\|\|\|\|\|\|\|\|\|\|\|\|\|\|\|\|\|\|\|\|\|\|\|\|\|\|\|\|\|\|\|\|\|\|\|\|\|\|\|\|\|\|\|\|\|\|\|\|\|\|\|\|\|\|\|\|\|\|\|\|\|\|\|\|\|\|\|\|\|\|\|\|\|\|\|\|\|\|\|\|\|\|\|\|\|\|\|\|\|\|\|\|\|\|\|\|\|\|\|\|\|\|\|\|\|\|\|\|\|\|\|\|\|\|\|\|\|\|\|\|\|\|\|\|\|\|\|\|\|\|\|\|\|\|\|\|\|\|\|\|\|\|\|\|\|\|\|\|\|\|\|\|\|\|\|\|\|\|\|\|\|\|\|\|\|\|\|\|\|\|\|\|\|\|\|\|\|\|\|\|\|\|\|\|\|\|\|\|\|\|\|\|\|\|\|\|\|\|\|\|\|\|\|\|\|\|\|\|\|\|\|\|\|\|\|\|\|\|\|\|\|\|\|\|\|\|\|\|\|\|\|\|\|\|\|\|\|\|\|\|\|\|\|\|\|\|\|\|\|\|\|\|\|\|\|\|\|\|\|\|\|\|\|\|\|\|\|\|\|\|\|\|\|\|\|\|\|\|\|\|\|\|\|\|\|\|\|\|\|\|\|\|\|\|\|\|\|\|\|\|\|\|\|\|\|\|\|\|\|\|\|\|\|\|\|\|\|\|\|\|\|\|\|\|\|\|\|\|\|\|\|\|\|\|\|\|\|\|\|\|\|\|\|\|\|\|\|\|\|\|\|\|\|\|\|\|\|\|\|\|\|\|\|\|\|\|\|\|\|\|\|\|\|\|\|\|\|\|\|\|\|\|\|\|\|\|\|\|\|\|\|\|\|\|\|\|\|\|\|\|\| \|\|\|\|\|\|\| \|\|\| \| \|\|\|\|\|\|\|\| \| \|\|\|\|\|\|\| \|\|\|\|\|\|\|\| \| \|\| \|\|\|\|\|\|\|\|\|\| \|\| \|\|\|\| \|\|\| \| \|\|\|\|\|\| \|\|\| \|\|\|\|\|\|\|\|\|\|\|\|\|\|\|\|\|\|\|\| \|\|\|\|\|\| \|\|\|\|\|\|\|\| \|\|\|\|\|\|\| 	GCGGCCCCTGCCCCGGCCTCAACGTCGCCGCAAACCACGGCTTCCTCGCCCGCGACGGCATAACCACCTTCGCCGAGCTCGTCAACGCCCAACAAAACATCTACAACGTCGGCTACGACCTCTCCCTCCTCCTCGCCTTCCTCGGCCTGCAAGCCGACGGAGACCTCGTAACAACCAAACTCTCCATCGGCTGCGACGCCACAACCCGCACCTCCATCGCGCCCTTACTCACCGGCTCACAACCCGGTCTTGCCGGCCACAACAAATTCGAAGGCGACACCTCCCTCACCCGCAACGACTTCTTCACCGGCGGCGGCGACAACTTCTCTTTCAACCGCACCCTGTTCGACATGATGTCGTCCACCACAGGCGGGAATTTCAACCGCGAAAACCTCGCGGTCTACCGCAAACAGCGCTACGACCAGTCCGTCGCGGAAAACGCCAACTTCTTCTTCGGACCCCTCGCGCTCCTCCTCTTCGGCGCCGCCTCGTTCCTCTACGAACTCATGCCATCGGGATCGAGGAACTACGCTCCCGACCAGGAAACCATGAATTCCTTCTTCATCGACGA-CGCTTGC-GTC-A-CTGGATCA-C-GCGTCGA-CCCTACTC-A-CG-CGACGTCGTC-GG-AGAT-CTG-C-ATGTAC-TCG-AGCGCCCGTTCTCTTCGGTG-CAACAC-GCTCAAGG-AAGTTTT ```      ``` 4	ATGAAGTCCCTCACCCTCCTCTCCCTCCTCGGCACGACCGCCCAAGCCTTCCCATGGGTCCAAAATGTCCCCGGCGTGGACAACTCCATGCTCTCCTCTCTCAATCGGCGACAGCAAAATCCTGCACCAGGTCCCGGCTCCGCTCAAACCTGCCCTTTCAACCCAAACCATGTCAACGCTCCCGAGGTGACAGACCAATTCCCCTACAACAACGCCAGAAAAGGTGGACGGGGAAACGAAAAGGGCGGCTACCTTGTCCCAGCTCCCTTCGACAACGTGCATCGCTACATCCCTCCGACAGCACGAGACATTCG	317 	\|\|\|\|\|\|\|\|\|\|\|\|\|\|\|\|\|\|\|\|\|\|\|\|\|\|\|\|\|\|\|\|\|\|\|\|\|\|\|\|\|\|\|\|\|\|\|\|\|\|\|\|\|\|\|\|\|\|\|\|\|\|\|\|\|\|\|\|\|\|\|\|\|\|\|\|\|\|\|\|\|\|\|\|\|\|\|\|\|\|\|\|\|\|\|\|\|\|\|\|\|\|\|\|\|\|\|\|\|\|\|\|\|\|\|\|\|\|\|\|\|\|\|\|\|\|\|\|\|\|\|\|\|\|\|\|\|\|\|\|\|\|\|\|\|\|\|\|\|\|\|\|\|\|\|\|\|\|\|\|\|\|\|\|\|\|\|\|\|\|\|\|\|\|\|\|\|\|\|\|\|\|\|\|\|\|\|\|\|\|\|\|\|\|\|\|\|\|\|\|\|\|\|\|\|\|\|\|\|\|\|\|\|\|\|\|\|\|\|\|\|\|\|\|\|\|\|\|\|\|\|\|\|\|\|\|\|\|\|\|\|\|\|\|\|\|\|\|\|\|\|\|\|\|\|\|\|\|\|\|\|\|\|\|\|\|\|\|\|\|\|\|\|\|\|\|\|\|\|\|\|\|\|\|\|\|\|\|\|\|\|\|\|\|\|\|\|\|\|\|\|\|\|\|\|\|\|\|\|\|\|\|\|\| 	ATGAAGTCCCTCACCCTCCTCTCCCTCCTCGGCACGACCGCCCAAGCCTTCCCATGGGTCCAAAATGTCCCCGGCGTGGACAACTCCATGCTCTCCTCTCTCAATCGGCGACAGCAAAATCCTGCACCAGGTCCCGGCTCCGCTCAAACCTGCCCTTTCAACCCAAACCATGTCAACGCTCCCGAGGTGACAGACCAATTCCCCTACAACAACGCCAGAAAAGGTGGACGGGGAAACGAAAAGGGCGGCTACCTTGTCCCAGCTCCCTTCGACAACGTGCATCGCTACATCCCTCCGACAGCACGAGACATTCG ``` |
| 22 | ggggacaagtttgtacaaaaaagcaggcttgatgCAGTCCGTCCGCCAGGGGCT, ggggaccactttgtacaagaaagctgggtcTAGCAAATCCCGCTGAATGA | Mycgr3\|74730  JGI  Ensembl | KOG1255 Succinyl-CoA synthetase, alpha subunit; | catalytic activity; metabolic process;  succinate-CoA ligase (GDP-forming) activity;  succinate-CoA ligase complex;  succinate-CoA ligase complex (GDP-forming); | 100 % |  |  |
| 22 | ggggacaagtttgtacaaaaaagcaggcttgatgGCTATAAAGGCAGCATCAAG, ggggaccactttgtacaagaaagctgggtcCAAAGTACACTCCTTGCCCT | Mycgr3\|74861  JGI  Ensembl |  |  | 100 % |  |  |
| 22 | ggggacaagtttgtacaaaaaagcaggcttgatgTCTCGCATGACAGTCCGAAC, ggggaccactttgtacaagaaagctgggtcTATGTTCGCTCCCTTACCGG | Mycgr3\|74910  JGI  Ensembl | KOG4129 Exopolyphosphatases and related proteins; | cytoplasm; exopolyphosphatase activity;  pyrophosphatase activity; | 100 % |  |  |
| 22 | ggggacaagtttgtacaaaaaagcaggcttgatgTCGACCAAGAAACCCTTCCA, ggggaccactttgtacaagaaagctgggtcGCCCAAGGCACGAGACAGCT | Mycgr3\|75791  JGI  Ensembl | KOG2614 Kynurenine 3-monooxygenase and related flavoprotein monooxygenases; | metabolic process; oxidoreductase activity;  salicylate 1-monooxygenase activity; | 100 % |  |  |
| 22 | ggggacaagtttgtacaaaaaagcaggcttgatgGCCTCCGACTACGAGACGTA, ggggaccactttgtacaagaaagctgggtcCAACTTCAAGATCTCAGTCT | Mycgr3\|76141  JGI  Ensembl |  |  | 0 % |  |  |
| 22 | ggggacaagtttgtacaaaaaagcaggcttgatgTCCGGCGTGACAAGACTGCC, ggggaccactttgtacaagaaagctgggtcCCAATAATCTCCAAATTTGC | Mycgr3\|76471  JGI  Ensembl | KOG4366 Predicted thioesterase; |  | 100 % |  |  |
| 22 | ggggacaagtttgtacaaaaaagcaggcttgatgGCGACTTCCATCCGCCTCCG, ggggaccactttgtacaagaaagctgggtcCCCTTCCCCAAAATCACCAT | Mycgr3\|77021  JGI  Ensembl | KOG3288 OTU-like cysteine protease; |  | 100 % |  | ``` 258	ATGGCGACTTCCATCCGCCTCCGAGTCCGAGGCCCTCAAGGAGTGGCGACGGTGACCATCAACAATGACGCCTCCTGGAGCGACCTCATTCACGAGATCTCCGAGAAGAGCGGAGTCGCCGACTTTGACCTCAAATACGGATATCCACCTCAGCCTCTGAACACCGAGTCCATCAGTCCAGACATGAAGCTATCCGACTTCCCTCAGAAACTCAACGGCGAACAACTCATCGTCATGCCGCGAGCTATTCAAGCCAAACTCTCCAGTCCTCTTGAAGGAACAACAAAGAACCAACAAAACGTCGAAACGTTACCCAAAGCGAGTTCATCATCAGCACCATCCCACAAGTCCGGAGACTTCCCAGACAAACCACTAAACCTCACGAGAAAGAAGCAAAACCTCGACTCTGACCCTCCCGAAGTACAAGTCCCTCTCCTCGACGGAGTCATGGTTCTGCGCGTGATGCCCGACGACAACTCCTGCATGTTCCGAGCATTGTCCTCCGCTGTCCTCGGCTCCGCCCTCGACGGCATGACCGAACTTCGCTCTGTCGTCGCGCAGACCATCCAATCTCAACCAGAGCTCTACACCGAAGGCATGCTCGAGAAGAAACCCACAGACTACTGCCAGTGGATCCAACGCGAAGACAGCTGGGGAGGTGGCATCGAGCTTTCGATTCTCAGCCAGCACTTCGATATTGAGATTTGCAGTATCAATGTGCAAGATTGTCGAGTGGACAAGTTTAATGAGGGGAAGGCTACGAGATGTCTGTTGGTGTATTCGGGGATTCACTACGATGTGTGTGCGGTGGCGCCTTATAGTGGTGCAGATCCGGAGTTCGATCGCAAGGTGTTTGAAGTGGTTCGAGTTGATGGCGAGGAGGAGATGGATGGAGGGGCGTTGGCTGGAGCGCTGGAGCTGTGCAGAGGATTGCAGAAGCGGCATTACTTCACGGACACGCATGGGTTCAAGTTGAAGTGCAATCAGTGTGGTCAGGGAGGAAATGGGCAGCAGTGGGCTGTTCAGCATGCGAAGAGTACCGGGCATGGTGATTTTGGGGAAGG	1321 	\|\|\|\|\|\|\|\|\|\|\|\|\|\|\|\|\|\|\|\|\|\|\|\|\|\|\|\|\|\|\|\|\|\|\|\|\|\|\|\|\|\|\|\|\|\|\|\|\|\|\|\|\|\|\|\|\|\|\|\|\|\|\|\|\|\|\|\|\|\|\|\|\|\|\|\|\|\|\|\|\|\|\|\|\|\|\|\|\|\|\|\|\|\|\|\|\|\|\|\|\|\|\|\|\|\|\|\|\|\|\|\|\|\|\|\|\|\|\|\|\|\|\|\|\|\|\|\|\|\|\|\|\|\|\|\|\|\|\|\|\|\|\|\|\|\|\|\|\|\|\|\|\|\|\|\|\|\|\|\|\|\|\|\|\|\|\|\|\|\|\|\|\|\|\|\|\|\|\|\|\|\|\|\|\|\|\|\|\|\|\|\|\|\|\|\|\|\|\|\|\|\|\|\|\|\|\|\|\|\|\|\|\|\|\|\|\|\|\|\|\|\|\|\|\|\|\|\|\|\|\|\|\|\|\|\|\|\|\|\|\|\|\|\|\|\|\|\|\|\|\|\|\|\|\|\|\|\|\|\|\|\|\|\|\|\|\|\|\|\|\|\|\|\|\|\|\|\|\|\|\|\|\|\|\|\|\|\|\|\|\|\|\|\|\|\|\|\|\|\|\|\|\|\|\|\|\|\|\|\|\|\|\|\|\|\|\|\|\|\|\|\|\|\|\|\|\|\|\|\|\|\|\|\|\|\|\|\|\|\|\|\|\|\|\|\|\|\|\|\|\|\|\|\|\|\|\|\|\|\|\|\|\|\|\|\|\|\|\|\|\|\|\|\|\|\|\|\|\|\|\|\|\|\|\|\|\|\|\|\|\|\|\|\|\|\|\|\|\|\|\|\|\|\|\|\|\|\|\|\|\|\|\|\|\|\|\|\|\|\|\|\|\|\|\|\|\|\|\|\|\|\|\|\|\|\|\|\|\|\|\|\|\|\|\|\|\|\|\|\|\|\|\|\|\|\|\|\|\|\|\|\|\|\|\|\|\|\|\|\|\|\|\|\|\|\|\|\|\|\|\|\|\|\|\|\|\|\|\|\|\|\|\|\|\|\|\|\|\|\|\|\|\|\|\|\|\|\|\|\|\|\|\|\|\|\|\|\|\|\|\|\|\|\|\|\|\|\|\|\|\|\|\|\|\|\|\|\|\|\|\|\|\|\|\|\|\|\|\|\|\|\|\|\|\|\|\|\|\|\|\|\|\|\|\|\|\|\|\|\|\|\|\|\|\|\|\|\|\|\|\|\|\|\|\|\|\|\|\|\|\|\|\|\|\|\|\|\|\|\|\|\|\|\|\|\|\|\|\|\|\|\|\|\|\|\|\|\|\|\|\|\|\|\|\|\|\|\|\|\|\|\|\|\|\|\|\|\|\|\|\|\|\|\|\|\|\|\|\|\|\|\|\|\|\|\|\|\|\|\|\|\|\|\|\|\|\|\|\|\|\|\|\|\|\|\|\|\|\|\|\|\|\|\|\|\|\|\|\|\|\|\|\|\|\|\|\|\|\|\|\|\|\|\|\|\|\|\|\|\|\|\|\|\|\|\|\|\|\|\|\|\|\|\|\|\|\|\|\|\|\|\|\|\|\|\|\|\|\|\|\|\|\|\|\|\|\|\|\|\|\|\|\|\|\|\|\|\|\|\|\|\|\|\|\|\|\|\|\|\|\|\|\|\|\|\|\|\|\|\|\|\|\|\|\|\|\|\|\|\|\|\|\|\|\|\|\|\|\|\|\|\|\|\|\|\|\|\|\|\|\|\|\|\|\|\|\|\|\|\|\|\|\|\|\|\|\|\|\|\|\|\|\|\|\|\|\|\|\|\|\|\|\|\|\|\|\|\|\|\|\|\|\|\|\|\|\|\|\|\|\|\|\|\|\|\|\|\|\|\|\|\|\|\|\|\|\|\|\|\|\|\|\|\|\|\|\|\|\|\|\|\|\|\|\|\|\|\|\|\|\|\|\|\|\|\|\|\|\|\|\|\|\|\|\|\|\|\|\|\|\|\|\|\|\|\|\|\|\|\|\|\|\|\|\|\| \|\|\|\|\|\|\|\|\|\|\|\|\|\|\|\|\|\|\|\|\|\|\|\|\|\|\|\|\|\|\|\|\|\|\|\|\|\|\|\|\|\|\|\|\|\|\|\|\|\|\|\|\|\| \|\|\|\|\|\|\| \|\|\|\|\|\|\|\|\|\|\|\|\|\|\|\|\|\|\|\|\|\|\|\|\|\|\|\|\|\|\|\| \|\|\|\|\|\|\|\|\|\|\|\|\|\| \|\|\|\| \|\|\|\|\|\|\|\|\|\|\| 	ATGGCGACTTCCATCCGCCTCCGAGTCCGAGGCCCTCAAGGAGTGGCGACGGTGACCATCAACAATGACGCCTCCTGGAGCGACCTCATTCACGAGATCTCCGAGAAGAGCGGAGTCGCCGACTTTGACCTCAAATACGGATATCCACCTCAGCCTCTGAACACCGAGTCCATCAGTCCAGACATGAAGCTATCCGACTTCCCTCAGAAACTCAACGGCGAACAACTCATCGTCATGCCGCGAGCTATTCAAGCCAAACTCTCCAGTCCTCTTGAAGGAACAACAAAGAACCAACAAAACGTCGAAACGTTACCCAAAGCGAGTTCATCATCAGCACCATCCCACAAGTCCGGAGACTTCCCAGACAAACCACTAAACCTCACGAGAAAGAAGCAAAACCTCGACTCTGACCCTCCCGAAGTACAAGTCCCTCTCCTCGACGGAGTCATGGTTCTGCGCGTGATGCCCGACGACAACTCCTGCATGTTCCGAGCATTGTCCTCCGCTGTCCTCGGCTCCGCCCTCGACGGCATGACCGAACTTCGCTCTGTCGTCGCGCAGACCATCCAATCTCAACCAGAGCTCTACACCGAAGGCATGCTCGAGAAGAAACCCACAGACTACTGCCAGTGGATCCAACGCGAAGACAGCTGGGGAGGTGGCATCGAGCTTTCGATTCTCAGCCAGCACTTCGATATTGAGATTTGCAGTATCAATGTGCAAGATTGTCGAGTGGACAAGTTTAATGAGGGGAAGGCTACGAGATGTCTGTTGGTGTATTCGGGGATTCACTACGATGTGTGTGCGGTGGCGCCTTATAGTGGTGCAGATCCGGAGTTCGATCGCAAGGTGTTTGAAGTGGTTCGAGTTGATGGCGAGGAGGAGATGGATGGAGGGGCGTTGGCTGGAGCGCTGGAGCTGTGCAGAGGATTGCAG-AGCGGCATTACTTCACGGACACGCATGGGTTCAAGTTGAAGTGCAATCAGTGTG-TCAGGGA-GAAATGGGCAGCAGTGGGCTGTTCAGCATGCG-AGAGTACCGGGCAT-GTGAATTTGGGGAAGG ```      ``` 258	ATGGCGACTTCCATCCGCCTCCGAGTCCGAGGCCCTCAAGGAGTGGCGACGGTGACCATCAACAATGACGCCTCCTGGAGCGACCTCATTCACGAGATCTCCGAGAAGAGCGGAGTCGCCGACTTTGACCTCAAATACGGATATCCACCTCAGCCTCTGAACACCGAGTCCATCAGTCCAGACATGAAGCTATCCGACTTCCCTCAGAAACTCAACGGCGAACAACTCATCGTCATGCCGCGAGCTATTCAAGCCAAACTCTCCAGTCCTCTTGAAGGAACAACAAAGAACCAACAAAACGTCGAAACGTTACCCAAAGCGAGTTCATCATCAGCACCATCCCACAAGTCCGGAGACTTCCCAGACAAACCACTAAACCTCACGAGAAAGAAGCAAAACCTCGACTCTGACCCTCCCGAAGTACAAGTCCCTCTCCTCGACGGAGTCATGGTTCTGCGCGTGATGCCCGACGACAACTCCTGCATGTTCCGAGCATTGTCCTCCGCTGTCCTCGGCTCCGCCCTCGACGGCATGACCGAACTTCGCTCTGTCGTCGCGCAGACCATCCAATCTCAACCAGAGCTCTACACCGAAGGCATGCTCGAGAAGAAACCCACAGACTACTGCCAGTGGATCCAACGCGAAGACAGCTGGGGAGGTGGCATCGAGCTTTCGATTCTCAGCCAGCACTTCGATATTGAGATTTGCAGTATCAATGTGCAAGATTGTCGAGTGGACAAGTTTAATGAGGGGAAGGCTACGAGATGTCTGTTGGTGTATTCGGGGATTCACTACGATGTGTGTGCGGTGGCGCCTTATAGTGGTGCAGATCCGGAGTTCGATCGCAAGGTGTTTGAAGTGG-TTCGAGTTGATGGCGAGGAGGAGATGGATGGAGGGGCGTTGGCTGGAGCGCTGGAGCTGTGCAGAGGATTGCAGAAGCGGCATTACTTCACGGACACGCATGGGTTCAAGTTGAAGTGCAATCAGTGTGGTCAGGGAGGAAATGGGCAGCAGTGGGCTGTTCAGCATGCGAAGAGTACCGGGCATGGTGATTTTGG-GGAA	1319 	\|\|\|\|\|\|\|\|\|\|\|\|\|\|\|\|\|\|\|\|\|\|\|\|\|\|\|\|\|\|\|\|\|\|\|\|\|\|\|\|\|\|\|\|\|\|\|\|\|\|\|\|\|\|\|\|\|\|\|\|\|\|\|\|\|\|\|\|\|\|\|\|\|\|\|\|\|\|\|\|\|\|\|\|\|\|\|\|\|\|\|\|\|\|\|\|\|\|\|\|\|\|\|\|\|\|\|\|\|\|\|\|\|\|\|\|\|\|\|\|\|\|\|\|\|\|\|\|\|\|\|\|\|\|\|\|\|\|\|\|\|\|\|\|\|\|\|\|\|\|\|\|\|\|\|\|\|\|\|\|\|\|\|\|\|\|\|\|\|\|\|\|\|\|\|\|\|\|\|\|\|\|\|\|\|\|\|\|\|\|\|\|\|\|\|\|\|\|\|\|\|\|\|\|\|\|\|\|\|\|\|\|\|\|\|\|\|\|\|\|\|\|\|\|\|\|\|\|\|\|\|\|\|\|\|\|\|\|\|\|\|\|\|\|\|\|\|\|\|\|\|\|\|\|\|\|\|\|\|\|\|\|\|\|\|\|\|\|\|\|\|\|\|\|\|\|\|\|\|\|\|\|\|\|\|\|\|\|\|\|\|\|\|\|\|\|\|\|\|\|\|\|\|\|\|\|\|\|\|\|\|\|\|\|\|\|\|\|\|\|\|\|\|\|\|\|\|\|\|\|\|\|\|\|\|\|\|\|\|\|\|\|\|\|\|\|\|\|\|\|\|\|\|\|\|\|\|\|\|\|\|\|\|\|\|\|\|\|\|\|\|\|\|\|\|\|\|\|\|\|\|\|\|\|\|\|\|\|\|\|\|\|\|\|\|\|\|\|\|\|\|\|\|\|\|\|\|\|\|\|\|\|\|\|\|\|\|\|\|\|\|\|\|\|\|\|\|\|\|\|\|\|\|\|\|\|\|\|\|\|\|\|\|\|\|\|\|\|\|\|\|\|\|\|\|\|\|\|\|\|\|\|\|\|\|\|\|\|\|\|\|\|\|\|\|\|\|\|\|\|\|\|\|\|\|\|\|\|\|\|\|\|\|\|\|\|\|\|\|\|\|\|\|\|\|\|\|\|\|\|\|\|\|\|\|\|\|\|\|\|\|\|\|\|\|\|\|\|\|\|\|\|\|\|\|\|\|\|\|\|\|\|\|\|\|\|\|\|\|\|\|\|\|\|\|\|\|\|\|\|\|\|\|\|\|\|\|\|\|\|\|\|\|\|\|\|\|\|\|\|\|\|\|\|\|\|\|\|\|\|\|\|\|\|\|\|\|\|\|\|\|\|\|\|\|\|\|\|\|\|\|\|\|\|\|\|\|\|\|\|\|\|\|\|\|\|\|\|\|\|\|\|\|\|\|\|\|\|\|\|\|\|\|\|\|\|\|\|\|\|\|\|\|\|\|\|\|\|\|\|\|\|\|\|\|\|\|\|\|\|\|\|\|\|\|\|\|\|\|\|\|\|\|\|\|\|\|\|\|\|\|\|\|\|\|\|\|\|\|\|\|\|\|\|\|\|\|\|\|\|\|\|\|\|\|\|\|\|\|\|\|\|\|\|\|\|\|\|\|\|\|\|\|\|\|\|\|\|\|\|\|\|\|\|\|\|\|\|\|\|\|\|\|\|\|\|\|\|\|\|\|\|\|\|\|\|\|\|\|\|\|\|\|\|\|\|\|\|\|\|\|\|\|\|\|\|\|\|\|\|\|\|\|\|\|\|\|\|\|\|\|\|\|\|\|\|\|\|\|\|\|\|\|\|\|\|\|\|\|\|\|\|\|\|\|\|\|\|\|\|\|\|\|\|\|\|\|\|\|\|\|\|\|\|\|\|\|\|\|\|\|\|\|\|\|\|\|\|\|\|\|\| \|\|\|\|\|\|\|\|\|\|\|\|\|\|\|\|\|\|\|\|\|\|\|\|\|\|\|\|\|\|\|\| \|\|\|\|\|\|\|\|\|\|\|\|\|\|\|\|\|\|\|\|\|\|\|\|\|\|\|\|\|\|\|\| \|\|\|\|\|\|\|\|\|\|\|\|\|\|\|\|\|\|\|\|\|\|\|\|\|\|\|\|\|\|\|\|\|\|\|\|\|\|\| \|\| \|\| \|\| \|\|\|\|\| \|\|\|\|\|\|\|\| \|\|\|\|  \|\| \|\|\|\|\|\| \|\|\|\|\|\|\|\|\| \|\|\|\| \|\|\|\|\|\|\|\|\|\| \|\|\|\|\|\|  \|\|\|\|\|\| \|\|\| \|\|\|\|\| \|\|\|\| 	ATGGCGACTTCCATCCGCCTCCGAGTCCGAGGCCCTCAAGGAGTGGCGACGGTGACCATCAACAATGACGCCTCCTGGAGCGACCTCATTCACGAGATCTCCGAGAAGAGCGGAGTCGCCGACTTTGACCTCAAATACGGATATCCACCTCAGCCTCTGAACACCGAGTCCATCAGTCCAGACATGAAGCTATCCGACTTCCCTCAGAAACTCAACGGCGAACAACTCATCGTCATGCCGCGAGCTATTCAAGCCAAACTCTCCAGTCCTCTTGAAGGAACAACAAAGAACCAACAAAACGTCGAAACGTTACCCAAAGCGAGTTCATCATCAGCACCATCCCACAAGTCCGGAGACTTCCCAGACAAACCACTAAACCTCACGAGAAAGAAGCAAAACCTCGACTCTGACCCTCCCGAAGTACAAGTCCCTCTCCTCGACGGAGTCATGGTTCTGCGCGTGATGCCCGACGACAACTCCTGCATGTTCCGAGCATTGTCCTCCGCTGTCCTCGGCTCCGCCCTCGACGGCATGACCGAACTTCGCTCTGTCGTCGCGCAGACCATCCAATCTCAACCAGAGCTCTACACCGAAGGCATGCTCGAGAAGAAACCCACAGACTACTGCCAGTGGATCCAACGCGAAGACAGCTGGGGAGGTGGCATCGAGCTTTCGATTCTCAGCCAGCACTTCGATATTGAGATTTGCAGTATCAATGTGCAAGATTGTCGAGTGGACAAGTTTAATGAGGGGAAGGCTACGAGATGTCTGTTGGTGTATTCGGGGATTCACTACGATGTGTGTGCGGTGGCGCCTTATAGTGGTGCAGATCCGGAGTTCGATCGCAAGGTGTTTGAAGTGGTTTCGAGTTGATGGCGAGGAGGAGATGGATGGA-GGGCGTTGGCTGGAGCGCTGGAGCTGTGCAGA-GATTGCAGAAGCGGCATTACTTCACGGACACGCATGGGT-CA-GT-GA-GTGCA-TCAGTGTG-TCAGC-AG-AAATGG-CAGCAGTGG-CTGT-CAGCATGCGA-GAGTAC--GGCATGCTGAATTTGGAGGAA ``` |
| 22 | ggggacaagtttgtacaaaaaagcaggcttgatgTCTCTCGCACGAACCCTCCT, ggggaccactttgtacaagaaagctgggtcCAACCTCCCATACCCACTCC | Mycgr3\|77127  JGI  Ensembl |  |  | 100 % | chr\_11:1521345 G->A; | ``` 75	GATGTCTCTCGCACGAACCCTCCTCTCCGCCCTCCTCCTCGCCACCACCACCACCTTCGCCCAAACCCACACCGAATGCAGCCCCCTCAACTCCACCTGTCCTCCAAACCCCGCCCTGGGAACCGAACTCCTCGAAGTCTTCAACTCCTCCTCCACCGAACTAAACTCCAAATACTGGCACTTTGACGGCGGTCAAGAGCTCGTTTCGTTCGGCAAAGATGGCGTCCACCTATCGCTCAAGGAGAAAGGCGATACCGTTACCATCAAGTCGAACTTTTACATCTTCTTCGGGACACTGGAGTTTATCATGAAAGCCGCTCCCGGGAATGGGATTATCAGTACGGTCATTCTGTTGAGTGATGATCTGGATGAGATTGATTGGGAGATCAAGGGTGGGAATACGACGGCTGTGTCGAATAATTACTATGG	503 	\|\|\|\|\|\|\|\|\|\|\|\|\|\|\|\|\|\|\|\|\|\|\|\|\|\|\|\|\|\|\|\|\|\|\|\|\|\|\|\|\|\|\|\|\|\|\|\|\|\|\|\|\|\|\|\|\|\|\|\|\|\|\|\|\|\|\|\|\|\|\|\|\|\|\|\|\|\|\|\|\|\|\|\|\|\|\|\|\|\|\|\|\|\|\|\|\|\|\|\|\|\|\|\|\|\|\|\|\|\|\|\|\|\|\|\|\|\|\|\|\|\|\|\|\|\|\|\|\|\|\|\|\|\|\|\|\|\|\|\|\|\|\|\|\|\|\|\|\|\|\|\|\|\|\|\|\|\|\|\|\|\|\|\|\|\|\|\|\|\|\|\|\|\|\|\|\|\|\|\|\|\|\|\|\|\|\|\|\|\|\|\|\|\|\|\|\|\|\|\|\|\|\|\|\|\|\|\|\|\|\|\|\|\|\|\|\|\|\|\|\|\|\|\|\|\|\|\|\|\|\|\|\|\|\|\|\|\|\|\|\|\|\|\|\|\|\|\|\|\|\|\|\|\|\|\|\|\|\|\|\|\|\|\|\|\|\|\|\|\|\|\|\|\|\|\|\|\|\|\|\|\|\|\|\|\|\|\|\|\|\|\|\|\|\|\|\|\|\|\|\|\|\|\|\|\|\|\|\|\|\|\|\|\|\|\|\|\|\|\|\|\|\|\|\|\|\|\|\|\|\|\|\|\|\|\|\|\|\|\|\|\|\|\|\|\|\|\|\|\|\|\|\|\|\|\|\|\|\|\|\|\|\|\|\|\|\|\|\|\|\|\|\|\|\|\|\|\|\|\|\|\|\|\|\|\|\|\|\|\|\|\|\|\|\|\|\|\|\|\|\|\|\|\|\|\|\|\|\|\|\|\|\|\|\|\|\|\|\|\|\|\|\|\|\|\|\|\|\| 	GATGTCTCTCGCACGAACCCTCCTCTCCGCCCTCCTCCTCGCCACCACCACCACCTTCGCCCAAACCCACACCGAATGCAGCCCCCTCAACTCCACCTGTCCTCCAAACCCCGCCCTGGGAACCGAACTCCTCGAAGTCTTCAACTCCTCCTCCACCGAACTAAACTCCAAATACTGGCACTTTGACGGCGGTCAAGAGCTCGTTTCGTTCGGCAAAGATGGCGTCCACCTATCGCTCAAGGAGAAAGGCGATACCGTTACCATCAAGTCGAACTTTTACATCTTCTTCGGGACACTGGAGTTTATCATGAAAGCCGCTCCCGGGAATGGGATTATCAGTACGGTCATTCTGTTGAGTGATGATCTGGATGAGATTGATTGGGAGATCAAGGGTGGGAATACGACGGCTGTGTCGAATAATTACTATGG ```      ``` 502	GGATGGGGCAATCAAGACCAACGATTCGCTGAAGATCCCGCTCTCGACGGTCCGCAAAATGACTACCACAACTACACGATCGACTGGACGCAGGAACGCATCGAGTACCTGATTGACAGCAACGTCGTGCGGACAGTACACTACGACGAACCAGGTCTCTACCCGCAAACTCCTTCGCGCGTACAATTCGGCGTCTGGTGTGGTGGATGTTCCAACCAACCAGGAACAGTCGAGTGGGCAGGTGGAGCGCCAGACTTCAGCAAAGG	767 	\|\|\|\|\|\|\|\|\|\|\|\|\|\|\|\|\|\|\|\|\|\|\|\|\|\|\|\|\|\|\|\|\|\|\|\|\|\|\|\|\|\|\|\| \|\|\|\|\|\|\|\|\|\|\|\|\|\|\|\|\|\|\|\|\|\|\|\|\|\|\|\|\|\|\|\|\|\|\|\|\|\|\|\|\|\|\|\|\|\|\|\|\|\|\|\|\|\|\|\|\|\|\|\|\|\|\|\|\|\|\|\|\|\|\|\|\|\|\|\|\|\|\|\|\|\|\|\|\|\|\|\|\|\|\|\|\|\|\|\|\|\|\|\|\|\|\|\|\|\|\|\|\|\|\|\|\|\|\|\|\|\|\|\|\|\|\|\|\|\|\|\|\|\|\|\|\|\|\|\|\|\|\|\|\|\|\|\|\|\|\|\|\|\|\|\|\|\|\|\|\|\|\|\|\|\|\|\|\|\|\|\|\|\|\|\|\|\|\|\|\|\|\|\|\|\|\|\|\|\|\|\|\|\|\|\|\|\|\|\|\|\|\|\|\|\|\|\|\|\|\|\|\|\|\|\|\|\|\|\|\|\|\|\|\| 	GGATGGGGCAATCAAGACCAACGATTCGCTGAAGATCCCGCTCTTGACGGTCCGCAAAATGACTACCACAACTACACGATCGACTGGACGCAGGAACGCATCGAGTACCTGATTGACAGCAACGTCGTGCGGACAGTACACTACGACGAACCAGGTCTCTACCCGCAAACTCCTTCGCGCGTACAATTCGGCGTCTGGTGTGGTGGATGTTCCAACCAACCAGGAACAGTCGAGTGGGCAGGTGGAGCGCCAGACTTCAGCAAAGG ```      ``` 767	GCCCCTTCACAGCCTCCATCGCCTCCCTCCGCGTCACCGACGCCCACACCAACGCCTCCACCTACACCTGGTCCGACCGCACGGGCTCCTACCAATCCATCGCCGTCACGGCCGGAAAATCCGAAGCCTACAAAGCCATCCACCACGAATCCACCC-GCGCCAAACTCTCCTCCACCTGGGCGGGCATGTCCCCCGGAGCGAAAATcggcgtcgcggcggggatattgggcgccgtggcggtggc-ggcggtg	1017 	\|\|\|\|\|\|\|\|\|\|\|\|\|\|\|\|\|\|\|\|\|\|\|\|\|\|\|\|\|\|\|\|\|\|\|\|\|\|\|\|\|\|\|\|\|\|\|\|\|\|\|\|\|\|\| \|\|\|\|\|\|\|\|\|\| \|\|\|\|\|\|\|\|\|\|\|\|\|\|\|\|\|\|\|\|\| \|\|\| \|\|\|\|\|\|\|\|\| \|\|\|\|\|\|\|\| \|\|\| \|\|\|\|\|\|\|\| \|\| \|\|\|\|\| \|\| \|\|\| \|\| \|\|\|\| \|\| \|\|\|\| \|\|\|\|\|\|\|\|\|\|\|\|\| \|\|\|\| \|\|\|\|\|\|\|\|\|\|\|\|\|\| \|\|\|\|\|\|\|\|\|\|\|\|\|\|\|\|\| \|\|\|  \|\|\|\|\|\|\| \|\|\|\| \|\|\|\|\|\|  \|\|\|\| \|\|\|\|\| \|\|\|\|\|\|\| 	GCCCCTTCACAGCCTCCATCGCCTCCCTCCGCGTCACCGACGCCCACACCAACGC-TCCACCTACA-CTGGTCCGACCGCACGGGCTC-TAC-AATCCATCG-CGTCACGGGCGG-AAATCCGAGGCTTACAA-GC-ATC-AC-ACGA-TC-ACCCGGCGCCAAACTCTC-TCCA-CTGGGCGGGCATGT-CCCCGGAGCGAAAATCG-CGTTCCGGCGGG-ATAT-GGGCGCA-TGGCCGTGGCCGGCGGTG ``` |
| 22 | ggggacaagtttgtacaaaaaagcaggcttgatgAGGGCTGGTACTTGGATAAC, ggggaccactttgtacaagaaagctgggtcTATGTGGAGCGGGACATACC | Mycgr3\|78715  JGI  Ensembl |  |  | 100 % |  |  |
| 22 | ggggacaagtttgtacaaaaaagcaggcttgatgTCCTCATTCTCCCAGACCCT, ggggaccactttgtacaagaaagctgggtcCCTCTTCCCCATCCAAGTCA | Mycgr3\|88401  JGI  Ensembl |  |  | 100 % |  |  |
| 22 | ggggacaagtttgtacaaaaaagcaggcttgatgGTTTCTCTCAAAGCCGCCGT, ggggaccactttgtacaagaaagctgggtcGTTCGGCGGCGACGGAGGCA | Mycgr3\|88944  JGI  Ensembl | KOG4701 Chitinase; |  | 100 % |  |  |
| 22 | ggggacaagtttgtacaaaaaagcaggcttgatgTCGCATTACTGCGACACCTG, ggggaccactttgtacaagaaagctgggtcGAGGAAACGTGCCGATTTTC | Mycgr3\|89702  JGI  Ensembl |  |  | 0 % |  |  |
| 22 | ggggacaagtttgtacaaaaaagcaggcttgatgGCTCTGTCCTTGCTATTCGC, ggggaccactttgtacaagaaagctgggtcGAGGGGGATCATGTCTTCCA | Mycgr3\|89737  JGI  Ensembl |  |  | 100 % | chr\_1:3971828 C->T; |  |
| 22 | ggggacaagtttgtacaaaaaagcaggcttgatgTCCGCCTGCGATATGGAATC, ggggaccactttgtacaagaaagctgggtcGAACTCCTTGCGAGTTCGAC | Mycgr3\|90036  JGI  Ensembl |  |  | 1.4383 % |  |  |
| 22 | ggggacaagtttgtacaaaaaagcaggcttgatgCAGCGATACCTTCAAGATCC, ggggaccactttgtacaagaaagctgggtcAGTCAAGCTCTTTCGCCTGC | Mycgr3\|90147  JGI  Ensembl |  |  | 100 % |  |  |
| 22 | ggggacaagtttgtacaaaaaagcaggcttgatgCAGCTCACGACCTCCATCCT, ggggaccactttgtacaagaaagctgggtcTTCCCACTTTTCATCAAATG | Mycgr3\|90676  JGI  Ensembl |  |  | 100 % |  |  |
| 22 | ggggacaagtttgtacaaaaaagcaggcttgatgGATTCAACCTGGGAACTCAC, ggggaccactttgtacaagaaagctgggtcTGCGCCCGGTGCCTCAGGGA | Mycgr3\|90837  JGI  Ensembl |  |  | 100 % |  | ``` 1	ATGGATTCAACCTGGGAACTCACATCCTCGATGGCGGCCTTCAAGCCCTACGACAGGCCCTCCAGCGCTCGTCCGCTCCTTGTATGCGGCATCCACCGCCGTCTTTCCGACAAGCAAATCGAGAAGCGTACATCCAAAGTCATCTACCACCGCTCTCATGGCAGCAATGAGAAGCCAGGATCCTGTATCTGCGACTACACCGAATTCCCGGAACTTCTCTCGGAACGTCTCAACTCTCAGAAAATG	246 	\|\|\|\|\|\|\|\|\|\|\|\|\|\|\|\|\|\|\|\|\|\|\|\|\|\|\|\|\|\|\|\|\|\|\|\|\|\|\|\|\|\|\|\|\|\|\|\|\|\|\|\|\|\|\|\|\|\|\|\|\|\|\|\|\|\|\|\|\|\|\|\|\|\|\|\|\|\|\|\|\|\|\|\|\|\|\|\|\|\|\|\|\|\|\|\|\|\|\|\|\|\|\|\|\|\|\|\|\|\|\|\|\|\|\|\|\|\|\|\|\|\|\|\|\|\|\|\|\|\|\|\|\|\|\|\|\|\|\|\|\|\|\|\|\|\|\|\|\|\|\|\|\|\|\|\|\|\|\|\|\|\|\|\|\|\|\|\|\|\|\|\|\|\|\|\|\|\|\|\|\|\|\|\|\|\|\|\|\|\|\|\|\|\|\|\|\|\|\|\|\|\|\|\|\|\|\|\|\|\|\|\|\|\|\|\|\|\|\|\|\|\|\|\|\|\|\|\|\|\|\|\|\|\|\|\|\|\|\|\|\|\|\|\|\|\| 	ATGGATTCAACCTGGGAACTCACATCCTCGATGGCGGCCTTCAAGCCCTACGACAGGCCCTCCAGCGCTCGTCCGCTCCTTGTATGCGGCATCCACCGCCGTCTTTCCGACAAGCAAATCGAGAAGCGTACATCCAAAGTCATCTACCACCGCTCTCATGGCAGCAATGAGAAGCCAGGATCCTGTATCTGCGACTACACCGAATTCCCGGAACTTCTCTCGGAACGTCTCAACTCTCAGAAAATG ```      ``` 246	GCCCTGCTCCATTCCCTCCCATGGAAAGTCCTTCATGGATCTCCCCACCGAGATGAGGATAAAATGTTACGAAGCCGCTCTGATCTCTCCTGACTCAATCGAATTCTGCCCCATGCCCTTCAACTTCACCGGTCGCGCTCCGTCCGGGAAAGAAGCTCAGGATGCCTTTGACGACGCGTGGTGCCCAGTCGAAGTCTATCACACTCAACTCATCGCCCGATTTTGCGCCCGCGCCAAACTCCTCCGTGTCAGCAAGACCATCAACGCCGAAGCGACTCCGGTTTTCTACGGTGGGAACCACTTCGCCTTCAGCAATATCTCCGGTTGGGTCGGATTGCTATTGTTCATGTACCGCATTGGGATGCACAACTGCCGACTGCTCCGGCGAATGACCGTTTGTCACCCCCTCTTGATGTTGAAGCCGCAGTCGCTCAAAGCAGACGACCATTTCCGCCGCAGTATCGTTGCATTGTTCTTCCCTTCCTCAGCGCATTCGTGGGGCTTCCTACCAGCGAGATGGTTTGATCACGGAAGACCAATGGATGGCAAACTCGTATTGGAGAAGATCTCGAAGCTAAAGGAACTAAACATCATCGTTGACTGTCCCCAAACTATGGGCATGTGGGACGGAGAAGCGGAGAGGGATTTAGTGGAGGGATTTGCGAGGCCTTTCCTCGACGAGAACAAGTTTGTGGACCTTCAGGTTACGGTGTTGGAGATGTGG	969 	\|\|\|\|\|\|\|\|\|\|\|\|\|\|\|\|\|\|\|\|\|\|\|\|\|\|\|\|\|\|\|\|\|\|\|\|\|\|\|\|\|\|\|\|\|\|\|\|\|\|\|\|\|\|\|\|\|\|\|\|\|\|\|\|\|\|\|\|\|\|\|\|\|\|\|\|\|\|\|\|\|\|\|\|\|\|\|\|\|\|\|\|\|\|\|\|\|\|\|\|\|\|\|\|\|\|\|\|\|\|\|\|\|\|\|\|\|\|\|\|\|\|\|\|\|\|\|\|\|\|\|\|\|\|\|\|\|\|\|\|\|\|\|\|\|\|\|\|\|\|\|\|\|\|\|\|\|\|\|\|\|\|\|\|\|\|\|\|\|\|\|\|\|\|\|\|\|\|\|\|\|\|\|\|\|\|\|\|\|\|\|\|\|\|\|\|\|\|\|\|\|\|\|\|\|\|\|\|\|\|\|\|\|\|\|\|\|\|\|\|\|\|\|\|\|\|\|\|\|\|\|\|\|\|\|\|\|\|\|\|\|\|\|\|\|\|\|\|\|\|\|\|\|\|\|\|\|\|\|\|\|\|\|\|\|\|\|\|\|\|\|\|\|\|\|\|\|\|\|\|\|\|\|\|\|\|\|\|\|\|\|\|\|\|\|\|\|\|\|\|\|\|\|\|\|\|\|\|\|\|\|\|\|\|\|\|\|\|\|\|\|\|\|\|\|\|\|\|\|\|\|\|\|\|\|\|\|\|\|\|\|\|\|\|\|\|\|\|\|\|\|\|\|\|\|\|\|\|\|\|\|\|\|\|\|\|\|\|\|\|\|\|\|\|\|\|\|\|\|\|\|\|\|\|\|\|\|\|\|\|\|\|\|\|\|\|\|\|\|\|\|\|\|\|\|\|\|\|\|\|\|\|\|\|\|\|\|\|\|\|\|\|\|\|\|\|\|\|\|\|\|\|\|\|\|\|\|\|\|\|\|\|\|\|\|\|\|\|\|\|\|\|\|\|\|\|\|\|\|\|\|\|\|\|\|\|\|\|\|\|\|\|\|\|\|\|\|\|\|\|\|\|\|\|\|\|\|\|\|\|\|\|\|\|\|\|\|\|\|\|\|\|\|\|\|\|\|\|\|\|\|\|\|\|\|\|\|\|\|\|\|\|\|\|\|\|\|\|\|\|\|\|\|\|\|\|\|\|\|\|\|\|\|\|\|\|\|\|\|\|\|\|\|\|\|\|\|\|\|\|\|\|\|\|\|\|\|\|\|\|\|\|\|\|\|\|\|\|\|\|\|\|\|\|\|\|\|\|\|\|\|\|\|\|\|\|\|\|\|\|\|\|\|\|\|\|\|\|\|\|\|\|\|\|\|\|\|\|\|\|\|\|\|\|\|\|\|\|\|\|\|\| \|\|\|\|\|\|\|\|\|\|\|\|\|\|\|\|\|\| \|\|\|\|\|\|\|\|\|\|\|\|\|\| \| \|\|\| \|\|\|\|\|\|\|\|\|\|\|\|\|\|\| \|\|\|\|\|\|\|\|\|\|\|\|\|\| \| \|\|\| \|\|\| \| \| \|\|\|\|\|\| 	GCCCTGCTCCATTCCCTCCCATGGAAAGTCCTTCATGGATCTCCCCACCGAGATGAGGATAAAATGTTACGAAGCCGCTCTGATCTCTCCTGACTCAATCGAATTCTGCCCCATGCCCTTCAACTTCACCGGTCGCGCTCCGTCCGGGAAAGAAGCTCAGGATGCCTTTGACGACGCGTGGTGCCCAGTCGAAGTCTATCACACTCAACTCATCGCCCGATTTTGCGCCCGCGCCAAACTCCTCCGTGTCAGCAAGACCATCAACGCCGAAGCGACTCCGGTTTTCTACGGTGGGAACCACTTCGCCTTCAGCAATATCTCCGGTTGGGTCGGATTGCTATTGTTCATGTACCGCATTGGGATGCACAACTGCCGACTGCTCCGGCGAATGACCGTTTGTCACCCCCTCTTGATGTTGAAGCCGCAGTCGCTCAAAGCAGACGACCATTTCCGCCGCAGTATCGTTGCATTGTTCTTCCCTTCCTCAGCGCATTCGTGGGGCTTCCTACCAGCGAGATGGTTTGATCACGGAAGACCAATGGATGGCAAACTCGTATTGGAGAAGATCTCGAAGCTAAAGGAACTAAACATCATCGTTGACTGTCCCCAAACTATGGGCATGTGGGACGGAG-AGCGGAGAGGGATTTAGT-GAGGGATTTGCGAG-C-TTT-CTCGACGAGAACAAG-TTGTGGACCTTCAG-T-ACG-TGT-GAA-ATGTGG ``` |
| 22 | ggggacaagtttgtacaaaaaagcaggcttgatgAAACTCTCTCTCCTAACCAT, ggggaccactttgtacaagaaagctgggtcAGCACCTCTCGTCTTCCTCG | Mycgr3\|91425  JGI  Ensembl | KOG1808 AAA ATPase containing von Willebrand factor type A (vWA) domain; |  | 100 % |  |  |
| 22 | ggggacaagtttgtacaaaaaagcaggcttgatgGCCATCTCTTGCTTCACGCT, ggggaccactttgtacaagaaagctgggtcGACCGTCCTCGCACTATCCC | Mycgr3\|91524  JGI  Ensembl |  |  | 100 % | chr\_3:1089186 G->A; |  |
| 22 | ggggacaagtttgtacaaaaaagcaggcttgatgAAAAGCATCCAGGAATGGCT, ggggaccactttgtacaagaaagctgggtcATGATACATCAACCGAGTGA | Mycgr3\|91638  JGI  Ensembl |  |  | 1.42965 % |  |  |
| 22 | ggggacaagtttgtacaaaaaagcaggcttgatgAAAACGCTCGTCTTGACTCA, ggggaccactttgtacaagaaagctgggtcCTGCACCTCCCTCCTGCCAG | Mycgr3\|92796  JGI  Ensembl |  |  | 100 % |  | ``` 1	ATGAAAACGCTCGTCTTGACTCACCGCGACCGCCGTACGCACGCAAATTTCCTGGCGTTGCCGCCCGAGGTGAGGCTGCGTTGCTACGAGTACATGTTGATCCCTCCATTACCCATACACGCCCTCCGCCACCAACCCACCTGCCACAACCACAATCCTCTCGCCGCCTCAATTCCGCCGCCATACATACCCGCCCTCCAAAATCGTCCCAAACCAGCCTCATGGCGCTCCCTCCTCCCCCGGGAAGTCTTATCCGAAAAGTGTCGCATCTCCTGTATCCAGCAATCCGGCCAAAGACGGAATCTCGCCATCCTTCGCGTAAGTCGATTGGTGTACAAGGAAGCCATAGAGATCTTGTGGGAAGATGAAAAATGGGGCTTCACTTTCACCGGTCTGCAGGATTTCGTGGATAGCATGAAGACCATGCCGCTGGAAGGTAGGTCGAGAGTGCGGAAGTTGGTGGTTAGAGAGGACAGGAATATGGATGTGCAGCGGAGGGGGTTTGGGTTGAAGGAGTGGTTGCATGCTGAGTTCTGGAGGTTCGTGGGAGGATGTCCGAGATTGAAGTATTTGGAGGTGCCGAGTACGATGGGTGGGGTGCTGGCGGGAGAGGTCGTGAAGCTGGGAAGCCTGGAGGTAGTGAGTGTGAGCGTGGTTGGGTGGATGACGTTCGTGGAGGGTATGGATGTGCACAGGGTGGAAATTGGAGTTAAAAAGGGGTTTAAAATTCGGAGCGGTGAGATCGCGAGGAATAGGAGGGAGATGAAGATGGAGGAGTGGTT-GAGAATAGCGGATGGCAGGCGTGGCGGGAGACGGGAGACGCCCGAGTGGATGCAAAGGGTTGATGCGGCGTTGGAAAGAGATCGAGATGTGATGGTCCATCAGGAAGGTGGGCGGGATTGGGTGACTGTGACGATGGCGGAGGGAATGCGGGTCGAAGTGGAGATATGGGG-ACTGCCAACGCATTCAAAAGCGGAGAGGAGGAAATTGTTTCTGC-AGGAGGCAAGGAAGCCGAAGACGTTGAAGGTTTCGAGGCGAACAGTGCCTGATCTTCCAGAACTGGGAGGTGAAAACGACGAGGAATTGGCGATGATGGCGAAAAAGTCGGAT	1110 	\|\|\|\|\|\|\|\|\|\|\|\|\|\|\|\|\|\|\|\|\|\|\|\|\|\|\|\|\|\|\|\|\|\|\|\|\|\|\|\|\|\|\|\|\|\|\|\|\|\|\|\|\|\|\|\|\|\|\|\|\|\|\|\|\|\|\|\|\|\|\|\|\|\|\|\|\|\|\|\|\|\|\|\|\|\|\|\|\|\|\|\|\|\|\|\|\|\|\|\|\|\|\|\|\|\|\|\|\|\|\|\|\|\|\|\|\|\|\|\|\|\|\|\|\|\|\|\|\|\|\|\|\|\|\|\|\|\|\|\|\|\|\|\|\|\|\|\|\|\|\|\|\|\|\|\|\|\|\|\|\|\|\|\|\|\|\|\|\|\|\|\|\|\|\|\|\|\|\|\|\|\|\|\|\|\|\|\|\|\|\|\|\|\|\|\|\|\|\|\|\|\|\|\|\|\|\|\|\|\|\|\|\|\|\|\|\|\|\|\|\|\|\|\|\|\|\|\|\|\|\|\|\|\|\|\|\|\|\|\|\|\|\|\|\|\|\|\|\|\|\|\|\|\|\|\|\|\|\|\|\|\|\|\|\|\|\|\|\|\|\|\|\|\|\|\|\|\|\|\|\|\|\|\|\|\|\|\|\|\|\|\|\|\|\|\|\|\|\|\|\|\|\|\|\|\|\|\|\|\|\|\|\|\|\|\|\|\|\|\|\|\|\|\|\|\|\|\|\|\|\|\|\|\|\|\|\|\|\|\|\|\|\|\|\|\|\|\|\|\|\|\|\|\|\|\|\|\|\|\|\|\|\|\|\|\|\|\|\|\|\|\|\|\|\|\|\|\|\|\|\|\|\|\|\|\|\|\|\|\|\|\|\|\|\|\|\|\|\|\|\|\|\|\|\|\|\|\|\|\|\|\|\|\|\|\|\|\|\|\|\|\|\|\|\|\|\|\|\|\|\|\|\|\|\|\|\|\|\|\|\|\|\|\|\|\|\|\|\|\|\|\|\|\|\|\|\|\|\|\|\|\|\|\|\|\|\|\|\|\|\|\|\|\|\|\|\|\|\|\|\|\|\|\|\|\|\|\|\|\|\|\|\|\|\|\|\|\|\|\|\|\|\|\|\|\|\|\|\|\|\|\|\|\|\|\|\|\|\|\|\|\|\|\|\|\|\|\|\|\|\|\|\|\|\|\|\|\|\|\|\|\|\|\|\|\|\|\|\|\|\|\|\|\|\|\|\|\|\|\|\|\|\|\|\|\|\|\|\|\|\|\|\|\|\|\|\|\|\|\|\|\|\|\|\|\|\|\|\|\|\|\|\|\|\|\|\|\|\|\|\|\|\|\|\|\|\|\|\|\|\|\|\|\|\|\|\|\|\|\|\|\|\|\|\|\|\|\|\|\|\|\|\|\|\|\|\|\|\|\|\|\|\|\|\|\|\|\|\|\|\|\|\|\|\|\|\|\|\|\|\|\|\|\|\|\|\|\|\|\|\|\|\|\|\|\|\|\|\|\|\|\|\|\|\|\|\|\|\|\|\|\|\|\|\|\|\|\|\|\|\|\|\|\|\|\|\|\|\|\|\|\|\|\|\|\|\|\|\|\|\|\|\|\|\|\|\|\|\|\|\|\|\|\|\|\|\|\|\|\|\|\|\|\|\|\|\|\|\|\|\|\|\|\|\|\|\|\|\|\|\|\|\|\|\|\|\|\|\|\|\|\|\|\|\|\|\|\|\|\|\|\| \|\|\| \|\|\|\|\|\|\|\|\|\|\|\|\|\|\|\|\|\|\| \|\|\|\|\|\|\|\|\|\|\|\|\|\|\|\|\|\|\|\|\|\|\|\|\|\|\|\|\|\|\|\|\|\|\|\|\|\|\|\|\|\|\|\|\|\|\|\|\|\|\|\|\|\|\|\|\|\|\|\|\|\|\|\|\|\|\|\|\|\|\|\|\|\|\|\|\|\| \|\|\|\|\|\|\|\|\|\|\|\|\|\|\|\| \|\|\|\|\|\|\|\|\|\|\|\|\|\|\|\|\|\|\|\|\| \|\|\|\|\|\|\|\|\| \|\|\|\|\|\|\|\| \|\|\|\|\|\|\|\|\|\| \|\|\|\|\|\|\|\|\|\|\|\|\|\|\|\|\|\| \|\|\|\|\| \|\|\|\|\| \|\| \|\|\|\|\|\|\|\|\|\| \|\|\|\| \|\|\| \| \|\|\|\| \|\| \|\|\|\|\|\|\|\|\|\| \|\| \| \|\|\|\| \| \|\| \|\| \| \|\|\|\|\| \|  \|\|  \|\|\| \|\|\|\|\|\|\|  \|\|\|\| \|\| \| \| \|\|\| \|\|\|  \|\|\|\|\|\|\|\|  \|\|\|\|\|\| 	ATGAAAACGCTCGTCTTGACTCACCGCGACCGCCGTACGCACGCAAATTTCCTGGCGTTGCCGCCCGAGGTGAGGCTGCGTTGCTACGAGTACATGTTGATCCCTCCATTACCCATACACGCCCTCCGCCACCAACCCACCTGCCACAACCACAATCCTCTCGCCGCCTCAATTCCGCCGCCATACATACCCGCCCTCCAAAATCGTCCCAAACCAGCCTCATGGCGCTCCCTCCTCCCCCGGGAAGTCTTATCCGAAAAGTGTCGCATCTCCTGTATCCAGCAATCCGGCCAAAGACGGAATCTCGCCATCCTTCGCGTAAGTCGATTGGTGTACAAGGAAGCCATAGAGATCTTGTGGGAAGATGAAAAATGGGGCTTCACTTTCACCGGTCTGCAGGATTTCGTGGATAGCATGAAGACCATGCCGCTGGAAGGTAGGTCGAGAGTGCGGAAGTTGGTGGTTAGAGAGGACAGGAATATGGATGTGCAGCGGAGGGGGTTTGGGTTGAAGGAGTGGTTGCATGCTGAGTTCTGGAGGTTCGTGGGAGGATGTCCGAGATTGAAGTATTTGGAGGTGCCGAGTACGATGGGTGGGGTGCTGGCGGGAGAGGTCGTGAAGCTGGGAAGCCTGGAGGTAGTGAGTGTGAGCGTGGTTGGGTGGATGACGTTCGTGGAGGGTATGGATGTGCACAGGGTGGAAATTGGAGTTAAAAAGGGGTTTAAAATTCGGAGCGGTGAGATCGCGAGGAATAGGAGGGAGATGAAGATGGAGGAGTGGTTTGAG-ATAGCGGATGGCAGGCGTGTCGGGAGACGGGAGACGCCCGAGTGGATGCAAAGGGTTGATGCGGCGTTGGAAAGAGATCGAGATGTGATGGTCCATCA-GAAGGTGGGCGGGATT-GGTGACTGTGACGATGGCGGAAGGAATGCGG-TCGAAGTG-AGATATGGGGCACTGCCAACGCATTCAAA-GCGGAAAGGAG-AA-TTGTTTCTGCTAGGAAGCA-G-AAGC-GA-GACGTTGAAG-TT-C-AGGCTA-CA-TGGC-GATCT-CGTGACTTGGTAGGTGAA--CGACCAGTA-TGGGC-ATGTAGGCGAAAAGTTCGGAT ``` |
| 22 | ggggacaagtttgtacaaaaaagcaggcttgatgATTCAAGTCATTAAGTTCTT, ggggaccactttgtacaagaaagctgggtcGTAACCAATCATCACATCGC | Mycgr3\|92926  JGI  Ensembl | KOG0157 Cytochrome P450 CYP4/CYP19/CYP26 subfamilies; |  | 100 % |  |  |
| 22 | ggggacaagtttgtacaaaaaagcaggcttgatgGCGACGATCCACGACCCACA, ggggaccactttgtacaagaaagctgggtcCTTCCCCTCAACTTCTTCCC | Mycgr3\|92956  JGI  Ensembl | KOG1307 K+-dependent Ca2+/Na+ exchanger NCKX1 and related proteins; |  | 100 % |  |  |
| 22 | ggggacaagtttgtacaaaaaagcaggcttgatgGATCGCAGGGAGCTGGAACG, ggggaccactttgtacaagaaagctgggtcATCCGCCTCTTTCCCGAACG | Mycgr3\|93078  JGI  Ensembl |  |  | 100 % | chr\_5:731120 G->A; |  |
| 22 | ggggacaagtttgtacaaaaaagcaggcttgatgCTCTTCCAATACGCAAACCT, ggggaccactttgtacaagaaagctgggtcACCGTTCTTGCCGCCCTCCT | Mycgr3\|93835  JGI  Ensembl |  |  | 100 % |  |  |
| 22 | ggggacaagtttgtacaaaaaagcaggcttgatgGCTTCTCCTCCATCCGGCGC, ggggaccactttgtacaagaaagctgggtcCAATGCAGCTTTCTTCGCAT | Mycgr3\|94670  JGI  Ensembl | KOG0800 FOG: Predicted E3 ubiquitin ligase; | protein binding; zinc ion binding; | 100 % | chr\_7:1935856 C->A; |  |
| 22 | ggggacaagtttgtacaaaaaagcaggcttgatgTCGTCCTCCAACTCCGCCAC, ggggaccactttgtacaagaaagctgggtcCCCCTCCGCCCTCCCACCCG | Mycgr3\|94835  JGI  Ensembl |  |  | 1.5897 % |  |  |
| 22 | ggggacaagtttgtacaaaaaagcaggcttgatgGCCACTCATCCACTCGAGAA, ggggaccactttgtacaagaaagctgggtcGACAGCGGGCTCCGCCTCAT | Mycgr3\|95531  JGI  Ensembl |  |  | 100 % |  |  |
| 22 | ggggacaagtttgtacaaaaaagcaggcttgatgCACTCCACCACCACTCTTTT, ggggaccactttgtacaagaaagctgggtcTCCTCCACTCCCTCCTCCTC | Mycgr3\|95797  JGI  Ensembl | KOG1187 Serine/threonine protein kinase; | UDP-N-acetylmuramoylalanyl-D-glutamyl-2,6-diaminopimelate-D-alanyl-D-alanine ligase activity; coenzyme F420-0 gamma-glutamyl ligase activity;  coenzyme F420-2 alpha-glutamyl ligase activity;  ribosomal S6-glutamic acid ligase activity; | 0 % |  |  |
| 22 | ggggacaagtttgtacaaaaaagcaggcttgatgCCTACCTCTCCCTCTCAAAA, ggggaccactttgtacaagaaagctgggtcTAATCCGGGTCCAATTGGCC | Mycgr3\|95850  JGI  Ensembl |  |  | 100 % |  |  |
| 22 | ggggacaagtttgtacaaaaaagcaggcttgatgGAACACAAATTGGCCGCCGG, ggggaccactttgtacaagaaagctgggtcCTGCCTCATCGCTCTTGTCA | Mycgr3\|96232  JGI  Ensembl |  |  | 100 % |  |  |
| 22 | ggggacaagtttgtacaaaaaagcaggcttgatgAACTCCATCATCATCGGCGC, ggggaccactttgtacaagaaagctgggtcGTCCTCCCCAACTGCGCCAC | Mycgr3\|97554  JGI  Ensembl |  |  | 100 % |  |  |
| 22 | ggggacaagtttgtacaaaaaagcaggcttgatgTTTCGTACTAACATACTTTC, ggggaccactttgtacaagaaagctgggtcCTTCTCTTCCAGCTCTCGGA | Mycgr3\|97584  JGI  Ensembl |  |  | 100 % |  |  |
| 22 | ggggacaagtttgtacaaaaaagcaggcttgatgGAGAATCATTCACGCTTCAA, ggggaccactttgtacaagaaagctgggtcAGCTGAAGTCAAACCAGCCT | Mycgr3\|97941  JGI  Ensembl | KOG4297 C-type lectin; |  | 100 % | chr\_19:194577 G->A; |  |
| 22 | ggggacaagtttgtacaaaaaagcaggcttgatgATCTCCTCGGTGCTCTCAGT, ggggaccactttgtacaagaaagctgggtcCTCCACAAGACCGAAAGAGA | Mycgr3\|98175  JGI  Ensembl | KOG1604 Predicted mutarotase; | aldose 1-epimerase activity; carbohydrate metabolic process;  isomerase activity; | 100 % |  |  |
| 22 | ggggacaagtttgtacaaaaaagcaggcttgatgTCCACCCAACTCGAGTCAGT, ggggaccactttgtacaagaaagctgggtcCTTCGTTTCGCCCTCTACAA | Mycgr3\|98377  JGI  Ensembl | KOG2757 Mannose-6-phosphate isomerase; | carbohydrate metabolic process; mannose-6-phosphate isomerase activity;  zinc ion binding; | 100 % |  |  |
| 22 | ggggacaagtttgtacaaaaaagcaggcttgatgGCAGCACAATCTCGGAAGCC, ggggaccactttgtacaagaaagctgggtcGTTCGCTCCCACCGTCGGAC | Mycgr3\|99779  JGI  Ensembl |  |  | 100 % |  |  |
| 22 | ggggacaagtttgtacaaaaaagcaggcttgatgAACGGCCAATCCCAAAAGCC, ggggaccactttgtacaagaaagctgggtcTTGATATGCTTTGCCACCGA | Mycgr3\|99841  JGI  Ensembl | KOG1260 Isocitrate lyase; | carboxyvinyl-carboxyphosphonate phosphorylmutase activity; catalytic activity;  metabolic process; | 100 % |  |  |
| 22 | ggggacaagtttgtacaaaaaagcaggcttgatgGCGGACAACTACATACTATC, ggggaccactttgtacaagaaagctgggtcGTCAAGGTCATCTCCTGGAG | Mycgr3\|99929  JGI  Ensembl |  |  | 100 % |  |  |
| 22 | ggggacaagtttgtacaaaaaagcaggcttgatgTCCACTTTCAAGAGTGTCCA, ggggaccactttgtacaagaaagctgggtcCAGCTTCGACTGTGCCCCTG | Mycgr3\|99937  JGI  Ensembl | KOG1196 Predicted NAD-dependent oxidoreductase; | 2-alkenal reductase activity; oxidoreductase activity;  zinc ion binding; | 100 % |  |  |
| 22 | ggggacaagtttgtacaaaaaagcaggcttgatgTCGCGATCCATGGCGCAACG, ggggaccactttgtacaagaaagctgggtcAAAGGTATCCTCATCCCAAT | Mycgr3\|101448  JGI  Ensembl |  |  | 100 % |  |  |
| 22 | ggggacaagtttgtacaaaaaagcaggcttgatgTCGACCTTCGGCCAGTACTA, ggggaccactttgtacaagaaagctgggtcCTCCTTCGACCCATTCGTCA | Mycgr3\|101785  JGI  Ensembl | KOG4492 Chorismate synthase; | aromatic amino acid family biosynthetic process; chorismate synthase activity;  extracellular region;  neurohypophyseal hormone activity; | 100 % |  |  |
| 22 | ggggacaagtttgtacaaaaaagcaggcttgatgACGTACACTGTCAATTACAA, ggggaccactttgtacaagaaagctgggtcACGCCCCCTGTGAGCACGCC | Mycgr3\|102798  JGI  Ensembl |  |  | 100 % |  |  |
| 22 | ggggacaagtttgtacaaaaaagcaggcttgatgGCCACCACCGACCCGAAACC, ggggaccactttgtacaagaaagctgggtcCTGCAACAACCTCAAAACCT | Mycgr3\|103255  JGI  Ensembl | KOG0455 Homoserine dehydrogenase; | amino acid biosynthetic process; homoserine dehydrogenase activity; | 1.95035 % |  |  |
| 22 | ggggacaagtttgtacaaaaaagcaggcttgatgCGTTCCTTCATCGTTGTCGC, ggggaccactttgtacaagaaagctgggtcCAGAGCCAAGCCGAAAATGG | Mycgr3\|105265  JGI  Ensembl | KOG2177 Predicted E3 ubiquitin ligase; |  | 100 % |  |  |
| 22 | ggggacaagtttgtacaaaaaagcaggcttgatgGACTCCTCCTCCCACCCGAG, ggggaccactttgtacaagaaagctgggtcGGTCCCAGCATTGAGCCACC | Mycgr3\|105910  JGI  Ensembl |  |  | 100 % |  |  |
| 22 | ggggacaagtttgtacaaaaaagcaggcttgatgTTGCTAGACCATCTAACAGA, ggggaccactttgtacaagaaagctgggtcGTCCAGCTCCATAGCGTGAG | Mycgr3\|106918  JGI  Ensembl |  |  | 100 % |  |  |
| 22 | ggggacaagtttgtacaaaaaagcaggcttgatgAACGCCGGCGGCGTCATCGC, ggggaccactttgtacaagaaagctgggtcATTTTGACCCGTGAATCGAG | Mycgr3\|108073  JGI  Ensembl | KOG1995 Conserved Zn-finger protein; |  | 2.24609 % |  |  |
| 22 | ggggacaagtttgtacaaaaaagcaggcttgatgGCTCCTACACCATCAACGGA, ggggaccactttgtacaagaaagctgggtcGATGAACCACAGCACCACTC | Mycgr3\|108268  JGI  Ensembl |  |  | 100 % |  |  |
| 22 | ggggacaagtttgtacaaaaaagcaggcttgatgTCGGTCGAATGCGATACAAT, ggggaccactttgtacaagaaagctgggtcTAAGCTCTGTACGAAAACCG | Mycgr3\|108309  JGI  Ensembl | KOG1515 Arylacetamide deacetylase; |  | 100 % |  |  |
| 22 | ggggacaagtttgtacaaaaaagcaggcttgatgTCTGGCCCATCCGTTGTCTG, ggggaccactttgtacaagaaagctgggtcTGCAAGTTCTTTCTCTTCCT | Mycgr3\|108533  JGI  Ensembl | KOG4852 Uncharacterized conserved protein; |  | 100 % |  |  |
| 22 | ggggacaagtttgtacaaaaaagcaggcttgatgAACTCCATGGCCCGTCCATT, ggggaccactttgtacaagaaagctgggtcTTCCCAGACCTTCGGCTGCT | Mycgr3\|110017  JGI  Ensembl |  |  | 100 % |  |  |
| 22 | ggggacaagtttgtacaaaaaagcaggcttgatgTCCGCCAGCAGCACAGATGA, ggggaccactttgtacaagaaagctgggtcACAAGGATGACTCCACGCTC | Mycgr3\|110274  JGI  Ensembl | KOG3671 Actin regulatory protein (Wiskott-Aldrich syndrome protein); | UDP-N-acetylmuramoylalanyl-D-glutamyl-2,6-diaminopimelate-D-alanyl-D-alanine ligase activity; coenzyme F420-0 gamma-glutamyl ligase activity;  coenzyme F420-2 alpha-glutamyl ligase activity;  ribosomal S6-glutamic acid ligase activity; | 100 % |  |  |
| 22 | ggggacaagtttgtacaaaaaagcaggcttgatgGTCCTCGAAGAAGCCACCAC, ggggaccactttgtacaagaaagctgggtcCCACCTCTTACCTCCACCTC | Mycgr3\|110767  JGI  Ensembl |  |  | 100 % |  | ``` 17	ATGGTCCTCGAAGAAGCCACCACACCTGAAGCTCCTTTCAGCGACTCTTCAACTCCCGACGACGACAATGACCTCTACTCCGGCGACCGCTCCCCCACCTACCTCGAGACCCGCTCGCCGACACGATGGCCCATCACCCGTCCAGGCCTCGAACCAATCTACATCAATCTAGATCAACTCCCGAAAGCATTCCTCATCTCAGGCGAGAACTCAGCATATAATAAAATCATCGGCGATGGGACGCAAAAGTACTGCGATGAGACAATTAAGCTACTCAACCGGCCTCTGCGGGAGGACGAAGCAACTGCTGCTGCCTACCATCTTGCGAAGACAATGCGGATCGCATCATACGGACCGCCTTTGGGAATACTAGCTGCGGCATTCATATGGAGGCGTGGAAAGGCGAAATTCCGCTTTCCTGGCTGGACACCGGGCGAGAAGTTCAACCCGGATCGCTTTGCGGGTGGACTTCTCAAGGGCAAGCCGGCACGATTCGCGTGGCATTTCCTGCGGATTAATGCGTACGGCGTGGTTGGGTCATTTGCAGGACAGGTCTTCTTCGGAAGCTATGCGATTACAGTGGGCAACGTTGGACGATTCCAAGATCCGCGCTTGAAGGACGTCACAGAGGCATTGAAAGCACGGCTAGAAGCGAATCGTGCAGCGGGCAAGCTGCCTGGAGGAAGAGAGGTAGATCAGACGGCAGGGAAGAGGGATGGTGAGACGTTTGACATGGCGAGGCAGAGGCAAAGAGTGCAAGAGCAGACAAGGAGGACAGCGGGACAGCAACAAGCACAGCGACAAAAGCCAGTTGCCGATGATATGAGCCCTACGGGTGGATCGTTCTCCGATGATTTCATGAATGAGGGCGCGGCGAGCGATGGCGGCATGTTGAGCGATTCGCAGATTCAGTCACTGCAGTCGCAGCAACAGAAGAGGGAGGAGAGATCGGAGGC-ACA-GACCGAAGCCGCACGAGCACGTCCAGTACAGCCGAACACGACAACA-CCCTCAACGCCATCACGGTCCGAACCTCCGACAAAGCAGTCGGGAAGTGCAT-GGGAGAG-AC-TACGGCAACAGGC-TGCTC-CAG	1103 	\|\|\|\|\|\|\|\|\|\|\|\|\|\|\|\|\|\|\|\|\|\|\|\|\|\|\|\|\|\|\|\|\|\|\|\|\|\|\|\|\|\|\|\|\|\|\|\|\|\|\|\|\|\|\|\|\|\|\|\|\|\|\|\|\|\|\|\|\|\|\|\|\|\|\|\|\|\|\|\|\|\|\|\|\|\|\|\|\|\|\|\|\|\|\|\|\|\|\|\|\|\|\|\|\|\|\|\|\|\|\|\|\|\|\|\|\|\|\|\|\|\|\|\|\|\|\|\|\|\|\|\|\|\|\|\|\|\|\|\|\|\|\|\|\|\|\|\|\|\|\|\|\|\|\|\|\|\|\|\|\|\|\|\|\|\|\|\|\|\|\|\|\|\|\|\|\|\|\|\|\|\|\|\|\|\|\|\|\|\|\|\|\|\|\|\|\|\|\|\|\|\|\|\|\|\|\|\|\|\|\|\|\|\|\|\|\|\|\|\|\|\|\|\|\|\|\|\|\|\|\|\|\|\|\|\|\|\|\|\|\|\|\|\|\|\|\|\|\|\|\|\|\|\|\|\|\|\|\|\|\|\|\|\|\|\|\|\|\|\|\|\|\|\|\|\|\|\|\|\|\|\|\|\|\|\|\|\|\|\|\|\|\|\|\|\|\|\|\|\|\|\|\|\|\|\|\|\|\|\|\|\|\|\|\|\|\|\|\|\|\|\|\|\|\|\|\|\|\|\|\|\|\|\|\|\|\|\|\|\|\|\|\|\|\|\|\|\|\|\|\|\|\|\|\|\|\|\|\|\|\|\|\|\|\|\|\|\|\|\|\|\|\|\|\|\|\|\|\|\|\|\|\|\|\|\|\|\|\|\|\|\|\|\|\|\|\|\|\|\|\|\|\|\|\|\|\|\|\|\|\|\|\|\|\|\|\|\|\|\|\|\|\|\|\|\|\|\|\|\|\|\|\|\|\|\|\|\|\|\|\|\|\|\|\|\|\|\|\|\|\|\|\|\|\|\|\|\|\|\|\|\|\|\|\|\|\|\|\|\|\|\|\|\|\|\|\|\|\|\|\|\|\|\|\|\|\|\|\|\|\|\|\|\|\|\|\|\|\|\|\|\|\|\|\|\|\|\|\|\|\|\|\|\|\|\|\|\|\|\|\|\|\|\|\|\|\|\|\|\|\|\|\|\|\|\|\|\|\|\|\|\|\|\|\|\|\|\|\|\|\|\|\|\|\|\|\|\|\|\|\|\|\|\|\|\|\|\|\|\|\|\|\|\|\|\|\|\|\|\|\|\|\|\|\|\|\|\|\|\|\|\|\|\|\|\|\|\|\|\|\|\|\|\|\|\|\|\|\|\|\|\|\|\|\|\|\|\|\|\|\|\|\|\|\|\|\|\|\|\|\|\|\|\|\|\|\|\|\|\|\|\|\|\|\|\|\|\|\|\|\|\|\|\|\|\|\|\|\|\|\|\|\|\|\|\|\|\|\|\|\|\|\|\|\|\|\|\|\|\|\|\|\|\|\|\|\|\|\|\|\|\|\|\|\|\|\|\|\|\|\|\|\|\|\|\|\|\|\|\|\|\|\|\|\|\|\|\|\|\|\|\|\|\|\|\|\|\|\|\|\|\|\|\|\|\|\|\|\|\|\|\|\|\|\|\|\|\|\|\|\|\|\|\|\|\|\|\|\|\|\|\|\|\|\|\|\|\|\|\|\|\|\|\|\|\|\|\|\|\|\|\|\|\|\|\|\|\|\|\|\|\|\|\|\|\|\|\|\|\|\|\|\|\|\|\|\|\|\|\|\|\|\|\|\|\|\|\|\|\|\|\|\|\|\|\|\|\|\|\|\|\|\|\|\|\|\|\|\|\|\|\|\|\|\|\|\|\|\|\|\|\|\|\|\|\|\|\|\|\|\|\|\|\|\|\| \|\|\|\|\|\|\|\|\|\|\|\|\|\|\|\|\|\|\|\|\|\|\|\|\|\|\|\|\|\|\|\|\|\|\|\|\|\|\|\|\|\|\|\|\|\|\|\|\|\|\|\|\|\|\|\|\|\|\|\|\|\|\|\|\|\|\|\|\|\|\|\|\|\|\|\|\|\|\|\|\|\|\|\|\|\|\|\|\| \|\|\| \|\|\|\|\|\|\|\|\|\|\|\|\|\|\|\|\|\|\|\|\|\|\|\|\|\|\|\|\|\|\|\|\|\|\|\|\|\|\|\|\|\|\|\|\|\| \|\|\|\|\|\|\|\|\|\|\|\|\|\|\|\|\|\|\| \|\|\|\|\|\|\|\|\|\|\|\|\|\| \|\|\|\|\|\|\|\| \|\|\|\|\|\|\|\| \|\|\|\|\|\|\| \|\| \|\|\|\| \|\|\|\| \| \| \|\|\|\|\| \|\|\| 	ATGGTCCTCGAAGAAGCCACCACACCTGAAGCTCCTTTCAGCGACTCTTCAACTCCCGACGACGACAATGACCTCTACTCCGGCGACCGCTCCCCCACCTACCTCGAGACCCGCTCGCCGACACGATGGCCCATCACCCGTCCAGGCCTCGAACCAATCTACATCAATCTAGATCAACTCCCGAAAGCATTCCTCATCTCAGGCGAGAACTCAGCATATAATAAAATCATCGGCGATGGGACGCAAAAGTACTGCGATGAGACAATTAAGCTACTCAACCGGCCTCTGCGGGAGGACGAAGCAACTGCTGCTGCCTACCATCTTGCGAAGACAATGCGGATCGCATCATACGGACCGCCTTTGGGAATACTAGCTGCGGCATTCATATGGAGGCGTGGAAAGGCGAAATTCCGCTTTCCTGGCTGGACACCGGGCGAGAAGTTCAACCCGGATCGCTTTGCGGGTGGACTTCTCAAGGGCAAGCCGGCACGATTCGCGTGGCATTTCCTGCGGATTAATGCGTACGGCGTGGTTGGGTCATTTGCAGGACAGGTCTTCTTCGGAAGCTATGCGATTACAGTGGGCAACGTTGGACGATTCCAAGATCCGCGCTTGAAGGACGTCACAGAGGCATTGAAAGCACGGCTAGAAGCGAATCGTGCAGCGGGCAAGCTGCCTGGAGGAAGAGAGGTAGATCAGACGGCAGGGAAGAGGGATGGTGAGACGTTTGACATGGCGAGGCAGAGGCAAAGAGTGCAAGAGCAGACAAGGAGGACAGCGGGACAGCAACAAGCACAGCGACAAAAGCCAGTTGCCGATGATATGAGCCCTACGGGTGGATCGTTCTCCGATGATTTCATGAATGA-GGCGCGGCGAGCGATGGCGGCATGTTGAGCGATTCGCAGATTCAGTCACTGCAGTCGCAGCAACAGAAGAGGGAGGAGAGATCGGAGGCCACAGGACCGAAGCCGCACGAGCACGTCCAGTACAGCCGAACACGACAACACCCCTCAACGCCATCACGGT-CGAACCTCCGACAA-GCAGTCGG-AAGTGCATTGGGAGAGGACCTACG-CAACGGCCCTGCTCTCAG ``` |
| 22 | ggggacaagtttgtacaaaaaagcaggcttgatgCCGAACATACGGTTGGCAGT, ggggaccactttgtacaagaaagctgggtcTTCATCCGTCAACTTATCCA | Mycgr3\|111281  JGI  Ensembl |  |  | 100 % |  | ``` 462	CCCTCCTCCTAGCAGGATGTTCGTCATCCTCGCCGTTAATACCGGGGATCTTCCTCATATCCTTCTATTACGAAATGTACACCCCGCGTTACGATCCCACACAAGTCAACCCCGGCGTCACAGCGGCCATCACCAACATCGTCGGCGGCGCGAGTTTGGAAGTAAGAGTGGGATATTTTGGACTCTGCATACAACCCGATGATGGAAGCTTCCTCTGCTCGAACAACGCCACGCTGCTGGCGCAACAAATCTCCATTGACGCGGATCCGCTCAATCTGATCTGGGTGGCCAATACGTTCAAGAACAGCGTTGTCTTTCCATACCTCCT	789 	\|\|\|\|\|\|\|\|\|\|\|\|\|\|\|\|\|\|\|\|\|\|\|\|\|\|\|\|\|\|\|\|\|\|\|\|\|\|\|\|\|\|\|\|\|\|\|\|\|\|\|\|\|\|\|\|\|\|\|\|\|\|\|\|\|\|\|\|\|\|\|\|\|\|\|\|\|\|\|\|\|\|\|\|\|\|\|\|\|\|\|\|\|\|\|\|\|\|\|\|\|\|\|\|\|\|\|\|\|\|\|\|\|\|\|\|\|\|\|\|\|\|\|\|\|\|\|\|\|\|\|\|\|\|\|\|\|\|\|\|\|\|\|\|\|\|\|\|\|\|\|\|\|\|\|\|\|\|\|\|\|\|\|\|\|\|\|\|\|\|\|\|\|\|\|\|\|\|\|\|\|\|\|\|\|\|\|\|\|\|\|\|\|\|\|\|\|\|\|\|\|\|\|\|\|\|\|\|\|\|\|\|\|\|\|\|\|\|\|\|\|\|\|\|\|\|\|\|\|\|\|\|\|\|\|\|\|\|\|\|\|\|\|\|\|\|\|\|\|\|\|\|\|\|\|\|\|\|\|\|\|\|\|\|\|\|\|\|\|\|\|\|\|\|\|\|\|\|\|\|\|\|\|\|\|\|\|\|\|\|\|\|\|\|\|\|\|\|\|\|\|\|\|\|\|\|\|\|\|\|\|\|\|\|\|\|\|\|\|\|\|\|\|\|\|\|\|\| 	CCCTCCTCCTAGCAGGATGTTCGTCATCCTCGCCGTTAATACCGGGGATCTTCCTCATATCCTTCTATTACGAAATGTACACCCCGCGTTACGATCCCACACAAGTCAACCCCGGCGTCACAGCGGCCATCACCAACATCGTCGGCGGCGCGAGTTTGGAAGTAAGAGTGGGATATTTTGGACTCTGCATACAACCCGATGATGGAAGCTTCCTCTGCTCGAACAACGCCACGCTGCTGGCGCAACAAATCTCCATTGACGCGGATCCGCTCAATCTGATCTGGGTGGCCAATACGTTCAAGAACAGCGTTGTCTTTCCATACCTCCT ```      ``` 359	ATGCCGAACATACGGTTGGCAGTCGGCATTCAGCGGCTGATACCCTTTCTTGGATACCACCATGTGTTGATGATCCTCATTGCACTGGCAATCATATTACTCT	461 	\|\|\|\|\|\|\|\|\|\|\|\|\|\|\|\|\|\|\|\|\|\|\|\|\|\|\|\|\|\|\|\|\|\|\|\|\|\|\|\|\|\|\|\|\|\|\|\|\|\|\|\|\|\|\|\|\|\|\|\|\|\|\|\|\|\|\|\|\|\|\|\|\|\|\|\|\|\|\|\|\|\|\|\|\|\|\|\|\|\|\|\|\|\|\|\|\|\|\|\|\|\|\| 	ATGCCGAACATACGGTTGGCAGTCGGCATTCAGCGGCTGATACCCTTTCTTGGATACCACCATGTGTTGATGATCCTCATTGCACTGGCAATCATATTACTCT ```      ``` 790	AATCGTCGCCATAATCCTCGCCTTCGGCACCTTCCTCCTCCTCGCCACCTTCCCGGGCTGGCACGAAGAACACGACGCCCGCACCGGGTCCGACATCGACATTAAACCCTTCCCCTCACGACCAGTCTCCCAAGTCGCCCTCGCGCTCATCTTCATCTCGTCCATATTCGTGCTCGTCTCCGTCCTCTGGCAACACACGGCCTCCGTCGCCGCGGCGCAGGTCGCTCAAGATTTCGGGAATGG-AAGCGTCAAGAGTGGAGTCGGTACGGCGGCGATGGTGTTGGGTTGGTTTGGATTTGCGCTGCTCATTATTGTGACGGTGGGATTGTTGGTTATGATTTTGAGTATCCATCTGTTGGATAAGTTGACGGATGA	1164 	\|\|\|\|\|\|\|\|\|\|\|\|\|\|\|\|\|\|\|\|\|\|\|\|\|\|\|\|\|\|\|\|\|\|\|\|\|\|\|\|\|\|\|\|\|\|\|\|\|\|\|\|\|\|\|\|\|\|\|\|\|\|\|\|\|\|\|\|\|\|\|\|\|\|\|\|\|\|\|\|\|\|\|\|\|\|\|\|\|\|\|\|\|\|\|\|\|\|\|\|\|\|\|\|\|\|\|\|\|\|\|\|\|\|\|\|\|\|\|\|\|\|\|\|\|\|\|\|\|\|\|\|\|\|\|\|\|\|\|\|\|\|\|\|\|\|\|\|\|\|\|\|\|\|\|\|\|\|\|\|\|\|\|\|\|\|\|\|\|\|\|\|\|\|\|\|\|\|\|\|\|\|\|\|\|\|\|\|\|\|\|\|\|\|\|\|\|\|\|\|\|\|\|\|\|\|\|\|\|\|\|\|\|\|\|\|\|\|\|\|\|\|\|\|\|\|\|\|\|\|\|\|\|\|\|\|\|\|\|\|\|\|\| \|\|\|\|\|\|\|\|\|\|\|\|\|\|\|\|\|\|\|\|\|\|\|\|\|\|\|\|\|\|\|\|\|\|\|\|\|\|\|\|\|\|\|\|\|\|\|\|\|\|\|\|\|\|\|\|\|\|\|\|\|\|\|\|\|\|\|\|\|\|\|\|\|\|\|\|\|\|\|\|\|\|\|\|\|\|\|\|\| \|\|\|\|\|\|\|\|\|\|\|\|\|\|\|\|\|\|\|\|\|\|\|\|\|\|\|\|\|\|\|\|\|\|\|\|\|\|\|\|\|\| 	AATCGTCGCCATAATCCTCGCCTTCGGCACCTTCCTCCTCCTCGCCACCTTCCCGGGCTGGCACGAAGAACACGACGCCCGCACCGGGTCCGACATCGACATTAAACCCTTCCCCTCACGACCAGTCTCCCAAGTCGCCCTCGCGCTCATCTTCATCTCGTCCATATTCGTGCTCGTCTCCGTCCTCTGGCAACACACGGCCTCCGTCGCCGCGGCGCAGGTCGCTCAAGATTTCGGGAATGGAAAGCGTCAAGAGTGGAGTCGGTACGGCGGCGATGGTGTTGGGTTGGTTTGGATTTGCGCTGCTCATTATTGTGACGGTGGGATTGTTGG-TATGATTTTGAGTATCCATCTGTTGGATAAGTTGACGGATGA ``` |
| 22 | ggggacaagtttgtacaaaaaagcaggcttgatgGCGCCCGCAGCCTCGAAGCA, ggggaccactttgtacaagaaagctgggtcAGCCTTCACTTGCTCCTTGA | Mycgr3\|111324  JGI  Ensembl | KOG0769 Predicted mitochondrial carrier protein; | binding; membrane; mitochondrial inner membrane;  transport;  transporter activity; | 100 % |  |  |
| 22 | ggggacaagtttgtacaaaaaagcaggcttgatgGTCTGTATTCGAGCAATCCT, ggggaccactttgtacaagaaagctgggtcGTCATTTTCAGTATCCGGTC | Mycgr3\|111350  JGI  Ensembl |  |  | 100 % | chr\_11:1427006 C->T; |  |
| 22 | ggggacaagtttgtacaaaaaagcaggcttgatgCCCGATTCCTTATTCTACAC, ggggaccactttgtacaagaaagctgggtcATCCTCCCCTGCCTCGGGCT | Mycgr3\|111762  JGI  Ensembl |  |  | 100 % |  |  |
| 23 | ggggacaagtttgtacaaaaaagcaggcttgatgTGTGGAAGATATGTTCTCAG, ggggaccactttgtacaagaaagctgggtcCTTCCCAAAGAAACTGGTAA | Mycgr3\|32208  JGI  Ensembl | KOG2618 Uncharacterized conserved protein; |  | 1.4782 % |  |  |
| 23 | ggggacaagtttgtacaaaaaagcaggcttgatgCGTTCAACTCTACTTCTCCT, ggggaccactttgtacaagaaagctgggtcCAAAGCCGGACACCCGAAAT | Mycgr3\|33131  JGI  Ensembl |  |  | 100 % |  |  |
| 23 | ggggacaagtttgtacaaaaaagcaggcttgatgAACTTCGAGGTCTCCGACGA, ggggaccactttgtacaagaaagctgggtcCAGCTTCGTCTCCTTTTTCG | Mycgr3\|36755  JGI  Ensembl | KOG0140 Medium-chain acyl-CoA dehydrogenase; | acyl-CoA dehydrogenase activity; electron transport;  oxidoreductase activity, acting on the CH-CH group of donors; | 100 % |  |  |
| 23 | ggggacaagtttgtacaaaaaagcaggcttgatgAATGACTCCACGATGGACCG, ggggaccactttgtacaagaaagctgggtcTCCCCACATCCCACCACTCC | Mycgr3\|37187  JGI  Ensembl |  |  | 100 % |  |  |
| 23 | ggggacaagtttgtacaaaaaagcaggcttgatgCCATCCCTGAATGGCCTGAT, ggggaccactttgtacaagaaagctgggtcTAGCAGAGAATGCAGCAAGA | Mycgr3\|37732  JGI  Ensembl | KOG0565 Inositol polyphosphate 5-phosphatase and related proteins; | inositol or phosphatidylinositol phosphatase activity; inositol-polyphosphate 5-phosphatase activity; | 100 % |  | ``` 1	ATGCCATCCCTGAATGGCCTGATCACAACCTTCAACTGCGGCCGGCAAGAAGTTCAAGTCGACTATTTCGCATCATCCCTGTATACAAGTCTCAAGTCCGGCTCCCTTCCTCCCGACCTCATTGTCCTCGCGCTCCAAGAAATCTCTCCGATCGGATTCAGCTTCATCGGAGGTTCCTTCATCGCGCCCTACTTTTCCCGCCTCGGCGAAGCTGTACTCCTCGCGACAGCCTGGGCGTTTTCGAAAGATGTGGTGTACGAGCATGTGGTCACACGAAATGCGGGCATGACGGGGATTATGGTATTCGCGAGGAAGGAGATGAAGGAGCGAATAAGAGGCATGAAAGAGGCAGGGACGGGAGTTGGATTCTGGGAGATGGGTAATAAAGGCGCGGTGGGCGTGAGGCTGGACTTGGAAGAGACGGCCATGACATTTGTGGCGGCACATTTGGCGCCGGCGGAGGATGCGTGCGAGAGGAGGAACAAGGATTGGAAGACTATATGTGAGAACCTGGTATTCGAGGATGGGGCCAAAAGCTTTCAGCCAAAGAGGTCGATAGCTGGCGACACATCAGAGAGCGAACCTCTGCTGGCGGAGGACAGGACAACTGGGCTAGGAAGAAACGGACCACATGGTATCTTTGGTCCTGAGCCGTCACACATCTTTTTCGCCGGTGACCTCAACTACCGTACATCCGACACACCACCCGACGAAAAGGACCATGAGAGCTGGCCGCAACCGATGGCCTCAGAATCCGACCCATCACATTTCAACCAACTCCTCTCAAAAGATCAATTGACCCGCGAGCTAAACAACGGTCGAACGTTGCATCACCTCGCCGAAGCAAACATCTCATTCCCTCCCACTTACAAGTACTCTACCCGCGCACAACAACTCGCCCGAGACAACGCACGGCAGTATACATCAGCCTCACAAGCGAATAAAGAACCGGGT-CCGACGATCAATCTGGAGGGCCGACAAGAACAGGTCTGGCTCTGGGCAAAGCATAGAGTACCGAGCTGGTGCGATCGCATATTGTATCTGTCGGACGCACAGCCGCAAGT-GCATTGG-TATACCGCG-CTGCCGG	1087 	\|\|\|\|\|\|\|\|\|\|\|\|\|\|\|\|\|\|\|\|\|\|\|\|\|\|\|\|\|\|\|\|\|\|\|\|\|\|\|\|\|\|\|\|\|\|\|\|\|\|\|\|\|\|\|\|\|\|\|\|\|\|\|\|\|\|\|\|\|\|\|\|\|\|\|\|\|\|\|\|\|\|\|\|\|\|\|\|\|\|\|\|\|\|\|\|\|\|\|\|\|\|\|\|\|\|\|\|\|\|\|\|\|\|\|\|\|\|\|\|\|\|\|\|\|\|\|\|\|\|\|\|\|\|\|\|\|\|\|\|\|\|\|\|\|\|\|\|\|\|\|\|\|\|\|\|\|\|\|\|\|\|\|\|\|\|\|\|\|\|\|\|\|\|\|\|\|\|\|\|\|\|\|\|\|\|\|\|\|\|\|\|\|\|\|\|\|\|\|\|\|\|\|\|\|\|\|\|\|\|\|\|\|\|\|\|\|\|\|\|\|\|\|\|\|\|\|\|\|\|\|\|\|\|\|\|\|\|\|\|\|\|\|\|\|\|\|\|\|\|\|\|\|\|\|\|\|\|\|\|\|\|\|\|\|\|\|\|\|\|\|\|\|\|\|\|\|\|\|\|\|\|\|\|\|\|\|\|\|\|\|\|\|\|\|\|\|\|\|\|\|\|\|\|\|\|\|\|\|\|\|\|\|\|\|\|\|\|\|\|\|\|\|\|\|\|\|\|\|\|\|\|\|\|\|\|\|\|\|\|\|\|\|\|\|\|\|\|\|\|\|\|\|\|\|\|\|\|\|\|\|\|\|\|\|\|\|\|\|\|\|\|\|\|\|\|\|\|\|\|\|\|\|\|\|\|\|\|\|\|\|\|\|\|\|\|\|\|\|\|\|\|\|\|\|\|\|\|\|\|\|\|\|\|\|\|\|\|\|\|\|\|\|\|\|\|\|\|\|\|\|\|\|\|\|\|\|\|\|\|\|\|\|\|\|\|\|\|\|\|\|\|\|\|\|\|\|\|\|\|\|\|\|\|\|\|\|\|\|\|\|\|\|\|\|\|\|\|\|\|\|\|\|\|\|\|\|\|\|\|\|\|\|\|\|\|\|\|\|\|\|\|\|\|\|\|\|\|\|\|\|\|\|\|\|\|\|\|\|\|\|\|\|\|\|\|\|\|\|\|\|\|\|\|\|\|\|\|\|\|\|\|\|\|\|\|\|\|\|\|\|\|\|\|\|\|\|\|\|\|\|\|\|\|\|\|\|\|\|\|\|\|\|\|\|\|\|\|\|\|\|\|\|\|\|\|\|\|\|\|\|\|\|\|\|\|\|\|\|\|\|\|\|\|\|\|\|\|\|\|\|\|\|\|\|\|\|\|\|\|\|\|\|\|\|\|\|\|\|\|\|\|\|\|\|\|\|\|\|\|\|\|\|\|\|\|\|\|\|\|\|\|\|\|\|\|\|\|\|\|\|\|\|\|\|\|\|\|\|\|\|\|\|\|\|\|\|\|\|\|\|\|\|\|\|\|\|\|\|\|\|\|\|\|\|\|\|\|\|\|\|\|\|\|\|\|\|\|\|\|\|\|\|\|\|\|\|\|\|\|\|\|\|\|\|\|\|\|\|\|\|\|\|\|\|\|\|\|\|\|\|\|\|\|\|\|\|\|\|\|\|\|\|\|\|\|\|\|\|\|\|\|\|\|\|\|\|\|\|\|\|\|\|\|\|\|\|\|\|\|\|\|\|\|\| \|\|\|\|\|\|\|\|\|\|\|\|\|\|\|\|\|\|\|\|\|\|\|\|\|\|\|\|\|\|\|\|\|\|\|\|\|\|\|\|\|\|\|\|\|\|\|\|\|\|\|\|\|\|\|\|\|\|\|\|\|\|\|\|\|\|\|\|\|\|\|\|\|\|\|\|\|\|\|\|\|\|\|\| \|\|\|\|\|\|\|\|\|\|\|\|\|\|\|\|\|\|\|\|\|\|\|\|\|\|\|\|\|\|\|\|\|\|\| \|\|\|\|\|\|\|\|\|\|\|\|\|\|\|\|\|\|\|\|\|\|\|\|\|\|\|\|\|\|\|\|\|\|\|\|\| \|\|\|\|\|\|\|\|\| \|\|\|\|\|\|\|\|\| \|\|\|\|\|\|\|\|\|\|\|\|\|\|\|\| \|\| \|\|\| \|\|\|\| \|\|\|\|\|\| \|\|\| \|\|\|\|\| \|\|\|\|\|\|\|  \|\|\|\|\|\|\| \|\|\|\|\|\|\|\|\|\|\| \|\|\|\|\|\|\| \|\|\|\|\| \|\|\| \|\|\|\|\|\|\| \|\|\|\|\|\|  \|\|\|\|\| \|\|\| \|\|\|\|\|\|\| 	ATGCCATCCCTGAATGGCCTGATCACAACCTTCAACTGCGGCCGGCAAGAAGTTCAAGTCGACTATTTCGCATCATCCCTGTATACAAGTCTCAAGTCCGGCTCCCTTCCTCCCGACCTCATTGTCCTCGCGCTCCAAGAAATCTCTCCGATCGGATTCAGCTTCATCGGAGGTTCCTTCATCGCGCCCTACTTTTCCCGCCTCGGCGAAGCTGTACTCCTCGCGACAGCCTGGGCGTTTTCGAAAGATGTGGTGTACGAGCATGTGGTCACACGAAATGCGGGCATGACGGGGATTATGGTATTCGCGAGGAAGGAGATGAAGGAGCGAATAAGAGGCATGAAAGAGGCAGGGACGGGAGTTGGATTCTGGGAGATGGGTAATAAAGGCGCGGTGGGCGTGAGGCTGGACTTGGAAGAGACGGCCATGACATTTGTGGCGGCACATTTGGCGCCGGCGGAGGATGCGTGCGAGAGGAGGAACAAGGATTGGAAGACTATATGTGAGAACCTGGTATTCGAGGATGGGGCCAAAAGCTTTCAGCCAAAGAGGTCGATAGCTGGCGACACATCAGAGAGCGAACCTCTGCTGGCGGAGGACAGGACAACTGGGCTAGGAAGAAACGGACCACATGGTATCTTTGGTCCTGAGCCGTCACACATCTTTTTCGCCGGTGACCTCAACTACCGTACATCCGACACACCACCCGACGAAAAGGACCATGAGAGCTGGCCGCAACCGATGGCCTCAGAATCCGACCCATCACATTTCAACCAACTCCTCTC-AAAGATCAATTGACCCGCGAGCTAAACAACGGTCGAACGTTGCATCACCTCGCCGAAGCAAACATCTCATTCCCTCCCACTTAC-AGTACTCTACCCGCGCACAACAACTCGCCCGAGAC-ACGCACGGCAGTATACATCAGCCTCACAAGCGAATAATGAACCGGGTTCCGACGATC-ATCTGGAGGGCCGACA-GA-CAG-TCTG-CTCTGG-CAA-GCATA-AGTACCGGCCTGGTGC-ATCGCATATTGGATCTGTC-GACGCCCAGGCGCAAGTTGCATTGTATATAC-GCGGCTGCCGG ``` |
| 23 | ggggacaagtttgtacaaaaaagcaggcttgatgTCGCAGGACCCGCTCACCCA, ggggaccactttgtacaagaaagctgggtcTAGCTTCGCACCACCCTCCT | Mycgr3\|40691  JGI  Ensembl | KOG0141 Isovaleryl-CoA dehydrogenase; | acyl-CoA dehydrogenase activity; electron transport;  oxidoreductase activity, acting on the CH-CH group of donors; | 100 % |  |  |
| 23 | ggggacaagtttgtacaaaaaagcaggcttgatgGAGTTGCTCAAATCAAGCCG, ggggaccactttgtacaagaaagctgggtcTGGAATGCCAGACTTCGGAT | Mycgr3\|40961  JGI  Ensembl | KOG1592 Asparaginase; | asparaginase activity; glycoprotein catabolic process; | 100 % |  |  |
| 23 | ggggacaagtttgtacaaaaaagcaggcttgatgGACTCCTCGATCCTCATCAT, ggggaccactttgtacaagaaagctgggtcGAGCCGCGCCGATGAGGTCC | Mycgr3\|43740  JGI  Ensembl | KOG2820 FAD-dependent oxidoreductase; | sarcosine oxidase activity; | 0 % |  |  |
| 23 | ggggacaagtttgtacaaaaaagcaggcttgatgCCTTACCGCAAAACAACAAC, ggggaccactttgtacaagaaagctgggtcTTCCCTCCCCCTCACCCTCC | Mycgr3\|45149  JGI  Ensembl |  |  | 100 % |  |  |
| 23 | ggggacaagtttgtacaaaaaagcaggcttgatgTCTCCCTTTCACAGAACAGC, ggggaccactttgtacaagaaagctgggtcATTCGCCTGACTCTCCGCCC | Mycgr3\|45731  JGI  Ensembl |  |  | 100 % |  |  |
| 23 | ggggacaagtttgtacaaaaaagcaggcttgatgGCTCAACCATCATCAGCAGC, ggggaccactttgtacaagaaagctgggtcGCCAACGCTAAGGACTGAAT | Mycgr3\|46807  JGI  Ensembl | KOG2950 Uncharacterized protein involved in protein-protein interaction, contains polyproline-binding GYF domain; |  | 100 % |  |  |
| 23 | ggggacaagtttgtacaaaaaagcaggcttgatgCTTTCCAAATTCATCCTTGC, ggggaccactttgtacaagaaagctgggtcGTAGCTGCGGACATGGTGGA | Mycgr3\|46866  JGI  Ensembl | KOG3544 Collagens (type IV and type XIII), and related proteins; |  | 100 % |  |  |
| 23 | ggggacaagtttgtacaaaaaagcaggcttgatgGCACTCTTCAGGCAGGCGTG, ggggaccactttgtacaagaaagctgggtcCTCTGCTTTTGTATGCTCAG | Mycgr3\|48084  JGI  Ensembl | KOG2762 Mannosyltransferase; | dolichyl-phosphate-mannose-glycolipid alpha-mannosyltransferase activity; endoplasmic reticulum;  integral to membrane;  mannosyltransferase activity; | 100 % |  |  |
| 23 | ggggacaagtttgtacaaaaaagcaggcttgatgGAGTTCATGACGGCGCTGCA, ggggaccactttgtacaagaaagctgggtcGACCATGACCCGCCTCAATC | Mycgr3\|48243  JGI  Ensembl | KOG0826 Predicted E3 ubiquitin ligase involved in peroxisome organization; | peroxisomal membrane; peroxisome organization and biogenesis; | 100 % |  |  |
| 23 | ggggacaagtttgtacaaaaaagcaggcttgatgGGCAGTTTCGAAGTTCTGAC, ggggaccactttgtacaagaaagctgggtcCTCCAGCTTTTGCTCCTTGC | Mycgr3\|48782  JGI  Ensembl | KOG3888 Gamma-butyrobetaine,2-oxoglutarate dioxygenase; |  | 0 % |  |  |
| 23 | ggggacaagtttgtacaaaaaagcaggcttgatgCACTGCACCATCTGCGGCGT, ggggaccactttgtacaagaaagctgggtcAGCCCATTTTGCCGCCTCCT | Mycgr3\|49969  JGI  Ensembl |  |  | 100 % |  |  |
| 23 | ggggacaagtttgtacaaaaaagcaggcttgatgAAAAGACTATCGTCCAGCTG, ggggaccactttgtacaagaaagctgggtcAGACAGGATGCCCTGAGCTC | Mycgr3\|50017  JGI  Ensembl | KOG1290 Serine/threonine protein kinase; | ATP binding; protein amino acid phosphorylation;  protein kinase activity;  protein serine/threonine kinase activity; | 100 % |  |  |
| 23 | ggggacaagtttgtacaaaaaagcaggcttgatgGCTCCCAAGTCCATCTTTTT, ggggaccactttgtacaagaaagctgggtcAGAGCTGTTGGCGCCAGTAG | Mycgr3\|50742  JGI  Ensembl |  |  | 100 % |  | ``` 621	GGACGGAGACAACTACCATTTCAACACCACTCTCTTCACGCAGATGATGTCCGTCTGCAACGGCAATTGCGGCCGTGAAGAACTCGCCCTCTACCGCCTGCAGCGCTACGAGGCCAGCAAGGCCGACAACGGCAACTTCTTCTTCGGCCCTGGCTCTCTGCTCCTCTTCGGCGCTGCGTCCTTTCTCTACGAGTTGATGCCAGGTAGTAGCCACAAGCCCGACGAGGCCACCATGATGTCGTTCTTCGGCGCCAGCAAGGACGCCTCAGGAAACTACGTCAACAACGGCCAAGAACGCCTTCCTGAGAA-CTGGTACAACCGTGTCGAGCACTACTCCAACGTCTCCGAGGAGATCCTCGCCATGTACGCGCTCCACCCCGTGCCCTTCGGCGGCAATGTTGGCAAGGGCAACTTCAACGGCCTCAACTCCGGAACCGACATCGTCAACGGCAAGATCAACCCCGATCTCCCTTACAACGCTCTCTGCCTCATGTACCAGGCCCTCACCGGTGCCCAGCCTGGTCTGTTGGGCGTGGCTGTCAACCTGCCTGTGAACAGCCTCAGTGCTATCACAAAGGGTCTTGCGCCGATGGCGGC-GAACTTGGGATGCCCCACAAACCACAACGATGGCAGCC	1255 	\|\|\|\|\|\|\|\|\|\|\|\| \|\| \|\| \|\|\|\| \| \|\|\|\|\|\|\|\|\|\|\|\|\|\|\|\|\|\|\|\|\|\|\|\|\| \|\| \|\|\|\|\|\|\|\|\|\|\|  \|\|\| \| \|\|\|\|\|\|\|\|\|\|\|\|\| \|\| \|\|\|\|\|\|\|\|\|\|\|\|\|\| \|\|\|\|\|\|\|\|\|\|\|\|\|\|\|\|\| \|\|\|\|\|\|\|\|\|\|\| \|\| \|\| \|\| \|\|\| \|\|\|\|\|\| \| \|\| \|\| \| \|\| \|\|\|\|\|\|\|\|\|\|\| \|\|\|\|\|\|\|  \|\|\|\| \|\|\|\| \|\|\|\|\|\|\|\|\|\|\|\|\|\|\|\|\|\|\|\|\|\| \|\|\|\|\|\|\|\|\|\|\|\|\|\|\|\| \|\|\|  \|\|\|\|\|\|\|\|\|\|\|\|\|\|\|\|\|\|\|\|\|\|\|\|\|\|\|\| \|\|\|\|\|\|\|\|\|\|\|\|\| \|\|\|\|\|\|\|\|\|\|\|\|\|\| \|\|\|\|\|\| \|\|\|\|\|\|\|\|\|\|\|\|\|\|\|\|\|\|\|\|\|\|\|\| \|\|\|\|\|\|\|\|\|\|\|\|\|\|\|\|\|\|\|\|\|\|\|\|\|\|\|\|\|\|\|\|\|\|\|\|\|\|\|\|\|\|\|\|\|\|\|\|\|\| \|\|\|\|\| \|\|\|\|\|\|\|\|\|\|\|\|\|\|\|\|\|\|\|\|\|\|\|\|\|\| \|\|\| \|\|\|\| \|\| \|\|\| \| \|\|\|\|\|\|\|\|\|\|\|\|\| \|\|\|\|\|\|\|\|\|\| \|\|\|\|\| \|\|\|\|\|\|\|\|\|\|\|\|\|\|\|\| \|\|\|\|\|\|\|\|\|\|\|\|\|\|\|\| \|\|\|\|\|\|\|\| \|\| \|\| \|\| \|\| \|\|\|\|\| \|\|\|\|\|  \| \|\|\|\|\|\|\|\|\|\|  \|\|\|\|\|\|\|\|\|\| \|\|\|\|\|\| \|\|\|\|\|\|\|\|\|\|\|\|\|\|  \| \|\| \|\|\|\|\| \|\|\| \| \|\| \|\|\|\|\|\|\|\|\|\|\|\|\| \|\| \|\|\|\|\| \| \|\|\|\|\|\|\|\| 	GGACGGAGACAATTAGCAGTTCAGCTCCACTCTCTTCACGCAGATGATGTCAGTATGCAACGGCAACAGCGTCGGTGAAGAACTCGCTCTGTACCGCCTGCAGCGATACGAGGCCAGCAAGGCTGACAACGGCAATTTGTTGTTTGGCGCTGGCTATATGATCGTTTTAGGCGCTGCGTCGTTTCTCTGAGAGTAGATGGCAGGTAGTAGCCACAAGCCCGATGAGGCCACCATGATGTAGTTAATCGGCGCCAGCAAGGACGCCTCAGGAAATTACGTCAACAACGCCCAAGAACGCCTTCTTGAGAAGCTGGTACAACCGTGTCGAGCACTAGTCCAACGTCTCCGAGGAGATCCTCGCCATGTACGCGCTCCACCCCGTGCCATTCGGGGGCAATGTTGGCAAGGGCAACTTCAATGGCGTCAAATCGGGACCTGACATCGTCAACGACAAGATCAACACCGATTTCCCTTACAACGCTCTTTGCCTCATGTACCAGGTCCTCACCGTTGTCCTGCTTGCTCTGTCGGGCGCAGATGTCAACCTGATTGTGAACAGCATCAGTGATATCACAAAGGGTCCGGAGCGGATGGTGGCAGCAC-TGGGATGCCCCACGAAGCACAAGGTTGGCAGCC ``` |
| 23 | ggggacaagtttgtacaaaaaagcaggcttgatgGTCAACATGGACTCTCCACA, ggggaccactttgtacaagaaagctgggtcCCAATCAAGTCCAGTATTCG | Mycgr3\|51997  JGI  Ensembl | KOG2876 Molybdenum cofactor biosynthesis pathway protein; | catalytic activity; iron-sulfur cluster binding; | 100 % |  |  |
| 23 | ggggacaagtttgtacaaaaaagcaggcttgatgAGCGATCTTCACAACACTCC, ggggaccactttgtacaagaaagctgggtcTGTGAACTTTTGATGGATGA | Mycgr3\|65218  JGI  Ensembl | KOG0676 Actin and related proteins; | protein binding; | 100 % |  |  |
| 23 | ggggacaagtttgtacaaaaaagcaggcttgatgGCCGACCAAGATCAAAGCAA, ggggaccactttgtacaagaaagctgggtcCATACTACTTCTCCCTCCAT | Mycgr3\|66245  JGI  Ensembl | KOG0725 Reductases with broad range of substrate specificities; | carbonyl reductase (NADPH) activity; metabolic process;  oxidoreductase activity; | 100 % |  |  |
| 23 | ggggacaagtttgtacaaaaaagcaggcttgatgGCGACCTCATTACGAACGCT, ggggaccactttgtacaagaaagctgggtcCCTCCACTCATGCTCTTCCT | Mycgr3\|66888  JGI  Ensembl | KOG0975 Branched chain aminotransferase BCAT1, pyridoxal phosphate enzymes type IV superfamily; | branched chain family amino acid metabolic process; branched-chain-amino-acid transaminase activity;  catalytic activity;  metabolic process; | 100 % |  |  |
| 23 | ggggacaagtttgtacaaaaaagcaggcttgatgCCCTCACAGACCTTCTTCCT, ggggaccactttgtacaagaaagctgggtcTACGAGCTCAAACTCGTAGA | Mycgr3\|67138  JGI  Ensembl | KOG2909 Vacuolar H+-ATPase V1 sector, subunit C; | ATP binding; hydrogen ion transporting ATP synthase activity, rotational mechanism;  hydrogen ion transporting ATPase activity, rotational mechanism;  hydrogen-transporting two-sector ATPase activity;  hydrolase activity, acting on acid anhydrides, catalyzing transmembrane movement of substances; proton-transporting ATP synthase complex;  proton-transporting two-sector ATPase complex; | 100 % |  | ``` 19	GATGCCCTCACAGACCTTCTTCCTCGTCTCGCTGCCCAACTCAATCAATCCCTCCGGAAACCGCGATGAAGCTCTTACCACTCTGCGCTCCGCCGTGAGCGCCGACGCAGCGACGACATATCCCTTTGCGATACCAGATTTCAAGGTCGGAACTCTCGACGCATTGGTGCAGCAAGCAGATGAGTTGGCGAAGCTGGAGCAGGGCTGCAAGGGCGTGGCGGACAAAGTCGCCGACTCATTAAGATCACTGTTGGAGGGCGACGAGGAGAAACTACAGGAGCAGAAGGTGGTCAATGACCG	318 	\|\|\|\|\|\|\|\|\|\|\|\|\|\|\|\|\|\|\|\|\|\|\|\|\|\|\|\|\|\|\|\|\|\|\|\|\|\|\|\|\|\|\|\|\|\|\|\|\|\|\|\|\|\|\|\|\|\|\|\|\|\|\|\|\|\|\|\|\|\|\|\|\|\|\|\|\|\|\|\|\|\|\|\|\|\|\|\|\|\|\|\|\|\|\|\|\|\|\|\|\|\|\|\|\|\|\|\|\|\|\|\|\|\|\|\|\|\|\|\|\|\|\|\|\|\|\|\|\|\|\|\|\|\|\|\|\|\|\|\|\|\|\|\|\|\|\|\|\|\|\|\|\|\|\|\|\|\|\|\|\|\|\|\|\|\|\|\|\|\|\|\|\|\|\|\|\|\|\|\|\|\|\|\|\|\|\|\|\|\|\|\|\|\|\|\|\|\|\|\|\|\|\|\|\|\|\|\|\|\|\|\|\|\|\|\|\|\|\|\|\|\|\|\|\|\|\|\|\|\|\|\|\|\|\|\|\|\|\|\|\|\|\|\|\|\|\|\|\|\|\|\|\|\|\|\|\|\|\|\|\|\|\|\|\|\|\|\|\|\|\|\|\|\|\|\|\|\|\|\|\|\|\|\|\|\|\|\|\|\|\|\|\|\|\|\|\|\|\|\| 	GATGCCCTCACAGACCTTCTTCCTCGTCTCGCTGCCCAACTCAATCAATCCCTCCGGAAACCGCGATGAAGCTCTTACCACTCTGCGCTCCGCCGTGAGCGCCGACGCAGCGACGACATATCCCTTTGCGATACCAGATTTCAAGGTCGGAACTCTCGACGCATTGGTGCAGCAAGCAGATGAGTTGGCGAAGCTGGAGCAGGGCTGCAAGGGCGTGGCGGACAAAGTCGCCGACTCATTAAGATCACTGTTGGAGGGCGACGAGGAGAAACTACAGGAGCAGAAGGTGGTCAATGACCG ```      ``` 309	TCAATGAC-CGACCGGTCGAGAACTACCTCCAATCTTTCCAATGGAACAAAGTCAAATATCGCGCCGACAAGCCCATCGCCGAGCTCATCGACACCCTGCAGAAAGAGATCGCAGCCGTGGACAACGACGTCAAGGCGAAATACAACCAATACAACACCTCCAAAACCGCACTCGCCAGCATGCAACGTAGCCATACCGGCAACCTCTCGCAAAAATCCCTAGCCTCCGTCGTCAACCCAGCGACTCTCCTGCAACCCGATGCATCAGAATACCTCGCGCAGCACATCATCGCGGTCCCCTCAAACCTTGTCAAGGATTTTCTCAAGTCATATGAGTCCCTCACACCAATGGTCGTTCCTCGCAGCGCGCAACTGCTGGCCAAAGATGACGAATTTCAGCTGTATGCGGTGACGGTGTTCAAGAAGCATTCGCAGGAGTTTGTGCATAAGTGTCGCGAGCAGCGGTGGACG-CCGCGGGAGATGAAATTCACAGAGGGAGGTCGGGAAGCGGAGGAGAAGGAATTGACCAAGTTGGAAAAGGAGGAGAGGAGGCTTTGGGGCGAGGCGTTACGATTGGGCAGGACGGGATACAGTGATGCTGTGCAAGGGTGGATACACGTGCTCACCTTGAGAGTGTTTGTGGAGTCT-GTGCTGCGA-TATGGGTTGCCTTTGGACTTTGTGTG---CGGTTTGCTCAAGA	1004 	\|\|\|\|\| \|\|  \|\|\|\|\|\|\|\|\|\|\|\|\|\|\|\|\|\|\|\|\|\|\|\|\|\|\|\|\|\|\|\|\|\|\|\|\|\|\|\|\|\|\|\|\|\|\|\|\|\|\|\|\|\|\|\|\|\|\|\|\|\|\|\|\|\|\|\|\|\|\|\|\|\|\|\|\|\|\|\|\|\|\|\|\|\|\|\|\|\|\|\|\|\|\|\|\|\|\|\|\|\|\|\|\|\|\|\|\|\|\|\|\|\|\|\|\|\|\|\|\|\|\|\|\|\|\|\|\|\|\|\|\|\|\|\|\|\|\|\|\|\|\|\|\|\|\|\|\|\|\|\|\|\|\|\|\|\|\|\|\|\|\|\|\|\|\|\|\|\|\|\|\|\|\|\|\|\|\|\|\|\|\|\|\|\|\|\|\|\|\|\|\|\|\|\|\|\|\|\|\|\|\|\|\|\|\|\|\|\|\|\|\|\|\|\|\|\|\|\|\|\|\|\|\|\|\|\|\|\|\|\|\|\|\|\|\|\|\|\|\|\|\|\|\|\|\|\|\|\|\|\|\|\|\|\|\|\|\|\|\|\|\|\|\|\|\|\|\|\|\|\|\|\|\|\|\|\|\|\|\|\|\|\|\|\|\|\|\|\|\|\|\|\|\|\|\|\|\|\|\|\|\|\|\|\|\|\|\|\|\|\|\|\|\|\|\|\|\|\|\|\|\|\|\|\|\|\|\|\|\|\|\|\|\|\|\|\|\|\|\|\|\|\|\|\|\|\|\|\|\|\|\|\|\|\|\|\|\|\|\|\|\|\|\|\|\|\|\|\|\|\|\|\|\|\|\|\|\|\|\|\|\|\|\|\|\|\|\|\|\|\|\|\|\|\|\|\|\|\|\|\|\|\|\|\|\|\|\|\|\|\|\|\|\|\|\|\|\|\|\|\|\|\|\|\|\|\|\|\|\|\|\|\|\|\|\|\|\|\|\|\|\|\|\|\|\|\|\|\|\|\|\|\|\|\|\|\|\|\|\| \|\|\|\|\|\|\|\|\|\|\|\|\|\|\|\|\|\|\|\|\|\|\|\|\|\|\|\| \|\|\|\|\|\| \|\|\| \|\| \|\|\| \| \| \| \|\|\| \| \|\| \|\|\|\|\| \|\|\|\| \|\|\|\|\|\| \|\|\|\|\|\|\|\|\|\|\|\|\| \|\|\| \|\|\|\|\| \|\|\|\|\| \|\|\|\|\|\|\|\|\|\|\|\|\|\|\|\|\|\|\|\|\|\|\|\|  \|\|\|\|\|\|\|\|\|\|\|\|\|\|\|\|\|\|\|\|\| \|\| \|\|\|\|\|\|\|\|\|\|\|\|\|\|\|\| \|\|\|\|\|\|\|\|\| \|\|\|\|\|\|\| \| \|\|\|\|\|\|\|\|\|\|\|\|\|\|\|\|   \|\|\|\|\|  \|\|\|\|\|\|\| 	TCAATCACAGGACCGGTCGAGAACTACCTCCAATCTTTCCAATGGAACAAAGTCAAATATCGCGCCGACAAGCCCATCGCCGAGCTCATCGACACCCTGCAGAAAGAGATCGCAGCCGTGGACAACGACGTCAAGGCGAAATACAACCAATACAACACCTCCAAAACCGCACTCGCCAGCATGCAACGTAGCCATACCGGCAACCTCTCGCAAAAATCCCTAGCCTCCGTCGTCAACCCAGCGACTCTCCTGCAACCCGATGCATCAGAATACCTCGCGCAGCACATCATCGCGGTCCCCTCAAACCTTGTCAAGGATTTTCTCAAGTCATATGAGTCCCTCACACCAATGGTCGTTCCTCGCAGCGCGCAACTGCTGGCCAAAGATGACGAATTTCAGCTGTATGCGGTGACGGTGTTCAAGAAGCATTCGCAGGAGTTTGTGCATAAGTGTCGCGAGCAGCGGTGGACGCCCGCGGGAGATGAAATTCACAGAGGGAG-TCGGGA-GCG-AG-AGA-G-A-T-GAC-A-GT-GGAAA-GGAG-AGAGGA-GCTTTGGGGCGAG-CGT-ACGAT-GGGCA-GACGGGATACAGTGATGCTGTGCA--GGTGGATACACGTGCTCACCT-GA-AGTGTTTGTGGAGTCTTGTGCTGCGAATATGGGT-G-CTTTGGACTTTGTGTGTGGCGGTTGCCTCAAGA ``` |
| 23 | ggggacaagtttgtacaaaaaagcaggcttgatgAGAACCCCATCAGGCTTCCT, ggggaccactttgtacaagaaagctgggtcATCCCAATTCTTGACCGCCT | Mycgr3\|68144  JGI  Ensembl | KOG4129 Exopolyphosphatases and related proteins; | exopolyphosphatase activity; | 100 % |  |  |
| 23 | ggggacaagtttgtacaaaaaagcaggcttgatgTCGCGCTACGGGAAATGGGA, ggggaccactttgtacaagaaagctgggtcTGTGTATGTACTCCCTGCAG | Mycgr3\|68796  JGI  Ensembl |  |  | 100 % |  | ``` 394	GGGTCAATACTCATCGCGTTCGTCGCGATTCTAGTCTCGCTATTCCTGCTATGGAAAACGGATCGCAAAAAGGCTGCGGTTGGAAGAAGG	483 	\|\|\|\|\|\|\|\|\|\|\|\|\|\|\|\|\|\|\|\|\|\|\|\|\|\|\|\|\|\|\|\|\|\|\|\|\|\|\|\| \|\|\|\|\|\|\|\|\|\|\|\|\|\|\|\|\|\|\|\|\|\|\|\|\|\|\|\|\|\|\|\|\|\|\|\|\|\|\|\|\|\|\|\|\|\|\|\|\| 	GGGTCAATACTCATCGCGTTCGTCGCGATTCTAGTCTCGCCATTCCTGCTATGGAAAACGGATCGCAAAAAGGCTGCGGTTGGAAGAAGG ```      ``` 217	ATGTCGCGCTACGGGAAATGGGACGACTTTTGTCGAGACAGTGGCAATGGGCATGCTACGATACCGGTCTGCAATCTGTTCGACCAGTCAAGGGCACGAGGAGGCAGTGAGCCGTACTGGGGCGGATGCGAGCTTACGGGCATTTCATTGGGAGGGAATTCATACCTCGCGAACTTGGG	395 	\|\|\|\|\|\|\|\|\|\|\|\|\|\|\|\|\|\|\|\|\|\|\|\|\|\|\|\|\|\|\|\|\|\|\|\|\|\|\|\|\|\|\|\|\|\|\|\|\|\|\|\|\|\|\|\|\|\|\|\|\|\|\|\|\|\|\|\|\|\|\|\|\|\|\|\|\|\|\|\|\|\|\|\|\|\|\|\|\|\|\|\|\|\|\|\|\|\|\|\|\|\|\|\|\|\|\|\|\|\|\|\|\|\|\|\|\|\|\|\|\|\|\|\|\|\|\|\|\|\|\|\|\|\|\|\|\|\|\|\|\|\|\|\|\|\|\|\|\|\|\|\|\|\|\|\|\|\|\|\|\|\|\|\|\|\|\|\|\|\|\|\|\|\|\|\|\|\|\| 	ATGTCGCGCTACGGGAAATGGGACGACTTTTGTCGAGACAGTGGCAATGGGCATGCTACGATACCGGTCTGCAATCTGTTCGACCAGTCAAGGGCACGAGGAGGCAGTGAGCCGTACTGGGGCGGATGCGAGCTTACGGGCATTTCATTGGGAGGGAATTCATACCTCGCGAACTTGGG ```      ``` 911	TTTTCCTCATCGGAGCCGCTCTGCTTTTCGCCATTGGTCAAGTCTTCCAATACGTGATTAGCGTCCACATCTGCAACGGCACGAAAGGAGCGATCAACGGCGGTCTCTTCTCAACACTGTTCACCCTACTTGCCGTCGCTGCGACATGGGTATTCTGGTCCAGCATCACAGAGGACGACTGGCCGATGCCCGCCACATCTGCAGGGAGTACATACAC	1127 	\|\|\|\|\|\|\|\|\|\|\|\|\|\|\|\|\|\|\|\|\|\|\|\|\|\|\|\|\|\|\|\|\|\|\|\|\|\|\|\|\|\|\|\|\|\|\|\|\|\|\|\|\|\|\|\|\|\|\|\|\|\|\|\|\|\|\|\|\|\|\|\|\|\|\|\|\|\|\|\|\|\|\|\|\|\|\|\|\|\|\|\|\|\|\|\| \|\|\|\|\|\|\|\|\|\|\| \|\|\|\| \|\|\|\|\|\| \|\|\|\|\|\|\|\| \|\|\|\|\|\|\|\|\|\|\|\|\|\|\|\|\|\|\| \|\|\| \|\|\| \| \|\|\|\|\|\|\|\|\|\|\|\|\| \|\|\|\|\|\|\|  \|\|\|\|\|\|\|\|\|\|\|\|\|\|\|\|\|\|\|\|\|\|\|\|\|\|\|\|\|\|\|\|\|\| 	TTTTCCTCATCGGAGCCGCTCTGCTTTTCGCCATTGGTCAAGTCTTCCAATACGTGATTAGCGTCCACATCTGCAACGGCACGAAAGGAGCGATCA-CGGCGGTCTCT-CTCA-CACTGT-CACCCTAC-TGCCGTCGCTGCGACATGG-TAT-CTG-T-CAGCATCACAGAG-ACGACTGA-CGATGCCCGCCACATCTGCAGGGAGTACATACAC ```      ``` 481	AGGGAAATGCAAATATTCCTCATCGGATACATAATCATTGAGATCTGTGAGATCTTCACGATCGGAGGCTTTCCGCTGGATGCCGCGGTTAGACGCGCATTTTCTGCCGCCCACATCGCTGCTGTTGTTGCTACACTATGGATCCTCATGATGAACGGCGCAGTCGGATACCAGCTCATCGACGACGGCACACCACTTTCGATTGGACTTGTCTTTGGCTCTGCGGCGGCGCTGTTCGTTGGCACTGGCTACATCGCACTCGACACAGCATTCAGCTGGACCGGCTTCTTCGACGATACGCTCCCACTACCCAATCAGGCATACGCGCTGTACACGTTGTACCAGCTCGTACCACTCGTCTTCTTGGTGGTGTACTTCCTGTTGGAGACATTCTTGGTTCTCAAGGTCCTAGGGGAGATCCGACCAATGC	910 	\|\|\|\|\|\|\|\|\|\|\|\|\|\|\|\|\|\|\|\|\|\|\|\|\|\|\|\|\|\|\|\|\|\|\|\|\|\|\|\|\|\|\|\|\|\|\|\|\|\|\|\|\|\|\|\|\|\|\|\|\|\|\|\|\|\|\|\|\|\|\|\|\|\|\|\|\|\|\|\|\|\|\|\|\|\|\|\|\|\|\|\|\|\|\|\|\|\|\|\|\|\|\|\|\|\|\|\|\|\|\|\|\|\|\|\|\|\|\|\|\|\|\|\|\|\|\|\|\|\|\|\|\|\|\|\|\|\|\|\|\|\|\|\|\|\|\|\|\|\|\|\|\|\|\|\|\|\|\|\|\|\|\|\|\|\|\|\|\|\|\|\|\|\|\|\|\|\|\|\|\|\|\|\|\|\|\|\|\|\|\|\|\|\|\|\|\|\|\|\|\|\|\|\|\|\|\|\|\|\|\|\|\|\|\|\|\|\|\|\|\|\|\|\|\|\|\|\|\|\|\|\|\|\|\|\|\|\|\|\|\|\|\|\|\|\|\|\|\|\|\|\|\|\|\|\|\|\|\|\|\|\|\|\|\|\|\|\|\|\|\|\|\|\|\|\|\|\|\|\|\|\|\|\|\|\|\|\|\|\|\|\|\|\|\|\|\|\|\|\|\|\|\|\|\|\|\|\|\|\|\|\|\|\|\|\|\|\|\|\|\|\|\|\|\|\|\|\|\|\|\|\|\|\|\|\|\|\|\|\|\|\|\|\|\|\|\|\|\|\|\|\|\|\|\|\|\|\|\|\|\|\|\|\|\|\|\|\|\|\|\|\|\|\|\|\|\|\|\|\|\|\|\|\|\|\|\|\|\|\|\|\|\|\|\|\|\|\|\|\|\|\|\|\|\|\|\|\|\|\|\|\|\|\|\|\|\|\|\|\|\|\|\|\|\|\|\|\|\|\| 	AGGGAAATGCAAATATTCCTCATCGGATACATAATCATTGAGATCTGTGAGATCTTCACGATCGGAGGCTTTCCGCTGGATGCCGCGGTTAGACGCGCATTTTCTGCCGCCCACATCGCTGCTGTTGTTGCTACACTATGGATCCTCATGATGAACGGCGCAGTCGGATACCAGCTCATCGACGACGGCACACCACTTTCGATTGGACTTGTCTTTGGCTCTGCGGCGGCGCTGTTCGTTGGCACTGGCTACATCGCACTCGACACAGCATTCAGCTGGACCGGCTTCTTCGACGATACGCTCCCACTACCCAATCAGGCATACGCGCTGTACACGTTGTACCAGCTCGTACCACTCGTCTTCTTGGTGGTGTACTTCCTGTTGGAGACATTCTTGGTTCTCAAGGTCCTAGGGGAGATCCGACCAATGC ``` |
| 23 | ggggacaagtttgtacaaaaaagcaggcttgatgCTGGACTCCGCCGGCGCGGG, ggggaccactttgtacaagaaagctgggtcATCCGCAACAATGACTTGAT | Mycgr3\|69046  JGI  Ensembl |  |  | 100 % |  |  |
| 23 | ggggacaagtttgtacaaaaaagcaggcttgatgGCGAAACCAAAAGCGGCGGC, ggggaccactttgtacaagaaagctgggtcGCGACCGACTTTGCCCGCAA | Mycgr3\|69260  JGI  Ensembl | KOG0680 Actin-related protein - Arp6p; | protein binding; | 100 % |  |  |
| 23 | ggggacaagtttgtacaaaaaagcaggcttgatgGCCACTCAAGCGTCAGGACT, ggggaccactttgtacaagaaagctgggtcTGCGGTCTTAATTCCCTTTG | Mycgr3\|69448  JGI  Ensembl | KOG0134 NADH:flavin oxidoreductase/12-oxophytodienoate reductase; | FMN binding; oxidoreductase activity; | 100 % |  |  |
| 23 | ggggacaagtttgtacaaaaaagcaggcttgatgGTCCTCCACAACCCAAACAA, ggggaccactttgtacaagaaagctgggtcCAGAATCGTACCAAATCCGA | Mycgr3\|70000  JGI  Ensembl | KOG2936 Uncharacterized conserved protein; |  | 100 % |  |  |
| 23 | ggggacaagtttgtacaaaaaagcaggcttgatgGCGGGCTTCGTCCAAAAAGG, ggggaccactttgtacaagaaagctgggtcCTCTCTGACAATCAAACTCG | Mycgr3\|70038  JGI  Ensembl | KOG1389 3-oxoacyl CoA thiolase; | acetyl-CoA C-acyltransferase activity; | 100 % |  |  |
| 23 | ggggacaagtttgtacaaaaaagcaggcttgatgGCTCTGACCACATCCATTAT, ggggaccactttgtacaagaaagctgggtcCACATACTCCGCACATATCG | Mycgr3\|70129  JGI  Ensembl |  |  | 100 % |  |  |
| 23 | ggggacaagtttgtacaaaaaagcaggcttgatgCTCACCTTCCGGCGTGCCCT, ggggaccactttgtacaagaaagctgggtcGTCGTCGTTCTCCGTTTGCT | Mycgr3\|70208  JGI  Ensembl |  |  | 100 % |  |  |
| 23 | ggggacaagtttgtacaaaaaagcaggcttgatgGCCGACGATACCTCCTCCTG, ggggaccactttgtacaagaaagctgggtcCTCACGATCCGTCTGCGTCC | Mycgr3\|70572  JGI  Ensembl | KOG2903 Predicted glutathione S-transferase; |  | 100 % |  | ``` 901	CTACGCTTCCTTAATCCGCTTCGACACAGTCTACGTCCAGCACTTCAAGTGCAACCTCGGCATGATTCGCTACAGCTACCCCATCCTTCACAA	993 	\|\|\|\|\|\|\|\|\|\|\|\|\|\|\|\|\|\|\|\|\|\|\|\|\|\|\|\|\|\|\|\|\|\|\|\|\|\|\|\|\|\|\|\|\|\|\|\|\|\|\|\|\|\|\|\|\|\|\|\|\|\|\|\|\|\|\|\|\|\|\|\|\|\|\|\|\|\|\|\|\|\|\|\|\|\|\|\|\|\|\|\|\| 	CTACGCTTCCTTAATCCGCTTCGACACAGTCTACGTCCAGCACTTCAAGTGCAACCTCGGCATGATTCGCTACAGCTACCCCATCCTTCACAA ```      ``` 191	ATGGCCGACGATACCTCCTCCTGGCACGCTGGGCCGGACGACTCCTTCCACGGCAAGATCACTCTCAATGGCCGCTTCGCACCCGAAGCCAACCGATACCACCTCTACGTCGGATACTTCTGCCCATTCGCACATCGAGCAATGATCGTGTACAAGCTGAAGCAGCTCGAGAAATACGCCGGCATCGAAATGAGCATCATGAGACCATACCCGAAAGGCGACGAGAACGGATGGCCGGGCTGGAGATTTAACGTCAAGGAGGAGGAAGATCAGTACGAGGGAGCTACAGTGGACAAACTCTTCGGCAGCAAGTACCTGCACGAGGTGTACTTCAAAGCAGACAAGGAATACAAGGGTCGCTACAGCGTGCCGGTGTTGTGGGACAAGAAGCTGAGTACGATCGTCAACAACGAATCGCATGAGTTGCTGAGAGATCTGCAGACGGAGTTCAATCCGCTTCTTCCGCGAGAGCTGCAAGATATCACGCTGTATCCTGAAGATCTTCGGGAGGAGGTCGATACCCTTGGTCAGCAGCTGCAACGAGATCTCAACACTCGTGTCTACAAGACTGGAAACGCAACGACACAGGAGGACTATGAGGAGAATCTTCCTATTGTGTTCGCCATGCTGAACAAGCTGGAGAAAGTAGCTGCAAAGTCCGGCGGTCCGTATATCCTTGGCAAGCATATGACCGAGGTGGACGTTCGCAC	900 	\|\|\|\|\|\|\|\|\|\|\|\|\|\|\|\|\|\|\|\|\|\|\|\|\|\|\|\|\|\|\|\|\|\|\|\|\|\|\|\|\|\|\|\|\|\|\|\|\|\|\|\|\|\|\|\|\|\|\|\|\|\|\|\|\|\|\|\|\|\|\|\|\|\|\|\|\|\|\|\|\|\|\|\|\|\|\|\|\|\|\|\|\|\|\|\|\|\|\|\|\|\|\|\|\|\|\|\|\|\|\|\|\|\|\|\|\|\|\|\|\|\|\|\|\|\|\|\|\|\|\|\|\|\|\|\|\|\|\|\|\|\|\|\|\|\|\|\|\|\|\|\|\|\|\|\|\|\|\|\|\|\|\|\|\|\|\|\|\|\|\|\|\|\|\|\|\|\|\|\|\|\|\|\|\|\|\|\|\|\|\|\|\|\|\|\|\|\|\|\|\|\|\|\|\|\|\|\|\|\|\|\|\|\|\|\|\|\|\|\|\|\|\|\|\|\|\|\|\|\|\|\|\|\|\|\|\|\|\|\|\|\|\|\|\|\|\|\|\|\|\|\|\|\|\|\|\|\|\|\|\|\|\|\|\|\|\|\|\|\|\|\|\|\|\|\|\|\|\|\|\|\|\|\|\|\|\|\|\|\|\|\|\|\|\|\|\|\|\|\|\|\|\|\|\|\|\|\|\|\|\|\|\|\|\|\|\|\|\|\|\|\|\|\|\|\|\|\|\|\|\|\|\|\|\|\|\|\|\|\|\|\|\|\|\|\|\|\|\|\|\|\|\|\|\|\|\|\|\|\|\|\|\|\|\|\|\|\|\|\|\|\|\|\|\|\|\|\|\|\|\|\|\|\|\|\|\|\|\|\|\|\|\|\|\|\|\|\|\|\|\|\|\|\|\|\|\|\|\|\|\|\|\|\|\|\|\|\|\|\|\|\|\|\|\|\|\|\|\|\|\|\|\|\|\|\|\|\|\|\|\|\|\|\|\|\|\|\|\|\|\|\|\|\|\|\|\|\|\|\|\|\|\|\|\|\|\|\|\|\|\|\|\|\|\|\|\|\|\|\|\|\|\|\|\|\|\|\|\|\|\|\|\|\|\|\|\|\|\|\|\|\|\|\|\|\|\|\|\|\|\|\|\|\|\|\|\|\|\|\|\|\|\|\|\|\|\|\|\|\|\|\|\|\|\|\|\|\|\|\|\|\|\|\|\|\|\|\|\|\|\|\|\|\|\|\|\|\|\|\|\|\|\|\|\|\|\|\|\|\|\|\|\|\|\|\|\|\|\|\|\|\|\|\|\|\|\|\|\|\|\|\|\|\|\|\|\|\|\|\|\|\|\|\|\|\|\|\|\|\|\|\|\|\|\|\|\|\|\|\|\|\|\|\|\|\|\|\|\|\|\|\|\|\|\|\|\|\|\|\|\|\|\|\|\|\|\|\|\|\|\|\|\|\|\|\|\|\|\|\|\|\|\|\|\|\|\|\|\|\|\|\|\|\|\|\|\|\|\|\|\|\|\|\|\|\|\|\|\|\|\|\|\|\|\|\|\|\|\|\|\|\|\|\|\|\|\|\|\|\| 	ATGGCCGACGATACCTCCTCCTGGCACGCTGGGCCGGACGACTCCTTCCACGGCAAGATCACTCTCAATGGCCGCTTCGCACCCGAAGCCAACCGATACCACCTCTACGTCGGATACTTCTGCCCATTCGCACATCGAGCAATGATCGTGTACAAGCTGAAGCAGCTCGAGAAATACGCCGGCATCGAAATGAGCATCATGAGACCATACCCGAAAGGCGACGAGAACGGATGGCCGGGCTGGAGATTTAACGTCAAGGAGGAGGAAGATCAGTACGAGGGAGCTACAGTGGACAAACTCTTCGGCAGCAAGTACCTGCACGAGGTGTACTTCAAAGCAGACAAGGAATACAAGGGTCGCTACAGCGTGCCGGTGTTGTGGGACAAGAAGCTGAGTACGATCGTCAACAACGAATCGCATGAGTTGCTGAGAGATCTGCAGACGGAGTTCAATCCGCTTCTTCCGCGAGAGCTGCAAGATATCACGCTGTATCCTGAAGATCTTCGGGAGGAGGTCGATACCCTTGGTCAGCAGCTGCAACGAGATCTCAACACTCGTGTCTACAAGACTGGAAACGCAACGACACAGGAGGACTATGAGGAGAATCTTCCTATTGTGTTCGCCATGCTGAACAAGCTGGAGAAAGTAGCTGCAAAGTCCGGCGGTCCGTATATCCTTGGCAAGCATATGACCGAGGTGGACGTTCGCAC ``` |
| 23 | ggggacaagtttgtacaaaaaagcaggcttgatgCCCTCCCTCCCTCGTCTCCG, ggggaccactttgtacaagaaagctgggtcAGCCTCCCTGGGAACATGGC | Mycgr3\|71206  JGI  Ensembl |  |  | 100 % |  |  |
| 23 | ggggacaagtttgtacaaaaaagcaggcttgatgAAGGCTATTCAATTCCACGG, ggggaccactttgtacaagaaagctgggtcCTCTTTAGACATGTCAACCA | Mycgr3\|71679  JGI  Ensembl | KOG0024 Sorbitol dehydrogenase; | (R,R)-butanediol dehydrogenase activity; oxidoreductase activity;  zinc ion binding; | 100 % |  |  |
| 23 | ggggacaagtttgtacaaaaaagcaggcttgatgGGTGTCTACACTGTCGCGGC, ggggaccactttgtacaagaaagctgggtcGTACGAAGGAGCGGCGAAGA | Mycgr3\|72582  JGI  Ensembl |  |  | 100 % |  |  |
| 23 | ggggacaagtttgtacaaaaaagcaggcttgatgTCGGTCCAACGCAACCCCAA, ggggaccactttgtacaagaaagctgggtcGAACAGAGGATAGGCAAGCG | Mycgr3\|72602  JGI  Ensembl | KOG0873 C-4 sterol methyl oxidase; |  | 100 % |  | ``` 232	ATGTCGGTCCAACGCAACCCCAAGGACAGCATGAAGTCCACATGGCGCACCGCCGACAAGTCCACTTGGAAATACCACCACTGGCTCTACGATATCCTCAATGTCCACCCAACCGACATCAACAAGCCGGTTCCCGTATTCCAGAAGTCCGACAAGATGCCCTACCTCGCTCACTGGTCAACACATCGCTGGATCCTCACGCATTCCCTGTGGCCCATCGCCGTGGCCCAGCTCTACACATATTACTTCGGCAAGAACATGCCAGTCATCGGAGCATTTGCATTCTACACCATCGCGTTCCACATCAACGCCATCAACCACATCTACATCCTCCGCCGCCTCGGACACAAATGGGGTTTCCTCGACGGCGACAAGCACGAGCGCGACCAGGTTCCTGACGTGAGCGTCAAGAAGGTTTTCGACTCGCTGTCTTTGACCGCCGTCGTCCGACCATTGATCACCATCTTCCTGGCGTATCGCAAAAACGTCGCCCCGATCGAATCTCTGAGCTGGTGGCTCCCGGTCGAAATTGGAGTCTATGGCATCGTCCTTGATTTCTTCTTCTACTGGTATCACCGCGTCATGCACGAGTACGAAGGACTGTGGAAGTTCCACCGCACGCACCACTTGACCAAGCACCCCAACCCATTGCTCTCCTTGTACGCCGACACCGAGCAGGAAATCTTCGACATTGCCATCGTGCCGATTGCGACCTGGGCAACGATGAAGTTGATGGGTTTCCCAATGGGTTTCGGAGACTGGTGGGTGTGCCATCACTACTTG	1014 	\|\|\|\|\|\|\|\|\|\|\|\|\|\|\|\|\|\|\|\|\|\|\|\|\|\|\|\|\|\|\|\|\|\|\|\|\|\|\|\|\|\|\|\|\|\|\|\|\|\|\|\|\|\|\|\|\|\|\|\|\|\|\|\|\|\|\|\|\|\|\|\|\|\|\|\|\|\|\|\|\|\|\|\|\|\|\|\|\|\|\|\|\|\|\|\|\|\|\|\|\|\|\|\|\|\|\|\|\|\|\|\|\|\|\|\|\|\|\|\|\|\|\|\|\|\|\|\|\|\|\|\|\|\|\|\|\|\|\|\|\|\|\|\|\|\|\|\|\|\|\|\|\|\|\|\|\|\|\|\|\|\|\|\|\|\|\|\|\|\|\|\|\|\|\|\|\|\|\|\|\|\|\|\|\|\|\|\|\|\|\|\|\|\|\|\|\|\|\|\|\|\|\|\|\|\|\|\|\|\|\|\|\|\|\|\|\|\|\|\|\|\|\|\|\|\|\|\|\|\|\|\|\|\|\|\|\|\|\|\|\|\|\|\|\|\|\|\|\|\|\|\|\|\|\|\|\|\|\|\|\|\|\|\|\|\|\|\|\|\|\|\|\|\|\|\|\|\|\|\|\|\|\|\|\|\|\|\|\|\|\|\|\|\|\|\|\|\|\|\|\|\|\|\|\|\|\|\|\|\|\|\|\|\|\|\|\|\|\|\|\|\|\|\|\|\|\|\|\|\|\|\|\|\|\|\|\|\|\|\|\|\|\|\|\|\|\|\|\|\|\|\|\|\|\|\|\|\|\|\|\|\|\|\|\|\|\|\|\|\|\|\|\|\|\|\|\|\|\|\|\|\|\|\|\|\|\|\|\|\|\|\|\|\|\|\|\|\|\|\|\|\|\|\|\|\|\|\|\|\|\|\|\|\|\|\|\|\|\|\|\|\|\|\|\|\|\|\|\|\|\|\|\|\|\|\|\|\|\|\|\|\|\|\|\|\|\|\|\|\|\|\|\|\|\|\|\|\|\|\|\|\|\|\|\|\|\|\|\|\|\|\|\|\|\|\|\|\|\|\|\|\|\|\|\|\|\|\|\|\|\|\|\|\|\|\|\|\|\|\|\|\|\|\|\|\|\|\|\|\|\|\|\|\|\|\|\|\|\|\|\|\|\|\|\|\|\|\|\|\|\|\|\|\|\|\|\|\|\|\|\|\|\|\|\|\|\|\|\|\|\|\|\|\|\|\|\|\|\|\|\|\|\|\|\|\|\|\|\|\|\|\|\|\|\|\|\|\|\|\|\|\|\|\|\|\|\|\|\|\|\|\|\|\|\|\|\|\|\|\|\|\|\|\|\|\|\|\|\|\|\|\|\|\|\|\|\|\|\|\|\|\|\|\|\|\|\|\|\|\|\|\|\|\|\|\|\|\|\|\|\|\|\|\|\|\|\|\|\|\|\|\|\|\|\|\|\|\|\|\|\|\|\|\|\|\|\|\|\|\|\|\|\|\|\|\|\|\|\|\|\|\|\|\|\|\|\|\|\|\|\|\|\|\|\|\|\|\|\|\|\|\|\|\|\|\|\|\|\|\|\|\|\|\|\|\|\|\|\|\|\|\|\|\|\|\|\|\|\|\|\|\|\|\|\|\|\|\|\|\|\|\|\|\|\|\|\|\|\|\|\|\|\|\|\|\|\|\|\|\|\|\|\|\|\|\|\|\|\|\|\|\|\|\|\|\|\|\|\|\|\|\|\| 	ATGTCGGTCCAACGCAACCCCAAGGACAGCATGAAGTCCACATGGCGCACCGCCGACAAGTCCACTTGGAAATACCACCACTGGCTCTACGATATCCTCAATGTCCACCCAACCGACATCAACAAGCCGGTTCCCGTATTCCAGAAGTCCGACAAGATGCCCTACCTCGCTCACTGGTCAACACATCGCTGGATCCTCACGCATTCCCTGTGGCCCATCGCCGTGGCCCAGCTCTACACATATTACTTCGGCAAGAACATGCCAGTCATCGGAGCATTTGCATTCTACACCATCGCGTTCCACATCAACGCCATCAACCACATCTACATCCTCCGCCGCCTCGGACACAAATGGGGTTTCCTCGACGGCGACAAGCACGAGCGCGACCAGGTTCCTGACGTGAGCGTCAAGAAGGTTTTCGACTCGCTGTCTTTGACCGCCGTCGTCCGACCATTGATCACCATCTTCCTGGCGTATCGCAAAAACGTCGCCCCGATCGAATCTCTGAGCTGGTGGCTCCCGGTCGAAATTGGAGTCTATGGCATCGTCCTTGATTTCTTCTTCTACTGGTATCACCGCGTCATGCACGAGTACGAAGGACTGTGGAAGTTCCACCGCACGCACCACTTGACCAAGCACCCCAACCCATTGCTCTCCTTGTACGCCGACACCGAGCAGGAAATCTTCGACATTGCCATCGTGCCGATTGCGACCTGGGCAACGATGAAGTTGATGGGTTTCCCAATGGGTTTCGGAGACTGGTGGGTGTGCCATCACTACTTG ``` |
| 23 | ggggacaagtttgtacaaaaaagcaggcttgatgTCCAAAGGCAAAGTCCTCAT, ggggaccactttgtacaagaaagctgggtcCTCATCCTTCCCACTAAACT | Mycgr3\|72813  JGI  Ensembl |  |  | 100 % |  |  |
| 23 | ggggacaagtttgtacaaaaaagcaggcttgatgAGCGAAATAGTCCGACGACG, ggggaccactttgtacaagaaagctgggtcCTTCCCAAACATCCGATCCA | Mycgr3\|74465  JGI  Ensembl | KOG1505 Lysophosphatidic acid acyltransferase LPAAT and related acyltransferases; | acyltransferase activity; glycerol-3-phosphate O-acyltransferase activity;  metabolic process;  protein binding; | 100 % |  |  |
| 23 | ggggacaagtttgtacaaaaaagcaggcttgatgTTCGGCAGCACACTCCGGCG, ggggaccactttgtacaagaaagctgggtcAGTAGCCTGAGGCGACAGTG | Mycgr3\|75418  JGI  Ensembl | KOG3042 Panthothenate synthetase; | pantoate-beta-alanine ligase activity; pantothenate biosynthetic process; | 100 % |  |  |
| 23 | ggggacaagtttgtacaaaaaagcaggcttgatgTCCGCCTCAATCCAAAACCT, ggggaccactttgtacaagaaagctgggtcCTCCTTCAACGGCGGTGTCG | Mycgr3\|77172  JGI  Ensembl | KOG1575 Voltage-gated shaker-like K+ channel, subunit beta/KCNAB; | oxidoreductase activity; pyridoxine 4-dehydrogenase activity; | 100 % |  |  |
| 23 | ggggacaagtttgtacaaaaaagcaggcttgatgCCGTCCTCCGTGGCCACGCA, ggggaccactttgtacaagaaagctgggtcCAGATCCACATCCTCCCCTC | Mycgr3\|77243  JGI  Ensembl |  |  | 100 % |  |  |
| 23 | ggggacaagtttgtacaaaaaagcaggcttgatgCTTGCTCGTGTTGCGCCGAT, ggggaccactttgtacaagaaagctgggtcTGGGTGGGACTTTTCCTGTT | Mycgr3\|77528  JGI  Ensembl | KOG1381 Para-hydroxybenzoate-polyprenyl transferase; | (S)-2,3-di-O-geranylgeranylglyceryl phosphate synthase activity; 1,4-dihydroxy-2-naphthoate octaprenyltransferase activity;  4-hydroxybenzoate octaprenyltransferase activity;  Sep-tRNA:Cys-tRNA synthase activity;  biosynthetic process;  integral to membrane;  prenyltransferase activity;  protein geranylgeranyltransferase activity;  protein prenyltransferase activity;  protoheme IX farnesyltransferase activity; | 100 % |  |  |
| 23 | ggggacaagtttgtacaaaaaagcaggcttgatgGCCACCTCGACCTTACCAGC, ggggaccactttgtacaagaaagctgggtcCAAAACCTCCAAAACCAGCC | Mycgr3\|79635  JGI  Ensembl |  |  | 100 % |  | ``` 156	ATGGCCACCTCGACCTTACCAGCTCCGACCGCATTCCCTTCTCCATTCCCAGTACCCTCAAAATCAACCAGCGGCCACATCAACGGAGTCCTCACCGAAGTCTCATCCATGGTCTTCTCCGACAAAATACTCATCACAATCTCTCAAGCCGGCAAGCTCGCACACTGGGTCCACGTGCCACTCGCGACAGCGACGGCCGATCCGCTTTCCGGACCAATTGCAAACTCGTCTTCCGAGAACGAGTTCCTGCCAATGAGCCATCTGACAGCGACCACTGTTCTGGGCGGAACCAGACGGGACGACGAGATCATAGGTCAGACTTTGGCTACTACGATTGCTAGTGCCCTTTTGGTGAAGAGACCTCAGGAGGAGCGTCTGCTTGTGATCGGCGCGGGGTGGGTGAACGCTGCCGAGACTTTGACGGGGAGGGCTCAGTTTGAAGAGGTTCTTGGGCTGGTTTTGGAGGTTTTG	626 	\|\|\|\|\|\|\|\|\|\|\|\|\|\|\|\|\|\|\|\|\|\|\|\|\|\|\|\|\|\|\|\|\|\|\|\|\|\|\|\|\|\|\|\|\|\|\|\|\|\|\|\|\|\|\|\|\|\|\|\|\|\|\|\|\|\|\|\|\|\|\|\|\|\|\|\|\|\|\|\|\|\|\|\|\|\|\|\|\|\|\|\|\|\|\|\|\|\|\|\|\|\|\|\|\|\|\|\|\|\|\|\|\|\|\|\|\|\|\|\|\|\|\|\|\|\|\|\|\|\|\|\|\|\|\|\|\|\|\|\|\|\|\|\|\|\|\|\|\|\|\|\|\|\|\|\|\|\|\|\|\|\|\|\|\|\|\|\|\|\|\|\|\|\|\|\|\|\|\|\|\|\|\|\|\|\|\|\|\|\|\|\|\|\|\|\|\|\|\|\|\|\|\|\|\|\|\|\|\|\|\|\|\|\|\|\|\|\|\|\|\|\|\|\|\|\|\|\|\|\|\|\|\|\|\|\|\|\|\|\|\|\|\|\|\|\|\|\|\|\|\|\|\|\|\|\|\|\|\|\|\|\|\|\|\|\|\|\|\|\|\|\|\|\|\|\|\|\|\|\|\|\|\|\|\|\|\|\|\|\|\|\|\|\|\|\|\|\|\|\|\|\|\|\|\|\|\|\|\|\|\|\|\|\|\|\|\|\|\|\|\|\|\|\|\|\|\|\|\|\|\|\|\|\|\|\|\|\|\|\|\|\|\|\|\|\|\|\|\|\|\|\|\|\|\|\|\|\|\|\|\|\|\|\|\|\|\|\|\|\|\|\|\|\|\|\|\|\|\|\|\|\|\|\|\|\|\|\|\|\|\|\|\|\|\|\|\|\|\|\|\|\|\|\|\|\|\|\|\|\|\|\|\|\|\|\|\|\|\|\|\|\|\|\|\|\|\|\|\|\|\|\|\|\|\|\|\|\|\|\|\|\|\|\|\|\|\|\|\|\|\|\|\|\|\|\|\|\|\|\|\|\|\|\|\|\|\|\|\|\|\| 	ATGGCCACCTCGACCTTACCAGCTCCGACCGCATTCCCTTCTCCATTCCCAGTACCCTCAAAATCAACCAGCGGCCACATCAACGGAGTCCTCACCGAAGTCTCATCCATGGTCTTCTCCGACAAAATACTCATCACAATCTCTCAAGCCGGCAAGCTCGCACACTGGGTCCACGTGCCACTCGCGACAGCGACGGCCGATCCGCTTTCCGGACCAATTGCAAACTCGTCTTCCGAGAACGAGTTCCTGCCAATGAGCCATCTGACAGCGACCACTGTTCTGGGCGGAACCAGACGGGACGACGAGATCATAGGTCAGACTTTGGCTACTACGATTGCTAGTGCCCTTTTGGTGAAGAGACCTCAGGAGGAGCGTCTGCTTGTGATCGGCGCGGGGTGGGTGAACGCTGCCGAGACTTTGACGGGGAGGGCTCAGTTTGAAGAGGTTCTTGGGCTGGTTTTGGAGGTTTTG ``` |
| 23 | ggggacaagtttgtacaaaaaagcaggcttgatgGGTCCGAAGAGCTGGCCCAA, ggggaccactttgtacaagaaagctgggtcAGAACGGGGACACCTTCCAT | Mycgr3\|86032  JGI  Ensembl |  |  | 100 % |  |  |
| 23 | ggggacaagtttgtacaaaaaagcaggcttgatgCCCATCTCAGCCCTCGACTC, ggggaccactttgtacaagaaagctgggtcCGCAATCCCCTTCTTCTCAC | Mycgr3\|86202  JGI  Ensembl | KOG2700 Adenylosuccinate lyase; | 3-carboxy-cis,cis-muconate cycloisomerase activity; catalytic activity; | 100 % |  |  |
| 23 | ggggacaagtttgtacaaaaaagcaggcttgatgGCTAATGTTGACATACACAA, ggggaccactttgtacaagaaagctgggtcTGGCACTTCACCATTGACCA | Mycgr3\|89091  JGI  Ensembl |  |  | 100 % |  |  |
| 23 | ggggacaagtttgtacaaaaaagcaggcttgatgGCCGGCGAGAGGCAACCAAG, ggggaccactttgtacaagaaagctgggtcCTCCAACCAAGTGTCCCTCT | Mycgr3\|90661  JGI  Ensembl |  |  | 1.61646 % |  |  |
| 23 | ggggacaagtttgtacaaaaaagcaggcttgatgGCCAGCGGATCAGCATATCT, ggggaccactttgtacaagaaagctgggtcGACGCTGGTTCGCATTCGCT | Mycgr3\|90710  JGI  Ensembl |  |  | 100 % |  |  |
| 23 | ggggacaagtttgtacaaaaaagcaggcttgatgAATCGAGGAAGCATCCCGGG, ggggaccactttgtacaagaaagctgggtcGTCAGCCTCCGTCCAAGGTT | Mycgr3\|91245  JGI  Ensembl |  |  | 100 % |  |  |
| 23 | ggggacaagtttgtacaaaaaagcaggcttgatgGTGCCTGGACCTGATTCCAT, ggggaccactttgtacaagaaagctgggtcTGCGCTGGCGGAAATGATGG | Mycgr3\|92288  JGI  Ensembl | KOG1721 FOG: Zn-finger; | intracellular; nucleic acid binding;  zinc ion binding; | 100 % |  | ``` 1	ATGGTGCCTGGACCTGATTCCATGGGCGCTCCGACCAGCTTCACTGCTCGGAGACCCAACGCTTCACATCTGCCATCGTTCGAACTGCCACCGCCTCCGCTTGCCGGACTGCAGCAAAAGTACCATCAACCAAGCTACTCCCACCACACATCGCAAGCACCCGCAACGACTCTGACGAGTGTTGGCAACCTGTTGACTCCTCCAAGCAACGGTTCCGCCGACGGCTTGAGTCCCGGCAATACTTCACACAGCACCGGTCCGGCATACAGCCCGAACGGCAGTGGAATGTGGCCACCACCATCGAACGGCGGCTCTACTCCGTCGTATGGGTTTCAACAAAGCAGCGCATCGCAGTATCAGCAGGGTCGCGGATTGTACTCGCCCTCTCTGAACTCGATTGTTCGCGGGAGCAACTCTCCCAGCGCAAGTGAACTTCCTCCTCCACCTTACGAAATGCCACACTACGCCAACTCGACCATGTCAATGTCGGCTCCTAATTTGCCCGCCATGCCAAACCACCAGCCGCAGATGATGGCTCCAGGACCTCAGCCATCGCCAGCAAACAGTCACGAGTCCTTCTCACGACCTCCACCGACGCCTTCCTATTCATATTCGTCGCAATCATCCGCGCCTCAGCAAAGTAGCTACTCGTACTCGGGACCTTCTCCCGTACAACAGAGTTCTAGCACTCAGTCCATGCCGAGAATGTCTCCAGTCAGCCAACAAGGCCAAGCGCCACCTGCTCAGCACCAGTCTcagcaacaacaacaacatcatcaacagcaacaccaccaccagtcccagcagcagcaacagTATGCGCAACATCGACCGTACAGCTACTCGATGCCTGTTCTCACCAACGTGAATAACCCGGGTGGACAGATGTCCATGGT-GGGAGGTGGAGTGCCTCACAGTATGATGGCACCCT-ACAACAGTGG-TCAAGCAGCCCACTACTCCCAGATGTAC-GCCCACCACCCACAGCAAGCCAACCCACCAAACGACAGACCTTTCCGATGCGACCAGTGCCCT-CAGAGCTTCAATCGCAACCACGACTTG	1059 	\|\|\|\|\|\|\|\|\|\|\|\|\|\|\|\|\|\|\|\|\|\|\|\|\|\|\|\|\|\|\|\|\|\|\|\|\|\|\|\|\|\|\|\|\|\|\|\|\|\|\|\|\|\|\|\|\|\|\|\|\|\|\|\|\|\|\|\|\|\|\|\|\|\|\|\|\|\|\|\|\|\|\|\|\|\|\|\|\|\|\|\|\|\|\|\|\|\|\|\|\|\|\|\|\|\|\|\|\|\|\|\|\|\|\|\|\|\|\|\|\|\|\|\|\|\|\|\|\|\|\|\|\|\|\|\|\|\|\|\|\|\|\|\|\|\|\|\|\|\|\|\|\|\|\|\|\|\|\|\|\|\|\|\|\|\|\|\|\|\|\|\|\|\|\|\|\|\|\|\|\|\|\|\|\|\|\|\|\|\|\|\|\|\|\|\|\|\|\|\|\|\|\|\|\|\|\|\|\|\|\|\|\|\|\|\|\|\|\|\|\|\|\|\|\|\|\|\|\|\|\|\|\|\|\|\|\|\|\|\|\|\|\|\|\|\|\|\|\|\|\|\|\|\|\|\|\|\|\|\|\|\|\|\|\|\|\|\|\|\|\|\|\|\|\|\|\|\|\|\|\|\|\|\|\|\|\|\|\|\|\|\|\|\|\|\|\|\|\|\|\|\|\|\|\|\|\|\|\|\|\|\|\|\|\|\|\|\|\|\|\|\|\|\|\|\|\|\|\|\|\|\|\|\|\|\|\|\|\|\|\|\|\|\|\|\|\|\|\|\|\|\|\|\|\|\|\|\|\|\|\|\|\|\|\|\|\|\|\|\|\|\|\|\|\|\|\|\|\|\|\|\|\|\|\|\|\|\|\|\|\|\|\|\|\|\|\|\|\|\|\|\|\|\|\|\|\|\|\|\|\|\|\|\|\|\|\|\|\|\|\|\|\|\|\|\|\|\|\|\|\|\|\|\|\|\|\|\|\|\|\|\|\|\|\|\|\|\|\|\|\|\|\|\|\|\|\|\|\|\|\|\|\|\|\|\|\|\|\|\|\|\|\|\|\|\|\|\|\|\|\|\|\|\|\|\|\|\|\|\|\|\|\|\|\|\|\|\|\|\|\|\|\|\|\|\|\|\|\|\|\|\|\|\|\|\|\|\|\|\|\|\|\|\|\|\|\|\|\|\|\|\|\|\|\|\|\|\|\|\|\|\|\|\|\|\|\|\|\|\|\|\|\|\|\|\|\|\|\|\|\|\|\|\|\|\|\|\|\|\|\|\|\|\|\|\|\|\|\|\|\|\|\|\|\|\|\|\|\|\|\|\|\|\|\|\|\|\|\|\|\|\|\|\|\|\|\|\|\|\|\|\|\|\|\|\|\|\|\|\|\|\|\|\|\|\|\|\|\|\|\|\|\|\|\|\|\|\|\|\|\|\|\|\|\|\|\|\|\|\|\|\|\|\|\|\|\|\|\|\|\|\|\|\|\|\|\|\|\|\|\|\|\|\|\|\|\|\|\|\|\|\|\|\|\|\|\|\|\|\|\|\|\|\|\|\|\|\|\|\|\|\|\|\|\|\|\|\|\|\|\|\|\|\|\|\|\|\|\|\|\|\|\|\|\|\|\|\|\|\|\|\|\|\|\|\|\|\|\|\|\|\|\|\|\|\|\|\|\|\|\|\|\|\|\|\|\|\|\|\|\|\|\|\|\|\|\|\|\|\|\|\|\|\|\|\|\|\|\|\|\|\|\|\|\|\|\|\|\|\|\|\|\|\|\|\|\|\|\|\|\|\|\|\|\|\|\|\|\|\|\|\|\|\|\|\|\|\|\|\|\|\|\| \|\|\|\|\|\|\| \|\|\|\|\|\|\|\|\|\|\|\|\|\|\|\|\|\|\|\|\|\|\|\|\|\|\|\|\|\|\|\|\|\|\|\|\|\| \|\|\|\|\|\|\|\| \|\|\|\|\|\|\|\|\|\|\|\|\|\|\|\| \|\|\|\|\|\|\|\|\|\|\|\|\|\|\|\|\|\|\|\|\|\|\|\|\|\|\|\|\|\|\|\|\|\|\| \|\|\| \|\|  \|\| \|\| \|\|\|\|\|\|\|\|\|\|\|\|\|\|\|\|\| \|\|\|\|\|\|\| \|\|\|\|\|\|\|\|\|\|\|\|\|\|\|\|\| \|\|\|\| \|\| \|\|\|\|\| \|\|\|\|  \|\|\|\|\|\|\|  \|\|\|\|\|\| \|\|\| \| \|\|\| \|\|\|\|\|\|\|\|\|\| \|\|\|\|\| \|\|  \|\|\|\|\|\| 	ATGGTGCCTGGACCTGATTCCATGGGCGCTCCGACCAGCTTCACTGCTCGGAGACCCAACGCTTCACATCTGCCATCGTTCGAACTGCCACCGCCTCCGCTTGCCGGACTGCAGCAAAAGTACCATCAACCAAGCTACTCCCACCACACATCGCAAGCACCCGCAACGACTCTGACGAGTGTTGGCAACCTGTTGACTCCTCCAAGCAACGGTTCCGCCGACGGCTTGAGTCCCGGCAATACTTCACACAGCACCGGTCCGGCATACAGCCCGAACGGCAGTGGAATGTGGCCACCACCATCGAACGGCGGCTCTACTCCGTCGTATGGGTTTCAACAAAGCAGCGCATCGCAGTATCAGCAGGGTCGCGGATTGTACTCGCCCTCTCTGAACTCGATTGTTCGCGGGAGCAACTCTCCCAGCGCAAGTGAACTTCCTCCTCCACCTTACGAAATGCCACACTACGCCAACTCGACCATGTCAATGTCGGCTCCTAATTTGCCCGCCATGCCAAACCACCAGCCGCAGATGATGGCTCCAGGACCTCAGCCATCGCCAGCAAACAGTCACGAGTCCTTCTCACGACCTCCACCGACGCCTTCCTATTCATATTCGTCGCAATCATCCGCGCCTCAGCAAAGTAGCTACTCGTACTCGGGACCTTCTCCCGTACAACAGAGTTCTAGCACTCAGTCCATGCCGAGAATGTCTCCAGTCAGCCAACAAGGCCAAGCGCCACCTGCTCAGCACCAGTCTCAGCAACAACAACAACATCATCAACAGCAACACCACCACCAGTCCCAGCAGCAGCAACAGTATGCGC-ACATCGATCGTACAGCTACTCGATGCCTGTTCTCACCAACGTGAATTACCCGGGTTGACAGATGTCCATGGTGGGGAGGTGGAGTGCCTCACAGTATGATGGCACCCTTACA-CATGGGATC-AGCAGCCCACTACTCCCTGATGTACGGCCCACCACCCACAGCAGGCCATCC-ACCAA-CGACGAACCTTTCGAATGCGAACAGGGTCCTTCAGAGCTTCA-TCGCA-CCCGGACTTG ``` |
| 23 | ggggacaagtttgtacaaaaaagcaggcttgatgGCCTCGGTCCGTTCTCGAGT, ggggaccactttgtacaagaaagctgggtcGCGTGAAGAAAGGTAGGAGA | Mycgr3\|92489  JGI  Ensembl | KOG2146 Splicing coactivator SRm160/300, subunit SRm160 (contains PWI domain); | poly(A)-specific ribonuclease activity; | 100 % |  |  |
| 23 | ggggacaagtttgtacaaaaaagcaggcttgatgCATTCCAGTCGCGGCAGATA, ggggaccactttgtacaagaaagctgggtcGTACTTGCAGCGGCCGCAAT | Mycgr3\|92938  JGI  Ensembl |  |  | 100 % |  |  |
| 23 | ggggacaagtttgtacaaaaaagcaggcttgatgGCCGTCTCACCATGGGCCTG, ggggaccactttgtacaagaaagctgggtcTCCCCTCGAAAGACCGTCTC | Mycgr3\|93322  JGI  Ensembl |  |  | 100 % |  |  |
| 23 | ggggacaagtttgtacaaaaaagcaggcttgatgACTTCTCCCACTTCCACTTC, ggggaccactttgtacaagaaagctgggtcCTTCTTCCCCGAATGCAACT | Mycgr3\|93440  JGI  Ensembl | KOG3599 Ca2+-modulated nonselective cation channel polycystin; |  | 100 % |  |  |
| 23 | ggggacaagtttgtacaaaaaagcaggcttgatgAACACACAAGGGACCTACCG, ggggaccactttgtacaagaaagctgggtcTTCATCCATGTCCAAGCTCT | Mycgr3\|93587  JGI  Ensembl |  |  | 100 % |  |  |
| 23 | ggggacaagtttgtacaaaaaagcaggcttgatgTCAGATACTCTATGGAACTT, ggggaccactttgtacaagaaagctgggtcGTCGTGGCGACGGGACCGGG | Mycgr3\|94224  JGI  Ensembl |  |  | 100 % |  |  |
| 23 | ggggacaagtttgtacaaaaaagcaggcttgatgAAGTACTCTCAGTCAACAGC, ggggaccactttgtacaagaaagctgggtcAGCGTTCGATCGCTTATAAC | Mycgr3\|94368  JGI  Ensembl |  |  | 100 % |  |  |
| 23 | ggggacaagtttgtacaaaaaagcaggcttgatgTCCGACCCGCACCGGAGAAA, ggggaccactttgtacaagaaagctgggtcAAAGCGATCGATCCCGAGAA | Mycgr3\|94676  JGI  Ensembl |  |  | 100 % |  |  |
| 23 | ggggacaagtttgtacaaaaaagcaggcttgatgCGCCTCTCGTCACCAGCAAC, ggggaccactttgtacaagaaagctgggtcTACACATGACGACGTGGAGG | Mycgr3\|94787  JGI  Ensembl |  |  | 100 % |  |  |
| 23 | ggggacaagtttgtacaaaaaagcaggcttgatgCCGGCCCAGAACGGTGCAAA, ggggaccactttgtacaagaaagctgggtcGCGATCAAGCTCCATTTCAT | Mycgr3\|94908  JGI  Ensembl |  |  | 100 % |  |  |
| 23 | ggggacaagtttgtacaaaaaagcaggcttgatgGACATCCACTCTCTCGCTAA, ggggaccactttgtacaagaaagctgggtcAGCGTAGTGGATATTGACGC | Mycgr3\|95041  JGI  Ensembl |  |  | 100 % |  |  |
| 23 | ggggacaagtttgtacaaaaaagcaggcttgatgGCATCGCTTCGCGCCGTGCG, ggggaccactttgtacaagaaagctgggtcAAGTTTTGCCATTTTGCGGC | Mycgr3\|95435  JGI  Ensembl | KOG2249 3'-5' exonuclease; |  | 100 % |  |  |
| 23 | ggggacaagtttgtacaaaaaagcaggcttgatgTCAATCACCACGGCCACTTC, ggggaccactttgtacaagaaagctgggtcCTTGCGGCCTGCAACCTGCG | Mycgr3\|95436  JGI  Ensembl | KOG2249 3'-5' exonuclease; | exonuclease activity; intracellular; | 100 % |  |  |
| 23 | ggggacaagtttgtacaaaaaagcaggcttgatgGATGACACTCTCGGACAGGT, ggggaccactttgtacaagaaagctgggtcTTTTGTGTCTTTGTATCGTG | Mycgr3\|96015  JGI  Ensembl |  |  | 100 % |  |  |
| 23 | ggggacaagtttgtacaaaaaagcaggcttgatgAGTCGGCCATCCAAGTCCCG, ggggaccactttgtacaagaaagctgggtcTATGTCAACCCTCTTCTTCA | Mycgr3\|97053  JGI  Ensembl | KOG2765 Predicted membrane protein; |  | 100 % |  |  |
| 23 | ggggacaagtttgtacaaaaaagcaggcttgatgGCGCCTCCACACACCTGGAC, ggggaccactttgtacaagaaagctgggtcTTGAACACGAGTCGATCGCT | Mycgr3\|97064  JGI  Ensembl | KOG2806 Chitinase; |  | 100 % |  |  |
| 23 | ggggacaagtttgtacaaaaaagcaggcttgatgTCTTCATATACATCCTGCTG, ggggaccactttgtacaagaaagctgggtcCGGAACCACCCGTGCGGCAT | Mycgr3\|97825  JGI  Ensembl |  |  | 100 % |  |  |
| 23 | ggggacaagtttgtacaaaaaagcaggcttgatgGCTGCCAGCGACTCTGCTGC, ggggaccactttgtacaagaaagctgggtcACAGCCCTCTGTGAGTGCGT | Mycgr3\|97956  JGI  Ensembl |  |  | 100 % |  | ``` 224	AGGGCCGGACGTGCGTGATGTACGGAGGATTGAGTGCAGGTAGGGAGCTGGTGATGCCGAAAGCCATGACCACGCAGGCGACGGTCTTTTGGCCCACCGTCGTGGGCTCTCGCGTCTCAGATTTGGGCCTCAGACAGCAAATCGTCTGTGTGGTACATGCATGTGGCTATCG	395 	\|\|\|\|\|\|\|\|\|\|\|\|\|\|\|\|\|\|\|\|\|\|\|\|\|\|\|\|\|\|\|\|\|\|\|\|\|\|\|\|\|\|\|\|\|\|\|\|\|\|\|\|\|\|\|\|\|\|\|\|\|\|\|\|\|\|\|\|\|\|\|\|\|\|\|\|\|\|\|\|\|\|\|\|\|\|\|\|\|\|\|\|\|\|\|\|\|\|\|\|\|\|\|\|\|\|\|\|\|\|\|\|\|\|\|\|\|\|\|\|\|\|\|\|\|\|\|\|\|\|\|\|\|\|\|\|\|\|\|\|\|\|\|\|\|\|\|\|\|\|\|\|\|\|\|\|\|\|\|\|\|\|\|\|\|\|\|\|\|\|\|\| 	AGGGCCGGACGTGCGTGATGTACGGAGGATTGAGTGCAGGTAGGGAGCTGGTGATGCCGAAAGCCATGACCACGCAGGCGACGGTCTTTTGGCCCACCGTCGTGGGCTCTCGCGTCTCAGATTTGGGCCTCAGACAGCAAATCGTCTGTGTGGTACATGCATGTGGCTATCG ```      ``` 1	ATGGCTGCCAGCGACTCTGCTGCCGCTCCTGCCGTGTCTTGTACTGCTGCGGCATCCCTCCTCTTCTTCATCGTGGCTTTGATATTGGCATGCTTCGCCGGCACTTCCATCGCTACCTTCTTCACTTCCGTCGTCGGCGGCATCATCGTCGCCGCTGTCTGGCTGGTCCTCTTCTTCGTCGTCGATCATGTGATCCTGTATCAGATCGGTGAATTCTACCAGAAGG	226 	\|\|\|\|\|\|\|\|\|\|\|\|\|\|\|\|\|\|\|\|\|\|\|\|\|\|\|\|\|\|\|\|\|\|\|\|\|\|\|\|\|\|\|\|\|\|\|\|\|\|\|\|\|\|\|\|\|\|\|\|\|\|\|\|\|\|\|\|\|\|\|\|\|\|\|\|\|\|\|\|\|\|\|\|\|\|\|\|\|\|\|\|\|\|\|\|\|\|\|\|\|\|\|\|\|\|\|\|\|\|\|\|\|\|\|\|\|\|\|\|\|\|\|\|\|\|\|\|\|\|\|\|\|\|\|\|\|\|\|\|\|\|\|\|\|\|\|\|\|\|\|\|\|\|\|\|\|\|\|\|\|\|\|\|\|\|\|\|\|\|\|\|\|\|\|\|\|\|\|\|\|\|\|\|\|\|\|\|\|\|\|\|\|\|\|\|\|\|\|\|\|\|\|\|\|\|\|\|\|\|\|\|\|\|\|\|\|\|\|\|\|\|\|\|\|\| 	ATGGCTGCCAGCGACTCTGCTGCCGCTCCTGCCGTGTCTTGTACTGCTGCGGCATCCCTCCTCTTCTTCATCGTGGCTTTGATATTGGCATGCTTCGCCGGCACTTCCATCGCTACCTTCTTCACTTCCGTCGTCGGCGGCATCATCGTCGCCGCTGTCTGGCTGGTCCTCTTCTTCGTCGTCGATCATGTGATCCTGTATCAGATCGGTGAATTCTACCAGAAGG ``` |
| 23 | ggggacaagtttgtacaaaaaagcaggcttgatgGCCGACCCAGCTCCACCCAC, ggggaccactttgtacaagaaagctgggtcGTCCTCCACACTCTTCCCAA | Mycgr3\|98669  JGI  Ensembl | KOG1181 FOG: Low-complexity; |  | 100 % |  |  |
| 23 | ggggacaagtttgtacaaaaaagcaggcttgatgCCTCCACTCAAGAGCTGGCT, ggggaccactttgtacaagaaagctgggtcGAACTTCATCTCCACCGCCC | Mycgr3\|98885  JGI  Ensembl | KOG2843 Fumarylacetoacetase; | aromatic amino acid family metabolic process; catalytic activity;  fumarylacetoacetase activity;  metabolic process; | 100 % |  |  |
| 23 | ggggacaagtttgtacaaaaaagcaggcttgatgACTGAGCCCGAGCTTGCAGA, ggggaccactttgtacaagaaagctgggtcCGGACGCACACCTACAAGAC | Mycgr3\|99133  JGI  Ensembl |  |  | 100 % |  |  |
| 23 | ggggacaagtttgtacaaaaaagcaggcttgatgGACGCCCTCATCGCCGCCCA, ggggaccactttgtacaagaaagctgggtcAATATACACCCTCTTCGCCT | Mycgr3\|99852  JGI  Ensembl | KOG2817 Predicted E3 ubiquitin ligase; |  | 100 % |  |  |
| 23 | ggggacaagtttgtacaaaaaagcaggcttgatgCCGCCGCGATACTTCAAGAA, ggggaccactttgtacaagaaagctgggtcAAGATCATTGTAGCCCAGAA | Mycgr3\|100022  JGI  Ensembl | KOG2787 Lanthionine synthetase C-like protein 1; |  | 100 % |  | ``` 145	ATGCCGCCGCGATACTTCAAGAACGACGCACCGCTCGCACGTCGAG	190 	\|\|\|\|\|\|\|\|\|\|\|\|\|\|\|\|\|\|\|\|\|\|\|\|\|\|\|\|\|\|\|\|\|\|\|\|\|\|\|\|\|\|\|\|\|\| 	ATGCCGCCGCGATACTTCAAGAACGACGCACCGCTCGCACGTCGAG ```      ``` 189	AGACCCCCGCCGCCAGCTCGAAGCCTCCCTCACCCGCCTCGTCCTCGAACACCCACCCACCTCGATCAAACCCGGCGGCGGGCTCTTCAAAGGCCCCATCTCCGTCGCCTACCTCTTCCTCGTCCTCCAGCAAATCTACTCCGACCTCCTCGTCGAAGACACACCCCTCGGTACCTGGTCCGCCGCCTACCTCCACCACGCCCAAGAACACATCGCCAGCTACGCCGGTCCTTCACCGGGAAAATGCGGCATCACCGACGACATTCTCGCGCTGCTGGCGATTGGCGCGGCGAGTAGTAAAGATGCGGACATGGCGGCGAATTTGTGTGATTATTCTGCTGAGGTGATTGATCCCGAGACGGAGAATGAGCTCATGTTTGGACGGGCGGGATACCTTTACCTCCTGCGGTTGGTGAAGGCAAGTTTTATGGATGATGCCAAGACGTTGCAGCTCATCACGGATACGCAGGAGGATGTCATTGATGCGATTCTCGACTCCCCACGGCCGTGGAAGTGGCAGGGGAAGACGTATATCGGAGCGATTCACGGCGCGATTGGGATTCTGACGCAAGTGGTGCTGACGAATCCGGGAAAGTACGCGGAGAAGTTGCAGGCGGAGTTGGCGGTGTTGTTGACGTATCAGTATGAAGGAGGGAATTTCCCGGCGAGTGTGCCGCCGGAGAAGGATCGTTTGGTGCAGGTTTGTCACGGGGCGCCCGGGGTGGTGTGTAGTTTGATTAGTGTGGTGGAGTTTTATGACGGGCCGTTGAGGGAGAAGATTGAGAAGGCGATTGTGAAGGGTAGGGAGTGTGTTTTGGAGAGGGGACTATTGACTAAGGAGCCGTGTGTTTGTCATG	1045 	\|\|\|\|\|\|\|\|\|\|\|\|\|\|\|\|\|\|\|\|\|\|\|\|\|\|\|\|\|\|\|\|\|\|\|\|\|\|\|\|\|\|\|\|\|\|\|\|\|\|\|\|\|\|\|\|\|\|\|\|\|\|\|\|\|\|\|\|\|\|\|\|\|\|\|\|\|\|\|\|\|\|\|\|\|\|\|\|\|\|\|\|\|\|\|\|\|\|\|\|\|\|\|\|\|\|\|\|\|\|\|\|\|\|\|\|\|\|\|\|\|\|\|\|\|\|\|\|\|\|\|\|\|\|\|\|\|\|\|\|\|\|\|\|\|\|\|\|\|\|\|\|\|\|\|\|\|\|\|\|\|\|\|\|\|\|\|\|\|\|\|\|\|\|\|\|\|\|\|\|\|\|\|\|\|\|\|\|\|\|\|\|\|\|\|\|\|\|\|\|\|\|\|\|\|\|\|\|\|\|\|\|\|\|\|\|\|\|\|\|\|\|\|\|\|\|\|\|\|\|\|\|\|\|\|\|\|\|\|\|\|\|\|\|\|\|\|\|\|\|\|\|\|\|\|\|\|\|\|\|\|\|\|\|\|\|\|\|\|\|\|\|\|\|\|\|\|\|\|\|\|\|\|\|\|\|\|\|\|\|\|\|\|\|\|\|\|\|\|\|\|\|\|\|\|\|\|\|\|\|\|\|\|\|\|\|\|\|\|\|\|\|\|\|\|\|\|\|\|\|\|\|\|\|\|\|\|\|\|\|\|\|\|\|\|\|\|\|\|\|\|\|\|\|\|\|\|\|\|\|\|\|\|\|\|\|\|\|\|\|\|\|\|\|\|\|\|\|\|\|\|\|\|\|\|\|\|\|\|\|\|\|\|\|\|\|\|\|\|\|\|\|\|\|\|\|\|\|\|\|\|\|\|\|\|\|\|\|\|\|\|\|\|\|\|\|\|\|\|\|\|\|\|\|\|\|\|\|\|\|\|\|\|\|\|\|\|\|\|\|\|\|\|\|\|\|\|\|\|\|\|\|\|\|\|\|\|\|\|\|\|\|\|\|\|\|\|\|\|\|\|\|\|\|\|\|\|\|\|\|\|\|\|\|\|\|\|\|\|\|\|\|\|\|\|\|\|\|\|\|\|\|\|\|\|\|\|\|\|\|\|\|\|\|\|\|\|\|\|\|\|\|\|\|\|\|\|\|\|\|\|\|\|\|\|\|\|\|\|\|\|\|\|\|\|\|\|\|\|\|\|\|\|\|\|\|\|\|\|\|\|\|\|\|\|\|\|\|\|\|\|\|\|\|\|\|\|\|\|\|\|\|\|\|\|\|\|\|\|\|\|\|\|\|\|\|\|\|\|\|\|\|\|\|\|\|\|\|\|\|\|\|\|\|\|\|\|\|\|\|\|\|\|\|\|\|\|\|\|\|\|\|\|\|\|\|\|\|\|\|\|\|\|\|\|\|\|\|\|\|\|\|\|\|\|\|\|\|\|\|\|\|\|\|\|\|\|\|\|\|\|\|\|\|\|\|\|\|\|\|\|\|\|\|\|\|\|\|\|\|\|\|\|\|\|\|\|\|\|\|\|\|\|\|\|\|\|\|\|\|\|\|\|\|\|\|\|\|\|\|\|\|\|\|\|\|\|\|\|\|\|\|\|\|\|\|\|\|\|\|\|\|\|\|\|\|\|\|\|\|\|\|\|\|\|\|\|\|\|\|\|\|\|\|\|\|\|\|\|\|\|\|\|\|\|\|\|\|\|\|\|\|\|\|\|\|\|\|\|\|\|\|\|\|\|\|\|\|\|\|\|\|\|\|\|\|\|\|\|\|\|\|\|\|\|\|\|\|\|\|\|\|\|\|\|\|\|\| \|\|\| \|\|\|\|\|\|\| \|\|\|\|\|\| 	AGACCCCCGCCGCCAGCTCGAAGCCTCCCTCACCCGCCTCGTCCTCGAACACCCACCCACCTCGATCAAACCCGGCGGCGGGCTCTTCAAAGGCCCCATCTCCGTCGCCTACCTCTTCCTCGTCCTCCAGCAAATCTACTCCGACCTCCTCGTCGAAGACACACCCCTCGGTACCTGGTCCGCCGCCTACCTCCACCACGCCCAAGAACACATCGCCAGCTACGCCGGTCCTTCACCGGGAAAATGCGGCATCACCGACGACATTCTCGCGCTGCTGGCGATTGGCGCGGCGAGTAGTAAAGATGCGGACATGGCGGCGAATTTGTGTGATTATTCTGCTGAGGTGATTGATCCCGAGACGGAGAATGAGCTCATGTTTGGACGGGCGGGATACCTTTACCTCCTGCGGTTGGTGAAGGCAAGTTTTATGGATGATGCCAAGACGTTGCAGCTCATCACGGATACGCAGGAGGATGTCATTGATGCGATTCTCGACTCCCCACGGCCGTGGAAGTGGCAGGGGAAGACGTATATCGGAGCGATTCACGGCGCGATTGGGATTCTGACGCAAGTGGTGCTGACGAATCCGGGAAAGTACGCGGAGAAGTTGCAGGCGGAGTTGGCGGTGTTGTTGACGTATCAGTATGAAGGAGGGAATTTCCCGGCGAGTGTGCCGCCGGAGAAGGATCGTTTGGTGCAGGTTTGTCACGGGGCGCCCGGGGTGGTGTGTAGTTTGATTAGTGTGGTGGAGTTTTATGACGGGCCGTTGAGGGAGAAGATTGAGAAGGCGATTGTGAAGGGTAGGGAGTGTGTTTTGGAGAGGGGACTATTGACTAAG-AGC-GTGTGTT-GTCATG ``` |
| 23 | ggggacaagtttgtacaaaaaagcaggcttgatgGCAGCTCCAGGAGGCAAATT, ggggaccactttgtacaagaaagctgggtcAAGGTACTTCGGCCTCGACG | Mycgr3\|100584  JGI  Ensembl |  |  | 100 % |  |  |
| 23 | ggggacaagtttgtacaaaaaagcaggcttgatgTCGGAAGCGATTCAAGACCC, ggggaccactttgtacaagaaagctgggtcTCCAATAAACGTGCAGGTCG | Mycgr3\|100619  JGI  Ensembl | KOG1709 Guanidinoacetate methyltransferase and related proteins; | guanidinoacetate N-methyltransferase activity; | 100 % |  |  |
| 23 | ggggacaagtttgtacaaaaaagcaggcttgatgCGCTTCAGGGCGAGCATTCA, ggggaccactttgtacaagaaagctgggtcCTGAGCGTAGGAAGAGATGT | Mycgr3\|101485  JGI  Ensembl | KOG3999 Checkpoint 9-1-1 complex, HUS1 component; |  | 100 % |  |  |
| 23 | ggggacaagtttgtacaaaaaagcaggcttgatgCCTGATCACGAAAGTGAACT, ggggaccactttgtacaagaaagctgggtcTGTTTTACAAGTTGGAGCAG | Mycgr3\|101931  JGI  Ensembl |  |  | 100 % |  |  |
| 23 | ggggacaagtttgtacaaaaaagcaggcttgatgGCCCACCCAACTCCATTCGA, ggggaccactttgtacaagaaagctgggtcCTCTGCGTGGAACTTTACAG | Mycgr3\|102520  JGI  Ensembl |  |  | 100 % |  |  |
| 23 | ggggacaagtttgtacaaaaaagcaggcttgatgTTCGGCTTCATAGCCGACAT, ggggaccactttgtacaagaaagctgggtcATCATCCTTCTTCGTAGGAG | Mycgr3\|103256  JGI  Ensembl | KOG1726 HVA22/DP1 gene product-related proteins; |  | 100 % |  | ``` 113	ATGTTCGGCTTCATAGCCGACATCCTCACCTCCCTCATCTCCATCCTCCTCCCCGTCTTCTTCTCCTACAAAGCGCTCCGCACCTCCGATCCCGCCGTCCTCACGCCATGGTTGATGTATTGGACGACGTTCTCACTCTTCCTCCTCGTCGAGAACCAATTTTACTTCATCCTATACTGGATCCCCTTCTACAGCTGGATGCGTCTCTTCATCCACATCTACCTCGTTCTACCGGGATCTCAAGGTAGCGTCTACCTTTACAAGCAATATCTGGGACCCTTTCTCGAAGACCACGAGCGTCAACTCGACCGTCTGATCAGTAATGTACACGCCAAGGGTCAGGCGGCAGGAATGGACGCTTTCAGACGAGCATGGGACTATATCCGAGTGCAATTCATGGGAGGAGAACCACGTTGTCCGACACCACCGCAGAGTCGCAATGTCAGCTATACGACCTCTCTCTTCAACCGCTTCGCCATGCCCAGCGCACGAGACGGACTCGCAGCAGCAGGTACATCGGACATTTTCAACCTCATCGGCAATGCTCTCAATCAGCATCCATCATCATCTTCCCGCACTGAACAAGCTCGGGATCTCGCTAGCTCAGGCACACTCGTCCCACCAGACCTCGCACCGGCCGAACGCGCGACTTTCGTGGACACCCAACGCGAGCGTCTTCGTACCCTCTTGCAAGCCTACGAGTCCGAAGCTCTGGGTAGCCACGTCGACACTCCACCAGGTGGGCTCTCACCTCGAGTCGGAACTCCTCGCCAGCCATCAAGTCGCAAACCTGTGCCCTCTTCGCAAGACGCTCTATCGCGGTCTCGATCAGAGTCGGAGTTTGAGGACCTAGGCTACGAACCTATGCCTGACCCGGAGCACTTTCTTCCACGAGAGGACGTGGCCTCAACGCCAAAGGAGGGAGGTGCAGGGTGGAGTAATTGGATTTGGGGCAATTATGGGGAGAGGGACTCTGCTGCTCCTACGA-AGAAGGATGA	1110 	\|\|\|\|\|\|\|\|\|\|\|\|\|\|\|\|\|\|\|\|\|\|\|\|\|\|\|\|\|\|\|\|\|\|\|\|\|\|\|\|\|\|\|\|\|\|\|\|\|\|\|\|\|\|\|\|\|\|\|\|\|\|\|\|\|\|\|\|\|\|\|\|\|\|\|\|\|\|\|\|\|\|\|\|\|\|\|\|\|\|\|\|\|\|\|\|\|\|\|\|\|\|\|\|\|\|\|\|\|\|\|\|\|\|\|\|\|\|\|\|\|\|\|\|\|\|\|\|\|\|\|\|\|\|\|\|\|\|\|\|\|\|\|\|\|\|\|\|\|\|\|\|\|\|\|\|\|\|\|\|\|\|\|\|\|\|\|\|\|\|\|\|\|\|\|\|\|\|\|\|\|\|\|\|\|\|\|\|\|\|\|\|\|\|\|\|\|\|\|\|\|\|\|\|\|\|\|\|\|\|\|\|\|\|\|\|\|\|\|\|\|\|\|\|\|\|\|\|\|\|\|\|\|\|\|\|\|\|\|\|\|\|\|\|\|\|\|\|\|\|\|\|\|\|\|\|\|\|\|\|\|\|\|\|\|\|\|\|\|\|\|\|\|\|\|\|\|\|\|\|\|\|\|\|\|\|\|\|\|\|\|\|\|\|\|\|\|\|\|\|\|\|\|\|\|\|\|\|\|\|\|\|\|\|\|\|\|\|\|\|\|\|\|\|\|\|\|\|\|\|\|\|\|\|\|\|\|\|\|\|\|\|\|\|\|\|\|\|\|\|\|\|\|\|\|\|\|\|\|\|\|\|\|\|\|\|\|\|\|\|\|\|\|\|\|\|\|\|\|\|\|\|\|\|\|\|\|\|\|\|\|\|\|\|\|\|\|\|\|\|\|\|\|\|\|\|\|\|\|\|\|\|\|\|\|\|\|\|\|\|\|\|\|\|\|\|\|\|\|\|\|\|\|\|\|\|\|\|\|\|\|\|\|\|\|\|\|\|\|\|\|\|\|\|\|\|\|\|\|\|\|\|\|\|\|\|\|\|\|\|\|\|\|\|\|\|\|\|\|\|\|\|\|\|\|\|\|\|\|\|\|\|\|\|\|\|\|\|\|\|\|\|\|\|\|\|\|\|\|\|\|\|\|\|\|\|\|\|\|\|\|\|\|\|\|\|\|\|\|\|\|\|\|\|\|\|\|\|\|\|\|\|\|\|\|\|\|\|\|\|\|\|\|\|\|\|\|\|\|\|\|\|\|\|\|\|\|\|\|\|\|\|\|\|\|\|\|\|\|\|\|\|\|\|\|\|\|\|\|\|\|\|\|\|\|\|\|\|\|\|\|\|\|\|\|\|\|\|\|\|\|\|\|\|\|\|\|\|\|\|\|\|\|\|\|\|\|\|\|\|\|\|\|\|\|\|\|\|\|\|\|\|\|\|\|\|\|\|\|\|\|\|\|\|\|\|\|\|\|\|\|\|\|\|\|\|\|\|\|\|\|\|\|\|\|\|\|\|\|\|\|\|\|\|\|\|\|\|\|\|\|\|\|\|\|\|\|\|\|\|\|\|\|\|\|\|\|\|\|\|\|\|\|\|\|\|\|\|\|\|\|\|\|\|\|\|\|\|\|\|\|\|\|\|\|\|\|\|\|\|\|\|\|\|\|\|\|\|\|\|\|\|\|\|\|\|\|\|\|\|\|\|\|\|\|\|\|\|\|\|\|\|\|\|\|\|\|\|\|\|\|\|\|\|\|\| \|\|\|\|\|\|\|\|\|\|\|\|\|\|\|\|\|\| \|\|\|\|\|\|\|\|\|\|\|\|\|\|\|\|\|\|\|\|\|\|\|\|\|\|\|\|\|\|\|\|\|\|\|\|\|\|\|\| \|\|\|\|\| \|\|\|\|\|\|\|\|\|\|\|\|\|\| \|\|\|\|\|\|\|\|\|\|\|\|\|\|\|\|\|\|\|\| \|\|\|\|\|\|\|\|\|\|\|\|\|\|\|\| \|\|\| \|\|\| \|\|\|\| \|\|  \|\| \|\|\|\|\|  \|\| \|\|\|\| \| \| \|\|\| \|\|\| \| \|\|\|\|\|\|\|\|\|\|\|\| \|\|\|\|\|\|\|\|\|\|\|\|\|\| \|\|\|\|\| \|\|\| \| \|\|\|\| 	ATGTTCGGCTTCATAGCCGACATCCTCACCTCCCTCATCTCCATCCTCCTCCCCGTCTTCTTCTCCTACAAAGCGCTCCGCACCTCCGATCCCGCCGTCCTCACGCCATGGTTGATGTATTGGACGACGTTCTCACTCTTCCTCCTCGTCGAGAACCAATTTTACTTCATCCTATACTGGATCCCCTTCTACAGCTGGATGCGTCTCTTCATCCACATCTACCTCGTTCTACCGGGATCTCAAGGTAGCGTCTACCTTTACAAGCAATATCTGGGACCCTTTCTCGAAGACCACGAGCGTCAACTCGACCGTCTGATCAGTAATGTACACGCCAAGGGTCAGGCGGCAGGAATGGACGCTTTCAGACGAGCATGGGACTATATCCGAGTGCAATTCATGGGAGGAGAACCACGTTGTCCGACACCACCGCAGAGTCGCAATGTCAGCTATACGACCTCTCTCTTCAACCGCTTCGCCATGCCCAGCGCACGAGACGGACTCGCAGCAGCAGGTACATCGGACATTTTCAACCTCATCGGCAATGCTCTCAATCAGCATCCATCATCATCTTCCCGCACTGAACAAGCTCGGGATCTCGCTAGCTCAGGCACACTCGTCCCACCAGACCTCGCACCGGCCGAACGCGCGACTTTCGTGGACACCCAACGCGAGCGTCTTCGTACCCTCTTGCAAGCCTACGAGTCCGAAGCTCTGGGTAGCCACGTCGACACTCCACCAGGTGGGCTCTCACCTCGAGTCGGAACTCCTCGCCAGCCATCAAGTCGC-AACCTGTGCCCTCTTCGC-AGACGCTCTATCGCGGTCTCGATCAGAGTCGGAGTTTGAG-ACCTA-GCTACGAACCTATG-CTGACCCGGAGCACTTTCTT-CACGAGAGGACGTGGC-TCA-CGC-AAAG-AG--AG-TGCAG--TG-AGTA-T-G-ATT-GGG-C-ATTATGGGGAGA-GGACTCTGCTGCTC-TACGAGAGATG-ATGA ``` |
| 23 | ggggacaagtttgtacaaaaaagcaggcttgatgGTTACCGCCGCTCCACAAGA, ggggaccactttgtacaagaaagctgggtcAATCTGCCTCTCGAGAACGA | Mycgr3\|105747  JGI  Ensembl | KOG1541 Predicted protein carboxyl methylase; |  | 100 % |  |  |
| 23 | ggggacaagtttgtacaaaaaagcaggcttgatgAAGGTCAACATCGCGTTCGT, ggggaccactttgtacaagaaagctgggtcCCAGCGGTCGTATTCTCCCA | Mycgr3\|107601  JGI  Ensembl |  |  | 100 % |  |  |
| 23 | ggggacaagtttgtacaaaaaagcaggcttgatgCCAGGCGCCGGCAAGCAACC, ggggaccactttgtacaagaaagctgggtcTGCCATCGCAAAGACCAGCA | Mycgr3\|107867  JGI  Ensembl | KOG1721 FOG: Zn-finger; |  | 100 % | chr\_2:586903 A->T; | ``` 55	ATGCCAGGCGCCGGCAAGCAACCGCCTCAGATCCAGGATCAAATCAAGGAATTGCTCGAGGCCGGCTTCGATCCCACTACGATCCATCGTCGTCTCAAAGTCGGCCGCAGTTCGATTTATCGCATGAAGCGATGCCTCGAACAGCATGGCACCAACTATATGCCGAAGGAGATGAACAAGAAGAATGGACGGCCGAAAGTGTTGACCCCGGAGCAGGAACTTG-AAGT	281 	\|\|\|\|\|\|\|\|\|\|\|\|\|\|\|\|\|\|\|\|\|\|\|\|\|\|\|\|\|\|\|\|\|\|\|\|\|\|\|\|\|\|\|\|\|\|\|\|\|\|\|\|\|\|\|\|\|\|\|\|\|\|\|\|\|\|\|\|\|\|\|\|\|\|\|\|\|\|\|\|\|\|\|\|\|\| \|\|\|\|\|\|\|\|\|\|\|\|\|\|\|\|\|\|\|\|\|\|\|\|\|\|\|\|\|\|\|\|\|\|\|\|\|\|\|\|\|\|\|\|\|\|\|\|\|\|\|\|\|\|\|\|\|\|\|\|\|\|\|\|\|\|\|\|\|\|\|\|\|\|\|\|\|\|\|\|\|\|\|\|\|\|\|\|\|\|\|\|\|\|\|\|\|\|\|\|\|\|\|\|\|\|\|\|\|\|\|\|\|\|\|\|\|\|\|\|\|\|\|\|\|\|\|\|\|\|\|\|\|\|\|\| \|\|\|\| 	ATGCCAGGCGCCGGCAAGCAACCGCCTCAGATCCAGGATCAAATCAAGGAATTGCTCGAGGCCGGCTTCGATCCCACTACGATCCAACGTCGTCTCAAAGTCGGCCGCAGTTCGATTTATCGCATGAAGCGATGCCTCGAACAGCATGGCACCAACTATATGCCGAAGGAGATGAACAAGAAGAATGGACGGCCGAAAGTGTTGACCCCGGAGCAGGAACTTGTAAGT ```      ``` 277	GAAGTCAGTGCCTGGCTCCGCGACCCCAAAAACCGGACCCGCTTCCTCGATGACCTAGTCTGGCTGATACACGACCGTTTCGGCATCGTCTGCAGTACAACGACCATGTCCAAAATGAAGCGAAAATGGCTCCGAGTGATCGACTACGAAGAGAACGGCACGCCCATCGACGACACCACTCGAGCACAATTGGAGGAGACGCATCCAGACCTCCCACTTCTCCACCAGCCGCACGCACCTGTGGCAGAGATGGGGAGCATGGCGTCCGGACTGGAATCCGATGGCATGGGCATGCAGCAACCCCTGCCGCCACAGATGGAGCAGCAACATCCCCATCCGCAACTGGAGCAACAACACCAAGACCCTCTGATCGAACAGCACCAGCTCGACGATCACTCTCACCTCGACCATACCCAGCACGCGGACTTTAGTCAACCCCAGCAGCAACATCTGCACGCAGACCTCTCGCAGCAACAACATCTCCACCCACACCTGGACCAGCACTCTCAACTGGACCAGCAGCAGCATCAACAACATCCCTACCCCGACCACCACCTCCAACCTCAACAACAACACGACGACATCTTATCCCTCCCTCCCCATCAACAACAGCACCACACTCAACATCCACTGGAAGATCCGTTGAACGTCTACCACCCACCTCCGGCTCATGATCCACAACTCGATGCGCAGCTAGAGCAGCAAATACGACATTCCGCGGAG-ATGGGCGTCATGGGTGGACATCACGGGCCGCATCTGG	1036 	\|\|\|\|\|\|\|\|\|\|\|\|\|\|\|\|\|\|\|\|\|\|\|\|\|\|\|\|\|\|\|\|\|\|\|\|\|\|\|\|\|\|\|\|\|\|\|\|\|\|\|\|\|\|\|\|\|\|\|\|\|\|\|\|\|\|\|\|\|\|\|\|\|\|\|\|\|\|\|\|\|\|\|\|\|\|\|\|\|\|\|\|\|\|\|\|\|\|\|\|\|\|\|\|\|\|\|\|\|\|\|\|\|\|\|\|\|\|\|\|\|\|\|\|\|\|\|\|\|\|\|\|\|\|\|\|\|\|\|\|\|\|\|\|\|\|\|\|\|\|\|\|\|\|\|\|\|\|\|\|\|\|\|\|\|\|\|\|\|\|\|\|\|\|\|\|\|\|\|\|\|\|\|\|\|\|\|\|\|\|\|\|\|\|\|\|\|\|\|\|\|\|\|\|\|\|\|\|\|\|\|\|\|\|\|\|\|\|\|\|\|\|\|\|\|\|\|\|\|\|\|\|\|\|\|\|\|\|\|\|\|\|\|\|\|\|\|\|\|\|\|\|\|\|\|\|\|\|\|\|\|\|\|\|\|\|\|\|\|\|\|\|\|\|\|\|\|\|\|\|\|\|\|\|\|\|\|\|\|\|\|\|\|\|\|\|\|\|\|\|\|\|\|\|\|\|\|\|\|\|\|\|\|\|\|\|\|\|\|\|\|\|\|\|\|\|\|\|\|\|\|\|\|\|\|\|\|\|\|\|\|\|\|\|\|\|\|\|\|\|\|\|\|\|\|\|\|\|\|\|\|\|\|\|\|\|\|\|\|\|\|\|\|\|\|\|\|\|\|\|\|\|\|\|\|\|\|\|\|\|\|\|\|\|\|\|\|\|\|\|\|\|\|\|\|\|\|\|\|\|\|\|\|\|\|\|\|\|\|\|\|\|\|\|\|\|\|\|\|\|\|\|\|\|\|\|\|\|\|\|\|\|\|\|\|\|\|\|\|\|\|\|\|\|\|\|\|\|\|\|\|\|\|\|\|\|\|\|\|\|\|\|\|\|\|\|\|\|\|\|\|\|\|\|\|\|\|\|\|\|\|\|\|\|\|\|\|\|\|\|\|\|\|\|\|\|\|\|\|\|\|\|\|\|\|\|\|\|\|\|\|\|\|\|\|\|\|\|\|\|\|\|\|\|\|\|\|\|\|\|\|\|\|\|\|\|\|\|\|\|\|\|\|\|\|\|\|\|\|\|\|\|\|\|\|\|\|\|\|\|\|\|\|\|\|\|\|\|\|\|\|\|\|\|\|\|\|\|\|\|\|\|\|\|\|\|\|\|\|\|\|\|\|\|\|\|\|\|\|\|\|\|\|\|\|\|\|\|\|\|\|\|\| \|\|\| \|\|\|\|\|\|\|\|\|\|\|\|\|\|\|\|\|\|\|\|\|\|\|\|\|\|\|\|\|\|\|\|\|\|\| \|\|\|\|\|\|\|\|\|\|\|\|\|\|\| \|\|\|\|\|\|\|\|\|\|\|\|\|\|\|\|\|\|\|\|\|\|\|\|\| \|\|\|\|\|\|\|\| \| \|\|\| \|\| \|\|\|\|\|\|\|\|\|\|\|\|\|\|\|\|\|\|\|\|\|\|\|\|\|\| \|\|\|\|\|\|\|\|\|\| 	GAAGTCAGTGCCTGGCTCCGCGACCCCAAAAACCGGACCCGCTTCCTCGATGACCTAGTCTGGCTGATACACGACCGTTTCGGCATCGTCTGCAGTACAACGACCATGTCCAAAATGAAGCGAAAATGGCTCCGAGTGATCGACTACGAAGAGAACGGCACGCCCATCGACGACACCACTCGAGCACAATTGGAGGAGACGCATCCAGACCTCCCACTTCTCCACCAGCCGCACGCACCTGTGGCAGAGATGGGGAGCATGGCGTCCGGACTGGAATCCGATGGCATGGGCATGCAGCAACCCCTGCCGCCACAGATGGAGCAGCAACATCCCCATCCGCAACTGGAGCAACAACACCAAGACCCTCTGATCGAACAGCACCAGCTCGACGATCACTCTCACCTCGACCATACCCAGCACGCGGACTTTAGTCAACCCCAGCAGCAACATCTGCACGCAGACCTCTCGCAGCAACAACATCTCCACCCACACCTGGACCAGCACTCTCAACTGGACCAGCAGCAGCATCAACAACATCCCTACCCCGACCACCACCTCCAACCTCAACAACAACACGACGACATCTTATCCCTCCCTCCCCATCAACAACAGCACCACACTCA-CAT-CACTGGAAGATCCGTTGAACGTCTACCACCCACCT-CGGCTCATGATCCAC-ACTCGATGCGCAGCTAGAGCAGCAA-TACGACAT-C-GCG-AGAATGGGCGTCATGGGTGGACATCACGG-CCGCATCTGG ``` |
| 23 | ggggacaagtttgtacaaaaaagcaggcttgatgGCACCCATCACTCGTTCCTT, ggggaccactttgtacaagaaagctgggtcTCTACTCGCCGTTTCGTCAT | Mycgr3\|108147  JGI  Ensembl | KOG4157 beta-1,6-N-acetylglucosaminyltransferase, contains WSC domain; |  | 100 % |  |  |
| 23 | ggggacaagtttgtacaaaaaagcaggcttgatgACCGACAGCAAGCAACAAAC, ggggaccactttgtacaagaaagctgggtcTTGGCCCTTCTCGCTCTTCA | Mycgr3\|108814  JGI  Ensembl | KOG0134 NADH:flavin oxidoreductase/12-oxophytodienoate reductase; | FMN binding; oxidoreductase activity; | 100 % |  |  |
| 23 | ggggacaagtttgtacaaaaaagcaggcttgatgGCGGACGCGAAGTTAGGCAG, ggggaccactttgtacaagaaagctgggtcCCCGTCCACGGCCTTCGTGA | Mycgr3\|109580  JGI  Ensembl |  |  | 0 % |  |  |
| 23 | ggggacaagtttgtacaaaaaagcaggcttgatgTTCGCCGTCAGGGCGATACT, ggggaccactttgtacaagaaagctgggtcCAGAGCGCTCCCTTTTGCGC | Mycgr3\|109582  JGI  Ensembl |  |  | 100 % |  |  |
| 23 | ggggacaagtttgtacaaaaaagcaggcttgatgGCTTCGCTGGGATTTACCAG, ggggaccactttgtacaagaaagctgggtcGCCGGCACAGGTCGACAGAA | Mycgr3\|109708  JGI  Ensembl |  |  | 100 % |  |  |
| 23 | ggggacaagtttgtacaaaaaagcaggcttgatgTTCTCTCAACGCGTTTCAAC, ggggaccactttgtacaagaaagctgggtcACGATATAGCAATCTATGCA | Mycgr3\|111010  JGI  Ensembl | KOG2665 Predicted FAD-dependent oxidoreductase; | aminobutyraldehyde dehydrogenase activity; | 100 % |  |  |
| 23 | ggggacaagtttgtacaaaaaagcaggcttgatgGCGACTGATCTGGACTTAGA, ggggaccactttgtacaagaaagctgggtcATCGTCAATTGTGATTGACG | Mycgr3\|111263  JGI  Ensembl |  |  | 1.96657 % |  |  |
| 23 | ggggacaagtttgtacaaaaaagcaggcttgatgGACTGCTTTCAAGACTTCTG, ggggaccactttgtacaagaaagctgggtcGGTGCTCCTAGTGCTGGACC | Mycgr3\|111502  JGI  Ensembl |  |  | 100 % |  | ``` 433	ATGGACTGCTTTCAAGACTTCTGCCTCTTCTGTGACCGGGACTCACCTGACGGTCCCTACTGCTCACAACGATGCAAACTCGCCGACCTTGAGAAGGCGAGCTCGTCCAACCCTGGATCACCCACATCATCTCGACACGACCACCGCGACACTCGATACGTACTGTCGCCAGCATACAAGTTCCACAGCGAGcaacaacaacaacaacaCCAACGAACACCAGGCTACGACGCTCCAGCGGCGAACAACTCTCGACGACCTCGATCATCCTACTTCATGTGGACAGCATCGCCCACAACCCAGGACCGATCACTCATCGAAGACCGCTCGCTATCACCCTCATCATCACGCACATCACTCTCATCAACGTCAAGCGGCTCCAGCACGACAGACGCCTTCTCCGCGCAAGCTAAGCAGCAACTTCAGGAGTATTTTAGCTCCTTCGACCAAGCTCGAGCTGCAAAGCGGAGGTCCAGCACTAGGAGCACCTGAC	925 	\|\|\|\|\|\|\|\|\|\|\|\|\|\|\|\|\|\|\|\|\|\|\|\|\|\|\|\|\|\|\|\|\|\|\|\|\|\|\|\|\|\|\|\|\|\|\|\|\|\|\|\|\|\|\|\|\|\|\|\|\|\|\|\|\|\|\|\|\|\|\|\|\|\|\|\|\|\|\|\|\|\|\|\|\|\|\|\|\|\|\|\|\|\|\|\|\|\|\|\|\|\|\|\|\|\|\|\|\|\|\|\|\|\|\|\|\|\|\|\|\|\|\|\|\|\|\|\|\|\|\|\|\|\|\|\|\|\|\|\|\|\|\|\|\|\|\|\|\|\|\|\|\|\|\|\|\|\|\|\|\|\|\|\|\|\|\|\|\|\|\|\|\|\|\|\|\|\|\|\|\|\|\|\|\|\|\|\|\|\|\|\|\|\|\|\|\|\|\|\|\|\|\|\|\|\|\|\|\|\|\|\|\|\|\|\|\|\|\|\|\|\|\|\|\|\|\|\|\|\|\|\|\|\|\|\|\|\|\|\|\|\|\|\|\|\|\|\|\|\|\|\|\|\|\|\|\|\|\|\|\|\|\|\|\|\|\|\|\|\|\|\|\|\|\|\|\|\|\|\|\|\|\|\|\|\|\|\|\|\|\|\|\|\|\|\|\|\|\|\|\|\|\|\|\|\|\|\|\|\|\|\|\|\|\|\|\|\|\|\|\|\|\|\|\|\|\|\|\|\|\|\|\|\|\|\|\|\|\|\|\|\|\|\|\|\|\|\|\|\|\|\|\|\|\|\|\|\|\|\|\|\|\|\|\|\|\|\|\|\|\|\|\|\|\|\|\|\|\|\|\|\|\|\|\|\|\|\|\|\|\|\|\|\|\|\|\|\|\|\|\|\|\|\|\|\|\|\|\|\|\|\|\|\|\|\|\|\|\|\|\|\|\|\|\|\|\|\|\|\|\|\|\|\|\|\|\|\|\|\|\|\|\|\|\|\|\|\|\|\|\|\|\|\|\|\|\|\|\|\|\|\|\|\|\|\|\|\|\|\|\|\|\|\|\|\|\|\|\|\|\|\|\|\|\|\|\|\|\| \|\|\| 	ATGGACTGCTTTCAAGACTTCTGCCTCTTCTGTGACCGGGACTCACCTGACGGTCCCTACTGCTCACAACGATGCAAACTCGCCGACCTTGAGAAGGCGAGCTCGTCCAACCCTGGATCACCCACATCATCTCGACACGACCACCGCGACACTCGATACGTACTGTCGCCAGCATACAAGTTCCACAGCGAGCAACAACAACAACAACACCAACGAACACCAGGCTACGACGCTCCAGCGGCGAACAACTCTCGACGACCTCGATCATCCTACTTCATGTGGACAGCATCGCCCACAACCCAGGACCGATCACTCATCGAAGACCGCTCGCTATCACCCTCATCATCACGCACATCACTCTCATCAACGTCAAGCGGCTCCAGCACGACAGACGCCTTCTCCGCGCAAGCTAAGCAGCAACTTCAGGAGTATTTTAGCTCCTTCGACCAAGCTCGAGCTGCAAAGCGGAGGTCCAGCACTAGGAGCACC-GAC ``` |
| 23 | ggggacaagtttgtacaaaaaagcaggcttgatgTCGAGCTCCTTCGGATTTCC, ggggaccactttgtacaagaaagctgggtcGAAACCGATGATCTCATCCG | Mycgr3\|111731  JGI  Ensembl |  |  | 100 % |  |  |
| 23 | ggggacaagtttgtacaaaaaagcaggcttgatgAAACGCTGCCGTCCAGCGGA, ggggaccactttgtacaagaaagctgggtcGACATCGGGGAACAGCTCGC | Mycgr3\|111776  JGI  Ensembl |  |  | 100 % |  |  |
| 24 | ggggacaagtttgtacaaaaaagcaggcttgatgAGTGCCCTACGATCCTCGGT, ggggaccactttgtacaagaaagctgggtcTCGCATCTCCAGCATCATAG | Mycgr3\|31803  JGI  Ensembl | KOG0558 Dihydrolipoamide transacylase (alpha-keto acid dehydrogenase E2 subunit); | 3-hydroxybutyryl-CoA thiolase activity; 3-ketopimelyl-CoA thiolase activity;  C-acyltransferase activity;  C-palmitoyltransferase activity;  H2A/H2B histone acetyltransferase activity;  H3/H4 histone acetyltransferase activity;  L-2-aminoadipate N-acetyltransferase activity;  N-acetyltransferase activity;  N-acyltransferase activity;  N-palmitoyltransferase activity;  N-succinyltransferase activity;  O-acetyltransferase activity;  O-acyltransferase activity;  O-octanoyltransferase activity;  O-palmitoyltransferase activity;  O-sinapoyltransferase activity;  O-succinyltransferase activity;  Ras palmitoyltransferase activity;  S-acetyltransferase activity;  S-acyltransferase activity;  S-malonyltransferase activity;  S-succinyltransferase activity;  UDP-3-O-[3-hydroxymyristoyl] glucosamine N-acyltransferase activity;  acetyltransferase activity;  acyl-CoA N-acyltransferase activity;  acylglycerol O-acyltransferase activity;  acyltransferase activity;  azetidine-2-carboxylic acid acetyltransferase activity;  benzoyl acetate-CoA thiolase activity;  carnitine O-acyltransferase activity;  dihydrolipoamide branched chain acyltransferase activity;  glucosaminyl-phosphotidylinositol O-acyltransferase activity;  keto acid formate lyase activity;  malonyltransferase activity;  metabolic process;  myristoyltransferase activity;  octanoyltransferase activity;  palmitoleoyl [acyl-carrier-protein]-dependent acyltransferase activity;  palmitoyltransferase activity;  peptidyl-lysine N6-myristoyltransferase activity;  peptidyl-lysine N6-palmitoyltransferase activity;  protein binding;  protein-cysteine S-acyltransferase activity;  protein-cysteine S-myristoyltransferase activity;  protein-cysteine S-palmitoleyltransferase activity;  serine O-acyltransferase activity;  sinapoyltransferase activity;  succinyltransferase activity; | 100 % |  | ``` 1	ATGAGTGCCCTACGATCCTCGGTCGCGAGAATCCTCCGCCTTCAACAACAACACGCTACACGTCCATCCGGTCTTTCTCGGCCACTCTTCTACCGACAATTTCACGCCTCTCAACGATCACACGTCGTGAAGCCTTTCCTGCTCGCAGACATTGGCGAGGGCATCACAGAATGCCAGCTGATACAATGGTTCGTCCAACCCGGAGCTCGCGTGGAGCAGTTCGACAAGCTGTGCGAGGTGCAGTCCGATAAAGCCAGCGTGGAGATTACCAGTCCATTCGATGGTGTAATCAAGAAGCTGCACTATGAGCCAGACGATATGGCGATCACGGGAAAAGCGTTGGTGGATATCGATATTCAAGGCGAGTTGAGCGAGGCGGATGAGGCGAAGCTCGGAGGAGGTGAAGGAGAGGGAAGCTCCAGCACAGATACGAAGGCGGAGACTTCAGCGGACGCAGAGGATGCAGTGAAACAGCAGGAGCAAGAAGCGGAAGGGTCACAGCAAGAGAGAGTACCAGAATCGAGGACAAATGGCAAACCTGAGCATTCATCACGCCCGAGCAAGGGCAACATGGGTGCTCTCGCAACGCCGGCGGTCAGGCATTTGACGAAGGAGTTCAAGGTCGACATCGCAAATATCGAAGGCACAGGGAAGGATGGCAGAGTACTGAAAGAGGATGTTCATAAGTTTGTCTCGGAGAGGGAGGAGAAGCAACAGCAGGCAACCACCATTGCGACGACAGCAAGCACAGAAGAGGACCGCAAAATCGCCCTCACGCCGGTTCAAAACCAAATGTTCAAGACCATGACCCGATCGCTCAACATCCCTCACTTCCTCTACACCTGCTCAGCAGACATGACGGCTGTCACGACAATGCGAAAGAGGATCAAGAAGACCTCCGACCAGAAAA-TCAC-ACACC-TC-GCATTCATC	930 	\|\|\|\|\|\|\|\|\|\|\|\|\|\|\|\|\|\|\|\|\|\|\|\|\|\|\|\|\|\|\|\|\|\|\|\|\|\|\|\|\|\|\|\|\|\|\|\|\|\|\|\|\|\|\|\|\|\|\|\|\|\|\|\|\|\|\|\|\|\|\|\|\|\|\|\|\|\|\|\|\|\|\|\|\|\|\|\|\|\|\|\|\|\|\|\|\|\|\|\|\|\|\|\|\|\|\|\|\|\|\|\|\|\|\|\|\|\|\|\|\|\|\|\|\|\|\|\|\|\|\|\|\|\|\|\|\|\|\|\|\|\|\|\|\|\|\|\|\|\|\|\|\|\|\|\|\|\|\|\|\|\|\|\|\|\|\|\|\|\|\|\|\|\|\|\|\|\|\|\|\|\|\|\|\|\|\|\|\|\|\|\|\|\|\|\|\|\|\|\|\|\|\|\|\|\|\|\|\|\|\|\|\|\|\|\|\|\|\|\|\|\|\|\|\|\|\|\|\|\|\|\|\|\|\|\|\|\|\|\|\|\|\|\|\|\|\|\|\|\|\|\|\|\|\|\|\|\|\|\|\|\|\|\|\|\|\|\|\|\|\|\|\|\|\|\|\|\|\|\|\|\|\|\|\|\|\|\|\|\|\|\|\|\|\|\|\|\|\|\|\|\|\|\|\|\|\|\|\|\|\|\|\|\|\|\|\|\|\|\|\|\|\|\|\|\|\|\|\|\|\|\|\|\|\|\|\|\|\|\|\|\|\|\|\|\|\|\|\|\|\|\|\|\|\|\|\|\|\|\|\|\|\|\|\|\|\|\|\|\|\|\|\|\|\|\|\|\|\|\|\|\|\|\|\|\|\|\|\|\|\|\|\|\|\|\|\|\|\|\|\|\|\|\|\|\|\|\|\|\|\|\|\|\|\|\|\|\|\|\|\|\|\|\|\|\|\|\|\|\|\|\|\|\|\|\|\|\|\|\|\|\|\|\|\|\|\|\|\|\|\|\|\|\|\|\|\|\|\|\|\|\|\|\|\|\|\|\|\|\|\|\|\|\|\|\|\|\|\|\|\|\|\|\|\|\|\|\|\|\|\|\|\|\|\|\|\|\|\|\|\|\|\|\|\|\|\|\|\|\|\|\|\|\|\|\|\|\|\|\|\|\|\|\|\|\|\|\|\|\|\|\|\|\|\|\|\|\|\|\|\|\|\|\|\|\|\|\|\|\|\|\|\|\|\|\|\|\|\|\|\|\|\|\|\|\|\|\|\|\|\|\|\|\|\|\|\|\|\|\|\|\|\|\|\|\|\|\|\|\|\|\|\|\|\|\|\|\|\|\|\|\|\|\|\|\|\|\|\|\|\|\|\|\|\|\|\|\|\|\|\|\|\|\|\|\|\|\|\|\|\|\|\|\|\|\|\|\|\|\|\|\|\|\|\|\|\|\|\|\|\|\|\|\|\|\|\|\|\|\|\|\|\|\|\|\|\|\|\|\|\|\|\|\|\|\|\|\|\|\|\|\|\|\|\|\|\|\|\|\|\|\|\|\|\|\|\|\|\|\|\|\|\|\|\|\|\|\|\|\|\|\|\|\|\|\|\|\|\|\|\|\|\|\|\|\|\|\|\|\|\|\|\|\|\|\|\|\|\|\|\|\|\|\|\|\|\|\|\|\|\|\|\|\|\|\|\|\|\|\|\|\|\|\|\|\|\|\|\|\|\|\|\|\|\|\|\|\|\|\|\|\|\|\|\|\|\|\|\|\|\|\|\|\|\|\|\|\|\|\|\|\|\|\|\|\|\|\|\|\|\|\|\|\|\|\|\|\|\|\|\|\|\|\|\|\|\|\|\|\|\|\|\|\|\|\|\|\|\|\|\|\|\|\|\|\|\|\|\|\|\|\|\|\|\|\|\|\|\|\|\|\|\|\|\|\|\|\|\|\|\|\| \|\|\|\|\|\|\|\|\|\|\|\|\|\|\|\|\|\|\|\|\|\|\|\|\|\|\|\|\|\|\|\|\|\|\|\|\| \|\|\|\| \|\|\|\|\| \|\| \|\|\|\|\|\|\|\|\| 	ATGAGTGCCCTACGATCCTCGGTCGCGAGAATCCTCCGCCTTCAACAACAACACGCTACACGTCCATCCGGTCTTTCTCGGCCACTCTTCTACCGACAATTTCACGCCTCTCAACGATCACACGTCGTGAAGCCTTTCCTGCTCGCAGACATTGGCGAGGGCATCACAGAATGCCAGCTGATACAATGGTTCGTCCAACCCGGAGCTCGCGTGGAGCAGTTCGACAAGCTGTGCGAGGTGCAGTCCGATAAAGCCAGCGTGGAGATTACCAGTCCATTCGATGGTGTAATCAAGAAGCTGCACTATGAGCCAGACGATATGGCGATCACGGGAAAAGCGTTGGTGGATATCGATATTCAAGGCGAGTTGAGCGAGGCGGATGAGGCGAAGCTCGGAGGAGGTGAAGGAGAGGGAAGCTCCAGCACAGATACGAAGGCGGAGACTTCAGCGGACGCAGAGGATGCAGTGAAACAGCAGGAGCAAGAAGCGGAAGGGTCACAGCAAGAGAGAGTACCAGAATCGAGGACAAATGGCAAACCTGAGCATTCATCACGCCCGAGCAAGGGCAACATGGGTGCTCTCGCAACGCCGGCGGTCAGGCATTTGACGAAGGAGTTCAAGGTCGACATCGCAAATATCGAAGGCACAGGGAAGGATGGCAGAGTACTGAAAGAGGATGTTCATAAGTTTGTCTCGGAGAGGGAGGAGAAGCAACAGCAGGCAACCACCATTGCGACGACAGCAAGCACAGAAGAGGACCGCAAAATCGCCCTCACGCCGGTTCAAAACCAAATGTTCAAGACCATGACCCGATCGCTCAACATCCCTCACTTCCTCTACACCTGCTCAGCAGACATGACGGCTGTCACGAC-ATGCGAAAGAGGATCAAGAAGACCTCCGACCAGAAAAATCACCACACCCTCCGCATTCATC ``` |
| 24 | ggggacaagtttgtacaaaaaagcaggcttgatgGCCGATACATCCTCCGCTCC, ggggaccactttgtacaagaaagctgggtcTTTACATTGAGGAGACATGG | Mycgr3\|35430  JGI  Ensembl | KOG0118 FOG: RRM domain; | nucleic acid binding; | 100 % | chr\_2:2832185 T->G; | ``` 1	ATGGCCGATACATCCTCCGCTCCCTCCTCGCTGCCTCAGCCACCGACTGGTGGTGCTTTCGAGAATGGCTCCCATATGCCGCCGCCGCTTAATATCCCTCAAAATACGAATCCGCTGCCCACCTCTATGAACAGTGCGTTCGAAATGGGCTCTCCGACCAGCGCTGGGGGCCTCGTCAGGCGTGCGGCCCCGGAGCCGAACAAAAGAGCTTTGTATGTGGGTGGACTCGATCCTCGCGTAACAGAGGACGTCTTGAAACAGATCTTCGAAACGACGGGCCATGTGCAGAATGTAAAGATCATTCCGGACAAAAAT	315 	\|\|\|\|\|\|\|\|\|\|\|\|\|\|\|\|\|\|\|\|\|\|\|\|\|\|\|\|\|\|\|\|\|\|\|\|\|\|\|\|\|\|\|\|\|\|\|\|\|\|\|\|\|\|\|\|\|\|\|\|\|\|\|\|\|\|\|\|\|\|\|\|\|\|\|\|\|\|\|\|\|\|\|\|\|\|\|\|\|\|\|\|\|\|\|\|\|\|\|\|\|\|\|\|\|\|\|\|\|\|\|\|\|\|\|\|\|\|\|\|\|\|\|\|\|\|\|\|\|\|\|\|\|\|\|\|\|\|\|\|\|\|\|\|\|\|\|\|\|\|\|\|\|\|\|\|\|\|\|\|\|\|\|\|\|\|\|\|\|\|\|\|\|\|\|\|\|\|\|\|\|\|\|\|\|\|\|\|\|\|\|\|\|\|\|\|\|\|\|\|\|\|\|\|\|\|\|\|\|\|\|\|\|\|\|\|\|\|\|\|\|\|\|\|\|\|\|\|\|\|\|\|\|\|\|\|\|\|\|\|\|\|\|\|\|\|\|\|\|\|\|\|\|\|\|\|\|\|\|\|\|\|\|\|\|\|\|\|\|\|\|\|\|\|\|\|\|\|\|\|\|\|\|\|\|\|\|\|\|\|\|\|\|\|\|\|\|\|\|\|\|\|\|\|\|\|\|\|\|\|\|\|\|\|\| 	ATGGCCGATACATCCTCCGCTCCCTCCTCGCTGCCTCAGCCACCGACTGGTGGTGCTTTCGAGAATGGCTCCCATATGCCGCCGCCGCTTAATATCCCTCAAAATACGAATCCGCTGCCCACCTCTATGAACAGTGCGTTCGAAATGGGCTCTCCGACCAGCGCTGGGGGCCTCGTCAGGCGTGCGGCCCCGGAGCCGAACAAAAGAGCTTTGTATGTGGGTGGACTCGATCCTCGCGTAACAGAGGACGTCTTGAAACAGATCTTCGAAACGACGGGCCATGTGCAGAATGTAAAGATCATTCCGGACAAAAAT ```      ``` 316	TTCCAGTCAAAGGGGTTCAACTACGGCTTTGTTGAATACGACGACCCCGGTGCCGCCGAAAGAGCCATGCAGACCCTCAACGGAAGGCGCGTGCATCAGCAGGAAATTCGGGTCAATTGGGCTTACCAGTCCAACACCACCTCCAAAGAGGACACTTCCAACCACTTCCACATTTTCGTCGGTGATCTCTCCAATGAGGTGAACGATGAGGTCCTCTTGCAGGCATTCTCAGCCTTTGGGACAGTTTCGGAGGCGCGCGTGATGTGGGACATGAAGACTGGCCGTTCGCGCGGATATGGTTTCGTAGCTTTCCGCGACCGCGCCGATGCTGAGAAGGCTCTGAGTTCAATGGATGGCGAGTGGCTCGGATCGCGCGCGATTCGCTGCAACTGGGCCAACCAGAAGGGTCAGCCATCATTTTCGCAGCAGCAGGCCATGGTTCAGATGGGCATGACTCCGACCAC-TCCATACG	787 	\|\|\|\|\|\|\|\|\|\|\|\|\|\|\|\|\|\|\|\|\|\|\|\|\|\|\|\|\|\|\|\|\|\|\|\|\|\|\|\|\|\|\|\|\|\|\|\|\|\|\|\|\|\|\|\|\|\|\|\|\|\|\|\|\|\|\|\|\|\|\|\|\|\|\|\|\|\|\|\|\|\|\|\|\|\|\|\|\|\|\|\|\|\|\|\|\|\|\|\|\|\|\|\|\|\|\|\|\|\|\|\|\|\|\|\|\|\|\|\|\|\|\|\|\|\|\|\|\|\|\|\|\|\|\|\|\|\|\|\|\|\|\|\|\|\|\|\|\|\|\|\|\|\|\|\|\|\|\|\|\|\|\|\|\|\|\|\|\|\|\|\|\|\|\|\|\|\|\|\|\|\|\|\|\|\|\|\|\|\|\|\|\|\|\|\|\|\|\|\|\|\|\|\|\|\|\|\|\|\|\|\|\|\|\|\|\|\|\|\|\|\|\|\|\|\|\|\|\|\|\|\|\|\|\|\|\|\|\|\|\|\|\|\|\|\|\|\|\|\|\|\|\|\|\|\|\|\|\|\|\|\|\|\|\|\|\|\|\|\|\|\|\|\|\|\|\|\|\|\|\|\|\|\|\|\|\|\|\|\|\|\|\|\|\|\|\|\|\|\|\|\|\|\|\|\|\|\|\|\|\|\|\|\|\|\|\|\|\|\|\|\|\|\|\|\|\|\|\|\|\|\|\|\| \| \|\|\|\|\|\|\|\|\|\|\| \|\| \|\|\|\|\|\|\|\|\| \|\|\|\|\|\|\|\|\|\|\|\|\|\|\|\|\|\|\|\|\|\|\|\|\|\| \|\|\|\|\|\| \| \| \|\|\| \|  \|\|\|\| \|\|\|\|\|\|\|\|\|\|\|\|\|\|\|\|\|\|\|\| \|\|\|\|\| \|\|\|\|\|\|\| \|\|\|\|\|\|\|\|\|\|\|\|\|\|\|\|\| \|\|\|\|\|\|\|\| 	TTCCAGTCAAAGGGGTTCAACTACGGCTTTGTTGAATACGACGACCCCGGTGCCGCCGAAAGAGCCATGCAGACCCTCAACGGAAGGCGCGTGCATCAGCAGGAAATTCGGGTCAATTGGGCTTACCAGTCCAACACCACCTCCAAAGAGGACACTTCCAACCACTTCCACATTTTCGTCGGTGATCTCTCCAATGAGGTGAACGATGAGGTCCTCTTGCAGGCATTCTCAGCCTTTGGGACAGTTTCGGAGGCGCGCGTGATGTGGGACATGAAGACTGGCCGTTCGCGCGGATATGGTTTCGTAGCTTTCCGCGACCGCGCCGATGCTGAGA-G-CTCTGAGTTCA-TG-ATGGCGAGT-GCTCGGATCGCGCGCGATTCGCTGCA-CTGGGC-A-C-AGA-G--TCAG-CATCATTTTCGCAGCAGCAG-CCATG-TTCAGAT-GGCATGACTCCGACCACCTCCATACG ``` |
| 24 | ggggacaagtttgtacaaaaaagcaggcttgatgGAAACCGCAAACTCAGCCGC, ggggaccactttgtacaagaaagctgggtcATACCTGGCGTCCATGTCTT | Mycgr3\|39611  JGI  Ensembl | KOG3937 mRNA splicing factor; |  | 0 % |  |  |
| 24 | ggggacaagtttgtacaaaaaagcaggcttgatgTCGAGAAGTCGCGCCCTTGT, ggggaccactttgtacaagaaagctgggtcCTCCTCCAGCGCAATCGCTC | Mycgr3\|40500  JGI  Ensembl | KOG3912 Predicted integral membrane protein; |  | 100 % |  |  |
| 24 | ggggacaagtttgtacaaaaaagcaggcttgatgGCTGAAGCGATAAAGTCCAT, ggggaccactttgtacaagaaagctgggtcTAACTCCACATCCCAAACAC | Mycgr3\|43113  JGI  Ensembl | KOG0280 Uncharacterized conserved protein; |  | 100 % |  |  |
| 24 | ggggacaagtttgtacaaaaaagcaggcttgatgACACCGAGCGCCATCAGCAT, ggggaccactttgtacaagaaagctgggtcAACATCGACCAAGTCCCTCG | Mycgr3\|44913  JGI  Ensembl | KOG0622 Ornithine decarboxylase; | catalytic activity; ornithine decarboxylase activity;  polyamine biosynthetic process; | 100 % |  |  |
| 24 | ggggacaagtttgtacaaaaaagcaggcttgatgAAGACCTCACTTCGGGCCAA, ggggaccactttgtacaagaaagctgggtcCAACGCCCCCAAAACCTCCC | Mycgr3\|48401  JGI  Ensembl | KOG0029 Amine oxidase; | amine oxidase activity; electron transport;  oxidoreductase activity; | 100 % |  | ``` 1	ATGAAGACCTCACTTCGGGCCAACGTCGGGATTCTCTCACTCCTCTGCATTCCAGCAGTCCTCGCCGACAACAACACACCTCAAACCGTCGACGTCGCAATAGTAGGCGGCGGCCTCTCCGGTCTCGCCGCCGCCAAAGTCCTCCTCGATGCCGGCAAAACTTTCACAATCCTCGAAGCCCGCGACCGTGTCGGCGGCAGAGTCCAAAACCAAGACCTCAAAAACGGCGGCGTCACCGAACTCGGCGCCGCATTCGTCGGCCCAACCCAAGACCGCGTGCTCGCCCTCGCCGCCTCCCTGAATCTTACTCTCATCCCCGAATATAATGCCGGCGACGTCGTCGCAGTCTTCGCCAACTCCACTTTGAAATACTCCACCGCGAGCCCGATCCCGCCGATCGACGACGAATCGTCCAAACTGCTGCTGCAAACTATCGCATTTCTTGAGACGACTGCTGGGACGATTAATACCACCGCGCCGTGGACGCATCCTAATGCGACGGTCTGGGATAGTCAGACGGTTGCCACTTTTTTCGATGGGCAGAGTCATCCTGTTGGTACCGTGCGCCGTTCGCTGTTGGATGTGTCGATTACGAGTATTTTCTCTGCTGAATCGGCTGAACTTTCTCTCTTCTACGCTCTTTCCTACATTGCGTCCGCAGGAAACGCTACCACACCGGGCTCCTTCATCCGCCTCATCTCCGTCGACAATGGAGCGCAACAAGCCCGCATCACAGGCGGCACTGCCCTTCTCGCTACCGGCCTGGCCAGTCAAGTCGGCTTATCCAACATCCAACTCAACTCCCCCGTCCGCTCCATCGCCCAGCAACCAGACGGCACCTACCTCCTCACCACCACCTCCGGCTGCACCACTTCCGCCCGCAAAGTCATAGTAGCCATGTcccccccTCTCGCCTCCCGCATCGAATACACCCCTCCCCTCCCGGCTTCCCGCGACCAACTCACTCAACGCATG-TTCATGGGTTCCCTCGCCAA	995 	\|\|\|\|\|\|\|\|\|\|\|\|\|\|\|\|\|\|\|\|\|\|\|\|\|\|\|\|\|\|\|\|\|\|\|\|\|\|\|\|\|\|\|\|\|\|\|\|\|\|\|\|\|\|\|\|\|\|\|\|\|\|\|\|\|\|\|\|\|\|\|\|\|\|\|\|\|\|\|\|\|\|\|\|\|\|\|\|\|\|\|\|\|\|\|\|\|\|\|\|\|\|\|\|\|\|\|\|\|\|\|\|\|\|\|\|\|\|\|\|\|\|\|\|\|\|\|\|\|\|\|\|\|\|\|\|\|\|\|\|\|\|\|\|\|\|\|\|\|\|\|\|\|\|\|\|\|\|\|\|\|\|\|\|\|\|\|\|\|\|\|\|\|\|\|\|\|\|\|\|\|\|\|\|\|\|\|\|\|\|\|\|\|\|\|\|\|\|\|\|\|\|\|\|\|\|\|\|\|\|\|\|\|\|\|\|\|\|\|\|\|\|\|\|\|\|\|\|\|\|\|\|\|\|\|\|\|\|\|\|\|\|\|\|\|\|\|\|\|\|\|\|\|\|\|\|\|\|\|\|\|\|\|\|\|\|\|\|\|\|\|\|\|\|\|\|\|\|\|\|\|\|\|\|\|\|\|\|\|\|\|\|\|\|\|\|\|\|\|\|\|\|\|\|\|\|\|\|\|\|\|\|\|\|\|\|\|\|\|\|\|\|\|\|\|\|\|\|\|\|\|\|\|\|\|\|\|\|\|\|\|\|\|\|\|\|\|\|\|\|\|\|\|\|\|\|\|\|\|\|\|\|\|\|\|\|\|\|\|\|\|\|\|\|\|\|\|\|\|\|\|\|\|\|\|\|\|\|\|\|\|\|\|\|\|\|\|\|\|\|\|\|\|\|\|\|\|\|\|\|\|\|\|\|\|\|\|\|\|\|\|\|\|\|\|\|\|\|\|\|\|\|\|\|\|\|\|\|\|\|\|\|\|\|\|\|\|\|\|\|\|\|\|\|\|\|\|\|\|\|\|\|\|\|\|\|\|\|\|\|\|\|\|\|\|\|\|\|\|\|\|\|\|\|\|\|\|\|\|\|\|\|\|\|\|\|\|\|\|\|\|\|\|\|\|\|\|\|\|\|\|\|\|\|\|\|\|\|\|\|\|\|\|\|\|\|\|\|\|\|\|\|\|\|\|\|\|\|\|\|\|\|\|\|\|\|\|\|\|\|\|\|\|\|\|\|\|\|\|\|\|\|\|\|\|\|\|\|\|\|\|\|\|\|\|\|\|\|\|\|\|\|\|\|\|\|\|\|\|\|\|\|\|\|\|\|\|\|\|\|\|\|\|\|\|\|\|\|\|\|\|\|\|\|\|\|\|\|\|\|\|\|\|\|\|\|\|\|\|\|\|\|\|\|\|\|\|\|\|\|\|\|\|\|\|\|\|\|\|\|\|\|\|\|\|\|\|\|\|\|\|\|\|\|\|\|\|\|\|\|\|\|\|\|\|\|\|\|\|\|\|\|\|\|\|\|\|\|\|\|\|\|\|\|\|\|\|\|\|\|\|\|\|\|\|\|\|\|\|\|\|\|\|\|\|\|\|\|\|\|\|\|\|\|\|\|\|\|\|\|\|\|\|\|\|\|\|\|\|\|\|\|\|\|\|\|\|\|\|\|\|\|\|\|\|\|\|\|\|\|\|\|\|\|\|\|\|\|\|\|\|\|\|\|\|\|\|\|\|\|\|\|\|\|\|\|\|\|\|\|\|\|\|\|\|\|\|\|\|\|\|\|\|\|\|\|\|\|\|\|\|\|\|\|\|\|\|\|\|\|\|\|\|\|\|\|\|\|\|\|\|\|\|\|\|\|\|\|\|\|\|\|\|\|\|\| \|\|\|\| \| \|\|\|\|\|\|\|\|\|\|\|\|\|\|\|\|\|\|\|\|\|\|\|\|\|\|\|\|\|\|\|\|\|\|\|\|\|\|\|\|\|\|\|\|\|\|\|\|\| \|   \|\|\|\|\|\| \|  \|\|\|\|\|\|\|\|\|\|\|\|\|\|\|\| \|\|\|\| \|\|\|\|\| \|\|\|\|\| \|\|\|\|\| \| \|\|\|\|\|\|\|\| \|\|\|\|\|\| \|\|\|\|\|\|\|\|\|\|\|\|\|\|\|\|\|\|\|\| 	ATGAAGACCTCACTTCGGGCCAACGTCGGGATTCTCTCACTCCTCTGCATTCCAGCAGTCCTCGCCGACAACAACACACCTCAAACCGTCGACGTCGCAATAGTAGGCGGCGGCCTCTCCGGTCTCGCCGCCGCCAAAGTCCTCCTCGATGCCGGCAAAACTTTCACAATCCTCGAAGCCCGCGACCGTGTCGGCGGCAGAGTCCAAAACCAAGACCTCAAAAACGGCGGCGTCACCGAACTCGGCGCCGCATTCGTCGGCCCAACCCAAGACCGCGTGCTCGCCCTCGCCGCCTCCCTGAATCTTACTCTCATCCCCGAATATAATGCCGGCGACGTCGTCGCAGTCTTCGCCAACTCCACTTTGAAATACTCCACCGCGAGCCCGATCCCGCCGATCGACGACGAATCGTCCAAACTGCTGCTGCAAACTATCGCATTTCTTGAGACGACTGCTGGGACGATTAATACCACCGCGCCGTGGACGCATCCTAATGCGACGGTCTGGGATAGTCAGACGGTTGCCACTTTTTTCGATGGGCAGAGTCATCCTGTTGGTACCGTGCGCCGTTCGCTGTTGGATGTGTCGATTACGAGTATTTTCTCTGCTGAATCGGCTGAACTTTCTCTCTTCTACGCTCTTTCCTACATTGCGTCCGCAGGAAACGCTACCACACCGGGCTCCTTCATCCGCCTCATCTCCGTCGACAATGGAGCGCAACAAGCCCGCATCACAGGCGGCACTGCCCTTCTCGCTACCGGCCTGGCCAGTCAAGTCGGCTTATCCAACATCCAACTCAACTCCCCCGTCCGCTCCATCGCCCAGCAACCAGACGGCACCTACCTC-TCAC-A-CACCTCCGGCTGCACCACTTCCGCCCGCAAAGTCATAGTAGCCATGTCCTC---TCTCGC-T--CGCATCGAATACACCC-TCCC-TCCCG-CTTCC-GCGAC-A-CTCACTCA-CGCATGTTTCATGGGTTCCCTCGCCAA ``` |
| 24 | ggggacaagtttgtacaaaaaagcaggcttgatgAAGGCAGTCATCGGACGCAG, ggggaccactttgtacaagaaagctgggtcCAACACAGCAGCAAAAGCAA | Mycgr3\|51025  JGI  Ensembl |  |  | 100 % |  |  |
| 24 | ggggacaagtttgtacaaaaaagcaggcttgatgTCCGAAGAACTGCTGGATGA, ggggaccactttgtacaagaaagctgggtcTCGTTTGCCTTTCTTCTTGC | Mycgr3\|61514  JGI  Ensembl | KOG3299 Uncharacterized conserved protein; |  | 100 % |  |  |
| 24 | ggggacaagtttgtacaaaaaagcaggcttgatgTATCTGCCCAGCACAAAATG, ggggaccactttgtacaagaaagctgggtcATCAATCGTCATGCGCTGCG | Mycgr3\|65650  JGI  Ensembl |  |  | 100 % |  |  |
| 24 | ggggacaagtttgtacaaaaaagcaggcttgatgGCCGGTCCTCTCCGCGACCT, ggggaccactttgtacaagaaagctgggtcATCCTCCAGCATGCCCGCCT | Mycgr3\|65916  JGI  Ensembl | KOG0569 Permease of the major facilitator superfamily; | L-arabinose isomerase activity; carbohydrate transport;  integral to membrane;  membrane;  sugar:hydrogen symporter activity;  transport;  transporter activity; | 100 % |  | ``` 1	ATGGCCGGTCCTCTCCGCGACCTCACGCCCTACTTCATCTACCTCCTCCTCACCGCAACCCTCGGCCCTCTTCTCTTCGGCTACCATCTCGCCGAACTCAACGCTCCCGCCGAAGTCATCACCTGCGCCAAAGAGAGCATCCACGGCACCACATCCACAAGGTTCTCACAATGCATCCGCATGAATCCCTCCCAGTTCGGCCTGGTGTCCTCTTCCTTCACTCTTGGAGGTCTCATAGGAGCTCTATCCGCAGGACCCATCACGACGAAATATGGTCGACTTCGAGCGATGTTGATATCGGGGATATTCGCCGTGGTCGGACCTGTCTTCGAAGCAGCAGCATCTAGCATTGGTGTCATGACCGTGGGAAGGCTCTTGTCAGGAGTAGGAGCCGGAATGGCGACCGTGGTGGTACCTATATACATCAGCGAGATCTCACCACCGGACAAGAGAGGACTATTTGGAGCTTTCACTCAGATCATGACCAACATGGGCATCCTCATTACACAGGCATTGGGACTCTTCCTGAGCAGAGGACAAATGTGGAGGATCATCTTGGGAGTCGGCGGAGCTATCGCTCTCGTGCAGACCATATCCCTCGGCCTTGCTGGACAAGAAAGCCCACAGTGGCTTGCAGACCATGGCAAAACTTCTCAAGCAAAGAGGATTCTACAAAAGCTCCGTGGGCATAAAGTCGATATCTCAGACGAGACCTCCACCTGGACATCTGCCCACAACGCCAATGCAAACGAAGAAGAAGAGACCCTTCTCGCCTCCTCCTCTACCCATGAAGCACCACCCACCATCGGCCCACTCGCCGTCCTCTCTCACCCAGACCACCGCTCCGCCGTCATCGCTGTCGTAACAATCATGATCGCCCAACAATTCACCGGCATCAACAGCATAGTCATGTACGGCGTCTCCCTCCTTTCCGACCTCCTAACCACCA	949 	\|\|\|\|\|\|\|\|\|\|\|\|\|\|\|\|\|\|\|\|\|\|\|\|\|\|\|\|\|\|\|\|\|\|\|\|\|\|\|\|\|\|\|\|\|\|\|\|\|\|\|\|\|\|\|\|\|\|\|\|\|\|\|\|\|\|\|\|\|\|\|\|\|\|\|\|\|\|\|\|\|\|\|\|\|\|\|\|\|\|\|\|\|\|\|\|\|\|\|\|\|\|\|\|\|\|\|\|\|\|\|\|\|\|\|\|\|\|\|\|\|\|\|\|\|\|\|\|\|\|\|\|\|\|\|\|\|\|\|\|\|\|\|\|\|\|\|\|\|\|\|\|\|\|\|\|\|\|\|\|\|\|\|\|\|\|\|\|\|\|\|\|\|\|\|\|\|\|\|\|\|\|\|\|\|\|\|\|\|\|\|\|\|\|\|\|\|\|\|\|\|\|\|\|\|\|\|\|\|\|\|\|\|\|\|\|\|\|\|\|\|\|\|\|\|\|\|\|\|\|\|\|\|\|\|\|\|\|\|\|\|\|\|\|\|\|\|\|\|\|\|\|\|\|\|\|\|\|\|\|\|\|\|\|\|\|\|\|\|\|\|\|\|\|\|\|\|\|\|\|\|\|\|\|\|\|\|\|\|\|\|\|\|\|\|\|\|\|\|\|\|\|\|\|\|\|\|\|\|\|\|\|\|\|\|\|\|\|\|\|\|\|\|\|\|\|\|\|\|\|\|\|\|\|\|\|\|\|\|\|\|\|\|\|\|\|\|\|\|\|\|\|\|\|\|\|\|\|\|\|\|\|\|\|\|\|\|\|\|\|\|\|\|\|\|\|\|\|\|\|\|\|\|\|\|\|\|\|\|\|\|\|\|\|\|\|\|\|\|\|\|\|\|\|\|\|\|\|\|\|\|\|\|\|\|\|\|\|\|\|\|\|\|\|\|\|\|\|\|\|\|\|\|\|\|\|\|\|\|\|\|\|\|\|\|\|\|\|\|\|\|\|\|\|\|\|\|\|\|\|\|\|\|\|\|\|\|\|\|\|\|\|\|\|\|\|\|\|\|\|\|\|\|\|\|\|\|\|\|\|\|\|\|\|\|\|\|\|\|\|\|\|\|\|\|\|\|\|\|\|\|\|\|\|\|\|\|\|\|\|\|\|\|\|\|\|\|\|\|\|\|\|\|\|\|\|\|\|\|\|\|\|\|\|\|\|\|\|\|\|\|\|\|\|\|\|\|\|\|\|\|\|\|\|\|\|\|\|\|\|\|\|\|\|\|\|\|\|\|\|\|\|\|\|\|\|\|\|\|\|\|\|\|\|\|\|\|\|\|\|\|\|\|\|\|\|\|\|\|\|\|\|\|\|\|\|\|\|\|\|\|\|\|\|\|\|\|\|\|\|\|\|\|\|\|\|\|\|\|\|\|\|\|\|\|\|\|\|\|\|\|\|\|\|\|\|\|\|\|\|\|\|\|\|\|\|\|\|\|\|\|\|\|\|\|\|\|\|\|\|\|\|\|\|\|\|\|\|\|\|\|\|\|\|\|\|\|\|\|\|\|\|\|\|\|\|\|\|\|\|\|\|\|\|\|\|\|\|\|\|\|\|\|\|\|\|\|\|\|\|\|\|\|\|\|\|\|\|\|\|\|\|\|\|\|\|\|\|\|\|\|\|\|\|\|\|\| \|\|\|\|\|\|\|\|\|\|\|\|\|\|\|\|\|\|\|\|\|\|\|\|\|\|\|\|\|\|\|\|\|\|\|\|\|\|\|\|\|\|\|\|\|\|\|\|\|\|\|\|\|\|\|\|\|\|\| \|\|\|\|\|\|\|\|\|\|\|\|\|\|\|\|\|\|\| \|\| \|\|\|\| \| \|\|\|\|\|\|\|\|\|\|\|\|\|\|\|\| \| \|\|\|\|\|\|\|\|\|\|\|\|\|\| \|\| \| \|\|\| \|\|\|\|\|\| \|\|\|\|\|\|\|\|\|\|\|\|\|\|\|\|\|\|\|\|\|\|\|\|\|\|\|\|\|\|\|\| \| \|\|\|\|\| \|\|\| \|\|\|\|\|\| 	ATGGCCGGTCCTCTCCGCGACCTCACGCCCTACTTCATCTACCTCCTCCTCACCGCAACCCTCGGCCCTCTTCTCTTCGGCTACCATCTCGCCGAACTCAACGCTCCCGCCGAAGTCATCACCTGCGCCAAAGAGAGCATCCACGGCACCACATCCACAAGGTTCTCACAATGCATCCGCATGAATCCCTCCCAGTTCGGCCTGGTGTCCTCTTCCTTCACTCTTGGAGGTCTCATAGGAGCTCTATCCGCAGGACCCATCACGACGAAATATGGTCGACTTCGAGCGATGTTGATATCGGGGATATTCGCCGTGGTCGGACCTGTCTTCGAAGCAGCAGCATCTAGCATTGGTGTCATGACCGTGGGAAGGCTCTTGTCAGGAGTAGGAGCCGGAATGGCGACCGTGGTGGTACCTATATACATCAGCGAGATCTCACCACCGGACAAGAGAGGACTATTTGGAGCTTTCACTCAGATCATGACCAACATGGGCATCCTCATTACACAGGCATTGGGACTCTTCCTGAGCAGAGGACAAATGTGGAGGATCATCTTGGGAGTCGGCGGAGCTATCGCTCTCGTGCAGACCATATCCCTCGGCCTTGCTGGACAAGAAAGCCCACAGTGGCTTGCAGACCATGGCAAAACTTCTCAAGCAAAGAGGATTCTACAAAAGCTCCGTGGGCATAAAGTCGATATCTCAGACGAGACCTCCACCTGGACATCTGCCCACAACGCCAATGCAAACGAAGAAG-AGAGACCCTTCTCGCCTCCTCCTCTACCCATGAAGCACCACCCACCATCGGCCCACTCG-CGTCCTCTCTCACCCAGAC-AC-GCTC-G-CGTCATCGCTGTCGTA-C-ATCATGATCGCCCA-CA-T-CAC-GGCATC-ACAGCATAGTCATGTACGGCGTCTCCCTCCTT-C-GACCTTCTA-CCACCA ``` |
| 24 | ggggacaagtttgtacaaaaaagcaggcttgatgGATTCCGTCAGTGCCGTCAC, ggggaccactttgtacaagaaagctgggtcAAAGCCTTCCCAGAACTCCA | Mycgr3\|66079  JGI  Ensembl |  |  | 100 % |  |  |
| 24 | ggggacaagtttgtacaaaaaagcaggcttgatgAGAGTCTTCTTCGCCTTCCT, ggggaccactttgtacaagaaagctgggtcTCTTGCGTCCGGCACATTGC | Mycgr3\|66368  JGI  Ensembl |  |  | 100 % |  |  |
| 24 | ggggacaagtttgtacaaaaaagcaggcttgatgCGGTCGCTTCTGACAAGGAG, ggggaccactttgtacaagaaagctgggtcTGGAGTCTTAGGGTTGTCGG | Mycgr3\|66375  JGI  Ensembl | KOG2707 Predicted metalloprotease with chaperone activity (RNAse H/HSP70 fold); | O-sialoglycoprotein endopeptidase activity; collagenase activity;  matrilysin-2 activity;  prenyl protein specific endopeptidase activity;  proteolysis;  stromelysin 3 activity; | 1.31853 % |  |  |
| 24 | ggggacaagtttgtacaaaaaagcaggcttgatgGCCCACCGAACGGAGACCAC, ggggaccactttgtacaagaaagctgggtcGATCCTCTCGGTAATCCAAC | Mycgr3\|66454  JGI  Ensembl | KOG1515 Arylacetamide deacetylase; |  | 100 % |  | ``` 1	ATGGCCCACCGAACGGAGACCACCCTCCTGGACAAGCTTCTCCTCCTACCACTGCTCGCAAAAGTCCTCATTGCCGTCTTCTTCCGCGCAATTTTGACACCGTTCAACACCGGCCCCAAAGGCAATGGAACCAAAAAAGACCTCATCTTCGTCGCCCTGCGAACTTTTCTCGGCGGCGTCAACTCCGCACAAGAACACTGGTTGAATACCCTCCAGATCACGGACCAGAACTACTTGGACTTTGCCAAAAAGCAGAATTTCGAGCCGGAGACGAGCACGCTGGAGTCTGGTCTGAAGCTACACTGGCTGGGACCAAGGACGGCGAGCAGAGTCCTGCTGTATTTCCACGGAGGTGGCTATGCCATGTCATGTTCGCCGGGGCATTTCCAATGGCTGTTCGACTTGCAGAACGACATTTCCAAGGACATCAGTATCGTTGTGCCATCGTATGCTGTCACTCCTGCGGCTCAGTATCCTACTCAGCTGAAGCAGGCGGCCGAGGCGTTACATTGGTTGATCGAAAAGCAAGGTAGAAGTCCACGGGAG	546 	\|\|\|\|\|\|\|\|\|\|\|\|\|\|\|\|\|\|\|\|\|\|\|\|\|\|\|\|\|\|\|\|\|\|\|\|\|\|\|\|\|\|\|\|\|\|\|\|\|\|\|\|\|\|\|\|\|\|\|\|\|\|\|\|\|\|\|\|\|\|\|\|\|\|\|\|\|\|\|\|\|\|\|\|\|\|\|\|\|\|\|\|\|\|\|\|\|\|\|\|\|\|\|\|\|\|\|\|\|\|\|\|\|\|\|\|\|\|\|\|\|\|\|\|\|\|\|\|\|\|\|\|\|\|\|\|\|\|\|\|\|\|\|\|\|\|\|\|\|\|\|\|\|\|\|\|\|\|\|\|\|\|\|\|\|\|\|\|\|\|\|\|\|\|\|\|\|\|\|\|\|\|\|\|\|\|\|\|\|\|\|\|\|\|\|\|\|\|\|\|\|\|\|\|\|\|\|\|\|\|\|\|\|\|\|\|\|\|\|\|\|\|\|\|\|\|\|\|\|\|\|\|\|\|\|\|\|\|\|\|\|\|\|\|\|\|\|\|\|\|\|\|\|\|\|\|\|\|\|\|\|\|\|\|\|\|\|\|\|\|\|\|\|\|\|\|\|\|\|\|\|\|\|\|\|\|\|\|\|\|\|\|\|\|\|\|\|\|\|\|\|\|\|\|\|\|\|\|\|\|\|\|\|\|\|\|\|\|\|\|\|\|\|\|\|\|\|\|\|\|\|\|\|\|\|\|\|\|\|\|\|\|\|\|\|\|\|\|\|\|\|\|\|\|\|\|\|\|\|\|\|\|\|\|\|\|\|\|\|\|\|\|\|\|\|\|\|\|\|\|\|\|\|\|\|\|\|\|\|\|\|\|\|\|\|\|\|\|\|\|\|\|\|\|\|\|\|\|\|\|\|\|\|\|\|\|\|\|\|\|\|\|\|\|\|\|\|\|\|\|\|\|\|\|\|\|\|\|\|\|\|\|\|\|\|\|\|\|\|\|\|\|\|\|\|\|\|\|\|\|\|\|\|\|\|\|\|\|\|\|\|\|\|\|\|\|\|\|\|\|\|\|\|\|\|\|\|\|\|\|\|\|\|\|\|\|\|\|\|\|\|\|\|\|\|\|\|\|\|\|\|\|\|\|\|\|\|\|\|\|\|\|\|\|\|\|\|\|\|\|\|\|\|\|\|\|\|\|\|\|\|\|\|\|\|\| 	ATGGCCCACCGAACGGAGACCACCCTCCTGGACAAGCTTCTCCTCCTACCACTGCTCGCAAAAGTCCTCATTGCCGTCTTCTTCCGCGCAATTTTGACACCGTTCAACACCGGCCCCAAAGGCAATGGAACCAAAAAAGACCTCATCTTCGTCGCCCTGCGAACTTTTCTCGGCGGCGTCAACTCCGCACAAGAACACTGGTTGAATACCCTCCAGATCACGGACCAGAACTACTTGGACTTTGCCAAAAAGCAGAATTTCGAGCCGGAGACGAGCACGCTGGAGTCTGGTCTGAAGCTACACTGGCTGGGACCAAGGACGGCGAGCAGAGTCCTGCTGTATTTCCACGGAGGTGGCTATGCCATGTCATGTTCGCCGGGGCATTTCCAATGGCTGTTCGACTTGCAGAACGACATTTCCAAGGACATCAGTATCGTTGTGCCATCGTATGCTGTCACTCCTGCGGCTCAGTATCCTACTCAGCTGAAGCAGGCGGCCGAGGCGTTACATTGGTTGATCGAAAAGCAAGGTAGAAGTCCACGGGAG ```      ``` 545	AGATCGTCCTCGGGGGCGACTCGGCTGGAGGTAACCTGTGCATGGCCCTTCTTAGCCACATTCTCCACCCCCACCCGGACGCTATCAAGATCCACCTGCGACAACCCCTCGCCAGTGCAATCCTGATCAGTCCATGGGTCAAATTCGCTACCGATGACCCCAGCTGGACGCGGAACGCAAGCACGGACATGTTTGTACCCACAGCCGGACATCGCTGGACGGATTTATTCCTCGG	779 	\|\|\|\|\|\|\|\|\|\|\|\|\|\|\|\|\|\|\|\|\|\|\|\|\|\|\|\|\|\|\|\|\|\|\|\|\|\|\|\|\|\|\|\|\|\|\|\|\|\|\|\|\|\|\|\|\|\|\|\|\|\|\|\|\|\|\|\|\|\|\|\|\|\|\|\|\|\|\|\|\|\|\|\|\|\|\|\|\|\|\|\|\|\|\|\|\|\|\|\|\|\|\|\|\|\|\|\|\|\|\|\|\|\|\|\|\|\|\|\|\|\|\|\|\|\|\|\|\|\|\|\|\|\|\|\|\|\|\|\|\|\|\|\|\|\|\|\|\|\|\|\|\|\|\|\|\|\|\|\|\|\|\|\|\|\|\|\|\|\|\|\|\|\|\|\|\|\|\|\|\|\|\|\|\|\|\|\|\|\|\| \|\|\|\|\|\|\|\|\|\|\|\|\|\|\|\|\|\|\|\|\|\|\|\|\|\|\|\|\|\|\|\|\|\|\|\|\|\|\|\|\|\|\| 	AGATCGTCCTCGGGGGCGACTCGGCTGGAGGTAACCTGTGCATGGCCCTTCTTAGCCACATTCTCCACCCCCACCCGGACGCTATCAAGATCCACCTGCGACAACCCCTCGCCAGTGCAATCCTGATCAGTCCATGGGTCAAATTCGCTACCGATGACCCCAGCTGGACGCGGAACGCAAGCACGGACATG-TTGTACCCACAGCCGGACATCGCTGGACGGATTTATTCCTCGG ``` |
| 24 | ggggacaagtttgtacaaaaaagcaggcttgatgAAGCTGCTGTTGTCTACCGG, ggggaccactttgtacaagaaagctgggtcGTCTACCAGCTCGGCCACTG | Mycgr3\|66652  JGI  Ensembl | KOG3023 Glutamate-cysteine ligase regulatory subunit; | DNA binding; glutamate-cysteine ligase activity;  glutamate-cysteine ligase complex;  oxidoreductase activity; | 100 % |  |  |
| 24 | ggggacaagtttgtacaaaaaagcaggcttgatgTCGTACAATCCCAGCAACGC, ggggaccactttgtacaagaaagctgggtcTCTCCATCCGCCAACCCTGC | Mycgr3\|66947  JGI  Ensembl | KOG0608 Warts/lats-like serine threonine kinases; |  | 100 % |  |  |
| 24 | ggggacaagtttgtacaaaaaagcaggcttgatgCCATTTCCTTTTGTGCTCCC, ggggaccactttgtacaagaaagctgggtcGTAATAGCTATTGCTCGTTG | Mycgr3\|67112  JGI  Ensembl |  |  | 100 % |  |  |
| 24 | ggggacaagtttgtacaaaaaagcaggcttgatgAAGATCATCATAGTCGGCGC, ggggaccactttgtacaagaaagctgggtcGTATGCTGCATCAGGGAATG | Mycgr3\|67335  JGI  Ensembl | KOG2614 Kynurenine 3-monooxygenase and related flavoprotein monooxygenases; | metabolic process; oxidoreductase activity; | 100 % |  |  |
| 24 | ggggacaagtttgtacaaaaaagcaggcttgatgCAGCACGTATTCTCGCACCA, ggggaccactttgtacaagaaagctgggtcTGACATCGTCGGCAGAAGGC | Mycgr3\|67501  JGI  Ensembl | KOG3036 Protein involved in cell differentiation/sexual development; |  | 100 % |  | ``` 701	GCACACATCCCACTCTTCCTCTACCCGTTCCTCAACACAACCTCCAAATCCCGACCCTTCGAATACCTCCGCCTGACATCCCTCGGCGTGATTGGCGCGCTGGTCAAAAACGACAGTTCCGATGTCATCAACTTCCTCCTCACCACCGAGATTATCCCGCTCTGCCTCCGTATCATGGAGACTGGATCGGAGCTGAGCAAGACGGTGGCCATCTTCATTGTGCAGAAGATCCTCCTCGACGACATGGGACTGCAGTATA-TCTGCCAGACATACGAGCGA	979 	\|\|\|\|\|\|\|\|\|\|\|\|\|\|\|\|\|\|\|\|\|\|\|\|\|\|\|\|\|\|\|\|\|\|\|\|\|\|\|\|\|\|\|\|\|\|\|\|\|\|\|\|\|\|\|\|\|\|\|\|\|\|\|\|\|\|\|\|\|\|\|\|\|\|\|\|\|\|\|\|\|\|\|\|\|\|\|\|\|\|\|\|\|\|\|\|\|\|\|\|\|\|\|\|\|\|\|\|\|\|\|\|\|\|\|\|\|\|\|\|\|\|\|\|\|\|\|\|\|\|\|\|\|\|\|\|\|\|\|\|\|\|\|\|\|\|\|\|\|\|\|\|\|\|\|\|\|\|\|\|\|\|\|\|\|\|\|\|\|\|\|\|\|\|\|\|\|\|\|\|\|\|\|\|\|\|\|\|\|\|\|\|\|\|\|\|\|\|\| \|\|\| \|\|\|\|\|\|\|\|\|\|\|\|\|\| \|\|\|\|\|\|\| \|\|\|\|\|\| \|\|\|\|\|\|\|\|\|\|\|\|\|\|\|\|\|\|\|\|\|\|\|\|\| \|\|\|\|\|\|\|\|\|\|\|\|\|\|\|\|\|\|\|\| 	GCACACATCCCACTCTTCCTCTACCCGTTCCTCAACACAACCTCCAAATCCCGACCCTTCGAATACCTCCGCCTGACATCCCTCGGCGTGATTGGCGCGCTGGTCAAAAACGACAGTTCCGATGTCATCAACTTCCTCCTCACCACCGAGATTATCCCGCTCTGCCTCCGTATCATGGAGACTGGATCGGAGCTGAGCA-GAC-GTGGCCATCTTCAT-GTGCAGA-GATCCT-CTCGACGACATGGGACTGCAGTATATTCTGCCAGACATACGAGCGA ```      ``` 158	ATGCAGCACGTATTCTCGCACCAGAGTCCATACCACCAGCAACCGGACTGGCAGCATCATCCGCAAGCACATCACCAATACGCCGCGCAATCACAAGCCGCCGCCCAGCAGCAACAACAGCAGCAGTCTCAGCAGCAACACTACGCTCGTCCCGCTCAGAACGGCGCAAGTAATGGTGGAGGTGGAAACGGCGTCGGACAGCTAGGAGGGGATCACAACGGTGGTGGTATCACGAGCGCGAGCGACAACATGAGTGAAGAGAACAGACGGGTATTGGAGTGGATCGCGCAGACGCTCAATGCGAATACGCGCGAGGCAGCGCTGTTGGAGCTGAGCAAGAAGAGAGAGCAGGTACCAGAGTTGGCGCTGATATTATGGCATTCTTTTGG	546 	\|\|\|\|\|\|\|\|\|\|\|\|\|\|\|\|\|\|\|\|\|\|\|\|\|\|\|\|\|\|\|\|\|\|\|\|\|\|\|\|\|\|\|\|\|\|\|\|\|\|\|\|\|\|\|\|\|\|\|\|\|\|\|\|\|\|\|\|\|\|\|\|\|\|\|\|\|\|\|\|\|\|\|\|\|\|\|\|\|\|\|\|\|\|\|\|\|\|\|\|\|\|\|\|\|\|\|\|\|\|\|\|\|\|\|\|\|\|\|\|\|\|\|\|\|\|\|\|\|\|\|\|\|\|\|\|\|\|\|\|\|\|\|\|\|\|\|\|\|\|\|\|\|\|\|\|\|\|\|\|\|\|\|\|\|\|\|\|\|\|\|\|\|\|\|\|\|\|\|\|\|\|\|\|\|\|\|\|\|\|\|\|\|\|\|\|\|\|\|\|\|\|\|\|\|\|\|\|\|\|\|\|\|\|\|\|\|\|\|\|\|\|\|\|\|\|\|\|\|\|\|\|\|\|\|\|\|\|\|\|\|\|\|\|\|\|\|\|\|\|\|\|\|\|\|\|\|\|\|\|\|\|\|\|\|\|\|\|\|\|\|\|\|\|\|\|\|\|\|\|\|\|\|\|\|\|\|\|\|\|\|\|\|\|\|\|\|\|\|\|\|\|\|\|\|\|\|\|\|\|\|\|\|\|\|\|\|\|\|\|\|\|\|\|\|\|\|\|\|\|\|\|\|\|\|\|\|\|\|\|\|\|\|\|\|\|\|\|\|\|\|\|\|\|\|\|\|\|\|\|\|\|\|\|\|\|\|\|\|\|\|\|\|\|\|\|\|\|\|\|\|\|\|\|\|\|\|\|\| 	ATGCAGCACGTATTCTCGCACCAGAGTCCATACCACCAGCAACCGGACTGGCAGCATCATCCGCAAGCACATCACCAATACGCCGCGCAATCACAAGCCGCCGCCCAGCAGCAACAACAGCAGCAGTCTCAGCAGCAACACTACGCTCGTCCCGCTCAGAACGGCGCAAGTAATGGTGGAGGTGGAAACGGCGTCGGACAGCTAGGAGGGGATCACAACGGTGGTGGTATCACGAGCGCGAGCGACAACATGAGTGAAGAGAACAGACGGGTATTGGAGTGGATCGCGCAGACGCTCAATGCGAATACGCGCGAGGCAGCGCTGTTGGAGCTGAGCAAGAAGAGAGAGCAGGTACCAGAGTTGGCGCTGATATTATGGCATTCTTTTGG ```      ``` 545	GGCGTCATGACGTCCCTTCTCCAGGAGATCATCTCGGTCTACCCGCTGCTCAATCCGTCACAGCTGACAGCGGCGGCTTCGAATCGGGTCTGCAATGCTTTGGCGCTGCTCCAGTGCGTTGCTAGTCACAGCGAGACACGCGGACTGTTCCTGAATG	701 	\|\|\|\|\|\|\|\|\|\|\|\|\|\|\|\|\|\|\|\|\|\|\|\|\|\|\|\|\|\|\|\|\|\|\|\|\|\|\|\|\|\|\|\|\|\|\|\|\|\|\|\|\|\|\|\|\|\|\|\|\|\|\|\|\|\|\|\|\|\|\|\|\|\|\|\|\|\|\|\|\|\|\|\|\|\|\|\|\|\|\|\|\|\|\|\|\|\|\|\|\|\|\|\|\|\|\|\|\|\|\|\|\|\|\|\|\|\|\|\|\|\|\|\|\|\|\|\|\|\|\|\|\|\|\|\|\|\|\|\|\|\|\|\|\|\|\|\|\|\|\|\|\|\|\|\|\| 	GGCGTCATGACGTCCCTTCTCCAGGAGATCATCTCGGTCTACCCGCTGCTCAATCCGTCACAGCTGACAGCGGCGGCTTCGAATCGGGTCTGCAATGCTTTGGCGCTGCTCCAGTGCGTTGCTAGTCACAGCGAGACACGCGGACTGTTCCTGAATG ``` |
| 24 | ggggacaagtttgtacaaaaaagcaggcttgatgCCCGTCACACACTTCACCAC, ggggaccactttgtacaagaaagctgggtcCTTTGCATAATGCGGCAACG | Mycgr3\|67509  JGI  Ensembl | KOG1417 Homogentisate 1,2-dioxygenase; | L-phenylalanine catabolic process; homogentisate 1,2-dioxygenase activity;  tyrosine metabolic process; | 100 % |  | ``` 29	ATGCCCGTCACACACTTCACCACCCCGGAGAAATACACCTACCTCCACGGCTTCTCCACCCACCACGAATCCTCCGCCCTCCCCAACGCCCTCCCCGTCGGCAGCAACAACCCCCAAAAAGCCCCCTACGGCCTGTACAACGAAAAACTCTCCGGAACAGCCTTCACCGCCCCCCGCCACGAAAACCTCCAAACATGGCTCTACCGCATCCTCCCGGCGTCCTCCCACTCCCCTTTCCAACCCCGTGAGACCTCCACCTACGACCTCAACCCGGGTGGTCTCCGCGACGCGAAATGGCATCAAATCCCCAACCAACTCCGCTGGGATCCCTTCGACTTGGACGAAAGTGTCGACTGGATCCACTCCCTCCGCCTCGTCGCGGGAGCCGGTGATCCGACCATGAAATCCGGACTAGGAATCCTCATCTTCGCAGCTGGGAAATCCATGGATGCCAACCACGCCTTCTACTCCGCTGACGGCGACTTGCTCATCGTGCTGCAACACGGCGTCCTCGACATCCGCACCGAACTCGGCAGCTTACTCGTCCGACCGAATGAAATCTGCGTGATACCACGGGGGATTCGACACCACGTCTCCCTCCCGTCAGGCCCAGCACGAGGGTACATCCTCGAACTCTACCAAGGCCACTTCACGCTCCCCGAACTCGGACCCATCGGATCCTCAGGCCTCGCTTCCCCGAGAGACTTCCAAGCCCCCGTGGCGAGCTACATCAACGACACCACCACCGAACACACCCTCTTCGCCAAATTCTCGGGTACCCTCTTCTCCGCGACTCAATCTCACTCCCCCTTCGACGTCGTCGCTTGGCATGGGAATTACTATCCTTACAAATACGACCTGGGACGGTTCGCCACGATCGGCAGCATTTCCCACGACCA-CCCCGATCCCTCCATATTCACAGTCCTCACAGCCCAGTCCCCCGTTG-CGGGAA	980 	\|\|\|\|\|\|\|\|\|\|\|\|\|\|\|\|\|\|\|\|\|\|\|\|\|\|\|\|\|\|\|\|\|\|\|\|\|\|\|\|\|\|\|\|\|\|\|\|\|\|\|\|\|\|\|\|\|\|\|\|\|\|\|\|\|\|\|\|\|\|\|\|\|\|\|\|\|\|\|\|\|\|\|\|\|\|\|\|\|\|\|\|\|\|\|\|\|\|\|\|\|\|\|\|\|\|\|\|\|\|\|\|\|\|\|\|\|\|\|\|\|\|\|\|\|\|\|\|\|\|\|\|\|\|\|\|\|\|\|\|\|\|\|\|\|\|\|\|\|\|\|\|\|\|\|\|\|\|\|\|\|\|\|\|\|\|\|\|\|\|\|\|\|\|\|\|\|\|\|\|\|\|\|\|\|\|\|\|\|\|\|\|\|\|\|\|\|\|\|\|\|\|\|\|\|\|\|\|\|\|\|\|\|\|\|\|\|\|\|\|\|\|\|\|\|\|\|\|\|\|\|\|\|\|\|\|\|\|\|\|\|\|\|\|\|\|\|\|\|\|\|\|\|\|\|\|\|\|\|\|\|\|\|\|\|\|\|\|\|\|\|\|\|\|\|\|\|\|\|\|\|\|\|\|\|\|\|\|\|\|\|\|\|\|\|\|\|\|\|\|\|\|\|\|\|\|\|\|\|\|\|\|\|\|\|\|\|\|\|\|\|\|\|\|\|\|\|\|\|\|\|\|\|\|\|\|\|\|\|\|\|\|\|\|\|\|\|\|\|\|\|\|\|\|\|\|\|\|\|\|\|\|\|\|\|\|\|\|\|\|\|\|\|\|\|\|\|\|\|\|\|\|\|\|\|\|\|\|\|\|\|\|\|\|\|\|\|\|\|\|\|\|\|\|\|\|\|\|\|\|\|\|\|\|\|\|\|\|\|\|\|\|\|\|\|\|\|\|\|\|\|\|\|\|\|\|\|\|\|\|\|\|\|\|\|\|\|\|\|\|\|\|\|\|\|\|\|\|\|\|\|\|\|\|\|\|\|\|\|\|\|\|\|\|\|\|\|\|\|\|\|\|\|\|\|\|\|\|\|\|\|\|\|\|\|\|\|\|\|\|\|\|\|\|\|\|\|\|\|\|\|\|\|\|\|\|\|\|\|\|\|\|\|\|\|\|\|\|\|\|\|\|\|\|\|\|\|\|\|\|\|\|\|\|\|\|\|\|\|\|\|\|\|\|\|\|\|\|\|\|\|\|\|\|\|\|\|\|\|\|\|\|\|\|\|\|\|\|\|\|\|\|\|\|\|\|\|\|\|\|\|\|\|\|\|\|\|\|\|\|\|\|\|\|\|\|\|\|\|\|\|\|\|\|\|\|\|\|\|\|\|\|\|\|\|\|\|\|\|\|\|\|\|\|\|\|\|\|\|\|\|\|\|\|\|\|\|\|\|\|\|\|\|\|\|\|\|\|\|\|\|\|\|\|\|\|\|\|\|\|\|\|\|\|\|\|\|\|\|\|\|\|\|\| \|\|\|\|\|\|\|\|\|\|\|\|\|\|\|\|\|\|\|\|\|\|\|\|\|\|\|\|\|\|\|\|\|\|\|\|\|\|\|\|\|\|\|\|\|\|\|\|\|\|\|\|\|\|\|\|\|\|\|\|\|\|\|\|\|\|\|\|\|\|\|\|\|\|\|\|\|\|\|\|\|\|\|\|\|\|\|\|\|\|\|\|\|\|\|\|\|\|\|\|\|\|\|\|\|\|\|\|\|\|\|\|\|\|\|\|\|\|\|\|\|\|\|\|\|\|\|\|\|\|\|\|\|\|\|\|\|\|\|\|\|\|\|\|\|\|\|\|\|\|\|\|\|\|\|\|\|\|\|\|\|\|\|\|\|\|\|\|\|\|\|\|\|\|\|\|\| \|\|\|\| \|\|\|\|\|\|\|\|\|\|\|\|\|\|\|\|\|\|\|\|\|\|\|\|\|\|\|\|\|\|\| \|\|\|\|\|\|\|\|\|\|\|\|\|\|\|\|\|\|\|\|\|\|\|\|\|\|\|\|\|\|\|\|\|\|\|\|\|\|\|\|\|\|\|\|\|\|\| \|\|\|\|\|\| 	ATGCCCGTCACACACTTCACCACCCCGGAGAAATACACCTACCTCCACGGCTTCTCCACCCACCACGAATCCTCCGCCCTCCCCAACGCCCTCCCCGTCGGCAGCAACAACCCCCAAAAAGCCCCCTACGGCCTGTACAACGAAAAACTCTCCGGAACAGCCTTCACCGCCCCCCGCCACGAAAACCTCCAAACATGGCTCTACCGCATCCTCCCGGCGTCCTCCCACTCCCCTTTCCAACCCCGTGAGACCTCCACCTACGACCTCAACCCGGGTGGTCTCCGCGACGCGAAATGGCATCAAATCCCCAACCAACTCCGCTGGGATCCCTTCGACTTGGACGAAAGTGTCGACTGGATCCACTCCCTCCGCCTCGTCGCGGGAGCCGGTGATCCGACCATGAAATCCGGACTAGGAATCCTCATCTTCGCAGCTGGGAAATCCATGGATGCCAACCACGCCTTCTACTCCGCTGACGGCGACTTGCTCATCGTGCTGCAACACGGCGTCCTCGACATCCGCACCGAACTCGGCAGCTTACTCGTCCGACCGAATGAAATCTGCGTGATACCACGGGGGATTCGACACCACGTCTCCCTCCCGTCAGGCCCAGCACGAGGGTACATCCTCGAACTCTACCAAGGCCACTTCACGCTCCCCGAACTCGGACCCATCGGATCCTCA-GCCTCGCTTCCCCGAGAGACTTCCAAGCCCCCGTGGCGAGCTACATCAACGACACCACCACCGAACACACCCTCTTCGCCAAATTCTCGGGTACCCTCTTCTCCGCGACTCAATCTCACTCCCCCTTCGACGTCGTCGCTTGGCATGGGAATTACTATCCTTACAAATACGACCTGG-ACGG-TCGCCACGATCGGCAGCATTTCCCACGACCACCCCCGATCCCTCCATATTCACAGTCCTCACAGCCCAGTCCCCCGTTGGCGGGAA ``` |
| 24 | ggggacaagtttgtacaaaaaagcaggcttgatgTCTTCCGAAGAGAGCTCCGC, ggggaccactttgtacaagaaagctgggtcCTTCCATCCGCCCTGAAGCT | Mycgr3\|67718  JGI  Ensembl | KOG2210 Oxysterol-binding protein; | steroid metabolic process; | 100 % |  |  |
| 24 | ggggacaagtttgtacaaaaaagcaggcttgatgTCCTTCGCTCGGAATCTCCT, ggggaccactttgtacaagaaagctgggtcATCCAACAGACGGGTGATCA | Mycgr3\|68887  JGI  Ensembl | KOG0460 Mitochondrial translation elongation factor Tu; | GTP binding; GTPase activity;  intracellular;  protein-synthesizing GTPase activity;  translation;  translation elongation factor activity;  translational elongation; | 100 % |  | ``` 247	AGACCACCCTCACAGCAGCCATCACCAAGCGACAAGCAGAAAAGGGCTTTGCCAACTTCCTCGAATATGGCGCCATTGACAAGGCTCCCGAGGAGCGCAAGCGCGGTATCACCATCTCCACCGCCCACATCGAATACCAGACCGACAACCGCCACTACGCCCACGTCGACTGCCCCGGTCACGCCGATTACATCAAGAACATGATCACAGGAGCTGCGTCCATGGACGGAGCCATCATCGTGGTGGCCGCCAGCGACGGACAAATGCCTCAGACTCGCGAGCACTTGCTCCTCGCCCGCCAAGTTGGTGTGAAGAGAATCGTAGTCTTCGTCAACAAGGTCGATGCGATCGAGGACAAGGAGATGTTGGAGCTCGTCGAGATGGAGATGCGCGAGCTGCTCTCCTCGTACGGCTTCGAGGGCGACGAGACTCCCATCATCATGGGTTCCGCTCTTTGCGCTCTCGACGGCCGCCAGCCAGACATTGGCGTTACCCAAATCGACAACCTCCTCGAAGCCGTTGACACCTGGATTCCCACACCCGAGCGAGACACCGACAAGCCCTTCCTCATGTCCGTCGAGGACGTCTTCTCCATTCCCGGCCGTGGCACAGTCGCCTCTGGCCGTGTCGAGCGTGGAGTGCTCAAGCGCGACTCCGAAGTTGAGTTGGTCGGCAAGAACGCCGCGCCCATCA	939 	\|\|\|\|\|\|\|\|\|\|\|\|\|\|\|\|\|\|\|\|\|\|\|\|\|\|\|\|\|\|\|\|\|\|\|\|\|\|\|\|\|\|\|\|\|\|\|\|\|\|\|\|\|\|\|\|\|\|\|\|\|\|\|\|\|\|\|\|\|\|\|\|\|\|\|\|\|\|\|\|\|\|\|\|\|\|\|\|\|\|\|\|\|\|\|\|\|\|\|\|\|\|\|\|\|\|\|\|\|\|\|\|\|\|\|\|\|\|\|\|\|\|\|\|\|\|\|\|\|\|\|\|\|\|\|\|\|\|\|\|\|\|\|\|\|\|\|\|\|\|\|\|\|\|\|\|\|\|\|\|\|\|\|\|\|\|\|\|\|\|\|\|\|\|\|\|\|\|\|\|\|\|\|\|\|\|\|\|\|\|\|\|\|\|\|\|\|\|\|\|\|\|\|\|\|\|\|\|\|\|\|\|\|\|\|\|\|\|\|\|\|\|\|\|\|\|\|\|\|\|\|\|\|\|\|\|\|\|\|\|\|\|\|\|\|\|\|\|\|\|\|\|\|\|\|\|\|\|\|\|\|\|\|\|\|\|\|\|\|\|\|\|\|\|\|\|\|\|\|\|\|\|\|\|\|\|\|\|\|\|\|\|\|\|\|\|\|\|\|\|\|\|\|\|\|\|\|\|\|\|\|\|\|\|\|\|\|\|\|\|\|\|\|\|\|\|\|\|\|\|\|\|\|\|\|\|\|\|\|\|\|\|\|\|\|\|\|\|\|\|\|\|\|\|\|\|\|\|\|\|\|\|\|\|\|\|\|\|\|\|\|\|\|\|\|\|\|\|\|\|\|\|\|\|\|\|\|\|\|\|\|\|\|\|\|\|\|\|\|\|\|\|\|\|\|\|\|\|\|\|\|\|\|\|\|\|\|\|\|\|\|\|\|\|\|\|\|\|\|\|\|\|\|\|\|\|\|\|\|\|\|\|\|\|\|\|\|\|\|\|\|\|\|\|\|\|\|\|\|\|\|\|\|\|\|\|\|\|\|\|\|\|\|\|\|\|\|\|\|\|\|\|\|\|\|\|\|\|\|\|\|\|\|\|\|\|\|\|\|\|\|\|\|\|\|\|\|\|\|\|\|\|\|\|\|\|\|\|\|\|\|\|\|\|\|\|\|\|\|\|\|\|\|\|\|\|\|\|\|\|\|\|\|\|\|\|\|\|\|\|\|\|\|\|\|\|\| \|\|\|\|\|\|\|\|\|\|\|\|\|\|\|\|\|\|\|\|\|\|\|\|\|\|\|\|\|\|\|\|\|\|\|\| \|\|\|\|\|\|\|\|\|\|\|\|\|\|\|\|\|\|\|\|\| \|\|\|\|  \|\|\|\|\|\|\|\|\|\|\|\|\| \|\|\|\|\|\|\|\| \|\|\|\|\|\|\|\|\| \|\| \|\| \|\|\|\| \| \|\|\|\|\|\| \| \|\|\|\|\|\|\|\|\|\|\|\|\|\|\| 	AGACCACCCTCACAGCAGCCATCACCAAGCGACAAGCAGAAAAGGGCTTTGCCAACTTCCTCGAATATGGCGCCATTGACAAGGCTCCCGAGGAGCGCAAGCGCGGTATCACCATCTCCACCGCCCACATCGAATACCAGACCGACAACCGCCACTACGCCCACGTCGACTGCCCCGGTCACGCCGATTACATCAAGAACATGATCACAGGAGCTGCGTCCATGGACGGAGCCATCATCGTGGTGGCCGCCAGCGACGGACAAATGCCTCAGACTCGCGAGCACTTGCTCCTCGCCCGCCAAGTTGGTGTGAAGAGAATCGTAGTCTTCGTCAACAAGGTCGATGCGATCGAGGACAAGGAGATGTTGGAGCTCGTCGAGATGGAGATGCGCGAGCTGCTCTCCTCGTACGGCTTCGAGGGCGACGAGACTCCCATCATCATGGGTTCCGCTCTTTGCGCTCTCGACGGCCGCCAGCCAGACATTGGCGTTACCCAAATCGACAACCTCCTCGAAGCCGTTGACACCTGGATTCCCACACCCGAGCGAGACACCGAC-AGCCCTTCCTCATGTCCGTCGAGGACGTCTTCTCCAGTCCCGGCCGTGGCACAGTCGC-TCTGT-CGTGTCGAGCGTG-AGTGCTCA-GCGCGACTC-GA-GT-GAGT-G-TCGGCA-G-ACGCCGCGCCCATCA ```      ``` 48	ATGTCCTTCGCTCGGAATCTCCTGCGCACCACGCGCGCCCTCCGCACCACCCAGAGCCCTCTCCAGTCCGCACTCTCCACTCGAGGACAGCAACAGTTCGCCGCCACCGCCGCATACACAGCACGAGGATATGCCACCGTATTTGAGCGCAGCAAGCCTCACGTCAATATTGGAACCATTGGCCACGTCGATCACGGAAAG	248 	\|\|\|\|\|\|\|\|\|\|\|\|\|\|\|\|\|\|\|\|\|\|\|\|\|\|\|\|\|\|\|\|\|\|\|\|\|\|\|\|\|\|\|\|\|\|\|\|\|\|\|\|\|\|\|\|\|\|\|\|\|\|\|\|\|\|\|\|\|\|\|\|\|\|\|\|\|\|\|\|\|\|\|\|\|\|\|\|\|\|\|\|\|\|\|\|\|\|\|\|\|\|\|\|\|\|\|\|\|\|\|\|\|\|\|\|\|\|\|\|\|\|\|\|\|\|\|\|\|\|\|\|\|\|\|\|\|\|\|\|\|\|\|\|\|\|\|\|\|\|\|\|\|\|\|\|\|\|\|\|\|\|\|\|\|\|\|\|\|\|\|\|\|\|\|\|\|\|\|\|\|\|\|\|\|\|\|\|\|\|\|\|\|\|\|\|\|\|\|\|\| 	ATGTCCTTCGCTCGGAATCTCCTGCGCACCACGCGCGCCCTCCGCACCACCCAGAGCCCTCTCCAGTCCGCACTCTCCACTCGAGGACAGCAACAGTTCGCCGCCACCGCCGCATACACAGCACGAGGATATGCCACCGTATTTGAGCGCAGCAAGCCTCACGTCAATATTGGAACCATTGGCCACGTCGATCACGGAAAG ``` |
| 24 | ggggacaagtttgtacaaaaaagcaggcttgatgGCCTCCGAACTTCCCAAGAC, ggggaccactttgtacaagaaagctgggtcATCCATCGTCAAAACAGCCC | Mycgr3\|69016  JGI  Ensembl | KOG0023 Alcohol dehydrogenase, class V; | alcohol dehydrogenase activity; alcohol dehydrogenase activity, iron-dependent;  alcohol dehydrogenase activity, metal ion-independent;  alcohol dehydrogenase activity, zinc-dependent;  oxidoreductase activity;  zinc ion binding; | 100 % |  |  |
| 24 | ggggacaagtttgtacaaaaaagcaggcttgatgCCTGGCCGCACACTTCCGAC, ggggaccactttgtacaagaaagctgggtcAAGCGACTTGAGCACCTTCG | Mycgr3\|69093  JGI  Ensembl | KOG0539 Sphingolipid fatty acid hydroxylase; | F420H2 dehydrogenase activity; heme binding;  malolactic enzyme activity;  regulation of oxidoreductase activity;  sulfur oxygenase reductase activity;  transition metal ion binding; | 100 % |  |  |
| 24 | ggggacaagtttgtacaaaaaagcaggcttgatgTCTTCCTCCCCACCATTTCA, ggggaccactttgtacaagaaagctgggtcGGCACTAGTCTCGTACTTTC | Mycgr3\|69990  JGI  Ensembl | KOG0255 Synaptic vesicle transporter SVOP and related transporters (major facilitator superfamily); | integral to membrane; membrane; transport; transporter activity; | 100 % |  |  |
| 24 | ggggacaagtttgtacaaaaaagcaggcttgatgGCGTTCGCTACCTTCGCCGT, ggggaccactttgtacaagaaagctgggtcCTTCTTCCCCACCGTCTCGC | Mycgr3\|70454  JGI  Ensembl | KOG1382 Multiple inositol polyphosphate phosphatase; | acid phosphatase activity; | 100 % |  |  |
| 24 | ggggacaagtttgtacaaaaaagcaggcttgatgGCCTCCAAACAGCGAACCGT, ggggaccactttgtacaagaaagctgggtcAAGCTTTGCTGCCGTCCGAA | Mycgr3\|72975  JGI  Ensembl |  |  | 100 % |  |  |
| 24 | ggggacaagtttgtacaaaaaagcaggcttgatgGGATCCGCTGGCCTTCCTCT, ggggaccactttgtacaagaaagctgggtcGATCAGCGGATTGGCTTTAT | Mycgr3\|73191  JGI  Ensembl |  |  | 0 % |  |  |
| 24 | ggggacaagtttgtacaaaaaagcaggcttgatgCCTGCACCAAAATTACAGCA, ggggaccactttgtacaagaaagctgggtcCTTCCTCCTCGCCATGACCT | Mycgr3\|73426  JGI  Ensembl | KOG0776 Geranylgeranyl pyrophosphate synthase/Polyprenyl synthetase; | (S)-2,3-di-O-geranylgeranylglyceryl phosphate synthase activity; 1,4-dihydroxy-2-naphthoate octaprenyltransferase activity;  4-hydroxybenzoate octaprenyltransferase activity;  Sep-tRNA:Cys-tRNA synthase activity;  isoprenoid biosynthetic process;  prenyltransferase activity;  protein geranylgeranyltransferase activity;  protein prenyltransferase activity;  protoheme IX farnesyltransferase activity; | 100 % |  | ``` 38	ATGCCTGCACCAAAATTACAGCACTCGCGTGCTTTGGTTGGGAAAGTCGAGACCTTAGCCACATCCGCATTCCGACCACGATTACGATGTCCGCAATGCCTCCTACCCATCTCCAACCGCACATTCCATAATGACCGACGGAAAGAGTATCATGATAGTACGCGACGGAGTAACGCTGGTGTTGCAGCAGCGGTGTCGGCAGCGAAGCAGATGTTCAATGGCCCGCCACCTTCAGGACCCATGGGCATTGATCCGTTGAGGAGTGTAGCGAAAGAGATGAAGTTCATGACGGGCAACATACAACAATTATTGGGATCCGGACATCCCATGCTGGACACTGTTGCGAAGTACTACACACAATGCGAAGGGAAATATGTCCGGCCAATGCTAGTGTTACTCATATCGCAAGCAACGGCCCTCCTCCCACGAGCACCACGACCTGCCTCTCCCGATGCCACAAATACACCCTTCGCTTCGGCATCCGTCCTCAACGACGCGAATCCTTCATCACCACTGGTGACAGCAGATACCGTCGAGGCTGACGCAGCTGCGATCGAAAGCGGTGTCCTACCAACACAACGACGACTGGCAGAAATAACCGAACTCATCCACACCGCTTCCCTCCTTCACGACGATGTTATCGACACCTCATTAGCACGGCGAGGCAATCCTTCGGCGAATGTCGCATTTGGCAACAAGATGGCGGTTTTGGCTGGGGACTTCCTGCTCGGGAGAGCATCAGTAGCGATGGCACGGCTACGCGACCCGGAAGTGACGGAGCTGCTCGCGACAGTCATTGCAAATCTGGTCGAGGGCGAGTTCATGCAATTGAAAAACACGGCTCTTGACGAGAAGCACCCAACATGGAGTGAGGACACAATAAGCTATTACCTACAAAAGACATACCTCAAGTCCGCCAG	957 	\|\|\|\|\|\|\|\|\|\|\|\|\|\|\|\|\|\|\|\|\|\|\|\|\|\|\|\|\|\|\|\|\|\|\|\|\|\|\|\|\|\|\|\|\|\|\|\|\|\|\|\|\|\|\|\|\|\|\|\|\|\|\|\|\|\|\|\|\|\|\|\|\|\|\|\|\|\|\|\|\|\|\|\|\|\|\|\|\|\|\|\|\|\|\|\|\|\|\|\|\|\|\|\|\|\|\|\|\|\|\|\|\|\|\|\|\|\|\|\|\|\|\|\|\|\|\|\|\|\|\|\|\|\|\|\|\|\|\|\|\|\|\|\|\|\|\|\|\|\|\|\|\|\|\|\|\|\|\|\|\|\|\|\|\|\|\|\|\|\|\|\|\|\|\|\|\|\|\|\|\|\|\|\|\|\|\|\|\|\|\|\|\|\|\|\|\|\|\|\|\|\|\|\|\|\|\|\|\|\|\|\|\|\|\|\|\|\|\|\|\|\|\|\|\|\|\|\|\|\|\|\|\|\|\|\|\|\|\|\|\|\|\|\|\|\|\|\|\|\|\|\|\|\|\|\|\|\|\|\|\|\|\|\|\|\|\|\|\|\|\|\|\|\|\|\|\|\|\|\|\|\|\|\|\|\|\|\|\|\|\|\|\|\|\|\|\|\|\|\|\|\|\|\|\|\|\|\|\|\|\|\|\|\|\|\|\|\|\|\|\|\|\|\|\|\|\|\|\|\|\|\|\|\|\|\|\|\|\|\|\|\|\|\|\|\|\|\|\|\|\|\|\|\|\|\|\|\|\|\|\|\|\|\|\|\|\|\|\|\|\|\|\|\|\|\|\|\|\|\|\|\|\|\|\|\|\|\|\|\|\|\|\|\|\|\|\|\|\|\|\|\|\|\|\|\|\|\|\|\|\|\|\|\|\|\|\|\|\|\|\|\|\|\|\|\|\|\|\|\|\|\|\|\|\|\|\|\|\|\|\|\|\|\|\|\|\|\|\|\|\|\|\|\|\|\|\|\|\|\|\|\|\|\|\|\|\|\|\|\|\|\|\|\|\|\|\|\|\|\|\|\|\|\|\|\|\|\|\|\|\|\|\|\|\|\|\|\|\|\|\|\|\|\|\|\|\|\|\|\|\|\|\|\|\|\|\|\|\|\|\|\|\|\|\|\|\|\|\|\|\|\|\|\|\|\|\|\|\|\|\|\|\|\|\|\|\|\|\|\|\|\|\|\|\|\|\|\|\|\|\|\|\|\|\|\|\|\|\|\|\|\|\|\|\|\|\|\|\|\|\|\|\|\|\|\|\|\|\|\|\|\|\|\|\|\|\|\|\|\|\|\|\|\|\|\|\|\|\|\|\|\|\|\|\|\|\|\|\|\|\|\|\|\|\|\|\|\|\|\|\|\|\|\|\|\|\|\|\|\|\|\|\|\|\|\|\|\|\|\|\|\|\|\|\|\|\|\|\|\|\|\|\|\|\|\|\|\|\|\|\|\|\|\|\|\|\|\|\|\|\|\|\|\|\|\|\|\|\|\|\|\|\|\|\|\|\|\|\|\|\|\|\|\|\|\|\|\|\|\|\|\|\|\|\|\|\|\|\|\|\|\|\|\|\|\|\|\|\|\|\|\|\|\|\|\|\|\|\|\|\|\|\|\|\|\|\|\|\|\|\|\|\|\|\|\|\|\|\|\|\|\|\|\|\|\|\|\|\|\|\|\|\|\|\|\|\|\|\|\|\|\|\|\|\|\|\|\|\|\|\|\|\|\|\|\|\|\|\|\|\|\|\|\|\|\|\|\|\|\|\|\|\|\|\|\|\|\|\|\|\|\|\|\|\|\|\|\|\|\|\| \|\|\|\|\|\|\|\|\|\|\|\|\|\|\|\|\|\|\|\|\| \|\|\|\|\|\|\| \|\|\|\|\|\|\|\|\|\| \|\|\|\|\|\|\|\|\|\|\|\|\|\|\|\|\|\|\|\|\|\|\|\|\|\|\|\|\|\|\|\|\|\|\|\|\|\|\|\|\|\|\|\|\|\|\| 	ATGCCTGCACCAAAATTACAGCACTCGCGTGCTTTGGTTGGGAAAGTCGAGACCTTAGCCACATCCGCATTCCGACCACGATTACGATGTCCGCAATGCCTCCTACCCATCTCCAACCGCACATTCCATAATGACCGACGGAAAGAGTATCATGATAGTACGCGACGGAGTAACGCTGGTGTTGCAGCAGCGGTGTCGGCAGCGAAGCAGATGTTCAATGGCCCGCCACCTTCAGGACCCATGGGCATTGATCCGTTGAGGAGTGTAGCGAAAGAGATGAAGTTCATGACGGGCAACATACAACAATTATTGGGATCCGGACATCCCATGCTGGACACTGTTGCGAAGTACTACACACAATGCGAAGGGAAATATGTCCGGCCAATGCTAGTGTTACTCATATCGCAAGCAACGGCCCTCCTCCCACGAGCACCACGACCTGCCTCTCCCGATGCCACAAATACACCCTTCGCTTCGGCATCCGTCCTCAACGACGCGAATCCTTCATCACCACTGGTGACAGCAGATACCGTCGAGGCTGACGCAGCTGCGATCGAAAGCGGTGTCCTACCAACACAACGACGACTGGCAGAAATAACCGAACTCATCCACACCGCTTCCCTCCTTCACGACGATGTTATCGACACCTCATTAGCACGGCGAGGCAATCCTTCGGCGAATGTCGCATTTGGCAACAAGATGGCGGTTTTGGCTGGGGACTTCCTGCTCGGGAGAGCATCAGTAGCGATGGCACGGCTACGCGACCCGGAAGTGACGGAGCTGCTCGCGACAGTCATTGCAAATCTGGTCGAGGGCGAGTTCATGCAATTG-AAAACACGGCTCTTGACGAGA-GCACCCA-CATGGAGTGA-GACACAATAAGCTATTACCTACAAAAGACATACCTCAAGTCCGCCAG ``` |
| 24 | ggggacaagtttgtacaaaaaagcaggcttgatgGACAATCGGGCGCACACCTC, ggggaccactttgtacaagaaagctgggtcCACCAACCCATCATTCTTCT | Mycgr3\|74263  JGI  Ensembl | KOG0224 Aquaporin (major intrinsic protein family); | membrane; transport; transporter activity; | 100 % |  |  |
| 24 | ggggacaagtttgtacaaaaaagcaggcttgatgGCACCGCCACTCCAGGAATC, ggggaccactttgtacaagaaagctgggtcACCCAGCAAAAAGGCGTTGA | Mycgr3\|74367  JGI  Ensembl | KOG2634 Initiator tRNA phosphoribosyl-transferase; | 7-cyano-7-deazaguanine tRNA-ribosyltransferase activity; NAD(P)-asparagine ADP-ribosyltransferase activity;  NAD(P)-cysteine ADP-ribosyltransferase activity;  NAD(P)-serine ADP-ribosyltransferase activity;  imidazoleglycerol-phosphate synthase activity;  transferase activity, transferring pentosyl groups; | 100 % |  |  |
| 24 | ggggacaagtttgtacaaaaaagcaggcttgatgCCTCGAGTATACCCACCCCT, ggggaccactttgtacaagaaagctgggtcTATCTTCTCCTCCTCCACCG | Mycgr3\|74485  JGI  Ensembl | KOG0764 Mitochondrial FAD carrier protein; | binding; membrane; mitochondrial inner membrane;  transport; | 100 % |  |  |
| 24 | ggggacaagtttgtacaaaaaagcaggcttgatgACGACCACCCTGTCTTACAC, ggggaccactttgtacaagaaagctgggtcCTGACCCTCCTTCTTCTCCC | Mycgr3\|75366  JGI  Ensembl | KOG4179 Lysyl hydrolase/glycosyltransferase family 25; | lipopolysaccharide biosynthetic process; | 100 % |  | ``` 475	CATCGTGGACAAAGGTATCTCCTCGGCAATCATCATGGAAGACGACGTGGACTGGGACATCCGCGTCAAATCCCAAATGCGCACCTACGCCAGAGCCTCAAGACTCCTCATGCAACCTCTTGAGGGTACAACAGACGAATTCCTCGACCCCACCTACCCTCATCCCAGCCCTGGAAATGAAGAAA-CCAAAAACTTCCACCTCGACGAAGAAACCGTCAGCGAACCCACCTCCTCCCCCTACGGCGACCTCCAATACTGGGACGTCCTCTGGCTCGGCCACTGCGGCGTGCGCTTCCCCTGGAAAGACGACAAATCCGTCCCTCTCGGTCGCGTCGTCATCCCCAACGACCCCACCGTCCCGGCGAAAAAGCACATCGACGTCGAATTCGGCAGCCGCCA	873 	\|\|\|\|\|\|\|\|\|\|\|\|\|\|\|\|\|\|\|\|\|\|\|\|\|\|\|\|\|\|\|\|\|\|\|\|\|\|\|\|\|\|\|\|\|\|\|\|\|\|\|\|\|\|\|\|\|\|\|\|\|\|\|\|\|\|\|\|\|\|\|\|\|\|\|\|\|\|\|\|\|\|\|\|\|\|\|\|\|\|\|\|\|\|\|\|\|\|\|\|\|\|\|\|\|\|\|\|\|\|\|\|\|\|\|\|\|\|\|\|\|\|\|\|\|\|\|\|\|\|\|\|\|\|\|\|\|\|\|\|\|\|\|\|\|\|\|\|\|\|\|\|\|\|\|\|\|\|\|\|\|\|\|\|\|\|\|\|\|\|\|\|\|\|\|\|\|\|\|\|\|\|\|\|\| \|\|\|\|\|\|\|\|\|\|\|\|\|\|\|\|\|\|\|\|\|\|\|\|\|\|\|\|\|\|\|\|\|\|\|\|\|\|\|\|\|\|\|\|\|\|\|\|\|\|\|\|\|\|\|\|\|\|\|\|\|\|\|\|\|\|\|\|\|\|\|\|\|\|\|\|\|\|\|\|\|\|\|\|\|\|\|\|\|\|\|\|\|\|\|\|\|\| \|\|\|\|\|\|\|\|\|\|\|\|\|\|\|\| \|\| \|\|\|\|\|\|\|\| \| \|\|\|\|\|\|\|\|\|\|\|\|\|\|\|\|\|\|\|\|\|\|\|\|\|\|\| \| \|\|\|\|\|\|\|\| \|\|\|\|\|\|\|\|\|\|\|\|\|\|  \|\|\|\|\|\|\|\|\|\|\|\|\| \|\|\|\|\|\|\|\|\|\|\|\|\|\|\| 	CATCGTGGACAAAGGTATCTCCTCGGCAATCATCATGGAAGACGACGTGGACTGGGACATCCGCGTCAAATCCCAAATGCGCACCTACGCCAGAGCCTCAAGACTCCTCATGCAACCTCTTGAGGGTACAACAGACGAATTCCTCGACCCCACCTACCCTCATCCCAGCCCTGGAAATGAAGAAACCCAAAAACTTCCACCTCGACGAAGAAACCGTCAGCGAACCCACCTCCTCCCCCTACGGCGACCTCCAATACTGGGACGTCCTCTGGCTCGGCCACTGC-GCGTGCGCTTCCCCTG-AA-GACGACAA-T-CGTCCCTCTCGGTCGCGTCGTCATCCC-A-CGACCCCA-CGTCCCGGCGAAAAGCCACATCGACGTCG-ATTCGGCAGCCGCCA ```      ``` 5	ATGACGACCACCCTGTCTTACACTCTCACGAGGTTCAGCTTTCTTTTCACCGGCCTCATTATCATTGTTCTCTGTTATTTTGGATACGAAAATGTTGCAGCCACGGTGGGAGATCCCTCGCCGCTGGCGAAGATCGCGCACACGAGCAATACACAGACACACAGAGTACAGTCGGAGAGAGTACAGATGGACAGAGTAACGCCAGAGCCGGTACAGATCAAGCACAGTCTCCTTGATGACACTATGAATGCAACACTGGGG	265 	\|\|\|\|\|\|\|\|\|\|\|\|\|\|\|\|\|\|\|\|\|\|\|\|\|\|\|\|\|\|\|\|\|\|\|\|\|\|\|\|\|\|\|\|\|\|\|\|\|\|\|\|\|\|\|\|\|\|\|\|\|\|\|\|\|\|\|\|\|\|\|\|\|\|\|\|\|\|\|\|\|\|\|\|\|\|\|\|\|\|\|\|\|\|\|\|\|\|\|\|\|\|\|\|\|\|\|\|\|\|\|\|\|\|\|\|\|\|\|\|\|\|\|\|\|\|\|\|\|\|\|\|\|\|\|\|\|\|\|\|\|\|\|\|\|\|\|\|\|\|\|\|\|\|\|\|\|\|\|\|\|\|\|\|\|\|\|\|\|\|\|\|\|\|\|\|\|\|\|\|\|\|\|\|\|\|\|\|\|\|\|\|\|\|\|\|\|\|\|\|\|\|\|\|\|\|\|\|\|\|\|\|\|\|\|\|\|\|\|\|\|\|\|\|\|\|\|\|\|\|\|\|\|\|\|\|\|\|\|\|\|\|\|\|\|\|\|\|\|\|\|\|\|\|\|\|\|\|\|\|\| 	ATGACGACCACCCTGTCTTACACTCTCACGAGGTTCAGCTTTCTTTTCACCGGCCTCATTATCATTGTTCTCTGTTATTTTGGATACGAAAATGTTGCAGCCACGGTGGGAGATCCCTCGCCGCTGGCGAAGATCGCGCACACGAGCAATACACAGACACACAGAGTACAGTCGGAGAGAGTACAGATGGACAGAGTAACGCCAGAGCCGGTACAGATCAAGCACAGTCTCCTTGATGACACTATGAATGCAACACTGGGG ```      ``` 265	GTTTGAGAAGATCTTCATCATCAATCTGGCGACCCGGACAGACCGCAGAGATGCCTTCACTCTCGCTGCTGCTTTCGTCGGACTGCAAGTCGAGGCCGTTGATGCTGCCACTAAAGTTGACGACAAGGCATGGCCTCCGGGTCCCGAGGAGTACCGTCCAAACCCCGGCGGAGCAGGCGCTTGGAGGTCTCATATGAATGTTTTAAGAGA	474 	\|\|\|\|\|\|\|\|\|\|\|\|\|\|\|\|\|\|\|\|\|\|\|\|\|\|\|\|\|\|\|\|\|\|\|\|\|\|\|\|\|\|\|\|\|\|\|\|\|\|\|\|\|\|\|\|\|\|\|\|\|\|\|\|\|\|\|\|\|\|\|\|\|\|\|\|\|\|\|\|\|\|\|\|\|\|\|\|\|\|\|\|\|\|\|\|\|\|\|\|\|\|\|\|\|\|\|\|\|\|\|\|\|\|\|\|\|\|\|\|\|\|\|\|\|\|\|\|\|\|\|\|\|\|\|\|\|\|\|\|\|\|\|\|\|\|\|\|\|\|\|\|\|\|\|\|\|\|\|\|\|\|\|\|\|\|\|\|\|\|\|\|\|\|\|\|\|\|\|\|\|\|\|\|\|\|\|\|\|\|\|\|\|\|\|\|\|\|\|\|\|\|\|\|\|\|\|\|\|\| 	GTTTGAGAAGATCTTCATCATCAATCTGGCGACCCGGACAGACCGCAGAGATGCCTTCACTCTCGCTGCTGCTTTCGTCGGACTGCAAGTCGAGGCCGTTGATGCTGCCACTAAAGTTGACGACAAGGCATGGCCTCCGGGTCCCGAGGAGTACCGTCCAAACCCCGGCGGAGCAGGCGCTTGGAGGTCTCATATGAATGTTTTAAGAGA ``` |
| 24 | ggggacaagtttgtacaaaaaagcaggcttgatgAGTGAGCTTCTGGAGTTCCT, ggggaccactttgtacaagaaagctgggtcAGCAGTGCTCTGCTCAGCCT | Mycgr3\|75652  JGI  Ensembl | KOG2911 Uncharacterized conserved protein; | protein transport; | 100 % |  | ``` 19	ATGAGTGAGCTTCTGGAGTTCCTGCGAACTCACGAAGAAGCTTTCCGTAG	68 	\|\|\|\|\|\|\|\|\|\|\|\|\|\|\|\|\|\|\|\|\|\|\|\|\|\|\|\|\|\|\|\|\|\|\|\|\|\|\|\|\|\|\|\|\|\|\|\|\|\| 	ATGAGTGAGCTTCTGGAGTTCCTGCGAACTCACGAAGAAGCTTTCCGTAG ```      ``` 61	TTCCGTAGTCGTTCACGTCTAGCTTCTCTCTACTCCGACTTTCGATCTCAGCGCCAAACGAATCCCGATGGCTACACAGCAAACTCATCCGCGTGGCTCCGAGCCTTGACCTCCGCAGCGAAGGCAGGTTTACTTCCTTCACCGAGTGGAGCTCAGAACGATCGCTTCGTGTTGAATGGTGGAGAGGAGCTCTTGAGGAACTTGGAAGGGGAATATGGCAGACCGCTGGCTTTGAGTGCTGTGCTGCAAGATGCCATACAAGCTGGAGCGCTCGTGGAACTGAGGACATTTCTGAACAAAAAAGAGAGCTTATACAAGAAGAGATGGGTGCCCACTGTAGGAGAAGTTGTAGGGTGGGGTTTGAGGCAACTGGGCGTGATTGGAGACAGCGAAGACAGACTTGTGAAGGGGAACTTTGTGGTGGTTGCAAACGTGGAGGCGGCGGCGAAAGCAGTACTGGACACGATTTCGAAGACGGCCACGAGTCCTAGCTCACGGATATTTTCACGAGAGATGTTCGCAAATACCTTTGGACCTCACCTCGGAGCACAGGACGGACTAAGTTCGAACGACCTGTCCGTCCTCATGGTGCACCTGGCGCGCGATCGACAGGCAATATCCTACCATGATGCCTCCGGTACCATCAAGTTCGCATCGCCCACCGAACCAGTTCCGTCGCCAGTTACAGAAGAAGATATATCGATAGCTTCACTCCGCTCTCTGATAGGTTCTCTGAGTCCTCAAATCGACCTCCT-CACGGATCAAATAGCAGCTTCAGACAAAGCTGCTCGGGAAGCAGTGAGCAAAAAGCAGCTCCACTCGGCGAAGATTGCGCTACGTCAAAAGAAGGCCGCAGAG-ACAAAGCTCAGCCAGCGAACAG-CGACACTC	948 	\|\|\|\| \|\|\|\|\|\|\|\|\|\|\|\|\|\|\|\|\|\|\|\|\|\|\|\|\|\|\|\|\|\|\|\|\|\|\|\|\|\|\|\|\|\|\|\|\|\|\|\|\|\|\|\|\|\|\|\|\|\|\|\|\|\|\|\|\|\|\|\|\|\|\|\|\|\|\|\|\|\|\|\|\|\|\|\|\|\|\|\|\|\|\|\|\|\|\|\|\|\|\|\|\|\|\|\|\|\|\|\|\|\|\|\|\|\|\|\|\|\|\|\|\|\|\|\|\|\|\|\|\|\|\|\|\|\|\|\|\|\|\|\|\|\|\|\|\|\|\|\|\|\|\|\|\|\|\|\|\|\|\|\|\|\|\|\|\|\|\|\|\|\|\|\|\|\|\|\|\|\|\|\|\|\|\|\|\|\|\|\|\|\|\|\|\|\|\|\|\|\|\|\|\|\|\|\|\|\|\|\|\|\|\|\|\|\|\|\|\|\|\|\|\|\|\|\|\|\|\|\|\|\|\|\|\|\|\|\|\|\|\|\|\|\|\|\|\|\|\|\|\|\|\|\|\|\|\|\|\|\|\|\|\|\|\|\|\|\|\|\|\|\|\|\|\|\|\|\|\|\|\|\|\|\|\|\|\|\|\|\|\|\|\|\|\|\|\|\|\|\|\|\|\|\|\|\|\|\|\|\|\|\|\|\|\|\|\|\|\|\|\|\|\|\|\|\|\|\|\|\|\|\|\|\|\|\|\|\|\|\|\|\|\|\|\|\|\|\|\|\|\|\|\|\|\|\|\|\|\|\|\|\|\|\|\|\|\|\|\|\|\|\|\|\|\|\|\|\|\|\|\|\|\|\|\|\|\|\|\|\|\|\|\|\|\|\|\|\|\|\|\|\|\|\|\|\|\|\|\|\|\|\|\|\|\|\|\|\|\|\|\|\|\|\|\|\|\|\|\|\|\|\|\|\|\|\|\|\|\|\|\|\|\|\|\|\|\|\|\|\|\|\|\|\|\|\|\|\|\|\|\|\|\|\|\|\|\|\|\|\|\|\|\|\|\|\|\|\|\|\|\|\|\|\|\|\|\|\|\|\|\|\|\|\|\|\|\|\|\|\|\|\|\|\|\|\|\|\|\|\|\|\|\|\|\|\|\|\|\|\|\|\|\|\|\|\|\|\|\|\|\|\|\|\|\|\|\|\|\|\|\|\|\|\|\|\|\|\|\|\|\|\|\|\|\|\|\|\|\|\|\|\|\|\|\|\|\|\|\|\|\|\|\|\|\|\|\|\|\|\|\|\|\|\|\|\|\|\|\|\|\|\|\|\|\|\|\|\|\|\|\|\|\|\|\|\|\|\|\|\|\|\|\|\|\|\|\|\|\|\|\|\|\|\|\|\|\|\|\|\|\|\|\|\|\|\|\|\|\|\|\|\|\|\|\|\|\|\|\|\|\|\|\|\|\|\|\|\|\|\|\|\|\|\|\|\|\|\|\|\|\|\|\|\|\|\|\|\|\|\|\|\|\|\|\|\|\|\|\|\|\|\|\|\|\|\|\|\|\|\|\|\|\|\|\|\|\|\|\|\|\|\|\|\|\|\|\|\|\|\|\|\|\|\|\|\|\|\|\|\|\|\|\|\|\|\|\|\|\|\|\|\|\|\|\|\|\|\|\|\|\|\| \|\|\|\|\|\|\|\|\|\|\|\|\|\|\|\|\|\|\|\|\|\|\|\|\|\|\|\|\|\|\|\|\|\|\|\|\|\| \|\|\|\|\|\|\|\|\|\|\|\|\|\| \|\|\|\|\|\|\| \|\|\|\|\|\|\|\| \|\|\|\|\|\|\|\|\|\|\|\|\|\|\|\|\|\| \|\|\|\| \|\|\|\|\|\|\|\| \|\|\|\|\|\|\|\|\|\|\|\|\|\|\|\|\|\|\|\|\|\| \|\|\|\|\|\|\|\| 	TTCCATAGTCGTTCACGTCTAGCTTCTCTCTACTCCGACTTTCGATCTCAGCGCCAAACGAATCCCGATGGCTACACAGCAAACTCATCCGCGTGGCTCCGAGCCTTGACCTCCGCAGCGAAGGCAGGTTTACTTCCTTCACCGAGTGGAGCTCAGAACGATCGCTTCGTGTTGAATGGTGGAGAGGAGCTCTTGAGGAACTTGGAAGGGGAATATGGCAGACCGCTGGCTTTGAGTGCTGTGCTGCAAGATGCCATACAAGCTGGAGCGCTCGTGGAACTGAGGACATTTCTGAACAAAAAAGAGAGCTTATACAAGAAGAGATGGGTGCCCACTGTAGGAGAAGTTGTAGGGTGGGGTTTGAGGCAACTGGGCGTGATTGGAGACAGCGAAGACAGACTTGTGAAGGGGAACTTTGTGGTGGTTGCAAACGTGGAGGCGGCGGCGAAAGCAGTACTGGACACGATTTCGAAGACGGCCACGAGTCCTAGCTCACGGATATTTTCACGAGAGATGTTCGCAAATACCTTTGGACCTCACCTCGGAGCACAGGACGGACTAAGTTCGAACGACCTGTCCGTCCTCATGGTGCACCTGGCGCGCGATCGACAGGCAATATCCTACCATGATGCCTCCGGTACCATCAAGTTCGCATCGCCCACCGAACCAGTTCCGTCGCCAGTTACAGAAGAAGATATATCGATAGCTTCACTCCGCTCTCTGATAGGTTCTCTGAGTCCTCAAATCGACCTCCTCCACGGATCAAATAGCAGCTTCAGACAAAGCTGCTCGGGGAGCAGTGAGCAAAA-GCAGCTC-ACTCGGCG-AGATTGCGCTACGTCAAA-GAAG-CCGCAGAGAACAAAGCTCAGCCAGCGAACAGGCGACACTC ``` |
| 24 | ggggacaagtttgtacaaaaaagcaggcttgatgGCGGAGTCGACTTCCACCGC, ggggaccactttgtacaagaaagctgggtcGATCAGACCAAACTCCTTGA | Mycgr3\|77663  JGI  Ensembl | KOG1502 Flavonol reductase/cinnamoyl-CoA reductase; |  | 100 % |  |  |
| 24 | ggggacaagtttgtacaaaaaagcaggcttgatgTCCACCCTCATATGGCCTCC, ggggaccactttgtacaagaaagctgggtcCGCATCGTCTGTATCTCCGT | Mycgr3\|77725  JGI  Ensembl |  |  | 100 % |  |  |
| 24 | ggggacaagtttgtacaaaaaagcaggcttgatgCTCTTTGTTCATCAGGCGCT, ggggaccactttgtacaagaaagctgggtcGTCCCACAGGACACCTCCAA | Mycgr3\|77835  JGI  Ensembl | KOG1339 Aspartyl protease; | pepsin A activity; proteolysis; | 100 % |  |  |
| 24 | ggggacaagtttgtacaaaaaagcaggcttgatgATCTCCAAAGCCGCAGCGAC, ggggaccactttgtacaagaaagctgggtcCAAGGATACATTCTTCGTAT | Mycgr3\|86584  JGI  Ensembl | KOG2614 Kynurenine 3-monooxygenase and related flavoprotein monooxygenases; | metabolic process; oxidoreductase activity;  salicylate 1-monooxygenase activity; | 100 % |  | ``` 1	ATGATCTCCAAAGCCGCAGCGACCGAGCCAAGAATCGCGATTGTCGGAGCCGGAGTGGGTGGTCTGGTCACTGCTCTTCATCTCCACGCACATGGATTCACTGCCATCGACATCTTCGAAGCCGCTGGACAACTGGTCTCTCTTGGAGTCGGCATCAATGTTCAACCGCATGCGGTCCTTGTCTTGAGAAATCTCGGTCTGCTGCCCAGGTTGGAGGCTATCGGCGTCGAGACCCAGG	238 	\|\|\|\|\|\|\|\|\|\|\|\|\|\|\|\|\|\|\|\|\|\|\|\|\|\|\|\|\|\|\|\|\|\|\|\|\|\|\|\|\|\|\|\|\|\|\|\|\|\|\|\|\|\|\|\|\|\|\|\|\|\|\|\|\|\|\|\|\|\|\|\|\|\|\|\|\|\|\|\|\|\|\|\|\|\|\|\|\|\|\|\|\|\|\|\|\|\|\|\|\|\|\|\|\|\|\|\|\|\|\|\|\|\|\|\|\|\|\|\|\|\|\|\|\|\|\|\|\|\|\|\|\|\|\|\|\|\|\|\|\|\|\|\|\|\|\|\|\|\|\|\|\|\|\|\|\|\|\|\|\|\|\|\|\|\|\|\|\|\|\|\|\|\|\|\|\|\|\|\|\|\|\|\|\|\|\|\|\|\|\|\|\|\|\|\|\|\|\|\|\|\|\|\|\|\|\|\|\|\|\|\|\|\|\|\|\|\|\|\|\|\|\|\|\|\|\|\|\|\|\|\|\|\|\|\|\|\| 	ATGATCTCCAAAGCCGCAGCGACCGAGCCAAGAATCGCGATTGTCGGAGCCGGAGTGGGTGGTCTGGTCACTGCTCTTCATCTCCACGCACATGGATTCACTGCCATCGACATCTTCGAAGCCGCTGGACAACTGGTCTCTCTTGGAGTCGGCATCAATGTTCAACCGCATGCGGTCCTTGTCTTGAGAAATCTCGGTCTGCTGCCCAGGTTGGAGGCTATCGGCGTCGAGACCCAGG ```      ``` 236	AGGAACTCAACTACTACGACAGACATGGCAACCCAATCATCAGCGAACCCAGGGGCAAATTTGCAGGGTACAAAGTGCCGCAGTTCTCCATCCACCGAGGAGAGCTACAACTCATGCTTCTTGATGCTGTGAAAGAACGACTTGGAGATGATAAAGTCCATCTCAATCATTCCATAGACAGTTTTGAACATCAAGAGAACGGGCAGGTGAAGCTCAGCTTCGTTCAGAAACGTTCCGGCGAGAACGCCGCTATTCCGGAGGTTGTCGTCCGCTTATGTGTGGCCGCCGACGGCATCAATTCGACAGTTCGCCGGATCCTGTATCCCAAAGAAGGACCTCCCAATTTCTCCGGACGCATGCTGTGGCGAGGCTGTGTTGAACGGAAACCGTATTTGACTTCAGCCAGCATGGTATGGACTGGCCATGCCAACCAGAAGTTCATTGCGTATCCGATTCGCAACTACGGGGAGGACGCTGGCTCGAAGTCTCTTGTCAACTGGATAGCCGAGCTGAGAGTTCGAGATGACGATGATCCGGATACCACTCCTCCCGAGAAAACGGACTGGCTGAACGACGTTCCGAAAGAGAGATTTGCAGGCCAGTTCGAGAAGTGGACCTTCGGCTTTCTCGACATTCCGCAGCTCATCGCTGAGAC	890 	\|\|\|\|\|\|\|\|\|\|\|\|\|\|\|\|\|\|\|\|\|\|\|\|\|\|\|\|\|\|\|\|\|\|\|\|\|\|\|\|\|\|\|\|\|\|\|\|\|\|\|\|\|\|\|\|\|\|\|\|\|\|\|\|\|\|\|\|\|\|\|\|\|\|\|\|\|\|\|\|\|\|\|\|\|\|\|\|\|\|\|\|\|\|\|\|\|\|\|\|\|\|\|\|\|\|\|\|\|\|\|\|\|\|\|\|\|\|\|\|\|\|\|\|\|\|\|\|\|\|\|\|\|\|\|\|\|\|\|\|\|\|\|\|\|\|\|\|\|\|\|\|\|\|\|\|\|\|\|\|\|\|\|\|\|\|\|\|\|\|\|\|\|\|\|\|\|\|\|\|\|\|\|\|\|\|\|\|\|\|\|\|\|\|\|\|\|\|\|\|\|\|\|\|\|\|\|\|\|\|\|\|\|\|\|\|\|\|\|\|\|\|\|\|\|\|\|\|\|\|\|\|\|\|\|\|\|\|\|\|\|\|\|\|\|\|\|\|\|\|\|\|\|\|\|\|\|\|\|\|\|\|\|\|\|\|\|\|\|\|\|\|\|\|\|\|\|\|\|\|\|\|\|\|\|\|\|\|\|\|\|\|\|\|\|\|\|\|\|\|\|\|\|\|\|\|\|\|\|\|\|\|\|\|\|\|\|\|\|\|\|\|\|\|\|\|\|\|\|\|\|\|\|\|\|\|\|\|\|\|\|\|\|\|\|\|\|\|\|\|\|\|\|\|\|\|\|\|\|\|\|\|\|\|\|\|\|\|\|\|\|\|\|\|\|\|\|\|\|\|\|\|\|\|\|\|\|\|\|\|\|\|\|\|\|\|\|\|\|\|\|\|\|\|\|\|\|\|\|\|\|\|\|\|\|\|\|\|\|\| \|\|\|\|\|\|\|\|\|\|\|\|\|\|\|\|\|\|\|\|\|\|\|\|\|\|\|\|\|\|\|\|\|\|\|\|\|\|\|\|\|\|\|\|\|\|\|\|\|\|\|\|\|\|\|\|\|\|\|\|\|\|\|\|\|\|\|\|\|\|\|\|\|\|\|\|\|\|\|\|\|\|\|\|\|\|\|\|\|\|\|\| \|\|\|\|\|\|\|\|\|\|\|\|\|\|\|\|\|\|\|\|\|\|\|\|\|\| \|\|\|\|\| \|\|\|\|\|\|\|\|\|\| \|\|\|\|\|\|\|  \|\|\| \|\|\|\|\|\| \| \|\|\| \|\| \|\|\|\|\| \|\|\| \|\| \|\| \|\|\|\|\|\| \|\|\| \|\| \|\|\|\|\|\| \|\|\|\|\|\|\|\|\|\|\|\|\|\|\|\|\|\|\|\|\|\|\|\|\|\|\|\|\|\|\| 	AGGAACTCAACTACTACGACAGACATGGCAACCCAATCATCAGCGAACCCAGGGGCAAATTTGCAGGGTACAAAGTGCCGCAGTTCTCCATCCACCGAGGAGAGCTACAACTCATGCTTCTTGATGCTGTGAAAGAACGACTTGGAGATGATAAAGTCCATCTCAATCATTCCATAGACAGTTTTGAACATCAAGAGAACGGGCAGGTGAAGCTCAGCTTCGTTCAGAAACGTTCCGGCGAGAACGCCGCTATTCCGGAGGTTGTCGTCCGCTTATGTGTGGCCGCCGACGGCATCAATTCGACAGTTCGCCGGATCCTGTATCCCAAAGAAGGACCTCCCAATTTCTCCGGACGCATGCTGTGGCGAGGCTGTGTTGAACGGAAACCGTATTTGACTTCAGCCAGCATGGTATGGACTGTCCATGCCAACCAGAAGTTCATTGCGTATCCGATTCGCAACTACGGGGAGGACGCTGGCTCGAAGTCTCTTGTCAACTGGATAGCCGAGCTGACAGTTCGAGATGACGATGATCCGGATA-CACTCGTCCCGAGAAA-CGGACTGA-TGA-CGACGT-CAGAA-GATAGATT-GCATGC-AGCTCGAGA-GTG-AC-TTCGGC-TTCTCGACATTCCGCAGCTCATCGCTGAGAC ``` |
| 24 | ggggacaagtttgtacaaaaaagcaggcttgatgTCGTGGCTTAGCCCGAAGCG, ggggaccactttgtacaagaaagctgggtcACGATCATACACAAAACCCA | Mycgr3\|88597  JGI  Ensembl | KOG4568 Cytoskeleton-associated protein and related proteins; |  | 100 % |  |  |
| 24 | ggggacaagtttgtacaaaaaagcaggcttgatgTCTCACTCTTCCGGGGAGGC, ggggaccactttgtacaagaaagctgggtcACCAATCCAGACTCGTCTTG | Mycgr3\|88868  JGI  Ensembl | KOG0039 Ferric reductase, NADH/NADPH oxidase and related proteins; |  | 100 % |  |  |
| 24 | ggggacaagtttgtacaaaaaagcaggcttgatgCCAGCGGACGCACGACTTCT, ggggaccactttgtacaagaaagctgggtcGCCAGCCGATCGATGAACAG | Mycgr3\|90148  JGI  Ensembl |  |  | 100 % |  |  |
| 24 | ggggacaagtttgtacaaaaaagcaggcttgatgACTCTGCCAATGTACTCGAT, ggggaccactttgtacaagaaagctgggtcTCCAGTGACCCGTCTGACCC | Mycgr3\|90485  JGI  Ensembl |  |  | 100 % |  |  |
| 24 | ggggacaagtttgtacaaaaaagcaggcttgatgTCTTTGCCGCCACCTAGCCA, ggggaccactttgtacaagaaagctgggtcAGATGTCGTGGTCAGCAATT | Mycgr3\|90527  JGI  Ensembl |  |  | 100 % |  |  |
| 24 | ggggacaagtttgtacaaaaaagcaggcttgatgGCCGAAGCACTGTTGCTGGA, ggggaccactttgtacaagaaagctgggtcCGTGTCCACGGAGCTCTCCG | Mycgr3\|91167  JGI  Ensembl |  |  | 100 % |  |  |
| 24 | ggggacaagtttgtacaaaaaagcaggcttgatgGGCCGCGAACTCAATAGTCT, ggggaccactttgtacaagaaagctgggtcTCCCTTGTAGTTCTTACAGC | Mycgr3\|92002  JGI  Ensembl | KOG0371 Serine/threonine protein phosphatase 2A, catalytic subunit; | bis(5'-nucleosyl)-tetraphosphatase (symmetrical) activity; hydrolase activity; | 100 % |  |  |
| 24 | ggggacaagtttgtacaaaaaagcaggcttgatgACAGCCATCACTGACTCTCA, ggggaccactttgtacaagaaagctgggtcCTTGTCCGTCTCCTCCAGCA | Mycgr3\|92100  JGI  Ensembl | KOG2700 Adenylosuccinate lyase; | 3-carboxy-cis,cis-muconate cycloisomerase activity; catalytic activity; | 100 % |  | ``` 331	CAGGACCTCACCGACACAGCCACAGTCCTCCAACTCCGAGACACCATCGTCCTCCTCGAGAAGTCCCTCGACACCATCGCCAACGCTCTAACCAAGCTCTGCAAAGACCACAAAACCACCCCAATGGCCGCCCGCAGCAACCTCCAACAAGCGGTCCCCATCTCCTTCGGCTTCAAAATGGCCCGTCTCCTCGCAACTTTCAACCGCCACCGTCACCGCCTCCTCGAACTCAAACCCCGTCTCCTCGTCCTCGAATTCTCCGGCGCAGCAGGGACCCTCGCGACCATCACGCCTTCCACATCCTACTCCCCCGCTGCGGAAACATCCACCCCTCTCGCTCTCCAATGCCAAACTCTCCTCGCCAGACACCTCGACCTCGCCGTTCCCAGTATAGCCTGGCACACCGAACGCGACACCTTCGCCGAGTTCTCCACTTTCCTCTCCCTCCTCACCGCCACCTGCGCGAAATTCGCAACGGATTTGAAATTGATGATGCAATCCGAAGTCAACGAAGCGCGGGAACCGTACGTCGCTCACCGCGGCTCCTCGTCCACCATGCCGCAAAAGCGAAATCCAATCGGATGCGCGTACATCTGCTCCATGGCGAGTTCAGTCC	946 	\|\|\|\|\|\|\|\|\|\|\|\|\|\|\|\|\|\|\|\|\|\|\|\|\|\|\|\|\|\|\|\|\|\|\|\|\|\|\|\|\|\|\|\|\|\|\|\|\|\|\|\|\|\|\|\|\|\|\|\|\|\|\|\|\|\|\|\|\|\|\|\|\|\|\|\|\|\|\|\|\|\|\|\|\|\|\|\|\|\|\|\|\|\|\|\|\|\|\|\|\|\|\|\|\|\|\|\|\|\|\|\|\|\|\|\|\|\|\|\|\|\|\|\|\|\|\|\|\|\|\|\|\|\|\|\|\|\|\|\|\|\|\|\|\|\|\|\|\|\|\|\|\|\|\|\|\|\|\|\|\|\|\|\|\|\|\|\|\|\|\|\|\|\|\|\|\|\|\|\|\|\|\|\|\|\|\|\|\|\|\|\|\|\|\|\|\|\|\|\|\|\|\|\|\|\|\|\|\|\|\|\|\|\|\|\|\|\|\|\|\|\|\|\|\|\|\|\|\|\|\|\|\|\|\|\|\|\|\|\|\|\|\|\|\|\|\|\|\|\|\|\|\|\|\|\|\|\|\|\|\|\|\|\|\|\|\|\|\|\| \|\|\|\|\|\|\|\|\|\|\|\|\|\|\|\|\|\|\|\|\|\|\|\|\|\|\|\|\|\|\|\|\|\|\|\|\|\|\|\|\|\|\|\|\|\|\|\|\|\|\|\|\|\|\|\|\|\|\|\|\|\|\|\|\|\|\|\|\|\|\|\|\|\|\|\|\|\|\|\|\|\|\|\|\|\|\|\|\|\|\|\|\|\|\|\|\|\|\|\|\|\|\|\|\|\|\|\|\|\|\|\|\|\|\|\|\|\|\|\|\|\|\|\|\|\|\|\|\|\|\|\|\|\|\|\|\|\|\|\|\|\|\|\|\|\|\|\|\|\|\|\|\|\|\| \|\|\|\|\|\|\|\|\|\|\|\|\|\|\|\|\|\|\|\|\|\|\|\|\|\|\|\|\|\|\|\|\|\|\|\|\|\|\|\|\| \|\|\|\|\|\|\|\|\|\|\|\|\|\|\|\|\| \|\|\|\|\|\|\|\|\|\|\|\|\|\|\|\| \|\|\|\| \|\|\| \|\|\|\| \|\|\|\|\|\|\|\|\|\|\|\|\|\|\|\|\|\|\| \|\|\| \|\| \|\|\|\|\|\|\|\|\|\|\|\|\|\| \|\|\|\|\| \|\|\|\|\|\|\|\|\| \|\|\|\|\|\|\|\|\|\|\|\|\|\|\|\|\|\|\|\|\|\|\|\|\|\|\|\|\|\|\|\|\| \|\|\|\|\|\| 	CAGGACCTCACCGACACAGCCACAGTCCTCCAACTCCGAGACACCATCGTCCTCCTCGAGAAGTCCCTCGACACCATCGCCAACGCTCTAACCAAGCTCTGCAAAGACCACAAAACCACCCCAATGGCCGCCCGCAGCAACCTCCAACAAGCGGTCCCCATCTCCTTCGGCTTCAAAATGGCCCGTCTCCTCGCAACTTTCAACCGCCACCGTCACCGCCTCCTCGAACTCAAACCCCGTCTCCTCGTCCTCGAATTCTCCGGCGCAGCA-GGACCCTCGCGACCATCACGCCTTCCACATCCTACTCCCCCGCTGCGGAAACATCCACCCCTCTCGCTCTCCAATGCCAAACTCTCCTCGCCAGACACCTCGACCTCGCCGTTCCCAGTATAGCCTGGCACACCGAACGCGACACCTTCGCCGAG-TCTCCACTTTCCTCTCCCTCCTCACCGCCACCTGCGCGAAA-TCGCAACGGATTTGAAA-TGATGATGCAATCCGA-GTCA-CGA-GCGC-GGAACCGTACGTCGCTCAC-GCG-CT-CTCGTCCACCATGC-GCAAA-GCGAAATCC-ATCGGATGCGCGTACATCTGCTCCATGGCGAGT-CAGTCC ```      ``` 1	ATGACAGCCATCACTGACTCTCAAATCTTCGGCAATGTCTTCTCCACGCCGGAGATAGCGAAGATCTGGTCTGACCGCCAGCGAACTCAATATTATCTCGACTTTGAAGCCGCATTGGCAAAAGCACAAGCGGAACTGGGCATTATTCCACAAAAAGCATGCGATGAGATTGTCAAGCATTGCACTTTGGATCAATTGGATTTTGATCTTTTGAGGCAGCAGACGGAATTGATCGGCTATCCGGTCTTGCCGGTGGTTCAGCAGCTGGTCAAGAAGATCAATGCGGTGGAAGATCGGCTGGGAGAATGGATTCATTGGGGGACTACGACGCAGG	334 	\|\|\|\|\|\|\|\|\|\|\|\|\|\|\|\|\|\|\|\|\|\|\|\|\|\|\|\|\|\|\|\|\|\|\|\|\|\|\|\|\|\|\|\|\|\|\|\|\|\|\|\|\|\|\|\|\|\|\|\|\|\|\|\|\|\|\|\|\|\|\|\|\|\|\|\|\|\|\|\|\|\|\|\|\|\|\|\|\|\|\|\|\|\|\|\|\|\|\|\|\|\|\|\|\|\|\|\|\|\|\|\|\|\|\|\|\|\|\|\|\|\|\|\|\|\|\|\|\|\|\|\|\|\|\|\|\|\|\|\|\|\|\|\|\|\|\|\|\|\|\|\|\|\|\|\|\|\|\|\|\|\|\|\|\|\|\|\|\|\|\|\|\|\|\|\|\|\|\|\|\|\|\|\|\|\|\|\|\|\|\|\|\|\|\|\|\|\|\|\|\|\|\|\|\|\|\|\|\|\|\|\|\|\|\|\|\|\|\|\|\|\|\|\|\|\|\|\|\|\|\|\|\|\|\|\|\|\|\|\|\|\|\|\|\|\|\|\|\|\|\|\|\|\|\|\|\|\|\|\|\|\|\|\|\|\|\|\|\|\|\|\|\|\|\|\|\|\|\|\|\|\|\|\|\|\|\|\|\|\|\|\|\|\|\|\|\|\|\|\|\|\|\|\|\|\|\|\|\|\|\|\|\|\|\|\|\|\|\|\|\|\|\|\|\|\|\|\|\|\|\|\|\|\| 	ATGACAGCCATCACTGACTCTCAAATCTTCGGCAATGTCTTCTCCACGCCGGAGATAGCGAAGATCTGGTCTGACCGCCAGCGAACTCAATATTATCTCGACTTTGAAGCCGCATTGGCAAAAGCACAAGCGGAACTGGGCATTATTCCACAAAAAGCATGCGATGAGATTGTCAAGCATTGCACTTTGGATCAATTGGATTTTGATCTTTTGAGGCAGCAGACGGAATTGATCGGCTATCCGGTCTTGCCGGTGGTTCAGCAGCTGGTCAAGAAGATCAATGCGGTGGAAGATCGGCTGGGAGAATGGATTCATTGGGGGACTACGACGCAGG ``` |
| 24 | ggggacaagtttgtacaaaaaagcaggcttgatgTCAACAGAATATACTCTACC, ggggaccactttgtacaagaaagctgggtcAGACTTTGCCTTCAGCCGGA | Mycgr3\|92214  JGI  Ensembl | KOG2064 Poly(ADP-ribose) glycohydrolase; |  | 100 % |  |  |
| 24 | ggggacaagtttgtacaaaaaagcaggcttgatgCGCTTCGCAGGCGCCAAAGC, ggggaccactttgtacaagaaagctgggtcCGAATGATAAATGAGGTGGC | Mycgr3\|92614  JGI  Ensembl |  |  | 100 % |  |  |
| 24 | ggggacaagtttgtacaaaaaagcaggcttgatgCCCAAGGAAGCAAAATCGTC, ggggaccactttgtacaagaaagctgggtcCGGTGCTCCAGGACCTGGGT | Mycgr3\|93101  JGI  Ensembl | KOG1812 Predicted E3 ubiquitin ligase; |  | 0 % |  |  |
| 24 | ggggacaagtttgtacaaaaaagcaggcttgatgATTCAATTTACCGCGCTGAG, ggggaccactttgtacaagaaagctgggtcCGACTGCTTCTCCTCTGTGC | Mycgr3\|93355  JGI  Ensembl | KOG1289 Amino acid transporters; | amino acid transmembrane transporter activity; amino acid transport;  membrane;  transport; | 100 % |  | ``` 1	ATGATTCAATTTACCGCGCTGAGCATGGCAGAACTGTGTTCTAGCATGCCGACTGTAGGAGGACTACACTACGCCGCCGCGATGTTGGCCCCAGAGGGTTGGGGACCGGCATGCAGCTGGTTCGTGGGATGGTCTAACATCTTCGGCTACTTGACCGCGCCATGCTCGTTGAATTACGCTTTGG	184 	\|\|\|\|\|\|\|\|\|\|\|\|\|\|\|\|\|\|\|\|\|\|\|\|\|\|\|\|\|\|\|\|\|\|\|\|\|\|\|\|\|\|\|\|\|\|\|\|\|\|\|\|\|\|\|\|\|\|\|\|\|\|\|\|\|\|\|\|\|\|\|\|\|\|\|\|\|\|\|\|\|\|\|\|\|\|\|\|\|\|\|\|\|\|\|\|\|\|\|\|\|\|\|\|\|\|\|\|\|\|\|\|\|\|\|\|\|\|\|\|\|\|\|\|\|\|\|\|\|\|\|\|\|\|\|\|\|\|\|\|\|\|\|\|\|\|\|\|\|\|\|\|\|\|\|\|\|\|\|\|\|\|\|\|\|\|\|\|\|\|\|\|\|\|\|\|\|\|\|\|\|\|\|\| 	ATGATTCAATTTACCGCGCTGAGCATGGCAGAACTGTGTTCTAGCATGCCGACTGTAGGAGGACTACACTACGCCGCCGCGATGTTGGCCCCAGAGGGTTGGGGACCGGCATGCAGCTGGTTCGTGGGATGGTCTAACATCTTCGGCTACTTGACCGCGCCATGCTCGTTGAATTACGCTTTGG ```      ``` 184	GCCTCGATGATCCTGACCTGCGGCTCCATCGCTCATCCATCCTACGTACCCACGAATTGGCACATCTACCTCCTCTTCCTCCTGCTCCTTACCATCGAGGGTCTCGTCGCAATGCAATCGACCAGATTCATCGGCAGATTCAACATCTTCGGCACAGTCTTCAATCTTCTCATCGTCATCATTTTCGTCATCTGGTTCCCGCTCGGTTCCATCAACATACCGAAGACCAACAGCTCTCACGACGTCTGGACCAATTTCACCAATGGTACTGCCTGGCCTACCCCATGGGCGACCATAATGGGCTTCCTCACTGCTGCGTATACCCTAGCAGGTTTCGACGCTCCTGTACACCTCGCAGAAGAATGCAGCAACTCAGCCATCGCCAGCCCGCGAGCGATCGTCATGGCCGCACAGTCGGGCTTGTATCTCGGCTGGGCCATTATCCTGGTCATCGCATACACAGTCAAGGACATTCCCGATGTCGTAAGTGGACAATACGGTCAACCGTTTGCCAGCCTGTGTCTGCAAGTCCTCGGATCAAAAGCTAGTCTCGCCTTGATCTGCCTCTCCATCATCGCGCAGTTCTCCGTGGCTACTGGATGTGCCATCACTGCGACGAGGGTCTTGTTTGCATTCTCGAGAGATGGTGCGCTGCCCGGTTCAAGCTGGTGGG-CGAAGA-TCGACTGTC	871 	\|\|\|\|\|\|\|\|\|\|\|\|\|\|\|\|\|\|\|\|\|\|\|\|\|\|\|\|\|\|\|\|\|\|\|\|\|\|\|\|\|\|\|\|\|\|\|\|\|\|\|\|\|\|\|\|\|\|\|\|\|\|\|\|\|\|\|\|\|\|\|\|\|\|\|\|\|\|\|\|\|\|\|\|\|\|\|\|\|\|\|\|\|\|\|\|\|\|\|\|\|\|\|\|\|\|\|\|\|\|\|\|\|\|\|\|\|\|\|\|\|\|\|\|\|\|\|\|\|\|\|\|\|\|\|\|\|\|\|\|\|\|\|\|\|\|\|\|\|\|\|\|\|\|\|\|\|\|\|\|\|\|\|\|\|\|\|\|\|\|\|\|\|\|\|\|\|\|\|\|\|\|\|\|\|\|\|\|\|\|\|\|\|\|\|\|\|\|\|\|\|\|\|\|\|\|\|\|\|\|\|\|\|\|\|\|\|\|\|\|\|\|\|\|\|\|\|\|\|\|\|\|\|\|\|\|\|\|\|\|\|\|\|\|\|\|\|\|\|\|\|\|\|\|\|\|\|\|\|\|\|\|\|\|\|\|\|\|\|\|\|\|\|\|\|\|\|\|\|\|\|\|\|\|\|\|\|\|\|\|\|\|\|\|\|\|\|\|\|\|\|\|\|\|\|\|\|\|\|\|\|\|\|\|\|\|\|\|\|\|\|\|\|\|\|\|\|\|\|\|\|\|\|\|\|\|\|\|\|\|\|\|\|\|\|\|\|\|\|\|\|\|\|\|\|\|\|\|\|\|\|\|\|\|\|\|\|\|\|\|\|\|\|\|\|\|\|\|\|\|\|\|\|\|\|\|\|\|\|\|\|\|\|\|\|\|\|\|\|\|\|\|\|\|\|\|\|\|\|\|\|\|\|\|\|\|\|\|\|\|\|\|\|\|\|\|\|\|\|\|\|\|\|\|\|\|\|\|\|\|\|\|\|\|\|\|\|\|\|\|\|\|\|\|\|\|\|\|\|\|\|\|\|\|\|\|\|\|\|\|\|\|\|\|\|\|\|\|\|\|\|\|\|\|\|\|\|\|\|\|\|\|\|\|\|\|\|\|\|\|\|\|\|\|\|\|\|\|\|\|\|\|\|\|\|\|\|\|\|\|\|\|\|\|\|\| \|\|\|\|\|\|\|\|\|\|\|\|\|\|\|\|\|\|\|\|\|\|\|\|\|\|\|\|\|\|\|\|\|\|\|\|\|\|\|\|\|\|\|\|\|\|\|\|\|\|\|\|\|\|\|\|\|\|\|\|\|\|\|\|\|\|\|\|\|\|\|\|\|\|\|\|\|\|\|\|\|\|\|\|\|\|\|\|\|\|\|\|\|\|\|\|\|\|\|\|\|\|\|\|\|\|\|\|\|\|\|\|\|\|\|\|\|\|\|\|\|\|\|\|\|\|\|\|\|\|\|\|\|\|\|\|\|\|\|\|\|\|\|\|\|\| \|\|\|\|\|\| \|\|\|\|\|\|\|\|\| 	GCCTCGATGATCCTGACCTGCGGCTCCATCGCTCATCCATCCTACGTACCCACGAATTGGCACATCTACCTCCTCTTCCTCCTGCTCCTTACCATCGAGGGTCTCGTCGCAATGCAATCGACCAGATTCATCGGCAGATTCAACATCTTCGGCACAGTCTTCAATCTTCTCATCGTCATCATTTTCGTCATCTGGTTCCCGCTCGGTTCCATCAACATACCGAAGACCAACAGCTCTCACGACGTCTGGACCAATTTCACCAATGGTACTGCCTGGCCTACCCCATGGGCGACCATAATGGGCTTCCTCACTGCTGCGTATACCCTAGCAGGTTTCGACGCTCCTGTACACCTCGCAGAAGAATGCAGCAACTCAGCCATCGCCAGCCCGCGAGCGATCGTCATGGCCGCACAGTCGGGCTTGTATCTCGGCTGGGCCATTATCCTGGTCATCGCATACACAGTCAAGGACATTCCCGATGTCGTAAGTGGACAATACGGTCAACCGTTTGCCAGCCTGTGTCTGC-AGTCCTCGGATCAAAAGCTAGTCTCGCCTTGATCTGCCTCTCCATCATCGCGCAGTTCTCCGTGGCTACTGGATGTGCCATCACTGCGACGAGGGTCTTGTTTGCATTCTCGAGAGATGGTGCGCTGCCCGGTTCAAGCTGGTGGGGCGAAGAATCGACTGTC ``` |
| 24 | ggggacaagtttgtacaaaaaagcaggcttgatgACGCGATCTGGTGAGCACGT, ggggaccactttgtacaagaaagctgggtcCATGATCGTGCAGCGAGCGG | Mycgr3\|93506  JGI  Ensembl | KOG2992 Nucleolar GTPase/ATPase p130; |  | 1.43836 % |  |  |
| 24 | ggggacaagtttgtacaaaaaagcaggcttgatgAACGCATTTGACTCTCGACA, ggggaccactttgtacaagaaagctgggtcCTTGTCTTGATCTCGACGCT | Mycgr3\|94769  JGI  Ensembl | KOG4297 C-type lectin; |  | 100 % |  |  |
| 24 | ggggacaagtttgtacaaaaaagcaggcttgatgTCCAACCAGCAGAATCAGAC, ggggaccactttgtacaagaaagctgggtcTTCATCAAAATCCATTTCAT | Mycgr3\|94818  JGI  Ensembl |  |  | 100 % |  |  |
| 24 | ggggacaagtttgtacaaaaaagcaggcttgatgGTCAATGTGTACAACTGTAC, ggggaccactttgtacaagaaagctgggtcGCGTCCATCACGAGGCGGAC | Mycgr3\|95082  JGI  Ensembl | KOG2992 Nucleolar GTPase/ATPase p130; |  | 100 % |  | ``` 1	ATGGTCAATGTGTACAACTGTACCCTGTGCTGCTTCAGGACCGCCTCCCTCGACGACATGAACACACATATCGCCGACTCCAACAACGGCCTTGTCCTACTCAAGTGCGCTGGTGAAGGCTGCAATACGCGCTTCTGTCTCGAGTCGCAGACCCTTAGCCACTTCCCGCACTGCGCGGCCATGCGGAGGTGGAAAGCGAAGACGAAGTATGGCCTTCAGATTACCGAGAAGCGCGTGAGACGCAAGTGCGATGTTTCGGAAGAGAGGAACGGACAGCACGCGACCTTCGCGACGAGTGTTGCTCTTGTAGCTCCTCCTACTCTGACAGCAGCGCCACCATCAACCAAGTCACCAAAGAAGAAGATGACGGTTGCGAAGAAGGCTCGAGCTTCGAAGCGGAAGGCTGAGGAGGACGACGACGTTGGCGACCAGGAAGAGCTGTCCAACGAGGATGACAACGCGCCGCCATCGGCCAAGCCATCCAAGAAAAAGGCGACGCCTGCGAAGAAGGGCGTGGCTCGGAAGCGGAAGGCTGTAGTGTTGGACCTTGATAACGAGGAAGAGCCGATCAGCGAGGAGGATGACTTGCGCATCCAGGACAAACCCGCGAGTGGTGAAGAACTCGATGATGAGCACGCCAACAACACCATCGGTGCACGTACTCAACGCGGCAACAAGAAGCCGCGTGTAGCCAACACCCAGGAAGCGAGCAACATCGCGGGCTCGTTTCCATCCAGACGTGCCAGGCGACCAGCC-GGCCTTACTTACGCGCCTTCGTGGTACGCCCGCCTTGAGGCGAAAGGCCACGATCTGAGGGAAACCAAGCGTAATTTCAAAGAGGCCGATACGATTGCGTGGAAAGCCGGTTTAGGGCCGCATCCATCGATTATACAACAAGTCTTCGATGAGTGGGCCTTGGAGGAGCAAGTGGGCAAGAACAGCGTGG	946 	\|\|\|\|\|\|\|\|\|\|\|\|\|\|\|\|\|\|\|\|\|\|\|\|\|\|\|\|\|\|\|\|\|\|\|\|\|\|\|\|\|\|\|\|\|\|\|\|\|\|\|\|\|\|\|\|\|\|\|\|\|\|\|\|\|\|\|\|\|\|\|\|\|\|\|\|\|\|\|\|\|\|\|\|\|\|\|\|\|\|\|\|\|\|\|\|\|\|\|\|\|\|\|\|\|\|\|\|\|\|\|\|\|\|\|\|\|\|\|\|\|\|\|\|\|\|\|\|\|\|\|\|\|\|\|\|\|\|\|\|\|\|\|\|\|\|\|\|\|\|\|\|\|\|\|\|\|\|\|\|\|\|\|\|\|\|\|\|\|\|\|\|\|\|\|\|\|\|\|\|\|\|\|\|\|\|\|\|\|\|\|\|\|\|\|\|\|\|\|\|\|\|\|\|\|\|\|\|\|\|\|\|\|\|\|\|\|\|\|\|\|\|\|\|\|\|\|\|\|\|\|\|\|\|\|\|\|\|\|\|\|\|\|\|\|\|\|\|\|\|\|\|\|\|\|\|\|\|\|\|\|\|\|\|\|\|\|\|\|\|\|\|\|\|\|\|\|\|\|\|\|\|\|\|\|\|\|\|\|\|\|\|\|\|\|\|\|\|\|\|\|\|\|\|\|\|\|\|\|\|\|\|\|\|\|\|\|\|\|\|\|\|\|\|\|\|\|\|\|\|\|\|\|\|\|\|\|\|\|\|\|\|\|\|\|\|\|\|\|\|\|\|\|\|\|\|\|\|\|\|\|\|\|\|\|\|\|\|\|\|\|\|\|\|\|\|\|\|\|\|\|\|\|\|\|\|\|\|\|\|\|\|\|\|\|\|\|\|\|\|\|\|\|\|\|\|\|\|\|\|\|\|\|\|\|\|\|\|\|\|\|\|\|\|\|\|\|\|\|\|\|\|\|\|\|\|\|\|\|\|\|\|\|\|\|\|\|\|\|\|\|\|\|\|\|\|\|\|\|\|\|\|\|\|\|\|\|\|\|\|\|\|\|\|\|\|\|\|\|\|\|\|\|\|\|\|\|\|\|\|\|\|\|\|\|\|\|\|\|\|\|\|\|\|\|\|\|\|\|\|\|\|\|\|\|\|\|\|\|\|\|\|\|\|\|\|\|\|\|\|\|\|\|\|\|\|\|\|\|\|\|\|\|\|\|\|\|\|\|\|\|\|\|\|\|\|\|\|\|\|\|\|\|\|\|\|\|\|\|\|\|\|\|\|\|\|\|\|\|\|\|\|\|\|\|\|\|\|\|\|\|\|\|\|\|\|\|\|\|\|\|\|\|\|\|\|\|\|\|\|\|\|\|\|\|\|\|\|\|\|\|\|\|\|\|\|\|\|\|\|\|\|\|\|\|\|\|\|\|\|\|\|\|\|\|\|\|\|\|\|\|\|\|\|\|\|\|\|\|\|\|\|\|\|\|\|\|\|\|\|\|\|\|\|\|\|\|\|\|\|\|\|\|\|\|\|\|\|\|\|\|\|\|\|\|\|\|\|\|\|\|\|\|\|\|\|\|\|\|\|\|\|\|\|\|\|\|\|\|\|\|\|\|\|\|\|\|\|\|\|\|\|\|\|\|\|\|\|\|\|\|\|\|\|\|\|\|\|\|\|\|\|\|\|\|\| \|\|\|\|\|\|\|\|\|\|\|\|\|\|\|\|\|\|\|\|\|\|\|\|\|\|\|\|\|\|\|\|\|\|\|\|\|\|\|\|\|\|\|\|\|\|\|\|\|\|\|\|\|\|\|\|\|\|\|\|\|\|\|\|\|\| \|\|\|\|\|\|\|\|\|\|\|\|\|\|\|\|\|\|\|\|\|\|\|\|\|\|\|\| \|\|\|\|\| \|\|\|\|\| \| \|\|\|\|\|\| \| \|\|\|\| \|\|\|\|\|\|\|\|\|\|\|\|\| \| \|\|\|\|\|\|\|\|\|\|\|\|\|\|\|\|\|\|\|\|\| \|\|\|\|\|\|\|\|\|\|\|\|\|\|\|\|\|\|\|\|\|\|\|\|\|\|\|\| 	ATGGTCAATGTGTACAACTGTACCCTGTGCTGCTTCAGGACCGCCTCCCTCGACGACATGAACACACATATCGCCGACTCCAACAACGGCCTTGTCCTACTCAAGTGCGCTGGTGAAGGCTGCAATACGCGCTTCTGTCTCGAGTCGCAGACCCTTAGCCACTTCCCGCACTGCGCGGCCATGCGGAGGTGGAAAGCGAAGACGAAGTATGGCCTTCAGATTACCGAGAAGCGCGTGAGACGCAAGTGCGATGTTTCGGAAGAGAGGAACGGACAGCACGCGACCTTCGCGACGAGTGTTGCTCTTGTAGCTCCTCCTACTCTGACAGCAGCGCCACCATCAACCAAGTCACCAAAGAAGAAGATGACGGTTGCGAAGAAGGCTCGAGCTTCGAAGCGGAAGGCTGAGGAGGACGACGACGTTGGCGACCAGGAAGAGCTGTCCAACGAGGATGACAACGCGCCGCCATCGGCCAAGCCATCCAAGAAAAAGGCGACGCCTGCGAAGAAGGGCGTGGCTCGGAAGCGGAAGGCTGTAGTGTTGGACCTTGATAACGAGGAAGAGCCGATCAGCGAGGAGGATGACTTGCGCATCCAGGACAAACCCGCGAGTGGTGAAGAACTCGATGATGAGCACGCCAACAACACCATCGGTGCACGTACTCAACGCGGCAACAAGAAGCCGCGTGTAGCCAACACCCAGGAAGCGAGCAACATCGCGGGCTCGTTTCCATCCAGACGTGCCAGGCGACCAGCCGGGCCTTACTTACGCGCCTTCGTGGTACGCCCGCCTTGAGGCGAAAGGCCACGATCTGAGGGAAACC-AGCGTAATTTCAAAGAGGCCGATACGAT-GCGTG-AAAGC-G-TTTAGG-C-GCAT-CATCGATTATACA-C-AGTCTTCGATGAGTGGGCCTT-GAGGAGCAAGTGGGCAAGAACAGCGTGG ``` |
| 24 | ggggacaagtttgtacaaaaaagcaggcttgatgCCCCGCACCGCCCGTGGCCC, ggggaccactttgtacaagaaagctgggtcGCGATTCGCCTTCCACCACT | Mycgr3\|95217  JGI  Ensembl |  |  | 100 % |  |  |
| 24 | ggggacaagtttgtacaaaaaagcaggcttgatgGTTTCTCACGCTGGCCGAGA, ggggaccactttgtacaagaaagctgggtcGCTGTTCGAATGCGTACGCT | Mycgr3\|95378  JGI  Ensembl |  |  | 100 % |  |  |
| 24 | ggggacaagtttgtacaaaaaagcaggcttgatgAAGCCAACGCATGGTCTCTC, ggggaccactttgtacaagaaagctgggtcCTGCGGCTGCCTATATCGGA | Mycgr3\|95545  JGI  Ensembl |  |  | 99.64689 % |  |  |
| 24 | ggggacaagtttgtacaaaaaagcaggcttgatgCAAGAAGAGTTCGATATCAT, ggggaccactttgtacaagaaagctgggtcGTCCTGAGACACTCTTTGTC | Mycgr3\|95960  JGI  Ensembl |  |  | 0 % |  |  |
| 24 | ggggacaagtttgtacaaaaaagcaggcttgatgGACCAATGGAGCGAAAAAGA, ggggaccactttgtacaagaaagctgggtcCGCCTCGATCTCGCCATTCT | Mycgr3\|96123  JGI  Ensembl |  |  | 100 % |  |  |
| 24 | ggggacaagtttgtacaaaaaagcaggcttgatgGTCATATCACTGAGCTCACA, ggggaccactttgtacaagaaagctgggtcTGTGTCACGCTTTTGGCCGA | Mycgr3\|96588  JGI  Ensembl |  |  | 100 % |  |  |
| 24 | ggggacaagtttgtacaaaaaagcaggcttgatgGCACCCCGCCAGATGGAGCC, ggggaccactttgtacaagaaagctgggtcCTGAGGAAAAAGTTCCTCCA | Mycgr3\|97868  JGI  Ensembl |  |  | 100 % |  |  |
| 24 | ggggacaagtttgtacaaaaaagcaggcttgatgCCTTCCCCCAAACTCAACCT, ggggaccactttgtacaagaaagctgggtcAACTCTCTTCCCACCAACAC | Mycgr3\|97907  JGI  Ensembl |  |  | 100 % |  |  |
| 24 | ggggacaagtttgtacaaaaaagcaggcttgatgTCGAGCGTCAAAGTCACCCG, ggggaccactttgtacaagaaagctgggtcGGCAAGACGATGCAGATCGC | Mycgr3\|98007  JGI  Ensembl |  |  | 100 % |  |  |
| 24 | ggggacaagtttgtacaaaaaagcaggcttgatgCCTGCATTCGACGAACGAGA, ggggaccactttgtacaagaaagctgggtcCTCAACCGCCACAGCAGCCG | Mycgr3\|99007  JGI  Ensembl | KOG1618 Predicted phosphatase; | hydrolase activity; metabolic process; | 100 % |  |  |
| 24 | ggggacaagtttgtacaaaaaagcaggcttgatgGCAGATCCACGGACAGGCAG, ggggaccactttgtacaagaaagctgggtcCAGTCCATTCATAACCTGCT | Mycgr3\|99683  JGI  Ensembl |  |  | 100 % |  |  |
| 24 | ggggacaagtttgtacaaaaaagcaggcttgatgAATCTCCCTCTAGCGGATCC, ggggaccactttgtacaagaaagctgggtcATCTCCACGTCGTCGCTTCG | Mycgr3\|99872  JGI  Ensembl | KOG1273 WD40 repeat protein; | 1-alkyl-2-acetylglycerophosphocholine esterase activity; | 100 % | chr\_4:1076205 T->C; | ``` 1	ATGAATCTCCCTCTAGCGGATCCGATCCTCCTCGCACAGGACATACCCGAGACTCTGACCGCGCGACTGCGGTCTTCCGGTCAGGCAGTATGCATGCGGTTCTCCCACCGCGGTGACCTGCTCGCTTCCGGTACGGCCAAAGGAACGATCGCGATCTTTGACCTCGAAACGAACGGAGTCGCGCGTAAATTGAGAGGTCACACAGCTGGACGCACAGTACAGTCGCTGTCATGGGAGAAGAGCGGCCGATATCTGCTCAGTAGCAGCGTAGATTGGAAGGTCATACTATGGGATCTCAACGATGGCAGTCGGTTGCGGACCATCAATCTGGGTGCGCCGGTGTACATTGCGGAGCTACATCCGTCAAACTTCAAAATGTGCGTGGCCGCATTGTATGAGTACAGGCCAGTGCTGGCCGACTTCACGAACGCCGCGAACGTCAAGCAGATGGCTCTACCCAATTTGCCCAAGCGTGCACCACACGAGGAGAGCGGCGAGAAGGCCGACGCGAAGCACTTTACAACAGTCGCAGCCTTTACACCTACTGGCTCGCACATCATCACTGGTACGACCAAGGGCTGGTTGAACGTGATCAGCACGACGACGCAGCAAACAGTCTACAGCACGCGACTCTGCAGCAAACCAATTCTCTTGATTCGGTTGAGTGGCAGTGGCCGCGACCTGCTCGTTAACGCATCCGACACAATCATCCGCACGATCAAACTGCCTGATCTCTCGAACCCGAAACTGCAGGACGATGACATTCATCTGGAAGTCGAGCATAAGTTTCAGGATGTCGTCAATCGTCTGTCATGGAATCACGTCGCCTTCAGTAGCAACGCCGACTATGTTATGGCGTCGACCTTGATGAATCACGACATCTACATCTGGGAGCGAGGTCATGGATCGTTGGTCAAGATTCTGGAAGGGCCGAGAGAAGAGCTAGGTGCAGTGG	955 	\|\|\|\|\|\|\|\|\|\|\|\|\|\| \|\|\|\|\|\|\|\|\|\|\|\|\|\|\|\|\|\|\|\|\|\|\|\|\|\|\|\|\|\|\|\|\|\|\|\|\|\|\|\|\|\|\|\|\|\|\|\|\|\|\|\|\|\|\|\|\|\|\|\|\|\|\|\|\|\|\|\|\|\|\|\|\|\|\|\|\|\|\|\|\|\|\|\|\|\|\|\|\|\|\|\|\|\|\|\|\|\|\|\|\|\|\|\|\|\|\|\|\|\|\|\|\|\|\|\|\|\|\|\|\|\|\|\|\|\|\|\|\|\|\|\|\|\|\|\|\|\|\|\|\|\|\|\|\|\|\|\|\|\|\|\|\|\|\|\|\|\|\|\|\|\|\|\|\|\|\|\|\|\|\|\|\|\|\|\|\|\|\|\|\|\|\|\|\|\|\|\|\|\|\|\|\|\|\|\|\|\|\|\|\|\|\|\|\|\|\|\|\|\|\|\|\|\|\|\|\|\|\|\|\|\|\|\|\|\|\|\|\|\|\|\|\|\|\|\|\|\|\|\|\|\|\|\|\|\|\|\|\|\|\|\|\|\|\|\|\|\|\|\|\|\|\|\|\|\|\|\|\|\|\|\|\|\|\|\|\|\|\|\|\|\|\|\|\|\|\|\|\|\|\|\|\|\|\|\|\|\|\|\|\|\|\|\|\|\|\|\|\|\|\|\|\|\|\|\|\|\|\|\|\|\|\|\|\|\|\|\|\|\|\|\|\|\|\|\|\|\|\|\|\|\|\|\|\|\|\|\|\|\|\|\|\|\|\|\|\|\|\|\|\|\|\|\|\|\|\|\|\|\|\|\|\|\|\|\|\|\|\|\|\|\|\|\|\|\|\|\|\|\|\|\|\|\|\|\|\|\|\|\|\|\|\|\|\|\|\|\|\|\|\|\|\|\|\|\|\|\|\|\|\|\|\|\|\|\|\|\|\|\|\|\|\|\|\|\|\|\|\|\|\|\|\|\|\|\|\|\|\|\|\|\|\|\|\|\|\|\|\|\|\|\|\|\|\|\|\|\|\|\|\|\|\|\|\|\|\|\|\|\|\|\|\|\|\|\|\|\|\|\|\|\|\|\|\|\|\|\|\|\|\|\|\|\|\|\|\|\|\|\|\|\|\|\|\|\|\|\|\|\|\|\|\|\|\|\|\|\|\|\|\|\|\|\|\|\|\|\|\|\|\|\|\|\|\|\|\|\|\|\|\|\|\|\|\|\|\|\|\|\|\|\|\|\|\|\|\|\|\|\|\|\|\|\|\|\|\|\|\|\|\|\|\|\|\|\|\|\|\|\|\|\|\|\|\|\|\|\|\|\|\|\|\|\|\|\|\|\|\|\|\|\|\|\|\|\|\|\|\|\|\|\|\|\|\|\|\|\|\|\|\|\|\|\|\|\|\|\|\|\|\|\|\|\|\|\|\|\|\|\|\|\|\|\|\|\|\|\|\|\|\|\|\|\|\|\|\|\|\|\|\|\|\|\|\|\|\|\|\|\|\|\|\|\|\|\|\|\|\|\|\|\|\|\|\|\|\|\|\|\|\|\|\|\|\|\|\|\|\|\|\|\|\|\|\|\|\|\|\|\|\|\|\|\|\|\|\|\|\|\|\|\|\|\|\|\|\|\|\|\|\|\|\|\|\|\|\|\|\|\|\|\|\|\|\|\|\|\|\|\|\|\|\|\|\|\|\|\|\|\|\|\|\|\|\|\| \|\|\|\|\|\|\|\|\|\|\|\|\|\|\|\|\|\|\|\|\|\| \|\|\|\|\|\|\|\|\|\|\|\|\|\|\|\|\|\|\|\|\|\|\|\|\|\|\|\|\|\|\|\|\|\|\| \|\|\|\|\|\|\|\|\|\|\|\|\|\|\|\|\|\|\| \|\|\|\|\|\|\|\|\|\|\|\|\|\|\|\|\|\|\|\|\|\|\|\|\|\|\| \|\|\|\|\| \|\|\|\|\| \| \|\| \|\|\|\|\|\|\|\|\|\|\|\|\|\|\|\|\|\|\|\|\|\|\|\|\|\|\|\|\|\|\|\|\|\|\|\|\|\|\| 	ATGAATCTCCCTCTGGCGGATCCGATCCTCCTCGCACAGGACATACCCGAGACTCTGACCGCGCGACTGCGGTCTTCCGGTCAGGCAGTATGCATGCGGTTCTCCCACCGCGGTGACCTGCTCGCTTCCGGTACGGCCAAAGGAACGATCGCGATCTTTGACCTCGAAACGAACGGAGTCGCGCGTAAATTGAGAGGTCACACAGCTGGACGCACAGTACAGTCGCTGTCATGGGAGAAGAGCGGCCGATATCTGCTCAGTAGCAGCGTAGATTGGAAGGTCATACTATGGGATCTCAACGATGGCAGTCGGTTGCGGACCATCAATCTGGGTGCGCCGGTGTACATTGCGGAGCTACATCCGTCAAACTTCAAAATGTGCGTGGCCGCATTGTATGAGTACAGGCCAGTGCTGGCCGACTTCACGAACGCCGCGAACGTCAAGCAGATGGCTCTACCCAATTTGCCCAAGCGTGCACCACACGAGGAGAGCGGCGAGAAGGCCGACGCGAAGCACTTTACAACAGTCGCAGCCTTTACACCTACTGGCTCGCACATCATCACTGGTACGACCAAGGGCTGGTTGAACGTGATCAGCACGACGACGCAGCAAACAGTCTACAGCACGCGACTCTGCAGCAAACCAATTCTCTTGATTCGGTTGAGTGGCAGTGGCCGCGACCTGCTCGTTAACGCATCCGACACAATCATCCGCACGATCAAACTGCCTGATCTCTCGAACCCGAAACTGCAGGACGATGACATTCATCTGGAAGTCGAGCATAAGTTTCA-GATGTCGTCAATCGTCTGTCAT-GAATCACGTCGCCTTCAGTAGCAACGCCGACTATG-TATGGCGTCGACCTTGATG-ATCACGACATCTACATCTGGGAGCGAG-TCATG-ATCGT-G-TC-AGATTCTGGAAGGGCCGAGAGAAGAGCTAGGTGCAGTGG ``` |
| 24 | ggggacaagtttgtacaaaaaagcaggcttgatgGACGTAGCCTACGATCACAT, ggggaccactttgtacaagaaagctgggtcCTCCCAATCATCATCATCAT | Mycgr3\|99952  JGI  Ensembl | KOG2690 Uncharacterized conserved protein, contains BSD domain; |  | 100 % |  |  |
| 24 | ggggacaagtttgtacaaaaaagcaggcttgatgCCACCCCCTCCTCCCCCTCG, ggggaccactttgtacaagaaagctgggtcTGTTGTTTCCTGCTCCTCCG | Mycgr3\|101591  JGI  Ensembl | KOG2614 Kynurenine 3-monooxygenase and related flavoprotein monooxygenases; | electron transport; metabolic process;  oxidoreductase activity; | 100 % |  | ``` 743	AGGGGAGGATGTGTTCCTGCTACCTCGTCAAAACATCCCACTCCCCCGAACTAGCGAAGTCCGCCACCAAACCCCGCCAACAAGCCCTTCAAAAAGAAATCTTCAcccgcctcttcgcctccttccccggctccctccgctcccgcgtcctctccggcctccgcACAACCTCCGACTTCTACGTCGCGCAAACCGCTCAAATCCACCTCCCGACTTGGTCGTCCGGTCGAGTCGCCCTCGTTGGGGACGCGGCGTATGCTCCCT-CACCGGTGACGGGGCAGGGGACTGCGATTGCGATTGTGG	1045 	\|\|\|\|\|\|\|\|\|\|\|\|\|\|\|\|\|\|\|\|\|\|\|\|\|\|\|\|\|\|\|\|\|\|\|\|\|\|\|\|\|\|\|\|\|\|\|\|\|\|\|\|\|\|\|\|\|\|\|\|\|\|\|\|\|\|\|\|\|\|\|\|\|\|\|\|\|\|\|\|\|\|\|\|\|\|\|\|\|\| \|\|\|\|\|\|\|\|\|\|\|\|\|\|\|\|\|\|\|\|\|\|\|\|\|\|\| \| \|\|\|\|\|\|\|\|\|\|\|\|\|\|\|\| \|\|\|\|\|\|\|\|\|\|\| \|\|\|\| \|\|\| \|\| \|\|\|\|\|  \|\|\|\|\|\|\|\|\|\|\|\|\|\|\|\|\|\|\|\|\|\|\| \|\|\|\|\|\|\|\|\|\|\|\|\| \|\|\|\|\|\|\|\| \|\|\|\|\|\|\| \| \|\|\|\|\|\|\|\|\|  \|\|\|\| \|\|\|\|\|\|\|\| \|\|\|\|\|\|\|\|\|\|\|\|\| \|\|\|\|\|\|\|\|\|\|\|\|\| \|\|\|\|\| \|\|\|\|\| \|\|\|\|\| \|\|\|\|\|\|\| 	AGGGGAGGATGTGTTCCTGCTACCTCGTCAAAACATCCCACTCCCCCGAACTAGCGAAGTCCGCCACCAAACCCCGCCAACAAGCCCTTC-AAAAGAAATCTTCACCCGCCTCTTCGC-T-CTTCCCCGGCTCCCTC-GCTCCCGCGTC-TCTC-GGC-TC-GCACA--CTCCGACTTCTACGTCGCGCAAA-CGCTCAAATCCAC-TCCCGACT-GGTCGTC-G-TCGAGTCGC--TCGT-GGGGACGC-GCGTATGCTCCCTCCACCGGTGACGGG-CAGGG-ACTGC-ATTGCAATTGTGG ```      ``` 1	ATGCCACCCCCTCCTCCCCCTCGAAGAATCCTGATCGTCGGCGGCGGCCCAGCCGGCGCAGTAGCAGCCTACCACCTCGCCAAATCCGGCCTCCAAGTGCTAATCCTCGAACGCTCCCCCTCTCCCACCACCTCCACCACGCGGGGTCAAGGAATCGACATCACTGGCCCGGCGATCTCCGTGATCCGACGAATGGAATTGTACGACGTCGTGCGAGCGAATTGTACCGGCGAGAAAGGCTTCTCCATGCTCGACGACTCGGGAAAGGTCATCGGGCGTCTTCCAGCGGCAGACCAAACAGATGGAGGCAAAGGTGGACTCCAATTGACGCAAGAACTGGAAATCATGCGCGGCAACATAACCCGCATCCTCGCGTCCGCCGCGCAAAGCCTTTCCTCGTCGCAGTCGTCGGCAGGTGATGGAAATGGAAAGGGAGGAGTAGAATACCGCTACGGCTCGTGGTTAACCTCCCTCTCGCAATCCTCCTCGTCCGTAACCGCCACGCTCAGCGACCTAGATGGCGAAGAGACGACGGTGTCCTTCCTCGCCGTGATTGGCGCCGACGGCATGAACTCCACCGTCCGGCGGCTAGCTTTCAGTCCGGAAGTGACTAGGAATTGCTACCGCCCTATCGGATCGCAATGGACGGCATTCTTCTCCATCCCGGCGGAGGAGAGAGACGGGGAGGATTCCGTGTTGCAGCAGGGAGTGGGATGTCGGACGGTATTAATCCGGCCAATCC	742 	\|\|\|\|\|\|\|\|\|\|\|\|\|\|\|\|\|\|\|\|\|\|\|\|\|\|\|\|\|\|\|\|\|\|\|\|\|\|\|\|\|\|\|\|\|\|\|\|\|\|\|\|\|\|\|\|\|\|\|\|\|\|\|\|\|\|\|\|\|\|\|\|\|\|\|\|\|\|\|\|\|\|\|\|\|\|\|\|\|\|\|\|\|\|\|\|\|\|\|\|\|\|\|\|\|\|\|\|\|\|\|\|\|\|\|\|\|\|\|\|\|\|\|\|\|\|\|\|\|\|\|\|\|\|\|\|\|\|\|\|\|\|\|\|\|\|\|\|\|\|\|\|\|\|\|\|\|\|\|\|\|\|\|\|\|\|\|\|\|\|\|\|\|\|\|\|\|\|\|\|\|\|\|\|\|\|\|\|\|\|\|\|\|\|\|\|\|\|\|\|\|\|\|\|\|\|\|\|\|\|\|\|\|\|\|\|\|\|\|\|\|\|\|\|\|\|\|\|\|\|\|\|\|\|\|\|\|\|\|\|\|\|\|\|\|\|\|\|\|\|\|\|\|\|\|\|\|\|\|\|\|\|\|\|\|\|\|\|\|\|\|\|\|\|\|\|\|\|\|\|\|\|\|\|\|\|\|\|\|\|\|\|\|\|\|\|\|\|\|\|\|\|\|\|\|\|\|\|\|\|\|\|\|\|\|\|\|\|\|\|\|\|\|\|\|\|\|\|\|\|\|\|\|\|\|\|\|\|\|\|\|\|\|\|\|\|\|\|\|\|\|\|\|\|\|\|\|\|\|\|\|\|\|\|\|\|\|\|\|\|\|\|\|\|\|\|\|\|\|\|\|\|\|\|\|\|\|\|\|\|\|\|\|\|\|\|\|\|\|\|\|\|\|\|\|\|\|\|\|\|\|\|\|\|\|\|\|\|\|\|\|\|\|\|\|\|\|\|\|\|\|\|\|\|\|\|\|\|\|\|\|\|\|\|\|\|\|\|\|\|\|\|\|\|\|\|\|\|\|\|\|\|\|\|\|\|\|\|\|\|\|\|\|\|\|\|\|\|\|\|\|\|\|\|\|\|\|\|\|\|\|\|\|\|\|\|\|\|\|\|\|\|\|\|\|\|\|\|\|\|\|\|\|\|\|\|\|\|\|\|\|\|\|\|\|\|\|\|\|\|\|\|\|\|\|\|\|\|\|\|\|\|\|\|\|\|\|\|\|\|\|\|\|\|\|\|\|\|\|\|\|\|\|\|\| \|\|\|\|\|\|\|\|\|\|\|\|\|\|\|\|\|\|\|\|\|\|\|\|\|\|\|\|\|\|\|\|\|\|\|\|\|\|\|\|\|\|\|\|\|\|\|\|\|\|\|\|\|\|\|\|\|\|\|\|\|\|\|\|\|\|\|\|\|\|\|\|\|\|\|\|\|\|\|\|\|\|\|\|\|\|\|\|\|\|\|\|\|\|\|\|\|\|\|\|\|\|\|\|\|\|\|\|\|\|\|\|\|\|\|\|\|\|\|\|\|\|\|\|\|\|\|\|\|\|\|\|\|\|\|\|\|\|\|\|\|\|\|\|\|\|\|\|\|\|\|\|\|\|\|\|\|\|\|\|\|\|\|\|\|\|\|\|\|\|\|\|\|\|\|\| 	ATGCCACCCCCTCCTCCCCCTCGAAGAATCCTGATCGTCGGCGGCGGCCCAGCCGGCGCAGTAGCAGCCTACCACCTCGCCAAATCCGGCCTCCAAGTGCTAATCCTCGAACGCTCCCCCTCTCCCACCACCTCCACCACGCGGGGTCAAGGAATCGACATCACTGGCCCGGCGATCTCCGTGATCCGACGAATGGAATTGTACGACGTCGTGCGAGCGAATTGTACCGGCGAGAAAGGCTTCTCCATGCTCGACGACTCGGGAAAGGTCATCGGGCGTCTTCCAGCGGCAGACCAAACAGATGGAGGCAAAGGTGGACTCCAATTGACGCAAGAACTGGAAATCATGCGCGGCAACATAACCCGCATCCTCGCGTCCGCCGCGCAAAGCCTTTCCTCGTCGCAGTCGTCGGCAGGTGATGGAAATGGAAAGGGAGGAGTAGAATACCGCTACGGCTCGTGGTTAACCTCCCTCTCGCAATCCTCCTCGTCCGTAACCGCCACGCTCAGCGACCTAGATGGCGAAGAGACGACGGTGTCCTTCCTCGCCGTGATTGGCGCCGACGACATGAACTCCACCGTCCGGCGGCTAGCTTTCAGTCCGGAAGTGACTAGGAATTGCTACCGCCCTATCGGATCGCAATGGACGGCATTCTTCTCCATCCCGGCGGAGGAGAGAGACGGGGAGGATTCCGTGTTGCAGCAGGGAGTGGGATGTCGGACGGTATTAATCCGGCCAATCC ``` |
| 24 | ggggacaagtttgtacaaaaaagcaggcttgatgCCCGTGCAACTGCTTCCTGC, ggggaccactttgtacaagaaagctgggtcAAACGTAGTCGCGTACTGCG | Mycgr3\|102441  JGI  Ensembl |  |  | 100 % |  | ``` 359	GATGCCCGTGCAACTGCTTCCTGCCTCGGCGCAGGCCTTTGCACCGCGTTCCGCTCCGGAGATCGTCCTCAACTCCAAGGTCGAGTCTTGGCTCACACAAACACTGAAGCGCATCAACCGGGTAAAGCGCCCACTCAACAGTGTTCCACAGCACCACCGATGTCTCACAGAAACTCTCGGCGGCACCCTCGCCATCTGGACACTTGCATCGCTCATGCTGCCCAAAGCACCCGATGCCGACCTGCGCAAGGATGACAATCCATTAACAGAAGCTCTGTTCAACCATCAGCTGATCCACATCGAGGCTTACGTCGTACACGTCGACATGGTGTCGCAACATGAAGTCGCCTTCAAGCTCACAACGGACACGATCGAGTTGCTTATCGAATACCACAAGGACATCTACTCTGTCGATGCTTCGGCAAGCACATGGTCCTGGCCCGAGAAGGAGTCGCACGTCAAGAAAATGCAAGATGAGTTCGTCCAAGCCGCCAACAAGTTTGTCTTTCGCACTCATGTCCGCGCGCTGGAGGGTTTGGAAGAGGACGGCGCCGGCGAGCTTCTCGAAGGTCGCTCCGAAGACGTCAAGAACGCCATCATGAACATGTTCGTCCCGCTACTCCCACCACCACCGCGAATCGTCGATGTCATCCACCCAGCACCCATCCTCCCGACATCATCGATGGCCGGTCACTGGTGGACACCATCGCCGCAGCAGCAGTTGGCGCCCGCGCATCCTGCGCCCGTAGACACATGGCGAGTGCTCCCATCGACACCGTCCCCGACGGCCACCACGTGCAGCGACTCCCGGTCTAACAGTTTCTGGCAAGACATGCAGTACGG-TCAGCAGGTCCAACTTCCGTCGCCCACGCCTTCGTTTAGCCAGCCCATTGCA-T-ACACAACGGCACCTGCGCAGTTCT-ATAGCTC-GCCACAGA-GCATCG	1299 	\|\|\|\|\|\|\|\|\|\|\|\|\|\|\|\|\|\|\|\|\|\|\|\|\|\|\|\|\|\|\|\|\|\|\|\|\|\|\|\|\|\|\|\|\|\|\|\|\|\|\|\|\|\|\|\|\|\|\|\|\|\|\|\|\|\|\|\|\|\|\|\|\|\|\|\|\|\|\|\|\|\|\|\|\|\|\|\|\|\|\|\|\|\|\|\|\|\|\|\|\|\|\|\|\|\|\|\|\|\|\|\|\|\|\|\|\|\|\|\|\|\|\|\|\|\|\|\|\|\|\|\|\|\|\|\|\|\|\|\|\|\|\|\|\|\|\|\|\|\|\|\|\|\|\|\|\|\|\|\|\|\|\|\|\|\|\|\|\|\|\|\|\|\|\|\|\|\|\|\|\|\|\|\|\|\|\|\|\|\|\|\|\|\|\|\|\|\|\|\|\|\|\|\|\|\|\|\|\|\|\|\|\|\|\|\|\|\|\|\|\|\|\|\|\|\|\|\|\|\|\|\|\|\|\|\|\|\|\|\|\|\|\|\|\|\|\|\|\|\|\|\|\|\|\|\|\|\|\|\|\|\|\|\|\|\|\|\|\|\|\|\|\|\|\|\|\|\|\|\|\|\|\|\|\|\|\|\|\|\|\|\|\|\|\|\|\|\|\|\|\|\|\|\|\|\|\|\|\|\|\|\|\|\|\|\|\|\|\|\|\|\|\|\|\|\|\|\|\|\|\|\|\|\|\|\|\|\|\|\|\|\|\|\|\|\|\|\|\|\|\|\|\|\|\|\|\|\|\|\|\|\|\|\|\|\|\|\|\|\|\|\|\|\|\|\|\|\|\|\|\|\|\|\|\|\|\|\|\|\|\|\|\|\|\|\|\|\|\|\|\|\|\|\|\|\|\|\|\|\|\|\|\|\|\|\|\|\|\|\|\|\|\|\|\|\|\|\|\|\|\|\|\|\|\|\|\|\|\|\|\|\|\|\|\|\|\|\|\|\|\|\|\|\|\|\|\|\|\|\|\|\|\|\|\|\|\|\|\|\|\|\|\|\|\|\|\|\|\|\|\|\|\|\|\|\|\|\|\|\|\|\|\|\|\|\|\|\|\|\|\|\|\|\|\|\|\|\|\|\|\|\|\|\|\|\|\|\|\|\|\|\|\|\|\|\|\|\|\|\|\|\|\|\|\|\|\|\|\|\|\|\|\|\|\|\|\|\|\|\|\|\|\|\|\|\|\|\|\|\|\|\|\|\|\|\|\|\|\|\|\|\|\|\|\|\|\|\|\|\|\|\|\|\|\|\|\|\|\|\|\|\|\|\|\|\|\|\|\|\|\|\|\|\|\|\|\|\|\|\|\|\|\|\|\|\|\|\|\|\|\|\|\|\|\|\|\|\|\|\|\|\|\|\|\|\|\|\|\|\|\|\|\|\|\|\|\|\|\|\|\|\|\|\|\|\|\|\|\|\|\|\|\|\|\|\|\|\|\|\|\|\|\|\|\|\|\|\|\|\|\|\|\|\|\|\|\|\|\|\|\|\|\|\|\|\|\|\|\|\|\|\|\|\|\|\|\|\|\|\|\|\|\|\|\|\|\|\|\|\|\|\|\|\|\|\|\|\|\|\|\|\|\|\|\|\|\|\|\|\|\|\|\|\|\|\|\|\|\|\|\|\|\|\|\|\|\|\|\|\|\|\|\|\|\|\|\|\|\|\|\|\|\|\|\|\|\|\|\|\|\|\|\|\|\|\|\|\|\|\|\|\|\|\|\|\|\|\|\|\|\|\|\|\|\|\|\|\|\|\|\|\|\|\|\|\|\|\|\|\|\|\|\|\|\|\|\|\|\|\|\|\|\|\|\|\|\|\|\|\|\|\|\| \|\|\|\|\|\|\| \|\|\|\|\|\| \|\|\|\|\|\|\|\|\|\|\|\|\| \|\|\|\|\|\|\|\|\|\|\|\|\|\|\|\|\|\|\|\|\|\|\| \| \|\|\|\|\|\|\|\|\|\|\|\|\|\|\|\|\|\|\|\|\|\|\|\| \|\|\|\|\|\|\| \|\|\|\|\|\|\|\| \|\|\|\|\|\| 	GATGCCCGTGCAACTGCTTCCTGCCTCGGCGCAGGCCTTTGCACCGCGTTCCGCTCCGGAGATCGTCCTCAACTCCAAGGTCGAGTCTTGGCTCACACAAACACTGAAGCGCATCAACCGGGTAAAGCGCCCACTCAACAGTGTTCCACAGCACCACCGATGTCTCACAGAAACTCTCGGCGGCACCCTCGCCATCTGGACACTTGCATCGCTCATGCTGCCCAAAGCACCCGATGCCGACCTGCGCAAGGATGACAATCCATTAACAGAAGCTCTGTTCAACCATCAGCTGATCCACATCGAGGCTTACGTCGTACACGTCGACATGGTGTCGCAACATGAAGTCGCCTTCAAGCTCACAACGGACACGATCGAGTTGCTTATCGAATACCACAAGGACATCTACTCTGTCGATGCTTCGGCAAGCACATGGTCCTGGCCCGAGAAGGAGTCGCACGTCAAGAAAATGCAAGATGAGTTCGTCCAAGCCGCCAACAAGTTTGTCTTTCGCACTCATGTCCGCGCGCTGGAGGGTTTGGAAGAGGACGGCGCCGGCGAGCTTCTCGAAGGTCGCTCCGAAGACGTCAAGAACGCCATCATGAACATGTTCGTCCCGCTACTCCCACCACCACCGCGAATCGTCGATGTCATCCACCCAGCACCCATCCTCCCGACATCATCGATGGCCGGTCACTGGTGGACACCATCGCCGCAGCAGCAGTTGGCGCCCGCGCATCCTGCGCCCGTAGACACATGGCGAGTGCTCCCATCGACACCGTCCCCGACGGCCACCACGTGCAGCGACTCCCGGTCTAACAGTTTCTGGCAAGACATGCAGTACGGGTCAGCAG-TCCAAC-TCCGTCGCCCACG-CTTCGTTTAGCCAGCCCATTGCAATAACACAACGGCACCTGCGCAGTTCTTATAGCTCCGCCACAGAAGCATCG ``` |
| 24 | ggggacaagtttgtacaaaaaagcaggcttgatgGCCGAAGTTTCAGCACCCTC, ggggaccactttgtacaagaaagctgggtcTACAGCAGTCGGCGTCTTCT | Mycgr3\|102488  JGI  Ensembl | KOG3239 Density-regulated protein related to translation initiation factor 1 (eIF-1/SUI1); | translation initiation factor activity; translational initiation; | 100 % |  |  |
| 24 | ggggacaagtttgtacaaaaaagcaggcttgatgTCGGCAGACATGGATCAGCT, ggggaccactttgtacaagaaagctgggtcATTGCCTCCCCTCCAGCTCC | Mycgr3\|103966  JGI  Ensembl |  |  | 100 % |  |  |
| 24 | ggggacaagtttgtacaaaaaagcaggcttgatgTCGCTGACGCTCCTTTCCAG, ggggaccactttgtacaagaaagctgggtcCGCCCTGACAGCAGATTCAA | Mycgr3\|104260  JGI  Ensembl | KOG2900 Biotin synthase; | biotin biosynthetic process; biotin synthase activity;  catalytic activity;  iron-sulfur cluster binding;  nucleic acid binding; | 100 % |  |  |
| 24 | ggggacaagtttgtacaaaaaagcaggcttgatgGAAGATGGCGTCTCCCAGAC, ggggaccactttgtacaagaaagctgggtcCTTGATGCCTTTCACTCGCA | Mycgr3\|104550  JGI  Ensembl | KOG1674 Cyclin; |  | 100 % |  |  |
| 24 | ggggacaagtttgtacaaaaaagcaggcttgatgGCCTCGTCAGGCTCTGGACT, ggggaccactttgtacaagaaagctgggtcGAGCAGGGAGTCTAGGCGAG | Mycgr3\|104882  JGI  Ensembl | KOG3983 Golgi protein; |  | 100 % |  | ``` 257	ATGGCCTCGTCAGGCTCTGGACTGACCCGACGCCGGGGTGCAGGCGGCGGAGGAGGAGCTGGCGACAATATCGGTGACGACCCCAGCCGCGTGTCCTCGCCTGCACCGAAACCCAACAACAACAACAACCAGTCCGGGCCCGAGACATCATACGAGTCCAGCGAAAACGGCCACAAAATCGCCTTCGACCCTCGCGACATCTCCGAGAATGAAGAGCGCTCCAAGCAGCCCAAACTCACGCTTATGGAGGAATGCTTGCTCATGGGTCTCAAGGACAAGCAGGGTTACTTGAGCTTCTGGAACGACAACATCAGCTATGCGCTGCGTGGGTGCATTGTTATTGAGTTGGCGTTTCGAGGGAGGATCAGCATGCAGAAAGACGCGAGTAGGCGGCGATTCCCCCTGGCGGACAGGCTGATTGAGGTGATTGACGACACATTGACGGGAGAAGTGCTACTGGACGAAGCGTTGAAGATGATGAAGAGCTCGGAGAAAATGAGTGTCAGTTCGTGGATTGATCTCATGAGCGGCGAG	790 	\|\|\|\|\|\|\|\|\|\|\|\|\|\|\|\|\|\|\|\|\|\|\|\|\|\|\|\|\|\|\|\|\|\|\|\|\|\|\|\|\|\|\|\|\|\|\|\|\|\|\|\|\|\|\|\|\|\|\|\|\|\|\|\|\|\|\|\|\|\|\|\|\|\|\|\|\|\|\|\|\|\|\|\|\|\|\|\|\|\|\|\|\|\|\|\|\|\|\|\|\|\|\|\|\|\|\|\|\|\|\|\|\|\|\|\|\|\|\|\|\|\|\|\|\|\|\|\|\|\|\|\|\|\|\|\|\|\|\|\|\|\|\|\|\|\|\|\|\|\|\|\|\|\|\|\|\|\|\|\|\|\|\|\|\|\|\|\|\|\|\|\|\|\|\|\|\|\|\|\|\|\|\|\|\|\|\|\|\|\|\|\|\|\|\|\|\|\|\|\|\|\|\|\|\|\|\|\|\|\|\|\|\|\|\|\|\|\|\|\|\|\|\|\|\|\|\|\|\|\|\|\|\|\|\|\|\|\|\|\|\|\|\|\|\|\|\|\|\|\|\|\|\|\|\|\|\|\|\|\|\|\|\|\|\|\|\|\|\|\|\|\|\|\|\|\|\|\|\|\|\|\|\|\|\|\|\|\|\|\|\|\|\|\|\|\|\|\|\|\|\|\|\|\|\|\|\|\|\|\|\|\|\|\|\|\|\|\|\|\|\|\|\|\|\|\|\|\|\|\|\|\|\|\|\|\|\|\|\|\|\|\|\|\|\|\|\|\|\|\|\|\|\|\|\|\|\|\|\|\|\|\|\|\|\|\|\|\|\|\|\|\|\|\|\|\|\|\|\|\|\|\|\|\|\|\|\|\|\|\|\|\|\|\|\|\|\|\|\|\|\|\|\|\|\|\|\|\|\|\|\|\|\|\|\|\|\|\|\|\|\|\|\|\|\|\|\|\|\|\|\|\|\|\|\|\|\|\|\|\|\|\|\|\|\|\|\|\|\|\|\|\|\|\|\|\|\|\|\|\|\|\|\|\|\|\|\|\|\|\|\|\|\|\|\|\|\|\|\|\|\|\|\|\|\|\|\|\|\|\|\|\|\|\|\|\|\|\|\|\|\|\|\|\|\|\|\|\|\|\|\|\|\|\|\|\|\|\|\|\|\|\|\|\|\|\|\|\|\|\| \|\|\| 	ATGGCCTCGTCAGGCTCTGGACTGACCCGACGCCGGGGTGCAGGCGGCGGAGGAGGAGCTGGCGACAATATCGGTGACGACCCCAGCCGCGTGTCCTCGCCTGCACCGAAACCCAACAACAACAACAACCAGTCCGGGCCCGAGACATCATACGAGTCCAGCGAAAACGGCCACAAAATCGCCTTCGACCCTCGCGACATCTCCGAGAATGAAGAGCGCTCCAAGCAGCCCAAACTCACGCTTATGGAGGAATGCTTGCTCATGGGTCTCAAGGACAAGCAGGGTTACTTGAGCTTCTGGAACGACAACATCAGCTATGCGCTGCGTGGGTGCATTGTTATTGAGTTGGCGTTTCGAGGGAGGATCAGCATGCAGAAAGACGCGAGTAGGCGGCGATTCCCCCTGGCGGACAGGCTGATTGAGGTGATTGACGACACATTGACGGGAGAAGTGCTACTGGACGAAGCGTTGAAGATGATGAAGAGCTCGGAGAAAATGAGTGTCAGTTCGTGGATTGATCTCATGAGCGGTGAG ```      ``` 785	GGCGAGACGTGGAATCTCATGAAGATCGGATACCAACTGAAGCAAGTACGCGAGCGATTGGCCAAGGGACTGGTCGACAAGGGAATTCTGCGAACAGAAAAGCGCAACTTCTTGCTTTTTGACATGGCTACACACCCAGTCGCGGATGGCGGTGCGAAAGACGAGATCAAAAGACGAGTGCGAGCTGTCCTGACAAACAGGACCGTTGTTCTGCCGGCCTCTCCTTTCCTGCCCGAAGGCATGGAGTTCCGGTATCTCAGGACAATCGCAATGCTCTGCGCTGCGTATGCCGCGAATGTGCTCGAGAACGCTCTGGTGACGCTCGATCATGAAGCACGTGAGCGGGCTTTCGCGCAAGTGGACGAACTGCTGGC-TG-AGTA	1164 	\|\|\|\|\|\|\|\|\|\|\|\|\|\|\|\|\|\|\|\|\|\|\|\|\|\|\|\|\|\|\|\|\|\|\|\|\|\|\|\|\|\|\|\|\|\|\|\|\|\|\|\|\|\|\|\|\|\|\|\|\|\|\|\|\|\|\|\|\|\|\|\|\|\|\|\|\|\|\|\|\|\|\|\|\|\|\|\|\|\|\|\|\|\|\|\|\|\|\|\|\|\|\|\|\|\|\|\|\|\|\|\|\|\|\|\|\|\|\|\|\|\|\|\|\|\|\|\|\|\|\|\|\|\|\|\|\|\|\|\|\|\|\|\|\|\|\|\|\|\|\|\|\|\|\|\|\|\|\|\|\|\|\|\|\|\|\|\|\|\|\|\|\|\|\|\|\|\|\|\|\|\|\|\|\|\|\|\|\|\|\|\|\|\|\|\|\|\|\|\|\|\|\|\|\|\|\|\|\|\|\|\|\|\|\|\|\|\|\|\|\|\|\|\|\|\|\|\|\|\|\|\|\|\|\|\|\|\|\|\|\|\|\|\|\|\|\|\|\|\|\|\|\|\|\|\|\|\|\|\|\|\|\|\|\|\|\|\|\|\|\|\|\|\|\|\|\|\|\|\|\|\|\|\|\|\|\|\|\|\|\|\|\|\|\|\|\|\|\|\|\|\|\|\|\|\| \|\|\|\|\|\|\|\|\|\|\|\|\|\|\|\|\|\|\|\|\|\|\|\|\|\|\|\|\|\|\|\|\|\|\|\|\|\|\|\|\|\|\|\|\|\|\|\|\|\|\|\|\|\|\|\|\|\|\|\|\|\|\|\|\|\|\| \|\| \|\|\|\| 	GGCGAGACGTGGAATCTCATGAAGATCGGATACCAACTGAAGCAAGTACGCGAGCGATTGGCCAAGGGACTGGTCGACAAGGGAATTCTGCGAACAGAAAAGCGCAACTTCTTGCTTTTTGACATGGCTACACACCCAGTCGCGGATGGCGGTGCGAAAGACGAGATCAAAAGACGAGTGCGAGCTGTCCTGACAAACAGGACCGTTGTTCTGCCGGCCTCTCCTTTCCTGCCCGAAGGCATGGAGTTCCGGTATCTCAGGACAATCGCAATGCTCTGCGCTGCGTATGCCGCGAATGTGCTCGAG-ACGCTCTGGTGACGCTCGATCATGAAGCACGTGAGCGGGCTTTCGCGCAAGTGGACGAACTGCTGGCCTGGAGTA ``` |
| 24 | ggggacaagtttgtacaaaaaagcaggcttgatgTCCACACAAGGACCTTCCCT, ggggaccactttgtacaagaaagctgggtcCCACGGACCCCAATTACCCC | Mycgr3\|105643  JGI  Ensembl |  |  | 100 % | chr\_9:211509 C->T; |  |
| 24 | ggggacaagtttgtacaaaaaagcaggcttgatgATCTCCAACGTTGTCGCTCT, ggggaccactttgtacaagaaagctgggtcCAAGAAGAAAGCGAGGACGG | Mycgr3\|106452  JGI  Ensembl | KOG1187 Serine/threonine protein kinase; | UDP-N-acetylmuramoylalanyl-D-glutamyl-2,6-diaminopimelate-D-alanyl-D-alanine ligase activity; coenzyme F420-0 gamma-glutamyl ligase activity;  coenzyme F420-2 alpha-glutamyl ligase activity;  ribosomal S6-glutamic acid ligase activity; | 1.85185 % |  |  |
| 24 | ggggacaagtttgtacaaaaaagcaggcttgatgGCCTATCCAGACGACAGACC, ggggaccactttgtacaagaaagctgggtcCGCAGGCCTTCCATTCTTGA | Mycgr3\|107165  JGI  Ensembl | KOG1874 KEKE-like motif-containing transcription regulator (Rlr1)/suppressor of sin4; | FK506-sensitive peptidyl-prolyl cis-trans isomerase; cyclophilin; cyclophilin-type peptidyl-prolyl cis-trans isomerase activity;  peptidyl-prolyl cis-trans isomerase activity; | 100 % |  |  |
| 24 | ggggacaagtttgtacaaaaaagcaggcttgatgGAGTCGCATCCGCCACCGTT, ggggaccactttgtacaagaaagctgggtcGCCACGCCTCATCGTTGCGG | Mycgr3\|107215  JGI  Ensembl | KOG1869 Splicing coactivator SRm160/300, subunit SRm300; |  | 100 % |  |  |
| 24 | ggggacaagtttgtacaaaaaagcaggcttgatgCCAGAAGCGATTGCCACCAC, ggggaccactttgtacaagaaagctgggtcTCCTACCGTCTTCCCTATCG | Mycgr3\|107300  JGI  Ensembl | KOG4765 Uncharacterized conserved protein; |  | 100 % |  |  |
| 24 | ggggacaagtttgtacaaaaaagcaggcttgatgGCCGACCGCACTCCAGTCTC, ggggaccactttgtacaagaaagctgggtcTGACTTTTTCCCTGTAAGAA | Mycgr3\|107425  JGI  Ensembl | KOG1181 FOG: Low-complexity; |  | 100 % |  |  |
| 24 | ggggacaagtttgtacaaaaaagcaggcttgatgTCGACTTACTTCACACTACC, ggggaccactttgtacaagaaagctgggtcAACCACCTTCTCGGTCTCCT | Mycgr3\|107818  JGI  Ensembl | KOG2992 Nucleolar GTPase/ATPase p130; |  | 100 % |  | ``` 1	ATGTCGACTTACTTCACACTACCCTCCTTCGCAAGGGCGAAGAAACACCAGGACGAGCTTGAGAAGAGCAATCCGCAGGAACCTGTGCTCAAAGAGGAGGATGAGAAGTTCTTGGAGAGACATTTGTCAATCGACGAGTCAGCCGGCGCCGCGACAAATGCAGCGACATCAGCGCCGAAGCCGAAGAAGGAGAAAGAGAAGACCCTAGACCTCCCAAGCCAAGAAGAGGCCGAGGCTGCGACAAGATCATGGAATGCGCAGAACACCTCAAGTGGAGGTACGGATACTGGAGATCACAAGCGGACATGGGCATCCTACATTCCGGCGGCTCTGGTGCCTTCGAAAGGCGACAGCTCCAAGCAGGCGGAAGAGAAGCAGTCCAGTGAAGGCGAGACTGGCCAGAAGCAGACATGGGCCATGTATGCATCGCAATATGTACCATCGTCAGCTACGACCCTGCCATCTCTCCCTGCGATGCCGGCAATCCCTGCTCTGAGCTCGTGGTATCGATCCAAGGACAAGGACGCCACACCGGAGCTGGTCTACAAAGAAGACGGAACTGTTGACGAGGCCGCCACGAAAGAGAAGCAAGAAAAGGAAGTCTCGGTACTTCTTGACAACCTCAACATGAGCAGCATCAACAACCGCGTCTTTGCACTCTCTGGTGAAACGCACAAGATCTACGAGCGCTTCGCTTTGGTGCTCAAGGACACCATTGAAGGAGGACCGACTGCATACGAAGACATGGAGAAGCTTATGAAGGATGCTGGACCCACGCTGGAGAAGCAGTTCCAGTCGATGCCACCCTTCGTACAGACTTTGGTCAAGAGTCTCCCCGCCAAGATGGGACTTGGCTTAGCTCCTGAGTTTCTCGCAACTCTCTCGGGCGACAAGCCCGATGACCATCTCAAAGCCCAAGGAGCGACTGCTTCCGCCAGCACTGTTTCTACAGACgaaaaggaaaagaagaagaaaaagaggaagaTACCTGGATTGAAGAAGTTGGTCAGCCAGCAGGGTGCCGTTGCGACGATGTTGCGCAATACTGTCTCCTTCATCACCACGCGATTCCCATTCCTGGCTTCGACGA-CAAATGTTGTCATGTCATTGTCGG	1114 	\|\|\|\|\|\|\|\|\|\|\|\|\|\|\|\|\|\|\|\|\|\|\|\|\|\|\|\|\|\|\|\|\|\|\|\|\|\|\|\|\|\|\|\|\|\|\|\|\|\|\|\|\|\|\|\|\|\|\|\|\|\|\|\|\|\|\|\|\|\|\|\|\|\|\|\|\|\|\|\|\|\|\|\|\|\|\|\|\|\|\|\|\|\|\|\|\|\|\|\|\|\|\|\|\|\|\|\|\|\|\|\|\|\|\|\|\|\|\|\|\|\|\|\|\|\|\|\|\|\|\|\|\|\|\|\|\|\|\|\|\|\|\|\|\|\|\|\|\|\|\|\|\|\|\|\|\|\|\|\|\|\|\|\|\|\|\|\|\|\|\|\|\|\|\|\|\|\|\|\|\|\|\|\|\|\|\|\|\|\|\|\|\|\|\|\|\|\|\|\|\|\|\|\|\|\|\|\|\|\|\|\|\|\|\|\|\|\|\|\|\|\|\|\|\|\|\|\|\|\|\|\|\|\|\|\|\|\|\|\|\|\|\|\|\|\|\|\|\|\|\|\|\|\|\|\|\|\|\|\|\|\|\|\|\|\|\|\|\|\|\|\|\|\|\|\|\|\|\|\|\|\|\|\|\|\|\|\|\|\|\|\|\|\|\|\|\|\|\|\|\|\|\|\|\|\|\|\|\|\|\|\|\|\|\|\|\|\|\|\|\|\|\|\|\|\|\|\|\|\|\|\|\|\|\|\|\|\|\|\|\|\|\|\|\|\|\|\|\|\|\|\|\|\|\|\|\|\|\|\|\|\|\|\|\|\|\|\|\|\|\|\|\|\|\|\|\|\|\|\|\|\|\|\|\|\|\|\|\|\|\|\|\|\|\|\|\|\|\|\|\|\|\|\|\|\|\|\|\|\|\|\|\|\|\|\|\|\|\|\|\|\|\|\|\|\|\|\|\|\|\|\|\|\|\|\|\|\|\|\|\|\|\|\|\|\|\|\|\|\|\|\|\|\|\|\|\|\|\|\|\|\|\|\|\|\|\|\|\|\|\|\|\|\|\|\|\|\|\|\|\|\|\|\|\|\|\|\|\|\|\|\|\|\|\|\|\|\|\|\|\|\|\|\|\|\|\|\|\|\|\|\|\|\|\|\|\|\|\|\|\|\|\|\|\|\|\|\|\|\|\|\|\|\|\|\|\|\|\|\|\|\|\|\|\|\|\|\|\|\|\|\|\|\|\|\|\|\|\|\|\|\|\|\|\|\|\|\|\|\|\|\|\|\|\|\|\|\|\|\|\|\|\|\|\|\|\|\|\|\|\|\|\|\|\|\|\|\|\|\|\|\|\|\|\|\|\|\|\|\|\|\|\|\|\|\|\|\|\|\|\|\|\|\|\|\|\|\|\|\|\|\|\|\|\|\|\|\|\|\|\|\|\|\|\|\|\|\|\|\|\|\|\|\|\|\|\|\|\|\|\|\|\|\|\|\|\|\|\|\|\|\|\|\|\|\|\|\|\|\|\|\|\|\|\|\|\|\|\|\|\|\|\|\|\|\|\|\|\|\|\|\|\|\|\|\|\|\|\|\|\|\|\|\|\|\|\|\|\|\|\|\|\|\|\|\|\|\|\|\|\|\|\|\|\|\|\|\|\|\|\|\|\|\|\|\|\|\|\|\|\|\|\|\|\|\|\|\|\|\|\|\|\|\|\|\|\|\|\|\|\|\|\|\|\|\|\|\|\|\|\|\|\|\|\|\|\|\|\|\|\|\|\|\|\|\|\|\|\|\|\|\|\|\|\|\|\|\|\|\|\|\|\|\|\|\|\|\|\|\|\|\|\|\|\|\|\|\|\|\|\|\|\|\|\|\|\|\|\|\|\|\|\|\|\|\|\|\|\|\|\|\|\|\|\|\|\|\|\|\|\|\|\|\|\|\|\|\|\|\|\|\|\|\|\|\|\|\|\|\|\|\|\|\|\|\|\|\|\|\|\|\|\|\|\|\|\|\|\|\|\|\|\|\|\|\|\|\|\|\|\|\|\|\|\|\|\|\|\|\|\|\|\|\|\|\|\|\|\|\|\|\|\|\|\|\|\|\|\|\|\|\|\|\|\|\|\|\|\|\|\|\|\|\|\|\|\|\|\|\|\|\|\|\|\|\|\| \|\| \|\|\|\|\| \|\|\| \| \|\|\|\|\|\|\|\| \|\|\|\|\| \|\| \|\| \|\|\|\|\|\|\| \|\|\|\|\|\| \|\|\| \|\| \|\|\|\|\|\|\|\|\| \|\|\|\|\|\|\|\|\|\|\|\|\|\|\|\|  \|\|\|\|\|\|\|\|\|\|\|\|\|\|\| \|\|\|\|\| \| \|\| \|\| \|\|\|\|\|\| \|\|\|\|\|\| \|\| \|\|\|\|\|\|\|\|\|\|\|\|\|\| 	ATGTCGACTTACTTCACACTACCCTCCTTCGCAAGGGCGAAGAAACACCAGGACGAGCTTGAGAAGAGCAATCCGCAGGAACCTGTGCTCAAAGAGGAGGATGAGAAGTTCTTGGAGAGACATTTGTCAATCGACGAGTCAGCCGGCGCCGCGACAAATGCAGCGACATCAGCGCCGAAGCCGAAGAAGGAGAAAGAGAAGACCCTAGACCTCCCAAGCCAAGAAGAGGCCGAGGCTGCGACAAGATCATGGAATGCGCAGAACACCTCAAGTGGAGGTACGGATACTGGAGATCACAAGCGGACATGGGCATCCTACATTCCGGCGGCTCTGGTGCCTTCGAAAGGCGACAGCTCCAAGCAGGCGGAAGAGAAGCAGTCCAGTGAAGGCGAGACTGGCCAGAAGCAGACATGGGCCATGTATGCATCGCAATATGTACCATCGTCAGCTACGACCCTGCCATCTCTCCCTGCGATGCCGGCAATCCCTGCTCTGAGCTCGTGGTATCGATCCAAGGACAAGGACGCCACACCGGAGCTGGTCTACAAAGAAGACGGAACTGTTGACGAGGCCGCCACGAAAGAGAAGCAAGAAAAGGAAGTCTCGGTACTTCTTGACAACCTCAACATGAGCAGCATCAACAACCGCGTCTTTGCACTCTCTGGTGAAACGCACAAGATCTACGAGCGCTTCGCTTTGGTGCTCAAGGACACCATTGAAGGAGGACCGACTGCATACGAAGACATGGAGAAGCTTATGAAGGATGCTGGACCCACGCTGGAGAAGCAGTTCCAGTCGATGCCACCCTTCGTACAGACTTTGGTCAAGAGTCTCCCCGCCAAGATGGGACTTGGCTTAGCTCCTGAGTTTCTCGCAACTCTCTCGGGCGACAAGCCCGATGACCATCTCAAAGCCCAAGGAGCGACTGCTTCCGCCAGCACTGTTTCTACAGACGAAAAGGAAAAGA-GA-GAAAAGGAG-A-GATACCTG-ATTGA-GA-GT-GGTCAGC-AGCAGG-TGC-GT-GCGACGATG-TGCGCAATACTGTCTC--TCATCACCACGCGAT-CCCAT-C-TGCCT-CGACGAACAAATG-TGCCATGTCATTGTCGG ``` |
| 24 | ggggacaagtttgtacaaaaaagcaggcttgatgTCGTACTTTTCTCAGCCGAC, ggggaccactttgtacaagaaagctgggtcTAGAAGGTTAGACAGCGAAC | Mycgr3\|108148  JGI  Ensembl |  |  | 100 % |  |  |
| 24 | ggggacaagtttgtacaaaaaagcaggcttgatgTGCCGCTTCCTTATCTACTC, ggggaccactttgtacaagaaagctgggtcAGCTGCACCAACCACAGCAC | Mycgr3\|108197  JGI  Ensembl | KOG1268 Glucosamine 6-phosphate synthetases, contain amidotransferase and phosphosugar isomerase domains; | metabolic process; | 100 % |  | ``` 255	ATGTGCCGCTTCCTTATCTACTCCGGACAATCGCCCATCCTCCTCTCCCACCTCATCACCTCTCCCACCCACAGCATCCTCACCCAGTCCTACGACTCCCGCCTCCGCATCGACCACACCCGGCCGCATAATGGCGACGGCTTCGGCGTGGGTTACTACACCACCACAGGCGTCCTCCCCACTCGTCGACGTTCCAGTCTAAGCAATGGCACGACCGACATCTTCACCGAGAGGCCCGCAGCCTCGCGTCAAGCTCTCCTCGACTTGGGTCCCGAACCGTGCATCTTCACCTCTACCATCCCCGCCTGGAACTGTCGCAACCTCGAACGTCTCGCCAGCAAGACTGTCAGTCCGCTCATCTTCGCACATGTCCGGGCCAGCACGGAGGGCGCTTTGAGTGACAGCAACTGCCACCCATTCGCACGCGGAGCGTTGATGTGGATGCACAATGGCGGTATTGGAGGTTGGAAACTGGGCGTCAAGAGGAAGTTGGTATCGGATGTGAACGACCGATGGTTTGAATGCGTGCAGGGCGCGACGGACAGCGAGTGGGCGTTTGCTCTGTTCTTGGACTCATTGGAGAAGGCGGGCGTGGATGCTGACGATGCGGTGAAGCAGAGGGATGGATTCGGGCATACCGTGTTGAGG-AAGGCGATGTTGAAGACGATTGAGAGGATCAATGGATATATTAGGTCGTTGCCGGCGAGCGTGAGGGAGGAAGGGGATGTGAGGAGCTTGCTGAACTTTGCGATTACGGATGGCAAGAGCGTGGTGTGCACAAGATATGTGTCGTCGAAGACGGACGAGGCGGCGAGTCTGTTCTTTTCCTCTGGCACGAACTGGGCAGAGCAGCCATCTGCAACA-CCACAGACCGGGCGAGGAAAGATTGGCAAGGGCGAATACAAGA-TGGAGCGGA	1170 	\|\|\|\|\|\|\|\|\|\|\|\|\|\|\|\|\|\|\|\|\|\|\|\|\|\|\|\|\|\|\|\|\|\|\|\|\|\|\|\|\|\|\|\|\|\|\|\|\|\|\|\|\|\|\|\|\|\|\|\|\|\|\|\|\|\|\|\|\|\|\|\|\|\|\|\|\|\|\|\|\|\|\|\|\|\|\|\|\|\|\|\|\|\|\|\|\|\|\|\|\|\|\|\|\|\|\|\|\|\|\|\|\|\|\|\|\|\|\|\|\|\|\|\|\|\|\|\|\|\|\|\|\|\|\|\|\|\|\|\|\|\|\|\|\|\|\|\|\|\|\|\|\|\|\|\|\|\|\|\|\|\|\|\|\|\|\|\|\|\|\|\|\|\|\|\|\|\|\|\|\|\|\|\|\|\|\|\|\|\|\|\|\|\|\|\|\|\|\|\|\|\|\|\|\|\|\|\|\|\|\|\|\|\|\|\|\|\|\|\|\|\|\|\|\|\|\|\|\|\|\|\|\|\|\|\|\|\|\|\|\|\|\|\|\|\|\|\|\|\|\|\|\|\|\|\|\|\|\|\|\|\|\|\|\|\|\|\|\|\|\|\|\|\|\|\|\|\|\|\|\|\|\|\|\|\|\|\|\|\|\|\|\|\|\|\|\|\|\|\|\|\|\|\|\|\|\|\|\|\|\|\|\|\|\|\|\|\|\|\|\|\|\|\|\|\|\|\|\|\|\|\|\|\|\|\|\|\|\|\|\|\|\|\|\|\|\|\|\|\|\|\|\|\|\|\|\|\|\|\|\|\|\|\|\|\|\|\|\|\|\|\|\|\|\|\|\|\|\|\|\|\|\|\|\|\|\|\|\|\|\|\|\|\|\|\|\|\|\|\|\|\|\|\|\|\|\|\|\|\|\|\|\|\|\|\|\|\|\|\|\|\|\|\|\|\|\|\|\|\|\|\|\|\|\|\|\|\|\|\|\|\|\|\|\|\|\|\|\|\|\|\|\|\|\|\|\|\|\|\|\|\|\|\|\|\|\|\|\|\|\|\|\|\|\|\|\|\|\|\|\|\|\|\|\|\|\|\|\|\|\|\|\|\|\|\|\|\|\|\|\|\|\|\|\|\|\|\|\|\|\|\|\|\|\|\|\|\|\|\|\|\|\|\|\|\|\|\|\|\|\|\|\|\|\|\|\|\|\|\|\|\|\|\|\|\|\|\|\|\|\|\|\|\|\|\|\|\|\|\|\|\|\|\|\|\|\|\|\|\|\|\|\|\|\|\|\|\|\|\|\|\|\|\|\|\|\|\|\|\|\|\|\|\|\|\|\|\|\|\|\|\|\|\|\|\|\|\|\|\|\|\|\|\|\|\|\|\|\|\|\|\|\|\|\|\|\|\|\|\|\|\|\|\|\|\|\|\|\|\|\|\|\|\|\|\|\|\| \|\|\|\|\|\|\|\|\|\|\|\|\|\|\|\|\|\|\|\|\|\|\|\|\|\|\|\|\|\|\|\|\|\|\|\|\|\|\|\|\|\|\| \|\|\|\|\|\|\|\|\|\|\|\|\|\|\|\|\|\|\|\|\|\|\|\|\|\|\|\|\|\|\|\|\|\|\|\|\|\|\|\|\|\|\|\|\|\|\|\|\|\|\|\|\|\|\|\|\|\|\|\|\|\|\|\|\|\|\|\|\|\|\|\|\|\|\|\|\|\|\|\|\|\|\|\|\|\|\|\|\|\|\|\|\|\|\|\|\|\|\|\|\|\|\|\|\|\|\|\|\|\|\|\|\|\|\|\|\|\|\|\|\|\|\|\|\|\|\|\|\|\|\|\|\|\|\|\|\|\|\|\|\|\|\|\|\|\|\|\|\|\|\|\|\|\|\|\|\|\|\|\|\|\|\|\|\|\|\|\|\|\|\|\| \|\|\|\|\|\|\|\|\|\|\|\|\|\|\|\|\|\|\|\|\|\|\|\|\|\|\|\|\|\|\|\|\|\|\|\|\|\|\|\|\|\|\| \|\|\|\|\|\|\|\|\| 	ATGTGCCGCTTCCTTATCTACTCCGGACAATCGCCCATCCTCCTCTCCCACCTCATCACCTCTCCCACCCACAGCATCCTCACCCAGTCCTACGACTCCCGCCTCCGCATCGACCACACCCGGCCGCATAATGGCGACGGCTTCGGCGTGGGTTACTACACCACCACAGGCGTCCTCCCCACTCGTCGACGTTCCAGTCTAAGCAATGGCACGACCGACATCTTCACCGAGAGGCCCGCAGCCTCGCGTCAAGCTCTCCTCGACTTGGGTCCCGAACCGTGCATCTTCACCTCTACCATCCCCGCCTGGAACTGTCGCAACCTCGAACGTCTCGCCAGCAAGACTGTCAGTCCGCTCATCTTCGCACATGTCCGGGCCAGCACGGAGGGCGCTTTGAGTGACAGCAACTGCCACCCATTCGCACGCGGAGCGTTGATGTGGATGCACAATGGCGGTATTGGAGGTTGGAAACTGGGCGTCAAGAGGAAGTTGGTATCGGATGTGAACGACCGATGGTTTGAATGCGTGCAGGGCGCGACGGACAGCGAGTGGGCGTTTGCTCTGTTCTTGGACTCATTGGAGAAGGCGGGCGTGGATGCTGACGATGCGGTGAAGCAGAGGGATGGATTCGGGCATACCGTGTTGAGGAAAGGCGATGTTGAAGACGATTGAGAGGATCAATGGATATATTA-GTCGTTGCCGGCGAGCGTGAGGGAGGAAGGGGATGTGAGGAGCTTGCTGAACTTTGCGATTACGGATGGCAAGAGCGTGGTGTGCACAAGATATGTGTCGTCGAAGACGGACGAGGCGGCGAGTCTGTTCTTTTCCTCTGGCACGAACTGGGCAGAGCAGCCATCTGCAACACCCACAGACCGGGCGAGGAAAGATTGGCAAGGGCGAATACAAGAATGGAGCGGA ``` |
| 24 | ggggacaagtttgtacaaaaaagcaggcttgatgGCGGAAAAGATTGAGAAGAC, ggggaccactttgtacaagaaagctgggtcCCCCGTCCACCCTTCCCTCC | Mycgr3\|108426  JGI  Ensembl | KOG3968 Atrazine chlorohydrolase/guanine deaminase; | guanine deaminase activity; hydrolase activity; | 1.08225 % |  |  |
| 24 | ggggacaagtttgtacaaaaaagcaggcttgatgTTCAGTCAGGGATCAGCAGT, ggggaccactttgtacaagaaagctgggtcATGTGAGTGAAGGCTATTTG | Mycgr3\|108931  JGI  Ensembl |  |  | 100 % |  |  |
| 24 | ggggacaagtttgtacaaaaaagcaggcttgatgATGTCGAACAACAACAACAA, ggggaccactttgtacaagaaagctgggtcTTCTTCCAGCTCGTTCTTCT | Mycgr3\|110384  JGI  Ensembl |  |  | 100 % |  |  |
| 24 | ggggacaagtttgtacaaaaaagcaggcttgatgAAGCTTCCATCACAGTCTAC, ggggaccactttgtacaagaaagctgggtcACCCTCGTTCGTGGCATTAT | Mycgr3\|110592  JGI  Ensembl |  | hydrolase activity, acting on ester bonds; metabolic process; | 100 % |  | ``` 278	ATGAAGCTTCCATCACAGTCTACATTCGCCCTGGTCGCGTTTGCGGCATTCAGCTATGCTAAGACGATCTACACCGGAGACACGCTTCAAGGCTATCCGGTCATTGCCTCTCTGGACATCAACGATGTGCCCGCGAACCAAATCTCCCGATTCTGGCTATCGCCCGCGTCCGGTCAAGGCGGATTGCCATACTTTCTCCCCATCTTCGTGGCTCGCGGATCGGAAGAGTCACTCGAGACTGGTCGGAAGTTCTCACTCTCTGCCTCGATTCACGGCGACGAGCTGAATCCTGTGGCGGTGGTGCAGAAAGTATTCGCCAGGCTCAACGAGACGGTGGCATCGGGAGACTTCAATGGCACAGTCATTGGACTGCCTACGCAGAATCCTAACGGCAACCTTATGAATCAGCGATACTTCTATACGAGCTCATCTAATGGCTTCCTCACGGATCTTAATCGGAACTTTCCTGGGATGAGCATTGCCGAAGGAGGCAGCCTACCCATGTCGTATACTGCTGCTATATGGAACAATATCTGGGGCAATACGTCGAACGTGGACATTGCAGTTGACCTCC	851 	\|\|\|\|\|\|\|\|\|\|\|\|\|\|\|\|\|\|\|\|\|\|\|\|\|\|\|\|\|\|\|\|\|\|\|\|\|\|\|\|\|\|\|\|\|\|\|\|\|\|\|\|\|\|\|\|\|\|\|\|\|\|\|\|\|\|\|\|\|\|\|\|\|\|\|\|\|\|\|\|\|\|\|\|\|\|\|\|\|\|\|\|\|\|\|\|\|\|\|\|\|\|\|\|\|\|\|\|\|\|\|\|\|\|\|\|\|\|\|\|\|\|\|\|\|\|\|\|\|\|\|\|\|\|\|\|\|\|\|\|\|\|\|\|\|\|\|\|\|\|\|\|\|\|\|\|\|\|\|\|\|\|\|\|\|\|\|\|\|\|\|\|\|\|\|\|\|\|\|\|\|\|\|\|\|\|\|\|\|\|\|\|\|\|\|\|\|\|\|\|\|\|\|\|\|\|\|\|\|\|\|\|\|\|\|\|\|\|\|\|\|\|\|\|\|\|\|\|\|\|\|\|\|\|\|\|\|\|\|\|\|\|\|\|\|\|\|\|\|\|\|\|\|\|\|\|\|\|\|\|\|\|\|\|\|\|\|\|\|\|\|\|\|\|\|\|\|\|\|\|\|\|\|\|\|\|\|\|\|\|\|\|\|\|\|\|\|\|\|\|\|\|\|\|\|\|\|\|\|\|\|\|\|\|\|\|\|\|\|\|\|\|\|\|\|\|\|\|\|\|\|\|\|\|\|\|\|\|\|\|\|\|\|\|\|\|\|\|\|\|\|\|\|\|\|\|\|\|\|\|\|\|\|\|\|\|\|\|\|\|\|\|\|\|\|\|\|\|\|\|\|\|\|\|\|\|\|\|\|\|\|\|\|\|\|\|\|\|\|\|\|\|\|\|\|\|\|\|\|\|\|\|\|\|\|\|\|\|\|\|\|\|\|\|\|\|\|\|\|\|\|\|\|\|\|\|\|\|\|\|\|\|\|\|\|\|\|\|\|\|\|\|\|\|\|\|\|\|\|\|\|\|\|\|\|\|\|\|\|\|\|\|\|\|\|\|\|\|\|\|\|\|\|\|\|\|\|\|\|\|\|\|\|\|\|\|\|\|\|\|\|\|\|\|\|\|\|\|\|\|\|\|\|\|\|\|\|\|\|\|\|\|\|\|\|\|\|\|\|\|\|\|\|\|\|\|\|\|\|\|\|\|\|\|\|\|\|\|\|\|\|\|\|\|\|\|\|\|\|\|\|\|\|\|\|\|\|\|\|\|\|\|\|\| 	ATGAAGCTTCCATCACAGTCTACATTCGCCCTGGTCGCGTTTGCGGCATTCAGCTATGCTAAGACGATCTACACCGGAGACACGCTTCAAGGCTATCCGGTCATTGCCTCTCTGGACATCAACGATGTGCCCGCGAACCAAATCTCCCGATTCTGGCTATCGCCCGCGTCCGGTCAAGGCGGATTGCCATACTTTCTCCCCATCTTCGTGGCTCGCGGATCGGAAGAGTCACTCGAGACTGGTCGGAAGTTCTCACTCTCTGCCTCGATTCACGGCGACGAGCTGAATCCTGTGGCGGTGGTGCAGAAAGTATTCGCCAGGCTCAACGAGACGGTGGCATCGGGAGACTTCAATGGCACAGTCATTGGACTGCCTACGCAGAATCCTAACGGCAACCTTATGAATCAGCGATACTTCTATACGAGCTCATCTAATGGCTTCCTCACGGATCTTAATCGGAACTTTCCTGGGATGAGCATTGCCGAAGGAGGCAGCCTACCCATGTCGTATACTGCTGCTATATGGAACAATATCTGGGGCAATACGTCGAACGTGGACATTGCAGTTGACCTCC ```      ``` 852	ACACTCTATCAACAGGCTCAATTGGCCCTCTCTGGGTCTACGCCGACTACGCATTAG-AAGGCGTGCAGAGAATCGCCGAGCTCGCCCAGCCAGACATGATCAAAATCGACCCTGGCCAAAGTGGCACCATCGAGACCTCCTTCGTCGAAAGAGGGATCCCAGCCATCACAATGGAGATCGGACCAGCAAATAACTGGAACGGAACGCTCATCTCAAGAGCTGTCGACTTTGTCTTTCGCCTGATGGACGATTTGCAAATGACCACTGGCGAGACTCCGACTCCTGATCTTTCCAACACTTACATCGCCAACAACTTCTCGGATGTGACGGTATCGCACTCTGGCTGGGTTGAACTGGATG-TGACTAC	1218 	\|\|\|\|\|\|\|\|\|\|\|\|\|\|\|\|\|\|\|\|\|\|\|\|\|\|\|\|\|\|\|\|\|\|\|\|\|\|\|\|\|\|\|\|\|\|\|\|\|\|\|\|\|\|\|\|\| \|\|\|\|\|\|\|\|\|\|\|\|\|\|\|\|\|\|\|\|\|\|\|\|\|\|\|\|\|\|\|\|\|\|\|\|\|\|\|\|\|\|\|\|\|\|\|\|\|\|\|\|\|\|\|\|\|\|\|\|\|\|\|\|\|\|\|\|\|\|\|\|\|\|\|\|\|\|\|\|\|\|\|\|\|\|\|\|\|\|\|\|\|\|\|\|\|\|\|\|\|\|\|\|\|\|\|\|\|\|\|\|\|\|\|\|\|\|\|\|\|\|\|\|\|\|\|\|\|\|\|\|\|\|\|\|\|\|\|\|\|\|\|\|\|\|\|\|\|\|\|\|\|\|\|\|\|\|\|\|\|\|\|\|\|\|\|\|\|\|\|\|\|\|\|\|\|\|\|\|\|\|\|\|\|\|\|\|\|\|\|\|\|\|\|\|\|\|\|\|\|\|\|\|\|\|\|\|\|\|\|\|\|\|\|\|\|\|\|\|\|\|\|\|\|\|\|\|\|\|\|\|\| \| \| \|\|\|\| \|\|\|\|\|\|\| \| \|\| \|\| \|\|\|\| \|\|\|\|\|\|\| \|\|\|\|\|\|\|\|\|\|\|\|\|\|\|\|\|\| \| \|\|\|\|\|\|\|\|\|\| \|\|\|\|\|\|\| 	ACACTCTATCAACAGGCTCAATTGGCCCTCTCTGGGTCTACGCCGACTACGCATTAGAAAGGCGTGCAGAGAATCGCCGAGCTCGCCCAGCCAGACATGATCAAAATCGACCCTGGCCAAAGTGGCACCATCGAGACCTCCTTCGTCGAAAGAGGGATCCCAGCCATCACAATGGAGATCGGACCAGCAAATAACTGGAACGGAACGCTCATCTCAAGAGCTGTCGACTTTGTCTTTCGCCTGATGGACGATTTGCAAATGACCACTGGCGAGACTCCGACTCCTGATCTT-C-A-CACT-ACATCGC-A-CA-CT-CTCG-ATGTGAC-GTATCGCACTCTGGCTGG-T-GAACTGGATGGTGACTAC ``` |
| 24 | ggggacaagtttgtacaaaaaagcaggcttgatgTCACTCGTTCACCCGCTCGG, ggggaccactttgtacaagaaagctgggtcAAACTGCTCCCTCGGCTTCA | Mycgr3\|110790  JGI  Ensembl |  |  | 100 % |  |  |
| 24 | ggggacaagtttgtacaaaaaagcaggcttgatgGCCTTGGACGAGCCAGAGCT, ggggaccactttgtacaagaaagctgggtcAGCTTCCAAATCCCCACGCC | Mycgr3\|111491  JGI  Ensembl | KOG2793 Putative N2,N2-dimethylguanosine tRNA methyltransferase; |  | 100 % |  |  |
| 24 | ggggacaagtttgtacaaaaaagcaggcttgatgTGGGACGTCTCGGCCTGGTT, ggggaccactttgtacaagaaagctgggtcCAGATCCAGCAGGCTGTCCG | Mycgr3\|111784  JGI  Ensembl |  |  | 1.43266 % |  |  |
| 24 | ggggacaagtttgtacaaaaaagcaggcttgatgTCCGACTTCAACGTCACATC, ggggaccactttgtacaagaaagctgggtcCGAACATCCAAATGCTGTCG | Mycgr3\|111795  JGI  Ensembl |  |  | 100 % |  |  |
| 25 | ggggacaagtttgtacaaaaaagcaggcttgatgCCCTCCGCCCCACCCATCCC, ggggaccactttgtacaagaaagctgggtcCATCTTGGACTTCTGGCACT | Mycgr3\|31720  JGI  Ensembl | KOG0139 Short-chain acyl-CoA dehydrogenase; | acyl-CoA dehydrogenase activity; electron transport;  isovaleryl-CoA dehydrogenase activity;  oxidoreductase activity, acting on the CH-CH group of donors; | 100 % |  |  |
| 25 | ggggacaagtttgtacaaaaaagcaggcttgatgTCCCCCCACCTCTCCGGACC, ggggaccactttgtacaagaaagctgggtcCAAAAACCCCCCCAAAGTCT | Mycgr3\|32667  JGI  Ensembl |  |  | 100 % | chr\_1:5928391 A->C; |  |
| 25 | ggggacaagtttgtacaaaaaagcaggcttgatgGACGACACCGACGGCGAACC, ggggaccactttgtacaagaaagctgggtcTCCGCCTCGAGCGATCCATA | Mycgr3\|33998  JGI  Ensembl |  |  | 100 % |  |  |
| 25 | ggggacaagtttgtacaaaaaagcaggcttgatgAGGCCCCGACTATTGCTGGT, ggggaccactttgtacaagaaagctgggtcCACAAAAATGGCCAGCACAA | Mycgr3\|34014  JGI  Ensembl |  |  | 100 % |  |  |
| 25 | ggggacaagtttgtacaaaaaagcaggcttgatgGGTCGATTGATTGTTCACCA, ggggaccactttgtacaagaaagctgggtcATTCCAGTCGACACCTCCAA | Mycgr3\|35448  JGI  Ensembl | KOG1339 Aspartyl protease; | aspartic-type endopeptidase activity; pepsin A activity;  proteolysis; | 11.33912 % | chr\_2:3535973 A->G; chr\_2:3532482 G->A; chr\_2:3385914 C->T; chr\_2:3270146 G->A; chr\_2:3537221 G->A; chr\_2:3374911 C->T; chr\_2:3461331 T->\*; chr\_2:3536866 T->C; |  |
| 25 | ggggacaagtttgtacaaaaaagcaggcttgatgGCAGAGATCGAAGTCCAAAG, ggggaccactttgtacaagaaagctgggtcCAATACCAGCGCACCGATGA | Mycgr3\|37295  JGI  Ensembl |  |  | 0 % |  |  |
| 25 | ggggacaagtttgtacaaaaaagcaggcttgatgGCTCGAACAACAAGGGCCGC, ggggaccactttgtacaagaaagctgggtcCTCGTCCGCAAACTCCGATG | Mycgr3\|40778  JGI  Ensembl | KOG1921 Endonuclease III; | base-excision repair; | 0 % |  |  |
| 25 | ggggacaagtttgtacaaaaaagcaggcttgatgGCCTCCTCACCGGCGTTCCA, ggggaccactttgtacaagaaagctgggtcAGCGGCCTGGACCCGCCGAC | Mycgr3\|43851  JGI  Ensembl | KOG2614 Kynurenine 3-monooxygenase and related flavoprotein monooxygenases; | metabolic process; oxidoreductase activity;  salicylate 1-monooxygenase activity; | 0 % |  |  |
| 25 | ggggacaagtttgtacaaaaaagcaggcttgatgCTCTGCTGCTTGAAGCGCGC, ggggaccactttgtacaagaaagctgggtcTGAAGCAGATAAGTCCAGCC | Mycgr3\|46302  JGI  Ensembl | KOG1290 Serine/threonine protein kinase; | ATP binding; protein amino acid phosphorylation;  protein kinase activity;  protein serine/threonine kinase activity;  protein-tyrosine kinase activity; | 100 % |  |  |
| 25 | ggggacaagtttgtacaaaaaagcaggcttgatgAAGTACCTACTCTTCTCGGC, ggggaccactttgtacaagaaagctgggtcACCGCCCAGCGCAGAAATTA | Mycgr3\|47119  JGI  Ensembl | KOG0029 Amine oxidase; |  | 1.43443 % |  |  |
| 25 | ggggacaagtttgtacaaaaaagcaggcttgatgACTTCCGAACGAGCCAACGG, ggggaccactttgtacaagaaagctgggtcGTCCTTCTTCTTATTCATCC | Mycgr3\|47411  JGI  Ensembl |  |  | 100 % |  |  |
| 25 | ggggacaagtttgtacaaaaaagcaggcttgatgCCCAAACGCCTCGTCCTTGC, ggggaccactttgtacaagaaagctgggtcGGCAGCTCCCGTGACGAATG | Mycgr3\|48331  JGI  Ensembl |  |  | 100 % | chr\_10:1385803 A->G; |  |
| 25 | ggggacaagtttgtacaaaaaagcaggcttgatgTTTCGGGCAGCCAGCGTAAG, ggggaccactttgtacaagaaagctgggtcTGCAAGCCACTCATCCCGCA | Mycgr3\|49843  JGI  Ensembl | KOG1290 Serine/threonine protein kinase; | ATP binding; protein amino acid phosphorylation;  protein kinase activity;  protein serine/threonine kinase activity; | 100 % |  |  |
| 25 | ggggacaagtttgtacaaaaaagcaggcttgatgCAGCGAATTGCTTTGCCTCG, ggggaccactttgtacaagaaagctgggtcCTTGAGGAAGCCGAACCAGC | Mycgr3\|54676  JGI  Ensembl | KOG4300 Predicted methyltransferase; |  | 100 % |  |  |
| 25 | ggggacaagtttgtacaaaaaagcaggcttgatgACCACCCGCTACCGAGTAGA, ggggaccactttgtacaagaaagctgggtcGGACGCATCACTCAACTCCG | Mycgr3\|56200  JGI  Ensembl |  |  | 100 % |  |  |
| 25 | ggggacaagtttgtacaaaaaagcaggcttgatgGCCGGCACTACGGAAAATGC, ggggaccactttgtacaagaaagctgggtcTCGATAGAGCACGGCTTCGA | Mycgr3\|57534  JGI  Ensembl | KOG1613 Exosomal 3'-5' exoribonuclease complex, subunit Rrp43; | 3'-5'-exoribonuclease activity; 5'-3' exoribonuclease activity;  RNA binding;  RNA processing; | 100 % |  |  |
| 25 | ggggacaagtttgtacaaaaaagcaggcttgatgGTCGGATCAAAGGTTCTTTC, ggggaccactttgtacaagaaagctgggtcCAATCGTCCAGCAGATCCAC | Mycgr3\|63538  JGI  Ensembl |  |  | 100 % |  |  |
| 25 | ggggacaagtttgtacaaaaaagcaggcttgatgGCCACTTCAACACAACAGGA, ggggaccactttgtacaagaaagctgggtcTCTCGCCACCGCAGCGCTGA | Mycgr3\|63945  JGI  Ensembl | KOG2758 Translation initiation factor 3, subunit e (eIF-3e); |  | 0 % |  |  |
| 25 | ggggacaagtttgtacaaaaaagcaggcttgatgGATGCGAACGCCTACCTGAA, ggggaccactttgtacaagaaagctgggtcTTTCGAGGCTTTGCTTTTCT | Mycgr3\|65191  JGI  Ensembl | KOG2809 Telomerase elongation inhibitor/RNA maturation protein PINX1; |  | 0 % |  |  |
| 25 | ggggacaagtttgtacaaaaaagcaggcttgatgGCCTCCCTCGCTCTCCGCCG, ggggaccactttgtacaagaaagctgggtcCAAATCCTCCGCGTCCACCA | Mycgr3\|65302  JGI  Ensembl | KOG1390 Acetyl-CoA acetyltransferase; | acetyl-CoA C-acetyltransferase activity; | 1.50215 % |  |  |
| 25 | ggggacaagtttgtacaaaaaagcaggcttgatgTACGAAAAACATGAGCAAGA, ggggaccactttgtacaagaaagctgggtcACAATGCGGTCCCAGCTTCC | Mycgr3\|65725  JGI  Ensembl | KOG2504 Monocarboxylate transporter; |  | 0 % |  |  |
| 25 | ggggacaagtttgtacaaaaaagcaggcttgatgAATGGCTTTGCGGACCGCGG, ggggaccactttgtacaagaaagctgggtcACTCATCATCGCTGCCTTCT | Mycgr3\|66006  JGI  Ensembl | KOG2667 COPII vesicle protein; |  | 100 % |  |  |
| 25 | ggggacaagtttgtacaaaaaagcaggcttgatgGGCAGCACCGCACCTCAATC, ggggaccactttgtacaagaaagctgggtcCAGCTTACTCTGCTCCTCCT | Mycgr3\|66463  JGI  Ensembl | KOG0134 NADH:flavin oxidoreductase/12-oxophytodienoate reductase; | FMN binding; NADPH dehydrogenase activity;  oxidoreductase activity; | 100 % |  |  |
| 25 | ggggacaagtttgtacaaaaaagcaggcttgatgGAGGAGGACCGTGCCATTTC, ggggaccactttgtacaagaaagctgggtcCGCAACGGCCTCCACCACCT | Mycgr3\|66970  JGI  Ensembl | KOG2648 Diphthamide biosynthesis protein; |  | 0 % |  |  |
| 25 | ggggacaagtttgtacaaaaaagcaggcttgatgGCTCAAATCAGAGGCACCGC, ggggaccactttgtacaagaaagctgggtcGTAGACCTTCTTCTTCTCAT | Mycgr3\|67588  JGI  Ensembl | KOG3998 Putative cargo transport protein ERV29; |  | 100 % |  |  |
| 25 | ggggacaagtttgtacaaaaaagcaggcttgatgGACCAGAATGCAACGACCGT, ggggaccactttgtacaagaaagctgggtcCATCCCAGCACCCTTCTGCT | Mycgr3\|67763  JGI  Ensembl | KOG0743 AAA+-type ATPase; | ATP binding; nucleoside-triphosphatase activity;  nucleotide binding; | 100 % |  |  |
| 25 | ggggacaagtttgtacaaaaaagcaggcttgatgCGGTCGCCGCAATCGAAACC, ggggaccactttgtacaagaaagctgggtcGCAACTCCAATCCCTCTCGG | Mycgr3\|68318  JGI  Ensembl |  |  | 100 % |  |  |
| 25 | ggggacaagtttgtacaaaaaagcaggcttgatgGCATTTTCTTGGTTTCCGGG, ggggaccactttgtacaagaaagctgggtcCTGCTCACTCTCGGCTCGGA | Mycgr3\|68570  JGI  Ensembl | KOG1838 Alpha/beta hydrolase; |  | 0 % |  |  |
| 25 | ggggacaagtttgtacaaaaaagcaggcttgatgACATCCTTCAGTACCTTATC, ggggaccactttgtacaagaaagctgggtcACTTTTTGTATACCACCACG | Mycgr3\|70537  JGI  Ensembl | KOG1441 Glucose-6-phosphate/phosphate and phosphoenolpyruvate/phosphate antiporter; |  | 100 % |  |  |
| 25 | ggggacaagtttgtacaaaaaagcaggcttgatgCCGCGCCTCGTCCGCCGCAC, ggggaccactttgtacaagaaagctgggtcAAACCTCCCAGTCTCTCTGC | Mycgr3\|71656  JGI  Ensembl |  |  | 100 % |  |  |
| 25 | ggggacaagtttgtacaaaaaagcaggcttgatgGCCGAATCCGTCCCCATCTC, ggggaccactttgtacaagaaagctgggtcCTTGACGCTGTTCTCGGATG | Mycgr3\|71669  JGI  Ensembl | KOG3098 Uncharacterized conserved protein; |  | 1.32188 % |  |  |
| 25 | ggggacaagtttgtacaaaaaagcaggcttgatgGCCAACACAGCCAACGACAC, ggggaccactttgtacaagaaagctgggtcACCCTCATTCTTATCCCACC | Mycgr3\|71742  JGI  Ensembl | KOG2511 Nicotinic acid phosphoribosyltransferase; | nicotinate nucleotide biosynthetic process; nicotinate phosphoribosyltransferase activity;  pyridine nucleotide biosynthetic process; | 100 % |  | ``` 1	ATGGCCAACACAGCCAACGACACGCCGGAAGGCATCTTCTCGCTCCTCGATACCGACCTCTACAAGCTCACCATGCAATGCGCCGTCCTGAAATACTTCCCCGACACCCACGTCCGATACAAGTTCACCAACCGCACGCCTCAGATGAAGTTCACGAGACAAGCTTTCAATTGGTTACATGCTCAGATCAGGAAGCTGGAAATTGTCGAGGTGACAAAGGAGGAAATCGAATGGCTGAGAAGGACGTGCAAATACCTCACACCTGAGTACCTCGACTATCTACAGACATTCCGATTGCGGCCTTCAGAGCAGGTGAAGGCTTCATTCATGCCGGACAGTGATGCCACAAGCGACTCAGTCACAGGTGAGGTCTCCTTGGCGATCAGCGGTATCTGGGTCGAGACAATCCTCTACGAGATCCCACTGCTAGCCCTCACATCAGAAGCATACTTTAAGTTCGTCGACAAGGATTGGAACCACAATGGTCAAGTCGCCAAAGCAAAAGAGAAGGCCGAGAAGCTCTTCGAAGCAGGCTGTGTCTTCAGCGAGTTCGGCAGTCGACGCAGAAGAGACTACAAAACCCACGAGATGGTCATTCAAGGCATCCTCCAAGCACAAGAGGAAGCAAAGAACAACGGCAAGAACTGGAAAGGCAAACTCACCGGCACATCCAATGTCCACCTCGCCATGAAACACAACACCACACCCTCCGGCACAGTCGCCCACGAATGGTTCATGGGCATCGCAGCCATTACGCAAGACTACGCCAACGCGAACGAGAAAGCGATGGAATACTGGATCGGCACCTTCGGGCCAGGCGTGCTGGGAATCGCACTCACCGACACCTTCGGCACGCCATCTTTCCTCAAAGCATTCGCCAAACCCGCTCCTTCATCCGACTCCAGCAGAACCTCCTCCGCCTCACCCTCACCAACCTACGCACAAGTCTTCGCCGGCGTACGGCAAGACTCCGGCGATCCCCTCGAATACGTACAAATGATCCGGCAATTCTACTCCAGTCAAAACATCACCTCCCCCAAAACGGTCGTCTTC-TCCGA-CAGCCTCA	1066 	\|\|\|\|\|\|\|\|\|\|\|\|\|\|\|\|\|\|\|\|\|\|\|\|\|\|\|\|\|\|\|\|\|\|\|\|\|\|\|\|\|\|\|\|\|\|\|\|\|\|\|\|\|\|\|\|\|\|\|\|\|\|\|\|\|\|\|\|\|\|\|\|\|\|\|\|\|\|\|\|\|\|\|\|\|\|\|\|\|\|\|\|\|\|\|\|\|\|\|\|\|\|\|\|\|\|\|\|\|\|\|\|\|\|\|\|\|\|\|\|\|\|\|\|\|\|\|\|\|\|\|\|\|\|\|\|\|\|\|\|\|\|\|\|\|\|\|\|\|\|\|\|\|\|\|\|\|\|\|\|\|\|\|\|\|\|\|\|\|\|\|\|\|\|\|\|\|\|\|\|\|\|\|\|\|\|\|\|\|\|\|\|\|\|\|\|\|\|\|\|\|\|\|\|\|\|\|\|\|\|\|\|\|\|\|\|\|\|\|\|\|\|\|\|\|\|\|\|\|\|\|\|\|\|\|\|\|\|\|\|\|\|\|\|\|\|\|\|\|\|\|\|\|\|\|\|\|\|\|\|\|\|\|\|\|\|\|\|\|\|\|\|\|\|\|\|\|\|\|\|\|\|\|\|\|\|\|\|\|\|\|\|\|\|\|\|\|\|\|\|\|\|\|\|\|\|\|\|\|\|\|\|\|\|\|\|\|\|\|\|\|\|\|\|\|\|\|\|\|\|\|\|\|\|\|\|\|\|\|\|\|\|\|\|\|\|\|\|\|\|\|\|\|\|\|\|\|\|\|\|\|\|\|\|\|\|\|\|\|\|\|\|\|\|\|\|\|\|\|\|\|\|\|\|\|\|\|\|\|\|\|\|\|\|\|\|\|\|\|\|\|\|\|\|\|\|\|\|\|\|\|\|\|\|\|\|\|\|\|\|\|\|\|\|\|\|\|\|\|\|\|\|\|\|\|\|\|\|\|\|\|\|\|\|\|\|\|\|\|\|\|\|\|\|\|\|\|\|\|\|\|\|\|\|\|\|\|\|\|\|\|\|\|\|\|\|\|\|\|\|\|\|\|\|\|\|\|\|\|\|\|\|\|\|\|\|\|\|\|\|\|\|\|\|\|\|\|\|\|\|\|\|\|\|\|\|\|\|\|\|\|\|\|\|\|\|\|\|\|\|\|\|\|\|\|\|\|\|\|\|\|\|\|\|\|\|\|\|\|\|\|\|\|\|\|\|\|\|\|\|\|\|\|\|\|\|\|\|\|\|\|\|\|\|\|\|\|\|\|\|\|\|\|\|\|\|\|\|\|\|\|\|\|\|\|\|\|\|\|\|\|\|\|\|\|\|\|\|\|\|\|\|\|\|\|\|\|\|\|\|\|\|\|\|\|\|\|\|\|\|\|\|\|\|\|\|\|\|\|\|\|\|\|\|\|\|\|\|\|\|\|\|\|\|\|\|\|\|\|\|\|\|\|\|\|\|\|\|\|\|\|\|\|\|\|\|\|\|\|\|\|\|\|\|\|\|\|\|\|\|\|\|\|\|\|\|\|\|\|\|\|\|\|\|\|\|\|\|\|\|\|\|\|\|\|\|\|\|\|\|\|\|\|\|\|\|\|\|\|\|\|\|\|\|\|\|\|\|\|\|\|\|\|\|\|\|\|\|\|\|\|\|\|\|\|\|\|\|\|\|\|\|\|\|\|\|\|\|\|\|\|\|\|\|\|\|\|\|\|\|\|\|\|\|\|\|\|\|\|\|\|\|\|\|\|\|\|\|\|\|\|\|\|\|\|\|\|\|\|\|\|\|\|\|\|\|\|\|\|\|\|\|\|\|\|\|\|\|\|\|\|\|\|\|\|\|\|\|\|\|\|\|\|\|\|\|\|\|\|\|\|\|\|\|\|\|\|\|\|\|\|\|\|\|\|\|\|\|\|\|\|\|\|\|\|\|\|\|\|\|\|\|\|\|\|\|\|\|\|\|\|\|\|\|\|\|\|\|\|\|\|\|\|\|\|\|\|\|\| \|\|\|\|\|\|\| \| \|\|\|\|\|\|\|\|\|\|\|\|\|\|\|\|\|\|\|\|\|\|\|\|\|\|\|\|\|\|\|\|\| \|\|\|\|\|\|\|\|\|\| \|\|\|\|\|\|\|\|\|\|\|\|\| \|\|\|\|\|\|\|\|\|\|\|\|\|\|\|\|\| \|\|\|\|\| \|\|\|\| \|\|\|\|\|\|\| \|\|\|\|\|\|\|\|\|\|\|\|\|\|\|\|\|\|\|\|\|\|\|\|\| \|\|\|\|\|\|\|\|\|\|\| \|\|\|\|\| \|\|\|\|\|\|\|\| 	ATGGCCAACACAGCCAACGACACGCCGGAAGGCATCTTCTCGCTCCTCGATACCGACCTCTACAAGCTCACCATGCAATGCGCCGTCCTGAAATACTTCCCCGACACCCACGTCCGATACAAGTTCACCAACCGCACGCCTCAGATGAAGTTCACGAGACAAGCTTTCAATTGGTTACATGCTCAGATCAGGAAGCTGGAAATTGTCGAGGTGACAAAGGAGGAAATCGAATGGCTGAGAAGGACGTGCAAATACCTCACACCTGAGTACCTCGACTATCTACAGACATTCCGATTGCGGCCTTCAGAGCAGGTGAAGGCTTCATTCATGCCGGACAGTGATGCCACAAGCGACTCAGTCACAGGTGAGGTCTCCTTGGCGATCAGCGGTATCTGGGTCGAGACAATCCTCTACGAGATCCCACTGCTAGCCCTCACATCAGAAGCATACTTTAAGTTCGTCGACAAGGATTGGAACCACAATGGTCAAGTCGCCAAAGCAAAAGAGAAGGCCGAGAAGCTCTTCGAAGCAGGCTGTGTCTTCAGCGAGTTCGGCAGTCGACGCAGAAGAGACTACAAAACCCACGAGATGGTCATTCAAGGCATCCTCCAAGCACAAGAGGAAGCAAAGAACAACGGCAAGAACTGGAAAGGCAAACTCACCGGCACATCCAATGTCCACCTCGCCATGAAACACAACACCACACCCTCCGGCACAGTCGCCCACGAATGGTTCATGGGCATCGCAGCCATTACGCAAGACTACGCCAACGCGAACGAGAAAGCGATGGAATACTGGATCGGCACCTTCGGGCCAGGCGTGCTGGGAATCGCACTCACCGACACCTTCGGCACGCCATCTTTCCTCAAAGCATTCGCCAAACCCGCTCCTTCATCCGACTCCAGCAGA-CCTCCTC-G-CTCACCCTCACCAACCTACGCACAAGTCTTCGC-GGCGTACGGC-AGACTCCGGCGAT-CCCTCGAATACGTACAA-TGATC-GGCA-TTCTACT-CAGTCAAAACATCACCTCCCCCAAA-CGGTCGTCTTCCTCCGAACAGCCTCA ``` |
| 25 | ggggacaagtttgtacaaaaaagcaggcttgatgACCAACATCACCCATCTTCC, ggggaccactttgtacaagaaagctgggtcAAGGCCAACCCACCCCCTCC | Mycgr3\|72393  JGI  Ensembl |  |  | 100 % |  |  |
| 25 | ggggacaagtttgtacaaaaaagcaggcttgatgACCTCTTTCACCGACATCTG, ggggaccactttgtacaagaaagctgggtcCTCTTCACCCGCACCGCCCT | Mycgr3\|72623  JGI  Ensembl | KOG1481 Cysteine synthase; | catalytic activity; cysteine synthase activity;  metabolic process;  pyridoxal phosphate binding; | 100 % |  |  |
| 25 | ggggacaagtttgtacaaaaaagcaggcttgatgATACGGTCGAGCTTCAGGCG, ggggaccactttgtacaagaaagctgggtcTTGCTCTAGCTCAGGCAATG | Mycgr3\|72674  JGI  Ensembl | KOG1290 Serine/threonine protein kinase; | 3-phosphoinositide-dependent protein kinase activity; AMP-activated protein kinase activity;  ATP binding;  DNA-dependent protein kinase activity;  GTP-dependent protein kinase activity;  JUN kinase activity;  JUN kinase kinase activity;  JUN kinase kinase kinase activity;  Janus kinase activity;  MAP kinase 1 activity;  MAP kinase 2 activity;  MAP kinase kinase activity;  MAP kinase kinase kinase kinase activity;  MAP/ERK kinase kinase activity;  MP kinase activity;  NF-kappaB-inducing kinase activity;  SAP kinase activity;  atypical protein kinase C activity;  calcium-dependent protein kinase C activity;  casein kinase I activity;  casein kinase activity;  cyclic nucleotide-dependent protein kinase activity;  cyclin-dependent protein kinase activating kinase activity;  eukaryotic translation initiation factor 2alpha kinase activity;  galactosyltransferase-associated kinase activity;  glycogen synthase kinase 3 activity;  protein amino acid phosphorylation;  protein kinase activity;  protein serine/threonine kinase activity;  receptor signaling protein serine/threonine kinase activity;  ribosomal protein S6 kinase activity;  transmembrane receptor protein kinase activity;  transmembrane receptor protein serine/threonine kinase activity; | 2.73038 % |  |  |
| 25 | ggggacaagtttgtacaaaaaagcaggcttgatgGCTCAGACACGCTCATCTCA, ggggaccactttgtacaagaaagctgggtcCCAAGTCGTGGGCACTTTGG | Mycgr3\|73513  JGI  Ensembl | KOG2992 Nucleolar GTPase/ATPase p130; |  | 100 % |  |  |
| 25 | ggggacaagtttgtacaaaaaagcaggcttgatgCCTCGACGCAAGGCCTTCGA, ggggaccactttgtacaagaaagctgggtcCTTGGCTGCAGCGCTCTTCG | Mycgr3\|74695  JGI  Ensembl | KOG2637 Uncharacterized conserved protein; |  | 0 % |  |  |
| 25 | ggggacaagtttgtacaaaaaagcaggcttgatgTCGGGCGGTTGGAACACCAT, ggggaccactttgtacaagaaagctgggtcCGACATATCCATCTCATCCG | Mycgr3\|75993  JGI  Ensembl | KOG2778 Ubiquitin C-terminal hydrolase; | intracellular; ubiquitin thiolesterase activity;  ubiquitin-dependent protein catabolic process; | 8.26129 % |  |  |
| 25 | ggggacaagtttgtacaaaaaagcaggcttgatgCCCGCGACCGGACCCTCTCG, ggggaccactttgtacaagaaagctgggtcATCGCAATACGCCAGATCCG | Mycgr3\|76332  JGI  Ensembl |  |  | 100 % |  | ``` 131	AGCCGGACTGGGCCCACGCGAGTCACGATGGAGAAGAGCCGGCACGAAAGCCTCAGGCGAACGTGCCATATATACAAGAGGAGGTGAAGGAGATTGGTGAAGATACCACACACCTGCTGTCACCGGAACAACAAGCAGCAGCCGCCAAAATTGTCAACTCTGAACGCCTGGAGATCAAGTCCAAGATCACGAAGAATGGAAAATACCTG-AAAGTCGACTTCCTGGGCGCCCATCGCGGATTCAACCCCAGCGTTATCCCTCA	392 	\|\|\| \|\|\|\|\|\|\|\|\|\|\|\|\|\|\|\|\|\|\|\|\|\|\|\|\|\|\|\|\|\|\|\|\|\|\|\|\|\|\|\|\|\|\|\|\|\|\|\|\|\|\|\|\|\|\|\|\|\|\|\|\|\|\|\|\|\|\|\|\|\|\|\|\|\|\|\|\|\|\|\|\|\|\|\|\|\|\|\|\|\|\|\|\|\|\|\|\|\|\|\|\|\|\|\|\|\|\|\|\|\|\|\|\|\|\|\|\|\|\|\|\|\|\|\|\|\|\|\|\|\|\|\|\| \|\|\|\|\|\|\|\|\|\|\|\|\|\|\|\|\|\|\|\|\|\|\|\|\|\|\|\|\|\|\|\|\|\|\|\|\|\|\|\|\|\|\|\| \| \|\|\|\|\|\|\|\| \|\| \|\|\| \|\|\|\|\|\|\|\| \|\|\|\|\|\| \|\|\|\|\|\|\|\|\|\|\|\|\|\| \|\|\|\|\|\|\|\|\|\|\| \|  \|\|\| \|\| \| \|\|\|\|\|\|\| 	AGCTGGACTGGGCCCACGCGAGTCACGATGGAGAAGAGCCGGCACGAAAGCCTCAGGCGAACGTGCCATATATACAAGAGGAGGTGAAGGAGATTGGTGAAGATACCACACACCTGCTGTCACCGGAACAACAAGCANCAGCCGCCAAAATTGTCAACTCTGAACGCCTGGAGATCAAGTCCNANATCACGAANAANGGANAATACCTGNAAAGTCNACTTCCTGGGCGCCNATCGCGGATTCNANNCCANCGNTNTCCCTCA ``` |
| 25 | ggggacaagtttgtacaaaaaagcaggcttgatgGCCTGGAGCTCTGACAACCT, ggggaccactttgtacaagaaagctgggtcAACAGTAAACCACCCATCCA | Mycgr3\|76546  JGI  Ensembl |  |  | 100 % |  |  |
| 25 | ggggacaagtttgtacaaaaaagcaggcttgatgTCGACAACACACACACAAGG, ggggaccactttgtacaagaaagctgggtcCTCCCTCAACTCCTTCTTCT | Mycgr3\|77145  JGI  Ensembl | KOG0761 Mitochondrial carrier protein CGI-69; | binding; membrane; transport; | 100 % | chr\_11:1567428 G->A; |  |
| 25 | ggggacaagtttgtacaaaaaagcaggcttgatgGCAGATCCATTGTACGAATT, ggggaccactttgtacaagaaagctgggtcGACCGCCGTCAATGGTGTCG | Mycgr3\|83550  JGI  Ensembl | KOG2069 Golgi transport complex subunit; |  | 0 % |  |  |
| 25 | ggggacaagtttgtacaaaaaagcaggcttgatgGAGGCACCGCTGGCACCGCA, ggggaccactttgtacaagaaagctgggtcGTCATCCAGTTCCTCAAATC | Mycgr3\|85675  JGI  Ensembl | KOG4464 Signaling protein RIC-8/synembryn (regulates neurotransmitter secretion); |  | 0 % |  |  |
| 25 | ggggacaagtttgtacaaaaaagcaggcttgatgAGAGTCCTTTCAACTGCCGG, ggggaccactttgtacaagaaagctgggtcCTTGCGAAGAAAAGCCTCAT | Mycgr3\|90125  JGI  Ensembl |  |  | 100 % |  |  |
| 25 | ggggacaagtttgtacaaaaaagcaggcttgatgCTGCTCCCTCGACAGGACAA, ggggaccactttgtacaagaaagctgggtcTGTCCGCGTCTCTCTTTCTG | Mycgr3\|92321  JGI  Ensembl | KOG0401 Translation initiation factor 4F, ribosome/mRNA-bridging subunit (eIF-4G); |  | 100 % |  |  |
| 25 | ggggacaagtttgtacaaaaaagcaggcttgatgGCGGCTGCCAGAAAGAAGAC, ggggaccactttgtacaagaaagctgggtcAACCAACTTCACGCCGCCGT | Mycgr3\|92363  JGI  Ensembl |  |  | 0 % |  |  |
| 25 | ggggacaagtttgtacaaaaaagcaggcttgatgTTGCAATTCATCCAAGACTA, ggggaccactttgtacaagaaagctgggtcGTCATCCACAAGAAATCGTG | Mycgr3\|92405  JGI  Ensembl |  |  | 0 % |  |  |
| 25 | ggggacaagtttgtacaaaaaagcaggcttgatgCCGAGCAAGCGTCGCCCACA, ggggaccactttgtacaagaaagctgggtcCTCCTCATCCTCGAGCGCGA | Mycgr3\|92442  JGI  Ensembl |  |  | 0 % |  |  |
| 25 | ggggacaagtttgtacaaaaaagcaggcttgatgGCCTCGCTCGGACCCATTGA, ggggaccactttgtacaagaaagctgggtcAGGCGGCTTAAACTTCTCGC | Mycgr3\|94392  JGI  Ensembl |  |  | 2.62985 % |  |  |
| 25 | ggggacaagtttgtacaaaaaagcaggcttgatgCATCTCACTCAAATTCTCGT, ggggaccactttgtacaagaaagctgggtcGCATGTATTGCCGCTGCTCA | Mycgr3\|94557  JGI  Ensembl |  |  | 100 % |  |  |
| 25 | ggggacaagtttgtacaaaaaagcaggcttgatgGACACCTTCAATACTCCAAG, ggggaccactttgtacaagaaagctgggtcTTCTCCAGATGCAGATCGGC | Mycgr3\|94631  JGI  Ensembl |  |  | 100 % |  |  |
| 25 | ggggacaagtttgtacaaaaaagcaggcttgatgGACGATCTCCCGGATGAACT, ggggaccactttgtacaagaaagctgggtcATTCGCAATCTCCTCGTCTT | Mycgr3\|95556  JGI  Ensembl | KOG0732 AAA+-type ATPase containing the bromodomain; |  | 0 % |  |  |
| 25 | ggggacaagtttgtacaaaaaagcaggcttgatgACTACAAACTCTCTCGAGGA, ggggaccactttgtacaagaaagctgggtcAAGACAATCACCAAAGTCCT | Mycgr3\|95753  JGI  Ensembl |  |  | 100 % | chr\_9:1118669 A->G; |  |
| 25 | ggggacaagtttgtacaaaaaagcaggcttgatgCCGCCAAAGCGACTTGATCA, ggggaccactttgtacaagaaagctgggtcACGTCTCCACACACCGAACT | Mycgr3\|95754  JGI  Ensembl |  |  | 0 % |  |  |
| 25 | ggggacaagtttgtacaaaaaagcaggcttgatgGCCGAAAAGTCTGAAAAGAA, ggggaccactttgtacaagaaagctgggtcATCGTCGTCCTCGTCATTGA | Mycgr3\|96303  JGI  Ensembl |  |  | 100 % |  |  |
| 25 | ggggacaagtttgtacaaaaaagcaggcttgatgGCCGCCAAAGCAGGAAAGAA, ggggaccactttgtacaagaaagctgggtcGAATCTGCTCTCGACTTTCT | Mycgr3\|97115  JGI  Ensembl | KOG1406 Peroxisomal 3-ketoacyl-CoA-thiolase P-44/SCP2; | acetyl-CoA C-acyltransferase activity; | 100 % |  |  |
| 25 | ggggacaagtttgtacaaaaaagcaggcttgatgTTCAGCTTCGACTTGGAGCA, ggggaccactttgtacaagaaagctgggtcGAGACTGCGATCCTCAGGGT | Mycgr3\|97560  JGI  Ensembl |  |  | 100 % |  |  |
| 25 | ggggacaagtttgtacaaaaaagcaggcttgatgCTTGATGTCAAGACCAAGGA, ggggaccactttgtacaagaaagctgggtcGGTGATGGTTAGAGGACTGC | Mycgr3\|97869  JGI  Ensembl |  |  | 10.50167 % |  |  |
| 25 | ggggacaagtttgtacaaaaaagcaggcttgatgTTCGTCATCGGGGATTTCAT, ggggaccactttgtacaagaaagctgggtcAAGATCTTTCTTTTTGTTCC | Mycgr3\|97948  JGI  Ensembl | KOG1216 von Willebrand factor and related coagulation proteins; |  | 1.36612 % |  |  |
| 25 | ggggacaagtttgtacaaaaaagcaggcttgatgAACGCCGCTGCTGCTCCAGT, ggggaccactttgtacaagaaagctgggtcGCGGCTCCGCCATCGTTTGC | Mycgr3\|98057  JGI  Ensembl |  |  | 100 % | chr\_20:348066 G->A; |  |
| 25 | ggggacaagtttgtacaaaaaagcaggcttgatgGAGATCAAGAAGCCCCAGTC, ggggaccactttgtacaagaaagctgggtcCAACCTCGACTGAATCTTCT | Mycgr3\|98156  JGI  Ensembl | KOG1406 Peroxisomal 3-ketoacyl-CoA-thiolase P-44/SCP2; | acetyl-CoA C-acyltransferase activity; | 100 % |  |  |
| 25 | ggggacaagtttgtacaaaaaagcaggcttgatgTCGAATCCTATCGTCATTCG, ggggaccactttgtacaagaaagctgggtcCCTCCACCCAACCCCCCTCT | Mycgr3\|98925  JGI  Ensembl |  |  | 100 % |  |  |
| 25 | ggggacaagtttgtacaaaaaagcaggcttgatgGACCGCCTCAAGAAGCTCTT, ggggaccactttgtacaagaaagctgggtcAGTAATAAAAAAGCTCAACC | Mycgr3\|99524  JGI  Ensembl | KOG1479 Nucleoside transporter; | membrane; nucleoside transmembrane transporter activity;  transport; | 1.43678 % |  |  |
| 25 | ggggacaagtttgtacaaaaaagcaggcttgatgGACAGCCCAAAGGACCTCCT, ggggaccactttgtacaagaaagctgggtcCGATGACCCACCATGCCTCA | Mycgr3\|100575  JGI  Ensembl | KOG3873 Sphingomyelinase family protein; | sphingomyelin phosphodiesterase activity; | 100 % |  |  |
| 25 | ggggacaagtttgtacaaaaaagcaggcttgatgTCGTCGAGCCTCTATAACCT, ggggaccactttgtacaagaaagctgggtcTCTGGCCTTCGCATCTTTTC | Mycgr3\|101027  JGI  Ensembl | KOG0885 Peptidyl-prolyl cis-trans isomerase; | FK506-sensitive peptidyl-prolyl cis-trans isomerase; cyclophilin; cyclophilin-type peptidyl-prolyl cis-trans isomerase activity;  peptidyl-prolyl cis-trans isomerase activity;  protein folding; | 2.9703 % |  |  |
| 25 | ggggacaagtttgtacaaaaaagcaggcttgatgGGCGCCTCCCCCGACTGGCT, ggggaccactttgtacaagaaagctgggtcTGCCTTGCGGGACCGTGTCA | Mycgr3\|101073  JGI  Ensembl | KOG3071 Fatty acyl-CoA elongase/Polyunsaturated fatty acid specific elongation enzyme; | integral to membrane; | 100 % |  |  |
| 25 | ggggacaagtttgtacaaaaaagcaggcttgatgGCCACAACCACAGCACAACG, ggggaccactttgtacaagaaagctgggtcGATAAAGCCAGCCTTTCGAA | Mycgr3\|101079  JGI  Ensembl | KOG2504 Monocarboxylate transporter; | integral to membrane; transport; transporter activity; | 100 % |  |  |
| 25 | ggggacaagtttgtacaaaaaagcaggcttgatgGTCTCCGACACGAAACTCTA, ggggaccactttgtacaagaaagctgggtcCAAAATCTCCTTCAACTTCT | Mycgr3\|101103  JGI  Ensembl | KOG0714 Molecular chaperone (DnaJ superfamily); | heat shock protein binding; protein folding;  unfolded protein binding; | 0 % |  |  |
| 25 | ggggacaagtttgtacaaaaaagcaggcttgatgCCTGCCGAGAAGATCACGAC, ggggaccactttgtacaagaaagctgggtcATTCGGATAATCCTTCAAAT | Mycgr3\|101124  JGI  Ensembl | KOG2450 Aldehyde dehydrogenase; | aldehyde dehydrogenase (NAD) activity; | 0 % |  |  |
| 25 | ggggacaagtttgtacaaaaaagcaggcttgatgTTGAACCAATATGGCCTCGA, ggggaccactttgtacaagaaagctgggtcCGCATACAAAATCTCCCGCA | Mycgr3\|101149  JGI  Ensembl |  |  | 100 % |  |  |
| 25 | ggggacaagtttgtacaaaaaagcaggcttgatgTCGACTCCCAAGAAGAACAT, ggggaccactttgtacaagaaagctgggtcTCTGATGCGGACGTTGTCCA | Mycgr3\|101694  JGI  Ensembl | KOG3867 Sulfatase; | N-sulfoglucosamine sulfohydrolase activity; metabolic process;  sulfuric ester hydrolase activity; | 0 % |  |  |
| 25 | ggggacaagtttgtacaaaaaagcaggcttgatgTCGTCCAATTCCGATGGACG, ggggaccactttgtacaagaaagctgggtcGAAGTCATCTCCGCGTTCCT | Mycgr3\|101727  JGI  Ensembl | KOG0764 Mitochondrial FAD carrier protein; | binding; membrane; mitochondrial inner membrane;  transport; | 100 % |  |  |
| 25 | ggggacaagtttgtacaaaaaagcaggcttgatgGCTTCTGCAAATGATGCTCC, ggggaccactttgtacaagaaagctgggtcCGCATCCACCGCCAGGATCT | Mycgr3\|101803  JGI  Ensembl | KOG1316 Argininosuccinate lyase; | arginine biosynthetic process via ornithine; argininosuccinate lyase activity;  catalytic activity; | 100 % |  |  |
| 25 | ggggacaagtttgtacaaaaaagcaggcttgatgAGCGCTCCTCCCACGTCGAC, ggggaccactttgtacaagaaagctgggtcCTTCGCGATCTCGACACCAC | Mycgr3\|103492  JGI  Ensembl | KOG0767 Mitochondrial phosphate carrier protein; | binding; membrane; mitochondrial inner membrane;  transport; | 0 % |  |  |
| 25 | ggggacaagtttgtacaaaaaagcaggcttgatgATCTCAAGAACCACCACACG, ggggaccactttgtacaagaaagctgggtcCTTCTCCGCCTGGTTCTTCG | Mycgr3\|103528  JGI  Ensembl | KOG4674 Uncharacterized conserved coiled-coil protein; |  | 0 % |  |  |
| 25 | ggggacaagtttgtacaaaaaagcaggcttgatgTTCAAGTCCAGCCTGCTCCG, ggggaccactttgtacaagaaagctgggtcGGTGACGGTGTATCCACCGT | Mycgr3\|103760  JGI  Ensembl | KOG0725 Reductases with broad range of substrate specificities; | D-arabinitol 2-dehydrogenase activity; metabolic process;  oxidoreductase activity; | 0 % |  |  |
| 25 | ggggacaagtttgtacaaaaaagcaggcttgatgACTACACTGCTCGATCTCCC, ggggaccactttgtacaagaaagctgggtcTGTTCCACCATACACCTCTT | Mycgr3\|105666  JGI  Ensembl |  |  | 100 % |  |  |
| 25 | ggggacaagtttgtacaaaaaagcaggcttgatgTCTCCCTCCATCAACATCAT, ggggaccactttgtacaagaaagctgggtcAACCATCATCGCGACGGCAC | Mycgr3\|106780  JGI  Ensembl | KOG1216 von Willebrand factor and related coagulation proteins; |  | 0 % |  |  |
| 25 | ggggacaagtttgtacaaaaaagcaggcttgatgACCCCCAACACACTTGACGA, ggggaccactttgtacaagaaagctgggtcGGTCGGAGGCTGTAGGGGTT | Mycgr3\|107290  JGI  Ensembl | KOG1269 SAM-dependent methyltransferases; |  | 100 % | chr\_1:3172374 G->T; |  |
| 25 | ggggacaagtttgtacaaaaaagcaggcttgatgTCGGTCGATTCAGAGCAGCT, ggggaccactttgtacaagaaagctgggtcAGAAGCCTGTGCGATTTCCA | Mycgr3\|107297  JGI  Ensembl |  |  | 0 % |  |  |
| 25 | ggggacaagtttgtacaaaaaagcaggcttgatgGCGAGATGGCTGACTTTTAG, ggggaccactttgtacaagaaagctgggtcCGACTCAATATCCGCTCGCT | Mycgr3\|107675  JGI  Ensembl | KOG1579 Homocysteine S-methyltransferase; | homocysteine S-methyltransferase activity; | 0 % |  |  |
| 25 | ggggacaagtttgtacaaaaaagcaggcttgatgACCATGACAGCGAAGCTCCG, ggggaccactttgtacaagaaagctgggtcAGCAGACGAGGCAGTCTGGT | Mycgr3\|107729  JGI  Ensembl | KOG0161 Myosin class II heavy chain; |  | 1.31488 % |  |  |
| 25 | ggggacaagtttgtacaaaaaagcaggcttgatgTTACGGCCGCGAAGCACGGT, ggggaccactttgtacaagaaagctgggtcAGAATCGAGCATGGAGTCAA | Mycgr3\|108734  JGI  Ensembl | KOG2992 Nucleolar GTPase/ATPase p130; |  | 100 % |  |  |
| 25 | ggggacaagtttgtacaaaaaagcaggcttgatgGCGCCCGCCGTTGAACGAGC, ggggaccactttgtacaagaaagctgggtcTTCCAAGTCACTGTCCTGTC | Mycgr3\|108817  JGI  Ensembl | KOG2177 Predicted E3 ubiquitin ligase; |  | 100 % |  |  |
| 25 | ggggacaagtttgtacaaaaaagcaggcttgatgAATCAGTTCCCCGCCTACTC, ggggaccactttgtacaagaaagctgggtcAGTCTCCATCGCATCCTCAT | Mycgr3\|108886  JGI  Ensembl |  |  | 100 % |  |  |
| 25 | ggggacaagtttgtacaaaaaagcaggcttgatgGGCGTCTCCGGCTTCCTCTT, ggggaccactttgtacaagaaagctgggtcCCAGTCGTGGACAAAGTCAG | Mycgr3\|109235  JGI  Ensembl |  |  | 100 % |  |  |
| 25 | ggggacaagtttgtacaaaaaagcaggcttgatgTCCGCCGAAGACAACTCTCC, ggggaccactttgtacaagaaagctgggtcTGCCTGAGCCTCTGGGATCT | Mycgr3\|109444  JGI  Ensembl | KOG2190 PolyC-binding proteins alphaCP-1 and related KH domain proteins; | RNA binding; | 100 % | chr\_5:1039392 C->T; |  |
| 25 | ggggacaagtttgtacaaaaaagcaggcttgatgACAACAGCAAACCTCCTCGC, ggggaccactttgtacaagaaagctgggtcCGCCGGCTGTGCAGTCGCCG | Mycgr3\|109856  JGI  Ensembl |  |  | 100 % |  |  |
| 25 | ggggacaagtttgtacaaaaaagcaggcttgatgGCTGTCACCAAGGACTCGAC, ggggaccactttgtacaagaaagctgggtcGTCGTCGAACTTGATGCCCT | Mycgr3\|110267  JGI  Ensembl | KOG2992 Nucleolar GTPase/ATPase p130; | exo-alpha-sialidase activity; | 100 % |  |  |
| 25 | ggggacaagtttgtacaaaaaagcaggcttgatgGCGGAAGACCAAGACACCTC, ggggaccactttgtacaagaaagctgggtcTTCCACGACTTGAATCTTCT | Mycgr3\|110284  JGI  Ensembl |  |  | 1.43247 % |  |  |
| 25 | ggggacaagtttgtacaaaaaagcaggcttgatgGAGACAATAACGACCACCGC, ggggaccactttgtacaagaaagctgggtcTCGTCCCGCACTAACCAGAC | Mycgr3\|111105  JGI  Ensembl | KOG2806 Chitinase; |  | 100 % |  |  |
| 25 | ggggacaagtttgtacaaaaaagcaggcttgatgTCTTCCAAGTTCTCCTGGAG, ggggaccactttgtacaagaaagctgggtcGTAAAACCTCCCTCCAACCC | Mycgr3\|111414  JGI  Ensembl | KOG1216 von Willebrand factor and related coagulation proteins; |  | 0 % |  |  |
| 25 | ggggacaagtttgtacaaaaaagcaggcttgatgGGCTACATCCGACAAGAACA, ggggaccactttgtacaagaaagctgggtcCTTCTTCTTCTTCTCCTCCA | Mycgr3\|111641  JGI  Ensembl | KOG2877 sn-1,2-diacylglycerol ethanolamine- and cholinephosphotranferases; | ethanolaminephosphotransferase activity; membrane; phospholipid biosynthetic process;  phosphotransferase activity, for other substituted phosphate groups; | 100 % |  |  |
| 25 | ggggacaagtttgtacaaaaaagcaggcttgatgGCCGACCCATCCGACATTCC, ggggaccactttgtacaagaaagctgggtcTCCAGCTTTACACATCTCCT | Mycgr3\|111803  JGI  Ensembl | KOG0161 Myosin class II heavy chain; |  | 100 % |  |  |
| 26 | ggggacaagtttgtacaaaaaagcaggcttgatgAGCCAAGATGGGTCGCCGCT, ggggaccactttgtacaagaaagctgggtcCCATCGTTGTGACTCAATCC | Mycgr3\|35590  JGI  Ensembl | KOG4159 Predicted E3 ubiquitin ligase; | ATP-dependent peptidase activity; ATP-dependent proteolysis;  protein binding;  zinc ion binding; | 0 % |  |  |
| 26 | ggggacaagtttgtacaaaaaagcaggcttgatgCAGGGTGTCTTACGCAGGAA, ggggaccactttgtacaagaaagctgggtcTACTTGTCCCGGGCACACCG | Mycgr3\|36631  JGI  Ensembl | KOG0157 Cytochrome P450 CYP4/CYP19/CYP26 subfamilies; | electron transport; heme binding;  iron ion binding;  monooxygenase activity; | 0 % |  |  |
| 26 | ggggacaagtttgtacaaaaaagcaggcttgatgGCCGACTACAGCAGACCACC, ggggaccactttgtacaagaaagctgggtcCAGCTTACCACTAGGCGCTT | Mycgr3\|36820  JGI  Ensembl | KOG2565 Predicted hydrolases or acyltransferases (alpha/beta hydrolase superfamily); | catalytic activity; epoxide hydrolase activity;  membrane;  microsome;  response to toxin; | 0 % |  |  |
| 26 | ggggacaagtttgtacaaaaaagcaggcttgatgGCGGGCTCGAATATAAACCC, ggggaccactttgtacaagaaagctgggtcCAACTCCGGCTTCGGTAGCG | Mycgr3\|38753  JGI  Ensembl | KOG1397 Ca2+/H+ antiporter VCX1 and related proteins; | integral to membrane; | 0 % |  |  |
| 26 | ggggacaagtttgtacaaaaaagcaggcttgatgCTAGCCCACCTCTTCGGAAC, ggggaccactttgtacaagaaagctgggtcAACTCCTTCTCCCCTCCCTC | Mycgr3\|40588  JGI  Ensembl | KOG3764 Vesicular amine transporter; | integral to membrane; transport; transporter activity; | 0 % |  |  |
| 26 | ggggacaagtttgtacaaaaaagcaggcttgatgGCAGCTAAGACTCCCGTGAA, ggggaccactttgtacaagaaagctgggtcAGGCCTCCAGCCATTGACCA | Mycgr3\|40852  JGI  Ensembl | KOG0409 Predicted dehydrogenase; |  | 0 % |  |  |
| 26 | ggggacaagtttgtacaaaaaagcaggcttgatgGCCACGACAGAATCATTTGT, ggggaccactttgtacaagaaagctgggtcGACCGTCCAACTCATCGACG | Mycgr3\|43467  JGI  Ensembl | KOG2456 Aldehyde dehydrogenase; |  | 0 % |  |  |
| 26 | ggggacaagtttgtacaaaaaagcaggcttgatgGCACCATTCGCTACAGGAAC, ggggaccactttgtacaagaaagctgggtcCACCACCACATTCGTCCTCG | Mycgr3\|44026  JGI  Ensembl | KOG2584 Dihydroorotase and related enzymes; | allantoinase activity; aspartic-type endopeptidase activity;  endodeoxyribonuclease activity, producing 5'-phosphomonoesters;  hydrolase activity;  hydrolase activity, acting on carbon-nitrogen (but not peptide) bonds, in cyclic amides; proteolysis; | 0 % |  |  |
| 26 | ggggacaagtttgtacaaaaaagcaggcttgatgGCTTTCCACATCTCGACCTT, ggggaccactttgtacaagaaagctgggtcTTCGTGGGCCCCGGTGATCC | Mycgr3\|45271  JGI  Ensembl | KOG0159 Cytochrome P450 CYP11/CYP12/CYP24/CYP27 subfamilies; | electron transport; heme binding;  iron ion binding;  monooxygenase activity;  unspecific monooxygenase activity; | 0 % |  |  |
| 26 | ggggacaagtttgtacaaaaaagcaggcttgatgCCCAAATCTCGTGGCCTCAC, ggggaccactttgtacaagaaagctgggtcTCTGCCTCTTGCTCGAACCC | Mycgr3\|47688  JGI  Ensembl | KOG3764 Vesicular amine transporter; | integral to membrane; response to antibiotic;  tetracycline transport;  tetracycline:hydrogen antiporter activity;  transport;  transporter activity; | 0 % |  |  |
| 26 | ggggacaagtttgtacaaaaaagcaggcttgatgACTGCGATAAGTCTCGCCTC, ggggaccactttgtacaagaaagctgggtcGCCAGCACCACGACTCTTCT | Mycgr3\|47899  JGI  Ensembl |  |  | 0 % |  |  |
| 26 | ggggacaagtttgtacaaaaaagcaggcttgatgGACTTTGCTAGCAGCGTCTC, ggggaccactttgtacaagaaagctgggtcATCATCAGGTCCGGTCAAGG | Mycgr3\|48419  JGI  Ensembl |  |  | 0 % |  |  |
| 26 | ggggacaagtttgtacaaaaaagcaggcttgatgTCGAATCAAGTCAAGGGTCG, ggggaccactttgtacaagaaagctgggtcGGTATACCATGTGGGCTCCG | Mycgr3\|49129  JGI  Ensembl | KOG0139 Short-chain acyl-CoA dehydrogenase; |  | 1.36808 % |  |  |
| 26 | ggggacaagtttgtacaaaaaagcaggcttgatgGCTCTTTCACGGCATCATGC, ggggaccactttgtacaagaaagctgggtcAATCCACCCCCCTTCCATCT | Mycgr3\|50144  JGI  Ensembl | KOG2880 SMAD6 interacting protein AMSH, contains JAB/MPN/Mov34 domain; | ubiquitin thiolesterase activity; | 0 % |  |  |
| 26 | ggggacaagtttgtacaaaaaagcaggcttgatgAACAGCTTCAATCTCTTGTC, ggggaccactttgtacaagaaagctgggtcCGCTTGTTGCGCTTCTCCTC | Mycgr3\|50300  JGI  Ensembl |  |  | 0 % |  |  |
| 26 | ggggacaagtttgtacaaaaaagcaggcttgatgGCCTCCTCCTCATCCTCACC, ggggaccactttgtacaagaaagctgggtcCACCGGCACATAACAATGCA | Mycgr3\|61457  JGI  Ensembl |  |  | 0 % |  |  |
| 26 | ggggacaagtttgtacaaaaaagcaggcttgatgGTCTCCTGGGCCACGGTTCA, ggggaccactttgtacaagaaagctgggtcTCGACCTTGCTGCTGCTCTC | Mycgr3\|64401  JGI  Ensembl |  |  | 0 % |  |  |
| 26 | ggggacaagtttgtacaaaaaagcaggcttgatgGTCAAGGCACTGTCGTTCAA, ggggaccactttgtacaagaaagctgggtcAGCATACTTGTCATGCTTGC | Mycgr3\|65433  JGI  Ensembl | KOG3962 Predicted actin-bundling protein; |  | 0 % |  |  |
| 26 | ggggacaagtttgtacaaaaaagcaggcttgatgAAGGACTCCCACGACCTCCC, ggggaccactttgtacaagaaagctgggtcCACTTTCCTCCATCTCCACC | Mycgr3\|67327  JGI  Ensembl | KOG2504 Monocarboxylate transporter; |  | 0 % |  |  |
| 26 | ggggacaagtttgtacaaaaaagcaggcttgatgGACCTGAACAATGAGACCCT, ggggaccactttgtacaagaaagctgggtcTTGTTCCTCGTCACCCAACG | Mycgr3\|68009  JGI  Ensembl | KOG1764 5'-AMP-activated protein kinase, gamma subunit; |  | 0 % |  |  |
| 26 | ggggacaagtttgtacaaaaaagcaggcttgatgCCCGCCGCAATATCAACACC, ggggaccactttgtacaagaaagctgggtcCCTCTTCAACACCTTCCTCG | Mycgr3\|68326  JGI  Ensembl | KOG1377 Uridine 5'- monophosphate synthase/orotate phosphoribosyltransferase; | 'de novo' pyrimidine base biosynthetic process; orotidine-5'-phosphate decarboxylase activity; | 32.65966 % |  | ``` 634	GTCTCCG-T-CTCCACCACAATCAGCATGAAATCCGAAGCCATCAGCCCGCAACCTCACCACCGACCTTCAATCCCTCGTGTCGCCTCCGACGACTCCGAAGATGAAGGAGACGACCCTGAAACTCTCGACCATCTCGGTCCACCTCCATACTTCCGCTCTCTCCTACTACTCGCGCAAATGTCTTCGGCCAACAACTTCTTTACACCAGAATACACAACCGCCTGTCTACACCACGCGAGGGAGAACAAGGACTTTGTCATGGGCTTCATTGCGCAAAAAGGGATGAATGAAGGGTCGGAGGACAATTTCATCACGATGACACCGGGTGTGCAACTGCAAGCGGGAGGTGATGGTATGGGTCAGCAGTACAACACTCCGGCCAAGGTGGTGG-GCGAAGGTGGCACC	1038 	\|\|\|\|\|\|\| \| \| \|\|\|\|\|\|\|\|\|\|\|\|\|\|\|\|\|\|\|\|\|\|\|\|\|\|\|\|\|\|\|\|\|\|\|\|\|\|\|\|\|\|\|\|\|\|\|\|\|\|\|\|\|\|\|\|\|\|\|\|\|\|\|\|\|\|\|\|\|\|\|\|\|\|\|\|\|\|\|\|\|\|\|\|\|\|\|\|\|\|\|\|\|\|\|\|\|\|\|\|\|\|\|\|\|\|\|\|\|\|\|\|\|\|\|\|\|\|\|\|\|\|\|\|\|\|\|\|\|\|\|\|\|\|\|\|\|\|\|\|\|\|\|\|\|\|\|\|\|\|\|\|\|\|\|\|\|\|\|\|\|\|\|\|\|\|\|\|\|\|\|\|\|\|\|\|\|\|\|\|\|\|\|\|\|\|\|\|\|\|\|\|\|\|\|\|\|\|\|\|\|\|\|\|\|\|\|\|\|\|\|\|\|\|\|\|\|\|\|\|\|\|\|\|\|\|\|\|\|\|\|\|\|\|\|\|\|\|\|\|\|\|\|\|\|\|\|\|\|\|\|\|\|\|\|\|\|\|\|\|\|\|\|\|\|\|\|\|\|\|\|\|\|\|\|\|\|\|\|\|\|\|\|\|\|\|\|\|\|\|\|\|\|\|\|\|\|\|\|\|\|\|\|\|\|\|\|\|\|\|\|\|\|\|\|\|\|\|\|\|\|\|\|\|\|\|\|\|\|\|\|\|\|\|\|\|\|\|\|\|\|\|\|\|\|\|\|\|\|\|\|\|\|\|\|\|\|\|\|\|\|\|\|\|\|\|\|\|\|\|\|\|\|\|\|\|\|\|\|\|\| \|\|\|\|\| \|\|\|\| \|\|\| 	GTCTCCGCTCCNCCACCACAATCAGCATGAAATCCGAAGCCATCAGCCCGCAACCTCACCACCGACCTTCAATCCCTCGTGTCGCCTCCGACGACTCCGAAGATGAAGGAGACGACCCTGAAACTCTCGACCATCTCGGTCCACCTCCATACTTCCGCTCTCTCCTACTACTCGCGCAAATGTCTTCGGCCAACAACTTCTTTACACCAGAATACACAACCGCCTGTCTACACCACGCGAGGGAGAACAAGGACTTTGTCATGGGCTTCATTGCGCAAAAAGGGATGAATGAAGGGTCGGAGGACAATTTCATCACGATGACACCGGGTGTGCAACTGCAAGCGGGAGGTGATGGTATGGGTCAGCAGTACAACACTCCGGCCAAGGTGGTGGTGCGAATGTGG-ACC ``` |
| 26 | ggggacaagtttgtacaaaaaagcaggcttgatgTCCGCCGTCTCCTCCGACAA, ggggaccactttgtacaagaaagctgggtcGTATGTCGCCGTGAGTCGCT | Mycgr3\|68338  JGI  Ensembl | KOG0143 Iron/ascorbate family oxidoreductases; | iron ion binding; oxidoreductase activity; | 0 % |  |  |
| 26 | ggggacaagtttgtacaaaaaagcaggcttgatgTACACCTCGACACAGCATTT, ggggaccactttgtacaagaaagctgggtcCAATCCAAAGCAATTGCCAT | Mycgr3\|68702  JGI  Ensembl | KOG2258 Glycerophosphoryl diester phosphodiesterase; | glycerol metabolic process; glycerophosphodiester phosphodiesterase activity; | 0 % |  |  |
| 26 | ggggacaagtttgtacaaaaaagcaggcttgatgAAGATTCCTGTGTCCAATTC, ggggaccactttgtacaagaaagctgggtcGATGTCATGCCCACAGGTAG | Mycgr3\|69294  JGI  Ensembl | KOG2195 Transferrin receptor and related proteins containing the protease-associated (PA) domain; | alanine aminopeptidase activity; peptidase activity;  proteolysis; | 0 % |  |  |
| 26 | ggggacaagtttgtacaaaaaagcaggcttgatgGCAGTCGACTACTCAGTATA, ggggaccactttgtacaagaaagctgggtcCAAGACTTCAATCTTTGCAG | Mycgr3\|69865  JGI  Ensembl | KOG2598 Phosphomethylpyrimidine kinase; | hydroxyethylthiazole kinase activity; thiamin biosynthetic process;  thiamin-phosphate diphosphorylase activity; | 0 % |  |  |
| 26 | ggggacaagtttgtacaaaaaagcaggcttgatgGCAGGCACGTCTCAGATTCC, ggggaccactttgtacaagaaagctgggtcATTAGCGTAGTAATTCACCA | Mycgr3\|70334  JGI  Ensembl | KOG0143 Iron/ascorbate family oxidoreductases; | iron ion binding; naringenin 3-dioxygenase activity;  oxidoreductase activity; | 0 % |  |  |
| 26 | ggggacaagtttgtacaaaaaagcaggcttgatgCCAGAATTCGCCTACGGCTC, ggggaccactttgtacaagaaagctgggtcGACCTCCTTGGCCTCGAATT | Mycgr3\|70433  JGI  Ensembl |  |  | 0 % |  |  |
| 26 | ggggacaagtttgtacaaaaaagcaggcttgatgGCATCTTTGAAAGATAGAAT, ggggaccactttgtacaagaaagctgggtcGCTGGCCTTCGCATTGTTCC | Mycgr3\|71071  JGI  Ensembl |  |  | 0 % |  |  |
| 26 | ggggacaagtttgtacaaaaaagcaggcttgatgATCCTAGTGACTCCGTCGCC, ggggaccactttgtacaagaaagctgggtcAGACACTGACCATCCTTCCT | Mycgr3\|71407  JGI  Ensembl | KOG2246 Galactosyltransferases; |  | 0 % |  |  |
| 26 | ggggacaagtttgtacaaaaaagcaggcttgatgGCACCCCACGCGGACACAAT, ggggaccactttgtacaagaaagctgggtcCATCACACTCCTCGCAGCAT | Mycgr3\|71664  JGI  Ensembl |  |  | 0 % |  |  |
| 26 | ggggacaagtttgtacaaaaaagcaggcttgatgTTTGCGAACATCGCACTTTT, ggggaccactttgtacaagaaagctgggtcATAGCCTGGTGGAGGGTGGT | Mycgr3\|72002  JGI  Ensembl |  |  | 0 % |  |  |
| 26 | ggggacaagtttgtacaaaaaagcaggcttgatgCCCGTCAAATCACGATTCAC, ggggaccactttgtacaagaaagctgggtcATTCGCATGCTTCTTCACTC | Mycgr3\|72288  JGI  Ensembl | KOG2667 COPII vesicle protein; |  | 0 % |  |  |
| 26 | ggggacaagtttgtacaaaaaagcaggcttgatgGCGACTCCTGCATCTCCTTC, ggggaccactttgtacaagaaagctgggtcATCCGCAACACTCTTCCCCG | Mycgr3\|72326  JGI  Ensembl | KOG1852 Cell cycle-associated protein; | protein binding; | 0 % |  |  |
| 26 | ggggacaagtttgtacaaaaaagcaggcttgatgCAGTGCACCCGCCTCACAGC, ggggaccactttgtacaagaaagctgggtcAATCACCTCCTCTCTCTCCC | Mycgr3\|72918  JGI  Ensembl |  |  | 0 % |  |  |
| 26 | ggggacaagtttgtacaaaaaagcaggcttgatgTCCTCCTCCTCGCCCAAAGA, ggggaccactttgtacaagaaagctgggtcGAATGTCTTATCCATCAACT | Mycgr3\|72966  JGI  Ensembl | KOG0758 Mitochondrial carnitine-acylcarnitine carrier protein; | binding; membrane; mitochondrial inner membrane;  transport;  transporter activity; | 0 % |  |  |
| 26 | ggggacaagtttgtacaaaaaagcaggcttgatgGTCCGCCACTCGTTCGAGGA, ggggaccactttgtacaagaaagctgggtcCACCTGCGCATACAGATAAT | Mycgr3\|73068  JGI  Ensembl | KOG2504 Monocarboxylate transporter; |  | 0 % |  |  |
| 26 | ggggacaagtttgtacaaaaaagcaggcttgatgCCGGCATCGAACATAGCAGA, ggggaccactttgtacaagaaagctgggtcTAAAGAAGCCTTCTTCGCCG | Mycgr3\|73661  JGI  Ensembl | KOG3016 Acyl-CoA thioesterase; | CoA hydrolase activity; acyl-CoA thioesterase activity; | 0 % |  |  |
| 26 | ggggacaagtttgtacaaaaaagcaggcttgatgGGCAACAAGAGGAAACGCAA, ggggaccactttgtacaagaaagctgggtcCGCCGAAGCCTCAACATCAA | Mycgr3\|74280  JGI  Ensembl |  |  | 0 % |  |  |
| 26 | ggggacaagtttgtacaaaaaagcaggcttgatgGACGTCGTCCTCGAGGTGTT, ggggaccactttgtacaagaaagctgggtcTTGCACCTTCTTCACCACGA | Mycgr3\|74506  JGI  Ensembl | KOG0872 Sterol C5 desaturase; | 1-chloro-2,2-bis(4-chlorophenyl)ethane dehydrogenase activity; 2,6-dihydroxycyclohexane-1-carboxyl-CoA dehydrogenase activity;  2-hydroxycyclohexane-1-carboxyl-CoA dehydrogenase activity;  carvone reductase activity;  catalytic activity;  metabolic process;  peptidyl-oxazoline dehydrogenase activity;  peptidyl-thiazoline dehydrogenase activity;  phytoene dehydrogenase activity; | 1.86667 % |  |  |
| 26 | ggggacaagtttgtacaaaaaagcaggcttgatgGCCTCCGCCCTCTTCTTCCT, ggggaccactttgtacaagaaagctgggtcTTGCACATCAGGCAGCCTTA | Mycgr3\|75112  JGI  Ensembl | KOG0937 Adaptor complexes medium subunit family; | clathrin adaptor complex; intracellular protein transport;  protein binding;  protein complex assembly;  vesicle-mediated transport; | 0 % |  |  |
| 26 | ggggacaagtttgtacaaaaaagcaggcttgatgGTGCGCGAGAGGGACTCGGA, ggggaccactttgtacaagaaagctgggtcCAAAAAGTACTCTCCTCTCC | Mycgr3\|75902  JGI  Ensembl | KOG0373 Serine/threonine specific protein phosphatase involved in cell cycle control, PP2A-related; | CTD phosphatase activity; calcium-dependent protein serine/threonine phosphatase activity;  calcium-dependent protein serine/threonine phosphatase regulator activity;  hydrolase activity;  magnesium-dependent protein serine/threonine phosphatase activity;  magnesium-dependent protein serine/threonine phosphatase complex;  myosin phosphatase activity;  myosin phosphatase complex;  myosin phosphatase regulator activity;  phosphoprotein phosphatase activity; | 1.35685 % |  |  |
| 26 | ggggacaagtttgtacaaaaaagcaggcttgatgAATTTCTGGACCAAATATGT, ggggaccactttgtacaagaaagctgggtcAACCTTCTGCAACCCCTTCC | Mycgr3\|76232  JGI  Ensembl |  |  | 0 % |  |  |
| 26 | ggggacaagtttgtacaaaaaagcaggcttgatgTCCAACACCGACTTCCTAGG, ggggaccactttgtacaagaaagctgggtcAGCACCCTCACTGCCAAACT | Mycgr3\|76672  JGI  Ensembl | KOG0739 AAA+-type ATPase; | ATP binding; microtubule-severing ATPase activity;  nucleoside-triphosphatase activity;  nucleotide binding; | 1.33992 % |  |  |
| 26 | ggggacaagtttgtacaaaaaagcaggcttgatgGTATCCTCTTTCACAGCCCC, ggggaccactttgtacaagaaagctgggtcCCACCCTCGCCACCTCACCC | Mycgr3\|76868  JGI  Ensembl |  |  | 0 % |  |  |
| 26 | ggggacaagtttgtacaaaaaagcaggcttgatgGCCTCCGGATTCATGCGCCT, ggggaccactttgtacaagaaagctgggtcATCCCATCTCAGGACTTCAG | Mycgr3\|76956  JGI  Ensembl | KOG4842 Protein involved in sister chromatid separation and/or segregation; |  | 0 % |  |  |
| 26 | ggggacaagtttgtacaaaaaagcaggcttgatgAGCACACCCGACTTCCCGCG, ggggaccactttgtacaagaaagctgggtcAGTTTCTTCGTCATCCTCAT | Mycgr3\|77261  JGI  Ensembl | KOG0255 Synaptic vesicle transporter SVOP and related transporters (major facilitator superfamily); | integral to membrane; transport; transporter activity; | 0 % |  |  |
| 26 | ggggacaagtttgtacaaaaaagcaggcttgatgGCGGTCCTCAAGCAAGCCCT, ggggaccactttgtacaagaaagctgggtcGATAAAGAGCTTATCTCCAT | Mycgr3\|77799  JGI  Ensembl | KOG2195 Transferrin receptor and related proteins containing the protease-associated (PA) domain; | alanine aminopeptidase activity; peptidase activity;  proteolysis; | 0 % |  |  |
| 26 | ggggacaagtttgtacaaaaaagcaggcttgatgCGTTTCGCTCTTTCCAGTCA, ggggaccactttgtacaagaaagctgggtcCGGCATTCCCGCCCTCACGT | Mycgr3\|86391  JGI  Ensembl | KOG2806 Chitinase; | carbohydrate metabolic process; chitin catabolic process;  chitinase activity;  endochitinase activity;  hydrolase activity, hydrolyzing O-glycosyl compounds; | 0 % |  |  |
| 26 | ggggacaagtttgtacaaaaaagcaggcttgatgACCTCCCGCGACGGATACAC, ggggaccactttgtacaagaaagctgggtcCGGCCTCGCCGTCGCCGCAG | Mycgr3\|88429  JGI  Ensembl | KOG0029 Amine oxidase; | FAD binding; amine oxidase activity;  electron transport;  oxidoreductase activity; | 0 % |  |  |
| 26 | ggggacaagtttgtacaaaaaagcaggcttgatgACCAATTTCGAATTCGTGAC, ggggaccactttgtacaagaaagctgggtcGTCTTGTTTGGAAGGTCCAT | Mycgr3\|89166  JGI  Ensembl |  |  | 0 % |  |  |
| 26 | ggggacaagtttgtacaaaaaagcaggcttgatgCTCGCCTTGAGACCTGTCGA, ggggaccactttgtacaagaaagctgggtcCTTCCTTCCAAACGTCCTGC | Mycgr3\|89839  JGI  Ensembl | KOG3599 Ca2+-modulated nonselective cation channel polycystin; |  | 0 % |  |  |
| 26 | ggggacaagtttgtacaaaaaagcaggcttgatgCCGGACCGCGACGAAGACAG, ggggaccactttgtacaagaaagctgggtcAAGCTGGATAGCAAACTTCT | Mycgr3\|91051  JGI  Ensembl | KOG0255 Synaptic vesicle transporter SVOP and related transporters (major facilitator superfamily); | integral to membrane; transport; transporter activity; | 0 % |  |  |
| 26 | ggggacaagtttgtacaaaaaagcaggcttgatgCTCCACCCCACGGGCGTCGA, ggggaccactttgtacaagaaagctgggtcTCTCGCAGACATCTTCCACT | Mycgr3\|92064  JGI  Ensembl |  |  | 0 % |  |  |
| 26 | ggggacaagtttgtacaaaaaagcaggcttgatgGCGCCGCCGATGGAAAACTT, ggggaccactttgtacaagaaagctgggtcGTCCTGGTTCTCCTCTTGAT | Mycgr3\|92126  JGI  Ensembl |  |  | 0 % |  |  |
| 26 | ggggacaagtttgtacaaaaaagcaggcttgatgTTTTTATTGCGCATTGACCT, ggggaccactttgtacaagaaagctgggtcAGAATCCTCAGATCCCCGAG | Mycgr3\|92645  JGI  Ensembl | KOG1339 Aspartyl protease; | pepsin A activity; proteolysis; | 1.50229 % |  |  |
| 26 | ggggacaagtttgtacaaaaaagcaggcttgatgTCTTATCATGAGCAGCATAA, ggggaccactttgtacaagaaagctgggtcGAGGTAGGGATAGATGCCTT | Mycgr3\|93095  JGI  Ensembl |  |  | 0 % |  |  |
| 26 | ggggacaagtttgtacaaaaaagcaggcttgatgCCATATCTGAACCGCAGGGG, ggggaccactttgtacaagaaagctgggtcCTCGTCATACTCGACCTCGA | Mycgr3\|93233  JGI  Ensembl | KOG1472 Histone acetyltransferase SAGA/ADA, catalytic subunit PCAF/GCN5 and related proteins; |  | 0 % |  |  |
| 26 | ggggacaagtttgtacaaaaaagcaggcttgatgCTCAAGCCGAGATCGCCCAG, ggggaccactttgtacaagaaagctgggtcGCATTTCTTTTTCGACGACG | Mycgr3\|94625  JGI  Ensembl | KOG1911 Heterochromatin-associated protein HP1 and related CHROMO domain proteins; |  | 0 % |  |  |
| 26 | ggggacaagtttgtacaaaaaagcaggcttgatgTCTCCACCACAAGGGAAATT, ggggaccactttgtacaagaaagctgggtcGCCCAGCCTTCCACGCTTCT | Mycgr3\|94683  JGI  Ensembl | KOG0824 Predicted E3 ubiquitin ligase; | protein binding; zinc ion binding; | 0 % |  |  |
| 26 | ggggacaagtttgtacaaaaaagcaggcttgatgGAAATCTACACCCGCGTTCC, ggggaccactttgtacaagaaagctgggtcACGCCTCATCAACACATCAC | Mycgr3\|94705  JGI  Ensembl | KOG4415 Uncharacterized conserved protein; |  | 0 % |  |  |
| 26 | ggggacaagtttgtacaaaaaagcaggcttgatgGGCTTCACAATCGACGCGGA, ggggaccactttgtacaagaaagctgggtcCTCAATGGTGAAGCTGCCTA | Mycgr3\|96354  JGI  Ensembl |  |  | 0 % |  |  |
| 26 | ggggacaagtttgtacaaaaaagcaggcttgatgCACTTCCTCCAGAGCCTCGT, ggggaccactttgtacaagaaagctgggtcGGATTGCACGAGGGGGGACT | Mycgr3\|96377  JGI  Ensembl |  |  | 0 % |  |  |
| 26 | ggggacaagtttgtacaaaaaagcaggcttgatgGCGCGAATCAAGACTCCAAA, ggggaccactttgtacaagaaagctgggtcCCGCCATGCTTTCCTTCCTT | Mycgr3\|96400  JGI  Ensembl | KOG1164 Casein kinase (serine/threonine/tyrosine protein kinase); |  | 0 % |  |  |
| 26 | ggggacaagtttgtacaaaaaagcaggcttgatgGCCGCGATGTACCCACCAAG, ggggaccactttgtacaagaaagctgggtcAAACATTCCTTGAAGTGCTT | Mycgr3\|96565  JGI  Ensembl |  |  | 1.46777 % |  |  |
| 26 | ggggacaagtttgtacaaaaaagcaggcttgatgATGTCTTCTCGCCACGCTCC, ggggaccactttgtacaagaaagctgggtcAGAGTCGTCTTCAGTCATTC | Mycgr3\|96676  JGI  Ensembl |  |  | 0 % |  |  |
| 26 | ggggacaagtttgtacaaaaaagcaggcttgatgTACCAATGTCGAAGCCATTG, ggggaccactttgtacaagaaagctgggtcGAAGGCAGCGCCGAAGCTGA | Mycgr3\|96946  JGI  Ensembl |  |  | 2.20065 % | chr\_12:326286 G->C; |  |
| 26 | ggggacaagtttgtacaaaaaagcaggcttgatgTTCTTCGTGCTCATGGCCAT, ggggaccactttgtacaagaaagctgggtcGCCAGACTTGCTCCTTTGCG | Mycgr3\|97166  JGI  Ensembl |  |  | 0 % |  |  |
| 26 | ggggacaagtttgtacaaaaaagcaggcttgatgGGTCAAGGCAGTGGACAGTC, ggggaccactttgtacaagaaagctgggtcGTGGCTCACACTGCTCACAC | Mycgr3\|97524  JGI  Ensembl | KOG2533 Permease of the major facilitator superfamily; |  | 0 % |  |  |
| 26 | ggggacaagtttgtacaaaaaagcaggcttgatgACCAAGGACCCGATGAGCGA, ggggaccactttgtacaagaaagctgggtcCCCATCCTCCCAACCCGCTA | Mycgr3\|97708  JGI  Ensembl | KOG1904 Transcription coactivator; |  | 0 % |  |  |
| 26 | ggggacaagtttgtacaaaaaagcaggcttgatgGACGGCTTCATCCAACCACA, ggggaccactttgtacaagaaagctgggtcGAGCTCCGGTGCATCATACC | Mycgr3\|97810  JGI  Ensembl |  |  | 0 % |  |  |
| 26 | ggggacaagtttgtacaaaaaagcaggcttgatgGACACCGCCAGCAACGACGA, ggggaccactttgtacaagaaagctgggtcCGACCCTTTTCCTGCATCGA | Mycgr3\|97946  JGI  Ensembl | KOG2748 Uncharacterized conserved protein, contains chromo domain; |  | 0 % |  |  |
| 26 | ggggacaagtttgtacaaaaaagcaggcttgatgTTCGAGCAGGAAGTTGTAGG, ggggaccactttgtacaagaaagctgggtcCCCATCTCGTACATCATCCT | Mycgr3\|98077  JGI  Ensembl |  |  | 0 % |  |  |
| 26 | ggggacaagtttgtacaaaaaagcaggcttgatgGCCCAGTCTCTCATCCAAGC, ggggaccactttgtacaagaaagctgggtcCGGGCGTGCGAGAGGGGAGG | Mycgr3\|98083  JGI  Ensembl |  |  | 0 % |  |  |
| 26 | ggggacaagtttgtacaaaaaagcaggcttgatgAACGCGGCAGTGACGATGGC, ggggaccactttgtacaagaaagctgggtcTTGTTGTCCTGTCGCTATAT | Mycgr3\|98124  JGI  Ensembl |  |  | 0 % |  |  |
| 26 | ggggacaagtttgtacaaaaaagcaggcttgatgTCGTCTTTGCGTACTTTTGC, ggggaccactttgtacaagaaagctgggtcCAACTTGCTAGACAGCACCG | Mycgr3\|98270  JGI  Ensembl | KOG1526 NADP-dependent isocitrate dehydrogenase; | isocitrate dehydrogenase (NADP+) activity; isocitrate dehydrogenase activity;  metabolic process;  oxidoreductase activity, acting on the CH-OH group of donors, NAD or NADP as acceptor; | 0 % |  |  |
| 26 | ggggacaagtttgtacaaaaaagcaggcttgatgCCGCGGATGCTCAGTCCGCC, ggggaccactttgtacaagaaagctgggtcGCCGGAAAAGAGTCTCGCAT | Mycgr3\|98361  JGI  Ensembl |  |  | 0 % |  |  |
| 26 | ggggacaagtttgtacaaaaaagcaggcttgatgTCGATCGATGCGCCCGCCTA, ggggaccactttgtacaagaaagctgggtcACCCTCCTTATCAAAACTAT | Mycgr3\|98901  JGI  Ensembl | KOG2759 Vacuolar H+-ATPase V1 sector, subunit H; | ATP binding; hydrogen ion transporting ATPase activity, rotational mechanism;  peripheral to membrane of membrane fraction;  vacuolar proton-transporting V-type ATPase, V1 domain; | 0 % |  |  |
| 26 | ggggacaagtttgtacaaaaaagcaggcttgatgCTTCGCGAAAGCCTCCTCGC, ggggaccactttgtacaagaaagctgggtcAGTCGACCCAATGGCTCCGA | Mycgr3\|100252  JGI  Ensembl |  |  | 0 % |  |  |
| 26 | ggggacaagtttgtacaaaaaagcaggcttgatgCTCTCTCAGCTGTTGAACCC, ggggaccactttgtacaagaaagctgggtcCATTCGATCGTGGATGTCTC | Mycgr3\|100278  JGI  Ensembl | KOG1721 FOG: Zn-finger; | intracellular; nucleic acid binding;  zinc ion binding; | 0 % |  |  |
| 26 | ggggacaagtttgtacaaaaaagcaggcttgatgTCCGAAAAAGAACGGCAATC, ggggaccactttgtacaagaaagctgggtcTCGATCGATTGATAGCAGCC | Mycgr3\|104744  JGI  Ensembl | KOG1441 Glucose-6-phosphate/phosphate and phosphoenolpyruvate/phosphate antiporter; |  | 0 % |  |  |
| 26 | ggggacaagtttgtacaaaaaagcaggcttgatgGCCTCCACCAAAACCCGTCC, ggggaccactttgtacaagaaagctgggtcCTCCGTACTCCACAACCCCA | Mycgr3\|106067  JGI  Ensembl | KOG1208 Dehydrogenases with different specificities (related to short-chain alcohol dehydrogenases); | (R)-2-hydroxyglutarate dehydrogenase activity; (R)-2-hydroxyisocaproate dehydrogenase activity;  2-hydroxytetrahydrofuran dehydrogenase activity;  3-keto sterol reductase activity;  3-ketoglucose-reductase activity;  5-exo-hydroxycamphor dehydrogenase activity;  C-3 sterol dehydrogenase (C-4 sterol decarboxylase) activity;  aldo-keto reductase activity;  epoxide dehydrogenase activity;  gluconate dehydrogenase activity;  isocitrate dehydrogenase activity;  membrane;  metabolic process;  mevaldate reductase activity;  oxidoreductase activity;  phenylcoumaran benzylic ether reductase activity;  steroid dehydrogenase activity;  steroid dehydrogenase activity, acting on the CH-CH group of donors;  steroid dehydrogenase activity, acting on the CH-OH group of donors, NAD or NADP as acceptor; transport; transporter activity; | 0 % |  |  |
| 26 | ggggacaagtttgtacaaaaaagcaggcttgatgGCCACCGCCACCATGTCTTC, ggggaccactttgtacaagaaagctgggtcCACGGCGCCAGGCGTAGCCA | Mycgr3\|107011  JGI  Ensembl | KOG2992 Nucleolar GTPase/ATPase p130; |  | 1.48675 % |  |  |
| 26 | ggggacaagtttgtacaaaaaagcaggcttgatgACGCCGCTTTTGCACTTCCC, ggggaccactttgtacaagaaagctgggtcGTCCTCCCAAATCGTCTCCA | Mycgr3\|107116  JGI  Ensembl | KOG1869 Splicing coactivator SRm160/300, subunit SRm300; | hydrogen ion transmembrane transporter activity; proton-transporting two-sector ATPase complex; | 0 % |  |  |
| 26 | ggggacaagtttgtacaaaaaagcaggcttgatgCGCGCTGCTACACTGGTCCA, ggggaccactttgtacaagaaagctgggtcCACCGAATGACTTCCTTCTG | Mycgr3\|107443  JGI  Ensembl |  |  | 0 % |  |  |
| 26 | ggggacaagtttgtacaaaaaagcaggcttgatgTCGTCGATACCACGTCGTGG, ggggaccactttgtacaagaaagctgggtcACCACTCAATAGATCCTTCA | Mycgr3\|107535  JGI  Ensembl | KOG2588 Predicted DNA-binding protein; | nucleus; regulation of transcription;  transcription regulator activity; | 0 % |  |  |
| 26 | ggggacaagtttgtacaaaaaagcaggcttgatgCAATCATGCTTGCAACGATG, ggggaccactttgtacaagaaagctgggtcCTCCTCCTTCAGCCATGGAT | Mycgr3\|107640  JGI  Ensembl | KOG1290 Serine/threonine protein kinase; | ATP binding; protein amino acid phosphorylation;  protein kinase activity;  protein serine/threonine kinase activity; | 1.27226 % |  |  |
| 26 | ggggacaagtttgtacaaaaaagcaggcttgatgGTCACGAAGAGATCTATCGA, ggggaccactttgtacaagaaagctgggtcGATATCGACATTCTGCCGCA | Mycgr3\|108055  JGI  Ensembl | KOG4210 Nuclear localization sequence binding protein; |  | 0 % |  |  |
| 26 | ggggacaagtttgtacaaaaaagcaggcttgatgTATTTTCAGCCCTTCCGGCC, ggggaccactttgtacaagaaagctgggtcCCAGAGAATCTCCGCTTGCG | Mycgr3\|108075  JGI  Ensembl |  |  | 0 % |  |  |
| 26 | ggggacaagtttgtacaaaaaagcaggcttgatgGGCGCAGCAGAGTCCAAAAA, ggggaccactttgtacaagaaagctgggtcTCCCTTCAAGATATTCCTCC | Mycgr3\|108571  JGI  Ensembl |  |  | 0 % |  |  |
| 26 | ggggacaagtttgtacaaaaaagcaggcttgatgGACCATCAGCCATCCACGCC, ggggaccactttgtacaagaaagctgggtcTCCTCTCTTGCTAAAGCGGC | Mycgr3\|109542  JGI  Ensembl |  |  | 0 % |  |  |
| 26 | ggggacaagtttgtacaaaaaagcaggcttgatgGCACCACGCGCAGCTGCCCT, ggggaccactttgtacaagaaagctgggtcATCATCTTCCACAATGATCC | Mycgr3\|109581  JGI  Ensembl | KOG0613 Projectin/twitchin and related proteins; |  | 1.26162 % |  |  |
| 26 | ggggacaagtttgtacaaaaaagcaggcttgatgGGCTTCTTCTCCCGCAAGAA, ggggaccactttgtacaagaaagctgggtcGAACACCGTGCCGTCGGACA | Mycgr3\|110387  JGI  Ensembl |  |  | 0 % |  |  |
| 26 | ggggacaagtttgtacaaaaaagcaggcttgatgCAGAGAGCTTCCAGAATGTC, ggggaccactttgtacaagaaagctgggtcCAATTTACACTTCTCCTCCA | Mycgr3\|111030  JGI  Ensembl | KOG2820 FAD-dependent oxidoreductase; | sarcosine oxidase activity; | 0 % |  |  |
| 26 | ggggacaagtttgtacaaaaaagcaggcttgatgGGCAACCAGGCCAACAGGAA, ggggaccactttgtacaagaaagctgggtcGTCTCTGACCTCGCGCAGGG | Mycgr3\|111706  JGI  Ensembl |  |  | 0 % |  |  |
| 27 | ggggacaagtttgtacaaaaaagcaggcttgatgGCTCAGTCTCCAGAACCGCA, ggggaccactttgtacaagaaagctgggtcCAAACTCCTCGTCCTCTCCT | Mycgr3\|32609  JGI  Ensembl | KOG0156 Cytochrome P450 CYP2 subfamily; | electron transport; heme binding;  iron ion binding;  monooxygenase activity; | 100 % |  |  |
| 27 | ggggacaagtttgtacaaaaaagcaggcttgatgCATCTCTTTTGGCTGCTCAT, ggggaccactttgtacaagaaagctgggtcCGATCCCGCCGTATCGATCC | Mycgr3\|35026  JGI  Ensembl | KOG2182 Hydrolytic enzymes of the alpha/beta hydrolase fold; | proteolysis; serine-type Pro-X carboxypeptidase activity;  serine-type peptidase activity; | 100 % |  |  |
| 27 | ggggacaagtttgtacaaaaaagcaggcttgatgTCACCACACGCTCTATCCGT, ggggaccactttgtacaagaaagctgggtcTCGCTTCTGCTCAAACGAGC | Mycgr3\|36273  JGI  Ensembl |  |  | 100 % |  |  |
| 27 | ggggacaagtttgtacaaaaaagcaggcttgatgGCCGCAAGCCCCATCAAGGA, ggggaccactttgtacaagaaagctgggtcCAATCGATATCTAAACTCTT | Mycgr3\|39937  JGI  Ensembl | KOG2533 Permease of the major facilitator superfamily; | integral to membrane; transport; transporter activity; | 0 % |  |  |
| 27 | ggggacaagtttgtacaaaaaagcaggcttgatgCCATTCTACCTAGCTGGACT, ggggaccactttgtacaagaaagctgggtcTATCGCCGAAGTGGAGTCCC | Mycgr3\|40824  JGI  Ensembl |  |  | 100 % |  |  |
| 27 | ggggacaagtttgtacaaaaaagcaggcttgatgGAGTCCACCGGCCGCGAAGC, ggggaccactttgtacaagaaagctgggtcAGCCTTCTGCACCACTTCCA | Mycgr3\|41244  JGI  Ensembl | KOG0255 Synaptic vesicle transporter SVOP and related transporters (major facilitator superfamily); | integral to membrane; transport; transporter activity; | 100 % |  |  |
| 27 | ggggacaagtttgtacaaaaaagcaggcttgatgTCATCCACCGACCCTCCCCT, ggggaccactttgtacaagaaagctgggtcGTACCCGCCCTGCTCCCTCC | Mycgr3\|44247  JGI  Ensembl | KOG2752 Uncharacterized conserved protein, contains N-recognin-type Zn-finger; | ubiquitin cycle; ubiquitin-protein ligase activity;  zinc ion binding; | 100 % | chr\_6:916019 C->T; |  |
| 27 | ggggacaagtttgtacaaaaaagcaggcttgatgCTCCGCCTCAAACCCTCGGT, ggggaccactttgtacaagaaagctgggtcGACTTCTCCAGTCATCAGCA | Mycgr3\|44636  JGI  Ensembl |  |  | 100 % |  |  |
| 27 | ggggacaagtttgtacaaaaaagcaggcttgatgGCCTTCCGAATCGTCTTTGC, ggggaccactttgtacaagaaagctgggtcAGAGAACGAGCGATAAATTG | Mycgr3\|46025  JGI  Ensembl | KOG2650 Zinc carboxypeptidase; |  | 0 % |  |  |
| 27 | ggggacaagtttgtacaaaaaagcaggcttgatgCGGACTCTCTCCGGCGGACT, ggggaccactttgtacaagaaagctgggtcAGCTTCCTTCCCACCCTCCT | Mycgr3\|47544  JGI  Ensembl |  |  | 100 % | chr\_9:372149 A->G; |  |
| 27 | ggggacaagtttgtacaaaaaagcaggcttgatgTCTCGCCACTCTTCCTTCCA, ggggaccactttgtacaagaaagctgggtcCACGGCCTTCATTACCTTCC | Mycgr3\|49090  JGI  Ensembl | KOG2504 Monocarboxylate transporter; | integral to membrane; transport; transporter activity; | 100 % |  |  |
| 27 | ggggacaagtttgtacaaaaaagcaggcttgatgAGAGTCTGGCGGTCCGCATC, ggggaccactttgtacaagaaagctgggtcACACTCACACCTCCCCACCC | Mycgr3\|49696  JGI  Ensembl | KOG0554 Asparaginyl-tRNA synthetase (mitochondrial); | ATP binding; aminoacyl-tRNA ligase activity;  asparagine-tRNA ligase activity;  asparaginyl-tRNA aminoacylation;  aspartate-tRNA ligase activity;  aspartyl-tRNA aminoacylation;  cytoplasm;  nucleic acid binding;  nucleotide binding;  tRNA aminoacylation for protein translation;  translation; | 100 % |  |  |
| 27 | ggggacaagtttgtacaaaaaagcaggcttgatgATCCTCACAGCGCTCTACGG, ggggaccactttgtacaagaaagctgggtcCCTCCTCTCCAACAGCACCT | Mycgr3\|50499  JGI  Ensembl | KOG0158 Cytochrome P450 CYP3/CYP5/CYP6/CYP9 subfamilies; | electron transport; heme binding;  iron ion binding;  monooxygenase activity; | 0 % |  |  |
| 27 | ggggacaagtttgtacaaaaaagcaggcttgatgTTTCCCATCCCAGTCCCCGA, ggggaccactttgtacaagaaagctgggtcCCAATAACACAAGCCTCCAT | Mycgr3\|52885  JGI  Ensembl |  |  | 100 % |  |  |
| 27 | ggggacaagtttgtacaaaaaagcaggcttgatgCTTGCAAAGTCCGCTCTTCA, ggggaccactttgtacaagaaagctgggtcACCAGTGATCTTGTAAATTG | Mycgr3\|53498  JGI  Ensembl | KOG1411 Aspartate aminotransferase/Glutamic oxaloacetic transaminase AAT1/GOT2; | amino acid metabolic process; aspartate transaminase activity;  biosynthetic process;  catalytic activity;  pyridoxal phosphate binding;  transaminase activity;  transferase activity, transferring nitrogenous groups; | 100 % |  |  |
| 27 | ggggacaagtttgtacaaaaaagcaggcttgatgCCACTCACTGTCCTCAGCGA, ggggaccactttgtacaagaaagctgggtcAAATCGCTCAATCGTAACTC | Mycgr3\|56546  JGI  Ensembl | KOG3007 Mu-crystallin; |  | 100 % |  |  |
| 27 | ggggacaagtttgtacaaaaaagcaggcttgatgGCTCTCTCACAAGCTGCTTA, ggggaccactttgtacaagaaagctgggtcATATCTCGTCAGCGCCGGAC | Mycgr3\|57881  JGI  Ensembl | KOG0024 Sorbitol dehydrogenase; | alcohol dehydrogenase activity; alcohol dehydrogenase activity, iron-dependent;  alcohol dehydrogenase activity, metal ion-independent;  alcohol dehydrogenase activity, zinc-dependent;  oxidoreductase activity;  zinc ion binding; | 0 % |  |  |
| 27 | ggggacaagtttgtacaaaaaagcaggcttgatgTTGTCCGGGATATTGCTCTT, ggggaccactttgtacaagaaagctgggtcAAATCGAATCTCATAGCTCC | Mycgr3\|61231  JGI  Ensembl | KOG0938 Adaptor complexes medium subunit family; | clathrin adaptor complex; intracellular protein transport;  protein binding;  protein complex assembly;  vesicle-mediated transport; | 100 % |  |  |
| 27 | ggggacaagtttgtacaaaaaagcaggcttgatgAGTCCCACGGAGATCAAGAA, ggggaccactttgtacaagaaagctgggtcTGCCGGCTTCCACTCATAAT | Mycgr3\|64902  JGI  Ensembl | KOG2653 6-phosphogluconate dehydrogenase; | NADP binding; pentose-phosphate shunt;  phosphogluconate dehydrogenase (decarboxylating) activity; | 100 % |  |  |
| 27 | ggggacaagtttgtacaaaaaagcaggcttgatgTTCGCCGCAGCCGTGACAGC, ggggaccactttgtacaagaaagctgggtcACGCTTGTTAGCGACCTCGA | Mycgr3\|65312  JGI  Ensembl | KOG0254 Predicted transporter (major facilitator superfamily); | L-arabinose isomerase activity; carbohydrate transport;  integral to membrane;  membrane;  sugar:hydrogen symporter activity;  transport;  transporter activity; | 100 % |  |  |
| 27 | ggggacaagtttgtacaaaaaagcaggcttgatgGGATCCGATTCGTTGGGAGG, ggggaccactttgtacaagaaagctgggtcCCATCCGAACTGCTCGGAGC | Mycgr3\|65806  JGI  Ensembl | KOG0039 Ferric reductase, NADH/NADPH oxidase and related proteins; | ferric-chelate reductase activity; | 1.15642 % |  |  |
| 27 | ggggacaagtttgtacaaaaaagcaggcttgatgCACAACCACGTTCACAATCC, ggggaccactttgtacaagaaagctgggtcCAGCGGAGCCAGACCACCAA | Mycgr3\|66426  JGI  Ensembl | KOG1116 Sphingosine kinase, involved in sphingolipid metabolism; | activation of protein kinase C activity; diacylglycerol kinase activity;  sphinganine kinase activity; | 100 % | chr\_1:3393051 G->A; chr\_1:3392984 C->A; |  |
| 27 | ggggacaagtttgtacaaaaaagcaggcttgatgGCCATTCTCCGCGTTGCGCC, ggggaccactttgtacaagaaagctgggtcATACTCCATAGCATCCTCAA | Mycgr3\|66658  JGI  Ensembl | KOG3857 Alcohol dehydrogenase, class IV; | alcohol dehydrogenase activity; alcohol dehydrogenase activity, iron-dependent;  alcohol dehydrogenase activity, metal ion-independent;  alcohol dehydrogenase activity, zinc-dependent;  metabolic process;  metal ion binding;  oxidoreductase activity; | 100 % |  |  |
| 27 | ggggacaagtttgtacaaaaaagcaggcttgatgCCTGGACCTCCATCGGCAAC, ggggaccactttgtacaagaaagctgggtcAAGCTTCCGACACCAATTTC | Mycgr3\|67202  JGI  Ensembl | KOG3880 Predicted small molecule transporter involved in cellular pH homeostasis (Batten disease protein in human); | membrane; | 100 % |  |  |
| 27 | ggggacaagtttgtacaaaaaagcaggcttgatgTCGAGGACCATCACCTACCA, ggggaccactttgtacaagaaagctgggtcTAATTTGCTACGAACCCACA | Mycgr3\|68078  JGI  Ensembl | KOG0538 Glycolate oxidase; | L-lactate dehydrogenase (cytochrome) activity; electron transport;  heme binding;  oxidoreductase activity;  transition metal ion binding; | 100 % |  |  |
| 27 | ggggacaagtttgtacaaaaaagcaggcttgatgACTTCTGGTAGCACGACCAC, ggggaccactttgtacaagaaagctgggtcTTCGATCAACTGGGCAGGAT | Mycgr3\|68348  JGI  Ensembl |  |  | 2.50313 % |  |  |
| 27 | ggggacaagtttgtacaaaaaagcaggcttgatgGTCGGTAAACGAGCTCTCCC, ggggaccactttgtacaagaaagctgggtcATTGATATTCGCATGCCCGG | Mycgr3\|68771  JGI  Ensembl | KOG1485 Mitochondrial Fe2+ transporter MMT1 and related transporters (cation diffusion facilitator superfamily); | cation transmembrane transporter activity; cation transport;  membrane; | 2.772 % |  |  |
| 27 | ggggacaagtttgtacaaaaaagcaggcttgatgACCACTCTCCAACCCAGACC, ggggaccactttgtacaagaaagctgggtcCTTCGCCTCCCACCCCGGCG | Mycgr3\|68972  JGI  Ensembl | KOG3720 Lysosomal & prostatic acid phosphatases; | acid phosphatase activity; | 100 % | chr\_2:3027622 A->G; |  |
| 27 | ggggacaagtttgtacaaaaaagcaggcttgatgCCTCCTCACACGTGCGAGAT, ggggaccactttgtacaagaaagctgggtcTCCTCCAGAAACTTCGATCT | Mycgr3\|69570  JGI  Ensembl | KOG2840 Uncharacterized conserved protein with similarity to predicted ATPase of the PP-loop superfamily; |  | 100 % |  |  |
| 27 | ggggacaagtttgtacaaaaaagcaggcttgatgCTCGGTGTGCTTCTGACGGC, ggggaccactttgtacaagaaagctgggtcACCCGATCCGTTCTTCCCAA | Mycgr3\|70510  JGI  Ensembl | KOG1231 Proteins containing the FAD binding domain; | FAD binding; oxidoreductase activity; | 100 % |  |  |
| 27 | ggggacaagtttgtacaaaaaagcaggcttgatgAGCGGCCAACCAATGCCCGT, ggggaccactttgtacaagaaagctgggtcGGAGGTCTTCTCCAGCGGAG | Mycgr3\|71033  JGI  Ensembl |  |  | 100 % |  |  |
| 27 | ggggacaagtttgtacaaaaaagcaggcttgatgGCGACTCTTGTACAGACATC, ggggaccactttgtacaagaaagctgggtcCTGCAGGAGATGGTCCCTGA | Mycgr3\|71065  JGI  Ensembl | KOG2785 C2H2-type Zn-finger protein; | intracellular; nucleic acid binding;  zinc ion binding; | 100 % | chr\_4:1301512 G->A; |  |
| 27 | ggggacaagtttgtacaaaaaagcaggcttgatgGCGCCAGCTTCGGAAGCAGG, ggggaccactttgtacaagaaagctgggtcATCGTATCCAATATCAGTCG | Mycgr3\|71081  JGI  Ensembl | KOG2614 Kynurenine 3-monooxygenase and related flavoprotein monooxygenases; | metabolic process; oxidoreductase activity;  salicylate 1-monooxygenase activity; | 0 % |  |  |
| 27 | ggggacaagtttgtacaaaaaagcaggcttgatgGTTCGGTCCTACATCTTCGT, ggggaccactttgtacaagaaagctgggtcCTCCGCCTCCGCTCGAAAGT | Mycgr3\|71198  JGI  Ensembl |  |  | 100 % |  |  |
| 27 | ggggacaagtttgtacaaaaaagcaggcttgatgCTCGACGAAAACCTTCCCGC, ggggaccactttgtacaagaaagctgggtcCTTCCCACTCATCAACCCAA | Mycgr3\|71228  JGI  Ensembl |  |  | 100 % |  |  |
| 27 | ggggacaagtttgtacaaaaaagcaggcttgatgGGTGTCCTGATAGAACCCAG, ggggaccactttgtacaagaaagctgggtcAATCTTCACCCCCTCCAAAA | Mycgr3\|71623  JGI  Ensembl |  |  | 100 % |  |  |
| 27 | ggggacaagtttgtacaaaaaagcaggcttgatgACCGAGACCAAAAAGTTGAA, ggggaccactttgtacaagaaagctgggtcCCAACGGAAGACCCTCCCAA | Mycgr3\|71908  JGI  Ensembl | KOG2614 Kynurenine 3-monooxygenase and related flavoprotein monooxygenases; | kynurenine 3-monooxygenase activity; metabolic process;  oxidoreductase activity; | 100 % |  |  |
| 27 | ggggacaagtttgtacaaaaaagcaggcttgatgGCCGCTCCCAAGCTTGCGAA, ggggaccactttgtacaagaaagctgggtcCTCTAACGCCTGTCCCTTCG | Mycgr3\|72635  JGI  Ensembl | KOG1401 Acetylornithine aminotransferase; | 4-aminobutyrate transaminase activity; pyridoxal phosphate binding;  transaminase activity; | 100 % |  |  |
| 27 | ggggacaagtttgtacaaaaaagcaggcttgatgTCTCCGTTCGGCGATATAAG, ggggaccactttgtacaagaaagctgggtcAGATATGGGCTCGGAACTCA | Mycgr3\|72843  JGI  Ensembl | KOG2533 Permease of the major facilitator superfamily; | integral to membrane; transport; transporter activity; | 0 % |  |  |
| 27 | ggggacaagtttgtacaaaaaagcaggcttgatgTCGGCCATGATCCCAATGGC, ggggaccactttgtacaagaaagctgggtcCTTCTTCTTGTTGATCTCCA | Mycgr3\|72961  JGI  Ensembl | KOG0552 FKBP-type peptidyl-prolyl cis-trans isomerase; | FK506-sensitive peptidyl-prolyl cis-trans isomerase; cyclophilin; cyclophilin-type peptidyl-prolyl cis-trans isomerase activity;  peptidyl-prolyl cis-trans isomerase activity;  protein folding; | 0 % |  |  |
| 27 | ggggacaagtttgtacaaaaaagcaggcttgatgGCGCCCAGCACCCCCCTAAT, ggggaccactttgtacaagaaagctgggtcTCGCTTCAACTGAGCGAATC | Mycgr3\|73382  JGI  Ensembl | KOG2027 Spindle pole body protein; |  | 100 % |  |  |
| 27 | ggggacaagtttgtacaaaaaagcaggcttgatgGCTCCCATCGCGATCGAGGA, ggggaccactttgtacaagaaagctgggtcGTTCTCAATCCCAATATGCA | Mycgr3\|74043  JGI  Ensembl | KOG1402 Ornithine aminotransferase; | ornithine-oxo-acid transaminase activity; pyridoxal phosphate binding;  transaminase activity; | 2.66667 % |  |  |
| 27 | ggggacaagtttgtacaaaaaagcaggcttgatgCCCATCAAGATCAGTAGTCT, ggggaccactttgtacaagaaagctgggtcGCTGACAGCTGTGTCCCGAG | Mycgr3\|75056  JGI  Ensembl |  |  | 100 % |  |  |
| 27 | ggggacaagtttgtacaaaaaagcaggcttgatgGCGTCCTCCAAAGCACCAAC, ggggaccactttgtacaagaaagctgggtcCAACTTCGCCTCCCCCCTAA | Mycgr3\|75580  JGI  Ensembl | KOG0538 Glycolate oxidase; | (S)-2-hydroxy-acid oxidase activity; electron transport;  glycolate oxidase activity;  glycolate oxidase complex;  oxidoreductase activity; | 100 % |  |  |
| 27 | ggggacaagtttgtacaaaaaagcaggcttgatgGACGATCTGACGGTGCAAGA, ggggaccactttgtacaagaaagctgggtcACTTGAGGGTCTCTCTTCGA | Mycgr3\|76570  JGI  Ensembl | KOG0254 Predicted transporter (major facilitator superfamily); | integral to membrane; response to antibiotic;  tetracycline transport;  tetracycline:hydrogen antiporter activity;  transport;  transporter activity; | 100 % | chr\_10:1440609 A->T; |  |
| 27 | ggggacaagtttgtacaaaaaagcaggcttgatgGCCACGCTCTCCAACTCGTT, ggggaccactttgtacaagaaagctgggtcTAACTTCGCCTTCGGGCCTT | Mycgr3\|77337  JGI  Ensembl | KOG1176 Acyl-CoA synthetase; | catalytic activity; cyanophycin synthetase activity;  metabolic process;  regulation of ligase activity; | 100 % |  |  |
| 27 | ggggacaagtttgtacaaaaaagcaggcttgatgCCAGCCGGAACAGCATCCCC, ggggaccactttgtacaagaaagctgggtcGACCTTCGCTTCCACTACAG | Mycgr3\|84494  JGI  Ensembl | KOG0254 Predicted transporter (major facilitator superfamily); | integral to membrane; response to antibiotic;  tetracycline transport;  tetracycline:hydrogen antiporter activity;  transport;  transporter activity; | 99.81741 % |  |  |
| 27 | ggggacaagtttgtacaaaaaagcaggcttgatgGCGGGCGACATTGAACTGCT, ggggaccactttgtacaagaaagctgggtcGGTCTTTGCGGTGCCGCCAT | Mycgr3\|86381  JGI  Ensembl |  |  | 1.22249 % |  |  |
| 27 | ggggacaagtttgtacaaaaaagcaggcttgatgAACAACGACTTCTTCCGCAA, ggggaccactttgtacaagaaagctgggtcTTTCTGAGCTGCCCTCCGCC | Mycgr3\|86496  JGI  Ensembl | KOG1911 Heterochromatin-associated protein HP1 and related CHROMO domain proteins; |  | 0 % |  |  |
| 27 | ggggacaagtttgtacaaaaaagcaggcttgatgGCCGGCCATGTCGCTTCGCT, ggggaccactttgtacaagaaagctgggtcGTAGTCCGGCCACTCTGGAT | Mycgr3\|86562  JGI  Ensembl | KOG2017 Molybdopterin synthase sulfurylase; | catalytic activity; negative regulation of ubiquitin-protein ligase activity;  positive regulation of ubiquitin-protein ligase activity;  regulation of ubiquitin-protein ligase activity;  regulation of ubiquitin-protein ligase activity during meiotic cell cycle;  ubiquitin-protein ligase activity; | 0 % |  |  |
| 27 | ggggacaagtttgtacaaaaaagcaggcttgatgGCGCAGCTGTCCAGCCTCTT, ggggaccactttgtacaagaaagctgggtcGTATCCACAAATCCCACTCC | Mycgr3\|86748  JGI  Ensembl | KOG0471 Alpha-amylase; | alpha-amylase activity; carbohydrate metabolic process;  catalytic activity;  cation binding; | 100 % |  |  |
| 27 | ggggacaagtttgtacaaaaaagcaggcttgatgGGCCCCAAAGGATCCGGCAA, ggggaccactttgtacaagaaagctgggtcGCTCCTCTCCTCCTGCTGCT | Mycgr3\|88791  JGI  Ensembl | KOG1187 Serine/threonine protein kinase; | UDP-N-acetylmuramoylalanyl-D-glutamyl-2,6-diaminopimelate-D-alanyl-D-alanine ligase activity; coenzyme F420-0 gamma-glutamyl ligase activity;  coenzyme F420-2 alpha-glutamyl ligase activity;  ribosomal S6-glutamic acid ligase activity; | 100 % |  |  |
| 27 | ggggacaagtttgtacaaaaaagcaggcttgatgTCAGTCCACCAATGTGATAC, ggggaccactttgtacaagaaagctgggtcTCGCCTGGAGACATGTCGTG | Mycgr3\|89921  JGI  Ensembl | KOG1325 Lysophospholipase; | proteolysis; subtilase activity; | 100 % |  |  |
| 27 | ggggacaagtttgtacaaaaaagcaggcttgatgGCTTCGCAGCACCATTTTGC, ggggaccactttgtacaagaaagctgggtcCTCCACGTTCAACCAAGTCC | Mycgr3\|90881  JGI  Ensembl | KOG1187 Serine/threonine protein kinase; |  | 100 % |  |  |
| 27 | ggggacaagtttgtacaaaaaagcaggcttgatgTACCGCATCCGAGGCAAGCC, ggggaccactttgtacaagaaagctgggtcTCCTCTTAGTCGGCCTCTGA | Mycgr3\|91207  JGI  Ensembl |  |  | 1.1772 % |  |  |
| 27 | ggggacaagtttgtacaaaaaagcaggcttgatgGCTTCCGCCTCCGCCCATCG, ggggaccactttgtacaagaaagctgggtcGTACTTGTTCTCCCAACCAC | Mycgr3\|92343  JGI  Ensembl |  |  | 100 % |  |  |
| 27 | ggggacaagtttgtacaaaaaagcaggcttgatgAATCATGAGAACCACTTCGA, ggggaccactttgtacaagaaagctgggtcCCTCCCCGCCGCCGCCATCA | Mycgr3\|93579  JGI  Ensembl | KOG3855 Monooxygenase involved in coenzyme Q (ubiquinone) biosynthesis; | 1-hydroxy-2-naphthoate hydroxylase activity; 1-hydroxy-2-oxolimonene 1,2-monooxygenase activity;  2-hydroxy-phenylacetate hydroxylase activity;  2-octaprenyl-3-methyl-6-methoxy-1,4-benzoquinol hydroxylase;  2-oxo-delta3-4,5,5-trimethylcyclopentenylacetyl-CoA 1,2-monooxygenase activity; 3,5-xylenol methylhydroxylase activity;  3-(3-hydroxy-phenyl)propionate hydroxylase activity;  3-methyl-2-oxo-1,2-dihydroquinoline 6-monooxygenase activity;  4-chlorobenzaldehyde oxidase activity;  4-nitrophenol 4-monooxygenase activity;  6-hydroxy-3-methyl-2-oxo-1,2-dihydroquinoline 6-monooxygenase activity;  6-hydroxynicotinate 3-monooxygenase activity;  FAD binding;  alpha-pinene monooxygenase [NADH] activity;  carbon disulfide oxygenase activity;  chlorophenol 4-monooxygenase activity;  dibenzothiophene monooxygenase activity;  dimethyl sulfide monooxygenase activity;  limonene 1,2-monooxygenase activity;  limonene monooxygenase activity;  metabolic process;  methanesulfonic acid monooxygenase activity;  oxidoreductase activity;  oxidoreductase activity, acting on paired donors, with incorporation or reduction of molecular oxygen, NADH or NADPH as one donor, and incorporation of one atom of oxygen; phenanthrene 1,2-monooxygenase activity;  phenanthrene 3,4-monooxygenase activity;  phenanthrene 9,10-monooxygenase activity;  phenylacetate hydroxylase activity;  styrene monooxygenase activity;  tetrahydrofuran hydroxylase activity;  toluene 2-monooxygenase activity;  toluene 3-monooxygenase activity;  toluene 4-monooxygenase activity;  toluene-4-sulfonate monooxygenase activity;  toluene-sulfonate methyl-monooxygenase activity;  ubiquinone biosynthetic process;  xylene monooxygenase activity; | 0 % |  |  |
| 27 | ggggacaagtttgtacaaaaaagcaggcttgatgACTTACGAAATGGCCACAAT, ggggaccactttgtacaagaaagctgggtcATACCTCTCATAAGCCCTCC | Mycgr3\|94197  JGI  Ensembl |  |  | 0 % |  |  |
| 27 | ggggacaagtttgtacaaaaaagcaggcttgatgTGCGGCGGCACTCAATTCCG, ggggaccactttgtacaagaaagctgggtcGCATCGCCCAGGTCCAGCCG | Mycgr3\|94909  JGI  Ensembl | KOG3544 Collagens (type IV and type XIII), and related proteins; |  | 0 % |  |  |
| 27 | ggggacaagtttgtacaaaaaagcaggcttgatgTCTTCTCCGCCACTCCACAC, ggggaccactttgtacaagaaagctgggtcCGACAGTGGTTCTCCCGCAA | Mycgr3\|95521  JGI  Ensembl |  |  | 100 % |  |  |
| 27 | ggggacaagtttgtacaaaaaagcaggcttgatgCCGTCTCTCCTCGACTTACC, ggggaccactttgtacaagaaagctgggtcACGACCGCCTCCTGTGGCCT | Mycgr3\|95582  JGI  Ensembl |  |  | 0 % |  |  |
| 27 | ggggacaagtttgtacaaaaaagcaggcttgatgAATAGCATCTCAAAAGTTGT, ggggaccactttgtacaagaaagctgggtcATCAATTTTCCGCCAGAATG | Mycgr3\|96097  JGI  Ensembl |  |  | 0 % |  |  |
| 27 | ggggacaagtttgtacaaaaaagcaggcttgatgCAGAGTCGGAGGAGTCGAGG, ggggaccactttgtacaagaaagctgggtcCCGCTGGAGAAAGACGATGA | Mycgr3\|97714  JGI  Ensembl |  |  | 2.34423 % |  |  |
| 27 | ggggacaagtttgtacaaaaaagcaggcttgatgAATCCATATAGTCTCTCCGC, ggggaccactttgtacaagaaagctgggtcTCTATCTCCGCGACTCTCGC | Mycgr3\|97801  JGI  Ensembl |  |  | 1.28993 % |  |  |
| 27 | ggggacaagtttgtacaaaaaagcaggcttgatgCCGACCAACTCCACCACCCG, ggggaccactttgtacaagaaagctgggtcCGCCGAACTTGCAACAAGTA | Mycgr3\|98036  JGI  Ensembl |  |  | 100 % |  |  |
| 27 | ggggacaagtttgtacaaaaaagcaggcttgatgCCGGAAAACGCCACTCTTCT, ggggaccactttgtacaagaaagctgggtcACATTCCTTGTAGATATCGC | Mycgr3\|98958  JGI  Ensembl | KOG0471 Alpha-amylase; | carbohydrate metabolic process; catalytic activity;  cation binding; | 100 % |  |  |
| 27 | ggggacaagtttgtacaaaaaagcaggcttgatgGCAGCACTGGAATCTGCCTC, ggggaccactttgtacaagaaagctgggtcCTCGATCTCCCACACGATCA | Mycgr3\|100157  JGI  Ensembl |  |  | 100 % |  |  |
| 27 | ggggacaagtttgtacaaaaaagcaggcttgatgGCGGCTTCTCAGTCGCAAGC, ggggaccactttgtacaagaaagctgggtcGCTTGGCCGCTTCACTTTGG | Mycgr3\|101417  JGI  Ensembl | KOG2627 Nuclear protein ES2; |  | 100 % |  |  |
| 27 | ggggacaagtttgtacaaaaaagcaggcttgatgCTCTCGCGCTCCGCTATCTC, ggggaccactttgtacaagaaagctgggtcCACCTTCAATCCCATTTCCT | Mycgr3\|101464  JGI  Ensembl | KOG2583 Ubiquinol cytochrome c reductase, subunit QCR2; | metalloendopeptidase activity; proteolysis; ubiquinol-cytochrome-c reductase activity;  zinc ion binding; | 100 % |  |  |
| 27 | ggggacaagtttgtacaaaaaagcaggcttgatgCGCTTCCACCAAGCGCTCCT, ggggaccactttgtacaagaaagctgggtcTCGTCGGGACATTGAGCGTC | Mycgr3\|102749  JGI  Ensembl | KOG4157 beta-1,6-N-acetylglucosaminyltransferase, contains WSC domain; |  | 100 % |  |  |
| 27 | ggggacaagtttgtacaaaaaagcaggcttgatgTCGCCATCCACCCGGCCGTC, ggggaccactttgtacaagaaagctgggtcATCTCGGGTCTTCCTCCTCT | Mycgr3\|103692  JGI  Ensembl | KOG1187 Serine/threonine protein kinase; | UDP-N-acetylmuramoylalanyl-D-glutamyl-2,6-diaminopimelate-D-alanyl-D-alanine ligase activity; coenzyme F420-0 gamma-glutamyl ligase activity;  coenzyme F420-2 alpha-glutamyl ligase activity;  ribosomal S6-glutamic acid ligase activity; | 100 % |  |  |
| 27 | ggggacaagtttgtacaaaaaagcaggcttgatgAAGTTCTCCACCATCCTGCC, ggggaccactttgtacaagaaagctgggtcAAGATCCATCTTCGCCACGA | Mycgr3\|104009  JGI  Ensembl | KOG1437 Fasciclin and related adhesion glycoproteins; |  | 0 % |  |  |
| 27 | ggggacaagtttgtacaaaaaagcaggcttgatgTACACTCTCTTCATTTTCTT, ggggaccactttgtacaagaaagctgggtcGCAGCCGCCGTTGCACCACG | Mycgr3\|107454  JGI  Ensembl |  |  | 0 % |  |  |
| 27 | ggggacaagtttgtacaaaaaagcaggcttgatgGCGGGATACTACCCCGACCC, ggggaccactttgtacaagaaagctgggtcAGACAACCTCAAAATCCCCA | Mycgr3\|107606  JGI  Ensembl | KOG4415 Uncharacterized conserved protein; |  | 100 % |  |  |
| 27 | ggggacaagtttgtacaaaaaagcaggcttgatgAGTGGGTTTCAGTTCCCTCC, ggggaccactttgtacaagaaagctgggtcCGCCTGATGTTTCTCTGCAA | Mycgr3\|107726  JGI  Ensembl | KOG1040 Polyadenylation factor I complex, subunit, Yth1 (CPSF subunit); | nucleic acid binding; zinc ion binding; | 100 % |  |  |
| 27 | ggggacaagtttgtacaaaaaagcaggcttgatgACCACAACGCGCCTGAACAC, ggggaccactttgtacaagaaagctgggtcCCCCGTCTGTTCCAACAGTG | Mycgr3\|107743  JGI  Ensembl |  |  | 100 % |  |  |
| 27 | ggggacaagtttgtacaaaaaagcaggcttgatgAAACAATGTATCCTCAAAGG, ggggaccactttgtacaagaaagctgggtcAGTGTGCGCCAGGTCGGTGC | Mycgr3\|107761  JGI  Ensembl | KOG4306 Glycosylphosphatidylinositol-specific phospholipase C; | intracellular signaling cascade; phospholipase C activity;  signal transduction; | 0 % |  |  |
| 27 | ggggacaagtttgtacaaaaaagcaggcttgatgGCAGGCGGGCAAGAGCAGCA, ggggaccactttgtacaagaaagctgggtcCAAGACATGTTGGATAGGCG | Mycgr3\|107958  JGI  Ensembl | KOG3069 Peroxisomal NUDIX hydrolase; | hydrolase activity; | 100 % |  |  |
| 27 | ggggacaagtttgtacaaaaaagcaggcttgatgTTTCGCTCCGGTGCTTCGCG, ggggaccactttgtacaagaaagctgggtcCTCCTTGTCCTCCTTCTCAT | Mycgr3\|108517  JGI  Ensembl | KOG3803 Transcription factor containing C2HC type Zn finger; |  | 100 % |  |  |
| 27 | ggggacaagtttgtacaaaaaagcaggcttgatgACTGATTACTCGAAGCTCAA, ggggaccactttgtacaagaaagctgggtcCCGTCTCCAGCTATCTCCTC | Mycgr3\|108780  JGI  Ensembl | KOG2992 Nucleolar GTPase/ATPase p130; |  | 100 % | chr\_3:2682945 C->T; |  |
| 27 | ggggacaagtttgtacaaaaaagcaggcttgatgGCCGACGATTTCGATTACGA, ggggaccactttgtacaagaaagctgggtcTTTTGCTCTCTTGGTGGCCC | Mycgr3\|108892  JGI  Ensembl | KOG2806 Chitinase; |  | 100 % |  |  |
| 27 | ggggacaagtttgtacaaaaaagcaggcttgatgACCACAACCACGCGGAAGCC, ggggaccactttgtacaagaaagctgggtcATCGTGCGGCAGATCGCGTG | Mycgr3\|109550  JGI  Ensembl |  |  | 0 % |  |  |
| 27 | ggggacaagtttgtacaaaaaagcaggcttgatgTCAAACCTTCCGTCACCCGC, ggggaccactttgtacaagaaagctgggtcCAGCTTTGAAAAGAGCGTCT | Mycgr3\|109741  JGI  Ensembl |  |  | 100 % |  |  |
| 27 | ggggacaagtttgtacaaaaaagcaggcttgatgGCGGACGAGCTACCGACGAT, ggggaccactttgtacaagaaagctgggtcGGTCATCAGATCCTCATCGT | Mycgr3\|109924  JGI  Ensembl | KOG2582 COP9 signalosome, subunit CSN3; |  | 1.69033 % |  |  |
| 27 | ggggacaagtttgtacaaaaaagcaggcttgatgCATCTCCTACGTCTAGGAGC, ggggaccactttgtacaagaaagctgggtcCTCAAAGCTAAAATACTTCG | Mycgr3\|110145  JGI  Ensembl | KOG1231 Proteins containing the FAD binding domain; | FAD binding; oxidoreductase activity; | 100 % |  |  |
| 27 | ggggacaagtttgtacaaaaaagcaggcttgatgTTTTCATTCCAGAGCCGCCT, ggggaccactttgtacaagaaagctgggtcTGGAGACGGGTTATACGTGC | Mycgr3\|110417  JGI  Ensembl | KOG1516 Carboxylesterase and related proteins; | carboxylesterase activity; | 100 % |  |  |
| 27 | ggggacaagtttgtacaaaaaagcaggcttgatgCTTCCTCCTGGATTGGAGAC, ggggaccactttgtacaagaaagctgggtcGAATGCAATGCCAGTGTCCC | Mycgr3\|110457  JGI  Ensembl |  |  | 100 % |  |  |
| 27 | ggggacaagtttgtacaaaaaagcaggcttgatgCCTGACTTTGCCTCGCACGA, ggggaccactttgtacaagaaagctgggtcGTCAGTGACCAGAGCTTCCG | Mycgr3\|111001  JGI  Ensembl |  |  | 0 % |  |  |
| 27 | ggggacaagtttgtacaaaaaagcaggcttgatgGGTGCAGACGCGACGGACGC, ggggaccactttgtacaagaaagctgggtcCTTCCACGACAAGAAGAACT | Mycgr3\|111045  JGI  Ensembl |  |  | 0 % |  |  |
| 27 | ggggacaagtttgtacaaaaaagcaggcttgatgGCGATTCGGCTCCAGGAGCC, ggggaccactttgtacaagaaagctgggtcATCGAGATGGACACCTGGTT | Mycgr3\|111745  JGI  Ensembl |  |  | 100 % |  |  |
| 28 | ggggacaagtttgtacaaaaaagcaggcttgatgGCGTACCTCTCCTTTCTCCG, ggggaccactttgtacaagaaagctgggtcATAGTTACCATCGCCGATGC | Mycgr3\|31615  JGI  Ensembl |  |  | 0 % |  |  |
| 28 | ggggacaagtttgtacaaaaaagcaggcttgatgTTCAAGACTGTCGAGAAAAA, ggggaccactttgtacaagaaagctgggtcGGGCAGCTGTAAAGCCTGCT | Mycgr3\|32867  JGI  Ensembl | KOG1212 Amidases; | amidase activity; carbon-nitrogen ligase activity, with glutamine as amido-N-donor; | 0 % |  |  |
| 28 | ggggacaagtttgtacaaaaaagcaggcttgatgCATCCCGTCATCATCGTATA, ggggaccactttgtacaagaaagctgggtcATTTTGTCTCCGTCTTAGCT | Mycgr3\|34026  JGI  Ensembl | KOG0158 Cytochrome P450 CYP3/CYP5/CYP6/CYP9 subfamilies; | 2-hydroxyisobutyrate 3-monooxygenase activity; 4-chlorophenoxyacetate monooxygenase activity;  4-nitrocatechol 4-monooxygenase activity;  alpha-pinene dehydrogenase activity;  alpha-pinene monooxygenase activity;  ammonia monooxygenase activity;  di-n-butyltin dioxygenase activity;  dimethylsilanediol hydroxylase activity;  electron transport;  fluorene oxygenase activity;  heme binding;  hydroxymethylsilanetriol oxidase activity;  iron ion binding;  limonene 8-monooxygenase activity;  lipid binding;  methyl tertiary butyl ether 3-monooxygenase activity;  methylsilanetriol hydroxylase activity;  mono-butyltin dioxygenase activity;  monooxygenase activity;  spheroidene monooxygenase activity;  tert-butyl alcohol 2-monooxygenase activity;  transport;  transporter activity;  tri-n-butyltin dioxygenase activity; | 1.11962 % |  |  |
| 28 | ggggacaagtttgtacaaaaaagcaggcttgatgGCCGGCTCACTGTCACTGCT, ggggaccactttgtacaagaaagctgggtcAGCTGATGCAAGACCACGCT | Mycgr3\|34547  JGI  Ensembl | KOG1383 Glutamate decarboxylase/sphingosine phosphate lyase; | carboxy-lyase activity; carboxylic acid metabolic process;  pyridoxal phosphate binding;  sphinganine-1-phosphate aldolase activity; | 0 % |  |  |
| 28 | ggggacaagtttgtacaaaaaagcaggcttgatgCCTATTATCCATCACATCAA, ggggaccactttgtacaagaaagctgggtcCAGCATCCTCTTCCTGAAAA | Mycgr3\|37024  JGI  Ensembl | KOG0157 Cytochrome P450 CYP4/CYP19/CYP26 subfamilies; | electron transport; heme binding;  iron ion binding;  monooxygenase activity; | 0 % |  |  |
| 28 | ggggacaagtttgtacaaaaaagcaggcttgatgGCTCCCTCTCCACCCACCAT, ggggaccactttgtacaagaaagctgggtcAATCGCCGGCCCCTCCCATC | Mycgr3\|38267  JGI  Ensembl |  |  | 0 % |  |  |
| 28 | ggggacaagtttgtacaaaaaagcaggcttgatgAAGTTTCAATACGCGACATG, ggggaccactttgtacaagaaagctgggtcCTTTGACTCAACGACATCCG | Mycgr3\|41422  JGI  Ensembl | KOG1211 Amidases; | carbon-nitrogen ligase activity, with glutamine as amido-N-donor; cobyrinate a,c-diamide synthase activity;  glutaminyl-tRNA synthase (glutamine-hydrolyzing) activity; | 1.24777 % |  |  |
| 28 | ggggacaagtttgtacaaaaaagcaggcttgatgGTTGTCTCACACAGCTTCGT, ggggaccactttgtacaagaaagctgggtcGTTGTAGCAGTACTGCGAGA | Mycgr3\|41969  JGI  Ensembl | KOG1325 Lysophospholipase; | lysophospholipase activity; phospholipase activity;  phospholipid catabolic process; | 0 % |  |  |
| 28 | ggggacaagtttgtacaaaaaagcaggcttgatgGGAGCACCTCATGCGAAGTA, ggggaccactttgtacaagaaagctgggtcCTTCTGATTTTGCCGTTCAT | Mycgr3\|42895  JGI  Ensembl | KOG2533 Permease of the major facilitator superfamily; | integral to membrane; transport; transporter activity; | 0 % |  |  |
| 28 | ggggacaagtttgtacaaaaaagcaggcttgatgTCTACTACGACTTATCCCAT, ggggaccactttgtacaagaaagctgggtcTTGCTCTCCGTCTCTTAGAT | Mycgr3\|43399  JGI  Ensembl | KOG0255 Synaptic vesicle transporter SVOP and related transporters (major facilitator superfamily); | integral to membrane; transport; transporter activity; | 0 % |  |  |
| 28 | ggggacaagtttgtacaaaaaagcaggcttgatgCCTGTCGTTCATTTGCTCGA, ggggaccactttgtacaagaaagctgggtcAACATCCTCTTCAAATCTCC | Mycgr3\|43769  JGI  Ensembl | KOG0623 Glutamine amidotransferase/cyclase; | 7-cyano-7-deazaguanine tRNA-ribosyltransferase activity; NAD(P)-asparagine ADP-ribosyltransferase activity;  NAD(P)-cysteine ADP-ribosyltransferase activity;  NAD(P)-serine ADP-ribosyltransferase activity;  catalytic activity;  cytoplasm;  histidine biosynthetic process;  imidazoleglycerol-phosphate synthase activity;  imidazoleglycerol-phosphate synthase complex;  transferase activity, transferring pentosyl groups; | 0 % |  |  |
| 28 | ggggacaagtttgtacaaaaaagcaggcttgatgAACTCTGAACGCTCATCGCA, ggggaccactttgtacaagaaagctgggtcCGCAGACTCCTTCTCTTGCG | Mycgr3\|44922  JGI  Ensembl | KOG0255 Synaptic vesicle transporter SVOP and related transporters (major facilitator superfamily); | integral to membrane; methylation; methyltransferase activity;  nucleic acid binding;  transport;  transporter activity; | 0 % |  |  |
| 28 | ggggacaagtttgtacaaaaaagcaggcttgatgCATCATTTGAACATTTTCTA, ggggaccactttgtacaagaaagctgggtcTACTGGATGCTTCGCTGCCA | Mycgr3\|47168  JGI  Ensembl | KOG2431 1, 2-alpha-mannosidase; | calcium ion binding; mannosyl-oligosaccharide 1,2-alpha-mannosidase activity;  membrane; | 0 % |  |  |
| 28 | ggggacaagtttgtacaaaaaagcaggcttgatgGCGTCACAGACGGGTGATGG, ggggaccactttgtacaagaaagctgggtcTATCGGCGCGTGGACCACTT | Mycgr3\|47254  JGI  Ensembl | KOG1287 Amino acid transporters; | amino acid transmembrane transporter activity; amino acid transport;  membrane;  transport; | 1.11633 % |  |  |
| 28 | ggggacaagtttgtacaaaaaagcaggcttgatgTTAAACACCAAATCTCCATC, ggggaccactttgtacaagaaagctgggtcACAGCTCTTTAACACTTTCC | Mycgr3\|48576  JGI  Ensembl | KOG1286 Amino acid transporters; | amino acid transmembrane transporter activity; amino acid transport;  integral to membrane;  membrane;  transport; | 0 % |  |  |
| 28 | ggggacaagtttgtacaaaaaagcaggcttgatgGACCAGTCATCGAGCGTCCC, ggggaccactttgtacaagaaagctgggtcCCGCTCCCTCCTCCGAAACT | Mycgr3\|48936  JGI  Ensembl | KOG0157 Cytochrome P450 CYP4/CYP19/CYP26 subfamilies; | electron transport; heme binding;  iron ion binding;  monooxygenase activity; | 0 % |  |  |
| 28 | ggggacaagtttgtacaaaaaagcaggcttgatgGCCGGAGCTGAGTTCATCTC, ggggaccactttgtacaagaaagctgggtcCATGGCCATCAAAGCATATG | Mycgr3\|48965  JGI  Ensembl | KOG1360 5-aminolevulinate synthase; | 8-amino-7-oxononanoate synthase activity; biosynthetic process;  electron transport;  heme binding;  peroxidase activity;  pyridoxal phosphate binding;  response to oxidative stress;  transferase activity, transferring nitrogenous groups; | 1.3658 % |  |  |
| 28 | ggggacaagtttgtacaaaaaagcaggcttgatgTACACTCACGCAGTATCAAT, ggggaccactttgtacaagaaagctgggtcAGCCATAACCCTATAAATTG | Mycgr3\|49571  JGI  Ensembl |  |  | 0 % |  |  |
| 28 | ggggacaagtttgtacaaaaaagcaggcttgatgCTCAAATCCCTCCCGCCTTC, ggggaccactttgtacaagaaagctgggtcCTTTCCACCAAGGCGAGATG | Mycgr3\|50896  JGI  Ensembl |  |  | 0 % |  |  |
| 28 | ggggacaagtttgtacaaaaaagcaggcttgatgTTCTCACGAATCAAAAATGT, ggggaccactttgtacaagaaagctgggtcGTCTGGAAAAGGCCAGACCT | Mycgr3\|51708  JGI  Ensembl |  |  | 0 % |  |  |
| 28 | ggggacaagtttgtacaaaaaagcaggcttgatgGCCACCGCTATTCTATCCCG, ggggaccactttgtacaagaaagctgggtcGCTCTGCCCTGGAGCAACAC | Mycgr3\|56782  JGI  Ensembl | KOG0370 Multifunctional pyrimidine synthesis protein CAD (includes carbamoyl-phophate synthetase, aspartate transcarbamylase, and glutamine amidotransferase); | biosynthetic process; carbamoyl-phosphate synthase activity;  catalytic activity;  glutamine metabolic process;  nitrogen compound metabolic process; | 0 % |  |  |
| 28 | ggggacaagtttgtacaaaaaagcaggcttgatgAAGAATACATTCAGCTTGCT, ggggaccactttgtacaagaaagctgggtcGAAGGTTCCAGTCTTCATAC | Mycgr3\|59905  JGI  Ensembl |  |  | 0 % |  |  |
| 28 | ggggacaagtttgtacaaaaaagcaggcttgatgAGCGGCACTCCCTACGCTCC, ggggaccactttgtacaagaaagctgggtcAATCTCCCTCGCCTTCTTCC | Mycgr3\|65914  JGI  Ensembl | KOG0666 Cyclin C-dependent kinase CDK8; | (S)-lactate 2-kinase activity; ATP binding;  ATP-dependent polynucleotide kinase activity;  GTP-dependent polynucleotide kinase activity;  LPPG:FO 2-phospho-L-lactate transferase activity;  cobinamide kinase activity;  cyclin-dependent protein kinase activating kinase regulator activity;  cyclin-dependent protein kinase regulator activity;  cytidine kinase activity;  eukaryotic elongation factor-2 kinase activator activity;  eukaryotic elongation factor-2 kinase regulator activity;  glycerate 2-kinase activity;  inositol tetrakisphosphate 2-kinase activity;  phenol kinase activity;  phosphatidylinositol phosphate kinase activity;  phosphofructokinase activity;  phosphoinositide 3-kinase activity;  phosphorylase kinase regulator activity;  polynucleotide kinase activity;  protein amino acid phosphorylation;  protein kinase activity;  protein serine/threonine kinase activity;  protein-tyrosine kinase activity; | 0 % |  |  |
| 28 | ggggacaagtttgtacaaaaaagcaggcttgatgCACCTCTCGTTTGCGCTGCT, ggggaccactttgtacaagaaagctgggtcGTCGGAGAACACGAACGTTG | Mycgr3\|66023  JGI  Ensembl | KOG2133 Transcriptional corepressor Atrophin-1/DRPLA; |  | 0 % |  |  |
| 28 | ggggacaagtttgtacaaaaaagcaggcttgatgCCGGGCATAATTTCAGATGA, ggggaccactttgtacaagaaagctgggtcCATGCTTTCCATAGCGTTTT | Mycgr3\|66209  JGI  Ensembl |  |  | 0 % |  |  |
| 28 | ggggacaagtttgtacaaaaaagcaggcttgatgTCTGCTCCGAATGGACCTTC, ggggaccactttgtacaagaaagctgggtcTGCTGAGGCAGCGGCCTCCA | Mycgr3\|66524  JGI  Ensembl | KOG1618 Predicted phosphatase; | hydrolase activity; metabolic process; | 0 % |  |  |
| 28 | ggggacaagtttgtacaaaaaagcaggcttgatgGAGCTGGTCACCTCGGCCGG, ggggaccactttgtacaagaaagctgggtcCAGTCCAGGCAATACCATCC | Mycgr3\|67350  JGI  Ensembl |  |  | 0 % |  |  |
| 28 | ggggacaagtttgtacaaaaaagcaggcttgatgATGAGCTTGTCTAGGACGGC, ggggaccactttgtacaagaaagctgggtcAAGCTTGATGTCACCACCGA | Mycgr3\|67764  JGI  Ensembl | KOG2901 Uncharacterized conserved protein; |  | 0 % |  |  |
| 28 | ggggacaagtttgtacaaaaaagcaggcttgatgACGACATACCGAGATGATGC, ggggaccactttgtacaagaaagctgggtcAAATCCTCCCTTCTCGAATT | Mycgr3\|69467  JGI  Ensembl | KOG0365 Beta subunit of farnesyltransferase; | (S)-2,3-di-O-geranylgeranylglyceryl phosphate synthase activity; 1,4-dihydroxy-2-naphthoate octaprenyltransferase activity;  4-hydroxybenzoate octaprenyltransferase activity;  Sep-tRNA:Cys-tRNA synthase activity;  catalytic activity;  prenyltransferase activity;  protein geranylgeranyltransferase activity;  protein prenyltransferase activity;  protoheme IX farnesyltransferase activity; | 0 % |  |  |
| 28 | ggggacaagtttgtacaaaaaagcaggcttgatgAAGCATACCGGTGTTCCATT, ggggaccactttgtacaagaaagctgggtcCTGCCGACGAGATCTAGGGT | Mycgr3\|69535  JGI  Ensembl | KOG0698 Serine/threonine protein phosphatase; | CTD phosphatase activity; calcium-dependent protein serine/threonine phosphatase activity;  calcium-dependent protein serine/threonine phosphatase regulator activity;  catalytic activity;  magnesium-dependent protein serine/threonine phosphatase activity;  magnesium-dependent protein serine/threonine phosphatase complex;  myosin phosphatase activity;  myosin phosphatase complex;  myosin phosphatase regulator activity;  phosphoprotein phosphatase activity;  protein amino acid dephosphorylation;  protein serine/threonine phosphatase activity;  protein serine/threonine phosphatase complex; | 0 % |  |  |
| 28 | ggggacaagtttgtacaaaaaagcaggcttgatgGATGTTCCAACGACTAAGTC, ggggaccactttgtacaagaaagctgggtcCAAAGAACTCCTCCCATCCA | Mycgr3\|70015  JGI  Ensembl | KOG0631 Galactokinase; | ATP binding; carbohydrate phosphorylation;  cytoplasm;  galactokinase activity;  galactose metabolic process;  kinase activity;  metabolic process;  phosphorylation;  phosphotransferase activity, alcohol group as acceptor; | 0 % |  |  |
| 28 | ggggacaagtttgtacaaaaaagcaggcttgatgAGCTCCGATTCCATCGAGGA, ggggaccactttgtacaagaaagctgggtcTGCGGCTTTGGCTTTCTTAT | Mycgr3\|70219  JGI  Ensembl | KOG2696 Histone acetyltransferase type b catalytic subunit; | histone acetyltransferase activity; | 0 % |  |  |
| 28 | ggggacaagtttgtacaaaaaagcaggcttgatgGCCGACCTCACCAAGCTTTC, ggggaccactttgtacaagaaagctgggtcACCCTCCCTCGGCGTCAATT | Mycgr3\|70822  JGI  Ensembl | KOG0157 Cytochrome P450 CYP4/CYP19/CYP26 subfamilies; | electron transport; heme binding;  iron ion binding;  monooxygenase activity;  oxidoreductase activity, acting on paired donors, with incorporation or reduction of molecular oxygen, reduced flavin or flavoprotein as one donor, and incorporation of one atom of oxygen; unspecific monooxygenase activity; | 0 % |  |  |
| 28 | ggggacaagtttgtacaaaaaagcaggcttgatgGACGACCATGGCGAGACCTC, ggggaccactttgtacaagaaagctgggtcCCGCTCTACAAACCCACTTA | Mycgr3\|72515  JGI  Ensembl | KOG1154 Gamma-glutamyl kinase; | RNA binding; amino acid biosynthetic process;  cytoplasm;  glutamate 5-kinase activity;  proline biosynthetic process; | 0 % |  |  |
| 28 | ggggacaagtttgtacaaaaaagcaggcttgatgGCTCTCTCCAAGTCCAGCCG, ggggaccactttgtacaagaaagctgggtcATGCCGATGCCCTTCATGCC | Mycgr3\|72682  JGI  Ensembl | KOG1483 Zn2+ transporter ZNT1 and related Cd2+/Zn2+ transporters (cation diffusion facilitator superfamily); | cation transmembrane transporter activity; cation transport;  membrane; | 0 % |  |  |
| 28 | ggggacaagtttgtacaaaaaagcaggcttgatgGCCGTGCCTCGCTCTCGGGA, ggggaccactttgtacaagaaagctgggtcGTGAGCGCCCACGGCGTCAG | Mycgr3\|73189  JGI  Ensembl | KOG4232 Delta 6-fatty acid desaturase/delta-8 sphingolipid desaturase; | heme binding; linoleoyl-CoA desaturase activity;  lipid metabolic process;  membrane;  oxidoreductase activity;  oxidoreductase activity, acting on paired donors, with oxidation of a pair of donors resulting in the reduction of molecular oxygen to two molecules of water; transition metal ion binding; | 0 % |  |  |
| 28 | ggggacaagtttgtacaaaaaagcaggcttgatgTCCTCCCAAAAGTCCGCCAT, ggggaccactttgtacaagaaagctgggtcGTCTCGTCCACCACCTACGA | Mycgr3\|74498  JGI  Ensembl |  |  | 0 % |  |  |
| 28 | ggggacaagtttgtacaaaaaagcaggcttgatgTTCGCCCGCAAAGCATCCTA, ggggaccactttgtacaagaaagctgggtcCTCGACACGCTTCTGCACCT | Mycgr3\|76417  JGI  Ensembl | KOG2929 Transcription factor, component of CCR4 transcriptional complex; | aminomethyltransferase activity; cytoplasm; glycine catabolic process; | 0 % |  |  |
| 28 | ggggacaagtttgtacaaaaaagcaggcttgatgAACTCGTCAACCCTTTCCAA, ggggaccactttgtacaagaaagctgggtcCCGCTGCCACTTCTTCGACT | Mycgr3\|76712  JGI  Ensembl | KOG1358 Serine palmitoyltransferase; | serine C-palmitoyltransferase activity; serine C-palmitoyltransferase complex; | 0 % |  |  |
| 28 | ggggacaagtttgtacaaaaaagcaggcttgatgGTGCAAGTCAAGCAAAAGGC, ggggaccactttgtacaagaaagctgggtcTTTCTTGACCAACAAACTCG | Mycgr3\|76815  JGI  Ensembl | KOG4513 Phosphoglycerate mutase; | 2,3-bisphosphoglycerate-dependent phosphoglycerate mutase activity; 2,3-bisphosphoglycerate-independent phosphoglycerate mutase activity;  catalytic activity;  cytoplasm;  glucose catabolic process;  manganese ion binding;  metal ion binding;  phosphoglycerate mutase activity; | 0 % |  |  |
| 28 | ggggacaagtttgtacaaaaaagcaggcttgatgCCAGCGATGCTCGACGATCC, ggggaccactttgtacaagaaagctgggtcCCTCCCCATGTGACCATTCT | Mycgr3\|76870  JGI  Ensembl |  |  | 0 % |  |  |
| 28 | ggggacaagtttgtacaaaaaagcaggcttgatgAAGTTCGGGCACGAATACTC, ggggaccactttgtacaagaaagctgggtcCATGACAGAGCAAGCCGGTC | Mycgr3\|77057  JGI  Ensembl | KOG4159 Predicted E3 ubiquitin ligase; | protein binding; zinc ion binding; | 0 % |  |  |
| 28 | ggggacaagtttgtacaaaaaagcaggcttgatgGTGCGCCACAAGAAAGACTT, ggggaccactttgtacaagaaagctgggtcCGGCGGCCTGTTCGGCGCAC | Mycgr3\|77085  JGI  Ensembl | KOG3154 Uncharacterized conserved protein; |  | 0 % |  |  |
| 28 | ggggacaagtttgtacaaaaaagcaggcttgatgCGGCACTCAAATGAAGACGA, ggggaccactttgtacaagaaagctgggtcTCGCTGTCCATCTTCGCTTT | Mycgr3\|77545  JGI  Ensembl | KOG0254 Predicted transporter (major facilitator superfamily); | L-arabinose isomerase activity; carbohydrate transport;  integral to membrane;  membrane;  sugar:hydrogen symporter activity;  transport;  transporter activity; | 0 % |  |  |
| 28 | ggggacaagtttgtacaaaaaagcaggcttgatgGCTAATGGTGGATTCCAACT, ggggaccactttgtacaagaaagctgggtcATCCCTCATCCTCTCATACA | Mycgr3\|77923  JGI  Ensembl | KOG1871 Ubiquitin-specific protease; | ubiquitin thiolesterase activity; ubiquitin-dependent protein catabolic process; | 0 % |  |  |
| 28 | ggggacaagtttgtacaaaaaagcaggcttgatgTCAACCACCACAACTACCAC, ggggaccactttgtacaagaaagctgggtcAACTGGTCCCAAAACCTCCT | Mycgr3\|77935  JGI  Ensembl | KOG1404 Alanine-glyoxylate aminotransferase AGT2; | adenosylmethionine-8-amino-7-oxononanoate transaminase activity; pyridoxal phosphate binding;  transaminase activity; | 0 % |  |  |
| 28 | ggggacaagtttgtacaaaaaagcaggcttgatgGCCGACAAGCAAGTTCCATT, ggggaccactttgtacaagaaagctgggtcCGAGCTGTCCTCCTTCGAAT | Mycgr3\|84245  JGI  Ensembl | KOG0252 Inorganic phosphate transporter; | integral to membrane; transport; transporter activity; | 0 % |  |  |
| 28 | ggggacaagtttgtacaaaaaagcaggcttgatgGGTGCTCATACCGCTCACAT, ggggaccactttgtacaagaaagctgggtcGGTGGTGAACATGTACCGCG | Mycgr3\|85019  JGI  Ensembl | KOG2533 Permease of the major facilitator superfamily; | integral to membrane; transport; transporter activity; | 0 % |  |  |
| 28 | ggggacaagtttgtacaaaaaagcaggcttgatgATCATACTAGCACTCGCGGC, ggggaccactttgtacaagaaagctgggtcCAAAACAGTCGCTTCTGTCC | Mycgr3\|87715  JGI  Ensembl | KOG0159 Cytochrome P450 CYP11/CYP12/CYP24/CYP27 subfamilies; | electron transport; heme binding;  iron ion binding;  monooxygenase activity; | 0 % |  |  |
| 28 | ggggacaagtttgtacaaaaaagcaggcttgatgGATATCTCGAAGACATCGAA, ggggaccactttgtacaagaaagctgggtcCAATGTATACCTGAATCCCA | Mycgr3\|88209  JGI  Ensembl | KOG2533 Permease of the major facilitator superfamily; | integral to membrane; transport; transporter activity; | 1.09763 % |  |  |
| 28 | ggggacaagtttgtacaaaaaagcaggcttgatgTCCTCATCAATGACCTCCCA, ggggaccactttgtacaagaaagctgggtcCCGTCTTCCCCTCCGATCCT | Mycgr3\|88700  JGI  Ensembl |  |  | 0 % |  |  |
| 28 | ggggacaagtttgtacaaaaaagcaggcttgatgACTGAAAATTCTAGCACTAC, ggggaccactttgtacaagaaagctgggtcAATCCTCCACCGCCAATTTG | Mycgr3\|89225  JGI  Ensembl |  |  | 0 % |  |  |
| 28 | ggggacaagtttgtacaaaaaagcaggcttgatgGCGGCGGCAGTGCAGGCTCA, ggggaccactttgtacaagaaagctgggtcCTCAAAGCAGCCTTTGCTTG | Mycgr3\|89384  JGI  Ensembl | KOG1869 Splicing coactivator SRm160/300, subunit SRm300; |  | 0 % |  |  |
| 28 | ggggacaagtttgtacaaaaaagcaggcttgatgACTGCTGTTGAAGGGTATCA, ggggaccactttgtacaagaaagctgggtcCAGGACGCAAGGACTTATGA | Mycgr3\|90338  JGI  Ensembl | KOG2614 Kynurenine 3-monooxygenase and related flavoprotein monooxygenases; | FAD binding; binding; electron transport;  oxidoreductase activity;  salicylate 1-monooxygenase activity; | 1.2761 % |  |  |
| 28 | ggggacaagtttgtacaaaaaagcaggcttgatgGACGACATAGTGTATTATAA, ggggaccactttgtacaagaaagctgggtcTGGCTTGTTAGCCTTGTGCT | Mycgr3\|90509  JGI  Ensembl | KOG1878 Nuclear receptor coregulator SMRT/SMRTER, contains Myb-like domains; |  | 0 % |  |  |
| 28 | ggggacaagtttgtacaaaaaagcaggcttgatgAAGCTGTCCGCACAACCGCC, ggggaccactttgtacaagaaagctgggtcCGCAAATCGCCCTCCCTCCG | Mycgr3\|91549  JGI  Ensembl |  |  | 0 % |  |  |
| 28 | ggggacaagtttgtacaaaaaagcaggcttgatgGATGATTCTTCGTCATACCC, ggggaccactttgtacaagaaagctgggtcCTCTTCCATCGCATCGCCAG | Mycgr3\|91791  JGI  Ensembl |  |  | 0 % |  |  |
| 28 | ggggacaagtttgtacaaaaaagcaggcttgatgTCGACTTTGCCCGGAAGGCA, ggggaccactttgtacaagaaagctgggtcTTCTGCTGAGGACTCGTCAA | Mycgr3\|94700  JGI  Ensembl |  |  | 0 % |  |  |
| 28 | ggggacaagtttgtacaaaaaagcaggcttgatgAAGAAGATCCCTGGGCAGAG, ggggaccactttgtacaagaaagctgggtcATTCGAAGCGCTTCCGCCGT | Mycgr3\|94725  JGI  Ensembl |  |  | 0 % |  |  |
| 28 | ggggacaagtttgtacaaaaaagcaggcttgatgCCGACTACATGCCCCAACAT, ggggaccactttgtacaagaaagctgggtcACACAACCCCAAACCAGCCT | Mycgr3\|95354  JGI  Ensembl | KOG1176 Acyl-CoA synthetase; | 4-coumarate-CoA ligase activity; catalytic activity;  metabolic process; | 0 % |  |  |
| 28 | ggggacaagtttgtacaaaaaagcaggcttgatgTTCTTCGCAAAAGCATTGCG, ggggaccactttgtacaagaaagctgggtcCCCAATCCATGGCCACGGCC | Mycgr3\|95555  JGI  Ensembl | KOG0672 Halotolerance protein HAL3 (contains flavoprotein domain); |  | 0 % |  |  |
| 28 | ggggacaagtttgtacaaaaaagcaggcttgatgAACACACTCGATGGTCTTCT, ggggaccactttgtacaagaaagctgggtcCAGGCCTTCCGAACGCTGGG | Mycgr3\|95590  JGI  Ensembl |  |  | 1.13095 % |  |  |
| 28 | ggggacaagtttgtacaaaaaagcaggcttgatgGAAGAGCGACAACGCAGCAG, ggggaccactttgtacaagaaagctgggtcGAAGTTGCGGCGCGAGCGTT | Mycgr3\|95982  JGI  Ensembl |  |  | 0 % |  |  |
| 28 | ggggacaagtttgtacaaaaaagcaggcttgatgCCGACGACCGTGGATTTCGA, ggggaccactttgtacaagaaagctgggtcCGAGGGCATCGCCAGCGTCG | Mycgr3\|96202  JGI  Ensembl |  |  | 0.69646 % |  |  |
| 28 | ggggacaagtttgtacaaaaaagcaggcttgatgGCTCCCACTACTCCCAAAGT, ggggaccactttgtacaagaaagctgggtcATAAGAATACTCATCCGGCG | Mycgr3\|96363  JGI  Ensembl |  |  | 0 % |  |  |
| 28 | ggggacaagtttgtacaaaaaagcaggcttgatgGCGACGATTCTGCAAAGCCT, ggggaccactttgtacaagaaagctgggtcGGATGTGCCTGCTCGGTCAC | Mycgr3\|96417  JGI  Ensembl |  |  | 0 % |  |  |
| 28 | ggggacaagtttgtacaaaaaagcaggcttgatgAAGTCTTCGCTTCTCTTCAT, ggggaccactttgtacaagaaagctgggtcCACTTGAAGCTGGGAGTAAG | Mycgr3\|96814  JGI  Ensembl |  |  | 0 % |  |  |
| 28 | ggggacaagtttgtacaaaaaagcaggcttgatgGAGCCGATGGCCGTGGGAGC, ggggaccactttgtacaagaaagctgggtcCGAAAATCGCAAAATCGCGA | Mycgr3\|97542  JGI  Ensembl |  |  | 1.11633 % |  |  |
| 28 | ggggacaagtttgtacaaaaaagcaggcttgatgCCGAAGCAAATGCCGACGGA, ggggaccactttgtacaagaaagctgggtcACATTCTCGCGGATGCGAGT | Mycgr3\|97989  JGI  Ensembl |  |  | 0 % |  |  |
| 28 | ggggacaagtttgtacaaaaaagcaggcttgatgGACTGGTCCTTTAACGGCTG, ggggaccactttgtacaagaaagctgggtcATCCCACCCATCCCAATCCC | Mycgr3\|98048  JGI  Ensembl |  |  | 0 % |  |  |
| 28 | ggggacaagtttgtacaaaaaagcaggcttgatgACCACCGAAGTCGCACTCTT, ggggaccactttgtacaagaaagctgggtcTTGAAAGTCCTGGTTTATGC | Mycgr3\|98129  JGI  Ensembl | KOG2395 Protein involved in vacuole import and degradation; |  | 0 % |  |  |
| 28 | ggggacaagtttgtacaaaaaagcaggcttgatgGCTATATCAACACTGATGGA, ggggaccactttgtacaagaaagctgggtcAAGTTGGACATCTATTGACG | Mycgr3\|98639  JGI  Ensembl | KOG0274 Cdc4 and related F-box and WD-40 proteins; |  | 0 % |  |  |
| 28 | ggggacaagtttgtacaaaaaagcaggcttgatgGGACAACATTCCGTCACTGG, ggggaccactttgtacaagaaagctgggtcGACTTTCTTACTATCCGCCA | Mycgr3\|99023  JGI  Ensembl | KOG1289 Amino acid transporters; | amino acid transmembrane transporter activity; amino acid transport;  integral to membrane;  membrane;  transport; | 0 % |  |  |
| 28 | ggggacaagtttgtacaaaaaagcaggcttgatgGCTCCTTCCTCGACAATACT, ggggaccactttgtacaagaaagctgggtcTTTGTGTCCGTTGCTGTCAT | Mycgr3\|99090  JGI  Ensembl | KOG1404 Alanine-glyoxylate aminotransferase AGT2; | pyridoxal phosphate binding; transaminase activity; | 0 % |  |  |
| 28 | ggggacaagtttgtacaaaaaagcaggcttgatgGCTTCCAAATACGCCCTCCC, ggggaccactttgtacaagaaagctgggtcCTGGAAGCTCTGCTGCAAGA | Mycgr3\|99257  JGI  Ensembl |  |  | 0 % |  |  |
| 28 | ggggacaagtttgtacaaaaaagcaggcttgatgGAGCGACCCTCGGGCGTATC, ggggaccactttgtacaagaaagctgggtcTTGCGGCTCGTCCTGAGTGA | Mycgr3\|100510  JGI  Ensembl | KOG0752 Mitochondrial solute carrier protein; | binding; membrane; mitochondrial inner membrane;  transport; | 0 % |  |  |
| 28 | ggggacaagtttgtacaaaaaagcaggcttgatgGCCACTGGACCAGTCAGCGA, ggggaccactttgtacaagaaagctgggtcCATCCACAAACATAGCTTCC | Mycgr3\|100589  JGI  Ensembl | KOG4328 WD40 protein; |  | 0 % |  |  |
| 28 | ggggacaagtttgtacaaaaaagcaggcttgatgGCGCCCAAAAACATCACAAT, ggggaccactttgtacaagaaagctgggtcCGTCCTGTAGTCACCATTGA | Mycgr3\|100901  JGI  Ensembl | KOG2786 Putative glutamate/ornithine acetyltransferase; | arginine biosynthetic process; glutamate N-acetyltransferase activity; | 1.29955 % |  |  |
| 28 | ggggacaagtttgtacaaaaaagcaggcttgatgCCAGAACAGTGGGAAGAAAT, ggggaccactttgtacaagaaagctgggtcAGCCTTAAGATCCTCAATGA | Mycgr3\|100958  JGI  Ensembl | KOG1212 Amidases; | amidase activity; carbon-nitrogen ligase activity, with glutamine as amido-N-donor; | 0 % |  |  |
| 28 | ggggacaagtttgtacaaaaaagcaggcttgatgGCATTCGGCAGCAGTAAGAA, ggggaccactttgtacaagaaagctgggtcGATAGCACCATACTCCGCTG | Mycgr3\|101642  JGI  Ensembl | KOG2210 Oxysterol-binding protein; | steroid metabolic process; | 1.4245 % |  |  |
| 28 | ggggacaagtttgtacaaaaaagcaggcttgatgTCTGTCAACACACCGACTGC, ggggaccactttgtacaagaaagctgggtcGTACTGCATCCCACCACTGC | Mycgr3\|101662  JGI  Ensembl | KOG1596 Fibrillarin and related nucleolar RNA-binding proteins; |  | 0 % |  |  |
| 28 | ggggacaagtttgtacaaaaaagcaggcttgatgGGTGACCTACAAGGCCGCAA, ggggaccactttgtacaagaaagctgggtcTCGCATGGCGTCCAAGCCGT | Mycgr3\|102121  JGI  Ensembl | KOG0660 Mitogen-activated protein kinase; | ATP binding; MAP kinase activity;  protein amino acid phosphorylation;  protein kinase activity;  protein serine/threonine kinase activity;  protein-tyrosine kinase activity; | 0 % |  |  |
| 28 | ggggacaagtttgtacaaaaaagcaggcttgatgGCCACCACTTTAACACCGGA, ggggaccactttgtacaagaaagctgggtcCAAGGCACTCGCTTTCCGCT | Mycgr3\|103523  JGI  Ensembl |  |  | 0 % |  |  |
| 28 | ggggacaagtttgtacaaaaaagcaggcttgatgGGTTTGAAGGGCGTCCACTT, ggggaccactttgtacaagaaagctgggtcCTTAGACTCGCCCTGCACCT | Mycgr3\|104277  JGI  Ensembl |  |  | 0 % |  |  |
| 28 | ggggacaagtttgtacaaaaaagcaggcttgatgCTCGCCCTAGCCATCACCGC, ggggaccactttgtacaagaaagctgggtcCGCCCTCCGAATCTCAAACT | Mycgr3\|105645  JGI  Ensembl | KOG1296 Uncharacterized conserved protein; |  | 0 % |  |  |
| 28 | ggggacaagtttgtacaaaaaagcaggcttgatgGCCGACAAAACAGCTCCTCG, ggggaccactttgtacaagaaagctgggtcTTCGCTGTCCGAGCTATCCT | Mycgr3\|106967  JGI  Ensembl | KOG2992 Nucleolar GTPase/ATPase p130; | calcium ion binding; exo-alpha-sialidase activity;  homophilic cell adhesion;  membrane; | 0 % |  |  |
| 28 | ggggacaagtttgtacaaaaaagcaggcttgatgGCTTACCAACATCAACAACC, ggggaccactttgtacaagaaagctgggtcTTGGACGAACCGTGGCTGAG | Mycgr3\|107198  JGI  Ensembl | KOG0653 Cyclin B and related kinase-activating proteins; | nucleus; | 0 % |  |  |
| 28 | ggggacaagtttgtacaaaaaagcaggcttgatgGCCGGCCGATCCGCACATGT, ggggaccactttgtacaagaaagctgggtcGAAGAAAGACTGACCATCCC | Mycgr3\|108163  JGI  Ensembl | KOG1124 FOG: TPR repeat; | nucleus; protein dimerization activity;  regulation of transcription, DNA-dependent;  sequence-specific DNA binding;  transcription factor activity; | 0 % |  |  |
| 28 | ggggacaagtttgtacaaaaaagcaggcttgatgGCTACCGAAACGCCCGTATT, ggggaccactttgtacaagaaagctgggtcGTAAATGCCTGTAGGACGGA | Mycgr3\|108816  JGI  Ensembl | KOG3947 Phosphoesterases; | hydrolase activity; | 1.24408 % |  |  |
| 28 | ggggacaagtttgtacaaaaaagcaggcttgatgGCGTCCTCCTCGCGCAATCC, ggggaccactttgtacaagaaagctgggtcCTCCACGCCGGCGTAAAGAT | Mycgr3\|108833  JGI  Ensembl | KOG2861 Uncharacterized conserved protein; |  | 0 % |  |  |
| 28 | ggggacaagtttgtacaaaaaagcaggcttgatgTCCACCTCAATCTTCCCACC, ggggaccactttgtacaagaaagctgggtcAAGTCTCGGCCTCTCCTGAT | Mycgr3\|109694  JGI  Ensembl | KOG2820 FAD-dependent oxidoreductase; | sarcosine oxidase activity; | 0 % |  |  |
| 28 | ggggacaagtttgtacaaaaaagcaggcttgatgGGAGGCAGCACCTTCACCAA, ggggaccactttgtacaagaaagctgggtcTCCCTCCTTCAATTCCTTCC | Mycgr3\|110490  JGI  Ensembl |  |  | 0 % |  |  |
| 28 | ggggacaagtttgtacaaaaaagcaggcttgatgGATCTACTCGTGATCAAGGC, ggggaccactttgtacaagaaagctgggtcCCGATGCAGCACATCATCTA | Mycgr3\|110804  JGI  Ensembl | KOG1187 Serine/threonine protein kinase; |  | 0 % |  |  |
| 29 | ggggacaagtttgtacaaaaaagcaggcttgatgGTTTCAGTGACTCCAGCCAT, ggggaccactttgtacaagaaagctgggtcAGAAGCTCCCCGAATTCGTG | Mycgr3\|31366  JGI  Ensembl | KOG0255 Synaptic vesicle transporter SVOP and related transporters (major facilitator superfamily); | integral to membrane; transport; transporter activity; | 100 % |  |  |
| 29 | ggggacaagtttgtacaaaaaagcaggcttgatgCGTGGAATCTCTGGCTTCGG, ggggaccactttgtacaagaaagctgggtcTGCAAGAACAAACAGTTCGA | Mycgr3\|33386  JGI  Ensembl | KOG3753 Circadian clock protein period; | mannan endo-1,6-alpha-mannosidase activity; | 100 % |  |  |
| 29 | ggggacaagtttgtacaaaaaagcaggcttgatgCCACGACCTGAGCTTGAGCA, ggggaccactttgtacaagaaagctgggtcGAGCCTCGACCGAACCCAAT | Mycgr3\|35091  JGI  Ensembl | KOG1516 Carboxylesterase and related proteins; | cholinesterase activity; | 0 % |  |  |
| 29 | ggggacaagtttgtacaaaaaagcaggcttgatgGCGTCTACCTCGGAACAAAC, ggggaccactttgtacaagaaagctgggtcCACTTCTCCCGTCTTCCTGC | Mycgr3\|39884  JGI  Ensembl |  |  | 0 % |  |  |
| 29 | ggggacaagtttgtacaaaaaagcaggcttgatgTCTGAAGAAAGCGCTCCGGT, ggggaccactttgtacaagaaagctgggtcACTCCCGCCGCCCTCCTTCC | Mycgr3\|40213  JGI  Ensembl | KOG2648 Diphthamide biosynthesis protein; |  | 100 % |  |  |
| 29 | ggggacaagtttgtacaaaaaagcaggcttgatgGAAAACAATCCAGCAGTAAC, ggggaccactttgtacaagaaagctgggtcCTTGTTGCCTTCAGCCTGAG | Mycgr3\|40431  JGI  Ensembl |  |  | 100 % |  | ``` 572	TGGATAGCATCACTGCGGCACAACTTCTACCCATCGCCGCCACGATCGTGGCATCAGCTGTTGGCTCGAATATCGCAAATGTGATGCCCAATCCACAGCTTGCTCTGGGCGCCATCATCGCTAGCTATGTGATGTGGGGGATGGCCGTTCCGATGGCGTTCTCAGTACTGGTCATCTACTACCAGCGCCTCGCATTGCACAAGATGCCACCGAGAGAGGTCATCGTGAGCGCCTTTCTGCCTCTGGGGCCCCTTGGCTACGGTGGCTATACAGTCTTGAACCTTGGCCGTGCCGCCGTCAAAGTCTTTCCGATGACGGGCTCTATCGATCCGATCGCTGGCAGAATCGCATATGTGATGGGTTTCTTCGTATCACTCGTGATGTGGGGATACGGACTGATCTGGTTTGCATTGGCTCTTGCAAGCATCTGGAAGGCGAGGCCATTCCCTTTCAATATGGGTTGGTGGGGAT	1042 	\|\|\|\|\|\|\|\|\|\|\|\|\|\|\|\|\|\|\|\|\|\|\|\|\|\|\|\|\|\|\|\|\|\|\|\|\|\|\|\|\|\|\|\|\|\|\|\|\|\|\|\|\|\|\|\|\|\|\|\|\|\|\|\|\|\|\|\|\|\|\|\|\|\|\|\|\|\|\|\|\|\|\|\|\|\|\|\|\|\|\|\|\|\|\|\|\|\|\|\|\|\|\|\|\|\|\|\|\|\|\|\|\|\|\|\|\|\|\|\|\|\|\|\|\|\|\|\|\|\|\|\|\|\|\|\|\|\|\|\|\|\|\|\|\|\|\|\|\|\|\|\|\|\|\|\|\|\|\|\|\|\|\|\|\|\|\|\|\|\|\|\|\|\|\|\|\|\|\|\|\|\|\|\|\|\|\|\|\|\|\|\|\|\|\|\|\|\|\|\|\|\|\|\|\|\|\|\|\|\|\|\|\|\|\|\|\|\|\|\|\|\|\|\|\|\|\|\|\|\|\|\|\|\|\|\|\|\|\|\|\|\|\|\|\|\|\|\|\|\|\|\|\|\|\|\|\|\|\|\|\|\|\|\|\|\|\|\|\|\|\|\|\|\|\|\|\|\|\|\|\|\|\|\|\|\|\|\|\|\|\|\|\|\|\|\|\|\|\|\|\|\|\|\|\|\|\|\|\|\|\|\|\|\|\|\|\|\|\|\|\|\|\|\|\|\|\|\|\|\|\|\|\|\|\|\|\|\|\|\|\|\|\|\|\|\|\|\|\|\|\|\|\|\|\|\|\|\|\|\|\|\|\|\|\|\|\|\|\|\|\|\|\|\|\|\|\|\|\|\|\|\|\|\|\|\|\|\|\|\|\|\|\|\|\|\|\|\|\|\|\|\|\|\|\|\|\|\|\|\|\|\|\|\|\|\|\|\|\|\|\|\|\|\|\|\|\|\|\|\|\|\|\|\|\|\|\|\|\|\|\|\|\|\|\|\|\|\|\|\|\|\|\|\|\|\|\|\|\|\|\|\|\|\|\|\|\|\|\|\|\| 	TGGATAGCATCACTGCGGCACAACTTCTACCCATCGCCGCCACGATCGTGGCATCAGCTGTTGGCTCGAATATCGCAAATGTGATGCCCAATCCACAGCTTGCTCTGGGCGCCATCATCGCTAGCTATGTGATGTGGGGGATGGCCGTTCCGATGGCGTTCTCAGTACTGGTCATCTACTACCAGCGCCTCGCATTGCACAAGATGCCACCGAGAGAGGTCATCGTGAGCGCCTTTCTGCCTCTGGGGCCCCTTGGCTACGGTGGCTATACAGTCTTGAACCTTGGCCGTGCCGCCGTCAAAGTCTTTCCGATGACGGGCTCTATCGATCCGATCGCTGGCAGAATCGCATATGTGATGGGTTTCTTCGTATCACTCGTGATGTGGGGATACGGACTGATCTGGTTTGCATTGGCTCTTGCAAGCATCTGGAAGGCGAGGCCATTCCCTTTCAATATGGGTTGGTGGGGAT ```      ``` 1043	TCACGTTCCCGCTCGGCGTATACTCCCTCAGCACAATCCTGATTGGGAAGGAACTGCCCTCGCTATTCTTCCGGGTTCTGGGCACTATATTTGCGACATGCGTCATTCTCCTCTGGATTCTCATCGCTGTACGGACCGGCAAGGGTGCATGGACTGGGAAAATGTTTCATGCGCCATGTCTTGCAAATTTGAAGAAACAACAAGAGCAGATGGAAGAAGAGAAGTCCATCGGGAGTACGAGTCAGAATACTCAGGCTGAAGGCAACAAG	1311 	\|\|\|\|\|\|\|\|\|\|\|\|\|\|\|\|\|\|\|\|\|\|\|\|\|\|\|\|\|\|\|\|\|\|\|\|\|\|\|\|\|\|\|\|\|\|\|\|\|\|\|\|\|\|\|\|\|\|\|\|\|\|\|\|\|\|\|\|\|\|\|\|\|\|\|\|\|\|\|\|\|\|\|\|\|\|\|\|\|\|\|\|\|\|\|\|\|\|\|\|\|\|\|\|\|\|\|\|\|\|\|\|\|\|\|\|\|\|\|\|\|\|\|\|\|\|\|\|\|\|\|\|\|\|\|\|\|\|\|\|\|\|\|\|\|\|\|\|\|\|\|\|\|\|\|\|\|\|\|\|\|\|\|\|\|\|\|\|\|\|\|\|\|\|\|\|\|\|\|\|\|\|\|\|\|\|\|\|\|\|\|\|\|\|\|\|\|\|\|\|\|\|\|\|\|\|\|\|\|\|\|\|\|\|\|\|\|\|\|\|\|\|\|\|\|\|\|\|\|\|\|\|\|\|\|\|\|\|\|\|\|\|\|\|\|\|\|\|\|\|\|\|\|\|\|\|\|\|\|\|\|\|\|\|\|\|\|\|\| 	TCACGTTCCCGCTCGGCGTATACTCCCTCAGCACAATCCTGATTGGGAAGGAACTGCCCTCGCTATTCTTCCGGGTTCTGGGCACTATATTTGCGACATGCGTCATTCTCCTCTGGATTCTCATCGCTGTACGGACCGGCAAGGGTGCATGGACTGGGAAAATGTTTCATGCGCCATGTCTTGCAAATTTGAAGAAACAACAAGAGCAGATGGAAGAAGAGAAGTCCATCGGGAGTACGAGTCAGAATACTCAGGCTGAAGGCAACAAG ``` |
| 29 | ggggacaagtttgtacaaaaaagcaggcttgatgTCGGAAGGAGGCCCCTCAAG, ggggaccactttgtacaagaaagctgggtcATATCTGATCGCTCGCTTGC | Mycgr3\|40740  JGI  Ensembl | KOG0255 Synaptic vesicle transporter SVOP and related transporters (major facilitator superfamily); | integral to membrane; transport; transporter activity; | 0 % |  |  |
| 29 | ggggacaagtttgtacaaaaaagcaggcttgatgTCGTACAATCCGTTCGGAAG, ggggaccactttgtacaagaaagctgggtcGGAACTCTCCTTCAACCTCC | Mycgr3\|40912  JGI  Ensembl | KOG1558 Fe2+/Zn2+ regulated transporter; | intracellular; metallopeptidase activity;  proteolysis;  zinc ion binding; | 100 % |  |  |
| 29 | ggggacaagtttgtacaaaaaagcaggcttgatgTGGAAGCTCGTCGACTTGAC, ggggaccactttgtacaagaaagctgggtcTACTCCGAATGATTCCTCCC | Mycgr3\|41234  JGI  Ensembl | KOG0039 Ferric reductase, NADH/NADPH oxidase and related proteins; | ferric-chelate reductase activity; | 100 % |  |  |
| 29 | ggggacaagtttgtacaaaaaagcaggcttgatgGCTACCGAGAAGCCGTTCTC, ggggaccactttgtacaagaaagctgggtcCTGGTTGTTTCCCGTTGCAA | Mycgr3\|41987  JGI  Ensembl | KOG4686 Predicted sugar transporter; | integral to membrane; transport; transporter activity; | 100 % |  |  |
| 29 | ggggacaagtttgtacaaaaaagcaggcttgatgCTGACTTGGTTTGATTACCA, ggggaccactttgtacaagaaagctgggtcGTACCACAGCGGGTTCTCCG | Mycgr3\|42328  JGI  Ensembl |  |  | 100 % |  |  |
| 29 | ggggacaagtttgtacaaaaaagcaggcttgatgGAGCAGCATCGTGTTTCATC, ggggaccactttgtacaagaaagctgggtcCTCAAGGACTGCCCGTCTGC | Mycgr3\|44362  JGI  Ensembl | KOG0159 Cytochrome P450 CYP11/CYP12/CYP24/CYP27 subfamilies; | electron transport; heme binding;  iron ion binding;  monooxygenase activity;  unspecific monooxygenase activity; | 100 % |  |  |
| 29 | ggggacaagtttgtacaaaaaagcaggcttgatgCAAGCTATACGCCAGTATAG, ggggaccactttgtacaagaaagctgggtcTCGCCAGCTTAAAGCCTTGC | Mycgr3\|45188  JGI  Ensembl | KOG0255 Synaptic vesicle transporter SVOP and related transporters (major facilitator superfamily); | integral to membrane; transport; transporter activity; | 0 % |  |  |
| 29 | ggggacaagtttgtacaaaaaagcaggcttgatgGCCTCCCAGATCCCCGAATG, ggggaccactttgtacaagaaagctgggtcTTGTGCCAGCTTCAGAGCTT | Mycgr3\|45386  JGI  Ensembl | KOG1212 Amidases; | amidase activity; carbon-nitrogen ligase activity, with glutamine as amido-N-donor; | 100 % |  |  |
| 29 | ggggacaagtttgtacaaaaaagcaggcttgatgGGCCCAGAACAACCTCTTCC, ggggaccactttgtacaagaaagctgggtcTTTCTTGATTCCAACAATGA | Mycgr3\|46277  JGI  Ensembl | KOG2651 rRNA adenine N-6-methyltransferase; |  | 0 % |  |  |
| 29 | ggggacaagtttgtacaaaaaagcaggcttgatgGGCATCCAAGAAGAGCCCTC, ggggaccactttgtacaagaaagctgggtcTAAAGCATATCTGAACCCCT | Mycgr3\|47026  JGI  Ensembl | KOG2533 Permease of the major facilitator superfamily; | integral to membrane; transport; transporter activity; | 0 % |  |  |
| 29 | ggggacaagtttgtacaaaaaagcaggcttgatgTGGGAGAAACTCGGTCGTCG, ggggaccactttgtacaagaaagctgggtcTCCGCAAACCACGCCCTCCT | Mycgr3\|48763  JGI  Ensembl | KOG2335 tRNA-dihydrouridine synthase; | F420H2 dehydrogenase activity; FAD binding;  malolactic enzyme activity;  regulation of oxidoreductase activity;  sulfur oxygenase reductase activity;  tRNA dihydrouridine synthase activity;  tRNA processing; | 100 % |  |  |
| 29 | ggggacaagtttgtacaaaaaagcaggcttgatgTTTGTCGGATCTTTCCTGGA, ggggaccactttgtacaagaaagctgggtcTTTAAGCTTTTCGATGACAG | Mycgr3\|50149  JGI  Ensembl | KOG2466 Uridine permease/thiamine transporter/allantoin transport; | membrane; nucleobase transmembrane transporter activity;  nucleobase, nucleoside, nucleotide and nucleic acid transport; | 1.12676 % |  |  |
| 29 | ggggacaagtttgtacaaaaaagcaggcttgatgGAGAAACACCCGCTTGATGC, ggggaccactttgtacaagaaagctgggtcGCTGTTCTCCTCGACGAACA | Mycgr3\|50352  JGI  Ensembl | KOG0254 Predicted transporter (major facilitator superfamily); | L-arabinose isomerase activity; carbohydrate transport;  catalytic activity;  integral to membrane;  membrane;  metal ion binding;  sugar:hydrogen symporter activity;  transport;  transporter activity; | 0 % |  |  |
| 29 | ggggacaagtttgtacaaaaaagcaggcttgatgATCGCAGCCAAGATACGGAC, ggggaccactttgtacaagaaagctgggtcCACTTGCGATTGCGCACGGT | Mycgr3\|51700  JGI  Ensembl |  |  | 0 % |  |  |
| 29 | ggggacaagtttgtacaaaaaagcaggcttgatgGCCGGCTCCTTCAATCCCCG, ggggaccactttgtacaagaaagctgggtcCCGCAACTCCCCCCTCCCAC | Mycgr3\|64892  JGI  Ensembl |  |  | 100 % |  |  |
| 29 | ggggacaagtttgtacaaaaaagcaggcttgatgGCCAACAAGCAGGACCCCCC, ggggaccactttgtacaagaaagctgggtcAAACAAATCCGTCATAGCTC | Mycgr3\|64976  JGI  Ensembl | KOG2448 Dihydroxy-acid dehydratase; | branched chain family amino acid biosynthetic process; catalytic activity;  dihydroxy-acid dehydratase activity;  metabolic process; | 0 % |  |  |
| 29 | ggggacaagtttgtacaaaaaagcaggcttgatgACACCTGCCATACCACCAGA, ggggaccactttgtacaagaaagctgggtcACGCAAGCCAGCGACATTGT | Mycgr3\|66489  JGI  Ensembl | KOG0255 Synaptic vesicle transporter SVOP and related transporters (major facilitator superfamily); | integral to membrane; transport; transporter activity; | 100 % |  |  |
| 29 | ggggacaagtttgtacaaaaaagcaggcttgatgTTTCGCTTCGTACAGCCATC, ggggaccactttgtacaagaaagctgggtcCTTCTCGTCCTCGGCCTTAC | Mycgr3\|66594  JGI  Ensembl |  |  | 100 % | chr\_1:3723227 T->A; |  |
| 29 | ggggacaagtttgtacaaaaaagcaggcttgatgGCCTCTTCCGCGCACTCCTA, ggggaccactttgtacaagaaagctgggtcAGTCGAACCAGAATTGTCAG | Mycgr3\|66885  JGI  Ensembl | KOG1304 Amino acid transporters; |  | 100 % |  |  |
| 29 | ggggacaagtttgtacaaaaaagcaggcttgatgTCGCTCAAGCAAGAAATCGA, ggggaccactttgtacaagaaagctgggtcCACCTCCTGCACCCAGACCT | Mycgr3\|67898  JGI  Ensembl | KOG4225 Sorbin and SH3 domain-containing protein; | nucleic acid binding; | 100 % |  |  |
| 29 | ggggacaagtttgtacaaaaaagcaggcttgatgGCCGGTGTAGAAGATGTCAT, ggggaccactttgtacaagaaagctgggtcCATTTCACGTTGGCTTTCAC | Mycgr3\|68030  JGI  Ensembl | KOG3021 Predicted kinase; | (S)-lactate 2-kinase activity; ATP-dependent polynucleotide kinase activity;  GTP-dependent polynucleotide kinase activity;  LPPG:FO 2-phospho-L-lactate transferase activity;  cobinamide kinase activity;  cyclin-dependent protein kinase activating kinase regulator activity;  cyclin-dependent protein kinase regulator activity;  cytidine kinase activity;  eukaryotic elongation factor-2 kinase activator activity;  eukaryotic elongation factor-2 kinase regulator activity;  glycerate 2-kinase activity;  inositol tetrakisphosphate 2-kinase activity;  kinase activity;  phenol kinase activity;  phosphatidylinositol phosphate kinase activity;  phosphofructokinase activity;  phosphoinositide 3-kinase activity;  phosphorylase kinase regulator activity;  polynucleotide kinase activity; | 100 % |  |  |
| 29 | ggggacaagtttgtacaaaaaagcaggcttgatgACCCCGTCACGGAGGACAGC, ggggaccactttgtacaagaaagctgggtcGCCGCTTCCATTCGATGAGA | Mycgr3\|68253  JGI  Ensembl |  |  | 100 % |  |  |
| 29 | ggggacaagtttgtacaaaaaagcaggcttgatgACGCACGACAATGTTCTCGA, ggggaccactttgtacaagaaagctgggtcGGACGCTGGGCTCTTCAGAG | Mycgr3\|68751  JGI  Ensembl |  |  | 100 % |  |  |
| 29 | ggggacaagtttgtacaaaaaagcaggcttgatgCCCTCCACCAGCCGAAAGAA, ggggaccactttgtacaagaaagctgggtcCACTTTGTCACCCACGATCT | Mycgr3\|68899  JGI  Ensembl | KOG0254 Predicted transporter (major facilitator superfamily); | L-arabinose isomerase activity; carbohydrate transport;  integral to membrane;  membrane;  sugar:hydrogen symporter activity;  transport;  transporter activity; | 100 % |  |  |
| 29 | ggggacaagtttgtacaaaaaagcaggcttgatgAAAAGTGCATTCGCGGCTCT, ggggaccactttgtacaagaaagctgggtcCAACTCATCCTTCCCCCCAC | Mycgr3\|69270  JGI  Ensembl | KOG2397 Protein kinase C substrate, 80 KD protein, heavy chain; | alpha-glucosidase activity; endoplasmic reticulum; | 100 % |  | ``` 1034	TTTACGCCTTCGAAAACTACCTCCCTCTCACCCTCCGCACCTGGCTCGACACCAAACTCCGCGACCTCCGCCTCCTACTCATCGAAAACGGCATCCTCGCCTCCAACCCCACCTCGGGCGAATCCACCACCCTCACCTCCGCCCGCACCCGCCTCACCACCGCGGAAAAATCCCTCGAAGACCTCCGTACTACCCTCACCTCCTCCCAATCCGACCTCACCACCTCCTTCGGCCCCGACGACGTCTTCCGCGCCCTCAAAGGCACTTGCATCAATACCGACAGCGGGGAGTACACCTACACCCTCTGCTTCCTCGACCGCACCGCGCAGAAACCGAAAAAGGGCGGTGGGGAGACGAATATGGGGAAATATACGGGTTTGGAAATGGTCACGGTGGATGAGGACTTGCCCGTTGATGGGAAAGGGTTGGGATCAGGGGAGAGGTGGGCGATGAAGTTTGAGAATGGACAGCATTGCTGGAATGGACCGAATCGGAGGACGACGGTTGTTTTGGGGTGTGCGGAGAAGGATGAGATTTGGAGGATTAGGGAGGAGGAGAAGTGTGTTTATCGGATGGAGGTGGGGACGCCAGCGGTTTGTGAggggagcggggggaagaaagaggggggtggggggAAGGATGAGTTG	1680 	\|\|\|\|\|\|\|\| \|\|\|\|\|\|\|\|\|\|\|\|\|\|\|\|\|\|\|\|\|\|\|\|\|\|\|\|\|\|\|\|\|\|\|\|\|\|\|\|\|\|\|\|\|\|\|\|\|\|\|\|\|\|\|\|\|\|\|\|\|\|\|\|\|\|\|\|\|\|\|\|\|\|\|\|\|\|\|\|\|\|\|\|\|\|\|\|\|\|\|\|\|\|\|\|\|\|\|\|\|\|\|\|\|\|\|\|\|\|\|\|\|\|\|\|\|\|\|\|\|\|\|\|\|\|\|\|\|\|\|\|\|\|\|\|\|\|\|\|\|\|\|\|\|\|\|\|\|\|\|\|\|\|\|\|\|\|\|\|\|\|\|\|\|\|\|\|\|\|\|\|\|\|\|\|\|\|\|\|\|\|\|\|\|\|\|\|\|\|\|\|\|\|\|\|\|\|\|\|\|\|\|\|\|\|\|\|\|\|\|\|\|\|\|\|\|\|\|\|\|\|\|\|\|\|\|\|\|\|\|\|\|\|\|\|\|\|\|\|\|\|\|\|\|\|\|\|\|\|\|\|\|\|\|\|\|\|\|\|\|\|\|\|\|\|\|\|\|\|\|\|\|\|\|\|\|\|\|\|\|\|\|\|\|\|\|\|\|\|\|\|\|\|\|\|\|\|\|\|\|\|\|\|\|\|\|\|\|\|\|\|\|\|\|\|\|\|\|\|\|\|\|\|\|\|\|\|\|\|\|\|\|\|\|\|\|\|\|\|\|\|\|\|\|\|\|\|\|\|\|\|\|\|\|\|\|\|\|\|\|\|\|\|\|\|\|\|\|\|\|\|\|\|\|\|\|\|\|\|\|\|\|\|\|\|\|\|\|\|\|\|\|\|\|\|\|\|\|\|\|\|\|\|\|\|\|\|\|\|\|\|\|\|\|\|\|\|\|\|\|\|\|\|\|\|\|\|\|\|\|\|\|\|\|\|\|\|\|\|\|\|\|\|\|\|\|\|\|\|\|\|\|\|\|\|\|\|\|\|\|\|\|\|\|\|\|\|\|\|\|\|\|\|\|\|\|\|\|\|\|\|\|\|\|\|\|\|\|\|\|\|\|\|\|\|\|\|\|\|\|\|\|\|\|\|\|\|\|\|\|\|\|\|\|\|\|\|\|\|\|\|\|\|\|\|\|\|\|\|\|\|\|\|\|\|\|\|\|\|\|\|\|\|\|\|\|\|\|\|\|\|\|\|\|\|\|\|\|\|\|\|\|\|\|\|\|\|\|\|\|\|\|\|\|\|\|\|\|\|\|\|\|\|\|\|\|\|\|\|\|\|\|\|\|\|\|\|\|\|\|\|\|\|\|\|\|\|\|\|\|\|\|\|\|\|\|\|\|\|\|\|\|\|\|\|\|\|\|\|\|\|\|\|\|\|\|\| 	TTTACGCC-TCGAAAACTACCTCCCTCTCACCCTCCGCACCTGGCTCGACACCAAACTCCGCGACCTCCGCCTCCTACTCATCGAAAACGGCATCCTCGCCTCCAACCCCACCTCGGGCGAATCCACCACCCTCACCTCCGCCCGCACCCGCCTCACCACCGCGGAAAAATCCCTCGAAGACCTCCGTACTACCCTCACCTCCTCCCAATCCGACCTCACCACCTCCTTCGGCCCCGACGACGTCTTCCGCGCCCTCAAAGGCACTTGCATCAATACCGACAGCGGGGAGTACACCTACACCCTCTGCTTCCTCGACCGCACCGCGCAGAAACCGAAAAAGGGCGGTGGGGAGACGAATATGGGGAAATATACGGGTTTGGAAATGGTCACGGTGGATGAGGACTTGCCCGTTGATGGGAAAGGGTTGGGATCAGGGGAGAGGTGGGCGATGAAGTTTGAGAATGGACAGCATTGCTGGAATGGACCGAATCGGAGGACGACGGTTGTTTTGGGGTGTGCGGAGAAGGATGAGATTTGGAGGATTAGGGAGGAGGAGAAGTGTGTTTATCGGATGGAGGTGGGGACGCCAGCGGTTTGTGAGGGGAGCGGGGGGAAGAAAGAGGGGGGTGGGGGGAAGGATGAGTTG ``` |
| 29 | ggggacaagtttgtacaaaaaagcaggcttgatgGCGCCCAAAATCCTTCGTCT, ggggaccactttgtacaagaaagctgggtcTGGCGGCGGGTTCAGGTCCA | Mycgr3\|69881  JGI  Ensembl | KOG0805 Carbon-nitrogen hydrolase; | hydrolase activity, acting on carbon-nitrogen (but not peptide) bonds; nitrilase activity;  nitrogen compound metabolic process; | 100 % |  |  |
| 29 | ggggacaagtttgtacaaaaaagcaggcttgatgAGGACGGCATTCCTCACATT, ggggaccactttgtacaagaaagctgggtcTCGTCTCCGCCGCCGCTTGA | Mycgr3\|69894  JGI  Ensembl | KOG2650 Zinc carboxypeptidase; | carboxypeptidase A activity; proteolysis; zinc ion binding; | 0 % |  |  |
| 29 | ggggacaagtttgtacaaaaaagcaggcttgatgGCCTTCAACTTCAACTGGTC, ggggaccactttgtacaagaaagctgggtcAGAGACGTATGCCGGCGGCG | Mycgr3\|70491  JGI  Ensembl | KOG2806 Chitinase; |  | 100 % |  |  |
| 29 | ggggacaagtttgtacaaaaaagcaggcttgatgTCGACCTTTGTTCTCACACG, ggggaccactttgtacaagaaagctgggtcCCTCGCCTTCTCTCCCCTCA | Mycgr3\|70812  JGI  Ensembl |  |  | 100 % |  |  |
| 29 | ggggacaagtttgtacaaaaaagcaggcttgatgAACCCTGCTATTACGAAAGA, ggggaccactttgtacaagaaagctgggtcCTTCGCCTTTCCGTTCGCCT | Mycgr3\|71278  JGI  Ensembl |  |  | 100 % | chr\_4:1816675 G->A; |  |
| 29 | ggggacaagtttgtacaaaaaagcaggcttgatgGCGTCACCCGCACCTCTCCT, ggggaccactttgtacaagaaagctgggtcATCCTTCCAATCCCACACCG | Mycgr3\|71299  JGI  Ensembl | KOG0581 Mitogen-activated protein kinase kinase (MAP2K); | (S)-lactate 2-kinase activity; ATP binding;  ATP-dependent polynucleotide kinase activity;  GTP-dependent polynucleotide kinase activity;  LPPG:FO 2-phospho-L-lactate transferase activity;  cobinamide kinase activity;  cyclin-dependent protein kinase activating kinase regulator activity;  cyclin-dependent protein kinase regulator activity;  cytidine kinase activity;  eukaryotic elongation factor-2 kinase activator activity;  eukaryotic elongation factor-2 kinase regulator activity;  glycerate 2-kinase activity;  inositol tetrakisphosphate 2-kinase activity;  phenol kinase activity;  phosphatidylinositol phosphate kinase activity;  phosphofructokinase activity;  phosphoinositide 3-kinase activity;  phosphorylase kinase regulator activity;  polynucleotide kinase activity;  protein amino acid phosphorylation;  protein kinase activity;  protein serine/threonine kinase activity;  protein-tyrosine kinase activity; | 100 % |  |  |
| 29 | ggggacaagtttgtacaaaaaagcaggcttgatgGCTCCTGGACTTGTTGAGAA, ggggaccactttgtacaagaaagctgggtcGTTGCTCTCGAATGGAGTCT | Mycgr3\|71416  JGI  Ensembl |  |  | 100 % |  | ``` 984	AGGAAGAGTCTGATTTCCTACTGAAGTTCCTGTTCGACCATGTGGCTCTTGGTGCAGACTTCCACGCTCGCGTGAAGTGGGAGAAGGGCACTGTGGTCGTTTGGGACAACCGCGTCACCGCACACACAGCGCTCTTGGACTGGCAGGATGGACAGAGGCGCCACCTTGCTAGGCTGACGCCTCGTGCCGAAAGG	1177 	\|\|\|\|\|\|\|\|\|\|\|\|\|\|\|\|\|\|\|\|\|\|\|\|\|\|\|\|\|\|\|\|\|\|\|\|\|\|\|\|\|\|\|\|\|\|\|\|\|\|\|\|\|\|\|\|\|\|\|\|\|\|\|\|\|\|\|\|\|\|\|\|\|\|\|\|\|\|\|\|\|\|\|\|\|\|\|\|\|\|\|\|\|\|\|\|\|\|\|\|\|\|\|\|\|\|\|\|\|\|\|\|\|\|\|\|\|\|\|\|\|\|\|\|\|\|\|\|\|\|\|\|\|\|\|\|\|\|\|\|\|\|\|\|\|\|\|\|\|\|\|\|\|\|\|\|\|\|\|\|\|\|\|\|\|\|\|\|\|\|\|\|\|\|\|\|\|\|\|\|\|\|\|\|\|\|\|\|\|\|\|\|\|\| 	AGGAAGAGTCTGATTTCCTACTGAAGTTCCTGTTCGACCATGTGGCTCTTGGTGCAGACTTCCACGCTCGCGTGAAGTGGGAGAAGGGCACTGTGGTCGTTTGGGACAACCGCGTCACCGCACACACAGCGCTCTTGGACTGGCAGGATGGACAGAGGCGCCACCTTGCTAGGCTGACGCCTCGTGCCGAAAGG ```      ``` 643	CAATTCGATCACATGGCACTCCGACGTTACGTACGAGATGCAGCCTCCGTCCACAACGTTCTTGTATCGTCTCGACGGACCTGATGCGGGAG	734 	\|\|\|\|\|\|\|\|\|\|\|\|\|\|\|\|\|\|\|\|\|\|\|\|\|\|\| \|\|\|\|\|\|\|\|\|\|\|\|\|\|\|\| \|\|\|\|\|\|\|\|\|\|\|\|\|\|\|\|\|\|\|\|\|\|\|\|\|\|\|\|\|\|\|\|\|\|\|\|\|\|\|\|\|\|\|\|\|\|\| 	CAATTCGATCACATGGCACTCCGACGT-ACGTACGAGATGCAGCTTCCGTCCACAACGTTCTTGTATCGTCTCGACGGACCTGATGCGGGAG ```      ``` 1175	AGGCCATTTGAGACTCCATTCGAGAGCAAC	1204 	\|\|\|\|\|\|\|\|\|\|\|\|\|\|\|\|\|\|\|\|\|\|\|\|\|\|\|\|\|\| 	AGGCCATTTGAGACTCCATTCGAGAGCAAC ```      ``` 735	GTGACACACTGTTCGCAAA-TATGGCAGAGGCGTACAAGAGGCTGTCACCCGCCTTTCAGG-AGCGCCTTCATGGCCTCAAAGCGGTCCACTCAGGCCACGAGCAAGCCCAGCAAGCCCTCGCACGCGGTGGCATTG	869 	\|\|\|\|\|\|\|\|\|\|\|\|\|\|\|\|\|\|\| \|\|\|\|\|\|\|\|\|\|\|\|\|\|\|\|\|\|\|\|\|\|\|\|\|\|\|\|\|\|\|\|\|\|\|\|\|\|\|\|\| \|\|\|\|\|\|\|\|\|\|\|\|\|\|\|\|\|\|\|\|\|\|\|\|\|\|\|\|\|\|\|\|\|\|\|\|\|\|\|\|\|\|\|\|\|\|\|\|\|\|\|\|\|\|\|\|\|\|\|\|\|\|\|\|\|\|\|\|\|\|\|\|\|\|\| 	GTGACACACTGTTCGCAAAATATGGCAGAGGCGTACAAGAGGCTGTCACCCGCCTTTCAGGGAGCGCCTTCATGGCCTCAAAGCGGTCCACTCAGGCCACGAGCAAGCCCAGCAAGCCCTCGCACGCGGTGGCATTG ```      ``` 869	GTCCGACGTGATCCAGTCTCGAACGTACATCCTATCGTCCGAACGCACCCAGTAACTGGCGAGAAGGCGTTGTTCGTCAACACTCAGTTCACCAAAAGAATAGTAGGGTTCAAGAAGG-AAG	989 	\|\|\|\|\|\|\|\|\|\|\|\|\|\|\|\|\|\|\|\|\|\|\|\|\|\|\|\|\|\|\|\|\|\|\|\|\|\|\|\|\|\|\|\|\|\|\|\|\|\|\|\|\|\|\|\|\|\|\|\|\|\|\|\|\|\|\|\|\|\|\|\|\|\|\|\|\|\|\|\|\|\|\|\|\|\|\|\|\|\|\|\|\|\|\|\|\|\|\|\|\|\|\|\|\|\|\|\|\|\|\|\|\|\|\|\|\|\| \|\|\| 	GTCCGACGTGATCCAGTCTCGAACGTACATCCTATCGTCCGAACGCACCCAGTAACTGGCGAGAAGGCGTTGTTCGTCAACACTCAGTTCACCAAAAGAATAGTAGGGTTCAAGAAGGTAAG ``` |
| 29 | ggggacaagtttgtacaaaaaagcaggcttgatgAGCGATAGCGCCCCGCCAGA, ggggaccactttgtacaagaaagctgggtcAGGCCACCACTTCATCGTCC | Mycgr3\|72103  JGI  Ensembl | KOG1430 C-3 sterol dehydrogenase/3-beta-hydroxysteroid dehydrogenase and related dehydrogenases; |  | 100 % |  |  |
| 29 | ggggacaagtttgtacaaaaaagcaggcttgatgGCATCCGAGAAACGAGTCGG, ggggaccactttgtacaagaaagctgggtcCTTCTTCCCCAACACCTCCA | Mycgr3\|73277  JGI  Ensembl | KOG2835 Phosphoribosylamidoimidazole-succinocarboxamide synthase; | 'de novo' IMP biosynthetic process; phosphoribosylaminoimidazole carboxylase activity;  phosphoribosylaminoimidazole carboxylase complex; | 100 % | chr\_6:1700992 G->A; |  |
| 29 | ggggacaagtttgtacaaaaaagcaggcttgatgTTGAGCTCTAGAAGAGCTTT, ggggaccactttgtacaagaaagctgggtcAAAGCGAGAGATGTCTCTTC | Mycgr3\|74173  JGI  Ensembl | KOG2495 NADH-dehydrogenase (ubiquinone); | FAD binding; NADH dehydrogenase activity;  electron transport;  oxidoreductase activity; | 0 % |  |  |
| 29 | ggggacaagtttgtacaaaaaagcaggcttgatgTCGTCGATACACCGCTGTAT, ggggaccactttgtacaagaaagctgggtcTATGTACTTGTAAGGGTTCA | Mycgr3\|74344  JGI  Ensembl | KOG1232 Proteins containing the FAD binding domain; | FAD binding; catalytic activity;  oxidoreductase activity; | 1.09133 % |  |  |
| 29 | ggggacaagtttgtacaaaaaagcaggcttgatgCCTTCCGCCAAACTCCTTCA, ggggaccactttgtacaagaaagctgggtcTGCCTTGGACAGGTAGATGC | Mycgr3\|75326  JGI  Ensembl | KOG2450 Aldehyde dehydrogenase; | aldehyde dehydrogenase (NAD) activity; | 100 % |  |  |
| 29 | ggggacaagtttgtacaaaaaagcaggcttgatgAACTCGACCCCTGTACCAAC, ggggaccactttgtacaagaaagctgggtcCGAGACCACATACAACTCCT | Mycgr3\|75353  JGI  Ensembl |  |  | 100 % |  |  |
| 29 | ggggacaagtttgtacaaaaaagcaggcttgatgGCTCGACTTGAAGATGACGA, ggggaccactttgtacaagaaagctgggtcGACTTTCAGCTCGGTACATG | Mycgr3\|76446  JGI  Ensembl | KOG0254 Predicted transporter (major facilitator superfamily); | integral to membrane; transport; transporter activity; | 100 % |  | ``` 667	ACAG-TCGTGGCG-TTCTTCTGC-GTACCCGCCGACTTCACAGCCAAGGAGCCCTTAACTTTCGAG-ACGCTCAAGAAATTCGACTTTTTGGGAA-CAGTCC-TGACGATTGCCGGCATCGGCATGTTCT-CTGCC-GCCTTGACGATGGGATCCACGGATGGCTGGACTG-CACCCTATGTGCTATCACTCCTCCTCATCGGGCTGGCTTTGATCGTCACC-TTCATATTCTGGGAAATGTGGTTTGAGTACCCACTCATGCCGATGGGAATCTGGAAAGACCGG-AACTTCTCCCTCGTTCTGGCGATTCTGGTATTGGGATTCTCGGCATTTACCATGTCCAGCTTCTTCCTCTCTCTCTACTTCCAAGATGTCTGGCACCTGTCCGCGCTCATGGTAGCCGTCTACCTTTTACCCATGGCCACCAATGGCGTCCTGGTGAACATCTTCGCCGGCTGGGCGCTCCACCGCATCTCGAACAAGATCCTGATGTGTGCAGCGACCCTCTCCTTCGTGCTGGCGTTCCTGCTCTTCGCACTAGCCACCGATGGCCAAGAATACTGGAAATTCTACTTCCCAGGTCTGCTACTTGTCGTCGTTGGAGCAGACATGGAGTTCAACGTGACAAATATGTATGTCATGTCTTCCATGCCGCCGGAACAGCAGAGCACCGCAGGAGGCATCTTTCAGACTGTGACGAAGCTTTGTATGACCATCGGCCTCGCACTAGCCACGGCTGTGTTCAACTCGGTCCAGAAGAGGCCGGCGTTGTCTGGCTACTGGGAGGAAGAGAGGCAGCCTTACGCGGCAGTGTACTGGTCGTCCTTTGCTTTTGCGATGGTGAGTCTGCTTCTCGTACCGTTGCTCACGATTGGAACGCAGGGAGGGAAGGAGAGGTGCAAAGATGGGGTCGTGGACGATGGAGACGTGGGCGGACAGACCGCGGTCGATGTGACAGAGGTTCAGACACATCCCAAGCAGGATGTAGTCGATGGTACACGGCCATGTACCGAGCTGAAAGTC	1680 	\|\|\|\| \|\|\| \|\|\|\| \|\|\|\|\|\|\|\|\| \|\|\|\|\|\|\|\|\|\|\|\|\| \|\|\|\|\|\|\|\|\|\| \|\|\|\|\|\|\|\|\|\|\|\|\| \|\|\| \|\|\|\|\|\|\|\|\| \|\| \|\|\|\|\|\|\|\|\|\|\|\|\|\|\| \|\|\|\|\|\| \|\|\|\|\|\|\|\|\|\|\|\|\|\|\|\|\|\|\|\|\|\|\|\|\|\|\| \|\|\|\|\| \|\|\|\|\|\|\|\|\|\|\|\|\| \|\|\|\|\|\|\| \|\| \|\|\|\|\|\|\|\|\| \|\|\|\|\|\|\|\|\|\|\|\|\|\|\|\|\|\|\|\|\|\|\|\|\|\|\|\|\|\|\|\|\|\|\|\|\|\|\|\|\|\|\|\|\|\|\|\|\|\| \|\|\|\|\|\|\|\|\|\|\|\|\|\|\|\|\|\|\|\|\|\|\|\|\|\|\|\|\|\|\|\|\|\|\|\|\|\|\|\|\|\|\|\|\|\|\|\|\|\|\|\|\|\|\|\|\|\|\|\|\|\|\| \|\|\|\|\|\|\|\|\|\|\|\|\|\|\|\|\|\|\|\|\|\|\|\|\|\|\|\|\|\|\|\|\|\|\|\|\|\|\|\|\|\|\|\|\|\|\|\|\|\|\|\|\|\|\|\|\|\|\|\|\|\|\|\|\|\|\|\|\|\|\|\|\|\|\|\|\|\|\|\|\|\|\|\|\|\|\|\|\|\|\|\|\|\|\|\|\|\|\|\|\|\|\|\|\|\|\|\|\|\|\|\|\|\|\|\|\|\|\|\|\|\|\|\|\|\|\|\|\|\|\|\|\|\|\|\|\|\|\|\|\|\|\|\|\|\|\|\|\|\|\|\|\|\|\|\|\|\|\|\|\|\|\|\|\|\|\|\|\|\|\|\|\|\|\|\|\|\|\|\|\|\|\|\|\|\|\|\|\|\|\|\|\|\|\|\|\|\|\|\|\|\|\|\|\|\|\|\|\|\|\|\|\|\|\|\|\|\|\|\|\|\|\|\|\|\|\|\|\|\|\|\|\|\|\|\|\|\|\|\|\|\|\|\|\|\|\|\|\|\|\|\|\|\|\|\|\|\|\|\|\|\|\|\|\|\|\|\|\|\|\|\|\|\|\|\|\|\|\|\|\|\|\|\|\|\|\|\|\|\|\|\|\|\|\|\|\|\|\|\|\|\|\|\|\|\|\|\|\|\|\|\|\|\|\|\|\|\|\|\|\|\|\|\|\|\|\|\|\|\|\|\|\|\|\|\|\|\|\|\|\|\|\|\|\|\|\|\|\|\|\|\|\|\|\|\|\|\|\|\|\|\|\|\|\|\|\|\|\|\|\|\|\|\|\|\|\|\|\|\|\|\|\|\|\|\|\|\|\|\|\|\|\|\|\|\|\|\|\|\|\|\|\|\|\|\|\|\|\|\|\|\|\|\|\|\|\|\|\|\|\|\|\|\|\|\|\|\|\|\|\|\|\|\|\|\|\|\|\|\|\|\|\|\|\|\|\|\|\|\|\|\|\|\|\|\|\|\|\|\|\|\|\|\|\|\|\|\|\|\|\|\|\|\|\|\|\|\|\|\|\|\|\|\|\|\|\|\|\|\|\|\|\|\|\|\|\|\|\|\|\|\|\|\|\|\|\|\|\|\|\|\|\|\|\|\|\|\|\|\|\|\|\|\|\|\|\|\|\|\|\|\|\|\|\|\|\|\|\|\|\|\|\|\|\|\|\|\|\|\|\|\|\|\|\|\|\|\|\|\|\|\|\|\|\|\|\|\|\|\|\|\|\|\|\|\|\|\|\|\|\|\|\|\|\|\|\|\|\|\|\|\|\|\|\|\|\|\|\|\|\|\|\|\|\|\|\|\|\|\|\|\|\|\|\|\|\|\|\|\|\|\|\|\|\|\|\|\|\|\|\|\|\|\|\|\|\|\|\|\|\|\|\|\|\|\|\|\|\|\|\|\|\|\|\|\|\|\|\|\|\|\|\|\|\|\|\|\|\|\|\|\|\|\|\|\|\|\|\|\|\|\|\|\|\|\|\|\|\|\|\|\|\|\|\|\|\|\|\|\|\|\|\|\|\|\|\|\|\|\|\|\|\|\|\|\|\|\|\|\|\|\|\|\|\|\|\|\|\|\|\|\|\|\|\|\|\|\| 	ACAGTTCGGGGCGTTTCTTCTGCTGTACCCGCCGACT-CACAGCCAAGAAGCCCTTAACTTTTGAGGACGCTCAAGGAAGTCGACTTTTTGGGAAACAGTCCATGACGATTGCCGGCATCGGCATGTTCTTCTGCCGGCCTTGACGATGGAATCCACG-ATTGCTGGACTGGCACCCTATGTGCTATCACTCCTCCTCATCGGGCTGGCTTTGATCGTCACCCTTCATATTCTGGGAAATGTGGTTTGAGTACCCACTCATGCCGATGGGAATCTGGAAAGACCGGGAACTTCTCCCTCGTTCTGGCGATTCTGGTATTGGGATTCTCGGCATTTACCATGTCCAGCTTCTTCCTCTCTCTCTACTTCCAAGATGTCTGGCACCTGTCCGCGCTCATGGTAGCCGTCTACCTTTTACCCATGGCCACCAATGGCGTCCTGGTGAACATCTTCGCCGGCTGGGCGCTCCACCGCATCTCGAACAAGATCCTGATGTGTGCAGCGACCCTCTCCTTCGTGCTGGCGTTCCTGCTCTTCGCACTAGCCACCGATGGCCAAGAATACTGGAAATTCTACTTCCCAGGTCTGCTACTTGTCGTCGTTGGAGCAGACATGGAGTTCAACGTGACAAATATGTATGTCATGTCTTCCATGCCGCCGGAACAGCAGAGCACCGCAGGAGGCATCTTTCAGACTGTGACGAAGCTTTGTATGACCATCGGCCTCGCACTAGCCACGGCTGTGTTCAACTCGGTCCAGAAGAGGCCGGCGTTGTCTGGCTACTGGGAGGAAGAGAGGCAGCCTTACGCGGCAGTGTACTGGTCGTCCTTTGCTTTTGCGATGGTGAGTCTGCTTCTCGTACCGTTGCTCACGATTGGAACGCAGGGAGGGAAGGAGAGGTGCAAAGATGGGGTCGTGGACGATGGAGACGTGGGCGGACAGACCGCGGTCGATGTGACAGAGGTTCAGACACATCCCAAGCAGGATGTAGTCGATGGTACACGGCCATGTACCGAGCTGAAAGTC ``` |
| 29 | ggggacaagtttgtacaaaaaagcaggcttgatgACCACCTCCAACGATACCGA, ggggaccactttgtacaagaaagctgggtcTCTCGACAGATCCCCTCCCT | Mycgr3\|76723  JGI  Ensembl |  |  | 100 % |  |  |
| 29 | ggggacaagtttgtacaaaaaagcaggcttgatgGCTTCCACTGCTCTGCTACT, ggggaccactttgtacaagaaagctgggtcGGTCGCGATCGGAGTGAATA | Mycgr3\|77280  JGI  Ensembl | KOG0157 Cytochrome P450 CYP4/CYP19/CYP26 subfamilies; | electron transport; heme binding;  iron ion binding;  monooxygenase activity;  unspecific monooxygenase activity; | 100 % |  |  |
| 29 | ggggacaagtttgtacaaaaaagcaggcttgatgAGCGCCGACGACAACACCCA, ggggaccactttgtacaagaaagctgggtcCGCAGCAGCCTCGACAGTCC | Mycgr3\|77678  JGI  Ensembl | KOG0101 Molecular chaperones HSP70/HSC70, HSP70 superfamily; | ATP binding; | 100 % |  |  |
| 29 | ggggacaagtttgtacaaaaaagcaggcttgatgTCCCGCCGACCGCCAGCAGG, ggggaccactttgtacaagaaagctgggtcGCCCCAAACATTCCCAAACA | Mycgr3\|85784  JGI  Ensembl | KOG0703 Predicted GTPase-activating protein; | ARF GTPase activator activity; regulation of ARF GTPase activity;  zinc ion binding; | 1.12423 % |  |  |
| 29 | ggggacaagtttgtacaaaaaagcaggcttgatgGAGCGACCATCCATAGCAGT, ggggaccactttgtacaagaaagctgggtcAATCGGCGAAGCAATCCACT | Mycgr3\|86441  JGI  Ensembl | KOG0156 Cytochrome P450 CYP2 subfamily; | dibenzothiophene-5,5-dioxide monooxygenase activity; electron transport;  heme binding;  iron ion binding;  monooxygenase activity; | 100 % |  |  |
| 29 | ggggacaagtttgtacaaaaaagcaggcttgatgGAGCCAATCGAGACTCTCGC, ggggaccactttgtacaagaaagctgggtcCGTCCCCGCCAGTCTCAACC | Mycgr3\|86654  JGI  Ensembl | KOG1220 Phosphoglucomutase/phosphomannomutase; | carbohydrate metabolic process; intramolecular transferase activity, phosphotransferases;  phosphomannomutase activity; | 1.0882 % |  |  |
| 29 | ggggacaagtttgtacaaaaaagcaggcttgatgCCCTCCTTCGACGAAGCTCC, ggggaccactttgtacaagaaagctgggtcGTAAATCTCGTCTCTCAAAA | Mycgr3\|86672  JGI  Ensembl | KOG1050 Trehalose-6-phosphate synthase component TPS1 and related subunits; | alpha,alpha-trehalose-phosphate synthase (UDP-forming) activity; catalytic activity;  trehalose biosynthetic process; | 0 % |  |  |
| 29 | ggggacaagtttgtacaaaaaagcaggcttgatgACTAGCGTAGACGATCTATT, ggggaccactttgtacaagaaagctgggtcGTCCAAGAAGTTGATCAAGG | Mycgr3\|87579  JGI  Ensembl | KOG2734 Uncharacterized conserved protein; |  | 2.16155 % |  |  |
| 29 | ggggacaagtttgtacaaaaaagcaggcttgatgTCCGACAAGTTCTACGTGGT, ggggaccactttgtacaagaaagctgggtcTTCAGAGCGCTTAAGCGGCG | Mycgr3\|88628  JGI  Ensembl |  |  | 0 % |  |  |
| 29 | ggggacaagtttgtacaaaaaagcaggcttgatgGCCATGGTCGGCCGGCAGAA, ggggaccactttgtacaagaaagctgggtcATCGCAATACGTCAAATCTC | Mycgr3\|88911  JGI  Ensembl | KOG0260 RNA polymerase II, large subunit; | DNA-directed RNA polymerase I activity; DNA-directed RNA polymerase II activity;  DNA-directed RNA polymerase III activity;  DNA-directed RNA polymerase activity;  RNA polymerase activity; | 0 % |  |  |
| 29 | ggggacaagtttgtacaaaaaagcaggcttgatgCCTGCCCAAACAATCCAGCA, ggggaccactttgtacaagaaagctgggtcCTCAAGGTTCTTGTAGAACT | Mycgr3\|89608  JGI  Ensembl | KOG1216 von Willebrand factor and related coagulation proteins; |  | 0 % |  |  |
| 29 | ggggacaagtttgtacaaaaaagcaggcttgatgTATACCAAGAAGATCCTCCT, ggggaccactttgtacaagaaagctgggtcGGTCCGCTCGTCCATATAAA | Mycgr3\|90561  JGI  Ensembl |  |  | 100 % |  |  |
| 29 | ggggacaagtttgtacaaaaaagcaggcttgatgTCTGTCATCGAAGTCGCAGG, ggggaccactttgtacaagaaagctgggtcACGGTACTTGGAGCCCTCTT | Mycgr3\|90850  JGI  Ensembl |  |  | 100 % |  |  |
| 29 | ggggacaagtttgtacaaaaaagcaggcttgatgGCTCCGAACAAGCTCGCCGA, ggggaccactttgtacaagaaagctgggtcGATGCCTGTATCAGCAGAGC | Mycgr3\|91619  JGI  Ensembl |  |  | 0 % |  |  |
| 29 | ggggacaagtttgtacaaaaaagcaggcttgatgGCAGGTCCCAGCGCACACAG, ggggaccactttgtacaagaaagctgggtcACCAATACTCTCGAACAGGG | Mycgr3\|91741  JGI  Ensembl |  |  | 100 % |  | ``` 1522	ATATATCAAATGCGGGAGGTCACAGGGCTTCAGAGTCGCCACCAGGGATCCACTCAGGACGATGAAGTGCTGCATGGTATACCTGCTCATTCCGAAATTGAGCAGGATGCCGGCACGACGGGTAACGACACTGTGGAAGGCGATGAGGATGGAGAACCGAAGGAGGACGAAGATGATGGGTGGGAACGTGTAATCTTCGTGCCTTTCGCTCGCGAGCTCCCGCAGTTGATTCGGGAGGCGCAGGCCCTGTTCGAGAGTATTGGT	1785 	\|\|\|\|\|\|\|\|\|\|\|\|\|\| \|\|\|\|\|\|\|\|\|\|\|\|\|\|\|\|\|\|\|\|\|\|\|\|\|\|\|\|\|\|\|\|\|\|\|\|\|\|\|\|\|\|\|\|\|\|\|\|\|\|\|\|\|\|\|\|\|\|\|\|\|\|\|\|\|\|\|\|\|\|\|\|\|\|\|\|\|\|\|\|\|\|\|\|\|\|\|\|\|\|\|\|\|\|\|\|\|\|\|\|\|\|\|\|\|\|\|\|\|\|\|\|\|\|\|\|\|\|\|\|\|\|\|\|\|\|\|\|\|\|\|\|\|\|\|\|\|\|\|\|\|\|\|\|\|\|\|\|\|\|\|\|\|\|\|\|\|\|\|\|\|\|\|\|\|\|\|\|\|\|\|\|\|\|\|\|\|\|\|\|\|\|\|\|\|\|\|\|\|\|\|\|\|\|\|\|\|\|\|\|\|\|\|\|\|\|\|\|\|\|\|\|\|\|\|\|\|\|\|\|\|\|\|\|\|\|\|\|\|\|\|\|\|\|\|\|\|\|\|\|\|\|\|\|\|\|\|\|\| 	ATATATCAAATGCGTGAGGTCACAGGGCTTCAGAGTCGCCACCAGGGATCCACTCAGGACGATGAAGTGCTGCATGGTATACCTGCTCATTCCGAAATTGAGCAGGATGCCGGCACGACGGGTAACGACACTGTGGAAGGCGATGAGGATGGAGAACCGAAGGAGGACGAAGATGATGGGTGGGAACGTGTAATCTTCGTGCCTTTCGCTCGCGAGCTCCCGCAGTTGATTCGGGAGGCGCAGGCCCTGTTCGAGAGTATTGGT ``` |
| 29 | ggggacaagtttgtacaaaaaagcaggcttgatgTCTGGGGAGAAGAAGGTGGC, ggggaccactttgtacaagaaagctgggtcCCTCGCAATCTCATATCCGG | Mycgr3\|92043  JGI  Ensembl | KOG1869 Splicing coactivator SRm160/300, subunit SRm300; | FK506-sensitive peptidyl-prolyl cis-trans isomerase; cyclophilin; cyclophilin-type peptidyl-prolyl cis-trans isomerase activity;  peptidyl-prolyl cis-trans isomerase activity; | 0 % |  |  |
| 29 | ggggacaagtttgtacaaaaaagcaggcttgatgGCGACTACCACGACGACATT, ggggaccactttgtacaagaaagctgggtcCGGGGCGCAGAAAAGGCGTC | Mycgr3\|92187  JGI  Ensembl | KOG0254 Predicted transporter (major facilitator superfamily); | integral to membrane; transport; transporter activity; | 1.11421 % |  |  |
| 29 | ggggacaagtttgtacaaaaaagcaggcttgatgGCGCAACCACAGGAGCCTCG, ggggaccactttgtacaagaaagctgggtcTACTTCGCTAATCCTCACAC | Mycgr3\|92638  JGI  Ensembl | KOG1181 FOG: Low-complexity; | UDP-N-acetylmuramoylalanyl-D-glutamyl-2,6-diaminopimelate-D-alanyl-D-alanine ligase activity; coenzyme F420-0 gamma-glutamyl ligase activity;  coenzyme F420-2 alpha-glutamyl ligase activity;  ribosomal S6-glutamic acid ligase activity; | 100 % |  |  |
| 29 | ggggacaagtttgtacaaaaaagcaggcttgatgACTACAACGCGACCAGAACC, ggggaccactttgtacaagaaagctgggtcGAAGCTGGCGGAGACGATTG | Mycgr3\|93251  JGI  Ensembl |  |  | 100 % |  |  |
| 29 | ggggacaagtttgtacaaaaaagcaggcttgatgTGTGTCATCGAGGAGAAGAT, ggggaccactttgtacaagaaagctgggtcTCGATAATACTGCGTCCTCT | Mycgr3\|93284  JGI  Ensembl | KOG4297 C-type lectin; | aspartic-type endopeptidase activity; proteolysis; | 0 % |  |  |
| 29 | ggggacaagtttgtacaaaaaagcaggcttgatgAGCAACTCAAGCTCAGAAGA, ggggaccactttgtacaagaaagctgggtcGAAGAAAAACCATTGCGAAA | Mycgr3\|94577  JGI  Ensembl | KOG1286 Amino acid transporters; | amino acid transmembrane transporter activity; amino acid transport;  membrane;  transport; | 100 % |  |  |
| 29 | ggggacaagtttgtacaaaaaagcaggcttgatgTCCAACGACTCCACCACTCG, ggggaccactttgtacaagaaagctgggtcCACGAACCGTATGAACGGAG | Mycgr3\|94724  JGI  Ensembl |  |  | 100 % |  |  |
| 29 | ggggacaagtttgtacaaaaaagcaggcttgatgTTCCGTGAATTGCAACCCAA, ggggaccactttgtacaagaaagctgggtcTCCCTCATCCCCAAGTTGTA | Mycgr3\|94745  JGI  Ensembl |  |  | 1.09447 % |  |  |
| 29 | ggggacaagtttgtacaaaaaagcaggcttgatgCCCGGCCGCCATGACGGACC, ggggaccactttgtacaagaaagctgggtcTGTCCGGAACGCAGGTTTCC | Mycgr3\|94768  JGI  Ensembl | KOG0431 Auxilin-like protein and related proteins containing DnaJ domain; |  | 100 % |  |  |
| 29 | ggggacaagtttgtacaaaaaagcaggcttgatgCCAAGTCTCCTCGACCTTCC, ggggaccactttgtacaagaaagctgggtcCTCGTCCGCATTCTTTCCGT | Mycgr3\|95591  JGI  Ensembl |  |  | 0 % |  |  |
| 29 | ggggacaagtttgtacaaaaaagcaggcttgatgCTTTTCCACACGCTGCTCGC, ggggaccactttgtacaagaaagctgggtcGGGGCAGGTGGTGGTGTTGT | Mycgr3\|95719  JGI  Ensembl |  |  | 0 % |  |  |
| 29 | ggggacaagtttgtacaaaaaagcaggcttgatgAACAATCCCTCACGCAGAAC, ggggaccactttgtacaagaaagctgggtcTTTGGCATTCTTCGATCCAA | Mycgr3\|96095  JGI  Ensembl |  |  | 100 % |  |  |
| 29 | ggggacaagtttgtacaaaaaagcaggcttgatgGCCAGCATCGCCCAACCCAT, ggggaccactttgtacaagaaagctgggtcGCATGCCCGCAGACGCCGCA | Mycgr3\|96361  JGI  Ensembl |  |  | 0 % |  |  |
| 29 | ggggacaagtttgtacaaaaaagcaggcttgatgGCGGAAGTAGAAACAAACGA, ggggaccactttgtacaagaaagctgggtcTCCCTCGTCCCACGTCTGCA | Mycgr3\|96962  JGI  Ensembl |  |  | 100 % | chr\_12:374087 C->A; |  |
| 29 | ggggacaagtttgtacaaaaaagcaggcttgatgCCAGGCCGAGCTATGGCGAG, ggggaccactttgtacaagaaagctgggtcCGAAGAGCAGAGATCATCGA | Mycgr3\|97134  JGI  Ensembl |  |  | 1.0951 % |  |  |
| 29 | ggggacaagtttgtacaaaaaagcaggcttgatgGCGAACAAAGCTGTCCTTCA, ggggaccactttgtacaagaaagctgggtcATCCTCACCTCCCACCTCAC | Mycgr3\|97180  JGI  Ensembl | KOG4308 LRR-containing protein; |  | 100 % |  |  |
| 29 | ggggacaagtttgtacaaaaaagcaggcttgatgGCGCACAACGAAGACACTCC, ggggaccactttgtacaagaaagctgggtcGGCCTCTAAGTTCCTCGTCG | Mycgr3\|97369  JGI  Ensembl | KOG0113 U1 small nuclear ribonucleoprotein (RRM superfamily); |  | 1.11483 % |  |  |
| 29 | ggggacaagtttgtacaaaaaagcaggcttgatgATTCTTCCGCAATTTGCTCG, ggggaccactttgtacaagaaagctgggtcTACGCAGACCAGATTCGACA | Mycgr3\|97518  JGI  Ensembl |  |  | 100 % |  |  |
| 29 | ggggacaagtttgtacaaaaaagcaggcttgatgAGCCAGGCCACGGGTGATGG, ggggaccactttgtacaagaaagctgggtcCTTCACCAGCTTGACAAGTA | Mycgr3\|97642  JGI  Ensembl |  |  | 1.07163 % |  |  |
| 29 | ggggacaagtttgtacaaaaaagcaggcttgatgGCTCCATCGAAGATCGACGC, ggggaccactttgtacaagaaagctgggtcGAGCGGCTCAAGCGCTCCCT | Mycgr3\|99833  JGI  Ensembl | KOG1285 Beta, beta-carotene 15,15'-dioxygenase and related enzymes; | beta-carotene 15,15'-monooxygenase activity; | 100 % |  | ``` 850	CGAACTGGCTCA-TCCTCTTTGTCTGGCCTTTCG-AGGCGTCTATCGAACGAATGAAGCGAGGCGGCCACCACTGGGCGTGGAGCAACGACCGCGGAATGACATTCATCGT-GGT-CCCTC-GCGACGCCGCTAAACCTGTCGCCCCAGGCTGGAAGCCAAACGAGGTCCGGTCCTACGGCTGGAAAACGGGCATGGCAATTCACACCGCC-GCCGCTTGGGAAGAT-CCATCTACG-AAGCAAATTTTCGTCGAAAGCTCTCGGGTCCACGACAACCCTTTCCCATTCTTTCCTCCTGACACTCCAGCTTCACATATGCCGGAGCCAGACACGACCGCCGATTATGTTCGCTGGACCATCGACCCAACGCAGCCCGATCGTAGCCAGCTACCAGACCCACAGGTGATCGTAGATGTGCCTGCGGAATTTCCACGTATCGACGAGCGATTCATGACGTCTCCTTACGACGTTGTGTGGATGAATGTCTTCGTCCCGCAGCATTCCGACGGGTCAAAGAATATTTACCATGGACTCAACGGGCTGGCCATGCATCGCCATTCCACGAACGAGACGAAATGGTTCTATGCTGGAGATGAGTCGTTGGTGCAGGAGCCGATCTTCGTTCCAAGGACGAAGGACGCACCGGAGGGAGATGGCTGGGTGATAGCGTTGGTGGAGAGAACCGGAAAAGCGGCGAGATGTGACTTGGTGGTCCTGGATACGAGGGAGTTCGAGAAGCCTGTCGCGATTGTGCAGTTGCCGTTCCACATGAAGGCGCAGGTTCACGGTAACTGGATCGAGGCGGATACATTGGGTGGGTACAGGTCAATCACCAGGGAAGTGAAGGACTTTGAGATTTCTGGCAAGGGAGCGCTTGAGCCGCTC	1727 	\|\|\|\|\|\|\|\| \|\|\| \|\| \|\| \|\|\|\|\|\|\|\|\|\| \|\|\|\| \|\|\|\| \| \|\|\|\|\|\|\|\|\|\|\|\|\|\|\|\|\|\|\|\|\|\|\|\|\|\|\|\|\|\|\|\|\|\|\|\|\|\|\|\|\|\|\|\|\|\|\|\|\|\|\|\|\|\|\| \|\|\|\|\|\|\|\|\|\|\|\|\| \|\|\| \|\|\|\|\| \| \|\|\|\|\|\|\|\|\| \|\|\|\|\|\|\|\|\|\|\|\|\|\|\|\|\|\|\|\|\|\|\|\|\|\|\|\|\|\|\|\|\|\|\|\|\|\|\|\|\|\|\|\|\|\|\|\|\|\|\|\|\|\|\|\|\|\|\|\|\|\|\|\|\|\|\|\|\|\|\|\|\|\|\|\|\| \|\|\|\|\|\|\|\|\|\|\|\|\|\|\| \|\|\|\|\|\|\|\|\| \|\|\|\|\|\|\|\|\|\|\|\|\|\|\|\|\|\|\|\|\|\|\|\|\|\|\|\|\|\|\|\|\|\|\|\|\|\|\|\|\|\|\|\|\|\|\|\|\|\|\|\|\|\|\|\|\|\|\|\|\|\|\|\|\|\|\|\|\|\|\|\|\|\|\|\|\|\|\|\|\|\|\|\|\|\|\|\|\|\|\|\|\|\|\|\|\|\|\|\|\|\|\|\|\|\|\|\|\|\|\|\|\|\|\|\|\|\|\|\|\|\|\|\|\|\|\|\|\|\|\|\|\|\|\|\|\|\|\|\|\|\|\|\|\|\|\|\|\|\|\|\|\|\|\|\|\|\|\|\|\|\|\|\|\|\|\|\|\|\|\|\|\|\|\|\|\|\|\|\|\|\|\|\|\|\|\|\|\|\|\|\|\|\|\|\|\|\|\|\|\|\|\|\|\|\|\|\|\|\|\|\|\|\|\|\|\|\|\|\|\|\|\|\|\|\|\|\|\|\|\|\|\|\|\|\|\|\|\|\|\|\|\|\|\|\|\|\|\|\|\|\|\|\|\|\|\|\|\|\|\|\|\|\|\|\|\|\|\|\|\|\|\|\|\|\|\|\|\|\|\|\|\|\|\|\|\|\|\|\|\|\|\|\|\|\|\|\|\|\|\|\|\|\|\|\|\|\|\|\|\|\|\|\|\|\|\|\|\|\|\|\|\|\|\|\|\|\|\|\|\|\|\|\|\|\|\|\|\|\|\|\|\|\|\|\|\|\|\|\|\|\|\|\|\|\|\|\|\|\|\|\|\|\|\|\|\|\|\|\|\|\|\|\|\|\|\|\|\|\|\|\|\|\|\|\|\|\|\|\|\|\|\|\|\|\|\|\|\|\|\|\|\|\|\|\|\|\|\|\|\|\|\|\|\|\|\|\|\|\|\|\|\|\|\|\|\|\|\|\|\|\|\|\|\|\|\|\|\|\|\|\|\|\|\|\|\|\|\|\|\|\|\|\|\|\|\|\|\|\|\|\|\|\|\|\|\|\|\|\|\|\|\|\|\|\|\|\|\|\|\|\|\|\|\|\|\|\|\|\|\|\|\|\|\|\|\|\|\|\|\|\|\|\|\|\|\|\|\|\|\|\|\|\|\|\|\|\|\|\|\|\|\|\|\|\|\|\|\|\|\|\|\|\|\|\|\|\|\|\|\|\|\|\|\|\|\|\|\|\|\|\|\|\|\|\|\|\|\|\|\|\|\|\|\|\|\|\|\|\|\|\|\|\|\|\|\|\|\|\|\|\|\|\|\|\|\|\|\|\|\|\|\|\|\|\|\|\|\|\|\|\|\|\|\|\|\|\|\|\|\|\|\|\|\|\|\|\|\|\|\|\|\|\|\|\|\|\|\|\|\|\|\|\|\|\|\|\|\|\|\|\|\|\|\|\|\|\| 	CGAACTGG-TCATTCATC-TTGTCTGGCC-TTCGAAGGC-TTTATCGAACGAATGAAGCGAGGCGGCCACCACTGGGCGTGGAGCAACGACCGCGGA-TGACATTCATCGTTGGTCCCCTCGGGGACGCCGCT-AACCTGTCGCCCCAGGCTGGAAGCCAAACGAGGTCCGGTCCTACGGCTGGAAAACGGGCATGGCAATTCACACCGCCCGCCGCTTGGGAAGATTCCATCTACGAAAGCAAATTTTCGTCGAAAGCTCTCGGGTCCACGACAACCCTTTCCCATTCTTTCCTCCTGACACTCCAGCTTCACATATGCCGGAGCCAGACACGACCGCCGATTATGTTCGCTGGACCATCGACCCAACGCAGCCCGATCGTAGCCAGCTACCAGACCCACAGGTGATCGTAGATGTGCCTGCGGAATTTCCACGTATCGACGAGCGATTCATGACGTCTCCTTACGACGTTGTGTGGATGAATGTCTTCGTCCCGCAGCATTCCGACGGGTCAAAGAATATTTACCATGGACTCAACGGGCTGGCCATGCATCGCCATTCCACGAACGAGACGAAATGGTTCTATGCTGGAGATGAGTCGTTGGTGCAGGAGCCGATCTTCGTTCCAAGGACGAAGGACGCACCGGAGGGAGATGGCTGGGTGATAGCGTTGGTGGAGAGAACCGGAAAAGCGGCGAGATGTGACTTGGTGGTCCTGGATACGAGGGAGTTCGAGAAGCCTGTCGCGATTGTGCAGTTGCCGTTCCACATGAAGGCGCAGGTTCACGGTAACTGGATCGAGGCGGATACATTGGGTGGGTACAGGTCAATCACCAGGGAAGTGAAGGACTTTGAGATTTCTGGCAAGGGAGCGCTTGAGCCGCTC ``` |
| 29 | ggggacaagtttgtacaaaaaagcaggcttgatgGCAACCATGGCCGTCGACAA, ggggaccactttgtacaagaaagctgggtcATCGTCATCTTCTCCGGGTG | Mycgr3\|100355  JGI  Ensembl | KOG1721 FOG: Zn-finger; | intracellular; nucleic acid binding;  zinc ion binding; | 100 % |  |  |
| 29 | ggggacaagtttgtacaaaaaagcaggcttgatgGCAGTCCGAGATCTCACGCT, ggggaccactttgtacaagaaagctgggtcAACGGCGAACAGCTTCCTTT | Mycgr3\|103968  JGI  Ensembl |  |  | 100 % |  | ``` 1206	TCGGTGGGCCCATGAGCGCCCCCGCGACGAACGACCCTCTGCCCTTTAGTCTTCTTGCATGGATCATCCCGCCAACATGGTCGGATGGTGCTAAAAGTCACTTGGCCCCAGCCACTGGGTCAGTCATGCTCATGTACTCCCTCGAGCTGTCGCCTCTACTCCAGCGTCCTTTTGTGGCTTCCTTTTCTCAGTATTTGGGCGAGCTCTCCTTCGGCATATACGCTATGCATAACACGGTCCGATGGATCGTCTGGGAAAAGTGGTTCGTTCCATGGCAAGCGGAGCATTTTGGACCGCCTTCCGAAATGTTGAATTGGGTACCAGGGTATCTTGTCATGACTGTCCTTGTGCTCTGGGCCGCTGAACTTTTCCGCATGGTGGATATACAATGCGTTCGGGTCGCGAGCCGCCTACAAAGGAAGCTGTTCGCCGTT	1639 	\|\|\|\|\|\|\| \|\|\|\|\|\|\|\|\|\|\|\|\|\|\|\|\|\|\|\|\|\|\|\|\|\|\|\|\|\|\|\|\|\|\|\|\|\|\|\|\|\|\|\|\|\|\|\|\|\|\|\|\|\|\|\|\|\|\|\|\|\|\|\|\|\|\|\|\|\|\|\|\|\|\|\|\|\|\|\|\|\|\|\|\|\|\|\|\|\|\|\|\|\|\|\|\|\|\|\|\|\|\|\|\|\|\|\|\|\|\|\|\|\|\|\|\|\|\|\|\|\|\|\|\|\|\|\|\|\|\|\|\|\|\|\|\|\|\|\|\|\|\|\|\|\|\|\|\|\|\|\|\|\|\|\|\|\|\|\|\|\|\|\|\|\|\|\|\|\|\|\|\|\|\|\|\|\| \|\|\|\|\|\|\|\|\|\|\|\|\|\|\|\|\|\|\|\|\|\|\|\|\|\|\|\|\|\|\|\|\|\|\|\|\|\|\|\|\|\|\|\|\|\|\|\|\|\|\|\|\|\|\|\|\|\|\|\|\|\|\|\|\|\|\|\|\|\|\|\|\|\|\|\|\|\|\|\|\|\|\|\|\|\|\|\|\|\|\|\|\|\|\|\|\|\|\|\|\|\|\|\|\|\|\|\|\|\|\|\|\|\|\|\|\|\|\|\|\|\|\|\|\|\|\|\|\|\|\|\|\|\|\|\|\|\|\|\|\|\|\|\|\|\|\|\|\|\|\|\|\|\|\|\|\|\|\|\|\|\|\|\|\|\|\|\|\|\|\|\|\|\|\|\|\|\|\|\|\|\|\|\|\|\|\|\|\|\|\|\|\|\|\|\|\|\|\|\|\|\|\|\|\|\|\|\|\|\|\|\|\|\|\|\|\|\|\|\|\|\|\|\|\|\|\|\|\|\|\|\|\|\|\|\|\|\|\|\|\|\|\|\|\|\|\| 	TCGGTGGACCCATGAGCGCCCCCGCGACGAACGACCCTCTGCCCTTTAGTCTTCTTGCATGGATCATCCCGCCAACATGGTCGGATGGTGCTAAAAGTCACTTGGCCCCAGCCACTGGGTCAGTCATGCTCATGTACTCCCTCGAGCTGTCGCCTCTACTCCAGCGTCCTTTTGTGGCTTCCTTTTGTCAGTATTTGGGCGAGCTCTCCTTCGGCATATACGCTATGCATAACACGGTCCGATGGATCGTCTGGGAAAAGTGGTTCGTTCCATGGCAAGCGGAGCATTTTGGACCGCCTTCCGAAATGTTGAATTGGGTACCAGGGTATCTTGTCATGACTGTCCTTGTGCTCTGGGCCGCTGAACTTTTCCGCATGGTGGATATACAATGCGTTCGGGTCGCGAGCCGCCTACAAAGGAAGCTGTTCGCCGTT ``` |
| 29 | ggggacaagtttgtacaaaaaagcaggcttgatgTCGCTCCTACGATCCTTTAC, ggggaccactttgtacaagaaagctgggtcGATCCAGATGAGCTGCTCGC | Mycgr3\|104538  JGI  Ensembl | KOG4157 beta-1,6-N-acetylglucosaminyltransferase, contains WSC domain; |  | 100 % |  |  |
| 29 | ggggacaagtttgtacaaaaaagcaggcttgatgGCCGACGAGAACAACACCAA, ggggaccactttgtacaagaaagctgggtcGTCCATGTAGAAGTCACCCT | Mycgr3\|105024  JGI  Ensembl | KOG3671 Actin regulatory protein (Wiskott-Aldrich syndrome protein); |  | 100 % |  | ``` 770	CTA-GTACCTCAGGCGAAGGCTTCAATTGGTCTGACAACTTCCGAAACCGCGCCGAACGCACCGCGTCCATCGCAGGAGGCATGACTGGAGTACCACCAGCCCCTCCTGCACATCTCCATCAATCTCGGGCTAAGAGCGTGGCTACAATGGAACCACCGGCGACGGTGATGCCCAAGCCAAAGGAACAAGTGCGCCCAGATCACTTTCAAGAGAGGATTCTGAAGGGTGACTTCTACATGGAC	1011 	\|\|\| \|\|\|\|\|\|\|\|\|\|\|\|\|\|\|\|\|\|\|\|\|\|\|\|\|\|\|\|\|\|\|\|\|\|\|\|\|\|\|\|\|\|\|\|\|\|\|\|\|\|\|\|\|\|\|\|\|\|\|\|\|\|\|\|\|\|\|\|\|\|\|\|\|\|\|\|\|\|\|\|\|\|\|\|\|\|\|\|\|\|\|\|\|\|\|\|\|\|\|\|\|\|\|\|\|\|\|\|\|\|\|\|\|\|\|\|\|\|\|\|\|\|\|\|\|\|\|\|\|\|\|\|\|\|\|\|\|\|\|\|\|\|\|\|\|\|\|\|\|\|\|\|\|\|\|\|\|\|\|\|\|\|\|\|\|\|\|\|\|\|\|\|\|\|\|\|\|\|\|\|\|\|\|\|\|\|\|\|\|\|\|\|\|\|\|\|\|\|\|\|\|\|\|\|\|\|\|\|\|\|\|\|\|\|\|\|\|\|\|\|\|\|\|\|\|\|\|\|\|\|\|\|\|\|\|\|\|\|\| 	CTAGGTACCTCAGGCGAAGGCTTCAATTGGTCTGACAACTTCCGAAACCGCGCCGAACGCACCGCGTCCATCGCAGGAGGCATGACTGGAGTACCACCAGCCCCTCCTGCACATCTCCATCAATCTCGGGCTAAGAGCGTGGCTACAATGGAACCACCGGCGACGGTGATGCCCAAGCCAAAGGAACAAGTGCGCCCAGATCACTTTCAAGAGAGGATTCTGAAGGGTGACTTCTACATGGAC ``` |
| 29 | ggggacaagtttgtacaaaaaagcaggcttgatgGACCTTCAGTCTCCAATTCG, ggggaccactttgtacaagaaagctgggtcCCTCTTCCGATCATCGTTGA | Mycgr3\|106678  JGI  Ensembl |  |  | 0 % |  |  |
| 29 | ggggacaagtttgtacaaaaaagcaggcttgatgGCTCCACACGCCAACGACTC, ggggaccactttgtacaagaaagctgggtcATAAAGCTTGTGCTCCAGTG | Mycgr3\|107427  JGI  Ensembl | KOG0693 Myo-inositol-1-phosphate synthase; | inositol biosynthetic process; inositol-3-phosphate synthase activity;  phospholipid biosynthetic process; | 100 % |  |  |
| 29 | ggggacaagtttgtacaaaaaagcaggcttgatgTCCGCCTCGGCGCCTCTTAG, ggggaccactttgtacaagaaagctgggtcACACTCCGACGCCCTCCGCT | Mycgr3\|107719  JGI  Ensembl | KOG1187 Serine/threonine protein kinase; |  | 0 % |  |  |
| 29 | ggggacaagtttgtacaaaaaagcaggcttgatgTCCTCCTCCATGTCCTACTC, ggggaccactttgtacaagaaagctgggtcCTTCTTCTCCGCGATCAACT | Mycgr3\|108116  JGI  Ensembl | KOG2806 Chitinase; |  | 100 % |  |  |
| 29 | ggggacaagtttgtacaaaaaagcaggcttgatgTTCGTCGCGCGATCCTTCAA, ggggaccactttgtacaagaaagctgggtcATAAGCACTAACACCTACCC | Mycgr3\|108179  JGI  Ensembl | KOG1869 Splicing coactivator SRm160/300, subunit SRm300; |  | 100 % |  | ``` 1250	CCGCAGCAGCAACACTCCACCTCAATTACCTGGCGGAGCAGCACCCTCAATCGACCACATGCTCTGGTCGCATCCCTTCCTCTTTCCACGTTTCAGTTCAGAAATTCGTCCGGCAACCGACAGCAATACATCCGAGGCTCAAAGACGATATCTCAATCCTGTTCCACTCACCTTTGAAGAATGGGAGGCGCCCTTCCAACAAGCGCTCCACGCACCTTATCTTGCGGAAGCCGTCGTCGACGCGCCCACCATCCAGATGCAGACGTACAACCGCAGCCGAGGTCGAGGCCGCGGAGGATCGAAACGAGGCAGTCCCAATCCCAGCTCGGAAGCCGGACGAGCGCTTCTTGGGCCCGAGAGTGCTCGGCAGAGGGAGGTGCGCGAGAATTCGGACTTCTTGAGAGTGGTTGTTTTGGAGATGAACATGCGGAGAGAAGGCAAGCTGGAGACGGGACGGGCGAAGATCTGGCTACCGCCGAGGGAGGTCAGTCCTGCTCCCTCCAAGACGGGAAGGGTGCCGAAGCGATGGGTAGGTGTTAGTGCTTAT	1796 	\|\|\|\|\|\|\|\|\|\|\|\|\|\|\|\|\|\|\|\|\|\|\|\|\|\|\|\|\|\|\|\|\|\|\|\|\|\|\| \|\|\|\|\|\|\|\|\|\|\|\|\|\|\|\|\|\|\|\|\|\|\|\|\|\|\|\|\|\|\|\|\|\|\|\| \|\|\|\|\|\|\|\|\|\|\|\|\|\|\|\|\|\|\|\|\|\|\|\|\|\|\|\|\|\|\|\|\|\|\|\|\|\|\|\|\|\|\|\|\|\|\|\|\|\|\|\|\|\|\|\|\|\|\|\|\|\|\|\|\|\|\|\|\|\|\|\|\|\|\|\|\|\|\|\|\|\|\|\|\|\|\|\|\|\|\|\|\|\|\|\|\|\|\|\|\|\|\|\|\|\|\|\|\|\|\|\|\|\|\|\|\|\|\|\|\|\|\|\|\|\|\|\|\|\|\|\|\|\|\|\|\|\|\|\|\|\|\|\|\|\|\|\|\|\|\|\|\|\|\|\|\|\|\|\|\|\|\|\|\|\|\|\|\|\|\|\|\|\|\|\|\|\|\|\|\|\|\|\|\|\|\|\|\|\|\|\|\|\|\|\|\|\|\|\|\|\|\|\|\|\|\|\|\|\|\|\|\|\|\|\|\|\|\|\|\|\|\|\|\|\|\|\|\|\|\|\|\|\|\|\|\|\|\|\|\|\|\|\|\|\|\|\|\|\|\|\|\|\|\|\|\|\|\|\|\|\|\|\|\|\|\|\|\|\|\|\|\|\|\|\|\|\|\|\|\|\|\|\|\|\|\|\|\|\|\|\|\|\|\|\|\|\|\|\|\|\|\|\|\|\|\|\|\|\|\|\|\|\|\|\|\|\|\|\|\|\|\|\|\|\|\|\|\|\|\|\|\|\|\|\|\|\|\|\|\|\|\|\|\|\|\|\|\|\|\|\|\|\|\|\|\|\|\|\|\|\|\|\|\|\|\|\|\|\|\|\|\|\|\|\|\|\|\|\|\|\|\|\|\|\|\|\|\|\|\|\|\|\|\|\|\|\|\|\|\|\|\|\|\|\|\|\|\|\|\|\|\|\|\|\|\|\|\|\|\|\|\|\|\|\|\|\|\|\|\|\|\|\|\|\|\|\|\|\|\|\|\|\|\|\|\|\|\|\|\|\|\|\|\|\|\|\|\|\|\|\|\|\|\|\|\|\|\|\| 	CCGCAGCAGCAACACTCCACCTCAATTACCTGGCGGAGCTGCACCCTCAATCGACCACATGCTCTGGTCGCATCCCGTCCTCTTTCCACGTTTCAGTTCAGAAATTCGTCCGGCAACCGACAGCAATACATCCGAGGCTCAAAGACGATATCTCAATCCTGTTCCACTCACCTTTGAAGAATGGGAGGCGCCCTTCCAACAAGCGCTCCACGCACCTTATCTTGCGGAAGCCGTCGTCGACGCGCCCACCATCCAGATGCAGACGTACAACCGCAGCCGAGGTCGAGGCCGCGGAGGATCGAAACGAGGCAGTCCCAATCCCAGCTCGGAAGCCGGACGAGCGCTTCTTGGGCCCGAGAGTGCTCGGCAGAGGGAGGTGCGCGAGAATTCGGACTTCTTGAGAGTGGTTGTTTTGGAGATGAACATGCGGAGAGAAGGCAAGCTGGAGACGGGACGGGCGAAGATCTGGCTACCGCCGAGGGAGGTCAGTCCTGCTCCCTCCAAGACGGGAAGGGTGCCGAAGCGATGGGTAGGTGTTAGTGCTTAT ``` |
| 29 | ggggacaagtttgtacaaaaaagcaggcttgatgAGGTTTGGTCTTGCTTTGCT, ggggaccactttgtacaagaaagctgggtcCTCCCTCCAATACCTTCCAT | Mycgr3\|108506  JGI  Ensembl | KOG2182 Hydrolytic enzymes of the alpha/beta hydrolase fold; | cysteine-type peptidase activity; proteolysis; serine-type peptidase activity; | 2.3571 % |  |  |
| 29 | ggggacaagtttgtacaaaaaagcaggcttgatgGCTCCCTCAAAACGTCAAAA, ggggaccactttgtacaagaaagctgggtcAGACCTCGTCGAGACAATAT | Mycgr3\|109293  JGI  Ensembl |  |  | 100 % |  |  |
| 29 | ggggacaagtttgtacaaaaaagcaggcttgatgGCGCCCCAAACCGTCACCGA, ggggaccactttgtacaagaaagctgggtcCAGATCCTTCGCCAACCTCG | Mycgr3\|109296  JGI  Ensembl | KOG1289 Amino acid transporters; | amino acid transmembrane transporter activity; amino acid transport;  membrane;  transport; | 100 % |  |  |
| 29 | ggggacaagtttgtacaaaaaagcaggcttgatgTCAATTGCCCCAAACTACAC, ggggaccactttgtacaagaaagctgggtcGGGCGGGGCTTCTCGCTTCA | Mycgr3\|109480  JGI  Ensembl | KOG2516 Protein involved in dolichol pathway for N-glycosylation (mannosyltransferase family); | dolichyl-phosphate-mannose-glycolipid alpha-mannosyltransferase activity; | 100 % |  | ``` 1344	CGAAGCAAAGCTTGTCAAGTCGACCTTCTGGAAGCAGTTCGACTACGTGCTGGCCGAGAAGCCGGAGAAAGTCATTGGGAATTGGAAGGAGATCGACACGATCAATGGCTTCGCTGGTGTGTCGATGAGGCCGGGAGACGAAGATGACATTCTTCCAATGCCGGCCGTGCTCGGACGTCCGGGCCGCATCGTACACGATGCGTACCAGCGAATGGCTCTTGTAGCTCGGCAGAAGTTCACGAAGGGATACTGGCCGGCCATACGTATGGCTCCGCGAATACATATTCTGAAGCGAGAAGCCCCGCCC	1650 	\|\|\|\|\|\|\|\|\|\|\|\|\|\|\|\|\|\|\|\|\|\|\|\|\|\|\|\|\|\|\|\|\|\|\|\|\|\|\|\|\|\|\|\|\|\|\|\|\|\|\|\|\|\|\|\|\|\|\|\|\|\|\|\|\|\|\|\|\|\|\|\|\|\|\|\|\|\|\|\|\|\|\|\|\|\|\|\|\|\|\|\|\|\|\|\|\|\|\|\|\|\|\|\|\|\|\|\|\|\|\|\|\|\|\|\|\|\|\|\|\|\|\|\|\|\|\|\|\|\|\|\|\|\|\|\|\|\|\|\|\|\|\|\|\|\|\|\|\|\|\|\|\|\|\|\|\|\|\|\|\|\|\|\|\|\|\|\|\|\|\|\|\|\|\|\|\|\|\|\|\|\|\|\|\|\|\|\|\|\|\|\|\|\|\|\|\|\|\|\|\|\|\|\|\|\|\|\|\|\|\|\|\|\|\|\|\|\|\|\|\|\|\|\|\|\|\|\|\|\|\|\|\|\|\|\|\|\|\|\|\|\|\|\|\|\|\|\|\|\|\|\|\|\|\|\|\|\|\|\|\|\|\|\|\|\|\|\|\|\|\|\|\|\|\|\|\|\|\|\|\|\|\|\|\|\|\|\|\|\|\|\|\|\|\|\|\|\|\|\|\|\|\|\|\|\|\| 	CGAAGCAAAGCTTGTCAAGTCGACCTTCTGGAAGCAGTTCGACTACGTGCTGGCCGAGAAGCCGGAGAAAGTCATTGGGAATTGGAAGGAGATCGACACGATCAATGGCTTCGCTGGTGTGTCGATGAGGCCGGGAGACGAAGATGACATTCTTCCAATGCCGGCCGTGCTCGGACGTCCGGGCCGCATCGTACACGATGCGTACCAGCGAATGGCTCTTGTAGCTCGGCAGAAGTTCACGAAGGGATACTGGCCGGCCATACGTATGGCTCCGCGAATACATATTCTGAAGCGAGAAGCCCCGCCC ``` |
| 29 | ggggacaagtttgtacaaaaaagcaggcttgatgACTCCACGACATTCCTTCAG, ggggaccactttgtacaagaaagctgggtcCAAGAACTTCTTGGGTGTGC | Mycgr3\|109961  JGI  Ensembl | KOG3839 Lectin VIP36, involved in the transport of glycoproteins carrying high mannose-type glycans; | membrane; | 100 % |  | ``` 712	CGTATACA-ATGT-GGAGAAGTTCGA-CGGACTGG-TATTGGTCG-TAGACC-AGTACGGA-GGCACTGGTG-GT-AAGATTCGTGGGTTCCTGAACGACGGATC-GCAGAATTATCGAGG--CCAGTCAGC-ACTAG-AGTCGCTAGCATTTGGACATTGCGATTATCACTACCGAAATCTGGGAAGGCCGAGCAAGTTGACAGTCA-GCTCG-AGCCAGTCCGGTCTCGATGTCAAGATCGACGACAAGACTTGTTTCTCGACAGCTACCGTCGCACTGCCCGAGGGCTACCACTTCGGAATTACCGCCTCTACGGGCGAGAGCAATCCCGACTCGATCGAGATATACAAGTTCCTGGTGAACTCCAAGCTCTCCCAGCACGAGCAAAAAATGAAGAATCCCCCCGCGCAGGCGGGAGATAAGAAACCTACACCTGGAACAACTCCTCGCAAGATGGACGCTTTTCCGGGTATCGTGGAGCTCATACCAGACCGTGAAGCAAAGGAATTCAAGTCTCAGACTGAGCAATTCTCGGACTTGCACGACCGATTGCAGGCCATGTCTCATCACATCGCCTACGTCTTCGGCGAGATCAAGACCCTCGCCGAGAGCTCGGAGACGCAACACAAGGAGTCGCTGGCCAAGAACACCGAGGTCCTCGATGCTGTCAAGAACATGAAAGGCCTGCCCCCTGACACGATTCGTCAGATCAGGGAGCTCTCTGAAAAGGTCGACAAGCTCGAAGCTTCTATGGCTCTCGTGTCGAAGTCTCTGGACGACGATCACTTCGGGAAGCTGTTCAAACAGATAGAGATGATGCACTTCGATGTGATGCCGGCTCAGATGAAGCATG	1550 	\|\|\|\|\|\|\|\| \| \|\| \|\|\|\|\|\|\|\|\|\|\|\| \|\|\|\| \|\|\| \|\|\|\|\|\|\|\|\| \|\|\|\|\|\| \|\|\|\|\|\|\|\| \|\|\|\|\|\| \|\|\| \|\| \|\|\|\|\|\|\|\|\|\|\|\|\|\|\|\|\|\|\|\|\|\|\|\|\|\|\|\|\| \|\|\|\|\|\|\|\|\|\|\|\|\|\|\|  \|\|\|\|\|\|\|\|\| \|\|\|\|\| \|\|\|\|\|\|\|\|\|\|\|\|\|\|\|\|\|\|\|\|\|\|\|\|\|\|\|\|\|\|\|\|\|\|\|\|\|\|\|\|\|\|\|\|\|\|\| \|\|\|\|\|\|\|\|\|\|\|\|\|\|\|\|\|\|\|\|\| \|\|\|\|\| \|\|\|\|\|\|\|\|\| \|\|\|\|\|\|\|\|\|\|\|\|\|\|\|\|\|\|\|\|\|\|\|\|\|\|\|\|\|\|\|\|\|\|\|\|\|\|\|\|\|\|\|\|\|\|\|\|\|\|\|\|\|\|\|\|\|\|\|\|\|\|\|\|\|\|\|\|\|\|\|\|\|\|\|\|\|\|\|\|\|\|\|\|\|\|\|\|\|\|\|\|\|\|\|\|\|\|\|\|\|\|\|\|\|\|\|\|\|\|\|\|\|\|\|\|\|\|\|\|\|\|\|\|\|\|\|\|\|\|\|\|\|\|\|\|\|\|\|\|\|\|\|\|\|\|\|\|\|\|\|\|\|\|\|\|\|\|\|\|\|\|\|\|\|\|\|\|\|\|\|\|\|\|\|\|\|\|\|\|\|\|\|\|\|\|\|\|\|\|\|\|\|\|\|\|\|\|\|\|\|\|\|\|\|\|\|\|\|\|\|\|\|\|\|\|\|\|\|\|\|\|\|\|\|\|\|\|\|\|\|\|\|\|\|\|\|\|\|\|\|\|\|\|\|\|\|\|\|\|\|\|\|\|\|\|\|\|\|\|\|\|\|\|\|\|\|\|\|\|\|\|\|\|\|\|\|\|\|\|\|\|\|\|\|\|\|\|\|\|\|\|\|\|\|\|\|\|\|\|\|\|\|\|\|\|\|\|\|\|\|\|\|\|\|\|\|\|\|\|\|\|\|\|\|\|\|\|\|\|\|\|\|\|\|\|\|\|\|\|\|\|\|\|\|\|\|\|\|\|\|\|\|\|\|\|\|\|\|\|\|\|\|\|\|\|\|\|\|\|\|\|\|\|\|\|\|\|\|\|\|\|\|\|\|\|\|\|\|\|\|\|\|\|\|\|\|\|\|\|\|\|\|\|\|\|\|\|\|\|\|\|\|\|\|\|\|\|\|\|\|\|\|\|\|\|\|\|\|\|\|\|\|\|\|\|\|\|\|\|\|\|\|\|\|\|\|\|\|\|\|\|\|\|\|\|\|\|\|\|\|\|\|\|\|\|\|\|\|\|\|\|\|\|\|\|\|\|\|\|\|\|\|\|\|\|\|\|\|\|\|\|\|\|\|\|\|\|\|\|\|\|\|\|\|\|\|\|\|\|\|\|\|\|\|\|\|\|\|\|\|\|\|\|\|\|\|\|\|\|\|\|\|\|\|\|\|\|\|\|\|\|\|\|\|\|\|\|\|\|\|\|\|\|\|\|\|\|\|\|\|\|\|\|\|\|\|\|\|\|\|\|\|\|\|\|\|\|\|\|\|\|\|\|\|\|\|\|\|\|\|\|\|\|\|\|\|\|\|\|\|\|\|\|\|\|\|\|\|\|\|\|\|\|\|\|\|\|\|\|\|\|\|\|\|\|\|\|\|\| 	CGTATACACAGGTGGGAGAAGTTCGAGCGGA-TGGTTATTGGTCGTTAGACCAAGTACGGAGGGCACT-GTGTGTAAAGATTCGTGGGTTCCTGAACGACGGATCTGCAGAATTATCGAGGGCCCAGTCAGCAACTAGAAGTCGCTAGCATTTGGACATTGCGATTATCACTACCGAAATCTGGGAGGGCCGAGCAAGTTGACAGTCAAGCTCGAAGCCAGTCCCGTCTCGATGTCAAGATCGACGACAAGACTTGTTTCTCGACAGCTACCGTCGCACTGCCCGAGGGCTACCACTTCGGAATTACCGCCTCTACGGGCGAGAGCAATCCCGACTCGATCGAGATATACAAGTTCCTGGTGAACTCCAAGCTCTCCCAGCACGAGCAAAAAATGAAGAATCCCCCCGCGCAGGCGGGAGATAAGAAACCTACACCTGGAACAACTCCTCGCAAGATGGACGCTTTTCCGGGTATCGTGGAGCTCATACCAGACCGTGAAGCAAAGGAATTCAAGTCTCAGACTGAGCAATTCTCGGACTTGCACGACCGATTGCAGGCCATGTCTCATCACATCGCCTACGTCTTCGGCGAGATCAAGACCCTCGCCGAGAGCTCGGAGACGCAACACAAGGAGTCGCTGGCCAAGAACACCGAGGTCCTCGATGCTGTCAAGAACATGAAAGGCCTGCCCCCTGACACGATTCGTCAGATCAGGGAGCTCTCTGAAAAGGTCGACAAGCTCGAAGCTTCTATGGCTCTCGTGTCGAAGTCTCTGGACGACGATCACTTCGGGAAGCTGTTCAAACAGATAGAGATGATGCACTTCGATGTGATGCCGGCTCAGATGAAGCATG ```      ``` 1550	GCTCTTCAGACCCATACCCCTAGCTTGGGGAAGTTGGCCTTGCTGCTCGTTGGAACTCAGGTCGTGCTCGCCGCTGCATACGTCGTGTACAAGCGGAGAAAGAACAGCACACCCAAGAAGTTCTTG	1675 	\|\|\|\|\|\|\|\|\|\|\|\|\|\|\|\|\|\|\|\|\|\|\|\|\|\|\|\|\|\|\|\|\|\|\|\|\|\|\|\|\|\|\|\|\|\|\|\|\|\|\|\|\|\|\|\|\|\|\|\|\|\|\|\|\|\|\|\|\|\|\|\|\|\|\|\|\|\|\|\|\|\|\|\|\|\|\|\|\|\|\|\|\|\|\|\|\|\|\|\|\|\|\|\|\|\|\|\|\|\|\|\|\|\|\|\|\|\|\|\|\|\|\|\|\|\| 	GCTCTTCAGACCCATACCCCTAGCTTGGGGAAGTTGGCCTTGCTGCTCGTTGGAACTCAGGTCGTGCTCGCCGCTGCATACGTCGTGTACAAGCGGAGAAAGAACAGCACACCCAAGAAGTTCTTG ``` |
| 29 | ggggacaagtttgtacaaaaaagcaggcttgatgGGCCGACGACACTATCTCAC, ggggaccactttgtacaagaaagctgggtcTTCCGCCTCAATTTCATCCT | Mycgr3\|110059  JGI  Ensembl |  |  | 100 % |  |  |
| 30 | ggggacaagtttgtacaaaaaagcaggcttgatgGGTATCCAACTTTGGCGCCG, ggggaccactttgtacaagaaagctgggtcAACTTTCTCCTCTGCGAATG | Mycgr3\|32616  JGI  Ensembl | KOG0254 Predicted transporter (major facilitator superfamily); | L-arabinose isomerase activity; carbohydrate transport;  glucose transmembrane transporter activity;  integral to membrane;  membrane;  sugar:hydrogen symporter activity;  transport;  transporter activity; | 0 % |  |  |
| 30 | ggggacaagtttgtacaaaaaagcaggcttgatgCCCTCCACAGGGCAAGACGC, ggggaccactttgtacaagaaagctgggtcAACTCCACAAACTGCACACA | Mycgr3\|33425  JGI  Ensembl | KOG4126 Alkaline phosphatase; | alkaline phosphatase activity; metabolic process; | 11.39924 % |  |  |
| 30 | ggggacaagtttgtacaaaaaagcaggcttgatgCCTAGCCTTCGAGATCTACT, ggggaccactttgtacaagaaagctgggtcCTTTTTCGGCGGGTTGTACT | Mycgr3\|33587  JGI  Ensembl | KOG1550 Extracellular protein SEL-1 and related proteins; |  | 0 % |  |  |
| 30 | ggggacaagtttgtacaaaaaagcaggcttgatgGAGACTAGAAGCGGCATTAT, ggggaccactttgtacaagaaagctgggtcACGTCCCCGCTCACGAGCTT | Mycgr3\|34052  JGI  Ensembl |  |  | 0 % |  |  |
| 30 | ggggacaagtttgtacaaaaaagcaggcttgatgGACGGCTTCAATGAAAAGGA, ggggaccactttgtacaagaaagctgggtcCTCAACATGCAAAACATTGC | Mycgr3\|36522  JGI  Ensembl | KOG0254 Predicted transporter (major facilitator superfamily); | L-arabinose isomerase activity; carbohydrate transport;  integral to membrane;  membrane;  sugar:hydrogen symporter activity;  transport;  transporter activity; | 0 % |  |  |
| 30 | ggggacaagtttgtacaaaaaagcaggcttgatgCGGTCCATCTTGCGACCCTT, ggggaccactttgtacaagaaagctgggtcCGCAGCAACCGCCTCAACTA | Mycgr3\|38294  JGI  Ensembl | KOG0255 Synaptic vesicle transporter SVOP and related transporters (major facilitator superfamily); | integral to membrane; transport; transporter activity; | 0 % |  |  |
| 30 | ggggacaagtttgtacaaaaaagcaggcttgatgCGGGCTGCACGGAGTTGGGC, ggggaccactttgtacaagaaagctgggtcAACTAAATCAGCATGAGCGC | Mycgr3\|40096  JGI  Ensembl | KOG3731 Sulfatases; | arylsulfatase activity; metabolic process;  sulfuric ester hydrolase activity; | 0 % |  |  |
| 30 | ggggacaagtttgtacaaaaaagcaggcttgatgGCCGAACAAACAGTGCCACT, ggggaccactttgtacaagaaagctgggtcCCCCTCCATCTCCTGCACTG | Mycgr3\|40465  JGI  Ensembl | KOG1184 Thiamine pyrophosphate-requiring enzyme; | magnesium ion binding; pyruvate decarboxylase activity;  thiamin pyrophosphate binding;  transferase activity; | 0 % |  |  |
| 30 | ggggacaagtttgtacaaaaaagcaggcttgatgTCGAACGGTGCGCCCAACGG, ggggaccactttgtacaagaaagctgggtcGATCAACACGTAAAGCGCAC | Mycgr3\|41169  JGI  Ensembl | KOG0380 Sterol O-acyltransferase/Diacylglycerol O-acyltransferase; | nucleic acid binding; sterol O-acyltransferase activity; | 0 % |  |  |
| 30 | ggggacaagtttgtacaaaaaagcaggcttgatgTCCAACTTTGAGAAGGACTA, ggggaccactttgtacaagaaagctgggtcTTGTGCTGTATCCTTCCTCC | Mycgr3\|41743  JGI  Ensembl | KOG0254 Predicted transporter (major facilitator superfamily); | integral to membrane; transport; transporter activity; | 0 % |  |  |
| 30 | ggggacaagtttgtacaaaaaagcaggcttgatgGGTCTCCCCAGCAAAACGAC, ggggaccactttgtacaagaaagctgggtcCTTCTCGTTCTTGTTCACCG | Mycgr3\|48350  JGI  Ensembl | KOG2466 Uridine permease/thiamine transporter/allantoin transport; | membrane; nucleobase transmembrane transporter activity;  nucleobase, nucleoside, nucleotide and nucleic acid transport; | 0 % |  |  |
| 30 | ggggacaagtttgtacaaaaaagcaggcttgatgATCACCCGGCGTCGATACGC, ggggaccactttgtacaagaaagctgggtcCGTCCCCGCCCTCCGAAACG | Mycgr3\|49195  JGI  Ensembl | KOG2431 1, 2-alpha-mannosidase; | calcium ion binding; mannosyl-oligosaccharide 1,2-alpha-mannosidase activity;  membrane; | 1.2419 % |  |  |
| 30 | ggggacaagtttgtacaaaaaagcaggcttgatgCGGCGCGCAATAATCCAAAA, ggggaccactttgtacaagaaagctgggtcAAGCTTCGCCTTCACCCCAT | Mycgr3\|50302  JGI  Ensembl | KOG0700 Protein phosphatase 2C/pyruvate dehydrogenase (lipoamide) phosphatase; | CTD phosphatase activity; calcium-dependent protein serine/threonine phosphatase activity;  calcium-dependent protein serine/threonine phosphatase regulator activity;  catalytic activity;  magnesium-dependent protein serine/threonine phosphatase activity;  magnesium-dependent protein serine/threonine phosphatase complex;  myosin phosphatase activity;  myosin phosphatase complex;  myosin phosphatase regulator activity;  phosphoprotein phosphatase activity; | 0 % |  |  |
| 30 | ggggacaagtttgtacaaaaaagcaggcttgatgCGCTCCCCGTTCAAGTCAAA, ggggaccactttgtacaagaaagctgggtcCAACAAATATCGATACCCCG | Mycgr3\|50651  JGI  Ensembl | KOG2533 Permease of the major facilitator superfamily; | integral to membrane; transport; transporter activity; | 0 % |  |  |
| 30 | ggggacaagtttgtacaaaaaagcaggcttgatgGCTGGGCACAACGATCCTAA, ggggaccactttgtacaagaaagctgggtcGCCATGCTTCTCCCTGGGAT | Mycgr3\|52118  JGI  Ensembl | KOG2709 Uncharacterized conserved protein; |  | 0 % |  |  |
| 30 | ggggacaagtttgtacaaaaaagcaggcttgatgGCCACCTCGACAACACCGGC, ggggaccactttgtacaagaaagctgggtcCAACGCATTGCCCAGGTCTT | Mycgr3\|52213  JGI  Ensembl | KOG0676 Actin and related proteins; | ATP binding; protein binding; | 1.40217 % |  |  |
| 30 | ggggacaagtttgtacaaaaaagcaggcttgatgGACGCTGCCAATCTCATAGA, ggggaccactttgtacaagaaagctgggtcTGACTGCGGCAGAACAAGCT | Mycgr3\|55724  JGI  Ensembl |  |  | 0 % |  |  |
| 30 | ggggacaagtttgtacaaaaaagcaggcttgatgGCGCGGCTCTCCTCCATACT, ggggaccactttgtacaagaaagctgggtcGTCGCTATCATCCAGCCCGC | Mycgr3\|58295  JGI  Ensembl | KOG3847 Phospholipase A2 (platelet-activating factor acetylhydrolase in humans); | 1-alkyl-2-acetylglycerophosphocholine esterase activity; 2-acetyl-1-alkylglycerophosphocholine esterase complex;  lipid catabolic process; | 0 % |  |  |
| 30 | ggggacaagtttgtacaaaaaagcaggcttgatgGCGTCTTTCTTCTCCAGCTG, ggggaccactttgtacaagaaagctgggtcGATCGACAAAGGCTCCAAGC | Mycgr3\|65177  JGI  Ensembl | KOG3847 Phospholipase A2 (platelet-activating factor acetylhydrolase in humans); | 1-alkyl-2-acetylglycerophosphocholine esterase activity; 2-acetyl-1-alkylglycerophosphocholine esterase complex;  ATP binding;  catalytic activity;  lipid catabolic process;  metabolic process; | 0 % |  |  |
| 30 | ggggacaagtttgtacaaaaaagcaggcttgatgTCTCCCAGCAAGGTGTCGTC, ggggaccactttgtacaagaaagctgggtcAGCCTTGTACGAGTGCATAA | Mycgr3\|65205  JGI  Ensembl | KOG2450 Aldehyde dehydrogenase; | aldehyde dehydrogenase [NAD(P)+] activity; | 0 % |  |  |
| 30 | ggggacaagtttgtacaaaaaagcaggcttgatgGCGGAAATTCCAATCAAAGT, ggggaccactttgtacaagaaagctgggtcTGCCGCCTTCTGCTCGGCCT | Mycgr3\|65843  JGI  Ensembl | KOG1872 Ubiquitin-specific protease; | protein modification process; ubiquitin thiolesterase activity;  ubiquitin-dependent protein catabolic process; | 0 % |  |  |
| 30 | ggggacaagtttgtacaaaaaagcaggcttgatgCCCGAGCGGCCCTTGACCAT, ggggaccactttgtacaagaaagctgggtcTTCAACCCTCTCATAAAACA | Mycgr3\|67691  JGI  Ensembl | KOG1867 Ubiquitin-specific protease; | ubiquitin thiolesterase activity; ubiquitin-dependent protein catabolic process; | 0 % |  |  |
| 30 | ggggacaagtttgtacaaaaaagcaggcttgatgGAACCCTGGTTGCACTCCAC, ggggaccactttgtacaagaaagctgggtcCAAGCCTCCATACCCACCCT | Mycgr3\|68020  JGI  Ensembl | KOG2913 Predicted membrane protein; |  | 0 % |  |  |
| 30 | ggggacaagtttgtacaaaaaagcaggcttgatgGACGCCGTCAAACGAGGCAT, ggggaccactttgtacaagaaagctgggtcCTTCAACTCAGGTTTCGCCT | Mycgr3\|68826  JGI  Ensembl | KOG1289 Amino acid transporters; | amino acid transmembrane transporter activity; amino acid transport;  membrane;  transport; | 0 % |  |  |
| 30 | ggggacaagtttgtacaaaaaagcaggcttgatgGCAGGCGACAACGAGAAAAG, ggggaccactttgtacaagaaagctgggtcCAACTTGCTCTCATCTTTGC | Mycgr3\|68968  JGI  Ensembl |  |  | 0 % |  |  |
| 30 | ggggacaagtttgtacaaaaaagcaggcttgatgGCCGCTCCACAGACAGAGTC, ggggaccactttgtacaagaaagctgggtcACTTTTCACCTCATCCTCTT | Mycgr3\|69159  JGI  Ensembl | KOG0255 Synaptic vesicle transporter SVOP and related transporters (major facilitator superfamily); | integral to membrane; transport; transporter activity; | 1.15291 % |  |  |
| 30 | ggggacaagtttgtacaaaaaagcaggcttgatgGTCAAAATGCTAGAGGATTC, ggggaccactttgtacaagaaagctgggtcTGTGCCTTTGCAGTACTCTA | Mycgr3\|70356  JGI  Ensembl |  |  | 0 % |  |  |
| 30 | ggggacaagtttgtacaaaaaagcaggcttgatgCCAAACCCAGTCAACACCGA, ggggaccactttgtacaagaaagctgggtcCAACTTAGAAGCCGCCGTCT | Mycgr3\|70677  JGI  Ensembl | KOG1261 Malate synthase; | glyoxylate cycle; malate synthase activity; | 0 % |  |  |
| 30 | ggggacaagtttgtacaaaaaagcaggcttgatgCCGAAAGGAATGCTCATTTT, ggggaccactttgtacaagaaagctgggtcGTCGTATTGGAACAATACGC | Mycgr3\|71222  JGI  Ensembl | KOG1448 Ribose-phosphate pyrophosphokinase; | magnesium ion binding; nucleoside metabolic process;  nucleotide biosynthetic process;  ribose phosphate diphosphokinase activity; | 0 % |  |  |
| 30 | ggggacaagtttgtacaaaaaagcaggcttgatgGATCCCAACAACCAGGGCTA, ggggaccactttgtacaagaaagctgggtcTGCGTGGATTGCTTGTTGTA | Mycgr3\|71484  JGI  Ensembl | KOG1362 Choline transporter-like protein; |  | 0 % |  |  |
| 30 | ggggacaagtttgtacaaaaaagcaggcttgatgGAGAAATCGACTCTTCCCAC, ggggaccactttgtacaagaaagctgggtcCTCCCCCTCAAACCTCGCCC | Mycgr3\|71524  JGI  Ensembl | KOG2275 Aminoacylase ACY1 and related metalloexopeptidases; | Gly-X carboxypeptidase activity; metallopeptidase activity;  proteolysis; | 0 % |  |  |
| 30 | ggggacaagtttgtacaaaaaagcaggcttgatgTCTCCGCCAATCACCGATGC, ggggaccactttgtacaagaaagctgggtcACCCACAGAACCCACACTCG | Mycgr3\|71654  JGI  Ensembl | KOG1289 Amino acid transporters; | amino acid transmembrane transporter activity; amino acid transport;  membrane;  transport; | 0 % |  |  |
| 30 | ggggacaagtttgtacaaaaaagcaggcttgatgGACATCGCCTTCCCGTATAC, ggggaccactttgtacaagaaagctgggtcACTCGCGATTCTCCCAACAC | Mycgr3\|72003  JGI  Ensembl | KOG3896 Dynactin, subunit p62; |  | 1.0983 % |  |  |
| 30 | ggggacaagtttgtacaaaaaagcaggcttgatgCTCCCACTCGTCCTCTCCCT, ggggaccactttgtacaagaaagctgggtcGATATCAAACTCGCATAGCC | Mycgr3\|72841  JGI  Ensembl |  |  | 1.27212 % |  |  |
| 30 | ggggacaagtttgtacaaaaaagcaggcttgatgTCCGCATCACACGTCTTCGT, ggggaccactttgtacaagaaagctgggtcCTTCTTCTTCCCGAACCCCA | Mycgr3\|73850  JGI  Ensembl | KOG2699 Predicted ubiquitin regulatory protein; |  | 0 % |  |  |
| 30 | ggggacaagtttgtacaaaaaagcaggcttgatgACAGCCCCTCGACAACACTC, ggggaccactttgtacaagaaagctgggtcATCGTGCATCGCCATCGAAG | Mycgr3\|74327  JGI  Ensembl | KOG0254 Predicted transporter (major facilitator superfamily); | integral to membrane; transport; transporter activity; | 0 % |  |  |
| 30 | ggggacaagtttgtacaaaaaagcaggcttgatgGCACACTCGTCGATGAGAGA, ggggaccactttgtacaagaaagctgggtcGACCAAGAGAAGAGCTCCTC | Mycgr3\|74641  JGI  Ensembl | KOG0255 Synaptic vesicle transporter SVOP and related transporters (major facilitator superfamily); | G-protein coupled receptor protein signaling pathway; integral to membrane;  rhodopsin-like receptor activity;  transport;  transporter activity; | 1.21107 % |  |  |
| 30 | ggggacaagtttgtacaaaaaagcaggcttgatgGCCGTAAATGCGACCGCCGG, ggggaccactttgtacaagaaagctgggtcCGGAGCTGTCGTCGGGAATT | Mycgr3\|75015  JGI  Ensembl | KOG0684 Cytochrome P450; | electron transport; heme binding;  iron ion binding;  monooxygenase activity; | 1.10926 % |  |  |
| 30 | ggggacaagtttgtacaaaaaagcaggcttgatgAGCGTACCCTATCTCGAGCT, ggggaccactttgtacaagaaagctgggtcGCAGCCCAAATACCTCTCCT | Mycgr3\|75274  JGI  Ensembl | KOG4203 Armadillo/beta-Catenin/plakoglobin; | ATP binding; carbohydrate metabolic process;  kinase activity;  metabolic process;  phosphoribulokinase activity;  phosphotransferase activity, alcohol group as acceptor;  uridine kinase activity; | 1.33721 % |  |  |
| 30 | ggggacaagtttgtacaaaaaagcaggcttgatgGCGAACGAAACTACTTCGGC, ggggaccactttgtacaagaaagctgggtcGATCTGACCTTGTTGAGCAT | Mycgr3\|76346  JGI  Ensembl |  |  | 1.31108 % |  |  |
| 30 | ggggacaagtttgtacaaaaaagcaggcttgatgTCCGACATGAGTGCCACCGT, ggggaccactttgtacaagaaagctgggtcGCAGCAATCGCCGTGGTCTC | Mycgr3\|76728  JGI  Ensembl | KOG0692 Pentafunctional AROM protein; | 3-dehydroquinate synthase activity; aromatic amino acid family biosynthetic process; | 0 % |  |  |
| 30 | ggggacaagtttgtacaaaaaagcaggcttgatgGCCAGAAACGACACCACCGT, ggggaccactttgtacaagaaagctgggtcGCTGCACGTTGCCATTGCGC | Mycgr3\|77474  JGI  Ensembl | KOG1305 Amino acid transporter protein; |  | 0 % |  |  |
| 30 | ggggacaagtttgtacaaaaaagcaggcttgatgGACCTCTCCGGCTTCGATCC, ggggaccactttgtacaagaaagctgggtcCTCCGCCCCTTGTTCCCTCT | Mycgr3\|84654  JGI  Ensembl | KOG1187 Serine/threonine protein kinase; |  | 0 % |  |  |
| 30 | ggggacaagtttgtacaaaaaagcaggcttgatgTCCTCGTCGCCTTTGCGCCC, ggggaccactttgtacaagaaagctgggtcATACGTCTCTCCCACATCGC | Mycgr3\|85339  JGI  Ensembl | KOG1853 LIS1-interacting protein NUDE; |  | 1.19435 % |  |  |
| 30 | ggggacaagtttgtacaaaaaagcaggcttgatgGAGATCGTCGAACAACCACT, ggggaccactttgtacaagaaagctgggtcCCCATCAAGGTAGTAGGCCA | Mycgr3\|86340  JGI  Ensembl | KOG1399 Flavin-containing monooxygenase; | cyclohexanone monooxygenase activity; | 0 % |  |  |
| 30 | ggggacaagtttgtacaaaaaagcaggcttgatgATCAAGCCTTTGCCGAAGCC, ggggaccactttgtacaagaaagctgggtcAACACCCTTCCCGAGCGCCG | Mycgr3\|88816  JGI  Ensembl | KOG0029 Amine oxidase; | amine oxidase activity; electron transport;  oxidoreductase activity; | 0 % |  |  |
| 30 | ggggacaagtttgtacaaaaaagcaggcttgatgGCAGACACGTCCACACAGCA, ggggaccactttgtacaagaaagctgggtcCTCCAGCACAAACTCATCGT | Mycgr3\|89474  JGI  Ensembl | KOG2992 Nucleolar GTPase/ATPase p130; | intracellular; | 0 % |  |  |
| 30 | ggggacaagtttgtacaaaaaagcaggcttgatgGACTCTGTCTCTGGACTCAA, ggggaccactttgtacaagaaagctgggtcCACAGCCTCTTCCCGGCGAG | Mycgr3\|90043  JGI  Ensembl | KOG0158 Cytochrome P450 CYP3/CYP5/CYP6/CYP9 subfamilies; | 2-hydroxyisobutyrate 3-monooxygenase activity; 4-chlorophenoxyacetate monooxygenase activity;  4-nitrocatechol 4-monooxygenase activity;  alpha-pinene dehydrogenase activity;  alpha-pinene monooxygenase activity;  ammonia monooxygenase activity;  di-n-butyltin dioxygenase activity;  dimethylsilanediol hydroxylase activity;  electron transport;  fluorene oxygenase activity;  heme binding;  hydroxymethylsilanetriol oxidase activity;  iron ion binding;  limonene 8-monooxygenase activity;  methyl tertiary butyl ether 3-monooxygenase activity;  methylsilanetriol hydroxylase activity;  mono-butyltin dioxygenase activity;  monooxygenase activity;  spheroidene monooxygenase activity;  tert-butyl alcohol 2-monooxygenase activity;  tri-n-butyltin dioxygenase activity; | 0 % |  |  |
| 30 | ggggacaagtttgtacaaaaaagcaggcttgatgTCCACCAGTGACGGAGAGCG, ggggaccactttgtacaagaaagctgggtcTGCGCCGTAGACAAGCATTG | Mycgr3\|90250  JGI  Ensembl |  |  | 0 % |  |  |
| 30 | ggggacaagtttgtacaaaaaagcaggcttgatgTACGAGGCGCTGGCAACATG, ggggaccactttgtacaagaaagctgggtcCTCGTCCTCGTCCGACGAAG | Mycgr3\|90594  JGI  Ensembl |  |  | 0 % |  |  |
| 30 | ggggacaagtttgtacaaaaaagcaggcttgatgGTCAGCCCACGACGCAGTAG, ggggaccactttgtacaagaaagctgggtcTCTTTCTGACTCGTCGGAGG | Mycgr3\|92210  JGI  Ensembl |  |  | 0 % |  |  |
| 30 | ggggacaagtttgtacaaaaaagcaggcttgatgCTGAAGGACAGCCGTCAGCA, ggggaccactttgtacaagaaagctgggtcCACTGATGTCGATTGTCCGT | Mycgr3\|92356  JGI  Ensembl | KOG1216 von Willebrand factor and related coagulation proteins; | exo-alpha-sialidase activity; | 1.0805 % |  |  |
| 30 | ggggacaagtttgtacaaaaaagcaggcttgatgTCCTCTGATCTGTTCGCCGC, ggggaccactttgtacaagaaagctgggtcCCGTCCCCTCAACCCCCCCT | Mycgr3\|92948  JGI  Ensembl | KOG1187 Serine/threonine protein kinase; |  | 0 % |  |  |
| 30 | ggggacaagtttgtacaaaaaagcaggcttgatgCGCGTTACTATCCTCCTGGC, ggggaccactttgtacaagaaagctgggtcCTGGACAAGTTCGTTGCTCG | Mycgr3\|93498  JGI  Ensembl |  |  | 0 % |  |  |
| 30 | ggggacaagtttgtacaaaaaagcaggcttgatgGGTGGCATGCAAACGACCGT, ggggaccactttgtacaagaaagctgggtcCTCCTCGTCGTCAGTGTACT | Mycgr3\|94803  JGI  Ensembl |  |  | 0 % |  |  |
| 30 | ggggacaagtttgtacaaaaaagcaggcttgatgTCTACAAATCATCTCCATAC, ggggaccactttgtacaagaaagctgggtcTTTGTGATTCATATTATCAC | Mycgr3\|94814  JGI  Ensembl |  |  | 0 % |  |  |
| 30 | ggggacaagtttgtacaaaaaagcaggcttgatgGCCGAGTCGACCGATGCGCA, ggggaccactttgtacaagaaagctgggtcGAAAGCCGTGACCTTGCTGC | Mycgr3\|94965  JGI  Ensembl |  |  | 0 % |  |  |
| 30 | ggggacaagtttgtacaaaaaagcaggcttgatgACCGGAGCCGACGATGACCG, ggggaccactttgtacaagaaagctgggtcTGACCAGAACCACCCGCACG | Mycgr3\|95389  JGI  Ensembl |  |  | 0 % |  |  |
| 30 | ggggacaagtttgtacaaaaaagcaggcttgatgGACGCGAACGCGAGCGCACT, ggggaccactttgtacaagaaagctgggtcCACTCCCCCATCCGCCTCGC | Mycgr3\|95588  JGI  Ensembl |  |  | 0 % |  |  |
| 30 | ggggacaagtttgtacaaaaaagcaggcttgatgGCGCCGCCAAATTTTGCCAT, ggggaccactttgtacaagaaagctgggtcGTGCTCGGAGTTGATCGTGA | Mycgr3\|96529  JGI  Ensembl | KOG0161 Myosin class II heavy chain; | ATP binding; catalytic activity;  metabolic process; | 0 % |  |  |
| 30 | ggggacaagtttgtacaaaaaagcaggcttgatgCCGACGAACCTTACTCTCAC, ggggaccactttgtacaagaaagctgggtcGAGAGGAGTTCTTACGTAAT | Mycgr3\|96649  JGI  Ensembl | KOG1181 FOG: Low-complexity; |  | 0 % |  |  |
| 30 | ggggacaagtttgtacaaaaaagcaggcttgatgGCTATCGAACCAAATCGTAC, ggggaccactttgtacaagaaagctgggtcCGCTTCCATCGGAATGCCCT | Mycgr3\|97138  JGI  Ensembl | KOG2510 SWI-SNF chromatin-remodeling complex protein; |  | 0 % |  |  |
| 30 | ggggacaagtttgtacaaaaaagcaggcttgatgGACATCGACCGAGCTGACAG, ggggaccactttgtacaagaaagctgggtcTCCTTCGACCGCCAGACTCT | Mycgr3\|97150  JGI  Ensembl |  |  | 0 % |  |  |
| 30 | ggggacaagtttgtacaaaaaagcaggcttgatgCATTACGCTTTCAACGACTC, ggggaccactttgtacaagaaagctgggtcTGCGCCACTGATAAAGCACG | Mycgr3\|97161  JGI  Ensembl |  |  | 0 % |  |  |
| 30 | ggggacaagtttgtacaaaaaagcaggcttgatgGACGACAATCCCGAGCGCTC, ggggaccactttgtacaagaaagctgggtcGCTCGGTGAGGTCCTCAATG | Mycgr3\|97539  JGI  Ensembl | KOG2992 Nucleolar GTPase/ATPase p130; |  | 0 % |  |  |
| 30 | ggggacaagtttgtacaaaaaagcaggcttgatgGCCTACGGACTTGGTCTTCC, ggggaccactttgtacaagaaagctgggtcAACCCTTTCGGCCTTCGATT | Mycgr3\|97675  JGI  Ensembl |  |  | 0 % |  |  |
| 30 | ggggacaagtttgtacaaaaaagcaggcttgatgGTCAGACAACGCCAGCGAAA, ggggaccactttgtacaagaaagctgggtcTCCATCCAGCCATGCCTGGG | Mycgr3\|97768  JGI  Ensembl |  |  | 0 % |  |  |
| 30 | ggggacaagtttgtacaaaaaagcaggcttgatgGCACGAACGAGAGCAATGGT, ggggaccactttgtacaagaaagctgggtcCGTATAGGCGACTTTGAAAT | Mycgr3\|98064  JGI  Ensembl | KOG3265 Histone chaperone involved in gene silencing; | 3-phosphoinositide-dependent protein kinase activity; AMP-activated protein kinase activity;  DNA-dependent protein kinase activity;  GTP-dependent protein kinase activity;  JUN kinase activity;  JUN kinase kinase activity;  JUN kinase kinase kinase activity;  Janus kinase activity;  MAP kinase 1 activity;  MAP kinase 2 activity;  MAP kinase kinase activity;  MAP kinase kinase kinase kinase activity;  MAP/ERK kinase kinase activity;  MP kinase activity;  NF-kappaB-inducing kinase activity;  SAP kinase activity;  atypical protein kinase C activity;  calcium-dependent protein kinase C activity;  casein kinase I activity;  casein kinase activity;  cyclic nucleotide-dependent protein kinase activity;  cyclin-dependent protein kinase activating kinase activity;  eukaryotic translation initiation factor 2alpha kinase activity;  galactosyltransferase-associated kinase activity;  glycogen synthase kinase 3 activity;  receptor signaling protein serine/threonine kinase activity;  ribosomal protein S6 kinase activity;  transmembrane receptor protein kinase activity;  transmembrane receptor protein serine/threonine kinase activity; | 0 % |  |  |
| 30 | ggggacaagtttgtacaaaaaagcaggcttgatgAGCGGGCGACCAGCGCCCCA, ggggaccactttgtacaagaaagctgggtcATACGTCCTTGCCAAATTCC | Mycgr3\|98959  JGI  Ensembl | KOG3677 RNA polymerase I-associated factor - PAF67; |  | 0 % |  |  |
| 30 | ggggacaagtttgtacaaaaaagcaggcttgatgCAGTACCAACTCGTCGCCGC, ggggaccactttgtacaagaaagctgggtcAAGGAAGAAAGCGGCCATGG | Mycgr3\|99917  JGI  Ensembl | KOG1187 Serine/threonine protein kinase; | DNA-directed RNA polymerase I activity; DNA-directed RNA polymerase II activity;  DNA-directed RNA polymerase III activity;  DNA-directed RNA polymerase activity;  RNA polymerase activity; | 0 % |  |  |
| 30 | ggggacaagtttgtacaaaaaagcaggcttgatgTCAACCGCAGCAGCGCCGCT, ggggaccactttgtacaagaaagctgggtcCCTCCTCCTCTCCCCACAAA | Mycgr3\|101114  JGI  Ensembl | KOG0156 Cytochrome P450 CYP2 subfamily; | electron transport; flavonoid 3'-monooxygenase activity;  heme binding;  iron ion binding;  monooxygenase activity; | 0 % |  |  |
| 30 | ggggacaagtttgtacaaaaaagcaggcttgatgGCATCTGCACGTTCGAGCAA, ggggaccactttgtacaagaaagctgggtcCGACTCATCTCCCTCGTCGA | Mycgr3\|101693  JGI  Ensembl |  |  | 0 % |  |  |
| 30 | ggggacaagtttgtacaaaaaagcaggcttgatgAGATACCTACTAGCACTCCC, ggggaccactttgtacaagaaagctgggtcCAGTTCATCCCTGAAGAATT | Mycgr3\|102744  JGI  Ensembl | KOG3394 Protein OS-9; | endoplasmic reticulum; | 16.49543 % |  |  |
| 30 | ggggacaagtttgtacaaaaaagcaggcttgatgCAAGAGGTGTGGGATTTGGA, ggggaccactttgtacaagaaagctgggtcGGCCGGAATCCTCGGCGAGG | Mycgr3\|106652  JGI  Ensembl |  |  | 0 % |  |  |
| 30 | ggggacaagtttgtacaaaaaagcaggcttgatgTCAGGTCTATGGAAGCGGTC, ggggaccactttgtacaagaaagctgggtcCACGGTCGGGCATCCATCAT | Mycgr3\|107001  JGI  Ensembl |  |  | 0 % |  |  |
| 30 | ggggacaagtttgtacaaaaaagcaggcttgatgGCGGAGGAGCAGCAAGTCGC, ggggaccactttgtacaagaaagctgggtcACCAACACTATCCTTGATCA | Mycgr3\|107462  JGI  Ensembl | KOG4166 Thiamine pyrophosphate-requiring enzyme; | acetolactate synthase activity; magnesium ion binding;  thiamin pyrophosphate binding;  transferase activity; | 0 % |  |  |
| 30 | ggggacaagtttgtacaaaaaagcaggcttgatgGAGGCAATACCATATTCCGA, ggggaccactttgtacaagaaagctgggtcGTGACCTGTGCCGTTCGCAG | Mycgr3\|107669  JGI  Ensembl | KOG0053 Cystathionine beta-lyases/cystathionine gamma-synthases; | O-acetylhomoserine aminocarboxypropyltransferase activity; amino acid metabolic process;  pyridoxal phosphate binding;  transferase activity, transferring alkyl or aryl (other than methyl) groups; | 0 % |  |  |
| 30 | ggggacaagtttgtacaaaaaagcaggcttgatgGATCCAACCACACTGGTGAC, ggggaccactttgtacaagaaagctgggtcAAACGAAACATCACCCGGAC | Mycgr3\|108013  JGI  Ensembl |  |  | 0 % |  |  |
| 30 | ggggacaagtttgtacaaaaaagcaggcttgatgGCGACTCTGGACAGACTACC, ggggaccactttgtacaagaaagctgggtcAAAGTCTTCCGCCCGCCACA | Mycgr3\|108391  JGI  Ensembl |  |  | 0 % |  |  |
| 30 | ggggacaagtttgtacaaaaaagcaggcttgatgGCCAAATCCACTATCGAGAA, ggggaccactttgtacaagaaagctgggtcGCCGAGCAAGACAACTGAGC | Mycgr3\|108486  JGI  Ensembl | KOG0471 Alpha-amylase; | alpha-glucosidase activity; carbohydrate metabolic process;  catalytic activity;  cation binding; | 0 % |  |  |
| 30 | ggggacaagtttgtacaaaaaagcaggcttgatgCCAGACGAGGCGGATCCGGC, ggggaccactttgtacaagaaagctgggtcACAACAGCTGAAGTCGGCGG | Mycgr3\|108606  JGI  Ensembl |  |  | 0 % |  |  |
| 30 | ggggacaagtttgtacaaaaaagcaggcttgatgCTCTCGCACGTAGCCGACCT, ggggaccactttgtacaagaaagctgggtcGAAGCCCTTCTCCGCCTCCA | Mycgr3\|109292  JGI  Ensembl |  |  | 1.54739 % |  |  |
| 30 | ggggacaagtttgtacaaaaaagcaggcttgatgGTTTACTCGCTGAACTACCG, ggggaccactttgtacaagaaagctgggtcGAAGAAATTGCCCTTCGGCA | Mycgr3\|109387  JGI  Ensembl | KOG3589 G protein signaling regulators; |  | 0 % |  |  |
| 30 | ggggacaagtttgtacaaaaaagcaggcttgatgGTACCCGCCAATGCCGCCAT, ggggaccactttgtacaagaaagctgggtcAGCTCTTCTCTGCGACTTGT | Mycgr3\|109439  JGI  Ensembl |  |  | 0 % |  |  |
| 30 | ggggacaagtttgtacaaaaaagcaggcttgatgACTCAATCGACAACCAGCAC, ggggaccactttgtacaagaaagctgggtcCCACAACCAGCCAAACCATC | Mycgr3\|109704  JGI  Ensembl | KOG2456 Aldehyde dehydrogenase; | aldehyde dehydrogenase (NAD) activity; | 0 % |  |  |
| 30 | ggggacaagtttgtacaaaaaagcaggcttgatgTCCACCGACGGCAGTGTTCA, ggggaccactttgtacaagaaagctgggtcGCATTGCTTGGGCTTCGACA | Mycgr3\|110075  JGI  Ensembl | KOG2177 Predicted E3 ubiquitin ligase; | protein binding; zinc ion binding; | 0 % |  |  |
| 30 | ggggacaagtttgtacaaaaaagcaggcttgatgGCCGACGGCATGCTTTCCGG, ggggaccactttgtacaagaaagctgggtcACTAGGCACTTTCCTCTTCT | Mycgr3\|110106  JGI  Ensembl | KOG1474 Transcription initiation factor TFIID, subunit BDF1 and related bromodomain proteins; | intracellular; nucleic acid binding;  zinc ion binding; | 0 % |  |  |
| 30 | ggggacaagtttgtacaaaaaagcaggcttgatgGACTTCCGAAAATTAGCCAA, ggggaccactttgtacaagaaagctgggtcTTCATCATCGTCATCGCTCT | Mycgr3\|110524  JGI  Ensembl | KOG0743 AAA+-type ATPase; | ATP binding; nucleoside-triphosphatase activity;  nucleotide binding; | 0 % |  |  |
| 30 | ggggacaagtttgtacaaaaaagcaggcttgatgCTGACTCTGATCATTTTTGC, ggggaccactttgtacaagaaagctgggtcATGCCGCTCGAACGTAATCA | Mycgr3\|110773  JGI  Ensembl | KOG0157 Cytochrome P450 CYP4/CYP19/CYP26 subfamilies; | electron transport; heme binding;  iron ion binding;  monooxygenase activity;  unspecific monooxygenase activity; | 0 % |  |  |
| 30 | ggggacaagtttgtacaaaaaagcaggcttgatgCAGGGATTTTTCGCGATAGC, ggggaccactttgtacaagaaagctgggtcTCCGCACGAGCGAACTCGAT | Mycgr3\|110888  JGI  Ensembl | KOG1339 Aspartyl protease; | pepsin A activity; phosphoenolpyruvate-dependent sugar phosphotransferase system;  proteolysis;  sugar:hydrogen symporter activity; | 0 % |  |  |
| 30 | ggggacaagtttgtacaaaaaagcaggcttgatgGCTGTCCTCATCACCCTGCT, ggggaccactttgtacaagaaagctgggtcGATGTCTTTCCTTTGTCCAT | Mycgr3\|111451  JGI  Ensembl | KOG1387 Glycosyltransferase; | 1,2-dihydroxy-phenanthrene glycosyltransferase activity; 1-phenanthrol glycosyltransferase activity;  9-phenanthrol UDP-glucuronosyltransferase activity;  9-phenanthrol glycosyltransferase activity;  O antigen polymerase activity;  UDP-N-acetylglucosamine-peptide N-acetylglucosaminyltransferase activity;  UDP-glucose:glycoprotein glucosyltransferase activity;  acetylgalactosaminyltransferase activity;  acetylglucosaminyltransferase activity;  alpha(1,3)-fucosyltransferase activity;  alpha(1,6)-fucosyltransferase activity;  alpha-1,2-galactosyltransferase activity;  alpha-1,2-mannosyltransferase activity;  alpha-1,3-galactosyltransferase activity;  alpha-1,3-mannosyltransferase activity;  alpha-1,6-mannosyltransferase activity;  beta-1,4-mannosyltransferase activity;  biosynthetic process;  cellulose synthase activity;  dolichyl pyrophosphate Glc1Man9GlcNAc2 alpha-1,3-glucosyltransferase activity; dolichyl pyrophosphate Man7GlcNAc2 alpha-1,3-glucosyltransferase activity;  dolichyl pyrophosphate Man9GlcNAc2 alpha-1,3-glucosyltransferase activity;  dolichyl-phosphate-glucose-glycolipid alpha-glucosyltransferase activity;  fucosyltransferase activity;  galactosyltransferase activity;  glucosyltransferase activity;  glycolipid mannosyltransferase activity;  inositol phosphoceramide synthase activity;  lipopolysaccharide-1,6-galactosyltransferase activity;  mannosyltransferase activity;  oligosaccharyl transferase activity;  phenanthrol glycosyltransferase activity; | 0 % |  |  |
| 30 | ggggacaagtttgtacaaaaaagcaggcttgatgCACTTGGCTGGGAGTCCCGT, ggggaccactttgtacaagaaagctgggtcATGGACGCCACAATCAAAGC | Mycgr3\|111799  JGI  Ensembl |  |  | 0 % |  |  |
| 31 | ggggacaagtttgtacaaaaaagcaggcttgatgGCCTTCTTTCAAGCCCTTCC, ggggaccactttgtacaagaaagctgggtcGAAATTGAAGTCCTCAAAAA | Mycgr3\|35159  JGI  Ensembl |  |  | 100 % |  |  |
| 31 | ggggacaagtttgtacaaaaaagcaggcttgatgGTCGCTCTCGCATACACGCC, ggggaccactttgtacaagaaagctgggtcCACGAATCCCGCCGCCGTTG | Mycgr3\|35402  JGI  Ensembl | KOG0745 Putative ATP-dependent Clp-type protease (AAA+ ATPase superfamily); | ATP binding; ATPase activity;  nucleoside-triphosphatase activity;  nucleotide binding;  protein binding;  protein transport; | 100 % |  |  |
| 31 | ggggacaagtttgtacaaaaaagcaggcttgatgCCATTTCCAGCGTTCTCCAT, ggggaccactttgtacaagaaagctgggtcTGAAGCCTTGCTACCCCGCA | Mycgr3\|39717  JGI  Ensembl | KOG2458 Endoplasmic reticulum protein EP58, contains filamin rod domain and KDEL motif; |  | 2.07006 % |  |  |
| 31 | ggggacaagtttgtacaaaaaagcaggcttgatgCCGGTCCCCGTCCAGATCGC, ggggaccactttgtacaagaaagctgggtcGAGCGGCACAAAAGGCAGAA | Mycgr3\|43263  JGI  Ensembl |  |  | 0 % |  |  |
| 31 | ggggacaagtttgtacaaaaaagcaggcttgatgGAGCCTGGGACCAACTGGAC, ggggaccactttgtacaagaaagctgggtcACGACCATTCGTCGTCACAC | Mycgr3\|43383  JGI  Ensembl | KOG2614 Kynurenine 3-monooxygenase and related flavoprotein monooxygenases; | metabolic process; oxidoreductase activity;  salicylate 1-monooxygenase activity; | 100 % |  |  |
| 31 | ggggacaagtttgtacaaaaaagcaggcttgatgCTTCACAGCCGATCATGCAC, ggggaccactttgtacaagaaagctgggtcTGATTCCTCAGTCTTTTCAC | Mycgr3\|43963  JGI  Ensembl | KOG1276 Protoporphyrinogen oxidase; | electron transport; oxidoreductase activity;  porphyrin biosynthetic process;  protoporphyrinogen oxidase activity;  sugar binding; | 100 % |  |  |
| 31 | ggggacaagtttgtacaaaaaagcaggcttgatgTCCGACGAGAAAGAGCAGCG, ggggaccactttgtacaagaaagctgggtcCCTTCCTCCACCACCCTCCC | Mycgr3\|45334  JGI  Ensembl | KOG2059 Ras GTPase-activating protein; |  | 100 % |  |  |
| 31 | ggggacaagtttgtacaaaaaagcaggcttgatgAAGGTTTCTGAGACAAGCGA, ggggaccactttgtacaagaaagctgggtcTGCGTTGCTTTTCTCCGATT | Mycgr3\|45546  JGI  Ensembl | KOG0254 Predicted transporter (major facilitator superfamily); | L-arabinose isomerase activity; carbohydrate transport;  integral to membrane;  membrane;  sugar:hydrogen symporter activity;  transport;  transporter activity; | 100 % |  |  |
| 31 | ggggacaagtttgtacaaaaaagcaggcttgatgTCCGACGACGAGGTTGATCC, ggggaccactttgtacaagaaagctgggtcGTACCATATTGATCGTGTCC | Mycgr3\|47506  JGI  Ensembl | KOG2524 Cobyrinic acid a,c-diamide synthase; |  | 0.97989 % |  |  |
| 31 | ggggacaagtttgtacaaaaaagcaggcttgatgCTGACTTCGTCACCCGCTGT, ggggaccactttgtacaagaaagctgggtcCGACAAGGTCTTCATGCCCG | Mycgr3\|48195  JGI  Ensembl |  |  | 2.05696 % |  |  |
| 31 | ggggacaagtttgtacaaaaaagcaggcttgatgAAGAACTATCTCCTGCTCGC, ggggaccactttgtacaagaaagctgggtcCCCACCCTCCGCCGTACACC | Mycgr3\|48841  JGI  Ensembl | KOG0602 Neutral trehalase; | alpha,alpha-trehalase activity; trehalose metabolic process; | 100 % |  |  |
| 31 | ggggacaagtttgtacaaaaaagcaggcttgatgCAGCATCCCCGCCTCACCTT, ggggaccactttgtacaagaaagctgggtcCAACTCATCATGCCCCTCCC | Mycgr3\|49510  JGI  Ensembl |  |  | 100 % |  |  |
| 31 | ggggacaagtttgtacaaaaaagcaggcttgatgGCGTACATACTTGCCCTCTC, ggggaccactttgtacaagaaagctgggtcTGCGGCATTCGGCATCCTCG | Mycgr3\|49899  JGI  Ensembl | KOG0626 Beta-glucosidase, lactase phlorizinhydrolase, and related proteins; | beta-glucosidase activity; carbohydrate metabolic process;  hydrolase activity, hydrolyzing O-glycosyl compounds; | 100 % |  |  |
| 31 | ggggacaagtttgtacaaaaaagcaggcttgatgGCTCCCTCACACCGACCTCG, ggggaccactttgtacaagaaagctgggtcCTGCGTCTTCTTTTCGCTCG | Mycgr3\|53203  JGI  Ensembl | KOG2575 Glucosyltransferase - Alg6p; | 1,2-dihydroxy-phenanthrene glycosyltransferase activity; 1-phenanthrol glycosyltransferase activity;  9-phenanthrol UDP-glucuronosyltransferase activity;  9-phenanthrol glycosyltransferase activity;  O antigen polymerase activity;  UDP-N-acetylglucosamine-peptide N-acetylglucosaminyltransferase activity;  UDP-glucose:glycoprotein glucosyltransferase activity;  acetylgalactosaminyltransferase activity;  acetylglucosaminyltransferase activity;  alpha(1,3)-fucosyltransferase activity;  alpha(1,6)-fucosyltransferase activity;  alpha-1,2-galactosyltransferase activity;  alpha-1,2-mannosyltransferase activity;  alpha-1,3-galactosyltransferase activity;  alpha-1,3-mannosyltransferase activity;  alpha-1,6-mannosyltransferase activity;  beta-1,4-mannosyltransferase activity;  cellulose synthase activity;  dolichyl pyrophosphate Glc1Man9GlcNAc2 alpha-1,3-glucosyltransferase activity; dolichyl pyrophosphate Man7GlcNAc2 alpha-1,3-glucosyltransferase activity;  dolichyl pyrophosphate Man9GlcNAc2 alpha-1,3-glucosyltransferase activity;  dolichyl-phosphate-glucose-glycolipid alpha-glucosyltransferase activity;  fucosyltransferase activity;  galactosyltransferase activity;  glucosyltransferase activity;  glycolipid mannosyltransferase activity;  inositol phosphoceramide synthase activity;  lipopolysaccharide-1,6-galactosyltransferase activity;  mannosyltransferase activity;  oligosaccharyl transferase activity;  phenanthrol glycosyltransferase activity; | 100 % |  |  |
| 31 | ggggacaagtttgtacaaaaaagcaggcttgatgGAGCTAGAGCACGACACACA, ggggaccactttgtacaagaaagctgggtcCAAAATGTACACCTTTCTCA | Mycgr3\|53585  JGI  Ensembl | KOG0396 Uncharacterized conserved protein; |  | 100 % |  |  |
| 31 | ggggacaagtttgtacaaaaaagcaggcttgatgGCCTACCCAGCCAACACGGG, ggggaccactttgtacaagaaagctgggtcATGATCCTCCTCAATATTAA | Mycgr3\|62049  JGI  Ensembl | KOG0254 Predicted transporter (major facilitator superfamily); | L-arabinose isomerase activity; carbohydrate transport;  integral to membrane;  membrane;  sugar:hydrogen symporter activity;  transport;  transporter activity; | 100 % |  |  |
| 31 | ggggacaagtttgtacaaaaaagcaggcttgatgAGCATCGAACTCCAAGAAAC, ggggaccactttgtacaagaaagctgggtcTCCGCGTCCCCTGCCTCGTC | Mycgr3\|62271  JGI  Ensembl | KOG2863 RNA lariat debranching enzyme; | T/G mismatch-specific endonuclease activity; hydrolase activity;  hydrolase activity, acting on ester bonds;  mRNA processing;  nuclease activity;  phosphoric ester hydrolase activity; | 100 % |  |  |
| 31 | ggggacaagtttgtacaaaaaagcaggcttgatgAGCGACGAATCGCAGACCGT, ggggaccactttgtacaagaaagctgggtcCTCCGGCGAGAGCACGCCCA | Mycgr3\|63492  JGI  Ensembl | KOG0628 Aromatic-L-amino-acid/L-histidine decarboxylase; | aromatic-L-amino-acid decarboxylase activity; carboxy-lyase activity;  carboxylic acid metabolic process;  pyridoxal phosphate binding; | 100 % |  |  |
| 31 | ggggacaagtttgtacaaaaaagcaggcttgatgGCCATGTTCGAACGCCGCTC, ggggaccactttgtacaagaaagctgggtcATCCGTCCCATACCCCTGAT | Mycgr3\|64910  JGI  Ensembl |  |  | 100 % |  |  |
| 31 | ggggacaagtttgtacaaaaaagcaggcttgatgCAGGTTCTGAACCTGATTGT, ggggaccactttgtacaagaaagctgggtcATAATCGAGGTGAAAGAGGC | Mycgr3\|65440  JGI  Ensembl | KOG0471 Alpha-amylase; | alpha-amylase activity; carbohydrate metabolic process;  catalytic activity;  cation binding; | 100 % | chr\_1:1621700 A->G; |  |
| 31 | ggggacaagtttgtacaaaaaagcaggcttgatgGCGATTCCACAAAGAAAGTC, ggggaccactttgtacaagaaagctgggtcGAAATGCTCCTTCCAGAACT | Mycgr3\|65626  JGI  Ensembl | KOG0039 Ferric reductase, NADH/NADPH oxidase and related proteins; |  | 100 % | chr\_1:1949662 C->T; chr\_1:1950073 G->T; |  |
| 31 | ggggacaagtttgtacaaaaaagcaggcttgatgGCGTTCATCCTGCTGCAACT, ggggaccactttgtacaagaaagctgggtcTTGCACATAGAACATTACAA | Mycgr3\|67381  JGI  Ensembl |  |  | 100 % |  |  |
| 31 | ggggacaagtttgtacaaaaaagcaggcttgatgCCAACCGCTCCCTCCCACGC, ggggaccactttgtacaagaaagctgggtcTTGCTGCTGTTGGTATGGCA | Mycgr3\|67865  JGI  Ensembl | KOG3227 Calcium-responsive transcription coactivator; | UDP-N-acetylmuramoylalanyl-D-glutamyl-2,6-diaminopimelate-D-alanyl-D-alanine ligase activity; coenzyme F420-0 gamma-glutamyl ligase activity;  coenzyme F420-2 alpha-glutamyl ligase activity;  intracellular;  nucleic acid binding;  ribosomal S6-glutamic acid ligase activity;  zinc ion binding; | 100 % |  |  |
| 31 | ggggacaagtttgtacaaaaaagcaggcttgatgAAAGAGCGGATATTCCTGAC, ggggaccactttgtacaagaaagctgggtcCATTTTATCAAAGACCTCCG | Mycgr3\|68116  JGI  Ensembl | KOG4730 D-arabinono-1, 4-lactone oxidase; | D-arabinono-1,4-lactone oxidase activity; FAD binding;  biosynthetic process;  electron transport;  membrane;  oxidoreductase activity;  oxidoreductase activity, acting on the CH-OH group of donors, oxygen as acceptor; | 100 % |  |  |
| 31 | ggggacaagtttgtacaaaaaagcaggcttgatgGCCGCCAACGACTACTACCA, ggggaccactttgtacaagaaagctgggtcCAGCTCGCGCTTAGACGTGG | Mycgr3\|68129  JGI  Ensembl | KOG2290 Rhomboid family proteins; | integral to membrane; | 100 % |  |  |
| 31 | ggggacaagtttgtacaaaaaagcaggcttgatgGTTCACCTGGGCAAAATTGC, ggggaccactttgtacaagaaagctgggtcACAAGGCGCGTGGGTCTTCC | Mycgr3\|68234  JGI  Ensembl | KOG1383 Glutamate decarboxylase/sphingosine phosphate lyase; | carboxy-lyase activity; carboxylic acid metabolic process;  glutamate decarboxylase activity;  glutamate metabolic process;  pyridoxal phosphate binding; | 100 % |  |  |
| 31 | ggggacaagtttgtacaaaaaagcaggcttgatgGCGTCGAGAAGCGCGACCAG, ggggaccactttgtacaagaaagctgggtcATTTCGTCCCCATCGTTTCC | Mycgr3\|68850  JGI  Ensembl | KOG1287 Amino acid transporters; | amino acid transmembrane transporter activity; amino acid transport;  membrane;  transport; | 100 % |  |  |
| 31 | ggggacaagtttgtacaaaaaagcaggcttgatgGAAGTCATCAAGGGCATTCA, ggggaccactttgtacaagaaagctgggtcACTCCCCGGCCCCTCCGGCG | Mycgr3\|69014  JGI  Ensembl | KOG4472 Glycolipid 2-alpha-mannosyltransferase (alpha-1,2-mannosyltransferase); | 1,2-dihydroxy-phenanthrene glycosyltransferase activity; 1-phenanthrol glycosyltransferase activity;  9-phenanthrol UDP-glucuronosyltransferase activity;  9-phenanthrol glycosyltransferase activity;  O antigen polymerase activity;  UDP-N-acetylglucosamine-peptide N-acetylglucosaminyltransferase activity;  UDP-glucose:glycoprotein glucosyltransferase activity;  acetylgalactosaminyltransferase activity;  acetylglucosaminyltransferase activity;  alpha(1,3)-fucosyltransferase activity;  alpha(1,6)-fucosyltransferase activity;  alpha-1,2-galactosyltransferase activity;  alpha-1,2-mannosyltransferase activity;  alpha-1,3-galactosyltransferase activity;  alpha-1,3-mannosyltransferase activity;  alpha-1,6-mannosyltransferase activity;  beta-1,4-mannosyltransferase activity;  cellulose synthase activity;  dolichyl pyrophosphate Glc1Man9GlcNAc2 alpha-1,3-glucosyltransferase activity; dolichyl pyrophosphate Man7GlcNAc2 alpha-1,3-glucosyltransferase activity;  dolichyl pyrophosphate Man9GlcNAc2 alpha-1,3-glucosyltransferase activity;  dolichyl-phosphate-glucose-glycolipid alpha-glucosyltransferase activity;  fucosyltransferase activity;  galactosyltransferase activity;  glucosyltransferase activity;  glycolipid 2-alpha-mannosyltransferase activity;  glycolipid mannosyltransferase activity;  inositol phosphoceramide synthase activity;  lipopolysaccharide-1,6-galactosyltransferase activity;  mannosyltransferase activity;  membrane;  oligosaccharyl transferase activity;  phenanthrol glycosyltransferase activity;  protein amino acid glycosylation; | 100 % |  |  |
| 31 | ggggacaagtttgtacaaaaaagcaggcttgatgGAATTGGAACGCCATACTCC, ggggaccactttgtacaagaaagctgggtcACCATCCAAATCAATAAAAC | Mycgr3\|69021  JGI  Ensembl | KOG0254 Predicted transporter (major facilitator superfamily); | L-arabinose isomerase activity; carbohydrate transport;  catalytic activity;  integral to membrane;  membrane;  sugar:hydrogen symporter activity;  transport;  transporter activity; | 0 % |  |  |
| 31 | ggggacaagtttgtacaaaaaagcaggcttgatgTCCGACCTCGAAAAGAAACC, ggggaccactttgtacaagaaagctgggtcCCAAATATGTCTCACCCGCG | Mycgr3\|69215  JGI  Ensembl | KOG2533 Permease of the major facilitator superfamily; | integral to membrane; transport; transporter activity; | 100 % |  |  |
| 31 | ggggacaagtttgtacaaaaaagcaggcttgatgCCATTTACCGCCCCATCATG, ggggaccactttgtacaagaaagctgggtcCAACTTCACCCCCGCCTTCC | Mycgr3\|69852  JGI  Ensembl | KOG1176 Acyl-CoA synthetase; | 4-coumarate-CoA ligase activity; catalytic activity;  metabolic process; | 100 % |  |  |
| 31 | ggggacaagtttgtacaaaaaagcaggcttgatgTTGCACTCATGGCAATCACT, ggggaccactttgtacaagaaagctgggtcTACCAGAAGGATTGAGAGCA | Mycgr3\|69907  JGI  Ensembl | KOG0471 Alpha-amylase; | alpha-amylase activity; carbohydrate metabolic process;  catalytic activity;  cation binding; | 3.70153 % |  |  |
| 31 | ggggacaagtttgtacaaaaaagcaggcttgatgGCGGTCGAAGTCGCATCGGA, ggggaccactttgtacaagaaagctgggtcCGCTGCTTCCGTGACCGCGG | Mycgr3\|70063  JGI  Ensembl | KOG2491 Nuclear matrix protein; |  | 100 % |  |  |
| 31 | ggggacaagtttgtacaaaaaagcaggcttgatgGTTCACACTCTACTCTTTGC, ggggaccactttgtacaagaaagctgggtcGTCGGCAAACAAATCTGGCG | Mycgr3\|70283  JGI  Ensembl | KOG0156 Cytochrome P450 CYP2 subfamily; | electron transport; heme binding;  iron ion binding;  monooxygenase activity; | 100 % |  |  |
| 31 | ggggacaagtttgtacaaaaaagcaggcttgatgAACACTCGCCAGGCGATCCA, ggggaccactttgtacaagaaagctgggtcCTCGACAAACTGCCTCGTCA | Mycgr3\|70550  JGI  Ensembl | KOG2111 Uncharacterized conserved protein, contains WD40 repeats; |  | 100 % | chr\_3:3367715 G->A; |  |
| 31 | ggggacaagtttgtacaaaaaagcaggcttgatgCCGGGCCCAAGTGTTCCCGG, ggggaccactttgtacaagaaagctgggtcAACCATGCCCGCCGGCCTCG | Mycgr3\|73051  JGI  Ensembl | KOG2688 Transcription-associated recombination protein - Thp1p; |  | 100 % |  |  |
| 31 | ggggacaagtttgtacaaaaaagcaggcttgatgGCATCAACAAAACAGCACTC, ggggaccactttgtacaagaaagctgggtcAATTTCGTGGCTCCCCCTCT | Mycgr3\|73238  JGI  Ensembl |  |  | 100 % |  |  |
| 31 | ggggacaagtttgtacaaaaaagcaggcttgatgTCAGATCCTCCCTCCCAAAG, ggggaccactttgtacaagaaagctgggtcCCGCTTCGCAAACCCAGGCG | Mycgr3\|74143  JGI  Ensembl |  |  | 100 % |  |  |
| 31 | ggggacaagtttgtacaaaaaagcaggcttgatgTCGTATTCAGAGAAAGCCGC, ggggaccactttgtacaagaaagctgggtcTAGACCGTCTCTTGATCTGT | Mycgr3\|74261  JGI  Ensembl | KOG0254 Predicted transporter (major facilitator superfamily); | L-arabinose isomerase activity; carbohydrate transport;  integral to membrane;  membrane;  sugar:hydrogen symporter activity;  transport;  transporter activity; | 1.23001 % |  |  |
| 31 | ggggacaagtttgtacaaaaaagcaggcttgatgATAAATCCCAACGATCTGAA, ggggaccactttgtacaagaaagctgggtcGGCGACGACACCTCGTTCCT | Mycgr3\|74334  JGI  Ensembl | KOG0157 Cytochrome P450 CYP4/CYP19/CYP26 subfamilies; | electron transport; heme binding;  iron ion binding;  monooxygenase activity; | 1.03359 % |  |  |
| 31 | ggggacaagtttgtacaaaaaagcaggcttgatgACAGCCAGAGCACAACCATC, ggggaccactttgtacaagaaagctgggtcCTGCATTGAGTTCTCCGACT | Mycgr3\|74399  JGI  Ensembl | KOG2614 Kynurenine 3-monooxygenase and related flavoprotein monooxygenases; | metabolic process; oxidoreductase activity; | 1.42566 % |  |  |
| 31 | ggggacaagtttgtacaaaaaagcaggcttgatgGACGTCAATGCAAAGCTCGA, ggggaccactttgtacaagaaagctgggtcGTGATCTACATCCTCCCATG | Mycgr3\|75137  JGI  Ensembl | KOG1284 Bifunctional GTP cyclohydrolase II/3,4-dihydroxy-2butanone-4-phosphate synthase; | GTP cyclohydrolase II activity; riboflavin biosynthetic process; | 100 % |  |  |
| 31 | ggggacaagtttgtacaaaaaagcaggcttgatgGAGAAGTCCTCCTTTCCCGC, ggggaccactttgtacaagaaagctgggtcGTCCTCCCACCTCTCACTCA | Mycgr3\|75537  JGI  Ensembl | KOG4389 Acetylcholinesterase/Butyrylcholinesterase; | carboxylesterase activity; | 100 % |  |  |
| 31 | ggggacaagtttgtacaaaaaagcaggcttgatgCGAGGCCACACCCACACCCA, ggggaccactttgtacaagaaagctgggtcCTTCCTCTTCCCTCGTCCCG | Mycgr3\|75924  JGI  Ensembl |  |  | 100 % |  |  |
| 31 | ggggacaagtttgtacaaaaaagcaggcttgatgTCTCAACGCGACCCCGATCT, ggggaccactttgtacaagaaagctgggtcCAACTCCTTCTTCAAAAAGT | Mycgr3\|76143  JGI  Ensembl | KOG1485 Mitochondrial Fe2+ transporter MMT1 and related transporters (cation diffusion facilitator superfamily); | cation transmembrane transporter activity; cation transport;  membrane; | 100 % |  |  |
| 31 | ggggacaagtttgtacaaaaaagcaggcttgatgAAGGGCCATATGGGTCACAT, ggggaccactttgtacaagaaagctgggtcCCATCCATATGTCTCGGTGA | Mycgr3\|76227  JGI  Ensembl |  |  | 1.16587 % |  |  |
| 31 | ggggacaagtttgtacaaaaaagcaggcttgatgGCCTGGTTGCTGGGGCTGGG, ggggaccactttgtacaagaaagctgggtcCACATGCATGCTCGCAAAGC | Mycgr3\|76449  JGI  Ensembl | KOG2771 Subunit of tRNA-specific adenosine-34 deaminase; | hydrolase activity; zinc ion binding; | 100 % | chr\_10:1022264 G->A; |  |
| 31 | ggggacaagtttgtacaaaaaagcaggcttgatgCATTTCTCGAAAACCCTCAT, ggggaccactttgtacaagaaagctgggtcGCCCTTCACCTTCTTCATCT | Mycgr3\|77203  JGI  Ensembl |  | beta-lactam antibiotic catabolic process; beta-lactamase activity;  response to antibiotic; | 100 % | chr\_12:200268 C->T; chr\_12:199980 G->A; |  |
| 31 | ggggacaagtttgtacaaaaaagcaggcttgatgTCGACCATGTCGTCCAACAG, ggggaccactttgtacaagaaagctgggtcTGCCAAATCCGACTTCGCTG | Mycgr3\|85426  JGI  Ensembl | KOG1289 Amino acid transporters; | amino acid transmembrane transporter activity; amino acid transport;  membrane;  transport; | 4.1365 % |  |  |
| 31 | ggggacaagtttgtacaaaaaagcaggcttgatgCCGCGACATGGCATGCCCGA, ggggaccactttgtacaagaaagctgggtcGATCGCAGTCTCGACAAACG | Mycgr3\|88642  JGI  Ensembl |  |  | 100 % |  |  |
| 31 | ggggacaagtttgtacaaaaaagcaggcttgatgCATTTGTTTTCGATCCTGGC, ggggaccactttgtacaagaaagctgggtcAGCTGTGAAGTGAGGTGTGA | Mycgr3\|88650  JGI  Ensembl | KOG2693 Putative zinc transporter; |  | 100 % |  |  |
| 31 | ggggacaagtttgtacaaaaaagcaggcttgatgGCTTCTCCCTTCGACGTCGA, ggggaccactttgtacaagaaagctgggtcGACGCTCGTTGAAAACTCAC | Mycgr3\|89242  JGI  Ensembl |  |  | 100 % |  |  |
| 31 | ggggacaagtttgtacaaaaaagcaggcttgatgGGAGCTTCAGCGCTGAAGGG, ggggaccactttgtacaagaaagctgggtcATAACGTTGCATGCTTTTGC | Mycgr3\|89301  JGI  Ensembl |  |  | 100 % |  |  |
| 31 | ggggacaagtttgtacaaaaaagcaggcttgatgCCTCCCTTCCTCTCCCTCCC, ggggaccactttgtacaagaaagctgggtcTCTCATTCGCTGGCGTTGCC | Mycgr3\|90345  JGI  Ensembl | KOG1738 Membrane-associated guanylate kinase-interacting protein/connector enhancer of KSR-like; |  | 100 % |  |  |
| 31 | ggggacaagtttgtacaaaaaagcaggcttgatgGAGAATTTGTTCAGCGTTGC, ggggaccactttgtacaagaaagctgggtcGCCCAAAGGTTGACCGTTGC | Mycgr3\|90360  JGI  Ensembl | KOG0725 Reductases with broad range of substrate specificities; | 3-oxoacyl-[acyl-carrier-protein] reductase activity; metabolic process;  oxidoreductase activity; | 100 % |  |  |
| 31 | ggggacaagtttgtacaaaaaagcaggcttgatgGCCTGGGCCACTTGGTCCAA, ggggaccactttgtacaagaaagctgggtcTCGTCCTCTTCCGGTAACAT | Mycgr3\|90760  JGI  Ensembl |  |  | 100 % |  |  |
| 31 | ggggacaagtttgtacaaaaaagcaggcttgatgACGCTATTCCGCGACACAAC, ggggaccactttgtacaagaaagctgggtcCATCGTCCTCTCCTCGGCTT | Mycgr3\|90853  JGI  Ensembl |  |  | 100 % | chr\_2:2191914 T->C; |  |
| 31 | ggggacaagtttgtacaaaaaagcaggcttgatgGCCGCGCCTCGAGACACGAT, ggggaccactttgtacaagaaagctgggtcATCCTCGGTGAAGGGAGAGT | Mycgr3\|91020  JGI  Ensembl |  |  | 100 % | chr\_2:2719954 G->A; |  |
| 31 | ggggacaagtttgtacaaaaaagcaggcttgatgCCCTCGACAGGTCAATTGCA, ggggaccactttgtacaagaaagctgggtcCGCCTCACGATGAACATAGA | Mycgr3\|91971  JGI  Ensembl | KOG1337 N-methyltransferase; |  | 100 % |  |  |
| 31 | ggggacaagtttgtacaaaaaagcaggcttgatgAAAGGGAGGGAGGATGTCGA, ggggaccactttgtacaagaaagctgggtcAATATCCATATCGATATCCC | Mycgr3\|92083  JGI  Ensembl | KOG4207 Predicted splicing factor, SR protein superfamily; | nucleic acid binding; | 2.08 % |  |  |
| 31 | ggggacaagtttgtacaaaaaagcaggcttgatgTCCAACTTCGTGCTCTTGCT, ggggaccactttgtacaagaaagctgggtcTCTCTCCATCATCCCCGCCG | Mycgr3\|92648  JGI  Ensembl | KOG1721 FOG: Zn-finger; | intracellular; nucleic acid binding;  zinc ion binding; | 100 % |  |  |
| 31 | ggggacaagtttgtacaaaaaagcaggcttgatgCTGGCTGACATCGGCGCTCA, ggggaccactttgtacaagaaagctgggtcCGTGCCCTCTGCATCAAATG | Mycgr3\|92654  JGI  Ensembl |  |  | 100 % |  |  |
| 31 | ggggacaagtttgtacaaaaaagcaggcttgatgCGTCAACGGATTGCTCCCAT, ggggaccactttgtacaagaaagctgggtcAAACAGACTTGTCAAACCCG | Mycgr3\|94266  JGI  Ensembl | KOG0436 Methionyl-tRNA synthetase; | ATP binding; aminoacyl-tRNA ligase activity;  cytoplasm;  methionine-tRNA ligase activity;  methionyl-tRNA aminoacylation;  nucleotide binding;  tRNA aminoacylation for protein translation;  translation; | 100 % |  |  |
| 31 | ggggacaagtttgtacaaaaaagcaggcttgatgTCAGAAATCGCTCCTGAAAG, ggggaccactttgtacaagaaagctgggtcGTGTCGCACTCTTCTCCCGC | Mycgr3\|94280  JGI  Ensembl | KOG0548 Molecular co-chaperone STI1; |  | 100 % |  |  |
| 31 | ggggacaagtttgtacaaaaaagcaggcttgatgGAAGACAATTTCCGCACCTG, ggggaccactttgtacaagaaagctgggtcGTCCGCGCTGTGCGCCAAAC | Mycgr3\|95742  JGI  Ensembl |  |  | 100 % |  |  |
| 31 | ggggacaagtttgtacaaaaaagcaggcttgatgGCGCCCGTTCTGCCCGGCCT, ggggaccactttgtacaagaaagctgggtcACTCCTCTCCAGTTCATGCC | Mycgr3\|96102  JGI  Ensembl |  |  | 100 % |  |  |
| 31 | ggggacaagtttgtacaaaaaagcaggcttgatgTCCTCATCCTCGACCCCACC, ggggaccactttgtacaagaaagctgggtcGCTGCTCATCTCGCTGAATG | Mycgr3\|96703  JGI  Ensembl |  |  | 26.66667 % |  |  |
| 31 | ggggacaagtttgtacaaaaaagcaggcttgatgGCCTCTGGTGTGTTTGCCTG, ggggaccactttgtacaagaaagctgggtcCCCAGGCGGCGTAACACACT | Mycgr3\|96877  JGI  Ensembl |  |  | 1.04603 % |  |  |
| 31 | ggggacaagtttgtacaaaaaagcaggcttgatgACAAGCCAGGCGCCAGTTCT, ggggaccactttgtacaagaaagctgggtcGCTGTGGAGACCAAGTGAAC | Mycgr3\|97748  JGI  Ensembl |  |  | 100 % |  |  |
| 31 | ggggacaagtttgtacaaaaaagcaggcttgatgATCGAAGAGCTTGAAGATGT, ggggaccactttgtacaagaaagctgggtcTGAAGAAGCGTAGTTGAAAG | Mycgr3\|97836  JGI  Ensembl |  |  | 100 % | chr\_17:453275 C->T; |  |
| 31 | ggggacaagtttgtacaaaaaagcaggcttgatgGCAGGACTCCATGTCTCCTT, ggggaccactttgtacaagaaagctgggtcGCTGTCGGATTCGAGGAGGA | Mycgr3\|98024  JGI  Ensembl | KOG1911 Heterochromatin-associated protein HP1 and related CHROMO domain proteins; |  | 100 % | chr\_20:136531 T->C; chr\_20:136946 A->G; |  |
| 31 | ggggacaagtttgtacaaaaaagcaggcttgatgGTGTCCCACCACGCGCCGCC, ggggaccactttgtacaagaaagctgggtcTATTTCTTCCCACGGCAAAT | Mycgr3\|98251  JGI  Ensembl |  |  | 100 % | chr\_1:1315067 C->A; |  |
| 31 | ggggacaagtttgtacaaaaaagcaggcttgatgTGCGGCATCTTCTGCTCGAT, ggggaccactttgtacaagaaagctgggtcTGTCAGCACGTGTGTGCCTT | Mycgr3\|98919  JGI  Ensembl | KOG0573 Asparagine synthase; |  | 100 % | chr\_1:5825232 C->T; |  |
| 31 | ggggacaagtttgtacaaaaaagcaggcttgatgGGCATTGACGAAACGGAAGT, ggggaccactttgtacaagaaagctgggtcCAGTGTGTATCGGAAGTCTG | Mycgr3\|100283  JGI  Ensembl | KOG2533 Permease of the major facilitator superfamily; |  | 100 % |  |  |
| 31 | ggggacaagtttgtacaaaaaagcaggcttgatgGCCGACATTGCGACTTCCAA, ggggaccactttgtacaagaaagctgggtcTGCGCTGGTAGTAGCCTCCG | Mycgr3\|100450  JGI  Ensembl | KOG4297 C-type lectin; |  | 100 % |  |  |
| 31 | ggggacaagtttgtacaaaaaagcaggcttgatgTTTTTGTTACGCCAATTGGC, ggggaccactttgtacaagaaagctgggtcGGCTTCGGGATACGCACACT | Mycgr3\|100496  JGI  Ensembl | KOG2499 Beta-N-acetylhexosaminidase; | beta-N-acetylglucosaminidase activity; beta-N-acetylhexosaminidase activity;  carbohydrate metabolic process; | 100 % | chr\_6:1249634 A->T; |  |
| 31 | ggggacaagtttgtacaaaaaagcaggcttgatgGATATCACACAGTTCATTGC, ggggaccactttgtacaagaaagctgggtcTCGCCCAAACCCAAACCAGC | Mycgr3\|101429  JGI  Ensembl | KOG2460 Signal recognition particle, subunit Srp68; | metallopeptidase activity; proteolysis; zinc ion binding; | 100 % |  |  |
| 31 | ggggacaagtttgtacaaaaaagcaggcttgatgTCAAACATTCCAGGCTTCCG, ggggaccactttgtacaagaaagctgggtcCTCGCCCTTCATACCACCCC | Mycgr3\|101579  JGI  Ensembl | KOG2806 Chitinase; | carbohydrate metabolic process; chitin catabolic process;  chitinase activity;  endochitinase activity;  hydrolase activity, hydrolyzing O-glycosyl compounds; | 100 % |  |  |
| 31 | ggggacaagtttgtacaaaaaagcaggcttgatgTCCTCCCTCGCCGCCCGCAG, ggggaccactttgtacaagaaagctgggtcCAGAATCTTCTCCTTGACGT | Mycgr3\|102306  JGI  Ensembl |  |  | 50.14409 % |  |  |
| 31 | ggggacaagtttgtacaaaaaagcaggcttgatgAAGGTCGCAGCGTCGGCCAT, ggggaccactttgtacaagaaagctgggtcAAACCACTCACCGCCCAACC | Mycgr3\|103135  JGI  Ensembl | KOG1282 Serine carboxypeptidases (lysosomal cathepsin A); | carboxypeptidase C activity; proteolysis; serine carboxypeptidase activity; | 100 % |  |  |
| 31 | ggggacaagtttgtacaaaaaagcaggcttgatgGACGTTCTCTCCGCATCTTC, ggggaccactttgtacaagaaagctgggtcAGTAGTATCACCACCATGAC | Mycgr3\|103568  JGI  Ensembl | KOG1414 Transcriptional activator FOSB/c-Fos and related bZIP transcription factors; | nucleus; protein dimerization activity;  regulation of transcription, DNA-dependent;  sequence-specific DNA binding;  transcription factor activity; | 100 % |  |  |
| 31 | ggggacaagtttgtacaaaaaagcaggcttgatgAAGTACGCTGGGATTCCCCT, ggggaccactttgtacaagaaagctgggtcCAAGGCAGCGAGAGCAATGG | Mycgr3\|105608  JGI  Ensembl | KOG3599 Ca2+-modulated nonselective cation channel polycystin; | diacylglycerol kinase activity; | 100 % |  |  |
| 31 | ggggacaagtttgtacaaaaaagcaggcttgatgAGCGCCAATGGCAACACCGA, ggggaccactttgtacaagaaagctgggtcGCTGTACAGCTTCTTCTCCA | Mycgr3\|106783  JGI  Ensembl | KOG2550 IMP dehydrogenase/GMP reductase; | GMP biosynthetic process; IMP dehydrogenase activity;  catalytic activity; | 100 % |  |  |
| 31 | ggggacaagtttgtacaaaaaagcaggcttgatgGAGTTGAGCGCGGCACCGAA, ggggaccactttgtacaagaaagctgggtcGTGGTAATCGGCAGATGGAA | Mycgr3\|107234  JGI  Ensembl |  |  | 100 % |  |  |
| 31 | ggggacaagtttgtacaaaaaagcaggcttgatgGATACCTCGGAGATCTTCTC, ggggaccactttgtacaagaaagctgggtcATCCGTCACTGATCCAGGCT | Mycgr3\|107825  JGI  Ensembl | KOG3846 L-kynurenine hydrolase; | NAD biosynthetic process; cytoplasm; kynureninase activity;  pyridoxal phosphate binding;  tryptophan catabolic process; | 0 % |  |  |
| 31 | ggggacaagtttgtacaaaaaagcaggcttgatgGCGTCCAGAGCTGCCGTCCG, ggggaccactttgtacaagaaagctgggtcCTCTCGTGACACATCGCGGC | Mycgr3\|107974  JGI  Ensembl | KOG2495 NADH-dehydrogenase (ubiquinone); | FAD binding; NADH dehydrogenase activity;  oxidoreductase activity; | 0 % |  |  |
| 31 | ggggacaagtttgtacaaaaaagcaggcttgatgACATCCGATTCTCTCTACGA, ggggaccactttgtacaagaaagctgggtcAGTGAATCGCTTAATCTTTC | Mycgr3\|108217  JGI  Ensembl |  |  | 1.15005 % |  |  |
| 31 | ggggacaagtttgtacaaaaaagcaggcttgatgCGCCGCTCAGCCGGCATCCT, ggggaccactttgtacaagaaagctgggtcCGGCTCGTTCACTGCTCCAT | Mycgr3\|108584  JGI  Ensembl |  |  | 100 % |  |  |
| 31 | ggggacaagtttgtacaaaaaagcaggcttgatgGAAGTTGCTGGAGTGGTACT, ggggaccactttgtacaagaaagctgggtcGAGTATTGCAGCGGTTCGGG | Mycgr3\|108587  JGI  Ensembl |  |  | 100 % |  |  |
| 31 | ggggacaagtttgtacaaaaaagcaggcttgatgGCAGTGCACTCGCTGTTAGC, ggggaccactttgtacaagaaagctgggtcAGATCCGCCTTTCTGCTTTC | Mycgr3\|109111  JGI  Ensembl | KOG1238 Glucose dehydrogenase/choline dehydrogenase/mandelonitrile lyase (GMC oxidoreductase family); | FAD binding; choline dehydrogenase activity;  oxidoreductase activity, acting on CH-OH group of donors; | 100 % |  |  |
| 31 | ggggacaagtttgtacaaaaaagcaggcttgatgGCCAAGCTCAAGGAGAGTAC, ggggaccactttgtacaagaaagctgggtcCATCCCATCAAACCCGTCCC | Mycgr3\|109156  JGI  Ensembl | KOG3696 Aspartyl beta-hydroxylase; |  | 100 % |  |  |
| 31 | ggggacaagtttgtacaaaaaagcaggcttgatgTTCCTCCGGCAGCTAGACTT, ggggaccactttgtacaagaaagctgggtcAGACACTATCTGAGGAGGAG | Mycgr3\|109253  JGI  Ensembl | KOG2760 Vacuolar sorting protein VPS36; | electron carrier activity; electron transport;  intracellular;  metal ion binding;  zinc ion binding; | 1.03655 % |  |  |
| 31 | ggggacaagtttgtacaaaaaagcaggcttgatgGCGGCGGTCGCGAATCAGCA, ggggaccactttgtacaagaaagctgggtcGTGACGTGTGCCGTTGATGT | Mycgr3\|110088  JGI  Ensembl | KOG1850 Myosin-like coiled-coil protein; |  | 100 % |  |  |
| 31 | ggggacaagtttgtacaaaaaagcaggcttgatgCTGTCCTTCCTCACCATCAT, ggggaccactttgtacaagaaagctgggtcCACCCTCTCATGCGGCACCA | Mycgr3\|110820  JGI  Ensembl | KOG4157 beta-1,6-N-acetylglucosaminyltransferase, contains WSC domain; |  | 100 % |  |  |
| 31 | ggggacaagtttgtacaaaaaagcaggcttgatgGAGTCGGTGATGCGGCCGAG, ggggaccactttgtacaagaaagctgggtcGGAAGCAAAGCCAACTTTCC | Mycgr3\|111361  JGI  Ensembl |  |  | 100 % |  |  |
| 31 | ggggacaagtttgtacaaaaaagcaggcttgatgGCTCGGATATTCGCAGAGGA, ggggaccactttgtacaagaaagctgggtcTTGCGCTGGAGTCACGCCGT | Mycgr3\|111656  JGI  Ensembl |  |  | 100 % |  |  |
| 32 | ggggacaagtttgtacaaaaaagcaggcttgatgGCAGACACATGCGACCAACC, ggggaccactttgtacaagaaagctgggtcGGACACCGGCGCATAGCGAC | Mycgr3\|32960  JGI  Ensembl | KOG0255 Synaptic vesicle transporter SVOP and related transporters (major facilitator superfamily); | integral to membrane; transport; transporter activity; | 99.9001 % |  |  |
| 32 | ggggacaagtttgtacaaaaaagcaggcttgatgGGTATCATCTCGATCCTTCT, ggggaccactttgtacaagaaagctgggtcCAACCTCGGATAAAGCCTCG | Mycgr3\|38151  JGI  Ensembl | KOG0157 Cytochrome P450 CYP4/CYP19/CYP26 subfamilies; | alkane 1-monooxygenase activity; electron transport;  heme binding;  iron ion binding;  monooxygenase activity; | 100 % |  |  |
| 32 | ggggacaagtttgtacaaaaaagcaggcttgatgAAACTCCTCACCAGTCTAGC, ggggaccactttgtacaagaaagctgggtcCAGCCCCGAATCATGCTTGG | Mycgr3\|38161  JGI  Ensembl | KOG1263 Multicopper oxidases; | copper ion binding; laccase activity;  oxidoreductase activity; | 100 % |  |  |
| 32 | ggggacaagtttgtacaaaaaagcaggcttgatgGAAGTTGACGAAAAGTCCGA, ggggaccactttgtacaagaaagctgggtcGCAGTGCCCTCGACGCTCGC | Mycgr3\|39188  JGI  Ensembl | KOG2533 Permease of the major facilitator superfamily; | integral to membrane; transport; transporter activity; | 2.134 % |  |  |
| 32 | ggggacaagtttgtacaaaaaagcaggcttgatgGATCCCTCAACACCACAGAC, ggggaccactttgtacaagaaagctgggtcTGCCGTCGGCAAGCTTGGGT | Mycgr3\|40797  JGI  Ensembl | KOG1721 FOG: Zn-finger; | intracellular; nucleic acid binding;  zinc ion binding; | 100 % |  | ``` 476	CATCCTACTACACCAACGGCCACGACAACTCCATGAGCAACCCATACCCGAGCTACCAGCCGAGCATGGCCAATGGCGCCGGGGCTGGAGGCTACGCCCAAACCGCTGGCAACGGTCAATACGCTGGATACGGCTCCGTCAGCTACCCCAACAGCTCTGCCATGGCCGACATACAGTCCATGGACACCCGCCGTCGTGCTATCGAGGCGCTCAACGACTTCCTCGGCGACATCAAGCGTCGTGCGATTGATCCTAGCAGCTACTACGACGTCGGTCACAGACTCCAAGGCAATGCGCTGCCTCTTCCCGTCCAATCTGGCACCGGCTACAACACTGGATACAACAACAGCGGATACACCGGCGGCAACTTCAACGCTGCGAGCCTCCTTGAGTCATTTAACCACGGCGGCATGAACGGAGGCGGTGGACAAGGTGGTGTTGCCACTGGAGGCTACTCTCTCCCACTTCCCAACGCCCGTACCAAGGGCGACCTCCAAGACATCGACCGCTTCCTCGAGCAGCTGCAAGCAACGGTCT-AC-GAGACCAGCAAGGGCTAC	1032 	\|\|\|\|\|\|\|\|\|\|\|\|\|\|\|\|\|\|\|\|\|\|\|\|\|\|\|\|\|\|\|\|\|\|\|\|\|\|\|\|\|\|\|\|\|\|\|\|\|\|\|\|\|\|\|\|\|\|\|\|\|\|\|\|\|\|\|\|\|\|\|\|\|\|\|\|\|\|\|\|\|\|\|\|\|\|\|\|\|\|\|\|\|\|\|\|\|\|\|\|\|\|\|\|\|\|\|\|\|\|\|\|\|\|\|\|\|\|\|\|\|\|\|\|\|\|\|\|\|\|\|\|\|\|\|\|\|\|\|\|\|\|\|\|\|\|\|\|\|\|\|\|\|\|\|\|\|\|\|\|\|\|\|\|\|\|\|\|\|\|\|\|\|\|\|\|\|\|\|\|\|\|\|\|\|\|\|\|\|\|\|\|\|\|\|\|\|\|\|\|\|\|\|\|\|\|\|\|\|\|\|\|\|\|\|\|\|\|\|\|\|\|\|\|\|\|\|\|\|\|\|\|\|\|\|\|\|\|\|\|\|\|\|\|\|\|\|\|\|\|\|\|\|\|\|\|\|\|\|\|\|\|\|\|\|\|\|\|\|\|\|\|\|\|\|\|\|\|\|\|\|\|\|\|\|\|\|\|\|\|\|\|\|\|\|\|\|\|\|\|\|\|\|\|\|\|\|\|\|\|\|\|\|\|\|\|\|\|\|\|\|\|\|\|\|\|\|\|\| \|\|\|\|\|\|\|\|\|\|\|\|\|\|\|\|\|\|\|\|\|\|\|\|\|\|\|\|\|\|\|\|\|\|\|\|\|\|\|\|\|\|\|\|\|\|\|\|\|\|\|\|\|\|\|\|\|\|\|\|\|\|\|\|\|\|\|\|\|\|\|\|\|\|\|\|\|\|\|\|\|\|\| \|\|\|\|\|\|\|\|\|\|\|\|\|\|\|\|\|\| \| \|\|\|\|\|\|\|\|\|\|\|\|\|\|\|\|\|\|\|\|\|\|\|\|\| \|\|\|\|\|\|\|\|\|\|\|\|\| \|\|\|\|\|\|\|\|\|\|\|\|\|\|\|\|\|\|\|\|\| \|\|\|\|\|\|\|\| \|\|\|\|\|\|\|\|\|\|\|\|\|\|\|\|\|\|\|\| \|\| \|\|\|\|\|\|\| \|\| \|\|\|\|\|\|\|\|\|\|\|\|\|\|\|\|\|\| 	CATCCTACTACACCAACGGCCACGACAACTCCATGAGCAACCCATACCCGAGCTACCAGCCGAGCATGGCCAATGGCGCCGGGGCTGGAGGCTACGCCCAAACCGCTGGCAACGGTCAATACGCTGGATACGGCTCCGTCAGCTACCCCAACAGCTCTGCCATGGCCGACATACAGTCCATGGACACCCGCCGTCGTGCTATCGAGGCGCTCAACGACTTCCTCGGCGACATCAAGCGTCGTGCGATTGATCCTAGCAGCTACTACGACGTCGGTCACAGACTCCAAGGCAATGCGCTGCCTCTTCCCGTCCAATCTGGCACCGGCTAC-ACACTGGATACAACAACAGCGGATACACCGGCGGCAACTTCAACGCTGCGAGCCTCCTTGAGTCATTTAACCACGGCGGCATG-ACGGAGGCGGTGGACAAG-T-GTGTTGCCACTGGAGGCTACTCTCT-CCACTTCCCAACG-CCGTACCAAGGGCGACCTCCA-GACATCGA-CGCTTCCTCGAGCAGCTGCA-GCCACGGTCTTACGGAGACCAGCAAGGGCTAC ```      ``` 1	ATGGATCCCTCAACACCACAGACGGTCGCGCAGGCCGTGCAAGCAGCCACGCAGGCCGCCGCGCATCAGCCCTCACCTCCCGTGCAGGCACCGTCGCACTCCACCTCGACCATCGACAACCTCACATGCCAATGGCAAGGTTGCGGAGAGCGATGCGACAACGCCGAAGCCCTCTATGACCACGTCTGCGAGCGTCACGTCGGTCGCAAGAGCACCAACAACCTCAACCTCACCTGCCAGTGGGGAGCCTGCCGAACCACCACCGTCAAACGCGACCACATCACCTCCCACATCCGCGTACATGTCCCGCTCAAACCACACAAGTGCGACTTTTGCGGCAAGGCCTTCAAGAGACCGCAGGATCTGAAGAAGCATGTCAAGACACACGCCGATGACAGCGTGTTGATGAACTCGCCGCATCAGGGACAAAATGGAAGAGGTGCCAGCGCAGGCATCGGACAACAGAATGGCAAGC	475 	\|\|\|\|\|\|\|\|\|\|\|\|\|\|\|\|\|\|\|\|\|\|\|\|\|\|\|\|\|\|\|\|\|\|\|\|\|\|\|\|\|\|\|\|\|\|\|\|\|\|\|\|\|\|\|\|\|\|\|\|\|\|\|\|\|\|\|\|\|\|\|\|\|\|\|\|\|\|\|\|\|\|\|\|\|\|\|\|\|\|\|\|\|\|\|\|\|\|\|\|\|\|\|\|\|\|\|\|\|\|\|\|\|\|\|\|\|\|\|\|\|\|\|\|\|\|\|\|\|\|\|\|\|\|\|\|\|\|\|\|\|\|\|\|\|\|\|\|\|\|\|\|\|\|\|\|\|\|\|\|\|\|\|\|\|\|\|\|\|\|\|\|\|\|\|\|\|\|\|\|\|\|\|\|\|\|\|\|\|\|\|\|\|\|\|\|\|\|\|\|\|\|\|\|\|\|\|\|\|\|\|\|\|\|\|\|\|\|\|\|\|\|\|\|\|\|\|\|\|\|\|\|\|\|\|\|\|\|\|\|\|\|\|\|\|\|\|\|\|\|\|\|\|\|\|\|\|\|\|\|\|\|\|\|\|\|\|\|\|\|\|\|\|\|\|\|\|\|\|\|\|\|\|\|\|\|\|\|\|\|\|\|\|\|\|\|\|\|\|\|\|\|\|\|\|\|\|\|\|\|\|\|\|\|\|\|\|\|\|\|\|\|\|\|\|\|\|\|\|\|\|\|\|\|\|\|\|\|\|\|\|\|\|\|\|\|\|\|\|\|\|\|\|\|\|\|\|\|\|\|\|\|\|\|\|\|\|\|\|\|\|\|\|\|\|\|\|\|\|\|\|\|\|\|\|\|\|\|\|\|\|\|\|\|\|\|\|\|\|\|\|\|\|\|\|\|\|\|\|\|\|\|\|\|\|\|\|\|\|\|\|\|\|\|\|\|\|\|\|\|\|\|\|\|\|\|\|\|\|\|\|\|\|\|\|\|\|\|\|\|\|\|\|\|\|\|\|\|\|\|\|\|\|\|\|\|\|\|\|\|\|\|\|\|\| 	ATGGATCCCTCAACACCACAGACGGTCGCGCAGGCCGTGCAAGCAGCCACGCAGGCCGCCGCGCATCAGCCCTCACCTCCCGTGCAGGCACCGTCGCACTCCACCTCGACCATCGACAACCTCACATGCCAATGGCAAGGTTGCGGAGAGCGATGCGACAACGCCGAAGCCCTCTATGACCACGTCTGCGAGCGTCACGTCGGTCGCAAGAGCACCAACAACCTCAACCTCACCTGCCAGTGGGGAGCCTGCCGAACCACCACCGTCAAACGCGACCACATCACCTCCCACATCCGCGTACATGTCCCGCTCAAACCACACAAGTGCGACTTTTGCGGCAAGGCCTTCAAGAGACCGCAGGATCTGAAGAAGCATGTCAAGACACACGCCGATGACAGCGTGTTGATGAACTCGCCGCATCAGGGACAAAATGGAAGAGGTGCCAGCGCAGGCATCGGACAACAGAATGGCAAGC ``` |
| 32 | ggggacaagtttgtacaaaaaagcaggcttgatgGCACCTCCAACTCAGTACCT, ggggaccactttgtacaagaaagctgggtcATCGAGATAAGTTGTGTTGA | Mycgr3\|42932  JGI  Ensembl |  |  | 0 % |  |  |
| 32 | ggggacaagtttgtacaaaaaagcaggcttgatgTCAGCCCGCCAGCGATCGAA, ggggaccactttgtacaagaaagctgggtcCTTCGAGGGGCGATAGTTGG | Mycgr3\|47406  JGI  Ensembl | KOG1185 Thiamine pyrophosphate-requiring enzyme; | acetolactate synthase activity; magnesium ion binding;  thiamin pyrophosphate binding;  transferase activity; | 100 % |  |  |
| 32 | ggggacaagtttgtacaaaaaagcaggcttgatgTATCTATTCATCACTGCTGA, ggggaccactttgtacaagaaagctgggtcCCGCCTTACCGCTTGACCCA | Mycgr3\|48051  JGI  Ensembl | KOG2084 Predicted histone tail methylase containing SET domain; | nucleus; | 100 % |  |  |
| 32 | ggggacaagtttgtacaaaaaagcaggcttgatgCGGCCATTGCGATGTACATA, ggggaccactttgtacaagaaagctgggtcCTTCCCCACACAGAACTTCT | Mycgr3\|48437  JGI  Ensembl | KOG1191 Mitochondrial GTPase; | GTP binding; intracellular; | 100 % |  |  |
| 32 | ggggacaagtttgtacaaaaaagcaggcttgatgGGAAAAATCGGTCGGGCTGA, ggggaccactttgtacaagaaagctgggtcCGTGATAATCTCTCCCGGAT | Mycgr3\|48880  JGI  Ensembl | KOG3769 Ribonuclease III domain proteins; | RNA binding; RNA processing;  double-stranded RNA binding;  intracellular;  ribonuclease III activity; | 100 % |  |  |
| 32 | ggggacaagtttgtacaaaaaagcaggcttgatgAGTTTTGTCGACGAGAAAGA, ggggaccactttgtacaagaaagctgggtcAGAATCATCCTCCAAGACCT | Mycgr3\|51069  JGI  Ensembl | KOG0254 Predicted transporter (major facilitator superfamily); | L-arabinose isomerase activity; carbohydrate transport;  integral to membrane;  membrane;  sugar:hydrogen symporter activity;  transport;  transporter activity; | 1.02407 % |  |  |
| 32 | ggggacaagtttgtacaaaaaagcaggcttgatgTGGTCGCGTCCGTTGCGGAC, ggggaccactttgtacaagaaagctgggtcTCCCGAGGTCACGACGGCAC | Mycgr3\|52355  JGI  Ensembl | KOG0700 Protein phosphatase 2C/pyruvate dehydrogenase (lipoamide) phosphatase; | CTD phosphatase activity; calcium-dependent protein serine/threonine phosphatase activity;  calcium-dependent protein serine/threonine phosphatase regulator activity;  catalytic activity;  magnesium-dependent protein serine/threonine phosphatase activity;  magnesium-dependent protein serine/threonine phosphatase complex;  myosin phosphatase activity;  myosin phosphatase complex;  myosin phosphatase regulator activity;  phosphoprotein phosphatase activity; | 0 % |  |  |
| 32 | ggggacaagtttgtacaaaaaagcaggcttgatgTTTCTATCCGTCCTTTTCAC, ggggaccactttgtacaagaaagctgggtcATGGTCTCGCTTCCGCATGC | Mycgr3\|53213  JGI  Ensembl | KOG0158 Cytochrome P450 CYP3/CYP5/CYP6/CYP9 subfamilies; | 2-hydroxyisobutyrate 3-monooxygenase activity; 4-chlorophenoxyacetate monooxygenase activity;  4-nitrocatechol 4-monooxygenase activity;  alpha-pinene dehydrogenase activity;  alpha-pinene monooxygenase activity;  ammonia monooxygenase activity;  di-n-butyltin dioxygenase activity;  dimethylsilanediol hydroxylase activity;  electron transport;  fluorene oxygenase activity;  heme binding;  hydroxymethylsilanetriol oxidase activity;  iron ion binding;  limonene 8-monooxygenase activity;  methyl tertiary butyl ether 3-monooxygenase activity;  methylsilanetriol hydroxylase activity;  mono-butyltin dioxygenase activity;  monooxygenase activity;  oxidoreductase activity, acting on paired donors, with incorporation or reduction of molecular oxygen; spheroidene monooxygenase activity;  tert-butyl alcohol 2-monooxygenase activity;  tri-n-butyltin dioxygenase activity; | 100 % |  |  |
| 32 | ggggacaagtttgtacaaaaaagcaggcttgatgAGCACCAAGCACTTCTTCCC, ggggaccactttgtacaagaaagctgggtcATTCAAGCCATCAAGAAGTC | Mycgr3\|54399  JGI  Ensembl | KOG2426 Dihydroxyacetone kinase/glycerone kinase; | glycerol metabolic process; glycerone kinase activity; | 100 % | chr\_1:5536812 C->T; chr\_1:5536794 C->T; |  |
| 32 | ggggacaagtttgtacaaaaaagcaggcttgatgTCTTTCCCGGTAGCCAATTC, ggggaccactttgtacaagaaagctgggtcCAACCTCGCCCTGTCCACTT | Mycgr3\|55571  JGI  Ensembl | KOG2844 Dimethylglycine dehydrogenase precursor; | metabolic process; oxidoreductase activity; | 100 % |  |  |
| 32 | ggggacaagtttgtacaaaaaagcaggcttgatgGCCTCCTCTACGAAGTATCT, ggggaccactttgtacaagaaagctgggtcTTGCTCTTTTTCGTAGCCGG | Mycgr3\|62211  JGI  Ensembl | KOG4472 Glycolipid 2-alpha-mannosyltransferase (alpha-1,2-mannosyltransferase); | 1,2-dihydroxy-phenanthrene glycosyltransferase activity; 1-phenanthrol glycosyltransferase activity;  9-phenanthrol UDP-glucuronosyltransferase activity;  9-phenanthrol glycosyltransferase activity;  O antigen polymerase activity;  UDP-N-acetylglucosamine-peptide N-acetylglucosaminyltransferase activity;  UDP-glucose:glycoprotein glucosyltransferase activity;  acetylgalactosaminyltransferase activity;  acetylglucosaminyltransferase activity;  alpha(1,3)-fucosyltransferase activity;  alpha(1,6)-fucosyltransferase activity;  alpha-1,2-galactosyltransferase activity;  alpha-1,2-mannosyltransferase activity;  alpha-1,3-galactosyltransferase activity;  alpha-1,3-mannosyltransferase activity;  alpha-1,6-mannosyltransferase activity;  beta-1,4-mannosyltransferase activity;  cellulose synthase activity;  dolichyl pyrophosphate Glc1Man9GlcNAc2 alpha-1,3-glucosyltransferase activity; dolichyl pyrophosphate Man7GlcNAc2 alpha-1,3-glucosyltransferase activity;  dolichyl pyrophosphate Man9GlcNAc2 alpha-1,3-glucosyltransferase activity;  dolichyl-phosphate-glucose-glycolipid alpha-glucosyltransferase activity;  fucosyltransferase activity;  galactosyltransferase activity;  glucosyltransferase activity;  glycolipid 2-alpha-mannosyltransferase activity;  glycolipid mannosyltransferase activity;  inositol phosphoceramide synthase activity;  lipopolysaccharide-1,6-galactosyltransferase activity;  mannosyltransferase activity;  membrane;  oligosaccharyl transferase activity;  phenanthrol glycosyltransferase activity;  protein amino acid glycosylation; | 0 % |  |  |
| 32 | ggggacaagtttgtacaaaaaagcaggcttgatgGAACCTCTTCTCTCCCGTCT, ggggaccactttgtacaagaaagctgggtcATCGTCCTTCTCATCGTCGT | Mycgr3\|62800  JGI  Ensembl | KOG1181 FOG: Low-complexity; |  | 100 % |  |  |
| 32 | ggggacaagtttgtacaaaaaagcaggcttgatgCCTCTCAACACGTCCGCCCT, ggggaccactttgtacaagaaagctgggtcCCAAGGCACAGCAGCACCGT | Mycgr3\|65045  JGI  Ensembl | KOG2410 Gamma-glutamyltransferase; | gamma-glutamyltransferase activity; | 3.14835 % |  |  |
| 32 | ggggacaagtttgtacaaaaaagcaggcttgatgGCCGACGTCGAGAAAAACAA, ggggaccactttgtacaagaaagctgggtcAGCGATCCAGAACCAGATGC | Mycgr3\|65946  JGI  Ensembl | KOG1286 Amino acid transporters; | amino acid transmembrane transporter activity; amino acid transport;  integral to membrane;  membrane;  transport; | 100 % |  |  |
| 32 | ggggacaagtttgtacaaaaaagcaggcttgatgAACACCCTCATCCGTCTCGC, ggggaccactttgtacaagaaagctgggtcCCATCCCTCCACCTCCTCCA | Mycgr3\|66283  JGI  Ensembl | KOG0157 Cytochrome P450 CYP4/CYP19/CYP26 subfamilies; | 2-hydroxyisobutyrate 3-monooxygenase activity; 4-chlorophenoxyacetate monooxygenase activity;  4-nitrocatechol 4-monooxygenase activity;  alpha-pinene dehydrogenase activity;  alpha-pinene monooxygenase activity;  ammonia monooxygenase activity;  di-n-butyltin dioxygenase activity;  dimethylsilanediol hydroxylase activity;  electron transport;  fluorene oxygenase activity;  heme binding;  hydroxymethylsilanetriol oxidase activity;  iron ion binding;  limonene 8-monooxygenase activity;  methyl tertiary butyl ether 3-monooxygenase activity;  methylsilanetriol hydroxylase activity;  mono-butyltin dioxygenase activity;  monooxygenase activity;  spheroidene monooxygenase activity;  tert-butyl alcohol 2-monooxygenase activity;  tri-n-butyltin dioxygenase activity; | 100 % |  |  |
| 32 | ggggacaagtttgtacaaaaaagcaggcttgatgGCCACCTTCGGTGCACGCCT, ggggaccactttgtacaagaaagctgggtcTCTCCCGCCAGTACTGTTCA | Mycgr3\|66773  JGI  Ensembl | KOG0255 Synaptic vesicle transporter SVOP and related transporters (major facilitator superfamily); | integral to membrane; transport; transporter activity; | 100 % |  |  |
| 32 | ggggacaagtttgtacaaaaaagcaggcttgatgGCTGGCGCTACCGAGACCAG, ggggaccactttgtacaagaaagctgggtcTACCATGCTGCCGTCAGATC | Mycgr3\|67791  JGI  Ensembl |  |  | 100 % |  |  |
| 32 | ggggacaagtttgtacaaaaaagcaggcttgatgTCAGAAACGTGGCGCTCCAT, ggggaccactttgtacaagaaagctgggtcGAGCCTGCTGTTGACTCCAC | Mycgr3\|68240  JGI  Ensembl | KOG1212 Amidases; | ATP binding; amidase activity;  aminoacyl-tRNA ligase activity;  carbon-nitrogen ligase activity, with glutamine as amido-N-donor;  cytoplasm;  nucleotide binding;  tRNA aminoacylation for protein translation;  translation; | 100 % |  |  |
| 32 | ggggacaagtttgtacaaaaaagcaggcttgatgGTCATGGCGGTTCCTAGCAT, ggggaccactttgtacaagaaagctgggtcCGTCCCAGCATACTTTGGCC | Mycgr3\|68390  JGI  Ensembl | KOG2415 Electron transfer flavoprotein ubiquinone oxidoreductase; | electron transport; electron-transferring-flavoprotein dehydrogenase activity;  oxidoreductase activity; | 100 % | chr\_2:1848155 G->A; |  |
| 32 | ggggacaagtttgtacaaaaaagcaggcttgatgGACTCCCTCAAGTCGCTCGG, ggggaccactttgtacaagaaagctgggtcACCACCAAACATCTTGTTGA | Mycgr3\|68415  JGI  Ensembl | KOG1286 Amino acid transporters; | amino acid transmembrane transporter activity; amino acid transport;  integral to membrane;  membrane;  oxidoreductase activity;  transport; | 100 % |  |  |
| 32 | ggggacaagtttgtacaaaaaagcaggcttgatgCTCCCAACCCCACCACCATC, ggggaccactttgtacaagaaagctgggtcGTAGGTCGCCGCCCAGGCGT | Mycgr3\|68429  JGI  Ensembl | KOG0583 Serine/threonine protein kinase; | (S)-lactate 2-kinase activity; ATP binding;  ATP-dependent polynucleotide kinase activity;  GTP-dependent polynucleotide kinase activity;  LPPG:FO 2-phospho-L-lactate transferase activity;  cobinamide kinase activity;  cyclin-dependent protein kinase activating kinase regulator activity;  cyclin-dependent protein kinase regulator activity;  cytidine kinase activity;  eukaryotic elongation factor-2 kinase activator activity;  eukaryotic elongation factor-2 kinase regulator activity;  glycerate 2-kinase activity;  inositol tetrakisphosphate 2-kinase activity;  phenol kinase activity;  phosphatidylinositol phosphate kinase activity;  phosphofructokinase activity;  phosphoinositide 3-kinase activity;  phosphorylase kinase regulator activity;  polynucleotide kinase activity;  protein amino acid phosphorylation;  protein kinase activity;  protein serine/threonine kinase activity;  protein-tyrosine kinase activity; | 100 % |  |  |
| 32 | ggggacaagtttgtacaaaaaagcaggcttgatgTCCGTCCCCGACCCCGGCAT, ggggaccactttgtacaagaaagctgggtcCTCTAAGCTCTCGGCACTCA | Mycgr3\|68456  JGI  Ensembl |  |  | 100 % |  |  |
| 32 | ggggacaagtttgtacaaaaaagcaggcttgatgCCGACCCTTTCGCTGGTCAA, ggggaccactttgtacaagaaagctgggtcCACACCAGACACACTCGCCG | Mycgr3\|68526  JGI  Ensembl | KOG4505 Na+/H+ antiporter; |  | 100 % |  |  |
| 32 | ggggacaagtttgtacaaaaaagcaggcttgatgGCCGACTCATTTGCCAACAC, ggggaccactttgtacaagaaagctgggtcTAGTCCCTCTTCCTCCTTCA | Mycgr3\|68686  JGI  Ensembl | KOG2626 Histone H3 (Lys4) methyltransferase complex, subunit CPS60/ASH2/BRE2; |  | 100 % |  | ``` 178	ATGGCCGACTCATTTGCCAACACGCCGCTCCAGGCCACCATGGACCAGGACCACGTGCCAAATGTTCCATCACCTCTCAATCCTGCCAGCTCGCGCGCCCCATCCAGGACCCCAGCACCCAATGCGACGCCGATCCAACGAGAACAACGAGAGAAGAAAGAGTCGCTGAAGAAGCGTGAGAATGCACTCAATGGCGGGACTCCTTCGTCGGCATCCAAGCGAAAAGCCAGTGTGGCACACTCCTATCCTTCACCCCAGCGATTTATGATCCCACCACCACGGGTGCAGGACTTCGAAAAGCCCAAGGACGATGTGATGATCCAGCACGAGCCTGCACCTCTTATGACACCCGGAAACGAGCGACAATTATACAAGCCGATTGATCTAGCGGAGAACAAGCGCGGCTATCGATACTCTCGCGCCATTGCGGATCCTCTGTTTCCTCACAAGCAGTTCTACCGATCGACTTCCCCTCCGCCGCATTATGCACGATTGAGCTTTGAAGATGCCGACAAGTGGATGCATTTCACGACGAATGCGTTGATAACGACAAACGAGAAGGGCTGGCGGATGGTGCGCTCCAATGTTTGTGCAAGAGAAGGAACATTGTACTACGAGCTCAAGATACATCGTGGAGTCCCGGTAGACGGACCAGATCCAGCTGCCAATGGACCGCAACCTCATGTGCGGTTCGGCTGGGCAAGACGAGAGGCACCTCTTGACGCACCTGTTGGTTTCGATGGCTACAGCTATGGCCTCACCGACATTCGGTTCGAGACAATGCACCGCAGCAGACCGGGCAAATTCTTCCACCCAAAGAAGACCAAGACCAAGACTGCCAAAGCCGCGCTCGCCGCTCCGACACCTGTTACACTGGCGCCAGAAGACCAACACGTCAAGGAAGGCGACGTGATCGGCGTCGAAATCCAACTTCCCGCGCTC-TCTCTGCAACAAAAAATCGTCAG-TGGAGTCTACAACCCCGCCGTGGACTCAGGAGACGGCTTCGACCAGTCCGCTGCAGCGTCTCTCGATGAAGAACCT-ATGGACG-TCATCCGCGACCG-TATTCCCG	1246 	\|\|\|\|\|\|\|\|\|\|\| \|\|\|\|\|\|\|\|\|\|\|\|\|\|\|\|\|\|\|\|\|\|\|\|\|\|\|\|\|\|\|\|\|\|\| \|\|\|\|\|\|\|\|\|\|\|\|\|\| \|\|\|\|\|\|\|\|\|\|\|\|\|\|\|\|\|\|\|\|\|\|\|\|\|\|\|\|\|\|\|\|\|\|\|\|\|\|\|\|\|\|\|\|\|\|\|\|\|\|\|\|\|\|\|\|\|\|\|\|\|\|\|\|\|\|\|\|\|\|\|\|\|\|\|\|\|\|\|\|\|\|\|\|\|\|\|\|\|\|\|\|\|\|\|\|\|\|\|\|\|\|\|\|\|\|\|\|\|\|\|\|\|\|\|\|\|\|\|\|\|\|\|\|\|\|\|\|\|\|\|\|\|\|\|\|\|\|\|\|\|\|\|\|\|\|\|\|\|\|\|\|\|\|\|\|\|\|\|\|\|\|\|\|\|\|\|\|\|\|\|\|\|\|\|\|\|\|\|\|\|\|\|\|\|\|\|\|\|\|\|\|\|\|\|\|\|\|\|\|\|\|\|\|\|\|\|\|\|\|\|\|\|\|\|\|\|\|\|\|\|\|\|\|\|\|\|\|\|\|\|\|\|\|\|\|\|\|\|\|\|\|\|\|\|\|\|\|\|\|\|\|\|\|\|\|\|\|\|\|\|\|\|\|\|\|\|\|\|\|\|\|\|\|\|\|\|\|\|\|\|\|\|\|\|\|\|\|\|\|\|\|\|\|\|\|\|\|\|\|\|\|\|\|\|\|\|\|\|\|\|\|\|\|\|\|\|\|\|\|\|\|\|\|\|\|\|\|\|\|\|\|\|\|\|\|\|\|\|\|\|\|\|\|\|\|\|\|\|\|\|\|\|\|\|\|\|\|\|\|\|\|\|\|\|\|\|\|\|\|\|\|\|\|\|\|\|\|\|\|\|\|\|\|\|\|\|\|\|\|\|\|\|\|\|\|\|\|\|\|\|\|\|\|\|\|\|\|\|\|\|\|\|\|\|\|\|\|\|\|\|\|\|\|\|\|\|\|\|\|\|\|\| \|\|\|\|\|\|\|\|\|\|\|\|\|\|\|\|\|\|\|\|\|\|\|\|\|\|\|\|\|\|\|\|\|\|\|\|\|\|\|\|\|\|\|\|\|\|\|\|\|\|\|\|\|\| \|\|\|\|\| \|\|\|\|\|\|\|\|\|\|\|\|\|\|\|\|\|\|\|\|\|\|\|\|\|\|\|\|\|\|\|\|\|\|\|\|\| \|\|\|\|\|\|\|\|\|\|\|\|\|\|\|\|\|\|\|\|\|\|\|\|\|\|\|\|\|\|\|\|\|\|\|\|\|\|\|\|\|\|\|\|\|\|\|\|\|\|\|\|\|\|\|\|\|\| \|\|\|\|\|\|\|\|\|\|\|\|\|\|\|\|\|\|\|\|\|\|\|\|\|\|\|\|\|\|\|\|\|\|\|\|\|\|\|\|\|\|\|\|\|\|\|\|\|\|\|\|\|\|\|\|\|\|\|\|\|\|\|\|\|\|\|\|\|\|\|\|\|\|\|\|\|\|\|\|\|\|\|\|\|\|\|\|\|\|\|\|\|\|\|\|\|\|\|\|\|\|\|\|\|\|\|\|\|\|\|\|\|\|\|\|\|\|\|\|\|\|\|\|\|\|\|\|\|\|\|\|\|\|\| \|\|\|\|\|\|\|\|\|\|\|\|\|\|\|\|\|\|\|\|\|\|\|\|\|\|\|\|\|\|\|\|\|\|\| \|\|\|\|\|\| \|\|\|\|\|\|\|\|\|\| \|\|\|\|\|\|\|\| \|\|\|\|  \|\|\| \|\|\|\|\|\|\|\|\|\|\|\|\|\|\| \|\|\|\| \|\|\|\| \|\|\|\|\|\|\|\|\|\|\|\|\| \|\|\|\|\|\|\|\|\|\|\|\|\|\|\|\|\|\|\|\|\|\| \|\|\|\|\|\|\|\|\| \|\|\|\|\| \|\|\|\|\|\|\|\|\|\| \|\|\|\|\|\|\|\|\|\|\|\| \|\|\|\|\|\|\|\|\|\|\|\| \| \| \| \| \|\|\|\|\|\|\|\|\|\|\|\|\|\|\|\|\|\|\|\|\|\|\|\|\|\|\| \|\|\|\|\|\| \|\|\|\|\|\|\|\|\|\|\|\| \| \|\| \|\| \|\|\|\|\|\|\| \|\|\|\|\| \|\| \| \|\| \|\|\|\|\|\|\|\| 	ATGGCCGACTCGTTTGCCAACACGCCGCTCCAGGCCACCATGGACCACGACCACGTGCCAAAGGTTCCATCACCTCTCAATCCTGCCAGCTCGCGCGCCCCATCCAGGACCCCAGCACCCAATGCGACGCCGATCCAACGAGAACAACGAGAGAAGAAAGAGTCGCTGAAGAAGCGTGAGAATGCACTCAATGGCGGGACTCCTTCGTCGGCATCCAAGCGAAAAGCCAGTGTGGCACACTCCTATCCTTCACCCCAGCGATTTATGATCCCACCACCACGGGTGCAGGACTTCGAAAAGCCCAAGGACGATGTGATGATCCAGCACGAGCCTGCACCTCTTATGACACCCGGAAACGAGCGACAATTATACAAGCCGATTGATCTAGCGGAGAACAAGCGCGGCTATCGATACTCTCGCGCCATTGCGGATCCTCTGTTTCCTCACAAGCAGTTCTACCGATCGACTTCCCCTCCGCCGCATTATGCACGATTGAACTTTGAAGATGCCGACAAGTGGATGCATTTCACGACGAATGCGTTGATAACGACCAACGACAAGGGCTGGCGGATGGTGCGCTCCAATGTTTGTGCAACAGAAGGAACATTGTACTACGAGCTCAAGATACATCGTGGAGTCCCGGTAGACGGACCACATCCAGCTGCCAATGGACCGCAACCTCATGTGCGGTTCGGCTGGGCAAGACGAGAGGCACCTCTTGACGCACCTGTTGGTTTCGATGGCTACAGCTATGGCCTCACCGACATTCGGTTCGAGACAATGCACCGCATCAGACCGGGCAAATTCTTCCACCCAAAGAAGACCATGACCAATACTGCCAAAGTCGCGCTCGTCGCTGAGAC-CCTGTTACACTGGCGTCAGATGACC-ACACGTCAAGGAATGCGACGTGATCGGCGTCGAAAT-CAACTTCCCCCGCTCGTCTCTGCAACGAAAAATCGTCAGATGGAGTCTACAAGCACAC-G-GGACTCAGGAGACGGCTTCGACCAGTCTGCTGCAACGTCTCTCGATGCAAAAGCTTATGGACGGTCATC-GCTAACGGTATTCCCG ``` |
| 32 | ggggacaagtttgtacaaaaaagcaggcttgatgTCCGACCGCGACTCCTCCGA, ggggaccactttgtacaagaaagctgggtcCTGCTTCGGACTAATCATCA | Mycgr3\|68689  JGI  Ensembl | KOG1237 H+/oligopeptide symporter; | membrane; oligopeptide transport;  transporter activity; | 100 % |  |  |
| 32 | ggggacaagtttgtacaaaaaagcaggcttgatgCCGCCCATTACACTCCCTTC, ggggaccactttgtacaagaaagctgggtcCTCCTTCCCCGATTCCGCGA | Mycgr3\|69451  JGI  Ensembl | KOG1270 Methyltransferases; | 2-polyprenyl-6-methoxy-1,4-benzoquinone methyltransferase activity; hexaprenyldihydroxybenzoate methyltransferase activity;  ubiquinone biosynthetic process; | 100 % |  |  |
| 32 | ggggacaagtttgtacaaaaaagcaggcttgatgAACTGCCCCTCACGAACCGA, ggggaccactttgtacaagaaagctgggtcGTTGACCCCAAGACCCGTCA | Mycgr3\|69657  JGI  Ensembl | KOG1291 Mn2+ and Fe2+ transporters of the NRAMP family; | membrane; transport; transporter activity; | 100 % |  |  |
| 32 | ggggacaagtttgtacaaaaaagcaggcttgatgTTCTCAACCACATTCCCGCT, ggggaccactttgtacaagaaagctgggtcCAAGTCAAACGTGTACTCCT | Mycgr3\|71051  JGI  Ensembl | KOG2121 Predicted metal-dependent hydrolase (beta-lactamase superfamily); | beta-lactam antibiotic catabolic process; beta-lactamase activity;  response to antibiotic; | 100 % |  |  |
| 32 | ggggacaagtttgtacaaaaaagcaggcttgatgGCAAGACTTGAAGCTGGAGA, ggggaccactttgtacaagaaagctgggtcGAAACAAATCCCCACGGGCA | Mycgr3\|71265  JGI  Ensembl |  |  | 100 % |  |  |
| 32 | ggggacaagtttgtacaaaaaagcaggcttgatgGGTCTCAAGCAATTCCTCCC, ggggaccactttgtacaagaaagctgggtcCCCATTAAGAATGGTATCCG | Mycgr3\|71439  JGI  Ensembl | KOG1238 Glucose dehydrogenase/choline dehydrogenase/mandelonitrile lyase (GMC oxidoreductase family); | FAD binding; choline dehydrogenase activity;  oxidoreductase activity, acting on CH-OH group of donors; | 100 % | chr\_4:2184117 G->T; |  |
| 32 | ggggacaagtttgtacaaaaaagcaggcttgatgGAATACGAAAAATGCTATGT, ggggaccactttgtacaagaaagctgggtcCAGCCCTTTCCAAGCGACCT | Mycgr3\|71542  JGI  Ensembl | KOG2538 Origin recognition complex, subunit 3; |  | 100 % |  |  |
| 32 | ggggacaagtttgtacaaaaaagcaggcttgatgGGCGTTTCCCGAAGAGCTGG, ggggaccactttgtacaagaaagctgggtcGTATATATATCTGAAGTTGA | Mycgr3\|72122  JGI  Ensembl | KOG2533 Permease of the major facilitator superfamily; | integral to membrane; transport; transporter activity; | 2.03868 % |  |  |
| 32 | ggggacaagtttgtacaaaaaagcaggcttgatgACAAGGGCAAGCGTCTTCAG, ggggaccactttgtacaagaaagctgggtcTATCAATTTCAGCATCCGCG | Mycgr3\|72248  JGI  Ensembl | KOG1397 Ca2+/H+ antiporter VCX1 and related proteins; | integral to membrane; | 100 % |  |  |
| 32 | ggggacaagtttgtacaaaaaagcaggcttgatgTATAGCAAATTGTTTCTGGC, ggggaccactttgtacaagaaagctgggtcAACTCTGAAGGATTCAACCT | Mycgr3\|72632  JGI  Ensembl | KOG4389 Acetylcholinesterase/Butyrylcholinesterase; | carboxylesterase activity; | 100 % |  |  |
| 32 | ggggacaagtttgtacaaaaaagcaggcttgatgGCTCTCATCGAGACGTCCCT, ggggaccactttgtacaagaaagctgggtcGTTGTCCGCGTTTCCAGACC | Mycgr3\|72642  JGI  Ensembl |  |  | 100 % |  |  |
| 32 | ggggacaagtttgtacaaaaaagcaggcttgatgTGGTACCACGTCTCGGAGCC, ggggaccactttgtacaagaaagctgggtcATGACCGTTAGCCTGTCCAT | Mycgr3\|73080  JGI  Ensembl | KOG2668 Flotillins; |  | 100 % |  |  |
| 32 | ggggacaagtttgtacaaaaaagcaggcttgatgGCGACGTTCACTCTTCGCGC, ggggaccactttgtacaagaaagctgggtcCAAGAACTGCCATCCTACAA | Mycgr3\|73109  JGI  Ensembl |  |  | 100 % |  |  |
| 32 | ggggacaagtttgtacaaaaaagcaggcttgatgGACTCAGACGATGAGTTCGG, ggggaccactttgtacaagaaagctgggtcGAGGTCGACATTGAACTCCC | Mycgr3\|73475  JGI  Ensembl | KOG1815 Predicted E3 ubiquitin ligase; | protein binding; zinc ion binding; | 100 % |  |  |
| 32 | ggggacaagtttgtacaaaaaagcaggcttgatgAGCACTGAACACAGCTACCT, ggggaccactttgtacaagaaagctgggtcGCCACTCAATTGCGGTCTGT | Mycgr3\|73819  JGI  Ensembl | KOG1212 Amidases; | ATP binding; amidase activity;  aminoacyl-tRNA ligase activity;  carbon-nitrogen ligase activity, with glutamine as amido-N-donor;  cytoplasm;  nucleotide binding;  tRNA aminoacylation for protein translation;  translation; | 100 % |  |  |
| 32 | ggggacaagtttgtacaaaaaagcaggcttgatgGACCTCATACGAAGAGCTGG, ggggaccactttgtacaagaaagctgggtcCAATTCCTTACGCGGCTCAT | Mycgr3\|74210  JGI  Ensembl | KOG2719 Metalloprotease; | membrane; metalloendopeptidase activity;  metallopeptidase activity;  proteolysis;  zinc ion binding; | 100 % | chr\_7:1611664 A->G; |  |
| 32 | ggggacaagtttgtacaaaaaagcaggcttgatgCGTGCCTTCTCGATCTCTTT, ggggaccactttgtacaagaaagctgggtcGTAAGCAAAGCCAGCCGCCA | Mycgr3\|75146  JGI  Ensembl | KOG1516 Carboxylesterase and related proteins; | carboxylesterase activity; cholinesterase activity; | 100 % |  |  |
| 32 | ggggacaagtttgtacaaaaaagcaggcttgatgGACAAAATCACCGACAAGAT, ggggaccactttgtacaagaaagctgggtcCGCCTCCAACAGATCCTTCC | Mycgr3\|75203  JGI  Ensembl | KOG0564 5,10-methylenetetrahydrofolate reductase; | methionine metabolic process; methylenetetrahydrofolate reductase (NADPH) activity; | 100 % |  |  |
| 32 | ggggacaagtttgtacaaaaaagcaggcttgatgGACCAGCTCTCCGGCTTCCG, ggggaccactttgtacaagaaagctgggtcCTCCGTTTTAGGCGACTCCC | Mycgr3\|75450  JGI  Ensembl | KOG0254 Predicted transporter (major facilitator superfamily); | L-arabinose isomerase activity; carbohydrate transport;  integral to membrane;  membrane;  sugar:hydrogen symporter activity;  transport;  transporter activity; | 100 % |  |  |
| 32 | ggggacaagtttgtacaaaaaagcaggcttgatgCCCTTTCCCAAAGCCTTTGG, ggggaccactttgtacaagaaagctgggtcAAACACCATCCCCTTGGTCT | Mycgr3\|76487  JGI  Ensembl | KOG0254 Predicted transporter (major facilitator superfamily); | integral to membrane; transport; transporter activity; | 100 % |  |  |
| 32 | ggggacaagtttgtacaaaaaagcaggcttgatgTCAATTCCAGCATTTGCACC, ggggaccactttgtacaagaaagctgggtcCCCAGGAAACCCACCCCTCG | Mycgr3\|77016  JGI  Ensembl | KOG4501 Transcription coactivator complex, P100 component; |  | 1.33745 % |  |  |
| 32 | ggggacaagtttgtacaaaaaagcaggcttgatgATACTGAAACCCTCGCCGCT, ggggaccactttgtacaagaaagctgggtcTTGTTCCACCCCGCCACCAC | Mycgr3\|77495  JGI  Ensembl | KOG1235 Predicted unusual protein kinase; |  | 100 % |  |  |
| 32 | ggggacaagtttgtacaaaaaagcaggcttgatgCGCCCACGCTCTCTACTTGC, ggggaccactttgtacaagaaagctgggtcCATCAACTCATACCTCCCAT | Mycgr3\|90244  JGI  Ensembl | KOG0260 RNA polymerase II, large subunit; |  | 100 % |  |  |
| 32 | ggggacaagtttgtacaaaaaagcaggcttgatgCGAGCATTGATCTTACTTCG, ggggaccactttgtacaagaaagctgggtcATCGCCCAAGCCCACAATCT | Mycgr3\|90471  JGI  Ensembl | KOG1282 Serine carboxypeptidases (lysosomal cathepsin A); | carboxypeptidase C activity; proteolysis; serine carboxypeptidase activity;  vacuolar carboxypeptidase Y; | 100 % |  |  |
| 32 | ggggacaagtttgtacaaaaaagcaggcttgatgTTCCGCACATTTGGGCTCTT, ggggaccactttgtacaagaaagctgggtcTTGCTTGTTGCACTCCTCGC | Mycgr3\|90758  JGI  Ensembl | KOG1516 Carboxylesterase and related proteins; | carboxylesterase activity; | 100 % |  |  |
| 32 | ggggacaagtttgtacaaaaaagcaggcttgatgCTATTCAAAATTCCGTTGGT, ggggaccactttgtacaagaaagctgggtcCAACGGGGTAACGGTAGGCG | Mycgr3\|91063  JGI  Ensembl |  |  | 100 % | chr\_2:2880568 A->G; chr\_2:2880821 A->C; |  |
| 32 | ggggacaagtttgtacaaaaaagcaggcttgatgCCGTCTCACAATCCAGGCGT, ggggaccactttgtacaagaaagctgggtcAGCCTCGCACTCGACAGTTC | Mycgr3\|91404  JGI  Ensembl |  |  | 100 % |  |  |
| 32 | ggggacaagtttgtacaaaaaagcaggcttgatgGCAGGACTCGCTCTCGATAA, ggggaccactttgtacaagaaagctgggtcTTCTTCCTCCTCCATATCAT | Mycgr3\|92384  JGI  Ensembl | KOG0591 NIMA (never in mitosis)-related G2-specific serine/threonine protein kinase; | ATP binding; carbohydrate metabolic process;  hydrolase activity, hydrolyzing O-glycosyl compounds;  protein amino acid phosphorylation;  protein kinase activity;  protein serine/threonine kinase activity; | 100 % |  |  |
| 32 | ggggacaagtttgtacaaaaaagcaggcttgatgGCTCGAATCGACCAGTCAGT, ggggaccactttgtacaagaaagctgggtcCACGATGTACCGGACCAGGA | Mycgr3\|93260  JGI  Ensembl |  |  | 100 % |  |  |
| 32 | ggggacaagtttgtacaaaaaagcaggcttgatgGATTACCGACCCGTAATATC, ggggaccactttgtacaagaaagctgggtcAACAAGACCTACGTCGTGCG | Mycgr3\|93512  JGI  Ensembl |  |  | 100 % |  |  |
| 32 | ggggacaagtttgtacaaaaaagcaggcttgatgGCTTCCTTCCTCGAGATCCT, ggggaccactttgtacaagaaagctgggtcGCTCAGGCGCATCTTTCGGC | Mycgr3\|94749  JGI  Ensembl |  |  | 100 % |  |  |
| 32 | ggggacaagtttgtacaaaaaagcaggcttgatgTCGCCCATGAAATCACTTCA, ggggaccactttgtacaagaaagctgggtcTTCGGAAACCACCGTATCAG | Mycgr3\|95611  JGI  Ensembl |  |  | 100 % |  |  |
| 32 | ggggacaagtttgtacaaaaaagcaggcttgatgGCCAAGCTGACCGAAAGCCC, ggggaccactttgtacaagaaagctgggtcTTCCCATACTCCAGTCTTGC | Mycgr3\|95744  JGI  Ensembl |  |  | 100 % |  |  |
| 32 | ggggacaagtttgtacaaaaaagcaggcttgatgGCCGAGAAGTACGATATCGT, ggggaccactttgtacaagaaagctgggtcCAACTGCGCTCGATCGCCAC | Mycgr3\|95830  JGI  Ensembl | KOG3855 Monooxygenase involved in coenzyme Q (ubiquinone) biosynthesis; | aromatic compound metabolic process; electron transport;  metabolic process;  monooxygenase activity;  oxidoreductase activity;  phenol 2-monooxygenase activity; | 100 % |  |  |
| 32 | ggggacaagtttgtacaaaaaagcaggcttgatgGCCGACATACAGCAATGGCG, ggggaccactttgtacaagaaagctgggtcGCATGGGCGGCTGTTCTCTC | Mycgr3\|96474  JGI  Ensembl |  |  | 100 % | chr\_11:184727 G->T; |  |
| 32 | ggggacaagtttgtacaaaaaagcaggcttgatgGCCTTCATCACCGAGATCTC, ggggaccactttgtacaagaaagctgggtcGACTCGACCATTGATGAAAT | Mycgr3\|97728  JGI  Ensembl |  |  | 100 % |  |  |
| 32 | ggggacaagtttgtacaaaaaagcaggcttgatgGCAGCAAACGGTGAAACCCA, ggggaccactttgtacaagaaagctgggtcTGTCCTCCCTAACACAGCCT | Mycgr3\|98679  JGI  Ensembl | KOG3855 Monooxygenase involved in coenzyme Q (ubiquinone) biosynthesis; | aromatic compound metabolic process; electron transport;  metabolic process;  monooxygenase activity;  oxidoreductase activity;  phenol 2-monooxygenase activity; | 100 % |  |  |
| 32 | ggggacaagtttgtacaaaaaagcaggcttgatgACTGAGCTCATCCGGGATAC, ggggaccactttgtacaagaaagctgggtcAATGTCCTTCCTCGCAAACT | Mycgr3\|98719  JGI  Ensembl | KOG0255 Synaptic vesicle transporter SVOP and related transporters (major facilitator superfamily); | integral to membrane; transport; transporter activity; | 100 % |  |  |
| 32 | ggggacaagtttgtacaaaaaagcaggcttgatgGCCGACACACCGGACTCGGA, ggggaccactttgtacaagaaagctgggtcCAGCGATCCATCCGCCCACT | Mycgr3\|99520  JGI  Ensembl | KOG1304 Amino acid transporters; |  | 100 % |  |  |
| 32 | ggggacaagtttgtacaaaaaagcaggcttgatgGCGCCTCCCGCCGCGACCAT, ggggaccactttgtacaagaaagctgggtcCAAACAGCCATTACCATTCT | Mycgr3\|100367  JGI  Ensembl | KOG0634 Aromatic amino acid aminotransferase and related proteins; | 6-aminohexanoate transaminase activity; N2-acetyl-L-lysine aminotransferase activity;  acetyldiaminopimelate aminotransferase activity;  aspartate-2-keto-4-methylthiobutyrate transaminase activity;  branched-chain amino acid:2-keto-4-methylthiobutyrate aminotransferase activity; catalytic activity;  methylation;  methyltransferase activity;  nucleic acid binding; | 100 % |  |  |
| 32 | ggggacaagtttgtacaaaaaagcaggcttgatgAAGCACACATTCAGCGCTGC, ggggaccactttgtacaagaaagctgggtcCAGCGTCAAAGCTGCGACGA | Mycgr3\|102047  JGI  Ensembl |  |  | 100 % |  |  |
| 32 | ggggacaagtttgtacaaaaaagcaggcttgatgAAGGGCTCAATCGCTATCCT, ggggaccactttgtacaagaaagctgggtcCAAAAGTGCCGCGCCGAACA | Mycgr3\|102341  JGI  Ensembl |  |  | 100 % |  |  |
| 32 | ggggacaagtttgtacaaaaaagcaggcttgatgGTCAACTCATTCGTCGCATC, ggggaccactttgtacaagaaagctgggtcTTGAGCGTGACCGTGGGGGT | Mycgr3\|102849  JGI  Ensembl | KOG3779 Homeobox transcription factor prospero; |  | 2.52193 % |  |  |
| 32 | ggggacaagtttgtacaaaaaagcaggcttgatgTTCAAGCGATCGTTCATCAC, ggggaccactttgtacaagaaagctgggtcCTTGGCAAGATCGGGGGTTC | Mycgr3\|103011  JGI  Ensembl |  |  | 100 % |  |  |
| 32 | ggggacaagtttgtacaaaaaagcaggcttgatgTCTCTCTGCTATCTCTCCGT, ggggaccactttgtacaagaaagctgggtcGAGCCATTGTTTGAACCATG | Mycgr3\|103387  JGI  Ensembl |  |  | 100 % |  |  |
| 32 | ggggacaagtttgtacaaaaaagcaggcttgatgCCTTCGTGGTTCAAACTGCT, ggggaccactttgtacaagaaagctgggtcCCTCTGCGCCAGCGCGATGT | Mycgr3\|107088  JGI  Ensembl |  |  | 100 % |  |  |
| 32 | ggggacaagtttgtacaaaaaagcaggcttgatgGAAGGCATCACGCGAATCCT, ggggaccactttgtacaagaaagctgggtcCTCCGCCTTCTCCAACCCCC | Mycgr3\|107914  JGI  Ensembl | KOG0255 Synaptic vesicle transporter SVOP and related transporters (major facilitator superfamily); | ATP binding; integral to membrane;  protein amino acid phosphorylation;  protein kinase activity;  transport;  transporter activity; | 100 % |  |  |
| 32 | ggggacaagtttgtacaaaaaagcaggcttgatgGCACCTCACGCGACCGAACC, ggggaccactttgtacaagaaagctgggtcTCGTACACTTGCACTCTGCT | Mycgr3\|107963  JGI  Ensembl | KOG1441 Glucose-6-phosphate/phosphate and phosphoenolpyruvate/phosphate antiporter; |  | 100 % |  |  |
| 32 | ggggacaagtttgtacaaaaaagcaggcttgatgGATCCTTCCACGGAGTCTCG, ggggaccactttgtacaagaaagctgggtcCGACGAACCACCTCCGCCAG | Mycgr3\|108003  JGI  Ensembl |  |  | 25.79449 % |  |  |
| 32 | ggggacaagtttgtacaaaaaagcaggcttgatgGCCTCTCTTCCTGCTCGTCA, ggggaccactttgtacaagaaagctgggtcTGAAGTCGTTATTGGTCCGA | Mycgr3\|108070  JGI  Ensembl | KOG3053 Uncharacterized conserved protein; |  | 100 % |  |  |
| 32 | ggggacaagtttgtacaaaaaagcaggcttgatgGCAGCGATTCGACCCGAGAG, ggggaccactttgtacaagaaagctgggtcCTTGCCGACGGACAGCGACA | Mycgr3\|108315  JGI  Ensembl |  |  | 100 % |  |  |
| 32 | ggggacaagtttgtacaaaaaagcaggcttgatgTCCTCCACACGAGTCTCGTT, ggggaccactttgtacaagaaagctgggtcCCGCCTTGGGAACCACGGAT | Mycgr3\|108793  JGI  Ensembl | KOG3671 Actin regulatory protein (Wiskott-Aldrich syndrome protein); | nucleus; regulation of transcription, DNA-dependent;  transcription factor activity;  zinc ion binding; | 100 % |  | ``` 5	ATGTCCTCCACACGAGTCTCGTTCGGCTCGCGTGCTGGCGGAGGCTCGGATTATCCAGGCGTCGGACAAGAAGTGTTCGTCAGACCCACCTCCCACTACGGCAGCTTCGACTCCCTCGTCGATCTCTCCAACGCCCTCCGAAACGAATGTGCCGACGCCGCTCTCAGCGATGCTCCCGCATGCACGACCAGCGAACTCTCCCTCAGCGGCAATGCGATCGTGCCGCTCATTCGACCCGGAGTCGGACCTTACTCTGACGGCGGGGAGCAGGACTTGCCCGCGGAATTCAAGACGAGCTATACGGTTCCGAATACAGACGGAAGCCCTGCTTCTAACGATATTGCATATCCGCAGGCTCCGATATCTGCAATCGTCAACGTCGCCTCGCCAAAGGAGCGTGCCATCGTCCAGCGCGCCGCCTCGCGAGGTATAATCCAAGCCCTCGAAGCCATCGACGGATTCCGGTACAGCTTCAACAACGCCTGGGCGGCCAAGGATGAAGAAGGGTCACGATTCTCGTACATCTGTCAGGACAGCATGCAGAACAAAGATCGGCACGCAAATGGATACCAGCGCACGGCGAAGCACTTGAAGGGTGGAGAATTCAGTCGTGGTCCACGCAAACCGACATATGATTGCAAGGGCAGTGTGAGCGTCAAGTTCAGCACGGCGAGGCAGAGGGTGGATGTATACTATCGACACTATGCGATTCATTCGACGGTCGCGGAGAGAAAGCCTCCGGAACGGCCGCCGCCTGGGTCAAGGAGGCTGTATAATAGTCGTGCCCCGGCTGTGAGTGCTGAGGCGCACGGGTTTGAGAATCCAGGTACGTTCTCGTTAGGCAACATATCACTACCGGTCGTTGCTTATTCGAAGTTTTATGCAGAAATGTACGACCAGCCCATGTACGGAGATGCTCCACCTTCCGGCTCCAACATTGAACGTCCATTGAAACGGAAGCGCGGTCAACCAAAGTCTTCAACACCCATCGCTTCAACAGCTAAACCCCTCTCCCTTGCCGACATCCTCCGCCAATCCTCATCTGCCAACGTCT-CTGATGATAAGTCCCCTCCTTTCCCGACCAGCTCCAACGC	1098 	\|\|\|\|\|\|\|\|\|\|\|\|\|\|\|\|\|\|\|\|\|\|\|\|\|\|\|\|\|\|\|\|\|\|\|\|\|\|\|\|\|\|\|\|\|\|\|\|\|\|\|\|\|\|\|\|\|\|\|\|\|\|\|\|\|\|\|\|\|\|\|\|\|\|\|\|\|\|\|\|\|\|\|\|\|\|\|\|\|\|\|\|\|\|\|\|\|\|\|\|\|\|\|\|\|\|\|\|\|\|\|\|\|\|\|\|\|\|\|\|\|\|\|\|\|\|\|\|\|\|\|\|\|\|\|\|\|\|\|\|\|\|\|\|\|\|\|\|\|\|\|\|\|\|\|\|\|\|\|\|\|\|\|\|\|\|\|\|\|\|\|\|\|\|\|\|\|\|\|\|\|\|\|\|\|\|\|\|\|\|\|\|\|\|\|\|\|\|\|\|\|\|\|\|\|\|\|\|\|\|\|\|\|\|\|\|\|\|\|\|\|\|\|\|\|\|\|\|\|\|\|\|\|\|\|\|\|\|\|\|\|\|\|\|\|\|\|\|\|\|\|\|\|\|\|\|\|\|\|\|\|\|\|\|\|\|\|\|\|\|\|\|\|\|\|\|\|\|\|\|\|\|\|\|\|\|\|\|\|\|\|\|\|\|\|\|\|\|\|\|\|\|\|\|\|\|\|\|\|\|\|\|\|\|\|\|\|\|\|\|\|\|\|\|\|\|\|\|\|\|\|\|\|\|\|\|\|\|\|\|\|\|\|\|\|\|\|\|\|\|\|\|\|\|\|\|\|\|\|\|\|\|\|\|\|\|\|\|\|\|\|\|\|\|\|\|\|\|\|\|\|\|\|\|\|\|\|\|\|\|\|\|\|\|\|\|\|\|\|\|\|\|\|\|\|\|\|\|\|\|\|\|\|\|\|\|\|\|\|\|\|\|\|\|\|\|\|\|\|\|\|\|\|\|\|\|\|\|\|\|\|\|\|\|\|\|\|\|\|\|\|\|\|\|\|\|\|\|\|\|\|\|\|\|\|\|\|\|\|\|\|\|\|\|\|\|\|\|\|\|\|\|\|\|\|\|\|\|\|\|\|\|\|\|\|\|\|\|\|\|\|\|\|\|\|\|\|\|\|\|\|\|\|\|\|\|\|\|\|\|\|\|\|\|\|\|\|\|\|\|\|\|\|\|\|\|\|\|\|\|\|\|\|\|\|\|\|\|\|\|\|\|\|\|\|\|\|\|\|\|\|\|\|\|\|\|\|\|\|\|\|\|\|\|\|\|\|\|\|\|\|\|\|\|\|\|\|\|\|\|\|\|\|\|\|\|\|\|\|\|\|\|\|\|\|\|\|\|\|\|\|\|\|\|\|\|\|\|\|\|\|\|\|\|\|\|\|\|\|\|\|\|\|\|\|\|\|\|\|\|\|\|\|\|\|\|\|\|\|\|\|\|\|\|\|\|\|\|\|\|\|\|\|\|\|\|\|\|\|\|\|\|\|\|\|\|\|\|\|\|\|\|\|\|\|\|\|\|\|\|\|\|\|\|\|\|\|\|\|\|\|\|\|\|\|\|\|\|\|\|\|\|\|\|\|\|\|\|\|\|\|\|\|\|\|\|\|\|\|\|\|\|\|\|\|\|\|\|\|\|\|\|\|\|\|\|\|\|\|\|\|\|\|\|\|\|\|\|\|\|\|\|\|\|\|\|\|\|\|\|\|\|\|\|\|\|\|\|\|\|\|\|\|\|\|\|\|\|\|\|\|\|\|\|\|\|\|\|\|\|\|\|\|\|\|\|\|\|\|\|\|\|\|\|\|\|\|\|\|\|\|\|\|\|\|\|\|\|\|\|\|\|\|\|\|\|\|\|\|\|\|\|\|\|\|\|\|\|\|\|\|\|\|\|\|\|\|\|\|\|\|\|\|\|\|\|\|\|\|\|\|\|\|\|\|\|\|\|\|\|\|\|\|\|\|\|\|\|\|\|\|\|\|\|\|\|\|\|\|\|\|\|\|\|\|\|\|\|\|\|\|\|\|\|\|\|\|\|\|\|\|\|\|\|\|\|\|\|\|\|\|\|\|\|\|\|\|\|\|\|\|\|\|\|\|\|\|\|\|\|\|\|\|\|\|\|\|\|\|\|\|\|\|\|\|\|\|\|\|\|\|\|\|\|\|\|\|\|\|\|\| \|\| \|\|\|\|\|\|\|\|\|\|\|\|\|\|\|\|\|\|\|\|\|\|\|\|\|\|\|\|\| \| \|\| \|\|\| \|\| \|\| \|\| \| \|\| \|\|\|\|\|\|\|\| \|\|\|\|\|\|\| \|\|\|\|\|\|\|\|\| \|\|\|\|\|\|\|\|\|\|\| \|\| \|\| \|\|\|\|  \|\|\|\|\|\| 	ATGTCCTCCACACGAGTCTCGTTCGGCTCGCGTGCTGGCGGAGGCTCGGATTATCCAGGCGTCGGACAAGAAGTGTTCGTCAGACCCACCTCCCACTACGGCAGCTTCGACTCCCTCGTCGATCTCTCCAACGCCCTCCGAAACGAATGTGCCGACGCCGCTCTCAGCGATGCTCCCGCATGCACGACCAGCGAACTCTCCCTCAGCGGCAATGCGATCGTGCCGCTCATTCGACCCGGAGTCGGACCTTACTCTGACGGCGGGGAGCAGGACTTGCCCGCGGAATTCAAGACGAGCTATACGGTTCCGAATACAGACGGAAGCCCTGCTTCTAACGATATTGCATATCCGCAGGCTCCGATATCTGCAATCGTCAACGTCGCCTCGCCAAAGGAGCGTGCCATCGTCCAGCGCGCCGCCTCGCGAGGTATAATCCAAGCCCTCGAAGCCATCGACGGATTCCGGTACAGCTTCAACAACGCCTGGGCGGCCAAGGATGAAGAAGGGTCACGATTCTCGTACATCTGTCAGGACAGCATGCAGAACAAAGATCGGCACGCAAATGGATACCAGCGCACGGCGAAGCACTTGAAGGGTGGAGAATTCAGTCGTGGTCCACGCAAACCGACATATGATTGCAAGGGCAGTGTGAGCGTCAAGTTCAGCACGGCGAGGCAGAGGGTGGATGTATACTATCGACACTATGCGATTCATTCGACGGTCGCGGAGAGAAAGCCTCCGGAACGGCCGCCGCCTGGGTCAAGGAGGCTGTATAATAGTCGTGCCCCGGCTGTGAGTGCTGAGGCGCACGGGTTTGAGAATCCAGGTACGTTCTCGTTAGGCAACATATCACTACCGGTCGTTGCTTATTCGAAGTTTTATGCAGAAATGTACGACCAGCCCATGTACGGAGATGCTCCACCTTCCGGCTCCAACATTGAACGTCCATTGAAACGGAAGCGCGGTCAACCAAAGTCTTCA-CA-CCATCGCTTCAACAGCTAAACCCCTCTCCAT-GC-GAC-TC-TC-GC-A-TC-TCATCTGC-AACGTCTTCTGATGATA-GTCCCCTCCTT-CC-GAACAGCCTCAACGC ``` |
| 32 | ggggacaagtttgtacaaaaaagcaggcttgatgAATACCGCCACGACGACAGC, ggggaccactttgtacaagaaagctgggtcCGTCTTCGTTGTGTTCCAAC | Mycgr3\|108832  JGI  Ensembl | KOG0380 Sterol O-acyltransferase/Diacylglycerol O-acyltransferase; | diacylglycerol O-acyltransferase activity; | 1.16066 % |  |  |
| 32 | ggggacaagtttgtacaaaaaagcaggcttgatgTGCTCAAACAACGAATCCAA, ggggaccactttgtacaagaaagctgggtcCCCGAGACTTGCCTGCCTGT | Mycgr3\|108869  JGI  Ensembl | KOG0161 Myosin class II heavy chain; |  | 100 % |  |  |
| 32 | ggggacaagtttgtacaaaaaagcaggcttgatgGCGTCCTTCCTCAATAAGGA, ggggaccactttgtacaagaaagctgggtcATTCCGATTTCCACAACTCA | Mycgr3\|108975  JGI  Ensembl | KOG0401 Translation initiation factor 4F, ribosome/mRNA-bridging subunit (eIF-4G); |  | 100 % |  |  |
| 32 | ggggacaagtttgtacaaaaaagcaggcttgatgTCGACAAACAACCAATCGAG, ggggaccactttgtacaagaaagctgggtcATACTGATAGAACGTCACAC | Mycgr3\|108989  JGI  Ensembl | KOG2304 3-hydroxyacyl-CoA dehydrogenase; | fatty acid metabolic process; oxidoreductase activity; | 100 % |  |  |
| 32 | ggggacaagtttgtacaaaaaagcaggcttgatgTCCGGCCCTTCATTGCCCCG, ggggaccactttgtacaagaaagctgggtcATACCGATATCGCGACCCGT | Mycgr3\|109004  JGI  Ensembl | KOG4246 Predicted DNA-binding protein, contains SAP domain; |  | 100 % |  |  |
| 32 | ggggacaagtttgtacaaaaaagcaggcttgatgTCCAGGAAATCAGTGTCGTC, ggggaccactttgtacaagaaagctgggtcAAAGTCATCTTCGTCATCAT | Mycgr3\|109371  JGI  Ensembl | KOG1869 Splicing coactivator SRm160/300, subunit SRm300; | DNA binding; DNA-directed RNA polymerase II, core complex;  hydrogen ion transmembrane transporter activity;  metabolic process;  oxidoreductase activity;  proton-transporting two-sector ATPase complex;  transcription from RNA polymerase II promoter; | 100 % |  |  |
| 32 | ggggacaagtttgtacaaaaaagcaggcttgatgCCCAACCTGATCTCGGGAGC, ggggaccactttgtacaagaaagctgggtcCCCCTTCCCTCTTGCCAATT | Mycgr3\|109426  JGI  Ensembl |  |  | 100 % |  |  |
| 32 | ggggacaagtttgtacaaaaaagcaggcttgatgGACCGCAACGAATGCCGTGG, ggggaccactttgtacaagaaagctgggtcGAGGTTTTGTACAACCTGCT | Mycgr3\|109632  JGI  Ensembl |  |  | 100 % |  |  |
| 32 | ggggacaagtttgtacaaaaaagcaggcttgatgGGTGACGAGTGTGCTAACTT, ggggaccactttgtacaagaaagctgggtcAACACCACAATTCACGACTC | Mycgr3\|110233  JGI  Ensembl |  |  | 100 % |  | ``` 1	ATGGGTGACGAGTGTGCTAACTTGCGTGTGTTCAACGCTCAGGTCAAGTGCGACGAGACTCACCCTCGCTGCGGTCCATGTTCGCGGCTCAATCGCGAGTGCGACTGGGAGCACCGATGGGCGTTCAACGATGCCACTCCTGTAACACAGGGCAAATACGCTAATATAAAGATTTCCGGAAATGTGGTATGGGATCCGTCCGTTCAAGCAAGTAGCTTTCCTCCGGCTAGTCGAAGACCCGATGATCTACCCGAGTTTGCTGCTCTTACAACCAATGAGGATCGCGAGCGAAAGGCGGAGACAAGACGACCAGGTACATTCTCTGTCGTAGCCACGCCTGAGAGCTTCTATGATCTCCCCGAATACGCCTGCGTTTCTCCCTCAAATCGACCAAGGGGCACTCATGGTTCATCTCGTGGCGCCCTAAGTGGCTCAAGACCTGGTTTTGGACGAGCTCAAACTTTTCCAGATCCCAACACCGTACTCCTCGATCGATTCGAAGAGACCTCGCCTACTTCTCTTAGTCCATCAATCGCAGTTGAAGAAAGTCGATCTGGCAGTGTTCCAGACATCCTCCAACATCTTTCACTGACAACACCGCCTCCACACGCTTCTACTGTCACGAGCGCACCTCCAACACCAGTGATCGGTCCCGACGATCACCTCCTTAACCACTTCCGCAACTACATTGCACCACGGCTGACCCAACCGCAACTCGATACTCCTTTCCCGACTCGACCCACCGACATATTCGAGGCAGAGTCCGTCCGCTTCCGACCTCTGTATCACGCGATCTGTGCCATCGCTGGTCTCAACCTCGCTTACAACGGGCGCGCTTCGATGGAGGACTCGATGCAACACTATCACCGAGCTCTGGCAGTGCAAAGTACTGCCACCGGAACGCCACAAGACCTTCTATCTGACGGTGTCTTCTTCCGTCATTTCCTCCTGTTCGTCTACGACATATGCATCCCCATGCAGACTTCATCATCTTCCGGCGAGCTCAGCGGTGCCGACATG-TGGGCCGAACACCTCTCGCACCTCCGCCTCCTCGCT-CTCCAG-CG-ATTCCATCG-CAACAACGCAT	1084 	\|\|\|\|\|\|\|\|\|\|\|\|\|\|\|\|\|\|\|\|\|\|\|\|\|\|\|\|\|\|\|\|\|\|\|\|\|\|\|\|\|\|\|\|\|\|\|\|\|\|\|\|\|\|\|\|\|\|\|\|\|\|\|\|\|\|\|\|\|\|\|\|\|\|\|\|\|\|\|\|\|\|\|\|\|\|\|\|\|\|\|\|\|\|\|\|\|\|\|\|\|\|\|\|\|\|\|\|\|\|\|\|\|\|\|\|\|\|\|\|\|\|\|\|\|\|\|\|\|\|\|\|\|\|\|\|\|\|\|\|\|\|\|\|\|\|\|\|\|\|\|\|\|\|\|\|\|\|\|\|\|\|\|\|\|\|\|\|\|\|\|\|\|\|\|\|\|\|\|\|\|\|\|\|\|\|\|\|\|\|\|\|\|\|\|\|\|\|\|\|\|\|\|\|\|\|\|\|\|\|\|\|\|\|\|\|\|\|\|\|\|\|\|\|\|\|\|\|\|\|\|\|\|\|\|\|\|\|\|\|\|\|\|\|\|\|\|\|\|\|\|\|\|\|\|\|\|\|\|\|\|\|\|\|\|\|\|\|\|\|\|\|\|\|\|\|\|\|\|\|\|\|\|\|\|\|\|\|\|\|\|\|\|\|\|\|\|\|\|\|\|\|\|\|\|\|\|\|\|\|\|\|\|\|\|\|\|\|\|\|\|\|\|\|\|\|\|\|\|\|\|\|\|\|\|\|\|\|\|\|\|\|\|\|\|\|\|\|\|\|\|\|\|\|\|\|\|\|\|\|\|\|\|\|\|\|\|\|\|\|\|\|\|\|\|\|\|\|\|\|\|\|\|\|\|\|\|\|\|\|\|\|\|\|\|\|\|\|\|\|\|\|\|\|\|\|\|\|\|\|\|\|\|\|\|\|\|\|\|\|\|\|\|\|\|\|\|\|\|\|\|\|\|\|\|\|\|\|\|\|\|\|\|\|\|\|\|\|\|\|\|\|\|\|\|\|\|\|\|\|\|\|\|\|\|\|\|\|\|\|\|\|\|\|\|\|\|\|\|\|\|\|\|\|\|\|\|\|\|\|\|\|\|\|\|\|\|\|\|\|\|\|\|\|\|\|\|\|\|\|\|\|\|\|\|\|\|\|\|\|\|\|\|\|\|\|\|\|\|\|\|\|\|\|\|\|\|\|\|\|\|\|\|\|\|\|\|\|\|\|\|\|\|\|\|\|\|\|\|\|\|\|\|\|\|\|\|\|\|\|\|\|\|\|\|\|\|\|\|\|\|\|\|\|\|\|\|\|\|\|\|\|\|\|\|\|\|\|\|\|\|\|\|\|\|\|\|\|\|\|\|\|\|\|\|\|\|\|\|\|\|\|\|\|\|\|\|\|\|\|\|\|\|\|\|\|\|\|\|\|\|\|\|\|\|\|\|\|\|\|\|\|\|\|\|\|\|\|\|\|\|\|\|\|\|\|\|\|\|\|\|\|\|\|\|\|\|\|\|\|\|\|\|\|\|\|\|\|\|\|\|\|\|\|\|\|\|\|\|\|\|\|\|\|\|\|\|\|\|\|\|\|\|\|\|\|\|\|\|\|\|\|\|\|\|\|\|\|\|\|\|\|\|\|\|\|\|\|\|\|\|\|\|\|\|\|\|\|\|\|\|\|\|\|\|\|\|\|\|\|\|\|\|\|\|\|\|\|\|\|\|\|\|\|\|\|\|\|\|\|\|\|\|\|\|\|\|\|\|\|\|\|\|\|\|\|\|\|\|\|\|\|\|\|\|\|\|\|\|\|\|\|\|\|\|\|\|\|\|\|\|\|\|\|\|\|\|\|\|\|\|\|\|\|\|\|\|\|\|\|\|\|\|\|\|\|\|\|\|\|\|\|\|\|\|\|\|\|\|\|\|\|\|\|\|\|\|\|\|\|\|\|\|\|\|\|\|\|\|\|\|\|\| \|\|\|\|\|\|\|\|\|\|\|\|\|\|\|\|\|\|\|\|\|\|\|\|\|\|\|\|\|\|\|\|\|\|\|\|\|\|\|\|\|\|\|\|\|\|\|\|\|\|\|\|\|\|\|\|\|\|\|\|\|\|\|\|\|\|\| \|\|\|\|\|\|\|\|\|\|\|\|\|\|\|\|\|\|\|\|\|\|\|\|\|\|\|\|\|\|\|\|\|\|\|\|\|\|\|\|\|\| \|\|\|\|\|\|\|\|\|\|\|\|\|\| \|\|\| \|\|\|\|\|\| \|\|\|\|\| \|\|\|\|\|\| \|\|\|\|\| \|\| \|\| \|\| \|\| \|\|\|\|\| \|\| \|\|\| \|\| \|\|\|\|\|\|\|\|\| \|\|\|\|\|\|\|\|\|\|\| 	ATGGGTGACGAGTGTGCTAACTTGCGTGTGTTCAACGCTCAGGTCAAGTGCGACGAGACTCACCCTCGCTGCGGTCCATGTTCGCGGCTCAATCGCGAGTGCGACTGGGAGCACCGATGGGCGTTCAACGATGCCACTCCTGTAACACAGGGCAAATACGCTAATATAAAGATTTCCGGAAATGTGGTATGGGATCCGTCCGTTCAAGCAAGTAGCTTTCCTCCGGCTAGTCGAAGACCCGATGATCTACCCGAGTTTGCTGCTCTTACAACCAATGAGGATCGCGAGCGAAAGGCGGAGACAAGACGACCAGGTACATTCTCTGTCGTAGCCACGCCTGAGAGCTTCTATGATCTCCCCGAATACGCCTGCGTTTCTCCCTCAAATCGACCAAGGGGCACTCATGGTTCATCTCGTGGCGCCCTAAGTGGCTCAAGACCTGGTTTTGGACGAGCTCAAACTTTTCCAGATCCCAACACCGTACTCCTCGATCGATTCGAAGAGACCTCGCCTACTTCTCTTAGTCCATCAATCGCAGTTGAAGAAAGTCGATCTGGCAGTGTTCCAGACATCCTCCAACATCTTTCACTGACAACACCGCCTCCACACGCTTCTACTGTCACGAGCGCACCTCCAACACCAGTGATCGGTCCCGACGATCACCTCCTTAACCACTTCCGCAACTACATTGCACCACGGCTGACCCAACCGCAACTCGATACTCCTTTCCCGACTCGACCCACCGACATATTCGAGGCAGAGTCCGTCCGCTTCCGACCTCTGTATCACGCGATCTGTGCCATCGCTGGTCTCAACCTCGCTTACAACGGGCGCGCTTCGATGGAGGACTCGATGCAACACTATCACCGAGCTCTGGCAGTGC-AAGTACTGCCACCGGAACGCCACAAGACCTTCTATCTGACGGTGTCTTCTTCCGTCATTTCCTCCTG-TCGTCTACGACATATGCATCCCCATGCAGACTTCATCATCTT-CGGCGAGCTCAGCG-TGC-GACATGGTGGGC-GAACAC-TCTCG-ACTTC-GCTTC-TCGCTTCTTCAGGCGGATTCCATCGGCAACAACGCAT ``` |
| 32 | ggggacaagtttgtacaaaaaagcaggcttgatgGCCCCTCGAACCACTCACTT, ggggaccactttgtacaagaaagctgggtcCCTCACTACGCTGTTGGACC | Mycgr3\|110270  JGI  Ensembl | KOG0255 Synaptic vesicle transporter SVOP and related transporters (major facilitator superfamily); | integral to membrane; transport; transporter activity; | 100 % |  |  |
| 32 | ggggacaagtttgtacaaaaaagcaggcttgatgCCCCGCATGATCGGTGGTTA, ggggaccactttgtacaagaaagctgggtcGTCTTCGAGGACAACCGCAG | Mycgr3\|110965  JGI  Ensembl | KOG0401 Translation initiation factor 4F, ribosome/mRNA-bridging subunit (eIF-4G); |  | 100 % |  |  |
| 32 | ggggacaagtttgtacaaaaaagcaggcttgatgGCGTGTCGCCCCGAAAGATC, ggggaccactttgtacaagaaagctgggtcAATGGTCCAGCCGTACTGAA | Mycgr3\|111728  JGI  Ensembl |  |  | 100 % |  |  |
| 32 | ggggacaagtttgtacaaaaaagcaggcttgatgTCGCGCGATTACCAGGCCGC, ggggaccactttgtacaagaaagctgggtcATCCATCAAACATATCAAGT | Mycgr3\|111788  JGI  Ensembl | KOG1029 Endocytic adaptor protein intersectin; |  | 100 % |  |  |
| 33 | ggggacaagtttgtacaaaaaagcaggcttgatgCATGTCGCACTGCATTTCCT, ggggaccactttgtacaagaaagctgggtcCGCATCGCTCAACAGACGAT | Mycgr3\|32893  JGI  Ensembl |  |  | 100 % |  |  |
| 33 | ggggacaagtttgtacaaaaaagcaggcttgatgGTCTGGACCACGACGAATCT, ggggaccactttgtacaagaaagctgggtcCCTTCTTTTAGGGTTCACAG | Mycgr3\|34630  JGI  Ensembl | KOG2248 3'-5' exonuclease; | exonuclease activity; intracellular; | 100 % |  |  |
| 33 | ggggacaagtttgtacaaaaaagcaggcttgatgTCCTCCTCTCAGGCCTTCCA, ggggaccactttgtacaagaaagctgggtcCCCGAATGCGAGCTTCCGCT | Mycgr3\|38315  JGI  Ensembl | KOG2095 DNA polymerase iota/DNA damage inducible protein; | DNA polymerase activity; DNA repair;  DNA-directed DNA polymerase activity; | 100 % |  |  |
| 33 | ggggacaagtttgtacaaaaaagcaggcttgatgGACGAACTCCTGTCGAGAGT, ggggaccactttgtacaagaaagctgggtcCTCAATTACAGATCCTCTCA | Mycgr3\|40143  JGI  Ensembl |  |  | 100 % |  |  |
| 33 | ggggacaagtttgtacaaaaaagcaggcttgatgCCGGACTTTTCGGACGACCT, ggggaccactttgtacaagaaagctgggtcCAGCTTCGCCAAAATGTTGT | Mycgr3\|40497  JGI  Ensembl | KOG0136 Acyl-CoA oxidase; | acyl-CoA oxidase activity; electron transport;  fatty acid beta-oxidation;  oxidoreductase activity, acting on the CH-CH group of donors;  peroxisome; | 100 % |  |  |
| 33 | ggggacaagtttgtacaaaaaagcaggcttgatgCGGCTTCTATACAGCATCCT, ggggaccactttgtacaagaaagctgggtcCCTCCAATTATCGGTGACTG | Mycgr3\|42503  JGI  Ensembl | KOG2421 Predicted starch-binding protein; | carbohydrate metabolic process; catalytic activity;  glucan 1,4-alpha-glucosidase activity;  polysaccharide catabolic process;  polysaccharide metabolic process; | 100 % |  |  |
| 33 | ggggacaagtttgtacaaaaaagcaggcttgatgGGCAGCGTCGGTGATCCAAA, ggggaccactttgtacaagaaagctgggtcATTATTACGTACCTCTGGAC | Mycgr3\|43176  JGI  Ensembl | KOG1399 Flavin-containing monooxygenase; | FAD binding; NADP binding;  flavin-containing monooxygenase activity;  oxidoreductase activity; | 100 % |  |  |
| 33 | ggggacaagtttgtacaaaaaagcaggcttgatgACACCCACCGACCAGCCGAA, ggggaccactttgtacaagaaagctgggtcTTGTGTGCGACCTTCTTGTT | Mycgr3\|46430  JGI  Ensembl | KOG1237 H+/oligopeptide symporter; | membrane; oligopeptide transport;  transporter activity; | 1.0355 % |  |  |
| 33 | ggggacaagtttgtacaaaaaagcaggcttgatgGCGTCCGTCATGCAGGCTCT, ggggaccactttgtacaagaaagctgggtcTGCTCCTGCTGCGACCATCG | Mycgr3\|55438  JGI  Ensembl | KOG0157 Cytochrome P450 CYP4/CYP19/CYP26 subfamilies; | dibenzothiophene-5,5-dioxide monooxygenase activity; electron transport;  heme binding;  iron ion binding;  monooxygenase activity; | 1.14811 % |  |  |
| 33 | ggggacaagtttgtacaaaaaagcaggcttgatgGCGGCTTCGGCATCTGCATC, ggggaccactttgtacaagaaagctgggtcTCCCTCTCGAAATCCTGCCA | Mycgr3\|62811  JGI  Ensembl | KOG2595 Predicted GTPase activator protein; | Rab GTPase activator activity; intracellular;  regulation of Rab GTPase activity; | 100 % |  |  |
| 33 | ggggacaagtttgtacaaaaaagcaggcttgatgAAATTTCGATTCAGTCGCAT, ggggaccactttgtacaagaaagctgggtcTGCACCAGAGAATCCGATCT | Mycgr3\|66386  JGI  Ensembl | KOG2458 Endoplasmic reticulum protein EP58, contains filamin rod domain and KDEL motif; |  | 100 % | chr\_1:3332006 G->A; |  |
| 33 | ggggacaagtttgtacaaaaaagcaggcttgatgACGGCGGCAGACGGCAATGG, ggggaccactttgtacaagaaagctgggtcCAGAACGACCAGCCCAGCCT | Mycgr3\|66656  JGI  Ensembl | KOG0068 D-3-phosphoglycerate dehydrogenase, D-isomer-specific 2-hydroxy acid dehydrogenase superfamily; | L-serine biosynthetic process; NAD binding;  cofactor binding;  metabolic process;  oxidoreductase activity, acting on the CH-OH group of donors, NAD or NADP as acceptor; phosphoglycerate dehydrogenase activity; | 100 % |  |  |
| 33 | ggggacaagtttgtacaaaaaagcaggcttgatgGGCGGTCTGAACCTTTCCAT, ggggaccactttgtacaagaaagctgggtcCGCCTTCTCTTCCGACTTGG | Mycgr3\|67469  JGI  Ensembl | KOG0254 Predicted transporter (major facilitator superfamily); | L-arabinose isomerase activity; integral to membrane;  transport;  transporter activity; | 0 % |  |  |
| 33 | ggggacaagtttgtacaaaaaagcaggcttgatgAAATGCGATGAAGTTAAGCC, ggggaccactttgtacaagaaagctgggtcCCGTGGGCGTGGTCTAGCCC | Mycgr3\|67482  JGI  Ensembl |  |  | 100 % | chr\_1:5565225 C->T; | ``` 812	TTCGACATATTTCAACGCTTCGTTACGGGCGTCAGTTTGTGGATCGATCTCTACGATCCTATGAAGCACTTCTCCACTTTAGTGCCACATCTGGCCTTGCATAACG	917 	\|\|\|\|\|\|\|\|\|\|\|\|\|\|\|\|\|\|\|\|\|\|\|\|\|\|\|\|\|\|\|\|\|\|\|\|\|\|\|\|\|\|\|\|\|\|\|\|\|\|\|\|\|\|\|\|\|\|\|\|\|\|\|\|\|\|\|\|\|\|\|\|\|\|\|\|\|\|\|\|\|\|\|\|\|\|\|\|\|\|\|\|\| \|\|\|\|\|\|\|\|\|\|\|\| 	TTCGACATATTTCAACGCTTCGTTACGGGCGTCAGTTTGTGGATCGATCTCTACGATCCTATGAAGCACTTCTCCACTTTAGTGCCACATCTGACCTTGCATAACG ```      ``` 194	ATGAAATGCGATGAAGTTAAGCCTGTCTGCGGTCCGTGTGGCAAAGCCAATCGAGTCTGTATCTATGGATCTCTTCCAGACAACGCAAGCGAAGACGGCAGCGTCGGCGCAACCCCCAAAGTTCATGCCTCCCCGATCAGCACTCCAAATGCTGCTCCAGTATCGAGTCACCGACGATGGCCCAGTGCTGCGGAATCGCTCGAAACGAGGCCGACTCCAAGAATGTACCCTGCCACGCCTGCTCCCACAGGCGACGAAGGCATGGTGCAACTCAGTCCGCAGAGTAGTTACAGCAATTCTACTGGATATGGCACGGAAGTCGCTCCGCTACGCTGGTTCGGACTACTTGCCGGAGACGCAGCTCAGAGCAATCTTGATACGCCAACGCTGGAAAGTCTGAGCGACGCAGCACTTGCTCGGCGGCAGGAGCTGTATCCCACGCAAGATGCGCGGCTTGCGTCCGCTGCGCAGCATGTGAACTTTGCGGAAGGTGTGGCGCTGCAGAACCCATCAAAGATTCTGCAAATATCGGCAATATCGCCAGCGCCGATGGATGGTAGCACGGATGAACGGCAGTTCTGGCAGTCTACAGAGCCAATACGACTCAAAGACCATGAA	811 	\|\|\|\|\|\|\|\|\|\|\|\|\|\|\|\|\|\|\|\|\|\|\|\|\|\|\|\|\|\|\|\|\|\|\|\|\|\|\|\|\|\|\|\|\|\|\|\|\|\|\|\|\|\|\|\|\|\|\|\|\|\|\|\|\|\|\|\|\|\|\|\|\|\|\|\|\|\|\|\|\|\|\|\|\|\|\|\|\|\|\|\|\|\|\|\|\|\|\|\|\|\|\|\|\|\|\|\|\|\|\|\|\|\|\|\|\|\|\|\|\|\|\|\|\|\|\|\|\|\|\|\|\|\|\|\|\|\|\|\|\|\|\|\|\|\|\|\|\|\|\|\|\|\|\|\|\|\|\|\|\|\|\|\|\|\|\|\|\|\|\|\|\|\|\|\|\|\|\|\|\|\|\|\|\|\|\|\|\|\|\|\|\|\|\|\|\|\|\|\|\|\|\|\|\|\|\|\|\|\|\|\|\|\|\|\|\|\|\|\|\|\|\|\|\|\|\|\|\|\|\|\|\|\|\|\|\|\|\|\|\|\|\|\|\|\|\|\|\|\|\|\|\|\|\|\|\|\|\|\|\|\|\|\|\|\|\|\|\|\|\|\|\|\|\|\|\|\|\|\|\|\|\|\|\|\|\|\|\|\|\|\|\|\|\|\|\|\|\|\|\|\|\|\|\|\|\|\|\|\|\|\|\|\|\|\|\|\|\|\|\|\|\|\|\|\|\|\|\|\|\|\|\|\|\|\|\|\|\|\|\|\|\|\|\|\|\|\|\|\|\|\|\|\|\|\|\|\|\|\|\|\|\|\|\|\|\|\|\|\|\|\|\|\|\|\|\|\|\|\|\|\|\|\|\|\|\|\|\|\|\|\|\|\|\|\|\|\|\|\|\|\|\|\|\|\|\|\|\|\|\|\|\|\|\|\|\|\|\|\|\|\|\|\|\|\|\|\|\|\|\|\|\|\|\|\|\|\|\|\|\|\|\|\|\|\|\|\|\|\|\|\|\|\|\|\|\|\|\|\|\|\|\|\|\|\|\|\|\|\|\|\|\|\|\|\|\|\|\|\|\|\|\|\|\|\|\|\|\|\|\|\|\|\|\|\|\|\|\|\|\|\|\|\|\|\|\|\|\|\|\|\|\|\|\|\|\|\|\|\|\|\|\|\|\|\|\|\|\|\|\|\|\|\|\|\|\|\|\|\|\|\|\|\|\|\|\|\|\|\|\|\|\|\|\|\|\|\|\|\|\|\|\|\|\|\|\|\|\|\|\|\|\|\|\|\|\|\|\|\|\|\|\|\|\|\|\|\|\|\|\|\|\|\|\|\|\|\|\|\|\|\|\|\|\|\|\|\|\|\|\|\|\|\|\|\|\|\| 	ATGAAATGCGATGAAGTTAAGCCTGTCTGCGGTCCGTGTGGCAAAGCCAATCGAGTCTGTATCTATGGATCTCTTCCAGACAACGCAAGCGAAGACGGCAGCGTCGGCGCAACCCCCAAAGTTCATGCCTCCCCGATCAGCACTCCAAATGCTGCTCCAGTATCGAGTCACCGACGATGGCCCAGTGCTGCGGAATCGCTCGAAACGAGGCCGACTCCAAGAATGTACCCTGCCACGCCTGCTCCCACAGGCGACGAAGGCATGGTGCAACTCAGTCCGCAGAGTAGTTACAGCAATTCTACTGGATATGGCACGGAAGTCGCTCCGCTACGCTGGTTCGGACTACTTGCCGGAGACGCAGCTCAGAGCAATCTTGATACGCCAACGCTGGAAAGTCTGAGCGACGCAGCACTTGCTCGGCGGCAGGAGCTGTATCCCACGCAAGATGCGCGGCTTGCGTCCGCTGCGCAGCATGTGAACTTTGCGGAAGGTGTGGCGCTGCAGAACCCATCAAAGATTCTGCAAATATCGGCAATATCGCCAGCGCCGATGGATGGTAGCACGGATGAACGGCAGTTCTGGCAGTCTACAGAGCCAATACGACTCAAAGACCATGAA ```      ``` 917	GAGGGTCTCATGAAGGCAGTCTTAGCACTAGGTGCACGACATTTGTCGATCAAGCCGCTAAATCGCGGAGAAGTGAATGTCGATCGCACGGCAGCCGTGCAATACTACTACGAAACGCTGCAATACCTGCAGTCCGCGATGCGCTTCACGTC-ATACAAGAACAGTCTC-GAGCTACT	1092 	\|\|\|\|\|\|\|\|\|\|\|\|\|\|\|\|\|\|\|\|\|\|\|\|\|\|\|\|\|\|\|\|\|\|\|\|\|\|\|\|\|\|\|\|\|\|\|\|\|\|\|\|\|\|\|\|\|\|\|\|\|\|\|\|\|\|\|\|\|\|\|\|\|\|\| \|\|\|\|\|\|\|\|\|\|\|\|\|\|\|\|\|\|\|\|\|\|\|\|\|\|\|\|\|\|\|\|\|\|\|\|\|\| \|\|\|\|\|\| \|\|\|\|\|\|\|\|\|\|\|\|\|\|\|\|\|\|\|\|\|\|\|\|\|\|\|\|\|\| \|\|\|\|\|\|\|\|\|\|\|\|\|\|\|\| \|\|\|\|\|\|\|\| 	GAGGGTCTCATGAAGGCAGTCTTAGCACTAGGTGCACGACATTTGTCGATCAAGCCGCTAAATCGCGGAGAAGTG-ATGTCGATCGCACGGCAGCCGTGCAATACTACTACGAA-CGCTGC-ATACCTGCAGTCCGCGATGCGCTTCACGTCAATACAAGAACAGTCTCCGAGCTACT ``` |
| 33 | ggggacaagtttgtacaaaaaagcaggcttgatgGCCGAGACCTACCCAGAACC, ggggaccactttgtacaagaaagctgggtcATGAGTCTCACTCCACTGCT | Mycgr3\|68224  JGI  Ensembl | KOG1175 Acyl-CoA synthetase; | acetoacetate-CoA ligase activity; catalytic activity;  metabolic process; | 1.91458 % |  |  |
| 33 | ggggacaagtttgtacaaaaaagcaggcttgatgGCGGATGAAGAGAGCCCCCT, ggggaccactttgtacaagaaagctgggtcCCCGAGACTCAAATTCCGCA | Mycgr3\|68741  JGI  Ensembl |  |  | 100 % | chr\_2:2567423 G->T; |  |
| 33 | ggggacaagtttgtacaaaaaagcaggcttgatgCATCTTTCATCAGCCTTTCT, ggggaccactttgtacaagaaagctgggtcAAAGGCCTTAAACTCCACCA | Mycgr3\|69100  JGI  Ensembl |  |  | 100 % |  |  |
| 33 | ggggacaagtttgtacaaaaaagcaggcttgatgCGTCCGCTGCGACGTGGAGT, ggggaccactttgtacaagaaagctgggtcCTTCGTTCGTAACAGAGGAT | Mycgr3\|69447  JGI  Ensembl | KOG2431 1, 2-alpha-mannosidase; | calcium ion binding; mannosyl-oligosaccharide 1,2-alpha-mannosidase activity;  membrane; | 1.03245 % |  |  |
| 33 | ggggacaagtttgtacaaaaaagcaggcttgatgCTACGCCTTGCATCGCCATC, ggggaccactttgtacaagaaagctgggtcAATCAAAGCCTGTCCCTTGA | Mycgr3\|70401  JGI  Ensembl | KOG0335 ATP-dependent RNA helicase; | 8-oxo-7,8-dihydroguanine triphosphatase activity; ATP binding;  ATP-dependent 5'-3' DNA helicase activity;  ATP-dependent helicase activity;  UDP-2,3-diacylglucosamine hydrolase activity;  bis(5'-nucleosyl)-tetraphosphatase activity;  dATP pyrophosphohydrolase activity;  diacylglycerol pyrophosphate phosphatase activity;  dihydroneopterin monophosphate phosphatase activity;  dihydroneopterin triphosphate pyrophosphohydrolase activity;  helicase activity;  nucleic acid binding;  pyrophosphatase activity;  thiamin-pyrophosphatase activity; | 100 % |  |  |
| 33 | ggggacaagtttgtacaaaaaagcaggcttgatgGCGGACACTGCTCAATATGG, ggggaccactttgtacaagaaagctgggtcACTAGGCGAGTATTTACTCA | Mycgr3\|70547  JGI  Ensembl | KOG0255 Synaptic vesicle transporter SVOP and related transporters (major facilitator superfamily); | integral to membrane; transport; transporter activity; | 100 % |  |  |
| 33 | ggggacaagtttgtacaaaaaagcaggcttgatgGCTCTCGCATCACACGTAGA, ggggaccactttgtacaagaaagctgggtcACAAACCGTCCGATGCACAC | Mycgr3\|70646  JGI  Ensembl | KOG1383 Glutamate decarboxylase/sphingosine phosphate lyase; | carboxy-lyase activity; carboxylic acid metabolic process;  glutamate decarboxylase activity;  glutamate metabolic process;  pyridoxal phosphate binding; | 100 % |  |  |
| 33 | ggggacaagtttgtacaaaaaagcaggcttgatgAGATACTCAGCTGTCTGGCT, ggggaccactttgtacaagaaagctgggtcGTCATCACTCAAGATGTGTC | Mycgr3\|71382  JGI  Ensembl | KOG1238 Glucose dehydrogenase/choline dehydrogenase/mandelonitrile lyase (GMC oxidoreductase family); | FAD binding; choline dehydrogenase activity;  oxidoreductase activity, acting on CH-OH group of donors; | 100 % |  |  |
| 33 | ggggacaagtttgtacaaaaaagcaggcttgatgGCCCTCCAACAAGCCGCCGA, ggggaccactttgtacaagaaagctgggtcCTCACCCCTCCCCTTCCCAT | Mycgr3\|71733  JGI  Ensembl | KOG1369 Hexokinase; | ATP binding; glucokinase activity;  glycolysis;  hexokinase activity; | 100 % | chr\_5:357285 G->A; |  |
| 33 | ggggacaagtttgtacaaaaaagcaggcttgatgAACGTGCTCAAGCTGCAGAA, ggggaccactttgtacaagaaagctgggtcACCCATCTTCTCCGACGCCG | Mycgr3\|72868  JGI  Ensembl | KOG0046 Ca2+-binding actin-bundling protein (fimbrin/plastin), EF-Hand protein superfamily; | actin binding; calcium ion binding; | 100 % |  |  |
| 33 | ggggacaagtttgtacaaaaaagcaggcttgatgCTCGTCGCCCTTATCGACCG, ggggaccactttgtacaagaaagctgggtcTGACCTGCTTGCCTTCTTGA | Mycgr3\|73230  JGI  Ensembl | KOG0157 Cytochrome P450 CYP4/CYP19/CYP26 subfamilies; | electron transport; heme binding;  iron ion binding;  monooxygenase activity;  oxidoreductase activity, acting on paired donors, with incorporation or reduction of molecular oxygen, reduced flavin or flavoprotein as one donor, and incorporation of one atom of oxygen; unspecific monooxygenase activity; | 20.77446 % |  |  |
| 33 | ggggacaagtttgtacaaaaaagcaggcttgatgTTCACACGCTCAGACTCCCC, ggggaccactttgtacaagaaagctgggtcATGGATGTCCACCGCAATCT | Mycgr3\|73782  JGI  Ensembl | KOG0254 Predicted transporter (major facilitator superfamily); | integral to membrane; transport; transporter activity; | 100 % |  |  |
| 33 | ggggacaagtttgtacaaaaaagcaggcttgatgGCGGCAACAACAACATGGCA, ggggaccactttgtacaagaaagctgggtcATCTTCCTTCAACCCCCTAC | Mycgr3\|74423  JGI  Ensembl | KOG0637 Sucrose transporter and related proteins; |  | 100 % |  |  |
| 33 | ggggacaagtttgtacaaaaaagcaggcttgatgCGGCCTGATCACACGGACGA, ggggaccactttgtacaagaaagctgggtcCATCCTCACCCTCCGATCAT | Mycgr3\|75087  JGI  Ensembl | KOG0293 WD40 repeat-containing protein; | nucleic acid binding; outer membrane-bounded periplasmic space;  transport;  transporter activity; | 100 % |  |  |
| 33 | ggggacaagtttgtacaaaaaagcaggcttgatgAGCATATGGCTGGAACGCTT, ggggaccactttgtacaagaaagctgggtcCGACTCTATCCCCAAGCCAT | Mycgr3\|76401  JGI  Ensembl | KOG1961 Vacuolar sorting protein VPS52/suppressor of actin Sac2; |  | 0 % |  |  |
| 33 | ggggacaagtttgtacaaaaaagcaggcttgatgCTGTCCACTCTGTTTCTCAT, ggggaccactttgtacaagaaagctgggtcCACCTCCCACGGCGGCCTCT | Mycgr3\|76675  JGI  Ensembl | KOG2182 Hydrolytic enzymes of the alpha/beta hydrolase fold; | cysteine-type peptidase activity; proteolysis; serine-type peptidase activity; | 100 % |  |  |
| 33 | ggggacaagtttgtacaaaaaagcaggcttgatgGAGGCTGCGAGTGCCCGCTA, ggggaccactttgtacaagaaagctgggtcAACCACTGATCCAAATCCAT | Mycgr3\|77089  JGI  Ensembl | KOG1087 Cytosolic sorting protein GGA2/TOM1; | clathrin adaptor complex; intracellular;  intracellular protein transport;  protein binding;  protein complex assembly;  protein transporter activity;  vesicle-mediated transport; | 100 % |  |  |
| 33 | ggggacaagtttgtacaaaaaagcaggcttgatgGCTCGCTTCTACCCACCGTC, ggggaccactttgtacaagaaagctgggtcACTACGTCCGTTCGCCCGGC | Mycgr3\|77233  JGI  Ensembl |  |  | 100 % |  |  |
| 33 | ggggacaagtttgtacaaaaaagcaggcttgatgTCGGAACGTGTAACGCGTTC, ggggaccactttgtacaagaaagctgggtcAGTAATATCGACAATCCATG | Mycgr3\|77785  JGI  Ensembl |  |  | 100 % |  |  |
| 33 | ggggacaagtttgtacaaaaaagcaggcttgatgTCCTTCATAACCGATACAGT, ggggaccactttgtacaagaaagctgggtcACTCAACCACCTCCCATACG | Mycgr3\|84646  JGI  Ensembl |  |  | 100 % |  |  |
| 33 | ggggacaagtttgtacaaaaaagcaggcttgatgTCACCCGAGGCCACTCTCCG, ggggaccactttgtacaagaaagctgggtcGGGCGAGTCGCGGCCGTTGG | Mycgr3\|89593  JGI  Ensembl |  |  | 2.13068 % |  |  |
| 33 | ggggacaagtttgtacaaaaaagcaggcttgatgGCATACGCCGACGCCAGATT, ggggaccactttgtacaagaaagctgggtcTGGTTCGCTTTTGTACGTGA | Mycgr3\|89777  JGI  Ensembl | KOG4415 Uncharacterized conserved protein; | nucleus; protein dimerization activity;  regulation of transcription, DNA-dependent;  sequence-specific DNA binding;  transcription factor activity; | 100 % |  | ``` 843	ACTAGCATCCCGACAGCGGAAGAAGAAGCACACGGAAGACTTGGAGGTCAAGGAGAAAAGCTACACTCAGCAGATTTCTTCTCTTCGAGCGCAAGTGGACGAGCTCACCCGTG-AGCGCGAGCACCGGGCACGAGACCAGCAAGCTACGCATCAGAGACTGCAGGAGGCTTTGCGGGCTATCGATACGATGCAGGATGAACGACGAGA-GCTCATCA	1057 	\|\|\|\|\|\|\|\|\|\|\|\|\|\|\|\|\|\|\|\| \| \|\|\|\|\|\|\|\|\|\|\|\|\|\|\|\|\|\|\|\|\|\|\| \|\|\|\|\|\|\|\| \|\|\|\|\|\|\|\|\|\|\|\|\|\|\|\|\|\|\|\|\|\|\|\|\|\|\|\|\|\|\|\|\|\|\|\|\| \|\| \|\|\|\|\|\|\|\|\|\|\|\|\|\|\|\| \|\|\|\|\|\|\|\|\|\|\| \|\|\|\|\|\|\|\|\|\| \|\|\|\| \|\|\|\|\|\|\|\|\|\|\|\|\|\|\|\|\|\|\|\|\|\| \|\| \|\|\|\|\|  \|\|\|\|\|\|\|\|\| \|\|\|\|\|\|\|\|\|\|\|\|\|\| \|\|\|\|\|\|\|\| \|\|\|\| \|\|\| 	ACTAGCATCCCGACAGCGGA-G-AGAAGCACACGGAAGACTTGGAG-TCAAGGAG-AAAGCTACACTCAGCAGATTTCTTCTCTTCGAGCGCA-GT-GACGAGCTCACCCGTGGAGCGCGAGCAC-GGGCACGAGA-CAGCCAGCTACGCATCAGAGACTGCAG-AGCCTTTGGCGGCTATCGAAACGATGCAGGATGA-CGACGAGAAGCTC-TCA ```      ``` 1	ATGGCATACGCCGACGCCAGATTCGGCTACCATCAGCCCCAGTCCGATTCCGTCTCCGCATCCGCCCTAGACAACAACAAACGAATCAACGCTTTTCTCGACGACGACTCTGTGGTGCTCGATCCCTCGGTGCTGGACTCTGACATCATGCAAAGTCCCTCTTCCGCCAACCCGTTCCGCAAGGACTCTCTCGCCAACAGCAATGGTGTGCTATCACCCGCCGACTCGCAAGCGGCATGGGACCATCCGTATGGTACTGGCATGTCTGCTGAACCTACCTCAGCAGGTGTTCCCCTTCCTTATCATGAAGACGGATCTGCCTACCTACAGCAACCGAATCCCATCGCGGTATCGTATGCTCATCAGCATCCACCACCTGCATCGTGGGCATTCGAACACGGTTCCGAGCACTGCGTGAGCAATGGTGCAACAGACTTCATGCCGCCGCCGCCTCAATTCGAGGTCCCTCGATACAGCCGTCACCGGACAGATAGCGTTCACGCCGGTTACAGTCAGCACCAACCGTCAATGCCCGCACACTTTATCGCGGCCCCTCAAGTTCAGACTCCCATGTCACCGCATAGTCATCAGGACTGGATGGGTATGGCCGAGCGTGAGATGGAGAATCGACCTTCGCACAAACGCATGCGGCCAGGCAGTCCTGCTCGTACGATGATCGACTTCCAACGACGCGATGGGATCCGGAAGAAGAACGGGCGCATCGACATTCCTCAGGAGAGGAGCATCGCGAACATTGATGAGCTTATTGAGAAAGAAACAGACGAGGATCAGCTGAAGGAATTGAAGCAACAGAAACGCCTGCTGAGGAATCGTGAGGCTGC	842 	\|\|\|\|\|\|\|\|\|\|\|\|\|\|\|\|\|\|\|\|\|\|\|\|\|\|\|\|\|\|\|\|\|\|\|\|\|\|\|\|\|\|\|\|\|\|\|\|\|\|\|\|\|\|\|\|\|\|\|\|\|\|\|\|\|\|\|\|\|\|\|\|\|\|\|\|\|\|\|\|\|\|\|\|\|\|\|\|\|\|\|\|\|\|\|\|\|\|\|\|\|\|\|\|\|\|\|\|\|\|\|\|\|\|\|\|\|\|\|\|\|\|\|\|\|\|\|\|\|\|\|\|\|\|\|\|\|\|\|\|\|\|\|\|\|\|\|\|\|\|\|\|\|\|\|\|\|\|\|\|\|\|\|\|\|\|\|\|\|\|\|\|\|\|\|\|\|\|\|\|\|\|\|\|\|\|\|\|\|\|\|\|\|\|\|\|\|\|\|\|\|\|\|\|\|\|\|\|\|\|\|\|\|\|\|\|\|\|\|\|\|\|\|\|\|\|\|\|\|\|\|\|\|\|\|\|\|\|\|\|\|\|\|\|\|\|\|\|\|\|\|\|\|\|\|\|\|\|\|\|\|\|\|\|\|\|\|\|\|\|\|\|\|\|\|\|\|\|\|\|\|\|\|\|\|\|\|\|\|\|\|\|\|\|\|\|\|\|\|\|\|\|\|\|\|\|\|\|\|\|\|\|\|\|\|\|\|\|\|\|\|\|\|\|\|\|\|\|\|\|\|\|\|\|\|\|\|\|\|\|\|\|\|\|\|\|\|\|\|\|\|\|\|\|\|\|\|\|\|\|\|\|\|\|\|\|\|\|\|\|\|\|\|\|\|\|\|\|\|\|\|\|\|\|\|\|\|\|\|\|\|\|\|\|\|\|\|\|\|\|\|\|\|\|\|\|\|\|\|\|\|\|\|\|\|\|\|\|\|\|\|\|\|\|\|\|\|\|\|\|\|\|\|\|\|\|\|\|\|\|\|\|\|\|\|\|\|\|\|\|\|\|\|\|\|\|\|\|\|\|\|\|\|\|\|\|\|\|\|\|\|\|\|\|\|\|\|\|\|\|\|\|\|\|\|\|\|\|\|\|\|\|\|\|\|\|\|\|\|\|\|\|\|\|\|\|\|\|\|\|\|\|\|\|\|\|\|\|\|\|\|\|\|\|\|\|\|\|\|\|\|\|\|\|\|\|\|\|\|\|\|\|\|\|\|\|\|\|\|\|\|\|\|\|\|\|\|\|\|\|\|\|\|\|\|\|\|\|\|\|\|\|\|\|\|\|\|\|\|\|\|\|\|\|\|\|\|\|\|\|\|\|\|\|\|\|\|\|\|\|\|\|\|\|\|\|\|\|\|\|\|\|\|\|\|\|\|\|\|\|\|\|\|\|\|\|\|\|\|\|\|\|\|\|\|\|\|\|\|\|\|\|\|\|\|\|\|\|\|\|\|\|\|\|\|\|\|\|\|\|\|\|\|\|\|\|\|\|\|\|\|\|\|\|\|\|\|\|\|\|\|\|\|\|\|\|\|\|\|\|\|\|\|\|\|\|\|\|\|\|\|\|\|\|\|\|\|\|\|\|\|\|\|\|\|\|\|\|\|\|\|\|\|\|\|\|\|\|\|\|\|\|\|\|\|\|\|\|\|\|\|\|\|\|\|\|\|\|\|\|\|\|\|\|\|\|\|\|\|\|\|\|\|\|\|\|\|\|\|\|\|\|\|\|\|\|\|\|\|\|\|\|\|\|\|\|\|\|\|\|\|\|\|\|\|\|\|\|\|\|\|\|\|\|\|\|\|\|\|\|\|\|\|\|\|\|\|\|\|\|\|\|\|\|\|\|\|\|\|\|\|\|\|\|\|\|\|\|\|\|\|\| 	ATGGCATACGCCGACGCCAGATTCGGCTACCATCAGCCCCAGTCCGATTCCGTCTCCGCATCCGCCCTAGACAACAACAAACGAATCAACGCTTTTCTCGACGACGACTCTGTGGTGCTCGATCCCTCGGTGCTGGACTCTGACATCATGCAAAGTCCCTCTTCCGCCAACCCGTTCCGCAAGGACTCTCTCGCCAACAGCAATGGTGTGCTATCACCCGCCGACTCGCAAGCGGCATGGGACCATCCGTATGGTACTGGCATGTCTGCTGAACCTACCTCAGCAGGTGTTCCCCTTCCTTATCATGAAGACGGATCTGCCTACCTACAGCAACCGAATCCCATCGCGGTATCGTATGCTCATCAGCATCCACCACCTGCATCGTGGGCATTCGAACACGGTTCCGAGCACTGCGTGAGCAATGGTGCAACAGACTTCATGCCGCCGCCGCCTCAATTCGAGGTCCCTCGATACAGCCGTCACCGGACAGATAGCGTTCACGCCGGTTACAGTCAGCACCAACCGTCAATGCCCGCACACTTTATCGCGGCCCCTCAAGTTCAGACTCCCATGTCACCGCATAGTCATCAGGACTGGATGGGTATGGCCGAGCGTGAGATGGAGAATCGACCTTCGCACAAACGCATGCGGCCAGGCAGTCCTGCTCGTACGATGATCGACTTCCAACGACGCGATGGGATCCGGAAGAAGAACGGGCGCATCGACATTCCTCAGGAGAGGAGCATCGCGAACATTGATGAGCTTATTGAGAAAGAAACAGACGAGGATCAGCTGAAGGAATTGAAGCAACAGAAACGCCTGCTGAGGAATCGTGAGGCTGC ``` |
| 33 | ggggacaagtttgtacaaaaaagcaggcttgatgCTCAGTTTCATCTCGACGCT, ggggaccactttgtacaagaaagctgggtcTCCCTCAACCGCAATCATCA | Mycgr3\|90331  JGI  Ensembl |  |  | 100 % |  |  |
| 33 | ggggacaagtttgtacaaaaaagcaggcttgatgTCCGACAACGACGAGCGCTC, ggggaccactttgtacaagaaagctgggtcGCCGTCAACCTCCATCTTCA | Mycgr3\|91745  JGI  Ensembl | KOG2897 DNA-binding protein YL1 and related proteins; |  | 100 % |  | ``` 1	ATGTCCGACAACGACGAGCGCTCGTGCTCGAGCTCCGAGAGTGATGGCGAAGATGTCACCACCACCGGACTCATCGCCACACGTGCGAAGCGATCCACAGCAGGAAATCTTTACGCGACTTTGCGCCAGAACCTCGACGACGAAGAACTACAGAAGGAACTACTCGCCGAAGAAGAGGAGGACGTCGGAGATTATGAGGGTACAGACAAGGATGACGACGATGATGCAGCGATGGACTCGTCCAGCGACGATGAAGAAGATGCCGGACCGCCAAAGGAGGGGGAGGTGGAGGATCTGGAGGGCGAGAAACAGTTGAAGAAAGCCGAGCGAGAGGCAGCAAGGAAGAAGCGAAAGATGGATGAAGCAAAGATGCGATTGCCCGCATGGCAGAAGAAGAAGAAGGTTAAATTGGCGGACGATGCAAAGACCGACGATGGGAGCGCTCCTAAACCCAAGAAGAAGAGCGAGCGGTCGAATTGGTTGCCCACAGCTGCCGACCTTCCTTCAAGACAGTCGACTCGTACATCGGCGGTCGAGTTCCGGACCCACGTGCATGAGAATCTCAAGCTGTCGAATGTGAGATCCGAGAATCAGAAGAAGGTCATGGCGAACCATATCCAGAAATCCAAGGCGAACAAGCGTGCGGATCTGACGCAGGAAGAACGGTTTGCCAAGGCTGCGAGGATCGAGAAAGAGACCGCGAGGGAATTTGGGAGATGGGAACGAGAAGAAGCTGAGAGACAACGCATACGAGAGGAGCAATTGGCGGCGAAGCGAAGGCGCACTATCGAGGGACCGGTGATTAGGTTCTGGAGCGGGAGTGTTATCTGGGAGGGCGATAAGATCAAAGTCAAGCGTGTACATCAAAATGGGAAGGTGGAGGAGGTGAAGGAGGAAGAAGAGGCCAAACTTGCGAAGGCCGCGACTGAGATGAATGGGAAGGACGAAGAGAAAGGAAACCGAAAGGACGAAGACGGAGATATCGCAATGGTCAACCCTGAACTGGATAAGCACGGTAACGCTGGGGAGACGGATAATACTGCTGTAAAAGCACCAG-TCGCTATCGCCGATTCGAAGCCT-CCAATG	1086 	\|\|\|\|\|\|\|\|\|\|\|\|\|\|\|\|\|\|\|\|\|\|\|\|\|\|\|\|\|\|\|\|\|\|\|\|\|\|\|\|\|\|\|\|\|\|\|\|\|\|\|\|\|\|\|\|\|\|\|\|\|\|\|\|\|\|\|\|\|\|\|\|\|\|\|\|\|\|\|\|\|\|\|\|\|\|\|\|\|\|\|\|\|\|\|\|\|\|\|\|\|\|\|\|\|\|\|\|\|\|\|\|\|\|\|\|\|\|\|\|\|\|\|\|\|\|\|\|\|\|\|\|\|\|\|\|\|\|\|\|\|\|\|\|\|\|\|\|\|\|\|\|\|\|\|\|\|\|\|\|\|\|\|\|\|\|\|\|\|\|\|\|\|\|\|\|\|\|\|\|\|\|\|\|\|\|\|\|\|\|\|\|\|\|\|\|\|\|\|\|\|\|\|\|\|\|\|\|\|\|\|\|\|\|\|\|\|\|\|\|\|\|\|\|\|\|\|\|\|\|\|\|\|\|\|\|\|\|\|\|\|\|\|\|\|\|\|\|\|\|\|\|\|\|\|\|\|\|\|\|\|\|\|\|\|\|\|\|\|\|\|\|\|\|\|\|\|\|\|\|\|\|\|\|\|\|\|\|\|\|\|\|\|\|\|\|\|\|\|\|\|\|\|\|\|\|\|\|\|\|\|\|\|\|\|\|\|\|\|\|\|\|\|\|\|\|\|\|\|\|\|\|\|\|\|\|\|\|\|\|\|\|\|\|\|\|\|\|\|\|\|\|\|\|\|\|\|\|\|\|\|\|\|\|\|\|\|\|\|\|\|\|\|\|\|\|\|\|\|\|\|\|\|\|\|\|\|\|\|\|\|\|\|\|\|\|\|\|\|\|\|\|\|\|\|\|\|\|\|\|\|\|\|\|\|\|\|\|\|\|\|\|\|\|\|\|\|\|\|\|\|\|\|\|\|\|\|\|\|\|\|\|\|\|\|\|\|\|\|\|\|\|\|\|\|\|\|\|\|\|\|\|\|\|\|\|\|\|\|\|\|\|\|\|\|\|\|\|\|\|\|\|\|\|\|\|\|\|\|\|\|\|\|\|\|\|\|\|\|\|\|\|\|\|\|\|\|\|\|\|\|\|\|\|\|\|\|\|\|\|\|\|\|\|\|\|\|\|\|\|\|\|\|\|\|\|\|\|\|\|\|\|\|\|\|\|\|\|\|\|\|\|\|\|\|\|\|\|\|\|\|\|\|\|\|\|\|\|\|\|\|\|\|\|\|\|\|\|\|\|\|\|\|\|\|\|\|\|\|\|\|\|\|\|\|\|\|\|\|\|\|\|\|\|\|\|\|\|\|\|\|\|\|\|\|\|\|\|\|\|\|\|\|\|\|\|\|\|\|\|\|\|\|\|\|\|\|\|\|\|\|\|\|\|\|\|\|\|\|\|\|\|\|\|\|\|\|\|\|\|\|\|\|\|\|\|\|\|\|\|\|\|\|\|\|\|\|\|\|\|\|\|\|\|\|\|\|\|\|\|\|\|\|\|\|\|\|\|\|\|\|\|\|\|\|\|\|\|\|\|\|\|\|\|\|\|\|\|\|\|\|\|\|\|\|\|\|\|\|\|\|\|\|\|\|\|\|\|\|\|\|\|\|\|\|\|\|\|\|\|\|\|\|\|\|\|\|\|\|\|\|\|\|\|\|\|\|\|\|\|\|\|\|\|\|\|\|\|\|\|\|\|\|\|\|\|\|\|\|\|\|\|\|\|\|\|\|\|\|\|\|\|\|\|\|\|\|\|\|\|\|\|\|\|\|\|\|\|\|\|\|\|\|\|\|\|\|\|\|\|\|\|\|\|\|\|\|\|\|\|\|\|\|\|\|\|\|\|\|\|\|\|\|\|\|\|\|\|\|\|\|\|\|\|\|\|\|\|\|\|\|\|\|\|\|\|\|\|\|\|\|\|\|\|\|\|\|\|\|\|\|\|\|\|\|\|\|\|\|\|\|\|\|\|\|\|\|\|\|\|\|\|\|\|\|\|\|\|\|\|\|\|\|\|\|\|\|\|\|\|\|\|\|\|\|\|\|\| \| \|\|\|\|\| \|\|\|\|\|\|\|\|\|\|\|\|\|\|\|\|\|\|\|\|\|\|\| \|\|\|\|\|\|\|\|\|\|\|\|\|\|\|\| \| \|\|\|\|\|\|\|\|\|\| \|\|\|\|\|\|\|\| \|\|\|\|\| \| \|\|\|\|\|\|\|\|\|\|\|\|\|\|\|\|\|\|\|\|\|\|\|\|\|\|\|\|\|\|\| \|\|\|\|\|\|\| \|\|\|\|\|\|\|\|\|\|\|\|\|\|\|\|\| \|\|\|\|\| \|\|\|\|\|\| 	ATGTCCGACAACGACGAGCGCTCGTGCTCGAGCTCCGAGAGTGATGGCGAAGATGTCACCACCACCGGACTCATCGCCACACGTGCGAAGCGATCCACAGCAGGAAATCTTTACGCGACTTTGCGCCAGAACCTCGACGACGAAGAACTACAGAAGGAACTACTCGCCGAAGAAGAGGAGGACGTCGGAGATTATGAGGGTACAGACAAGGATGACGACGATGATGCAGCGATGGACTCGTCCAGCGACGATGAAGAAGATGCCGGACCGCCAAAGGAGGGGGAGGTGGAGGATCTGGAGGGCGAGAAACAGTTGAAGAAAGCCGAGCGAGAGGCAGCAAGGAAGAAGCGAAAGATGGATGAAGCAAAGATGCGATTGCCCGCATGGCAGAAGAAGAAGAAGGTTAAATTGGCGGACGATGCAAAGACCGACGATGGGAGCGCTCCTAAACCCAAGAAGAAGAGCGAGCGGTCGAATTGGTTGCCCACAGCTGCCGACCTTCCTTCAAGACAGTCGACTCGTACATCGGCGGTCGAGTTCCGGACCCACGTGCATGAGAATCTCAAGCTGTCGAATGTGAGATCCGAGAATCAGAAGAAGGTCATGGCGAACCATATCCAGAAATCCAAGGCGAACAAGCGTGCGGATCTGACGCAGGAAGAACGGTTTGCCAAGGCTGCGAGGATCGAGAAAGAGACCGCGAGGGAATTTGGGAGATGGGAACGAGAAGAAGCTGAGAGACAACGCATACGAGAGGAGCAATTGGCGGCGAAGCGAAGGCGCACTATCGAGGGACCGGTGATTAGGTTCTGGAGCGGGAGTGTTATCTGGGAGGGCGATAAGATCAAAGTCAAGCGTGTACATCAAAATGGGAAGGTGGAGGAGGTGAAGGAGGAAGAAGAGGCCAAACTTGCGAAGGCCGCGACTGAGATGAATGGTA-GGACG-AGAGAAAGGAAACCGAAAGGACG-AGACGGAGATATCGCATT-GTCAACCCTGGACTGGATA-GCACG-T-ACGCTGGGGAGACGGATAATACTGCTGTAAAGGCACCAGTTCGCTATCGCCGATTCGGAGCCTTCCAATG ``` |
| 33 | ggggacaagtttgtacaaaaaagcaggcttgatgGGTGCTGAGATTTCAAAGAC, ggggaccactttgtacaagaaagctgggtcACCGAAGGTCGGAAACACCG | Mycgr3\|92250  JGI  Ensembl | KOG0516 Dystonin, GAS (Growth-arrest-specific protein), and related proteins; |  | 100 % |  |  |
| 33 | ggggacaagtttgtacaaaaaagcaggcttgatgTCTGTTCTCAATCTGAATCC, ggggaccactttgtacaagaaagctgggtcAGATCTGCCGTCAGGTACTG | Mycgr3\|92357  JGI  Ensembl |  |  | 100 % |  |  |
| 33 | ggggacaagtttgtacaaaaaagcaggcttgatgCCGGCGGAACTTCATCAACC, ggggaccactttgtacaagaaagctgggtcGACTTCGAAGACGCGGGAGC | Mycgr3\|93297  JGI  Ensembl |  |  | 100 % |  |  |
| 33 | ggggacaagtttgtacaaaaaagcaggcttgatgGCTACTGCTCAGAGTGACTT, ggggaccactttgtacaagaaagctgggtcGTCCGGCACAGGTGACAATT | Mycgr3\|93450  JGI  Ensembl | KOG0254 Predicted transporter (major facilitator superfamily); | L-arabinose isomerase activity; carbohydrate transport;  integral to membrane;  membrane;  sugar:hydrogen symporter activity;  transport;  transporter activity; | 100 % |  |  |
| 33 | ggggacaagtttgtacaaaaaagcaggcttgatgGCGACTTCATACTTCCCGGA, ggggaccactttgtacaagaaagctgggtcACTGAGAAGCACTCTGATCT | Mycgr3\|93836  JGI  Ensembl | KOG3758 Uncharacterized conserved protein; |  | 100 % |  |  |
| 33 | ggggacaagtttgtacaaaaaagcaggcttgatgAACATGTCATCCGCCACGGT, ggggaccactttgtacaagaaagctgggtcCTTCTGAAGCATCCTCTCTT | Mycgr3\|94837  JGI  Ensembl |  |  | 100 % |  |  |
| 33 | ggggacaagtttgtacaaaaaagcaggcttgatgGGGATCCTCGACTTGCCCAA, ggggaccactttgtacaagaaagctgggtcTGGTGTTGAAGCCCATGGCG | Mycgr3\|95089  JGI  Ensembl |  |  | 100 % |  |  |
| 33 | ggggacaagtttgtacaaaaaagcaggcttgatgAAGTCTCTCGTTCACGTCGC, ggggaccactttgtacaagaaagctgggtcGACCTTGTAGCCAACCAGAC | Mycgr3\|95144  JGI  Ensembl | KOG1216 von Willebrand factor and related coagulation proteins; |  | 0 % |  |  |
| 33 | ggggacaagtttgtacaaaaaagcaggcttgatgGTCCTTCGTCGAAAGTCCGT, ggggaccactttgtacaagaaagctgggtcAACAATAGTATTCCATGGCT | Mycgr3\|95304  JGI  Ensembl |  |  | 100 % |  |  |
| 33 | ggggacaagtttgtacaaaaaagcaggcttgatgGGAATCGCGGCGCCGATTCC, ggggaccactttgtacaagaaagctgggtcCTCGAGACCTTCGTCTGCTC | Mycgr3\|95643  JGI  Ensembl |  |  | 0.99385 % |  |  |
| 33 | ggggacaagtttgtacaaaaaagcaggcttgatgGCCACTCCTCGAGTCACACG, ggggaccactttgtacaagaaagctgggtcGATGAAGCGGCGGTCAACAG | Mycgr3\|95868  JGI  Ensembl | KOG4805 Uncharacterized conserved protein; |  | 100 % |  |  |
| 33 | ggggacaagtttgtacaaaaaagcaggcttgatgGACCCAGCAAACACTCCTCT, ggggaccactttgtacaagaaagctgggtcGACGTCTGGCGCCTGCGCCT | Mycgr3\|96010  JGI  Ensembl |  |  | 100 % |  |  |
| 33 | ggggacaagtttgtacaaaaaagcaggcttgatgTCAGATCTGGGCGGCGGTGC, ggggaccactttgtacaagaaagctgggtcATCCCCTCCAGTCCTCCGAC | Mycgr3\|97494  JGI  Ensembl | KOG4297 C-type lectin; |  | 100 % |  |  |
| 33 | ggggacaagtttgtacaaaaaagcaggcttgatgCAAGCGCAATCAAACGCACC, ggggaccactttgtacaagaaagctgggtcGGCTCGAGGATGTAGGGCGA | Mycgr3\|97777  JGI  Ensembl |  |  | 100 % |  |  |
| 33 | ggggacaagtttgtacaaaaaagcaggcttgatgGCGAAATTGCGCCGCAGTCC, ggggaccactttgtacaagaaagctgggtcCTCTTCGTGCACTGCTTCGA | Mycgr3\|97805  JGI  Ensembl | KOG2739 Leucine-rich acidic nuclear protein; |  | 0 % |  |  |
| 33 | ggggacaagtttgtacaaaaaagcaggcttgatgGCTTCAGTAACATCACTAGA, ggggaccactttgtacaagaaagctgggtcCGGCTTCGGCCTTGGCCCAA | Mycgr3\|98363  JGI  Ensembl | KOG2046 Calponin; |  | 100 % |  |  |
| 33 | ggggacaagtttgtacaaaaaagcaggcttgatgTCGTTCCTCGGCGGGCCAGA, ggggaccactttgtacaagaaagctgggtcGAAGTCGAATTGCCCTCGCA | Mycgr3\|98722  JGI  Ensembl | KOG1125 TPR repeat-containing protein; |  | 0 % |  |  |
| 33 | ggggacaagtttgtacaaaaaagcaggcttgatgGACGGCTGGCAGGCAGTGCA, ggggaccactttgtacaagaaagctgggtcCGCGTCCCAGTTGGAAGCGT | Mycgr3\|98908  JGI  Ensembl | KOG2992 Nucleolar GTPase/ATPase p130; | cytoplasm; inorganic diphosphatase activity;  magnesium ion binding;  phosphate metabolic process; | 100 % |  |  |
| 33 | ggggacaagtttgtacaaaaaagcaggcttgatgCCGCCAAAGGTCAATATCGA, ggggaccactttgtacaagaaagctgggtcGAGATCGCCCAACTCCTTCA | Mycgr3\|99913  JGI  Ensembl | KOG4389 Acetylcholinesterase/Butyrylcholinesterase; | carboxylesterase activity; | 100 % |  |  |
| 33 | ggggacaagtttgtacaaaaaagcaggcttgatgTCTGACGGGAGACTGCAGTC, ggggaccactttgtacaagaaagctgggtcCCATGCGTTGACGTCAAAGT | Mycgr3\|100024  JGI  Ensembl | KOG0120 Splicing factor U2AF, large subunit (RRM superfamily); | RNA binding; mRNA processing;  nucleic acid binding;  nucleus; | 100 % |  |  |
| 33 | ggggacaagtttgtacaaaaaagcaggcttgatgACCTACGGAGCCAAGATCAT, ggggaccactttgtacaagaaagctgggtcACGGTCTGCCACGGCGCTTG | Mycgr3\|100226  JGI  Ensembl | KOG1185 Thiamine pyrophosphate-requiring enzyme; | magnesium ion binding; thiamin pyrophosphate binding;  transferase activity; | 100 % |  |  |
| 33 | ggggacaagtttgtacaaaaaagcaggcttgatgTCGACTTCATCAGAGAACGA, ggggaccactttgtacaagaaagctgggtcGATAAGATCCTCAACCATGT | Mycgr3\|100227  JGI  Ensembl | KOG1177 Long chain fatty acid acyl-CoA ligase; | catalytic activity; fatty-acid ligase activity;  long-chain-fatty-acid-CoA ligase activity;  medium-chain-fatty-acid-CoA ligase activity;  metabolic process;  short-chain-fatty-acid-CoA ligase activity;  very-long-chain-fatty-acid-CoA ligase activity; | 100 % |  |  |
| 33 | ggggacaagtttgtacaaaaaagcaggcttgatgGCCACCGCACAGCACGCCGA, ggggaccactttgtacaagaaagctgggtcTGCATCCTCCGCCTCTACCC | Mycgr3\|100794  JGI  Ensembl | KOG0201 Serine/threonine protein kinase; | 3-phosphoinositide-dependent protein kinase activity; AMP-activated protein kinase activity;  ATP binding;  DNA-dependent protein kinase activity;  GTP-dependent protein kinase activity;  JUN kinase activity;  JUN kinase kinase activity;  JUN kinase kinase kinase activity;  Janus kinase activity;  MAP kinase 1 activity;  MAP kinase 2 activity;  MAP kinase kinase activity;  MAP kinase kinase kinase kinase activity;  MAP/ERK kinase kinase activity;  MP kinase activity;  NF-kappaB-inducing kinase activity;  SAP kinase activity;  atypical protein kinase C activity;  calcium-dependent protein kinase C activity;  casein kinase I activity;  casein kinase activity;  cyclic nucleotide-dependent protein kinase activity;  cyclin-dependent protein kinase activating kinase activity;  eukaryotic translation initiation factor 2alpha kinase activity;  galactosyltransferase-associated kinase activity;  glycogen synthase kinase 3 activity;  protein amino acid phosphorylation;  protein kinase activity;  protein serine/threonine kinase activity;  protein-tyrosine kinase activity;  receptor signaling protein serine/threonine kinase activity;  ribosomal protein S6 kinase activity;  transmembrane receptor protein kinase activity;  transmembrane receptor protein serine/threonine kinase activity; | 87.65957 % |  |  |
| 33 | ggggacaagtttgtacaaaaaagcaggcttgatgCCCGACCAACCCTCCACCAA, ggggaccactttgtacaagaaagctgggtcTCCCGCCAAAACGTGGACCC | Mycgr3\|100869  JGI  Ensembl | KOG4229 Myosin VII, myosin IXB and related myosins; |  | 100 % |  |  |
| 33 | ggggacaagtttgtacaaaaaagcaggcttgatgGACGGCCAGGCCTCTCAGGA, ggggaccactttgtacaagaaagctgggtcCATTGATATTCTCCGCGCCA | Mycgr3\|100967  JGI  Ensembl | KOG0375 Serine-threonine phosphatase 2B, catalytic subunit; | CTD phosphatase activity; calcium-dependent protein serine/threonine phosphatase activity;  calcium-dependent protein serine/threonine phosphatase regulator activity;  hydrolase activity;  magnesium-dependent protein serine/threonine phosphatase activity;  magnesium-dependent protein serine/threonine phosphatase complex;  myosin phosphatase activity;  myosin phosphatase complex;  myosin phosphatase regulator activity;  phosphoprotein phosphatase activity; | 100 % |  |  |
| 33 | ggggacaagtttgtacaaaaaagcaggcttgatgCGGAATCGGCTCGCAGTGGG, ggggaccactttgtacaagaaagctgggtcTCGCCAAATCTGCTCCTTCC | Mycgr3\|101252  JGI  Ensembl | KOG0927 Predicted transporter (ABC superfamily); | ATP binding; ATPase activity;  nucleoside-triphosphatase activity;  nucleotide binding; | 100 % |  |  |
| 33 | ggggacaagtttgtacaaaaaagcaggcttgatgGTGGGCGCAAGGCATGCCGC, ggggaccactttgtacaagaaagctgggtcACCCAGAGACGCCAGAATAG | Mycgr3\|101338  JGI  Ensembl | KOG2344 Exocyst component protein and related proteins; | exocyst; exocytosis; | 100 % | chr\_10:146052 G->A; |  |
| 33 | ggggacaagtttgtacaaaaaagcaggcttgatgGGATATTACGACGAAGACGG, ggggaccactttgtacaagaaagctgggtcCAAGCGGCTACCGTGGACGA | Mycgr3\|103731  JGI  Ensembl | KOG3271 Translation initiation factor 5A (eIF-5A); | translation initiation factor activity; translational initiation; | 100 % |  |  |
| 33 | ggggacaagtttgtacaaaaaagcaggcttgatgGTCCGATCAACCATCATCGC, ggggaccactttgtacaagaaagctgggtcCAATGACCGAGCGTGCGCAC | Mycgr3\|103950  JGI  Ensembl |  |  | 100 % |  |  |
| 33 | ggggacaagtttgtacaaaaaagcaggcttgatgGCGATCGCACGCCATGGCTC, ggggaccactttgtacaagaaagctgggtcGACCAGCTTTCCTCTTCCAC | Mycgr3\|106901  JGI  Ensembl | KOG4466 Component of histone deacetylase complex (breast carcinoma metastasis suppressor 1 protein in human); |  | 100 % | chr\_1:1316185 G->A; chr\_1:1316203 C->A; |  |
| 33 | ggggacaagtttgtacaaaaaagcaggcttgatgGCTGCCCAAATCTTCACCTG, ggggaccactttgtacaagaaagctgggtcATCACTCTCGTACGCAATGC | Mycgr3\|107169  JGI  Ensembl |  |  | 100 % |  |  |
| 33 | ggggacaagtttgtacaaaaaagcaggcttgatgTCGTCCGATCGCCCTGCCAG, ggggaccactttgtacaagaaagctgggtcAGGAACATGCTGGCGTCGGT | Mycgr3\|107196  JGI  Ensembl | KOG2992 Nucleolar GTPase/ATPase p130; |  | 0 % |  |  |
| 33 | ggggacaagtttgtacaaaaaagcaggcttgatgGCTCCCCGCGGCTCCGCATA, ggggaccactttgtacaagaaagctgggtcCTTCTCCATCAATCCCCTCA | Mycgr3\|107790  JGI  Ensembl |  |  | 100 % |  |  |
| 33 | ggggacaagtttgtacaaaaaagcaggcttgatgCTCCTCAGCTCCGCCCTCCT, ggggaccactttgtacaagaaagctgggtcGGAGGATGGGGAGCAGGAGC | Mycgr3\|108014  JGI  Ensembl |  |  | 0 % |  |  |
| 33 | ggggacaagtttgtacaaaaaagcaggcttgatgGCTTTCAACAACTTCAACGA, ggggaccactttgtacaagaaagctgggtcATACTCCTCATCACTCCCAT | Mycgr3\|108274  JGI  Ensembl |  |  | 100 % |  |  |
| 33 | ggggacaagtttgtacaaaaaagcaggcttgatgCATCTCCTCAAAGTCCTCTC, ggggaccactttgtacaagaaagctgggtcAGTGATTGGCTCGATCGCAA | Mycgr3\|108349  JGI  Ensembl | KOG4157 beta-1,6-N-acetylglucosaminyltransferase, contains WSC domain; |  | 100 % |  |  |
| 33 | ggggacaagtttgtacaaaaaagcaggcttgatgGCCACAGCAGACGTCCTTCA, ggggaccactttgtacaagaaagctgggtcCTTCTCAAAATACTGCATCT | Mycgr3\|108578  JGI  Ensembl | KOG1399 Flavin-containing monooxygenase; | FAD binding; electron transport;  oxidoreductase activity; | 100 % |  |  |
| 33 | ggggacaagtttgtacaaaaaagcaggcttgatgCCACCTGCATTGCCAGACAC, ggggaccactttgtacaagaaagctgggtcTTTCGTTAATCCCACCCGCT | Mycgr3\|108849  JGI  Ensembl | KOG2992 Nucleolar GTPase/ATPase p130; | chromatin; chromatin assembly or disassembly;  chromatin binding;  nucleus; | 1.89663 % |  |  |
| 33 | ggggacaagtttgtacaaaaaagcaggcttgatgAAATCTGCTGGTATCATCGC, ggggaccactttgtacaagaaagctgggtcCCTCATATACTCCTTCGCAT | Mycgr3\|108908  JGI  Ensembl | KOG4389 Acetylcholinesterase/Butyrylcholinesterase; | carboxylesterase activity; | 1.95408 % |  |  |
| 33 | ggggacaagtttgtacaaaaaagcaggcttgatgGCCGACACTCACTCCGATCG, ggggaccactttgtacaagaaagctgggtcTCCCGTCTCGACCTTGACCT | Mycgr3\|108995  JGI  Ensembl | KOG2827 Uncharacterized conserved protein; |  | 100 % | chr\_4:726115 C->T; |  |
| 33 | ggggacaagtttgtacaaaaaagcaggcttgatgGATCTCCTCAGGCTTGAACA, ggggaccactttgtacaagaaagctgggtcCGACTTGATCCTCGCAGCCC | Mycgr3\|109095  JGI  Ensembl | KOG2992 Nucleolar GTPase/ATPase p130; |  | 100 % |  |  |
| 33 | ggggacaagtttgtacaaaaaagcaggcttgatgGTCATCATCCTCCAAGTCGG, ggggaccactttgtacaagaaagctgggtcCGTCCCACGAATACCCCAGC | Mycgr3\|109136  JGI  Ensembl |  |  | 100 % |  |  |
| 33 | ggggacaagtttgtacaaaaaagcaggcttgatgAGCGCCAGCAATGTCCGCAC, ggggaccactttgtacaagaaagctgggtcATCCGCCTTCCCACTCTTGC | Mycgr3\|109266  JGI  Ensembl | KOG0940 Ubiquitin protein ligase RSP5/NEDD4; | cell wall catabolic process; | 100 % |  |  |
| 33 | ggggacaagtttgtacaaaaaagcaggcttgatgGCCGCCGTGATGACAGTCCC, ggggaccactttgtacaagaaagctgggtcCCCCTGCCTCCCATCCTCCT | Mycgr3\|109284  JGI  Ensembl | KOG3712 RFX family transcription factor; | DNA binding; nucleus; regulation of transcription, DNA-dependent;  transcription;  transcription factor activity;  zinc ion binding; | 1.85185 % |  |  |
| 33 | ggggacaagtttgtacaaaaaagcaggcttgatgACAATGGCTCGCTTGCGAAA, ggggaccactttgtacaagaaagctgggtcCCGCCAGCTTTGACTGGACC | Mycgr3\|109571  JGI  Ensembl | KOG1181 FOG: Low-complexity; |  | 100 % |  |  |
| 33 | ggggacaagtttgtacaaaaaagcaggcttgatgGAGATCCCCAAGACCGAGGC, ggggaccactttgtacaagaaagctgggtcTCCCGCGAAAAATTGTAAGC | Mycgr3\|109829  JGI  Ensembl |  |  | 100 % |  |  |
| 33 | ggggacaagtttgtacaaaaaagcaggcttgatgAAAATGCACGACGACCTTCC, ggggaccactttgtacaagaaagctgggtcTCCTCCATCATCATCGCCCT | Mycgr3\|109897  JGI  Ensembl |  |  | 100 % |  |  |
| 33 | ggggacaagtttgtacaaaaaagcaggcttgatgCCTTCCACTAGGGACCTCCT, ggggaccactttgtacaagaaagctgggtcATAGTAAGGCGCCGCGGACG | Mycgr3\|109922  JGI  Ensembl | KOG4701 Chitinase; | exo-alpha-sialidase activity; | 100 % |  |  |
| 33 | ggggacaagtttgtacaaaaaagcaggcttgatgTGCCCGACTCCAGATGATGC, ggggaccactttgtacaagaaagctgggtcTGACACCACCGCCGTCTTGG | Mycgr3\|110230  JGI  Ensembl | KOG2367 Alpha-isopropylmalate synthase/homocitrate synthase; | carboxylic acid metabolic process; catalytic activity;  homocitrate synthase activity;  transferase activity, transferring acyl groups, acyl groups converted into alkyl on transfer; | 100 % |  |  |
| 33 | ggggacaagtttgtacaaaaaagcaggcttgatgGGCTTCCTCAAGCACTTCCG, ggggaccactttgtacaagaaagctgggtcCCTCCACAAATCATGCTTGC | Mycgr3\|110511  JGI  Ensembl |  |  | 100 % | chr\_8:1377262 C->G; chr\_8:1377432 G->T; |  |
| 33 | ggggacaagtttgtacaaaaaagcaggcttgatgAGTGGCTTCGAGATCGCGGG, ggggaccactttgtacaagaaagctgggtcCAGAGCAATCTGCGTCGCCC | Mycgr3\|110759  JGI  Ensembl |  |  | 1.04767 % |  |  |
| 33 | ggggacaagtttgtacaaaaaagcaggcttgatgGCTGCTGTGGTCGAGAAAGC, ggggaccactttgtacaagaaagctgggtcCGAGCGTGGGGCAGCGGCAC | Mycgr3\|111025  JGI  Ensembl | KOG2567 Uncharacterized conserved protein; |  | 100 % |  |  |
| 33 | ggggacaagtttgtacaaaaaagcaggcttgatgTTTGAGACTCACCGATCTCG, ggggaccactttgtacaagaaagctgggtcTGCCGGCGAAGCACCGACTC | Mycgr3\|111085  JGI  Ensembl | KOG2419 Phosphatidylserine decarboxylase; | phosphatidylserine decarboxylase activity; phospholipid biosynthetic process; | 100 % |  |  |
| 33 | ggggacaagtttgtacaaaaaagcaggcttgatgCTGCTCGATATGAAGCTCCT, ggggaccactttgtacaagaaagctgggtcATACGAAGGCGGCGAGAGTG | Mycgr3\|111476  JGI  Ensembl |  |  | 0 % |  |  |
| 33 | ggggacaagtttgtacaaaaaagcaggcttgatgGGCCGAATGACGAGAGCGAA, ggggaccactttgtacaagaaagctgggtcGGATCCCGCAGCATTCGGGA | Mycgr3\|111542  JGI  Ensembl | KOG2992 Nucleolar GTPase/ATPase p130; |  | 0 % |  |  |
| 33 | ggggacaagtttgtacaaaaaagcaggcttgatgTCCACCTTCCTCGACATAGC, ggggaccactttgtacaagaaagctgggtcCTGGGGTACACAATTGTCGA | Mycgr3\|111708  JGI  Ensembl |  |  | 1.95704 % |  |  |
| 34 | ggggacaagtttgtacaaaaaagcaggcttgatgTCCTTCAAGCGCTCCAGGGC, ggggaccactttgtacaagaaagctgggtcGTCGTCTTCAGAATCACTCC | Mycgr3\|31345  JGI  Ensembl | KOG2138 Predicted RNA binding protein, contains G-patch domain; | intracellular; nucleic acid binding; | 100 % |  |  |
| 34 | ggggacaagtttgtacaaaaaagcaggcttgatgGGCCTGGAGGAGTTTGAGAA, ggggaccactttgtacaagaaagctgggtcCTGCGGCTCCATCAACGCCT | Mycgr3\|33196  JGI  Ensembl | KOG2477 Uncharacterized conserved protein; |  | 0 % |  |  |
| 34 | ggggacaagtttgtacaaaaaagcaggcttgatgTTTACGCATACCGCACCAGC, ggggaccactttgtacaagaaagctgggtcGAGATCACCATATTCCTCTC | Mycgr3\|34982  JGI  Ensembl | KOG1238 Glucose dehydrogenase/choline dehydrogenase/mandelonitrile lyase (GMC oxidoreductase family); | FAD binding; choline dehydrogenase activity;  oxidoreductase activity, acting on CH-OH group of donors; | 100 % |  |  |
| 34 | ggggacaagtttgtacaaaaaagcaggcttgatgGACGCTCCATCACAGCGGTC, ggggaccactttgtacaagaaagctgggtcCCAACTATCATTGTCAAACA | Mycgr3\|35076  JGI  Ensembl | KOG2806 Chitinase; | nucleus; protein dimerization activity;  regulation of transcription, DNA-dependent;  sequence-specific DNA binding;  transcription factor activity; | 100 % |  |  |
| 34 | ggggacaagtttgtacaaaaaagcaggcttgatgCCATGGTATAGCCGCCACAG, ggggaccactttgtacaagaaagctgggtcAACAGAGATGATGGCCTTGG | Mycgr3\|35889  JGI  Ensembl |  |  | 0 % |  |  |
| 34 | ggggacaagtttgtacaaaaaagcaggcttgatgCCAGCGAGCCGCACTGCGCG, ggggaccactttgtacaagaaagctgggtcCACAGCACCATAGAGATCCC | Mycgr3\|36302  JGI  Ensembl |  |  | 100 % |  | ``` 1	ATGCCAGCGAGCCGCACTGCGCGGCGAGGCCTCGCCGAGAAAGCCTGCAACGAATGTCAGAGACGGAAGACGAGATGCATACCAAGCGAAGACGATTCGCGCTGCGCCTTCTGCCAGCGCGCCGGGAAGGCGTGTGTCTTTGAGAAACCACCGGCCAGGACGCCGTTGACTCGGCAGAATCTTGATCAAGCGGAGGCACGATGTCGACAATTGGAAGAAGTCCTCACGCGGCGTGAGGCTGAGATGTACTCGAATCATGCCAATGGAGGGGAGAATCCGCAGCAGCCAGCGCACAAGGCACACGCTGCAGACCACAATGCTGGGGAGGGGAATGCGGACTACGAGTGGAACGAGGCCTCGGATCATGAAGCGAGAGAGCAGTCATCAGATAGCGATCATGCCAGAGATGGCATGGCAGCGCTGACGAGTCGAAATGCGGGCTCAGGCTATCTGGG	455 	\|\|\|\|\|\|\|\|\|\|\|\|\|\|\|\|\|\|\|\|\|\|\|\|\|\|\|\|\|\|\|\|\|\|\|\|\|\|\|\|\|\|\|\|\|\|\|\|\|\|\|\|\|\|\|\|\|\|\|\|\|\|\|\|\|\|\|\|\|\|\|\|\|\|\|\|\|\|\|\|\|\|\|\|\|\|\|\|\|\|\|\|\|\|\|\|\|\|\|\|\|\|\|\|\|\|\|\|\|\|\|\|\|\|\|\|\|\|\|\|\|\|\|\|\|\|\|\|\|\|\|\|\|\|\|\|\|\|\|\|\|\|\|\|\|\|\|\|\|\|\|\|\|\|\|\|\|\|\|\|\|\|\|\|\|\|\|\|\|\|\|\|\|\|\|\|\|\|\|\|\|\|\|\|\|\|\|\|\|\|\|\|\|\|\|\|\|\|\|\|\|\|\|\|\|\|\|\|\|\|\|\|\|\|\|\|\|\|\|\|\|\|\|\|\|\|\|\|\|\|\|\|\|\|\|\|\|\|\|\|\|\|\|\|\|\|\|\|\|\|\|\|\|\|\|\|\|\|\|\|\|\|\|\|\|\|\|\|\|\|\|\|\|\|\|\|\|\|\|\|\|\|\|\|\|\|\|\|\|\|\|\|\|\|\|\|\|\|\|\|\|\|\|\|\|\|\|\|\|\|\|\|\|\|\|\|\|\|\|\|\|\|\|\|\|\|\|\|\|\|\|\|\|\|\|\|\|\|\|\|\|\|\|\|\|\|\|\|\|\|\|\|\|\|\|\|\|\|\|\|\|\|\|\|\|\|\|\|\|\|\|\|\|\|\|\|\|\|\|\|\|\|\|\|\|\|\|\|\|\|\|\|\|\|\|\|\|\|\|\|\|\|\|\|\|\|\|\|\|\|\|\|\|\|\|\|\|\|\|\|\|\|\|\|\|\|\|\|\|\|\|\|\|\|\|\|\|\|\|\|\|\|\|\|\|\|\|\|\|\|\|\|\|\|\| 	ATGCCAGCGAGCCGCACTGCGCGGCGAGGCCTCGCCGAGAAAGCCTGCAACGAATGTCAGAGACGGAAGACGAGATGCATACCAAGCGAAGACGATTCGCGCTGCGCCTTCTGCCAGCGCGCCGGGAAGGCGTGTGTCTTTGAGAAACCACCGGCCAGGACGCCGTTGACTCGGCAGAATCTTGATCAAGCGGAGGCACGATGTCGACAATTGGAAGAAGTCCTCACGCGGCGTGAGGCTGAGATGTACTCGAATCATGCCAATGGAGGGGAGAATCCGCAGCAGCCAGCGCACAAGGCACACGCTGCAGACCACAATGCTGGGGAGGGGAATGCGGACTACGAGTGGAACGAGGCCTCGGATCATGAAGCGAGAGAGCAGTCATCAGATAGCGATCATGCCAGAGATGGCATGGCAGCGCTGACGAGTCGAAATGCGGGCTCAGGCTATCTGGG ```      ``` 454	GGCACGAGTTCCGGCCACGAACTGTTGCAATCAGTATCCTCTTTTCTCCCTCCAGCAACCGGCGCTGGTCGCCGCAGCAAGAGTGCCGCGCAGGAGACGCCGAATTCGAACGCTTCTGGGCACGGCTTACCGCCATCCTCGTATGCGAGCTCCTTGTCCGCCTCGTCGATCCAAGAAGTGCTCATATCCGCATACTTTACCCGGTATAACAGCAGCTATCCCATCCTCAACGAGAAGTCCTTTCGCGATCAGTGTGCTTCGAGGAAGTCGCTGCCTCGATCCTCGTCATGGCACATCACGTACTACGCTGTGCTCGCGATGGGCGAGTGGTTATCCGGCAGTGGTGCGGACGACAGCCCTTCTCTATACTACGAGGCCGCACGATCGTTATTGAAGCCCGAAGTCCTCGAATCCGGCTCCATCAACAATGTACAGGCATTTCTCATTCTTGGGAACTACCTGCAGAAAACGGATCGGCCCAACACAGGTTACAATTATCTCGGCATCGCTCTGAGACTCGCTATGGGTATTGGACTTCATCGTGAGCTTCCAAGTACGTCGACTG-CGGGACCAT-CGA	1030 	\|\|\|\|\|\|\|\|\|\|\|\|\|\|\|\|\|\|\|\|\|\|\|\|\|\|\|\|\|\|\|\|\|\|\|\|\|\|\|\|\|\|\|\|\|\|\|\|\|\|\|\|\|\|\|\|\|\|\|\|\|\|\|\|\|\|\|\|\|\|\|\|\|\|\|\|\|\|\|\|\|\|\|\|\|\|\|\|\|\|\|\|\|\|\|\|\|\|\|\|\|\|\|\|\|\|\|\|\|\|\|\|\|\|\|\|\|\|\|\|\|\|\|\|\|\|\|\|\|\|\|\|\|\|\|\|\|\|\|\|\|\|\|\|\|\|\|\|\|\|\|\|\|\|\|\|\|\|\|\|\|\|\|\|\|\|\|\|\|\|\|\|\|\|\|\|\|\|\|\|\|\|\|\|\|\|\|\|\|\|\|\|\|\|\|\|\|\|\|\|\|\|\|\|\|\|\|\|\|\|\|\|\|\|\|\|\|\|\|\|\|\|\|\|\|\|\|\|\|\|\|\|\|\|\|\|\|\|\|\|\|\|\|\|\|\|\|\|\|\|\|\|\|\|\|\|\|\|\|\|\|\|\|\|\|\|\|\|\|\|\|\|\|\|\|\|\|\|\|\|\|\|\|\|\|\|\|\|\|\|\|\|\|\|\|\|\|\|\|\|\|\|\|\|\|\|\|\|\|\|\|\|\|\|\|\|\|\|\|\|\|\|\|\|\|\|\|\|\|\|\|\|\|\|\|\|\|\|\|\|\|\|\|\|\|\|\|\|\|\|\|\|\|\|\|\|\|\|\|\|\|\|\|\|\|\|\|\|\|\|\|\|\|\|\|\|\|\|\|\|\|\|\|\|\|\|\|\|\|\|\|\|\|\|\|\|\|\|\|\|\|\|\|\|\|\|\|\|\|\|\|\|\|\|\|\|\|\|\|\|\|\|\|\|\|\|\|\|\|\|\|\|\|\|\|\|\|\|\|\|\|\|\|\|\|\|\|\|\|\|\|\|\|\|\|\|\|\|\|\|\|\|\|\|\|\|\|\| \| \|\|\|\|\|\|\|\|\|\|\|\|\|\|\|\| \| \|\|\| \| \|\|\|\|\| \|\|\|\|\|\|\|\|\|\|\|\|\|\|\|\|\|\|\|\|\|\|\|\| \|\|\| \| \|\| \|\|\|\|\|\|\|\|\|\|\|\|\|\|  \|\|\|\|\|\|\|\|\|\|\|\|\| \|\|\|\|\|\|\|\|\| \|\|\| 	GGCACGAGTTCCGGCCACGAACTGTTGCAATCAGTATCCTCTTTTCTCCCTCCAGCAACCGGCGCTGGTCGCCGCAGCAAGAGTGCCGCGCAGGAGACGCCGAATTCGAACGCTTCTGGGCACGGCTTACCGCCATCCTCGTATGCGAGCTCCTTGTCCGCCTCGTCGATCCAAGAAGTGCTCATATCCGCATACTTTACCCGGTATAACAGCAGCTATCCCATCCTCAACGAGAAGTCCTTTCGCGATCAGTGTGCTTCGAGGAAGTCGCTGCCTCGATCCTCGTCATGGCACATCACGTACTACGCTGTGCTCGCGATGGGCGAGTGGTTATCCGGCAGTGGTGCGGACGACAGCCCTTCTCTATACTACGAGGCCGCACGATCGTTATTGAAGCCCGAAGTCCTCGAATCCGGCTCCATCAACAATGTACAGGCATTTCTCATTCTTGGGAACTACCTGCAGAAA-C-GATCGGCCCAACACAG-T-ACA-T-ATCTC-GCATCGCTCTGAGACTCGCTATGG-TAT-G-AC-TCATCGTGAGCTTC--AGTACGTCGACTGGCGGGACCATTCGA ``` |
| 34 | ggggacaagtttgtacaaaaaagcaggcttgatgAAGGAAAGATCTGGCTTGTT, ggggaccactttgtacaagaaagctgggtcGGATCCGTGCCTGCCTCGTC | Mycgr3\|39012  JGI  Ensembl | KOG1172 Na+-independent Cl/HCO3 exchanger AE1 and related transporters (SLC4 family); | anion exchanger activity; anion transport;  integral to membrane; | 100 % |  |  |
| 34 | ggggacaagtttgtacaaaaaagcaggcttgatgGCTGCCCTTCCTCAATTCAC, ggggaccactttgtacaagaaagctgggtcGGCGGCAACGGCGGCGCCGG | Mycgr3\|42292  JGI  Ensembl | KOG2493 Na+/Pi symporter; | inorganic phosphate transmembrane transporter activity; membrane; phosphate transport; | 100 % |  |  |
| 34 | ggggacaagtttgtacaaaaaagcaggcttgatgGCATCGCGCAGGAGCTTGGG, ggggaccactttgtacaagaaagctgggtcTCCCAATCCCTCTTCATAAA | Mycgr3\|48386  JGI  Ensembl | KOG1842 FYVE finger-containing protein; | 1-phosphatidylinositol-3-phosphate 5-kinase activity; intracellular;  nucleic acid binding;  zinc ion binding; | 0.85934 % |  |  |
| 34 | ggggacaagtttgtacaaaaaagcaggcttgatgGCCAAGGTCCTCAATCGCAT, ggggaccactttgtacaagaaagctgggtcGACCCTCTGCTCCACCTCCT | Mycgr3\|49829  JGI  Ensembl | KOG0236 Sulfate/bicarbonate/oxalate exchanger SAT-1 and related transporters (SLC26 family); | integral to membrane; transport; transporter activity; | 100 % |  |  |
| 34 | ggggacaagtttgtacaaaaaagcaggcttgatgGAGAGCTACAAATACACACC, ggggaccactttgtacaagaaagctgggtcCGCAAGCAAGATGGCGGTGG | Mycgr3\|50442  JGI  Ensembl |  |  | 0.94251 % |  |  |
| 34 | ggggacaagtttgtacaaaaaagcaggcttgatgTCCAAGGACTACGAGAAAGG, ggggaccactttgtacaagaaagctgggtcGACAATATACCGATAGCCCT | Mycgr3\|60310  JGI  Ensembl | KOG2533 Permease of the major facilitator superfamily; | integral to membrane; transport; transporter activity; | 2.7551 % |  |  |
| 34 | ggggacaagtttgtacaaaaaagcaggcttgatgCGAACCACTGCTCCAGCCGT, ggggaccactttgtacaagaaagctgggtcCAAGAAGAGTGCGATGATCG | Mycgr3\|64715  JGI  Ensembl | KOG1325 Lysophospholipase; | lysophospholipase activity; phospholipase activity;  phospholipid catabolic process; | 100 % |  |  |
| 34 | ggggacaagtttgtacaaaaaagcaggcttgatgGTGCTGGGACATGCTCGAAA, ggggaccactttgtacaagaaagctgggtcTGACTCCGCGTCCACGAAGA | Mycgr3\|64917  JGI  Ensembl | KOG2210 Oxysterol-binding protein; | steroid metabolic process; | 100 % | chr\_1:348677 G->T; |  |
| 34 | ggggacaagtttgtacaaaaaagcaggcttgatgGCACCTAACGCGTCCAAGAA, ggggaccactttgtacaagaaagctgggtcCTCTCCCTCCACATCAGCCG | Mycgr3\|65589  JGI  Ensembl | KOG4191 Histone acetyltransferases PCAF/SAGA/ADA, subunit TADA3L/NGG1; |  | 100 % |  |  |
| 34 | ggggacaagtttgtacaaaaaagcaggcttgatgGCCGACACAGAAAATCTAGA, ggggaccactttgtacaagaaagctgggtcTTGCCCTTCCTTGACAGCCC | Mycgr3\|66265  JGI  Ensembl | KOG2624 Triglyceride lipase-cholesterol esterase; | lipid metabolic process; | 100 % |  |  |
| 34 | ggggacaagtttgtacaaaaaagcaggcttgatgGGAAAACGCCGCAAACGCTC, ggggaccactttgtacaagaaagctgggtcTTTAACAGCCACAAATCCCA | Mycgr3\|66834  JGI  Ensembl | KOG2248 3'-5' exonuclease; | DNA binding; exonuclease activity;  intracellular; | 100 % |  |  |
| 34 | ggggacaagtttgtacaaaaaagcaggcttgatgCCATACCACCACAGACTTCC, ggggaccactttgtacaagaaagctgggtcAGCCTGCACCGGCGAATCCT | Mycgr3\|67933  JGI  Ensembl | KOG1238 Glucose dehydrogenase/choline dehydrogenase/mandelonitrile lyase (GMC oxidoreductase family); | FAD binding; choline dehydrogenase activity;  oxidoreductase activity, acting on CH-OH group of donors; | 1.89639 % |  |  |
| 34 | ggggacaagtttgtacaaaaaagcaggcttgatgAGCGGCGTCGCGACCAAGCA, ggggaccactttgtacaagaaagctgggtcCTCATGAGCTTTGGATGCTT | Mycgr3\|68012  JGI  Ensembl | KOG0698 Serine/threonine protein phosphatase; | CTD phosphatase activity; calcium-dependent protein serine/threonine phosphatase activity;  calcium-dependent protein serine/threonine phosphatase regulator activity;  catalytic activity;  magnesium-dependent protein serine/threonine phosphatase activity;  magnesium-dependent protein serine/threonine phosphatase complex;  myosin phosphatase activity;  myosin phosphatase complex;  myosin phosphatase regulator activity;  phosphoprotein phosphatase activity;  protein amino acid dephosphorylation;  protein serine/threonine phosphatase activity;  protein serine/threonine phosphatase complex; | 0 % |  |  |
| 34 | ggggacaagtttgtacaaaaaagcaggcttgatgCGCGAAGTCAACTTCAGCAT, ggggaccactttgtacaagaaagctgggtcAGTATGACTCTGATTCTGCT | Mycgr3\|68176  JGI  Ensembl | KOG1869 Splicing coactivator SRm160/300, subunit SRm300; | zinc ion binding; | 100 % |  |  |
| 34 | ggggacaagtttgtacaaaaaagcaggcttgatgCGAACCGCCCTTCCCCTCGC, ggggaccactttgtacaagaaagctgggtcAAAGTTCGGCGACAACATAT | Mycgr3\|68458  JGI  Ensembl | KOG1236 Predicted unusual protein kinase; |  | 72.98294 % |  |  |
| 34 | ggggacaagtttgtacaaaaaagcaggcttgatgGCGGCCCTCGATGCTGTGCG, ggggaccactttgtacaagaaagctgggtcCTCCAAATCCTCAGGTGCAT | Mycgr3\|68790  JGI  Ensembl | KOG2331 Predicted glycosylhydrolase; |  | 0.96685 % |  |  |
| 34 | ggggacaagtttgtacaaaaaagcaggcttgatgTCTACCGCCGTCGCAACGCC, ggggaccactttgtacaagaaagctgggtcAGAATGAAATAAATCCCAGG | Mycgr3\|69186  JGI  Ensembl | KOG0391 SNF2 family DNA-dependent ATPase; |  | 100 % | chr\_2:3770505 G->A; |  |
| 34 | ggggacaagtttgtacaaaaaagcaggcttgatgAATCACGAAGACACAGCCGC, ggggaccactttgtacaagaaagctgggtcGATCTTCTCCTTCGGCTCTT | Mycgr3\|69338  JGI  Ensembl | KOG1237 H+/oligopeptide symporter; | membrane; oligopeptide transport;  transporter activity; | 100 % |  |  |
| 34 | ggggacaagtttgtacaaaaaagcaggcttgatgGCCAGCACCGCCGAGAAGCT, ggggaccactttgtacaagaaagctgggtcCTTCCCCGCCAACCTCTTCA | Mycgr3\|70436  JGI  Ensembl | KOG2505 Ankyrin repeat protein; | intracellular; nucleic acid binding;  zinc ion binding; | 1.08965 % |  |  |
| 34 | ggggacaagtttgtacaaaaaagcaggcttgatgCGTACAGTGTCTTCGGCCGC, ggggaccactttgtacaagaaagctgggtcCGACTCCGTCTCTCCCTCTT | Mycgr3\|70525  JGI  Ensembl | KOG0238 3-Methylcrotonyl-CoA carboxylase, biotin-containing subunit/Propionyl-CoA carboxylase, alpha chain/Acetyl-CoA carboxylase, biotin carboxylase subunit; | ATP binding; catalytic activity;  ligase activity;  metabolic process;  methylcrotonoyl-CoA carboxylase activity; | 100 % |  |  |
| 34 | ggggacaagtttgtacaaaaaagcaggcttgatgCGGACCTCCAGCGAGTCGCA, ggggaccactttgtacaagaaagctgggtcAGTCTCCTTATAATACCCCC | Mycgr3\|70577  JGI  Ensembl | KOG0254 Predicted transporter (major facilitator superfamily); |  | 100 % | chr\_4:34852 A->G; |  |
| 34 | ggggacaagtttgtacaaaaaagcaggcttgatgACCGACTGGAAGGCTGCCGG, ggggaccactttgtacaagaaagctgggtcGCCGTGCAGAGCCTTGCCCA | Mycgr3\|71143  JGI  Ensembl | KOG1212 Amidases; | amidase activity; carbon-nitrogen ligase activity, with glutamine as amido-N-donor; | 0 % |  |  |
| 34 | ggggacaagtttgtacaaaaaagcaggcttgatgTCTCAAGAATCCTTCGACGC, ggggaccactttgtacaagaaagctgggtcCACGAACGGCTTGAAAGCTG | Mycgr3\|71361  JGI  Ensembl | KOG1399 Flavin-containing monooxygenase; |  | 100 % |  |  |
| 34 | ggggacaagtttgtacaaaaaagcaggcttgatgGCCCAACGAGTATTCCTAGA, ggggaccactttgtacaagaaagctgggtcATAAGTCCTCTTCACCGGCG | Mycgr3\|71498  JGI  Ensembl | KOG2403 Succinate dehydrogenase, flavoprotein subunit; | FAD binding; electron transport;  oxidoreductase activity;  oxidoreductase activity, acting on the CH-CH group of donors;  succinate dehydrogenase (ubiquinone) activity;  tricarboxylic acid cycle; | 100 % |  |  |
| 34 | ggggacaagtttgtacaaaaaagcaggcttgatgTCGCGTTCACGAAATCTGCT, ggggaccactttgtacaagaaagctgggtcGTCGCCCATCGTATCGCTTC | Mycgr3\|72009  JGI  Ensembl | KOG1947 Leucine rich repeat proteins, some proteins contain F-box; | protein binding; | 100 % |  |  |
| 34 | ggggacaagtttgtacaaaaaagcaggcttgatgGGCGGCATCCACGACGACCA, ggggaccactttgtacaagaaagctgggtcCTCAATCCTCCTCGCACCAC | Mycgr3\|72701  JGI  Ensembl | KOG2614 Kynurenine 3-monooxygenase and related flavoprotein monooxygenases; | metabolic process; oxidoreductase activity; | 100 % |  |  |
| 34 | ggggacaagtttgtacaaaaaagcaggcttgatgTCCACCCTGCGACGGCGGAT, ggggaccactttgtacaagaaagctgggtcGTCTTCGGGATTTCCTGCAT | Mycgr3\|73528  JGI  Ensembl | KOG2369 Lecithin:cholesterol acyltransferase (LCAT)/Acyl-ceramide synthase; | lipid metabolic process; phosphatidylcholine-sterol O-acyltransferase activity;  phospholipid:diacylglycerol acyltransferase activity; | 100 % |  |  |
| 34 | ggggacaagtttgtacaaaaaagcaggcttgatgAAGGTCGCCTGTATTTCTAT, ggggaccactttgtacaagaaagctgggtcGATCCCTCCATCAACATAAT | Mycgr3\|74078  JGI  Ensembl | KOG1516 Carboxylesterase and related proteins; | acetylcholinesterase activity; | 100 % |  |  |
| 34 | ggggacaagtttgtacaaaaaagcaggcttgatgAACGCTCACAGCAATATGGA, ggggaccactttgtacaagaaagctgggtcAAACATCTTCCCCATAAACC | Mycgr3\|74471  JGI  Ensembl | KOG1286 Amino acid transporters; | amino acid transmembrane transporter activity; amino acid transport;  integral to membrane;  membrane;  transport; | 100 % |  |  |
| 34 | ggggacaagtttgtacaaaaaagcaggcttgatgCCTTCTTTTGGGCCAAGAAT, ggggaccactttgtacaagaaagctgggtcCTTGTGTGTCTCGAGCTTCG | Mycgr3\|75102  JGI  Ensembl |  |  | 0 % |  |  |
| 34 | ggggacaagtttgtacaaaaaagcaggcttgatgGAGCTCCCATCCGCGAGGCC, ggggaccactttgtacaagaaagctgggtcCGCCGTCCGCTCAAACTCCC | Mycgr3\|75341  JGI  Ensembl | KOG1211 Amidases; | carbon-nitrogen ligase activity, with glutamine as amido-N-donor; catalytic activity;  cobyrinate a,c-diamide synthase activity;  glutaminyl-tRNA synthase (glutamine-hydrolyzing) activity;  metabolic process; | 95.75672 % |  |  |
| 34 | ggggacaagtttgtacaaaaaagcaggcttgatgACGAGTTTGGTCAAAAGACA, ggggaccactttgtacaagaaagctgggtcGTACCGACCCACTCCAACAT | Mycgr3\|75467  JGI  Ensembl |  |  | 100 % |  |  |
| 34 | ggggacaagtttgtacaaaaaagcaggcttgatgGGCGACCAACAACCGAGCTC, ggggaccactttgtacaagaaagctgggtcACATCGCTCAAACACACGGT | Mycgr3\|75629  JGI  Ensembl | KOG1979 DNA mismatch repair protein - MLH1 family; | ATP binding; mismatch repair;  mismatched DNA binding;  nucleus; | 100 % |  |  |
| 34 | ggggacaagtttgtacaaaaaagcaggcttgatgGCAGCAACCCCCTTCACCCC, ggggaccactttgtacaagaaagctgggtcTGGCTCATCCTTGACCGCGT | Mycgr3\|76495  JGI  Ensembl | KOG3823 Uncharacterized conserved protein; |  | 100 % |  |  |
| 34 | ggggacaagtttgtacaaaaaagcaggcttgatgTCTAATATCATCGGCAACCG, ggggaccactttgtacaagaaagctgggtcAAGGCGGCCGTCTGCCTGAG | Mycgr3\|76944  JGI  Ensembl | KOG3791 Predicted RNA-binding protein involved in translational regulation; |  | 100 % |  |  |
| 34 | ggggacaagtttgtacaaaaaagcaggcttgatgGCCTCTCAGTCTGCGGAGAG, ggggaccactttgtacaagaaagctgggtcATCATCAGGATACTCCTCCT | Mycgr3\|77046  JGI  Ensembl |  |  | 100 % |  |  |
| 34 | ggggacaagtttgtacaaaaaagcaggcttgatgCTTCGCAATATCTCGCCAGT, ggggaccactttgtacaagaaagctgggtcTCCCTCGAACACAGAATGCT | Mycgr3\|78014  JGI  Ensembl | KOG1235 Predicted unusual protein kinase; |  | 100 % |  |  |
| 34 | ggggacaagtttgtacaaaaaagcaggcttgatgGTCCTCCCAGTCGAGAATCC, ggggaccactttgtacaagaaagctgggtcCCACTTCTGACAGAATTCCT | Mycgr3\|78024  JGI  Ensembl | KOG2852 Possible oxidoreductase; |  | 100 % |  |  |
| 34 | ggggacaagtttgtacaaaaaagcaggcttgatgCATTCTCTGACCGTGCTCCA, ggggaccactttgtacaagaaagctgggtcATGACTAGGCATGGAAGCAT | Mycgr3\|87705  JGI  Ensembl | KOG2024 Beta-Glucuronidase GUSB (glycosylhydrolase superfamily 2); | beta-glucuronidase activity; carbohydrate metabolic process;  hydrolase activity, hydrolyzing O-glycosyl compounds;  methylation;  methyltransferase activity;  nucleic acid binding; | 100 % |  |  |
| 34 | ggggacaagtttgtacaaaaaagcaggcttgatgGCGAAACCTGGAGATGACTG, ggggaccactttgtacaagaaagctgggtcGTGCTCGTAGAAAGGGGCAT | Mycgr3\|88879  JGI  Ensembl |  |  | 100 % | chr\_1:1203814 A->G; chr\_1:1203269 A->T; chr\_1:1203234 C->A; |  |
| 34 | ggggacaagtttgtacaaaaaagcaggcttgatgTCCATTACTCAAGCGCCTTT, ggggaccactttgtacaagaaagctgggtcCTGCAACGGCTGGCCCTTGC | Mycgr3\|89002  JGI  Ensembl |  |  | 100 % |  |  |
| 34 | ggggacaagtttgtacaaaaaagcaggcttgatgCCATCTCGGCCAAGAAGGCA, ggggaccactttgtacaagaaagctgggtcCCGCCTCGGAAAGTGAGCAT | Mycgr3\|89261  JGI  Ensembl | KOG1216 von Willebrand factor and related coagulation proteins; | exo-alpha-sialidase activity; | 100 % |  |  |
| 34 | ggggacaagtttgtacaaaaaagcaggcttgatgTCGTACAGCTCAAGAAAGTA, ggggaccactttgtacaagaaagctgggtcGATGGCACCTTTCTTCTCGT | Mycgr3\|89429  JGI  Ensembl |  |  | 100 % |  |  |
| 34 | ggggacaagtttgtacaaaaaagcaggcttgatgGACTGGACTGGAGGCACTCG, ggggaccactttgtacaagaaagctgggtcAAACGAGAGGCTGTCACTCA | Mycgr3\|89632  JGI  Ensembl |  |  | 100 % |  |  |
| 34 | ggggacaagtttgtacaaaaaagcaggcttgatgTTGGAGTGTCGTGCATGTGC, ggggaccactttgtacaagaaagctgggtcCTGCTCTCGCATCGCGATCT | Mycgr3\|90455  JGI  Ensembl | KOG4197 FOG: PPR repeat; |  | 100 % |  |  |
| 34 | ggggacaagtttgtacaaaaaagcaggcttgatgGCTTGGCCTCGACAAAACCT, ggggaccactttgtacaagaaagctgggtcCATATCGCCAAACACCATCT | Mycgr3\|91094  JGI  Ensembl |  |  | 23.31002 % |  |  |
| 34 | ggggacaagtttgtacaaaaaagcaggcttgatgACCTACTACCGCTTCCAGGC, ggggaccactttgtacaagaaagctgggtcTACTTCGGTCCCAGTAGTCG | Mycgr3\|91321  JGI  Ensembl | KOG3599 Ca2+-modulated nonselective cation channel polycystin; | chitinase activity; endochitinase activity; | 100 % |  |  |
| 34 | ggggacaagtttgtacaaaaaagcaggcttgatgAACTCAGCAACCTCCACTTT, ggggaccactttgtacaagaaagctgggtcGGGCTCAGAGATTCGCGCGG | Mycgr3\|94177  JGI  Ensembl |  |  | 0 % |  |  |
| 34 | ggggacaagtttgtacaaaaaagcaggcttgatgGCGACCGCGCCTTGCCCCAC, ggggaccactttgtacaagaaagctgggtcCTCCTCATCATCATCATCCA | Mycgr3\|94193  JGI  Ensembl | KOG2992 Nucleolar GTPase/ATPase p130; |  | 100 % | chr\_6:2640866 A->G; |  |
| 34 | ggggacaagtttgtacaaaaaagcaggcttgatgAAGGGGGGACACGAGTTGGG, ggggaccactttgtacaagaaagctgggtcCCGCCCCCACTTCACCCGCC | Mycgr3\|94827  JGI  Ensembl |  |  | 0 % |  |  |
| 34 | ggggacaagtttgtacaaaaaagcaggcttgatgGCATCTCATCTTCTCAAGCT, ggggaccactttgtacaagaaagctgggtcCGCCTTCAACACCTCTCCAG | Mycgr3\|95621  JGI  Ensembl | KOG2195 Transferrin receptor and related proteins containing the protease-associated (PA) domain; | glutamate carboxypeptidase II activity; peptidase activity;  proteolysis; | 100 % |  |  |
| 34 | ggggacaagtttgtacaaaaaagcaggcttgatgGTTCAAACTAGGAGCAAGAC, ggggaccactttgtacaagaaagctgggtcTAAAGTTGCCTCTCCCGTCA | Mycgr3\|96092  JGI  Ensembl | KOG3627 Trypsin; |  | 0.87236 % |  |  |
| 34 | ggggacaagtttgtacaaaaaagcaggcttgatgTCCGAAGGCATGTGCTACCG, ggggaccactttgtacaagaaagctgggtcTCTATCATACGCTTCATTCT | Mycgr3\|97162  JGI  Ensembl |  |  | 100 % |  |  |
| 34 | ggggacaagtttgtacaaaaaagcaggcttgatgTTTGCGGCCACTCAGTCTCG, ggggaccactttgtacaagaaagctgggtcTTGCGAAGGTTGGAAGGGTG | Mycgr3\|97727  JGI  Ensembl |  |  | 100 % |  |  |
| 34 | ggggacaagtttgtacaaaaaagcaggcttgatgTTCTTTCCGTCGAGTTCACT, ggggaccactttgtacaagaaagctgggtcCTTGTCATCGTCGTCCTCTG | Mycgr3\|97978  JGI  Ensembl |  |  | 0 % |  |  |
| 34 | ggggacaagtttgtacaaaaaagcaggcttgatgCTTATCGACGAAGTTTTGCT, ggggaccactttgtacaagaaagctgggtcCCAAACGCCCAACGGCAAGA | Mycgr3\|98411  JGI  Ensembl | KOG0158 Cytochrome P450 CYP3/CYP5/CYP6/CYP9 subfamilies; | electron transport; heme binding;  iron ion binding;  monooxygenase activity; | 100 % |  |  |
| 34 | ggggacaagtttgtacaaaaaagcaggcttgatgATCACGAACGAAATGGCACC, ggggaccactttgtacaagaaagctgggtcTAGCCTGACACGTCCAGCCT | Mycgr3\|98456  JGI  Ensembl | KOG2296 Integral membrane protein; |  | 100 % |  |  |
| 34 | ggggacaagtttgtacaaaaaagcaggcttgatgCAGCCAGCCCTGGTCAAGTC, ggggaccactttgtacaagaaagctgggtcAGTTGGCCAAGATCGCGGCG | Mycgr3\|98764  JGI  Ensembl |  |  | 100 % |  |  |
| 34 | ggggacaagtttgtacaaaaaagcaggcttgatgGCTCCCAATGATTCGTCCTC, ggggaccactttgtacaagaaagctgggtcATCATAGTCCTGACCAACAC | Mycgr3\|99247  JGI  Ensembl | KOG0254 Predicted transporter (major facilitator superfamily); | carbohydrate transport; integral to membrane;  membrane;  sugar:hydrogen symporter activity;  transport;  transporter activity; | 100 % |  |  |
| 34 | ggggacaagtttgtacaaaaaagcaggcttgatgCCTTGCTTCAAGGGCCTTGC, ggggaccactttgtacaagaaagctgggtcAAGCTCCTCATCTCCATCTT | Mycgr3\|99365  JGI  Ensembl | KOG2992 Nucleolar GTPase/ATPase p130; |  | 100 % |  |  |
| 34 | ggggacaagtttgtacaaaaaagcaggcttgatgCCTCCCGTGGCAGTGGGATT, ggggaccactttgtacaagaaagctgggtcCTTCTTCTTCCTTCCCTTAC | Mycgr3\|99680  JGI  Ensembl | KOG0804 Cytoplasmic Zn-finger protein BRAP2 (BRCA1 associated protein); | UDP-N-acetylmuramoylalanyl-D-glutamyl-2,6-diaminopimelate-D-alanyl-D-alanine ligase activity; coenzyme F420-0 gamma-glutamyl ligase activity;  coenzyme F420-2 alpha-glutamyl ligase activity;  protein binding;  ribosomal S6-glutamic acid ligase activity;  zinc ion binding; | 100 % |  |  |
| 34 | ggggacaagtttgtacaaaaaagcaggcttgatgACCTACCATGACCTTACACA, ggggaccactttgtacaagaaagctgggtcCAATTGCCTGCTCGAAGAAC | Mycgr3\|99750  JGI  Ensembl |  |  | 22.31813 % |  |  |
| 34 | ggggacaagtttgtacaaaaaagcaggcttgatgGTACTGTCACTCTCTGGAGC, ggggaccactttgtacaagaaagctgggtcCCAGTCTCCCAAGAGCGGAT | Mycgr3\|99840  JGI  Ensembl | KOG1282 Serine carboxypeptidases (lysosomal cathepsin A); | proteolysis; serine carboxypeptidase activity; | 100 % |  |  |
| 34 | ggggacaagtttgtacaaaaaagcaggcttgatgTCCACTCACAAGCAAGACGA, ggggaccactttgtacaagaaagctgggtcCAACTTCGCCTTCAACTTGC | Mycgr3\|100498  JGI  Ensembl | KOG1175 Acyl-CoA synthetase; | catalytic activity; metabolic process;  propionate-CoA ligase activity; | 1.94497 % |  |  |
| 34 | ggggacaagtttgtacaaaaaagcaggcttgatgTCCACCACCACTGCTCCCGT, ggggaccactttgtacaagaaagctgggtcATGGAACATCCCATGCACCA | Mycgr3\|101096  JGI  Ensembl | KOG1111 N-acetylglucosaminyltransferase complex, subunit PIG-A/SPT14, required for phosphatidylinositol biosynthesis/Sulfolipid synthase; | 1,2-dihydroxy-phenanthrene glycosyltransferase activity; 1-phenanthrol glycosyltransferase activity;  9-phenanthrol UDP-glucuronosyltransferase activity;  9-phenanthrol glycosyltransferase activity;  O antigen polymerase activity;  UDP-N-acetylglucosamine-peptide N-acetylglucosaminyltransferase activity;  UDP-glucose:glycoprotein glucosyltransferase activity;  acetylgalactosaminyltransferase activity;  acetylglucosaminyltransferase activity;  alpha(1,3)-fucosyltransferase activity;  alpha(1,6)-fucosyltransferase activity;  alpha-1,2-galactosyltransferase activity;  alpha-1,2-mannosyltransferase activity;  alpha-1,3-galactosyltransferase activity;  alpha-1,3-mannosyltransferase activity;  alpha-1,6-mannosyltransferase activity;  beta-1,4-mannosyltransferase activity;  biosynthetic process;  cellulose synthase activity;  dolichyl pyrophosphate Glc1Man9GlcNAc2 alpha-1,3-glucosyltransferase activity; dolichyl pyrophosphate Man7GlcNAc2 alpha-1,3-glucosyltransferase activity;  dolichyl pyrophosphate Man9GlcNAc2 alpha-1,3-glucosyltransferase activity;  dolichyl-phosphate-glucose-glycolipid alpha-glucosyltransferase activity;  fucosyltransferase activity;  galactosyltransferase activity;  glucosyltransferase activity;  glycolipid mannosyltransferase activity;  inositol phosphoceramide synthase activity;  lipopolysaccharide-1,6-galactosyltransferase activity;  mannosyltransferase activity;  oligosaccharyl transferase activity;  phenanthrol glycosyltransferase activity; | 100 % |  |  |
| 34 | ggggacaagtttgtacaaaaaagcaggcttgatgCGTTCCGTCTTTGCTGTCGC, ggggaccactttgtacaagaaagctgggtcGTAGCCGGAGATTGATGGGG | Mycgr3\|103393  JGI  Ensembl | KOG3017 Defense-related protein containing SCP domain; | extracellular region; | 100 % |  |  |
| 34 | ggggacaagtttgtacaaaaaagcaggcttgatgCGCTTCTTCACTTCCCTTGC, ggggaccactttgtacaagaaagctgggtcAACGATGCTGATCAATTTGA | Mycgr3\|104299  JGI  Ensembl |  |  | 100 % |  |  |
| 34 | ggggacaagtttgtacaaaaaagcaggcttgatgATGCGCCGAGCATTGACGAC, ggggaccactttgtacaagaaagctgggtcATTGAACGCGTTGCTCCTCG | Mycgr3\|106853  JGI  Ensembl | KOG0342 ATP-dependent RNA helicase pitchoune; | 8-oxo-7,8-dihydroguanine triphosphatase activity; ATP binding;  ATP-dependent 5'-3' DNA helicase activity;  ATP-dependent helicase activity;  UDP-2,3-diacylglucosamine hydrolase activity;  bis(5'-nucleosyl)-tetraphosphatase activity;  dATP pyrophosphohydrolase activity;  diacylglycerol pyrophosphate phosphatase activity;  dihydroneopterin monophosphate phosphatase activity;  dihydroneopterin triphosphate pyrophosphohydrolase activity;  helicase activity;  nucleic acid binding;  pyrophosphatase activity;  thiamin-pyrophosphatase activity; | 100 % |  |  |
| 34 | ggggacaagtttgtacaaaaaagcaggcttgatgAAGTCCAGACCATCCACGAC, ggggaccactttgtacaagaaagctgggtcCCGCTTCTCCATCGAAGGCC | Mycgr3\|107363  JGI  Ensembl | KOG1339 Aspartyl protease; |  | 100 % |  |  |
| 34 | ggggacaagtttgtacaaaaaagcaggcttgatgGAGCGTCCTTCGCCCATCAC, ggggaccactttgtacaagaaagctgggtcAAGCCGCGCATCAAGTCTTC | Mycgr3\|107923  JGI  Ensembl |  |  | 100 % |  |  |
| 34 | ggggacaagtttgtacaaaaaagcaggcttgatgTCGAGCGCCACAAAAGAGCC, ggggaccactttgtacaagaaagctgggtcTCCCGTCCCATTAACGCTAT | Mycgr3\|108131  JGI  Ensembl | KOG1922 Rho GTPase effector BNI1 and related formins; |  | 100 % | chr\_2:2239643 C->T; chr\_2:2239448 T->A; |  |
| 34 | ggggacaagtttgtacaaaaaagcaggcttgatgCTCATCCAATGCCCTAAGTT, ggggaccactttgtacaagaaagctgggtcACCAAACGCCAGCGCCCATG | Mycgr3\|108203  JGI  Ensembl |  |  | 100 % |  |  |
| 34 | ggggacaagtttgtacaaaaaagcaggcttgatgCCGCAACAGGCAGCGCCCGC, ggggaccactttgtacaagaaagctgggtcAGCCTTCCTTTTCCTCGCAG | Mycgr3\|108215  JGI  Ensembl | KOG2489 Transmembrane protein; |  | 100 % |  |  |
| 34 | ggggacaagtttgtacaaaaaagcaggcttgatgGCACGCCAACCAGAGTCAAC, ggggaccactttgtacaagaaagctgggtcCAGAGCTTTGACGTCGCTGC | Mycgr3\|108333  JGI  Ensembl | KOG2617 Citrate synthase; | cellular carbohydrate metabolic process; transferase activity, transferring acyl groups, acyl groups converted into alkyl on transfer; | 100 % |  |  |
| 34 | ggggacaagtttgtacaaaaaagcaggcttgatgCTTACTCAAGACTTCCTGCA, ggggaccactttgtacaagaaagctgggtcCTCCTGCCTGCCCGGCCACT | Mycgr3\|108509  JGI  Ensembl |  |  | 100 % |  |  |
| 34 | ggggacaagtttgtacaaaaaagcaggcttgatgTCATCGGAAACACTCCAGCG, ggggaccactttgtacaagaaagctgggtcTCGAGCTCGTGAGTCGCTCT | Mycgr3\|108605  JGI  Ensembl |  |  | 100 % |  |  |
| 34 | ggggacaagtttgtacaaaaaagcaggcttgatgGATCTGGCATCCATCGTCCC, ggggaccactttgtacaagaaagctgggtcTTGAACCATACGAACACGAC | Mycgr3\|109248  JGI  Ensembl |  |  | 0 % |  |  |
| 34 | ggggacaagtttgtacaaaaaagcaggcttgatgTCCCGCTCCGCCACCAGCAA, ggggaccactttgtacaagaaagctgggtcAGCAGTGACGGCGAACGCAT | Mycgr3\|109321  JGI  Ensembl |  |  | 100 % |  |  |
| 34 | ggggacaagtttgtacaaaaaagcaggcttgatgGTTCTCGGAGTCACCAACGT, ggggaccactttgtacaagaaagctgggtcGCTCATCTGCACCGAGATAT | Mycgr3\|109942  JGI  Ensembl | KOG4525 Jacalin-like lectin domain-containing protein; | collagenase activity; matrilysin-2 activity;  prenyl protein specific endopeptidase activity;  stromelysin 3 activity; | 0 % |  |  |
| 34 | ggggacaagtttgtacaaaaaagcaggcttgatgGACGAGGCTGGCAGAATACG, ggggaccactttgtacaagaaagctgggtcCGAAATCTGCGGTTTACTCC | Mycgr3\|110012  JGI  Ensembl | KOG4231 Intracellular membrane-bound Ca2+-independent phospholipase A2; | lipid metabolic process; | 100 % |  |  |
| 34 | ggggacaagtttgtacaaaaaagcaggcttgatgGCTATTTTCAAGACTCTCGT, ggggaccactttgtacaagaaagctgggtcCCATCCCGGTTGGGAAGGGT | Mycgr3\|110349  JGI  Ensembl |  |  | 100 % |  |  |
| 34 | ggggacaagtttgtacaaaaaagcaggcttgatgTTCACCGGCTCCCGGCCAAC, ggggaccactttgtacaagaaagctgggtcACGAGCGTTCTCCCCAATGC | Mycgr3\|110573  JGI  Ensembl |  |  | 100 % |  |  |
| 34 | ggggacaagtttgtacaaaaaagcaggcttgatgGCCCCAAAACACTCCAAGGA, ggggaccactttgtacaagaaagctgggtcCAATTTCGCATGTTCCCTCG | Mycgr3\|110645  JGI  Ensembl |  |  | 100 % |  |  |
| 34 | ggggacaagtttgtacaaaaaagcaggcttgatgTACCCAAATCAGGGTGGCTA, ggggaccactttgtacaagaaagctgggtcTCTCCCGCCCTTTCTTCCCA | Mycgr3\|110701  JGI  Ensembl | KOG2510 SWI-SNF chromatin-remodeling complex protein; | UDP-N-acetylmuramoylalanyl-D-glutamyl-2,6-diaminopimelate-D-alanyl-D-alanine ligase activity; coenzyme F420-0 gamma-glutamyl ligase activity;  coenzyme F420-2 alpha-glutamyl ligase activity;  ribosomal S6-glutamic acid ligase activity; | 0 % |  |  |
| 34 | ggggacaagtttgtacaaaaaagcaggcttgatgTTCAGCATTCGTCGCCTCCG, ggggaccactttgtacaagaaagctgggtcAGACTTGCCTTCGATGGAGG | Mycgr3\|110811  JGI  Ensembl | KOG2187 tRNA uracil-5-methyltransferase and related tRNA-modifying enzymes; | GPI-anchor transamidase activity; RNA binding;  RNA methyltransferase activity;  RNA processing;  cyanophycinase activity;  regulation of hydrolase activity; | 49.02326 % |  |  |
| 34 | ggggacaagtttgtacaaaaaagcaggcttgatgAGCGCCGGCAACTACGCCGC, ggggaccactttgtacaagaaagctgggtcTGACGTCCCAAACACCGGCG | Mycgr3\|110974  JGI  Ensembl | KOG2510 SWI-SNF chromatin-remodeling complex protein; |  | 1.1194 % |  |  |
| 34 | ggggacaagtttgtacaaaaaagcaggcttgatgGATGAAGAGATGCAGTCGGG, ggggaccactttgtacaagaaagctgggtcCCCAGCACTACTCAACCAAG | Mycgr3\|111096  JGI  Ensembl | KOG0115 RNA-binding protein p54nrb (RRM superfamily); | DNA binding; nucleus; regulation of transcription, DNA-dependent;  transcription;  transcription factor activity;  zinc ion binding; | 100 % |  | ``` 1	ATGGATGAAGAGATGCAGTCGGGACAACCCGAGTCGCAAGGCTCAGAGCAGCCAGACAATAGCAGTGGAAAACAGCCCACTGCAACCAAACAGCTGAGACGCGCCTGCGATTGTTGTCGCAAGCGCAAAGTCAGATGCGACGGTCAGGACCCATGCGGGCCATGCAAGAAGGCCTCGATCCGCTGTGCATATCTCACGCCACCGAAGAAGAAAGGCCCAAAAGGGTTGAGAAGTGCTAGAGTGCTGCATGCGTTACGCAGGATAGACGACGATGCTACTTCGGCTCCGACGAGTCCACACAGCCCCAACATGCCCTTTGGCAATTGGGGTTGGAACAGCAATGGAACATCCAATGCTGGACCAGCATCCTATGGAGGTCCTCCACCAACGGGTCTTCCAGGTCCGCACGAGCTGCTGTACGGACAACAACCTGGCGTAGCGCCTGACCAGCCATACTACCAGCAGCCGTCGAACCCGCCCACGTCCCGACCGCCATATCCGCCGCCCTACCGACAAGAGACAGCATATCAGCAAGGCTGGACGCCGAAGCAGCCGGAATCGGTGTCTACTGCGAGTGAGAGTATACAGCGTGTCTTTTCGCCTTCATCGTCCGAGCACCACACCCAGCGAGCATTGTCACCAGACAACTTCCTACCATATGTGAGGCTCTTCTTCCAAAACATGTACAGCATCATGCCGGTCATCGACCGAAACGTCTACCTTGACCCGGTAATCTACACGAACCGACCAATGAGCTCAGAGCTTTATGCATTCATGTGTGCATTGTGCGCCGCGACGATCGTTCAACTCGACGACTCGATTCCGCAACCACCTCCTTTGCATCCTGGAAAGCCCTCTGATTCTCTGTTTGCCGAGGAATGTTTGAGAGAGAGGAAGACGTTCGACTACGTCGACAATATGTCACCAATGAGCATAATGACCTCGTTTTTTCTATTCGCATATTACGGCAATTACGAGAAGCATACAAAGGCGTGGCACTATCTGCAAGAAAGCATCACATTCAGTG-AGAATCTGGAGATGGACGATGAAGACAGCTACG-CGAAAC-TCGATCCGACCGAGGC-TCAGTGGC	1090 	\|\|\|\|\|\|\|\|\|\|\|\|\|\|\|\|\|\|\|\|\|\|\|\|\|\|\|\|\|\|\|\|\|\|\|\|\|\|\|\|\|\|\|\|\|\|\|\|\|\|\|\|\|\|\|\|\|\|\|\|\|\|\|\|\|\|\|\|\|\|\|\|\|\|\|\|\|\|\|\|\|\|\|\|\|\|\|\|\|\|\|\|\|\|\|\|\|\|\|\|\|\|\|\|\|\|\|\|\|\|\|\|\|\|\|\|\|\|\|\|\|\|\|\|\|\|\|\|\|\|\|\|\|\|\|\|\|\|\|\|\|\|\|\|\|\|\|\|\|\|\|\|\|\|\|\|\|\|\|\|\|\|\|\|\|\|\|\|\|\|\|\|\|\|\|\|\|\|\|\|\|\|\|\|\|\|\|\|\|\|\|\|\|\|\|\|\|\|\|\|\|\|\|\|\|\|\|\|\|\|\|\|\|\|\|\|\|\|\|\|\|\|\|\|\|\|\|\|\|\|\|\|\|\|\|\|\|\|\|\|\|\|\|\|\|\|\|\|\|\|\|\|\|\|\|\|\|\|\|\|\|\|\|\|\|\|\|\|\|\|\|\|\|\|\|\|\|\|\|\|\|\|\|\|\|\|\|\|\|\|\|\|\|\|\|\|\|\|\|\|\|\|\|\|\|\|\|\|\|\|\|\|\|\|\|\|\|\|\|\|\|\|\|\|\|\|\|\|\|\|\|\|\|\|\|\|\|\|\|\|\|\|\|\|\|\|\|\|\|\|\|\|\|\|\|\|\|\|\|\|\|\|\|\|\|\|\|\|\|\|\|\|\|\|\|\|\|\|\|\|\|\|\|\|\|\|\|\|\|\|\|\|\|\|\|\|\|\|\|\|\|\|\|\|\|\|\|\|\|\|\|\|\|\|\|\|\|\|\|\|\|\|\|\|\|\|\|\|\|\|\|\|\|\|\|\|\|\|\|\|\|\|\|\|\|\|\|\|\|\|\|\|\|\|\|\|\|\|\|\|\|\|\|\|\|\|\|\|\|\|\|\|\|\|\|\|\|\|\|\|\|\|\|\|\|\|\|\|\|\|\|\|\|\|\|\|\|\|\|\|\|\|\|\|\|\|\|\|\|\|\|\|\|\|\|\|\|\|\|\|\|\|\|\|\|\|\|\|\|\|\|\|\|\|\|\|\|\|\|\|\|\|\|\|\|\|\|\|\|\|\|\|\|\|\|\|\|\|\|\|\|\|\|\|\|\|\|\|\|\|\|\|\|\|\|\|\|\|\|\|\|\|\|\|\|\|\|\|\|\|\|\|\|\|\|\|\|\|\|\|\|\|\|\|\|\|\|\|\|\|\|\|\|\|\|\|\|\|\|\|\|\|\|\|\|\|\|\|\|\|\|\|\|\|\|\|\|\|\|\|\|\|\|\|\|\|\|\|\|\|\|\|\|\|\|\|\|\|\|\|\|\|\|\|\|\|\|\|\|\|\|\|\|\|\|\|\|\|\|\|\|\|\|\|\|\|\|\|\|\|\|\|\|\|\|\|\|\|\|\|\|\|\|\|\|\|\|\|\|\|\|\|\|\|\|\|\|\|\|\|\|\|\|\|\|\|\|\|\|\|\|\|\|\|\|\|\|\|\|\|\|\|\|\|\|\|\|\|\|\|\|\|\|\|\|\|\|\|\|\|\|\|\|\|\|\|\|\|\|\|\|\|\|\|\|\|\|\|\|\|\|\|\|\|\|\|\|\|\|\|\|\|\|\|\|\|\|\|\|\|\|\|\|\|\|\|\|\|\|\|\|\|\|\|\|\|\|\|\|\|\|\|\|\|\|\|\|\|\|\|\|\|\|\|\|\|\|\|\|\|\|\|\|\|\|\|\|\|\|\|\|\|\|\|\|\|\|\|\|\|\|\|\|\|\|\|\|\|\|\|\|\|\|\|\|\|\|\|\|\|\|\|\|\|\|\|\|\|\|\|\|\|\|\|\|\|\|\|\|\|\|\|\|\|\|\|\|\|\|\|\|\|\|\|\| \|\|\|\|\|\|\|\|\|\|\|\|\|\|\|\|\|\|\| \|\|\|\|\|\|\|\|\|\|\|\|\|\|\|\|\|\|\|\|\|\|\|\|\|\|\|\|\|\|\|\|\|\|\|\|\|\|\|\|\|\|\| \|\|\|\|\|  \|\|\|\|\|\|\|\|\|\|\|\|\|\|\|\|\|\|\| \|\|\|\|\|\|\|\|\|\|\|\|\|\| \|\|\|\|\| \|\|\| \|\|\|\|\|\|\|\|\| \|\|\|\|\|\|\|\|\|\|\|\|\|\|\|\|\|\|\| \|\|\|\|\|\| \|\|\|\|\|\| \|\|\|\|\|\|\|\|\| \|\|\|\|\|\|\|\| 	ATGGATGAAGAGATGCAGTCGGGACAACCCGAGTCGCAAGGCTCAGAGCAGCCAGACAATAGCAGTGGAAAACAGCCCACTGCAACCAAACAGCTGAGACGCGCCTGCGATTGTTGTCGCAAGCGCAAAGTCAGATGCGACGGTCAGGACCCATGCGGGCCATGCAAGAAGGCCTCGATCCGCTGTGCATATCTCACGCCACCGAAGAAGAAAGGCCCAAAAGGGTTGAGAAGTGCTAGAGTGCTGCATGCGTTACGCAGGATAGACGACGATGCTACTTCGGCTCCGACGAGTCCACACAGCCCCAACATGCCCTTTGGCAATTGGGGTTGGAACAGCAATGGAACATCCAATGCTGGACCAGCATCCTATGGAGGTCCTCCACCAACGGGTCTTCCAGGTCCGCACGAGCTGCTGTACGGACAACAACCTGGCGTAGCGCCTGACCAGCCATACTACCAGCAGCCGTCGAACCCGCCCACGTCCCGACCGCCATATCCGCCGCCCTACCGACAAGAGACAGCATATCAGCAAGGCTGGACGCCGAAGCAGCCGGAATCGGTGTCTACTGCGAGTGAGAGTATACAGCGTGTCTTTTCGCCTTCATCGTCCGAGCACCACACCCAGCGAGCATTGTCACCAGACAACTTCCTACCATATGTGAGGCTCTTCTTCCAAAACATGTACAGCATCATGCCGGTCATCGACCGAAACGTCTACCTTGACCCGGTAATCTACACGAACCGACCAATGAGCTCAGAGCTTTATGCATTCATGTGTGCATTGTGCGCCGCGACGATCGTTCAACTCGACGACTCGATTCCGCAACCACCTCCTTTGCATCCTGGAAAGCCCTCTGATTCTCTGTTTGCCGAGGAATGTTTGAGAGAGAGGAAGACGTTCGACTACGTCGAC-ATATGTCACCAATGAGCAT-ATGACCTCGTTTTTTCTATTCGCATATTACGGCAATTACGAGA-GCATAACAAGGCGTGGCACTATCTGC-AGAAAGCATCACAT-CAGTGGAGA-TCTGGAGAT-GACGATGAAGACAGCTACGGCGAAACCTCGATC-GACCGAGGCCTCAGTGGC ``` |
| 34 | ggggacaagtttgtacaaaaaagcaggcttgatgAACATGGCACCTCCAGCAAT, ggggaccactttgtacaagaaagctgggtcAACCCTCACCTGACCCTGCA | Mycgr3\|111164  JGI  Ensembl | KOG1216 von Willebrand factor and related coagulation proteins; | ATP-dependent DNA helicase activity; ATP-dependent RNA helicase activity;  ATP-dependent helicase activity;  ATPase activity;  ATPase activity, coupled;  ATPase activity, coupled to transmembrane movement of ions;  ATPase activity, coupled to transmembrane movement of substances;  ATPase activity, uncoupled;  DNA-dependent ATPase activity;  RNA-dependent ATPase activity;  protein-transmembrane transporting ATPase activity;  single-stranded DNA-dependent ATP-dependent DNA helicase activity; | 100 % |  | ``` 3	GAACATGGCACCTCCAGCAATGGACGACGATGGCGATACATCTAGCATCTCCTCTCCACCGCAATCTCCGCAAAATCACCACGAAGACGAGACGATCAAAGTCGCCCCTCGTTATCTTGGTGAAGGTGTTGTTCACCACCCGGCGGCTTCGGCTTCGGCTTCCTCCTCTGCCACACCACTCCCCGGAACTGCACCAGTCAAACCCAAACGCACACGCAAACCAAAGTTGGATGCAAATGGCAATCCTGTCTTGCCAAAGCCGAGGAAAAAGGCCGAACCAAAAGTCAAGGCGGAAGGAGGTGCAACAACGACGGCTCCTGCGACGAAAAGGAGAAAGAAGACTGCACCGGAGGAAAAGCCTGCCACCAGTACTGCCATAAGTGGTGATGTCCGCCAAACAACCCTCACCGGCCTGCATAGTCACTTCCTCCCGCCTGGCAATCCCGCCTCGTCCGCCCCACCTCCACCAATGCAGCAGCAACAACCTCATCCACCACCGCCGCAAACATTCTCATCTTCATCACTCGCCAACACAACAACATCAACATCCGACACATTGATGCGCAGCAGTCTACCACCACCACTCCTCTCATCAACTCCCACACCACGACCATTCTCCAGCGGACAGAATTACGATCCTATTCGCGGACAAAATTACGATTCCATTCGTGGTGCTATCGACGCGACTTCATCACGACC-GCCTGTGCTATCGAACGGTGCGCAATCCGCACATG	736 	\|\|\|\|\|\|\|\| \|\|\|\|\|\|\| \|\|\|\|\|\|\|\|\|\|\|\|\|\|\|\|\|\|\|\|\|\|\|\|\|\|\|\|\|\|\|\|\|\|\|\|\|\|\|\|\|\|\|\|\|\|\|\|\|\|\|\| \|\|\|\|\|\|\|\|\|\|\|\| \|\|\|\|\|\|\|\|\|\| \|\|\|\|\|\| \|\|\|\| \|\|\|\|\|\|\|\|\|\|\|\|\|\|\|\|\|\|\|\|\|\|\|\|\|\|\|\|\|\|\|\| \|\| \|\|\|\|\|\|\|\|\|\|\|\|\|\|\|\| \|\|\|\|\|\|\|\|\|\|\|\|\|\|\|\|\|\|\|\|\|\|\|\|\|\|\|\|\|\|\|\|\|\|\|\| \|\|\|\|\|\|\|\|\|\|\|\|\|\|\|\|\|\|\|\|\|\|\| \|\|\|\|\|\|\|\|\|\|\|\|\|\|\|\|\|\|\|\|\|\|\|\|\|\|\|\|\|\|\|\|\|\|\|\|\|\| \|\|\|\|\|\|\|\|\|\|\|\|\|\|\|\|\|\|\|\|\|\|\|\|\|\|\|\|\|\|\|\|\|\|\|  \|\|\|\|\|\|\|\|\|\|\|\|\|\|\|\|\|\| \|\|\|\|\|\|\|\|\|\|\|\|\|\|\|\|\|\|\|\| \|\|\|\|\|\|\|\|\|\|\|\|\|\|\|\|\|\|\|\|\|\|\|\|\|\|\|\|\|\|\|\|\|\|\|\|\|\| \|\|\|\|\|\|\|\|\|\|\|\|\|\|\|\|\|\|\|\|\|\|\| \|\| \|\|\|\|\|\|\|\|\| \|\|\|\|\|\|\|  \|\|\|\|\| \|\|\|\|\| \|\|\|\|  \|\|\|\|\|\|\|\|\|\|\|\|\|\|\|\|\|\|\|\|\|\|\|\| \| \|\|\| \|\|   \|\| \|\|\|\|\|\| \|\|\|\|\|\|\|\|\|\|\|\|\|\|\|\|\| \|\|\|\|\|\|\|\|\|\|\|\|\|\|\|\|\|\|\|\|\|\|\|\|\|\|  \|\|\| \|\|\|\|\|\|\|\| \|\| \|\|\|\|\|\|\|\|\|\|\|\|\|\|\|\|\|\|\| \|\|\|\|\|\|\| \|  \|\|\|\|\|\|\|\|\|\|\|\|\|\|\|\|\| \|\|\|\|\|\|\| \|\|\|\| \|\|\| \|\|\|\|\|\|\|\|\|\|\|\|\|\|\|\|\|\|\|\|\|\|\|\|\|\|\|\|\|\|\|\|\|\| \|\|\|\| \|\|\|\|\|\|\|\|\|\| \|\|\|\|\| \| \|\|\|\|\|\|\| \|\|\|\|\|\|\|\|\|\| \|\|\|\| \|\|\| \|\|\|\|\|\|\|\|\|\|\| \|\|\|\|\|\|\|\|\|\|\|\| \| \|\|\|\|\|\|\|\| 	GAACATGGAACCTCCAACAATGGACGACGATGGCGATACATCTAGCATCTCCTCTCCACCGCAATCTCCACAAAATCACCACAAAGACGAGACAATCAAACTCGCACCTCGTTATCTTGGTGAAGGTGTTGTTCACCAACCAGCGGCTTCGGCTTCGGATTCCTCCTCTGCCACACCACTCCCCGGAACTGCACC-GTCAAACCCAAACGCACACGCAATCCAAAGTTGGATGCAAATGGCAATCCTGTCTTGCCAAATCCGAGGAAAAAGGCCGAACCAAAAGTCAAGGCGGATTGAGGTGCAACAACGACGGTTCCTGCGACGAAAAGGAGAAGGAAGACTGCACCGGAGGAAAAGCCTGCCACCAGTACTGACATAAGTGGTGATGTCCGCCAAATAATCCTCACCGGTCTGCATATACACTTACTCCCTCCTGTTAATCCCGCCTCGTCCGCCCCACCTTCTCCATTGTCTCAACAACAATCTCATCCACCACCGCCGTAAACATTCTCATCTTCATCACTCGCCG-CAC-ACAACATCTACTTCCGACACATTGATGCGCATCAGTCTAGCCGCACCACTCCTCTCATCACCTCCCACCCCACCACCTTTCTCCAGCGGACAGAATTACGATCCTATTCGCGCACAACATTACGATTCTATTCGCGTTGCTATC-ACGCGACTTCTTCACTACCTGCCTGTGCTATTGAACGGTGCGCACTACGCACATG ``` |
| 34 | ggggacaagtttgtacaaaaaagcaggcttgatgATTCTCCGACCATTGGCAAT, ggggaccactttgtacaagaaagctgggtcCTTCCGCAGCGCGTACAGCA | Mycgr3\|111223  JGI  Ensembl | KOG0624 dsRNA-activated protein kinase inhibitor P58, contains TPR and DnaJ domains; | heat shock protein binding; | 0.90304 % |  |  |
| 34 | ggggacaagtttgtacaaaaaagcaggcttgatgGGTATCATCCAAAAGATCGA, ggggaccactttgtacaagaaagctgggtcCAATCTCCTCGACTGCTCAG | Mycgr3\|111483  JGI  Ensembl | KOG2806 Chitinase; | exo-alpha-sialidase activity; | 1.77839 % |  |  |
| 35 | ggggacaagtttgtacaaaaaagcaggcttgatgTGTCCGCTGTACGCCGCGAC, ggggaccactttgtacaagaaagctgggtcAGAACCAGATGGCGCCGAAG | Mycgr3\|38282  JGI  Ensembl | KOG2089 Metalloendopeptidase family - saccharolysin & thimet oligopeptidase; | metalloendopeptidase activity; metallopeptidase activity;  proteolysis;  saccharolysin activity;  zinc ion binding; | 100 % |  |  |
| 35 | ggggacaagtttgtacaaaaaagcaggcttgatgCACATCCGGAATATATTTGC, ggggaccactttgtacaagaaagctgggtcCGGCAGACTATTGAACAGCT | Mycgr3\|38337  JGI  Ensembl |  |  | 100 % |  |  |
| 35 | ggggacaagtttgtacaaaaaagcaggcttgatgGCCGCTCCACCGTCTATCGA, ggggaccactttgtacaagaaagctgggtcAATACGAGAATTGCCATTGT | Mycgr3\|39064  JGI  Ensembl |  |  | 100 % |  |  |
| 35 | ggggacaagtttgtacaaaaaagcaggcttgatgGAGAACGAGCGTCTGAAAGC, ggggaccactttgtacaagaaagctgggtcCTTCCGCACCCCATCATTCA | Mycgr3\|40311  JGI  Ensembl | KOG1399 Flavin-containing monooxygenase; | FAD binding; oxidoreductase activity; | 100 % |  |  |
| 35 | ggggacaagtttgtacaaaaaagcaggcttgatgGCGCTCTCGAACATCCTCTC, ggggaccactttgtacaagaaagctgggtcCGACGACCCATCCAACACCA | Mycgr3\|43499  JGI  Ensembl | KOG2100 Dipeptidyl aminopeptidase; | dipeptidyl-peptidase IV activity; membrane; proteolysis; serine-type peptidase activity;  tripeptidyl-peptidase activity; | 100 % |  |  |
| 35 | ggggacaagtttgtacaaaaaagcaggcttgatgGACGCCGCCGGCCAACTCCT, ggggaccactttgtacaagaaagctgggtcAACAAACCTCCTACCATCTT | Mycgr3\|44328  JGI  Ensembl | KOG2922 Uncharacterized conserved protein; |  | 100 % |  |  |
| 35 | ggggacaagtttgtacaaaaaagcaggcttgatgGGGTCGCGACACCTGTCAGT, ggggaccactttgtacaagaaagctgggtcCCAAGTCAAATCGGGCAGAT | Mycgr3\|46813  JGI  Ensembl |  |  | 100 % |  |  |
| 35 | ggggacaagtttgtacaaaaaagcaggcttgatgGCCACCATCGTCCAGCGACC, ggggaccactttgtacaagaaagctgggtcCCCCTGCCTCGAATAAGGCC | Mycgr3\|48113  JGI  Ensembl | KOG1716 Dual specificity phosphatase; | dephosphorylation; phosphoric monoester hydrolase activity;  prenylated protein tyrosine phosphatase activity;  protein amino acid dephosphorylation;  protein tyrosine phosphatase activity;  protein tyrosine/serine/threonine phosphatase activity; | 100 % |  |  |
| 35 | ggggacaagtttgtacaaaaaagcaggcttgatgAGCCACATCGATCCCGGAGC, ggggaccactttgtacaagaaagctgggtcCTCCCTCCTCGCACCCCTCC | Mycgr3\|51359  JGI  Ensembl | KOG1418 Tandem pore domain K+ channel; | ion channel activity; ion transport;  membrane;  potassium channel activity;  potassium ion transport; | 100 % |  |  |
| 35 | ggggacaagtttgtacaaaaaagcaggcttgatgTTCTCCTCCACCGTGAGACA, ggggaccactttgtacaagaaagctgggtcATAAGTTCGCTTGAAAGGTG | Mycgr3\|53658  JGI  Ensembl | KOG2403 Succinate dehydrogenase, flavoprotein subunit; | FAD binding; electron transport;  oxidoreductase activity;  oxidoreductase activity, acting on the CH-CH group of donors;  succinate dehydrogenase (ubiquinone) activity;  tricarboxylic acid cycle; | 1.10999 % |  |  |
| 35 | ggggacaagtttgtacaaaaaagcaggcttgatgGCACGCATCTTTCCCAAACT, ggggaccactttgtacaagaaagctgggtcCTCTGCCCCGTTCCTCGACA | Mycgr3\|58167  JGI  Ensembl |  |  | 100 % |  |  |
| 35 | ggggacaagtttgtacaaaaaagcaggcttgatgTCGGACATTGAGAAGATGAT, ggggaccactttgtacaagaaagctgggtcGGTCTTGCGAAGGTAAAGCG | Mycgr3\|58200  JGI  Ensembl | KOG0061 Transporter, ABC superfamily (Breast cancer resistance protein); | ATP binding; ATPase activity;  nucleoside-triphosphatase activity;  nucleotide binding; | 100 % |  |  |
| 35 | ggggacaagtttgtacaaaaaagcaggcttgatgATGGGATCTGTAACTGGGAA, ggggaccactttgtacaagaaagctgggtcTGCTACCACCTCCACGCTCA | Mycgr3\|65466  JGI  Ensembl | KOG3957 Predicted L-carnitine dehydratase/alpha-methylacyl-CoA racemase; | catalytic activity; metabolic process; | 100 % |  |  |
| 35 | ggggacaagtttgtacaaaaaagcaggcttgatgTTCTCCAAAGTCCCCGGCTT, ggggaccactttgtacaagaaagctgggtcTGCCAACTTCTCTGGCGACT | Mycgr3\|65706  JGI  Ensembl | KOG2385 Uncharacterized conserved protein; | catalytic activity; | 100 % |  |  |
| 35 | ggggacaagtttgtacaaaaaagcaggcttgatgAATGAAGGCACGCTCGAGGA, ggggaccactttgtacaagaaagctgggtcCTTCACTTTGCTCATGACCG | Mycgr3\|65888  JGI  Ensembl | KOG2286 Exocyst complex subunit SEC6; |  | 100 % |  |  |
| 35 | ggggacaagtttgtacaaaaaagcaggcttgatgTCAGGAGACGAGCTCGTTGA, ggggaccactttgtacaagaaagctgggtcTAGCTCCACAGCCTTGGCAG | Mycgr3\|65952  JGI  Ensembl | KOG4201 Anthranilate synthase component II; | anthranilate synthase activity; biosynthetic process;  catalytic activity;  indole-3-glycerol-phosphate synthase activity;  metabolic process;  phosphoribosylanthranilate isomerase activity;  tryptophan metabolic process; | 100 % |  |  |
| 35 | ggggacaagtttgtacaaaaaagcaggcttgatgGCCAAGAAGGAGCCTTTGAA, ggggaccactttgtacaagaaagctgggtcCTTCCTCTCAAGGTTGCTGT | Mycgr3\|66582  JGI  Ensembl | KOG0254 Predicted transporter (major facilitator superfamily); | integral to membrane; transport; transporter activity; | 1.68365 % |  |  |
| 35 | ggggacaagtttgtacaaaaaagcaggcttgatgGCCTCAATGCGGAGCTCTCA, ggggaccactttgtacaagaaagctgggtcCTTCCCTCCATTAGCAACCT | Mycgr3\|67374  JGI  Ensembl | KOG1255 Succinyl-CoA synthetase, alpha subunit; | catalytic activity; metabolic process; | 100 % |  |  |
| 35 | ggggacaagtttgtacaaaaaagcaggcttgatgGGTATGTTCAGCAAGAGCGA, ggggaccactttgtacaagaaagctgggtcCTTCTCCCCAAACATCTCCT | Mycgr3\|67669  JGI  Ensembl | KOG1286 Amino acid transporters; | amino acid transmembrane transporter activity; amino acid transport;  membrane;  transport; | 100 % |  |  |
| 35 | ggggacaagtttgtacaaaaaagcaggcttgatgGCTTCCCTGATACTGGACAC, ggggaccactttgtacaagaaagctgggtcCCACTCCACCCGATCATCAG | Mycgr3\|67705  JGI  Ensembl | KOG0039 Ferric reductase, NADH/NADPH oxidase and related proteins; | ferric-chelate reductase activity; | 0 % |  |  |
| 35 | ggggacaagtttgtacaaaaaagcaggcttgatgCCTCACTACACCCTCCCCAT, ggggaccactttgtacaagaaagctgggtcCGAATACACATTCCCCAAAT | Mycgr3\|67759  JGI  Ensembl | KOG0681 Actin-related protein - Arp5p; | protein binding; | 100 % | chr\_2:341110 G->A; | ``` 1	ATGCCTCACTACACCCTCCCCATCCAGACCCAATCTGGCATCAACTCTCCCAAAGGACCACCTTCTCACATCTACAACGTCAGCGATCCTCCATACCAAGGCCAACTTCCCGCCGACCGACACGCCTACTCGCAAACCACCTCCGACACAGCAATAGTCATCGACAATGGCTCCTCTTCCGTCCGCGCCGGCTGGCAGACCGACGCGACTCCCCGCCTGCAATTCCCACCGCTCATGGCGCGCTATACCGACCGGAAATTGAATCGCAAACTCATGTTCATCGGAAGTGAGATCTACTTTGACGGCACGGCGCGTGGACAGGCGAAGAATGTGTATGAGCCGGGGAGCAACATTGTCAATAACTGGGATGTGCAGGAGGGGGTGTTGGACTACATTTTCTTGAAGCTGGGCATTGATGGGTCAAAGGGAGCGGTCGACCGGCCGGTGGTCATGACGGAGCCGCTGGCAAATTTGGGCTATACGAGGAAGGTCATGAGTGAGATTCTCTTTGAGCTGTATAATGTGCCGAGCGTGGCGTATGGTGTGGACTCGCTATTTAGTTATGACTACAATGGTGGGAACACGGGGTTGGTCGTTTCGAGCGCGAATATGAGCACGCATTTGATACCGGTGGTGGACAAGCAACCGCTCATCAGTCATGCGACACGCCTGGACTGGGGACGATCACATTGCGTGGAGTACCTGCAGAGGCTTCTGAAGACGAAGTACCCAGGACTTCTCACCTCTGGTAAGGTCACAGATACACAGATTGAGGATCTTGTGCGGCAACATTGCTACGTCAGCCAGAATTACGACCAGGAGACCGTGAAGATGCTGGAGTGGACAGGCTTGGAGGATCGTGATCACATCGTGCAGCTTCCATTCACCGAGAAGGAGGTGATCCAAAAGACGGAAGAGGAAATCAAGCGGGCAGAGGAGAAGAGACGAGAGGGTGGAAGAAGGCTACAAGAACAAGCAGCGAAGATGCGTCTGGAGAAGCTGGTCAGGAAAGAGCAAGAGCTTGAGTACTTCAAGCAGCTGCAAGGACAAGTACAAGAAGCAACGACAAAGAAGGAGATCAGATCATTACTGGA-AGA	1097 	\|\|\|\|\|\|\|\|\|\|\|\|\|\|\|\|\|\|\|\|\|\|\|\|\|\|\|\|\|\|\|\|\|\|\|\|\|\|\|\|\|\|\|\|\|\|\|\|\|\|\|\|\|\|\|\|\|\|\|\|\|\|\|\|\|\|\|\|\|\|\|\|\|\|\|\|\|\|\|\|\|\|\|\|\|\|\|\|\|\|\|\|\|\|\|\|\|\|\|\|\|\|\|\|\|\|\|\|\|\|\|\|\|\|\|\|\|\|\|\|\|\|\|\|\|\|\|\|\|\|\|\|\|\|\|\|\|\|\|\|\|\|\|\|\|\|\|\|\|\|\|\|\|\|\|\|\|\|\|\|\|\|\|\|\|\|\|\|\|\|\|\|\|\|\|\|\|\|\|\|\|\|\|\|\|\|\|\|\|\|\|\|\|\|\|\|\|\|\|\|\|\|\|\|\|\|\|\|\|\|\|\|\|\|\|\|\|\|\|\|\|\|\|\|\|\|\|\|\|\|\|\|\|\|\|\|\|\|\|\|\|\|\|\|\|\|\|\|\|\|\|\|\|\|\|\|\|\|\|\|\|\|\|\|\|\|\|\|\|\|\|\|\|\|\|\|\|\|\|\|\|\|\|\|\|\|\|\|\|\|\|\|\|\|\|\|\|\|\|\|\|\|\|\|\|\|\|\|\|\|\|\|\|\|\|\|\|\|\|\|\|\|\|\|\|\|\|\|\|\|\|\|\|\|\|\|\|\|\|\|\|\|\|\|\|\|\|\|\|\|\|\|\|\|\|\|\|\|\|\|\|\|\|\|\|\|\|\|\|\|\|\|\|\|\|\|\|\|\|\|\|\|\|\|\|\|\|\|\|\|\|\|\|\|\|\|\|\|\|\|\|\|\|\|\|\|\|\|\|\|\|\|\|\|\|\|\|\|\|\|\|\|\|\|\|\|\|\|\|\|\|\|\|\|\|\|\|\|\|\|\|\|\|\|\|\|\|\|\|\|\|\|\|\|\|\|\|\|\|\|\|\|\|\|\|\|\|\|\|\|\|\|\|\|\|\|\|\|\|\|\|\|\|\|\|\|\|\|\|\|\|\|\|\|\|\|\|\|\|\|\|\|\|\|\|\|\|\|\|\|\|\|\|\|\|\|\|\|\|\|\|\|\|\|\|\|\|\|\|\|\|\|\|\|\|\|\|\|\|\|\|\|\|\|\|\|\|\|\|\|\|\|\|\|\|\|\|\|\|\|\|\|\|\|\|\|\|\|\|\|\|\|\|\|\|\|\|\|\|\|\|\|\|\|\|\|\|\|\|\|\|\|\|\|\|\|\|\|\|\|\|\|\|\|\|\|\|\|\|\|\|\|\|\|\|\|\|\|\|\|\|\|\|\|\|\|\|\|\|\|\|\|\|\|\|\|\|\|\|\|\|\|\|\|\|\|\|\|\|\|\|\|\|\|\|\|\|\|\|\|\|\|\|\|\|\|\|\|\|\|\|\|\|\|\|\|\|\|\|\|\|\|\|\|\|\|\|\|\|\|\|\|\|\|\|\|\|\|\|\|\|\|\|\|\|\|\|\|\|\|\|\|\|\|\|\|\|\|\|\|\|\|\|\|\|\|\|\|\|\|\|\|\|\|\|\|\|\|\|\|\|\|\|\|\|\|\|\|\|\|\|\|\|\|\|\|\|\|\|\|\|\|\|\|\|\|\|\|\|\|\|\|\|\|\|\|\|\|\|\|\|\|\|\|\|\|\|\|\|\|\|\|\|\|\|\|\|\|\|\|\|\|\|\|\|\|\|\|\|\|\|\|\|\|\|\|\|\|\|\|\|\|\|\|\|\|\|\|\|\|\|\|\|\|\|\|\|\|\|\|\|\|\|\|\|\|\|\|\|\|\|\|\|\|\|\|\|\|\|\|\|\|\|\|\|\|\|\|\|\|\|\|\|\|\|\|\|\|\|\|\|\|\|\|\|\|\|\|\|\|\|\|\|\|\|\|\|\|\|\|\|\|\|\|\|\|\| \|\|\|\|\| \|\|\|\|\|\|\|\|\|\|\|\|\|\|\|\|\|\|\|\|\|\|\|\|\|\|\|\|\|\|\|\|\|\|\|\| \|\|\|\|\|\|\|\|\|\|\|\|\|\|\|\|\|\|\|\|\|\|\|\|\|\|\|\|\|\| \|\|\|\|\|\|\|\|\|\|\|\|\|\| \|\|\|\|\|\|\|\|\|\|\|\|\|\|\|\|\|\|\| \|\|\|\|\| \|\|\|\|\|\|\| \|\| \|\|\|\|\|\|\|\|\| \| \|\|\| \|\|\|\|\| \|\|\|\|\|\| \|\|\|\|\|\| \| \|\|\|\|\|\|\|\|\|\|\|\|\|\|\|\|\|\|\|\|\|\| \|\|\| 	ATGCCTCACTACACCCTCCCCATCCAGACCCAATCTGGCATCAACTCTCCCAAAGGACCACCTTCTCACATCTACAACGTCAGCGATCCTCCATACCAAGGCCAACTTCCCGCCGACCGACACGCCTACTCGCAAACCACCTCCGACACAGCAATAGTCATCGACAATGGCTCCTCTTCCGTCCGCGCCGGCTGGCAGACCGACGCGACTCCCCGCCTGCAATTCCCACCGCTCATGGCGCGCTATACCGACCGGAAATTGAATCGCAAACTCATGTTCATCGGAAGTGAGATCTACTTTGACGGCACGGCGCGTGGACAGGCGAAGAATGTGTATGAGCCGGGGAGCAACATTGTCAATAACTGGGATGTGCAGGAGGGGGTGTTGGACTACATTTTCTTGAAGCTGGGCATTGATGGGTCAAAGGGAGCGGTCGACCGGCCGGTGGTCATGACGGAGCCGCTGGCAAATTTGGGCTATACGAGGAAGGTCATGAGTGAGATTCTCTTTGAGCTGTATAATGTGCCGAGCGTGGCGTATGGTGTGGACTCGCTATTTAGTTATGACTACAATGGTGGGAACACGGGGTTGGTCGTTTCGAGCGCGAATATGAGCACGCATTTGATACCGGTGGTGGACAAGCAACCGCTCATCAGTCATGCGACACGCCTGGACTGGGGACGATCACATTGCGTGGAGTACCTGCAGAGGCTTCTGAAGACGAAGTACCCAGGACTTCTCACCTCTGGTAAGGTCACAGATACACAGATTGAGGATCTTGTGCGGCAACATTGCTACGTCAGCCAGAATTACGACCAGGAGACCGTGAAGATGCTGGAGTGGACAGGCTTGGAGGATCGTGATCACATCGTGCAGCTTCCATTCACCGAGAAGGAGGTGATCCAAA-GACGG-AGAGGAAATCAAGCGGGCAGAGGAGAAGAGACGAGA-GGTGGAAGAAGGCTACAAGAACAAGCAGCG-AGATGCGTCTGGAG-AGCTGGTCAGGAAAGAGCA-GAGCT-GAGTACT-CA-GCAGCTGCA-G-ACA-GTACA-GAAGCA-CGACAA-G-AGGAGATCAGATCATTACTGGAGAGA ``` |
| 35 | ggggacaagtttgtacaaaaaagcaggcttgatgCCAGACTCGCCGATACTAAA, ggggaccactttgtacaagaaagctgggtcCTGCTCTCCCTCATCATCCT | Mycgr3\|69058  JGI  Ensembl | KOG1426 FOG: RCC1 domain; | electron transport; heme binding;  iron ion binding;  monooxygenase activity; | 100 % | chr\_2:3374911 C->T; |  |
| 35 | ggggacaagtttgtacaaaaaagcaggcttgatgCGATCCCTCCACCTCACAGC, ggggaccactttgtacaagaaagctgggtcCAGCAACAGCGCGCAACCAA | Mycgr3\|69329  JGI  Ensembl |  |  | 100 % |  |  |
| 35 | ggggacaagtttgtacaaaaaagcaggcttgatgGCAGACAGCAAAGCCGAGCC, ggggaccactttgtacaagaaagctgggtcTCCAGTACAGCGCTCCCAAG | Mycgr3\|69859  JGI  Ensembl | KOG2534 DNA polymerase IV (family X); | DNA binding; DNA polymerase activity;  DNA repair;  DNA replication;  DNA-directed DNA polymerase activity; | 0 % |  |  |
| 35 | ggggacaagtttgtacaaaaaagcaggcttgatgGCTACAAAGACCCGTGGGGA, ggggaccactttgtacaagaaagctgggtcGTTGACCGGCACACCATTGG | Mycgr3\|69914  JGI  Ensembl | KOG1382 Multiple inositol polyphosphate phosphatase; | acid phosphatase activity; | 100 % |  |  |
| 35 | ggggacaagtttgtacaaaaaagcaggcttgatgGGAAACTCCAACTCCAACAT, ggggaccactttgtacaagaaagctgggtcACCATGGCTAATCACCACCC | Mycgr3\|70929  JGI  Ensembl | KOG0598 Ribosomal protein S6 kinase and related proteins; | ATP binding; protein amino acid phosphorylation;  protein kinase activity;  protein serine/threonine kinase activity;  protein-tyrosine kinase activity; | 100 % |  |  |
| 35 | ggggacaagtttgtacaaaaaagcaggcttgatgCGTTTCACCAAGTCCTTCCT, ggggaccactttgtacaagaaagctgggtcTCCCACAAGCGGCTGCGATC | Mycgr3\|70944  JGI  Ensembl |  |  | 100 % |  |  |
| 35 | ggggacaagtttgtacaaaaaagcaggcttgatgGCCTCCTCTGGCACAGCACT, ggggaccactttgtacaagaaagctgggtcTCCCTTCTCCACCAGCCCGG | Mycgr3\|71363  JGI  Ensembl | KOG1973 Chromatin remodeling protein, contains PHD Zn-finger; | protein binding; zinc ion binding; | 100 % |  |  |
| 35 | ggggacaagtttgtacaaaaaagcaggcttgatgGCGCTTCTGCACCACCGTCC, ggggaccactttgtacaagaaagctgggtcGCAATCATTGCTCATCTCTC | Mycgr3\|72330  JGI  Ensembl |  |  | 100 % |  |  |
| 35 | ggggacaagtttgtacaaaaaagcaggcttgatgGGCGAGTCACGCCCAACCGC, ggggaccactttgtacaagaaagctgggtcCTCTCGCACGACCATGAACG | Mycgr3\|72479  JGI  Ensembl | KOG3855 Monooxygenase involved in coenzyme Q (ubiquinone) biosynthesis; | FAD binding; aromatic compound metabolic process;  electron transport;  metabolic process;  monooxygenase activity;  oxidoreductase activity;  phenol 2-monooxygenase activity; | 100 % |  |  |
| 35 | ggggacaagtttgtacaaaaaagcaggcttgatgCGCACATCGACCTTGTTCCG, ggggaccactttgtacaagaaagctgggtcATCGACCTTTCCTTGTGCTG | Mycgr3\|72541  JGI  Ensembl | KOG0191 Thioredoxin/protein disulfide isomerase; | cell redox homeostasis; protein disulfide isomerase activity; | 100 % |  |  |
| 35 | ggggacaagtttgtacaaaaaagcaggcttgatgTCCTCTCCCACCACCACCCC, ggggaccactttgtacaagaaagctgggtcCAACTTACTCTTCAGCCCCA | Mycgr3\|74664  JGI  Ensembl | KOG1175 Acyl-CoA synthetase; | acetoacetate-CoA ligase activity; catalytic activity;  metabolic process; | 100 % |  |  |
| 35 | ggggacaagtttgtacaaaaaagcaggcttgatgCCTGCCCAGCGGCAGCCGAT, ggggaccactttgtacaagaaagctgggtcATCCAATCCAATCAACTCCT | Mycgr3\|74826  JGI  Ensembl | KOG1303 Amino acid transporters; |  | 100 % |  |  |
| 35 | ggggacaagtttgtacaaaaaagcaggcttgatgGCCGACGATGGGATGATGCT, ggggaccactttgtacaagaaagctgggtcTCCCAAGTTAAACTCCGAAG | Mycgr3\|75420  JGI  Ensembl | KOG0348 ATP-dependent RNA helicase; | 8-oxo-7,8-dihydroguanine triphosphatase activity; ATP binding;  ATP-dependent 5'-3' DNA helicase activity;  ATP-dependent helicase activity;  UDP-2,3-diacylglucosamine hydrolase activity;  bis(5'-nucleosyl)-tetraphosphatase activity;  dATP pyrophosphohydrolase activity;  diacylglycerol pyrophosphate phosphatase activity;  dihydroneopterin monophosphate phosphatase activity;  dihydroneopterin triphosphate pyrophosphohydrolase activity;  helicase activity;  nucleic acid binding;  pyrophosphatase activity;  thiamin-pyrophosphatase activity; | 100 % |  |  |
| 35 | ggggacaagtttgtacaaaaaagcaggcttgatgAGGGCTTCCTTCGCATCCAT, ggggaccactttgtacaagaaagctgggtcCGGCACCGTATCCGCCGTAC | Mycgr3\|75584  JGI  Ensembl |  |  | 100 % |  |  |
| 35 | ggggacaagtttgtacaaaaaagcaggcttgatgTCGGAGAGCAAGGGACTATC, ggggaccactttgtacaagaaagctgggtcAATGCCCAACCCAGGCACAG | Mycgr3\|75597  JGI  Ensembl | KOG2215 Exocyst complex subunit; |  | 100 % |  |  |
| 35 | ggggacaagtttgtacaaaaaagcaggcttgatgACCGAACGCCGCCCCATACA, ggggaccactttgtacaagaaagctgggtcATCCGCTGCATTTCTCCTCG | Mycgr3\|75806  JGI  Ensembl | KOG1347 Uncharacterized membrane protein, predicted efflux pump; | antiporter activity; drug transporter activity;  membrane;  multidrug transport; | 100 % |  |  |
| 35 | ggggacaagtttgtacaaaaaagcaggcttgatgCCCTCCAAACGCCCCGTGGC, ggggaccactttgtacaagaaagctgggtcACACCAACTGAACACCGGCG | Mycgr3\|75991  JGI  Ensembl | KOG2381 Phosphatidylinositol 4-kinase; | phosphotransferase activity, alcohol group as acceptor; | 12.18873 % |  |  |
| 35 | ggggacaagtttgtacaaaaaagcaggcttgatgGCCGAACTTGCGATCATGGA, ggggaccactttgtacaagaaagctgggtcACGGCTATCGTCGCCGCCGC | Mycgr3\|76222  JGI  Ensembl | KOG2641 Predicted seven transmembrane receptor - rhodopsin family; |  | 100 % |  |  |
| 35 | ggggacaagtttgtacaaaaaagcaggcttgatgGAAGACTTCCCTTTCCACCA, ggggaccactttgtacaagaaagctgggtcAGTGTAGTTCTTGCTAAGGT | Mycgr3\|82218  JGI  Ensembl |  |  | 100 % |  |  |
| 35 | ggggacaagtttgtacaaaaaagcaggcttgatgTCCGATACCGTGGATCTGGA, ggggaccactttgtacaagaaagctgggtcTGACATGCCAGTGTACGCCG | Mycgr3\|85078  JGI  Ensembl |  |  | 100 % |  |  |
| 35 | ggggacaagtttgtacaaaaaagcaggcttgatgATGATTCAGCCAACTCGAGT, ggggaccactttgtacaagaaagctgggtcCTCCCGTAACCTCTCAACAC | Mycgr3\|92404  JGI  Ensembl | KOG3841 TEF-1 and related transcription factor, TEAD family; | nucleus; regulation of transcription, DNA-dependent;  transcription factor activity; | 100 % |  | ``` 447	AGCGTTGGCCATCTTCCCACCTATGGGTCGCAACAAGCAAAGCCACAGAGACAAGCTACGAGGCCGCAATGAACTAATCTCAGACTACATCTACCAAGAGACACGAGTTTTACGCACCCGCAAGCAAGTCTCCAGTCATATTCAGGTCTTGAAGCCATTCACCAGAGGTGATCCGGCCATCACGAAGTACTTTGTAAAACAAAGAGGCCACGGCGATCACTACGGTGGCAGCAGATCGTCTCACGTCGATCGAAGGCACACTTCAAGATACCCAGTCAATGCACCGCCTCACGACATGTGCAACGGTCGGTTTCCGATGCCGCGGGTCGACACACGAGACTGGTCCAGCTCTCACATCCCGCTCGGGACATTTGAGCCCTTGAGCTTTGAGATGTTCGTTCAGCAGAAGTTCGTGCAGGACCACGACAAACCGGACATTGTGAAACGTCTACA-CACGTA-TACCGCATCGACCCATCATCCGTGTCTGGCAGATGAGC-ACTTCA-GAGACTGGCCCACTCTGGGCT-ATACATACCCACAGCTC-GCCAAGATGCATGCGAGGCGGCCGTTGAA-TTGCAA-CCTC-GTT-GC-TG-CGCAAGC	1040 	\|\|\|\|\|\|\|\|\|\|\|\|\|\|\|\|\|\|\|\|\|\|\|\|\|\|\|\|\|\|\|\|\|\|\|\|\|\|\|\|\|\|\|\|\|\|\|\|\|\|\|\|\|\|\|\|\|\|\|\|\|\|\|\|\|\|\|\|\|\|\|\|\|\|\|\|\|\|\|\|\|\|\|\|\|\|\|\|\|\|\|\|\|\|\|\|\|\|\|\|\|\|\|\|\|\|\|\|\|\|\|\|\|\|\|\|\|\|\|\|\|\|\|\|\|\|\|\|\|\|\|\|\|\|\|\|\|\|\|\|\|\|\|\|\|\|\|\|\|\|\|\|\|\|\|\|\|\|\|\|\|\|\|\|\|\|\|\|\|\|\|\|\|\|\|\|\|\|\|\|\|\|\|\|\|\|\|\|\|\|\|\|\|\|\|\|\|\|\|\|\|\|\|\|\|\|\|\|\|\|\|\|\|\|\|\|\|\|\|\|\|\|\|\|\|\|\|\|\|\|\|\|\|\|\|\|\|\|\|\|\|\|\|\|\|\|\|\|\|\|\|\|\|\|\|\|\|\|\|\|\|\|\|\|\|\|\|\|\|\|\|\|\|\|\|\|\|\|\|\|\|\|\|\|\|\|\|\|\|\|\|\|\|\|\|\|\|\|\|\|\|\|\|\|\|\|\|\|\|\|\|\|\|\|\|\|\|\|\|\|\|\|\|\|\|\|\|\|\|\|\|\|\|\|\|\|\|\|\|\|\|\|\|\|\|\|\|\|\|\|\|\|\|\|\|\|\|\|\|\|\|\|\|\|\|\|\|\|\|\|\|\|\|\|\|\|\|\|\|\|\|\|\|\|\|\|\|\|\|\|\|\|\|\|\|\|\|\|\|\|\|\|\|\|\|\|\|\|\|\|\|\|\|\|\|\|\|\|\|\|\|\|\|\|\|\|\|\|\|\|\|\|\|\|\|\|\|\|\|\|\|\|\|\|\|\|\|\|\|\|\|\|\| \|\|\|\|\|\| \|\|\|\|\|\|\|\|\|\|\|\|\| \|\|\|\|\| \|\|\|\|\|\|\|\|\|\|\|\|\|\|\|\|\|\| \|\|\|\|\|\| \|\|\|\|\|\|\| \|\|\|\|\|\|\|\| \|\|\|\| \|\|\|\|\|\|\|\|\|\|\|\|\|\|\|\|\| \|\|\| \|\|\|\|\|\|\|\|\|\|\|\|\|\|\|\| \|\|\|\|\|\|\|\| \|\|\|\|\|\| \|\|\|\| \|\|\| \|\| \|\| \|\|\|\|\|\|\| 	AGCGTTGGCCATCTTCCCACCTATGGGTCGCAACAAGCAAAGCCACAGAGACAAGCTACGAGGCCGCAATGAACTAATCTCAGACTACATCTACCAAGAGACACGAGTTTTACGCACCCGCAAGCAAGTCTCCAGTCATATTCAGGTCTTGAAGCCATTCACCAGAGGTGATCCGGCCATCACGAAGTACTTTGTAAAACAAAGAGGCCACGGCGATCACTACGGTGGCAGCAGATCGTCTCACGTCGATCGAAGGCACACTTCAAGATACCCAGTCAATGCACCGCCTCACGACATGTGCAACGGTCGGTTTCCGATGCCGCGGGTCGACACACGAGACTGGTCCAGCTCTCACATCCCGCTCGGGACATTTGAGCCCTTGAGCTTTGAGATGTTCGTTCAGCAGAAGTTCGTGCAGGACCACGACAAACCGGACATTGTGAAACGTCTACAACACGTATTACCGCATCGACC-ATCAT-CGTGTCTGGCAGATGAGCCACTTCAGGAGACTGACCCACTCTTGGCTAATACATACCCACAGCTCCGCC-AGATGCATGCGAGGCGTCCGTTGAAATTGCAAACCTCCGTTTGCCTGGCGCAAGC ```      ``` 1	ATGATGATTCAGCCAACTCGAGTCTTGCCCTCCAATGCCTCTCCTCTGCGTGACGCTGATAGTCTTCATGGCCCGAGGGTGTTGCAAGAACACTCTGGCAATCGCCAATACCACAACTACTCATACACTGAGCATTCACAACCGAAATATCCTCCGAGCCTGTCGACGGAGAACAGCTACATCGGACATCAGCCACTACATCACGAAGCTCACTATTACCAACAACACCCCGCACAACGCTACCATGGTGGAAGTAGACGGTCCCGGCCTCCCACGCGCGAAGAGCAGGATGCATGGATGCGAAAGTGTCTCAAGGAGGCGGACGTCCTCTACGCACGCTTCAGGGCTTCCGATGGCTACGCGAAATATCGTCAGCGTCAACAGAGGGAAGAGAAGCCGGGCACGGAACAGAAATGGCCAGAGCACCTCGAGGTAGCTTTCTTCCAAG	448 	\|\|\|\|\|\|\|\|\|\|\|\|\|\|\|\|\|\|\|\|\|\|\|\|\|\|\|\|\|\|\|\|\|\|\|\|\|\|\|\|\|\|\|\|\|\|\|\|\|\|\|\|\|\|\|\|\|\|\|\|\|\|\|\|\|\|\|\|\|\|\|\|\|\|\|\|\|\|\|\|\|\|\|\|\|\|\|\|\|\|\|\|\|\|\|\|\|\|\|\|\|\|\|\|\|\|\|\|\|\|\|\|\|\|\|\|\|\|\|\|\|\|\|\|\|\|\|\|\|\|\|\|\|\|\|\|\|\|\|\|\|\|\|\|\|\|\|\|\|\|\|\|\|\|\|\|\|\|\|\|\|\|\|\|\|\|\|\|\|\|\|\|\|\|\|\|\|\|\|\|\|\|\|\|\|\|\|\|\|\|\|\|\|\|\|\|\|\|\|\|\|\|\|\|\|\|\|\|\|\|\|\|\|\|\|\|\|\|\|\|\|\|\|\|\|\|\|\|\|\|\|\|\|\|\|\|\|\|\|\|\|\|\|\|\|\|\|\|\|\|\|\|\|\|\|\|\|\|\|\|\|\|\|\|\|\|\|\|\|\|\|\|\|\|\|\|\|\|\|\|\|\|\|\|\|\|\|\|\|\|\|\|\|\|\|\|\|\|\|\|\|\|\|\|\|\|\|\|\|\|\|\|\|\|\|\|\|\|\|\|\|\|\|\|\|\|\|\|\|\|\|\|\|\|\|\|\|\|\|\|\|\|\|\|\|\|\|\|\|\|\|\|\|\|\|\|\|\|\|\|\|\|\|\|\|\|\|\|\|\|\|\|\|\|\|\|\|\|\|\|\|\|\|\|\|\|\|\|\|\|\|\|\|\|\|\|\|\|\|\|\|\|\|\|\|\|\|\|\|\|\|\|\|\|\|\|\|\|\|\|\|\|\|\|\|\|\|\|\|\|\|\|\|\|\|\|\|\|\|\|\|\|\|\|\|\|\|\| 	ATGATGATTCAGCCAACTCGAGTCTTGCCCTCCAATGCCTCTCCTCTGCGTGACGCTGATAGTCTTCATGGCCCGAGGGTGTTGCAAGAACACTCTGGCAATCGCCAATACCACAACTACTCATACACTGAGCATTCACAACCGAAATATCCTCCGAGCCTGTCGACGGAGAACAGCTACATCGGACATCAGCCACTACATCACGAAGCTCACTATTACCAACAACACCCCGCACAACGCTACCATGGTGGAAGTAGACGGTCCCGGCCTCCCACGCGCGAAGAGCAGGATGCATGGATGCGAAAGTGTCTCAAGGAGGCGGACGTCCTCTACGCACGCTTCAGGGCTTCCGATGGCTACGCGAAATATCGTCAGCGTCAACAGAGGGAAGAGAAGCCGGGCACGGAACAGAAATGGCCAGAGCACCTCGAGGTAGCTTTCTTCCAAG ``` |
| 35 | ggggacaagtttgtacaaaaaagcaggcttgatgCCGACCTACAGCGCATGCGG, ggggaccactttgtacaagaaagctgggtcAACCACAACAAACCTCATCC | Mycgr3\|92420  JGI  Ensembl | KOG4475 FOG: Immunoglobin and related proteins; |  | 100 % |  |  |
| 35 | ggggacaagtttgtacaaaaaagcaggcttgatgCTGTCCTCCTATGCGACCCG, ggggaccactttgtacaagaaagctgggtcATGCTCCGCTTCCACAACTC | Mycgr3\|92641  JGI  Ensembl |  |  | 100 % |  |  |
| 35 | ggggacaagtttgtacaaaaaagcaggcttgatgGCTGCCGCGAAGAAGCAATC, ggggaccactttgtacaagaaagctgggtcACCCCGACTCCTCTTCGCCC | Mycgr3\|92720  JGI  Ensembl | KOG4475 FOG: Immunoglobin and related proteins; |  | 0 % |  |  |
| 35 | ggggacaagtttgtacaaaaaagcaggcttgatgGCGTTCCTCTCCCAGAATCG, ggggaccactttgtacaagaaagctgggtcCTCATCATCCTCATCGTCGT | Mycgr3\|92747  JGI  Ensembl | KOG1216 von Willebrand factor and related coagulation proteins; | exo-alpha-sialidase activity; | 100 % |  |  |
| 35 | ggggacaagtttgtacaaaaaagcaggcttgatgGATCGTGGATTTGGGCAGCC, ggggaccactttgtacaagaaagctgggtcAGCAGGGCTGGAGCAGTCAT | Mycgr3\|93009  JGI  Ensembl | KOG4678 FOG: Calponin homology domain; |  | 0.90293 % |  |  |
| 35 | ggggacaagtttgtacaaaaaagcaggcttgatgTCCGGCATCGAAGTCGCAGG, ggggaccactttgtacaagaaagctgggtcTTTGGACAACCAAACATTGT | Mycgr3\|93031  JGI  Ensembl |  |  | 100 % |  |  |
| 35 | ggggacaagtttgtacaaaaaagcaggcttgatgATCACCTCTGCCAGGCTACA, ggggaccactttgtacaagaaagctgggtcGCTGCCAGGTGTTGCGCCAG | Mycgr3\|93175  JGI  Ensembl |  |  | 100 % |  |  |
| 35 | ggggacaagtttgtacaaaaaagcaggcttgatgGCACGCCCTCGCAAGTCGCG, ggggaccactttgtacaagaaagctgggtcCGACTCGATCCTCTTTGGAT | Mycgr3\|93296  JGI  Ensembl |  |  | 100 % |  |  |
| 35 | ggggacaagtttgtacaaaaaagcaggcttgatgGACGACGACGGCGAGCCGCA, ggggaccactttgtacaagaaagctgggtcCATCTTGAACAGCACTACCT | Mycgr3\|93376  JGI  Ensembl |  |  | 1.71086 % |  |  |
| 35 | ggggacaagtttgtacaaaaaagcaggcttgatgAGGGCCACCACTCGCTGCTC, ggggaccactttgtacaagaaagctgggtcAAGCATAGCACTTAACCTCT | Mycgr3\|93697  JGI  Ensembl | KOG1257 NADP+-dependent malic enzyme; | malate dehydrogenase (oxaloacetate-decarboxylating) (NADP+) activity; malate metabolic process;  malic enzyme activity; | 1.84768 % |  |  |
| 35 | ggggacaagtttgtacaaaaaagcaggcttgatgGCGGTTAAAACACCAGTAGC, ggggaccactttgtacaagaaagctgggtcAGGGATTGCGCTGGTGAACT | Mycgr3\|94071  JGI  Ensembl |  |  | 0.837 % |  |  |
| 35 | ggggacaagtttgtacaaaaaagcaggcttgatgCTGCAACCCAGCCATCCCGC, ggggaccactttgtacaagaaagctgggtcACCCGCGTGGGATCTCTCTT | Mycgr3\|96439  JGI  Ensembl |  |  | 100 % |  |  |
| 35 | ggggacaagtttgtacaaaaaagcaggcttgatgGCCGACGTAATTGCGCTCCA, ggggaccactttgtacaagaaagctgggtcCCACCGGCCGTAGTATTCTT | Mycgr3\|96528  JGI  Ensembl | KOG0161 Myosin class II heavy chain; |  | 100 % |  |  |
| 35 | ggggacaagtttgtacaaaaaagcaggcttgatgGTCACTTTCATAAACGCTCT, ggggaccactttgtacaagaaagctgggtcCCGGGCGATTGATTCTCGGC | Mycgr3\|97167  JGI  Ensembl | KOG3599 Ca2+-modulated nonselective cation channel polycystin; | glucan 1,4-alpha-glucosidase activity; nutrient reservoir activity; | 100 % |  |  |
| 35 | ggggacaagtttgtacaaaaaagcaggcttgatgGACGATGAGGCGCTGGAGAA, ggggaccactttgtacaagaaagctgggtcGTATGTCAAGATTTTCAGAG | Mycgr3\|97753  JGI  Ensembl |  |  | 100 % |  |  |
| 35 | ggggacaagtttgtacaaaaaagcaggcttgatgATCGCAGACGTCGGAGACCG, ggggaccactttgtacaagaaagctgggtcTCGCGGCGTCTCAGCCGGCA | Mycgr3\|97852  JGI  Ensembl |  |  | 0 % |  |  |
| 35 | ggggacaagtttgtacaaaaaagcaggcttgatgGACGTGTCACATGCGCAATC, ggggaccactttgtacaagaaagctgggtcACGAATCACACCCCACTCGC | Mycgr3\|98409  JGI  Ensembl | KOG0305 Anaphase promoting complex, Cdc20, Cdh1, and Ama1 subunits; | hydrogen ion transmembrane transporter activity; proton-transporting two-sector ATPase complex; | 100 % |  |  |
| 35 | ggggacaagtttgtacaaaaaagcaggcttgatgGAGCAACCCAAGCCCTCGAC, ggggaccactttgtacaagaaagctgggtcCAAATCGTCAAGGCCCATGC | Mycgr3\|98983  JGI  Ensembl | KOG0242 Kinesin-like protein; | ATP binding; microtubule associated complex;  microtubule motor activity;  microtubule-based movement; | 100 % |  | ``` 1	ATGGAGCAACCCAAGCCCTCGACCGCGCTGTTCGATGTCTTTCTGAGACTCCGCCCTTCCACTGCCGTCGACAAGGAGCGATTTCTGGACGTCGAGCCCGGTGGCCTCTCAGCCCCAACCCATATCACGATCAAGCCTCCGACTGGCGATACAAGAAAGAGAGCAATTGAGAGATTCGCATTTACGAGAGTGTTTGAGGAGAATGCTGGACAGCG	215 	\|\|\|\|\|\|\|\|\|\|\|\|\|\|\|\|\|\|\|\|\|\|\|\|\|\|\|\|\|\|\|\| \|\|\|\|\|\|\|\|  \| \|\|\|\|\|\|\|\|\|\|\|\|\|\|\| \|\|\|\|\| \|\|\|\|\|\|\|\|\|\|\|\|\|\|\|\|\|\|\|\|\|\|\|\|\|\|\|\|\|\|\| \|\|\|\|\|\|\|\|\|\|\|\|\|\|\|\|\|\|\|\|\|\|\| \|\|\|\|\|\|\|\|\|\|\|\| \|\|\|\|\|\|\| \|\|\|\|\|\|\|\|\|\|\|\|\|\|\|\|\|\|\|\|\| \|\|\|\|\|\|\|\|\|\|\|\|\|\|\|\|\|\|\|\|\|\|\|\|\|\|\|\| \|\|\|\|\|\|\|\|\|\|\|\|\|\|\|\|\|\|\|\| 	ATGGAGCAACCCAAGCCCTCGACCGCGCTGTTTGATGTCTTCTTAAGACTCCGCCCTTCCGCTGCCATCGACAAGGAGCGATTTCTGGACGTCGAGCCGGGTGGCCTCTCAGCCCCAACCCACATCACGATCAAGTCTCCGACGGGCGATACAAGAAAGAGAGCAGTTGAGAGATTCGCATTTACGAGAGTGTTCGAGGAGAATGCTGGACAGCG ```      ``` 197	AGGAGAATGCTGGACAGCGCGAACTATTCGAGACCACGGGAGCCGTGGAGCTCATTC	253 	\|\|\|\|\|\|\|\|\|\|\|\|\|\|\|\|\|\|\|\|\|\|\|\|\|\|\|\|\| \|\|\| \|\|\|\|\|\|\|\|\|\|\|\|\|\|\| \|\|\|\|\|\|\| 	AGGAGAATGCTGGACAGCGCGAACTATTCTAGAACACGGGAGCCGTGGATCTCATTC ``` |
| 35 | ggggacaagtttgtacaaaaaagcaggcttgatgTCCCTCTTCGGCGACGACGA, ggggaccactttgtacaagaaagctgggtcATCCCCCAACGGCATCGCCT | Mycgr3\|99206  JGI  Ensembl | KOG2273 Membrane coat complex Retromer, subunit VPS5/SNX1, Sorting nexins, and related PX domain-containing proteins; | cell communication; phosphoinositide binding;  protein binding; | 0.89888 % |  |  |
| 35 | ggggacaagtttgtacaaaaaagcaggcttgatgGACGACCAAGAGTATCTCGA, ggggaccactttgtacaagaaagctgggtcATAATACGGCCGACCCTCCT | Mycgr3\|100473  JGI  Ensembl | KOG2146 Splicing coactivator SRm160/300, subunit SRm160 (contains PWI domain); |  | 100 % | chr\_6:1095135 A->G; |  |
| 35 | ggggacaagtttgtacaaaaaagcaggcttgatgGTCGGCCTCGGACCACGGCG, ggggaccactttgtacaagaaagctgggtcCATACCCTTGAGCATGGACT | Mycgr3\|100586  JGI  Ensembl | KOG1369 Hexokinase; | ATP binding; glycolysis; hexokinase activity; | 100 % |  |  |
| 35 | ggggacaagtttgtacaaaaaagcaggcttgatgCCATTCTTACAACTTTTGAA, ggggaccactttgtacaagaaagctgggtcCAGCCTTCTCCTCTGCTCCT | Mycgr3\|101182  JGI  Ensembl | KOG3660 Sodium-neurotransmitter symporter; | integral to plasma membrane; membrane; neurotransmitter transport;  neurotransmitter:sodium symporter activity; | 100 % | chr\_9:758950 G->T; |  |
| 35 | ggggacaagtttgtacaaaaaagcaggcttgatgTCTTTTCTGGCTCTTGAAGG, ggggaccactttgtacaagaaagctgggtcCAGGATGCCTTCGGGCTTTC | Mycgr3\|101763  JGI  Ensembl | KOG1200 Mitochondrial/plastidial beta-ketoacyl-ACP reductase; | carbohydrate metabolic process; hydrolase activity, acting on carbon-nitrogen (but not peptide) bonds;  metabolic process;  oxidoreductase activity; | 100 % |  |  |
| 35 | ggggacaagtttgtacaaaaaagcaggcttgatgGACACAAAATTCGACGACTC, ggggaccactttgtacaagaaagctgggtcATCCGAGATCAACTCCTCCG | Mycgr3\|101913  JGI  Ensembl | KOG1289 Amino acid transporters; | amino acid transmembrane transporter activity; amino acid transport;  membrane;  transport; | 100 % |  |  |
| 35 | ggggacaagtttgtacaaaaaagcaggcttgatgGGTGTCTTTGACAAGTTTAC, ggggaccactttgtacaagaaagctgggtcAGCAATCGATGCGCGGCGCT | Mycgr3\|102977  JGI  Ensembl | KOG1286 Amino acid transporters; | amino acid transmembrane transporter activity; amino acid transport;  membrane;  transport; | 100 % |  |  |
| 35 | ggggacaagtttgtacaaaaaagcaggcttgatgAGTGCTCCCATCGCTGATGA, ggggaccactttgtacaagaaagctgggtcAGGCCTACTCTCCTCCTCAT | Mycgr3\|104715  JGI  Ensembl | KOG1237 H+/oligopeptide symporter; | membrane; oligopeptide transport;  transporter activity; | 100 % |  |  |
| 35 | ggggacaagtttgtacaaaaaagcaggcttgatgAAGTTCTCCACCGTCCTCCC, ggggaccactttgtacaagaaagctgggtcGAAAGCGGCGACGGCCATCA | Mycgr3\|105310  JGI  Ensembl |  |  | 0 % |  |  |
| 35 | ggggacaagtttgtacaaaaaagcaggcttgatgGCTGGTGGCCTCGTCGCTAC, ggggaccactttgtacaagaaagctgggtcAAAGGCAGACGCATTACGCT | Mycgr3\|105594  JGI  Ensembl | KOG0254 Predicted transporter (major facilitator superfamily); | L-arabinose isomerase activity; carbohydrate transport;  integral to membrane;  membrane;  sugar:hydrogen symporter activity;  transport;  transporter activity; | 100 % |  |  |
| 35 | ggggacaagtttgtacaaaaaagcaggcttgatgTCCGCCAAATCCATCCTCGA, ggggaccactttgtacaagaaagctgggtcAGCCTCGAACTCCTTCACAC | Mycgr3\|105760  JGI  Ensembl | KOG1254 ATP-citrate lyase; | ATP citrate synthase activity; | 100 % | chr\_9:1471573 C->T; |  |
| 35 | ggggacaagtttgtacaaaaaagcaggcttgatgGCACCAGCAATCGGTATCGA, ggggaccactttgtacaagaaagctgggtcGTCAACCTCCTCGACGGTTG | Mycgr3\|105895  JGI  Ensembl | KOG0101 Molecular chaperones HSP70/HSC70, HSP70 superfamily; | ATP binding; cysteine-type endopeptidase activity;  proteolysis; | 100 % |  |  |
| 35 | ggggacaagtttgtacaaaaaagcaggcttgatgGCCTCTCTTTTCGCCCTACT, ggggaccactttgtacaagaaagctgggtcACCCGCGCTTTTCAAATGCA | Mycgr3\|106799  JGI  Ensembl | KOG4003 Pyrazinamidase/nicotinamidase PNC1; | catalytic activity; metabolic process; | 100 % |  |  |
| 35 | ggggacaagtttgtacaaaaaagcaggcttgatgCCACCACTGGTTCTCGCAAC, ggggaccactttgtacaagaaagctgggtcAGCCAACCATGAGCAGAGCT | Mycgr3\|106886  JGI  Ensembl | KOG2051 Nonsense-mediated mRNA decay 2 protein; |  | 100 % |  | ``` 998	ATGGCAGAACTCGATGACGAGGAGCTGGCTGACATGGGATGCTTTGCCCTTCGTCCCTTGGACGATGTCGATGAGGTTGACGGTTTCGAGTATGAGGAGGACGATGGCTT	1107 	\|\|\|\|\|\|\|\|\|\|\|\| \|\|\|\|\|\|\|\|\|\|\|\|\|\|\|\|\|\|\|\|\|\|\|\|\|\|\|\|\|\|\|\|\|\|\| \| \|\|\|\|\|\|\|\|\|\|\| \|\|\|\|\|\|\|\|\|\|\|\|\|\| \|\|\|\|\|\|\|\|\|\|\|\|\|\|\| \|\|\|\|\|\|\|\|\|\|\|\|\|\|\|\| 	ATGGCAGAACTCNATGACGAGGAGCTGGCTGACATGGGATGCTTTGCCTTGCGTCCCTTGGATGATGTCGATGAGGTCGACGGTTTCGAGTATAAGGAGGACGATGGCTT ``` |
| 35 | ggggacaagtttgtacaaaaaagcaggcttgatgGCGCCCGACAGAGGCTCATC, ggggaccactttgtacaagaaagctgggtcCGCATGTGGCACGCCAGCAC | Mycgr3\|106955  JGI  Ensembl | KOG2262 Sexual differentiation process protein ISP4; |  | 0 % |  |  |
| 35 | ggggacaagtttgtacaaaaaagcaggcttgatgGAGGAGAGGCAAACAGAACT, ggggaccactttgtacaagaaagctgggtcGGCGAAGAAGGCTCTCTTCG | Mycgr3\|107195  JGI  Ensembl | KOG2094 Predicted DNA damage inducible protein; | DNA polymerase activity; DNA repair;  DNA-directed DNA polymerase activity; | 0 % |  |  |
| 35 | ggggacaagtttgtacaaaaaagcaggcttgatgCAGGACGACAACAGTCCGCT, ggggaccactttgtacaagaaagctgggtcCACTTCTATCCACTCATCCT | Mycgr3\|107288  JGI  Ensembl | KOG0334 RNA helicase; |  | 100 % |  |  |
| 35 | ggggacaagtttgtacaaaaaagcaggcttgatgACTTCAACATACAGAACTTC, ggggaccactttgtacaagaaagctgggtcCGTCCACCAGCCCCCCTGTA | Mycgr3\|107384  JGI  Ensembl | KOG2412 Nuclear-export-signal (NES)-containing protein/polyadenylated-RNA export factor; |  | 0.9434 % |  |  |
| 35 | ggggacaagtttgtacaaaaaagcaggcttgatgCCAAAGGAGAACGCTGGAAG, ggggaccactttgtacaagaaagctgggtcTAGAAGGCCGTCTGGCCACA | Mycgr3\|107684  JGI  Ensembl | KOG1515 Arylacetamide deacetylase; |  | 100 % |  |  |
| 35 | ggggacaagtttgtacaaaaaagcaggcttgatgTCGCTTGACCGCTCACCGAG, ggggaccactttgtacaagaaagctgggtcTGCGTCTTGCGGTTGAGCTT | Mycgr3\|107800  JGI  Ensembl | KOG4748 Subunit of Golgi mannosyltransferase complex; | integral to membrane; transferase activity, transferring hexosyl groups; | 1.12474 % |  |  |
| 35 | ggggacaagtttgtacaaaaaagcaggcttgatgGCTAGGAGTCAGGCGCTGCA, ggggaccactttgtacaagaaagctgggtcAAGCTCCTTCTTCCCCTCTT | Mycgr3\|107801  JGI  Ensembl | KOG2513 Protein required for meiotic chromosome segregation; |  | 100 % |  |  |
| 35 | ggggacaagtttgtacaaaaaagcaggcttgatgCGAATCCGCCTGCGCATCGA, ggggaccactttgtacaagaaagctgggtcCGCCACCTCGACCACACCCT | Mycgr3\|109018  JGI  Ensembl | KOG2992 Nucleolar GTPase/ATPase p130; | binding; | 100 % |  |  |
| 35 | ggggacaagtttgtacaaaaaagcaggcttgatgTCCGCGTCTGACCAACAATC, ggggaccactttgtacaagaaagctgggtcTGTGTTCCTCAAAAACACCC | Mycgr3\|109078  JGI  Ensembl | KOG2133 Transcriptional corepressor Atrophin-1/DRPLA; | nucleus; regulation of transcription, DNA-dependent;  transcription factor activity;  zinc ion binding; | 100 % |  | ``` 1	ATGTCCGCGTCTGACCAACAATCGGAAGCGTACCCGTCGCCCTCCGCCGCATCGAACAATCATCCCTTCTACCAGCATCTCCAACAACAAAGCCTCGAATCCCGCACACCACAGCAGAATGGCAGGAACACCCAGTCGCCCGGTCCTGGCGACGACAATACGAACAGAAGAATGTCGGAAGATTCACAAATGCAGGCACATCTGGACGTAATGGGCAGCTTTGGCAATTTTGGCTCAGTGCCAGGACAATTACCATATCCTCCTCCCGGTGCAGCGTCCTCCTCACAAGTCCCTCACGGTTCCGTCGCATATGCACAAGCGAACGGCACGGCCGATGCTGGGCCTCCACCGCCGGACAAGAGCAAGCAGCGAACGAAAGTCAGTCGGGCTTGTGATGAGTGCAGACGGAAGAAGATACGATGTGACGCGGTCGATGAAGTGCCGAGCATGCCATGTAACAACTGTGCTCGGACGGCAGCGAAGTGCACATTCAGTCGACAGCCGATGAAGCGAGGACCGAGCAAGGGCTACATCAAAGAGCTGGCAGAACGTCTAAACTCCCTCGAACAGCAGGTGCAACCTCCACATCTGGAACAGCAGCATCGGTACGAGATACAGCAAGTGCAACAGGCGATGGCGGAGGGACTTCAAGCGTATACACATGATCCTCGTATCATGCCGGATCCACGAATAGGTGATAAGCGAACGCATGAGGTCGCGGAAGGGTTACATCCAGGACAGCAGGCGATATATTCAAATGTGGGCAATGCGCCGCCGCCGCAGCAGCACATGTACGCGCCGCATATGCAAACACCAGGACCGACACACCAGCAGCATAATGTCCAGCCCTTCTGGCGGTTGTCAGGGAGTGAGAACCGACACGATACAGTTGGCATGGCCTTTGAAGCGGAGAACAGCACGACGATCACGAACTTTGACTGGGACGAGGCGGTGATTGACGAGTACTACCGGATCATACATCAG-ACATTCCCTCTGCTCGCGAACAATAAAAATCGCCTACGACAGCGGCTAGCTGAATG-TCCCGCATCGTTACGAGA-GGCATTCCTCG	1069 	\|\|\|\|\|\|\|\|\|\|\|\|\|\|\|\|\|\|\|\|\|\|\|\|\|\|\|\|\|\|\|\|\|\|\|\|\|\|\|\|\|\|\|\|\|\|\|\|\|\|\|\|\|\|\|\|\|\|\|\|\|\|\|\|\|\|\|\|\|\|\|\|\|\|\|\|\|\|\|\|\|\|\|\|\|\|\|\|\|\|\|\|\|\|\|\|\|\|\|\|\|\|\|\|\|\|\|\|\|\|\|\|\|\|\|\|\|\|\|\|\|\|\|\|\|\|\|\|\|\|\|\|\|\|\|\|\|\|\|\|\|\|\|\|\|\|\|\|\|\|\|\|\|\|\|\|\|\|\|\|\|\|\|\|\|\|\|\|\|\|\|\|\|\|\|\|\|\|\|\|\|\|\|\|\|\|\|\|\|\|\|\|\|\|\|\|\|\|\|\|\|\|\|\|\|\|\|\|\|\|\|\|\|\|\|\|\|\|\|\|\|\|\|\|\|\|\|\|\|\|\|\|\|\|\|\|\|\|\|\|\|\|\|\|\|\|\|\|\|\|\|\|\|\|\|\|\|\|\|\|\|\|\|\|\|\|\|\|\|\|\|\|\|\|\|\|\|\|\|\|\|\|\|\|\|\|\|\|\|\|\|\|\|\|\|\|\|\|\|\|\|\|\|\|\|\|\|\|\|\|\|\|\|\|\|\|\|\|\|\|\|\|\|\|\|\|\|\|\|\|\|\|\|\|\|\|\|\|\|\|\|\|\|\|\|\|\|\|\|\|\|\|\|\|\|\|\|\|\|\|\|\|\|\|\|\|\|\|\|\|\|\|\|\|\|\|\|\|\|\|\|\|\|\|\|\|\|\|\|\|\|\|\|\|\|\|\|\|\|\|\|\|\|\|\|\|\|\|\|\|\|\|\|\|\|\|\|\|\|\|\|\|\|\|\|\|\|\|\|\|\|\|\|\|\|\|\|\|\|\|\|\|\|\|\|\|\|\|\|\|\|\|\|\|\|\|\|\|\|\|\|\|\|\|\|\|\|\|\|\|\|\|\|\|\|\|\|\|\|\|\|\|\|\|\|\|\|\|\|\|\|\|\|\|\|\|\|\|\|\|\|\|\|\|\|\|\|\|\|\|\|\|\|\|\|\|\|\|\|\|\|\|\|\|\|\|\|\|\|\|\|\|\|\|\|\|\|\|\|\|\|\|\|\|\|\|\|\|\|\|\|\|\|\|\|\|\|\|\|\|\|\|\|\|\|\|\|\|\|\|\|\|\|\|\|\|\|\|\|\|\|\|\|\|\|\|\|\|\|\|\|\|\|\|\|\|\|\|\|\|\|\|\|\|\|\|\|\|\|\|\|\|\|\|\|\|\|\|\|\|\|\|\|\|\|\|\|\|\|\|\|\|\|\|\|\|\|\|\|\|\|\|\|\|\|\|\|\|\|\|\|\|\|\|\|\|\|\|\|\|\|\|\|\|\|\|\|\|\|\|\|\|\|\|\|\|\|\|\|\|\|\|\|\|\|\|\|\|\|\|\|\|\|\|\|\|\|\|\|\|\|\|\|\|\|\|\|\|\|\|\|\|\|\|\|\|\|\|\|\|\|\|\|\|\|\|\|\|\|\|\|\|\|\|\|\|\|\|\|\|\|\|\|\|\|\|\|\|\|\|\|\|\|\|\|\|\|\|\|\|\|\|\|\|\|\|\|\|\|\|\|\|\|\|\|\|\|\|\|\|\|\|\|\|\|\|\|\|\|\|\|\|\|\|\|\|\|\|\|\|\|\|\|\|\|\|\|\|\|\|\|\|\|\|\|\|\|\|\|\|\|\|\|\|\|\|\|\|\|\|\|\|\|\|\|\|\|\|\|\|\|\|\|\|\|\|\|\|\|\|\|\|\|\|\|\|\|\|\|\|\|\|\|\| \|\|\|\|\|\|\|\|\|\|\|\|\|\|\|\|\|\|\|\|\|\|\|\|\|\|\|\|\|\|\|\|\|\|\|\|\|\|\|\|\|\|\|\|\|\|\| \|\|\|\|\|\|\|\|\|\|\|\|\|\|\|\|\|\|\|\|\|\|\|\|\|\|\|\|\|\|\|\|\|\|\|\|\| \|\|\|\|\|\|\|\|\|\|\|\|\|\|\|\|\|\|\|\|\|\|\|\|\|\|\|\|\|\|\|\|\| \|\|\|\|\|\|\|\|\|\|\|\|\|\|\|\|\|\| \|\|\|\|\|\|\|\|\|\|\|\|\| \|\|\|\|\|\|\|\|\|\|\|\|\|\|\|\|\|\|\|\|\|\|\| \|\|\|\|\|\|\|\|\|\|\|\|\|\|\|\|\|\| \|\|\|\|\|\|\| \|\|\| 	ATGTCCGCGTCTGACCAACAATCGGAAGCGTACCCGTCGCCCTCCGCCGCATCGAACAATCATCCCTTCTACCAGCATCTCCAACAACAAAGCCTCGAATCCCGCACACCACAGCAGAATGGCAGGAACACCCAGTCGCCCGGTCCTGGCGACGACAATACGAACAGAAGAATGTCGGAAGATTCACAAATGCAGGCACATCTGGACGTAATGGGCAGCTTTGGCAATTTTGGCTCAGTGCCAGGACAATTACCATATCCTCCTCCCGGTGCAGCGTCCTCCTCACAAGTCCCTCACGGTTCCGTCGCATATGCACAAGCGAACGGCACGGCCGATGCTGGGCCTCCACCGCCGGACAAGAGCAAGCAGCGAACGAAAGTCAGTCGGGCTTGTGATGAGTGCAGACGGAAGAAGATACGATGTGACGCGGTCGATGAAGTGCCGAGCATGCCATGTAACAACTGTGCTCGGACGGCAGCGAAGTGCACATTCAGTCGACAGCCGATGAAGCGAGGACCGAGCAAGGGCTACATCAAAGAGCTGGCAGAACGTCTAAACTCCCTCGAACAGCAGGTGCAACCTCCACATCTGGAACAGCAGCATCGGTACGAGATACAGCAAGTGCAACAGGCGATGGCGGAGGGACTTCAAGCGTATACACATGATCCTCGTATCATGCCGGATCCACGAATAGGTGATAAGCGAACGCATGAGGTCGCGGAAGGGTTACATCCAGGACAGCAGGCGATATATTCAAATGTGGGCAATGCGCCGCCGCCGCAGCAGCACATGTACGCGCCGCATATGCAAACACCAGGACCGACACACCAGCAGCATAATGTCCAGCCCTTCTGGCGGTTGTCA-GGAGTGAGAACCGACACGATACAGTTGGCATGGCCTTTGAAGCGGAG-ACAGCACGACGATCACGAACTTTGACTGGGACGAGGC-GTGATTGACGAGTACTACCGGATCATACATCAGAACATTCCCTCTGCTCGCG-ACAATAAAAATCG-CTACGACAGCGGCTAGCTGAATGTTCCCGCATCGTTACGAGAAGGCATTCTTCG ``` |
| 35 | ggggacaagtttgtacaaaaaagcaggcttgatgGACGACTTTGGCAAGCAACA, ggggaccactttgtacaagaaagctgggtcCAGCATCAACCTCTCCAGCT | Mycgr3\|109219  JGI  Ensembl | KOG1721 FOG: Zn-finger; | intracellular; nucleic acid binding;  zinc ion binding; | 100 % | chr\_4:2053459 G->C; |  |
| 35 | ggggacaagtttgtacaaaaaagcaggcttgatgTCGCGATACCGAGCTTCAGA, ggggaccactttgtacaagaaagctgggtcAGTCAACGTCGCCATCGCCG | Mycgr3\|109465  JGI  Ensembl | KOG2253 U1 snRNP complex, subunit SNU71 and related PWI-motif proteins; |  | 100 % |  |  |
| 35 | ggggacaagtttgtacaaaaaagcaggcttgatgGCGGGCATGTCCAGCAAACG, ggggaccactttgtacaagaaagctgggtcAATCTTACTCCTCGCCGCCT | Mycgr3\|109524  JGI  Ensembl | KOG3069 Peroxisomal NUDIX hydrolase; | hydrolase activity; post-translational protein modification;  regulation of protein metabolic process;  small conjugating protein ligase activity; | 0.87796 % |  |  |
| 35 | ggggacaagtttgtacaaaaaagcaggcttgatgTCGACCACGGCGCCCTTGAC, ggggaccactttgtacaagaaagctgggtcCTCCTCCCTACTCCTTTCCC | Mycgr3\|109848  JGI  Ensembl | KOG1869 Splicing coactivator SRm160/300, subunit SRm300; | G-protein coupled receptor protein signaling pathway; integral to membrane;  rhodopsin-like receptor activity; | 100 % |  |  |
| 35 | ggggacaagtttgtacaaaaaagcaggcttgatgAAGCTCTCCGCAGTCCTCGC, ggggaccactttgtacaagaaagctgggtcAGATGCAAAGACAGGTGCCG | Mycgr3\|109984  JGI  Ensembl |  |  | 100 % |  |  |
| 35 | ggggacaagtttgtacaaaaaagcaggcttgatgTCGACCATGATCATGTCCAG, ggggaccactttgtacaagaaagctgggtcATCGTAATCCGGGTTCGCAA | Mycgr3\|110399  JGI  Ensembl | KOG2806 Chitinase; |  | 100 % |  |  |
| 35 | ggggacaagtttgtacaaaaaagcaggcttgatgGTCAACATCGACATTCCGCA, ggggaccactttgtacaagaaagctgggtcAGGGCTGCTCTCGCCAATCC | Mycgr3\|110636  JGI  Ensembl | KOG0788 S-adenosylmethionine decarboxylase; | adenosylmethionine decarboxylase activity; catalytic activity;  spermidine biosynthetic process;  spermine biosynthetic process; | 100 % |  |  |
| 35 | ggggacaagtttgtacaaaaaagcaggcttgatgGTCGTGTTCTGGCGACGATA, ggggaccactttgtacaagaaagctgggtcTCCCTCCCGCCCCAGCCTCT | Mycgr3\|110937  JGI  Ensembl |  |  | 100 % |  |  |
| 35 | ggggacaagtttgtacaaaaaagcaggcttgatgTCCAAAGAAAGTTCCAATCA, ggggaccactttgtacaagaaagctgggtcCTGAGGGCTGTTCTCGCGAT | Mycgr3\|111433  JGI  Ensembl | KOG1981 SOK1 kinase belonging to the STE20/SPS1/GC kinase family; |  | 100 % |  |  |
| 35 | ggggacaagtttgtacaaaaaagcaggcttgatgCTCTCCAGTAACATGCAGAG, ggggaccactttgtacaagaaagctgggtcCTTCTTCTCCTCCCTACCTC | Mycgr3\|111557  JGI  Ensembl | KOG0391 SNF2 family DNA-dependent ATPase; | DNA-directed RNA polymerase I activity; DNA-directed RNA polymerase II activity;  DNA-directed RNA polymerase III activity;  DNA-directed RNA polymerase activity;  RNA polymerase activity; | 100 % |  |  |
| 35 | ggggacaagtttgtacaaaaaagcaggcttgatgGATTGCTTCGTTGCCACACG, ggggaccactttgtacaagaaagctgggtcCTGGCCTCTCCTAGGCAACG | Mycgr3\|111679  JGI  Ensembl | KOG3947 Phosphoesterases; | hydrolase activity; | 0.92389 % |  |  |
| 36 | ggggacaagtttgtacaaaaaagcaggcttgatgGTTCGAGGAACGGTGCTCGG, ggggaccactttgtacaagaaagctgggtcCTCTATCGAAAGTGCAGTAT | Mycgr3\|31465  JGI  Ensembl |  |  | 0.81666 % |  |  |
| 36 | ggggacaagtttgtacaaaaaagcaggcttgatgGCTGACGACGAGTTCTCAGC, ggggaccactttgtacaagaaagctgggtcACCACCAAGCACACTCTTCA | Mycgr3\|34911  JGI  Ensembl | KOG0161 Myosin class II heavy chain; |  | 0 % |  |  |
| 36 | ggggacaagtttgtacaaaaaagcaggcttgatgTCCATCCAATCCGTGGCGGT, ggggaccactttgtacaagaaagctgggtcCAACCTCGCCCTCCCCTCTC | Mycgr3\|37086  JGI  Ensembl | KOG1238 Glucose dehydrogenase/choline dehydrogenase/mandelonitrile lyase (GMC oxidoreductase family); | cholesterol oxidase activity; oxidoreductase activity, acting on CH-OH group of donors; | 0.8107 % |  |  |
| 36 | ggggacaagtttgtacaaaaaagcaggcttgatgGCCCTCTCGACAGAATATCC, ggggaccactttgtacaagaaagctgggtcTCCAGCTAGCAGCCCGAACA | Mycgr3\|37351  JGI  Ensembl |  |  | 100 % | chr\_2:3778962 T->C; |  |
| 36 | ggggacaagtttgtacaaaaaagcaggcttgatgGCAGAGTCCACGAAAGCACA, ggggaccactttgtacaagaaagctgggtcAGCCCCCCTAAGAGCCTCCC | Mycgr3\|39347  JGI  Ensembl | KOG2319 Vacuolar assembly/sorting protein VPS9; |  | 100 % |  |  |
| 36 | ggggacaagtttgtacaaaaaagcaggcttgatgCTGACTACAACTCAATGCGC, ggggaccactttgtacaagaaagctgggtcTAGCTCAGTGTTTGAACATA | Mycgr3\|46264  JGI  Ensembl | KOG2089 Metalloendopeptidase family - saccharolysin & thimet oligopeptidase; | metalloendopeptidase activity; metallopeptidase activity;  proteolysis;  saccharolysin activity;  zinc ion binding; | 100 % | chr\_8:2133632 G->A; chr\_8:2132470 C->T; |  |
| 36 | ggggacaagtttgtacaaaaaagcaggcttgatgAAGAGACTCGGGCGTGAGGC, ggggaccactttgtacaagaaagctgggtcAGTGTGCACTCCCCACTTGC | Mycgr3\|49589  JGI  Ensembl | KOG2090 Metalloendopeptidase family - mitochondrial intermediate peptidase; | metalloendopeptidase activity; metallopeptidase activity;  mitochondrial intermediate peptidase activity;  proteolysis;  zinc ion binding; | 100 % |  |  |
| 36 | ggggacaagtttgtacaaaaaagcaggcttgatgGACTCTATTCGCCCATTTCT, ggggaccactttgtacaagaaagctgggtcCCGCTGGACGACTCTATCAC | Mycgr3\|49727  JGI  Ensembl |  |  | 100 % |  |  |
| 36 | ggggacaagtttgtacaaaaaagcaggcttgatgCCAGGCAAGAAAGTCAAGAG, ggggaccactttgtacaagaaagctgggtcGACCACTTTCTTCACCTCTA | Mycgr3\|56708  JGI  Ensembl |  |  | 100 % |  |  |
| 36 | ggggacaagtttgtacaaaaaagcaggcttgatgGCATCGTGGTTCGGGGGCGC, ggggaccactttgtacaagaaagctgggtcCAGCTCGATCAAACTCTCCT | Mycgr3\|57915  JGI  Ensembl | KOG1818 Membrane trafficking and cell signaling protein HRS, contains VHS and FYVE domains; | intracellular protein transport; receptor signaling protein tyrosine kinase activity; | 0 % |  |  |
| 36 | ggggacaagtttgtacaaaaaagcaggcttgatgCTTCAACACAGTGGACGCGG, ggggaccactttgtacaagaaagctgggtcCTCCGCCTCCACGACACCCA | Mycgr3\|66729  JGI  Ensembl | KOG1051 Chaperone HSP104 and related ATP-dependent Clp proteases; | ATP binding; nucleoside-triphosphatase activity;  nucleotide binding; | 0 % |  |  |
| 36 | ggggacaagtttgtacaaaaaagcaggcttgatgGCAGCCACCGCAAACGGCCA, ggggaccactttgtacaagaaagctgggtcCCACTTCTCGCCTTGGCCCG | Mycgr3\|66768  JGI  Ensembl |  |  | 100 % |  |  |
| 36 | ggggacaagtttgtacaaaaaagcaggcttgatgACTAGCATCGCGGGAGCGAA, ggggaccactttgtacaagaaagctgggtcACCAGTGGGACTGCCACAAG | Mycgr3\|66912  JGI  Ensembl | KOG1303 Amino acid transporters; | membrane; transport; transporter activity; | 100 % |  |  |
| 36 | ggggacaagtttgtacaaaaaagcaggcttgatgCAAGCAGAGATCAGGACGCA, ggggaccactttgtacaagaaagctgggtcATGTTGTACCGCAACACCGC | Mycgr3\|67008  JGI  Ensembl | KOG0062 ATPase component of ABC transporters with duplicated ATPase domains/Translation elongation factor EF-3b; | ATP binding; ATPase activity;  nucleoside-triphosphatase activity;  nucleotide binding; | 100 % |  |  |
| 36 | ggggacaagtttgtacaaaaaagcaggcttgatgGCCGTAGCTCTAGAGAACGC, ggggaccactttgtacaagaaagctgggtcCAATCTGGCCTTGGGCGCCG | Mycgr3\|68560  JGI  Ensembl | KOG1256 Long-chain acyl-CoA synthetases (AMP-forming); | catalytic activity; fatty-acid ligase activity;  long-chain-fatty-acid-CoA ligase activity;  medium-chain-fatty-acid-CoA ligase activity;  metabolic process;  short-chain-fatty-acid-CoA ligase activity;  very-long-chain-fatty-acid-CoA ligase activity; | 0.92464 % |  |  |
| 36 | ggggacaagtttgtacaaaaaagcaggcttgatgGACAAGCTGAACGAGGCCTC, ggggaccactttgtacaagaaagctgggtcGAAACCAAGGGACACGTGCG | Mycgr3\|68779  JGI  Ensembl | KOG0470 1,4-alpha-glucan branching enzyme/starch branching enzyme II; | 1,4-alpha-glucan branching enzyme activity; carbohydrate metabolic process;  catalytic activity;  cation binding;  hydrolase activity, hydrolyzing O-glycosyl compounds; | 100 % |  |  |
| 36 | ggggacaagtttgtacaaaaaagcaggcttgatgGAGAACTTGAGTCACCCCGT, ggggaccactttgtacaagaaagctgggtcCAGTAGTCGAAAGTCATACG | Mycgr3\|69322  JGI  Ensembl |  |  | 100 % | chr\_3:451732 T->C; chr\_3:451723 C->T; |  |
| 36 | ggggacaagtttgtacaaaaaagcaggcttgatgGCTGCTCTCGACCCCAAAAG, ggggaccactttgtacaagaaagctgggtcCGAACTCCTCATCCCCATCG | Mycgr3\|69737  JGI  Ensembl | KOG0253 Synaptic vesicle transporter SV2 (major facilitator superfamily); | integral to membrane; transport; transporter activity; | 100 % | chr\_3:1468662 G->A; |  |
| 36 | ggggacaagtttgtacaaaaaagcaggcttgatgTCGGCCGCATTACAAGAGAG, ggggaccactttgtacaagaaagctgggtcCCAAGTCTCAGCAATATACT | Mycgr3\|69850  JGI  Ensembl | KOG1158 NADP/FAD dependent oxidoreductase; | [methionine synthase] reductase activity; electron transport;  oxidoreductase activity; | 100 % |  |  |
| 36 | ggggacaagtttgtacaaaaaagcaggcttgatgGCGAACACTTCTGTCGAGTC, ggggaccactttgtacaagaaagctgggtcCATGACATTGCATCCACCGC | Mycgr3\|70710  JGI  Ensembl | KOG4341 F-box protein containing LRR; | protein binding; | 100 % |  |  |
| 36 | ggggacaagtttgtacaaaaaagcaggcttgatgTCTGCGCTGGCTGCTCCTCG, ggggaccactttgtacaagaaagctgggtcCTCCCCGCCCTTCTTCTTTT | Mycgr3\|71233  JGI  Ensembl | KOG2118 Predicted membrane protein, contains two CBS domains; |  | 100 % | chr\_4:1740319 G->A; |  |
| 36 | ggggacaagtttgtacaaaaaagcaggcttgatgAATTCTTTGGCTGCTGGAAA, ggggaccactttgtacaagaaagctgggtcTTTCCTGACTGGCTCAAGGC | Mycgr3\|71564  JGI  Ensembl | KOG2436 Acetylglutamate kinase/acetylglutamate synthase; | acetylglutamate kinase activity; amino-acid N-acetyltransferase activity;  arginine biosynthetic process; | 100 % |  |  |
| 36 | ggggacaagtttgtacaaaaaagcaggcttgatgCGGTTCTCCTCCTTTGCCGC, ggggaccactttgtacaagaaagctgggtcCGACTGCGCCAAAAACAATT | Mycgr3\|72506  JGI  Ensembl |  |  | 100 % |  |  |
| 36 | ggggacaagtttgtacaaaaaagcaggcttgatgGCTACCGAAGCTGCCACCAC, ggggaccactttgtacaagaaagctgggtcCAACTTGCTGTCAACACAGC | Mycgr3\|73347  JGI  Ensembl | KOG1186 Copper amine oxidase; | amine metabolic process; amine oxidase activity;  copper ion binding;  quinone binding; | 100 % |  |  |
| 36 | ggggacaagtttgtacaaaaaagcaggcttgatgACCACGGCAGGTCTCAATGT, ggggaccactttgtacaagaaagctgggtcTGTCTGCAGAGACAGCGATA | Mycgr3\|73868  JGI  Ensembl | KOG2316 Predicted ATPase (PP-loop superfamily); |  | 1.06545 % |  |  |
| 36 | ggggacaagtttgtacaaaaaagcaggcttgatgCCGCCGGGTTACTCTGACCC, ggggaccactttgtacaagaaagctgggtcCGGACATGCATCCTGGTTTG | Mycgr3\|74077  JGI  Ensembl | KOG2089 Metalloendopeptidase family - saccharolysin & thimet oligopeptidase; | metalloendopeptidase activity; metallopeptidase activity;  proteolysis;  saccharolysin activity;  zinc ion binding; | 100 % | chr\_7:1316093 G->A; |  |
| 36 | ggggacaagtttgtacaaaaaagcaggcttgatgGCCGCCACCATGGACCTTCA, ggggaccactttgtacaagaaagctgggtcTGGCTCCAACCAAGTCTGCA | Mycgr3\|74083  JGI  Ensembl | KOG2142 Molybdenum cofactor sulfurase; | catalytic activity; molybdenum ion binding;  pyridoxal phosphate binding; | 1.59731 % |  |  |
| 36 | ggggacaagtttgtacaaaaaagcaggcttgatgTTCTTCAAACATGAGCTGCT, ggggaccactttgtacaagaaagctgggtcCAGACTATCCATCCCTCCAC | Mycgr3\|74711  JGI  Ensembl |  |  | 100 % |  |  |
| 36 | ggggacaagtttgtacaaaaaagcaggcttgatgCCTAACAGGCTCTCGAAACT, ggggaccactttgtacaagaaagctgggtcACAACATCCACCACCGCCAC | Mycgr3\|75159  JGI  Ensembl | KOG1283 Serine carboxypeptidases; |  | 1.66028 % |  |  |
| 36 | ggggacaagtttgtacaaaaaagcaggcttgatgCTGCACGAGGTCCTCCTAGC, ggggaccactttgtacaagaaagctgggtcCACGTCCAGTAACTTCTCCT | Mycgr3\|75424  JGI  Ensembl | KOG2065 Gamma-tubulin ring complex protein; | microtubule cytoskeleton organization and biogenesis; microtubule organizing center;  spindle pole; | 100 % |  |  |
| 36 | ggggacaagtttgtacaaaaaagcaggcttgatgGACGCCATTAAGAAGGCTTT, ggggaccactttgtacaagaaagctgggtcTGAGGCATTACTCGGCTGCA | Mycgr3\|75925  JGI  Ensembl | KOG0590 Checkpoint kinase and related serine/threonine protein kinases; | (S)-lactate 2-kinase activity; ATP binding;  ATP-dependent polynucleotide kinase activity;  GTP-dependent polynucleotide kinase activity;  LPPG:FO 2-phospho-L-lactate transferase activity;  cobinamide kinase activity;  cyclin-dependent protein kinase activating kinase regulator activity;  cyclin-dependent protein kinase regulator activity;  cytidine kinase activity;  eukaryotic elongation factor-2 kinase activator activity;  eukaryotic elongation factor-2 kinase regulator activity;  glycerate 2-kinase activity;  inositol tetrakisphosphate 2-kinase activity;  phenol kinase activity;  phosphatidylinositol phosphate kinase activity;  phosphofructokinase activity;  phosphoinositide 3-kinase activity;  phosphorylase kinase regulator activity;  polynucleotide kinase activity;  protein amino acid phosphorylation;  protein kinase activity;  protein serine/threonine kinase activity;  protein-tyrosine kinase activity; | 0 % |  |  |
| 36 | ggggacaagtttgtacaaaaaagcaggcttgatgGCCGCCACCGCCAGCCGCTC, ggggaccactttgtacaagaaagctgggtcATTATTCGCCGGACTATCCC | Mycgr3\|77377  JGI  Ensembl | KOG0042 Glycerol-3-phosphate dehydrogenase; | glycerol-3-phosphate dehydrogenase activity; glycerol-3-phosphate dehydrogenase complex;  glycerol-3-phosphate metabolic process;  oxidoreductase activity; | 100 % | chr\_12:594575 G->A; |  |
| 36 | ggggacaagtttgtacaaaaaagcaggcttgatgATGGTGACGGTCAAGTATAT, ggggaccactttgtacaagaaagctgggtcTACATCCTTCACGAAGCCAA | Mycgr3\|77642  JGI  Ensembl |  |  | 0 % |  |  |
| 36 | ggggacaagtttgtacaaaaaagcaggcttgatgAAGTCTCCAACCTTGATTCT, ggggaccactttgtacaagaaagctgggtcCTCTACCACCAAATGCCCCT | Mycgr3\|85505  JGI  Ensembl |  |  | 100 % | chr\_4:196614 G->A; chr\_4:196151 G->A; |  |
| 36 | ggggacaagtttgtacaaaaaagcaggcttgatgGCGCGAAAGGATTTCCACCA, ggggaccactttgtacaagaaagctgggtcCGTGAAGAAGGCCTCCAATC | Mycgr3\|86617  JGI  Ensembl | KOG1133 Helicase of the DEAD superfamily; | 8-oxo-7,8-dihydroguanine triphosphatase activity; ATP binding;  ATP-dependent 5'-3' DNA helicase activity;  ATP-dependent DNA helicase activity;  ATP-dependent helicase activity;  DNA binding;  UDP-2,3-diacylglucosamine hydrolase activity;  bis(5'-nucleosyl)-tetraphosphatase activity;  dATP pyrophosphohydrolase activity;  diacylglycerol pyrophosphate phosphatase activity;  dihydroneopterin monophosphate phosphatase activity;  dihydroneopterin triphosphate pyrophosphohydrolase activity;  hydrolase activity, acting on acid anhydrides, in phosphorus-containing anhydrides; nucleic acid binding;  nucleobase, nucleoside, nucleotide and nucleic acid metabolic process;  nucleotide-excision repair;  nucleus;  pyrophosphatase activity;  thiamin-pyrophosphatase activity; | 0 % |  |  |
| 36 | ggggacaagtttgtacaaaaaagcaggcttgatgCTTGCTGCGAGGAGAGTGGC, ggggaccactttgtacaagaaagctgggtcCGCCTTCTGCGCCCCCTTCT | Mycgr3\|88201  JGI  Ensembl | KOG0453 Aconitase/homoaconitase (aconitase superfamily); | 4 iron, 4 sulfur cluster binding; aconitate hydratase activity;  metabolic process;  tricarboxylic acid cycle; | 100 % |  |  |
| 36 | ggggacaagtttgtacaaaaaagcaggcttgatgCCGAAGTCACTGAGCCGTCG, ggggaccactttgtacaagaaagctgggtcGCAGCACTTGGACATTAGCA | Mycgr3\|88647  JGI  Ensembl |  |  | 100 % | chr\_1:344682 G->T; chr\_1:344655 A->C; |  |
| 36 | ggggacaagtttgtacaaaaaagcaggcttgatgGCCAACGAAGAAGTTCCTCA, ggggaccactttgtacaagaaagctgggtcATCCTCATCGCTGGCCGCGA | Mycgr3\|90406  JGI  Ensembl | KOG0151 Predicted splicing regulator, contains RRM, SWAP and RPR domains; |  | 0 % |  |  |
| 36 | ggggacaagtttgtacaaaaaagcaggcttgatgCGAGTATCAATCATTGCATA, ggggaccactttgtacaagaaagctgggtcCCCGTACCGAGCATCGATCT | Mycgr3\|90989  JGI  Ensembl |  |  | 0 % |  |  |
| 36 | ggggacaagtttgtacaaaaaagcaggcttgatgCCCACCAAGCCGAGATCGAC, ggggaccactttgtacaagaaagctgggtcGACGCTGGCCTCCGATCTCT | Mycgr3\|91019  JGI  Ensembl | KOG1869 Splicing coactivator SRm160/300, subunit SRm300; |  | 0.8589 % |  |  |
| 36 | ggggacaagtttgtacaaaaaagcaggcttgatgTCTGAACCGCTTCTGGCGCA, ggggaccactttgtacaagaaagctgggtcTGGACTGACTTGTGGGTATA | Mycgr3\|91137  JGI  Ensembl |  |  | 100 % |  |  |
| 36 | ggggacaagtttgtacaaaaaagcaggcttgatgGCGACGCCAGAGACGATCCC, ggggaccactttgtacaagaaagctgggtcTCTATACTTGGGGTCCTTCT | Mycgr3\|91287  JGI  Ensembl |  |  | 0 % |  |  |
| 36 | ggggacaagtttgtacaaaaaagcaggcttgatgGCGATATTCAACTTTCGCAA, ggggaccactttgtacaagaaagctgggtcCTTCTTCTTCCCAAACAGCC | Mycgr3\|92070  JGI  Ensembl | KOG1869 Splicing coactivator SRm160/300, subunit SRm300; |  | 100 % | chr\_3:2998109 G->A; chr\_3:2998324 A->G; chr\_3:2998381 G->A; |  |
| 36 | ggggacaagtttgtacaaaaaagcaggcttgatgGCCGCCACAGGAGGTTCCGC, ggggaccactttgtacaagaaagctgggtcCTTCTTGAAACCAGGCGGCG | Mycgr3\|92966  JGI  Ensembl | KOG4674 Uncharacterized conserved coiled-coil protein; |  | 0 % |  |  |
| 36 | ggggacaagtttgtacaaaaaagcaggcttgatgAGGGACGAGGACGAGCTGCG, ggggaccactttgtacaagaaagctgggtcATCGTCCAAGCTTTCCCAAT | Mycgr3\|93554  JGI  Ensembl | KOG2992 Nucleolar GTPase/ATPase p130; |  | 0 % |  |  |
| 36 | ggggacaagtttgtacaaaaaagcaggcttgatgCTCCCACAGTCGCTCTTCTC, ggggaccactttgtacaagaaagctgggtcTTCCTCCATCAGTAACGCCG | Mycgr3\|94952  JGI  Ensembl |  |  | 100 % |  |  |
| 36 | ggggacaagtttgtacaaaaaagcaggcttgatgGCCGACATTAAGACTGCCGC, ggggaccactttgtacaagaaagctgggtcCAGCGCTGCCAAATCGTCTT | Mycgr3\|95253  JGI  Ensembl | KOG1181 FOG: Low-complexity; | electron transport; | 100 % |  |  |
| 36 | ggggacaagtttgtacaaaaaagcaggcttgatgAAGGCCGCTGTACTTCTCCC, ggggaccactttgtacaagaaagctgggtcCAACGCCAAAACAGCAGCCA | Mycgr3\|95442  JGI  Ensembl | KOG4597 Serine proteinase inhibitor (KU family) with thrombospondin repeats; | exo-alpha-sialidase activity; | 0.80354 % |  |  |
| 36 | ggggacaagtttgtacaaaaaagcaggcttgatgCCGTCCCAACCACCCTTCCC, ggggaccactttgtacaagaaagctgggtcCAACTTCCCTTTCGCCTGCA | Mycgr3\|95543  JGI  Ensembl | KOG3730 Acyl-CoA:dihydroxyactetone-phosphate acyltransferase DHAPAT; | acyltransferase activity; glycerol-3-phosphate O-acyltransferase activity;  metabolic process; | 1.58924 % |  |  |
| 36 | ggggacaagtttgtacaaaaaagcaggcttgatgTCTCCACCGTCGCCTCCGCC, ggggaccactttgtacaagaaagctgggtcAATCGCCGTCCCAACCCCAA | Mycgr3\|95576  JGI  Ensembl |  |  | 100 % |  |  |
| 36 | ggggacaagtttgtacaaaaaagcaggcttgatgGACGCCCGCTATTCCAAGAC, ggggaccactttgtacaagaaagctgggtcGCCTTGTCCAGCTTGAGATG | Mycgr3\|96009  JGI  Ensembl |  |  | 100 % |  |  |
| 36 | ggggacaagtttgtacaaaaaagcaggcttgatgGACGGCAACGCACTGTCGCC, ggggaccactttgtacaagaaagctgggtcTGCTACCGGCACGAGTCGTT | Mycgr3\|97076  JGI  Ensembl |  |  | 0 % |  |  |
| 36 | ggggacaagtttgtacaaaaaagcaggcttgatgAGTAGACCGTACGATTATCG, ggggaccactttgtacaagaaagctgggtcCCCCGCCGTCGTCTCCCCAG | Mycgr3\|97342  JGI  Ensembl | KOG1187 Serine/threonine protein kinase; |  | 100 % |  |  |
| 36 | ggggacaagtttgtacaaaaaagcaggcttgatgCCAGCCCGAGCACCACAACA, ggggaccactttgtacaagaaagctgggtcTTTCTGCTTCTTCTTCCGGG | Mycgr3\|97580  JGI  Ensembl |  |  | 100 % |  |  |
| 36 | ggggacaagtttgtacaaaaaagcaggcttgatgACCTGCACAGGTTGTAGCAA, ggggaccactttgtacaagaaagctgggtcCGTCTTCTTCCTGCCAACGG | Mycgr3\|97582  JGI  Ensembl | KOG0207 Cation transport ATPase; | ATP binding; ATPase activity, coupled to transmembrane movement of ions, phosphorylative mechanism; catalytic activity;  copper-exporting ATPase activity;  hydrolase activity;  hydrolase activity, acting on acid anhydrides, catalyzing transmembrane movement of substances; integral to membrane;  membrane;  metabolic process;  metal ion binding;  metal ion transmembrane transporter activity;  metal ion transport;  transport; | 0 % |  |  |
| 36 | ggggacaagtttgtacaaaaaagcaggcttgatgATGAATTCGACCAATTACAG, ggggaccactttgtacaagaaagctgggtcCAGGCACGCGGAGGATTTCT | Mycgr3\|97610  JGI  Ensembl |  |  | 0 % |  |  |
| 36 | ggggacaagtttgtacaaaaaagcaggcttgatgCCACCGTCCACCACATGGGT, ggggaccactttgtacaagaaagctgggtcTTGATCGATTTCGGGGTCGC | Mycgr3\|97943  JGI  Ensembl |  |  | 100 % |  |  |
| 36 | ggggacaagtttgtacaaaaaagcaggcttgatgGCCGAGCAAGCACTGTCGAA, ggggaccactttgtacaagaaagctgggtcTGCTGCCCCTGCTTCGAGTT | Mycgr3\|98612  JGI  Ensembl | KOG0064 Peroxisomal long-chain acyl-CoA transporter, ABC superfamily; | ATP binding; ATPase activity;  membrane;  nucleoside-triphosphatase activity;  nucleotide binding;  transport; | 0 % |  |  |
| 36 | ggggacaagtttgtacaaaaaagcaggcttgatgCCACTCCTCCCCATCCTCTT, ggggaccactttgtacaagaaagctgggtcAATCAACGCACTCAGCACAC | Mycgr3\|99292  JGI  Ensembl | KOG1224 Para-aminobenzoate (PABA) synthase ABZ1; | anthranilate synthase activity; biosynthetic process;  catalytic activity;  glutamine metabolic process;  metabolic process;  oxo-acid-lyase activity; | 100 % | chr\_2:3073359 T->A; chr\_2:3074188 G->A; chr\_2:3072890 T->A; chr\_2:3072947 A->G; chr\_2:3072793 G->T; |  |
| 36 | ggggacaagtttgtacaaaaaagcaggcttgatgGTCCTCGAACGCCTCGCACA, ggggaccactttgtacaagaaagctgggtcCTTGCAGCAAGCGTCATTGC | Mycgr3\|100872  JGI  Ensembl | KOG1186 Copper amine oxidase; | amine metabolic process; amine oxidase activity;  copper ion binding;  protein binding;  quinone binding; | 100 % |  |  |
| 36 | ggggacaagtttgtacaaaaaagcaggcttgatgGCTGGGAAGAGGCTGCTCGA, ggggaccactttgtacaagaaagctgggtcCAGCTTGACAGACTTTGCCC | Mycgr3\|100915  JGI  Ensembl | KOG1234 ABC (ATP binding cassette) 1 protein; |  | 100 % | chr\_8:789451 A->C; chr\_8:790578 T->C; chr\_8:790454 G->A; chr\_8:789512 G->A; chr\_8:788907 G->A; chr\_8:789554 T->C; chr\_8:790480 G->A; chr\_8:788930 T->C; chr\_8:789215 T->C; |  |
| 36 | ggggacaagtttgtacaaaaaagcaggcttgatgCCGTTCTTCGACTTCCCGCA, ggggaccactttgtacaagaaagctgggtcAATCACAGCCCACTCCAACA | Mycgr3\|101408  JGI  Ensembl | KOG2722 Predicted membrane protein; | integral to membrane; | 100 % | chr\_10:649429 C->T; |  |
| 36 | ggggacaagtttgtacaaaaaagcaggcttgatgGGGCCTCCAGTCGGCGATCA, ggggaccactttgtacaagaaagctgggtcGAGAGAATGACCACCACGCT | Mycgr3\|102217  JGI  Ensembl | KOG1187 Serine/threonine protein kinase; | metallopeptidase activity; proteolysis; zinc ion binding; | 100 % |  |  |
| 36 | ggggacaagtttgtacaaaaaagcaggcttgatgATGAAGAACGCCGTCCTCAT, ggggaccactttgtacaagaaagctgggtcAGCGAAGAAAGCCAAGAAGG | Mycgr3\|103427  JGI  Ensembl | KOG1216 von Willebrand factor and related coagulation proteins; | UDP-N-acetylmuramoylalanyl-D-glutamyl-2,6-diaminopimelate-D-alanyl-D-alanine ligase activity; coenzyme F420-0 gamma-glutamyl ligase activity;  coenzyme F420-2 alpha-glutamyl ligase activity;  ribosomal S6-glutamic acid ligase activity; | 100 % | chr\_2:3019228 G->C; chr\_2:3020700 C->T; |  |
| 36 | ggggacaagtttgtacaaaaaagcaggcttgatgTCAGAACGCCCGGAACTCTT, ggggaccactttgtacaagaaagctgggtcATTTCGGACTTTCGGAGCAT | Mycgr3\|106807  JGI  Ensembl |  |  | 0 % |  |  |
| 36 | ggggacaagtttgtacaaaaaagcaggcttgatgGACAGTCGATCACAACGACT, ggggaccactttgtacaagaaagctgggtcAATCCAGCCACCCTCCACCG | Mycgr3\|107151  JGI  Ensembl | KOG1124 FOG: TPR repeat; |  | 100 % |  |  |
| 36 | ggggacaagtttgtacaaaaaagcaggcttgatgGAACACTCCGGAGACGGATT, ggggaccactttgtacaagaaagctgggtcATTGGGAGTTGGAGGTTTGC | Mycgr3\|107327  JGI  Ensembl |  |  | 0 % |  |  |
| 36 | ggggacaagtttgtacaaaaaagcaggcttgatgTCTTCGCCCACCGCGTCGAC, ggggaccactttgtacaagaaagctgggtcGAACGCCCAAGACCCTCTCC | Mycgr3\|107616  JGI  Ensembl | KOG2992 Nucleolar GTPase/ATPase p130; |  | 100 % |  |  |
| 36 | ggggacaagtttgtacaaaaaagcaggcttgatgCCTCCCAAGAATATCACAGC, ggggaccactttgtacaagaaagctgggtcCCAGGTCTCGTCCCACCAGT | Mycgr3\|107703  JGI  Ensembl |  |  | 100 % | chr\_1:5390600 G->A; |  |
| 36 | ggggacaagtttgtacaaaaaagcaggcttgatgTCTGAAGCACACAACACCGC, ggggaccactttgtacaagaaagctgggtcCGCGCTCGCGGTGACCGGAG | Mycgr3\|107781  JGI  Ensembl | KOG4475 FOG: Immunoglobin and related proteins; |  | 0 % |  |  |
| 36 | ggggacaagtttgtacaaaaaagcaggcttgatgGCTCCTCCACAGGACCCTCC, ggggaccactttgtacaagaaagctgggtcGGCAGAGAAGGCCTGCTGGG | Mycgr3\|107949  JGI  Ensembl | KOG1878 Nuclear receptor coregulator SMRT/SMRTER, contains Myb-like domains; | protein modification process; | 0 % |  |  |
| 36 | ggggacaagtttgtacaaaaaagcaggcttgatgGCGACCTTCTTGACTCAATC, ggggaccactttgtacaagaaagctgggtcTCCAACGGTCGTCGGATACT | Mycgr3\|108025  JGI  Ensembl |  |  | 100 % | chr\_2:1572039 G->A; |  |
| 36 | ggggacaagtttgtacaaaaaagcaggcttgatgGCTGCTGTTCCAAAGGACTT, ggggaccactttgtacaagaaagctgggtcCAAGATCGTCGCCCCAAACA | Mycgr3\|108245  JGI  Ensembl | KOG4757 Predicted telomere binding protein; | DNA binding; nuclear chromosome, telomeric region;  telomere maintenance; | 0 % |  |  |
| 36 | ggggacaagtttgtacaaaaaagcaggcttgatgAACGGACGGCTGGTGAATGT, ggggaccactttgtacaagaaagctgggtcCTCGCCATTCCACGATGGGA | Mycgr3\|108566  JGI  Ensembl | KOG0676 Actin and related proteins; |  | 2.49261 % |  |  |
| 36 | ggggacaagtttgtacaaaaaagcaggcttgatgGCAGAGGCCGTTGCTTTAAC, ggggaccactttgtacaagaaagctgggtcGTCGCTACCATCAATGGCCA | Mycgr3\|108872  JGI  Ensembl | KOG4177 Ankyrin; |  | 0 % |  |  |
| 36 | ggggacaagtttgtacaaaaaagcaggcttgatgCCGGGTTCCAAGCCATTCAC, ggggaccactttgtacaagaaagctgggtcCTCTTCTTTGCTGCTCAATG | Mycgr3\|109288  JGI  Ensembl | KOG0613 Projectin/twitchin and related proteins; |  | 100 % |  |  |
| 36 | ggggacaagtttgtacaaaaaagcaggcttgatgCATCTCACCAACCCGTTCGC, ggggaccactttgtacaagaaagctgggtcAACAGCAATCGTGGGAAGAG | Mycgr3\|109435  JGI  Ensembl | KOG1216 von Willebrand factor and related coagulation proteins; |  | 100 % |  |  |
| 36 | ggggacaagtttgtacaaaaaagcaggcttgatgGACTCCAGCCGAAAACGCAG, ggggaccactttgtacaagaaagctgggtcTCTCGTCAAGTTGGCAAAAA | Mycgr3\|109619  JGI  Ensembl | KOG1827 Chromatin remodeling complex RSC, subunit RSC1/Polybromo and related proteins; |  | 100 % |  |  |
| 36 | ggggacaagtttgtacaaaaaagcaggcttgatgCATATCACCAAGAAGCTGGC, ggggaccactttgtacaagaaagctgggtcCGCCTTCTTGTCCGCGCCGT | Mycgr3\|109648  JGI  Ensembl | KOG0564 5,10-methylenetetrahydrofolate reductase; | methionine metabolic process; methylenetetrahydrofolate reductase (NADPH) activity; | 0 % |  |  |
| 36 | ggggacaagtttgtacaaaaaagcaggcttgatgCGGTTACATTCCATCTTTAA, ggggaccactttgtacaagaaagctgggtcGACTCCGGAATCACTCTGCT | Mycgr3\|109666  JGI  Ensembl | KOG1263 Multicopper oxidases; | carbohydrate metabolic process; catalytic activity;  laccase activity;  oxidoreductase activity; | 100 % | chr\_5:2495001 C->T; chr\_5:2493451 G->A; |  |
| 36 | ggggacaagtttgtacaaaaaagcaggcttgatgGCAGTCAACAGACTGCGCAG, ggggaccactttgtacaagaaagctgggtcAAACAAGTCCAGCAGGTCTG | Mycgr3\|109857  JGI  Ensembl | KOG1061 Vesicle coat complex AP-1/AP-2/AP-4, beta subunit; | intracellular protein transport; membrane coat;  protein binding;  protein complex assembly;  vesicle-mediated transport; | 0.84071 % |  |  |
| 36 | ggggacaagtttgtacaaaaaagcaggcttgatgAAGCTCTTCAACCTCCTCGT, ggggaccactttgtacaagaaagctgggtcTCCAGAGCCAGTCCCGCGAA | Mycgr3\|110156  JGI  Ensembl | KOG1474 Transcription initiation factor TFIID, subunit BDF1 and related bromodomain proteins; |  | 100 % |  |  |
| 36 | ggggacaagtttgtacaaaaaagcaggcttgatgGCACGATATCCGACCTTTGA, ggggaccactttgtacaagaaagctgggtcCGGCAACCGGCCAACTTTCT | Mycgr3\|110212  JGI  Ensembl | KOG2053 Mitochondrial inheritance and actin cytoskeleton organization protein; |  | 100 % |  |  |
| 36 | ggggacaagtttgtacaaaaaagcaggcttgatgGGCTCGTTCCTCTCCGCACC, ggggaccactttgtacaagaaagctgggtcGGTAGCGATGGGCACAGCGT | Mycgr3\|110409  JGI  Ensembl | KOG0161 Myosin class II heavy chain; |  | 100 % |  |  |
| 36 | ggggacaagtttgtacaaaaaagcaggcttgatgCTTGCCATGCACCGGGCTCT, ggggaccactttgtacaagaaagctgggtcATTGGCTGATCTATGCGAGG | Mycgr3\|110654  JGI  Ensembl | KOG2495 NADH-dehydrogenase (ubiquinone); | FAD binding; NADH dehydrogenase activity;  calcium ion binding;  oxidoreductase activity; | 100 % |  |  |
| 36 | ggggacaagtttgtacaaaaaagcaggcttgatgGGATCAACGAACTCGGATTC, ggggaccactttgtacaagaaagctgggtcTAGGTCTGTCCCTCTAGCCT | Mycgr3\|111077  JGI  Ensembl | KOG2614 Kynurenine 3-monooxygenase and related flavoprotein monooxygenases; | salicylate 1-monooxygenase activity; | 100 % |  |  |
| 36 | ggggacaagtttgtacaaaaaagcaggcttgatgCTCAACCGAGCACTAACACG, ggggaccactttgtacaagaaagctgggtcGAGTCTCTCAACGGCACCCG | Mycgr3\|111104  JGI  Ensembl | KOG0787 Dehydrogenase kinase; | ATP binding; [3-methyl-2-oxobutanoate dehydrogenase (acetyl-transferring)] kinase activity; | 100 % |  |  |
| 36 | ggggacaagtttgtacaaaaaagcaggcttgatgTCAACCCTCCCATCCGAAGG, ggggaccactttgtacaagaaagctgggtcATCCCAATTGACCTCCTTCG | Mycgr3\|111178  JGI  Ensembl | KOG1060 Vesicle coat complex AP-3, beta subunit; | 1,2-dihydroxy-phenanthrene glycosyltransferase activity; 1-phenanthrol glycosyltransferase activity;  9-phenanthrol UDP-glucuronosyltransferase activity;  9-phenanthrol glycosyltransferase activity;  O antigen polymerase activity;  UDP-N-acetylglucosamine-peptide N-acetylglucosaminyltransferase activity;  UDP-glucose:glycoprotein glucosyltransferase activity;  acetylgalactosaminyltransferase activity;  acetylglucosaminyltransferase activity;  alpha(1,3)-fucosyltransferase activity;  alpha(1,6)-fucosyltransferase activity;  alpha-1,2-galactosyltransferase activity;  alpha-1,2-mannosyltransferase activity;  alpha-1,3-galactosyltransferase activity;  alpha-1,3-mannosyltransferase activity;  alpha-1,6-mannosyltransferase activity;  beta-1,4-mannosyltransferase activity;  cellulose synthase activity;  dolichyl pyrophosphate Glc1Man9GlcNAc2 alpha-1,3-glucosyltransferase activity; dolichyl pyrophosphate Man7GlcNAc2 alpha-1,3-glucosyltransferase activity;  dolichyl pyrophosphate Man9GlcNAc2 alpha-1,3-glucosyltransferase activity;  dolichyl-phosphate-glucose-glycolipid alpha-glucosyltransferase activity;  fucosyltransferase activity;  galactosyltransferase activity;  glucosyltransferase activity;  glycolipid mannosyltransferase activity;  inositol phosphoceramide synthase activity;  lipopolysaccharide-1,6-galactosyltransferase activity;  mannosyltransferase activity;  oligosaccharyl transferase activity;  phenanthrol glycosyltransferase activity; | 100 % | chr\_11:261930 C->A; |  |
| 36 | ggggacaagtttgtacaaaaaagcaggcttgatgCGTGGCAGTCAACTTGCCTC, ggggaccactttgtacaagaaagctgggtcCACCACACCGTCGCGTCGTC | Mycgr3\|111417  JGI  Ensembl | KOG2737 Putative metallopeptidase; | metalloendopeptidase activity; metallopeptidase activity;  proteolysis;  zinc ion binding; | 100 % |  |  |
| 36 | ggggacaagtttgtacaaaaaagcaggcttgatgGACAAGATGCGCCCTTTATC, ggggaccactttgtacaagaaagctgggtcCCTCCTCGCCGCGATATTCG | Mycgr3\|111479  JGI  Ensembl | KOG4701 Chitinase; |  | 100 % |  |  |
| 36 | ggggacaagtttgtacaaaaaagcaggcttgatgCCGATCCACGTAGTTTTGGC, ggggaccactttgtacaagaaagctgggtcGTAGGATCGTGCTCGTGCTT | Mycgr3\|111600  JGI  Ensembl | KOG1215 Low-density lipoprotein receptors containing Ca2+-binding EGF-like domains; |  | 1.75064 % |  |  |
| 36 | ggggacaagtttgtacaaaaaagcaggcttgatgCCAACCGACACCTCAACCCC, ggggaccactttgtacaagaaagctgggtcCTTCCTCCTCCACCGTTTCG | Mycgr3\|111736  JGI  Ensembl | KOG0161 Myosin class II heavy chain; |  | 0 % |  |  |
| 36 | ggggacaagtttgtacaaaaaagcaggcttgatgGCAAGACTAAACACGAGGAA, ggggaccactttgtacaagaaagctgggtcACCAGAGACAATGTCCCGCA | Mycgr3\|111796  JGI  Ensembl | KOG1181 FOG: Low-complexity; |  | 100 % |  |  |
| 37 | ggggacaagtttgtacaaaaaagcaggcttgatgGCCCCGGACCCGTTCGACCA, ggggaccactttgtacaagaaagctgggtcAATCTTCCCGATCCACCCGA | Mycgr3\|32527  JGI  Ensembl |  |  | 100 % | chr\_1:5274739 G->A; |  |
| 37 | ggggacaagtttgtacaaaaaagcaggcttgatgTACCACACCTTTCAAGCTGC, ggggaccactttgtacaagaaagctgggtcCGTGCCATCTTGTGGCCACT | Mycgr3\|36903  JGI  Ensembl |  |  | 100 % |  | ``` 1	ATGTACCACACCTTTCAAGCTGCGCCGTCCTTTCCTGCGAGTGGAGACGGTAGCAAAGGACCCCGACCACCTGGTAGCTCTTCATCGAGGCGAATCTCCACTACGAATGCATGCATCGAG	120 	\|\|\|\|\|\|\|\|\|\|\|\|\|\|\|\|\|\|\|\|\|\|\|\|\|\|\|\|\|\|\|\|\|\|\|\|\|\|\|\|\|\|\|\|\|\|\|\|\|\|\|\|\|\|\|\|\|\|\|\|\|\|\|\|\|\|\|\|\|\|\|\|\|\|\|\|\|\|\|\|\|\|\|\|\|\|\|\|\|\|\|\|\|\|\|\|\|\|\|\|\|\|\|\|\|\|\|\|\|\|\|\|\|\|\|\|\|\|\|\| 	ATGTACCACACCTTTCAAGCTGCGCCGTCCTTTCCTGCGAGTGGAGACGGTAGCAAAGGACCCCGACCACCTGGTAGCTCTTCATCGAGGCGAATCTCCACTACGAATGCATGCATCGAG ```      ``` 240	ACTGGTCGAGAGACTTCAATCCCAAGTCGAACAATACCACTCGCTCTTCGCTCGACTGTTCCCCGGCAAGGAACTGGAGAACATCCTTACGATGTCGAGAGAAGGCCTCGTCAACCTAGCTGTGACCCTGCCAGCGGCCAAACTCTCTCCACCTTCTGCAGAAGCCCGGGACCGGCATCAAGTGAAGTCAGAGACAGTCGAATCAGAGACAGCCGACAGTCTTGAAGCTCTCGAGCAGGCCCCCGAGCAGGACTCCACATCTGGCGAATCGAGAAAGCTCCACGCGAAGATACAGGCCATCTCTGACGATGTCAATGGACTGTCGCTGTCAATCGACAAAGCTTCATCATACACGGGCGTTTCCTCCATCAACGCGGCGATGAAGGTTATCCTCAAGACAGCGCCTTTCGCTTGGGCTTTCATTTCGCAGCCTTCCGCACAAACATCCAACCCAAGTCGCGCTACTACTCCACCACCACATGTGCGAGATCCGGACCCATATTACTTGCCTTCCGCGGACGTCGGCGAAAAGCTCATTGAATCATATTTCAAATATGTCCACGTACAGATGCCAATGGTGGACGAGGAACAGTTTCGACATGGCTATCTGTACGGAACACGTAGAGATCCGGCCTGGCTGTCACTGCTGAACATGGTGTTTGCTC-TCGGGTCCTTAGCTAGTTCGACATGCGACAATGAGGAACACA--TTGC-TTTC-TTCCAGCGAG-CGAGGAAA-C-ATCTC	978 	\|\|\|\|\|\|\|\|\|\|\|\|\|\|\|\|\|\|\|\|\|\|\|\|\|\|\|\|\|\|\|\|\|\|\|\|\|\|\|\|\|\|\|\|\|\|\|\|\|\|\|\|\|\|\|\|\|\|\|\|\|\|\|\|\|\|\|\|\|\|\|\|\|\|\|\|\|\|\|\|\|\|\|\|\|\|\|\|\|\|\|\|\|\|\|\|\|\|\|\|\|\|\|\|\|\|\|\|\|\|\|\|\|\|\|\|\|\|\|\|\|\|\|\|\|\|\|\|\|\|\|\|\|\|\|\|\|\|\|\|\|\|\|\|\|\|\|\|\|\|\|\|\|\|\|\|\|\|\|\|\|\|\|\|\|\|\|\|\|\|\|\|\|\|\|\|\|\|\|\|\|\|\|\|\|\|\|\|\|\|\|\|\|\|\|\|\|\|\|\|\|\|\|\|\|\|\|\|\|\|\|\|\|\|\|\|\|\|\|\|\|\|\|\|\|\|\|\|\|\|\|\|\|\|\|\|\|\|\|\|\|\|\|\|\|\|\|\|\|\|\|\|\|\|\|\|\|\|\|\|\|\|\|\|\|\|\|\|\|\|\|\|\|\|\|\|\|\|\|\|\|\|\|\|\|\|\|\|\|\|\|\|\|\|\|\|\|\|\|\|\|\|\|\|\|\|\|\|\|\|\|\|\|\|\|\|\|\|\|\|\|\|\|\|\|\|\|\|\|\|\|\|\|\|\|\|\|\|\|\|\|\|\|\|\|\|\|\|\|\|\|\|\|\|\|\|\|\|\|\|\|\|\|\|\|\|\|\|\|\|\|\|\|\|\|\|\|\|\|\|\|\|\|\|\|\|\|\|\|\|\|\|\|\|\|\|\|\|\|\|\|\|\|\|\|\|\|\|\|\|\|\|\|\|\|\|\|\|\|\|\|\|\|\|\|\|\|\|\|\|\|\|\|\|\|\|\|\|\|\|\|\|\|\|\|\|\|\|\|\|\|\|\|\|\|\|\|\|\|\|\|\|\|\|\|\|\|\|\|\|\|\|\|\|\|\|\|\|\|\|\|\|\|\|\|\|\|\|\|\|\|\|\|\|\|\|\|\|\|\|\|\|\|\|\|\|\|\|\|\|\|\|\|\|\|\|\|\|\|\|\|\|\|\|\|\|\|\|\|\|\|\|\|\|\|\|\|\|\|\|\|\|\|\|\|\|\|\|\|\|\|\|\|\|\|\|\|\|\|\|\|\|\|\|\|\|\|\|\|\|\|\|\|\|\|\|\|\|\|\|\|\|\|\|\|\|\| \|\|\|\|\|\|\|\|\|\|\|\|\|\|\|\|\|\|\|\|\|\|\|\|\|\|\|\|\|\|\|\|\|\|\|\|\|\|\|\|\|\|\|\|\|\|\| \|\|\|\|\|\|\|\|\|\|\|\|\|\| \|\|\|\|\|\|\|\|\| \|\|\|\| \|\|\|\|\|\|\|\|\| \|\|\|\|\|\| \|\|\|\|\|\|\|\|\|\|\|\|\|  \|\|\|\|\|\|\|\|\|\|  \|\|\|\| \|\|\|\| \|\|\|\|\|\|\|\|\|\| \|\|\|\|\|\|\|\| \| \|\|\|\|\| 	ACTGGTCGAGAGACTTCAATCCCAAGTCGAACAATACCACTCGCTCTTCGCTCGACTGTTCCCCGGCAAGGAACTGGAGAACATCCTTACGATGTCGAGAGAAGGCCTCGTCAACCTAGCTGTGACCCTGCCAGCGGCCAAACTCTCTCCACCTTCTGCAGAAGCCCGGGACCGGCATCAAGTGAAGTCAGAGACAGTCGAATCAGAGACAGCCGACAGTCTTGAAGCTCTCGAGCAGGCCCCCGAGCAGGACTCCACATCTGGCGAATCGAGAAAGCTCCACGCGAAGATACAGGCCATCTCTGACGATGTCAATGGACTGTCGCTGTCAATCGACAAAGCTTCATCATACACGGGCGTTTCCTCCATCAACGCGGCGATGAAGGTTATCCTCAAGACAGCGCCTTTCGCTTGGGCTTTCATTTCGCAGCCTTCCGCACAAACATCCAACCCAAGTCGCGCTACTACTCCACCACCACATGTGCGAGATCCGGACCCATATTACTTGCCTTCCGCGGACGTCGGCGAAAAGCTCATTGAATCATATTTCAAATATGTCCACGTACAGATGCCAATGGTGGACGAGG-ACAGTTTCGACATGGCTATCTGTACGGAACACGTAGAGATCCGGCCT-GCTGTCACTGCTGA-CATGGTGTT-GCTCCTCGGGTCCT-AGCTAG-TCGACATGCGACATGGAGGAACACACATTGCCTTTCCTTCCAGCGAGGCGAGGAAAACCATCTC ```      ``` 119	AGTGTCGACGACGTAAGATCCGCTGTGATGGCAATCAGCCTTGCGGCCAGTGTAATTGGTATCAGCATCCGGAGCTATGCGCGTACTCGAAGCCAGCACAAAGAGTCATCCCGTCGCGAAA	239 	\|\|\|\|\|\|\|\|\|\|\|\|\|\|\|\|\|\|\|\|\|\|\|\|\|\|\|\|\|\|\|\|\|\|\|\|\|\|\|\|\|\|\|\|\|\|\|\|\|\|\|\|\|\|\|\|\|\|\|\|\|\|\|\|\|\|\|\|\|\|\|\|\|\|\|\|\|\|\|\|\|\|\|\|\|\|\|\|\|\|\|\|\|\|\|\|\|\|\|\|\|\|\|\|\|\|\|\|\|\|\|\|\|\|\|\|\|\|\|\|\| 	AGTGTCGACGACGTAAGATCCGCTGTGATGGCAATCAGCCTTGCGGCCAGTGTAATTGGTATCAGCATCCGGAGCTATGCGCGTACTCGAAGCCAGCACAAAGAGTCATCCCGTCGCGAAA ``` |
| 37 | ggggacaagtttgtacaaaaaagcaggcttgatgGCGTCCTTTGTCTCTTCTCT, ggggaccactttgtacaagaaagctgggtcTGCCACACAGCTCTTAACCT | Mycgr3\|39597  JGI  Ensembl |  |  | 100 % |  |  |
| 37 | ggggacaagtttgtacaaaaaagcaggcttgatgCGTGCCGCTCCGTGGTTGGC, ggggaccactttgtacaagaaagctgggtcCAACTCAAATCCCCCCGTCA | Mycgr3\|42323  JGI  Ensembl |  |  | 100 % |  |  |
| 37 | ggggacaagtttgtacaaaaaagcaggcttgatgGCTTTCCAGCCGTCTGGCGG, ggggaccactttgtacaagaaagctgggtcAATGTTCTTAAAGAAGATTT | Mycgr3\|46105  JGI  Ensembl | KOG0642 Cell-cycle nuclear protein, contains WD-40 repeats; |  | 0 % |  |  |
| 37 | ggggacaagtttgtacaaaaaagcaggcttgatgAACCGCCGCACGAGCAGCTA, ggggaccactttgtacaagaaagctgggtcCGAAATCGCCTCCCTCCGAT | Mycgr3\|47177  JGI  Ensembl | KOG0475 Cl- channel CLC-3 and related proteins (CLC superfamily); | chloride transport; membrane; voltage-gated chloride channel activity; | 99.92203 % |  |  |
| 37 | ggggacaagtttgtacaaaaaagcaggcttgatgGAGCGTCACATCGACCGTGG, ggggaccactttgtacaagaaagctgggtcCTCATTTTTTCGCCATTGTA | Mycgr3\|48531  JGI  Ensembl | KOG2000 Gamma-tubulin complex, DGRIP91/SPC98 component; |  | 100 % | chr\_10:571191 G->A; |  |
| 37 | ggggacaagtttgtacaaaaaagcaggcttgatgGGCAACCATTGGGGTATCAG, ggggaccactttgtacaagaaagctgggtcCAATTCATCGTGATGGCCAC | Mycgr3\|48810  JGI  Ensembl | KOG2161 Glucosidase I; | endoplasmic reticulum; mannosyl-oligosaccharide glucosidase activity;  oligosaccharide metabolic process; | 0 % |  |  |
| 37 | ggggacaagtttgtacaaaaaagcaggcttgatgGGTAACCAGCCCTCCAAAGA, ggggaccactttgtacaagaaagctgggtcGTAACTCTGAGGGAACTGGC | Mycgr3\|52472  JGI  Ensembl | KOG0374 Serine/threonine specific protein phosphatase PP1, catalytic subunit; | CTD phosphatase activity; calcium-dependent protein serine/threonine phosphatase activity;  calcium-dependent protein serine/threonine phosphatase regulator activity;  hydrogen ion transmembrane transporter activity;  hydrolase activity;  magnesium-dependent protein serine/threonine phosphatase activity;  magnesium-dependent protein serine/threonine phosphatase complex;  myosin phosphatase activity;  myosin phosphatase complex;  myosin phosphatase regulator activity;  phosphoprotein phosphatase activity;  proton-transporting two-sector ATPase complex; | 0 % |  |  |
| 37 | ggggacaagtttgtacaaaaaagcaggcttgatgTTCGCCTTGAAATCGCTCTC, ggggaccactttgtacaagaaagctgggtcGTAAGGCGAGTTGACGATCT | Mycgr3\|59611  JGI  Ensembl | KOG2395 Protein involved in vacuole import and degradation; |  | 100 % |  |  |
| 37 | ggggacaagtttgtacaaaaaagcaggcttgatgGCGCGCTCGCCTTTCCCTGT, ggggaccactttgtacaagaaagctgggtcTGTGATGGTGAACTCCCCAC | Mycgr3\|65151  JGI  Ensembl | KOG2232 Ceramidases; |  | 100 % | chr\_1:1007387 A->G; |  |
| 37 | ggggacaagtttgtacaaaaaagcaggcttgatgAGCAATCAGAGCTCATCTCC, ggggaccactttgtacaagaaagctgggtcCATCTTCGACTCCTTCGGTT | Mycgr3\|66126  JGI  Ensembl | KOG1175 Acyl-CoA synthetase; | acetoacetate-CoA ligase activity; catalytic activity;  metabolic process; | 100 % | chr\_1:2857545 A->G; |  |
| 37 | ggggacaagtttgtacaaaaaagcaggcttgatgGCCACCACCGAAAAACACCA, ggggaccactttgtacaagaaagctgggtcCTGCCTTCCCCTCAGATCTC | Mycgr3\|66868  JGI  Ensembl | KOG0255 Synaptic vesicle transporter SVOP and related transporters (major facilitator superfamily); | integral to membrane; transport; transporter activity; | 100 % |  |  |
| 37 | ggggacaagtttgtacaaaaaagcaggcttgatgCCTCCATTCTCCAGGTCCAG, ggggaccactttgtacaagaaagctgggtcTCCCACAAACCCACCGCCCT | Mycgr3\|66965  JGI  Ensembl |  |  | 100 % |  |  |
| 37 | ggggacaagtttgtacaaaaaagcaggcttgatgCGTCTCCCTTTCCATCTCCT, ggggaccactttgtacaagaaagctgggtcAGCATACCGCGTACTCCCAT | Mycgr3\|67183  JGI  Ensembl | KOG3607 Meltrins, fertilins and related Zn-dependent metalloproteinases of the ADAMs family; | collagenase activity; matrilysin-2 activity;  metalloendopeptidase activity;  prenyl protein specific endopeptidase activity;  proteolysis;  stromelysin 3 activity; | 100 % | chr\_1:4837152 C->G; |  |
| 37 | ggggacaagtttgtacaaaaaagcaggcttgatgAAGGGTTGTCTCAATCATCT, ggggaccactttgtacaagaaagctgggtcCTGGACATCGTTCTGAGGAC | Mycgr3\|67250  JGI  Ensembl |  |  | 100 % |  |  |
| 37 | ggggacaagtttgtacaaaaaagcaggcttgatgGAGGAGGGCGACAGCGCTGC, ggggaccactttgtacaagaaagctgggtcACCACTGCGCTGAGACATCA | Mycgr3\|67446  JGI  Ensembl | KOG2043 Signaling protein SWIFT and related BRCT domain proteins; | intracellular; | 100 % |  |  |
| 37 | ggggacaagtttgtacaaaaaagcaggcttgatgGCCGATCGTCGTGGCGCTCT, ggggaccactttgtacaagaaagctgggtcAGTCGGCGGCAACCCCTCTC | Mycgr3\|67987  JGI  Ensembl | KOG0828 Predicted E3 ubiquitin ligase; | protein binding; zinc ion binding; | 0.85371 % |  |  |
| 37 | ggggacaagtttgtacaaaaaagcaggcttgatgAAAGACTCGGGAGATGAGGA, ggggaccactttgtacaagaaagctgggtcAAACCAATAACTCTCACCAC | Mycgr3\|69002  JGI  Ensembl |  |  | 100 % |  |  |
| 37 | ggggacaagtttgtacaaaaaagcaggcttgatgGTGGGCTGGAAACGCAACAA, ggggaccactttgtacaagaaagctgggtcGAAGATATATCCGAGCGAGT | Mycgr3\|69847  JGI  Ensembl | KOG1286 Amino acid transporters; | amino acid transmembrane transporter activity; amino acid transport;  integral to membrane;  membrane;  transport; | 100 % |  |  |
| 37 | ggggacaagtttgtacaaaaaagcaggcttgatgGTATTGCGGACATTGGGGTG, ggggaccactttgtacaagaaagctgggtcGTCTCCACCATGCAGTATCG | Mycgr3\|70294  JGI  Ensembl | KOG1238 Glucose dehydrogenase/choline dehydrogenase/mandelonitrile lyase (GMC oxidoreductase family); | FAD binding; choline dehydrogenase activity;  oxidoreductase activity, acting on CH-OH group of donors; | 100 % |  |  |
| 37 | ggggacaagtttgtacaaaaaagcaggcttgatgGCCTTGCGTGGGCCGAACAT, ggggaccactttgtacaagaaagctgggtcCCTGCCCTTCGCCTTGACCT | Mycgr3\|70532  JGI  Ensembl | KOG0057 Mitochondrial Fe/S cluster exporter, ABC superfamily; | ATP binding; ATP-dependent DNA helicase activity;  ATP-dependent RNA helicase activity;  ATP-dependent helicase activity;  ATPase activity;  ATPase activity, coupled;  ATPase activity, coupled to transmembrane movement of ions;  ATPase activity, coupled to transmembrane movement of substances;  ATPase activity, uncoupled;  DNA-dependent ATPase activity;  RNA-dependent ATPase activity;  integral to membrane;  nucleoside-triphosphatase activity;  nucleotide binding;  protein-transmembrane transporting ATPase activity;  single-stranded DNA-dependent ATP-dependent DNA helicase activity;  transport; | 100 % | chr\_3:3340752 C->T; |  |
| 37 | ggggacaagtttgtacaaaaaagcaggcttgatgACTGCGCTTCTGCTTACATC, ggggaccactttgtacaagaaagctgggtcCAAAGGCATCTCATATTCAC | Mycgr3\|71284  JGI  Ensembl |  |  | 100 % |  |  |
| 37 | ggggacaagtttgtacaaaaaagcaggcttgatgTACGCAGTCATAGCAGCCCT, ggggaccactttgtacaagaaagctgggtcCTCTTCGTAATCGAAAGCAG | Mycgr3\|71401  JGI  Ensembl |  |  | 100 % |  |  |
| 37 | ggggacaagtttgtacaaaaaagcaggcttgatgCCACAATCCGAATTCGTCCC, ggggaccactttgtacaagaaagctgggtcGGCGCAAGAGATCTTGAATC | Mycgr3\|71695  JGI  Ensembl | KOG1066 Glucosidase II catalytic (alpha) subunit and related enzymes, glycosyl hydrolase family 31; | carbohydrate metabolic process; glucan 1,3-alpha-glucosidase activity;  hydrolase activity, acting on acid anhydrides, catalyzing transmembrane movement of substances; hydrolase activity, hydrolyzing O-glycosyl compounds;  membrane;  proton transport;  proton-transporting two-sector ATPase complex; | 100 % |  |  |
| 37 | ggggacaagtttgtacaaaaaagcaggcttgatgTCCTCCTGCTTCGGATCCCG, ggggaccactttgtacaagaaagctgggtcAGTCATACTCCGAACATGCG | Mycgr3\|72810  JGI  Ensembl |  |  | 100 % |  |  |
| 37 | ggggacaagtttgtacaaaaaagcaggcttgatgCCGGCCCATCTCACCCGCCA, ggggaccactttgtacaagaaagctgggtcAGGACTCACGTCTCTCTGCA | Mycgr3\|72989  JGI  Ensembl | KOG4197 FOG: PPR repeat; |  | 100 % | chr\_6:1013776 C->T; chr\_6:1013444 T->C; |  |
| 37 | ggggacaagtttgtacaaaaaagcaggcttgatgGCGGCACTTTCATGGCTGTG, ggggaccactttgtacaagaaagctgggtcCAACTCTGCCCTCATCTTGC | Mycgr3\|73377  JGI  Ensembl | KOG3009 Predicted carbohydrate kinase, contains PfkB domain; |  | 100 % | chr\_6:1903027 G->A; |  |
| 37 | ggggacaagtttgtacaaaaaagcaggcttgatgGGTGAACCACAGCACCGCGA, ggggaccactttgtacaagaaagctgggtcCTGAATCGTACTCGCACTCG | Mycgr3\|74660  JGI  Ensembl | KOG3742 Glycogen synthase; | glycogen (starch) synthase activity; glycogen biosynthetic process; | 100 % | chr\_8:1001157 G->A; |  |
| 37 | ggggacaagtttgtacaaaaaagcaggcttgatgCAGGCCGGCAACATTCTTCA, ggggaccactttgtacaagaaagctgggtcCCTACCTCCCGGCGGCGCCC | Mycgr3\|77508  JGI  Ensembl | KOG2220 Predicted signal transduction protein; | prenylated protein tyrosine phosphatase activity; protein tyrosine phosphatase activity; | 0 % |  |  |
| 37 | ggggacaagtttgtacaaaaaagcaggcttgatgGCTTCGACATCTAGCAAGAT, ggggaccactttgtacaagaaagctgggtcCAGCGCCTTCAAGCCATCTC | Mycgr3\|86218  JGI  Ensembl | KOG0236 Sulfate/bicarbonate/oxalate exchanger SAT-1 and related transporters (SLC26 family); | integral to membrane; membrane; secondary active sulfate transmembrane transporter activity;  sulfate transport;  transport;  transporter activity; | 100 % | chr\_5:2080087 G->A; chr\_5:2082453 G->A; |  |
| 37 | ggggacaagtttgtacaaaaaagcaggcttgatgTCCTCGAGCGTCGACTGGGA, ggggaccactttgtacaagaaagctgggtcACCCACCCATCCTCTCCCAT | Mycgr3\|90261  JGI  Ensembl | KOG4805 Uncharacterized conserved protein; |  | 100 % | chr\_1:5786356 G->A; chr\_1:5787737 A->G; |  |
| 37 | ggggacaagtttgtacaaaaaagcaggcttgatgTCTCGTCCGCTCGCGGCCGA, ggggaccactttgtacaagaaagctgggtcACCCTCCTTGCCCTTCAGCT | Mycgr3\|90278  JGI  Ensembl |  |  | 100 % | chr\_1:5851422 C->T; chr\_1:5852089 T->C; |  |
| 37 | ggggacaagtttgtacaaaaaagcaggcttgatgGCTACTTCCCACGCCGGCAT, ggggaccactttgtacaagaaagctgggtcGCAAGTGTCTGAAAGTTTCG | Mycgr3\|90446  JGI  Ensembl | KOG0692 Pentafunctional AROM protein; | 3-dehydroquinate dehydratase activity; ATP binding;  aromatic amino acid family biosynthetic process;  cytoplasm;  shikimate 5-dehydrogenase activity;  shikimate kinase activity; | 1.45958 % |  |  |
| 37 | ggggacaagtttgtacaaaaaagcaggcttgatgGCCGCAGCCACCGATGCGCC, ggggaccactttgtacaagaaagctgggtcCTCCGGCTTCAGCTTCAACT | Mycgr3\|90574  JGI  Ensembl | KOG4475 FOG: Immunoglobin and related proteins; | glucan 1,4-alpha-glucosidase activity; | 100 % | chr\_2:1101832 C->G; |  |
| 37 | ggggacaagtttgtacaaaaaagcaggcttgatgGACACCAAGTCGCCAGCCAG, ggggaccactttgtacaagaaagctgggtcTCCAAGCCATCCATCATCAT | Mycgr3\|91635  JGI  Ensembl | KOG4475 FOG: Immunoglobin and related proteins; |  | 100 % | chr\_3:1464956 G->A; chr\_3:1464719 C->T; |  |
| 37 | ggggacaagtttgtacaaaaaagcaggcttgatgGCCACCTCATCCCGCGTCGT, ggggaccactttgtacaagaaagctgggtcCAACCTTGCTCTGAACTCCT | Mycgr3\|91876  JGI  Ensembl | KOG2844 Dimethylglycine dehydrogenase precursor; | aminomethyltransferase activity; catalytic activity;  cytoplasm;  dimethylglycine dehydrogenase activity;  glycine catabolic process; | 100 % | chr\_3:2309610 C->T; |  |
| 37 | ggggacaagtttgtacaaaaaagcaggcttgatgTCGACGGTGGCATGGCGTGG, ggggaccactttgtacaagaaagctgggtcCCTGCTTTGCCATTCCAGTC | Mycgr3\|93368  JGI  Ensembl | KOG4266 Subtilisin kexin isozyme-1/site 1 protease, subtilase superfamily; | proteolysis; subtilase activity; | 0 % |  |  |
| 37 | ggggacaagtttgtacaaaaaagcaggcttgatgTCCAACGACTCCAACAAGCC, ggggaccactttgtacaagaaagctgggtcCTCCTTGCCTTTGTTCTTCT | Mycgr3\|94462  JGI  Ensembl | KOG1187 Serine/threonine protein kinase; |  | 100 % |  |  |
| 37 | ggggacaagtttgtacaaaaaagcaggcttgatgGCGAAAACTAAAGGCATCGC, ggggaccactttgtacaagaaagctgggtcCCGTTTTGCACCAGTAGGCT | Mycgr3\|94916  JGI  Ensembl |  |  | 100 % |  |  |
| 37 | ggggacaagtttgtacaaaaaagcaggcttgatgCCCAAACAGCCAGCGGACTA, ggggaccactttgtacaagaaagctgggtcGTGCTTCATATCCCCACCCG | Mycgr3\|95105  JGI  Ensembl | KOG1803 DNA helicase; |  | 100 % |  |  |
| 37 | ggggacaagtttgtacaaaaaagcaggcttgatgCCAGAAGAGCTTCCGCTCAC, ggggaccactttgtacaagaaagctgggtcCCCAATCCACCTCGGCAGCT | Mycgr3\|95722  JGI  Ensembl |  |  | 0 % |  |  |
| 37 | ggggacaagtttgtacaaaaaagcaggcttgatgTCGAAATACAACGAAGAAGA, ggggaccactttgtacaagaaagctgggtcCACGCGCACGCTGCTTGCGT | Mycgr3\|95921  JGI  Ensembl | KOG3854 SPRT-like metalloprotease; | protein kinase CK2 activity; | 100 % |  |  |
| 37 | ggggacaagtttgtacaaaaaagcaggcttgatgTCCGATCCTCTCAACTCTTT, ggggaccactttgtacaagaaagctgggtcAAAGAACCTCCGATATTCTC | Mycgr3\|95993  JGI  Ensembl | KOG0384 Chromodomain-helicase DNA-binding protein; |  | 1.46287 % |  |  |
| 37 | ggggacaagtttgtacaaaaaagcaggcttgatgAACGCTGAAGACAGACACAA, ggggaccactttgtacaagaaagctgggtcCTGAGAGCAATCCTCACACG | Mycgr3\|96052  JGI  Ensembl | KOG1081 Transcription factor NSD1 and related SET domain proteins; | protein binding; zinc ion binding; | 100 % |  |  |
| 37 | ggggacaagtttgtacaaaaaagcaggcttgatgCCAAGATTTAGCGCTACTTG, ggggaccactttgtacaagaaagctgggtcGTTCCGAGTGAACGGATGGA | Mycgr3\|97139  JGI  Ensembl | KOG0101 Molecular chaperones HSP70/HSC70, HSP70 superfamily; | binding; | 100 % |  |  |
| 37 | ggggacaagtttgtacaaaaaagcaggcttgatgGCGTACGACGCCGAAGAAGC, ggggaccactttgtacaagaaagctgggtcCTCCTTCTTCTTGTCCTCCT | Mycgr3\|97179  JGI  Ensembl | KOG1216 von Willebrand factor and related coagulation proteins; |  | 100 % | chr\_12:1209303 T->C; |  |
| 37 | ggggacaagtttgtacaaaaaagcaggcttgatgCGGGTAACACATCATCTGGC, ggggaccactttgtacaagaaagctgggtcATTGGAGTTCTGAGTGTATG | Mycgr3\|97361  JGI  Ensembl | KOG2089 Metalloendopeptidase family - saccharolysin & thimet oligopeptidase; | metalloendopeptidase activity; proteolysis; saccharolysin activity; | 100 % | chr\_13:490249 G->A; |  |
| 37 | ggggacaagtttgtacaaaaaagcaggcttgatgAACACCTCCAACACTCAAAC, ggggaccactttgtacaagaaagctgggtcTAGGAAACACCGTGCCGGTA | Mycgr3\|97571  JGI  Ensembl | KOG1181 FOG: Low-complexity; |  | 1.50957 % |  |  |
| 37 | ggggacaagtttgtacaaaaaagcaggcttgatgCCACGGCAGATACTACGACA, ggggaccactttgtacaagaaagctgggtcCATGGCTGTCCGGTGGAATG | Mycgr3\|98008  JGI  Ensembl |  |  | 0.87786 % |  |  |
| 37 | ggggacaagtttgtacaaaaaagcaggcttgatgACTTTTGCGGGAAAACACAA, ggggaccactttgtacaagaaagctgggtcGCAGTGGAGAGCGATCAGAT | Mycgr3\|98099  JGI  Ensembl |  |  | 100 % |  |  |
| 37 | ggggacaagtttgtacaaaaaagcaggcttgatgAATGTACACCGATTCGCAAA, ggggaccactttgtacaagaaagctgggtcAGACTCTCCAGCCATGTATG | Mycgr3\|98120  JGI  Ensembl | KOG0613 Projectin/twitchin and related proteins; | myosin ATPase activity; | 1.43646 % |  |  |
| 37 | ggggacaagtttgtacaaaaaagcaggcttgatgGCTGCCACCGAAGAGCCGCC, ggggaccactttgtacaagaaagctgggtcGTAAGGCCACTTCCAATCCG | Mycgr3\|98369  JGI  Ensembl |  |  | 100 % | chr\_1:2058694 A->G; chr\_1:2058543 G->A; |  |
| 37 | ggggacaagtttgtacaaaaaagcaggcttgatgGCACAAATCACGCTCAACCC, ggggaccactttgtacaagaaagctgggtcTTTCGCCGTCCCATTCATGG | Mycgr3\|99393  JGI  Ensembl | KOG2440 Pyrophosphate-dependent phosphofructo-1-kinase; | 6-phosphofructokinase activity; 6-phosphofructokinase complex;  cytoplasm;  glycolysis; | 100 % | chr\_3:428607 G->A; |  |
| 37 | ggggacaagtttgtacaaaaaagcaggcttgatgCAACGAGTCAAAAACGTCGG, ggggaccactttgtacaagaaagctgggtcCGTCTCAACACTCTCAACCA | Mycgr3\|99982  JGI  Ensembl | KOG0458 Elongation factor 1 alpha; | GTP binding; GTPase activity;  protein-synthesizing GTPase activity;  translation; | 100 % |  |  |
| 37 | ggggacaagtttgtacaaaaaagcaggcttgatgACCGCCTCGGAAGACTATGT, ggggaccactttgtacaagaaagctgggtcTTGATGAAGTGGAAGGTCTC | Mycgr3\|100276  JGI  Ensembl | KOG1292 Xanthine/uracil transporters; | membrane; protein binding;  transport;  transporter activity; | 100 % |  |  |
| 37 | ggggacaagtttgtacaaaaaagcaggcttgatgCCTCCTCATCCACTCAAAAG, ggggaccactttgtacaagaaagctgggtcCTTGCAGCATGCATCGCCAT | Mycgr3\|100462  JGI  Ensembl | KOG1186 Copper amine oxidase; | amine metabolic process; amine oxidase activity;  copper ion binding;  quinone binding; | 100 % |  |  |
| 37 | ggggacaagtttgtacaaaaaagcaggcttgatgATGGCAGGTATGGAGGCGGA, ggggaccactttgtacaagaaagctgggtcCCCGCAGTAAATATCGAGCA | Mycgr3\|101025  JGI  Ensembl |  |  | 100 % | chr\_8:1444466 T->A; chr\_8:1444621 T->C; chr\_8:1443214 C->T; chr\_8:1444434 C->T; |  |
| 37 | ggggacaagtttgtacaaaaaagcaggcttgatgCGCATGCGTTGGGAGAGTAT, ggggaccactttgtacaagaaagctgggtcACTCGTTGCAGGCACTGTCG | Mycgr3\|101265  JGI  Ensembl | KOG2458 Endoplasmic reticulum protein EP58, contains filamin rod domain and KDEL motif; |  | 100 % | chr\_9:1361990 A->G; |  |
| 37 | ggggacaagtttgtacaaaaaagcaggcttgatgAGTACGCCGAACAAGCGGCG, ggggaccactttgtacaagaaagctgggtcTTCCTCTTCGTCCTCGTCCA | Mycgr3\|101535  JGI  Ensembl | KOG0967 ATP-dependent DNA ligase I; | ATP binding; DNA ligase (ATP) activity;  DNA recombination;  DNA repair;  DNA replication; | 0.70214 % |  |  |
| 37 | ggggacaagtttgtacaaaaaagcaggcttgatgCGCCTCGCATACTACGCCGG, ggggaccactttgtacaagaaagctgggtcAGCCTCCGGTTCAGGCACCT | Mycgr3\|101607  JGI  Ensembl | KOG0802 E3 ubiquitin ligase; | protein binding; zinc ion binding; | 100 % | chr\_11:998087 A->G; chr\_11:999739 G->A; |  |
| 37 | ggggacaagtttgtacaaaaaagcaggcttgatgCCTCCTCGCATTTCCTTCGC, ggggaccactttgtacaagaaagctgggtcCATAAAATACCGCATCCCCT | Mycgr3\|101742  JGI  Ensembl |  |  | 100 % |  |  |
| 37 | ggggacaagtttgtacaaaaaagcaggcttgatgGAATACTCTACGCATGCATC, ggggaccactttgtacaagaaagctgggtcCTCCTCTCGCTTGCGCTTCT | Mycgr3\|105116  JGI  Ensembl | KOG1721 FOG: Zn-finger; | intracellular; nucleic acid binding;  zinc ion binding; | 100 % |  |  |
| 37 | ggggacaagtttgtacaaaaaagcaggcttgatgGTGCAATCCTCCATCCTCGG, ggggaccactttgtacaagaaagctgggtcCTTGGAAGCGTGCTTCTCAC | Mycgr3\|106395  JGI  Ensembl | KOG2263 Methionine synthase II (cobalamin-independent); | 5-methyltetrahydropteroyltriglutamate-homocysteine S-methyltransferase activity; methionine biosynthetic process; | 100 % |  |  |
| 37 | ggggacaagtttgtacaaaaaagcaggcttgatgGAGTTTTCTGGATGGCCGAT, ggggaccactttgtacaagaaagctgggtcAGAAAATCCAGCACCCAGTC | Mycgr3\|106752  JGI  Ensembl | KOG1721 FOG: Zn-finger; | intracellular; nucleic acid binding;  zinc ion binding; | 100 % |  |  |
| 37 | ggggacaagtttgtacaaaaaagcaggcttgatgTCTTCTCTAGGCGTATTCAA, ggggaccactttgtacaagaaagctgggtcATTCCCAACCTCCGCAACCG | Mycgr3\|106793  JGI  Ensembl | KOG1721 FOG: Zn-finger; | intracellular; nucleic acid binding;  nucleus;  regulation of transcription, DNA-dependent;  transcription factor activity;  zinc ion binding; | 100 % | chr\_1:666083 G->A; chr\_1:664873 G->C; chr\_1:665431 C->T; | ``` 1	ATGTCTTCTCTAGGCGTATTCAAGTGCACCCAGTGCGACAAGGTCTACAAGACTCGGACCTCTCTGACTCGCCACGCCCACAACCACTCCCGCGGACCGCCTCAACATCAGTGCGATACTTGTGGCGTCCTATTCGCCCGGCGCGACATTCTGAACCGCCATGTGAGCAACGGCCATTGCGCGAATTCAAATGCTGCCAGACATCGGTGCCATACCGCGTGCGAGGCGTGTCGCGGAGCCAGGATCAAATGCGACGGCCAGGTTCCATGTCGCGCGTGTTTCGAGGGTCGCCGAGAATGTAAATATCCTACCACTACGGGCAGGATCAGCCATGCGAGCCGGGCACCTAGGAGCGTGAAAGCGGAAGGGAGTGGAGAAGTGGATGAGAAGTCAGTGGGCTCGCCCTACGCCGATGCTTCCGATGAAGGCATGCATGGATCTGAGGACCGAGAATCACCTGAAGAGAGTGTCCATGGACCACGTGCGAACGAGCGGACCCATAGCCAGGCCCTGATGACCCCGACTTCACTTTCTCCCGCGATCGGATCTGTAGCGCCATATAAGGAAATGCTGACTGCAGGCGCTGCCGACAGTCATAATTTGAGGCTGGCCGGAGATGTACATGAGGAGAAGGCCTGGCCATGGCTGCATGAGAATTTGTTTTTATTGAACGGCGGGAAATCACAAAGCCGGAAACCATCACTACATCACCAGAACGCACGACCAACATCGACAGCACAGGATGCCAAAGGCTCGAGCATCTCATCTCCAACCAGCAATAATACGCACTCTCCTCTCTCGAACACCATCGAAAATCTCGTTGCGTATGCGGCGAACACGGCCTTCTCGCCCGATGACCGCGGAATCAGACGGCGGTATTGGGAATCCGTTTCACTACAACT-GGCCGACATATTCCAAGACGCCAGTAAAAAATCTGTTGTTGATCCACATCGCACTTTGTACCAGCTCGCCGAATCCTTCCTCACCAACTTCAATTCTGGATGGCCCA-TATTGAACCACGAACAGCTGGATCCGGATACACATCATCCTGTGCTATTCCTGGTAGTTGGT-GCGATCGGAGCGATGTTCGGCGA-CGCA	1098 	\|\|\|\|\|\|\|\|\|\|\|\|\|\|\|\|\|\|\|\|\|\|\|\|\|\|\|\|\|\|\|\|\|\|\|\|\|\|\|\|\|\|\|\|\|\|\|\|\|\|\|\|\|\|\|\|\|\|\|\|\|\|\|\|\|\|\|\|\|\|\|\|\|\|\|\|\|\|\|\|\|\|\|\|\|\|\|\|\|\|\|\|\|\|\|\|\|\|\|\|\|\|\|\|\|\|\|\|\|\|\|\|\|\|\|\|\|\|\|\|\|\|\|\|\|\|\|\|\|\|\|\|\|\|\|\|\|\|\|\|\|\|\|\|\|\|\|\|\|\|\|\|\|\|\|\|\|\|\|\|\|\|\|\|\|\|\|\|\|\|\|\|\|\|\|\|\|\|\|\|\|\|\|\|\|\|\|\|\|\|\|\|\|\|\|\|\|\|\|\|\|\|\|\|\|\|\|\|\|\|\|\|\|\|\|\|\|\|\|\|\|\|\|\|\|\|\|\|\|\|\|\|\|\|\|\|\|\|\|\|\|\|\|\|\|\|\|\|\|\|\|\|\|\|\|\|\|\|\|\|\|\|\|\|\|\|\|\|\|\|\|\|\|\|\|\|\|\|\|\|\|\|\|\|\|\|\|\|\|\|\|\|\|\|\|\|\|\|\|\|\|\|\|\|\|\|\|\|\|\|\|\|\|\|\|\|\|\|\|\|\|\|\|\|\|\|\|\|\|\|\|\|\|\|\|\|\|\|\|\|\|\|\|\|\|\|\|\|\|\|\|\|\|\|\|\|\|\|\|\|\|\|\|\|\|\|\|\|\|\|\|\|\|\|\|\|\|\|\|\|\|\|\|\|\|\|\|\|\|\|\|\|\|\|\|\|\|\|\|\|\|\|\|\|\|\|\|\|\|\|\|\|\|\|\|\|\|\|\|\|\|\|\|\|\|\|\|\|\|\|\|\|\|\|\|\|\|\|\|\|\|\|\|\|\|\|\|\|\|\|\|\|\|\|\|\|\|\|\|\|\|\|\|\|\|\|\|\|\|\|\|\|\|\|\|\|\|\|\|\|\|\|\|\|\|\|\|\|\|\|\|\|\|\|\|\|\|\|\|\|\|\|\|\|\|\|\|\|\|\|\|\|\|\|\|\|\|\|\|\|\|\|\|\|\|\|\|\|\|\|\|\|\|\|\|\|\|\|\|\|\|\|\|\|\|\|\|\|\|\|\|\|\|\|\|\|\|\|\|\|\|\|\|\|\|\|\|\|\|\|\|\|\|\|\|\|\|\|\|\|\|\|\|\|\|\|\|\|\|\|\|\|\|\|\|\|\|\|\|\|\|\|\|\|\|\|\|\|\|\|\|\|\|\|\|\|\|\|\|\|\|\|\|\|\|\|\|\|\|\|\|\|\|\|\|\|\|\|\|\|\|\|\|\|\|\|\|\|\|\|\|\|\|\|\|\|\|\|\|\|\|\|\|\|\|\|\|\|\|\|\|\|\|\|\|\|\|\|\|\|\|\|\|\|\|\|\|\|\|\|\|\|\|\|\|\|\|\|\|\|\|\|\|\|\|\|\|\|\|\|\|\|\|\|\|\|\|\|\|\|\|\|\|\|\|\|\|\|\|\|\|\|\|\|\|\|\|\|\|\|\|\|\|\|\|\|\|\|\|\|\|\|\|\|\|\|\|\|\|\|\|\|\|\|\|\|\|\|\|\|\|\|\|\|\|\|\|\|\|\|\|\|\|\|\|\|\|\|\|\|\|\|\|\|\|\|\|\|\|\|\|\|\|\|\|\|\|\|\|\|\|\|\|\|\|\|\|\|\|\|\|\|\|\|\|\|\|\|\|\|\|\|\|\|\|\|\|\|\|\|\|\|\|\|\|\|\|\|\|\|\|\|\|\|\|\|\|\|\|\|\|\|\|\|\|\|\|\|\|\|\|\|\|\|\|\|\|\|\|\|\|\|\|\|\|\|\|\|\|\|\|\|\|\|\|\|\|\|\|\|\|\| \|\|\|\|\|\|\|\|\|\|\|\|\|\|\|\|\|\|\|\|\|\|\|\|\|\|\|\|\|\|\|\|\|\|\|\|\|\|\|\|\|\|\| \|\|\|\|\|\|\|\|\|\|\|\|\|\|\|\|\|\|\|\|\|\|\|\|\|\|\|\|\|\|\|\|\|\|\|\|\|\|\|\|\|\| \|\|\|\|\|  \|\|\|\|\|\|\|\|\|\|\|\|\| \|\|\| \|\| \| \|\|\| \|\|\|\|\|\|\|\|\|\|\|\|\| \|\|\|\| \|\|\|\|\|\| \|\|\|\|\|\|\|\| \|\|\|\|\| \|\|\|\| \|\|\| \|\|\|\|\|\|\|\|\|\|\|\|\|\|\|\|\|\|\|\|\|\|\| \|\|\|\| 	ATGTCTTCTCTAGGCGTATTCAAGTGCACCCAGTGCGACAAGGTCTACAAGACTCGGACCTCTCTGACTCGCCACGCCCACAACCACTCCCGCGGACCGCCTCAACATCAGTGCGATACTTGTGGCGTCCTATTCGCCCGGCGCGACATTCTGAACCGCCATGTGAGCAACGGCCATTGCGCGAATTCAAATGCTGCCAGACATCGGTGCCATACCGCGTGCGAGGCGTGTCGCGGAGCCAGGATCAAATGCGACGGCCAGGTTCCATGTCGCGCGTGTTTCGAGGGTCGCCGAGAATGTAAATATCCTACCACTACGGGCAGGATCAGCCATGCGAGCCGGGCACCTAGGAGCGTGAAAGCGGAAGGGAGTGGAGAAGTGGATGAGAAGTCAGTGGGCTCGCCCTACGCCGATGCTTCCGATGAAGGCATGCATGGATCTGAGGACCGAGAATCACCTGAAGAGAGTGTCCATGGACCACGTGCGAACGAGCGGACCCATAGCCAGGCCCTGATGACCCCGACTTCACTTTCTCCCGCGATCGGATCTGTAGCGCCATATAAGGAAATGCTGACTGCAGGCGCTGCCGACAGTCATAATTTGAGGCTGGCCGGAGATGTACATGAGGAGAAGGCCTGGCCATGGCTGCATGAGAATTTGTTTTTATTGAACGGCGGGAAATCACAAAGCCGGAAACCATCACTACATCACCAGAACGCACGACCAACATCGACAGCACAGGATGCCAAAGGCTCGAGCATCTCATCTCCAACCAGCAATAATACGCACTCTCCTCTCTCGAACACCATCGAAAATCTCGTTGCGTATGCGGCGAACACGGCCTTCTCGCCCGATGACCGCGGAATCAGACGGCGGTATTGGGAATCCGTTTCACTACAACTGGGCCGACATATTCCAAGACGCCAGTAAAAAATCTGTTGTTGAT-CACATCGCACTTTGTACCAGCTCGCCGAATCCTTCCTCACCA-CTTCA--TCTGGATGGCCCAATAT-GA-C-ACG-ACAGCTGGATCCG-ATAC-CATCAT-CTGTGCTAATCCTGTTAGT-GGTGGCGATCGGAGCGATGTTCGGCGAACGCA ``` |
| 37 | ggggacaagtttgtacaaaaaagcaggcttgatgGCGGCCCAGTCAACGCTCCG, ggggaccactttgtacaagaaagctgggtcTGATTCCGACTCTTGAGTTC | Mycgr3\|106868  JGI  Ensembl | KOG0064 Peroxisomal long-chain acyl-CoA transporter, ABC superfamily; | ATP binding; ATPase activity;  membrane;  nucleoside-triphosphatase activity;  nucleotide binding;  transport; | 100 % |  |  |
| 37 | ggggacaagtttgtacaaaaaagcaggcttgatgGACGATATTGCAGCAATGTC, ggggaccactttgtacaagaaagctgggtcATTGCTGACGATCACGCGCT | Mycgr3\|107069  JGI  Ensembl | KOG0039 Ferric reductase, NADH/NADPH oxidase and related proteins; | ferric-chelate reductase activity; | 1.89573 % |  |  |
| 37 | ggggacaagtttgtacaaaaaagcaggcttgatgTGCTCTCCCTTTCCGACAAT, ggggaccactttgtacaagaaagctgggtcATAAGGCGGCGCACTCATCA | Mycgr3\|107106  JGI  Ensembl | KOG1877 Putative transmembrane protein cmp44E; |  | 100 % |  |  |
| 37 | ggggacaagtttgtacaaaaaagcaggcttgatgTCATCAACACGCACCGCCAT, ggggaccactttgtacaagaaagctgggtcGATGAGTACCTCAAGCTGGT | Mycgr3\|107221  JGI  Ensembl |  |  | 100 % | chr\_1:2875111 G->A; |  |
| 37 | ggggacaagtttgtacaaaaaagcaggcttgatgAATGCCCATAGTGGAGTCGA, ggggaccactttgtacaagaaagctgggtcCCTACTCCAGCCGAACAGTC | Mycgr3\|107296  JGI  Ensembl | KOG2992 Nucleolar GTPase/ATPase p130; |  | 100 % | chr\_1:3217624 A->G; |  |
| 37 | ggggacaagtttgtacaaaaaagcaggcttgatgTCGAGCGACCAAGGCGCAAA, ggggaccactttgtacaagaaagctgggtcCTCCTCCTCGGACTCTTCGG | Mycgr3\|107338  JGI  Ensembl |  |  | 0.77864 % |  |  |
| 37 | ggggacaagtttgtacaaaaaagcaggcttgatgGACACCTCTGCCGGGCAATC, ggggaccactttgtacaagaaagctgggtcTTCCTCCACCGCGCTGACAG | Mycgr3\|107401  JGI  Ensembl | KOG0391 SNF2 family DNA-dependent ATPase; |  | 0 % |  |  |
| 37 | ggggacaagtttgtacaaaaaagcaggcttgatgACGACCATGGCGATGCCCAT, ggggaccactttgtacaagaaagctgggtcAATCACAATATACTCCCCAT | Mycgr3\|107474  JGI  Ensembl |  |  | 0.77882 % |  |  |
| 37 | ggggacaagtttgtacaaaaaagcaggcttgatgCGGCCCTCGGACGTCCTCGC, ggggaccactttgtacaagaaagctgggtcTCTGGCAGTCCGGCTTCTTG | Mycgr3\|107828  JGI  Ensembl | KOG1119 Mitochondrial Fe-S cluster biosynthesis protein ISA2 (contains a HesB-like domain); |  | 100 % | chr\_2:313145 G->A; |  |
| 37 | ggggacaagtttgtacaaaaaagcaggcttgatgCGTTTCGGCAAGACCCTGGA, ggggaccactttgtacaagaaagctgggtcAAACGTCGACGACGACAAAT | Mycgr3\|107959  JGI  Ensembl | KOG1161 Protein involved in vacuolar polyphosphate accumulation, contains SPX domain; |  | 100 % | chr\_2:1180833 T->A; |  |
| 37 | ggggacaagtttgtacaaaaaagcaggcttgatgTTCCTCGATGAACAACGTCT, ggggaccactttgtacaagaaagctgggtcCTCATTCGATGTGGACCGCC | Mycgr3\|108169  JGI  Ensembl | KOG2146 Splicing coactivator SRm160/300, subunit SRm160 (contains PWI domain); |  | 100 % |  |  |
| 37 | ggggacaagtttgtacaaaaaagcaggcttgatgTCCTCTCCTCCGCCGCCATC, ggggaccactttgtacaagaaagctgggtcCCCAAGCGAACCCGAGTGCA | Mycgr3\|108753  JGI  Ensembl | KOG0946 ER-Golgi vesicle-tethering protein p115; |  | 100 % |  |  |
| 37 | ggggacaagtttgtacaaaaaagcaggcttgatgCCGCCGTCGCCGCCGGCGCG, ggggaccactttgtacaagaaagctgggtcCGACACCAAATGCCCATACT | Mycgr3\|108842  JGI  Ensembl | KOG1844 PHD Zn-finger proteins; | exo-alpha-sialidase activity; protein binding;  zinc ion binding; | 0.9255 % |  |  |
| 37 | ggggacaagtttgtacaaaaaagcaggcttgatgGGAGACCACTCGACTTCATC, ggggaccactttgtacaagaaagctgggtcGTGCCCCGAAGCGCCGTACA | Mycgr3\|109112  JGI  Ensembl | KOG1187 Serine/threonine protein kinase; | hydrogen ion transmembrane transporter activity; proton-transporting two-sector ATPase complex; | 100 % |  |  |
| 37 | ggggacaagtttgtacaaaaaagcaggcttgatgGACCACGATGCGCCGGAAGT, ggggaccactttgtacaagaaagctgggtcCAGCAACCCCTTACACCCAT | Mycgr3\|109341  JGI  Ensembl |  |  | 0.77093 % |  |  |
| 37 | ggggacaagtttgtacaaaaaagcaggcttgatgTCTTCGACGGACCAGCACTC, ggggaccactttgtacaagaaagctgggtcCTGCATCAAAGCAAGACTCA | Mycgr3\|109342  JGI  Ensembl | KOG0401 Translation initiation factor 4F, ribosome/mRNA-bridging subunit (eIF-4G); |  | 100 % | chr\_5:274741 C->T; chr\_5:275099 C->G; chr\_5:275204 G->A; |  |
| 37 | ggggacaagtttgtacaaaaaagcaggcttgatgGCACAGACTCCAGCCCATCT, ggggaccactttgtacaagaaagctgggtcAAAGCTCGCAAACTCCAGCC | Mycgr3\|109427  JGI  Ensembl | KOG1426 FOG: RCC1 domain; | calcium ion binding; zinc ion binding; | 100 % | chr\_5:864548 T->A; |  |
| 37 | ggggacaagtttgtacaaaaaagcaggcttgatgGCAGGCTCTAGAGCTAAACA, ggggaccactttgtacaagaaagctgggtcGGCGTCAGACCTTCCAGACT | Mycgr3\|109451  JGI  Ensembl |  |  | 100 % |  |  |
| 37 | ggggacaagtttgtacaaaaaagcaggcttgatgGACAATGTCTTGGAAGACAT, ggggaccactttgtacaagaaagctgggtcAGCGATGCTGACGAAGTCGC | Mycgr3\|110027  JGI  Ensembl | KOG0029 Amine oxidase; | electron transport; oxidoreductase activity;  polyamine oxidase activity; | 0.86531 % |  |  |
| 37 | ggggacaagtttgtacaaaaaagcaggcttgatgGAAGCACCGAACTGGGCACA, ggggaccactttgtacaagaaagctgggtcTGGCCCACCAAGCAACTTGC | Mycgr3\|110516  JGI  Ensembl |  |  | 100 % |  |  |
| 37 | ggggacaagtttgtacaaaaaagcaggcttgatgGCCGACGATGGACCAGCCCT, ggggaccactttgtacaagaaagctgggtcCCGACTAAACTCAACCAACC | Mycgr3\|110574  JGI  Ensembl |  |  | 100 % |  |  |
| 37 | ggggacaagtttgtacaaaaaagcaggcttgatgTCAATACGACACTACACCGC, ggggaccactttgtacaagaaagctgggtcTCGTCTGCCTCCACCCTCAA | Mycgr3\|110760  JGI  Ensembl | KOG3227 Calcium-responsive transcription coactivator; | histone acetyltransferase activity; | 100 % |  |  |
| 37 | ggggacaagtttgtacaaaaaagcaggcttgatgTCGGCCGGCGGAAAACAGGA, ggggaccactttgtacaagaaagctgggtcTCCCATCAGCTTCTTCAACA | Mycgr3\|110810  JGI  Ensembl | KOG4151 Myosin assembly protein/sexual cycle protein and related proteins; |  | 100 % | chr\_9:1344466 G->A; |  |
| 37 | ggggacaagtttgtacaaaaaagcaggcttgatgAAGACGTTTGAATTGCCCGC, ggggaccactttgtacaagaaagctgggtcAGCTCCCAATAACGCTACAA | Mycgr3\|111591  JGI  Ensembl | KOG2806 Chitinase; | exo-alpha-sialidase activity; | 0.7965 % |  |  |
| 37 | ggggacaagtttgtacaaaaaagcaggcttgatgTCTTCGGACCCCCACTCCGC, ggggaccactttgtacaagaaagctgggtcGACCTCAACAAAACTCATCC | Mycgr3\|111756  JGI  Ensembl | KOG1216 von Willebrand factor and related coagulation proteins; |  | 100 % | chr\_16:93481 G->C; chr\_16:93239 C->T; |  |
| 38 | ggggacaagtttgtacaaaaaagcaggcttgatgCCGGTTCTATATGGTGAGGG, ggggaccactttgtacaagaaagctgggtcGTTTCGCGTCCACTGGCAAC | Mycgr3\|32494  JGI  Ensembl | KOG0504 FOG: Ankyrin repeat; |  | 100 % | chr\_1:5267350 C->T; |  |
| 38 | ggggacaagtttgtacaaaaaagcaggcttgatgGAAGTTCAAGATCTGAGTCC, ggggaccactttgtacaagaaagctgggtcTCTCTCAGGTGCGGGGGCAG | Mycgr3\|34265  JGI  Ensembl |  |  | 100 % | chr\_1:2608847 C->T; |  |
| 38 | ggggacaagtttgtacaaaaaagcaggcttgatgGCGCTCGCTCCTGCCAACGA, ggggaccactttgtacaagaaagctgggtcTCCTCCGCCTCCGCTTCCAC | Mycgr3\|38597  JGI  Ensembl | KOG2175 Protein predicted to be involved in carbohydrate metabolism; |  | 58.84732 % | chr\_3:2984319 G->C; |  |
| 38 | ggggacaagtttgtacaaaaaagcaggcttgatgCAATACGTACGAGCCGTCTC, ggggaccactttgtacaagaaagctgggtcTTTCTTCCACCCCGTCTTTC | Mycgr3\|41117  JGI  Ensembl | KOG2116 Protein involved in plasmid maintenance/nuclear protein involved in lipid metabolism; | hydrogen ion transmembrane transporter activity; proton-transporting two-sector ATPase complex; | 100 % |  |  |
| 38 | ggggacaagtttgtacaaaaaagcaggcttgatgCCGGAGGCTACTGACGTCGC, ggggaccactttgtacaagaaagctgggtcGTCACCAACCACATCATCCA | Mycgr3\|41772  JGI  Ensembl | KOG0442 Structure-specific endonuclease ERCC1-XPF, catalytic component XPF/ERCC4; | DNA binding; DNA metabolic process;  DNA repair;  T/G mismatch-specific endonuclease activity;  endonuclease activity;  nuclease activity;  nucleic acid binding;  nucleus;  phosphoric ester hydrolase activity;  protein binding; | 0 % |  |  |
| 38 | ggggacaagtttgtacaaaaaagcaggcttgatgAGTGGACGGACCACACCAAA, ggggaccactttgtacaagaaagctgggtcCATCGTAACACATCCCACGC | Mycgr3\|45396  JGI  Ensembl | KOG3745 Exocyst subunit - Sec10p; |  | 100 % |  |  |
| 38 | ggggacaagtttgtacaaaaaagcaggcttgatgGCCAGCTTCCAGACCTACAA, ggggaccactttgtacaagaaagctgggtcAGGTAAATGCAACCCCGTCG | Mycgr3\|48895  JGI  Ensembl |  |  | 100 % | chr\_10:1185159 T->A; |  |
| 38 | ggggacaagtttgtacaaaaaagcaggcttgatgGTGAGTACCTCCATCCCCCG, ggggaccactttgtacaagaaagctgggtcCCGTTCCCGATCAACCCCGA | Mycgr3\|51429  JGI  Ensembl |  |  | 100 % |  |  |
| 38 | ggggacaagtttgtacaaaaaagcaggcttgatgGCGAGTGCTGACCTCCTCCG, ggggaccactttgtacaagaaagctgggtcAAGGTCATCCATCCCATGGC | Mycgr3\|56270  JGI  Ensembl | KOG0598 Ribosomal protein S6 kinase and related proteins; | 3-phosphoinositide-dependent protein kinase activity; AMP-activated protein kinase activity;  ATP binding;  DNA-dependent protein kinase activity;  GTP-dependent protein kinase activity;  JUN kinase activity;  JUN kinase kinase activity;  JUN kinase kinase kinase activity;  Janus kinase activity;  MAP kinase 1 activity;  MAP kinase 2 activity;  MAP kinase kinase activity;  MAP kinase kinase kinase kinase activity;  MAP/ERK kinase kinase activity;  MP kinase activity;  NF-kappaB-inducing kinase activity;  SAP kinase activity;  atypical protein kinase C activity;  calcium-dependent protein kinase C activity;  casein kinase I activity;  casein kinase activity;  cyclic nucleotide-dependent protein kinase activity;  cyclin-dependent protein kinase activating kinase activity;  eukaryotic translation initiation factor 2alpha kinase activity;  galactosyltransferase-associated kinase activity;  glycogen synthase kinase 3 activity;  protein amino acid phosphorylation;  protein kinase activity;  protein serine/threonine kinase activity;  protein-tyrosine kinase activity;  receptor signaling protein serine/threonine kinase activity;  ribosomal protein S6 kinase activity;  transmembrane receptor protein kinase activity;  transmembrane receptor protein serine/threonine kinase activity; | 0.755 % |  |  |
| 38 | ggggacaagtttgtacaaaaaagcaggcttgatgAGCACGCAGAAATCCGAGCT, ggggaccactttgtacaagaaagctgggtcTTCATGGCCGCCATTAACGC | Mycgr3\|65376  JGI  Ensembl | KOG0602 Neutral trehalase; | alpha,alpha-trehalase activity; calcium ion binding;  cytoplasm;  trehalose catabolic process;  trehalose metabolic process; | 100 % |  |  |
| 38 | ggggacaagtttgtacaaaaaagcaggcttgatgGCAACCCCCACCGCCAATAC, ggggaccactttgtacaagaaagctgggtcTCCTCTGCTGAACCAGCTGG | Mycgr3\|65760  JGI  Ensembl | KOG0240 Kinesin (SMY1 subfamily); | ATP binding; microtubule associated complex;  microtubule motor activity;  microtubule-based movement;  plus-end-directed microtubule motor activity; | 0 % |  |  |
| 38 | ggggacaagtttgtacaaaaaagcaggcttgatgGCGGCACAGCAGTCTCCAAC, ggggaccactttgtacaagaaagctgggtcGGTCGCCTGCTCTGCGCTGC | Mycgr3\|66803  JGI  Ensembl | KOG1124 FOG: TPR repeat; | UDP-N-acetylmuramoylalanyl-D-glutamyl-2,6-diaminopimelate-D-alanyl-D-alanine ligase activity; coenzyme F420-0 gamma-glutamyl ligase activity;  coenzyme F420-2 alpha-glutamyl ligase activity;  ribosomal S6-glutamic acid ligase activity; | 1.36411 % |  |  |
| 38 | ggggacaagtttgtacaaaaaagcaggcttgatgGCCAGCAACCAAGTCCTCCC, ggggaccactttgtacaagaaagctgggtcCTTTGCCGCGGCCAACTCTC | Mycgr3\|67635  JGI  Ensembl | KOG1156 N-terminal acetyltransferase; | peptide alpha-N-acetyltransferase activity; | 0 % |  |  |
| 38 | ggggacaagtttgtacaaaaaagcaggcttgatgGCGGACGATCACAAATACCA, ggggaccactttgtacaagaaagctgggtcCTCAAACTTCAACCCCCCAA | Mycgr3\|67686  JGI  Ensembl | KOG2195 Transferrin receptor and related proteins containing the protease-associated (PA) domain; | glutamate carboxypeptidase II activity; peptidase activity;  proteolysis; | 100 % |  |  |
| 38 | ggggacaagtttgtacaaaaaagcaggcttgatgAGTTCGAGCAAACCGCACGG, ggggaccactttgtacaagaaagctgggtcCATGTCATAACTCGCCGCGA | Mycgr3\|68017  JGI  Ensembl | KOG0448 Mitofusin 1 GTPase, involved in mitochondrila biogenesis; | carboxylic acid metabolic process; hydrolase activity, acting on acid anhydrides, acting on GTP, involved in cellular and subcellular movement; transferase activity, transferring acyl groups, acyl groups converted into alkyl on transfer; | 100 % |  |  |
| 38 | ggggacaagtttgtacaaaaaagcaggcttgatgGCGATTGACCGGCGGCGCGC, ggggaccactttgtacaagaaagctgggtcCAACTCTTCATGCTCCACCT | Mycgr3\|68653  JGI  Ensembl | KOG0104 Molecular chaperones GRP170/SIL1, HSP70 superfamily; | ATP binding; endoplasmic reticulum; | 4.9545 % |  |  |
| 38 | ggggacaagtttgtacaaaaaagcaggcttgatgGCGGGGGCATCTCCTGGCGC, ggggaccactttgtacaagaaagctgggtcCTGCCACACTCCTCTCGACT | Mycgr3\|69529  JGI  Ensembl | KOG0588 Serine/threonine protein kinase; | ATP binding; protein amino acid phosphorylation;  protein kinase activity;  protein serine/threonine kinase activity;  protein-tyrosine kinase activity; | 100 % |  |  |
| 38 | ggggacaagtttgtacaaaaaagcaggcttgatgGCGCCCTCAGAGAGCTTGTT, ggggaccactttgtacaagaaagctgggtcCTTCAACTCCGTCAACTCGT | Mycgr3\|69748  JGI  Ensembl | KOG2189 Vacuolar H+-ATPase V0 sector, subunit a; | hydrogen ion transporting ATP synthase activity, rotational mechanism; hydrogen ion transporting ATPase activity, rotational mechanism;  hydrogen-transporting two-sector ATPase activity;  membrane;  proton transport;  proton-transporting ATP synthase complex; | 100 % |  |  |
| 38 | ggggacaagtttgtacaaaaaagcaggcttgatgCCTTCAGTCTCAGAGTTTTT, ggggaccactttgtacaagaaagctgggtcAAGCGCATAGTTCTCTGTAT | Mycgr3\|70729  JGI  Ensembl | KOG0039 Ferric reductase, NADH/NADPH oxidase and related proteins; | ferric-chelate reductase activity; | 100 % | chr\_4:485381 A->G; chr\_4:484598 A->G; |  |
| 38 | ggggacaagtttgtacaaaaaagcaggcttgatgACGACTCTCGCCGCTCGCTC, ggggaccactttgtacaagaaagctgggtcTGTCTTCGGGTTGACAAATT | Mycgr3\|71128  JGI  Ensembl | KOG0198 MEKK and related serine/threonine protein kinases; | 3-phosphoinositide-dependent protein kinase activity; AMP-activated protein kinase activity;  ATP binding;  DNA-dependent protein kinase activity;  GTP-dependent protein kinase activity;  JUN kinase activity;  JUN kinase kinase activity;  JUN kinase kinase kinase activity;  Janus kinase activity;  MAP kinase 1 activity;  MAP kinase 2 activity;  MAP kinase kinase activity;  MAP kinase kinase kinase kinase activity;  MAP/ERK kinase kinase activity;  MP kinase activity;  NF-kappaB-inducing kinase activity;  SAP kinase activity;  atypical protein kinase C activity;  calcium-dependent protein kinase C activity;  casein kinase I activity;  casein kinase activity;  cyclic nucleotide-dependent protein kinase activity;  cyclin-dependent protein kinase activating kinase activity;  eukaryotic translation initiation factor 2alpha kinase activity;  galactosyltransferase-associated kinase activity;  glycogen synthase kinase 3 activity;  protein amino acid phosphorylation;  protein kinase activity;  protein serine/threonine kinase activity;  protein-tyrosine kinase activity;  receptor signaling protein serine/threonine kinase activity;  ribosomal protein S6 kinase activity;  transmembrane receptor protein kinase activity;  transmembrane receptor protein serine/threonine kinase activity; | 59.52717 % |  |  |
| 38 | ggggacaagtttgtacaaaaaagcaggcttgatgGGATCCCGACCGGAGCTCAA, ggggaccactttgtacaagaaagctgggtcCAAAGCCAGCACTCCCTTCA | Mycgr3\|72798  JGI  Ensembl | KOG0219 Mismatch repair ATPase MSH2 (MutS family); | ATP binding; ATP synthesis coupled proton transport;  MutSalpha complex;  MutSbeta complex;  hydrogen ion transporting ATP synthase activity, rotational mechanism;  hydrogen ion transporting ATPase activity, rotational mechanism;  membrane;  mismatch repair;  mismatched DNA binding;  proton-transporting two-sector ATPase complex; | 100 % |  |  |
| 38 | ggggacaagtttgtacaaaaaagcaggcttgatgAAAGACACGCAAGCCATGGA, ggggaccactttgtacaagaaagctgggtcAGCCTCAATCGTCACGCCCT | Mycgr3\|73800  JGI  Ensembl | KOG0519 Sensory transduction histidine kinase; | ATP binding; histone arginine kinase activity;  membrane;  peptidyl-histidine phosphorylation;  phosphorylation;  protein histidine kinase activity;  regulation of transcription, DNA-dependent;  signal transduction;  transferase activity, transferring phosphorus-containing groups;  two-component response regulator activity;  two-component sensor activity;  two-component signal transduction system (phosphorelay); | 100 % | chr\_7:610006 T->A; |  |
| 38 | ggggacaagtttgtacaaaaaagcaggcttgatgGCAGACAAGAGCATTGACAT, ggggaccactttgtacaagaaagctgggtcAAACCGCACCACTGCCTTCG | Mycgr3\|74016  JGI  Ensembl | KOG2533 Permease of the major facilitator superfamily; | integral to membrane; transport; transporter activity; | 100 % |  |  |
| 38 | ggggacaagtttgtacaaaaaagcaggcttgatgAACCCAGGCCCCGCGCCTCG, ggggaccactttgtacaagaaagctgggtcATGCCCACCTCCCTTTCTGA | Mycgr3\|75043  JGI  Ensembl |  |  | 1.56125 % |  |  |
| 38 | ggggacaagtttgtacaaaaaagcaggcttgatgGACTTTGCCATGAATGGAGT, ggggaccactttgtacaagaaagctgggtcTTGATACCACTCATCCTCAT | Mycgr3\|75413  JGI  Ensembl |  |  | 7.25259 % |  |  |
| 38 | ggggacaagtttgtacaaaaaagcaggcttgatgTCGACCTCCCGCACCCTCCC, ggggaccactttgtacaagaaagctgggtcCAACCGACTCTCCACCCCAC | Mycgr3\|75552  JGI  Ensembl | KOG2004 Mitochondrial ATP-dependent protease PIM1/LON; | ATP binding; ATP-dependent peptidase activity;  ATP-dependent proteolysis;  endopeptidase La activity;  nucleoside-triphosphatase activity;  nucleotide binding;  proteolysis;  serine-type endopeptidase activity; | 1.40562 % |  |  |
| 38 | ggggacaagtttgtacaaaaaagcaggcttgatgCCCTCCACTCCGCCGCCGCT, ggggaccactttgtacaagaaagctgggtcGCTTCCAACGCCTCCACCAT | Mycgr3\|76044  JGI  Ensembl | KOG2197 Ypt/Rab-specific GTPase-activating protein GYP7 and related proteins; | Rab GTPase activator activity; intracellular;  regulation of Rab GTPase activity; | 0.72106 % |  |  |
| 38 | ggggacaagtttgtacaaaaaagcaggcttgatgACTGCCTCTGAGCCTGCAGT, ggggaccactttgtacaagaaagctgggtcCGACTTCATTGATCTTCGTG | Mycgr3\|76301  JGI  Ensembl |  |  | 100 % |  |  |
| 38 | ggggacaagtttgtacaaaaaagcaggcttgatgCGACCGCTACGCTTCCTGCT, ggggaccactttgtacaagaaagctgggtcTCGCCCACCGCCTCTCTCCA | Mycgr3\|77796  JGI  Ensembl | KOG4628 Predicted E3 ubiquitin ligase; | protein binding; zinc ion binding; | 1.5885 % |  |  |
| 38 | ggggacaagtttgtacaaaaaagcaggcttgatgGAAGCGGCTGTACGCTGGTC, ggggaccactttgtacaagaaagctgggtcCTTCGCAGTGCACAGGCATT | Mycgr3\|83759  JGI  Ensembl | KOG1008 Uncharacterized conserved protein, contains WD40 repeats; | membrane; transport; transporter activity; | 0.67736 % |  |  |
| 38 | ggggacaagtttgtacaaaaaagcaggcttgatgAGCTCGCCGCAATGGCAGAC, ggggaccactttgtacaagaaagctgggtcACGATTCAAACTCAACAACC | Mycgr3\|88345  JGI  Ensembl | KOG2622 Putative myrosinase precursor; |  | 6.26712 % |  |  |
| 38 | ggggacaagtttgtacaaaaaagcaggcttgatgTCGTCGTCATCCTGGGCCGG, ggggaccactttgtacaagaaagctgggtcTTTCCTCCTCTGCGCCGACC | Mycgr3\|88655  JGI  Ensembl |  |  | 1.32069 % |  |  |
| 38 | ggggacaagtttgtacaaaaaagcaggcttgatgGCTTCACTTCCAGCACCACC, ggggaccactttgtacaagaaagctgggtcTCCAGGCGCTCCATCACCCA | Mycgr3\|88695  JGI  Ensembl | KOG0516 Dystonin, GAS (Growth-arrest-specific protein), and related proteins; |  | 100 % |  |  |
| 38 | ggggacaagtttgtacaaaaaagcaggcttgatgGGCTACTCCACTTTGCTGCT, ggggaccactttgtacaagaaagctgggtcCCGCCAGACGGGATGTATGA | Mycgr3\|88714  JGI  Ensembl | KOG2806 Chitinase; | exo-alpha-sialidase activity; | 100 % |  |  |
| 38 | ggggacaagtttgtacaaaaaagcaggcttgatgCCATCACCACATATTCCCTC, ggggaccactttgtacaagaaagctgggtcGAGCTCCAAGTGCACCTCTT | Mycgr3\|89735  JGI  Ensembl | KOG0061 Transporter, ABC superfamily (Breast cancer resistance protein); | ATP binding; ATPase activity;  nucleoside-triphosphatase activity;  nucleotide binding; | 0 % |  |  |
| 38 | ggggacaagtttgtacaaaaaagcaggcttgatgCCCACCCTCAAGCAGATCAC, ggggaccactttgtacaagaaagctgggtcCCACCCGGCGTTGCCTTTTC | Mycgr3\|89755  JGI  Ensembl | KOG1999 RNA polymerase II transcription elongation factor DSIF/SUPT5H/SPT5; | histone acetyltransferase activity; | 100 % |  |  |
| 38 | ggggacaagtttgtacaaaaaagcaggcttgatgCCGGGTCCGACACAACCCAC, ggggaccactttgtacaagaaagctgggtcCAACATGCCGCCACCTCCGC | Mycgr3\|89814  JGI  Ensembl |  |  | 2.14977 % |  |  |
| 38 | ggggacaagtttgtacaaaaaagcaggcttgatgCAACCTCCCAAACTACTGCT, ggggaccactttgtacaagaaagctgggtcCTCCATGCGGTTCTTCTTGC | Mycgr3\|89827  JGI  Ensembl | KOG3599 Ca2+-modulated nonselective cation channel polycystin; |  | 100 % |  |  |
| 38 | ggggacaagtttgtacaaaaaagcaggcttgatgGGACTGATCAAGCTCACCAT, ggggaccactttgtacaagaaagctgggtcGCGCCGGGAAATCTCGATCT | Mycgr3\|90066  JGI  Ensembl |  |  | 0 % |  |  |
| 38 | ggggacaagtttgtacaaaaaagcaggcttgatgTCGACTCCTACCATACCGCC, ggggaccactttgtacaagaaagctgggtcCAACTCATCCTCCATCTTTG | Mycgr3\|90493  JGI  Ensembl | KOG1181 FOG: Low-complexity; | DNA binding; | 100 % |  |  |
| 38 | ggggacaagtttgtacaaaaaagcaggcttgatgAAGTCGCACTCTTCTCTCGC, ggggaccactttgtacaagaaagctgggtcCGCGAGCATGGATATCGTCC | Mycgr3\|90507  JGI  Ensembl |  |  | 100 % |  |  |
| 38 | ggggacaagtttgtacaaaaaagcaggcttgatgAACACAACCACTAATGATAA, ggggaccactttgtacaagaaagctgggtcGACATTGCTCTGCGAGCCTG | Mycgr3\|90544  JGI  Ensembl | KOG1869 Splicing coactivator SRm160/300, subunit SRm300; |  | 100 % |  |  |
| 38 | ggggacaagtttgtacaaaaaagcaggcttgatgCTTACATCGCTTTGCCATCG, ggggaccactttgtacaagaaagctgggtcATCGCTGTCCCCTGAGATCA | Mycgr3\|90854  JGI  Ensembl |  |  | 100 % |  |  |
| 38 | ggggacaagtttgtacaaaaaagcaggcttgatgGCGCCGGTGCGGAAGAAGAG, ggggaccactttgtacaagaaagctgggtcCAACATACTCCTCACCCCCT | Mycgr3\|91172  JGI  Ensembl |  |  | 0.66041 % |  |  |
| 38 | ggggacaagtttgtacaaaaaagcaggcttgatgTCGGCTGCCACCTCCCCCAC, ggggaccactttgtacaagaaagctgggtcCGGGCTTTTCGCTGCTAGAC | Mycgr3\|91248  JGI  Ensembl | KOG1759 Macrophage migration inhibitory factor; |  | 100 % |  |  |
| 38 | ggggacaagtttgtacaaaaaagcaggcttgatgACAGCCATTTCGAGGAAGCG, ggggaccactttgtacaagaaagctgggtcACGATCGCTGTAAGACTGGC | Mycgr3\|92872  JGI  Ensembl |  |  | 0.71429 % |  |  |
| 38 | ggggacaagtttgtacaaaaaagcaggcttgatgTCTTTCATACTGCAGCCTGG, ggggaccactttgtacaagaaagctgggtcCTGGATCGCACGCTTGAAGT | Mycgr3\|92972  JGI  Ensembl | KOG0516 Dystonin, GAS (Growth-arrest-specific protein), and related proteins; |  | 1.32248 % |  |  |
| 38 | ggggacaagtttgtacaaaaaagcaggcttgatgGCGACCCCGGCGAACGAATA, ggggaccactttgtacaagaaagctgggtcGGAGCCGCCTTTCTCGGAAC | Mycgr3\|93593  JGI  Ensembl | KOG0959 N-arginine dibasic convertase NRD1 and related Zn2+-dependent endopeptidases, insulinase superfamily; |  | 1.40642 % |  |  |
| 38 | ggggacaagtttgtacaaaaaagcaggcttgatgGTTGGCGGAGTTGCAAAGCT, ggggaccactttgtacaagaaagctgggtcTCTCTTCCACGACGCTCCCG | Mycgr3\|93735  JGI  Ensembl | KOG2992 Nucleolar GTPase/ATPase p130; |  | 0.76004 % |  |  |
| 38 | ggggacaagtttgtacaaaaaagcaggcttgatgGCATCAAAAGACGCCCAAGC, ggggaccactttgtacaagaaagctgggtcCGAGTCATCGAGCATATCAT | Mycgr3\|93972  JGI  Ensembl | KOG2992 Nucleolar GTPase/ATPase p130; |  | 100 % |  |  |
| 38 | ggggacaagtttgtacaaaaaagcaggcttgatgGCACCAACCAGTGCATCGCC, ggggaccactttgtacaagaaagctgggtcGTCAGCGGCCCCCTCGACCA | Mycgr3\|94223  JGI  Ensembl |  |  | 100 % | chr\_7:93017 G->A; |  |
| 38 | ggggacaagtttgtacaaaaaagcaggcttgatgGGCTGCAACACATCGGTGGC, ggggaccactttgtacaagaaagctgggtcGGACATTTCCCAGAACTCGC | Mycgr3\|94762  JGI  Ensembl |  |  | 100 % |  |  |
| 38 | ggggacaagtttgtacaaaaaagcaggcttgatgGATCGACCAGAGCTCGGCCG, ggggaccactttgtacaagaaagctgggtcTTTCAAGCTGCTGCAGTTCG | Mycgr3\|94915  JGI  Ensembl |  |  | 100 % |  |  |
| 38 | ggggacaagtttgtacaaaaaagcaggcttgatgACACCCCCTCGCCCGTCGGA, ggggaccactttgtacaagaaagctgggtcGATCCACCCACCGTCTTCCT | Mycgr3\|95064  JGI  Ensembl | KOG1216 von Willebrand factor and related coagulation proteins; | cyclin-dependent protein kinase activity; | 15.95197 % |  |  |
| 38 | ggggacaagtttgtacaaaaaagcaggcttgatgGAGGAAGCCAACACTGCTGC, ggggaccactttgtacaagaaagctgggtcCTCGTCAAGCCTCCTCGGCA | Mycgr3\|95214  JGI  Ensembl | KOG1633 F-box protein JEMMA and related proteins with JmjC, PHD, F-box and LRR domains; |  | 1.40499 % |  |  |
| 38 | ggggacaagtttgtacaaaaaagcaggcttgatgAAGCTATTCGCTGGACTCCT, ggggaccactttgtacaagaaagctgggtcTGTACTCGAAATCGCAAAGG | Mycgr3\|95663  JGI  Ensembl | KOG0251 Clathrin assembly protein AP180 and related proteins, contain ENTH domain; | calcium ion binding; lipid catabolic process;  phospholipase A2 activity; | 100 % |  |  |
| 38 | ggggacaagtttgtacaaaaaagcaggcttgatgACCTCGCTCGGGGAGCTTCC, ggggaccactttgtacaagaaagctgggtcGACGTTGGCGTCCCAATTGG | Mycgr3\|95752  JGI  Ensembl |  |  | 100 % |  |  |
| 38 | ggggacaagtttgtacaaaaaagcaggcttgatgGACGCGGACGACGCCATGGT, ggggaccactttgtacaagaaagctgggtcCATTTCCCCACCCCCACCAC | Mycgr3\|95845  JGI  Ensembl | KOG1187 Serine/threonine protein kinase; |  | 0 % |  |  |
| 38 | ggggacaagtttgtacaaaaaagcaggcttgatgGCCATGTGGCCCTTCAAACG, ggggaccactttgtacaagaaagctgggtcAACCTCCTCCGATTTCTCAG | Mycgr3\|95885  JGI  Ensembl | KOG1869 Splicing coactivator SRm160/300, subunit SRm300; |  | 100 % | chr\_9:1627489 G->A; |  |
| 38 | ggggacaagtttgtacaaaaaagcaggcttgatgGCGTCGAATAAATTCTTTAC, ggggaccactttgtacaagaaagctgggtcGATTTTCTGAGCGTCGATCA | Mycgr3\|96806  JGI  Ensembl | KOG0161 Myosin class II heavy chain; | acid phosphatase activity; | 100 % | chr\_11:1388508 A->T; |  |
| 38 | ggggacaagtttgtacaaaaaagcaggcttgatgGCTCACTCATCAGTACTAAA, ggggaccactttgtacaagaaagctgggtcCTGATGGTCACTCCGCCCTC | Mycgr3\|96947  JGI  Ensembl |  |  | 88.19444 % |  |  |
| 38 | ggggacaagtttgtacaaaaaagcaggcttgatgCATCTCCACCTCCTCCTCGC, ggggaccactttgtacaagaaagctgggtcACGGTCATCAGATTCCTTCT | Mycgr3\|97203  JGI  Ensembl |  |  | 1.34205 % |  |  |
| 38 | ggggacaagtttgtacaaaaaagcaggcttgatgGAGCAATCTCCCACCCTGCT, ggggaccactttgtacaagaaagctgggtcGTGGAACCCATTGAAGATCA | Mycgr3\|97321  JGI  Ensembl |  |  | 28.38352 % |  |  |
| 38 | ggggacaagtttgtacaaaaaagcaggcttgatgGACCGAGGAACTATGGCCCG, ggggaccactttgtacaagaaagctgggtcTCGAGGACTCTTATAATAGC | Mycgr3\|97511  JGI  Ensembl | KOG0800 FOG: Predicted E3 ubiquitin ligase; | protein binding; zinc ion binding; | 100 % |  |  |
| 38 | ggggacaagtttgtacaaaaaagcaggcttgatgGCCGCACCATCATCATCTCA, ggggaccactttgtacaagaaagctgggtcGGTTTCGAAATGCGGATCTA | Mycgr3\|97739  JGI  Ensembl | KOG2992 Nucleolar GTPase/ATPase p130; |  | 100 % |  |  |
| 38 | ggggacaagtttgtacaaaaaagcaggcttgatgGCACCCACTTTTAAAAATGT, ggggaccactttgtacaagaaagctgggtcCAGGCCGTTATACCCACAGC | Mycgr3\|98656  JGI  Ensembl |  |  | 100 % |  |  |
| 38 | ggggacaagtttgtacaaaaaagcaggcttgatgCGTTTCCCGCTCATAGCAGC, ggggaccactttgtacaagaaagctgggtcAGCAAACTGCCATCTGGCAT | Mycgr3\|99418  JGI  Ensembl | KOG2103 Uncharacterized conserved protein; | electron transport; | 100 % |  |  |
| 38 | ggggacaagtttgtacaaaaaagcaggcttgatgTGTGGGATCCTCGCCGTTAT, ggggaccactttgtacaagaaagctgggtcAATGTCTTCACATGCCTGCG | Mycgr3\|100095  JGI  Ensembl | KOG0572 Glutamine phosphoribosylpyrophosphate amidotransferase; | amidophosphoribosyltransferase activity; metabolic process;  nucleoside metabolic process;  purine base biosynthetic process;  transferase activity; | 0.85227 % |  |  |
| 38 | ggggacaagtttgtacaaaaaagcaggcttgatgGAGGCTGCGGCGCTGAAGGC, ggggaccactttgtacaagaaagctgggtcACCAGCTCGGACCAAGTGAG | Mycgr3\|101229  JGI  Ensembl | KOG0371 Serine/threonine protein phosphatase 2A, catalytic subunit; | CTD phosphatase activity; calcium-dependent protein serine/threonine phosphatase activity;  calcium-dependent protein serine/threonine phosphatase regulator activity;  hydrolase activity;  magnesium-dependent protein serine/threonine phosphatase activity;  magnesium-dependent protein serine/threonine phosphatase complex;  myosin phosphatase activity;  myosin phosphatase complex;  myosin phosphatase regulator activity;  phosphoprotein phosphatase activity; | 100 % |  |  |
| 38 | ggggacaagtttgtacaaaaaagcaggcttgatgATACGCTGCCCCTCGATCAA, ggggaccactttgtacaagaaagctgggtcCTCCAACGGGATGCCCTCGT | Mycgr3\|104147  JGI  Ensembl | KOG2436 Acetylglutamate kinase/acetylglutamate synthase; | N-acetyl-gamma-glutamyl-phosphate reductase activity; NAD binding;  acetylglutamate kinase activity;  amino acid biosynthetic process;  amino acid metabolic process;  arginine biosynthetic process;  cytoplasm;  oxidoreductase activity, acting on the aldehyde or oxo group of donors, NAD or NADP as acceptor; | 0.75075 % |  |  |
| 38 | ggggacaagtttgtacaaaaaagcaggcttgatgTCCATGTCACGTCTACCGCA, ggggaccactttgtacaagaaagctgggtcCCGGTACGGATCATAGTCAC | Mycgr3\|106740  JGI  Ensembl | KOG0161 Myosin class II heavy chain; | exo-alpha-sialidase activity; | 100 % |  |  |
| 38 | ggggacaagtttgtacaaaaaagcaggcttgatgTCTGTGGAGGAGATGGACCT, ggggaccactttgtacaagaaagctgggtcAGCTTCATGTACCCCAACAA | Mycgr3\|106788  JGI  Ensembl |  |  | 100 % | chr\_1:638223 A->G; |  |
| 38 | ggggacaagtttgtacaaaaaagcaggcttgatgTCTCAAGACTCGCTGGCTCG, ggggaccactttgtacaagaaagctgggtcTCTGGCCACTGTCAGGCGTA | Mycgr3\|106912  JGI  Ensembl | KOG2992 Nucleolar GTPase/ATPase p130; |  | 100 % |  |  |
| 38 | ggggacaagtttgtacaaaaaagcaggcttgatgCATGACCTCAGAAAGAAGGC, ggggaccactttgtacaagaaagctgggtcGCGTGTGAAAGTTTGGCGTG | Mycgr3\|107024  JGI  Ensembl | KOG2842 Interferon-related protein PC4 like; | cysteine protease inhibitor activity; | 100 % |  |  |
| 38 | ggggacaagtttgtacaaaaaagcaggcttgatgAGCATTGGCACTGGCGGTCT, ggggaccactttgtacaagaaagctgggtcAAAGACATAGGTCTTCCTCT | Mycgr3\|107136  JGI  Ensembl | KOG1878 Nuclear receptor coregulator SMRT/SMRTER, contains Myb-like domains; |  | 100 % |  |  |
| 38 | ggggacaagtttgtacaaaaaagcaggcttgatgGCTCTGGCCGTTGGATCATC, ggggaccactttgtacaagaaagctgggtcCGCCATCATCGGATTGATGA | Mycgr3\|107138  JGI  Ensembl | KOG1703 Adaptor protein Enigma and related PDZ-LIM proteins; | zinc ion binding; | 100 % | chr\_1:2520792 T->C; |  |
| 38 | ggggacaagtttgtacaaaaaagcaggcttgatgGCCACAAGTACTGCTTTCGC, ggggaccactttgtacaagaaagctgggtcTCCTGCGCGGGCTTTCTTTG | Mycgr3\|108024  JGI  Ensembl | KOG2992 Nucleolar GTPase/ATPase p130; |  | 0.81202 % |  |  |
| 38 | ggggacaagtttgtacaaaaaagcaggcttgatgTTCAAACGCTTCTCCACCCG, ggggaccactttgtacaagaaagctgggtcCAGTAGCGGGAAAGCCAAAA | Mycgr3\|108649  JGI  Ensembl | KOG0503 Asparaginase; | amino acid metabolic process; asparaginase activity; | 100 % |  |  |
| 38 | ggggacaagtttgtacaaaaaagcaggcttgatgCCCGCTGTGAAGATCGCGGC, ggggaccactttgtacaagaaagctgggtcACCCGCAGCAGTAACCGCCG | Mycgr3\|108745  JGI  Ensembl | KOG0161 Myosin class II heavy chain; |  | 100 % | chr\_3:2497139 C->T; chr\_3:2497447 G->A; |  |
| 38 | ggggacaagtttgtacaaaaaagcaggcttgatgGGCACACTGACGGGCGGAGA, ggggaccactttgtacaagaaagctgggtcCGACCGGAACACCTTCACCT | Mycgr3\|108796  JGI  Ensembl | KOG4172 Predicted E3 ubiquitin ligase; | protein binding; zinc ion binding; | 0.68799 % |  |  |
| 38 | ggggacaagtttgtacaaaaaagcaggcttgatgGACTTCTGGGCTCGCCTGCT, ggggaccactttgtacaagaaagctgggtcTGCGCTTCGAACCTCCCTGT | Mycgr3\|108906  JGI  Ensembl | KOG3695 Uncharacterized conserved protein; |  | 100 % |  |  |
| 38 | ggggacaagtttgtacaaaaaagcaggcttgatgGCGGCGAGCTATGGACAGTC, ggggaccactttgtacaagaaagctgggtcAACTTCCATCCCATCGTCCC | Mycgr3\|109766  JGI  Ensembl |  |  | 0.64233 % |  |  |
| 38 | ggggacaagtttgtacaaaaaagcaggcttgatgGCCAAACGAACGGAGACGAC, ggggaccactttgtacaagaaagctgggtcCACTTGACTTTTTGACCTTG | Mycgr3\|109813  JGI  Ensembl | KOG4760 Uncharacterized conserved protein; |  | 100 % |  |  |
| 38 | ggggacaagtttgtacaaaaaagcaggcttgatgGCACCCGCGGCCATGGCTTT, ggggaccactttgtacaagaaagctgggtcAGTCCTCGGCAAGCCAAGGA | Mycgr3\|110078  JGI  Ensembl |  |  | 1.31853 % |  |  |
| 38 | ggggacaagtttgtacaaaaaagcaggcttgatgTGCGCAGACGACGATGACTT, ggggaccactttgtacaagaaagctgggtcGGACATGGCCTCGTACGGGT | Mycgr3\|110344  JGI  Ensembl | KOG1104 Nuclear cap-binding complex, subunit NCBP1/CBP80; |  | 72.43263 % | chr\_8:395490 G->A; chr\_8:395196 C->T; chr\_8:397121 C->T; |  |
| 38 | ggggacaagtttgtacaaaaaagcaggcttgatgGAGACTGTCTCCTACGTACC, ggggaccactttgtacaagaaagctgggtcAAATTCGAAACCACCCCACC | Mycgr3\|110389  JGI  Ensembl | KOG1216 von Willebrand factor and related coagulation proteins; | DNA binding; glucan 1,4-alpha-glucosidase activity;  nucleus;  transcription;  zinc ion binding; | 100 % |  |  |
| 38 | ggggacaagtttgtacaaaaaagcaggcttgatgTCAACGACGCGCCGAAATCC, ggggaccactttgtacaagaaagctgggtcCCACCTGCTGCTAGCACTGC | Mycgr3\|110431  JGI  Ensembl | KOG2294 Transcription factor of the Forkhead/HNF3 family; | nucleus; regulation of transcription, DNA-dependent;  sequence-specific DNA binding;  transcription factor activity; | 40.3854 % |  |  |
| 38 | ggggacaagtttgtacaaaaaagcaggcttgatgGCCGAATTACCCACACCGCC, ggggaccactttgtacaagaaagctgggtcCAACCCCAACGGATTCCCCG | Mycgr3\|110638  JGI  Ensembl |  |  | 100 % |  |  |
| 38 | ggggacaagtttgtacaaaaaagcaggcttgatgCCCTCCATACCTCTTGTGGA, ggggaccactttgtacaagaaagctgggtcGCTCTTATGACTCTCCTCTT | Mycgr3\|110861  JGI  Ensembl |  |  | 100 % |  |  |
| 38 | ggggacaagtttgtacaaaaaagcaggcttgatgGCGACCACCACCTCCACTAC, ggggaccactttgtacaagaaagctgggtcGCTTTTCGCGGAGAAGGGCG | Mycgr3\|111056  JGI  Ensembl |  |  | 0.71225 % |  |  |
| 38 | ggggacaagtttgtacaaaaaagcaggcttgatgGATGCACGGGCCGAACTCAA, ggggaccactttgtacaagaaagctgggtcCTGATATCGGTACTCCTGCG | Mycgr3\|111065  JGI  Ensembl | KOG2510 SWI-SNF chromatin-remodeling complex protein; | DNA polymerase activity; DNA-directed DNA polymerase activity; | 11.88738 % |  |  |
| 38 | ggggacaagtttgtacaaaaaagcaggcttgatgGATAACCTTCTTAACGCCGA, ggggaccactttgtacaagaaagctgggtcCTCCCGGAAATAGTGCTGCA | Mycgr3\|111257  JGI  Ensembl | KOG1075 FOG: Reverse transcriptase; |  | 15.27399 % |  |  |
| 38 | ggggacaagtttgtacaaaaaagcaggcttgatgCAGCAGCGTAGTGACCTTCA, ggggaccactttgtacaagaaagctgggtcGGCCTCTTTGTCAGTCCCGC | Mycgr3\|111306  JGI  Ensembl | KOG2992 Nucleolar GTPase/ATPase p130; |  | 100 % |  |  |
| 38 | ggggacaagtttgtacaaaaaagcaggcttgatgCCGCCCAAACGGGCTCGCAG, ggggaccactttgtacaagaaagctgggtcCTGCACCAGCCTCGAATCCC | Mycgr3\|111322  JGI  Ensembl | KOG1827 Chromatin remodeling complex RSC, subunit RSC1/Polybromo and related proteins; | DNA binding; histone acetyltransferase activity; | 100 % |  |  |
| 38 | ggggacaagtttgtacaaaaaagcaggcttgatgCCGCCTGCCGCTGATATGAC, ggggaccactttgtacaagaaagctgggtcCATGGCTGTGACAGAGCAGA | Mycgr3\|111598  JGI  Ensembl | KOG2615 Permease of the major facilitator superfamily; | integral to membrane; response to antibiotic;  tetracycline transport;  tetracycline:hydrogen antiporter activity;  transport;  transporter activity; | 0 % |  |  |
| 38 | ggggacaagtttgtacaaaaaagcaggcttgatgGCTCCATCCGCATCAGACCA, ggggaccactttgtacaagaaagctgggtcCCCCCCGACTCCCACCTTCT | Mycgr3\|111752  JGI  Ensembl | KOG2654 Uncharacterized conserved protein; |  | 100 % |  |  |
| 39 | ggggacaagtttgtacaaaaaagcaggcttgatgGAGAAAGACGATCTTCCTAA, ggggaccactttgtacaagaaagctgggtcCAACATCCTGACCATGCCCC | Mycgr3\|31572  JGI  Ensembl |  |  | 100 % |  |  |
| 39 | ggggacaagtttgtacaaaaaagcaggcttgatgCTGTCCGCATTCAAGGCACA, ggggaccactttgtacaagaaagctgggtcACTCCATGCAGCAGACCTTC | Mycgr3\|34254  JGI  Ensembl | KOG2063 Vacuolar assembly/sorting proteins VPS39/VAM6/VPS3; | small GTPase regulator activity; | 0.65975 % |  |  |
| 39 | ggggacaagtttgtacaaaaaagcaggcttgatgGCAGCCGAAGCCGAGACCGA, ggggaccactttgtacaagaaagctgggtcCTTGGATTGCGCGTATCGTG | Mycgr3\|39819  JGI  Ensembl | KOG0255 Synaptic vesicle transporter SVOP and related transporters (major facilitator superfamily); | integral to membrane; membrane; transport; transporter activity; | 100 % |  |  |
| 39 | ggggacaagtttgtacaaaaaagcaggcttgatgAAGCAGCGATTCTCCTCCTT, ggggaccactttgtacaagaaagctgggtcCCTCTTCTTGCCCCTGCCTC | Mycgr3\|43818  JGI  Ensembl | KOG2030 Predicted RNA-binding protein; | ATP-dependent DNA helicase activity; ATP-dependent RNA helicase activity;  ATP-dependent helicase activity;  ATPase activity;  ATPase activity, coupled;  ATPase activity, coupled to transmembrane movement of ions;  ATPase activity, coupled to transmembrane movement of substances;  ATPase activity, uncoupled;  DNA-dependent ATPase activity;  RNA-dependent ATPase activity;  protein-transmembrane transporting ATPase activity;  single-stranded DNA-dependent ATP-dependent DNA helicase activity; | 0.60661 % |  |  |
| 39 | ggggacaagtttgtacaaaaaagcaggcttgatgCCGGCCTCCAAGGCTCAAGA, ggggaccactttgtacaagaaagctgggtcGTCCTTCATCTCCTCATCCT | Mycgr3\|44761  JGI  Ensembl | KOG0961 Predicted Zn2+-dependent endopeptidase, insulinase superfamily; | metalloendopeptidase activity; proteolysis; zinc ion binding; | 100 % | chr\_7:908384 G->A; |  |
| 39 | ggggacaagtttgtacaaaaaagcaggcttgatgGACCAGGGACCAATCGATGC, ggggaccactttgtacaagaaagctgggtcCACCCCAGCACTAAACTCCG | Mycgr3\|47228  JGI  Ensembl | KOG1888 Putative phosphoinositide phosphatase; |  | 0 % |  |  |
| 39 | ggggacaagtttgtacaaaaaagcaggcttgatgGAGCAGCAGCTAGTGACTTT, ggggaccactttgtacaagaaagctgggtcCTCGCTCATAGTTCGCAGCT | Mycgr3\|47930  JGI  Ensembl | KOG2274 Predicted importin 9; | cytoplasm; intracellular protein transport;  nuclear pore;  nucleus;  protein import into nucleus, docking;  protein transporter activity; | 100 % | chr\_9:709370 G->A; |  |
| 39 | ggggacaagtttgtacaaaaaagcaggcttgatgGCCACCATCAAGGCCATCGA, ggggaccactttgtacaagaaagctgggtcCGTCCTCGCCTTGAGCCATG | Mycgr3\|48403  JGI  Ensembl | KOG1978 DNA mismatch repair protein - MLH2/PMS1/Pms2 family; | ATP binding; mismatch repair;  mismatched DNA binding; | 0 % |  |  |
| 39 | ggggacaagtttgtacaaaaaagcaggcttgatgCCGATCTCCGCAGAAAGAGA, ggggaccactttgtacaagaaagctgggtcTCCCGCCTTCGGCACCCCCT | Mycgr3\|48999  JGI  Ensembl | KOG2347 Sec5 subunit of exocyst complex; |  | 100 % |  |  |
| 39 | ggggacaagtttgtacaaaaaagcaggcttgatgGCACAACAGCAAAAGACAGT, ggggaccactttgtacaagaaagctgggtcTTTCTGCGAGTACTCCTCGA | Mycgr3\|49587  JGI  Ensembl | KOG1939 Oxoprolinase; | hydrolase activity; | 100 % | chr\_11:1586888 A->C; chr\_11:1587529 T->C; |  |
| 39 | ggggacaagtttgtacaaaaaagcaggcttgatgGCGGAATCACAGCAGTCAAA, ggggaccactttgtacaagaaagctgggtcCCTCTGCCACTCCTTCAACC | Mycgr3\|50095  JGI  Ensembl | KOG1178 Non-ribosomal peptide synthetase/alpha-aminoadipate reductase and related enzymes; | catalytic activity; cofactor binding;  fatty-acid ligase activity;  long-chain-fatty-acid-CoA ligase activity;  medium-chain-fatty-acid-CoA ligase activity;  metabolic process;  short-chain-fatty-acid-CoA ligase activity;  very-long-chain-fatty-acid-CoA ligase activity; | 100 % |  |  |
| 39 | ggggacaagtttgtacaaaaaagcaggcttgatgCGCGCGTCGATCCGACGCCA, ggggaccactttgtacaagaaagctgggtcTTCGCCGGACTGGATGAATG | Mycgr3\|51580  JGI  Ensembl | KOG0243 Kinesin-like protein; | ATP binding; microtubule associated complex;  microtubule motor activity;  microtubule-based movement; | 100 % | chr\_14:573865 T->C; |  |
| 39 | ggggacaagtttgtacaaaaaagcaggcttgatgGCCCCGAGTAGAGGGAAAGG, ggggaccactttgtacaagaaagctgggtcGTCGCTTTCCAACCAGTCAG | Mycgr3\|57932  JGI  Ensembl | KOG2179 Nucleotide excision repair complex XPC-HR23B, subunit XPC/DPB11; | damaged DNA binding; nucleotide-excision repair;  nucleus; | 100 % |  |  |
| 39 | ggggacaagtttgtacaaaaaagcaggcttgatgCCTCCGAAGGGCAAGACGAA, ggggaccactttgtacaagaaagctgggtcTCTACCATTCGCTCCGACCT | Mycgr3\|63782  JGI  Ensembl | KOG0161 Myosin class II heavy chain; | exo-alpha-sialidase activity; | 100 % |  |  |
| 39 | ggggacaagtttgtacaaaaaagcaggcttgatgCGCCGCCATCCTTCGCGAGC, ggggaccactttgtacaagaaagctgggtcCGACTTCCCACCACCCCCTC | Mycgr3\|65060  JGI  Ensembl | KOG1947 Leucine rich repeat proteins, some proteins contain F-box; | protein binding; | 100 % |  |  |
| 39 | ggggacaagtttgtacaaaaaagcaggcttgatgGCGGAAGCCAACCAACAATG, ggggaccactttgtacaagaaagctgggtcAGGATGGCCAGGTGGTCGTC | Mycgr3\|65067  JGI  Ensembl | KOG0056 Heavy metal exporter HMT1, ABC superfamily; | ATP binding; ATPase activity;  ATPase activity, coupled to transmembrane movement of substances;  integral to membrane;  nucleoside-triphosphatase activity;  nucleotide binding;  transport; | 100 % | chr\_1:771220 C->T; |  |
| 39 | ggggacaagtttgtacaaaaaagcaggcttgatgGCTTACCAGGGAGCTGGAGG, ggggaccactttgtacaagaaagctgggtcCCTCTTATTACAGCATCCCA | Mycgr3\|65104  JGI  Ensembl | KOG2571 Chitin synthase/hyaluronan synthase (glycosyltransferases); | chitin biosynthetic process; chitin synthase activity; | 0 % |  |  |
| 39 | ggggacaagtttgtacaaaaaagcaggcttgatgCTTGAAAGCCTGGCATTGCC, ggggaccactttgtacaagaaagctgggtcCAGTTCCTCTTCCTCCCAGT | Mycgr3\|65527  JGI  Ensembl | KOG1380 Heme A farnesyltransferase; | (S)-2,3-di-O-geranylgeranylglyceryl phosphate synthase activity; 1,4-dihydroxy-2-naphthoate octaprenyltransferase activity;  4-hydroxybenzoate octaprenyltransferase activity;  Sep-tRNA:Cys-tRNA synthase activity;  heme biosynthetic process;  integral to membrane;  prenyltransferase activity;  protein geranylgeranyltransferase activity;  protein prenyltransferase activity;  protoheme IX farnesyltransferase activity; | 0.60606 % |  |  |
| 39 | ggggacaagtttgtacaaaaaagcaggcttgatgGCCAAGACACGCAAAGGGAG, ggggaccactttgtacaagaaagctgggtcAATCTCAAACTCCTTCCAGC | Mycgr3\|66060  JGI  Ensembl | KOG2194 Aminopeptidases of the M20 family; | peptidase activity; proteolysis; | 100 % |  |  |
| 39 | ggggacaagtttgtacaaaaaagcaggcttgatgCACCTCCAAACCAGCGAAGG, ggggaccactttgtacaagaaagctgggtcGTACACCTCAAGCACATACC | Mycgr3\|66305  JGI  Ensembl | KOG1158 NADP/FAD dependent oxidoreductase; | electron transport; oxidoreductase activity; | 100 % |  |  |
| 39 | ggggacaagtttgtacaaaaaagcaggcttgatgAAGGACTCATTGCTGAGTTC, ggggaccactttgtacaagaaagctgggtcATTGCAGTCCGGACAGCAAG | Mycgr3\|66635  JGI  Ensembl | KOG1310 WD40 repeat protein; |  | 0 % |  |  |
| 39 | ggggacaagtttgtacaaaaaagcaggcttgatgTCTGAAATCCAACTCTACCT, ggggaccactttgtacaagaaagctgggtcCTCTTCCTCCTCGTTGCCTT | Mycgr3\|66670  JGI  Ensembl | KOG1967 DNA repair/transcription protein Mms19; |  | 2.0462 % |  |  |
| 39 | ggggacaagtttgtacaaaaaagcaggcttgatgCGCTTCACCAAGCCATCGTG, ggggaccactttgtacaagaaagctgggtcCCCATCCGTGATCATGGCCT | Mycgr3\|67073  JGI  Ensembl | KOG0973 Histone transcription regulator HIRA, WD repeat superfamily; | nucleus; regulation of transcription;  transcription regulator activity; | 0 % |  |  |
| 39 | ggggacaagtttgtacaaaaaagcaggcttgatgCTGGCCGTTTCGCATCGCTC, ggggaccactttgtacaagaaagctgggtcCTGGTATCCGTAATTCTGCG | Mycgr3\|67427  JGI  Ensembl | KOG3227 Calcium-responsive transcription coactivator; | histone acetyltransferase activity; | 100 % | chr\_1:5456767 G->A; | ``` 143	TGATGCTGGCCGTTTCGCATCGCTCCTCCGCGTTACCTACATTATTACACCGCGGTGTCATGGACGAGTCTGGCACCATCAACCCGTCTGCTCTTCACGCGTCCG	247 	\|\|\|\|\|\|\|\|\|\|\|\|\|\|\|\|\|\|\|\|\|\|\|\|\|\|\|\|\|\|\|\|\|\|\|\|\|\|\|\|\|\|\|\|\|\|\|\|\|\|\|\|\|\|\|\|\|\|\|\|\|\|\|\|\|\|\|\|\|\|\|\|\|\|\|\|\|\|\|\|\|\|\|\|\|\|\|\|\|\|\|\|\|\|\|\|\|\|\|\|\|\|\|\|\| 	TGATGCTGGCCGTTTCGCATCGCTCCTCCGCGTTACCTACATTATTACACCGCGGTGTCATGGACGAGTCTGGCACCATCAACCCGTCTGCTCTTCACGCGTCCG ```      ``` 247	GCCAACACTATCGCCCCCAATCTTCTCTCGCAAACCCAACCGAGCCCGCGCGGCATCAAACGCAGTCGCTCGCCTGATGGTTATACCGCGAATGATAACGACGGCGCAG	355 	\|\|\|\|\|\|\|\|\|\|\|\|\|\|\|\|\|\|\|\|\|\|\|\|\|\|\|\|\|\|\|\|\|\|\|\|\|\|\|\|\|\|\|\|\|\|\|\|\|\|\|\|\|\|\|\|\|\|\|\|\|\|\|\|\|\|\|\|\|\|\|\|\|\|\|\|\|\|\|\|\|\|\|\|\|\|\|\|\|\|\|\|\|\|\|\|\|\|\|\|\|\|\|\|\|\|\|\|\| 	GCCAACACTATCGCCCCCAATCTTCTCTCGCAAACCCAACCGAGCCCGCGCGGCATCAAACGCAGTCGCTCGCCTGATGGTTATACCGCGAATGATAACGACGGCGCAG ```      ``` 354	AGATGACGCCAAGACTGGAAGCATCAACATCAACAAGGCTCGGAAACTAAAGGCCGGCGAAAGTCCCGCGCCATCGCAAATGACGGTCGCCGCCAACATGCCGCCGCACGCGGTGCAGACACCGCAGCAACGCACATCCTCGCTGCCGCAGCAGTCACCCAGCCAGGCCTCGCCGCAACGTTCGACACCTGGATCCGCGACGAAGCCAGCCACGATCAAAGCGCTGCCGACCGTACGCGACCACACGACCGATCAACTCGGCCCAGAAGGCGACGAATACCTTCCACGCGAAATCGACGATCATGGAGAGACAAAAGTGTCGCGCGATGGATACCCGCTGGACGGTCGCAAATACAAATGTCGAACCTTTCAGGTTCCCAATCGAGGAGAGAAGCTTTTCATGCTGGCCACGGAATGTGCGCGCGTGCTGGGATATCGCGACTCTTACCTATTGTTCAACAAGAATCGGTCGCTGTTCAAGATCATCGCCAATCAGACGGAAAAGGACAACCTCATACATCAAGAGATACTCCCATACTCGTATCGATCGCGACAAATAGCCATCGTCACGGCTC-GCTCAATGTTTCGACAATTCGGCAGTCGCTTGATCGAAGGTGGCAG	974 	\|\|\|\|\|\|\|\|\|\|\|\|\|\|\|\|\|\|\|\|\|\|\|\|\|\|\|\|\|\|\|\|\|\|\|\|\|\|\|\|\|\|\|\|\|\|\|\|\|\|\|\|\|\|\|\|\|\|\|\|\|\|\|\|\|\|\|\|\|\|\|\|\|\|\|\|\|\|\|\|\|\|\|\|\|\|\|\|\|\|\|\|\|\|\|\|\|\|\|\|\|\|\|\|\|\|\|\|\|\|\|\|\|\|\|\|\|\|\|\|\|\|\|\|\|\|\|\|\|\|\|\|\|\|\|\|\|\|\|\|\|\|\|\|\|\|\|\|\|\|\|\|\|\|\|\|\|\|\|\|\|\|\|\|\|\|\|\|\|\|\|\|\|\|\|\|\|\|\|\|\|\|\|\|\|\|\|\|\|\|\|\|\|\|\|\|\|\|\|\|\|\|\|\|\|\|\|\|\|\|\|\|\|\|\|\|\|\|\|\|\|\|\|\|\|\|\|\|\|\|\|\|\|\|\|\|\|\|\|\|\|\|\|\|\|\|\|\|\|\|\|\|\|\|\|\|\|\|\|\|\|\|\|\|\|\|\|\|\|\|\|\|\|\|\|\|\|\|\|\|\|\|\|\|\|\|\|\|\|\|\|\|\|\|\|\|\|\|\|\|\|\|\|\|\|\|\|\|\|\|\|\|\|\|\|\|\|\|\|\|\|\|\|\|\|\|\|\|\|\|\|\|\|\|\|\|\|\|\|\|\|\|\|\|\|\|\|\|\|\|\|\|\|\|\|\|\|\|\|\|\|\|\|\|\|\|\|\|\|\|\|\| \|\|\|\|\|\|\|\|\|\|\|\|\|\|\|\|\|\|\|\|\|\|\|\|\|\|\|\|\|\|\|\|\|\|\|\|\|\|\|\|\|\|\|\|\|\|\|\|\|\|\|\|\|\|\|\|\|\|\|\|\|\|\|\|\|\|\|\|\|\|\|\|\|\|\|\|\|\|\|\|\|\|\|\|\|\|\|\|\|\|\|\|\|\|\|\|\|\|\|\|\|\|\|\|\|\| \|\|\|\|\|\|\|\|\|\| \|\|\|\|\|\|\|\|\|\|\|\|\|\|\|\|\|\| \| \|\|\|\|\|\|\|\|\|\| \|\|\|\|\|\|\|\| \|\|\|\|\|\|\|\|\|\|\|\|\|\|\|\|\|\|\|\|\|\|\|\| \|\|\|\| \|\|\|\|\|\|\|\|\| \|\|\| \|\|\|\|\| \|\|\|\|\|\|\|\|\|\| \| \|\| \|\|\|\|\|\|\| \|\|\|\|\|\|\|\|\|\|\|\| \|\|\| 	AGATGACGCCAAGACTGGAAGCATCAACATCAACAAGGCTCGGAAACTAAAGGCCGGCGAAAGTCCCGCGCCATCGCAAATGACGGTCGCCGCCAACATGCCGCCGCACGCGGTGCAGACACCGCAGCAACGCACATCCTCGCTGCCGCAGCAGTCACCCAGCCAGGCCTCGCCGCAACGTTCGACACCTGGATCCGCGACGAAGCCAGCCACGATCAAAGCGCTGCCGACCGTACGCGACCACACGACCGATCAACTCGGCCCAGAAGGCGACGAATACCTTCCACGCGAAATCGACGATCATGGAGAGACAAAAGTGTCGCGCGATGGATACCCGCTGGACGGTCGCAAATACAAATGTCGAACCTTTCA-GTTCCCAATCGAGGAGAGAAGCTTTTCATGCTGGCCACGGAATGTGCGCGCGTGCTGGGATATCGCGACTCTTACCTATTGTTCAACAAGAATCGGTCGCTGTTCA-GATCATCGCC-ATCAGACGGAAAAGGACA-C-TCATACATCA-GAGATACT-CCATACTCGTATCGATCGCGACAA-TAGC-ATCGTCACG-CTCCGCTCA-TGTTTCGACA-T-CG-CAGTCGC-TGATCGAAGGTGCCAG ``` |
| 39 | ggggacaagtttgtacaaaaaagcaggcttgatgGCGGGTCTGCTCGCGAGACA, ggggaccactttgtacaagaaagctgggtcTCGTCCGGTCCTCCCTCGCC | Mycgr3\|67825  JGI  Ensembl | KOG1134 Uncharacterized conserved protein; | membrane; | 0.68404 % |  |  |
| 39 | ggggacaagtttgtacaaaaaagcaggcttgatgGAGCTGGCCGACTCTCTGGA, ggggaccactttgtacaagaaagctgggtcAGTACCTGAGATGCCCAGAT | Mycgr3\|68302  JGI  Ensembl | KOG2089 Metalloendopeptidase family - saccharolysin & thimet oligopeptidase; | metalloendopeptidase activity; metallopeptidase activity;  proteolysis;  saccharolysin activity;  zinc ion binding; | 0.66313 % |  |  |
| 39 | ggggacaagtttgtacaaaaaagcaggcttgatgGACGAAGTCGAGAACCCAGA, ggggaccactttgtacaagaaagctgggtcATACCCCTCAATCGGACCCT | Mycgr3\|69142  JGI  Ensembl | KOG3680 Uncharacterized conserved protein; |  | 100 % | chr\_2:3665273 C->T; |  |
| 39 | ggggacaagtttgtacaaaaaagcaggcttgatgTCCAGCAACCTCCATCCCTC, ggggaccactttgtacaagaaagctgggtcAACACCATTATAAGGCCTCA | Mycgr3\|69304  JGI  Ensembl |  |  | 100 % | chr\_3:414639 G->T; chr\_3:414587 C->A; |  |
| 39 | ggggacaagtttgtacaaaaaagcaggcttgatgCTCGCCCTCACCCTCGCTGC, ggggaccactttgtacaagaaagctgggtcAAGCATCCTCTTCACCCCTC | Mycgr3\|69330  JGI  Ensembl | KOG4125 Acid trehalase; | alpha,alpha-trehalase activity; carbohydrate binding;  carbohydrate metabolic process;  catalytic activity; | 100 % | chr\_3:504400 G->A; |  |
| 39 | ggggacaagtttgtacaaaaaagcaggcttgatgTCTGCTACAGCTCTTCGTTT, ggggaccactttgtacaagaaagctgggtcCGTAGGCATGGGAGCAGCAG | Mycgr3\|69333  JGI  Ensembl | KOG2040 Glycine dehydrogenase (decarboxylating); | glycine cleavage system; glycine dehydrogenase (decarboxylating) activity;  glycine metabolic process; | 3.70828 % |  |  |
| 39 | ggggacaagtttgtacaaaaaagcaggcttgatgCTCGCCTCGGGGATGGCACT, ggggaccactttgtacaagaaagctgggtcGCTCATACTGAACCCATCAG | Mycgr3\|70017  JGI  Ensembl | KOG0056 Heavy metal exporter HMT1, ABC superfamily; | ATP binding; ATPase activity;  ATPase activity, coupled to transmembrane movement of substances;  integral to membrane;  nucleoside-triphosphatase activity;  nucleotide binding;  transport; | 100 % |  |  |
| 39 | ggggacaagtttgtacaaaaaagcaggcttgatgGTGTCCACAAAGACCGTTCC, ggggaccactttgtacaagaaagctgggtcCGAAAAGATATCACTCCAAT | Mycgr3\|70343  JGI  Ensembl | KOG1158 NADP/FAD dependent oxidoreductase; | FMN binding; NADPH-hemoprotein reductase activity;  electron carrier activity;  electron transport;  heme binding;  iron ion binding;  monooxygenase activity;  oxidoreductase activity; | 100 % |  |  |
| 39 | ggggacaagtttgtacaaaaaagcaggcttgatgAAGTTCTCTCACAGCATCCA, ggggaccactttgtacaagaaagctgggtcAAACCCAGCCGCAAGCATCA | Mycgr3\|71423  JGI  Ensembl | KOG1281 Na+/dicarboxylate, Na+/tricarboxylate and phosphate transporters; | membrane; sodium ion transport;  transporter activity; | 100 % | chr\_4:2140883 G->A; |  |
| 39 | ggggacaagtttgtacaaaaaagcaggcttgatgGCTGGCGAACAGCTACGATG, ggggaccactttgtacaagaaagctgggtcCAACTTCGCCCTCCCCCCAT | Mycgr3\|72489  JGI  Ensembl | KOG1206 Peroxisomal multifunctional beta-oxidation protein and related enzymes; | metabolic process; oxidoreductase activity; | 100 % | chr\_5:2105136 A->G; chr\_5:2105227 C->T; |  |
| 39 | ggggacaagtttgtacaaaaaagcaggcttgatgGATGATACGACTTCCGACGC, ggggaccactttgtacaagaaagctgggtcGTCCGTTATCACCCTCAAAG | Mycgr3\|72999  JGI  Ensembl | KOG2101 Intermediate filament-like protein, sorting nexins, and related proteins containing PX (PhoX) domain(s); | intracellular signaling cascade; | 100 % |  |  |
| 39 | ggggacaagtttgtacaaaaaagcaggcttgatgCTATCACCCGCCGACCGCTC, ggggaccactttgtacaagaaagctgggtcTCGCTCATATCTTGGCAAAT | Mycgr3\|73804  JGI  Ensembl | KOG2806 Chitinase; |  | 100 % | chr\_7:615485 T->C; |  |
| 39 | ggggacaagtttgtacaaaaaagcaggcttgatgCGTGGCTCCAATGGCTCATG, ggggaccactttgtacaagaaagctgggtcCTCAAACTTCACCCTCCACC | Mycgr3\|74623  JGI  Ensembl | KOG1066 Glucosidase II catalytic (alpha) subunit and related enzymes, glycosyl hydrolase family 31; | alpha-glucosidase activity; carbohydrate metabolic process;  hydrolase activity, hydrolyzing O-glycosyl compounds; | 1.29054 % |  |  |
| 39 | ggggacaagtttgtacaaaaaagcaggcttgatgGCTCCTTCATTCACCTCACG, ggggaccactttgtacaagaaagctgggtcTTGACCATGCTCGGCAACAG | Mycgr3\|75953  JGI  Ensembl | KOG2127 Calmodulin-binding protein CRAG, contains DENN domain; |  | 1.33333 % |  |  |
| 39 | ggggacaagtttgtacaaaaaagcaggcttgatgGACGACTTCAACACCCTCGA, ggggaccactttgtacaagaaagctgggtcCTTCCGGAAAGTCATGCTCT | Mycgr3\|76687  JGI  Ensembl | KOG2206 Exosome 3'-5' exoribonuclease complex, subunit PM/SCL-100 (Rrp6); | 3'-5' exonuclease activity; 3'-5'-exoribonuclease activity;  5'-3' exoribonuclease activity;  intracellular;  nucleic acid binding;  nucleobase, nucleoside, nucleotide and nucleic acid metabolic process; | 100 % |  |  |
| 39 | ggggacaagtttgtacaaaaaagcaggcttgatgCGCCTCGAACAGATTCCCGG, ggggaccactttgtacaagaaagctgggtcCACATAATACCTCGGCTCAT | Mycgr3\|77121  JGI  Ensembl | KOG3979 FGF receptor activating protein 1; |  | 100 % | chr\_11:1500725 A->G; chr\_11:1502159 G->A; |  |
| 39 | ggggacaagtttgtacaaaaaagcaggcttgatgGCTCAAACGCCATCATTGAA, ggggaccactttgtacaagaaagctgggtcACCGTCAGAAGTGTCCATCC | Mycgr3\|77616  JGI  Ensembl | KOG2042 Ubiquitin fusion degradation protein-2; | protein ubiquitination; ubiquitin ligase complex;  ubiquitin-protein ligase activity; | 100 % | chr\_12:1357161 G->A; |  |
| 39 | ggggacaagtttgtacaaaaaagcaggcttgatgGCTGCCTTTGACGAAAAGTC, ggggaccactttgtacaagaaagctgggtcCAACTCATCCCTCCCCTGCT | Mycgr3\|77815  JGI  Ensembl |  |  | 2.0955 % |  |  |
| 39 | ggggacaagtttgtacaaaaaagcaggcttgatgCTGTTTCTCTCCCTTCTTCC, ggggaccactttgtacaagaaagctgggtcCTTCAACTTAAGGAAGTACG | Mycgr3\|83794  JGI  Ensembl |  |  | 0 % |  |  |
| 39 | ggggacaagtttgtacaaaaaagcaggcttgatgGCGCCATCAAGCACAAACAA, ggggaccactttgtacaagaaagctgggtcAGCCAGTAGCCGTTGGATGT | Mycgr3\|85666  JGI  Ensembl | KOG1001 Helicase-like transcription factor HLTF/DNA helicase RAD5, DEAD-box superfamily; | ATP binding; ATP-dependent DNA helicase activity;  ATP-dependent RNA helicase activity;  ATP-dependent helicase activity;  ATPase activity;  ATPase activity, coupled;  ATPase activity, coupled to transmembrane movement of ions;  ATPase activity, coupled to transmembrane movement of substances;  ATPase activity, uncoupled;  DNA binding;  DNA-dependent ATPase activity;  RNA-dependent ATPase activity;  helicase activity;  nucleic acid binding;  protein binding;  protein-transmembrane transporting ATPase activity;  single-stranded DNA-dependent ATP-dependent DNA helicase activity;  zinc ion binding; | 0 % |  |  |
| 39 | ggggacaagtttgtacaaaaaagcaggcttgatgTCGAGCCTCGGCTCTGCTGT, ggggaccactttgtacaagaaagctgggtcTATCCGTCGCAGCTCAAACG | Mycgr3\|86272  JGI  Ensembl | KOG2024 Beta-Glucuronidase GUSB (glycosylhydrolase superfamily 2); | beta-galactosidase activity; beta-galactosidase complex;  carbohydrate metabolic process;  hydrolase activity, hydrolyzing O-glycosyl compounds; | 100 % | chr\_5:2483621 T->C; chr\_5:2483545 T->C; chr\_5:2484493 T->G; |  |
| 39 | ggggacaagtttgtacaaaaaagcaggcttgatgTTCTCTCCTCTCGTCGTCGC, ggggaccactttgtacaagaaagctgggtcTACCTGCAGGGTCGCAGCCA | Mycgr3\|88691  JGI  Ensembl | KOG2806 Chitinase; | glucan 1,4-alpha-glucosidase activity; | 1.25274 % |  |  |
| 39 | ggggacaagtttgtacaaaaaagcaggcttgatgGCCAATTCGACGTGGCTCCC, ggggaccactttgtacaagaaagctgggtcCTTGCACCTCTCCACAAGCA | Mycgr3\|89108  JGI  Ensembl | KOG0575 Polo-like serine/threonine protein kinase; | ATP binding; MAP kinase kinase kinase activity;  protein amino acid phosphorylation;  protein kinase activity;  protein serine/threonine kinase activity;  protein-tyrosine kinase activity; | 100 % |  |  |
| 39 | ggggacaagtttgtacaaaaaagcaggcttgatgCGTTCGTTCACAGCATTGGC, ggggaccactttgtacaagaaagctgggtcTCCATTCATCTCAAACAAAT | Mycgr3\|89487  JGI  Ensembl | KOG3599 Ca2+-modulated nonselective cation channel polycystin; |  | 100 % | chr\_1:3119657 G->A; |  |
| 39 | ggggacaagtttgtacaaaaaagcaggcttgatgTCCTGCGTACATCAGGATGT, ggggaccactttgtacaagaaagctgggtcGTCATCAGGCGTGATGTCCA | Mycgr3\|89719  JGI  Ensembl |  |  | 0 % |  |  |
| 39 | ggggacaagtttgtacaaaaaagcaggcttgatgTCTCTCGAAGCATTGCCCGC, ggggaccactttgtacaagaaagctgggtcATTCAAGCGCAAGAACTGAT | Mycgr3\|90149  JGI  Ensembl |  |  | 100 % |  |  |
| 39 | ggggacaagtttgtacaaaaaagcaggcttgatgGCGAGCCATTTCTCCGAACT, ggggaccactttgtacaagaaagctgggtcTGAAAACGATTCTACCACTC | Mycgr3\|90373  JGI  Ensembl |  |  | 100 % |  |  |
| 39 | ggggacaagtttgtacaaaaaagcaggcttgatgCATTTGCTCGCGCACTGGTG, ggggaccactttgtacaagaaagctgggtcGGAAGATTGAAAGTCGCCGA | Mycgr3\|92805  JGI  Ensembl | KOG4307 RNA binding protein RBM12/SWAN; |  | 100 % |  |  |
| 39 | ggggacaagtttgtacaaaaaagcaggcttgatgGACCAGCCCCGCGAGCGTCC, ggggaccactttgtacaagaaagctgggtcACCGTACCCATTCGAAGCGC | Mycgr3\|92834  JGI  Ensembl |  |  | 0.5886 % |  |  |
| 39 | ggggacaagtttgtacaaaaaagcaggcttgatgTCTCGCGGCTTGGACGGCAT, ggggaccactttgtacaagaaagctgggtcTCTGATCTGTTGCTGCCTGG | Mycgr3\|92941  JGI  Ensembl | KOG4781 Uncharacterized conserved protein; |  | 100 % |  |  |
| 39 | ggggacaagtttgtacaaaaaagcaggcttgatgACTTCGCACATCGTCCAGAT, ggggaccactttgtacaagaaagctgggtcTCTCCTCCGACCAGAGTCTG | Mycgr3\|93083  JGI  Ensembl |  |  | 100 % | chr\_5:748012 T->A; chr\_5:748705 G->A; |  |
| 39 | ggggacaagtttgtacaaaaaagcaggcttgatgGCACGACCCAGGCAATTTGG, ggggaccactttgtacaagaaagctgggtcGACCTCGAATCCCAACTCAT | Mycgr3\|93373  JGI  Ensembl | KOG2458 Endoplasmic reticulum protein EP58, contains filamin rod domain and KDEL motif; |  | 0.672 % |  |  |
| 39 | ggggacaagtttgtacaaaaaagcaggcttgatgGACCTCCGCACATATGTTTC, ggggaccactttgtacaagaaagctgggtcGATCTTCGGCATCTTCCTAT | Mycgr3\|94085  JGI  Ensembl | KOG0923 mRNA splicing factor ATP-dependent RNA helicase; | ATP-dependent DNA helicase activity; ATP-dependent RNA helicase activity;  ATP-dependent helicase activity;  ATPase activity;  ATPase activity, coupled;  ATPase activity, coupled to transmembrane movement of ions;  ATPase activity, coupled to transmembrane movement of substances;  ATPase activity, uncoupled;  DNA-dependent ATPase activity;  RNA-dependent ATPase activity;  helicase activity;  protein-transmembrane transporting ATPase activity;  single-stranded DNA-dependent ATP-dependent DNA helicase activity; | 1.20594 % |  |  |
| 39 | ggggacaagtttgtacaaaaaagcaggcttgatgAAACGAAAGCGCCCAGACCG, ggggaccactttgtacaagaaagctgggtcAGCAGCGTAGTAACAGTCTG | Mycgr3\|94324  JGI  Ensembl |  |  | 100 % | chr\_7:521776 C->T; chr\_7:521629 T->G; |  |
| 39 | ggggacaagtttgtacaaaaaagcaggcttgatgTCACGAGCACTGAGCTCCGA, ggggaccactttgtacaagaaagctgggtcAGGGGCCCCGGCAGGGACAG | Mycgr3\|95261  JGI  Ensembl |  |  | 100 % | chr\_8:1427580 C->T; chr\_8:1428599 G->A; |  |
| 39 | ggggacaagtttgtacaaaaaagcaggcttgatgGGATTCATTTCTGCACAGTA, ggggaccactttgtacaagaaagctgggtcATCGCCTACGCTCGCAACGC | Mycgr3\|95274  JGI  Ensembl | KOG1965 Sodium/hydrogen exchanger protein; | integral to membrane; regulation of pH;  sodium ion transport;  sodium:hydrogen antiporter activity;  solute:hydrogen antiporter activity; | 100 % |  |  |
| 39 | ggggacaagtttgtacaaaaaagcaggcttgatgGCGGACGATGCTCTGAGCTG, ggggaccactttgtacaagaaagctgggtcCAGCTCGCTGCTTTGTTGAG | Mycgr3\|95893  JGI  Ensembl | KOG0613 Projectin/twitchin and related proteins; | biosynthetic process; catalytic activity;  pyridoxal phosphate binding; | 0 % |  |  |
| 39 | ggggacaagtttgtacaaaaaagcaggcttgatgGAGGGCGCTGACCATTGTTT, ggggaccactttgtacaagaaagctgggtcGGGCTTGGATCGACGATCCG | Mycgr3\|96362  JGI  Ensembl |  |  | 1.45358 % |  |  |
| 39 | ggggacaagtttgtacaaaaaagcaggcttgatgGACTCTCGCAGACGAATGAG, ggggaccactttgtacaagaaagctgggtcACGGCTCTTGTAGCATCTCG | Mycgr3\|97730  JGI  Ensembl |  |  | 100 % | chr\_16:245198 C->T; |  |
| 39 | ggggacaagtttgtacaaaaaagcaggcttgatgCGCACGAAATCCAGAGCACC, ggggaccactttgtacaagaaagctgggtcAGCCCAAGCAGCTCCGAGAG | Mycgr3\|97837  JGI  Ensembl | KOG0971 Microtubule-associated protein dynactin DCTN1/Glued; | chromosome, pericentric region; meiotic chromosome segregation;  nucleus; | 0 % |  |  |
| 39 | ggggacaagtttgtacaaaaaagcaggcttgatgGAAGTGGCGAACATGAGGCG, ggggaccactttgtacaagaaagctgggtcACGGGGGGCGTCGAGACGGT | Mycgr3\|98085  JGI  Ensembl |  |  | 100 % |  |  |
| 39 | ggggacaagtttgtacaaaaaagcaggcttgatgGCGCTCCCCAAACACAGCAG, ggggaccactttgtacaagaaagctgggtcATAGCGCCTCTCATATCCAC | Mycgr3\|98224  JGI  Ensembl | KOG1995 Conserved Zn-finger protein; |  | 100 % |  |  |
| 39 | ggggacaagtttgtacaaaaaagcaggcttgatgGCAGGGATTGAACAAATCAT, ggggaccactttgtacaagaaagctgggtcAAGCTCAAACACCTCCTGCA | Mycgr3\|98235  JGI  Ensembl | KOG1737 Oxysterol-binding protein; | steroid metabolic process; | 1.32013 % |  |  |
| 39 | ggggacaagtttgtacaaaaaagcaggcttgatgCAGGTCTCCACGATGGCCAA, ggggaccactttgtacaagaaagctgggtcGACGAACGGATCTGCTGCCA | Mycgr3\|99123  JGI  Ensembl | KOG0874 Sphingolipid hydroxylase; | catalytic activity; metabolic process; | 0 % |  |  |
| 39 | ggggacaagtttgtacaaaaaagcaggcttgatgGGCAACAAGCCCGACACTGC, ggggaccactttgtacaagaaagctgggtcGACCTTCTCCTTCTCTTCGT | Mycgr3\|100176  JGI  Ensembl | KOG0202 Ca2+ transporting ATPase; | ATP binding; ATPase activity, coupled to transmembrane movement of ions, phosphorylative mechanism; calcium-transporting ATPase activity;  catalytic activity;  cation transport;  hydrolase activity, acting on acid anhydrides, catalyzing transmembrane movement of substances; integral to membrane;  membrane;  metabolic process;  transport; | 0 % |  |  |
| 39 | ggggacaagtttgtacaaaaaagcaggcttgatgGCAGTCGGTCAGCTTAAGAG, ggggaccactttgtacaagaaagctgggtcCTCATCGTACTCAACAGTCA | Mycgr3\|100421  JGI  Ensembl | KOG2408 Peroxidase/oxygenase; | electron transport; heme binding;  peroxidase activity;  response to oxidative stress; | 100 % | chr\_6:540468 C->T; |  |
| 39 | ggggacaagtttgtacaaaaaagcaggcttgatgTCCCGCGCCAATGCCCCGAT, ggggaccactttgtacaagaaagctgggtcCACTCTCCGACTGTAACCGC | Mycgr3\|100515  JGI  Ensembl | KOG0202 Ca2+ transporting ATPase; | ATP binding; ATPase activity, coupled to transmembrane movement of ions, phosphorylative mechanism; calcium ion transport;  calcium-transporting ATPase activity;  catalytic activity;  cation transport;  hydrolase activity, acting on acid anhydrides, catalyzing transmembrane movement of substances; integral to membrane;  membrane;  metabolic process;  transport; | 100 % |  |  |
| 39 | ggggacaagtttgtacaaaaaagcaggcttgatgGCAGCCCAGGAGAGTGGTAA, ggggaccactttgtacaagaaagctgggtcCGCCTTCCCTCTGCCAGCAC | Mycgr3\|100579  JGI  Ensembl | KOG2308 Phosphatidic acid-preferring phospholipase A1, contains DDHD domain; | 1-oxa-2-oxocycloheptane lactonase activity; 3,4-dihydrocoumarin hydrolase activity;  butyrolactone hydrolase activity;  catalytic activity;  lipase activity;  metal ion binding;  phospholipase activity;  sulfolactone hydrolase activity; | 0.63898 % |  |  |
| 39 | ggggacaagtttgtacaaaaaagcaggcttgatgGGCGCCTCAGAGAGCAAGCT, ggggaccactttgtacaagaaagctgggtcCTTCATACTAATACTGCTGA | Mycgr3\|101041  JGI  Ensembl | KOG2226 Proteins containing regions of low-complexity; |  | 0 % |  |  |
| 39 | ggggacaagtttgtacaaaaaagcaggcttgatgGCCACTCCCACACAACCTCA, ggggaccactttgtacaagaaagctgggtcTTGAGGAGACTTGGCGCCTC | Mycgr3\|101108  JGI  Ensembl | KOG4198 RNA-binding Ran Zn-finger protein and related proteins; | intracellular; nucleic acid binding;  zinc ion binding; | 100 % |  |  |
| 39 | ggggacaagtttgtacaaaaaagcaggcttgatgGCCTTCCTCGCCGATCCCCC, ggggaccactttgtacaagaaagctgggtcAGCCTCGTTCCTCGGTCGTC | Mycgr3\|107149  JGI  Ensembl | KOG0113 U1 small nuclear ribonucleoprotein (RRM superfamily); | FK506-sensitive peptidyl-prolyl cis-trans isomerase; cyclophilin; cyclophilin-type peptidyl-prolyl cis-trans isomerase activity;  peptidyl-prolyl cis-trans isomerase activity; | 100 % | chr\_1:2570685 G->A; |  |
| 39 | ggggacaagtttgtacaaaaaagcaggcttgatgGAGGGTGCGTTCACCCATGT, ggggaccactttgtacaagaaagctgggtcATCATAGCCACCGACGCCGA | Mycgr3\|107419  JGI  Ensembl | KOG1802 RNA helicase nonsense mRNA reducing factor (pNORF1); |  | 0.64103 % |  |  |
| 39 | ggggacaagtttgtacaaaaaagcaggcttgatgCACACCATGCGTTGCATCGT, ggggaccactttgtacaagaaagctgggtcCTCCGACAACGTCTGGTTCC | Mycgr3\|107565  JGI  Ensembl | KOG2230 Predicted beta-mannosidase; | beta-mannosidase activity; carbohydrate metabolic process;  hydrolase activity, hydrolyzing O-glycosyl compounds; | 0 % |  |  |
| 39 | ggggacaagtttgtacaaaaaagcaggcttgatgGCCAAAGCACTCCCCTCCAG, ggggaccactttgtacaagaaagctgggtcCTGCGGTCTTCCCAAGACGT | Mycgr3\|107834  JGI  Ensembl | KOG2820 FAD-dependent oxidoreductase; |  | 100 % |  |  |
| 39 | ggggacaagtttgtacaaaaaagcaggcttgatgGACCGGAGATCTCTGCTGGA, ggggaccactttgtacaagaaagctgggtcCTTTTCCCGACGGACAGGGA | Mycgr3\|107859  JGI  Ensembl | KOG4157 beta-1,6-N-acetylglucosaminyltransferase, contains WSC domain; |  | 100 % |  |  |
| 39 | ggggacaagtttgtacaaaaaagcaggcttgatgGATGTGCCTGCTGCAGACTT, ggggaccactttgtacaagaaagctgggtcTGCTGCAGCGACCTCGATAC | Mycgr3\|107920  JGI  Ensembl | KOG1869 Splicing coactivator SRm160/300, subunit SRm300; |  | 100 % |  |  |
| 39 | ggggacaagtttgtacaaaaaagcaggcttgatgGCGTGGGAGCAACTCTCCAT, ggggaccactttgtacaagaaagctgggtcCCGTCCGACATTCGTGCCCC | Mycgr3\|108173  JGI  Ensembl | KOG4505 Na+/H+ antiporter; | integral to membrane; regulation of pH;  solute:hydrogen antiporter activity; | 100 % |  |  |
| 39 | ggggacaagtttgtacaaaaaagcaggcttgatgGCGACCGCCACGACTGCGCG, ggggaccactttgtacaagaaagctgggtcGTTGTCCGAGTCGTCCGAAT | Mycgr3\|108294  JGI  Ensembl | KOG2043 Signaling protein SWIFT and related BRCT domain proteins; |  | 0 % |  |  |
| 39 | ggggacaagtttgtacaaaaaagcaggcttgatgAAGCGAGAGAGGTCGAGCAG, ggggaccactttgtacaagaaagctgggtcCTCGGCGGTATTGTTCTGTG | Mycgr3\|108298  JGI  Ensembl |  |  | 100 % |  |  |
| 39 | ggggacaagtttgtacaaaaaagcaggcttgatgTCTCCCCCAAAATACGACCT, ggggaccactttgtacaagaaagctgggtcTGTTGAGATCATGACACTGT | Mycgr3\|108445  JGI  Ensembl | KOG2450 Aldehyde dehydrogenase; |  | 100 % |  |  |
| 39 | ggggacaagtttgtacaaaaaagcaggcttgatgGATCCCGACTCTTCTCAGAG, ggggaccactttgtacaagaaagctgggtcAAGCCGCCTCGTTCGCTTCT | Mycgr3\|108675  JGI  Ensembl | KOG0391 SNF2 family DNA-dependent ATPase; | glucan 1,4-alpha-glucosidase activity; | 1.8775 % |  |  |
| 39 | ggggacaagtttgtacaaaaaagcaggcttgatgGCCGACGAGCCGGAGCCACC, ggggaccactttgtacaagaaagctgggtcATGGTTCACCGCCGTGAAAA | Mycgr3\|108739  JGI  Ensembl | KOG0825 PHD Zn-finger protein; | UDP-N-acetylmuramoylalanyl-D-glutamyl-2,6-diaminopimelate-D-alanyl-D-alanine ligase activity; coenzyme F420-0 gamma-glutamyl ligase activity;  coenzyme F420-2 alpha-glutamyl ligase activity;  response to stress;  ribosomal S6-glutamic acid ligase activity; | 0 % |  |  |
| 39 | ggggacaagtttgtacaaaaaagcaggcttgatgCCGTCAACGCCCAGTCATCG, ggggaccactttgtacaagaaagctgggtcCTTCGGATTTGAGTTTTGGG | Mycgr3\|108985  JGI  Ensembl | KOG2276 Metalloexopeptidases; | metallopeptidase activity; proteolysis; | 1.23839 % |  |  |
| 39 | ggggacaagtttgtacaaaaaagcaggcttgatgTGGAGATGCGATGCTTCGTC, ggggaccactttgtacaagaaagctgggtcAATGCTCTCCACCCGATACG | Mycgr3\|109051  JGI  Ensembl |  |  | 100 % | chr\_4:1062030 G->A; chr\_4:1063928 A->G; chr\_4:1063584 C->T; |  |
| 39 | ggggacaagtttgtacaaaaaagcaggcttgatgCACAAGATCTCCAACTTCAC, ggggaccactttgtacaagaaagctgggtcATGCGGAGGCCCAATCGGGG | Mycgr3\|111066  JGI  Ensembl | KOG2208 Vigilin; | RNA binding; | 99.90945 % |  |  |
| 39 | ggggacaagtttgtacaaaaaagcaggcttgatgACTATGGCCGCGAACGGATA, ggggaccactttgtacaagaaagctgggtcGTGAAAGCCATGGGTCGTAT | Mycgr3\|111074  JGI  Ensembl |  |  | 100 % |  | ``` 348	GCGTGCATACCATGCAGACAACGCAAAGTCCGCTGCGACTTGGGCTCCGTCGACAACCCTCACGATCCACCATGCACACGCTGTCGTCGCGAGGCAAAGGAGTGCTACTTCTCCGCCACACGACGCAAGAAAAAGAATGCAGGCGACGAGAATACTGGTTCGGAGGCAGGCGACGATAATGTAACATACGAGATCCGCAGCAACAGAAAGCGACCGAGGACATCCGCGCCGCCTGAGGTGGCTGTCGAAGAGGACGACTACGACTACGACGATCCACCACGGACACCAGGCGGTTCCATCGGACGGACACAGCCTTTGCGCAGACCACCAGCGCCGAAGCCAATTCAGTATGGCGAAGAAGATGAGAAGCAAAGCGAACATACTACAGCACTGTTGCAGCAAGCCGAGGTGCATGGAGGTCACGATGCACTGAAGCTCTTGTACTCTGCAGCGGTGCACGGAAGAGCGGACAGCAGCACGAGCAGCGGTGTCGCGCGGCCCGCCTTCACTGGTTTCACTGCATCCTCTCTGCCGTCTGCAACTAGTCCGATCATGGAAAAGGCCGGTAATGTAAGCTATACCAATGGCCCGCTGGCCATGTTCCAGCACGATGGGACGTGGAAGACTCCCACTGACCAGATGTCGTCCAAGCCTGCGAACTTTGCGCGTATGCCGAGCAATACTCCCGATTGCTCCACTGCACTCAAAGCATGGACCC-GGTTCAAGTTCGTCCGCGCCGGCTGGTTCACCGCCCAAGAAGGCATCGATTACATCCTCTACTTCTACAAGCATTTGGCTCCG-CTCACGCCTGTTGCGCTTCCGGACTTTCGGCAGCCTTCGACAC-A-CAAGAAGATGCTGGAAGAAGAGCCGATGCTGACTGTATCCATGCTT-A-CGATTGCCTCACGTC-ATA	1257 	\|\|\|\|\|\|\|\|\|\|\|\|\|\|\|\|\|\|\|\|\|\|\|\|\|\|\|\|\|\|\|\|\|\|\|\|\|\|\|\|\|\|\|\|\|\|\|\|\|\|\|\|\|\|\|\|\|\|\|\|\|\|\|\|\|\|\|\|\|\|\|\|\|\|\|\|\|\|\|\|\|\|\|\|\|\|\|\|\|\|\|\|\|\|\|\|\|\|\|\|\|\|\|\|\|\|\|\|\|\|\|\|\|\|\|\|\|\|\|\|\|\|\|\|\|\|\|\|\|\|\|\|\|\|\|\|\|\|\|\|\|\|\|\|\|\|\|\|\|\|\|\|\|\|\|\|\|\|\|\|\|\|\|\|\|\|\|\|\|\|\|\|\|\|\|\|\|\|\|\|\|\|\|\|\|\|\|\|\|\|\|\|\|\|\|\|\|\|\|\|\|\|\|\|\|\|\|\|\|\|\|\|\|\|\|\|\|\|\|\|\|\|\|\|\|\|\|\|\|\|\|\|\|\|\|\|\|\|\|\|\|\|\|\|\|\|\|\|\|\|\|\|\|\|\|\|\|\|\|\|\|\|\|\|\|\|\|\|\|\|\|\|\|\|\|\|\|\|\|\|\|\|\|\|\|\|\|\|\|\|\|\|\|\|\|\|\|\|\|\|\|\|\|\|\|\|\|\|\|\|\|\|\|\|\|\|\|\|\|\|\|\|\|\|\|\|\|\|\|\|\|\|\|\|\|\|\|\|\|\|\|\|\|\|\|\|\|\|\|\|\|\|\|\|\|\|\|\|\|\|\|\|\|\|\|\|\|\|\|\|\|\|\|\|\|\|\|\|\|\|\|\|\|\|\|\|\|\|\|\|\|\|\|\|\|\|\|\|\|\|\|\|\|\|\|\|\|\|\|\|\|\|\|\|\|\|\|\|\|\|\|\|\|\|\|\|\|\|\|\|\|\|\|\|\|\|\|\|\|\|\|\|\|\|\|\|\|\|\|\|\|\|\|\|\|\|\|\|\|\|\|\|\|\|\|\|\|\|\|\|\|\|\|\|\|\|\|\|\|\|\|\|\|\|\|\|\|\|\|\|\|\|\|\|\|\|\|\|\|\|\|\|\|\|\|\|\|\|\|\|\|\|\|\|\|\|\|\|\|\|\|\|\|\|\|\|\|\|\|\|\|\|\|\|\|\|\|\|\|\|\|\|\|\|\|\|\|\|\|\|\|\|\|\|\|\|\|\|\|\|\|\|\|\|\|\|\|\|\|\|\|\|\|\|\|\|\|\|\|\|\|\|\|\|\|\|\|\|\|\|\|\|\|\|\|\|\|\|\|\|\|\|\|\|\|\|\|\|\|\|\|\|\|\|\|\|\|\|\|\|\|\|\|\|\|\|\|\|\|\|\|\|\|\|\|\|\|\|\|\|\|\|\|\|\|\|\|\|\|\|\|\|\|\|\|\|\|\|\|\|\|\|\|\|\|\|\|\|\|\|\|\|\|\|\|\|\|\|\|\|\|\|\|\|\|\|\|\|\|\|\|\|\|\|\|\|\|\|\|\|\|\|\|\|\|\|\|\|\|\|\|\|\|\|\|\|\|\| \|\|\|\|\|\|\|\|\|\|\|\|\|\|\|\|\|\|\|\|\|\|\|\|\|\|\|\|\|\|\|\|\|\|\|\|\|\|\|\|\|\|\|\|\|\|\|\|\|\|\|\|\|\|\|\|\|\|\|\|\|\|\|\|\|\|\|\| \|\|\|\|\|\|\|\|\|\|\|\|\|\| \|\|\|\|\|\|\|\|\|\|\|\|\|\|\|\|\|\|\|\|\|\|\|\|\|\|\|\|\|\|\|\|\|\| \|\|\|\|\| \|\| \| \|\|\|\|\|\|\|\|\|\|\|\|\| \| \| \|\|\|\|\| \|\|\|\|\|\|\|\|\|\|\|\|\|\|\|\|\|\|\|\|\|\| \| \|\|\|\|  \|\|\|\|\|\|\|\|\| \|\|\| 	GCGTGCATACCATGCAGACAACGCAAAGTCCGCTGCGACTTGGGCTCCGTCGACAACCCTCACGATCCACCATGCACACGCTGTCGTCGCGAGGCAAAGGAGTGCTACTTCTCCGCCACACGACGCAAGAAAAAGAATGCAGGCGACGAGAATACTGGTTCGGAGGCAGGCGACGATAATGTAACATACGAGATCCGCAGCAACAGAAAGCGACCGAGGACATCCGCGCCGCCTGAGGTGGCTGTCGAAGAGGACGACTACGACTACGACGATCCACCACGGACACCAGGCGGTTCCATCGGACGGACACAGCCTTTGCGCAGACCACCAGCGCCGAAGCCAATTCAGTATGGCGAAGAAGATGAGAAGCAAAGCGAACATACTACAGCACTGTTGCAGCAAGCCGAGGTGCATGGAGGTCACGATGCACTGAAGCTCTTGTACTCTGCAGCGGTGCACGGAAGAGCGGACAGCAGCACGAGCAGCGGTGTCGCGCGGCCCGCCTTCACTGGTTTCACTGCATCCTCTCTGCCGTCTGCAACTAGTCCGATCATGGAAAAGGCCGGTAATGTAAGCTATACCAATGGCCCGCTGGCCATGTTCCAGCACGATGGGACGTGGAAGACTCCCACTGACCAGATGTCGTCCAAGCCTGCGAACTTTGCGCGTATGCCGAGCAATACTCCCGATTGCTCCACTGCACTCAAAGCATGGACCCGGGTTCAAGTTCGTCCGCGCCGGCTGGTTCACCGCCCAAGAAGGCATCGATTACATCCTCTACTTCTAC-AGCATTTGGCTCCGCCTCACGCCTGTTGCGCTTCCGGACTTTCGGCAGC-TTCGAAACCACCAAGAAGATGCTG-A-G-AGAGCGGATGCTGACTGTATCCATGCTTTAACGATGCCCTCACGTCTATA ```      ``` 228	ATGACTATGGCCGCGAACGGATATGCGCTGGCACAAGCAGACCAGTATACTGCGCGATCTCAGTCGCCAAAGACGCAAGCCGCTGCTGGCCTTGCGCCGGGACATCAGCGCACATATCAGG	348 	\|\|\|\|\|\|\|\|\|\|\|\|\|\|\|\|\|\|\|\|\|\|\|\|\|\|\|\|\|\|\|\|\|\|\|\|\|\|\|\|\|\|\|\|\|\|\|\|\|\|\|\|\|\|\|\|\|\|\|\|\|\|\|\|\|\|\|\|\|\|\|\|\|\|\|\|\|\|\|\|\|\|\|\|\|\|\|\|\|\|\|\|\|\|\|\|\|\|\|\|\|\|\|\|\|\|\|\|\|\|\|\|\|\|\|\|\|\|\|\|\| 	ATGACTATGGCCGCGAACGGATATGCGCTGGCACAAGCAGACCAGTATACTGCGCGATCTCAGTCGCCAAAGACGCAAGCCGCTGCTGGCCTTGCGCCGGGACATCAGCGCACATATCAGG ``` |
| 39 | ggggacaagtttgtacaaaaaagcaggcttgatgGCTTCCATATTCACGTACGA, ggggaccactttgtacaagaaagctgggtcCTCGCAGACACGCGTGGTGA | Mycgr3\|111226  JGI  Ensembl | KOG1143 Predicted translation elongation factor; |  | 0.56697 % |  |  |
| 39 | ggggacaagtttgtacaaaaaagcaggcttgatgAGACAATATCGCCAACCATG, ggggaccactttgtacaagaaagctgggtcAACAGCCGTACTCCTCCCCT | Mycgr3\|111431  JGI  Ensembl | KOG3599 Ca2+-modulated nonselective cation channel polycystin; |  | 100 % |  |  |
| 39 | ggggacaagtttgtacaaaaaagcaggcttgatgACGCTGTTCGAAAAAAGTAT, ggggaccactttgtacaagaaagctgggtcCATCCCCTCCCCCAAACTCG | Mycgr3\|111677  JGI  Ensembl | KOG0161 Myosin class II heavy chain; |  | 100 % |  |  |
| 40 | ggggacaagtttgtacaaaaaagcaggcttgatgCCGATCGTTCACATCG, ggggaccactttgtacaagaaagctgggtcGAATTTGCCAGGCTCAAAGT | Mycgr3\|31813  JGI  Ensembl |  |  | 100 % |  |  |
| 40 | ggggacaagtttgtacaaaaaagcaggcttgatgTCCGACCCGTCCTCAT, ggggaccactttgtacaagaaagctgggtcTAGCGAGGCGGCCGCTTCAA | Mycgr3\|32106  JGI  Ensembl |  |  | 100 % |  |  |
| 40 | ggggacaagtttgtacaaaaaagcaggcttgatgACACGCGCCGCACAGA, ggggaccactttgtacaagaaagctgggtcATCCACACTAACACCCTTCG | Mycgr3\|37747  JGI  Ensembl | KOG4841 Dolichol-phosphate mannosyltransferase, subunit 3; |  | 100 % |  |  |
| 40 | ggggacaagtttgtacaaaaaagcaggcttgatgAACTTCTTCACCACCA, ggggaccactttgtacaagaaagctgggtcAAGCCAGACACATCCAGTTC | Mycgr3\|41491  JGI  Ensembl |  |  | 100 % |  |  |
| 40 | ggggacaagtttgtacaaaaaagcaggcttgatgCAGCGCGCACCACCCT, ggggaccactttgtacaagaaagctgggtcAGAATCCCGACTTCCCATGA | Mycgr3\|44402  JGI  Ensembl | KOG1142 Transcription initiation factor TFIID, subunit TAF12 (also component of histone acetyltransferase SAGA); | transcription factor TFIID complex; transcription initiation;  transcription initiation factor activity; | 100 % |  |  |
| 40 | ggggacaagtttgtacaaaaaagcaggcttgatgCAGTCCTTCAACCTGC, ggggaccactttgtacaagaaagctgggtcTAGAAGCCCTTTCCTCGCGA | Mycgr3\|46900  JGI  Ensembl |  |  | 100 % |  |  |
| 40 | ggggacaagtttgtacaaaaaagcaggcttgatgCTCGGCATCCTCAAAT, ggggaccactttgtacaagaaagctgggtcTTCCCTTAAGCTCGCAATTC | Mycgr3\|50731  JGI  Ensembl | KOG3808 Uncharacterized conserved protein; |  | 100 % |  |  |
| 40 | ggggacaagtttgtacaaaaaagcaggcttgatgGCGAGCACAACAGCAT, ggggaccactttgtacaagaaagctgggtcCCCAGTGTCCTTCTGCCGCA | Mycgr3\|51114  JGI  Ensembl | KOG3801 Uncharacterized conserved protein BCN92; |  | 100 % |  |  |
| 40 | ggggacaagtttgtacaaaaaagcaggcttgatgGACCAAGCCAAAGACC, ggggaccactttgtacaagaaagctgggtcTGCGCCGCCAACGAGGATGT | Mycgr3\|51258  JGI  Ensembl | KOG3498 Preprotein translocase, gamma subunit; | P-P-bond-hydrolysis-driven protein transmembrane transporter activity; intracellular protein transport;  membrane;  protein targeting;  protein transport; | 100 % |  |  |
| 40 | ggggacaagtttgtacaaaaaagcaggcttgatgAGCTTCACCTTTATCA, ggggaccactttgtacaagaaagctgggtcAGGCGTCGGAGCGAAGATAA | Mycgr3\|51607  JGI  Ensembl |  |  | 100 % |  |  |
| 40 | ggggacaagtttgtacaaaaaagcaggcttgatgGACAAGGACATCAATG, ggggaccactttgtacaagaaagctgggtcTTCGTCGTCGTCATCCTGGA | Mycgr3\|55886  JGI  Ensembl | KOG1781 Small Nuclear ribonucleoprotein splicing factor; | mRNA metabolic process; ribonucleoprotein complex; | 100 % |  |  |
| 40 | ggggacaagtttgtacaaaaaagcaggcttgatgGCGAAACTAGGCAACC, ggggaccactttgtacaagaaagctgggtcCTTGCTCGTCTTCGCCGCAG | Mycgr3\|72923  JGI  Ensembl |  |  | 100 % |  |  |
| 40 | ggggacaagtttgtacaaaaaagcaggcttgatgGCCCAATTTCGCCCCA, ggggaccactttgtacaagaaagctgggtcCCAGCTCGCCTTCTTCACTC | Mycgr3\|77845  JGI  Ensembl | KOG1741 Mitochondrial/chloroplast ribosomal protein S14/S29; | intracellular; ribosome; structural constituent of ribosome;  translation; | 100 % |  |  |
| 40 | ggggacaagtttgtacaaaaaagcaggcttgatgGGAAAAGTCCACGGAT, ggggaccactttgtacaagaaagctgggtcTGCCGCGCCTGGGTTGGGGT | Mycgr3\|78410  JGI  Ensembl | KOG0009 Ubiquitin-like/40S ribosomal S30 protein fusion; | intracellular; ribosome; structural constituent of ribosome;  translation; | 100 % |  |  |
| 40 | ggggacaagtttgtacaaaaaagcaggcttgatgTCGCGCGAAGCCTACA, ggggaccactttgtacaagaaagctgggtcACGAGCTTCAAATTGAATCA | Mycgr3\|78914  JGI  Ensembl | KOG3507 DNA-directed RNA polymerase, subunit RPB7.0; | DNA binding; DNA-directed RNA polymerase I activity;  DNA-directed RNA polymerase II activity;  DNA-directed RNA polymerase III activity;  DNA-directed RNA polymerase activity;  RNA polymerase activity;  transcription; | 100 % |  |  |
| 40 | ggggacaagtttgtacaaaaaagcaggcttgatgCACTTTTCCAAGCTCG, ggggaccactttgtacaagaaagctgggtcAATATTGCAGATGATAACGC | Mycgr3\|79161  JGI  Ensembl |  |  | 100 % |  |  |
| 40 | ggggacaagtttgtacaaaaaagcaggcttgatgCTCCTCAGCATGGCTC, ggggaccactttgtacaagaaagctgggtcGGCGCAAGCCGCTCCTCCTC | Mycgr3\|79286  JGI  Ensembl |  |  | 100 % |  |  |
| 40 | ggggacaagtttgtacaaaaaagcaggcttgatgTCTCGAATCATTCTTT, ggggaccactttgtacaagaaagctgggtcGCAATCGATCGGCCTTCGCT | Mycgr3\|79484  JGI  Ensembl |  |  | 100 % |  |  |
| 40 | ggggacaagtttgtacaaaaaagcaggcttgatgTACAAGTTCGCCTTCG, ggggaccactttgtacaagaaagctgggtcCGCATTCCTTGGCGCAGTCA | Mycgr3\|80321  JGI  Ensembl | KOG0260 RNA polymerase II, large subunit; |  | 100 % |  |  |
| 40 | ggggacaagtttgtacaaaaaagcaggcttgatgCAGATCTCCCAGATGT, ggggaccactttgtacaagaaagctgggtcCGATTGGTTGAAGCAGAGCG | Mycgr3\|80332  JGI  Ensembl |  |  | 100 % | chr\_4:637490 G->A; |  |
| 40 | ggggacaagtttgtacaaaaaagcaggcttgatgTCGATCCTCCTCGCGC, ggggaccactttgtacaagaaagctgggtcACTCGCCCTTGCAGCACTCG | Mycgr3\|81208  JGI  Ensembl |  |  | 100 % |  |  |
| 40 | ggggacaagtttgtacaaaaaagcaggcttgatgCAGCTCTTCCAGCTCG, ggggaccactttgtacaagaaagctgggtcATAGCAGAAGGCATATTGTT | Mycgr3\|82029  JGI  Ensembl |  |  | 100 % |  |  |
| 40 | ggggacaagtttgtacaaaaaagcaggcttgatgAGGGCGTGTGTGCGCG, ggggaccactttgtacaagaaagctgggtcGCAGAACTCCGCCATGCTGA | Mycgr3\|82038  JGI  Ensembl |  |  | 100 % |  |  |
| 40 | ggggacaagtttgtacaaaaaagcaggcttgatgATCACCGACGCCTCAC, ggggaccactttgtacaagaaagctgggtcTGACTTCTGGCTCTTCCCAC | Mycgr3\|82148  JGI  Ensembl |  |  | 100 % |  |  |
| 40 | ggggacaagtttgtacaaaaaagcaggcttgatgTTCTCCTCCTCGAACC, ggggaccactttgtacaagaaagctgggtcGCGACGAGACATCTTGTGGA | Mycgr3\|82659  JGI  Ensembl |  |  | 100 % |  |  |
| 40 | ggggacaagtttgtacaaaaaagcaggcttgatgAAGCTCACCATTCTTA, ggggaccactttgtacaagaaagctgggtcGGCCTTGTCGCCATCATCAA | Mycgr3\|82925  JGI  Ensembl |  |  | 100 % |  |  |
| 40 | ggggacaagtttgtacaaaaaagcaggcttgatgAAGCTCATCACCCTCG, ggggaccactttgtacaagaaagctgggtcAGTACAGACGCATGTGCCGG | Mycgr3\|83081  JGI  Ensembl |  |  | 100 % |  |  |
| 40 | ggggacaagtttgtacaaaaaagcaggcttgatgCAGGCAAGATCGCAAG, ggggaccactttgtacaagaaagctgggtcGACGCCGCGGAACTGCTCTA | Mycgr3\|83239  JGI  Ensembl |  |  | 100 % |  |  |
| 40 | ggggacaagtttgtacaaaaaagcaggcttgatgAAGATCTCGACTCTCC, ggggaccactttgtacaagaaagctgggtcATCCCACATAATCCCACGAC | Mycgr3\|84191  JGI  Ensembl |  |  | 100 % |  |  |
| 40 | ggggacaagtttgtacaaaaaagcaggcttgatgGCCGAGATCGAAGATG, ggggaccactttgtacaagaaagctgggtcTTGCTGTGGCGATATGAGGA | Mycgr3\|84746  JGI  Ensembl | KOG3460 Small nuclear ribonucleoprotein (snRNP) LSM3; | mRNA metabolic process; ribonucleoprotein complex; | 100 % |  |  |
| 40 | ggggacaagtttgtacaaaaaagcaggcttgatgATGGGATGGCTTGTGG, ggggaccactttgtacaagaaagctgggtcTTCGGCATCTTTCTTCAGCC | Mycgr3\|85399  JGI  Ensembl |  |  | 100 % |  |  |
| 40 | ggggacaagtttgtacaaaaaagcaggcttgatgAAGCTCTCCATCAACT, ggggaccactttgtacaagaaagctgggtcAGAAATGCAGGAGCCTCCAC | Mycgr3\|87205  JGI  Ensembl |  |  | 100 % |  |  |
| 40 | ggggacaagtttgtacaaaaaagcaggcttgatgCATCTCATGTACACCC, ggggaccactttgtacaagaaagctgggtcATGCGTACTCTGCTGGGTGA | Mycgr3\|87284  JGI  Ensembl | KOG0497 Oxidosqualene-lanosterol cyclase and related proteins; | lanosterol synthase activity; | 100 % |  |  |
| 40 | ggggacaagtttgtacaaaaaagcaggcttgatgTCTTTCCAAGACGCAA, ggggaccactttgtacaagaaagctgggtcAGCGGCAATGGTAACCTCAA | Mycgr3\|88095  JGI  Ensembl | KOG0900 40S ribosomal protein S20; | intracellular; ribosome; small ribosomal subunit;  structural constituent of ribosome;  translation; | 100 % |  |  |
| 40 | ggggacaagtttgtacaaaaaagcaggcttgatgCATCTCAGCCTACCAT, ggggaccactttgtacaagaaagctgggtcCCCGCACACACCGCTCGTAC | Mycgr3\|88664  JGI  Ensembl |  |  | 100 % |  |  |
| 40 | ggggacaagtttgtacaaaaaagcaggcttgatgCTCGTGCTCGCAGCAA, ggggaccactttgtacaagaaagctgggtcCAACTCCACCTTTCCACTGA | Mycgr3\|90092  JGI  Ensembl |  |  | 100 % |  |  |
| 40 | ggggacaagtttgtacaaaaaagcaggcttgatgAGGATCCACACTCTGC, ggggaccactttgtacaagaaagctgggtcGTACCACCTTTCGCAAGTAG | Mycgr3\|90236  JGI  Ensembl |  |  | 100 % |  |  |
| 40 | ggggacaagtttgtacaaaaaagcaggcttgatgAAATCTACCATCGTCT, ggggaccactttgtacaagaaagctgggtcGATAGTGTAGACAATCGAAT | Mycgr3\|90420  JGI  Ensembl |  |  | 100 % |  |  |
| 40 | ggggacaagtttgtacaaaaaagcaggcttgatgCAATTCTTCTACACCG, ggggaccactttgtacaagaaagctgggtcCTCGACCTTAGGCCAGGCCT | Mycgr3\|91285  JGI  Ensembl |  |  | 100 % |  |  |
| 40 | ggggacaagtttgtacaaaaaagcaggcttgatgCCCAAGGTGCCATGGA, ggggaccactttgtacaagaaagctgggtcGGTCCGACAGGGACCATCGA | Mycgr3\|91505  JGI  Ensembl |  |  | 100 % | chr\_3:947507 T->C; |  |
| 40 | ggggacaagtttgtacaaaaaagcaggcttgatgCTCGGACTACTCATCA, ggggaccactttgtacaagaaagctgggtcGTTTGGATTAGCAATGCAGG | Mycgr3\|92365  JGI  Ensembl |  |  | 100 % |  |  |
| 40 | ggggacaagtttgtacaaaaaagcaggcttgatgCAGATTTCTCAGCTCG, ggggaccactttgtacaagaaagctgggtcAGCATCACTCGGCGGGTAGT | Mycgr3\|94107  JGI  Ensembl |  |  | 100 % |  |  |
| 40 | ggggacaagtttgtacaaaaaagcaggcttgatgCAGCTCACCCCTCTCA, ggggaccactttgtacaagaaagctgggtcCCTCACCCTCTCCAACCTCC | Mycgr3\|94330  JGI  Ensembl |  |  | 100 % |  |  |
| 40 | ggggacaagtttgtacaaaaaagcaggcttgatgCAGATCCACATCTTCG, ggggaccactttgtacaagaaagctgggtcGGCTCCTGGGGGCGTTTGGC | Mycgr3\|95491  JGI  Ensembl |  |  | 100 % |  |  |
| 40 | ggggacaagtttgtacaaaaaagcaggcttgatgCAAATCAACCTCATCT, ggggaccactttgtacaagaaagctgggtcTCCCTCACCCATTCCCACCG | Mycgr3\|95665  JGI  Ensembl |  |  | 100 % |  |  |
| 40 | ggggacaagtttgtacaaaaaagcaggcttgatgTTCTCCTCGTTTCCGA, ggggaccactttgtacaagaaagctgggtcGTACTTCGTCGGCGTCACCT | Mycgr3\|95714  JGI  Ensembl |  |  | 100 % |  |  |
| 40 | ggggacaagtttgtacaaaaaagcaggcttgatgCAATTCTCAATCTTCC, ggggaccactttgtacaagaaagctgggtcCTGCCTAGGGCAATTGAACC | Mycgr3\|95958  JGI  Ensembl |  |  | 10.52632 % |  |  |
| 40 | ggggacaagtttgtacaaaaaagcaggcttgatgAAGGTCGCCATCCTCA, ggggaccactttgtacaagaaagctgggtcGTGGCTGTAGCCCTTTGGTG | Mycgr3\|96225  JGI  Ensembl |  |  | 100 % |  |  |
| 40 | ggggacaagtttgtacaaaaaagcaggcttgatgCTCATAGTGCTGCTAG, ggggaccactttgtacaagaaagctgggtcTGACTCTAGCACGGACGCGT | Mycgr3\|96731  JGI  Ensembl |  |  | 100 % |  |  |
| 40 | ggggacaagtttgtacaaaaaagcaggcttgatgAAGATCTCCATCCTCA, ggggaccactttgtacaagaaagctgggtcCCCTGATGCCACGCTTGCCG | Mycgr3\|96882  JGI  Ensembl |  |  | 100 % |  |  |
| 40 | ggggacaagtttgtacaaaaaagcaggcttgatgAAGACTTTAACTATCC, ggggaccactttgtacaagaaagctgggtcACAAGGGAAGACGTTGGGAC | Mycgr3\|99124  JGI  Ensembl |  |  | 100 % |  |  |
| 40 | ggggacaagtttgtacaaaaaagcaggcttgatgAAGTTCTCTCTTGCTC, ggggaccactttgtacaagaaagctgggtcATTGTTGATAGAAACAATCT | Mycgr3\|99350  JGI  Ensembl |  |  | 100 % |  |  |
| 40 | ggggacaagtttgtacaaaaaagcaggcttgatgAACATCTTCCAGCTTT, ggggaccactttgtacaagaaagctgggtcAGCGCAATCCGCCGAACCAT | Mycgr3\|100649  JGI  Ensembl |  |  | 100 % |  |  |
| 40 | ggggacaagtttgtacaaaaaagcaggcttgatgTCCAGTGATCTTCCGC, ggggaccactttgtacaagaaagctgggtcCAGAAAAGACGCCGAGTGAA | Mycgr3\|100838  JGI  Ensembl |  |  | 100 % |  |  |
| 40 | ggggacaagtttgtacaaaaaagcaggcttgatgCGAGTGTTCGGACTGA, ggggaccactttgtacaagaaagctgggtcGATCATGACGGAGACGTATG | Mycgr3\|101077  JGI  Ensembl |  |  | 100 % |  |  |
| 40 | ggggacaagtttgtacaaaaaagcaggcttgatgGCCTCCGCAGCTGCCC, ggggaccactttgtacaagaaagctgggtcACTCAACAGCACCAAAATCA | Mycgr3\|101457  JGI  Ensembl | KOG3459 Small nuclear ribonucleoprotein (snRNP) Sm core protein; | mRNA metabolic process; ribonucleoprotein complex; | 100 % |  |  |
| 40 | ggggacaagtttgtacaaaaaagcaggcttgatgCGCTTCGCTCACGTCG, ggggaccactttgtacaagaaagctgggtcGGGACACTTGCCCTTGCAAG | Mycgr3\|101652  JGI  Ensembl |  |  | 100 % |  |  |
| 40 | ggggacaagtttgtacaaaaaagcaggcttgatgGTGAACTACTACTTCG, ggggaccactttgtacaagaaagctgggtcCGCACCATACTCCGGCGGCT | Mycgr3\|102037  JGI  Ensembl |  |  | 100 % |  |  |
| 40 | ggggacaagtttgtacaaaaaagcaggcttgatgTTCTCCCTCTTCCTCC, ggggaccactttgtacaagaaagctgggtcCGCAATCGGCTCGTATCCAG | Mycgr3\|102119  JGI  Ensembl |  |  | 100 % |  |  |
| 40 | ggggacaagtttgtacaaaaaagcaggcttgatgCGGACACCCTGGACGA, ggggaccactttgtacaagaaagctgggtcCCAACTATACCACTGCCTCT | Mycgr3\|102256  JGI  Ensembl |  |  | 100 % |  |  |
| 40 | ggggacaagtttgtacaaaaaagcaggcttgatgAGTAAAGGTACTGTAC, ggggaccactttgtacaagaaagctgggtcTGTATCATGGATCCGGAAAT | Mycgr3\|102513  JGI  Ensembl |  |  | 100 % |  |  |
| 40 | ggggacaagtttgtacaaaaaagcaggcttgatgAAGCACCTCGCGGCAT, ggggaccactttgtacaagaaagctgggtcGTCGAAGAGACCGAAGCCCA | Mycgr3\|102737  JGI  Ensembl | KOG3449 60S acidic ribosomal protein P2; | intracellular; ribosome; structural constituent of ribosome;  translational elongation; | 100 % |  |  |
| 40 | ggggacaagtttgtacaaaaaagcaggcttgatgCCCGCTGTTTTTTTTT, ggggaccactttgtacaagaaagctgggtcACGGTATCGACTCTCGCAGG | Mycgr3\|102982  JGI  Ensembl |  |  | 100 % |  |  |
| 40 | ggggacaagtttgtacaaaaaagcaggcttgatgCAGCTCTCCATCATCC, ggggaccactttgtacaagaaagctgggtcGTACAGAGTCGGGCGCGCCG | Mycgr3\|102983  JGI  Ensembl |  |  | 100 % |  |  |
| 40 | ggggacaagtttgtacaaaaaagcaggcttgatgAATTTCTCCCTGCAAC, ggggaccactttgtacaagaaagctgggtcTCGCGATGTTAGCAATCGTG | Mycgr3\|103126  JGI  Ensembl |  |  | 100 % |  |  |
| 40 | ggggacaagtttgtacaaaaaagcaggcttgatgATCACCTCTTTCACCC, ggggaccactttgtacaagaaagctgggtcTAGTGACTGCACAAGCACCT | Mycgr3\|103519  JGI  Ensembl |  |  | 100 % |  |  |
| 40 | ggggacaagtttgtacaaaaaagcaggcttgatgAAGCTCTCCACCCTCT, ggggaccactttgtacaagaaagctgggtcATGCGTGTGGACGTGTGCAT | Mycgr3\|103555  JGI  Ensembl |  |  | 100 % |  |  |
| 40 | ggggacaagtttgtacaaaaaagcaggcttgatgCACGCCTTCAAGCTCT, ggggaccactttgtacaagaaagctgggtcAGAACACACCATCTCACCCT | Mycgr3\|103572  JGI  Ensembl |  |  | 100 % |  | ``` 32	ATGCACGCCTTCAAGCTCTGCGCCGTCTTCTCTATGATGGCTGCTACTGCTTTCGCTCATAAATGCAAACCCAACCCACCCGCCAGTGGTTACTGCTACTACAGTCCAACCGACATGCGAGCCTGCCACCACTATCACGAGTGCAAGACCCCGAACGGCGCTTGCGGCTACAACAACAAGGGTGAGATGGTGTGTTCT	229 	\|\|\|\|\|\|\|\|\|\|\|\|\|\|\|\|\|\|\|\|\|\|\|\|\|\|\|\|\|\|\|\|\|\|\|\|\|\|\|\|\|\|\|\|\|\|\|\|\|\|\|\|\|\|\|\|\|\|\|\|\|\|\|\|\|\|\|\|\|\|\|\|\|\|\|\|\|\|\|\|\|\|\|\|\|\|\|\|\|\|\|\|\|\|\|\|\|\|\|\|\|\|\|\|\|\|\|\|\|\|\|\|\|\|\|\|\|\|\|\|\|\|\|\|\|\|\|\|\|\|\|\|\|\|\|\|\|\|\|\|\|\|\|\|\|\|\|\|\|\|\|\|\|\|\|\|\|\|\|\|\|\|\|\|\|\|\|\|\|\|\|\|\|\|\|\|\|\|\|\|\|\|\|\|\|\|\|\|\|\|\|\|\|\|\|\|\|\| 	ATGCACGCCTTCAAGCTCTGCGCCGTCTTCTCTATGATGGCTGCTACTGCTTTCGCTCATAAATGCAAACCCAACCCACCCGCCAGTGGTTACTGCTACTACAGTCCAACCGACATGCGAGCCTGCCACCACTATCACGAGTGCAAGACCCCGAACGGCGCTTGCGGCTACAACAACAAGGGTGAGATGGTGTGTTCT ``` |
| 40 | ggggacaagtttgtacaaaaaagcaggcttgatgAACTCCATCCTCCATC, ggggaccactttgtacaagaaagctgggtcGCTGACCTCATTTGTTGGAG | Mycgr3\|103650  JGI  Ensembl |  |  | 100 % |  |  |
| 40 | ggggacaagtttgtacaaaaaagcaggcttgatgAGACACACAATATGCA, ggggaccactttgtacaagaaagctgggtcGCAGCAGGGAAAGGGGACGA | Mycgr3\|103716  JGI  Ensembl |  |  | 100 % |  |  |
| 40 | ggggacaagtttgtacaaaaaagcaggcttgatgTCCGCAGTGGTGGTGG, ggggaccactttgtacaagaaagctgggtcCGCCCGCGGTCGAACTTCAA | Mycgr3\|103758  JGI  Ensembl |  |  | 100 % |  |  |
| 40 | ggggacaagtttgtacaaaaaagcaggcttgatgCAGTTCTCCAACGTCG, ggggaccactttgtacaagaaagctgggtcGGAGGTCTGATCTGAGCAGA | Mycgr3\|104383  JGI  Ensembl |  |  | 100 % |  |  |
| 40 | ggggacaagtttgtacaaaaaagcaggcttgatgAACTTGGAAGTGAACC, ggggaccactttgtacaagaaagctgggtcCTGCTTAGGCTGTCGCATGA | Mycgr3\|104431  JGI  Ensembl |  |  | 100 % |  |  |
| 40 | ggggacaagtttgtacaaaaaagcaggcttgatgGCCGGAATCCTTGGTG, ggggaccactttgtacaagaaagctgggtcCTCGTCATCCGCCTCGATGT | Mycgr3\|104853  JGI  Ensembl | KOG3494 Ubiquinol cytochrome c oxidoreductase, subunit QCR9; | mitochondrial electron transport, ubiquinol to cytochrome c; mitochondrial envelope;  ubiquinol-cytochrome-c reductase activity; | 100 % |  |  |
| 40 | ggggacaagtttgtacaaaaaagcaggcttgatgTCGTCCTCCATCACCA, ggggaccactttgtacaagaaagctgggtcCTTCTTCCCCAACTTCTCCG | Mycgr3\|104965  JGI  Ensembl |  |  | 100 % |  |  |
| 40 | ggggacaagtttgtacaaaaaagcaggcttgatgCTCTCCAACTACCTGA, ggggaccactttgtacaagaaagctgggtcGACGGTAGCCCTGTTCTCCG | Mycgr3\|105182  JGI  Ensembl |  |  | 100 % |  |  |
| 40 | ggggacaagtttgtacaaaaaagcaggcttgatgCTGGTCCTCACCGCCG, ggggaccactttgtacaagaaagctgggtcTTCTGGAATACACCGCCCCG | Mycgr3\|105682  JGI  Ensembl |  |  | 100 % |  |  |
| 40 | ggggacaagtttgtacaaaaaagcaggcttgatgAAGCTTTCCACCATTA, ggggaccactttgtacaagaaagctgggtcGTTGGGAATGCATCCGCAAG | Mycgr3\|105825  JGI  Ensembl |  |  | 100 % |  |  |
| 40 | ggggacaagtttgtacaaaaaagcaggcttgatgCTCGGGACTACGGTGC, ggggaccactttgtacaagaaagctgggtcGCCATTGAACAGCTCGTCCT | Mycgr3\|106021  JGI  Ensembl |  |  | 100 % |  |  |
| 40 | ggggacaagtttgtacaaaaaagcaggcttgatgCACGCTTTCAAGGTCT, ggggaccactttgtacaagaaagctgggtcGGCTGGAGCGTCATCGTGGC | Mycgr3\|106125  JGI  Ensembl |  |  | 100 % |  |  |
| 40 | ggggacaagtttgtacaaaaaagcaggcttgatgAAGCTTCTGACAACCA, ggggaccactttgtacaagaaagctgggtcGCAACACGCAAGACGTAGAA | Mycgr3\|106127  JGI  Ensembl |  |  | 100 % |  |  |
| 40 | ggggacaagtttgtacaaaaaagcaggcttgatgCTCTGGATCTCAGGAG, ggggaccactttgtacaagaaagctgggtcCAGCCGATTTCGCTCGTCGT | Mycgr3\|106229  JGI  Ensembl |  |  | 100 % |  |  |
| 40 | ggggacaagtttgtacaaaaaagcaggcttgatgAAGGCCTTCGCAATCA, ggggaccactttgtacaagaaagctgggtcCTCGAACAGCAAGCTTCCAC | Mycgr3\|106436  JGI  Ensembl |  |  | 100 % | chr\_13:55702 C->T; |  |
| 40 | ggggacaagtttgtacaaaaaagcaggcttgatgCATTTTGTCCAGCTCG, ggggaccactttgtacaagaaagctgggtcTTCCCCAAGCGCCTGGAATG | Mycgr3\|106502  JGI  Ensembl |  |  | 100 % |  |  |
| 40 | ggggacaagtttgtacaaaaaagcaggcttgatgATGAGGATGGCGATTT, ggggaccactttgtacaagaaagctgggtcAAAGTCGCAGACATTGGGCG | Mycgr3\|106576  JGI  Ensembl |  |  | 100 % |  |  |
| 40 | ggggacaagtttgtacaaaaaagcaggcttgatgCAAAGCTCGAAACTGA, ggggaccactttgtacaagaaagctgggtcTCTCCGCCTCGCGATACGCA | Mycgr3\|106703  JGI  Ensembl |  |  | 100 % |  |  |
| 40 | ggggacaagtttgtacaaaaaagcaggcttgatgGCTGCCACCTCTGCGA, ggggaccactttgtacaagaaagctgggtcAGACTTCAGCATCTTGTTGC | Mycgr3\|107493  JGI  Ensembl | KOG4786 Ubinuclein, nuclear protein interacting with cellular and viral transcription factors; |  | 100 % |  |  |
| 40 | ggggacaagtttgtacaaaaaagcaggcttgatgAAGCTCTCCATGGCAT, ggggaccactttgtacaagaaagctgggtcCGCGTTTCCGGTTGCATCGT | Mycgr3\|108877  JGI  Ensembl |  |  | 100 % |  |  |
| 40 | ggggacaagtttgtacaaaaaagcaggcttgatgTACGCCTCCCTCAACT, ggggaccactttgtacaagaaagctgggtcAGGCACTCTCTTCACTTTGC | Mycgr3\|110918  JGI  Ensembl |  |  | 100 % |  |  |
| 41 | ggggacaagtttgtacaaaaaagcaggcttgatgTTCTTCGGAGGACTGG, ggggaccactttgtacaagaaagctgggtcCGCCGCCTTCTCTGCACTGA | Mycgr3\|18598  JGI  Ensembl |  |  | 100 % |  | ``` 1	ATGTTCTTCGGAGGACTGGCATTCACCTGCTTGACTGCTTTGGTGACTCGACGAGCATTGACGAAGAAGAAGCTGGCAATGTATCCGCATCTACAACCGACCGTTGGCAAGGACGCGAAACTGGAGTTGCCAAAGTTTACACCTTCAAATACACCGCCGAAAGCTGAGGGAGGTTTGGACGCTGCGGAAGCTCTTTTCCTGGCTACGCTGAATGTGGGAGCGGTGTTCATGGCTAGCGTGGGTGCGTTTATGAAGTTCGCCGATGTCGCGGATACGGAAGATTTGAGAGATTATGTGAGGAGTGGGATTGGCTATGATGTCTATGCTGGGGACACAGAGGCGGACAGAGAGATTGAGCAGTGGTTTGCGGAGGTGCTGAGCAGAAAGGATGGC	393 	\|\|\|\|\|\|\|\|\|\|\|\|\|\|\|\|\|\|\|\|\|\|\|\|\|\|\|\|\|\|\|\|\|\|\|\|\|\|\|\|\|\|\|\|\|\|\|\|\|\|\|\|\|\|\|\|\|\|\|\|\|\|\|\|\|\|\|\|\|\|\|\|\|\|\|\|\|\|\|\|\|\|\|\|\|\|\|\|\|\|\|\|\|\|\|\|\|\|\|\|\|\|\|\|\|\|\|\|\|\|\|\|\|\|\|\|\|\|\|\|\|\|\|\|\|\|\|\|\|\|\|\|\|\|\|\|\|\|\|\|\|\|\|\|\|\|\|\|\|\|\|\|\|\|\|\|\|\|\|\|\|\|\|\|\|\|\|\|\|\|\|\|\|\|\|\|\|\|\|\|\|\|\|\|\|\|\|\|\|\|\|\|\|\|\|\|\|\|\|\|\|\|\|\|\|\|\|\|\|\|\|\|\|\|\|\|\|\|\|\|\|\|\|\|\|\|\|\|\|\|\|\|\|\|\|\|\|\|\|\|\|\|\|\|\|\|\|\|\|\|\|\|\|\|\|\|\|\|\|\|\|\|\|\|\|\|\|\|\|\|\|\|\|\|\|\|\|\|\|\|\|\|\|\|\|\|\|\|\|\|\|\|\|\|\|\|\|\|\|\|\|\|\|\|\|\|\|\|\|\|\|\|\|\|\|\|\|\|\|\|\|\|\|\|\|\|\|\|\|\|\|\|\|\|\|\|\|\|\|\|\|\|\|\|\|\|\|\|\|\|\|\|\|\|\|\|\|\|\|\|\|\|\|\|\|\|\|\|\|\|\|\|\|\|\|\|\|\|\|\|\|\|\|\|\|\|\|\|\|\|\|\|\| 	ATGTTCTTCGGAGGACTGGCATTCACCTGCTTGACTGCTTTGGTGACTCGACGAGCATTGACGAAGAAGAAGCTGGCAATGTATCCGCATCTACAACCGACCGTTGGCAAGGACGCGAAACTGGAGTTGCCAAAGTTTACACCTTCAAATACACCGCCGAAAGCTGAGGGAGGTTTGGACGCTGCGGAAGCTCTTTTCCTGGCTACGCTGAATGTGGGAGCGGTGTTCATGGCTAGCGTGGGTGCGTTTATGAAGTTCGCCGATGTCGCGGATACGGAAGATTTGAGAGATTATGTGAGGAGTGGGATTGGCTATGATGTCTATGCTGGGGACACAGAGGCGGACAGAGAGATTGAGCAGTGGTTTGCGGAGGTGCTGAGCAGAAAGGATGGC ``` |
| 41 | ggggacaagtttgtacaaaaaagcaggcttgatgCCTCAAGCACAACCCG, ggggaccactttgtacaagaaagctgggtcATCCCTCCCTCCACCTCCAC | Mycgr3\|19663  JGI  Ensembl | KOG1780 Small Nuclear ribonucleoprotein G; | mRNA metabolic process; ribonucleoprotein complex; | 100 % |  | ``` 103	GTCTTCCTGAACATCGTCCTCGACGAAGCCACCGAGTCGAAACCCAACGCGGAAAAAGTGCGATTGGGAATGTGCGTCATTCGCGGGAATTCCGTCGTCATGATGGAGGCGCTGGATCGGATTGGAGGA	231 	\|\|\|\|\|\|\|\|\|\|\|\|\|\|\|\|\|\|\|\|\|\|\|\|\|\|\|\|\|\|\|\|\|\|\|\|\|\|\|\|\|\|\|\|\|\|\|\|\|\|\|\|\|\|\|\|\|\|\|\|\|\|\|\|\|\|\|\|\|\|\|\|\|\|\|\|\|\|\|\|\|\|\|\|\|\|\|\|\|\|\|\|\|\|\|\|\|\|\|\|\|\|\|\|\|\|\|\|\|\|\|\|\|\|\|\|\|\|\|\|\|\|\|\|\|\|\|\|\| 	GTCTTCCTGAACATCGTCCTCGACGAAGCCACCGAGTCGAAACCCAACGCGGAAAAAGTGCGATTGGGAATGTGCGTCATTCGCGGGAATTCCGTCGTCATGATGGAGGCGCTGGATCGGATTGGAGGA ```      ``` 1	ATGCCTCAAGCACAACCCGAACTCAAAAAG-TAC	33 	\|\|\|\|\|\|\|\|\|\|\|\|\|\|\|\|\|\|\|\|\|\|\|\|\|\|\|\|\|\| \|\|\| 	ATGCCTCAAGCACAACCCGAACTCAAAAAGGTAC ```      ``` 29	AGTACCTCGACAAACGCGTTGAAGTCCAACTCAACGGCTCCCGAAAAGTCATGGGTACTCTCCGTGGATACGACGT	104 	\|\|\|\|\|\|\|\|\|\|\|\|\|\|\|\|\|\|\|\|\|\|\|\|\|\|\|\|\|\|\|\|\|\|\|\|\|\|\|\|\|\|\|\|\|\|\|\|\|\|\|\|\|\|\|\|\|\|\|\|\|\|\|\|\|\|\|\|\|\|\|\|\|\|\|\| 	AGTACCTCGACAAACGCGTTGAAGTCCAACTCAACGGCTCCCGAAAAGTCATGGGTACTCTCCGTGGATACGACGT ``` |
| 41 | ggggacaagtttgtacaaaaaagcaggcttgatgGCCTCAGGCACATCCA, ggggaccactttgtacaagaaagctgggtcCTCCTCCGCCTCCGCCTTCG | Mycgr3\|28506  JGI  Ensembl | KOG3217 Protein tyrosine phosphatase; | protein amino acid dephosphorylation; protein tyrosine phosphatase activity; | 100 % |  | ``` 1	ATGGCCTCAGGCACATCCACCCCCAGCGGCCGCCAGGTCTCCGTCCTTTTCGTCTGCCTCGGAAACATCTGCCGATCTCCCATGGCAGAAGGCGTGTTCCGGCATCTCACCAACTTCGAAACCTCCGATCCACACCCTCTCATCGCCGCCATCGACTCCTGTGGCACGGGAGCCTACCATGCCGGCGATCGACCCGACTCTCGCACACTTTCCGTACTGAAGGATAACGGGCTCTCGAGCTACAAACATGCCGCGCGAAAAGTGCGGGTGCCGGAGGACTTTGAGAAGTTTGATTACCTGATCGCGATGGACGAGGATAATTACATGGATCTGAGGGATTTGGTGAAGAGGACGAAGAAGAACAAGGGGACGCTGGGAGATGAGCATTTGGACAAGGTGAGGTTGTATGGAGAGTTCGGCGGCAAGGACAAAAAGGAGGAGATTGGCGATCCTTATTATGGTGGGAGAGAGGGCTTCACGATCGCGTACGAGCAGGTGGTGAGGTGTGGGAAGGGGCTGTTGAAGGAGATTGAGGAG	537 	\|\|\|\|\|\|\|\|\|\|\|\|\|\|\|\|\|\|\|\|\|\|\|\|\|\|\|\|\|\|\|\|\|\|\|\|\|\|\|\|\|\|\|\|\|\|\|\|\|\|\|\|\|\|\|\|\|\|\|\|\|\|\|\|\|\| \|\|\|\|\|\|\|\|\|\|\|\|\|\|\|\|\|\|\|\| \| \|\|\|\|\|\|\|\|\|\|\|\|\|\|\|\|\|\|\|\|\|\|\|\|\|\|\|\|\|\|\|\|\|\|\|\|\|\|\|\|\|\|\|\|\|\|\|\|\|\|\|\|\|\|\|\|\|\|\|\| \|\| \|\|\|\|\|\|\|\|\|\|\|\|\|\|\|\|\|\|\|\|\|\|\|\|\|\|\|\|\|\|\|\|\|\|\|\|\|\|\|\|\|\|\|\|\|\|\|\|\|\|\|\|\|\|\|\|\|\|\|\|\|\|\|\|\|\|\|\|\|\|\|\|\|\|\|\|\|\|\|\|\|\|\|\|\|\|\|\|\|\|\|\|\|\|\|\|\|\|\|\|\|\|\|\|\|\|\|\|\|\|\|\|\|\|\|\|\|\|\|\|\|\|\|\|\|\|\|\|\|\|\|\|\|\|\|\|\|\|\|\|\|\|\|\|\|\|\|\|\|\|\|\|\|\|\|\|\|\|\|\|\|\|\|\|\|\|\|\|\|\|\|\|\|\|\|\|\|\|\|\|\|\|\|\|\|\|\|\|\|\|\|\|\|\|\|\|\|\|\|\|\|\|\|\|\|\|\|\|\|\|\|\|\|\|\|\|\|\|\|\|\|\|\|\|\|\|\|\|\|\|\|\|\|\|\|\|\|\|\|\|\|\|\|\|\|\|\|\|\|\|\|\|\|\|\|\|\|\|\|\|\|\|\|\|\|\|\|\|\|\|\|\|\|\|\|\|\|\|\|\|\|\|\|\|\|\|\|\|\|\|\|\|\|\|\|\|\|\|\|\|\|\|\|\|\|\|\|\|\|\|\|\|\|\|\|\|\|\|\|\|\|\|\|\|\|\|\|\|\|\|\| \|\|\|\|\|\|\|\|\|\|\|\|\|\|\|\|\|\|\|\|\|\|\|\|\|\|\|\|\|\|\|\|\|\|\|\|\|\|\|\|\|\|\|\|\|\|\|\|\|\|\| 	ATGGCCTCAGGCACATCCACCCCCAGCGGCCGCCAGGTCTCCGTCCTTTTCGTCTGCCTCGGAAACTTCTGCCGATCTCCCATGGCATACGGCGTGTTCCGGCATCTCACCAACTTCGAAACCTCCGATCCACACCCTCTCATCGCCGCCTTCTACTCCTGTGGCACGGGAGCCTACCATGCCGGCGATCGACCCGACTCTCGCACACTTTCCGTACTGAAGGATAACGGGCTCTCGAGCTACAAACATGCCGCGCGAAAAGTGCGGGTGCCGGAGGACTTTGAGAAGTTTGATTACCTGATCGCGATGGACGAGGATAATTACATGGATCTGAGGGATTTGGTGAAGAGGACGAAGAAGAACAAGGGGACGCTGGGAGATGAGCATTTGGACAAGGTGAGGTTGTATGGAGAGTTCGGCGGCAAGGACAAAAAGGAGGAGATTGGCGATCCTTATTATGGTGGGAGAGAGGGCTTCACGATCGCATACGAGCAGGTGGTGAGGTGTGGGAAGGGGCTGTTGAAGGAGATTGAGGAG ``` |
| 41 | ggggacaagtttgtacaaaaaagcaggcttgatgGTGAAGCCAATTACCG, ggggaccactttgtacaagaaagctgggtcCTGGCCCTCCGCCCGCTGGT | Mycgr3\|31400  JGI  Ensembl |  |  | 100 % |  | ``` 100	GGCTTCCACGTTCCATCGGTTCGCCGCCGCGATTTGTTCTACGCGAAGCTCGAGGACCAGCGGGCGGAGGGCCAG	174 	\|\|\|\|\|\|\|\|\|\|\|\|\|\|\|\|\|\|\|\|\|\|\|\|\|\|\|\|\|\|\|\|\|\|\|\|\|\|\|\|\|\|\|\|\|\|\|\|\|\|\|\|\|\|\|\|\|\|\|\|\|\|\|\|\|\|\|\|\|\|\|\|\|\|\| 	GGCTTCCACGTTCCATCGGTTCGCCGCCGCGATTTGTTCTACGCGAAGCTCGAGGACCAGCGGGCGGAGGGCCAG ```      ``` 22	ATGCTCCGAAGGGGCCTCGTTCTCGATCTCTCAGTCGCATTCGGT	66 	\|\|\|\|\|\|\|\|\|\|\|\|\|\|\|\|\|\|\|\|\|\|\|\|\|\|\|\|\|\|\|\|\|\|\|\|\|\|\|\|\|\|\|\|\| 	ATGCTCCGAAGGGGCCTCGTTCTCGATCTCTCAGTCGCATTCGGT ```      ``` 64	GGTCTCGGAACAGCCTTCGGATACGGCTGGTGGTACGG	101 	\|\|\|\|\|\|\|\|\|\|\|\|\|\|\|\|\|\|\|\|\|\|\|\|\|\|\|\|\|\|\|\|\|\|\|\|\|\| 	GGTCTCGGAACAGCCTTCGGATACGGCTGGTGGTACGG ``` |
| 41 | ggggacaagtttgtacaaaaaagcaggcttgatgCGTCCTCAAGTCCACA, ggggaccactttgtacaagaaagctgggtcCTCTTTCCTGAACTGCATCA | Mycgr3\|32491  JGI  Ensembl | KOG3302 TATA-box binding protein (TBP), component of TFIID and TFIIIB; | DNA binding; RNA polymerase II transcription factor activity;  nucleus;  regulation of transcription, DNA-dependent;  transcription factor TFIID complex;  transcription initiation from RNA polymerase II promoter; | 100 % |  |  |
| 41 | ggggacaagtttgtacaaaaaagcaggcttgatgTCCGACTTTGGCGATG, ggggaccactttgtacaagaaagctgggtcCTCCTTCGCCCCCTCCGCCG | Mycgr3\|32785  JGI  Ensembl | KOG4747 Two-component phosphorelay intermediate involved in MAP kinase cascade regulation; | histone arginine kinase activity; signal transducer activity;  signal transduction;  two-component sensor activity; | 100 % |  | ``` 44	AGATCCTCGAAATGGACGACGACGAAGAAGAGCGCGAATTCAGCCGATCGATCGTGTTTGACTTTTTCGAACAAGCCGCCAATACATTCAGCCAGATGGACAAGTCATTG	153 	\|\|\|\|\|\|\|\|\|\|\|\|\|\|\|\|\|\|\|\| \|\|\|\|\|\|\|\| \|\|\|\|\|\|\|\|\|\|\|\|\|\|\|\|\|\|\|\|\|\|\|\|\|\|\|\|\|\|\|\|\|\|\|\|\|\|\|\|\|\|\|\|\|\|\|\|\|\|\|\|\|\|\|\|\|\|\|\|\|\|\|\|\|\|\|\|\|\|\|\|\|\|\|\|\|\|\|\| 	AGATCCTCGAAATGGACGACTACGAAGAATAGCGCGAATTCAGCCGATCGATCGTGTTTGACTTTTTCGAACAAGCCGCCAATACATTCAGCCAGATGGACAAGTCATTG ```      ``` 1	ATGTCCGACTTTGGCGATGTTATTGACACGCTCACATTCGAGCAG	45 	\|\|\|\|\|\|\|\|\|\|\|\|\|\|\|\|\|\|\|\|\|\|\|\|\|\|\|\|\|\|\|\|\|\|\|\|\|\|\|\|\|\|\|\|\| 	ATGTCCGACTTTGGCGATGTTATTGACACGCTCACATTCGAGCAG ```      ``` 153	GGAAGCCAAAGACCTCGAAAACCTCTCCGCCCTAGGCCACTTCCTCAAAGGTTCCTCCGCCACTCTGGGACTGACCAAAGTCAAAGATTCGTGCGAGAAGATTCAACACTTTGGCGCGCGCAAGAATGAGACGGGCGATCAGGACGAGCCGGATGATGAGGTCAGTCTAAGCAAGTGTCGGACGACGATTGAGCAGACGAAGGTGGAGTTTGCCGAGGCGGAGAGGAAGTTGAGGAGGTTTTATCATGATGATGGACCGGCGGA-GGGGGCGAA-GGAG	429 	\|\|\|\|\|\|\|\|\|\|\|\|\|\|\|\|\|\|\|\|\|\|\|\|\|\|\|\|\|\|\|\|\|\|\|\|\|\|\|\|\|\|\|\|\|\|\|\|\|\|\|\|\|\|\|\|\|\|\|\|\|\|\|\|\|\|\|\|\|\|\|\|\|\|\|\|\|\|\|\|\|\|\|\|\|\|\|\|\|\|\|\|\|\|\|\|\|\|\|\|\|\|\|\|\|\|\|\|\|\|\|\|\|\|\|\|\|\|\|\|\|\|\|\|\|\|\|\|\|\|\|\|\|\|\|\|\|\|\|\|\|\|\|\|\|\|\|\|\|\|\|\|\|\|\|\|\|\|\|\|\|\|\|\|\|\|\|\|\|\|\|\|\|\|\|\|\|\|\|\|\|\|\|\|\|\|\|\|\|\|\|\|\|\|\|\|\|\|\|\|\|\|\|\|\|\|\|\|\|\|\|\|\|\|\|\|\|\|\|\|\|\|\|\|\|\|\|\|\|\|\|\|\|\|\|\|\|\|\|\|\|\|\|\|\|\|\|\|\|\|\|\|\|\|\|\|\|\|\|\|\|\|\|\| \|\|\|\|\|\|\|\|\| \|\|\|\| 	GGAAGCCAAAGACCTCGAAAACCTCTCCGCCCTAGGCCACTTCCTCAAAGGTTCCTCCGCCACTCTGGGACTGACCAAAGTCAAAGATTCGTGCGAGAAGATTCAACACTTTGGCGCGCGCAAGAATGAGACGGGCGATCAGGACGAGCCGGATGATGAGGTCAGTCTAAGCAAGTGTCGGACGACGATTGAGCAGACGAAGGTGGAGTTTGCCGAGGCGGAGAGGAAGTTGAGGAGGTTTTATCATGATGATGGACCGGCGGATGGGGGCGAAAGGAG ``` |
| 41 | ggggacaagtttgtacaaaaaagcaggcttgatgGACGTCGACGGCGCAT, ggggaccactttgtacaagaaagctgggtcGACCTTCACGTGCTGGCCCA | Mycgr3\|33739  JGI  Ensembl | KOG3167 Box H/ACA snoRNP component, involved in ribosomal RNA pseudouridinylation; | ribonucleoprotein complex; ribosome biogenesis and assembly; | 100 % |  | ``` 151	GCTGCCAAAGCCAAAGCCCTCAAACGCGGCGTAAAAGAATGCGTCAAATCCATCCGCAAATCCCCACCATCCACGCCCGCCAGCCTGGCCTCCTCCGCCCTTCCCAACGGCATCGTAATCCTCGCCGCCGACATCTCACCCATGGACGTCATCAGCCACATTCCAGTGCTATGCGAGGACCACGGAATCCCCTACATCTACGTCCCTTCGCGCGCGGAATTGGGCGCGGCGGGAAGCACGAAGAGGCCGACGAGTGTGGTCATGTTGACGCCGGCAGTGGGAAAGAAGGGCGAGGGCGCGGAGGAGTGGGCGGAGGCGTATAAGGAGCTTGCGAAGTTGGCGCAAAAGCTGGGCCAGCACGTGAAGGTC	519 	\|\|\|\|\|\|\|\|\|\|\|\|\|\|\|\|\|\|\|\|\|\|\|\|\|\|\|\|\|\|\|\|\|\|\|\|\|\|\|\|\|\|\|\|\|\|\|\|\|\|\|\|\|\|\|\|\|\|\|\|\|\|\|\|\|\|\|\|\|\|\|\|\|\|\|\|\|\|\|\|\|\|\|\|\|\|\|\|\|\|\|\|\|\|\|\|\|\|\|\|\|\|\|\|\|\|\|\|\|\|\|\|\|\|\|\|\|\|\|\|\|\|\|\|\|\|\|\|\|\|\|\|\|\|\|\|\|\|\|\|\|\|\|\|\|\|\|\|\|\|\|\|\|\|\|\|\|\|\|\|\|\|\|\|\|\|\|\|\|\|\|\|\|\|\|\|\|\|\|\|\|\|\|\|\|\|\|\|\|\|\|\|\|\|\|\|\|\|\|\|\|\|\|\|\|\|\|\|\|\|\|\|\|\|\|\|\|\|\|\|\|\|\|\|\|\|\|\|\|\|\|\|\|\|\|\|\|\|\|\|\|\|\|\|\|\|\|\|\|\|\|\|\|\|\|\|\|\|\|\|\|\|\|\|\|\|\|\|\|\|\|\|\|\|\|\|\|\|\|\|\|\|\|\|\|\|\|\|\|\|\|\|\|\|\|\|\|\|\|\|\|\|\|\|\|\|\|\|\|\|\|\|\|\|\|\|\|\|\|\|\|\|\|\|\|\|\|\|\|\|\|\|\|\|\|\|\|\|\|\|\|\|\|\|\|\|\|\|\|\|\|\|\|\|\|\|\|\|\|\|\|\|\|\|\|\|\|\|\| 	GCTGCCAAAGCCAAAGCCCTCAAACGCGGCGTAAAAGAATGCGTCAAATCCATCCGCAAATCCCCACCATCCACGCCCGCCAGCCTGGCCTCCTCCGCCCTTCCCAACGGCATCGTAATCCTCGCCGCCGACATCTCACCCATGGACGTCATCAGCCACATTCCAGTGCTATGCGAGGACCACGGAATCCCCTACATCTACGTCCCTTCGCGCGCGGAATTGGGCGCGGCGGGAAGCACGAAGAGGCCGACGAGTGTGGTCATGTTGACGCCGGCAGTGGGAAAGAAGGGCGAGGGCGCGGAGGAGTGGGCGGAGGCGTATAAGGAGCTTGCGAAGTTGGCGCAAAAGCTGGGCCAGCACGTGAAGGTC ```      ``` 1	ATGGACGTCGACGGCGCATCGCTCGTCCAGGTCGAGGACGGCGACGACAAGAAGACTGTTCTCGCAATTCCTCTCGCCGCATTGGTGCCTTTCGCGAACCCATTGGCCGACGACAAGAGCCAGAAGAAAGTTCTCAAGAGCGTCAAGAAGG	151 	\|\|\|\|\|\|\|\|\|\|\|\|\|\|\|\|\|\|\|\|\|\|\|\|\|\|\|\|\|\|\|\|\|\|\|\|\|\|\|\|\|\|\|\|\|\|\|\|\|\|\|\|\|\|\|\|\|\|\|\|\|\|\|\|\|\|\|\|\|\|\|\|\|\|\|\|\|\|\|\|\|\|\|\|\|\|\|\|\|\|\|\|\|\|\|\|\|\|\|\|\|\|\|\|\|\|\|\|\|\|\|\|\|\|\|\|\|\|\|\|\|\|\|\|\|\|\|\|\|\|\|\|\|\|\|\|\|\|\|\|\|\|\|\|\|\|\|\|\|\|\| 	ATGGACGTCGACGGCGCATCGCTCGTCCAGGTCGAGGACGGCGACGACAAGAAGACTGTTCTCGCAATTCCTCTCGCCGCATTGGTGCCTTTCGCGAACCCATTGGCCGACGACAAGAGCCAGAAGAAAGTTCTCAAGAGCGTCAAGAAGG ``` |
| 41 | ggggacaagtttgtacaaaaaagcaggcttgatgTTCTGCTTGAGATGTA, ggggaccactttgtacaagaaagctgggtcCAGCCGTATCGTTGCCCCAA | Mycgr3\|36494  JGI  Ensembl |  |  | 100 % |  | ``` 1	ATGTTCTGCTTGAGATGTAAGACAACCCGGCGTGACCTCGTGGAGAGCATTACTAACCTACCTGTTTCAGCCTGGGTGTCTCTTCTGGTCTTTCTCAGCACCGCGTCGCCTCTATACACATTGATATATCTTATCGGACACTTCTGGCTCAGTCGACCCTGCGT	164 	\|\|\|\|\|\|\|\|\|\|\|\|\|\|\|\|\|\|\|\|\|\|\|\|\|\|\|\|\|\|\|\|\|\|\|\|\|\|\|\|\|\|\|\|\|\|\|\|\|\|\|\|\|\|\|\|\|\|\|\|\|\|\|\|\|\|\|\|\|\|\|\|\|\|\|\|\|\|\|\|\|\|\|\|\|\|\|\|\|\|\|\|\|\|\|\|\|\|\|\|\|\|\|\|\|\|\|\|\|\|\|\|\|\|\|\|\|\|\|\|\|\|\|\|\|\|\|\|\|\|\|\|\|\|\|\|\|\|\|\|\|\|\|\|\|\|\|\|\|\|\|\|\|\|\|\|\|\|\|\|\|\|\|\| 	ATGTTCTGCTTGAGATGTAAGACAACCCGGCGTGACCTCGTGGAGAGCATTACTAACCTACCTGTTTCAGCCTGGGTGTCTCTTCTGGTCTTTCTCAGCACCGCGTCGCCTCTATACACATTGATATATCTTATCGGACACTTCTGGCTCAGTCGACCCTGCGT ```      ``` 165	ATACTGCACGTTCCTCCTCTGCTGCGTTGTGATTTCCCTACTCGACTTCTCCGCGGACTGGTTCGAACCAAGATGGACCGATGCCTCGACTTCGATCAACGAGACTGCGACGTTCCTCTTCAGCGGCAATGCGACGCTCACAGATGCCGTTATGGAGACTGCCAGCCTGGCACTGTCGGCTTTGAACGGGACAGGTGGATCGCTGGTCAGCTCGGTAATGGGTGGTGTAAAGAGCAATGCGGTGTCTGGCGAGCCTGTGCGAATGGTTGCCGCCAGCATACCAAATGGAAGCAACGTGTTTGAGTGGCTGCGCGGATTACTGGAGAAATTCCAATTCAGAATACCTTGTGTTGGGGCAACGATACGGCTG	534 	\|\|\|\|\|\|\|\|\|\|\|\|\|\|\|\|\|\|\|\|\|\|\|\|\|\|\|\|\|\|\|\|\|\|\|\|\|\|\|\|\|\|\|\|\|\|\|\|\|\|\|\|\|\|\|\|\|\|\|\|\|\|\|\|\|\|\|\|\|\|\|\|\|\|\|\|\|\|\|\|\|\|\|\|\|\|\|\|\|\|\|\|\|\|\|\|\|\|\|\|\|\|\|\|\|\|\|\|\|\|\|\|\|\|\|\|\|\|\|\|\|\|\|\|\|\|\|\|\|\|\|\|\|\|\|\|\|\|\|\|\|\|\|\|\|\|\|\|\|\|\|\|\|\|\|\|\|\|\|\|\|\|\|\|\|\|\|\|\|\|\|\|\|\|\|\|\|\|\|\|\|\|\|\|\|\|\|\|\|\|\|\|\|\|\|\|\|\|\|\|\|\|\|\|\|\|\|\|\|\|\|\|\|\|\|\|\|\|\|\|\|\|\|\|\|\|\|\|\|\|\|\|\|\|\|\|\|\|\|\|\|\|\|\|\|\|\|\|\|\|\|\|\|\|\|\|\|\|\|\|\|\|\|\|\|\|\|\|\|\|\|\|\|\|\|\|\|\|\|\|\|\|\|\|\|\|\|\|\|\|\|\|\|\|\|\|\|\|\|\|\|\|\|\|\|\|\|\|\|\|\|\|\|\|\|\|\|\|\|\|\|\|\|\|\|\|\|\|\|\|\|\|\|\|\|\|\|\|\|\|\|\|\|\|\|\|\|\|\|\|\|\|\|\|\|\|\|\|\|\|\|\|\|\|\|\|\|\|\|\| 	ATACTGCACGTTCCTCCTCTGCTGCGTTGTGATTTCCCTACTCGACTTCTCCGCGGACTGGTTCGAACCAAGATGGACCGATGCCTCGACTTCGATCAACGAGACTGCGACGTTCCTCTTCAGCGGCAATGCGACGCTCACAGATGCCGTTATGGAGACTGCCAGCCTGGCACTGTCGGCTTTGAACGGGACAGGTGGATCGCTGGTCAGCTCGGTAATGGGTGGTGTAAAGAGCAATGCGGTGTCTGGCGAGCCTGTGCGAATGGTTGCCGCCAGCATACCAAATGGAAGCAACGTGTTTGAGTGGCTGCGCGGATTACTGGAGAAATTCCAATTCAGAATACCTTGTGTTGGGGCAACGATACGGCTG ``` |
| 41 | ggggacaagtttgtacaaaaaagcaggcttgatgTTCACTCGACTCACAT, ggggaccactttgtacaagaaagctgggtcGCGAGTATCGCTTGCCGTTG | Mycgr3\|38680  JGI  Ensembl |  |  | 100 % |  | ``` 1	ATGTTCACTCGACTCACATCACCGCTAACAGTGAGCTTGCTGGCTTCTACAGTCGCAGCCAAATGCTCATCATTCGGCATTGACTTCCAAGGAGGCAACACCTACTTCCAAAACGTCTCGTCGACGGATCCCTTCACGTTTGTGTCGACGTTCGAGG	157 	\|\|\|\|\|\|\|\|\|\|\|\|\|\|\|\|\|\|\|\|\|\|\|\|\|\|\|\|\|\|\|\|\|\|\|\|\|\|\|\|\|\|\|\|\|\|\|\|\|\|\|\|\|\|\|\|\|\|\|\|\|\|\|\|\|\|\|\|\|\|\|\|\|\|\|\|\|\|\|\|\|\|\|\|\|\|\|\|\|\|\|\|\|\|\|\|\|\|\|\|\|\|\|\|\|\|\|\|\|\|\|\|\|\|\|\|\|\|\|\|\|\|\|\|\|\|\|\|\|\|\|\|\|\|\|\|\|\|\|\|\|\|\|\|\|\|\|\|\|\|\|\|\|\|\|\|\| 	ATGTTCACTCGACTCACATCACCGCTAACAGTGAGCTTGCTGGCTTCTACAGTCGCAGCCAAATGCTCATCATTCGGCATTGACTTCCAAGGAGGCAACACCTACTTCCAAAACGTCTCGTCGACGGATCCCTTCACGTTTGTGTCGACGTTCGAGG ```      ``` 157	GCTAACGAAGATATAGGCTGCCAGAAAGACATCACAAACAACATTCTCTACGATGAAGCCGGAGACGAGTGGGAGTGCACGGAGACGCCAACTCAACCAGATGATACCAAGCAGCTATCCACGTGTCCAGTGAACAAGAGCGATCTCCACACCGGCAGTTGGGGAGTCGTGATCATCTCAAACAATGGTGACGCTGAACCGATCGCCTTCGATCGCAGGTTCAGCCTTGTTGTTGGCGAGCCTGCTACAACAACGTTCACTCCTACTGTGACTGTAAGACCTTCATGTGAATCAACGGCAAGCGATACTCGC	468 	\|\|\|\|\|\|\|\|\|\|\|\|\|\|\|\|\|\|\|\|\|\|\|\|\|\|\|\|\|\|\|\|\|\|\|\|\|\|\|\|\|\|\|\|\|\|\|\|\|\|\|\|\|\|\|\|\|\|\|\|\|\|\|\|\|\|\|\|\|\|\|\|\|\|\|\|\|\|\|\|\|\|\|\|\|\|\|\|\|\|\|\|\|\|\|\|\|\|\|\|\|\|\|\|\|\|\|\|\|\|\|\|\|\|\|\|\|\|\|\|\|\|\|\|\|\|\|\|\|\|\|\|\|\|\|\|\|\|\|\|\|\|\|\|\|\|\|\|\|\|\|\|\|\|\|\|\|\|\|\|\|\|\|\|\|\|\|\|\|\|\|\|\|\|\|\|\|\|\|\|\|\|\|\|\|\|\|\|\|\|\|\|\|\|\|\|\|\|\|\|\|\|\|\|\|\|\|\|\|\|\|\|\|\|\|\|\|\|\|\|\|\|\|\|\|\|\|\|\|\|\|\|\|\|\|\|\|\|\|\|\|\|\|\|\|\|\|\|\|\|\|\|\|\|\|\|\|\|\|\|\|\|\|\|\|\|\|\|\|\|\|\|\|\|\|\|\|\|\|\|\|\|\|\|\|\|\|\|\|\|\|\|\|\|\|\|\|\|\|\|\|\|\|\|\|\|\|\|\|\|\|\| 	GCTAACGAAGATATAGGCTGCCAGAAAGACATCACAAACAACATTCTCTACGATGAAGCCGGAGACGAGTGGGAGTGCACGGAGACGCCAACTCAACCAGATGATACCAAGCAGCTATCCACGTGTCCAGTGAACAAGAGCGATCTCCACACCGGCAGTTGGGGAGTCGTGATCATCTCAAACAATGGTGACGCTGAACCGATCGCCTTCGATCGCAGGTTCAGCCTTGTTGTTGGCGAGCCTGCTACAACAACGTTCACTCCTACTGTGACTGTAAGACCTTCATGTGAATCAACGGCAAGCGATACTCGC ``` |
| 41 | ggggacaagtttgtacaaaaaagcaggcttgatgAGAATGATCTCGACAT, ggggaccactttgtacaagaaagctgggtcCTCTGTGGGATCCGTGCTAC | Mycgr3\|41229  JGI  Ensembl | KOG3413 Mitochondrial matrix protein frataxin, involved in Fe/S protein biosynthesis; |  | 100 % |  | ``` 219	CAGCGCTGGCGTCCTCTCAATAACATTTCCGCCCGCTGGCACGTACATCATCAACAAGCAGCCGCCGAACAAGCAGATCTGGCTGTCGTCACCAATCTCCGGTCCCAAGCGATACGATTGGGTTATCGAGGGCGAGGGCATGCATCAGAAGGAGGGCGGAGGAACGGGAGAATGGGTGTATTTGCGTGACGGCACGAATCTCACAGAGCTGTTGGGCGAAGAGCTGGGCATCCACATCTATGCACAGGCGGGGGAGGAGATGAAGGGTAGCACGGATCCCACAGAG	504 	\|\|\| \|\|\|\|\|\|\|\|\|\|\|\|\|\|\|\|\|\|\|\|\|\|\|\|\|\|\|\|\|\|\|\|\|\|\|\|\|\|\|\|\|\|\|\|\|\|\|\|\|\|\|\|\|\|\|\|\|\|\|\|\|\|\|\|\|\|\|\|\|\|\|\|\|\|\|\|\|\|\|\|\|\|\|\|\|\|\|\|\|\|\|\|\|\|\|\|\|\|\|\|\|\|\|\|\|\|\|\|\|\|\|\|\|\|\|\|\|\|\|\|\|\|\|\|\|\|\|\|\|\|\|\|\|\|\|\|\|\|\|\|\|\|\|\|\|\|\|\|\|\|\|\|\|\|\|\|\|\|\|\|\|\|\|\|\|\|\|\|\|\|\|\|\|\|\|\|\|\|\|\|\|\|\|\|\|\|\|\|\|\|\|\|\|\|\|\|\|\|\|\|\|\|\|\|\|\|\|\|\|\|\|\|\|\|\|\|\|\|\|\|\|\|\|\|\|\|\|\|\|\|\|\|\|\|\|\|\|\|\|\|\|\|\|\|\|\|\|\|\|\|\|\|\|\|\|\|\|\|\|\|\|\|\|\|\|\|\|\|\|\|\|\|\|\|\|\|\|\|\|\|\|\| 	CAG-GCTGGCGTCCTCTCAATAACATTTCCGCCCGCTGGCACGTACATCATCAACAAGCAGCCGCCGAACAAGCAGATCTGGCTGTCGTCACCAATCTCCGGTCCCAAGCGATACGATTGGGTTATCGAGGGCGAGGGCATGCATCAGAAGGAGGGCGGAGGAACGGGAGAATGGGTGTATTTGCGTGACGGCACGAATCTCACAGAGCTGTTGGGCGAAGAGCTGGGCATCCACATCTATGCACAGGCGGGGGAGGAGATGAAGGGTAGCACGGATCCCACAGAG ```      ``` 1	ATGAGAATGATCTCGACATCAAAGGCTTCAATGGCCGGACTCATGCCTGATTCAGAAGATCCCGAGCCGCCGACAGTCAAGGCGACAGAAGCTCCGCCCGCTCAACCAACAGAGCTCAGCGATGAAGCATACGCCACGGTTGCGGAGGAATACATGGAATTAGTTCATGAGAAGGCAGAACAGATCCAGGAGACACGGGAAGATGTTGAAGTCGAGTACAGCG	223 	\|\|\|\|\|\|\|\|\|\|\|\|\|\|\|\|\|\|\|\|\|\|\|\|\|\|\|\|\|\|\|\|\|\|\|\|\|\|\|\|\|\|\|\|\|\|\|\|\|\|\|\|\|\| \|\|\|\|\|\|\|\|\|\|\|\|\|\|\|\|\|\|\|\|\|\|\|\|\|\|\|\|\|\|\|\|\|\|\|\|\|\|\|\|\|\|\|\|\|\|\|\|\|\|\|\|\|\|\|\|\|\|\|\|\|\|\|\|\|\|\|\|\|\|\|\|\|\|\|\|\|\|\|\|\|\|\|\|\|\|\|\|\|\|\|\|\|\|\|\|\|\|\|\|\|\|\|\|\|\|\|\|\|\|\|\|\|\|\|\|\|\|\|\|\|\|\|\|\|\|\|\|\|\|\|\|\|\|\|\|\|\|\|\|\|\|\|\|\|\|\|\|\|\|\|\|\|\|\|\|\|\|\|\|\|\|\|\|\|\|\|\| 	ATGAGAATGATCTCGACATCAAAGGCTTCAATGGCCGGACTCATGCCTGATTCATAAGATCCCGAGCCGCCGACAGTCAAGGCGACAGAAGCTCCGCCCGCTCAACCAACAGAGCTCAGCGATGAAGCATACGCCACGGTTGCGGAGGAATACATGGAATTAGTTCATGAGAAGGCAGAACAGATCCAGGAGACACGGGAAGATGTTGAAGTCGAGTACAGCG ``` |
| 41 | ggggacaagtttgtacaaaaaagcaggcttgatgGCATCAGTACTCGCAA, ggggaccactttgtacaagaaagctgggtcTCCCGTCTTCTCCAGCAACT | Mycgr3\|42338  JGI  Ensembl | KOG0723 Molecular chaperone (DnaJ superfamily); | heat shock protein binding; | 100 % |  | ``` 52	GGCCGCGCTGGCCTAGTCGCATTACGAAGAGCACGAGGTGGCGGAACAGCACTCGGGAAATCCTACTACAAAGGCGGCTTCGAAAAGACGATGAACCGAAAAGAGGCAGCTTTGATCTTGGAGACATC	179 	\|\|\|\|\|\|\|\|\|\|\|\|\|\|\|\|\|\|\|\|\|\|\|\|\|\|\|\|\|\|\|\|\|\|\|\|\|\|\|\|\|\|\|\|\|\|\|\|\|\|\|\|\|\|\|\|\|\|\|\|\|\|\|\|\|\|\|\|\|\|\|\|\|\|\|\|\|\|\|\|\|\|\|\|\|\|\|\|\|\|\|\|\|\|\|\|\|\|\|\|\|\|\|\|\|\|\|\|\|\|\|\|\|\|\|\|\|\|\|\|\|\|\|\|\|\|\|\| 	GGCCGCGCTGGCCTAGTCGCATTACGAAGAGCACGAGGTGGCGGAACAGCACTCGGGAAATCCTACTACAAAGGCGGCTTCGAAAAGACGATGAACCGAAAAGAGGCAGCTTTGATCTTGGAGACATC ```      ``` 180	AGAACGAGGCATAACGAAAGAAATGATTCGAAAGAAGCACCGGCAAATGATGTTGTTAAACCACCCGGATCGAGGAGGAAGTCCATATCTGGCTACAAAGATCAACGAGGCGAAAGAGTTGCTGGAGAAGACGGGA	315 	\|\|\|\|\|\|\|\|\|\|\|\|\|\|\|\|\|\|\|\|\|\|\|\|\|\|\|\|\|\|\|\|\|\|\|\|\|\|\|\|\|\|\|\|\|\|\|\|\|\|\|\|\|\|\|\|\|\|\|\|\|\|\|\|\|\|\|\|\|\|\|\|\|\|\|\|\|\|\|\|\|\|\|\|\|\|\|\|\|\|\|\|\|\|\|\|\|\|\|\|\|\|\|\|\|\|\|\|\|\|\|\|\|\|\|\|\|\|\|\|\|\|\|\|\|\|\|\|\|\|\|\|\|\|\|\| 	AGAACGAGGCATAACGAAAGAAATGATTCGAAAGAAGCACCGGCAAATGATGTTGTTAAACCACCCGGATCGAGGAGGAAGTCCATATCTGGCTACAAAGATCAACGAGGCGAAAGAGTTGCTGGAGAAGACGGGA ```      ``` 1	ATGGCATCAGTACTCGCAATAGGCGTCGGCGTCGCAGCCGCAGCATTCTTCG	52 	\|\|\|\|\|\|\|\|\|\|\|\|\|\|\|\|\|\|\|\|\|\|\|\|\|\|\|\|\|\|\|\|\|\|\|\|\|\|\|\|\|\|\|\|\|\|\|\|\|\|\|\| 	ATGGCATCAGTACTCGCAATAGGCGTCGGCGTCGCAGCCGCAGCATTCTTCG ``` |
| 41 | ggggacaagtttgtacaaaaaagcaggcttgatgACATTTTCCAACCCAG, ggggaccactttgtacaagaaagctgggtcCCTCATCAAGTGTCCCTTCA | Mycgr3\|42731  JGI  Ensembl |  |  | 100 % |  | ``` 1	ATGACATTTTCCAACCCAGGCGGCATCCTCATGCTCCTTGTAAATCTTGCCTACTCCATCGGCGCATTCTTCTTCGACTACAGCGACACTCACGTCCTCAACCCACGATGGCCTCCACATGCAAAGTTCCACAACGGACAGACCATGTCTTTAGGAGCCATGCAGGCCGCGCTCTCGATCTACCTCATTCTGCGTCCGACTGCGAACAAGACGGAGAAGAGAGATAGTCTGTGGCTCGCGGCGCTGGTTGGGAGTCTGTACTGCGCGGCGGGACTGTCTGCGATTCTGTATCCGGGAACGAAATGGGCGGA	311 	\|\|\|\|\|\|\|\|\|\|\|\|\|\|\|\|\|\|\|\|\|\|\|\|\|\|\|\|\|\|\|\|\|\|\|\|\|\|\|\|\|\|\|\|\|\|\|\|\|\|\|\|\|\|\|\|\|\|\|\|\|\|\|\|\|\|\|\|\|\|\|\|\|\|\|\|\|\|\|\|\|\|\|\|\|\|\|\|\|\|\|\|\|\|\|\|\|\|\|\|\|\|\|\|\|\|\|\|\|\|\|\|\|\|\|\|\|\|\|\|\|\|\|\|\|\|\|\|\|\|\|\|\|\|\|\|\|\|\|\|\|\|\|\|\|\|\|\|\|\|\|\|\|\|\|\|\|\|\|\|\|\|\|\|\|\|\|\|\|\|\|\|\|\|\|\|\|\|\|\|\|\|\|\|\|\|\|\|\|\|\|\|\|\|\|\|\|\|\|\|\|\|\|\|\|\|\|\|\|\|\|\|\|\|\|\|\|\|\|\|\|\|\|\|\|\|\|\|\|\|\|\|\|\|\|\|\|\|\|\|\|\|\|\|\|\|\|\|\|\|\|\|\|\|\|\|\|\|\|\|\|\|\|\|\|\|\|\|\|\|\|\|\|\|\|\|\|\|\|\|\|\|\|\|\|\|\|\|\|\|\|\|\|\|\|\|\|\|\|\|\|\|\|\|\|\|\|\|\|\|\| 	ATGACATTTTCCAACCCAGGCGGCATCCTCATGCTCCTTGTAAATCTTGCCTACTCCATCGGCGCATTCTTCTTCGACTACAGCGACACTCACGTCCTCAACCCACGATGGCCTCCACATGCAAAGTTCCACAACGGACAGACCATGTCTTTAGGAGCCATGCAGGCCGCGCTCTCGATCTACCTCATTCTGCGTCCGACTGCGAACAAGACGGAGAAGAGAGATAGTCTGTGGCTCGCGGCGCTGGTTGGGAGTCTGTACTGCGCGGCGGGACTGTCTGCGATTCTGTATCCGGGAACGAAATGGGCGGA ```      ``` 312	TCCTGAATTCGCGGCCAATGGTGAGCAGAAGTTCATCTTCGGGGGATGCGTCGTTTTGATATGGGCCGGGTACTGGGTCGAAACGAGGAGGCTGCAGCATGTGAAGGGACACTTGATGAGG	432 	\|\|\|\|\|\|\|\|\|\|\|\|\|\|\|\|\|\|\|\|\|\|\|\|\|\|\|\|\|\|\|\|\|\|\|\|\|\|\|\|\|\|\|\|\|\|\|\|\|\|\|\|\|\|\|\|\|\|\|\|\|\|\|\|\|\|\|\|\|\|\|\|\|\|\|\|\|\|\|\|\|\|\|\|\|\|\|\|\|\|\|\|\|\|\|\|\|\|\|\|\|\|\|\|\|\|\|\|\|\|\|\|\|\|\|\|\|\|\|\|\| 	TCCTGAATTCGCGGCCAATGGTGAGCAGAAGTTCATCTTCGGGGGATGCGTCGTTTTGATATGGGCCGGGTACTGGGTCGAAACGAGGAGGCTGCAGCATGTGAAGGGACACTTGATGAGG ``` |
| 41 | ggggacaagtttgtacaaaaaagcaggcttgatgGCTGACGTCGACATGA, ggggaccactttgtacaagaaagctgggtcGCGACCATACTTCTGGAACT | Mycgr3\|46230  JGI  Ensembl | KOG2930 SCF ubiquitin ligase, Rbx1 component; | negative regulation of ubiquitin-protein ligase activity; positive regulation of ubiquitin-protein ligase activity;  protein binding;  regulation of ubiquitin-protein ligase activity;  regulation of ubiquitin-protein ligase activity during meiotic cell cycle;  ubiquitin-protein ligase activity;  zinc ion binding; | 100 % |  | ``` 107	AGTGGAACGCCGTCGCGCTCTGGGCATGGGACATCGTTGTGGATAACTGCGCCATTTGCAGAAACCACATCATGGACTTGTGTATCGACTGTCAGGCCAACCAAGCTTCGGCGACCAGCGAGGAGTGCACAGTCGCTTGGGGAATCTGCAAT	258 	\|\|\|\|\|\|\|\|\|\|\|\|\|\|\|\|\|\|\|\|\|\|\|\|\|\|\|\|\|\|\|\|\|\|\|\|\|\|\|\|\|\|\|\|\|\|\|\|\|\|\|\|\|\|\|\|\|\|\|\|\|\|\|\|\|\|\|\|\|\|\|\|\|\|\|\|\|\|\|\|\|\|\|\|\|\|\|\|\|\|\|\|\|\|\|\|\|\|\|\|\|\|\|\|\|\|\|\|\|\|\|\|\|\|\|\|\|\|\|\|\|\|\|\|\|\|\|\|\|\|\|\|\|\|\|\|\|\|\|\|\|\|\|\|\|\|\|\|\|\|\|\| 	AGTGGAACGCCGTCGCGCTCTGGGCATGGGACATCGTTGTGGATAACTGCGCCATTTGCAGAAACCACATCATGGACTTGTGTATCGACTGTCAGGCCAACCAAGCTTCGGCGACCAGCGAGGAGTGCACAGTCGCTTGGGGAATCTGCAAT ```      ``` 259	CACGCCTTCCACTTCCACTGCATCTCGAGATGGCTCAAGACACGCCAGGTCTGCCCGCTGGACAATCGAGACTGGGAGTTCCAGAAGTATGGTCGC	354 	\|\|\|\|\|\|\|\|\|\|\|\|\|\|\|\|\|\|\|\|\|\|\|\|\|\|\|\|\|\|\|\|\|\|\|\|\|\|\|\|\|\|\|\|\|\|\|\|\|\|\|\|\|\|\|\|\|\|\|\|\|\|\|\|\|\|\|\|\|\|\|\|\|\|\|\|\|\|\|\|\|\|\|\|\|\|\|\|\|\|\|\|\|\|\|\| 	CACGCCTTCCACTTCCACTGCATCTCGAGATGGCTCAAGACACGCCAGGTCTGCCCGCTGGACAATCGAGACTGGGAGTTCCAGAAGTATGGTCGC ```      ``` 1	ATGGCTGACGTCGACATGACCGATGCTCCACCCAAGGCGGTGGCGACGAGCAAGGCTGGTGCAAAAGGCGGCGAAGCGGATGGCAAGAAGCGATTCGAGGTCAAGAAG-TGG	111 	\|\|\|\|\|\|\|\|\|\|\|\|\|\|\|\|\|\|\|\|\|\|\|\|\|\|\|\|\|\|\|\|\|\|\|\|\|\|\|\|\|\|\|\|\|\|\|\|\|\|\|\|\|\|\|\|\|\|\|\|\|\|\|\|\|\|\|\|\|\|\|\|\|\|\|\|\|\|\|\|\|\|\|\|\|\|\|\|\|\|\|\|\|\|\|\|\|\|\|\|\|\|\|\|\|\|\|\| \|\|\| 	ATGGCTGACGTCGACATGACCGATGCTCCACCCAAGGCGGTGGCGACGAGCAAGGCTGGTGCAAAAGGCGGCGAAGCGGATGGCAAGAAGCGATTCGAGGTCAAGAAGGTGG ``` |
| 41 | ggggacaagtttgtacaaaaaagcaggcttgatgGCGAGATATCGCAACA, ggggaccactttgtacaagaaagctgggtcGCACGATACTAAAGAAGATG | Mycgr3\|47646  JGI  Ensembl | KOG3467 Histone H4; | DNA binding; nucleosome; nucleosome assembly;  nucleus; | 100 % |  | ``` 1	ATGGCGAGATATCGCAACAGTAACGCCATCATCCGTTCAGATGTGCCGCACGCTTCGTCGAGCTCTGCGCCGCGTCCGACCGCTTCCTCGACTGGCGGCAAAGGAAAGACCTCGTTCCACGGCGGAGTTGGATCCAAGGGACTCGGCGTCACTCCACACAAGCGGCACCG	170 	\|\|\|\|\|\|\|\|\|\|\|\|\|\|\|\|\|\|\|\|\|\|\|\|\|\|\|\|\|\|\|\|\|\|\|\|\|\|\|\|\|\|\|\|\|\|\|\|\|\|\|\|\|\|\|\|\|\|\|\|\|\|\|\|\|\|\|\|\|\|\|\|\|\|\|\|\|\|\|\|\|\|\|\|\|\|\|\|\|\|\|\|\|\|\|\|\|\|\|\|\|\|\|\|\|\|\|\|\|\|\|\|\|\|\|\|\|\|\|\|\|\|\|\|\|\|\|\|\|\|\|\|\|\|\|\|\|\|\|\|\|\|\|\|\|\|\|\|\|\|\|\|\|\|\|\|\|\|\|\|\|\|\|\|\|\|\|\|\|\| 	ATGGCGAGATATCGCAACAGTAACGCCATCATCCGTTCAGATGTGCCGCACGCTTCGTCGAGCTCTGCGCCGCGTCCGACCGCTTCCTCGACTGGCGGCAAAGGAAAGACCTCGTTCCACGGCGGAGTTGGATCCAAGGGACTCGGCGTCACTCCACACAAGCGGCACCG ```      ``` 307	CTTCTCAAGGATATCGCCGCGGTCGTCGAGCTCTCAGGACGGAAGACGGTCTGCGTCACAGATGTGGTGTTCGTACTCAACCGCGTAGGAAACCCGATCTACGTAAGTTTGATGCTCCGCACATCTTCTTTAGTATCGTGC	447 	\|\|\|\|\|\|\|\|\|\|\|\|\|\|\|\|\|\|\|\|\|\|\|\|\|\|\|\|\|\|\|\|\|\|\|\|\|\|\|\|\|\|\|\|\|\|\|\|\|\|\|\|\|\|\|\|\|\|\|\|\|\|\|\|\|\|\|\|\|\|\|\|\|\|\|\|\|\|\|\|\|\|\|\|\|\|\|\|\|\|\|\|\|\|\|\|\|\|\|\|\|\|\|\|\|\|\|\|\|\|\|\|\|\|\|\|\|\|\|\|\|\|\|\|\|\|\|\|\|\|\|\|\|\|\|\|\|\|\|\|\| 	CTTCTCAAGGATATCGCCGCGGTCGTCGAGCTCTCAGGACGGAAGACGGTCTGCGTCACAGATGTGGTGTTCGTACTCAACCGCGTAGGAAACCCGATCTACGTAAGTTTGATGCTCCGCACATCTTCTTTAGTATCGTGC ```      ``` 170	GCAAGATCTTACGCGACAATATTCAAGGCGTCACAAAAGGCGACATCCGTCGAATGGCCCGCCGTGGTGGAATCAAGCGTATTGCAGGCGGGATCTACGAGGAGACTCGTCTCGTCTTGAGAAAGCGTCTGGAAGTT	306 	\|\|\|\|\|\|\|\|\|\|\|\|\|\|\|\|\|\|\|\|\|\|\|\|\|\|\|\|\|\|\|\|\|\|\|\|\|\|\|\|\|\|\|\|\|\|\|\|\|\|\|\|\|\|\|\|\|\|\|\|\|\|\|\|\|\|\|\|\|\|\|\|\|\|\|\|\|\|\|\|\|\|\|\|\|\|\|\|\|\|\|\|\|\|\|\|\|\|\|\|\|\|\|\|\|\|\|\|\|\|\|\|\|\|\|\|\|\|\|\|\|\|\|\|\|\|\|\|\|\|\|\|\|\|\|\|\| 	GCAAGATCTTACGCGACAATATTCAAGGCGTCACAAAAGGCGACATCCGTCGAATGGCCCGCCGTGGTGGAATCAAGCGTATTGCAGGCGGGATCTACGAGGAGACTCGTCTCGTCTTGAGAAAGCGTCTGGAAGTT ``` |
| 41 | ggggacaagtttgtacaaaaaagcaggcttgatgAACCGCCATCTCCGCG, ggggaccactttgtacaagaaagctgggtcCCTCCCACCCACAGCCGCAT | Mycgr3\|56988  JGI  Ensembl |  |  | 3.39286 % |  |  |
| 41 | ggggacaagtttgtacaaaaaagcaggcttgatgGCGCAGCCAGCAGCGC, ggggaccactttgtacaagaaagctgggtcCGTCGCCTTTTCCTTCGTCT | Mycgr3\|62604  JGI  Ensembl | KOG3463 Transcription initiation factor IIA, gamma subunit; | RNA polymerase II transcription factor activity; transcription factor TFIIA complex;  transcription initiation from RNA polymerase II promoter; | 100 % |  | ``` 171	CATCGGTCTCGCGCTGGCGGATACGCTCGATGATCTGATCAGCAGCCGTCGCATCGAGCCGCAGCTTGCGATGCGCATCATGGCCAACTTCGATCAGCATATTGCTTCGGTGCTTGGGGAGAAGGTTAAGGCGAGGATGAATTTCAAGG	319 	\|\|\|\|\|\|\|\|\|\|\|\|\|\|\|\|\|\|\|\|\|\|\|\|\|\|\|\|\|\|\|\|\|\|\|\|\|\|\|\|\|\|\|\|\|\|\|\|\|\|\|\|\|\|\|\|\|\|\|\|\|\|\|\|\|\|\|\|\|\|\|\|\|\|\|\|\|\|\|\|\|\|\|\|\|\|\|\|\|\|\|\|\|\|\|\|\|\|\|\|\|\|\|\|\|\|\|\|\|\|\|\|\|\|\|\|\|\|\|\|\|\|\|\|\|\|\|\|\|\|\|\|\|\|\|\|\|\|\|\|\|\|\|\|\|\|\|\|\| 	CATCGGTCTCGCGCTGGCGGATACGCTCGATGATCTGATCAGCAGCCGTCGCATCGAGCCGCAGCTTGCGATGCGCATCATGGCCAACTTCGATCAGCATATTGCTTCGGTGCTTGGGGAGAAGGTTAAGGCGAGGATGAATTTCAAGG ```      ``` 317	AGGGCCACTTGGACGTCTACCGATTCTGCGACGAAGTCTGGACATTTGTGATCAAAGACGTCAAGTTCAAGATGGATAACAACACACAACAATTGGATGCGGAGAAGGTCAAGATCGTGGCTTGCCAGACGAAGGAAAAGGCGACG	462 	\|\|\|\|\|\|\|\|\|\|\|\|\|\|\|\|\|\|\|\|\|\|\|\|\|\|\|\|\|\|\|\|\|\|\|\|\|\|\|\|\|\|\|\|\|\|\|\|\|\|\|\|\|\|\|\|\|\|\|\|\|\|\|\|\|\|\|\|\|\|\|\|\|\|\|\|\|\|\|\|\|\|\|\|\|\|\|\|\|\|\|\|\|\|\|\|\|\|\|\|\|\|\|\|\|\|\|\|\|\|\|\|\|\|\|\|\|\|\|\|\|\|\|\|\|\|\|\|\|\|\|\|\|\|\|\|\|\|\|\|\|\|\|\|\|\| 	AGGGCCACTTGGACGTCTACCGATTCTGCGACGAAGTCTGGACATTTGTGATCAAAGACGTCAAGTTCAAGATGGATAACAACACACAACAATTGGATGCGGAGAAGGTCAAGATCGTGGCTTGCCAGACGAAGGAAAAGGCGACG ```      ``` 121	ATGGCGCAGCCAGCAGCGCAGGGCTTCTACGAGCTTTACAGAGGCACATC	170 	\|\|\|\|\|\|\|\|\|\|\|\|\|\|\|\|\|\|\|\|\|\|\|\|\|\|\|\|\|\|\|\|\|\|\|\|\|\|\|\|\|\|\|\|\|\|\|\|\|\| 	ATGGCGCAGCCAGCAGCGCAGGGCTTCTACGAGCTTTACAGAGGCACATC ``` |
| 41 | ggggacaagtttgtacaaaaaagcaggcttgatgGCCGGTGGAACCCAGC, ggggaccactttgtacaagaaagctgggtcCCACAACTTATACCACGGCT | Mycgr3\|63422  JGI  Ensembl |  |  | 3.76238 % |  |  |
| 41 | ggggacaagtttgtacaaaaaagcaggcttgatgGCTCACCGCATAATAA, ggggaccactttgtacaagaaagctgggtcATCTCTGTACAGCTTCGGCC | Mycgr3\|70050  JGI  Ensembl | KOG3442 Uncharacterized conserved protein; |  | 100 % |  | ``` 1	ATGGCTCACCGCATAATAACTCAAATGGCCTTTACCGGCGCCCGGGTACTTGGTCGAGCATTTACTGAAGCATACAAGCAAGCATCCGCCTCTCAG	96 	\|\|\|\|\|\|\|\|\|\|\|\|\|\|\|\|\|\|\|\|\|\|\|\|\|\|\|\|\|\|\|\|\|\|\|\|\|\|\|\|\|\|\|\|\|\|\|\|\|\|\|\|\|\|\|\|\|\|\|\|\|\|\|\|\|\|\|\|\|\|\|\|\|\|\|\|\|\|\|\|\|\|\|\|\|\|\|\|\|\|\|\|\|\|\|\| 	ATGGCTCACCGCATAATAACTCAAATGGCCTTTACCGGCGCCCGGGTACTTGGTCGAGCATTTACTGAAGCATACAAGCAAGCATCCGCCTCTCAG ```      ``` 94	CAGAAATACGCGCAAGCAGGCCAGACCGGCGCAAACACCCTATCATCAGCAGGCTTGACCCTCAGCGAAGCCTGCCAAATCCTCAACGTTCCTCCTCCGAAAGCCGGCACGACAGATCTCGTCAAAGTCCACAACAACTTCAAGCGACTATTCGACCTGAACGACCCGAAGAAAGGCGGCTCATTTTACATCCAAAGTAAAGTGCTACGCGCGCGGGAAAGATTAGAGCTGGAGGTACGTGAAGCGGAAGCGGCGAAAGAACGAGAAGACGAGAAGAAGCACGGCTGGAGGCCGAAGCTGTACAGAGAT	402 	\|\|\|\|\|\|\|\|\|\|\|\|\|\|\|\|\|\|\|\|\|\|\|\|\|\|\|\|\|\|\|\|\|\|\|\|\|\|\|\|\|\|\|\|\|\|\|\|\|\|\|\|\|\|\|\|\|\|\|\|\|\|\|\|\|\|\|\|\|\|\|\|\|\|\|\|\|\|\|\|\|\|\|\|\|\|\|\|\|\|\|\|\|\|\|\|\|\|\|\|\|\|\|\|\|\|\|\|\|\|\|\|\|\|\|\|\|\|\|\|\|\|\|\|\|\|\|\|\|\|\|\|\|\|\|\|\|\|\|\|\|\|\|\|\|\|\|\|\|\|\|\|\|\|\|\|\|\|\|\|\|\|\|\|\|\|\|\|\|\|\|\|\|\|\|\|\|\|\|\|\|\|\|\|\|\|\|\|\|\|\|\|\|\|\|\|\|\|\|\|\|\|\|\|\|\|\|\|\|\|\|\|\|\|\|\|\|\|\|\|\|\|\|\|\|\|\|\|\|\|\|\|\|\|\|\|\|\|\|\|\|\|\|\|\|\|\|\|\|\|\|\|\|\|\|\|\|\|\|\|\|\|\|\|\|\|\|\|\|\|\|\|\|\|\|\|\|\|\|\|\|\|\|\|\|\|\|\|\|\|\|\|\|\|\|\|\|\|\|\|\|\|\|\|\|\|\|\|\| 	CAGAAATACGCGCAAGCAGGCCAGACCGGCGCAAACACCCTATCATCAGCAGGCTTGACCCTCAGCGAAGCCTGCCAAATCCTCAACGTTCCTCCTCCGAAAGCCGGCACGACAGATCTCGTCAAAGTCCACAACAACTTCAAGCGACTATTCGACCTGAACGACCCGAAGAAAGGCGGCTCATTTTACATCCAAAGTAAAGTGCTACGCGCGCGGGAAAGATTAGAGCTGGAGGTACGTGAAGCGGAAGCGGCGAAAGAACGAGAAGACGAGAAGAAGCACGGCTGGAGGCCGAAGCTGTACAGAGAT ``` |
| 41 | ggggacaagtttgtacaaaaaagcaggcttgatgAACTCACCATTCTGGC, ggggaccactttgtacaagaaagctgggtcCATCATAATCCTCCCCCGAC | Mycgr3\|72452  JGI  Ensembl | KOG3472 Predicted small membrane protein; |  | 100 % |  | ``` 1	ATGAACTCACCATTCTGGCTCGTCGGCGCCCTGTCCGCTGCCTCATCAGTCGGCATCGGAGCTTTCGGAGCTCATGGCTTGAAGAATCGCGGCCTGGAGCCGGCGAAGATCGCTTCATTTCAGACGGCGGCGCATTACCAG	141 	\|\|\|\|\|\|\|\|\|\|\|\|\|\|\|\|\|\|\|\|\|\|\|\|\|\|\|\|\|\|\|\|\|\|\|\|\|\|\|\|\|\|\|\|\|\|\|\|\|\|\|\|\|\|\|\|\|\|\|\|\|\|\|\|\|\|\|\|\|\|\|\|\|\|\|\|\|\|\|\|\|\|\|\|\|\|\|\|\|\|\|\|\|\|\|\|\|\|\|\|\|\|\|\|\|\|\|\|\|\|\|\|\|\|\|\|\|\|\|\|\|\|\|\|\|\|\|\|\|\|\|\|\|\|\|\|\|\|\|\|\| 	ATGAACTCACCATTCTGGCTCGTCGGCGCCCTGTCCGCTGCCTCATCAGTCGGCATCGGAGCTTTCGGAGCTCATGGCTTGAAGAATCGCGGCCTGGAGCCGGCGAAGATCGCTTCATTTCAGACGGCGGCGCATTACCAG ```      ``` 138	CCAGCTCGTCCACTCCATCGCCCTACTCGTAGCCGAACACGCTGCTCCAAAGAACGTCTGGGCCAAGAGCTTGTTCACAGCCGGGATTACGATGTTTTCTGGCTCGATCTATCTTCTCGTTCTGGATGGACAGACGTTCAAGTCTCTTGGTCCGGTCACGCCGTTGGGAGGGATGTGTTTGATGGGAGGATGGATTGCTCTGGCTTTTGGGAGTCGGGGGAGGATTATGATG	369 	\|\|\|\|\|\|\|\|\|\|\|\|\|\|\|\|\|\|\|\|\|\|\|\|\|\|\|\|\|\|\|\|\|\|\|\|\|\|\|\|\|\|\|\|\|\|\|\|\|\|\|\|\|\|\|\|\|\|\|\|\|\|\|\|\|\|\|\|\|\|\|\|\|\|\|\|\|\|\|\|\|\|\|\|\|\|\|\|\|\|\|\|\|\|\|\|\|\|\|\|\|\|\|\|\|\|\|\|\|\|\|\|\|\|\|\|\|\|\|\|\|\|\|\|\|\|\|\|\|\|\|\|\|\|\|\|\|\|\|\|\|\|\|\|\|\|\|\|\|\|\|\|\|\|\|\|\|\|\|\|\|\|\|\|\|\|\|\|\|\|\|\|\|\|\|\|\|\|\|\|\|\|\|\|\|\|\|\|\|\|\|\|\|\|\|\|\|\|\|\|\|\|\|\|\|\|\|\|\|\|\|\|\|\|\|\|\|\|\|\|\|\|\|\|\|\|\|\|\|\|\|\| 	CCAGCTCGTCCACTCCATCGCCCTACTCGTAGCCGAACACGCTGCTCCAAAGAACGTCTGGGCCAAGAGCTTGTTCACAGCCGGGATTACGATGTTTTCTGGCTCGATCTATCTTCTCGTTCTGGATGGACAGACGTTCAAGTCTCTTGGTCCGGTCACGCCGTTGGGAGGGATGTGTTTGATGGGAGGATGGATTGCTCTGGCTTTTGGGAGTCGGGGGAGGATTATGATG ``` |
| 41 | ggggacaagtttgtacaaaaaagcaggcttgatgTCTGAGTCTCAAGGTC, ggggaccactttgtacaagaaagctgggtcTATCGCGCTGGGGTAGAGCA | Mycgr3\|74429  JGI  Ensembl | KOG0887 60S ribosomal protein L35A/L37; | intracellular; ribosome; structural constituent of ribosome;  translation; | 100 % |  | ``` 60	GGGCAAGCACCTTTCGTACCAACGTGGCAAGCGCAACACCAACCCCGACACCTCTCTCATCCAGATTGAGGGTGTCGAGAACACCAAGGACGCAACCTTCTACCTGGGCAAGCGCATCGCATACGTCTACCGCGCCAAGAAGGAGATTCGAGGAAGCAAGATCCGTGTGATCTGGGGCAAGGTCACGAGGACACACGG	257 	\|\|\|\|\|\|\|\|\|\|\|\|\|\|\|\|\|\|\|\|\|\|\|\|\|\|\|\|\|\|\|\|\|\|\|\|\|\|\|\|\|\|\|\|\|\|\|\|\|\|\|\|\|\|\|\|\|\|\|\|\|\|\|\|\|\|\|\|\|\|\|\|\|\|\|\|\|\|\|\|\|\|\|\|\|\|\|\|\|\|\|\|\|\|\|\|\|\|\|\|\|\|\|\|\|\|\|\|\|\|\|\|\|\|\|\|\|\|\|\|\|\|\|\|\|\|\|\|\|\|\|\|\|\|\|\|\|\|\|\|\|\|\|\|\|\|\|\|\|\|\|\|\|\|\|\|\|\|\|\|\|\|\|\|\|\|\|\|\|\|\|\|\|\|\|\|\|\|\|\|\|\|\|\|\|\|\|\|\|\|\|\|\|\|\|\|\|\| 	GGGCAAGCACCTTTCGTACCAACGTGGCAAGCGCAACACCAACCCCGACACCTCTCTCATCCAGATTGAGGGTGTCGAGAACACCAAGGACGCAACCTTCTACCTGGGCAAGCGCATCGCATACGTCTACCGCGCCAAGAAGGAGATTCGAGGAAGCAAGATCCGTGTGATCTGGGGCAAGGTCACGAGGACACACGG ```      ``` 251	CACACGGCAACTCCGGCGTCGTGCGAGCACAGTTCAGGCACAACCTGCCAGCCAAGTCCTTCGGCGCGATGGTCCGAATCATGCTCTACCCCAGCGCGATA	351 	\|\|\|\| \|\|\|\|\|\|\|\|\|\|\|\|\|\|\|\|\|\|\|\|\|\|\|\|\|\|\|\|\|\|\|\|\|\|\|\|\|\|\|\|\|\|\|\|\|\|\|\|\|\|\|\|\|\|\|\|\|\|\|\|\|\|\|\|\|\|\|\|\|\|\|\|\|\|\|\|\|\|\|\|\|\|\|\|\|\|\|\|\|\|\|\|\|\|\|\| 	CACA-GGCAACTCCGGCGTCGTGCGAGCACAGTTCAGGCACAACCTGCCAGCCAAGTCCTTCGGCGCGATGGTCCGAATCATGCTCTACCCCAGCGCGATA ```      ``` 25	ATGTCTGAGTCTCAAGGTCACCGGTTATACGTTAAG	60 	\|\|\|\|\|\|\|\|\|\|\|\|\|\|\|\|\|\|\|\|\|\|\|\|\|\|\|\|\|\|\|\|\|\|\|\| 	ATGTCTGAGTCTCAAGGTCACCGGTTATACGTTAAG ``` |
| 41 | ggggacaagtttgtacaaaaaagcaggcttgatgTCCGCCGAAGCCTCCG, ggggaccactttgtacaagaaagctgggtcGATCATGAGTCTCTCAACCT | Mycgr3\|75672  JGI  Ensembl | KOG3387 60S ribosomal protein 15.5kD/SNU13, NHP2/L7A family (includes ribonuclease P subunit p38), involved in splicing; | intracellular; ribonucleoprotein complex;  ribosome;  ribosome biogenesis and assembly;  structural constituent of ribosome;  translation; | 100 % |  | ``` 570	AGCTACCAAGACCCTCAACCGTGGTGTTGCTGAGTTGATCATTCTCGCCGCCGACACGACTCCACTGGCCATCCTCCTCCATCTCCCACTCCTCTGTGAAGACAAGAACGTGCCATACGTCTACGTGCCGAGCAAGATGGCTCTCGGCCGCGCTTGCGGTGTCAGCCGCGCTGTCATTGCCGCGAGCATCACCACCAACGAGGCCAGTGACTTGATGGGACAGATTCGCGCGACCAAGGATCAGGTTGAGAGACTCATGATC	831 	\|\|\|\|\|\|\|\|\|\|\|\|\|\|\|\|\|\|\|\|\|\|\|\|\|\|\|\|\|\|\|\|\|\|\|\|\|\|\|\|\|\|\|\|\|\|\|\|\|\|\|\|\|\|\|\|\|\|\|\|\|\|\|\|\|\|\|\|\|\|\|\|\|\|\|\|\|\|\|\|\|\|\|\|\|\|\|\|\|\|\|\|\|\|\|\|\|\|\|\|\|\|\|\|\|\|\|\|\|\|\|\|\|\|\|\|\|\|\|\|\|\|\|\|\|\|\|\|\|\|\|\|\|\|\|\|\|\|\|\|\|\|\|\|\|\|\|\|\|\|\|\|\|\|\|\|\|\|\|\|\|\|\|\|\|\|\|\|\|\|\|\|\|\|\|\|\|\|\|\|\|\|\|\|\|\|\|\|\|\|\|\|\|\|\|\|\|\|\|\|\|\|\|\|\|\|\|\|\|\|\|\|\|\|\|\|\|\|\|\|\|\|\|\|\|\|\|\|\|\|\|\|\|\|\|\|\|\|\|\|\|\|\|\|\|\|\|\|\|\|\|\|\|\|\|\|\|\|\|\|\|\| 	AGCTACCAAGACCCTCAACCGTGGTGTTGCTGAGTTGATCATTCTCGCCGCCGACACGACTCCACTGGCCATCCTCCTCCATCTCCCACTCCTCTGTGAAGACAAGAACGTGCCATACGTCTACGTGCCGAGCAAGATGGCTCTCGGCCGCGCTTGCGGTGTCAGCCGCGCTGTCATTGCCGCGAGCATCACCACCAACGAGGCCAGTGACTTGATGGGACAGATTCGCGCGACCAAGGATCAGGTTGAGAGACTCATGATC ```      ``` 454	ATGTCCGCCGAAGCCTCCGCCGCATGGCCCGTGGCCGACCAGGCCCTCACCCAGCAGCTGTTGGATCTCCTCCAGCAGGCTACCCACTACCGCCAGGCCAAGAAGGGAGCCAACGAAG	571 	\|\|\|\|\|\|\|\|\|\|\|\|\|\|\|\|\|\|\|\|\|\|\|\|\|\|\|\|\|\|\|\|\|\|\|\|\|\|\|\|\|\|\|\|\|\|\|\|\|\|\|\|\|\|\|\|\|\|\|\|\|\|\|\|\|\|\|\|\|\|\|\|\|\|\|\|\|\|\|\|\|\|\|\|\|\|\|\|\|\|\|\|\|\|\|\|\|\|\|\|\|\|\|\|\|\|\|\|\|\|\|\|\|\|\|\|\|\| 	ATGTCCGCCGAAGCCTCCGCCGCATGGCCCGTGGCCGACCAGGCCCTCACCCAGCAGCTGTTGGATCTCCTCCAGCAGGCTACCCACTACCGCCAGGCCAAGAAGGGAGCCAACGAAG ``` |
| 41 | ggggacaagtttgtacaaaaaagcaggcttgatgTCGTTGACCACCGCGC, ggggaccactttgtacaagaaagctgggtcCGGCGCCGGGTTATCCTTCA | Mycgr3\|77491  JGI  Ensembl | KOG0424 Ubiquitin-protein ligase; | negative regulation of ubiquitin-protein ligase activity; positive regulation of ubiquitin-protein ligase activity;  post-translational protein modification;  regulation of protein metabolic process;  regulation of ubiquitin-protein ligase activity;  regulation of ubiquitin-protein ligase activity during meiotic cell cycle;  small conjugating protein ligase activity;  ubiquitin-protein ligase activity; | 100 % |  | ``` 122	TGGCGAAAGGACCATCCTTTCGGATTCGTCGCAAGGCCAATGAAGACACCACAAGGCGCCCTCGACCTCAAACGCTGGGACTGTGCGATTCCAGGCAAAGACAAGACTATCTGGGAGGGCGGACTCTTCAAACTCGAAGTGATCTTTCCGGATGAGTA	279 	\|\|\|\| \|\|\|\|\|\|\|\|\|\|\|\|\|\|\|\|\|\|\|\|\|\|\|\|\|\|\|\|\|\|\|\|\|\|\|\|\|\|\| \|\| \|\| \|\|\|\|\| \|\|\|\|\|\|\|\|\|\|\|\|\|\|\|\|\|\|\|\|\|\|\|\|\|\|\|\|\| \|\|\|\|\|\|\| \|\|\|\|\|\|\|\|\|\|\|\|\|\|\|\|\|\|\|\|\|\|\|\|\|\|\|\|\|\|\|\|\|\|\|\|\|\|\|\|\|\|\|\|\|\|\|\|\|\|\|\|\|\|\|\|\|\|\| \|\|\| 	TGGCTAAAGGACCATCCTTTCGGATTCGTCGCAAGGCCAATGAATACTCCTCAAGGGGCCCTCGACCTCAAACGCTGGGACTGTGCTATTCCAGTCAAAGACAAGACTATCTGGGAGGGCGGACTCTTCAAACTCGAAGTGATCTTTCCGGATG-GTA ```      ``` 275	GAGTACCCCACGAAACCACCAAAGTGCAAATTCACGCCTCCACTCTTCCACCCAAACGTCTACCCATCCGGCACCGTCTGCCTCAGCATCCTCAACGAAGAAGAAGGCTGGAAGCCCGCGATCACGATCAAGGAAATATTGCTGGGCATTCAGATGCTGTTGGACGAGGTCAACCCGGACTCCCCGGCGCAAGCGGAGGCCTACAACCTGTTCAAGAAGGATCGCGCGCAGTATGAGAAGCGCATCAAGTCGGTGGTGAAGGATAACCCGGC	546 	\|\|\|\|\|\|\|\| \|\|\|\|\|\|\|\|\|\|\|\|\|\|\|\|\|\|\|\|\|\|\|\|\|\|\|\|\|\|\|\|\|\|\|\|\|\|\|\|\|\|\|\|\|\|\|\|\|\|\|\|\|\|\|\|\|\|\|\|\|\|\|\|\|\|\|\|\|\|\|\|\|\|\|\|\|\|\|\|\|\|\|\|\|\|\|\|\|  \|  \|\|\|\|\|\|\|\|\|\|\|\|\|\|\|\|\|\|\|\|\|\|\|\|\|\|\|\|\|\|\|\|\|\| \|\|\|\|\|\|\|\|\|\|\|\|\| \|\|\|\|\|\|\|\|\|\|\|\|\|\|\|\|\|\|\|\|\|\|\|\|\|\|\|\|\|\|\|\|\|\|\|\|\|\|\|\|\|\|\|\|\|\|\|\|\|\|\|\|\|\|\|\|\|\|\|\|\|\|\|\|\| \|\|\|\|\|\|\|\|\|\|\|\|\|\|\|\|\|\|\|\|\| \| \|\|\|\|\|\|\|\|\|\|\|\|\|\|\|\| \|\|\|\|\|\|\| \|\|\|\|\| 	GAGTACCCAACGAAACCACCAAAGTGCAAATTCACGCCTCCACTCTTCCACCCAAACGTCTACCCATCCGGCACCGTCTGCCTCAGCATCCTCAACGACCATTAAGGCTGGAAGCCCGCGATCACGATCAAGGAAATTTTGCTGGGCATTCTGATGCTGTTGGACGAGGTCAACCCGGACTCCCCGGCGCAAGCGGAGGCCTACAACCTGTTCAAGATGGATCGCGCGCAGTATGAGAACCACATCAAGTCGGTGGTGTAGGATAATCCGGC ``` |
| 41 | ggggacaagtttgtacaaaaaagcaggcttgatgAGTCCCGGCGCCATCA, ggggaccactttgtacaagaaagctgggtcTGCATTCCCTTTGAAGAGCG | Mycgr3\|80419  JGI  Ensembl |  |  | 100 % |  | ``` 140	ATGAGTCCCGGCGCCATCATGGCGATGAAGAGCGCCTTTCCCGCTATACTCGCTCTGCTCGCGAGTCCGCAATTCGCCATCGCTGCTGGTGTCGGCGTCGGCGTCACGATCATCGCCTTCGGCGGCTACAAGATCATCAAGAAGATCAAAGCAAGGGATGAGAAGAAGTTGCTTGAGGATGGTGTTGCGATCCCGGAAAGTCCGACCGAGGAGATGGACGAGCTGCGCGAACTTGATCGAATCGAAATGTGGCGTCGTGGTATCGCGGACGAGCAGGCTGCAAGCTCGGTCGGAACCAGCGTCGAGGGCAATTTTGTCACCCCTGGCGCGACGCGGACATTGATTGAGGAGGGAAAGATCACAGAGGCGGACCTTAAGCCGAAGGAGAAGAAGCATCGCAAAAAGAAGTCGCGGAGTGAGGCGAGTGGATCTACAAGCAAGACGAGTAAGACGAAGAAGACGAAAGCTTCGAAGGAACCGTCAATGTTCCGGTCGCTCTTCAAAGGGAATGCA	652 	\|\|\|\|\|\|\|\|\|\|\|\|\|\|\|\|\|\|\|\|\|\|\|\|\|\|\|\|\|\|\|\|\|\|\|\|\|\|\|\|\|\|\|\|\|\|\|\|\|\|\|\|\|\|\|\|\|\|\|\|\|\|\|\|\|\|\|\|\|\|\|\|\|\|\|\|\|\|\|\|\|\|\|\|\|\|\|\|\|\|\|\|\|\|\|\|\|\|\|\|\|\|\|\|\|\|\|\|\|\|\|\|\|\|\|\|\|\|\|\|\|\|\|\|\|\|\|\|\|\|\|\|\|\|\|\|\|\|\|\|\|\|\|\|\|\|\|\|\|\|\|\|\|\|\|\|\|\|\|\|\|\|\|\|\|\|\|\|\|\|\|\|\|\|\|\|\|\|\|\|\|\|\|\|\|\|\|\|\|\|\|\|\|\|\|\|\|\|\|\|\|\|\|\|\|\|\|\|\|\|\|\|\|\|\|\|\|\|\|\|\|\|\|\|\|\|\|\|\|\|\|\|\|\|\|\|\|\|\|\|\|\|\|\|\|\|\|\|\|\|\|\|\|\|\|\|\|\|\|\|\|\|\|\|\|\|\|\|\|\|\|\|\|\|\|\|\|\|\|\|\|\|\|\|\|\|\|\|\|\|\|\|\|\|\|\|\|\|\|\|\|\|\|\|\|\|\|\|\|\|\|\|\|\|\|\|\|\|\|\|\|\|\|\|\|\|\|\|\|\|\|\|\|\|\|\|\|\|\|\|\|\|\|\|\|\|\|\|\|\|\|\|\|\|\|\|\|\|\|\|\|\|\|\|\|\|\|\|\|\|\|\|\|\|\|\|\|\|\|\|\|\|\|\|\|\|\|\|\|\|\|\|\|\|\|\|\|\|\|\|\|\|\|\|\|\|\|\|\|\|\|\|\|\|\|\|\|\|\|\|\|\|\|\|\|\|\|\|\|\|\|\|\|\|\|\|\|\|\|\|\|\|\|\|\|\|\|\|\|\|\|\|\|\|\|\|\|\|\|\|\|\|\|\|\|\|\|\|\|\|\|\|\|\|\|\|\|\|\|\|\|\|\|\|\|\|\|\|\|\|\|\|\|\|\|\|\|\|\|\|\|\|\|\|\|\|\|\|\|\|\|\|\| 	ATGAGTCCCGGCGCCATCATGGCGATGAAGAGCGCCTTTCCCGCTATACTCGCTCTGCTCGCGAGTCCGCAATTCGCCATCGCTGCTGGTGTCGGCGTCGGCGTCACGATCATCGCCTTCGGCGGCTACAAGATCATCAAGAAGATCAAAGCAAGGGATGAGAAGAAGTTGCTTGAGGATGGTGTTGCGATCCCGGAAAGTCCGACCGAGGAGATGGACGAGCTGCGCGAACTTGATCGAATCGAAATGTGGCGTCGTGGTATCGCGGACGAGCAGGCTGCAAGCTCGGTCGGAACCAGCGTCGAGGGCAATTTTGTCACCCCTGGCGCGACGCGGACATTGATTGAGGAGGGAAAGATCACAGAGGCGGACCTTAAGCCGAAGGAGAAGAAGCATCGCAAAAAGAAGTCGCGGAGTGAGGCGAGTGGATCTACAAGCAAGACGAGTAAGACGAAGAAGACGAAAGCTTCGAAGGAACCGTCAATGTTCCGGTCGCTCTTCAAAGGGAATGCA ``` |
| 41 | ggggacaagtttgtacaaaaaagcaggcttgatgCCTACCAGACTCACCA, ggggaccactttgtacaagaaagctgggtcAGCGACGAGCTGAACAACAC | Mycgr3\|81123  JGI  Ensembl | KOG1742 60s ribosomal protein L15/L27; | intracellular; ribosome; structural constituent of ribosome;  translation; | 100 % |  | ``` 58	TCACGTTTCGGCCGGTCATGGTCGTGTCGGAAAGGCGAGAAAGCATCCCGGTGGTCGCGGTATGGCTGGTGGTCAGCACCACCACCGCACCAACTTGGACAAGTACCATCCCGGTTACTTCGGAAAGGTCGGTATGAGATACTTCCACAAGCTCCAGAACCAGTTCTGGAAGCCGGTCATCAACCTCGACAAG	250 	\|\|\|\|\| \|\|\|\|\|\|\|\|\|\|\| \|\| \|\|\|\|\|\|\|\|\|\| \|\|\|\| \|       \|    \|\|\|\|\|\|\|\|\|\|\|\|\|\|\|\|\|\|\|    \|  \|  \| \|\|  \|\|\| \|\| \|\|\|\|\|\|\|\|\|\|\|\|    \|\|\|\|\|\|\|\|\|\|\|\|\|\|\|\|\|\|\|\|\|\|\|\|\|\|\|\|\|\|\|\|\|\|\|\|\|\|\|\| \|\|\|\|\|\|\|\|\|\|\|\|\|\|\|\|\|\|\|\|\| \| \|\|\|\|\| \|   \|\|\|\|\|\|\|\|\|\| 	TCACGGTTCGGCCGGTCGTGTTCGTGTCGGATAGGCCATTTTTTTTTTTTGTGGTCGCGGTATGGCTGG-CCCCTCCTTCTCCTTCGCTCCTACTTGGACAAGT-TTTTCCCGGTTACTTCGGAAAGGTCGGTATGAGATACTTCCACGAGCTCCAGAACCAGTTCTGGATGACGGTCTTTG-CCTCGACAAG ```      ``` 249	AGCTCTGGTCTCTCGTCCCAATCGAGCAGCGCGAGAAGTACCTCGCCAACAAGTCCGCCGACACCGCTCCAGTCCTCGACCTCCTCCCTCTCGGCTACAGCAAGGTCCTCGGAAAGGGTCGTCTCCCTCAGGTCCCGATCATCGTCCGCGCCCGCTACGTCAGCAAGGACGCCGAGCGCAAGATCAAGGAGGCCGGCGGTGTTGTTCAGCTCGTCGCT	466 	\|\|\|\|\|\|\|\|\|\|\|\|\|\|\|\|\|\|\|\|\|\|\|\|\|\|\|\|\|\|\|\|\|\|\|\|\|\|\|\|\|\|\|\|\|\|\|\|\|\|  \|\|\|\|\|\|\|\|\|\|\|\|\|\|\|\|\|\|  \|\|\| \| \|\|\| \| \|\|\|\|\|\|\|\|\|\|\|\|\|\|\|\|\|\|\|\|\|\|\|\|\|\|\|\|\| \|\|\|\|\|\|\|\|\|\|\|\|\|\|\|\|\|\|\|\|\|\|\|\|\|\|\|\|\|\|\| \|\|\|\|\|\| \|\|\|\|\|\|\|\|     \|\|\|\| \|\| \|\|\|\|\| \|\|\|\|\|\|\|\|\|\| \|\|\|\|\|\| \| \| \|\|\|\|\|\|\| \|\|\|\|\|\|\|\| 	AGCTCTGGTCTCTCGTCCCAATCGAGCAGCGCGAGAAGTACCTCGCCAACCTGTCCGCCGACACCGCTCCTTTCCACTACCACATCCCTCTCGGCTACAGCAAGGTCCTCGGACAGGGTCGTCTCCCTCAGGTCCCGATCATCGT-CGCGCCTGCTACGTCTTTC-GGACCCCTAGCGCCAGATCAAGGACGCCGGCCGCGCTGTTCAGATCGTCGCT ``` |
| 41 | ggggacaagtttgtacaaaaaagcaggcttgatgTCCTCGCAACTGCTCC, ggggaccactttgtacaagaaagctgggtcGTTCTGGTCTGGCATGCCTC | Mycgr3\|81468  JGI  Ensembl | KOG1775 U6 snRNA-associated Sm-like protein; | mRNA metabolic process; ribonucleoprotein complex; | 100 % |  | ``` 246	GCTTATTCCGGGAGGCATGCCAGACCAGAAC	276 	\|\|\|\|\|\|\|\|\|\|\|\|\|\|\|\|\|\|\|\|\|\|\|\|\|\|\|\|\|\|\| 	GCTTATTCCGGGAGGCATGCCAGACCAGAAC ```      ``` 183	CGACTACTCTGGCGCTCAAACGAAGCTTCCAAAGATTCTCTTGAATGGCAATAATGTGGCGATG	246 	\|\|\|\|\|\|\|\|\|\|\|\|\|\|\|\|\|\|\|\|\|\|\|\|\|\|\|\|\|\|\|\|\|\|\|\|\|\|\|\|\|\|\|\|\|\|\|\|\|\|\|\|\|\|\|\|\|\|\|\|\|\|\|\| 	CGACTACTCTGGCGCTCAAACGAAGCTTCCAAAGATTCTCTTGAATGGCAATAATGTGGCGATG ```      ``` 115	GAATTCAGTGGCACTCTACTCGGTTTCGATGACTTCGTCAATATG	159 	\|\|\|\|\|\|\|\|\|\|\|\|\|\|\|\|\|\|\|\|\|\|\|\|\|\|\|\|\|\|\|\|\|\|\|\|\|\|\|\| \|\|\|\| 	GAATTCAGTGGCACTCTACTCGGTTTCGATGACTTCGTCAGTATG ```      ``` 155	ATATGGTGCTTGAAGACGTGACGGAATA	182 	\|\|\|\|\|\|\|\|\|\|\|\|\|\|\|\|\|\|\|\|\|\|\|\|\|\|\|\| 	ATATGGTGCTTGAAGACGTGACGGAATA ```      ``` 33	GATGTCCTCGCAACTGCTCCCGCTAGAGCTCATTGACAGATGTGTCGGCTCGCCGATATGGGTGATCATGAAGGGCGACAAGG	115 	\|\|\|\|\|\|\|\|\|\|\|\|\|\|\|\|\|\|\|\|\|\|\|\|\|\|\|\|\|\|\|\|\|\|\|\|\|\|\|\|\|\|\|\|\|\|\|\|\|\|\|\|\|\|\|\|\|\|\|\|\|\|\|\|\|\|\|\|\|\|\|\|\|\|\|\|\|\|\|\|\|\|\| 	GATGTCCTCGCAACTGCTCCCGCTAGAGCTCATTGACAGATGTGTCGGCTCGCCGATATGGGTGATCATGAAGGGCGACAAGG ``` |
| 41 | ggggacaagtttgtacaaaaaagcaggcttgatgGGAGGCGGAGCATCAG, ggggaccactttgtacaagaaagctgggtcTATAATCGGATCCGCGGCCA | Mycgr3\|81942  JGI  Ensembl | KOG3225 Mitochondrial import inner membrane translocase, subunit TIM22; | mitochondrial inner membrane; protein transport;  protein transporter activity; | 100 % |  | ``` 9	ATGGGAGGCGGAGCATCAGCCGGCATGGATCCTCAGAAAATCCAAGAGCAGCAGATGATCAAGCTGATGC-AATC	82 	\|\|\|\|\|\|\|\|\|\|\|\|\|\|\|\|\|\|\|\|\|\|\|\|\|\|\|\|\|\|\|\|\|\|\|\|\|\|\|\|\|\|\|\|\|\|\|\|\|\|\|\|\|\|\|\|\|\|\|\|\|\|\|\|\|\| \|\|\| \|\|\|\| 	ATGGGAGGCGGAGCATCAGCCGGCATGGATCCTCAGAAAATCCAAGAGCAGCAGATGATCAAGCTGGTGCGAATC ```      ``` 74	GATGCAATCCGGCATGGAATCCTGCCCCGTAAAAATGGCCCTCGCCGGCGGAATGGGCTTCGGCCTCGGCGGTCTCTTCGGCCTCTTCATGGCCTCGATGCGCTACGACACGCCCATGTCCTCCGCCATCCCCGGCAGCGCCGCCGCAGAAATCGAAAAGATCCCCATGCGCCAACAGATCAAAGCGGGATTCAAAGACATGGGAAAATCTTCGTGGTCGTCGGCCAAGAATTTCGGATACATCGGCGGAGTGTACTCGGGCAGTGAGTGCGTGATTGAAGGGTTCCGTGGGAAAAATGATCTGGCGAATGGAGTCGCGGCGGGGTGTTTTACGGGAGGGTTTTTGGCGAAGAGTGCGGGGCCGCAGGCGATGGCGATTGGGTGTGCGGGTTTTGCAGCGTTTTCGGCGGCGATTGATGCGTATATGAGGATGGAGAGTAATGATG	519 	\|\|\|\|\|\|\|\|\|\|\|\|\|\|\|\|\|\|\|\|\|\|\|\|\|\|\|\|\|\|\|\|\|\|\|\|\|\|\|\|\|\|\|\|\|\|\|\|\|\|\|\|\|\|\|\|\|\|\|\|\|\|\|\|\|\|\|\|\|\|\|\|\|\|\|\|\|\|\|\|\|\|\|\|\|\|\|\|\|\|\|\|\|\|\|\|\|\|\|\|\|\|\|\|\|\|\|\|\|\|\|\|\|\|\|\|\|\|\|\|\|\|\|\|\|\|\|\|\|\|\|\|\|\|\|\|\|\|\|\|\|\|\|\|\|\|\|\|\|\|\|\|\|\|\|\|\|\|\|\|\|\|\|\|\|\|\|\|\|\|\|\|\|\|\|\|\|\|\|\|\|\|\|\|\|\|\|\|\|\|\|\|\|\|\|\|\|\|\|\|\|\|\|\|\|\|\|\|\|\|\|\|\|\|\|\|\|\|\|\|\|\|\|\|\|\|\|\|\|\|\|\|\|\|\|\|\|\|\|\|\|\|\|\|\|\|\|\|\|\|\|\|\|\|\|\|\|\|\|\|\|\|\|\|\|\|\|\|\|\|\|\|\|\|\|\|\|\|\|\|\|\|\|\|\|\|\|\|\|\|\|\|\|\|\|\|\|\|\|\|\|\|\|\|\|\|\|\|\|\|\|\|\|\|\|\|\|\|\|\|\|\|\|\|\|\|\|\|\|\|\|\|\|\|\|\|\|\|\|\|\|\|\|\|\|\|\|\|\|\|\|\|\|\|\|\|\|\|\|\|\|\|\|\|\|\|\|\|\|\|\|\|\|\|\|\|\|\|\|\|\|\|\|\|\|\|\|\|\|\|\|\|\|\|\|\|\|\|\|\|\|\|\|\|\|\|\|\|\|\|\|\|\|\|\|\|\|\|\|\|\|\|\|\|\|\|\|\|\|\|\|\|\|\|\|\|\|\|\|\|\|\|\|\|\|\| 	GATGCAATCCGGCATGGAATCCTGCCCCGTAAAAATGGCCCTCGCCGGCGGAATGGGCTTCGGCCTCGGCGGTCTCTTCGGCCTCTTCATGGCCTCGATGCGCTACGACACGCCCATGTCCTCCGCCATCCCCGGCAGCGCCGCCGCAGAAATCGAAAAGATCCCCATGCGCCAACAGATCAAAGCGGGATTCAAAGACATGGGAAAATCTTCGTGGTCGTCGGCCAAGAATTTCGGATACATCGGCGGAGTGTACTCGGGCAGTGAGTGCGTGATTGAAGGGTTCCGTGGGAAAAATGATCTGGCGAATGGAGTCGCGGCGGGGTGTTTTACGGGAGGGTTTTTGGCGAAGAGTGCGGGGCCGCAGGCGATGGCGATTGGGTGTGCGGGTTTTGCAGCGTTTTCGGCGGCGATTGATGCGTATATGAGGATGGAGAGTAATGATG ``` |
| 41 | ggggacaagtttgtacaaaaaagcaggcttgatgAAGCCCGTCGTTTCCG, ggggaccactttgtacaagaaagctgggtcTCGCAGTGCAATCTCTCCTC | Mycgr3\|81999  JGI  Ensembl |  |  | 100 % |  | ``` 336	GGCTTTCCTACTATTTTGCGGATGCCAGGCGATCTTGCACAAACGAGCGAGTCGGAGAGGAGAGATTGCACTGCGA	411 	\|\|\|\|\|\|\|\|\|\|\|\|\|\|\|\|\|\|\|\|\|\|\|\|\|\|\|\|\|\|\|\|\|\|\|\|\|\|\|\|\|\|\|\|\|\|\|\|\|\|\|\|\|\|\|\|\|\|\|\|\|\|\|\|\|\|\|\|\|\|\|\|\|\|\|\| 	GGCTTTCCTACTATTTTGCGGATGCCAGGCGATCTTGCACAAACGAGCGAGTCGGAGAGGAGAGATTGCACTGCGA ```      ``` 232	GGCATGTTCGCCAACAACAACCACTCCCTGATGGGATCGAAAGAGGACCCGAAGGACGGAGCCAAAGTCGCTACGGCCGTGTTTAGTGCTGTGGCTGTATATGGGG	337 	\|\|\|\|\|\|\|\|\|\|\|\|\|\|\|\|\|\|\|\|\|\|\|\|\|\|\|\|\|\|\|\|\|\|\|\|\|\|\|\|\|\|\|\|\|\|\|\|\|\|\|\|\|\|\|\|\|\|\|\|\|\|\|\|\|\|\|\|\|\|\|\|\|\|\|\|\|\|\|\|\|\|\|\|\|\|\|\|\|\|\|\|\|\|\|\|\|\|\|\|\|\|\|\|\|\| 	GGCATGTTCGCCAACAACAACCACTCCCTGATGGGATCGAAAGAGGACCCGAAGGACGGAGCCAAAGTCGCTACGGCCGTGTTTAGTGCTGTGGCTGTATATGGGG ```      ``` 185	GCATCGTCATCTCCATCTTTGCGATCGTGATCCTCAGCACAATTGGCGG	233 	\|\|\|\|\|\|\|\|\|\|\|\|\|\|\|\|\|\|\|\|\|\|\|\|\|\|\|\|\|\|\|\|\|\|\|\|\|\|\|\|\|\|\|\|\|\|\|\|\| 	GCATCGTCATCTCCATCTTTGCGATCGTGATCCTCAGCACAATTGGCGG ```      ``` 148	ATGAAGCCCGTCGTTTCCGCCTTCAACGCATGGACATG	185 	\|\|\|\|\|\|\|\|\|\|\|\|\|\|\|\|\|\|\|\|\|\|\|\|\|\|\|\|\|\|\|\|\|\|\|\|\|\| 	ATGAAGCCCGTCGTTTCCGCCTTCAACGCATGGACATG ``` |
| 41 | ggggacaagtttgtacaaaaaagcaggcttgatgGCCATGGACACCACAC, ggggaccactttgtacaagaaagctgggtcTCGATAGAATTCACCCCTCT | Mycgr3\|87549  JGI  Ensembl | KOG3423 Transcription initiation factor TFIID, subunit TAF10 (also component of histone acetyltransferase SAGA); | RNA polymerase II transcription factor activity; nucleus; transcription initiation; | 100 % |  | ``` 1	ATGGCCATGGACACCACACAGGATGCGCCAGCACCTAACTCAGCGCCCGAGACCAATGGTCTACCAGCAATACCTACTGTTCTCGAACCACGGATACCTCAAAAGAAAGATACCTCTCTGCGCGAATTCCTAGGAAAGATGGACGACTACGCTCCTCTCATTCCCGACGCCGTCACAGACTACTACCTCACACTCGCGGGTCTCCCACCACCACCCGAGACCGACCGTCGTCTCGCCCGACTCCTCGCTCTTGCGACGCAGAAGTTCATTGCAGACATTGCAGCCGACGCATACCAATACAGCCGGATCCGTAGCACGAACACCACGTCCAACAATCCTCTAACGGGTCTTGGAGGTGCCGCGGGATTTGGAATGAATATGCCGCAAGGCGGTGAAGGCGCGGATGGCGCTGGAAAGGGTGGGAAGAAAGATGTTGGTAAGGCGGCACCTTTAGGCGGACCACGTGCGGGATATGGAGGTGGAGGTGCAGGCGCCGGTCAAGGTCGGACTGTGCTCACCATGGAAGATTTGGGTATGGCGGTGGGCGAGTACGGTGTCAATGTCAAGAGGGGTGAATTCTATCGA	585 	\|\|\|\|\|\|\|\|\|\|\|\|\|\|\|\|\|\|\|\|\|\|\|\|\|\|\|\|\|\|\|\|\|\|\|\|\|\|\|\|\|\|\|\|\|\|\|\|\|\|\|\|\|\|\|\|\|\|\|\|\|\|\|\|\|\|\|\|\|\|\|\|\|\|\|\|\|\|\|\|\|\|\|\|\|\|\|\|\|\|\|\|\|\|\|\|\|\|\|\|\|\|\|\|\|\|\|\|\|\|\|\|\|\|\|\|\|\|\|\|\|\|\|\|\|\|\|\|\|\|\|\|\|\|\|\|\|\|\|\|\|\|\|\|\|\|\|\|\|\|\|\|\|\|\|\|\|\|\|\|\|\|\|\|\|\|\|\|\|\|\|\|\|\|\|\|\|\|\|\|\|\|\|\|\|\|\|\|\|\|\|\|\|\|\|\|\|\|\|\|\|\|\|\|\|\|\|\|\|\|\|\|\|\|\|\|\|\|\|\|\|\|\|\|\|\|\|\|\|\|\|\|\|\|\|\|\|\|\|\|\|\|\|\|\|\|\|\|\|\|\|\|\|\|\|\|\|\|\|\|\|\|\|\|\|\|\|\|\|\|\|\|\|\|\|\|\|\|\|\|\|\|\|\|\|\|\|\|\|\|\|\|\|\|\|\|\|\|\|\|\|\|\|\|\|\|\|\|\|\|\|\|\|\|\|\|\|\|\|\|\|\|\|\|\|\|\|\|\|\|\|\|\|\|\|\|\|\|\|\|\|\|\|\|\|\|\|\|\|\|\|\|\|\|\|\|\|\|\|\|\|\|\|\|\|\|\|\|\|\|\|\|\|\|\|\|\|\|\|\|\|\|\|\|\|\|\|\|\|\|\|\|\|\|\|\|\|\|\|\|\|\|\|\|\|\|\|\|\|\|\|\|\|\|\|\|\|\|\|\|\|\|\|\|\|\|\|\|\|\|\|\|\|\|\|\|\|\|\|\|\|\|\|\|\|\|\|\|\|\|\|\|\|\|\|\|\|\|\|\|\|\|\|\|\|\|\|\|\|\|\|\|\|\|\|\|\|\|\|\|\|\|\|\|\|\|\|\|\|\|\|\|\|\|\|\|\|\|\|\|\|\|\|\|\|\|\|\|\|\|\|\|\|\|\|\|\|\|\|\|\|\|\|\|\|\|\|\|\|\|\|\|\|\|\|\|\|\|\|\|\|\|\|\|\|\|\|\|\|\|\|\|\|\|\|\|\|\|\|\|\|\|\|\|\|\|\|\|\|\|\|\|\|\|\|\|\|\|\|\|\|\|\|\|\| 	ATGGCCATGGACACCACACAGGATGCGCCAGCACCTAACTCAGCGCCCGAGACCAATGGTCTACCAGCAATACCTACTGTTCTCGAACCACGGATACCTCAAAAGAAAGATACCTCTCTGCGCGAATTCCTAGGAAAGATGGACGACTACGCTCCTCTCATTCCCGACGCCGTCACAGACTACTACCTCACACTCGCGGGTCTCCCACCACCACCCGAGACCGACCGTCGTCTCGCCCGACTCCTCGCTCTTGCGACGCAGAAGTTCATTGCAGACATTGCAGCCGACGCATACCAATACAGCCGGATCCGTAGCACGAACACCACGTCCAACAATCCTCTAACGGGTCTTGGAGGTGCCGCGGGATTTGGAATGAATATGCCGCAAGGCGGTGAAGGCGCGGATGGCGCTGGAAAGGGTGGGAAGAAAGATGTTGGTAAGGCGGCACCTTTAGGCGGACCACGTGCGGGATATGGAGGTGGAGGTGCAGGCGCCGGTCAAGGTCGGACTGTGCTCACCATGGAAGATTTGGGTATGGCGGTGGGCGAGTACGGTGTCAATGTCAAGAGGGGTGAATTCTATCGA ``` |
| 41 | ggggacaagtttgtacaaaaaagcaggcttgatgCTCGAATGCATCGGCG, ggggaccactttgtacaagaaagctgggtcCCCCTTGCGAACCCCAACAT | Mycgr3\|88098  JGI  Ensembl | KOG2698 GTP cyclohydrolase I; | GTP cyclohydrolase I activity; aromatic compound biosynthetic process;  cytoplasm; | 100 % |  | ``` 221	AGATCCACATCGCCTACATACCCAACAGCACCGTCATCGGCCTCTCCAAAGTCCCGCGCATCGTGCAAATGTACTCTCACAGACTGCAAATCCAAGAGCGTCTGACGTCGCAAGTGGCTTATGCCGTTCAAGATGTCGTGAAGGCGAAGGGAGTCGCGGTCGTGATGGAGGCTTCGCACTTGTGTATGAAGATGCGGGGGGTTGAGCAGACTGACGCTACTACTGTCACGAGCTGCTTCATGGGGTGTTTCGATGAGAAGCCAGAGCTTCGAAGCGAGTTTTTGAGGTATGTTGGGGTTCGCAAGGGG	528 	\|\|\|\|\|\|\|\|\|\|\|\|\|\|\|\|\|\|\|\|\|\|\|\|\|\|\|\|\|\|\|\|\|\|\|\|\|\|\|\|\|\|\|\|\|\|\|\|\|\|\|\|\|\|\|\|\|\|\|\|\|\|\|\|\|\|\|\|\|\|\|\|\|\|\|\|\|\|\|\|\|\|\|\|\|\|\|\|\|\|\|\|\|\|\|\|\|\|\|\|\|\|\|\|\|\|\|\|\|\|\|\|\|\|\|\|\|\|\|\|\|\|\|\|\|\|\|\|\|\|\|\|\|\|\|\|\|\|\|\|\|\|\|\|\|\|\|\|\|\|\|\|\|\|\|\|\|\|\|\|\|\|\|\|\|\|\|\|\|\|\|\|\|\|\|\|\|\|\|\|\|\|\|\|\|\|\|\|\|\|\|\|\|\|\|\|\|\|\|\|\|\|\|\|\|\|\|\|\|\|\|\|\|\|\|\|\|\|\|\|\|\|\|\|\|\|\|\|\|\|\|\|\|\|\|\|\|\|\|\|\|\|\|\|\|\|\|\|\|\|\|\|\|\|\|\|\|\|\|\|\|\|\|\|\|\|\|\|\|\|\|\|\|\|\|\|\|\|\|\|\|\|\|\|\|\|\|\|\|\|\|\|\|\|\|\|\|\|\|\|\|\|\|\|\|\|\|\| 	AGATCCACATCGCCTACATACCCAACAGCACCGTCATCGGCCTCTCCAAAGTCCCGCGCATCGTGCAAATGTACTCTCACAGACTGCAAATCCAAGAGCGTCTGACGTCGCAAGTGGCTTATGCCGTTCAAGATGTCGTGAAGGCGAAGGGAGTCGCGGTCGTGATGGAGGCTTCGCACTTGTGTATGAAGATGCGGGGGGTTGAGCAGACTGACGCTACTACTGTCACGAGCTGCTTCATGGGGTGTTTCGATGAGAAGCCAGAGCTTCGAAGCGAGTTTTTGAGGTATGTTGGGGTTCGCAAGGGG ```      ``` 1	ATGCTCGAATGCATCGGCGAAGATCCCAATCGAAAAGACCTCCTCGAAACACCACGTCGTTACGCCCAAGCCCTCATCGATCTCACCTCCGGCTACAAGCTCGACACTCACGCCATCGTTGGCGAAGCTCTTTTCCATGGGGCATATACTGGTATGGTCGTTGTCAAGGACATCGAGGTCCACAGTCTTTGCGAACATCATCTTCTTCCCTTTTTAGGCAAG	222 	\|\|\|\|\|\|\|\|\|\|\|\|\|\|\|\|\|\|\|\|\|\|\|\|\|\|\|\|\|\|\|\|\|\|\|\|\|\|\|\|\|\|\|\|\|\|\|\|\|\|\|\|\|\|\|\|\|\|\|\|\|\|\|\|\|\|\|\|\|\|\|\|\|\|\|\|\|\|\|\|\|\|\|\|\|\|\|\|\|\|\|\|\|\|\|\|\|\|\|\|\|\|\|\|\|\|\|\|\|\|\|\|\|\|\|\|\|\|\|\|\|\|\|\|\|\|\|\|\|\|\|\|\|\|\|\|\|\|\|\|\|\|\|\|\|\|\|\|\|\|\|\|\|\|\|\|\|\|\|\|\|\|\|\|\|\|\|\|\|\|\|\|\|\|\|\|\|\|\|\|\|\|\|\|\|\|\|\|\|\|\|\|\|\|\|\|\|\|\|\|\|\|\|\|\|\|\|\|\|\|\|\|\|\|\|\|\|\|\|\|\|\| 	ATGCTCGAATGCATCGGCGAAGATCCCAATCGAAAAGACCTCCTCGAAACACCACGTCGTTACGCCCAAGCCCTCATCGATCTCACCTCCGGCTACAAGCTCGACACTCACGCCATCGTTGGCGAAGCTCTTTTCCATGGGGCATATACTGGTATGGTCGTTGTCAAGGACATCGAGGTCCACAGTCTTTGCGAACATCATCTTCTTCCCTTTTTAGGCAAG ``` |
| 41 | ggggacaagtttgtacaaaaaagcaggcttgatgACCACCCCGGTTTACG, ggggaccactttgtacaagaaagctgggtcGAGCACATACATGGAATCAA | Mycgr3\|89156  JGI  Ensembl | KOG2533 Permease of the major facilitator superfamily; |  | 100 % |  | ``` 1	ATGACCACCCCGGTTTACGTGGTCGGCTGCATCTCGCTCATCGTCCAGCTCTGCTATTCCGACAAGGTCAAGAAGCGTGGTGTTTTCATCATCTCCTGCGGTGTCCCGGGCTCGGTCGGCTACCTGATCTGTGTCGGGAGCTCGAATCCTGGTGCTGGATATGCCGGAATGTTTGTGCTTGTCCTAGG	188 	\|\|\|\|\|\|\|\|\|\|\|\|\|\|\|\|\|\|\|\|\|\|\|\|\|\|\|\|\|\|\|\|\|\|\|\|\|\|\|\|\|\|\|\|\|\|\|\|\|\|\|\|\|\|\|\|\|\|\|\|\|\|\|\|\|\|\|\|\|\|\|\|\|\|\|\|\|\|\|\|\|\|\|\|\|\|\|\|\|\|\|\|\|\|\|\|\|\|\|\|\|\|\|\|\|\|\|\|\|\|\|\|\|\|\|\|\|\|\|\|\|\|\|\|\|\|\|\|\|\|\|\|\|\|\|\|\|\|\|\|\|\|\|\|\|\|\|\|\|\|\|\|\|\|\|\|\|\|\|\|\|\|\|\|\|\|\|\|\|\|\|\|\|\|\|\|\|\|\|\|\|\|\|\|\|\|\|\| 	ATGACCACCCCGGTTTACGTGGTCGGCTGCATCTCGCTCATCGTCCAGCTCTGCTATTCCGACAAGGTCAAGAAGCGTGGTGTTTTCATCATCTCCTGCGGTGTCCCGGGCTCGGTCGGCTACCTGATCTGTGTCGGGAGCTCGAATCCTGGTGCTGGATATGCCGGAATGTTTGTGCTTGTCCTAGG ```      ``` 185	TAGGGCTCTACCCTATTTCCACACTTGCCGTAATCTGGATCACCAATAACCTCTCGCCCGACTCCAAGCGCGCGGTCGGCATGCCCTTCGCGTACAGTATCGCGAACTTGTCCAATCTCGTGTCGTCGCAGTTGTATCCCAGCCAGCAAGGACCACGATACGTACAGGGTAATGCTGTGTCCGCTGGTCTGACGGTGGTCGCCGGGTTTCTGTATGATGCATGCTGGGTCCTGCTGAGGCGCAGGAACATGAAGAAGGCGAAGCTGATCGCCGAGGGTGCAACGAACAATGGGCTAGAAGGTGATCAGAGTCTTGATTCCATGTATGTGCTC	516 	\|\|\|\|\|\|\|\|\|\|\|\|\|\|\|\|\|\|\|\|\|\|\|\|\|\|\|\|\|\|\|\|\|\|\|\|\|\|\|\|\|\|\|\|\|\|\|\|\|\|\|\|\|\|\|\|\|\|\|\|\|\|\|\|\|\|\|\|\|\|\|\|\|\|\|\|\|\|\|\|\|\|\|\|\|\|\|\|\|\|\|\|\|\|\|\|\|\|\|\|\|\|\|\|\|\|\|\|\|\|\|\|\|\|\|\|\|\|\|\|\|\|\|\|\|\|\|\|\|\|\|\|\|\|\|\|\|\|\|\|\|\|\|\|\|\|\|\|\|\|\|\|\|\|\|\|\|\|\|\|\|\|\|\|\|\|\|\|\|\|\|\|\|\|\|\|\|\|\|\|\|\|\|\|\|\|\|\|\|\|\|\|\|\|\|\|\|\|\|\|\|\|\|\|\|\|\|\|\|\|\|\|\|\|\|\|\|\|\|\|\|\|\|\|\|\|\|\|\|\|\|\|\|\|\|\|\|\|\|\|\|\|\|\|\|\|\|\|\|\|\|\|\|\|\|\|\|\|\|\|\|\|\|\|\|\|\|\|\|\|\|\|\|\|\|\|\|\|\|\|\|\|\|\|\|\|\|\|\|\|\|\|\|\|\|\|\|\|\|\|\|\|\|\|\|\|\|\|\|\|\|\|\|\|\|\|\|\|\|\|\|\|\|\|\|\|\|\|\|\|\|\| 	TAGGGCTCTACCCTATTTCCACACTTGCCGTAATCTGGATCACCAATAACCTCTCGCCCGACTCCAAGCGCGCGGTCGGCATGCCCTTCGCGTACAGTATCGCGAACTTGTCCAATCTCGTGTCGTCGCAGTTGTATCCCAGCCAGCAAGGACCACGATACGTACAGGGTAATGCTGTGTCCGCTGGTCTGACGGTGGTCGCCGGGTTTCTGTATGATGCATGCTGGGTCCTGCTGAGGCGCAGGAACATGAAGAAGGCGAAGCTGATCGCCGAGGGTGCAACGAACAATGGGCTAGAAGGTGATCAGAGTCTTGATTCCATGTATGTGCTC ``` |
| 41 | ggggacaagtttgtacaaaaaagcaggcttgatgCAGCCAACCTCAATGC, ggggaccactttgtacaagaaagctgggtcCTCAAAGCAAGGGTTGAACG | Mycgr3\|89878  JGI  Ensembl |  |  | 40.51724 % |  |  |
| 41 | ggggacaagtttgtacaaaaaagcaggcttgatgAAGATCACCTTCTTCT, ggggaccactttgtacaagaaagctgggtcTGGCTGGCGGATGCGATGTG | Mycgr3\|90520  JGI  Ensembl |  |  | 100 % |  | ``` 261	CACGCGGGAGATCCAACTAGTGGTGCATGCACATCGCATCCGCCAGCCA	309 	\|\|\|\|\|\|\|\|\|\|\|\|\|\|\|\|\|\|\|\|\|\|\|\|\|\|\|\|\|\|\|\|\|\|\|\|\|\|\|\|\|\|\|\|\|\|\|\|\| 	CACGCGGGAGATCCAACTAGTGGTGCATGCACATCGCATCCGCCAGCCA ```      ``` 117	GTACCACGGCAATGGAGCTATATGGTGCCCTCCGACACCTCAGCCACCCACAACTCAAATCCGATCTGAGGAACCAAAAGTTCCCAATGCTCCCAATACTGTCAATGCTCCAAGAGACATCCTTGATGACCACAATGGAAATCC	260 	\|\|\|\|\|\|\|\|\|\|\|\|\|\|\|\|\|\|\|\|\|\|\|\|\|\|\|\|\|\|\|\|\|\|\|\|\|\|\|\|\|\|\|\|\|\|\|\|\|\|\|\|\|\|\|\|\|\|\|\|\|\|\|\|\|\|\|\|\|\|\|\|\|\|\|\|\|\|\|\|\|\|\|\|\|\|\|\|\|\|\|\|\|\|\|\|\|\|\|\|\|\|\|\|\|\|\|\|\|\|\|\|\|\|\|\|\|\|\|\|\|\|\|\|\|\|\|\|\|\|\|\|\|\|\|\|\|\|\|\|\|\|\|\| 	GTACCACGGCAATGGAGCTATATGGTGCCCTCCGACACCTCAGCCACCCACAACTCAAATCCGATCTGAGGAACCAAAAGTTCCCAATGCTCCCAATACTGTCAATGCTCCAAGAGACATCCTTGATGACCACAATGGAAATCC ```      ``` 64	CAGCAGCGAGACAAGGTCACTTCCAAAGATGACTACGAAGGGTGCGACAAGAGG	117 	\|\|\|\|\|\|\|\|\|\|\|\|\|\|\|\|\|\|\|\|\|\|\|\|\|\|\|\|\|\|\|\|\|\|\|\|\|\|\|\|\|\|\|\|\|\|\|\|\|\|\|\|\|\| 	CAGCAGCGAGACAAGGTCACTTCCAAAGATGACTACGAAGGGTGCGACAAGAGG ```      ``` 1	ATGAAGATCACCTTCTTCTCCACCCTCGCCATGCTGTCCGCAGCTGTGCTCGCAGCTCCTGTC	63 	\|\|\|\|\|\|\|\|\|\|\|\|\|\|\|\|\|\|\|\|\|\|\|\|\|\|\|\|\|\|\|\|\|\|\|\|\|\|\|\|\|\|\|\|\|\|\|\|\|\|\|\|\|\|\|\|\|\|\|\|\|\|\| 	ATGAAGATCACCTTCTTCTCCACCCTCGCCATGCTGTCCGCAGCTGTGCTCGCAGCTCCTGTC ``` |
| 41 | ggggacaagtttgtacaaaaaagcaggcttgatgCCCTCCTTCACCACCA, ggggaccactttgtacaagaaagctgggtcATCCGCACGAACCATTCCAC | Mycgr3\|90699  JGI  Ensembl |  |  | 100 % |  | ``` 213	TGCCGGCGTGACGCAAACCTTCCTTCCCGACACTGGATTCGACTGTTTGTTGATGCAG	270 	\|\|\|\|\|\|\|\|\|\|\|\|\|\|\|\|\|\|\|\|\|\|\|\|\|\|\|\|\|\|\|\|\|\|\|\|\|\|\|\|\|\|\|\|\|\|\|\|\|\|\|\|\|\|\|\|\|\| 	TGCCGGCGTGACGCAAACCTTCCTTCCCGACACTGGATTCGACTGTTTGTTGATGCAG ```      ``` 270	GAATAAATGTGACGCCTCCGTCCCCTTCGCCGACATCTCCTCGGATGAGACCGAAATCGCCCGCGACCTCTCTGGCCTCAAGTGGATCGGCAACGCGACGAGCTACATTTGCTTCTCGGAGGTCATGGCCGCCAAGCGATTCATTCGCATCTCGAGACTCAAAGCTTCGGGTGGAATGGTTCGTGCGGAT	459 	\|\|\|\|\|\|\|\|\|\|\|\|\|\|\|\|\|\|\|\|\|\|\|\|\|\|\|\|\|\|\|\|\|\|\|\|\|\|\|\|\|\|\|\|\|\|\|\|\|\|\|\|\|\|\|\|\|\|\|\|\|\|\|\|\|\|\|\|\|\|\|\|\|\|\|\|\|\|\|\|\|\|\|\|\|\|\|\|\|\|\|\|\|\|\|\|\|\|\|\|\|\|\|\|\|\|\|\|\|\|\|\|\|\|\|\|\|\|\|\|\|\|\|\|\|\|\|\|\|\|\|\|\|\|\|\|\|\|\|\|\|\|\|\|\|\|\|\|\|\|\|\|\|\|\|\|\|\|\|\|\|\|\|\|\|\|\|\|\|\|\|\|\|\|\|\|\|\|\|\|\|\|\| \|\|\|\|\|\| 	GAATAAATGTGACGCCTCCGTCCCCTTCGCCGACATCTCCTCGGATGAGACCGAAATCGCCCGCGACCTCTCTGGCCTCAAGTGGATCGGCAACGCGACGAGCTACATTTGCTTCTCGGAGGTCATGGCCGCCAAGCGATTCATTCGCATCTCGAGACTCAAAGCTTCGGGTGGAATGGTTCGAGCGGAT ```      ``` 1	ATGCCCTCCTTCACCACCATCCTCACCCTCCTCGCCGCCACCGCCATCACTTTCACCACCGCCCTCCCATCTCCCCACCTGTTCACGCCACGCGCACCCACCAACATCACCGACCTCAATGCCGGCGTGAATATCTGCACCGAGCCCGACTTCGGGGGCGAGTGCTACCATTCCTCGTGGCCGAAGAACACCTGCATCGATTTGAGAGGATA	212 	\|\|\|\|\|\|\|\|\|\|\|\|\|\|\|\|\|\|\|\|\|\|\|\|\|\|\|\|\|\|\|\|\|\|\|\|\|\|\|\|\|\|\|\|\|\|\|\|\|\|\|\|\|\|\|\|\|\|\|\|\|\|\|\|\|\|\|\|\|\|\|\|\|\|\|\|\|\|\|\|\|\|\|\|\|\|\|\|\|\|\|\|\|\|\|\|\|\|\|\|\|\|\|\|\|\|\|\|\|\|\|\|\|\|\|\|\|\|\|\|\|\|\|\|\|\|\|\|\|\|\|\|\|\|\|\|\|\|\|\|\|\|\|\|\|\|\|\|\|\|\|\|\|\|\|\|\|\|\|\|\|\|\|\|\|\|\|\|\|\|\|\|\|\|\|\|\|\|\|\|\|\|\|\|\|\|\|\|\|\|\|\|\|\|\|\|\|\|\|\|\|\|\|\|\|\|\|\|\|\|\|\| 	ATGCCCTCCTTCACCACCATCCTCACCCTCCTCGCCGCCACCGCCATCACTTTCACCACCGCCCTCCCATCTCCCCACCTGTTCACGCCACGCGCACCCACCAACATCACCGACCTCAATGCCGGCGTGAATATCTGCACCGAGCCCGACTTCGGGGGCGAGTGCTACCATTCCTCGTGGCCGAAGAACACCTGCATCGATTTGAGAGGATA ``` |
| 41 | ggggacaagtttgtacaaaaaagcaggcttgatgGCCATGATGTTCTTCC, ggggaccactttgtacaagaaagctgggtcATCGCGCTGGGTGCCGGCAG | Mycgr3\|91252  JGI  Ensembl |  |  | 100 % |  | ``` 1	ATGGCCATGATGTTCTTCCTCATCTCTGCGATGTTGCACCTTGGACACGCTTCCGCCCTGCCCCATCTTCACAGCCGACAGGCTACCTTTGAGAGACCTGCACAAGCTCCTTACGCCTTCAACGATACGGTCGAATATGTCAAGCAGGTGGTGGCTGCAACTGTCGCAGCTGAACAGGACCTCATCCAGGACATTTTGAAGTGTGTGGATATCACCAATGCCACCTTTGAGAACACGATTCTGCCATATCTTCAGTTTGACAGCGTCCGAAATGCACAATTGCAACCTTTCTACGTCTATGATGATGCCTCGGACATCGGGGGTATCGCGAGCGACATCGACGCCAACGCCACGAGTGAGACGACGAAGACATTTACGAACGCCGATTATTTCCTTTTAGTCGACCATATCTACCAGGCCAACTTGGACGACAAATCAGCCCTGAGCGAAGAGAACCAACTCATGCTGGTTTCATTCTGGAATGCATTCGCCAGCAGTGGATTAAACATTCCTGCCGGCACCCAGCGCGAT	531 	\|\|\|\|\|\|\|\|\|\|\|\|\|\|\|\|\|\|\|\|\|\|\|\|\|\|\|\|\|\|\|\|\|\|\|\|\|\|\|\|\|\|\|\|\|\|\|\|\|\|\|\|\|\|\|\|\|\|\|\|\|\|\|\|\|\|\|\|\|\|\|\|\|\|\|\|\|\|\|\|\|\|\|\|\|\|\|\|\|\|\|\|\|\|\|\|\|\|\|\|\|\|\|\|\|\|\|\|\|\|\|\|\|\|\|\|\|\|\|\|\|\|\|\|\|\|\|\|\|\|\|\|\|\|\|\|\|\|\|\|\|\|\|\|\|\|\|\|\|\|\|\|\|\|\|\|\|\|\|\|\|\|\|\|\|\|\|\|\|\|\|\|\|\|\|\|\|\|\|\|\|\|\|\|\|\|\|\|\|\|\|\|\|\|\|\|\|\|\|\|\|\|\|\|\|\|\|\|\|\|\|\|\|\|\|\|\|\|\|\|\|\|\|\|\|\|\|\|\|\|\|\|\|\|\|\|\|\|\|\|\|\|\|\|\|\|\|\|\|\|\|\|\|\|\|\|\|\|\|\|\|\|\|\|\|\|\|\|\|\|\|\|\|\|\|\|\|\|\|\|\|\|\|\|\|\|\|\|\|\|\|\|\|\|\|\|\|\|\|\|\|\|\|\|\|\|\|\|\|\|\|\|\|\|\|\|\|\|\|\|\|\|\|\|\|\|\|\|\|\|\|\|\|\|\|\|\|\|\|\|\|\|\|\|\|\|\|\|\|\|\|\|\|\|\|\|\|\|\|\|\|\|\|\|\|\|\|\|\|\|\|\|\|\|\|\|\|\|\|\|\|\|\|\|\|\|\|\|\|\|\|\|\|\|\|\|\|\|\|\|\|\|\|\|\|\|\|\|\|\|\|\|\|\|\|\|\|\|\|\|\|\|\|\|\|\|\|\|\|\|\|\|\|\|\|\|\|\|\|\|\|\|\|\|\|\|\|\|\|\|\|\|\|\|\|\|\|\|\|\|\|\|\|\|\|\|\|\|\|\|\|\|\|\|\|\|\|\|\|\|\|\|\|\|\|\|\|\|\|\|\|\|\|\|\|\|\|\|\|\|\|\|\|\|\|\|\|\|\|\|\|\|\|\|\|\|\|\|\|\|\|\|\|\|\|\|\|\|\|\|\| 	ATGGCCATGATGTTCTTCCTCATCTCTGCGATGTTGCACCTTGGACACGCTTCCGCCCTGCCCCATCTTCACAGCCGACAGGCTACCTTTGAGAGACCTGCACAAGCTCCTTACGCCTTCAACGATACGGTCGAATATGTCAAGCAGGTGGTGGCTGCAACTGTCGCAGCTGAACAGGACCTCATCCAGGACATTTTGAAGTGTGTGGATATCACCAATGCCACCTTTGAGAACACGATTCTGCCATATCTTCAGTTTGACAGCGTCCGAAATGCACAATTGCAACCTTTCTACGTCTATGATGATGCCTCGGACATCGGGGGTATCGCGAGCGACATCGACGCCAACGCCACGAGTGAGACGACGAAGACATTTACGAACGCCGATTATTTCCTTTTAGTCGACCATATCTACCAGGCCAACTTGGACGACAAATCAGCCCTGAGCGAAGAGAACCAACTCATGCTGGTTTCATTCTGGAATGCATTCGCCAGCAGTGGATTAAACATTCCTGCCGGCACCCAGCGCGAT ``` |
| 41 | ggggacaagtttgtacaaaaaagcaggcttgatgCAGTTTACTCAGATTT, ggggaccactttgtacaagaaagctgggtcGTAGTGAAAACAAAATGCTG | Mycgr3\|91471  JGI  Ensembl |  |  | 6.65499 % |  |  |
| 41 | ggggacaagtttgtacaaaaaagcaggcttgatgCACTCATCCACCATCA, ggggaccactttgtacaagaaagctgggtcGTTCTTTATTTTCTCCACTG | Mycgr3\|91704  JGI  Ensembl |  |  | 100 % |  | ``` 177	CGCCGCGGACAGCGCCAACATTCCAGTGGAGAAGAGAGGCGAATGGGACTACGAGCCCACCCCTCACTACTACCCACCCGCCGCGGACAGCGCCAACATTCCAGTGGAGAAGAGAGGCGAGTGGGACTACGAGCCCACCCCTCACTACTACCCACCCGCCGCGGACAGCGCCAACATTCCAGTGGAGAAGAGAGGCGAGTGGGACTACGAGCCCACCTCTCACTACTACCCACCCGCCGCGGACAGCGCCAACATTCCAGTGGAGAA	443 	\|\|\| \|\|\|\|\| \| \|\| \|\|\|\|\|\|\|\|\|\|\|\|\|\|\|\|\|\|\|\|\|\|\|\|\|\|\| \|\|\|\|\|\|\|\|\|\|\|\|\|\|\|\|\|\|\|\|\|\|\|\|\|\|\|\|\|\|\|\|\|\|\|\|\|\|\|\|\|\|\|\|\|\|\|\|\|\|\|\|\|\|\|\|\|\|\|\|\|\|\|\|\|\|\|\|\|\|\|\|\|\|\|\|\| \|\|\|\|\|\|\|\|\|\|\|\|\|\|\|\|\|\|\|\|\|\|\|\|\|\|\|\|\|\|\|\|\|\|\|\|\|\|\|\|\|\|\|\|\|\|\|\|\|\|\|\|\|\|\|\|\|\|\|\|\|\|\|\|\|\|\|\|\|\|\|\|\|\|\|\|\|\|\|\|\|\|\|\|\|\|\|\|\|\|\|\|\|\|\|\| \|\|\|\|\|\|\|\|\|\|\|\|\|\|\|\|\|\|\|\|\|\|\|\|\|\|\|\|\|\|\|\|\|\|\|\|\|\|\|\|\|\|\|\|\|\|\|\|\| 	CGCGGCGGAGAACGTCAACATTCCAGTGGAGAAGAGAGGCGAGTGGGACTACGAGCCCACCCCTCACTACTACCCACCCGCCGCGGACAGCGCCAACATTCCAGTGGAGAAGAGAGGCGAATGGGACTACGAGCCCACCCCTCACTACTACCCACCCGCCGCGGACAGCGCCAACATTCCAGTGGAGAAGAGAGGCGAGTGGGACTACGAGCCCACCCCTCACTACTACCCACCCGCCGCGGACAGCGCCAACATTCCAGTGGAGAA ```      ``` 255	CGCCGCGGACAGCGCCAACATTCCAGTGGAGAAGAGAGGCGAGTGGGACTACGAGCCCACCCCTCACTACTACCCACCCGCCGCGGACAGCGCCAACATTCCAGTGGAGAAGAGAGGCGAGTGGGACTACGAGCCCACCTCTCACTACTACCCACCCGCCGCGGACAGCGCCAACATTCCAGTGGAGAA	443 	\|\|\| \|\|\|\|\| \| \|\| \|\|\|\|\|\|\|\|\|\|\|\|\|\|\|\|\|\|\|\|\|\|\|\|\|\|\|\|\|\|\|\|\|\|\|\|\|\|\|\|\|\|\|\|\|\|\|\|\|\|\|\|\|\|\|\|\|\|\|\|\|\|\|\|\|\|\|\|\|\|\|\|\|\|\|\|\|\|\|\|\|\|\|\|\|\|\|\|\|\|\|\|\|\|\|\|\|\|\|\|\|\|\|\|\| \|\|\|\|\|\|\|\|\|\|\|\|\|\|\|\|\|\| \|\|\|\|\|\|\|\|\|\|\|\|\|\|\|\|\|\|\|\|\|\|\|\|\|\|\|\|\|\|\|\|\|\|\|\|\|\|\|\|\|\|\|\|\|\|\|\|\| 	CGCGGCGGAGAACGTCAACATTCCAGTGGAGAAGAGAGGCGAGTGGGACTACGAGCCCACCCCTCACTACTACCCACCCGCCGCGGACAGCGCCAACATTCCAGTGGAGAAGAGAGGCGAATGGGACTACGAGCCCACCCCTCACTACTACCCACCCGCCGCGGACAGCGCCAACATTCCAGTGGAGAA ```      ``` 333	CGCCGCGGACAGCGCCAACATTCCAGTGGAGAAGAGAGGCGAGTGGGACTACGAGCCCACCTCTCACTACTACCCACCCGCCGCGGACAGCGCCAACATTCCAGTGGAGAA	443 	\|\|\| \|\|\|\|\| \| \|\| \|\|\|\|\|\|\|\|\|\|\|\|\|\|\|\|\|\|\|\|\|\|\|\|\|\|\|\|\|\|\|\|\|\|\|\|\|\|\|\|\|\|\|\|\|\| \|\|\|\|\|\|\|\|\|\|\|\|\|\|\|\|\|\|\|\|\|\|\|\|\|\|\|\|\|\|\|\|\|\|\|\|\|\|\|\|\|\|\|\|\|\|\|\|\| 	CGCGGCGGAGAACGTCAACATTCCAGTGGAGAAGAGAGGCGAGTGGGACTACGAGCCCACCCCTCACTACTACCCACCCGCCGCGGACAGCGCCAACATTCCAGTGGAGAA ```      ``` 99	CGCGGCGGAGAACGTCAACATTCCAGTGGAGAAGAGAGGCGAGTGGGACTACGAGCCCACCCCTCACTACTACCCACCCGCCGCGGACAGCGCCAACATTCCAGTGGAGAAGAGAGGCGAATGGGACTACGAGCCCACCCCTCACTACTACCCACCCGCCGCGGACAGCGCCAACATTCCAGTGGAGAA	287 	\|\|\| \|\|\|\|\| \| \|\| \|\|\|\|\|\|\|\|\|\|\|\|\|\|\|\|\|\|\|\|\|\|\|\|\|\|\|\|\|\|\|\|\|\|\|\|\|\|\|\|\|\|\|\|\|\|\|\|\|\|\|\|\|\|\|\|\|\|\|\|\|\|\|\|\|\|\|\|\|\|\|\|\|\|\|\|\|\|\|\|\|\|\|\|\|\|\|\|\|\|\|\|\|\|\|\|\|\|\|\|\|\|\|\|\| \|\|\|\|\|\|\|\|\|\|\|\|\|\|\|\|\|\| \|\|\|\|\|\|\|\|\|\|\|\|\|\|\|\|\|\|\|\|\|\|\|\|\|\|\|\|\|\|\|\|\|\|\|\|\|\|\|\|\|\|\|\|\|\|\|\|\| 	CGCCGCGGACAGCGCCAACATTCCAGTGGAGAAGAGAGGCGAGTGGGACTACGAGCCCACCCCTCACTACTACCCACCCGCCGCGGACAGCGCCAACATTCCAGTGGAGAAGAGAGGCGAGTGGGACTACGAGCCCACCTCTCACTACTACCCACCCGCCGCGGACAGCGCCAACATTCCAGTGGAGAA ```      ``` 99	CGCGGCGGAGAACGTCAACATTCCAGTGGAGAAGAGAGGCGAGTGGGACTACGAGCCCACCCCTCACTACTACCCACCCGCCGCGGACAGCGCCAACATTCCAGTGGAGAAGAGAGGCGAATGGGACTACGAGCCCACCCCTCACTACTACCCACCCGCCGCGGACAGCGCCAACATTCCAGTGGAGAAGAGAGGCGAGTGGGACTACGAGCCCACCCCTCACTACTACCCACCCGCCGCGGACAGCGCCAACATTCCAGTGGAGAA	365 	\|\|\| \|\|\|\|\| \| \|\| \|\|\|\|\|\|\|\|\|\|\|\|\|\|\|\|\|\|\|\|\|\|\|\|\|\|\| \|\|\|\|\|\|\|\|\|\|\|\|\|\|\|\|\|\|\|\|\|\|\|\|\|\|\|\|\|\|\|\|\|\|\|\|\|\|\|\|\|\|\|\|\|\|\|\|\|\|\|\|\|\|\|\|\|\|\|\|\|\|\|\|\|\|\|\|\|\|\|\|\|\|\|\|\| \|\|\|\|\|\|\|\|\|\|\|\|\|\|\|\|\|\|\|\|\|\|\|\|\|\|\|\|\|\|\|\|\|\|\|\|\|\|\|\|\|\|\|\|\|\|\|\|\|\|\|\|\|\|\|\|\|\|\|\|\|\|\|\|\|\|\|\|\|\|\|\|\|\|\|\|\|\|\|\|\|\|\|\|\|\|\|\|\|\|\|\|\|\|\|\| \|\|\|\|\|\|\|\|\|\|\|\|\|\|\|\|\|\|\|\|\|\|\|\|\|\|\|\|\|\|\|\|\|\|\|\|\|\|\|\|\|\|\|\|\|\|\|\|\| 	CGCCGCGGACAGCGCCAACATTCCAGTGGAGAAGAGAGGCGAATGGGACTACGAGCCCACCCCTCACTACTACCCACCCGCCGCGGACAGCGCCAACATTCCAGTGGAGAAGAGAGGCGAGTGGGACTACGAGCCCACCCCTCACTACTACCCACCCGCCGCGGACAGCGCCAACATTCCAGTGGAGAAGAGAGGCGAGTGGGACTACGAGCCCACCTCTCACTACTACCCACCCGCCGCGGACAGCGCCAACATTCCAGTGGAGAA ```      ``` 99	CGCGGCGGAGAACGTCAACATTCCAGTGGAGAAGAGAGGCGAGTGGGACTACGAGCCCACCCCTCACTACTACCCACCCGCCGCGGACAGCGCCAACATTCCAGTGGAGAA	209 	\|\|\| \|\|\|\|\| \| \|\| \|\|\|\|\|\|\|\|\|\|\|\|\|\|\|\|\|\|\|\|\|\|\|\|\|\|\|\|\|\|\|\|\|\|\|\|\|\|\|\|\|\|\|\|\|\| \|\|\|\|\|\|\|\|\|\|\|\|\|\|\|\|\|\|\|\|\|\|\|\|\|\|\|\|\|\|\|\|\|\|\|\|\|\|\|\|\|\|\|\|\|\|\|\|\| 	CGCCGCGGACAGCGCCAACATTCCAGTGGAGAAGAGAGGCGAGTGGGACTACGAGCCCACCTCTCACTACTACCCACCCGCCGCGGACAGCGCCAACATTCCAGTGGAGAA ```      ``` 1	ATGCACTCATCCACCATCATTGCCTTGGCGTCTGTCCTTGCTACTGGCATCGTCCACGCCCTTCCCCAGGCTGGAGTGCGCGGTGACCGGTATGCGCGCGCGGCGGAGAACGTCAACATTCCAGTGGAGAAGAGAGGCGAGTGGGACTACGAGCCCACCCCTCACTACTACCCACCCGCCGCGGACAGCGCCAACATTCCAGTGGAGAAGAGAGGCGAATGGGACTACGAGCCCACCCCTCACTACTACCCACCCGCCGCGGACAGCGCCAACATTCCAGTGGAGAAGAGAGGCGAGTGGGACTACGAGCCCACCCCTCACTACTACCCACCCGCCGCGGACAGCGCCAACATTCCAGTGGAGAAGAGAGGCGAGTGGGACTACGAGCCCACCTCTCACTACTACCCACCCGCCGCGGACAGCGCCAACATTCCAGTGGAGAAAATAAAGAAC	453 	\|\|\|\|\|\|\|\|\|\|\|\|\|\|\|\|\|\|\|\|\|\|\|\|\|\|\|\|\|\|\|\|\|\|\|\|\|\|\|\|\|\|\|\|\|\|\|\|\|\|\|\|\|\|\|\|\|\|\|\|\|\|\|\|\|\|\|\|\|\|\|\|\|\|\|\|\|\|\|\|\|\|\|\|\|\|\|\|\|\|\|\|\|\|\|\|\|\|\|\|\|\|\|\|\|\|\|\|\|\|\|\|\|\|\|\|\|\|\|\|\|\|\|\|\|\|\|\|\|\|\|\|\|\|\|\|\|\|\|\|\|\|\|\|\|\|\|\|\|\|\|\|\|\|\|\|\|\|\|\|\|\|\|\|\|\|\|\|\|\|\|\|\|\|\|\|\|\|\|\|\|\|\|\|\|\|\|\|\|\|\|\|\|\|\|\|\|\|\|\|\|\|\|\|\|\|\|\|\|\|\|\|\|\|\|\|\|\|\|\|\|\|\|\|\|\|\|\|\|\|\|\|\|\|\|\|\|\|\|\|\|\|\|\|\|\|\|\|\|\|\|\|\|\|\|\|\|\|\|\|\|\|\|\|\|\|\|\|\|\|\|\|\|\|\|\|\|\|\|\|\|\|\|\|\|\|\|\|\|\|\|\|\|\|\|\|\|\|\|\|\|\|\|\|\|\|\|\|\|\|\|\|\|\|\|\|\|\|\|\|\|\|\|\|\|\|\|\|\|\|\|\|\|\|\|\|\|\|\|\|\|\|\|\|\|\|\|\|\|\|\|\|\|\|\|\|\|\|\|\|\|\|\|\|\|\|\|\|\|\|\|\|\|\|\|\|\|\|\|\|\|\|\|\|\|\|\|\|\|\|\|\|\|\|\|\|\|\|\|\|\|\|\|\|\|\|\|\|\|\|\|\|\|\|\|\|\|\|\|\|\|\|\|\|\|\|\|\|\|\|\|\|\|\|\|\|\|\|\|\|\|\|\|\|\|\|\|\|\|\|\|\|\| 	ATGCACTCATCCACCATCATTGCCTTGGCGTCTGTCCTTGCTACTGGCATCGTCCACGCCCTTCCCCAGGCTGGAGTGCGCGGTGACCGGTATGCGCGCGCGGCGGAGAACGTCAACATTCCAGTGGAGAAGAGAGGCGAGTGGGACTACGAGCCCACCCCTCACTACTACCCACCCGCCGCGGACAGCGCCAACATTCCAGTGGAGAAGAGAGGCGAATGGGACTACGAGCCCACCCCTCACTACTACCCACCCGCCGCGGACAGCGCCAACATTCCAGTGGAGAAGAGAGGCGAGTGGGACTACGAGCCCACCCCTCACTACTACCCACCCGCCGCGGACAGCGCCAACATTCCAGTGGAGAAGAGAGGCGAGTGGGACTACGAGCCCACCTCTCACTACTACCCACCCGCCGCGGACAGCGCCAACATTCCAGTGGAGAAAATAAAGAAC ``` |
| 41 | ggggacaagtttgtacaaaaaagcaggcttgatgAATCGCTTGACATTCG, ggggaccactttgtacaagaaagctgggtcACACCGCGGACTCGGGTACT | Mycgr3\|91995  JGI  Ensembl |  |  | 100 % |  | ``` 1	ATGAATCGCTTGACATTCGCCACGATAGTGGCATCCCTCTTCGCTACCACCACTGCCCAAGCTCCACCCTTCCACATCGAGAACTTCGCCGCTCAAGACATTCGCATTCCAAGCGAGTCCCACAACGTCTCCTTCACGGTCAAACACCTCGGAGCAGCTCCTCAACAGGGAGGAGACGCGCCATTCGACCTCTCCATCGCCTGGGAAG	208 	\|\|\|\|\|\|\|\|\|\|\|\|\|\|\|\|\|\|\|\|\|\|\|\|\|\|\|\|\|\|\|\|\|\|\|\|\|\|\|\|\|\|\|\|\|\|\|\|\|\|\|\|\|\|\|\|\|\|\|\|\|\|\|\|\|\|\|\|\|\|\|\|\|\|\|\|\|\|\|\|\|\|\|\|\|\|\|\|\|\|\|\|\|\|\|\|\|\|\|\|\|\|\|\|\|\|\|\|\|\|\|\|\|\|\|\|\|\|\|\|\|\|\|\|\|\|\|\|\|\|\|\|\|\|\|\|\|\|\|\|\|\|\|\|\|\|\|\|\|\|\|\|\|\|\|\|\|\|\|\|\|\|\|\|\|\|\|\|\|\|\|\|\|\|\|\|\|\|\|\|\|\|\|\|\|\|\|\|\|\|\|\|\|\|\|\|\|\|\|\|\|\|\|\| \|\|\| 	ATGAATCGCTTGACATTCGCCACGATAGTGGCATCCCTCTTCGCTACCACCACTGCCCAAGCTCCACCCTTCCACATCGAGAACTTCGCCGCTCAAGACATTCGCATTCCAAGCGAGTCCCACAACGTCTCCTTCACGGTCAAACACCTCGGAGCAGCTCCTCAACAGGGAGGAGACGCGCCATTCGACCTCTCCATCGCCTGGTAAG ```      ``` 194	CCATCGC--CTGGGAAGGCTGCGACCCACCCCCCTGCTACATCAATAGCGAAGGGACAGGATGGTACACTCGCATCGTTCCAGGCTCCTACAAAGGCGCCACCGACTTCACGCTCGAGATCGTCAAGCATAATATTTACAAGGCTGCTACGACGTGGACGGCCAGGATCCCCTACGTGGAGGGAAGCGCAGGGTATGACTGCTCGCGAGGTGGACAGAGTACGCTTTGTGGGTTCCAGGACGAGAGCGATTCGACATATGAGACTGTGTATGTCGGTGGTTTGGAAGCGGAGTACCCGAGTCCGCGGTGT	501 	\|\|\|\|\|\|\|  \| \|\|\|\|\|\|\|\|\|\|\|\|\|\|\|\|\|\|\|\|\|\|\|\|\|\|\|\|\|\|\|\|\|\|\|\|\|\|\|\|\|\|\|\|\|\|\|\|\|\|\|\|\|\|\|\|\|\|\|\|\|\|\|\|\|\|\|\|\|\|\|\|\|\|\|\|\|\|\|\|\|\|\|\|\|\|\|\|\|\|\|\|\|\|\|\|\|\|\|\|\|\|\|\|\|\|\|\|\|\|\|\|\|\|\|\|\|\|\|\|\|\|\|\|\|\|\|\|\|\|\|\|\|\|\|\|\|\|\|\|\|\|\|\|\|\|\|\|\|\|\|\|\|\|\|\|\|\|\|\|\|\|\|\|\|\|\|\|\|\|\|\|\|\|\|\|\|\|\|\|\|\|\|\|\|\|\|\|\|\|\|\|\|\|\|\|\|\|\|\|\|\|\|\|\|\|\|\|\|\|\|\|\|\|\|\|\|\|\|\|\|\|\|\|\|\|\|\|\|\|\|\|\|\|\|\|\|\|\|\|\|\|\|\|\|\|\|\|\|\|\|\|\|\|\|\|\|\|\|\|\|\|\|\|\|\|\|\|\|\|\|\|\|\|\|\|\|\|\|\|\|\|\|\|\|\|\|\|\|\|\|\|\|\|\|\|\|\|\| 	CCATCGCAACAGGGAAGGCTGCGACCCACCCCCCTGCTACATCAATAGCGAAGGGACAGGATGGTACACTCGCATCGTTCCAGGCTCCTACAAAGGCGCCACCGACTTCACGCTCGAGATCGTCAAGCATAATATTTACAAGGCTGCTACGACGTGGACGGCCAGGATCCCCTACGTGGAGGGAAGCGCAGGGTATGACTGCTCGCGAGGTGGACAGAGTACGCTTTGTGGGTTCCAGGACGAGAGCGATTCGACATATGAGACTGTGTATGTCGGTGGTTTGGAAGCGGAGTACCCGAGTCCGCGGTGT ``` |
| 41 | ggggacaagtttgtacaaaaaagcaggcttgatgCAGATCTTCAAGCTCT, ggggaccactttgtacaagaaagctgggtcTCGTCCAGTCATGAGCATCG | Mycgr3\|92640  JGI  Ensembl |  |  | 100 % |  | ``` 1	ATGCAGATCTTCAAGCTCTCCATCATTCTCTTCATGACCTCGGCGGCCATGGCGGAATTT	60 	\|\|\|\|\|\|\|\|\|\|\|\|\|\|\|\|\|\|\|\|\|\|\|\|\|\|\|\|\|\|\|\|\|\|\|\|\|\|\|\|\|\|\|\|\|\|\|\|\|\|\|\|\|\|\|\|\|\|\|\| 	ATGCAGATCTTCAAGCTCTCCATCATTCTCTTCATGACCTCGGCGGCCATGGCGGAATTT ```      ``` 145	ACCCAAACACCAGGCTGGGCAAGTACGGAGCAGACTGCAGTTAAAGGCGAACGTGCGCGGAAGAGCCGCAGGTCTCCGAGCTTTGGATGCCCCATCTCGGGAGACAGCGCGATCTCGAATGACGGGATGAAGGTGGAGGACAGCGATGGAAGACAATCGACGGGTACCGTCCGGGAAAGGTTGGACGCTGACAATCCGATGCTCATGACTGGACGA	360 	\|\|\|\|\|\|\|\|\|\|\|\|\|\|\|\|\|\|\|\|\|\|\|\|\|\|\|\|\|\|\|\|\|\|\|\|\|\|\|\|\|\|\|\|\|\|\|\|\|\|\|\|\|\|\|\|\|\|\|\|\|\|\|\|\|\|\|\|\|\|\|\|\|\|\|\|\|\|\|\|\|\|\|\|\|\|\|\|\|\|\|\|\|\|\|\|\|\|\|\|\|\|\|\|\|\|\|\|\|\|\|\|\|\|\|\|\|\|\|\|\|\|\|\|\|\|\|\|\|\|\|\|\|\|\|\|\|\|\|\|\|\|\|\|\|\|\|\|\|\|\|\|\|\|\|\|\|\|\|\|\|\|\|\|\|\|\|\|\|\|\|\|\|\|\|\|\|\|\|\|\|\|\| \|\|\|\|\|\|\|\|\|\|\|\|\|\|\|\|\|\|\|\|\|\|\|\|\|\|\|\|\|\|\|\| 	ACCCAAACACCAGGCTGGGCAAGTACGGAGCAGACTGCAGTTAAAGGCGAACGTGCGCGGAAGAGCCGCAGGTCTCCGAGCTTTGGATGCCCCATCTCGGGAGACAGCGCGATCTCGAATGACGGGATGAAGGTGGAGGACAGCGATGGAAGACAATCGACGGGTACCGTCCGGGAAAGGTTGAACGCTGACAATCCGATGCTCATGACTGGACGA ```      ``` 61	ATCTGCGAAGTCATGTCGGAAGAACACCCGGAGGGTCGTTGCATCGGCAGAAAAGGAGACAGGGGCATCTACGATTGCGGCAGC	144 	\|\|\|\|\|\|\|\|\|\|\|\|\|\|\|\|\|\|\|\|\|\|\|\|\|\|\|\|\|\|\|\|\|\|\|\|\|\|\|\|\|\|\|\|\|\|\|\|\|\|\|\|\|\|\|\|\|\|\|\|\|\|\|\|\|\|\|\|\|\|\|\|\|\|\|\|\|\|\|\|\|\|\|\| 	ATCTGCGAAGTCATGTCGGAAGAACACCCGGAGGGTCGTTGCATCGGCAGAAAAGGAGACAGGGGCATCTACGATTGCGGCAGC ``` |
| 41 | ggggacaagtttgtacaaaaaagcaggcttgatgAAGTCCAGCCTCACCG, ggggaccactttgtacaagaaagctgggtcCGACTTGACCACTTGTTGAG | Mycgr3\|92826  JGI  Ensembl |  |  | 100 % |  | ``` 274	GATGCTGGCGGCAGGTTATTGTTGATGCGCATCGGGACTCAACAAGTGGTCAAGTCG	330 	\|\|\|\|\|\|\|\|\|\|\|\|\|\|\|\|\|\|\|\|\|\|\|\|\|\|\|\|\|\|\|\|\|\|\|\|\|\|\|\|\|\|\|\|\|\|\|\|\|\|\|\|\|\|\|\|\| 	GATGCTGGCGGCAGGTTATTGTTGATGCGCATCGGGACTCAACAAGTGGTCAAGTCG ```      ``` 1	ATGAAGTCCAGCCTCACCGTCCTTACTCTCGCGGCAGCAGGTTTTCTCTGCGCAGCAGCCTCCCCAACGCTCAAGGTTCGCCAGAACGATGTTCCCAAATGTGCCGACGGCCGCGGAGATGGGCTGCGCAGCTACGATAGCCG	143 	\|\|\|\|\|\|\|\|\|\|\|\|\|\|\|\|\|\|\|\|\|\|\|\|\|\|\|\|\|\|\|\|\|\|\|\|\|\|\|\|\|\|\|\|\|\|\|\|\|\|\|\|\|\|\|\|\|\|\|\|\|\|\|\|\|\|\|\|\|\|\|\|\|\|\|\|\|\|\|\|\|\|\|\|\|\|\|\|\|\|\|\|\|\|\|\|\|\|\|\|\|\|\|\|\|\|\|\|\|\|\|\|\|\|\|\|\|\|\|\|\|\|\|\|\|\|\|\|\|\|\|\|\|\|\|\|\|\|\|\|\|\|\| 	ATGAAGTCCAGCCTCACCGTCCTTACTCTCGCGGCAGCAGGTTTTCTCTGCGCAGCAGCCTCCCCAACGCTCAAGGTTCGCCAGAACGATGTTCCCAAATGTGCCGACGGCCGCGGAGATGGGCTGCGCAGCTACGATAGCCG ```      ``` 143	GAAGCAAGTGCGAAGACGGCTGCAAGTCCTTGAAAGAGGAAGCCGAAGGCGACAACAAAGGTGGGATCTGCAGTGGATTCTGTACCACCTACCCCGGACCCCAGAGTTTTGGAATCATCATCCGCTGCAATG	274 	\|\|\|\|\|\|\|\|\|\|\|\|\|\|\|\|\|\|\|\|\|\|\|\|\|\|\|\|\|\|\|\|\|\|\|\|\|\|\|\|\|\|\|\|\|\|\|\|\|\|\|\|\|\|\|\|\|\|\|\|\|\|\|\|\|\|\|\|\|\|\|\|\|\|\|\|\|\|\|\|\|\|\|\|\|\|\|\|\|\|\|\|\|\|\|\|\|\|\|\|\|\|\|\|\|\|\|\|\|\|\|\|\|\|\|\|\|\|\|\|\|\|\|\|\|\|\|\|\|\|\|\| 	GAAGCAAGTGCGAAGACGGCTGCAAGTCCTTGAAAGAGGAAGCCGAAGGCGACAACAAAGGTGGGATCTGCAGTGGATTCTGTACCACCTACCCCGGACCCCAGAGTTTTGGAATCATCATCCGCTGCAATG ``` |
| 41 | ggggacaagtttgtacaaaaaagcaggcttgatgCCTTCGACTACCACCC, ggggaccactttgtacaagaaagctgggtcGCGGCCAGTGAAGCCTTTAG | Mycgr3\|93501  JGI  Ensembl |  |  | 100 % |  |  |
